# Predicting Hosts Based on Early SARS-CoV-2 Samples and Analyzing Later World-wide Pandemic in 2020

**DOI:** 10.1101/2021.03.21.436312

**Authors:** Qian Guo, Mo Li, Chunhui Wang, Jinyuan Guo, Xiaoqing Jiang, Jie Tan, Shufang Wu, Peihong Wang, Tingting Xiao, Man Zhou, Zhencheng Fang, Yonghong Xiao, Huaiqiu Zhu

## Abstract

The SARS-CoV-2 pandemic has raised the concern for identifying hosts of the virus since the early-stage outbreak. To address this problem, we proposed a deep learning method, DeepHoF, based on extracting the viral genomic features automatically, to predict host likelihood scores on five host types, including plant, germ, invertebrate, non-human vertebrate and human, for novel viruses. DeepHoF made up for the lack of an accurate tool applicable to any novel virus and overcame the limitation of the sequence similarity-based methods, reaching a satisfactory AUC of 0.987 on the five-classification. Additionally, to fill the gap in the efficient inference of host species for SARS-CoV-2 using existed tools, we conducted a deep analysis on the host likelihood profile calculated by DeepHoF. Using the isolates sequenced in the earliest stage of COVID-19, we inferred minks, bats, dogs and cats were potential hosts of SARS-CoV-2, while minks might be one of the most noteworthy hosts. Several genes of SARS-CoV-2 demonstrated their significance in determining the host range. Furthermore, the large-scale genome analysis, based on DeepHoF’s computation for the later world-wide pandemic in 2020, disclosed the uniformity of host range among SARS-CoV-2 samples and the strong association of SARS-CoV-2 between humans and minks.

## Introduction

The global COVID-19 pandemic caused by severe acute respiratory syndrome coronavirus 2 (SARS-CoV-2) has raised the long-lasting quest for hosts of the virus since the pandemic outbreak, meanwhile the majority view is that the virus probably originated from bats [1]. So far there have been many discussions for the potential hosts despite an initial pointer to *Manis javanica* (pangolins) [2, 3], most of the suppositions were based on the increasing cases of animal infection, such as dogs, cats, tigers, lions, and minks [4, 5], *etc*. Several studies performed experiments to investigate the susceptibility of a limited number of model animals [6-8]. At the same time, some studies attempted to reveal the range of hosts based on analysis of molecular sequence or structural information [9, 10]. For instance, Damas *et al*., [10] conducted a computational analysis based on host receptor similarity using the angiotensin-converting enzyme 2 (ACE2) protein and evaluated the infection risks for a broad range of animals. As the pandemic spreads, minks, which were even not referred to as high infection animal in above peer-review articles, have been frequently reported massively infected with COVID-19 over the world [5], and were the only known animal reported to transmit SARS-CoV-2 to humans [11, 12]. It is worth mentioning that, in January, 2020, we have reported in the form of a preprint archive with predicting minks as a potential host based on the six earliest sequenced SARS-CoV-2 isolates [13]. However, the later complication of pandemic prompts peoples again to have a full review of the issue of host determination for SARS-CoV-2. This raises a new challenge, which is how to implement and improve the capability of computational methods to predict the hosts of a novel virus like SARS-CoV-2, especially when we have relatively small amounts of samples of sequencing viral data at the early stage of the pandemic outbreak. It is certainly constructive for similar pandemic caused by novel viruses in the future.

Generally, the host range of viruses is dependent on molecular interactions between viruses and host cells including receptor recognition, adaptions to the host cellular machinery and evading innate immune recognition [14]. Of these, receptor recognition that facilitates the attachment of viruses to the host cells is the most primary step. Thus, the glycoproteins that viruses use to recognize the host receptor as well as the whole genome sequences are widely used in identifying the potential hosts of viruses [1]. To detect the potential host and pathogenicity of novel viruses, the conventional computational methods are almost based on similarity of either virus genome composition or host receptor. Limitations of the both strategies lie in that they assume phylogeny may reflect host association. However, this assumption is untenable from the perspective of epidemiology and evolution. On the one hand, viruses occasionally shift between distantly related host species. On the other hand, owing to the long-term adaptation to the hosts, the viral genomic characteristics acquired from hosts can be quite incompatible with the virus phylogenetic groups [15]. The specificity of recognition between viruses and host species also involves structural information in some key domains of both viral proteins and host receptor proteins, such as the receptor-binding domain, that sequence similarity is insufficient to explain. For example, the civet-specific K479 and S487 residues of SARS-CoV spike glycoprotein can efficiently bind to civet ACE2 but have much less affinity to human ACE2 [16, 17]. This is also the reason that the similarity-based method of host ACE2 proteins sequences fails to predict minks as host of high and very high risk for SARS-CoV-2 infection [10].

Until now, several published tools aimed to identify the hosts of viruses exceeded the limitation of sequence-similarity-based strategies by machine learning methods with viral sequences or their genomic traits related to virus-host interactions, such as ViralHostPredictor [15], HostPhinder [18], WIsH [19], Host Taxon Predictor [20], and VIDHOP [21]. While these tools performed well under some conditions, they are actually not considered feasible to be applied to a novel virus without the knowledge of host range, like SARS-CoV-2. HostPhinder and WIsH predict hosts for only bacteriophages and they are inappropriate for non-phage viruses. Host Taxon Predictor focuses on distinguish bacteriophages and eukaryotic viruses. ViralHostPredictor predicts hosts and the existence and identity of arthropod vectors for human-infecting RNA viruses by Gradient boosting machines with the features of selected evolutionary genomic traits and phylogenetic information. It also illustrated the better ability of machine learning methods to predict virus hosts compared to the way of sequence similarity comparison. However, ViralHostPredictor cannot determine whether human is the host of a novel virus. With the utilization of evolutionary signatures, ViralHostPredictor lacks power to predict incidental hosts which do not maintain long-term circulation of new viruses. Moreover, the predictive abilities of the methods above rely on the handcrafted features like codon pair scores, *k*-mer frequencies and amino acid biases, which might neglect other important information encoded in the virus genomes. VIDHOP, a deep-learning-based tool, is designed to predict potential hosts of viruses, but its application was limited into three viral species: influenza A, rabies lyssavirus and rotavirus A.

To address the challenge of predicting probable hosts of a novel virus like SARS-CoV-2, we proposed the host prediction algorithm DeepHoF (**Deep** learning-based **Ho**st **F**inder) in the current study. Developed based on BiPath Convolutional Neural Network (BiPathCNN), DeepHoF automatically extracts the genomic features from the input viral sequences. The model finally outputs five host likelihood scores and their *p*-values on five host types, including plant, germ, invertebrate, non-human vertebrate (refers to other vertebrates except humans) and human, where all the living organism hosts are covered. DeepHoF was designed as a five-class classifier containing five independent nodes in the output layer with sigmoid activation and binary cross-entropy loss function for each node, corresponding to five independent binary classifications on the five host types individually. DeepHoF made up for the lack of efficient method applicable for any novel virus and significantly outperformed the Basic Local Alignment Search Tool (BLAST)-based strategy with the evidently high AUC of 0.987 on the classification of five host types. In January 2020, we have reported the host prediction for six earliest sequenced SARS-CoV-2 isolates employing our algorithm [13]. In this study, we furthered the work using the 17 earliest sampled SARS-CoV-2 isolates, which provides essential information in the early epidemic of the virus. DeepHoF evaluated the host likelihood scores on humans and non-human vertebrates for the earliest samples and characterized the isolates with their host likelihood score profiles. As there existed a blank in the inference of host species for SARS-CoV-2 using the tools which were state of the art, we conducted a deep analysis on the host likelihood score profile predicted by DeepHoF to find the detailed hosts, including both reservoirs and susceptible hosts which are not discriminated in this study. We inferred minks, bats, dogs and cats were the probable hosts, while minks maybe one of the most noteworthy hosts. The inference was supported by the infection facts or animal experiments in the later pandemic. Based on our model, several genes of SARS-CoV-2 were further investigated and demonstrated their significance in determining the host likelihood scores on human or the host range for SARS-CoV-2, respectively. With a large-scale genome analysis based on DeepHoF’s computation for the later world-wide pandemic, the uniformity of host inference among a large number of SARS-CoV-2 samples was verified, and the association of SARS-CoV-2 between humans and minks was disclosed. Supported by the satisfactory performance on five host type classification and the successful application in SARS-CoV-2, DeepHoF has the capability to provide reliable host information of novel virus, and is expected to narrow the time lag between novel virus discovery and prevention at the early-stage of epidemic prevention.

## Results

### Performance of the DeepHoF algorithm

The DeepHoF algorithm is designed as a five-class classifier using the deep learning method of BiPathCNN (see Methods). Herein five likelihood scores on five host types, including plants, germs, invertebrates, non-human vertebrates, and humans, are calculated by DeepHoF. The host likelihood score profile consisting of five predicted scores, is then analysed in depth to find the specific hosts of a novel virus such as SARS-CoV-2 in this study. As mentioned above, the existed bioinformatics tools [15, 18-21] were not designed to perform the prediction of the host likelihood scores on the five host types for any given virus, and thus cannot be compared with DeepHoF directly. And therefore, we compared the performance of DeepHoF model with BLAST (details of finding host using BLAST are described in Supplementary Methods), adopting six classification metrics: true-positive rate (TPR), false-positive rate (FPR), area under the curve (AUC), precision, accuracy and F1-score. To assess the performance of predicting novel viruses, we used training and test datasets divided in chronological order [22] (Methods). There is no overlap of virus species in training and test sets. With an evident higher AUC of 0.987, DeepHoF can significantly outperform BLAST (with the average AUC of 0.833) as shown in **Figure 1A** and **Table 1** (a detailed comparison on each host type is illustrated in Supplemental Figure S1 and Table S1).

**Figure 1.**
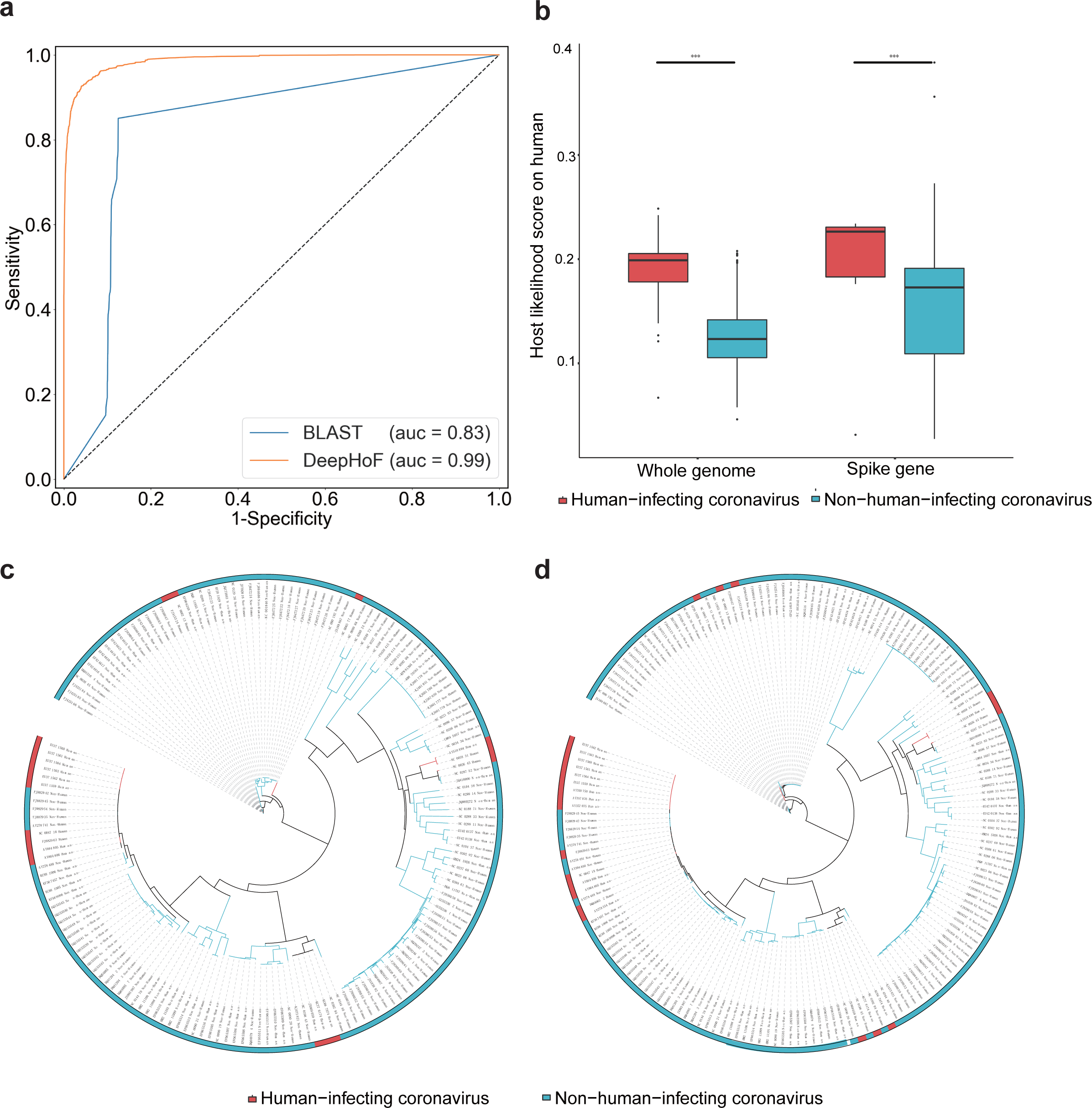
DeepHoF outperforms BLAST and well learns the information of virus hosts. **A**. Average ROC curves and AUC values of DeepHoF and BLAST. DeepHoF performs better than BLAST on average AUC of five host types. **B**. Comparison of host likelihood scores predicted by DeepHoF between human-infecting and non-human-infecting coronaviruses on human. The former performed higher probabilities than the latter (two-sided unpaired Welch Two Sample *t*-test, *t*_(43.843)_ = 8.265 and *t*_(38.016)_ = 4.674, *p*-values = 1.732×10^−10^ and 3.657×10^−5^. *** *p*-value < 0.0001, *t*-values and degrees of freedom were presented as *t*_(df)_). **C**. Phylogenetic analyses of whole genomes of coronaviruses. **D**. Phylogenetic analyses of S genes of coronaviruses. Maximum-likelihood phylogenic trees were built by RAxML [41] with 1,000 bootstrap replicates and visualized with iTOL [43]. The whole genomes and the S genes of the human-infecting coronaviruses could not be distinguished from the non-human-infecting ones. (Red: human-infecting coronaviruses; Blue: non-human-infecting coronaviruses).

**Table 1.** Performance metrics of DeepHoF and BLAST.

| Methods | Precision | Accuracy | TPR | FPR | AUC | F1-score |
| --- | --- | --- | --- | --- | --- | --- |
| BLAST | 0.699 | 0.892 | 0.888 | 0.107 | 0.833 | 0.896 |
| DeepHoF | 0.968 | 0.964 | 0.865 | 0.008 | 0.987 | 0.963 |
| TPR: true-positive rate; FPR: false-positive rate; AUC: area under the curve |  |  |  |  |  |  |

In addition, we compared the utility of DeepHoF and a phylogenetic tree to discriminate the human-infecting and non-human-infecting coronaviruses using their whole genome sequences. As shown in Figure 1B (the left), DeepHoF could identify evidently higher probabilities of human-infecting coronaviruses to infect humans (two-sided unpaired Welch Two Sample *t*-test, *p*-value = 1.732×10^−10^). However, the phylogenetic analysis result was not satisfactory owing to the weak homology among the human-infecting coronaviruses, which were scattered around the phylogenetic tree of coronaviruses (Figure 1C). The comparison was similar for the inferences using their spike glycoprotein coding genes (S genes) as shown in Figure 1B (the right), and D (two-sided unpaired Welch Two Sample *t*-test, *p*-value=3.657×10^−5^). This result is nontrivial because S genes are essential in coronavirus-host interaction [23]. Clearly, DeepHoF can overcome the limitation of sequence similarity-based method and shows superior predictive ability especially for novel viruses.

### Host prediction of SARS-CoV-2

The accurate prediction of hosts of earliest detected isolates can undoubtedly assist the public health system to take more appropriate preventive measures at the early stage of the pandemic outbreak. In view of this, we focused on the prediction with SARS-CoV-2 isolates sequenced in the earliest stage of COVID-19 detection, which is closer to the most recent common ancestor of SARS-CoV-2. Previous to this paper, we have reported the prediction for the six earliest sequenced SARS-CoV-2 isolates using our algorithm on 21 January, 2020 [13]. In this study, we further strengthened the prediction of hosts of SARS-CoV-2 with all 17 earliest detected isolates (including the six earliest ones) sequenced in December, 2019. Herein we take NC_045512 (complete genome of SARS-CoV-2 isolate, Wuhan-Hu-1, collected on 31 December 2019 in Wuhan, China, and used as the representative genome of SARS-CoV-2 in most studies) as an example to illustrate the workflow of DeepHoF on SARS-CoV-2 isolates (**Figure 2**).

**Figure 2.**
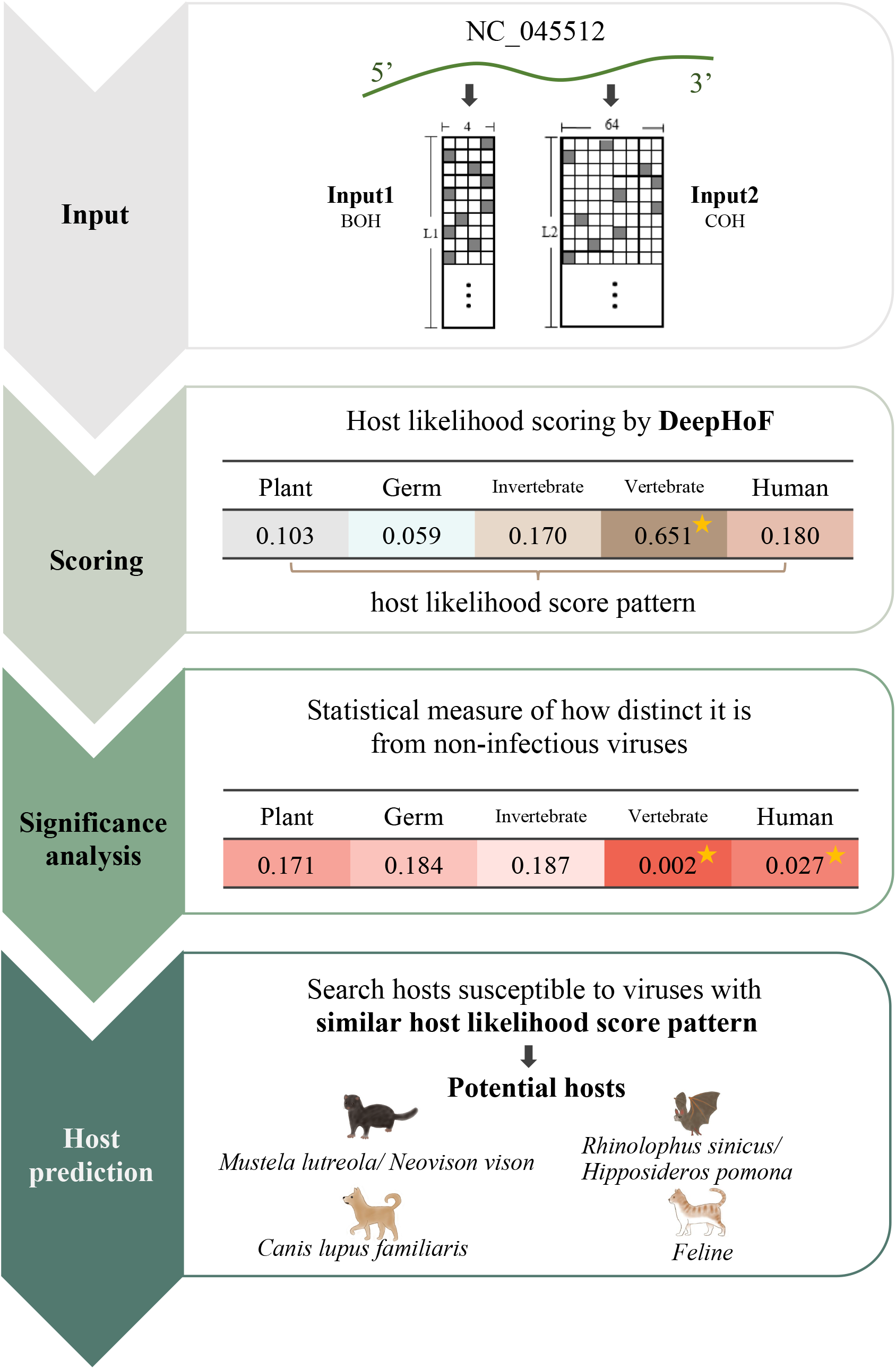
The workflow of application of DeepHoF on NC_045512. In the application of DeepHoF on SARS-CoV-2 NC_045512, the whole genome of NC_045512 was the only input required by the pre-trained DeepHoF model and coded into BOH and COH matrix for BiPathCNN network. DeepHoF output the host likelihood scores of NC_045512 on five host types respectively and the corresponding significance. The hosts of NC_045512 were predicted to be non-human vertebrates and humans with *p*-values less than 0.05. Simultaneously, NC_045512 was characterized by its host likelihood score profile. Susceptible to viruses with similar profile, *Mustela lutreola/ Neovison vison, Rhinolophus sinicus, Canis lupus familiaris, Hipposideros pomona* and Feline were output as the probable hosts of NC_045512. BOH: base one-hot matrix, COH: codon one-hot matrix.

For all the 17 SARS-CoV-2 isolates listed in **Figure 3A**, the host likelihood scores on non-human vertebrates and humans were assigned *p*-values less than 0.05 (0.002 and 0.027 respectively), illustrating a high possibility of non-human vertebrates and humans (Methods) to be the hosts of SARS-CoV-2. Besides, compared to other coronaviruses released on RefSeq [24], the high similarity of human and non-human vertebrate host likelihood scores among SARS-CoV-2, SARS-CoV and MERS-CoV (Figure 3B), would raise an alarm when the infection capabilities of SARS-CoV-2 was uncertain in the early stage of pandemic.

**Figure 3.**
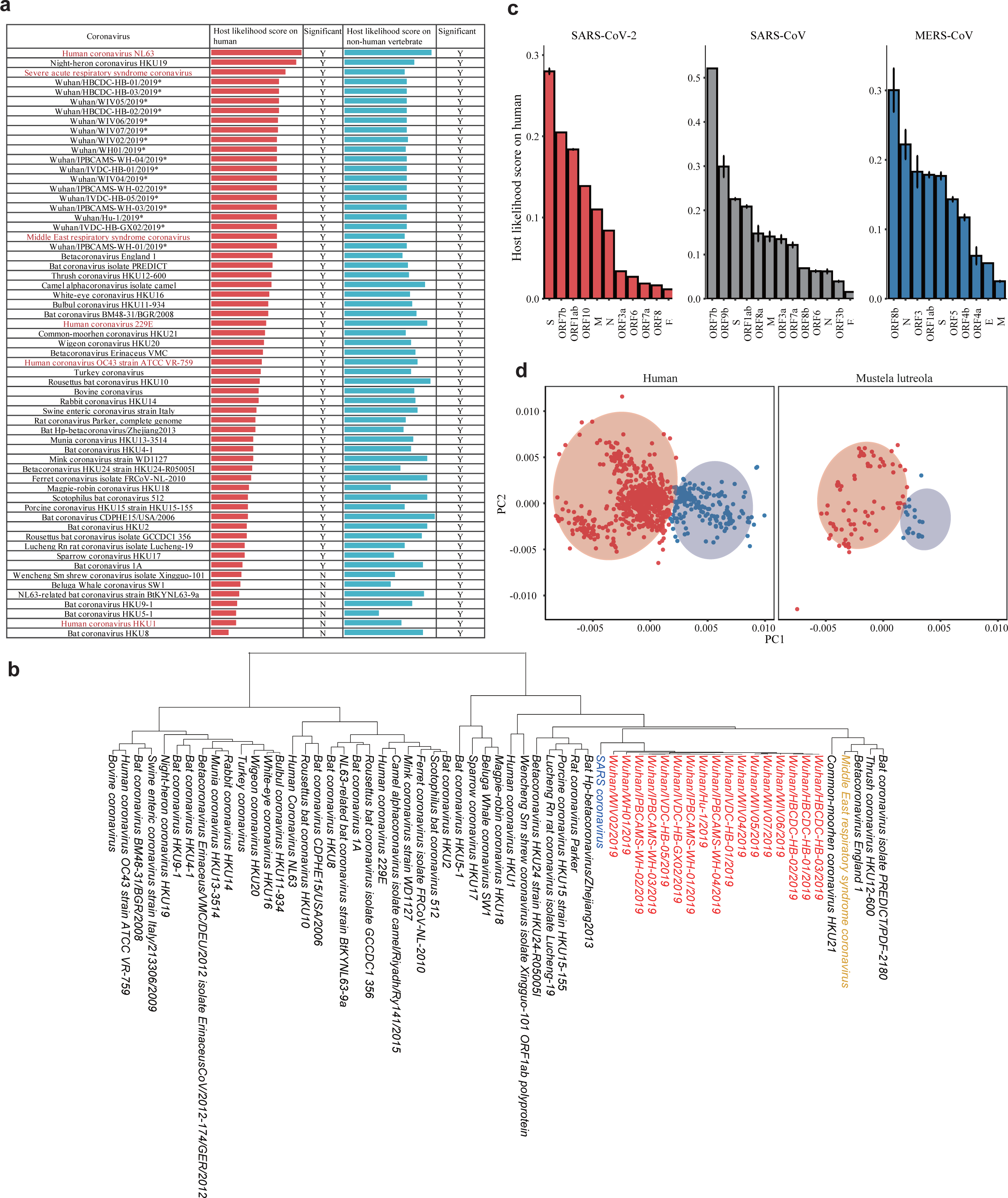
Evaluation of host likelihood scores of SARS-CoV-2. The contribution of each gene in the prediction and the visualization of host likelihood score profiles of SARS-CoV-2 isolates sampled in Netherlands. **A**. Host likelihood scores of 17 earliest detected SARS-CoV-2 isolates and other coronaviruses on humans and non-human vertebrates. SARS-CoV-2 showed high host likelihood scores on both humans and non-human vertebrates with *p*-values less than 0.05. In addition, SARS-CoV-2 was predicted lower score than SARS-CoV and comparable score to MERS-CoV on human. As for host likelihood scores on non-human vertebrates, SARS-CoV-2, SARS-CoV and MERS-CoV were close to each other. Host likelihood scores have *p*-values less than 0.05 are marked ‘Y (yes)’. (Red: human-infecting coronaviruses; *: the 17 earliest collected SARS-CoV-2 isolates). **B**. Hierarchical clustering of early-stage SARS-CoV-2 and other coronaviruses using five-dimensional host likelihood score profiles given by DeepHoF. The profile of SARS-CoV-2 was close to that of SARS-CoV and MERS-CoV (Red: SARS-CoV-2; Blue: SARS-CoV; Yellow: MERS-CoV). **C**. Contributions of the protein coding genes on determining the host likelihood scores of SARS-CoV-2, SARS-CoV and MERS-CoV on human. The structural genes, ORF1ab and group-specific genes contributed differently in the three coronaviruses (two-sided unpaired Welch Two Sample t-test, *p*-value < 0.05, see in Supplemental Figure S3). S, ORF7b and ORF1ab were the most pivotal in SARS-CoV-2. ORF7b, ORF9b and S were the most considerable in SARS-CoV. ORF8b, N and ORF3 contributed the most in MERS-CoV (S: spike glycoprotein coding gene; M: membrane/matrix glycoprotein coding gene; N: nucleocapsid phosphoprotein coding gene; E: envelope coding gene). **D**. Principal component analysis (PCA) of host likelihood score profiles of SARS-CoV-2 detected on humans and minks in Netherlands. The host likelihood score profiles of mink-derived and human-derived SARS-CoV-2 isolates in Netherlands are distributed in a consistent mode, containing a major cluster and divergence. The host likelihood score profiles of human-derived (left) and mink-derived (right) SARS-CoV-2 isolates in Netherlands distributed in a consistent mode, both containing a major cluster (red) and divergence (blue). The major cluster and the divergence were divided by the pam function of R package cluster.

To describe the contribution of each gene in the determination of the host likelihood scores of SARS-CoV-2 isolates (use NC_045512 as a representation), we used each gene sequence of SARS-CoV-2 as the input of DeepHoF and predicted the host likelihood scores for each gene. We found that the S gene, ORF1ab and ORF7b indeed acquired high likelihood scores on human host type and thus playing important roles in determining human as the host (Figure 3C). The fact that several domains on S gene and ORF1ab are essential for the coronavirus-host fusion process, host survival or viral replication [25-27] suggests the rationality of our findings. It is noteworthy that the linear correlation between the lengths and the host likelihood scores for genes is not tenable (Supplemental Figure S2). This shows that the importance of ORF1ab is not due to the remarkable length of the gene. Additionally, our prediction proposes the necessity of further experimental research on the function of ORF7b in SARS-CoV-2. Furthermore, we explored how each gene functioning on coronavirus life circle [25-28] contributed to the human host likelihood scores of SARS-CoV-2, SAR-CoV and MERS-CoV using the earliest sequenced samples, including 12 SARS-CoV isolates, 9 MERS-CoV isolates and 17 SARS-CoV-2 isolates released in NCBI in 2003, 2012 and 2019, respectively (Supplemental Table S2). The contributions of these genes were represented by their host likelihood scores on human. We found that ORF1ab was relatively important in the prediction for all these viruses, which was possibly due to its functions in viral replication and host survival [27]. The structural genes (S, M, N, and E genes) in these three viruses contributed differently on the human host type, illustrating these genes functioned inconsistently in these viruses. Specifically, S gene, participating in virus-host fusion process, contributed more in SARS-CoV-2 and SARS-CoV, while N gene, eliciting the strong specific antibody responses, played the most important role in MERS-CoV. Two equivalent genes, ORF9b, attaching membrane in virion assembly of SARS-CoV, and ORF8b, related to or immune evasion of MERS-CoV, made high contributions on human host likelihood scores for the two viruses. Moreover, two group-specific genes, ORF7b with unclear function in SARS-CoV, and ORF3 associated with virial replication and pathogenesis in MERS-CoV contributed significantly in the two viruses (Figure 3C, Supplemental Figure S3). These discrepancies might indicate the different significance of these genes among the three coronaviruses in the interaction with human and give hints to the target of drug design. It is disappointed that host determination for SARS-CoV-2 is extremely difficult due to the limited knowledge of the virus world. Therefore, the sequences and host information of viruses contained in the public database should be valued and fully utilized. To fill the gap in the efficient inference of host species for SARS-CoV-2 using the tools which were state of the art, we deeply analyzed the host likelihood profiles of viruses output by DeepHoF to seek specific vertebrate hosts of the early-stage SARS-CoV-2 isolates. In this study, we proposed that viruses with the same host species possessed the host likelihood score profiles close in the five-dimensional space. Based on this assumption, we compared the host likelihood score profile of SARS-CoV-2 with those of the non-human vertebrate viruses released in GenBank [29] before the pandemic outbreak of SARS-CoV-2 (Methods). We found that minks (*Mustela lutreola/Neovison vison*) were the most probable host, followed by Chinese rufous horseshoe bats (*Rhinolophus sinicus*), dogs (*Canis lupus familiaris*), Pomona roundleaf bats (*Hipposideros Pomona*) and cat family (*Felidae*) (**Table 2**, Supplemental Table S3). In contrast, minks, Chinese rufous horseshoe bats, dogs and cat family were respectively classified into very low, low or medium groups by Damas *et al*., [10], who divided 410 vertebrate species into five categories from very high to very low depending on the susceptibility to SARS-CoV-2 based on the analysis of sequence similarity of ACE2 and protein structure of ACE2/SARS-CoV-2 S-binding interface from the vertebrates. In the later world-wide pandemic, it should be pointed out that all the probable hosts we predicted were proved by animal experiments or the infection events [5], which illustrated the usefulness of such analysis for the host inference of SARS-CoV-2. Remarkably, SARS-CoV-2 has been reported largely to infect farmed minks in Netherlands, Denmark, Spain, the United States, Sweden, Italy, Greece, France, Lithuania, Canada, and Poland from April to Febrary, 2021. As of Febrary, 2021, SARS-CoV-2 had been reported to sweep 69 and 207 mink farms in Netherlands and Denmark, respectively, which accelerated the cull of minks and killed the fur industry in the two countries. On 9 October, 2020, at least 10,000 minks were reported dead at Utah and Wisconsin mink farms in the USA, and they were believed infected by SARS-CoV-2 [5] (Table 2).

**Table 2.**
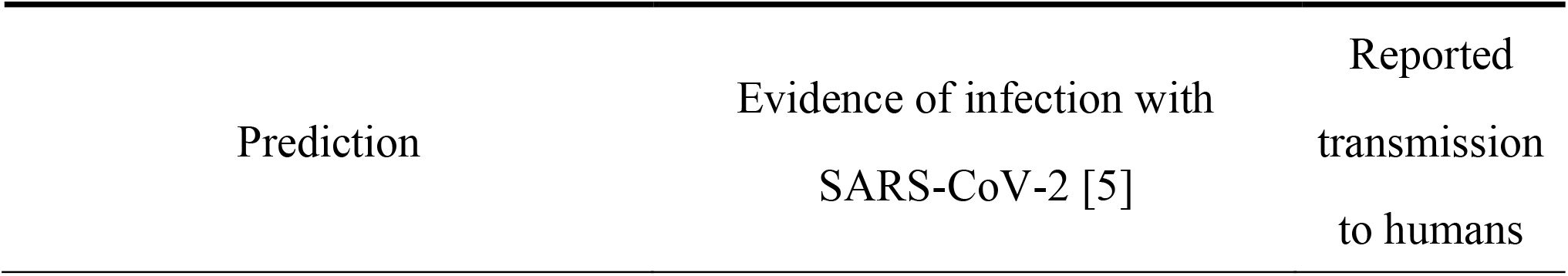
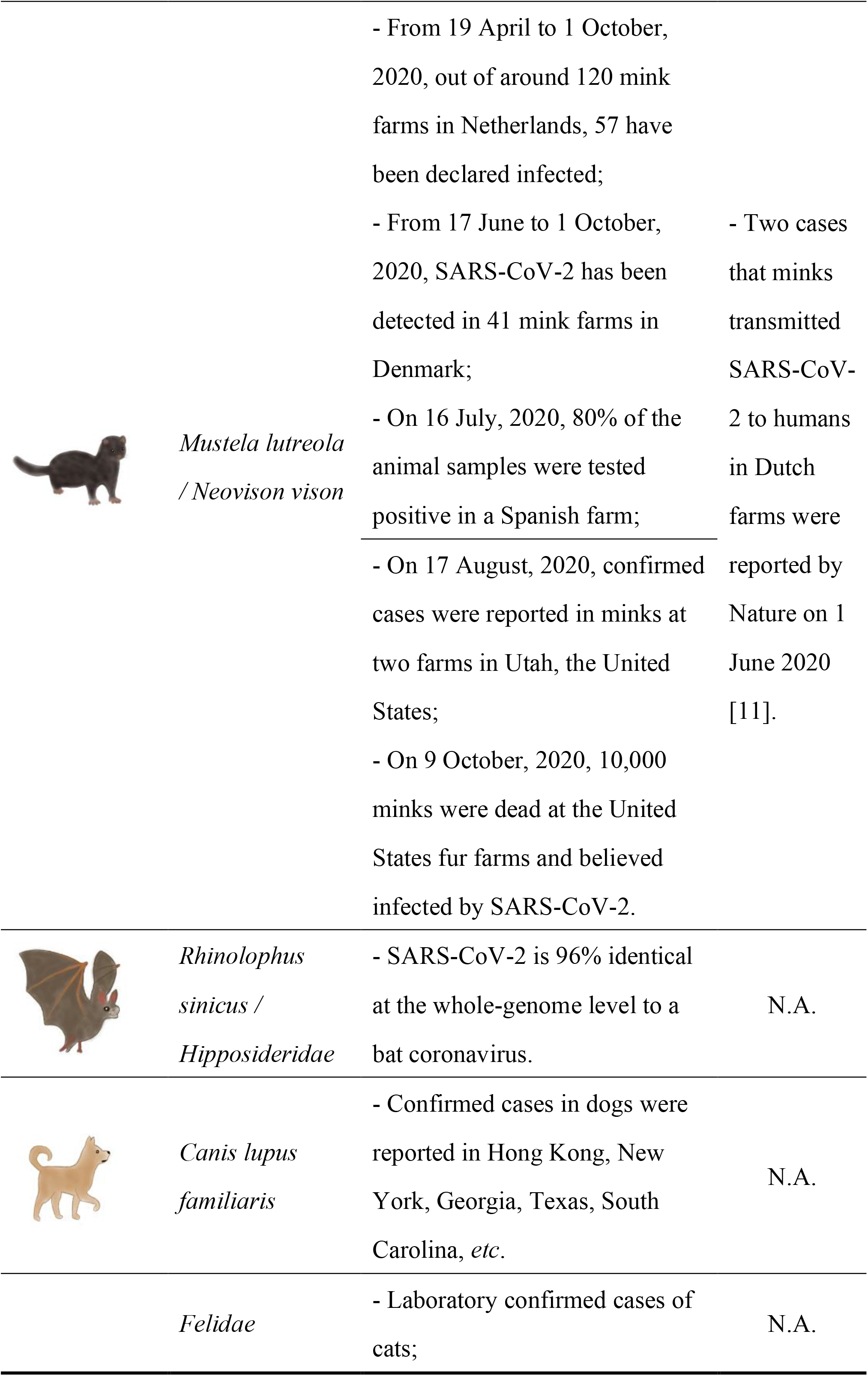
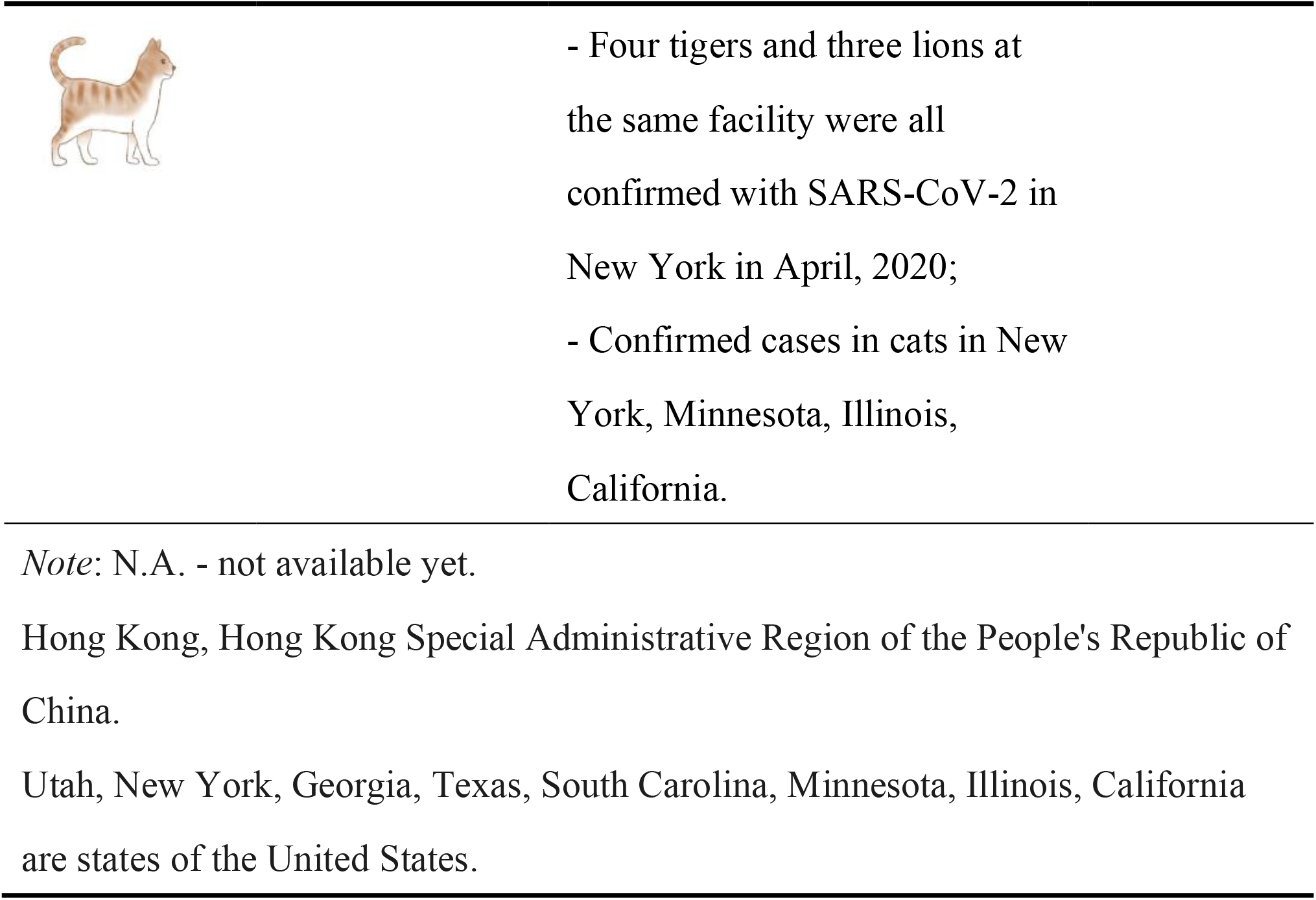
Host prediction results of SARS-CoV-2.

When evaluating the contributions of 11 genes of SARS-CoV-2 in determining mink as the most probable host, we found ORF1ab and ORF8 contributed the most (Supplemental Table S4), which suggesting that genes show different contributions when determining different hosts. The rationality of this result is supported by the roles of ORF1ab in viral replication and host survival [27], and the roles of ORF8 related to immune evasion [30]. However, the interaction between the two genes and the mink cell should merit the further attention and investigation.

Additionally, novel coronaviruses, which possess high sequence similarity with SARS-CoV-2, were found on pangolin [2, 3] in China. Even though these pangolin-associated coronaviruses were assigned similar host likelihood score profiles with early-stage SARS-CoV-2 isolates, our analysis demonstrated that the similarity of profiles between SARS-CoV-2 and pangolin-associated coronaviruses was lower than those between SARS-CoV-2 and certain viruses of mink and Chinese rufous horseshoe bat.

### Association of SARS-CoV-2 between humans and minks

In April 2020, farmed minks in Netherlands were noticed to be infected by SARS-CoV-2 because of the abnormal mortality [4]. Even though all the mink farms in Netherlands have been screened mandatorily since 28 May 2020, the transmission of coronavirus among the mink population did not seem to cease. Thus, a million farmed minks were culled in Netherlands, and followed by a plan to cull 2.5 million farmed minks in Denmark.

Characterizing SARS-CoV-2 by their host likelihood score profiles, we found the isolates detected on humans and minks in Netherlands distributed in a consistent mode, where both groups were divided into a major cluster and a divergence (Figure 3D, 1,746 SARS-CoV-2 samples collected from humans in Netherlands as of September 15 and 153 SARS-CoV-2 samples collected from farmed minks in Netherlands as of October 15 were used respectively, Methods). For SARS-CoV-2, as the host likelihood score on susceptible hosts such as human and mink can also indicate the likelihood to infect these animals, the mode of host likelihood score profile can reflect its property of viral infection. Consequently, the consistency mentioned above hinted the close infection-related behaviors of SARS-CoV-2 on humans and minks in Netherlands and thus illustrated the association of SARS-CoV-2 isolates collected from the two populations. Furthermore, nine of 14 high-frequency variants in human-derived SARS-CoV-2 genomes sequenced in Netherlands were absent in the genomes detected in other countries. Herein we used NC_045512 as the reference for variant calling, regarded the variants with ≥5% frequency as high-frequency ones and filtered out the synonymous single nucleotide polymorphisms (SNPs) (Supplemental Table S5). Among these unique high-frequency variants in Dutch human-derived SARS-CoV-2, two were found in Dutch mink-derived SARS-CoV-2, thus proved the circulation of SARS-CoV-2 between humans and minks in Netherlands. It was remarkable that our findings could be supported by the conclusions from a research team in Netherland, who utilized more detailed information about patients and related mink farms [12]. In the 2020 world-wide pandemic, minks are the only animal that has been reported to transmit SARS-CoV-2 to humans [11, 12]. We further compared the high-frequency variants of SARS-CoV-2 isolates in humans and minks in Netherlands. Except for four common variants, SARS-CoV-2 isolates derived from minks still had 23 unique high-frequency variants and six were found on S protein that is related to virus-host fusion process. This result indicated that the virus might have gained higher diversity after the intra-species circulation among mink herd and inter-species circulation between minks and human. As the mink infections are expanding worldwide, the association and circulation of SARS-CoV-2 between humans and minks in Netherlands notifies us of the importance to take precautions of the bidirectional transmission in other regions.

### Retrospective analysis of the world-wide pandemic

To verify the stability and uniformity of the host inference among SARS-CoV-2 samples, retrospective analysis of more isolates in the lasting pandemic was required. As the surge in variants of SARS-CoV-2 complicated the host prediction of the novel virus, we utilized 102,804 SARS-CoV-2 genomes released on GISAID EpiCoV Database (https://www.gisaid.org/) [31] as of 15 September 2020, before the rapid accumulation of mutations in SARS-CoV-2. We picked out 53,759 genomes which met the quality standard given by Chinese Academy of Sciences [32] and trimmed their varied-length 5′-and 3′-untranslated regions (UTR) based on the annotation of NC_045512 (Methods). We calculated the host likelihood score profiles of the 53,759 isolates (Supplemental Table S5) and conducted principal component analysis (PCA) on the profiles. As shown in **Figure 4A**, we found a clear cluster of all SARS-CoV-2 isolates with 17 earliest ones locating in the center. The kernel density estimation curves displayed on the first two principal components were approximately normally distributed. As the profiles of the 53,759 isolates are under the normal distribution mentioned above, the host range of SARS-CoV-2 isolates keep consistent throughout the pandemic and it is therefore reasonable that the validity of the host inference using the earliest 17 isolates would be efficient in the later pandemic.

**Figure 4.**
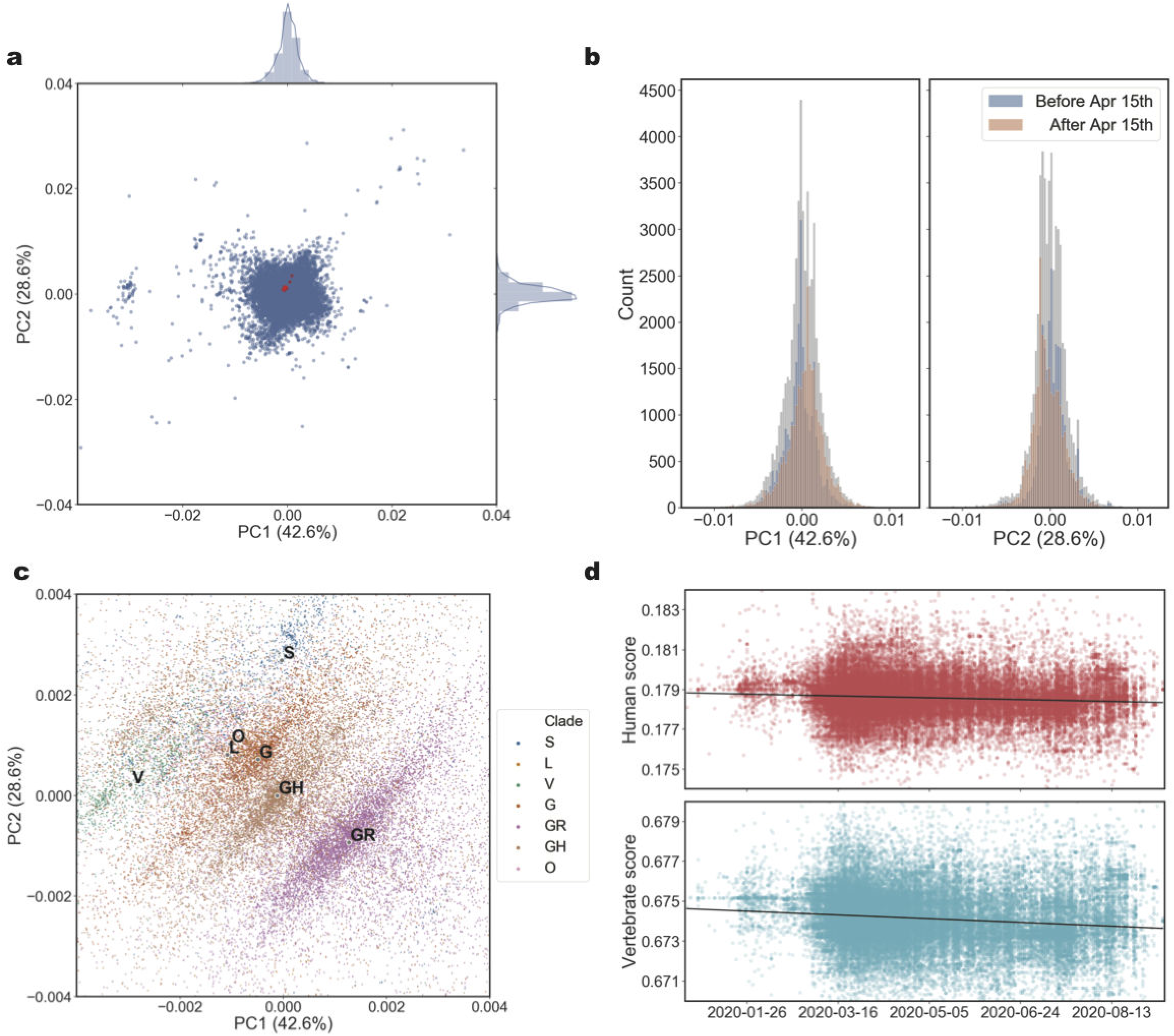
Entirety and divergence in the host likelihood score profiles of 53,759 SARS-CoV-2 isolates in the later world-wide pandemic. **A**. PCA of host likelihood score profiles of 53,759 SARS-CoV-2 isolates and the distribution on each principal component. All the host s likelihood core profiles of 53,759 SARS-CoV-2 isolates were clustered with 17 earliest sequenced isolates located in the center and the density curves displayed on each principal component were approximate normal distribution. **B**. Distributions of host likelihood score profiles of 53,759 SARS-CoV-2 isolates collected before and after 15 April 2020. When the SARS-CoV-2 isolates were divided chronologically using 15 April 2020 as the split date, which divided the 53,759 isolates into two parts more evenly than other dates. The host likelihood score profiles of SARS-CoV-2 before and after 15 April 2020 had divergent distributions on each principal component (two-sided two-sample Kolmogorov-Smirnov test, *p*-value = 0, n_isolates_ = 26,167 before 15 April 2020 and 27,592 after 15 April 2020. Blue, 26,167 isolates collected before 15 April 2020; Red, 27,592 isolates collected after 15 April 2020; Grey, all the 53,759 isolates). **C**. GISAID clades represented in PCA of host likelihood score profiles of 53,759 SARS-CoV-2 genomes. All the 53,759 samples representing 53,759 host likelihood score profiles were painted with six different colours corresponding to six different GISAID clades of SARS-CoV-2. SARS-CoV-2 isolates fell into several clear fusiform clusters with different colours according to their clades. **D**. Time series of the host likelihood scores on humans and non-human vertebrates for SARS-CoV-2 in the later world-wide pandemic. The host likelihood scores on humans and non-human vertebrates descend gradually with time (linear regression model analysis, *R*-squared = 6.806×10^−3^ and 1.431 × 10 ^-2^, *t*_(53,757)_ = −19.22 and *t*_(53,757)_ = −27.96, *p*-values = 5.543 × 10^−84^ and 3.292×10^−272^, slopes = −1.853×10^−6^ and −3.768×10^−6^).

However, when the SARS-CoV-2 isolates were divided chronologically using 15 April 2020 as the split date, which divided 53,759 isolates into two parts more evenly than other dates, we found that the two subsets have divergent distributions in each of the two dimensions of PCA (two-sided two-sample Kolmogorov-Smirnov test, *p*-value = 0, n_isolates_ = 26,167 before 15 April 2020 and 27,592 after 15 April 2020) (Figure 4B). The approximately normal distribution of SARS-CoV-2 genomes and their time-dependent feature indicate the overall consistency and a certain extent of divergence in the host likelihood score profiles of SARS-CoV-2 isolates.

To explain the divergence among host likelihood score profiles, we identified all variants in 53,759 genomes (Supplemental Table S5). The 13 high-frequency variants were located on S gene, N gene, ORF1ab, ORF8 and ORF3a, some of which are related to virus-host fusion process [22, 33]. Furthermore, we annotated our PCA result with the GISAID nomenclature system [31] which divides all SARS-CoV-2 genomes into six major clades based on marker variants that appeared over time. Most of the marker variants were recognized as high-frequency variants in the variant calling. As we can see in Figure 4C, SARS-CoV-2 isolates fell into several clear fusiform clusters according to their clades. This indicated that those marker variants might explain the divergence among host likelihood score profiles. When we manually mutated the 17 earliest sequenced genomes with those marker variants, we found the variants marking each clade drove the earliest sequenced SARS-CoV-2 to the corresponding cluster of the clade (Supplemental Figure S4), which verified our previous speculation and demonstrated the efficacy of DeepHoF to identify the important variants emerging in the virus’s evolution. However, as the consistency of the distribution of host likelihood score profiles were not disturbed, it hinted that these mutations did not change the host range of SARS-CoV-2.

Furthermore, to explore the trend of host likelihood of the SARS-CoV-2 over time, we finally examined the relationships between sampling time and the host likelihood scores on non-human vertebrates and humans (Figure 4D). We found that both scores gradually descended. As the host likelihood scores on susceptible hosts also indicate the likelihood to be infected by SARS-CoV-2 from a computational point of view, the trends might indicate the gradually descending infectiousness to human and other vertebrates from the outbreak to 15 September 2020. Those trends may not be so pronounced, but they should arouse our attention.

## Discussion

In summary, we proposed a deep learning method, DeepHoF, based on extracting the viral genomic features, to calculate the host likelihood scores on five host types. DeepHoF made up for the vacancy of a universal tool feasible to any novel virus. For the identification of five host types, our model can significantly outperform BLAST and well discriminate the human-infecting and non-human-infecting viruses like coronaviruses. Overcoming the limitation of sequence similarity-based methods to disclose the host information of novel viruses, DeepHoF demonstrated the practicality to SARS-CoV-2 in the 2020 pandemic. Using 17 SARS-CoV-2 isolates sequenced in the earliest stage of COVID-19 detection, DeepHoF evaluated the host likelihood scores on humans and non-human vertebrates for SARS-CoV-2. Filling the gap in predicting the host species for any novel virus that remained unsolved using the tools which were state of the art, we further analyzed the host likelihood score profile to further infer the specific hosts of SARS-CoV-2. The hosts determined by DeepHoF can be either reservoirs or susceptible middle hosts, which are not discriminated in this study. We found minks, bats, dogs and cats could be potential hosts of SARS-CoV-2, while minks might be one of the most noteworthy animal hosts. Due to mutations, the host likelihood score profiles of the isolates in the long period of the later pandemic had slightly varied, but followed normal distribution where those of the early 17 isolates locate in the center. As a consequence, the host range inferred with the profiles of the isolates during the pandemic was consistent with the inference using the early samples. Additionally, based on the model, we further found three genes (S gene, ORF7b and ORF1ab) and two genes (ORF1ab and ORF8) were significant in determining the host likelihood score on human and the host range for SARS-CoV-2, respectively. The genes involving virus-host fusion process (S gene), viral replication (ORF1ab) and host survival (ORF1ab) played a significant role in determining human as the host, while the genes related to viral replication (ORF1ab), host survival (ORF1ab) and immune evasion (ORF8) were significant to determine the host range for SARS-CoV-2. For the prevention and control of a novel epidemic disease such as COVID-19, the prediction of probable hosts is essential at the early stage of the epidemic outbreak. In view of this, our study is expected to play a potentially effective role in support of those efforts.

Furthermore, according to the analysis results of host likelihood score profiles of humans and minks in Netherlands, we found a strong association of SARS-CoV-2 isolates collected from the two populations and disclosed the contribution of mink on higher divergence in SARS-CoV-2. The phenomenon coincided with the analysis result of variant calling and could be explained by characteristics of minks in virus circulation. As reported by previous studies about avian-derived influenza A virus, minks serve as a significant node in the viral transmission network, connecting animals from different families and acting as domesticators for viral adaptation to mammals [34]. As the only one animal that has been reported to transmit SARS-CoV-2 to humans, the role of minks in the evolution of SARS-CoV-2 should be studied in depth. Therefore, with a large-scale genome analysis based on DeepHoF’s computation for the later world-wide pandemic, it should not be slighted for the relationship of SARS-CoV-2 between humans and minks.

Although we have applied DeepHoF to SARS-CoV-2 in the current study, the application of DeepHoF is not limited to this virus. DeepHoF is also feasible to determine the host ranges for many other novel viruses, such as the small circular rep-encoding ssDNA viruses newly discovered on wild animals and domestic animals or in the environment. However, limitations of DeepHoF lie in that it does not consider the host sequence information, which can be improved in the future. DeepHoF also does not discriminate between reservoir hosts, vector hosts and other susceptible hosts. Meanwhile, the present study is expected to be further confirmed with both the ongoing events of pandemic and additional experimental findings, and the interpretation of our analysis should be still kept a certain caution.

Represented by SARS-CoV-2, more complex and larger numbers of viral genome data will be produced in similar epidemics in the future. In addition, the metagenome and the metavirome can also be used in the prevention and control of the epidemic. The United States Agency for International Development launched the Global Virus Program in 2018 to reduce possible epidemiological threats by studying metaviromic samples from more than 35 countries around the world [35]. It is estimated that there are about 1.67 million novel viruses in mammals, birds and other important hosts of zoonotic viruses. Among them, 631,000-827,000 have the potential to cause zoonotic diseases [35]. However, only 263 viruses from 25 virus families have been confirmed to infect humans [36]. Newly emerged infectious viruses keep threatening our health and well-being. Under the circumstances, using computational methods to discover pathogenetic viruses and acquire knowledge, including the host range, about novel viruses can provide timely response in the prevention of epidemics and pandemics. In the future, the detection of novel viruses will rely more heavily on high-throughput sequencing technologies such as metagenomics and metaviromics. Thus, more robust tools designed for metagenomes and metaviromes are required.

## Materials and Methods

### Datasets construction for training and test

We downloaded 63,049 whole viral genomes from GenBank by 9 July, 2019, and tagged them with five host labels (plant, germ, invertebrate, non-human vertebrate and human), which were integrated from the host metadata provided by GenBank (Supplemental Table S6). The five host types covered all the living organism hosts. For viruses infecting multiple host types, multiple labels were given. Following the data collection procedure, short fragments were generated randomly from those tagged whole genomes because of the computational cost in long sequence processing. The training set was constructed with short fragments from 55,283 genomes released before 1 January, 2018, and the test set was constructed with the rest (the Accession list and the host information of the genomes used for training and test are in Supplemental Table S7). There is non-overlap of virus species in the training and test sets.

### Mathematical representation of viral whole genomes

Due to the long-term adaptation to natural reservoirs, viruses share some evolutionary signatures in nucleotide sequences, such as codon pair, dinucleotide, codon, and amino acid biases, with their natural reservoirs [15]. Besides, viral proteins, especially the receptors that are effectively attached to the host cell membrane, are crucial factors for viruses to invade and infect the host cells [37]. In brief, the genome compositions of viruses can inform host-virus correlation.

Herein, we represent a given viral sequence with a base one-hot matrix (BOH) and a codon one-hot matrix (COH), digitizing the genetic information of the virus on nucleotide and codon level respectively. To start with, bases and codons are encoded with one-hot format to work with deep learning algorithms. In the coding of BOH, each consecutive base of a query sequence linked by its complementary strand is encoded by one-hot. For COH, we do not extract ORFs since coding sequences make up most of the viral genome. Instead, we directly concatenate the six phases of the input sequence (Supplemental Figure S5), and then each consecutive codon of the joined sequences is encoded by one-hot. Consequently, for an input sequence of length L, it will be transformed to a BOH matrix, with the size of 2L×4, and a COH matrix, with the size of 2L×64.

### BiPathCNN Model descriptions

In building the framework of DeepHoF, we firstly utilize a BiPathCNN [38], containing two CNN paths, digging information from the BOH matrix and COH matrix respectively. The information is naturally corresponding to the viral genomic features for the viruses which infect the same kind of hosts. After independent convolution and pooling operations at the beginning, the two paths are combined by a concatenation layer. Following a normalization layer, five prediction scores will be provided by five sub-paths, containing five independent nodes, corresponding to five independent binary classifications on plant, germ, invertebrate, non-human vertebrate and human individually, in the output layer with sigmoid activation and binary cross-entropy loss function for each node. The architecture of DeepHoF is shown in Supplemental Figure S6 and the details of each layer in BiPathCNN are described in Supplementary Information.

### Implementation of DeepHoF

In the practical application, viral nucleotide sequence is the only input required by DeepHoF. For a viral whole genome sequence (or a partial genome sequence), a cut window moves along the long sequence without overlapping to separate it into suitable fragments for the pre-trained BiPathCNN model. DeepHoF firstly predicts the host infection scores for each fragment. Then it calculates the final score by weighting and summing the predicted scores of each fragment. For example, a 2,000 bp query sequence is separated into three consecutive fragments, corresponding to the first 800 bp, the middle 800 bp and the last 400 bp of the query sequence. Then DeepHoF predicts the three fragments independently and calculates the weighted average of the three predicted score vectors with the weights of 800/2,000, 800/2,000, and 400/2,000 respectively. For each input sequence, DeepHoF outputs five scores on five host types, respectively. Besides, DeepHoF provides the *p*-values of each score, statistically measuring of how distinct the scores are compared with those of non-infectious viruses [22]. For example, if an input virus has a score of 0.4 on human, we compare 0.4 with the scores of non-human viruses in our dataset and provide the *p*-value as a judgment basis. If the *p*-value is less than 0.05, we conclude that human is the probable host of the input virus with a significantly higher score on human host type than non-human viruses.

As the host likelihood score profile of a virus, consisting of the five predicted scores given by DeepHoF, can be regarded as a host-related feature vector extracted by DeepHoF, we utilize it to characterize the virus. It is logistical to regard the viruses with the same host species possess the similar host likelihood score profiles. Based on this assumption, the potential host species of a virus can be inferred by the analysis of the profiles. To quantitatively compare host likelihood score profiles between viruses, we calculated the Euclidean distance between the profiles. In the case of SARS-CoV-2, we searched the detailed vertebrate host of the earliest detected isolates, which are closer to the most recent common ancestor of SARS-CoV-2. To start with, we added the host annotations provided by Virus-Host DB [39] to the vertebrate viruses included in GenBank. Here, the average of host likelihood score profiles of 17 earliest sequenced isolates was used as the representation of SARS-CoV-2. We calculated the Euclidean distance between the profile of SARS-CoV-2 and that of each non-human vertebrate virus (discovered before the outbreak of SARS-CoV-2). We regarded the vertebrate infected by a virus possessing profile close to that of SARS-CoV-2 was the probable host of SARS-CoV-2.

### Data filtering and trimming for SARS-CoV-2 genome sequences

There were 102,804 SARS-CoV-2 genomes released on GISAID EpiCoV Database as of 15th September 2020. We downloaded all the sequences and filtered them with the quality standard given by the Chinese Academy of Sciences [32]. Because the UTRs were not taken as seriously as the protein-coding regions and the lengths of sequenced UTRs varied a lot in different SARS-CoV-2 genomes, we trimmed the 5′- and 3′-UTR according to the annotation of NC_045512 to get rid of noises. Thus, we finally got 53,759 clean sequences.

### Phylogenetic analysis and single nucleotide polymorphisms analysis

In this study, we applied Clustal Omega software [40] (version 1.2.4) for multiple sequence alignment and RAxML software [41] (version 8.2.12) for phylogenetic tree building using maximum likelihood methods with 1000 bootstrap replicates. Snippy [42] (version 4.4.3) was utilized for variant calling, using NC_045512 as the reference genome. In this study, we filtered out the synonymous SNPs and regarded the variants with ≥ 5% frequency as high-frequency ones. Commands of the three tools are included in Supplementary Information.

## Authors’ contributions

HQZ and YHX co-supervised the study. QG, ML, CHW, JYG, XQJ and HQZ developed the DeepHoF model, conducted the analyses and wrote the manuscript. HQZ and ZCF helped with designing the model. MZ calculated performance metrics of DeepHoF and BLAST. PHW helped with phylogeny analysis and SNP analysis. JT, SFW and TTX made plots and table for the results. All authors read and approved the final manuscript.

## Acknowledgements

We acknowledge GISAID (https://www.gisaid.org/) for facilitating open data sharing. This work was supported by the National Key Research and Development Program of China (2017YFC1200205) and the National Natural Science Foundation of China (32070667, 31671366). Part of the analysis was performed on the High Performance Computing Platform of the Center for Life Science of Peking University.

## Supplementary material

**Supplementary material Supplemental Figure S1-S6, Supplemental Table S1, S3 and S6 and Supplemental Methods**

**Supplemental Figure S1 ROC curves and AUC values of DeepHoF and BLAST on five host types**

DeepHoF performs better than BLAST on AUC of each host type.

**Supplemental Figure S2 The untenable linear correlations between the lengths and the host likelihood scores for genes of SARS-CoV-2**

For the genes of SARS-CoV-2, there is no statistical significance in the linear correlations between the lengths and the host likelihood scores on plant (**A**), germ (**B**), invertebrate (**C**), vertebrate (**D**) and human (**E**).

**Supplemental Figure S3 Human host likelihood scores of 5 genes of SARS-CoV-2, SARS-CoV and MERS-CoV**

Although all the three coronaviruses possess ORF1ab and four structural genes (S, M, N, E), these genes made different contributions on human host likelihood scores in these three viruses (two-sided unpaired Welch Two Sample *t*-test, *p*-value < 0.05). S gene and M gene contributed more in SARS-CoV-2 and SARS-CoV, while N gene and E gene were more significant in MERS-CoV.

**Supplemental Figure S4 Visualization of the host likelihood score profiles of SARS-CoV-2 isolates from different GISAID clades and the manually mutated SARS-CoV-2 isolates on two-dimensional PCA**

SARS-CoV-2 isolates fall into several clear fusiform clusters with different colors according to their clades. Manually mutated with specific marker variants, the 17 earliest sequenced isolates move to the corresponding fusiform cluster of the clade that is represented by the specific marker variants.

**Supplemental Figure S5 Six phases of an input sequence**

For coding the COH matrix of a given sequence, we represented it with the direct conjunction of its six phases, generated from its complementary strand and itself. **Supplemental Figure S6 Structure of BiPathCNN in DeepHoF**

BOH matrix and COH matrix are input into two paths independently and transformed by the convolution and pooling layers at the beginning. A concatenation layer and a normalization layer combine the output of the two paths. Five sub-paths process the combined intermediate output individually. Each sub-path contains a full connection layer, a normalization layer and an output layer with sigmoid activation and binary cross-entropy loss function. The five sub-paths output the host likelihood scores on five host types respectively.

**Supplemental Table S1 Comparison of performance of DeepHoF and BLAST on each host type classification**

**Supplemental Table S3 Top 20 hosts predicted by DeepHoF on SARS-CoV-2 Supplemental Table S6 Subtypes in five host types**

**Other supplementary material for this manuscript includes the following: Supplemental Table S2 Metadata and host likelihood scores of genes for SARS-CoV, MERS-CoV and SARS-COV-2 isolates**

**Supplemental Table S4 Contributions of 11 genes in the determination of hosts for SARS-CoV-2**

**Supplemental Table S5 Metadata, host likelihood score profiles, and high frequency SNPs on 53759 SARS-CoV-2 isolates**

**Supplemental Table S7 Host information of the viral genomes in training and test sets of DeepHoF**

**Supplemental Table S8 Acknowledge of sequence data of SARS-CoV-2 in GISAID**

## Data statement

Data utilized in the analysis of SARS-CoV-2, including the host likelihood score profiles and the metadata of 53,759 SARS-CoV-2 isolates, are available in the main text and Supplementary Information. The trimmed sequences of 53,759 isolates and the training and test sets of DeepHoF have been deposited on our lab homepage http://cqb.pku.edu.cn/ZhuLab/DeepHoF/.

The open source code utilized in this study has been deposited on GitHub https://github.com/PKUbioinfo-ZhuLab/DeepHoF and our lab homepage http://cqb.pku.edu.cn/ZhuLab/DeepHoF/

## References

[1] Zhou P, Yang X-L, Wang X-G, Hu B, Zhang L, Zhang W, et al. A pneumonia outbreak associated with a new coronavirus of probable bat origin. nature 2020;579:270–3.

[2] Lam TT-Y, Jia N, Zhang Y-W, Shum MH-H, Jiang J-F, Zhu H-C, et al. Identifying SARS-CoV-2-related coronaviruses in Malayan pangolins. Nature 2020;583:282–5.

[3] Xiao K, Zhai J, Feng Y, Zhou N, Zhang X, Zou J-J, et al. Isolation of SARS-CoV-2-related coronavirus from Malayan pangolins. Nature 2020;583:286–9.

[4] Oreshkova N, Molenaar RJ, Vreman S, Harders F, Munnink BBO, Hakze-van Der Honing RW, et al. SARS-CoV-2 infection in farmed minks, the Netherlands, April and May 2020. Eurosurveillance 2020;25:2001005.

[5] OIE. COVID-19 Portal: Events in Animals. https://www.oie.int/en/scientific-expertise/specific-information-and-recommendations/questions-and-answers-on-2019novel-coronavirus/events-in-animals/ (Oct 25 2020, date last accessed).

[6] Shi J, Wen Z, Zhong G, Yang H, Wang C, Huang B, et al. Susceptibility of ferrets, cats, dogs, and other domesticated animals to SARS–coronavirus 2. Science 2020;368:1016–20.

[7] Sia SF, Yan L-M, Chin AW, Fung K, Choy K-T, Wong AY, et al. Pathogenesis and transmission of SARS-CoV-2 in golden hamsters. Nature 2020;583:834–8.

[8] Munster VJ, Feldmann F, Williamson BN, Van Doremalen N, Pérez-Pérez L, Schulz J, et al. Respiratory disease in rhesus macaques inoculated with SARS-CoV-2. Nature 2020;585:268–72.

[9] Santini JM, Edwards SJ. Host range of SARS-CoV-2 and implications for public health. The Lancet Microbe 2020;1:e141–e2.

[10] Damas J, Hughes GM, Keough KC, Painter CA, Persky NS, Corbo M, et al. Broad host range of SARS-CoV-2 predicted by comparative and structural analysis of ACE2 in vertebrates. Proceedings of the National Academy of Sciences 2020;117:22311–22.

[11] Mallapaty S. What’s the risk that animals will spread the coronavirus. Nature 2020.

[12] Munnink BBO, Sikkema RS, Nieuwenhuijse DF, Molenaar RJ, Munger E, Molenkamp R, et al. Transmission of SARS-CoV-2 on mink farms between humans and mink and back to humans. Science 2021;371:172–7.

[13] Guo Q, Li M, Wang C, Wang P, Fang Z, tan J, et al. Host and infectivity prediction of Wuhan 2019 novel coronavirus using deep learning algorithm. bioRxiv 2020:2020.01.21.914044.

[14] Rothenburg S, Brennan G. Species-specific host–virus interactions: implications for viral host range and virulence. Trends in microbiology 2020;28:46–56.

[15] Babayan SA, Orton RJ, Streicker DG. Predicting reservoir hosts and arthropod vectors from evolutionary signatures in RNA virus genomes. Science 2018;362:577–80.

[16] Lu G, Wang Q, Gao GF. Bat-to-human: spike features determining ‘host jump’of coronaviruses SARS-CoV, MERS-CoV, and beyond. Trends in microbiology 2015;23:468–78.

[17] Li W, Zhang C, Sui J, Kuhn JH, Moore MJ, Luo S, et al. Receptor and viral determinants of SARS-coronavirus adaptation to human ACE2. The EMBO journal 2005;24:1634–43.

[18] Villarroel J, Kleinheinz KA, Jurtz VI, Zschach H, Lund O, Nielsen M, et al. HostPhinder: a phage host prediction tool. Viruses 2016;8:116.

[19] Galiez C, Siebert M, Enault F, Vincent J, Söding J. WIsH: who is the host? Predicting prokaryotic hosts from metagenomic phage contigs. Bioinformatics 2017;33:3113–4.

[20] Gałan W, Bąk M, Jakubowska M. Host taxon predictor-a tool for predicting taxon of the host of a newly discovered virus. Scientific reports 2019;9:1–13.

[21] Mock F, Viehweger A, Barth E, Marz M. VIDHOP, viral host prediction with deep learning. Bioinformatics 2020.

[22] Ren J, Ahlgren NA, Lu YY, Fuhrman JA, Sun F. VirFinder: a novel k-mer based tool for identifying viral sequences from assembled metagenomic data. Microbiome 2017;5:1–20.

[23] Belouzard S, Millet JK, Licitra BN, Whittaker GR. Mechanisms of coronavirus cell entry mediated by the viral spike protein. Viruses 2012;4:1011–33.

[24] Brister JR, Ako-Adjei D, Bao Y, Blinkova O. NCBI viral genomes resource. Nucleic acids research 2015;43:D571–D7.

[25] Li Y-H, Hu C-Y, Wu N-P, Yao H-P, Li L-J. Molecular characteristics, functions, and related pathogenicity of MERS-CoV proteins. Engineering 2019;5:940–7.

[26] Cheng VC, Lau SK, Woo PC, Yuen KY. Severe acute respiratory syndrome coronavirus as an agent of emerging and reemerging infection. Clinical microbiology reviews 2007;20:660–94.

[27] Hu B, Guo H, Zhou P, Shi Z-L. Characteristics of SARS-CoV-2 and COVID-19. Nature Reviews Microbiology 2020:1–14.

[28] Wong L-YR, Ye Z-W, Lui P-Y, Zheng X, Yuan S, Zhu L, et al. Middle East Respiratory Syndrome Coronavirus ORF8b Accessory Protein Suppresses Type I IFN Expression by Impeding HSP70-Dependent Activation of IRF3 Kinase IKKε. The Journal of Immunology 2020;205:1564–79.

[29] Sayers EW, Cavanaugh M, Clark K, Pruitt KD, Schoch CL, Sherry ST, et al. GenBank. Nucleic acids research 2021;49:D92–D6.

[30] Young BE, Fong S-W, Chan Y-H, Mak T-M, Ang LW, Anderson DE, et al. Effects of a major deletion in the SARS-CoV-2 genome on the severity of infection and the inflammatory response: an observational cohort study. The Lancet 2020;396:603–11.

[31] Shu Y, McCauley J. GISAID: Global initiative on sharing all influenza data–from vision to reality. Eurosurveillance 2017;22:30494.

[32] Zhao W-M, Song S-H, Chen M-L, Zou D, Ma L-N, Ma Y-K, et al. The 2019 novel coronavirus resource. Yi chuan = Hereditas 2020;42:212–21.

[33] Lan J, Ge J, Yu J, Shan S, Zhou H, Fan S, et al. Structure of the SARS-CoV-2 spike receptor-binding domain bound to the ACE2 receptor. Nature 2020;581:215–20.

[34] Xue R, Tian Y, Hou T, Bao D, Chen H, Teng Q, et al. H9N2 influenza virus isolated from minks has enhanced virulence in mice. Transboundary and emerging diseases 2018;65:904–10.

[35] Carroll D, Daszak P, Wolfe ND, Gao GF, Morel CM, Morzaria S, et al. The global virome project. Science 2018;359:872–4.

[36] King AM, Lefkowitz E, Adams MJ, Carstens EB. Virus taxonomy: ninth report of the International Committee on Taxonomy of Viruses. Elsevier, 2011.

[37] Dimitrov DS. Virus entry: molecular mechanisms and biomedical applications. Nature Reviews Microbiology 2004;2:109–22.

[38] Fang Z, Tan J, Wu S, Li M, Xu C, Xie Z, et al. PPR-Meta: a tool for identifying phages and plasmids from metagenomic fragments using deep learning. Gigascience 2019;8:giz066.

[39] Mihara T, Nishimura Y, Shimizu Y, Nishiyama H, Yoshikawa G, Uehara H, et al. Linking virus genomes with host taxonomy. Viruses 2016;8:66.

[40] Sievers F, Wilm A, Dineen D, Gibson TJ, Karplus K, Li W, et al. Fast, scalable generation of high-quality protein multiple sequence alignments using Clustal Omega. Molecular systems biology 2011;7:539.

[41] Stamatakis A. RAxML-VI-HPC: maximum likelihood-based phylogenetic analyses with thousands of taxa and mixed models. Bioinformatics 2006;22:2688–90.

[42] Seemann T. Snippy: rapid bacterial SNP calling and core genome alignments. https://github.com/tseemann/snippy.git (Oct 25 2020, date last accessed).

[43] Letunic I, Bork P. Interactive Tree Of Life (iTOL) v4: recent updates and new developments. Nucleic acids research 2019;47:W256–W9.

