## Supplementary material for "Predicting Hosts Based on Early SARS-CoV-2 Samples and Analyzing Later World-wide Pandemic in 2020"

### Supplemantary Meterial for

**The following sections include:**

Supplementary Methods

Supplemental Tables 1, 3 and 6

Supplemental Figs. 1 to 6

**Other Supplementary Material for this manuscript includes the following:**

Supplemental Table S2 Metadata and host likelihood scores of genes for SARS-CoV, MERS-CoV and SARS-COV-2 isolates

Supplemental Table S4 Contributions of 11 genes in the determination of hosts for SARS-CoV-2

Supplemental Table S5 Metadata, host likelihood score profiles, and high frequency SNPs on 53759 SARS-CoV-2 isolates

Supplemental Table S7 Host information of the viral genomes in training and test sets of DeepHoF

Supplemental Table S8 Acknowledge of sequence data of SARS-CoV-2 in GISAID

#### Supplementary Methods

##### Host prediction using BLAST

As there are no bioinformatics tools for comparison, we compare evaluation metrics of BLAST and DeepHoF. The host prediction using BLAST are divided into the following steps:

(1) Make BLAST database with the viral whole genomes used in the training dataset of DeepHoF. Besides, all the viral genomes are annotated with the corresponding host types.

Command: makeBLASTdb -in input_file -dbtype nucl -out database_name

(2) Align the whole genome sequences in test dataset of DeepHoF to the BLAST database made at the step (1).

Command: BLASTn -query input_file -task BLASTn -db database_name -out output_file -evalue 1e-1 -outfmt 6 -qcov_hsp_perc 0.5 -num_threads 30 -max_target_seqs 1

(3) Assign host prediction result for each query sequence. For example, for query sequence A, if virus B, which is annotated with host type "human, invertebrate", is the best hit for it, then the hosts of virus B (human and invertebrate) will be regarded as the predicted host types of viral sequence A. Meanwhile, the identity between sequence A and virus B is assigned as the prediction scores on both "human" and "invertebrate" host types, while the scores on other host types are assigned with zero. Besides, if there are no hit for viral sequence A, then the prediction scores are assigned with the difference between [1,1,1,1,1] and the true label vector. For example, if the viral A has host types of plant and germ, its label vector is [1, 1, 0, 0, 0], and if there are no hit for viral sequence A, the predicted score vector will be provided with [0, 0, 1, 1, 1] by

BLAST. (Note: the five labels in the label vector are corresponding to plants, germs, invertebrates, vertebrates except human, and human, respectively)

**The details of each layer of DeepHoF**

Input: BOH(COH)

Layer b1(c1): 1D convolutional layer, including 512 convolution kernels with the length of six. Activation function: ReLU (Rectified Linear Unit, $y = \max(0, x)$).

Layer b2(c2): global average pooling layer.

Layer 3: the concatenation layer, combining the output of the "base path" and "codon path".

Layer 4: batch normalization layers with the dropout operation, used to facilitate the convergence and prevent the overfitting.

Layers 5a-5e, 6a-6e and 7a-7e: full connection layers, followed by batch normalization layers with the dropout operation. After the dropout operation, five sigmoid layers calculate five prediction scores, each corresponding to host likelihood scores on plants, germs, invertebrates, vertebrates and human, respectively.

**Commands of software used in phylogenetic analysis and protein annotation**

Clustalo: clustalo --full -force -i Multiple_sequence_input_file --distmat-out=name.mat --guidetree-out name.guide.nwk -o name.aln --threads 30 --outfmt=a2m

RAxML: raxmlHPC-HYBRID -s multi-alignmentfile -w outputdir -n coronavirus -m GTRGAMMA -T 40 -N 100 -p 20170808 -f a -x 20170808

Snippy: snippy --outdir outidr --ctgs fastafile –ref NC_045512.gbk --cpus 50

**Supplementary Tables**

**Supplemental Table S1 Comparison of performance of DeepHoF and BLAST on each host type classification**

| Metrics | Plant | | Germ | | Invertebrate | |
|---|---|---|---|---|---|---|
| | BLAST | DeepHoF | BLAST | DeepHoF | BLAST | DeepHoF |
| TPR | 0.72807 | 0.412281 | 0.901219 | 0.904577 | 0.808696 | 0.913043 |
| FPR | 0.101413 | 0.002748 | 0.12494 | 0.011922 | 0.112158 | 0.020024 |
| AUC | 0.776833 | 0.973326 | 0.833512 | 0.994894 | 0.809932 | 0.993924 |
| Precision | 0.096737 | 0.691176 | 0.951138 | 0.99514 | 0.097895 | 0.406977 |
| Accuracy | 0.89608 | 0.988654 | 0.894146 | 0.927153 | 0.886668 | 0.978984 |

| Metrics | Vertebrate | | Human | | Average | |
|---|---|---|---|---|---|---|
| | BLAST | DeepHoF | BLAST | DeepHoF | BLAST | DeepHoF |
| TPR | 0.840225 | 0.808065 | 0.95873 | 0.75873 | 0.887814 | 0.864929 |
| FPR | 0.096774 | 0.003623 | 0.109599 | 0.003087 | 0.106788 | 0.007986 |
| AUC | 0.832527 | 0.976409 | 0.873414 | 0.996917 | 0.832533 | 0.987131 |
| Precision | 0.745809 | 0.986908 | 0.436101 | 0.956 | 0.699312 | 0.968049 |
| Accuracy | 0.887313 | 0.948814 | 0.895952 | 0.977566 | 0.892032 | 0.964234 |

**Supplemental Table S3  Top 20 hosts predicted by DeepHoF on SARS-CoV-2**

| Rank | Host name | Euclidean distance |
|---|---|---|
| 1 | Neovison vison/ Mustela lutreola | 0.01886 |
| 2 | Rhinolophus sinicus | 0.02647 |
| 3 | Canis lupus familiaris | 0.03508 |
| 4 | Hipposideros pomona | 0.03878 |
| 5 | Rhinolophus affinis | 0.04301 |
| 6 | Feliformia/ Felidae | 0.05125 |
| 7 | Ailuropoda melanoleuca | 0.05456 |
| 8 | Meleagris gallopavo | 0.06631 |
| 9 | Homo sapiens/ Paguma larvata | 0.07209 |
| 10 | Sus scrofa | 0.07290 |
| 11 | Plecotus auritus | 0.07410 |
| 12 | Enhydra lutris kenyoni | 0.07444 |
| 13 | Gorilla gorilla | 0.07839 |
| 14 | Galliformes/ Gallus gallus | 0.07906 |
| 15 | Canis lupus signatus | 0.08073 |
| 16 | Pan troglodytes | 0.08151 |
| 17 | Gallus gallus | 0.08597 |
| 18 | Pipistrellus | 0.08734 |
| 19 | Chelonia mydas | 0.08987 |
| 20 | Zosteropidae | 0.09105 |

**Supplemental Table S6 Subtypes in five host types**

| Host type | Host subtypes |
|---|---|
| plant | algae, diatom, plant |
| germ | Archaea, bacteria, fungi, protozoa |
| invertebrate | invertebrate |
| vertebrate | vertebrates except human |
| human | human |

#### Supplementary Figures

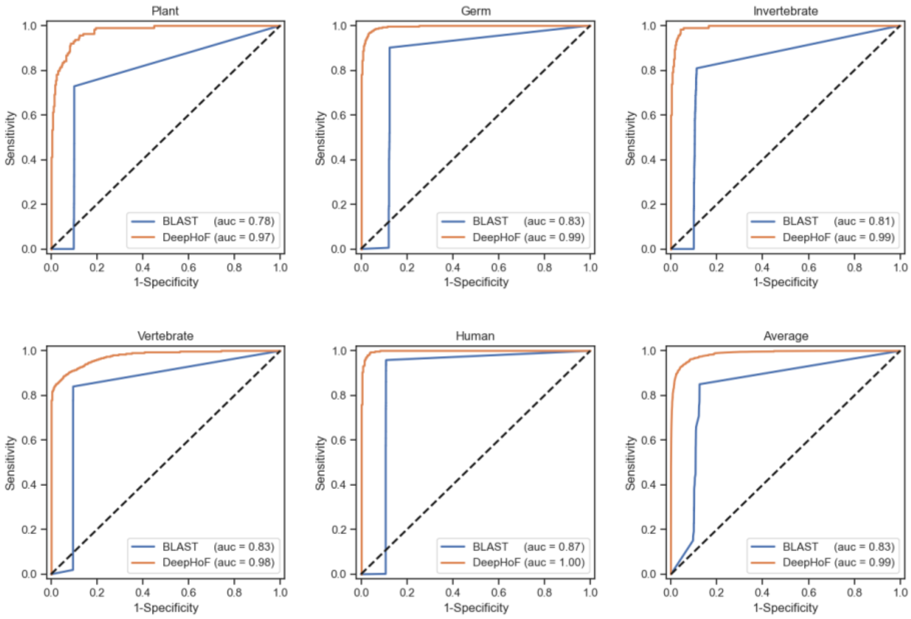


**Supplemental Figure 1 ROC curves and AUC values of DeepHoF and BLAST on five host types**

DeepHoF performs better than BLAST on AUC of each host type.

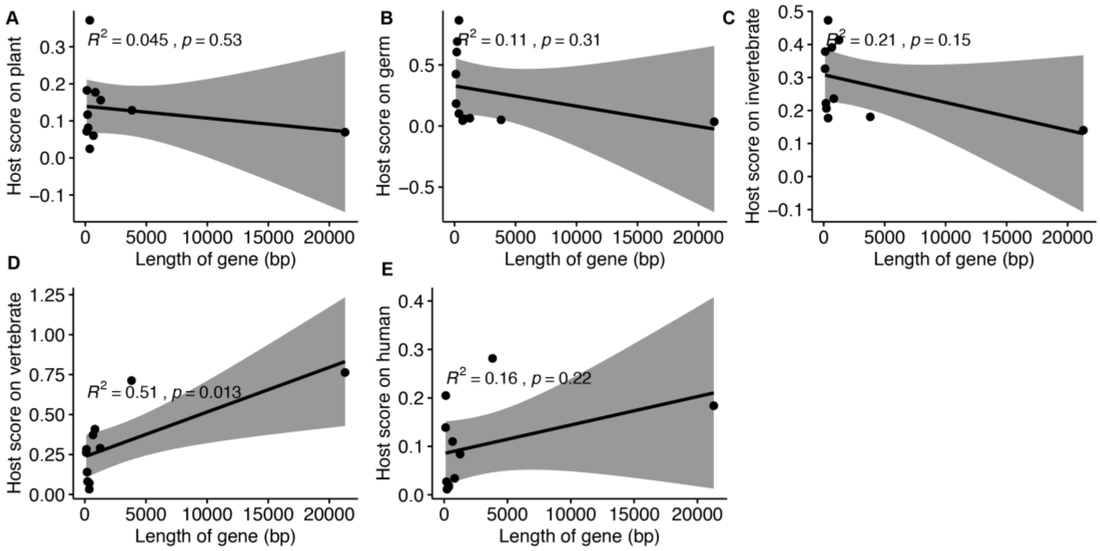


**Supplemental Figure S2 The untenable linear correlations between the lengths and the host likelihood scores for genes of SARS-CoV-2**

For the genes of SARS-CoV-2, there is no statistical significance in the linear correlations between the lengths and the host likelihood scores on plant (**A**), germ (**B**), invertebrate (**C**), vertebrate (**D**) and human (**E**).

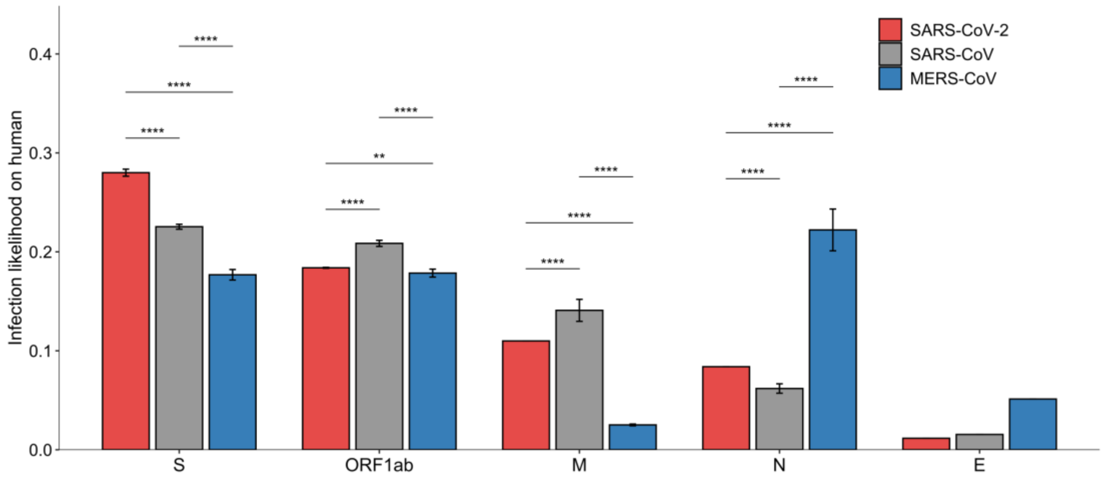


**Supplemental Figure 3 Human host likelihood scores of 5 genes of SARS-CoV-2, SARS-CoV and MERS-CoV**

Although all the three coronaviruses possess ORF1ab and four structural genes (S, M, N, E), these genes made different contributions on human host likelihood scores in these three viruses (two-sided unpaired Welch Two Sample *t*-test, *p*-value $< 0.05$). S gene and M gene contributed more in SARS-CoV-2 and SARS-CoV, while N gene and E gene were more significant in MERS-CoV.

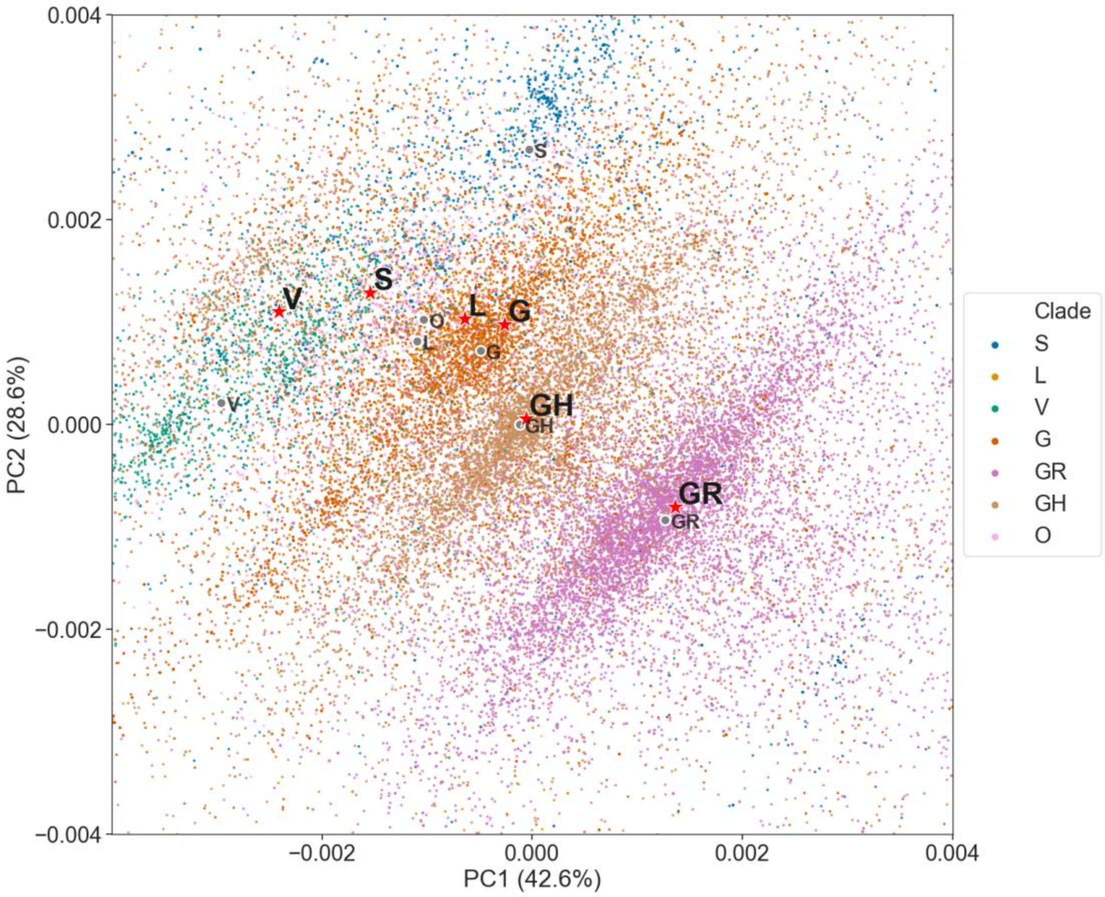


**Supplemental Figure S4 Visualization of the host likelihood score profiles of SARS-CoV-2 isolates from different GISAID clades and the manually mutated SARS-CoV-2 isolates on two-dimensional PCA**

SARS-CoV-2 isolates fall into several clear fusiform clusters with different colors according to their clades. Manually mutated with specific marker variants, the 17 earliest sequenced isolates move to the corresponding fusiform cluster of the clade that is represented by the specific marker variants.

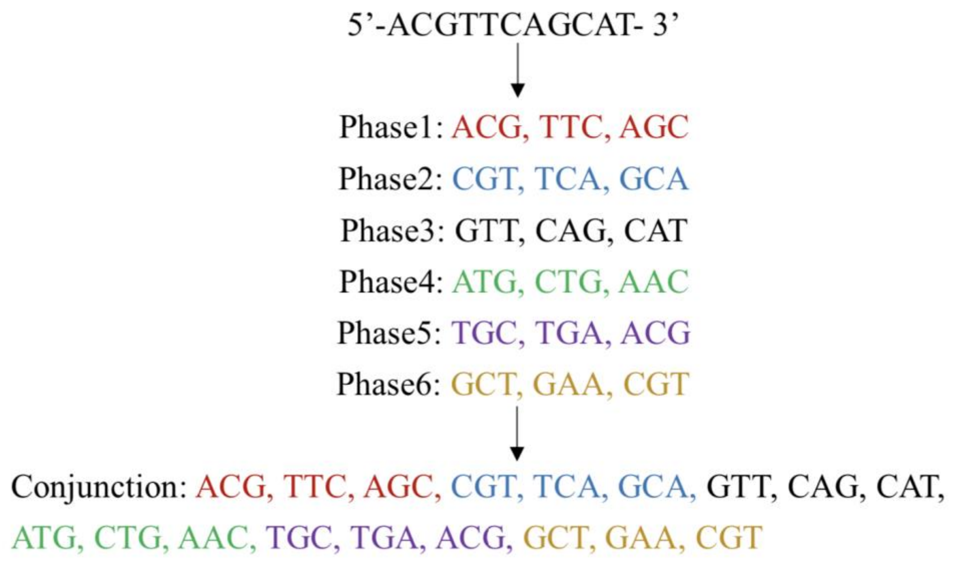


**Supplemental Figure 5  Six phases of an input sequence**

For coding the COH matrix of a given sequence, we represented it with the direct conjunction of its six phases, generated from its complementary strand and itself.

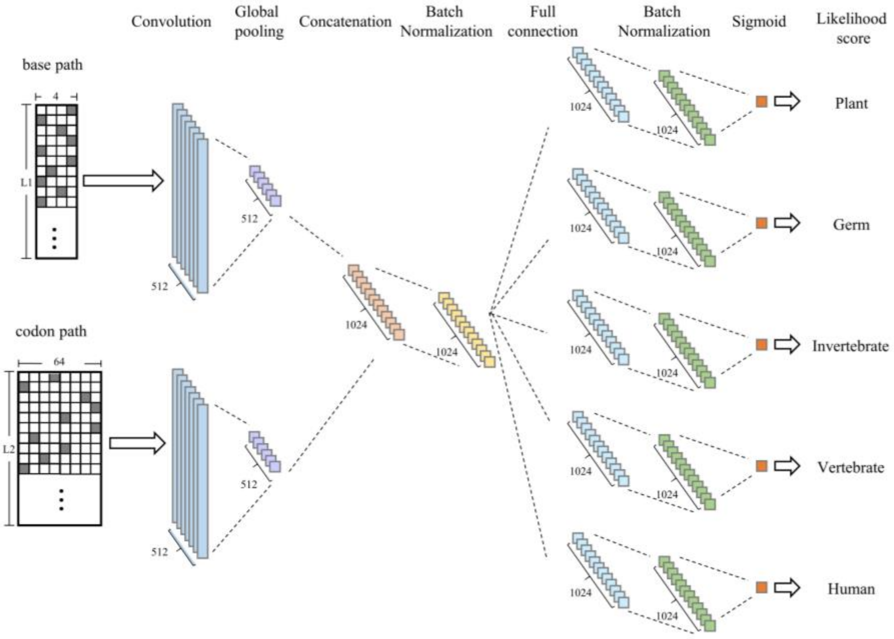


**Supplemental Figure S6 Structure of BiPathCNN in DeepHoF**

BOH matrix and COH matrix are input into two paths independently and transformed by the convolution and pooling layers at the beginning. A concatenation layer and a normalization layer combine the output of the two paths. Five sub-paths process the combined intermediate output individually. Each sub-path contains a full connection layer, a normalization layer and an output layer with sigmoid activation and binary cross-entropy loss function. The five sub-paths output the host likelihood scores on five host types respectively.
