## Supplemental Figure S8 for "Predicting Hosts Based on Early SARS-CoV-2 Samples and Analyzing Later World-wide Pandemic in 2020"

We gratefully acknowledge the following Authors from the Originating laboratories responsible for obtaining the specimens, as well as the Submitting laboratories where the genome data were generated and shared via GISAID, on which this research is based.

All Submitters of data may be contacted directly via www.gisaid.org

| Accession ID | Originating Laboratory | Submitting Laboratory | Authors |
|---|---|---|---|
| EPI_ISL_547877 | Maximum Containment Laboratory, National Institute of Virology | Maximum Containment Laboratory, National Institute of Virology | Yadav,P.D., Shete-aich,A., Nyayanit,D.A., Abraham,P. |

We gratefully acknowledge the following Authors from the Originating laboratories responsible for obtaining the specimens, as well as the Submitting laboratories where the genome data were generated and shared via GISAID, on which this research is based.

All Submitters of data may be contacted directly via www.gisaid.org

| Accession ID | Originating Laboratory | Submitting Laboratory | Authors |
|---|---|---|---|
| EPI_ISL_402119 | National Institute for Viral Disease Control and Prevention, China CDC | National Institute for Viral Disease Control and Prevention, China CDC | Wenjie TanXiang ZhaoWenling WangXuejun MaYongzhong JiangRoujian Lu, Ji Wang, Weimin ZhouPeihua NiuPeipei LiuFaxian ZhanWeifeng ShiBaoying HuangJun LiuLi ZhaoYao MengXiaozhou HeFei YeNa ZhuYang LiJing ChenWenbo XuGeorge F. GaoGuizhen Wu |
| EPI_ISL_402120 | National Institute for Viral Disease Control and Prevention, China CDC | National Institute for Viral Disease Control and Prevention, China CDC | Wenjie TanXiang ZhaoWenling WangXuejun MaYongzhong JiangRoujian LuJi WangWeimin ZhouPeihua NiuPeipei LiuFaxian ZhanWeifeng ShiBaoying HuangJun LiuLi ZhaoYao MengXiaozhou HeFei YeNa ZhuYang LiJing ChenWenbo XuGeorge F. GaoGuizhen Wu |
| EPI_ISL_402121 | National Institute for Viral Disease Control and Prevention, China CDC | National Institute for Viral Disease Control and Prevention, China CDC | Wenjie TanXuejun MaXiang ZhaoWenling WangYongzhong JiangRoujian LuJi WangPeihua Niu, Weimin Zhou, Faxian ZhanWeifeng ShiBaoying HuangJun LiuLi ZhaoYao MengFei YeNa Zhu, Xiaozhou HePeipei Liu, Yang LiJing ChenWenbo Xu George F. GaoGuizhen Wu |
| EPI_ISL_402123 | Institute of Pathogen Biology, Chinese Academy of Medical Sciences & Peking Union Medical College | Institute of Pathogen Biology, Chinese Academy of Medical Sciences & Peking Union Medical College | Lili Ren, Jianwei Wang, Qi Jin, Zichun Xiang, Zhiqiang Wu, Chao Wu, Yiwei Liu |
| EPI_ISL_402124 | Wuhan Jinyintan Hospital | Wuhan Institute of Virology, Chinese Academy of Sciences | Peng Zhou, Xing-Lou Yang, Ding-Yu Zhang, Lei Zhang, Yan Zhu, Hao-Rui Si, Zhengli Shi |
| EPI_ISL_402125 | National Institute for Communicable Disease Control and Prevention (ICDC) Chinese Center for Disease Control and Prevention (China CDC) | National Institute for Communicable Disease Control and Prevention (ICDC) Chinese Center for Disease Control and Prevention (China CDC) | Zhang,Y.-Z., Wu,F., Chen,Y.-M., Pei,Y.-Y., Xu,L., Wang,W., Zhao,S., Yu,B., Hu,Y., Tao,Z.-W., Song,Z.-G., Tian,J.-H., Zhang,Y.-L., Liu,Y., Zheng,J.-J., Dai,F.-H., Wang,Q.-M., She,J.-L. and Zhu,T.-Y. |
| EPI_ISL_402126 | Dept. of Virology III, National Institute of Infectious Diseases | Dept. of Virology III, National Institute of Infectious Diseases | Naganori Nao, Kazuya Shirato, Shutoku Matsuyama, Makoto Takeda |
| EPI_ISL_402127, EPI_ISL_402128, EPI_ISL_402129, EPI_ISL_402130 | Wuhan Jinyintan Hospital | Wuhan Institute of Virology, Chinese Academy of Sciences | Peng Zhou, Xing-Lou Yang, Ding-Yu Zhang, Lei Zhang, Yan Zhu, Hao-Rui Si, Zhengli Shi |
| EPI_ISL_402131 | Wuhan Institute of Virology, Chinese Academy of Sciences | Wuhan Institute of Virology, Chinese Academy of Sciences | Yan Zhu, Ping Yu, Bei Li, Ben Hu, Hao-Rui Si, Xing-Lou Yang, Peng Zhou, Zheng-Li Shi |
| EPI_ISL_402132 | Wuhan Jinyintan Hospital | Hubei Provincial Center for Disease Control and Prevention | Bin Fang, Xiang Li, Xiao Yu, Linlin Liu, Bo Yang, Faxian Zhan, Guojun Ye, Xixiang Huo, Junqiang Xu, Bo Yu, Kun Cai, Jing Li, Yongzhong Jiang. |
| EPI_ISL_403928, EPI_ISL_403929, EPI_ISL_403930, EPI_ISL_403931 | Institute of Pathogen Biology, Chinese Academy of Medical Sciences & Peking Union Medical College | Institute of Pathogen Biology, Chinese Academy of Medical Sciences & Peking Union Medical College | Lili Ren, Jianwei Wang, Qi Jin, Zichun Xiang, Zhiqiang Wu, Chao Wu, Yiwei Liu |
| EPI_ISL_403932, EPI_ISL_403933, EPI_ISL_403934, EPI_ISL_403935, EPI_ISL_403936, EPI_ISL_403937 | Guangdong Provincial Center for Diseases Control and Prevention; Guangdong Provincial Public Health | Department of Microbiology, Guangdong Provincial Center for Diseases Control and Prevention | Min Kang, Jie Wu, Jing Lu, Tao Liu, Baisheng Li, Shujiang Mei, Feng Ruan, Lifeng Lin, Changwen Ke, Haojie Zhong, Yingtao Zhang, Lirong Zou, Xuguang Chen, Qi Zhu, Jianpeng Xiao, Jianxiang Geng, Zhe Liu, Jianxiong Hu, Weilin Zeng, Xing Li, Yuhuang Liao, Xiujuan Tang, Songjian Xiao, Ying Wang, Yingchao Song, Xue Zhuang, Lijun Liang, Guanhao He, Huihong Deng, Tie Song, Jianfeng He, Wenjun Ma |
| EPI_ISL_403962, EPI_ISL_403963 | Bamrasnaradura Hospital | 1. Department of Medical Sciences, Ministry of Public Health, Thailand 2. Thai Red Cross Emerging Infectious Diseases - Health Science Centre 3. Department of Disease Control, Ministry of Public Health, Thailand | Pilailuk,Okada; Siripaporn,Phuygun; Thanutsapa,Thanadachakul; Supaporn,Wacharapluesadee; Sittiporn,Parnmen; Warawan,Wongboot; Sunthareeya,Waicharoen; Rome,Buathong; Malinee,Chittaganpitch; Nanthawan,Mekha |
| EPI_ISL_404227 | Zhejiang Provincial Center for Disease Control and Prevention | Department of Microbiology, Zhejiang Provincial Center for Disease Control and Prevention | Yin Chen, Yanjun Zhang, Haiyan Mao, Junhang Pan, Xiuyu Lou, Yiyu Lu, Juying Yan, Hanping Zhu, Jian Gao, Yan Feng, Yi Sun, Hao Yan, Zhen Li, Yisheng Sun, Liming Gong, Qiong Ge, Wen Shi, Xinying Wang, Wenwu Yao, Zhangnv Yang, Fang Xu, Chen Chen, Enfu Chen, Zhen Wang, Zhiping Chen, Jianmin Jiang, Chonggao Hu |
| EPI_ISL_404228 | Zhejiang Provincial Center for Disease Control and Prevention | Department of Microbiology, Zhejiang Provincial Center for Disease Control and Prevention | Yanjun Zhang, Yin Chen, Haiyan Mao, Junhang Pan, Xiuyu Lou, Yiyu Lu, Juying Yan, Hanping Zhu, Jian Gao, Yan Feng, Yi Sun, Hao Yan, Zhen Li, Yisheng Sun, Liming Gong, Qiong Ge, Wen Shi, Xinying Wang, Wenwu Yao, Zhangnv Yang, Fang Xu, Chen Chen, Enfu Chen, Zhen Wang, Zhiping Chen, Jianmin Jiang, Chonggao Hu |
| EPI_ISL_404253 | IL Department of Public Health Chicago Laboratory | Pathogen Discovery, Respiratory Viruses Branch, Division of Viral Diseases, Centers for Dieases Control and Prevention | Ying Tao, Krista Queen, Clinton R. Paden, Jing Zhang, Yan Li, Anna Uehara, Xiaoyan Lu, Brian Lynch, Senthil Kumar K. Sakthivel, Brett L. Whitaker, Shifaq Kamili, Lijuan Wang, Janna' R. Murray, Susan I. Gerber, Stephen Lindstrom, Suxiang Tong |
| EPI_ISL_404895 | Providence Regional Medical Center | Division of Viral Diseases, Centers for Disease Control and Prevention | Queen,K., Tao,Y., Li,Y., Paden,C.R., Lu,X., Zhang,J., Gerber,S.I., Lindstrom,S., Tong,S. |
| EPI_ISL_405839, EPI_ISL_406030 | The University of Hong Kong - Shenzhen Hospital | Li Ka Shing Faculty of Medicine, The University of Hong Kong | Chan,J.F.-W., Yuan,S., Kok,K.H., To,K.K.-W., Chu,H., Yang,J., Xing,F., Liu,J., Yip,C.C.-Y., Poon,R.W.-S., Tsai,H.W., Lo,S.K.-F., Chan,K.H., Poon,V.K.-M., Chan,W.M., Ip,J.D., Cai,J.P., Cheng,V.C.-C., Chen,H., Hui,C.K.-M. and Yuen,K.Y. |
| EPI_ISL_406031 | Centers for Disease Control, R.O.C. (Taiwan) | Centers for Disease Control, R.O.C. (Taiwan) | Ji-Rong Yang, Yu-Chi Lin, Jung-Jung Mu, Ming-Tsan Liu, Shu-Ying Li |
| EPI_ISL_406034 | California Department of Public Health | Pathogen Discovery, Respiratory Viruses Branch, Division of Viral Diseases, Centers for Dieases Control and Prevention | Anna Uehara, Krista Queen, Ying Tao, Yan Li, Clinton R. Paden, Jing Zhang, Xiaoyan Lu, Brian Lynch, Senthil Kumar K. Sakthivel, Brett L. Whitaker, Shifaq Kamili, Lijuan Wang, Janna' R. Murray, Susan I. Gerber, Stephen Lindstrom, Suxiang Tong |
| EPI_ISL_406036 | California Department of Public Health | Pathogen Discovery, Respiratory Viruses Branch, Division of Viral Diseases, Centers for Dieases Control and Prevention | Anna Uehara, Krista Queen, Ying Tao, Yan Li, Clinton R. Paden, Jing Zhang, Xiaoyan Lu, Brian Lynch, Senthil Kumar K. Sakthivel, Brett L. Whitaker, Shifaq Kamili, Lijuan Wang, Janna' R. Murray, Susan I. Gerber, Stephen Lindstrom, Suxiang Tong |
| EPI_ISL_406223 | Arizona Department of Health Services | Pathogen Discovery, Respiratory Viruses Branch, Division of Viral Diseases, Centers for Dieases Control and Prevention | Ying Tao, Clinton R. Paden, Krista Queen, Anna Uehara, Yan Li, Jing Zhang, Xiaoyan Lu, Brian Lynch, Senthil Kumar K. Sakthivel, Brett L. Whitaker, Shifaq Kamili, Lijuan Wang, Janna' R. Murray, Susan I. Gerber, Stephen Lindstrom, Suxiang Tong |
| EPI_ISL_406531 | Guangdong Provincial Center for Diseases Contorl and Prevention; Guangdong Provinical Public Health | Guangdong Provincial Center for Disease Control and Prevention | Min Kang, Jie Wu, Jing Lu, Tao Liu, Baisheng Li, Shujiang Mei, Feng Ruan, Lifeng Lin, Changwen Ke, Haojie Zhong, Yingtao Zhang, Lirong Zou, Xuguang Chen, Qi Zhu, Jianpeng Xiao, Jianxiang Geng, Zhe Liu, Jianxiong Hu, Weilin Zeng, Xing Li, Yuhuang Liao, Xiujuan Tang, Songjian Xiao, Ying Wang, Yingchao Song, Xue Zhuang, Lijun Liang, Guanhao He, Huihong Deng, Tie Song, Jianfeng He, Wenjun Ma |
| EPI_ISL_406533 | Guangdong Provincial Center for Diseases Contorl and Prevention; Guangdong Provinical Public Health | Guangdong Provincial Center for Diseases Control and Prevention | Min Kang, Jie Wu, Jing Lu, Tao Liu, Baisheng Li, Shujiang Mei, Feng Ruan, Lifeng Lin, Changwen Ke, Haojie Zhong, Yingtao Zhang, Lirong Zou, Xuguang Chen, Qi Zhu, Jianpeng Xiao, Jianxiang Geng, Zhe Liu, Jianxiong Hu, Weilin Zeng, Xing Li, Yuhuang Liao, Xiujuan Tang, Songjian Xiao, Ying Wang, Yingchao Song, Xue Zhuang, Lijun Liang, Guanhao He, Huihong Deng, Tie Song, Jianfeng He, Wenjun Ma |
| EPI_ISL_406534, EPI_ISL_406535, EPI_ISL_406536 | Guangdong Provincial Center for Diseases Control and Prevention; Guangdong Provinical Public Health | Guangdong Provincial Center for Diseases Control and Prevention | Min Kang, Jie Wu, Jing Lu, Tao Liu, Baisheng Li, Shujiang Mei, Feng Ruan, Lifeng Lin, Changwen Ke, Haojie Zhong, Yingtao Zhang, Lirong Zou, Xuguang Chen, Qi Zhu, Jianpeng Xiao, Jianxiang Geng, Zhe Liu, Jianxiong Hu, Weilin Zeng, Xing Li, Yuhuang Liao, Xiujuan Tang, Songjian Xiao, Ying Wang, Yingchao Song, Xue Zhuang, Lijun Liang, Guanhao He, Huihong Deng, Tie Song, Jianfeng He, Wenjun Ma |
| EPI_ISL_406538 | Guangdong Provincial Center for Diseases Control and Prevention;Guangdong Provincial Institute of Public Health | Guangdong Provincial Center for Diseases Control and Prevention | Min Kang, Jie Wu, Jing Lu, Tao Liu, Baisheng Li, Shujiang Mei, Feng Ruan, Lifeng Lin, Changwen Ke, Haojie Zhong, Yingtao Zhang, Lirong Zou, Xuguang Chen, Qi Zhu, Jianpeng Xiao, Jianxiang Geng, Zhe Liu, Jianxiong Hu, Weilin Zeng, Xing Li, Yuhuang Liao, Xiujuan Tang, Songjian Xiao, Ying Wang, Yingchao Song, Xue Zhuang, Lijun Liang, Guanhao He, Huihong Deng, Tie Song, Jianfeng He, Wenjun Ma |
| EPI_ISL_406592 | Shenzhen Third People's Hospital | Shenzhen Key Laboratory of Pathogen and Immunity, National Clinical Research Center for Infectious Disease,Shenzhen Third People's Hospital | Yang Yang, Chenguang Shen, Li Xing, Zhixiang Xu, Haixia Zheng, Yingxia Liu |
| EPI_ISL_406593, EPI_ISL_406594, EPI_ISL_406595 | Shenzhen Key Laboratory of Pathogen and Immunity, National Clinical Research Center for Infectious Disease, Shenzhen Third People's Hospital | Shenzhen Key Laboratory of Pathogen and Immunity, National Clinical Research Center for Infectious Disease, Shenzhen Third People's Hospital | Yang Yang, Chenguang Shen, Li Xing, Zhixiang Xu, Haixia Zheng, Yingxia Liu |
| EPI_ISL_406596, EPI_ISL_406597 | Department of Infectious and Tropical Diseases, Bichat Claude Bernard Hospital, Paris | National Reference Center for Viruses of Respiratory Infections, Institut Pasteur, Paris | Mélanie Albert, Marion Barbet, Sylvie Behillil, Méline Bizard, Angela Brisebarre, Flora Donati, Vincent Enouf, Maud Vanpeene, Sylvie van der Werf, Yazdan Yazdanpanah, Xavier Lescure. |
| EPI_ISL_406716, EPI_ISL_406717 | State Key Laboratory of Virology, Wuhan University | State Key Laboratory of Virology, Wuhan University | Chen,L., Liu,W., Zhang,Q., Xu,K., Ye,G., Wu,W., Sun,Z., Liu,F., Wu,K., Mei,Y., Zhang,W., Chen,Y., Li,Y., Shi,M., Lan,K. and Liu,Y. |
| EPI_ISL_406798, EPI_ISL_406799, EPI_ISL_406800, EPI_ISL_406801 | General Hospital of Central Theater Command of People's Liberation Army of China | BGI & Institute of Microbiology, Chinese Academy of Sciences & Shandong First Medical University & Shandong Academy of Medical Sciences & General Hospital of Central Theater Command of People's Liberation Army of China | Weijun Chen, Yuhai Bi, Weifeng Shi and Zhenhong Hu |
| EPI_ISL_406844 | Monash Medical Centre | Collaboration between the University of Melbourne at The Peter Doherty Institute for Infection and Immunity, and the Victorian Infectious Disease Reference Laboratory | Caly,L., Seemann,T., Schultz,M., Druce,J. and Taiaroa,G |
| EPI_ISL_406862 | Charité Universitätsmedizin Berlin, Institute of Virology; Institut für Mikrobiologie der Bundeswehr, Munich | Charité Universitätsmedizin Berlin, Institute of Virology | Victor M Corman, Julia Schneider, Talitha Veith, Barbara Mühlemann, Markus Antwerpen, Christian Drosten, Roman Wölfel |
| EPI_ISL_406959 | Virology Laboratory, INMI L. Spallanzani | Virology Laboratory, INMI L. Spallanzani | Capobianchi,M.R., Carletti,F., Lalle,E., Bordi,L., Marsella,P., Colavita,F., Matusali,G., Nicastri,E., Ippolito,G. and Castilletti,C. |
| EPI_ISL_406960 | Virology Laboratory, INMI L. Spallanzani | Virology Laboratory, INMI L. Spallanzani | Capobianchi,M.R., Lalle,E., Carletti,F., Bordi,L., Marsella,P., Colavita,F., Matusali,G., Nicastri,E., Ippolito,G. and Castilletti,C. |
| EPI_ISL_406970 | Hangzhou Center for Disease and Control Microbiology Lab | Hangzhou Center for Disease and Control Microbiology Lab | Yu Hua, Wang Haoqiu, Li Jun, Yu Xinfeng |
| EPI_ISL_406973 | Singapore General Hospital | National Public Health Laboratory | Mak, TM; Octavia S; Chavatte JM; Zhou, ZY; Cui, L; Lin, RTP |
| EPI_ISL_407071 | Respiratory Virus Unit, Microbiology Services Colindale, Public Health England | Respiratory Virus Unit, Microbiology Services Colindale, Public Health England | Monica Galiano, Shahjahan Miah, Richard Myers, Angie Lackenby, Omolola Akinbami, Tiina Talts, Leena Bhaw, Kirstin Edwards, Jonathan Hubb, Joanna Ellis, Maria Zambon |
| EPI_ISL_407073 | Respiratory Virus Unit, Microbiology Services | Respiratory Virus Unit, Microbiology Services Colindale, Public | Monica Galiano, Shahjahan Miah, Richard Myers, Angie Lackenby, Omolola Akinbami, Tiina Talts, Leena Bhaw, Kirstin Edwards, Jonathan Hubb, Joanna Ellis, Maria Zambon. |

| | | | |
|---|---|---|---|
| | Colindale, Public Health England | Health England | |
| EPI_ISL_407079 | Lapland Central Hospital | Department of Virology, University of Helsinki and Helsinki University Hospital, Helsinki, Finland | Teemu Smura, Suvi Kuivanen, Hannimari Kallio-Kokko, Olli Vapalahti |
| EPI_ISL_407084 | Department of Virology III, National Institute of Infectious Diseases | Pathogen Genomics Center, National Institute of Infectious Diseases | Tsuyoshi Sekizuka, Shutoku Matsuyama, Naganori Nao, Kazuya Shirato, Shinji Watanabe, Makoto Takeda, Makoto Kuroda |
| EPI_ISL_407193 | Korea Centers for Disease Control & Prevention (KCDC) Center for Laboratory Control of Infectious Diseases Division of Viral Diseases | Korea Centers for Disease Control & Prevention (KCDC) Center for Laboratory Control of Infectious Diseases Division of Viral Diseases | Jeong-Min Kim, Yoon-Seok Chung, Namjoo Lee, Mi-Seon Kim, SangHee Woo, Hye-Joon Jo, Sehee Park, Heui Man Kim, Myung Guk Han |
| EPI_ISL_407214, EPI_ISL_407215 | Washington State Department of Health | Pathogen Discovery, Respiratory Viruses Branch, Division of Viral Diseases, Centers for Dieases Control and Prevention | Krista Queen, Azaibi Tamin, Jennifer Harcourt, Ying Tao, Clinton R. Paden, Jing Zhang, Yan Li, Anna Uehara, Xiaoyan Lu, Shifaq Kamili, Rashi Gautam, Haibin Wang, Janna' R. Murray, Susan I. Gerber, Stephen Lindstrom, Natalie Thornburg, Suxiang Tong |
| EPI_ISL_407313 | Hangzhou Center for Disease Control and Prevention | Hangzhou Center for Disease Control and Prevention | Jun Li, Haoqiu Wang, Hua Yu, Lingfeng Mao, Xinfen Yu, Zhou Sun, Qingxin Kong, Xin Qian, Shuchang Chen, Xuchu Wang |
| EPI_ISL_407893 | Centre for Infectious Diseases and Microbiology Laboratory Services | NSW Health Pathology - Institute of Clinical Pathology and Medical Research; Westmead Hospital; University of Sydney | Eden J-S, Carter I, Rahman H, Holmes EC, Rockett R, O'Sullivan MV, Sintchenko V, Chen SC, Maddocks S, Kok J and Dwyer DE for the 2019-nCoV Study Group |
| EPI_ISL_407894, EPI_ISL_407896 | Pathology Queensland | Public Health Virology Laboratory | Ben Huang, Alyssa Pyke, Amanda De Jong, Andrew Van Den Hurk, Carmel Taylor, David Warrilow, Doris Genge, Elisabeth Gamez, Glen Hewitson, Ian Maxwell Mackay, Inga Sultana, Jamie McMahon, Jean Barcelon, Judy Northill, Mitchell Finger, Natalie Simpson, Neelima Nair, Peter Burtonclay, Peter Moore, Sarah Wheatley, Sean Moody, Sonja Hall-Mendelin, Timothy Gardam, and Frederick Moore. |
| EPI_ISL_407976 | KU Leuven, Clinical and Epidemiological Virology | KU Leuven, Clinical and Epidemiological Virology | Bert Vanmechelen, Elke Wollants, Annabel Rector, Els Keyaerts, Lies Laenen, Marc Van Ranst, and Piet Maes |
| EPI_ISL_407987 | Singapore General Hospital | Programme in Emerging Infectious Diseases, Duke-NUS Medical School | Danielle E Anderson, Martin Linster, Yan Zhuang, Jayanthi Jayakumar, Kian Sing Chan, Lynette LE Oon, Jenny GH Low, Yvonne CF Su, Linfa Wang, Gavin JD Smith |
| EPI_ISL_407988 | National Centre for Infectious Diseases | Programme in Emerging Infectious Diseases, Duke-NUS Medical School | Danielle E Anderson, Martin Linster, Yan Zhuang, Jayanthi Jayakumar, David CB Lye, Yee Sin Leo, Barnaby E Young, Yvonne CF Su, Linfa Wang, Gavin JD Smith |
| EPI_ISL_408008 | California Department of Health | Pathogen Discovery, Respiratory Viruses Branch, Division of Viral Diseases, Centers for Dieases Control and Prevention | Krista Queen, Jing Zhang, Yan Li, Ying Tao, Anna Uehara, Clinton Paden, Xiaoyan Lu, Brian Lynch, Senthil Kumar K. Sakthivel, Brett L. Whitaker, Shifaq Kamili, Lijuan Wang, Janna' R. Murray, Susan I. Gerber, Stephen Lindstrom, Suxiang Tong |
| EPI_ISL_408009 | California Department of Health | Pathogen Discovery, Respiratory Viruses Branch, Division of Viral Diseases, Centers for Dieases Control and Prevention | Krista Queen, Jing Zhang, Yan Li, Ying Tao, Anna Uehara, Clinton Paden, Xiaoyan Lu, Brian Lynch, Senthil Kumar K. Sakthivel, Brett L. Whitaker, Shifaq Kamili, Lijuan Wang, Janna' R. Murray, Susan I. Gerber, Stephen Lindstrom, Suxiang Tong |
| EPI_ISL_408010 | California Department of Health | Pathogen Discovery, Respiratory Viruses Branch, Division of Viral Diseases, Centers for Dieases Control and Prevention | Ying Tao, Krista Queen, Jing Zhang, Yan Li, Anna Uehara, Clinton Paden, Xiaoyan Lu, Brian Lynch, Senthil Kumar K. Sakthivel, Brett L. Whitaker, Shifaq Kamili, Lijuan Wang, Janna' R. Murray, Susan I. Gerber, Stephen Lindstrom, Suxiang Tong |
| EPI_ISL_408068 | Virology Laboratory National Institute for Infectious Diseases 'Lazzaro Spallanzani' IRCCS | Virology Laboratory National Institute for Infectious Diseases 'Lazzaro Spallanzani' IRCCS | Capobianchi,M.R., Carletti,F., Lalle,E., Bordi,L., Marsella,P.,Colavita,F., Matusali,G., Nicastri,E., Ippolito,G. and Castilletti,C. |
| EPI_ISL_408430 | Department of Infectious and Tropical Diseases, Bichat Claude Bernard Hospital, Paris | National Reference Center for Viruses of Respiratory Infections, Institut Pasteur, Paris | Mélanie Albert, Marion Barbet, Sylvie Behillil, Méline Bizard, Angela Brisebarre, Flora Donati, Vincent Enouf, Maud Vanpeene, Sylvie van der Werf, Yazdan Yazdanpanah, Xavier Lescure |
| EPI_ISL_408431 | Sorbonne Université, Inserm et Assistance Publique-Hôpitaux de Paris (Pitié Salpétrière) | National Reference Center for Viruses of Respiratory Infections, Institut Pasteur, Paris | Mélanie Albert, Marion Barbet, Sylvie Behillil, Méline Bizard, Angela Brisebarre, Flora Donati, Vincent Enouf, Maud Vanpeene, Sylvie van der Werf, Sonia Burrel, Anne-Geneviève Marcelin, Vincent Calvez, David Boutolleau, Elise Klément, Valérie Pourcher, Eric Caumes. |
| EPI_ISL_408478 | Yongchuan District Center for Disease Control and Prevention | Chongqing Municipal Center for Disease Control and Prevention | Ye Sheng, Tang Yun, Ling Hua,Yu zhen,Chen Shuang,Tan ZhangPing, Su Kun, Li Qing, Tang Wenge, Rong Rong |
| EPI_ISL_408479 | Zhongxian Center for Disease Control and Prevention | Chongqing Municipal Center for Disease Control and Prevention | Ye Sheng, Tang Yun, Ling Hua, Zhang Hong, Yu zhen,Chen Shuang,Tan ZhangPing, Su Kun, Li Qin, Tang Wenge, Rong Rong |
| EPI_ISL_408480 | National Institute for Viral Disease Control and Prevention, China CDC | National Institute for Viral Disease Control & Prevention, CCDC | Wenjie TanXiaoqing FuXiang ZhaoWenling Wang Peihua NiuRoujian Lu,Yanhong SunBaoying HuangLi ZhaoFei YeWenbo XuGeorge F. GaoGuizhen Wu |
| EPI_ISL_408481 | National Institute for Viral Disease Control and Prevention, China CDC | National Institute for Viral Disease Control & Prevention, CCDC | Wenjie Tan, Hengqin Wang, Xiang Zhao, Wenling Wang, Peihua Niu, Roujian Lu, Sheng Ye, Baoying Huang, Li Zhao, Fei Ye, Wenbo Xu, George F. Gao, Guizhen Wu |
| EPI_ISL_408482 | National Institute for Viral Disease Control and Prevention, China CDC | National Institute for Viral Disease Control & Prevention, CCDC | Wenjie Tan, Zhaoguo Wang, Xiang Zhao, Wenling Wang, Peihua Niu, Roujian Lu, Ti Liu, Baoying Huang, Li Zhao, Fei Ye, Wenbo Xu, George F. Gao, Guizhen Wu |
| EPI_ISL_408483 | National Institute for Viral Disease Control and Prevention, China CDC | National Institute for Viral Disease Control & Prevention, CCDC | Wenjie TanZhen Teng,Xiang ZhaoWenling Wang Peihua NiuRoujian Lu,Chongshan Li,Baoying HuangLi ZhaoFei YeWenbo XuGeorge F. GaoGuizhen Wu |
| EPI_ISL_408484 | National Institute for Viral Disease Control and Prevention, China CDC | National Institute for Viral Disease Control & Prevention, CCDC | Wenjie Tan, Jianan Xu, Wenling Wang, Peihua Niu, Roujian Lu, Huiping Yang, Xiang Zhao, Baoying Huang, Li Zhao, Fei Ye, Wenbo Xu, George F. Gao, Guizhen Wu |
| EPI_ISL_408485 | National Institute for Viral Disease Control and Prevention, China CDC | National Institute for Viral Disease Control & Prevention, CCDC | Wenjie Tan,Quanyi Wang,Wenling Wang, Peihua Niu,Roujian Lu,Yang Pan,Xiang Zhao,Baoying Huang,Li Zhao,Fei Ye,Wenbo Xu,George F. Gao,Guizhen Wu |
| EPI_ISL_408486 | National Institute for Viral Disease Control and Prevention, China CDC | National Institute for Viral Disease Control & Prevention, CCDC | Wenjie Tan, Yong Shi, Wenling Wang, Peihua Niu, Roujian Lu, Jianxiong Li, Xiang Zhao, Baoying Huang, Li Zhao, Fei Ye, Wenbo Xu, George F. Gao, Guizhen Wu |
| EPI_ISL_408487 | National Institute for Viral Disease Control and Prevention, China CDC | National Institute for Viral Disease Control & Prevention, China CDC | Wenjie Tan, Jin Xu, Wenling Wang, Peihua Niu, Roujian Lu, Xueyong Huang, Xiang Zhao, Baoying Huang, Li Zhao, Fei Ye, Wenbo Xu, George F. Gao, Guizhen Wu |
| EPI_ISL_408488 | National Institute for Viral Disease Control and Prevention, China CDC | National Institute for Viral Disease Control & Prevention, CCDC | Wenjie Tan, Shenjiao Wang, Wenling Wang, Peihua Niu, Roujian Lu, Kangchen Zhao, Xiang Zhao, Baoying Huang, Li Zhao, Fei Ye, Wenbo Xu, George F. Gao, Guizhen Wu |
| EPI_ISL_408489 | Department of Laboratory Medicine, National Taiwan University Hospital | Microbial Genomics Core Lab, National Taiwan University Centers of Genomic and Precision Medicine | Shiou-Hwei Yeh, You-Yu Lin, Ya-Yun Lai, Chiao-Ling Li, Shan-Chwen Chang, Pei-Jer Chen, Sui-Yuan Chang |
| EPI_ISL_408511, EPI_ISL_408512, EPI_ISL_408513 | Institute of Viral Disease Control and Prevention, China CDC | Institute of Viral Disease Control and Prevention, China CDC | William J. Liu, Peipei Liu, Xiang Zhao, Peihua Niu, Yingze Zhao, Wenwen Lei, Ziqian Xu, Beiwei Ye, Weifeng Shi, Roujian Lu, Wenjie Tan, Zhixiao Chen, Yuchao Wu, Juan Song, Dayan Wang, Jun Han, Wenbo Xu, George F. Gao, Guizhen Wu |
| EPI_ISL_408514, EPI_ISL_408515 | Institute of Viral Disease Control and Prevention, China CDC | Institute of Viral Disease Control and Prevention, China CDC | William J. Liu, Peipei Liu, Xiang Zhao, Peihua Niu, Yingze Zhao, Wenwen Lei, Ziqian Xu, Shumei Zou, Wei Zhen, Beiwei Ye, Mengjie Yang, Weifeng Shi, Roujian Lu, Wenjie Tan, Zhixiao Chen, Yuchao Wu, Juan Song, Weimin Zhou, Dayan Wang, Jun Han, Wenbo Xu, George F. Gao, Guizhen Wu |
| EPI_ISL_408665, EPI_ISL_408666, EPI_ISL_408667 | Dept. of Virology III, National Institute of Infectious Diseases | Pathogen Genomics Center, National Institute of Infectious Diseases | Tsuyoshi Sekizuka, Shutoku Matsuyama, Naganori Nao, Kazuya Shirato, Makoto Takeda, Makoto Kuroda |
| EPI_ISL_408668 | National Influenza Center - National Institute of Hygiene and Epidemiology (NIHE) | National Influenza Center - National Institute of Hygiene and Epidemiology (NIHE) | Ung Thi Hong Trang, Hoang Vu Mai Phuong, Nguyen Le Khanh Hang, Nguyen Vu Son, Le Thi Thanh, Vuong Duc Cuong, Nguyen Phuong Anh, Pham Thi Hien, Tran Thu Huong, Le Thi Quynh Mai, |
| EPI_ISL_408669 | Dept. of Virology III, National Institute of Infectious Diseases | Pathogen Genomics Center, National Institute of Infectious Diseases | Tsuyoshi Sekizuka, Shutoku Matsuyama, Naganori Nao, Kazuya Shirato, Makoto Takeda, Makoto Kuroda |
| EPI_ISL_408670 | Wisconsin Department of Health Services | Pathogen Discovery, Respiratory Viruses Branch, Division of Viral Diseases, Centers for Dieases Control and Prevention | Jing Zhang, Anna Uehara, Krista Queen, Yan Li, Ying Tao, Clinton R. Paden, Xiaoyan Lu, Brian Lynch, Senthil Kumar K. Sakthivel, Brett L. Whitaker, Shifaq Kamili, Lijuan Wang, Janna' R. Murray, Susan I. Gerber, Stephen Lindstrom, Suxiang Tong |
| EPI_ISL_408975 | Queen Elizabeth Hospital | Hong Kong Department of Health | Mak Gannon C.K., Cheng Peter K.C., Lam Edman T.K., Chan Rickjason C.W., Tsang Dominic N.C. |
| EPI_ISL_408976 | Centre for Infectious Diseases and Microbiology Laboratory Services | NSW Health Pathology - Institute of Clinical Pathology and Medical Research; Westmead Hospital; University of Sydney | Rockett R, Sadsad R, Eden J-S, Carter I, Rahman H, Holmes EC, O'Sullivan MV, Sintchenko V, Chen SC, Maddocks S, Kok J and Dwyer DE for the 2019-nCoV Study Group* |
| EPI_ISL_408977 | Serology, Virology and OTDS Laboratories (SAViD), NSW Health Pathology Randwick | NSW Health Pathology - Institute of Clinical Pathology and Medical Research; Centre for Infectious Diseases and Microbiology Laboratory Services; Westmead Hospital; University of Sydney | Eden J-S, Carter I, Rahman H, Rawlinson W, Holmes EC, Rockett R, O'Sullivan MV, Sintchenko V, Chen SC, Maddocks S, Kok J and Dwyer DE for the 2019-nCoV Study Group* |
| EPI_ISL_408978 | Wuhan Fourth Hospital | Beijing Genomics Institute (BGI) | Weijun Chen |
| EPI_ISL_408994 | Prince of Wales Hospital | Hong Kong Department of Health | Mak Gannon C.K., Cheng Peter K.C., Lam Edman T.K., Chan Rickjason C.W., Tsang Dominic N.C. |
| EPI_ISL_408995 | Tuen Mun Hospital | Hong Kong Department of Health | Mak Gannon C.K., Cheng Peter K.C., Lam Edman T.K., Chan Rickjason C.W., Tsang Dominic N.C. |
| EPI_ISL_408996 | Prince of Wales Hospital | Hong Kong Department of Health | Mak Gannon C.K., Cheng Peter K.C., Lam Edman T.K., Chan Rickjason C.W., Tsang Dominic N.C. |
| EPI_ISL_408997 | Prince of Wales Hospital | Hong Kong Department of Health | Mak Gannon C.K., Cheng Peter K.C., Lam Edman T.K., Chan Rickjason C.W., Tsang Dominic N.C. |
| EPI_ISL_408998 | Ruttonjee Hospital | Hong Kong Department of Health | Mak Gannon C.K., Cheng Peter K.C., Lam Edman T.K., Chan Rickjason C.W., Tsang Dominic N.C. |
| EPI_ISL_408999 | Prince of Wales Hospital | Hong Kong Department of Health | Mak Gannon C.K., Cheng Peter K.C., Lam Edman T.K., Chan Rickjason C.W., Tsang Dominic N.C. |
| EPI_ISL_409000 | Tuen Mun Hospital | Hong Kong Department of Health | Mak Gannon C.K., Cheng Peter K.C., Lam Edman T.K., Chan Rickjason C.W., Tsang Dominic N.C. |
| EPI_ISL_409001 | Queen Mary Hospital | Hong Kong Department of Health | Mak Gannon C.K., Cheng Peter K.C., Lam Edman T.K., Chan Rickjason C.W., Tsang Dominic N.C. |
| EPI_ISL_409002 | Queen Mary Hospital | Hong Kong Department of Health | Mak Gannon C.K., Cheng Peter K.C., Lam Edman T.K., Chan Rickjason C.W., Tsang Dominic N.C. |
| EPI_ISL_409020 | Queen Elizabeth Hospital | Hong Kong Department of Health | Mak Gannon C.K., Cheng Peter K.C., Lam Edman T.K., Chan Rickjason C.W., Tsang Dominic N.C. |
| EPI_ISL_409022 | Princess Margaret Hospital | Hong Kong Department of Health | Mak Gannon C.K., Cheng Peter K.C., Lam Edman T.K., Chan Rickjason C.W., Tsang Dominic N.C. |
| EPI_ISL_409023 | Queen Elizabeth Hospital | Hong Kong Department of Health | Mak Gannon C.K., Cheng Peter K.C., Lam Edman T.K., Chan Rickjason C.W., Tsang Dominic N.C. |

| | | | |
|---|---|---|---|
| EPI_ISL_409024 | Caritas Medical Centre | Hong Kong Department of Health | Mak Gannon C.K., Cheng Peter K.C., Lam Edman T.K., Chan Rickjason C.W., Tsang Dominic N.C. |
| EPI_ISL_409025 | Queen Elizabeth Hospital | Hong Kong Department of Health | Mak Gannon C.K., Cheng Peter K.C., Lam Edman T.K., Chan Rickjason C.W., Tsang Dominic N.C. |
| EPI_ISL_409026 | Queen Elizabeth Hospital | Hong Kong Department of Health | Mak Gannon C.K., Cheng Peter K.C., Lam Edman T.K., Chan Rickjason C.W., Tsang Dominic N.C. |
| EPI_ISL_409027 | Tseung Kwan O Hospital | Hong Kong Department of Health | Mak Gannon C.K., Cheng Peter K.C., Lam Edman T.K., Chan Rickjason C.W., Tsang Dominic N.C. |
| EPI_ISL_409067 | Massachusetts Department of Public Health | Pathogen Discovery, Respiratory Viruses Branch, Division of Viral Diseases, Centers for Dieases Control and Prevention | Clinton R. Paden, Jing Zhang, Krista Queen, Yan Li, Ying Tao, Anna Uehara, Xiaoyan Lu, Brian Lynch, Senthil Kumar K. Sakthivel, Brett L. Whitaker, Shifaq Kamili, Lijuan Wang, Janna' R. Murray, Susan I. Gerber, Stephen Lindstrom, Suxiang Tong |
| EPI_ISL_410044 | California Department of Public Health | Pathogen Discovery, Respiratory Viruses Branch, Division of Viral Diseases, Centers for Dieases Control and Prevention | Jing Zhang, Krista Queen, Yan Li, Ying Tao, Anna Uehara, Clinton R. Paden, Xiaoyan Lu, Brian Lynch, Senthil Kumar K. Sakthivel, Brett L. Whitaker, Shifaq Kamili, Lijuan Wang, Janna' R. Murray, Susan I. Gerber, Stephen Lindstrom, Suxiang Tong |
| EPI_ISL_410045 | IL Department of Public Health Chicago Laboratory | Pathogen Discovery, Respiratory Viruses Branch, Division of Viral Diseases, Centers for Dieases Control and Prevention | Yan Li, Jing Zhang, Krista Queen, Ying Tao, Anna Uehara, Clinton R. Paden, Xiaoyan Lu, Brian Lynch, Senthil Kumar K. Sakthivel, Brett L. Whitaker, Shifaq Kamili, Lijuan Wang, Janna' R. Murray, Susan I. Gerber, Stephen Lindstrom, Suxiang Tong |
| EPI_ISL_410218 | Department of Laboratory Medicine, National Taiwan University Hospital | Microbial Genomics Core Lab, National Taiwan University Centers of Genomic and Precision Medicine | Shiou-Hwei Yeh, You-Yu Lin, Ya-Yun Lai, Chiao-Ling Li, Shan-Chwen Chang, Pei-Jer Chen, Sui-Yuan Chang |
| EPI_ISL_410301 | National Influenza Centre, National Public Health Laboratory, Kathmandu, Nepal | The University of Hong Kong | Ranjit Sah , Runa Jha, Daniel Chu, Haogao Gu, Malik Peiris, Anup Bastola, Alfonso J. Rodriguez-Morales, Bibek Kumar Lal, Basu Dev Pandey, Leo Poon |
| EPI_ISL_410302 | Amalea Dulcene Nicolasora Research Institute for Tropical Medicine, Molecular Biology Laboratory | Amalea Dulcene Nicolasora Research Institute for Tropical Medicine, Molecular Biology Laboratory | Nicolasora,A.D., Mercado,E.S., Polotan,F.M., Manalo,J.G., Medado,I.P., Tujan,M.A., Onza,O.T. and Cruz,K.M. |
| EPI_ISL_410314 | Joanna Ina Manalo Research Institute for Tropical Medicine, Molecular Biology Laboratory | Joanna Ina Manalo Research Institute for Tropical Medicine, Molecular Biology Laboratory | Manalo,J.I. and Nicolasora,A.D. |
| EPI_ISL_410344, EPI_ISL_410345 | Joanna Ina Manalo Research Institute for Tropical Medicine, Molecular Biology Laboratory | Joanna Ina Manalo Research Institute for Tropical Medicine, Molecular Biology Laboratory | Mercado,E.S., Manalo,J.I., Nicolasora,A.D., Medado,I.P., Tujan,M.A., Onza,O.T., Cruz,K.M. and Polotan,F.M. |
| EPI_ISL_410486 | CNR Virus des Infections Respiratoires - France SUD | CNR Virus des Infections Respiratoires - France SUD | Bal, Antonin; Destras, Gregory; Gaymard, Alexandre; Bouscambert-Duchamp, Maude; Cheynet, Valérie; Brengel-Pesce, Karen; Morfin-Sherpa, Florence; Valette, Martine; Josset, Laurence; Lina, Bruno. |
| EPI_ISL_410487, EPI_ISL_410488, EPI_ISL_410489 | National Public Health Laboratory | National Public Health Laboratory | Yu Kie,C., Norazimah,T., Rehan Shuhada,A.B., Selvanesan,S., Noorliza,M.N. and Hani,M.H. |
| EPI_ISL_410531, EPI_ISL_410532 | Dept. of Pathology, National Institute of Infectious Diseases | Pathogen Genomics Center, National Institute of Infectious Diseases | Tsuyoshi Sekizuka, Harutaka Katano, Shutoku Matsuyama, Naganori Nao, Kazuya Shirato, Motoi Suzuki, Hideki Hasegawa, Takaji Wakita, Makoto Takeda, Tadaki Suzuki, Makoto Kuroda |
| EPI_ISL_410535 | National Centre for Infectious Diseases | Programme in Emerging Infectious Diseases, Duke-NUS Medical School | Danielle E Anderson, Martin Linster, Yan Zhuang, Jayanthi Jayakumar, David CB Lye, Yee Sin Leo, Barnaby E Young, Yvonne CF Su, Gavin JD Smith |
| EPI_ISL_410536, EPI_ISL_410537 | Singapore General Hospital, Molecular Laboratory, Division of Pathology | Programme in Emerging Infectious Diseases, Duke-NUS Medical School | Danielle E Anderson, Martin Linster, Yan Zhuang, Jayanthi Jayakumar, Kian Sing Chan, Lynette LE Oon, Shirin Kalimuddin, Jenny GH Low, Yvonne CF Su, Gavin JD Smith |
| EPI_ISL_410538, EPI_ISL_410539, EPI_ISL_410540, EPI_ISL_410541, EPI_ISL_410542, EPI_ISL_410543, EPI_ISL_410544 | Beijing Institute of Microbiology and Epidemiology | Beijing Institute of Microbiology and Epidemiology | Wu-Chun Cao; Tommy Tsan-Yuk Lam; Na Jia; Ya-Wei Zhang; Jia-Fu Jiang; Bao-Gui Jiang |
| EPI_ISL_410545 | INMI Lazzaro Spallanzani IRCCS | Laboratory of Virology, INMI Lazzaro Spallanzani IRCCS | Maria R. Capobianchi, Cesare E. M. Gruber, Martina Rueca, Barbara Bartolini, Francesco Messina, Emanuela Giombini, Francesca Colavita, Concetta Castilletti, Eleonora Lalle, Fabrizio Carletti, Emanuele Nicastri, Giuseppe Ippolito. |
| EPI_ISL_410546 | INMI Lazzaro Spallanzani IRCCS | Laboratory of Virology, INMI Lazzaro Spallanzani IRCCS | Maria R. Capobianchi, Cesare E. M. Gruber, Martina Rueca, Fabrizio Carletti, Barbara Bartolini, Francesco Messina, Emanuela Giombini, Francesca Colavita, Concetta Castilletti, Eleonora Lalle, Emanuele Nicastri, Giuseppe Ippolito. |
| EPI_ISL_410713, EPI_ISL_410714 | National Public Health Laboratory, National Centre for Infectious Diseases | National Public Health Laboratory, National Centre for Infectious Diseases | Octavia S, Mak TM, Cui L, Lin RTP |
| EPI_ISL_410715 | National Public Health Laboratory, National Centre for Infectious Diseases | National Public Health Laboratory, National Centre for Infectious Diseases | Octavia S, Mak TM, Cui L, Lin RTP |
| EPI_ISL_410716 | National Public Health Laboratory, National Centre for Infectious Diseases | National Centre for Infectious Diseases, National Centre for Infectious Diseases | Octavia S, Mak TM, Cui L, Lin RTP |
| EPI_ISL_410717, EPI_ISL_410718 | Pathology Queensland | Public Health Virology Laboratory | Ben Huang, Alyssa Pyke, Amanda De Jong, Andrew Van Den Hurk, Carmel Taylor, David Warrilow, Doris Genge, Elisabeth Gamez, Glen Hewitson, Ian Maxwell Mackay, Inga Sultana, Jamie McMahon, Jean Barcelon, Judy Northill, Mitchell Finger, Natalie Simpson, Neelima Nair, Peter Burtonclay, Peter Moore, Sarah Wheatley, Sean Moody, Sonja Hall-Mendelin, Timothy Gardam, and Frederick Moore. |
| EPI_ISL_410719 | National Public Health Laboratory | National Public Health Laboratory | Octavia S, Mak TM, Cui L, Lin RTP |
| EPI_ISL_410720 | Department of Infectious and Tropical Diseases, Bichat Claude Bernard Hospital, Paris | National Reference Center for Viruses of Respiratory Infections, Institut Pasteur, Paris | Mélanie Albert, Marion Barbet, Sylvie Behillil, Méline Bizard, Angela Brisebarre, Flora Donati, Vincent Enouf, Maud Vanpeene, Sylvie van der Werf, Yazdan Yazdanpanah, Xavier Lescure. |
| EPI_ISL_410721 | South China Agricultural University | South China Agricultural University | Yongyi Shen, Lihua Xiao, Wu Chen |
| EPI_ISL_410984 | Department of Infectious and Tropical Diseases, Bichat Claude Bernard Hospital, Paris | National Reference Center for Viruses of Respiratory Infections, Institut Pasteur, Paris | Mélanie Albert, Marion Barbet, Sylvie Behillil, Méline Bizard, Angela Brisebarre, Flora Donati, Vincent Enouf, Maud Vanpeene, Sylvie van der Werf, Yazdan Yazdanpanah, Xavier Lescure |
| EPI_ISL_411060, EPI_ISL_411066 | Fujian Center for Disease Control and Prevention | Fujian Center for Disease Control and Prevention | Chen Wei, Zhang Yanhua, He Wenxiang, Weng Yuwei |
| EPI_ISL_411218 | Department of Infectious and Tropical Diseases, Bichat Claude Bernard Hospital, Paris | Laboratoire Virpath, CIRI U111, UCBL1, INSERM, CNRS, ENS Lyon | Olivier Terrier, Aurélien Traversier, Julien Fouret, Yazdan Yazdanpanah, Xavier Lescure, Catherine Legras-Lachuer, Alexandre Gaymard, Bruno Lina, Manuel Rosa-Calatrava |
| EPI_ISL_411219 | Department of Infectious and Tropical Diseases, Bichat Claude Bernard Hospital, Paris | Laboratoire Virpath, CIRI U111, UCBL1, INSERM, CNRS, ENS Lyon | Olivier Terrier, Aurélien Traversier, Julien Fouret, Yazdan Yazdanpanah, Xavier Lescure, Alexandre Gaymard, Bruno Lina, Manuel Rosa-Calatrava |
| EPI_ISL_411220 | Department of Infectious and Tropical Diseases, Bichat Claude Bernard Hospital, Paris | Laboratoire Virpath, CIRI U111, UCBL1, INSERM, CNRS, ENS Lyon | Olivier Terrier, Aurélien Traversier, Julien Fouret, Yazdan Yazdanpanah, Xavier Lescure, Alexandre Gaymard, Bruno Lina, Manuel Rosa-Calatrava |
| EPI_ISL_411902 | Virology Unit, Institut Pasteur du Cambodge. | Virology Unit, Institut Pasteur du Cambodge (Sequencing done by: Jessica E Manning/Jennifer A Bohl at Malaria and Vector Research Research Laboratory, National Institute of Allergy and Infectious Diseases and Vida Ahyong from Chan-Zuckerberg Biohub) | Erik A Karlsson, Jennifer A Bohl, Vida Ahyong, Veasna Duong, Philippe Dussart, Jessica E Manning. |
| EPI_ISL_411915 | Laboratory Medicine | Department of Laboratory Medicine, Lin-Kou Chang Gung Memorial Hospital, Taoyuan, Taiwan. | Kuo-Chien Tsao, Yu-Nong Gong, Shu-Li Yang, Yi-Chun Li, Chung-Guei Huang, Yhu-Chering Huang, Shin-Ru Shih |
| EPI_ISL_411926, EPI_ISL_411927 | Taiwan Centers for Disease Control | Taiwan Centers for Disease Control | Ji-Rong Yang, Yu-Chi-Lin, Jung-Jung Mu, Ming-Tsan-Liu |
| EPI_ISL_411929 | Department of Clinical Diagnostics | Department of Clinical Diagnostics | Park,W.B., Kwon,N.-J., Choi,S.-J., Kang,C.K., Choe,P.G., Kim,J.Y., Yun,J., Lee,G.-W., Seong,M.-W., Kim,N., Seo,J.-S. and Oh,M.-D. |
| EPI_ISL_411949 | unknown | Pathogenic microbiology laboratoryHuashan Hospital, Fudan University | Jing-Wen Ai, Yi Zhang, Hao-Cheng Zhang, Teng Xu, Wen-Hong Zhang |
| EPI_ISL_411950 | NHC Key laboratory of Enteric Pathogenic Microbiology, Institute of Pathogenic Microbiology | Jiangsu Provincial Center for Disease Control & Prevention | Lunbiao Cui,Kangchen Zhao,Xiaojuan Zhu,Yiyue Ge,Tao Wu,Bin Wu,Yin Chen,Fengcai Zhu,Baoli Zhu,Ming Wu |
| EPI_ISL_411951 | Unit for Laboratory Development and Technology Transfer, Public Health Agency of Sweden | Unit for Laboratory Development and Technology Transfer, Public Health Agency of Sweden | Bengner,M., Palmerus,M., Lindsjo,O., Lind Karlberg,M., Monteil,V., Appelberg,S., Brave,A., Muradrasoli,S. and Tegmark-Wisell,K. |
| EPI_ISL_411952, EPI_ISL_411953 | NHC Key laboratory of Enteric Pathogenic Microbiology, Institute of Pathogenic Microbiology | Jiangsu Provincial Center for Disease Control & Prevention | Kangchen Zhao, Xiaojuan Zhu, Lunbiao Cui, Tao Wu, Yiyue Ge, Bin Wu, Yin Chen, Fengcai Zhu, Baoli Zhu, Ming Wu |
| EPI_ISL_411954, EPI_ISL_411955 | California Department of Public Health | Pathogen Discovery, Respiratory Viruses Branch, Division of Viral Diseases, Centers for Dieases Control and Prevention | Krista Queen, Anna Uehara, Jing Zhang, Yan Li, Ying Tao, Clinton R. Paden, Haibin Wang, Shifaq Kamili, Xiaoyan Lu, Brian Lynch, Senthil Kumar K. Sakthivel, Brett L. Whitaker, Lijuan Wang, Janna' R. Murray, Susan I. Gerber, Stephen Lindstrom, Suxiang Tong |
| EPI_ISL_411956 | Texas Department of State Health Services | Pathogen Discovery, Respiratory Viruses Branch, Division of Viral Diseases, Centers for Dieases Control and Prevention | Krista Queen, Anna Uehara, Jing Zhang, Yan Li, Ying Tao, Clinton R. Paden, Haibin Wang, Shifaq Kamili, Xiaoyan Lu, Brian Lynch, Senthil Kumar K. Sakthivel, Brett L. Whitaker, Lijuan Wang, Janna' R. Murray, Susan I. Gerber, Stephen Lindstrom, Suxiang Tong |
| EPI_ISL_411957 | Key Laboratory of Human Diseases, Comparative Medicine, Institute of Laboratory Animal Science | Key Laboratory of Human Diseases, Comparative Medicine, Institute of Laboratory Animal Science | Linlin,B., Lili,R., Shuran,G., Jiangning,L., Feifei,Q., Qi,L., Fengdi,L., Jing,X., Wei,D., Pin,Y., Yanfeng,X., Yajin,Q., Hong,G., Qiang,W., Mingya,L., Guanpeng,W., Shunyi,W., Zhiqi,S., Li,G., Lan,C., Conghui,W., Ying,W., Xinming,W., Yan,X., Qi,J. and Chuan,Q. |
| EPI_ISL_411958 | Bioinfo, Vision Medicals, Lianhe, | Bioinfo, Vision Medicals, Lianhe, | Zhang,W.H |
| EPI_ISL_411959, EPI_ISL_411960, EPI_ISL_411961, EPI_ISL_411962, EPI_ISL_411963, EPI_ISL_411964, EPI_ISL_411965, EPI_ISL_411966, EPI_ISL_411967 | Bioinfo, Vision Medicals, Lianhe | Bioinfo, Vision Medicals, Lianhe | Zhang,W.H |
| EPI_ISL_412026 | Second Hospital of Anhui Medical University | Second Hospital of Anhui Medical University | Changtai Wang, Zhongping Liua, Zixiang Chen, Xin Huang, Mengyuan Xua, Tengfei He, Mengji Lu, Zhenhua Zhang |
| EPI_ISL_412028 | Hong Kong Department of Health | School of Public Health, The University of Hon g Kong | Dominic N.C. Tsang, Daniel K.W. Chu, Leo L.M. Poon, Malik Peiris |
| EPI_ISL_412029 | Hong Kong Department of Health | The University of Hong Kong | Dominic N.C. Tsang, Daniel K.W. Chu, Leo L.M. Poon, Malik Peiris |

| | | | |
|---|---|---|---|
| EPI_ISL_412030 | Hong Kong Department of Health | School of Public Health, The University of Hon g Kong | Dominic N.C. Tsang, Daniel K.W. Chu, Leo L.M. Poon, Malik Peiris |
| EPI_ISL_412034, EPI_ISL_412035, EPI_ISL_412036, EPI_ISL_412037, EPI_ISL_412038, EPI_ISL_412039 | Department of Clinical Laboratory, Tongji Medical College, Huazhong University of Science and Technology | Department of Clinical Laboratory, Tongji Medical College, Huazhong University of Science and Technology | Liu,W., Zhang,Q., Song,H., Xiang,R., Sun,Z. and Liu,Y |
| EPI_ISL_412041 | University of Hong Kong-Shenzhen Hospital | University of Hong Kong-Shenzhen Hospital | Chan,J.F.-W., Yuan,S., Kok,K.H., To,K.K.-W., Chu,H., Yang,J., Xing,F., Liu,J., Yip,C.C.-Y., Poon,R.W.-S., Tsai,H.W., Lo,S.K.-F., Chan,K.H., Poon,V.K.-M., Chan,W.M., Ip,J.D., Cai,J.P., Cheng,V.C.-C., Chen,H., Hui,C.K.-M. and Yuen,K.Y |
| EPI_ISL_412042 | University of Hong Kong-Shenzhen Hospital | University of Hong Kong-Shenzhen Hospital | Chan,J.F.-W., Yuan,S., Kok,K.H., To,K.K.-W., Chu,H., Yang,J., Xing,F., Liu,J., Yip,C.C.-Y., Poon,R.W.-S., Tsai,H.W., Lo,S.K.-F., Chan,K.H., Poon,V.K.-M., Chan,W.M., Ip,J.D., Cai,J.P., Cheng,V.C.-C., Chen,H., Hui,C.K.-M. and Yuen,K.Y. |
| EPI_ISL_412043 | University of Hong Kong-Shenzhen Hospital | Li Ka Shing Faculty of Medicine, The University of Hong Kong | Chan,J.F.-W., Yuan,S., Kok,K.H., To,K.K.-W., Chu,H., Yang,J., Xing,F., Liu,J., Yip,C.C.-Y., Poon,R.W.-S., Tsai,H.W., Lo,S.K.-F., Chan,K.H., Poon,V.K.-M., Chan,W.M., Ip,J.D., Cai,J.P., Cheng,V.C.-C., Chen,H., Hui,C.K.-M. and Yuen,K.Y. |
| EPI_ISL_412044 | unknown | University of Hong Kong-Shenzhen Hospital | Chan,J.F.-W., Yuan,S., Kok,K.H., To,K.K.-W., Chu,H., Yang,J., Xing,F., Liu,J., Yip,C.C.-Y., Poon,R.W.-S., Tsai,H.W., Lo,S.K.-F., Chan,K.H., Poon,V.K.-M., Chan,W.M., Ip,J.D., Cai,J.P., Cheng,V.C.-C., Chen,H., Hui,C.K.-M. and Yuen,K.Y. |
| EPI_ISL_412045 | University of Hong Kong-Shenzhen Hospital | Li Ka Shing Faculty of Medicine, The University of Hong Kong | Chan,J.F.-W., Yuan,S., Kok,K.H., To,K.K.-W., Chu,H., Yang,J., Xing,F., Liu,J., Yip,C.C.-Y., Poon,R.W.-S., Tsai,H.W., Lo,S.K.-F., Chan,K.H., Poon,V.K.-M., Chan,W.M., Ip,J.D., Cai,J.P., Cheng,V.C.-C., Chen,H., Hui,C.K.-M. and Yuen,K.Y. |
| EPI_ISL_412046 | unknown | University of Hong Kong-Shenzhen Hospital | Chan,J.F.-W., Yuan,S., Kok,K.H., To,K.K.-W., Chu,H., Yang,J., Xing,F., Liu,J., Yip,C.C.-Y., Poon,R.W.-S., Tsai,H.W., Lo,S.K.-F., Chan,K.H., Poon,V.K.-M., Chan,W.M., Ip,J.D., Cai,J.P., Cheng,V.C.-C., Chen,H., Hui,C.K.-M. and Yuen,K.Y. |
| EPI_ISL_412049 | University of Hong Kong- Shenzhen Hospital | University of Hong Kong- Shenzhen Hospital | Chan,J.F.-W., Yuan,S., Kok,K.H., To,K.K.-W., Chu,H., Yang,J., Xing,F., Liu,J., Yip,C.C.-Y., Poon,R.W.-S., Tsai,H.W., Lo,S.K.-F., Chan,K.H., Poon,V.K.-M., Chan,W.M., Ip,J.D., Cai,J.P., Cheng,V.C.-C., Chen,H., Hui,C.K.-M. and Yuen,K.Y. |
| EPI_ISL_412050, EPI_ISL_412051, EPI_ISL_412052, EPI_ISL_412053, EPI_ISL_412054 | The University of Hong Kong- Shenzhen Hospital | The University of Hong Kong- Shenzhen Hospital | Chan,J.F.-W., Yuan,S., Kok,K.H., To,K.K.-W., Chu,H., Yang,J., Xing,F., Liu,J., Yip,C.C.-Y., Poon,R.W.-S., Tsai,H.W., Lo,S.K.-F., Chan,K.H., Poon,V.K.-M., Chan,W.M., Ip,J.D., Cai,J.P., Cheng,V.C.-C., Chen,H., Hui,C.K.-M. and Yuen,K.Y. |
| EPI_ISL_412116 | Respiratory Virus Unit, Microbiology Services Colindale, Public Health England | Respiratory Virus Unit, Microbiology Services Colindale, Public Health England | Monica Galiano, Shahjahan Miah, Angie Lackenby, Omolola Akinbami, Tiina Talts, Leena Bhaw, Richard Myers, Steven Platt, Kirstin Edwards, Jonathan Hubb, Joanna Ellis, Maria Zambon |
| EPI_ISL_412387 | Shandong Provincial Center for Disease Control and Prevention | Beijing Institute of Microbiology and Epidemiology | Xiao-Lin Jiang, Wen-Kui Sun, Xiang-Na Zhao, Yang Hang, Zeng-Qiang Kou, Lin-Yao, Li-Jun Duan, Xiao Wei, Mai-Juan Ma, Dian-Ming Kang |
| EPI_ISL_412418, EPI_ISL_412419, EPI_ISL_412420, EPI_ISL_412421, EPI_ISL_412422, EPI_ISL_412423, EPI_ISL_412424, EPI_ISL_412425, EPI_ISL_412426 | Shandong Provincial Center for Disease Control and Prevention | Beijing Institute of Microbiology and Epidemiology | Xiao-Lin Jiang, Wen-Kui Sun, Xiang-Na Zhao, Yang Hang, Zeng-Qiang Kou, Lin-Yao, Li-Jun Duan, Xiao Wei, Dian-Ming Kang, Mai-Juan Ma |
| EPI_ISL_412459 | Jingzhou Center for Disease Control and Prevention | Hubei Provincial Center for Disease Control and Prevention | Bin Fang, Xiang Li, Xiao Yu, Linlin Liu, Bo Yang, Faxian Zhan, Guojun Ye, Xixiang Huo, Junqiang Xu, Bo Yu, Kun Cai, Jing Li, Maoyi Chen,Jie Hu, Chunlin Mao, Yongzhong Jiang. |
| EPI_ISL_412860 | SCSFRI, South China Sea Fisheries Research Institute, Chinese Academy of Fishery Sciences (SCSFRI, CAFS) | SCSFRI, South China Sea Fisheries Research Institute, Chinese Academy of Fishery Sciences (SCSFRI, CAFS) | Jiang,J.-Z., Liu,P. and Chen,J.-P. |
| EPI_ISL_412862 | California Department of Public Health | Pathogen Discovery, Respiratory Viruses Branch, Division of Viral Diseases, Centers for Disease Control and Prevention | Krista Queen, Anna Uehara, Jing Zhang, Yan Li, Ying Tao, Clinton R. Paden, Haibin Wang, Shifaq Kamili, Xiaoyan Lu, Brian Lynch, Senthil Kumar K. Sakthivel, Brett L. Whitaker, Lijuan Wang, Janna' R. Murray, Jasmine Padilla, Justin Lee, Susan I. Gerber, Stephen Lindstrom, Suxiang Tong |
| EPI_ISL_412869 | Division of Viral Diseases, Center for Laboratory Control of Infectious Diseases, Korea Centers for Diseases Control and Prevention | Division of Viral Diseases, Center for Laboratory Control of Infectious Diseases, Korea Centers for Diseases Control and Prevention | Jeong-Min Kim, Yoon-Seok Chung, Namjoo Lee, Mi-Seon Kim, Sang Hee Woo, Hye-Jun Jo, Sehee Park, Heui Man Kim, Myung Guk Han |
| EPI_ISL_412870 | Division of Viral Diseases, Center for Laboratory Control of Infectious Diseases, Korea Centers for Diseases Control and Prevention | Division of Viral Diseases, Center for Laboratory Control of Infectious Diseases, Korea Centers for Diseases Control and Prevention | Jeong-Min Kim, Yoon-Seok Chung, Namjoo Lee, Mi-Seon Kim, Sang Hee Woo, Hye-Jun Jo, Sehee Park, Heui Man Kim, Myung Guk Han |
| EPI_ISL_412871 | Division of Viral Diseases, Center for Laboratory Control of Infectious Diseases, Korea Centers for Diseases Control and Prevention | Division of Viral Diseases, Center for Laboratory Control of Infectious Diseases, Korea Centers for Diseases Control and Prevention | Jeong-Min Kim, Yoon-Seok Chung, Namjoo Lee, Mi-Seon Kim, Sang Hee Woo, Hye-Jun Jo, Sehee Park, Heui Man Kim, Myung Guk Han |
| EPI_ISL_412872 | Division of Viral Diseases, Center for Laboratory Control of Infectious Diseases, Korea Centers for Diseases Control and Prevention | Division of Viral Diseases, Center for Laboratory Control of Infectious Diseases, Korea Centers for Diseases Control and Prevention | Jeong-Min Kim, Yoon-Seok Chung, Namjoo Lee, Mi-Seon Kim, Sang Hee Woo, Hye-Jun Jo, Sehee Park, Heui Man Kim, Myung Guk Han |
| EPI_ISL_412873 | Division of Viral Diseases, Center for Laboratory Control of Infectious Diseases, Korea Centers for Diseases Control and Prevention | Division of Viral Diseases, Center for Laboratory Control of Infectious Diseases, Korea Centers for Diseases Control and Prevention | Jeong-Min Kim, Yoon-Seok Chung, Namjoo Lee, Mi-Seon Kim, Sang Hee Woo, Hye-Jun Jo, Sehee Park, Heui Man Kim, Myung Guk Han |
| EPI_ISL_412898, EPI_ISL_412899, EPI_ISL_412900 | Wuhan Jinyintan Hospital | Hubei Provincial Center for Disease Control and Prevention | Bin Fang, Xiang Li, Xiao Yu, Linlin Liu, Bo Yang, Faxian Zhan, Guojun Ye, Xixiang Huo, Junqiang Xu, Bo Yu, Kun Cai, Jing Li, Yongzhong Jiang. |
| EPI_ISL_412912 | State Health Office Baden-Württemberg | Charité Universitätsmedizin Berlin, Institute of Virology | Victor M Corman, Julia Schneider, Barbara Mühlemann, Talitha Veith, Jörn Beheim-Schwarzbach, Terry Jones, Rainer Oehme, Silke Fischer, Christian Drosten |
| EPI_ISL_412964 | Hospital Israelita Albert Einstein | Instituto Adolfo Lutz Interdisciplinary Procedures Center Strategic Laboratory | Jaqueline Goes de Jesus, Claudio Tavares Sacchi, Daniela Bernardes Borges da Silva, Ingra Morales Claro, Flávia Cristina da Silva Sales, Claudia Regina Gonçalves, Joshua Quick, Maria do Carmo, Sampaio Tavares Timenetsky, Nicholas James Loman, Andrew Rambaut, Ester Cerdeira Sabino, Nuno Rodrigues Faria |
| EPI_ISL_412965 | BCCDC Public Health Laboratory | BCCDC Public Health Laboratory | Harrigan, Prystajecky, Krajden, Lee, Kamelian, Lapointe, Choi, Hoang, Sekirov, Levett, Tyson, Loman, Quick, Li, Gilmour |
| EPI_ISL_412966 | Technology Centre, Guangzhou Customs | Technology Centre, Guangzhou Customs | Shi,Y., Sun,J., Zheng,K., Huang,J. and Zhao,J. |
| EPI_ISL_412967 | Technology Centre, Guangzhou Customs | Technology Centre, Guangzhou Customs | Shi,Y., Zheng,K., Sun,J., Huang,J., Zhu,A., Zhuang,Z., Dai,J., Chen,Z., Sun,F., Zhang,Z., Li,X. and Wang,Y. |
| EPI_ISL_412968, EPI_ISL_412969 | Takayuki Hishiki Kanagawa Prefectural Institute of Public Health | Takayuki Hishiki Kanagawa Prefectural Institute of Public Health | Hishiki,T., Suzuki,R., Sakuragi,J., Usui,K., Tanaka,Y., Kawai,J., Kogo,Y., Matsuki,Y., An,T., Hayashizaki,Y. and Takasaki,T. |
| EPI_ISL_412970 | Washington State Department of Health | Seattle Flu Study | Helen Chu, Michael Boeckh, Janet Englund, Michael Famulare, Barry Lutz, Deborah Nickerson, Mark Rieder, Lea Starita, Matthew Thompson, Jay Shendure, and Trevor Bedford |
| EPI_ISL_412971 | HUS Diagnostiikkakeskus, Hallinto | Department of Virology Faculty of Medicine, Medicum University of Helsinki | Teemu Smura, Suvi Kuivanen, Hannimari Kallio-Kokko, Olli Vapalahti |
| EPI_ISL_412972 | Instituto Nacional de Enfermedades Respiratorias | Instituto de Diagnostico y Referencia Epidemiologicos (INDRE) | Ramirez-Gonzalez Ernesto, Garces-Ayala Fabiola, Araiza-Rodriguez Adnan, Mendieta-Condado Edgar, Rodriguez-Maldonado Abril, Wong-Arambula Claudia, Vazquez-Perez Joel, Martinez Arturo, Boukadida Celia, Munoz-Medina Esteban, Sanchez Alejandro, Isa Pavel, Taboada Blanca, Lopez Susana, Arias Carlos, Barrera-Badillo Gisela, Hernandez-Rivas Lucia, Lopez-Martinez Irma |
| EPI_ISL_412973 | Department of Infectious Diseases, Istituto Superiore di Sanità, Roma , Italy | Virology Laboratory, Scientific Department, Army Medical Center | Paola Stefanelli, Stefano Fiore, Antonella Marchi, Eleonora Benedetti, Concetta Fabiani, Giovanni Faggioni, Antonella Fortunato, Riccardo De Santis, Silvia Fillo, Anna Anselmo, Andrea Ciammaruconi, Stefano Palomba, Florigio Lista |
| EPI_ISL_412974 | Department of Infectious Diseases, Istituto Superiore di Sanità, Rome, Italy | Virology Laboratory, Scientific Department, Army Medical Center | Paola Stefanelli, Stefano Fiore, Antonella Marchi, Eleonora Benedetti, Concetta Fabiani, Giovanni Faggioni, Antonella Fortunato, Silvia Fillo, Riccardo De Santis, Andrea Ciammaruconi, Giancarlo Petralito, Filippo Molinari, Florigio Lista |
| EPI_ISL_412975 | Centre for Infectious Diseases and Microbiology Laboratory Services | NSW Health Pathology - Institute of Clinical Pathology and Medical Research; Westmead Hospital; University of Sydney | Eden J-S, Carter I, Rahman H, Holmes EC, Rockett R, O'Sullivan MV, Sintchenko V, Chen SC, Maddocks S, Kok J and Dwyer DE for the 2019-nCoV Study Group |
| EPI_ISL_412976, EPI_ISL_412977 | Shandong First Medical University & Shandong Academy of Medical Sciences | Institute of Microbiology, Chinese Academy of Sciences | Weifeng Shi, Tao Hu, Hong Zhou, Juan Li, Xing Chen, Alice Catherine Hughes, Yuhai Bi |
| EPI_ISL_412978 | The Central Hospital Of Wuhan | Hubei Provincial Center for Disease Control and Prevention | Bin Fang, Xiang Li, Xiao Yu, Linlin Liu, Bo Yang, Faxian Zhan, Guojun Ye, Xixiang Huo, Junqiang Xu, Bo Yu, Kun Cai, Jing Li, Yongzhong Jiang. |
| EPI_ISL_412979, EPI_ISL_412980 | Union Hospital of Tongji Medical College, Huazhong University of Science and Technology | Hubei Provincial Center for Disease Control and Prevention | Bin Fang, Xiang Li, Xiao Yu, Linlin Liu, Bo Yang, Faxian Zhan, Guojun Ye, Xixiang Huo, Junqiang Xu, Bo Yu, Kun Cai, Jing Li, Yongzhong Jiang. |
| EPI_ISL_412981 | CR&WISCO GENERAL HOSPITAL | Hubei Provincial Center for Disease Control and Prevention | Bin Fang, Xiang Li, Xiao Yu, Linlin Liu, Bo Yang, Faxian Zhan, Guojun Ye, Xixiang Huo, Junqiang Xu, Bo Yu, Kun Cai, Jing Li, Yongzhong Jiang. |
| EPI_ISL_412982 | Wuhan Lung Hospital | Hubei Provincial Center for Disease Control and Prevention | Bin Fang, Xiang Li, Xiao Yu, Linlin Liu, Bo Yang, Faxian Zhan, Guojun Ye, Xixiang Huo, Junqiang Xu, Bo Yu, Kun Cai, Jing Li, Yongzhong Jiang. |
| EPI_ISL_412983 | Tianmen Center for Disease Control and Prevention | Hubei Provincial Center for Disease Control and Prevention | Bin Fang, Xiang Li, Xiao Yu, Linlin Liu, Bo Yang, Faxian Zhan, Guojun Ye, Xixiang Huo, Junqiang Xu, Bo Yu, Kun Cai, Jing Li, YiFa Zhu, Yangyang Tao,Xierong Li,Yongzhong Jiang. |
| EPI_ISL_413014 | Public Health Ontario Laboratory | Ontario Agency for Health Protection and Promotion (OAHPP) | Alireza Eshaghi, Samir N Patel, Jonathan B Gubbay, Vanessa G Allen, Christine Frantz, Aimin Li, Sandeep Nagra |
| EPI_ISL_413015 | Public Health Ontario Laboratory | National Microbiology Laboratory | Shari Tyson, Anna Majer, Erika Landry, Morag Graham, Grace Seo, Philip Mabon, Natalie Knox, Adrian Zetner, Samira Mubareka, Rob Kozak, Jocelyne Lew, Darryl Falzarano, Gerdts Volker, Jonathan Gubbay, Stephanie Booth, Guillaume Poliquin, Tom Graefenhan, Matthew Gilmour, Nathalie Bastien, Yan Li, Timothy Booth |
| EPI_ISL_413016 | Hospital Israelita Albert Einstein | Instituto Adolfo Lutz, Interdisciplinary Procedures Center, Strategic Laboratory | Jaqueline Goes de Jesus, Claudio Tavares Sacchi, Fabiana Cristina Pereira dos Santos, Ingra Morales Claro, Flávia Cristina da Silva Sales, Claudia Regina Gonçalves, Joshua Quick, Maria do Carmo Sampaio Tavares Timenetsky, Nicholas James Loman, Andrew Rambaut, Ester Cerdeira Sabino, Nuno Rodrigues Faria |
| EPI_ISL_413017, EPI_ISL_413018 | Department of Microbiology, Institute for Viral Diseases, College of Medicine, Korea University | Department of Microbiology, Institute for Viral Diseases, College of Medicine, Korea University | Changmin Kang, Joon-Yong Bae, Jungmin Lee, Heedo Park, Juyoung Cho, Jeonghun Kim, Gee eun Lee, Cui Chunguang, Kyeong-ryeol Shin, Dong Min Kim, Jin Il Kim, Man-Seong Park |
| EPI_ISL_413019, EPI_ISL_413020 | Department of Internal Medicine, Triemli Hospital | Institute of Medical Virology, University of Zurich | Stefan Schmutz, Maryam Zaheri, Verena Kufner, Patrick Redli, Fiona Steiner, Jon Huder, Riccarda Capaul, Andrea Zbinden, Jürg Böni, Michael Huber, Gerhard Eich, Alexandra Trkola |
| EPI_ISL_413021 | Klinik Hirslanden Zurich | Institute of Medical Virology, University of Zurich | Stefan Schmutz, Maryam Zaheri, Verena Kufner, Gabriela Ziltener, Patrick Redli, Fiona Steiner, Jon Huder, Riccarda Capaul, Andrea Zbinden, Jürg Böni, Michael Huber, Christian Ruef, Alexandra Trkola |
| EPI_ISL_413022, EPI_ISL_413023, EPI_ISL_413024 | Division of Infectious Diseases, University Hospital Zurich | Institute of Medical Virology, University of Zurich | Stefan Schmutz, Maryam Zaheri, Verena Kufner, Gabriela Ziltener, Patrick Redli, Fiona Steiner, Jon Huder, Riccarda Capaul, Andrea Zbinden, Jürg Böni, Michael Huber, Roberto Speck, Alexandra Trkola |
| EPI_ISL_413025 | Harborview Medical Center | UW Virology Lab | Pavitra Roychoudhury, Arun Nalla, Hong Xie, Keith Jerome, Alexander Greninger |
| EPI_ISL_413213, EPI_ISL_413214 | Centre for Infectious Diseases and Microbiology Laboratory Services | NSW Health Pathology - Institute of Clinical Pathology and Medical Research; Westmead Hospital; University of Sydney | Eden J-S, Carter I, Rahman H, Holmes EC, Rockett R, O'Sullivan MV, Sintchenko V, Chen SC, Maddocks S, Kok J and Dwyer DE for the 2019-nCoV Study Group* |
| EPI_ISL_413219 | National Institute of Health Research and Development | National Institute of Health Research and Development | Setiawaty,V; Subangkit; Pawestri,HA; Puspa,KD; Ikawati,HD; Nugraha,AA; Hariastuti,NI; Ramadhany,R; Susilarini,NK; Pratiwi,E; Agustiningsih; Kurniawati,J; Siswanto |

| | | | |
|---|---|---|---|
| EPI_ISL_413221 | West of Scotland Specialist Virology Centre, NHSGGC | MRC-University of Glasgow Centre for Virus Research | Emma Thomson, Antonia Ho; James Shephard, Shirin Ashraf; Kathy Smollett, Daniel Mair, Stephen Carmichael, Ana da Silva Filipe; Richard Orton, Josh Singer, David L Robertson; Andrew Rambaut; Alasdair MacLean, Rory Gunson. |
| EPI_ISL_413455 | Washington State Public Health Lab | University of Washington Virology Lab | Pavitra Roychoudhury, Arun Nalla, Hong Xie, Keith Jerome, Alexander Greninger |
| EPI_ISL_413456 | Seattle Flu Study, University of Washington Medical Center | Seattle Flu Study, University of Washington Medical Center | Chu et al |
| EPI_ISL_413457, EPI_ISL_413458 | Washington State Public Health Lab | UW Virology Lab | Pavitra Roychoudhury, Arun Nalla, Hong Xie, Keith Jerome, Alexander Greninger |
| EPI_ISL_413459 | Department of Pathology, Toshima Hospital | Pathogen Genomics Center, National Institute of Infectious Diseases | Tsuyoshi Sekizuka, Kentaro Itokawa, Takuya Adachi, Masahiro Sano, Jun Yamazaki, Ippei Miyamoto, Haruka Nishioka, Ja-Mun Chong, Noriko Nakajima, Yuko Sato, Minoru Tobiume, Harutaka Katano, Tadaki Suzuki, Makoto Kuroda |
| EPI_ISL_413485 | Department of microbiology laboratory,Anhui Provincial Center for Disease Control and Prevention | Department of microbiology laboratory,Anhui Provincial Center for Disease Control and Prevention | Weiwei Li,Jun He,Yong Sun,Junling Yu,Qingqing Chen,Yuan Yuan,Yonglin Shi,Zhuhui Zhang,Yinglu Ge,Weidong Li,Bin Su,Zhirong Liu |
| EPI_ISL_413486 | Valley Medical Center | University of Washington Virology Lab | Pavitra Roychoudhury, Arun Nalla, Hong Xie, Keith Jerome, Alexander Greninger |
| EPI_ISL_413487 | Harborview Medical Center | University of Washington Virology Lab | Pavitra Roychoudhury, Arun Nalla, Hong Xie, Keith Jerome, Alexander Greninger |
| EPI_ISL_413488 | Center of Medical Microbiology, Virology, and Hospital Hygiene, University of Duesseldorf | Center of Medical Microbiology, Virology, and Hospital Hygiene, University of Duesseldorf | Ortwin Adams, Marcel Andree, Alexander Dilthey, Torsten Feldt, Sandra Hauka, Torsten Houwaart, Björn-Erik Jensen, Detlef Kindgen-Milles, Malte Kohns Vasconcelos, Klaus Pfeffer, Tina Senff, Daniel Strelow, Jörg Timm, Andreas Walker, Tobias Wienemann |
| EPI_ISL_413489 | Laboratorio di Microbiologia e Virologia, Università Vita-Salute San Raffaele, Milano | Laboratorio di Microbiologia e Virologia, Università Vita-Salute San Raffaele, Milano | R.A Diotti, E. Criscuolo, M. Castelli, V. Caputo, R. Ferrarese, M. Sampaolo, E. Boeri, I. Negri, V. Amato, G. Lo Raso, C. Di Resta, R. Burioni, M. Clementi, N. Mancini & N. Clementi |
| EPI_ISL_413490 | Auckland Hospital | Institute of Environmental Science and Research (ESR) | Matt Storey, Xiaoyun Ren, Gary McAuliffe, Sally Roberts, Matthew Blakiston, Erasmus Smit, Lauren Jelly, Joep de Ligt |
| EPI_ISL_413491 | Princess Margaret Hospital | Hong Kong Department of Health | Mak Gannon C.K., Cheng Peter K.C., Lam Edman T.K., Chan Rickjason C.W., Tsang Dominic N.C. |
| EPI_ISL_413492 | Queen Mary Hospital | Hong Kong Department of Health | Mak Gannon C.K., Cheng Peter K.C., Lam Edman T.K., Chan Rickjason C.W., Tsang Dominic N.C. |
| EPI_ISL_413493, EPI_ISL_413494 | Princess Margaret Hospital | Hong Kong Department of Health | Mak Gannon C.K., Cheng Peter K.C., Lam Edman T.K., Chan Rickjason C.W., Tsang Dominic N.C. |
| EPI_ISL_413495, EPI_ISL_413496 | Ruttonjee Hospital | Hong Kong Department of Health | Mak Gannon C.K., Cheng Peter K.C., Lam Edman T.K., Chan Rickjason C.W., Tsang Dominic N.C. |
| EPI_ISL_413497 | Ruttonjee Hospital | Hong Kong Department of Health | Mak Gannon C.K., Cheng Peter K.C., Lam Edman T.K., Chan Rickjason C.W., Tsang Dominic N.C. |
| EPI_ISL_413498 | Pamela Youde Nethersole Eastern Hospital | Hong Kong Department of Health | Mak Gannon C.K., Cheng Peter K.C., Lam Edman T.K., Chan Rickjason C.W., Tsang Dominic N.C. |
| EPI_ISL_413513 | Division of Infectious Diseases, Department of Internal Medicine, Korea University College of Medicine | Department of Microbiology, Institute for Viral Diseases, College of Medicine, Korea University | Changmin Kang, Joon-Yong Bae, Jungmin Lee, Jin Gu Yoon, Heedo Park, Juyoung Cho, Jeonghun Kim, Gee Eun Lee, Cui Chunguang, Kyeong-ryeol Shin, Ji Yun Noh, Joon Young Song, Hee Jin Cheong, Woo Joo Kim, Jin Il Kim, Man-Seong Park |
| EPI_ISL_413514 | Department of Microbiology, Institute for Viral Diseases, College of Medicine, Korea University | Department of Microbiology, Institute for Viral Diseases, College of Medicine, Korea University | Changmin Kang, Joon-Yong Bae, Jungmin Lee, Jin Gu Yoon, Heedo Park, Juyoung Cho, Jeonghun Kim, Gee Eun Lee, Cui Chunguang, Kyeong-ryeol Shin, Ji Yun Noh, Joon Young Song, Hee Jin Cheong, Woo Joo Kim, Jin Il Kim, Man-Seong Park |
| EPI_ISL_413515 | Division of Infectious Diseases, Department of Internal Medicine, Korea University College of Medicine | Department of Microbiology, Institute for Viral Diseases, College of Medicine, Korea University | Changmin Kang, Joon-Yong Bae, Jungmin Lee, Jin Gu Yoon, Heedo Park, Juyoung Cho, Jeonghun Kim, Gee Eun Lee, Cui Chunguang, Kyeong-ryeol Shin, Ji Yun Noh, Joon Young Song, Hee Jin Cheong, Woo Joo Kim, Jin Il Kim, Man-Seong Park |
| EPI_ISL_413516 | Department of Microbiology, Institute for Viral Diseases, College of Medicine, Korea University | Department of Microbiology, Institute for Viral Diseases, College of Medicine, Korea University | Changmin Kang, Joon-Yong Bae, Jungmin Lee, Jin Gu Yoon, Heedo Park, Juyoung Cho, Jeonghun Kim, Gee Eun Lee, Cui Chunguang, Kyeong-ryeol Shin, Ji Yun Noh, Joon Young Song, Hee Jin Cheong, Woo Joo Kim, Jin Il Kim, Man-Seong Park |
| EPI_ISL_413517 | Gastrointestinal and Liver Diseases Research Center, Iran University of Medical Sciences | Gastrointestinal and Liver Diseases Research Center, Iran University of Medical Sciences | Karbalaie Niya,M.H., Laali,A., Tabibzadeh,A., Safarnezhad Tameshkel,F., Zamani,F., Sohrabi,M.R., Ranjbar,M., Savaj,S., Rezaie,N., Ajdarkosh,H., Keyvani,H., Khoonsari,M., Ameli,M., Nikkhah,M., Ghanbari,B., Faraji,A., Jamshidi Makiani,M. and Rohani,M. |
| EPI_ISL_413518, EPI_ISL_413519, EPI_ISL_413520, EPI_ISL_413521 | Infectious Disease Control Center, Center for Disease Control and Prevention of PLA | Infectious Disease Control Center, Center for Disease Control and Prevention of PLA | Li,J., Li,L., Li,Z., Qiu,S., Song,H., Li,P. and Li,P. |
| EPI_ISL_413522 | Indian Council of Medical Research - National Institute of Virology | National Influenza Center, Indian Council of Medical Research - National Institute of Virology | Potdar V, Yadav PD, Choudhary ML, Shete-Aich A |
| EPI_ISL_413523 | Indian Council of Medical Research-National Institute of Virology | National Influenza Center, Indian Council of Medical Research-National Institute of Virology | Potdar V, Yadav PD, Choudhary ML, Shete-Aich A |
| EPI_ISL_413550 | Centre for Human and Zoonotic Virology (CHAZVY), College of Medicine University of Lagos/Lagos University Teaching Hospital (LUTH), part of the Laboratory Network of the Nigeria Centre for Disease Control (NCDC) | African Centre of Excellence for Genomics of Infectious Diseases (ACEGID), Redeemer's University, Ede, Osun State, Nigeria | Oluniyi P.E., Ajogbasile F.V., Kayode A., Oguzie J., Folarin O.A., Ihekweazu C. Happi C.T. |
| EPI_ISL_413553 | Iran National Influenza Center | Iran National Influenza Center | Jila Yavarian, Nazanin Zahra Shafiei Jandaghi, Kaveh Sadeghi, Fatemeh Ajaminejad, Nastaran Ghavvami and Talat Mokhtari Azad |
| EPI_ISL_413554 | Iran National Influenza Center | Iran National Influenza Center | Jila Yavarian, Nazanin Zahra Shafiei Jandaghi, Kaveh Sadeghi, Nastaran Ghavvami, Fatemeh Ajami Nejad, Fatemeh Saadatmand and Talat Mokhtari Azad |
| EPI_ISL_413555 | Wales Specialist Virology Centre | Public Health Wales Microbiology Cardiff | Catherine Moore, Cen Sabu, Joanne Watkins, Sally Corden, Tom Connor |
| EPI_ISL_413556 | Wales Specialist Virology Centre | Public Health Wales Microbiology Cardiff | Catherine Moore, Tim Jones, Joanne Watkins, Sally Corden, Tom Connor |
| EPI_ISL_413557 | California Department of Public Health | Chiu Laboratory, University of California, San Francisco | Xianding Deng, Scot Federman, Chao-Yang Pan, Hugo Guevara,Wei Gu, Debra A. Wadford, and Charles Y. Chiu |
| EPI_ISL_413558, EPI_ISL_413559 | California Department of Public Health | Chiu Laboratory, University of California, San Francisco | Xianding Deng, Scot Federman, Chao-Yang Pan, Hugo Guevara,Wei Gu, Debra A. Wadford, and Charles Y. Chiu |
| EPI_ISL_413560 | Seattle Flu Study | Seattle Flu Study | Chu et al |
| EPI_ISL_413561 | California Department of Public Health | Chiu Laboratory, University of California, San Francisco | Xianding Deng, Scot Federman, Chao-Yang Pan, Hugo Guevara,Wei Gu, Debra A. Wadford, and Charles Y. Chiu |
| EPI_ISL_413562, EPI_ISL_413563 | UW Virology Lab | UW Virology Lab | Pavitra Roychoudhury, Hong Xie, Keith Jerome, Alexander Greninger |
| EPI_ISL_413564 | MHC West-Brabant | Erasmus Medical Center | David Nieuwenhuijse, Bas Oude Munnink, Reina Sikkema, Claudia Schapendonk, Irina Chestakova, Anne van der Linden, Mark Pronk, Pascal Lexmond, Corien Swaan, Manon Haverkate, Madelief Mollers, Mart Stein, Sandra Kengne Kamga Mobou, Jeroen van Kampen, Jolanda Voermans, Aura Timen, Corine GeurtsvanKessel, Annemiek van der Eijk, Richard Molenkamp, Marion Koopmans, on behalf of the Dutch national COVID-19 response team. |
| EPI_ISL_413565 | Foundation Pamm | Erasmus Medical Center | David Nieuwenhuijse, Bas Oude Munnink, Reina Sikkema, Claudia Schapendonk, Irina Chestakova, Anne van der Linden, Mark Pronk, Pascal Lexmond, Corien Swaan, Manon Haverkate, Madelief Mollers, Mart Stein, Sandra Kengne Kamga Mobou, Jeroen van Kampen, Jolanda Voermans, Aura Timen, Corine GeurtsvanKessel, Annemiek van der Eijk, Richard Molenkamp, Marion Koopmans, on behalf of the Dutch national COVID-19 response team. |
| EPI_ISL_413566 | MHC Gooi & Vechtstreek | Erasmus Medical Center | David Nieuwenhuijse, Bas Oude Munnink, Reina Sikkema, Claudia Schapendonk, Irina Chestakova, Anne van der Linden, Mark Pronk, Pascal Lexmond, Corien Swaan, Manon Haverkate, Madelief Mollers, Mart Stein, Sandra Kengne Kamga Mobou, Jeroen van Kampen, Jolanda Voermans, Aura Timen, Corine GeurtsvanKessel, Annemiek van der Eijk, Richard Molenkamp, Marion Koopmans, on behalf of the Dutch national COVID-19 response team. |
| EPI_ISL_413567 | Unknown | Erasmus Medical Center | David Nieuwenhuijse, Bas Oude Munnink, Reina Sikkema, Claudia Schapendonk, Irina Chestakova, Anne van der Linden, Mark Pronk, Pascal Lexmond, Corien Swaan, Manon Haverkate, Madelief Mollers, Mart Stein, Sandra Kengne Kamga Mobou, Jeroen van Kampen, Jolanda Voermans, Aura Timen, Corine GeurtsvanKessel, Annemiek van der Eijk, Richard Molenkamp, Marion Koopmans, on behalf of the Dutch national COVID-19 response team. |
| EPI_ISL_413568 | MHC Drente | Erasmus Medical Center | David Nieuwenhuijse, Bas Oude Munnink, Reina Sikkema, Claudia Schapendonk, Irina Chestakova, Anne van der Linden, Mark Pronk, Pascal Lexmond, Corien Swaan, Manon Haverkate, Madelief Mollers, Mart Stein, Sandra Kengne Kamga Mobou, Jeroen van Kampen, Jolanda Voermans, Aura Timen, Corine GeurtsvanKessel, Annemiek van der Eijk, Richard Molenkamp, Marion Koopmans, on behalf of the Dutch national COVID-19 response team. |
| EPI_ISL_413569, EPI_ISL_413570 | RIVM | Erasmus Medical Center | David Nieuwenhuijse, Bas Oude Munnink, Reina Sikkema, Claudia Schapendonk, Irina Chestakova, Anne van der Linden, Mark Pronk, Pascal Lexmond, Corien Swaan, Manon Haverkate, Madelief Mollers, Mart Stein, Sandra Kengne Kamga Mobou, Jeroen van Kampen, Jolanda Voermans, Aura Timen, Corine GeurtsvanKessel, Annemiek van der Eijk, Richard Molenkamp, Marion Koopmans, on behalf of the Dutch national COVID-19 response team. |
| EPI_ISL_413571 | MHC Brabant Zuidoost | Erasmus Medical Center | David Nieuwenhuijse, Bas Oude Munnink, Reina Sikkema, Claudia Schapendonk, Irina Chestakova, Anne van der Linden, Mark Pronk, Pascal Lexmond, Corien Swaan, Manon Haverkate, Madelief Mollers, Mart Stein, Sandra Kengne Kamga Mobou, Jeroen van Kampen, Jolanda Voermans, Aura Timen, Corine GeurtsvanKessel, Annemiek van der Eijk, Richard Molenkamp, Marion Koopmans, on behalf of the Dutch national COVID-19 response team. |
| EPI_ISL_413572 | MHC Kennemerland | Erasmus Medical Center | David Nieuwenhuijse, Bas Oude Munnink, Reina Sikkema, Claudia Schapendonk, Irina Chestakova, Anne van der Linden, Mark Pronk, Pascal Lexmond, Corien Swaan, Manon Haverkate, Madelief Mollers, Mart Stein, Sandra Kengne Kamga Mobou, Jeroen van Kampen, Jolanda Voermans, Aura Timen, Corine GeurtsvanKessel, Annemiek van der Eijk, Richard Molenkamp, Marion Koopmans, on behalf of the Dutch national COVID-19 response team. |
| EPI_ISL_413573 | Dienst Gezondheid & Jeugd Zuid-Holland Zuid | Erasmus Medical Center | David Nieuwenhuijse, Bas Oude Munnink, Reina Sikkema, Claudia Schapendonk, Irina Chestakova, Anne van der Linden, Mark Pronk, Pascal Lexmond, Corien Swaan, Manon Haverkate, Madelief Mollers, Mart Stein, Sandra Kengne Kamga Mobou, Jeroen van Kampen, Jolanda Voermans, Aura Timen, Corine GeurtsvanKessel, Annemiek van der Eijk, Richard Molenkamp, Marion Koopmans, on behalf of the Dutch national COVID-19 response team. |
| EPI_ISL_413574 | MHC West-Brabant | Erasmus Medical Center | David Nieuwenhuijse, Bas Oude Munnink, Reina Sikkema, Claudia Schapendonk, Irina Chestakova, Anne van der Linden, Mark Pronk, Pascal Lexmond, Corien Swaan, Manon Haverkate, Madelief Mollers, Mart Stein, Sandra Kengne Kamga Mobou, Jeroen van Kampen, Jolanda Voermans, Aura Timen, Corine GeurtsvanKessel, Annemiek van der Eijk, Richard Molenkamp, Marion Koopmans, on behalf of the Dutch national COVID-19 response team. |
| EPI_ISL_413575, EPI_ISL_413576 | RIVM | Erasmus Medical Center | David Nieuwenhuijse, Bas Oude Munnink, Reina Sikkema, Claudia Schapendonk, Irina Chestakova, Anne van der Linden, Mark Pronk, Pascal Lexmond, Corien Swaan, Manon Haverkate, Madelief Mollers, Mart Stein, Sandra Kengne Kamga Mobou, Jeroen van Kampen, Jolanda Voermans, Aura Timen, Corine GeurtsvanKessel, Annemiek van der Eijk, Richard Molenkamp, Marion Koopmans, on behalf of the Dutch national COVID-19 response team. |
| EPI_ISL_413577 | MHC Gooi & Vechtstreek | Erasmus Medical Center | David Nieuwenhuijse, Bas Oude Munnink, Reina Sikkema, Claudia Schapendonk, Irina Chestakova, Anne van der Linden, Mark Pronk, Pascal Lexmond, Corien Swaan, Manon Haverkate, Madelief Mollers, Mart Stein, Sandra Kengne Kamga Mobou, Jeroen van Kampen, Jolanda Voermans, Aura Timen, Corine GeurtsvanKessel, Annemiek van der Eijk, Richard Molenkamp, Marion Koopmans, on behalf of the Dutch national COVID-19 response team. |
| EPI_ISL_413578 | ErasmusMC | Erasmus Medical Center | David Nieuwenhuijse, Bas Oude Munnink, Reina Sikkema, Claudia Schapendonk, Irina Chestakova, Anne van der Linden, Mark Pronk, Pascal Lexmond, Corien Swaan, Manon Haverkate, Madelief Mollers, Mart Stein, Sandra Kengne Kamga Mobou, Jeroen van Kampen, Jolanda Voermans, Aura Timen, Corine GeurtsvanKessel, Annemiek van der Eijk, Richard Molenkamp, Marion Koopmans, on behalf of the Dutch national COVID-19 response team. |
| EPI_ISL_413579 | MHC Haaglanden | Erasmus Medical Center | David Nieuwenhuijse, Bas Oude Munnink, Reina Sikkema, Claudia Schapendonk, Irina Chestakova, Anne van der Linden, Mark Pronk, Pascal Lexmond, Corien Swaan, Manon Haverkate, Madelief Mollers, Mart Stein, Sandra Kengne Kamga Mobou, Jeroen van Kampen, Jolanda Voermans, Aura Timen, Corine GeurtsvanKessel, Annemiek van der Eijk, Richard Molenkamp, Marion Koopmans, on behalf of the Dutch national COVID-19 response team. |
| EPI_ISL_413580 | MHC Hart voor Brabant | Erasmus Medical Center | David Nieuwenhuijse, Bas Oude Munnink, Reina Sikkema, Claudia Schapendonk, Irina Chestakova, Anne van der Linden, Mark Pronk, Pascal Lexmond, Corien Swaan, Manon Haverkate, Madelief Mollers, Mart Stein, Sandra Kengne Kamga Mobou, Jeroen van Kampen, Jolanda Voermans, Aura Timen, Corine GeurtsvanKessel, Annemiek van der Eijk, Richard Molenkamp, Marion Koopmans, on behalf of the Dutch national COVID-19 response team. |
| EPI_ISL_413581 | RIVM | Erasmus Medical Center | David Nieuwenhuijse, Bas Oude Munnink, Reina Sikkema, Claudia Schapendonk, Irina Chestakova, Anne van der Linden, Mark Pronk, Pascal Lexmond, Corien Swaan, Manon Haverkate, Madelief Mollers, Mart Stein, Sandra Kengne Kamga Mobou, Jeroen van Kampen, Jolanda Voermans, Aura Timen, Corine GeurtsvanKessel, Annemiek van der Eijk, Richard Molenkamp, Marion Koopmans, on behalf of the Dutch national COVID-19 response team. |

| | | | |
|---|---|---|---|
| EPI_ISL_413582 | ErasmusMC | Erasmus Medical Center | David Nieuwenhuijse, Bas Oude Munnink, Reina Sikkema, Claudia Schapendonk, Irina Chestakova, Anne van der Linden, Mark Pronk, Pascal Lexmond, Corien Swaan, Manon Haverkate, Madelief Mollers, Mart Stein, Sandra Kengne Kamga Mobou, Jeroen van Kampen, Jolanda Voermans, Aura Timen, Corine GeurtsvanKessel, Annemiek van der Eijk, Richard Molenkamp, Marion Koopmans, on behalf of the Dutch national COVID-19 response team. |
| EPI_ISL_413583 | MHC Rotterdam-Rijnmond | Erasmus Medical Center | David Nieuwenhuijse, Bas Oude Munnink, Reina Sikkema, Claudia Schapendonk, Irina Chestakova, Anne van der Linden, Mark Pronk, Pascal Lexmond, Corien Swaan, Manon Haverkate, Madelief Mollers, Mart Stein, Sandra Kengne Kamga Mobou, Jeroen van Kampen, Jolanda Voermans, Aura Timen, Corine GeurtsvanKessel, Annemiek van der Eijk, Richard Molenkamp, Marion Koopmans, on behalf of the Dutch national COVID-19 response team. |
| EPI_ISL_413584, EPI_ISL_413585 | unknown | Erasmus Medical Center | David Nieuwenhuijse, Bas Oude Munnink, Reina Sikkema, Claudia Schapendonk, Irina Chestakova, Anne van der Linden, Mark Pronk, Pascal Lexmond, Corien Swaan, Manon Haverkate, Madelief Mollers, Mart Stein, Sandra Kengne Kamga Mobou, Jeroen van Kampen, Jolanda Voermans, Aura Timen, Corine GeurtsvanKessel, Annemiek van der Eijk, Richard Molenkamp, Marion Koopmans, on behalf of the Dutch national COVID-19 response team. |
| EPI_ISL_413586, EPI_ISL_413587 | Foundation Elisabeth-Tweesteden Ziekenhuis | Erasmus Medical Center | David Nieuwenhuijse, Bas Oude Munnink, Reina Sikkema, Claudia Schapendonk, Irina Chestakova, Anne van der Linden, Mark Pronk, Pascal Lexmond, Corien Swaan, Manon Haverkate, Madelief Mollers, Mart Stein, Sandra Kengne Kamga Mobou, Jeroen van Kampen, Jolanda Voermans, Aura Timen, Corine GeurtsvanKessel, Annemiek van der Eijk, Richard Molenkamp, Marion Koopmans, on behalf of the Dutch national COVID-19 response team. |
| EPI_ISL_413588, EPI_ISL_413589, EPI_ISL_413590 | MHC Utrecht | Erasmus Medical Center | David Nieuwenhuijse, Bas Oude Munnink, Reina Sikkema, Claudia Schapendonk, Irina Chestakova, Anne van der Linden, Mark Pronk, Pascal Lexmond, Corien Swaan, Manon Haverkate, Madelief Mollers, Mart Stein, Sandra Kengne Kamga Mobou, Jeroen van Kampen, Jolanda Voermans, Aura Timen, Corine GeurtsvanKessel, Annemiek van der Eijk, Richard Molenkamp, Marion Koopmans, on behalf of the Dutch national COVID-19 response team. |
| EPI_ISL_413591 | MHC Flevoland | Erasmus Medical Center | David Nieuwenhuijse, Bas Oude Munnink, Reina Sikkema, Claudia Schapendonk, Irina Chestakova, Anne van der Linden, Mark Pronk, Pascal Lexmond, Corien Swaan, Manon Haverkate, Madelief Mollers, Mart Stein, Sandra Kengne Kamga Mobou, Jeroen van Kampen, Jolanda Voermans, Aura Timen, Corine GeurtsvanKessel, Annemiek van der Eijk, Richard Molenkamp, Marion Koopmans, on behalf of the Dutch national COVID-19 response team. |
| EPI_ISL_413592 | Department of Laboratory Medicine, National Taiwan University Hospital | Microbial Genomics Core Lab, National Taiwan University Centers of Genomic and Precision Medicine | Shiou-Hwei Yeh, You-Yu Lin, Ya-Yun Lai, Chiao-Ling Li, Shan-Chwen Chang, Pei-Jer Chen, Sui-Yuan Chang |
| EPI_ISL_413593 | Laboratoire National de Santé | Erasmus Medical Center | David Nieuwenhuijse, Bas Oude Munnink, Reina Sikkema, Claudia Schapendonk, Irina Chestakova, Anne van der Linden, Mark Pronk, Pascal Lexmond, T. Abdelrahman, G. Fournier, J. Mossong, T. Nguyen, Jeroen van Kampen, Jolanda Voermans, Corine GeurtsvanKessel, Annemiek van der Eijk, Richard Molenkamp, Marion Koopmans, on behalf of the Dutch national COVID-19 response team. |
| EPI_ISL_413594 | Centre for Infectious Diseases and Microbiology Laboratory Services | NSW Health Pathology - Institute of Clinical Pathology and Medical Research; Westmead Hospital; University of Sydney | Rockett R, Eden J-S, Lam C, Gray K, Timms, V, Gall, M, Alicia, A, Carter I, Rahman H, Holmes EC, , O'Sullivan MV, Sintchenko V, Chen SC, Maddocks S, Kok J and Dwyer DE for the 2019-nCoV Study Group* |
| EPI_ISL_413595 | Centre for Infectious Diseases and Microbiology Laboratory Services | NSW Health Pathology - Institute of Clinical Pathology and Medical Research; Westmead Hospital; University of Sydney | Rockett R, Eden J-S, Lam C, Gray K, Timms, V, Gall, M, Carter I, Rahman H, Holmes EC, O'Sullivan MV, Sintchenko V, Chen SC, Maddocks S, Kok J and Dwyer DE for the 2019-nCoV Study Group* |
| EPI_ISL_413596 | Centre for Infectious Diseases and Microbiology - Public Health | NSW Health Pathology - Institute of Clinical Pathology and Medical Research; Westmead Hospital; University of Sydney | Rockett R, Eden J-S, Lam C, Gray K, Timms, V, Gall, M, Carter I, Rahman H, Holmes EC, O'Sullivan MV, Sintchenko V, Chen SC, Maddocks S, Kok J and Dwyer DE for the 2019-nCoV Study Group* |
| EPI_ISL_413597 | Centre for Infectious Diseases and Microbiology- Public Health | NSW Health Pathology - Institute of Clinical Pathology and Medical Research; Westmead Hospital; University of Sydney | Lam C, Eden J-S, Rockett R, Gray K, Timms, V, Gall, M, Carter I, Rahman H, Holmes EC, O'Sullivan MV, Sintchenko V, Chen SC, Maddocks S, Kok J and Dwyer DE for the 2019-nCoV Study Group* |
| EPI_ISL_413598 | Centre for Infectious Diseases and Microbiology - Public Health | NSW Health Pathology - Institute of Clinical Pathology and Medical Research; Westmead Hospital; University of Sydney | Gray K, Eden J-S, Lam C, Rockett R, Timms, V, Gall, M, Carter I, Rahman H, Holmes EC, O'Sullivan MV, Sintchenko V, Chen SC, Maddocks S, Kok J and Dwyer DE for the 2019-nCoV Study Group* |
| EPI_ISL_413599 | Centre for Infectious Diseases and Microbiology - Public Health | NSW Health Pathology - Institute of Clinical Pathology and Medical Research; Westmead Hospital; University of Sydney | Timms, V, Eden J-S, Lam C, Gray K, Rockett R, Gall, M, Carter I, Rahman H, Holmes EC, O'Sullivan MV, Sintchenko V, Chen SC, Maddocks S, Kok J and Dwyer DE for the 2019-nCoV Study Group* |
| EPI_ISL_413600 | Centre for Infectious Diseases and Microbiology - Public Health | NSW Health Pathology - Institute of Clinical Pathology and Medical Research; Westmead Hospital; University of Sydney | Gall, M, Eden J-S, Lam C, Gray K, Timms, V, Rockett R, Carter I, Rahman H, Holmes EC, O'Sullivan MV, Sintchenko V, Chen SC, Maddocks S, Kok J and Dwyer DE for the 2019-nCoV Study Group* |
| EPI_ISL_413601 | UW Virology Lab | UW Virology Lab | Pavitra Roychoudhury, Hong Xie, Keith Jerome, Alexander Greninger |
| EPI_ISL_413602, EPI_ISL_413603 | Department of Virology and Immunology, University of Helsinki and Helsinki University Hospital, Huslab Finland | Department of Virology, Faculty of Medicine, University of Helsinki, Helsinki, Finland | Teemu Smura, Hannimari Kallio-Kokko, Olli Vapalahti |
| EPI_ISL_413604 | Department of Virology and Immunology, University of Helsinki and Helsinki University Hospital, Huslab Finland | Department of Virology, Faculty of Medicine, University of Helsinki, Helsinki, Finland | Teemu Smura, Hannimari Kallio-Kokko, Olli Vapalahti |
| EPI_ISL_413605 | Department of Virology and Immunology, University of Helsinki and Helsinki University Hospital, Huslab Finland | Department of Virology, Faculty of Medicine, University of Helsinki, Helsinki, Finland | Teemu Smura, Hannimari Kallio-Kokko, Olli Vapalahti |
| EPI_ISL_413606, EPI_ISL_413607, EPI_ISL_413608, EPI_ISL_413609, EPI_ISL_413610, EPI_ISL_413611 | unknown | Pathogen Discovery, Respiratory Viruses Branch, Division of Viral Diseases, Centers for Dieases Control and Prevention | Anna Uehara, Ying Tao, Clinton R. Paden, Krista Queen, Jing Zhang, Yan Li, Mary S. Keckler, Alison S Laufer Halpin, Haibin Wang, Jasmine Padilla, Justin Lee, Christopher A. Elkins, Susan I. Gerber, Suxiang Tong |
| EPI_ISL_413612, EPI_ISL_413613, EPI_ISL_413614, EPI_ISL_413615, EPI_ISL_413616, EPI_ISL_413617 | unknown | Pathogen Discovery, Respiratory Viruses Branch, Division of Viral Diseases, Centers for Dieases Control and Prevention | Ying Tao, Clinton R. Paden, Krista Queen, Anna Uehara, Jing Zhang, Yan Li, Haibin Wang, Shifaq Kamili, Xiaoyan Lu, Brian Lynch, Senthil Kumar K. Sakthivel, Brett L. Whitaker, Lijuan Wang, Janna' R. Murray, Jasmine Padilla, Justin Lee, Susan I. Gerber, Stephen Lindstrom, Suxiang Tong |
| EPI_ISL_413618, EPI_ISL_413619, EPI_ISL_413620, EPI_ISL_413621, EPI_ISL_413622, EPI_ISL_413623 | unknown | Pathogen Discovery, Respiratory Viruses Branch, Division of Viral Diseases, Centers for Dieases Control and Prevention | Clinton R. Paden, Ying Tao, Krista Queen, Anna Uehara, Jing Zhang, Yan Li, Haibin Wang, Shifaq Kamili, Xiaoyan Lu, Brian Lynch, Senthil Kumar K. Sakthivel, Brett L. Whitaker, Lijuan Wang, Janna' R. Murray, Jasmine Padilla, Justin Lee, Susan I. Gerber, Stephen Lindstrom, Suxiang Tong |
| EPI_ISL_413647 | Centro Hospital do Porto, E.P.E. - H. Geral de Santo Antonio | Instituto Nacional de Saude (INSA) | Raquel Guiomar, Inês Costa, Pedro Pechirra, Joana Mendonça, Luís Vieira, Helena Ramos, Joana Isidro, Vítor Borges, João Paulo Gomes |
| EPI_ISL_413648 | Centro Hospitalar e Universitário de Sao Joao, Porto | Instituto Nacional de Saude (INSA) | Raquel Guiomar, Inês Costa, Pedro Pechirra, Joana Mendonça, Luís Vieira, João Tiago Guimarães, Joana Isidro, Vítor Borges, João Paulo Gomes |
| EPI_ISL_413649, EPI_ISL_413650, EPI_ISL_413651, EPI_ISL_413652, EPI_ISL_413653 | UW Virology Lab | UW Virology Lab | Pavitra Roychoudhury, Hong Xie, Keith Jerome, Alexander Greninger |
| EPI_ISL_413691, EPI_ISL_413692, EPI_ISL_413693, EPI_ISL_413694, EPI_ISL_413695, EPI_ISL_413696, EPI_ISL_413697, EPI_ISL_413711, EPI_ISL_413729, EPI_ISL_413746, EPI_ISL_413747, EPI_ISL_413748, EPI_ISL_413749, EPI_ISL_413750, EPI_ISL_413751, EPI_ISL_413752, EPI_ISL_413753, EPI_ISL_413761, EPI_ISL_413791, EPI_ISL_413809 | | | |
| see above | Weifang Center for Disease Control and Prevention | Weifang Center for Disease Control and Prevention & BGI-Shenzhen | Qing Nie, Xingguang Li, Erik M Volz, Han Fu, Haowei Wang, Xiaoyue Xi, Wei Chen, Dehui Liu, Yingying Chen, Mengmeng Tian, Wei Tan, Junjie Zai, Wanying Sun, Jiandong Li, Junhua Li |
| EPI_ISL_413850, EPI_ISL_413851, EPI_ISL_413852, EPI_ISL_413853, EPI_ISL_413854, EPI_ISL_413855, EPI_ISL_413856, EPI_ISL_413857, EPI_ISL_413858, EPI_ISL_413859, EPI_ISL_413860, EPI_ISL_413861, EPI_ISL_413862, EPI_ISL_413863, EPI_ISL_413864 | | | |
| see above | Guangdong Provincial Institution of Public Health, Guangdong Provinical Center for Disease Control and Prevention | Guangdong Provincial Institution of Public Health | Jing Lu, Louis du Plessis, Liu Zhe, Jiufeng Sun, Sarah François, Huifang Lin, Moritz Kraemer, Jingju Peng, Qianlin Xiong, Runyu Yuan, Lilian Zeng, Pingping Zhou, Chuming Liang, Tao Liu, Wei Li, Juan Su, Huanying Zheng, Kang Min, Song Tie, Bo Peng, Shisong Fang, Wenzhe Su, Kuibiao Li, Ruilin Sun, Ru bai, Xi Tang, Minfeng Liang, Nuno Faria, Josh Quick, Andrew Rambaut, Verity Hill, Wenjun Ma, Nick Loman, Oliver Pybus, Changwen Ke |
| EPI_ISL_413865 | Guangdong Provincial Institution of Public Health, Guangdong Provinical Center for Disease Control and Prevention | Guangdong Provincial Institution of Public Health | Jing Lu, Louis du Plessis, Liu Zhe, Jiufeng Sun, Sarah François, Huifang Lin, Moritz Kraemer, Jingju Peng, Qianlin Xiong, Runyu Yuan, Lilian Zeng, Pingping Zhou, Chuming Liang, Tao Liu, Wei Li, Juan Su, Huanying Zheng, Kang Min, Song Tie, Bo Peng, Shisong Fang, Wenzhe Su, Kuibiao Li, Ruilin Sun, Ru bai, Xi Tang, Minfeng Liang, Nuno Faria, Josh Quick, Andrew Rambaut, Verity Hill, Wenjun Ma, Nick Loman, Oliver Pybus, Changwen Ke |
| EPI_ISL_413866 | Guangdong Provincial Institution of Public Health, Guangdong Provinical Center for Disease Control and Prevention | Guangdong Provincial Institution of Public Health | Jing Lu, Louis du Plessis, Liu Zhe, Jiufeng Sun, Sarah François, Huifang Lin, Moritz Kraemer, Jingju Peng, Qianlin Xiong, Runyu Yuan, Lilian Zeng, Pingping Zhou, Chuming Liang, Tao Liu, Wei Li, Juan Su, Huanying Zheng, Kang Min, Song Tie, Bo Peng, Shisong Fang, Wenzhe Su, Kuibiao Li, Ruilin Sun, Ru bai, Xi Tang, Minfeng Liang, Nuno Faria, Josh Quick, Andrew Rambaut, Verity Hill, Wenjun Ma, Nick Loman, Oliver Pybus, Changwen Ke |
| EPI_ISL_413867 | Guangdong Provincial Institution of Public Health, Guangdong Provinical Center for Disease Control and Prevention | Guangdong Provincial Institution of Public Health | Jing Lu, Louis du Plessis, Liu Zhe, Jiufeng Sun, Sarah François, Huifang Lin, Moritz Kraemer, Jingju Peng, Qianlin Xiong, Runyu Yuan, Lilian Zeng, Pingping Zhou, Chuming Liang, Tao Liu, Wei Li, Juan Su, Huanying Zheng, Kang Min, Song Tie, Bo Peng, Shisong Fang, Wenzhe Su, Kuibiao Li, Ruilin Sun, Ru bai, Xi Tang, Minfeng Liang, Nuno Faria, Josh Quick, Andrew Rambaut, Verity Hill, Wenjun Ma, Nick Loman, Oliver Pybus, Changwen Ke |
| EPI_ISL_413868 | Guangdong Provincial Institution of Public Health, Guangdong Provinical Center for Disease Control and Prevention | Guangdong Provincial Institution of Public Health | Jing Lu, Louis du Plessis, Liu Zhe, Jiufeng Sun, Sarah François, Huifang Lin, Moritz Kraemer, Jingju Peng, Qianlin Xiong, Runyu Yuan, Lilian Zeng, Pingping Zhou, Chuming Liang, Tao Liu, Wei Li, Juan Su, Huanying Zheng, Kang Min, Song Tie, Bo Peng, Shisong Fang, Wenzhe Su, Kuibiao Li, Ruilin Sun, Ru bai, Xi Tang, Minfeng Liang, Nuno Faria, Josh Quick, Andrew Rambaut, Verity Hill, Wenjun Ma, Nick Loman, Oliver Pybus, Changwen Ke |
| EPI_ISL_413869 | Guangdong Provincial Institution of Public Health, Guangdong Provinical Center for Disease Control and Prevention | Guangdong Provincial Institution of Public Health | Jing Lu, Louis du Plessis, Liu Zhe, Jiufeng Sun, Sarah François, Huifang Lin, Moritz Kraemer, Jingju Peng, Qianlin Xiong, Runyu Yuan, Lilian Zeng, Pingping Zhou, Chuming Liang, Tao Liu, Wei Li, Juan Su, Huanying Zheng, Kang Min, Song Tie, Bo Peng, Shisong Fang, Wenzhe Su, Kuibiao Li, Ruilin Sun, Ru bai, Xi Tang, Minfeng Liang, Nuno Faria, Josh Quick, Andrew Rambaut, Verity Hill, Wenjun Ma, Nick Loman, Oliver Pybus, Changwen Ke |
| EPI_ISL_413870, EPI_ISL_413871, EPI_ISL_413872, EPI_ISL_413873, EPI_ISL_413874, EPI_ISL_413875, EPI_ISL_413876, EPI_ISL_413877, EPI_ISL_413878, EPI_ISL_413879, EPI_ISL_413880, EPI_ISL_413881, EPI_ISL_413882, EPI_ISL_413883, EPI_ISL_413884, EPI_ISL_413885, EPI_ISL_413886, EPI_ISL_413887, EPI_ISL_413888, EPI_ISL_413889, EPI_ISL_413890, EPI_ISL_413891, EPI_ISL_413892, EPI_ISL_413893, EPI_ISL_413894, EPI_ISL_413895, EPI_ISL_413896, EPI_ISL_413897, EPI_ISL_413898, EPI_ISL_413899, EPI_ISL_413900, EPI_ISL_413901, EPI_ISL_413902 | | | |
| see above | Guangdong Provincial Institution of Public Health, Guangdong Provinical Center for Disease Control and Prevention | Guangdong Provincial Institution of Public Health | Jing Lu, Louis du Plessis, Liu Zhe, Jiufeng Sun, Sarah François, Huifang Lin, Moritz Kraemer, Jingju Peng, Qianlin Xiong, Runyu Yuan, Lilian Zeng, Pingping Zhou, Chuming Liang, Tao Liu, Wei Li, Juan Su, Huanying Zheng, Kang Min, Song Tie, Bo Peng, Shisong Fang, Wenzhe Su, Kuibiao Li, Ruilin Sun, Ru bai, Xi Tang, Minfeng Liang, Nuno Faria, Josh Quick, Andrew Rambaut, Verity Hill, Wenjun Ma, Nick Loman, Oliver Pybus, Changwen Ke |
| EPI_ISL_413904 | Iran National Influenza Center | Iran National Influenza Center | Nazanin Zahra Shafiei Jandaghi,Jila Yavarian,Fatemeh AjamiNejad, Nastaran Ghavvami,Kaveh Sadeghi and Talat Mokhtari Azad |
| EPI_ISL_413905 | Iran National Influenza Center | Iran National Influenza Center | Nazanin Zahra Shafiei Jandaghi, Jila Yavarian, Vahid Salimi, Kaveh Sadeghi and Talat Mokhtari Azad |
| EPI_ISL_413906 | Iran National Influenza Center | Iran National Influenza Center | Jila Yavarian, Nazanin Zahra Shafiei Jandaghi, Kaveh Sadeghi, Nastaran Ghavvami, Fatemeh Ajaminejad and Talat Mokhtari Azad |
| EPI_ISL_413922, EPI_ISL_413924, EPI_ISL_413925, EPI_ISL_413926, EPI_ISL_413928 | California Department of Public Health | Chiu Laboratory, University of California, San Francisco | Xianding Deng, Scot Federman, Chao-Yang Pan, Hugo Guevara,Wei Gu, Debra A. Wadford, and Charles Y. Chiu |
| EPI_ISL_413930 | California Department of Public Health | Chiu Laboratory UCSF-Abbott Viral Diagnostics and Discovery Center University of California, San Francisco | Xianding Deng, Scot Federman, Guixia Yu, Chao-Yang Pan, Hugo Guevara, Alicia Sotomayor-Gonzalez, Allan Gopez, Wei Gu, Steve Miller, Debra A. Wadford, and Charles Y. Chiu |

| | | | |
|---|---|---|---|
| EPI_ISL_413931 | California Department of Public Health | Chiu Laboratory, University of California, San Francisco | Xianding Deng, Scot Federman, Chao-Yang Pan, Hugo Guevara,Wei Gu, Debra A. Wadford, and Charles Y. Chiu |
| EPI_ISL_413949 | Iran National Influenza Center | Iran National Influenza Center | Jila Yavarian, Nazanin Zahra Shafiei Jandaghi,Kaveh Sadeghi, Saeedeh Mahfozi and Talat Mokhtari Azad |
| EPI_ISL_413955 | Iran National Influenza Center | Iran National Influenza Center | Nazanin Zahra Shafiei Jandaghi, Jila Yavarian, Kaveh Sadeghi, Vahid Salimi, Simin Abbasi and Talat Mokhtari Azad |
| EPI_ISL_413996, EPI_ISL_413997, EPI_ISL_413999 | Laboratoire de Virologie, HUG | Swiss National Reference Centre for Influenza | LAUBSCHER Florian et al. |
| EPI_ISL_414005, EPI_ISL_414006, EPI_ISL_414007, EPI_ISL_414008, EPI_ISL_414009, EPI_ISL_414010, EPI_ISL_414011, EPI_ISL_414012, EPI_ISL_414013 | Respiratory Virus Unit, Microbiology Services Colindale, Public Health England | Respiratory Virus Unit, Microbiology Services Colindale, Public Health England | Monica Galiano, Shahjahan Miah, Angie Lackenby, Omolola Akinbami, Tiina Talts, Leena Bhaw, Richard Myers, Steven Platt, Kirstin Edwards, Jonathan Hubb, Joanna Ellis, Maria Zambon |
| EPI_ISL_414014 | Hospital Israelita Albert Einstein | Instituto Adolfo Lutz, Interdiciplinary Procedures Center, Strategic Laboratory | Claudio Tavares Sacchi, Claudia Regina Gonçalves, Katia Correia dos Santos, Carlos Henrique Camargo, Maria do Carmo Sampaito Tavares Timenetsky, Terezinha Maria de Paiva, Ester Cerdeira Sabino |
| EPI_ISL_414015 | Hospital São Joaquim Beneficencia Portuguesa | Instituto Adolfo Lutz, Interdiciplinary Procedures Center, Strategic Laboratory | Claudio Tavares Sacchi, Claudia Regina Gonçalves, SimoneGuadagnucci Morillo, Carlos Henrique Camargo, Maria do Carmo Sampaito Tavares Timenetsky, Fabiana Cristina Pereira dos Santos Terezinha Maria de Paiva, Ester Cerdeira Sabino |
| EPI_ISL_414016 | Hospital São Joaquim Beneficencia Portuguesa | Instituto Adolfo Lutz, Interdiciplinary Procedures Center, Strategic Laboratory | Claudio Tavares Sacchi, Claudia Regina Gonçalves, Audrey Cilli, Carlos Henrique Camargo, Maria do Carmo Sampaito Tavares Timenetsky, Daniela Bernardes Borges da Silva, Terezinha Maria de Paiva, Ester Cerdeira Sabino |
| EPI_ISL_414017 | Hospital São Joaquim Beneficencia Portuguesa | Instituto Adolfo Lutz, Interdiciplinary Procedures Center, Strategic Laboratory | Claudio Tavares Sacchi, Claudia Regina Gonçalves, Fabiana Cristina Pereira dos Santos, Carlos Henrique Camargo, Maria do Carmo Sampaito Tavares Timenetsky, Daniela Bernardes Borges da Silva, Terezinha Maria de Paiva, Ester Cerdeira Sabino |
| EPI_ISL_414019, EPI_ISL_414020, EPI_ISL_414021, EPI_ISL_414022, EPI_ISL_414023 | Laboratoire de Virologie, HUG | Swiss National Reference Centre for Influenza | LAUBSCHER Florian et al. |
| EPI_ISL_414024, EPI_ISL_414025, EPI_ISL_414026 | West of Scotland Specialist Virology Centre, NHSGGC | MRC-University of Glasgow Centre for Virus Research | Emma Thomson, Antonia Ho; Kathy Smollett, Daniel Mair, Stephen Carmichael, Ana da Silva Filipe; Richard Orton, David L Robertson; Alasdair MacLean, Rory Gunson. |
| EPI_ISL_414027 | West of Scotland Specialist Virology Centre, NHSGGC | MRC-University of Glasgow Centre for Virus Research | Emma Thomson, Antonia Ho; Kathy Smollett, Daniel Mair, Stephen Carmichael, Ana da Silva Filipe; Richard Orton, David L Robertson; Alasdair MacLean, Rory Gunson. |
| EPI_ISL_414040, EPI_ISL_414041, EPI_ISL_414042, EPI_ISL_414043, EPI_ISL_414044 | Respiratory Virus Unit, Microbiology Services Colindale, Public Health England | Respiratory Virus Unit, Microbiology Services Colindale, Public Health England | Monica Galiano, Shahjahan Miah, Angie Lackenby, Omolola Akinbami, Tiina Talts, Leena Bhaw, Richard Myers, Steven Platt, Kirstin Edwards, Jonathan Hubb, Joanna Ellis, Maria Zambon |
| EPI_ISL_414045 | LACEN RJ - Laboratório Central de Saúde Pública Noel Nutels | Instituto Oswaldo Cruz FIOCRUZ - Laboratory of Respiratory Viruses and Measles (LVRS) | Paola Resende, Alisson Fabri, Joilson Xavier, Sunando Roy, Fernando Motta, Aline Mattos, Milene Miranda, Cristiana Garcia, Braulia Caetano, Maria Ogrzewalska, Jonathan Lopes, Luciana Appolinario, Maria Nóbrega, Marilda Siqueira |
| EPI_ISL_414363, EPI_ISL_414364, EPI_ISL_414365, EPI_ISL_414366, EPI_ISL_414367, EPI_ISL_414368, EPI_ISL_414369 | UW Virology Lab | UW Virology Lab | Pavitra Roychoudhury, Hong Xie, Keith Jerome, Alexander Greninger |
| EPI_ISL_414371 | Iran National Influenza Center | Iran National Influenza Center | Nazanin Zahra Shafiei Jandaghi, Jila Yavarian, Kaveh Sadeghi, Nastaran Ghavvami and Talat Mokhtari Azad |
| EPI_ISL_414372 | Iran National Influenza Center | Iran National Influenza Center | Jila Yavarian, Nazanin Zahra Shafiei Jandaghi, Kaveh Sadeghi, Fatemeh Ajaminejad, Fatemeh Saadatmand and Talat Mokhtari Azad |
| EPI_ISL_414373 | Iran National Influenza Center | Iran National Influenza Center | Kaveh Sadeghi, Jila Yavarian, Nazanin Zahra Shafiei Jandaghi, Ahmad Nejati, Najmeh Parhizghari, Soad Ghabeshi and Talat Mokhtari Azad |
| EPI_ISL_414374 | Iran National Influenza Center | Iran National Influenza Center | Jila Yavarian, Nazanin Zahra Shafiei Jandaghi, Kaveh Sadeghi, Saeedeh Mahfozi, Simin Abbasi and Talat Mokhtari Azad |
| EPI_ISL_414375, EPI_ISL_414376, EPI_ISL_414377 | National Institute of Health Research and Development | National Institute of Health Research and Development | Setiawaty,V; Subangkit; Puspa,KD; Ikawati,HD; Nugraha, AA; Susilarini, NK; Agustiningsih; Ramadhany,R; Pratiwi,E; Hariastuti,NI; Kurniawati,J; Pawestri,HA; Siswanto |
| EPI_ISL_414378 | National Centre for Infectious Diseases | Programme in Emerging Infectious Diseases, Duke-NUS Medical School | Danielle E Anderson, Martin Linster, Yan Zhuang, Jayanthi Jayakumar, Louisa Sun, David CB Lye, Yee Sin Leo, Barnaby E Young, Yvonne CF Su, Gavin JD Smith |
| EPI_ISL_414379, EPI_ISL_414380 | National Centre for Infectious Diseases | Programme in Emerging Infectious Diseases, Duke-NUS Medical School | Danielle E Anderson, Martin Linster, Yan Zhuang, Jayanthi Jayakumar, David CB Lye, Yee Sin Leo, Barnaby E Young, Yvonne CF Su, Gavin JD Smith |
| EPI_ISL_414414 | Pathology Queensland | Public Health Virology Laboratory | Bixing Huang, Alyssa Pyke, Amanda De Jong, Andrew Van Den Hurk, Carmel Taylor, David Warrilow, Doris Genge, Elisabeth Gamez, Glen Hewitson, Ian Maxwell Mackay, Inga Sultana, Jamie McMahon, Jean Barcelon, Judy Northill, Mitchell Finger, Natalie Simpson, Neelima Nair, Peter Burtonclay, Peter Moore, Sarah Wheatley, Sean Moody, Sonja Hall-Mendelin, Timothy Gardam, and Frederick Moore |
| EPI_ISL_414423, EPI_ISL_414424, EPI_ISL_414425, EPI_ISL_414426, EPI_ISL_414427, EPI_ISL_414428, EPI_ISL_414429, EPI_ISL_414430, EPI_ISL_414431, EPI_ISL_414432, EPI_ISL_414433, EPI_ISL_414434, EPI_ISL_414435, EPI_ISL_414436, EPI_ISL_414437, EPI_ISL_414438, EPI_ISL_414439, EPI_ISL_414440, EPI_ISL_414441, EPI_ISL_414442, EPI_ISL_414443, EPI_ISL_414444, EPI_ISL_414445, EPI_ISL_414446, EPI_ISL_414448, EPI_ISL_414449, EPI_ISL_414450, EPI_ISL_414451, EPI_ISL_414452, EPI_ISL_414453, EPI_ISL_414454, EPI_ISL_414455, EPI_ISL_414456, EPI_ISL_414457, EPI_ISL_414458, EPI_ISL_414459, EPI_ISL_414460, EPI_ISL_414461, EPI_ISL_414462, EPI_ISL_414463, EPI_ISL_414464, EPI_ISL_414465, EPI_ISL_414466, EPI_ISL_414467, EPI_ISL_414468, EPI_ISL_414469, EPI_ISL_414470, EPI_ISL_414471 | | | |
| see above | Dutch COVID-19 response team | Erasmus Medical Center | David Nieuwenhuijse, Bas Oude Munnink, Reina Sikkema, Claudia Schapendonk, Irina Chestakova, Anne van der Linden, Mark Pronk, Pascal Lexmond, Corien Swaan, Manon Haverkate, Madelief Mollers, Mart Stein, Sandra Kengne Kamga Mobou, Jeroen van Kampen, Jolanda Voermans, Aura Timen, Corine GeurtsvanKessel, Annemiek van der Eijk, Richard Molenkamp, Marion Koopmans, on behalf of the Dutch national COVID-19 response team. |
| EPI_ISL_414475 | Iran National Influenza Center | Iran National Influenza Center | Jila Yavarian, Nazanin Zahra Shafiei Jandaghi, Ahmad Nejati, Simin Abbasi and Talat Mokhtari Azad |
| EPI_ISL_414476 | MSHS Clinical Microbiology Laboratories | MSHS Pathogen Surveillance Program | Gopi Patel, Emilia Sordillo, Melissa Gitman, Alberto Paniz-mondolfi, Matthew Hernandez, Shelcie Fabre, Jose Polanco, Ana Sylvia Gonzalez-Reiche, Zenab Khan, Nancy Francoeur, Melissa Smith, Robert Sebra, Lisa Miorin, Wen-chun Liu, Randy Albrecht, Judith Aberg, Florian Krammer, Adolfo Garcia-Sarstre, Viviana Simon, Harm van Bakel |
| EPI_ISL_414477 | The National Institute of Public Health Center for Epidemiology and Microbiology | State Veterinary Institute Prague | Alexander Nagy, Oldrich Bartos, Helena Jirincova, Klara Labska, Ludmila Novakova, Olga Storkanova, Dusan Trnka, Jaromira Vecerova |
| EPI_ISL_414479, EPI_ISL_414480, EPI_ISL_414481 | unknown | Pathogen Discovery, Respiratory Viruses Branch, Division of Viral Diseases, Centers for Disease Control and Prevention | Ying Tao, Krista Queen, Clinton R. Paden, Anna Uehara, Jing Zhang, Yan Li, Mary S. Keckler, Alison S. Laufer Halpin, Haibin Wang, Jasmine Padilla, Justin Lee, Christopher A. Elkins, Susan I. Gerber, Suxiang Tong |
| EPI_ISL_414482, EPI_ISL_414483, EPI_ISL_414484, EPI_ISL_414485 | unknown | Pathogen Discovery, Respiratory Viruses Branch, Division of Viral Diseases, Centers for Disease Control and Prevention | Krista Queen, Anna Uehara, Ying Tao, Clinton R. Paden, Jing Zhang, Yan Li, Haibin Wang, Shifaq Kamili, Xiaoyan Lu, Brian Lynch, Senthil Kumar K. Sakthivel, Brett L. Whitaker, Lijuan Wang, Janna' R. Murray, Jasmine Padilla, Justin Lee, Susan I. Gerber, Stephen Lindstrom, Suxiang Tong |
| EPI_ISL_414486 | Wales Specialist Virology Centre | Public Health Wales Microbiology Cardiff | Catherine Moore, Joanne Watkins, Sally Corden, Tom Connor |
| EPI_ISL_414487 | UCD National Virus Reference Laboratory | UCD National Virus Reference Laboratory | Michael Carr, Gabriel Gonzalez, Jonathan Dean, Suzie Coughlan, Alison Murphy, Kevin Byrne, Ken Wolfe, Jeff Connell, Brendan Loftus, Cillian F De Gascun |
| EPI_ISL_414488 | Wales Specialist Virology Centre | Public Health Wales Microbiology Cardiff | Catherine Moore, Joanne Watkins, Sally Corden, Tom Connor |
| EPI_ISL_414495 | Servicio Microbiología. Hospital Clínico Universitario. Valencia. | Sequencing and Bioinformatics Service. Molecular Epidemiology Laboratory. FISABIO-Public Health | David Navarro, Maria Alma Bracho, Giuseppe D'Auria, Griselda De Marco, Neris Garcia-Gonzalez, Fernando Gonzalez-Candelas |
| EPI_ISL_414496 | Servicio Microbiologia. Hospital Clinico Universitario. Valencia. | Sequencing and Bioinformatics Service. Molecular Epidemiology Laboratory. FISABIO-Public Health | David Navarro, María Alma Bracho, Giuseppe D'Auria, Griselda De Marco, Neris Garcia-Gonzalez, Fernando Gonzalez-Candelas |
| EPI_ISL_414497, EPI_ISL_414498, EPI_ISL_414499 | Center of Medical Microbiology, Virology, and Hospital Hygiene, University of Duesseldorf | Center of Medical Microbiology, Virology, and Hospital Hygiene, University of Duesseldorf | Ortwin Adams, Marcel Andree, Alexander Dilthey, Torsten Feldt, Sandra Hauka, Torsten Houwaart, Björn-Erik Jensen, Detlef Kindgen-Milles, Malte Kohns Vasconcelos, Klaus Pfeffer, Tina Senff, Daniel Strelow, Jörg Timm, Andreas Walker, Tobias Wienemann |
| EPI_ISL_414500, EPI_ISL_414501 | Virology Department, Sheffield Teaching Hospitals NHS Foundation Trust | Department of Infection, Immunity and Cardiovascular Disease, The Florey Institute, The Medical School, University of Sheffield | Thushan de Silva, Matthew Parker, Matthew Wyles, Mehmet Yavuz, Mohammad Raza, Cariad Evans |
| EPI_ISL_414504, EPI_ISL_414505, EPI_ISL_414506, EPI_ISL_414507, EPI_ISL_414508, EPI_ISL_414509 | Center of Medical Microbiology, Virology, and Hospital Hygiene, University of Duesseldorf | Center of Medical Microbiology, Virology, and Hospital Hygiene, University of Duesseldorf | Ortwin Adams, Marcel Andree, Alexander Dilthey, Torsten Feldt, Sandra Hauka, Torsten Houwaart, Björn-Erik Jensen, Detlef Kindgen-Milles, Malte Kohns Vasconcelos, Klaus Pfeffer, Tina Senff, Daniel Strelow, Jörg Timm, Andreas Walker, Tobias Wienemann |
| EPI_ISL_414510 | Key Laboratory of Medical Molecular Virology (MOE/NHC/CAMS), School of Basic Medicine, Shanghai Medical College, Fudan University | Key Laboratory of Medical Molecular Virology (MOE/NHC/CAMS), School of Basic Medicine, Shanghai Medical College, Fudan University | Zhang,R., Yi,Z., Wang,Y., Teng,Z., Xu,W., Song,W., Cai,X., Sun,Z., Gu,C., Zhou,Y., Chen,H., Ye,R., Han,W., Zhu,Y., Feng,F., Fang,F., Li,C., Zhang,X., Qu,D., Fu,C., Xie,Y. and Yuan,Z. |
| EPI_ISL_414511 | Department of Microbiology; Ryota Kumagai Tokyo Metropolitan Institute of Public Health | Tokyo Metoropolitan Institute of Public Health | Kumagai,R., Yoshida,I., Nagashima,M., Chiba,T. and Sadamasu,K. |
| EPI_ISL_414512 | Dipartimento di Medicina e Chirurgia, University of Insubria and Ospedale di Circolo e Fondazione Macchi | Dipartimento di Medicina e Chirurgia, University of Insubria and Ospedale di Circolo e Fondazione Macchi | Genoni,A.P., Baj,A. and Rossi,A. |
| EPI_ISL_414513 | Gastrointestinal and Liver Diseases Research Center, Iran University of Medical Sciences | Gastrointestinal and Liver Diseases Research Center, Iran University of Medical Sciences | Karbalaie Niya,M.H., Laali,A., Tabibzadeh,A., Safarnezhad Tameshkel,F., Zamani,F., Sohrabi,M.R., Ranjbar,M., Savaj,S., Rezaie,N., Ajdarkosh,H., Keyvani,H., Khoonsari,M., Ameli,M., Nikkhah,M., Ghanbari,B., Faraji,A., Jamshidi Makiani,M. and Roham,M. |
| EPI_ISL_414514 | Gastrointestinal and Liver Diseases Research Center, Iran University of Medical Sciences | Gastrointestinal and Liver Diseases Research Center, Iran University of Medical Sciences | Karbalaie Niya,M.H., Laali,A., Tabibzadeh,A., Safarnezhad Tameshkel,F., Zamani,F., Sohrabi,M.R., Ranjbar,M., Savaj,S., Rezaie,N., Ajdarkosh,H., Keyvani,H., Khoonsari,M., Ameli,M., Nikkhah,M., Ghanbari,B., Faraji,A., Jamshidi Makiani,M. and Roham,M. |
| EPI_ISL_414515 | Biotechnology, National Centre for Disease Control | Biotechnology, National Centre for Disease Control | Kumar,P., Dhar,M., Vashistha,H., Singh,P., Sharma,U., Singh,S., Saini,N., Lall,H., Bala,M., Singh,S.K. and Rakshit,P. |
| EPI_ISL_414516 | Molecular Pathology, Mehr Pathobiology Lab | Molecular Pathology, Mehr Pathobiology Lab | Hamedi Asl,D., Soleimani,M., Mirzapour,M., Shabadori,A. and Kamali,M. |
| EPI_ISL_414517 | Hong Kong Department of Health | School of Public Health, The University of Hong Kong | Dominic N.C. Tsang, Daniel K.W. Chu, Leo L.M. Poon, Malik Peiris |
| EPI_ISL_414518 | Tai Lung Veterinary Laboratory, Agriculture, Fisheries and Conservation Department | School of Public Health, The University of Hong Kong | Thomas S.H. Chung, Christopher J Brackman, Daniel K.W. Chu, Leo L.M. Poon, Malik Peiris |

| | | | |
|---|---|---|---|
| EPI_ISL_414519 | Hong Kong Department of Health | School of Public Health, The University of Hong Kong | Dominic N.C. Tsang, Daniel K.W. Chu, Leo L.M. Poon, Malik Peiris |
| EPI_ISL_414520, EPI_ISL_414521 | Bundeswehr Institute of Microbiology | Bundeswehr Institute of Microbiology | Mathias C Walter, Markus H Antwerpen and Roman Wölfel |
| EPI_ISL_414522, EPI_ISL_414523, EPI_ISL_414524, EPI_ISL_414525, EPI_ISL_414526 | Respiratory Virus Unit, Microbiology Services Colindale, Public Health England | Respiratory Virus Unit, Microbiology Services Colindale, Public Health England | Monica Galiano, Shahjahan Miah, Angie Lackenby, Omolola Akinbami, Tiina Talts, Leena Bhaw, Richard Myers, Steven Platt, Kirstin Edwards, Jonathan Hubb, Joanna Ellis, Maria Zambon |
| EPI_ISL_414527, EPI_ISL_414528 | Hong Kong Department of Health | School of Public Health, The University of Hong Kong | Dominic N.C. Tsang, Daniel K.W. Chu, Leo L.M. Poon, Malik Peiris |
| EPI_ISL_414529, EPI_ISL_414530, EPI_ISL_414531, EPI_ISL_414532, EPI_ISL_414533, EPI_ISL_414534, EPI_ISL_414535, EPI_ISL_414536, EPI_ISL_414537, EPI_ISL_414538, EPI_ISL_414539, EPI_ISL_414540, EPI_ISL_414541, EPI_ISL_414542, EPI_ISL_414543, EPI_ISL_414544, EPI_ISL_414545, EPI_ISL_414546, EPI_ISL_414547, EPI_ISL_414548, EPI_ISL_414549, EPI_ISL_414550, EPI_ISL_414551, EPI_ISL_414552, EPI_ISL_414553, EPI_ISL_414554, EPI_ISL_414555, EPI_ISL_414556, EPI_ISL_414557, EPI_ISL_414558, EPI_ISL_414559, EPI_ISL_414560, EPI_ISL_414561, EPI_ISL_414562, EPI_ISL_414563, EPI_ISL_414564, EPI_ISL_414565, EPI_ISL_414566 | | | |
| see above | Dutch COVID-19 response team | Erasmus Medical Center | David Nieuwenhuijse, Bas Oude Munnink, Reina Sikkema, Claudia Schapendonk, Irina Chestakova, Anne van der Linden, Mark Pronk, Pascal Lexmond, Corien Swaan, Manon Haverkate, Madelief Mollers, Mart Stein, Sandra Kengne Kamga Mobou, Jeroen van Kampen, Jolanda Voermans, Aura Timen, Corine GeurtsvanKessel, Annemiek van der Eijk, Richard Molenkamp, Marion Koopmans, on behalf of the Dutch national COVID-19 response team. |
| EPI_ISL_414567 | Hong Kong Department of Health | School of Public Health, The University of Hong Kong | Dominic N.C. Tsang, Daniel K.W. Chu, Leo L.M. Poon, Malik Peiris |
| EPI_ISL_414568 | Pasteur Institute of Iran | Pasteur Institute of Iran | Arash Arashkia, Kayhan Azadmanesh, Mohammad Hassan Pouriayevali, Tahmineh Jalali, Zahra Ahmadi, Mohammad Sadegh Shams Nosrati, Ali Maleki, Zabihollah Shoja, Sanam Azad-Mazjiri, Neda Amin, Mehdi Rohani, Saber Esmaeili, Ahmad Ghasemi, Amir Hesam Nemati, Ahmad Mahmoudi, Zahra Fereydouni, Mahsa Tavakolirad, Tahereh Mohammadi, Sahar Khakifirouz, Mehdi Fazlalipour, Maryam Rostamtabar, Arezoo Parikhani, Hesam Karimi, Kazem Baesi, Seyed Dawood Mousavi Nasab, Mahmood Barati, Mohammad Reza Asadi Karam, Mehri Habibi, Fatemeh Fotouhi-Chahooki, Neda Afzali, Ali Torabi, Azita Eshratkhah mohammadnejad, Seyedeh Sahar Bathaeian, Mina Bahri, Mohamad Mahdi Mortazavipour, Seyedeh Atefe Hosseini, Farideh Niknam, Parastoo Yekta Sanati, Hadiseh Shokouhi, Azam Amirian, Afsaneh zokaei, Hajarossadat Ghaderi, Mahboobeh Rafigh, Elmira Vadaye kheiri, Akram Agharezaei, Akram Abouie Mehrizi, seyedeh Zahra Moravej, Mostafa Salehi-Vaziri |
| EPI_ISL_414569, EPI_ISL_414571 | Hong Kong Department of Health | School of Public Health, The University of Hong Kong | Dominic N.C. Tsang, Daniel K.W. Chu, Leo L.M. Poon, Malik Peiris |
| EPI_ISL_414572 | Pasteur Institute of Iran | Pasteur Institute of Iran | Arash Arashkia, Kayhan Azadmanesh, Mohammad Hassan Pouriayevali, Tahmineh Jalali, Zahra Ahmadi, Mohammad Sadegh Shams Nosrati, Ali Maleki, Zabihollah Shoja, Sanam Azad-Mazjiri, Neda Amin, Mehdi Rohani, Saber Esmaeili, Ahmad Ghasemi, Amir Hesam Nemati, Ahmad Mahmoudi, Zahra Fereydouni, Mahsa Tavakolirad, Tahereh Mohammadi, Sahar Khakifirouz, Mehdi Fazlalipour, Maryam Rostamtabar, Arezoo Parikhani, Hesam Karimi, Kazem Baesi, Seyed Dawood Mousavi Nasab, Mahmood Barati, Mohammad Reza Asadi Karam, Mehri Habibi, Fatemeh Fotouhi-Chahooki, Neda Afzali, Ali Torabi, Azita Eshratkhah mohammadnejad, Seyedeh Sahar Bathaeian, Mina Bahri, Mohamad Mahdi Mortazavipour, Seyedeh Atefe Hosseini, Farideh Niknam, Parastoo Yekta Sanati, Hadiseh Shokouhi, Azam Amirian, Afsaneh zokaei, Hajarossadat Ghaderi, Mahboobeh Rafigh, Elmira Vadaye kheiri, Akram Agharezaei, Akram Abouie Mehrizi, seyedeh Zahra Moravej, Mostafa Salehi-Vaziri |
| EPI_ISL_414573 | Pasteur Institute of Iran | Pasteur Institute of Iran | Arash Arashkia, Kayhan Azadmanesh, Mohammad Hassan Pouriayevali, Tahmineh Jalali, Zahra Ahmadi, Mohammad Sadegh Shams Nosrati, Ali Maleki, Zabihollah Shoja, Sanam Azad-Mazjiri, Neda Amin, Mehdi Rohani, Saber Esmaeili, Ahmad Ghasemi, Amir Hesam Nemati, Ahmad Mahmoudi, Zahra Fereydouni, Mahsa Tavakolirad, Tahereh Mohammadi, Sahar Khakifirouz, Mehdi Fazlalipour, Maryam Rostamtabar, Arezoo Parikhani, Hesam Karimi, Kazem Baesi, Seyed Dawood Mousavi Nasab, Mahmood Barati, Mohammad Reza Asadi Karam, Mehri Habibi, Fatemeh Fotouhi-Chahooki, Neda Afzali, Ali Torabi, Azita Eshratkhah mohammadnejad, Seyedeh Sahar Bathaeian, Mina Bahri, Mohamad Mahdi Mortazavipour, Seyedeh Atefe Hosseini, Farideh Niknam, Parastoo Yekta Sanati, Hadiseh Shokouhi, Azam Amirian, Afsaneh zokaei, Hajarossadat Ghaderi, Mahboobeh Rafigh, Elmira Vadaye kheiri, Akram Agharezaei, Akram Abouie Mehrizi, seyedeh Zahra Moravej, Mostafa Salehi-Vaziri |
| EPI_ISL_414574 | Center of Medical Microbiology, Virology, and Hospital Hygiene, University of Duesseldorf | Center of Medical Microbiology, Virology, and Hospital Hygiene, University of Duesseldorf | Ortwin Adams, Marcel Andree, Alexander Dilthey, Torsten Feldt, Sandra Hauka, Torsten Houwaart, Björn-Erik Jensen, Detlef Kindgen-Milles, Malte Kohns Vasconcelos, Klaus Pfeffer, Tina Senff, Daniel Strelow, Jörg Timm, Andreas Walker, Tobias Wienemann |
| EPI_ISL_414575 | Pasteur Institute of Iran | Pasteur Institute of Iran | Arash Arashkia, Kayhan Azadmanesh, Mohammad Hassan Pouriayevali, Tahmineh Jalali, Zahra Ahmadi, Mohammad Sadegh Shams Nosrati, Ali Maleki, Zabihollah Shoja, Sanam Azad-Mazjiri, Neda Amin, Mehdi Rohani, Saber Esmaeili, Ahmad Ghasemi, Amir Hesam Nemati, Ahmad Mahmoudi, Zahra Fereydouni, Mahsa Tavakolirad, Tahereh Mohammadi, Sahar Khakifirouz, Mehdi Fazlalipour, Maryam Rostamtabar, Arezoo Parikhani, Hesam Karimi, Kazem Baesi, Seyed Dawood Mousavi Nasab, Mahmood Barati, Mohammad Reza Asadi Karam, Mehri Habibi, Fatemeh Fotouhi-Chahooki, Neda Afzali, Ali Torabi, Azita Eshratkhah mohammadnejad, Seyedeh Sahar Bathaeian, Mina Bahri, Mohamad Mahdi Mortazavipour, Seyedeh Atefe Hosseini, Farideh Niknam, Parastoo Yekta Sanati, Hadiseh Shokouhi, Azam Amirian, Afsaneh zokaei, Hajarossadat Ghaderi, Mahboobeh Rafigh, Elmira Vadaye kheiri, Mina Agharezaei, Akram Abouie Mehrizi, seyedeh Zahra Moravej, Mostafa Salehi-Vaziri |
| EPI_ISL_414577, EPI_ISL_414578 | Hospital de Talca, Chile | Instituto de Salud Publica de Chile | Andrés E. Castillo, Bárbara Parra, Paz Tapia, Alejandra Acevedo, Jaime Lagos, Winston Andrade, Loredana Arata, Gabriel Leal, Gisselle Barra, Carolina Tambley, Javier Tognarelli, Patricia Bustos, Soledad Ulloa, Rodrigo Fasce, Jorge Fernández. |
| EPI_ISL_414579 | Clinica Alemana de Santiago, Chile | Instituto de Salud Publica de Chile | Andrés E. Castillo, Bárbara Parra, Paz Tapia, Alejandra Acevedo, Jaime Lagos, Winston Andrade, Loredana Arata, Gabriel Leal, Gisselle Barra, Carolina Tambley, Javier Tognarelli, Patricia Bustos, Soledad Ulloa, Rodrigo Fasce, Jorge Fernández. |
| EPI_ISL_414580 | Clinica Santa Maria, Santiago, Chile | Instituto de Salud Publica de Chile | Andrés E. Castillo, Bárbara Parra, Paz Tapia, Alejandra Acevedo, Jaime Lagos, Winston Andrade, Loredana Arata, Gabriel Leal, Gisselle Barra, Carolina Tambley, Javier Tognarelli, Patricia Bustos, Soledad Ulloa, Rodrigo Fasce, Jorge Fernández. |
| EPI_ISL_414584, EPI_ISL_414585, EPI_ISL_414586, EPI_ISL_414587 | UCD National Virus Reference Laboratory | UCD National Virus Reference Laboratory | Michael Carr, Gabriel Gonzalez, Jonathan Dean, Suzie Coughlan, Alison Murphy, Kevin Byrne, Ken Wolfe, Jeff Connell, Brendan Loftus, Cillian F De Gascun |
| EPI_ISL_414588, EPI_ISL_414589, EPI_ISL_414590 | Minnesota Department of Health, Public Health Laboratory | Minnesota Department of Health, Public Health Laboratory | Matt Plumb, Jake Garfin and Xiong Wang |
| EPI_ISL_414591, EPI_ISL_414592, EPI_ISL_414593, EPI_ISL_414594, EPI_ISL_414595, EPI_ISL_414596, EPI_ISL_414597 | UW Virology Lab | UW Virology Lab | Pavitra Roychoudhury, Hong Xie, Keith Jerome, Alexander Greninger |
| EPI_ISL_414598 | Servicio Microbiologia, Hospital Clinico Universitario, Valencia | Sequencing and Bioinformatics Service and Molecular Epidemiology Research Group. FISABIO-Public Health. | David Navarro, Maria Alma Bracho, Giuseppe D'Auria, Griselda De Marco, Neris Garcia-Gonzalez, Fernando Gonzalez-Candelas |
| EPI_ISL_414599 | Department of Surgical Sciences, University of Cagliari | Department of Surgical Sciences, University of Cagliari | Scano,A., Fais,S., Loddo,M., Palmieri,G., Scioscia,R., DelRio,N.M.C., Coghe,F. and Orru,G. |
| EPI_ISL_414600 | Laboratoire de Virologie Institut de Virologie - INSERM U 1109 Hôpitaux Universitaires de Strasbourg | National Reference Center for Viruses of Respiratory Infections, Institut Pasteur, Paris | Mélnie Albert, Marion Barbet, Sylvie Behillil, Méline Bizard, Angela Brisebarre, Flora Donati Vincent Enouf, Maud Vanpeene, Sylvie van der Werf, Samira Fafi-Kremer |
| EPI_ISL_414616, EPI_ISL_414617, EPI_ISL_414618, EPI_ISL_414619, EPI_ISL_414620, EPI_ISL_414621, EPI_ISL_414622 | UW Virology Lab | UW Virology Lab | Pavitra Roychoudhury, Hong Xie, Keith Jerome, Alexander Greninger |
| EPI_ISL_414623 | Laboratoire de Virologie Institut de Virologie - INSERM U 1109 Hôpitaux Universitaires de Strasbourg | National Reference Center for Viruses of Respiratory Infections, Institut Pasteur, Paris | Mélnie Albert, Marion Barbet, Sylvie Behillil, Méline Bizard, Angela Brisebarre, Flora Donati Vincent Enouf, Maud Vanpeene, Sylvie van der Werf, Samira Fafi-Kremer |
| EPI_ISL_414624 | Centre Hositalier Universitaire de Rouen Laboratoire de Virologie | National Reference Center for Viruses of Respiratory Infections, Institut Pasteur, Paris | Mélnie Albert, Marion Barbet, Sylvie Behillil, Méline Bizard, Angela Brisebarre, Flora Donati Vincent Enouf, Maud Vanpeene, Sylvie van der Werf, Jean-Christophe Plantier |
| EPI_ISL_414625 | Centre Hospitalier Régional Universitaire de Nantes Laboratoire de Virologie | National Reference Center for Viruses of Respiratory Infections, Institut Pasteur, Paris | Mélnie Albert, Marion Barbet, Sylvie Behillil, Méline Bizard, Angela Brisebarre, Flora Donati Vincent Enouf, Maud Vanpeene, Sylvie van der Werf, Marianne Coste-Burel |
| EPI_ISL_414626 | unknown | National Reference Center for Viruses of Respiratory Infections, Institut Pasteur, Paris | Mélnie Albert, Marion Barbet, Sylvie Behillil, Méline Bizard, Angela Brisebarre, Flora Donati Vincent Enouf, Maud Vanpeene, Sylvie van der Werf |
| EPI_ISL_414627, EPI_ISL_414628, EPI_ISL_414629, EPI_ISL_414630 | Centre Hospitalier Compiègne Laboratoire de Biologie | National Reference Center for Viruses of Respiratory Infections, Institut Pasteur, Paris | Mélnie Albert, Marion Barbet, Sylvie Behillil, Méline Bizard, Angela Brisebarre, Flora Donati Vincent Enouf, Maud Vanpeene, Sylvie van der Werf, Raulin Olivia |
| EPI_ISL_414631, EPI_ISL_414632 | Hôpital Robert Debré Laboratoire de Virologie | National Reference Center for Viruses of Respiratory Infections, Institut Pasteur, Paris | Mélnie Albert, Marion Barbet, Sylvie Behillil, Méline Bizard, Angela Brisebarre, Flora Donati Vincent Enouf, Maud Vanpeene, Sylvie van der Werf, Laurent Andreoletti |
| EPI_ISL_414633 | Centre Hospitalier René Dubois Laboratoire de Microbiologie - Bât A | National Reference Center for Viruses of Respiratory Infections, Institut Pasteur, Paris | Mélnie Albert, Marion Barbet, Sylvie Behillil, Méline Bizard, Angela Brisebarre, Flora Donati Vincent Enouf, Maud Vanpeene, Sylvie van der Werf, Pascale Martres |
| EPI_ISL_414634, EPI_ISL_414635, EPI_ISL_414636, EPI_ISL_414637, EPI_ISL_414638 | Centre Hospitalier Compiègne Laboratoire de Biologie | National Reference Center for Viruses of Respiratory Infections, Institut Pasteur, Paris | Mélnie Albert, Marion Barbet, Sylvie Behillil, Méline Bizard, Angela Brisebarre, Flora Donati Vincent Enouf, Maud Vanpeene, Sylvie van der Werf, Raulin Olivia |
| EPI_ISL_414639 | NYU Langone Health | Departments of Pathology and Medicine, New York University School of Medicine | John Chen, Dacia Dimartino, Xiaojun Feng, Adriana Heguy, Megan Hogan, Emily Huang, George Jour, Christian Marier, Matthew T. Maurano, Mark J. Mulligan, Peter Meyn, Marie Samanovic-Golden, Amy Rapkiewicz, Guomiao Shen, Matija Snuderl, Gael Westby, Paul Zappile |
| EPI_ISL_414640, EPI_ISL_414641, EPI_ISL_414642, EPI_ISL_414643, EPI_ISL_414644, EPI_ISL_414645, EPI_ISL_414646 | Department of Virology and Immunology, University of Helsinki and Helsinki University Hospital, Huslab Finland | Department of Virology, Faculty of Medicine, University of Helsinki, Helsinki, Finland | Teemu Smura, Hannimari Kallio-Kokko, Olli Vapalahti |
| EPI_ISL_414647 | Viral Respiratory Lab, National Institute for Biomedical Research (INRB) | Pathogen Sequencing Lab, National Institute for Biomedical Research (INRB) | Placide Mbala-Kingebeni, Edith Nkwembe, Eddy Kinganda-Lusamaki, Amuri Aziza, Catherine Pratt, Matthias Pauthner, Josh Quick, Allison Black, James Hadfield, Trevor Bedford, Ian Goodfellow, Nick Loman, Kristian Andersen, Michael Wiley, Steve Ahuka-Mundeke, Jean-Jacques Muyembe Tamfum |
| EPI_ISL_414648 | Andersen Lab, The Scripps Research Institute | Andersen Lab, The Scripps Research Institute | Mark Zeller, Catie Anderson, Emily Spender, Sarah Topol, Raphaelle Klitting, Refugio Robles-Sikisaka, Karthik Gangavarapu, Laura Nicholson, Kristian Andersen |
| EPI_ISL_414663, EPI_ISL_414686 | State Key Laboratory of Respiratory Disease, National Clinical Research Center for Respiratory Disease, Guangzhou Institute of Respiratory Health, the First Affiliated Hospital of Guangzhou Medical University | The First Affiliated Hospital of Guangzhou Medical University & BGI-Shenzhen | Zhao et al |
| EPI_ISL_414687 | State Key Laboratory of Respiratory Disease, National Clinical Research Center for Respiratory Disease, Guangzhou Institute of Respiratory Health, the First Affiliated Hospital of Guangzhou Medical University | the First Affiliated Hospital of Guangzhou Medical University & BGI-Shenzhen | Zhao et al |
| EPI_ISL_414688, EPI_ISL_414689, EPI_ISL_414690, EPI_ISL_414691, EPI_ISL_414692 | State Key Laboratory of Respiratory Disease, National Clinical Research Center for Respiratory Disease, Guangzhou Institute of Respiratory Health, the First Affiliated Hospital of Guangzhou Medical University | The First Affiliated Hospital of Guangzhou Medical University & BGI-Shenzhen | Zhao et al |

| | | | |
|---|---|---|---|
| EPI_ISL_414934, EPI_ISL_414935, EPI_ISL_414936, EPI_ISL_414937, EPI_ISL_414938, EPI_ISL_414939, EPI_ISL_414940, EPI_ISL_414941 | Shandong Provincial Center for Disease Control and Prevention | Beijing Institute of Microbiology and Epidemiology | Xiao-Lin Jiang, Xiao-Li Zhang, Xiang-Na Zhao, Cun-Bao Li, Jie Lei, Zeng-Qiang Kou, Wen-Kui Sun, Yang Hang, Feng Gao, Sheng-Xiang Ji, Can-Fang Lin, Bo Pang, Ming-Xiao Yao, Guo-Lin Wang, Lin Yao, Li-Jun Duan, Xiao Wei, Dian-Ming Kang, Mai-Juan Ma |
| EPI_ISL_414945 | Iran National Influenza Center | Iran National Influenza Center | Nazanin Zahra Shafiei Jandaghi, Jila Yavarian,Kaveh Sadeghi, Vahid Salimi, Simin Abbasi, Saeedeh Mahfozi and Talat Mokhtari Azad |
| EPI_ISL_414946 | Iran National Influenza Center | Iran National Influenza Center | Jila Yavarian,Nazanin Zahra Shafiei Jandaghi,Kaveh Sadeghi, Nastaran Ghavvami, Fatemeh Ajaminejad, Fatemeh Saadatmand and Talat Mokhtari Azad |
| EPI_ISL_414948 | Iran National Influenza Center | Iran National Influenza Center | Jila Yavarian, Nazanin Zahra Shafiei Jandaghi and Talat Mokhtari Azad |
| EPI_ISL_414949 | Regional Virus Laboratory, Belfast | Public Health Wales Microbiology Cardiff | Tanya Curran, Conall McCaughey, Catherine Moore, Joanne Watkins, Sally Corden, Tom Connor |
| EPI_ISL_415041 | Wales Specialist Virology Centre | Public Health Wales Microbiology Cardiff | Catherine Moore, Joanne Watkins, Sally Corden, Tom Connor |
| EPI_ISL_415105 | Laboratório Central de Saúde Pública Professor Gonçalo Moniz – LACEN/BA | Instituto Oswaldo Cruz FIOCRUZ - Laboratory of Respiratory Viruses and Measles (LVRS) | Paola Resende, Allison Fabri, Joilson Xavier, Sunando Roy, Fernando Motta, Aline Mattos, Milene Miranda, Cristiana Garcia, Braulia Caetano, Maria Ogrzewalska, Jonathan Lopes, Luciana Appolinario, Maria Nóbrega, Marilda Siqueira |
| EPI_ISL_415128 | LACEN/ES - Laboratório Central de Saúde Pública do Espírito Santo | Instituto Oswaldo Cruz FIOCRUZ - Laboratory of Respiratory Viruses and Measles (LVRS) | Paola Resende, Allison Fabri, Joilson Xavier, Sunando Roy, Fernando Motta, Aline Mattos, Milene Miranda, Cristiana Garcia, Braulia Caetano, Maria Ogrzewalska, Jonathan Lopes, Luciana Appolinario, Maria Nóbrega, Marilda Siqueira |
| EPI_ISL_415129, EPI_ISL_415130, EPI_ISL_415131, EPI_ISL_415132, EPI_ISL_415133, EPI_ISL_415134, EPI_ISL_415135, EPI_ISL_415136, EPI_ISL_415137, EPI_ISL_415138, EPI_ISL_415139, EPI_ISL_415140, EPI_ISL_415141, EPI_ISL_415142, EPI_ISL_415143, EPI_ISL_415144, EPI_ISL_415145, EPI_ISL_415146, EPI_ISL_415147, EPI_ISL_415148, EPI_ISL_415149, EPI_ISL_415150 | | | |
| see above | Respiratory Virus Unit, Microbiology Services Colindale, Public Health England | Respiratory Virus Unit, Microbiology Services Colindale, Public Health England | Monica Galiano, Shahjahan Miah, Angie Lackenby, Omolola Akinbami, Tiina Talts, Leena Bhaw, Richard Myers, Steven Platt, Kirstin Edwards, Jonathan Hubb, Joanna Ellis, Maria Zambon |
| EPI_ISL_415151 | MSHS Clinical Microbiology Laboratories | MSHS Pathogen Surveillance Program | Gopi Patel, Emilia Sordillo, Melissa Gitman, Alberto Paniz-mondolfi, Matthew Hernandez, Shelcie Fabre, Jose Polanco, Ana Silvia Gonzalez-Reiche, Zenab Khan, Nancy Francoeur, Melissa Smith, Robert Sebra, Lisa Miorin, Wen-chun Liu, Randy Albrecht, Judith Aberg, Florian Krammer, Adolfo Garcia-Sarstre, Viviana Simon, Harm van Bakel |
| EPI_ISL_415152 | Gorgas Memorial Institute for Health Studies | Gorgas Memorial Institute for Health Studies | Danilo Franco, Sandra Lopez-Verges, Elimelec Valdespino, Claudia Gonzalez, Oris Chavarria, Ambar Moreno, Yamilka Diaz, Leyda Abrego, Juan M. Pascale, Alexander A. Martinez. |
| EPI_ISL_415153 | KU Leuven, Clinical and Epidemiological Virology | KU Leuven, Clinical and Epidemiological Virology | Bert Vanmechelen, Joan Marti-Carreras, Tony Wawina, Marc Van Ranst, Piet Maes |
| EPI_ISL_415154 | KU Leuven, Clincal and Epidemiological Virology | KU Leuven, Clincal and Epidemiological Virology | Bert Vanmechelen, Joan Marti-Carreras, Tony Wawina, Marc Van Ranst, Piet Maes. |
| EPI_ISL_415155 | KU Leuven, Clinical and Epidemiological Virology | KU Leuven, Clinical and Epidemiological Virology | Bert Vanmechelen, Joan Marti-Carreras, Tony Wawina, Marc Van Ranst, Piet Maes |
| EPI_ISL_415156, EPI_ISL_415157, EPI_ISL_415158, EPI_ISL_415159 | KU Leuven, Clinical and Epidemiological Virology | KU Leuven, Clinical and Epidemiological Virology | Bert Vanmechelen, Joan Marti-Carreras, Tony Wawina, Piet Maes |
| EPI_ISL_415435, EPI_ISL_415453 | Wales Specialist Virology Centre | Public Health Wales Microbiology Cardiff | Catherine Moore, Joanne Watkins, Sally Corden, Tom Connor |
| EPI_ISL_415454, EPI_ISL_415455, EPI_ISL_415456, EPI_ISL_415457, EPI_ISL_415458, EPI_ISL_415459 | University Hospitals of Geneva Laboratory of Virology | University Hospitals of Geneva Laboratory of Virology | Laubscher F. |
| EPI_ISL_415460, EPI_ISL_415461, EPI_ISL_415462, EPI_ISL_415463, EPI_ISL_415464, EPI_ISL_415465, EPI_ISL_415466, EPI_ISL_415467, EPI_ISL_415468, EPI_ISL_415469, EPI_ISL_415470, EPI_ISL_415471, EPI_ISL_415472, EPI_ISL_415473, EPI_ISL_415474, EPI_ISL_415475, EPI_ISL_415476, EPI_ISL_415477, EPI_ISL_415478, EPI_ISL_415479, EPI_ISL_415480, EPI_ISL_415481, EPI_ISL_415482, EPI_ISL_415483, EPI_ISL_415484, EPI_ISL_415485, EPI_ISL_415486, EPI_ISL_415487, EPI_ISL_415488, EPI_ISL_415489, EPI_ISL_415490, EPI_ISL_415491, EPI_ISL_415492, EPI_ISL_415493, EPI_ISL_415494, EPI_ISL_415495, EPI_ISL_415496, EPI_ISL_415497, EPI_ISL_415498, EPI_ISL_415499, EPI_ISL_415500, EPI_ISL_415501, EPI_ISL_415502, EPI_ISL_415503, EPI_ISL_415504, EPI_ISL_415505, EPI_ISL_415506, EPI_ISL_415507, EPI_ISL_415508, EPI_ISL_415509, EPI_ISL_415510, EPI_ISL_415511, EPI_ISL_415512, EPI_ISL_415513, EPI_ISL_415514, EPI_ISL_415515, EPI_ISL_415516, EPI_ISL_415517, EPI_ISL_415518, EPI_ISL_415519, EPI_ISL_415520, EPI_ISL_415521, EPI_ISL_415522, EPI_ISL_415523, EPI_ISL_415524, EPI_ISL_415525, EPI_ISL_415526, EPI_ISL_415527, EPI_ISL_415528, EPI_ISL_415529, EPI_ISL_415530, EPI_ISL_415531, EPI_ISL_415532, EPI_ISL_415533, EPI_ISL_415534, EPI_ISL_415535 | | | |
| see above | Dutch COVID-19 response team | Erasmus Medical Center | David Nieuwenhuijse, Bas Oude Munnink, Reina Sikkema, Claudia Schapendonk, Irina Chestakova, Anne van der Linden, Mark Pronk, Pascal Lexmond, Corien Swaan, Manon Haverkate, Madelief Mollers, Mart Stein, Sandra Kengne Kamga Mobou, Jeroen van Kampen, Jolanda Voermans, Aura Timen, Corine GeurtsvanKessel, Annemiek van der Eijk, Richard Molenkamp, Marion Koopmans, on behalf of the Dutch national COVID-19 response team. |
| EPI_ISL_415536, EPI_ISL_415537, EPI_ISL_415538 | Wales Specialist Virology Centre | Public Health Wales Microbiology Cardiff | Catherine Moore, Joanne Watkins, Sally Corden, Tom Connor |
| EPI_ISL_415539, EPI_ISL_415541, EPI_ISL_415542, EPI_ISL_415543, EPI_ISL_415544 | Utah Public Health Laboratory | Utah Public Health Laboratory | Erin Young, Kelly Oakeson |
| EPI_ISL_415577, EPI_ISL_415578, EPI_ISL_415579, EPI_ISL_415580, EPI_ISL_415581, EPI_ISL_415582, EPI_ISL_415583, EPI_ISL_415584, EPI_ISL_415585, EPI_ISL_415586, EPI_ISL_415587, EPI_ISL_415588, EPI_ISL_415589, EPI_ISL_415590 | | | |
| see above | BCCDC Public Health Laboratory | BCCDC Public Health Laboratory | Harrigan, Prystajecky, Krajden, Lee, Kamelian, Lapointe, Choi, Hoang, Sekirov, Levett, Tyson, Snutch, Loman, Quick, Li, Gilmour |
| EPI_ISL_415591, EPI_ISL_415592, EPI_ISL_415593, EPI_ISL_415594, EPI_ISL_415595, EPI_ISL_415596, EPI_ISL_415597, EPI_ISL_415598, EPI_ISL_415599, EPI_ISL_415600, EPI_ISL_415601, EPI_ISL_415602, EPI_ISL_415603, EPI_ISL_415604, EPI_ISL_415605, EPI_ISL_415606, EPI_ISL_415607, EPI_ISL_415608, EPI_ISL_415609, EPI_ISL_415610, EPI_ISL_415611, EPI_ISL_415612, EPI_ISL_415613, EPI_ISL_415614, EPI_ISL_415615, EPI_ISL_415616, EPI_ISL_415617, EPI_ISL_415618, EPI_ISL_415619, EPI_ISL_415620, EPI_ISL_415621, EPI_ISL_415622, EPI_ISL_415623, EPI_ISL_415624, EPI_ISL_415625, EPI_ISL_415626, EPI_ISL_415627 | | | |
| see above | UW Virology Lab | UW Virology Lab | Pavitra Roychoudhury, Hong Xie, Keith Jerome, Alexander Greninger |
| EPI_ISL_415628 | West of Scotland Specialist Virology Centre, NHSGGC | MRC-University of Glasgow Centre for Virus Research | Kathy Smollett, Daniel Mair, Stephen Carmichael, Ana da Silva Filipe; Richard Orton, David L Robertson; Alasdair MacLean, Rory Gunson; Natasha Jesudason, Kathy Li, Antonia Ho; Emma Thomson. |
| EPI_ISL_415629 | Virology Department, Royal Infirmary of Edinburgh, NHS Lothian | Virology Department, Royal Infirmary of Edinburgh, NHS Lothian | McHugh M, Dewar R, O'Toole Á, Rambaut A, Williams TC, Templeton K |
| EPI_ISL_415630, EPI_ISL_415631 | West of Scotland Specialist Virology Centre, NHSGGC | MRC-University of Glasgow Centre for Virus Research | Kathy Smollett, Daniel Mair, Stephen Carmichael, Ana da Silva Filipe; Richard Orton, David L Robertson; Alasdair MacLean, Rory Gunson; Natasha Jesudason, Kathy Li, Antonia Ho; Emma Thomson. |
| EPI_ISL_415632, EPI_ISL_415633, EPI_ISL_415634, EPI_ISL_415635, EPI_ISL_415636, EPI_ISL_415637, EPI_ISL_415638, EPI_ISL_415639, EPI_ISL_415640 | Virology Department, Royal Infirmary of Edinburgh, NHS Lothian | Virology Department, Royal Infirmary of Edinburgh, NHS Lothian | McHugh M, Dewar R, O'Toole Á, Rambaut A, Williams TC, Templeton K |
| EPI_ISL_415641, EPI_ISL_415642, EPI_ISL_415643, EPI_ISL_415644 | R. G. Lugar Center for Public Health Research, National Center for Disease Control and Public Health (NCDC) of Georgia. | R. G. Lugar Center for Public Health Research, National Center for Disease Control and Public Health (NCDC) of Georgia. | Nato Kotaria, Marine Murtskhvaladze, Ann Machablishvili, Lela Sabadze, Mari Gavashelidze, Ana Papkiauri, Meri Pantsulaia, Gvantsa Brachveli, Tata Imnadze, Tamar Jashiashvili, Tea Tevdoradze, Ketevan Sidamonidze, Ekaterine Khmaladze, Ekaterine Zhghenti, Roena Sukhiashvili, Mariam Zakalashvili, Lela Urushadze, Magda Dgebuadze, Giorgi Tomashvili, Davit Tsaguria, Ekaterine Zangaladze, Nino Berishvili, Gvantsa Chanturia, Adam Kotorashvili, Maia Alkhazashvili, Irma Burjanadze, Anna Kasradze, Khatuna Zakhashvili, Paata Imnadze, Amiran Gamkrelidze. |
| EPI_ISL_415646 | Department of Virus and Microbiological Special diagnostics, Statens Serum Institut, Copenhagen, Denmark. | ViFU | Morten Rasmussen, Maiken Worsoe Rosenstierne , Anders Fomsgaard |
| EPI_ISL_415647 | Department of Virus and Microbiological Special diagnostics, Statens Serum Institut, Copenhagen, Denmark. | Statens Serum Institute | Morten Rasmussen, Maiken Worsoe Rosenstierne , Anders Fomsgaard |
| EPI_ISL_415648 | Department of Virus and Microbiological Special diagnostics, Statens Serum Institut, Copenhagen, Denmark. | ViFU | Morten Rasmussen, Maiken Worsoe Rosenstierne , Anders Fomsgaard |
| EPI_ISL_415649 | unknown | National Reference Center for Viruses of Respiratory Infections, Institut Pasteur, Paris | Mélnie Albert, Marion Barbet, Sylvie Behillil, Méline Bizard, Angela Brisebarre, Flora Donati Vincent Enouf, Maud Vanpeene, Sylvie van der Werf |
| EPI_ISL_415650 | Hôpital Instruction des Armées - BEGIN | National Reference Center for Viruses of Respiratory Infections, Institut Pasteur, Paris | Mélnie Albert, Marion Barbet, Sylvie Behillil, Méline Bizard, Angela Brisebarre, Flora Donati Vincent Enouf, Maud Vanpeene, Sylvie van der Werf, Christine Bigaillon |
| EPI_ISL_415651 | Unknown | National Reference Center for Viruses of Respiratory Infections, Institut Pasteur, Paris | Mélnie Albert, Marion Barbet, Sylvie Behillil, Méline Bizard, Angela Brisebarre, Flora Donati Vincent Enouf, Maud Vanpeene, Sylvie van der Werf |
| EPI_ISL_415652 | unknown | National Reference Center for Viruses of Respiratory Infections, Institut Pasteur, Paris | Mélnie Albert, Marion Barbet, Sylvie Behillil, Méline Bizard, Angela Brisebarre, Flora Donati Vincent Enouf, Maud Vanpeene, Sylvie van der Werf |
| EPI_ISL_415653, EPI_ISL_415654 | Centre Hospitalier Compiègne Laboratoire de Biologie | National Reference Center for Viruses of Respiratory Infections, Institut Pasteur, Paris | Mélnie Albert, Marion Barbet, Sylvie Behillil, Méline Bizard, Angela Brisebarre, Flora Donati Vincent Enouf, Maud Vanpeene, Sylvie van der Werf, Raulin Olivia |
| EPI_ISL_415655, EPI_ISL_415656 | Wales Specialist Virology Centre | Public Health Wales Microbiology Cardiff | Catherine Moore, Joanne Watkins, Sally Corden, Tom Connor |
| EPI_ISL_415657 | Wales Specialist Virology Centre | Public Health Wales Microbiology Cardiff | Catherine Moore, Joanne watkins, Sally Corden, Tom Connor |
| EPI_ISL_415658 | Laboratory of Molecular Virology, Pontificia Universidad Católica de Chile | MSHS Pathogen Surveillance Program | Rafael A. Medina, Pablo Vial, Tamara Garcia, Eileen Serrano, Ana Silvia Gonzalez-Reiche, Zenab Khan, Mitchell Sullivan, Ajay Obla, Matthew Hernandez, Hala Alshammary, Juan Soto, Shwetha Sridhar Hara, Ying-Chih Wang, Melissa Smith, Robert Sebra, Viviana Simon, Harm van Bakel |
| EPI_ISL_415659 | Wales Specialist Virology Centre | Public Health Wales Microbiology Cardiff | Catherine Moore, Joanne Watkins, Sally Corden, Tom Connor |
| EPI_ISL_415660, EPI_ISL_415661 | Laboratory of Molecular Virology, Pontificia Universidad Católica de Chile | MSHS Pathogen Surveillance Program | Rafael A. Medina, Pablo Vial, Tamara Garcia, Eileen Serrano, Ana Silvia Gonzalez-Reiche, Zenab Khan, Mitchell Sullivan, Ajay Obla, Matthew Hernandez, Hala Alshammary, Juan Soto, Shwetha Sridhar Hara, Ying-Chih Wang, Melissa Smith, Robert Sebra, Viviana Simon, Harm van Bakel |
| EPI_ISL_415698, EPI_ISL_415699, EPI_ISL_415700, EPI_ISL_415701, EPI_ISL_415702, EPI_ISL_415703 | University Hospitals of Geneva Laboratory of Virology | University Hospitals of Geneva Laboratory of Virology | Laubscher F. |
| EPI_ISL_415704, EPI_ISL_415705 | University Hospitals of Geneva Laboratory of Virology | University Hospitals of Geneva Laboratory of Virology | Laubscher F. |
| EPI_ISL_415706, EPI_ISL_415707, EPI_ISL_415708 | University Hospitals of Geneva Laboratory of Virology | University Hospitals of Geneva Laboratory of Virology | Laubscher F. |
| EPI_ISL_415709 | State Key Laboratory for Diagnosis and Treatment of | State Key Laboratory for Diagnosis and Treatment of | Hangping Yao, Nanping Wu, Chao Jiang, Xiangyun Lu, Linfang Cheng, Fumin Liu, Zhigang Wu, Haibo Wu, Changzhong Jin, Min Zheng, Lanjuan Li |

| | | | |
|---|---|---|---|
| | Infectious Diseases, National Clinical Research Center for Infectious Diseases, First Affiliated Hospital, Zhejiang University School of Medicine, Hangzhou, China. 310003 | Infectious Diseases, National Clinical Research Center for Infectious Diseases, First Affiliated Hospital, Zhejiang University School of Medicine, Hangzhou, China. 310003 | |
| EPI_ISL_415710 | WHO National Influenza Centre Russian Federation | WHO National Influenza Centre Russian Federation | Andrey Komissarov, Artem Fadeev, Anna Ivanova, Daria Danilenko |
| EPI_ISL_415711 | State Key Laboratory for Diagnosis and Treatment of Infectious Diseases, National Clinical Research Center for Infectious Diseases, First Affiliated Hospital, Zhejiang University School of Medicine, Hangzhou, China. 310003 | State Key Laboratory for Diagnosis and Treatment of Infectious Diseases, National Clinical Research Center for Infectious Diseases, First Affiliated Hospital, Zhejiang University School of Medicine, Hangzhou, China. 310003 | Hangping Yao, Nanping Wu, Chao Jiang, Xiangyun Lu, Linfang Cheng, Fumin Liu, Zhigang Wu, Haibo Wu, Changzhong Jin, Min Zheng, Lanjuan Li |
| EPI_ISL_415741, EPI_ISL_415742, EPI_ISL_415743 | Laboratory Medicine | Department of Laboratory Medicine, Lin-Kou Chang Gung Memorial Hospital, Taoyuan, Taiwan | Kuo-Chien Tsao, Yu-Nong Gong, Shu-Li Yang, Yi-Chun Liu, Chung-Guei Huang, Po-Wei Huang, Mei-Jen Hsiao, Cheng-Ta Yang, Cheng-Hsun Chiu, Chi-Hsien Huang, Kuang-Tso Le, Shu-Min Lin, Peng-Nien Huang, Kuo-Ming Lee, Guang-Wu Chen, Shin-Ru Shih |
| EPI_ISL_415787 | Laboratorio de Referencia Nacional de Virus Respiratorio. Instituto Nacional de Salud. Peru | Laboratorio de Referencia Nacional de Biotecnologia y Biologia Molecular.Instituto Nacional de Salud.Peru | Carlos Padilla Rojas, Priscila Lope Pari, Karolyn Vega Chozo, Johanna Balbuena Torres, Omar Caceres Rey, Hemri Bailon Calderon, Maribel Huaringa Nuñez, Nancy Rojas Serrano |
| EPI_ISL_415916, EPI_ISL_415918, EPI_ISL_415919, EPI_ISL_415920, EPI_ISL_415977, EPI_ISL_415978, EPI_ISL_415991, EPI_ISL_416024 | Wales Specialist Virology Centre | Public Health Wales Microbiology Cardiff | Catherine Moore, Joanne Watkins, Sally Corden, Tom Connor |
| EPI_ISL_416025 | Laboratorio de Referencia Nacional de Virus Respiratorios. Instituto Nacional de Salud Peru | Laboratorio de Referencia Nacional de Biotecnologia y Biologia Molecular. Instituto Nacional de Salud Peru | Carlos Padilla Rojas, Priscila Lope Pari, Karolyn Vega Chozo, Johanna Balbuena Torres, Omar Caceres Rey, Henri Bailon Calderon, Maribel Huaringa Nuñez, Nancy Rojas Serrano |
| EPI_ISL_416026, EPI_ISL_416027 | Wales Specialist Virology Centre | Public Health Wales Microbiology Cardiff | Catherine Moore, Joanne Watkins, Sally Corden, Tom Connor |
| EPI_ISL_416028 | National Influenza Center - Instituto Adolfo Lutz | Instituto Adolfo Lutz, Interdiciplinary Procedures Center, Strategic Laboratory | Claudio Tavares Sacchi, Claudia Regina Gonçalves, Carlos Henrique Camargo, Fabiana Cristina Pereira dos Santos, Daniela Bernardes Borges da Silva, Simone Guadagnucci Morillo, Adriano Abbud, Adriana Bugno, Maria do Carmo Sampaio Tavares Timenetsky, Terezinha Maria de Paiva |
| EPI_ISL_416029 | Laboiratório Fleury | Instituto Adolfo Lutz, Interdiciplinary Procedures Center, Strategic Laboratory | Claudio Tavares Sacchi, Claudia Regina Gonçalves, Carlos Henrique Camargo, Fabiana Cristina Pereira dos Santos, Daniela Bernardes Borges da Silva, Simone Guadagnucci Morillo, Adriano Abbud, Adriana Bugno, Maria do Carmo Sampaio Tavares Timenetsky, Terezinha Maria de Paiva |
| EPI_ISL_416031, EPI_ISL_416032 | National Influenza Center - Instituto Adolfo Lutz | Instituto Adolfo Lutz, Interdiciplinary Procedures Center, Strategic Laboratory | Claudio Tavares Sacchi, Claudia Regina Gonçalves, Carlos Henrique Camargo, Fabiana Cristina Pereira dos Santos, Daniela Bernardes Borges da Silva, Simone Guadagnucci Morillo, Adriano Abbud, Adriana Bugno, Maria do Carmo Sampaio Tavares Timenetsky, Terezinha Maria de Paiva |
| EPI_ISL_416033, EPI_ISL_416034 | Hospital Israelita Albert Einstein | Instituto Adolfo Lutz, Interdiciplinary Procedures Center, Strategic Laboratory | Claudio Tavares Sacchi, Claudia Regina Gonçalves, Carlos Henrique Camargo, Erica Valessa Ramos Gomes, Fabiana Cristina Pereira dos Santos, Daniela Bernardes Borges da Silva, Simone Guadagnucci Morillo, Adriano Abbud, Adriana Bugno, Maria do Carmo Sampaio Tavares Timenetsky, Terezinha Maria de Paiva |
| EPI_ISL_416035, EPI_ISL_416036 | National Influenza Center - Instituto Adolfo Lutz | Instituto Adolfo Lutz, Interdiciplinary Procedures Center, Strategic Laboratory | Claudio Tavares Sacchi, Claudia Regina Gonçalves, Carlos Henrique Camargo, Erica Valessa Ramos Gomes, Fabiana Cristina Pereira dos Santos, Daniela Bernardes Borges da Silva, Simone Guadagnucci Morillo, Adriano Abbud, Adriana Bugno, Maria do Carmo Sampaio Tavares Timenetsky, Terezinha Maria de Paiva |
| EPI_ISL_416042 | State Key Laboratory for Diagnosis and Treatment of Infectious Diseases, National Clinical Research Center for Infectious Diseases, First Affiliated Hospital, Zhejiang University School of Medicine, Hangzhou, China. 310003 | State Key Laboratory for Diagnosis and Treatment of Infectious Diseases, National Clinical Research Center for Infectious Diseases, First Affiliated Hospital, Zhejiang University School of Medicine, Hangzhou, China. 310003 | Hangping Yao, Nanping Wu, Chao Jiang, Xiangyun Lu, Linfang Cheng, Fumin Liu, Zhigang Wu, Haibo Wu, Changzhong Jin, Min Zheng, Lanjuan Li |
| EPI_ISL_416044, EPI_ISL_416046, EPI_ISL_416047 | State Key Laboratory for Diagnosis and Treatment of Infectious Diseases, National Clinical Research Center for Infectious Diseases, First Affiliated Hospital, Zhejiang University School of Medicine, Hangzhou, China 310003 | State Key Laboratory for Diagnosis and Treatment of Infectious Diseases, National Clinical Research Center for Infectious Diseases, First Affiliated Hospital, Zhejiang University School of Medicine, Hangzhou, China 310003 | Hangping Yao, Nanping Wu, Chao Jiang, Xiangyun Lu, Linfang Cheng, Fumin Liu, Zhigang Wu, Haibo Wu, Changzhong Jin, Min Zheng, Lanjuan Li |
| EPI_ISL_416140 | Department of Virus and Microbiological Special diagnostics, Statens Serum Institut, Copenhagen, Denmark. | Statens Serum Institute | Morten Rasmussen, Maiken Worsoe Rosenstierne , Anders Fomsgaard |
| EPI_ISL_416141, EPI_ISL_416142 | Department of Virus and Microbiological Special diagnostics, Statens Serum Institut, Copenhagen, Denmark. | Statens Serum Institute | Morten Rasmussen, Maiken Worsoe Rosenstierne , Anders Fomsgaard |
| EPI_ISL_416143, EPI_ISL_416144, EPI_ISL_416153 | Department of Virus and Microbiological Special diagnostics, Statens Serum Institut, Copenhagen, Denmark. | ViFU | Morten Rasmussen, Maiken Worsoe Rosenstierne , Anders Fomsgaard |
| EPI_ISL_416314, EPI_ISL_416315 | Department of Microbiology, Faculty of Medicine, The Chinese University of Hong Kong, Hong Kong SAR, China | Department of Microbiology, Faculty of Medicine, Chinese University of Hong Kong, Hong Kong SAR, China | Zigui Chen, Paul KS Chan |
| EPI_ISL_416316, EPI_ISL_416317, EPI_ISL_416318, EPI_ISL_416319, EPI_ISL_416320, EPI_ISL_416321, EPI_ISL_416322, EPI_ISL_416323, EPI_ISL_416324, EPI_ISL_416325, EPI_ISL_416326, EPI_ISL_416327, EPI_ISL_416328, EPI_ISL_416329, EPI_ISL_416330, EPI_ISL_416331, EPI_ISL_416332, EPI_ISL_416333, EPI_ISL_416334, EPI_ISL_416335, EPI_ISL_416336, EPI_ISL_416337, EPI_ISL_416338, EPI_ISL_416339, EPI_ISL_416340, EPI_ISL_416341, EPI_ISL_416342, EPI_ISL_416343, EPI_ISL_416344, EPI_ISL_416345, EPI_ISL_416346, EPI_ISL_416347, EPI_ISL_416348, EPI_ISL_416349, EPI_ISL_416350, EPI_ISL_416351, EPI_ISL_416352, EPI_ISL_416353, EPI_ISL_416354, EPI_ISL_416355, EPI_ISL_416356, EPI_ISL_416357, EPI_ISL_416358, EPI_ISL_416359, EPI_ISL_416360, EPI_ISL_416361, EPI_ISL_416362, EPI_ISL_416363, EPI_ISL_416364, EPI_ISL_416365, EPI_ISL_416366, EPI_ISL_416367, EPI_ISL_416368, EPI_ISL_416369, EPI_ISL_416370, EPI_ISL_416371, EPI_ISL_416372, EPI_ISL_416373, EPI_ISL_416374, EPI_ISL_416375, EPI_ISL_416376, EPI_ISL_416377, EPI_ISL_416378, EPI_ISL_416379, EPI_ISL_416380, EPI_ISL_416381, EPI_ISL_416382, EPI_ISL_416383, EPI_ISL_416384, EPI_ISL_416385, EPI_ISL_416386, EPI_ISL_416387, EPI_ISL_416388, EPI_ISL_416389, EPI_ISL_416390, EPI_ISL_416391, EPI_ISL_416392, EPI_ISL_416393, EPI_ISL_416394, EPI_ISL_416395, EPI_ISL_416396, EPI_ISL_416397, EPI_ISL_416398, EPI_ISL_416399, EPI_ISL_416400, EPI_ISL_416401, EPI_ISL_416402, EPI_ISL_416403, EPI_ISL_416404, EPI_ISL_416405, EPI_ISL_416406, EPI_ISL_416407, EPI_ISL_416408, EPI_ISL_416409 | | | |
| see above | Shanghai Public Health Clinical Center, Shanghai Medical College, Fudan University | National Research Center for Translational Medicine (Shanghai), Ruijin Hospital affiliated to Shanghai Jiao Tong University School of Medicine & Shanghai Public Health Clinical Center | Shengyue Wang, Xiaonan Zhang, Gang Lu, Yun Tan, Yun Ling, Hongzhou Lu, Saijuan Chen |
| EPI_ISL_416410, EPI_ISL_416411, EPI_ISL_416412, EPI_ISL_416413, EPI_ISL_416414, EPI_ISL_416415 | Victorian Infectious Diseases Reference Laboratory (VIDRL) | Victorian Infectious Diseases Reference Laboratory and Microbiological Diagnostic Unit Public Health Laboratory, Doherty Institute | Caly L., Seemann T., Schultz M., Druce J., Taiaroa, G. |
| EPI_ISL_416416, EPI_ISL_416417, EPI_ISL_416418, EPI_ISL_416419 | Connecticut State Department of Public Health | Grubaugh Lab - Yale School of Public Health | Joseph Fauver, Chantal Vogels, Anderson Brito, Tara Alpert, Nagarjuna Cheemarla, Ellen Foxman, Anthony Muyombwe, Jafar Razeq, Richard Martinello, Albert Ko, Marie-Louise Landry, Nathan Grubaugh |
| EPI_ISL_416420, EPI_ISL_416421, EPI_ISL_416422, EPI_ISL_416423, EPI_ISL_416424 | Yale Clinical Virology Laboratory | Grubaugh Lab - Yale School of Public Health | Joseph Fauver, Chantal Vogels, Anderson Brito, Tara Alpert, Nagarjuna Cheemarla, Ellen Foxman, Anthony Muyombwe, Jafar Razeq, Richard Martinello, Albert Ko, Marie-Louise Landry, Nathan Grubaugh |
| EPI_ISL_416425 | State Key Laboratory for Diagnosis and Treatment of Infectious Diseases, National Clinical Research Center for Infectious Diseases, First Affiliated Hospital, Zhejiang University School of Medicine, Hangzhou, China 310003 | State Key Laboratory for Diagnosis and Treatment of Infectious Diseases, National Clinical Research Center for Infectious Diseases, First Affiliated Hospital, Zhejiang University School of Medicine, Hangzhou, China 310003 | Hangping Yao, Nanping Wu, Chao Jiang, Xiangyun Lu, Linfang Cheng, Fumin Liu, Zhigang Wu, Haibo Wu, Changzhong Jin, Min Zheng, Lanjuan Li |
| EPI_ISL_416426 | Virological Research Group, Szentágothai Research Centre, University of Pécs | Bioinformatics Research Group, Szentágothai Research Centre, University of Pécs | Péter Urbán, Endre Gábor Tóth, Gábor Kemenesi, Róbert Herczeg, Attila Gyenesei, Ferenc Jakab |
| EPI_ISL_416427, EPI_ISL_416428, EPI_ISL_416429, EPI_ISL_416430, EPI_ISL_416431 | National Influenza Center, National Institute of Hygiene and Epidemiology (NIHE) | National Influenza Center, National Institute of Hygiene and Epidemiology (NIHE) | Le Quynh Mai, Taichiro Takemura, Meng Ling Moi, Takeshi Nabeshima, Nguyen Le Khanh Hang, Hoang Vu Mai Phuong, Ung Thi Hong Trang, Le Thi Thanh, Nguyen Vu Son, Vuong Duc Cuong, Pham Thi Hien, Tran Thu Huong, Nguyen Phuong Anh, Pham Hong Quynh Anh, Kouichi Morita, Futoshi Hasebe, Dang Duc Anh |
| EPI_ISL_416432 | Clinical Microbiology Lab | Infectious Disease Research Department, King Abdullah International Medical Research Center (KAIMRC) | Majed Alghoribi, Sadeem Alhayli, Abdulrahman Alswaji, Liliane Okdah, Sameera Al Johani, Michel Doumith |
| EPI_ISL_416433, EPI_ISL_416434, EPI_ISL_416435, EPI_ISL_416436, EPI_ISL_416437, EPI_ISL_416438, EPI_ISL_416439, EPI_ISL_416440, EPI_ISL_416441, EPI_ISL_416442, EPI_ISL_416443, EPI_ISL_416444, EPI_ISL_416445, EPI_ISL_416446, EPI_ISL_416447, EPI_ISL_416448, EPI_ISL_416449, EPI_ISL_416450, EPI_ISL_416451, EPI_ISL_416452, EPI_ISL_416453, EPI_ISL_416454, EPI_ISL_416455, EPI_ISL_416456 | | | |
| see above | UW Virology Lab | UW Virology Lab | Pavitra Roychoudhury, Hong Xie, Keith Jerome, Alexander Greninger |
| EPI_ISL_416457 | Andersen Lab, The Scripps Research Institute | Andersen Lab, The Scripps Research Institute | Mark Zeller, Catie Anderson, Emily Spender, Sarah Topol, Raphaelle Klitting, Refugio Robles-Sikisaka, Karthik Gangavarapu, Laura Nicholson, Kristian Andersen |
| EPI_ISL_416458 | Virology laboratory Ministry of Health Kuwait sequenced at Dasman Diabetes Institute | Dasman Diabetes Institute | Fahd Al-Mulla, Sumi John, Sara Alqabandi, Rasheeba iqbal, Motasem Melhem, Ebaa alOzairi, Qais Al-Duwairi |
| EPI_ISL_416459, EPI_ISL_416460, EPI_ISL_416461, EPI_ISL_416462, EPI_ISL_416463, EPI_ISL_416464, EPI_ISL_416465, EPI_ISL_416466 | Seattle Flu Study | Seattle Flu Study | Chu et al |
| EPI_ISL_416467, EPI_ISL_416468, EPI_ISL_416469, EPI_ISL_416470, EPI_ISL_416471, EPI_ISL_416472 | KU Leuven, Clinical and Epidemiological Virology | KU Leuven, Clinical and Epidemiological Virology | Bert Vanmechelen, Tony Wawina, Joan Marti-Carreras, Piet Maes |
| EPI_ISL_416473, EPI_ISL_416474 | State Key Laboratory for Diagnosis and Treatment of Infectious Diseases, National Clinical Research Center | State Key Laboratory for Diagnosis and Treatment of Infectious Diseases, National Clinical Research Center for | Hangping Yao, Nanping Wu, Chao Jiang, Xiangyun Lu, Linfang Cheng, Fumin Liu, Zhigang Wu, Haibo Wu, Changzhong Jin, Min Zheng, Lanjuan Li |

| | | | |
|---|---|---|---|
| | for Infectious Diseases, First Affiliated Hospital, Zhejiang University School of Medicine, Hangzhou, China 310003 | Infectious Diseases, First Affiliated Hospital, Zhejiang University School of Medicine, Hangzhou, China 310003 | |
| EPI_ISL_416475, EPI_ISL_416476 | KU Leuven, Clinical and Epidemiological Virology | KU Leuven, Clinical and Epidemiological Virology | Bert Vanmechelen, Tony Wawina, Joan Marti-Carreras, Piet Maes |
| EPI_ISL_416477, EPI_ISL_416478, EPI_ISL_416479 | R. G. Lugar Center for Public Health Research, National Center for Disease Control and Public Health (NCDC) of Georgia. | R. G. Lugar Center for Public Health Research, National Center for Disease Control and Public Health (NCDC) of Georgia. | Marine Murtskhvaladze, Nato Kotaria, Ann Machablishvili, Lela Sabadze, Mari Gavashelidze, Ana Papkiauri, Meri Pantsulaia, Gvantsa Brachveli, Tata Imnadze, Tamar Jashiashvili, Tea Tevdoradze, Ketevan Sidamonidze, Ekaterine Khmaladze, Ekaterine Zhghenti, Roena Sukhiashvili, Mariam Zakalashvili, Lela Urushadze, Magda Dgebuadze, Giorgi Tomashvili, Davit Tsaguria, Ekaterine Zangaladze, Nino Berishvili, Gvantsa Chanturia, Adam Kotorashvili, Maia Alkhazashvili, Irma Burjanadze, Anna Kasradze, Khatuna Zakhashvili, Paata Imnadze, Amiran Gamkrelidze. |
| EPI_ISL_416480 | R. G. Lugar Center for Public Health Research, National Center for Disease Control and Public Health (NCDC) of Georgia. | R. G. Lugar Center for Public Health Research, National Center for Disease Control and Public Health (NCDC) of Georgia. | Ann Machablishvili, Nato Kotaria, Marine Murtskhvaladze, Lela Sabadze, Mari Gavashelidze, Ana Papkiauri, Meri Pantsulaia, Gvantsa Brachveli, Tata Imnadze, Tamar Jashiashvili, Tea Tevdoradze, Ketevan Sidamonidze, Ekaterine Khmaladze, Ekaterine Zhghenti, Roena Sukhiashvili, Mariam Zakalashvili, Lela Urushadze, Magda Dgebuadze, Giorgi Tomashvili, Davit Tsaguria, Ekaterine Zangaladze, Nino Berishvili, Gvantsa Chanturia, Adam Kotorashvili, Maia Alkhazashvili, Irma Burjanadze, Anna Kasradze, Khatuna Zakhashvili, Paata Imnadze, Amiran Gamkrelidze. |
| EPI_ISL_416481 | R. G. Lugar Center for Public Health Research, National Center for Disease Control and Public Health (NCDC) of Georgia. | R. G. Lugar Center for Public Health Research, National Center for Disease Control and Public Health (NCDC) of Georgia. | Gvantsa Chanturia, Marine Murtskhvaladze, Nato Kotaria, Ann Machablishvili, Lela Sabadze, Mari Gavashelidze, Ana Papkiauri, Meri Pantsulaia, Gvantsa Brachveli, Tata Imnadze, Tamar Jashiashvili, Tea Tevdoradze, Ketevan Sidamonidze, Ekaterine Khmaladze, Ekaterine Zhghenti, Roena Sukhiashvili, Mariam Zakalashvili, Lela Urushadze, Magda Dgebuadze, Giorgi Tomashvili, Davit Tsaguria, Ekaterine Zangaladze, Nino Berishvili, Adam Kotorashvili, Maia Alkhazashvili, Irma Burjanadze, Anna Kasradze, Khatuna Zakhashvili, Paata Imnadze, Amiran Gamkrelidze. |
| EPI_ISL_416482 | R. G. Lugar Center for Public Health Research, National Center for Disease Control and Public Health (NCDC) of Georgia. | R. G. Lugar Center for Public Health Research, National Center for Disease Control and Public Health (NCDC) of Georgia. | Adam Kotorashvili, Marine Murtskhvaladze, Nato Kotaria, Ann Machablishvili, Lela Sabadze, Mari Gavashelidze, Ana Papkiauri, Meri Pantsulaia, Gvantsa Brachveli, Tata Imnadze, Tamar Jashiashvili, Tea Tevdoradze, Ketevan Sidamonidze, Ekaterine Khmaladze, Ekaterine Zhghenti, Roena Sukhiashvili, Mariam Zakalashvili, Lela Urushadze, Magda Dgebuadze, Giorgi Tomashvili, Davit Tsaguria, Ekaterine Zangaladze, Nino Berishvili, Gvantsa Chanturia, Maia Alkhazashvili, Irma Burjanadze, Anna Kasradze, Khatuna Zakhashvili, Paata Imnadze, Amiran Gamkrelidze. |
| EPI_ISL_416483 | Servicio de Microbiología. Consorcio Hospital General Universitario de Valencia | Sequencing and Bioinformatics Service and Molecular Epidemiology Research Group. FISABIO-Public Health | Maria Alma Bracho, Maria Dolores Ocete, Concepcion Gimeno, Giuseppe D'Auria, Griselda De Marco, Neris Garcia-Gonzalez, Fernando Gonzalez-Candelas |
| EPI_ISL_416484 | Servicio de Microbiología. Consorcio Hospital General Universitario de Valencia | Sequencing and Bioinformatics Service and Molecular Epidemiology Research Group. FISABIO-Public Health | Maria Dolores Ocete, Concepcion Gimeno, Giuseppe D'Auria, Griselda De Marco, Neris Garcia-Gonzalez, Maria Alma Bracho, Fernando Gonzalez-Candelas |
| EPI_ISL_416485 | Servicio de Microbiologia. Consorcio Hospital General Universitario de Valencia | Sequencing and Bioinformatics Service and Molecular Epidemiology Research Group. FISABIO-Public Health | Griselda De Marco, Neris Garcia-Gonzalez, Maria Alma Bracho, Maria Dolores Ocete, Concepcion Gimeno, Giuseppe D'Auria, Fernando Gonzalez-Candelas |
| EPI_ISL_416486 | Servicio de Microbiología. Consorcio Hospital General Universitario de Valencia | Sequencing and Bioinformatics Service and Molecular Epidemiology Research Group. FISABIO-Public Health | Neris Garcia-Gonzalez, Maria Alma Bracho, Maria Dolores Ocete, Concepcion Gimeno, Giuseppe D'Auria, Griselda De Marco, Fernando Gonzalez-Candelas |
| EPI_ISL_416487 | Servicio de Microbiología. Consorcio Hospital General Universitario de Valencia | Sequencing and Bioinformatics Service and Molecular Epidemiology Research Group. FISABIO-Public Health | Giuseppe D'Auria, Griselda De Marco, Neris Garcia-Gonzalez, Maria Alma Bracho, Maria Dolores Ocete, Concepcion Gimeno, Fernando Gonzalez-Candelas |
| EPI_ISL_416488 | ViroGenetics - BSL3 Laboratory of Virology; Human Genome Variation Research Group & Genomics Centre MCB; Bioinformatics Research Group Department of Virology | ViroGenetics - BSL3 Laboratory of Virology; Human Genome Variation Research Group & Genomics Centre MCB; Bioinformatics Research Group Department of Virology | Aleksandra Milewska, Ewelina Pośpiech, Agata Jarosz, Adrianna Klajmon, Kamila Marszałek, Katarzyna Pancer, Magdalena Rzeczkowska, Tomasz Wołkowicz, Katarzyna Zacharczuk, Agnieszka Kołakowska-Kulesza, Natalia Wolaniuk, Ewelina Hallman-Szelińska, Paweł P Łabaj, Wojciech Branicki, Krzysztof Pyrć |
| EPI_ISL_416489, EPI_ISL_416491, EPI_ISL_416492 | University of Wisconsin-Madison AIDS Vaccine Research Laboratories | University of Wisconsin-Madison AIDS Vaccine Research Laboratories | Gage Moreno, Katarina Braun, et al. AIDS Vaccine Research Laboratories |
| EPI_ISL_416493 | CH Jean de Navarre Laboratoire de Biologie | National Reference Center for Viruses of Respiratory Infections, Institut Pasteur, Paris | Mélnie Albert, Marion Barbet, Sylvie Behillil, Méline Bizard, Angela Brisebarre, Flora Donati, Etienne Simon-Lorière, Vincent Enouf, Maud Vanpeene, Sylvie van der Werf |
| EPI_ISL_416494 | Centre Hositalier Universitaire de Rouen Laboratoire de Virologie | National Reference Center for Viruses of Respiratory Infections, Institut Pasteur, Paris | Mélnie Albert, Marion Barbet, Sylvie Behillil, Méline Bizard, Angela Brisebarre, Flora Donati, Etienne Simon-Lorière, Vincent Enouf, Maud Vanpeene, Sylvie van der Werf, Jean-Christophe Plantier |
| EPI_ISL_416495, EPI_ISL_416496, EPI_ISL_416497 | Centre Hospitalier Compiègne Laboratoire de Biologie | National Reference Center for Viruses of Respiratory Infections, Institut Pasteur, Paris | Mélnie Albert, Marion Barbet, Sylvie Behillil, Méline Bizard, Angela Brisebarre, Flora Donati, Etienne Simon-Lorière, Vincent Enouf, Maud Vanpeene, Sylvie van der Werf, Raulin Olivia |
| EPI_ISL_416498 | Institut Médico légal- Hop R. Poincaré | National Reference Center for Viruses of Respiratory Infections, Institut Pasteur, Paris | Mélnie Albert, Marion Barbet, Sylvie Behillil, Méline Bizard, Angela Brisebarre, Flora Donati, Etienne Simon-Lorière, Vincent Enouf, Maud Vanpeene, Sylvie van der Werf |
| EPI_ISL_416499, EPI_ISL_416500 | LABM GH nord Essonne | National Reference Center for Viruses of Respiratory Infections, Institut Pasteur, Paris | Mélnie Albert, Marion Barbet, Sylvie Behillil, Méline Bizard, Angela Brisebarre, Flora Donati, Etienne Simon-Lorière, Vincent Enouf, Maud Vanpeene, Sylvie van der Werf |
| EPI_ISL_416501 | Hopital franco britannique - Service des Urgences | National Reference Center for Viruses of Respiratory Infections, Institut Pasteur, Paris | Mélnie Albert, Marion Barbet, Sylvie Behillil, Méline Bizard, Angela Brisebarre, Flora Donati, Etienne Simon-Lorière, Vincent Enouf, Maud Vanpeene, Sylvie van der Werf |
| EPI_ISL_416502, EPI_ISL_416503, EPI_ISL_416504, EPI_ISL_416505, EPI_ISL_416506, EPI_ISL_416507, EPI_ISL_416508, EPI_ISL_416509, EPI_ISL_416510, EPI_ISL_416511, EPI_ISL_416512, EPI_ISL_416513 | | | |
| see above | CHRU Pontchaillou - Laboratoire de Virologie | National Reference Center for Viruses of Respiratory Infections, Institut Pasteur, Paris | Mélnie Albert, Marion Barbet, Sylvie Behillil, Méline Bizard, Angela Brisebarre, Flora Donati, Etienne Simon-Lorière, Vincent Enouf, Maud Vanpeene, Sylvie van der Werf, Gisèle Lagathu |
| EPI_ISL_416514, EPI_ISL_416515, EPI_ISL_416516, EPI_ISL_416517, EPI_ISL_416518 | Victorian Infectious Diseases Reference Laboratory (VIDRL) | Victorian Infectious Diseases Reference Laboratory and Microbiological Diagnostic Unit Public Health Laboratory, Doherty Institute | Caly L., Seemann T., Schultz M., Taiaroa, G., Druce J. |
| EPI_ISL_416519, EPI_ISL_416520 | Auckland Hospital | Institute of Environmental Science and Research (ESR) | Matt Storey, Xiaoyun Ren, Gary McAuliffe, Sally Roberts, Matthew Blakiston, Erasmus Smit, Lauren Jelly, Joep de Ligt |
| EPI_ISL_416521 | Public Health Laboratory | Public Health Laboratory, Saudi CDC | Albarrag, A |
| EPI_ISL_416522 | Public Health Laboratory, Saudi CDC | Public Health Laboratory, Saudi CDC | Albarrag,A |
| EPI_ISL_416523 | University of Wisconsin-Madison AIDS Vaccine Research Laboratories | University of Wisconsin-Madison AIDS Vaccine Research Laboratories | Gage Moreno, Katarina Braun, et al. AIDS Vaccine Research Laboratories |
| EPI_ISL_416524 | Saitama Medical University Hospital | Saitama Medical University | Kazuo Imai |
| EPI_ISL_416525 | Saitama Medical University | Saitama Medical University | Kazuo Imai |
| EPI_ISL_416526 | Auckland Hospital | Institute of Environmental Science and Research (ESR) | Matt Storey, Xiaoyun Ren, Gary McAuliffe, Sally Roberts, Matthew Blakiston, Erasmus Smit, Lauren Jelly, Joep de Ligt |
| EPI_ISL_416538 | Wellington Hospital | Institute of Environmental Science and Research (ESR) | Wellington SCL, Wellington Hospital, Riddiford Street, Newtown, Wellington 6021, New Zealand |
| EPI_ISL_416539 | Wellington Hospital | Institute of Environmental Science and Research (ESR) | Matt Storey, Xiaoyun Ren, Craig Thornley, Maxim Bloomfield, Erasmus Smit, Lauren Jelly, Joep de Ligt |
| EPI_ISL_416541 | Dasman Diabetes Institute and Virology Laboratory Ministry of Health | Dasman Diabetes Institute | Fahd Al-Mulla, Sumi John, Rasheeba Iqbal, Motasem Melhem, Ebaa AlOzairi, Sara Al-Qabandi, Qais Al-Duwairi |
| EPI_ISL_416542 | Dasman Diabetes Institute | Dasman Diabetes Institute | Fahd Al-Mulla, Sumi John, Rasheeba Iqbal, Motasem Melhem, Ebaa AlOzairi, Sara Al-Qabandi, Qais Al-Duwairi |
| EPI_ISL_416543 | Dasman Diabetes Institute | Dasman Diabetes Institute | Fahd Al-Mulla, Rasheeba Iqbal, Sumi John, Motasem Melhem, Ebaa AlOzairi, Sara Al-Qabandi, Qais Al-Duwairi |
| EPI_ISL_416565, EPI_ISL_416566, EPI_ISL_416567, EPI_ISL_416568, EPI_ISL_416569, EPI_ISL_416570, EPI_ISL_416571, EPI_ISL_416572, EPI_ISL_416573, EPI_ISL_416574, EPI_ISL_416575, EPI_ISL_416576, EPI_ISL_416577, EPI_ISL_416578, EPI_ISL_416579, EPI_ISL_416580, EPI_ISL_416581, EPI_ISL_416582, EPI_ISL_416583, EPI_ISL_416584, EPI_ISL_416585, EPI_ISL_416586, EPI_ISL_416587, EPI_ISL_416588, EPI_ISL_416589, EPI_ISL_416590, EPI_ISL_416591, EPI_ISL_416592, EPI_ISL_416593, EPI_ISL_416594, EPI_ISL_416595, EPI_ISL_416596, EPI_ISL_416597, EPI_ISL_416598, EPI_ISL_416599, EPI_ISL_416600, EPI_ISL_416601, EPI_ISL_416602, EPI_ISL_416603, EPI_ISL_416604, EPI_ISL_416605, EPI_ISL_416606, EPI_ISL_416607, EPI_ISL_416608, EPI_ISL_416609, EPI_ISL_416610, EPI_ISL_416611, EPI_ISL_416612, EPI_ISL_416613, EPI_ISL_416614, EPI_ISL_416615, EPI_ISL_416616, EPI_ISL_416617, EPI_ISL_416618, EPI_ISL_416619, EPI_ISL_416620, EPI_ISL_416621, EPI_ISL_416622, EPI_ISL_416623, EPI_ISL_416624, EPI_ISL_416625, EPI_ISL_416626, EPI_ISL_416627, EPI_ISL_416628, EPI_ISL_416629, EPI_ISL_416630, EPI_ISL_416631, EPI_ISL_416632, EPI_ISL_416633, EPI_ISL_416634 | | | |
| see above | Japanese Quarantine Stations | Pathogen Genomics Center, National Institute of Infectious Diseases | Tsuyoshi Sekizuka, Kentaro Itokawa, Rina Tanaka, Masanori Hashino, Tsutomu Kageyama, Shinji Saito, Ikuyo Takayama, Hideki Hasegawa, Takuri Takahashi, Hajime Kamiya, Takuya Yamagishi, Motoi Suzuki, Takaji Wakita, Makoto Kuroda |
| EPI_ISL_416635, EPI_ISL_416636, EPI_ISL_416637, EPI_ISL_416638, EPI_ISL_416639, EPI_ISL_416640, EPI_ISL_416641, EPI_ISL_416642, EPI_ISL_416643, EPI_ISL_416644, EPI_ISL_416645, EPI_ISL_416646, EPI_ISL_416647, EPI_ISL_416648, EPI_ISL_416649, EPI_ISL_416650, EPI_ISL_416651, EPI_ISL_416652, EPI_ISL_416653, EPI_ISL_416654, EPI_ISL_416655, EPI_ISL_416656, EPI_ISL_416657, EPI_ISL_416658, EPI_ISL_416659, EPI_ISL_416660, EPI_ISL_416661, EPI_ISL_416662, EPI_ISL_416663, EPI_ISL_416664, EPI_ISL_416665, EPI_ISL_416666, EPI_ISL_416667, EPI_ISL_416668, EPI_ISL_416669, EPI_ISL_416670, EPI_ISL_416671, EPI_ISL_416672, EPI_ISL_416673, EPI_ISL_416674, EPI_ISL_416675, EPI_ISL_416676, EPI_ISL_416677, EPI_ISL_416678, EPI_ISL_416679, EPI_ISL_416680, EPI_ISL_416681, EPI_ISL_416682, EPI_ISL_416683, EPI_ISL_416684, EPI_ISL_416685, EPI_ISL_416686, EPI_ISL_416687, EPI_ISL_416688, EPI_ISL_416689, EPI_ISL_416690, EPI_ISL_416691, EPI_ISL_416692, EPI_ISL_416693, EPI_ISL_416694, EPI_ISL_416695, EPI_ISL_416696, EPI_ISL_416697, EPI_ISL_416698, EPI_ISL_416699, EPI_ISL_416700, EPI_ISL_416701, EPI_ISL_416702, EPI_ISL_416703, EPI_ISL_416704, EPI_ISL_416705, EPI_ISL_416706, EPI_ISL_416707, EPI_ISL_416708, EPI_ISL_416709, EPI_ISL_416710, EPI_ISL_416711, EPI_ISL_416712, EPI_ISL_416713, EPI_ISL_416714, EPI_ISL_416715, EPI_ISL_416716, EPI_ISL_416717, EPI_ISL_416718, EPI_ISL_416719, EPI_ISL_416720, EPI_ISL_416721, EPI_ISL_416722, EPI_ISL_416723, EPI_ISL_416724, EPI_ISL_416725, EPI_ISL_416726, EPI_ISL_416727, EPI_ISL_416728, EPI_ISL_416729 | | | |
| see above | UW Virology Lab | UW Virology Lab | Pavitra Roychoudhury, Hong Xie, Keith Jerome, Alexander Greninger |
| EPI_ISL_416730, EPI_ISL_416731, EPI_ISL_416732, EPI_ISL_416733, EPI_ISL_416734, EPI_ISL_416735, EPI_ISL_416736, EPI_ISL_416737, EPI_ISL_416738, EPI_ISL_416739, EPI_ISL_416740 | | | |
| see above | Virology Department, Sheffield Teaching Hospitals NHS Foundation Trust | Department of Infection, Immunity and Cardiovascular Disease, The Florey Institute, The Medical School, University of Sheffield | Thushan de Silva, Matthew Parker, Adri Angyal, Rebecca Brown, Matthew Wyles, Mehmet Yavuz, Mohammad Raza, Cariad Evans |
| EPI_ISL_416741 | National Public Health Surveillance Laboratory, Vilnius, Lithuania | Charite Universitaetsmedizin Berlin, Institute of Virology | Victor M Corman, Julia Schneider, Jörn Beheim-Schwarzbach, Talitha Veith, Barbara Muehlemann, Terry Jones, Ana Steponkiene, Christian Drosten |
| EPI_ISL_416742, EPI_ISL_416743 | NRL for Influenza, Centrum Epidemiology and Microbiology of National Institute of Public Health, Czech Republic | Charite Universitaetsmedizin Berlin, Institute of Virology | Victor M Corman, Julia Schneider, Jörn Beheim-Schwarzbach, Talitha Veith, Barbara Muehlemann, Terry Jones, Akexander Nagy, Jaromira Vecerova, Dusan Trnka, Ludmila Novakova, Helena Jirincova, Christian Drosten |
| EPI_ISL_416744 | Virological Research Group, Szentágothai Research Centre | Bioinformatics Research Group, Szentágothai Research Centre | Péter Urbán, Endre Gábor Tóth, Gábor Kemenesi, Róbert Herczeg, Attila Gyenesei, Ferenc Jakab |
| EPI_ISL_416745, EPI_ISL_416746 | CNR Virus des Infections Respiratoires - France SUD | CNR Virus des Infections Respiratoires - France SUD | Bal, Antonin; Destras, Gregory; Gaymard, Alexandre; Bouscambert-Duchamp, Maude; Cheynet, Valérie; Brengel-Pesce, Karen; Morfin-Sherpa, Florence; Valette, Martine; Josset, Laurence; Lina, Bruno. |
| EPI_ISL_416747, EPI_ISL_416748 | Institut des Agents Infectieux (IAI) Hospices Civils de Lyon | CNR Virus des Infections Respiratoires - France SUD | Bal, Antonin; Destras, Gregory; Gaymard, Alexandre; Bouscambert-Duchamp, Maude; Cheynet, Valérie; Brengel-Pesce, Karen; Morfin-Sherpa, Florence; Valette, Martine; Josset, Laurence; Lina, Bruno. |
| EPI_ISL_416749 | Centre Hospitalier de Valence | CNR Virus des Infections Respiratoires - France SUD | Bal, Antonin; Destras, Gregory; Gaymard, Alexandre; Bouscambert-Duchamp, Maude; Cheynet, Valérie; Brengel-Pesce, Karen; Morfin-Sherpa, Florence; Valette, Martine; Josset, Laurence; Lina, Bruno. |
| EPI_ISL_416750 | Institut des Agents Infectieux (IAI) Hospices Civils de | CNR Virus des Infections Respiratoires - France SUD | Bal, Antonin; Destras, Gregory; Gaymard, Alexandre; Bouscambert-Duchamp, Maude; Cheynet, Valérie; Brengel-Pesce, Karen; Morfin-Sherpa, Florence; Valette, Martine; Josset, Laurence; Lina, Bruno. |

| | | | |
|---|---|---|---|
| | Lyon | | |
| EPI_ISL_416751, EPI_ISL_416752 | CHU Gabriel Montpied | CNR Virus des Infections Respiratoires - France SUD | Bal, Antonin; Destras, Gregory; Gaymard, Alexandre; Bouscambert-Duchamp, Maude; Cheynet, Valérie; Brengel-Pesce, Karen; Morfin-Sherpa, Florence; Valette, Martine; Josset, Laurence; Lina, Bruno. |
| EPI_ISL_416753, EPI_ISL_416754, EPI_ISL_416756 | Institut des Agents Infectieux (IAI) Hospices Civils de Lyon | CNR Virus des Infections Respiratoires - France SUD | Bal, Antonin; Destras, Gregory; Gaymard, Alexandre; Bouscambert-Duchamp, Maude; Cheynet, Valérie; Brengel-Pesce, Karen; Morfin-Sherpa, Florence; Valette, Martine; Josset, Laurence; Lina, Bruno. |
| EPI_ISL_416757 | Centre Hospitalier de Bourg en Bresse | CNR Virus des Infections Respiratoires - France SUD | Bal, Antonin; Destras, Gregory; Gaymard, Alexandre; Bouscambert-Duchamp, Maude; Cheynet, Valérie; Brengel-Pesce, Karen; Morfin-Sherpa, Florence; Valette, Martine; Josset, Laurence; Lina, Bruno. |
| EPI_ISL_416758 | Institut des Agents Infectieux (IAI) Hospices Civils de Lyon | CNR Virus des Infections Respiratoires - France SUD | Bal, Antonin; Destras, Gregory; Gaymard, Alexandre; Bouscambert-Duchamp, Maude; Cheynet, Valérie; Brengel-Pesce, Karen; Morfin-Sherpa, Florence; Valette, Martine; Josset, Laurence; Lina, Bruno. |
| EPI_ISL_416829 | National Public Health Laboratory | Malaysia Genome Institute | Mohd Noor Mat Isa, Irni Suhayu Sapian, Yusuf Muhammad Noor, Nurhezreen Md Iqbal, Mohd Faizal Abu Bakar, Enizza Kasim, Shamsidar Sopie, Siti Noraini Othman, Azrin Ahmad, Nor Azfa Johari, Norazimah Tajudin, Noorliza Mohamad Noordin, W Afiza W Mohd Arifin, Rehan Shuhada Abu Bakar, Yu Kie Chem, Selvanesan Sengol, Hani Mat Hussin, Shahrul Hisham Zainal Ariffin |
| EPI_ISL_416830, EPI_ISL_416831, EPI_ISL_416832 | NYU Langone Health | Department of Pathology and Medicine, New York University School of Medicine | John Chen, Dacia Dimartino, Xiaojun Feng, Adriana Heguy, Megan Hogan, Emily Huang, George Jour, Christian Marier, Matt Maurano, Mark Mulligan, Peter Meyn, Marie Samanovic-Golden, Amy Rapkiewicz, Guomiao Shen, Matija Snuderl, Gael Westby, Paul Zappile |
| EPI_ISL_416866, EPI_ISL_416884 | National Public Health Laboratory | Malaysia Genome Institute | Mohd Noor Mat Isa, Irni Suhayu Sapian, Yusuf Muhammad Noor, Nurhezreen Md Iqbal, Mohd Faizal Abu Bakar, Enizza Kasim, Shamsidar Sopie, Siti Noraini Othman, Azrin Ahmad, Nor Azfa Johari, Norazimah Tajudin, Noorliza Mohamad Noordin, W Afiza W Mohd Arifin, Rehan Shuhada Abu Bakar, Yu Kie Chem, Selvanesan Sengol, Hani Mat Hussin, Shahrul Hisham Zainal Ariffin |
| EPI_ISL_416885, EPI_ISL_416886 | National Public Health Laboratory | Malaysia Genome Institute | Mohd Noor Mat Isa, Irni Suhayu Sapian, Yusuf Muhammad Noor, Nurhezreen Md Iqbal, Mohd Faizal Abu Bakar, Enizza Kasim, Shamsidar Sopie, Siti Noraini Othman, Azrin Ahmad, Nor Azfa Johari, Norazimah Tajudin, Noorliza Mohamad Noordin, W Afiza W Mohd Arifin, Rehan Shuhada Abu Bakar, Yu Kie Chem, Selvanesan Sengol, Hani Mat Hussin, Shahrul Hisham Zainal Ariffin |
| EPI_ISL_416907 | National Public Health Laboratory | Malaysia Genome Institute | Mohd Noor Mat Isa, Irni Suhayu Sapian, Yusuf Muhammad Noor, Nurhezreen Md Iqbal, Mohd Faizal Abu Bakar, Enizza Kasim, Shamsidar Sopie, Siti Noraini Othman, Azrin Ahmad, Nor Azfa Johari, Norazimah Tajudin, Noorliza Mohamad Noordin, W Afiza W Mohd Arifin, Rehan Shuhada Abu Bakar, Yu Kie Chem, Selvanesan Sengol, Hani Mat Hussin, Shahrul Hisham Zainal Ariffin |
| EPI_ISL_416994 | COMPLEJO ASISTENCIAL UNIVERSITARIO DE BURGOS | Instituto de Salud Carlos III | Iglesias-Caballero, M. Molinero Calamita, M. González-Esguevillas, M. Camarero S. Pozo F. Casas I. Jiménez P. Jiménez M. Zaballos A. Monzón, S. Varona, S. Juliá M. Cuesta I. Megias Lobón, G. Hospital: ------ |
| EPI_ISL_416997, EPI_ISL_417004 | Department of Clinical Microbiology | GIGA Medical Genomics | Durkin Keith, Artesi Maria, Bontems Sébastien, Boreux Raphaël, Meex Cécile, Melin Pierrette, Hayette Marie-Pierre, Bours Vincent. |
| EPI_ISL_417005 | Department of Clinical Microbiology | GIGA Medical Genomics | Durkin Keith, Artesi Maria, Bontems Sébastien, Boreux Raphaël, Meex Cécile, Melin Pierrette, Hayette Marie-Pierre, Bours Vincent. |
| EPI_ISL_417006 | Department of Clinical Microbiology | GIGA Medical Genomics | Durkin Keith, Artesi Maria, Bontems Sébastien, Boreux Raphaël, Meex Cécile, Melin Pierrette, Hayette Marie-Pierre, Bours Vincent. |
| EPI_ISL_417007 | HOSPITAL SANTA MARIA NAI | Instituto de Salud Carlos III | Iglesias-Caballero, M. Molinero Calamita, M. González-Esguevillas, M. Camarero S. Pozo F. Casas I. Jiménez, P. Jiménez, M. Zaballos, A. Monzón, S. Varona, S. Juliá, M. Cuesta, I. García Costa, J. |
| EPI_ISL_417008, EPI_ISL_417009 | Department of Clinical Microbiology | GIGA Medical Genomics | Durkin Keith, Artesi Maria, Bontems Sébastien, Boreux Raphaël, Meex Cécile, Melin Pierrette, Hayette Marie-Pierre, Bours Vincent. |
| EPI_ISL_417010 | FUNDACION JIMENEZ DIAZ | Instituto de Salud Carlos III | Iglesias-Caballero, M. Molinero Calamita, M. González-Esguevillas, M. Camarero, S. Pozo, F. Casas, I. Jiménez, P. Jiménez, M. Zaballos, A. Monzón, S. Varona, S. Juliá, M. Cuesta, I. Fernández Roblas, R. |
| EPI_ISL_417011 | Department of Clinical Microbiology | GIGA Medical Genomics | Durkin Keith, Artesi Maria, Bontems Sébastien, Boreux Raphaël, Meex Cécile, Melin Pierrette, Hayette Marie-Pierre, Bours Vincent. |
| EPI_ISL_417012, EPI_ISL_417013, EPI_ISL_417014, EPI_ISL_417015, EPI_ISL_417016, EPI_ISL_417017, EPI_ISL_417018, EPI_ISL_417019, EPI_ISL_417020 | Department of Clinical Microbiology | GIGA Medical Genomics | Durkin Keith, Artesi Maria, Bontems Sébastien, Boreux Raphaël, Meex Cécile, Melin Pierrette, Hayette Marie-Pierre, Bours Vincent. |
| EPI_ISL_417021, EPI_ISL_417022, EPI_ISL_417023, EPI_ISL_417025 | Department of Clinical Microbiology | GIGA Medical Genomics | Durkin Keith, Artesi Maria, Bontems Sébastien, Boreux Raphaël, Meex Cécile, Melin Pierrette, Hayette Marie-Pierre, Bours Vincent. |
| EPI_ISL_417026, EPI_ISL_417027, EPI_ISL_417028 | Utah Public Health Laboratory | Utah Public Health Laboratory | Erin Young, Kelly Oakeson |
| EPI_ISL_417030 | Centre for Infectious Diseases and Microbiology Laboratory Services | NSW Health Pathology - Institute of Clinical Pathology and Medical Research; Westmead Hospital; University of Sydney | Eden J-S, Rockett R, Carter I, Rahman H, Holmes EC, O'Sullivan MV, Sintchenko V, Chen SC, Maddocks S, Kok J and Dwyer DE for the 2019-nCoV Study Group* |
| EPI_ISL_417031 | Pathology Queensland | Public Health Virology Laboratory | Bixing Huang, Alyssa Pyke, Amanda De Jong, Andrew Van Den Hurk, Carmel Taylor, David Warrilow, Doris Genge, Elisabeth Gamez, Glen Hewitson, Ian Maxwell Mackay, Inga Sultana, Jamie McMahon, Jean Barcelon, Judy Northill, Mitchell Finger, Natalie Simpson, Neelima Nair, Peter Burtonclay, Peter Moore, Sarah Wheatley, Sean Moody, Sonja Hall-Mendelin, Timothy Gardam, and Frederick Moore |
| EPI_ISL_417032 | Rockhampton Base Hospital | Public Health Virology Laboratory | Bixing Huang, Alyssa Pyke, Amanda De Jong, Andrew Van Den Hurk, Carmel Taylor, David Warrilow, Doris Genge, Elisabeth Gamez, Glen Hewitson, Ian Maxwell Mackay, Inga Sultana, Jamie McMahon, Jean Barcelon, Judy Northill, Mitchell Finger, Natalie Simpson, Neelima Nair, Peter Burtonclay, Peter Moore, Sarah Wheatley, Sean Moody, Sonja Hall-Mendelin, Timothy Gardam, and Frederick Moore |
| EPI_ISL_417033 | Sullivan Nicolaides Pathology | Public Health Virology Laboratory | Bixing Huang, Alyssa Pyke, Amanda De Jong, Andrew Van Den Hurk, Carmel Taylor, David Warrilow, Doris Genge, Elisabeth Gamez, Glen Hewitson, Ian Maxwell Mackay, Inga Sultana, Jamie McMahon, Jean Barcelon, Judy Northill, Mitchell Finger, Natalie Simpson, Neelima Nair, Peter Burtonclay, Peter Moore, Sarah Wheatley, Sean Moody, Sonja Hall-Mendelin, Timothy Gardam, and Frederick Moore |
| EPI_ISL_417034 | Laboratorio de Ecologia de Doencas Transmissiveis na Amazonia, Instituto Leonidas e Maria Deane - Fiocruz Amazonia | Laboratorio de Ecologia de Doencas Transmissiveis na Amazonia, Instituto Leonidas e Maria Deane - Fiocruz Amazonia | Valdinete Nascimento, André Corado, Fernanda Nascimento, Ágatha Costa, Debora Duarte, Luciana Gonçalves, Michele Jesus, Sérgio Luz, Felipe Naveca |
| EPI_ISL_417064 | Prince of Wales Hospital | Hong Kong Department of Health | Alan K.L. Tsang, Peter C.W. Yip, Edman T.K. Lam, Rickjason C.W. Chan, Dominic N.C. Tsang |
| EPI_ISL_417065, EPI_ISL_417066, EPI_ISL_417067, EPI_ISL_417068, EPI_ISL_417069, EPI_ISL_417070, EPI_ISL_417071, EPI_ISL_417072, EPI_ISL_417073, EPI_ISL_417074, EPI_ISL_417075, EPI_ISL_417076, EPI_ISL_417077, EPI_ISL_417078, EPI_ISL_417079, EPI_ISL_417080, EPI_ISL_417081, EPI_ISL_417082, EPI_ISL_417083, EPI_ISL_417084, EPI_ISL_417085, EPI_ISL_417086, EPI_ISL_417087, EPI_ISL_417088, EPI_ISL_417089, EPI_ISL_417090, EPI_ISL_417091, EPI_ISL_417092, EPI_ISL_417093, EPI_ISL_417094, EPI_ISL_417095, EPI_ISL_417096, EPI_ISL_417097, EPI_ISL_417098, EPI_ISL_417099, EPI_ISL_417100, EPI_ISL_417101, EPI_ISL_417102, EPI_ISL_417103, EPI_ISL_417104, EPI_ISL_417105, EPI_ISL_417106, EPI_ISL_417107, EPI_ISL_417108, EPI_ISL_417109, EPI_ISL_417110, EPI_ISL_417111, EPI_ISL_417112, EPI_ISL_417113, EPI_ISL_417114, EPI_ISL_417115, EPI_ISL_417116, EPI_ISL_417117, EPI_ISL_417118, EPI_ISL_417119, EPI_ISL_417120, EPI_ISL_417121, EPI_ISL_417122, EPI_ISL_417123, EPI_ISL_417124, EPI_ISL_417125, EPI_ISL_417126, EPI_ISL_417127, EPI_ISL_417128, EPI_ISL_417129, EPI_ISL_417130, EPI_ISL_417131, EPI_ISL_417132, EPI_ISL_417133, EPI_ISL_417134, EPI_ISL_417135, EPI_ISL_417136, EPI_ISL_417137, EPI_ISL_417138, EPI_ISL_417139, EPI_ISL_417140, EPI_ISL_417141, EPI_ISL_417142, EPI_ISL_417143, EPI_ISL_417144, EPI_ISL_417145, EPI_ISL_417146, EPI_ISL_417147, EPI_ISL_417148, EPI_ISL_417149, EPI_ISL_417150, EPI_ISL_417151, EPI_ISL_417152, EPI_ISL_417153, EPI_ISL_417154, EPI_ISL_417155, EPI_ISL_417156, EPI_ISL_417157, EPI_ISL_417158, EPI_ISL_417159, EPI_ISL_417160, EPI_ISL_417161, EPI_ISL_417162 | | | |
| see above | Washington State Department of Health | Seattle Flu Study | Chu etl al |
| EPI_ISL_417163, EPI_ISL_417164, EPI_ISL_417165 | Seattle Flu Study | Seattle Flu Study | Chu etl al |
| EPI_ISL_417166, EPI_ISL_417167, EPI_ISL_417168, EPI_ISL_417169, EPI_ISL_417170, EPI_ISL_417171, EPI_ISL_417172, EPI_ISL_417173, EPI_ISL_417174, EPI_ISL_417175 | Washington State Department of Health | Seattle Flu Study | Chu etl al |
| EPI_ISL_417176, EPI_ISL_417177, EPI_ISL_417178, EPI_ISL_417179 | Department of Pathology, Princess Margaret Hospital | Department of Health Technology and Informatics, Faculty of Health and Social Science, The Hong Kong Polytechnic University | Kenneth Siu-Sing LEUNG, Timothy Ting-Leung NG, Alan Ka-Lun WU, Miranda Chong-Yee YAU, Hiu-Yin LAO, Ming-Pan CHOI, Kingsley King-Gee TAM, Lam-Kwong LEE, Barry Kin-Chung WONG, Alex Yat-Man HO, Kam-Tong Yip, Kwok-Cheung LUNG, Raymond Wai-To LIU, Eugene Yuk-Keung TSO, Wai-Shing LEUNG, Man-Chun CHAN, Yuk-Yung NG, Kit-Man SIN, Kitty Sau-Chun FUNG, Sandy Ka-Yee CHAU, Wing-Kin TO, Tak-Lun Que, David Ho-Keung SHUM, Shea Ping YIP, Wing Cheong YAM, Gilman Kit-Hang SIU |
| EPI_ISL_417180, EPI_ISL_417181, EPI_ISL_417182 | Department of Pathology, United Christian Hospital | Department of Health Technology and Informatics, Faculty of Health and Social Science, The Hong Kong Polytechnic University | Kenneth Siu-Sing LEUNG, Timothy Ting-Leung NG, Alan Ka-Lun WU, Miranda Chong-Yee YAU, Hiu-Yin LAO, Ming-Pan CHOI, Kingsley King-Gee TAM, Lam-Kwong LEE, Barry Kin-Chung WONG, Alex Yat-Man HO, Kam-Tong Yip, Kwok-Cheung LUNG, Raymond Wai-To LIU, Eugene Yuk-Keung TSO, Wai-Shing LEUNG, Man-Chun CHAN, Yuk-Yung NG, Kit-Man SIN, Kitty Sau-Chun FUNG, Sandy Ka-Yee CHAU, Wing-Kin TO, Tak-Lun Que, David Ho-Keung SHUM, Shea Ping YIP, Wing Cheong YAM, Gilman Kit-Hang SIU |
| EPI_ISL_417183, EPI_ISL_417184 | Department of Clinical Pathology, Pamela Youde Nethersole Eastern Hospital | Department of Health Technology and Informatics, Faculty of Health and Social Science, The Hong Kong Polytechnic University | Kenneth Siu-Sing LEUNG, Timothy Ting-Leung NG, Alan Ka-Lun WU, Miranda Chong-Yee YAU, Hiu-Yin LAO, Ming-Pan CHOI, Kingsley King-Gee TAM, Lam-Kwong LEE, Barry Kin-Chung WONG, Alex Yat-Man HO, Kam-Tong Yip, Kwok-Cheung LUNG, Raymond Wai-To LIU, Eugene Yuk-Keung TSO, Wai-Shing LEUNG, Man-Chun CHAN, Yuk-Yung NG, Kit-Man SIN, Kitty Sau-Chun FUNG, Sandy Ka-Yee CHAU, Wing-Kin TO, Tak-Lun Que, David Ho-Keung SHUM, Shea Ping YIP, Wing Cheong YAM, Gilman Kit-Hang SIU |
| EPI_ISL_417185 | Department of Pathology, United Christian Hospital | Department of Health Technology and Informatics, Faculty of Health and Social Science, The Hong Kong Polytechnic University | Kenneth Siu-Sing LEUNG, Timothy Ting-Leung NG, Alan Ka-Lun WU, Miranda Chong-Yee YAU, Hiu-Yin LAO, Ming-Pan CHOI, Kingsley King-Gee TAM, Lam-Kwong LEE, Barry Kin-Chung WONG, Alex Yat-Man HO, Kam-Tong Yip, Kwok-Cheung LUNG, Raymond Wai-To LIU, Eugene Yuk-Keung TSO, Wai-Shing LEUNG, Man-Chun CHAN, Yuk-Yung NG, Kit-Man SIN, Kitty Sau-Chun FUNG, Sandy Ka-Yee CHAU, Wing-Kin TO, Tak-Lun Que, David Ho-Keung SHUM, Shea Ping YIP, Wing Cheong YAM, Gilman Kit-Hang SIU |
| EPI_ISL_417186 | National Institute for Communicable Diseases of the National Health Laboratory Service | National Institute for Communicable Diseases of the National Health Laboratory Service | Allam M, Kwenda S, van Heusden P, Khumalo Z, Mohale T, Subramoney K, von Gottberg, A, Ismail A, Bhiman JN |
| EPI_ISL_417187, EPI_ISL_417188 | Department of Clinical Pathology, Pamela Youde Nethersole Eastern Hospital | Department of Health Technology and Informatics, Faculty of Health and Social Science, The Hong Kong Polytechnic University | Kenneth Siu-Sing LEUNG, Timothy Ting-Leung NG, Alan Ka-Lun WU, Miranda Chong-Yee YAU, Hiu-Yin LAO, Ming-Pan CHOI, Kingsley King-Gee TAM, Lam-Kwong LEE, Barry Kin-Chung WONG, Alex Yat-Man HO, Kam-Tong Yip, Kwok-Cheung LUNG, Raymond Wai-To LIU, Eugene Yuk-Keung TSO, Wai-Shing LEUNG, Man-Chun CHAN, Yuk-Yung NG, Kit-Man SIN, Kitty Sau-Chun FUNG, Sandy Ka-Yee CHAU, Wing-Kin TO, Tak-Lun Que, David Ho-Keung SHUM, Shea Ping YIP, Wing Cheong YAM, Gilman Kit-Hang SIU |
| EPI_ISL_417189 | Minnesota Department of Health, Public Health Laboratory | Minnesota Department of Health, Public Health Laboratory | Matt Plumb, Jake Garfin and Xiong Wang |
| EPI_ISL_417190 | Department of Clinical Pathology, Pamela Youde Nethersole Eastern Hospital | Department of Health Technology and Informatics, Faculty of Health and Social Science, The Hong Kong Polytechnic University | Kenneth Siu-Sing LEUNG, Timothy Ting-Leung NG, Alan Ka-Lun WU, Miranda Chong-Yee YAU, Hiu-Yin LAO, Ming-Pan CHOI, Kingsley King-Gee TAM, Lam-Kwong LEE, Barry Kin-Chung WONG, Alex Yat-Man HO, Kam-Tong Yip, Kwok-Cheung LUNG, Raymond Wai-To LIU, Eugene Yuk-Keung TSO, Wai-Shing LEUNG, Man-Chun CHAN, Yuk-Yung NG, Kit-Man SIN, Kitty Sau-Chun FUNG, Sandy Ka-Yee CHAU, Wing-Kin TO, Tak-Lun Que, David Ho-Keung SHUM, Shea Ping YIP, Wing Cheong YAM, Gilman Kit-Hang SIU |
| EPI_ISL_417191, EPI_ISL_417192 | Minnesota Department of Health, Public Health Laboratory | Minnesota Department of Health, Public Health Laboratory | Matt Plumb, Jake Garfin and Xiong Wang |
| EPI_ISL_417193 | Department of Clinical Pathology, Pamela Youde Nethersole Eastern Hospital | Department of Health Technology and Informatics, Faculty of Health and Social Science, The Hong Kong Polytechnic University | Kenneth Siu-Sing LEUNG, Timothy Ting-Leung NG, Alan Ka-Lun WU, Miranda Chong-Yee YAU, Hiu-Yin LAO, Ming-Pan CHOI, Kingsley King-Gee TAM, Lam-Kwong LEE, Barry Kin-Chung WONG, Alex Yat-Man HO, Kam-Tong Yip, Kwok-Cheung LUNG, Raymond Wai-To LIU, Eugene Yuk-Keung TSO, Wai-Shing LEUNG, Man-Chun CHAN, Yuk-Yung NG, Kit-Man SIN, Kitty Sau-Chun FUNG, Sandy Ka-Yee CHAU, Wing-Kin TO, Tak-Lun Que, David Ho-Keung SHUM, Shea Ping YIP, Wing Cheong YAM, Gilman Kit-Hang SIU |
| EPI_ISL_417194 | Minnesota Department of Health, Public Health Laboratory | Minnesota Department of Health, Public Health Laboratory | Matt Plumb, Jake Garfin and Xiong Wang |
| EPI_ISL_417195 | Department of Clinical Pathology, Pamela Youde Nethersole Eastern Hospital | Department of Health Technology and Informatics, Faculty of Health and Social Science, The Hong Kong Polytechnic | Kenneth Siu-Sing LEUNG, Timothy Ting-Leung NG, Alan Ka-Lun WU, Miranda Chong-Yee YAU, Hiu-Yin LAO, Ming-Pan CHOI, Kingsley King-Gee TAM, Lam-Kwong LEE, Barry Kin-Chung WONG, Alex Yat-Man HO, Kam-Tong Yip, Kwok-Cheung LUNG, Raymond Wai-To LIU, Eugene Yuk-Keung TSO, Wai-Shing LEUNG, Man-Chun CHAN, Yuk-Yung NG, Kit-Man SIN, Kitty Sau-Chun FUNG, Sandy Ka-Yee CHAU, Wing-Kin TO, Tak-Lun Que, David Ho-Keung SHUM, Shea Ping YIP, Wing Cheong YAM, |

| | | | |
|---|---|---|---|
| | | University | Gilman Kit-Hang SIU |
| EPI_ISL_417196 | Minnesota Department of Health, Public Health Laboratory | Minnesota Department of Health, Public Health Laboratory | Matt Plumb, Jake Garfin and Xiong Wang |
| EPI_ISL_417197 | Department of Clinical Pathology, Pamela Youde Nethersole Eastern Hospital | Department of Health Technology and Informatics, Faculty of Health and Social Science, The Hong Kong Polytechnic University | Kenneth Siu-Sing LEUNG, Timothy Ting-Leung NG, Alan Ka-Lun WU, Miranda Chong-Yee YAU, Hiu-Yin LAO, Ming-Pan CHOI, Kingsley King-Gee TAM, Lam-Kwong LEE, Barry Kin-Chung WONG, Alex Yat-Man HO, Kam-Tong Yip, Kwok-Cheung LUNG, Raymond Wai-To LIU, Eugene Yuk-Keung TSO, Wai-Shing LEUNG, Man-Chun CHAN, Yuk-Yung NG, Kit-Man SIN, Kitty Sau-Chun FUNG, Sandy Ka-Yee CHAU, Wing-Kin TO, Tak-Lun Que, David Ho-Keung SHUM, Shea Ping YIP, Wing Cheong YAM, Gilman Kit-Hang SIU |
| EPI_ISL_417198 | Minnesota Department of Health, Public Health Laboratory | Minnesota Department of Health, Public Health Laboratory | Matt Plumb, Jake Garfin and Xiong Wang |
| EPI_ISL_417199 | Department of Clinical Pathology, Pamela Youde Nethersole Eastern Hospital | Department of Health Technology and Informatics, Faculty of Health and Social Science, The Hong Kong Polytechnic University | Chong-Yee YAU, Hiu-Yin LAO, Ming-Pan CHOI, Kingsley King-Gee TAM, Lam-Kwong LEE, Barry Kin-Chung WONG, Alex Yat-Man HO, Kam-Tong Yip, Kwok-Cheung LUNG, Raymond Wai-To LIU, Eugene Yuk-Keung TSO, Wai-Shing LEUNG, Man-Chun CHAN, Yuk-Yung NG, Kit-Man SIN, Kitty Sau-Chun FUNG, Sandy Ka-Yee CHAU, Wing-Kin TO, Tak-Lun Que, David Ho-Keung SHUM, Shea Ping YIP, Wing Cheong YAM, Gilman Kit-Hang SIU |
| EPI_ISL_417200, EPI_ISL_417201, EPI_ISL_417202, EPI_ISL_417203, EPI_ISL_417204 | University of Wisconsin-Madison AIDS Vaccine Research Laboratories | University of Wisconsin-Madison AIDS Vaccine Research Laboratories | Gage Moreno, Katarina Braun, et al. AIDS Vaccine Research Laboratories |
| EPI_ISL_417205 | Servicio de Microbiología. Consorcio Hospital General Universitario de Valencia | Sequencing and Bioinformatics Service and Molecular Epidemiology Research Group. FISABIO-Public Health | Maria Alma Bracho, Maria Dolores Ocete, Concepcion Gimeno, Giuseppe D'Auria, Griselda De Marco, Neris Garcia-Gonzalez, Fernando Gonzalez-Candelas |
| EPI_ISL_417206 | Servicio de Microbiologia. Consorcio Hospital General Universitario de Valencia | Sequencing and Bioinformatics Service and Molecular Epidemiology Research Group. FISABIO-Public Health | Maria Alma Bracho, Maria Dolores Ocete, Concepcion Gimeno, Giuseppe D'Auria, Griselda De Marco, Neris Garcia-Gonzalez, Fernando Gonzalez-Candelas |
| EPI_ISL_417211, EPI_ISL_417212 | Dunedin Hospital | University of Otago | M.E. Quiñones-Mateu, B. Lawley, J. Grant, R. Harfoot, J. Ussher |
| EPI_ISL_417213, EPI_ISL_417214, EPI_ISL_417215, EPI_ISL_417216, EPI_ISL_417217, EPI_ISL_417218, EPI_ISL_417219, EPI_ISL_417220, EPI_ISL_417221, EPI_ISL_417222, EPI_ISL_417223, EPI_ISL_417224, EPI_ISL_417225, EPI_ISL_417226, EPI_ISL_417227, EPI_ISL_417228, EPI_ISL_417229, EPI_ISL_417230, EPI_ISL_417231, EPI_ISL_417232, EPI_ISL_417233, EPI_ISL_417234, EPI_ISL_417235, EPI_ISL_417236, EPI_ISL_417237, EPI_ISL_417238, EPI_ISL_417239, EPI_ISL_417240, EPI_ISL_417241, EPI_ISL_417242, EPI_ISL_417243, EPI_ISL_417244, EPI_ISL_417245, EPI_ISL_417246, EPI_ISL_417247, EPI_ISL_417248, EPI_ISL_417249, EPI_ISL_417250, EPI_ISL_417251, EPI_ISL_417252, EPI_ISL_417253, EPI_ISL_417254, EPI_ISL_417255, EPI_ISL_417256, EPI_ISL_417257, EPI_ISL_417258, EPI_ISL_417259, EPI_ISL_417260, EPI_ISL_417261, EPI_ISL_417262, EPI_ISL_417263, EPI_ISL_417264, EPI_ISL_417265, EPI_ISL_417266, EPI_ISL_417267, EPI_ISL_417268, EPI_ISL_417269, EPI_ISL_417270, EPI_ISL_417271, EPI_ISL_417272, EPI_ISL_417273, EPI_ISL_417274, EPI_ISL_417275, EPI_ISL_417276, EPI_ISL_417277, EPI_ISL_417278, EPI_ISL_417279, EPI_ISL_417280, EPI_ISL_417281, EPI_ISL_417282, EPI_ISL_417283, EPI_ISL_417284, EPI_ISL_417285, EPI_ISL_417286, EPI_ISL_417287, EPI_ISL_417288, EPI_ISL_417289, EPI_ISL_417290, EPI_ISL_417291, EPI_ISL_417292, EPI_ISL_417293, EPI_ISL_417294, EPI_ISL_417295, EPI_ISL_417296, EPI_ISL_417297, EPI_ISL_417298, EPI_ISL_417299, EPI_ISL_417300, EPI_ISL_417301, EPI_ISL_417302, EPI_ISL_417303, EPI_ISL_417304, EPI_ISL_417305, EPI_ISL_417306, EPI_ISL_417307, EPI_ISL_417308, EPI_ISL_417309, EPI_ISL_417310, EPI_ISL_417311, EPI_ISL_417312, EPI_ISL_417313, EPI_ISL_417314, EPI_ISL_417315 | | | |
| see above | Respiratory Virus Unit, Microbiology Services Colindale, Public Health England | Respiratory Virus Unit, Microbiology Services Colindale, Public Health England | Monica Galiano, Shahjahan Miah, Angie Lackenby, Omolola Akinbami, Tiina Talts, Leena Bhaw, Richard Myers, Steven Platt, Kirstin Edwards, Jonathan Hubb, Joanna Ellis, Maria Zambon |
| EPI_ISL_417316 | Chiu Laboratory, University of California, San Francisco | Chiu Laboratory, University of California, San Francisco | Xianding Deng, Scot Federman, Wei Gu, and Charles Y. Chiu |
| EPI_ISL_417317, EPI_ISL_417318, EPI_ISL_417319, EPI_ISL_417320, EPI_ISL_417321 | Santa Clara County Public Health Department | Chiu Laboratory, University of California, San Francisco | Xianding Deng, Scot Federman, Wei Gu, Elsa Villarino, Brandon Bonin, Debra A. Wadford, and Charles Y. Chiu |
| EPI_ISL_417322, EPI_ISL_417323, EPI_ISL_417324, EPI_ISL_417325, EPI_ISL_417326, EPI_ISL_417327, EPI_ISL_417328, EPI_ISL_417329 | California Department of Public Health | Chiu Laboratory, University of California, San Francisco | Xianding Deng, Scot Federman, Chao-Yang Pan, Hugo Guevara,Wei Gu, Debra A. Wadford, and Charles Y. Chiu |
| EPI_ISL_417330, EPI_ISL_417331, EPI_ISL_417332 | Chiu Laboratory, University of California, San Francisco | Chiu Laboratory, University of California, San Francisco | Xianding Deng, Scot Federman, Wei Gu, and Charles Y. Chiu |
| EPI_ISL_417333, EPI_ISL_417334, EPI_ISL_417335, EPI_ISL_417336, EPI_ISL_417337 | Institut des Agents Infectieux (IAI), Hospices Civils de Lyon | CNR Virus des Infections Respiratoires - France SUD | Antonin Bal, Gregory Destras, Gwendolyne Burfin, Solenne Brun, Carine Moustaud, Raphaelle Lamy, Alexandre Gaymard, Maude Bouscambert-Duchamp, Florence Morfin-Sherpa, Martine Valette, Laurence Josset, Bruno Lina |
| EPI_ISL_417338 | Centre Hospitalier de Macon | CNR Virus des Infections Respiratoires - France SUD | Antonin Bal, Gregory Destras, Gwendolyne Burfin, Solenne Brun, Carine Moustaud, Raphaelle Lamy, Alexandre Gaymard, Maude Bouscambert-Duchamp, Florence Morfin-Sherpa, Martine Valette, Laurence Josset, Bruno Lina |
| EPI_ISL_417339 | Institut des Agents Infectieux (IAI), Hospices Civils de Lyon | CNR Virus des Infections Respiratoires - France SUD | Antonin Bal, Gregory Destras, Gwendolyne Burfin, Solenne Brun, Carine Moustaud, Raphaelle Lamy, Alexandre Gaymard, Maude Bouscambert-Duchamp, Florence Morfin-Sherpa, Martine Valette, Laurence Josset, Bruno Lina |
| EPI_ISL_417340 | Centre Hospitalier de Bourg en Bresse | CNR Virus des Infections Respiratoires - France SUD | Antonin Bal, Gregory Destras, Gwendolyne Burfin, Solenne Brun, Carine Moustaud, Raphaelle Lamy, Alexandre Gaymard, Maude Bouscambert-Duchamp, Florence Morfin-Sherpa, Martine Valette, Laurence Josset, Bruno Lina |
| EPI_ISL_417341, EPI_ISL_417342, EPI_ISL_417343, EPI_ISL_417344, EPI_ISL_417345, EPI_ISL_417346, EPI_ISL_417347, EPI_ISL_417348, EPI_ISL_417349, EPI_ISL_417350, EPI_ISL_417351, EPI_ISL_417352, EPI_ISL_417353, EPI_ISL_417354, EPI_ISL_417355, EPI_ISL_417356, EPI_ISL_417357, EPI_ISL_417358, EPI_ISL_417359, EPI_ISL_417360, EPI_ISL_417361, EPI_ISL_417362, EPI_ISL_417363, EPI_ISL_417364, EPI_ISL_417365, EPI_ISL_417366, EPI_ISL_417367, EPI_ISL_417368, EPI_ISL_417369, EPI_ISL_417370, EPI_ISL_417371, EPI_ISL_417372, EPI_ISL_417373, EPI_ISL_417374, EPI_ISL_417375, EPI_ISL_417376, EPI_ISL_417377, EPI_ISL_417378, EPI_ISL_417379, EPI_ISL_417380, EPI_ISL_417381, EPI_ISL_417382 | | | |
| see above | UW Virology Lab | UW Virology Lab | Pavitra Roychoudhury, Hong Xie, Keith Jerome, Alexander Greninger |
| EPI_ISL_417383 | Centre for Infectious Diseases and Microbiology Public Health | NSW Health Pathology - Institute of Clinical Pathology and Medical Research; Westmead Hospital; University of Sydney | Rockett R, Eden J-S, Lam C, Gray K, Timms V, Gall M, Arnott A, Sadsad R, Carter I, Rahman H, Holmes EC, O'Sullivan MV, Sintchenko V, Chen SC, Maddocks S, Kok J and Dwyer DE for the 2019-nCoV Study Group |
| EPI_ISL_417384 | Centre for Infectious Diseases and Microbiology Public Health | NSW Health Pathology - Institute of Clinical Pathology and Medical Research; Westmead Hospital; University of Sydney | Eden J-S, Lam C, Gray K, Timms V, Gall M, Arnott A, Sadsad R, Carter I, Rahman H, Holmes EC, O'Sullivan MV, Sintchenko V, Chen SC, Maddocks S, Kok J, Dwyer DE and Rockett R for the 2019-nCoV Study Group |
| EPI_ISL_417385 | Centre for Infectious Diseases and Microbiology Public Health | NSW Health Pathology - Institute of Clinical Pathology and Medical Research; Westmead Hospital; University of Sydney | Lam C, Gray K, Timms V, Gall M, Arnott A, Sadsad R, Carter I, Rahman H, Holmes EC, O'Sullivan MV, Sintchenko V, Chen SC, Maddocks S, Kok J, Dwyer DE, Rockett R and Eden J-S for the 2019-nCoV Study Group |
| EPI_ISL_417386 | Centre for Infectious Diseases and Microbiology Public Health | NSW Health Pathology - Institute of Clinical Pathology and Medical Research; Westmead Hospital; University of Sydney | Gray K, Timms V, Gall M, Arnott A, Sadsad R, Carter I, Rahman H, Holmes EC, O'Sullivan MV, Sintchenko V, Chen SC, Maddocks S, Kok J, Dwyer DE, Rockett R, Eden J-S and Lam C for the 2019-nCoV Study Group |
| EPI_ISL_417387 | Centre for Infectious Diseases and Microbiology Public Health | NSW Health Pathology - Institute of Clinical Pathology and Medical Research; Westmead Hospital; University of Sydney | Timms V, Gall M, Arnott A, Sadsad R, Carter I, Rahman H, Holmes EC, O'Sullivan MV, Sintchenko V, Chen SC, Maddocks S, Kok J, Dwyer DE, Rockett R, Eden J-S, Lam C and Gray K for the 2019-nCoV Study Group |
| EPI_ISL_417388 | Centre for Infectious Diseases and Microbiology Public Health | NSW Health Pathology - Institute of Clinical Pathology and Medical Research; Westmead Hospital; University of Sydney | Gall M, Arnott A, Sadsad R, Carter I, Rahman H, Holmes EC, O'Sullivan MV, Sintchenko V, Chen SC, Maddocks S, Kok J, Dwyer DE, Rockett R, Eden J-S, Lam C, Gray K and Timms V for the 2019-nCoV Study Group |
| EPI_ISL_417389 | Centre for Infectious Diseases and Microbiology Public Health | NSW Health Pathology - Institute of Clinical Pathology and Medical Research; Westmead Hospital; University of Sydney | Arnott A, Sadsad R, Carter I, Rahman H, Holmes EC, O'Sullivan MV, Sintchenko V, Chen SC, Maddocks S, Kok J, Dwyer DE, Rockett R, Eden J-S, Lam C, Gray K, Timms V and Gall M for the 2019-nCoV Study Group |
| EPI_ISL_417390 | Centre for Infectious Diseases and Microbiology Public Health | NSW Health Pathology - Institute of Clinical Pathology and Medical Research; Westmead Hospital; University of Sydney | Sadsad R, Carter I, Rahman H, Holmes EC, O'Sullivan MV, Sintchenko V, Chen SC, Maddocks S, Kok J, Dwyer DE, Rockett R, Eden J-S, Lam C, Gray K, Timms V, Gall M and Arnott A for the 2019-nCoV Study Group |
| EPI_ISL_417391 | Centre for Infectious Diseases and Microbiology Public Health | NSW Health Pathology - Institute of Clinical Pathology and Medical Research; Westmead Hospital; University of Sydney | Carter I, Rahman H, Holmes EC, O'Sullivan MV, Sintchenko V, Chen SC, Maddocks S, Kok J, Dwyer DE, Rockett R, Eden J-S, Lam C, Gray K, Timms V, Gall M, Arnott A and Sadsad R for the 2019-nCoV Study Group |
| EPI_ISL_417392 | Centre for Infectious Diseases and Microbiology Public Health | NSW Health Pathology - Institute of Clinical Pathology and Medical Research; Westmead Hospital; University of Sydney | Rahman H, Holmes EC, O'Sullivan MV, Sintchenko V, Chen SC, Maddocks S, Kok J, Dwyer DE, Rockett R, Eden J-S, Lam C, Gray K, Timms V, Gall M, Arnott A, Sadsad R and Carter I for the 2019-nCoV Study Group |
| EPI_ISL_417393 | Centre for Infectious Diseases and Microbiology Public Health | NSW Health Pathology - Institute of Clinical Pathology and Medical Research; Westmead Hospital; University of Sydney | Holmes EC, O'Sullivan MV, Sintchenko V, Chen SC, Maddocks S, Kok J, Dwyer DE, Rockett R, Eden J-S, Lam C, Gray K, Timms V, Gall M, Arnott A, Sadsad R, Carter I and Rahman H for the 2019-nCoV Study Group |
| EPI_ISL_417394 | Centre for Infectious Diseases and Microbiology Public Health | NSW Health Pathology - Institute of Clinical Pathology and Medical Research; Westmead Hospital; University of Sydney | O'Sullivan MV, Sintchenko V, Chen SC, Maddocks S, Kok J, Dwyer DE, Rockett R, Eden J-S, Lam C, Gray K, Timms V, Gall M, Arnott A, Sadsad R, Carter I, Rahman H and Holmes EC for the 2019-nCoV Study Group |
| EPI_ISL_417395 | Centre for Infectious Diseases and Microbiology Public Health | NSW Health Pathology - Institute of Clinical Pathology and Medical Research; Westmead Hospital; University of Sydney | Sintchenko V, Chen SC, Maddocks S, Kok J, Dwyer DE, Rockett R, Eden J-S, Lam C, Gray K, Timms V, Gall M, Arnott A, Sadsad R, Carter I, Rahman H, Holmes EC and O'Sullivan MV for the 2019-nCoV Study Group |
| EPI_ISL_417396 | Centre for Infectious Diseases and Microbiology Public Health | NSW Health Pathology - Institute of Clinical Pathology and Medical Research; Westmead Hospital; University of Sydney | Chen SC, Maddocks S, Kok J, Dwyer DE, Rockett R, Eden J-S, Lam C, Gray K, Timms V, Gall M, Arnott A, Sadsad R, Carter I, Rahman H, Holmes EC, O'Sullivan MV and Sintchenko V for the 2019-nCoV Study Group |
| EPI_ISL_417397 | Centre for Infectious Diseases and Microbiology Public Health | NSW Health Pathology - Institute of Clinical Pathology and Medical Research; Westmead Hospital; University of Sydney | Maddocks S, Kok J, Dwyer DE, Rockett R, Eden J-S, Lam C, Gray K, Timms V, Gall M, Arnott A, Sadsad R, Carter I, Rahman H, Holmes EC, O'Sullivan MV, Sintchenko V and Chen SC for the 2019-nCoV Study Group |
| EPI_ISL_417398 | Centre for Infectious Diseases and Microbiology Public Health | NSW Health Pathology - Institute of Clinical Pathology and Medical Research; Westmead Hospital; University of Sydney | Kok J, Dwyer DE, Rockett R, Eden J-S, Lam C, Gray K, Timms V, Gall M, Arnott A, Sadsad R, Carter I, Rahman H, Holmes EC, O'Sullivan MV, Sintchenko V, Chen SC and Maddocks S for the 2019-nCoV Study Group |
| EPI_ISL_417399 | Centre for Infectious Diseases and Microbiology Public Health | NSW Health Pathology - Institute of Clinical Pathology and Medical Research; Westmead Hospital; University of Sydney | Dwyer DE, Rockett R, Eden J-S, Lam C, Gray K, Timms V, Gall M, Arnott A, Sadsad R, Carter I, Rahman H, Holmes EC, O'Sullivan MV, Sintchenko V, Chen SC, Maddocks S and Kok J for the 2019-nCoV Study Group |
| EPI_ISL_417400 | Centre for Infectious Diseases and Microbiology Public Health | NSW Health Pathology - Institute of Clinical Pathology and Medical Research; Westmead Hospital; University of Sydney | Rockett R, Eden J-S, Lam C, Gray K, Timms V, Gall M, Arnott A, Sadsad R, Carter I, Rahman H, Holmes EC, O'Sullivan MV, Sintchenko V, Chen SC, Maddocks S, Kok J and Dwyer DE for the 2019-nCoV Study Group |
| EPI_ISL_417401 | Centre for Infectious Diseases and Microbiology Public Health | NSW Health Pathology - Institute of Clinical Pathology and Medical Research; Westmead Hospital; University of Sydney | Eden J-S, Lam C, Gray K, Timms V, Gall M, Arnott A, Sadsad R, Carter I, Rahman H, Holmes EC, O'Sullivan MV, Sintchenko V, Chen SC, Maddocks S, Kok J, Dwyer DE and Rockett R for the 2019-nCoV Study Group |
| EPI_ISL_417402 | Centre for Infectious Diseases and Microbiology Public Health | NSW Health Pathology - Institute of Clinical Pathology and Medical Research; Westmead Hospital; University of Sydney | Lam C, Gray K, Timms V, Gall M, Arnott A, Sadsad R, Carter I, Rahman H, Holmes EC, O'Sullivan MV, Sintchenko V, Chen SC, Maddocks S, Kok J, Dwyer DE, Rockett R and Eden J-S for the 2019-nCoV Study Group |
| EPI_ISL_417403 | Centre for Infectious Diseases and Microbiology Public Health | NSW Health Pathology - Institute of Clinical Pathology and Medical Research; Westmead Hospital; University of Sydney | Gray K, Timms V, Gall M, Arnott A, Sadsad R, Carter I, Rahman H, Holmes EC, O'Sullivan MV, Sintchenko V, Chen SC, Maddocks S, Kok J, Dwyer DE, Rockett R, Eden J-S and Lam C for the 2019-nCoV Study Group |
| EPI_ISL_417404 | Centre for Infectious Diseases and Microbiology Public Health | NSW Health Pathology - Institute of Clinical Pathology and Medical Research; Westmead Hospital; University of Sydney | Timms V, Gall M, Arnott A, Sadsad R, Carter I, Rahman H, Holmes EC, O'Sullivan MV, Sintchenko V, Chen SC, Maddocks S, Kok J, Dwyer DE, Rockett R, Eden J-S, Lam C and Gray K for the 2019-nCoV Study Group |

| | | | |
|---|---|---|---|
| EPI_ISL_417405 | Centre for Infectious Diseases and Microbiology Public Health | NSW Health Pathology - Institute of Clinical Pathology and Medical Research; Westmead Hospital; University of Sydney | Gall M, Arnott A, Sadsad R, Carter I, Rahman H, Holmes EC, O'Sullivan MV, Sintchenko V, Chen SC, Maddocks S, Kok J, Dwyer DE, Rockett R, Eden J-S, Lam C, Gray K and Timms V for the 2019-nCoV Study Group |
| EPI_ISL_417406 | Centre for Infectious Diseases and Microbiology Public Health | NSW Health Pathology - Institute of Clinical Pathology and Medical Research; Westmead Hospital; University of Sydney | Arnott A, Sadsad R, Carter I, Rahman H, Holmes EC, O'Sullivan MV, Sintchenko V, Chen SC, Maddocks S, Kok J, Dwyer DE, Rockett R, Eden J-S, Lam C, Gray K, Timms V and Gall M for the 2019-nCoV Study Group |
| EPI_ISL_417407 | Centre for Infectious Diseases and Microbiology Public Health | NSW Health Pathology - Institute of Clinical Pathology and Medical Research; Westmead Hospital; University of Sydney | Sadsad R, Carter I, Rahman H, Holmes EC, O'Sullivan MV, Sintchenko V, Chen SC, Maddocks S, Kok J, Dwyer DE, Rockett R, Eden J-S, Lam C, Gray K, Timms V, Gall M and Arnott A for the 2019-nCoV Study Group |
| EPI_ISL_417408 | Centre for Infectious Diseases and Microbiology Public Health | NSW Health Pathology - Institute of Clinical Pathology and Medical Research; Westmead Hospital; University of Sydney | Carter I, Rahman H, Holmes EC, O'Sullivan MV, Sintchenko V, Chen SC, Maddocks S, Kok J, Dwyer DE, Rockett R, Eden J-S, Lam C, Gray K, Timms V, Gall M, Arnott A and Sadsad R for the 2019-nCoV Study Group |
| EPI_ISL_417409 | Centre for Infectious Diseases and Microbiology Public Health | NSW Health Pathology - Institute of Clinical Pathology and Medical Research; Westmead Hospital; University of Sydney | Rahman H, Holmes EC, O'Sullivan MV, Sintchenko V, Chen SC, Maddocks S, Kok J, Dwyer DE, Rockett R, Eden J-S, Lam C, Gray K, Timms V, Gall M, Arnott A, Sadsad R and Carter I for the 2019-nCoV Study Group |
| EPI_ISL_417410 | Centre for Infectious Diseases and Microbiology Public Health | NSW Health Pathology - Institute of Clinical Pathology and Medical Research; Westmead Hospital; University of Sydney | Holmes EC, O'Sullivan MV, Sintchenko V, Chen SC, Maddocks S, Kok J, Dwyer DE, Rockett R, Eden J-S, Lam C, Gray K, Timms V, Gall M, Arnott A, Sadsad R, Carter I and Rahman H for the 2019-nCoV Study Group |
| EPI_ISL_417411 | Centre for Infectious Diseases and Microbiology Public Health | NSW Health Pathology - Institute of Clinical Pathology and Medical Research; Westmead Hospital; University of Sydney | O'Sullivan MV, Sintchenko V, Chen SC, Maddocks S, Kok J, Dwyer DE, Rockett R, Eden J-S, Lam C, Gray K, Timms V, Gall M, Arnott A, Sadsad R, Carter I, Rahman H and Holmes EC for the 2019-nCoV Study Group |
| EPI_ISL_417412 | Centre for Infectious Diseases and Microbiology Public Health | NSW Health Pathology - Institute of Clinical Pathology and Medical Research; Westmead Hospital; University of Sydney | Sintchenko V, Chen SC, Maddocks S, Kok J, Dwyer DE, Rockett R, Eden J-S, Lam C, Gray K, Timms V, Gall M, Arnott A, Sadsad R, Carter I, Rahman H, Holmes EC and O'Sullivan MV for the 2019-nCoV Study Group |
| EPI_ISL_417413 | Ministry of Health Turkey | Ministry of Health Turkey | Fatma Bayrakdar,Ayşe Başak Altaş,Yasemin Coşgun,Gülay Korukluoğlu,Selçuk Kılıç |
| EPI_ISL_417418 | Laboratory of Molecular Virology International Center fro Genetic Engineering and Biotechnology (ICGEB) | ARGO Open Lab Platform for Genome sequencing | Licastro D, Rajasekharan S, Dal Monego S, Segat L, D'Agaro P, Marcello A |
| EPI_ISL_417419 | Laboratory of Molecular Virology International Center for Genetic Engineering and Biotechnology (ICGEB) | ARGO Open Lab Platform for Genome sequencing | Licastro D, Rajasekharan S, Dal Monego S, Segat L, D'Agaro P, Marcello A |
| EPI_ISL_417420 | Jiangxi province Center for Disease Control and Prevention | Jiangxi province Center for Disease Control and Prevention | Li jian Xiong |
| EPI_ISL_417421 | Laboratory of Molecular Virology International Center for Genetic Engineering and Biotechnology (ICGEB) | ARGO Open Lab Platform for Genome sequencing | Licastro D, Rajasekharan S, Dal Monego S, Segat L, D'Agaro P, Marcello A |
| EPI_ISL_417422 | KU Leuven, Clinical and Epidemiological Virology | KU Leuven, Clinical and Epidemiological Virology | Joan Marti-Carerras, Tony Wawina, Bert Vanmechelen, Piet Maes |
| EPI_ISL_417423 | Laboratory of Molecular Virology International Center for Genetic Engineering and Biotechnology (ICGEB) | ARGO Open Lab Platform for Genome sequencing | Licastro D, Rajasekharan, Dal Monego S, Segat L, D'Agaro P, Marcello A |
| EPI_ISL_417424, EPI_ISL_417425, EPI_ISL_417426, EPI_ISL_417427, EPI_ISL_417428, EPI_ISL_417429, EPI_ISL_417430 | KU Leuven, Clinical and Epidemiological Virology | KU Leuven, Clinical and Epidemiological Virology | Joan Marti-Carerras, Tony Wawina, Bert Vanmechelen, Piet Maes |
| EPI_ISL_417432 | UCD National Virus Reference Laboratory | Public Health Wales Microbiology Cardiff | Catherine Dempsey, Olive Murphy, Michael Prentice, Cillian F De Gascun, |
| EPI_ISL_417433, EPI_ISL_417434, EPI_ISL_417435, EPI_ISL_417436, EPI_ISL_417437, EPI_ISL_417438, EPI_ISL_417439, EPI_ISL_417440, EPI_ISL_417441, EPI_ISL_417442 | Viral Respiratory Lab, National Institute for Biomedical Research (INRB) | Pathogen Sequencing Lab, National Institute for Biomedical Research (INRB) | Placide Mbala-Kingebeni, Edith Nkwembe, Eddy Kinganda-Lusamaki, Amuri Aziza, Catherine Pratt, Matthias Pauthner, Josh Quick, Allison Black, James Hadfield, Trevor Bedford, Ian Goodfellow, Nick Loman, Kristian Andersen, Michael Wiley, Steve Ahuka-Mundeke, Jean-Jacques Muyembe Tamfum |
| EPI_ISL_417443 | State Key Laboratory for Emerging Infectious Diseases Department of Microbiology Li Ka Shing Faculty of Medicine The University of Hong Kong | State Key Laboratory for Emerging Infectious Diseases Department of Microbiology Li Ka Shing Faculty of Medicine The University of Hong Kong | Pui Wang, Siu-Ying Lau, Shaofeng Deng, Bobo Wing-Yee Mok, Wenjun Song, Kwok-Yung Yuen, Honglin Chen |
| EPI_ISL_417444 | Department of Healthcare Biotechnology, National University of Sciences and Technology (NUST) | Department of Healthcare Biotechnology, National University of Sciences and Technology (NUST) | Javed,A., Niazi,S.K., Ghani,E., Saqib,M., Janjua,H.A., Corman,V.M. and Zohaib,A. |
| EPI_ISL_417445, EPI_ISL_417446, EPI_ISL_417447 | Laboratory of Infectious Diseases, Department of Biomedical and Clinical Sciences L. Sacco, University of Milan | Laboratory of Infectious Diseases, Department of Biomedical and Clinical Sciences L. Sacco, University of Milan | Gianguglielmo Zehender, Alessia Lai, Annalisa Bergna, Luca Meroni, Agostino Riva, Claudia Balotta, Maciej Tarkowski, Arianna Gabrieli, Dario Bernacchia, Stefano Rusconi, Giuliano Rizzardini, Spinello Antinori, Massimo Galli |
| EPI_ISL_417448, EPI_ISL_417449, EPI_ISL_417450, EPI_ISL_417451, EPI_ISL_417452, EPI_ISL_417453, EPI_ISL_417454, EPI_ISL_417455, EPI_ISL_417456 | UW Virology Lab | UW Virology Lab | Pavitra Roychoudhury, Hong Xie, Keith Jerome, Alexander Greninger |
| EPI_ISL_417457, EPI_ISL_417458, EPI_ISL_417459, EPI_ISL_417460, EPI_ISL_417461, EPI_ISL_417462, EPI_ISL_417463, EPI_ISL_417464, EPI_ISL_417465, EPI_ISL_417466, EPI_ISL_417467, EPI_ISL_417468 | | | |
| see above | Center of Medical Microbiology, Virology, and Hospital Hygiene, University of Duesseldorf | Center of Medical Microbiology, Virology, and Hospital Hygiene, University of Duesseldorf | Ortwin Adams, Marcel Andree, Alexander Dilthey, Torsten Feldt, Sandra Hauka, Torsten Houwaart, Björn-Erik Jensen, Detlef Kindgen-Milles, Malte Kohns Vasconcelos, Klaus Pfeffer, Tina Senff, Daniel Strelow, Jörg Timm, Andreas Walker, Tobias Wienemann |
| EPI_ISL_417469, EPI_ISL_417470, EPI_ISL_417471, EPI_ISL_417472, EPI_ISL_417473, EPI_ISL_417474, EPI_ISL_417475, EPI_ISL_417476, EPI_ISL_417477, EPI_ISL_417478, EPI_ISL_417479, EPI_ISL_417480 | | | |
| see above | Minnesota Department of Health, Public Health Laboratory | Minnesota Department of Health, Public Health Laboratory | Matt Plumb, Jake Garfin and Xiong Wang |
| EPI_ISL_417481 | deCODE genetics | deCODE genetics | Daniel F Gudbjartsson, Agnar Helgason, Hakon Jonsson, Olafur T Magnusson, Pall Melsted, Gudmundur L Norddahl, Jona Saemundsdottir, Asgeir Sigurdsson, Patrick Sulem, Arna B Agustsdottir, Berglind Eiriksdottir, Run Fridriksdottir, Elisabet E Gardarsdottir, Gudmundur Georgsson, Olafia S Gretarsdottir, Kjartan R Gudmundsson, Thora R Gunnarsdottir, Arnaldur Gylfason, Hilma Holm, Brynjar O Jensson, Aslaug Jonasdottir, Kamilla S Josefsdottir, Thordur Kristjansson, Droplaug N Magnusdottir, Louise le Roux, Gudrun Sigmundsdottir, Gardar Sveinbjornsson, Kristin E Sveinsdottir, Maney Sveinsdottir, Emil A Thorarensen, Bjarni Thorbjornsson, Gisli Masson, Ingileif Jonsdottir, Alma Moller, Thorolfur Gudnason, Karl G Kristinsson, Unnur Thorsteinsdottir, Kari Stefansson |
| EPI_ISL_417482 | Institute of Microbiology, Universidad San Francisco de Quito | Institute of Microbiology, Universidad San Francisco de Quito | Sully Márquez, Belén Prado-Vivar, Juan José Guadalupe, Bernardo Gutiérrez, Manuel Jibaja, Milton Tobar, Verónica Barragán, Patricio Rojas-Silva, Gabriel Trueba, Michelle Grunauer, Paúl Cárdenas |
| EPI_ISL_417483 | Oslo University Hospital, Department of Medical Microbiology | Norwegian Institute of Public Health | Kathrine Stene-Johansen, Kamilla Heddeland Instefjord, Hilde Elshaug, Karoline Bragstad, Olav Hungnes |
| EPI_ISL_417484 | Oslo University Hospital, Department of Medical Microbiology | Norwegian Institute of Public Health, Department of Virology | Kathrine Stene-Johansen, Kamilla Heddeland Instefjord, Hilde Elshaug, Karoline Bragstad, Olav Hungnes |
| EPI_ISL_417485 | University Hospital of Northern Norway, Department for Microbiology and Infectious Disease Control | Norwegian Institute of Public Health, Department of Virology | Kathrine Stene-Johansen, Kamilla Heddeland Instefjord, Hilde Elshaug, Karoline Bragstad, Olav Hungnes |
| EPI_ISL_417486 | Hospital of Southern Norway - Kristiansand, Department of Medical Microbiology | Norwegian Institute of Public Health, Department of Virology | Kathrine Stene-Johansen, Kamilla Heddeland Instefjord, Hilde Elshaug, Karoline Bragstad, Olav Hungnes |
| EPI_ISL_417487 | Hospital of Southern Norway - Kristiansand, Department of Medical Microbiology | Norwegian Institute of Public Health, Department of Virology | Kathrine Stene-Johansen, Kamilla Heddeland Instefjord, Hilde Elshaug, Karoline Bragstad, Olav Hungnes |
| EPI_ISL_417488, EPI_ISL_417489, EPI_ISL_417490 | Oslo University Hospital, Department of Medical Microbiology | Norwegian Institute of Public Health, Department of Virology | Kathrine Stene-Johansen, Kamilla Heddeland Instefjord, Hilde Elshaug, Karoline Bragstad, Olav Hungnes |
| EPI_ISL_417491 | Virology Laboratory, Department of Biomedical Sciences and Public Health, University Politecnica delle Marche | Virology and Legal Medicine Laboratories, Department of Biomedical Sciences and Public Health, University Politecnica delle Marche | Bagnarelli,P., Caucci,S., Di Sante,L., Menzo,S., Alessandrini,F., Onofri,V., Turchi,C., Tagliabracci,A. |
| EPI_ISL_417492, EPI_ISL_417493, EPI_ISL_417494, EPI_ISL_417495, EPI_ISL_417496, EPI_ISL_417497, EPI_ISL_417498, EPI_ISL_417499, EPI_ISL_417500, EPI_ISL_417501, EPI_ISL_417502, EPI_ISL_417503 | | | |
| see above | Minnesota Department of Health, Public Health Laboratory | Minnesota Department of Health, Public Health Laboratory | Matt Plumb, Jake Garfin and Xiong Wang |
| EPI_ISL_417504, EPI_ISL_417505, EPI_ISL_417506, EPI_ISL_417507, EPI_ISL_417508, EPI_ISL_417509, EPI_ISL_417510, EPI_ISL_417512, EPI_ISL_417513, EPI_ISL_417514, EPI_ISL_417515, EPI_ISL_417516, EPI_ISL_417517 | | | |
| see above | University of Wisconsin-Madison AIDS Vaccine Research Laboratories | University of Wisconsin-Madison AIDS Vaccine Research Laboratories | Gage Moreno, Katarina Braun, et al. AIDS Vaccine Research Laboratories |
| EPI_ISL_417518, EPI_ISL_417519, EPI_ISL_417520, EPI_ISL_417521, EPI_ISL_417522, EPI_ISL_417523, EPI_ISL_417524, EPI_ISL_417525 | Laboratory Medicine | Department of Laboratory Medicine, Lin-Kou Chang Gung Memorial Hospital, Taoyuan, Taiwan | Kuo-Chien Tsao, Yu-Nong Gong, Shu-Li Yang, Yi-Chun Liu, Chung-Guei Huang, Po-Wei Huang, Mei-Jen Hsiao, Cheng-Ta Yang, Cheng-Hsun Chiu, Peng-Nien Huang, Kuo-Ming Lee, Guang-Wu Chen , Shin-Ru Shih |
| EPI_ISL_417526, EPI_ISL_417527, EPI_ISL_417528, EPI_ISL_417529, EPI_ISL_417530, EPI_ISL_417531, EPI_ISL_417532, EPI_ISL_417533, EPI_ISL_417534 | Laboratoire Nationale de Santé, Microbiology, Virology | Laboratoire Nationale de Santé, Microbiology, Epidemiology and Microbial Genomics | Anke Wienecke-Baldacchino, Ardashel Latsuzbaia, Jessica Tapp, Catherine Ragimbeau, Guillaume Fournier, Tamir Abdelrahman, Trung Nguyen Nguyen, Joel Mossong |

| | | | |
|---|---|---|---|
| EPI_ISL_417535, EPI_ISL_417536, EPI_ISL_417537, EPI_ISL_417538, EPI_ISL_417539, EPI_ISL_417540, EPI_ISL_417541, EPI_ISL_417542, EPI_ISL_417543, EPI_ISL_417544, EPI_ISL_417545, EPI_ISL_417546, EPI_ISL_417547, EPI_ISL_417548, EPI_ISL_417549 | | | |
| see above | deCODE genetics | deCODE genetics | Daniel F Gudbjartsson; Agnar Helgason; Hakon Jonsson; Olafur T Magnusson; Pall Melsted; Gudmundur L Norddahl; Jona Saemundsdottir; Asgeir Sigurdsson; Patrick Sulem; Arna B Agustsdottir; Berglind Eiriksdottir; Run Fridriksdottir; Elisabet E Gardarsdottir; Gudmundur Georgsson; Olafia S Gretarsdottir; Kjartan R Gudmundsson; Thora R Gunnarsdottir; Arnaldur Gylfason; Hilma Holm; Brynjar O Jensson; Aslaug Jonasdottir; Kamilla S Josefsdottir; Thordur Kristjansson; Droplaug N Magnusdottir; Louise le Roux; Gudrun Sigmundsdottir; Gardar Sveinbjornsson; Kristin E Sveinsdottir; Maney Sveinsdottir; Emil A Thorarensen; Bjarni Thorbjornsson; Gisli Masson; Ingileif Jonsdottir; Alma Moller; Thorolfur Gudnason; Karl G Kristinsson; Unnur Thorsteinsdottir; Kari Stefansson |
| EPI_ISL_417550, EPI_ISL_417551 | The National University Hospital of Iceland | deCODE genetics | Daniel F Gudbjartsson; Agnar Helgason; Hakon Jonsson; Olafur T Magnusson; Pall Melsted; Gudmundur L Norddahl; Jona Saemundsdottir; Asgeir Sigurdsson; Patrick Sulem; Arna B Agustsdottir; Berglind Eiriksdottir; Run Fridriksdottir; Elisabet E Gardarsdottir; Gudmundur Georgsson; Olafia S Gretarsdottir; Kjartan R Gudmundsson; Thora R Gunnarsdottir; Arnaldur Gylfason; Hilma Holm; Brynjar O Jensson; Aslaug Jonasdottir; Kamilla S Josefsdottir; Thordur Kristjansson; Droplaug N Magnusdottir; Louise le Roux; Gudrun Sigmundsdottir; Gardar Sveinbjornsson; Kristin E Sveinsdottir; Maney Sveinsdottir; Emil A Thorarensen; Bjarni Thorbjornsson; Gisli Masson; Ingileif Jonsdottir; Alma Moller; Thorolfur Gudnason; Karl G Kristinsson; Unnur Thorsteinsdottir; Kari Stefansson |
| EPI_ISL_417552 | deCODE genetics | deCODE genetics | Daniel F Gudbjartsson; Agnar Helgason; Hakon Jonsson; Olafur T Magnusson; Pall Melsted; Gudmundur L Norddahl; Jona Saemundsdottir; Asgeir Sigurdsson; Patrick Sulem; Arna B Agustsdottir; Berglind Eiriksdottir; Run Fridriksdottir; Elisabet E Gardarsdottir; Gudmundur Georgsson; Olafia S Gretarsdottir; Kjartan R Gudmundsson; Thora R Gunnarsdottir; Arnaldur Gylfason; Hilma Holm; Brynjar O Jensson; Aslaug Jonasdottir; Kamilla S Josefsdottir; Thordur Kristjansson; Droplaug N Magnusdottir; Louise le Roux; Gudrun Sigmundsdottir; Gardar Sveinbjornsson; Kristin E Sveinsdottir; Maney Sveinsdottir; Emil A Thorarensen; Bjarni Thorbjornsson; Gisli Masson; Ingileif Jonsdottir; Alma Moller; Thorolfur Gudnason; Karl G Kristinsson; Unnur Thorsteinsdottir; Kari Stefansson |
| EPI_ISL_417553, EPI_ISL_417554, EPI_ISL_417555, EPI_ISL_417556, EPI_ISL_417557, EPI_ISL_417558, EPI_ISL_417559, EPI_ISL_417560, EPI_ISL_417561, EPI_ISL_417562, EPI_ISL_417563, EPI_ISL_417564, EPI_ISL_417565, EPI_ISL_417566, EPI_ISL_417567, EPI_ISL_417568, EPI_ISL_417569, EPI_ISL_417570, EPI_ISL_417571, EPI_ISL_417572, EPI_ISL_417573, EPI_ISL_417574, EPI_ISL_417575, EPI_ISL_417576, EPI_ISL_417577, EPI_ISL_417578, EPI_ISL_417579, EPI_ISL_417580, EPI_ISL_417581, EPI_ISL_417582, EPI_ISL_417583, EPI_ISL_417584, EPI_ISL_417585, EPI_ISL_417586, EPI_ISL_417587, EPI_ISL_417588, EPI_ISL_417589 | | | |
| see above | The National University Hospital of Iceland | deCODE genetics | Daniel F Gudbjartsson; Agnar Helgason; Hakon Jonsson; Olafur T Magnusson; Pall Melsted; Gudmundur L Norddahl; Jona Saemundsdottir; Asgeir Sigurdsson; Patrick Sulem; Arna B Agustsdottir; Berglind Eiriksdottir; Run Fridriksdottir; Elisabet E Gardarsdottir; Gudmundur Georgsson; Olafia S Gretarsdottir; Kjartan R Gudmundsson; Thora R Gunnarsdottir; Arnaldur Gylfason; Hilma Holm; Brynjar O Jensson; Aslaug Jonasdottir; Kamilla S Josefsdottir; Thordur Kristjansson; Droplaug N Magnusdottir; Louise le Roux; Gudrun Sigmundsdottir; Gardar Sveinbjornsson; Kristin E Sveinsdottir; Maney Sveinsdottir; Emil A Thorarensen; Bjarni Thorbjornsson; Gisli Masson; Ingileif Jonsdottir; Alma Moller; Thorolfur Gudnason; Karl G Kristinsson; Unnur Thorsteinsdottir; Kari Stefansson |
| EPI_ISL_417590 | deCODE genetics | deCODE genetics | Daniel F Gudbjartsson; Agnar Helgason; Hakon Jonsson; Olafur T Magnusson; Pall Melsted; Gudmundur L Norddahl; Jona Saemundsdottir; Asgeir Sigurdsson; Patrick Sulem; Arna B Agustsdottir; Berglind Eiriksdottir; Run Fridriksdottir; Elisabet E Gardarsdottir; Gudmundur Georgsson; Olafia S Gretarsdottir; Kjartan R Gudmundsson; Thora R Gunnarsdottir; Arnaldur Gylfason; Hilma Holm; Brynjar O Jensson; Aslaug Jonasdottir; Kamilla S Josefsdottir; Thordur Kristjansson; Droplaug N Magnusdottir; Louise le Roux; Gudrun Sigmundsdottir; Gardar Sveinbjornsson; Kristin E Sveinsdottir; Maney Sveinsdottir; Emil A Thorarensen; Bjarni Thorbjornsson; Gisli Masson; Ingileif Jonsdottir; Alma Moller; Thorolfur Gudnason; Karl G Kristinsson; Unnur Thorsteinsdottir; Kari Stefansson |
| EPI_ISL_417591, EPI_ISL_417592, EPI_ISL_417593, EPI_ISL_417594, EPI_ISL_417595, EPI_ISL_417596, EPI_ISL_417597, EPI_ISL_417598, EPI_ISL_417599, EPI_ISL_417600, EPI_ISL_417601, EPI_ISL_417602, EPI_ISL_417603, EPI_ISL_417604, EPI_ISL_417605, EPI_ISL_417606, EPI_ISL_417607, EPI_ISL_417608, EPI_ISL_417609, EPI_ISL_417610, EPI_ISL_417611, EPI_ISL_417612, EPI_ISL_417613, EPI_ISL_417614, EPI_ISL_417615, EPI_ISL_417616, EPI_ISL_417617 | | | |
| see above | The National University Hospital of Iceland | deCODE genetics | Daniel F Gudbjartsson; Agnar Helgason; Hakon Jonsson; Olafur T Magnusson; Pall Melsted; Gudmundur L Norddahl; Jona Saemundsdottir; Asgeir Sigurdsson; Patrick Sulem; Arna B Agustsdottir; Berglind Eiriksdottir; Run Fridriksdottir; Elisabet E Gardarsdottir; Gudmundur Georgsson; Olafia S Gretarsdottir; Kjartan R Gudmundsson; Thora R Gunnarsdottir; Arnaldur Gylfason; Hilma Holm; Brynjar O Jensson; Aslaug Jonasdottir; Kamilla S Josefsdottir; Thordur Kristjansson; Droplaug N Magnusdottir; Louise le Roux; Gudrun Sigmundsdottir; Gardar Sveinbjornsson; Kristin E Sveinsdottir; Maney Sveinsdottir; Emil A Thorarensen; Bjarni Thorbjornsson; Gisli Masson; Ingileif Jonsdottir; Alma Moller; Thorolfur Gudnason; Karl G Kristinsson; Unnur Thorsteinsdottir; Kari Stefansson |
| EPI_ISL_417618 | deCODE genetics | deCODE genetics | Daniel F Gudbjartsson; Agnar Helgason; Hakon Jonsson; Olafur T Magnusson; Pall Melsted; Gudmundur L Norddahl; Jona Saemundsdottir; Asgeir Sigurdsson; Patrick Sulem; Arna B Agustsdottir; Berglind Eiriksdottir; Run Fridriksdottir; Elisabet E Gardarsdottir; Gudmundur Georgsson; Olafia S Gretarsdottir; Kjartan R Gudmundsson; Thora R Gunnarsdottir; Arnaldur Gylfason; Hilma Holm; Brynjar O Jensson; Aslaug Jonasdottir; Kamilla S Josefsdottir; Thordur Kristjansson; Droplaug N Magnusdottir; Louise le Roux; Gudrun Sigmundsdottir; Gardar Sveinbjornsson; Kristin E Sveinsdottir; Maney Sveinsdottir; Emil A Thorarensen; Bjarni Thorbjornsson; Gisli Masson; Ingileif Jonsdottir; Alma Moller; Thorolfur Gudnason; Karl G Kristinsson; Unnur Thorsteinsdottir; Kari Stefansson |
| EPI_ISL_417619, EPI_ISL_417620, EPI_ISL_417621, EPI_ISL_417622, EPI_ISL_417623, EPI_ISL_417624, EPI_ISL_417625, EPI_ISL_417626, EPI_ISL_417627, EPI_ISL_417628, EPI_ISL_417629, EPI_ISL_417630, EPI_ISL_417631, EPI_ISL_417632, EPI_ISL_417633, EPI_ISL_417634, EPI_ISL_417635, EPI_ISL_417636, EPI_ISL_417637, EPI_ISL_417638, EPI_ISL_417639, EPI_ISL_417640, EPI_ISL_417641, EPI_ISL_417642, EPI_ISL_417643, EPI_ISL_417644, EPI_ISL_417645, EPI_ISL_417646, EPI_ISL_417647, EPI_ISL_417648, EPI_ISL_417649, EPI_ISL_417650, EPI_ISL_417651, EPI_ISL_417652, EPI_ISL_417653, EPI_ISL_417654 | | | |
| see above | The National University Hospital of Iceland | deCODE genetics | Daniel F Gudbjartsson; Agnar Helgason; Hakon Jonsson; Olafur T Magnusson; Pall Melsted; Gudmundur L Norddahl; Jona Saemundsdottir; Asgeir Sigurdsson; Patrick Sulem; Arna B Agustsdottir; Berglind Eiriksdottir; Run Fridriksdottir; Elisabet E Gardarsdottir; Gudmundur Georgsson; Olafia S Gretarsdottir; Kjartan R Gudmundsson; Thora R Gunnarsdottir; Arnaldur Gylfason; Hilma Holm; Brynjar O Jensson; Aslaug Jonasdottir; Kamilla S Josefsdottir; Thordur Kristjansson; Droplaug N Magnusdottir; Louise le Roux; Gudrun Sigmundsdottir; Gardar Sveinbjornsson; Kristin E Sveinsdottir; Maney Sveinsdottir; Emil A Thorarensen; Bjarni Thorbjornsson; Gisli Masson; Ingileif Jonsdottir; Alma Moller; Thorolfur Gudnason; Karl G Kristinsson; Unnur Thorsteinsdottir; Kari Stefansson |
| EPI_ISL_417655, EPI_ISL_417656, EPI_ISL_417657, EPI_ISL_417658, EPI_ISL_417659, EPI_ISL_417660, EPI_ISL_417661, EPI_ISL_417662, EPI_ISL_417663, EPI_ISL_417664, EPI_ISL_417665, EPI_ISL_417666, EPI_ISL_417667, EPI_ISL_417668, EPI_ISL_417669, EPI_ISL_417670, EPI_ISL_417671, EPI_ISL_417672 | | | |
| see above | deCODE genetics | deCODE genetics | Daniel F Gudbjartsson; Agnar Helgason; Hakon Jonsson; Olafur T Magnusson; Pall Melsted; Gudmundur L Norddahl; Jona Saemundsdottir; Asgeir Sigurdsson; Patrick Sulem; Arna B Agustsdottir; Berglind Eiriksdottir; Run Fridriksdottir; Elisabet E Gardarsdottir; Gudmundur Georgsson; Olafia S Gretarsdottir; Kjartan R Gudmundsson; Thora R Gunnarsdottir; Arnaldur Gylfason; Hilma Holm; Brynjar O Jensson; Aslaug Jonasdottir; Kamilla S Josefsdottir; Thordur Kristjansson; Droplaug N Magnusdottir; Louise le Roux; Gudrun Sigmundsdottir; Gardar Sveinbjornsson; Kristin E Sveinsdottir; Maney Sveinsdottir; Emil A Thorarensen; Bjarni Thorbjornsson; Gisli Masson; Ingileif Jonsdottir; Alma Moller; Thorolfur Gudnason; Karl G Kristinsson; Unnur Thorsteinsdottir; Kari Stefansson |
| EPI_ISL_417673, EPI_ISL_417674, EPI_ISL_417675 | The National University Hospital of Iceland | deCODE genetics | Daniel F Gudbjartsson; Agnar Helgason; Hakon Jonsson; Olafur T Magnusson; Pall Melsted; Gudmundur L Norddahl; Jona Saemundsdottir; Asgeir Sigurdsson; Patrick Sulem; Arna B Agustsdottir; Berglind Eiriksdottir; Run Fridriksdottir; Elisabet E Gardarsdottir; Gudmundur Georgsson; Olafia S Gretarsdottir; Kjartan R Gudmundsson; Thora R Gunnarsdottir; Arnaldur Gylfason; Hilma Holm; Brynjar O Jensson; Aslaug Jonasdottir; Kamilla S Josefsdottir; Thordur Kristjansson; Droplaug N Magnusdottir; Louise le Roux; Gudrun Sigmundsdottir; Gardar Sveinbjornsson; Kristin E Sveinsdottir; Maney Sveinsdottir; Emil A Thorarensen; Bjarni Thorbjornsson; Gisli Masson; Ingileif Jonsdottir; Alma Moller; Thorolfur Gudnason; Karl G Kristinsson; Unnur Thorsteinsdottir; Kari Stefansson |
| EPI_ISL_417676 | deCODE genetics | deCODE genetics | Daniel F Gudbjartsson; Agnar Helgason; Hakon Jonsson; Olafur T Magnusson; Pall Melsted; Gudmundur L Norddahl; Jona Saemundsdottir; Asgeir Sigurdsson; Patrick Sulem; Arna B Agustsdottir; Berglind Eiriksdottir; Run Fridriksdottir; Elisabet E Gardarsdottir; Gudmundur Georgsson; Olafia S Gretarsdottir; Kjartan R Gudmundsson; Thora R Gunnarsdottir; Arnaldur Gylfason; Hilma Holm; Brynjar O Jensson; Aslaug Jonasdottir; Kamilla S Josefsdottir; Thordur Kristjansson; Droplaug N Magnusdottir; Louise le Roux; Gudrun Sigmundsdottir; Gardar Sveinbjornsson; Kristin E Sveinsdottir; Maney Sveinsdottir; Emil A Thorarensen; Bjarni Thorbjornsson; Gisli Masson; Ingileif Jonsdottir; Alma Moller; Thorolfur Gudnason; Karl G Kristinsson; Unnur Thorsteinsdottir; Kari Stefansson |
| EPI_ISL_417677, EPI_ISL_417678, EPI_ISL_417679, EPI_ISL_417680, EPI_ISL_417681, EPI_ISL_417682, EPI_ISL_417683, EPI_ISL_417684, EPI_ISL_417685, EPI_ISL_417686, EPI_ISL_417687, EPI_ISL_417688, EPI_ISL_417689, EPI_ISL_417690, EPI_ISL_417691, EPI_ISL_417692, EPI_ISL_417693, EPI_ISL_417694, EPI_ISL_417695, EPI_ISL_417696, EPI_ISL_417697, EPI_ISL_417698, EPI_ISL_417699, EPI_ISL_417700, EPI_ISL_417701, EPI_ISL_417702, EPI_ISL_417703, EPI_ISL_417704, EPI_ISL_417705, EPI_ISL_417706, EPI_ISL_417707, EPI_ISL_417708, EPI_ISL_417709, EPI_ISL_417710, EPI_ISL_417711, EPI_ISL_417712, EPI_ISL_417713, EPI_ISL_417714, EPI_ISL_417715, EPI_ISL_417716, EPI_ISL_417717, EPI_ISL_417718, EPI_ISL_417719, EPI_ISL_417720, EPI_ISL_417721, EPI_ISL_417722, EPI_ISL_417723, EPI_ISL_417724, EPI_ISL_417725, EPI_ISL_417726, EPI_ISL_417727, EPI_ISL_417728, EPI_ISL_417729, EPI_ISL_417730, EPI_ISL_417731, EPI_ISL_417732, EPI_ISL_417733, EPI_ISL_417734, EPI_ISL_417735, EPI_ISL_417736, EPI_ISL_417737, EPI_ISL_417738, EPI_ISL_417739, EPI_ISL_417740, EPI_ISL_417741, EPI_ISL_417742, EPI_ISL_417743, EPI_ISL_417744, EPI_ISL_417745, EPI_ISL_417746, EPI_ISL_417747, EPI_ISL_417748, EPI_ISL_417749, EPI_ISL_417750, EPI_ISL_417751, EPI_ISL_417752, EPI_ISL_417753, EPI_ISL_417754, EPI_ISL_417755, EPI_ISL_417756, EPI_ISL_417757, EPI_ISL_417758, EPI_ISL_417759, EPI_ISL_417760, EPI_ISL_417761, EPI_ISL_417762, EPI_ISL_417763, EPI_ISL_417764, EPI_ISL_417765, EPI_ISL_417766, EPI_ISL_417767, EPI_ISL_417768, EPI_ISL_417769, EPI_ISL_417770, EPI_ISL_417771, EPI_ISL_417772, EPI_ISL_417773, EPI_ISL_417774, EPI_ISL_417775, EPI_ISL_417776, EPI_ISL_417777, EPI_ISL_417778, EPI_ISL_417779, EPI_ISL_417780, EPI_ISL_417781, EPI_ISL_417782, EPI_ISL_417783, EPI_ISL_417784, EPI_ISL_417785, EPI_ISL_417786, EPI_ISL_417787, EPI_ISL_417788, EPI_ISL_417789, EPI_ISL_417790, EPI_ISL_417791, EPI_ISL_417792, EPI_ISL_417793, EPI_ISL_417794, EPI_ISL_417795, EPI_ISL_417796, EPI_ISL_417797, EPI_ISL_417798, EPI_ISL_417799, EPI_ISL_417800, EPI_ISL_417801, EPI_ISL_417802, EPI_ISL_417803, EPI_ISL_417804, EPI_ISL_417805, EPI_ISL_417806, EPI_ISL_417807, EPI_ISL_417808, EPI_ISL_417809, EPI_ISL_417810, EPI_ISL_417811, EPI_ISL_417812, EPI_ISL_417813, EPI_ISL_417814, EPI_ISL_417815, EPI_ISL_417816, EPI_ISL_417817, EPI_ISL_417818, EPI_ISL_417819, EPI_ISL_417820, EPI_ISL_417821, EPI_ISL_417822, EPI_ISL_417823, EPI_ISL_417824, EPI_ISL_417825, EPI_ISL_417826, EPI_ISL_417827, EPI_ISL_417828, EPI_ISL_417829, EPI_ISL_417830, EPI_ISL_417831, EPI_ISL_417832, EPI_ISL_417833, EPI_ISL_417834, EPI_ISL_417835, EPI_ISL_417836, EPI_ISL_417837, EPI_ISL_417838, EPI_ISL_417839, EPI_ISL_417840, EPI_ISL_417841, EPI_ISL_417842, EPI_ISL_417843, EPI_ISL_417844, EPI_ISL_417845, EPI_ISL_417846, EPI_ISL_417847, EPI_ISL_417848, EPI_ISL_417849, EPI_ISL_417850, EPI_ISL_417851, EPI_ISL_417852, EPI_ISL_417853, EPI_ISL_417854, EPI_ISL_417855, EPI_ISL_417856, EPI_ISL_417857, EPI_ISL_417858, EPI_ISL_417859, EPI_ISL_417860, EPI_ISL_417861, EPI_ISL_417862, EPI_ISL_417863, EPI_ISL_417864, EPI_ISL_417865, EPI_ISL_417866, EPI_ISL_417867, EPI_ISL_417868, EPI_ISL_417869, EPI_ISL_417870, EPI_ISL_417871, EPI_ISL_417872, EPI_ISL_417873, EPI_ISL_417874, EPI_ISL_417875, EPI_ISL_417876 | | | |
| see above | The National University Hospital of Iceland | deCODE genetics | Daniel F Gudbjartsson; Agnar Helgason; Hakon Jonsson; Olafur T Magnusson; Pall Melsted; Gudmundur L Norddahl; Jona Saemundsdottir; Asgeir Sigurdsson; Patrick Sulem; Arna B Agustsdottir; Berglind Eiriksdottir; Run Fridriksdottir; Elisabet E Gardarsdottir; Gudmundur Georgsson; Olafia S Gretarsdottir; Kjartan R Gudmundsson; Thora R Gunnarsdottir; Arnaldur Gylfason; Hilma Holm; Brynjar O Jensson; Aslaug Jonasdottir; Kamilla S Josefsdottir; Thordur Kristjansson; Droplaug N Magnusdottir; Louise le Roux; Gudrun Sigmundsdottir; Gardar Sveinbjornsson; Kristin E Sveinsdottir; Maney Sveinsdottir; Emil A Thorarensen; Bjarni Thorbjornsson; Gisli Masson; Ingileif Jonsdottir; Alma Moller; Thorolfur Gudnason; Karl G Kristinsson; Unnur Thorsteinsdottir; Kari Stefansson |
| EPI_ISL_417877 | Institute of Virology, Biomedical Research Center of the Slovak Academy of Sciences, Bratislava; Public Health Authority of the Slovak Republic, Bratislava | Institute of Virology, Biomedical Research Center of the Slovak Academy of Sciences, Bratislava; Comenius University Science Park, Bratislava | Monika Slávikova, Martina Ličková, Sabina Fumačová Havlíková, Juraj Koči, Juraj Kopáček, Elena Tichá, Edita Staroňová, Jaroslav Budiš, Werner Krampl, Miroslav Böhmer, Diana Rusňáková, Tomáš Szemeš, Boris Klempa |
| EPI_ISL_417878 | Institute of Virology, Biomedical Research Center of the Slovak Academy of Sciences, Bratislava; Public Health Authority of the Slovak Republic, Bratislava | Institute of Virology, Biomedical Research Center of the Slovak Academy of Sciences, Bratislava; Comenius University Science Park, Bratislava | Monika Slávikova, Martina Ličková, Sabina Fumačová Havlíková, Juraj Koči, Juraj Kopáček, Elena Tichá, Edita Staroňová, Jaroslav Budiš, Werner Krampl, Miroslav Böhmer, Diana Rusňáková, Tomáš Szemeš, Boris Klempa |
| EPI_ISL_417879 | Institute of Virology, Biomedical Research Center of the Slovak Academy of Sciences, Bratislava; Public Health Authority of the Slovak Republic, Bratislava | Institute of Virology, Biomedical Research Center of the Slovak Academy of Sciences, Bratislava; Comenius University Science Park, Bratislava | Monika Slávikova, Martina Ličková, Sabina Fumačová Havlíková, Juraj Koči, Juraj Kopáček, Elena Tichá, Edita Staroňová, Jaroslav Budiš, Werner Krampl, Miroslav Böhmer, Diana Rusňáková, Tomáš Szemeš, Boris Klempa |
| EPI_ISL_417880 | Institute of Virology, Biomedical Research Center of the Slovak Academy of Sciences, Bratislava; Public Health Authority of the Slovak Republic, Bratislava | Institute of Virology, Biomedical Research Center of the Slovak Academy of Sciences, Bratislava; Comenius University Science Park, Bratislava | Monika Slávikova, Martina Ličková, Sabina Fumačová Havlíková, Juraj Koči, Juraj Kopáček, Elena Tichá, Edita Staroňová, Jaroslav Budiš, Werner Krampl, Miroslav Böhmer, Diana Rusňáková, Tomáš Szemeš, Boris Klempa |
| EPI_ISL_417917 | Department of Medical Microbiology, University Malaya Medical Centre | Department of Medical Microbiology | Yoong Min CHONG, Sasheela PONNAMPALAVANAR, Sharifah Faridah SYED OMAR, Adeeba KAMARULZAMAN,Vijayan MUNUSAMY, Chee Kuan WONG, Cindy Shuan Ju TEH, I-Ching SAM, Yoke Fun Chan, University Malaya Medical Centre COVID Team |
| EPI_ISL_417918 | Department of Medical Microbiology, University Malaya Medical Centre | Department of Medical Microbiology, Faculty of Medicine, University of Malaya | Yoong Min CHONG, Sasheela PONNAMPALAVANAR, Sharifah Faridah SYED OMAR, Adeeba KAMARULZAMAN,Vijayan MUNUSAMY, Chee Kuan WONG, Cindy Shuan Ju TEH, I-Ching SAM, Yoke Fun Chan, University Malaya Medical Centre COVID Team |
| EPI_ISL_417919 | Department of Medical Microbiology, University Malaya Medical Centre | Department of Medical Microbiology, Faculty of Medicine, University of Malaya | Yoong Min CHONG, Sasheela PONNAMPALAVANAR, Sharifah Faridah SYED OMAR, Adeeba KAMARULZAMAN,Vijayan MUNUSAMY, Chee Kuan WONG, Fadhil Hadi JAMALUDDIN, Han Ming GAN, Cindy Shuan Ju TEH, I-Ching SAM, Yoke Fun CHAN, University Malaya Medical Centre COVID Team |
| EPI_ISL_417920 | Department of Medical Microbiology, University Malaya Medical Centre | Department of Medical Microbiology, Faculty of Medicine, University of Malaya | Yoong Min CHONG, Sasheela PONNAMPALAVANAR, Sharifah Faridah SYED OMAR, Adeeba KAMARULZAMAN,Vijayan MUNUSAMY, Chee Kuan WONG, Fadhil Hadi JAMALUDDIN, Cindy Shuan Ju TEH, I-Ching SAM, Yoke Fun Chan, University Malaya Medical Centre COVID Team |
| EPI_ISL_417921 | INMI Lazzaro Spallanzani IRCCS | Laboratory of Virology, INMI Lazzaro Spallanzani IRCCS | Martina Rueca, Barbara Bartolini, Francesco Messina, Cesare E. M. Gruber, Emanuela Giombini, Maria R. Capobianchi, Fabrizio Carletti, Francesca Colavita, Concetta Castilletti, Eleonora Lalle, Daniele Lapa, Giuseppe Ippolito. |
| EPI_ISL_417922 | INMI Lazzaro Spallanzani IRCCS | Laboratory of Virology, INMI Lazzaro Spallanzani IRCCS | Cesare E. M. Gruber, Martina Rueca, Barbara Bartolini, Francesco Messina, Emanuela Giombini, Maria R. Capobianchi, Fabrizio Carletti, Francesca Colavita, Concetta Castilletti, Eleonora Lalle, Daniele Lapa, Giuseppe Ippolito. |
| EPI_ISL_417923 | INMI Lazzaro Spallanzani IRCCS | Laboratory of Virology, INMI Lazzaro Spallanzani IRCCS | Francesco Messina, Barbara Bartolini, Martina Rueca, Cesare E. M. Gruber, Emanuela Giombini, Maria R. Capobianchi, Fabrizio Carletti, Francesca Colavita, Concetta Castilletti, Eleonora Lalle, Daniele Lapa, Giuseppe Ippolito. |
| EPI_ISL_417924 | Secretaría de Salud Medellín | Instituto Nacional de Salud, Universidad Cooperativa de Colombia, Instituto Alexander von Humboldt, Imperial College-London, London School of Hygiene & Tropical Medicine | Marcela Mercado-Reyes, Katherine Laiton-Donato, Diego A. Álvarez-Díaz, Carlos Franco-Muñoz, Jose A. Usme-Ciro, Gloria Puerto, Nicolás D. Franco-Sierra, Mailyn A. Gonzalez, Zulma M. Cucunubá, Christian Julian Villabona-Arenas, Liz Villabona-Arenas, Sussy Echeverria-Londoño, Astrid C. Flórez, Sergio Gomez Rangel, Luz Dary Rodriguez, Juliana Barbosa, Erika Ospitia, Diana Marcela Walteros-Acero, Martha Lucia Ospina Martinez |
| EPI_ISL_417925 | Laboratório Simili | Bioinformatics Laboratory / LNCC | Filipe Romero, Ana Paula Guimarães, Mariane Talon, Luiz Gonzaga Paula de Almeida, Ronaldo da Silva, Francisco Junior, Diana Mariani, Lídia Boullosa, Alexandra Gerber, Jaqueline Goes de Jesus, Ingra Morales Claro, Ester Cerdeira Sabino, Nuno Rodrigues Faria, Terezinha Marta Pereira, Pinto Castiñeiras, Isabela de Carvalho Leitão, Rafael de Mello Galliez, Cássia Cristina Alves Gonçalves, Érica Ramos dos Santos Nascimento, Richard Araújo Maia, Mauro Teixeira,Cristiano Xavier Lima, Orlando Ferreira Jr., Rodrigo Brindeiro, Luciana Jesus Costa e André Felipe Santos, Laboratorio Hermes Pardini, Laboratorio Simile, Amilcar Tanuri, Renato Santana Aguiar e Ana Tereza Vasconcelos |
| EPI_ISL_417926 | Laboratório Simili | Bioinformatics Laboratory / LNCC | Filipe Romero, Ana Paula Guimarães, Mariane Talon, Luiz Gonzaga Paula de Almeida, Ronaldo da Silva Francisco Junior, Diana Mariani, Lídia Boullosa, Alexandra Gerber, Jaqueline Goes de Jesus, Ingra Morales Claro, Ester Cerdeira Sabino, Nuno Rodrigues Faria, Terezinha Marta Pereira, Pinto Castiñeiras, Isabela de Carvalho Leitão, Rafael de Mello Galliez, Cássia Cristina Alves Gonçalves, Érica Ramos dos Santos Nascimento, Richard Araújo Maia, Mauro Teixeira,Cristiano Xavier Lima, |

| | | | |
|---|---|---|---|
| | | | Orlando Ferreira Jr., Rodrigo Brindeiro, Luciana Jesus Costa e André Felipe Santos, Laboratorio Hermes Pardini, Laboratorio Simile, Amilcar Tanuri, Renato Santana Aguiar e Ana Tereza Vasconcelos |
| EPI_ISL_417928, EPI_ISL_417929, EPI_ISL_417930 | Laboratório Hermes Pardini | Bioinformatics Laboratory - LNCC | Filipe Romero, Ana Paula Guimarães, Mariane Talon, Luiz Gonzaga Paula de Almeida, Ronaldo da Silva Francisco Junior, Diana Mariani, Lídia Boullosa, Alexandra Gerber, Jaqueline Goes de Jesus, Ingra Morales Claro, Ester Cerdeira Sabino, Nuno Rodrigues Faria, Terezinha Marta Pereira, Pinto Castiñeiras, Isabela de Carvalho Leitão, Rafael de Mello Galliez, Cássia Cristina Alves Gonçalves, Érica Ramos dos Santos Nascimento, Richard Araújo Maia, Mauro Teixeira,Cristiano Xavier Lima, Orlando Ferreira Jr., Rodrigo Brindeiro, Luciana Jesus Costa e André Felipe Santos, Laboratorio Hermes Pardini, Laboratorio Simile, Amilcar Tanuri, Renato Santana Aguiar e Ana Tereza Vasconcelos |
| EPI_ISL_417931 | UCSF Clinical Microbiology Laboratory | Chan-Zuckerberg Biohub | Shaun Arevalo, Josh Batson, Olga Botvinnik, Gloria Castaneda, Angela Detweiler, David Dynerman, Samantha Hao, Jack Kamm, Amy Kistler, G. Renuka Kumar, Chaz Langelier, Lucy Li, Steve Miller, Lusajo Mwakibete, Norma Neff, Angela Pisco, Maira Phelps, Michelle Tan, Chunyu Zhao |
| EPI_ISL_417932, EPI_ISL_417933 | UCSF Clinical Microbiology Laboratory | Chan-Zuckerberg Biohub | Shaun Arevalo, Josh Batson, Olga Botvinnik, Gloria Castaneda, Angela Detweiler, David Dynerman, Samantha Hao, Jack Kamm, Amy Kistler, G. Renuka Kumar, Chaz Langelier, Lucy Li, Steve Miller, Lusajo Mwakibete, Norma Neff, Angela Pisco, Maira Phelps, Michelle Tan, Chunyu Zhao |
| EPI_ISL_417934 | Laboratório Hermes Pardini | Bioinformatics Laboratory - LNCC | Filipe Romero, Ana Paula Guimarães, Mariane Talon, Luiz Gonzaga Paula de Almeida, Ronaldo da Silva Francisco Junior, Diana Mariani, Lídia Boullosa, Alexandra Gerber, Jaqueline Goes de Jesus, Ingra Morales Claro, Ester Cerdeira Sabino, Nuno Rodrigues Faria, Terezinha Marta Pereira, Pinto Castiñeiras, Isabela de Carvalho Leitão, Rafael de Mello Galliez, Cássia Cristina Alves Gonçalves, Érica Ramos dos Santos Nascimento, Richard Araújo Maia, Mauro Teixeira,Cristiano Xavier Lima, Orlando Ferreira Jr., Rodrigo Brindeiro, Luciana Jesus Costa e André Felipe Santos, Laboratorio Hermes Pardini, Laboratorio Simile, Amilcar Tanuri, Renato Santana Aguiar e Ana Tereza Vasconcelos |
| EPI_ISL_417935 | UCSF Clinical Microbiology Laboratory | Chan-Zuckerberg Biohub | Shaun Arevalo, Josh Batson, Olga Botvinnik, Gloria Castaneda, Angela Detweiler, David Dynerman, Samantha Hao, Jack Kamm, Amy Kistler, G. Renuka Kumar, Chaz Langelier, Lucy Li, Steve Miller, Lusajo Mwakibete, Norma Neff, Angela Pisco, Maira Phelps, Michelle Tan, Chunyu Zhao |
| EPI_ISL_417936 | Laboratório Hermes Pardini | Bioinformatics Laboratory - LNCC | Filipe Romero, Ana Paula Guimarães, Mariane Talon, Luiz Gonzaga Paula de Almeida, Ronaldo da Silva Francisco Junior, Diana Mariani, Lídia Boullosa, Alexandra Gerber, Jaqueline Goes de Jesus, Ingra Morales Claro, Ester Cerdeira Sabino, Nuno Rodrigues Faria, Terezinha Marta Pereira, Pinto Castiñeiras, Isabela de Carvalho Leitão, Rafael de Mello Galliez, Cássia Cristina Alves Gonçalves, Érica Ramos dos Santos Nascimento, Richard Araújo Maia, Mauro Teixeira,Cristiano Xavier Lima, Orlando Ferreira Jr., Rodrigo Brindeiro, Luciana Jesus Costa e André Felipe Santos, Laboratorio Hermes Pardini, Laboratorio Simile, Amilcar Tanuri, Renato Santana Aguiar e Ana Tereza Vasconcelos |
| EPI_ISL_417937, EPI_ISL_417938, EPI_ISL_417939 | UCSF Clinical Microbiology Laboratory | Chan-Zuckerberg Biohub | Shaun Arevalo, Josh Batson, Olga Botvinnik, Gloria Castaneda, Angela Detweiler, David Dynerman, Samantha Hao, Jack Kamm, Amy Kistler, G. Renuka Kumar, Chaz Langelier, Lucy Li, Steve Miller, Lusajo Mwakibete, Norma Neff, Angela Pisco, Maira Phelps, Michelle Tan, Chunyu Zhao |
| EPI_ISL_417940 | Laboratório Hermes Pardini | Bioinformatics Laboratory - LNCC | Filipe Romero, Ana Paula Guimarães, Mariane Talon, Luiz Gonzaga Paula de Almeida, Ronaldo da Silva Francisco Junior, Diana Mariani, Lídia Boullosa,Alexandra Gerber, Jaqueline Goes de Jesus, Ingra Morales Claro, Ester Cerdeira Sabino, Nuno Rodrigues Faria, Terezinha Marta Pereira, Pinto Castiñeiras, Isabela de Carvalho Leitão, Rafael de Mello Galliez, Cássia Alves Gonçalves, Érica Ramos dos Santos Nascimento, Richard Araújo Maia, Mauro Teixeira,Cristiano Xavier Lima, Orlando Ferreira Jr., Rodrigo Brindeiro, Luciana Jesus Costa e André Felipe Santos, Laboratorio Hermes Pardini, Laboratorio Simile, Amilcar Tanuri, Renato Santana Aguiar e Ana Tereza Vasconcelos |
| EPI_ISL_417941, EPI_ISL_417942 | Viral Respiratory Lab, National Institute for Biomedical Research (INRB) | Pathogen Sequencing Lab, National Institute for Biomedical Research (INRB) | Placide Mbala-Kingebeni, Edith Nkwembe, Eddy Kinganda-Lusamaki, Amuri Aziza, Catherine Pratt, Matthias Pauthner, Josh Quick, Allison Black, James Hadfield, Trevor Bedford, Ian Goodfellow, Nick Loman, Kristian Andersen, Michael Wiley, Steve Ahuka-Mundeke, Jean-Jacques Muyembe Tamfum |
| EPI_ISL_417943 | Laboratório Hermes Pardini | Bioinformatics Laboratory | Filipe Romero, Ana Paula Guimarães, Mariane Talon, Luiz Gonzaga Paula de Almeida, Ronaldo da Silva Francisco Junior, Diana Mariani, Lídia Boullosa,Alexandra Gerber, Jaqueline Goes de Jesus, Ingra Morales Claro, Ester Cerdeira Sabino, Nuno Rodrigues Faria, Terezinha Marta Pereira, Pinto Castiñeiras, Isabela de Carvalho Leitão, Rafael de Mello Galliez, Cássia Alves Gonçalves, Érica Ramos dos Santos Nascimento, Richard Araújo Maia, Mauro Teixeira,Cristiano Xavier Lima, Orlando Ferreira Jr., Rodrigo Brindeiro, Luciana Jesus Costa e André Felipe Santos, Laboratorio Hermes Pardini, Laboratorio Simile, Amilcar Tanuri, Renato Santana Aguiar e Ana Tereza Vasconcelos |
| EPI_ISL_417944 | Viral Respiratory Lab, National Institute for Biomedical Research (INRB) | Pathogen Sequencing Lab, National Institute for Biomedical Research (INRB) | Placide Mbala-Kingebeni, Edith Nkwembe, Eddy Kinganda-Lusamaki, Amuri Aziza, Catherine Pratt, Matthias Pauthner, Josh Quick, Allison Black, James Hadfield, Trevor Bedford, Ian Goodfellow, Nick Loman, Kristian Andersen, Michael Wiley, Steve Ahuka-Mundeke, Jean-Jacques Muyembe Tamfum |
| EPI_ISL_417945 | Laboratório Hermes Pardini | Bioinformatics Laboratory - LNCC | Filipe Romero, Ana Paula Guimarães, Mariane Talon, Luiz Gonzaga Paula de Almeida, Ronaldo da Silva Francisco Junior, Diana Mariani, Lídia Boullosa,Alexandra Gerber, Jaqueline Goes de Jesus, Ingra Morales Claro, Ester Cerdeira Sabino, Nuno Rodrigues Faria, Terezinha Marta Pereira, Pinto Castiñeiras, Isabela de Carvalho Leitão, Rafael de Mello Galliez, Cássia Alves Gonçalves, Érica Ramos dos Santos Nascimento, Richard Araújo Maia, Mauro Teixeira,Cristiano Xavier Lima, Orlando Ferreira Jr., Rodrigo Brindeiro, Luciana Jesus Costa e André Felipe Santos, Laboratorio Hermes Pardini, Laboratorio Simile, Amilcar Tanuri, Renato Santana Aguiar e Ana Tereza Vasconcelos |
| EPI_ISL_417946, EPI_ISL_417947, EPI_ISL_417948 | Viral Respiratory Lab, National Institute for Biomedical Research (INRB) | Pathogen Sequencing Lab, National Institute for Biomedical Research (INRB) | Placide Mbala-Kingebeni, Edith Nkwembe, Eddy Kinganda-Lusamaki, Amuri Aziza, Catherine Pratt, Matthias Pauthner, Josh Quick, Allison Black, James Hadfield, Trevor Bedford, Ian Goodfellow, Nick Loman, Kristian Andersen, Michael Wiley, Steve Ahuka-Mundeke, Jean-Jacques Muyembe Tamfum |
| EPI_ISL_417949 | Laboratório Hermes Pardini | Bioinformatics Laboratory - LNCC | Filipe Romero, Ana Paula Guimarães, Mariane Talon, Luiz Gonzaga Paula de Almeida, Ronaldo da Silva Francisco Junior, Diana Mariani, Lídia Boullosa,Alexandra Gerber, Jaqueline Goes de Jesus, Ingra Morales Claro, Ester Cerdeira Sabino, Nuno Rodrigues Faria, Terezinha Marta Pereira, Pinto Castiñeiras, Isabela de Carvalho Leitão, Rafael de Mello Galliez, Cássia Alves Gonçalves, Érica Ramos dos Santos Nascimento, Richard Araújo Maia, Mauro Teixeira,Cristiano Xavier Lima, Orlando Ferreira Jr., Rodrigo Brindeiro, Luciana Jesus Costa e André Felipe Santos, Laboratorio Hermes Pardini, Laboratorio Simile, Amilcar Tanuri, Renato Santana Aguiar e Ana Tereza Vasconcelos |
| EPI_ISL_417950 | Viral Respiratory Lab, National Institute for Biomedical Research (INRB) | Pathogen Sequencing Lab, National Institute for Biomedical Research (INRB) | Placide Mbala-Kingebeni, Edith Nkwembe, Eddy Kinganda-Lusamaki, Amuri Aziza, Catherine Pratt, Matthias Pauthner, Josh Quick, Allison Black, James Hadfield, Trevor Bedford, Ian Goodfellow, Nick Loman, Kristian Andersen, Michael Wiley, Steve Ahuka-Mundeke, Jean-Jacques Muyembe Tamfum |
| EPI_ISL_417951 | Universidade Federal do Rio de Janeiro | Bioinformatics Laboratory - LNCC | Filipe Romero, Ana Paula Guimarães, Mariane Talon, Luiz Gonzaga Paula de Almeida, Ronaldo da Silva Francisco Junior, Diana Mariani, Lídia Boullosa,Alexandra Gerber, Jaqueline Goes de Jesus, Ingra Morales Claro, Ester Cerdeira Sabino, Nuno Rodrigues Faria, Terezinha Marta Pereira, Pinto Castiñeiras, Isabela de Carvalho Leitão, Rafael de Mello Galliez, Cássia Alves Gonçalves, Érica Ramos dos Santos Nascimento, Richard Araújo Maia, Mauro Teixeira,Cristiano Xavier Lima, Orlando Ferreira Jr., Rodrigo Brindeiro, Luciana Jesus Costa e André Felipe Santos, Laboratorio Hermes Pardini, Laboratorio Simile, Amilcar Tanuri, Renato Santana Aguiar e Ana Tereza Vasconcelos |
| EPI_ISL_417952 | Hospital Universitario 12 de Octubre | Hospital Universitario La Paz | Elias Dahdouh, Sara González, Fernando Lázaro, Esther Viedma, Natalia Stella, Julio García, Juan Carlos Galán, Rafael Cantón, Mª Dolores Folgueira, Rafael Delgado, Jesús Mingorance |
| EPI_ISL_417953 | Universidade Federal do Rio de Janeiro | Bioinformatics Laboratory - LNCC | Filipe Romero, Ana Paula Guimarães, Mariane Talon, Luiz Gonzaga Paula de Almeida, Ronaldo da Silva Francisco Junior, Diana Mariani, Lídia Boullosa,Alexandra Gerber, Jaqueline Goes de Jesus, Ingra Morales Claro, Ester Cerdeira Sabino, Nuno Rodrigues Faria, Terezinha Marta Pereira, Pinto Castiñeiras, Isabela de Carvalho Leitão, Rafael de Mello Galliez, Cássia Alves Gonçalves, Érica Ramos dos Santos Nascimento, Richard Araújo Maia, Mauro Teixeira,Cristiano Xavier Lima, Orlando Ferreira Jr., Rodrigo Brindeiro, Luciana Jesus Costa e André Felipe Santos, Laboratorio Hermes Pardini, Laboratorio Simile, Amilcar Tanuri, Renato Santana Aguiar e Ana Tereza Vasconcelos |
| EPI_ISL_417954 | Hospital Universitario 12 de Octubre | Hospital Universitario La Paz | Elias Dahdouh, Sara González, Fernando Lázaro, Esther Viedma, Natalia Stella, Julio García, Juan Carlos Galán, Rafael Cantón, Mª Dolores Folgueira, Rafael Delgado, Jesús Mingorance |
| EPI_ISL_417955 | Viral Respiratory Lab, National Institute for Biomedical Research (INRB) | Pathogen Sequencing Lab, National Institute for Biomedical Research (INRB) | Placide Mbala-Kingebeni, Edith Nkwembe, Eddy Kinganda-Lusamaki, Amuri Aziza, Catherine Pratt, Matthias Pauthner, Josh Quick, Allison Black, James Hadfield, Trevor Bedford, Ian Goodfellow, Nick Loman, Kristian Andersen, Michael Wiley, Steve Ahuka-Mundeke, Jean-Jacques Muyembe Tamfum |
| EPI_ISL_417956 | Hospital Universitario 12 de Octubre | Hospital Universitario La Paz | Elias Dahdouh, Sara González, Fernando Lázaro, Esther Viedma, Natalia Stella, Julio García, Juan Carlos Galán, Rafael Cantón, Mª Dolores Folgueira, Rafael Delgado, Jesús Mingorance |
| EPI_ISL_417957 | Hospital Universitario 12 de Octubre | Hospital Universitario La Paz | Elias Dahdouh, Sara González, Fernando Lázaro, Esther Viedma, Natalia Stella, Julio García, Juan Carlos Galán, Rafael Cantón, Mª Dolores Folgueira, Rafael Delgado, Jesús Mingorance |
| EPI_ISL_417958, EPI_ISL_417959, EPI_ISL_417960 | Utah Public Health Laboratory | Utah Public Health Laboratory | Erin Young, Kelly Oakeson |
| EPI_ISL_417961, EPI_ISL_417963 | Hospital Universitario 12 de Octubre | Hospital Universitario La Paz | Elias Dahdouh, Sara González, Fernando Lázaro, Esther Viedma, Natalia Stella, Julio García, Juan Carlos Galán, Rafael Cantón, Mª Dolores Folgueira, Rafael Delgado, Jesús Mingorance |
| EPI_ISL_417964, EPI_ISL_417966 | Utah Public Health Laboratory | Utah Public Health Laboratory | Erin Young, Kelly Oakeson |
| EPI_ISL_417967 | Hospital Universitario 12 de Octubre | Hospital Universitario La Paz | Elias Dahdouh, Sara González, Fernando Lázaro, Esther Viedma, Natalia Stella, Julio García, Juan Carlos Galán, Rafael Cantón, Mª Dolores Folgueira, Rafael Delgado, Jesús Mingorance |
| EPI_ISL_417969 | Hospital Universitario La Paz | Hospital Universitario La Paz | Elias Dahdouh, Sara González, Fernando Lázaro, Esther Viedma, Natalia Stella, Julio García, Juan Carlos Galán, Rafael Cantón, Mª Dolores Folgueira, Rafael Delgado, Jesús Mingorance |
| EPI_ISL_417970, EPI_ISL_417971 | Utah Public Health Laboratory | Utah Public Health Laboratory | Erin Young, Kelly Oakeson |
| EPI_ISL_417972 | Hospital Universitario La Paz | Hospital Universitario La Paz | Elias Dahdouh, Sara González, Fernando Lázaro, Esther Viedma, Natalia Stella, Julio García, Juan Carlos Galán, Rafael Cantón, Mª Dolores Folgueira, Rafael Delgado, Jesús Mingorance |
| EPI_ISL_417973, EPI_ISL_417974 | Utah Public Health Laboratory | Utah Public Health Laboratory | Erin Young, Kelly Oakeson |
| EPI_ISL_417975 | Hospital Universitario La Paz | Hospital Universitario La Paz | Elias Dahdouh, Sara González, Fernando Lázaro, Esther Viedma, Natalia Stella, Julio García, Juan Carlos Galán, Rafael Cantón, Mª Dolores Folgueira, Rafael Delgado, Jesús Mingorance |
| EPI_ISL_417976, EPI_ISL_417977 | Utah Public Health Laboratory | Utah Public Health Laboratory | Erin Young, Kelly Oakeson |
| EPI_ISL_417978 | Hospital Universitario La Paz | Hospital Universitario La Paz | Elias Dahdouh, Sara González, Fernando Lázaro, Esther Viedma, Natalia Stella, Julio García, Juan Carlos Galán, Rafael Cantón, Mª Dolores Folgueira, Rafael Delgado, Jesús Mingorance |
| EPI_ISL_417979, EPI_ISL_417980, EPI_ISL_417981 | Hospital Universitario Ramón y Cajal | Hospital Universitario La Paz | Elias Dahdouh, Sara González, Fernando Lázaro, Esther Viedma, Natalia Stella, Julio García, Juan Carlos Galán, Rafael Cantón, Mª Dolores Folgueira, Rafael Delgado, Jesús Mingorance |
| EPI_ISL_417982, EPI_ISL_417983, EPI_ISL_417984 | Universidade Federal do Rio de Janeiro | Bioinformatics Laboratory - LNCC | Filipe Romero, Ana Paula Guimarães, Mariane Talon, Luiz Gonzaga Paula de Almeida, Ronaldo da Silva Francisco Junior, Diana Mariani, Lídia Boullosa,Alexandra Gerber, Jaqueline Goes de Jesus, Ingra Morales Claro, Ester Cerdeira Sabino, Nuno Rodrigues Faria, Terezinha Marta Pereira, Pinto Castiñeiras, Isabela de Carvalho Leitão, Rafael de Mello Galliez, Cássia Alves Gonçalves, Érica Ramos dos Santos Nascimento, Richard Araújo Maia, Mauro Teixeira,Cristiano Xavier Lima, Orlando Ferreira Jr., Rodrigo Brindeiro, Luciana Jesus Costa e André Felipe Santos, Laboratorio Hermes Pardini, Laboratorio Simile, Amilcar Tanuri, Renato Santana Aguiar e Ana Tereza Vasconcelos |
| EPI_ISL_417985 | Universidade Federal do Rio de Janeiro | Bioinformatics Laboratory - LNCC | Filipe Romero, Ana Paula Guimarães, Mariane Talon, Luiz Gonzaga Paula de Almeida, Ronaldo da Silva Francisco Junior, Diana Mariani, Lídia Boullosa,Alexandra Gerber, Jaqueline Goes de Jesus, Ingra Morales Claro, Ester Cerdeira Sabino, Nuno Rodrigues Faria, Terezinha Marta Pereira, Pinto Castiñeiras, Isabela de Carvalho Leitão, Rafael de Mello Galliez, Cássia Alves Gonçalves, Érica Ramos dos Santos Nascimento, Richard Araújo Maia, Mauro Teixeira,Cristiano Xavier Lima, Orlando Ferreira Jr., Rodrigo Brindeiro, Luciana Jesus Costa e André Felipe Santos, Laboratorio Hermes Pardini, Laboratorio Simile, Amilcar Tanuri, Renato Santana Aguiar e Ana Tereza Vasconcelos |
| EPI_ISL_417986, EPI_ISL_417987 | Centro Hospitalar e Universitario de Sao Joao, Porto | Instituto Nacional de Saude (INSA) | Guiomar et al |
| EPI_ISL_417988 | CHULC - H Curry Cabral | Instituto Nacional de Saude (INSA) | Guiomar et al |
| EPI_ISL_417989 | Centro Hospitalar e Universitario de Sao Joao, Porto | Instituto Nacional de Saude (INSA) | Guiomar et al |
| EPI_ISL_417990, EPI_ISL_417991 | CHULC - H Curry Cabral | Instituto Nacional de Saude (INSA) | Guiomar et al |
| EPI_ISL_417992, EPI_ISL_417993 | CHULC - H D Estefania | Instituto Nacional de Saude (INSA) | Guiomar et al |
| EPI_ISL_417994, EPI_ISL_417995, EPI_ISL_417996 | CHULC - H Curry Cabral | Instituto Nacional de Saude (INSA) | Guiomar et al |
| EPI_ISL_417997, EPI_ISL_417998, EPI_ISL_417999 | Centro Hospital do Porto, E.P.E. - H. Geral de Santo Antonio | Instituto Nacional de Saude (INSA) | Guiomar et al |
| EPI_ISL_418000, EPI_ISL_418001 | ARS Algarve - Laboratório Laura Ayres | Instituto Nacional de Saude (INSA) | Guiomar et al |
| EPI_ISL_418002 | CHU Coimbra | Instituto Nacional de Saude (INSA) | Guiomar et al |
| EPI_ISL_418003 | H Braga | Instituto Nacional de Saude (INSA) | Guiomar et al |
| EPI_ISL_418004 | ARS Algarve - Laboratório Laura Ayres | Instituto Nacional de Saude (INSA) | Guiomar et al |
| EPI_ISL_418005 | CHU Coimbra - Pediátrico | Instituto Nacional de Saude (INSA) | Guiomar et al |
| EPI_ISL_418006 | CHBarreiro Montijo | Instituto Nacional de Saude (INSA) | Guiomar et al |
| EPI_ISL_418007, EPI_ISL_418008 | H Braga | Instituto Nacional de Saude (INSA) | Guiomar et al |
| EPI_ISL_418009 | HSE Ilha Terceira - Angra do Heroismo | Instituto Nacional de Saude (INSA) | Guiomar et al |

| | | | |
|---|---|---|---|
| EPI_ISL_418010, EPI_ISL_418011, EPI_ISL_418012, EPI_ISL_418013, EPI_ISL_418014, EPI_ISL_418015, EPI_ISL_418016 | CHULC - H Curry Cabral | Instituto Nacional de Saude (INSA) | Guiomar et al |
| EPI_ISL_418017 | CHMT | Instituto Nacional de Saude (INSA) | Guiomar et al |
| EPI_ISL_418018 | H Garcia de Orta | Instituto Nacional de Saude (INSA) | Guiomar et al |
| EPI_ISL_418019, EPI_ISL_418020, EPI_ISL_418021, EPI_ISL_418022 | H Braga | Instituto Nacional de Saude (INSA) | Guiomar et al |
| EPI_ISL_418023 | H Evora | Instituto Nacional de Saude (INSA) | Guiomar et al |
| EPI_ISL_418024 | CHUA - Faro | Instituto Nacional de Saude (INSA) | Guiomar et al |
| EPI_ISL_418025 | H Santarem | Instituto Nacional de Saude (INSA) | Guiomar et al |
| EPI_ISL_418026 | H Dr. Nelio Mendonca - Funchal | Instituto Nacional de Saude (INSA) | Guiomar et al |
| EPI_ISL_418027 | CHTMAD | Instituto Nacional de Saude (INSA) | Guiomar et al |
| EPI_ISL_418028, EPI_ISL_418029, EPI_ISL_418030, EPI_ISL_418031, EPI_ISL_418032, EPI_ISL_418033, EPI_ISL_418034, EPI_ISL_418035, EPI_ISL_418036, EPI_ISL_418037, EPI_ISL_418038, EPI_ISL_418039, EPI_ISL_418040, EPI_ISL_418041, EPI_ISL_418042, EPI_ISL_418043, EPI_ISL_418044, EPI_ISL_418045, EPI_ISL_418046, EPI_ISL_418047, EPI_ISL_418048, EPI_ISL_418049, EPI_ISL_418050, EPI_ISL_418051, EPI_ISL_418052, EPI_ISL_418053, EPI_ISL_418054, EPI_ISL_418055, EPI_ISL_418056, EPI_ISL_418057, EPI_ISL_418058, EPI_ISL_418059, EPI_ISL_418060, EPI_ISL_418061, EPI_ISL_418062, EPI_ISL_418063, EPI_ISL_418064, EPI_ISL_418065, EPI_ISL_418066, EPI_ISL_418067, EPI_ISL_418068, EPI_ISL_418069, EPI_ISL_418070, EPI_ISL_418071, EPI_ISL_418072, EPI_ISL_418073, EPI_ISL_418074, EPI_ISL_418075, EPI_ISL_418076, EPI_ISL_418077, EPI_ISL_418078, EPI_ISL_418079, EPI_ISL_418080, EPI_ISL_418081, EPI_ISL_418082 | | | |
| see above | UW Virology Lab | UW Virology Lab | Pavitra Roychoudhury, Hong Xie, Keith Jerome, Alexander Greninger |
| EPI_ISL_418083, EPI_ISL_418084, EPI_ISL_418085, EPI_ISL_418086, EPI_ISL_418087, EPI_ISL_418088, EPI_ISL_418089, EPI_ISL_418090, EPI_ISL_418091, EPI_ISL_418092, EPI_ISL_418093, EPI_ISL_418094, EPI_ISL_418095, EPI_ISL_418096, EPI_ISL_418097, EPI_ISL_418098, EPI_ISL_418099, EPI_ISL_418100, EPI_ISL_418101, EPI_ISL_418102, EPI_ISL_418103, EPI_ISL_418104, EPI_ISL_418105, EPI_ISL_418106, EPI_ISL_418107, EPI_ISL_418108, EPI_ISL_418109, EPI_ISL_418110, EPI_ISL_418111, EPI_ISL_418112, EPI_ISL_418113, EPI_ISL_418114, EPI_ISL_418115, EPI_ISL_418116, EPI_ISL_418117, EPI_ISL_418118, EPI_ISL_418119, EPI_ISL_418120, EPI_ISL_418121, EPI_ISL_418122, EPI_ISL_418123, EPI_ISL_418124, EPI_ISL_418125, EPI_ISL_418126, EPI_ISL_418127, EPI_ISL_418128, EPI_ISL_418129, EPI_ISL_418130, EPI_ISL_418131, EPI_ISL_418132, EPI_ISL_418133, EPI_ISL_418134, EPI_ISL_418135, EPI_ISL_418136, EPI_ISL_418137, EPI_ISL_418138, EPI_ISL_418139, EPI_ISL_418140, EPI_ISL_418141, EPI_ISL_418142, EPI_ISL_418143, EPI_ISL_418144, EPI_ISL_418145, EPI_ISL_418146, EPI_ISL_418147, EPI_ISL_418148, EPI_ISL_418149, EPI_ISL_418150, EPI_ISL_418151, EPI_ISL_418152, EPI_ISL_418153, EPI_ISL_418154, EPI_ISL_418155, EPI_ISL_418156, EPI_ISL_418157, EPI_ISL_418158, EPI_ISL_418159, EPI_ISL_418160, EPI_ISL_418161, EPI_ISL_418162, EPI_ISL_418163, EPI_ISL_418164, EPI_ISL_418165 | | | |
| see above | Wales Specialist Virology Centre | Public Health Wales Microbiology Cardiff | Catherine Moore, Joanne Watkins, Sally Corden, Sara Rey, Matt Bull, Tom Connor |
| EPI_ISL_418182 | Hospital Universitario 12 de Octubre | Hospital Universitario La Paz | Elias Dahdouh, Sara González, Fernando Lázaro, Esther Viedma, Natalia Stella, Julio García, Juan Carlos Galán, Rafael Cantón, Mª Dolores Folgueira, Rafael Delgado, Jesús Mingorance |
| EPI_ISL_418183 | Virological Research Group, Szentágothai Research Centre | Bioinformatics Research Group, Szentágothai Research Centre | Péter Urbán, Endre Gábor Tóth, Gábor Kemenesi, Róbert Herczeg, Attila Gyenesei, Ferenc Jakab |
| EPI_ISL_418184 | Gundersen Molecular Diagnostics Laboratory | Kabara Cancer Research Institute | Craig S. Richmond & Paraic A. Kenny |
| EPI_ISL_418185, EPI_ISL_418186 | Gundersen Molecular Diagnostic Laboratory | Kabara Cancer Research Institute | Craig S. Richmond & Paraic A. Kenny |
| EPI_ISL_418187, EPI_ISL_418188, EPI_ISL_418189 | Gundersen Molecular Diagnostics Laboratory | Kabara Cancer Research Institute | Craig S. Richmond & Paraic A. Kenny |
| EPI_ISL_418190, EPI_ISL_418191, EPI_ISL_418192, EPI_ISL_418193, EPI_ISL_418194, EPI_ISL_418195, EPI_ISL_418196, EPI_ISL_418197, EPI_ISL_418198, EPI_ISL_418199, EPI_ISL_418200, EPI_ISL_418201, EPI_ISL_418202, EPI_ISL_418203, EPI_ISL_418204, EPI_ISL_418205 | | | |
| see above | NYU Langone Health | Department of Pathology and Medicine, New York University School of Medicine | Margaret Black, John Cadley, Paolo Cotzia, John Chen, Dacia Dimartino, Xiaojun Feng, Adriana Heguy, Megan Hogan, Emily Huang, George Jour, Christian Marier, Matthew T. Maurano, Mark J. Mulligan, Peter Meyn, Jared Pinnell, Amy Rapkiewicz, Marie Samanovic-Golden, Antonio Serrano, Guomiao Shen, Matija Snuderl, Nick Vulpescu, Gael Westby, Paul Zappile |
| EPI_ISL_418206, EPI_ISL_418207, EPI_ISL_418208, EPI_ISL_418209, EPI_ISL_418210, EPI_ISL_418211 | Institut Pasteur Dakar | Institut Pasteur de Dakar | Ndongo Dia, Ousmane Faye, Amadou Alpha Sall |
| EPI_ISL_418212 | Institut Pasteur Dakar | Institut Pasteur de Dakar | Ndongo Dia, Ousmane Faye, Amadou Alpha sall |
| EPI_ISL_418213, EPI_ISL_418214 | Institut Pasteur Dakar | Institut Pasteur de Dakar | Ndongo Dia, Ousmane Faye, Amadou Alpha Sall |
| EPI_ISL_418215 | Instirut Pasteur Dakar | Institut Pasteur de Dakar | Ndongo Dia, Ousmane Faye, Amadou Alpha Sall |
| EPI_ISL_418216, EPI_ISL_418217 | Institut Pasteur Dakar | Institut Pasteur de Dakar | Ndongo Dia, Ousmane Faye, Amadou Alpha Sall |
| EPI_ISL_418218 | Centre Hospitalier Compiègne Laboratoire de Biologie | National Reference Center for Viruses of Respiratory Infections, Institut Pasteur, Paris | Mélanie Albert, Marion Barbet, Sylvie Behillil, Méline Bizard, Angela Brisebarre, Flora Donati, Fabiana Gambaro, Etienne Simon-Lorière, Vincent Enouf, Maud Vanpeene, Sylvie van der Werf, Raulin Olivia |
| EPI_ISL_418219 | CHU - Hôpital Cavale Blanche - Labo. de Virologie | National Reference Center for Viruses of Respiratory Infections, Institut Pasteur, Paris | Mélanie Albert, Marion Barbet, Sylvie Behillil, Méline Bizard, Angela Brisebarre, Flora Donati, Fabiana Gambaro, Etienne Simon-Lorière, Vincent Enouf, Maud Vanpeene, Sylvie van der Werf, Léa Pilorge |
| EPI_ISL_418220, EPI_ISL_418221 | Centre Hospitalier Compiègne Laboratoire de Biologie | National Reference Center for Viruses of Respiratory Infections, Institut Pasteur, Paris | Mélanie Albert, Marion Barbet, Sylvie Behillil, Méline Bizard, Angela Brisebarre, Flora Donati, Fabiana Gambaro, Etienne Simon-Lorière, Vincent Enouf, Maud Vanpeene, Sylvie van der Werf, Raulin Olivia |
| EPI_ISL_418222 | CHRU Bretonneau - Serv. Bacterio-Virol. | National Reference Center for Viruses of Respiratory Infections, Institut Pasteur, Paris | Mélanie Albert, Marion Barbet, Sylvie Behillil, Méline Bizard, Angela Brisebarre, Flora Donati, Fabiana Gambaro, Etienne Simon-Lorière, Vincent Enouf, Maud Vanpeene, Sylvie van der Werf, Julien Marlet |
| EPI_ISL_418223, EPI_ISL_418224, EPI_ISL_418225 | Centre Hospitalier Compiègne Laboratoire de Biologie | National Reference Center for Viruses of Respiratory Infections, Institut Pasteur, Paris | Mélanie Albert, Marion Barbet, Sylvie Behillil, Méline Bizard, Angela Brisebarre, Flora Donati, Fabiana Gambaro, Etienne Simon-Lorière, Vincent Enouf, Maud Vanpeene, Sylvie van der Werf, Raulin Olivia |
| EPI_ISL_418226 | EHPAD - Résidences les Cèdres | National Reference Center for Viruses of Respiratory Infections, Institut Pasteur, Paris | Mélanie Albert, Marion Barbet, Sylvie Behillil, Méline Bizard, Angela Brisebarre, Flora Donati, Etienne Simon-Lorière, Vincent Enouf, Maud Vanpeene, Sylvie van der Werf |
| EPI_ISL_418227, EPI_ISL_418228 | Centre Hospitalier Compiègne Laboratoire de Biologie | National Reference Center for Viruses of Respiratory Infections, Institut Pasteur, Paris | Mélanie Albert, Marion Barbet, Sylvie Behillil, Méline Bizard, Angela Brisebarre, Flora Donati, Etienne Simon-Lorière, Vincent Enouf, Maud Vanpeene, Sylvie van der Werf, Raulin Olivia |
| EPI_ISL_418229 | Hopital franco britannique - Laboratoire | National Reference Center for Viruses of Respiratory Infections, Institut Pasteur, Paris | Mélanie Albert, Marion Barbet, Sylvie Behillil, Méline Bizard, Angela Brisebarre, Flora Donati, Etienne Simon-Lorière, Vincent Enouf, Maud Vanpeene, Sylvie van der Werf, Marianne Asso Bonnet |
| EPI_ISL_418230 | Clinique AVERAY LA BROUSTE, Med. Polyvalente | National Reference Center for Viruses of Respiratory Infections, Institut Pasteur, Paris | Mélanie Albert, Marion Barbet, Sylvie Behillil, Méline Bizard, Angela Brisebarre, Flora Donati, Etienne Simon-Lorière, Vincent Enouf, Maud Vanpeene, Sylvie van der Werf, Elsa Ngwem |
| EPI_ISL_418231 | Centre Hospitalier Compiègne Laboratoire de Biologie | National Reference Center for Viruses of Respiratory Infections, Institut Pasteur, Paris | Mélanie Albert, Marion Barbet, Sylvie Behillil, Méline Bizard, Angela Brisebarre, Flora Donati, Etienne Simon-Lorière, Vincent Enouf, Maud Vanpeene, Sylvie van der Werf, Raulin Olivia |
| EPI_ISL_418232, EPI_ISL_418233 | Service des Urgences | National Reference Center for Viruses of Respiratory Infections, Institut Pasteur, Paris | Mélanie Albert, Marion Barbet, Sylvie Behillil, Méline Bizard, Angela Brisebarre, Flora Donati, Etienne Simon-Lorière, Vincent Enouf, Maud Vanpeene, Sylvie van der Werf, Boubkeur |
| EPI_ISL_418234 | LABM GH nord Essonne | National Reference Center for Viruses of Respiratory Infections, Institut Pasteur, Paris | Mélanie Albert, Marion Barbet, Sylvie Behillil, Méline Bizard, Angela Brisebarre, Flora Donati, Etienne Simon-Lorière, Vincent Enouf, Maud Vanpeene, Sylvie van der Werf, Christine Lambert |
| EPI_ISL_418235 | Cabinet médical | National Reference Center for Viruses of Respiratory Infections, Institut Pasteur, Paris | Mélanie Albert, Marion Barbet, Sylvie Behillil, Méline Bizard, Angela Brisebarre, Flora Donati, Etienne Simon-Lorière, Vincent Enouf, Maud Vanpeene, Sylvie van der Werf |
| EPI_ISL_418236, EPI_ISL_418237, EPI_ISL_418238, EPI_ISL_418239 | Centre Hospitalier Compiègne Laboratoire de Biologie | National Reference Center for Viruses of Respiratory Infections, Institut Pasteur, Paris | Mélanie Albert, Marion Barbet, Sylvie Behillil, Méline Bizard, Angela Brisebarre, Flora Donati, Etienne Simon-Lorière, Vincent Enouf, Maud Vanpeene, Sylvie van der Werf, Raulin Olivia |
| EPI_ISL_418240 | LABM GH nord Essonne | National Reference Center for Viruses of Respiratory Infections, Institut Pasteur, Paris | Mélanie Albert, Marion Barbet, Sylvie Behillil, Méline Bizard, Angela Brisebarre, Flora Donati, Etienne Simon-Lorière, Vincent Enouf, Maud Vanpeene, Sylvie van der Werf, Christine Lambert |
| EPI_ISL_418241, EPI_ISL_418242 | NIC Viral Respiratory Unit - Institut Pasteur of Algeria | National Reference Center for Viruses of Respiratory Infections, Institut Pasteur, Paris | Mélanie Albert, Marion Barbet, Sylvie Behillil, Méline Bizard, Angela Brisebarre, Flora Donati, Etienne Simon-Lorière, Vincent Enouf, Maud Vanpeene, Sylvie van der Werf, Fawzi Derrar |
| EPI_ISL_418243, EPI_ISL_418244 | HOSPITAL UNIVERSITARIO VIRGEN DE LAS NIEVES | Instituto de Salud Carlos III | Iglesias-Caballero, M. Molinero Calamita, M. González-Esguevillas, M. Camarero, S. Pozo, F. Casas, I. Jiménez, P. Jiménez, M. Zaballos, A. Monzón, S. Varona, S. Juliá, M. Cuesta, I. Sanbonmatsu S. |
| EPI_ISL_418245, EPI_ISL_418246 | Hospital General y Universitario de Guadalajara | Instituto de Salud Carlos III | Iglesias-Caballero, M. Molinero Calamita, M. González-Esguevillas, M. Camarero, S. Pozo, F. Casas, I. Jiménez, P. Jiménez, M. Zaballos, A. Monzón, S. Varona, S. Juliá, M. Cuesta, I. Gonzalez-Praetorius A. |
| EPI_ISL_418247 | HOSPITAL GENERAL DE SEGOVIA | Instituto de Salud Carlos III | Iglesias-Caballero, M. Molinero Calamita, M. González-Esguevillas, M. Camarero, S. Pozo, F. Casas, I. Jiménez, P. Jiménez, M. Zaballos, A. Monzón, S. Varona, S. Juliá, M. Cuesta, I. Hernando-Real S. |
| EPI_ISL_418248, EPI_ISL_418249 | COMPLEJO ASISTENCIAL UNIVERSITARIO DE BURGOS | Instituto de Salud Carlos III | Iglesias-Caballero, M. Molinero Calamita, M. González-Esguevillas, M. Camarero, S. Pozo, F. Casas, I. Jiménez, P. Jiménez, M. Zaballos, A. Monzón, S. Varona, S. Juliá, M. Cuesta, I. Megias-Lobon G. |
| EPI_ISL_418250 | HOSPITAL CLINIC | Instituto de Salud Carlos III | Iglesias-Caballero, M. Molinero Calamita, M. González-Esguevillas, M. Camarero, S. Pozo, F. Casas, I. Jiménez, P. Jiménez, M. Zaballos, A. Monzón, S. Varona, S. Juliá, M. Cuesta, I. Marcos M.A |
| EPI_ISL_418251 | HOSPITAL UNIVERSITARIO LA PAZ | Instituto de Salud Carlos III | Iglesias-Caballero, M. Molinero Calamita, M. González-Esguevillas, M. Camarero, S. Pozo, F. Casas, I. Jiménez, P. Jiménez, M. Zaballos, A. Monzón, S. Varona, S. Juliá, M. Cuesta, I. Romero P. |
| EPI_ISL_418252 | FUNDACION JIMENEZ DIAZ | Instituto de Salud Carlos III | Iglesias-Caballero, M. Molinero Calamita, M. González-Esguevillas, M. Camarero, S. Pozo, F. Casas, I. Jiménez, P. Jiménez, M. Zaballos, A. Monzón, S. Varona, S. Juliá, M. Cuesta, I. Fernández Roblas, R. |
| EPI_ISL_418253 | HOSPITAL TXAGORRITXU | Instituto de Salud Carlos III | Iglesias-Caballero, M. Molinero Calamita, M. González-Esguevillas, M. Camarero, S. Pozo, F. Casas, I. Jiménez, P. Jiménez, M. Zaballos, A. Monzón, S. Varona, S. Juliá, M. Cuesta, I. Gomez-Gonzalez C. |
| EPI_ISL_418254 | NYU Langone Health | Department of Pathology and Medicine, New York University School of Medicine | Margaret Black, John Cadley, Paolo Cotzia, John Chen, Dacia Dimartino, Xiaojun Feng, Adriana Heguy, Megan Hogan, Emily Huang, George Jour, Christian Marier, Matthew T. Maurano, Mark J. Mulligan, Peter Meyn, Jared Pinnell, Amy Rapkiewicz, Marie Samanovic-Golden, Antonio Serrano, Guomiao Shen, Matija Snuderl, Nick Vulpescu, Gael Westby, Paul Zappile |
| EPI_ISL_418255 | Presidio Ospedaliero "S. Spirito" - PESCARA | Istituto Zooprofilattico Sperimentale dell'Abruzzo e Molise "G. Caporale" | Lorusso A, Marcacci M, Cammà C, Monaco F, Puglia I, Di Pasquale A, Rinaldi A, Mangone I, Savini G |
| EPI_ISL_418256 | Ospedale “San Liberatore” di Atri | Istituto Zooprofilattico Sperimentale dell'Abruzzo e Molise "G. Caporale" | Lorusso A, Marcacci M, Di Domenico M, Puglia I, Curini V, Ancora M, Di Pasquale A, Rinaldi A, Mangone I, Cammà C, Savini G. |
| EPI_ISL_418257 | Ospedale Civile Giuseppe Mazzini, Teramo | Istituto Zooprofilattico Sperimentale dell'Abruzzo e Molise "G.Caporale" | Lorusso A, Marcacci M, Di Domenico M, Puglia I, Curini V, Ancora M, Di Pasquale A, Rinaldi A, Mangone I, Cammà C, Savini G. |

| | | | |
|---|---|---|---|
| EPI_ISL_418258, EPI_ISL_418259 | Presidio ospedaliero "Santo Spirito" | Istituto Zooprofilattico Sperimentale dell'Abruzzo e Molise "G. Caporale" | Lorusso A, Marcacci M, Di Domenico M, Puglia I, Curini V, Ancora M, Di Pasquale A, Rinaldi A, Mangone I, Cammà C, Savini G. |
| EPI_ISL_418260, EPI_ISL_418261 | Ospedale Civile Giuseppe Mazzini | Istituto Zooprofilattico Sperimentale dell'Abruzzo e Molise "G. Caporale" | Lorusso A, Marcacci M, Di Domenico M, Puglia I, Curini V, Ancora M, Di Pasquale A, Rinaldi A, Mangone I, Cammà C, Savini G. |
| EPI_ISL_418262 | Instituto Nacional de Salud | Instituto Nacional de Salud Universidad Cooperativa de Colombia Instituto Alexander von Humboldt Imperial College-London London School of Hygiene & Tropical Medicine | Marcela Mercado-Reyes, Katherine Laiton-Donato, Diego A. Álvarez-Díaz, Carlos Franco-Muñoz, Jose A. Usme-Ciro, Gloria Puerto, Nicolas D. Franco-Sierra, Mailyn A.Gonzalez, Zulma M. Cucunubá, Christian Julian Villabona-Arenas, Liz Villabona-Arenas, Sussy Echeverria, Astrid C. Flórez, Sergio Gomez Rangel, Luz Dary Rodriguez, Juliana Barbosa, Erika Ospitia, Diana Marcela Walteros-Acero, Nuno Rodrigues Faria, Martha Lucia Ospina Martinez |
| EPI_ISL_418263, EPI_ISL_418264, EPI_ISL_418265 | Laboratory of Microbiology, Department of Medicine, National and Kapodistrian University of Athens, Greece | Laboratory of Biology, Department of Medicine, Democritus University of Thrace, Greece | Maria Bampali, Elisavet Gatzidou, Nikolaos Dovrolis, Stavroula Veletza, Nikolaos Spanakis, Ioannis Karakasiliotis |
| EPI_ISL_418267 | Microbiology and Immunology department, Pasteur institute in Ho Chi Minh city | Microbiology and Immunology department, Pasteur institute in Ho Chi Minh city | Nguyen,H.T., Cao,T.M., Pham,H.T.T., Vu,N.P.H., Dao,M.H., Huynh,L.T.K., Nguyen,L.T., Nguyen,N.T., Nguyen,T.T.N., Nguyen,A.H., Luong,Q.C., Nguyen,T.V., Tran,K.C., Pham,Q.D., Tran,T., Hoang,C.Q., Nguyen,T.T., Le,H.Q., Phung,T.M., Vo,T.N.A., Nguyen,S.N., Pham,D.T., Nguyen,T.V. and Phan,L.T. |
| EPI_ISL_418268 | Hospital Universitari Germans Trias i Pujol(HUGTiP)/Fundació Lluita contra la SIDA (FLSida)/IRTA-CReSA | IrsiCaixa AIDS Research Lab | Pilar Armengol, Marc Noguera-Julian, Jordi Rodón, Julia Vergara, Lidia Ruiz, Nuria Izquierdo, Jorge Carrillo, Roger Paredes, Albert Bensaid, Julia Blanco, Joaquim Segalés, Bonaventura Clotet |
| EPI_ISL_418269 | Microbiology and Immunology department, Pasteur institute in Ho Chi Minh city | Microbiology and Immunology department, Pasteur institute in Ho Chi Minh city | Cao,T.M., Nguyen,H.T., Pham,H.T.T., Vu,N.P.H., Dao,M.H., Huynh,L.T.K., Nguyen,L.T., Nguyen,N.T., Nguyen,T.T.N., Nguyen,A.H., Luong,Q.C., Nguyen,T.V., Tran,K.C., Pham,Q.D., Tran,T., Hoang,C.Q., Nguyen,T.T., Le,H.Q., Phung,T.M., Vo,T.N.A., Nguyen,S.N., Pham,D.T., Phan,L.T. and Nguyen,T.V. |
| EPI_ISL_418270 | KU Leuven, Clinical and Epidemiological Virology | KU Leuven, Clinical and Epidemiological Virology | Tony Wawina, Joan Marti-Carreras, Bert Vanmechelen, Piet Maes |
| EPI_ISL_418271, EPI_ISL_418273 | University Hospital Basel, Clinical Virology | University Hospital Basel, Labormedizin | Hirsch, H., Leuzinger, K., Seth-Smith, H., Mari, A., Roloff, T., Egli, A. |
| EPI_ISL_418274 | University Hospital Basel, Clinical Virology | University Hospital Basel, Clinical Bacteriology | Hirsch, H., Leuzinger, K., Seth-Smith, H., Mari, A., Roloff, T., Egli, A.University Hospital Basel, Clinical Bacteriology |
| EPI_ISL_418275, EPI_ISL_418277, EPI_ISL_418278, EPI_ISL_418279, EPI_ISL_418280, EPI_ISL_418281, EPI_ISL_418282, EPI_ISL_418283, EPI_ISL_418284, EPI_ISL_418285 | University Hospital Basel, Clinical Virology | University Hospital Basel, Clinical Bacteriology | Hirsch, H., Leuzinger, K., Seth-Smith, H., Mari, A., Roloff, T., Egli, A. |
| EPI_ISL_418286, EPI_ISL_418287, EPI_ISL_418288, EPI_ISL_418289, EPI_ISL_418290, EPI_ISL_418291, EPI_ISL_418292, EPI_ISL_418293, EPI_ISL_418294, EPI_ISL_418295, EPI_ISL_418296, EPI_ISL_418297, EPI_ISL_418298, EPI_ISL_418299, EPI_ISL_418300, EPI_ISL_418301, EPI_ISL_418302, EPI_ISL_418303, EPI_ISL_418304, EPI_ISL_418305, EPI_ISL_418306, EPI_ISL_418307, EPI_ISL_418308, EPI_ISL_418309, EPI_ISL_418310, EPI_ISL_418311, EPI_ISL_418312, EPI_ISL_418313, EPI_ISL_418314, EPI_ISL_418315, EPI_ISL_418316, EPI_ISL_418317, EPI_ISL_418318, EPI_ISL_418319, EPI_ISL_418320, EPI_ISL_418321 | | | |
| see above | Virology Department, Sheffield Teaching Hospitals NHS Foundation Trust | Department of Infection, Immunity and Cardiovascular Disease, The Florey Institute, The Medical School, University of Sheffield | Thushan de Silva, Matthew Parker, Adri Angyal, Rebecca Brown, Rachel Tucker, Paul Parsons, Danielle Groves, Alex Keeley, Dave Partridge, Matthew Wyles, Benjamin Lindsey, Mehmet Yavuz, Mohammad Raza, Cariad Evans |
| EPI_ISL_418322, EPI_ISL_418323, EPI_ISL_418324, EPI_ISL_418325, EPI_ISL_418326, EPI_ISL_418327, EPI_ISL_418328, EPI_ISL_418329, EPI_ISL_418330, EPI_ISL_418331, EPI_ISL_418332, EPI_ISL_418333, EPI_ISL_418334, EPI_ISL_418335, EPI_ISL_418336, EPI_ISL_418337, EPI_ISL_418338, EPI_ISL_418339, EPI_ISL_418340, EPI_ISL_418341, EPI_ISL_418342, EPI_ISL_418343, EPI_ISL_418344, EPI_ISL_418345, EPI_ISL_418346, EPI_ISL_418347, EPI_ISL_418348, EPI_ISL_418349, EPI_ISL_418350, EPI_ISL_418351, EPI_ISL_418352, EPI_ISL_418353, EPI_ISL_418354, EPI_ISL_418355, EPI_ISL_418356, EPI_ISL_418357, EPI_ISL_418358, EPI_ISL_418359, EPI_ISL_418360, EPI_ISL_418361, EPI_ISL_418362, EPI_ISL_418363, EPI_ISL_418364, EPI_ISL_418365, EPI_ISL_418366, EPI_ISL_418367, EPI_ISL_418368, EPI_ISL_418369, EPI_ISL_418370, EPI_ISL_418371, EPI_ISL_418372, EPI_ISL_418373, EPI_ISL_418374, EPI_ISL_418375, EPI_ISL_418376, EPI_ISL_418377, EPI_ISL_418378, EPI_ISL_418379, EPI_ISL_418380, EPI_ISL_418381, EPI_ISL_418382, EPI_ISL_418383, EPI_ISL_418384 | | | |
| see above | Public Health Ontario Laboratories | Public Health Ontario Laboratories | Alireza Eshaghi, Samir N Patel, Jonathan B Gubbay, Vanessa G Allen, Christine Frantz, Aimin Li, Sandeep Nagra |
| EPI_ISL_418385, EPI_ISL_418386, EPI_ISL_418387, EPI_ISL_418388, EPI_ISL_418389, EPI_ISL_418390, EPI_ISL_418391, EPI_ISL_418392, EPI_ISL_418393, EPI_ISL_418394, EPI_ISL_418395, EPI_ISL_418396, EPI_ISL_418397, EPI_ISL_418398, EPI_ISL_418399, EPI_ISL_418400, EPI_ISL_418401, EPI_ISL_418402, EPI_ISL_418403, EPI_ISL_418404, EPI_ISL_418405, EPI_ISL_418406, EPI_ISL_418407, EPI_ISL_418408, EPI_ISL_418409, EPI_ISL_418410, EPI_ISL_418411 | | | |
| see above | Department of Virology and Immunology, University of Helsinki and Helsinki University Hospital, Huslab Finland | Department of Virology, Faculty of Medicine, University of Helsinki, Helsinki, Finland | Teemu Smura, Hannimari Kallio-Kokko, Olli Vapalahti |
| EPI_ISL_418412 | Centre Hospitalier des Vals d'Ardeche | CNR Virus des Infections Respiratoires - France SUD | Antonin Bal, Gregory Destras, Gwendolyne Burfin, Solenne Brun, Carine Moustaud, Raphaelle Lamy, Alexandre Gaymard, Maude Bouscambert-Duchamp, Florence Morfin-Sherpa, Martine Valette, Bruno Lina, Laurence Josset |
| EPI_ISL_418413 | Centre Hospitalier de Macon | CNR Virus des Infections Respiratoires - France SUD | Antonin Bal, Gregory Destras, Gwendolyne Burfin, Solenne Brun, Carine Moustaud, Raphaelle Lamy, Alexandre Gaymard, Maude Bouscambert-Duchamp, Florence Morfin-Sherpa, Martine Valette, Bruno Lina, Laurence Josset |
| EPI_ISL_418414, EPI_ISL_418415 | Centre Hospitalier de Valence | CNR Virus des Infections Respiratoires - France SUD | Antonin Bal, Gregory Destras, Gwendolyne Burfin, Solenne Brun, Carine Moustaud, Raphaelle Lamy, Alexandre Gaymard, Maude Bouscambert-Duchamp, Florence Morfin-Sherpa, Martine Valette, Bruno Lina, Laurence Josset |
| EPI_ISL_418416 | GH Les Portes du Sud | CNR Virus des Infections Respiratoires - France SUD | Antonin Bal, Gregory Destras, Gwendolyne Burfin, Solenne Brun, Carine Moustaud, Raphaelle Lamy, Alexandre Gaymard, Maude Bouscambert-Duchamp, Florence Morfin-Sherpa, Martine Valette, Bruno Lina, Laurence Josset |
| EPI_ISL_418417 | Centre Hospitalier de Valence | CNR Virus des Infections Respiratoires - France SUD | Antonin Bal, Gregory Destras, Gwendolyne Burfin, Solenne Brun, Carine Moustaud, Raphaelle Lamy, Alexandre Gaymard, Maude Bouscambert-Duchamp, Florence Morfin-Sherpa, Martine Valette, Bruno Lina, Laurence Josset |
| EPI_ISL_418418, EPI_ISL_418419 | Centre Hospitalier Saint Joseph Saint Luc | CNR Virus des Infections Respiratoires - France SUD | Antonin Bal, Gregory Destras, Gwendolyne Burfin, Solenne Brun, Carine Moustaud, Raphaelle Lamy, Alexandre Gaymard, Maude Bouscambert-Duchamp, Florence Morfin-Sherpa, Martine Valette, Bruno Lina, Laurence Josset |
| EPI_ISL_418420, EPI_ISL_418421, EPI_ISL_418422, EPI_ISL_418423, EPI_ISL_418424, EPI_ISL_418425 | Institut des Agents Infectieux (IAI), Hospices Civils de Lyon | CNR Virus des Infections Respiratoires - France SUD | Antonin Bal, Gregory Destras, Gwendolyne Burfin, Solenne Brun, Carine Moustaud, Raphaelle Lamy, Alexandre Gaymard, Maude Bouscambert-Duchamp, Florence Morfin-Sherpa, Martine Valette, Bruno Lina, Laurence Josset |
| EPI_ISL_418426 | Centre Hospitalier de Bourg en Bresse | CNR Virus des Infections Respiratoires - France SUD | Antonin Bal, Gregory Destras, Gwendolyne Burfin, Solenne Brun, Carine Moustaud, Raphaelle Lamy, Alexandre Gaymard, Maude Bouscambert-Duchamp, Florence Morfin-Sherpa, Martine Valette, Bruno Lina, Laurence Josset |
| EPI_ISL_418427 | Hopital Privé de l'Est Lyonnais | CNR Virus des Infections Respiratoires - France SUD | Antonin Bal, Gregory Destras, Gwendolyne Burfin, Solenne Brun, Carine Moustaud, Raphaelle Lamy, Alexandre Gaymard, Maude Bouscambert-Duchamp, Florence Morfin-Sherpa, Martine Valette, Bruno Lina, Laurence Josset |
| EPI_ISL_418428 | Centre Hospitalier Lucien Hussel | CNR Virus des Infections Respiratoires - France SUD | Antonin Bal, Gregory Destras, Gwendolyne Burfin, Solenne Brun, Carine Moustaud, Raphaelle Lamy, Alexandre Gaymard, Maude Bouscambert-Duchamp, Florence Morfin-Sherpa, Martine Valette, Bruno Lina, Laurence Josset |
| EPI_ISL_418429, EPI_ISL_418430, EPI_ISL_418431, EPI_ISL_418432 | Institut des Agents Infectieux (IAI), Hospices Civils de Lyon | CNR Virus des Infections Respiratoires - France SUD | Antonin Bal, Gregory Destras, Gwendolyne Burfin, Solenne Brun, Carine Moustaud, Raphaelle Lamy, Alexandre Gaymard, Maude Bouscambert-Duchamp, Florence Morfin-Sherpa, Martine Valette, Bruno Lina, Laurence Josset |
| EPI_ISL_418433, EPI_ISL_418434, EPI_ISL_418435 | University Hospital Basel, Clinical Virology | University Hospital Basel, Clinical Bacteriology | Hirsch, H., Leuzinger, K., Seth-Smith, H., Mari, A., Roloff, T., Egli, A. |
| EPI_ISL_418436 | University Hospital Basel, Clinical Virology | University Hospital Basel, Clinical Virology | Hirsch, H., Leuzinger, K., Seth-Smith, H., Mari, A., Roloff, T., Egli, A. |
| EPI_ISL_418437, EPI_ISL_418438, EPI_ISL_418439, EPI_ISL_418440 | University Hospital Basel, Clinical Virology | University Hospital Basel, Clinical Bacteriology | Hirsch, H., Leuzinger, K., Seth-Smith, H., Mari, A., Roloff, T., Egli, A. |
| EPI_ISL_418441, EPI_ISL_418442, EPI_ISL_418502, EPI_ISL_418503, EPI_ISL_418504 | Hangzhou Center for Disease Control and Prevention | Inspection Center of Hangzhou Center for Disease Control and Prevention | Yu hua, Wang haoqiu, Li jun, Yu xinfeng, Pan jingcao |
| EPI_ISL_418506 | Hangzhou Center for Disease Control and Prevention | Inspection Center of Hanghzou Center for Disease Control and Prevention | Yu hua, Wang haoqiu, Li jun, Yu xinfeng, Pan jingcao |
| EPI_ISL_418507, EPI_ISL_418508, EPI_ISL_418509 | Hangzhou Center for Disease Control and Prevention | Inspection Center of Hangzhou Center for Disease Control and Prevention | Yu hua, Wang haoqiu, Li jun, Yu xinfeng, Pan jingcao |
| EPI_ISL_418510 | Hangzhou Center for Disease Control and Prevention | Insepction Center of Hangzhou Center for Disease Control and Prevention | Yu hua, Wang haoqiu, Li jun, Yu xinfeng, Pan jingcao |
| EPI_ISL_418511, EPI_ISL_418512, EPI_ISL_418513, EPI_ISL_418514, EPI_ISL_418515 | Hangzhou Center for Disease Control and Prevention | Inspection Center of Hangzhou Center for Disease Control and Prevention | Yu hua, Wang haoqiu, Li jun, Yu xinfeng, Pan jingcao |
| EPI_ISL_418516, EPI_ISL_418548, EPI_ISL_418580, EPI_ISL_418581, EPI_ISL_418582, EPI_ISL_418583, EPI_ISL_418584 | UCD National Virus Reference Laboratory | UCD National Virus Reference Laboratory | Michael Carr, Gabriel Gonzalez, Jonathan Dean, Suzie Coughlan, Alison Murphy, Kevin Byrne, Ken Wolfe, Jeff Connell, Brendan Loftus, Cillian F De Gascun |
| EPI_ISL_418623, EPI_ISL_418624, EPI_ISL_418625, EPI_ISL_418626, EPI_ISL_418627, EPI_ISL_418628, EPI_ISL_418629, EPI_ISL_418630, EPI_ISL_418631, EPI_ISL_418632, EPI_ISL_418633, EPI_ISL_418634, EPI_ISL_418635, EPI_ISL_418636, EPI_ISL_418637, EPI_ISL_418638, EPI_ISL_418639, EPI_ISL_418640, EPI_ISL_418641, EPI_ISL_418642, EPI_ISL_418643, EPI_ISL_418644, EPI_ISL_418645, EPI_ISL_418646, EPI_ISL_418647, EPI_ISL_418648, EPI_ISL_418649, EPI_ISL_418650, EPI_ISL_418651, EPI_ISL_418652, EPI_ISL_418653, EPI_ISL_418654, EPI_ISL_418655, EPI_ISL_418656, EPI_ISL_418657, EPI_ISL_418658, EPI_ISL_418659, EPI_ISL_418660, EPI_ISL_418661, EPI_ISL_418662, EPI_ISL_418663, EPI_ISL_418664, EPI_ISL_418665, EPI_ISL_418666 | | | |
| see above | Department of Clinical Microbiology | GIGA Medical Genomics | Keith Durkin, Maria Artesi, Sébastien Bontems, Raphaël Boreux, Cécile Meex, Pierrette Melin, Marie-Pierre Hayette, Vincent Bours. |
| EPI_ISL_418667, EPI_ISL_418668, EPI_ISL_418669, EPI_ISL_418670, EPI_ISL_418671, EPI_ISL_418672, EPI_ISL_418673, EPI_ISL_418674, EPI_ISL_418675, EPI_ISL_418676, EPI_ISL_418677, EPI_ISL_418678, EPI_ISL_418679, EPI_ISL_418680, EPI_ISL_418681, EPI_ISL_418682, EPI_ISL_418683, EPI_ISL_418684, EPI_ISL_418685, EPI_ISL_418686, EPI_ISL_418687, EPI_ISL_418688, EPI_ISL_418689, EPI_ISL_418690, EPI_ISL_418691, EPI_ISL_418692, EPI_ISL_418693, EPI_ISL_418694, EPI_ISL_418695, EPI_ISL_418696, EPI_ISL_418697, EPI_ISL_418698, EPI_ISL_418699, EPI_ISL_418700, EPI_ISL_418701, EPI_ISL_418702, EPI_ISL_418703, EPI_ISL_418704, EPI_ISL_418705, EPI_ISL_418706, EPI_ISL_418707, EPI_ISL_418708, EPI_ISL_418709, EPI_ISL_418710, EPI_ISL_418711, EPI_ISL_418712, EPI_ISL_418713, EPI_ISL_418714, EPI_ISL_418715, EPI_ISL_418716, EPI_ISL_418717, EPI_ISL_418718, EPI_ISL_418719, EPI_ISL_418720, EPI_ISL_418721, EPI_ISL_418722, EPI_ISL_418723, EPI_ISL_418724, EPI_ISL_418725, EPI_ISL_418726, EPI_ISL_418727, EPI_ISL_418728, EPI_ISL_418729, EPI_ISL_418730, EPI_ISL_418731, EPI_ISL_418732, EPI_ISL_418733, EPI_ISL_418734, EPI_ISL_418735, EPI_ISL_418736, EPI_ISL_418737, EPI_ISL_418738, EPI_ISL_418739, EPI_ISL_418740, EPI_ISL_418741, EPI_ISL_418742, EPI_ISL_418743, EPI_ISL_418744, EPI_ISL_418745, EPI_ISL_418746, EPI_ISL_418747, EPI_ISL_418748, EPI_ISL_418749, EPI_ISL_418750, EPI_ISL_418751, EPI_ISL_418752, EPI_ISL_418753, EPI_ISL_418754, EPI_ISL_418755, EPI_ISL_418756, EPI_ISL_418757, EPI_ISL_418758, EPI_ISL_418759, EPI_ISL_418760, EPI_ISL_418761, EPI_ISL_418762, EPI_ISL_418763, EPI_ISL_418764, EPI_ISL_418765, EPI_ISL_418766, EPI_ISL_418767, EPI_ISL_418768, EPI_ISL_418769, EPI_ISL_418770 | | | |
| see above | Respiratory Virus Unit, Microbiology Services Colindale, Public Health England | Respiratory Virus Unit, Microbiology Services Colindale, Public Health England | Monica Galiano, Shahjahan Miah, Angie Lackenby, Omolola Akinbami, Tiina Talts, Leena Bhaw, Richard Myers, Steven Platt, Kirstin Edwards, Jonathan Hubb, Joanna Ellis, Maria Zambon |
| EPI_ISL_418771, EPI_ISL_418772, EPI_ISL_418773, EPI_ISL_418774, EPI_ISL_418775, EPI_ISL_418776, EPI_ISL_418777 | WA State Department of Health | Pathogen Discovery, Respiratory Viruses Branch, Division of Viral Diseases, Centers for Disease Control and Prevention | Jing Zhang, Ying Tao, Clinton R. Paden, Krista Queen, Anna Uehara, Yan Li, Haibin Wang, Jesica Jacobs, Denny Russell, Brian Hiatt, Jessica Gant, Suxiang Tong |
| EPI_ISL_418778, EPI_ISL_418779, EPI_ISL_418780, EPI_ISL_418781, EPI_ISL_418782, EPI_ISL_418783, EPI_ISL_418784, EPI_ISL_418785, EPI_ISL_418786, EPI_ISL_418787, EPI_ISL_418788, EPI_ISL_418789, EPI_ISL_418790, EPI_ISL_418791 | | | |
| see above | WA State Department of Health | Pathogen Discovery, Respiratory Viruses Branch, Division of Viral Diseases, Centers for Disease Control and Prevention | Ying Tao, Jing Zhang, Clinton R. Paden, Krista Queen, Anna Uehara, Yan Li, Haibin Wang, Jesica Jacobs, Denny Russell, Brian Hiatt, Jessica Gant, Suxiang Tong |

| | | | |
|---|---|---|---|
| EPI_ISL_418792, EPI_ISL_418793 | KU Leuven, Clinical and Epidemiological Virology | KU Leuven, Clinical and Epidemiological Virology | Bert Vanmechelen, Tony Wawina, Joan Marti-Carreras, Piet Maes |
| EPI_ISL_418794, EPI_ISL_418795, EPI_ISL_418796, EPI_ISL_418797, EPI_ISL_418798 | KU Leuven, Clinical and Epidemiological Virology | KU Leuven, Clinical and Epidemiological Virology | Bert Vanmechelen, Joan Marti-Carreras, Tony Wawina, Piet Maes |
| EPI_ISL_418799 | Mater Pathology | Public Health Virology Laboratory | Bixing Huang, Alyssa Pyke, Amanda De Jong, Andrew Van Den Hurk, Carmel Taylor, David Warrilow, Doris Genge, Elisabeth Gamez, Glen Hewitson, Ian Maxwell Mackay, Inga Sultana, Jamie McMahon, Jean Barcelon, Judy Northill, Mitchell Finger, Natalie Simpson, Neelima Nair, Peter Burtonclay, Peter Moore, Sarah Wheatley, Sean Moody, Sonja Hall-Mendelin, Timothy Gardam, and Frederick Moore |
| EPI_ISL_418800 | KU Leuven, Clinical and Epidemiological Virology | KU Leuven, Clinical and Epidemiological Virology | Bert Vanmechelen, Joan Marti-Carreras, Tony Wawina, Piet Maes |
| EPI_ISL_418801 | Mater Pathology | Public Health Virology Laboratory | Bixing Huang, Alyssa Pyke, Amanda De Jong, Andrew Van Den Hurk, Carmel Taylor, David Warrilow, Doris Genge, Elisabeth Gamez, Glen Hewitson, Ian Maxwell Mackay, Inga Sultana, Jamie McMahon, Jean Barcelon, Judy Northill, Mitchell Finger, Natalie Simpson, Neelima Nair, Peter Burtonclay, Peter Moore, Sarah Wheatley, Sean Moody, Sonja Hall-Mendelin, Timothy Gardam, and Frederick Moore |
| EPI_ISL_418802, EPI_ISL_418803, EPI_ISL_418804 | Pathology Queensland | Public Health Virology Laboratory | Bixing Huang, Alyssa Pyke, Amanda De Jong, Andrew Van Den Hurk, Carmel Taylor, David Warrilow, Doris Genge, Elisabeth Gamez, Glen Hewitson, Ian Maxwell Mackay, Inga Sultana, Jamie McMahon, Jean Barcelon, Judy Northill, Mitchell Finger, Natalie Simpson, Neelima Nair, Peter Burtonclay, Peter Moore, Sarah Wheatley, Sean Moody, Sonja Hall-Mendelin, Timothy Gardam, and Frederick Moore |
| EPI_ISL_418805, EPI_ISL_418806 | KU Leuven, Clinical and Epidemiological Virology | KU Leuven, Clinical and Epidemiological Virology | Bert Vanmechelen, Joan Marti-Carreras, Tony Wawina, Piet Maes |
| EPI_ISL_418807, EPI_ISL_418808 | Pathology Queensland | Public Health Virology Laboratory | Bixing Huang, Alyssa Pyke, Amanda De Jong, Andrew Van Den Hurk, Carmel Taylor, David Warrilow, Doris Genge, Elisabeth Gamez, Glen Hewitson, Ian Maxwell Mackay, Inga Sultana, Jamie McMahon, Jean Barcelon, Judy Northill, Mitchell Finger, Natalie Simpson, Neelima Nair, Peter Burtonclay, Peter Moore, Sarah Wheatley, Sean Moody, Sonja Hall-Mendelin, Timothy Gardam, and Frederick Moore |
| EPI_ISL_418809 | University of Wisconsin - Madison: Influenza Research Institute | University of Wisconsin Madison, AIDS Vaccine Research Laboratories | Katarina Braun, Gage Moreno, Peter Halfmann, et al. |
| EPI_ISL_418811 | Dr. Georges-L.-Dumont University Hospital Centre | National Microbiology Laboratory | Anna Majer, Shari Tyson, Grace Seo, Philip Mabon, Natalie Knox, Morag Graham, Richard Garceau, Guillaume Desnoyers, Nathalie Bastien, Yan Li, Matthew Gilmour, Timothy Booth |
| EPI_ISL_418812, EPI_ISL_418813 | Cadham Provincial Laboratory | National Microbiology Laboratory | Anna Majer, Shari Tyson, Grace Seo, Philip Mabon, Natalie Knox, Morag Graham, Paul Van Caeseele, Jared Bullard, David Alexander, Kerry Dust, Nathalie Bastien, Yan Li, Matthew Gilmour, Timothy Booth |
| EPI_ISL_418814 | Queen Elizabeth II Health Science Centre | National Microbiology Laboratory | Anna Majer, Shari Tyson, Grace Seo, Philip Mabon, Natalie Knox, Morag Graham, Todd Hatchette, Jason LeBlanc, Nathalie Bastien, Yan Li, Matthew Gilmour, Timothy Booth |
| EPI_ISL_418815 | Department of Clinical Pathology, Pamela Youde Nethersole Eastern Hospital | Department of Health Technology and Informatics, Faculty of Health and Social Science, The Hong Kong Polytechnic University | Kenneth Siu-Sing LEUNG, Timothy Ting-Leung NG, Alan Ka-Lun WU, Miranda Chong-Yee YAU, Hiu-Yin LAO, Ming-Pan CHOI, Kingsley King-Gee TAM, Lam-Kwong LEE, Barry Kin-Chung WONG, Alex Yat-Man HO, Kam-Tong YIP, Kwok-Cheung LUNG, Raymond Wai-To LIU, Eugene Yuk-Keung TSO, Wai-Shing LEUNG, Man-Chun CHAN, Yuk-Yung NG, Kit-Man SIN, Kitty Sau-Chun FUNG, Sandy Ka-Yee CHAU, Wing-Kin TO, Tak-Lun QUE, David Ho-Keung SHUM, Shea Ping YIP, Wing Cheong YAM, Gilman Kit-Hang SIU |
| EPI_ISL_418816, EPI_ISL_418817, EPI_ISL_418818, EPI_ISL_418819, EPI_ISL_418820, EPI_ISL_418821, EPI_ISL_418822, EPI_ISL_418823, EPI_ISL_418824, EPI_ISL_418825, EPI_ISL_418826, EPI_ISL_418827, EPI_ISL_418828, EPI_ISL_418829, EPI_ISL_418830, EPI_ISL_418831, EPI_ISL_418832, EPI_ISL_418833, EPI_ISL_418834, EPI_ISL_418835, EPI_ISL_418836, EPI_ISL_418837, EPI_ISL_418838, EPI_ISL_418839, EPI_ISL_418840, EPI_ISL_418841, EPI_ISL_418842, EPI_ISL_418843, EPI_ISL_418844, EPI_ISL_418845, EPI_ISL_418846, EPI_ISL_418847, EPI_ISL_418848, EPI_ISL_418849, EPI_ISL_418850, EPI_ISL_418851, EPI_ISL_418852, EPI_ISL_418853, EPI_ISL_418854, EPI_ISL_418855, EPI_ISL_418856, EPI_ISL_418857, EPI_ISL_418858, EPI_ISL_418859 | | | |
| see above | BCCDC Public Health Laboratory | BCCDC Public Health Laboratory | Harrigan, Prystajecky, Krajden, Lee, Kamelian, Lapointe, Choi, Hoang, Sekirov, Levett, Tyson, Snutch, Loman, Quick, Li, Gilmour |
| EPI_ISL_418860, EPI_ISL_418861 | Hospital Universitari Vall d'Hebron (HUVH) - Vall d'Hebron Research Institute (VHIR) | Hospital Universitari Vall d'Hebron (HUVH) - Vall d'Hebron Research Institute (VHIR) | Cristina Andrés, Dàmir Garcia-Cehic, Maria Piñana, Mercedes Guerrero-Murillo, Ariadna Rando, Tomàs Pumarola, Maria Gema Codina, Andrés Antón, Josep Quer |
| EPI_ISL_418863 | KU Leuven, Clinical and Epidemiological Virology | KU Leuven, Clinical and Epidemiological Virology | Bert Vanmechelen, Joan Marti-Carreras, Tony Wawina, Piet Maes |
| EPI_ISL_418864 | Virginia DCLS | Virginia DCLS | Virginia DCLS |
| EPI_ISL_418865 | California Department of Public Health | University of California, San Francisco | Xianding Deng, Scot Federman, Chao-Yang Pan, Hugo Guevara,Wei Gu, Debra A. Wadford, and Charles Y. Chiu |
| EPI_ISL_418866, EPI_ISL_418867, EPI_ISL_418868, EPI_ISL_418869, EPI_ISL_418870, EPI_ISL_418871, EPI_ISL_418872, EPI_ISL_418873, EPI_ISL_418874, EPI_ISL_418875, EPI_ISL_418876, EPI_ISL_418877, EPI_ISL_418878, EPI_ISL_418879, EPI_ISL_418880, EPI_ISL_418881, EPI_ISL_418882, EPI_ISL_418883, EPI_ISL_418884, EPI_ISL_418885, EPI_ISL_418886, EPI_ISL_418887, EPI_ISL_418888, EPI_ISL_418889, EPI_ISL_418890, EPI_ISL_418891, EPI_ISL_418892, EPI_ISL_418893, EPI_ISL_418894, EPI_ISL_418895, EPI_ISL_418896, EPI_ISL_418897, EPI_ISL_418898, EPI_ISL_418899, EPI_ISL_418900, EPI_ISL_418901, EPI_ISL_418902, EPI_ISL_418903, EPI_ISL_418904, EPI_ISL_418905, EPI_ISL_418906, EPI_ISL_418907, EPI_ISL_418908, EPI_ISL_418909, EPI_ISL_418910, EPI_ISL_418911, EPI_ISL_418912, EPI_ISL_418913, EPI_ISL_418914, EPI_ISL_418915, EPI_ISL_418916, EPI_ISL_418917, EPI_ISL_418918, EPI_ISL_418919, EPI_ISL_418920, EPI_ISL_418921, EPI_ISL_418922, EPI_ISL_418923, EPI_ISL_418924, EPI_ISL_418925, EPI_ISL_418926, EPI_ISL_418927, EPI_ISL_418928, EPI_ISL_418929, EPI_ISL_418930, EPI_ISL_418931, EPI_ISL_418932, EPI_ISL_418933, EPI_ISL_418934, EPI_ISL_418935, EPI_ISL_418936, EPI_ISL_418937, EPI_ISL_418938, EPI_ISL_418939, EPI_ISL_418940, EPI_ISL_418941, EPI_ISL_418942, EPI_ISL_418943, EPI_ISL_418944, EPI_ISL_418945, EPI_ISL_418946, EPI_ISL_418947, EPI_ISL_418948, EPI_ISL_418949, EPI_ISL_418950, EPI_ISL_418951, EPI_ISL_418952, EPI_ISL_418953, EPI_ISL_418954, EPI_ISL_418955 | | | |
| see above | UW Virology Lab | UW Virology Lab | Pavitra Roychoudhury, Hong Xie, Keith Jerome, Alexander Greninger |
| EPI_ISL_418956, EPI_ISL_418957, EPI_ISL_418958 | Virginia DCLS | Virginia DCLS | Virginia DCLS |
| EPI_ISL_418959 | Universidade Federal do Rio de Janeiro - UFRJ | Bioinformatics Laboratory - LNCC | Filipe Romero, Ana Paula Guimarães, Mariane Talon, Luiz Gonzaga Paula de Almeida, Ronaldo da Silva Francisco Junior, Diana Mariani, Lídia Boullosa, Alexandra Gerber, Jaqueline Goes de Jesus, Ingra Morales Claro, Ester Cerdeira Sabino, Nuno Rodrigues Faria, Terezinha Marta Pereira, Pinto Castiñeiras, Isabela de Carvalho Leitão, Rafael de Mello Galliez, Cássia Cristina Alves Gonçalves, Érica Ramos dos Santos Nascimento, Richard Araújo Maia, Mauro Teixeira,Cristiano Xavier Lima, Orlando Ferreira Jr., Rodrigo Brindeiro, Luciana Jesus Costa e André Felipe Santos, Laboratorio Hermes Pardini, Laboratorio Simile, Amilcar Tanuri, Renato Santana Aguiar, e Ana Tereza Vasconcelos |
| EPI_ISL_418961, EPI_ISL_418962, EPI_ISL_418963, EPI_ISL_418964, EPI_ISL_418965, EPI_ISL_418966, EPI_ISL_418967 | Utah Public Health Laboratory | Utah Public Health Laboratory | Erin Young, Kelly Oakeson |
| EPI_ISL_418968, EPI_ISL_418969, EPI_ISL_418970, EPI_ISL_418971, EPI_ISL_418972, EPI_ISL_418973, EPI_ISL_418974, EPI_ISL_418975, EPI_ISL_418976, EPI_ISL_418977, EPI_ISL_418978, EPI_ISL_418979, EPI_ISL_418980 | | | |
| see above | NYU Langone Health | Department of Pathology and Medicine, New York University School of Medicine | Maria Aguero-Rosenfeld, Margaret Black, John Cadley, Paolo Cotzia, John Chen, Dacia Dimartino, Xiaojun Feng, Adriana Heguy, Megan Hogan, Emily Huang, George Jour, Christian Marier, Matthew T. Maurano, Mark J. Mulligan, Peter Meyn, Jared Pinnell, Sitharam Ramaswami, Amy Rapkiewicz, Marie Samanovic-Golden, Antonio Serrano, Guomiao Shen, Matija Snuderl, Nick Vulpescu, Gael Westby, Paul Zappile, Yutong Zhang |
| EPI_ISL_418981, EPI_ISL_418982, EPI_ISL_418983, EPI_ISL_418984, EPI_ISL_418985, EPI_ISL_418986, EPI_ISL_418987 | KU Leuven, Clinical and Epidemiological Virology | KU Leuven, Clinical and Epidemiological Virology | Bert Vanmechelen, Joan Marti-Carreras, Tony Wawina, Piet Maes |
| EPI_ISL_418988 | Institute information KU Leuven, Clinical and Epidemiological Virology | Institute information KU Leuven, Clinical and Epidemiological Virology | Bert Vanmechelen, Joan Marti-Carreras, Tony Wawina, Piet Maes |
| EPI_ISL_418989 | KU Leuven, Clinical and Epidemiological Virology | KU Leuven, Clinical and Epidemiological Virology | Bert Vanmechelen, Joan Marti-Carreras, Tony Wawina, Piet Maes |
| EPI_ISL_418990, EPI_ISL_418991 | State Key Laboratory for Diagnosis and Treatment of Infectious Diseases, National Clinical Research Center for Infectious Diseases, First Affiliated Hospital, Zhejiang University School of Medicine, Hangzhou, China 310003 | State Key Laboratory for Diagnosis and Treatment of Infectious Diseases, National Clinical Research Center for Infectious Diseases, First Affiliated Hospital, Zhejiang University School of Medicine, Hangzhou, China 310003 | Hangping Yao, Nanping Wu, Chao Jiang, Xiangyun Lu, Linfang Cheng, Fumin Liu, Zhigang Wu, Haibo Wu, Changzhong Jin, Min Zheng, Lanjuan Li |
| EPI_ISL_418992, EPI_ISL_418993, EPI_ISL_418994, EPI_ISL_418995, EPI_ISL_418996, EPI_ISL_418997, EPI_ISL_418998, EPI_ISL_418999, EPI_ISL_419000, EPI_ISL_419001 | National Public Health Laboratory, National Centre for Infectious Diseases | National Public Health Laboratory, National Centre for Infectious Diseases | Mak TM, Octavia S, Cui L, Lin RTP |
| EPI_ISL_419168 | Centre Hospitalier de Valence | CNR Virus des Infections Respiratoires - France SUD | Antonin Bal, Gregory Destras, Gwendolyne Burfin, Solenne Brun, Carine Moustaud, Raphaelle Lamy, Alexandre Gaymard, Maude Bouscambert-Duchamp, Florence Morfin-Sherpa, Martine Valette, Bruno Lina, Laurence Josset |
| EPI_ISL_419169, EPI_ISL_419170, EPI_ISL_419171, EPI_ISL_419172, EPI_ISL_419173 | Institut des Agents Infectieux (IAI), Hospices Civils de Lyon | CNR Virus des Infections Respiratoires - France SUD | Antonin Bal, Gregory Destras, Gwendolyne Burfin, Solenne Brun, Carine Moustaud, Raphaelle Lamy, Alexandre Gaymard, Maude Bouscambert-Duchamp, Florence Morfin-Sherpa, Martine Valette, Bruno Lina, Laurence Josset |
| EPI_ISL_419174, EPI_ISL_419175, EPI_ISL_419176 | Centre Hospitalier de Macon | CNR Virus des Infections Respiratoires - France SUD | Antonin Bal, Gregory Destras, Gwendolyne Burfin, Solenne Brun, Carine Moustaud, Raphaelle Lamy, Alexandre Gaymard, Maude Bouscambert-Duchamp, Florence Morfin-Sherpa, Martine Valette, Bruno Lina, Laurence Josset |
| EPI_ISL_419177, EPI_ISL_419178, EPI_ISL_419179, EPI_ISL_419180, EPI_ISL_419181, EPI_ISL_419182 | Institut des Agents Infectieux (IAI), Hospices Civils de Lyon | CNR Virus des Infections Respiratoires - France SUD | Antonin Bal, Gregory Destras, Gwendolyne Burfin, Solenne Brun, Carine Moustaud, Raphaelle Lamy, Alexandre Gaymard, Maude Bouscambert-Duchamp, Florence Morfin-Sherpa, Martine Valette, Bruno Lina, Laurence Josset |
| EPI_ISL_419183 | Centre Hospitalier de Bourg en Bresse | CNR Virus des Infections Respiratoires - France SUD | Antonin Bal, Gregory Destras, Gwendolyne Burfin, Solenne Brun, Carine Moustaud, Raphaelle Lamy, Alexandre Gaymard, Maude Bouscambert-Duchamp, Florence Morfin-Sherpa, Martine Valette, Bruno Lina, Laurence Josset |
| EPI_ISL_419184 | Institut des Agents Infectieux (IAI), Hospices Civils de Lyon | CNR Virus des Infections Respiratoires - France SUD | Antonin Bal, Gregory Destras, Gwendolyne Burfin, Solenne Brun, Carine Moustaud, Raphaelle Lamy, Alexandre Gaymard, Maude Bouscambert-Duchamp, Florence Morfin-Sherpa, Martine Valette, Bruno Lina, Laurence Josset |
| EPI_ISL_419185, EPI_ISL_419186 | Centre Hospitalier de Bourg en Bresse | CNR Virus des Infections Respiratoires - France SUD | Antonin Bal, Gregory Destras, Gwendolyne Burfin, Solenne Brun, Carine Moustaud, Raphaelle Lamy, Alexandre Gaymard, Maude Bouscambert-Duchamp, Florence Morfin-Sherpa, Martine Valette, Bruno Lina, Laurence Josset |
| EPI_ISL_419187, EPI_ISL_419188 | Centre Hospitalier de Macon | CNR Virus des Infections Respiratoires - France SUD | Antonin Bal, Gregory Destras, Gwendolyne Burfin, Solenne Brun, Carine Moustaud, Raphaelle Lamy, Alexandre Gaymard, Maude Bouscambert-Duchamp, Florence Morfin-Sherpa, Martine Valette, Bruno Lina, Laurence Josset |
| EPI_ISL_419210 | The Chaim Sheba Medical Center | Israel Institute for Biological Research | Inbar Cohen-Gihon, Ofir Israeli, Ohad Shifman, Dana Stein, Sharon Melamed, Nir Paran, Tomer Israely, Hagit Achdout, Yfat Yahalom Ronen, Hadas Tamir, Boaz Politi, Lilach Cherry, Einat Vitner, Orly Laskar, Shay Weiss, Michal Mandelboim, Oran Erster, Gili Regev-Yochay, Gadi Segal, Shmuel Yitzhaki, Shmuel C. Shapira, Adi Beth-Din, Anat Zvi |
| EPI_ISL_419211 | Central Virology Laboratory | Israel Institute for Biological Research | Inbar Cohen-Gihon, Ofir Israeli, Ohad Shifman, Dana Stein, Sharon Melamed, Nir Paran, Tomer Israely, Hagit Achdout, Yfat Yahalom Ronen, Hadas Tamir, Boaz Politi, Lilach Cherry, Einat Vitner, Orly Laskar, Shay Weiss, Michal Mandelboim, Oran Erster, Gili Regev-Yochay, Gadi Segal, Shmuel Yitzhaki, Shmuel C. Shapira, Adi Beth-Din, Anat Zvi |
| EPI_ISL_419213 | Department of Clinical Pathology, Pamela Youde Nethersole Eastern Hospital | Department of Health Technology and Informatics, Faculty of Health and Social Science, The Hong Kong Polytechnic University | Kenneth Siu-Sing LEUNG, Timothy Ting-Leung NG, Alan Ka-Lun WU, Miranda Chong-Yee YAU, Hiu-Yin LAO, Ming-Pan CHOI, Kingsley King-Gee TAM, Lam-Kwong LEE, Barry Kin-Chung WONG, Alex Yat-Man HO, Kam-Tong YIP, Kwok-Cheung LUNG, Raymond Wai-To LIU, Eugene Yuk-Keung TSO, Wai-Shing LEUNG, Man-Chun CHAN, Yuk-Yung NG, Kit-Man SIN, Kitty Sau-Chun FUNG, Sandy Ka-Yee CHAU, Wing-Kin TO, Tak-Lun QUE, David Ho-Keung SHUM, Shea Ping YIP, Wing Cheong YAM, Gilman Kit-Hang SIU |
| EPI_ISL_419214, EPI_ISL_419215, EPI_ISL_419216 | Department of Clinical Pathology, Pamela Youde Nethersole Eastern Hospital | Department of Health Technology and Informatics, Faculty of Health and Social Science, The Hong Kong Polytechnic University | Kenneth Siu-Sing LEUNG, Timothy Ting-Leung NG, Alan Ka-Lun WU, Miranda Chong-Yee YAU, Hiu-Yin LAO, Ming-Pan CHOI, Kingsley King-Gee TAM, Lam-Kwong LEE, Barry Kin-Chung WONG, Alex Yat-Man HO, Kam-Tong YIP, Kwok-Cheung LUNG, Raymond Wai-To LIU, Eugene Yuk-Keung TSO, Wai-Shing LEUNG, Man-Chun CHAN, Yuk-Yung NG, Kit-Man SIN, Kitty Sau-Chun FUNG, Sandy Ka-Yee CHAU, Wing-Kin TO, Tak-Lun QUE, David Ho-Keung SHUM, Shea Ping YIP, Wing Cheong YAM, Gilman Kit-Hang SIU |
| EPI_ISL_419217 | Department of Pathology, Princess Margaret Hospital | Department of Health Technology and Informatics, Faculty of | Kenneth Siu-Sing LEUNG, Timothy Ting-Leung NG, Alan Ka-Lun WU, Miranda Chong-Yee YAU, Hiu-Yin LAO, Ming-Pan CHOI, Kingsley King-Gee TAM, Lam-Kwong LEE, Barry Kin-Chung WONG, Alex Yat-Man HO, Kam-Tong YIP, Kwok-Cheung LUNG, |

| | | | |
|---|---|---|---|
| | | Health and Social Science, The Hong Kong Polytechnic University | Raymond Wai-To LIU, Eugene Yuk-Keung TSO, Wai-Shing LEUNG, Man-Chun CHAN, Yuk-Yung NG, Kit-Man SIN, Kitty Sau-Chun FUNG, Sandy Ka-Yee CHAU, Wing-Kin TO, Tak-Lun QUE, David Ho-Keung SHUM, Shea Ping YIP, Wing Cheong YAM, Gilman Kit-Hang SIU |
| EPI_ISL_419218, EPI_ISL_419219 | Department of Clinical Pathology, Pamela Youde Nethersole Eastern Hospital | Department of Health Technology and Informatics, Faculty of Health and Social Science, The Hong Kong Polytechnic University | Kenneth Siu-Sing LEUNG, Timothy Ting-Leung NG, Alan Ka-Lun WU, Miranda Chong-Yee YAU, Hiu-Yin LAO, Ming-Pan CHOI, Kingsley King-Gee TAM, Lam-Kwong LEE, Barry Kin-Chung WONG, Alex Yat-Man HO, Kam-Tong YIP, Kwok-Cheung LUNG, Raymond Wai-To LIU, Eugene Yuk-Keung TSO, Wai-Shing LEUNG, Man-Chun CHAN, Yuk-Yung NG, Kit-Man SIN, Kitty Sau-Chun FUNG, Sandy Ka-Yee CHAU, Wing-Kin TO, Tak-Lun QUE, David Ho-Keung SHUM, Shea Ping YIP, Wing Cheong YAM, Gilman Kit-Hang SIU |
| EPI_ISL_419221 | Department of Pathology, United Christian Hospital | Department of Health Technology and Informatics, Faculty of Health and Social Science, The Hong Kong Polytechnic University | Kenneth Siu-Sing LEUNG, Timothy Ting-Leung NG, Alan Ka-Lun WU, Miranda Chong-Yee YAU, Hiu-Yin LAO, Ming-Pan CHOI, Kingsley King-Gee TAM, Lam-Kwong LEE, Barry Kin-Chung WONG, Alex Yat-Man HO, Kam-Tong YIP, Kwok-Cheung LUNG, Raymond Wai-To LIU, Eugene Yuk-Keung TSO, Wai-Shing LEUNG, Man-Chun CHAN, Yuk-Yung NG, Kit-Man SIN, Kitty Sau-Chun FUNG, Sandy Ka-Yee CHAU, Wing-Kin TO, Tak-Lun QUE, David Ho-Keung SHUM, Shea Ping YIP, Wing Cheong YAM, Gilman Kit-Hang SIU |
| EPI_ISL_419222, EPI_ISL_419223 | Department of Pathology, Princess Margaret Hospital | Department of Health Technology and Informatics, Faculty of Health and Social Science, The Hong Kong Polytechnic University | Kenneth Siu-Sing LEUNG, Timothy Ting-Leung NG, Alan Ka-Lun WU, Miranda Chong-Yee YAU, Hiu-Yin LAO, Ming-Pan CHOI, Kingsley King-Gee TAM, Lam-Kwong LEE, Barry Kin-Chung WONG, Alex Yat-Man HO, Kam-Tong YIP, Kwok-Cheung LUNG, Raymond Wai-To LIU, Eugene Yuk-Keung TSO, Wai-Shing LEUNG, Man-Chun CHAN, Yuk-Yung NG, Kit-Man SIN, Kitty Sau-Chun FUNG, Sandy Ka-Yee CHAU, Wing-Kin TO, Tak-Lun QUE, David Ho-Keung SHUM, Shea Ping YIP, Wing Cheong YAM, Gilman Kit-Hang SIU |
| EPI_ISL_419224, EPI_ISL_419225, EPI_ISL_419226, EPI_ISL_419227, EPI_ISL_419228, EPI_ISL_419229 | Department of Clinical Pathology, Pamela Youde Nethersole Eastern Hospital | Department of Health Technology and Informatics, Faculty of Health and Social Science, The Hong Kong Polytechnic University | Kenneth Siu-Sing LEUNG, Timothy Ting-Leung NG, Alan Ka-Lun WU, Miranda Chong-Yee YAU, Hiu-Yin LAO, Ming-Pan CHOI, Kingsley King-Gee TAM, Lam-Kwong LEE, Barry Kin-Chung WONG, Alex Yat-Man HO, Kam-Tong YIP, Kwok-Cheung LUNG, Raymond Wai-To LIU, Eugene Yuk-Keung TSO, Wai-Shing LEUNG, Man-Chun CHAN, Yuk-Yung NG, Kit-Man SIN, Kitty Sau-Chun FUNG, Sandy Ka-Yee CHAU, Wing-Kin TO, Tak-Lun QUE, David Ho-Keung SHUM, Shea Ping YIP, Wing Cheong YAM, Gilman Kit-Hang SIU |
| EPI_ISL_419230 | Hospital Universitario Virgen de las Nieves | Instituto de Salud Carlos III | Iglesias-Caballero, M.; Molinero Calamita, M.; González-Esguevillas, M.; Camarero, S.; Pozo, F.; Casas, I.; Jiménez, P.; Jiménez, M.; Zaballos, A.; Monzón, S.; Varona, S.; Juliá, M.; Cuesta, I.; Sanbonmatsu, S. |
| EPI_ISL_419231 | Department of Clinical Pathology, Tuen Mun Hospital, 23 Tsing Chung Koon Road, Tuen Mun, N.T. | Department of Health Technology and Informatics, Faculty of Health and Social Science, The Hong Kong Polytechnic University | Kenneth Siu-Sing LEUNG, Timothy Ting-Leung NG, Alan Ka-Lun WU, Miranda Chong-Yee YAU, Hiu-Yin LAO, Ming-Pan CHOI, Kingsley King-Gee TAM, Lam-Kwong LEE, Barry Kin-Chung WONG, Alex Yat-Man HO, Kam-Tong YIP, Kwok-Cheung LUNG, Raymond Wai-To LIU, Eugene Yuk-Keung TSO, Wai-Shing LEUNG, Man-Chun CHAN, Yuk-Yung NG, Kit-Man SIN, Kitty Sau-Chun FUNG, Sandy Ka-Yee CHAU, Wing-Kin TO, Tak-Lun QUE, David Ho-Keung SHUM, Shea Ping YIP, Wing Cheong YAM, Gilman Kit-Hang SIU |
| EPI_ISL_419232 | Department of Clinical Pathology, Pamela Youde Nethersole Eastern Hospital | Department of Health Technology and Informatics, Faculty of Health and Social Science, The Hong Kong Polytechnic University | Kenneth Siu-Sing LEUNG, Timothy Ting-Leung NG, Alan Ka-Lun WU, Miranda Chong-Yee YAU, Hiu-Yin LAO, Ming-Pan CHOI, Kingsley King-Gee TAM, Lam-Kwong LEE, Barry Kin-Chung WONG, Alex Yat-Man HO, Kam-Tong YIP, Kwok-Cheung LUNG, Raymond Wai-To LIU, Eugene Yuk-Keung TSO, Wai-Shing LEUNG, Man-Chun CHAN, Yuk-Yung NG, Kit-Man SIN, Kitty Sau-Chun FUNG, Sandy Ka-Yee CHAU, Wing-Kin TO, Tak-Lun QUE, David Ho-Keung SHUM, Shea Ping YIP, Wing Cheong YAM, Gilman Kit-Hang SIU |
| EPI_ISL_419233 | Hospital Universitario de Canarias | Instituto de Salud Carlos III | Iglesias-Caballero, M.; Molinero Calamita, M.; González-Esguevillas, M.; Camarero, S.; Pozo, F.; Casas, I.; Jiménez, P.; Jiménez, M.; Zaballos, A.; Monzón, S.; Varona, S.; Juliá, M.; Cuesta, I.; Castro, B. |
| EPI_ISL_419234 | Hospital San Pedro | Instituto de Salud Carlos III | Iglesias-Caballero, M.; Molinero Calamita, M.; González-Esguevillas, M.; Camarero, S.; Pozo, F.; Casas, I.; Jiménez, P.; Jiménez, M.; Zaballos, A.; Monzón, S.; Varona, S.; Juliá, M.; Cuesta, I.; Alonso, C. |
| EPI_ISL_419235, EPI_ISL_419236, EPI_ISL_419237 | Fundacion Jimenez Diaz | Instituto de Salud Carlos III | Iglesias-Caballero, M.; Molinero Calamita, M.; González-Esguevillas, M.; Camarero, S.; Pozo, F.; Casas, I.; Jiménez, P.; Jiménez, M.; Zaballos, A.; Monzón, S.; Varona, S.; Juliá, M.; Cuesta, I.; Fernández, R. |
| EPI_ISL_419238 | HOSPITAL DE CRUCES. | Instituto de Salud Carlos III | Iglesias-Caballero, M. Molinero Calamita, M. González-Esguevillas, M. Camarero, S. Pozo, F. Casas, I. Jiménez, P. Jiménez, M. Zaballos, A. Monzón, S. Varona, S. Juliá, M. Cuesta, I. Aranzamendi, M. |
| EPI_ISL_419240 | HOSPITAL TXAGORRITXU | Instituto de Salud Carlos III | Iglesias-Caballero, M. Molinero Calamita, M. González-Esguevillas, M. Camarero, S. Pozo, F. Casas, I. Jiménez, P. Jiménez, M. Zaballos, A. Monzón, S. Varona, S. Juliá, M. Cuesta, I. Gómez, C |
| EPI_ISL_419241 | Department of Clinical Pathology, Pamela Youde Nethersole Eastern Hospital | Department of Health Technology and Informatics, Faculty of Health and Social Science, The Hong Kong Polytechnic University | Kenneth Siu-Sing LEUNG, Timothy Ting-Leung NG, Alan Ka-Lun WU, Miranda Chong-Yee YAU, Hiu-Yin LAO, Ming-Pan CHOI, Kingsley King-Gee TAM, Lam-Kwong LEE, Barry Kin-Chung WONG, Alex Yat-Man HO, Kam-Tong YIP, Kwok-Cheung LUNG, Raymond Wai-To LIU, Eugene Yuk-Keung TSO, Wai-Shing LEUNG, Man-Chun CHAN, Yuk-Yung NG, Kit-Man SIN, Kitty Sau-Chun FUNG, Sandy Ka-Yee CHAU, Wing-Kin TO, Tak-Lun QUE, David Ho-Keung SHUM, Shea Ping YIP, Wing Cheong YAM, Gilman Kit-Hang SIU |
| EPI_ISL_419242 | Department of Clinical Pathology, Tuen Mun Hospital | Department of Health Technology and Informatics, Faculty of Health and Social Science, The Hong Kong Polytechnic University | Kenneth Siu-Sing LEUNG, Timothy Ting-Leung NG, Alan Ka-Lun WU, Miranda Chong-Yee YAU, Hiu-Yin LAO, Ming-Pan CHOI, Kingsley King-Gee TAM, Lam-Kwong LEE, Barry Kin-Chung WONG, Alex Yat-Man HO, Kam-Tong YIP, Kwok-Cheung LUNG, Raymond Wai-To LIU, Eugene Yuk-Keung TSO, Wai-Shing LEUNG, Man-Chun CHAN, Yuk-Yung NG, Kit-Man SIN, Kitty Sau-Chun FUNG, Sandy Ka-Yee CHAU, Wing-Kin TO, Tak-Lun QUE, David Ho-Keung SHUM, Shea Ping YIP, Wing Cheong YAM, Gilman Kit-Hang SIU |
| EPI_ISL_419243, EPI_ISL_419244, EPI_ISL_419245, EPI_ISL_419246, EPI_ISL_419247, EPI_ISL_419248, EPI_ISL_419249, EPI_ISL_419250, EPI_ISL_419251, EPI_ISL_419252 | Department of Clinical Pathology, Pamela Youde Nethersole Eastern Hospital | Department of Health Technology and Informatics, Faculty of Health and Social Science, The Hong Kong Polytechnic University | Kenneth Siu-Sing LEUNG, Timothy Ting-Leung NG, Alan Ka-Lun WU, Miranda Chong-Yee YAU, Hiu-Yin LAO, Ming-Pan CHOI, Kingsley King-Gee TAM, Lam-Kwong LEE, Barry Kin-Chung WONG, Alex Yat-Man HO, Kam-Tong YIP, Kwok-Cheung LUNG, Raymond Wai-To LIU, Eugene Yuk-Keung TSO, Wai-Shing LEUNG, Man-Chun CHAN, Yuk-Yung NG, Kit-Man SIN, Kitty Sau-Chun FUNG, Sandy Ka-Yee CHAU, Wing-Kin TO, Tak-Lun QUE, David Ho-Keung SHUM, Shea Ping YIP, Wing Cheong YAM, Gilman Kit-Hang SIU |
| EPI_ISL_419253 | Department of Pathology, United Christian Hospital | Department of Health Technology and Informatics, Faculty of Health and Social Science, The Hong Kong Polytechnic University | Kenneth Siu-Sing LEUNG, Timothy Ting-Leung NG, Alan Ka-Lun WU, Miranda Chong-Yee YAU, Hiu-Yin LAO, Ming-Pan CHOI, Kingsley King-Gee TAM, Lam-Kwong LEE, Barry Kin-Chung WONG, Alex Yat-Man HO, Kam-Tong YIP, Kwok-Cheung LUNG, Raymond Wai-To LIU, Eugene Yuk-Keung TSO, Wai-Shing LEUNG, Man-Chun CHAN, Yuk-Yung NG, Kit-Man SIN, Kitty Sau-Chun FUNG, Sandy Ka-Yee CHAU, Wing-Kin TO, Tak-Lun QUE, David Ho-Keung SHUM, Shea Ping YIP, Wing Cheong YAM, Gilman Kit-Hang SIU |
| EPI_ISL_419254 | INMI Lazzaro Spallanzani IRCCS | Laboratory of Virology, INMI Lazzaro Spallanzani IRCCS | Barbara Bartolini, Martina Rueca, Francesco Messina, Cesare E. M. Gruber, Emanuela Giombini, Maria R. Capobianchi, Fabrizio Carletti, Francesca Colavita, Concetta Castilletti, Eleonora Lalle, Daniele Lapa, Giuseppe Ippolito. |
| EPI_ISL_419255 | INMI Lazzaro Spallanzani IRCCS | INMI Lazzaro Spallanzani IRCCS | Antonino Di Caro, Cesare E. M. Gruber, Martina Rueca, Barbara Bartolini, Francesco Messina, Emanuela Giombini, Maria R. Capobianchi, Fabrizio Carletti, Francesca Colavita, Concetta Castilletti, Eleonora Lalle, Daniele Lapa, Giuseppe Ippolito. |
| EPI_ISL_419256, EPI_ISL_419257, EPI_ISL_419258 | Virginia DCLS | Virginia DCLS | Virginia DCLS |
| EPI_ISL_419259 | Lab voor klinische biologie | Onderzoeksgroep Virologie | Laurens Lambrechts, Nick Vereecke, Marthe Pauwels, Basiel Cole, Bruno Verhasselt, Linos Vandekerckhove, Hans Nauwynck, Sebastiaan Theuns |
| EPI_ISL_419260, EPI_ISL_419261, EPI_ISL_419262, EPI_ISL_419263 | Virginia DCLS | Virginia DCLS | Virginia DCLS |
| EPI_ISL_419264 | Lab voor klinische biologie | Onderzoeksgroep Virologie | Nick Vereecke, Laurens Lambrechts, Marthe Pauwels, Basiel Cole, Bruno Verhasselt, Linos Vandekerckhove, Hans Nauwynck, Sebastiaan Theuns |
| EPI_ISL_419265 | Lab voor klinische biologie | Onderzoeksgroep Virologie | Laurens Lambrechts, Nick Vereecke, Marthe Pauwels, Basiel Cole, Bruno Verhasselt, Linos Vandekerckhove, Hans Nauwynck, Sebastiaan Theuns |
| EPI_ISL_419266 | Lab voor klinische biologie | Onderzoeksgroep Virologie | Nick Vereecke, Laurens Lambrechts, Marthe Pauwels, Basiel Cole, Bruno Verhasselt, Linos Vandekerckhove, Hans Nauwynck, Sebastiaan Theuns |
| EPI_ISL_419296 | Kochi Prefectural Institute of Public Health | Pathogen Genomics Center, National Institute of Infectious Diseases | Tsuyoshi Sekizuka, Akihiko Tokaji, Kentaro Itokawa, Rina Tanaka, Masanori Hashino, Hajime Kamiya, Motoi Suzuki, Makoto Kuroda |
| EPI_ISL_419297, EPI_ISL_419298 | Chiba Prefectural Institute of Public Health | Pathogen Genomics Center, National Institute of Infectious Diseases | Tsuyoshi Sekizuka, Masakatsu Taira, Yushi Hachisu, Kentaro Itokawa, Rina Tanaka, Masanori Hashino, Hajime Kamiya, Motoi Suzuki, Makoto Kuroda |
| EPI_ISL_419299, EPI_ISL_419300 | Ishikawa Prefectural Institute of Public Health and Environmental Science | Pathogen Genomics Center, National Institute of Infectious Diseases | Tsuyoshi Sekizuka, Sanae Kuramoto, Eri Nariai, Kentaro Itokawa, Rina Tanaka, Masanori Hashino, Hajime Kamiya, Motoi Suzuki, Makoto Kuroda |
| EPI_ISL_419301, EPI_ISL_419302, EPI_ISL_419303, EPI_ISL_419304, EPI_ISL_419305, EPI_ISL_419306, EPI_ISL_419307, EPI_ISL_419308 | Saitama Prefectural Institute of Public Health | Pathogen Genomics Center, National Institute of Infectious Diseases | Tsuyoshi Sekizuka, Michiyo Shinohara, Tsuyoshi Kishimoto, Kentaro Itokawa, Rina Tanaka, Masanori Hashino, Hajime Kamiya, Motoi Suzuki, Makoto Kuroda |
| EPI_ISL_419309, EPI_ISL_419310, EPI_ISL_419311 | Chiba Prefectural Institute of Public Health | Pathogen Genomics Center, National Institute of Infectious Diseases | Tsuyoshi Sekizuka, Masakatsu Taira, Yushi Hachisu, Kentaro Itokawa, Rina Tanaka, Masanori Hashino, Hajime Kamiya, Motoi Suzuki, Makoto Kuroda |
| EPI_ISL_419313 | Molecular Biology and Biotechnology Lab II | Molecular Biology and Biotechnology Lab II | Tayyaba Zainab, Sana Shamshad, Azka Noureen, Aimen Malik, Muhammad Javaid Asad, Kumail Ali Rizvi |
| EPI_ISL_419386 | Hospital Prof. Doutor Fernando Fonseca, EPE | Instituto Gulbenkian de Ciência | João Costa, Cathy Paulino, Joao Sobral, Susana Ladeiro, Ricardo Leite |
| EPI_ISL_419387 | Hospital Prof. Doutor Fernando Fonseca, EPE | Instituto Gulbenkian de Ciência | João Costa, Cathy Paulino, Joao Sobral, Susana Ladeiro, Ricardo Leite |
| EPI_ISL_419388, EPI_ISL_419390, EPI_ISL_419391, EPI_ISL_419392, EPI_ISL_419393, EPI_ISL_419394, EPI_ISL_419395, EPI_ISL_419396, EPI_ISL_419397 | Minnesota Department of Health, Public Health Laboratory | Minnesota Department of Health, Public Health Laboratory | Matt Plumb, Jake Garfin and Xiong Wang |
| EPI_ISL_419398, EPI_ISL_419399, EPI_ISL_419400, EPI_ISL_419401, EPI_ISL_419402, EPI_ISL_419403, EPI_ISL_419404, EPI_ISL_419405, EPI_ISL_419406, EPI_ISL_419407, EPI_ISL_419408, EPI_ISL_419409, EPI_ISL_419410, EPI_ISL_419411, EPI_ISL_419412, EPI_ISL_419413, EPI_ISL_419414, EPI_ISL_419415, EPI_ISL_419416, EPI_ISL_419417, EPI_ISL_419418, EPI_ISL_419419, EPI_ISL_419420, EPI_ISL_419421, EPI_ISL_419422, EPI_ISL_419423, EPI_ISL_419424, EPI_ISL_419425, EPI_ISL_419426, EPI_ISL_419427, EPI_ISL_419428, EPI_ISL_419429, EPI_ISL_419430, EPI_ISL_419431, EPI_ISL_419432, EPI_ISL_419433, EPI_ISL_419434, EPI_ISL_419435, EPI_ISL_419436, EPI_ISL_419437, EPI_ISL_419438, EPI_ISL_419439, EPI_ISL_419440, EPI_ISL_419441, EPI_ISL_419442, EPI_ISL_419443, EPI_ISL_419444, EPI_ISL_419445, EPI_ISL_419446, EPI_ISL_419447, EPI_ISL_419448, EPI_ISL_419449, EPI_ISL_419450, EPI_ISL_419451, EPI_ISL_419452, EPI_ISL_419453, EPI_ISL_419454, EPI_ISL_419455, EPI_ISL_419456, EPI_ISL_419457, EPI_ISL_419458, EPI_ISL_419459, EPI_ISL_419460, EPI_ISL_419461, EPI_ISL_419462, EPI_ISL_419463, EPI_ISL_419464, EPI_ISL_419465, EPI_ISL_419466, EPI_ISL_419467, EPI_ISL_419468, EPI_ISL_419469, EPI_ISL_419470, EPI_ISL_419471, EPI_ISL_419472, EPI_ISL_419473, EPI_ISL_419474, EPI_ISL_419475, EPI_ISL_419476, EPI_ISL_419477, EPI_ISL_419478, EPI_ISL_419479, EPI_ISL_419480, EPI_ISL_419481, EPI_ISL_419482, EPI_ISL_419483, EPI_ISL_419484, EPI_ISL_419485, EPI_ISL_419486, EPI_ISL_419487, EPI_ISL_419488, EPI_ISL_419489, EPI_ISL_419490, EPI_ISL_419491, EPI_ISL_419492, EPI_ISL_419493, EPI_ISL_419494, EPI_ISL_419495, EPI_ISL_419496, EPI_ISL_419497, EPI_ISL_419498, EPI_ISL_419499, EPI_ISL_419500, EPI_ISL_419501, EPI_ISL_419502, EPI_ISL_419503, EPI_ISL_419504, EPI_ISL_419505, EPI_ISL_419506, EPI_ISL_419507, EPI_ISL_419508, EPI_ISL_419509, EPI_ISL_419510, EPI_ISL_419511 | | | |
| see above | Wales Specialist Virology Centre | Public Health Wales Microbiology Cardiff | Catherine Moore, Joanne Watkins, Sally Corden, Sara Rey, Matt Bull, Tom Connor |
| EPI_ISL_419512, EPI_ISL_419513, EPI_ISL_419514, EPI_ISL_419515, EPI_ISL_419516 | Yale COVID-19 Biorepository | Grubaugh Lab - Yale School of Public Health | Joseph Fauver, Tara Alpert, Anderson Brito, Anne Wyllie, Chantal Vogels, Mary Petrone, Chaney Kalinich, Isabel Ott, Arnau Casanovas, Catherine Muenker, Adam Moore, Alice Lu, Maria Tokuyama, Patrick Wong, Peiwen Lu, Saad Omer, Richard Martinello, Allison Nelson, Shelli Farhadian, Akiko Iwasaki, Charlese Dela Cruz, Albert Ko, Nathan Grubaugh |
| EPI_ISL_419517, EPI_ISL_419518, EPI_ISL_419519, EPI_ISL_419520, EPI_ISL_419521, EPI_ISL_419522, EPI_ISL_419523, EPI_ISL_419524, EPI_ISL_419525, EPI_ISL_419526, EPI_ISL_419527, EPI_ISL_419528 | | | |
| see above | Yale Clinical Virology Laboratory | Grubaugh Lab - Yale School of Public Health | Joseph Fauver, Anderson Brito, Tara Alpert, Chantal Vogels, Ellen Foxman, Albert Ko, Marie Landry, Nathan Grubaugh |
| EPI_ISL_419541, EPI_ISL_419542, EPI_ISL_419543, EPI_ISL_419544, EPI_ISL_419545, EPI_ISL_419546, EPI_ISL_419547, EPI_ISL_419548, EPI_ISL_419549, EPI_ISL_419550, EPI_ISL_419551, EPI_ISL_419552 | | | |
| see above | Center of Medical Microbiology, Virology, and Hospital Hygiene, University of Duesseldorf | Center of Medical Microbiology, Virology, and Hospital Hygiene, University of Duesseldorf | Ortwin Adams, Marcel Andree, Alexander Dilthey, Torsten Feldt, Sandra Hauka, Torsten Houwaart, Björn-Erik Jensen, Detlef Kindgen-Milles, Malte Kohns Vasconcelos, Klaus Pfeffer, Tina Senff, Daniel Strelow, Jörg Timm, Andreas Walker, Tobias Wienemann |
| EPI_ISL_419553 | RI State Health Laboratories | Pathogen Discovery, Respiratory Viruses Branch, Division of Viral Diseases, Centers for Disease Control and Prevention | Ying Tao, Jing Zhang, Krista Queen, Anna Uehara, Clinton R. Paden, Yan Li, Haibin Wang, Jasmine Padilla, Justin Lee, Suxiang Tong |
| EPI_ISL_419554 | California Department of Public Health | Pathogen Discovery, Respiratory Viruses Branch, Division of Viral Diseases, Centers for Disease Control and Prevention | Ying Tao, Jing Zhang, Krista Queen, Anna Uehara, Clinton R. Paden, Yan Li, Haibin Wang, Jasmine Padilla, Justin Lee, Suxiang Tong |
| EPI_ISL_419555 | WA State Department of Health | Pathogen Discovery, Respiratory Viruses Branch, Division of | Ying Tao, Jing Zhang, Krista Queen, Anna Uehara, Clinton R. Paden, Yan Li, Haibin Wang, Jasmine Padilla, Justin Lee, Suxiang Tong |

| | | | |
|---|---|---|---|
| | | Viral Diseases, Centers for Disease Control and Prevention | |
| EPI_ISL_419556, EPI_ISL_419557 | GA Department of Public Health Laboratory | Pathogen Discovery, Respiratory Viruses Branch, Division of Viral Diseases, Centers for Disease Control and Prevention | Ying Tao, Jing Zhang, Krista Queen, Anna Uehara, Clinton R. Paden, Yan Li, Haibin Wang, Jasmine Padilla, Justin Lee, Suxiang Tong |
| EPI_ISL_419558 | OR State PHL-Virology/Immunology Section | Pathogen Discovery, Respiratory Viruses Branch, Division of Viral Diseases, Centers for Disease Control and Prevention | Ying Tao, Jing Zhang, Krista Queen, Anna Uehara, Clinton R. Paden, Yan Li, Haibin Wang, Jasmine Padilla, Justin Lee, Suxiang Tong |
| EPI_ISL_419559, EPI_ISL_419560 | FL Bureau of Public Health Laboratories-Tampa | Pathogen Discovery, Respiratory Viruses Branch, Division of Viral Diseases, Centers for Disease Control and Prevention | Anna Uehara, Ying Tao, Jing Zhang, Krista Queen, Clinton R. Paden, Yan Li, Haibin Wang, Jasmine Padilla, Justin Lee, Suxiang Tong |
| EPI_ISL_419562, EPI_ISL_419563, EPI_ISL_419564, EPI_ISL_419565, EPI_ISL_419566, EPI_ISL_419567, EPI_ISL_419568, EPI_ISL_419569, EPI_ISL_419570, EPI_ISL_419571, EPI_ISL_419572, EPI_ISL_419573, EPI_ISL_419574, EPI_ISL_419575, EPI_ISL_419576, EPI_ISL_419577, EPI_ISL_419578, EPI_ISL_419579, EPI_ISL_419580, EPI_ISL_419581, EPI_ISL_419582, EPI_ISL_419583, EPI_ISL_419584, EPI_ISL_419585, EPI_ISL_419586, EPI_ISL_419587, EPI_ISL_419588, EPI_ISL_419589, EPI_ISL_419590, EPI_ISL_419591, EPI_ISL_419592, EPI_ISL_419593, EPI_ISL_419594, EPI_ISL_419595, EPI_ISL_419596, EPI_ISL_419597, EPI_ISL_419598, EPI_ISL_419599, EPI_ISL_419600, EPI_ISL_419601, EPI_ISL_419602, EPI_ISL_419603, EPI_ISL_419604, EPI_ISL_419605, EPI_ISL_419606, EPI_ISL_419607 | | | |
| see above | Laboratoire National de Santé, Microbiology, Virology | Laboratoire National de Santé, Microbiology, Epidemiology and Microbial Genomics | Anke Wienecke-Baldacchino, Ardashel Latsuzbaia, Jessica Tapp, Catherine Ragimbeau, Guillaume Fournier, Tamir Abdelrahman, Trung Nguyen Nguyen, Joel Mossong |
| EPI_ISL_419651, EPI_ISL_419652 | Gundersen Molecular Diagnostics Laboratory | Kabara Cancer Research Institute | Craig S. Richmond & Paraic A. Kenny |
| EPI_ISL_419654, EPI_ISL_419655, EPI_ISL_419656, EPI_ISL_419657, EPI_ISL_419658, EPI_ISL_419659, EPI_ISL_419660, EPI_ISL_419661, EPI_ISL_419662, EPI_ISL_419663, EPI_ISL_419664, EPI_ISL_419665, EPI_ISL_419666, EPI_ISL_419667, EPI_ISL_419668, EPI_ISL_419669, EPI_ISL_419670, EPI_ISL_419671, EPI_ISL_419672, EPI_ISL_419673, EPI_ISL_419674 | | | |
| see above | Center for Virology, Medical University of Vienna | Bergthaler laboratory, CeMM Research Center for Molecular Medicine of the Austrian Academy of Sciences | Alexandra Popa, Benedikt Agerer, Henrique Colaco, Lukas Endler, Jakob-Wendelin Genger, Alexander Lercher, Mark Smyth, Thomas Penz, Michael Schuster, Judith Aberle, Stephan Aberle, Elisabeth Puchhammer-Stöckl, Christoph Bock, Andreas Bergthaler |
| EPI_ISL_419675 | Servicio de Microbiología. Consorcio Hospital General Universitario de Valencia | Sequencing and Bioinformatics Service and Molecular Epidemiology Research Group. FISABIO-Public Health | Maria Alma Bracho, Maria Dolores Ocete, Giuseppe D'Auria, Griselda De Marco, Neris Garcia-Gonzalez, Concepcion Gimeno, Fernando Gonzalez-Candelas |
| EPI_ISL_419676 | Servicio de Microbiología. Consorcio Hospital General Universitario de Valencia | Sequencing and Bioinformatics Service and Molecular Epidemiology Research Group. FISABIO-Public Health | Maria Dolores Ocete, Giuseppe D'Auria, Griselda De Marco, Neris Garcia-Gonzalez, Maria Alma Bracho, Concepcion Gimeno, Fernando Gonzalez-Candelas |
| EPI_ISL_419677 | Servicio de Microbiología. Consorcio Hospital General Universitario de Valencia | Sequencing and Bioinformatics Service and Molecular Epidemiology Research Group. FISABIO-Public Health | Giuseppe D'Auria, Griselda De Marco, Neris Garcia-Gonzalez, Maria Alma Bracho, Maria Dolores Ocete, Concepcion Gimeno, Fernando Gonzalez-Candelas |
| EPI_ISL_419678 | Servicio de Microbiología. Consorcio Hospital General Universitario de Valencia | Sequencing and Bioinformatics Service and Molecular Epidemiology Research Group. FISABIO-Public Health | Griselda De Marco, Neris Garcia-Gonzalez, Maria Alma Bracho, Maria Dolores Ocete, Giuseppe D'Auria, Concepcion Gimeno, Fernando Gonzalez-Candelas |
| EPI_ISL_419679 | Servicio de Microbiología. Consorcio Hospital General Universitario de Valencia | Sequencing and Bioinformatics Service and Molecular Epidemiology Research Group. FISABIO-Public Health | Neris Garcia-Gonzalez, Maria Alma Bracho, Maria Dolores Ocete, Giuseppe D'Auria, Griselda De Marco, Concepcion Gimeno, Fernando Gonzalez-Candelas |
| EPI_ISL_419680 | Servicio de Microbiología. Consorcio Hospital General Universitario de Valencia | Sequencing and Bioinformatics Service and Molecular Epidemiology Research Group. FISABIO-Public Health | Maria Alma Bracho, Maria Dolores Ocete, Giuseppe D'Auria, Griselda De Marco, Neris Garcia-Gonzalez, Concepcion Gimeno, Fernando Gonzalez-Candelas |
| EPI_ISL_419681 | Servicio de Microbiología. Consorcio Hospital General Universitario de Valencia | Sequencing and Bioinformatics Service and Molecular Epidemiology Research Group. FISABIO-Public Health | Maria Dolores Ocete, Giuseppe D'Auria, Griselda De Marco, Neris Garcia-Gonzalez, Maria Alma Bracho, Concepcion Gimeno, Fernando Gonzalez-Candelas |
| EPI_ISL_419682 | Servicio de Microbiología. Consorcio Hospital General Universitario de Valencia | Sequencing and Bioinformatics Service and Molecular Epidemiology Research Group. FISABIO-Public Health | Giuseppe D'Auria, Griselda De Marco, Neris Garcia-Gonzalez, Maria Alma Bracho, Maria Dolores Ocete, Concepcion Gimeno, Fernando Gonzalez-Candelas |
| EPI_ISL_419683 | Servicio de Microbiología. Consorcio Hospital General Universitario de Valencia | Sequencing and Bioinformatics Service and Molecular Epidemiology Research Group. FISABIO-Public Health | Griselda De Marco, Neris Garcia-Gonzalez, Maria Alma Bracho, Maria Dolores Ocete, Giuseppe D'Auria, Concepcion Gimeno, Fernando Gonzalez-Candelas |
| EPI_ISL_419684 | Servicio de Microbiología. Consorcio Hospital General Universitario de Valencia | Sequencing and Bioinformatics Service and Molecular Epidemiology Research Group. FISABIO-Public Health | Neris Garcia-Gonzalez, Maria Alma Bracho, Maria Dolores Ocete, Giuseppe D'Auria, Griselda De Marco, Concepcion Gimeno, Fernando Gonzalez-Candelas |
| EPI_ISL_419685 | Servicio de Microbiología. Consorcio Hospital General Universitario de Valencia | Sequencing and Bioinformatics Service and Molecular Epidemiology Research Group. FISABIO-Public Health | Maria Alma Bracho, Maria Dolores Ocete, Giuseppe D'Auria, Griselda De Marco, Neris Garcia-Gonzalez, Concepcion Gimeno, Fernando Gonzalez-Candelas |
| EPI_ISL_419686 | Servicio de Microbiología. Consorcio Hospital General Universitario de Valencia | Sequencing and Bioinformatics Service and Molecular Epidemiology Research Group. FISABIO-Public Health | Maria Dolores Ocete, Giuseppe D'Auria, Griselda De Marco, Neris Garcia-Gonzalez, Maria Alma Bracho, Concepcion Gimeno, Fernando Gonzalez-Candelas |
| EPI_ISL_419687 | Servicio de Microbiología. Consorcio Hospital General Universitario de Valencia | Sequencing and Bioinformatics Service and Molecular Epidemiology Research Group. FISABIO-Public Health | Giuseppe D'Auria, Griselda De Marco, Neris Garcia-Gonzalez, Maria Alma Bracho, Maria Dolores Ocete, Concepcion Gimeno, Fernando Gonzalez-Candelas |
| EPI_ISL_419688 | Servicio de Microbiología. Consorcio Hospital General Universitario de Valencia | Sequencing and Bioinformatics Service and Molecular Epidemiology Research Group. FISABIO-Public Health | Griselda De Marco, Neris Garcia-Gonzalez, Maria Alma Bracho, Maria Dolores Ocete, Giuseppe D'Auria, Concepcion Gimeno, Fernando Gonzalez-Candelas |
| EPI_ISL_419689 | Servicio de Microbiología. Consorcio Hospital General Universitario de Valencia | Sequencing and Bioinformatics Service and Molecular Epidemiology Research Group. FISABIO-Public Health | Neris Garcia-Gonzalez, Maria Alma Bracho, Maria Dolores Ocete, Giuseppe D'Auria, Griselda De Marco, Concepcion Gimeno, Fernando Gonzalez-Candelas |
| EPI_ISL_419690 | Servicio de Microbiología. Consorcio Hospital General Universitario de Valencia | Sequencing and Bioinformatics Service and Molecular Epidemiology Research Group. FISABIO-Public Health | Maria Alma Bracho, Maria Dolores Ocete, Giuseppe D'Auria, Griselda De Marco, Neris Garcia-Gonzalez, Concepcion Gimeno, Fernando Gonzalez-Candelas |
| EPI_ISL_419691 | E. Gulbja Laboratorija | Charité Universitätsmedizin Berlin, Institute of Virology | Victor M Corman, Julia Schneider, Barbara Mühlemann, Talitha Veith, Jörn Beheim-Schwarzbach, Terry Jones, Dr. Didzis Gavars, Mikus Gavars, Dmitrijs Perminovs, Christian Drosten |
| EPI_ISL_419692 | The Republican Research and Practical Center for Epidemiology and Microbiology | Charité Universitätsmedizin Berlin, Institute of Virology | Victor M Corman, Julia Schneider, Barbara Mühlemann, Talitha Veith, Jörn Beheim-Schwarzbach, Terry Jones, Natallia Shmialiova, Natallia Sivets, Christian Drosten |
| EPI_ISL_419693 | The Republican Research and Practical Center for Epidemiology and Microbiology | Charité Universitätsmedizin Berlin, Institute of Virology | Victor M Corman, Julia Schneider, Barbara Mühlemann, Talitha Veith, Jörn Beheim-Schwarzbach, Terry Jones, Natallia Shmialiova, Natallia Sivets, Christian Drosten |
| EPI_ISL_419696, EPI_ISL_419697, EPI_ISL_419698, EPI_ISL_419699, EPI_ISL_419700, EPI_ISL_419701, EPI_ISL_419702, EPI_ISL_419703, EPI_ISL_419704, EPI_ISL_419705 | NYU Langone Health | Departments of Pathology and Medicine, New York University School of Medicine | Maria Aguero-Rosenfeld, Margaret Black, John Cadley, Paolo Cotzia, John Chen, Dacia Dimartino, Xiaojun Feng, Adriana Heguy, Megan Hogan, Emily Huang, George Jour, Christian Marier, Matthew T. Maurano, Mark J. Mulligan, Peter Meyn, Jared Pinnell, Sitharam Ramaswami, Amy Rapkiewicz, Marie Samanovic-Golden, Antonio Serrano, Guomiao Shen, Matija Snuderl, Nick Vulpescu, Gael Westby, Paul Zappile, Yutong Zhang |
| EPI_ISL_419706 | Virginia DCLS | Virginia DCLS | Virginia DCLS |
| EPI_ISL_419707 | HOSPITAL CLINIC | Instituto de Salud Carlos III | Iglesias-Caballero, M. Molinero Calamita, M. González-Esguevillas, M. Camarero S. Pozo F. Casas I. Jiménez, P. Jiménez, M. Zaballos, A. Monzón, S. Varona, S. Juliá, M. Cuesta, I. Marcos, M.A |
| EPI_ISL_419708 | Virginia DCLS | Virginia DCLS | Virginia DCLS |
| EPI_ISL_419709 | HOSPITAL TXAGORRITXU | Instituto de Salud Carlos III | Iglesias-Caballero, M. Molinero Calamita, M. González-Esguevillas, M. Camarero S. Pozo F. Casas I. Jiménez, P. Jiménez, M. Zaballos, A. Monzón, S. Varona, S. Juliá, M. Cuesta, I. Gómez, C. |
| EPI_ISL_419710, EPI_ISL_419711, EPI_ISL_419712, EPI_ISL_419713 | Virginia DCLS | Virginia DCLS | Virginia DCLS |
| EPI_ISL_419714, EPI_ISL_419715, EPI_ISL_419716, EPI_ISL_419717, EPI_ISL_419718, EPI_ISL_419719, EPI_ISL_419720, EPI_ISL_419721, EPI_ISL_419722, EPI_ISL_419723, EPI_ISL_419724, EPI_ISL_419725, EPI_ISL_419726, EPI_ISL_419727, EPI_ISL_419728, EPI_ISL_419729, EPI_ISL_419730, EPI_ISL_419731, EPI_ISL_419732 | | | |
| see above | Microbiological Diagnostic Unit Public Health Laboratory | Microbiological Diagnostic Unit Public Health Laboratory | Seemann T., Schultz M., Sait, M., Sherry, N. |
| EPI_ISL_419733, EPI_ISL_419734, EPI_ISL_419735, EPI_ISL_419736, EPI_ISL_419737, EPI_ISL_419738, EPI_ISL_419739, EPI_ISL_419740, EPI_ISL_419741, EPI_ISL_419742, EPI_ISL_419743, EPI_ISL_419744, EPI_ISL_419745, EPI_ISL_419746, EPI_ISL_419747, EPI_ISL_419748, EPI_ISL_419749, EPI_ISL_419750, EPI_ISL_419751, EPI_ISL_419752, EPI_ISL_419753, EPI_ISL_419754, EPI_ISL_419755, EPI_ISL_419756, EPI_ISL_419757, EPI_ISL_419758, EPI_ISL_419759, EPI_ISL_419760, EPI_ISL_419761, EPI_ISL_419762, EPI_ISL_419763, EPI_ISL_419764, EPI_ISL_419765, EPI_ISL_419766, EPI_ISL_419767, EPI_ISL_419768, EPI_ISL_419769, EPI_ISL_419770, EPI_ISL_419771, EPI_ISL_419772, EPI_ISL_419773, EPI_ISL_419774, EPI_ISL_419775, EPI_ISL_419776, EPI_ISL_419777, EPI_ISL_419778, EPI_ISL_419779, EPI_ISL_419780, EPI_ISL_419781, EPI_ISL_419782, EPI_ISL_419783, EPI_ISL_419784, EPI_ISL_419785, EPI_ISL_419786, EPI_ISL_419787, EPI_ISL_419788, EPI_ISL_419789, EPI_ISL_419790, EPI_ISL_419791, EPI_ISL_419792, EPI_ISL_419793, EPI_ISL_419794, EPI_ISL_419795, EPI_ISL_419796, EPI_ISL_419797, EPI_ISL_419798, EPI_ISL_419799, EPI_ISL_419800, EPI_ISL_419801, EPI_ISL_419802, EPI_ISL_419803, EPI_ISL_419804, EPI_ISL_419805, EPI_ISL_419806, EPI_ISL_419807, EPI_ISL_419808, EPI_ISL_419809, EPI_ISL_419810, EPI_ISL_419811, EPI_ISL_419812, EPI_ISL_419813, EPI_ISL_419814, EPI_ISL_419815, EPI_ISL_419816, EPI_ISL_419817, EPI_ISL_419818, EPI_ISL_419819, EPI_ISL_419820, EPI_ISL_419821, EPI_ISL_419822 | | | |
| see above | Victorian Infectious Diseases Reference Laboratory (VIDRL) | Victorian Infectious Diseases Reference Laboratory and Microbiological Diagnostic Unit Public Health Laboratory, Doherty Institute | Caly L., Seemann T., Sait, M., Schultz M., Druce J., Sherry, N. |
| EPI_ISL_419823, EPI_ISL_419824, EPI_ISL_419825 | Microbiological Diagnostic Unit Public Health Laboratory | Microbiological Diagnostic Unit Public Health Laboratory | Seemann T., Schultz M., Sait, M., Sherry, N. |
| EPI_ISL_419826 | Victorian Infectious Diseases Reference Laboratory (VIDRL) | Victorian Infectious Diseases Reference Laboratory and Microbiological Diagnostic Unit Public Health Laboratory, Doherty Institute | Caly L., Seemann T., Sait, M., Schultz M., Druce J., Sherry, N. |
| EPI_ISL_419827, EPI_ISL_419828, EPI_ISL_419829, EPI_ISL_419830 | Microbiological Diagnostic Unit Public Health Laboratory | Microbiological Diagnostic Unit Public Health Laboratory | Seemann T., Schultz M., Sait, M., Sherry, N. |
| EPI_ISL_419831, EPI_ISL_419832, EPI_ISL_419833 | Royal Darwin Hospital | Victorian Infectious Diseases Reference Laboratory and Microbiological Diagnostic Unit Public Health Laboratory, Doherty Institute | Meumann, E., Seemann T., Sait, M., Schultz M., Caly L., Druce J. |
| EPI_ISL_419834 | Victorian Infectious Diseases Reference Laboratory (VIDRL) | Victorian Infectious Diseases Reference Laboratory and Microbiological Diagnostic Unit Public Health Laboratory, Doherty Institute | Caly L., Seemann T., Sait, M., Schultz M., Druce J., Sherry, N. |
| EPI_ISL_419835 | Royal Darwin Hospital | Victorian Infectious Diseases Reference Laboratory and Microbiological Diagnostic Unit Public Health Laboratory, Doherty Institute | Meumann, E., Seemann T., Sait, M., Schultz M., Caly L., Druce J. |

| | | | |
|---|---|---|---|
| EPI_ISL_419836, EPI_ISL_419837, EPI_ISL_419838, EPI_ISL_419839, EPI_ISL_419840, EPI_ISL_419841, EPI_ISL_419842, EPI_ISL_419843, EPI_ISL_419844, EPI_ISL_419845, EPI_ISL_419846, EPI_ISL_419847, EPI_ISL_419848, EPI_ISL_419849, EPI_ISL_419850, EPI_ISL_419851, EPI_ISL_419852, EPI_ISL_419853, EPI_ISL_419854, EPI_ISL_419855, EPI_ISL_419856, EPI_ISL_419857, EPI_ISL_419858, EPI_ISL_419859, EPI_ISL_419860, EPI_ISL_419861, EPI_ISL_419862, EPI_ISL_419863, EPI_ISL_419864, EPI_ISL_419865, EPI_ISL_419866, EPI_ISL_419867, EPI_ISL_419868, EPI_ISL_419869, EPI_ISL_419870, EPI_ISL_419871, EPI_ISL_419872, EPI_ISL_419873, EPI_ISL_419874, EPI_ISL_419875, EPI_ISL_419876, EPI_ISL_419877, EPI_ISL_419878, EPI_ISL_419879, EPI_ISL_419880, EPI_ISL_419881, EPI_ISL_419882, EPI_ISL_419883, EPI_ISL_419884, EPI_ISL_419885, EPI_ISL_419886, EPI_ISL_419887, EPI_ISL_419888, EPI_ISL_419889, EPI_ISL_419890, EPI_ISL_419891, EPI_ISL_419892, EPI_ISL_419893, EPI_ISL_419894, EPI_ISL_419895, EPI_ISL_419896, EPI_ISL_419897, EPI_ISL_419898, EPI_ISL_419899, EPI_ISL_419900, EPI_ISL_419901, EPI_ISL_419902, EPI_ISL_419903, EPI_ISL_419904, EPI_ISL_419905, EPI_ISL_419906, EPI_ISL_419907, EPI_ISL_419908, EPI_ISL_419909, EPI_ISL_419910, EPI_ISL_419911, EPI_ISL_419912, EPI_ISL_419913, EPI_ISL_419914, EPI_ISL_419915, EPI_ISL_419916, EPI_ISL_419917, EPI_ISL_419918, EPI_ISL_419919, EPI_ISL_419920, EPI_ISL_419921, EPI_ISL_419922, EPI_ISL_419923, EPI_ISL_419924, EPI_ISL_419925, EPI_ISL_419926, EPI_ISL_419927, EPI_ISL_419928, EPI_ISL_419929, EPI_ISL_419930, EPI_ISL_419931, EPI_ISL_419932, EPI_ISL_419933, EPI_ISL_419934, EPI_ISL_419935, EPI_ISL_419936, EPI_ISL_419937, EPI_ISL_419938, EPI_ISL_419939, EPI_ISL_419940, EPI_ISL_419941, EPI_ISL_419942, EPI_ISL_419943, EPI_ISL_419944, EPI_ISL_419945, EPI_ISL_419946, EPI_ISL_419947, EPI_ISL_419948, EPI_ISL_419949, EPI_ISL_419950, EPI_ISL_419951, EPI_ISL_419952, EPI_ISL_419953, EPI_ISL_419954, EPI_ISL_419955, EPI_ISL_419956, EPI_ISL_419957, EPI_ISL_419958, EPI_ISL_419959, EPI_ISL_419960, EPI_ISL_419961, EPI_ISL_419962, EPI_ISL_419963, EPI_ISL_419964, EPI_ISL_419965, EPI_ISL_419966, EPI_ISL_419967, EPI_ISL_419968, EPI_ISL_419969, EPI_ISL_419970, EPI_ISL_419971, EPI_ISL_419972, EPI_ISL_419973, EPI_ISL_419974, EPI_ISL_419975, EPI_ISL_419976, EPI_ISL_419977, EPI_ISL_419978, EPI_ISL_419979 | | | |
| see above | Victorian Infectious Diseases Reference Laboratory (VIDRL) | Victorian Infectious Diseases Reference Laboratory and Microbiological Diagnostic Unit Public Health Laboratory, Doherty Institute | Caly L., Seemann T., Sait, M., Schultz M., Druce J., Sherry, N. |
| EPI_ISL_419980, EPI_ISL_419981 | Microbiological Diagnostic Unit Public Health Laboratory | Microbiological Diagnostic Unit Public Health Laboratory | Seemann T., Schultz M., Sait, M., Sherry, N. |
| EPI_ISL_419982, EPI_ISL_419983, EPI_ISL_419984, EPI_ISL_419985, EPI_ISL_419986, EPI_ISL_419987, EPI_ISL_419988, EPI_ISL_419989, EPI_ISL_419990, EPI_ISL_419991, EPI_ISL_419992, EPI_ISL_419993, EPI_ISL_419994, EPI_ISL_419995, EPI_ISL_419996, EPI_ISL_419997, EPI_ISL_419998 | | | |
| see above | Victorian Infectious Diseases Reference Laboratory (VIDRL) | Victorian Infectious Diseases Reference Laboratory and Microbiological Diagnostic Unit Public Health Laboratory, Doherty Institute | Caly L., Seemann T., Sait, M., Schultz M., Druce J., Sherry, N. |
| EPI_ISL_419999, EPI_ISL_420000, EPI_ISL_420001, EPI_ISL_420002, EPI_ISL_420003 | Microbiological Diagnostic Unit Public Health Laboratory | Microbiological Diagnostic Unit Public Health Laboratory | Seemann T., Schultz M., Sait, M., Sherry, N. |
| EPI_ISL_420004, EPI_ISL_420005 | Victorian Infectious Diseases Reference Laboratory (VIDRL) | Victorian Infectious Diseases Reference Laboratory and Microbiological Diagnostic Unit Public Health Laboratory, Doherty Institute | Caly L., Seemann T., Sait, M., Schultz M., Druce J., Sherry, N. |
| EPI_ISL_420006, EPI_ISL_420007, EPI_ISL_420008, EPI_ISL_420009, EPI_ISL_420010, EPI_ISL_420011, EPI_ISL_420012, EPI_ISL_420013, EPI_ISL_420014, EPI_ISL_420015, EPI_ISL_420016, EPI_ISL_420017 | | | |
| see above | Microbiological Diagnostic Unit Public Health Laboratory | Microbiological Diagnostic Unit Public Health Laboratory | Seemann T., Schultz M., Sait, M., Sherry, N. |
| EPI_ISL_420018, EPI_ISL_420019, EPI_ISL_420020, EPI_ISL_420021, EPI_ISL_420022, EPI_ISL_420023, EPI_ISL_420024, EPI_ISL_420025, EPI_ISL_420026, EPI_ISL_420027, EPI_ISL_420028, EPI_ISL_420029 | | | |
| see above | Virginia DCLS | Virginia DCLS | Virginia DCLS |
| EPI_ISL_420030, EPI_ISL_420031, EPI_ISL_420032, EPI_ISL_420033, EPI_ISL_420034, EPI_ISL_420035 | Viral Respiratory Lab, National Institute for Biomedical Research (INRB) | Pathogen Sequencing Lab, National Institute for Biomedical Research (INRB) | Placide Mbala-Kingebeni, Edith Nkwembe, Eddy Kinganda-Lusamaki, Amuri Aziza, Catherine Pratt, Matthias Pauthner, Josh Quick, Allison Black, James Hadfield, Trevor Bedford, Ian Goodfellow, Nick Loman, Kristian Andersen, Michael Wiley, Steve Ahuka-Mundeke, Jean-Jacques Muyembe Tamfum |
| EPI_ISL_420036 | Victorian Infectious Diseases Reference Laboratory (VIDRL) | Victorian Infectious Diseases Reference Laboratory and Microbiological Diagnostic Unit Public Health Laboratory, Doherty Institute | Caly L., Seemann T., Sait, M., Schultz M., Druce J., Sherry, N. |
| EPI_ISL_420037 | NIC Viral Respiratory Unit - Institut Pasteur of Algeria | National Reference Center for Viruses of Respiratory Infections, Institut Pasteur, Paris | Mélanie Albert, Marion Barbet, Sylvie Behillil, Méline Bizard, Angela Brisebarre, Flora Donati, Etienne Simon-Lorière, Vincent Enouf, Maud Vanpeene, Sylvie van der Werf, Fawzi Derrar |
| EPI_ISL_420038 | Sentinelles network | National Reference Center for Viruses of Respiratory Infections, Institut Pasteur, Paris | Mélanie Albert, Marion Barbet, Sylvie Behillil, Méline Bizard, Angela Brisebarre, Flora Donati, Etienne Simon-Lorière, Vincent Enouf, Maud Vanpeene, Sylvie van der Werf |
| EPI_ISL_420039, EPI_ISL_420040 | L'Air du Temps | National Reference Center for Viruses of Respiratory Infections, Institut Pasteur, Paris | Mélanie Albert, Marion Barbet, Sylvie Behillil, Méline Bizard, Angela Brisebarre, Flora Donati, Etienne Simon-Lorière, Vincent Enouf, Maud Vanpeene, Sylvie van der Werf |
| EPI_ISL_420041 | CH Compiègne Laboratoire de Biologie | National Reference Center for Viruses of Respiratory Infections, Institut Pasteur, Paris | Mélanie Albert, Marion Barbet, Sylvie Behillil, Méline Bizard, Angela Brisebarre, Flora Donati, Etienne Simon-Lorière, Vincent Enouf, Maud Vanpeene, Sylvie van der Werf, Raulin Olivia |
| EPI_ISL_420042 | Service de Biologie clinique | National Reference Center for Viruses of Respiratory Infections, Institut Pasteur, Paris | Mélanie Albert, Marion Barbet, Sylvie Behillil, Méline Bizard, Angela Brisebarre, Flora Donati, Etienne Simon-Lorière, Vincent Enouf, Maud Vanpeene, Sylvie van der Werf |
| EPI_ISL_420043 | CMIP | National Reference Center for Viruses of Respiratory Infections, Institut Pasteur, Paris | Mélanie Albert, Marion Barbet, Sylvie Behillil, Méline Bizard, Angela Brisebarre, Flora Donati, Etienne Simon-Lorière, Vincent Enouf, Maud Vanpeene, Sylvie van der Werf |
| EPI_ISL_420044 | CH Jean de Navarre Laboratoire de Biologie | National Reference Center for Viruses of Respiratory Infections, Institut Pasteur, Paris | Mélanie Albert, Marion Barbet, Sylvie Behillil, Méline Bizard, Angela Brisebarre, Flora Donati, Etienne Simon-Lorière, Vincent Enouf, Maud Vanpeene, Sylvie van der Werf |
| EPI_ISL_420045 | Sentinelles network | National Reference Center for Viruses of Respiratory Infections, Institut Pasteur, Paris | Mélanie Albert, Marion Barbet, Sylvie Behillil, Méline Bizard, Angela Brisebarre, Flora Donati, Etienne Simon-Lorière, Vincent Enouf, Maud Vanpeene, Sylvie van der Werf |
| EPI_ISL_420046, EPI_ISL_420047 | Résidence Villa Caroline | National Reference Center for Viruses of Respiratory Infections, Institut Pasteur, Paris | Mélanie Albert, Marion Barbet, Sylvie Behillil, Méline Bizard, Angela Brisebarre, Flora Donati, Etienne Simon-Lorière, Vincent Enouf, Maud Vanpeene, Sylvie van der Werf |
| EPI_ISL_420048 | Service de Biologie Médicale - BP 125 | National Reference Center for Viruses of Respiratory Infections, Institut Pasteur, Paris | Mélanie Albert, Marion Barbet, Sylvie Behillil, Méline Bizard, Angela Brisebarre, Flora Donati, Etienne Simon-Lorière, Vincent Enouf, Maud Vanpeene, Sylvie van der Werf, Christine Lambert |
| EPI_ISL_420049, EPI_ISL_420050 | CH Compiègne Laboratoire de Biologie | National Reference Center for Viruses of Respiratory Infections, Institut Pasteur, Paris | Mélanie Albert, Marion Barbet, Sylvie Behillil, Méline Bizard, Angela Brisebarre, Flora Donati, Etienne Simon-Lorière, Vincent Enouf, Maud Vanpeene, Sylvie van der Werf, Raulin Olivia |
| EPI_ISL_420051 | Résidence Eleusis | National Reference Center for Viruses of Respiratory Infections, Institut Pasteur, Paris | Mélanie Albert, Marion Barbet, Sylvie Behillil, Méline Bizard, Angela Brisebarre, Flora Donati, Etienne Simon-Lorière, Vincent Enouf, Maud Vanpeene, Sylvie van der Werf |
| EPI_ISL_420052 | Résidence les Marines | National Reference Center for Viruses of Respiratory Infections, Institut Pasteur, Paris | Mélanie Albert, Marion Barbet, Sylvie Behillil, Méline Bizard, Angela Brisebarre, Flora Donati, Etienne Simon-Lorière, Vincent Enouf, Maud Vanpeene, Sylvie van der Werf |
| EPI_ISL_420053 | CH Jean de Navarre Laboratoire de Biologie | National Reference Center for Viruses of Respiratory Infections, Institut Pasteur, Paris | Mélanie Albert, Marion Barbet, Sylvie Behillil, Méline Bizard, Angela Brisebarre, Flora Donati, Etienne Simon-Lorière, Vincent Enouf, Maud Vanpeene, Sylvie van der Werf |
| EPI_ISL_420054 | Résidence de maintenon | National Reference Center for Viruses of Respiratory Infections, Institut Pasteur, Paris | Mélanie Albert, Marion Barbet, Sylvie Behillil, Méline Bizard, Angela Brisebarre, Flora Donati, Etienne Simon-Lorière, Vincent Enouf, Maud Vanpeene, Sylvie van der Werf |
| EPI_ISL_420055 | Sentinelles network | National Reference Center for Viruses of Respiratory Infections, Institut Pasteur, Paris | Mélanie Albert, Marion Barbet, Sylvie Behillil, Méline Bizard, Angela Brisebarre, Flora Donati, Etienne Simon-Lorière, Vincent Enouf, Maud Vanpeene, Sylvie van der Werf |
| EPI_ISL_420056, EPI_ISL_420057 | CH Compiègne Laboratoire de Biologie | National Reference Center for Viruses of Respiratory Infections, Institut Pasteur, Paris | Mélanie Albert, Marion Barbet, Sylvie Behillil, Méline Bizard, Angela Brisebarre, Flora Donati, Etienne Simon-Lorière, Vincent Enouf, Maud Vanpeene, Sylvie van der Werf, Raulin Olivia |
| EPI_ISL_420058, EPI_ISL_420059, EPI_ISL_420060 | Service de Biologie Médicale - BP 125 | National Reference Center for Viruses of Respiratory Infections, Institut Pasteur, Paris | Mélanie Albert, Marion Barbet, Sylvie Behillil, Méline Bizard, Angela Brisebarre, Flora Donati, Etienne Simon-Lorière, Vincent Enouf, Maud Vanpeene, Sylvie van der Werf, Christine Lambert |
| EPI_ISL_420061 | CMIP | National Reference Center for Viruses of Respiratory Infections, Institut Pasteur, Paris | Mélanie Albert, Marion Barbet, Sylvie Behillil, Méline Bizard, Angela Brisebarre, Flora Donati, Etienne Simon-Lorière, Vincent Enouf, Maud Vanpeene, Sylvie van der Werf |
| EPI_ISL_420062 | Service de Biologie Médicale - BP 125 | National Reference Center for Viruses of Respiratory Infections, Institut Pasteur, Paris | Mélanie Albert, Marion Barbet, Sylvie Behillil, Méline Bizard, Angela Brisebarre, Flora Donati, Etienne Simon-Lorière, Vincent Enouf, Maud Vanpeene, Sylvie van der Werf, Christine Lambert |
| EPI_ISL_420063 | Labo BM - Site de Juvisy - Hopital Général | National Reference Center for Viruses of Respiratory Infections, Institut Pasteur, Paris | Mélanie Albert, Marion Barbet, Sylvie Behillil, Méline Bizard, Angela Brisebarre, Flora Donati, Etienne Simon-Lorière, Vincent Enouf, Maud Vanpeene, Sylvie van der Werf |
| EPI_ISL_420064 | Service de Biologie Médicale - BP 125 | National Reference Center for Viruses of Respiratory Infections, Institut Pasteur, Paris | Mélanie Albert, Marion Barbet, Sylvie Behillil, Méline Bizard, Angela Brisebarre, Flora Donati, Etienne Simon-Lorière, Vincent Enouf, Maud Vanpeene, Sylvie van der Werf, Christine Lambert |
| EPI_ISL_420065, EPI_ISL_420066, EPI_ISL_420067 | Health Board Laboratory of Communicable Diseases | Charité Universitätsmedizin Berlin, Institute of Virology | Victor M Corman, Jörn Beheim-Schwarzbach, Barbara Mühlemann, Talitha Veith, Julia Schneider, Liidia Dotsenko, Natalja Kuznetsova, Terry Jones, Christian Drosten |
| EPI_ISL_420069, EPI_ISL_420070, EPI_ISL_420071 | Institut Pasteur Dakar | Institut Pasteur de Dakar | Ndongo Dia, Moussa Moise Diagne, Mamadou Diop, Ousmane Faye, Amadou Alpha Sall |
| EPI_ISL_420072, EPI_ISL_420073, EPI_ISL_420074 | Institut Pasteur Dakar | Institut Pasteur de Dakar | Ndongo Dia, Moussa Moise Diagne, Mamadou Diop, Ousmane Faye , Amadou Alpha Sall |
| EPI_ISL_420075 | Institut pasteur Dakar | Institut Pasteur de Dakar | Ndongo Dia, Moussa Moise Diagne, Mamadou Diop, Ousmane Faye , Amadou Alpha Sall |
| EPI_ISL_420076 | Institut Pasteur Dakar | Institut Pasteur de Dakar | Ndongo Dia, Moussa Moise Diagne, Mamadou Diop, Ousmane Faye , Ndongo Dia |
| EPI_ISL_420077, EPI_ISL_420078, EPI_ISL_420079 | Institut Pasteur Dakar | Institut Pasteur de Dakar | Ndongo Dia, Moussa Moise Diagne, Mamadou Diop, Ousmane Faye , Amadou Alpha Sall |
| EPI_ISL_420080 | WHO National Influenza Centre Russian Federation | WHO National Influenza Centre Russian Federation | Andrey Komissarov, Artem Fadeev, Anna Ivanova, Daria Danilenko |
| EPI_ISL_420081 | WHO National Influenza Centre Russian Federation | WHO National Influenza Centre Russian Federation | Andrey Komissarov, Artem Fadeev, Mariia Sergeeva, Anna Ivanova, Daria Danilenko |
| EPI_ISL_420082 | Centers for Disease Control, R.O.C. (Taiwan) | Centers for Disease Control, R.O.C. (Taiwan) | Ji-Rong Yang, Yu-Chi-Lin, Jung-Jung Mu, Ming-Tsan Liu |
| EPI_ISL_420083, EPI_ISL_420084, EPI_ISL_420085 | Centers for Disease Control, R.O.C. (Taiwan) | Centers for Disease Control, R.O.C. (Taiwan) | Ji-Rong Yang, Yu-Chi Lin, Jung-Jung Mu, Ming-Tsan Liu |

| | | | |
|---|---|---|---|
| EPI_ISL_420086, EPI_ISL_420087, EPI_ISL_420088, EPI_ISL_420089, EPI_ISL_420090, EPI_ISL_420091, EPI_ISL_420092, EPI_ISL_420093, EPI_ISL_420094, EPI_ISL_420095 | Yale Clinical Virology Laboratory | Grubaugh Lab - Yale School of Public Health | Joseph Fauver, Anderson Brito, Tara Alpert, Chantal Vogels, Ellen Foxman, Albert Ko, Marie Landry, Nathan Grubaugh |
| EPI_ISL_420096 | Yale Clinical Virology Laboratory | Grubaugh Lab - Yale School of Public Health | Joseph Fauver, Anderson Brito, Tara Alpert, Chantal Vogels, Ellen Foxman, Albert Ko, Marie Landry, Nathan Grubaugh |
| EPI_ISL_420097, EPI_ISL_420098 | Yale Clinical Virology Laboratory | Grubaugh Lab - Yale School of Public Health | Joseph Fauver, Anderson Brito, Tara Alpert, Chantal Vogels, Ellen Foxman, Albert Ko, Marie Landry, Nathan Grubaugh |
| EPI_ISL_420099, EPI_ISL_420100, EPI_ISL_420101, EPI_ISL_420102, EPI_ISL_420103, EPI_ISL_420104, EPI_ISL_420105, EPI_ISL_420106, EPI_ISL_420107, EPI_ISL_420108, EPI_ISL_420109, EPI_ISL_420110, EPI_ISL_420111 | | | |
| see above | National Centre for Infectious Diseases | Programme in Emerging Infectious Diseases, Duke-NUS Medical School | Danielle E Anderson, Martin Linster, Yan Zhuang, Jayanthi Jayakumar, David CB Lye, Yee Sin Leo, Barnaby E Young, Yvonne CF Su, Gavin JD Smith |
| EPI_ISL_420112 | Servicio de Microbiología. Consorcio Hospital General Universitario de Valencia | Sequencing and Bioinformatics Service and Molecular Epidemiology Research Group. FISABIO-Public Health | Lidia Ruiz Roldan, Marta Pla Diaz, Neris Garcia-Gonzalez, Loreto Ferrús Abad, Inma Galán Vendrell, Paula Ruiz-Hueso, Mariana Reyes-Prieto, Vicente Soriano Chirona, Maria Alma Bracho, Griselda De Marco, Beatriz Beamud, Maria Dolores Ocete, Llúcia Martínez-Priego, Concepcion Gimeno, Giuseppe D'Auria, Fernando Gonzalez-Candelas |
| EPI_ISL_420113 | Servicio de Microbiología. Consorcio Hospital General Universitario de Valencia | Sequencing and Bioinformatics Service and Molecular Epidemiology Research Group. FISABIO-Public Health | Beatriz Beamud, Lidia Ruiz Roldan, Marta Pla Diaz, Neris Garcia-Gonzalez, Loreto Ferrús Abad, Inma Galán Vendrell, Paula Ruiz-Hueso, Mariana Reyes-Prieto, Vicente Soriano Chirona, Maria Alma Bracho, Maria Dolores Ocete, Llúcia Martínez-PriegoGriselda De Marco, , Concepcion Gimeno, Giuseppe D'Auria, Fernando Gonzalez-Candelas |
| EPI_ISL_420114 | Servicio de Microbiología. Consorcio Hospital General Universitario de Valencia | Sequencing and Bioinformatics Service and Molecular Epidemiology Research Group. FISABIO-Public Health | Griselda De Marco, Beatriz Beamud, Lidia Ruiz Roldan, Marta Pla Diaz, Neris Garcia-Gonzalez, Loreto Ferrús Abad, Inma Galán Vendrell, Paula Ruiz-Hueso, Mariana Reyes-Prieto, Vicente Soriano Chirona, Maria Alma Bracho, Maria Dolores Ocete, Llúcia Martínez-Priego, Concepcion Gimeno, Giuseppe D'Auria, Fernando Gonzalez-Candelas |
| EPI_ISL_420115 | Servicio de Microbiología. Consorcio Hospital General Universitario de Valencia | Sequencing and Bioinformatics Service and Molecular Epidemiology Research Group. FISABIO-Public Health | Marta Pla Diaz, Neris Garcia-Gonzalez, Loreto Ferrús Abad, Inma Galán Vendrell, Paula Ruiz-Hueso, Mariana Reyes-Prieto, Vicente Soriano Chirona, Maria Alma Bracho, Griselda De Marco, Beatriz Beamud, Lidia Ruiz Roldan, Maria Dolores Ocete, Llúcia Martínez-Priego, Concepcion Gimeno, Giuseppe D'Auria, Fernando Gonzalez-Candelas |
| EPI_ISL_420116 | Servicio de Microbiología. Consorcio Hospital General Universitario de Valencia | Sequencing and Bioinformatics Service and Molecular Epidemiology Research Group. FISABIO-Public Health | Neris Garcia-Gonzalez, Loreto Ferrús Abad, Inma Galán Vendrell, Paula Ruiz-Hueso, Mariana Reyes-Prieto, Vicente Soriano Chirona, Maria Alma Bracho, Griselda De Marco, Beatriz Beamud, Lidia Ruiz Roldan, Marta Pla Diaz, Maria Dolores Ocete, Llúcia Martínez-Priego, Concepcion Gimeno, Giuseppe D'Auria, Fernando Gonzalez-Candelas |
| EPI_ISL_420117 | Servicio de Microbiología. Consorcio Hospital General Universitario de Valencia | Sequencing and Bioinformatics Service and Molecular Epidemiology Research Group. FISABIO-Public Health | Loreto Ferrús Abad, Inma Galán Vendrell, Paula Ruiz-Hueso, Mariana Reyes-Prieto, Vicente Soriano Chirona, Maria Alma Bracho, Griselda De Marco, Beatriz Beamud, Lidia Ruiz Roldan, Marta Pla Diaz,Neris Garcia-Gonzalez, Maria Dolores Ocete, Llúcia Martínez-Priego, Concepcion Gimeno, Giuseppe D'Auria, Fernando Gonzalez-Candelas |
| EPI_ISL_420118 | Servicio de Microbiología. Consorcio Hospital General Universitario de Valencia | Sequencing and Bioinformatics Service and Molecular Epidemiology Research Group. FISABIO-Public Health | Inma Galán Vendrell, Paula Ruiz-Hueso, Mariana Reyes-Prieto, Vicente Soriano Chirona, Maria Alma Bracho, Griselda De Marco, Beatriz Beamud, Lidia Ruiz Roldan, Marta Pla Diaz,Neris Garcia-Gonzalez, Loreto Ferrús Abad, Maria Dolores Ocete, Llúcia Martínez-Priego, Concepcion Gimeno, Giuseppe D'Auria, Fernando Gonzalez-Candelas |
| EPI_ISL_420119 | Servicio de Microbiología. Consorcio Hospital General Universitario de Valencia | Sequencing and Bioinformatics Service and Molecular Epidemiology Research Group. FISABIO-Public Health | Paula Ruiz-Hueso, Mariana Reyes-Prieto, Vicente Soriano Chirona, Maria Alma Bracho, Griselda De Marco, Beatriz Beamud, Lidia Ruiz Roldan, Marta Pla Diaz,Neris Garcia-Gonzalez, Loreto Ferrús Abad, Inma Galán Vendrell, Maria Dolores Ocete, Llúcia Martínez-Priego, Concepcion Gimeno, Giuseppe D'Auria, Fernando Gonzalez-Candelas |
| EPI_ISL_420120 | Servicio de Microbiología. Consorcio Hospital General Universitario de Valencia | Sequencing and Bioinformatics Service and Molecular Epidemiology Research Group. FISABIO-Public Health | Mariana Reyes-Prieto, Vicente Soriano Chirona, Maria Alma Bracho, Griselda De Marco, Beatriz Beamud, Lidia Ruiz Roldan, Marta Pla Diaz,Neris Garcia-Gonzalez, Loreto Ferrús Abad, Inma Galán Vendrell, Paula Ruiz-Hueso, Maria Dolores Ocete, Llúcia Martínez-Priego, Concepcion Gimeno, Giuseppe D'Auria, Fernando Gonzalez-Candelas |
| EPI_ISL_420121 | Servicio de Microbiología. Consorcio Hospital General Universitario de Valencia | Sequencing and Bioinformatics Service and Molecular Epidemiology Research Group. FISABIO-Public Health | Vicente Soriano Chirona, Maria Alma Bracho, Griselda De Marco, Beatriz Beamud, Lidia Ruiz Roldan, Marta Pla Diaz,Neris Garcia-Gonzalez, Loreto Ferrús Abad, Inma Galán Vendrell, Paula Ruiz-Hueso, Mariana Reyes-Prieto, Maria Dolores Ocete, Llúcia Martínez-Priego, Concepcion Gimeno, Giuseppe D'Auria, Fernando Gonzalez-Candelas |
| EPI_ISL_420122 | Servicio de Microbiología. Consorcio Hospital General Universitario de Valencia | Sequencing and Bioinformatics Service and Molecular Epidemiology Research Group. FISABIO-Public Health | Maria Alma Bracho, Griselda De Marco, Beatriz Beamud, Lidia Ruiz Roldan, Marta Pla Diaz, Neris Garcia-Gonzalez, Loreto Ferrús Abad, Inma Galán Vendrell, Paula Ruiz-Hueso, Mariana Reyes-Prieto, Vicente Soriano Chirona, Maria Dolores Ocete, Llúcia Martínez-Priego, Concepcion Gimeno, Giuseppe D'Auria, Fernando Gonzalez-Candelas |
| EPI_ISL_420123 | Servicio de Microbiología. Consorcio Hospital General Universitario de Valencia | Sequencing and Bioinformatics Service and Molecular Epidemiology Research Group. FISABIO-Public Health | Maria Dolores Ocete, Maria Alma Bracho, Griselda De Marco, Beatriz Beamud, Lidia Ruiz Roldan, Marta Pla Diaz, Neris Garcia-Gonzalez, Loreto Ferrús Abad, Inma Galán Vendrell, Paula Ruiz-Hueso, Mariana Reyes-Prieto, Vicente Soriano Chirona, Llúcia Martínez-Priego, Concepcion Gimeno, Giuseppe D'Auria, Fernando Gonzalez-Candelas |
| EPI_ISL_420124 | Servicio de Microbiología. Consorcio Hospital General Universitario de Valencia | Sequencing and Bioinformatics Service and Molecular Epidemiology Research Group. FISABIO-Public Health | Concepcion Gimeno, Maria Alma Bracho, Griselda De Marco, Beatriz Beamud, Lidia Ruiz Roldan, Marta Pla Diaz, Neris Garcia-Gonzalez, Loreto Ferrús Abad, Inma Galán Vendrell, Paula Ruiz-Hueso, Mariana Reyes-Prieto, Vicente Soriano Chirona, Maria Dolores Ocete, Llúcia Martínez-Priego, Giuseppe D'Auria, Fernando Gonzalez-Candelas |
| EPI_ISL_420125 | Servicio de Microbiologia. Hospital Clinico Universitario de Valencia | Sequencing and Bioinformatics Service and Molecular Epidemiology Research Group. FISABIO-Public Health | David Navarro, Giuseppe D'Auria, Llúcia Martínez-Priego, Maria Alma Bracho, Griselda De Marco, Beatriz Beamud, Lidia Ruiz Roldan, Marta Pla Diaz, Neris Garcia-Gonzalez, Loreto Ferrús Abad, Inma Galán Vendrell, Paula Ruiz-Hueso, Mariana Reyes-Prieto, Vicente Soriano Chirona, Fernando Gonzalez-Candelas |
| EPI_ISL_420126 | Servicio de Microbiologia. Hospital Clinico Universitario de Valencia | Sequencing and Bioinformatics Service and Molecular Epidemiology Research Group. FISABIO-Public Health | Llúcia Martínez-Priego, Maria Alma Bracho, Griselda De Marco, Beatriz Beamud, Lidia Ruiz Roldan, Marta Pla Diaz, Neris Garcia-Gonzalez, Loreto Ferrús Abad, Inma Galán Vendrell, Paula Ruiz-Hueso, Mariana Reyes-Prieto, Vicente Soriano Chirona, David Navarro, Giuseppe D'Auria, Fernando Gonzalez-Candelas |
| EPI_ISL_420127 | Servicio de Microbiologia. Hospital Clinico Universitario de Valencia | Sequencing and Bioinformatics Service and Molecular Epidemiology Research Group. FISABIO-Public Health | Maria Alma Bracho, Griselda De Marco, Beatriz Beamud, Lidia Ruiz Roldan, Marta Pla Diaz, Neris Garcia-Gonzalez, Loreto Ferrús Abad, Inma Galán Vendrell, Paula Ruiz-Hueso, Mariana Reyes-Prieto, Vicente Soriano Chirona, David Navarro, Llúcia Martínez-Priego, Giuseppe D'Auria, Fernando Gonzalez-Candelas |
| EPI_ISL_420128 | Servicio de Microbiologia. Hospital Clinico Universitario de Valencia | Sequencing and Bioinformatics Service and Molecular Epidemiology Research Group. FISABIO-Public Health | Griselda De Marco, Beatriz Beamud, Lidia Ruiz Roldan, Marta Pla Diaz, Neris Garcia-Gonzalez, Loreto Ferrús Abad, Inma Galán Vendrell, Paula Ruiz-Hueso, Mariana Reyes-Prieto, Vicente Soriano Chirona, David Navarro, Maria Alma Bracho, Llúcia Martínez-Priego, Giuseppe D'Auria, Fernando Gonzalez-Candelas |
| EPI_ISL_420129 | Servicio de Microbiologia. Hospital Clinico Universitario de Valencia | Sequencing and Bioinformatics Service and Molecular Epidemiology Research Group. FISABIO-Public Health | Inma Galán Vendrell, Loreto Ferrús Abad, Maria Alma Bracho, Griselda De Marco, Beatriz Beamud, Lidia Ruiz Roldan, Marta Pla Diaz, Neris Garcia-Gonzalez, Paula Ruiz-Hueso, Mariana Reyes-Prieto, Vicente Soriano Chirona, David Navarro, Llúcia Martínez-Priego, Giuseppe D'Auria, Fernando Gonzalez-Candelas |
| EPI_ISL_420130 | Servicio de Microbiologia. Hospital Clinico Universitario de Valencia | Sequencing and Bioinformatics Service and Molecular Epidemiology Research Group. FISABIO-Public Health | David Navarro, Loreto Ferrús Abad, Maria Alma Bracho, Griselda De Marco, Beatriz Beamud, Lidia Ruiz Roldan, Marta Pla Diaz, Neris Garcia-Gonzalez, Inma Galán Vendrell, Paula Ruiz-Hueso, Mariana Reyes-Prieto, Vicente Soriano Chirona, Sandra Carbo, Ivan Ansari, Llúcia Martínez-Priego, Giuseppe D'Auria, Fernando Gonzalez-Candelas |
| EPI_ISL_420131 | Servicio de Microbiologia. Hospital Clinico Universitario de Valencia | Sequencing and Bioinformatics Service and Molecular Epidemiology Research Group. FISABIO-Public Health | Paula Ruiz-Hueso,Loreto Ferrús Abad, Maria Alma Bracho, Griselda De Marco, Beatriz Beamud, Sandra Carbo, Lidia Ruiz Roldan, Marta Pla Diaz, Neris Garcia-Gonzalez, Inma Galán Vendrell, Mariana Reyes-Prieto, Vicente Soriano Chirona, Ivan Ansari, David Navarro, Llúcia Martínez-Priego, Giuseppe D'Auria, Fernando Gonzalez-Candelas |
| EPI_ISL_420132 | Servicio de Microbiologia. Hospital Clinico Universitario de Valencia | Sequencing and Bioinformatics Service and Molecular Epidemiology Research Group. FISABIO-Public Health | Giuseppe D'Auria,Sandra Carbo, Loreto Ferrús Abad, Maria Alma Bracho, Griselda De Marco, Beatriz Beamud, Lidia Ruiz Roldan, Marta Pla Diaz, Neris Garcia-Gonzalez, Inma Galán Vendrell, Paula Ruiz-Hueso, Mariana Reyes-Prieto, Vicente Soriano Chirona, Ivan Ansari, David Navarro, Llúcia Martínez-Priego, Fernando Gonzalez-Candelas |
| EPI_ISL_420134 | Akershus University Hospital, Department for Microbiology and Infectious Disease Control | Norwegian Institute of Public Health, Department of Virology | Kathrine Stene-Johansen, Kamilla Heddeland Instefjord, Hilde Elshaug, Karoline Bragstad, Olav Hungnes |
| EPI_ISL_420135 | Oslo University Hospital, Department of Medical Microbiology | Norwegian Institute of Public Health, Department of Virology | Kathrine Stene-Johansen, Kamilla Heddeland Instefjord, Hilde Elshaug, Karoline Bragstad, Olav Hungnes |
| EPI_ISL_420136 | Akershus University Hospital, Department for Microbiology and Infectious Disease Control | Norwegian Institute of Public Health, Department of Virology | Kathrine Stene-Johansen, Kamilla Heddeland Instefjord, Hilde Elshaug, Karoline Bragstad, Olav Hungnes |
| EPI_ISL_420137 | Vestfold Hospital, Tonsberg Department of Microbiology | Norwegian Institute of Public Health, Department of Virology | Kathrine Stene-Johansen, Kamilla Heddeland Instefjord, Hilde Elshaug, Karoline Bragstad, Olav Hungnes |
| EPI_ISL_420138 | University Hospital of Northern Norway, Department for Microbiology and Infectious Disease Control | Norwegian Institute of Public Health, Department of Virology | Kathrine Stene-Johansen, Kamilla Heddeland Instefjord, Hilde Elshaug, Karoline Bragstad, Olav Hungnes |
| EPI_ISL_420139 | Akershus University Hospital, Department for Microbiology and Infectious Disease Control | Norwegian Institute of Public Health, Department of Virology | Kathrine Stene-Johansen, Kamilla Heddeland Instefjord, Hilde Elshaug, Karoline Bragstad, Olav Hungnes |
| EPI_ISL_420140 | Department for Virology, Molecular Biology and Genome Research, R. G. Lugar Center for Public Health Research, National Center for Disease Control and Public Health (NCDC) of Georgia. | Department for Virology, Molecular Biology and Genome Research, R. G. Lugar Center for Public Health Research, National Center for Disease Control and Public Health (NCDC) of Georgia. | Nato Kotaria, Marine Murtskhvaladze, Ann Machablishvili, Lela Sabadze, Mari Gavashelidze, Ana Papkiauri, Meri Pantsulaia, Gvantsa Brachveli, Tata Imnadze, Tamar Jashiashvili, Tea Tevdoradze, Ketevan Sidamonidze, Ekaterine Khmaladze, Ekaterine Zhghenti, Roena Sukhiashvili, Mariam Zakalashvili, Lela Urushadze, Magda Dgebuadze, Giorgi Tomashvili, Davit Tsaguria, Ekaterine Zangaladze, Nino Berishvili, Gvantsa Chanturia, Adam Kotorashvili, Maia Alkhazashvili, Irma Burjanadze, Anna Kasradze, Khatuna Zakhashvili, Paata Imnadze, Amiran Gamkrelidze |
| EPI_ISL_420141 | Furst Medical Laboratory | Norwegian Institute of Public Health, Department of Virology | Kathrine Stene-Johansen, Kamilla Heddeland Instefjord, Hilde Elshaug, Karoline Bragstad, Olav Hungnes |
| EPI_ISL_420142 | Department for Virology, Molecular Biology and Genome Research, R. G. Lugar Center for Public Health Research, National Center for Disease Control and Public Health (NCDC) of Georgia. | Department for Virology, Molecular Biology and Genome Research, R. G. Lugar Center for Public Health Research, National Center for Disease Control and Public Health (NCDC) of Georgia. | Marine Murtskhvaladze, Ann Machablishvili, Lela Sabadze, Mari Gavashelidze, Ana Papkiauri, Meri Pantsulaia, Gvantsa Brachveli, Tata Imnadze, Tamar Jashiashvili, Tea Tevdoradze, Ketevan Sidamonidze, Ekaterine Khmaladze, Ekaterine Zhghenti, Roena Sukhiashvili, Mariam Zakalashvili, Lela Urushadze, Magda Dgebuadze, Giorgi Tomashvili, Davit Tsaguria, Ekaterine Zangaladze, Nino Berishvili, Gvantsa Chanturia, Adam Kotorashvili, Maia Alkhazashvili, Irma Burjanadze, Anna Kasradze, Khatuna Zakhashvili, Paata Imnadze, Amiran Gamkrelidze. |
| EPI_ISL_420143 | Unilabs Laboratory Medicine | Norwegian Institute of Public Health, Department of Virology | Kathrine Stene-Johansen, Kamilla Heddeland Instefjord, Hilde Elshaug, Karoline Bragstad, Olav Hungnes |
| EPI_ISL_420144 | Department for Virology, Molecular Biology and Genome Research, R. G. Lugar Center for Public Health Research, National Center for Disease Control and Public Health (NCDC) of Georgia. | Department for Virology, Molecular Biology and Genome Research, R. G. Lugar Center for Public Health Research, National Center for Disease Control and Public Health (NCDC) of Georgia. | Gvantsa Chanturia, Ann Machablishvili, Nato Kotaria, Marine Murtskhvaladze, Lela Sabadze, Mari Gavashelidze, Ana Papkiauri, Meri Pantsulaia, Gvantsa Brachveli, Tata Imnadze, Tamar Jashiashvili, Tea Tevdoradze, Ketevan Sidamonidze, Ekaterine Khmaladze, Ekaterine Zhghenti, Roena Sukhiashvili, Mariam Zakalashvili, Lela Urushadze, Magda Dgebuadze, Giorgi Tomashvili, Davit Tsaguria, Ekaterine Zangaladze, Nino Berishvili, Adam Kotorashvili, Maia Alkhazashvili, Irma Burjanadze, Anna Kasradze, Khatuna Zakhashvili, Paata Imnadze, Amiran Gamkrelidze. |
| EPI_ISL_420145 | Forde Hospital Department of Microbiology | Norwegian Institute of Public Health, Department of Virology | Kathrine Stene-Johansen, Kamilla Heddeland Instefjord, Hilde Elshaug, Karoline Bragstad, Olav Hungnes |
| EPI_ISL_420146 | Furst Medical Laboratory | Norwegian Institute of Public Health, Department of Virology | Kathrine Stene-Johansen, Kamilla Heddeland Instefjord, Hilde Elshaug, Karoline Bragstad, Olav Hungnes |
| EPI_ISL_420147 | Hospital of Southern Norway - Kristiansand, Department of Medical Microbiology | Norwegian Institute of Public Health, Department of Virology | Kathrine Stene-Johansen, Kamilla Heddeland Instefjord, Hilde Elshaug, Karoline Bragstad, Olav Hungnes |
| EPI_ISL_420148 | Ostfold Hospital Trust -Kalnes Centre for Laboratory Medicine Section for gene technology and infection serology | Norwegian Institute of Public Health, Department of Virology | Kathrine Stene-Johansen, Kamilla Heddeland Instefjord, Hilde Elshaug, Karoline Bragstad, Olav Hungnes |
| EPI_ISL_420149 | Stavanger University Hospital, Department of Medical Microbiology | Norwegian Institute of Public Health, Department of Virology | Kathrine Stene-Johansen, Kamilla Heddeland Instefjord, Hilde Elshaug, Karoline Bragstad, Olav Hungnes |
| EPI_ISL_420150 | Oslo University Hospital, Department of Medical Microbiology | Norwegian Institute of Public Health, Department of Virology | Kathrine Stene-Johansen, Kamilla Heddeland Instefjord, Hilde Elshaug, Karoline Bragstad, Olav Hungnes |
| EPI_ISL_420151 | Nordland Hospital - Bodo, Laboratory Department, Molecular Biology Unit | Norwegian Institute of Public Health, Department of Virology | Kathrine Stene-Johansen, Kamilla Heddeland Instefjord, Hilde Elshaug, Karoline Bragstad, Olav Hungnes |

| | | | |
|---|---|---|---|
| EPI_ISL_420152, EPI_ISL_420153 | University Hospital of Northern Norway, Department for Microbiology and Infectious Disease Control | Norwegian Institute of Public Health, Department of Virology | Kathrine Stene-Johansen, Kamilla Heddeland Instefjord, Hilde Elshaug, Karoline Bragstad, Olav Hungnes |
| EPI_ISL_420154, EPI_ISL_420155, EPI_ISL_420156, EPI_ISL_420157, EPI_ISL_420158, EPI_ISL_420159, EPI_ISL_420160, EPI_ISL_420161, EPI_ISL_420162, EPI_ISL_420163, EPI_ISL_420164, EPI_ISL_420165, EPI_ISL_420166, EPI_ISL_420167, EPI_ISL_420168, EPI_ISL_420169, EPI_ISL_420170, EPI_ISL_420171, EPI_ISL_420172, EPI_ISL_420173, EPI_ISL_420174, EPI_ISL_420175, EPI_ISL_420176, EPI_ISL_420177, EPI_ISL_420178, EPI_ISL_420179, EPI_ISL_420180, EPI_ISL_420181, EPI_ISL_420182, EPI_ISL_420183, EPI_ISL_420184, EPI_ISL_420185, EPI_ISL_420186, EPI_ISL_420187, EPI_ISL_420188, EPI_ISL_420189, EPI_ISL_420190, EPI_ISL_420191, EPI_ISL_420192, EPI_ISL_420193, EPI_ISL_420194, EPI_ISL_420195, EPI_ISL_420196, EPI_ISL_420197, EPI_ISL_420198, EPI_ISL_420199, EPI_ISL_420200, EPI_ISL_420201, EPI_ISL_420202, EPI_ISL_420203, EPI_ISL_420204, EPI_ISL_420205, EPI_ISL_420206, EPI_ISL_420207, EPI_ISL_420208, EPI_ISL_420209, EPI_ISL_420210, EPI_ISL_420211, EPI_ISL_420212, EPI_ISL_420213, EPI_ISL_420214, EPI_ISL_420215, EPI_ISL_420216, EPI_ISL_420217, EPI_ISL_420218, EPI_ISL_420219, EPI_ISL_420220, EPI_ISL_420221, EPI_ISL_420222, EPI_ISL_420223, EPI_ISL_420224, EPI_ISL_420225, EPI_ISL_420226, EPI_ISL_420227, EPI_ISL_420228, EPI_ISL_420229, EPI_ISL_420230, EPI_ISL_420231, EPI_ISL_420232, EPI_ISL_420233, EPI_ISL_420234, EPI_ISL_420235, EPI_ISL_420236, EPI_ISL_420237, EPI_ISL_420238, EPI_ISL_420239, EPI_ISL_420240, EPI_ISL_420241, EPI_ISL_420242, EPI_ISL_420243, EPI_ISL_420244, EPI_ISL_420245, EPI_ISL_420246, EPI_ISL_420247, EPI_ISL_420248, EPI_ISL_420249, EPI_ISL_420250, EPI_ISL_420251, EPI_ISL_420252, EPI_ISL_420253, EPI_ISL_420254, EPI_ISL_420255, EPI_ISL_420256, EPI_ISL_420257, EPI_ISL_420258, EPI_ISL_420259, EPI_ISL_420260, EPI_ISL_420261, EPI_ISL_420262, EPI_ISL_420263, EPI_ISL_420264, EPI_ISL_420265, EPI_ISL_420266, EPI_ISL_420267, EPI_ISL_420268, EPI_ISL_420269, EPI_ISL_420270, EPI_ISL_420271, EPI_ISL_420272, EPI_ISL_420273, EPI_ISL_420274, EPI_ISL_420275, EPI_ISL_420276, EPI_ISL_420277, EPI_ISL_420278, EPI_ISL_420279, EPI_ISL_420280, EPI_ISL_420281, EPI_ISL_420282, EPI_ISL_420283, EPI_ISL_420284, EPI_ISL_420285, EPI_ISL_420286, EPI_ISL_420287, EPI_ISL_420288, EPI_ISL_420289, EPI_ISL_420290, EPI_ISL_420291, EPI_ISL_420292 | | | |
| see above | Virology Department, Sheffield Teaching Hospitals NHS Foundation Trust | Department of Infection, Immunity and Cardiovascular Disease, The Florey Institute, The Medical School, University of Sheffield | Thushan de Silva, Matthew Parker, Adri Angyal, Rebecca Brown, Rachel Tucker, Paul Parsons, Luke Green, Danielle Groves, Alex Keeley, Dave Partridge, Matthew Wyles, Benjamin Lindsey, Mehmet Yavuz, Mohammad Raza, Cariad Evans |
| EPI_ISL_420293 | Wildlife Conservation Society, Bronx Zoo | Diagnostic Virology Laboratory, United States Department of Agriculture, National Veterinary Services Laboratories | Patrick K. Mitchell, Renee R. Anderson, Brittany Chilson, Roopa Venugopalan, D. G. Diel, Laura B. Goodman, L. Wang, F. Yuan, Y. Fang, Mary Lea Killian, Kerrie Franzen, Nichole Hines Bergeson, Ivan Kuzmin, Melinda Jenkins-Moore, Tod P. Stuber |
| EPI_ISL_420294, EPI_ISL_420295 | Institute of Microbiology and Immunology, Faculty of Medicine, University of Ljubljana | Institute of Microbiology and Immunology, Faculty of Medicine, University of Ljubljana | Samo Zakotnik, Tomaž Mark Zorec, Lucijan Skubic, Miša Korva, Mario Poljak, Tatjana Avšič - Županc |
| EPI_ISL_420296, EPI_ISL_420297, EPI_ISL_420298, EPI_ISL_420299, EPI_ISL_420300, EPI_ISL_420301, EPI_ISL_420302 | NYU Langone Health | Departments of Pathology and Medicine, New York University School of Medicine | Maria Aguero-Rosenfeld, Margaret Black, John Cadley, Paolo Cotzia, John Chen, Dacia Dimartino, Xiaojun Feng, Adriana Heguy, Megan Hogan, Emily Huang, George Jour, Christian Marier, Matthew T. Maurano, Mark J. Mulligan, Peter Meyn, Jared Pinnell, Sitharam Ramaswami, Amy Rapkiewicz, Marie Samanovic-Golden, Antonio Serrano, Guomiao Shen, Matija Snuderl, Nick Vulpescu, Gael Westby, Paul Zappile, Yutong Zhang |
| EPI_ISL_420303 | Alaska State Virology Laboratory | Alaska State Virology Laboratory | Chen, J |
| EPI_ISL_420304, EPI_ISL_420305, EPI_ISL_420306 | Alaska State Virology Laboratory | Alaska State Virology Laboratory | Chen, J. |
| EPI_ISL_420307, EPI_ISL_420308, EPI_ISL_420309 | NYU Langone Health | Departments of Pathology and Medicine, New York University School of Medicine | Maria Aguero-Rosenfeld, Margaret Black, John Cadley, Paolo Cotzia, John Chen, Dacia Dimartino, Xiaojun Feng, Adriana Heguy, Megan Hogan, Emily Huang, George Jour, Christian Marier, Matthew T. Maurano, Mark J. Mulligan, Peter Meyn, Jared Pinnell, Sitharam Ramaswami, Amy Rapkiewicz, Marie Samanovic-Golden, Antonio Serrano, Guomiao Shen, Matija Snuderl, Nick Vulpescu, Gael Westby, Paul Zappile, Yutong Zhang |
| EPI_ISL_420310 | University Hospital of Northern Norway, Department for Microbiology and Infectious Disease Control | Norwegian Institute of Public Health, Department of Virology | Kathrine Stene-Johansen, Kamilla Heddeland Instefjord, Hilde Elshaug, Karoline Bragstad, Olav Hungnes |
| EPI_ISL_420311, EPI_ISL_420312 | Akershus University Hospital, Department for Microbiology and Infectious Disease Control | Norwegian Institute of Public Health, Department of Virology | Kathrine Stene-Johansen, Kamilla Heddeland Instefjord, Hilde Elshaug, Karoline Bragstad, Olav Hungnes |
| EPI_ISL_420313 | Furst Medical Laboratory | Norwegian Institute of Public Health, Department of Virology | Kathrine Stene-Johansen, Kamilla Heddeland Instefjord, Hilde Elshaug, Karoline Bragstad, Olav Hungnes |
| EPI_ISL_420314, EPI_ISL_420315, EPI_ISL_420316, EPI_ISL_420317, EPI_ISL_420318, EPI_ISL_420319, EPI_ISL_420320, EPI_ISL_420321, EPI_ISL_420322, EPI_ISL_420323, EPI_ISL_420324, EPI_ISL_420325, EPI_ISL_420326, EPI_ISL_420327, EPI_ISL_420328, EPI_ISL_420329, EPI_ISL_420330, EPI_ISL_420331, EPI_ISL_420332, EPI_ISL_420333, EPI_ISL_420334, EPI_ISL_420335, EPI_ISL_420336, EPI_ISL_420337, EPI_ISL_420338, EPI_ISL_420339, EPI_ISL_420340, EPI_ISL_420341, EPI_ISL_420342, EPI_ISL_420343, EPI_ISL_420344, EPI_ISL_420345, EPI_ISL_420346, EPI_ISL_420347, EPI_ISL_420348, EPI_ISL_420349, EPI_ISL_420350, EPI_ISL_420351, EPI_ISL_420352, EPI_ISL_420353, EPI_ISL_420354, EPI_ISL_420355, EPI_ISL_420356, EPI_ISL_420357, EPI_ISL_420358, EPI_ISL_420359, EPI_ISL_420360, EPI_ISL_420361, EPI_ISL_420362, EPI_ISL_420363, EPI_ISL_420364, EPI_ISL_420365, EPI_ISL_420366, EPI_ISL_420367, EPI_ISL_420368, EPI_ISL_420369, EPI_ISL_420370, EPI_ISL_420371, EPI_ISL_420372, EPI_ISL_420373, EPI_ISL_420374, EPI_ISL_420375, EPI_ISL_420376, EPI_ISL_420377, EPI_ISL_420378, EPI_ISL_420379, EPI_ISL_420380, EPI_ISL_420381, EPI_ISL_420382, EPI_ISL_420383, EPI_ISL_420384, EPI_ISL_420385, EPI_ISL_420386, EPI_ISL_420387, EPI_ISL_420388, EPI_ISL_420389, EPI_ISL_420390, EPI_ISL_420391, EPI_ISL_420392, EPI_ISL_420393, EPI_ISL_420394, EPI_ISL_420395, EPI_ISL_420396, EPI_ISL_420397, EPI_ISL_420398, EPI_ISL_420399, EPI_ISL_420400, EPI_ISL_420401, EPI_ISL_420402, EPI_ISL_420403, EPI_ISL_420404, EPI_ISL_420405, EPI_ISL_420406, EPI_ISL_420407, EPI_ISL_420408, EPI_ISL_420409, EPI_ISL_420410, EPI_ISL_420411, EPI_ISL_420412, EPI_ISL_420413, EPI_ISL_420414, EPI_ISL_420415, EPI_ISL_420416, EPI_ISL_420417, EPI_ISL_420418, EPI_ISL_420419, EPI_ISL_420420, EPI_ISL_420421, EPI_ISL_420422, EPI_ISL_420423, EPI_ISL_420424, EPI_ISL_420425, EPI_ISL_420426, EPI_ISL_420427, EPI_ISL_420428, EPI_ISL_420429, EPI_ISL_420430, EPI_ISL_420431, EPI_ISL_420432, EPI_ISL_420433, EPI_ISL_420434, EPI_ISL_420435, EPI_ISL_420436, EPI_ISL_420437, EPI_ISL_420438, EPI_ISL_420439, EPI_ISL_420440, EPI_ISL_420441, EPI_ISL_420442, EPI_ISL_420443, EPI_ISL_420444, EPI_ISL_420445, EPI_ISL_420446, EPI_ISL_420447, EPI_ISL_420448, EPI_ISL_420449, EPI_ISL_420450, EPI_ISL_420451, EPI_ISL_420452, EPI_ISL_420453, EPI_ISL_420454 | | | |
| see above | KU Leuven, Clinical and Epidemiological Virology | KU Leuven, Clinical and Epidemiological Virology | Joan Marti-Carreras, Bert Vanmechelen, Tony Wawina, Piet Maes |
| EPI_ISL_420455 | Department of Clinical Pathology, Pamela Youde Nethersole Eastern Hospital | Department of Health Technology and Informatics, Faculty of Health and Social Science, The Hong Kong Polytechnic University | Kenneth Siu-Sing LEUNG, Timothy Ting-Leung NG, Alan Ka-Lun WU, Miranda Chong-Yee YAU, Hiu-Yin LAO, Ming-Pan CHOI, Kingsley King-Gee TAM, Lam-Kwong LEE, Barry Kin-Chung WONG, Alex Yat-Man HO, Kam-Tong YIP, Kwok-Cheung LUNG, Raymond Wai-To LIU, Eugene Yuk-Keung TSO, Wai-Shing LEUNG, Man-Chun CHAN, Yuk-Yung NG, Kit-Man SIN, Kitty Sau-Chun FUNG, Sandy Ka-Yee CHAU, Wing-Kin TO, Tak-Lun QUE, David Ho-Keung SHUM, Shea Ping YIP, Wing Cheong YAM, Gilman Kit-Hang SIU |
| EPI_ISL_420456 | PathWest Laboratory Medicine WA | PathWest Laboratory Medicine WA | Chisha Sikazwe, Jurissa Lang, Avram Levy, David Smith and David Speers |
| EPI_ISL_420457, EPI_ISL_420458, EPI_ISL_420459, EPI_ISL_420460, EPI_ISL_420461, EPI_ISL_420462, EPI_ISL_420463, EPI_ISL_420464, EPI_ISL_420465, EPI_ISL_420466, EPI_ISL_420467, EPI_ISL_420468, EPI_ISL_420469, EPI_ISL_420470, EPI_ISL_420471, EPI_ISL_420472, EPI_ISL_420473, EPI_ISL_420474, EPI_ISL_420475, EPI_ISL_420476, EPI_ISL_420477, EPI_ISL_420478, EPI_ISL_420479, EPI_ISL_420480, EPI_ISL_420481, EPI_ISL_420482, EPI_ISL_420483, EPI_ISL_420484, EPI_ISL_420485, EPI_ISL_420486, EPI_ISL_420487, EPI_ISL_420488, EPI_ISL_420489, EPI_ISL_420490, EPI_ISL_420491, EPI_ISL_420492, EPI_ISL_420493, EPI_ISL_420494, EPI_ISL_420495, EPI_ISL_420496, EPI_ISL_420497, EPI_ISL_420498, EPI_ISL_420499, EPI_ISL_420500, EPI_ISL_420501, EPI_ISL_420502, EPI_ISL_420503, EPI_ISL_420504, EPI_ISL_420505, EPI_ISL_420506, EPI_ISL_420507, EPI_ISL_420508, EPI_ISL_420509, EPI_ISL_420510, EPI_ISL_420511, EPI_ISL_420512, EPI_ISL_420513, EPI_ISL_420514, EPI_ISL_420515, EPI_ISL_420516, EPI_ISL_420517, EPI_ISL_420518, EPI_ISL_420519, EPI_ISL_420520, EPI_ISL_420521, EPI_ISL_420522, EPI_ISL_420523, EPI_ISL_420524, EPI_ISL_420525, EPI_ISL_420526, EPI_ISL_420527, EPI_ISL_420528, EPI_ISL_420529, EPI_ISL_420530 | | | |
| see above | Respiratory Virus Unit, Microbiology Services Colindale, Public Health England | Respiratory Virus Unit, Microbiology Services Colindale, Public Health England | Monica Galiano, Shahjahan Miah, Angie Lackenby, Omolola Akinbami, Tiina Talts, Leena Bhaw, Richard Myers, Steven Platt, Kirstin Edwards, Jonathan Hubb, Joanna Ellis, Maria Zambon |
| EPI_ISL_420531 | Department of Microbiology, PathWest QEII Medical Centre | Department of Microbiology, PathWest QEII Medical Centre | Chisha Sikazwe, Jurissa Lang, Avram Levy, David Speers and David Smith |
| EPI_ISL_420532 | Department of Microbiology, PathWest QEII Medical Centre | Department of Microbiology, PathWest QEII Medical Centre | Chisha Sikazwe, Jurissa Lang, Avram Levy, David Speers and David Smith |
| EPI_ISL_420533 | Department of Microbiology, PathWest QEII Medical Centre | Department of Microbiology, PathWest QEII Medical Centre | Chisha Sikazwe, Jurissa Lang, Avram Levy, David Speers and David Smith |
| EPI_ISL_420534 | Department of Microbiology, PathWest QEII Medical Centre | Department of Microbiology, PathWest QEII Medical Centre | Chisha Sikazwe, Jurissa Lang, Avram Levy, David Speers and David Smith |
| EPI_ISL_420536, EPI_ISL_420537, EPI_ISL_420538 | Department of Microbiology, PathWest QEII Medical Centre | Department of Microbiology, PathWest QEII Medical Centre | Chisha Sikazwe, Jurissa Lang, Avram Levy, David Speers and David Smith |
| EPI_ISL_420539 | Department of Microbiology, PathWest QEII Medical Centre | Department of Microbiology, PathWest QEII Medical Centre | Chisha Sikazwe, Jurissa Lang, Avram Levy, David Speers and David Smith |
| EPI_ISL_420540 | SYNLAB Eesti OÜ | Charité Universitätsmedizin Berlin, Institute of Virology | Victor M Corman, Jörn Beheim-Schwarzbach, Barbara Mühlemann, Talitha Veith, Julia Schneider, Paul Naaber, Terry Jones, Christian Drosten |
| EPI_ISL_420541 | Institute of Microbiology and Immunology, Faculty of Medicine, University of Ljubljana | Institute of Microbiology and Immunology, Faculty of Medicine, University of Ljubljana | Tomaž Mark Zorec, Samo Zakotnik, Lucijan Skubic, Miša Korva, Tatjana Avšič - Županc, Mario Poljak |
| EPI_ISL_420542 | Respiratory Virus Unit, Microbiology Services Colindale, Public Health England | Respiratory Virus Unit, Microbiology Services Colindale, Public Health England | Monica Galiano, Shahjahan Miah, Angie Lackenby, Omolola Akinbami, Tiina Talts, Leena Bhaw, Richard Myers, Steven Platt, Kirstin Edwards, Jonathan Hubb, Joanna Ellis, Maria Zambon |
| EPI_ISL_420543 | National Influenza Center, Indian Council of Medical Research - National Institute of Virology | Indian Council of Medical Research-National Institute of Virology, Microbial Containment Complex | Pragya D. Yadav. Savita Patil, Varsha Potdar, Prasad Sarkale, Dimpal A. Nyayanit, Gajanan Sapkal, Anita M. Shete, Atanu Basu, Lalit Dar, M Choudhary, Amita Jain, Bharati Malhotra, Pranita Gawande, Sarah Cherian, Priya Abraham |
| EPI_ISL_420544 | Indian Council of Medical Research-National Institute of Virology, Microbial Containment Complex | Indian Council of Medical Research-National Institute of Virology, Microbial Containment Complex | Pragya D. Yadav. Savita Patil, Varsha Potdar, Prasad Sarkale, Dimpal A. Nyayanit, Gajanan Sapkal, Anita M. Shete, Atanu Basu, Lalit Dar, M Choudhary, Amita Jain, Bharati Malhotra, Pranita Gawande, Sarah Cherian, Priya Abraham |
| EPI_ISL_420545 | National Influenza Center, Indian Council of Medical Research - National Institute of Virology | Indian Council of Medical Research-National Institute of Virology, Microbial Containment Complex | Pragya D. Yadav. Savita Patil, Varsha Potdar, Prasad Sarkale, Dimpal A. Nyayanit, Gajanan Sapkal, Anita M. Shete, Atanu Basu, Lalit Dar, M Choudhary, Amita Jain, Bharati Malhotra, Pranita Gawande, Sarah Cherian, Priya Abraham |
| EPI_ISL_420546 | Indian Council of Medical Research-National Institute of Virology, Microbial Containment Complex | Indian Council of Medical Research-National Institute of Virology, Microbial Containment Complex | Pragya D. Yadav. Savita Patil, Varsha Potdar, Prasad Sarkale, Dimpal A. Nyayanit, Gajanan Sapkal, Anita M. Shete, Atanu Basu, Lalit Dar, M Choudhary, Amita Jain, Bharati Malhotra, Pranita Gawande, Sarah Cherian, Priya Abraham |
| EPI_ISL_420547 | National Influenza Center, Indian Council of Medical Research - National Institute of Virology | Indian Council of Medical Research-National Institute of Virology, Microbial Containment Complex | Pragya D. Yadav. Savita Patil, Varsha Potdar, Prasad Sarkale, Dimpal A. Nyayanit, Gajanan Sapkal, Anita M. Shete, Atanu Basu, Lalit Dar, M Choudhary, Amita Jain, Bharati Malhotra, Pranita Gawande, Sarah Cherian, Priya Abraham |
| EPI_ISL_420548 | Indian Council of Medical Research-National Institute of Virology, Microbial Containment Complex | Indian Council of Medical Research-National Institute of Virology, Microbial Containment Complex | Pragya D. Yadav. Savita Patil, Varsha Potdar, Prasad Sarkale, Dimpal A. Nyayanit, Gajanan Sapkal, Anita M. Shete, Atanu Basu, Lalit Dar, M Choudhary, Amita Jain, Bharati Malhotra, Pranita Gawande, Sarah Cherian, Priya Abraham |
| EPI_ISL_420549 | National Influenza Center, Indian Council of Medical Research - National Institute of Virology | Indian Council of Medical Research-National Institute of Virology, Microbial Containment Complex | Pragya D. Yadav. Savita Patil, Varsha Potdar, Prasad Sarkale, Dimpal A. Nyayanit, Gajanan Sapkal, Anita M. Shete, Atanu Basu, Lalit Dar, M Choudhary, Amita Jain, Bharati Malhotra, Pranita Gawande, Sarah Cherian, Priya Abraham |
| EPI_ISL_420550 | Indian Council of Medical Research-National Institute of Virology, Microbial Containment Complex | Indian Council of Medical Research-National Institute of Virology, Microbial Containment Complex | Pragya D. Yadav. Savita Patil, Varsha Potdar, Prasad Sarkale, Dimpal A. Nyayanit, Gajanan Sapkal, Anita M. Shete, Atanu Basu, Lalit Dar, M Choudhary, Amita Jain, Bharati Malhotra, Pranita Gawande, Sarah Cherian, Priya Abraham |
| EPI_ISL_420551 | National Influenza Center, Indian Council of Medical Research - National Institute of Virology | Indian Council of Medical Research-National Institute of Virology, Microbial Containment Complex | Pragya D. Yadav. Savita Patil, Varsha Potdar, Prasad Sarkale, Dimpal A. Nyayanit, Gajanan Sapkal, Anita M. Shete, Atanu Basu, Lalit Dar, M Choudhary, Amita Jain, Bharati Malhotra, Pranita Gawande, Sarah Cherian, Priya Abraham |
| EPI_ISL_420552 | Indian Council of Medical Research-National Institute of Virology, Microbial Containment Complex | Indian Council of Medical Research-National Institute of Virology, Microbial Containment Complex | Pragya D. Yadav. Savita Patil, Varsha Potdar, Prasad Sarkale, Dimpal A. Nyayanit, Gajanan Sapkal, Anita M. Shete, Atanu Basu, Lalit Dar, M Choudhary, Amita Jain, Bharati Malhotra, Pranita Gawande, Sarah Cherian, Priya Abraham |
| EPI_ISL_420553 | National Influenza Center, Indian Council of Medical Research - National Institute of Virology | Indian Council of Medical Research-National Institute of Virology, Microbial Containment Complex | Pragya D. Yadav. Savita Patil, Varsha Potdar, Prasad Sarkale, Dimpal A. Nyayanit, Gajanan Sapkal, Anita M. Shete, Atanu Basu, Lalit Dar, M Choudhary, Amita Jain, Bharati Malhotra, Pranita Gawande, Sarah Cherian, Priya Abraham |
| EPI_ISL_420554 | Indian Council of Medical Research-National Institute of Virology, Microbial Containment Complex | Indian Council of Medical Research-National Institute of Virology, Microbial Containment Complex | Pragya D. Yadav. Savita Patil, Varsha Potdar, Prasad Sarkale, Dimpal A. Nyayanit, Gajanan Sapkal, Anita M. Shete, Atanu Basu, Lalit Dar, M Choudhary, Amita Jain, Bharati Malhotra, Pranita Gawande, Sarah Cherian, Priya Abraham |
| EPI_ISL_420555 | National Influenza Center, Indian Council of Medical Research - National Institute of Virology | Indian Council of Medical Research-National Institute of Virology, Microbial Containment Complex | Pragya D. Yadav. Savita Patil, Varsha Potdar, Prasad Sarkale, Dimpal A. Nyayanit, Gajanan Sapkal, Anita M. Shete, Atanu Basu, Lalit Dar, M Choudhary, Amita Jain, Bharati Malhotra, Pranita Gawande, Sarah Cherian, Priya Abraham |
| EPI_ISL_420556 | Indian Council of Medical Research-National Institute of Virology, Microbial Containment Complex | Indian Council of Medical Research-National Institute of Virology, Microbial Containment Complex | Pragya D. Yadav. Savita Patil, Varsha Potdar, Prasad Sarkale, Dimpal A. Nyayanit, Gajanan Sapkal, Anita M. Shete, Atanu Basu, Lalit Dar, M Choudhary, Amita Jain, Bharati Malhotra, Pranita Gawande, Sarah Cherian, Priya Abraham |
| EPI_ISL_420563 | Ospedale Civile Giuseppe Mazzini | Istituto Zooprofilattico Sperimentale dell'Abruzzo e Molise "G. | Lorusso A, Marcacci M, Di Domenico M, Ancora M, Curini V, Mangone I, Rinaldi A, Di Pasquale A, Cammà C, Puglia I, Savini G |

| | | | |
|---|---|---|---|
| | | Caporale" | |
| EPI_ISL_420564 | Ospedale Civile Castel Di Sangro | Istituto Zooprofilattico Sperimentale dell'Abruzzo e Molise "G.Caporale" | Lorusso A, Marcacci M, Di Domenico M, Ancora M, Curini V, Mangone I, Rinaldi A, Di Pasquale A, Cammà C, Puglia I, Savini G |
| EPI_ISL_420565 | Ospedale Civile Giuseppe Mazzini | Istituto Zooprofilattico Sperimentale dell'Abruzzo e Molise "G.Caporale" | Lorusso A, Marcacci M, Di Domenico M, Ancora M, Curini V, Mangone I, Rinaldi A, Di Pasquale A, Cammà C, Puglia I, Savini G |
| EPI_ISL_420566, EPI_ISL_420567 | Ospedale Regionale San Salvatore | Istituto Zooprofilattico Sperimentale dell'Abruzzo e Molise "G.Caporale" | Lorusso A, Marcacci M, Di Domenico M, Ancora M, Curini V, Mangone I, Rinaldi A, Di Pasquale A, Cammà C, Puglia I, Savini G |
| EPI_ISL_420568, EPI_ISL_420569 | Ospedale Civile Giuseppe Mazzini | Istituto Zooprofilattico Sperimentale dell'Abruzzo e Molise "G.Caporale" | Lorusso A, Marcacci M, Di Domenico M, Ancora M, Curini V, Mangone I, Rinaldi A, Di Pasquale A, Cammà C, Puglia I, Savini G |
| EPI_ISL_420570, EPI_ISL_420571, EPI_ISL_420572, EPI_ISL_420573, EPI_ISL_420574, EPI_ISL_420575, EPI_ISL_420576, EPI_ISL_420577, EPI_ISL_420578, EPI_ISL_420579, EPI_ISL_420580, EPI_ISL_420581, EPI_ISL_420582 | | | |
| see above | NYU Langone Health | Departments of Pathology and Medicine, New York University School of Medicine | Maria Aguero-Rosenfeld, Brendan Belovarac, Margaret Black, Ludovic Boytard, John Cadley, Paolo Cotzia, John Chen, Dacia Dimartino, Xiaojun Feng, Tatyana Gindin, Adriana Heguy, Megan Hogan, Emily Huang, George Jour, Andrew Lytle, Christian Marier, Matthew T. Maurano, Mark J. Mulligan, Peter Meyn, Iman Osman, Jared Pinnell, Sitharam Ramaswami, Amy Rapkiewicz, Marie Samanovic-Golden, Antonio Serrano, Guomiao Shen, Matija Snuderl, Theodore Vougiouklakis, Nick Vulpescu, Gael Westby, Paul Zappile, Yutong Zhang |
| EPI_ISL_420583 | Ospedale Civile Giuseppe Mazzini | Istituto Zooprofilattico Sperimentale dell'Abruzzo e Molise "G.Caporale" | Lorusso A, Marcacci M, Di Domenico M, Ancora M, Curini V, Mangone I, Rinaldi A, Di Pasquale A, Cammà C, Puglia I, Savini G |
| EPI_ISL_420584, EPI_ISL_420585, EPI_ISL_420586, EPI_ISL_420587, EPI_ISL_420588, EPI_ISL_420589, EPI_ISL_420590, EPI_ISL_420591 | NYU Langone Health | Departments of Pathology and Medicine, New York University School of Medicine | Maria Aguero-Rosenfeld, Brendan Belovarac, Margaret Black, Ludovic Boytard, John Cadley, Paolo Cotzia, John Chen, Dacia Dimartino, Xiaojun Feng, Tatyana Gindin, Adriana Heguy, Megan Hogan, Emily Huang, George Jour, Andrew Lytle, Christian Marier, Matthew T. Maurano, Mark J. Mulligan, Peter Meyn, Iman Osman, Jared Pinnell, Sitharam Ramaswami, Amy Rapkiewicz, Marie Samanovic-Golden, Antonio Serrano, Guomiao Shen, Matija Snuderl, Theodore Vougiouklakis, Nick Vulpescu, Gael Westby, Paul Zappile, Yutong Zhang |
| EPI_ISL_420592 | Ospedale Civile Giuseppe Mazzini | Istituto Zooprofilattico Sperimentale dell'Abruzzo e Molise "G.Caporale" | Lorusso A, Marcacci M, Di Domenico M, Ancora M, Curini V, Mangone I, Rinaldi A, Di Pasquale A, Cammà C, Puglia I, Savini G |
| EPI_ISL_420598, EPI_ISL_420599, EPI_ISL_420600 | Servicio Virosis Respiratorias-Departamento Virología-INEI | Instituto Nacional Enfermedades Infecciosas C.G.Malbran | Baumeister E., Avaro M., Benedetti E., Russo M., Dattero ME, Pontoriero A., Cisterna D., Molina V., Perandones C., Tuduri E., Lorenzo F., Poklepovich T., Campos J. |
| EPI_ISL_420604, EPI_ISL_420605, EPI_ISL_420606, EPI_ISL_420607, EPI_ISL_420608, EPI_ISL_420609, EPI_ISL_420610, EPI_ISL_420611 | Institut des Agents Infectieux (IAI), Hospices Civils de Lyon | CNR Virus des Infections Respiratoires - France SUD | Antonin Bal, Gregory Destras, Gwendolyne Burfin, Solenne Brun, Carine Moustaud, Raphaelle Lamy, Alexandre Gaymard, Maude Bouscambert-Duchamp, Florence Morfin-Sherpa, Martine Valette, Bruno Lina, Laurence Josset |
| EPI_ISL_420612, EPI_ISL_420613, EPI_ISL_420614 | Centre Hospitalier de Macon | CNR Virus des Infections Respiratoires - France SUD | Antonin Bal, Gregory Destras, Gwendolyne Burfin, Solenne Brun, Carine Moustaud, Raphaelle Lamy, Alexandre Gaymard, Maude Bouscambert-Duchamp, Florence Morfin-Sherpa, Martine Valette, Bruno Lina, Laurence Josset |
| EPI_ISL_420615, EPI_ISL_420616 | Institut des Agents Infectieux (IAI), Hospices Civils de Lyon | CNR Virus des Infections Respiratoires - France SUD | Antonin Bal, Gregory Destras, Gwendolyne Burfin, Solenne Brun, Carine Moustaud, Raphaelle Lamy, Alexandre Gaymard, Maude Bouscambert-Duchamp, Florence Morfin-Sherpa, Martine Valette, Bruno Lina, Laurence Josset |
| EPI_ISL_420617 | Centre Hospitalier Saint Joseph Saint Luc | CNR Virus des Infections Respiratoires - France SUD | Antonin Bal, Gregory Destras, Gwendolyne Burfin, Solenne Brun, Carine Moustaud, Raphaelle Lamy, Alexandre Gaymard, Maude Bouscambert-Duchamp, Florence Morfin-Sherpa, Martine Valette, Bruno Lina, Laurence Josset |
| EPI_ISL_420618, EPI_ISL_420619 | Institut des Agents Infectieux (IAI), Hospices Civils de Lyon | CNR Virus des Infections Respiratoires - France SUD | Antonin Bal, Gregory Destras, Gwendolyne Burfin, Solenne Brun, Carine Moustaud, Raphaelle Lamy, Alexandre Gaymard, Maude Bouscambert-Duchamp, Florence Morfin-Sherpa, Martine Valette, Bruno Lina, Laurence Josset |
| EPI_ISL_420620 | Centre Hospitalier de Bourg en Bresse | CNR Virus des Infections Respiratoires - France SUD | Antonin Bal, Gregory Destras, Gwendolyne Burfin, Solenne Brun, Carine Moustaud, Raphaelle Lamy, Alexandre Gaymard, Maude Bouscambert-Duchamp, Florence Morfin-Sherpa, Martine Valette, Bruno Lina, Laurence Josset |
| EPI_ISL_420621, EPI_ISL_420622, EPI_ISL_420623, EPI_ISL_420624, EPI_ISL_420625 | Institut des Agents Infectieux (IAI), Hospices Civils de Lyon | CNR Virus des Infections Respiratoires - France SUD | Antonin Bal, Gregory Destras, Gwendolyne Burfin, Solenne Brun, Carine Moustaud, Raphaelle Lamy, Alexandre Gaymard, Maude Bouscambert-Duchamp, Florence Morfin-Sherpa, Martine Valette, Bruno Lina, Laurence Josset |
| EPI_ISL_420628, EPI_ISL_420629, EPI_ISL_420630 | Virginia DCLS | Virginia DCLS | Virginia DCLS |
| EPI_ISL_420631, EPI_ISL_420632, EPI_ISL_420633, EPI_ISL_420634, EPI_ISL_420635, EPI_ISL_420636, EPI_ISL_420637, EPI_ISL_420638, EPI_ISL_420639, EPI_ISL_420640, EPI_ISL_420641, EPI_ISL_420642, EPI_ISL_420643, EPI_ISL_420644, EPI_ISL_420645, EPI_ISL_420646, EPI_ISL_420647, EPI_ISL_420648, EPI_ISL_420649, EPI_ISL_420650, EPI_ISL_420651, EPI_ISL_420652, EPI_ISL_420653, EPI_ISL_420654, EPI_ISL_420655, EPI_ISL_420656, EPI_ISL_420657, EPI_ISL_420658, EPI_ISL_420659, EPI_ISL_420660, EPI_ISL_420661, EPI_ISL_420662, EPI_ISL_420663, EPI_ISL_420664, EPI_ISL_420665, EPI_ISL_420666, EPI_ISL_420667, EPI_ISL_420668, EPI_ISL_420669, EPI_ISL_420670, EPI_ISL_420671, EPI_ISL_420672, EPI_ISL_420673, EPI_ISL_420674, EPI_ISL_420675, EPI_ISL_420676, EPI_ISL_420677, EPI_ISL_420678, EPI_ISL_420679, EPI_ISL_420680, EPI_ISL_420681, EPI_ISL_420682, EPI_ISL_420683, EPI_ISL_420684, EPI_ISL_420685, EPI_ISL_420686, EPI_ISL_420687, EPI_ISL_420688, EPI_ISL_420689, EPI_ISL_420690, EPI_ISL_420691, EPI_ISL_420692, EPI_ISL_420693, EPI_ISL_420694, EPI_ISL_420695, EPI_ISL_420696, EPI_ISL_420697, EPI_ISL_420698, EPI_ISL_420699, EPI_ISL_420700, EPI_ISL_420701, EPI_ISL_420702, EPI_ISL_420703, EPI_ISL_420704, EPI_ISL_420705, EPI_ISL_420706, EPI_ISL_420707, EPI_ISL_420708, EPI_ISL_420709, EPI_ISL_420710, EPI_ISL_420711, EPI_ISL_420712, EPI_ISL_420713, EPI_ISL_420714, EPI_ISL_420715, EPI_ISL_420716, EPI_ISL_420717, EPI_ISL_420718, EPI_ISL_420719, EPI_ISL_420720, EPI_ISL_420721, EPI_ISL_420722, EPI_ISL_420723, EPI_ISL_420724, EPI_ISL_420725, EPI_ISL_420726, EPI_ISL_420727, EPI_ISL_420728, EPI_ISL_420729, EPI_ISL_420730, EPI_ISL_420731, EPI_ISL_420732, EPI_ISL_420733, EPI_ISL_420734, EPI_ISL_420735, EPI_ISL_420736, EPI_ISL_420737, EPI_ISL_420738, EPI_ISL_420739, EPI_ISL_420740, EPI_ISL_420741, EPI_ISL_420742, EPI_ISL_420743, EPI_ISL_420744, EPI_ISL_420745, EPI_ISL_420746, EPI_ISL_420747, EPI_ISL_420748, EPI_ISL_420749, EPI_ISL_420750, EPI_ISL_420751, EPI_ISL_420752, EPI_ISL_420753, EPI_ISL_420754, EPI_ISL_420755, EPI_ISL_420756, EPI_ISL_420757, EPI_ISL_420758, EPI_ISL_420759, EPI_ISL_420760, EPI_ISL_420761, EPI_ISL_420762, EPI_ISL_420763, EPI_ISL_420764, EPI_ISL_420765, EPI_ISL_420766, EPI_ISL_420767, EPI_ISL_420768, EPI_ISL_420769, EPI_ISL_420770, EPI_ISL_420771, EPI_ISL_420772, EPI_ISL_420773, EPI_ISL_420774, EPI_ISL_420775, EPI_ISL_420776, EPI_ISL_420777 | | | |
| see above | Respiratory Virus Unit, Microbiology Services Colindale, Public Health England | Respiratory Virus Unit, Microbiology Services Colindale, Public Health England | Monica Galiano, Shahjahan Miah, Angie Lackenby, Omolola Akinbami, Tiina Talts, Leena Bhaw, Richard Myers, Steven Platt, Kirstin Edwards, Jonathan Hubb, Joanna Ellis, Maria Zambon |
| EPI_ISL_420784 | AZ Department of Health Services | Pathogen Discovery, Respiratory Viruses Branch, Division of Viral Diseases, Centers for Disease Control and Prevention | Krista Queen, Yan Li, Ying Tao, Jing Zhang, Anne Uehara, Clinton R. Paden, Haibin Wang, Rachel Marine, Mary S. Keckler, Alison S. Laufer Halpin, Jasmine Padilla, Justin Lee, Christopher A. Elkins, Suxiang Tong |
| EPI_ISL_420785 | FL Bureau of Health Laboratories Tampa | Pathogen Discovery, Respiratory Viruses Branch, Division of Viral Diseases, Centers for Disease Control and Prevention | Krista Queen, Yan Li, Ying Tao, Jing Zhang, Anne Uehara, Clinton R. Paden, Haibin Wang, Rachel Marine, Mary S. Keckler, Alison S. Laufer Halpin, Jasmine Padilla, Justin Lee, Christopher A. Elkins, Suxiang Tong |
| EPI_ISL_420786, EPI_ISL_420787, EPI_ISL_420788 | GA Department of Public Health | Pathogen Discovery, Respiratory Viruses Branch, Division of Viral Diseases, Centers for Disease Control and Prevention | Krista Queen, Yan Li, Ying Tao, Jing Zhang, Anne Uehara, Clinton R. Paden, Haibin Wang, Rachel Marine, Mary S. Keckler, Alison S. Laufer Halpin, Jasmine Padilla, Justin Lee, Christopher A. Elkins, Suxiang Tong |
| EPI_ISL_420789, EPI_ISL_420790 | Illinois Department of Public Health Chicago Laboratory | Pathogen Discovery, Respiratory Viruses Branch, Division of Viral Diseases, Centers for Disease Control and Prevention | Krista Queen, Yan Li, Ying Tao, Jing Zhang, Anne Uehara, Clinton R. Paden, Haibin Wang, Rachel Marine, Mary S. Keckler, Alison S. Laufer Halpin, Jasmine Padilla, Justin Lee, Christopher A. Elkins, Suxiang Tong |
| EPI_ISL_420791, EPI_ISL_420792 | NH Department of Health and Human Services Public Health Labs | Pathogen Discovery, Respiratory Viruses Branch, Division of Viral Diseases, Centers for Disease Control and Prevention | Krista Queen, Yan Li, Ying Tao, Jing Zhang, Anne Uehara, Clinton R. Paden, Haibin Wang, Rachel Marine, Mary S. Keckler, Alison S. Laufer Halpin, Jasmine Padilla, Justin Lee, Christopher A. Elkins, Suxiang Tong |
| EPI_ISL_420793 | NYC Department of Health and Mental Hygiene | Pathogen Discovery, Respiratory Viruses Branch, Division of Viral Diseases, Centers for Disease Control and Prevention | Krista Queen, Yan Li, Ying Tao, Jing Zhang, Anne Uehara, Clinton R. Paden, Haibin Wang, Rachel Marine, Mary S. Keckler, Alison S. Laufer Halpin, Jasmine Padilla, Justin Lee, Christopher A. Elkins, Suxiang Tong |
| EPI_ISL_420794 | Oregon State Public Health- Virology section | Pathogen Discovery, Respiratory Viruses Branch, Division of Viral Diseases, Centers for Disease Control and Prevention | Krista Queen, Yan Li, Ying Tao, Jing Zhang, Anne Uehara, Clinton R. Paden, Haibin Wang, Rachel Marine, Mary S. Keckler, Alison S. Laufer Halpin, Jasmine Padilla, Justin Lee, Christopher A. Elkins, Suxiang Tong |
| EPI_ISL_420795 | RI State Health Laboratory | Pathogen Discovery, Respiratory Viruses Branch, Division of Viral Diseases, Centers for Disease Control and Prevention | Krista Queen, Yan Li, Ying Tao, Jing Zhang, Anne Uehara, Clinton R. Paden, Haibin Wang, Rachel Marine, Mary S. Keckler, Alison S. Laufer Halpin, Jasmine Padilla, Justin Lee, Christopher A. Elkins, Suxiang Tong |
| EPI_ISL_420796, EPI_ISL_420797, EPI_ISL_420798 | Texas DSHS Lab Services | Pathogen Discovery, Respiratory Viruses Branch, Division of Viral Diseases, Centers for Disease Control and Prevention | Krista Queen, Yan Li, Ying Tao, Jing Zhang, Anne Uehara, Clinton R. Paden, Haibin Wang, Rachel Marine, Mary S. Keckler, Alison S. Laufer Halpin, Jasmine Padilla, Justin Lee, Christopher A. Elkins, Suxiang Tong |
| EPI_ISL_420799, EPI_ISL_420800, EPI_ISL_420801 | Brian D. Allgood Army Community Hospital | Pathogen Discovery, Respiratory Viruses Branch, Division of Viral Diseases, Centers for Disease Control and Prevention | Krista Queen, Yan Li, Ying Tao, Jing Zhang, Anne Uehara, Clinton R. Paden, Haibin Wang, Rachel Marine, Mary S. Keckler, Alison S. Laufer Halpin, Jasmine Padilla, Justin Lee, Christopher A. Elkins, Suxiang Tong |
| EPI_ISL_420803, EPI_ISL_420804, EPI_ISL_420805, EPI_ISL_420806, EPI_ISL_420807, EPI_ISL_420808, EPI_ISL_420809, EPI_ISL_420810, EPI_ISL_420811, EPI_ISL_420812, EPI_ISL_420813, EPI_ISL_420814, EPI_ISL_420815, EPI_ISL_420816, EPI_ISL_420817, EPI_ISL_420818, EPI_ISL_420819, EPI_ISL_420820, EPI_ISL_420821, EPI_ISL_420822, EPI_ISL_420823, EPI_ISL_420824, EPI_ISL_420825 | | | |
| see above | Utah Public Health Laboratory | Utah Public Health Laboratory | Erin Young, Kelly Oakeson |
| EPI_ISL_420838, EPI_ISL_420839, EPI_ISL_420840, EPI_ISL_420841, EPI_ISL_420842, EPI_ISL_420843, EPI_ISL_420844, EPI_ISL_420845, EPI_ISL_420846, EPI_ISL_420847, EPI_ISL_420848, EPI_ISL_420849, EPI_ISL_420850, EPI_ISL_420851, EPI_ISL_420852, EPI_ISL_420853, EPI_ISL_420854 | | | |
| see above | Viral Respiratory Lab, National Institute for Biomedical Research (INRB) | Pathogen Sequencing Lab, National Institute for Biomedical Research (INRB) | Placide Mbala-Kingebeni, Edith Nkwembe, Eddy Kinganda-Lusamaki, Amuri Aziza, Catherine Pratt, Matthias Pauthner, Josh Quick, Allison Black, James Hadfield, Trevor Bedford, Ian Goodfellow, Nick Loman, Kristian Andersen, Michael Wiley, Steve Ahuka-Mundeke, Jean-Jacques Muyembe Tamfum |
| EPI_ISL_420855, EPI_ISL_420876, EPI_ISL_420877 | Geelong Centre for Emerging Infectious Diseases | Geelong Centre for Emerging Infectious Diseases | Chamings,A., Raj Bhatta T., Alexandersen S. |
| EPI_ISL_420878, EPI_ISL_420879 | Mater Pathology | Public Health Virology Laboratory | Bixing Huang, Alyssa Pyke, Amanda De Jong, Andrew Van Den Hurk, Carmel Taylor, David Warrilow, Doris Genge, Elisabeth Gamez, Glen Hewitson, Ian Maxwell Mackay, Inga Sultana, Jamie McMahon, Jean Barcelon, Judy Northill, Mitchell Finger, Natalie Simpson, Neelima Nair, Peter Burtonclay, Peter Moore, Sarah Wheatley, Sean Moody, Sonja Hall-Mendelin, Timothy Gardam, and Frederick Moore |
| EPI_ISL_420889, EPI_ISL_420890 | Takayuki Hishiki Kanagawa Prefectural Institute of Public Health | Takayuki Hishiki Kanagawa Prefectural Institute of Public Health | Hishiki,T., Suzuki,R., Sakuragi,J., Usui,K., Tanaka,Y., Kawai,J., Kogo,Y., Matsuki,Y., An,T., Hayashizaki,Y. and Takasaki,T. |
| EPI_ISL_420898 | Max von Pettenkofer Institute, Virology, National Reference Center for Retroviruses, LMU Munich | Laboratory for Functional Genome Analysis, Dept. Genomics, Gene Center of the LMU Munich | Max Muenchhoff, Stefan Krebs, Alexander Graf, Ashok Varadharajan, Oliver Keppler, Helmut Blum |
| EPI_ISL_420899 | Max von Pettenkofer Institute, Virology, National Reference Center for Retroviruses, LMU Munich | Laboratory for Functional Genome Analysis, Dept. Genomics, Gene Center of the LMU Munich | Max Muenchhoff, Stefan Krebs, Alexander Graf, Ashok Varadharajan, Oliver Keppler, Helmut Blum |
| EPI_ISL_420900 | Max von Pettenkofer Institute, Virology, National Reference Center for Retroviruses, LMU Munich | Laboratory for Functional Genome Analysis, Dept. Genomics, Gene Center of the LMU Munich | Max Muenchhoff, Stefan Krebs, Alexander Graf, Ashok Varadharajan, Oliver Keppler, Helmut Blum |
| EPI_ISL_420901, EPI_ISL_420902 | Max von Pettenkofer Institute, Virology, National Reference Center for Retroviruses, LMU Munich | Laboratory for Functional Genome Analysis, Dept. Genomics, Gene Center of the LMU Munich | Max Muenchhoff, Stefan Krebs, Alexander Graf, Ashok Varadharajan, Oliver Keppler, Helmut Blum |
| EPI_ISL_420903, EPI_ISL_420904 | Max von Pettenkofer Institute, Virology, National Reference Center for Retroviruses, LMU Munich | Laboratory for Functional Genome Analysis, Dept. Genomics, Gene Center of the LMU Munich | Max Muenchhoff, Stefan Krebs, Alexander Graf, Ashok Varadharajan, Oliver Keppler, Helmut Blum |

| | | | |
|---|---|---|---|
| EPI_ISL_420905, EPI_ISL_420906 | Max von Pettenkofer Institute, Virology, National Reference Center for Retroviruses, LMU Munich | Laboratory for Functional Genome Analysis, Dept. Genomics, Gene Center of the LMU Munich | Max Muenchhoff, Stefan Krebs, Alexander Graf, Ashok Varadharajan, Oliver Keppler, Helmut Blum |
| EPI_ISL_420907 | Max von Pettenkofer Institute, Virology, National Reference Center for Retroviruses, LMU Munich | Laboratory for Functional Genome Analysis, Dept. Genomics, Gene Center of the LMU Munich | Max Muenchhoff, Stefan Krebs, Alexander Graf, Ashok Varadharajan, Oliver Keppler, Helmut Blum |
| EPI_ISL_420908, EPI_ISL_420909 | Max von Pettenkofer Institute, Virology, National Reference Center for Retroviruses, LMU Munich | Laboratory for Functional Genome Analysis, Dept. Genomics, Gene Center of the LMU Munich | Max Muenchhoff, Stefan Krebs, Alexander Graf, Ashok Varadharajan, Oliver Keppler, Helmut Blum |
| EPI_ISL_420910 | Respiratory Virus Unit, Microbiology Services Colindale, Public Health England | Respiratory Virus Unit, Microbiology Services Colindale, Public Health England | Monica Galiano, Shahjahan Miah, Angie Lackenby, Omolola Akinbami, Tiina Talts, Leena Bhaw, Richard Myers, Steven Platt, Kirstin Edwards, Jonathan Hubb, Joanna Ellis, Maria Zambon |
| EPI_ISL_420911, EPI_ISL_420912 | Max von Pettenkofer Institute, Virology, National Reference Center for Retroviruses, LMU Munich | Laboratory for Functional Genome Analysis, Dept. Genomics, Gene Center of the LMU Munich | Max Muenchhoff, Stefan Krebs, Alexander Graf, Ashok Varadharajan, Oliver Keppler, Helmut Blum |
| EPI_ISL_420913, EPI_ISL_420914, EPI_ISL_420915, EPI_ISL_420916, EPI_ISL_420917, EPI_ISL_420918, EPI_ISL_420919, EPI_ISL_420920, EPI_ISL_420921, EPI_ISL_420922, EPI_ISL_420923, EPI_ISL_420924, EPI_ISL_420925, EPI_ISL_420926, EPI_ISL_420927, EPI_ISL_420928, EPI_ISL_420929, EPI_ISL_420930, EPI_ISL_420931, EPI_ISL_420932, EPI_ISL_420933, EPI_ISL_420934, EPI_ISL_420935, EPI_ISL_420936, EPI_ISL_420937, EPI_ISL_420938, EPI_ISL_420939, EPI_ISL_420940, EPI_ISL_420941, EPI_ISL_420942, EPI_ISL_420943, EPI_ISL_420944, EPI_ISL_420945, EPI_ISL_420946, EPI_ISL_420947, EPI_ISL_420948, EPI_ISL_420949, EPI_ISL_420950, EPI_ISL_420951, EPI_ISL_420952, EPI_ISL_420953, EPI_ISL_420954, EPI_ISL_420955, EPI_ISL_420956, EPI_ISL_420957, EPI_ISL_420958, EPI_ISL_420959, EPI_ISL_420960, EPI_ISL_420961, EPI_ISL_420962, EPI_ISL_420963, EPI_ISL_420964, EPI_ISL_420965, EPI_ISL_420966, EPI_ISL_420967, EPI_ISL_420968, EPI_ISL_420969, EPI_ISL_420970, EPI_ISL_420971, EPI_ISL_420972, EPI_ISL_420973, EPI_ISL_420974, EPI_ISL_420975, EPI_ISL_420976, EPI_ISL_420977, EPI_ISL_420978, EPI_ISL_420979, EPI_ISL_420980, EPI_ISL_420981, EPI_ISL_420982, EPI_ISL_420983, EPI_ISL_420984, EPI_ISL_420985, EPI_ISL_420986, EPI_ISL_420987, EPI_ISL_420988, EPI_ISL_420989, EPI_ISL_420990, EPI_ISL_420991, EPI_ISL_420992, EPI_ISL_420993, EPI_ISL_420994, EPI_ISL_420995, EPI_ISL_420996, EPI_ISL_420997, EPI_ISL_420998, EPI_ISL_420999, EPI_ISL_421000, EPI_ISL_421001, EPI_ISL_421002, EPI_ISL_421003, EPI_ISL_421004, EPI_ISL_421005, EPI_ISL_421006, EPI_ISL_421007, EPI_ISL_421008, EPI_ISL_421009, EPI_ISL_421010, EPI_ISL_421011 | | | |
| see above | Wales Specialist Virology Centre | Public Health Wales Microbiology Cardiff | Catherine Moore, Joanne Watkins, Sally Corden, Malorie Perry, Simon Cottrell Sara Rey, Matt Bull, Tom Connor |
| EPI_ISL_421171 | Hospital Universitario 12 de Octubre | Hospital Universitario 12 de Octubre | Esther Viedma, Sara González, Elias Dahdouh, Raúl Recio, Fernando Lázaro, Julio García, Mª Dolores Folgueira, Jesús Mingorance, Rafael Delgado |
| EPI_ISL_421172, EPI_ISL_421173, EPI_ISL_421174, EPI_ISL_421175 | Hospital Universitario 12 de Octubre | Hospital Universitario 12 de Octubre | Esther Viedma, Sara González, Elias Dahdouh, Raúl Recio, Fernando Lázaro, Julio García, Mª Dolores Folgueira, Jesús Mingorance, Rafael Delgado |
| EPI_ISL_421176 | Hospital Universitario 12 de Octubre | Hospital Universitario 12 de Octubre | Esther Viedma, Sara González, Elias Dahdouh, Raúl Recio, Fernando Lázaro, Julio García, Mª Dolores Folgueira, Jesús Mingorance, Rafael Delgado |
| EPI_ISL_421177, EPI_ISL_421178, EPI_ISL_421179, EPI_ISL_421180 | Hospital Universitario 12 de Octubre | Hospital Universitario 12 de Octubre | Esther Viedma, Sara González, Elias Dahdouh, Raúl Recio, Fernando Lázaro, Julio García, Mª Dolores Folgueira, Jesús Mingorance, Rafael Delgado |
| EPI_ISL_421182, EPI_ISL_421183, EPI_ISL_421184, EPI_ISL_421185, EPI_ISL_421186, EPI_ISL_421187, EPI_ISL_421188, EPI_ISL_421189, EPI_ISL_421190, EPI_ISL_421191, EPI_ISL_421192, EPI_ISL_421193, EPI_ISL_421194, EPI_ISL_421195, EPI_ISL_421196, EPI_ISL_421197, EPI_ISL_421198, EPI_ISL_421199, EPI_ISL_421200, EPI_ISL_421201, EPI_ISL_421202, EPI_ISL_421203, EPI_ISL_421204, EPI_ISL_421205, EPI_ISL_421206, EPI_ISL_421207, EPI_ISL_421208, EPI_ISL_421209, EPI_ISL_421210, EPI_ISL_421211, EPI_ISL_421212, EPI_ISL_421213, EPI_ISL_421214 | | | |
| see above | Department of Clinical Microbiology | GIGA Medical Genomics | Keith Durkin, Maria Artesi, Sébastien Bontems, Raphaël Boreux, Cécile Meex, Pierrette Melin, Marie-Pierre Hayette, Vincent Bours. |
| EPI_ISL_421221, EPI_ISL_421222, EPI_ISL_421223, EPI_ISL_421224, EPI_ISL_421225, EPI_ISL_421226, EPI_ISL_421227, EPI_ISL_421228, EPI_ISL_421229, EPI_ISL_421230, EPI_ISL_421231, EPI_ISL_421232, EPI_ISL_421233, EPI_ISL_421234, EPI_ISL_421235, EPI_ISL_421236 | | | |
| see above | Hangzhou Center for Diseases Control and Prevention | Hangzhou Center for Diseases Control and Prevention | Jun Li, Haoqiu Wang, Lingfeng Mao, Hua Yu, Xinfen Yu, Zhou Sun, Xin Qian, Shuchang Chen, Junfang Chen, Xuchu Wang |
| EPI_ISL_421237, EPI_ISL_421238, EPI_ISL_421239, EPI_ISL_421240, EPI_ISL_421241, EPI_ISL_421242, EPI_ISL_421243, EPI_ISL_421244, EPI_ISL_421245, EPI_ISL_421246, EPI_ISL_421247, EPI_ISL_421248, EPI_ISL_421249, EPI_ISL_421250, EPI_ISL_421251, EPI_ISL_421252, EPI_ISL_421253, EPI_ISL_421254, EPI_ISL_421256, EPI_ISL_421257, EPI_ISL_421258, EPI_ISL_421259, EPI_ISL_421260, EPI_ISL_421261, EPI_ISL_421262 | | | |
| see above | Jiangxi Province Center for Disease Control and Prevention | Jiangxi Province Center for Disease Control and Prevention | JianXiong Li,Ying Xiong,Tian Gong,Yong Shi,Jun Zhou,Fang Xiao,ShiWen Liu,XiaoQing Liu,Gang Xu,DaJin Xiao,Xin Ran,YanNi Zhang |
| EPI_ISL_421272 | Wyoming Public Health Laboratory | Center for Global Health, University of New Mexico Health Sciences Center | Daryl Domman, Kurt Schwalm, Rob Christensen, Wanda Manley, Cari Sloma, Noah Hull, Darrell Dinwiddie |
| EPI_ISL_421275 | Russian State Collection of Viruses | Pathogenic Microorganisms Variability Laboratory | Alexey Shchetinin, Maria Nikiforova, Nadezhda Kuznetsova, Ekaterina Aksenova, Marina Kunda, Natalia Ryzhova, Olga Voronina, Inna Dolzhikova, Daria Grousova, Andrey Botikov, Denis Logunov, Alexander Gintsburg, Vladimir Gushchin |
| EPI_ISL_421279 | Clinical Diagnostics Laboratory, Diagnostic & Experimental Pathology, Lilly Research Laboratories | Clinical Diagnostics Laboratory, Diagnostic & Experimental Pathology, Lilly Research Laboratories | Tim Holzer, Mayuri Vaidya, Angie Fulford, Sam McNeely, Rachael Redmond, Phil Ebert, John Calley, Leslie O'Neill Reising, Pat Finnegan, Erin Wray, John McElwee, Jeff Fill, Joe Oakley, Andrew Schade |
| EPI_ISL_421281 | Clinical Diagnostics Laboratory, Diagnostic & Experimental Pathology, Lilly Research Laboratories | Clinical Diagnostics Laboratory, Diagnostic & Experimental Pathology, Lilly Research Laboratories | Tim Holzer, Mayuri Vaidya, Angie Fulford, Sam McNeely, Rachael Redmond, Phil Ebert, John Calley, Leslie O'Neill Reising, Pat Finnegan, Erin Wray, John McElwee, Jeff Fill, Joe Oakley, Andrew Schade |
| EPI_ISL_421283, EPI_ISL_421284, EPI_ISL_421285, EPI_ISL_421286, EPI_ISL_421287, EPI_ISL_421288, EPI_ISL_421289, EPI_ISL_421290, EPI_ISL_421291, EPI_ISL_421292, EPI_ISL_421293, EPI_ISL_421294, EPI_ISL_421295, EPI_ISL_421296, EPI_ISL_421297, EPI_ISL_421298, EPI_ISL_421299, EPI_ISL_421300, EPI_ISL_421301, EPI_ISL_421302, EPI_ISL_421303, EPI_ISL_421304, EPI_ISL_421305, EPI_ISL_421306, EPI_ISL_421307, EPI_ISL_421308, EPI_ISL_421309, EPI_ISL_421310, EPI_ISL_421311, EPI_ISL_421312, EPI_ISL_421313, EPI_ISL_421314, EPI_ISL_421315, EPI_ISL_421316, EPI_ISL_421317, EPI_ISL_421318, EPI_ISL_421319, EPI_ISL_421320, EPI_ISL_421321, EPI_ISL_421322, EPI_ISL_421323, EPI_ISL_421324, EPI_ISL_421325, EPI_ISL_421326, EPI_ISL_421327, EPI_ISL_421328, EPI_ISL_421329, EPI_ISL_421330, EPI_ISL_421331, EPI_ISL_421332, EPI_ISL_421333, EPI_ISL_421334, EPI_ISL_421335, EPI_ISL_421336, EPI_ISL_421338, EPI_ISL_421339, EPI_ISL_421340, EPI_ISL_421341, EPI_ISL_421342, EPI_ISL_421343 | | | |
| see above | University of Wisconsin-Madison AIDS Vaccine Research Laboratories | University of Wisconsin-Madison AIDS Vaccine Research Laboratories | Gage Moreno, Katarina Braun, et al. AIDS Vaccine Research Laboratories |
| EPI_ISL_421345, EPI_ISL_421346, EPI_ISL_421347 | Wyoming Public Health Laboratory | Center for Global Health, University of New Mexico Health Sciences Center | Daryl Domman, Kurt Schwalm, Rob Christensen, Wanda Manley, Cari Sloma, Noah Hull, Darrell Dinwiddie |
| EPI_ISL_421348, EPI_ISL_421349, EPI_ISL_421350, EPI_ISL_421351, EPI_ISL_421352, EPI_ISL_421353, EPI_ISL_421354, EPI_ISL_421355, EPI_ISL_421356, EPI_ISL_421357, EPI_ISL_421358, EPI_ISL_421359, EPI_ISL_421360, EPI_ISL_421361, EPI_ISL_421362, EPI_ISL_421363, EPI_ISL_421364, EPI_ISL_421365, EPI_ISL_421366, EPI_ISL_421367, EPI_ISL_421368, EPI_ISL_421369, EPI_ISL_421370, EPI_ISL_421371, EPI_ISL_421372, EPI_ISL_421373, EPI_ISL_421374, EPI_ISL_421375, EPI_ISL_421376, EPI_ISL_421377, EPI_ISL_421378, EPI_ISL_421379, EPI_ISL_421380, EPI_ISL_421381, EPI_ISL_421382, EPI_ISL_421383, EPI_ISL_421384, EPI_ISL_421385, EPI_ISL_421386, EPI_ISL_421387, EPI_ISL_421388, EPI_ISL_421389, EPI_ISL_421390, EPI_ISL_421391, EPI_ISL_421392, EPI_ISL_421393, EPI_ISL_421394, EPI_ISL_421395, EPI_ISL_421396, EPI_ISL_421397, EPI_ISL_421398, EPI_ISL_421399, EPI_ISL_421400, EPI_ISL_421401, EPI_ISL_421402, EPI_ISL_421403, EPI_ISL_421404, EPI_ISL_421405, EPI_ISL_421406, EPI_ISL_421407, EPI_ISL_421408, EPI_ISL_421409, EPI_ISL_421410, EPI_ISL_421411, EPI_ISL_421412, EPI_ISL_421413, EPI_ISL_421414, EPI_ISL_421415, EPI_ISL_421416, EPI_ISL_421417, EPI_ISL_421418, EPI_ISL_421419, EPI_ISL_421420, EPI_ISL_421421, EPI_ISL_421422, EPI_ISL_421423, EPI_ISL_421424, EPI_ISL_421425, EPI_ISL_421426, EPI_ISL_421427, EPI_ISL_421428, EPI_ISL_421429, EPI_ISL_421430, EPI_ISL_421431, EPI_ISL_421432, EPI_ISL_421433, EPI_ISL_421434, EPI_ISL_421435 | | | |
| see above | MSHS Clinical Microbiology Laboratories | MSHS Pathogen Surveillance Program | Ana S. Gonzalez-Reiche, Mitchell Sullivan, Ajay Obla, Gopi Patel, Emilia Sordillo, Melissa Gitman, Alberto Paniz-mondolfi, Matthew Hernandez, Shelcie Fabre, Jose Polanco, Zenab Khan, Bremy Albuquerque, Jayeeta Dutta, Juan Soto, Shwetha Sridhar Hara, Ying-Chih Wang, Melissa Smith, Robert Sebra, Lisa Miorin, Wen-chun Liu, Randy Albrecht, Judith Aberg, Florian Krammer, Adolfo Garcia-Sarstre, Viviana Simon, Harm van Bakel |
| EPI_ISL_421446, EPI_ISL_421447, EPI_ISL_421448 | H Guimaraes | Instituto Nacional de Saude (INSA) | Guiomar et al |
| EPI_ISL_421449 | H Dr. Nelio Mendonca - Funchal | Instituto Nacional de Saude (INSA) | Guiomar et al |
| EPI_ISL_421450, EPI_ISL_421451, EPI_ISL_421452 | Instituto Nacional de Saude (INSA) | Instituto Nacional de Saude (INSA) | Guiomar et al |
| EPI_ISL_421453 | CHTMAD | Instituto Nacional de Saude (INSA) | Guiomar et al |
| EPI_ISL_421454 | H Beatriz Angelo | Instituto Nacional de Saude (INSA) | Guiomar et al |
| EPI_ISL_421455 | CH Barreiro Montijo | Instituto Nacional de Saude (INSA) | Guiomar et al |
| EPI_ISL_421456 | Instituto Nacional de Saude (INSA) | Instituto Nacional de Saude (INSA) | Guiomar et al |
| EPI_ISL_421457 | H Dr Nelio Mendonca - Funchal | Instituto Nacional de Saude (INSA) | Guiomar et al |
| EPI_ISL_421458 | H Beatriz Angelo | Instituto Nacional de Saude (INSA) | Guiomar et al |
| EPI_ISL_421459, EPI_ISL_421460, EPI_ISL_421461 | H Dr. Nelio Mendonca - Funchal | Instituto Nacional de Saude (INSA) | Guiomar et al |
| EPI_ISL_421462 | H Santarem | Instituto Nacional de Saude (INSA) | Guiomar et al |
| EPI_ISL_421463 | HSE Ilha Terceira - Angra do Heroismo | Instituto Nacional de Saude (INSA) | Guiomar et al |
| EPI_ISL_421464, EPI_ISL_421465 | CHTMAD | Instituto Nacional de Saude (INSA) | Guiomar et al |
| EPI_ISL_421466, EPI_ISL_421467 | H Evora | Instituto Nacional de Saude (INSA) | Guiomar et al |
| EPI_ISL_421468, EPI_ISL_421469, EPI_ISL_421470, EPI_ISL_421471 | H Santarem | Instituto Nacional de Saude (INSA) | Guiomar et al |
| EPI_ISL_421472, EPI_ISL_421473, EPI_ISL_421474, EPI_ISL_421475, EPI_ISL_421476, EPI_ISL_421477, EPI_ISL_421478 | Instituto Nacional de Saude (INSA) | Instituto Nacional de Saude (INSA) | Guiomar et al |
| EPI_ISL_421479, EPI_ISL_421480 | CH Barreiro Montijo | Instituto Nacional de Saude (INSA) | Guiomar et al |
| EPI_ISL_421481 | H Beatriz Angelo | Instituto Nacional de Saude (INSA) | Guiomar et al |
| EPI_ISL_421482, EPI_ISL_421483, EPI_ISL_421484 | CH VN Gaia - Espinho | Instituto Nacional de Saude (INSA) | Guiomar et al |
| EPI_ISL_421485, EPI_ISL_421486 | CH Barreiro Montijo | Instituto Nacional de Saude (INSA) | Guiomar et al |
| EPI_ISL_421487 | H Beatriz Angelo | Instituto Nacional de Saude (INSA) | Guiomar et al |
| EPI_ISL_421488 | H Santarem | Instituto Nacional de Saude (INSA) | Guiomar et al |
| EPI_ISL_421489 | HSE Ilha Terceira - Angra do Heroismo | Instituto Nacional de Saude (INSA) | Guiomar et al |
| EPI_ISL_421490 | H Santarem | Instituto Nacional de Saude (INSA) | Guiomar et al |
| EPI_ISL_421491 | H Beatriz Angelo | Instituto Nacional de Saude (INSA) | Guiomar et al |
| EPI_ISL_421492 | H Santarem | Instituto Nacional de Saude (INSA) | Guiomar et al |
| EPI_ISL_421493, EPI_ISL_421494 | HSE Ilha Terceira - Angra do Heroismo | Instituto Nacional de Saude (INSA) | Guiomar et al |
| EPI_ISL_421495 | H Santarem | Instituto Nacional de Saude (INSA) | Guiomar et al |

| | | | |
|---|---|---|---|
| EPI_ISL_421496, EPI_ISL_421497, EPI_ISL_421498, EPI_ISL_421499 | Instituto Nacional de Saude (INSA) | Instituto Nacional de Saude (INSA) | Guiomar et al |
| EPI_ISL_421500 | CH Compiègne Laboratoire de Biologie | National Reference Center for Viruses of Respiratory Infections, Institut Pasteur, Paris | Mélanie Albert, Marion Barbet, Sylvie Behillil, Méline Bizard, Angela Brisebarre, Flora Donati, Etienne Simon-Lorière, Vincent Enouf, Maud Vanpeene, Sylvie van der Werf, Raulin Olivia |
| EPI_ISL_421501 | Service de Biologie Médicale - BP 125 | National Reference Center for Viruses of Respiratory Infections, Institut Pasteur, Paris | Mélanie Albert, Marion Barbet, Sylvie Behillil, Méline Bizard, Angela Brisebarre, Flora Donati, Etienne Simon-Lorière, Vincent Enouf, Maud Vanpeene, Sylvie van der Werf, Christine Lambert |
| EPI_ISL_421502, EPI_ISL_421503 | Parc des Dames | National Reference Center for Viruses of Respiratory Infections, Institut Pasteur, Paris | Mélanie Albert, Marion Barbet, Sylvie Behillil, Méline Bizard, Angela Brisebarre, Flora Donati, Etienne Simon-Lorière, Vincent Enouf, Maud Vanpeene, Sylvie van der Werf |
| EPI_ISL_421504, EPI_ISL_421505, EPI_ISL_421506 | Service de Biologie Médicale - BP 125 | National Reference Center for Viruses of Respiratory Infections, Institut Pasteur, Paris | Mélanie Albert, Marion Barbet, Sylvie Behillil, Méline Bizard, Angela Brisebarre, Flora Donati, Etienne Simon-Lorière, Vincent Enouf, Maud Vanpeene, Sylvie van der Werf, Christine Lambert |
| EPI_ISL_421507, EPI_ISL_421508 | Le Château de Seine-Port | National Reference Center for Viruses of Respiratory Infections, Institut Pasteur, Paris | Mélanie Albert, Marion Barbet, Sylvie Behillil, Méline Bizard, Angela Brisebarre, Flora Donati, Etienne Simon-Lorière, Vincent Enouf, Maud Vanpeene, Sylvie van der Werf |
| EPI_ISL_421509, EPI_ISL_421510, EPI_ISL_421511 | CH Compiègne Laboratoire de Biologie | National Reference Center for Viruses of Respiratory Infections, Institut Pasteur, Paris | Mélanie Albert, Marion Barbet, Sylvie Behillil, Méline Bizard, Angela Brisebarre, Flora Donati, Etienne Simon-Lorière, Vincent Enouf, Maud Vanpeene, Sylvie van der Werf, Raulin Olivia |
| EPI_ISL_421512 | Service de Biologie Médicale - BP 125 | National Reference Center for Viruses of Respiratory Infections, Institut Pasteur, Paris | Mélanie Albert, Marion Barbet, Sylvie Behillil, Méline Bizard, Angela Brisebarre, Flora Donati, Etienne Simon-Lorière, Vincent Enouf, Maud Vanpeene, Sylvie van der Werf, Christine Lambert |
| EPI_ISL_421513 | Service de Biologie clinique | National Reference Center for Viruses of Respiratory Infections, Institut Pasteur, Paris | Mélanie Albert, Marion Barbet, Sylvie Behillil, Méline Bizard, Angela Brisebarre, Flora Donati, Etienne Simon-Lorière, Vincent Enouf, Maud Vanpeene, Sylvie van der Werf, Christine Lambert |
| EPI_ISL_421514 | Sentinelles network | National Reference Center for Viruses of Respiratory Infections, Institut Pasteur, Paris | Mélanie Albert, Marion Barbet, Sylvie Behillil, Méline Bizard, Angela Brisebarre, Flora Donati, Etienne Simon-Lorière, Vincent Enouf, Maud Vanpeene, Sylvie van der Werf |
| EPI_ISL_421515 | Servicio de Microbiologia. Hospital Clinico Universitario de Valencia | Sequencing and Bioinformatics Service and Molecular Epidemiology Research Group. FISABIO-Public Health | Giuseppe D'Auria, Llúcia Martínez-Priego, Maria Alma Bracho, Griselda De Marco, Beatriz Beamud, Lidia Ruiz Roldan, Marta Pla Diaz, Neris Garcia-Gonzalez, Loreto Ferrús Abad, Inma Galán Vendrell, Paula Ruiz-Hueso, Mariana Reyes-Prieto, Vicente Soriano Chirona, David Navarro, Fernando Gonzalez-Candelas |
| EPI_ISL_421516 | Servicio de Microbiologia. Hospital Clinico Universitario de Valencia | Sequencing and Bioinformatics Service and Molecular Epidemiology Research Group. FISABIO-Public Health | Loreto Ferrús Abad, Maria Alma Bracho, Griselda De Marco, Beatriz Beamud, Lidia Ruiz Roldan, Marta Pla Diaz, Neris Garcia-Gonzalez, Inma Galán Vendrell, Paula Ruiz-Hueso, Mariana Reyes-Prieto, Vicente Soriano Chirona, David Navarro, Llúcia Martínez-Priego, Giuseppe D'Auria, Fernando Gonzalez-Candelas |
| EPI_ISL_421517 | Servicio de Microbiologia. Hospital Clinico Universitario de Valencia | Sequencing and Bioinformatics Service and Molecular Epidemiology Research Group. FISABIO-Public Health | Sandra Carbo, Loreto Ferrús Abad, Maria Alma Bracho, Griselda De Marco, Beatriz Beamud, Lidia Ruiz Roldan, Marta Pla Diaz, Neris Garcia-Gonzalez, Inma Galán Vendrell, Paula Ruiz-Hueso, Mariana Reyes-Prieto, Vicente Soriano Chirona, Ivan Ansari, David Navarro, Llúcia Martínez-Priego, Giuseppe D'Auria, Fernando Gonzalez-Candelas |
| EPI_ISL_421518 | Servicio de Microbiologia. Hospital Clinico Universitario de Valencia | Sequencing and Bioinformatics Service and Molecular Epidemiology Research Group. FISABIO-Public Health | Inma Galán Vendrell, Paula Ruiz-Hueso, Sandra Carbo, Loreto Ferrús Abad, Maria Alma Bracho, Griselda De Marco, Beatriz Beamud, Lidia Ruiz Roldan, Marta Pla Diaz, Neris Garcia-Gonzalez, Mariana Reyes-Prieto, Vicente Soriano Chirona, Ivan Ansari, David Navarro, Llúcia Martínez-Priego, Giuseppe D'Auria, Fernando Gonzalez-Candelas |
| EPI_ISL_421519 | Servicio de Microbiologia. Hospital Clinico Universitario de Valencia | Sequencing and Bioinformatics Service and Molecular Epidemiology Research Group. FISABIO-Public Health | David Navarro, Maria Alma Bracho, Griselda De Marco, Beatriz Beamud, Lidia Ruiz Roldan, Marta Pla Diaz, Neris Garcia-Gonzalez, Inma Galán Vendrell, Sandra Carbo, Loreto Ferrús Abad, Paula Ruiz-Hueso, Mariana Reyes-Prieto, Vicente Soriano Chirona, Ivan Ansari, Llúcia Martínez-Priego, Giuseppe D'Auria, Fernando Gonzalez-Candelas |
| EPI_ISL_421520 | Servicio de Microbiologia. Hospital Clinico Universitario de Valencia | Sequencing and Bioinformatics Service and Molecular Epidemiology Research Group. FISABIO-Public Health | Loreto Ferrús Abad, Maria Alma Bracho, Griselda De Marco, Sandra Carbo, Beatriz Beamud, Lidia Ruiz Roldan, Marta Pla Diaz, Neris Garcia-Gonzalez, Inma Galán Vendrell, Paula Ruiz-Hueso, Mariana Reyes-Prieto, Vicente Soriano Chirona, Ivan Ansari, David Navarro, Llúcia Martínez-Priego, Giuseppe D'Auria, Fernando Gonzalez-Candelas |
| EPI_ISL_421531 | State Key Laboratory of Agricultural Microbiology | State Key Laboratory of Agricultural Microbiology | Meilin Jin |
| EPI_ISL_421543, EPI_ISL_421544, EPI_ISL_421545, EPI_ISL_421546, EPI_ISL_421547, EPI_ISL_421548, EPI_ISL_421549, EPI_ISL_421550, EPI_ISL_421551, EPI_ISL_421552, EPI_ISL_421553, EPI_ISL_421554 | | | |
| see above | Wyoming Public Health Laboratory | Center for Global Health, University of New Mexico Health Sciences Center | Daryl Domman, Kurt Schwalm, Rob Christensen, Wanda Manley, Cari Sloma, Noah Hull, Darrell Dinwiddie |
| EPI_ISL_421560, EPI_ISL_421561, EPI_ISL_421562, EPI_ISL_421563 | Utah Public Health Laboratory | Utah Public Health Laboratory | Erin Young, Kelly Oakeson |
| EPI_ISL_421572 | Molecular Diagnostic Services and FLowpath | KRISP, KZN Research Innovation and Sequencing Platform | Giandhari J, Pillay S, Ngcapu S, Samsunder N, Lessells R, Chimukangara B, Deforche K, Tegally H, Wilkinson E, de Oliveira T |
| EPI_ISL_421573 | Molecular Diagnostic Services | KRISP, KZN Research Innovation and Sequencing Platform | Giandhari J, Pillay S, Ngcapu S, Samsunder N, Lessells R, Chimukangara B, Deforche K, Tegally H, Wilkinson E, de Oliveira T |
| EPI_ISL_421574, EPI_ISL_421575 | Molecular Diagnostic Services | KRISP, KZN Research Innovation and Sequencing Platform | Giandhari J, Pillay S, Ngcapu S, Samsunder N, Lessells R, Chimukangara B, Deforche K, Tegally H, Wilkinson E, de Oliveira T |
| EPI_ISL_421576 | Molecular Diagnostic Services | KRISP, KZN Research Innovation and Sequencing Platform | Giandhari J, Pillay S, Ngcapu S, Samsunder N, Lessells R, Chimukangara B, Deforche K, Tegally H, Wilkinson E, de Oliveira T |
| EPI_ISL_421577, EPI_ISL_421578, EPI_ISL_421579, EPI_ISL_421580, EPI_ISL_421581, EPI_ISL_421582, EPI_ISL_421583, EPI_ISL_421584, EPI_ISL_421585, EPI_ISL_421586, EPI_ISL_421587, EPI_ISL_421588, EPI_ISL_421589, EPI_ISL_421590, EPI_ISL_421591, EPI_ISL_421592 | | | |
| see above | NYU Langone Health | Departments of Pathology and Medicine, New York University School of Medicine | Maria Aguero-Rosenfeld, Brendan Belovarac, Margaret Black, Ludovic Boytard, John Cadley, Paolo Cotzia, John Chen, Dacia Dimartino, Xiaojun Feng, Tatyana Gindin, Adriana Heguy, Megan Hogan, Emily Huang, George Jour, Andrew Lytle, Christian Marier, Matthew T. Maurano, Mark J. Mulligan, Peter Meyn, Iman Osman, Jared Pinnell, Sitharam Ramaswami, Amy Rapkiewicz, Marie Samanovic-Golden, Antonio Serrano, Guomiao Shen, Matija Snuderl, Theodore Vougiouklakis, Nick Vulpescu, Gael Westby, Paul Zappile, Yutong Zhang |
| EPI_ISL_421593, EPI_ISL_421594, EPI_ISL_421595, EPI_ISL_421596, EPI_ISL_421597, EPI_ISL_421598, EPI_ISL_421599, EPI_ISL_421600, EPI_ISL_421601, EPI_ISL_421602, EPI_ISL_421603, EPI_ISL_421604, EPI_ISL_421605, EPI_ISL_421606, EPI_ISL_421607, EPI_ISL_421608, EPI_ISL_421609, EPI_ISL_421610, EPI_ISL_421611, EPI_ISL_421612, EPI_ISL_421613, EPI_ISL_421614, EPI_ISL_421615, EPI_ISL_421616, EPI_ISL_421617, EPI_ISL_421618, EPI_ISL_421619, EPI_ISL_421620, EPI_ISL_421621, EPI_ISL_421622, EPI_ISL_421623, EPI_ISL_421624, EPI_ISL_421625, EPI_ISL_421626, EPI_ISL_421627, EPI_ISL_421628, EPI_ISL_421629, EPI_ISL_421630, EPI_ISL_421631, EPI_ISL_421632, EPI_ISL_421633, EPI_ISL_421634, EPI_ISL_421635 | | | |
| see above | MSHS Clinical Microbiology Laboratories | MSHS Pathogen Surveillance Program | Ana S. Gonzalez-Reiche, Mitchell Sullivan, Ajay Obla, Gopi Patel, Emilia Sordillo, Melissa Gitman, Alberto Paniz-mondolfi, Matthew Hernandez, Shelcie Fabre, Jose Polanco, Zenab Khan, Bremy Albuquerque, Jayeeta Dutta, Juan Soto, Shwetha Sridhar Hara, Ying-Chih Wang, Melissa Smith, Robert Sebra, Lisa Miorin, Wen-chun Liu, Randy Albrecht, Judith Aberg, Florian Krammer, Adolfo Garcia-Sarstre, Viviana Simon, Harm van Bakel |
| EPI_ISL_421636 | Pathology North | Public Health Virology Laboratory | Bixing Huang, Alyssa Pyke, Amanda De Jong, Andrew Van Den Hurk, Carmel Taylor, David Warrilow, Doris Genge, Elisabeth Gamez, Glen Hewitson, Ian Maxwell Mackay, Inga Sultana, Jamie McMahon, Jean Barcelon, Judy Northill, Mitchell Finger, Natalie Simpson, Neelima Nair, Peter Burtonclay, Peter Moore, Sarah Wheatley, Sean Moody, Sonja Hall-Mendelin, Timothy Gardam, and Frederick Moore |
| EPI_ISL_421641, EPI_ISL_421651 | Centers for Disease Control, R.O.C. (Taiwan) | Centers for Disease Control, R.O.C. (Taiwan) | Ji-Rong Yang, Yu-Chi Lin, Jung-Jung Mu, Ming-Tsan Liu |
| EPI_ISL_421652 | Dasman Diabetes Institute | Dasman Diabetes Institute | Fahd Al-Mulla, Rasheeba Iqbal, Sumi John, Ebaa Al-Ozairi, Qais Al-Duwairi |
| EPI_ISL_421653, EPI_ISL_421654, EPI_ISL_421655, EPI_ISL_421656 | E. Gulbja Laboratorija | Latvian Biomedical Research and Study Centre | Ivars Silamiķelis, Kaspars Megnis, Monta Ustinova, Ņikita Zrelovs, Vita Rovīte, Mikus Gavars, Dmitrijs Perminovs, Uga Dumpis, Jānis Kloviņš |
| EPI_ISL_421660 | The Ohio State University | The Ohio State University-James Molecular Lab at Polaris | Huolin Tu, Matthew Avenarius, Preeti Panchioli, Sean Caruthers, Joan-Miquel Balada-Llasat, Jason Garee, Matt Hunt, Xiaokang Pan, Dan Jones |
| EPI_ISL_421662, EPI_ISL_421663, EPI_ISL_421664, EPI_ISL_421665, EPI_ISL_421666, EPI_ISL_421667, EPI_ISL_421668, EPI_ISL_421669, EPI_ISL_421670, EPI_ISL_421671, EPI_ISL_421672 | | | |
| see above | National Influenza Center, Indian Council of Medical Research - National Institute of Virology | Indian Council of Medical Research-National Institute of Virology, Microbial Containment Complex | Pragya D. Yadav, Varsha Potdar, Savita Patil, Dimpal A. Nyayanit, Triparna Majumdar, Manohar. L. Chaudhary, Gururaj Deshpande, Padinjaremattathil Thankappan Ullas, Anita Shete-Aich, Hitesh Dighe, Sreelekshmy Mohandas, Gajanan Sapkal, Atanu Basu, Amita Jain, Bharti Malhotra, Deepika Chaudhary, Sarah Cherian, Priya Abraham |
| EPI_ISL_421675 | The Ohio State University Wexner Medical Center | The Ohio State University James Molecular lab | Huolin Tu, Sean Caruthers, Matthew Avenarius, Joan-Miquel Balada-Llasat, Matthew Hunt, Preeti Panchioli, Xiaokang Pen, Jason Garee, Pam Snyder, Dan Jones |
| EPI_ISL_421683, EPI_ISL_421684, EPI_ISL_421685, EPI_ISL_421686, EPI_ISL_421687, EPI_ISL_421688, EPI_ISL_421689, EPI_ISL_421690, EPI_ISL_421691, EPI_ISL_421692 | Minnesota Department of Health, Public Health Laboratory | Minnesota Department of Health, Public Health Laboratory | Matt Plumb, Jacob Garfin, Xiong Wang |
| EPI_ISL_421704, EPI_ISL_421705, EPI_ISL_421706, EPI_ISL_421707, EPI_ISL_421708, EPI_ISL_421709, EPI_ISL_421710, EPI_ISL_421711, EPI_ISL_421712, EPI_ISL_421713, EPI_ISL_421714, EPI_ISL_421715, EPI_ISL_421716, EPI_ISL_421717, EPI_ISL_421718, EPI_ISL_421719, EPI_ISL_421720, EPI_ISL_421721, EPI_ISL_421722, EPI_ISL_421723, EPI_ISL_421724, EPI_ISL_421725, EPI_ISL_421726, EPI_ISL_421727, EPI_ISL_421728, EPI_ISL_421729, EPI_ISL_421730, EPI_ISL_421731, EPI_ISL_421732, EPI_ISL_421733 | | | |
| see above | NYU Langone Health | Departments of Pathology and Medicine, New York University School of Medicine | Maria Aguero-Rosenfeld, Brendan Belovarac, Margaret Black, Ludovic Boytard, John Cadley, Paolo Cotzia, John Chen, Dacia Dimartino, Xiaojun Feng, Tatyana Gindin, Adriana Heguy, Megan Hogan, Emily Huang, George Jour, Andrew Lytle, Christian Marier, Matthew T. Maurano, Mark J. Mulligan, Peter Meyn, Iman Osman, Jared Pinnell, Sitharam Ramaswami, Amy Rapkiewicz, Marie Samanovic-Golden, Antonio Serrano, Guomiao Shen, Matija Snuderl, Theodore Vougiouklakis, Nick Vulpescu, Gael Westby, Paul Zappile, Yutong Zhang |
| EPI_ISL_421734, EPI_ISL_421735, EPI_ISL_421736, EPI_ISL_421737, EPI_ISL_421738, EPI_ISL_421739, EPI_ISL_421740, EPI_ISL_421741, EPI_ISL_421742, EPI_ISL_421743, EPI_ISL_421744, EPI_ISL_421745, EPI_ISL_421746, EPI_ISL_421747, EPI_ISL_421748, EPI_ISL_421749, EPI_ISL_421750, EPI_ISL_421751, EPI_ISL_421752, EPI_ISL_421753, EPI_ISL_421754, EPI_ISL_421755, EPI_ISL_421756, EPI_ISL_421757, EPI_ISL_421758, EPI_ISL_421759, EPI_ISL_421760, EPI_ISL_421761, EPI_ISL_421762, EPI_ISL_421763 | | | |
| see above | Laboratoire National de Sante, Microbiology, Virology | Laboratoire National de Sante, Microbiology, Epidemiology and Microbial Genomics | Anke Wienecke-Baldacchino, Ardashel Latsuzbaia, Jessica Tapp, Catherine Ragimbeau, Guillaume Fournier, Tamir Abdelrahman, Trung Nguyen Nguyen, Joel Mossong |
| EPI_ISL_421768, EPI_ISL_421769, EPI_ISL_421770, EPI_ISL_421771, EPI_ISL_421772, EPI_ISL_421773, EPI_ISL_421774, EPI_ISL_421775, EPI_ISL_421776, EPI_ISL_421777, EPI_ISL_421778, EPI_ISL_421779, EPI_ISL_421780, EPI_ISL_421781, EPI_ISL_421782, EPI_ISL_421783, EPI_ISL_421784, EPI_ISL_421785, EPI_ISL_421786, EPI_ISL_421787, EPI_ISL_421788, EPI_ISL_421789, EPI_ISL_421790, EPI_ISL_421791, EPI_ISL_421792, EPI_ISL_421793, EPI_ISL_421794, EPI_ISL_421795, EPI_ISL_421796, EPI_ISL_421797, EPI_ISL_421798, EPI_ISL_421799, EPI_ISL_421800, EPI_ISL_421801, EPI_ISL_421802, EPI_ISL_421803, EPI_ISL_421804, EPI_ISL_421805, EPI_ISL_421806, EPI_ISL_421807, EPI_ISL_421808, EPI_ISL_421809, EPI_ISL_421810, EPI_ISL_421811, EPI_ISL_421812, EPI_ISL_421813, EPI_ISL_421814, EPI_ISL_421815, EPI_ISL_421816, EPI_ISL_421817, EPI_ISL_421818, EPI_ISL_421819, EPI_ISL_421820, EPI_ISL_421821, EPI_ISL_421822, EPI_ISL_421823, EPI_ISL_421824, EPI_ISL_421825, EPI_ISL_421826, EPI_ISL_421827, EPI_ISL_421828, EPI_ISL_421829, EPI_ISL_421830, EPI_ISL_421831, EPI_ISL_421832, EPI_ISL_421833, EPI_ISL_421834, EPI_ISL_421835, EPI_ISL_421836, EPI_ISL_421837, EPI_ISL_421838, EPI_ISL_421839, EPI_ISL_421840, EPI_ISL_421841, EPI_ISL_421842, EPI_ISL_421843, EPI_ISL_421844, EPI_ISL_421845, EPI_ISL_421846, EPI_ISL_421847, EPI_ISL_421848, EPI_ISL_421849, EPI_ISL_421850, EPI_ISL_421851, EPI_ISL_421852, EPI_ISL_421853, EPI_ISL_421854, EPI_ISL_421855, EPI_ISL_421856, EPI_ISL_421857, EPI_ISL_421858, EPI_ISL_421859, EPI_ISL_421860, EPI_ISL_421861, EPI_ISL_421862, EPI_ISL_421863, EPI_ISL_421864, EPI_ISL_421865, EPI_ISL_421866, EPI_ISL_421867, EPI_ISL_421868, EPI_ISL_421869, EPI_ISL_421870, EPI_ISL_421871, EPI_ISL_421872, EPI_ISL_421873, EPI_ISL_421874, EPI_ISL_421875, EPI_ISL_421876, EPI_ISL_421877, EPI_ISL_421878, EPI_ISL_421879, EPI_ISL_421880, EPI_ISL_421881, EPI_ISL_421882, EPI_ISL_421883, EPI_ISL_421884, EPI_ISL_421885, EPI_ISL_421886, EPI_ISL_421887, EPI_ISL_421888, EPI_ISL_421889, EPI_ISL_421890, EPI_ISL_421891, EPI_ISL_421892, EPI_ISL_421893, EPI_ISL_421894, EPI_ISL_421895, EPI_ISL_421896, EPI_ISL_421897, EPI_ISL_421898, EPI_ISL_421899, EPI_ISL_421900, EPI_ISL_421901, EPI_ISL_421902, EPI_ISL_421903, EPI_ISL_421904, EPI_ISL_421905, EPI_ISL_421906, EPI_ISL_421907, EPI_ISL_421908, EPI_ISL_421909, EPI_ISL_421910, EPI_ISL_421911, EPI_ISL_421912, EPI_ISL_421913, EPI_ISL_421914, EPI_ISL_421915, EPI_ISL_421916, EPI_ISL_421917, EPI_ISL_421918, EPI_ISL_421919, EPI_ISL_421920, EPI_ISL_421921, EPI_ISL_421922, EPI_ISL_421923, EPI_ISL_421924, EPI_ISL_421925, EPI_ISL_421926, EPI_ISL_421927, EPI_ISL_421928, EPI_ISL_421929, EPI_ISL_421930, EPI_ISL_421931, EPI_ISL_421932, EPI_ISL_421933, EPI_ISL_421934, EPI_ISL_421935, EPI_ISL_421936, EPI_ISL_421937, EPI_ISL_421938, EPI_ISL_421939, EPI_ISL_421940, EPI_ISL_421941, EPI_ISL_421942, EPI_ISL_421943, EPI_ISL_421944, EPI_ISL_421945, EPI_ISL_421946, EPI_ISL_421947, EPI_ISL_421948, EPI_ISL_421949, EPI_ISL_421951, EPI_ISL_421952, EPI_ISL_421953, EPI_ISL_421954, EPI_ISL_421955, EPI_ISL_421956, EPI_ISL_421957, EPI_ISL_421958, EPI_ISL_421959, EPI_ISL_421960, EPI_ISL_421961, EPI_ISL_421962, EPI_ISL_421963, EPI_ISL_421964, EPI_ISL_421965, EPI_ISL_421966, EPI_ISL_421967, EPI_ISL_421968, EPI_ISL_421969, EPI_ISL_421970, EPI_ISL_421971, EPI_ISL_421972, EPI_ISL_421973, EPI_ISL_421974, EPI_ISL_421975, EPI_ISL_421976, EPI_ISL_421977, EPI_ISL_421978, EPI_ISL_421979, EPI_ISL_421980, EPI_ISL_421981, EPI_ISL_421982, EPI_ISL_421983, EPI_ISL_421984, EPI_ISL_421985, EPI_ISL_421986, EPI_ISL_421987, EPI_ISL_421988, EPI_ISL_421989, EPI_ISL_421990, EPI_ISL_421991, EPI_ISL_421992, EPI_ISL_421993, EPI_ISL_421994, EPI_ISL_421995, EPI_ISL_421996, EPI_ISL_421997, EPI_ISL_421998, EPI_ISL_421999, EPI_ISL_422000, EPI_ISL_422001, EPI_ISL_422002, EPI_ISL_422003, EPI_ISL_422004, EPI_ISL_422005, EPI_ISL_422006, EPI_ISL_422007, EPI_ISL_422008, EPI_ISL_422009, EPI_ISL_422010, EPI_ISL_422011, EPI_ISL_422012 | | | |
| see above | Respiratory Virus Unit, Microbiology Services Colindale, Public Health England | Respiratory Virus Unit, Microbiology Services Colindale, Public Health England | Monica Galiano, Shahjahan Miah, Angie Lackenby, Omolola Akinbami, Tiina Talts, Leena Bhaw, Richard Myers, Steven Platt, Kirstin Edwards, Jonathan Hubb, Joanna Ellis, Maria Zambon |
| EPI_ISL_422016, EPI_ISL_422017, EPI_ISL_422018, EPI_ISL_422019, EPI_ISL_422020, EPI_ISL_422021, EPI_ISL_422022, EPI_ISL_422023, EPI_ISL_422024, EPI_ISL_422025, EPI_ISL_422026, EPI_ISL_422027, EPI_ISL_422028, EPI_ISL_422029, EPI_ISL_422030, EPI_ISL_422031, EPI_ISL_422032, EPI_ISL_422033, EPI_ISL_422034, EPI_ISL_422035, EPI_ISL_422036, EPI_ISL_422037, EPI_ISL_422038, EPI_ISL_422039, EPI_ISL_422040, EPI_ISL_422041, EPI_ISL_422042, EPI_ISL_422043, EPI_ISL_422044, EPI_ISL_422045, EPI_ISL_422046, EPI_ISL_422047, EPI_ISL_422048, EPI_ISL_422049, EPI_ISL_422050, EPI_ISL_422051, EPI_ISL_422052, EPI_ISL_422053, EPI_ISL_422054, EPI_ISL_422055, EPI_ISL_422056, EPI_ISL_422057, EPI_ISL_422058, EPI_ISL_422059, EPI_ISL_422060, EPI_ISL_422061, EPI_ISL_422062, EPI_ISL_422063, | | | |

| | | | |
|---|---|---|---|
| EPI_ISL_422064, EPI_ISL_422065, EPI_ISL_422066, EPI_ISL_422067, EPI_ISL_422068, EPI_ISL_422069, EPI_ISL_422070, EPI_ISL_422071, EPI_ISL_422072, EPI_ISL_422073, EPI_ISL_422074, EPI_ISL_422075, EPI_ISL_422076, EPI_ISL_422077, EPI_ISL_422078, EPI_ISL_422079, EPI_ISL_422080, EPI_ISL_422081, EPI_ISL_422082, EPI_ISL_422083, EPI_ISL_422084, EPI_ISL_422085, EPI_ISL_422086, EPI_ISL_422087, EPI_ISL_422088, EPI_ISL_422089, EPI_ISL_422090, EPI_ISL_422091, EPI_ISL_422092, EPI_ISL_422093, EPI_ISL_422094, EPI_ISL_422095, EPI_ISL_422096, EPI_ISL_422097, EPI_ISL_422098, EPI_ISL_422099, EPI_ISL_422100, EPI_ISL_422102, EPI_ISL_422103, EPI_ISL_422104, EPI_ISL_422105, EPI_ISL_422106, EPI_ISL_422107, EPI_ISL_422108, EPI_ISL_422109, EPI_ISL_422110, EPI_ISL_422111, EPI_ISL_422112, EPI_ISL_422113, EPI_ISL_422114, EPI_ISL_422115, EPI_ISL_422116, EPI_ISL_422117, EPI_ISL_422118, EPI_ISL_422119, EPI_ISL_422120, EPI_ISL_422121, EPI_ISL_422122, EPI_ISL_422123, EPI_ISL_422124, EPI_ISL_422125, EPI_ISL_422126, EPI_ISL_422127, EPI_ISL_422128, EPI_ISL_422129, EPI_ISL_422130, EPI_ISL_422131, EPI_ISL_422132, EPI_ISL_422133, EPI_ISL_422134, EPI_ISL_422135, EPI_ISL_422136, EPI_ISL_422137, EPI_ISL_422138, EPI_ISL_422139, EPI_ISL_422140, EPI_ISL_422141, EPI_ISL_422142, EPI_ISL_422143, EPI_ISL_422144, EPI_ISL_422145, EPI_ISL_422146, EPI_ISL_422147, EPI_ISL_422148, EPI_ISL_422149, EPI_ISL_422150, EPI_ISL_422151, EPI_ISL_422152, EPI_ISL_422153, EPI_ISL_422154, EPI_ISL_422155, EPI_ISL_422156, EPI_ISL_422157, EPI_ISL_422158, EPI_ISL_422159, EPI_ISL_422160, EPI_ISL_422161, EPI_ISL_422162, EPI_ISL_422163, EPI_ISL_422164, EPI_ISL_422165, EPI_ISL_422166, EPI_ISL_422167, EPI_ISL_422168, EPI_ISL_422169, EPI_ISL_422170, EPI_ISL_422171, EPI_ISL_422172, EPI_ISL_422173, EPI_ISL_422174, EPI_ISL_422175, EPI_ISL_422176, EPI_ISL_422177, EPI_ISL_422178, EPI_ISL_422179, EPI_ISL_422180, EPI_ISL_422181, EPI_ISL_422182, EPI_ISL_422183, EPI_ISL_422184, EPI_ISL_422185, EPI_ISL_422186, EPI_ISL_422187, EPI_ISL_422188, EPI_ISL_422189, EPI_ISL_422190, EPI_ISL_422191, EPI_ISL_422192, EPI_ISL_422193, EPI_ISL_422194, EPI_ISL_422195, EPI_ISL_422196, EPI_ISL_422197, EPI_ISL_422198, EPI_ISL_422199, EPI_ISL_422200, EPI_ISL_422201, EPI_ISL_422202, EPI_ISL_422203, EPI_ISL_422204, EPI_ISL_422205, EPI_ISL_422206, EPI_ISL_422207, EPI_ISL_422208, EPI_ISL_422209, EPI_ISL_422210, EPI_ISL_422211, EPI_ISL_422212, EPI_ISL_422213, EPI_ISL_422214, EPI_ISL_422215, EPI_ISL_422216, EPI_ISL_422217, EPI_ISL_422218, EPI_ISL_422219, EPI_ISL_422220, EPI_ISL_422221, EPI_ISL_422222, EPI_ISL_422223, EPI_ISL_422224, EPI_ISL_422225, EPI_ISL_422226, EPI_ISL_422227, EPI_ISL_422228, EPI_ISL_422229, EPI_ISL_422230, EPI_ISL_422231, EPI_ISL_422232, EPI_ISL_422233, EPI_ISL_422234, EPI_ISL_422235, EPI_ISL_422236, EPI_ISL_422237, EPI_ISL_422238, EPI_ISL_422239, EPI_ISL_422240, EPI_ISL_422241, EPI_ISL_422242, EPI_ISL_422243, EPI_ISL_422244, EPI_ISL_422245, EPI_ISL_422246, EPI_ISL_422247, EPI_ISL_422248, EPI_ISL_422249, EPI_ISL_422250, EPI_ISL_422251, EPI_ISL_422252, EPI_ISL_422253, EPI_ISL_422254, EPI_ISL_422255, EPI_ISL_422256, EPI_ISL_422257, EPI_ISL_422258, EPI_ISL_422259, EPI_ISL_422260, EPI_ISL_422261, EPI_ISL_422262, EPI_ISL_422263, EPI_ISL_422264, EPI_ISL_422265, EPI_ISL_422266, EPI_ISL_422267, EPI_ISL_422268, EPI_ISL_422269, EPI_ISL_422270, EPI_ISL_422271, EPI_ISL_422272, EPI_ISL_422273, EPI_ISL_422274, EPI_ISL_422275, EPI_ISL_422276, EPI_ISL_422277, EPI_ISL_422278, EPI_ISL_422279, EPI_ISL_422280, EPI_ISL_422281, EPI_ISL_422282, EPI_ISL_422283, EPI_ISL_422284, EPI_ISL_422285, EPI_ISL_422286, EPI_ISL_422287, EPI_ISL_422288, EPI_ISL_422289, EPI_ISL_422290, EPI_ISL_422291, EPI_ISL_422292, EPI_ISL_422293, EPI_ISL_422294, EPI_ISL_422295, EPI_ISL_422296, EPI_ISL_422297, EPI_ISL_422298, EPI_ISL_422299, EPI_ISL_422300, EPI_ISL_422301, EPI_ISL_422302, EPI_ISL_422303, EPI_ISL_422304, EPI_ISL_422305, EPI_ISL_422306, EPI_ISL_422307, EPI_ISL_422308, EPI_ISL_422309, EPI_ISL_422310, EPI_ISL_422311, EPI_ISL_422312, EPI_ISL_422313, EPI_ISL_422314, EPI_ISL_422315, EPI_ISL_422316, EPI_ISL_422317, EPI_ISL_422318, EPI_ISL_422319, EPI_ISL_422320, EPI_ISL_422321, EPI_ISL_422322, EPI_ISL_422323, EPI_ISL_422324, EPI_ISL_422325, EPI_ISL_422326, EPI_ISL_422327, EPI_ISL_422328, EPI_ISL_422329, EPI_ISL_422330, EPI_ISL_422331, EPI_ISL_422332, EPI_ISL_422333, EPI_ISL_422334, EPI_ISL_422335, EPI_ISL_422336, EPI_ISL_422337, EPI_ISL_422338, EPI_ISL_422339, EPI_ISL_422340, EPI_ISL_422341, EPI_ISL_422342, EPI_ISL_422343, EPI_ISL_422344, EPI_ISL_422345, EPI_ISL_422346, EPI_ISL_422347, EPI_ISL_422348, EPI_ISL_422349, EPI_ISL_422350, EPI_ISL_422351, EPI_ISL_422352, EPI_ISL_422353, EPI_ISL_422354, EPI_ISL_422355, EPI_ISL_422356, EPI_ISL_422357, EPI_ISL_422358, EPI_ISL_422359, EPI_ISL_422360, EPI_ISL_422361, EPI_ISL_422362, EPI_ISL_422363, EPI_ISL_422364, EPI_ISL_422365, EPI_ISL_422366, EPI_ISL_422368, EPI_ISL_422369, EPI_ISL_422370, EPI_ISL_422371, EPI_ISL_422372, EPI_ISL_422373, EPI_ISL_422374, EPI_ISL_422375, EPI_ISL_422376, EPI_ISL_422377 | | | |
| see above | Wales Specialist Virology Centre | Public Health Wales Microbiology Cardiff | Catherine Moore, Johnathan Evans, Malorie Perry, Simon Cottrell, Alec Birchley, Alexander Adams, Amy Gaskin, Bree Gatica-Wilcox, Jason Coombes, Lauren Gilbert, Lee Graham, Nicole Pacchiarini, Sara Kumziene-Summerhayes, Sarah Taylor, Sophie Jones, Sara Rey, Matthew Bull, Joanne Watkins, Sally Corden, Tom Connor |
| EPI_ISL_422382 | NMIMR, Department of Virology | WACCBIP, University of Ghana | Joyce M. Ngoi, Bright Adu, Collins M. Misita, Selassie Kumordjie, Miriam Eshun, Linda Boatemaa, Vanessa Magnussen, Erasmus Kotey, Fred Tei-Maya, Dominic S. Y. Amuzu, Peter Quashie, Augustina Arjaquah, Ivy Asante, Evelyn Bonney, George B. Kyei, Kofi Bonney, Gordon A. Awandare, William Ampofo |
| EPI_ISL_422384, EPI_ISL_422387, EPI_ISL_422390, EPI_ISL_422394, EPI_ISL_422397, EPI_ISL_422398, EPI_ISL_422399, EPI_ISL_422400, EPI_ISL_422401, EPI_ISL_422402, EPI_ISL_422403, EPI_ISL_422404, EPI_ISL_422405, EPI_ISL_422406 | | | |
| see above | NMIMR, Department of Virology | WACCBIP, University of Ghana | Joyce M. Ngoi, Bright Adu, Collins M. Morang'a, Selassie Kumordjie, Miriam Eshun, Linda Boatemaa, Vanessa Magnussen, Erasmus Kotey, Fred Tei-Maya, Dominic S. Y. Amuzu, Peter Quashie, Augustina Arjarquah, Ivy Asante, Evelyn Bonney, George B. Kyei, Kofi Bonney, Abraham Kwabena Anang, Gordon A. Awandare, William Ampofo |
| EPI_ISL_422407, EPI_ISL_422408, EPI_ISL_422409, EPI_ISL_422410, EPI_ISL_422411, EPI_ISL_422412, EPI_ISL_422413, EPI_ISL_422414, EPI_ISL_422415, EPI_ISL_422416, EPI_ISL_422417, EPI_ISL_422418, EPI_ISL_422419, EPI_ISL_422420, EPI_ISL_422421, EPI_ISL_422422 | | | |
| see above | Department of Laboratory Medicine, National Taiwan University Hospital | Microbial Genomics Core Lab, National Taiwan University Centers of Genomic and Precision Medicine | Shiou-Hwei Yeh, You-Yu Lin, Ya-Yun Lai, Chiao-Ling Li, Shan-Chwen Chang, Pei-Jer Chen, Sui-Yuan Chang |
| EPI_ISL_422424 | Jaber Al Ahmad Al Sabah Hospital | Dasman diabetes Institute | Fahd Al-Mulla, Rasheeba Iqbal, Sumi John, Ebaa Al-Ozairi, Qais Al-Duwairi |
| EPI_ISL_422425 | Zhejiang Provincial Center for Disease Control and Prevention | Zhejiang Provincial Center for Disease Control and Prevention | Yanjun Zhang, Yi Sun |
| EPI_ISL_422426, EPI_ISL_422427 | JABER AL AHMAD AL SABAH HOSPITAL – KUWAIT CITY | Dasman Diabetes Institute | Fahd Al-Mulla, Rasheeba Iqbal, Sumi John, Ebaa Al-Ozairi, Qais Al-Duwairi |
| EPI_ISL_422428, EPI_ISL_422429, EPI_ISL_422430, EPI_ISL_422431, EPI_ISL_422432, EPI_ISL_422433, EPI_ISL_422434, EPI_ISL_422435 | National Public Health Laboratory, National Centre for Infectious Diseases | National Public Health Laboratory, National Centre for Infectious Diseases | Mak TM, Octavia S, Cui L, Lin RTP |
| EPI_ISL_422436 | Respiratory Virus Unit, Microbiology Services Colindale, Public Health England | Respiratory Virus Unit, Microbiology Services Colindale, Public Health England | Monica Galiano, Shahjahan Miah, Angie Lackenby, Omolola Akinbami, Tiina Talts, Leena Bhaw, Richard Myers, Steven Platt, Kirstin Edwards, Jonathan Hubb, Joanna Ellis, Maria Zambon |
| EPI_ISL_422437 | ULSS9 Distretto di Bussolengo | Istituto Zooprofilattico Sperimentale delle Venezie | Adelaide Milani, Alessia Schivo, Annalisa Salviato, Erika Giorgia Quaranta, Gianpiero Zamperin, Ambra Pastori, Bianca Zecchin, Alice Fusaro, Calogero Terregino, Antonia Ricci |
| EPI_ISL_422438 | ULSS9 Distretto di Bussolengo | Istituto Zooprofilattico Sperimentale delle Venezie | Adelaide Milani, Alessia Schivo, Annalisa Salviato, Erika Giorgia Quaranta, Gianpiero Zamperin, Ambra Pastori, Bianca Zecchin, Alice Fusaro, Calogero Terregino, Antonia Ricci |
| EPI_ISL_422453, EPI_ISL_422459 | Gundersen Molecular Diagnostics Laboratory | Kabara Cancer Research Institute | Craig S. Richmond & Paraic A. Kenny |
| EPI_ISL_422461, EPI_ISL_422462, EPI_ISL_422463 | Gundersen Molecular Diagnostics Laboratory | Kabara Cancer Research Institute | Craig S. Richmond, Paraic A. Kenny |
| EPI_ISL_422464 | Gundersen Molecular Diagnostics Laboratory | Kabara Cancer Research Institute | Craig S. Richmond; Paraic A. Kenny |
| EPI_ISL_422465 | Gundersen Molecular Diagnostics Laboratory | Kabara Cancer Research Institute | Craig S. Richmond, Paraic A. Kenny |
| EPI_ISL_422488, EPI_ISL_422489, EPI_ISL_422490, EPI_ISL_422491, EPI_ISL_422492, EPI_ISL_422493, EPI_ISL_422494, EPI_ISL_422495, EPI_ISL_422496, EPI_ISL_422497, EPI_ISL_422498, EPI_ISL_422499, EPI_ISL_422500, EPI_ISL_422501, EPI_ISL_422502, EPI_ISL_422503, EPI_ISL_422504, EPI_ISL_422505, EPI_ISL_422506, EPI_ISL_422507, EPI_ISL_422508, EPI_ISL_422509, EPI_ISL_422510, EPI_ISL_422511, EPI_ISL_422512, EPI_ISL_422513, EPI_ISL_422514, EPI_ISL_422515, EPI_ISL_422516, EPI_ISL_422517, EPI_ISL_422518, EPI_ISL_422519, EPI_ISL_422520, EPI_ISL_422521, EPI_ISL_422522, EPI_ISL_422523, EPI_ISL_422524, EPI_ISL_422525, EPI_ISL_422526, EPI_ISL_422527, EPI_ISL_422528, EPI_ISL_422529, EPI_ISL_422530, EPI_ISL_422531, EPI_ISL_422532, EPI_ISL_422533, EPI_ISL_422534, EPI_ISL_422535, EPI_ISL_422536, EPI_ISL_422537, EPI_ISL_422538, EPI_ISL_422539, EPI_ISL_422540, EPI_ISL_422541, EPI_ISL_422542, EPI_ISL_422543, EPI_ISL_422544, EPI_ISL_422545, EPI_ISL_422546, EPI_ISL_422547, EPI_ISL_422548, EPI_ISL_422549, EPI_ISL_422550, EPI_ISL_422551, EPI_ISL_422552, EPI_ISL_422553, EPI_ISL_422554, EPI_ISL_422555, EPI_ISL_422556, EPI_ISL_422557, EPI_ISL_422558, EPI_ISL_422559, EPI_ISL_422560, EPI_ISL_422561, EPI_ISL_422562 | | | |
| see above | MSHS Clinical Microbiology Laboratories | MSHS Pathogen Surveillance Program | Ana S. Gonzalez-Reiche, Mitchell Sullivan, Ajay Obla, Gopi Patel, Emilia Sordillo, Melissa Gitman, Alberto Paniz-mondolfi, Matthew Hernandez, Shelcie Fabre, Jose Polanco, Zenab Khan, Bremy Albuquerque, Jayeeta Dutta, Juan Soto, Shwetha Sridhar Hara, Ying-Chih Wang, Melissa Smith, Robert Sebra, Lisa Miorin, Wen-chun Liu, Randy Albrecht, Judith Aberg, Florian Krammer, Adolfo Garcia-Sarstre, Viviana Simon, Harm van Bakel |
| EPI_ISL_422563 | Institute of Microbiology Universidad San Francisco de Quito | Institute of Microbiology Universidad San Francisco de Quito | Belen Prado-Vivar, Sully Marquez, Juan Jose Guadalupe, Bernardo Gutierrez, Francisco Mora, Juan Gaviria, Alejandra Ramones, Franklin Espinoza, Edison Ligña, Jorge Reyes, Patricio Rojas-Silva, Veronica Barragan, Gabriel Trueba, Michelle Grunauer, Paul Cardenas |
| EPI_ISL_422564 | Institute of Microbiology Universidad San Francisco de Quito | Institute of Microbiology Universidad San Francisco de Quito | Juan Jose Guadalupe, Belen Prado-Vivar, Sully Marquez, Bernardo Gutierrez, Francisco Mora, Juan Gaviria, Alejandra Ramones, Franklin Espinoza, Edison Ligña, Jorge Reyes, Patricio Rojas-Silva, Veronica Barragan, Gabriel Trueba, Michelle Grunauer, Paul Cardenas |
| EPI_ISL_422565 | Institute of Microbiology Universidad San Francisco de Quito | Institute of Microbiology Universidad San Francisco de Quito | Sully Marquez, Belen Prado-Vivar, Juan Jose Guadalupe, Bernardo Gutierrez, Francisco Mora, Juan Gaviria, Alejandra Ramones, Franklin Espinoza, Edison Ligña, Jorge Reyes, Patricio Rojas-Silva, Veronica Barragan, Gabriel Trueba, Michelle Grunauer, Paul Cardenas |
| EPI_ISL_422566, EPI_ISL_422567, EPI_ISL_422568, EPI_ISL_422569, EPI_ISL_422570, EPI_ISL_422571, EPI_ISL_422572, EPI_ISL_422573, EPI_ISL_422574, EPI_ISL_422575, EPI_ISL_422576, EPI_ISL_422577, EPI_ISL_422578, EPI_ISL_422579, EPI_ISL_422580, EPI_ISL_422581, EPI_ISL_422582, EPI_ISL_422583, EPI_ISL_422584, EPI_ISL_422585, EPI_ISL_422586, EPI_ISL_422587, EPI_ISL_422588, EPI_ISL_422589, EPI_ISL_422590, EPI_ISL_422591, EPI_ISL_422592, EPI_ISL_422593, EPI_ISL_422594, EPI_ISL_422595, EPI_ISL_422596, EPI_ISL_422597, EPI_ISL_422598, EPI_ISL_422599, EPI_ISL_422600, EPI_ISL_422601, EPI_ISL_422602, EPI_ISL_422603, EPI_ISL_422604, EPI_ISL_422605, EPI_ISL_422606, EPI_ISL_422607, EPI_ISL_422608, EPI_ISL_422609, EPI_ISL_422610, EPI_ISL_422611, EPI_ISL_422612, EPI_ISL_422613, EPI_ISL_422614, EPI_ISL_422615, EPI_ISL_422616, EPI_ISL_422617, EPI_ISL_422618, EPI_ISL_422619, EPI_ISL_422620, EPI_ISL_422621, EPI_ISL_422622, EPI_ISL_422623, EPI_ISL_422624, EPI_ISL_422625, EPI_ISL_422626, EPI_ISL_422627, EPI_ISL_422628, EPI_ISL_422629, EPI_ISL_422630, EPI_ISL_422631, EPI_ISL_422632, EPI_ISL_422633, EPI_ISL_422634, EPI_ISL_422635 | | | |
| see above | Dutch COVID-19 response team | Erasmus Medical Center | Bas Oude Munnink, David Nieuwenhuijse, Reina Sikkema, Claudia Schapendonk, Irina Chestakova, Anne van der Linden, Theo Bestebroer, Stefan van Nieuwkoop, Mark Pronk, Pascal Lexmond, Corien Swaan, Manon Haverkate, Madelief Mollers, Mart Stein, Sandra Kengne Kamga Mobou, Jeroen van Kampen, Jolanda Voermans, Aura Timen, Corine GeurtsvanKessel, Annemiek van der Eijk, Richard Molenkamp, Marion Koopmans, on behalf of the Dutch national COVID-19 response team. |
| EPI_ISL_422636 | The National Institute of Public Health Center for Epidemiology and Microbiology | State Veterinary Institute Prague | Alexander Nagy, Helena Jirincova, Klara Labska, Ludmila Novakova, Olga Storkanova, Dusan Trnka, Jaromira Vecerova |
| EPI_ISL_422637, EPI_ISL_422638, EPI_ISL_422639, EPI_ISL_422640, EPI_ISL_422641, EPI_ISL_422642, EPI_ISL_422643, EPI_ISL_422644, EPI_ISL_422645, EPI_ISL_422646, EPI_ISL_422647, EPI_ISL_422648, EPI_ISL_422649, EPI_ISL_422650, EPI_ISL_422651, EPI_ISL_422652, EPI_ISL_422653, EPI_ISL_422654, EPI_ISL_422655, EPI_ISL_422656, EPI_ISL_422657, EPI_ISL_422658, EPI_ISL_422659, EPI_ISL_422660, EPI_ISL_422661, EPI_ISL_422662, EPI_ISL_422663, EPI_ISL_422664, EPI_ISL_422665, EPI_ISL_422666, EPI_ISL_422667, EPI_ISL_422668, EPI_ISL_422669, EPI_ISL_422670, EPI_ISL_422671, EPI_ISL_422672, EPI_ISL_422673, EPI_ISL_422674, EPI_ISL_422675, EPI_ISL_422676, EPI_ISL_422677, EPI_ISL_422678, EPI_ISL_422679, EPI_ISL_422680, EPI_ISL_422681, EPI_ISL_422682, EPI_ISL_422683, EPI_ISL_422684, EPI_ISL_422685, EPI_ISL_422686, EPI_ISL_422687, EPI_ISL_422688, EPI_ISL_422689, EPI_ISL_422690, EPI_ISL_422691, EPI_ISL_422692, EPI_ISL_422693, EPI_ISL_422694, EPI_ISL_422695, EPI_ISL_422696, EPI_ISL_422697, EPI_ISL_422698, EPI_ISL_422699, EPI_ISL_422700, EPI_ISL_422701, EPI_ISL_422702, EPI_ISL_422703, EPI_ISL_422704, EPI_ISL_422705, EPI_ISL_422706, EPI_ISL_422707, EPI_ISL_422708, EPI_ISL_422709, EPI_ISL_422710, EPI_ISL_422711, EPI_ISL_422712, EPI_ISL_422713, EPI_ISL_422714, EPI_ISL_422715, EPI_ISL_422716, EPI_ISL_422717, EPI_ISL_422718, EPI_ISL_422719, EPI_ISL_422720, EPI_ISL_422721, EPI_ISL_422722, EPI_ISL_422723, EPI_ISL_422724, EPI_ISL_422725, EPI_ISL_422726, EPI_ISL_422727, EPI_ISL_422728, EPI_ISL_422729, EPI_ISL_422730, EPI_ISL_422731, EPI_ISL_422732, EPI_ISL_422733, EPI_ISL_422734, EPI_ISL_422735, EPI_ISL_422736, EPI_ISL_422737, EPI_ISL_422738, EPI_ISL_422739, EPI_ISL_422740, EPI_ISL_422741, EPI_ISL_422742, EPI_ISL_422743, EPI_ISL_422744, EPI_ISL_422745, EPI_ISL_422746, EPI_ISL_422747, EPI_ISL_422748, EPI_ISL_422749, EPI_ISL_422750, EPI_ISL_422751, EPI_ISL_422752, EPI_ISL_422753, EPI_ISL_422754, EPI_ISL_422755, EPI_ISL_422756, EPI_ISL_422757, EPI_ISL_422758, EPI_ISL_422759, EPI_ISL_422760, EPI_ISL_422761, EPI_ISL_422762, EPI_ISL_422763, EPI_ISL_422764, EPI_ISL_422765, EPI_ISL_422766, EPI_ISL_422767, EPI_ISL_422768, EPI_ISL_422769, EPI_ISL_422770, EPI_ISL_422771, EPI_ISL_422772, EPI_ISL_422773, EPI_ISL_422774, EPI_ISL_422775, EPI_ISL_422776, EPI_ISL_422777, EPI_ISL_422778, EPI_ISL_422779, EPI_ISL_422780, EPI_ISL_422781, EPI_ISL_422782, EPI_ISL_422783, EPI_ISL_422784, EPI_ISL_422785, EPI_ISL_422786, EPI_ISL_422787, EPI_ISL_422788, EPI_ISL_422789, EPI_ISL_422790, EPI_ISL_422791, EPI_ISL_422792, EPI_ISL_422793, EPI_ISL_422794, EPI_ISL_422795, EPI_ISL_422796, EPI_ISL_422797, EPI_ISL_422798, EPI_ISL_422799, EPI_ISL_422800, EPI_ISL_422801, EPI_ISL_422802, EPI_ISL_422803, EPI_ISL_422804, EPI_ISL_422805, EPI_ISL_422806, EPI_ISL_422807, EPI_ISL_422808, EPI_ISL_422809, EPI_ISL_422810, EPI_ISL_422811, EPI_ISL_422812, EPI_ISL_422813, EPI_ISL_422814, EPI_ISL_422815, EPI_ISL_422816, EPI_ISL_422817, EPI_ISL_422818, EPI_ISL_422819, EPI_ISL_422820, EPI_ISL_422821, EPI_ISL_422822, EPI_ISL_422823, EPI_ISL_422824, EPI_ISL_422825, EPI_ISL_422826, EPI_ISL_422827, EPI_ISL_422828, EPI_ISL_422829, EPI_ISL_422830, EPI_ISL_422831, EPI_ISL_422832, EPI_ISL_422833, EPI_ISL_422834, EPI_ISL_422835, EPI_ISL_422836, EPI_ISL_422837, EPI_ISL_422838, EPI_ISL_422839, EPI_ISL_422840, EPI_ISL_422841, EPI_ISL_422842, EPI_ISL_422843, EPI_ISL_422844, EPI_ISL_422845, EPI_ISL_422846, EPI_ISL_422847, EPI_ISL_422848, EPI_ISL_422849, EPI_ISL_422850, EPI_ISL_422851, EPI_ISL_422852, EPI_ISL_422853, EPI_ISL_422854, EPI_ISL_422855, EPI_ISL_422856, EPI_ISL_422857, EPI_ISL_422858, EPI_ISL_422859, EPI_ISL_422860, EPI_ISL_422861, EPI_ISL_422862, EPI_ISL_422863, EPI_ISL_422864, EPI_ISL_422865, EPI_ISL_422866, EPI_ISL_422867, EPI_ISL_422868, EPI_ISL_422869, EPI_ISL_422870, EPI_ISL_422871, EPI_ISL_422872, EPI_ISL_422873, EPI_ISL_422874, EPI_ISL_422875, EPI_ISL_422876, EPI_ISL_422877, EPI_ISL_422878, EPI_ISL_422879, EPI_ISL_422880, EPI_ISL_422881, EPI_ISL_422882, EPI_ISL_422883, EPI_ISL_422884, EPI_ISL_422885, EPI_ISL_422886, EPI_ISL_422887, EPI_ISL_422888, EPI_ISL_422889, EPI_ISL_422890, EPI_ISL_422891, EPI_ISL_422892, EPI_ISL_422893, EPI_ISL_422894, EPI_ISL_422895, EPI_ISL_422896, EPI_ISL_422897, EPI_ISL_422898, EPI_ISL_422899, EPI_ISL_422900, EPI_ISL_422901, EPI_ISL_422902, EPI_ISL_422903, EPI_ISL_422904, EPI_ISL_422905, EPI_ISL_422906, EPI_ISL_422907, EPI_ISL_422908, EPI_ISL_422909, EPI_ISL_422910, EPI_ISL_422911, EPI_ISL_422912, EPI_ISL_422913, EPI_ISL_422914, EPI_ISL_422915, EPI_ISL_422916, EPI_ISL_422917, EPI_ISL_422918, EPI_ISL_422919, EPI_ISL_422920, EPI_ISL_422921, EPI_ISL_422922, EPI_ISL_422923, EPI_ISL_422924, EPI_ISL_422925, EPI_ISL_422926, EPI_ISL_422927, EPI_ISL_422928, EPI_ISL_422929, EPI_ISL_422930, EPI_ISL_422931, EPI_ISL_422932, EPI_ISL_422933, EPI_ISL_422934, EPI_ISL_422935, EPI_ISL_422936, EPI_ISL_422937, EPI_ISL_422938, EPI_ISL_422939, EPI_ISL_422940, EPI_ISL_422941, EPI_ISL_422942, EPI_ISL_422943, EPI_ISL_422944, EPI_ISL_422945, EPI_ISL_422946, EPI_ISL_422947, EPI_ISL_422948, EPI_ISL_422949, EPI_ISL_422950, EPI_ISL_422951, EPI_ISL_422952, EPI_ISL_422953, EPI_ISL_422954, EPI_ISL_422955, EPI_ISL_422956, EPI_ISL_422957, EPI_ISL_422958, EPI_ISL_422959, EPI_ISL_422960 | | | |
| see above | Dutch COVID-19 response team | Erasmus Medical Center | Bas Oude Munnink, David Nieuwenhuijse, Reina Sikkema, Claudia Schapendonk, Irina Chestakova, Anne van der Linden, Theo Bestebroer, Stefan van Nieuwkoop, Mark Pronk, Pascal Lexmond, Corien Swaan, Manon Haverkate, Madelief Mollers, Mart Stein, Sandra Kengne Kamga Mobou, Jeroen van Kampen, Jolanda Voermans, Aura Timen, Corine GeurtsvanKessel, Annemiek van der Eijk, Richard Molenkamp, Marion Koopmans, on behalf of the Dutch national COVID-19 response team. |
| EPI_ISL_422961, EPI_ISL_422962, EPI_ISL_422963, EPI_ISL_422964, EPI_ISL_422965, EPI_ISL_422966, EPI_ISL_422967, EPI_ISL_422968, EPI_ISL_422969, EPI_ISL_422970, EPI_ISL_422971, EPI_ISL_422972, EPI_ISL_422973, EPI_ISL_422974, EPI_ISL_422975, EPI_ISL_422976, EPI_ISL_422977, EPI_ISL_422978, EPI_ISL_422979, EPI_ISL_422980, EPI_ISL_422981, EPI_ISL_422982, EPI_ISL_422983, EPI_ISL_422984, EPI_ISL_422985, EPI_ISL_422986, EPI_ISL_422987, EPI_ISL_422988, EPI_ISL_422989, EPI_ISL_422990, EPI_ISL_422991, EPI_ISL_422992, EPI_ISL_422993, EPI_ISL_422994, EPI_ISL_422995, EPI_ISL_422996, EPI_ISL_422997, EPI_ISL_422998, EPI_ISL_422999, EPI_ISL_423000, EPI_ISL_423001, EPI_ISL_423002, EPI_ISL_423003, EPI_ISL_423004, EPI_ISL_423005, EPI_ISL_423006, EPI_ISL_423007, EPI_ISL_423008, EPI_ISL_423009, EPI_ISL_423010, EPI_ISL_423011, EPI_ISL_423012, EPI_ISL_423013, EPI_ISL_423014, EPI_ISL_423015, EPI_ISL_423016, EPI_ISL_423017, EPI_ISL_423018, EPI_ISL_423019, EPI_ISL_423020, EPI_ISL_423021, EPI_ISL_423022, EPI_ISL_423023, EPI_ISL_423024, EPI_ISL_423025, EPI_ISL_423026, EPI_ISL_423027, EPI_ISL_423028, EPI_ISL_423029, EPI_ISL_423030, EPI_ISL_423031, EPI_ISL_423032, EPI_ISL_423033 | | | |
| see above | UW Virology Lab | UW Virology Lab | Pavitra Roychoudhury, Hong Xie, Keith Jerome, Alexander Greninger |
| EPI_ISL_423034 | Dutch COVID-19 response team | Erasmus Medical Center | Bas Oude Munnink, David Nieuwenhuijse, Reina Sikkema, Claudia Schapendonk, Irina Chestakova, Anne van der Linden, Theo Bestebroer, Stefan van Nieuwkoop, Mark Pronk, Pascal Lexmond, Corien Swaan, Manon Haverkate, Madelief Mollers, Mart Stein, Sandra Kengne Kamga Mobou, Jeroen van Kampen, Jolanda Voermans, Aura Timen, Corine GeurtsvanKessel, Annemiek van der Eijk, Richard Molenkamp, Marion Koopmans, on behalf of the Dutch national COVID-19 response team. |
| EPI_ISL_423039, EPI_ISL_423040, EPI_ISL_423041, EPI_ISL_423042, EPI_ISL_423043 | Ramathibodi Hospital | COVID-19 Network Investigations (CONI) Alliance | Elizabeth Batty, Wasun Chantratita, Thanat Chookajorn, Stefan Fernandez, Angkana Huang, Anthony R. Jones, Khajohn Joonsalak, Chonticha Klungtong, Theerarat Kochakarn, Namfon Kotanan, Krittikorn Kumpornsin, Wudtichai Manasatienkij, Bhakbhoom Panthan, Ekawat Pasomsub, Insee Sensorn, Arporn Wangwiwatsin |
| EPI_ISL_423044, EPI_ISL_423045, EPI_ISL_423046, EPI_ISL_423047, EPI_ISL_423048, EPI_ISL_423049, EPI_ISL_423050, EPI_ISL_423051, EPI_ISL_423052, EPI_ISL_423053, EPI_ISL_423054, EPI_ISL_423055, EPI_ISL_423056, EPI_ISL_423057, EPI_ISL_423058, EPI_ISL_423059, EPI_ISL_423060, EPI_ISL_423061, EPI_ISL_423062, EPI_ISL_423063, EPI_ISL_423064, EPI_ISL_423065, EPI_ISL_423066, EPI_ISL_423067, EPI_ISL_423068, EPI_ISL_423069, EPI_ISL_423070, EPI_ISL_423071, EPI_ISL_423072, EPI_ISL_423073, EPI_ISL_423074, EPI_ISL_423075, EPI_ISL_423076, EPI_ISL_423077, EPI_ISL_423078, EPI_ISL_423079, EPI_ISL_423080, EPI_ISL_423081, EPI_ISL_423082, EPI_ISL_423083, EPI_ISL_423084, EPI_ISL_423085, EPI_ISL_423086, EPI_ISL_423087, EPI_ISL_423088, EPI_ISL_423089, EPI_ISL_423090, EPI_ISL_423091, EPI_ISL_423092, EPI_ISL_423093, EPI_ISL_423094, EPI_ISL_423095, EPI_ISL_423096, EPI_ISL_423097, EPI_ISL_423098, EPI_ISL_423099, EPI_ISL_423100, EPI_ISL_423101, EPI_ISL_423102, EPI_ISL_423103, EPI_ISL_423104, EPI_ISL_423105, EPI_ISL_423106, EPI_ISL_423107, EPI_ISL_423108, EPI_ISL_423109, EPI_ISL_423110, EPI_ISL_423111, EPI_ISL_423112, EPI_ISL_423113, EPI_ISL_423114, EPI_ISL_423115, EPI_ISL_423116, EPI_ISL_423117, EPI_ISL_423118, EPI_ISL_423119, EPI_ISL_423120, EPI_ISL_423121, EPI_ISL_423122, EPI_ISL_423123, EPI_ISL_423124, EPI_ISL_423125, EPI_ISL_423126, EPI_ISL_423127, EPI_ISL_423128, EPI_ISL_423129, EPI_ISL_423130, EPI_ISL_423131, EPI_ISL_423132, EPI_ISL_423133, EPI_ISL_423134, EPI_ISL_423135, EPI_ISL_423136, EPI_ISL_423137, EPI_ISL_423138, EPI_ISL_423139, EPI_ISL_423140, EPI_ISL_423141, EPI_ISL_423142, EPI_ISL_423143, EPI_ISL_423144, EPI_ISL_423145, EPI_ISL_423146, EPI_ISL_423147, EPI_ISL_423148, EPI_ISL_423149, EPI_ISL_423150, EPI_ISL_423151, EPI_ISL_423152, EPI_ISL_423153, EPI_ISL_423154, EPI_ISL_423155, EPI_ISL_423156, EPI_ISL_423157, EPI_ISL_423158, EPI_ISL_423159, EPI_ISL_423160, EPI_ISL_423161, EPI_ISL_423162, EPI_ISL_423163, EPI_ISL_423164, EPI_ISL_423165, EPI_ISL_423166, EPI_ISL_423167, EPI_ISL_423168, EPI_ISL_423169, EPI_ISL_423170, EPI_ISL_423171, EPI_ISL_423172, EPI_ISL_423173, EPI_ISL_423174, EPI_ISL_423175, EPI_ISL_423176, EPI_ISL_423177, EPI_ISL_423178, EPI_ISL_423179, EPI_ISL_423180, EPI_ISL_423181, EPI_ISL_423182, EPI_ISL_423183, EPI_ISL_423184, EPI_ISL_423185, EPI_ISL_423186, EPI_ISL_423187, EPI_ISL_423188, EPI_ISL_423189, EPI_ISL_423190, EPI_ISL_423191, EPI_ISL_423192, EPI_ISL_423193, EPI_ISL_423194, EPI_ISL_423195, EPI_ISL_423196, EPI_ISL_423197, EPI_ISL_423198, EPI_ISL_423199, EPI_ISL_423200, EPI_ISL_423201, EPI_ISL_423202, EPI_ISL_423203, EPI_ISL_423204, EPI_ISL_423205, EPI_ISL_423206, EPI_ISL_423207, EPI_ISL_423208, EPI_ISL_423209, EPI_ISL_423210, EPI_ISL_423211, EPI_ISL_423212, EPI_ISL_423213, EPI_ISL_423214, EPI_ISL_423215, EPI_ISL_423216, EPI_ISL_423217, EPI_ISL_423218, EPI_ISL_423219, EPI_ISL_423220, EPI_ISL_423221, EPI_ISL_423222, EPI_ISL_423223, EPI_ISL_423224, EPI_ISL_423225, EPI_ISL_423226, EPI_ISL_423227, EPI_ISL_423228, EPI_ISL_423229, EPI_ISL_423230, EPI_ISL_423231, EPI_ISL_423232, EPI_ISL_423233, EPI_ISL_423234, EPI_ISL_423235, EPI_ISL_423236, EPI_ISL_423237, EPI_ISL_423238, EPI_ISL_423239, EPI_ISL_423240, EPI_ISL_423241, EPI_ISL_423242, EPI_ISL_423243, EPI_ISL_423244, EPI_ISL_423245, EPI_ISL_423246, EPI_ISL_423247, EPI_ISL_423248, EPI_ISL_423249, EPI_ISL_423250, EPI_ISL_423251, EPI_ISL_423252, EPI_ISL_423253, EPI_ISL_423254, EPI_ISL_423255, EPI_ISL_423256, EPI_ISL_423257, EPI_ISL_423258, EPI_ISL_423259, EPI_ISL_423260, EPI_ISL_423261, EPI_ISL_423262, EPI_ISL_423263, EPI_ISL_423264, EPI_ISL_423265, EPI_ISL_423266, EPI_ISL_423267, EPI_ISL_423268, EPI_ISL_423269, EPI_ISL_423270, EPI_ISL_423271, EPI_ISL_423272, EPI_ISL_423273, EPI_ISL_423274, EPI_ISL_423275, EPI_ISL_423276, EPI_ISL_423277, EPI_ISL_423278, EPI_ISL_423279, EPI_ISL_423280, EPI_ISL_423281, EPI_ISL_423282, EPI_ISL_423283, | | | |

| | | | |
|---|---|---|---|
| EPI_ISL_423284, EPI_ISL_423285, EPI_ISL_423286, EPI_ISL_423287, EPI_ISL_423288, EPI_ISL_423289, EPI_ISL_423290, EPI_ISL_423291, EPI_ISL_423292, EPI_ISL_423293, EPI_ISL_423294, EPI_ISL_423295, EPI_ISL_423296, EPI_ISL_423297, EPI_ISL_423298, EPI_ISL_423299, EPI_ISL_423300, EPI_ISL_423301, EPI_ISL_423302, EPI_ISL_423303, EPI_ISL_423304, EPI_ISL_423305, EPI_ISL_423306, EPI_ISL_423307, EPI_ISL_423308, EPI_ISL_423309, EPI_ISL_423310, EPI_ISL_423311, EPI_ISL_423312, EPI_ISL_423313, EPI_ISL_423314, EPI_ISL_423315, EPI_ISL_423316, EPI_ISL_423317, EPI_ISL_423318, EPI_ISL_423319, EPI_ISL_423320, EPI_ISL_423321, EPI_ISL_423322, EPI_ISL_423323, EPI_ISL_423324, EPI_ISL_423325, EPI_ISL_423326, EPI_ISL_423327, EPI_ISL_423328, EPI_ISL_423329, EPI_ISL_423330, EPI_ISL_423331, EPI_ISL_423332, EPI_ISL_423333, EPI_ISL_423334, EPI_ISL_423335, EPI_ISL_423336, EPI_ISL_423337, EPI_ISL_423338, EPI_ISL_423339, EPI_ISL_423340, EPI_ISL_423341, EPI_ISL_423342, EPI_ISL_423343, EPI_ISL_423344, EPI_ISL_423345, EPI_ISL_423346, EPI_ISL_423347, EPI_ISL_423348, EPI_ISL_423349, EPI_ISL_423350, EPI_ISL_423351, EPI_ISL_423352, EPI_ISL_423353, EPI_ISL_423354, EPI_ISL_423355, EPI_ISL_423356, EPI_ISL_423357, EPI_ISL_423358, EPI_ISL_423359, EPI_ISL_423360, EPI_ISL_423361, EPI_ISL_423362, EPI_ISL_423363, EPI_ISL_423364, EPI_ISL_423365, EPI_ISL_423366, EPI_ISL_423367, EPI_ISL_423368, EPI_ISL_423369, EPI_ISL_423370, EPI_ISL_423371, EPI_ISL_423372, EPI_ISL_423373, EPI_ISL_423374, EPI_ISL_423375, EPI_ISL_423376, EPI_ISL_423377, EPI_ISL_423378, EPI_ISL_423379, EPI_ISL_423380, EPI_ISL_423381, EPI_ISL_423382, EPI_ISL_423383, EPI_ISL_423384, EPI_ISL_423385, EPI_ISL_423386, EPI_ISL_423387, EPI_ISL_423388, EPI_ISL_423389, EPI_ISL_423390, EPI_ISL_423391, EPI_ISL_423392, EPI_ISL_423393, EPI_ISL_423394, EPI_ISL_423395, EPI_ISL_423396, EPI_ISL_423397, EPI_ISL_423398, EPI_ISL_423399, EPI_ISL_423400, EPI_ISL_423401, EPI_ISL_423402, EPI_ISL_423403, EPI_ISL_423404, EPI_ISL_423405, EPI_ISL_423406, EPI_ISL_423407, EPI_ISL_423408, EPI_ISL_423409, EPI_ISL_423410, EPI_ISL_423411, EPI_ISL_423412, EPI_ISL_423413, EPI_ISL_423414, EPI_ISL_423415, EPI_ISL_423416, EPI_ISL_423417, EPI_ISL_423418, EPI_ISL_423419, EPI_ISL_423420, EPI_ISL_423421, EPI_ISL_423422, EPI_ISL_423423, EPI_ISL_423424, EPI_ISL_423425, EPI_ISL_423426, EPI_ISL_423427, EPI_ISL_423428, EPI_ISL_423429, EPI_ISL_423430, EPI_ISL_423431, EPI_ISL_423432, EPI_ISL_423433, EPI_ISL_423434, EPI_ISL_423435, EPI_ISL_423436, EPI_ISL_423437, EPI_ISL_423438, EPI_ISL_423439, EPI_ISL_423440, EPI_ISL_423441, EPI_ISL_423442, EPI_ISL_423443, EPI_ISL_423444, EPI_ISL_423445, EPI_ISL_423446, EPI_ISL_423447, EPI_ISL_423448, EPI_ISL_423449, EPI_ISL_423450, EPI_ISL_423451, EPI_ISL_423452, EPI_ISL_423453, EPI_ISL_423454, EPI_ISL_423455, EPI_ISL_423456, EPI_ISL_423457, EPI_ISL_423458, EPI_ISL_423459, EPI_ISL_423460, EPI_ISL_423461, EPI_ISL_423462, EPI_ISL_423463, EPI_ISL_423464, EPI_ISL_423465, EPI_ISL_423466, EPI_ISL_423467, EPI_ISL_423468, EPI_ISL_423469, EPI_ISL_423470, EPI_ISL_423471, EPI_ISL_423472, EPI_ISL_423473, EPI_ISL_423474, EPI_ISL_423475, EPI_ISL_423476, EPI_ISL_423477, EPI_ISL_423478, EPI_ISL_423479, EPI_ISL_423480, EPI_ISL_423481, EPI_ISL_423482, EPI_ISL_423483, EPI_ISL_423484, EPI_ISL_423485, EPI_ISL_423486, EPI_ISL_423487, EPI_ISL_423488, EPI_ISL_423489, EPI_ISL_423490, EPI_ISL_423491, EPI_ISL_423492, EPI_ISL_423493, EPI_ISL_423494, EPI_ISL_423495, EPI_ISL_423496, EPI_ISL_423497, EPI_ISL_423498, EPI_ISL_423499, EPI_ISL_423500, EPI_ISL_423501, EPI_ISL_423502, EPI_ISL_423503, EPI_ISL_423504, EPI_ISL_423505, EPI_ISL_423506, EPI_ISL_423507, EPI_ISL_423508, EPI_ISL_423509, EPI_ISL_423510, EPI_ISL_423511, EPI_ISL_423512, EPI_ISL_423513, EPI_ISL_423514, EPI_ISL_423515, EPI_ISL_423516, EPI_ISL_423517, EPI_ISL_423518, EPI_ISL_423519, EPI_ISL_423520, EPI_ISL_423521, EPI_ISL_423522, EPI_ISL_423523, EPI_ISL_423524, EPI_ISL_423525, EPI_ISL_423526, EPI_ISL_423527, EPI_ISL_423528, EPI_ISL_423529, EPI_ISL_423530, EPI_ISL_423531, EPI_ISL_423532, EPI_ISL_423533, EPI_ISL_423534, EPI_ISL_423535, EPI_ISL_423536, EPI_ISL_423537, EPI_ISL_423538, EPI_ISL_423539, EPI_ISL_423540, EPI_ISL_423541, EPI_ISL_423542, EPI_ISL_423543, EPI_ISL_423544, EPI_ISL_423545, EPI_ISL_423546, EPI_ISL_423547, EPI_ISL_423548, EPI_ISL_423549, EPI_ISL_423550, EPI_ISL_423551, EPI_ISL_423552, EPI_ISL_423553, EPI_ISL_423554, EPI_ISL_423555, EPI_ISL_423556, EPI_ISL_423557, EPI_ISL_423558, EPI_ISL_423559, EPI_ISL_423560, EPI_ISL_423561, EPI_ISL_423562, EPI_ISL_423563, EPI_ISL_423564, EPI_ISL_423565, EPI_ISL_423566, EPI_ISL_423567, EPI_ISL_423568, EPI_ISL_423569, EPI_ISL_423570, EPI_ISL_423571, EPI_ISL_423572, EPI_ISL_423573, EPI_ISL_423574, EPI_ISL_423575, EPI_ISL_423576, EPI_ISL_423577, EPI_ISL_423578, EPI_ISL_423579, EPI_ISL_423580, EPI_ISL_423581, EPI_ISL_423582, EPI_ISL_423583, EPI_ISL_423584, EPI_ISL_423585, EPI_ISL_423586, EPI_ISL_423587, EPI_ISL_423588, EPI_ISL_423589, EPI_ISL_423590, EPI_ISL_423591, EPI_ISL_423592, EPI_ISL_423593, EPI_ISL_423594, EPI_ISL_423595, EPI_ISL_423596, EPI_ISL_423597, EPI_ISL_423598, EPI_ISL_423599, EPI_ISL_423600, EPI_ISL_423601, EPI_ISL_423602, EPI_ISL_423603, EPI_ISL_423604, EPI_ISL_423605, EPI_ISL_423606, EPI_ISL_423607, EPI_ISL_423608, EPI_ISL_423609, EPI_ISL_423610, EPI_ISL_423611, EPI_ISL_423612, EPI_ISL_423613, EPI_ISL_423614, EPI_ISL_423615, EPI_ISL_423616, EPI_ISL_423617, EPI_ISL_423618, EPI_ISL_423619, EPI_ISL_423620, EPI_ISL_423621, EPI_ISL_423622, EPI_ISL_423623, EPI_ISL_423624, EPI_ISL_423625, EPI_ISL_423626, EPI_ISL_423627, EPI_ISL_423628, EPI_ISL_423629, EPI_ISL_423630, EPI_ISL_423631, EPI_ISL_423632, EPI_ISL_423633, EPI_ISL_423634, EPI_ISL_423635, EPI_ISL_423636, EPI_ISL_423637, EPI_ISL_423638, EPI_ISL_423639, EPI_ISL_423640, EPI_ISL_423641, EPI_ISL_423642, EPI_ISL_423643, EPI_ISL_423644, EPI_ISL_423645, EPI_ISL_423646, EPI_ISL_423647, EPI_ISL_423648, EPI_ISL_423649, EPI_ISL_423650, EPI_ISL_423651, EPI_ISL_423652, EPI_ISL_423653, EPI_ISL_423654, EPI_ISL_423655, EPI_ISL_423656, EPI_ISL_423657, EPI_ISL_423658, EPI_ISL_423659, EPI_ISL_423660, EPI_ISL_423661, EPI_ISL_423662, EPI_ISL_423663, EPI_ISL_423664, EPI_ISL_423665, EPI_ISL_423666, EPI_ISL_423667, EPI_ISL_423668, EPI_ISL_423669, EPI_ISL_423670, EPI_ISL_423671, EPI_ISL_423672, EPI_ISL_423673, EPI_ISL_423674, EPI_ISL_423675, EPI_ISL_423676, EPI_ISL_423677, EPI_ISL_423678, EPI_ISL_423679, EPI_ISL_423680, EPI_ISL_423681, EPI_ISL_423682, EPI_ISL_423683, EPI_ISL_423684, EPI_ISL_423685, EPI_ISL_423686, EPI_ISL_423687, EPI_ISL_423688, EPI_ISL_423689, EPI_ISL_423690, EPI_ISL_423691, EPI_ISL_423692, EPI_ISL_423693, EPI_ISL_423694, EPI_ISL_423695, EPI_ISL_423696, EPI_ISL_423697, EPI_ISL_423698, EPI_ISL_423699, EPI_ISL_423700, EPI_ISL_423701, EPI_ISL_423702, EPI_ISL_423703, EPI_ISL_423704, EPI_ISL_423705, EPI_ISL_423706, EPI_ISL_423707, EPI_ISL_423708, EPI_ISL_423709, EPI_ISL_423710, EPI_ISL_423711, EPI_ISL_423712, EPI_ISL_423713, EPI_ISL_423714, EPI_ISL_423715, EPI_ISL_423716, EPI_ISL_423717, EPI_ISL_423718, EPI_ISL_423719, EPI_ISL_423720, EPI_ISL_423721, EPI_ISL_423722, EPI_ISL_423723, EPI_ISL_423724, EPI_ISL_423725, EPI_ISL_423726, EPI_ISL_423727, EPI_ISL_423728, EPI_ISL_423729, EPI_ISL_423730, EPI_ISL_423731, EPI_ISL_423732, EPI_ISL_423733, EPI_ISL_423734, EPI_ISL_423735, EPI_ISL_423736, EPI_ISL_423737, EPI_ISL_423738, EPI_ISL_423739, EPI_ISL_423740, EPI_ISL_423741, EPI_ISL_423742, EPI_ISL_423743, EPI_ISL_423744, EPI_ISL_423745, EPI_ISL_423746, EPI_ISL_423747, EPI_ISL_423748, EPI_ISL_423749, EPI_ISL_423750, EPI_ISL_423751, EPI_ISL_423752, EPI_ISL_423753, EPI_ISL_423754, EPI_ISL_423755, EPI_ISL_423756, EPI_ISL_423757, EPI_ISL_423758, EPI_ISL_423759, EPI_ISL_423760, EPI_ISL_423761, EPI_ISL_423762, EPI_ISL_423763, EPI_ISL_423764, EPI_ISL_423765, EPI_ISL_423766, EPI_ISL_423767, EPI_ISL_423768, EPI_ISL_423769, EPI_ISL_423770, EPI_ISL_423771, EPI_ISL_423772, EPI_ISL_423773, EPI_ISL_423774, EPI_ISL_423775, EPI_ISL_423776, EPI_ISL_423777, EPI_ISL_423778, EPI_ISL_423779, EPI_ISL_423780, EPI_ISL_423781, EPI_ISL_423782, EPI_ISL_423783, EPI_ISL_423784, EPI_ISL_423785, EPI_ISL_423786, EPI_ISL_423787, EPI_ISL_423788, EPI_ISL_423789, EPI_ISL_423790, EPI_ISL_423791, EPI_ISL_423792, EPI_ISL_423793, EPI_ISL_423794, EPI_ISL_423795, EPI_ISL_423796, EPI_ISL_423797, EPI_ISL_423798, EPI_ISL_423799, EPI_ISL_423800, EPI_ISL_423801, EPI_ISL_423802, EPI_ISL_423803, EPI_ISL_423804, EPI_ISL_423805, EPI_ISL_423806, EPI_ISL_423807, EPI_ISL_423808, EPI_ISL_423809, EPI_ISL_423810, EPI_ISL_423811, EPI_ISL_423812, EPI_ISL_423813, EPI_ISL_423814, EPI_ISL_423815, EPI_ISL_423816, EPI_ISL_423817, EPI_ISL_423818, EPI_ISL_423819, EPI_ISL_423820, EPI_ISL_423821, EPI_ISL_423822, EPI_ISL_423823, EPI_ISL_423824, EPI_ISL_423825, EPI_ISL_423826, EPI_ISL_423827, EPI_ISL_423828, EPI_ISL_423829, EPI_ISL_423830, EPI_ISL_423831, EPI_ISL_423832, EPI_ISL_423833, EPI_ISL_423834, EPI_ISL_423835, EPI_ISL_423836, EPI_ISL_423837, EPI_ISL_423838, EPI_ISL_423839, EPI_ISL_423840, EPI_ISL_423841, EPI_ISL_423842, EPI_ISL_423843, EPI_ISL_423844, EPI_ISL_423845, EPI_ISL_423846, EPI_ISL_423847, EPI_ISL_423848, EPI_ISL_423849, EPI_ISL_423850, EPI_ISL_423851, EPI_ISL_423852, EPI_ISL_423853, EPI_ISL_423854, EPI_ISL_423855, EPI_ISL_423856, EPI_ISL_423857, EPI_ISL_423858, EPI_ISL_423859, EPI_ISL_423860, EPI_ISL_423861, EPI_ISL_423862, EPI_ISL_423863, EPI_ISL_423864, EPI_ISL_423865, EPI_ISL_423866, EPI_ISL_423867, EPI_ISL_423868, EPI_ISL_423869, EPI_ISL_423870, EPI_ISL_423871, EPI_ISL_423872, EPI_ISL_423873, EPI_ISL_423874, EPI_ISL_423875, EPI_ISL_423876, EPI_ISL_423877, EPI_ISL_423878, EPI_ISL_423879, EPI_ISL_423880, EPI_ISL_423881, EPI_ISL_423882, EPI_ISL_423883, EPI_ISL_423884, EPI_ISL_423885, EPI_ISL_423886, EPI_ISL_423887, EPI_ISL_423888, EPI_ISL_423889, EPI_ISL_423890, EPI_ISL_423891, EPI_ISL_423892, EPI_ISL_423893, EPI_ISL_423894, EPI_ISL_423895, EPI_ISL_423896, EPI_ISL_423897, EPI_ISL_423898, EPI_ISL_423899, EPI_ISL_423900, EPI_ISL_423901, EPI_ISL_423902, EPI_ISL_423903, EPI_ISL_423904, EPI_ISL_423905, EPI_ISL_423906, EPI_ISL_423907, EPI_ISL_423908, EPI_ISL_423909, EPI_ISL_423910, EPI_ISL_423911, EPI_ISL_423912, EPI_ISL_423913, EPI_ISL_423914, EPI_ISL_423915, EPI_ISL_423916, EPI_ISL_423917, EPI_ISL_423918, EPI_ISL_423919, EPI_ISL_423920, EPI_ISL_423921, EPI_ISL_423922, EPI_ISL_423923, EPI_ISL_423924, EPI_ISL_423925, EPI_ISL_423926, EPI_ISL_423927, EPI_ISL_423928, EPI_ISL_423929, EPI_ISL_423930, EPI_ISL_423931, EPI_ISL_423932, EPI_ISL_423933, EPI_ISL_423934, EPI_ISL_423935, EPI_ISL_423936, EPI_ISL_423937, EPI_ISL_423938, EPI_ISL_423939, EPI_ISL_423940, EPI_ISL_423941, EPI_ISL_423942, EPI_ISL_423943, EPI_ISL_423944, EPI_ISL_423945, EPI_ISL_423946, EPI_ISL_423947, EPI_ISL_423948, EPI_ISL_423949, EPI_ISL_423950, EPI_ISL_423951, EPI_ISL_423952, EPI_ISL_423953, EPI_ISL_423954, EPI_ISL_423955, EPI_ISL_423956, EPI_ISL_423957, EPI_ISL_423958, EPI_ISL_423959, EPI_ISL_423960, EPI_ISL_423961, EPI_ISL_423962, EPI_ISL_423963, EPI_ISL_423964, EPI_ISL_423965, EPI_ISL_423966, EPI_ISL_423967, EPI_ISL_423968, EPI_ISL_423969, EPI_ISL_423970, EPI_ISL_423971, EPI_ISL_423972, EPI_ISL_423973, EPI_ISL_423974, EPI_ISL_423975, EPI_ISL_423976, EPI_ISL_423977, EPI_ISL_423978, EPI_ISL_423979, EPI_ISL_423980, EPI_ISL_423981, EPI_ISL_423982, EPI_ISL_423983, EPI_ISL_423984, EPI_ISL_423985, EPI_ISL_423986, EPI_ISL_423987, EPI_ISL_423988, EPI_ISL_423989, EPI_ISL_423990, EPI_ISL_423991, EPI_ISL_423992, EPI_ISL_423993, EPI_ISL_423994, EPI_ISL_423995, EPI_ISL_423996, EPI_ISL_423997, EPI_ISL_423998, EPI_ISL_423999, EPI_ISL_424000, EPI_ISL_424001, EPI_ISL_424002, EPI_ISL_424003, EPI_ISL_424004, EPI_ISL_424005, EPI_ISL_424006, EPI_ISL_424007, EPI_ISL_424008, EPI_ISL_424009, EPI_ISL_424010, EPI_ISL_424011, EPI_ISL_424012, EPI_ISL_424013, EPI_ISL_424014, EPI_ISL_424015, EPI_ISL_424016, EPI_ISL_424017, EPI_ISL_424018, EPI_ISL_424019, EPI_ISL_424020, EPI_ISL_424021, EPI_ISL_424022, EPI_ISL_424023, EPI_ISL_424024, EPI_ISL_424025, EPI_ISL_424026, EPI_ISL_424027, EPI_ISL_424028, EPI_ISL_424029, EPI_ISL_424030, EPI_ISL_424031, EPI_ISL_424032, EPI_ISL_424033, EPI_ISL_424034, EPI_ISL_424035, EPI_ISL_424036, EPI_ISL_424037, EPI_ISL_424038, EPI_ISL_424039, EPI_ISL_424040, EPI_ISL_424041, EPI_ISL_424042, EPI_ISL_424043, EPI_ISL_424044, EPI_ISL_424045, EPI_ISL_424046, EPI_ISL_424047, EPI_ISL_424048, EPI_ISL_424049, EPI_ISL_424050, EPI_ISL_424051, EPI_ISL_424052, EPI_ISL_424053, EPI_ISL_424054, EPI_ISL_424055, EPI_ISL_424056, EPI_ISL_424057, EPI_ISL_424058, EPI_ISL_424059, EPI_ISL_424060, EPI_ISL_424061, EPI_ISL_424062, EPI_ISL_424063, EPI_ISL_424064, EPI_ISL_424065, EPI_ISL_424066, EPI_ISL_424067, EPI_ISL_424068, EPI_ISL_424069, EPI_ISL_424070, EPI_ISL_424071, EPI_ISL_424072, EPI_ISL_424073, EPI_ISL_424074, EPI_ISL_424075, EPI_ISL_424076, EPI_ISL_424077, EPI_ISL_424078, EPI_ISL_424079, EPI_ISL_424080, EPI_ISL_424081, EPI_ISL_424082, EPI_ISL_424083, EPI_ISL_424084, EPI_ISL_424085, EPI_ISL_424086, EPI_ISL_424087, EPI_ISL_424088, EPI_ISL_424089, EPI_ISL_424090, EPI_ISL_424091, EPI_ISL_424092, EPI_ISL_424093, EPI_ISL_424094, EPI_ISL_424095, EPI_ISL_424096, EPI_ISL_424097, EPI_ISL_424098, EPI_ISL_424099, EPI_ISL_424100, EPI_ISL_424101, EPI_ISL_424102, EPI_ISL_424103, EPI_ISL_424104, EPI_ISL_424105, EPI_ISL_424106, EPI_ISL_424107, EPI_ISL_424108, EPI_ISL_424109, EPI_ISL_424110, EPI_ISL_424111, EPI_ISL_424112, EPI_ISL_424113, EPI_ISL_424114, EPI_ISL_424115, EPI_ISL_424116, EPI_ISL_424117, EPI_ISL_424118, EPI_ISL_424119, EPI_ISL_424120, EPI_ISL_424121, EPI_ISL_424122, EPI_ISL_424123, EPI_ISL_424124, EPI_ISL_424125, EPI_ISL_424126, EPI_ISL_424127, EPI_ISL_424128, EPI_ISL_424129, EPI_ISL_424130, EPI_ISL_424131, EPI_ISL_424132, EPI_ISL_424133, EPI_ISL_424134, EPI_ISL_424135, EPI_ISL_424136, EPI_ISL_424137, EPI_ISL_424138, EPI_ISL_424139, EPI_ISL_424140, EPI_ISL_424141, EPI_ISL_424142, EPI_ISL_424143, EPI_ISL_424144, EPI_ISL_424145, EPI_ISL_424146, EPI_ISL_424147, EPI_ISL_424148, EPI_ISL_424149, EPI_ISL_424150, EPI_ISL_424151, EPI_ISL_424152, EPI_ISL_424153, EPI_ISL_424154, EPI_ISL_424155, EPI_ISL_424156, EPI_ISL_424157, EPI_ISL_424158, EPI_ISL_424159, EPI_ISL_424160, EPI_ISL_424161, EPI_ISL_424162, EPI_ISL_424163, EPI_ISL_424164, EPI_ISL_424165 | | | |
| see above | Respiratory Virus Unit, Microbiology Services Colindale, Public Health England | Respiratory Virus Unit, Microbiology Services Colindale, Public Health England | Monica Galiano, Shahjahan Miah, Angie Lackenby, Omolola Akinbami, Tiina Talts, Leena Bhaw, Richard Myers, Steven Platt, Kirstin Edwards, Jonathan Hubb, Joanna Ellis, Maria Zambon |
| EPI_ISL_424166, EPI_ISL_424167, EPI_ISL_424168, EPI_ISL_424169, EPI_ISL_424170, EPI_ISL_424171, EPI_ISL_424172, EPI_ISL_424173, EPI_ISL_424174, EPI_ISL_424175, EPI_ISL_424176, EPI_ISL_424177, EPI_ISL_424178, EPI_ISL_424179, EPI_ISL_424180, EPI_ISL_424181, EPI_ISL_424182, EPI_ISL_424183, EPI_ISL_424184, EPI_ISL_424185, EPI_ISL_424186, EPI_ISL_424187, EPI_ISL_424188, EPI_ISL_424189, EPI_ISL_424190, EPI_ISL_424191, EPI_ISL_424192, EPI_ISL_424193, EPI_ISL_424194, EPI_ISL_424195, EPI_ISL_424196, EPI_ISL_424197, EPI_ISL_424198, EPI_ISL_424199, EPI_ISL_424200, EPI_ISL_424201, EPI_ISL_424202, EPI_ISL_424203, EPI_ISL_424204, EPI_ISL_424205, EPI_ISL_424206, EPI_ISL_424207, EPI_ISL_424208, EPI_ISL_424209, EPI_ISL_424210, EPI_ISL_424211, EPI_ISL_424212, EPI_ISL_424213, EPI_ISL_424214, EPI_ISL_424215, EPI_ISL_424216, EPI_ISL_424217, EPI_ISL_424218, EPI_ISL_424219, EPI_ISL_424220, EPI_ISL_424221, EPI_ISL_424222, EPI_ISL_424223, EPI_ISL_424224, EPI_ISL_424225, EPI_ISL_424226, EPI_ISL_424227, EPI_ISL_424228, EPI_ISL_424229, EPI_ISL_424230, EPI_ISL_424231, EPI_ISL_424232, EPI_ISL_424233, EPI_ISL_424234, EPI_ISL_424235, EPI_ISL_424236, EPI_ISL_424237, EPI_ISL_424238, EPI_ISL_424239, EPI_ISL_424240, EPI_ISL_424241, EPI_ISL_424242, EPI_ISL_424243, EPI_ISL_424244, EPI_ISL_424245, EPI_ISL_424246, EPI_ISL_424247, EPI_ISL_424248, EPI_ISL_424249, EPI_ISL_424250, EPI_ISL_424251, EPI_ISL_424252, EPI_ISL_424253, EPI_ISL_424254, EPI_ISL_424255, EPI_ISL_424256, EPI_ISL_424257, EPI_ISL_424258, EPI_ISL_424259, EPI_ISL_424260, EPI_ISL_424261, EPI_ISL_424262, EPI_ISL_424263, EPI_ISL_424264, EPI_ISL_424265, EPI_ISL_424266, EPI_ISL_424267, EPI_ISL_424268, EPI_ISL_424269, EPI_ISL_424270, EPI_ISL_424271, EPI_ISL_424272, EPI_ISL_424273, EPI_ISL_424274, EPI_ISL_424275, EPI_ISL_424276, EPI_ISL_424277, EPI_ISL_424278, EPI_ISL_424279, EPI_ISL_424280, EPI_ISL_424281, EPI_ISL_424282, EPI_ISL_424283, EPI_ISL_424284, EPI_ISL_424285, EPI_ISL_424286, EPI_ISL_424287, EPI_ISL_424288, EPI_ISL_424289, EPI_ISL_424290, EPI_ISL_424291, EPI_ISL_424292, EPI_ISL_424293, EPI_ISL_424294, EPI_ISL_424295, EPI_ISL_424296, EPI_ISL_424297, EPI_ISL_424298, EPI_ISL_424299, EPI_ISL_424300, EPI_ISL_424301, EPI_ISL_424302, EPI_ISL_424303, EPI_ISL_424304, EPI_ISL_424305, EPI_ISL_424306, EPI_ISL_424307, EPI_ISL_424308, EPI_ISL_424309, EPI_ISL_424310, EPI_ISL_424311, EPI_ISL_424312, EPI_ISL_424313, EPI_ISL_424314, EPI_ISL_424315, EPI_ISL_424316, EPI_ISL_424317, EPI_ISL_424318, EPI_ISL_424319, EPI_ISL_424320, EPI_ISL_424321, EPI_ISL_424322, EPI_ISL_424323, EPI_ISL_424324, EPI_ISL_424325, EPI_ISL_424326, EPI_ISL_424327, EPI_ISL_424328, EPI_ISL_424329, EPI_ISL_424330, EPI_ISL_424331, EPI_ISL_424332, EPI_ISL_424333, EPI_ISL_424334, EPI_ISL_424335, EPI_ISL_424336, EPI_ISL_424337, EPI_ISL_424338, EPI_ISL_424339, EPI_ISL_424340, EPI_ISL_424341 | | | |
| see above | UW Virology Lab | UW Virology Lab | Pavitra Roychoudhury, Hong Xie, Keith Jerome, Alexander Greninger |
| EPI_ISL_424342 | INMI Lazzaro Spallanzani IRCCS | Laboratory of Virology, INMI Lazzaro Spallanzani IRCCS | Concetta Castilletti, Barbara Bartolini, Martina Rueca, Cesare Ernesto Maria Gruber, Francesco Messina, Fabrizio Carletti, Eleonora Lalle, Licia Bordi, Giulia Matusali, Francesca Colavita, Maria Rosaria Capobianchi, Francesco Vairo, Giuseppe Ippolito, Antonino Di Caro |
| EPI_ISL_424343 | INMI Lazzaro Spallanzani IRCCS | Laboratory of Virology, INMI Lazzaro Spallanzani IRCCS | Fabrizio Carletti, Barbara Bartolini, Martina Rueca, Cesare Ernesto Maria Gruber, Francesco Messina, Eleonora Lalle, Licia Bordi, Giulia Matusali, Francesca Colavita, Maria Rosaria Capobianchi, Concetta Castilletti, Francesco Vairo, Giuseppe Ippolito, Antonino Di Caro |
| EPI_ISL_424344 | INMI Lazzaro Spallanzani IRCCS | Laboratory of Virology, INMI Lazzaro Spallanzani IRCCS | Eleonora Lalle, Barbara Bartolini, Martina Rueca, Cesare Ernesto Maria Gruber, Francesco Messina, Fabrizio Carletti, Licia Bordi, Giulia Matusali, Francesca Colavita, Maria Rosaria Capobianchi, Concetta Castilletti, Francesco Vairo, Giuseppe Ippolito, Antonino Di Caro |
| EPI_ISL_424345 | Instituto Nacional de Enfermedades Respiratorias | Instituto Nacional de Enfermedades Respiratorias | Joel Armando Vázquez Pérez, Celia Boukadida, Santiago Avila Ríos, Mario Mújica Sánchez, José Arturo Martínez Orozco, Eduardo Becerril Vargas, Jorge Salas Hernández, Irma López Martínez, Lucia Hernández Rivas, Gisela Barrera Badillo, Edgar Mendieta Condado, Fabiola Garcés Ayala, Adnan Araiza Rodríguez, José Ernesto Ramírez González, Victor Hugo Borja Aburto, Concepción Grajales Muñiz, Cesar Raúl González Bonilla, Carolina González Torres, Francisco Javier Gaytán Cervantes, José Esteban Muñoz Medina, Guillermo M. Ruiz-Palacios, Pilar Ramos Cervantes, Violeta Ibarra Gonzalez, Fernando Ledesma Barrientos, Luis Alberto García Andrade, Alfredo Ponce de León Garduño, Blanca Taboada, Alejandro Sánchez, Pavel Isa, Ricardo Grande, Gloria Vázquez, Francisco Pulido, Carlos F. Arias. |
| EPI_ISL_424346, EPI_ISL_424347 | Alaska State Virology Laboratory | Alaska State Virology Laboratory | Chen, J. |
| EPI_ISL_424348 | Instituto Nacional de Enfermedades Respiratorias | Instituto Nacional de Enfermedades Respiratorias | Joel Armando Vázquez Pérez, Celia Boukadida, Santiago Avila Ríos, Mario Mújica Sánchez, José Arturo Martínez Orozco, Eduardo Becerril Vargas, Jorge Salas Hernández, Irma López Martínez, Lucia Hernández Rivas, Gisela Barrera Badillo, Edgar Mendieta Condado, Fabiola Garcés Ayala, Adnan Araiza Rodríguez, José Ernesto Ramírez González, Victor Hugo Borja Aburto, Concepción Grajales Muñiz, Cesar Raúl González Bonilla, Carolina González Torres, Francisco Javier Gaytán Cervantes, José Esteban Muñoz Medina, Guillermo M. Ruiz-Palacios, Pilar Ramos Cervantes, Violeta Ibarra Gonzalez, Fernando Ledesma Barrientos, Luis Alberto García Andrade, Alfredo Ponce de León Garduño, Blanca Taboada, Alejandro Sánchez, Pavel Isa, Ricardo Grande, Gloria Vázquez, Francisco Pulido, Carlos F. Arias. |
| EPI_ISL_424349 | Human Genetic Research Center, Kawsar Biotech Company | Human Genetic Research Center, Kawsar Biotech Company | Zeinali,S., Khosravi,M.A., Abbasalipour Bashash,M., Mostafavi Jabbari,S., Firoozi,M., Pourtavakoli,S., Khateri,E., Zeinali,R. and Hoseini,F. |
| EPI_ISL_424352 | Clinical Laboratory, Fuyang City Center for Disease Control and Prevention | Clinical Laboratory, Fuyang City Center for Disease Control and Prevention | Ge,B. |
| EPI_ISL_424353 | Dirk Dittmer | Dirk Dittmer | Bailey,A.G., Caro-Vegas,C., Thompson,C., Dittmer,D., Eason,A.B., Juarez,A., Landis,J.T., McNamara,R.P., Miller,M.B., Moorad,R., Pluta,L.J., Seltzer,T.A., Villamor,F. and Vahrson,W. |
| EPI_ISL_424354 | Dirk Dittmer | Dirk Dittmer | Bailey,A.G., Caro-Vegas,C.P., Dittmer,D., Eason,A.B., Juarez,A., Landis,J.T., McNamara,R.P., Miller,M.B., Moorad,R., Pluta,L.J., Seltzer,T.A., Thompson,C., Vahrson,W. and Villamor,F. |
| EPI_ISL_424355, EPI_ISL_424356, EPI_ISL_424357, EPI_ISL_424358, EPI_ISL_424359, EPI_ISL_424360 | Beijing Institute of Microbiology and Epidemiology | Beijing Institute of Microbiology and Epidemiology | Fan,H., Qin,E., Wu,Y., Guo,Y., Zhang,X., Yong,Y., Hou,J., Xu,Z., Mu,J., Teng,Y., Mi,Z., Yang,R., Song,Y., Li,B. and Cui,Y. |
| EPI_ISL_424361, EPI_ISL_424362, EPI_ISL_424363, EPI_ISL_424364, EPI_ISL_424365 | National Influenza Center, Indian Council of Medical Research - National Institute of Virology | Indian Council of Medical Research-National Institute of Virology, Microbial Containment Complex | Pragya D. Yadav, Varsha Potdar, Savita Patil, Dimpal A. Nyayanit, Triparna Majumdar, Manohar. L. Chaudhary, Gururaj Deshpande, Padinjaremattathil Thankappan Ullas, Anita Shete-Aich, Hitesh Dighe, Sreelekshmy Mohandas, Gajanan Sapkal, Atanu Basu, Amita Jain, Bharti Malhotra, Deepika Chaudhary, Sarah Cherian, Priya Abraham |
| EPI_ISL_424366 | Vaccine Research, Development and Application Center, Erciyes University | Gen Era Diagnostics Inc. | Shaikh Terkis Islam Pavel, Hazel Yetiskin, Gunsu Aydin, Can Holyavkin, Muhammet Ali Uygut, Zehra B Dursun, İlhami Celik, Alper Iseri, Aykut Ozdarendeli |
| EPI_ISL_424367, EPI_ISL_424368, EPI_ISL_424369, EPI_ISL_424370, EPI_ISL_424371, EPI_ISL_424372, EPI_ISL_424373, EPI_ISL_424374, EPI_ISL_424375, EPI_ISL_424376 | The National University Hospital of Iceland | deCODE genetics | Daniel F Gudbjartsson; Agnar Helgason; Hakon Jonsson; Olafur T Magnusson; Pall Melsted; Gudmundur L Norddahl; Jona Saemundsdottir; Asgeir Sigurdsson; Patrick Sulem; Arna B Agustsdottir; Berglind Eiriksdottir; Run Fridriksdottir; Elisabet E Gardarsdottir; Gudmundur Georgsson; Olafia S Gretarsdottir; Kjartan R Gudmundsson; Thora R Gunnarsdottir; Arnaldur Gylfason; Hilma Holm; Brynjar O Jensson; Aslaug Jonasdottir; Kamilla S Josefsdottir; Thordur Kristjansson; Droplaug N Magnusdottir; Louise le Roux; Gudrun Sigmundsdottir; Gardar Sveinbjornsson; Kristin E Sveinsdottir; Maney Sveinsdottir; Emil A Thorarensen; Bjarni Thorbjornsson; Gisli Masson; Ingileif Jonsdottir; Alma Moller; Thorolfur Gudnason; Karl G Kristinsson; Unnur Thorsteinsdottir; Kari Stefansson |
| EPI_ISL_424377, EPI_ISL_424378 | deCODE genetics | deCODE genetics | Daniel F Gudbjartsson; Agnar Helgason; Hakon Jonsson; Olafur T Magnusson; Pall Melsted; Gudmundur L Norddahl; Jona Saemundsdottir; Asgeir Sigurdsson; Patrick Sulem; Arna B Agustsdottir; Berglind Eiriksdottir; Run Fridriksdottir; Elisabet E Gardarsdottir; Gudmundur Georgsson; Olafia S Gretarsdottir; Kjartan R Gudmundsson; Thora R Gunnarsdottir; Arnaldur Gylfason; Hilma Holm; Brynjar O Jensson; Aslaug Jonasdottir; Kamilla S Josefsdottir; Thordur Kristjansson; Droplaug N Magnusdottir; Louise le Roux; Gudrun Sigmundsdottir; Gardar Sveinbjornsson; Kristin E Sveinsdottir; Maney Sveinsdottir; Emil A Thorarensen; Bjarni Thorbjornsson; Gisli Masson; Ingileif Jonsdottir; Alma Moller; Thorolfur Gudnason; Karl G Kristinsson; Unnur Thorsteinsdottir; Kari Stefansson |
| EPI_ISL_424379, EPI_ISL_424380, EPI_ISL_424381, EPI_ISL_424382, EPI_ISL_424383, EPI_ISL_424384, EPI_ISL_424385, EPI_ISL_424386, EPI_ISL_424387, EPI_ISL_424388, EPI_ISL_424389, EPI_ISL_424390, EPI_ISL_424391, EPI_ISL_424392, EPI_ISL_424393, EPI_ISL_424394, EPI_ISL_424395, EPI_ISL_424396, EPI_ISL_424397, EPI_ISL_424398, EPI_ISL_424399, EPI_ISL_424400, EPI_ISL_424401, EPI_ISL_424402, EPI_ISL_424403, EPI_ISL_424404, EPI_ISL_424405, EPI_ISL_424406, EPI_ISL_424407, EPI_ISL_424408, EPI_ISL_424409, EPI_ISL_424410, EPI_ISL_424411, EPI_ISL_424412, EPI_ISL_424413, EPI_ISL_424414, EPI_ISL_424415, EPI_ISL_424416, EPI_ISL_424417, EPI_ISL_424418, EPI_ISL_424419, EPI_ISL_424420, EPI_ISL_424421, EPI_ISL_424422, EPI_ISL_424423, EPI_ISL_424424, EPI_ISL_424425, EPI_ISL_424426, EPI_ISL_424427, EPI_ISL_424428, EPI_ISL_424429, EPI_ISL_424430, EPI_ISL_424431, EPI_ISL_424432, EPI_ISL_424433, EPI_ISL_424434, EPI_ISL_424435, EPI_ISL_424436, EPI_ISL_424437, EPI_ISL_424438, EPI_ISL_424439, EPI_ISL_424440, EPI_ISL_424441, EPI_ISL_424442, EPI_ISL_424443, EPI_ISL_424444, EPI_ISL_424445, EPI_ISL_424446, EPI_ISL_424447, EPI_ISL_424448, EPI_ISL_424449, EPI_ISL_424450, EPI_ISL_424451, EPI_ISL_424452, EPI_ISL_424453, EPI_ISL_424454, EPI_ISL_424455, EPI_ISL_424456, EPI_ISL_424457, EPI_ISL_424458, EPI_ISL_424459, EPI_ISL_424460, EPI_ISL_424461, EPI_ISL_424462, EPI_ISL_424463, EPI_ISL_424464, EPI_ISL_424465, EPI_ISL_424466, EPI_ISL_424467, EPI_ISL_424468, EPI_ISL_424469, EPI_ISL_424470, EPI_ISL_424471, EPI_ISL_424472, EPI_ISL_424473, EPI_ISL_424474, EPI_ISL_424475, EPI_ISL_424476, EPI_ISL_424477, EPI_ISL_424478, EPI_ISL_424479, EPI_ISL_424480, EPI_ISL_424481, EPI_ISL_424482, EPI_ISL_424483, EPI_ISL_424484, EPI_ISL_424485, EPI_ISL_424486, EPI_ISL_424487, EPI_ISL_424488, EPI_ISL_424489, EPI_ISL_424490, EPI_ISL_424491, EPI_ISL_424492, EPI_ISL_424493, EPI_ISL_424494, EPI_ISL_424495, EPI_ISL_424496, EPI_ISL_424497, EPI_ISL_424498, EPI_ISL_424499, EPI_ISL_424500, EPI_ISL_424501, EPI_ISL_424502, EPI_ISL_424503, EPI_ISL_424504, EPI_ISL_424505, EPI_ISL_424506, EPI_ISL_424507, EPI_ISL_424508, EPI_ISL_424509, EPI_ISL_424510, EPI_ISL_424511, EPI_ISL_424512, EPI_ISL_424513, EPI_ISL_424514, EPI_ISL_424515, EPI_ISL_424516, EPI_ISL_424517, EPI_ISL_424518, EPI_ISL_424519, EPI_ISL_424520, EPI_ISL_424521, EPI_ISL_424522, EPI_ISL_424523, EPI_ISL_424524, EPI_ISL_424525, EPI_ISL_424526, EPI_ISL_424527, EPI_ISL_424528, EPI_ISL_424529, EPI_ISL_424530, EPI_ISL_424531, EPI_ISL_424532 | | | |
| see above | The National University Hospital of Iceland | deCODE genetics | Daniel F Gudbjartsson; Agnar Helgason; Hakon Jonsson; Olafur T Magnusson; Pall Melsted; Gudmundur L Norddahl; Jona Saemundsdottir; Asgeir Sigurdsson; Patrick Sulem; Arna B Agustsdottir; Berglind Eiriksdottir; Run Fridriksdottir; Elisabet E Gardarsdottir; Gudmundur Georgsson; Olafia S Gretarsdottir; Kjartan R Gudmundsson; Thora R Gunnarsdottir; Arnaldur Gylfason; Hilma Holm; Brynjar O Jensson; Aslaug Jonasdottir; Kamilla S Josefsdottir; Thordur Kristjansson; Droplaug N Magnusdottir; Louise le Roux; Gudrun Sigmundsdottir; Gardar Sveinbjornsson; Kristin E Sveinsdottir; Maney Sveinsdottir; Emil A Thorarensen; Bjarni Thorbjornsson; Gisli Masson; Ingileif Jonsdottir; Alma Moller; Thorolfur Gudnason; Karl G Kristinsson; Unnur Thorsteinsdottir; Kari Stefansson |
| EPI_ISL_424533, EPI_ISL_424534, EPI_ISL_424535, EPI_ISL_424536, EPI_ISL_424537, EPI_ISL_424538, EPI_ISL_424539, EPI_ISL_424540, EPI_ISL_424541, EPI_ISL_424542, EPI_ISL_424543, EPI_ISL_424544, EPI_ISL_424545, EPI_ISL_424546, EPI_ISL_424547, EPI_ISL_424548, EPI_ISL_424549, EPI_ISL_424550, EPI_ISL_424551 | | | |
| see above | deCODE genetics | deCODE genetics | Daniel F Gudbjartsson; Agnar Helgason; Hakon Jonsson; Olafur T Magnusson; Pall Melsted; Gudmundur L Norddahl; Jona Saemundsdottir; Asgeir Sigurdsson; Patrick Sulem; Arna B Agustsdottir; Berglind Eiriksdottir; Run Fridriksdottir; Elisabet E Gardarsdottir; Gudmundur Georgsson; Olafia S Gretarsdottir; Kjartan R Gudmundsson; Thora R Gunnarsdottir; Arnaldur Gylfason; Hilma Holm; Brynjar O Jensson; Aslaug Jonasdottir; Kamilla S Josefsdottir; Thordur Kristjansson; Droplaug N |

| | | | |
|---|---|---|---|
| | | | Magnusdottir; Louise le Roux; Gudrun Sigmundsdottir; Gardar Sveinbjornsson; Kristin E Sveinsdottir; Maney Sveinsdottir; Emil A Thorarensen; Bjarni Thorbjornsson; Gisli Masson; Ingileif Jonsdottir; Alma Moller; Thorolfur Gudnason; Karl G Kristinsson; Unnur Thorsteinsdottir; Kari Stefansson |
| EPI_ISL_424552 | The National University Hospital of Iceland | deCODE genetics | Daniel F Gudbjartsson; Agnar Helgason; Hakon Jonsson; Olafur T Magnusson; Pall Melsted; Gudmundur L Norddahl; Jona Saemundsdottir; Asgeir Sigurdsson; Patrick Sulem; Arna B Agustsdottir; Berglind Eiriksdottir; Run Fridriksdottir; Elisabet E Gardarsdottir; Gudmundur Georgsson; Olafia S Gretarsdottir; Kjartan R Gudmundsson; Thora R Gunnarsdottir; Arnaldur Gylfason; Hilma Holm; Brynjar O Jensson; Aslaug Jonasdottir; Kamilla S Josefsdottir; Thordur Kristjansson; Droplaug N Magnusdottir; Louise le Roux; Gudrun Sigmundsdottir; Gardar Sveinbjornsson; Kristin E Sveinsdottir; Maney Sveinsdottir; Emil A Thorarensen; Bjarni Thorbjornsson; Gisli Masson; Ingileif Jonsdottir; Alma Moller; Thorolfur Gudnason; Karl G Kristinsson; Unnur Thorsteinsdottir; Kari Stefansson |
| EPI_ISL_424553 | deCODE genetics | deCODE genetics | Daniel F Gudbjartsson; Agnar Helgason; Hakon Jonsson; Olafur T Magnusson; Pall Melsted; Gudmundur L Norddahl; Jona Saemundsdottir; Asgeir Sigurdsson; Patrick Sulem; Arna B Agustsdottir; Berglind Eiriksdottir; Run Fridriksdottir; Elisabet E Gardarsdottir; Gudmundur Georgsson; Olafia S Gretarsdottir; Kjartan R Gudmundsson; Thora R Gunnarsdottir; Arnaldur Gylfason; Hilma Holm; Brynjar O Jensson; Aslaug Jonasdottir; Kamilla S Josefsdottir; Thordur Kristjansson; Droplaug N Magnusdottir; Louise le Roux; Gudrun Sigmundsdottir; Gardar Sveinbjornsson; Kristin E Sveinsdottir; Maney Sveinsdottir; Emil A Thorarensen; Bjarni Thorbjornsson; Gisli Masson; Ingileif Jonsdottir; Alma Moller; Thorolfur Gudnason; Karl G Kristinsson; Unnur Thorsteinsdottir; Kari Stefansson |
| EPI_ISL_424554, EPI_ISL_424555, EPI_ISL_424556, EPI_ISL_424557, EPI_ISL_424558, EPI_ISL_424559 | The National University Hospital of Iceland | deCODE genetics | Daniel F Gudbjartsson; Agnar Helgason; Hakon Jonsson; Olafur T Magnusson; Pall Melsted; Gudmundur L Norddahl; Jona Saemundsdottir; Asgeir Sigurdsson; Patrick Sulem; Arna B Agustsdottir; Berglind Eiriksdottir; Run Fridriksdottir; Elisabet E Gardarsdottir; Gudmundur Georgsson; Olafia S Gretarsdottir; Kjartan R Gudmundsson; Thora R Gunnarsdottir; Arnaldur Gylfason; Hilma Holm; Brynjar O Jensson; Aslaug Jonasdottir; Kamilla S Josefsdottir; Thordur Kristjansson; Droplaug N Magnusdottir; Louise le Roux; Gudrun Sigmundsdottir; Gardar Sveinbjornsson; Kristin E Sveinsdottir; Maney Sveinsdottir; Emil A Thorarensen; Bjarni Thorbjornsson; Gisli Masson; Ingileif Jonsdottir; Alma Moller; Thorolfur Gudnason; Karl G Kristinsson; Unnur Thorsteinsdottir; Kari Stefansson |
| EPI_ISL_424560, EPI_ISL_424561, EPI_ISL_424562 | deCODE genetics | deCODE genetics | Daniel F Gudbjartsson; Agnar Helgason; Hakon Jonsson; Olafur T Magnusson; Pall Melsted; Gudmundur L Norddahl; Jona Saemundsdottir; Asgeir Sigurdsson; Patrick Sulem; Arna B Agustsdottir; Berglind Eiriksdottir; Run Fridriksdottir; Elisabet E Gardarsdottir; Gudmundur Georgsson; Olafia S Gretarsdottir; Kjartan R Gudmundsson; Thora R Gunnarsdottir; Arnaldur Gylfason; Hilma Holm; Brynjar O Jensson; Aslaug Jonasdottir; Kamilla S Josefsdottir; Thordur Kristjansson; Droplaug N Magnusdottir; Louise le Roux; Gudrun Sigmundsdottir; Gardar Sveinbjornsson; Kristin E Sveinsdottir; Maney Sveinsdottir; Emil A Thorarensen; Bjarni Thorbjornsson; Gisli Masson; Ingileif Jonsdottir; Alma Moller; Thorolfur Gudnason; Karl G Kristinsson; Unnur Thorsteinsdottir; Kari Stefansson |
| EPI_ISL_424563, EPI_ISL_424564 | The National University Hospital of Iceland | deCODE genetics | Daniel F Gudbjartsson; Agnar Helgason; Hakon Jonsson; Olafur T Magnusson; Pall Melsted; Gudmundur L Norddahl; Jona Saemundsdottir; Asgeir Sigurdsson; Patrick Sulem; Arna B Agustsdottir; Berglind Eiriksdottir; Run Fridriksdottir; Elisabet E Gardarsdottir; Gudmundur Georgsson; Olafia S Gretarsdottir; Kjartan R Gudmundsson; Thora R Gunnarsdottir; Arnaldur Gylfason; Hilma Holm; Brynjar O Jensson; Aslaug Jonasdottir; Kamilla S Josefsdottir; Thordur Kristjansson; Droplaug N Magnusdottir; Louise le Roux; Gudrun Sigmundsdottir; Gardar Sveinbjornsson; Kristin E Sveinsdottir; Maney Sveinsdottir; Emil A Thorarensen; Bjarni Thorbjornsson; Gisli Masson; Ingileif Jonsdottir; Alma Moller; Thorolfur Gudnason; Karl G Kristinsson; Unnur Thorsteinsdottir; Kari Stefansson |
| EPI_ISL_424565, EPI_ISL_424566, EPI_ISL_424567, EPI_ISL_424568, EPI_ISL_424569, EPI_ISL_424570 | deCODE genetics | deCODE genetics | Daniel F Gudbjartsson; Agnar Helgason; Hakon Jonsson; Olafur T Magnusson; Pall Melsted; Gudmundur L Norddahl; Jona Saemundsdottir; Asgeir Sigurdsson; Patrick Sulem; Arna B Agustsdottir; Berglind Eiriksdottir; Run Fridriksdottir; Elisabet E Gardarsdottir; Gudmundur Georgsson; Olafia S Gretarsdottir; Kjartan R Gudmundsson; Thora R Gunnarsdottir; Arnaldur Gylfason; Hilma Holm; Brynjar O Jensson; Aslaug Jonasdottir; Kamilla S Josefsdottir; Thordur Kristjansson; Droplaug N Magnusdottir; Louise le Roux; Gudrun Sigmundsdottir; Gardar Sveinbjornsson; Kristin E Sveinsdottir; Maney Sveinsdottir; Emil A Thorarensen; Bjarni Thorbjornsson; Gisli Masson; Ingileif Jonsdottir; Alma Moller; Thorolfur Gudnason; Karl G Kristinsson; Unnur Thorsteinsdottir; Kari Stefansson |
| EPI_ISL_424571, EPI_ISL_424572, EPI_ISL_424573, EPI_ISL_424574, EPI_ISL_424575, EPI_ISL_424576, EPI_ISL_424577, EPI_ISL_424578, EPI_ISL_424579, EPI_ISL_424580, EPI_ISL_424581, EPI_ISL_424582, EPI_ISL_424583, EPI_ISL_424584, EPI_ISL_424585, EPI_ISL_424586, EPI_ISL_424587, EPI_ISL_424588, EPI_ISL_424589, EPI_ISL_424590, EPI_ISL_424591, EPI_ISL_424592, EPI_ISL_424593, EPI_ISL_424594, EPI_ISL_424595, EPI_ISL_424596, EPI_ISL_424597, EPI_ISL_424598, EPI_ISL_424599, EPI_ISL_424600, EPI_ISL_424601, EPI_ISL_424602, EPI_ISL_424603, EPI_ISL_424604, EPI_ISL_424605, EPI_ISL_424606, EPI_ISL_424607 | | | |
| see above | The National University Hospital of Iceland | deCODE genetics | Daniel F Gudbjartsson; Agnar Helgason; Hakon Jonsson; Olafur T Magnusson; Pall Melsted; Gudmundur L Norddahl; Jona Saemundsdottir; Asgeir Sigurdsson; Patrick Sulem; Arna B Agustsdottir; Berglind Eiriksdottir; Run Fridriksdottir; Elisabet E Gardarsdottir; Gudmundur Georgsson; Olafia S Gretarsdottir; Kjartan R Gudmundsson; Thora R Gunnarsdottir; Arnaldur Gylfason; Hilma Holm; Brynjar O Jensson; Aslaug Jonasdottir; Kamilla S Josefsdottir; Thordur Kristjansson; Droplaug N Magnusdottir; Louise le Roux; Gudrun Sigmundsdottir; Gardar Sveinbjornsson; Kristin E Sveinsdottir; Maney Sveinsdottir; Emil A Thorarensen; Bjarni Thorbjornsson; Gisli Masson; Ingileif Jonsdottir; Alma Moller; Thorolfur Gudnason; Karl G Kristinsson; Unnur Thorsteinsdottir; Kari Stefansson |
| EPI_ISL_424608, EPI_ISL_424609 | deCODE genetics | deCODE genetics | Daniel F Gudbjartsson; Agnar Helgason; Hakon Jonsson; Olafur T Magnusson; Pall Melsted; Gudmundur L Norddahl; Jona Saemundsdottir; Asgeir Sigurdsson; Patrick Sulem; Arna B Agustsdottir; Berglind Eiriksdottir; Run Fridriksdottir; Elisabet E Gardarsdottir; Gudmundur Georgsson; Olafia S Gretarsdottir; Kjartan R Gudmundsson; Thora R Gunnarsdottir; Arnaldur Gylfason; Hilma Holm; Brynjar O Jensson; Aslaug Jonasdottir; Kamilla S Josefsdottir; Thordur Kristjansson; Droplaug N Magnusdottir; Louise le Roux; Gudrun Sigmundsdottir; Gardar Sveinbjornsson; Kristin E Sveinsdottir; Maney Sveinsdottir; Emil A Thorarensen; Bjarni Thorbjornsson; Gisli Masson; Ingileif Jonsdottir; Alma Moller; Thorolfur Gudnason; Karl G Kristinsson; Unnur Thorsteinsdottir; Kari Stefansson |
| EPI_ISL_424610, EPI_ISL_424611, EPI_ISL_424612, EPI_ISL_424613, EPI_ISL_424614, EPI_ISL_424615, EPI_ISL_424616, EPI_ISL_424617, EPI_ISL_424618, EPI_ISL_424619, EPI_ISL_424620, EPI_ISL_424621, EPI_ISL_424622, EPI_ISL_424623, EPI_ISL_424624 | | | |
| see above | The National University Hospital of Iceland | deCODE genetics | Daniel F Gudbjartsson; Agnar Helgason; Hakon Jonsson; Olafur T Magnusson; Pall Melsted; Gudmundur L Norddahl; Jona Saemundsdottir; Asgeir Sigurdsson; Patrick Sulem; Arna B Agustsdottir; Berglind Eiriksdottir; Run Fridriksdottir; Elisabet E Gardarsdottir; Gudmundur Georgsson; Olafia S Gretarsdottir; Kjartan R Gudmundsson; Thora R Gunnarsdottir; Arnaldur Gylfason; Hilma Holm; Brynjar O Jensson; Aslaug Jonasdottir; Kamilla S Josefsdottir; Thordur Kristjansson; Droplaug N Magnusdottir; Louise le Roux; Gudrun Sigmundsdottir; Gardar Sveinbjornsson; Kristin E Sveinsdottir; Maney Sveinsdottir; Emil A Thorarensen; Bjarni Thorbjornsson; Gisli Masson; Ingileif Jonsdottir; Alma Moller; Thorolfur Gudnason; Karl G Kristinsson; Unnur Thorsteinsdottir; Kari Stefansson |
| EPI_ISL_424625, EPI_ISL_424626, EPI_ISL_424627 | Instituto Nacional de Enfermedades Respiratorias | Instituto Nacional de Enfermedades Respiratorias | Joel Armando Vázquez Pérez, Celia Boukadida, Santiago Avila Ríos, Mario Mújica Sánchez, José Arturo Martínez Orozco, Eduardo Becerril Vargas, Jorge Salas Hernández, Irma López Martínez, Lucia Hernández Rivas, Gisela Barrera Badillo, Edgar Mendieta Condado, Fabiola Garcés Ayala, Adnan Araiza Rodríguez, José Ernesto Ramírez González, Victor Hugo Borja Aburto, Concepción Grajales Muñiz, Cesar Raúl González Bonilla, Carolina González Torres, Francisco Javier Gaytán Cervantes, José Esteban Muñoz Medina, Guillermo M. Ruiz-Palacios, Pilar Ramos Cervantes, Violeta Ibarra Gonzalez, Fernando Ledesma Barrientos, Luis Alberto García Andrade, Alfredo Ponce de León Garduño, Blanca Taboada, Alejandro Sánchez, Pavel Isa, Ricardo Grande, Gloria Vázquez, Francisco Pulido, Carlos F. Arias. |
| EPI_ISL_424628, EPI_ISL_424629, EPI_ISL_424630, EPI_ISL_424631, EPI_ISL_424632, EPI_ISL_424633, EPI_ISL_424634, EPI_ISL_424635, EPI_ISL_424636, EPI_ISL_424637, EPI_ISL_424638, EPI_ISL_424639, EPI_ISL_424640, EPI_ISL_424641, EPI_ISL_424642, EPI_ISL_424643, EPI_ISL_424644, EPI_ISL_424645, EPI_ISL_424646, EPI_ISL_424647, EPI_ISL_424648, EPI_ISL_424649, EPI_ISL_424650, EPI_ISL_424651, EPI_ISL_424652, EPI_ISL_424653, EPI_ISL_424654, EPI_ISL_424655, EPI_ISL_424656, EPI_ISL_424657, EPI_ISL_424658, EPI_ISL_424659, EPI_ISL_424660, EPI_ISL_424661, EPI_ISL_424662, EPI_ISL_424663, EPI_ISL_424664, EPI_ISL_424665 | | | |
| see above | Department of Clinical Microbiology | GIGA Medical Genomics | Keith Durkin, Maria Artesi, Sébastien Bontems, Raphaël Boreux, Cécile Meex, Pierrette Melin, Marie-Pierre Hayette, Vincent Bours. |
| EPI_ISL_424666 | Instituto de Diagnóstico y Referencia Epidemiológicos | Instituto de Diagnóstico y Referencia Epidemiológicos | José Ernesto Ramírez González, Irma López Martínez, Lucia Hernández Rivas, Gisela Barrera Badillo, Edgar Mendieta Condado, Fabiola Garcés Ayala, Adnan Araiza Rodríguez, Celia Boukadida, Santiago Avila Ríos, Mario Mújica Sánchez, José Arturo Martínez Orozco,Eduardo Becerril Vargas, Joel Armando Vázquez Pérez, Victor Hugo Borja Aburto, Concepción Grajales Muñiz, Cesar Raúl González Bonilla, Carolina González Torres, Francisco Javier Gaytán Cervantes, José Esteban Muñoz Medina, Guillermo M. Ruiz-Palacios, Pilar Ramos Cervantes, Violeta Ibarra Gonzalez, Fernando Ledesma Barrientos, Luis Alberto García Andrade, Alfredo Ponce de León Garduño, Blanca Taboada, Alejandro Sánchez, Pavel Isa, Ricardo Grande, Gloria Vázquez, Francisco Pulido, Carlos F. Arias. |
| EPI_ISL_424667 | Laboratorio Estatal de Salud Publica del Estado de México | Instituto de Diagnóstico y Referencia Epidemiológicos | Irma López Martínez, José Ernesto Ramírez González, Lucia Hernández Rivas, Gisela Barrera Badillo, Edgar Mendieta Condado, Fabiola Garcés Ayala, Adnan Araiza Rodríguez, Celia Boukadida, Santiago Avila Ríos, Mario Mújica Sánchez, José Arturo Martínez Orozco,Eduardo Becerril Vargas, Joel Armando Vázquez Pérez, Victor Hugo Borja Aburto, Concepción Grajales Muñiz, Cesar Raúl González Bonilla, Carolina González Torres, Francisco Javier Gaytán Cervantes, José Esteban Muñoz Medina, Guillermo M. Ruiz-Palacios, Pilar Ramos Cervantes, Violeta Ibarra Gonzalez, Fernando Ledesma Barrientos, Luis Alberto García Andrade, Alfredo Ponce de León Garduño, Blanca Taboada, Alejandro Sánchez, Pavel Isa, Ricardo Grande, Gloria Vázquez, Francisco Pulido, Carlos F. Arias. |
| EPI_ISL_424668, EPI_ISL_424669 | Arizona State University Health Services | Arizona State University | Rabia Maqsood, LaRinda A. Holland, Emily A. Kaelin, Bereket Estifanos, Nicholas J. Mellor, Jason Steel, Lily I. Wu, Arvind Varsani, Rolf U. Halden, Brenda G. Hogue, Matthew Scotch, Efrem S. Lim |
| EPI_ISL_424670 | Laboratorio Estatal de Salud Publica del Estado de Queretaro | Instituto de Diagnóstico y Referencia Epidemiologicos | Gisela Barrera Badillo, Irma López Martínez, Lucia Hernández Rivas, Edgar Mendieta Condado, Fabiola Garcés Ayala, Adnan Araiza Rodríguez, Celia Boukadida, Santiago Avila Ríos, Mario Mújica Sánchez, José Arturo Martínez Orozco,Eduardo Becerril Vargas, Joel Armando Vázquez Pérez, Victor Hugo Borja Aburto, Concepción Grajales Muñiz, Cesar Raúl González Bonilla, Carolina González Torres, Francisco Javier Gaytán Cervantes, José Esteban Muñoz Medina, Guillermo M. Ruiz-Palacios, Pilar Ramos Cervantes, Violeta Ibarra Gonzalez, Fernando Ledesma Barrientos, Luis Alberto García Andrade, Alfredo Ponce de León Garduño, Blanca Taboada, Alejandro Sánchez, Pavel Isa, Ricardo Grande, Gloria Vázquez, Francisco Pulido, Carlos F. Arias,José Ernesto Ramírez González |
| EPI_ISL_424671 | Arizona State University Health Services | Arizona State University | Rabia Maqsood, LaRinda A. Holland, Emily A. Kaelin, Bereket Estifanos, Nicholas J. Mellor, Jason Steel, Lily I. Wu, Arvind Varsani, Rolf U. Halden, Brenda G. Hogue, Matthew Scotch, Efrem S. Lim |
| EPI_ISL_424672 | Laboratorio Estatal de Salud Publica del Estado de Puebla | Instituto de Diagnostico y Referencia Epidemiologicos | Fabiola Garcés Ayala, Gisela Barrera Badillo, Irma López Martínez, Lucia Hernández Rivas, Edgar Mendieta Condado, Adnan Araiza Rodríguez, Celia Boukadida, Santiago Avila Ríos, Mario Mújica Sánchez, José Arturo Martínez Orozco,Eduardo Becerril Vargas, Joel Armando Vázquez Pérez, Victor Hugo Borja Aburto, Concepción Grajales Muñiz, Cesar Raúl González Bonilla, Carolina González Torres, Francisco Javier Gaytán Cervantes, José Esteban Muñoz Medina, Guillermo M. Ruiz-Palacios, Pilar Ramos Cervantes, Violeta Ibarra Gonzalez, Fernando Ledesma Barrientos, Luis Alberto García Andrade, Alfredo Ponce de León Garduño, Blanca Taboada, Alejandro Sánchez, Pavel Isa, Ricardo Grande, Gloria Vázquez, Francisco Pulido, Carlos F. Arias,José Ernesto Ramírez González |
| EPI_ISL_424673 | Instituto de Diagnostico y Referencia Epidemiologicos | Instituto de Diagnostico y Referencia Epidemiologicos | Adnan Araiza Rodríguez, Edgar Mendieta Condado,Fabiola Garcés Ayala, Gisela Barrera Badillo, Irma López Martínez, Lucia Hernández Rivas, Celia Boukadida, Santiago Avila Ríos, Mario Mújica Sánchez, José Arturo Martínez Orozco,Eduardo Becerril Vargas, Joel Armando Vázquez Pérez, Victor Hugo Borja Aburto, Concepción Grajales Muñiz, Cesar Raúl González Bonilla, Carolina González Torres, Francisco Javier Gaytán Cervantes, José Esteban Muñoz Medina, Guillermo M. Ruiz-Palacios, Pilar Ramos Cervantes, Violeta Ibarra Gonzalez, Fernando Ledesma Barrientos, Luis Alberto García Andrade, Alfredo Ponce de León Garduño, Blanca Taboada, Alejandro Sánchez, Pavel Isa, Ricardo Grande, Gloria Vázquez, Francisco Pulido, Carlos F. Arias,José Ernesto Ramírez González |
| EPI_ISL_424703 | Klinisk mikrobiologi, Region Västerbotten | Unit for Biological Agents, Department for CBRN Defence and Security, Swedish Defence Research Agency | FOI Bioinformatics team |
| EPI_ISL_424731 | Hospital General Regional No.66, Ciudad Juárez, Chihuahua. | Laboratorio Central de Epidemiología-DLVIE / Laboratorio de Secuenciación-Centro de Instrumentos. Instituto Mexicano del Seguro Social | Muñoz-Medina JE, González-Torres C, Gaytán-Cervantes FJ, López-Martínez I, Hernández-Rivas L, Barrera-Badillo G, Mendieta-Condado E, Garcés-Ayala F, Araiza-Rodríguez A, Ramírez-González JE, Boukadida C, Ávila-Ríos S, Mújica-Sánchez M, Martínez-Orozco JA, Becerril-Vargas E, Vázquez-Pérez JA, Taboada B, Sánchez A, Isa P, Grande R, Vázquez G, Pulido F. López S, Arias C, Ramos-Cervantes P, Ruiz-Palacios GM, Ibarra-González V, Ledesma-Barrientos F, García-Andrade LA, de León-Garduño AP, González-Bonilla CR, Grajales-Muñiz C, Borja-Aburto VH. |
| EPI_ISL_424841, EPI_ISL_424842 | SC Dept of Health and Env. Control-Bureau of Laboratories | Pathogen Discovery, Respiratory Viruses Branch, Division of Viral Diseases, Centers for Disease Control and Prevention | Yan Li, Krista Queen, Clinton R. Paden, Rachel Marine, Anna Uehara, Ying Tao, Jing Zhang, Haibin Wang, Mary S. Keckler, Alison S. Laufer Halpin, Christopher A. Elkins, Suxiang Tong |
| EPI_ISL_424843, EPI_ISL_424844, EPI_ISL_424845, EPI_ISL_424846, EPI_ISL_424847 | MA State Public Health Laboratory | Pathogen Discovery, Respiratory Viruses Branch, Division of Viral Diseases, Centers for Disease Control and Prevention | Yan Li, Krista Queen, Clinton R. Paden, Rachel Marine, Anna Uehara, Ying Tao, Jing Zhang, Haibin Wang, Mary S. Keckler, Alison S. Laufer Halpin, Christopher A. Elkins, Suxiang Tong |
| EPI_ISL_424848, EPI_ISL_424849 | AZ SPHL, Arizona Department of Health Services | Pathogen Discovery, Respiratory Viruses Branch, Division of Viral Diseases, Centers for Disease Control and Prevention | Yan Li, Krista Queen, Clinton R. Paden, Rachel Marine, Anna Uehara, Ying Tao, Jing Zhang, Haibin Wang, Mary S. Keckler, Alison S. Laufer Halpin, Christopher A. Elkins, Suxiang Tong |
| EPI_ISL_424850, EPI_ISL_424851 | IL Department of Public Health Chicago Laboratory | Pathogen Discovery, Respiratory Viruses Branch, Division of Viral Diseases, Centers for Disease Control and Prevention | Yan Li, Krista Queen, Clinton R. Paden, Rachel Marine, Anna Uehara, Ying Tao, Jing Zhang, Haibin Wang, Mary S. Keckler, Alison S. Laufer Halpin, Christopher A. Elkins, Suxiang Tong |
| EPI_ISL_424852 | DC Public Health Lab/ Dept. of Forensic Sciences | Pathogen Discovery, Respiratory Viruses Branch, Division of Viral Diseases, Centers for Disease Control and Prevention | Yan Li, Krista Queen, Clinton R. Paden, Rachel Marine, Anna Uehara, Ying Tao, Jing Zhang, Haibin Wang, Mary S. Keckler, Alison S. Laufer Halpin, Christopher A. Elkins, Suxiang Tong |
| EPI_ISL_424853, EPI_ISL_424854 | FL Bureau of Public Health Laboratories-Miami | Pathogen Discovery, Respiratory Viruses Branch, Division of Viral Diseases, Centers for Disease Control and Prevention | Yan Li, Krista Queen, Clinton R. Paden, Rachel Marine, Anna Uehara, Ying Tao, Jing Zhang, Haibin Wang, Mary S. Keckler, Alison S. Laufer Halpin, Christopher A. Elkins, Suxiang Tong |
| EPI_ISL_424855 | FL Bureau of Public Health Laboratories-Tampa | Pathogen Discovery, Respiratory Viruses Branch, Division of Viral Diseases, Centers for Disease Control and Prevention | Yan Li, Krista Queen, Clinton R. Paden, Rachel Marine, Anna Uehara, Ying Tao, Jing Zhang, Haibin Wang, Mary S. Keckler, Alison S. Laufer Halpin, Christopher A. Elkins, Suxiang Tong |
| EPI_ISL_424856, EPI_ISL_424857 | FL Bur. of Public Health Laboratories-Jacksonville | Pathogen Discovery, Respiratory Viruses Branch, Division of Viral Diseases, Centers for Disease Control and Prevention | Yan Li, Krista Queen, Clinton R. Paden, Rachel Marine, Anna Uehara, Ying Tao, Jing Zhang, Haibin Wang, Mary S. Keckler, Alison S. Laufer Halpin, Christopher A. Elkins, Suxiang Tong |
| EPI_ISL_424858, EPI_ISL_424859, EPI_ISL_424860, EPI_ISL_424861, EPI_ISL_424862, EPI_ISL_424863, EPI_ISL_424864 | GA Department of Public Health Laboratory | Pathogen Discovery, Respiratory Viruses Branch, Division of Viral Diseases, Centers for Disease Control and Prevention | Yan Li, Krista Queen, Clinton R. Paden, Rachel Marine, Anna Uehara, Ying Tao, Jing Zhang, Haibin Wang, Mary S. Keckler, Alison S. Laufer Halpin, Christopher A. Elkins, Suxiang Tong |

| | | | |
|---|---|---|---|
| EPI_ISL_424865 | IA State Hygienic Laboratory | Pathogen Discovery, Respiratory Viruses Branch, Division of Viral Diseases, Centers for Disease Control and Prevention | Yan Li, Krista Queen, Clinton R. Paden, Rachel Marine, Anna Uehara, Ying Tao, Jing Zhang, Haibin Wang, Mary S. Keckler, Alison S. Laufer Halpin, Christopher A. Elkins, Suxiang Tong |
| EPI_ISL_424866 | IN State Department of Health Laboratory Services | Pathogen Discovery, Respiratory Viruses Branch, Division of Viral Diseases, Centers for Disease Control and Prevention | Yan Li, Krista Queen, Clinton R. Paden, Rachel Marine, Anna Uehara, Ying Tao, Jing Zhang, Haibin Wang, Mary S. Keckler, Alison S. Laufer Halpin, Christopher A. Elkins, Suxiang Tong |
| EPI_ISL_424867 | KS Health and Environmental Laboratories | Pathogen Discovery, Respiratory Viruses Branch, Division of Viral Diseases, Centers for Disease Control and Prevention | Yan Li, Krista Queen, Clinton R. Paden, Rachel Marine, Anna Uehara, Ying Tao, Jing Zhang, Haibin Wang, Mary S. Keckler, Alison S. Laufer Halpin, Christopher A. Elkins, Suxiang Tong |
| EPI_ISL_424868 | LA Office of Public Health Laboratories | Pathogen Discovery, Respiratory Viruses Branch, Division of Viral Diseases, Centers for Disease Control and Prevention | Yan Li, Krista Queen, Clinton R. Paden, Rachel Marine, Anna Uehara, Ying Tao, Jing Zhang, Haibin Wang, Mary S. Keckler, Alison S. Laufer Halpin, Christopher A. Elkins, Suxiang Tong |
| EPI_ISL_424869 | MD DOH Laboratories Administration | Pathogen Discovery, Respiratory Viruses Branch, Division of Viral Diseases, Centers for Disease Control and Prevention | Yan Li, Krista Queen, Clinton R. Paden, Rachel Marine, Anna Uehara, Ying Tao, Jing Zhang, Haibin Wang, Mary S. Keckler, Alison S. Laufer Halpin, Christopher A. Elkins, Suxiang Tong |
| EPI_ISL_424870 | MO State Public Health Laboratory | Pathogen Discovery, Respiratory Viruses Branch, Division of Viral Diseases, Centers for Disease Control and Prevention | Yan Li, Krista Queen, Clinton R. Paden, Rachel Marine, Anna Uehara, Ying Tao, Jing Zhang, Haibin Wang, Mary S. Keckler, Alison S. Laufer Halpin, Christopher A. Elkins, Suxiang Tong |
| EPI_ISL_424871, EPI_ISL_424872, EPI_ISL_424873 | NC State Laboratory of Public Health | Pathogen Discovery, Respiratory Viruses Branch, Division of Viral Diseases, Centers for Disease Control and Prevention | Yan Li, Krista Queen, Clinton R. Paden, Rachel Marine, Anna Uehara, Ying Tao, Jing Zhang, Haibin Wang, Mary S. Keckler, Alison S. Laufer Halpin, Christopher A. Elkins, Suxiang Tong |
| EPI_ISL_424874, EPI_ISL_424875 | NE Public Health Laboratory | Pathogen Discovery, Respiratory Viruses Branch, Division of Viral Diseases, Centers for Disease Control and Prevention | Yan Li, Krista Queen, Clinton R. Paden, Rachel Marine, Anna Uehara, Ying Tao, Jing Zhang, Haibin Wang, Mary S. Keckler, Alison S. Laufer Halpin, Christopher A. Elkins, Suxiang Tong |
| EPI_ISL_424876, EPI_ISL_424877 | NH Dept. of Health and Human Services Public Health Labs | Pathogen Discovery, Respiratory Viruses Branch, Division of Viral Diseases, Centers for Disease Control and Prevention | Yan Li, Krista Queen, Clinton R. Paden, Rachel Marine, Anna Uehara, Ying Tao, Jing Zhang, Haibin Wang, Mary S. Keckler, Alison S. Laufer Halpin, Christopher A. Elkins, Suxiang Tong |
| EPI_ISL_424878 | NJ Public Health and Environmental Laboratories | Pathogen Discovery, Respiratory Viruses Branch, Division of Viral Diseases, Centers for Disease Control and Prevention | Yan Li, Krista Queen, Clinton R. Paden, Rachel Marine, Anna Uehara, Ying Tao, Jing Zhang, Haibin Wang, Mary S. Keckler, Alison S. Laufer Halpin, Christopher A. Elkins, Suxiang Tong |
| EPI_ISL_424879 | NV-Southern Nevada Public Health Laboratory | Pathogen Discovery, Respiratory Viruses Branch, Division of Viral Diseases, Centers for Disease Control and Prevention | Yan Li, Krista Queen, Clinton R. Paden, Rachel Marine, Anna Uehara, Ying Tao, Jing Zhang, Haibin Wang, Mary S. Keckler, Alison S. Laufer Halpin, Christopher A. Elkins, Suxiang Tong |
| EPI_ISL_424880 | OH Department of Health Laboratory | Pathogen Discovery, Respiratory Viruses Branch, Division of Viral Diseases, Centers for Disease Control and Prevention | Yan Li, Krista Queen, Clinton R. Paden, Rachel Marine, Anna Uehara, Ying Tao, Jing Zhang, Haibin Wang, Mary S. Keckler, Alison S. Laufer Halpin, Christopher A. Elkins, Suxiang Tong |
| EPI_ISL_424881, EPI_ISL_424882, EPI_ISL_424883, EPI_ISL_424884, EPI_ISL_424885, EPI_ISL_424886 | PA Department of Health, Bureau of Laboratories | Pathogen Discovery, Respiratory Viruses Branch, Division of Viral Diseases, Centers for Disease Control and Prevention | Yan Li, Krista Queen, Clinton R. Paden, Rachel Marine, Anna Uehara, Ying Tao, Jing Zhang, Haibin Wang, Mary S. Keckler, Alison S. Laufer Halpin, Christopher A. Elkins, Suxiang Tong |
| EPI_ISL_424887 | RI State Health Laboratories | Pathogen Discovery, Respiratory Viruses Branch, Division of Viral Diseases, Centers for Disease Control and Prevention | Yan Li, Krista Queen, Clinton R. Paden, Rachel Marine, Anna Uehara, Ying Tao, Jing Zhang, Haibin Wang, Mary S. Keckler, Alison S. Laufer Halpin, Christopher A. Elkins, Suxiang Tong |
| EPI_ISL_424888 | SC Dept of Health and Env. Control-Bureau of Laboratories | Pathogen Discovery, Respiratory Viruses Branch, Division of Viral Diseases, Centers for Disease Control and Prevention | Yan Li, Krista Queen, Clinton R. Paden, Rachel Marine, Anna Uehara, Ying Tao, Jing Zhang, Haibin Wang, Mary S. Keckler, Alison S. Laufer Halpin, Christopher A. Elkins, Suxiang Tong |
| EPI_ISL_424889, EPI_ISL_424890 | UT-Unified State Labs: Public Health Utah DOH | Pathogen Discovery, Respiratory Viruses Branch, Division of Viral Diseases, Centers for Disease Control and Prevention | Yan Li, Krista Queen, Clinton R. Paden, Rachel Marine, Anna Uehara, Ying Tao, Jing Zhang, Haibin Wang, Mary S. Keckler, Alison S. Laufer Halpin, Christopher A. Elkins, Suxiang Tong |
| EPI_ISL_424891, EPI_ISL_424892, EPI_ISL_424893, EPI_ISL_424894 | VA-Division of Consolidated Laboratory Services | Pathogen Discovery, Respiratory Viruses Branch, Division of Viral Diseases, Centers for Disease Control and Prevention | Yan Li, Krista Queen, Clinton R. Paden, Rachel Marine, Anna Uehara, Ying Tao, Jing Zhang, Haibin Wang, Mary S. Keckler, Alison S. Laufer Halpin, Christopher A. Elkins, Suxiang Tong |
| EPI_ISL_424895, EPI_ISL_424896, EPI_ISL_424897, EPI_ISL_424898, EPI_ISL_424899, EPI_ISL_424900 | IA State Hygienic Laboratory | Pathogen Discovery, Respiratory Viruses Branch, Division of Viral Diseases, Centers for Disease Control and Prevention | Ying Tao, Clinton R. Paden, Jing Zhang, Krista Queen, Anna Uehara, Yan Li, Haibin Wang, Mary S. Keckler, Alison S. Laufer Halpin, Christopher A. Elkins, Suxiang Tong |
| EPI_ISL_424901 | NJ Public Health and Environmental Laboratories | Pathogen Discovery, Respiratory Viruses Branch, Division of Viral Diseases, Centers for Disease Control and Prevention | Ying Tao, Clinton R. Paden, Jing Zhang, Krista Queen, Anna Uehara, Yan Li, Haibin Wang, Mary S. Keckler, Alison S. Laufer Halpin, Christopher A. Elkins, Suxiang Tong |
| EPI_ISL_424902, EPI_ISL_424903, EPI_ISL_424904, EPI_ISL_424905 | SC Dept of Health and Env. Control-Bureau of Laboratories | Pathogen Discovery, Respiratory Viruses Branch, Division of Viral Diseases, Centers for Disease Control and Prevention | Ying Tao, Clinton R. Paden, Jing Zhang, Krista Queen, Anna Uehara, Yan Li, Haibin Wang, Mary S. Keckler, Alison S. Laufer Halpin, Christopher A. Elkins, Suxiang Tong |
| EPI_ISL_424906 | NYC Department of Health and Mental Hygiene | Pathogen Discovery, Respiratory Viruses Branch, Division of Viral Diseases, Centers for Disease Control and Prevention | Ying Tao, Clinton R. Paden, Jing Zhang, Krista Queen, Anna Uehara, Yan Li, Haibin Wang, Mary S. Keckler, Alison S. Laufer Halpin, Christopher A. Elkins, Suxiang Tong |
| EPI_ISL_424907 | VA-Division of Consolidated Laboratory Services | Pathogen Discovery, Respiratory Viruses Branch, Division of Viral Diseases, Centers for Disease Control and Prevention | Ying Tao, Clinton R. Paden, Jing Zhang, Krista Queen, Anna Uehara, Yan Li, Haibin Wang, Mary S. Keckler, Alison S. Laufer Halpin, Christopher A. Elkins, Suxiang Tong |
| EPI_ISL_424908, EPI_ISL_424909, EPI_ISL_424910, EPI_ISL_424911, EPI_ISL_424912, EPI_ISL_424913, EPI_ISL_424914, EPI_ISL_424915, EPI_ISL_424916, EPI_ISL_424917, EPI_ISL_424918, EPI_ISL_424919, EPI_ISL_424920 | | | |
| see above | MA State Public Health Laboratory | Pathogen Discovery, Respiratory Viruses Branch, Division of Viral Diseases, Centers for Disease Control and Prevention | Ying Tao, Clinton R. Paden, Jing Zhang, Krista Queen, Anna Uehara, Yan Li, Haibin Wang, Mary S. Keckler, Alison S. Laufer Halpin, Christopher A. Elkins, Suxiang Tong |
| EPI_ISL_424929, EPI_ISL_424930, EPI_ISL_424931, EPI_ISL_424932, EPI_ISL_424933, EPI_ISL_424934, EPI_ISL_424935, EPI_ISL_424936, EPI_ISL_424937, EPI_ISL_424938, EPI_ISL_424939, EPI_ISL_424940, EPI_ISL_424941, EPI_ISL_424942, EPI_ISL_424943, EPI_ISL_424944, EPI_ISL_424945, EPI_ISL_424946, EPI_ISL_424947, EPI_ISL_424948, EPI_ISL_424949, EPI_ISL_424950, EPI_ISL_424951, EPI_ISL_424952, EPI_ISL_424953, EPI_ISL_424954, EPI_ISL_424955, EPI_ISL_424956, EPI_ISL_424957, EPI_ISL_424958, EPI_ISL_424959, EPI_ISL_424960, EPI_ISL_424961, EPI_ISL_424962, EPI_ISL_424963, EPI_ISL_424964, EPI_ISL_424965, EPI_ISL_424966, EPI_ISL_424967, EPI_ISL_424968 | | | |
| see above | NYU Langone Health | Departments of Pathology and Medicine, New York University School of Medicine | Maria Aguero-Rosenfeld, Brendan Belovarac, Margaret Black, Ludovic Boytard, John Cadley, Paolo Cotzia, John Chen, Dacia Dimartino, Xiaojun Feng, Tatyana Gindin, Adriana Heguy, Megan Hogan, Emily Huang, George Jour, Andrew Lytle, Christian Marier, Matthew T. Maurano, Mark J. Mulligan, Peter Meyn, Iman Osman, Jared Pinnell, Sitharam Ramaswami, Amy Rapkiewicz, Marie Samanovic-Golden, Antonio Serrano, Guomiao Shen, Matija Snuderl, Theodore Vougiouklakis, Nick Vulpescu, Gael Westby, Paul Zappile, Yutong Zhang |
| EPI_ISL_424969, EPI_ISL_424970, EPI_ISL_424971, EPI_ISL_424972, EPI_ISL_424973, EPI_ISL_424974, EPI_ISL_424975, EPI_ISL_424978 | Laboratory Medicine | Department of Laboratory Medicine, Lin-Kou Chang Gung Memorial Hospital, Taoyuan, Taiwan | Kuo-Chien Tsao, Yu-Nong Gong, Shu-Li Yang, Yi-Chun Liu, Chung-Guei Huang, Mei-Jen Hsiao, Po-Wei Huang, Cheng-Ta Yang, Cheng-Hsun Chiu, Peng-Nien Huang, Kuo-Ming Lee, Guang-Wu Chen, Shin-Ru Shih |
| EPI_ISL_424981, EPI_ISL_424982 | Bureau of Laboratories, Michigan Department of Health and Human Services | Michigan Department of Health and Human Services | Blankenship HM, Riner D, Soehnlen MK |
| EPI_ISL_424983 | Dirk Dittmer | Dirk Dittmer | Bailey,A.G., Caro-Vegas,C., Dittmer,D., Eason,A.B., Juarez,A., Landis,J.T., McNamara,R.P., Miller,M.B., Moorad,R., Pluta,L.J., Seltzer,T.A., Thompson,C., Vahrson,W. and Villamor,F. |
| EPI_ISL_424984, EPI_ISL_424985, EPI_ISL_424986 | Bureau of Laboratories, Michigan Department of Health and Human Services | Michigan Department of Health and Human Services | Blankenship HM, Riner D, Soehnlen MK |
| EPI_ISL_424987 | Dirk Dittmer | Dirk Dittmer | Bailey,A.G., Caro-Vegas,C.P., Dittmer,D., Eason,A.B., Juarez,A., Landis,J.T., McNamara,R.P., Miller,M.B., Moorad,R., Pluta,L.J., Seltzer,T.A., Thompson,C., Vahrson,W. and Villamor,F. |
| EPI_ISL_424988, EPI_ISL_424989, EPI_ISL_424990, EPI_ISL_424992 | Bureau of Laboratories, Michigan Department of Health and Human Services | Michigan Department of Health and Human Services | Blankenship HM, Riner D, Soehnlen MK |
| EPI_ISL_424993 | CHU Purpan - Laboratoire de Virologie - Institut Fédératif de Biologie | Laboratoire de virologie - École Nationale Vétérinaire de Toulouse | Croville,G., Guerin,J.-L. and Izopet,J. |
| EPI_ISL_425023, EPI_ISL_425024 | Respiratory Virus Unit, Microbiology Services Colindale, Public Health England | Respiratory Virus Unit, Microbiology Services Colindale, Public Health England | Steven Platt, Shahjahan Miah, Angie Lackenby, Omolola Akinbami, Tiina Talts, Leena Bhaw, Richard Myers, Monica Galiano, Kirstin Edwards, Jonathan Hubb, Joanna Ellis, Maria Zambon |
| EPI_ISL_425048, EPI_ISL_425050 | Lab voor klinische biologie | Onderzoeksgroep Virologie | Laurens Lambrechts, Nick Vereecke, Marthe Pauwels, Basiel Cole, Bruno Verhasselt, Linos Vandekerckhove, Hans Nauwynck, Sebastiaan Theuns |
| EPI_ISL_425051, EPI_ISL_425052, EPI_ISL_425053, EPI_ISL_425054, EPI_ISL_425055 | Lab voor klinische biologie | Onderzoeksgroep Virologie | Nick Vereecke, Laurens Lambrechts, Marthe Pauwels, Basiel Cole, Bruno Verhasselt, Linos Vandekerckhove, Hans Nauwynck, Sebastiaan Theuns |
| EPI_ISL_425056, EPI_ISL_425057, EPI_ISL_425058 | Lab voor klinische biologie | Onderzoeksgroep Virologie | Laurens Lambrechts, Nick Vereecke, Marthe Pauwels, Basiel Cole, Bruno Verhasselt, Linos Vandekerckhove, Hans Nauwynck, Sebastiaan Theuns |
| EPI_ISL_425059, EPI_ISL_425060, EPI_ISL_425061 | Lab voor klinische biologie | Onderzoeksgroep Virologie | Nick Vereecke, Laurens Lambrechts, Marthe Pauwels, Basiel Cole, Bruno Verhasselt, Linos Vandekerckhove, Hans Nauwynck, Sebastiaan Theuns |
| EPI_ISL_425062, EPI_ISL_425063, EPI_ISL_425064 | Lab voor klinische biologie | Onderzoeksgroep Virologie | Laurens Lambrechts, Nick Vereecke, Marthe Pauwels, Jozefien De Clercq, Bruno Verhasselt, Linos Vandekerckhove, Hans Nauwynck, Sebastiaan Theuns |
| EPI_ISL_425117 | Division of Viral Diseases, Center for Laboratory Control of Infectious Diseases, Korea Centers for Diseases Control and Prevention | Division of Viral Diseases, Center for Laboratory Control of Infectious Diseases, Korea Centers for Diseases Control and Prevention | Jeong-Min Kim, Yoon-Seok Chung, Namjoo Lee, Mi-Seon Kim, Sang Hee Woo, Hye-Jun Jo, Sehee Park, Heui Man Kim, Jun-Sub Kim, Junhyeong Jang, Dong Hyun Song, Daesang Lee, Seong Tae Jeong, Myung Guk Han |
| EPI_ISL_425118 | Division of Viral Diseases, Center for Laboratory Control of Infectious Diseases, Korea Centers for Diseases Control and Prevention | Division of Viral Diseases, Center for Laboratory Control of Infectious Diseases, Korea Centers for Diseases Control and Prevention | Jeong-Min Kim, Yoon-Seok Chung, Namjoo Lee, Mi-Seon Kim, Sang Hee Woo, Hye-Jun Jo, Sehee Park, Heui Man Kim, Jun-Sub Kim, Junhyeong Jang, Dong Hyun Song, Daesang Lee, Seong Tae Jeong, Myung Guk Han |
| EPI_ISL_425120, EPI_ISL_425121, EPI_ISL_425122, EPI_ISL_425123, EPI_ISL_425124, EPI_ISL_425125, EPI_ISL_425126, EPI_ISL_425127, EPI_ISL_425128 | Center of Medical Microbiology, Virology, and Hospital Hygiene, University of Duesseldorf | Center of Medical Microbiology, Virology, and Hospital Hygiene, University of Duesseldorf | Ortwin Adams, Marcel Andree, Alexander Dilthey, Torsten Feldt, Sandra Hauka, Torsten Houwaart, Björn-Erik Jensen, Detlef Kindgen-Milles, Malte Kohns Vasconcelos, Klaus Pfeffer, Tina Senff, Daniel Strelow, Jörg Timm, Andreas Walker, Tobias Wienemann |
| EPI_ISL_425129, EPI_ISL_425130, | Center of Medical Microbiology, Virology, and Hospital | Center of Medical Microbiology, Virology, and Hospital | Ortwin Adams, Marcel Andree, Alexander Dilthey, Torsten Feldt, Sandra Hauka, Torsten Houwaart, Bjorn-Erik Jensen, Detlef Kindgen-Milles, Malte Kohns Vasconcelos, Klaus Pfeffer, Tina Senff, Daniel Strelow, Jorg Timm, Andreas Walker, Tobias |

| | | | |
|---|---|---|---|
| EPI_ISL_425131, EPI_ISL_425132, EPI_ISL_425133, EPI_ISL_425134, EPI_ISL_425135, EPI_ISL_425136, EPI_ISL_425137 | Hygiene, University of Duesseldorf | Hygiene, University of Duesseldorf | Wienemann |
| EPI_ISL_425138, EPI_ISL_425139 | Center of Medical Microbiology, Virology, and Hospital Hygiene, University of Duesseldorf | Center of Medical Microbiology, Virology, and Hospital Hygiene, University of Duesseldorf | Ortwin Adams, Marcel Andree, Alexander Dilthey, Torsten Feldt, Sandra Hauka, Torsten Houwaart, Björn-Erik Jensen, Detlef Kindgen-Milles, Malte Kohns Vasconcelos, Klaus Pfeffer, Tina Senff, Daniel Strelow, Jörg Timm,Andreas Walker, Tobias Wienemann |
| EPI_ISL_425140 | Center of Medical Microbiology, Virology, and Hospital Hygiene, University of Duesseldorf | Center of Medical Microbiology, Virology, and Hospital Hygiene, University of Duesseldorf | Ortwin Adams, Marcel Andree, Alexander Dilthey, Torsten Feldt, Sandra Hauka, Torsten Houwaart, Björn-Erik Jensen, Detlef Kindgen-Milles, Malte Kohns Vasconcelos, Klaus Pfeffer, Tina Senff, Daniel Strelow, Jörg Timm,Andreas Walker, Tobias Wienemann |
| EPI_ISL_425142, EPI_ISL_425143, EPI_ISL_425144, EPI_ISL_425145, EPI_ISL_425146, EPI_ISL_425147, EPI_ISL_425148, EPI_ISL_425149, EPI_ISL_425150, EPI_ISL_425151, EPI_ISL_425152, EPI_ISL_425153, EPI_ISL_425154, EPI_ISL_425155, EPI_ISL_425156, EPI_ISL_425157, EPI_ISL_425158, EPI_ISL_425159, EPI_ISL_425160, EPI_ISL_425161, EPI_ISL_425162, EPI_ISL_425163, EPI_ISL_425164, EPI_ISL_425165, EPI_ISL_425166, EPI_ISL_425167, EPI_ISL_425168, EPI_ISL_425169, EPI_ISL_425170, EPI_ISL_425171, EPI_ISL_425172, EPI_ISL_425173, EPI_ISL_425174, EPI_ISL_425175, EPI_ISL_425176 | | | |
| see above | University of Wisconsin-Madison AIDS Vaccine Research Laboratories | University of Wisconsin-Madison AIDS Vaccine Research Laboratories | Gage Moreno, Katarina Braun, et al. AIDS Vaccine Research Laboratories |
| EPI_ISL_425177 | Public Health Ontario | Public Health Agency of Canada - National Microbiology Laboratory | Amrit S. Boese, Nikesh Tailor, Anders Leung, Joshua Quick, Shari Tyson, Morag Graham, Jonathan Audet, Natalie Knox, Darwyn Kobasa |
| EPI_ISL_425178 | Servicio de Microbiología. Consorcio Hospital General Universitario de Valencia | Sequencing and Bioinformatics Service and Molecular Epidemiology Research Group. FISABIO-Public Health | David Navarro, Maria Alma Bracho, Griselda De Marco, Beatriz Beamud, Lidia Ruiz Roldan, Marta Pla Diaz, Neris Garcia-Gonzalez, Inma Galán Vendrell, Sandra Carbo, Loreto Ferrús Abad, Paula Ruiz-Hueso, Mariana Reyes-Prieto, Vicente Soriano Chirona, Ivan Ansari, Llúcia Martínez-Priego, Giuseppe D'Auria, Fernando Gonzalez-Candelas |
| EPI_ISL_425179 | Servicio de Microbiología. Consorcio Hospital General Universitario de Valencia | Sequencing and Bioinformatics Service and Molecular Epidemiology Research Group. FISABIO-Public Health | David Navarro, Maria Alma Bracho, Griselda De Marco, Beatriz Beamud, Lidia Ruiz Roldan, Marta Pla Diaz, Neris Garcia-Gonzalez, Inma Galán Vendrell, Sandra Carbo, Loreto Ferrús Abad, Paula Ruiz-Hueso, Mariana Reyes-Prieto, Vicente Soriano Chirona, Ivan Ansari, David Navarro, Llúcia Martínez-Priego, Giuseppe D'Auria, Fernando Gonzalez-Candelas |
| EPI_ISL_425180 | Servicio de Microbiología. Consorcio Hospital General Universitario de Valencia | Sequencing and Bioinformatics Service and Molecular Epidemiology Research Group. FISABIO-Public Health | Griselda De Marco, Beatriz Beamud, Lidia Ruiz Roldan, Marta Pla Diaz, Neris Garcia-Gonzalez, Inma Galán Vendrell, Sandra Carbo, Loreto Ferrús Abad, Paula Ruiz-Hueso, Mariana Reyes-Prieto, Vicente Soriano Chirona, Ivan Ansari, David Navarro, Maria Alma Bracho, Llúcia Martínez-Priego, Giuseppe D'Auria, Fernando Gonzalez-Candelas |
| EPI_ISL_425181 | Servicio de Microbiología. Consorcio Hospital General Universitario de Valencia | Sequencing and Bioinformatics Service and Molecular Epidemiology Research Group. FISABIO-Public Health | Beatriz Beamud, Lidia Ruiz Roldan, Marta Pla Diaz, Neris Garcia-Gonzalez, Inma Galán Vendrell, Sandra Carbo, Loreto Ferrús Abad, Paula Ruiz-Hueso, Mariana Reyes-Prieto, Vicente Soriano Chirona, Ivan Ansari, David Navarro, Maria Alma Bracho, Griselda De Marco, Llúcia Martínez-Priego, Giuseppe D'Auria, Fernando Gonzalez-Candelas |
| EPI_ISL_425182 | Servicio de Microbiología. Consorcio Hospital General Universitario de Valencia | Sequencing and Bioinformatics Service and Molecular Epidemiology Research Group. FISABIO-Public Health | Lidia Ruiz Roldan, Marta Pla Diaz, Neris Garcia-Gonzalez, Inma Galán Vendrell, Sandra Carbo, Loreto Ferrús Abad, Paula Ruiz-Hueso, Mariana Reyes-Prieto, Vicente Soriano Chirona, Ivan Ansari, David Navarro, Maria Alma Bracho, Griselda De Marco, Beatriz Beamud, Llúcia Martínez-Priego, Giuseppe D'Auria, Fernando Gonzalez-Candelas |
| EPI_ISL_425183 | Servicio de Microbiología. Consorcio Hospital General Universitario de Valencia | Sequencing and Bioinformatics Service and Molecular Epidemiology Research Group. FISABIO-Public Health | Marta Pla Diaz, Neris Garcia-Gonzalez, Inma Galán Vendrell, Sandra Carbo, Loreto Ferrús Abad, Paula Ruiz-Hueso, Mariana Reyes-Prieto, Vicente Soriano Chirona, Ivan Ansari, David Navarro, Maria Alma Bracho, Griselda De Marco, Beatriz Beamud, Lidia Ruiz Roldan, Llúcia Martínez-Priego, Giuseppe D'Auria, Fernando Gonzalez-Candelas |
| EPI_ISL_425184 | Servicio de Microbiología. Consorcio Hospital General Universitario de Valencia | Sequencing and Bioinformatics Service and Molecular Epidemiology Research Group. FISABIO-Public Health | Neris Garcia-Gonzalez, Inma Galán Vendrell, Sandra Carbo, Loreto Ferrús Abad, Paula Ruiz-Hueso, Mariana Reyes-Prieto, Vicente Soriano Chirona, Ivan Ansari, David Navarro, Maria Alma Bracho, Griselda De Marco, Beatriz Beamud, Lidia Ruiz Roldan, Marta Pla Diaz, Llúcia Martínez-Priego, Giuseppe D'Auria, Fernando Gonzalez-Candelas |
| EPI_ISL_425185 | Servicio de Microbiología. Consorcio Hospital General Universitario de Valencia | Sequencing and Bioinformatics Service and Molecular Epidemiology Research Group. FISABIO-Public Health | Inma Galán Vendrell, Sandra Carbo, Loreto Ferrús Abad, Paula Ruiz-Hueso, Mariana Reyes-Prieto, Vicente Soriano Chirona, Ivan Ansari, David Navarro, Maria Alma Bracho, Griselda De Marco, Beatriz Beamud, Lidia Ruiz Roldan, Marta Pla Diaz, Neris Garcia-Gonzalez, Llúcia Martínez-Priego, Giuseppe D'Auria, Fernando Gonzalez-Candelas |
| EPI_ISL_425186 | Servicio de Microbiología. Consorcio Hospital General Universitario de Valencia | Sequencing and Bioinformatics Service and Molecular Epidemiology Research Group. FISABIO-Public Health | Sandra Carbo, Loreto Ferrús Abad, Paula Ruiz-Hueso, Mariana Reyes-Prieto, Vicente Soriano Chirona, Ivan Ansari, David Navarro, Maria Alma Bracho, Griselda De Marco, Beatriz Beamud, Lidia Ruiz Roldan, Marta Pla Diaz, Neris Garcia-Gonzalez, Inma Galán Vendrell, Llúcia Martínez-Priego, Giuseppe D'Auria, Fernando Gonzalez-Candelas |
| EPI_ISL_425187 | Servicio de Microbiología. Consorcio Hospital General Universitario de Valencia | Sequencing and Bioinformatics Service and Molecular Epidemiology Research Group. FISABIO-Public Health | Loreto Ferrús Abad, Paula Ruiz-Hueso, Mariana Reyes-Prieto, Vicente Soriano Chirona, Ivan Ansari, David Navarro, Maria Alma Bracho, Griselda De Marco, Beatriz Beamud, Lidia Ruiz Roldan, Marta Pla Diaz, Neris Garcia-Gonzalez, Inma Galán Vendrell, Sandra Carbo, Llúcia Martínez-Priego, Giuseppe D'Auria, Fernando Gonzalez-Candelas |
| EPI_ISL_425188 | Servicio de Microbiología. Consorcio Hospital General Universitario de Valencia | Sequencing and Bioinformatics Service and Molecular Epidemiology Research Group. FISABIO-Public Health | Paula Ruiz-Hueso, Mariana Reyes-Prieto, Vicente Soriano Chirona, Ivan Ansari, David Navarro, Maria Alma Bracho, Griselda De Marco, Beatriz Beamud, Lidia Ruiz Roldan, Marta Pla Diaz, Neris Garcia-Gonzalez, Inma Galán Vendrell, Sandra Carbo, Loreto Ferrús Abad, Llúcia Martínez-Priego, Giuseppe D'Auria, Fernando Gonzalez-Candelas |
| EPI_ISL_425189 | Servicio de Microbiología. Consorcio Hospital General Universitario de Valencia | Sequencing and Bioinformatics Service and Molecular Epidemiology Research Group. FISABIO-Public Health | Mariana Reyes-Prieto, Vicente Soriano Chirona, Ivan Ansari, David Navarro, Maria Alma Bracho, Griselda De Marco, Beatriz Beamud, Lidia Ruiz Roldan, Marta Pla Diaz, Neris Garcia-Gonzalez, Inma Galán Vendrell, Sandra Carbo, Loreto Ferrús Abad, Paula Ruiz-Hueso, Llúcia Martínez-Priego, Giuseppe D'Auria, Fernando Gonzalez-Candelas |
| EPI_ISL_425190 | Servicio de Microbiología. Consorcio Hospital General Universitario de Valencia | Sequencing and Bioinformatics Service and Molecular Epidemiology Research Group. FISABIO-Public Health | Vicente Soriano Chirona, Ivan Ansari, David Navarro, Maria Alma Bracho, Griselda De Marco, Beatriz Beamud, Lidia Ruiz Roldan, Marta Pla Diaz, Neris Garcia-Gonzalez, Inma Galán Vendrell, Sandra Carbo, Loreto Ferrús Abad, Paula Ruiz-Hueso, Mariana Reyes-Prieto, Llúcia Martínez-Priego, Giuseppe D'Auria, Fernando Gonzalez-Candelas |
| EPI_ISL_425191 | Servicio de Microbiología. Consorcio Hospital General Universitario de Valencia | Sequencing and Bioinformatics Service and Molecular Epidemiology Research Group. FISABIO-Public Health | Ivan Ansari, David Navarro, Maria Alma Bracho, Griselda De Marco, Beatriz Beamud, Lidia Ruiz Roldan, Marta Pla Diaz, Neris Garcia-Gonzalez, Inma Galán Vendrell, Sandra Carbo, Loreto Ferrús Abad, Paula Ruiz-Hueso, Mariana Reyes-Prieto, Vicente Soriano Chirona, Llúcia Martínez-Priego, Giuseppe D'Auria, Fernando Gonzalez-Candelas |
| EPI_ISL_425192 | Servicio de Microbiología. Consorcio Hospital General Universitario de Valencia | Sequencing and Bioinformatics Service and Molecular Epidemiology Research Group. FISABIO-Public Health | David Navarro, Maria Alma Bracho, Griselda De Marco, Beatriz Beamud, Lidia Ruiz Roldan, Marta Pla Diaz, Neris Garcia-Gonzalez, Inma Galán Vendrell, Sandra Carbo, Loreto Ferrús Abad, Paula Ruiz-Hueso, Mariana Reyes-Prieto, Vicente Soriano Chirona, Ivan Ansari, Llúcia Martínez-Priego, Giuseppe D'Auria, Fernando Gonzalez-Candelas |
| EPI_ISL_425193 | Servicio de Microbiología. Consorcio Hospital General Universitario de Valencia | Sequencing and Bioinformatics Service and Molecular Epidemiology Research Group. FISABIO-Public Health | Maria Alma Bracho, Griselda De Marco, Beatriz Beamud, Lidia Ruiz Roldan, Marta Pla Diaz, Neris Garcia-Gonzalez, Inma Galán Vendrell, Sandra Carbo, Loreto Ferrús Abad, Paula Ruiz-Hueso, Mariana Reyes-Prieto, Vicente Soriano Chirona, Ivan Ansari, David Navarro, Llúcia Martínez-Priego, Giuseppe D'Auria, Fernando Gonzalez-Candelas |
| EPI_ISL_425194 | Servicio de Microbiología. Consorcio Hospital General Universitario de Valencia | Sequencing and Bioinformatics Service and Molecular Epidemiology Research Group. FISABIO-Public Health | Griselda De Marco, Beatriz Beamud, Lidia Ruiz Roldan, Marta Pla Diaz, Neris Garcia-Gonzalez, Inma Galán Vendrell, Sandra Carbo, Loreto Ferrús Abad, Paula Ruiz-Hueso, Mariana Reyes-Prieto, Vicente Soriano Chirona, Ivan Ansari, David Navarro, Maria Alma Bracho, Llúcia Martínez-Priego, Giuseppe D'Auria, Fernando Gonzalez-Candelas |
| EPI_ISL_425195 | Servicio de Microbiología. Consorcio Hospital General Universitario de Valencia | Sequencing and Bioinformatics Service and Molecular Epidemiology Research Group. FISABIO-Public Health | Beatriz Beamud, Lidia Ruiz Roldan, Marta Pla Diaz, Neris Garcia-Gonzalez, Inma Galán Vendrell, Sandra Carbo, Loreto Ferrús Abad, Paula Ruiz-Hueso, Mariana Reyes-Prieto, Vicente Soriano Chirona, Ivan Ansari, David Navarro, Maria Alma Bracho, Griselda De Marco, Llúcia Martínez-Priego, Giuseppe D'Auria, Fernando Gonzalez-Candelas |
| EPI_ISL_425196 | Servicio de Microbiología. Consorcio Hospital General Universitario de Valencia | Sequencing and Bioinformatics Service and Molecular Epidemiology Research Group. FISABIO-Public Health | Lidia Ruiz Roldan, Marta Pla Diaz, Neris Garcia-Gonzalez, Inma Galán Vendrell, Sandra Carbo, Loreto Ferrús Abad, Paula Ruiz-Hueso, Mariana Reyes-Prieto, Vicente Soriano Chirona, Ivan Ansari, David Navarro, Maria Alma Bracho, Griselda De Marco, Beatriz Beamud, Llúcia Martínez-Priego, Giuseppe D'Auria, Fernando Gonzalez-Candelas |
| EPI_ISL_425197 | Servicio de Microbiología. Consorcio Hospital General Universitario de Valencia | Sequencing and Bioinformatics Service and Molecular Epidemiology Research Group. FISABIO-Public Health | Marta Pla Diaz, Neris Garcia-Gonzalez, Inma Galán Vendrell, Sandra Carbo, Loreto Ferrús Abad, Paula Ruiz-Hueso, Mariana Reyes-Prieto, Vicente Soriano Chirona, Ivan Ansari, David Navarro, Maria Alma Bracho, Griselda De Marco, Beatriz Beamud, Lidia Ruiz Roldan, Llúcia Martínez-Priego, Giuseppe D'Auria, Fernando Gonzalez-Candelas |
| EPI_ISL_425198 | Servicio de Microbiología. Consorcio Hospital General Universitario de Valencia | Sequencing and Bioinformatics Service and Molecular Epidemiology Research Group. FISABIO-Public Health | Neris Garcia-Gonzalez, Inma Galán Vendrell, Sandra Carbo, Loreto Ferrús Abad, Paula Ruiz-Hueso, Mariana Reyes-Prieto, Vicente Soriano Chirona, Ivan Ansari, David Navarro, Maria Alma Bracho, Griselda De Marco, Beatriz Beamud, Lidia Ruiz Roldan, Marta Pla Diaz, Llúcia Martínez-Priego, Giuseppe D'Auria, Fernando Gonzalez-Candelas |
| EPI_ISL_425199 | Servicio de Microbiologia. Hospital Clinico Universitario de Valencia | Sequencing and Bioinformatics Service and Molecular Epidemiology Research Group. FISABIO-Public Health | Inma Galán Vendrell, Paula Ruiz-Hueso, Mariana Reyes-Prieto, Vicente Soriano Chirona, Maria Alma Bracho, Griselda De Marco, Beatriz Beamud, Lidia Ruiz Roldan, Marta Pla Diaz,Neris Garcia-Gonzalez, Loreto Ferrús Abad, Maria Dolores Ocete, Llúcia Martínez-Priego, Concepcion Gimeno, Giuseppe D'Auria, Fernando Gonzalez-Candelas |
| EPI_ISL_425200 | Servicio de Microbiologia. Hospital Clinico Universitario de Valencia | Sequencing and Bioinformatics Service and Molecular Epidemiology Research Group. FISABIO-Public Health | Paula Ruiz-Hueso, Mariana Reyes-Prieto, Vicente Soriano Chirona, Maria Alma Bracho, Griselda De Marco, Beatriz Beamud, Lidia Ruiz Roldan, Marta Pla Diaz,Neris Garcia-Gonzalez, Loreto Ferrús Abad, Maria Dolores Ocete, Llúcia Martínez-Priego, Inma Galán Vendrell, Concepcion Gimeno, Giuseppe D'Auria, Fernando Gonzalez-Candelas |
| EPI_ISL_425201 | Servicio de Microbiologia. Hospital Clinico Universitario de Valencia | Sequencing and Bioinformatics Service and Molecular Epidemiology Research Group. FISABIO-Public Health | Mariana Reyes-Prieto, Vicente Soriano Chirona, Maria Alma Bracho, Griselda De Marco, Beatriz Beamud, Lidia Ruiz Roldan, Marta Pla Diaz,Neris Garcia-Gonzalez, Loreto Ferrús Abad, Maria Dolores Ocete, Llúcia Martínez-Priego, Inma Galán Vendrell, Paula Ruiz-Hueso,Concepcion Gimeno, Giuseppe D'Auria, Fernando Gonzalez-Candelas |
| EPI_ISL_425202 | Servicio de Microbiologia. Hospital Clinico Universitario de Valencia | Sequencing and Bioinformatics Service and Molecular Epidemiology Research Group. FISABIO-Public Health | Vicente Soriano Chirona, Maria Alma Bracho, Griselda De Marco, Beatriz Beamud, Lidia Ruiz Roldan, Marta Pla Diaz,Neris Garcia-Gonzalez, Loreto Ferrús Abad, Maria Dolores Ocete, Inma Galán Vendrell, Paula Ruiz-Hueso, Mariana Reyes-Prieto, Llúcia Martínez-Priego, Concepcion Gimeno, Giuseppe D'Auria, Fernando Gonzalez-Candelas |
| EPI_ISL_425203 | Servicio de Microbiologia. Hospital Clinico Universitario de Valencia | Sequencing and Bioinformatics Service and Molecular Epidemiology Research Group. FISABIO-Public Health | Maria Alma Bracho, Griselda De Marco, Beatriz Beamud, Lidia Ruiz Roldan, Marta Pla Diaz,Neris Garcia-Gonzalez, Loreto Ferrús Abad, Maria Dolores Ocete, Inma Galán Vendrell, Paula Ruiz-Hueso, Mariana Reyes-Prieto, Vicente Soriano Chirona, Llúcia Martínez-Priego, Concepcion Gimeno, Giuseppe D'Auria, Fernando Gonzalez-Candelas |
| EPI_ISL_425204 | Servicio de Microbiologia. Hospital Clinico Universitario de Valencia | Sequencing and Bioinformatics Service and Molecular Epidemiology Research Group. FISABIO-Public Health | Maria Dolores Ocete, Inma Galán Vendrell, Paula Ruiz-Hueso, Mariana Reyes-Prieto, Vicente Soriano Chirona, Maria Alma Bracho, Griselda De Marco, Beatriz Beamud, Lidia Ruiz Roldan, Marta Pla Diaz,Neris Garcia-Gonzalez, Loreto Ferrús Abad, Llúcia Martínez-Priego, Concepcion Gimeno, Giuseppe D'Auria, Fernando Gonzalez-Candelas |
| EPI_ISL_425205 | Servicio de Microbiologia. Hospital Clinico Universitario de Valencia | Sequencing and Bioinformatics Service and Molecular Epidemiology Research Group. FISABIO-Public Health | Griselda De Marco, Beatriz Beamud, Lidia Ruiz Roldan, Marta Pla Diaz,Neris Garcia-Gonzalez, Loreto Ferrús Abad, Maria Dolores Ocete, Inma Galán Vendrell, Paula Ruiz-Hueso, Mariana Reyes-Prieto, Vicente Soriano Chirona, Maria Alma Bracho, Llúcia Martínez-Priego, Concepcion Gimeno, Giuseppe D'Auria, Fernando Gonzalez-Candelas |
| EPI_ISL_425206 | Servicio de Microbiologia. Hospital Clinico Universitario de Valencia | Sequencing and Bioinformatics Service and Molecular Epidemiology Research Group. FISABIO-Public Health | Beatriz Beamud, Lidia Ruiz Roldan, Marta Pla Diaz,Neris Garcia-Gonzalez, Loreto Ferrús Abad, Maria Dolores Ocete, Inma Galán Vendrell, Paula Ruiz-Hueso, Mariana Reyes-Prieto, Vicente Soriano Chirona, Maria Alma Bracho, Griselda De Marco, Llúcia Martínez-Priego, Concepcion Gimeno, Giuseppe D'Auria, Fernando Gonzalez-Candelas |
| EPI_ISL_425207 | Servicio de Microbiologia. Hospital Clinico Universitario de Valencia | Sequencing and Bioinformatics Service and Molecular Epidemiology Research Group. FISABIO-Public Health | Lidia Ruiz Roldan, Marta Pla Diaz,Neris Garcia-Gonzalez, Loreto Ferrús Abad, Maria Dolores Ocete, Inma Galán Vendrell, Paula Ruiz-Hueso, Mariana Reyes-Prieto, Vicente Soriano Chirona, Maria Alma Bracho, Griselda De Marco, Beatriz Beamud, Llúcia Martínez-Priego, Concepcion Gimeno, Giuseppe D'Auria, Fernando Gonzalez-Candelas |
| EPI_ISL_425208 | Servicio de Microbiologia. Hospital Clinico Universitario de Valencia | Sequencing and Bioinformatics Service and Molecular Epidemiology Research Group. FISABIO-Public Health | Marta Pla Diaz,Neris Garcia-Gonzalez, Loreto Ferrús Abad, Maria Dolores Ocete, Inma Galán Vendrell, Paula Ruiz-Hueso, Mariana Reyes-Prieto, Vicente Soriano Chirona, Maria Alma Bracho, Griselda De Marco, Beatriz Beamud, Lidia Ruiz Roldan, Llúcia Martínez-Priego, Concepcion Gimeno, Giuseppe D'Auria, Fernando Gonzalez-Candelas |
| EPI_ISL_425209 | Servicio de Microbiologia. Hospital Clinico Universitario de Valencia | Sequencing and Bioinformatics Service and Molecular Epidemiology Research Group. FISABIO-Public Health | Neris Garcia-Gonzalez, Loreto Ferrús Abad, Maria Dolores Ocete, Inma Galán Vendrell, Paula Ruiz-Hueso, Mariana Reyes-Prieto, Vicente Soriano Chirona, Maria Alma Bracho, Griselda De Marco, Beatriz Beamud, Lidia Ruiz Roldan, Marta Pla Diaz, Llúcia Martínez-Priego, Concepcion Gimeno, Giuseppe D'Auria, Fernando Gonzalez-Candelas |
| EPI_ISL_425210 | Servicio de Microbiologia. Hospital Clinico Universitario de Valencia | Sequencing and Bioinformatics Service and Molecular Epidemiology Research Group. FISABIO-Public Health | Loreto Ferrús Abad, Maria Dolores Ocete, Inma Galán Vendrell, Paula Ruiz-Hueso, Mariana Reyes-Prieto, Vicente Soriano Chirona, Maria Alma Bracho, Griselda De Marco, Beatriz Beamud, Lidia Ruiz Roldan, Marta Pla Diaz,Neris Garcia-Gonzalez, Llúcia Martínez-Priego, Concepcion Gimeno, Giuseppe D'Auria, Fernando Gonzalez-Candelas |
| EPI_ISL_425211 | Servicio de Microbiologia. Hospital Clinico Universitario de Valencia | Sequencing and Bioinformatics Service and Molecular Epidemiology Research Group. FISABIO-Public Health | Loreto Ferrús Abad, Maria Dolores Ocete,Inma Galán Vendrell, Paula Ruiz-Hueso, Mariana Reyes-Prieto, Vicente Soriano Chirona, Maria Alma Bracho, Griselda De Marco, Beatriz Beamud, Lidia Ruiz Roldan, Marta Pla Diaz,Neris Garcia-Gonzalez, Llúcia Martínez-Priego, Concepcion Gimeno, Giuseppe D'Auria, Fernando Gonzalez-Candelas |
| EPI_ISL_425212 | Servicio de Microbiología. Consorcio Hospital General Universitario de Valencia | Sequencing and Bioinformatics Service and Molecular Epidemiology Research Group. FISABIO-Public Health | David Navarro, Maria Alma Bracho, Griselda De Marco, Beatriz Beamud, Lidia Ruiz Roldan, Marta Pla Diaz, Neris Garcia-Gonzalez, Inma Galán Vendrell, Sandra Carbo, Loreto Ferrús Abad, Paula Ruiz-Hueso, Mariana Reyes-Prieto, Vicente Soriano Chirona, Ivan Ansari, Llúcia Martínez-Priego, Giuseppe D'Auria, Fernando Gonzalez-Candelas |
| EPI_ISL_425213 | Servicio de Microbiología. Consorcio Hospital General Universitario de Valencia | Sequencing and Bioinformatics Service and Molecular Epidemiology Research Group. FISABIO-Public Health | Maria Alma Bracho, Griselda De Marco, Beatriz Beamud, Lidia Ruiz Roldan, Marta Pla Diaz, Neris Garcia-Gonzalez, Inma Galán Vendrell, Sandra Carbo, Loreto Ferrús Abad, Paula Ruiz-Hueso, Mariana Reyes-Prieto, Vicente Soriano Chirona, Ivan Ansari, David Navarro, Llúcia Martínez-Priego, Giuseppe D'Auria, Fernando Gonzalez-Candelas |
| EPI_ISL_425214 | Servicio de Microbiología. Consorcio Hospital General Universitario de Valencia | Sequencing and Bioinformatics Service and Molecular Epidemiology Research Group. FISABIO-Public Health | Griselda De Marco, Beatriz Beamud, Lidia Ruiz Roldan, Marta Pla Diaz, Neris Garcia-Gonzalez, Inma Galán Vendrell, Sandra Carbo, Loreto Ferrús Abad, Paula Ruiz-Hueso, Mariana Reyes-Prieto, Vicente Soriano Chirona, Ivan Ansari, David Navarro, Maria Alma Bracho, Llúcia Martínez-Priego, Giuseppe D'Auria, Fernando Gonzalez-Candelas |
| EPI_ISL_425215 | Servicio de Microbiología. Consorcio Hospital General Universitario de Valencia | Sequencing and Bioinformatics Service and Molecular Epidemiology Research Group. FISABIO-Public Health | Beatriz Beamud, Lidia Ruiz Roldan, Marta Pla Diaz, Neris Garcia-Gonzalez, Inma Galán Vendrell, Sandra Carbo, Loreto Ferrús Abad, Paula Ruiz-Hueso, Mariana Reyes-Prieto, Vicente Soriano Chirona, Ivan Ansari, David Navarro, Maria Alma Bracho, Griselda De Marco, Llúcia Martínez-Priego, Giuseppe D'Auria, Fernando Gonzalez-Candelas |
| EPI_ISL_425216 | Servicio de Microbiología. Consorcio Hospital General | Sequencing and Bioinformatics Service and Molecular | Lidia Ruiz Roldan, Marta Pla Diaz, Neris Garcia-Gonzalez, Inma Galán Vendrell, Sandra Carbo, Loreto Ferrús Abad, Paula Ruiz-Hueso, Mariana Reyes-Prieto, Vicente Soriano Chirona, Ivan Ansari, David Navarro, Maria Alma Bracho, Griselda De |

| | | | |
|---|---|---|---|
| | Universitario de Valencia | Epidemiology Research Group. FISABIO-Public Health | Marco, Beatriz Beamud, Llúcia Martínez-Priego, Giuseppe D'Auria, Fernando Gonzalez-Candelas |
| EPI_ISL_425217 | Servicio de Microbiología. Consorcio Hospital General Universitario de Valencia | Sequencing and Bioinformatics Service and Molecular Epidemiology Research Group. FISABIO-Public Health | Marta Pla Diaz, Neris Garcia-Gonzalez, Inma Galán Vendrell, Sandra Carbo, Loreto Ferrús Abad, Paula Ruiz-Hueso, Mariana Reyes-Prieto, Vicente Soriano Chirona, Ivan Ansari, David Navarro, Maria Alma Bracho, Griselda De Marco, Beatriz Beamud, Lidia Ruiz Roldan, Llúcia Martínez-Priego, Giuseppe D'Auria, Fernando Gonzalez-Candelas |
| EPI_ISL_425218, EPI_ISL_425219 | Servicio de Microbiología. Consorcio Hospital General Universitario de Valencia | Sequencing and Bioinformatics Service and Molecular Epidemiology Research Group. FISABIO-Public Health | Neris Garcia-Gonzalez, Inma Galán Vendrell, Sandra Carbo, Loreto Ferrús Abad, Paula Ruiz-Hueso, Mariana Reyes-Prieto, Vicente Soriano Chirona, Ivan Ansari, David Navarro, Maria Alma Bracho, Griselda De Marco, Beatriz Beamud, Lidia Ruiz Roldan, Marta Pla Diaz, Llúcia Martínez-Priego, Giuseppe D'Auria, Fernando Gonzalez-Candelas |
| EPI_ISL_425220 | Servicio de Microbiología. Consorcio Hospital General Universitario de Valencia | Sequencing and Bioinformatics Service and Molecular Epidemiology Research Group. FISABIO-Public Health | Inma Galán Vendrell, Sandra Carbo, Loreto Ferrús Abad, Paula Ruiz-Hueso, Mariana Reyes-Prieto, Vicente Soriano Chirona, Ivan Ansari, David Navarro, Maria Alma Bracho, Griselda De Marco, Beatriz Beamud, Lidia Ruiz Roldan, Marta Pla Diaz, Neris Garcia-Gonzalez, Llúcia Martínez-Priego, Giuseppe D'Auria, Fernando Gonzalez-Candelas |
| EPI_ISL_425221 | Servicio de Microbiología. Consorcio Hospital General Universitario de Valencia | Sequencing and Bioinformatics Service and Molecular Epidemiology Research Group. FISABIO-Public Health | Sandra Carbo, Loreto Ferrús Abad, Paula Ruiz-Hueso, Mariana Reyes-Prieto, Vicente Soriano Chirona, Ivan Ansari, David Navarro, Maria Alma Bracho, Griselda De Marco, Beatriz Beamud, Lidia Ruiz Roldan, Marta Pla Diaz, Neris Garcia-Gonzalez, Inma Galán Vendrell, Llúcia Martínez-Priego, Giuseppe D'Auria, Fernando Gonzalez-Candelas |
| EPI_ISL_425222 | Servicio de Microbiología. Consorcio Hospital General Universitario de Valencia | Sequencing and Bioinformatics Service and Molecular Epidemiology Research Group. FISABIO-Public Health | Loreto Ferrús Abad, Paula Ruiz-Hueso, Mariana Reyes-Prieto, Vicente Soriano Chirona, Ivan Ansari, David Navarro, Maria Alma Bracho, Griselda De Marco, Beatriz Beamud, Lidia Ruiz Roldan, Marta Pla Diaz, Neris Garcia-Gonzalez, Inma Galán Vendrell, Sandra Carbo, Llúcia Martínez-Priego, Giuseppe D'Auria, Fernando Gonzalez-Candelas |
| EPI_ISL_425223 | Servicio de Microbiología. Consorcio Hospital General Universitario de Valencia | Sequencing and Bioinformatics Service and Molecular Epidemiology Research Group. FISABIO-Public Health | Paula Ruiz-Hueso, Mariana Reyes-Prieto, Vicente Soriano Chirona, Ivan Ansari, David Navarro, Maria Alma Bracho, Griselda De Marco, Beatriz Beamud, Lidia Ruiz Roldan, Marta Pla Diaz, Neris Garcia-Gonzalez, Inma Galán Vendrell, Sandra Carbo, Loreto Ferrús Abad, Llúcia Martínez-Priego, Giuseppe D'Auria, Fernando Gonzalez-Candelas |
| EPI_ISL_425224 | Wadsworth Center, New York State Department of Health | Wadsworth Center, New York State Department of Health | Kirsten St. George, Daryl M. Lamson, Sara Griesemer, Jonathan Plitnick, Navjot Singh, Matthew D. Shudt, Erica Lasek-Nesselquist |
| EPI_ISL_425228, EPI_ISL_425229 | Laboratory of Molecular Genetics, 2nd Faculty of Medicine, Charles University in Prague, Prague, Czech Republic | Laboratory of Molecular Genetics, 2nd Faculty of Medicine, Charles University in Prague, Prague, Czech Republic | Lenka Kramná, Kateřina Poláčková, Ondřej Cinek |
| EPI_ISL_425231, EPI_ISL_425232, EPI_ISL_425233, EPI_ISL_425234, EPI_ISL_425235, EPI_ISL_425236, EPI_ISL_425237, EPI_ISL_425238, EPI_ISL_425239, EPI_ISL_425240, EPI_ISL_425241, EPI_ISL_425242, EPI_ISL_425243, EPI_ISL_425244, EPI_ISL_425245, EPI_ISL_425246, EPI_ISL_425247, EPI_ISL_425248, EPI_ISL_425249, EPI_ISL_425250, EPI_ISL_425251, EPI_ISL_425252, EPI_ISL_425253, EPI_ISL_425254, EPI_ISL_425255, EPI_ISL_425256, EPI_ISL_425257, EPI_ISL_425258, EPI_ISL_425259, EPI_ISL_425260, EPI_ISL_425261, EPI_ISL_425262, EPI_ISL_425263, EPI_ISL_425264, EPI_ISL_425265, EPI_ISL_425266, EPI_ISL_425267, EPI_ISL_425268, EPI_ISL_425269, EPI_ISL_425270, EPI_ISL_425271, EPI_ISL_425272, EPI_ISL_425273, EPI_ISL_425274, EPI_ISL_425275, EPI_ISL_425276, EPI_ISL_425277, EPI_ISL_425278, EPI_ISL_425279, EPI_ISL_425280, EPI_ISL_425281, EPI_ISL_425282, EPI_ISL_425283, EPI_ISL_425284, EPI_ISL_425285, EPI_ISL_425286, EPI_ISL_425287, EPI_ISL_425288, EPI_ISL_425289, EPI_ISL_425290, EPI_ISL_425291, EPI_ISL_425292, EPI_ISL_425293, EPI_ISL_425294, EPI_ISL_425295, EPI_ISL_425296, EPI_ISL_425297, EPI_ISL_425298, EPI_ISL_425299, EPI_ISL_425300, EPI_ISL_425301, EPI_ISL_425302, EPI_ISL_425303, EPI_ISL_425304, EPI_ISL_425305, EPI_ISL_425306, EPI_ISL_425307, EPI_ISL_425308, EPI_ISL_425309, EPI_ISL_425310, EPI_ISL_425311, EPI_ISL_425312, EPI_ISL_425313, EPI_ISL_425314, EPI_ISL_425315, EPI_ISL_425316, EPI_ISL_425317, EPI_ISL_425318, EPI_ISL_425319, EPI_ISL_425320, EPI_ISL_425321, EPI_ISL_425322, EPI_ISL_425323, EPI_ISL_425324, EPI_ISL_425325, EPI_ISL_425326, EPI_ISL_425327, EPI_ISL_425328, EPI_ISL_425329, EPI_ISL_425330, EPI_ISL_425331, EPI_ISL_425332, EPI_ISL_425333, EPI_ISL_425334, EPI_ISL_425335, EPI_ISL_425336, EPI_ISL_425337, EPI_ISL_425338, EPI_ISL_425339, EPI_ISL_425340, EPI_ISL_425341, EPI_ISL_425342, EPI_ISL_425343, EPI_ISL_425344, EPI_ISL_425345, EPI_ISL_425346, EPI_ISL_425347, EPI_ISL_425348, EPI_ISL_425349, EPI_ISL_425350, EPI_ISL_425351, EPI_ISL_425352, EPI_ISL_425353, EPI_ISL_425354, EPI_ISL_425355, EPI_ISL_425356, EPI_ISL_425357, EPI_ISL_425358, EPI_ISL_425359, EPI_ISL_425360, EPI_ISL_425361, EPI_ISL_425362, EPI_ISL_425363, EPI_ISL_425364, EPI_ISL_425365, EPI_ISL_425366, EPI_ISL_425367, EPI_ISL_425368, EPI_ISL_425369, EPI_ISL_425370, EPI_ISL_425371, EPI_ISL_425373, EPI_ISL_425374, EPI_ISL_425375, EPI_ISL_425376, EPI_ISL_425377, EPI_ISL_425378, EPI_ISL_425379, EPI_ISL_425380, EPI_ISL_425381, EPI_ISL_425382, EPI_ISL_425383, EPI_ISL_425384, EPI_ISL_425385, EPI_ISL_425386, EPI_ISL_425387, EPI_ISL_425388, EPI_ISL_425389, EPI_ISL_425390, EPI_ISL_425391, EPI_ISL_425392, EPI_ISL_425393, EPI_ISL_425394, EPI_ISL_425395, EPI_ISL_425396, EPI_ISL_425397, EPI_ISL_425398, EPI_ISL_425399, EPI_ISL_425400, EPI_ISL_425401, EPI_ISL_425402, EPI_ISL_425403, EPI_ISL_425404, EPI_ISL_425405, EPI_ISL_425406, EPI_ISL_425407, EPI_ISL_425408, EPI_ISL_425409, EPI_ISL_425410, EPI_ISL_425411, EPI_ISL_425412, EPI_ISL_425413, EPI_ISL_425414, EPI_ISL_425415, EPI_ISL_425416, EPI_ISL_425417, EPI_ISL_425418, EPI_ISL_425419, EPI_ISL_425420, EPI_ISL_425421, EPI_ISL_425422, EPI_ISL_425423, EPI_ISL_425424, EPI_ISL_425425, EPI_ISL_425426, EPI_ISL_425427, EPI_ISL_425428, EPI_ISL_425429, EPI_ISL_425430, EPI_ISL_425431, EPI_ISL_425432, EPI_ISL_425433, EPI_ISL_425434, EPI_ISL_425435, EPI_ISL_425436, EPI_ISL_425437, EPI_ISL_425438, EPI_ISL_425439, EPI_ISL_425440, EPI_ISL_425441, EPI_ISL_425442, EPI_ISL_425443, EPI_ISL_425444, EPI_ISL_425445, EPI_ISL_425446, EPI_ISL_425447, EPI_ISL_425448, EPI_ISL_425449, EPI_ISL_425450, EPI_ISL_425451, EPI_ISL_425452, EPI_ISL_425453, EPI_ISL_425454, EPI_ISL_425455, EPI_ISL_425456, EPI_ISL_425457, EPI_ISL_425458, EPI_ISL_425459, EPI_ISL_425460, EPI_ISL_425461 | | | |
| see above | Department of Pathology, University of Cambridge | COVID-19 Genomics UK (COG-UK) Consortium | Luke W Meredith, M. Estee Torok , Myra Hosmillo, William L. Hamilton, Martin D. Curran, Theresa Feltwell, Anna Yakovleva, Charlotte J. Houldcroft, Aminu S. Jahun, Sarah L. Caddy, Ian Goodfellow |
| EPI_ISL_425462, EPI_ISL_425463, EPI_ISL_425464, EPI_ISL_425465, EPI_ISL_425466, EPI_ISL_425467, EPI_ISL_425468, EPI_ISL_425469, EPI_ISL_425470, EPI_ISL_425471, EPI_ISL_425472, EPI_ISL_425473, EPI_ISL_425474, EPI_ISL_425475, EPI_ISL_425476, EPI_ISL_425477, EPI_ISL_425478, EPI_ISL_425479, EPI_ISL_425480, EPI_ISL_425481, EPI_ISL_425482, EPI_ISL_425483, EPI_ISL_425484, EPI_ISL_425485, EPI_ISL_425486, EPI_ISL_425487, EPI_ISL_425488, EPI_ISL_425489, EPI_ISL_425490, EPI_ISL_425491, EPI_ISL_425492, EPI_ISL_425493, EPI_ISL_425494, EPI_ISL_425495, EPI_ISL_425496, EPI_ISL_425497, EPI_ISL_425498, EPI_ISL_425499, EPI_ISL_425500, EPI_ISL_425501, EPI_ISL_425502, EPI_ISL_425503, EPI_ISL_425504, EPI_ISL_425505, EPI_ISL_425506, EPI_ISL_425507, EPI_ISL_425508, EPI_ISL_425509, EPI_ISL_425510, EPI_ISL_425511, EPI_ISL_425512, EPI_ISL_425513, EPI_ISL_425514, EPI_ISL_425515, EPI_ISL_425516, EPI_ISL_425517, EPI_ISL_425518, EPI_ISL_425519, EPI_ISL_425520, EPI_ISL_425521, EPI_ISL_425522, EPI_ISL_425523, EPI_ISL_425524, EPI_ISL_425525, EPI_ISL_425526, EPI_ISL_425527, EPI_ISL_425528, EPI_ISL_425529, EPI_ISL_425530, EPI_ISL_425531, EPI_ISL_425532, EPI_ISL_425533, EPI_ISL_425534, EPI_ISL_425535, EPI_ISL_425536, EPI_ISL_425537, EPI_ISL_425538, EPI_ISL_425539, EPI_ISL_425540, EPI_ISL_425541, EPI_ISL_425542, EPI_ISL_425543, EPI_ISL_425544, EPI_ISL_425545, EPI_ISL_425546, EPI_ISL_425547, EPI_ISL_425548, EPI_ISL_425549, EPI_ISL_425550, EPI_ISL_425551, EPI_ISL_425552, EPI_ISL_425553, EPI_ISL_425554, EPI_ISL_425555, EPI_ISL_425556, EPI_ISL_425557, EPI_ISL_425558, EPI_ISL_425559, EPI_ISL_425560, EPI_ISL_425561, EPI_ISL_425562, EPI_ISL_425563, EPI_ISL_425564, EPI_ISL_425565, EPI_ISL_425566, EPI_ISL_425567, EPI_ISL_425568, EPI_ISL_425569, EPI_ISL_425570, EPI_ISL_425571, EPI_ISL_425572, EPI_ISL_425573, EPI_ISL_425574, EPI_ISL_425575, EPI_ISL_425576, EPI_ISL_425577, EPI_ISL_425579, EPI_ISL_425580, EPI_ISL_425581, EPI_ISL_425582, EPI_ISL_425583, EPI_ISL_425584, EPI_ISL_425585, EPI_ISL_425586, EPI_ISL_425587, EPI_ISL_425588, EPI_ISL_425589, EPI_ISL_425590, EPI_ISL_425591, EPI_ISL_425592, EPI_ISL_425593, EPI_ISL_425594, EPI_ISL_425595, EPI_ISL_425596, EPI_ISL_425597, EPI_ISL_425598, EPI_ISL_425599, EPI_ISL_425600, EPI_ISL_425601, EPI_ISL_425602, EPI_ISL_425603, EPI_ISL_425604, EPI_ISL_425605, EPI_ISL_425606, EPI_ISL_425607, EPI_ISL_425608, EPI_ISL_425609, EPI_ISL_425610, EPI_ISL_425611, EPI_ISL_425612, EPI_ISL_425613, EPI_ISL_425614, EPI_ISL_425615, EPI_ISL_425616, EPI_ISL_425617, EPI_ISL_425618, EPI_ISL_425619, EPI_ISL_425620, EPI_ISL_425621, EPI_ISL_425622, EPI_ISL_425623, EPI_ISL_425624, EPI_ISL_425625, EPI_ISL_425626, EPI_ISL_425627, EPI_ISL_425628, EPI_ISL_425629, EPI_ISL_425630, EPI_ISL_425631, EPI_ISL_425632, EPI_ISL_425633, EPI_ISL_425634, EPI_ISL_425635, EPI_ISL_425636, EPI_ISL_425637, EPI_ISL_425638, EPI_ISL_425639, EPI_ISL_425640, EPI_ISL_425641, EPI_ISL_425642, EPI_ISL_425643, EPI_ISL_425644, EPI_ISL_425645, EPI_ISL_425646 | | | |
| see above | Queens Medical Centre, Clinical Microbiology Department / DeepSeq Nottingham | COVID-19 Genomics UK (COG-UK) Consortium | Gemma Clark, Wendy Smith, Manjinder Khakh, Hannah Howson-Wells, Jonathan Ball, Patrick McClure, Joseph Chappell, Theocharis Tsoleridis, Nadine Holmes, Matthew Carlisle, Christopher Moore, Fei Sang, Johnny Debebe, Victoria Wright, Matthew Loose |
| EPI_ISL_425647, EPI_ISL_425648, EPI_ISL_425649, EPI_ISL_425650, EPI_ISL_425651, EPI_ISL_425652, EPI_ISL_425653, EPI_ISL_425654, EPI_ISL_425655, EPI_ISL_425656, EPI_ISL_425657, EPI_ISL_425658, EPI_ISL_425659, EPI_ISL_425660, EPI_ISL_425661, EPI_ISL_425662, EPI_ISL_425663, EPI_ISL_425664, EPI_ISL_425665, EPI_ISL_425666, EPI_ISL_425667, EPI_ISL_425668, EPI_ISL_425669, EPI_ISL_425670, EPI_ISL_425671, EPI_ISL_425672, EPI_ISL_425673, EPI_ISL_425674, EPI_ISL_425675, EPI_ISL_425676, EPI_ISL_425677, EPI_ISL_425678, EPI_ISL_425679, EPI_ISL_425680, EPI_ISL_425681, EPI_ISL_425682, EPI_ISL_425683, EPI_ISL_425684, EPI_ISL_425685, EPI_ISL_425686, EPI_ISL_425687, EPI_ISL_425688, EPI_ISL_425689, EPI_ISL_425690, EPI_ISL_425691, EPI_ISL_425692, EPI_ISL_425693, EPI_ISL_425694, EPI_ISL_425695, EPI_ISL_425696, EPI_ISL_425697, EPI_ISL_425698, EPI_ISL_425699, EPI_ISL_425700, EPI_ISL_425701, EPI_ISL_425702, EPI_ISL_425703, EPI_ISL_425704, EPI_ISL_425705, EPI_ISL_425706, EPI_ISL_425707, EPI_ISL_425708, EPI_ISL_425709, EPI_ISL_425710, EPI_ISL_425711, EPI_ISL_425712, EPI_ISL_425713, EPI_ISL_425714, EPI_ISL_425715, EPI_ISL_425716, EPI_ISL_425717, EPI_ISL_425718, EPI_ISL_425719, EPI_ISL_425720, EPI_ISL_425721, EPI_ISL_425722, EPI_ISL_425723, EPI_ISL_425724, EPI_ISL_425725, EPI_ISL_425726, EPI_ISL_425727, EPI_ISL_425728, EPI_ISL_425729, EPI_ISL_425730, EPI_ISL_425731, EPI_ISL_425732, EPI_ISL_425733, EPI_ISL_425734, EPI_ISL_425735, EPI_ISL_425736, EPI_ISL_425737, EPI_ISL_425738, EPI_ISL_425739, EPI_ISL_425740, EPI_ISL_425741, EPI_ISL_425742, EPI_ISL_425743, EPI_ISL_425744, EPI_ISL_425745, EPI_ISL_425746, EPI_ISL_425747, EPI_ISL_425748, EPI_ISL_425749, EPI_ISL_425750, EPI_ISL_425751, EPI_ISL_425752, EPI_ISL_425753, EPI_ISL_425754, EPI_ISL_425755, EPI_ISL_425756, EPI_ISL_425757, EPI_ISL_425758, EPI_ISL_425759, EPI_ISL_425760, EPI_ISL_425761, EPI_ISL_425762, EPI_ISL_425763, EPI_ISL_425764, EPI_ISL_425765, EPI_ISL_425766, EPI_ISL_425767, EPI_ISL_425768, EPI_ISL_425769, EPI_ISL_425770, EPI_ISL_425771, EPI_ISL_425772, EPI_ISL_425773, EPI_ISL_425774, EPI_ISL_425775, EPI_ISL_425776, EPI_ISL_425777, EPI_ISL_425778, EPI_ISL_425779, EPI_ISL_425780, EPI_ISL_425781, EPI_ISL_425782, EPI_ISL_425783, EPI_ISL_425784, EPI_ISL_425785, EPI_ISL_425786, EPI_ISL_425787, EPI_ISL_425788, EPI_ISL_425789, EPI_ISL_425790, EPI_ISL_425791, EPI_ISL_425792, EPI_ISL_425793, EPI_ISL_425794, EPI_ISL_425795, EPI_ISL_425796, EPI_ISL_425797, EPI_ISL_425798, EPI_ISL_425799, EPI_ISL_425800, EPI_ISL_425801, EPI_ISL_425802, EPI_ISL_425803, EPI_ISL_425804, EPI_ISL_425805, EPI_ISL_425806, EPI_ISL_425807, EPI_ISL_425808, EPI_ISL_425809, EPI_ISL_425810, EPI_ISL_425811, EPI_ISL_425812, EPI_ISL_425813, EPI_ISL_425814, EPI_ISL_425815, EPI_ISL_425816, EPI_ISL_425817, EPI_ISL_425818 | | | |
| see above | West of Scotland Specialist Virology Centre, NHSGGC / MRC-University of Glasgow Centre for Virus Research | COVID-19 Genomics UK (COG-UK) Consortium | Ana da Silva Filipe, Kathy Smollett, Stephen Carmichael, Natasha Johnson, Daniel Mair, Lily Tong, Jenna Nichols; Sarah McDonald; Richard Orton, Joseph Hughes, Sreenu Vattipally, David L Robertson; Kathy Li, Natasha Jesudason, Rajiv Shah, James Shepherd, Antonia Ho, Emma Thomson; Alasdair MacLean, Rory Gunson. |
| EPI_ISL_425819, EPI_ISL_425820, EPI_ISL_425821, EPI_ISL_425822, EPI_ISL_425823, EPI_ISL_425824, EPI_ISL_425825, EPI_ISL_425826, EPI_ISL_425827, EPI_ISL_425828, EPI_ISL_425829, EPI_ISL_425830, EPI_ISL_425831, EPI_ISL_425832, EPI_ISL_425833, EPI_ISL_425834, EPI_ISL_425835, EPI_ISL_425836, EPI_ISL_425837, EPI_ISL_425838, EPI_ISL_425839, EPI_ISL_425840, EPI_ISL_425841, EPI_ISL_425842, EPI_ISL_425843, EPI_ISL_425844, EPI_ISL_425845, EPI_ISL_425846, EPI_ISL_425847, EPI_ISL_425848, EPI_ISL_425849, EPI_ISL_425850, EPI_ISL_425851, EPI_ISL_425852, EPI_ISL_425853, EPI_ISL_425854, EPI_ISL_425855, EPI_ISL_425856, EPI_ISL_425857, EPI_ISL_425858, EPI_ISL_425859, EPI_ISL_425860, EPI_ISL_425861, EPI_ISL_425862, EPI_ISL_425863, EPI_ISL_425864, EPI_ISL_425865, EPI_ISL_425866, EPI_ISL_425867, EPI_ISL_425868, EPI_ISL_425869, EPI_ISL_425870, EPI_ISL_425871, EPI_ISL_425872, EPI_ISL_425873, EPI_ISL_425874, EPI_ISL_425875, EPI_ISL_425876, EPI_ISL_425877, EPI_ISL_425878, EPI_ISL_425879, EPI_ISL_425880, EPI_ISL_425881, EPI_ISL_425882, EPI_ISL_425883, EPI_ISL_425884, EPI_ISL_425885, EPI_ISL_425886, EPI_ISL_425887, EPI_ISL_425888, EPI_ISL_425889, EPI_ISL_425890, EPI_ISL_425891, EPI_ISL_425892, EPI_ISL_425893, EPI_ISL_425894, EPI_ISL_425895, EPI_ISL_425896, EPI_ISL_425897, EPI_ISL_425898, EPI_ISL_425899, EPI_ISL_425900, EPI_ISL_425901, EPI_ISL_425902, EPI_ISL_425903, EPI_ISL_425904, EPI_ISL_425905, EPI_ISL_425906, EPI_ISL_425907, EPI_ISL_425908, EPI_ISL_425909, EPI_ISL_425910, EPI_ISL_425911, EPI_ISL_425912, EPI_ISL_425913, EPI_ISL_425914, EPI_ISL_425915, EPI_ISL_425916, EPI_ISL_425917, EPI_ISL_425918, EPI_ISL_425919, EPI_ISL_425920, EPI_ISL_425921, EPI_ISL_425922, EPI_ISL_425923, EPI_ISL_425924, EPI_ISL_425925, EPI_ISL_425926, EPI_ISL_425927, EPI_ISL_425928, EPI_ISL_425929, EPI_ISL_425930, EPI_ISL_425931, EPI_ISL_425932, EPI_ISL_425933, EPI_ISL_425934, EPI_ISL_425935, EPI_ISL_425936, EPI_ISL_425937, EPI_ISL_425938, EPI_ISL_425939, EPI_ISL_425940, EPI_ISL_425941, EPI_ISL_425942, EPI_ISL_425943, EPI_ISL_425944, EPI_ISL_425945, EPI_ISL_425946, EPI_ISL_425947, EPI_ISL_425948, EPI_ISL_425949, EPI_ISL_425950, EPI_ISL_425951, EPI_ISL_425952, EPI_ISL_425953, EPI_ISL_425954, EPI_ISL_425955, EPI_ISL_425956, EPI_ISL_425957, EPI_ISL_425958, EPI_ISL_425959, EPI_ISL_425960, EPI_ISL_425961, EPI_ISL_425962, EPI_ISL_425963, EPI_ISL_425964, EPI_ISL_425965, EPI_ISL_425966, EPI_ISL_425967, EPI_ISL_425968, EPI_ISL_425969, EPI_ISL_425970, EPI_ISL_425971, EPI_ISL_425972, EPI_ISL_425973, EPI_ISL_425974, EPI_ISL_425975, EPI_ISL_425976, EPI_ISL_425977, EPI_ISL_425978, EPI_ISL_425979, EPI_ISL_425980, EPI_ISL_425981, EPI_ISL_425982, EPI_ISL_425983, EPI_ISL_425984, EPI_ISL_425985, EPI_ISL_425986, EPI_ISL_425987, EPI_ISL_425988, EPI_ISL_425989, EPI_ISL_425990, EPI_ISL_425991, EPI_ISL_425992, EPI_ISL_425993, EPI_ISL_425994, EPI_ISL_425995, EPI_ISL_425996, EPI_ISL_425997, EPI_ISL_425998, EPI_ISL_425999, EPI_ISL_426000, EPI_ISL_426001, EPI_ISL_426002, EPI_ISL_426003, EPI_ISL_426004, EPI_ISL_426005, EPI_ISL_426006, EPI_ISL_426007, EPI_ISL_426008, EPI_ISL_426009, EPI_ISL_426010, EPI_ISL_426011, EPI_ISL_426012, EPI_ISL_426013, EPI_ISL_426014, EPI_ISL_426015, EPI_ISL_426016, EPI_ISL_426017, EPI_ISL_426018, EPI_ISL_426019, EPI_ISL_426020, EPI_ISL_426021, EPI_ISL_426022, EPI_ISL_426023, EPI_ISL_426024 | | | |
| see above | Virology Department, Royal Infirmary of Edinburgh, NHS Lothian / School of Biological Sciences, University of Edinburgh / Institute of Genetics and Molecular Medicine, University of Edinburgh | COVID-19 Genomics UK (COG-UK) Consortium | McHugh M, Dewar R, Rooke S, Gallagher M, Balcaza C, O'Toole A, Hill V, McCrone JT, Colquhoun R, Yu X, Jackson B, Scher E, Rambaut A, Williams TC, Templeton K |
| EPI_ISL_426025, EPI_ISL_426026, EPI_ISL_426027, EPI_ISL_426028, EPI_ISL_426029, EPI_ISL_426030, EPI_ISL_426031, EPI_ISL_426032, EPI_ISL_426033, EPI_ISL_426034, EPI_ISL_426035, EPI_ISL_426036, EPI_ISL_426037, EPI_ISL_426038, EPI_ISL_426039, EPI_ISL_426040, EPI_ISL_426041, EPI_ISL_426042, EPI_ISL_426043, EPI_ISL_426044, EPI_ISL_426045, EPI_ISL_426046, EPI_ISL_426047, EPI_ISL_426048, EPI_ISL_426049, EPI_ISL_426050 | | | |
| see above | Wadsworth Center, New York State Department.of Health | Wadsworth Center, New York State Department.of Health | Kirsten St. George, Daryl M. Lamson, Sara Griesemer, Jonathan Plitnick, Navjot Singh, Matthew D. Shudt, Erica Lasek-Nesselquist |
| EPI_ISL_426051 | Laboratory of Molecular Genetics, 2nd Faculty of Medicine, Charles University in Prague, Prague, Czech Republic | Laboratory of Molecular Genetics, 2nd Faculty of Medicine, Charles University in Prague, Prague, Czech Republic | Lenka Kramná, Kateřina Poláčková, Ondřej Cinek |
| EPI_ISL_426052, EPI_ISL_426053, EPI_ISL_426054, EPI_ISL_426055, EPI_ISL_426056, EPI_ISL_426057, EPI_ISL_426058, EPI_ISL_426059, EPI_ISL_426060, EPI_ISL_426061, EPI_ISL_426062, EPI_ISL_426063, EPI_ISL_426064, EPI_ISL_426065, EPI_ISL_426066, EPI_ISL_426067, EPI_ISL_426068, EPI_ISL_426069, EPI_ISL_426070, EPI_ISL_426071, EPI_ISL_426072, EPI_ISL_426073, EPI_ISL_426074, EPI_ISL_426075, EPI_ISL_426076, EPI_ISL_426077, EPI_ISL_426078, EPI_ISL_426079, EPI_ISL_426080, EPI_ISL_426081, EPI_ISL_426082, EPI_ISL_426083, EPI_ISL_426084, EPI_ISL_426085, EPI_ISL_426086, EPI_ISL_426087, EPI_ISL_426088, EPI_ISL_426089, EPI_ISL_426090, EPI_ISL_426091, EPI_ISL_426092, EPI_ISL_426093, EPI_ISL_426094, EPI_ISL_426095, EPI_ISL_426096, EPI_ISL_426097, EPI_ISL_426098, EPI_ISL_426099, EPI_ISL_426100, EPI_ISL_426101, EPI_ISL_426102, EPI_ISL_426103, EPI_ISL_426104, EPI_ISL_426105, EPI_ISL_426106, EPI_ISL_426107, EPI_ISL_426108, EPI_ISL_426109, EPI_ISL_426110, EPI_ISL_426111, EPI_ISL_426112, EPI_ISL_426113, EPI_ISL_426114, EPI_ISL_426115, EPI_ISL_426116, EPI_ISL_426117, EPI_ISL_426118, EPI_ISL_426119, EPI_ISL_426120, EPI_ISL_426121, EPI_ISL_426122, EPI_ISL_426123, EPI_ISL_426124, EPI_ISL_426125, EPI_ISL_426126, EPI_ISL_426127, EPI_ISL_426128, EPI_ISL_426129, EPI_ISL_426130, EPI_ISL_426131, EPI_ISL_426132, EPI_ISL_426133, EPI_ISL_426134, EPI_ISL_426135, EPI_ISL_426136, EPI_ISL_426137 | | | |
| see above | UW Virology Lab | UW Virology Lab | Pavitra Roychoudhury, Hong Xie, Keith Jerome, Alexander Greninger |
| EPI_ISL_426159, EPI_ISL_426160, EPI_ISL_426161 | Gundersen Molecular Diagnostics Laboratory | Kabara Cancer Research Institute | Craig S. Richmond, Paraic A. Kenny |
| EPI_ISL_426163 | Division of Viral Diseases, Center for Laboratory Control of Infectious Diseases, Korea Centers for Diseases Control and Prevention | Division of Viral Diseases, Center for Laboratory Control of Infectious Diseases, Korea Centers for Diseases Control and Prevention | Jeong-Min Kim, Yoon-Seok Chung, Namjoo Lee, Mi-Seon Kim, Sang Hee Woo, Hye-Jun Jo, Sehee Park, Heui Man Kim, Jun-Sub Kim, Junhyeong Jang, Myung Guk Han |
| EPI_ISL_426164 | Division of Viral Diseases, Center for Laboratory Control of Infectious Diseases, Korea Centers for Diseases Control and Prevention | Division of Viral Diseases, Center for Laboratory Control of Infectious Diseases, Korea Centers for Diseases Control and Prevention | Jeong-Min Kim, Yoon-Seok Chung, Namjoo Lee, Mi-Seon Kim, Sang Hee Woo, Hye-Jun Jo, Sehee Park, Heui Man Kim, Jun-Sub Kim, Junhyeong Jang, Dong Hyun Song, Daesang Lee, Seong Tae Jeong, Myung Guk Han |
| EPI_ISL_426166, EPI_ISL_426167, EPI_ISL_426168 | Division of Viral Diseases, Center for Laboratory Control of Infectious Diseases, Korea Centers for Diseases Control and Prevention | Division of Viral Diseases, Center for Laboratory Control of Infectious Diseases, Korea Centers for Diseases Control and Prevention | Jeong-Min Kim, Yoon-Seok Chung, Namjoo Lee, Mi-Seon Kim, Sang Hee Woo, Hye-Jun Jo, Sehee Park, Heui Man Kim, Jun-Sub Kim, Junhyeong Jang, Myung Guk Han |
| EPI_ISL_426169, EPI_ISL_426171 | Division of Viral Diseases, Center for Laboratory Control of Infectious Diseases, Korea Centers for Diseases Control and Prevention | Division of Viral Diseases, Center for Laboratory Control of Infectious Diseases, Korea Centers for Diseases Control and Prevention | Jeong-Min Kim, Yoon-Seok Chung, Namjoo Lee, Mi-Seon Kim, Sang Hee Woo, Hye-Jun Jo, Sehee Park, Heui Man Kim, Jun-Sub Kim, Junhyeong Jang, Dong Hyun Song, Daesang Lee, Seong Tae Jeong, Myung Guk Han |
| EPI_ISL_426173 | Division of Viral Diseases, Center for Laboratory Control of Infectious Diseases, Korea Centers for Diseases Control and Prevention | Division of Viral Diseases, Center for Laboratory Control of Infectious Diseases, Korea Centers for Diseases Control and Prevention | Jeong-Min Kim, Yoon-Seok Chung, Namjoo Lee, Mi-Seon Kim, Sang Hee Woo, Hye-Jun Jo, Sehee Park, Heui Man Kim, Jun-Sub Kim, Junhyeong Jang, Myung Guk Han |
| EPI_ISL_426179 | National Influenza Center, Indian Council of Medical Research - National Institute of Virology | Indian Council of Medical Research-National Institute of Virology, Microbial Containment Complex | Pragya D. Yadav, Varsha Potdar, Savita Patil, Dimpal A. Nyayanit, Triparna Majumdar, Manohar. L. Chaudhary, Gururaj Deshpande, Padinjaremattathil Thankappan Ullas, Anita Shete-Aich, Hitesh Dighe, Sreelekshmy Mohandas, Gajanan Sapkal, Atanu Basu, Amita Jain, Bharti Malhotra, Deepika Chaudhary, Sarah Cherian, Priya Abraham |
| EPI_ISL_426180, EPI_ISL_426181, EPI_ISL_426182, EPI_ISL_426183, EPI_ISL_426187 | Division of Viral Diseases, Center for Laboratory Control of Infectious Diseases, Korea Centers for Diseases Control and Prevention | Division of Viral Diseases, Center for Laboratory Control of Infectious Diseases, Korea Centers for Diseases Control and Prevention | Jeong-Min Kim, Yoon-Seok Chung, Namjoo Lee, Mi-Seon Kim, Sang Hee Woo, Hye-Jun Jo, Sehee Park, Heui Man Kim, Jun-Sub Kim, Junhyeong Jang, Myung Guk Han |

| | | | |
|---|---|---|---|
| EPI_ISL_426285, EPI_ISL_426286, EPI_ISL_426287, EPI_ISL_426288, EPI_ISL_426289 | E. Gulbja Laboratorija | Latvian Biomedical Research and Study Centre | Ivars Silamiķelis, Kaspars Megnis, Monta Ustinova, Ņikita Zrelovs, Vita Rovīte, Mikus Gavars, Dmitrijs Perminovs, Uga Dumpis, Jānis Kloviņš |
| EPI_ISL_426290 | Wadsworth Center, New York State Department of Health | Wadsworth Center, New York State Department.of Health | Kirsten St. George, Daryl M. Lamson, Sara Griesemer, Jonathan Plitnick, Navjot Singh, Matthew D. Shudt, Erica Lasek-Nesselquist |
| EPI_ISL_426291, EPI_ISL_426292, EPI_ISL_426293, EPI_ISL_426294, EPI_ISL_426295, EPI_ISL_426296, EPI_ISL_426297, EPI_ISL_426298, EPI_ISL_426299, EPI_ISL_426300, EPI_ISL_426301, EPI_ISL_426302, EPI_ISL_426303, EPI_ISL_426304, EPI_ISL_426305, EPI_ISL_426306, EPI_ISL_426307, EPI_ISL_426308, EPI_ISL_426309, EPI_ISL_426310, EPI_ISL_426311, EPI_ISL_426312, EPI_ISL_426313, EPI_ISL_426314, EPI_ISL_426315, EPI_ISL_426316, EPI_ISL_426317, EPI_ISL_426318, EPI_ISL_426319, EPI_ISL_426320, EPI_ISL_426321, EPI_ISL_426322, EPI_ISL_426323, EPI_ISL_426324, EPI_ISL_426325, EPI_ISL_426326, EPI_ISL_426327, EPI_ISL_426328 | | | |
| see above | Wadsworth Center, New York State Department.of Health | Wadsworth Center, New York State Department.of Health | Kirsten St. George, Daryl M. Lamson, Sara Griesemer, Jonathan Plitnick, Navjot Singh, Matthew D. Shudt, Erica Lasek-Nesselquist |
| EPI_ISL_426356, EPI_ISL_426357, EPI_ISL_426358, EPI_ISL_426359, EPI_ISL_426360 | Laboratory of Molecular Genetics, 2nd Faculty of Medicine, Charles University in Prague, Prague, Czech Republic | Laboratory of Molecular Genetics, 2nd Faculty of Medicine, Charles University in Prague, Prague, Czech Republic | Lenka Kramna, Katerina Polackova, Ondrej Cinek |
| EPI_ISL_426361, EPI_ISL_426362, EPI_ISL_426363 | Instituto Nacional de Ciencias Medicas y Nutricion Salvador Zubiran | Instituto Nacional de Ciencias Medicas y Nutricion Salvador Zubiran | Guillermo M. Ruiz-Palacios, Pilar Ramos Cervantes, Violeta Ibarra Gonzalez, Fernando Ledesma Barrientos, Luis Alberto García Andrade, Alfredo Ponce de León Garduño, Irma López Martínez, Lucia Hernández Rivas, Gisela Barrera Badillo, Edgar Mendieta Condado, Fabiola Garcés Ayala, Adnan Araiza Rodríguez, José Ernesto Ramírez González, Celia Boukadida, Santiago Avila Ríos, Mario Mújica Sánchez, José Arturo Martínez Orozco, Eduardo Becerril Vargas, Joel Armando Vázquez Pérez, Victor Hugo Borja Aburto, Concepción Grajales Muñiz, Cesar Raúl González Bonilla, Carolina González Torres, Francisco Javier Gaytán Cervantes, José Esteban Muñoz Medina, Blanca Taboada, Alejandro Sánchez, Pavel Isa, Ricardo Grande, Gloria Vázquez, Francisco Pulido, Carlos F. Arias |
| EPI_ISL_426364 | Instituto Nacional de Ciencias Medicas y Nutricion Salvador Zubiran | Instituto Nacional de Ciencias Medicas y Nutricion | Guillermo M. Ruiz-Palacios, Pilar Ramos Cervantes, Violeta Ibarra Gonzalez, Fernando Ledesma Barrientos, Luis Alberto García Andrade, Alfredo Ponce de León Garduño, Irma López Martínez, Lucia Hernández Rivas, Gisela Barrera Badillo, Edgar Mendieta Condado, Fabiola Garcés Ayala, Adnan Araiza Rodríguez, José Ernesto Ramírez González, Celia Boukadida, Santiago Avila Ríos, Mario Mújica Sánchez, José Arturo Martínez Orozco, Eduardo Becerril Vargas, Joel Armando Vázquez Pérez, Victor Hugo Borja Aburto, Concepción Grajales Muñiz, Cesar Raúl González Bonilla, Carolina González Torres, Francisco Javier Gaytán Cervantes, José Esteban Muñoz Medina, Blanca Taboada, Alejandro Sánchez, Pavel Isa, Ricardo Grande, Gloria Vázquez, Francisco Pulido, Carlos F. Arias |
| EPI_ISL_426365 | Instituto Nacional de Ciencias Medicas y Nutricion Salvador Zubiran | Instituto Nacional de Ciencias Medicas y Nutricion Salvador Zubiran | Guillermo M. Ruiz-Palacios, Pilar Ramos Cervantes, Violeta Ibarra Gonzalez, Fernando Ledesma Barrientos, Luis Alberto García Andrade, Alfredo Ponce de León Garduño, Irma López Martínez, Lucia Hernández Rivas, Gisela Barrera Badillo, Edgar Mendieta Condado, Fabiola Garcés Ayala, Adnan Araiza Rodríguez, José Ernesto Ramírez González, Celia Boukadida, Santiago Avila Ríos, Mario Mújica Sánchez, José Arturo Martínez Orozco, Eduardo Becerril Vargas, Joel Armando Vázquez Pérez, Victor Hugo Borja Aburto, Concepción Grajales Muñiz, Cesar Raúl González Bonilla, Carolina González Torres, Francisco Javier Gaytán Cervantes, José Esteban Muñoz Medina, Blanca Taboada, Alejandro Sánchez, Pavel Isa, Ricardo Grande, Gloria Vázquez, Francisco Pulido, Carlos F. Arias |
| EPI_ISL_426379 | The National Laboratory of Health, Environment and Food, Maribor, Slovenia | The National Laboratory of Health, Environment and Food, Maribor, Slovenia | Mahnic A., Hedzet S., Janezic S., Duh D., Zavrsnik J., Blazun Vosner H., Rupnik M. |
| EPI_ISL_426380 | Hong Kong Sanatorium & Hospital | Hong Kong Department of Health | Mak Gannon C.K., Cheng Peter K.C., Lam Edman T.K., Chan Rickjason C.W., Tsang Dominic N.C. |
| EPI_ISL_426381 | Ruttonjee Hospital | Hong Kong Department of Health | Mak Gannon C.K., Cheng Peter K.C., Lam Edman T.K., Chan Rickjason C.W., Tsang Dominic N.C. |
| EPI_ISL_426382 | Ruttonjee Hospital | Hong Kong Department of Health | Mak Gannon C.K., Cheng Peter K.C., Lam Edman T.K., Chan Rickjason C.W., Tsang Dominic N.C. |
| EPI_ISL_426383 | Pamela Youde Nethersole Eastern Hospital | Hong Kong Department of Health | Mak Gannon C.K., Cheng Peter K.C., Lam Edman T.K., Chan Rickjason C.W., Tsang Dominic N.C. |
| EPI_ISL_426384 | Ruttonjee Hospital | Hong Kong Department of Health | Mak Gannon C.K., Cheng Peter K.C., Lam Edman T.K., Chan Rickjason C.W., Tsang Dominic N.C. |
| EPI_ISL_426385 | Pamela Youde Nethersole Eastern Hospital | Hong Kong Department of Health | Mak Gannon C.K., Cheng Peter K.C., Lam Edman T.K., Chan Rickjason C.W., Tsang Dominic N.C. |
| EPI_ISL_426386 | Ruttonjee Hospital | Hong Kong Department of Health | Mak Gannon C.K., Cheng Peter K.C., Lam Edman T.K., Chan Rickjason C.W., Tsang Dominic N.C. |
| EPI_ISL_426387, EPI_ISL_426388 | Pamela Youde Nethersole Eastern Hospital | Hong Kong Department of Health | Mak Gannon C.K., Cheng Peter K.C., Lam Edman T.K., Chan Rickjason C.W., Tsang Dominic N.C. |
| EPI_ISL_426389 | Ruttonjee Hospital | Hong Kong Department of Health | Mak Gannon C.K., Cheng Peter K.C., Lam Edman T.K., Chan Rickjason C.W., Tsang Dominic N.C. |
| EPI_ISL_426390 | Prince of Wales Hospital | Hong Kong Department of Health | Mak Gannon C.K., Cheng Peter K.C., Lam Edman T.K., Chan Rickjason C.W., Tsang Dominic N.C. |
| EPI_ISL_426391 | Pamela Youde Nethersole Eastern Hospital | Hong Kong Department of Health | Mak Gannon C.K., Cheng Peter K.C., Lam Edman T.K., Chan Rickjason C.W., Tsang Dominic N.C. |
| EPI_ISL_426392 | Ruttonjee Hospital | Hong Kong Department of Health | Mak Gannon C.K., Cheng Peter K.C., Lam Edman T.K., Chan Rickjason C.W., Tsang Dominic N.C. |
| EPI_ISL_426393 | Kwong Wah Hospital | Hong Kong Department of Health | Mak Gannon C.K., Cheng Peter K.C., Lam Edman T.K., Chan Rickjason C.W., Tsang Dominic N.C. |
| EPI_ISL_426394 | Queen Elizabeth Hospital | Hong Kong Department of Health | Mak Gannon C.K., Cheng Peter K.C., Lam Edman T.K., Chan Rickjason C.W., Tsang Dominic N.C. |
| EPI_ISL_426395, EPI_ISL_426396 | Pamela Youde Nethersole Eastern Hospital | Hong Kong Department of Health | Mak Gannon C.K., Cheng Peter K.C., Lam Edman T.K., Chan Rickjason C.W., Tsang Dominic N.C. |
| EPI_ISL_426397 | Queen Mary Hospital | Hong Kong Department of Health | Mak Gannon C.K., Cheng Peter K.C., Lam Edman T.K., Chan Rickjason C.W., Tsang Dominic N.C. |
| EPI_ISL_426398, EPI_ISL_426399 | Ruttonjee Hospital | Hong Kong Department of Health | Mak Gannon C.K., Cheng Peter K.C., Lam Edman T.K., Chan Rickjason C.W., Tsang Dominic N.C. |
| EPI_ISL_426400 | Pamela Youde Nethersole Eastern Hospital | Hong Kong Department of Health | Mak Gannon C.K., Cheng Peter K.C., Lam Edman T.K., Chan Rickjason C.W., Tsang Dominic N.C. |
| EPI_ISL_426401 | United Christian Hospital | Hong Kong Department of Health | Mak Gannon C.K., Cheng Peter K.C., Lam Edman T.K., Chan Rickjason C.W., Tsang Dominic N.C. |
| EPI_ISL_426402 | Queen Mary Hospital | Hong Kong Department of Health | Mak Gannon C.K., Cheng Peter K.C., Lam Edman T.K., Chan Rickjason C.W., Tsang Dominic N.C. |
| EPI_ISL_426403 | United Christian Hospital | Hong Kong Department of Health | Mak Gannon C.K., Cheng Peter K.C., Lam Edman T.K., Chan Rickjason C.W., Tsang Dominic N.C. |
| EPI_ISL_426404 | New Territories Families Clinic | Hong Kong Department of Health | Mak Gannon C.K., Cheng Peter K.C., Lam Edman T.K., Chan Rickjason C.W., Tsang Dominic N.C. |
| EPI_ISL_426405 | Queen Mary Hospital | Hong Kong Department of Health | Mak Gannon C.K., Cheng Peter K.C., Lam Edman T.K., Chan Rickjason C.W., Tsang Dominic N.C. |
| EPI_ISL_426406 | Queen Elizabeth Hospital | Hong Kong Department of Health | Mak Gannon C.K., Cheng Peter K.C., Lam Edman T.K., Chan Rickjason C.W., Tsang Dominic N.C. |
| EPI_ISL_426407 | Caritas Medical Centre | Hong Kong Department of Health | Mak Gannon C.K., Cheng Peter K.C., Lam Edman T.K., Chan Rickjason C.W., Tsang Dominic N.C. |
| EPI_ISL_426408 | Ruttonjee Hospital | Hong Kong Department of Health | Mak Gannon C.K., Cheng Peter K.C., Lam Edman T.K., Chan Rickjason C.W., Tsang Dominic N.C. |
| EPI_ISL_426409 | Queen Mary Hospital | Hong Kong Department of Health | Mak Gannon C.K., Cheng Peter K.C., Lam Edman T.K., Chan Rickjason C.W., Tsang Dominic N.C. |
| EPI_ISL_426410 | Central Kowloon Health Centre | Hong Kong Department of Health | Mak Gannon C.K., Cheng Peter K.C., Lam Edman T.K., Chan Rickjason C.W., Tsang Dominic N.C. |
| EPI_ISL_426411, EPI_ISL_426412 | Pok Oi Hospital | Hong Kong Department of Health | Mak Gannon C.K., Cheng Peter K.C., Lam Edman T.K., Chan Rickjason C.W., Tsang Dominic N.C. |
| EPI_ISL_426414 | Sir M P Shah Government Medical College | Gujarat Biotechnology Research Centre | Ramesh Pandit, Tejas Shah, Ankit Hinsu, Pritesh Sabara, Apurvasinh Puvar, Janvi Raval, Monika Gandhi, Pinal Trivedi, Maharshi Pandya, Amit Kanani, Akanksha Verma, Nitin Savaliya, Raghawendra Kumar, Dinesh Kumar, Zubair Saiyed, Dipa Kinariwala, Disha Patel, Binita Aring, Geeta Vaghela, Sonia Barve, Bhavesh Modi, Kairavi Joshi, Nidhi Sood, Pranay Shah, R D Dixit, Snehal Bagatharia, Madhvi Joshi, Chaitanya Joshi |
| EPI_ISL_426415 | Sir M P Shah Government Medical College, Jamnagar | Gujarat Biotechnology Research Centre, Gandhinagar | Ramesh Pandit, Tejas Shah, Ankit Hinsu, Pritesh Sabara, Apurvasinh Puvar, Janvi Raval, Monika Gandhi, Pinal Trivedi, Maharshi Pandya, Amit Kanani, Akanksha Verma, Nitin Savaliya, Raghvendra Kumar, Dinesh Kumar, Zuber Saiyed, Dipa Kinariwala, Disha Patel, Binita Aring, Geeta Vaghela, Sonia Barve, Bhavesh Modi, Kairavi Joshi, Nidhi Sood, Pranay Shah, R D Dixit, Snehal Bagatharia, Madhvi Joshi, Chaitanya Joshi |
| EPI_ISL_426416 | CT-Dr. Katherine A. Kelley State Public Health Lab | Pathogen Discovery, Respiratory Viruses Branch, Division of Viral Diseases, Centers for Disease Control and Prevention | Anna Uehara, Yan Li, Krista Queen, Clinton R. Paden, Rachel Marine, Ying Tao, Jing Zhang, Haibin Wang, Mary S. Keckler, Alison S. Laufer Halpin, Christopher A. Elkins, Suxiang Tong |
| EPI_ISL_426417, EPI_ISL_426418, EPI_ISL_426419 | GA Department of Public Health Laboratory | Pathogen Discovery, Respiratory Viruses Branch, Division of Viral Diseases, Centers for Disease Control and Prevention | Anna Uehara, Yan Li, Krista Queen, Clinton R. Paden, Rachel Marine, Ying Tao, Jing Zhang, Haibin Wang, Mary S. Keckler, Alison S. Laufer Halpin, Christopher A. Elkins, Suxiang Tong |
| EPI_ISL_426420, EPI_ISL_426421 | HI Dept. of Health, State Laboratories Division | Pathogen Discovery, Respiratory Viruses Branch, Division of Viral Diseases, Centers for Disease Control and Prevention | Anna Uehara, Yan Li, Krista Queen, Clinton R. Paden, Rachel Marine, Ying Tao, Jing Zhang, Haibin Wang, Mary S. Keckler, Alison S. Laufer Halpin, Christopher A. Elkins, Suxiang Tong |
| EPI_ISL_426422, EPI_ISL_426423, EPI_ISL_426424 | IN State Department of Health Laboratory Services | Pathogen Discovery, Respiratory Viruses Branch, Division of Viral Diseases, Centers for Disease Control and Prevention | Krista Queen, Yan Li, Anna Uehara, Clinton R. Paden, Rachel Marine, Ying Tao, Jing Zhang, Haibin Wang, Mary S. Keckler, Alison S. Laufer Halpin, Christopher A. Elkins, Suxiang Tong |
| EPI_ISL_426425 | MD DOH Laboratories Administration | Pathogen Discovery, Respiratory Viruses Branch, Division of Viral Diseases, Centers for Disease Control and Prevention | Krista Queen, Yan Li, Anna Uehara, Clinton R. Paden, Rachel Marine, Ying Tao, Jing Zhang, Haibin Wang, Mary S. Keckler, Alison S. Laufer Halpin, Christopher A. Elkins, Suxiang Tong |
| EPI_ISL_426426, EPI_ISL_426427 | MN PHL Division, Minnesota Department of Health | Pathogen Discovery, Respiratory Viruses Branch, Division of Viral Diseases, Centers for Disease Control and Prevention | Krista Queen, Yan Li, Anna Uehara, Clinton R. Paden, Rachel Marine, Ying Tao, Jing Zhang, Haibin Wang, Mary S. Keckler, Alison S. Laufer Halpin, Christopher A. Elkins, Suxiang Tong |
| EPI_ISL_426428 | NC State Laboratory of Public Health | Pathogen Discovery, Respiratory Viruses Branch, Division of Viral Diseases, Centers for Disease Control and Prevention | Krista Queen, Yan Li, Anna Uehara, Clinton R. Paden, Rachel Marine, Ying Tao, Jing Zhang, Haibin Wang, Mary S. Keckler, Alison S. Laufer Halpin, Christopher A. Elkins, Suxiang Tong |
| EPI_ISL_426429 | NV State Public Health Laboratory | Pathogen Discovery, Respiratory Viruses Branch, Division of Viral Diseases, Centers for Disease Control and Prevention | Krista Queen, Yan Li, Anna Uehara, Clinton R. Paden, Rachel Marine, Ying Tao, Jing Zhang, Haibin Wang, Mary S. Keckler, Alison S. Laufer Halpin, Christopher A. Elkins, Suxiang Tong |
| EPI_ISL_426430, EPI_ISL_426431 | OH Department of Health Laboratory | Pathogen Discovery, Respiratory Viruses Branch, Division of Viral Diseases, Centers for Disease Control and Prevention | Krista Queen, Yan Li, Anna Uehara, Clinton R. Paden, Rachel Marine, Ying Tao, Jing Zhang, Haibin Wang, Mary S. Keckler, Alison S. Laufer Halpin, Christopher A. Elkins, Suxiang Tong |
| EPI_ISL_426432, EPI_ISL_426433, EPI_ISL_426434 | PA Department of Health, Bureau of Laboratories | Pathogen Discovery, Respiratory Viruses Branch, Division of Viral Diseases, Centers for Disease Control and Prevention | Krista Queen, Yan Li, Anna Uehara, Clinton R. Paden, Rachel Marine, Ying Tao, Jing Zhang, Haibin Wang, Mary S. Keckler, Alison S. Laufer Halpin, Christopher A. Elkins, Suxiang Tong |
| EPI_ISL_426435 | RI State Health Laboratories | Pathogen Discovery, Respiratory Viruses Branch, Division of Viral Diseases, Centers for Disease Control and Prevention | Krista Queen, Yan Li, Anna Uehara, Clinton R. Paden, Rachel Marine, Ying Tao, Jing Zhang, Haibin Wang, Mary S. Keckler, Alison S. Laufer Halpin, Christopher A. Elkins, Suxiang Tong |
| EPI_ISL_426436 | WA State Department of Health | Pathogen Discovery, Respiratory Viruses Branch, Division of Viral Diseases, Centers for Disease Control and Prevention | Jing Zhang, Ying Tao, Clinton R. Paden, Krista Queen, Anna Uehara, Yan Li, Haibin Wang, Jesica Jacobs, Denny Russell, Brian Hiatt, Jessica Gant, Suxiang Tong |
| EPI_ISL_426437 | WA State Department of Health | Pathogen Discovery, Respiratory Viruses Branch, Division of Viral Diseases, Centers for Disease Control and Prevention | Ying Tao, Jing Zhang, Clinton R. Paden, Krista Queen, Anna Uehara, Yan Li, Haibin Wang, Jesica Jacobs, Denny Russell, Brian Hiatt, Jessica Gant, Suxiang Tong |
| EPI_ISL_426438, EPI_ISL_426439 | WA State Department of Health | Pathogen Discovery, Respiratory Viruses Branch, Division of | Jing Zhang, Ying Tao, Clinton R. Paden, Krista Queen, Anna Uehara, Yan Li, Haibin Wang, Jesica Jacobs, Denny Russell, Brian Hiatt, Jessica Gant, Suxiang Tong |

| | | | |
|---|---|---|---|
| | | Viral Diseases, Centers for Disease Control and Prevention | |
| EPI_ISL_426440, EPI_ISL_426441 | WA State Department of Health | Pathogen Discovery, Respiratory Viruses Branch, Division of Viral Diseases, Centers for Disease Control and Prevention | Ying Tao, Jing Zhang, Clinton R. Paden, Krista Queen, Anna Uehara, Yan Li, Haibin Wang, Jesica Jacobs, Denny Russell, Brian Hiatt, Jessica Gant, Suxiang Tong |
| EPI_ISL_426442 | WA State Department of Health | Pathogen Discovery, Respiratory Viruses Branch, Division of Viral Diseases, Centers for Disease Control and Prevention | Jing Zhang, Ying Tao, Clinton R. Paden, Krista Queen, Anna Uehara, Yan Li, Haibin Wang, Jesica Jacobs, Denny Russell, Brian Hiatt, Jessica Gant, Suxiang Tong |
| EPI_ISL_426443, EPI_ISL_426444 | WA State Department of Health | Pathogen Discovery, Respiratory Viruses Branch, Division of Viral Diseases, Centers for Disease Control and Prevention | Ying Tao, Jing Zhang, Clinton R. Paden, Krista Queen, Anna Uehara, Yan Li, Haibin Wang, Jesica Jacobs, Denny Russell, Brian Hiatt, Jessica Gant, Suxiang Tong |
| EPI_ISL_426445 | WA State Department of Health | Pathogen Discovery, Respiratory Viruses Branch, Division of Viral Diseases, Centers for Disease Control and Prevention | Jing Zhang, Ying Tao, Clinton R. Paden, Krista Queen, Anna Uehara, Yan Li, Haibin Wang, Jesica Jacobs, Denny Russell, Brian Hiatt, Jessica Gant, Suxiang Tong |
| EPI_ISL_426446, EPI_ISL_426447, EPI_ISL_426448, EPI_ISL_426449 | WA State Department of Health | Pathogen Discovery, Respiratory Viruses Branch, Division of Viral Diseases, Centers for Disease Control and Prevention | Ying Tao, Jing Zhang, Clinton R. Paden, Krista Queen, Anna Uehara, Yan Li, Haibin Wang, Jesica Jacobs, Denny Russell, Brian Hiatt, Jessica Gant, Suxiang Tong |
| EPI_ISL_426450 | WA State Department of Health | Pathogen Discovery, Respiratory Viruses Branch, Division of Viral Diseases, Centers for Disease Control and Prevention | Jing Zhang, Ying Tao, Clinton R. Paden, Krista Queen, Anna Uehara, Yan Li, Haibin Wang, Jesica Jacobs, Denny Russell, Brian Hiatt, Jessica Gant, Suxiang Tong |
| EPI_ISL_426451, EPI_ISL_426452, EPI_ISL_426453 | WA State Department of Health | Pathogen Discovery, Respiratory Viruses Branch, Division of Viral Diseases, Centers for Disease Control and Prevention | Ying Tao, Jing Zhang, Clinton R. Paden, Krista Queen, Anna Uehara, Yan Li, Haibin Wang, Jesica Jacobs, Denny Russell, Brian Hiatt, Jessica Gant, Suxiang Tong |
| EPI_ISL_434560 | Department of Microbiology, The University of Hong Kong | Department of Microbiology, The University of Hong Kong | Lau,S.K.P., Luk,H.K.H., Wong,A.C.P., Li,K.S.M., Zhu,L., He,Z., Fung,J., Chan,T.T.Y., Fung,K.S.C. and Woo,P.C.Y. |
| EPI_ISL_434563, EPI_ISL_434564, EPI_ISL_434565, EPI_ISL_434566, EPI_ISL_434567, EPI_ISL_434568, EPI_ISL_434569 | unknown | Microbiology, The University of Hong Kong | To,K.K.W. and Yuen,K.-Y. |
| EPI_ISL_434571 | Microbiology, The University of Hong Kong | Microbiology, The University of Hong Kong | Chan,J.F.W. and Yuen,K.-Y. |
| EPI_ISL_450198, EPI_ISL_450199, EPI_ISL_450200, EPI_ISL_450201, EPI_ISL_450202, EPI_ISL_450203, EPI_ISL_450204, EPI_ISL_450205, EPI_ISL_450206, EPI_ISL_450207, EPI_ISL_450208, EPI_ISL_450209, EPI_ISL_450210, EPI_ISL_450211 | | | |
| see above | Department of Virology | Department of Virology | Boehmer,M.M., Buchholz,U., Corman,V.M., Hoch,M., Katz,K., Marosevic,D.V., Boehm,S., Woudenberg,T., Ackermann,N., Konrad,R., Eberle,U., Treis,B., Dangel,A., Bengs,K., Fingerle,V., Berger,A., Hoermansdorfer,S., Ippisch,S., Wicklein,B., Grahl,A., Poertner,K., Muller,N., Zeitlmann,N., Boender,T.S., Cai,W., Reich,A., an der Heiden,M., Rexroth,U., Hamouda,O., Schneider,J., Veith,T., Muehlemann,B., Woelfel,R., Antwerpen,M., Walter,M., Protzer,U., Liebl,B., Haas,W., Sing,A., Drosten,C., Zapf,A., Jones,T.C. |
| EPI_ISL_450403 | School of Public Health, The University of Hong Kong | School of Public Health, The University of Hong Kong | Sit,T.H.S., Brackman,C.J., Sims,L.D., Tsang,D.N.C., Chu,D.K.W., Perera,R.A.P.M., Poon,L.L.M. and Peiris,M. |
| EPI_ISL_450404, EPI_ISL_450405 | unknown | School of Public Health, The University of Hong Kong | Sit,T.H.S., Brackman,C.J., Sims,L.D., Tsang,D.N.C., Chu,D.K.W., Perera,R.A.P.M., Poon,L.L.M. and Peiris,M. |
| EPI_ISL_450413 | Center for Diagnostics, Institute of Medical Microbiology, Virology and Hygiene | University Medical Center Hamburg-Eppendorf | Huang,J., Pfefferle,S. and Fischer,N. |
| EPI_ISL_450444 | 20 Dongda Street, Fengtai District, Beijing, Beijing 100071, China | Dept. OPA, Beijing Institute of Microbiology and Epidemiology | Zhang,X.A., Fan,H., Qi,R.Z., Zheng,W., Zheng,K., Gong,J.H., Fang,L.Q. and Liu,W. |
| EPI_ISL_450482 | National Influenza and other Respiratory Viruses Centre-Tunisia, Virology Unit, Microbiology Laboratory, Charles Nicolle Hospital | National Influenza and other Respiratory Viruses Centre-Tunisia, Virology Unit, Microbiology Laboratory, Charles Nicolle Hospital | El Moussi,A., Abid,S., Ben Nasr,M., Landolsi,I., Charaa,L., Enigrou,D. and Boutiba,I. |
| EPI_ISL_450483 | Molecular Genetic, Immuno Gene Center | Molecular Genetic, Immuno Gene Center | Dlovan,M.F., Haval,F.M., Hazha,H.J. and Ariamand,A. |
| EPI_ISL_450488 | Institute of Biomedical & Genetic Engineering | Institute of Biomedical & Genetic Engineering | Hashmi,A.H., Ajmal,M. and Ahmad,N. |
| EPI_ISL_450489 | Wuhan Institute of Virology, Chinese Academy of Sciences | Wuhan Institute of Virology, Chinese Academy of Sciences | Si,H., Zhu,Y., Lin,H., Xie,S., Shi,Z. and Zhou,P. |
| EPI_ISL_450490, EPI_ISL_450491, EPI_ISL_450492, EPI_ISL_450493, EPI_ISL_450494 | unknown | National Influenza and other Respiratory Viruses Centre-Tunisia | El Moussi,A., Abid,S., Ben Nasr,M., Landolsi,I., Charaa,L., Ferjeni,A., Arab Ennigrou,D., Boutiba,I. |
| EPI_ISL_450500, EPI_ISL_450501, EPI_ISL_450502, EPI_ISL_450503, EPI_ISL_450504 | unknown | CAS Key Laboratory of Special Pathogens and Biosafety and Center for Emerging Infectious Diseases | Si,H., Zhu,Y., Lin,H., Xie,S., Shi,Z., Zhou,P. |
| EPI_ISL_450506 | Clinical Laboratory, Hospital Israelita Albert Einstein | Clinical Laboratory, Hospital Israelita Albert Einstein | Malta,F., Amgarten,D., de Oliveira,D.B.L., Araujo,D.B., Machado,R.R.G., Santana,R.A.F., Mangueira,C.L.P., Durigon,E.L. and Pinho,J.R.R. |
| EPI_ISL_450507 | Nigerian Institute of Medical Research | Nigerian Institute of Medical Research | Shaibu,J.O., Onwuamah,C.K., James,A.B., Okwuraiwe,A.P., Amoo,O.S., Salu,O.B., Ige,F.A., Okoli,L.C., Ahmed,R.A., Sokei,J., Oyefolu,A.O., Omilabu,S.A., Salako,B.L. and Audu,R.A. |
| EPI_ISL_514752 | Infectious Disease Control Center, Center for Disease Control and Prevention of PLA | Infectious Disease Control Center, Center for Disease Control and Prevention of PLA | Li, P. |
| EPI_ISL_524437, EPI_ISL_524438, EPI_ISL_524439, EPI_ISL_524440, EPI_ISL_524441, EPI_ISL_524442, EPI_ISL_524443 | University of Washington Virology Lab | Laboratory Medicine, University of Washington | Roychoudhury,P., Greninger,A., Jerome,K. |
| EPI_ISL_524444 | University of Washington Virology Lab | Fred Hutchinson Cancer Research Center | Roychoudhury,P., Greninger,A., Jerome,K. |
| EPI_ISL_527871 | Institute of Medical Biology, Chinese Academy of Medical Sciences and Peking Union Medical College | Institute of Medical Biology, Chinese Academy of Medical Sciences and Peking Union Medical College | Xu,X., Liao,Y., Wang,L., Zhou,X., Xie,Z., Chen,H., Fan,S., Liu,L.,Zheng,H., Jiang,G. and Li,Q. |
| EPI_ISL_529149, EPI_ISL_529150 | Technology Centre, Guangzhou Customs | Technology Centre, Guangzhou Customs | Huang, J., Shi, Y., Sun, J., Zheng, K., Zhu, ., Sun, F., Zhuang, Z., Dai, J., Zhang, Z., Huang, S., Wang, Y., Li, X. |
| EPI_ISL_529155, EPI_ISL_529156, EPI_ISL_529157, EPI_ISL_529158, EPI_ISL_529159 | University of Washington, Laboratory Medicine | University of Washington, Laboratory Medicine | Roychoudhury,P., Greninger,A., Jerome,K. |
| EPI_ISL_529160, EPI_ISL_529161 | University of Washington, Laboratory Medicine | University of Washington, Laboratory Medicine | Roychoudhury, P., Greninger, A., Jerome, K. |
| EPI_ISL_529169 | Michigan Department of Health and Human Services, Bureau of Laboratories | Michigan Department of Health and Human Services, Bureau of Laboratories | Blankenship,H.M., Riner,D., Soehnlen,M. |
| EPI_ISL_529170, EPI_ISL_529200 | University of Washington, Laboratory Medicine | University of Washington, Laboratory Medicine | Roychoudhury,P., Greninger,A., Jerome,K. |
| EPI_ISL_529201, EPI_ISL_529204 | Utah Public Health Laboratory | Utah Public Health Laboratory | Erin Young, Kelly Oakeson |
| EPI_ISL_529208, EPI_ISL_529209, EPI_ISL_529210, EPI_ISL_529211, EPI_ISL_529212 | Laboratory Medicine, University of Washington | University of Washington, Laboratory Medicine | Roychoudhury,P., Greninger,A., Jerome,K. |
| EPI_ISL_529213, EPI_ISL_529214, EPI_ISL_529215, EPI_ISL_529216, EPI_ISL_529217 | Beijing Institute of Microbiology and Epidemiology | Beijing Institute of Microbiology and Epidemiology | Fan, Hang; Qin, E.; Wu, Y.; Guo, Y.; Zhang, X.; Yong, Y.; Hou, J.; Xu, Z.; Mu, J.; Teng, Yue; Mi, Z.; Yang, R.; Song, Yajun.; Li, B.; Cui, Y. |
| EPI_ISL_547870, EPI_ISL_547871, EPI_ISL_547872, EPI_ISL_547873, EPI_ISL_547874, EPI_ISL_547875, EPI_ISL_547876, EPI_ISL_547877 | Maximum Containment Laboratory, National Institute of Virology | Maximum Containment Laboratory, National Institute of Virology | Yadav,P.D., Shete-aich,A., Nyayanit,D.A., Abraham,P. |

We gratefully acknowledge the following Authors from the Originating laboratories responsible for obtaining the specimens, as well as the Submitting laboratories where the genome data were generated and shared via GISAID, on which this research is based.

All Submitters of data may be contacted directly via www.gisaid.org

| Accession ID | Originating Laboratory | Submitting Laboratory | Authors |
|---|---|---|---|
| EPI_ISL_426454, EPI_ISL_426455, EPI_ISL_426456, EPI_ISL_426457, EPI_ISL_426458, EPI_ISL_426459, EPI_ISL_426460, EPI_ISL_426461, EPI_ISL_426462, EPI_ISL_426463, EPI_ISL_426464, EPI_ISL_426465, EPI_ISL_426466, EPI_ISL_426467, EPI_ISL_426468, EPI_ISL_426469, EPI_ISL_426470, EPI_ISL_426471, EPI_ISL_426472, EPI_ISL_426473, EPI_ISL_426474, EPI_ISL_426475 | | | |
| see above | Virginia DCLS | Virginia DCLS | Virginia DCLS |
| EPI_ISL_426476, EPI_ISL_426477, EPI_ISL_426478 | Microbial Genomics Laboratory, Institut Pasteur Montevideo | Microbial Genomics Laboratory, Institut Pasteur Montevideo, Uruguay | Cecilia Salazar, Florencia Díaz-Viraqué, Marianoel Pereira, Pilar Moreno, Gonzalo Moratorio, Gregorio Iraola |
| EPI_ISL_426479, EPI_ISL_426480 | Microbial Genomics Laboratory, Institut Pasteur Montevideo | Microbial Genomics Laboratory, Institut Pasteur Montevideo | Cecilia Salazar, Florencia Díaz-Viraqué, Marianoel Pereira, Pilar Moreno, Gonzalo Moratorio, Gregorio Iraola |
| EPI_ISL_426481, EPI_ISL_426482 | Microbial Genomics Laboratory, Institut Pasteur Montevideo, Uruguay | Microbial Genomics Laboratory, Institut Pasteur Montevideo | Cecilia Salazar, Florencia Díaz-Viraqué, Marianoel Pereira, Pilar Moreno, Gonzalo Moratorio, Gregorio Iraola |
| EPI_ISL_426483 | AZ SPHL, Arizona Department of Health Services | TGen North | Jolene Bowers, Megan Folkerts, Darrin Lemmer, Dave Engelthaler |
| EPI_ISL_426484 | n/a | Worobey Lab on behalf of the Arizona COVID-19 Genomics Union | Brendan B. Larsen, Megan Folkerts, Krystal Sheridan, Ashlyn Pfeiffer, Danielle Yasquez, Hayley Yaglom, Darrin Lemmer, Jolene Bowers,Evan Bolyen, Jason W. Sahl, Nicholas A. Bokulich, J. Gregory Caporaso, Crystal Hepp, Jason Ladner,David M. Engelthaler, Paul Keim, Michael Worobey |
| EPI_ISL_426485 | AZ SPHL, Arizona Department of Health Services | TGen North | Jolene Bowers, Megan Folkerts, Darrin Lemmer, Dave Engelthaler |
| EPI_ISL_426499, EPI_ISL_426500, EPI_ISL_426501, EPI_ISL_426502, EPI_ISL_426503, EPI_ISL_426504, EPI_ISL_426505, EPI_ISL_426506, EPI_ISL_426507, EPI_ISL_426508, EPI_ISL_426509, EPI_ISL_426510, EPI_ISL_426511 | | | |
| see above | TGen North | TGen North | Jolene Bowers, Megan Folkerts, Darrin Lemmer, Dave Engelthaler |
| EPI_ISL_426512, EPI_ISL_426513, EPI_ISL_426514, EPI_ISL_426515, EPI_ISL_426516, EPI_ISL_426517, EPI_ISL_426518, EPI_ISL_426519 | AZ SPHL, Arizona Department of Health Services | TGen North | Jolene Bowers, Megan Folkerts, Darrin Lemmer, Dave Engelthaler |
| EPI_ISL_426520, EPI_ISL_426521, EPI_ISL_426522, EPI_ISL_426523, EPI_ISL_426524, EPI_ISL_426525, EPI_ISL_426526 | TGen North | TGen North | Jolene Bowers, Megan Folkerts, Darrin Lemmer, Dave Engelthaler |
| EPI_ISL_426527, EPI_ISL_426528, EPI_ISL_426529, EPI_ISL_426530, EPI_ISL_426531 | AZ SPHL, Arizona Department of Health Services | TGen North | Jolene Bowers, Megan Folkerts, Darrin Lemmer, Dave Engelthaler |
| EPI_ISL_426532, EPI_ISL_426533, EPI_ISL_426534, EPI_ISL_426535, EPI_ISL_426536 | TGen North | TGen North | Jolene Bowers, Megan Folkerts, Darrin Lemmer, Dave Engelthaler |
| EPI_ISL_426537, EPI_ISL_426538, EPI_ISL_426539, EPI_ISL_426540, EPI_ISL_426541, EPI_ISL_426542, EPI_ISL_426543, EPI_ISL_426544, EPI_ISL_426545, EPI_ISL_426546, EPI_ISL_426547, EPI_ISL_426548, EPI_ISL_426549, EPI_ISL_426550, EPI_ISL_426551, EPI_ISL_426552, EPI_ISL_426553, EPI_ISL_426554, EPI_ISL_426555 | | | |
| see above | AZ SPHL, Arizona Department of Health Services | TGen North | Jolene Bowers, Megan Folkerts, Darrin Lemmer, Dave Engelthaler |
| EPI_ISL_426556, EPI_ISL_426557 | TGen North | TGen North | Jolene Bowers, Megan Folkerts, Darrin Lemmer, Dave Engelthaler |
| EPI_ISL_426558, EPI_ISL_426559, EPI_ISL_426560, EPI_ISL_426561, EPI_ISL_426562, EPI_ISL_426563, EPI_ISL_426564, EPI_ISL_426565, EPI_ISL_426566, EPI_ISL_426567, EPI_ISL_426568, EPI_ISL_426569 | | | |
| see above | AZ SPHL, Arizona Department of Health Services | TGen North | Jolene Bowers, Megan Folkerts, Darrin Lemmer, Dave Engelthaler |
| EPI_ISL_426580 | Instituto Sabin | Laboratory of Virology | Fernando L Melo,Gustavo Barra, Ticiane H Santa-Rita, Pedro G Mesquita, Ikaro A Andrade, Tatsuya Nagata, Bergmann M Ribeiro |
| EPI_ISL_426581 | Motol University Hospital | Institute of Applied Biotechnologies a.s. | Petr Brož, Jan Geryk, Petr Klempt, Martin Kašný, Adam Novotný, Kateřina Kvapilová, Pavel Dřevínek, Petr Kvapil, Milan Macek |
| EPI_ISL_426583 | Microbial Genomics Laboratory, Institut Pasteur Monteivdeo | Microbial Genomics Laboratory, Institut Pasteur Montevideo, Uruguay | Cecilia Salazar, Florencia Díaz-Viraqué, Marianoel Pereira, Pilar Moreno, Gonzalo Moratorio, Gregorio Iraola |
| EPI_ISL_426584 | Microbial Genomics Laboratory, Institut Pasteur Montevideo, Uruguay | Microbial Genomics Laboratory, Institut Pasteur Montevideo, Uruguay | Cecilia Salazar, Florencia Díaz-Viraqué, Marianoel Pereira, Pilar Moreno, Gonzalo Moratorio, Gregorio Iraola |
| EPI_ISL_426617, EPI_ISL_426618, EPI_ISL_426619, EPI_ISL_426620, EPI_ISL_426621, EPI_ISL_426622, EPI_ISL_426623, EPI_ISL_426624, EPI_ISL_426625, EPI_ISL_426626 | NYU Langone Health | Departments of Pathology and Medicine, New York University School of Medicine | Maria Aguero-Rosenfeld, Brendan Belovarac, Margaret Black, Ludovic Boytard, John Cadley, Paolo Cotzia, John Chen, Dacia Dimartino, Xiaojun Feng, Tatyana Gindin, Emily Guzman, Adriana Heguy, Megan Hogan, Emily Huang, George Jour, Andrew Lytle, Christian Marier, Matthew T. Maurano, Mark J. Mulligan, Peter Meyn, Iman Osman, Jared Pinnell, Vanessa Raabe, Sitharam Ramaswami, Amy Rapkiewicz, Marie Samanovic-Golden, Antonio Serrano, Guomiao Shen, Matija Snuderl, Theodore Vougiouklakis, Nick Vulpescu, Gael Westby, Paul Zappile, Yutong Zhang |
| EPI_ISL_426627, EPI_ISL_426628 | Ochsner Health | BioInfoExperts, LLC | Amy Feehan, David Nolan, Rebecca Rose, Susanna Lamers, Sissy Cross, Julia-Garcia-Diaz, Tong Yang, Luke Caruso, David Moraga Amador, Wayra Navia, Lydia Von Borstel, Xiao Hui Zhou |
| EPI_ISL_426629 | TSGH-CP molecular lab | TSGH-CP molecular lab | Cherng-Lih Perng, Ming-Jr Jian, Chih-Kai Chang, Jung-Chung Lin, Kuo-Ming Yeh, Chien-Wen Chen, Sheng-Kang Chiu, Hsing-Yi Chung, Shih-Hung Tsai, Kuo-Sheng Hung, Feng-Yee Chang, Hung-Sheng Shang |
| EPI_ISL_426630 | TSGH-CP molecular lab | TSGH-CP molecular lab | Cherng-Lih Perng, Ming-Jr Jian, Chih-Kai Chang, Jung-Chung Lin, Kuo-Ming Yeh, Chien-Wen Chen, Sheng-Kang Chiu, Hsing-Yi Chung, Shih-Hung Tsai, Kuo-Sheng Hung, Feng-Yee Chang, Hung-Sheng Shang |
| EPI_ISL_426631, EPI_ISL_426632 | TSGH-CP molecular lab | TSGH-CP molecular lab | Cherng-Lih Perng, Ming-Jr Jian, Chih-Kai Chang, Jung-Chung Lin, Kuo-Ming Yeh, Chien-Wen Chen, Sheng-Kang Chiu, Hsing-Yi Chung, Shih-Hung Tsai, Kuo-Sheng Hung, Tien-Yao Chang, Feng-Yee Chang, Hung-Sheng Shang |
| EPI_ISL_426633, EPI_ISL_426634, EPI_ISL_426635, EPI_ISL_426636 | Royal Darwin Hospital Pathology | Microbiological Diagnostic Unit Public Health Laboratory and Victorian Infectious Diseases Reference Laboratory, Doherty Institute | Meumann, E., Caly L., Seemann T., Sait, M., Schultz M., Druce J., Sherry, N. |
| EPI_ISL_426637, EPI_ISL_426638, EPI_ISL_426639, EPI_ISL_426640, EPI_ISL_426641, EPI_ISL_426642, EPI_ISL_426643, EPI_ISL_426644, EPI_ISL_426645, EPI_ISL_426646, EPI_ISL_426647, EPI_ISL_426648, EPI_ISL_426649, EPI_ISL_426650, EPI_ISL_426651, EPI_ISL_426652, EPI_ISL_426653, EPI_ISL_426654, EPI_ISL_426655, EPI_ISL_426656, EPI_ISL_426657, EPI_ISL_426658, EPI_ISL_426659, EPI_ISL_426660, EPI_ISL_426661, EPI_ISL_426662, EPI_ISL_426663, EPI_ISL_426664, EPI_ISL_426665, EPI_ISL_426666, EPI_ISL_426667, EPI_ISL_426668, EPI_ISL_426669, EPI_ISL_426670, EPI_ISL_426671, EPI_ISL_426672, EPI_ISL_426673, EPI_ISL_426674, EPI_ISL_426675, EPI_ISL_426676, EPI_ISL_426677, EPI_ISL_426678, EPI_ISL_426679, EPI_ISL_426680, EPI_ISL_426681, EPI_ISL_426682, EPI_ISL_426683, EPI_ISL_426684, EPI_ISL_426685, EPI_ISL_426686, EPI_ISL_426687, EPI_ISL_426688, EPI_ISL_426689, EPI_ISL_426690, EPI_ISL_426691, EPI_ISL_426692, EPI_ISL_426693, EPI_ISL_426694, EPI_ISL_426695, EPI_ISL_426696, EPI_ISL_426697, EPI_ISL_426698, EPI_ISL_426699, EPI_ISL_426700, EPI_ISL_426701, EPI_ISL_426702, EPI_ISL_426703, EPI_ISL_426704, EPI_ISL_426705, EPI_ISL_426706, EPI_ISL_426707, EPI_ISL_426708, EPI_ISL_426709, EPI_ISL_426710, EPI_ISL_426711, EPI_ISL_426712, EPI_ISL_426713, EPI_ISL_426714, EPI_ISL_426715, EPI_ISL_426716, EPI_ISL_426717, EPI_ISL_426718, EPI_ISL_426719, EPI_ISL_426720, EPI_ISL_426721, EPI_ISL_426722, EPI_ISL_426723, EPI_ISL_426724, EPI_ISL_426725, EPI_ISL_426726, EPI_ISL_426727, EPI_ISL_426728, EPI_ISL_426729, EPI_ISL_426730, EPI_ISL_426731, EPI_ISL_426732, EPI_ISL_426733, EPI_ISL_426734, EPI_ISL_426735, EPI_ISL_426736, EPI_ISL_426737, EPI_ISL_426738, EPI_ISL_426739, EPI_ISL_426740, EPI_ISL_426741, EPI_ISL_426742, EPI_ISL_426743, EPI_ISL_426744, EPI_ISL_426745, EPI_ISL_426746, EPI_ISL_426747, EPI_ISL_426748, EPI_ISL_426749, EPI_ISL_426750, EPI_ISL_426751, EPI_ISL_426752, EPI_ISL_426753, EPI_ISL_426754, EPI_ISL_426755, EPI_ISL_426756, EPI_ISL_426757, EPI_ISL_426758, EPI_ISL_426759, EPI_ISL_426760, EPI_ISL_426761, EPI_ISL_426762, EPI_ISL_426763, EPI_ISL_426764, EPI_ISL_426765, EPI_ISL_426766, EPI_ISL_426767, EPI_ISL_426768, EPI_ISL_426769, EPI_ISL_426770, EPI_ISL_426771, EPI_ISL_426772, EPI_ISL_426773, EPI_ISL_426774, EPI_ISL_426775, EPI_ISL_426776, EPI_ISL_426777, EPI_ISL_426778, EPI_ISL_426779, EPI_ISL_426780, EPI_ISL_426781, EPI_ISL_426782, EPI_ISL_426783, EPI_ISL_426784, EPI_ISL_426785, EPI_ISL_426786, EPI_ISL_426787, EPI_ISL_426788, EPI_ISL_426789, EPI_ISL_426790, EPI_ISL_426791, EPI_ISL_426792, EPI_ISL_426793, EPI_ISL_426794, EPI_ISL_426795, EPI_ISL_426796, EPI_ISL_426797, EPI_ISL_426798, EPI_ISL_426799, EPI_ISL_426800, EPI_ISL_426801, EPI_ISL_426802, EPI_ISL_426803, EPI_ISL_426804, EPI_ISL_426805, EPI_ISL_426806, EPI_ISL_426807, EPI_ISL_426808, EPI_ISL_426809, EPI_ISL_426810, EPI_ISL_426811, EPI_ISL_426812, EPI_ISL_426813, EPI_ISL_426814, EPI_ISL_426815, EPI_ISL_426816, EPI_ISL_426817, EPI_ISL_426818, EPI_ISL_426819, EPI_ISL_426820, EPI_ISL_426821, EPI_ISL_426822, EPI_ISL_426823, EPI_ISL_426824, EPI_ISL_426825, EPI_ISL_426826, EPI_ISL_426827, EPI_ISL_426828, EPI_ISL_426829, EPI_ISL_426830, EPI_ISL_426831, EPI_ISL_426832, EPI_ISL_426833, EPI_ISL_426834, EPI_ISL_426835, EPI_ISL_426836, EPI_ISL_426837, EPI_ISL_426838, EPI_ISL_426839, EPI_ISL_426840, EPI_ISL_426841, EPI_ISL_426842, EPI_ISL_426843, EPI_ISL_426844, EPI_ISL_426845, EPI_ISL_426846, EPI_ISL_426847, EPI_ISL_426848, EPI_ISL_426849, EPI_ISL_426850, EPI_ISL_426851, EPI_ISL_426852, EPI_ISL_426853, EPI_ISL_426854, EPI_ISL_426855, EPI_ISL_426856, EPI_ISL_426857, EPI_ISL_426858, EPI_ISL_426859, EPI_ISL_426860, EPI_ISL_426861, EPI_ISL_426862, EPI_ISL_426863, EPI_ISL_426864, EPI_ISL_426865, EPI_ISL_426866, EPI_ISL_426867, EPI_ISL_426869, EPI_ISL_426870, EPI_ISL_426871, EPI_ISL_426872, EPI_ISL_426873, EPI_ISL_426874, EPI_ISL_426875, EPI_ISL_426876, EPI_ISL_426877, EPI_ISL_426878, EPI_ISL_426879, EPI_ISL_426880, EPI_ISL_426881, EPI_ISL_426882 | | | |
| see above | Victorian Infectious Diseases Reference Laboratory (VIDRL) | Microbiological Diagnostic Unit Public Health Laboratory and Victorian Infectious Diseases Reference Laboratory, Doherty Institute | Caly L., Seemann T., Sait, M., Schultz M., Druce J., Sherry, N. |
| EPI_ISL_426883, EPI_ISL_426884, EPI_ISL_426885, EPI_ISL_426886, EPI_ISL_426887, EPI_ISL_426888, EPI_ISL_426889, EPI_ISL_426890, EPI_ISL_426891, EPI_ISL_426892, EPI_ISL_426893, EPI_ISL_426894, EPI_ISL_426895, EPI_ISL_426896, EPI_ISL_426897 | | | |
| see above | Motol University Hospital | Institute of Applied Biotechnologies a.s. | Petr Brož, Jan Geryk, Petr Klempt, Martin Kašný, Adam Novotný, Kateřina Kvapilová, Pavel Dřevínek, Petr Kvapil, Milan Macek |
| EPI_ISL_426898, EPI_ISL_426899, EPI_ISL_426900, EPI_ISL_426901, EPI_ISL_426902, EPI_ISL_426903, EPI_ISL_426904 | Royal Darwin Hospital Pathology | Microbiological Diagnostic Unit Public Health Laboratory and Victorian Infectious Diseases Reference Laboratory, Doherty Institute | Meumann, E., Caly L., Seemann T., Sait, M., Schultz M., Druce J., Sherry, N. |
| EPI_ISL_426905, EPI_ISL_426906, EPI_ISL_426907, EPI_ISL_426908, EPI_ISL_426909, EPI_ISL_426910, EPI_ISL_426911, EPI_ISL_426912, EPI_ISL_426913, EPI_ISL_426914, EPI_ISL_426915, EPI_ISL_426916, EPI_ISL_426917, EPI_ISL_426918, EPI_ISL_426919, EPI_ISL_426920, EPI_ISL_426921, EPI_ISL_426922 | | | |
| see above | Microbiological Diagnostic Unit Public Health Laboratory | Microbiological Diagnostic Unit Public Health Laboratory | Seemann T., Schultz M., Sait, M., Sherry, N. |
| EPI_ISL_426923, EPI_ISL_426924, EPI_ISL_426925, EPI_ISL_426926, EPI_ISL_426927, EPI_ISL_426928, EPI_ISL_426929, EPI_ISL_426930, EPI_ISL_426931, EPI_ISL_426932, EPI_ISL_426933, EPI_ISL_426934, EPI_ISL_426935, EPI_ISL_426936, EPI_ISL_426937, EPI_ISL_426938, EPI_ISL_426939, EPI_ISL_426940, EPI_ISL_426941, EPI_ISL_426942, EPI_ISL_426943, EPI_ISL_426944, EPI_ISL_426945, EPI_ISL_426946, EPI_ISL_426947, EPI_ISL_426948, EPI_ISL_426949, EPI_ISL_426950, EPI_ISL_426951, EPI_ISL_426952, EPI_ISL_426953, EPI_ISL_426954, EPI_ISL_426955, EPI_ISL_426956, EPI_ISL_426957, EPI_ISL_426958, EPI_ISL_426959, EPI_ISL_426960, EPI_ISL_426961, EPI_ISL_426962, EPI_ISL_426963, EPI_ISL_426964, EPI_ISL_426965, EPI_ISL_426966, EPI_ISL_426967, EPI_ISL_426968, EPI_ISL_426969, EPI_ISL_426970, EPI_ISL_426971, EPI_ISL_426972, EPI_ISL_426973, EPI_ISL_426974, EPI_ISL_426975, EPI_ISL_426976, EPI_ISL_426977, EPI_ISL_426978, EPI_ISL_426979, EPI_ISL_426980, EPI_ISL_426981, EPI_ISL_426982, EPI_ISL_426983, EPI_ISL_426984, EPI_ISL_426985, EPI_ISL_426986, EPI_ISL_426987, EPI_ISL_426988, EPI_ISL_426989, EPI_ISL_426990, EPI_ISL_426991, EPI_ISL_426992, EPI_ISL_426993, EPI_ISL_426994, EPI_ISL_426995, EPI_ISL_426996, EPI_ISL_426997, EPI_ISL_426998, EPI_ISL_426999, EPI_ISL_427000, EPI_ISL_427001, EPI_ISL_427002, EPI_ISL_427003, EPI_ISL_427004, EPI_ISL_427005, EPI_ISL_427006, EPI_ISL_427007, EPI_ISL_427008, EPI_ISL_427009, EPI_ISL_427010, EPI_ISL_427011, EPI_ISL_427012, EPI_ISL_427013, EPI_ISL_427014, EPI_ISL_427015, EPI_ISL_427016, EPI_ISL_427017, EPI_ISL_427018, EPI_ISL_427019, EPI_ISL_427020, EPI_ISL_427021, EPI_ISL_427022, EPI_ISL_427023, EPI_ISL_427024, EPI_ISL_427025, EPI_ISL_427026, EPI_ISL_427027, EPI_ISL_427028, EPI_ISL_427029, EPI_ISL_427030, EPI_ISL_427031, EPI_ISL_427032, EPI_ISL_427033, EPI_ISL_427034, EPI_ISL_427035, EPI_ISL_427036, EPI_ISL_427037, EPI_ISL_427038, EPI_ISL_427039, EPI_ISL_427040, EPI_ISL_427041, EPI_ISL_427042 | | | |
| see above | Victorian Infectious Diseases Reference Laboratory (VIDRL) | Microbiological Diagnostic Unit Public Health Laboratory and Victorian Infectious Diseases Reference Laboratory, Doherty Institute | Caly L., Seemann T., Sait, M., Schultz M., Druce J., Sherry, N. |
| EPI_ISL_427043 | Laboratory of Microbiology, Medical School, National and Kapodistrian University of Athens | Laboratory of Biology, Department of Medicine, Democritus University of Thrace | Bampali,M., Dovrolis,N., Gatzidou,E., Froukala,E., Stavropoulou,A., Veletza,S., Tsakris,A., Spanakis,N. and Karakasiliotis,I. |
| EPI_ISL_427044, EPI_ISL_427045, EPI_ISL_427046, EPI_ISL_427047, EPI_ISL_427048, EPI_ISL_427049, | Victorian Infectious Diseases Reference Laboratory (VIDRL) | Microbiological Diagnostic Unit Public Health Laboratory and Victorian Infectious Diseases Reference Laboratory, Doherty | Caly L., Seemann T., Sait, M., Schultz M., Druce J., Sherry, N. |

| | | | |
|---|---|---|---|
| EPI_ISL_427050, EPI_ISL_427051, EPI_ISL_427052, EPI_ISL_427053 | | Institute | |
| EPI_ISL_427054, EPI_ISL_427055, EPI_ISL_427056, EPI_ISL_427057, EPI_ISL_427058, EPI_ISL_427059, EPI_ISL_427060, EPI_ISL_427061, EPI_ISL_427062, EPI_ISL_427063, EPI_ISL_427064, EPI_ISL_427065, EPI_ISL_427066, EPI_ISL_427067, EPI_ISL_427068, EPI_ISL_427069, EPI_ISL_427070, EPI_ISL_427071, EPI_ISL_427072, EPI_ISL_427073, EPI_ISL_427074, EPI_ISL_427075, EPI_ISL_427076, EPI_ISL_427077, EPI_ISL_427078 | | | |
| see above | Microbiological Diagnostic Unit Public Health Laboratory | Microbiological Diagnostic Unit Public Health Laboratory | Seemann T., Schultz M., Sait, M., Sherry, N. |
| EPI_ISL_427079, EPI_ISL_427080, EPI_ISL_427081, EPI_ISL_427082, EPI_ISL_427083, EPI_ISL_427084, EPI_ISL_427085, EPI_ISL_427086, EPI_ISL_427087, EPI_ISL_427088, EPI_ISL_427089, EPI_ISL_427090, EPI_ISL_427091, EPI_ISL_427092, EPI_ISL_427093, EPI_ISL_427094, EPI_ISL_427095, EPI_ISL_427096, EPI_ISL_427097, EPI_ISL_427098, EPI_ISL_427099, EPI_ISL_427100, EPI_ISL_427101, EPI_ISL_427102, EPI_ISL_427103, EPI_ISL_427104, EPI_ISL_427105, EPI_ISL_427106, EPI_ISL_427107, EPI_ISL_427108, EPI_ISL_427109, EPI_ISL_427110, EPI_ISL_427111, EPI_ISL_427112, EPI_ISL_427113, EPI_ISL_427114, EPI_ISL_427115, EPI_ISL_427116, EPI_ISL_427117, EPI_ISL_427118, EPI_ISL_427119, EPI_ISL_427120, EPI_ISL_427121, EPI_ISL_427122, EPI_ISL_427123, EPI_ISL_427124, EPI_ISL_427125, EPI_ISL_427126, EPI_ISL_427127, EPI_ISL_427128, EPI_ISL_427129, EPI_ISL_427130, EPI_ISL_427131, EPI_ISL_427132 | | | |
| see above | Victorian Infectious Diseases Reference Laboratory (VIDRL) | Microbiological Diagnostic Unit Public Health Laboratory and Victorian Infectious Diseases Reference Laboratory, Doherty Institute | Caly L., Seemann T., Sait, M., Schultz M., Druce J., Sherry, N. |
| EPI_ISL_427133 | Microbiological Diagnostic Unit Public Health Laboratory | Microbiological Diagnostic Unit Public Health Laboratory | Seemann T., Schultz M., Sait, M., Sherry, N. |
| EPI_ISL_427134, EPI_ISL_427135, EPI_ISL_427136, EPI_ISL_427137, EPI_ISL_427138, EPI_ISL_427139, EPI_ISL_427140, EPI_ISL_427141, EPI_ISL_427142, EPI_ISL_427143, EPI_ISL_427144, EPI_ISL_427145, EPI_ISL_427146, EPI_ISL_427147 | | | |
| see above | Victorian Infectious Diseases Reference Laboratory (VIDRL) | Microbiological Diagnostic Unit Public Health Laboratory and Victorian Infectious Diseases Reference Laboratory, Doherty Institute | Caly L., Seemann T., Sait, M., Schultz M., Druce J., Sherry, N. |
| EPI_ISL_427148, EPI_ISL_427149 | Microbiological Diagnostic Unit Public Health Laboratory | Microbiological Diagnostic Unit Public Health Laboratory | Seemann T., Schultz M., Sait, M., Sherry, N. |
| EPI_ISL_427150, EPI_ISL_427151, EPI_ISL_427152, EPI_ISL_427153, EPI_ISL_427154, EPI_ISL_427155, EPI_ISL_427156, EPI_ISL_427157, EPI_ISL_427158, EPI_ISL_427159, EPI_ISL_427160 | | | |
| see above | Victorian Infectious Diseases Reference Laboratory (VIDRL) | Microbiological Diagnostic Unit Public Health Laboratory and Victorian Infectious Diseases Reference Laboratory, Doherty Institute | Caly L., Seemann T., Sait, M., Schultz M., Druce J., Sherry, N. |
| EPI_ISL_427161, EPI_ISL_427162, EPI_ISL_427163, EPI_ISL_427164, EPI_ISL_427165, EPI_ISL_427166, EPI_ISL_427167, EPI_ISL_427168, EPI_ISL_427169, EPI_ISL_427170, EPI_ISL_427171, EPI_ISL_427172, EPI_ISL_427173, EPI_ISL_427174, EPI_ISL_427175, EPI_ISL_427176, EPI_ISL_427177, EPI_ISL_427178, EPI_ISL_427179, EPI_ISL_427180, EPI_ISL_427181, EPI_ISL_427182, EPI_ISL_427183, EPI_ISL_427184, EPI_ISL_427185, EPI_ISL_427186, EPI_ISL_427187, EPI_ISL_427188, EPI_ISL_427189, EPI_ISL_427190, EPI_ISL_427191, EPI_ISL_427192, EPI_ISL_427193, EPI_ISL_427194, EPI_ISL_427195, EPI_ISL_427196, EPI_ISL_427197, EPI_ISL_427198, EPI_ISL_427199, EPI_ISL_427200, EPI_ISL_427201, EPI_ISL_427202, EPI_ISL_427203, EPI_ISL_427204, EPI_ISL_427205, EPI_ISL_427206, EPI_ISL_427207, EPI_ISL_427208, EPI_ISL_427209, EPI_ISL_427210, EPI_ISL_427211, EPI_ISL_427212, EPI_ISL_427213, EPI_ISL_427214, EPI_ISL_427215, EPI_ISL_427216, EPI_ISL_427217, EPI_ISL_427218, EPI_ISL_427219, EPI_ISL_427220, EPI_ISL_427221, EPI_ISL_427222, EPI_ISL_427223, EPI_ISL_427224, EPI_ISL_427225, EPI_ISL_427226, EPI_ISL_427227, EPI_ISL_427228, EPI_ISL_427229, EPI_ISL_427230, EPI_ISL_427231, EPI_ISL_427232, EPI_ISL_427233, EPI_ISL_427234, EPI_ISL_427235, EPI_ISL_427236, EPI_ISL_427237, EPI_ISL_427238, EPI_ISL_427239, EPI_ISL_427240, EPI_ISL_427241, EPI_ISL_427242, EPI_ISL_427243, EPI_ISL_427244, EPI_ISL_427245, EPI_ISL_427246, EPI_ISL_427247, EPI_ISL_427248, EPI_ISL_427249, EPI_ISL_427250, EPI_ISL_427251, EPI_ISL_427252, EPI_ISL_427253, EPI_ISL_427254, EPI_ISL_427255, EPI_ISL_427256, EPI_ISL_427257, EPI_ISL_427258, EPI_ISL_427259, EPI_ISL_427260, EPI_ISL_427261, EPI_ISL_427262, EPI_ISL_427263, EPI_ISL_427264, EPI_ISL_427265, EPI_ISL_427266, EPI_ISL_427267, EPI_ISL_427268, EPI_ISL_427269, EPI_ISL_427270 | | | |
| see above | UW Virology Lab | UW Virology Lab | Pavitra Roychoudhury, Hong Xie, Keith Jerome, Alexander Greninger |
| EPI_ISL_427271, EPI_ISL_427272 | AZ SPHL, Arizona Department of Health Services | TGen North | Jolene Bowers, Megan Folkerts, Darrin Lemmer, Dave Engelthaler |
| EPI_ISL_427273, EPI_ISL_427274, EPI_ISL_427275, EPI_ISL_427276, EPI_ISL_427277, EPI_ISL_427278, EPI_ISL_427279, EPI_ISL_427280, EPI_ISL_427281, EPI_ISL_427282, EPI_ISL_427283, EPI_ISL_427284, EPI_ISL_427285, EPI_ISL_427286, EPI_ISL_427287 | | | |
| see above | Minnesota Department of Health, Public Health Laboratory | Minnesota Department of Health, Public Health Laboratory | Matt Plumb, Jacob Garfin and Xiong Wang |
| EPI_ISL_427288 | The Ohio State University | The Ohio State University-James Molecular Lab at Polaris | Huolin Tu, Preeti Panchioli, Jason Garee, Matthew Hunt, Joan-Miquel Balada-Llasat, Erica Vincent, Weiqiang Zhao, Dan Jones |
| EPI_ISL_427289 | The Ohio State University | The Ohio State University-James Molecular Lab at Polaris | Huolin Tu, Joan-Miquel Balada-Llasat, Jason Garee, Matthew Hunt, Preeti Pancholi, Erica Vincent, Xiaokang Zhao, Dan Jones |
| EPI_ISL_427290 | The Ohio State University | The Ohio State University-James Molecular Lab at Polaris | Huolin Tu, Jason Garee, Matthew Hunt, Joan-Miquel Balada-Llasat, Preeti Pancholi, Erica Vincent, Rongqin Ren, Dan Jones |
| EPI_ISL_427291 | The Ohio State University | The Ohio State University-James Molecular Lab at Polaris | Huolin Tu, Matthew Hunt, Preeti Pancholi, Jason Garee, Joan-Miquel Balada-Llasat, Erica Vincent, Weiqiang Zhao, Dan Jones |
| EPI_ISL_427292 | LACEN-AL - Laboratorio Central de Alagoas | Instituto Oswaldo Cruz FIOCRUZ - Laboratory of Respiratory Viruses and Measles (LVRS) | Paola Resende, Fernando Motta, Luciana Appolinario, Sunando Roy, Aline Mattos, Milene Miranda, Cristiana Garcia, Braulia Caetano, Maria Ogrzewalska, Priscila Born, Jonathan Lopes, Marilda Siqueira |
| EPI_ISL_427293 | LACEN-BA - Laboratório Central de Saúde Pública Professor Gonçalo Moniz | Instituto Oswaldo Cruz FIOCRUZ - Laboratory of Respiratory Viruses and Measles (LVRS) | Paola Resende, Fernando Motta, Luciana Appolinario, Sunando Roy, Aline Mattos, Milene Miranda, Cristiana Garcia, Braulia Caetano, Maria Ogrzewalska, Priscila Born, Jonathan Lopes, Marilda Siqueira |
| EPI_ISL_427294, EPI_ISL_427295, EPI_ISL_427296, EPI_ISL_427297, EPI_ISL_427298, EPI_ISL_427299, EPI_ISL_427300, EPI_ISL_427301, EPI_ISL_427302, EPI_ISL_427303, EPI_ISL_427304 | | | |
| see above | Instituto Oswaldo Cruz FIOCRUZ - Laboratory of Respiratory Viruses and Measles (LVRS) | Instituto Oswaldo Cruz FIOCRUZ - Laboratory of Respiratory Viruses and Measles (LVRS) | Paola Resende, Fernando Motta, Luciana Appolinario, Sunando Roy, Aline Mattos, Milene Miranda, Cristiana Garcia, Braulia Caetano, Maria Ogrzewalska, Priscila Born, Jonathan Lopes, Marilda Siqueira |
| EPI_ISL_427305, EPI_ISL_427306 | LACEN-SC - Laboratorio Central de Santa Catarina | Instituto Oswaldo Cruz FIOCRUZ - Laboratory of Respiratory Viruses and Measles (LVRS) | Paola Resende, Fernando Motta, Luciana Appolinario, Sunando Roy, Aline Mattos, Milene Miranda, Cristiana Garcia, Braulia Caetano, Maria Ogrzewalska, Priscila Born, Jonathan Lopes, Marilda Siqueira |
| EPI_ISL_427307, EPI_ISL_427308, EPI_ISL_427309, EPI_ISL_427310, EPI_ISL_427311, EPI_ISL_427312, EPI_ISL_427313, EPI_ISL_427314, EPI_ISL_427315, EPI_ISL_427316, EPI_ISL_427317, EPI_ISL_427318, EPI_ISL_427319, EPI_ISL_427320, EPI_ISL_427321, EPI_ISL_427322, EPI_ISL_427323, EPI_ISL_427324, EPI_ISL_427325, EPI_ISL_427326, EPI_ISL_427327, EPI_ISL_427328, EPI_ISL_427329, EPI_ISL_427330, EPI_ISL_427331, EPI_ISL_427332, EPI_ISL_427333, EPI_ISL_427334, EPI_ISL_427335, EPI_ISL_427336, EPI_ISL_427337, EPI_ISL_427338, EPI_ISL_427339 | | | |
| see above | WHO National Influenza Centre Russian Federation | WHO National Influenza Centre Russian Federation | Andrey Komissarov, Artem Fadeev, Mariia Sergeeva, Anna Ivanova, Daria Danilenko |
| EPI_ISL_427340, EPI_ISL_427341, EPI_ISL_427342, EPI_ISL_427343, EPI_ISL_427344, EPI_ISL_427345, EPI_ISL_427346, EPI_ISL_427347, EPI_ISL_427348, EPI_ISL_427349, EPI_ISL_427350, EPI_ISL_427351, EPI_ISL_427352, EPI_ISL_427353, EPI_ISL_427354, EPI_ISL_427355, EPI_ISL_427356, EPI_ISL_427357, EPI_ISL_427358, EPI_ISL_427359, EPI_ISL_427360, EPI_ISL_427361, EPI_ISL_427362, EPI_ISL_427363, EPI_ISL_427364, EPI_ISL_427365, EPI_ISL_427366, EPI_ISL_427367, EPI_ISL_427368, EPI_ISL_427369, EPI_ISL_427370, EPI_ISL_427371, EPI_ISL_427372, EPI_ISL_427373, EPI_ISL_427374, EPI_ISL_427375, EPI_ISL_427376, EPI_ISL_427377, EPI_ISL_427378, EPI_ISL_427379, EPI_ISL_427380, EPI_ISL_427381, EPI_ISL_427382, EPI_ISL_427383, EPI_ISL_427384, EPI_ISL_427385, EPI_ISL_427386, EPI_ISL_427387, EPI_ISL_427388, EPI_ISL_427389, EPI_ISL_427390 | | | |
| see above | Department of Clinical Microbiology | GIGA Medical Genomics | Keith Durkin, Maria Artesi, Sébastien Bontems, Raphaël Boreux, Cécile Meex, Pierrette Melin, Marie-Pierre Hayette, Vincent Bours. |
| EPI_ISL_427391 | Genomic Laboratory (GLAB) (Conjoint lab of Health Directorate of Istanbul and Istanbul Technical University) | Genomic Laboratory (GLAB), Istanbul Technical University | Ilker Karacan, Tugba Kizilboga Akgun, Bugra Agaoglu, Gizem Alkurt, Jale Yildiz, Betsi Köse, Elifnaz Çelik, Mehtap Aydın, Levent Doganay, Gizem Dinler Doganay |
| EPI_ISL_427392, EPI_ISL_427393, EPI_ISL_427394, EPI_ISL_427395, EPI_ISL_427396, EPI_ISL_427397, EPI_ISL_427398 | TSGH-CP molecular lab | TSGH-CP molecular lab | Cherng-Lih Perng, Ming-Jr Jian, Chih-Kai Chang, Jung-Chung Lin, Kuo-Ming Yeh, Chien-Wen Chen, Sheng-Kang Chiu, Hsing-Yi Chung, Shih-Hung Tsai, Kuo-Sheng Hung, Tien-Yao Chang, Feng-Yee Chang, Hung-Sheng Shang |
| EPI_ISL_427404, EPI_ISL_427405, EPI_ISL_427406, EPI_ISL_427407, EPI_ISL_427408, EPI_ISL_427409, EPI_ISL_427410, EPI_ISL_427411, EPI_ISL_427412, EPI_ISL_427414, EPI_ISL_427415, EPI_ISL_427416, EPI_ISL_427417, EPI_ISL_427418, EPI_ISL_427419, EPI_ISL_427420 | | | |
| see above | Ministry of Public Health (MoPH) | Biomedical Research Center (BRC) | Abdullatif Al-Khal, Muna A. S. Al-Maslamani, Ajaeb D. M. H. Al-Nabet, Peter V. Coyle, Einas A. E. Al-Kuwari, Nourah B. M. Younes, Hamad E. Al-Romaihi, Salih Al-Marri, Mohammed Al-Thani, Fatiha M. Benslimane, Heba A. Al-Khatib, Sonia Boughattas, Hadi M. Yassine, Asmaa A. Al-Thani. |
| EPI_ISL_427427, EPI_ISL_427428, EPI_ISL_427429, EPI_ISL_427430, EPI_ISL_427431, EPI_ISL_427432, EPI_ISL_427433, EPI_ISL_427434, EPI_ISL_427435, EPI_ISL_427436, EPI_ISL_427437, EPI_ISL_427438, EPI_ISL_427439, EPI_ISL_427440, EPI_ISL_427441, EPI_ISL_427442, EPI_ISL_427443, EPI_ISL_427444, EPI_ISL_427445, EPI_ISL_427446, EPI_ISL_427447, EPI_ISL_427448, EPI_ISL_427449, EPI_ISL_427450, EPI_ISL_427451, EPI_ISL_427452, EPI_ISL_427453, EPI_ISL_427454, EPI_ISL_427455, EPI_ISL_427456, EPI_ISL_427457, EPI_ISL_427458, EPI_ISL_427459, EPI_ISL_427460, EPI_ISL_427461, EPI_ISL_427462 | | | |
| see above | University of Wisconsin-Madison AIDS Vaccine Research Laboratories | University of Wisconsin-Madison AIDS Vaccine Research Laboratories | Gage Moreno, Katarina Braun, et al. AIDS Vaccine Research Laboratories |
| EPI_ISL_427469, EPI_ISL_427470, EPI_ISL_427471, EPI_ISL_427472, EPI_ISL_427473, EPI_ISL_427474, EPI_ISL_427475, EPI_ISL_427476, EPI_ISL_427477, EPI_ISL_427478, EPI_ISL_427479, EPI_ISL_427480, EPI_ISL_427481, EPI_ISL_427482, EPI_ISL_427483, EPI_ISL_427484, EPI_ISL_427485, EPI_ISL_427486, EPI_ISL_427487, EPI_ISL_427488, EPI_ISL_427489, EPI_ISL_427490, EPI_ISL_427491, EPI_ISL_427492, EPI_ISL_427493, EPI_ISL_427494, EPI_ISL_427495, EPI_ISL_427496, EPI_ISL_427497, EPI_ISL_427498, EPI_ISL_427499, EPI_ISL_427500, EPI_ISL_427501, EPI_ISL_427502, EPI_ISL_427503, EPI_ISL_427504, EPI_ISL_427505, EPI_ISL_427506, EPI_ISL_427507, EPI_ISL_427508, EPI_ISL_427509, EPI_ISL_427510, EPI_ISL_427511, EPI_ISL_427512, EPI_ISL_427513, EPI_ISL_427514, EPI_ISL_427515, EPI_ISL_427516, EPI_ISL_427517, EPI_ISL_427518, EPI_ISL_427519, EPI_ISL_427520, EPI_ISL_427521, EPI_ISL_427524, EPI_ISL_427525 | | | |
| see above | NYU Langone Health | Departments of Pathology and Medicine, New York University School of Medicine | Maria Aguero-Rosenfeld, Brendan Belovarac, Margaret Black, Ludovic Boytard, John Cadley, Paolo Cotzia, John Chen, Dacia Dimartino, Xiaojun Feng, Tatyana Gindin, Emily Guzman, Adriana Heguy, Megan Hogan, Emily Huang, George Jour, Andrew Lytle, Christian Marier, Matthew T. Maurano, Mark J. Mulligan, Peter Meyn, Iman Osman, Jared Pinnell, Vanessa Raabe, Sitharam Ramaswami, Amy Rapkiewicz, Marie Samanovic-Golden, Antonio Serrano, Guomiao Shen, Matija Snuderl, Theodore Vougiouklakis, Nick Vulpescu, Gael Westby, Paul Zappile, Yutong Zhang |
| EPI_ISL_427526, EPI_ISL_427527, EPI_ISL_427528, EPI_ISL_427529, EPI_ISL_427530, EPI_ISL_427531, EPI_ISL_427532, EPI_ISL_427533, EPI_ISL_427534, EPI_ISL_427535, EPI_ISL_427536, EPI_ISL_427537, EPI_ISL_427538, EPI_ISL_427539, EPI_ISL_427540, EPI_ISL_427541, EPI_ISL_427542, EPI_ISL_427543, EPI_ISL_427544, EPI_ISL_427545, EPI_ISL_427546, EPI_ISL_427547, EPI_ISL_427548, EPI_ISL_427549, EPI_ISL_427550, EPI_ISL_427551, EPI_ISL_427552, EPI_ISL_427553, EPI_ISL_427554, EPI_ISL_427555, EPI_ISL_427556, EPI_ISL_427557, EPI_ISL_427558, EPI_ISL_427559, EPI_ISL_427560, EPI_ISL_427561, EPI_ISL_427562, EPI_ISL_427563, EPI_ISL_427564, EPI_ISL_427565, EPI_ISL_427566, EPI_ISL_427567, EPI_ISL_427568, EPI_ISL_427569, EPI_ISL_427570, EPI_ISL_427571, EPI_ISL_427572, EPI_ISL_427573, EPI_ISL_427574, EPI_ISL_427575, EPI_ISL_427576, EPI_ISL_427577, EPI_ISL_427578, EPI_ISL_427579, EPI_ISL_427580, EPI_ISL_427581, EPI_ISL_427582, EPI_ISL_427583, EPI_ISL_427584, EPI_ISL_427585, EPI_ISL_427586, EPI_ISL_427587, EPI_ISL_427588, EPI_ISL_427589, EPI_ISL_427590, EPI_ISL_427591, EPI_ISL_427592, EPI_ISL_427593, EPI_ISL_427594, EPI_ISL_427595, EPI_ISL_427596, EPI_ISL_427597, EPI_ISL_427598, EPI_ISL_427599, EPI_ISL_427600, EPI_ISL_427601, EPI_ISL_427602, EPI_ISL_427603, EPI_ISL_427604, EPI_ISL_427605, EPI_ISL_427606, EPI_ISL_427607, EPI_ISL_427608, EPI_ISL_427609, EPI_ISL_427610, EPI_ISL_427611, EPI_ISL_427612, EPI_ISL_427613, EPI_ISL_427614, EPI_ISL_427615, EPI_ISL_427616, EPI_ISL_427617, EPI_ISL_427618 | | | |
| see above | NewYork-Presbyterian & Mason Lab | Mason Lab | Daniel J. Butler, Christopher Mozsary, Cem Meydan, David Danko, Jonathan Foox, Joel Rosiene, Alon Shaiber, Matthew MacKay, Ebrahim Afshinnekoo, Fritz J. Sedlazeck, Nikolay A. Ivanov, Maria Sierra, Craig D. Westover, Krista Ryon, Benjamin Young, Chandrima Bhattacharya, Phyllis Ruggiero, Justyna Gawrys, Iman Hajirasouliha, Dmitry Meleshko, Mirella Salvatore, Dong Xu, Jenny Xiang, John Sipley, Lin Cong, Arryn Craney, Priya Velu, Lars F. Westblade, Massimo Loda, Shawn Levy, Melissa Cushing, Marcin Imielinski, Hanna Rennert, Christopher E. Mason |
| EPI_ISL_427619, EPI_ISL_427620, EPI_ISL_427621, EPI_ISL_427622 | Alaska State Virology Laboratory | Alaska State Virology Laboratory | Chen, J. |
| EPI_ISL_427627, EPI_ISL_427628, EPI_ISL_427629, EPI_ISL_427630, EPI_ISL_427631, EPI_ISL_427632, EPI_ISL_427633, EPI_ISL_427634, EPI_ISL_427635, EPI_ISL_427636, EPI_ISL_427637, EPI_ISL_427638, EPI_ISL_427639, EPI_ISL_427640, EPI_ISL_427641, EPI_ISL_427642 | | | |
| see above | NYU Langone Health | Departments of Pathology and Medicine, New York University School of Medicine | Maria Aguero-Rosenfeld, Brendan Belovarac, Margaret Black, Ludovic Boytard, John Cadley, Paolo Cotzia, John Chen, Dacia Dimartino, Xiaojun Feng, Tatyana Gindin, Emily Guzman, Adriana Heguy, Megan Hogan, Emily Huang, George Jour, Andrew Lytle, Christian Marier, Matthew T. Maurano, Mark J. Mulligan, Peter Meyn, Iman Osman, Jared Pinnell, Vanessa Raabe, Sitharam Ramaswami, Amy Rapkiewicz, Marie Samanovic-Golden, Antonio Serrano, Guomiao Shen, Matija Snuderl, Theodore Vougiouklakis, Nick Vulpescu, Gael Westby, Paul Zappile, Yutong Zhang |
| EPI_ISL_427643 | Centre for Infectious Diseases and Microbiology Public Health | NSW Health Pathology - Institute of Clinical Pathology and Medical Research; Westmead Hospital; University of Sydney | Timms V, Gall M, Arnott A, Sadsad R, Draper J, Sim E, Bachmann N, Rockett R, Lam C, Gray K, Carter I, Holmes EC, O'Sullivan MV, Byun R, Sintchenko V, Chen SC, Eden JS, Maddocks S, Kok J, Propenko M, Sorrell T, Chang S, Basile K, Dwyer DE for the 2019-nCoV Study Group |
| EPI_ISL_427644 | Centre for Infectious Diseases and Microbiology Public Health | NSW Health Pathology - Institute of Clinical Pathology and Medical Research; Westmead Hospital; University of Sydney | Rockett R, Lam C, Gray K, Timms V, Gall M, Arnott A, Sadsad R, Draper J, Sim E, Bachmann N, Carter I, Holmes EC, O'Sullivan MV, Byun R, Sintchenko V, Chen SC, Eden JS, Maddocks S, Kok J, Propenko M, Sorrell T, Chang S, Basile K, Dwyer DE for the 2019-nCoV Study Group |
| EPI_ISL_427645 | Centre for Infectious Diseases and Microbiology Public Health | NSW Health Pathology - Institute of Clinical Pathology and Medical Research; Westmead Hospital; University of Sydney | Gray K, Timms V, Gall M, Arnott A, Sadsad R, Draper J, Sim E, Bachmann N, Rockett R, Lam C, Carter I, Holmes EC, O'Sullivan MV, Byun R, Sintchenko V, Chen SC, Eden JS, Maddocks S, Kok J, Propenko M, Sorrell T, Chang S, Basile K, Dwyer DE for the 2019-nCoV Study Group |
| EPI_ISL_427646 | Centre for Infectious Diseases and Microbiology Public Health | NSW Health Pathology - Institute of Clinical Pathology and Medical Research; Westmead Hospital; University of Sydney | Timms V, Gall M, Arnott A, Sadsad R, Draper J, Sim E, Bachmann N, Rockett R, Lam C, Gray K, Carter I, Holmes EC, O'Sullivan MV, Byun R, Sintchenko V, Chen SC, Eden JS, Maddocks S, Kok J, Propenko M, Sorrell T, Chang S, Basile K, Dwyer DE for the 2019-nCoV Study Group |
| EPI_ISL_427647 | Centre for Infectious Diseases and Microbiology Public Health | NSW Health Pathology - Institute of Clinical Pathology and | Gray K, Timms V, Gall M, Arnott A, Sadsad R, Draper J, Sim E, Bachmann N, Rockett R, Lam C, Carter I, Holmes EC, O'Sullivan MV, Byun R, Sintchenko V, Chen SC, Eden JS, Maddocks S, Kok J, Propenko M, Sorrell T, |

| | | | |
|---|---|---|---|
| | | Medical Research; Westmead Hospital; University of Sydney | Chang S, Basile K, Dwyer DE for the 2019-nCoV Study Group |
| EPI_ISL_427648 | Centre for Infectious Diseases and Microbiology Public Health | NSW Health Pathology - Institute of Clinical Pathology and Medical Research; Westmead Hospital; University of Sydney | Bachmann N, Rockett R, Lam C, Gray K, Timms V, Gall M, Arnott A, Sadsad R, Draper J, Sim E, Carter I, Holmes EC, O’Sullivan MV, Byun R, Sintchenko V, Chen SC, Eden JS, Maddocks S, Kok J, Propenko M, Sorrell T, Chang S, Basile K, Dwyer DE for the 2019-nCoV Study Group |
| EPI_ISL_427649 | Centre for Infectious Diseases and Microbiology Public Health | NSW Health Pathology - Institute of Clinical Pathology and Medical Research; Westmead Hospital; University of Sydney | Timms V, Gall M, Arnott A, Sadsad R, Draper J, Sim E, Bachmann N, Rockett R, Lam C, Gray K, Carter I, Holmes EC, O’Sullivan MV, Byun R, Sintchenko V, Chen SC, Eden JS, Maddocks S, Kok J, Propenko M, Sorrell T, Chang S, Basile K, Dwyer DE for the 2019-nCoV Study Group |
| EPI_ISL_427650 | Centre for Infectious Diseases and Microbiology Public Health | NSW Health Pathology - Institute of Clinical Pathology and Medical Research; Westmead Hospital; University of Sydney | Gall M, Arnott A, Sadsad R, Draper J, Sim E, Bachmann N, Rockett R, Lam C, Gray K, Timms V, Carter I, Holmes EC, O’Sullivan MV, Byun R, Sintchenko V, Chen SC, Eden JS, Maddocks S, Kok J, Propenko M, Sorrell T, Chang S, Basile K, Dwyer DE for the 2019-nCoV Study Group |
| EPI_ISL_427651 | Centre for Infectious Diseases and Microbiology Public Health | NSW Health Pathology - Institute of Clinical Pathology and Medical Research; Westmead Hospital; University of Sydney | Arnott A, Sadsad R, Draper J, Sim E, Bachmann N, Rockett R, Lam C, Gray K, Timms V, Gall M, Carter I, Holmes EC, O’Sullivan MV, Byun R, Sintchenko V, Chen SC, Eden JS, Maddocks S, Kok J, Propenko M, Sorrell T, Chang S, Basile K, Dwyer DE for the 2019-nCoV Study Group |
| EPI_ISL_427652 | Centre for Infectious Diseases and Microbiology Public Health | NSW Health Pathology - Institute of Clinical Pathology and Medical Research; Westmead Hospital; University of Sydney | Lam C, Gray K, Timms, V, Gall M, Arnott A, Sadsad R, Draper J, Sim E, Bachmann N, Rockett R, Carter I, Holmes EC, O’Sullivan MV, Byun R, Sintchenko V, Chen SC, Eden JS, Maddocks S, Kok J, Propenko M, Sorrell T, Chang S, Basile K, Dwyer DE for the 2019-nCoV Study Group |
| EPI_ISL_427653 | Centre for Infectious Diseases and Microbiology Public Health | NSW Health Pathology - Institute of Clinical Pathology and Medical Research; Westmead Hospital; University of Sydney | Rockett R, Lam C, Gray K, Timms V, Gall M, Arnott A, Sadsad R, Draper J, Sim E, Bachmann N, Carter I, Holmes EC, O’Sullivan MV, Byun R, Sintchenko V, Chen SC, Eden JS, Maddocks S, Kok J, Propenko M, Sorrell T, Chang S, Basile K, Dwyer DE for the 2019-nCoV Study Group |
| EPI_ISL_427654 | Centre for Infectious Diseases and Microbiology Public Health | NSW Health Pathology - Institute of Clinical Pathology and Medical Research; Westmead Hospital; University of Sydney | Sadsad R, Draper J, Sim E, Bachmann N, Rockett R, Lam C, Gray K, Timms V, Gall M, Arnott A, Carter I, Holmes EC, O’Sullivan MV, Byun R, Sintchenko V, Chen SC, Eden JS, Maddocks S, Kok J, Propenko M, Sorrell T, Chang S, Basile K, Dwyer DE for the 2019-nCoV Study Group |
| EPI_ISL_427655 | Centre for Infectious Diseases and Microbiology Public Health | NSW Health Pathology - Institute of Clinical Pathology and Medical Research; Westmead Hospital; University of Sydney | Draper J, Sim E, Bachmann N, Rockett R, Lam C, Gray K, Timms V, Gall M, Arnott A, Sadsad R, Carter I, Holmes EC, O’Sullivan MV, Byun R, Sintchenko V, Chen SC, Eden JS, Maddocks S, Kok J, Propenko M, Sorrell T, Chang S, Basile K, Dwyer DE for the 2019-nCoV Study Group |
| EPI_ISL_427656 | Centre for Infectious Diseases and Microbiology Public Health | NSW Health Pathology - Institute of Clinical Pathology and Medical Research; Westmead Hospital; University of Sydney | Bachmann N, Rockett R, Lam C, Gray K, Timms V, Gall M, Arnott A, Sadsad R, Draper J, Sim E, Carter I, Holmes EC, O’Sullivan MV, Byun R, Sintchenko V, Chen SC, Eden JS, Maddocks S, Kok J, Propenko M, Sorrell T, Chang S, Basile K, Dwyer DE for the 2019-nCoV Study Group |
| EPI_ISL_427657 | Centre for Infectious Diseases and Microbiology Public Health | NSW Health Pathology - Institute of Clinical Pathology and Medical Research; Westmead Hospital; University of Sydney | Rockett R, Lam C, Gray K, Timms V, Gall M, Arnott A, Sadsad R, Draper J, Sim E, Bachmann N, Carter I, Holmes EC, O’Sullivan MV, Byun R, Sintchenko V, Chen SC, Eden JS, Maddocks S, Kok J, Propenko M, Sorrell T, Chang S, Basile K, Dwyer DE for the 2019-nCoV Study Group |
| EPI_ISL_427658 | Centre for Infectious Diseases and Microbiology Public Health | NSW Health Pathology - Institute of Clinical Pathology and Medical Research; Westmead Hospital; University of Sydney | Sim E, Bachmann N, Rockett R, Lam C, Gray K, Timms V, Gall M, Arnott A, Sadsad R, Draper J, Carter I, Holmes EC, O’Sullivan MV, Byun R, Sintchenko V, Chen SC, Eden JS, Maddocks S, Kok J, Propenko M, Sorrell T, Chang S, Basile K, Dwyer DE for the 2019-nCoV Study Group |
| EPI_ISL_427659 | Centre for Infectious Diseases and Microbiology Public Health | NSW Health Pathology - Institute of Clinical Pathology and Medical Research; Westmead Hospital; University of Sydney | Bachmann N, Rockett R, Lam C, Gray K, Timms V, Gall M, Arnott A, Sadsad R, Draper J, Sim E, Carter I, Holmes EC, O’Sullivan MV, Byun R, Sintchenko V, Chen SC, Eden JS, Maddocks S, Kok J, Propenko M, Sorrell T, Chang S, Basile K, Dwyer DE for the 2019-nCoV Study Group |
| EPI_ISL_427660 | Centre for Infectious Diseases and Microbiology Public Health | NSW Health Pathology - Institute of Clinical Pathology and Medical Research; Westmead Hospital; University of Sydney | Rockett R, Lam C, Gray K, Timms V, Gall M, Arnott A, Sadsad R, Draper J, Sim E, Bachmann N, Carter I, Holmes EC, O’Sullivan MV, Byun R, Sintchenko V, Chen SC, Eden JS, Maddocks S, Kok J, Propenko M, Sorrell T, Chang S, Basile K, Dwyer DE for the 2019-nCoV Study Group |
| EPI_ISL_427661 | Centre for Infectious Diseases and Microbiology Public Health | NSW Health Pathology - Institute of Clinical Pathology and Medical Research; Westmead Hospital; University of Sydney | Sim E, Bachmann N, Rockett R, Lam C, Gray K, Timms V, Gall M, Arnott A, Sadsad R, Draper J, Carter I, Holmes EC, O’Sullivan MV, Byun R, Sintchenko V, Chen SC, Eden JS, Maddocks S, Kok J, Propenko M, Sorrell T, Chang S, Basile K, Dwyer DE for the 2019-nCoV Study Group |
| EPI_ISL_427662 | Centre for Infectious Diseases and Microbiology Public Health | NSW Health Pathology - Institute of Clinical Pathology and Medical Research; Westmead Hospital; University of Sydney | Lam C, Gray K, Timms, V, Gall M, Arnott A, Sadsad R, Draper J, Sim E, Bachmann N, Rockett R, Carter I, Holmes EC, O’Sullivan MV, Byun R, Sintchenko V, Chen SC, Eden JS, Maddocks S, Kok J, Propenko M, Sorrell T, Chang S, Basile K, Dwyer DE for the 2019-nCoV Study Group |
| EPI_ISL_427663, EPI_ISL_427664, EPI_ISL_427665 | Centre for Infectious Diseases and Microbiology Public Health | NSW Health Pathology - Institute of Clinical Pathology and Medical Research; Westmead Hospital; University of Sydney | Timms V, Gall M, Arnott A, Sadsad R, Draper J, Sim E, Bachmann N, Rockett R, Lam C, Gray K, Carter I, Holmes EC, O’Sullivan MV, Byun R, Sintchenko V, Chen SC, Eden JS, Maddocks S, Kok J, Propenko M, Sorrell T, Chang S, Basile K, Dwyer DE for the 2019-nCoV Study Group |
| EPI_ISL_427666 | Centre for Infectious Diseases and Microbiology Public Health | NSW Health Pathology - Institute of Clinical Pathology and Medical Research; Westmead Hospital; University of Sydney | Bachmann N, Rockett R, Lam C, Gray K, Timms V, Gall M, Arnott A, Sadsad R, Draper J, Sim E, Carter I, Holmes EC, O’Sullivan MV, Byun R, Sintchenko V, Chen SC, Eden JS, Maddocks S, Kok J, Propenko M, Sorrell T, Chang S, Basile K, Dwyer DE for the 2019-nCoV Study Group |
| EPI_ISL_427667 | Centre for Infectious Diseases and Microbiology Public Health | NSW Health Pathology - Institute of Clinical Pathology and Medical Research; Westmead Hospital; University of Sydney | Gall M, Arnott A, Sadsad R, Draper J, Sim E, Bachmann N, Rockett R, Lam C, Gray K, Timms V, Carter I, Holmes EC, O’Sullivan MV, Byun R, Sintchenko V, Chen SC, Eden JS, Maddocks S, Kok J, Propenko M, Sorrell T, Chang S, Basile K, Dwyer DE for the 2019-nCoV Study Group |
| EPI_ISL_427668 | Centre for Infectious Diseases and Microbiology Public Health | NSW Health Pathology - Institute of Clinical Pathology and Medical Research; Westmead Hospital; University of Sydney | Lam C, Gray K, Timms, V, Gall M, Arnott A, Sadsad R, Draper J, Sim E, Bachmann N, Rockett R, Carter I, Holmes EC, O’Sullivan MV, Byun R, Sintchenko V, Chen SC, Eden JS, Maddocks S, Kok J, Propenko M, Sorrell T, Chang S, Basile K, Dwyer DE for the 2019-nCoV Study Group |
| EPI_ISL_427669 | Centre for Infectious Diseases and Microbiology Public Health | NSW Health Pathology - Institute of Clinical Pathology and Medical Research; Westmead Hospital; University of Sydney | Sadsad R, Draper J, Sim E, Bachmann N, Rockett R, Lam C, Gray K, Timms V, Gall M, Arnott A, Carter I, Holmes EC, O’Sullivan MV, Byun R, Sintchenko V, Chen SC, Eden JS, Maddocks S, Kok J, Propenko M, Sorrell T, Chang S, Basile K, Dwyer DE for the 2019-nCoV Study Group |
| EPI_ISL_427670 | Centre for Infectious Diseases and Microbiology Public Health | NSW Health Pathology - Institute of Clinical Pathology and Medical Research; Westmead Hospital; University of Sydney | Arnott A, Sadsad R, Draper J, Sim E, Bachmann N, Rockett R, Lam C, Gray K, Timms V, Gall M, Carter I, Holmes EC, O’Sullivan MV, Byun R, Sintchenko V, Chen SC, Eden JS, Maddocks S, Kok J, Propenko M, Sorrell T, Chang S, Basile K, Dwyer DE for the 2019-nCoV Study Group |
| EPI_ISL_427671 | Centre for Infectious Diseases and Microbiology Public Health | NSW Health Pathology - Institute of Clinical Pathology and Medical Research; Westmead Hospital; University of Sydney | Bachmann N, Rockett R, Lam C, Gray K, Timms V, Gall M, Arnott A, Sadsad R, Draper J, Sim E, Carter I, Holmes EC, O’Sullivan MV, Byun R, Sintchenko V, Chen SC, Eden JS, Maddocks S, Kok J, Propenko M, Sorrell T, Chang S, Basile K, Dwyer DE for the 2019-nCoV Study Group |
| EPI_ISL_427672, EPI_ISL_427673 | Centre for Infectious Diseases and Microbiology Public Health | NSW Health Pathology - Institute of Clinical Pathology and Medical Research; Westmead Hospital; University of Sydney | Gray K, Timms V, Gall M, Arnott A, Sadsad R, Draper J, Sim E, Bachmann N, Rockett R, Lam C, Carter I, Holmes EC, O’Sullivan MV, Byun R, Sintchenko V, Chen SC, Eden JS, Maddocks S, Kok J, Propenko M, Sorrell T, Chang S, Basile K, Dwyer DE for the 2019-nCoV Study Group |
| EPI_ISL_427674 | Centre for Infectious Diseases and Microbiology Public Health | NSW Health Pathology - Institute of Clinical Pathology and Medical Research; Westmead Hospital; University of Sydney | Gall M, Arnott A, Sadsad R, Draper J, Sim E, Bachmann N, Rockett R, Lam C, Gray K, Timms V, Carter I, Holmes EC, O’Sullivan MV, Byun R, Sintchenko V, Chen SC, Eden JS, Maddocks S, Kok J, Propenko M, Sorrell T, Chang S, Basile K, Dwyer DE for the 2019-nCoV Study Group |
| EPI_ISL_427675 | Centre for Infectious Diseases and Microbiology Public Health | NSW Health Pathology - Institute of Clinical Pathology and Medical Research; Westmead Hospital; University of Sydney | Bachmann N, Rockett R, Lam C, Gray K, Timms V, Gall M, Arnott A, Sadsad R, Draper J, Sim E, Carter I, Holmes EC, O’Sullivan MV, Byun R, Sintchenko V, Chen SC, Eden JS, Maddocks S, Kok J, Propenko M, Sorrell T, Chang S, Basile K, Dwyer DE for the 2019-nCoV Study Group |
| EPI_ISL_427676 | Centre for Infectious Diseases and Microbiology Public Health | NSW Health Pathology - Institute of Clinical Pathology and Medical Research; Westmead Hospital; University of Sydney | Draper J, Sim E, Bachmann N, Rockett R, Lam C, Gray K, Timms V, Gall M, Arnott A, Sadsad R, Carter I, Holmes EC, O’Sullivan MV, Byun R, Sintchenko V, Chen SC, Eden JS, Maddocks S, Kok J, Propenko M, Sorrell T, Chang S, Basile K, Dwyer DE for the 2019-nCoV Study Group |
| EPI_ISL_427677 | Centre for Infectious Diseases and Microbiology Public Health | NSW Health Pathology - Institute of Clinical Pathology and Medical Research; Westmead Hospital; University of Sydney | Lam C, Gray K, Timms, V, Gall M, Arnott A, Sadsad R, Draper J, Sim E, Bachmann N, Rockett R, Carter I, Holmes EC, O’Sullivan MV, Byun R, Sintchenko V, Chen SC, Eden JS, Maddocks S, Kok J, Propenko M, Sorrell T, Chang S, Basile K, Dwyer DE for the 2019-nCoV Study Group |
| EPI_ISL_427678 | Centre for Infectious Diseases and Microbiology Public Health | NSW Health Pathology - Institute of Clinical Pathology and Medical Research; Westmead Hospital; University of Sydney | Timms V, Gall M, Arnott A, Sadsad R, Draper J, Sim E, Bachmann N, Rockett R, Lam C, Gray K, Carter I, Holmes EC, O’Sullivan MV, Byun R, Sintchenko V, Chen SC, Eden JS, Maddocks S, Kok J, Propenko M, Sorrell T, Chang S, Basile K, Dwyer DE for the 2019-nCoV Study Group |
| EPI_ISL_427679, EPI_ISL_427680 | Centre for Infectious Diseases and Microbiology Public Health | NSW Health Pathology - Institute of Clinical Pathology and Medical Research; Westmead Hospital; University of Sydney | Draper J, Sim E, Bachmann N, Rockett R, Lam C, Gray K, Timms V, Gall M, Arnott A, Sadsad R, Carter I, Holmes EC, O’Sullivan MV, Byun R, Sintchenko V, Chen SC, Eden JS, Maddocks S, Kok J, Propenko M, Sorrell T, Chang S, Basile K, Dwyer DE for the 2019-nCoV Study Group |
| EPI_ISL_427681 | Centre for Infectious Diseases and Microbiology Public Health | NSW Health Pathology - Institute of Clinical Pathology and Medical Research; Westmead Hospital; University of Sydney | Rockett R, Lam C, Gray K, Timms V, Gall M, Arnott A, Sadsad R, Draper J, Sim E, Bachmann N, Carter I, Holmes EC, O’Sullivan MV, Byun R, Sintchenko V, Chen SC, Eden JS, Maddocks S, Kok J, Propenko M, Sorrell T, Chang S, Basile K, Dwyer DE for the 2019-nCoV Study Group |
| EPI_ISL_427682 | Centre for Infectious Diseases and Microbiology Public Health | NSW Health Pathology - Institute of Clinical Pathology and Medical Research; Westmead Hospital; University of Sydney | Timms V, Gall M, Arnott A, Sadsad R, Draper J, Sim E, Bachmann N, Rockett R, Lam C, Gray K, Carter I, Holmes EC, O’Sullivan MV, Byun R, Sintchenko V, Chen SC, Eden JS, Maddocks S, Kok J, Propenko M, Sorrell T, Chang S, Basile K, Dwyer DE for the 2019-nCoV Study Group |
| EPI_ISL_427683 | Centre for Infectious Diseases and Microbiology Public Health | NSW Health Pathology - Institute of Clinical Pathology and Medical Research; Westmead Hospital; University of Sydney | Arnott A, Sadsad R, Draper J, Sim E, Bachmann N, Rockett R, Lam C, Gray K, Timms V, Gall M, Carter I, Holmes EC, O’Sullivan MV, Byun R, Sintchenko V, Chen SC, Eden JS, Maddocks S, Kok J, Propenko M, Sorrell T, Chang S, Basile K, Dwyer DE for the 2019-nCoV Study Group |
| EPI_ISL_427684 | Centre for Infectious Diseases and Microbiology Public Health | NSW Health Pathology - Institute of Clinical Pathology and Medical Research; Westmead Hospital; University of Sydney | Draper J, Sim E, Bachmann N, Rockett R, Lam C, Gray K, Timms V, Gall M, Arnott A, Sadsad R, Carter I, Holmes EC, O’Sullivan MV, Byun R, Sintchenko V, Chen SC, Eden JS, Maddocks S, Kok J, Propenko M, Sorrell T, Chang S, Basile K, Dwyer DE for the 2019-nCoV Study Group |
| EPI_ISL_427685 | Centre for Infectious Diseases and Microbiology Public Health | NSW Health Pathology - Institute of Clinical Pathology and Medical Research; Westmead Hospital; University of Sydney | Sadsad R, Draper J, Sim E, Bachmann N, Rockett R, Lam C, Gray K, Timms V, Gall M, Arnott A, Carter I, Holmes EC, O’Sullivan MV, Byun R, Sintchenko V, Chen SC, Eden JS, Maddocks S, Kok J, Propenko M, Sorrell T, Chang S, Basile K, Dwyer DE for the 2019-nCoV Study Group |
| EPI_ISL_427686, EPI_ISL_427687 | Centre for Infectious Diseases and Microbiology Public Health | NSW Health Pathology - Institute of Clinical Pathology and Medical Research; Westmead Hospital; University of Sydney | Gray K, Timms V, Gall M, Arnott A, Sadsad R, Draper J, Sim E, Bachmann N, Rockett R, Lam C, Carter I, Holmes EC, O’Sullivan MV, Byun R, Sintchenko V, Chen SC, Eden JS, Maddocks S, Kok J, Propenko M, Sorrell T, Chang S, Basile K, Dwyer DE for the 2019-nCoV Study Group |
| EPI_ISL_427688 | Centre for Infectious Diseases and Microbiology Public Health | NSW Health Pathology - Institute of Clinical Pathology and Medical Research; Westmead Hospital; University of Sydney | Sim E, Bachmann N, Rockett R, Lam C, Gray K, Timms V, Gall M, Arnott A, Sadsad R, Draper J, Carter I, Holmes EC, O’Sullivan MV, Byun R, Sintchenko V, Chen SC, Eden JS, Maddocks S, Kok J, Propenko M, Sorrell T, Chang S, Basile K, Dwyer DE for the 2019-nCoV Study Group |
| EPI_ISL_427689 | Centre for Infectious Diseases and Microbiology Public Health | NSW Health Pathology - Institute of Clinical Pathology and Medical Research; Westmead Hospital; University of Sydney | Arnott A, Sadsad R, Draper J, Sim E, Bachmann N, Rockett R, Lam C, Gray K, Timms V, Gall M, Carter I, Holmes EC, O’Sullivan MV, Byun R, Sintchenko V, Chen SC, Eden JS, Maddocks S, Kok J, Propenko M, Sorrell T, Chang S, Basile K, Dwyer DE for the 2019-nCoV Study Group |
| EPI_ISL_427690 | Centre for Infectious Diseases and Microbiology Public Health | NSW Health Pathology - Institute of Clinical Pathology and Medical Research; Westmead Hospital; University of Sydney | Draper J, Sim E, Bachmann N, Rockett R, Lam C, Gray K, Timms V, Gall M, Arnott A, Sadsad R, Carter I, Holmes EC, O’Sullivan MV, Byun R, Sintchenko V, Chen SC, Eden JS, Maddocks S, Kok J, Propenko M, Sorrell T, Chang S, Basile K, Dwyer DE for the 2019-nCoV Study Group |
| EPI_ISL_427691 | Centre for Infectious Diseases and Microbiology Public Health | NSW Health Pathology - Institute of Clinical Pathology and Medical Research; Westmead Hospital; University of Sydney | Gall M, Arnott A, Sadsad R, Draper J, Sim E, Bachmann N, Rockett R, Lam C, Gray K, Timms V, Carter I, Holmes EC, O’Sullivan MV, Byun R, Sintchenko V, Chen SC, Eden JS, Maddocks S, Kok J, Propenko M, Sorrell T, Chang S, Basile K, Dwyer DE for the 2019-nCoV Study Group |
| EPI_ISL_427692 | Centre for Infectious Diseases and Microbiology Public Health | NSW Health Pathology - Institute of Clinical Pathology and Medical Research; Westmead Hospital; University of Sydney | Timms V, Gall M, Arnott A, Sadsad R, Draper J, Sim E, Bachmann N, Rockett R, Lam C, Gray K, Carter I, Holmes EC, O’Sullivan MV, Byun R, Sintchenko V, Chen SC, Eden JS, Maddocks S, Kok J, Propenko M, Sorrell T, Chang S, Basile K, Dwyer DE for the 2019-nCoV Study Group |
| EPI_ISL_427693, EPI_ISL_427694 | Centre for Infectious Diseases and Microbiology Public Health | NSW Health Pathology - Institute of Clinical Pathology and Medical Research; Westmead Hospital; University of Sydney | Arnott A, Sadsad R, Draper J, Sim E, Bachmann N, Rockett R, Lam C, Gray K, Timms V, Gall M, Carter I, Holmes EC, O’Sullivan MV, Byun R, Sintchenko V, Chen SC, Eden JS, Maddocks S, Kok J, Propenko M, Sorrell T, Chang S, Basile K, Dwyer DE for the 2019-nCoV Study Group |
| EPI_ISL_427695 | Centre for Infectious Diseases and Microbiology Public Health | NSW Health Pathology - Institute of Clinical Pathology and Medical Research; Westmead Hospital; University of Sydney | Timms V, Gall M, Arnott A, Sadsad R, Draper J, Sim E, Bachmann N, Rockett R, Lam C, Gray K, Carter I, Holmes EC, O’Sullivan MV, Byun R, Sintchenko V, Chen SC, Eden JS, Maddocks S, Kok J, Propenko M, Sorrell T, Chang S, Basile K, Dwyer DE for the 2019-nCoV Study Group |
| EPI_ISL_427696 | Centre for Infectious Diseases and Microbiology Public Health | NSW Health Pathology - Institute of Clinical Pathology and Medical Research; Westmead Hospital; University of Sydney | Gray K, Timms V, Gall M, Arnott A, Sadsad R, Draper J, Sim E, Bachmann N, Rockett R, Lam C, Carter I, Holmes EC, O’Sullivan MV, Byun R, Sintchenko V, Chen SC, Eden JS, Maddocks S, Kok J, Propenko M, Sorrell T, Chang S, Basile K, Dwyer DE for the 2019-nCoV Study Group |
| EPI_ISL_427697 | Centre for Infectious Diseases and Microbiology Public Health | NSW Health Pathology - Institute of Clinical Pathology and Medical Research; Westmead Hospital; University of Sydney | Gall M, Arnott A, Sadsad R, Draper J, Sim E, Bachmann N, Rockett R, Lam C, Gray K, Timms V, Carter I, Holmes EC, O’Sullivan MV, Byun R, Sintchenko V, Chen SC, Eden JS, Maddocks S, Kok J, Propenko M, Sorrell T, Chang S, Basile K, Dwyer DE for the 2019-nCoV Study Group |
| EPI_ISL_427698 | Centre for Infectious Diseases and Microbiology Public Health | NSW Health Pathology - Institute of Clinical Pathology and | Timms V, Gall M, Arnott A, Sadsad R, Draper J, Sim E, Bachmann N, Rockett R, Lam C, Gray K, Carter I, Holmes EC, O’Sullivan MV, Byun R, Sintchenko V, Chen SC, Eden JS, Maddocks S, Kok J, Propenko M, Sorrell T, |

| | | | |
|---|---|---|---|
| | | Medical Research; Westmead Hospital; University of Sydney | Chang S, Basile K, Dwyer DE for the 2019-nCoV Study Group |
| EPI_ISL_427699 | Centre for Infectious Diseases and Microbiology Public Health | NSW Health Pathology - Institute of Clinical Pathology and Medical Research; Westmead Hospital; University of Sydney | Sim E, Bachmann N, Rockett R, Lam C, Gray K, Timms V, Gall M, Arnott A, Sadsad R, Draper J, Carter I, Holmes EC, O’Sullivan MV, Byun R, Sintchenko V, Chen SC, Eden JS, Maddocks S, Kok J, Propenko M, Sorrell T, Chang S, Basile K, Dwyer DE for the 2019-nCoV Study Group |
| EPI_ISL_427700 | Centre for Infectious Diseases and Microbiology Public Health | NSW Health Pathology - Institute of Clinical Pathology and Medical Research; Westmead Hospital; University of Sydney | Gall M, Arnott A, Sadsad R, Draper J, Sim E, Bachmann N, Rockett R, Lam C, Gray K, Timms V, Carter I, Holmes EC, O’Sullivan MV, Byun R, Sintchenko V, Chen SC, Eden JS, Maddocks S, Kok J, Propenko M, Sorrell T, Chang S, Basile K, Dwyer DE for the 2019-nCoV Study Group |
| EPI_ISL_427701 | Centre for Infectious Diseases and Microbiology Public Health | NSW Health Pathology - Institute of Clinical Pathology and Medical Research; Westmead Hospital; University of Sydney | Draper J, Sim E, Bachmann N, Rockett R, Lam C, Gray K, Timms V, Gall M, Arnott A, Sadsad R, Carter I, Holmes EC, O’Sullivan MV, Byun R, Sintchenko V, Chen SC, Eden JS, Maddocks S, Kok J, Propenko M, Sorrell T, Chang S, Basile K, Dwyer DE for the 2019-nCoV Study Group |
| EPI_ISL_427702 | Centre for Infectious Diseases and Microbiology Public Health | NSW Health Pathology - Institute of Clinical Pathology and Medical Research; Westmead Hospital; University of Sydney | Arnott A, Sadsad R, Draper J, Sim E, Bachmann N, Rockett R, Lam C, Gray K, Timms V, Gall M, Carter I, Holmes EC, O’Sullivan MV, Byun R, Sintchenko V, Chen SC, Eden JS, Maddocks S, Kok J, Propenko M, Sorrell T, Chang S, Basile K, Dwyer DE for the 2019-nCoV Study Group |
| EPI_ISL_427703 | Centre for Infectious Diseases and Microbiology Public Health | NSW Health Pathology - Institute of Clinical Pathology and Medical Research; Westmead Hospital; University of Sydney | Gray K, Timms V, Gall M, Arnott A, Sadsad R, Draper J, Sim E, Bachmann N, Rockett R, Lam C, Carter I, Holmes EC, O’Sullivan MV, Byun R, Sintchenko V, Chen SC, Eden JS, Maddocks S, Kok J, Propenko M, Sorrell T, Chang S, Basile K, Dwyer DE for the 2019-nCoV Study Group |
| EPI_ISL_427704 | Centre for Infectious Diseases and Microbiology Public Health | NSW Health Pathology - Institute of Clinical Pathology and Medical Research; Westmead Hospital; University of Sydney | Timms V, Gall M, Arnott A, Sadsad R, Draper J, Sim E, Bachmann N, Rockett R, Lam C, Gray K, Carter I, Holmes EC, O’Sullivan MV, Byun R, Sintchenko V, Chen SC, Eden JS, Maddocks S, Kok J, Propenko M, Sorrell T, Chang S, Basile K, Dwyer DE for the 2019-nCoV Study Group |
| EPI_ISL_427705 | Centre for Infectious Diseases and Microbiology Public Health | NSW Health Pathology - Institute of Clinical Pathology and Medical Research; Westmead Hospital; University of Sydney | Sadsad R, Draper J, Sim E, Bachmann N, Rockett R, Lam C, Gray K, Timms V, Gall M, Arnott A, Carter I, Holmes EC, O’Sullivan MV, Byun R, Sintchenko V, Chen SC, Eden JS, Maddocks S, Kok J, Propenko M, Sorrell T, Chang S, Basile K, Dwyer DE for the 2019-nCoV Study Group |
| EPI_ISL_427706 | Centre for Infectious Diseases and Microbiology Public Health | NSW Health Pathology - Institute of Clinical Pathology and Medical Research; Westmead Hospital; University of Sydney | Lam C, Gray K, Timms, V, Gall M, Arnott A, Sadsad R, Draper J, Sim E, Bachmann N, Rockett R, Carter I, Holmes EC, O’Sullivan MV, Byun R, Sintchenko V, Chen SC, Eden JS, Maddocks S, Kok J, Propenko M, Sorrell T, Chang S, Basile K, Dwyer DE for the 2019-nCoV Study Group |
| EPI_ISL_427707 | Centre for Infectious Diseases and Microbiology Public Health | NSW Health Pathology - Institute of Clinical Pathology and Medical Research; Westmead Hospital; University of Sydney | Arnott A, Sadsad R, Draper J, Sim E, Bachmann N, Rockett R, Lam C, Gray K, Timms V, Gall M, Carter I, Holmes EC, O’Sullivan MV, Byun R, Sintchenko V, Chen SC, Eden JS, Maddocks S, Kok J, Propenko M, Sorrell T, Chang S, Basile K, Dwyer DE for the 2019-nCoV Study Group |
| EPI_ISL_427708 | Centre for Infectious Diseases and Microbiology Public Health | NSW Health Pathology - Institute of Clinical Pathology and Medical Research; Westmead Hospital; University of Sydney | Gray K, Timms V, Gall M, Arnott A, Sadsad R, Draper J, Sim E, Bachmann N, Rockett R, Lam C, Carter I, Holmes EC, O’Sullivan MV, Byun R, Sintchenko V, Chen SC, Eden JS, Maddocks S, Kok J, Propenko M, Sorrell T, Chang S, Basile K, Dwyer DE for the 2019-nCoV Study Group |
| EPI_ISL_427709, EPI_ISL_427710 | Centre for Infectious Diseases and Microbiology Public Health | NSW Health Pathology - Institute of Clinical Pathology and Medical Research; Westmead Hospital; University of Sydney | Sim E, Bachmann N, Rockett R, Lam C, Gray K, Timms V, Gall M, Arnott A, Sadsad R, Draper J, Carter I, Holmes EC, O’Sullivan MV, Byun R, Sintchenko V, Chen SC, Eden JS, Maddocks S, Kok J, Propenko M, Sorrell T, Chang S, Basile K, Dwyer DE for the 2019-nCoV Study Group |
| EPI_ISL_427711 | ACT Pathology, The Canberra Hospital | NSW Health Pathology - Institute of Clinical Pathology and Medical Research; Westmead Hospital; University of Sydney | Rockett R, Lam C, Gray K, Timms V, Gall M, Arnott A, Sadsad R, Draper J, Sim E, Bachmann N, Carter I, Holmes EC, O’Sullivan MV, Byun R, Sintchenko V, Chen SC, Eden JS, Maddocks S, Kok J, Propenko M, Sorrell T, Chang S, Basile K, Dwyer DE for the 2019-nCoV Study Group |
| EPI_ISL_427712 | Centre for Infectious Diseases and Microbiology Public Health | NSW Health Pathology - Institute of Clinical Pathology and Medical Research; Westmead Hospital; University of Sydney | Sim E, Bachmann N, Rockett R, Lam C, Gray K, Timms V, Gall M, Arnott A, Sadsad R, Draper J, Carter I, Holmes EC, O’Sullivan MV, Byun R, Sintchenko V, Chen SC, Eden JS, Maddocks S, Kok J, Propenko M, Sorrell T, Chang S, Basile K, Dwyer DE for the 2019-nCoV Study Group |
| EPI_ISL_427713 | Centre for Infectious Diseases and Microbiology Public Health | NSW Health Pathology - Institute of Clinical Pathology and Medical Research; Westmead Hospital; University of Sydney | Bachmann N, Rockett R, Lam C, Gray K, Timms V, Gall M, Arnott A, Sadsad R, Draper J, Sim E, Carter I, Holmes EC, O’Sullivan MV, Byun R, Sintchenko V, Chen SC, Eden JS, Maddocks S, Kok J, Propenko M, Sorrell T, Chang S, Basile K, Dwyer DE for the 2019-nCoV Study Group |
| EPI_ISL_427714 | Centre for Infectious Diseases and Microbiology Public Health | NSW Health Pathology - Institute of Clinical Pathology and Medical Research; Westmead Hospital; University of Sydney | Rockett R, Lam C, Gray K, Timms V, Gall M, Arnott A, Sadsad R, Draper J, Sim E, Bachmann N, Carter I, Holmes EC, O’Sullivan MV, Byun R, Sintchenko V, Chen SC, Eden JS, Maddocks S, Kok J, Propenko M, Sorrell T, Chang S, Basile K, Dwyer DE for the 2019-nCoV Study Group |
| EPI_ISL_427715, EPI_ISL_427716 | Centre for Infectious Diseases and Microbiology Public Health | NSW Health Pathology - Institute of Clinical Pathology and Medical Research; Westmead Hospital; University of Sydney | Lam C, Gray K, Timms, V, Gall M, Arnott A, Sadsad R, Draper J, Sim E, Bachmann N, Rockett R, Carter I, Holmes EC, O’Sullivan MV, Byun R, Sintchenko V, Chen SC, Eden JS, Maddocks S, Kok J, Propenko M, Sorrell T, Chang S, Basile K, Dwyer DE for the 2019-nCoV Study Group |
| EPI_ISL_427717 | ACT Pathology, The Canberra Hospital | NSW Health Pathology - Institute of Clinical Pathology and Medical Research; Westmead Hospital; University of Sydney | Sim E, Bachmann N, Rockett R, Lam C, Gray K, Timms V, Gall M, Arnott A, Sadsad R, Draper J, Carter I, Holmes EC, O’Sullivan MV, Byun R, Sintchenko V, Chen SC, Eden JS, Maddocks S, Kok J, Propenko M, Sorrell T, Chang S, Basile K, Dwyer DE for the 2019-nCoV Study Group |
| EPI_ISL_427718 | ACT Pathology, The Canberra Hospital | NSW Health Pathology - Institute of Clinical Pathology and Medical Research; Westmead Hospital; University of Sydney | Gray K, Timms V, Gall M, Arnott A, Sadsad R, Draper J, Sim E, Bachmann N, Rockett R, Lam C, Carter I, Holmes EC, O’Sullivan MV, Byun R, Sintchenko V, Chen SC, Eden JS, Maddocks S, Kok J, Propenko M, Sorrell T, Chang S, Basile K, Dwyer DE for the 2019-nCoV Study Group |
| EPI_ISL_427719 | ACT Pathology, The Canberra Hospital | NSW Health Pathology - Institute of Clinical Pathology and Medical Research; Westmead Hospital; University of Sydney | Timms V, Gall M, Arnott A, Sadsad R, Draper J, Sim E, Bachmann N, Rockett R, Lam C, Gray K, Carter I, Holmes EC, O’Sullivan MV, Byun R, Sintchenko V, Chen SC, Eden JS, Maddocks S, Kok J, Propenko M, Sorrell T, Chang S, Basile K, Dwyer DE for the 2019-nCoV Study Group |
| EPI_ISL_427720 | Centre for Infectious Diseases and Microbiology Public Health | NSW Health Pathology - Institute of Clinical Pathology and Medical Research; Westmead Hospital; University of Sydney | Sim E, Bachmann N, Rockett R, Lam C, Gray K, Timms V, Gall M, Arnott A, Sadsad R, Draper J, Carter I, Holmes EC, O’Sullivan MV, Byun R, Sintchenko V, Chen SC, Eden JS, Maddocks S, Kok J, Propenko M, Sorrell T, Chang S, Basile K, Dwyer DE for the 2019-nCoV Study Group |
| EPI_ISL_427721 | ACT Pathology, The Canberra Hospital | NSW Health Pathology - Institute of Clinical Pathology and Medical Research; Westmead Hospital; University of Sydney | Sim E, Bachmann N, Rockett R, Lam C, Gray K, Timms V, Gall M, Arnott A, Sadsad R, Draper J, Carter I, Holmes EC, O’Sullivan MV, Byun R, Sintchenko V, Chen SC, Eden JS, Maddocks S, Kok J, Propenko M, Sorrell T, Chang S, Basile K, Dwyer DE for the 2019-nCoV Study Group |
| EPI_ISL_427722 | Centre for Infectious Diseases and Microbiology Public Health | NSW Health Pathology - Institute of Clinical Pathology and Medical Research; Westmead Hospital; University of Sydney | Draper J, Sim E, Bachmann N, Rockett R, Lam C, Gray K, Timms V, Gall M, Arnott A, Sadsad R, Carter I, Holmes EC, O’Sullivan MV, Byun R, Sintchenko V, Chen SC, Eden JS, Maddocks S, Kok J, Propenko M, Sorrell T, Chang S, Basile K, Dwyer DE for the 2019-nCoV Study Group |
| EPI_ISL_427723, EPI_ISL_427724 | Centre for Infectious Diseases and Microbiology Public Health | NSW Health Pathology - Institute of Clinical Pathology and Medical Research; Westmead Hospital; University of Sydney | Lam C, Gray K, Timms, V, Gall M, Arnott A, Sadsad R, Draper J, Sim E, Bachmann N, Rockett R, Carter I, Holmes EC, O’Sullivan MV, Byun R, Sintchenko V, Chen SC, Eden JS, Maddocks S, Kok J, Propenko M, Sorrell T, Chang S, Basile K, Dwyer DE for the 2019-nCoV Study Group |
| EPI_ISL_427725 | Centre for Infectious Diseases and Microbiology Public Health | NSW Health Pathology - Institute of Clinical Pathology and Medical Research; Westmead Hospital; University of Sydney | Timms V, Gall M, Arnott A, Sadsad R, Draper J, Sim E, Bachmann N, Rockett R, Lam C, Gray K, Carter I, Holmes EC, O’Sullivan MV, Byun R, Sintchenko V, Chen SC, Eden JS, Maddocks S, Kok J, Propenko M, Sorrell T, Chang S, Basile K, Dwyer DE for the 2019-nCoV Study Group |
| EPI_ISL_427726 | Centre for Infectious Diseases and Microbiology Public Health | NSW Health Pathology - Institute of Clinical Pathology and Medical Research; Westmead Hospital; University of Sydney | Rockett R, Lam C, Gray K, Timms V, Gall M, Arnott A, Sadsad R, Draper J, Sim E, Bachmann N, Carter I, Holmes EC, O’Sullivan MV, Byun R, Sintchenko V, Chen SC, Eden JS, Maddocks S, Kok J, Propenko M, Sorrell T, Chang S, Basile K, Dwyer DE for the 2019-nCoV Study Group |
| EPI_ISL_427727 | Centre for Infectious Diseases and Microbiology Public Health | NSW Health Pathology - Institute of Clinical Pathology and Medical Research; Westmead Hospital; University of Sydney | Lam C, Gray K, Timms, V, Gall M, Arnott A, Sadsad R, Draper J, Sim E, Bachmann N, Rockett R, Carter I, Holmes EC, O’Sullivan MV, Byun R, Sintchenko V, Chen SC, Eden JS, Maddocks S, Kok J, Propenko M, Sorrell T, Chang S, Basile K, Dwyer DE for the 2019-nCoV Study Group |
| EPI_ISL_427728 | Centre for Infectious Diseases and Microbiology Public Health | NSW Health Pathology - Institute of Clinical Pathology and Medical Research; Westmead Hospital; University of Sydney | Rockett R, Lam C, Gray K, Timms V, Gall M, Arnott A, Sadsad R, Draper J, Sim E, Bachmann N, Carter I, Holmes EC, O’Sullivan MV, Byun R, Sintchenko V, Chen SC, Eden JS, Maddocks S, Kok J, Propenko M, Sorrell T, Chang S, Basile K, Dwyer DE for the 2019-nCoV Study Group |
| EPI_ISL_427729 | Centre for Infectious Diseases and Microbiology Public Health | NSW Health Pathology - Institute of Clinical Pathology and Medical Research; Westmead Hospital; University of Sydney | Gall M, Arnott A, Sadsad R, Draper J, Sim E, Bachmann N, Rockett R, Lam C, Gray K, Timms V, Carter I, Holmes EC, O’Sullivan MV, Byun R, Sintchenko V, Chen SC, Eden JS, Maddocks S, Kok J, Propenko M, Sorrell T, Chang S, Basile K, Dwyer DE for the 2019-nCoV Study Group |
| EPI_ISL_427730, EPI_ISL_427731, EPI_ISL_427732 | Centre for Infectious Diseases and Microbiology Public Health | NSW Health Pathology - Institute of Clinical Pathology and Medical Research; Westmead Hospital; University of Sydney | Gray K, Timms V, Gall M, Arnott A, Sadsad R, Draper J, Sim E, Bachmann N, Rockett R, Lam C, Carter I, Holmes EC, O’Sullivan MV, Byun R, Sintchenko V, Chen SC, Eden JS, Maddocks S, Kok J, Propenko M, Sorrell T, Chang S, Basile K, Dwyer DE for the 2019-nCoV Study Group |
| EPI_ISL_427733 | Centre for Infectious Diseases and Microbiology Public Health | NSW Health Pathology - Institute of Clinical Pathology and Medical Research; Westmead Hospital; University of Sydney | Gall M, Arnott A, Sadsad R, Draper J, Sim E, Bachmann N, Rockett R, Lam C, Gray K, Timms V, Carter I, Holmes EC, O’Sullivan MV, Byun R, Sintchenko V, Chen SC, Eden JS, Maddocks S, Kok J, Propenko M, Sorrell T, Chang S, Basile K, Dwyer DE for the 2019-nCoV Study Group |
| EPI_ISL_427734 | Centre for Infectious Diseases and Microbiology Public Health | NSW Health Pathology - Institute of Clinical Pathology and Medical Research; Westmead Hospital; University of Sydney | Timms V, Gall M, Arnott A, Sadsad R, Draper J, Sim E, Bachmann N, Rockett R, Lam C, Gray K, Carter I, Holmes EC, O’Sullivan MV, Byun R, Sintchenko V, Chen SC, Eden JS, Maddocks S, Kok J, Propenko M, Sorrell T, Chang S, Basile K, Dwyer DE for the 2019-nCoV Study Group |
| EPI_ISL_427735 | Centre for Infectious Diseases and Microbiology Public Health | NSW Health Pathology - Institute of Clinical Pathology and Medical Research; Westmead Hospital; University of Sydney | Arnott A, Sadsad R, Draper J, Sim E, Bachmann N, Rockett R, Lam C, Gray K, Timms V, Gall M, Carter I, Holmes EC, O’Sullivan MV, Byun R, Sintchenko V, Chen SC, Eden JS, Maddocks S, Kok J, Propenko M, Sorrell T, Chang S, Basile K, Dwyer DE for the 2019-nCoV Study Group |
| EPI_ISL_427736 | Centre for Infectious Diseases and Microbiology Public Health | NSW Health Pathology - Institute of Clinical Pathology and Medical Research; Westmead Hospital; University of Sydney | Draper J, Sim E, Bachmann N, Rockett R, Lam C, Gray K, Timms V, Gall M, Arnott A, Sadsad R, Carter I, Holmes EC, O’Sullivan MV, Byun R, Sintchenko V, Chen SC, Eden JS, Maddocks S, Kok J, Propenko M, Sorrell T, Chang S, Basile K, Dwyer DE for the 2019-nCoV Study Group |
| EPI_ISL_427737 | Centre for Infectious Diseases and Microbiology Public Health | NSW Health Pathology - Institute of Clinical Pathology and Medical Research; Westmead Hospital; University of Sydney | Rockett R, Lam C, Gray K, Timms V, Gall M, Arnott A, Sadsad R, Draper J, Sim E, Bachmann N, Carter I, Holmes EC, O’Sullivan MV, Byun R, Sintchenko V, Chen SC, Eden JS, Maddocks S, Kok J, Propenko M, Sorrell T, Chang S, Basile K, Dwyer DE for the 2019-nCoV Study Group |
| EPI_ISL_427738 | Centre for Infectious Diseases and Microbiology Public Health | NSW Health Pathology - Institute of Clinical Pathology and Medical Research; Westmead Hospital; University of Sydney | Lam C, Gray K, Timms, V, Gall M, Arnott A, Sadsad R, Draper J, Sim E, Bachmann N, Rockett R, Carter I, Holmes EC, O’Sullivan MV, Byun R, Sintchenko V, Chen SC, Eden JS, Maddocks S, Kok J, Propenko M, Sorrell T, Chang S, Basile K, Dwyer DE for the 2019-nCoV Study Group |
| EPI_ISL_427739 | Centre for Infectious Diseases and Microbiology Public Health | NSW Health Pathology - Institute of Clinical Pathology and Medical Research; Westmead Hospital; University of Sydney | Arnott A, Sadsad R, Draper J, Sim E, Bachmann N, Rockett R, Lam C, Gray K, Timms V, Gall M, Carter I, Holmes EC, O’Sullivan MV, Byun R, Sintchenko V, Chen SC, Eden JS, Maddocks S, Kok J, Propenko M, Sorrell T, Chang S, Basile K, Dwyer DE for the 2019-nCoV Study Group |
| EPI_ISL_427740 | Centre for Infectious Diseases and Microbiology Public Health | NSW Health Pathology - Institute of Clinical Pathology and Medical Research; Westmead Hospital; University of Sydney | Sadsad R, Draper J, Sim E, Bachmann N, Rockett R, Lam C, Gray K, Timms V, Gall M, Arnott A, Carter I, Holmes EC, O’Sullivan MV, Byun R, Sintchenko V, Chen SC, Eden JS, Maddocks S, Kok J, Propenko M, Sorrell T, Chang S, Basile K, Dwyer DE for the 2019-nCoV Study Group |
| EPI_ISL_427741 | Centre for Infectious Diseases and Microbiology Public Health | NSW Health Pathology - Institute of Clinical Pathology and Medical Research; Westmead Hospital; University of Sydney | Draper J, Sim E, Bachmann N, Rockett R, Lam C, Gray K, Timms V, Gall M, Arnott A, Sadsad R, Carter I, Holmes EC, O’Sullivan MV, Byun R, Sintchenko V, Chen SC, Eden JS, Maddocks S, Kok J, Propenko M, Sorrell T, Chang S, Basile K, Dwyer DE for the 2019-nCoV Study Group |
| EPI_ISL_427742 | Centre for Infectious Diseases and Microbiology Public Health | NSW Health Pathology - Institute of Clinical Pathology and Medical Research; Westmead Hospital; University of Sydney | Lam C, Gray K, Timms, V, Gall M, Arnott A, Sadsad R, Draper J, Sim E, Bachmann N, Rockett R, Carter I, Holmes EC, O’Sullivan MV, Byun R, Sintchenko V, Chen SC, Eden JS, Maddocks S, Kok J, Propenko M, Sorrell T, Chang S, Basile K, Dwyer DE for the 2019-nCoV Study Group |
| EPI_ISL_427743 | Centre for Infectious Diseases and Microbiology Public Health | NSW Health Pathology - Institute of Clinical Pathology and Medical Research; Westmead Hospital; University of Sydney | Sim E, Bachmann N, Rockett R, Lam C, Gray K, Timms V, Gall M, Arnott A, Sadsad R, Draper J, Carter I, Holmes EC, O’Sullivan MV, Byun R, Sintchenko V, Chen SC, Eden JS, Maddocks S, Kok J, Propenko M, Sorrell T, Chang S, Basile K, Dwyer DE for the 2019-nCoV Study Group |
| EPI_ISL_427744 | Centre for Infectious Diseases and Microbiology Public Health | NSW Health Pathology - Institute of Clinical Pathology and Medical Research; Westmead Hospital; University of Sydney | Bachmann N, Rockett R, Lam C, Gray K, Timms V, Gall M, Arnott A, Sadsad R, Draper J, Sim E, Carter I, Holmes EC, O’Sullivan MV, Byun R, Sintchenko V, Chen SC, Eden JS, Maddocks S, Kok J, Propenko M, Sorrell T, Chang S, Basile K, Dwyer DE for the 2019-nCoV Study Group |
| EPI_ISL_427745 | Centre for Infectious Diseases and Microbiology Public Health | NSW Health Pathology - Institute of Clinical Pathology and Medical Research; Westmead Hospital; University of Sydney | Gray K, Timms V, Gall M, Arnott A, Sadsad R, Draper J, Sim E, Bachmann N, Rockett R, Lam C, Carter I, Holmes EC, O’Sullivan MV, Byun R, Sintchenko V, Chen SC, Eden JS, Maddocks S, Kok J, Propenko M, Sorrell T, Chang S, Basile K, Dwyer DE for the 2019-nCoV Study Group |
| EPI_ISL_427746 | Centre for Infectious Diseases and Microbiology Public Health | NSW Health Pathology - Institute of Clinical Pathology and Medical Research; Westmead Hospital; University of Sydney | Gall M, Arnott A, Sadsad R, Draper J, Sim E, Bachmann N, Rockett R, Lam C, Gray K, Timms V, Carter I, Holmes EC, O’Sullivan MV, Byun R, Sintchenko V, Chen SC, Eden JS, Maddocks S, Kok J, Propenko M, Sorrell T, Chang S, Basile K, Dwyer DE for the 2019-nCoV Study Group |
| EPI_ISL_427747 | Centre for Infectious Diseases and Microbiology Public Health | NSW Health Pathology - Institute of Clinical Pathology and Medical Research; Westmead Hospital; University of Sydney | Sadsad R, Draper J, Sim E, Bachmann N, Rockett R, Lam C, Gray K, Timms V, Gall M, Arnott A, Carter I, Holmes EC, O’Sullivan MV, Byun R, Sintchenko V, Chen SC, Eden JS, Maddocks S, Kok J, Propenko M, Sorrell T, Chang S, Basile K, Dwyer DE for the 2019-nCoV Study Group |
| EPI_ISL_427748 | Centre for Infectious Diseases and Microbiology Public Health | NSW Health Pathology - Institute of Clinical Pathology and | Timms V, Gall M, Arnott A, Sadsad R, Draper J, Sim E, Bachmann N, Rockett R, Lam C, Gray K, Carter I, Holmes EC, O’Sullivan MV, Byun R, Sintchenko V, Chen SC, Eden JS, Maddocks S, Kok J, Propenko M, Sorrell T, |

| | | | |
|---|---|---|---|
| | | Medical Research; Westmead Hospital; University of Sydney | Chang S, Basile K, Dwyer DE for the 2019-nCoV Study Group |
| EPI_ISL_427749 | Centre for Infectious Diseases and Microbiology Public Health | NSW Health Pathology - Institute of Clinical Pathology and Medical Research; Westmead Hospital; University of Sydney | Bachmann N, Rockett R, Lam C, Gray K, Timms V, Gall M, Arnott A, Sadsad R, Draper J, Sim E, Carter I, Holmes EC, O’Sullivan MV, Byun R, Sintchenko V, Chen SC, Eden JS, Maddocks S, Kok J, Propenko M, Sorrell T, Chang S, Basile K, Dwyer DE for the 2019-nCoV Study Group |
| EPI_ISL_427750 | Centre for Infectious Diseases and Microbiology Public Health | NSW Health Pathology - Institute of Clinical Pathology and Medical Research; Westmead Hospital; University of Sydney | Rockett R, Lam C, Gray K, Timms V, Gall M, Arnott A, Sadsad R, Draper J, Sim E, Bachmann N, Carter I, Holmes EC, O’Sullivan MV, Byun R, Sintchenko V, Chen SC, Eden JS, Maddocks S, Kok J, Propenko M, Sorrell T, Chang S, Basile K, Dwyer DE for the 2019-nCoV Study Group |
| EPI_ISL_427751 | Centre for Infectious Diseases and Microbiology Public Health | NSW Health Pathology - Institute of Clinical Pathology and Medical Research; Westmead Hospital; University of Sydney | Arnott A, Sadsad R, Draper J, Sim E, Bachmann N, Rockett R, Lam C, Gray K, Timms V, Gall M, Carter I, Holmes EC, O’Sullivan MV, Byun R, Sintchenko V, Chen SC, Eden JS, Maddocks S, Kok J, Propenko M, Sorrell T, Chang S, Basile K, Dwyer DE for the 2019-nCoV Study Group |
| EPI_ISL_427752 | Centre for Infectious Diseases and Microbiology Public Health | NSW Health Pathology - Institute of Clinical Pathology and Medical Research; Westmead Hospital; University of Sydney | Lam C, Gray K, Timms, V, Gall M, Arnott A, Sadsad R, Draper J, Sim E, Bachmann N, Rockett R, Carter I, Holmes EC, O’Sullivan MV, Byun R, Sintchenko V, Chen SC, Eden JS, Maddocks S, Kok J, Propenko M, Sorrell T, Chang S, Basile K, Dwyer DE for the 2019-nCoV Study Group |
| EPI_ISL_427753 | Centre for Infectious Diseases and Microbiology Public Health | NSW Health Pathology - Institute of Clinical Pathology and Medical Research; Westmead Hospital; University of Sydney | Arnott A, Sadsad R, Draper J, Sim E, Bachmann N, Rockett R, Lam C, Gray K, Timms V, Gall M, Carter I, Holmes EC, O’Sullivan MV, Byun R, Sintchenko V, Chen SC, Eden JS, Maddocks S, Kok J, Propenko M, Sorrell T, Chang S, Basile K, Dwyer DE for the 2019-nCoV Study Group |
| EPI_ISL_427754, EPI_ISL_427755 | Centre for Infectious Diseases and Microbiology Public Health | NSW Health Pathology - Institute of Clinical Pathology and Medical Research; Westmead Hospital; University of Sydney | Sadsad R, Draper J, Sim E, Bachmann N, Rockett R, Lam C, Gray K, Timms V, Gall M, Arnott A, Carter I, Holmes EC, O’Sullivan MV, Byun R, Sintchenko V, Chen SC, Eden JS, Maddocks S, Kok J, Propenko M, Sorrell T, Chang S, Basile K, Dwyer DE for the 2019-nCoV Study Group |
| EPI_ISL_427756 | Centre for Infectious Diseases and Microbiology Public Health | NSW Health Pathology - Institute of Clinical Pathology and Medical Research; Westmead Hospital; University of Sydney | Gall M, Arnott A, Sadsad R, Draper J, Sim E, Bachmann N, Rockett R, Lam C, Gray K, Timms V, Carter I, Holmes EC, O’Sullivan MV, Byun R, Sintchenko V, Chen SC, Eden JS, Maddocks S, Kok J, Propenko M, Sorrell T, Chang S, Basile K, Dwyer DE for the 2019-nCoV Study Group |
| EPI_ISL_427757 | Centre for Infectious Diseases and Microbiology Public Health | NSW Health Pathology - Institute of Clinical Pathology and Medical Research; Westmead Hospital; University of Sydney | Draper J, Sim E, Bachmann N, Rockett R, Lam C, Gray K, Timms V, Gall M, Arnott A, Sadsad R, Carter I, Holmes EC, O’Sullivan MV, Byun R, Sintchenko V, Chen SC, Eden JS, Maddocks S, Kok J, Propenko M, Sorrell T, Chang S, Basile K, Dwyer DE for the 2019-nCoV Study Group |
| EPI_ISL_427758 | Centre for Infectious Diseases and Microbiology Public Health | NSW Health Pathology - Institute of Clinical Pathology and Medical Research; Westmead Hospital; University of Sydney | Bachmann N, Rockett R, Lam C, Gray K, Timms V, Gall M, Arnott A, Sadsad R, Draper J, Sim E, Carter I, Holmes EC, O’Sullivan MV, Byun R, Sintchenko V, Chen SC, Eden JS, Maddocks S, Kok J, Propenko M, Sorrell T, Chang S, Basile K, Dwyer DE for the 2019-nCoV Study Group |
| EPI_ISL_427759 | Centre for Infectious Diseases and Microbiology Public Health | NSW Health Pathology - Institute of Clinical Pathology and Medical Research; Westmead Hospital; University of Sydney | Sim E, Bachmann N, Rockett R, Lam C, Gray K, Timms V, Gall M, Arnott A, Sadsad R, Draper J, Carter I, Holmes EC, O’Sullivan MV, Byun R, Sintchenko V, Chen SC, Eden JS, Maddocks S, Kok J, Propenko M, Sorrell T, Chang S, Basile K, Dwyer DE for the 2019-nCoV Study Group |
| EPI_ISL_427760 | Centre for Infectious Diseases and Microbiology Public Health | NSW Health Pathology - Institute of Clinical Pathology and Medical Research; Westmead Hospital; University of Sydney | Sadsad R, Draper J, Sim E, Bachmann N, Rockett R, Lam C, Gray K, Timms V, Gall M, Arnott A, Carter I, Holmes EC, O’Sullivan MV, Byun R, Sintchenko V, Chen SC, Eden JS, Maddocks S, Kok J, Propenko M, Sorrell T, Chang S, Basile K, Dwyer DE for the 2019-nCoV Study Group |
| EPI_ISL_427761 | Centre for Infectious Diseases and Microbiology Public Health | NSW Health Pathology - Institute of Clinical Pathology and Medical Research; Westmead Hospital; University of Sydney | Arnott A, Sadsad R, Draper J, Sim E, Bachmann N, Rockett R, Lam C, Gray K, Timms V, Gall M, Carter I, Holmes EC, O’Sullivan MV, Byun R, Sintchenko V, Chen SC, Eden JS, Maddocks S, Kok J, Propenko M, Sorrell T, Chang S, Basile K, Dwyer DE for the 2019-nCoV Study Group |
| EPI_ISL_427762 | Centre for Infectious Diseases and Microbiology Public Health | NSW Health Pathology - Institute of Clinical Pathology and Medical Research; Westmead Hospital; University of Sydney | Bachmann N, Rockett R, Lam C, Gray K, Timms V, Gall M, Arnott A, Sadsad R, Draper J, Sim E, Carter I, Holmes EC, O’Sullivan MV, Byun R, Sintchenko V, Chen SC, Eden JS, Maddocks S, Kok J, Propenko M, Sorrell T, Chang S, Basile K, Dwyer DE for the 2019-nCoV Study Group |
| EPI_ISL_427763 | Centre for Infectious Diseases and Microbiology Public Health | NSW Health Pathology - Institute of Clinical Pathology and Medical Research; Westmead Hospital; University of Sydney | Lam C, Gray K, Timms, V, Gall M, Arnott A, Sadsad R, Draper J, Sim E, Bachmann N, Rockett R, Carter I, Holmes EC, O’Sullivan MV, Byun R, Sintchenko V, Chen SC, Eden JS, Maddocks S, Kok J, Propenko M, Sorrell T, Chang S, Basile K, Dwyer DE for the 2019-nCoV Study Group |
| EPI_ISL_427764 | Centre for Infectious Diseases and Microbiology Public Health | NSW Health Pathology - Institute of Clinical Pathology and Medical Research; Westmead Hospital; University of Sydney | Gall M, Arnott A, Sadsad R, Draper J, Sim E, Bachmann N, Rockett R, Lam C, Gray K, Timms V, Carter I, Holmes EC, O’Sullivan MV, Byun R, Sintchenko V, Chen SC, Eden JS, Maddocks S, Kok J, Propenko M, Sorrell T, Chang S, Basile K, Dwyer DE for the 2019-nCoV Study Group |
| EPI_ISL_427765 | Centre for Infectious Diseases and Microbiology Public Health | NSW Health Pathology - Institute of Clinical Pathology and Medical Research; Westmead Hospital; University of Sydney | Rockett R, Lam C, Gray K, Timms V, Gall M, Arnott A, Sadsad R, Draper J, Sim E, Bachmann N, Carter I, Holmes EC, O’Sullivan MV, Byun R, Sintchenko V, Chen SC, Eden JS, Maddocks S, Kok J, Propenko M, Sorrell T, Chang S, Basile K, Dwyer DE for the 2019-nCoV Study Group |
| EPI_ISL_427766 | Centre for Infectious Diseases and Microbiology Public Health | NSW Health Pathology - Institute of Clinical Pathology and Medical Research; Westmead Hospital; University of Sydney | Draper J, Sim E, Bachmann N, Rockett R, Lam C, Gray K, Timms V, Gall M, Arnott A, Sadsad R, Carter I, Holmes EC, O’Sullivan MV, Byun R, Sintchenko V, Chen SC, Eden JS, Maddocks S, Kok J, Propenko M, Sorrell T, Chang S, Basile K, Dwyer DE for the 2019-nCoV Study Group |
| EPI_ISL_427767 | Centre for Infectious Diseases and Microbiology Public Health | NSW Health Pathology - Institute of Clinical Pathology and Medical Research; Westmead Hospital; University of Sydney | Gray K, Timms V, Gall M, Arnott A, Sadsad R, Draper J, Sim E, Bachmann N, Rockett R, Lam C, Carter I, Holmes EC, O’Sullivan MV, Byun R, Sintchenko V, Chen SC, Eden JS, Maddocks S, Kok J, Propenko M, Sorrell T, Chang S, Basile K, Dwyer DE for the 2019-nCoV Study Group |
| EPI_ISL_427768 | Centre for Infectious Diseases and Microbiology Public Health | NSW Health Pathology - Institute of Clinical Pathology and Medical Research; Westmead Hospital; University of Sydney | Draper J, Sim E, Bachmann N, Rockett R, Lam C, Gray K, Timms V, Gall M, Arnott A, Sadsad R, Carter I, Holmes EC, O’Sullivan MV, Byun R, Sintchenko V, Chen SC, Eden JS, Maddocks S, Kok J, Propenko M, Sorrell T, Chang S, Basile K, Dwyer DE for the 2019-nCoV Study Group |
| EPI_ISL_427769 | Centre for Infectious Diseases and Microbiology Public Health | NSW Health Pathology - Institute of Clinical Pathology and Medical Research; Westmead Hospital; University of Sydney | Gall M, Arnott A, Sadsad R, Draper J, Sim E, Bachmann N, Rockett R, Lam C, Gray K, Timms V, Carter I, Holmes EC, O’Sullivan MV, Byun R, Sintchenko V, Chen SC, Eden JS, Maddocks S, Kok J, Propenko M, Sorrell T, Chang S, Basile K, Dwyer DE for the 2019-nCoV Study Group |
| EPI_ISL_427770 | Centre for Infectious Diseases and Microbiology Public Health | NSW Health Pathology - Institute of Clinical Pathology and Medical Research; Westmead Hospital; University of Sydney | Arnott A, Sadsad R, Draper J, Sim E, Bachmann N, Rockett R, Lam C, Gray K, Timms V, Gall M, Carter I, Holmes EC, O’Sullivan MV, Byun R, Sintchenko V, Chen SC, Eden JS, Maddocks S, Kok J, Propenko M, Sorrell T, Chang S, Basile K, Dwyer DE for the 2019-nCoV Study Group |
| EPI_ISL_427771 | Centre for Infectious Diseases and Microbiology Public Health | NSW Health Pathology - Institute of Clinical Pathology and Medical Research; Westmead Hospital; University of Sydney | Timms V, Gall M, Arnott A, Sadsad R, Draper J, Sim E, Bachmann N, Rockett R, Lam C, Gray K, Carter I, Holmes EC, O’Sullivan MV, Byun R, Sintchenko V, Chen SC, Eden JS, Maddocks S, Kok J, Propenko M, Sorrell T, Chang S, Basile K, Dwyer DE for the 2019-nCoV Study Group |
| EPI_ISL_427772 | Centre for Infectious Diseases and Microbiology Public Health | NSW Health Pathology - Institute of Clinical Pathology and Medical Research; Westmead Hospital; University of Sydney | Gall M, Arnott A, Sadsad R, Draper J, Sim E, Bachmann N, Rockett R, Lam C, Gray K, Timms V, Carter I, Holmes EC, O’Sullivan MV, Byun R, Sintchenko V, Chen SC, Eden JS, Maddocks S, Kok J, Propenko M, Sorrell T, Chang S, Basile K, Dwyer DE for the 2019-nCoV Study Group |
| EPI_ISL_427773 | Centre for Infectious Diseases and Microbiology Public Health | NSW Health Pathology - Institute of Clinical Pathology and Medical Research; Westmead Hospital; University of Sydney | Lam C, Gray K, Timms, V, Gall M, Arnott A, Sadsad R, Draper J, Sim E, Bachmann N, Rockett R, Carter I, Holmes EC, O’Sullivan MV, Byun R, Sintchenko V, Chen SC, Eden JS, Maddocks S, Kok J, Propenko M, Sorrell T, Chang S, Basile K, Dwyer DE for the 2019-nCoV Study Group |
| EPI_ISL_427774 | Centre for Infectious Diseases and Microbiology Public Health | NSW Health Pathology - Institute of Clinical Pathology and Medical Research; Westmead Hospital; University of Sydney | Arnott A, Sadsad R, Draper J, Sim E, Bachmann N, Rockett R, Lam C, Gray K, Timms V, Gall M, Carter I, Holmes EC, O’Sullivan MV, Byun R, Sintchenko V, Chen SC, Eden JS, Maddocks S, Kok J, Propenko M, Sorrell T, Chang S, Basile K, Dwyer DE for the 2019-nCoV Study Group |
| EPI_ISL_427775, EPI_ISL_427776, EPI_ISL_427777 | Centre for Infectious Diseases and Microbiology Public Health | NSW Health Pathology - Institute of Clinical Pathology and Medical Research; Westmead Hospital; University of Sydney | Sadsad R, Draper J, Sim E, Bachmann N, Rockett R, Lam C, Gray K, Timms V, Gall M, Arnott A, Carter I, Holmes EC, O’Sullivan MV, Byun R, Sintchenko V, Chen SC, Eden JS, Maddocks S, Kok J, Propenko M, Sorrell T, Chang S, Basile K, Dwyer DE for the 2019-nCoV Study Group |
| EPI_ISL_427778 | Centre for Infectious Diseases and Microbiology Public Health | NSW Health Pathology - Institute of Clinical Pathology and Medical Research; Westmead Hospital; University of Sydney | Bachmann N, Rockett R, Lam C, Gray K, Timms V, Gall M, Arnott A, Sadsad R, Draper J, Sim E, Carter I, Holmes EC, O’Sullivan MV, Byun R, Sintchenko V, Chen SC, Eden JS, Maddocks S, Kok J, Propenko M, Sorrell T, Chang S, Basile K, Dwyer DE for the 2019-nCoV Study Group |
| EPI_ISL_427779 | Centre for Infectious Diseases and Microbiology Public Health | NSW Health Pathology - Institute of Clinical Pathology and Medical Research; Westmead Hospital; University of Sydney | Sadsad R, Draper J, Sim E, Bachmann N, Rockett R, Lam C, Gray K, Timms V, Gall M, Arnott A, Carter I, Holmes EC, O’Sullivan MV, Byun R, Sintchenko V, Chen SC, Eden JS, Maddocks S, Kok J, Propenko M, Sorrell T, Chang S, Basile K, Dwyer DE for the 2019-nCoV Study Group |
| EPI_ISL_427780 | Centre for Infectious Diseases and Microbiology Public Health | NSW Health Pathology - Institute of Clinical Pathology and Medical Research; Westmead Hospital; University of Sydney | Rockett R, Lam C, Gray K, Timms V, Gall M, Arnott A, Sadsad R, Draper J, Sim E, Bachmann N, Carter I, Holmes EC, O’Sullivan MV, Byun R, Sintchenko V, Chen SC, Eden JS, Maddocks S, Kok J, Propenko M, Sorrell T, Chang S, Basile K, Dwyer DE for the 2019-nCoV Study Group |
| EPI_ISL_427781 | Centre for Infectious Diseases and Microbiology Public Health | NSW Health Pathology - Institute of Clinical Pathology and Medical Research; Westmead Hospital; University of Sydney | Sadsad R, Draper J, Sim E, Bachmann N, Rockett R, Lam C, Gray K, Timms V, Gall M, Arnott A, Carter I, Holmes EC, O’Sullivan MV, Byun R, Sintchenko V, Chen SC, Eden JS, Maddocks S, Kok J, Propenko M, Sorrell T, Chang S, Basile K, Dwyer DE for the 2019-nCoV Study Group |
| EPI_ISL_427782 | Centre for Infectious Diseases and Microbiology Public Health | NSW Health Pathology - Institute of Clinical Pathology and Medical Research; Westmead Hospital; University of Sydney | Lam C, Gray K, Timms, V, Gall M, Arnott A, Sadsad R, Draper J, Sim E, Bachmann N, Rockett R, Carter I, Holmes EC, O’Sullivan MV, Byun R, Sintchenko V, Chen SC, Eden JS, Maddocks S, Kok J, Propenko M, Sorrell T, Chang S, Basile K, Dwyer DE for the 2019-nCoV Study Group |
| EPI_ISL_427783 | Centre for Infectious Diseases and Microbiology Public Health | NSW Health Pathology - Institute of Clinical Pathology and Medical Research; Westmead Hospital; University of Sydney | Sim E, Bachmann N, Rockett R, Lam C, Gray K, Timms V, Gall M, Arnott A, Sadsad R, Draper J, Carter I, Holmes EC, O’Sullivan MV, Byun R, Sintchenko V, Chen SC, Eden JS, Maddocks S, Kok J, Propenko M, Sorrell T, Chang S, Basile K, Dwyer DE for the 2019-nCoV Study Group |
| EPI_ISL_427784 | Centre for Infectious Diseases and Microbiology Public Health | NSW Health Pathology - Institute of Clinical Pathology and Medical Research; Westmead Hospital; University of Sydney | Draper J, Sim E, Bachmann N, Rockett R, Lam C, Gray K, Timms V, Gall M, Arnott A, Sadsad R, Carter I, Holmes EC, O’Sullivan MV, Byun R, Sintchenko V, Chen SC, Eden JS, Maddocks S, Kok J, Propenko M, Sorrell T, Chang S, Basile K, Dwyer DE for the 2019-nCoV Study Group |
| EPI_ISL_427785 | Centre for Infectious Diseases and Microbiology Public Health | NSW Health Pathology - Institute of Clinical Pathology and Medical Research; Westmead Hospital; University of Sydney | Gray K, Timms V, Gall M, Arnott A, Sadsad R, Draper J, Sim E, Bachmann N, Rockett R, Lam C, Carter I, Holmes EC, O’Sullivan MV, Byun R, Sintchenko V, Chen SC, Eden JS, Maddocks S, Kok J, Propenko M, Sorrell T, Chang S, Basile K, Dwyer DE for the 2019-nCoV Study Group |
| EPI_ISL_427786 | Centre for Infectious Diseases and Microbiology Public Health | NSW Health Pathology - Institute of Clinical Pathology and Medical Research; Westmead Hospital; University of Sydney | Sim E, Bachmann N, Rockett R, Lam C, Gray K, Timms V, Gall M, Arnott A, Sadsad R, Draper J, Carter I, Holmes EC, O’Sullivan MV, Byun R, Sintchenko V, Chen SC, Eden JS, Maddocks S, Kok J, Propenko M, Sorrell T, Chang S, Basile K, Dwyer DE for the 2019-nCoV Study Group |
| EPI_ISL_427787 | Centre for Infectious Diseases and Microbiology Public Health | NSW Health Pathology - Institute of Clinical Pathology and Medical Research; Westmead Hospital; University of Sydney | Bachmann N, Rockett R, Lam C, Gray K, Timms V, Gall M, Arnott A, Sadsad R, Draper J, Sim E, Carter I, Holmes EC, O’Sullivan MV, Byun R, Sintchenko V, Chen SC, Eden JS, Maddocks S, Kok J, Propenko M, Sorrell T, Chang S, Basile K, Dwyer DE for the 2019-nCoV Study Group |
| EPI_ISL_427788 | Centre for Infectious Diseases and Microbiology Public Health | NSW Health Pathology - Institute of Clinical Pathology and Medical Research; Westmead Hospital; University of Sydney | Sim E, Bachmann N, Rockett R, Lam C, Gray K, Timms V, Gall M, Arnott A, Sadsad R, Draper J, Carter I, Holmes EC, O’Sullivan MV, Byun R, Sintchenko V, Chen SC, Eden JS, Maddocks S, Kok J, Propenko M, Sorrell T, Chang S, Basile K, Dwyer DE for the 2019-nCoV Study Group |
| EPI_ISL_427789 | Centre for Infectious Diseases and Microbiology Public Health | NSW Health Pathology - Institute of Clinical Pathology and Medical Research; Westmead Hospital; University of Sydney | Rockett R, Lam C, Gray K, Timms V, Gall M, Arnott A, Sadsad R, Draper J, Sim E, Bachmann N, Carter I, Holmes EC, O’Sullivan MV, Byun R, Sintchenko V, Chen SC, Eden JS, Maddocks S, Kok J, Propenko M, Sorrell T, Chang S, Basile K, Dwyer DE for the 2019-nCoV Study Group |
| EPI_ISL_427790 | Centre for Infectious Diseases and Microbiology Public Health | NSW Health Pathology - Institute of Clinical Pathology and Medical Research; Westmead Hospital; University of Sydney | Sadsad R, Draper J, Sim E, Bachmann N, Rockett R, Lam C, Gray K, Timms V, Gall M, Arnott A, Carter I, Holmes EC, O’Sullivan MV, Byun R, Sintchenko V, Chen SC, Eden JS, Maddocks S, Kok J, Propenko M, Sorrell T, Chang S, Basile K, Dwyer DE for the 2019-nCoV Study Group |
| EPI_ISL_427791 | Centre for Infectious Diseases and Microbiology Public Health | NSW Health Pathology - Institute of Clinical Pathology and Medical Research; Westmead Hospital; University of Sydney | Rockett R, Lam C, Gray K, Timms V, Gall M, Arnott A, Sadsad R, Draper J, Sim E, Bachmann N, Carter I, Holmes EC, O’Sullivan MV, Byun R, Sintchenko V, Chen SC, Eden JS, Maddocks S, Kok J, Propenko M, Sorrell T, Chang S, Basile K, Dwyer DE for the 2019-nCoV Study Group |
| EPI_ISL_427792 | Centre for Infectious Diseases and Microbiology Public Health | NSW Health Pathology - Institute of Clinical Pathology and Medical Research; Westmead Hospital; University of Sydney | Lam C, Gray K, Timms, V, Gall M, Arnott A, Sadsad R, Draper J, Sim E, Bachmann N, Rockett R, Carter I, Holmes EC, O’Sullivan MV, Byun R, Sintchenko V, Chen SC, Eden JS, Maddocks S, Kok J, Propenko M, Sorrell T, Chang S, Basile K, Dwyer DE for the 2019-nCoV Study Group |
| EPI_ISL_427793 | Centre for Infectious Diseases and Microbiology Public Health | NSW Health Pathology - Institute of Clinical Pathology and Medical Research; Westmead Hospital; University of Sydney | Gray K, Timms V, Gall M, Arnott A, Sadsad R, Draper J, Sim E, Bachmann N, Rockett R, Lam C, Carter I, Holmes EC, O’Sullivan MV, Byun R, Sintchenko V, Chen SC, Eden JS, Maddocks S, Kok J, Propenko M, Sorrell T, Chang S, Basile K, Dwyer DE for the 2019-nCoV Study Group |
| EPI_ISL_427794 | Centre for Infectious Diseases and Microbiology Public Health | NSW Health Pathology - Institute of Clinical Pathology and Medical Research; Westmead Hospital; University of Sydney | Timms V, Gall M, Arnott A, Sadsad R, Draper J, Sim E, Bachmann N, Rockett R, Lam C, Gray K, Carter I, Holmes EC, O’Sullivan MV, Byun R, Sintchenko V, Chen SC, Eden JS, Maddocks S, Kok J, Propenko M, Sorrell T, Chang S, Basile K, Dwyer DE for the 2019-nCoV Study Group |
| EPI_ISL_427795 | Centre for Infectious Diseases and Microbiology Public Health | NSW Health Pathology - Institute of Clinical Pathology and Medical Research; Westmead Hospital; University of Sydney | Sadsad R, Draper J, Sim E, Bachmann N, Rockett R, Lam C, Gray K, Timms V, Gall M, Arnott A, Carter I, Holmes EC, O’Sullivan MV, Byun R, Sintchenko V, Chen SC, Eden JS, Maddocks S, Kok J, Propenko M, Sorrell T, Chang S, Basile K, Dwyer DE for the 2019-nCoV Study Group |
| EPI_ISL_427796 | Centre for Infectious Diseases and Microbiology Public Health | NSW Health Pathology - Institute of Clinical Pathology and | Draper J, Sim E, Bachmann N, Rockett R, Lam C, Gray K, Timms V, Gall M, Arnott A, Sadsad R, Carter I, Holmes EC, O’Sullivan MV, Byun R, Sintchenko V, Chen SC, Eden JS, Maddocks S, Kok J, Propenko M, Sorrell T, |

| | | | |
|---|---|---|---|
| | | Medical Research; Westmead Hospital; University of Sydney | Chang S, Basile K, Dwyer DE for the 2019-nCoV Study Group |
| EPI_ISL_427797 | Centre for Infectious Diseases and Microbiology Public Health | NSW Health Pathology - Institute of Clinical Pathology and Medical Research; Westmead Hospital; University of Sydney | Bachmann N, Rockett R, Lam C, Gray K, Timms V, Gall M, Arnott A, Sadsad R, Draper J, Sim E, Carter I, Holmes EC, O’Sullivan MV, Byun R, Sintchenko V, Chen SC, Eden JS, Maddocks S, Kok J, Propenko M, Sorrell T, Chang S, Basile K, Dwyer DE for the 2019-nCoV Study Group |
| EPI_ISL_427798 | Centre for Infectious Diseases and Microbiology Public Health | NSW Health Pathology - Institute of Clinical Pathology and Medical Research; Westmead Hospital; University of Sydney | Rockett R, Lam C, Gray K, Timms V, Gall M, Arnott A, Sadsad R, Draper J, Sim E, Bachmann N, Carter I, Holmes EC, O’Sullivan MV, Byun R, Sintchenko V, Chen SC, Eden JS, Maddocks S, Kok J, Propenko M, Sorrell T, Chang S, Basile K, Dwyer DE for the 2019-nCoV Study Group |
| EPI_ISL_427799 | Centre for Infectious Diseases and Microbiology Public Health | NSW Health Pathology - Institute of Clinical Pathology and Medical Research; Westmead Hospital; University of Sydney | Draper J, Sim E, Bachmann N, Rockett R, Lam C, Gray K, Timms V, Gall M, Arnott A, Sadsad R, Carter I, Holmes EC, O’Sullivan MV, Byun R, Sintchenko V, Chen SC, Eden JS, Maddocks S, Kok J, Propenko M, Sorrell T, Chang S, Basile K, Dwyer DE for the 2019-nCoV Study Group |
| EPI_ISL_427800 | Centre for Infectious Diseases and Microbiology Public Health | NSW Health Pathology - Institute of Clinical Pathology and Medical Research; Westmead Hospital; University of Sydney | Gall M, Arnott A, Sadsad R, Draper J, Sim E, Bachmann N, Rockett R, Lam C, Gray K, Timms V, Carter I, Holmes EC, O’Sullivan MV, Byun R, Sintchenko V, Chen SC, Eden JS, Maddocks S, Kok J, Propenko M, Sorrell T, Chang S, Basile K, Dwyer DE for the 2019-nCoV Study Group |
| EPI_ISL_427801 | Centre for Infectious Diseases and Microbiology Public Health | NSW Health Pathology - Institute of Clinical Pathology and Medical Research; Westmead Hospital; University of Sydney | Rockett R, Lam C, Gray K, Timms V, Gall M, Arnott A, Sadsad R, Draper J, Sim E, Bachmann N, Carter I, Holmes EC, O’Sullivan MV, Byun R, Sintchenko V, Chen SC, Eden JS, Maddocks S, Kok J, Propenko M, Sorrell T, Chang S, Basile K, Dwyer DE for the 2019-nCoV Study Group |
| EPI_ISL_427802 | Centre for Infectious Diseases and Microbiology Public Health | NSW Health Pathology - Institute of Clinical Pathology and Medical Research; Westmead Hospital; University of Sydney | Sim E, Bachmann N, Rockett R, Lam C, Gray K, Timms V, Gall M, Arnott A, Sadsad R, Draper J, Carter I, Holmes EC, O’Sullivan MV, Byun R, Sintchenko V, Chen SC, Eden JS, Maddocks S, Kok J, Propenko M, Sorrell T, Chang S, Basile K, Dwyer DE for the 2019-nCoV Study Group |
| EPI_ISL_427803 | Centre for Infectious Diseases and Microbiology Public Health | NSW Health Pathology - Institute of Clinical Pathology and Medical Research; Westmead Hospital; University of Sydney | Timms V, Gall M, Arnott A, Sadsad R, Draper J, Sim E, Bachmann N, Rockett R, Lam C, Gray K, Carter I, Holmes EC, O’Sullivan MV, Byun R, Sintchenko V, Chen SC, Eden JS, Maddocks S, Kok J, Propenko M, Sorrell T, Chang S, Basile K, Dwyer DE for the 2019-nCoV Study Group |
| EPI_ISL_427804 | Centre for Infectious Diseases and Microbiology Public Health | NSW Health Pathology - Institute of Clinical Pathology and Medical Research; Westmead Hospital; University of Sydney | Arnott A, Sadsad R, Draper J, Sim E, Bachmann N, Rockett R, Lam C, Gray K, Timms V, Gall M, Carter I, Holmes EC, O’Sullivan MV, Byun R, Sintchenko V, Chen SC, Eden JS, Maddocks S, Kok J, Propenko M, Sorrell T, Chang S, Basile K, Dwyer DE for the 2019-nCoV Study Group |
| EPI_ISL_427805 | Centre for Infectious Diseases and Microbiology Public Health | NSW Health Pathology - Institute of Clinical Pathology and Medical Research; Westmead Hospital; University of Sydney | Bachmann N, Rockett R, Lam C, Gray K, Timms V, Gall M, Arnott A, Sadsad R, Draper J, Sim E, Carter I, Holmes EC, O’Sullivan MV, Byun R, Sintchenko V, Chen SC, Eden JS, Maddocks S, Kok J, Propenko M, Sorrell T, Chang S, Basile K, Dwyer DE for the 2019-nCoV Study Group |
| EPI_ISL_427806 | Centre for Infectious Diseases and Microbiology Public Health | NSW Health Pathology - Institute of Clinical Pathology and Medical Research; Westmead Hospital; University of Sydney | Timms V, Gall M, Arnott A, Sadsad R, Draper J, Sim E, Bachmann N, Rockett R, Lam C, Gray K, Carter I, Holmes EC, O’Sullivan MV, Byun R, Sintchenko V, Chen SC, Eden JS, Maddocks S, Kok J, Propenko M, Sorrell T, Chang S, Basile K, Dwyer DE for the 2019-nCoV Study Group |
| EPI_ISL_427807, EPI_ISL_427808 | Centre for Infectious Diseases and Microbiology Public Health | NSW Health Pathology - Institute of Clinical Pathology and Medical Research; Westmead Hospital; University of Sydney | Gall M, Arnott A, Sadsad R, Draper J, Sim E, Bachmann N, Rockett R, Lam C, Gray K, Timms V, Carter I, Holmes EC, O’Sullivan MV, Byun R, Sintchenko V, Chen SC, Eden JS, Maddocks S, Kok J, Propenko M, Sorrell T, Chang S, Basile K, Dwyer DE for the 2019-nCoV Study Group |
| EPI_ISL_427809 | Division of Viral Diseases, Center for Laboratory Control of Infectious Diseases, Korea Centers for Diseases Control and Prevention | Division of Viral Diseases, Center for Laboratory Control of Infectious Diseases, Korea Centers for Diseases Control and Prevention | Jeong-Min Kim, Yoon-Seok Chung, Namjoo Lee, Mi-Seon Kim, Sang Hee Woo, Hye-Jun Jo, Sehee Park, Heui Man Kim, Jun-Sub Kim, Junhyeong Jang, Myung Guk Han |
| EPI_ISL_427810, EPI_ISL_427811, EPI_ISL_427812, EPI_ISL_427813 | Division of Viral Diseases, Center for Laboratory Control of Infectious Diseases, Korea Centers for Diseases Control and Prevention | Division of Viral Diseases, Center for Laboratory Control of Infectious Diseases, Korea Centers for Diseases Control and Prevention | Jeong-Min Kim, Yoon-Seok Chung, Namjoo Lee, Mi-Seon Kim, Sang Hee Woo, Hye-Jun Jo, Sehee Park, Heui Man Kim, Jun-Sub Kim, Junhyeong Jang, Dong Hyun Song, Daesang Lee, Seong Tae Jeong, Myung Guk Han |
| EPI_ISL_427815 | WHO National Influenza Centre Russian Federation | WHO National Influenza Centre Russian Federation | Andrey Komissarov, Artem Fadeev, Anna Ivanova, Daria Danilenko |
| EPI_ISL_428148 | Klinisk mikrobiologi, Region Västerbotten | Unit for Biological Agents, Department for CBRN Defence and Security, Swedish Defence Research Agency | FOI bioinformatics team |
| EPI_ISL_428201 | Klinisk mikrobiologi, Region Västerbotten | Unit for Biological Agents, Department for CBRN Defence and Security, Swedish Defence Research Agency | FOI Bioinformatics team |
| EPI_ISL_428202, EPI_ISL_428203, EPI_ISL_428204, EPI_ISL_428205, EPI_ISL_428206 | Nebraska Public Health Laboratory | UNMC COVID-19 Response Team | UNMC COVID-19 Response Team |
| EPI_ISL_428207, EPI_ISL_428208 | National Institute of Health Research and Development | National Institute of Health Research and Development | Setiawaty,V;Subangkit;Puspa,KD;Ikawati,HD;Nugraha,AA;Hariastuti,NI;Ramadhany,R;Susilarini,NK;Pratiwi,E;Agustiningsih;Kurniawati,J;Pawestri,HA;Siswanto |
| EPI_ISL_428209 | Laboratory of Molecular Biology, Diagnostyka sp. z o.o. | Laboratory of Recombinant Vaccines | Lukasz Rabalski, Anna Piotrowska-Mietelska, Boguslaw Szewczyk, Krystyna Bienkowska-Szewczyk |
| EPI_ISL_428229 | TSGH-CP molecular lab | TSGH-CP molecular lab | Cherng-Lih Perng, Ming-Jr Jian, Chih-Kai Chang, Jung-Chung Lin, Kuo-Ming Yeh, Chien-Wen Chen, Sheng-Kang Chiu, Hsing-Yi Chung, Shih-Hung Tsai, Kuo-Sheng Hung, Tien-Yao Chang, Feng-Yee Chang, Hung-Sheng Shang |
| EPI_ISL_428230 | TSGH-CP molecular lab | TSGH-CP molecular lab | Cherng-Lih Perng, Ming-Jr Jian, Chih-Kai Chang, Jung-Chung Lin, Kuo-Ming Yeh, Chien-Wen Chen, Sheng-Kang Chiu, Hsing-Yi Chung, Shih-Hung Tsai, Kuo-Sheng Hung, Tien-Yao Chang, Feng-Yee Chang, Hung-Sheng Shang |
| EPI_ISL_428231 | TSGH-CP molecular lab | TSGH-CP molecular lab | Cherng-Lih Perng, Ming-Jr JIAN, Chih-Kai Chang, Jung-Chung Lin, Kuo-Ming Yeh, Chien-Wen Chen, Sheng-Kang Chiu, Hsing-Yi Chung, Shih-Hung Tsai, Kuo-Sheng Hung, Tien-Yao Chang, Feng-Yee Chang, Hung-Sheng Shang |
| EPI_ISL_428232, EPI_ISL_428233, EPI_ISL_428234, EPI_ISL_428235, EPI_ISL_428236 | Hematology Laboratory, Section of Molecular Diagnostics, University Clinical Centre, Medical University of Gdansk | Department of Virology, Faculty of Medicine, University of Helsinki, Helsinki, Finland | Marlena Robakowska, Aneta Szulc, Maciej Grzybek, Olli Vapalahti, Teemu Smura |
| EPI_ISL_428250 | Yale Clinical Virology Laboratory | Grubaugh Lab - Yale School of Public Health | Joseph Fauver, Anderson Brito, Tara Alpert, Chantal Vogels, Ellen Foxman, Albert Ko, Marie Landry, Nathan Grubaugh |
| EPI_ISL_428252, EPI_ISL_428253, EPI_ISL_428254, EPI_ISL_428255, EPI_ISL_428256, EPI_ISL_428257, EPI_ISL_428258, EPI_ISL_428259, EPI_ISL_428260, EPI_ISL_428261, EPI_ISL_428262, EPI_ISL_428263, EPI_ISL_428264, EPI_ISL_428265, EPI_ISL_428266, EPI_ISL_428267, EPI_ISL_428268, EPI_ISL_428269, EPI_ISL_428270, EPI_ISL_428271, EPI_ISL_428272, EPI_ISL_428273, EPI_ISL_428274, EPI_ISL_428275, EPI_ISL_428276, EPI_ISL_428277, EPI_ISL_428278, EPI_ISL_428279, EPI_ISL_428280, EPI_ISL_428281, EPI_ISL_428282, EPI_ISL_428283, EPI_ISL_428284, EPI_ISL_428285, EPI_ISL_428286, EPI_ISL_428287, EPI_ISL_428288, EPI_ISL_428289, EPI_ISL_428290, EPI_ISL_428291, EPI_ISL_428292, EPI_ISL_428293, EPI_ISL_428294, EPI_ISL_428295, EPI_ISL_428296, EPI_ISL_428297, EPI_ISL_428298, EPI_ISL_428299, EPI_ISL_428300, EPI_ISL_428301, EPI_ISL_428302, EPI_ISL_428303, EPI_ISL_428304, EPI_ISL_428305, EPI_ISL_428306, EPI_ISL_428307, EPI_ISL_428308, EPI_ISL_428309, EPI_ISL_428310, EPI_ISL_428311, EPI_ISL_428312, EPI_ISL_428313, EPI_ISL_428314, EPI_ISL_428315, EPI_ISL_428316, EPI_ISL_428317, EPI_ISL_428318, EPI_ISL_428319, EPI_ISL_428320, EPI_ISL_428321, EPI_ISL_428322, EPI_ISL_428323, EPI_ISL_428324, EPI_ISL_428325, EPI_ISL_428326, EPI_ISL_428327, EPI_ISL_428328, EPI_ISL_428329, EPI_ISL_428330, EPI_ISL_428331, EPI_ISL_428332, EPI_ISL_428333, EPI_ISL_428334, EPI_ISL_428335, EPI_ISL_428336, EPI_ISL_428337, EPI_ISL_428338, EPI_ISL_428339, EPI_ISL_428340, EPI_ISL_428341, EPI_ISL_428342, EPI_ISL_428343, EPI_ISL_428344, EPI_ISL_428345 | | | |
| see above | University of Wisconsin-Madison AIDS Vaccine Research Laboratories | University of Wisconsin-Madison AIDS Vaccine Research Laboratories | Gage Moreno, Katarina Braun, et al. AIDS Vaccine Research Laboratories |
| EPI_ISL_428346 | Genomic Laboratory (GLAB) (Conjoint lab of Health Directorate of Istanbul and Istanbul Technical University) | Genomic Laboratory (GLAB), Istanbul Technical University | Ilker Karacan, Tugba Kizilboga Akgun, Bugra Agaoglu, Gizem Alkurt, Jale Yildiz, Betsi Köse, Elifnaz Çelik, Arzu Irvem, Yasemin Kendir Demirkol, Ozlem Akgun Dogan, Mehtap Aydın, Levent Doganay, Gizem Dinler Doganay |
| EPI_ISL_428347 | Service de Biologie Médicale - BP 125 | National Reference Center for Viruses of Respiratory Infections, Institut Pasteur, Paris | Mélanie Albert, Marion Barbet, Sylvie Behillil, Méline Bizard, Angela Brisebarre, Flora Donati, Etienne Simon-Lorière, Vincent Enouf, Maud Vanpeene, Sylvie van der Werf |
| EPI_ISL_428348 | Maison de Santé du Val d'Ormois | National Reference Center for Viruses of Respiratory Infections, Institut Pasteur, Paris | Mélanie Albert, Marion Barbet, Sylvie Behillil, Méline Bizard, Angela Brisebarre, Flora Donati, Etienne Simon-Lorière, Vincent Enouf, Maud Vanpeene, Sylvie van der Werf |
| EPI_ISL_428349 | Service de Biologie Médicale - BP 125 | National Reference Center for Viruses of Respiratory Infections, Institut Pasteur, Paris | Mélanie Albert, Marion Barbet, Sylvie Behillil, Méline Bizard, Angela Brisebarre, Flora Donati, Etienne Simon-Lorière, Vincent Enouf, Maud Vanpeene, Sylvie van der Werf |
| EPI_ISL_428350 | CH Jean de Navarre Laboratoire de Biologie | National Reference Center for Viruses of Respiratory Infections, Institut Pasteur, Paris | Mélanie Albert, Marion Barbet, Sylvie Behillil, Méline Bizard, Angela Brisebarre, Flora Donati, Etienne Simon-Lorière, Vincent Enouf, Maud Vanpeene, Sylvie van der Werf |
| EPI_ISL_428351, EPI_ISL_428352 | GH Nord Essonne Service de Biologie clinique | National Reference Center for Viruses of Respiratory Infections, Institut Pasteur, Paris | Mélanie Albert, Marion Barbet, Sylvie Behillil, Méline Bizard, Angela Brisebarre, Flora Donati, Etienne Simon-Lorière, Vincent Enouf, Maud Vanpeene, Sylvie van der Werf |
| EPI_ISL_428353 | CH Compiègne Laboratoire de Biologie | National Reference Center for Viruses of Respiratory Infections, Institut Pasteur, Paris | Mélanie Albert, Marion Barbet, Sylvie Behillil, Méline Bizard, Angela Brisebarre, Flora Donati, Etienne Simon-Lorière, Vincent Enouf, Maud Vanpeene, Sylvie van der Werf |
| EPI_ISL_428354 | LABM GH nord Essonne de Longjumeau - BP 125 | National Reference Center for Viruses of Respiratory Infections, Institut Pasteur, Paris | Mélanie Albert, Marion Barbet, Sylvie Behillil, Méline Bizard, Angela Brisebarre, Flora Donati, Etienne Simon-Lorière, Vincent Enouf, Maud Vanpeene, Sylvie van der Werf |
| EPI_ISL_428355, EPI_ISL_428356, EPI_ISL_428357 | Institut Médico légal- Hop R. Poincaré | National Reference Center for Viruses of Respiratory Infections, Institut Pasteur, Paris | Mélanie Albert, Marion Barbet, Sylvie Behillil, Méline Bizard, Angela Brisebarre, Flora Donati, Etienne Simon-Lorière, Vincent Enouf, Maud Vanpeene, Sylvie van der Werf |
| EPI_ISL_428358 | CH Jeanne de Navarre Laboratoire de Biologie | National Reference Center for Viruses of Respiratory Infections, Institut Pasteur, Paris | Mélanie Albert, Marion Barbet, Sylvie Behillil, Méline Bizard, Angela Brisebarre, Flora Donati, Etienne Simon-Lorière, Vincent Enouf, Maud Vanpeene, Sylvie van der Werf |
| EPI_ISL_428359, EPI_ISL_428360 | CH Compiègne Laboratoire de Biologie | National Reference Center for Viruses of Respiratory Infections, Institut Pasteur, Paris | Mélanie Albert, Marion Barbet, Sylvie Behillil, Méline Bizard, Angela Brisebarre, Flora Donati, Etienne Simon-Lorière, Vincent Enouf, Maud Vanpeene, Sylvie van der Werf |
| EPI_ISL_428361, EPI_ISL_428362 | LABM GH nord Essonne de Longjumeau - BP 125 | National Reference Center for Viruses of Respiratory Infections, Institut Pasteur, Paris | Mélanie Albert, Marion Barbet, Sylvie Behillil, Méline Bizard, Angela Brisebarre, Flora Donati, Etienne Simon-Lorière, Vincent Enouf, Maud Vanpeene, Sylvie van der Werf |
| EPI_ISL_428363 | GH Nord Essonne Service de Biologie clinique | National Reference Center for Viruses of Respiratory Infections, Institut Pasteur, Paris | Mélanie Albert, Marion Barbet, Sylvie Behillil, Méline Bizard, Angela Brisebarre, Flora Donati, Etienne Simon-Lorière, Vincent Enouf, Maud Vanpeene, Sylvie van der Werf |
| EPI_ISL_428364 | Cabinet Médical | National Reference Center for Viruses of Respiratory Infections, Institut Pasteur, Paris | Mélanie Albert, Marion Barbet, Sylvie Behillil, Méline Bizard, Angela Brisebarre, Flora Donati, Etienne Simon-Lorière, Vincent Enouf, Maud Vanpeene, Sylvie van der Werf |
| EPI_ISL_428365 | LABM GH nord Essonne de Longjumeau - BP 125 | National Reference Center for Viruses of Respiratory Infections, Institut Pasteur, Paris | Mélanie Albert, Marion Barbet, Sylvie Behillil, Méline Bizard, Angela Brisebarre, Flora Donati, Etienne Simon-Lorière, Vincent Enouf, Maud Vanpeene, Sylvie van der Werf |
| EPI_ISL_428366 | CH Jeanne de Navarre Laboratoire de Biologie | National Reference Center for Viruses of Respiratory Infections, Institut Pasteur, Paris | Mélanie Albert, Marion Barbet, Sylvie Behillil, Méline Bizard, Angela Brisebarre, Flora Donati, Etienne Simon-Lorière, Vincent Enouf, Maud Vanpeene, Sylvie van der Werf |
| EPI_ISL_428367 | Cabinet Médical | National Reference Center for Viruses of Respiratory Infections, Institut Pasteur, Paris | Mélanie Albert, Marion Barbet, Sylvie Behillil, Méline Bizard, Angela Brisebarre, Flora Donati, Etienne Simon-Lorière, Vincent Enouf, Maud Vanpeene, Sylvie van der Werf |
| EPI_ISL_428368 | Genomic Laboratory (GLAB) (Conjoint lab of Health Directorate of Istanbul and Istanbul Technical University) | Genomic Laboratory (GLAB), Istanbul Technical University | Ilker Karacan, Tugba Kizilboga Akgun, Bugra Agaoglu, Gizem Alkurt, Jale Yildiz, Betsi Köse, Elifnaz Çelik, Arzu Irvem, Yasemin Kendir Demirkol, Ozlem Akgun Dogan, Mehtap Aydın, Levent Doganay, Gizem Dinler Doganay |
| EPI_ISL_428369 | Yale Clinical Virology Laboratory | Grubaugh Lab - Yale School of Public Health | Joseph Fauver, Anderson Brito, Tara Alpert, Chantal Vogels, Ellen Foxman, Albert Ko, Marie Landry, Nathan Grubaugh |

| | | | |
|---|---|---|---|
| EPI_ISL_428370, EPI_ISL_428371, EPI_ISL_428372 | Yale Clinical Virology Laboratory | Grubaugh Lab - Yale School of Public Health | Joseph Fauver, Anderson Brito, Tara Alpert, Chantal Vogels, Ellen Foxman, Albert Ko, Marie Landry, Nathan Grubaugh |
| EPI_ISL_428373, EPI_ISL_428374, EPI_ISL_428375, EPI_ISL_428376, EPI_ISL_428377, EPI_ISL_428378, EPI_ISL_428379, EPI_ISL_428380 | Yale COVID-19 Biorepository | Grubaugh Lab - Yale School of Public Health | Joseph Fauver, Tara Alpert, Anderson Brito, Anne Wyllie, Chantal Vogels, Mary Petrone, Chaney Kalinich, Isabel Ott, Arnau Casanovas, Catherine Muenker, Adam Moore, Alice Lu, Maria Tokuyama, Patrick Wong, Peiwen Lu, Saad Omer, Richard Martinello, Allison Nelson, Shelli Farhadian, Akiko Iwasaki, Charlese Dela Cruz, Albert Ko, Nathan Grubaugh |
| EPI_ISL_428381, EPI_ISL_428382 | Yale Clinical Virology Laboratory | Grubaugh Lab - Yale School of Public Health | Joseph Fauver, Anderson Brito, Tara Alpert, Chantal Vogels, Ellen Foxman, Albert Ko, Marie Landry, Nathan Grubaugh |
| EPI_ISL_428384, EPI_ISL_428385, EPI_ISL_428386, EPI_ISL_428387, EPI_ISL_428388, EPI_ISL_428389, EPI_ISL_428390, EPI_ISL_428391, EPI_ISL_428392, EPI_ISL_428393, EPI_ISL_428394, EPI_ISL_428395, EPI_ISL_428396, EPI_ISL_428397, EPI_ISL_428398 | | | |
| see above | Yale COVID-19 Biorepository | Grubaugh Lab - Yale School of Public Health | Joseph Fauver, Tara Alpert, Anderson Brito, Anne Wyllie, Chantal Vogels, Mary Petrone, Chaney Kalinich, Isabel Ott, Arnau Casanovas, Catherine Muenker, Adam Moore, Alice Lu, Maria Tokuyama, Patrick Wong, Peiwen Lu, Saad Omer, Richard Martinello, Allison Nelson, Shelli Farhadian, Akiko Iwasaki, Charlese Dela Cruz, Albert Ko, Nathan Grubaugh |
| EPI_ISL_428399 | Yale COVID-19 Biorepository | Grubaugh Lab - Yale School of Public Health | Joseph Fauver, Tara Alpert, Anderson Brito, Anne Wyllie, Chantal Vogels, Mary Petrone, Cole Jensen, Chaney Kalinich, Isabel Ott, Arnau Casanovas, Catherine Muenker, Adam Moore, Alice Lu, Maria Tokuyama, Patrick Wong, Peiwen Lu, Saad Omer, Richard Martinello, Allison Nelson, Shelli Farhadian, Akiko Iwasaki, Charlese Dela Cruz, Albert Ko, Nathan Grubaugh |
| EPI_ISL_428400, EPI_ISL_428401, EPI_ISL_428402, EPI_ISL_428403, EPI_ISL_428404, EPI_ISL_428405 | Yale COVID-19 Biorepository | Grubaugh Lab - Yale School of Public Health | Joseph Fauver, Tara Alpert, Anderson Brito, Anne Wyllie, Chantal Vogels, Mary Petrone, Cole Jensen, Chaney Kalinich, Isabel Ott, Arnau Casanovas, Catherine Muenker, Adam Moore, Alice Lu, Maria Tokuyama, Patrick Wong, Peiwen Lu, Saad Omer, Richard Martinello, Allison Nelson, Shelli Farhadian, Akiko Iwasaki, Charlese Dela Cruz, Albert Ko, Nathan Grubaugh |
| EPI_ISL_428440, EPI_ISL_428441, EPI_ISL_428442, EPI_ISL_428443, EPI_ISL_428444, EPI_ISL_428445, EPI_ISL_428446, EPI_ISL_428447, EPI_ISL_428448, EPI_ISL_428449, EPI_ISL_428450, EPI_ISL_428451, EPI_ISL_428452, EPI_ISL_428453, EPI_ISL_428454, EPI_ISL_428455, EPI_ISL_428456, EPI_ISL_428457, EPI_ISL_428458, EPI_ISL_428459, EPI_ISL_428460, EPI_ISL_428461, EPI_ISL_428462, EPI_ISL_428463, EPI_ISL_428464, EPI_ISL_428465, EPI_ISL_428466, EPI_ISL_428467, EPI_ISL_428468, EPI_ISL_428469, EPI_ISL_428470, EPI_ISL_428471, EPI_ISL_428472, EPI_ISL_428473, EPI_ISL_428474, EPI_ISL_428475, EPI_ISL_428476, EPI_ISL_428477, EPI_ISL_428478 | | | |
| see above | Guangdong Provincial Center for Diseases Control and Prevention;Guangdong Provincial Institute of Public Health | School of Public Health, The University of Hong Kong | Bosheng Li, Haogao Gu, Lijun Liang, Zhencui Li, Hui-Ling Yen, Yao Hu, Yingchao Song , Hanri Zeng, Tie Song, Jie Wu, Leo L.M. Poon |
| EPI_ISL_428479, EPI_ISL_428480, EPI_ISL_428481, EPI_ISL_428482, EPI_ISL_428483, EPI_ISL_428484, EPI_ISL_428485, EPI_ISL_428486, EPI_ISL_428487 | District Surveillence Unit | Department of Neurovirology, National Institute of Mental Health and Neuroscience (NIMHANS) | Chitra Pattabiraman, Vijayalakshmi Reddy, Harsha PK, Risha Rasheed, Shafeeq S Hameed, Manjunatha Venkataswamy, Anita Desai, Ravi Vasanthapuram |
| EPI_ISL_428488, EPI_ISL_428489, EPI_ISL_428490, EPI_ISL_428491 | Centers for Disease Control, R.O.C. (Taiwan) | Centers for Disease Control, R.O.C. (Taiwan) | Ji-Rong Yang, Yu-Chi Lin, Jung-Jung Mu, Ming-Tsan Liu |
| EPI_ISL_428670 | Centre for Dengue Research | Centre for Dengue Research | Chandima Jeewandara, Dinuka Ariyaratne, Laksiri Gomes, Deshni Jayathilaka, Ananda Wijewickrama, Eranga Narangoda, Damayanthi Idampitiya, Neelika Malaige |
| EPI_ISL_428671 | Centre for Dengue Research | Centre for Dengue Research | Chandima Jeewandara, Dinuka Ariyatane, Laksiri Gomes, Deshni Jayathilaka, Diyanath Ranasinghe, Ananda Wijewickrama, Eranga Narangoda, Damayanthi Tdampitiya, Neelika Malavige |
| EPI_ISL_428672, EPI_ISL_428673 | Centre for Dengue Research | Centre for Dengue Research | Chandima Jeewandara, Dinuka Ariyaratne, Laksiri Gomes, Deshni Jayathilaka, Diyanath Ranasinghe, Ananda Wijewickrama, Eranga Narangoda, Damayanthi Idampitiya, Neelika Malavige |
| EPI_ISL_428674, EPI_ISL_428675, EPI_ISL_428676, EPI_ISL_428677, EPI_ISL_428678, EPI_ISL_428679, EPI_ISL_428680, EPI_ISL_428681, EPI_ISL_428682 | Hospital Universitario La Paz | Hospital Universitario 12 de Octubre | Elias Dahdouh, Sara González, Raúl Recio, Fernando Lázaro, Esther Viedma, Natalia Stella, Julio García, Juan Carlos Galán, Rafael Cantón, Mª Dolores Folgueira, Rafael Delgado, Jesús Mingorance |
| EPI_ISL_428683, EPI_ISL_428684, EPI_ISL_428685, EPI_ISL_428686, EPI_ISL_428687, EPI_ISL_428688, EPI_ISL_428689, EPI_ISL_428690, EPI_ISL_428691, EPI_ISL_428692, EPI_ISL_428693, EPI_ISL_428694 | | | |
| see above | Hospital Universitario 12 de Octubre | Hospital Universitario 12 de Octubre | Sara González, Raúl Recio,Elias Dahdouh, Fernando Lázaro, Esther Viedma, Natalia Stella, Julio García, Juan Carlos Galán, Rafael Cantón, Mª Dolores Folgueira, Rafael Delgado, Jesús Mingorance |
| EPI_ISL_428695, EPI_ISL_428696, EPI_ISL_428697, EPI_ISL_428698, EPI_ISL_428699 | Hospital Universitario 12 de Octubre | Hospital Universitario 12 de Octubre | Raúl Recio, Sara González, Elias Dahdouh, Fernando Lázaro, Esther Viedma, Natalia Stella, Julio García, Juan Carlos Galán, Rafael Cantón, Mª Dolores Folgueira, Rafael Delgado, Jesús Mingorance |
| EPI_ISL_428700, EPI_ISL_428701, EPI_ISL_428702, EPI_ISL_428703, EPI_ISL_428704, EPI_ISL_428705, EPI_ISL_428706, EPI_ISL_428707, EPI_ISL_428708, EPI_ISL_428709, EPI_ISL_428710, EPI_ISL_428711 | | | |
| see above | Hospital Universitario 12 de Octubre | Hospital Universitario 12 de Octubre | Esther Viedma, Sara González, Raúl Recio, Elias Dahdouh, Fernando Lázaro, Julio García, Mª Dolores Folgueira, Jesús Mingorance, Rafael Delgado |
| EPI_ISL_428712, EPI_ISL_428713, EPI_ISL_428714, EPI_ISL_428715, EPI_ISL_428716, EPI_ISL_428717, EPI_ISL_428718, EPI_ISL_428719, EPI_ISL_428720, EPI_ISL_428721, EPI_ISL_428722, EPI_ISL_428723 | | | |
| see above | Ministry of Health Turkey | Ministry of Health Turkey | Fatma Bayrakdar,Ayşe Başak Altaş,Yasemin Coşgun,Gülay Korukluoğlu,Selçuk Kılıç |
| EPI_ISL_428724, EPI_ISL_428725, EPI_ISL_428726, EPI_ISL_428727, EPI_ISL_428728, EPI_ISL_428729, EPI_ISL_428730, EPI_ISL_428731, EPI_ISL_428732 | University of Wisconsin-Madison AIDS Vaccine Research Laboratories | University of Wisconsin-Madison AIDS Vaccine Research Laboratories | Gage Moreno, Katarina Braun, et al. AIDS Vaccine Research Laboratories |
| EPI_ISL_428740 | Yale Clinical Virology Laboratory | Grubaugh Lab - Yale School of Public Health | Joseph Fauver, Anderson Brito, Tara Alpert, Chantal Vogels, Ellen Foxman, Albert Ko, Marie Landry, Nathan Grubaugh |
| EPI_ISL_428745, EPI_ISL_428746, EPI_ISL_428747, EPI_ISL_428748, EPI_ISL_428749, EPI_ISL_428750, EPI_ISL_428751, EPI_ISL_428752 | Yale COVID-19 Biorepository | Grubaugh Lab - Yale School of Public Health | Joseph Fauver, Tara Alpert, Anderson Brito, Anne Wyllie, Chantal Vogels, Mary Petrone, Cole Jensen, Chaney Kalinich, Isabel Ott, Arnau Casanovas, Catherine Muenker, Adam Moore, Alice Lu, Maria Tokuyama, Patrick Wong, Peiwen Lu, Saad Omer, Richard Martinello, Allison Nelson, Shelli Farhadian, Akiko Iwasaki, Charlese Dela Cruz, Albert Ko, Nathan Grubaugh |
| EPI_ISL_428757, EPI_ISL_428758, EPI_ISL_428759, EPI_ISL_428760, EPI_ISL_428761, EPI_ISL_428762, EPI_ISL_428763, EPI_ISL_428764, EPI_ISL_428765, EPI_ISL_428766, EPI_ISL_428767, EPI_ISL_428768, EPI_ISL_428769, EPI_ISL_428770, EPI_ISL_428771, EPI_ISL_428772, EPI_ISL_428773, EPI_ISL_428774, EPI_ISL_428775, EPI_ISL_428776, EPI_ISL_428777, EPI_ISL_428778, EPI_ISL_428779, EPI_ISL_428780, EPI_ISL_428781, EPI_ISL_428782, EPI_ISL_428783, EPI_ISL_428784, EPI_ISL_428785, EPI_ISL_428786, EPI_ISL_428787, EPI_ISL_428788, EPI_ISL_428789, EPI_ISL_428790, EPI_ISL_428791, EPI_ISL_428792, EPI_ISL_428793, EPI_ISL_428794, EPI_ISL_428795, EPI_ISL_428796, EPI_ISL_428797, EPI_ISL_428798, EPI_ISL_428799, EPI_ISL_428800, EPI_ISL_428801, EPI_ISL_428802, EPI_ISL_428803, EPI_ISL_428804, EPI_ISL_428805 | | | |
| see above | NYU Langone Health | Departments of Pathology and Medicine, New York University School of Medicine | Maria Aguero-Rosenfeld, Brendan Belovarac, Margaret Black, Ludovic Boytard, John Cadley, Paolo Cotzia, John Chen, Dacia Dimartino, Xiaojun Feng, Tatyana Gindin, Emily Guzman, Adriana Heguy, Megan Hogan, Emily Huang, George Jour, Andrew Lytle, Christian Marier, Matthew T. Maurano, Mark J. Mulligan, Peter Meyn, Iman Osman, Jared Pinnell, Vanessa Raabe, Sitharam Ramaswami, Amy Rapkiewicz, Marie Samanovic-Golden, Antonio Serrano, Guomiao Shen, Matija Snuderl, Theodore Vougiouklakis, Nick Vulpescu, Gael Westby, Paul Zappile, Yutong Zhang |
| EPI_ISL_428822, EPI_ISL_428823, EPI_ISL_428824, EPI_ISL_428825, EPI_ISL_428826, EPI_ISL_428827, EPI_ISL_428828, EPI_ISL_428829, EPI_ISL_428830, EPI_ISL_428831, EPI_ISL_428832, EPI_ISL_428833, EPI_ISL_428834, EPI_ISL_428835, EPI_ISL_428836, EPI_ISL_428837, EPI_ISL_428838, EPI_ISL_428839, EPI_ISL_428840, EPI_ISL_428841, EPI_ISL_428842, EPI_ISL_428843, EPI_ISL_428844, EPI_ISL_428845, EPI_ISL_428846, EPI_ISL_428847, EPI_ISL_428848, EPI_ISL_428849, EPI_ISL_428850 | | | |
| see above | National Public Health Laboratory, National Centre for Infectious Diseases | National Public Health Laboratory, National Centre for Infectious Diseases | Mak TM, Octavia S, Chavatte JM, Cui L, Lin RTP |
| EPI_ISL_428851, EPI_ISL_428852 | FSBSI "Chumakov Federal Scientific Center for Research and Development of Immune-and-Biological Products of Russian Academy of Sciences" | FSBSI "Chumakov Federal Scientific Center for Research and Development of Immune-and-Biological Products of Russian Academy of Sciences" & NRC "Kurchatov institute" | Liubov Kozlovskaya, Anastasia Piniaeva, Georgy Ignatyev, Anna Shishova, Aydar Ishmukhametov, Mikhail Rychev, Egor Prokhorchuk, Denis Protsenko, Anastasia Berestovskaya |
| EPI_ISL_428853 | Laboratory of Molecular Virology International Center for Genetic Engineering and Biotechnology (ICGEB) | ARGO Open Lab Platform for Genome Sequencing | Licastro D, Rajasekharan S, Dal Monego S, Segat L, D'Agaro P, Marcello A |
| EPI_ISL_428854 | Laboratory of Molecular Virology International Center for Genetic Engineering and Biotechnology (ICGEB) | ARGO Open Lab Platform for Genome sequencing | Licastro D, Rajasekharan S, Dal Monego S, Segat L, D'Agaro P, Marcello A |
| EPI_ISL_428855 | MRCG at LSHTM Geomics lab | MRCG at LSHTM Genomics lab | Sesay et al |
| EPI_ISL_428856 | MRCG at LSHTM Genomics Lab | MRCG at LSHTM Genomics lab | Sesay et al |
| EPI_ISL_428857 | MRCG at LSHTM Genomics lab | MRCG at LSHTM Genomics lab | Sesay et al |
| EPI_ISL_428860, EPI_ISL_428861 | State Research Center of Virology and Biotechnology VECTOR, Department of Collection of Microorganisms | State Research Center of Virology and Biotechnology VECTOR, Department of Collection of Microorganisms | Oleg V. Pyankov, Sergey A. Bodnev, Tatyana V. Tregubchak, Alexander N. Shvalov, Elena V. Gavrilova, Rinat A. Maksyutov |
| EPI_ISL_428862, EPI_ISL_428863 | State Research Center of Virology and Biotechnology VECTOR, Department of Collection of Microorganisms | State Research Center of Virology and Biotechnology VECTOR, Department of Collection of Microorganisms | Oleg V. Pyankov, Sergey A. Bodnev, Anastasiya M. Smirnova, Anastasiya A. Nazarenko, Tatyana V. Tregubchak, Alexander N. Shvalov, Elena V. Gavrilova, Rinat A. Maksyutov |
| EPI_ISL_428864, EPI_ISL_428865, EPI_ISL_428866, EPI_ISL_428867, EPI_ISL_428868, EPI_ISL_428869, EPI_ISL_428870, EPI_ISL_428871 | State Research Center of Virology and Biotechnology VECTOR, Department of Collection of Microorganisms | State Research Center of Virology and Biotechnology VECTOR, Department of Collection of Microorganisms | Oleg V. Pyankov, Sergey A. Bodnev, Tatyana V. Tregubchak, Alexander N. Shvalov, Elena V. Gavrilova, Rinat A. Maksyutov |
| EPI_ISL_428872, EPI_ISL_428873, EPI_ISL_428874, EPI_ISL_428875 | State Research Center of Virology and Biotechnology VECTOR, Department of Collection of Microorganisms | State Research Center of Virology and Biotechnology VECTOR, Department of Collection of Microorganisms | Sergey A. Bodnev, Oleg V. Pyankov, Tatyana V. Tregubchak, Alexander N. Shvalov, Elena V. Gavrilova, Rinat A. Maksyutov |
| EPI_ISL_428876, EPI_ISL_428877 | State Research Center of Virology and Biotechnology VECTOR, Department of Collection of Microorganisms | State Research Center of Virology and Biotechnology VECTOR, Department of Collection of Microorganisms | Sergey A. Bodnev, Oleg V. Pyankov, Anastasiya M. Smirnova, Anastasiya A. Nazarenko, Tatyana V. Tregubchak, Alexander N. Shvalov, Elena V. Gavrilova, Rinat A. Maksyutov |
| EPI_ISL_428878, EPI_ISL_428879, EPI_ISL_428880, EPI_ISL_428881 | State Research Center of Virology and Biotechnology VECTOR, Department of Collection of Microorganisms | State Research Center of Virology and Biotechnology VECTOR, Department of Collection of Microorganisms | Sergey A. Bodnev, Oleg V. Pyankov, Tatyana V. Tregubchak, Alexander N. Shvalov, Elena V. Gavrilova, Rinat A. Maksyutov |
| EPI_ISL_428882, EPI_ISL_428883, EPI_ISL_428884, EPI_ISL_428885, EPI_ISL_428886, EPI_ISL_428887, EPI_ISL_428888, EPI_ISL_428889, EPI_ISL_428890, EPI_ISL_428891, EPI_ISL_428892, EPI_ISL_428893, EPI_ISL_428894 | | | |
| see above | State Research Center of Virology and Biotechnology VECTOR, Department of Collection of Microorganisms | State Research Center of Virology and Biotechnology VECTOR, Department of Collection of Microorganisms | Oleg V. Pyankov, Sergey A. Bodnev, Tatyana V. Tregubchak, Alexander N. Shvalov, Elena V. Gavrilova, Rinat A. Maksyutov |
| EPI_ISL_428895, EPI_ISL_428896, EPI_ISL_428897, EPI_ISL_428898, EPI_ISL_428899, EPI_ISL_428900, EPI_ISL_428901, EPI_ISL_428902, EPI_ISL_428903, EPI_ISL_428904, EPI_ISL_428905 | | | |
| see above | State Research Center of Virology and Biotechnology VECTOR, Department of Collection of Microorganisms | State Research Center of Virology and Biotechnology VECTOR, Department of Collection of Microorganisms | Sergey A. Bodnev, Oleg V. Pyankov, Tatyana V. Tregubchak, Alexander N. Shvalov, Elena V. Gavrilova, Rinat A. Maksyutov |
| EPI_ISL_428906, EPI_ISL_428907, EPI_ISL_428908, EPI_ISL_428909, EPI_ISL_428910, EPI_ISL_428911, EPI_ISL_428912, EPI_ISL_428913, EPI_ISL_428914, EPI_ISL_428915, EPI_ISL_428916 | | | |
| see above | State Research Center of Virology and Biotechnology VECTOR, Department of Collection of Microorganisms | State Research Center of Virology and Biotechnology VECTOR, Department of Collection of Microorganisms | Oleg V. Pyankov, Sergey A. Bodnev, Tatyana V. Tregubchak, Alexander N. Shvalov, Elena V. Gavrilova, Rinat A. Maksyutov |
| EPI_ISL_428917, EPI_ISL_428918, EPI_ISL_428919, EPI_ISL_428920, EPI_ISL_428921, EPI_ISL_428922, EPI_ISL_428923, EPI_ISL_428924 | State Research Center of Virology and Biotechnology VECTOR, Department of Collection of Microorganisms | State Research Center of Virology and Biotechnology VECTOR, Department of Collection of Microorganisms | Sergey A. Bodnev, Oleg V. Pyankov, Tatyana V. Tregubchak, Alexander N. Shvalov, Elena V. Gavrilova, Rinat A. Maksyutov |
| EPI_ISL_428925, EPI_ISL_428926, EPI_ISL_428927, EPI_ISL_428928, EPI_ISL_428929, EPI_ISL_428930, | ViroGenetics - BSL3 Laboratory of Virology; Human Genome Variation Research Group & Genomics Centre MCB; | ViroGenetics - BSL3 Laboratory of Virology; Human Genome Variation Research Group & Genomics Centre MCB; | Wojciech Branicki, Ewelina Pośpiech, Michał Kowalski, Agnieszka Starowicz, Adrianna Klajmon, Aleksandra Pisarek, Danuta Piniewska-Róg, Kamila Marszałek, Tomasz Gromowski, Katarzyna Kopera, Katarzyna Dudek, Inga Drebot, Katarzyna Guła, Magda Pachota, Aleksandra Synowiec, Marek Sanak, Jarosław Foremny, Paweł P Łabaj, Krzysztof Pyrć |

| | | | |
|---|---|---|---|
| EPI_ISL_428931, EPI_ISL_428932 | Bioinformatics Research Group; Wojewódzka Stacja Sanitarno-Epidemiologiczna w Krakowie | Bioinformatics Research Group | |
| EPI_ISL_428935, EPI_ISL_428936 | University of Wisconsin-Madison AIDS Vaccine Research Laboratories | University of Wisconsin-Madison AIDS Vaccine Research Laboratories | Gage Moreno, Katarina Braun, et al. AIDS Vaccine Research Laboratories |
| EPI_ISL_428939, EPI_ISL_428940, EPI_ISL_428941, EPI_ISL_428942, EPI_ISL_428943, EPI_ISL_428944, EPI_ISL_428945, EPI_ISL_428946, EPI_ISL_428947, EPI_ISL_428948, EPI_ISL_428949, EPI_ISL_428950, EPI_ISL_428951, EPI_ISL_428952, EPI_ISL_428953, EPI_ISL_428954, EPI_ISL_428955, EPI_ISL_428956, EPI_ISL_428957, EPI_ISL_428958, EPI_ISL_428959, EPI_ISL_428960, EPI_ISL_428961, EPI_ISL_428962 | | | |
| see above | Laboratoire National de Sante, Microbiology, Virology | Laboratoire National de Sante, Microbiology, Epidemiology and Microbial Genomics | Anke Wienecke-Baldacchino, Ardashel Latsuzbaia, Jessica Tapp, Catherine Ragimbeau, Guillaume Fournier, Tamir Abdelrahman, Trung Nguyen Nguyen, Joel Mossong |
| EPI_ISL_428990, EPI_ISL_428991, EPI_ISL_428992, EPI_ISL_428993, EPI_ISL_428994, EPI_ISL_428995, EPI_ISL_428996, EPI_ISL_428997, EPI_ISL_428998, EPI_ISL_428999, EPI_ISL_429000, EPI_ISL_429001, EPI_ISL_429002, EPI_ISL_429003, EPI_ISL_429004, EPI_ISL_429005, EPI_ISL_429006, EPI_ISL_429007, EPI_ISL_429008, EPI_ISL_429009, EPI_ISL_429010, EPI_ISL_429011, EPI_ISL_429012, EPI_ISL_429013, EPI_ISL_429014, EPI_ISL_429015, EPI_ISL_429016, EPI_ISL_429017, EPI_ISL_429018, EPI_ISL_429019, EPI_ISL_429020, EPI_ISL_429021, EPI_ISL_429022, EPI_ISL_429023, EPI_ISL_429024, EPI_ISL_429025, EPI_ISL_429026, EPI_ISL_429027, EPI_ISL_429028, EPI_ISL_429029, EPI_ISL_429030, EPI_ISL_429031, EPI_ISL_429032, EPI_ISL_429033, EPI_ISL_429034, EPI_ISL_429035, EPI_ISL_429036, EPI_ISL_429037, EPI_ISL_429038, EPI_ISL_429039, EPI_ISL_429040, EPI_ISL_429041, EPI_ISL_429042, EPI_ISL_429043, EPI_ISL_429044, EPI_ISL_429045, EPI_ISL_429046, EPI_ISL_429047, EPI_ISL_429048, EPI_ISL_429049, EPI_ISL_429050, EPI_ISL_429051, EPI_ISL_429052, EPI_ISL_429053, EPI_ISL_429054, EPI_ISL_429055, EPI_ISL_429056, EPI_ISL_429057, EPI_ISL_429058, EPI_ISL_429059, EPI_ISL_429060, EPI_ISL_429061, EPI_ISL_429062, EPI_ISL_429063, EPI_ISL_429064, EPI_ISL_429065, EPI_ISL_429066, EPI_ISL_429067, EPI_ISL_429068, EPI_ISL_429069, EPI_ISL_429070, EPI_ISL_429071, EPI_ISL_429072, EPI_ISL_429073 | | | |
| see above | UCSF Clinical Microbiology Laboratory | Chan-Zuckerberg Biohub | CZB Cliahub Consortium |
| EPI_ISL_429074, EPI_ISL_429075 | The First Affiliated Hospital of Guangzhou Medical University | BGI-shenzhen & The First Affiliated Hospital of Guangzhou Medical University | Yanqun Wang, Daxi Wang, Lu Zhang, Wanying Sun, Zhaoyong Zhang et al. |
| EPI_ISL_429076 | The First Affiliated Hospital of Guangzhou Medical University | BGI-shenzhen & The First Affiliated Hospital of Guangzhou Medical University | |
| EPI_ISL_429077 | The First Affiliated Hospital of Guangzhou Medical University | BGI-shenzhen & The First Affiliated Hospital of Guangzhou Medical University | Yanqun Wang, Daxi Wang, Lu Zhang, Wanying Sun, Zhaoyong Zhang et al. |
| EPI_ISL_429078, EPI_ISL_429079, EPI_ISL_429080, EPI_ISL_429081 | The First Affiliated Hospital of Guangzhou Medical University | BGI-shenzhen & The First Affiliated Hospital of Guangzhou Medical University | |
| EPI_ISL_429082, EPI_ISL_429083 | The First Affiliated Hospital of Guangzhou Medical University | BGI-shenzhen & The First Affiliated Hospital of Guangzhou Medical University | Yanqun Wang, Daxi Wang, Lu Zhang, Wanying Sun, Zhaoyong Zhang et al. |
| EPI_ISL_429084 | The First Affiliated Hospital of Guangzhou Medical University | BGI-shenzhen & The First Affiliated Hospital of Guangzhou Medical University | |
| EPI_ISL_429085 | The First Affiliated Hospital of Guangzhou Medical University | BGI-shenzhen & The First Affiliated Hospital of Guangzhou Medical University | Yanqun Wang, Daxi Wang, Lu Zhang, Wanying Sun, Zhaoyong Zhang et al. |
| EPI_ISL_429086 | The First Affiliated Hospital of Guangzhou Medical University | BGI-shenzhen & The First Affiliated Hospital of Guangzhou Medical University | |
| EPI_ISL_429088, EPI_ISL_429089, EPI_ISL_429090, EPI_ISL_429091, EPI_ISL_429092, EPI_ISL_429093 | The First Affiliated Hospital of Guangzhou Medical University | BGI-shenzhen & The First Affiliated Hospital of Guangzhou Medical University | Yanqun Wang, Daxi Wang, Lu Zhang, Wanying Sun, Zhaoyong Zhang et al. |
| EPI_ISL_429094, EPI_ISL_429095 | The First Affiliated Hospital of Guangzhou Medical University | BGI-shenzhen & The First Affiliated Hospital of Guangzhou Medical University | |
| EPI_ISL_429096, EPI_ISL_429097, EPI_ISL_429098 | The First Affiliated Hospital of Guangzhou Medical University | BGI-shenzhen & The First Affiliated Hospital of Guangzhou Medical University | Yanqun Wang, Daxi Wang, Lu Zhang, Wanying Sun, Zhaoyong Zhang et al. |
| EPI_ISL_429099, EPI_ISL_429100, EPI_ISL_429101, EPI_ISL_429102, EPI_ISL_429103, EPI_ISL_429104, EPI_ISL_429105 | The First Affiliated Hospital of Guangzhou Medical University | BGI-shenzhen & The First Affiliated Hospital of Guangzhou Medical University | |
| EPI_ISL_429115, EPI_ISL_429116, EPI_ISL_429117, EPI_ISL_429118, EPI_ISL_429119, EPI_ISL_429120, EPI_ISL_429121, EPI_ISL_429122, EPI_ISL_429123, EPI_ISL_429124, EPI_ISL_429125 | | | |
| see above | Klinisk mikrobiologi och vardhygien Halmstad | The Public Health Agency of Sweden | Arne Kotz, Olov Svartstrom, Maria Lind Karlberg, Anna-Malin Linde, Oskar Karlsson Lindsjo, Anna Risberg, Shaman Muradrasoli, Karin Tegmark-Wisell |
| EPI_ISL_429126, EPI_ISL_429127 | Unilabs Skovde | The Public Health Agency of Sweden | Tobias Kollberg, Helena Enroth, Olov Svartstrom, Maria Lind Karlberg, Anna-Malin Linde, Oskar Karlsson Lindsjo, Anna Risberg, Shaman Muradrasoli, Karin Tegmark-Wisell |
| EPI_ISL_429128, EPI_ISL_429129, EPI_ISL_429130, EPI_ISL_429131, EPI_ISL_429132, EPI_ISL_429133, EPI_ISL_429134, EPI_ISL_429135 | Laboratoriemedicin | The Public Health Agency of Sweden | Olov Svartstrom, Maria Lind Karlberg, Anna-Malin Linde, Oskar Karlsson Lindsjo, Anna Risberg, Shaman Muradrasoli, Karin Tegmark-Wisell |
| EPI_ISL_429136, EPI_ISL_429137, EPI_ISL_429138, EPI_ISL_429139, EPI_ISL_429140, EPI_ISL_429141, EPI_ISL_429142, EPI_ISL_429143, EPI_ISL_429144, EPI_ISL_429145, EPI_ISL_429146, EPI_ISL_429147, EPI_ISL_429148, EPI_ISL_429150, EPI_ISL_429151, EPI_ISL_429152, EPI_ISL_429153, EPI_ISL_429154, EPI_ISL_429155, EPI_ISL_429156, EPI_ISL_429157, EPI_ISL_429158, EPI_ISL_429160 | | | |
| see above | Klinisk mikrobiologi Orebro | The Public Health Agency of Sweden | Martin Sundqvist, Olov Svartstrom, Maria Lind Karlberg, Anna-Malin Linde, Oskar Karlsson Lindsjo, Anna Risberg, Shaman Muradrasoli, Karin Tegmark-Wisell |
| EPI_ISL_429161, EPI_ISL_429162, EPI_ISL_429163 | The Public Health Agency of Sweden | The Public Health Agency of Sweden | Olov Svartstrom, Maria Lind Karlberg, Anna-Malin Linde, Oskar Karlsson Lindsjo, Anna Risberg, Shaman Muradrasoli, Karin Tegmark-Wisell |
| EPI_ISL_429164, EPI_ISL_429165, EPI_ISL_429166, EPI_ISL_429167, EPI_ISL_429168, EPI_ISL_429169, EPI_ISL_429170, EPI_ISL_429171, EPI_ISL_429172, EPI_ISL_429173, EPI_ISL_429174, EPI_ISL_429175, EPI_ISL_429176, EPI_ISL_429177, EPI_ISL_429178, EPI_ISL_429179, EPI_ISL_429180, EPI_ISL_429181, EPI_ISL_429182, EPI_ISL_429183, EPI_ISL_429184 | | | |
| see above | Ramathibodi Hospital | COVID-19 Network Investigations (CONI) Alliance | Elizabeth Batty, Wasun Chantratita, Thanat Chookajorn, Stefan Fernandez, Angkana Huang, Poramate Jiaranai, Anthony R. Jones, Khajohn Joonsalak, Chonticha Klungtong, Theerarat Kochakarn, Namfon Kotanan, Krittikorn Kumpornsin, Wudtichai Manasatienkij, Bhakbhoom Panthan, Ekawat Pasomsub, Kingkan Rakmanee, Insee Sensorn, Janjira Thaipadungpanit, Arporn Wangwiwatsin,Treewat Watthanachockchai |
| EPI_ISL_429196, EPI_ISL_429197, EPI_ISL_429199, EPI_ISL_429200, EPI_ISL_429201, EPI_ISL_429203, EPI_ISL_429204, EPI_ISL_429206, EPI_ISL_429207, EPI_ISL_429208, EPI_ISL_429209 | | | |
| see above | University Hospitals of Geneva Laboratory of Virology | University Hospitals of Geneva Laboratory of Virology | Laubscher F. |
| EPI_ISL_429210 | University Hospitals of Geneva Laboratory of Virology | University Hospitals of Geneva Laboratory of Virology | Laubscher F. |
| EPI_ISL_429211, EPI_ISL_429212, EPI_ISL_429213, EPI_ISL_429214, EPI_ISL_429215, EPI_ISL_429217, EPI_ISL_429218 | University Hospitals of Geneva Laboratory of Virology | University Hospitals of Geneva Laboratory of Virology | Laubscher F. |
| EPI_ISL_429219 | University Hospitals of Geneva Laboratory of Virology | University Hospitals of Geneva Laboratory of Virology | Laubscher F. |
| EPI_ISL_429220, EPI_ISL_429221, EPI_ISL_429222, EPI_ISL_429223 | University Hospitals of Geneva Laboratory of Virology | University Hospitals of Geneva Laboratory of Virology | Laubscher F. |
| EPI_ISL_429226, EPI_ISL_429227 | Presidio Ospedaliero Santo Spirito | Istituto Zooprofilattico Sperimentale dell'Abruzzo e Molise "G. Caporale" | Lorusso A, Marcacci M, Di Domenico M, Ancora M, Curini V, Mangone I, Rinaldi A, Di Pasquale A, Camma C, Puglia I, Savini G |
| EPI_ISL_429228 | Ospedale Civile Giuseppe Mazzini | Istituto Zooprofilattico Sperimentale dell'Abruzzo e Molise "G. Caporale" | Lorusso A, Marcacci M, Di Domenico M, Ancora M, Curini V, Mangone I, Rinaldi A, Di Pasquale A, Camma C, Puglia I, Savini G |
| EPI_ISL_429229 | Ospedale Regionale San Salvatore | Istituto Zooprofilattico Sperimentale dell'Abruzzo e Molise "G. Caporale" | Lorusso A, Marcacci M, Di Domenico M, Ancora M, Curini V, Mangone I, Rinaldi A, Di Pasquale A, Camma C, Puglia I, Savini G |
| EPI_ISL_429230, EPI_ISL_429231, EPI_ISL_429232, EPI_ISL_429233, EPI_ISL_429234, EPI_ISL_429235 | Ospedale Civile Giuseppe Mazzini | Istituto Zooprofilattico Sperimentale dell'Abruzzo e Molise "G. Caporale" | Lorusso A, Marcacci M, Di Domenico M, Ancora M, Curini V, Mangone I, Rinaldi A, Di Pasquale A, Camma C, Puglia I, Savini G |
| EPI_ISL_429236 | Ospedale Civile S. Liberatore di Atri | Istituto Zooprofilattico Sperimentale dell'Abruzzo e Molise "G. Caporale" | Lorusso A, Marcacci M, Di Domenico M, Ancora M, Curini V, Mangone I, Rinaldi A, Di Pasquale A, Camma C, Puglia I, Savini G |
| EPI_ISL_429239 | Department of Clinical Laboratory, the First People's Hospital of Yunnan Province | Department of Clinical Laboratory, the First People's Hospital of Yunnan Province | Yi Sun,Ziqin Dian,Ya Xu,Guiqian Zhang,Xin Fan,Yu Zhang |
| EPI_ISL_429254, EPI_ISL_429255 | Viral Respiratory Lab, National Institute for Biomedical Research (INRB) | Pathogen Sequencing Lab, National Institute for Biomedical Research (INRB) | Placide Mbala-Kingebeni, Edith Nkwembe, Eddy Kinganda-Lusamaki, Amuri Aziza, Catherine Pratt, Matthias Pauthner, Josh Quick, Allison Black, James Hadfield, Trevor Bedford, Ian Goodfellow, Nick Loman, Kristian Andersen, Michael Wiley, Steve Ahuka-Mundeke, Jean-Jacques Muyembe Tamfum |
| EPI_ISL_429256 | Health Sciences Technology Park, Avicena, 8, 18016 Granada. Spain | Sequencing and Bioinformatics Service FISABIO-Public Health | Joaquín Mendoza, Almudena Rojas, Pablo Mendoza |
| EPI_ISL_429257 | Microbial Genomics Laboratory, Institut Pasteur Montevideo | Microbial Genomics Laboratory, Institut Pasteur Montevideo | Cecilia Salazar, Florencia Díaz-Viraqué, Marianoel Pereira, Pilar Moreno, Gonzalo Moratorio, Gregorio Iraola |
| EPI_ISL_429258, EPI_ISL_429259 | Viral Respiratory Lab, National Institute for Biomedical Research (INRB) | Pathogen Sequencing Lab, National Institute for Biomedical Research (INRB) | Placide Mbala-Kingebeni, Edith Nkwembe, Eddy Kinganda-Lusamaki, Amuri Aziza, Catherine Pratt, Matthias Pauthner, Josh Quick, Allison Black, James Hadfield, Trevor Bedford, Ian Goodfellow, Nick Loman, Kristian Andersen, Michael Wiley, Steve Ahuka-Mundeke, Jean-Jacques Muyembe Tamfum |
| EPI_ISL_429262, EPI_ISL_429263, EPI_ISL_429264, EPI_ISL_429265, EPI_ISL_429266, EPI_ISL_429267, EPI_ISL_429268, EPI_ISL_429269, EPI_ISL_429270, EPI_ISL_429271, EPI_ISL_429272, EPI_ISL_429273, EPI_ISL_429274, EPI_ISL_429275, EPI_ISL_429276, EPI_ISL_429277, EPI_ISL_429278, EPI_ISL_429279, EPI_ISL_429280, EPI_ISL_429281, EPI_ISL_429282, EPI_ISL_429283, EPI_ISL_429284, EPI_ISL_429285, EPI_ISL_429286, EPI_ISL_429287, EPI_ISL_429288, EPI_ISL_429289, EPI_ISL_429290, EPI_ISL_429291, EPI_ISL_429292, EPI_ISL_429293, EPI_ISL_429294, EPI_ISL_429295, EPI_ISL_429296, EPI_ISL_429297, EPI_ISL_429298, EPI_ISL_429299, EPI_ISL_429300, EPI_ISL_429301, EPI_ISL_429302, EPI_ISL_429303, EPI_ISL_429304, EPI_ISL_429305, EPI_ISL_429306, EPI_ISL_429307, EPI_ISL_429308, EPI_ISL_429309, EPI_ISL_429310, EPI_ISL_429311, EPI_ISL_429312, EPI_ISL_429313, EPI_ISL_429314, EPI_ISL_429315, EPI_ISL_429316, EPI_ISL_429317, EPI_ISL_429318, EPI_ISL_429320, EPI_ISL_429321, EPI_ISL_429322, EPI_ISL_429323, EPI_ISL_429324, EPI_ISL_429325, EPI_ISL_429326, EPI_ISL_429327, EPI_ISL_429328, EPI_ISL_429329, EPI_ISL_429330, EPI_ISL_429331, EPI_ISL_429332 | | | |
| see above | Department of Clinical Microbiology, Copenhagen University Hospital, Hvidovre, Kettegaard Alle 30, 2650 Hvidovre. | Albertsen lab, Department of Chemistry and Bioscience, Aalborg University, Denmark | Rasmus Kirkegaard |
| EPI_ISL_429333, EPI_ISL_429334, EPI_ISL_429335, EPI_ISL_429336, EPI_ISL_429337, EPI_ISL_429338, EPI_ISL_429339, EPI_ISL_429340, EPI_ISL_429341, EPI_ISL_429342, EPI_ISL_429343, EPI_ISL_429344, EPI_ISL_429345, EPI_ISL_429346, EPI_ISL_429347, EPI_ISL_429348, EPI_ISL_429349, EPI_ISL_429350, EPI_ISL_429351, EPI_ISL_429352, EPI_ISL_429353, EPI_ISL_429354, EPI_ISL_429355, EPI_ISL_429356, EPI_ISL_429357, EPI_ISL_429358, EPI_ISL_429359, EPI_ISL_429360, EPI_ISL_429361, EPI_ISL_429362, EPI_ISL_429363, EPI_ISL_429364, EPI_ISL_429365, EPI_ISL_429366, EPI_ISL_429367, EPI_ISL_429368, EPI_ISL_429369, EPI_ISL_429370, EPI_ISL_429371, EPI_ISL_429372, EPI_ISL_429373, EPI_ISL_429374, EPI_ISL_429375, EPI_ISL_429376, EPI_ISL_429377, EPI_ISL_429378, EPI_ISL_429379, EPI_ISL_429380, EPI_ISL_429381, EPI_ISL_429382, EPI_ISL_429383, EPI_ISL_429384, EPI_ISL_429385, EPI_ISL_429386, EPI_ISL_429387, EPI_ISL_429388, EPI_ISL_429389, EPI_ISL_429390, EPI_ISL_429391, EPI_ISL_429392, EPI_ISL_429393, EPI_ISL_429394, EPI_ISL_429395, EPI_ISL_429396, EPI_ISL_429397, EPI_ISL_429398, EPI_ISL_429399, EPI_ISL_429400, EPI_ISL_429401, EPI_ISL_429402, EPI_ISL_429403, EPI_ISL_429404, EPI_ISL_429405, EPI_ISL_429406, EPI_ISL_429407, EPI_ISL_429408, EPI_ISL_429409, EPI_ISL_429410, EPI_ISL_429411, EPI_ISL_429412, EPI_ISL_429413, EPI_ISL_429414, EPI_ISL_429415, EPI_ISL_429416, EPI_ISL_429417, EPI_ISL_429418, EPI_ISL_429419, EPI_ISL_429420, EPI_ISL_429421, EPI_ISL_429422, EPI_ISL_429423, EPI_ISL_429424, EPI_ISL_429425, EPI_ISL_429426, EPI_ISL_429427, EPI_ISL_429428, EPI_ISL_429429, EPI_ISL_429430, EPI_ISL_429431, EPI_ISL_429432, EPI_ISL_429433, EPI_ISL_429434, EPI_ISL_429435, EPI_ISL_429436, EPI_ISL_429437, EPI_ISL_429438, EPI_ISL_429439, EPI_ISL_429440, EPI_ISL_429441, EPI_ISL_429442, EPI_ISL_429443, EPI_ISL_429444, EPI_ISL_429445, EPI_ISL_429446, EPI_ISL_429447, EPI_ISL_429448, EPI_ISL_429449, EPI_ISL_429450, EPI_ISL_429451, EPI_ISL_429452, EPI_ISL_429453, EPI_ISL_429454, EPI_ISL_429455, EPI_ISL_429456, EPI_ISL_429457, EPI_ISL_429458, EPI_ISL_429459, EPI_ISL_429460, EPI_ISL_429461, EPI_ISL_429462, EPI_ISL_429463, EPI_ISL_429464, EPI_ISL_429465, EPI_ISL_429466, EPI_ISL_429467, EPI_ISL_429468, EPI_ISL_429469, EPI_ISL_429470, EPI_ISL_429471, EPI_ISL_429472, EPI_ISL_429473, EPI_ISL_429474, EPI_ISL_429475, EPI_ISL_429476, EPI_ISL_429477, EPI_ISL_429478, EPI_ISL_429479, EPI_ISL_429480, EPI_ISL_429481, EPI_ISL_429482, EPI_ISL_429483, EPI_ISL_429484, EPI_ISL_429485, EPI_ISL_429486, EPI_ISL_429487, EPI_ISL_429488, EPI_ISL_429489, EPI_ISL_429490, EPI_ISL_429491, EPI_ISL_429492, EPI_ISL_429493, EPI_ISL_429494, EPI_ISL_429495, EPI_ISL_429496, EPI_ISL_429497, EPI_ISL_429498, EPI_ISL_429499, EPI_ISL_429500, EPI_ISL_429501, EPI_ISL_429502, EPI_ISL_429503, EPI_ISL_429504, EPI_ISL_429505, EPI_ISL_429506, EPI_ISL_429507, EPI_ISL_429508, EPI_ISL_429509, EPI_ISL_429510, EPI_ISL_429511, EPI_ISL_429512, EPI_ISL_429513, EPI_ISL_429514, EPI_ISL_429515, EPI_ISL_429516, EPI_ISL_429517, EPI_ISL_429518, EPI_ISL_429519, EPI_ISL_429520, EPI_ISL_429521, EPI_ISL_429522, EPI_ISL_429523, EPI_ISL_429524, | | | |

| | | | |
|---|---|---|---|
| EPI_ISL_429525, EPI_ISL_429526, EPI_ISL_429527, EPI_ISL_429528, EPI_ISL_429529, EPI_ISL_429530, EPI_ISL_429531, EPI_ISL_429532, EPI_ISL_429533, EPI_ISL_429534, EPI_ISL_429535, EPI_ISL_429536, EPI_ISL_429537, EPI_ISL_429538, EPI_ISL_429539, EPI_ISL_429540, EPI_ISL_429541, EPI_ISL_429542, EPI_ISL_429543, EPI_ISL_429544, EPI_ISL_429545, EPI_ISL_429546, EPI_ISL_429547, EPI_ISL_429548, EPI_ISL_429549, EPI_ISL_429550, EPI_ISL_429551, EPI_ISL_429552, EPI_ISL_429553, EPI_ISL_429554, EPI_ISL_429555, EPI_ISL_429556, EPI_ISL_429557, EPI_ISL_429558, EPI_ISL_429559, EPI_ISL_429560, EPI_ISL_429561, EPI_ISL_429562, EPI_ISL_429563, EPI_ISL_429564, EPI_ISL_429565, EPI_ISL_429566, EPI_ISL_429567, EPI_ISL_429568, EPI_ISL_429569, EPI_ISL_429570, EPI_ISL_429571, EPI_ISL_429572, EPI_ISL_429573, EPI_ISL_429574, EPI_ISL_429575, EPI_ISL_429576, EPI_ISL_429577, EPI_ISL_429578, EPI_ISL_429579, EPI_ISL_429580, EPI_ISL_429581, EPI_ISL_429582, EPI_ISL_429583, EPI_ISL_429584, EPI_ISL_429585, EPI_ISL_429586, EPI_ISL_429587, EPI_ISL_429588, EPI_ISL_429589, EPI_ISL_429590 | | | |
| see above | Department of Virus and Microbiological Special Diagnostics, Statens Serum Institut, Copenhagen, Denmark, Artillerivej 5, 2300 Copenahgen S | Albertsen lab, Department of Chemistry and Bioscience, Aalborg University, Denmark | Rasmus Kirkegaard |
| EPI_ISL_429597, EPI_ISL_429598, EPI_ISL_429599, EPI_ISL_429600, EPI_ISL_429601, EPI_ISL_429602, EPI_ISL_429603, EPI_ISL_429604, EPI_ISL_429605, EPI_ISL_429606, EPI_ISL_429607, EPI_ISL_429608, EPI_ISL_429609, EPI_ISL_429610, EPI_ISL_429611, EPI_ISL_429612, EPI_ISL_429613, EPI_ISL_429614, EPI_ISL_429615, EPI_ISL_429616, EPI_ISL_429617, EPI_ISL_429618, EPI_ISL_429619, EPI_ISL_429620, EPI_ISL_429621, EPI_ISL_429622, EPI_ISL_429623, EPI_ISL_429624, EPI_ISL_429625, EPI_ISL_429626, EPI_ISL_429627, EPI_ISL_429628, EPI_ISL_429629, EPI_ISL_429630, EPI_ISL_429631, EPI_ISL_429632, EPI_ISL_429633, EPI_ISL_429634, EPI_ISL_429635, EPI_ISL_429636, EPI_ISL_429637, EPI_ISL_429638, EPI_ISL_429639, EPI_ISL_429640, EPI_ISL_429641, EPI_ISL_429642, EPI_ISL_429643, EPI_ISL_429644, EPI_ISL_429645, EPI_ISL_429646, EPI_ISL_429647, EPI_ISL_429648, EPI_ISL_429649, EPI_ISL_429650, EPI_ISL_429651, EPI_ISL_429652, EPI_ISL_429653, EPI_ISL_429654, EPI_ISL_429655, EPI_ISL_429656 | | | |
| see above | UW Virology Lab | UW Virology Lab | Pavitra Roychoudhury, Hong Xie, Keith Jerome, Alexander Greninger |
| EPI_ISL_429659, EPI_ISL_429663 | Institute for Public Health | Laboratory for advanced genomics | Filip Rokić, Lovro Trgovec-Greif, Neven Sučić, Tomislav Rukavina, Igor Jurak, Oliver Vugrek |
| EPI_ISL_429664, EPI_ISL_429665, EPI_ISL_429666, EPI_ISL_429667, EPI_ISL_429668, EPI_ISL_429669, EPI_ISL_429670, EPI_ISL_429671, EPI_ISL_429672, EPI_ISL_429673, EPI_ISL_429674, EPI_ISL_429675, EPI_ISL_429676, EPI_ISL_429677, EPI_ISL_429678, EPI_ISL_429679, EPI_ISL_429680, EPI_ISL_429681, EPI_ISL_429682, EPI_ISL_429683, EPI_ISL_429684, EPI_ISL_429685, EPI_ISL_429686, EPI_ISL_429687, EPI_ISL_429688, EPI_ISL_429689, EPI_ISL_429690, EPI_ISL_429691, EPI_ISL_429692, EPI_ISL_429693, EPI_ISL_429694, EPI_ISL_429695, EPI_ISL_429696, EPI_ISL_429697, EPI_ISL_429698, EPI_ISL_429699, EPI_ISL_429700, EPI_ISL_429701, EPI_ISL_429702, EPI_ISL_429703 | | | |
| see above | Central Public Health Laboratory/Octávio Magalhães Institute (IOM) from the Ezequiel Dias Foundation (FUNED) | Instituto Octávio Magalhães / Fundação Ezequiel Dias (IOM/Funed) | Talita Adelino, Joilson Xavier, Marta Giovanetti, Vagner Fonseca, Marcos Vinícius Silva, Luiz Carlos Junior Alcantara, Marluce Aparecida Assunção Oliveira |
| EPI_ISL_429705 | Institute for Public Health | Laboratory for advanced genomics | Filip Rokić, Lovro Trgovec-Greif, Neven Sučić, Tomislav Rukavina, Igor Jurak, Oliver Vugrek |
| EPI_ISL_429706, EPI_ISL_429707, EPI_ISL_429708, EPI_ISL_429709, EPI_ISL_429710, EPI_ISL_429711, EPI_ISL_429712, EPI_ISL_429713, EPI_ISL_429714, EPI_ISL_429715, EPI_ISL_429716, EPI_ISL_429717, EPI_ISL_429718, EPI_ISL_429719, EPI_ISL_429720, EPI_ISL_429721, EPI_ISL_429722, EPI_ISL_429723, EPI_ISL_429724, EPI_ISL_429725, EPI_ISL_429726, EPI_ISL_429727, EPI_ISL_429728, EPI_ISL_429729, EPI_ISL_429730, EPI_ISL_429731, EPI_ISL_429732, EPI_ISL_429733, EPI_ISL_429734, EPI_ISL_429735, EPI_ISL_429736, EPI_ISL_429737, EPI_ISL_429738, EPI_ISL_429739, EPI_ISL_429740, EPI_ISL_429741, EPI_ISL_429742, EPI_ISL_429743, EPI_ISL_429744, EPI_ISL_429745, EPI_ISL_429746, EPI_ISL_429747, EPI_ISL_429748, EPI_ISL_429749, EPI_ISL_429750, EPI_ISL_429751, EPI_ISL_429752, EPI_ISL_429753, EPI_ISL_429754, EPI_ISL_429755, EPI_ISL_429756, EPI_ISL_429757, EPI_ISL_429758, EPI_ISL_429759, EPI_ISL_429760, EPI_ISL_429761, EPI_ISL_429762, EPI_ISL_429763, EPI_ISL_429764, EPI_ISL_429765, EPI_ISL_429766, EPI_ISL_429767, EPI_ISL_429768, EPI_ISL_429769, EPI_ISL_429770, EPI_ISL_429771, EPI_ISL_429772, EPI_ISL_429773, EPI_ISL_429774, EPI_ISL_429775, EPI_ISL_429776, EPI_ISL_429777, EPI_ISL_429778, EPI_ISL_429779, EPI_ISL_429780, EPI_ISL_429781, EPI_ISL_429782, EPI_ISL_429783, EPI_ISL_429784, EPI_ISL_429785, EPI_ISL_429786, EPI_ISL_429787, EPI_ISL_429788, EPI_ISL_429789, EPI_ISL_429790, EPI_ISL_429791 | | | |
| see above | Laboratoire National de Sante, Microbiology, Virology | Laboratoire National de Sante, Microbiology, Epidemiology and Microbial Genomics | Anke Wienecke-Baldacchino, Ardashel Latsuzbaia, Jessica Tapp, Catherine Ragimbeau, Guillaume Fournier, Tamir Abdelrahman, Trung Nguyen Nguyen, Joel Mossong |
| EPI_ISL_429792 | Institute for Public Health | Laboratory for advanced genomics | Filip Rokić, Lovro Trgovec-Greif, Neven Sučić, Tomislav Rukavina, Igor Jurak, Oliver Vugrek |
| EPI_ISL_429793, EPI_ISL_429794, EPI_ISL_429795, EPI_ISL_429796, EPI_ISL_429797, EPI_ISL_429798, EPI_ISL_429799, EPI_ISL_429800 | Laboratoire National de Sante, Microbiology, Virology | Laboratoire National de Sante, Microbiology, Epidemiology and Microbial Genomics | Anke Wienecke-Baldacchino, Ardashel Latsuzbaia, Jessica Tapp, Catherine Ragimbeau, Guillaume Fournier, Tamir Abdelrahman, Trung Nguyen Nguyen, Joel Mossong |
| EPI_ISL_429802, EPI_ISL_429803, EPI_ISL_429805 | Institute for Public Health | Laboratory for advanced genomics | Filip Rokić, Lovro Trgovec-Greif, Neven Sučić, Tomislav Rukavina, Igor Jurak, Oliver Vugrek |
| EPI_ISL_429806 | Dr. Georges-L.-Dumont University Hospital Centre | National Microbiology Laboratory | Anna Majer, Shari Tyson, Grace Seo, Kristyn Burak, Philip Mabon, Elsie Grudeski, Rhiannon Huzarewich, Russell Mandes, Jennifer Tanner, Natalie Knox, Morag Graham, Gary Van Domselaar, Richard Garceau, Guillaume Desnoyers, Nathalie Bastien, Yan Li, Timothy Booth, Matthew Gilmour |
| EPI_ISL_429807 | Cadham Provincial Laboratory | National Microbiology Laboratory | Anna Majer, Shari Tyson, Grace Seo, Kristyn Burak, Philip Mabon, Elsie Grudeski, Rhiannon Huzarewich, Russell Mandes, Jennifer Tanner, Natalie Knox, Morag Graham, Gary Van Domselaar, Paul Van Caeseele, Jared Bullard, David Alexander, Kerry Dust, Nathalie Bastien, Yan Li, Timothy Booth, Matthew Gilmour |
| EPI_ISL_429811 | Dr. Georges-L.-Dumont University Hospital Centre | National Microbiology Laboratory | Anna Majer, Shari Tyson, Grace Seo, Kristyn Burak, Philip Mabon, Elsie Grudeski, Rhiannon Huzarewich, Russell Mandes, Jennifer Tanner, Natalie Knox, Morag Graham, Gary Van Domselaar, Richard Garceau, Guillaume Desnoyers, Nathalie Bastien, Yan Li, Timothy Booth, Matthew Gilmour |
| EPI_ISL_429812, EPI_ISL_429813, EPI_ISL_429814 | Queen Elizabeth II Health Science Centre | National Microbiology Laboratory | Anna Majer, Shari Tyson, Grace Seo, Kristyn Burak, Philip Mabon, Elsie Grudeski, Rhiannon Huzarewich, Russell Mandes, Jennifer Tanner, Natalie Knox, Morag Graham, Gary Van Domselaar, Todd Hatchette, Jason LeBlanc, Nathalie Bastien, Yan Li, Timothy Booth, Matthew Gilmour |
| EPI_ISL_429815 | Public Health Laboratory | National Microbiology Laboratory | Anna Majer, Shari Tyson, Grace Seo, Kristyn Burak, Philip Mabon, Elsie Grudeski, Rhiannon Huzarewich, Russell Mandes, Jennifer Tanner, Natalie Knox, Morag Graham, Gary Van Domselaar, Robert Needle, Yang Yu, Adel Malek, Laura Gilbert, George Zahariadis, Nathalie Bastien, Yan Li, Timothy Booth, Matthew Gilmour |
| EPI_ISL_429816 | Dr. Georges-L.-Dumont University Hospital Centre | National Microbiology Laboratory | Anna Majer, Shari Tyson, Grace Seo, Kristyn Burak, Philip Mabon, Elsie Grudeski, Rhiannon Huzarewich, Russell Mandes, Jennifer Tanner, Natalie Knox, Morag Graham, Gary Van Domselaar, Richard Garceau, Guillaume Desnoyers, Nathalie Bastien, Yan Li, Timothy Booth, Matthew Gilmour |
| EPI_ISL_429817, EPI_ISL_429818, EPI_ISL_429819 | Cadham Provincial Laboratory | National Microbiology Laboratory | Anna Majer, Shari Tyson, Grace Seo, Kristyn Burak, Philip Mabon, Elsie Grudeski, Rhiannon Huzarewich, Russell Mandes, Jennifer Tanner, Natalie Knox, Morag Graham, Gary Van Domselaar, Paul Van Caeseele, Jared Bullard, David Alexander, Kerry Dust, Nathalie Bastien, Yan Li, Timothy Booth, Matthew Gilmour |
| EPI_ISL_429820 | Cadham Provincial Laboratory | National Microbiology Laboratory | Anna Majer, Shari Tyson, Grace Seo, Kristyn Burak, Philip Mabon, Elsie Grudeski, Rhiannon Huzarewich, Russell Mandez, Jennifer Tanner, Natalie Knox, Morag Graham, Gary Van Domselaar, Paul Van Caeseele, Jared Bullard, David Alexander, Kerry Dust, Nathalie Bastien, Yan Li, Timothy Booth, Matthew Gilmour |
| EPI_ISL_429843 | Gundersen Molecular Diagnostics Laboratory | Kabara Cancer Research Institute | Craig S. Richmond, Paraic A. Kenny |
| EPI_ISL_429844 | Gundersen Molecular Diagnostics Laboratory | Kabara Cancer Research Institute | Craig S. Richmond; Paraic A. Kenny |
| EPI_ISL_429845, EPI_ISL_429846, EPI_ISL_429847, EPI_ISL_429848 | Gundersen Molecular Diagnostics Laboratory | Kabara Cancer Research Institute | Craig S. Richmond, Paraic A. Kenny |
| EPI_ISL_429852 | Centers for Disease Control and Prevention of Lishui | Department of InspectionCenters for Disease Control and Prevention of Lishui | Wang Xiaoguang,Ji Qiaoying,Ji Jiansong,Ye Bifeng,Ye Ling |
| EPI_ISL_429853 | Centers for Disease Control and Prevention of Lishui | Department of InspectionCenters for Disease Control and Prevention of Lishui | Wang Xiaoguang,Ji Qiaoying,Ji Jiansong,Ye Bifeng,Ye Ling |
| EPI_ISL_429854, EPI_ISL_429855 | Centers for Disease Control and Prevention of Lishui | Department of InspectionCenters for Disease Control and Prevention of Lishui | Wang Xiaoguang,Ji Qiaoying,Ji Jiansong,Ye Bifeng,Ye Ling |
| EPI_ISL_429861, EPI_ISL_429862, EPI_ISL_429863, EPI_ISL_429864, EPI_ISL_429865, EPI_ISL_429866, EPI_ISL_429867, EPI_ISL_429868, EPI_ISL_429869, EPI_ISL_429870, EPI_ISL_429871, EPI_ISL_429872, EPI_ISL_429873 | | | |
| see above | Ministry of Health Turkey | Ministry of Health Turkey | Fatma Bayrakdar,Ayşe Başak Altaş,Yasemin Coşgun,Gülay Korukluoğlu,Selçuk Kılıç |
| EPI_ISL_429874 | Microbiology, Virology and Biemergency Laboratory-ASST FBF Sacco | Microbiology, Virology and Biemergency Laboratory-ASST FBF Sacco | Rimoldi SG, Stefani F |
| EPI_ISL_429875, EPI_ISL_429876 | California Department of Public Health | Chiu Laboratory, University of California, San Francisco | Xianding Deng, Scot Federman, Chao-Yang Pan, Hugo Guevara,Wei Gu, Debra A. Wadford, and Charles Y. Chiu |
| EPI_ISL_429877, EPI_ISL_429878 | Chiu Laboratory, University of California, San Francisco | Chiu Laboratory, University of California, San Francisco | Xianding Deng, Scot Federman, Wei Gu, and Charles Y. Chiu |
| EPI_ISL_429879, EPI_ISL_429880 | Santa Clara County Public Health Department | Chiu Laboratory, University of California, San Francisco | Xianding Deng, Scot Federman, Wei Gu, Elsa Villarino, Brandon Bonin, Debra A. Wadford, and Charles Y. Chiu |
| EPI_ISL_429881 | California Department of Public Health | Chiu Laboratory, University of California, San Francisco | Xianding Deng, Scot Federman, Wei Gu, and Charles Y. Chiu |
| EPI_ISL_429882 | Centers for Disease Control, R.O.C. (Taiwan) | Centers for Disease Control, R.O.C. (Taiwan) | Ji-Rong Yang, Yu-Chi Lin, Jung-Jung Mu, Ming-Tsan Liu |
| EPI_ISL_429883 | Centers for Disease Control, R.O.C. (Taiwan) | Centers for Disease Control, R.O.C. (Taiwan) | Ji-Rong Yang, Yu-Chi Lin, Jung-Jung Mu, Ming-Tsan-Liu |
| EPI_ISL_429884 | Centers for Disease Control, R.O.C. (Taiwan) | Centers for Disease Control, R.O.C. (Taiwan) | Ji-Rong Yang, Yu-Chi Lin, Jung-Jung Mu, Ming-Tsan Liu |
| EPI_ISL_429968 | Centre Hospitalier Compiègne Laboratoire de Biologie | National Reference Center for Viruses of Respiratory Infections, Institut Pasteur, Paris | Mélanie Albert, Marion Barbet, Sylvie Behillil, Méline Bizard, Angela Brisebarre, Flora Donati, Fabiana Gambaro, Etienne Simon-Lorière, Vincent Enouf, Maud Vanpeene, Sylvie van der Werf, Raulin Olivia |
| EPI_ISL_429969, EPI_ISL_429970, EPI_ISL_429971, EPI_ISL_429972, EPI_ISL_429973, EPI_ISL_429974, EPI_ISL_429975, EPI_ISL_429976, EPI_ISL_429977, EPI_ISL_429978, EPI_ISL_429979, EPI_ISL_429980, EPI_ISL_429981, EPI_ISL_429982, EPI_ISL_429983, EPI_ISL_429984, EPI_ISL_429985, EPI_ISL_429986, EPI_ISL_429987, EPI_ISL_429988, EPI_ISL_429989 | | | |
| see above | Virginia DCLS | Virginia DCLS | Virginia DCLS |
| EPI_ISL_429990 | Rady's Childrens Hospital | Andersen lab at Scripps Research | SEARCH Alliance San Diego with Christina Clarke, Michelle Vanderpool, Teresa Mueller, Denise Malicki |
| EPI_ISL_429991 | Andersen lab at Scripps Research | Andersen lab at Scripps Research | SEARCH Alliance San Diego |
| EPI_ISL_429992, EPI_ISL_429993, EPI_ISL_429994, EPI_ISL_429995, EPI_ISL_429996, EPI_ISL_429997, EPI_ISL_429998, EPI_ISL_429999, EPI_ISL_430000, EPI_ISL_430001, EPI_ISL_430002, EPI_ISL_430003, EPI_ISL_430004, EPI_ISL_430005, EPI_ISL_430006, EPI_ISL_430007, EPI_ISL_430008, EPI_ISL_430009 | | | |
| see above | Biolab Diagnostic Laboratories | Andersen lab at Scripps Research | Issa Abu-Dayyeh, Ahmad Tibi, Lama Hussein, Lina Mohammad, Zein Naber, Amid Abdelnour with SEARCH Alliance San Diego |
| EPI_ISL_430010 | OSU Wexner Medical Center | James Molecular Lab - OSUWMC | Huolin Tu, Preeti Pancholi, Matt Avenarius, Erica Vincent, Matt Hunt, Dan Jones |
| EPI_ISL_430011, EPI_ISL_430012, EPI_ISL_430013, EPI_ISL_430014, EPI_ISL_430015 | Biolab Diagnostic Laboratories | Andersen lab at Scripps Research | Issa Abu-Dayyeh, Ahmad Tibi, Lama Hussein, Lina Mohammad, Zein Naber, Amid Abdelnour with SEARCH Alliance San Diego |
| EPI_ISL_430016 | Andersen lab at Scripps Research | Andersen lab at Scripps Research | SEARCH Alliance San Diego |
| EPI_ISL_430018 | Molecular Pathology Division, Department of Pathology, Hong Kong Sanatorium & Hospital | Molecular Pathology Division, Department of Pathology, Hong Kong Sanatorium & Hospital | Chun Hang AU, Wai Sing CHAN, Ho Yin LAM, Dona N. HO, Simon Y.M. LAM, Jonpaul S.T. ZEE, Tsun Leung CHAN, Edmond S.K. MA |
| EPI_ISL_430019, EPI_ISL_430020, EPI_ISL_430021, EPI_ISL_430022, EPI_ISL_430023, EPI_ISL_430024, EPI_ISL_430025, EPI_ISL_430026, EPI_ISL_430027, EPI_ISL_430028, EPI_ISL_430029, EPI_ISL_430030, EPI_ISL_430031, EPI_ISL_430032, EPI_ISL_430033, EPI_ISL_430034, EPI_ISL_430035, EPI_ISL_430036, EPI_ISL_430037, EPI_ISL_430038, EPI_ISL_430039, EPI_ISL_430040, EPI_ISL_430041, EPI_ISL_430042, EPI_ISL_430043, EPI_ISL_430044, EPI_ISL_430045, EPI_ISL_430046, EPI_ISL_430047, EPI_ISL_430048, EPI_ISL_430049, EPI_ISL_430050, EPI_ISL_430051, EPI_ISL_430052, EPI_ISL_430053, EPI_ISL_430054, EPI_ISL_430055, EPI_ISL_430056, EPI_ISL_430057, EPI_ISL_430058, EPI_ISL_430059, EPI_ISL_430060, EPI_ISL_430061, EPI_ISL_430062 | | | |
| see above | Utah Public Health Laboratory | Utah Public Health Laboratory | Erin Young, Kelly Oakeson |
| EPI_ISL_430063 | Molecular Pathology Division, Department of Pathology, Hong Kong Sanatorium & Hospital | Molecular Pathology Division, Department of Pathology, Hong Kong Sanatorium & Hospital | Chun Hang AU, Wai Sing CHAN, Ho Yin LAM, Dona N. HO, Simon Y.M. LAM, Jonpaul S.T. ZEE, Tsun Leung CHAN, Edmond S.K. MA |
| EPI_ISL_430064, EPI_ISL_430065, EPI_ISL_430066 | Geelong Centre for Emerging Infectious Diseases | Geelong Centre for Emerging Infectious Diseases | Chamings A., Bhatta T.R., Alexandersen S. |
| EPI_ISL_430067, EPI_ISL_430068, EPI_ISL_430069, EPI_ISL_430070, EPI_ISL_430071, EPI_ISL_430072, EPI_ISL_430073, EPI_ISL_430074, EPI_ISL_430075, EPI_ISL_430076, EPI_ISL_430077, EPI_ISL_430078, EPI_ISL_430079, EPI_ISL_430080, EPI_ISL_430081, EPI_ISL_430082, EPI_ISL_430083, EPI_ISL_430084, EPI_ISL_430085, EPI_ISL_430086, EPI_ISL_430087, EPI_ISL_430088, EPI_ISL_430089, EPI_ISL_430090, EPI_ISL_430091, EPI_ISL_430092, EPI_ISL_430093, EPI_ISL_430094, EPI_ISL_430095, EPI_ISL_430096, EPI_ISL_430097, EPI_ISL_430098, EPI_ISL_430099, EPI_ISL_430100, EPI_ISL_430101, EPI_ISL_430102, EPI_ISL_430103, EPI_ISL_430104, EPI_ISL_430105, EPI_ISL_430106, EPI_ISL_430107, EPI_ISL_430108, EPI_ISL_430109, EPI_ISL_430110, EPI_ISL_430111, EPI_ISL_430112 | | | |
| see above | WHO National Influenza Centre Russian Federation | WHO National Influenza Centre Russian Federation | Andrey Komissarov, Artem Fadeev, Mariia Sergeeva, Anna Ivanova, Daria Danilenko |

| | | | |
|---|---|---|---|
| EPI_ISL_430113, EPI_ISL_430114, EPI_ISL_430115, EPI_ISL_430116, EPI_ISL_430117, EPI_ISL_430118, EPI_ISL_430119, EPI_ISL_430120, EPI_ISL_430121, EPI_ISL_430122, EPI_ISL_430123, EPI_ISL_430124, EPI_ISL_430125, EPI_ISL_430126, EPI_ISL_430127, EPI_ISL_430128, EPI_ISL_430129, EPI_ISL_430130, EPI_ISL_430131, EPI_ISL_430132, EPI_ISL_430133, EPI_ISL_430134, EPI_ISL_430135, EPI_ISL_430136, EPI_ISL_430137, EPI_ISL_430138, EPI_ISL_430139, EPI_ISL_430140, EPI_ISL_430141, EPI_ISL_430142, EPI_ISL_430143, EPI_ISL_430144, EPI_ISL_430145, EPI_ISL_430146, EPI_ISL_430147, EPI_ISL_430148, EPI_ISL_430149, EPI_ISL_430150, EPI_ISL_430151, EPI_ISL_430152, EPI_ISL_430153, EPI_ISL_430154, EPI_ISL_430155, EPI_ISL_430156 | | | |
| see above | Seattle Flu Study | Seattle Flu Study | Chu et al |
| EPI_ISL_430160, EPI_ISL_430161, EPI_ISL_430162, EPI_ISL_430163, EPI_ISL_430164, EPI_ISL_430165, EPI_ISL_430166, EPI_ISL_430167, EPI_ISL_430168, EPI_ISL_430169, EPI_ISL_430170, EPI_ISL_430171, EPI_ISL_430172, EPI_ISL_430173, EPI_ISL_430174, EPI_ISL_430175, EPI_ISL_430176, EPI_ISL_430177, EPI_ISL_430178, EPI_ISL_430179, EPI_ISL_430180, EPI_ISL_430181, EPI_ISL_430182, EPI_ISL_430183, EPI_ISL_430184, EPI_ISL_430185, EPI_ISL_430186, EPI_ISL_430187, EPI_ISL_430188, EPI_ISL_430189, EPI_ISL_430190, EPI_ISL_430191, EPI_ISL_430192, EPI_ISL_430193, EPI_ISL_430194, EPI_ISL_430196, EPI_ISL_430197, EPI_ISL_430198, EPI_ISL_430199, EPI_ISL_430200, EPI_ISL_430201, EPI_ISL_430202, EPI_ISL_430203, EPI_ISL_430204, EPI_ISL_430205, EPI_ISL_430206, EPI_ISL_430207, EPI_ISL_430208, EPI_ISL_430209, EPI_ISL_430210, EPI_ISL_430211, EPI_ISL_430212, EPI_ISL_430213, EPI_ISL_430214, EPI_ISL_430215, EPI_ISL_430216, EPI_ISL_430217, EPI_ISL_430218, EPI_ISL_430219, EPI_ISL_430220, EPI_ISL_430221, EPI_ISL_430222, EPI_ISL_430223, EPI_ISL_430224, EPI_ISL_430225, EPI_ISL_430226, EPI_ISL_430227, EPI_ISL_430228, EPI_ISL_430229, EPI_ISL_430230, EPI_ISL_430231, EPI_ISL_430232, EPI_ISL_430233, EPI_ISL_430234, EPI_ISL_430235, EPI_ISL_430236, EPI_ISL_430237, EPI_ISL_430238, EPI_ISL_430239, EPI_ISL_430240, EPI_ISL_430241, EPI_ISL_430242, EPI_ISL_430243, EPI_ISL_430244, EPI_ISL_430245, EPI_ISL_430246, EPI_ISL_430247, EPI_ISL_430248, EPI_ISL_430249, EPI_ISL_430250, EPI_ISL_430251, EPI_ISL_430252, EPI_ISL_430253, EPI_ISL_430254, EPI_ISL_430255, EPI_ISL_430256, EPI_ISL_430257, EPI_ISL_430258, EPI_ISL_430259, EPI_ISL_430260, EPI_ISL_430261, EPI_ISL_430262, EPI_ISL_430263, EPI_ISL_430264, EPI_ISL_430265, EPI_ISL_430266, EPI_ISL_430267, EPI_ISL_430268, EPI_ISL_430269, EPI_ISL_430270, EPI_ISL_430271, EPI_ISL_430272, EPI_ISL_430273, EPI_ISL_430274, EPI_ISL_430275, EPI_ISL_430276, EPI_ISL_430277, EPI_ISL_430278, EPI_ISL_430279, EPI_ISL_430280, EPI_ISL_430281, EPI_ISL_430282, EPI_ISL_430283, EPI_ISL_430284, EPI_ISL_430285, EPI_ISL_430286, EPI_ISL_430287, EPI_ISL_430288, EPI_ISL_430289, EPI_ISL_430290, EPI_ISL_430291, EPI_ISL_430292, EPI_ISL_430293, EPI_ISL_430294, EPI_ISL_430295, EPI_ISL_430296 | | | |
| see above | Washington State Department of Health | Seattle Flu Study | Chu et al |
| EPI_ISL_430297 | National Institute for Communicable Diseases of the National Health Laboratory Service | National Institute for Communicable Diseases of the National Health Laboratory Service | Allam M, Kwenda S, van Heusden P, Khumalo Z, Mohale T, Subramoney K, von Gottberg, A, Ismail A, Bhiman JN |
| EPI_ISL_430319, EPI_ISL_430320, EPI_ISL_430321, EPI_ISL_430322, EPI_ISL_430323, EPI_ISL_430324, EPI_ISL_430325, EPI_ISL_430326, EPI_ISL_430327, EPI_ISL_430328, EPI_ISL_430329, EPI_ISL_430330, EPI_ISL_430331, EPI_ISL_430332, EPI_ISL_430333, EPI_ISL_430334, EPI_ISL_430335, EPI_ISL_430336, EPI_ISL_430337, EPI_ISL_430338, EPI_ISL_430339, EPI_ISL_430340, EPI_ISL_430341, EPI_ISL_430342, EPI_ISL_430343, EPI_ISL_430344, EPI_ISL_430345, EPI_ISL_430346, EPI_ISL_430347, EPI_ISL_430348, EPI_ISL_430349, EPI_ISL_430350, EPI_ISL_430351, EPI_ISL_430352, EPI_ISL_430353, EPI_ISL_430354, EPI_ISL_430355, EPI_ISL_430356, EPI_ISL_430357, EPI_ISL_430358, EPI_ISL_430359, EPI_ISL_430360, EPI_ISL_430361, EPI_ISL_430362, EPI_ISL_430363, EPI_ISL_430364, EPI_ISL_430365, EPI_ISL_430366, EPI_ISL_430367, EPI_ISL_430368, EPI_ISL_430369, EPI_ISL_430370, EPI_ISL_430371, EPI_ISL_430372, EPI_ISL_430373, EPI_ISL_430374, EPI_ISL_430375, EPI_ISL_430376, EPI_ISL_430377, EPI_ISL_430378, EPI_ISL_430379, EPI_ISL_430380, EPI_ISL_430381, EPI_ISL_430382, EPI_ISL_430383, EPI_ISL_430384, EPI_ISL_430385, EPI_ISL_430386, EPI_ISL_430387, EPI_ISL_430388, EPI_ISL_430389, EPI_ISL_430390, EPI_ISL_430391, EPI_ISL_430392, EPI_ISL_430393, EPI_ISL_430394, EPI_ISL_430395, EPI_ISL_430396, EPI_ISL_430397, EPI_ISL_430398, EPI_ISL_430399, EPI_ISL_430400, EPI_ISL_430401, EPI_ISL_430402, EPI_ISL_430403, EPI_ISL_430404, EPI_ISL_430405, EPI_ISL_430406, EPI_ISL_430407, EPI_ISL_430408, EPI_ISL_430409, EPI_ISL_430410, EPI_ISL_430411, EPI_ISL_430412, EPI_ISL_430413, EPI_ISL_430414, EPI_ISL_430415, EPI_ISL_430416, EPI_ISL_430417, EPI_ISL_430418, EPI_ISL_430419, EPI_ISL_430420, EPI_ISL_430421, EPI_ISL_430422, EPI_ISL_430423, EPI_ISL_430424, EPI_ISL_430425, EPI_ISL_430426, EPI_ISL_430427, EPI_ISL_430428, EPI_ISL_430429, EPI_ISL_430430, EPI_ISL_430431, EPI_ISL_430432, EPI_ISL_430433, EPI_ISL_430434 | | | |
| see above | NYU Langone Health | Departments of Pathology and Medicine, New York University School of Medicine | Maria Aguero-Rosenfeld, Brendan Belovarac, Margaret Black, Ludovic Boytard, John Cadley, Paolo Cotzia, John Chen, Dacia Dimartino, Xiaojun Feng, Tatyana Gindin, Emily Guzman, Adriana Heguy, Megan Hogan, Emily Huang, George Jour, Lawrence H. Lin, Raven Luther, Andrew Lytle, Christian Marier, Matthew T. Maurano, Mark J. Mulligan, Peter Meyn, Raquel Ordonez Ciriza, Iman Osman, Jared Pinnell, Vanessa Raabe, Sitharam Ramaswami, Amy Rapkiewicz, Andre M. Ribeiro-dos-Santos, Marie Samanovic-Golden, Antonio Serrano, Guomiao Shen, Matija Snuderl, Theodore Vougiouklakis, Nick Vulpescu, Gael Westby, Paul Zappile, Yutong Zhang |
| EPI_ISL_430439 | Institute for Medical Research, Infectious Disease Research Centre, National Institutes of Health, Ministry of Health Malaysia | Institute for Medical Research Infectious Disease Research Centre, National Institutes of Health, Ministry of Health Malaysia | Suppiah.J, Mohd-Zawawi.Z, Kalyanasundram.J, Azizan.M-A, Mat-Sharani.S, Hisham.H-A, Tan.L-P, Abdul-Wahid.M-Z, Mohd-Zain.R, Ahmad.N, Thayan.R |
| EPI_ISL_430440 | Institute for Medical Research, Infectious Disease Research Centre, National Institutes of Health, Ministry of Health Malaysia | Institute for Medical Research, Infectious Disease Research Centre, National Institutes of Health, Ministry of Health Malaysia | Suppiah.J, Mohd-Zawawi.Z, Kalyanasundram.J, Azizan.M-A, Mat-Sharani.S, Hisham.H-A, Tan.L-P, Abdul-Wahid.M-Z, Tengku-Abd-Rashid.T-R, Mohd-Zain.R, Ahmad.N, Thayan.R |
| EPI_ISL_430441, EPI_ISL_430442, EPI_ISL_430443, EPI_ISL_430444 | Institute for Medical Research, Infectious Disease Research Centre, National Institutes of Health, Ministry of Health Malaysia | Institute for Medical Research, Infectious Disease Research Centre, National Institutes of Health, Ministry of Health Malaysia | Suppiah.J, Mohd-Zawawi.Z, Kalyanasundram.J, Azizan.M-A, Mat-Sharani.S, Hisham.H-A, Tan.L-P, Abdul-Wahid.M-Z, Tengku-Rogayah.TAR, Mohd-Zain.R, Ahmad.N, Thayan.R |
| EPI_ISL_430456 | Rizal Medical Center | Research Institute for Tropical Medicine | Medado,I.A.P., Bautista,C.T., Onza,O.J.T., Polotan,F.G.M., Brunker, K., Mercado,E.S., Manalo, D.L., Demetria, C.S. |
| EPI_ISL_430464, EPI_ISL_430465, EPI_ISL_430466, EPI_ISL_430467, EPI_ISL_430468 | ICMR-National Institute of Cholera and Enteric Diseases | National Institute of Biomedical Genomics | Arindam Maitra, Mamta Chawla Sarkar, Sreedhar Chinnaswamy, Hasina Banu, Ananya Chatterjee, Shanta Dutta, Saumitra Das |
| EPI_ISL_430469 | Hellenic Pasteur Institute, Public Health Laboratories | Hellenic Pasteur Institute, Public Health Laboratories, Unit of Bioinformatics and Applied Genomics | Vasiliki Pogka, Timokratis Karamitros, Athanasios Kossyvakis, Antonios Kalliaropoulos, Horefti Elina, Evangelidou Maria, Androniki Voulgari-Kokota, Aspasia Kontou, Andreas Mentis |
| EPI_ISL_430470, EPI_ISL_430471, EPI_ISL_430472 | Microbiological Diagnostic Unit Public Health Laboratory | Microbiological Diagnostic Unit Public Health Laboratory | Seemann T., Schultz M., Sait, M., Sherry, N. |
| EPI_ISL_430473, EPI_ISL_430474, EPI_ISL_430475, EPI_ISL_430476, EPI_ISL_430477, EPI_ISL_430478, EPI_ISL_430479, EPI_ISL_430480, EPI_ISL_430481, EPI_ISL_430482, EPI_ISL_430483, EPI_ISL_430484, EPI_ISL_430485, EPI_ISL_430486, EPI_ISL_430487, EPI_ISL_430488, EPI_ISL_430489, EPI_ISL_430490, EPI_ISL_430491, EPI_ISL_430492, EPI_ISL_430493 | | | |
| see above | Victorian Infectious Diseases Reference Laboratory (VIDRL) | Microbiological Diagnostic Unit Public Health Laboratory and Victorian Infectious Diseases Reference Laboratory, The Peter Doherty Institute for Infection and Immunity | Caly L., Seemann T., Sait, M., Schultz M., Druce J., Sherry, N. |
| EPI_ISL_430494, EPI_ISL_430495, EPI_ISL_430496, EPI_ISL_430497 | Royal Darwin Hospital Pathology | Microbiological Diagnostic Unit Public Health Laboratory and Victorian Infectious Diseases Reference Laboratory, The Peter Doherty Institute for Infection and Immunity | Meumann, E., Caly L., Seemann T., Sait, M., Schultz M., Druce J., Sherry, N. |
| EPI_ISL_430498, EPI_ISL_430499, EPI_ISL_430500, EPI_ISL_430501, EPI_ISL_430502, EPI_ISL_430503, EPI_ISL_430504, EPI_ISL_430505, EPI_ISL_430506, EPI_ISL_430507, EPI_ISL_430508, EPI_ISL_430509, EPI_ISL_430510, EPI_ISL_430511, EPI_ISL_430512, EPI_ISL_430513, EPI_ISL_430514, EPI_ISL_430515, EPI_ISL_430516, EPI_ISL_430517, EPI_ISL_430518, EPI_ISL_430519, EPI_ISL_430520, EPI_ISL_430521, EPI_ISL_430522, EPI_ISL_430523, EPI_ISL_430524, EPI_ISL_430525, EPI_ISL_430526, EPI_ISL_430527, EPI_ISL_430528, EPI_ISL_430529, EPI_ISL_430530, EPI_ISL_430531, EPI_ISL_430532, EPI_ISL_430533, EPI_ISL_430534, EPI_ISL_430535, EPI_ISL_430536, EPI_ISL_430537, EPI_ISL_430538, EPI_ISL_430539, EPI_ISL_430540, EPI_ISL_430541, EPI_ISL_430542, EPI_ISL_430543, EPI_ISL_430544, EPI_ISL_430545, EPI_ISL_430546, EPI_ISL_430547, EPI_ISL_430548, EPI_ISL_430549, EPI_ISL_430550, EPI_ISL_430551, EPI_ISL_430552, EPI_ISL_430553, EPI_ISL_430554, EPI_ISL_430555, EPI_ISL_430556, EPI_ISL_430557, EPI_ISL_430558, EPI_ISL_430559, EPI_ISL_430560, EPI_ISL_430561, EPI_ISL_430562, EPI_ISL_430563, EPI_ISL_430564, EPI_ISL_430565, EPI_ISL_430566, EPI_ISL_430567, EPI_ISL_430568, EPI_ISL_430569, EPI_ISL_430570, EPI_ISL_430571, EPI_ISL_430572, EPI_ISL_430573, EPI_ISL_430574, EPI_ISL_430575, EPI_ISL_430576, EPI_ISL_430577, EPI_ISL_430578, EPI_ISL_430579, EPI_ISL_430580, EPI_ISL_430581, EPI_ISL_430582, EPI_ISL_430583, EPI_ISL_430584, EPI_ISL_430585, EPI_ISL_430586, EPI_ISL_430587, EPI_ISL_430588, EPI_ISL_430589, EPI_ISL_430590, EPI_ISL_430591, EPI_ISL_430592, EPI_ISL_430593, EPI_ISL_430594, EPI_ISL_430595, EPI_ISL_430596, EPI_ISL_430597, EPI_ISL_430598, EPI_ISL_430599, EPI_ISL_430600, EPI_ISL_430601, EPI_ISL_430602, EPI_ISL_430603, EPI_ISL_430604, EPI_ISL_430605, EPI_ISL_430606, EPI_ISL_430607, EPI_ISL_430608, EPI_ISL_430609, EPI_ISL_430610, EPI_ISL_430611, EPI_ISL_430612, EPI_ISL_430613, EPI_ISL_430614, EPI_ISL_430615, EPI_ISL_430616, EPI_ISL_430617, EPI_ISL_430618, EPI_ISL_430619, EPI_ISL_430620, EPI_ISL_430621, EPI_ISL_430622, EPI_ISL_430623, EPI_ISL_430624, EPI_ISL_430625, EPI_ISL_430626, EPI_ISL_430627, EPI_ISL_430628, EPI_ISL_430629, EPI_ISL_430630 | | | |
| see above | Victorian Infectious Diseases Reference Laboratory (VIDRL) | Microbiological Diagnostic Unit Public Health Laboratory and Victorian Infectious Diseases Reference Laboratory, The Peter Doherty Institute for Infection and Immunity | Caly L., Seemann T., Sait, M., Schultz M., Druce J., Sherry, N. |
| EPI_ISL_430631, EPI_ISL_430632, EPI_ISL_430633, EPI_ISL_430634, EPI_ISL_430635, EPI_ISL_430636 | Royal Darwin Hospital Pathology | Microbiological Diagnostic Unit Public Health Laboratory and Victorian Infectious Diseases Reference Laboratory, The Peter Doherty Institute for Infection and Immunity | Meumann, E., Caly L., Seemann T., Sait, M., Schultz M., Druce J., Sherry, N. |
| EPI_ISL_430637, EPI_ISL_430638 | Victorian Infectious Diseases Reference Laboratory (VIDRL) | Microbiological Diagnostic Unit Public Health Laboratory and Victorian Infectious Diseases Reference Laboratory, The Peter Doherty Institute for Infection and Immunity | Caly L., Seemann T., Sait, M., Schultz M., Druce J., Sherry, N. |
| EPI_ISL_430639, EPI_ISL_430640, EPI_ISL_430641, EPI_ISL_430642, EPI_ISL_430643, EPI_ISL_430644, EPI_ISL_430645, EPI_ISL_430646, EPI_ISL_430647, EPI_ISL_430648, EPI_ISL_430649, EPI_ISL_430650, EPI_ISL_430651, EPI_ISL_430652, EPI_ISL_430653, EPI_ISL_430654, EPI_ISL_430655, EPI_ISL_430656, EPI_ISL_430657, EPI_ISL_430658, EPI_ISL_430659, EPI_ISL_430660, EPI_ISL_430661, EPI_ISL_430662, EPI_ISL_430663, EPI_ISL_430664, EPI_ISL_430665, EPI_ISL_430666, EPI_ISL_430667, EPI_ISL_430668, EPI_ISL_430669, EPI_ISL_430670, EPI_ISL_430671, EPI_ISL_430672, EPI_ISL_430673, EPI_ISL_430674, EPI_ISL_430675, EPI_ISL_430676, EPI_ISL_430677, EPI_ISL_430678, EPI_ISL_430679, EPI_ISL_430680, EPI_ISL_430681, EPI_ISL_430682, EPI_ISL_430683, EPI_ISL_430684, EPI_ISL_430685, EPI_ISL_430686, EPI_ISL_430687 | | | |
| see above | Microbiological Diagnostic Unit Public Health Laboratory | Microbiological Diagnostic Unit Public Health Laboratory | Seemann T., Schultz M., Sait, M., Sherry, N. |
| EPI_ISL_430688, EPI_ISL_430689, EPI_ISL_430690, EPI_ISL_430691, EPI_ISL_430692, EPI_ISL_430693, EPI_ISL_430694, EPI_ISL_430695, EPI_ISL_430696, EPI_ISL_430697, EPI_ISL_430698, EPI_ISL_430699, EPI_ISL_430700, EPI_ISL_430701, EPI_ISL_430702, EPI_ISL_430703, EPI_ISL_430704, EPI_ISL_430705, EPI_ISL_430706, EPI_ISL_430707, EPI_ISL_430708, EPI_ISL_430709, EPI_ISL_430710, EPI_ISL_430711, EPI_ISL_430712, EPI_ISL_430713, EPI_ISL_430714 | | | |
| see above | Victorian Infectious Diseases Reference Laboratory (VIDRL) | Microbiological Diagnostic Unit Public Health Laboratory and Victorian Infectious Diseases Reference Laboratory, The Peter Doherty Institute for Infection and Immunity | Caly L., Seemann T., Sait, M., Schultz M., Druce J., Sherry, N. |
| EPI_ISL_430715, EPI_ISL_430716, EPI_ISL_430717 | Microbiological Diagnostic Unit Public Health Laboratory | Microbiological Diagnostic Unit Public Health Laboratory | Seemann T., Schultz M., Sait, M., Sherry, N. |
| EPI_ISL_430718 | Hospital Universitario 12 de Octubre | Hospital Universitario 12 de Octubre | Sara González, Raúl Recio,Elias Dahdouh, Fernando Lázaro, Esther Viedma, Natalia Stella, Julio García, Juan Carlos Galán, Rafael Cantón, Mª Dolores Folgueira, Rafael Delgado, Jesús Mingorance |
| EPI_ISL_430719, EPI_ISL_430720, EPI_ISL_430721 | Hospital Universitario La Paz | Hospital Universitario 12 de Octubre | Elias Dahdouh, Sara González, Raúl Recio, Fernando Lázaro, Esther Viedma, Natalia Stella, Julio García, Juan Carlos Galán, Rafael Cantón, Mª Dolores Folgueira, Rafael Delgado, Jesús Mingorance |
| EPI_ISL_430722, EPI_ISL_430723, EPI_ISL_430724, EPI_ISL_430725, EPI_ISL_430726, EPI_ISL_430727, EPI_ISL_430728, EPI_ISL_430729, EPI_ISL_430730, EPI_ISL_430731, EPI_ISL_430732, EPI_ISL_430733, EPI_ISL_430734, EPI_ISL_430735, EPI_ISL_430736, EPI_ISL_430737, EPI_ISL_430738, EPI_ISL_430739, EPI_ISL_430740, EPI_ISL_430741, EPI_ISL_430742, EPI_ISL_430743, EPI_ISL_430744, EPI_ISL_430745, EPI_ISL_430746 | | | |
| see above | Chinese PLA Institute for Disease Control and Prevention | Chinese PLA Institute for Disease Control and Prevention | Peng LiJinhui Li, Lizhong Li |
| EPI_ISL_430791, EPI_ISL_430792 | UCSF Clinical Microbiology Laboratory | Chan-Zuckerberg Biohub | CZB Cliahub Consortium |
| EPI_ISL_430793, EPI_ISL_430794 | Laboratorio Análisis Clínicos, Unidad de Servicios Diagnósticos, Swiss Medical Group | Área de Secuenciación del Laboratorio de Virología del Hospital de Niños Dr. Ricardo Gutierrez | Nabaes Jodar, MS; Goya, S; Natale, MI; Lusso, S; Sanchez, O; Guevara, D; Vicario, SM; Mistchenko, AS; Valinotto, LE; Viegas, M. |
| EPI_ISL_430795 | Laboratorio de Virología del Hospital de Niños Dr. Ricardo Gutierrez | Área de Secuenciación del Laboratorio de Virología del Hospital de Niños Dr. Ricardo Gutierrez | Nabaes Jodar, MS; Goya, S; Natale, MI; Lusso, S; Gravis, E; Mistchenko, AS; Valinotto, LE; Viegas, M. |
| EPI_ISL_430796, EPI_ISL_430797, EPI_ISL_430798 | Departamento de Biología y genética molecular, IACA Laboratorios. | Área de Secuenciación del Laboratorio de Virología del Hospital de Niños Dr. Ricardo Gutierrez | Nabaes Jodar, MS; Goya, S; Natale, MI; Lusso, S; Tittarelli, E; Suárez, A; Masciovecchio MV; Streitenberger ER; Mistchenko, AS; Valinotto, LE; Viegas, M. |
| EPI_ISL_430799, EPI_ISL_430800, EPI_ISL_430801 | Laboratorio de Virología del Hospital de Niños Dr. Ricardo Gutierrez | Área de Secuenciación del Laboratorio de Virología del Hospital de Niños Dr. Ricardo Gutierrez | Nabaes Jodar, MS; Goya, S; Natale, MI; Lusso, S; Gravis, E; Mistchenko, AS; Valinotto, LE; Viegas, M. |
| EPI_ISL_430802 | Departamento de Biología y genética molecular, IACA Laboratorios. | Área de Secuenciación del Laboratorio de Virología del Hospital de Niños Dr. Ricardo Gutierrez | Nabaes Jodar, MS; Goya, S; Natale, MI; Lusso, S; Tittarelli, E; Suárez, A; Masciovecchio MV; Streitenberger ER; Mistchenko, AS; Valinotto, LE; Viegas, M. |
| EPI_ISL_430803, EPI_ISL_430804 | Laboratorio de Virología del Hospital de Niños Dr. Ricardo Gutierrez | Área de Secuenciación del Laboratorio de Virología del Hospital de Niños Dr. Ricardo Gutierrez | Nabaes Jodar, MS; Goya, S; Natale, MI; Lusso, S; Gravis, E; Mistchenko, AS; Valinotto, LE; Viegas, M. |
| EPI_ISL_430805, EPI_ISL_430806 | Departamento de Biología y genética molecular, IACA Laboratorios. | Área de Secuenciación del Laboratorio de Virología del Hospital de Niños Dr. Ricardo Gutierrez | Nabaes Jodar, MS; Goya, S; Natale, MI; Lusso, S; Tittarelli, E; Suárez, A; Masciovecchio MV; Streitenberger ER; Mistchenko, AS; Valinotto, LE; Viegas, M. |
| EPI_ISL_430807 | Laboratorio de Virología del Hospital de Niños Dr. Ricardo Gutierrez | Área de Secuenciación del Laboratorio de Virología del Hospital de Niños Dr. Ricardo Gutierrez | Nabaes Jodar, MS; Goya, S; Natale, MI; Lusso, S; Gravis, E; Mistchenko, AS; Valinotto, LE; Viegas, M. |
| EPI_ISL_430808 | Departamento de Biología y genética molecular, IACA Laboratorios. | Área de Secuenciación del Laboratorio de Virología del Hospital de Niños Dr. Ricardo Gutierrez | Nabaes Jodar, MS; Goya, S; Natale, MI; Lusso, S; Tittarelli, E; Suárez, A; Masciovecchio MV; Streitenberger ER; Mistchenko, AS; Valinotto, LE; Viegas, M. |
| EPI_ISL_430809, EPI_ISL_430810, EPI_ISL_430811, EPI_ISL_430812, EPI_ISL_430813, EPI_ISL_430814, EPI_ISL_430815, EPI_ISL_430816, EPI_ISL_430817, | Laboratorio de Virología del Hospital de Niños Dr. Ricardo Gutierrez | Área de Secuenciación del Laboratorio de Virología del Hospital de Niños Dr. Ricardo Gutierrez | Nabaes Jodar, MS; Goya, S; Natale, MI; Lusso, S; Gravis, E; Mistchenko, AS; Valinotto, LE; Viegas, M. |

| | | | |
|---|---|---|---|
| EPI_ISL_430818 | | | |
| EPI_ISL_430819 | Center of Scientific Excellence for Influenza Viruses,National Research Centre (NRC), Egypt. | Center of Scientific Excellence for Influenza Viruses,National Research Centre (NRC), Egypt. | Mohamed Ahmed Ali, Ahmed Kandeil, Ahmed Mostafa, Rabeh El-Shesheny, Mahmoud Shehata, Wael Roshdy, Shymaa Showky Ahmed , Amal Naguib, Nancy M. El Guindy, Mokhtar Gomaa, Ahmed El-Taweel, Ahmed E Kayed, Yassmin Moatasim, Omnia Kutkat, Sara Mahmoud, Mina Kamel, Abo Shama, M Noura, Mohamed El Sayes |
| EPI_ISL_430820 | Center of Scientific Excellence for Influenza Viruses, National Research Centre (NRC), Egypt. | Center of Scientific Excellence for Influenza Viruses, National Research Centre (NRC), Egypt. | Mohamed Ahmed Ali, Ahmed Kandeil, Ahmed Mostafa, Rabeh El-Shesheny, Mahmoud Shehata, Wael Roshdy, Shymaa Showky Ahmed , Amal Naguib, Mokhtar Gomaa, Ahmed El-Taweel, Ahmed E Kayed, Yassmin Moatasim, Omnia Kutkat, Sara Mahmoud, Mina Kamel, Abo Shama, M Noura, Mohamed El Sayes, Nancy M. El Guindy |
| EPI_ISL_430837 | n/a | Thai National Influenza Center, Department of medical Science, Ministry of Public Health, Thailand | Pilailuk,Okada; Siripaporn,Phuygun; Thanutsapa,Thanadachakul;Sittiporn,Parnmen;Warawan,Wongboot;Sunthareeya,Waicharoen; Malinee,Chittaganpitch |
| EPI_ISL_430838 | Makati Medical Center | Research Institute for Tropical Medicine | Medado,I.A.P., Bautista,C.T., Onza,O.J.T., Polotan,F.G.M., Brunker, K., Mercado,E.S., Manalo, D.L., Demetria, C.S. |
| EPI_ISL_430839 | Research Institute for Tropical Medicine | Research Institute for Tropical Medicine | Medado,I.A.P., Bautista,C.T., Onza,O.J.T., Polotan,F.G.M., Brunker, K., Mercado,E.S., Manalo, D.L., Demetria, C.S. |
| EPI_ISL_430840 | Veterans Memorial Medical Center | Research Institute for Tropical Medicine | Medado,I.A.P., Bautista,C.T., Onza,O.J.T., Polotan,F.G.M., Brunker, K., Mercado,E.S., Manalo, D.L., Demetria, C.S. |
| EPI_ISL_430841 | Praram 9 Hospital | National Institute of Health. Department of medical Sciences, Ministry of Public Health, Thailand | Pilailuk,Okada; Siripaporn,Phuygun; Thanutsapa,Thanadachakul; Sittiporn,Parnmen;Warawan,Wongboot; Sunthareeya,Waicharoen; Malinee,Chittaganpitch |
| EPI_ISL_430842 | Central chest Institute of Thailand | National Institute of Health. Department of medical Sciences, Ministry of Public Health, Thailand | Pilailuk,Okada; Siripaporn,Phuygun; Thanutsapa,Thanadachakul; Sittiporn,Parnmen;Warawan,Wongboot; Sunthareeya,Waicharoen; Malinee,Chittaganpitch |
| EPI_ISL_430843 | Bethany Hospital | Research Institute for Tropical Medicine | Medado,I.A.P., Bautista,C.T., Onza,O.J.T., Polotan,F.G.M., Brunker, K., Mercado,E.S., Manalo, D.L., Demetria, C.S. |
| EPI_ISL_430844 | Lung Center of the Philippines | Research Institute for Tropical Medicine | Medado,I.A.P., Bautista,C.T., Onza,O.J.T., Polotan,F.G.M., Brunker, K., Mercado,E.S., Manalo, D.L., Demetria, C.S. |
| EPI_ISL_430845 | Pasig City General Hospital | Research Institute for Tropical Medicine | Medado,I.A.P., Bautista,C.T., Onza,O.J.T., Polotan,F.G.M., Brunker, K., Mercado,E.S., Manalo, D.L., Demetria, C.S. |
| EPI_ISL_430846 | General Intensive Care Unit, Raymond Poincaré Hospital (AP-HP), Lab Inflammation & Infection, U1173 University Paris Saclay-UVSQ/INSERM, Garches, France. | Institut Pasteur, Laboratory for Urgent Response to biological Threats | Annane Djillali, Vanhomwegen Jessica, Caro Valérie, Manuguerra Jean-Claude |
| EPI_ISL_430847 | HS mikrobiologi virus | The Public Health Agency of Sweden | Zhibing Yun, Oskar Karlsson Lindsjo, Maria Lind Karlberg, Anna-Malin Linde, Olov Svartstrom, Anna Risberg, Shaman Muradrasoli, Karin Tegmark-Wisell |
| EPI_ISL_430848, EPI_ISL_430849, EPI_ISL_430850, EPI_ISL_430851, EPI_ISL_430852, EPI_ISL_430853, EPI_ISL_430854, EPI_ISL_430855 | Klinisk mikrobiologi och vardhygien Halmstad | The Public Health Agency of Sweden | Arne Kotz, Oskar Karlsson Lindsjo, Maria Lind Karlberg, Anna-Malin Linde, Olov Svartstrom, Anna Risberg, Shaman Muradrasoli, Karin Tegmark-Wisell |
| EPI_ISL_430856, EPI_ISL_430857, EPI_ISL_430858, EPI_ISL_430859 | Laboratoriemedicin | The Public Health Agency of Sweden | Oskar Karlsson Lindsjo, Maria Lind Karlberg, Anna-Malin Linde, Olov Svartstrom, Anna Risberg, Shaman Muradrasoli, Karin Tegmark-Wisell |
| EPI_ISL_430860, EPI_ISL_430861 | Klinisk mikrobiologi Orebro | The Public Health Agency of Sweden | Martin Sundqvist, Oskar Karlsson Lindsjo, Maria Lind Karlberg, Anna-Malin Linde, Olov Svartstrom, Anna Risberg, Shaman Muradrasoli, Karin Tegmark-Wisell |
| EPI_ISL_430862 | The Public Health Agency of Sweden | The Public Health Agency of Sweden | Oskar Karlsson Lindsjo, Maria Lind Karlberg, Anna-Malin Linde, Olov Svartstrom, Anna Risberg, Shaman Muradrasoli, Karin Tegmark-Wisell |
| EPI_ISL_430863 | Klinisk mikrobiologi Orebro | The Public Health Agency of Sweden | Martin Sundqvist, Oskar Karlsson Lindsjo, Maria Lind Karlberg, Anna-Malin Linde, Olov Svartstrom, Anna Risberg, Shaman Muradrasoli, Karin Tegmark-Wisell |
| EPI_ISL_430864 | The Public Health Agency of Sweden | The Public Health Agency of Sweden | Oskar Karlsson Lindsjo, Maria Lind Karlberg, Anna-Malin Linde, Olov Svartstrom, Anna Risberg, Shaman Muradrasoli, Karin Tegmark-Wisell |
| EPI_ISL_430867, EPI_ISL_430868, EPI_ISL_430869, EPI_ISL_430870, EPI_ISL_430871, EPI_ISL_430872, EPI_ISL_430873, EPI_ISL_430874, EPI_ISL_430875, EPI_ISL_430876, EPI_ISL_430877, EPI_ISL_430878, EPI_ISL_430879, EPI_ISL_430880, EPI_ISL_430881, EPI_ISL_430882, EPI_ISL_430883, EPI_ISL_430884, EPI_ISL_430885, EPI_ISL_430886, EPI_ISL_430887, EPI_ISL_430888, EPI_ISL_430889, EPI_ISL_430890, EPI_ISL_430891, EPI_ISL_430892, EPI_ISL_430893, EPI_ISL_430894, EPI_ISL_430895, EPI_ISL_430896, EPI_ISL_430897, EPI_ISL_430898, EPI_ISL_430899, EPI_ISL_430900, EPI_ISL_430901, EPI_ISL_430902, EPI_ISL_430903, EPI_ISL_430904, EPI_ISL_430905, EPI_ISL_430906, EPI_ISL_430907, EPI_ISL_430908, EPI_ISL_430909, EPI_ISL_430910, EPI_ISL_430911, EPI_ISL_430912, EPI_ISL_430913, EPI_ISL_430914, EPI_ISL_430915, EPI_ISL_430916, EPI_ISL_430917, EPI_ISL_430918, EPI_ISL_430919, EPI_ISL_430920, EPI_ISL_430921, EPI_ISL_430922, EPI_ISL_430923, EPI_ISL_430924, EPI_ISL_430925, EPI_ISL_430926, EPI_ISL_430927, EPI_ISL_430928, EPI_ISL_430929, EPI_ISL_430930, EPI_ISL_430931, EPI_ISL_430932, EPI_ISL_430933, EPI_ISL_430934, EPI_ISL_430935, EPI_ISL_430936, EPI_ISL_430937, EPI_ISL_430938, EPI_ISL_430939, EPI_ISL_430940, EPI_ISL_430941, EPI_ISL_430942, EPI_ISL_430943, EPI_ISL_430944, EPI_ISL_430945, EPI_ISL_430946, EPI_ISL_430947, EPI_ISL_430948, EPI_ISL_430949, EPI_ISL_430950, EPI_ISL_430951, EPI_ISL_430952, EPI_ISL_430953, EPI_ISL_430954, EPI_ISL_430955, EPI_ISL_430956, EPI_ISL_430957, EPI_ISL_430958, EPI_ISL_430959, EPI_ISL_430960, EPI_ISL_430961, EPI_ISL_430962, EPI_ISL_430963, EPI_ISL_430964, EPI_ISL_430965, EPI_ISL_430966, EPI_ISL_430967, EPI_ISL_430968, EPI_ISL_430969, EPI_ISL_430970, EPI_ISL_430971, EPI_ISL_430972, EPI_ISL_430973, EPI_ISL_430974, EPI_ISL_430975, EPI_ISL_430976, EPI_ISL_430977, EPI_ISL_430978, EPI_ISL_430979, EPI_ISL_430980 | | | |
| see above | UW Virology Lab | UW Virology Lab | Pavitra Roychoudhury, Hong Xie, Keith Jerome, Alexander Greninger |
| EPI_ISL_431011, EPI_ISL_431012 | Viral Respiratory Lab, National Institute for Biomedical Research (INRB) | Pathogen Sequencing Lab, National Institute for Biomedical Research (INRB) | Placide Mbala-Kingebeni, Edith Nkwembe, Eddy Kinganda-Lusamaki, Amuri Aziza, Francisca Muyembe Mawete, Catherine Pratt, Matthias Pauthner, Josh Quick, Allison Black, James Hadfield, Trevor Bedford, Ian Goodfellow, Andrew Rambaut, Nick Loman, Kristian Andersen, Michael Wiley, Steve Ahuka-Mundeke, Jean-Jacques Muyembe Tamfum |
| EPI_ISL_431013 | Alaska State Virology Laboratory | Alaska State Virology Laboratory | Jack Chen |
| EPI_ISL_431014, EPI_ISL_431015, EPI_ISL_431016, EPI_ISL_431017, EPI_ISL_431018, EPI_ISL_431019 | Alaska State Virology Laboratory | Alaska State Virology Laboratory | Jack Chen, Ph.D. |
| EPI_ISL_431080, EPI_ISL_431081, EPI_ISL_431082, EPI_ISL_431083, EPI_ISL_431084, EPI_ISL_431085, EPI_ISL_431086, EPI_ISL_431087, EPI_ISL_431088, EPI_ISL_431089, EPI_ISL_431090, EPI_ISL_431091, EPI_ISL_431092, EPI_ISL_431093, EPI_ISL_431094, EPI_ISL_431095, EPI_ISL_431096 | | | |
| see above | Yale COVID-19 Biorepository | Grubaugh Lab - Yale School of Public Health | Joseph Fauver, Tara Alpert, Anderson Brito, Anne Wyllie, Chantal Vogels, Mary Petrone, Cole Jensen, Chaney Kalinich, Isabel Ott, Arnau Casanovas, Catherine Muenker, Adam Moore, Alice Lu, Maria Tokuyama, Patrick Wong, Peiwen Lu, Saad Omer, Richard Martinello, Allison Nelson, Shelli Farhadian, Akiko Iwasaki, Charlese Dela Cruz, Albert Ko, Nathan Grubaugh |
| EPI_ISL_431101 | Department of Microbiology,Gandhi Medical College and Hospital | Virus Research Laboratory, Department of Zoology, Osmania University,Hyderabad,India | Muttineni Radhakrishna, Nagamani K, Thrilok Chander B, Raja Rao M, Kalyani Putty, Ravikumar P, Sunitha P, Pankaj Singh D, Anand Kumar K, Amit A. Upadhyay Steven E. Bosinger, Rama Amara |
| EPI_ISL_431102 | Department of MicroBiology,Gandhi Medical College and Hospital,Secendrabad,Hyderabad,India | Department of Microbiology, Gandhi Medical College and Hospital, Secendrabad, Hyderabad | Nagamani K, Muttineni Radhakrishna, Thrilok Chander B, Raja Rao M, Kalyani Putty, Ravikumar P, Sunitha P, Pankaj Singh D, Anand Kumar K, Amit A. Upadhyay, Steven E. Bosinger, Rama Amara |
| EPI_ISL_431103 | Department of Microbiology, Gandhi Medical College and Hospital, Secendrabad, Hyderabad, India | Department of Microbiology, Gandhi Medical College and Hospital, Secendrabad, Hyderabad, India | Nagamani K, Muttineni Radhakrishna, Thrilok Chander B, Raja Rao M, Kalyani Putty, Ravikumar P, Sunitha P, Pankaj Singh D, Anand Kumar K, Amit A. Upadhyay, Steven E. Bosinger, Rama Amara |
| EPI_ISL_431117 | Department of Microbiology, Gandhi Medical College and Hospital, Secendrabad, Hyderabad, India | Department of Microbiology, Gandhi Medical College and Hospital, Secendrabad, Hyderabad, India | Thrilok Chander B, Muttineni Radhakrishna, Nagamani K, Raja Rao M, Kalyani Putty, Ravikumar P, Sunitha P, Pankaj Singh D, Anand Kumar K, Amit A. Upadhyay, Steven E.Bosinger, Rama Amara |
| EPI_ISL_431118, EPI_ISL_431180, EPI_ISL_431240, EPI_ISL_431292 | Fujian Center for Disease Control and Prevention | Fujian Center for Disease Control and Prevention | Lin Qi, Huang Zhimiao, Zhang Yanhua, Weng Yuwei |
| EPI_ISL_431778 | Virology, Wageningen Bioveterinary Research | Virology, Wageningen Bioveterinary Research | Oreshkova,N., Vreman,S., Molenaar,R.J., Harders,F., Hakze van der Honing,R.W., Gerhards,N., Bouwstra,R., Hissink,H., Smit,L., Tacken,M., Weesendorp,E., Stegeman,A. and van der Poel,W.H.M. |
| EPI_ISL_431779, EPI_ISL_431780 | Fujian Center for Disease Control and Prevention | Fujian Center for Disease Control and Prevention | Lin Qi, Huang Zhimiao, Zhang Yanhua, Weng Yuwei |
| EPI_ISL_431781, EPI_ISL_431782, EPI_ISL_431783, EPI_ISL_431784, EPI_ISL_431785 | Fujian Center for Disease Control and Prevention | Fujian Center for Disease Control and Prevention | Lin Qi, Huang Zhimiao, Zhang Yanhua, Weng Yuwei |
| EPI_ISL_431833 | National Institutes of Health, University of the Philippines Manila | Philippine Genome Center, University of the Philippines System | Carlo M. Lapid, Francis A. Tablizo, Benedict A. Maralit, Jan Michael C. Yap, Raul V. Destura, Marissa M. Alejandria, El King D. Morado, Joshua Gregor A. Dizon, Jo-Hannah S. Llames, Shiela Mae M. Araiza, Kris P. Punayan, Kristianne Arielle D. Gabriel, Shebna Rose D. Fabilloren, Shana F. Genavia, Jarvin E. Nipales, Alessandra C. Sanchez, Haifa L.Gaza, Joy Ann Petronio-Santos, Julius Aaron Mejia, Maribell Dollete, Sonia Salamat, Christina Tan, Bernard Demot, John Mark Velasco, Eva Maria Cutiongco-de la Paz, and Cynthia P. Saloma |
| EPI_ISL_431901, EPI_ISL_431902, EPI_ISL_431903, EPI_ISL_431904, EPI_ISL_431905, EPI_ISL_431906, EPI_ISL_431907, EPI_ISL_431908, EPI_ISL_431909, EPI_ISL_431910, EPI_ISL_431911, EPI_ISL_431912, EPI_ISL_431913, EPI_ISL_431914, EPI_ISL_431915, EPI_ISL_431916, EPI_ISL_431917, EPI_ISL_431918, EPI_ISL_431919, EPI_ISL_431920, EPI_ISL_431921, EPI_ISL_431922, EPI_ISL_431923, EPI_ISL_431924, EPI_ISL_431925, EPI_ISL_431926, EPI_ISL_431927, EPI_ISL_431928, EPI_ISL_431929, EPI_ISL_431930, EPI_ISL_431931, EPI_ISL_431932, EPI_ISL_431933, EPI_ISL_431934, EPI_ISL_431935, EPI_ISL_431936, EPI_ISL_431937, EPI_ISL_431938, EPI_ISL_431939, EPI_ISL_431940, EPI_ISL_431941, EPI_ISL_431942, EPI_ISL_431943, EPI_ISL_431944, EPI_ISL_431945, EPI_ISL_431946, EPI_ISL_431947, EPI_ISL_431948, EPI_ISL_431949, EPI_ISL_431950, EPI_ISL_431951, EPI_ISL_431952, EPI_ISL_431953, EPI_ISL_431954, EPI_ISL_431955, EPI_ISL_431956, EPI_ISL_431957, EPI_ISL_431958, EPI_ISL_431959, EPI_ISL_431960, EPI_ISL_431961, EPI_ISL_431962, EPI_ISL_431963, EPI_ISL_431964, EPI_ISL_431965, EPI_ISL_431966, EPI_ISL_431967, EPI_ISL_431968, EPI_ISL_431969, EPI_ISL_431970, EPI_ISL_431971, EPI_ISL_431972, EPI_ISL_431973, EPI_ISL_431974, EPI_ISL_431975, EPI_ISL_431976, EPI_ISL_431977, EPI_ISL_431978, EPI_ISL_431979, EPI_ISL_431980, EPI_ISL_431981, EPI_ISL_431982, EPI_ISL_431983, EPI_ISL_431984, EPI_ISL_431985, EPI_ISL_431986, EPI_ISL_431987, EPI_ISL_431988, EPI_ISL_431989, EPI_ISL_431990, EPI_ISL_431991, EPI_ISL_431992, EPI_ISL_431993, EPI_ISL_431994, EPI_ISL_431995, EPI_ISL_431996, EPI_ISL_431997, EPI_ISL_431998, EPI_ISL_431999, EPI_ISL_432000, EPI_ISL_432001, EPI_ISL_432002, EPI_ISL_432003, EPI_ISL_432004, EPI_ISL_432005, EPI_ISL_432006, EPI_ISL_432007, EPI_ISL_432008, EPI_ISL_432009, EPI_ISL_432010, EPI_ISL_432011, EPI_ISL_432012, EPI_ISL_432013, EPI_ISL_432014, EPI_ISL_432015, EPI_ISL_432016, EPI_ISL_432017, EPI_ISL_432018, EPI_ISL_432019, EPI_ISL_432020, EPI_ISL_432021, EPI_ISL_432022, EPI_ISL_432023, EPI_ISL_432024, EPI_ISL_432025, EPI_ISL_432026, EPI_ISL_432027, EPI_ISL_432028, EPI_ISL_432029, EPI_ISL_432030, EPI_ISL_432031, EPI_ISL_432032, EPI_ISL_432033, EPI_ISL_432034, EPI_ISL_432035, EPI_ISL_432036, EPI_ISL_432037, EPI_ISL_432038, EPI_ISL_432039, EPI_ISL_432040, EPI_ISL_432041, EPI_ISL_432042, EPI_ISL_432043, EPI_ISL_432044, EPI_ISL_432045, EPI_ISL_432046, EPI_ISL_432047, EPI_ISL_432048, EPI_ISL_432049, EPI_ISL_432050, EPI_ISL_432051, EPI_ISL_432052, EPI_ISL_432053, EPI_ISL_432054, EPI_ISL_432055, EPI_ISL_432056, EPI_ISL_432057, EPI_ISL_432058, EPI_ISL_432059, EPI_ISL_432060, EPI_ISL_432061, EPI_ISL_432062, EPI_ISL_432063, EPI_ISL_432064, EPI_ISL_432065, EPI_ISL_432066, EPI_ISL_432067, EPI_ISL_432068, EPI_ISL_432069, EPI_ISL_432070, EPI_ISL_432071, EPI_ISL_432072, EPI_ISL_432073, EPI_ISL_432074, EPI_ISL_432075, EPI_ISL_432076, EPI_ISL_432077, EPI_ISL_432078, EPI_ISL_432079, EPI_ISL_432080, EPI_ISL_432081, EPI_ISL_432082, EPI_ISL_432083, EPI_ISL_432084, EPI_ISL_432085, EPI_ISL_432086, EPI_ISL_432087, EPI_ISL_432088, EPI_ISL_432089, EPI_ISL_432090, EPI_ISL_432091, EPI_ISL_432092, EPI_ISL_432093, EPI_ISL_432094, EPI_ISL_432095, EPI_ISL_432096, EPI_ISL_432097, EPI_ISL_432098, EPI_ISL_432099, EPI_ISL_432100, EPI_ISL_432101, EPI_ISL_432102, EPI_ISL_432103, EPI_ISL_432104, EPI_ISL_432105, EPI_ISL_432106, EPI_ISL_432107, EPI_ISL_432108, EPI_ISL_432109, EPI_ISL_432110, EPI_ISL_432111, EPI_ISL_432112, EPI_ISL_432113, EPI_ISL_432114, EPI_ISL_432115, EPI_ISL_432116, EPI_ISL_432117, EPI_ISL_432118, EPI_ISL_432119, EPI_ISL_432120, EPI_ISL_432121, EPI_ISL_432122, EPI_ISL_432123, EPI_ISL_432124, EPI_ISL_432125, EPI_ISL_432126, EPI_ISL_432127, EPI_ISL_432128, EPI_ISL_432129, EPI_ISL_432130, EPI_ISL_432131, EPI_ISL_432132, EPI_ISL_432133, EPI_ISL_432134, EPI_ISL_432135, EPI_ISL_432136, EPI_ISL_432137, EPI_ISL_432138, EPI_ISL_432139, EPI_ISL_432140, EPI_ISL_432141, EPI_ISL_432142, EPI_ISL_432143, EPI_ISL_432144, EPI_ISL_432145, EPI_ISL_432146, EPI_ISL_432147, EPI_ISL_432148, EPI_ISL_432149, EPI_ISL_432150, EPI_ISL_432151, EPI_ISL_432152, EPI_ISL_432153, EPI_ISL_432154, EPI_ISL_432155, EPI_ISL_432156, EPI_ISL_432157, EPI_ISL_432158, EPI_ISL_432159, EPI_ISL_432160, EPI_ISL_432161, EPI_ISL_432162, EPI_ISL_432163, EPI_ISL_432164, EPI_ISL_432165, EPI_ISL_432166, EPI_ISL_432167, EPI_ISL_432168, EPI_ISL_432169, EPI_ISL_432170, EPI_ISL_432171, EPI_ISL_432172, EPI_ISL_432173, EPI_ISL_432174, EPI_ISL_432175, EPI_ISL_432176, EPI_ISL_432177, EPI_ISL_432178, EPI_ISL_432179, EPI_ISL_432180, EPI_ISL_432181, EPI_ISL_432182, EPI_ISL_432183, EPI_ISL_432184, EPI_ISL_432185, EPI_ISL_432186, EPI_ISL_432187, EPI_ISL_432188, EPI_ISL_432189, EPI_ISL_432190, EPI_ISL_432191, EPI_ISL_432192, EPI_ISL_432193, EPI_ISL_432194, EPI_ISL_432195, EPI_ISL_432196, EPI_ISL_432197, EPI_ISL_432198, EPI_ISL_432199, EPI_ISL_432200, EPI_ISL_432201, EPI_ISL_432202, EPI_ISL_432203, EPI_ISL_432204, EPI_ISL_432205, EPI_ISL_432206, EPI_ISL_432207, EPI_ISL_432208, EPI_ISL_432209, EPI_ISL_432210, EPI_ISL_432211, EPI_ISL_432212, EPI_ISL_432213, EPI_ISL_432214, EPI_ISL_432215, EPI_ISL_432216, EPI_ISL_432217, EPI_ISL_432218, EPI_ISL_432219, EPI_ISL_432220, EPI_ISL_432221, EPI_ISL_432222, EPI_ISL_432223, EPI_ISL_432224, EPI_ISL_432225, EPI_ISL_432226, EPI_ISL_432227, EPI_ISL_432228, EPI_ISL_432229, EPI_ISL_432230, EPI_ISL_432231, EPI_ISL_432232, EPI_ISL_432233, EPI_ISL_432234, EPI_ISL_432235, EPI_ISL_432236, EPI_ISL_432237, EPI_ISL_432238, EPI_ISL_432239, EPI_ISL_432240, EPI_ISL_432241, EPI_ISL_432242, EPI_ISL_432243, EPI_ISL_432244, EPI_ISL_432245, EPI_ISL_432246, EPI_ISL_432247, EPI_ISL_432248, EPI_ISL_432249, EPI_ISL_432250, EPI_ISL_432251, EPI_ISL_432252, EPI_ISL_432253, EPI_ISL_432254, EPI_ISL_432255, EPI_ISL_432256, EPI_ISL_432257, EPI_ISL_432258, EPI_ISL_432259, EPI_ISL_432260, EPI_ISL_432261, EPI_ISL_432262, EPI_ISL_432263, EPI_ISL_432264, EPI_ISL_432265, EPI_ISL_432266, EPI_ISL_432267, EPI_ISL_432268, EPI_ISL_432269, EPI_ISL_432270, EPI_ISL_432271, EPI_ISL_432272, EPI_ISL_432273, EPI_ISL_432274, EPI_ISL_432275, EPI_ISL_432276, EPI_ISL_432277, EPI_ISL_432278, EPI_ISL_432279, EPI_ISL_432280, EPI_ISL_432281, EPI_ISL_432282, EPI_ISL_432283, EPI_ISL_432284, EPI_ISL_432285, EPI_ISL_432286, EPI_ISL_432287, EPI_ISL_432288, EPI_ISL_432289, EPI_ISL_432290, EPI_ISL_432291, EPI_ISL_432292, EPI_ISL_432293, EPI_ISL_432294, EPI_ISL_432295, EPI_ISL_432296, EPI_ISL_432297, EPI_ISL_432298, EPI_ISL_432299, EPI_ISL_432300, EPI_ISL_432301, EPI_ISL_432302, EPI_ISL_432303, EPI_ISL_432304, EPI_ISL_432305, EPI_ISL_432306, EPI_ISL_432307, EPI_ISL_432308, EPI_ISL_432309, EPI_ISL_432310, EPI_ISL_432311, EPI_ISL_432312, EPI_ISL_432313, EPI_ISL_432314, EPI_ISL_432315, EPI_ISL_432316, EPI_ISL_432317, EPI_ISL_432318, EPI_ISL_432319, EPI_ISL_432320, EPI_ISL_432321, EPI_ISL_432322, EPI_ISL_432323, EPI_ISL_432324, EPI_ISL_432325, EPI_ISL_432326, EPI_ISL_432327, EPI_ISL_432328, EPI_ISL_432329, EPI_ISL_432330, EPI_ISL_432331, EPI_ISL_432332, EPI_ISL_432333, EPI_ISL_432334, EPI_ISL_432335, EPI_ISL_432336, EPI_ISL_432337, EPI_ISL_432338, EPI_ISL_432339, EPI_ISL_432340, EPI_ISL_432341, EPI_ISL_432342, EPI_ISL_432343, EPI_ISL_432344, EPI_ISL_432345, EPI_ISL_432346, EPI_ISL_432347, EPI_ISL_432348, EPI_ISL_432349, EPI_ISL_432350, EPI_ISL_432351, EPI_ISL_432352, EPI_ISL_432353, EPI_ISL_432354, EPI_ISL_432355, EPI_ISL_432356, EPI_ISL_432357, EPI_ISL_432358, EPI_ISL_432359, EPI_ISL_432360, EPI_ISL_432361, EPI_ISL_432362, EPI_ISL_432363, EPI_ISL_432364, EPI_ISL_432365, EPI_ISL_432366, EPI_ISL_432367, EPI_ISL_432368, EPI_ISL_432369, EPI_ISL_432370, EPI_ISL_432371, EPI_ISL_432372, EPI_ISL_432373, EPI_ISL_432374, EPI_ISL_432375, EPI_ISL_432376, EPI_ISL_432377, EPI_ISL_432378, EPI_ISL_432379, EPI_ISL_432380, EPI_ISL_432381, EPI_ISL_432382, EPI_ISL_432383, EPI_ISL_432384, EPI_ISL_432385, EPI_ISL_432386, EPI_ISL_432387, EPI_ISL_432388, EPI_ISL_432389, EPI_ISL_432390, EPI_ISL_432391, EPI_ISL_432392, EPI_ISL_432393, EPI_ISL_432394, EPI_ISL_432395, EPI_ISL_432396, EPI_ISL_432397, EPI_ISL_432398, EPI_ISL_432399, EPI_ISL_432400, EPI_ISL_432401, EPI_ISL_432402, EPI_ISL_432403, EPI_ISL_432404, EPI_ISL_432405, EPI_ISL_432406, EPI_ISL_432407, EPI_ISL_432408, EPI_ISL_432409, EPI_ISL_432410, EPI_ISL_432411, EPI_ISL_432412, EPI_ISL_432413, EPI_ISL_432414, EPI_ISL_432415, EPI_ISL_432416, EPI_ISL_432417, EPI_ISL_432418, EPI_ISL_432419, EPI_ISL_432420, EPI_ISL_432421, EPI_ISL_432422, EPI_ISL_432423, EPI_ISL_432424, EPI_ISL_432425, EPI_ISL_432426, EPI_ISL_432427, EPI_ISL_432428, EPI_ISL_432429, EPI_ISL_432430, EPI_ISL_432431, EPI_ISL_432432, EPI_ISL_432433, EPI_ISL_432434, EPI_ISL_432435, EPI_ISL_432436, EPI_ISL_432437, EPI_ISL_432438, EPI_ISL_432439, EPI_ISL_432440, EPI_ISL_432441, EPI_ISL_432442, EPI_ISL_432443, EPI_ISL_432444, EPI_ISL_432445, EPI_ISL_432446, EPI_ISL_432447, EPI_ISL_432448, EPI_ISL_432449, EPI_ISL_432450 | | | |
| see above | Wales Specialist Virology Centre | Public Health Wales Microbiology Cardiff | Catherine Moore, Johnathan Evans, Malorie Perry, Simon Cottrell, Alec Birchley, Alexander Adams, Amy Gaskin, Bree Gatica-Wilcox, Jason Coombes, Lauren Gilbert, Lee Graham, Nicole Pacchiarini, Sara Kumziene-Summerhayes, Sarah Taylor, Sophie Jones, Sara Rey, Matthew Bull, Joanne Watkins, Sally Corden, Tom Connor |
| EPI_ISL_432451, EPI_ISL_432452, EPI_ISL_432453, EPI_ISL_432454, EPI_ISL_432455, EPI_ISL_432456, EPI_ISL_432457, EPI_ISL_432458, EPI_ISL_432459, EPI_ISL_432460, EPI_ISL_432461, EPI_ISL_432462, EPI_ISL_432463, EPI_ISL_432464, EPI_ISL_432465, EPI_ISL_432466, EPI_ISL_432467, EPI_ISL_432468, EPI_ISL_432469, EPI_ISL_432470, EPI_ISL_432471, EPI_ISL_432472, EPI_ISL_432473, EPI_ISL_432474, EPI_ISL_432475, EPI_ISL_432476, EPI_ISL_432477, EPI_ISL_432478, EPI_ISL_432479, EPI_ISL_432480, EPI_ISL_432481, EPI_ISL_432482, EPI_ISL_432483, EPI_ISL_432484, EPI_ISL_432485, EPI_ISL_432486, EPI_ISL_432487, EPI_ISL_432488, EPI_ISL_432489, EPI_ISL_432490, EPI_ISL_432491, EPI_ISL_432492, EPI_ISL_432493, EPI_ISL_432494, EPI_ISL_432495, EPI_ISL_432496, EPI_ISL_432497, EPI_ISL_432498, EPI_ISL_432499, EPI_ISL_432500, EPI_ISL_432501, EPI_ISL_432502, EPI_ISL_432503, EPI_ISL_432504, EPI_ISL_432505, EPI_ISL_432506, EPI_ISL_432507, EPI_ISL_432508, EPI_ISL_432509, EPI_ISL_432510, EPI_ISL_432511, EPI_ISL_432512, EPI_ISL_432513, EPI_ISL_432514, EPI_ISL_432515, EPI_ISL_432516, EPI_ISL_432517, EPI_ISL_432518, EPI_ISL_432519, EPI_ISL_432520, EPI_ISL_432521, EPI_ISL_432522, EPI_ISL_432523, EPI_ISL_432524, EPI_ISL_432525, EPI_ISL_432526, EPI_ISL_432527, EPI_ISL_432528, EPI_ISL_432529, EPI_ISL_432530, EPI_ISL_432531, EPI_ISL_432532, EPI_ISL_432533, EPI_ISL_432534, EPI_ISL_432535, EPI_ISL_432536, EPI_ISL_432537, EPI_ISL_432538, EPI_ISL_432539, EPI_ISL_432540, EPI_ISL_432541, EPI_ISL_432542, EPI_ISL_432543, EPI_ISL_432544, EPI_ISL_432545, EPI_ISL_432546, EPI_ISL_432547, EPI_ISL_432548, EPI_ISL_432549, EPI_ISL_432550, EPI_ISL_432551, EPI_ISL_432552, EPI_ISL_432553, EPI_ISL_432554, EPI_ISL_432555, EPI_ISL_432556, EPI_ISL_432557, EPI_ISL_432558, EPI_ISL_432559, EPI_ISL_432560, EPI_ISL_432561, EPI_ISL_432562, EPI_ISL_432563, EPI_ISL_432564, EPI_ISL_432565, EPI_ISL_432566, EPI_ISL_432567, EPI_ISL_432568, EPI_ISL_432569, EPI_ISL_432570, EPI_ISL_432571, EPI_ISL_432572, EPI_ISL_432573, EPI_ISL_432574, EPI_ISL_432575, EPI_ISL_432576, EPI_ISL_432577, EPI_ISL_432578, EPI_ISL_432579, EPI_ISL_432580, EPI_ISL_432581, EPI_ISL_432582, EPI_ISL_432583, EPI_ISL_432584, EPI_ISL_432585, EPI_ISL_432586, EPI_ISL_432587, EPI_ISL_432588, EPI_ISL_432589, EPI_ISL_432590, EPI_ISL_432591, EPI_ISL_432592, EPI_ISL_432593, EPI_ISL_432594, EPI_ISL_432595, EPI_ISL_432596, EPI_ISL_432597, EPI_ISL_432598, EPI_ISL_432599, EPI_ISL_432600, EPI_ISL_432601, EPI_ISL_432602, EPI_ISL_432603, EPI_ISL_432604, EPI_ISL_432605, EPI_ISL_432606, EPI_ISL_432607, EPI_ISL_432608, EPI_ISL_432609, EPI_ISL_432610, EPI_ISL_432611, EPI_ISL_432612, EPI_ISL_432613, EPI_ISL_432614, EPI_ISL_432615, EPI_ISL_432616, EPI_ISL_432617, EPI_ISL_432618, | | | |

EPI_ISL_432619, EPI_ISL_432620, EPI_ISL_432621, EPI_ISL_432622, EPI_ISL_432623, EPI_ISL_432624, EPI_ISL_432625, EPI_ISL_432626, EPI_ISL_432627, EPI_ISL_432628, EPI_ISL_432629, EPI_ISL_432630, EPI_ISL_432631, EPI_ISL_432632, EPI_ISL_432633, EPI_ISL_432634, EPI_ISL_432635, EPI_ISL_432636, EPI_ISL_432637, EPI_ISL_432638, EPI_ISL_432639, EPI_ISL_432640, EPI_ISL_432641, EPI_ISL_432642, EPI_ISL_432643, EPI_ISL_432644, EPI_ISL_432645, EPI_ISL_432646, EPI_ISL_432647, EPI_ISL_432648, EPI_ISL_432649, EPI_ISL_432650, EPI_ISL_432651, EPI_ISL_432652, EPI_ISL_432653, EPI_ISL_432654, EPI_ISL_432655, EPI_ISL_432656, EPI_ISL_432657, EPI_ISL_432658, EPI_ISL_432659, EPI_ISL_432660, EPI_ISL_432661, EPI_ISL_432662, EPI_ISL_432663, EPI_ISL_432664, EPI_ISL_432665, EPI_ISL_432666, EPI_ISL_432667

| | | | |
|---|---|---|---|
| see above | Virology Department, Sheffield Teaching Hospitals NHS Foundation Trust / Virology Department, Sheffield Teaching Hospitals NHS Foundation Trust | COVID-19 Genomics UK (COG-UK) Consortium | Thushan de Silva, Matthew Parker,Adri Angyal, Rebecca Brown, Luke Green, Rachel Tucker, Paul Parsons, Danielle Groves, Alex Keeley, Dave Partridge, Matthew Wyles, Benjamin Lindsey, Mehmet Yavuz, Mohammad Raza, Cariad Evans |

EPI_ISL_432668, EPI_ISL_432669, EPI_ISL_432670, EPI_ISL_432671, EPI_ISL_432672, EPI_ISL_432673, EPI_ISL_432674, EPI_ISL_432675, EPI_ISL_432676, EPI_ISL_432677, EPI_ISL_432678, EPI_ISL_432679, EPI_ISL_432680, EPI_ISL_432681, EPI_ISL_432682, EPI_ISL_432683, EPI_ISL_432684, EPI_ISL_432685, EPI_ISL_432686, EPI_ISL_432687, EPI_ISL_432688, EPI_ISL_432689, EPI_ISL_432690, EPI_ISL_432691, EPI_ISL_432692, EPI_ISL_432693, EPI_ISL_432694, EPI_ISL_432695, EPI_ISL_432696, EPI_ISL_432697, EPI_ISL_432698, EPI_ISL_432699, EPI_ISL_432700, EPI_ISL_432701, EPI_ISL_432702, EPI_ISL_432703, EPI_ISL_432704, EPI_ISL_432705, EPI_ISL_432706, EPI_ISL_432707, EPI_ISL_432708, EPI_ISL_432709

| | | | |
|---|---|---|---|
| see above | Queens Medical Centre, Clinical Microbiology Department / DeepSeq Nottingham | COVID-19 Genomics UK (COG-UK) Consortium | Gemma Clark, Wendy Smith, Manjinder Khakh, Hannah Howson-Wells, Jonathan Ball, Patrick McClure, Joseph Chappell, Theocharis Tsoleridis, Nadine Holmes, Matthew Carlisle, Christopher Moore, Fei Sang, Johnny Debebe, Victoria Wright, Matthew Loose |

EPI_ISL_432710, EPI_ISL_432711, EPI_ISL_432712, EPI_ISL_432713, EPI_ISL_432714, EPI_ISL_432715, EPI_ISL_432716, EPI_ISL_432717, EPI_ISL_432718, EPI_ISL_432719, EPI_ISL_432720, EPI_ISL_432721, EPI_ISL_432722, EPI_ISL_432723, EPI_ISL_432724, EPI_ISL_432725, EPI_ISL_432726, EPI_ISL_432727, EPI_ISL_432728, EPI_ISL_432729, EPI_ISL_432730, EPI_ISL_432731, EPI_ISL_432732, EPI_ISL_432733, EPI_ISL_432734, EPI_ISL_432735, EPI_ISL_432736, EPI_ISL_432737, EPI_ISL_432738, EPI_ISL_432739, EPI_ISL_432740, EPI_ISL_432741, EPI_ISL_432742, EPI_ISL_432743, EPI_ISL_432744, EPI_ISL_432745, EPI_ISL_432746, EPI_ISL_432747, EPI_ISL_432748, EPI_ISL_432749, EPI_ISL_432750, EPI_ISL_432751, EPI_ISL_432752, EPI_ISL_432753, EPI_ISL_432754, EPI_ISL_432755, EPI_ISL_432756, EPI_ISL_432757, EPI_ISL_432758, EPI_ISL_432759, EPI_ISL_432760, EPI_ISL_432761, EPI_ISL_432762, EPI_ISL_432763, EPI_ISL_432764, EPI_ISL_432765, EPI_ISL_432766, EPI_ISL_432767, EPI_ISL_432768, EPI_ISL_432769, EPI_ISL_432770, EPI_ISL_432771, EPI_ISL_432772, EPI_ISL_432773, EPI_ISL_432774, EPI_ISL_432775, EPI_ISL_432776, EPI_ISL_432777, EPI_ISL_432778, EPI_ISL_432779, EPI_ISL_432780, EPI_ISL_432781, EPI_ISL_432782, EPI_ISL_432783, EPI_ISL_432784, EPI_ISL_432785, EPI_ISL_432786, EPI_ISL_432787, EPI_ISL_432788, EPI_ISL_432789, EPI_ISL_432790, EPI_ISL_432791, EPI_ISL_432792, EPI_ISL_432793, EPI_ISL_432794, EPI_ISL_432795, EPI_ISL_432796, EPI_ISL_432797, EPI_ISL_432798, EPI_ISL_432799, EPI_ISL_432800, EPI_ISL_432801, EPI_ISL_432802, EPI_ISL_432803, EPI_ISL_432804, EPI_ISL_432805, EPI_ISL_432806, EPI_ISL_432807, EPI_ISL_432808, EPI_ISL_432809, EPI_ISL_432810, EPI_ISL_432811, EPI_ISL_432812, EPI_ISL_432813, EPI_ISL_432814, EPI_ISL_432815, EPI_ISL_432816, EPI_ISL_432817, EPI_ISL_432818, EPI_ISL_432819, EPI_ISL_432820, EPI_ISL_432821, EPI_ISL_432822, EPI_ISL_432823, EPI_ISL_432824, EPI_ISL_432825, EPI_ISL_432826, EPI_ISL_432827, EPI_ISL_432828, EPI_ISL_432829, EPI_ISL_432830, EPI_ISL_432831, EPI_ISL_432832, EPI_ISL_432833, EPI_ISL_432834, EPI_ISL_432835, EPI_ISL_432836, EPI_ISL_432837, EPI_ISL_432838, EPI_ISL_432839, EPI_ISL_432840, EPI_ISL_432841, EPI_ISL_432842, EPI_ISL_432843, EPI_ISL_432844, EPI_ISL_432845, EPI_ISL_432846, EPI_ISL_432847, EPI_ISL_432848, EPI_ISL_432849, EPI_ISL_432850, EPI_ISL_432851, EPI_ISL_432852, EPI_ISL_432853, EPI_ISL_432854, EPI_ISL_432855, EPI_ISL_432856, EPI_ISL_432857, EPI_ISL_432858, EPI_ISL_432859, EPI_ISL_432860, EPI_ISL_432861, EPI_ISL_432862, EPI_ISL_432863, EPI_ISL_432864, EPI_ISL_432865, EPI_ISL_432866, EPI_ISL_432867

| | | | |
|---|---|---|---|
| see above | Virology Department, Sheffield Teaching Hospitals NHS Foundation Trust / Virology Department, Sheffield Teaching Hospitals NHS Foundation Trust | COVID-19 Genomics UK (COG-UK) Consortium | Thushan de Silva, Matthew Parker,Adri Angyal, Rebecca Brown, Luke Green, Rachel Tucker, Paul Parsons, Danielle Groves, Alex Keeley, Dave Partridge, Matthew Wyles, Benjamin Lindsey, Mehmet Yavuz, Mohammad Raza, Cariad Evans |

EPI_ISL_432868, EPI_ISL_432869, EPI_ISL_432870, EPI_ISL_432871, EPI_ISL_432872, EPI_ISL_432873, EPI_ISL_432874, EPI_ISL_432875, EPI_ISL_432876, EPI_ISL_432877, EPI_ISL_432878, EPI_ISL_432879, EPI_ISL_432880, EPI_ISL_432881, EPI_ISL_432882, EPI_ISL_432883, EPI_ISL_432884, EPI_ISL_432885, EPI_ISL_432886, EPI_ISL_432887, EPI_ISL_432888, EPI_ISL_432889, EPI_ISL_432890, EPI_ISL_432891, EPI_ISL_432892, EPI_ISL_432893, EPI_ISL_432894, EPI_ISL_432895, EPI_ISL_432896, EPI_ISL_432897, EPI_ISL_432898, EPI_ISL_432899

| | | | |
|---|---|---|---|
| see above | Virology Department, Royal Infirmary of Edinburgh, NHS Lothian / School of Biological Sciences, University of Edinburgh / Institute of Genetics and Molecular Medicine, University of Edinburgh | COVID-19 Genomics UK (COG-UK) Consortium | McHugh M, Dewar R, Rooke S, Gallagher M, Balcaza C, O'Toole A, Hill V, McCrone JT, Colquhoun R, Yu X, Jackson B, Rambaut A, Williams TC, Templeton K |

EPI_ISL_432900, EPI_ISL_432901, EPI_ISL_432902, EPI_ISL_432903, EPI_ISL_432904, EPI_ISL_432905, EPI_ISL_432906, EPI_ISL_432907, EPI_ISL_432908, EPI_ISL_432909, EPI_ISL_432910, EPI_ISL_432911, EPI_ISL_432912, EPI_ISL_432913, EPI_ISL_432914, EPI_ISL_432915, EPI_ISL_432916, EPI_ISL_432917, EPI_ISL_432918, EPI_ISL_432919, EPI_ISL_432920, EPI_ISL_432921, EPI_ISL_432922, EPI_ISL_432923, EPI_ISL_432924, EPI_ISL_432925, EPI_ISL_432926, EPI_ISL_432927, EPI_ISL_432928, EPI_ISL_432929, EPI_ISL_432930, EPI_ISL_432931, EPI_ISL_432932, EPI_ISL_432933, EPI_ISL_432934, EPI_ISL_432935, EPI_ISL_432936, EPI_ISL_432937, EPI_ISL_432938, EPI_ISL_432939, EPI_ISL_432940, EPI_ISL_432941, EPI_ISL_432942, EPI_ISL_432943, EPI_ISL_432944, EPI_ISL_432945, EPI_ISL_432946, EPI_ISL_432947, EPI_ISL_432948, EPI_ISL_432949, EPI_ISL_432950, EPI_ISL_432951, EPI_ISL_432952, EPI_ISL_432953, EPI_ISL_432954, EPI_ISL_432955, EPI_ISL_432956, EPI_ISL_432957, EPI_ISL_432958, EPI_ISL_432959, EPI_ISL_432960, EPI_ISL_432961, EPI_ISL_432962, EPI_ISL_432963, EPI_ISL_432964, EPI_ISL_432965, EPI_ISL_432966, EPI_ISL_432967, EPI_ISL_432968, EPI_ISL_432969, EPI_ISL_432970, EPI_ISL_432971, EPI_ISL_432972, EPI_ISL_432973, EPI_ISL_432974, EPI_ISL_432975, EPI_ISL_432976, EPI_ISL_432977, EPI_ISL_432978, EPI_ISL_432979, EPI_ISL_432980, EPI_ISL_432981, EPI_ISL_432982, EPI_ISL_432983, EPI_ISL_432984, EPI_ISL_432985, EPI_ISL_432986, EPI_ISL_432987, EPI_ISL_432988, EPI_ISL_432989, EPI_ISL_432990, EPI_ISL_432991, EPI_ISL_432992, EPI_ISL_432993, EPI_ISL_432994, EPI_ISL_432995, EPI_ISL_432996, EPI_ISL_432997, EPI_ISL_432998, EPI_ISL_432999, EPI_ISL_433000, EPI_ISL_433001, EPI_ISL_433002, EPI_ISL_433003, EPI_ISL_433004, EPI_ISL_433005, EPI_ISL_433006, EPI_ISL_433007, EPI_ISL_433008, EPI_ISL_433009, EPI_ISL_433010, EPI_ISL_433011, EPI_ISL_433012, EPI_ISL_433013, EPI_ISL_433014, EPI_ISL_433015, EPI_ISL_433016, EPI_ISL_433017, EPI_ISL_433018, EPI_ISL_433019, EPI_ISL_433020, EPI_ISL_433021, EPI_ISL_433022, EPI_ISL_433023, EPI_ISL_433024, EPI_ISL_433025, EPI_ISL_433026, EPI_ISL_433027, EPI_ISL_433028, EPI_ISL_433029, EPI_ISL_433030, EPI_ISL_433031, EPI_ISL_433032, EPI_ISL_433033, EPI_ISL_433034, EPI_ISL_433035, EPI_ISL_433036, EPI_ISL_433037, EPI_ISL_433038, EPI_ISL_433039, EPI_ISL_433040, EPI_ISL_433041, EPI_ISL_433042, EPI_ISL_433043, EPI_ISL_433044, EPI_ISL_433045, EPI_ISL_433046, EPI_ISL_433047, EPI_ISL_433048, EPI_ISL_433049, EPI_ISL_433050, EPI_ISL_433051, EPI_ISL_433052, EPI_ISL_433053, EPI_ISL_433054, EPI_ISL_433055, EPI_ISL_433056, EPI_ISL_433057, EPI_ISL_433058, EPI_ISL_433059, EPI_ISL_433060, EPI_ISL_433061, EPI_ISL_433062, EPI_ISL_433063, EPI_ISL_433064, EPI_ISL_433065, EPI_ISL_433066, EPI_ISL_433067

| | | | |
|---|---|---|---|
| see above | Queens Medical Centre, Clinical Microbiology Department / DeepSeq Nottingham | COVID-19 Genomics UK (COG-UK) Consortium | Gemma Clark, Wendy Smith, Manjinder Khakh, Hannah Howson-Wells, Jonathan Ball, Patrick McClure, Joseph Chappell, Theocharis Tsoleridis, Nadine Holmes, Matthew Carlisle, Christopher Moore, Fei Sang, Johnny Debebe, Victoria Wright, Matthew Loose |

EPI_ISL_433068, EPI_ISL_433069, EPI_ISL_433070, EPI_ISL_433071, EPI_ISL_433072, EPI_ISL_433073, EPI_ISL_433074, EPI_ISL_433075, EPI_ISL_433076, EPI_ISL_433077, EPI_ISL_433078, EPI_ISL_433079, EPI_ISL_433080, EPI_ISL_433081, EPI_ISL_433082, EPI_ISL_433083, EPI_ISL_433084, EPI_ISL_433085, EPI_ISL_433086, EPI_ISL_433087, EPI_ISL_433088, EPI_ISL_433089, EPI_ISL_433090, EPI_ISL_433091, EPI_ISL_433092, EPI_ISL_433093, EPI_ISL_433094, EPI_ISL_433095, EPI_ISL_433096, EPI_ISL_433097, EPI_ISL_433098, EPI_ISL_433099, EPI_ISL_433100, EPI_ISL_433101, EPI_ISL_433102, EPI_ISL_433103, EPI_ISL_433104, EPI_ISL_433105, EPI_ISL_433106, EPI_ISL_433107, EPI_ISL_433108, EPI_ISL_433109, EPI_ISL_433110, EPI_ISL_433111, EPI_ISL_433112, EPI_ISL_433113, EPI_ISL_433114, EPI_ISL_433115, EPI_ISL_433116, EPI_ISL_433117, EPI_ISL_433118, EPI_ISL_433119, EPI_ISL_433120, EPI_ISL_433121, EPI_ISL_433122, EPI_ISL_433123, EPI_ISL_433124, EPI_ISL_433125, EPI_ISL_433126, EPI_ISL_433127, EPI_ISL_433128, EPI_ISL_433129, EPI_ISL_433130, EPI_ISL_433131, EPI_ISL_433132, EPI_ISL_433133, EPI_ISL_433134, EPI_ISL_433135, EPI_ISL_433136, EPI_ISL_433137, EPI_ISL_433138, EPI_ISL_433139, EPI_ISL_433140, EPI_ISL_433141, EPI_ISL_433142, EPI_ISL_433143, EPI_ISL_433144, EPI_ISL_433145, EPI_ISL_433146, EPI_ISL_433147, EPI_ISL_433148, EPI_ISL_433149, EPI_ISL_433150, EPI_ISL_433151, EPI_ISL_433152, EPI_ISL_433153, EPI_ISL_433154, EPI_ISL_433155, EPI_ISL_433156, EPI_ISL_433157, EPI_ISL_433158, EPI_ISL_433159, EPI_ISL_433160, EPI_ISL_433161, EPI_ISL_433162, EPI_ISL_433163, EPI_ISL_433164, EPI_ISL_433165, EPI_ISL_433166, EPI_ISL_433167, EPI_ISL_433168, EPI_ISL_433169, EPI_ISL_433170, EPI_ISL_433171, EPI_ISL_433172, EPI_ISL_433173, EPI_ISL_433174, EPI_ISL_433175, EPI_ISL_433176, EPI_ISL_433177, EPI_ISL_433178, EPI_ISL_433179, EPI_ISL_433180, EPI_ISL_433181, EPI_ISL_433182, EPI_ISL_433183, EPI_ISL_433184, EPI_ISL_433185, EPI_ISL_433186, EPI_ISL_433187, EPI_ISL_433188, EPI_ISL_433189, EPI_ISL_433190, EPI_ISL_433191, EPI_ISL_433192, EPI_ISL_433193, EPI_ISL_433194, EPI_ISL_433195, EPI_ISL_433196, EPI_ISL_433197, EPI_ISL_433198, EPI_ISL_433199, EPI_ISL_433200, EPI_ISL_433201, EPI_ISL_433202, EPI_ISL_433203, EPI_ISL_433204, EPI_ISL_433205, EPI_ISL_433206, EPI_ISL_433207, EPI_ISL_433208, EPI_ISL_433209, EPI_ISL_433210, EPI_ISL_433211, EPI_ISL_433212, EPI_ISL_433213, EPI_ISL_433214, EPI_ISL_433215, EPI_ISL_433216, EPI_ISL_433217, EPI_ISL_433218, EPI_ISL_433219, EPI_ISL_433220, EPI_ISL_433221, EPI_ISL_433222, EPI_ISL_433223, EPI_ISL_433224, EPI_ISL_433225, EPI_ISL_433226, EPI_ISL_433227, EPI_ISL_433228, EPI_ISL_433229, EPI_ISL_433230, EPI_ISL_433231, EPI_ISL_433232, EPI_ISL_433233, EPI_ISL_433234, EPI_ISL_433235, EPI_ISL_433236, EPI_ISL_433237, EPI_ISL_433238, EPI_ISL_433239, EPI_ISL_433240, EPI_ISL_433241, EPI_ISL_433242, EPI_ISL_433243, EPI_ISL_433244, EPI_ISL_433245, EPI_ISL_433246, EPI_ISL_433247, EPI_ISL_433248, EPI_ISL_433249, EPI_ISL_433250, EPI_ISL_433251, EPI_ISL_433252, EPI_ISL_433253, EPI_ISL_433254, EPI_ISL_433255, EPI_ISL_433256, EPI_ISL_433257, EPI_ISL_433258, EPI_ISL_433259, EPI_ISL_433260, EPI_ISL_433261, EPI_ISL_433262, EPI_ISL_433263, EPI_ISL_433264, EPI_ISL_433265, EPI_ISL_433266, EPI_ISL_433267

| | | | |
|---|---|---|---|
| see above | Virology Department, Royal Infirmary of Edinburgh, NHS Lothian / School of Biological Sciences, University of Edinburgh / Institute of Genetics and Molecular Medicine, University of Edinburgh | COVID-19 Genomics UK (COG-UK) Consortium | McHugh M, Dewar R, Rooke S, Gallagher M, Balcaza C, O'Toole A, Hill V, McCrone JT, Colquhoun R, Yu X, Jackson B, Rambaut A, Williams TC, Templeton K |

EPI_ISL_433268, EPI_ISL_433269, EPI_ISL_433270, EPI_ISL_433271, EPI_ISL_433272, EPI_ISL_433273, EPI_ISL_433274, EPI_ISL_433275, EPI_ISL_433276, EPI_ISL_433277, EPI_ISL_433278, EPI_ISL_433279, EPI_ISL_433280, EPI_ISL_433281, EPI_ISL_433282, EPI_ISL_433283, EPI_ISL_433284, EPI_ISL_433285, EPI_ISL_433286, EPI_ISL_433287, EPI_ISL_433288, EPI_ISL_433289, EPI_ISL_433290, EPI_ISL_433291, EPI_ISL_433292, EPI_ISL_433293, EPI_ISL_433294, EPI_ISL_433295, EPI_ISL_433296, EPI_ISL_433297, EPI_ISL_433298, EPI_ISL_433299, EPI_ISL_433300, EPI_ISL_433301, EPI_ISL_433302, EPI_ISL_433303, EPI_ISL_433304, EPI_ISL_433305, EPI_ISL_433306, EPI_ISL_433307, EPI_ISL_433308, EPI_ISL_433309, EPI_ISL_433310, EPI_ISL_433311, EPI_ISL_433312, EPI_ISL_433313, EPI_ISL_433314, EPI_ISL_433315, EPI_ISL_433316, EPI_ISL_433317, EPI_ISL_433318, EPI_ISL_433319, EPI_ISL_433320, EPI_ISL_433321, EPI_ISL_433322, EPI_ISL_433323, EPI_ISL_433324, EPI_ISL_433325, EPI_ISL_433326, EPI_ISL_433327, EPI_ISL_433328, EPI_ISL_433329, EPI_ISL_433330, EPI_ISL_433331, EPI_ISL_433332, EPI_ISL_433333, EPI_ISL_433334, EPI_ISL_433335, EPI_ISL_433336, EPI_ISL_433337, EPI_ISL_433338, EPI_ISL_433339, EPI_ISL_433340, EPI_ISL_433341, EPI_ISL_433342, EPI_ISL_433343, EPI_ISL_433344, EPI_ISL_433345, EPI_ISL_433346, EPI_ISL_433347, EPI_ISL_433348, EPI_ISL_433349, EPI_ISL_433350, EPI_ISL_433351, EPI_ISL_433352, EPI_ISL_433353, EPI_ISL_433354, EPI_ISL_433355, EPI_ISL_433356, EPI_ISL_433357, EPI_ISL_433358, EPI_ISL_433359, EPI_ISL_433360, EPI_ISL_433361, EPI_ISL_433362, EPI_ISL_433363, EPI_ISL_433364, EPI_ISL_433365, EPI_ISL_433366, EPI_ISL_433367, EPI_ISL_433368, EPI_ISL_433369, EPI_ISL_433370, EPI_ISL_433371, EPI_ISL_433372, EPI_ISL_433373

| | | | |
|---|---|---|---|
| see above | West of Scotland Specialist Virology Centre, NHSGGC / MRC-University of Glasgow Centre for Virus Research | COVID-19 Genomics UK (COG-UK) Consortium | Ana da Silva Filipe, Natasha Johnson, Kathy Smollett, Daniel Mair, Stephen Carmichael, Lily Tong, Jenna Nichols, Elihu Aranday-Cortes, Kirstyn Brunker, Yasmin Parr, Kyriaki Nomikou; Sarah McDonald, Marc Niebel, Patawee Asamaphan; Richard Orton, Joseph Hughes, Sreenu Vattipally, David L Robertson; Alasdair MacLean, Rory Gunson; Kathy Li, Natasha Jesudason, Rajiv Shah, James Shepherd, Antonia Ho, Emma Thomson |

EPI_ISL_433374, EPI_ISL_433375, EPI_ISL_433376, EPI_ISL_433377, EPI_ISL_433378, EPI_ISL_433379, EPI_ISL_433380, EPI_ISL_433381, EPI_ISL_433382, EPI_ISL_433383, EPI_ISL_433384, EPI_ISL_433385, EPI_ISL_433386, EPI_ISL_433387, EPI_ISL_433388, EPI_ISL_433389, EPI_ISL_433390, EPI_ISL_433391, EPI_ISL_433392, EPI_ISL_433393, EPI_ISL_433394, EPI_ISL_433395, EPI_ISL_433396, EPI_ISL_433397, EPI_ISL_433398, EPI_ISL_433399, EPI_ISL_433400, EPI_ISL_433401, EPI_ISL_433402, EPI_ISL_433403, EPI_ISL_433404, EPI_ISL_433405, EPI_ISL_433406, EPI_ISL_433407, EPI_ISL_433408, EPI_ISL_433409, EPI_ISL_433410, EPI_ISL_433411, EPI_ISL_433412, EPI_ISL_433413, EPI_ISL_433414, EPI_ISL_433415, EPI_ISL_433416, EPI_ISL_433417, EPI_ISL_433418, EPI_ISL_433419, EPI_ISL_433420, EPI_ISL_433421, EPI_ISL_433422, EPI_ISL_433423, EPI_ISL_433424, EPI_ISL_433425, EPI_ISL_433426, EPI_ISL_433427, EPI_ISL_433428, EPI_ISL_433429, EPI_ISL_433430, EPI_ISL_433431, EPI_ISL_433432, EPI_ISL_433433, EPI_ISL_433434, EPI_ISL_433435, EPI_ISL_433436, EPI_ISL_433437, EPI_ISL_433438, EPI_ISL_433439, EPI_ISL_433440, EPI_ISL_433441, EPI_ISL_433442, EPI_ISL_433443, EPI_ISL_433444, EPI_ISL_433445, EPI_ISL_433446, EPI_ISL_433447, EPI_ISL_433448, EPI_ISL_433449, EPI_ISL_433450, EPI_ISL_433451, EPI_ISL_433452, EPI_ISL_433453, EPI_ISL_433454, EPI_ISL_433455, EPI_ISL_433456, EPI_ISL_433457, EPI_ISL_433458, EPI_ISL_433459, EPI_ISL_433460, EPI_ISL_433461, EPI_ISL_433462, EPI_ISL_433463, EPI_ISL_433464, EPI_ISL_433465

| | | | |
|---|---|---|---|
| see above | Virology Department, Royal Infirmary of Edinburgh, NHS Lothian / School of Biological Sciences, University of Edinburgh / Institute of Genetics and Molecular Medicine, University of Edinburgh | COVID-19 Genomics UK (COG-UK) Consortium | McHugh M, Dewar R, Rooke S, Gallagher M, Balcaza C, O'Toole A, Hill V, McCrone JT, Colquhoun R, Yu X, Jackson B, Rambaut A, Williams TC, Templeton K |

EPI_ISL_433466, EPI_ISL_433467, EPI_ISL_433468, EPI_ISL_433469, EPI_ISL_433470, EPI_ISL_433471, EPI_ISL_433472, EPI_ISL_433473, EPI_ISL_433474, EPI_ISL_433475, EPI_ISL_433476, EPI_ISL_433477, EPI_ISL_433478, EPI_ISL_433479, EPI_ISL_433480, EPI_ISL_433481, EPI_ISL_433482, EPI_ISL_433483, EPI_ISL_433484, EPI_ISL_433485, EPI_ISL_433486, EPI_ISL_433487, EPI_ISL_433488, EPI_ISL_433489, EPI_ISL_433490, EPI_ISL_433491, EPI_ISL_433492, EPI_ISL_433493, EPI_ISL_433494, EPI_ISL_433495, EPI_ISL_433496, EPI_ISL_433497

| | | | |
|---|---|---|---|
| see above | Department of Pathology, University of Cambridge | COVID-19 Genomics UK (COG-UK) Consortium | Luke W Meredith, M. Estee Torok , Myra Hosmillo, William L. Hamilton, Martin D. Curran, Theresa Feltwell, Grant Hall, Anna Yakovleva, Fahad A Khokhar, Charlotte J. Houldcroft, Laura G Caller, Aminu S. Jahun, Sarah L. Caddy, Ian Goodfellow |

EPI_ISL_433498, EPI_ISL_433499, EPI_ISL_433500, EPI_ISL_433501, EPI_ISL_433502, EPI_ISL_433503, EPI_ISL_433504, EPI_ISL_433505, EPI_ISL_433506, EPI_ISL_433507, EPI_ISL_433508, EPI_ISL_433509, EPI_ISL_433510, EPI_ISL_433511, EPI_ISL_433512, EPI_ISL_433513, EPI_ISL_433514, EPI_ISL_433515, EPI_ISL_433516, EPI_ISL_433517, EPI_ISL_433518, EPI_ISL_433519, EPI_ISL_433520, EPI_ISL_433521, EPI_ISL_433522, EPI_ISL_433523, EPI_ISL_433524, EPI_ISL_433525, EPI_ISL_433526, EPI_ISL_433527, EPI_ISL_433528, EPI_ISL_433529, EPI_ISL_433530, EPI_ISL_433531, EPI_ISL_433532, EPI_ISL_433533, EPI_ISL_433534, EPI_ISL_433535, EPI_ISL_433536, EPI_ISL_433537, EPI_ISL_433538, EPI_ISL_433539, EPI_ISL_433540, EPI_ISL_433541, EPI_ISL_433542, EPI_ISL_433543, EPI_ISL_433544, EPI_ISL_433545, EPI_ISL_433546, EPI_ISL_433547, EPI_ISL_433548, EPI_ISL_433549, EPI_ISL_433550, EPI_ISL_433551, EPI_ISL_433552, EPI_ISL_433553, EPI_ISL_433554, EPI_ISL_433555, EPI_ISL_433556, EPI_ISL_433557, EPI_ISL_433558, EPI_ISL_433559, EPI_ISL_433560, EPI_ISL_433561, EPI_ISL_433562, EPI_ISL_433563, EPI_ISL_433564, EPI_ISL_433565, EPI_ISL_433566, EPI_ISL_433567, EPI_ISL_433568, EPI_ISL_433569, EPI_ISL_433570, EPI_ISL_433571, EPI_ISL_433572, EPI_ISL_433573, EPI_ISL_433574, EPI_ISL_433575, EPI_ISL_433576, EPI_ISL_433577, EPI_ISL_433578, EPI_ISL_433579, EPI_ISL_433580, EPI_ISL_433581, EPI_ISL_433582, EPI_ISL_433583, EPI_ISL_433584, EPI_ISL_433585, EPI_ISL_433586, EPI_ISL_433587, EPI_ISL_433588, EPI_ISL_433589, EPI_ISL_433590, EPI_ISL_433591, EPI_ISL_433592, EPI_ISL_433593, EPI_ISL_433594, EPI_ISL_433595, EPI_ISL_433596, EPI_ISL_433597, EPI_ISL_433598, EPI_ISL_433599, EPI_ISL_433600, EPI_ISL_433601, EPI_ISL_433602, EPI_ISL_433603, EPI_ISL_433604, EPI_ISL_433605, EPI_ISL_433606, EPI_ISL_433607, EPI_ISL_433608, EPI_ISL_433609, EPI_ISL_433610, EPI_ISL_433611, EPI_ISL_433612, EPI_ISL_433613, EPI_ISL_433614, EPI_ISL_433615, EPI_ISL_433616, EPI_ISL_433617, EPI_ISL_433618, EPI_ISL_433619, EPI_ISL_433620, EPI_ISL_433621, EPI_ISL_433622, EPI_ISL_433623, EPI_ISL_433624, EPI_ISL_433625, EPI_ISL_433626, EPI_ISL_433627, EPI_ISL_433628, EPI_ISL_433629, EPI_ISL_433630, EPI_ISL_433631, EPI_ISL_433632, EPI_ISL_433633, EPI_ISL_433634, EPI_ISL_433635, EPI_ISL_433636, EPI_ISL_433637, EPI_ISL_433638, EPI_ISL_433639, EPI_ISL_433640, EPI_ISL_433641, EPI_ISL_433642, EPI_ISL_433643, EPI_ISL_433644, EPI_ISL_433645, EPI_ISL_433646, EPI_ISL_433647, EPI_ISL_433648, EPI_ISL_433649, EPI_ISL_433650, EPI_ISL_433651, EPI_ISL_433652, EPI_ISL_433653, EPI_ISL_433654, EPI_ISL_433655, EPI_ISL_433656, EPI_ISL_433657, EPI_ISL_433658, EPI_ISL_433659, EPI_ISL_433660, EPI_ISL_433661, EPI_ISL_433662, EPI_ISL_433663, EPI_ISL_433664, EPI_ISL_433665

| | | | |
|---|---|---|---|
| see above | West of Scotland Specialist Virology Centre, NHSGGC / MRC-University of Glasgow Centre for Virus Research | COVID-19 Genomics UK (COG-UK) Consortium | Ana da Silva Filipe, Natasha Johnson, Kathy Smollett, Daniel Mair, Stephen Carmichael, Lily Tong, Jenna Nichols, Elihu Aranday-Cortes, Kirstyn Brunker, Yasmin Parr, Kyriaki Nomikou; Sarah McDonald, Marc Niebel, Patawee Asamaphan; Richard Orton, Joseph Hughes, Sreenu Vattipally, David L Robertson; Alasdair MacLean, Rory Gunson; Kathy Li, Natasha Jesudason, Rajiv Shah, James Shepherd, Antonia Ho, Emma Thomson |

EPI_ISL_433666, EPI_ISL_433667, EPI_ISL_433668, EPI_ISL_433669, EPI_ISL_433670, EPI_ISL_433671, EPI_ISL_433672, EPI_ISL_433673, EPI_ISL_433674, EPI_ISL_433675, EPI_ISL_433676, EPI_ISL_433677, EPI_ISL_433678, EPI_ISL_433679, EPI_ISL_433680, EPI_ISL_433681, EPI_ISL_433682, EPI_ISL_433683, EPI_ISL_433684, EPI_ISL_433685, EPI_ISL_433686, EPI_ISL_433687, EPI_ISL_433688, EPI_ISL_433689, EPI_ISL_433690, EPI_ISL_433691, EPI_ISL_433692, EPI_ISL_433693, EPI_ISL_433694, EPI_ISL_433695, EPI_ISL_433696, EPI_ISL_433697, EPI_ISL_433698, EPI_ISL_433699, EPI_ISL_433700, EPI_ISL_433701, EPI_ISL_433702, EPI_ISL_433703, EPI_ISL_433704, EPI_ISL_433705, EPI_ISL_433706, EPI_ISL_433707, EPI_ISL_433708, EPI_ISL_433709, EPI_ISL_433710, EPI_ISL_433711, EPI_ISL_433712, EPI_ISL_433713, EPI_ISL_433714, EPI_ISL_433715, EPI_ISL_433716, EPI_ISL_433717, EPI_ISL_433718, EPI_ISL_433719, EPI_ISL_433720, EPI_ISL_433721, EPI_ISL_433722, EPI_ISL_433723, EPI_ISL_433724, EPI_ISL_433725, EPI_ISL_433726, EPI_ISL_433727, EPI_ISL_433728, EPI_ISL_433729, EPI_ISL_433730, EPI_ISL_433731, EPI_ISL_433732, EPI_ISL_433733, EPI_ISL_433734, EPI_ISL_433735, EPI_ISL_433736, EPI_ISL_433737, EPI_ISL_433738, EPI_ISL_433739, EPI_ISL_433740, EPI_ISL_433741, EPI_ISL_433742, EPI_ISL_433743, EPI_ISL_433744, EPI_ISL_433745, EPI_ISL_433746, EPI_ISL_433747, EPI_ISL_433748, EPI_ISL_433749, EPI_ISL_433750, EPI_ISL_433751, EPI_ISL_433752, EPI_ISL_433753, EPI_ISL_433754, EPI_ISL_433755, EPI_ISL_433756, EPI_ISL_433757, EPI_ISL_433758, EPI_ISL_433759, EPI_ISL_433760, EPI_ISL_433761, EPI_ISL_433762, EPI_ISL_433763, EPI_ISL_433764, EPI_ISL_433765, EPI_ISL_433766, EPI_ISL_433767, EPI_ISL_433768, EPI_ISL_433769, EPI_ISL_433770, EPI_ISL_433771, EPI_ISL_433772, EPI_ISL_433773, EPI_ISL_433774, EPI_ISL_433775, EPI_ISL_433776, EPI_ISL_433777, EPI_ISL_433778, EPI_ISL_433779, EPI_ISL_433780, EPI_ISL_433781, EPI_ISL_433782, EPI_ISL_433783, EPI_ISL_433784, EPI_ISL_433785, EPI_ISL_433786, EPI_ISL_433787, EPI_ISL_433788, EPI_ISL_433789, EPI_ISL_433790, EPI_ISL_433791, EPI_ISL_433792, EPI_ISL_433793, EPI_ISL_433794, EPI_ISL_433795, EPI_ISL_433796, EPI_ISL_433797, EPI_ISL_433798, EPI_ISL_433799, EPI_ISL_433800, EPI_ISL_433801, EPI_ISL_433802, EPI_ISL_433803, EPI_ISL_433804, EPI_ISL_433805, EPI_ISL_433806, EPI_ISL_433807, EPI_ISL_433808, EPI_ISL_433809, EPI_ISL_433810, EPI_ISL_433811, EPI_ISL_433812, EPI_ISL_433813, EPI_ISL_433814, EPI_ISL_433815, EPI_ISL_433816, EPI_ISL_433817, EPI_ISL_433818, EPI_ISL_433819, EPI_ISL_433820, EPI_ISL_433821, EPI_ISL_433822, EPI_ISL_433823, EPI_ISL_433824, EPI_ISL_433825, EPI_ISL_433826, EPI_ISL_433827, EPI_ISL_433828, EPI_ISL_433829, EPI_ISL_433830, EPI_ISL_433831, EPI_ISL_433832, EPI_ISL_433833, EPI_ISL_433834, EPI_ISL_433835, EPI_ISL_433836, EPI_ISL_433837, EPI_ISL_433838, EPI_ISL_433839, EPI_ISL_433840, EPI_ISL_433841, EPI_ISL_433842, EPI_ISL_433843, EPI_ISL_433844, EPI_ISL_433845, EPI_ISL_433846, EPI_ISL_433847, EPI_ISL_433848, EPI_ISL_433849, EPI_ISL_433850, EPI_ISL_433851, EPI_ISL_433852, EPI_ISL_433853, EPI_ISL_433854, EPI_ISL_433855, EPI_ISL_433856, EPI_ISL_433857, EPI_ISL_433858, EPI_ISL_433859, EPI_ISL_433860, EPI_ISL_433861, EPI_ISL_433862, EPI_ISL_433863, EPI_ISL_433864, EPI_ISL_433865, EPI_ISL_433866, EPI_ISL_433867, EPI_ISL_433868, EPI_ISL_433869, EPI_ISL_433870, EPI_ISL_433871, EPI_ISL_433872, EPI_ISL_433873, EPI_ISL_433874, EPI_ISL_433875, EPI_ISL_433876, EPI_ISL_433877, EPI_ISL_433878, EPI_ISL_433879, EPI_ISL_433880, EPI_ISL_433881, EPI_ISL_433882, EPI_ISL_433883, EPI_ISL_433884, EPI_ISL_433885, EPI_ISL_433886, EPI_ISL_433887, EPI_ISL_433888, EPI_ISL_433889, EPI_ISL_433890, EPI_ISL_433891, EPI_ISL_433892, EPI_ISL_433893, EPI_ISL_433894, EPI_ISL_433895, EPI_ISL_433896, EPI_ISL_433897, EPI_ISL_433898, EPI_ISL_433899, EPI_ISL_433900, EPI_ISL_433901, EPI_ISL_433902, EPI_ISL_433903, EPI_ISL_433904, EPI_ISL_433905, EPI_ISL_433906, EPI_ISL_433907, EPI_ISL_433908, EPI_ISL_433909, EPI_ISL_433910, EPI_ISL_433911, EPI_ISL_433912, EPI_ISL_433913, EPI_ISL_433914, EPI_ISL_433915, EPI_ISL_433916, EPI_ISL_433917, EPI_ISL_433918, EPI_ISL_433919, EPI_ISL_433920, EPI_ISL_433921, EPI_ISL_433922, EPI_ISL_433923, EPI_ISL_433924, EPI_ISL_433925, EPI_ISL_433926, EPI_ISL_433927, EPI_ISL_433928, EPI_ISL_433929, EPI_ISL_433930, EPI_ISL_433931, EPI_ISL_433932, EPI_ISL_433933, EPI_ISL_433934, EPI_ISL_433935, EPI_ISL_433936, EPI_ISL_433937, EPI_ISL_433938, EPI_ISL_433939, EPI_ISL_433940, EPI_ISL_433941, EPI_ISL_433942, EPI_ISL_433943, EPI_ISL_433944, EPI_ISL_433945, EPI_ISL_433946, EPI_ISL_433947, EPI_ISL_433948, EPI_ISL_433949, EPI_ISL_433950, EPI_ISL_433951, EPI_ISL_433952, EPI_ISL_433953, EPI_ISL_433954, EPI_ISL_433955, EPI_ISL_433956, EPI_ISL_433957, EPI_ISL_433958, EPI_ISL_433959, EPI_ISL_433960, EPI_ISL_433961, EPI_ISL_433962, EPI_ISL_433963, EPI_ISL_433964, EPI_ISL_433965, EPI_ISL_433966, EPI_ISL_433967, EPI_ISL_433968, EPI_ISL_433969, EPI_ISL_433970, EPI_ISL_433971, EPI_ISL_433972, EPI_ISL_433973, EPI_ISL_433974, EPI_ISL_433975, EPI_ISL_433976, EPI_ISL_433977, EPI_ISL_433978, EPI_ISL_433979, EPI_ISL_433980, EPI_ISL_433981, EPI_ISL_433982, EPI_ISL_433983, EPI_ISL_433984, EPI_ISL_433985, EPI_ISL_433986, EPI_ISL_433987, EPI_ISL_433988, EPI_ISL_433989, EPI_ISL_433990, EPI_ISL_433991, EPI_ISL_433992, EPI_ISL_433993, EPI_ISL_433994, EPI_ISL_433995, EPI_ISL_433996, EPI_ISL_433997, EPI_ISL_433998, EPI_ISL_433999, EPI_ISL_434000, EPI_ISL_434001, EPI_ISL_434002, EPI_ISL_434003, EPI_ISL_434004, EPI_ISL_434005, EPI_ISL_434006, EPI_ISL_434007, EPI_ISL_434008, EPI_ISL_434009, EPI_ISL_434010, EPI_ISL_434011, EPI_ISL_434012, EPI_ISL_434013, EPI_ISL_434014, EPI_ISL_434015, EPI_ISL_434016, EPI_ISL_434017, EPI_ISL_434018, EPI_ISL_434019, EPI_ISL_434020, EPI_ISL_434021, EPI_ISL_434022, EPI_ISL_434023, EPI_ISL_434024, EPI_ISL_434025, EPI_ISL_434026, EPI_ISL_434027, EPI_ISL_434028, EPI_ISL_434029, EPI_ISL_434030, EPI_ISL_434031, EPI_ISL_434032, EPI_ISL_434033, EPI_ISL_434034, EPI_ISL_434035, EPI_ISL_434036, EPI_ISL_434037, EPI_ISL_434038, EPI_ISL_434039, EPI_ISL_434040, EPI_ISL_434041, EPI_ISL_434042, EPI_ISL_434043, EPI_ISL_434044, EPI_ISL_434045, EPI_ISL_434046, EPI_ISL_434047, EPI_ISL_434048, EPI_ISL_434049, EPI_ISL_434050, EPI_ISL_434051, EPI_ISL_434052, EPI_ISL_434053, EPI_ISL_434054, EPI_ISL_434055, EPI_ISL_434056, EPI_ISL_434057, EPI_ISL_434058, EPI_ISL_434059, EPI_ISL_434060, EPI_ISL_434061, EPI_ISL_434062

| | | | |
|---|---|---|---|
| see above | Department of Pathology, University of Cambridge | COVID-19 Genomics UK (COG-UK) Consortium | Luke W Meredith, M. Estee Torok , Myra Hosmillo, William L. Hamilton, Martin D. Curran, Theresa Feltwell, Grant Hall, Anna Yakovleva, Fahad A Khokhar, Charlotte J. Houldcroft, Laura G Caller, Aminu S. Jahun, Sarah L. Caddy, Ian Goodfellow |

EPI_ISL_434063, EPI_ISL_434064, EPI_ISL_434065, EPI_ISL_434066, EPI_ISL_434067, EPI_ISL_434068, EPI_ISL_434069, EPI_ISL_434070, EPI_ISL_434071, EPI_ISL_434072, EPI_ISL_434073, EPI_ISL_434074, EPI_ISL_434075, EPI_ISL_434076, EPI_ISL_434077, EPI_ISL_434078, EPI_ISL_434079, EPI_ISL_434080, EPI_ISL_434081, EPI_ISL_434082, EPI_ISL_434083, EPI_ISL_434084, EPI_ISL_434085, EPI_ISL_434086, EPI_ISL_434087, EPI_ISL_434088, EPI_ISL_434089, EPI_ISL_434090, EPI_ISL_434091, EPI_ISL_434092, EPI_ISL_434093, EPI_ISL_434094, EPI_ISL_434095, EPI_ISL_434096, EPI_ISL_434097, EPI_ISL_434098, EPI_ISL_434099, EPI_ISL_434100, EPI_ISL_434101, EPI_ISL_434102, EPI_ISL_434103, EPI_ISL_434104, EPI_ISL_434105, EPI_ISL_434106, EPI_ISL_434107, EPI_ISL_434108, EPI_ISL_434109, EPI_ISL_434110, EPI_ISL_434111, EPI_ISL_434112, EPI_ISL_434113, EPI_ISL_434114, EPI_ISL_434115, EPI_ISL_434116, EPI_ISL_434117, EPI_ISL_434118, EPI_ISL_434119, EPI_ISL_434120, EPI_ISL_434121, EPI_ISL_434122, EPI_ISL_434123, EPI_ISL_434124, EPI_ISL_434125, EPI_ISL_434126, EPI_ISL_434127, EPI_ISL_434128, EPI_ISL_434129, EPI_ISL_434130, EPI_ISL_434131, EPI_ISL_434132, EPI_ISL_434133, EPI_ISL_434134, EPI_ISL_434135, EPI_ISL_434136, EPI_ISL_434137, EPI_ISL_434138, EPI_ISL_434139, EPI_ISL_434140, EPI_ISL_434141, EPI_ISL_434142, EPI_ISL_434143, EPI_ISL_434144, EPI_ISL_434145, EPI_ISL_434146, EPI_ISL_434147, EPI_ISL_434148, EPI_ISL_434149, EPI_ISL_434150, EPI_ISL_434151, EPI_ISL_434152, EPI_ISL_434153, EPI_ISL_434154, EPI_ISL_434155, EPI_ISL_434156, EPI_ISL_434157, EPI_ISL_434158, EPI_ISL_434159, EPI_ISL_434160, EPI_ISL_434161, EPI_ISL_434162, EPI_ISL_434163, EPI_ISL_434164, EPI_ISL_434165, EPI_ISL_434166, EPI_ISL_434167, EPI_ISL_434168, EPI_ISL_434169, EPI_ISL_434170, EPI_ISL_434171, EPI_ISL_434172, EPI_ISL_434173, EPI_ISL_434174, EPI_ISL_434175, EPI_ISL_434176, EPI_ISL_434177, EPI_ISL_434178, EPI_ISL_434179, EPI_ISL_434180, EPI_ISL_434181, EPI_ISL_434182, EPI_ISL_434183, EPI_ISL_434184, EPI_ISL_434185, EPI_ISL_434186, EPI_ISL_434187, EPI_ISL_434188, EPI_ISL_434189, EPI_ISL_434190, EPI_ISL_434191, EPI_ISL_434192, EPI_ISL_434193, EPI_ISL_434194, EPI_ISL_434195, EPI_ISL_434196, EPI_ISL_434197, EPI_ISL_434198, EPI_ISL_434199, EPI_ISL_434200, EPI_ISL_434201, EPI_ISL_434202, EPI_ISL_434203, EPI_ISL_434204, EPI_ISL_434205, EPI_ISL_434206, EPI_ISL_434207, EPI_ISL_434208, EPI_ISL_434209, EPI_ISL_434210, EPI_ISL_434211, EPI_ISL_434212, EPI_ISL_434213, EPI_ISL_434214, EPI_ISL_434215, EPI_ISL_434216, EPI_ISL_434217, EPI_ISL_434218, EPI_ISL_434219, EPI_ISL_434220, EPI_ISL_434221, EPI_ISL_434222, EPI_ISL_434223, EPI_ISL_434224, EPI_ISL_434225, EPI_ISL_434226, EPI_ISL_434227, EPI_ISL_434228, EPI_ISL_434229, EPI_ISL_434230, EPI_ISL_434231, EPI_ISL_434232, EPI_ISL_434233, EPI_ISL_434234, EPI_ISL_434235, EPI_ISL_434236, EPI_ISL_434237, EPI_ISL_434238, EPI_ISL_434239, EPI_ISL_434240, EPI_ISL_434241, EPI_ISL_434242, EPI_ISL_434243, EPI_ISL_434244, EPI_ISL_434245, EPI_ISL_434246, EPI_ISL_434247, EPI_ISL_434248, EPI_ISL_434249, EPI_ISL_434250, EPI_ISL_434251, EPI_ISL_434252, EPI_ISL_434253, EPI_ISL_434254, EPI_ISL_434255, EPI_ISL_434256, EPI_ISL_434257, EPI_ISL_434258, EPI_ISL_434259, EPI_ISL_434260, EPI_ISL_434261, EPI_ISL_434262, EPI_ISL_434263, EPI_ISL_434264, EPI_ISL_434265, EPI_ISL_434266, EPI_ISL_434267, EPI_ISL_434268, EPI_ISL_434269, EPI_ISL_434270, EPI_ISL_434271, EPI_ISL_434272, EPI_ISL_434273, EPI_ISL_434274, EPI_ISL_434275, EPI_ISL_434276, EPI_ISL_434277, EPI_ISL_434278,

| | | | |
|---|---|---|---|
| EPI_ISL_434279, EPI_ISL_434280, EPI_ISL_434281, EPI_ISL_434282, EPI_ISL_434283, EPI_ISL_434284, EPI_ISL_434285, EPI_ISL_434286, EPI_ISL_434287, EPI_ISL_434288, EPI_ISL_434289, EPI_ISL_434290, EPI_ISL_434291, EPI_ISL_434292, EPI_ISL_434293, EPI_ISL_434294, EPI_ISL_434295, EPI_ISL_434296, EPI_ISL_434297, EPI_ISL_434298, EPI_ISL_434299, EPI_ISL_434300, EPI_ISL_434301, EPI_ISL_434302, EPI_ISL_434303, EPI_ISL_434304, EPI_ISL_434305, EPI_ISL_434306, EPI_ISL_434307, EPI_ISL_434308, EPI_ISL_434309, EPI_ISL_434310, EPI_ISL_434311, EPI_ISL_434312, EPI_ISL_434313, EPI_ISL_434314, EPI_ISL_434315, EPI_ISL_434316, EPI_ISL_434317, EPI_ISL_434318, EPI_ISL_434319, EPI_ISL_434320, EPI_ISL_434321, EPI_ISL_434322, EPI_ISL_434323, EPI_ISL_434324, EPI_ISL_434325, EPI_ISL_434326, EPI_ISL_434327, EPI_ISL_434328, EPI_ISL_434329, EPI_ISL_434330, EPI_ISL_434331, EPI_ISL_434332, EPI_ISL_434333, EPI_ISL_434334, EPI_ISL_434335, EPI_ISL_434336, EPI_ISL_434337, EPI_ISL_434338, EPI_ISL_434339, EPI_ISL_434340, EPI_ISL_434341, EPI_ISL_434342, EPI_ISL_434343, EPI_ISL_434344, EPI_ISL_434345, EPI_ISL_434346 | | | |
| see above | Washington State Department of Health | Seattle Flu Study | Chu et al |
| EPI_ISL_434347, EPI_ISL_434348, EPI_ISL_434349, EPI_ISL_434350, EPI_ISL_434351, EPI_ISL_434352, EPI_ISL_434353, EPI_ISL_434354, EPI_ISL_434355 | Lab voor klinische biologie | Onderzoeksgroep Virologie | Laurens Lambrechts, Nick Vereecke, Marthe Pauwels, Jozefien De Clercq, Bruno Verhasselt, Linos Vandekerckhove, Hans Nauwynck, Sebastiaan Theuns |
| EPI_ISL_434356, EPI_ISL_434357, EPI_ISL_434358, EPI_ISL_434359, EPI_ISL_434360, EPI_ISL_434361, EPI_ISL_434362, EPI_ISL_434363, EPI_ISL_434364, EPI_ISL_434365 | Lab voor klinische biologie | Onderzoeksgroep Virologie | Nick Vereecke, Laurens Lambrechts, Marthe Pauwels, Jozefien De Clercq, Bruno Verhasselt, Linos Vandekerckhove, Hans Nauwynck, Sebastiaan Theuns |
| EPI_ISL_434366, EPI_ISL_434367, EPI_ISL_434368, EPI_ISL_434369, EPI_ISL_434370, EPI_ISL_434371 | Hospital AZ Rivierenland | Institute of Tropical Medicine | Philippe Selhorst, Colin Anthony |
| EPI_ISL_434372, EPI_ISL_434373, EPI_ISL_434374, EPI_ISL_434375, EPI_ISL_434376, EPI_ISL_434377, EPI_ISL_434378, EPI_ISL_434379, EPI_ISL_434380, EPI_ISL_434381, EPI_ISL_434382, EPI_ISL_434383 | | | |
| see above | Hospital AZ Rivierenland | Institute of Tropical Medicine | Philippe Selhorst, Colin Anthony, |
| EPI_ISL_434384, EPI_ISL_434385, EPI_ISL_434386 | Hospital AZ Rivierenland | Institute of Tropical Medicine | Philippe Selhorst, Colin Anthony |
| EPI_ISL_434455, EPI_ISL_434456, EPI_ISL_434457, EPI_ISL_434458, EPI_ISL_434459, EPI_ISL_434460, EPI_ISL_434461, EPI_ISL_434462, EPI_ISL_434463, EPI_ISL_434464, EPI_ISL_434465, EPI_ISL_434466, EPI_ISL_434467, EPI_ISL_434468, EPI_ISL_434469, EPI_ISL_434470, EPI_ISL_434471, EPI_ISL_434472, EPI_ISL_434473, EPI_ISL_434474, EPI_ISL_434475, EPI_ISL_434476, EPI_ISL_434477, EPI_ISL_434478, EPI_ISL_434479, EPI_ISL_434480, EPI_ISL_434481, EPI_ISL_434482, EPI_ISL_434483, EPI_ISL_434484, EPI_ISL_434485, EPI_ISL_434486 | | | |
| see above | Laboratory of Microbiology, Medical School, National and Kapodistrian University of Athens | Laboratory of Biology, Department of Medicine, Democritus University of Thrace | Kassela K., Bampali,M., Dovrolis,N., Gatzidou,E., Froukala,E., Stavropoulou,A., Veletza,S., Tsakris,A., Spanakis,N. and Karakasiliotis,I. |
| EPI_ISL_434487, EPI_ISL_434488, EPI_ISL_434489, EPI_ISL_434490, EPI_ISL_434491, EPI_ISL_434492, EPI_ISL_434493, EPI_ISL_434494, EPI_ISL_434495, EPI_ISL_434496, EPI_ISL_434497, EPI_ISL_434498, EPI_ISL_434499, EPI_ISL_434500, EPI_ISL_434501, EPI_ISL_434502, EPI_ISL_434503, EPI_ISL_434504, EPI_ISL_434505, EPI_ISL_434506, EPI_ISL_434507, EPI_ISL_434508, EPI_ISL_434509, EPI_ISL_434510, EPI_ISL_434511, EPI_ISL_434512, EPI_ISL_434513, EPI_ISL_434514, EPI_ISL_434515 | | | |
| see above | Laboratoire National de Sante, Microbiology, Virology | Laboratoire National de Sante, Microbiology, Epidemiology and Microbial Genomics | Anke Wienecke-Baldacchino, Ardashel Latsuzbaia, Jessica Tapp, Catherine Ragimbeau, Guillaume Fournier, Tamir Abdelrahman, Trung Nguyen Nguyen, Joel Mossong |
| EPI_ISL_434516 | Biolab Diagnostic Laboratories | Andersen lab at Scripps Research | Issa Abu-Dayyeh, Ahmad Tibi, Lama Hussein, Lina Mohammad, Zein Naber, Amid Abdelnour with SEARCH Alliance San Diego |
| EPI_ISL_434517, EPI_ISL_434518, EPI_ISL_434519, EPI_ISL_434520, EPI_ISL_434521, EPI_ISL_434522, EPI_ISL_434523, EPI_ISL_434524, EPI_ISL_434525, EPI_ISL_434526, EPI_ISL_434527, EPI_ISL_434528, EPI_ISL_434529, EPI_ISL_434530, EPI_ISL_434531, EPI_ISL_434532 | | | |
| see above | Robert Garry lab | Andersen lab at Scripps Research | Allison Smither, Gilberto Sabino-Santos, Patricia Snarski, Lilia Melnik, Antoinette Bell, Kaylynn Genemaras, Arnaud Drouin, Dahlene Fusco, Robert Garry with SEARCH Alliance San Diego |
| EPI_ISL_434533 | Area de Salud Alajuela Sur | Inciensa, Instituto Costarricense de Investigación y Enseñanza en Nutrición y Salud | Francisco Duarte, Hebleen Porras, Claudio Soto-Garita, Estela Cordero, Adriana Godinez & Melany Calderon |
| EPI_ISL_434534 | National Institute for Viral Disease Control and Prevention, China CDC | National Institute for Viral Disease Control and Prevention, China CDC, Yunnan Provincial CDC | Wenjie Tan, Roujian Lu, Wenling Wang, Peihua Niu, Huijuan Wang, Baoying Huang, Li Zhao, Fei Ye, Guizhen Wu |
| EPI_ISL_434535 | Area de Salud Alajuela Sur | Inciensa, Instituto Costarricense de Investigación y Enseñanza en Nutrición y Salud | Francisco Duarte, Hebleen Porras, Claudio Soto-Garita, Estela Cordero, Adriana Godinez & Melany Calderon |
| EPI_ISL_434536 | Hospital San Vicente de Paul | Inciensa, Instituto Costarricense de Investigación y Enseñanza en Nutrición y Salud | Francisco Duarte, Hebleen Porras, Claudio Soto-Garita, Estela Cordero, Adriana Godinez & Melany Calderon |
| EPI_ISL_434538 | COOPESAIN | Inciensa, Instituto Costarricense de Investigación y Enseñanza en Nutrición y Salud | Francisco Duarte, Hebleen Porras, Claudio Soto-Garita, Estela Cordero, Adriana Godinez & Melany Calderon |
| EPI_ISL_434539 | Area de Salud Orotina | Inciensa, Instituto Costarricense de Investigación y Enseñanza en Nutrición y Salud | Francisco Duarte, Hebleen Porras, Claudio Soto-Garita, Estela Cordero, Adriana Godinez & Melany Calderon |
| EPI_ISL_434540 | EBAIS Concepción Norte | Inciensa, Instituto Costarricense de Investigación y Enseñanza en Nutrición y Salud | Francisco Duarte, Hebleen Porras, Claudio Soto-Garita, Estela Cordero, Adriana Godinez & Melany Calderon |
| EPI_ISL_434541, EPI_ISL_434542, EPI_ISL_434543, EPI_ISL_434544, EPI_ISL_434545, EPI_ISL_434546, EPI_ISL_434547, EPI_ISL_434548, EPI_ISL_434549, EPI_ISL_434550, EPI_ISL_434551, EPI_ISL_434552, EPI_ISL_434553 | | | |
| see above | Puerto Rico Department of Health | Centers for Disease Control and Prevention, Dengue Branch | Gilberto A. Santiago, Glenda Gonzalez, Betzabel Flores, Keyla Charriez, Fabiola Cruz, Chaney Kalinich, Joseph Fauver, Jessica I. Falcon, Nathan Grubaugh, Jorge L. Munoz-Jordan |
| EPI_ISL_434554, EPI_ISL_434555, EPI_ISL_434556, EPI_ISL_434557, EPI_ISL_434558 | National Institutes of Health, University of the Philippines Manila | Philippine Genome Center | Carlo M. Lapid, Francis A. Tablizo, Benedict A. Maralit, Jan Michael C. Yap, Raul V. Destura, Marissa M. Alejandria, El King D. Morado, Joshua Gregor A. Dizon, Jo-Hannah S. Llames, Shiela Mae M. Araiza, Kris P. Punayan, Kristianne Arielle D. Gabriel, Shebna Rose D. Fabilloren, Shana F. Genavia, Jarvin E. Nipales, Alessandra C. Sanchez, Haifa L.Gaza, Joy Ann Petronio-Santos, Julius Aaron Mejia, Maribell Dollete, Sonia Salamat, Christina Tan, Bernard Demot, John Mark Velasco, Eva Maria Cutiongco-de la Paz, and Cynthia P. Saloma |
| EPI_ISL_434561, EPI_ISL_434562 | Department of Microbiology; Ryota Kumagai Tokyo Metropolitan Institute of Public Health | Department of Microbiology; Ryota Kumagai Tokyo Metropolitan Institute of Public Health | Kumagai,R., Yoshida,I., Asakura,H., Nagashima,M., Chiba,T. and Sadamasu,K. |
| EPI_ISL_434570 | unknown | Microbiology | To,K.K.W. and Yuen,K.-Y. |
| EPI_ISL_434572 | The National Institute of Public Health Center for Epidemiology and Microbiology | The National Institute of Public Health Center for Epidemiology and Microbiology | Alexander Nagy, Helena Jirincova, Ludmila Novakova, Dusan Trnka, Jaromira Vecerova |
| EPI_ISL_434586, EPI_ISL_434587, EPI_ISL_434588 | Johns Hopkins Hospital Department of Pathology | Johns Hopkins Hospital Department of Pathology | Peter M. Thielen, Thomas Mehoke, Shirlee Wohl, Srividya Ramakrishnan, Oluwaseun Nwulia-Falade, Amanda Ernlund, Melanie Kirsche, Paul Morris, Norah Sadowski, Nídiá Trovao, Victoria Gniazdowski, Michael Schatz, Stuart C. Ray, Winston Timp, Heba Mostafa |
| EPI_ISL_434590, EPI_ISL_434591, EPI_ISL_434592, EPI_ISL_434593, EPI_ISL_434594, EPI_ISL_434595, EPI_ISL_434596, EPI_ISL_434597, EPI_ISL_434598, EPI_ISL_434599, EPI_ISL_434600, EPI_ISL_434601, EPI_ISL_434602, EPI_ISL_434603, EPI_ISL_434604, EPI_ISL_434605, EPI_ISL_434606 | | | |
| see above | Virginia DCLS | Virginia DCLS | Virginia DCLS |
| EPI_ISL_434607, EPI_ISL_434608, EPI_ISL_434609, EPI_ISL_434610, EPI_ISL_434611, EPI_ISL_434612, EPI_ISL_434613, EPI_ISL_434614, EPI_ISL_434615 | University of Wisconsin-Madison AIDS Vaccine Research Laboratories | University of Wisconsin-Madison AIDS Vaccine Research Laboratories | Gage Moreno, Katarina Braun, et al. AIDS Vaccine Research Laboratories |
| EPI_ISL_434616, EPI_ISL_434617, EPI_ISL_434618, EPI_ISL_434619, EPI_ISL_434620, EPI_ISL_434621, EPI_ISL_434622, EPI_ISL_434623, EPI_ISL_434624, EPI_ISL_434625, EPI_ISL_434626, EPI_ISL_434627, EPI_ISL_434628, EPI_ISL_434629, EPI_ISL_434630, EPI_ISL_434631, EPI_ISL_434632, EPI_ISL_434633, EPI_ISL_434634, EPI_ISL_434635 | | | |
| see above | CHU Purpan - Laboratoire de Virologie - Institut Fédératif de Biologie | Laboratoire de virologie - École Nationale Vétérinaire de Toulouse | Guillaume Croville, Jean-Luc Guérin, Jacques Izopet |
| EPI_ISL_434636 | Lednicky Laboratory, Emerging Pathogens Institute, University of Florida | Lednicky Laboratory at Emerging Pathogens Institute, University of Florida | Elbadry,M.A., Subramaniam,K., Waltzek,T.B., Stephenson,C.J.,Gibson,J.C., Alam,M., Morris,J.G. Jr. and Lednicky,J.A. |
| EPI_ISL_434637 | Lednicky Laboratory, Emerging Pathogens Institute, University of Florida. | Lednicky Laboratory, Emerging Pathogens Institute, University of Florida. | Elbadry,M.A.; Subramaniam,K.; Waltzek,T.B.; Gibson,J.C.; Stephenson,C.J.; Morris,J.G. Jr. and Lednicky,J.A. |
| EPI_ISL_434638, EPI_ISL_434639, EPI_ISL_434640 | Johns Hopkins Hospital Department of Pathology | Johns Hopkins Hospital Department of Pathology | Peter M. Thielen, Thomas Mehoke, Shirlee Wohl, Srividya Ramakrishnan, Oluwaseun Nwulia-Falade, Amanda Ernlund, Melanie Kirsche, Paul Morris, Norah Sadowski, Nídiá Trovao, Victoria Gniazdowski, Michael Schatz, Stuart C. Ray, Winston Timp, Heba Mostafa |
| EPI_ISL_434641, EPI_ISL_434642 | Laboratoriemedicin | The Public Health Agency of Sweden | Oskar Karlsson Lindsjo, Maria Lind Karlberg, Anna-Malin Linde, Olov Svartstrom, Anna Risberg, Shaman Muradrasoli, Karin Tegmark-Wisell |
| EPI_ISL_434643 | Uppsala Narakut Aleris | The Public Health Agency of Sweden | Annika Nilsson, Oskar Karlsson Lindsjo, Maria Lind Karlberg, Anna-Malin Linde, Olov Svartstrom, Anna Risberg, Theresa Enkirch, Mia Brytting, Karin Tegmark-Wisell |
| EPI_ISL_434644 | Kungsholmsdoktorn | The Public Health Agency of Sweden | Linus Hammar, Oskar Karlsson Lindsjo, Maria Lind Karlberg, Anna-Malin Linde, Olov Svartstrom, Anna Risberg, Theresa Enkirch, Mia Brytting, Karin Tegmark-Wisell |
| EPI_ISL_434645 | Svardsjo VC | The Public Health Agency of Sweden | Tommy Janers, Oskar Karlsson Lindsjo, Maria Lind Karlberg, Anna-Malin Linde, Olov Svartstrom, Anna Risberg, Theresa Enkirch, Mia Brytting, Karin Tegmark-Wisell |
| EPI_ISL_434646 | Ulltuna Vardcentral | The Public Health Agency of Sweden | Heidi Lindback, Oskar Karlsson Lindsjo, Maria Lind Karlberg, Anna-Malin Linde, Olov Svartstrom, Anna Risberg, Theresa Enkirch, Mia Brytting, Karin Tegmark-Wisell |
| EPI_ISL_434647, EPI_ISL_434648 | Victoria Vard och Halsa | The Public Health Agency of Sweden | Sarah Henriksson, Oskar Karlsson Lindsjo, Maria Lind Karlberg, Anna-Malin Linde, Olov Svartstrom, Anna Risberg, Theresa Enkirch, Mia Brytting, Karin Tegmark-Wisell |
| EPI_ISL_434649 | Svardsjo VC | The Public Health Agency of Sweden | Tommy Janers, Oskar Karlsson Lindsjo, Maria Lind Karlberg, Anna-Malin Linde, Olov Svartstrom, Anna Risberg, Theresa Enkirch, Mia Brytting, Karin Tegmark-Wisell |
| EPI_ISL_434650 | Follinge Halsocentral | The Public Health Agency of Sweden | Kerstin Persson Moberg, Oskar Karlsson Lindsjo, Maria Lind Karlberg, Anna-Malin Linde, Olov Svartstrom, Anna Risberg, Theresa Enkirch, Mia Brytting, Karin Tegmark-Wisell |
| EPI_ISL_434651 | Krokoms Halsocentral | The Public Health Agency of Sweden | Martin Ersson, Oskar Karlsson Lindsjo, Maria Lind Karlberg, Anna-Malin Linde, Olov Svartstrom, Anna Risberg, Theresa Enkirch, Mia Brytting, Karin Tegmark-Wisell |
| EPI_ISL_434652 | Ektorps Vardcentral | The Public Health Agency of Sweden | Eva Espmark, Oskar Karlsson Lindsjo, Maria Lind Karlberg, Anna-Malin Linde, Olov Svartstrom, Anna Risberg, Theresa Enkirch, Mia Brytting, Karin Tegmark-Wisell |
| EPI_ISL_434653 | Trollbackens VC | The Public Health Agency of Sweden | Amelie Holmqvist, Oskar Karlsson Lindsjo, Maria Lind Karlberg, Anna-Malin Linde, Olov Svartstrom, Anna Risberg, Theresa Enkirch, Mia Brytting, Karin Tegmark-Wisell |
| EPI_ISL_434654 | Saroledens Familjelakare | The Public Health Agency of Sweden | Katarina Jarbur, Oskar Karlsson Lindsjo, Maria Lind Karlberg, Anna-Malin Linde, Olov Svartstrom, Anna Risberg, Theresa Enkirch, Mia Brytting, Karin Tegmark-Wisell |
| EPI_ISL_434655 | Uppsala Narakut Aleris | The Public Health Agency of Sweden | Annika Nilsson, Oskar Karlsson Lindsjo, Maria Lind Karlberg, Anna-Malin Linde, Olov Svartstrom, Anna Risberg, Theresa Enkirch, Mia Brytting, Karin Tegmark-Wisell |
| EPI_ISL_434656 | Trollbackens VC | The Public Health Agency of Sweden | Amelie Holmqvist, Oskar Karlsson Lindsjo, Maria Lind Karlberg, Anna-Malin Linde, Olov Svartstrom, Anna Risberg, Theresa Enkirch, Mia Brytting, Karin Tegmark-Wisell |
| EPI_ISL_434657 | Kristianstadkliniken | The Public Health Agency of Sweden | Mia Settergren Hammer, Oskar Karlsson Lindsjo, Maria Lind Karlberg, Anna-Malin Linde, Olov Svartstrom, Anna Risberg, Theresa Enkirch, Mia Brytting, Karin Tegmark-Wisell |
| EPI_ISL_434658 | Lundens VC | The Public Health Agency of Sweden | Marita Dagner, Oskar Karlsson Lindsjo, Maria Lind Karlberg, Anna-Malin Linde, Olov Svartstrom, Anna Risberg, Theresa Enkirch, Mia Brytting, Karin Tegmark-Wisell |
| EPI_ISL_434659 | Knivsta VC | The Public Health Agency of Sweden | Johanna Carlson, Oskar Karlsson Lindsjo, Maria Lind Karlberg, Anna-Malin Linde, Olov Svartstrom, Anna Risberg, Theresa Enkirch, Mia Brytting, Karin Tegmark-Wisell |
| EPI_ISL_434660 | Narhalsan Backa vardcentral | The Public Health Agency of Sweden | Mats Olsson, Oskar Karlsson Lindsjo, Maria Lind Karlberg, Anna-Malin Linde, Olov Svartstrom, Anna Risberg, Theresa Enkirch, Mia Brytting, Karin Tegmark-Wisell |

| | | | |
|---|---|---|---|
| EPI_ISL_434661, EPI_ISL_434662 | Omtanken Grimmered | The Public Health Agency of Sweden | Bernd Sengpiel, Oskar Karlsson Lindsjo, Maria Lind Karlberg, Anna-Malin Linde, Olov Svartstrom, Anna Risberg, Theresa Enkirch, Mia Brytting, Karin Tegmark-Wisell |
| EPI_ISL_434663 | Surbrunns VC | The Public Health Agency of Sweden | Erik Embring, Oskar Karlsson Lindsjo, Maria Lind Karlberg, Anna-Malin Linde, Olov Svartstrom, Anna Risberg, Theresa Enkirch, Mia Brytting, Karin Tegmark-Wisell |
| EPI_ISL_434664 | Omtanken Grimmered | The Public Health Agency of Sweden | Bernd Sengpiel, Oskar Karlsson Lindsjo, Maria Lind Karlberg, Anna-Malin Linde, Olov Svartstrom, Anna Risberg, Theresa Enkirch, Mia Brytting, Karin Tegmark-Wisell |
| EPI_ISL_434665 | Uppsala Narakut Aleris | The Public Health Agency of Sweden | Annika Nilsson, Oskar Karlsson Lindsjo, Maria Lind Karlberg, Anna-Malin Linde, Olov Svartstrom, Anna Risberg, Theresa Enkirch, Mia Brytting, Karin Tegmark-Wisell |
| EPI_ISL_434666 | Hornefors Halsocentral | The Public Health Agency of Sweden | Camilla Eiback, Oskar Karlsson Lindsjo, Maria Lind Karlberg, Anna-Malin Linde, Olov Svartstrom, Anna Risberg, Theresa Enkirch, Mia Brytting, Karin Tegmark-Wisell |
| EPI_ISL_434667 | Ulltuna Vardcentral | The Public Health Agency of Sweden | Heidi Lindback, Oskar Karlsson Lindsjo, Maria Lind Karlberg, Anna-Malin Linde, Olov Svartstrom, Anna Risberg, Theresa Enkirch, Mia Brytting, Karin Tegmark-Wisell |
| EPI_ISL_434668 | Kungsors VC | The Public Health Agency of Sweden | Jessica Karlsson, Oskar Karlsson Lindsjo, Maria Lind Karlberg, Anna-Malin Linde, Olov Svartstrom, Anna Risberg, Theresa Enkirch, Mia Brytting, Karin Tegmark-Wisell |
| EPI_ISL_434669 | Lakargruppen | The Public Health Agency of Sweden | Boris Klanger, Oskar Karlsson Lindsjo, Maria Lind Karlberg, Anna-Malin Linde, Olov Svartstrom, Anna Risberg, Theresa Enkirch, Mia Brytting, Karin Tegmark-Wisell |
| EPI_ISL_434670 | Narhalsan Molnlycke, Barn och ungdomsmedicin | The Public Health Agency of Sweden | Mats Reimer, Oskar Karlsson Lindsjo, Maria Lind Karlberg, Anna-Malin Linde, Olov Svartstrom, Anna Risberg, Theresa Enkirch, Mia Brytting, Karin Tegmark-Wisell |
| EPI_ISL_434671 | Narhalsan Sjobo vardcentral | The Public Health Agency of Sweden | Lovisa Hjerten, Oskar Karlsson Lindsjo, Maria Lind Karlberg, Anna-Malin Linde, Olov Svartstrom, Anna Risberg, Theresa Enkirch, Mia Brytting, Karin Tegmark-Wisell |
| EPI_ISL_434672 | Surbrunns VC | The Public Health Agency of Sweden | Erik Embring, Oskar Karlsson Lindsjo, Maria Lind Karlberg, Anna-Malin Linde, Olov Svartstrom, Anna Risberg, Theresa Enkirch, Mia Brytting, Karin Tegmark-Wisell |
| EPI_ISL_434673 | Omtanken Grimmered | The Public Health Agency of Sweden | Bernd Sengpiel, Oskar Karlsson Lindsjo, Maria Lind Karlberg, Anna-Malin Linde, Olov Svartstrom, Anna Risberg, Theresa Enkirch, Mia Brytting, Karin Tegmark-Wisell |
| EPI_ISL_434674 | Narhalsan Backa vardcentral | The Public Health Agency of Sweden | Mats Olsson, Oskar Karlsson Lindsjo, Maria Lind Karlberg, Anna-Malin Linde, Olov Svartstrom, Anna Risberg, Theresa Enkirch, Mia Brytting, Karin Tegmark-Wisell |
| EPI_ISL_434675, EPI_ISL_434676 | Surbrunns VC | The Public Health Agency of Sweden | Erik Embring, Oskar Karlsson Lindsjo, Maria Lind Karlberg, Anna-Malin Linde, Olov Svartstrom, Anna Risberg, Theresa Enkirch, Mia Brytting, Karin Tegmark-Wisell |
| EPI_ISL_434677 | Lednicky Laboratory at Emerging Pathogens Institute | Lednicky Laboratory at Emerging Pathogens Institute | Shankar,S.N., Wu,C.-Y., Clugston,J.R., Elbadry,M.A., Morris,J.G. Jr. and Lednicky,J.A. |
| EPI_ISL_434678, EPI_ISL_434679, EPI_ISL_434680, EPI_ISL_434681 | Viral Respiratory Lab, National Institute for Biomedical Research (INRB) | Pathogen Sequencing Lab, National Institute for Biomedical Research (INRB) | Placide Mbala-Kingebeni; Edith Nkwembe; Eddy Kinganda-Lusamaki; Amuri Aziza; Francisca Muyembe Mawete; Catherine Pratt; Matthias Pauthner; Josh Quick; Allison Black; James Hadfield; Trevor Bedford; Ian Goodfellow; Andrew Rambaut; Nick Loman; Kristian Andersen; Michael Wiley; Steve Ahuka-Mundeke; Jean-Jacques Muyembe Tamfum |
| EPI_ISL_434682, EPI_ISL_434683, EPI_ISL_434684, EPI_ISL_434685, EPI_ISL_434686, EPI_ISL_434687, EPI_ISL_434688, EPI_ISL_434689, EPI_ISL_434690, EPI_ISL_434691 | Johns Hopkins Hospital Department of Pathology | Johns Hopkins Hospital Department of Pathology | Peter M. Thielen, Thomas Mehoke, Shirlee Wohl, Srividya Ramakrishnan, Melanie Kirsche, Amanda Ernlund, Oluwaseun Falade-Nwulia, Timothy Gilpatrick, Paul Morris, Norah Sadowski, Nídiá Trovao, Victoria Gniazdowski, Michael Schatz, Stuart C. Ray, Winston Timp, Heba Mostafa |
| EPI_ISL_434692, EPI_ISL_434693, EPI_ISL_434694 | Bamrasnaradura hospital | National Institute of Health. Department of medical Sciences, Ministry of Public Health, Thailand | Pilailuk,Okada; Siripaporn,Phuygun; Thanutsapa,Thanadachakul; Sittiporn,Parnmen;Warawan,Wongboot; Sunthareeya,Waicharoen; Malinee,Chittaganpitch |
| EPI_ISL_434695 | unknown | National Institute of Health. Department of medical Sciences, Ministry of Public Health, Thailand | Pilailuk,Okada; Siripaporn,Phuygun; Thanutsapa,Thanadachakul; Sittiporn,Parnmen;Warawan,Wongboot; Sunthareeya,Waicharoen; Malinee,Chittaganpitch |
| EPI_ISL_434696 | Bamrasnaradura hospital | National Institute of Health. Department of medical Sciences, Ministry of Public Health, Thailand | Pilailuk,Okada; Siripaporn,Phuygun; Thanutsapa,Thanadachakul; Sittiporn,Parnmen;Warawan,Wongboot; Sunthareeya,Waicharoen; Malinee,Chittaganpitch |
| EPI_ISL_434697 | unknown | National Institute of Health. Department of medical Sciences, Ministry of Public Health, Thailand | Pilailuk,Okada; Siripaporn,Phuygun; Thanutsapa,Thanadachakul; Sittiporn,Parnmen;Warawan,Wongboot; Sunthareeya,Waicharoen; Malinee,Chittaganpitch |
| EPI_ISL_434698, EPI_ISL_434699 | Praram 9 Hospital | National Institute of Health. Department of medical Sciences, Ministry of Public Health, Thailand | Pilailuk,Okada; Siripaporn,Phuygun; Thanutsapa,Thanadachakul; Sittiporn,Parnmen;Warawan,Wongboot; Sunthareeya,Waicharoen; Malinee,Chittaganpitch |
| EPI_ISL_434700 | Ramkhamhaeng Hospital | National Institute of Health. Department of medical Sciences, Ministry of Public Health, Thailand | Pilailuk,Okada; Siripaporn,Phuygun; Thanutsapa,Thanadachakul; Sittiporn,Parnmen;Warawan,Wongboot; Sunthareeya,Waicharoen; Malinee,Chittaganpitch |
| EPI_ISL_434701 | Panyanunthaphikkhu Chonprathan Medical Center (PCMC) | National Institute of Health. Department of medical Sciences, Ministry of Public Health, Thailand | Pilailuk,Okada; Siripaporn,Phuygun; Thanutsapa,Thanadachakul; Sittiporn,Parnmen;Warawan,Wongboot; Sunthareeya,Waicharoen; Malinee,Chittaganpitch |
| EPI_ISL_434702, EPI_ISL_434703, EPI_ISL_434704, EPI_ISL_434705, EPI_ISL_434706 | Praram 9 Hospital | National Institute of Health. Department of medical Sciences, Ministry of Public Health, Thailand | Pilailuk,Okada; Siripaporn,Phuygun; Thanutsapa,Thanadachakul; Sittiporn,Parnmen;Warawan,Wongboot; Sunthareeya,Waicharoen; Malinee,Chittaganpitch |
| EPI_ISL_434707 | Thammasat University Hospital | National Institute of Health. Department of medical Sciences, Ministry of Public Health, Thailand | Pilailuk,Okada; Siripaporn,Phuygun; Thanutsapa,Thanadachakul; Sittiporn,Parnmen;Warawan,Wongboot; Sunthareeya,Waicharoen; Malinee,Chittaganpitch |
| EPI_ISL_434708, EPI_ISL_434709 | unknown | National Institute of Health. Department of medical Sciences, Ministry of Public Health, Thailand | Pilailuk,Okada; Siripaporn,Phuygun; Thanutsapa,Thanadachakul; Sittiporn,Parnmen;Warawan,Wongboot; Sunthareeya,Waicharoen; Malinee,Chittaganpitch |
| EPI_ISL_434710, EPI_ISL_434711 | Viral Respiratory Lab, National Institute for Biomedical Research (INRB) | Pathogen Sequencing Lab, National Institute for Biomedical Research (INRB) | Placide Mbala-Kingebeni, Edith Nkwembe, Eddy Kinganda-Lusamaki, Adrienne Amuri Aziza, Francisca Muyembe Mawete, Catherine Pratt, Matthias Pauthner, Josh Quick, Allison Black, James Hadfield, Trevor Bedford, Ian Goodfellow, Andrew Rambaut, Nick Loman, Kristian Andersen, Michael Wiley, Steve Ahuka-Mundeke, Jean-Jacques Muyembe Tamfum |
| EPI_ISL_434712, EPI_ISL_434713, EPI_ISL_434714, EPI_ISL_434715, EPI_ISL_434716, EPI_ISL_434717, EPI_ISL_434718, EPI_ISL_434719, EPI_ISL_434720, EPI_ISL_434721, EPI_ISL_434722, EPI_ISL_434723, EPI_ISL_434724, EPI_ISL_434725, EPI_ISL_434726, EPI_ISL_434727, EPI_ISL_434728, EPI_ISL_434729, EPI_ISL_434730, EPI_ISL_434731, EPI_ISL_434732, EPI_ISL_434733, EPI_ISL_434734, EPI_ISL_434735, EPI_ISL_434736, EPI_ISL_434737, EPI_ISL_434738, EPI_ISL_434739, EPI_ISL_434740, EPI_ISL_434741, EPI_ISL_434742, EPI_ISL_434743, EPI_ISL_434744, EPI_ISL_434745, EPI_ISL_434746, EPI_ISL_434747, EPI_ISL_434748, EPI_ISL_434749, EPI_ISL_434750, EPI_ISL_434751, EPI_ISL_434752, EPI_ISL_434753, EPI_ISL_434754, EPI_ISL_434755, EPI_ISL_434756, EPI_ISL_434757, EPI_ISL_434758, EPI_ISL_434759, EPI_ISL_434760, EPI_ISL_434761, EPI_ISL_434762, EPI_ISL_434763, EPI_ISL_434764, EPI_ISL_434765, EPI_ISL_434766, EPI_ISL_434767, EPI_ISL_434768, EPI_ISL_434769, EPI_ISL_434770, EPI_ISL_434771, EPI_ISL_434772, EPI_ISL_434773, EPI_ISL_434774, EPI_ISL_434775, EPI_ISL_434776, EPI_ISL_434777, EPI_ISL_434778, EPI_ISL_434779, EPI_ISL_434780, EPI_ISL_434781, EPI_ISL_434782, EPI_ISL_434783, EPI_ISL_434784, EPI_ISL_434785, EPI_ISL_434786, EPI_ISL_434787, EPI_ISL_434788, EPI_ISL_434789, EPI_ISL_434790, EPI_ISL_434791, EPI_ISL_434792, EPI_ISL_434793, EPI_ISL_434794, EPI_ISL_434795, EPI_ISL_434796, EPI_ISL_434797, EPI_ISL_434798, EPI_ISL_434799, EPI_ISL_434800, EPI_ISL_434801, EPI_ISL_434802, EPI_ISL_434803, EPI_ISL_434804, EPI_ISL_434805, EPI_ISL_434806, EPI_ISL_434807, EPI_ISL_434808, EPI_ISL_434809, EPI_ISL_434810, EPI_ISL_434811, EPI_ISL_434812, EPI_ISL_434813, EPI_ISL_434814, EPI_ISL_434815, EPI_ISL_434816, EPI_ISL_434817, EPI_ISL_434818, EPI_ISL_434819, EPI_ISL_434820, EPI_ISL_434821, EPI_ISL_434822, EPI_ISL_434823, EPI_ISL_434824, EPI_ISL_434825, EPI_ISL_434826, EPI_ISL_434827, EPI_ISL_434828, EPI_ISL_434829, EPI_ISL_434830, EPI_ISL_434831, EPI_ISL_434832, EPI_ISL_434833, EPI_ISL_434834, EPI_ISL_434835, EPI_ISL_434836, EPI_ISL_434837, EPI_ISL_434838, EPI_ISL_434839, EPI_ISL_434840, EPI_ISL_434841, EPI_ISL_434842, EPI_ISL_434843, EPI_ISL_434844, EPI_ISL_434845, EPI_ISL_434846, EPI_ISL_434847, EPI_ISL_434848, EPI_ISL_434849, EPI_ISL_434850, EPI_ISL_434851, EPI_ISL_434852, EPI_ISL_434853, EPI_ISL_434854, EPI_ISL_434855, EPI_ISL_434856, EPI_ISL_434857, EPI_ISL_434858, EPI_ISL_434859, EPI_ISL_434860, EPI_ISL_434861, EPI_ISL_434862, EPI_ISL_434863, EPI_ISL_434864, EPI_ISL_434865, EPI_ISL_434866, EPI_ISL_434867, EPI_ISL_434868, EPI_ISL_434869, EPI_ISL_434870, EPI_ISL_434871, EPI_ISL_434872, EPI_ISL_434873, EPI_ISL_434874, EPI_ISL_434875, EPI_ISL_434876, EPI_ISL_434877, EPI_ISL_434878, EPI_ISL_434879, EPI_ISL_434880, EPI_ISL_434881, EPI_ISL_434882, EPI_ISL_434883, EPI_ISL_434884, EPI_ISL_434885, EPI_ISL_434886, EPI_ISL_434887, EPI_ISL_434888, EPI_ISL_434889, EPI_ISL_434890, EPI_ISL_434891, EPI_ISL_434892, EPI_ISL_434893, EPI_ISL_434894, EPI_ISL_434895, EPI_ISL_434896, EPI_ISL_434897, EPI_ISL_434898, EPI_ISL_434899, EPI_ISL_434900, EPI_ISL_434901, EPI_ISL_434902, EPI_ISL_434903, EPI_ISL_434904, EPI_ISL_434905, EPI_ISL_434906, EPI_ISL_434907, EPI_ISL_434908, EPI_ISL_434909, EPI_ISL_434910, EPI_ISL_434911, EPI_ISL_434912, EPI_ISL_434913, EPI_ISL_434914, EPI_ISL_434915, EPI_ISL_434916, EPI_ISL_434917, EPI_ISL_434918, EPI_ISL_434919, EPI_ISL_434920, EPI_ISL_434921, EPI_ISL_434922, EPI_ISL_434923, EPI_ISL_434924, EPI_ISL_434925, EPI_ISL_434926, EPI_ISL_434927, EPI_ISL_434928, EPI_ISL_434929, EPI_ISL_434930, EPI_ISL_434931, EPI_ISL_434932, EPI_ISL_434933, EPI_ISL_434934, EPI_ISL_434935, EPI_ISL_434936, EPI_ISL_434937, EPI_ISL_434938, EPI_ISL_434939, EPI_ISL_434940, EPI_ISL_434941, EPI_ISL_434942, EPI_ISL_434943, EPI_ISL_434944, EPI_ISL_434945, EPI_ISL_434946, EPI_ISL_434947, EPI_ISL_434948, EPI_ISL_434949, EPI_ISL_434950, EPI_ISL_434951, EPI_ISL_434952, EPI_ISL_434953, EPI_ISL_434954, EPI_ISL_434955, EPI_ISL_434956, EPI_ISL_434957, EPI_ISL_434958, EPI_ISL_434959, EPI_ISL_434960, EPI_ISL_434961, EPI_ISL_434962, EPI_ISL_434963, EPI_ISL_434964, EPI_ISL_434965, EPI_ISL_434966, EPI_ISL_434967, EPI_ISL_434968, EPI_ISL_434969, EPI_ISL_434970, EPI_ISL_434971, EPI_ISL_434972, EPI_ISL_434973, EPI_ISL_434974, EPI_ISL_434975, EPI_ISL_434976, EPI_ISL_434977, EPI_ISL_434978, EPI_ISL_434979, EPI_ISL_434980, EPI_ISL_434981, EPI_ISL_434982, EPI_ISL_434983, EPI_ISL_434984, EPI_ISL_434985, EPI_ISL_434986, EPI_ISL_434987, EPI_ISL_434988, EPI_ISL_434989, EPI_ISL_434990, EPI_ISL_434991, EPI_ISL_434992, EPI_ISL_434993, EPI_ISL_434994, EPI_ISL_434995, EPI_ISL_434996, EPI_ISL_434997, EPI_ISL_434998, EPI_ISL_434999, EPI_ISL_435000, EPI_ISL_435001, EPI_ISL_435002, EPI_ISL_435003, EPI_ISL_435004, EPI_ISL_435005, EPI_ISL_435006, EPI_ISL_435007, EPI_ISL_435008, EPI_ISL_435009, EPI_ISL_435010, EPI_ISL_435011, EPI_ISL_435012, EPI_ISL_435013, EPI_ISL_435014, EPI_ISL_435015, EPI_ISL_435016, EPI_ISL_435017, EPI_ISL_435018, EPI_ISL_435019, EPI_ISL_435020, EPI_ISL_435021, EPI_ISL_435022, EPI_ISL_435023, EPI_ISL_435024, EPI_ISL_435025, EPI_ISL_435026, EPI_ISL_435027, EPI_ISL_435028, EPI_ISL_435029, EPI_ISL_435030, EPI_ISL_435031 | | | |
| see above | Houston Methodist Hospital | Houston Methodist Hospital | S. Wesley Long, Randall J. Olsen, Paul A. Christensen, David W. Bernard, James J. Davis, Maulik Shukla, Marcus Nguyen, Matthew Ojeda Saavedra, Concepcion C. Cantu, Prasanti Yerramilli, Layne Pruitt, Sishir Subedi, Heather Hendrickson, Ghazaleh Eskandari, Muthiah Kumaraswami, Jason S. McLellan, Hakon Jonsson, Kari Stefansson, and James M. Musser |
| EPI_ISL_435032, EPI_ISL_435033 | Viral Respiratory Lab, National Institute for Biomedical Research (INRB) | Pathogen Sequencing Lab, National Institute for Biomedical Research (INRB) | Placide Mbala-Kingebeni, Edith Nkwembe, Eddy Kinganda-Lusamaki, Adrienne Amuri Aziza, Francisca Muyembe Mawete, Catherine Pratt, Matthias Pauthner, Josh Quick, Allison Black, James Hadfield, Trevor Bedford, Ian Goodfellow, Andrew Rambaut, Nick Loman, Kristian Andersen, Michael Wiley, Steve Ahuka-Mundeke, Jean-Jacques Muyembe Tamfum |
| EPI_ISL_435034 | LSUHS Emerging Viral Threat Laboratory | Microbial Genome Sequencing Center | Jeremy P. Kamil, John A. Vanchiere, Rona S. Scott, Camille F. Abshire, Abida Siddiqa, Byeong-Jae Lee, Chan-ki Min, Md Maksudul Alam, Monica Gestal-Cartele, Edna Ondari, Adam Greer, Malgorzata Bienkowska-Haba, Katarzyna Zwolinska, Michelle M. Arnold, Jason M. Bodily, Andrew D. Yurochko, Paul M. Weinberger, Christopher G. Kevil, Martin J. Sapp, Daniel J. Snyder, Vaughn S. Cooper |
| EPI_ISL_435035 | LSUHS Emerging Viral Threat Laboratory | Microbial Genome Sequencing Center | Jeremy P. Kamil, John A. Vanchiere, Rona S. Scott, Camille F. Abshire, Abida Siddiqa, Byeong-Jae Lee, Chan-ki Min, Md Maksudul Alam, Monica Gestal-Cartele, Edna Ondari, Adam Greer, Malgorzata Bienkowska-Haba, Katarzyna Zwolinska, Jason M. Bodily, Andrew D. Yurochko, Paul M. Weinberger, Christopher G. Kevil, Martin J. Sapp, Daniel J. Snyder, Vaughn S. Cooper |
| EPI_ISL_435036 | LSUHS Emerging Viral Threat Laboratory | Microbial Genome Sequencing Center | Jeremy P. Kamil, John A. Vanchiere, Rona S. Scott, Camille F. Abshire, Abida Siddiqa, Byeong-Jae Lee, Chan-ki Min, Md Maksudul Alam, Monica Gestal-Cartele, Edna Ondari, Adam Greer, Malgorzata Bienkowska-Haba, Katarzyna Zwolinska, Jason M. Bodily, Andrew D. Yurochko, Paul M. Weinberger, Christopher G. Kevil, Martin J. Sapp, Daniel J. Snyder, Vaughn S. Cooper |
| EPI_ISL_435037, EPI_ISL_435038, EPI_ISL_435039, EPI_ISL_435040, EPI_ISL_435041, EPI_ISL_435042, EPI_ISL_435043, EPI_ISL_435044 | LSUHS Emerging Viral Threat Laboratory | Microbial Genome Sequencing Center | Jeremy P. Kamil, John A. Vanchiere, Rona S. Scott, Camille F. Abshire, Abida Siddiqa, Byeong-Jae Lee, Chan-ki Min, Md Maksudul Alam, Monica Gestal-Cartele, Edna Ondari, Adam Greer, Malgorzata Bienkowska-Haba, Katarzyna Zwolinska, Jason M. Bodily, Andrew D. Yurochko, Paul M. Weinberger, Christopher G. Kevil, Martin J. Sapp, Daniel J. Snyder, Vaughn S. Cooper |
| EPI_ISL_435045, EPI_ISL_435046, EPI_ISL_435047, EPI_ISL_435048 | Laboratory of Applied Genetics | RSE "National Center for Biotechnology" | Alexandr Shevtsov, Ilyas Akhmetollayev, Viktoriya Lutsay, Asylulan Amirgazin, Ruslan Kalendar, Yerlan Ramanculov |
| EPI_ISL_435049 | B.J. Medical College and Civil hospital | Gujarat Biotechnology Research Centre | Pinal Trivedi, Maharshi Pandya, Amit Kanani, Akanksha Verma, Nitin Savaliya, Raghawendra Kumar, Dinesh Kumar, Zuber Saiyed, Dipa Kinariwala, Disha Patel, Binita Aring, Geeta Vaghela, Sonia Barve, Bhavesh Modi, Kairavi Joshi, Gaurishankar Shrimali, Nidhi Sood, Pranay Shah, R D Dixit, Snehal Bagatharia, Kamlesh J Upadhyay, Ramesh Pandit, Tejas Shah, Ankit Hinsu, Pritesh Sabara, Apurvasinh Puvar, Janvi Raval, Monika Gandhi, Neha Rajpara, Chaitanya Joshi, Madhvi Joshi |
| EPI_ISL_435050 | B.J. Medical College and Civil hospital | Gujarat Biotechnology Research Centre | Ankit Hinsu, Pritesh Sabara, Apurvasinh Puvar, Janvi Raval, Monika Gandhi, Pinal Trivedi, Maharshi Pandya, Amit Kanani, Akanksha Verma, Nitin Savaliya, Raghawendra Kumar, Dinesh Kumar, Zuber Saiyed, Dipa Kinariwala, Disha Patel, Binita Aring, Geeta Vaghela, Sonia Barve, Bhavesh Modi, Kairavi Joshi, Gaurishankar Shrimali, Nidhi Sood, Pranay Shah, R D Dixit, Snehal Bagatharia, Kamlesh J Upadhyay, Ramesh Pandit, Tejas Shah, Dipeshwari Shewale, Chaitanya Joshi, Madhvi Joshi |
| EPI_ISL_435051 | B.J. Medical College and Civil hospital | Gujarat Biotechnology Research Centre | Pritesh Sabara, Apurvasinh Puvar, Janvi Raval, Monika Gandhi, Pinal Trivedi, Maharshi Pandya, Amit Kanani, Akanksha Verma, Nitin Savaliya, Raghawendra Kumar, Dinesh Kumar, Zuber Saiyed, Dipa Kinariwala, Disha Patel, Binita Aring, Geeta Vaghela, Sonia Barve, Bhavesh Modi, Kairavi Joshi, Gaurishankar Shrimali, Nidhi Sood, Pranay Shah, R D Dixit, Snehal Bagatharia, Kamlesh J Upadhyay, Ramesh Pandit, Tejas Shah, Ankit Hinsu, Vasudha Sharma, Chaitanya Joshi, Madhvi Joshi |
| EPI_ISL_435052 | B.J. Medical College and Civil hospital | Gujarat Biotechnology Research Centre | Apurvasinh Puvar, Janvi Raval, Monika Gandhi, Pinal Trivedi, Maharshi Pandya, Amit Kanani, Akanksha Verma, Nitin Savaliya, Raghawendra Kumar, Dinesh Kumar, Zuber Saiyed, Dipa Kinariwala, Disha Patel, Binita Aring, Geeta Vaghela, Sonia Barve, Bhavesh Modi, Kairavi Joshi, Gaurishankar Shrimali, Nidhi Sood, Pranay Shah, R D Dixit, Snehal Bagatharia, Kamlesh J Upadhyay, Ramesh Pandit, Tejas Shah, Ankit Hinsu, Pritesh Sabara, Pooja P Doshi, Chaitanya Joshi, Madhvi Joshi |
| EPI_ISL_435053 | B.J. Medical College and Civil hospital | Gujarat Biotechnology Research Centre | Janvi Raval, Monika Gandhi, Pinal Trivedi, Maharshi Pandya, Amit Kanani, Akanksha Verma, Nitin Savaliya, Raghawendra Kumar, Dinesh Kumar, Zuber Saiyed, Dipa Kinariwala, Disha Patel, Binita Aring, Geeta Vaghela, Sonia Barve, Bhavesh Modi, Kairavi Joshi, Gaurishankar Shrimali, Nidhi Sood, Pranay Shah, R D Dixit, Snehal Bagatharia, Kamlesh J Upadhyay, Ramesh Pandit, Tejas Shah, Ankit Hinsu, Pritesh Sabara, Apurvasinh Puvar, Nidhi Patel, Chaitanya Joshi, Madhvi Joshi |
| EPI_ISL_435054 | B.J. Medical College and Civil hospital | Gujarat Biotechnology Research Centre | Monika Gandhi, Pinal Trivedi, Maharshi Pandya, Amit Kanani, Akanksha Verma, Nitin Savaliya, Raghawendra Kumar, Dinesh Kumar, Zuber Saiyed, Dipa Kinariwala, Disha Patel, Binita Aring, Geeta Vaghela, Sonia Barve, Bhavesh Modi, Kairavi Joshi, Gaurishankar Shrimali, Nidhi Sood, Pranay Shah, R D Dixit, Snehal Bagatharia, Kamlesh J Upadhyay, Ramesh Pandit, Tejas Shah, Ankit Hinsu, Pritesh Sabara, Apurvasinh Puvar, Janvi Raval, Priti Pandita, Chaitanya Joshi, Madhvi Joshi |
| EPI_ISL_435055 | Gujarat Biotechnology Research Centre | Gujarat Biotechnology Research Centre | Tejas Shah, Ankit Hinsu, Pritesh Sabara, Apurvasinh Puvar, Janvi Raval, Monika Gandhi, Pinal Trivedi, Maharshi Pandya, Amit Kanani, Akanksha Verma, Nitin Savaliya, Raghawendra Kumar, Dinesh Kumar, Zuber |

| | | | |
|---|---|---|---|
| | | | Saiyed, Dipa Kinariwala, Disha Patel, Binita Aring, Geeta Vaghela, Sonia Barve, Bhavesh Modi, Kairavi Joshi, Gaurishankar Shrimali, Nidhi Sood, Pranay Shah, R D Dixit, Snehal Bagatharia, Kamlesh J Upadhyay, Ramesh Pandit, Anjali Rajwal, Chaitanya Joshi, Madhvi Joshi |
| EPI_ISL_435056 | Gujarat Biotechnology Research Centre | Gujarat Biotechnology Research Centre | Maharshi Pandya, Amit Kanani, Akanksha Verma, Nitin Savaliya, Raghawendra Kumar, Dinesh Kumar, Zuber Saiyed, Dipa Kinariwala, Disha Patel, Binita Aring, Geeta Vaghela, Sonia Barve, Bhavesh Modi, Kairavi Joshi, Gaurishankar Shrimali, Nidhi Sood, Pranay Shah, R D Dixit, Snehal Bagatharia, Kamlesh J Upadhyay, Ramesh Pandit, Tejas Shah, Ankit Hinsu, Pritesh Sabara, Apurvasinh Puvar, Janvi Raval, Monika Gandhi, Pinal Trivedi, Afzal Ansari, Chaitanya Joshi, Madhvi Joshi |
| EPI_ISL_435057 | T.C. Sağlık Bakanlığı Adıyaman İl Sağlık Müdürlüğü Adıyaman Eğitim Ve Araştırma Hastanesi | VETAL Animal Health Products Company, BSL3+ Production Laboratuary, Turkey | Fatma Nilay Tutak, Haluk Uluca, Fethiye Sevimli, O. Ugur Sezerman |
| EPI_ISL_435058, EPI_ISL_435059 | National Institute for Communicable Diseases of the National Health Laboratory Service | National Institute for Communicable Diseases of the National Health Laboratory Service | Allam M, Kwenda S, van Heusden P, Khumalo Z, Mohale T, Subramoney K, von Gottberg, A, Ismail A, Bhiman JN |
| EPI_ISL_435060, EPI_ISL_435061, EPI_ISL_435062, EPI_ISL_435063, EPI_ISL_435064, EPI_ISL_435065, EPI_ISL_435066, EPI_ISL_435067, EPI_ISL_435068, EPI_ISL_435069, EPI_ISL_435070, EPI_ISL_435071, EPI_ISL_435072, EPI_ISL_435073, EPI_ISL_435074, EPI_ISL_435075, EPI_ISL_435076, EPI_ISL_435077, EPI_ISL_435078, EPI_ISL_435079, EPI_ISL_435080, EPI_ISL_435081, EPI_ISL_435082, EPI_ISL_435083, EPI_ISL_435084, EPI_ISL_435085, EPI_ISL_435086, EPI_ISL_435087, EPI_ISL_435088, EPI_ISL_435089, EPI_ISL_435090, EPI_ISL_435091, EPI_ISL_435092, EPI_ISL_435093, EPI_ISL_435094, EPI_ISL_435095, EPI_ISL_435096, EPI_ISL_435097, EPI_ISL_435098, EPI_ISL_435099, EPI_ISL_435100, EPI_ISL_435101, EPI_ISL_435102, EPI_ISL_435103, EPI_ISL_435104, EPI_ISL_435105, EPI_ISL_435106, EPI_ISL_435107, EPI_ISL_435108, EPI_ISL_435109, EPI_ISL_435110, EPI_ISL_435111, EPI_ISL_435112 | | | |
| see above | National Centre for Disease control (NCDC), CSIR-Institute of Genomics and Integrative Biology (CSIR-IGIB) | NCDC/CSIR-IGIB | Pramod Kumar, Rajesh Pandey, Pooja Sharma, Mahesh Dhar, Vivekanand A, Bharathram Uppili, Himanshu Vashisht, Saruchi Wadhwa, Nishu Tyagi, Uma Sharma, Priyanka Singh, Hemlata Lall, Meena Datta, Poonam Gupta, Nidhi Saini, Aarti Tewari, Bibhash Nandi, Dhirendra Kumar, Satyabrata Bag, Varun Jaiswal, Hema Gogia, Preeti Madan, Simrita Singh, Prateek Singh, Debasis Dash, Mitali Mukerji, Manju Bala, Sandhya Kabra, Sujeet Singh, Mohammed Faruq, Anurag Agrawal, Partha Rakshit |
| EPI_ISL_435113, EPI_ISL_435114, EPI_ISL_435116, EPI_ISL_435117, EPI_ISL_435118 | Viral Respiratory Lab, National Institute for Biomedical Research (INRB) | Pathogen Sequencing Lab, National Institute for Biomedical Research (INRB) | Placide Mbala-Kingebeni, Edith Nkwembe, Eddy Kinganda-Lusamaki, Adrienne Amuri Aziza, Francisca Muyembe Mawete, Catherine Pratt, Matthias Pauthner, Josh Quick, Allison Black, James Hadfield, Trevor Bedford, Ian Goodfellow, Andrew Rambaut, Nick Loman, Kristian Andersen, Michael Wiley, Steve Ahuka-Mundeke, Jean-Jacques Muyembe Tamfum |
| EPI_ISL_435119 | Mohammed Bin Rashid University of Medicine and Health Sciences | Al Jalila Children's Hospital | Ahmad Abou Tayoun, Tom Loney, Hamda Khansaheb, Sathishkumar Ramaswamy, Divinlal Harilal, Zulfa Omar Deesi, Rupa Murthy Varghese, Hanan Al Suwaidi, Abdulmajeed Alkhaja, Mohammed Uddin, Rifat Hamoudi, Rabih Halwani, Abiola Catherine Senok, Qutayba Hamid, Norbert Nowotny, Alawi Alsheikh-Ali |
| EPI_ISL_435120, EPI_ISL_435121, EPI_ISL_435122, EPI_ISL_435123, EPI_ISL_435124, EPI_ISL_435125, EPI_ISL_435126, EPI_ISL_435127, EPI_ISL_435128, EPI_ISL_435129, EPI_ISL_435130, EPI_ISL_435131, EPI_ISL_435132, EPI_ISL_435133, EPI_ISL_435134, EPI_ISL_435135, EPI_ISL_435136, EPI_ISL_435137, EPI_ISL_435138, EPI_ISL_435139, EPI_ISL_435140, EPI_ISL_435141, EPI_ISL_435142, EPI_ISL_435143 | | | |
| see above | Mohammed Bin Rashid University of Medicine and Health Sciences | Al Jalila Genomics Center | Ahmad Abou Tayoun, Tom Loney, Hamda Khansaheb, Sathishkumar Ramaswamy, Divinlal Harilal, Zulfa Omar Deesi, Rupa Murthy Varghese, Hanan Al Suwaidi, Abdulmajeed Alkhaja, Mohammed Uddin, Rifat Hamoudi, Rabih Halwani, Abiola Catherine Senok, Qutayba Hamid, Norbert Nowotny, Alawi Alsheikh-Ali |
| EPI_ISL_435144 | Hospital Universitario La Paz | Hospital Universitario 12 de Octubre | Elias Dahdouh, Sara González, Raúl Recio, Fernando Lázaro, Esther Viedma, Natalia Stella, Julio García, Juan Carlos Galán, Rafael Cantón, Mª Dolores Folgueira, Rafael Delgado, Jesús Mingorance |
| EPI_ISL_435145 | Ospedale Civile Giuseppe Mazzini | Istituto Zooprofilattico Sperimentale dell'Abruzzo e Molise "G.Caporale" | Lorusso A, Marcacci M, Di Domenico M, Ancora M, Curini V, Mangone I, Rinaldi A, Di Pasquale A, Cammà C, Puglia I, Savini G |
| EPI_ISL_435146, EPI_ISL_435147 | Villa Serena del Dr. Leonardo Petruzzi | Istituto Zooprofilattico Sperimentale dell'Abruzzo e Molise "G.Caporale" | Lorusso A, Marcacci M, Di Domenico M, Ancora M, Curini V, Mangone I, Rinaldi A, Di Pasquale A, Cammà C, Puglia I, Savini G |
| EPI_ISL_435148 | Ospedale SS Annunziata | Istituto Zooprofilattico Sperimentale dell'Abruzzo e Molise "G.Caporale" | Lorusso A, Marcacci M, Di Domenico M, Ancora M, Curini V, Mangone I, Rinaldi A, Di Pasquale A, Cammà C, Puglia I, Savini G |
| EPI_ISL_435149 | SERVIZIO DI IGIENE E SANITÀ PUBBLICA ASL Teramo | Istituto Zooprofilattico Sperimentale dell'Abruzzo e Molise "G.Caporale" | Lorusso A, Marcacci M, Di Domenico M, Ancora M, Curini V, Mangone I, Rinaldi A, Di Pasquale A, Cammà C, Puglia I, Savini G |
| EPI_ISL_435150, EPI_ISL_435151 | Ospedale SS Annunziata | Istituto Zooprofilattico Sperimentale dell'Abruzzo e Molise "G.Caporale" | Lorusso A, Marcacci M, Di Domenico M, Ancora M, Curini V, Mangone I, Rinaldi A, Di Pasquale A, Cammà C, Puglia I, Savini G |
| EPI_ISL_435152 | Servizio di Igiene, Epidemiologia e Sanità Pubblica (SIESP) Avezzano | Istituto Zooprofilattico Sperimentale dell'Abruzzo e Molise "G.Caporale" | Lorusso A, Marcacci M, Di Domenico M, Ancora M, Curini V, Mangone I, Rinaldi A, Di Pasquale A, Cammà C, Puglia I, Savini G |
| EPI_ISL_435153, EPI_ISL_435154, EPI_ISL_435155 | SERVIZIO DI IGIENE E SANITÀ PUBBLICA ASL Teramo | Istituto Zooprofilattico Sperimentale dell'Abruzzo e Molise "G.Caporale" | Lorusso A, Marcacci M, Di Domenico M, Ancora M, Curini V, Mangone I, Rinaldi A, Di Pasquale A, Cammà C, Puglia I, Savini G |
| EPI_ISL_435156, EPI_ISL_435157, EPI_ISL_435158, EPI_ISL_435159, EPI_ISL_435160, EPI_ISL_435161, EPI_ISL_435162, EPI_ISL_435163, EPI_ISL_435164, EPI_ISL_435165, EPI_ISL_435166, EPI_ISL_435167, EPI_ISL_435168 | | | |
| see above | Viral Respiratory Lab, National Institute for Biomedical Research (INRB) | Pathogen Sequencing Lab, National Institute for Biomedical Research (INRB) | Placide Mbala-Kingebeni, Edith Nkwembe, Eddy Kinganda-Lusamaki, Amuri Aziza, Francisca Muyembe Mawete, Catherine Pratt, Matthias Pauthner, Josh Quick, Allison Black, James Hadfield, Trevor Bedford, Ian Goodfellow, Andrew Rambaut, Nick Loman, Kristian Andersen, Michael Wiley, Steve Ahuka-Mundeke, Jean-Jacques Muyembe Tamfum |
| EPI_ISL_435281 | Medistra Hospital Jakarta | Eijkman Institute for Molecular Biology, Ministry of Research and Technology/National Agency for Research and Innovation | Edison Johar, Frilasita A Yudhaputri, Hidayat Trimarsanto, David H Muljono, Safarina G Malik, Khin Saw Myint, Amin Soebandrio |
| EPI_ISL_435282, EPI_ISL_435283 | RS Pondok Indah Hospital – Pondok Indah | Eijkman Institute for Molecular Biology, Ministry of Research and Technology/National Agency for Research and Innovation | Edison Johar, Frilasita A Yudhaputri, Hidayat Trimarsanto, David H Muljono, Safarina G Malik, Khin Saw Myint, Amin Soebandrio |
| EPI_ISL_435284 | Central Virology Laboratory, Israel Ministry of Health | Central Virology Laboratory, Israel Ministry of Health | Neta Zuckerman, Efrat Bucris, Oran Erster, Danit Sofer, Orna Mor, Ella Mendelson, Michal Mandelboim |
| EPI_ISL_435286 | Central Virology Laboratory, Israel Ministry of Health | Central Virology Laboratory, Israel Ministry of Health | eta Zuckerman, Efrat Bucris, Oran Erster, Orna Mor, Ella Mendelson, Michal Mandelboim, Danit Sofer |
| EPI_ISL_435287 | Central Virology Laboratory, Israel Ministry of Health | Central Virology Laboratory, Israel Ministry of Health | Neta Zuckerman, Efrat Bucris, Oran Erster, Danit Sofer, Orna Mor, Ella Mendelson, Michal Mandelboim |
| EPI_ISL_435289 | Central Virology Laboratory, Israel Ministry of Health | Central Virology Laboratory, Israel Ministry of Health | Neta Zuckerman, Efrat Bucris, Oran Erster, Danit Sofer, Orna Mor, Ella Mendelson, Michal Mandelboim |
| EPI_ISL_435291 | Central Virology Laboratory, Israel Ministry of Health | Central Virology Laboratory, Israel Ministry of Health | Neta Zuckerman, Efrat Bucris, Oran Erster, Danit Sofer, Orna Mor, Ella Mendelson, Michal Mandelboim |
| EPI_ISL_435292 | Central Virology Laboratory, Israel Ministry of Health | Central Virology Laboratory, Israel Ministry of Health | Neta Zuckerman, Efrat Bucris, Oran Erster, Danit Sofer, Ella Mendelson, Michal Mandelboim, Orna Mor |
| EPI_ISL_435303 | National Hospital of Tropical Diseases | Oxford University Clinical Research Unit, Hanoi, Vietnam | Nguyen Thi Tam, Van Dinh Trang, Nguyen Thu Trang, Nguyen Thi Ngoc Diep, Le Nguyen Minh Hoa, Pham Ngoc Thach, H.Rogier van Doorn, on behalf of the OUCRU COVID-19 research group |
| EPI_ISL_435305 | National Hospital of Tropical Diseases | Oxford University Clinical Research Unit, Hanoi, Vietnam | Nguyen Thi Tam, Van Dinh Trang, Nguyen Thu Trang, Nguyen Thi Ngoc Diep, Le Nguyen Minh Hoa, Pham Ngoc Thach, H. Rogier van Doorn, on behalf of the OUCRU COVID-19 research group |
| EPI_ISL_435308 | National Hospital of Tropical Diseases | Oxford University Clinical Research Unit, Hanoi, Vietnam | Nguyen Thi Tam, Van Dinh Trang, Nguyen Thu Trang, Nguyen Thi Ngoc Diep, Le Nguyen Minh Hoa, Pham Ngoc Thach, H. Rogier van Doorn, on behalf of the OUCRU COVID-19 research group |
| EPI_ISL_435310 | National Hospital of Tropical Diseases | Oxford University Clinical Research Unit, Hanoi, Vietnam | Nguyen Thi Tam, Van Dinh Trang, Nguyen Thu Trang, Nguyen Thi Ngoc Diep, Le Nguyen Minh Hoa, Pham Ngoc Thach, H. Rogier van Doorn, on behalf of the OUCRU COVID-19 research group |
| EPI_ISL_435311 | National Hospital of Tropical Diseases | Oxford University Clinical Research Unit, Hanoi, Vietnam | Nguyen Thi Tam, Van Dinh Trang, Nguyen Thu Trang, Nguyen Thi Ngoc Diep, Le Nguyen Minh Hoa, Pham Ngoc Thach, H. Rogier van Doorn, on behalf of the OUCRU COVID-19 research group |
| EPI_ISL_435312, EPI_ISL_435313 | National Hospital of Tropical Diseases | Oxford University Clinical Research Unit, Hanoi, Vietnam | Nguyen Thi Tam, Van Dinh Trang, Nguyen Thu Trang, Nguyen Thi Ngoc Diep, Le Nguyen Minh Hoa, Pham Ngoc Thach, H. Rogier van Doorn, on behalf of the OUCRU COVID-19 research group |
| EPI_ISL_435314 | National Hospital of Tropical Diseases | Oxford University Clinical Research Unit, Hanoi, Vietnam | Nguyen Thi Tam, Van Dinh Trang, Nguyen Thu Trang, Nguyen Thi Ngoc Diep, Le Nguyen Minh Hoa, Pham Ngoc Thach, H. Rogier van Doorn, on behalf of the OUCRU COVID-19 research group |
| EPI_ISL_435315, EPI_ISL_435316, EPI_ISL_435317 | National Hospital of Tropical Diseases | Oxford University Clinical Research Unit, Hanoi, Vietnam | Nguyen Thi Tam, Van Dinh Trang, Nguyen Thu Trang, Nguyen Thi Ngoc Diep, Le Nguyen Minh Hoa, Pham Ngoc Thach, H. Rogier van Doorn, on behalf of the OUCRU COVID-19 research group |
| EPI_ISL_435343, EPI_ISL_435344 | Laboratoire de microbiologie, Hopital de Verdun | Smith Laboratory, Centre de Recherche CHU Sainte-Justine | Martin Smith, Marieke Rozendaal, Ivan Pavlov |
| EPI_ISL_435345, EPI_ISL_435346, EPI_ISL_435347 | Laboratoire de microbiologie, Hopital de Verdun | Smith Laboratory, Centre de Recherche CHU Sainte-Justine | Martin Smith, Marieke Rozendaal, Ivan Pavlov |
| EPI_ISL_435348, EPI_ISL_435349, EPI_ISL_435350, EPI_ISL_435351, EPI_ISL_435352, EPI_ISL_435353, EPI_ISL_435354, EPI_ISL_435355, EPI_ISL_435356, EPI_ISL_435357, EPI_ISL_435358, EPI_ISL_435359, EPI_ISL_435360, EPI_ISL_435361, EPI_ISL_435362, EPI_ISL_435363, EPI_ISL_435364, EPI_ISL_435365, EPI_ISL_435366, EPI_ISL_435367, EPI_ISL_435368, EPI_ISL_435369, EPI_ISL_435370, EPI_ISL_435371, EPI_ISL_435372, EPI_ISL_435373, EPI_ISL_435374, EPI_ISL_435375, EPI_ISL_435376, EPI_ISL_435377, EPI_ISL_435378, EPI_ISL_435379, EPI_ISL_435380, EPI_ISL_435381, EPI_ISL_435382, EPI_ISL_435383, EPI_ISL_435384, EPI_ISL_435385, EPI_ISL_435386, EPI_ISL_435387, EPI_ISL_435388, EPI_ISL_435389, EPI_ISL_435390, EPI_ISL_435391, EPI_ISL_435392, EPI_ISL_435393 | | | |
| see above | Utah Public Health Laboratory | Utah Public Health Laboratory | Erin Young, Kelly Oakeson |
| EPI_ISL_435394, EPI_ISL_435395, EPI_ISL_435396, EPI_ISL_435397, EPI_ISL_435398, EPI_ISL_435399, EPI_ISL_435400, EPI_ISL_435401, EPI_ISL_435402 | Gundersen Molecular Diagnostics Laboratory | Kabara Cancer Research Institute | Craig S. Richmond, Paraic A. Kenny |
| EPI_ISL_435403, EPI_ISL_435404, EPI_ISL_435405, EPI_ISL_435406, EPI_ISL_435407, EPI_ISL_435408, EPI_ISL_435409, EPI_ISL_435410, EPI_ISL_435411, EPI_ISL_435412, EPI_ISL_435413, EPI_ISL_435414, EPI_ISL_435415, EPI_ISL_435416, EPI_ISL_435417, EPI_ISL_435418, EPI_ISL_435419, EPI_ISL_435420, EPI_ISL_435421, EPI_ISL_435422, EPI_ISL_435423, EPI_ISL_435424, EPI_ISL_435425, EPI_ISL_435426, EPI_ISL_435427, EPI_ISL_435428, EPI_ISL_435429, EPI_ISL_435430, EPI_ISL_435431 | | | |
| see above | Virological Research Group, Szentágothai Research Centre | Bioinformatics Research Group, Szentágothai Research Centre | Péter Urbán, Endre Gábor Tóth, Gábor Kemenesi, Róbert Herczeg, Attila Gyenesei, Ferenc Jakab |
| EPI_ISL_435441, EPI_ISL_435442, EPI_ISL_435443, EPI_ISL_435444 | Alaska State Virology Laboratory | Alaska State Virology Laboratory | Jack Chen, Ph.D. |
| EPI_ISL_435445, EPI_ISL_435446, EPI_ISL_435447, EPI_ISL_435448, EPI_ISL_435449, EPI_ISL_435450, EPI_ISL_435451, EPI_ISL_435452, EPI_ISL_435453, EPI_ISL_435454, EPI_ISL_435455, EPI_ISL_435456, EPI_ISL_435457, EPI_ISL_435458, EPI_ISL_435459, EPI_ISL_435460, EPI_ISL_435461, EPI_ISL_435462, EPI_ISL_435463, EPI_ISL_435464, EPI_ISL_435465, EPI_ISL_435466, EPI_ISL_435467, EPI_ISL_435468, EPI_ISL_435469, EPI_ISL_435470, EPI_ISL_435471, EPI_ISL_435472 | | | |
| see above | Robert Garry lab | Andersen lab at Scripps Research | Allison Smither, Gilberto Sabino-Santos, Patricia Snarski, Lilia Melnik, Antoinette Bell, Kaylynn Genemaras, Arnaud Drouin, Dahlene Fusco, Robert Garry with SEARCH Alliance San Diego |
| EPI_ISL_435473, EPI_ISL_435474 | Rady's Childrens Hospital | Andersen lab at Scripps Research | SEARCH Alliance San Diego |
| EPI_ISL_435475, EPI_ISL_435476, EPI_ISL_435477, EPI_ISL_435478, EPI_ISL_435479, EPI_ISL_435480, EPI_ISL_435481, EPI_ISL_435482, EPI_ISL_435483, EPI_ISL_435484, EPI_ISL_435485, EPI_ISL_435486, EPI_ISL_435487, EPI_ISL_435488, EPI_ISL_435489, EPI_ISL_435490, EPI_ISL_435491, EPI_ISL_435492, EPI_ISL_435493, EPI_ISL_435494, EPI_ISL_435495, EPI_ISL_435496, EPI_ISL_435497, EPI_ISL_435498, EPI_ISL_435499, EPI_ISL_435500, EPI_ISL_435501, EPI_ISL_435502, EPI_ISL_435503, EPI_ISL_435504, EPI_ISL_435505, EPI_ISL_435506, EPI_ISL_435507, EPI_ISL_435508, EPI_ISL_435509, EPI_ISL_435510, EPI_ISL_435511, EPI_ISL_435512, EPI_ISL_435513, EPI_ISL_435514, EPI_ISL_435515, EPI_ISL_435516, EPI_ISL_435517, EPI_ISL_435518, EPI_ISL_435519, EPI_ISL_435520, EPI_ISL_435521, EPI_ISL_435522, EPI_ISL_435523, EPI_ISL_435524, EPI_ISL_435525, EPI_ISL_435526, EPI_ISL_435527, EPI_ISL_435528, EPI_ISL_435529, EPI_ISL_435530, EPI_ISL_435531, EPI_ISL_435532, EPI_ISL_435533, EPI_ISL_435534, EPI_ISL_435535, EPI_ISL_435536, EPI_ISL_435537, EPI_ISL_435538, EPI_ISL_435539, EPI_ISL_435540, EPI_ISL_435541, EPI_ISL_435542, EPI_ISL_435543, EPI_ISL_435544, EPI_ISL_435545, EPI_ISL_435546, EPI_ISL_435547, EPI_ISL_435548, EPI_ISL_435549 | | | |
| see above | NYU Langone Health | Departments of Pathology and Medicine, New York University School of Medicine | Maria Aguero-Rosenfeld, Brendan Belovarac, Margaret Black, Ludovic Boytard, John Cadley, Paolo Cotzia, John Chen, Dacia Dimartino, Xiaojun Feng, Tatyana Gindin, Emily Guzman, Adriana Heguy, Megan Hogan, Emily Huang, George Jour, Lawrence H. Lin, Raven Luther, Andrew Lytle, Christian Marier, Matthew T. Maurano, Mark J. Mulligan, Peter Meyn, Raquel Ordonez Ciriza, Iman Osman, Jared Pinnell, Vanessa Raabe, Sitharam Ramaswami, Amy Rapkiewicz, Andre M. Ribeiro-dos-Santos, Marie Samanovic-Golden, Antonio Serrano, Guomiao Shen, Matija Snuderl, Theodore Vougiouklakis, Nick Vulpescu, Gael Westby, Paul Zappile, Yutong Zhang |
| EPI_ISL_435550, EPI_ISL_435551, EPI_ISL_435552, EPI_ISL_435553, EPI_ISL_435554 | LSUHS Emerging Viral Threat Laboratory | Microbial Genome Sequencing Center | Rona S. Scott, Jeremy P. Kamil, John A. Vanchiere, Camille F. Abshire, Abida Siddiqa, Byeong-Jae Lee, Chan-ki Min, Md Maksudul Alam, Monica Gestal-Cartele, Edna Ondari, Adam Greer, Malgorzata Bienkowska-Haba, Katarzyna Zwolinska, Jason M. Bodily, Andrew D. Yurochko, Paul M. Weinberger, Christopher G. Kevil, Martin J. Sapp, Daniel J. Snyder, Vaughn S. Cooper |
| EPI_ISL_435555, EPI_ISL_435556, EPI_ISL_435557, EPI_ISL_435558, EPI_ISL_435559, EPI_ISL_435560, EPI_ISL_435561, EPI_ISL_435562, EPI_ISL_435563, EPI_ISL_435564, EPI_ISL_435565, EPI_ISL_435566, EPI_ISL_435567, EPI_ISL_435568 | | | |
| see above | LSUHS Emerging Viral Threat Laboratory | Microbial Genome Sequencing Center | John A. Vanchiere, Jeremy P. Kamil, Rona S. Scott, Camille F. Abshire, Abida Siddiqa, Byeong-Jae Lee, Chan-ki Min, Md Maksudul Alam, Monica Gestal-Cartele, Edna Ondari, Adam Greer, Malgorzata Bienkowska-Haba, |

| | | | |
|---|---|---|---|
| | | | Katarzyna Zwolinska, Jason M. Bodily, Andrew D. Yurochko, Paul M. Weinberger, Christopher G. Kevil, Martin J. Sapp, Daniel J. Snyder, Vaughn S. Cooper |
| EPI_ISL_435569, EPI_ISL_435570, EPI_ISL_435571, EPI_ISL_435572, EPI_ISL_435573, EPI_ISL_435574, EPI_ISL_435575, EPI_ISL_435576, EPI_ISL_435577, EPI_ISL_435578, EPI_ISL_435579 | | | |
| see above | LSUHS Emerging Viral Threat Laboratory | Microbial Genome Sequencing Center | Jeremy P. Kamil, John A. Vanchiere, Rona S. Scott, Camille F. Abshire, Abida Siddiqa, Byeong-Jae Lee, Chan-ki Min, Md Maksudul Alam, Monica Gestal-Cartele, Edna Ondari, Adam Greer, Malgorzata Bienkowska-Haba, Katarzyna Zwolinska, Jason M. Bodily, Andrew D. Yurochko, Paul M. Weinberger, Christopher G. Kevil, Martin J. Sapp, Daniel J. Snyder, Vaughn S. Cooper |
| EPI_ISL_435580, EPI_ISL_435581, EPI_ISL_435582, EPI_ISL_435583, EPI_ISL_435584, EPI_ISL_435585, EPI_ISL_435586, EPI_ISL_435587, EPI_ISL_435588, EPI_ISL_435589, EPI_ISL_435590, EPI_ISL_435591, EPI_ISL_435592, EPI_ISL_435593, EPI_ISL_435594, EPI_ISL_435595, EPI_ISL_435596, EPI_ISL_435597, EPI_ISL_435598, EPI_ISL_435599, EPI_ISL_435600, EPI_ISL_435601, EPI_ISL_435602, EPI_ISL_435603, EPI_ISL_435604, EPI_ISL_435605, EPI_ISL_435606, EPI_ISL_435607, EPI_ISL_435608, EPI_ISL_435609, EPI_ISL_435610, EPI_ISL_435611, EPI_ISL_435612, EPI_ISL_435613, EPI_ISL_435614, EPI_ISL_435615, EPI_ISL_435616, EPI_ISL_435617, EPI_ISL_435618, EPI_ISL_435619, EPI_ISL_435620, EPI_ISL_435621, EPI_ISL_435622, EPI_ISL_435623, EPI_ISL_435624, EPI_ISL_435625, EPI_ISL_435626, EPI_ISL_435627, EPI_ISL_435628, EPI_ISL_435629, EPI_ISL_435630, EPI_ISL_435631, EPI_ISL_435632, EPI_ISL_435633, EPI_ISL_435634, EPI_ISL_435635, EPI_ISL_435636, EPI_ISL_435637, EPI_ISL_435638, EPI_ISL_435639, EPI_ISL_435640, EPI_ISL_435641, EPI_ISL_435642, EPI_ISL_435643, EPI_ISL_435644, EPI_ISL_435645, EPI_ISL_435646, EPI_ISL_435647, EPI_ISL_435648, EPI_ISL_435649, EPI_ISL_435650, EPI_ISL_435651, EPI_ISL_435652, EPI_ISL_435653, EPI_ISL_435654, EPI_ISL_435655, EPI_ISL_435656, EPI_ISL_435657, EPI_ISL_435658, EPI_ISL_435659, EPI_ISL_435660, EPI_ISL_435661, EPI_ISL_435662, EPI_ISL_435663, EPI_ISL_435664, EPI_ISL_435665, EPI_ISL_435666, EPI_ISL_435667, EPI_ISL_435668, EPI_ISL_435669, EPI_ISL_435670, EPI_ISL_435671, EPI_ISL_435672, EPI_ISL_435673 | | | |
| see above | Santa Clara County Public Health Department | Chiu Laboratory, University of California, San Francisco | Xianding Deng, Scot Federman, Wei Gu, Elsa Villarino, Brandon Bonin, Debra A. Wadford, and Charles Y. Chiu |
| EPI_ISL_435674, EPI_ISL_435675, EPI_ISL_435676, EPI_ISL_435677 | National Virology Reference Laboratory | National Public Health Laboratory, National Centre for Infectious Diseases | Mak Tze Minn, Octavia Sophie, Chavatte Jean-Marc, Zaini Zainun, Taib Surita, Cui Lin, Lin Raymond Tzer Pin |
| EPI_ISL_435678, EPI_ISL_435679, EPI_ISL_435680, EPI_ISL_435681, EPI_ISL_435682, EPI_ISL_435683, EPI_ISL_435684, EPI_ISL_435685, EPI_ISL_435686, EPI_ISL_435687, EPI_ISL_435688, EPI_ISL_435689, EPI_ISL_435690, EPI_ISL_435691, EPI_ISL_435692, EPI_ISL_435693, EPI_ISL_435694, EPI_ISL_435695, EPI_ISL_435696, EPI_ISL_435697, EPI_ISL_435698, EPI_ISL_435699, EPI_ISL_435700 | | | |
| see above | National Public Health Laboratory, National Centre for Infectious Diseases | National Public Health Laboratory, National Centre for Infectious Diseases | Mak Tze Minn, Octavia Sophie, Chavatte Jean-Marc, Cui Lin, Lin Raymond Tzer Pin |
| EPI_ISL_435702, EPI_ISL_435703, EPI_ISL_435704, EPI_ISL_435705, EPI_ISL_435706, EPI_ISL_435707, EPI_ISL_435708, EPI_ISL_435709 | Yale COVID-19 Biorepository | Grubaugh Lab - Yale School of Public Health | Joseph Fauver, Tara Alpert, Anderson Brito, Anne Wyllie, Chantal Vogels, Mary Petrone, Cole Jensen, Chaney Kalinich, Isabel Ott, Arnau Casanovas, Catherine Muenker, Adam Moore, Alice Lu, Maria Tokuyama, Patrick Wong, Peiwen Lu, Saad Omer, Richard Martinello, Allison Nelson, Shelli Farhadian, Akiko Iwasaki, Charlese Dela Cruz, Albert Ko, Nathan Grubaugh |
| EPI_ISL_435710, EPI_ISL_435711, EPI_ISL_435712, EPI_ISL_435713, EPI_ISL_435714, EPI_ISL_435715, EPI_ISL_435716, EPI_ISL_435717, EPI_ISL_435718, EPI_ISL_435719 | Connecticut State Department of Public Health | Grubaugh Lab - Yale School of Public Health | Joseph Fauver, Tara Alpert, Anderson Brito, Anne Wyllie, Chantal Vogels, Mary Petrone, Cole Jensen, Chaney Kalinich, Isabel Ott, Arnau Casanovas, Catherine Muenker, Adam Moore, Alice Lu, Maria Tokuyama, Patrick Wong, Peiwen Lu, Saad Omer, Richard Martinello, Allison Nelson, Shelli Farhadian, Akiko Iwasaki, Charlese Dela Cruz, Albert Ko, Nathan Grubaugh |
| EPI_ISL_435720 | Yale Clinical virology | Grubaugh Lab - Yale School of Public Health | Joseph Fauver, Tara Alpert, Anderson Brito, Anne Wyllie, Chantal Vogels, Mary Petrone, Cole Jensen, Chaney Kalinich, Isabel Ott, Arnau Casanovas, Catherine Muenker, Adam Moore, Alice Lu, Maria Tokuyama, Patrick Wong, Peiwen Lu, Saad Omer, Richard Martinello, Allison Nelson, Shelli Farhadian, Akiko Iwasaki, Charlese Dela Cruz, Albert Ko, Nathan Grubaugh |
| EPI_ISL_435721, EPI_ISL_435722 | NYU Langone Health | Departments of Pathology and Medicine, New York University School of Medicine | Maria Aguero-Rosenfeld, Brendan Belovarac, Margaret Black, Ludovic Boytard, John Cadley, Paolo Cotzia, John Chen, Dacia Dimartino, Xiaojun Feng, Tatyana Gindin, Emily Guzman, Adriana Heguy, Megan Hogan, Emily Huang, George Jour, Lawrence H. Lin, Raven Luther, Andrew Lytle, Christian Marier, Matthew T. Maurano, Mark J. Mulligan, Peter Meyn, Raquel Ordonez Ciriza, Iman Osman, Jared Pinnell, Vanessa Raabe, Sitharam Ramaswami, Amy Rapkiewicz, Andre M. Ribeiro-dos-Santos, Marie Samanovic-Golden, Antonio Serrano, Guomiao Shen, Matija Snuderl, Theodore Vougiouklakis, Nick Vulpescu, Gael Westby, Paul Zappile, Yutong Zhang |
| EPI_ISL_435723 | Laboratory of Genomics & Bioinformatics, Institute of Immunology and Experimental Therapy, Polish Academy of Sciences Oddział Mikrobiologii Wojewódzkiej Stacji Sanitarno Epidemiologiczna | Laboratory of Genomics & Bioinformatics, Institute of Immunology and Experimental Therapy, Polish Academy of Sciences | Aleksandra Herud, Dorota Kujawa, Dariusz Martynowski, Krzysztof Jakub Pawlik, Joanna Sikorska, Paulina Żebrowska, Grażyna Zalewska, Oskar Karpiński and Łukasz Łaczmański |
| EPI_ISL_436040, EPI_ISL_436041, EPI_ISL_436042, EPI_ISL_436043 | DC Public Health Lab Dept of Forensic Science | Pathogen Discovery, Respiratory Viruses Branch, Division of Viral Diseases, Centers for Disease Control and Prevention | Ying Tao, Jing Zhang, Krista Queen, Yan Li, Anna Uehara, Clinton R. Paden, Haibin Wang, Zachary Weiner, Bettina Bankamp, Suxiang Tong |
| EPI_ISL_436044 | Louisiana Office of Public Health Laboratories | Pathogen Discovery, Respiratory Viruses Branch, Division of Viral Diseases, Centers for Disease Control and Prevention | Ying Tao, Jing Zhang, Krista Queen, Yan Li, Anna Uehara, Clinton R. Paden, Haibin Wang, Zachary Weiner, Bettina Bankamp, Suxiang Tong |
| EPI_ISL_436045, EPI_ISL_436046 | US VI Department of Health | Pathogen Discovery, Respiratory Viruses Branch, Division of Viral Diseases, Centers for Disease Control and Prevention | Ying Tao, Jing Zhang, Krista Queen, Yan Li, Anna Uehara, Clinton R. Paden, Haibin Wang, Zachary Weiner, Bettina Bankamp, Suxiang Tong |
| EPI_ISL_436047, EPI_ISL_436048, EPI_ISL_436049, EPI_ISL_436050, EPI_ISL_436051, EPI_ISL_436052, EPI_ISL_436053, EPI_ISL_436054, EPI_ISL_436055, EPI_ISL_436056, EPI_ISL_436057, EPI_ISL_436058, EPI_ISL_436059, EPI_ISL_436060, EPI_ISL_436061, EPI_ISL_436062, EPI_ISL_436063, EPI_ISL_436064, EPI_ISL_436065, EPI_ISL_436066, EPI_ISL_436067, EPI_ISL_436068, EPI_ISL_436069, EPI_ISL_436070, EPI_ISL_436071, EPI_ISL_436072, EPI_ISL_436073, EPI_ISL_436074, EPI_ISL_436075, EPI_ISL_436076, EPI_ISL_436077, EPI_ISL_436078, EPI_ISL_436079, EPI_ISL_436080, EPI_ISL_436081, EPI_ISL_436082 | | | |
| see above | NYC Department of Health and Mental Hygiene | Pathogen Discovery, Respiratory Viruses Branch, Division of Viral Diseases, Centers for Disease Control and Prevention | Ying Tao, Krista Queen, Christy Harrison, Jennifer Rakeman, Clinton R. Paden, Jing Zhang, Anna Uehara, Yan Li, Haibin Wang, Jasmine Padilla, Justin Lee, Bettina Bankamp, Zachary Weiner, Suxiang Tong |
| EPI_ISL_436097 | Prince Charles Hospital | Public Health Virology Laboratory, Forensics and Scientific Services, Queensland Health | Alyssa Pyke, Neelima Nair, Natalie Simpson, Lisa Leckie, Jamie McMahon, Jean Barcelon, Amanda De Jong, Sean Moody, Doris Genge, Glen Hewitson, Peter Burtonclay, Judy Northill, Ian Maxwell Mackay, Carmel Taylor, Bixing Huang, David Warrilow, Mitchell Finger, Peter Moore, Sarah Wheatley, Sonja Hall-Mendelin, Andrew Van Den Hurk, Elisabeth Gamez, Inga Sultana and Frederick Moore |
| EPI_ISL_436098 | Royal Brisbane and Women's Hospital | Public Health Virology Laboratory, Forensic and Scientific Services, Queensland Health | Alyssa Pyke, Neelima Nair, Natalie Simpson, Lisa Leckie, Jamie McMahon, Jean Barcelon, Amanda De Jong, Sean Moody, Doris Genge, Glen Hewitson, Peter Burtonclay, Judy Northill, Ian Maxwell Mackay, Carmel Taylor, Bixing Huang, David Warrilow, Mitchell Finger, Peter Moore, Sarah Wheatley, Sonja Hall-Mendelin, Andrew Van Den Hurk, Elisabeth Gamez, Inga Sultana and Frederick Moore |
| EPI_ISL_436099 | TSGH-CP molecular lab | TSGH-CP molecular lab | Cherng-Lih Perng, Ming-Jr JIAN, Chih-Kai Chang, Jung-Chung Lin, Kuo-Ming Yeh, Chien-Wen Chen, Sheng-Kang Chiu, Hsing-Yi Chung, Shih-Hung Tsai, Kuo-Sheng Hung, Tien-Yao Chang, Feng-Yee Chang, Hung-Sheng Shang |
| EPI_ISL_436100 | TSGH-CP molecular lab | TSGH-CP molecular lab | "No. 325, Sec.2, Chenggong Road, Neihu District, Taipei City, Taiwan Postal code11490 Division of Clinical Pathology, Department of Pathology" |
| EPI_ISL_436101, EPI_ISL_436102, EPI_ISL_436103, EPI_ISL_436104 | TSGH-CP molecular lab | TSGH-CP molecular lab | Cherng-Lih Perng, Ming-Jr JIAN, Chih-Kai Chang, Jung-Chung Lin, Kuo-Ming Yeh, Chien-Wen Chen, Sheng-Kang Chiu, Hsing-Yi Chung, Shih-Hung Tsai, Kuo-Sheng Hung, Tien-Yao Chang, Feng-Yee Chang, Hung-Sheng Shang |
| EPI_ISL_436105 | TSGH-CP molecular lab | TSGH-CP molecular lab | Cherng-Lih Perng, Ming-Jr Jian, Chih-Kai Chang, Jung-Chung Lin, Kuo-Ming Yeh, Chien-Wen Chen, Sheng-Kang Chiu, Hsing-Yi Chung, Shih-Hung Tsai, Kuo-Sheng Hung, Tien-Yao Chang, Feng-Yee Chang, Hung-Sheng Shang |
| EPI_ISL_436106, EPI_ISL_436107, EPI_ISL_436108 | TSGH-CP molecular lab | TSGH-CP molecular lab | Cherng-Lih Perng, Ming-Jr JIAN, Chih-Kai Chang, Jung-Chung Lin, Kuo-Ming Yeh, Chien-Wen Chen, Sheng-Kang Chiu, Hsing-Yi Chung, Shih-Hung Tsai, Kuo-Sheng Hung, Tien-Yao Chang, Feng-Yee Chang, Hung-Sheng Shang |
| EPI_ISL_436111, EPI_ISL_436112, EPI_ISL_436113, EPI_ISL_436114, EPI_ISL_436115, EPI_ISL_436116, EPI_ISL_436117, EPI_ISL_436118, EPI_ISL_436119, EPI_ISL_436120, EPI_ISL_436121, EPI_ISL_436122, EPI_ISL_436123, EPI_ISL_436124, EPI_ISL_436125, EPI_ISL_436126, EPI_ISL_436127, EPI_ISL_436128, EPI_ISL_436129, EPI_ISL_436130, EPI_ISL_436131, EPI_ISL_436132 | | | |
| see above | Victorian Infectious Diseases Reference Laboratory (VIDRL) | Microbiological Diagnostic Unit Public Health Laboratory and Victorian Infectious Diseases Reference Laboratory, Doherty Institute | Caly L., Seemann T., Sait, M., Schultz M., Druce J., Sherry, N. |
| EPI_ISL_436137, EPI_ISL_436138, EPI_ISL_436139, EPI_ISL_436140, EPI_ISL_436141, EPI_ISL_436156, EPI_ISL_436157 | District Surveillance Unit | Department of Neurovirology, National Institute of Mental Health and Neuroscience (NIMHANS) | Chitra Pattabiraman, Vijayalakshmi Reddy, Harsha PK, Risha Rasheed, Shafeeq S Hameed, Manjunatha Venkataswamy, Anita Desai, Ravi Vasanthapuram |
| EPI_ISL_436194 | Viral Respiratory Lab, National Institute for Biomedical Research (INRB) | Pathogen Sequencing Lab, National Institute for Biomedical Research (INRB) | Placide Mbala-Kingebeni, Edith Nkwembe, Eddy Kinganda-Lusamaki, Amuri Aziza, Francisca Muyembe Mawete, Catherine Pratt, Matthias Pauthner, Josh Quick, Allison Black, James Hadfield, Trevor Bedford, Ian Goodfellow, Andrew Rambaut, Nick Loman, Kristian Andersen, Michael Wiley, Steve Ahuka-Mundeke, Jean-Jacques Muyembe Tamfum |
| EPI_ISL_436195 | Servicio de Microbiología. Consorcio Hospital General Universitario de Valencia | Sequencing and Bioinformatics Service and Molecular Epidemiology Research Group. FISABIO-Public Health | Maria Dolores Ocete, Inma Galán Vendrell, Paula Ruiz-Hueso, Mariana Reyes-Prieto, Vicente Soriano Chirona, Maria Alma Bracho, Griselda De Marco, Beatriz Beamud, Lidia Ruiz Roldan, Marta Pla Diaz,Neris Garcia-Gonzalez, Loreto Ferrús Abad, Llúcia Martínez-Priego, Concepcion Gimeno, Giuseppe D'Auria, Fernando Gonzalez-Candelas |
| EPI_ISL_436196 | Servicio de Microbiología. Consorcio Hospital General Universitario de Valencia | Sequencing and Bioinformatics Service and Molecular Epidemiology Research Group. FISABIO-Public Health | Griselda De Marco, Beatriz Beamud, Lidia Ruiz Roldan, Marta Pla Diaz,Neris Garcia-Gonzalez, Loreto Ferrús Abad, Maria Dolores Ocete, Inma Galán Vendrell, Paula Ruiz-Hueso, Mariana Reyes-Prieto, Vicente Soriano Chirona, Maria Alma Bracho, Llúcia Martínez-Priego, Concepcion Gimeno, Giuseppe D'Auria, Fernando Gonzalez-Candelas |
| EPI_ISL_436197 | Servicio de Microbiología. Consorcio Hospital General Universitario de Valencia | Sequencing and Bioinformatics Service and Molecular Epidemiology Research Group. FISABIO-Public Health | Beatriz Beamud, Lidia Ruiz Roldan, Marta Pla Diaz,Neris Garcia-Gonzalez, Loreto Ferrús Abad, Maria Dolores Ocete, Inma Galán Vendrell, Paula Ruiz-Hueso, Mariana Reyes-Prieto, Vicente Soriano Chirona, Maria Alma Bracho, Griselda De Marco, Llúcia Martínez-Priego, Concepcion Gimeno, Giuseppe D'Auria, Fernando Gonzalez-Candelas |
| EPI_ISL_436198 | Servicio de Microbiología. Consorcio Hospital General Universitario de Valencia | Sequencing and Bioinformatics Service and Molecular Epidemiology Research Group. FISABIO-Public Health | Lidia Ruiz Roldan, Marta Pla Diaz,Neris Garcia-Gonzalez, Loreto Ferrús Abad, Maria Dolores Ocete, Inma Galán Vendrell, Paula Ruiz-Hueso, Mariana Reyes-Prieto, Vicente Soriano Chirona, Maria Alma Bracho, Griselda De Marco, Beatriz Beamud, Llúcia Martínez-Priego, Concepcion Gimeno, Giuseppe D'Auria, Fernando Gonzalez-Candelas |
| EPI_ISL_436199 | Servicio de Microbiología. Consorcio Hospital General Universitario de Valencia | Sequencing and Bioinformatics Service and Molecular Epidemiology Research Group. FISABIO-Public Health | Marta Pla Diaz,Neris Garcia-Gonzalez, Loreto Ferrús Abad, Maria Dolores Ocete, Inma Galán Vendrell, Paula Ruiz-Hueso, Mariana Reyes-Prieto, Vicente Soriano Chirona, Maria Alma Bracho, Griselda De Marco, Beatriz Beamud, Lidia Ruiz Roldan, Llúcia Martínez-Priego, Concepcion Gimeno, Giuseppe D'Auria, Fernando Gonzalez-Candelas |
| EPI_ISL_436200 | Servicio de Microbiología. Consorcio Hospital General Universitario de Valencia | Sequencing and Bioinformatics Service and Molecular Epidemiology Research Group. FISABIO-Public Health | Neris Garcia-Gonzalez, Loreto Ferrús Abad, Maria Dolores Ocete, Inma Galán Vendrell, Paula Ruiz-Hueso, Mariana Reyes-Prieto, Vicente Soriano Chirona, Maria Alma Bracho, Griselda De Marco, Beatriz Beamud, Lidia Ruiz Roldan, Marta Pla Diaz, Llúcia Martínez-Priego, Concepcion Gimeno, Giuseppe D'Auria, Fernando Gonzalez-Candelas |
| EPI_ISL_436201 | Servicio de Microbiología. Consorcio Hospital General Universitario de Valencia | Sequencing and Bioinformatics Service and Molecular Epidemiology Research Group. FISABIO-Public Health | Loreto Ferrús Abad, Maria Dolores Ocete, Inma Galán Vendrell, Paula Ruiz-Hueso, Mariana Reyes-Prieto, Vicente Soriano Chirona, Maria Alma Bracho, Griselda De Marco, Beatriz Beamud, Lidia Ruiz Roldan, Marta Pla Diaz,Neris Garcia-Gonzalez, Llúcia Martínez-Priego, Concepcion Gimeno, Giuseppe D'Auria, Fernando Gonzalez-Candelas |
| EPI_ISL_436202 | Servicio de Microbiología. Consorcio Hospital General Universitario de Valencia | Sequencing and Bioinformatics Service and Molecular Epidemiology Research Group. FISABIO-Public Health | Loreto Ferrús Abad, Maria Dolores Ocete,Inma Galán Vendrell, Paula Ruiz-Hueso, Mariana Reyes-Prieto, Vicente Soriano Chirona, Maria Alma Bracho, Griselda De Marco, Beatriz Beamud, Lidia Ruiz Roldan, Marta Pla Diaz,Neris Garcia-Gonzalez, Llúcia Martínez-Priego, Concepcion Gimeno, Giuseppe D'Auria, Fernando Gonzalez-Candelas |
| EPI_ISL_436203 | Servicio de Microbiología. Consorcio Hospital General Universitario de Valencia | Sequencing and Bioinformatics Service and Molecular Epidemiology Research Group. FISABIO-Public Health | Maria Dolores Ocete, Inma Galán Vendrell, Paula Ruiz-Hueso, Mariana Reyes-Prieto, Vicente Soriano Chirona, Maria Alma Bracho, Griselda De Marco, Beatriz Beamud, Lidia Ruiz Roldan, Marta Pla Diaz,Neris Garcia-Gonzalez, Loreto Ferrús Abad, Llúcia Martínez-Priego, Concepcion Gimeno, Giuseppe D'Auria, Fernando Gonzalez-Candelas |
| EPI_ISL_436204 | Servicio de Microbiología. Consorcio Hospital General Universitario de Valencia | Sequencing and Bioinformatics Service and Molecular Epidemiology Research Group. FISABIO-Public Health | Griselda De Marco, Beatriz Beamud, Lidia Ruiz Roldan, Marta Pla Diaz,Neris Garcia-Gonzalez, Loreto Ferrús Abad, Maria Dolores Ocete, Inma Galán Vendrell, Paula Ruiz-Hueso, Mariana Reyes-Prieto, Vicente Soriano Chirona, Maria Alma Bracho, Llúcia Martínez-Priego, Concepcion Gimeno, Giuseppe D'Auria, Fernando Gonzalez-Candelas |
| EPI_ISL_436205 | Servicio de Microbiología. Consorcio Hospital General Universitario de Valencia | Sequencing and Bioinformatics Service and Molecular Epidemiology Research Group. FISABIO-Public Health | Beatriz Beamud, Lidia Ruiz Roldan, Marta Pla Diaz,Neris Garcia-Gonzalez, Loreto Ferrús Abad, Maria Dolores Ocete, Inma Galán Vendrell, Paula Ruiz-Hueso, Mariana Reyes-Prieto, Vicente Soriano Chirona, Maria Alma Bracho, Griselda De Marco, Llúcia Martínez-Priego, Concepcion Gimeno, Giuseppe D'Auria, Fernando Gonzalez-Candelas |
| EPI_ISL_436206 | Servicio de Microbiología. Consorcio Hospital General Universitario de Valencia | Sequencing and Bioinformatics Service and Molecular Epidemiology Research Group. FISABIO-Public Health | Lidia Ruiz Roldan, Marta Pla Diaz,Neris Garcia-Gonzalez, Loreto Ferrús Abad, Maria Dolores Ocete, Inma Galán Vendrell, Paula Ruiz-Hueso, Mariana Reyes-Prieto, Vicente Soriano Chirona, Maria Alma Bracho, Griselda De Marco, Beatriz Beamud, Llúcia Martínez-Priego, Concepcion Gimeno, Giuseppe D'Auria, Fernando Gonzalez-Candelas |
| EPI_ISL_436207 | Servicio de Microbiología. Consorcio Hospital General Universitario de Valencia | Sequencing and Bioinformatics Service and Molecular Epidemiology Research Group. FISABIO-Public Health | Marta Pla Diaz,Neris Garcia-Gonzalez, Loreto Ferrús Abad, Maria Dolores Ocete, Inma Galán Vendrell, Paula Ruiz-Hueso, Mariana Reyes-Prieto, Vicente Soriano Chirona, Maria Alma Bracho, Griselda De Marco, Beatriz Beamud, Lidia Ruiz Roldan, Llúcia Martínez-Priego, Concepcion Gimeno, Giuseppe D'Auria, Fernando Gonzalez-Candelas |
| EPI_ISL_436208 | Servicio de Microbiología. Consorcio Hospital General Universitario de Valencia | Sequencing and Bioinformatics Service and Molecular Epidemiology Research Group. FISABIO-Public Health | Neris Garcia-Gonzalez, Loreto Ferrús Abad, Maria Dolores Ocete, Inma Galán Vendrell, Paula Ruiz-Hueso, Mariana Reyes-Prieto, Vicente Soriano Chirona, Maria Alma Bracho, Griselda De Marco, Beatriz Beamud, Lidia Ruiz Roldan, Marta Pla Diaz, Llúcia Martínez-Priego, Concepcion Gimeno, Giuseppe D'Auria, Fernando Gonzalez-Candelas |

| | | | |
|---|---|---|---|
| EPI_ISL_436209 | Servicio de Microbiología. Consorcio Hospital General Universitario de Valencia | Sequencing and Bioinformatics Service and Molecular Epidemiology Research Group. FISABIO-Public Health | Loreto Ferrús Abad, Maria Dolores Ocete, Inma Galán Vendrell, Paula Ruiz-Hueso, Mariana Reyes-Prieto, Vicente Soriano Chirona, Maria Alma Bracho, Griselda De Marco, Beatriz Beamud, Lidia Ruiz Roldan, Marta Pla Diaz,Neris Garcia-Gonzalez, Llúcia Martínez-Priego, Concepcion Gimeno, Giuseppe D'Auria, Fernando Gonzalez-Candelas |
| EPI_ISL_436210 | Servicio de Microbiología. Consorcio Hospital General Universitario de Valencia | Sequencing and Bioinformatics Service and Molecular Epidemiology Research Group. FISABIO-Public Health | Loreto Ferrús Abad, Maria Dolores Ocete,Inma Galán Vendrell, Paula Ruiz-Hueso, Mariana Reyes-Prieto, Vicente Soriano Chirona, Maria Alma Bracho, Griselda De Marco, Beatriz Beamud, Lidia Ruiz Roldan, Marta Pla Diaz,Neris Garcia-Gonzalez, Llúcia Martínez-Priego, Concepcion Gimeno, Giuseppe D'Auria, Fernando Gonzalez-Candelas |
| EPI_ISL_436211 | Servicio de Microbiología. Consorcio Hospital General Universitario de Valencia | Sequencing and Bioinformatics Service and Molecular Epidemiology Research Group. FISABIO-Public Health | Maria Dolores Ocete, Inma Galán Vendrell, Paula Ruiz-Hueso, Mariana Reyes-Prieto, Vicente Soriano Chirona, Maria Alma Bracho, Griselda De Marco, Beatriz Beamud, Lidia Ruiz Roldan, Marta Pla Diaz,Neris Garcia-Gonzalez, Loreto Ferrús Abad, Llúcia Martínez-Priego, Concepcion Gimeno, Giuseppe D'Auria, Fernando Gonzalez-Candelas |
| EPI_ISL_436212 | Servicio de Microbiología. Consorcio Hospital General Universitario de Valencia | Sequencing and Bioinformatics Service and Molecular Epidemiology Research Group. FISABIO-Public Health | Griselda De Marco, Beatriz Beamud, Lidia Ruiz Roldan, Marta Pla Diaz,Neris Garcia-Gonzalez, Loreto Ferrús Abad, Maria Dolores Ocete, Inma Galán Vendrell, Paula Ruiz-Hueso, Mariana Reyes-Prieto, Vicente Soriano Chirona, Maria Alma Bracho, Llúcia Martínez-Priego, Concepcion Gimeno, Giuseppe D'Auria, Fernando Gonzalez-Candelas |
| EPI_ISL_436213 | Servicio de Microbiología. Consorcio Hospital General Universitario de Valencia | Sequencing and Bioinformatics Service and Molecular Epidemiology Research Group. FISABIO-Public Health | Beatriz Beamud, Lidia Ruiz Roldan, Marta Pla Diaz,Neris Garcia-Gonzalez, Loreto Ferrús Abad, Maria Dolores Ocete, Inma Galán Vendrell, Paula Ruiz-Hueso, Mariana Reyes-Prieto, Vicente Soriano Chirona, Maria Alma Bracho, Griselda De Marco, Llúcia Martínez-Priego, Concepcion Gimeno, Giuseppe D'Auria, Fernando Gonzalez-Candelas |
| EPI_ISL_436214 | Servicio de Microbiología. Consorcio Hospital General Universitario de Valencia | Sequencing and Bioinformatics Service and Molecular Epidemiology Research Group. FISABIO-Public Health | Lidia Ruiz Roldan, Marta Pla Diaz,Neris Garcia-Gonzalez, Loreto Ferrús Abad, Maria Dolores Ocete, Inma Galán Vendrell, Paula Ruiz-Hueso, Mariana Reyes-Prieto, Vicente Soriano Chirona, Maria Alma Bracho, Griselda De Marco, Beatriz Beamud, Llúcia Martínez-Priego, Concepcion Gimeno, Giuseppe D'Auria, Fernando Gonzalez-Candelas |
| EPI_ISL_436215 | Servicio de Microbiología. Consorcio Hospital General Universitario de Valencia | Sequencing and Bioinformatics Service and Molecular Epidemiology Research Group. FISABIO-Public Health | Marta Pla Diaz,Neris Garcia-Gonzalez, Loreto Ferrús Abad, Maria Dolores Ocete, Inma Galán Vendrell, Paula Ruiz-Hueso, Mariana Reyes-Prieto, Vicente Soriano Chirona, Maria Alma Bracho, Griselda De Marco, Beatriz Beamud, Lidia Ruiz Roldan, Llúcia Martínez-Priego, Concepcion Gimeno, Giuseppe D'Auria, Fernando Gonzalez-Candelas |
| EPI_ISL_436216 | Servicio de Microbiología. Consorcio Hospital General Universitario de Valencia | Sequencing and Bioinformatics Service and Molecular Epidemiology Research Group. FISABIO-Public Health | Neris Garcia-Gonzalez, Loreto Ferrús Abad, Maria Dolores Ocete, Inma Galán Vendrell, Paula Ruiz-Hueso, Mariana Reyes-Prieto, Vicente Soriano Chirona, Maria Alma Bracho, Griselda De Marco, Beatriz Beamud, Lidia Ruiz Roldan, Marta Pla Diaz, Llúcia Martínez-Priego, Concepcion Gimeno, Giuseppe D'Auria, Fernando Gonzalez-Candelas |
| EPI_ISL_436217 | Servicio de Microbiología. Consorcio Hospital General Universitario de Valencia | Sequencing and Bioinformatics Service and Molecular Epidemiology Research Group. FISABIO-Public Health | Loreto Ferrús Abad, Maria Dolores Ocete, Inma Galán Vendrell, Paula Ruiz-Hueso, Mariana Reyes-Prieto, Vicente Soriano Chirona, Maria Alma Bracho, Griselda De Marco, Beatriz Beamud, Lidia Ruiz Roldan, Marta Pla Diaz,Neris Garcia-Gonzalez, Llúcia Martínez-Priego, Concepcion Gimeno, Giuseppe D'Auria, Fernando Gonzalez-Candelas |
| EPI_ISL_436218 | Servicio de Microbiología. Consorcio Hospital General Universitario de Valencia | Sequencing and Bioinformatics Service and Molecular Epidemiology Research Group. FISABIO-Public Health | Loreto Ferrús Abad, Maria Dolores Ocete,Inma Galán Vendrell, Paula Ruiz-Hueso, Mariana Reyes-Prieto, Vicente Soriano Chirona, Maria Alma Bracho, Griselda De Marco, Beatriz Beamud, Lidia Ruiz Roldan, Marta Pla Diaz,Neris Garcia-Gonzalez, Llúcia Martínez-Priego, Concepcion Gimeno, Giuseppe D'Auria, Fernando Gonzalez-Candelas |
| EPI_ISL_436219 | Servicio de Microbiología. Consorcio Hospital General Universitario de Valencia | Sequencing and Bioinformatics Service and Molecular Epidemiology Research Group. FISABIO-Public Health | Maria Dolores Ocete, Inma Galán Vendrell, Paula Ruiz-Hueso, Mariana Reyes-Prieto, Vicente Soriano Chirona, Maria Alma Bracho, Griselda De Marco, Beatriz Beamud, Lidia Ruiz Roldan, Marta Pla Diaz,Neris Garcia-Gonzalez, Loreto Ferrús Abad, Llúcia Martínez-Priego, Concepcion Gimeno, Giuseppe D'Auria, Fernando Gonzalez-Candelas |
| EPI_ISL_436220 | Servicio de Microbiología. Consorcio Hospital General Universitario de Valencia | Sequencing and Bioinformatics Service and Molecular Epidemiology Research Group. FISABIO-Public Health | Griselda De Marco, Beatriz Beamud, Lidia Ruiz Roldan, Marta Pla Diaz,Neris Garcia-Gonzalez, Loreto Ferrús Abad, Maria Dolores Ocete, Inma Galán Vendrell, Paula Ruiz-Hueso, Mariana Reyes-Prieto, Vicente Soriano Chirona, Maria Alma Bracho, Llúcia Martínez-Priego, Concepcion Gimeno, Giuseppe D'Auria, Fernando Gonzalez-Candelas |
| EPI_ISL_436221 | Servicio de Microbiología. Consorcio Hospital General Universitario de Valencia | Sequencing and Bioinformatics Service and Molecular Epidemiology Research Group. FISABIO-Public Health | Beatriz Beamud, Lidia Ruiz Roldan, Marta Pla Diaz,Neris Garcia-Gonzalez, Loreto Ferrús Abad, Maria Dolores Ocete, Inma Galán Vendrell, Paula Ruiz-Hueso, Mariana Reyes-Prieto, Vicente Soriano Chirona, Maria Alma Bracho, Griselda De Marco, Llúcia Martínez-Priego, Concepcion Gimeno, Giuseppe D'Auria, Fernando Gonzalez-Candelas |
| EPI_ISL_436222 | Servicio de Microbiología. Consorcio Hospital General Universitario de Valencia | Sequencing and Bioinformatics Service and Molecular Epidemiology Research Group. FISABIO-Public Health | Lidia Ruiz Roldan, Marta Pla Diaz,Neris Garcia-Gonzalez, Loreto Ferrús Abad, Maria Dolores Ocete, Inma Galán Vendrell, Paula Ruiz-Hueso, Mariana Reyes-Prieto, Vicente Soriano Chirona, Maria Alma Bracho, Griselda De Marco, Beatriz Beamud, Llúcia Martínez-Priego, Concepcion Gimeno, Giuseppe D'Auria, Fernando Gonzalez-Candelas |
| EPI_ISL_436223 | Servicio de Microbiología. Consorcio Hospital General Universitario de Valencia | Sequencing and Bioinformatics Service and Molecular Epidemiology Research Group. FISABIO-Public Health | Marta Pla Diaz,Neris Garcia-Gonzalez, Loreto Ferrús Abad, Maria Dolores Ocete, Inma Galán Vendrell, Paula Ruiz-Hueso, Mariana Reyes-Prieto, Vicente Soriano Chirona, Maria Alma Bracho, Griselda De Marco, Beatriz Beamud, Lidia Ruiz Roldan, Llúcia Martínez-Priego, Concepcion Gimeno, Giuseppe D'Auria, Fernando Gonzalez-Candelas |
| EPI_ISL_436224 | Servicio de Microbiología. Consorcio Hospital General Universitario de Valencia | Sequencing and Bioinformatics Service and Molecular Epidemiology Research Group. FISABIO-Public Health | Neris Garcia-Gonzalez, Loreto Ferrús Abad, Maria Dolores Ocete, Inma Galán Vendrell, Paula Ruiz-Hueso, Mariana Reyes-Prieto, Vicente Soriano Chirona, Maria Alma Bracho, Griselda De Marco, Beatriz Beamud, Lidia Ruiz Roldan, Marta Pla Diaz, Llúcia Martínez-Priego, Concepcion Gimeno, Giuseppe D'Auria, Fernando Gonzalez-Candelas |
| EPI_ISL_436225 | Servicio de Microbiología. Consorcio Hospital General Universitario de Valencia | Sequencing and Bioinformatics Service and Molecular Epidemiology Research Group. FISABIO-Public Health | Loreto Ferrús Abad, Maria Dolores Ocete, Inma Galán Vendrell, Paula Ruiz-Hueso, Mariana Reyes-Prieto, Vicente Soriano Chirona, Maria Alma Bracho, Griselda De Marco, Beatriz Beamud, Lidia Ruiz Roldan, Marta Pla Diaz,Neris Garcia-Gonzalez, Llúcia Martínez-Priego, Concepcion Gimeno, Giuseppe D'Auria, Fernando Gonzalez-Candelas |
| EPI_ISL_436226 | Servicio de Microbiología. Consorcio Hospital General Universitario de Valencia | Sequencing and Bioinformatics Service and Molecular Epidemiology Research Group. FISABIO-Public Health | Maria Dolores Ocete, Inma Galán Vendrell, Paula Ruiz-Hueso, Mariana Reyes-Prieto, Vicente Soriano Chirona, Maria Alma Bracho, Griselda De Marco, Beatriz Beamud, Lidia Ruiz Roldan, Marta Pla Diaz,Neris Garcia-Gonzalez, Loreto Ferrús Abad, Llúcia Martínez-Priego, Concepcion Gimeno, Giuseppe D'Auria, Fernando Gonzalez-Candelas |
| EPI_ISL_436227 | Servicio de Microbiología. Consorcio Hospital General Universitario de Valencia | Sequencing and Bioinformatics Service and Molecular Epidemiology Research Group. FISABIO-Public Health | Griselda De Marco, Beatriz Beamud, Lidia Ruiz Roldan, Marta Pla Diaz,Neris Garcia-Gonzalez, Loreto Ferrús Abad, Maria Dolores Ocete, Inma Galán Vendrell, Paula Ruiz-Hueso, Mariana Reyes-Prieto, Vicente Soriano Chirona, Maria Alma Bracho, Llúcia Martínez-Priego, Concepcion Gimeno, Giuseppe D'Auria, Fernando Gonzalez-Candelas |
| EPI_ISL_436228 | Servicio de Microbiología. Consorcio Hospital General Universitario de Valencia | Sequencing and Bioinformatics Service and Molecular Epidemiology Research Group. FISABIO-Public Health | Beatriz Beamud, Lidia Ruiz Roldan, Marta Pla Diaz,Neris Garcia-Gonzalez, Loreto Ferrús Abad, Maria Dolores Ocete, Inma Galán Vendrell, Paula Ruiz-Hueso, Mariana Reyes-Prieto, Vicente Soriano Chirona, Maria Alma Bracho, Griselda De Marco, Llúcia Martínez-Priego, Concepcion Gimeno, Giuseppe D'Auria, Fernando Gonzalez-Candelas |
| EPI_ISL_436229 | Servicio de Microbiología. Consorcio Hospital General Universitario de Valencia | Sequencing and Bioinformatics Service and Molecular Epidemiology Research Group. FISABIO-Public Health | Lidia Ruiz Roldan, Marta Pla Diaz,Neris Garcia-Gonzalez, Loreto Ferrús Abad, Maria Dolores Ocete, Inma Galán Vendrell, Paula Ruiz-Hueso, Mariana Reyes-Prieto, Vicente Soriano Chirona, Maria Alma Bracho, Griselda De Marco, Beatriz Beamud, Llúcia Martínez-Priego, Concepcion Gimeno, Giuseppe D'Auria, Fernando Gonzalez-Candelas |
| EPI_ISL_436230 | Servicio de Microbiología. Consorcio Hospital General Universitario de Valencia | Sequencing and Bioinformatics Service and Molecular Epidemiology Research Group. FISABIO-Public Health | Marta Pla Diaz,Neris Garcia-Gonzalez, Loreto Ferrús Abad, Maria Dolores Ocete, Inma Galán Vendrell, Paula Ruiz-Hueso, Mariana Reyes-Prieto, Vicente Soriano Chirona, Maria Alma Bracho, Griselda De Marco, Beatriz Beamud, Lidia Ruiz Roldan, Llúcia Martínez-Priego, Concepcion Gimeno, Giuseppe D'Auria, Fernando Gonzalez-Candelas |
| EPI_ISL_436231 | Servicio de Microbiología. Consorcio Hospital General Universitario de Valencia | Sequencing and Bioinformatics Service and Molecular Epidemiology Research Group. FISABIO-Public Health | Neris Garcia-Gonzalez, Loreto Ferrús Abad, Maria Dolores Ocete, Inma Galán Vendrell, Paula Ruiz-Hueso, Mariana Reyes-Prieto, Vicente Soriano Chirona, Maria Alma Bracho, Griselda De Marco, Beatriz Beamud, Lidia Ruiz Roldan, Marta Pla Diaz, Llúcia Martínez-Priego, Concepcion Gimeno, Giuseppe D'Auria, Fernando Gonzalez-Candelas |
| EPI_ISL_436232 | Servicio de Microbiología. Consorcio Hospital General Universitario de Valencia | Sequencing and Bioinformatics Service and Molecular Epidemiology Research Group. FISABIO-Public Health | Loreto Ferrús Abad, Maria Dolores Ocete, Inma Galán Vendrell, Paula Ruiz-Hueso, Mariana Reyes-Prieto, Vicente Soriano Chirona, Maria Alma Bracho, Griselda De Marco, Beatriz Beamud, Lidia Ruiz Roldan, Marta Pla Diaz,Neris Garcia-Gonzalez, Llúcia Martínez-Priego, Concepcion Gimeno, Giuseppe D'Auria, Fernando Gonzalez-Candelas |
| EPI_ISL_436233 | Servicio de Microbiología. Consorcio Hospital General Universitario de Valencia | Sequencing and Bioinformatics Service and Molecular Epidemiology Research Group. FISABIO-Public Health | Loreto Ferrús Abad, Maria Dolores Ocete,Inma Galán Vendrell, Paula Ruiz-Hueso, Mariana Reyes-Prieto, Vicente Soriano Chirona, Maria Alma Bracho, Griselda De Marco, Beatriz Beamud, Lidia Ruiz Roldan, Marta Pla Diaz,Neris Garcia-Gonzalez, Llúcia Martínez-Priego, Concepcion Gimeno, Giuseppe D'Auria, Fernando Gonzalez-Candelas |
| EPI_ISL_436234 | Servicio de Microbiología. Consorcio Hospital General Universitario de Valencia | Sequencing and Bioinformatics Service and Molecular Epidemiology Research Group. FISABIO-Public Health | Maria Dolores Ocete, Inma Galán Vendrell, Paula Ruiz-Hueso, Mariana Reyes-Prieto, Vicente Soriano Chirona, Maria Alma Bracho, Griselda De Marco, Beatriz Beamud, Lidia Ruiz Roldan, Marta Pla Diaz,Neris Garcia-Gonzalez, Loreto Ferrús Abad, Llúcia Martínez-Priego, Concepcion Gimeno, Giuseppe D'Auria, Fernando Gonzalez-Candelas |
| EPI_ISL_436235, EPI_ISL_436236, EPI_ISL_436237, EPI_ISL_436238, EPI_ISL_436239, EPI_ISL_436240, EPI_ISL_436241, EPI_ISL_436242, EPI_ISL_436243, EPI_ISL_436244, EPI_ISL_436245, EPI_ISL_436246, EPI_ISL_436247, EPI_ISL_436248, EPI_ISL_436249, EPI_ISL_436250, EPI_ISL_436251, EPI_ISL_436252, EPI_ISL_436253, EPI_ISL_436254, EPI_ISL_436255, EPI_ISL_436256, EPI_ISL_436257, EPI_ISL_436258, EPI_ISL_436259, EPI_ISL_436260, EPI_ISL_436261, EPI_ISL_436262, EPI_ISL_436263, EPI_ISL_436264, EPI_ISL_436265, EPI_ISL_436266, EPI_ISL_436267, EPI_ISL_436268, EPI_ISL_436269, EPI_ISL_436270 | | | |
| see above | Servicio de Microbiologia. Hospital Universitario Doctor Peset | Sequencing and Bioinformatics Service and Molecular Epidemiology Research Group. FISABIO-Public Health | Juan Alberola Enguídanos, Juan Jose Camarena Miñana, Rosa González Pellicer, Neris Garcia-Gonzalez, Inma Galán Vendrell, Sandra Carbo, Loreto Ferrús Abad, Paula Ruiz-Hueso, Mariana Reyes-Prieto, Vicente Soriano Chirona, Ivan Ansari, Maria Alma Bracho, Griselda De Marco, Beatriz Beamud, Lidia Ruiz Roldan, Marta Pla Diaz, Llúcia Martínez-Priego, Giuseppe D'Auria, Jose Miguel Nogueira Coito, Fernando Gonzalez-Candelas |
| EPI_ISL_436271 | Servicio de Microbiología. Consorcio Hospital General Universitario de Valencia | Sequencing and Bioinformatics Service and Molecular Epidemiology Research Group. FISABIO-Public Health | Griselda De Marco, Beatriz Beamud, Lidia Ruiz Roldan, Marta Pla Diaz,Neris Garcia-Gonzalez, Loreto Ferrús Abad, Maria Dolores Ocete, Inma Galán Vendrell, Paula Ruiz-Hueso, Mariana Reyes-Prieto, Vicente Soriano Chirona, Maria Alma Bracho, Llúcia Martínez-Priego, Concepcion Gimeno, Giuseppe D'Auria, Fernando Gonzalez-Candelas |
| EPI_ISL_436272 | Servicio de Microbiología. Consorcio Hospital General Universitario de Valencia | Sequencing and Bioinformatics Service and Molecular Epidemiology Research Group. FISABIO-Public Health | Beatriz Beamud, Lidia Ruiz Roldan, Marta Pla Diaz,Neris Garcia-Gonzalez, Loreto Ferrús Abad, Maria Dolores Ocete, Inma Galán Vendrell, Paula Ruiz-Hueso, Mariana Reyes-Prieto, Vicente Soriano Chirona, Maria Alma Bracho, Griselda De Marco, Llúcia Martínez-Priego, Concepcion Gimeno, Giuseppe D'Auria, Fernando Gonzalez-Candelas |
| EPI_ISL_436273 | Servicio de Microbiología. Consorcio Hospital General Universitario de Valencia | Sequencing and Bioinformatics Service and Molecular Epidemiology Research Group. FISABIO-Public Health | Lidia Ruiz Roldan, Marta Pla Diaz,Neris Garcia-Gonzalez, Loreto Ferrús Abad, Maria Dolores Ocete, Inma Galán Vendrell, Paula Ruiz-Hueso, Mariana Reyes-Prieto, Vicente Soriano Chirona, Maria Alma Bracho, Griselda De Marco, Beatriz Beamud, Llúcia Martínez-Priego, Concepcion Gimeno, Giuseppe D'Auria, Fernando Gonzalez-Candelas |
| EPI_ISL_436274 | Servicio de Microbiología. Consorcio Hospital General Universitario de Valencia | Sequencing and Bioinformatics Service and Molecular Epidemiology Research Group. FISABIO-Public Health | Marta Pla Diaz,Neris Garcia-Gonzalez, Loreto Ferrús Abad, Maria Dolores Ocete, Inma Galán Vendrell, Paula Ruiz-Hueso, Mariana Reyes-Prieto, Vicente Soriano Chirona, Maria Alma Bracho, Griselda De Marco, Beatriz Beamud, Lidia Ruiz Roldan, Llúcia Martínez-Priego, Concepcion Gimeno, Giuseppe D'Auria, Fernando Gonzalez-Candelas |
| EPI_ISL_436275 | Servicio de Microbiología. Consorcio Hospital General Universitario de Valencia | Sequencing and Bioinformatics Service and Molecular Epidemiology Research Group. FISABIO-Public Health | Neris Garcia-Gonzalez, Loreto Ferrús Abad, Maria Dolores Ocete, Inma Galán Vendrell, Paula Ruiz-Hueso, Mariana Reyes-Prieto, Vicente Soriano Chirona, Maria Alma Bracho, Griselda De Marco, Beatriz Beamud, Lidia Ruiz Roldan, Marta Pla Diaz, Llúcia Martínez-Priego, Concepcion Gimeno, Giuseppe D'Auria, Fernando Gonzalez-Candelas |
| EPI_ISL_436276 | Servicio de Microbiología. Consorcio Hospital General Universitario de Valencia | Sequencing and Bioinformatics Service and Molecular Epidemiology Research Group. FISABIO-Public Health | Loreto Ferrús Abad, Maria Dolores Ocete, Inma Galán Vendrell, Paula Ruiz-Hueso, Mariana Reyes-Prieto, Vicente Soriano Chirona, Maria Alma Bracho, Griselda De Marco, Beatriz Beamud, Lidia Ruiz Roldan, Marta Pla Diaz,Neris Garcia-Gonzalez, Llúcia Martínez-Priego, Concepcion Gimeno, Giuseppe D'Auria, Fernando Gonzalez-Candelas |
| EPI_ISL_436277 | Servicio de Microbiología. Consorcio Hospital General Universitario de Valencia | Sequencing and Bioinformatics Service and Molecular Epidemiology Research Group. FISABIO-Public Health | Neris Garcia-Gonzalez, Loreto Ferrús Abad, Maria Dolores Ocete, Inma Galán Vendrell, Paula Ruiz-Hueso, Mariana Reyes-Prieto, Vicente Soriano Chirona, Maria Alma Bracho, Griselda De Marco, Beatriz Beamud, Lidia Ruiz Roldan, Marta Pla Diaz, Llúcia Martínez-Priego, Concepcion Gimeno, Giuseppe D'Auria, Fernando Gonzalez-Candelas |
| EPI_ISL_436278 | Servicio de Microbiologia. Hospital Clinico Universitario de Valencia | Sequencing and Bioinformatics Service and Molecular Epidemiology Research Group. FISABIO-Public Health | David Navarro, Maria Alma Bracho, Griselda De Marco, Beatriz Beamud, Lidia Ruiz Roldan, Marta Pla Diaz, Neris Garcia-Gonzalez, Inma Galán Vendrell, Sandra Carbo, Loreto Ferrús Abad, Paula Ruiz-Hueso, Mariana Reyes-Prieto, Vicente Soriano Chirona, Ivan Ansari, David Navarro, Llúcia Martínez-Priego, Giuseppe D'Auria, Fernando Gonzalez-Candelas |
| EPI_ISL_436279 | Servicio de Microbiologia. Hospital Clinico Universitario de Valencia | Sequencing and Bioinformatics Service and Molecular Epidemiology Research Group. FISABIO-Public Health | Griselda De Marco, Beatriz Beamud, Lidia Ruiz Roldan, Marta Pla Diaz, Neris Garcia-Gonzalez, Inma Galán Vendrell, Sandra Carbo, Loreto Ferrús Abad, Paula Ruiz-Hueso, Mariana Reyes-Prieto, Vicente Soriano Chirona, Ivan Ansari, David Navarro, Maria Alma Bracho, Llúcia Martínez-Priego, Giuseppe D'Auria, Fernando Gonzalez-Candelas |
| EPI_ISL_436280 | Servicio de Microbiologia. Hospital Clinico Universitario de Valencia | Sequencing and Bioinformatics Service and Molecular Epidemiology Research Group. FISABIO-Public Health | Beatriz Beamud, Lidia Ruiz Roldan, Marta Pla Diaz, Neris Garcia-Gonzalez, Inma Galán Vendrell, Sandra Carbo, Loreto Ferrús Abad, Paula Ruiz-Hueso, Mariana Reyes-Prieto, Vicente Soriano Chirona, Ivan Ansari, David Navarro, Maria Alma Bracho, Griselda De Marco, Llúcia Martínez-Priego, Giuseppe D'Auria, Fernando Gonzalez-Candelas |
| EPI_ISL_436281 | Servicio de Microbiologia. Hospital Clinico Universitario de Valencia | Sequencing and Bioinformatics Service and Molecular Epidemiology Research Group. FISABIO-Public Health | Lidia Ruiz Roldan, Marta Pla Diaz, Neris Garcia-Gonzalez, Inma Galán Vendrell, Sandra Carbo, Loreto Ferrús Abad, Paula Ruiz-Hueso, Mariana Reyes-Prieto, Vicente Soriano Chirona, Ivan Ansari, David Navarro, Maria Alma Bracho, Griselda De Marco, Beatriz Beamud, Llúcia Martínez-Priego, Giuseppe D'Auria, Fernando Gonzalez-Candelas |
| EPI_ISL_436282 | Servicio de Microbiologia. Hospital Clinico Universitario de Valencia | Sequencing and Bioinformatics Service and Molecular Epidemiology Research Group. FISABIO-Public Health | Marta Pla Diaz, Neris Garcia-Gonzalez, Inma Galán Vendrell, Sandra Carbo, Loreto Ferrús Abad, Paula Ruiz-Hueso, Mariana Reyes-Prieto, Vicente Soriano Chirona, Ivan Ansari, David Navarro, Maria Alma Bracho, Griselda De Marco, Beatriz Beamud, Lidia Ruiz Roldan, Llúcia Martínez-Priego, Giuseppe D'Auria, Fernando Gonzalez-Candelas |
| EPI_ISL_436283 | Servicio de Microbiologia. Hospital Clinico Universitario de Valencia | Sequencing and Bioinformatics Service and Molecular Epidemiology Research Group. FISABIO-Public Health | Neris Garcia-Gonzalez, Inma Galán Vendrell, Sandra Carbo, Loreto Ferrús Abad, Paula Ruiz-Hueso, Mariana Reyes-Prieto, Vicente Soriano Chirona, Ivan Ansari, David Navarro, Maria Alma Bracho, Griselda De Marco, Beatriz Beamud, Lidia Ruiz Roldan, Marta Pla Diaz, Llúcia Martínez-Priego, Giuseppe D'Auria, Fernando Gonzalez-Candelas |
| EPI_ISL_436284 | Servicio de Microbiologia. Hospital Clinico Universitario de Valencia | Sequencing and Bioinformatics Service and Molecular Epidemiology Research Group. FISABIO-Public Health | Inma Galán Vendrell, Sandra Carbo, Loreto Ferrús Abad, Paula Ruiz-Hueso, Mariana Reyes-Prieto, Vicente Soriano Chirona, Ivan Ansari, David Navarro, Maria Alma Bracho, Griselda De Marco, Beatriz Beamud, Lidia Ruiz Roldan, Marta Pla Diaz, Neris Garcia-Gonzalez, Llúcia Martínez-Priego, Giuseppe D'Auria, Fernando Gonzalez-Candelas |
| EPI_ISL_436285 | Servicio de Microbiologia. Hospital Clinico Universitario de Valencia | Sequencing and Bioinformatics Service and Molecular Epidemiology Research Group. FISABIO-Public Health | Sandra Carbo, Loreto Ferrús Abad, Paula Ruiz-Hueso, Mariana Reyes-Prieto, Vicente Soriano Chirona, Ivan Ansari, David Navarro, Maria Alma Bracho, Griselda De Marco, Beatriz Beamud, Lidia Ruiz Roldan, Marta Pla Diaz, Neris Garcia-Gonzalez, Inma Galán Vendrell, Llúcia Martínez-Priego, Giuseppe D'Auria, Fernando Gonzalez-Candelas |
| EPI_ISL_436286 | Servicio de Microbiologia. Hospital Clinico Universitario de Valencia | Sequencing and Bioinformatics Service and Molecular Epidemiology Research Group. FISABIO-Public Health | Loreto Ferrús Abad, Paula Ruiz-Hueso, Mariana Reyes-Prieto, Vicente Soriano Chirona, Ivan Ansari, David Navarro, Maria Alma Bracho, Griselda De Marco, Beatriz Beamud, Lidia Ruiz Roldan, Marta Pla Diaz, Neris Garcia-Gonzalez, Inma Galán Vendrell, Sandra Carbo, Llúcia Martínez-Priego, Giuseppe D'Auria, Fernando Gonzalez-Candelas |
| EPI_ISL_436287 | Servicio de Microbiologia. Hospital Clinico Universitario de Valencia | Sequencing and Bioinformatics Service and Molecular Epidemiology Research Group. FISABIO-Public Health | Paula Ruiz-Hueso, Mariana Reyes-Prieto, Vicente Soriano Chirona, Ivan Ansari, David Navarro, Maria Alma Bracho, Griselda De Marco, Beatriz Beamud, Lidia Ruiz Roldan, Marta Pla Diaz, Neris Garcia-Gonzalez, Inma Galán Vendrell, Sandra Carbo, Loreto Ferrús Abad, Llúcia Martínez-Priego, Giuseppe D'Auria, Fernando Gonzalez-Candelas |

| | | | |
|---|---|---|---|
| EPI_ISL_436288 | Servicio de Microbiologia. Hospital Clinico Universitario de Valencia | Sequencing and Bioinformatics Service and Molecular Epidemiology Research Group. FISABIO-Public Health | Mariana Reyes-Prieto, Vicente Soriano Chirona, Ivan Ansari, David Navarro, Maria Alma Bracho, Griselda De Marco, Beatriz Beamud, Lidia Ruiz Roldan, Marta Pla Diaz, Neris Garcia-Gonzalez, Inma Galán Vendrell, Sandra Carbo, Loreto Ferrús Abad, Paula Ruiz-Hueso, Llúcia Martínez-Priego, Giuseppe D'Auria, Fernando Gonzalez-Candelas |
| EPI_ISL_436289 | Servicio de Microbiologia. Hospital Clinico Universitario de Valencia | Sequencing and Bioinformatics Service and Molecular Epidemiology Research Group. FISABIO-Public Health | Vicente Soriano Chirona, Ivan Ansari, David Navarro, Maria Alma Bracho, Griselda De Marco, Beatriz Beamud, Lidia Ruiz Roldan, Marta Pla Diaz, Neris Garcia-Gonzalez, Inma Galán Vendrell, Sandra Carbo, Loreto Ferrús Abad, Paula Ruiz-Hueso, Mariana Reyes-Prieto, Llúcia Martínez-Priego, Giuseppe D'Auria, Fernando Gonzalez-Candelas |
| EPI_ISL_436290 | Servicio de Microbiologia. Hospital Clinico Universitario de Valencia | Sequencing and Bioinformatics Service and Molecular Epidemiology Research Group. FISABIO-Public Health | Ivan Ansari, David Navarro, Maria Alma Bracho, Griselda De Marco, Beatriz Beamud, Lidia Ruiz Roldan, Marta Pla Diaz, Neris Garcia-Gonzalez, Inma Galán Vendrell, Sandra Carbo, Loreto Ferrús Abad, Paula Ruiz-Hueso, Mariana Reyes-Prieto, Vicente Soriano Chirona, Llúcia Martínez-Priego, Giuseppe D'Auria, Fernando Gonzalez-Candelas |
| EPI_ISL_436291 | Servicio de Microbiologia. Hospital Clinico Universitario de Valencia | Sequencing and Bioinformatics Service and Molecular Epidemiology Research Group. FISABIO-Public Health | David Navarro, Maria Alma Bracho, Griselda De Marco, Beatriz Beamud, Lidia Ruiz Roldan, Marta Pla Diaz, Neris Garcia-Gonzalez, Inma Galán Vendrell, Sandra Carbo, Loreto Ferrús Abad, Paula Ruiz-Hueso, Mariana Reyes-Prieto, Vicente Soriano Chirona, Ivan Ansari, Llúcia Martínez-Priego, Giuseppe D'Auria, Fernando Gonzalez-Candelas |
| EPI_ISL_436292 | Servicio de Microbiologia. Hospital Clinico Universitario de Valencia | Sequencing and Bioinformatics Service and Molecular Epidemiology Research Group. FISABIO-Public Health | Maria Alma Bracho, Griselda De Marco, Beatriz Beamud, Lidia Ruiz Roldan, Marta Pla Diaz, Neris Garcia-Gonzalez, Inma Galán Vendrell, Sandra Carbo, Loreto Ferrús Abad, Paula Ruiz-Hueso, Mariana Reyes-Prieto, Vicente Soriano Chirona, Ivan Ansari, David Navarro, Llúcia Martínez-Priego, Giuseppe D'Auria, Fernando Gonzalez-Candelas |
| EPI_ISL_436293 | Servicio de Microbiologia. Hospital Clinico Universitario de Valencia | Sequencing and Bioinformatics Service and Molecular Epidemiology Research Group. FISABIO-Public Health | Griselda De Marco, Beatriz Beamud, Lidia Ruiz Roldan, Marta Pla Diaz, Neris Garcia-Gonzalez, Inma Galán Vendrell, Sandra Carbo, Loreto Ferrús Abad, Paula Ruiz-Hueso, Mariana Reyes-Prieto, Vicente Soriano Chirona, Ivan Ansari, David Navarro, Maria Alma Bracho, Llúcia Martínez-Priego, Giuseppe D'Auria, Fernando Gonzalez-Candelas |
| EPI_ISL_436294 | Servicio de Microbiologia. Hospital Clinico Universitario de Valencia | Sequencing and Bioinformatics Service and Molecular Epidemiology Research Group. FISABIO-Public Health | Beatriz Beamud, Lidia Ruiz Roldan, Marta Pla Diaz, Neris Garcia-Gonzalez, Inma Galán Vendrell, Sandra Carbo, Loreto Ferrús Abad, Paula Ruiz-Hueso, Mariana Reyes-Prieto, Vicente Soriano Chirona, Ivan Ansari, David Navarro, Maria Alma Bracho, Griselda De Marco, Llúcia Martínez-Priego, Giuseppe D'Auria, Fernando Gonzalez-Candelas |
| EPI_ISL_436295 | Servicio de Microbiologia. Hospital Clinico Universitario de Valencia | Sequencing and Bioinformatics Service and Molecular Epidemiology Research Group. FISABIO-Public Health | Lidia Ruiz Roldan, Marta Pla Diaz, Neris Garcia-Gonzalez, Inma Galán Vendrell, Sandra Carbo, Loreto Ferrús Abad, Paula Ruiz-Hueso, Mariana Reyes-Prieto, Vicente Soriano Chirona, Ivan Ansari, David Navarro, Maria Alma Bracho, Griselda De Marco, Beatriz Beamud, Llúcia Martínez-Priego, Giuseppe D'Auria, Fernando Gonzalez-Candelas |
| EPI_ISL_436296 | Servicio de Microbiologia. Hospital Clinico Universitario de Valencia | Sequencing and Bioinformatics Service and Molecular Epidemiology Research Group. FISABIO-Public Health | Marta Pla Diaz, Neris Garcia-Gonzalez, Inma Galán Vendrell, Sandra Carbo, Loreto Ferrús Abad, Paula Ruiz-Hueso, Mariana Reyes-Prieto, Vicente Soriano Chirona, Ivan Ansari, David Navarro, Maria Alma Bracho, Griselda De Marco, Beatriz Beamud, Lidia Ruiz Roldan, Llúcia Martínez-Priego, Giuseppe D'Auria, Fernando Gonzalez-Candelas |
| EPI_ISL_436297 | Servicio de Microbiologia. Hospital Clinico Universitario de Valencia | Sequencing and Bioinformatics Service and Molecular Epidemiology Research Group. FISABIO-Public Health | Neris Garcia-Gonzalez, Inma Galán Vendrell, Sandra Carbo, Loreto Ferrús Abad, Paula Ruiz-Hueso, Mariana Reyes-Prieto, Vicente Soriano Chirona, Ivan Ansari, David Navarro, Maria Alma Bracho, Griselda De Marco, Beatriz Beamud, Lidia Ruiz Roldan, Marta Pla Diaz, Llúcia Martínez-Priego, Giuseppe D'Auria, Fernando Gonzalez-Candelas |
| EPI_ISL_436298 | Servicio de Microbiologia. Hospital Clinico Universitario de Valencia | Sequencing and Bioinformatics Service and Molecular Epidemiology Research Group. FISABIO-Public Health | David Navarro, Maria Alma Bracho, Griselda De Marco, Beatriz Beamud, Lidia Ruiz Roldan, Marta Pla Diaz, Neris Garcia-Gonzalez, Inma Galán Vendrell, Sandra Carbo, Loreto Ferrús Abad, Paula Ruiz-Hueso, Mariana Reyes-Prieto, Vicente Soriano Chirona, Ivan Ansari, David Navarro, Llúcia Martínez-Priego, Giuseppe D'Auria, Fernando Gonzalez-Candelas |
| EPI_ISL_436299 | Servicio de Microbiologia. Hospital Clinico Universitario de Valencia | Sequencing and Bioinformatics Service and Molecular Epidemiology Research Group. FISABIO-Public Health | Griselda De Marco, Beatriz Beamud, Lidia Ruiz Roldan, Marta Pla Diaz, Neris Garcia-Gonzalez, Inma Galán Vendrell, Sandra Carbo, Loreto Ferrús Abad, Paula Ruiz-Hueso, Mariana Reyes-Prieto, Vicente Soriano Chirona, Ivan Ansari, David Navarro, Maria Alma Bracho, Llúcia Martínez-Priego, Giuseppe D'Auria, Fernando Gonzalez-Candelas |
| EPI_ISL_436300 | Servicio de Microbiologia. Hospital Clinico Universitario de Valencia | Sequencing and Bioinformatics Service and Molecular Epidemiology Research Group. FISABIO-Public Health | Beatriz Beamud, Lidia Ruiz Roldan, Marta Pla Diaz, Neris Garcia-Gonzalez, Inma Galán Vendrell, Sandra Carbo, Loreto Ferrús Abad, Paula Ruiz-Hueso, Mariana Reyes-Prieto, Vicente Soriano Chirona, Ivan Ansari, David Navarro, Maria Alma Bracho, Griselda De Marco, Llúcia Martínez-Priego, Giuseppe D'Auria, Fernando Gonzalez-Candelas |
| EPI_ISL_436301 | Servicio de Microbiologia. Hospital Clinico Universitario de Valencia | Sequencing and Bioinformatics Service and Molecular Epidemiology Research Group. FISABIO-Public Health | Lidia Ruiz Roldan, Marta Pla Diaz, Neris Garcia-Gonzalez, Inma Galán Vendrell, Sandra Carbo, Loreto Ferrús Abad, Paula Ruiz-Hueso, Mariana Reyes-Prieto, Vicente Soriano Chirona, Ivan Ansari, David Navarro, Maria Alma Bracho, Griselda De Marco, Beatriz Beamud, Llúcia Martínez-Priego, Giuseppe D'Auria, Fernando Gonzalez-Candelas |
| EPI_ISL_436302 | Servicio de Microbiologia. Hospital Clinico Universitario de Valencia | Sequencing and Bioinformatics Service and Molecular Epidemiology Research Group. FISABIO-Public Health | Marta Pla Diaz, Neris Garcia-Gonzalez, Inma Galán Vendrell, Sandra Carbo, Loreto Ferrús Abad, Paula Ruiz-Hueso, Mariana Reyes-Prieto, Vicente Soriano Chirona, Ivan Ansari, David Navarro, Maria Alma Bracho, Griselda De Marco, Beatriz Beamud, Lidia Ruiz Roldan, Llúcia Martínez-Priego, Giuseppe D'Auria, Fernando Gonzalez-Candelas |
| EPI_ISL_436303 | Servicio de Microbiologia. Hospital Clinico Universitario de Valencia | Sequencing and Bioinformatics Service and Molecular Epidemiology Research Group. FISABIO-Public Health | Neris Garcia-Gonzalez, Inma Galán Vendrell, Sandra Carbo, Loreto Ferrús Abad, Paula Ruiz-Hueso, Mariana Reyes-Prieto, Vicente Soriano Chirona, Ivan Ansari, David Navarro, Maria Alma Bracho, Griselda De Marco, Beatriz Beamud, Lidia Ruiz Roldan, Marta Pla Diaz, Llúcia Martínez-Priego, Giuseppe D'Auria, Fernando Gonzalez-Candelas |
| EPI_ISL_436304 | Servicio de Microbiologia. Hospital Clinico Universitario de Valencia | Sequencing and Bioinformatics Service and Molecular Epidemiology Research Group. FISABIO-Public Health | Inma Galán Vendrell, Sandra Carbo, Loreto Ferrús Abad, Paula Ruiz-Hueso, Mariana Reyes-Prieto, Vicente Soriano Chirona, Ivan Ansari, David Navarro, Maria Alma Bracho, Griselda De Marco, Beatriz Beamud, Lidia Ruiz Roldan, Marta Pla Diaz, Neris Garcia-Gonzalez, Llúcia Martínez-Priego, Giuseppe D'Auria, Fernando Gonzalez-Candelas |
| EPI_ISL_436305 | Servicio de Microbiologia. Hospital Clinico Universitario de Valencia | Sequencing and Bioinformatics Service and Molecular Epidemiology Research Group. FISABIO-Public Health | Sandra Carbo, Loreto Ferrús Abad, Paula Ruiz-Hueso, Mariana Reyes-Prieto, Vicente Soriano Chirona, Ivan Ansari, David Navarro, Maria Alma Bracho, Griselda De Marco, Beatriz Beamud, Lidia Ruiz Roldan, Marta Pla Diaz, Neris Garcia-Gonzalez, Inma Galán Vendrell, Llúcia Martínez-Priego, Giuseppe D'Auria, Fernando Gonzalez-Candelas |
| EPI_ISL_436306 | Servicio de Microbiologia. Hospital Clinico Universitario de Valencia | Sequencing and Bioinformatics Service and Molecular Epidemiology Research Group. FISABIO-Public Health | Loreto Ferrús Abad, Paula Ruiz-Hueso, Mariana Reyes-Prieto, Vicente Soriano Chirona, Ivan Ansari, David Navarro, Maria Alma Bracho, Griselda De Marco, Beatriz Beamud, Lidia Ruiz Roldan, Marta Pla Diaz, Neris Garcia-Gonzalez, Inma Galán Vendrell, Sandra Carbo, Llúcia Martínez-Priego, Giuseppe D'Auria, Fernando Gonzalez-Candelas |
| EPI_ISL_436307 | Servicio de Microbiologia. Hospital Clinico Universitario de Valencia | Sequencing and Bioinformatics Service and Molecular Epidemiology Research Group. FISABIO-Public Health | Paula Ruiz-Hueso, Mariana Reyes-Prieto, Vicente Soriano Chirona, Ivan Ansari, David Navarro, Maria Alma Bracho, Griselda De Marco, Beatriz Beamud, Lidia Ruiz Roldan, Marta Pla Diaz, Neris Garcia-Gonzalez, Inma Galán Vendrell, Sandra Carbo, Loreto Ferrús Abad, Llúcia Martínez-Priego, Giuseppe D'Auria, Fernando Gonzalez-Candelas |
| EPI_ISL_436308 | Servicio de Microbiologia. Hospital Clinico Universitario de Valencia | Sequencing and Bioinformatics Service and Molecular Epidemiology Research Group. FISABIO-Public Health | Mariana Reyes-Prieto, Vicente Soriano Chirona, Ivan Ansari, David Navarro, Maria Alma Bracho, Griselda De Marco, Beatriz Beamud, Lidia Ruiz Roldan, Marta Pla Diaz, Neris Garcia-Gonzalez, Inma Galán Vendrell, Sandra Carbo, Loreto Ferrús Abad, Paula Ruiz-Hueso, Llúcia Martínez-Priego, Giuseppe D'Auria, Fernando Gonzalez-Candelas |
| EPI_ISL_436309 | Servicio de Microbiologia. Hospital Clinico Universitario de Valencia | Sequencing and Bioinformatics Service and Molecular Epidemiology Research Group. FISABIO-Public Health | Vicente Soriano Chirona, Ivan Ansari, David Navarro, Maria Alma Bracho, Griselda De Marco, Beatriz Beamud, Lidia Ruiz Roldan, Marta Pla Diaz, Neris Garcia-Gonzalez, Inma Galán Vendrell, Sandra Carbo, Loreto Ferrús Abad, Paula Ruiz-Hueso, Mariana Reyes-Prieto, Llúcia Martínez-Priego, Giuseppe D'Auria, Fernando Gonzalez-Candelas |
| EPI_ISL_436310 | Servicio de Microbiologia. Hospital Clinico Universitario de Valencia | Sequencing and Bioinformatics Service and Molecular Epidemiology Research Group. FISABIO-Public Health | Ivan Ansari, David Navarro, Maria Alma Bracho, Griselda De Marco, Beatriz Beamud, Lidia Ruiz Roldan, Marta Pla Diaz, Neris Garcia-Gonzalez, Inma Galán Vendrell, Sandra Carbo, Loreto Ferrús Abad, Paula Ruiz-Hueso, Mariana Reyes-Prieto, Vicente Soriano Chirona, Llúcia Martínez-Priego, Giuseppe D'Auria, Fernando Gonzalez-Candelas |
| EPI_ISL_436311 | Servicio de Microbiologia. Hospital Clinico Universitario de Valencia | Sequencing and Bioinformatics Service and Molecular Epidemiology Research Group. FISABIO-Public Health | David Navarro, Maria Alma Bracho, Griselda De Marco, Beatriz Beamud, Lidia Ruiz Roldan, Marta Pla Diaz, Neris Garcia-Gonzalez, Inma Galán Vendrell, Sandra Carbo, Loreto Ferrús Abad, Paula Ruiz-Hueso, Mariana Reyes-Prieto, Vicente Soriano Chirona, Ivan Ansari, Llúcia Martínez-Priego, Giuseppe D'Auria, Fernando Gonzalez-Candelas |
| EPI_ISL_436312 | Servicio de Microbiologia. Hospital Clinico Universitario de Valencia | Sequencing and Bioinformatics Service and Molecular Epidemiology Research Group. FISABIO-Public Health | Maria Alma Bracho, Griselda De Marco, Beatriz Beamud, Lidia Ruiz Roldan, Marta Pla Diaz, Neris Garcia-Gonzalez, Inma Galán Vendrell, Sandra Carbo, Loreto Ferrús Abad, Paula Ruiz-Hueso, Mariana Reyes-Prieto, Vicente Soriano Chirona, Ivan Ansari, David Navarro, Llúcia Martínez-Priego, Giuseppe D'Auria, Fernando Gonzalez-Candelas |
| EPI_ISL_436313 | Servicio de Microbiologia. Hospital Clinico Universitario de Valencia | Sequencing and Bioinformatics Service and Molecular Epidemiology Research Group. FISABIO-Public Health | Griselda De Marco, Beatriz Beamud, Lidia Ruiz Roldan, Marta Pla Diaz, Neris Garcia-Gonzalez, Inma Galán Vendrell, Sandra Carbo, Loreto Ferrús Abad, Paula Ruiz-Hueso, Mariana Reyes-Prieto, Vicente Soriano Chirona, Ivan Ansari, David Navarro, Maria Alma Bracho, Llúcia Martínez-Priego, Giuseppe D'Auria, Fernando Gonzalez-Candelas |
| EPI_ISL_436314 | Servicio de Microbiologia. Hospital Clinico Universitario de Valencia | Sequencing and Bioinformatics Service and Molecular Epidemiology Research Group. FISABIO-Public Health | Beatriz Beamud, Lidia Ruiz Roldan, Marta Pla Diaz, Neris Garcia-Gonzalez, Inma Galán Vendrell, Sandra Carbo, Loreto Ferrús Abad, Paula Ruiz-Hueso, Mariana Reyes-Prieto, Vicente Soriano Chirona, Ivan Ansari, David Navarro, Maria Alma Bracho, Griselda De Marco, Llúcia Martínez-Priego, Giuseppe D'Auria, Fernando Gonzalez-Candelas |
| EPI_ISL_436315 | Servicio de Microbiologia. Hospital Clinico Universitario de Valencia | Sequencing and Bioinformatics Service and Molecular Epidemiology Research Group. FISABIO-Public Health | Lidia Ruiz Roldan, Marta Pla Diaz, Neris Garcia-Gonzalez, Inma Galán Vendrell, Sandra Carbo, Loreto Ferrús Abad, Paula Ruiz-Hueso, Mariana Reyes-Prieto, Vicente Soriano Chirona, Ivan Ansari, David Navarro, Maria Alma Bracho, Griselda De Marco, Beatriz Beamud, Llúcia Martínez-Priego, Giuseppe D'Auria, Fernando Gonzalez-Candelas |
| EPI_ISL_436316 | Servicio de Microbiologia. Hospital Clinico Universitario de Valencia | Sequencing and Bioinformatics Service and Molecular Epidemiology Research Group. FISABIO-Public Health | Marta Pla Diaz, Neris Garcia-Gonzalez, Inma Galán Vendrell, Sandra Carbo, Loreto Ferrús Abad, Paula Ruiz-Hueso, Mariana Reyes-Prieto, Vicente Soriano Chirona, Ivan Ansari, David Navarro, Maria Alma Bracho, Griselda De Marco, Beatriz Beamud, Lidia Ruiz Roldan, Llúcia Martínez-Priego, Giuseppe D'Auria, Fernando Gonzalez-Candelas |
| EPI_ISL_436317 | Servicio de Microbiologia. Hospital Clinico Universitario de Valencia | Sequencing and Bioinformatics Service and Molecular Epidemiology Research Group. FISABIO-Public Health | Neris Garcia-Gonzalez, Inma Galán Vendrell, Sandra Carbo, Loreto Ferrús Abad, Paula Ruiz-Hueso, Mariana Reyes-Prieto, Vicente Soriano Chirona, Ivan Ansari, David Navarro, Maria Alma Bracho, Griselda De Marco, Beatriz Beamud, Lidia Ruiz Roldan, Marta Pla Diaz, Llúcia Martínez-Priego, Giuseppe D'Auria, Fernando Gonzalez-Candelas |
| EPI_ISL_436318 | Servicio de Microbiologia. Hospital Clinico Universitario de Valencia | Sequencing and Bioinformatics Service and Molecular Epidemiology Research Group. FISABIO-Public Health | David Navarro, Maria Alma Bracho, Griselda De Marco, Beatriz Beamud, Lidia Ruiz Roldan, Marta Pla Diaz, Neris Garcia-Gonzalez, Inma Galán Vendrell, Sandra Carbo, Loreto Ferrús Abad, Paula Ruiz-Hueso, Mariana Reyes-Prieto, Vicente Soriano Chirona, Ivan Ansari, David Navarro, Llúcia Martínez-Priego, Giuseppe D'Auria, Fernando Gonzalez-Candelas |
| EPI_ISL_436319 | Servicio de Microbiologia. Hospital Clinico Universitario de Valencia | Sequencing and Bioinformatics Service and Molecular Epidemiology Research Group. FISABIO-Public Health | Griselda De Marco, Beatriz Beamud, Lidia Ruiz Roldan, Marta Pla Diaz, Neris Garcia-Gonzalez, Inma Galán Vendrell, Sandra Carbo, Loreto Ferrús Abad, Paula Ruiz-Hueso, Mariana Reyes-Prieto, Vicente Soriano Chirona, Ivan Ansari, David Navarro, Maria Alma Bracho, Llúcia Martínez-Priego, Giuseppe D'Auria, Fernando Gonzalez-Candelas |
| EPI_ISL_436320 | Servicio de Microbiologia. Hospital Clinico Universitario de Valencia | Sequencing and Bioinformatics Service and Molecular Epidemiology Research Group. FISABIO-Public Health | Beatriz Beamud, Lidia Ruiz Roldan, Marta Pla Diaz, Neris Garcia-Gonzalez, Inma Galán Vendrell, Sandra Carbo, Loreto Ferrús Abad, Paula Ruiz-Hueso, Mariana Reyes-Prieto, Vicente Soriano Chirona, Ivan Ansari, David Navarro, Maria Alma Bracho, Griselda De Marco, Llúcia Martínez-Priego, Giuseppe D'Auria, Fernando Gonzalez-Candelas |
| EPI_ISL_436321 | Servicio de Microbiologia. Hospital Clinico Universitario de Valencia | Sequencing and Bioinformatics Service and Molecular Epidemiology Research Group. FISABIO-Public Health | Lidia Ruiz Roldan, Marta Pla Diaz, Neris Garcia-Gonzalez, Inma Galán Vendrell, Sandra Carbo, Loreto Ferrús Abad, Paula Ruiz-Hueso, Mariana Reyes-Prieto, Vicente Soriano Chirona, Ivan Ansari, David Navarro, Maria Alma Bracho, Griselda De Marco, Beatriz Beamud, Llúcia Martínez-Priego, Giuseppe D'Auria, Fernando Gonzalez-Candelas |
| EPI_ISL_436322 | Servicio de Microbiologia. Hospital Clinico Universitario de Valencia | Sequencing and Bioinformatics Service and Molecular Epidemiology Research Group. FISABIO-Public Health | Marta Pla Diaz, Neris Garcia-Gonzalez, Inma Galán Vendrell, Sandra Carbo, Loreto Ferrús Abad, Paula Ruiz-Hueso, Mariana Reyes-Prieto, Vicente Soriano Chirona, Ivan Ansari, David Navarro, Maria Alma Bracho, Griselda De Marco, Beatriz Beamud, Lidia Ruiz Roldan, Llúcia Martínez-Priego, Giuseppe D'Auria, Fernando Gonzalez-Candelas |
| EPI_ISL_436323 | Servicio de Microbiologia. Hospital Clinico Universitario de Valencia | Sequencing and Bioinformatics Service and Molecular Epidemiology Research Group. FISABIO-Public Health | Neris Garcia-Gonzalez, Inma Galán Vendrell, Sandra Carbo, Loreto Ferrús Abad, Paula Ruiz-Hueso, Mariana Reyes-Prieto, Vicente Soriano Chirona, Ivan Ansari, David Navarro, Maria Alma Bracho, Griselda De Marco, Beatriz Beamud, Lidia Ruiz Roldan, Marta Pla Diaz, Llúcia Martínez-Priego, Giuseppe D'Auria, Fernando Gonzalez-Candelas |
| EPI_ISL_436324 | Servicio de Microbiologia. Hospital Clinico Universitario de Valencia | Sequencing and Bioinformatics Service and Molecular Epidemiology Research Group. FISABIO-Public Health | Inma Galán Vendrell, Sandra Carbo, Loreto Ferrús Abad, Paula Ruiz-Hueso, Mariana Reyes-Prieto, Vicente Soriano Chirona, Ivan Ansari, David Navarro, Maria Alma Bracho, Griselda De Marco, Beatriz Beamud, Lidia Ruiz Roldan, Marta Pla Diaz, Neris Garcia-Gonzalez, Llúcia Martínez-Priego, Giuseppe D'Auria, Fernando Gonzalez-Candelas |
| EPI_ISL_436325 | Servicio de Microbiologia. Hospital Clinico Universitario de Valencia | Sequencing and Bioinformatics Service and Molecular Epidemiology Research Group. FISABIO-Public Health | Sandra Carbo, Loreto Ferrús Abad, Paula Ruiz-Hueso, Mariana Reyes-Prieto, Vicente Soriano Chirona, Ivan Ansari, David Navarro, Maria Alma Bracho, Griselda De Marco, Beatriz Beamud, Lidia Ruiz Roldan, Marta Pla Diaz, Neris Garcia-Gonzalez, Inma Galán Vendrell, Llúcia Martínez-Priego, Giuseppe D'Auria, Fernando Gonzalez-Candelas |
| EPI_ISL_436326 | Servicio de Microbiologia. Hospital Clinico Universitario de Valencia | Sequencing and Bioinformatics Service and Molecular Epidemiology Research Group. FISABIO-Public Health | Loreto Ferrús Abad, Paula Ruiz-Hueso, Mariana Reyes-Prieto, Vicente Soriano Chirona, Ivan Ansari, David Navarro, Maria Alma Bracho, Griselda De Marco, Beatriz Beamud, Lidia Ruiz Roldan, Marta Pla Diaz, Neris Garcia-Gonzalez, Inma Galán Vendrell, Sandra Carbo, Llúcia Martínez-Priego, Giuseppe D'Auria, Fernando Gonzalez-Candelas |
| EPI_ISL_436327 | Servicio de Microbiologia. Hospital Clinico Universitario de Valencia | Sequencing and Bioinformatics Service and Molecular Epidemiology Research Group. FISABIO-Public Health | Paula Ruiz-Hueso, Mariana Reyes-Prieto, Vicente Soriano Chirona, Ivan Ansari, David Navarro, Maria Alma Bracho, Griselda De Marco, Beatriz Beamud, Lidia Ruiz Roldan, Marta Pla Diaz, Neris Garcia-Gonzalez, Inma Galán Vendrell, Sandra Carbo, Loreto Ferrús Abad, Llúcia Martínez-Priego, Giuseppe D'Auria, Fernando Gonzalez-Candelas |
| EPI_ISL_436328 | Servicio de Microbiologia. Hospital Clinico Universitario de Valencia | Sequencing and Bioinformatics Service and Molecular Epidemiology Research Group. FISABIO-Public Health | Mariana Reyes-Prieto, Vicente Soriano Chirona, Ivan Ansari, David Navarro, Maria Alma Bracho, Griselda De Marco, Beatriz Beamud, Lidia Ruiz Roldan, Marta Pla Diaz, Neris Garcia-Gonzalez, Inma Galán Vendrell, Sandra Carbo, Loreto Ferrús Abad, Paula Ruiz-Hueso, Llúcia Martínez-Priego, Giuseppe D'Auria, Fernando Gonzalez-Candelas |
| EPI_ISL_436329 | Servicio de Microbiologia. Hospital Clinico Universitario de Valencia | Sequencing and Bioinformatics Service and Molecular Epidemiology Research Group. FISABIO-Public Health | Vicente Soriano Chirona, Ivan Ansari, David Navarro, Maria Alma Bracho, Griselda De Marco, Beatriz Beamud, Lidia Ruiz Roldan, Marta Pla Diaz, Neris Garcia-Gonzalez, Inma Galán Vendrell, Sandra Carbo, Loreto Ferrús Abad, Paula Ruiz-Hueso, Mariana Reyes-Prieto, Llúcia Martínez-Priego, Giuseppe D'Auria, Fernando Gonzalez-Candelas |
| EPI_ISL_436330 | Servicio de Microbiologia. Hospital Clinico Universitario de Valencia | Sequencing and Bioinformatics Service and Molecular Epidemiology Research Group. FISABIO-Public Health | Ivan Ansari, David Navarro, Maria Alma Bracho, Griselda De Marco, Beatriz Beamud, Lidia Ruiz Roldan, Marta Pla Diaz, Neris Garcia-Gonzalez, Inma Galán Vendrell, Sandra Carbo, Loreto Ferrús Abad, Paula Ruiz-Hueso, Mariana Reyes-Prieto, Vicente Soriano Chirona, Llúcia Martínez-Priego, Giuseppe D'Auria, Fernando Gonzalez-Candelas |
| EPI_ISL_436331 | Servicio de Microbiologia. Hospital Clinico Universitario de Valencia | Sequencing and Bioinformatics Service and Molecular Epidemiology Research Group. FISABIO-Public Health | David Navarro, Maria Alma Bracho, Griselda De Marco, Beatriz Beamud, Lidia Ruiz Roldan, Marta Pla Diaz, Neris Garcia-Gonzalez, Inma Galán Vendrell, Sandra Carbo, Loreto Ferrús Abad, Paula Ruiz-Hueso, Mariana Reyes-Prieto, Vicente Soriano Chirona, Ivan Ansari, Llúcia Martínez-Priego, Giuseppe D'Auria, Fernando Gonzalez-Candelas |
| EPI_ISL_436332 | Servicio de Microbiologia. Hospital Clinico Universitario de Valencia | Sequencing and Bioinformatics Service and Molecular Epidemiology Research Group. FISABIO-Public Health | Maria Alma Bracho, Griselda De Marco, Beatriz Beamud, Lidia Ruiz Roldan, Marta Pla Diaz, Neris Garcia-Gonzalez, Inma Galán Vendrell, Sandra Carbo, Loreto Ferrús Abad, Paula Ruiz-Hueso, Mariana Reyes-Prieto, Vicente Soriano Chirona, Ivan Ansari, David Navarro, Llúcia Martínez-Priego, Giuseppe D'Auria, Fernando Gonzalez-Candelas |

| | | | |
|---|---|---|---|
| EPI_ISL_436333 | Servicio de Microbiologia. Hospital Clinico Universitario de Valencia | Sequencing and Bioinformatics Service and Molecular Epidemiology Research Group. FISABIO-Public Health | Griselda De Marco, Beatriz Beamud, Lidia Ruiz Roldan, Marta Pla Diaz, Neris Garcia-Gonzalez, Inma Galán Vendrell, Sandra Carbo, Loreto Ferrús Abad, Paula Ruiz-Hueso, Mariana Reyes-Prieto, Vicente Soriano Chirona, Ivan Ansari, David Navarro, Maria Alma Bracho, Llúcia Martínez-Priego, Giuseppe D'Auria, Fernando Gonzalez-Candelas |
| EPI_ISL_436334 | Servicio de Microbiologia. Hospital Clinico Universitario de Valencia | Sequencing and Bioinformatics Service and Molecular Epidemiology Research Group. FISABIO-Public Health | Beatriz Beamud, Lidia Ruiz Roldan, Marta Pla Diaz, Neris Garcia-Gonzalez, Inma Galán Vendrell, Sandra Carbo, Loreto Ferrús Abad, Paula Ruiz-Hueso, Mariana Reyes-Prieto, Vicente Soriano Chirona, Ivan Ansari, David Navarro, Maria Alma Bracho, Griselda De Marco, Llúcia Martínez-Priego, Giuseppe D'Auria, Fernando Gonzalez-Candelas |
| EPI_ISL_436335 | Servicio de Microbiologia. Hospital Clinico Universitario de Valencia | Sequencing and Bioinformatics Service and Molecular Epidemiology Research Group. FISABIO-Public Health | Lidia Ruiz Roldan, Marta Pla Diaz, Neris Garcia-Gonzalez, Inma Galán Vendrell, Sandra Carbo, Loreto Ferrús Abad, Paula Ruiz-Hueso, Mariana Reyes-Prieto, Vicente Soriano Chirona, Ivan Ansari, David Navarro, Maria Alma Bracho, Griselda De Marco, Beatriz Beamud, Llúcia Martínez-Priego, Giuseppe D'Auria, Fernando Gonzalez-Candelas |
| EPI_ISL_436336 | Servicio de Microbiologia. Hospital Clinico Universitario de Valencia | Sequencing and Bioinformatics Service and Molecular Epidemiology Research Group. FISABIO-Public Health | Marta Pla Diaz, Neris Garcia-Gonzalez, Inma Galán Vendrell, Sandra Carbo, Loreto Ferrús Abad, Paula Ruiz-Hueso, Mariana Reyes-Prieto, Vicente Soriano Chirona, Ivan Ansari, David Navarro, Maria Alma Bracho, Griselda De Marco, Beatriz Beamud, Lidia Ruiz Roldan, Llúcia Martínez-Priego, Giuseppe D'Auria, Fernando Gonzalez-Candelas |
| EPI_ISL_436337 | Servicio de Microbiologia. Hospital Clinico Universitario de Valencia | Sequencing and Bioinformatics Service and Molecular Epidemiology Research Group. FISABIO-Public Health | Neris Garcia-Gonzalez, Inma Galán Vendrell, Sandra Carbo, Loreto Ferrús Abad, Paula Ruiz-Hueso, Mariana Reyes-Prieto, Vicente Soriano Chirona, Ivan Ansari, David Navarro, Maria Alma Bracho, Griselda De Marco, Beatriz Beamud, Lidia Ruiz Roldan, Marta Pla Diaz, Llúcia Martínez-Priego, Giuseppe D'Auria, Fernando Gonzalez-Candelas |
| EPI_ISL_436338 | Servicio de Microbiologia. Hospital Clinico Universitario de Valencia | Sequencing and Bioinformatics Service and Molecular Epidemiology Research Group. FISABIO-Public Health | David Navarro, Maria Alma Bracho, Griselda De Marco, Beatriz Beamud, Lidia Ruiz Roldan, Marta Pla Diaz, Neris Garcia-Gonzalez, Inma Galán Vendrell, Sandra Carbo, Loreto Ferrús Abad, Paula Ruiz-Hueso, Mariana Reyes-Prieto, Vicente Soriano Chirona, Ivan Ansari, David Navarro, Llúcia Martínez-Priego, Giuseppe D'Auria, Fernando Gonzalez-Candelas |
| EPI_ISL_436339 | Servicio de Microbiologia. Hospital Clinico Universitario de Valencia | Sequencing and Bioinformatics Service and Molecular Epidemiology Research Group. FISABIO-Public Health | Griselda De Marco, Beatriz Beamud, Lidia Ruiz Roldan, Marta Pla Diaz, Neris Garcia-Gonzalez, Inma Galán Vendrell, Sandra Carbo, Loreto Ferrús Abad, Paula Ruiz-Hueso, Mariana Reyes-Prieto, Vicente Soriano Chirona, Ivan Ansari, David Navarro, Maria Alma Bracho, Llúcia Martínez-Priego, Giuseppe D'Auria, Fernando Gonzalez-Candelas |
| EPI_ISL_436340 | Servicio de Microbiologia. Hospital Clinico Universitario de Valencia | Sequencing and Bioinformatics Service and Molecular Epidemiology Research Group. FISABIO-Public Health | Beatriz Beamud, Lidia Ruiz Roldan, Marta Pla Diaz, Neris Garcia-Gonzalez, Inma Galán Vendrell, Sandra Carbo, Loreto Ferrús Abad, Paula Ruiz-Hueso, Mariana Reyes-Prieto, Vicente Soriano Chirona, Ivan Ansari, David Navarro, Maria Alma Bracho, Griselda De Marco, Llúcia Martínez-Priego, Giuseppe D'Auria, Fernando Gonzalez-Candelas |
| EPI_ISL_436341 | Servicio de Microbiologia. Hospital Clinico Universitario de Valencia | Sequencing and Bioinformatics Service and Molecular Epidemiology Research Group. FISABIO-Public Health | Lidia Ruiz Roldan, Marta Pla Diaz, Neris Garcia-Gonzalez, Inma Galán Vendrell, Sandra Carbo, Loreto Ferrús Abad, Paula Ruiz-Hueso, Mariana Reyes-Prieto, Vicente Soriano Chirona, Ivan Ansari, David Navarro, Maria Alma Bracho, Griselda De Marco, Beatriz Beamud, Llúcia Martínez-Priego, Giuseppe D'Auria, Fernando Gonzalez-Candelas |
| EPI_ISL_436342 | Servicio de Microbiologia. Hospital Clinico Universitario de Valencia | Sequencing and Bioinformatics Service and Molecular Epidemiology Research Group. FISABIO-Public Health | Marta Pla Diaz, Neris Garcia-Gonzalez, Inma Galán Vendrell, Sandra Carbo, Loreto Ferrús Abad, Paula Ruiz-Hueso, Mariana Reyes-Prieto, Vicente Soriano Chirona, Ivan Ansari, David Navarro, Maria Alma Bracho, Griselda De Marco, Beatriz Beamud, Lidia Ruiz Roldan, Llúcia Martínez-Priego, Giuseppe D'Auria, Fernando Gonzalez-Candelas |
| EPI_ISL_436343 | Servicio de Microbiologia. Hospital Clinico Universitario de Valencia | Sequencing and Bioinformatics Service and Molecular Epidemiology Research Group. FISABIO-Public Health | Neris Garcia-Gonzalez, Inma Galán Vendrell, Sandra Carbo, Loreto Ferrús Abad, Paula Ruiz-Hueso, Mariana Reyes-Prieto, Vicente Soriano Chirona, Ivan Ansari, David Navarro, Maria Alma Bracho, Griselda De Marco, Beatriz Beamud, Lidia Ruiz Roldan, Marta Pla Diaz, Llúcia Martínez-Priego, Giuseppe D'Auria, Fernando Gonzalez-Candelas |
| EPI_ISL_436344 | Servicio de Microbiologia. Hospital Clinico Universitario de Valencia | Sequencing and Bioinformatics Service and Molecular Epidemiology Research Group. FISABIO-Public Health | Inma Galán Vendrell, Sandra Carbo, Loreto Ferrús Abad, Paula Ruiz-Hueso, Mariana Reyes-Prieto, Vicente Soriano Chirona, Ivan Ansari, David Navarro, Maria Alma Bracho, Griselda De Marco, Beatriz Beamud, Lidia Ruiz Roldan, Marta Pla Diaz, Neris Garcia-Gonzalez, Llúcia Martínez-Priego, Giuseppe D'Auria, Fernando Gonzalez-Candelas |
| EPI_ISL_436345 | Servicio de Microbiologia. Hospital Clinico Universitario de Valencia | Sequencing and Bioinformatics Service and Molecular Epidemiology Research Group. FISABIO-Public Health | Sandra Carbo, Loreto Ferrús Abad, Paula Ruiz-Hueso, Mariana Reyes-Prieto, Vicente Soriano Chirona, Ivan Ansari, David Navarro, Maria Alma Bracho, Griselda De Marco, Beatriz Beamud, Lidia Ruiz Roldan, Marta Pla Diaz, Neris Garcia-Gonzalez, Inma Galán Vendrell, Llúcia Martínez-Priego, Giuseppe D'Auria, Fernando Gonzalez-Candelas |
| EPI_ISL_436346 | Servicio de Microbiologia. Hospital Clinico Universitario de Valencia | Sequencing and Bioinformatics Service and Molecular Epidemiology Research Group. FISABIO-Public Health | Loreto Ferrús Abad, Paula Ruiz-Hueso, Mariana Reyes-Prieto, Vicente Soriano Chirona, Ivan Ansari, David Navarro, Maria Alma Bracho, Griselda De Marco, Beatriz Beamud, Lidia Ruiz Roldan, Marta Pla Diaz, Neris Garcia-Gonzalez, Inma Galán Vendrell, Sandra Carbo, Llúcia Martínez-Priego, Giuseppe D'Auria, Fernando Gonzalez-Candelas |
| EPI_ISL_436347 | Servicio de Microbiologia. Hospital Clinico Universitario de Valencia | Sequencing and Bioinformatics Service and Molecular Epidemiology Research Group. FISABIO-Public Health | Paula Ruiz-Hueso, Mariana Reyes-Prieto, Vicente Soriano Chirona, Ivan Ansari, David Navarro, Maria Alma Bracho, Griselda De Marco, Beatriz Beamud, Lidia Ruiz Roldan, Marta Pla Diaz, Neris Garcia-Gonzalez, Inma Galán Vendrell, Sandra Carbo, Loreto Ferrús Abad, Llúcia Martínez-Priego, Giuseppe D'Auria, Fernando Gonzalez-Candelas |
| EPI_ISL_436348 | Servicio de Microbiologia. Hospital Clinico Universitario de Valencia | Sequencing and Bioinformatics Service and Molecular Epidemiology Research Group. FISABIO-Public Health | Mariana Reyes-Prieto, Vicente Soriano Chirona, Ivan Ansari, David Navarro, Maria Alma Bracho, Griselda De Marco, Beatriz Beamud, Lidia Ruiz Roldan, Marta Pla Diaz, Neris Garcia-Gonzalez, Inma Galán Vendrell, Sandra Carbo, Loreto Ferrús Abad, Paula Ruiz-Hueso, Llúcia Martínez-Priego, Giuseppe D'Auria, Fernando Gonzalez-Candelas |
| EPI_ISL_436349 | Servicio de Microbiologia. Hospital Clinico Universitario de Valencia | Sequencing and Bioinformatics Service and Molecular Epidemiology Research Group. FISABIO-Public Health | Vicente Soriano Chirona, Ivan Ansari, David Navarro, Maria Alma Bracho, Griselda De Marco, Beatriz Beamud, Lidia Ruiz Roldan, Marta Pla Diaz, Neris Garcia-Gonzalez, Inma Galán Vendrell, Sandra Carbo, Loreto Ferrús Abad, Paula Ruiz-Hueso, Mariana Reyes-Prieto, Llúcia Martínez-Priego, Giuseppe D'Auria, Fernando Gonzalez-Candelas |
| EPI_ISL_436350 | Servicio de Microbiologia. Hospital Clinico Universitario de Valencia | Sequencing and Bioinformatics Service and Molecular Epidemiology Research Group. FISABIO-Public Health | Ivan Ansari, David Navarro, Maria Alma Bracho, Griselda De Marco, Beatriz Beamud, Lidia Ruiz Roldan, Marta Pla Diaz, Neris Garcia-Gonzalez, Inma Galán Vendrell, Sandra Carbo, Loreto Ferrús Abad, Paula Ruiz-Hueso, Mariana Reyes-Prieto, Vicente Soriano Chirona, Llúcia Martínez-Priego, Giuseppe D'Auria, Fernando Gonzalez-Candelas |
| EPI_ISL_436351 | Servicio de Microbiologia. Hospital Clinico Universitario de Valencia | Sequencing and Bioinformatics Service and Molecular Epidemiology Research Group. FISABIO-Public Health | David Navarro, Maria Alma Bracho, Griselda De Marco, Beatriz Beamud, Lidia Ruiz Roldan, Marta Pla Diaz, Neris Garcia-Gonzalez, Inma Galán Vendrell, Sandra Carbo, Loreto Ferrús Abad, Paula Ruiz-Hueso, Mariana Reyes-Prieto, Vicente Soriano Chirona, Ivan Ansari, Llúcia Martínez-Priego, Giuseppe D'Auria, Fernando Gonzalez-Candelas |
| EPI_ISL_436352 | Servicio de Microbiologia. Hospital Clinico Universitario de Valencia | Sequencing and Bioinformatics Service and Molecular Epidemiology Research Group. FISABIO-Public Health | Maria Alma Bracho, Griselda De Marco, Beatriz Beamud, Lidia Ruiz Roldan, Marta Pla Diaz, Neris Garcia-Gonzalez, Inma Galán Vendrell, Sandra Carbo, Loreto Ferrús Abad, Paula Ruiz-Hueso, Mariana Reyes-Prieto, Vicente Soriano Chirona, Ivan Ansari, David Navarro, Llúcia Martínez-Priego, Giuseppe D'Auria, Fernando Gonzalez-Candelas |
| EPI_ISL_436353 | Servicio de Microbiologia. Hospital Clinico Universitario de Valencia | Sequencing and Bioinformatics Service and Molecular Epidemiology Research Group. FISABIO-Public Health | Griselda De Marco, Beatriz Beamud, Lidia Ruiz Roldan, Marta Pla Diaz, Neris Garcia-Gonzalez, Inma Galán Vendrell, Sandra Carbo, Loreto Ferrús Abad, Paula Ruiz-Hueso, Mariana Reyes-Prieto, Vicente Soriano Chirona, Ivan Ansari, David Navarro, Maria Alma Bracho, Llúcia Martínez-Priego, Giuseppe D'Auria, Fernando Gonzalez-Candelas |
| EPI_ISL_436354 | Servicio de Microbiologia. Hospital Clinico Universitario de Valencia | Sequencing and Bioinformatics Service and Molecular Epidemiology Research Group. FISABIO-Public Health | Beatriz Beamud, Lidia Ruiz Roldan, Marta Pla Diaz, Neris Garcia-Gonzalez, Inma Galán Vendrell, Sandra Carbo, Loreto Ferrús Abad, Paula Ruiz-Hueso, Mariana Reyes-Prieto, Vicente Soriano Chirona, Ivan Ansari, David Navarro, Maria Alma Bracho, Griselda De Marco, Llúcia Martínez-Priego, Giuseppe D'Auria, Fernando Gonzalez-Candelas |
| EPI_ISL_436355 | Servicio de Microbiologia. Hospital Clinico Universitario de Valencia | Sequencing and Bioinformatics Service and Molecular Epidemiology Research Group. FISABIO-Public Health | Lidia Ruiz Roldan, Marta Pla Diaz, Neris Garcia-Gonzalez, Inma Galán Vendrell, Sandra Carbo, Loreto Ferrús Abad, Paula Ruiz-Hueso, Mariana Reyes-Prieto, Vicente Soriano Chirona, Ivan Ansari, David Navarro, Maria Alma Bracho, Griselda De Marco, Beatriz Beamud, Llúcia Martínez-Priego, Giuseppe D'Auria, Fernando Gonzalez-Candelas |
| EPI_ISL_436356 | Servicio de Microbiologia. Hospital Clinico Universitario de Valencia | Sequencing and Bioinformatics Service and Molecular Epidemiology Research Group. FISABIO-Public Health | Marta Pla Diaz, Neris Garcia-Gonzalez, Inma Galán Vendrell, Sandra Carbo, Loreto Ferrús Abad, Paula Ruiz-Hueso, Mariana Reyes-Prieto, Vicente Soriano Chirona, Ivan Ansari, David Navarro, Maria Alma Bracho, Griselda De Marco, Beatriz Beamud, Lidia Ruiz Roldan, Llúcia Martínez-Priego, Giuseppe D'Auria, Fernando Gonzalez-Candelas |
| EPI_ISL_436357 | Servicio de Microbiologia. Hospital Clinico Universitario de Valencia | Sequencing and Bioinformatics Service and Molecular Epidemiology Research Group. FISABIO-Public Health | Neris Garcia-Gonzalez, Inma Galán Vendrell, Sandra Carbo, Loreto Ferrús Abad, Paula Ruiz-Hueso, Mariana Reyes-Prieto, Vicente Soriano Chirona, Ivan Ansari, David Navarro, Maria Alma Bracho, Griselda De Marco, Beatriz Beamud, Lidia Ruiz Roldan, Marta Pla Diaz, Llúcia Martínez-Priego, Giuseppe D'Auria, Fernando Gonzalez-Candelas |
| EPI_ISL_436358 | Servicio de Microbiologia. Hospital Clinico Universitario de Valencia | Sequencing and Bioinformatics Service and Molecular Epidemiology Research Group. FISABIO-Public Health | Lidia Ruiz Roldan, Marta Pla Diaz, Neris Garcia-Gonzalez, Inma Galán Vendrell, Sandra Carbo, Loreto Ferrús Abad, Paula Ruiz-Hueso, Mariana Reyes-Prieto, Vicente Soriano Chirona, Ivan Ansari, David Navarro, Maria Alma Bracho, Griselda De Marco, Beatriz Beamud, Llúcia Martínez-Priego, Giuseppe D'Auria, Fernando Gonzalez-Candelas |
| EPI_ISL_436359 | Servicio de Microbiologia. Hospital Clinico Universitario de Valencia | Sequencing and Bioinformatics Service and Molecular Epidemiology Research Group. FISABIO-Public Health | Marta Pla Diaz, Neris Garcia-Gonzalez, Inma Galán Vendrell, Sandra Carbo, Loreto Ferrús Abad, Paula Ruiz-Hueso, Mariana Reyes-Prieto, Vicente Soriano Chirona, Ivan Ansari, David Navarro, Maria Alma Bracho, Griselda De Marco, Beatriz Beamud, Lidia Ruiz Roldan, Llúcia Martínez-Priego, Giuseppe D'Auria, Fernando Gonzalez-Candelas |
| EPI_ISL_436360 | Servicio de Microbiologia. Hospital Clinico Universitario de Valencia | Sequencing and Bioinformatics Service and Molecular Epidemiology Research Group. FISABIO-Public Health | Neris Garcia-Gonzalez, Inma Galán Vendrell, Sandra Carbo, Loreto Ferrús Abad, Paula Ruiz-Hueso, Mariana Reyes-Prieto, Vicente Soriano Chirona, Ivan Ansari, David Navarro, Maria Alma Bracho, Griselda De Marco, Beatriz Beamud, Lidia Ruiz Roldan, Marta Pla Diaz, Llúcia Martínez-Priego, Giuseppe D'Auria, Fernando Gonzalez-Candelas |
| EPI_ISL_436361 | Servicio de Microbiología. Consorcio Hospital General Universitario de Valencia | Sequencing and Bioinformatics Service and Molecular Epidemiology Research Group. FISABIO-Public Health | Neris Garcia-Gonzalez, Loreto Ferrús Abad, Maria Dolores Ocete, Inma Galán Vendrell, Paula Ruiz-Hueso, Mariana Reyes-Prieto, Vicente Soriano Chirona, Maria Alma Bracho, Griselda De Marco, Beatriz Beamud, Lidia Ruiz Roldan, Marta Pla Diaz, Llúcia Martínez-Priego, Concepcion Gimeno, Giuseppe D'Auria, Fernando Gonzalez-Candelas |
| EPI_ISL_436362 | Servicio de Microbiologia. Hospital Clinico Universitario de Valencia | Sequencing and Bioinformatics Service and Molecular Epidemiology Research Group. FISABIO-Public Health | David Navarro, Maria Alma Bracho, Griselda De Marco, Beatriz Beamud, Lidia Ruiz Roldan, Marta Pla Diaz, Neris Garcia-Gonzalez, Inma Galán Vendrell, Sandra Carbo, Loreto Ferrús Abad, Paula Ruiz-Hueso, Mariana Reyes-Prieto, Vicente Soriano Chirona, Ivan Ansari, David Navarro, Llúcia Martínez-Priego, Giuseppe D'Auria, Fernando Gonzalez-Candelas |
| EPI_ISL_436363 | Servicio de Microbiologia. Hospital Clinico Universitario de Valencia | Sequencing and Bioinformatics Service and Molecular Epidemiology Research Group. FISABIO-Public Health | Griselda De Marco, Beatriz Beamud, Lidia Ruiz Roldan, Marta Pla Diaz, Neris Garcia-Gonzalez, Inma Galán Vendrell, Sandra Carbo, Loreto Ferrús Abad, Paula Ruiz-Hueso, Mariana Reyes-Prieto, Vicente Soriano Chirona, Ivan Ansari, David Navarro, Maria Alma Bracho, Llúcia Martínez-Priego, Giuseppe D'Auria, Fernando Gonzalez-Candelas |
| EPI_ISL_436364 | Servicio de Microbiologia. Hospital Clinico Universitario de Valencia | Sequencing and Bioinformatics Service and Molecular Epidemiology Research Group. FISABIO-Public Health | Beatriz Beamud, Lidia Ruiz Roldan, Marta Pla Diaz, Neris Garcia-Gonzalez, Inma Galán Vendrell, Sandra Carbo, Loreto Ferrús Abad, Paula Ruiz-Hueso, Mariana Reyes-Prieto, Vicente Soriano Chirona, Ivan Ansari, David Navarro, Maria Alma Bracho, Griselda De Marco, Llúcia Martínez-Priego, Giuseppe D'Auria, Fernando Gonzalez-Candelas |
| EPI_ISL_436365 | Servicio de Microbiologia. Hospital Clinico Universitario de Valencia | Sequencing and Bioinformatics Service and Molecular Epidemiology Research Group. FISABIO-Public Health | Lidia Ruiz Roldan, Marta Pla Diaz, Neris Garcia-Gonzalez, Inma Galán Vendrell, Sandra Carbo, Loreto Ferrús Abad, Paula Ruiz-Hueso, Mariana Reyes-Prieto, Vicente Soriano Chirona, Ivan Ansari, David Navarro, Maria Alma Bracho, Griselda De Marco, Beatriz Beamud, Llúcia Martínez-Priego, Giuseppe D'Auria, Fernando Gonzalez-Candelas |
| EPI_ISL_436366 | Servicio de Microbiologia. Hospital Clinico Universitario de Valencia | Sequencing and Bioinformatics Service and Molecular Epidemiology Research Group. FISABIO-Public Health | Marta Pla Diaz, Neris Garcia-Gonzalez, Inma Galán Vendrell, Sandra Carbo, Loreto Ferrús Abad, Paula Ruiz-Hueso, Mariana Reyes-Prieto, Vicente Soriano Chirona, Ivan Ansari, David Navarro, Maria Alma Bracho, Griselda De Marco, Beatriz Beamud, Lidia Ruiz Roldan, Llúcia Martínez-Priego, Giuseppe D'Auria, Fernando Gonzalez-Candelas |
| EPI_ISL_436367 | Servicio de Microbiologia. Hospital Clinico Universitario de Valencia | Sequencing and Bioinformatics Service and Molecular Epidemiology Research Group. FISABIO-Public Health | Neris Garcia-Gonzalez, Inma Galán Vendrell, Sandra Carbo, Loreto Ferrús Abad, Paula Ruiz-Hueso, Mariana Reyes-Prieto, Vicente Soriano Chirona, Ivan Ansari, David Navarro, Maria Alma Bracho, Griselda De Marco, Beatriz Beamud, Lidia Ruiz Roldan, Marta Pla Diaz, Llúcia Martínez-Priego, Giuseppe D'Auria, Fernando Gonzalez-Candelas |
| EPI_ISL_436368 | Servicio de Microbiologia. Hospital Clinico Universitario de Valencia | Sequencing and Bioinformatics Service and Molecular Epidemiology Research Group. FISABIO-Public Health | Inma Galán Vendrell, Sandra Carbo, Loreto Ferrús Abad, Paula Ruiz-Hueso, Mariana Reyes-Prieto, Vicente Soriano Chirona, Ivan Ansari, David Navarro, Maria Alma Bracho, Griselda De Marco, Beatriz Beamud, Lidia Ruiz Roldan, Marta Pla Diaz, Neris Garcia-Gonzalez, Llúcia Martínez-Priego, Giuseppe D'Auria, Fernando Gonzalez-Candelas |
| EPI_ISL_436369 | Servicio de Microbiologia. Hospital Clinico Universitario de Valencia | Sequencing and Bioinformatics Service and Molecular Epidemiology Research Group. FISABIO-Public Health | Sandra Carbo, Loreto Ferrús Abad, Paula Ruiz-Hueso, Mariana Reyes-Prieto, Vicente Soriano Chirona, Ivan Ansari, David Navarro, Maria Alma Bracho, Griselda De Marco, Beatriz Beamud, Lidia Ruiz Roldan, Marta Pla Diaz, Neris Garcia-Gonzalez, Inma Galán Vendrell, Llúcia Martínez-Priego, Giuseppe D'Auria, Fernando Gonzalez-Candelas |
| EPI_ISL_436370 | Servicio de Microbiologia. Hospital Clinico Universitario de Valencia | Sequencing and Bioinformatics Service and Molecular Epidemiology Research Group. FISABIO-Public Health | Loreto Ferrús Abad, Paula Ruiz-Hueso, Mariana Reyes-Prieto, Vicente Soriano Chirona, Ivan Ansari, David Navarro, Maria Alma Bracho, Griselda De Marco, Beatriz Beamud, Lidia Ruiz Roldan, Marta Pla Diaz, Neris Garcia-Gonzalez, Inma Galán Vendrell, Sandra Carbo, Llúcia Martínez-Priego, Giuseppe D'Auria, Fernando Gonzalez-Candelas |
| EPI_ISL_436371 | Servicio de Microbiologia. Hospital Clinico Universitario de Valencia | Sequencing and Bioinformatics Service and Molecular Epidemiology Research Group. FISABIO-Public Health | Paula Ruiz-Hueso, Mariana Reyes-Prieto, Vicente Soriano Chirona, Ivan Ansari, David Navarro, Maria Alma Bracho, Griselda De Marco, Beatriz Beamud, Lidia Ruiz Roldan, Marta Pla Diaz, Neris Garcia-Gonzalez, Inma Galán Vendrell, Sandra Carbo, Loreto Ferrús Abad, Llúcia Martínez-Priego, Giuseppe D'Auria, Fernando Gonzalez-Candelas |
| EPI_ISL_436372, EPI_ISL_436373, EPI_ISL_436374, EPI_ISL_436375, EPI_ISL_436376, EPI_ISL_436377, EPI_ISL_436378, EPI_ISL_436379, EPI_ISL_436380, EPI_ISL_436381 | Servicio de Microbiologia. Hospital Universitario Doctor Peset | Sequencing and Bioinformatics Service and Molecular Epidemiology Research Group. FISABIO-Public Health | Juan Alberola Enguídanos, Juan Jose Camarena Miñana, Rosa González Pellicer, Neris Garcia-Gonzalez, Inma Galán Vendrell, Sandra Carbo, Loreto Ferrús Abad, Paula Ruiz-Hueso, Mariana Reyes-Prieto, Vicente Soriano Chirona, Ivan Ansari, Maria Alma Bracho, Griselda De Marco, Beatriz Beamud, Lidia Ruiz Roldan, Marta Pla Diaz, Llúcia Martínez-Priego, Giuseppe D'Auria, Jose Miguel Nogueira Coito, Fernando Gonzalez-Candelas |
| EPI_ISL_436382 | Servicio de Microbiología. Consorcio Hospital General Universitario de Valencia | Sequencing and Bioinformatics Service and Molecular Epidemiology Research Group. FISABIO-Public Health | Loreto Ferrús Abad, Maria Dolores Ocete,Inma Galán Vendrell, Paula Ruiz-Hueso, Mariana Reyes-Prieto, Vicente Soriano Chirona, Maria Alma Bracho, Griselda De Marco, Beatriz Beamud, Lidia Ruiz Roldan, Marta Pla Diaz,Neris Garcia-Gonzalez, Llúcia Martínez-Priego, Concepcion Gimeno, Giuseppe D'Auria, Fernando Gonzalez-Candelas |
| EPI_ISL_436383 | Servicio de Microbiología. Consorcio Hospital General Universitario de Valencia | Sequencing and Bioinformatics Service and Molecular Epidemiology Research Group. FISABIO-Public Health | Maria Dolores Ocete, Inma Galán Vendrell, Paula Ruiz-Hueso, Mariana Reyes-Prieto, Vicente Soriano Chirona, Maria Alma Bracho, Griselda De Marco, Beatriz Beamud, Lidia Ruiz Roldan, Marta Pla Diaz,Neris Garcia-Gonzalez, Loreto Ferrús Abad, Llúcia Martínez-Priego, Concepcion Gimeno, Giuseppe D'Auria, Fernando Gonzalez-Candelas |
| EPI_ISL_436384 | Servicio de Microbiología. Consorcio Hospital General Universitario de Valencia | Sequencing and Bioinformatics Service and Molecular Epidemiology Research Group. FISABIO-Public Health | Griselda De Marco, Beatriz Beamud, Lidia Ruiz Roldan, Marta Pla Diaz,Neris Garcia-Gonzalez, Loreto Ferrús Abad, Maria Dolores Ocete, Inma Galán Vendrell, Paula Ruiz-Hueso, Mariana Reyes-Prieto, Vicente Soriano Chirona, Maria Alma Bracho, Llúcia Martínez-Priego, Concepcion Gimeno, Giuseppe D'Auria, Fernando Gonzalez-Candelas |
| EPI_ISL_436385 | Servicio de Microbiología. Consorcio Hospital General Universitario de Valencia | Sequencing and Bioinformatics Service and Molecular Epidemiology Research Group. FISABIO-Public Health | Beatriz Beamud, Lidia Ruiz Roldan, Marta Pla Diaz,Neris Garcia-Gonzalez, Loreto Ferrús Abad, Maria Dolores Ocete, Inma Galán Vendrell, Paula Ruiz-Hueso, Mariana Reyes-Prieto, Vicente Soriano Chirona, Maria Alma Bracho, Griselda De Marco, Llúcia Martínez-Priego, Concepcion Gimeno, Giuseppe D'Auria, Fernando Gonzalez-Candelas |
| EPI_ISL_436386 | Servicio de Microbiología. Consorcio Hospital General | Sequencing and Bioinformatics Service and Molecular | Lidia Ruiz Roldan, Marta Pla Diaz,Neris Garcia-Gonzalez, Loreto Ferrús Abad, Maria Dolores Ocete, Inma Galán Vendrell, Paula Ruiz-Hueso, Mariana Reyes-Prieto, Vicente Soriano Chirona, Maria Alma Bracho, Griselda |

| | | | |
|---|---|---|---|
| | Universitario de Valencia | Epidemiology Research Group. FISABIO-Public Health | De Marco, Beatriz Beamud, Llúcia Martínez-Priego, Concepcion Gimeno, Giuseppe D'Auria, Fernando Gonzalez-Candelas |
| EPI_ISL_436387 | Servicio de Microbiología. Consorcio Hospital General Universitario de Valencia | Sequencing and Bioinformatics Service and Molecular Epidemiology Research Group. FISABIO-Public Health | Marta Pla Diaz,Neris Garcia-Gonzalez, Loreto Ferrús Abad, Maria Dolores Ocete, Inma Galán Vendrell, Paula Ruiz-Hueso, Mariana Reyes-Prieto, Vicente Soriano Chirona, Maria Alma Bracho, Griselda De Marco, Beatriz Beamud, Lidia Ruiz Roldan, Llúcia Martínez-Priego, Concepcion Gimeno, Giuseppe D'Auria, Fernando Gonzalez-Candelas |
| EPI_ISL_436388 | Servicio de Microbiología. Consorcio Hospital General Universitario de Valencia | Sequencing and Bioinformatics Service and Molecular Epidemiology Research Group. FISABIO-Public Health | Neris Garcia-Gonzalez, Loreto Ferrús Abad, Maria Dolores Ocete, Inma Galán Vendrell, Paula Ruiz-Hueso, Mariana Reyes-Prieto, Vicente Soriano Chirona, Maria Alma Bracho, Griselda De Marco, Beatriz Beamud, Lidia Ruiz Roldan, Marta Pla Diaz, Llúcia Martínez-Priego, Concepcion Gimeno, Giuseppe D'Auria, Fernando Gonzalez-Candelas |
| EPI_ISL_436389 | Servicio de Microbiología. Consorcio Hospital General Universitario de Valencia | Sequencing and Bioinformatics Service and Molecular Epidemiology Research Group. FISABIO-Public Health | Loreto Ferrús Abad, Maria Dolores Ocete, Inma Galán Vendrell, Paula Ruiz-Hueso, Mariana Reyes-Prieto, Vicente Soriano Chirona, Maria Alma Bracho, Griselda De Marco, Beatriz Beamud, Lidia Ruiz Roldan, Marta Pla Diaz,Neris Garcia-Gonzalez, Llúcia Martínez-Priego, Concepcion Gimeno, Giuseppe D'Auria, Fernando Gonzalez-Candelas |
| EPI_ISL_436390 | Servicio de Microbiología. Consorcio Hospital General Universitario de Valencia | Sequencing and Bioinformatics Service and Molecular Epidemiology Research Group. FISABIO-Public Health | Maria Dolores Ocete, Inma Galán Vendrell, Paula Ruiz-Hueso, Mariana Reyes-Prieto, Vicente Soriano Chirona, Maria Alma Bracho, Griselda De Marco, Beatriz Beamud, Lidia Ruiz Roldan, Marta Pla Diaz,Neris Garcia-Gonzalez, Loreto Ferrús Abad, Llúcia Martínez-Priego, Concepcion Gimeno, Giuseppe D'Auria, Fernando Gonzalez-Candelas |
| EPI_ISL_436391 | Servicio de Microbiología. Consorcio Hospital General Universitario de Valencia | Sequencing and Bioinformatics Service and Molecular Epidemiology Research Group. FISABIO-Public Health | Griselda De Marco, Beatriz Beamud, Lidia Ruiz Roldan, Marta Pla Diaz,Neris Garcia-Gonzalez, Loreto Ferrús Abad, Maria Dolores Ocete, Inma Galán Vendrell, Paula Ruiz-Hueso, Mariana Reyes-Prieto, Vicente Soriano Chirona, Maria Alma Bracho, Llúcia Martínez-Priego, Concepcion Gimeno, Giuseppe D'Auria, Fernando Gonzalez-Candelas |
| EPI_ISL_436392 | Servicio de Microbiología. Consorcio Hospital General Universitario de Valencia | Sequencing and Bioinformatics Service and Molecular Epidemiology Research Group. FISABIO-Public Health | Beatriz Beamud, Lidia Ruiz Roldan, Marta Pla Diaz,Neris Garcia-Gonzalez, Loreto Ferrús Abad, Maria Dolores Ocete, Inma Galán Vendrell, Paula Ruiz-Hueso, Mariana Reyes-Prieto, Vicente Soriano Chirona, Maria Alma Bracho, Griselda De Marco, Llúcia Martínez-Priego, Concepcion Gimeno, Giuseppe D'Auria, Fernando Gonzalez-Candelas |
| EPI_ISL_436393 | Servicio de Microbiología. Consorcio Hospital General Universitario de Valencia | Sequencing and Bioinformatics Service and Molecular Epidemiology Research Group. FISABIO-Public Health | Lidia Ruiz Roldan, Marta Pla Diaz,Neris Garcia-Gonzalez, Loreto Ferrús Abad, Maria Dolores Ocete, Inma Galán Vendrell, Paula Ruiz-Hueso, Mariana Reyes-Prieto, Vicente Soriano Chirona, Maria Alma Bracho, Griselda De Marco, Beatriz Beamud, Llúcia Martínez-Priego, Concepcion Gimeno, Giuseppe D'Auria, Fernando Gonzalez-Candelas |
| EPI_ISL_436394 | Servicio de Microbiología. Consorcio Hospital General Universitario de Valencia | Sequencing and Bioinformatics Service and Molecular Epidemiology Research Group. FISABIO-Public Health | Marta Pla Diaz,Neris Garcia-Gonzalez, Loreto Ferrús Abad, Maria Dolores Ocete, Inma Galán Vendrell, Paula Ruiz-Hueso, Mariana Reyes-Prieto, Vicente Soriano Chirona, Maria Alma Bracho, Griselda De Marco, Beatriz Beamud, Lidia Ruiz Roldan, Llúcia Martínez-Priego, Concepcion Gimeno, Giuseppe D'Auria, Fernando Gonzalez-Candelas |
| EPI_ISL_436395 | Servicio de Microbiología. Consorcio Hospital General Universitario de Valencia | Sequencing and Bioinformatics Service and Molecular Epidemiology Research Group. FISABIO-Public Health | Neris Garcia-Gonzalez, Loreto Ferrús Abad, Maria Dolores Ocete, Inma Galán Vendrell, Paula Ruiz-Hueso, Mariana Reyes-Prieto, Vicente Soriano Chirona, Maria Alma Bracho, Griselda De Marco, Beatriz Beamud, Lidia Ruiz Roldan, Marta Pla Diaz, Llúcia Martínez-Priego, Concepcion Gimeno, Giuseppe D'Auria, Fernando Gonzalez-Candelas |
| EPI_ISL_436396 | Servicio de Microbiologia. Hospital Clinico Universitario de Valencia | Sequencing and Bioinformatics Service and Molecular Epidemiology Research Group. FISABIO-Public Health | Sandra Carbo, Loreto Ferrús Abad, Paula Ruiz-Hueso, Mariana Reyes-Prieto, Vicente Soriano Chirona, Ivan Ansari, David Navarro, Maria Alma Bracho, Griselda De Marco, Beatriz Beamud, Lidia Ruiz Roldan, Marta Pla Diaz, Neris Garcia-Gonzalez, Inma Galán Vendrell, Llúcia Martínez-Priego, Giuseppe D'Auria, Fernando Gonzalez-Candelas |
| EPI_ISL_436397 | Servicio de Microbiologia. Hospital Clinico Universitario de Valencia | Sequencing and Bioinformatics Service and Molecular Epidemiology Research Group. FISABIO-Public Health | Loreto Ferrús Abad, Paula Ruiz-Hueso, Mariana Reyes-Prieto, Vicente Soriano Chirona, Ivan Ansari, David Navarro, Maria Alma Bracho, Griselda De Marco, Beatriz Beamud, Lidia Ruiz Roldan, Marta Pla Diaz, Neris Garcia-Gonzalez, Inma Galán Vendrell, Sandra Carbo, Llúcia Martínez-Priego, Giuseppe D'Auria, Fernando Gonzalez-Candelas |
| EPI_ISL_436398 | Servicio de Microbiologia. Hospital Clinico Universitario de Valencia | Sequencing and Bioinformatics Service and Molecular Epidemiology Research Group. FISABIO-Public Health | Paula Ruiz-Hueso, Mariana Reyes-Prieto, Vicente Soriano Chirona, Ivan Ansari, David Navarro, Maria Alma Bracho, Griselda De Marco, Beatriz Beamud, Lidia Ruiz Roldan, Marta Pla Diaz, Neris Garcia-Gonzalez, Inma Galán Vendrell, Sandra Carbo, Loreto Ferrús Abad, Llúcia Martínez-Priego, Giuseppe D'Auria, Fernando Gonzalez-Candelas |
| EPI_ISL_436399 | Servicio de Microbiologia. Hospital Clinico Universitario de Valencia | Sequencing and Bioinformatics Service and Molecular Epidemiology Research Group. FISABIO-Public Health | Mariana Reyes-Prieto, Vicente Soriano Chirona, Ivan Ansari, David Navarro, Maria Alma Bracho, Griselda De Marco, Beatriz Beamud, Lidia Ruiz Roldan, Marta Pla Diaz, Neris Garcia-Gonzalez, Inma Galán Vendrell, Sandra Carbo, Loreto Ferrús Abad, Paula Ruiz-Hueso, Llúcia Martínez-Priego, Giuseppe D'Auria, Fernando Gonzalez-Candelas |
| EPI_ISL_436400 | Servicio de Microbiologia. Hospital Clinico Universitario de Valencia | Sequencing and Bioinformatics Service and Molecular Epidemiology Research Group. FISABIO-Public Health | Vicente Soriano Chirona, Ivan Ansari, David Navarro, Maria Alma Bracho, Griselda De Marco, Beatriz Beamud, Lidia Ruiz Roldan, Marta Pla Diaz, Neris Garcia-Gonzalez, Inma Galán Vendrell, Sandra Carbo, Loreto Ferrús Abad, Paula Ruiz-Hueso, Mariana Reyes-Prieto, Llúcia Martínez-Priego, Giuseppe D'Auria, Fernando Gonzalez-Candelas |
| EPI_ISL_436401 | Servicio de Microbiologia. Hospital Clinico Universitario de Valencia | Sequencing and Bioinformatics Service and Molecular Epidemiology Research Group. FISABIO-Public Health | Ivan Ansari, David Navarro, Maria Alma Bracho, Griselda De Marco, Beatriz Beamud, Lidia Ruiz Roldan, Marta Pla Diaz, Neris Garcia-Gonzalez, Inma Galán Vendrell, Sandra Carbo, Loreto Ferrús Abad, Paula Ruiz-Hueso, Mariana Reyes-Prieto, Vicente Soriano Chirona, Llúcia Martínez-Priego, Giuseppe D'Auria, Fernando Gonzalez-Candelas |
| EPI_ISL_436402 | Servicio de Microbiologia. Hospital Clinico Universitario de Valencia | Sequencing and Bioinformatics Service and Molecular Epidemiology Research Group. FISABIO-Public Health | David Navarro, Maria Alma Bracho, Griselda De Marco, Beatriz Beamud, Lidia Ruiz Roldan, Marta Pla Diaz, Neris Garcia-Gonzalez, Inma Galán Vendrell, Sandra Carbo, Loreto Ferrús Abad, Paula Ruiz-Hueso, Mariana Reyes-Prieto, Vicente Soriano Chirona, Ivan Ansari, Llúcia Martínez-Priego, Giuseppe D'Auria, Fernando Gonzalez-Candelas |
| EPI_ISL_436403 | Servicio de Microbiologia. Hospital Clinico Universitario de Valencia | Sequencing and Bioinformatics Service and Molecular Epidemiology Research Group. FISABIO-Public Health | Maria Alma Bracho, Griselda De Marco, Beatriz Beamud, Lidia Ruiz Roldan, Marta Pla Diaz, Neris Garcia-Gonzalez, Inma Galán Vendrell, Sandra Carbo, Loreto Ferrús Abad, Paula Ruiz-Hueso, Mariana Reyes-Prieto, Vicente Soriano Chirona, Ivan Ansari, David Navarro, Llúcia Martínez-Priego, Giuseppe D'Auria, Fernando Gonzalez-Candelas |
| EPI_ISL_436404 | Servicio de Microbiologia. Hospital Clinico Universitario de Valencia | Sequencing and Bioinformatics Service and Molecular Epidemiology Research Group. FISABIO-Public Health | Griselda De Marco, Beatriz Beamud, Lidia Ruiz Roldan, Marta Pla Diaz, Neris Garcia-Gonzalez, Inma Galán Vendrell, Sandra Carbo, Loreto Ferrús Abad, Paula Ruiz-Hueso, Mariana Reyes-Prieto, Vicente Soriano Chirona, Ivan Ansari, David Navarro, Maria Alma Bracho, Llúcia Martínez-Priego, Giuseppe D'Auria, Fernando Gonzalez-Candelas |
| EPI_ISL_436405 | Servicio de Microbiologia. Hospital Clinico Universitario de Valencia | Sequencing and Bioinformatics Service and Molecular Epidemiology Research Group. FISABIO-Public Health | Beatriz Beamud, Lidia Ruiz Roldan, Marta Pla Diaz, Neris Garcia-Gonzalez, Inma Galán Vendrell, Sandra Carbo, Loreto Ferrús Abad, Paula Ruiz-Hueso, Mariana Reyes-Prieto, Vicente Soriano Chirona, Ivan Ansari, David Navarro, Maria Alma Bracho, Griselda De Marco, Llúcia Martínez-Priego, Giuseppe D'Auria, Fernando Gonzalez-Candelas |
| EPI_ISL_436406 | Servicio de Microbiologia. Hospital Clinico Universitario de Valencia | Sequencing and Bioinformatics Service and Molecular Epidemiology Research Group. FISABIO-Public Health | Lidia Ruiz Roldan, Marta Pla Diaz, Neris Garcia-Gonzalez, Inma Galán Vendrell, Sandra Carbo, Loreto Ferrús Abad, Paula Ruiz-Hueso, Mariana Reyes-Prieto, Vicente Soriano Chirona, Ivan Ansari, David Navarro, Maria Alma Bracho, Griselda De Marco, Beatriz Beamud, Llúcia Martínez-Priego, Giuseppe D'Auria, Fernando Gonzalez-Candelas |
| EPI_ISL_436407 | Servicio de Microbiologia. Hospital Clinico Universitario de Valencia | Sequencing and Bioinformatics Service and Molecular Epidemiology Research Group. FISABIO-Public Health | Marta Pla Diaz, Neris Garcia-Gonzalez, Inma Galán Vendrell, Sandra Carbo, Loreto Ferrús Abad, Paula Ruiz-Hueso, Mariana Reyes-Prieto, Vicente Soriano Chirona, Ivan Ansari, David Navarro, Maria Alma Bracho, Griselda De Marco, Beatriz Beamud, Lidia Ruiz Roldan, Llúcia Martínez-Priego, Giuseppe D'Auria, Fernando Gonzalez-Candelas |
| EPI_ISL_436408 | Servicio de Microbiologia. Hospital Clinico Universitario de Valencia | Sequencing and Bioinformatics Service and Molecular Epidemiology Research Group. FISABIO-Public Health | Neris Garcia-Gonzalez, Inma Galán Vendrell, Sandra Carbo, Loreto Ferrús Abad, Paula Ruiz-Hueso, Mariana Reyes-Prieto, Vicente Soriano Chirona, Ivan Ansari, David Navarro, Maria Alma Bracho, Griselda De Marco, Beatriz Beamud, Lidia Ruiz Roldan, Marta Pla Diaz, Llúcia Martínez-Priego, Giuseppe D'Auria, Fernando Gonzalez-Candelas |
| EPI_ISL_436409 | Servicio de Microbiologia. Hospital Clinico Universitario de Valencia | Sequencing and Bioinformatics Service and Molecular Epidemiology Research Group. FISABIO-Public Health | Lidia Ruiz Roldan, Marta Pla Diaz, Neris Garcia-Gonzalez, Inma Galán Vendrell, Sandra Carbo, Loreto Ferrús Abad, Paula Ruiz-Hueso, Mariana Reyes-Prieto, Vicente Soriano Chirona, Ivan Ansari, David Navarro, Maria Alma Bracho, Griselda De Marco, Beatriz Beamud, Llúcia Martínez-Priego, Giuseppe D'Auria, Fernando Gonzalez-Candelas |
| EPI_ISL_436410 | Servicio de Microbiologia. Hospital Clinico Universitario de Valencia | Sequencing and Bioinformatics Service and Molecular Epidemiology Research Group. FISABIO-Public Health | Marta Pla Diaz, Neris Garcia-Gonzalez, Inma Galán Vendrell, Sandra Carbo, Loreto Ferrús Abad, Paula Ruiz-Hueso, Mariana Reyes-Prieto, Vicente Soriano Chirona, Ivan Ansari, David Navarro, Maria Alma Bracho, Griselda De Marco, Beatriz Beamud, Lidia Ruiz Roldan, Llúcia Martínez-Priego, Giuseppe D'Auria, Fernando Gonzalez-Candelas |
| EPI_ISL_436411 | Servicio de Microbiologia. Hospital Clinico Universitario de Valencia | Sequencing and Bioinformatics Service and Molecular Epidemiology Research Group. FISABIO-Public Health | Neris Garcia-Gonzalez, Inma Galán Vendrell, Sandra Carbo, Loreto Ferrús Abad, Paula Ruiz-Hueso, Mariana Reyes-Prieto, Vicente Soriano Chirona, Ivan Ansari, David Navarro, Maria Alma Bracho, Griselda De Marco, Beatriz Beamud, Lidia Ruiz Roldan, Marta Pla Diaz, Llúcia Martínez-Priego, Giuseppe D'Auria, Fernando Gonzalez-Candelas |
| EPI_ISL_436412 | Viral Respiratory Lab, National Institute for Biomedical Research (INRB) | Pathogen Sequencing Lab, National Institute for Biomedical Research (INRB) | Placide Mbala-Kingebeni, Edith Nkwembe, Eddy Kinganda-Lusamaki, Amuri Aziza, Francisca Muyembe Mawete, Catherine Pratt, Matthias Pauthner, Josh Quick, Allison Black, James Hadfield, Trevor Bedford, Ian Goodfellow, Andrew Rambaut, Nick Loman, Kristian Andersen, Michael Wiley, Steve Ahuka-Mundeke, Jean-Jacques Muyembe Tamfum |
| EPI_ISL_436413, EPI_ISL_436414, EPI_ISL_436415, EPI_ISL_436416, EPI_ISL_436417, EPI_ISL_436418, EPI_ISL_436419, EPI_ISL_436420, EPI_ISL_436421, EPI_ISL_436422, EPI_ISL_436423, EPI_ISL_436424, EPI_ISL_436425, EPI_ISL_436426, EPI_ISL_436427, EPI_ISL_436428, EPI_ISL_436429, EPI_ISL_436430, EPI_ISL_436431, EPI_ISL_436432, EPI_ISL_436433, EPI_ISL_436434, EPI_ISL_436435, EPI_ISL_436436, EPI_ISL_436437, EPI_ISL_436438, EPI_ISL_436439, EPI_ISL_436440, EPI_ISL_436441, EPI_ISL_436442, EPI_ISL_436443, EPI_ISL_436444, EPI_ISL_436445, EPI_ISL_436446, EPI_ISL_436447, EPI_ISL_436448, EPI_ISL_436449, EPI_ISL_436450, EPI_ISL_436451, EPI_ISL_436452, EPI_ISL_436453, EPI_ISL_436454, EPI_ISL_436455, EPI_ISL_436456, EPI_ISL_436457, EPI_ISL_436458, EPI_ISL_436459, EPI_ISL_436460, EPI_ISL_436461, EPI_ISL_436462, EPI_ISL_436463 | | | |
| see above | National Centre for Disease control (NCDC) | NCDC/CSIR-IGIB | Pramod Kumar#, Rajesh Pandey#, Pooja Sharma, Mahesh S Dhar, Vivekanand A, Bharathram Uppili, Himanshu Vashisht, Saruchi Wadhwa, Nishu Tyagi, Uma Sharma, Priyanka Singh, Hemlata Lall, Meena Datta, Poonam Gupta, Nidhi Saini, Aarti Tewari, Bibhash Nandi, Dhirendra Kumar, Satyabrata Bag, Varun Jaiswal, Hema Gogia, Preeti Madan, Simrita Singh, Prateek Singh, Debasis Dash, Mitali Mukerji, Manju Bala, Sandhya Kabra, Sujeet Singh, Mohammed Faruq, Anurag Agrawal*, Partha Rakshit* |
| EPI_ISL_436464 | Alaska State Virology Laboratory | Alaska State Virology Laboratory | Jack Chen |
| EPI_ISL_436466, EPI_ISL_436467, EPI_ISL_436468, EPI_ISL_436469, EPI_ISL_436470, EPI_ISL_436471, EPI_ISL_436472, EPI_ISL_436473, EPI_ISL_436474, EPI_ISL_436475, EPI_ISL_436476, EPI_ISL_436477, EPI_ISL_436478, EPI_ISL_436479, EPI_ISL_436480, EPI_ISL_436481, EPI_ISL_436482, EPI_ISL_436483, EPI_ISL_436484, EPI_ISL_436485, EPI_ISL_436486, EPI_ISL_436487, EPI_ISL_436488, EPI_ISL_436489, EPI_ISL_436490, EPI_ISL_436491, EPI_ISL_436492, EPI_ISL_436493, EPI_ISL_436494, EPI_ISL_436495, EPI_ISL_436496, EPI_ISL_436497, EPI_ISL_436498, EPI_ISL_436499, EPI_ISL_436500, EPI_ISL_436501, EPI_ISL_436502, EPI_ISL_436503, EPI_ISL_436504 | | | |
| see above | UPMC Clinical Laboratory | Microbial Genome Sequencing Center, Microbial Genomic Epidemiological Laboratory | Dan Snyder, Stephanie L Mitchell, Mustapha M Mustapha, Marissa P Griffith, Vatsala R Srinivasa, Kady D Waggle, Chinelo Ezeonwuku, Jane W. Marsh, Lee H. Harrison, Vaughn S. Cooper |
| EPI_ISL_436505, EPI_ISL_436506, EPI_ISL_436507, EPI_ISL_436508, EPI_ISL_436509, EPI_ISL_436510, EPI_ISL_436511, EPI_ISL_436512, EPI_ISL_436513, EPI_ISL_436514, EPI_ISL_436515, EPI_ISL_436516, EPI_ISL_436517, EPI_ISL_436518, EPI_ISL_436519, EPI_ISL_436520, EPI_ISL_436521, EPI_ISL_436522, EPI_ISL_436523, EPI_ISL_436524, EPI_ISL_436525, EPI_ISL_436526, EPI_ISL_436527, EPI_ISL_436528, EPI_ISL_436529, EPI_ISL_436530, EPI_ISL_436531, EPI_ISL_436532, EPI_ISL_436533, EPI_ISL_436534, EPI_ISL_436535, EPI_ISL_436536, EPI_ISL_436537, EPI_ISL_436538, EPI_ISL_436539, EPI_ISL_436540, EPI_ISL_436541, EPI_ISL_436542, EPI_ISL_436543, EPI_ISL_436544, EPI_ISL_436545, EPI_ISL_436546, EPI_ISL_436547, EPI_ISL_436548, EPI_ISL_436549, EPI_ISL_436550, EPI_ISL_436551, EPI_ISL_436552, EPI_ISL_436553, EPI_ISL_436554, EPI_ISL_436555, EPI_ISL_436556, EPI_ISL_436557, EPI_ISL_436558, EPI_ISL_436559, EPI_ISL_436560, EPI_ISL_436561, EPI_ISL_436562, EPI_ISL_436563 | | | |
| see above | Florida Bureau of Public Health Laboratories | Florida Bureau of Public Health Laboratories | Sarah Schmedes, Jason Blanton |
| EPI_ISL_436564, EPI_ISL_436565, EPI_ISL_436566, EPI_ISL_436567, EPI_ISL_436568, EPI_ISL_436569, EPI_ISL_436570, EPI_ISL_436571, EPI_ISL_436572, EPI_ISL_436573, EPI_ISL_436574, EPI_ISL_436575, EPI_ISL_436576, EPI_ISL_436577, EPI_ISL_436578, EPI_ISL_436579, EPI_ISL_436580, EPI_ISL_436581, EPI_ISL_436582, EPI_ISL_436583, EPI_ISL_436584, EPI_ISL_436585, EPI_ISL_436586, EPI_ISL_436587, EPI_ISL_436588, EPI_ISL_436589, EPI_ISL_436590, EPI_ISL_436591, EPI_ISL_436592, EPI_ISL_436593, EPI_ISL_436594, EPI_ISL_436595, EPI_ISL_436596, EPI_ISL_436597, EPI_ISL_436598, EPI_ISL_436599, EPI_ISL_436600, EPI_ISL_436601, EPI_ISL_436602, EPI_ISL_436603, EPI_ISL_436604, EPI_ISL_436605, EPI_ISL_436606, EPI_ISL_436607, EPI_ISL_436608, EPI_ISL_436609, EPI_ISL_436610, EPI_ISL_436611, EPI_ISL_436612, EPI_ISL_436613, EPI_ISL_436614, EPI_ISL_436615, EPI_ISL_436616, EPI_ISL_436617, EPI_ISL_436618, EPI_ISL_436619, EPI_ISL_436620, EPI_ISL_436621, EPI_ISL_436622, EPI_ISL_436623, EPI_ISL_436624, EPI_ISL_436625, EPI_ISL_436626, EPI_ISL_436627, EPI_ISL_436628, EPI_ISL_436629, EPI_ISL_436630, EPI_ISL_436631, EPI_ISL_436632, EPI_ISL_436633, EPI_ISL_436634, EPI_ISL_436635, EPI_ISL_436636, EPI_ISL_436637, EPI_ISL_436638, EPI_ISL_436639, EPI_ISL_436640 | | | |
| see above | University of Wisconsin-Madison AIDS Vaccine Research Laboratories | University of Wisconsin-Madison AIDS Vaccine Research Laboratories | Gage Moreno, Katarina Braun, et al. AIDS Vaccine Research Laboratories |
| EPI_ISL_436641, EPI_ISL_436642, EPI_ISL_436643, EPI_ISL_436644, EPI_ISL_436645, EPI_ISL_436646, EPI_ISL_436647, EPI_ISL_436648, EPI_ISL_436649, EPI_ISL_436650, EPI_ISL_436651, EPI_ISL_436652, EPI_ISL_436653, EPI_ISL_436654, EPI_ISL_436655, EPI_ISL_436656, EPI_ISL_436657, EPI_ISL_436658, EPI_ISL_436659, EPI_ISL_436660, EPI_ISL_436661, EPI_ISL_436662, EPI_ISL_436663, EPI_ISL_436664, EPI_ISL_436665, EPI_ISL_436666, EPI_ISL_436667, EPI_ISL_436668, EPI_ISL_436669, EPI_ISL_436670, EPI_ISL_436671, EPI_ISL_436672, EPI_ISL_436673, EPI_ISL_436674, EPI_ISL_436675, EPI_ISL_436676, EPI_ISL_436677, EPI_ISL_436678, EPI_ISL_436679, EPI_ISL_436680, EPI_ISL_436681, EPI_ISL_436682, EPI_ISL_436683 | | | |
| see above | County of Santa Clara Public Health Department | Chan-Zuckerberg Biohub | CZB Cliahub Consortium |
| EPI_ISL_436684, EPI_ISL_436685, EPI_ISL_436686, EPI_ISL_436687 | KRISP, KZN Research Innovation and Sequencing Platform | KRISP, KZN Research Innovation and Sequencing Platform | Giandhari J, Pillay S, Lessells R, Chimukangara B, Deforche K, Tegally H, Wilkinson E, de Oliveira T |
| EPI_ISL_436688, EPI_ISL_436689 | Victorian Infectious Diseases Reference Laboratory (VIDRL) | Microbiological Diagnostic Unit Public Health Laboratory and Victorian Infectious Diseases Reference Laboratory, The Peter Doherty Institute for Infection & Immunity | Caly L., Seemann T., Sait, M., Schultz M., Druce J., Sherry, N. |
| EPI_ISL_436715 | Genomics and Computational Biology Lab, Scientific Research Institute of Physical-Chemical Medicine, FMBA of Russia | Genomics and Computational Biology Lab, Scientific Research Institute of Physical-Chemical Medicine, FMBA of Russia | A. Pavlenko, O. Guskova, K. Klimina, V. Veselovsky, A. Manolov, D. Fedorov, V. Govorun and E. Ilina |
| EPI_ISL_436716 | Genomics and Computational Biology Lab, Scientific Research Institute of Physical-Chemical Medicine, FMBA of Russia | Genomics and Computational Biology Lab, Scientific Research Institute of Physical-Chemical Medicine, FMBA of Russia | A. Pavlenko, O. Guskova, K. Klimina, V. Veselovsky, A. Manolov, D. Fedorov, V. Govorun and E. Ilina |
| EPI_ISL_436717 | Genomics and Computational Biology Lab, Scientific Research Institute of Physical-Chemical Medicine, FMBA of Russia | Genomics and Computational Biology Lab, Scientific Research Institute of Physical-Chemical Medicine, FMBA of Russia | A. Pavlenko, O. Guskova, K. Klimina, V. Veselovsky, A. Manolov, D. Fedorov, V. Govorun and E. Ilina |
| EPI_ISL_436718 | Ospedale Regionale San Salvatore | Istituto Zooprofilattico Sperimentale dell'Abruzzo e Molise "G.Caporale" | Lorusso A, Marcacci M, Di Domenico M, Ancora M, Curini V, Mangone I, Rinaldi A, Di Pasquale A, Cammà C, Puglia I, Savini G |

| | | | |
|---|---|---|---|
| EPI_ISL_436719, EPI_ISL_436720, EPI_ISL_436721, EPI_ISL_436722 | Ospedale Civile S. Liberatore di Atri | Istituto Zooprofilattico Sperimentale dell'Abruzzo e Molise "G.Caporale" | Lorusso A, Marcacci M, Di Domenico M, Ancora M, Curini V, Mangone I, Rinaldi A, Di Pasquale A, Cammà C, Puglia I, Savini G |
| EPI_ISL_436723 | Ospedale Civile Giuseppe Mazzini | Istituto Zooprofilattico Sperimentale dell'Abruzzo e Molise "G.Caporale" | Lorusso A, Marcacci M, Di Domenico M, Ancora M, Curini V, Mangone I, Rinaldi A, Di Pasquale A, Cammà C, Puglia I, Savini G |
| EPI_ISL_436724 | Ospedale Civile S. Liberatore di Atri | Istituto Zooprofilattico Sperimentale dell'Abruzzo e Molise "G.Caporale" | Lorusso A, Marcacci M, Di Domenico M, Ancora M, Curini V, Mangone I, Rinaldi A, Di Pasquale A, Cammà C, Puglia I, Savini G |
| EPI_ISL_436725 | RSA/RP Villa San Giovanni - Gruppo Edos | Istituto Zooprofilattico Sperimentale dell'Abruzzo e Molise "G.Caporale" | Lorusso A, Marcacci M, Di Domenico M, Ancora M, Curini V, Mangone I, Rinaldi A, Di Pasquale A, Cammà C, Puglia I, Savini G |
| EPI_ISL_436726, EPI_ISL_436727, EPI_ISL_436728, EPI_ISL_436729 | SERVIZIO DI IGIENE E SANITÀ PUBBLICA ASL Teramo | Istituto Zooprofilattico Sperimentale dell'Abruzzo e Molise "G.Caporale" | Lorusso A, Marcacci M, Di Domenico M, Ancora M, Curini V, Mangone I, Rinaldi A, Di Pasquale A, Cammà C, Puglia I, Savini G |
| EPI_ISL_436730 | Servizio di igiene epidemiologia e sanità pubblica (Siesp) Chieti | Istituto Zooprofilattico Sperimentale dell'Abruzzo e Molise "G.Caporale" | Lorusso A, Marcacci M, Di Domenico M, Ancora M, Curini V, Mangone I, Rinaldi A, Di Pasquale A, Cammà C, Puglia I, Savini G |
| EPI_ISL_436731, EPI_ISL_436732 | Ospedale Civile S. Liberatore di Atri | Istituto Zooprofilattico Sperimentale dell'Abruzzo e Molise "G.Caporale" | Lorusso A, Marcacci M, Di Domenico M, Ancora M, Curini V, Mangone I, Rinaldi A, Di Pasquale A, Cammà C, Puglia I, Savini G |
| EPI_ISL_436733, EPI_ISL_436734, EPI_ISL_436735, EPI_ISL_436736, EPI_ISL_436737, EPI_ISL_436738, EPI_ISL_436739, EPI_ISL_436740, EPI_ISL_436741, EPI_ISL_436742, EPI_ISL_436743 | | | |
| see above | NYU Langone Health | Departments of Pathology and Medicine, New York University School of Medicine | Maria Aguero-Rosenfeld, Brendan Belovarac, Margaret Black, Ludovic Boytard, John Cadley, Paolo Cotzia, John Chen, Dacia Dimartino, Xiaojun Feng, Tatyana Gindin, Emily Guzman, Adriana Heguy, Megan Hogan, Emily Huang, George Jour, Alireza Khodadadi-Jamayran, Lawrence H. Lin, Raven Luther, Andrew Lytle, Christian Marier, Matthew T. Maurano, Mark J. Mulligan, Peter Meyn, Raquel Ordonez Ciriza, Iman Osman, Jared Pinnell, Vanessa Raabe, Sitharam Ramaswami, Amy Rapkiewicz, Andre M. Ribeiro-dos-Santos, Marie Samanovic-Golden, Antonio Serrano, Guomiao Shen, Matija Snuderl, Theodore Vougiouklakis, Nick Vulpescu, Gael Westby, Paul Zappile, Yutong Zhang |
| EPI_ISL_436800, EPI_ISL_436801, EPI_ISL_436802, EPI_ISL_436803, EPI_ISL_436804, EPI_ISL_436805, EPI_ISL_436806, EPI_ISL_436807, EPI_ISL_436808, EPI_ISL_436809, EPI_ISL_436810, EPI_ISL_436811, EPI_ISL_436812, EPI_ISL_436813, EPI_ISL_436814, EPI_ISL_436815, EPI_ISL_436816, EPI_ISL_436817, EPI_ISL_436818, EPI_ISL_436819, EPI_ISL_436820, EPI_ISL_436821, EPI_ISL_436822, EPI_ISL_436823, EPI_ISL_436824, EPI_ISL_436825, EPI_ISL_436826, EPI_ISL_436827, EPI_ISL_436828, EPI_ISL_436829, EPI_ISL_436830, EPI_ISL_436831, EPI_ISL_436832, EPI_ISL_436833, EPI_ISL_436834, EPI_ISL_436835, EPI_ISL_436836, EPI_ISL_436837, EPI_ISL_436838, EPI_ISL_436839, EPI_ISL_436840, EPI_ISL_436841, EPI_ISL_436842, EPI_ISL_436843, EPI_ISL_436844, EPI_ISL_436845, EPI_ISL_436846, EPI_ISL_436847, EPI_ISL_436848, EPI_ISL_436849, EPI_ISL_436850, EPI_ISL_436851, EPI_ISL_436852, EPI_ISL_436853, EPI_ISL_436854, EPI_ISL_436855, EPI_ISL_436856, EPI_ISL_436857, EPI_ISL_436858, EPI_ISL_436859, EPI_ISL_436860, EPI_ISL_436861, EPI_ISL_436862, EPI_ISL_436863, EPI_ISL_436864, EPI_ISL_436865, EPI_ISL_436866, EPI_ISL_436867, EPI_ISL_436868, EPI_ISL_436869, EPI_ISL_436870, EPI_ISL_436871, EPI_ISL_436872, EPI_ISL_436873, EPI_ISL_436874, EPI_ISL_436875, EPI_ISL_436876, EPI_ISL_436877, EPI_ISL_436878, EPI_ISL_436879, EPI_ISL_436880, EPI_ISL_436881, EPI_ISL_436882, EPI_ISL_436883, EPI_ISL_436884, EPI_ISL_436885, EPI_ISL_436886, EPI_ISL_436887, EPI_ISL_436888, EPI_ISL_436889, EPI_ISL_436890 | | | |
| see above | Michigan Department of Health and Human Services, Bureau of Laboratories | Michigan Department of Health and Human Services, Bureau of Laboratories | Blankenship HM, Riner D, Soehnlen MK |
| EPI_ISL_436891, EPI_ISL_436892, EPI_ISL_436893, EPI_ISL_436894, EPI_ISL_436895, EPI_ISL_436896, EPI_ISL_436897, EPI_ISL_436898, EPI_ISL_436899, EPI_ISL_436900 | Gundersen Molecular Diagnostics Laboratory | Kabara Cancer Research Institute | Craig S. Richmond, Paraic A. Kenny |
| EPI_ISL_436901, EPI_ISL_436902, EPI_ISL_436903, EPI_ISL_436904, EPI_ISL_436905, EPI_ISL_436906, EPI_ISL_436907, EPI_ISL_436908, EPI_ISL_436909, EPI_ISL_436910, EPI_ISL_436911, EPI_ISL_436912, EPI_ISL_436913, EPI_ISL_436914, EPI_ISL_436915, EPI_ISL_436916, EPI_ISL_436917, EPI_ISL_436918, EPI_ISL_436919, EPI_ISL_436920, EPI_ISL_436921, EPI_ISL_436922, EPI_ISL_436923, EPI_ISL_436924, EPI_ISL_436925 | | | |
| see above | Utah Public Health Laboratory | Utah Public Health Laboratory | Erin Young, Kelly Oakeson |
| EPI_ISL_436926 | $x^2$ | Utah Public Health Laboratory | Erin Young, Kelly Oakeson |
| EPI_ISL_436927, EPI_ISL_436928, EPI_ISL_436929, EPI_ISL_436930, EPI_ISL_436931, EPI_ISL_436932, EPI_ISL_436933, EPI_ISL_436934, EPI_ISL_436935, EPI_ISL_436936, EPI_ISL_436937, EPI_ISL_436938 | | | |
| see above | Utah Public Health Laboratory | Utah Public Health Laboratory | Erin Young, Kelly Oakeson |
| EPI_ISL_436939, EPI_ISL_436940, EPI_ISL_436941, EPI_ISL_436942, EPI_ISL_436943, EPI_ISL_436944, EPI_ISL_436945, EPI_ISL_436946, EPI_ISL_436947, EPI_ISL_436948, EPI_ISL_436949, EPI_ISL_436950, EPI_ISL_436951, EPI_ISL_436952, EPI_ISL_436953, EPI_ISL_436954, EPI_ISL_436955, EPI_ISL_436956, EPI_ISL_436957, EPI_ISL_436958, EPI_ISL_436959, EPI_ISL_436960, EPI_ISL_436961 | | | |
| see above | Ochsner Health | Bioinfoexperts, LLC | Amy Feehan, David J. Nolan, Rebecca Rose, Sissy Cross, David Moraga Amador, Tong Yang, Luke Caruso, Wayra Navia, Lydia Von Borstel, Xiao Hui Zhou, Julia-Garcia-Diaz, Susanna L. Lamers |
| EPI_ISL_436962, EPI_ISL_436963, EPI_ISL_436964, EPI_ISL_436965, EPI_ISL_436966, EPI_ISL_436967, EPI_ISL_436968, EPI_ISL_436969, EPI_ISL_436970, EPI_ISL_436971, EPI_ISL_436972, EPI_ISL_436973, EPI_ISL_436974, EPI_ISL_436975, EPI_ISL_436976, EPI_ISL_436977, EPI_ISL_436978, EPI_ISL_436979, EPI_ISL_436980, EPI_ISL_436981, EPI_ISL_436982, EPI_ISL_436983, EPI_ISL_436984, EPI_ISL_436985, EPI_ISL_436986, EPI_ISL_436987, EPI_ISL_436988, EPI_ISL_436989, EPI_ISL_436990, EPI_ISL_436991, EPI_ISL_436992, EPI_ISL_436993, EPI_ISL_436994, EPI_ISL_436995, EPI_ISL_436996, EPI_ISL_436997, EPI_ISL_436998, EPI_ISL_436999, EPI_ISL_437000, EPI_ISL_437001, EPI_ISL_437002, EPI_ISL_437003, EPI_ISL_437004, EPI_ISL_437005, EPI_ISL_437006, EPI_ISL_437007, EPI_ISL_437008, EPI_ISL_437009, EPI_ISL_437010, EPI_ISL_437011, EPI_ISL_437012, EPI_ISL_437013, EPI_ISL_437014, EPI_ISL_437015, EPI_ISL_437016, EPI_ISL_437017, EPI_ISL_437018, EPI_ISL_437019, EPI_ISL_437020, EPI_ISL_437021, EPI_ISL_437022, EPI_ISL_437023, EPI_ISL_437024, EPI_ISL_437025, EPI_ISL_437026, EPI_ISL_437027, EPI_ISL_437028, EPI_ISL_437029, EPI_ISL_437030, EPI_ISL_437031, EPI_ISL_437032, EPI_ISL_437033, EPI_ISL_437034, EPI_ISL_437035, EPI_ISL_437036, EPI_ISL_437037, EPI_ISL_437038, EPI_ISL_437039, EPI_ISL_437040, EPI_ISL_437041, EPI_ISL_437042 | | | |
| see above | Department of Virus and Microbiological Special Diagnostics, Statens Serum Institut, Copenhagen, Denmark, Artillerivej 5, 2300 Copenahgen S | Albertsen lab, Department of Chemistry and Bioscience, Aalborg University, Denmark | Rasmus Kirkegaard |
| EPI_ISL_437043, EPI_ISL_437044, EPI_ISL_437045, EPI_ISL_437046, EPI_ISL_437047, EPI_ISL_437048, EPI_ISL_437049, EPI_ISL_437050, EPI_ISL_437051, EPI_ISL_437052, EPI_ISL_437053, EPI_ISL_437054, EPI_ISL_437055, EPI_ISL_437056, EPI_ISL_437057, EPI_ISL_437058, EPI_ISL_437059, EPI_ISL_437060, EPI_ISL_437061, EPI_ISL_437062, EPI_ISL_437063, EPI_ISL_437064, EPI_ISL_437065, EPI_ISL_437066, EPI_ISL_437067, EPI_ISL_437068, EPI_ISL_437069, EPI_ISL_437070, EPI_ISL_437071, EPI_ISL_437072, EPI_ISL_437073, EPI_ISL_437074, EPI_ISL_437075, EPI_ISL_437076, EPI_ISL_437077, EPI_ISL_437078, EPI_ISL_437079, EPI_ISL_437080, EPI_ISL_437081, EPI_ISL_437082, EPI_ISL_437083, EPI_ISL_437084, EPI_ISL_437085, EPI_ISL_437086, EPI_ISL_437087, EPI_ISL_437088 | | | |
| see above | County of Santa Clara Public Health | Chan-Zuckerberg Biohub | CZB Cliahub Consortium |
| EPI_ISL_437089, EPI_ISL_437090, EPI_ISL_437091, EPI_ISL_437092, EPI_ISL_437093, EPI_ISL_437094, EPI_ISL_437095, EPI_ISL_437096 | Latvijas Infektoloģijas centrs | Latvian Biomedical Research and Study Centre | Ivars Silamiķelis, Kaspars Megnis, Monta Ustinova, Ņikita Zrelovs, Vita Rovīte, Jeļena Storoženko, Tatjana Kolupajeva, Oksana Savicka, Uga Dumpis, Jānis Kloviņš |
| EPI_ISL_437097, EPI_ISL_437098, EPI_ISL_437099, EPI_ISL_437100, EPI_ISL_437101, EPI_ISL_437102, EPI_ISL_437103, EPI_ISL_437104, EPI_ISL_437105, EPI_ISL_437106, EPI_ISL_437107, EPI_ISL_437108, EPI_ISL_437109, EPI_ISL_437110, EPI_ISL_437111, EPI_ISL_437112, EPI_ISL_437113, EPI_ISL_437114, EPI_ISL_437115, EPI_ISL_437116, EPI_ISL_437117, EPI_ISL_437118, EPI_ISL_437119, EPI_ISL_437120, EPI_ISL_437121, EPI_ISL_437122, EPI_ISL_437123, EPI_ISL_437124, EPI_ISL_437125, EPI_ISL_437126, EPI_ISL_437127, EPI_ISL_437128, EPI_ISL_437129, EPI_ISL_437130, EPI_ISL_437131, EPI_ISL_437132, EPI_ISL_437133, EPI_ISL_437134, EPI_ISL_437135, EPI_ISL_437136, EPI_ISL_437137, EPI_ISL_437138, EPI_ISL_437139, EPI_ISL_437140, EPI_ISL_437141, EPI_ISL_437142, EPI_ISL_437143, EPI_ISL_437144, EPI_ISL_437145, EPI_ISL_437146, EPI_ISL_437147, EPI_ISL_437148, EPI_ISL_437149, EPI_ISL_437150, EPI_ISL_437151, EPI_ISL_437152, EPI_ISL_437153, EPI_ISL_437154, EPI_ISL_437155, EPI_ISL_437156, EPI_ISL_437157, EPI_ISL_437158, EPI_ISL_437159, EPI_ISL_437160, EPI_ISL_437161, EPI_ISL_437162, EPI_ISL_437163, EPI_ISL_437164, EPI_ISL_437165, EPI_ISL_437166, EPI_ISL_437167, EPI_ISL_437168, EPI_ISL_437169, EPI_ISL_437170, EPI_ISL_437171, EPI_ISL_437172, EPI_ISL_437173, EPI_ISL_437174, EPI_ISL_437175, EPI_ISL_437176, EPI_ISL_437177, EPI_ISL_437178, EPI_ISL_437179, EPI_ISL_437180, EPI_ISL_437181, EPI_ISL_437182, EPI_ISL_437183, EPI_ISL_437184, EPI_ISL_437185, EPI_ISL_437186 | | | |
| see above | Michigan Department of Health and Human Services, Bureau of Laboratories | Michigan Department of Health and Human Services, Bureau of Laboratories | Blankenship HM, Riner D, Soehnlen MK |
| EPI_ISL_437187 | Siloam Hospitals | Institute of Tropical Disease, Universitas Airlangga | Kazufumi Shimizu, Krisnoadi Rahardjo, Aldise M Nastri, Jezzy R Dewantari, Rima R Prasetya, Maria M Padmidewi, Gatot Soegiarto, Laksmi Wulandari, Retno A Setyoningrum, Resti Y Meliana, Yohko K Shimizu, Mitsuhiro Nishimura, Yasuko Mori, Soetjipto, Maria I Lusida |
| EPI_ISL_437188 | RSUD Dr. Soetomo | Institute of Tropical Disease, Universitas Airlangga | Krisnoadi Rahardjo, Aldise M Nastri, Jezzy R Dewantari, Rima R Prasetya, Joni Wahyuhadi, Gatot Soegiarto, Laksmi Wulandari, Retno A Setyoningrum, Resti Y Meliana, Yohko K Shimizu, Mitsuhiro Nishimura, Yasuko Mori, Soetjipto, Kazufumi Shimizu, Maria I Lusida |
| EPI_ISL_437189 | Pusat Pertamina Hospital | Eijkman Institute for Molecular Biology, Ministry of Research and Technology/National Agency for Research and Innovation | Edison Johar, Frilasita A Yudhaputri, Hidayat Trimarsanto, David H Muljono, Safarina G Malik, Khin Saw Myint, Amin Soebandrio |
| EPI_ISL_437190, EPI_ISL_437191 | RS Pondok Indah Hospital – Pondok Indah | Eijkman Institute for Molecular Biology, Ministry of Research and Technology/National Agency for Research and Innovation | Edison Johar, Frilasita A Yudhaputri, Hidayat Trimarsanto, David H Muljono, Safarina G Malik, Khin Saw Myint, Amin Soebandrio |
| EPI_ISL_437192 | Mitra Keluarga Kelapa Gading Hospital | Eijkman Institute for Molecular Biology, Ministry of Research and Technology/National Agency for Research and Innovation | Edison Johar, Frilasita A Yudhaputri, Hidayat Trimarsanto, David H Muljono, Safarina G Malik, Khin Saw Myint, Amin Soebandrio |
| EPI_ISL_437193, EPI_ISL_437194, EPI_ISL_437195, EPI_ISL_437196 | Viral Respiratory Lab, National Institute for Biomedical Research (INRB) | Pathogen Sequencing Lab, National Institute for Biomedical Research (INRB) | Placide Mbala-Kingebeni, Edith Nkwembe, Eddy Kinganda-Lusamaki, Amuri Aziza, Francisca Muyembe Mawete, Catherine Pratt, Matthias Pauthner, Josh Quick, Allison Black, James Hadfield, Trevor Bedford, Ian Goodfellow, Andrew Rambaut, Nick Loman, Kristian Andersen, Michael Wiley, Steve Ahuka-Mundeke, Jean-Jacques Muyembe Tamfum |
| EPI_ISL_437197 | Diagnostic- and Research Institute of Pathology, Medical University of Graz | Diagnostic- and Research Institute of Pathology, Medical University of Graz | Karl Kashofer, Peter Regitnig, Martin Zacharias, Gregor Gorkiewicz |
| EPI_ISL_437198, EPI_ISL_437199, EPI_ISL_437200, EPI_ISL_437201, EPI_ISL_437202 | Diagnostic- and Research Institute of Pathology, Medical University of Graz | Diagnostic- and Research Institute of Pathology, Medical University of Graz | Karl Kashofer, Peter Regitnig, Martin Zacharias, Gregor Gorkiewicz |
| EPI_ISL_437203 | Diagnostic- and Research Institute of Pathology, Medical University of Graz | Diagnostic- and Research Institute of Pathology, Medical University of Graz | Karl Kashofer, Peter Regitnig, Martin Zacharias, Gregor Gorkiewicz |
| EPI_ISL_437204, EPI_ISL_437205, EPI_ISL_437206, EPI_ISL_437207, EPI_ISL_437208, EPI_ISL_437209, EPI_ISL_437210, EPI_ISL_437211, EPI_ISL_437212, EPI_ISL_437213, EPI_ISL_437214, EPI_ISL_437215, EPI_ISL_437216, EPI_ISL_437217, EPI_ISL_437218, EPI_ISL_437219, EPI_ISL_437220, EPI_ISL_437221, EPI_ISL_437222, EPI_ISL_437223, EPI_ISL_437224, EPI_ISL_437225, EPI_ISL_437226, EPI_ISL_437227, EPI_ISL_437228, EPI_ISL_437229, EPI_ISL_437230, EPI_ISL_437231, EPI_ISL_437232, EPI_ISL_437233, EPI_ISL_437234, EPI_ISL_437235, EPI_ISL_437236, EPI_ISL_437237, EPI_ISL_437238, EPI_ISL_437239, EPI_ISL_437240, EPI_ISL_437241, EPI_ISL_437242, EPI_ISL_437243, EPI_ISL_437244, EPI_ISL_437245, EPI_ISL_437246, EPI_ISL_437247, EPI_ISL_437248, EPI_ISL_437249, EPI_ISL_437250, EPI_ISL_437251, EPI_ISL_437252, EPI_ISL_437253, EPI_ISL_437254, EPI_ISL_437255, EPI_ISL_437256, EPI_ISL_437257, EPI_ISL_437258, EPI_ISL_437259, EPI_ISL_437260, EPI_ISL_437261, EPI_ISL_437262, EPI_ISL_437263, EPI_ISL_437264, EPI_ISL_437265, EPI_ISL_437266, EPI_ISL_437267, EPI_ISL_437268, EPI_ISL_437269, EPI_ISL_437270, EPI_ISL_437271, EPI_ISL_437272, EPI_ISL_437273, EPI_ISL_437274, EPI_ISL_437275, EPI_ISL_437276, EPI_ISL_437277, EPI_ISL_437278, EPI_ISL_437279, EPI_ISL_437280, EPI_ISL_437281, EPI_ISL_437282, EPI_ISL_437283, EPI_ISL_437284, EPI_ISL_437285, EPI_ISL_437286, EPI_ISL_437287, EPI_ISL_437288, EPI_ISL_437289, EPI_ISL_437290, EPI_ISL_437291, EPI_ISL_437292, EPI_ISL_437293, EPI_ISL_437294, EPI_ISL_437295, EPI_ISL_437296, EPI_ISL_437297 | | | |
| see above | Max von Pettenkofer Institute, Virology, National Reference Center for Retroviruses, LMU München | Laboratory for Functional Genome Analysis, Dept. Genomics, Gene Center of the LMU Munich | Max Muenchhoff, Stefan Krebs, Alexander Graf, Oliver Keppler, Helmut Blum |
| EPI_ISL_437298, EPI_ISL_437299, EPI_ISL_437300, EPI_ISL_437301, EPI_ISL_437302, EPI_ISL_437303 | Diagnostic- and Research Institute of Pathology, Medical University of Graz | Diagnostic- and Research Institute of Pathology, Medical University of Graz | Karl Kashofer, Peter Regitnig, Martin Zacharias, Gregor Gorkiewicz |
| EPI_ISL_437304, EPI_ISL_437305, EPI_ISL_437306, EPI_ISL_437307, EPI_ISL_437308, EPI_ISL_437309, EPI_ISL_437310, EPI_ISL_437311, EPI_ISL_437312, EPI_ISL_437313, EPI_ISL_437314, EPI_ISL_437315, EPI_ISL_437316, EPI_ISL_437317, EPI_ISL_437318 | | | |
| see above | Ministry of Health Turkey | Ministry of Health Turkey | Fatma Bayrakdar,Tülin Demir,Süleyman Yalçın, Selçuk Kılıç |
| EPI_ISL_437319, EPI_ISL_437320, EPI_ISL_437321 | Ministry of Health Turkey | Ministry of Health Turkey | Fatma Bayrakdar,Ayşe Başak Altaş,Yasemin Coşgun,Süleyman Yalçın, Gülay Korukluoğlu,Selçuk Kılıç |
| EPI_ISL_437322 | Ministry of Health Turkey | Ministry of Health Turkey | Fatma Bayrakdar,Tülin Demir,Süleyman Yalçın, Selçuk Kılıç |
| EPI_ISL_437323, EPI_ISL_437324, EPI_ISL_437325, EPI_ISL_437326, EPI_ISL_437327, EPI_ISL_437328, EPI_ISL_437329, EPI_ISL_437330 | Ministry of Health Turkey | Ministry of Health Turkey | Fatma Bayrakdar,Ayşe Başak Altaş,Yasemin Coşgun,Süleyman Yalçın, Gülay Korukluoğlu,Selçuk Kılıç |
| EPI_ISL_437331 | Ministry of Health Turkey | Ministry of Health Turkey | Fatma Bayrakdar,Tülin Demir,Süleyman Yalçın, Selçuk Kılıç |
| EPI_ISL_437332, EPI_ISL_437333, EPI_ISL_437334, | Ministry of Health Turkey | Ministry of Health Turkey | Fatma Bayrakdar,Ayşe Başak Altaş,Yasemin Coşgun,Süleyman Yalçın, Gülay Korukluoğlu,Selçuk Kılıç |

| | | | |
|---|---|---|---|
| EPI_ISL_437335 | | | |
| EPI_ISL_437336 | TSGH-CP molecular lab, Division of Clinical Pathology, Department of Pathology | TSGH-CP molecular lab, Division of Clinical Pathology, Department of Pathology | Cherng-Lih Perng, Ming-Jr JIAN, Chih-Kai Chang, Jung-Chung Lin, Kuo-Ming Yeh, Chien-Wen Chen, Sheng-Kang Chiu, Hsing-Yi Chung, Shih-Hung Tsai, Kuo-Sheng Hung, Tien-Yao Chang, Feng-Yee Chang, Hung-Sheng Shang |
| EPI_ISL_437337, EPI_ISL_437338, EPI_ISL_437339, EPI_ISL_437340, EPI_ISL_437341, EPI_ISL_437342, EPI_ISL_437343, EPI_ISL_437344, EPI_ISL_437345, EPI_ISL_437346, EPI_ISL_437347, EPI_ISL_437348 | | | |
| see above | Viral Respiratory Lab, National Institute for Biomedical Research (INRB) | Pathogen Sequencing Lab, National Institute for Biomedical Research (INRB) | Placide Mbala-Kingebeni, Edith Nkwembe, Eddy Kinganda-Lusamaki, Amuri Aziza, Francisca Muyembe Mawete, Catherine Pratt, Matthias Pauthner, Josh Quick, Allison Black, James Hadfield, Trevor Bedford, Ian Goodfellow, Andrew Rambaut, Nick Loman, Kristian Andersen, Michael Wiley, Steve Ahuka-Mundeke, Jean-Jacques Muyembe Tamfum |
| EPI_ISL_437349 | Ecole nationale vétérinaire d'Alfort-laboratoire de santé animale Anses UMR 1161 de virologie ENVA-Anses-INRAE | Institut Pasteur CIBU-ERI | Sophie Le Poder, Corinne Sailleau, Marine Dumarest, Bernard Klonjkowski, Stéphan Zientara |
| EPI_ISL_437350, EPI_ISL_437351, EPI_ISL_437352, EPI_ISL_437353, EPI_ISL_437354, EPI_ISL_437355, EPI_ISL_437356, EPI_ISL_437357, EPI_ISL_437358 | Viral Respiratory Lab, National Institute for Biomedical Research (INRB) | Pathogen Sequencing Lab, National Institute for Biomedical Research (INRB) | Placide Mbala-Kingebeni, Edith Nkwembe, Eddy Kinganda-Lusamaki, Amuri Aziza, Francisca Muyembe Mawete, Catherine Pratt, Matthias Pauthner, Josh Quick, Allison Black, James Hadfield, Trevor Bedford, Ian Goodfellow, Andrew Rambaut, Nick Loman, Kristian Andersen, Michael Wiley, Steve Ahuka-Mundeke, Jean-Jacques Muyembe Tamfum |
| EPI_ISL_437359 | Max von Pettenkofer Institute, Virology, National Reference Center for Retroviruses, LMU München | Laboratory for Functional Genome Analysis, Dept. Genomics, Gene Center of the LMU Munich | Max Muenchhoff, Stefan Krebs, Alexander Graf, Oliver Keppler, Helmut Blum |
| EPI_ISL_437360, EPI_ISL_437361, EPI_ISL_437362, EPI_ISL_437363, EPI_ISL_437364, EPI_ISL_437365, EPI_ISL_437366, EPI_ISL_437367, EPI_ISL_437368, EPI_ISL_437369, EPI_ISL_437370, EPI_ISL_437371, EPI_ISL_437372, EPI_ISL_437373, EPI_ISL_437374, EPI_ISL_437375, EPI_ISL_437376, EPI_ISL_437377, EPI_ISL_437378, EPI_ISL_437379, EPI_ISL_437380, EPI_ISL_437381, EPI_ISL_437382, EPI_ISL_437383, EPI_ISL_437384, EPI_ISL_437385, EPI_ISL_437386 | | | |
| see above | Minnesota Department of Health, Public Health Laboratory | Minnesota Department of Health, Public Health Laboratory | Matt Plumb, Jacob Garfin, and Xiong Wang |
| EPI_ISL_437387, EPI_ISL_437388, EPI_ISL_437389, EPI_ISL_437390, EPI_ISL_437391, EPI_ISL_437392, EPI_ISL_437393, EPI_ISL_437394, EPI_ISL_437395, EPI_ISL_437396, EPI_ISL_437397, EPI_ISL_437398, EPI_ISL_437399, EPI_ISL_437400, EPI_ISL_437401, EPI_ISL_437402, EPI_ISL_437403, EPI_ISL_437404, EPI_ISL_437405, EPI_ISL_437406, EPI_ISL_437407, EPI_ISL_437408, EPI_ISL_437409, EPI_ISL_437410, EPI_ISL_437411, EPI_ISL_437412, EPI_ISL_437413, EPI_ISL_437414, EPI_ISL_437415, EPI_ISL_437416, EPI_ISL_437417, EPI_ISL_437418, EPI_ISL_437419, EPI_ISL_437420, EPI_ISL_437421, EPI_ISL_437422, EPI_ISL_437423, EPI_ISL_437424, EPI_ISL_437425, EPI_ISL_437426, EPI_ISL_437427, EPI_ISL_437428, EPI_ISL_437429, EPI_ISL_437430, EPI_ISL_437431, EPI_ISL_437432 | | | |
| see above | Virginia DCLS | Virginia DCLS | Virginia DCLS |
| EPI_ISL_437433 | Bozeman Health Deaconess Hospital | Wiedenheft lab, Montana State University | Artem Nemudryi, Anna Nemudraia, Kevin Surya, Tanner Wiegand, Murat Buyukyoruk, Royce Wilkinson, Blake Wiedenheft |
| EPI_ISL_437434 | Bozeman Water Reclamation Facility | Wiedenheft lab, Montana State University | Artem Nemudryi, Anna Nemudraia, Kevin Surya, Tanner Wiegand, Murat Buyukyoruk, Royce Wilkinson, Blake Wiedenheft |
| EPI_ISL_437435, EPI_ISL_437436 | Veterinary Specialized Institue Kraljevo | Veterinary Specialized Institue Kraljevo | Dejan Vidanovic. Bojana Tesovic, Milanko Sekler, Marko Dmitric, Kazimir Matovic, Zoran Debeljak, Nikola Vaskovic, Tamas Petrovic, Jeremy Volkening, Claudio L Afonso |
| EPI_ISL_437437 | Alaska State Virology Laboratory | Alaska State Virology Laboratory | Jack Chen, Ph.D. |
| EPI_ISL_437438 | Department of MicroBiology, Government Medical College, Surat | Gujarat Biotechnology Research Centre | Amit Kanani, Akanksha Verma, Nitin Savaliya, Raghawendra Kumar, Dinesh Kumar, Zuber Saiyed, Dipa Kinariwala, Disha Patel, Binita Aring, Neeta Khandelwal, Geeta Vaghela, Sonia Barve, Bhavesh Modi, Kairavi Joshi, Gaurishankar Shrimali, Nidhi Sood, Pranay Shah, R D Dixit, Snehal Bagatharia, Kamlesh J Upadhyay, Ramesh Pandit, Tejas Shah, Ankit Hinsu, Pritesh Sabara, Apurvasinh Puvar, Janvi Raval, Monika Gandhi, Pinal Trivedi, Maharshi Pandya, Neelam Nathani, Chaitanya Joshi, Madhvi Joshi |
| EPI_ISL_437439 | Department of MicroBiology, Government Medical College, Surat | Gujarat Biotechnology Research Centre | Akanksha Verma, Nitin Savaliya, Raghawendra Kumar, Dinesh Kumar, Zuber Saiyed, Dipa Kinariwala, Disha Patel, Binita Aring, Neeta Khandelwal, Geeta Vaghela, Sonia Barve, Bhavesh Modi, Kairavi Joshi, Gaurishankar Shrimali, Nidhi Sood, Pranay Shah, R D Dixit, Snehal Bagatharia, Kamlesh J Upadhyay, Ramesh Pandit, Tejas Shah, Ankit Hinsu, Pritesh Sabara, Apurvasinh Puvar, Janvi Raval, Monika Gandhi, Pinal Trivedi, Maharshi Pandya, Amit Kanani, Armi Chaudhari, Chaitanya Joshi, Madhvi Joshi |
| EPI_ISL_437440 | Department of MicroBiology, Government Medical College, Surat | Gujarat Biotechnology Research Centre | Nitin Savaliya, Raghawendra Kumar, Dinesh Kumar, Zuber Saiyed, Dipa Kinariwala, Disha Patel, Binita Aring, Neeta Khandelwal, Geeta Vaghela, Sonia Barve, Bhavesh Modi, Kairavi Joshi, Gaurishankar Shrimali, Nidhi Sood, Pranay Shah, R D Dixit, Snehal Bagatharia, Kamlesh J Upadhyay, Ramesh Pandit, Tejas Shah, Ankit Hinsu, Pritesh Sabara, Apurvasinh Puvar, Janvi Raval, Monika Gandhi, Pinal Trivedi, Maharshi Pandya, Amit Kanani, Akanksha Verma, Bhavya Jindal, Chaitanya Joshi, Madhvi Joshi |
| EPI_ISL_437441 | Department of MicroBiology, Government Medical College, Surat | Gujarat Biotechnology Research Centre | Raghawendra Kumar, Dinesh Kumar, Zuber Saiyed, Dipa Kinariwala, Disha Patel, Binita Aring, Neeta Khandelwal, Geeta Vaghela, Sonia Barve, Bhavesh Modi, Kairavi Joshi, Gaurishankar Shrimali, Nidhi Sood, Pranay Shah, R D Dixit, Snehal Bagatharia, Kamlesh J Upadhyay, Ramesh Pandit, Tejas Shah, Ankit Hinsu, Pritesh Sabara, Apurvasinh Puvar, Janvi Raval, Monika Gandhi, Pinal Trivedi, Maharshi Pandya, Amit Kanani, Akanksha Verma, Nitin Savaliya, Anjali Rajwar, Chaitanya Joshi, Madhvi Joshi |
| EPI_ISL_437442 | Department of MicroBiology, Government Medical College, Surat | Gujarat Biotechnology Research Centre | Dinesh Kumar, Zuber Saiyed, Dipa Kinariwala, Disha Patel, Binita Aring, Neeta Khandelwal, Geeta Vaghela, Sonia Barve, Bhavesh Modi, Kairavi Joshi, Gaurishankar Shrimali, Nidhi Sood, Pranay Shah, R D Dixit, Snehal Bagatharia, Kamlesh J Upadhyay, Ramesh Pandit, Tejas Shah, Ankit Hinsu, Pritesh Sabara, Apurvasinh Puvar, Janvi Raval, Monika Gandhi, Pinal Trivedi, Maharshi Pandya, Amit Kanani, Akanksha Verma, Nitin Savaliya, Raghawendra Kumar, Dipeshwari Shewale, Chaitanya Joshi, Madhvi Joshi |
| EPI_ISL_437443 | Department of MicroBiology, Government Medical College, Surat | Gujarat Biotechnology Research Centre | Zuber Saiyed, Dipa Kinariwala, Disha Patel, Binita Aring, Neeta Khandelwal, Geeta Vaghela, Sonia Barve, Bhavesh Modi, Kairavi Joshi, Gaurishankar Shrimali, Nidhi Sood, Pranay Shah, R D Dixit, Snehal Bagatharia, Kamlesh J Upadhyay, Ramesh Pandit, Tejas Shah, Ankit Hinsu, Pritesh Sabara, Apurvasinh Puvar, Janvi Raval, Monika Gandhi, Pinal Trivedi, Maharshi Pandya, Amit Kanani, Akanksha Verma, Nitin Savaliya, Raghawendra Kumar, Dinesh Kumar, Sharmistha Majumdar, Chaitanya Joshi, Madhvi Joshi |
| EPI_ISL_437444 | Department of MicroBiology, Government Medical College, Surat | Gujarat Biotechnology Research Centre | Dipa Kinariwala, Disha Patel, Binita Aring, Neeta Khandelwal, Geeta Vaghela, Sonia Barve, Bhavesh Modi, Kairavi Joshi, Gaurishankar Shrimali, Nidhi Sood, Pranay Shah, R D Dixit, Snehal Bagatharia, Kamlesh J Upadhyay, Ramesh Pandit, Tejas Shah, Ankit Hinsu, Pritesh Sabara, Apurvasinh Puvar, Janvi Raval, Monika Gandhi, Pinal Trivedi, Maharshi Pandya, Amit Kanani, Akanksha Verma, Nitin Savaliya, Raghawendra Kumar, Dinesh Kumar, Zuber Saiyed, Pooja P Doshi, Chaitanya Joshi, Madhvi Joshi |
| EPI_ISL_437445 | B.J. Medical College and Civil hospital | Gujarat Biotechnology Research Centre | Disha Patel, Binita Aring, Neeta Khandelwal, Geeta Vaghela, Sonia Barve, Bhavesh Modi, Kairavi Joshi, Gaurishankar Shrimali, Nidhi Sood, Pranay Shah, R D Dixit, Snehal Bagatharia, Kamlesh J Upadhyay, Ramesh Pandit, Tejas Shah, Ankit Hinsu, Pritesh Sabara, Apurvasinh Puvar, Janvi Raval, Monika Gandhi, Pinal Trivedi, Maharshi Pandya, Amit Kanani, Akanksha Verma, Nitin Savaliya, Raghawendra Kumar, Dinesh Kumar, Zuber Saiyed, Dipa Kinariwala, Nidhi Patel, Chaitanya Joshi, Madhvi Joshi |
| EPI_ISL_437446 | B.J. Medical College and Civil hospital | Gujarat Biotechnology Research Centre | Binita Aring, Neeta Khandelwal, Geeta Vaghela, Sonia Barve, Bhavesh Modi, Kairavi Joshi, Gaurishankar Shrimali, Nidhi Sood, Pranay Shah, R D Dixit, Snehal Bagatharia, Kamlesh J Upadhyay, Ramesh Pandit, Tejas Shah, Ankit Hinsu, Pritesh Sabara, Apurvasinh Puvar, Janvi Raval, Monika Gandhi, Pinal Trivedi, Maharshi Pandya, Amit Kanani, Akanksha Verma, Nitin Savaliya, Raghawendra Kumar, Dinesh Kumar, Zuber Saiyed, Dipa Kinariwala, Disha Patel, Priti Pandita, Chaitanya Joshi, Madhvi Joshi |
| EPI_ISL_437447 | B.J. Medical College and Civil hospital | Gujarat Biotechnology Research Centre | Neeta Khandelwal, Geeta Vaghela, Sonia Barve, Bhavesh Modi, Kairavi Joshi, Gaurishankar Shrimali, Nidhi Sood, Pranay Shah, R D Dixit, Snehal Bagatharia, Kamlesh J Upadhyay, Ramesh Pandit, Tejas Shah, Ankit Hinsu, Pritesh Sabara, Apurvasinh Puvar, Janvi Raval, Monika Gandhi, Pinal Trivedi, Maharshi Pandya, Amit Kanani, Akanksha Verma, Nitin Savaliya, Raghawendra Kumar, Dinesh Kumar, Zuber Saiyed, Dipa Kinariwala, Disha Patel, Binita Aring, Neha Rajpara, Chaitanya Joshi, Madhvi Joshi |
| EPI_ISL_437448 | B.J. Medical College and Civil hospital | Gujarat Biotechnology Research Centre | Geeta Vaghela, Sonia Barve, Bhavesh Modi, Kairavi Joshi, Gaurishankar Shrimali, Nidhi Sood, Pranay Shah, R D Dixit, Snehal Bagatharia, Kamlesh J Upadhyay, Ramesh Pandit, Tejas Shah, Ankit Hinsu, Pritesh Sabara, Apurvasinh Puvar, Janvi Raval, Monika Gandhi, Pinal Trivedi, Maharshi Pandya, Amit Kanani, Akanksha Verma, Nitin Savaliya, Raghawendra Kumar, Dinesh Kumar, Zuber Saiyed, Dipa Kinariwala, Disha Patel, Binita Aring, Neeta Khandelwal, Afzal Ansari, Chaitanya Joshi, Madhvi Joshi |
| EPI_ISL_437449 | B.J. Medical College and Civil hospital | Gujarat Biotechnology Research Centre | Sonia Barve, Bhavesh Modi, Kairavi Joshi, Gaurishankar Shrimali, Nidhi Sood, Pranay Shah, R D Dixit, Snehal Bagatharia, Kamlesh J Upadhyay, Ramesh Pandit, Tejas Shah, Ankit Hinsu, Pritesh Sabara, Apurvasinh Puvar, Janvi Raval, Monika Gandhi, Pinal Trivedi, Maharshi Pandya, Amit Kanani, Akanksha Verma, Nitin Savaliya, Raghawendra Kumar, Dinesh Kumar, Zuber Saiyed, Dipa Kinariwala, Disha Patel, Binita Aring, Neeta Khandelwal, Geeta Vaghela, Neelam Nathani, Chaitanya Joshi, Madhvi Joshi |
| EPI_ISL_437450 | B.J. Medical College and Civil hospital | Gujarat Biotechnology Research Centre | Bhavesh Modi, Kairavi Joshi, Gaurishankar Shrimali, Nidhi Sood, Pranay Shah, R D Dixit, Snehal Bagatharia, Kamlesh J Upadhyay, Ramesh Pandit, Tejas Shah, Ankit Hinsu, Pritesh Sabara, Apurvasinh Puvar, Janvi Raval, Monika Gandhi, Pinal Trivedi, Maharshi Pandya, Amit Kanani, Akanksha Verma, Nitin Savaliya, Raghawendra Kumar, Dinesh Kumar, Zuber Saiyed, Dipa Kinariwala, Disha Patel, Binita Aring, Neeta Khandelwal, Geeta Vaghela, Sonia Barve, Armi Chaudhari, Chaitanya Joshi, Madhvi Joshi |
| EPI_ISL_437451 | B.J. Medical College and Civil hospital | Gujarat Biotechnology Research Centre | Kairavi Joshi, Gaurishankar Shrimali, Nidhi Sood, Pranay Shah, R D Dixit, Snehal Bagatharia, Kamlesh J Upadhyay, Ramesh Pandit, Tejas Shah, Ankit Hinsu, Pritesh Sabara, Apurvasinh Puvar, Janvi Raval, Monika Gandhi, Pinal Trivedi, Maharshi Pandya, Amit Kanani, Akanksha Verma, Nitin Savaliya, Raghawendra Kumar, Dinesh Kumar, Zuber Saiyed, Dipa Kinariwala, Disha Patel, Binita Aring, Neeta Khandelwal, Geeta Vaghela, Sonia Barve, Bhavesh Modi, Bhavya Jindal, Chaitanya Joshi, Madhvi Joshi |
| EPI_ISL_437452 | B.J. Medical College and Civil hospital | Gujarat Biotechnology Research Centre | Gaurishankar Shrimali, Nidhi Sood, Pranay Shah, R D Dixit, Snehal Bagatharia, Kamlesh J Upadhyay, Ramesh Pandit, Tejas Shah, Ankit Hinsu, Pritesh Sabara, Apurvasinh Puvar, Janvi Raval, Monika Gandhi, Pinal Trivedi, Maharshi Pandya, Amit Kanani, Akanksha Verma, Nitin Savaliya, Raghawendra Kumar, Dinesh Kumar, Zuber Saiyed, Dipa Kinariwala, Disha Patel, Binita Aring, Neeta Khandelwal, Geeta Vaghela, Sonia Barve, Bhavesh Modi, Kairavi Joshi, Chaitanya Joshi, Anjali Rajwar, Madhvi Joshi |
| EPI_ISL_437453 | B.J. Medical College and Civil hospital | Gujarat Biotechnology Research Centre | Nidhi Sood, Pranay Shah, R D Dixit, Snehal Bagatharia, Kamlesh J Upadhyay, Ramesh Pandit, Tejas Shah, Ankit Hinsu, Pritesh Sabara, Apurvasinh Puvar, Janvi Raval, Monika Gandhi, Pinal Trivedi, Maharshi Pandya, Amit Kanani, Akanksha Verma, Nitin Savaliya, Raghawendra Kumar, Dinesh Kumar, Zuber Saiyed, Dipa Kinariwala, Disha Patel, Binita Aring, Neeta Khandelwal, Geeta Vaghela, Sonia Barve, Bhavesh Modi, Kairavi Joshi, Gaurishankar Shrimali, Chaitanya Joshi, Dipeshwari Shewale, Madhvi Joshi |
| EPI_ISL_437454 | B.J. Medical College and Civil hospital | Gujarat Biotechnology Research Centre | Pranay Shah, R D Dixit, Snehal Bagatharia, Kamlesh J Upadhyay, Ramesh Pandit, Tejas Shah, Ankit Hinsu, Pritesh Sabara, Apurvasinh Puvar, Janvi Raval, Monika Gandhi, Pinal Trivedi, Maharshi Pandya, Amit Kanani, Akanksha Verma, Nitin Savaliya, Raghawendra Kumar, Dinesh Kumar, Zuber Saiyed, Dipa Kinariwala, Disha Patel, Binita Aring, Neeta Khandelwal, Geeta Vaghela, Sonia Barve, Bhavesh Modi, Kairavi Joshi, Gaurishankar Shrimali, Nidhi Sood, Chaitanya Joshi, Sharmistha Majumdar, Madhvi Joshi |
| EPI_ISL_437455, EPI_ISL_437456, EPI_ISL_437457, EPI_ISL_437458 | Clinical Diagnostics Laboratory, Diagnostic & Experimental Pathology, Lilly Research Laboratories | Clinical Diagnostics Laboratory, Diagnostic & Experimental Pathology, Lilly Research Laboratories | Tim Holzer, Mayuri Vaidya, Angie Fulford, Sam McNeely, Rachael Redmond, Phil Ebert, John Calley, Leslie ONeill Reising, Pat Finnegan, Erin Wray, John McElwee, Jeff Fill, Joe Oakley, Andrew Schade |
| EPI_ISL_437459, EPI_ISL_437460, EPI_ISL_437461, EPI_ISL_437462, EPI_ISL_437463, EPI_ISL_437464, EPI_ISL_437465, EPI_ISL_437466, EPI_ISL_437467, EPI_ISL_437468, EPI_ISL_437469, EPI_ISL_437470, EPI_ISL_437471, EPI_ISL_437472, EPI_ISL_437473, EPI_ISL_437474, EPI_ISL_437475, EPI_ISL_437476, EPI_ISL_437477, EPI_ISL_437478, EPI_ISL_437479 | | | |
| see above | Pathogen Genomics Lab King Abdullah University of Science and Technology(KAUST) | Pathogen Genomics Lab King Abdullah University of Science and Technology(KAUST) | Sharif Hala,Raeece Naeem,Sara Mfarrej,Arnab Pain |
| EPI_ISL_437481, EPI_ISL_437482, EPI_ISL_437483, EPI_ISL_437484, EPI_ISL_437485, EPI_ISL_437486, EPI_ISL_437487, EPI_ISL_437488, EPI_ISL_437489, EPI_ISL_437490, EPI_ISL_437491, EPI_ISL_437492, EPI_ISL_437493, EPI_ISL_437494, EPI_ISL_437495, EPI_ISL_437496, EPI_ISL_437497 | | | |
| see above | Pathogen Genomics Lab King Abdullah University of Science and Technology(KAUST) | Pathogen Genomics Lab King Abdullah University of Science and Technology(KAUST) | Sara Mfarrej,Raeece Naeem,Sharif Hala,Amit Subudhi,Fathia Rached,Arnab Pain |
| EPI_ISL_437498, EPI_ISL_437499 | Biotechnology Center for Advanced Technologies | Biotechnology Center for Advanced Technologies | Abdullaev,A., Abdurakhimov,A., Muminov,M., Nuriddinov,S., Dalimova,D., Tsoy,V., Tsay,E., Bozorov,S., Charishnikova,O., Dalimova,D. and Turdikulova,S. |
| EPI_ISL_437512 | Human Genetic Research Center, Kawsar Biotech Company | Human Genetic Research Center, Kawsar Biotech Company | Khosravi,M.A., Abbasalipour,M., Zeinali,S., Sabeghi,S., Kehsvar,Y., Hosseini,F. and Haghdoost,Y. |
| EPI_ISL_437513, EPI_ISL_437514, EPI_ISL_437515, EPI_ISL_437516, EPI_ISL_437517, EPI_ISL_437518 | Alaska State Virology Laboratory | Alaska State Virology Laboratory | Jack Chen, Ph.D. |
| EPI_ISL_437519 | The National Institute of Public Health Center for Epidemiology and Microbiology | The National Institute of Public Health Center for Epidemiology and Microbiology | Alexander Nagy, Helena Jirincova, Ludmila Novakova, Dusan Trnka, Jaromira Vecerova |
| EPI_ISL_437520, EPI_ISL_437521, EPI_ISL_437522, EPI_ISL_437523, EPI_ISL_437524, EPI_ISL_437525, EPI_ISL_437526, EPI_ISL_437527, EPI_ISL_437528, EPI_ISL_437529, EPI_ISL_437530, EPI_ISL_437531, EPI_ISL_437532, EPI_ISL_437533, EPI_ISL_437534, EPI_ISL_437535 | | | |
| see above | OHSU Lab Services Molecular Microbiology Lab | Oregon SARS-CoV-2 Genome Sequencing Center | Brendan L. O'Connell, Ruth V. Nichols, Alec J. Hirsch, Guang Fan, Daniel N. Streblow, William B. Messer, Andrew C. Adey, Benjamin N. Bimber, Brian J. O'Roak |
| EPI_ISL_437536, EPI_ISL_437537, EPI_ISL_437538, | ICMR-National Institute of Cholera and Enteric Diseases | National Institute of Biomedical Genomics | Arindam Maitra, Mamta Chawla Sarkar, Sreedhar Chinnaswamy, Hasina Banu, Ananya Chatterjee, Shanta Dutta, Saumitra Das |

| | | | |
|---|---|---|---|
| EPI_ISL_437539 | | | |
| EPI_ISL_437540, EPI_ISL_437541, EPI_ISL_437542, EPI_ISL_437543, EPI_ISL_437544, EPI_ISL_437545, EPI_ISL_437546, EPI_ISL_437547, EPI_ISL_437548 | Robert Garry lab | Andersen lab at Scripps Research | Allison Smither, Gilberto Sabino-Santos, Patricia Snarski, Lilia Melnik, Antoinette Bell, Kaylynn Genemaras, Arnaud Drouin, Dahlene Fusco, Robert Garry with SEARCH Alliance San Diego |
| EPI_ISL_437549, EPI_ISL_437550, EPI_ISL_437551, EPI_ISL_437552, EPI_ISL_437553, EPI_ISL_437554, EPI_ISL_437555, EPI_ISL_437556, EPI_ISL_437557, EPI_ISL_437558, EPI_ISL_437559, EPI_ISL_437560, EPI_ISL_437561, EPI_ISL_437562, EPI_ISL_437563, EPI_ISL_437564, EPI_ISL_437565, EPI_ISL_437566, EPI_ISL_437567, EPI_ISL_437568, EPI_ISL_437569, EPI_ISL_437570, EPI_ISL_437571, EPI_ISL_437572, EPI_ISL_437573, EPI_ISL_437574, EPI_ISL_437575, EPI_ISL_437576, EPI_ISL_437577, EPI_ISL_437578, EPI_ISL_437579, EPI_ISL_437580, EPI_ISL_437581, EPI_ISL_437582, EPI_ISL_437583, EPI_ISL_437584, EPI_ISL_437585, EPI_ISL_437586, EPI_ISL_437587, EPI_ISL_437588, EPI_ISL_437589, EPI_ISL_437590, EPI_ISL_437591, EPI_ISL_437592, EPI_ISL_437593, EPI_ISL_437594, EPI_ISL_437595, EPI_ISL_437596, EPI_ISL_437597, EPI_ISL_437598, EPI_ISL_437599, EPI_ISL_437600 | | | |
| see above | Scripps Medical Laboratory | Andersen lab at Scripps Research | SEARCH Alliance San Diego with Michael Quigley, Ellen Stefanski, Ian Mchardy |
| EPI_ISL_437601 | Keio University School of Medicine | Keio University School of Medicine | Kenjiro Kosaki, Yuka Iwasaki, Toshiki Takenouchi, Haruhiko Siomi, |
| EPI_ISL_437602, EPI_ISL_437603, EPI_ISL_437604, EPI_ISL_437605, EPI_ISL_437606, EPI_ISL_437607, EPI_ISL_437608, EPI_ISL_437609, EPI_ISL_437610, EPI_ISL_437611, EPI_ISL_437612, EPI_ISL_437613, EPI_ISL_437614, EPI_ISL_437615, EPI_ISL_437616, EPI_ISL_437617, EPI_ISL_437618, EPI_ISL_437619, EPI_ISL_437620, EPI_ISL_437621, EPI_ISL_437622, EPI_ISL_437623, EPI_ISL_437624 | | | |
| see above | unknown | Faculty of Medicine | Rodpan,A., Joyjinda,Y., Wacharapluesadee,S., Buathong,R., Ghai,S., Petcharat,S., Bunprakob,S., Sirichan,N., Prasithsirikul,W., Mungaomklang,A., Plipat,T. and Hemachudha,T. |
| EPI_ISL_437625 | Laboratory of Genomics & Bioinformatics, Institute of Immunology and Experimental Therapy, Polish Academy of Sciences Oddzial Mikrobiologii Wojewodzkiej Stacji Sanitarno-Epidemiologicznej. | Laboratory of Genomics & Bioinformatics, Institute of Immunology and Experimental Therapy, Polish Academy of Sciences | Dorota Kujawa, Aleksandra Herud, Dariusz Martynowski, Krzysztof Jakub Pawlik, Joanna Sikorska, Paulina Zebrowska, Grazyna Zalewska, Oskar Karpinski and Lukasz Laczmanski |
| EPI_ISL_437626 | Department of Microbiology,Gandhi Medical College and Hospital | Department of Veterinary Biotechnalogy, College of Veterinary Science, Rajendranagar, PV Narsimha Rao Telengana Veterinary University | Kalyani Putty, Muttineni Radhakrishna, Nagamani K, Thrilok Chander B, Raja Rao M, Ravikumar P, Sunitha P, Pankaj Singh D, Anand Kumar K, Amit A. Upadhyay, Steven Bosinger, Rama Amara |
| EPI_ISL_437627, EPI_ISL_437628, EPI_ISL_437629, EPI_ISL_437630, EPI_ISL_437631, EPI_ISL_437632, EPI_ISL_437633, EPI_ISL_437634, EPI_ISL_437635, EPI_ISL_437636, EPI_ISL_437637, EPI_ISL_437638, EPI_ISL_437639, EPI_ISL_437640, EPI_ISL_437641, EPI_ISL_437642, EPI_ISL_437643, EPI_ISL_437644, EPI_ISL_437645, EPI_ISL_437646, EPI_ISL_437647, EPI_ISL_437648, EPI_ISL_437649, EPI_ISL_437650, EPI_ISL_437651, EPI_ISL_437652, EPI_ISL_437653, EPI_ISL_437654, EPI_ISL_437655, EPI_ISL_437656, EPI_ISL_437657, EPI_ISL_437658, EPI_ISL_437659, EPI_ISL_437660, EPI_ISL_437661, EPI_ISL_437662, EPI_ISL_437663, EPI_ISL_437664, EPI_ISL_437665, EPI_ISL_437666, EPI_ISL_437667, EPI_ISL_437668, EPI_ISL_437669, EPI_ISL_437670, EPI_ISL_437671, EPI_ISL_437672, EPI_ISL_437673, EPI_ISL_437674, EPI_ISL_437675, EPI_ISL_437676, EPI_ISL_437677, EPI_ISL_437678, EPI_ISL_437679, EPI_ISL_437680, EPI_ISL_437681, EPI_ISL_437682, EPI_ISL_437683 | | | |
| see above | Department of Virus and Microbiological Special Diagnostics, Statens Serum Institut, Copenhagen, Denmark, Artillerivej 5, 2300 Copenahgen S | Albertsen lab, Department of Chemistry and Bioscience, Aalborg University, Denmark | Rasmus Kirkegaard |
| EPI_ISL_437684, EPI_ISL_437685, EPI_ISL_437686, EPI_ISL_437687, EPI_ISL_437688 | UCD National Virus Reference Laboratory | UCD National Virus Reference Laboratory | Michael J. Carr, Gabriel Gonzalez, Brendan Crowley, Cillian F De Gascun |
| EPI_ISL_437689 | Laboratory for Urgent Response to Biological Threats | Institut Pasteur CIBU / ERI | V. Caro, A. Kwasiborski, V. Hourdel, C. Balière, J. Vanhomwegen, C. Batéjat, JC. Manuguerra |
| EPI_ISL_437690 | Laboratory for Urgent Response to Biological Threats | Institut Pasteur CIBU /ERI | V. Caro, A. Kwasiborski, H. Hourdel, C. Balière, J. Vanhomwegen, C. Batéjat, JC. Manuguerra |
| EPI_ISL_437691, EPI_ISL_437692, EPI_ISL_437693, EPI_ISL_437694, EPI_ISL_437695, EPI_ISL_437696, EPI_ISL_437697, EPI_ISL_437698, EPI_ISL_437699, EPI_ISL_437700, EPI_ISL_437701, EPI_ISL_437702, EPI_ISL_437703, EPI_ISL_437704, EPI_ISL_437705, EPI_ISL_437706, EPI_ISL_437707, EPI_ISL_437708, EPI_ISL_437709, EPI_ISL_437710, EPI_ISL_437711, EPI_ISL_437712, EPI_ISL_437713, EPI_ISL_437714, EPI_ISL_437715, EPI_ISL_437716, EPI_ISL_437717, EPI_ISL_437718, EPI_ISL_437719, EPI_ISL_437720, EPI_ISL_437721, EPI_ISL_437722, EPI_ISL_437723, EPI_ISL_437724, EPI_ISL_437725, EPI_ISL_437726, EPI_ISL_437727, EPI_ISL_437728, EPI_ISL_437729, EPI_ISL_437730, EPI_ISL_437731, EPI_ISL_437732, EPI_ISL_437733, EPI_ISL_437734, EPI_ISL_437735, EPI_ISL_437736, EPI_ISL_437737, EPI_ISL_437738, EPI_ISL_437739, EPI_ISL_437740, EPI_ISL_437741, EPI_ISL_437742, EPI_ISL_437743, EPI_ISL_437744, EPI_ISL_437745, EPI_ISL_437746, EPI_ISL_437747, EPI_ISL_437748, EPI_ISL_437749, EPI_ISL_437750, EPI_ISL_437751, EPI_ISL_437752, EPI_ISL_437753, EPI_ISL_437754, EPI_ISL_437755, EPI_ISL_437756, EPI_ISL_437757, EPI_ISL_437758, EPI_ISL_437759, EPI_ISL_437760, EPI_ISL_437761, EPI_ISL_437762 | | | |
| see above | Pathogen Genomics Lab King Abdullah University of Science and Technology(KAUST) | Pathogen Genomics Lab King Abdullah University of Science and Technology(KAUST) | Sharif Hala,Fadwa Alofi,Afrah Alsomali, Asim Khogeer, Sara Mfarrej, Khaled Algithami,Raeece Naeem, Amit Kumar Subudhi,Fathia Ben-Rached, Rahul Salunke, Anwar Hashem, Naif Almontashiri, Arnab Pain |
| EPI_ISL_437763, EPI_ISL_437764, EPI_ISL_437765, EPI_ISL_437766, EPI_ISL_437767, EPI_ISL_437768, EPI_ISL_437769, EPI_ISL_437770, EPI_ISL_437771, EPI_ISL_437772, EPI_ISL_437773, EPI_ISL_437774, EPI_ISL_437775, EPI_ISL_437776, EPI_ISL_437777, EPI_ISL_437778, EPI_ISL_437779, EPI_ISL_437780, EPI_ISL_437781, EPI_ISL_437782, EPI_ISL_437783, EPI_ISL_437784, EPI_ISL_437785, EPI_ISL_437786, EPI_ISL_437787, EPI_ISL_437788, EPI_ISL_437789, EPI_ISL_437790, EPI_ISL_437791, EPI_ISL_437792, EPI_ISL_437793, EPI_ISL_437794, EPI_ISL_437795, EPI_ISL_437796, EPI_ISL_437797, EPI_ISL_437798, EPI_ISL_437799, EPI_ISL_437800, EPI_ISL_437801, EPI_ISL_437802 | | | |
| see above | Virginia DCLS | Virginia DCLS | Virginia DCLS |
| EPI_ISL_437803, EPI_ISL_437804, EPI_ISL_437805, EPI_ISL_437806, EPI_ISL_437807, EPI_ISL_437808, EPI_ISL_437809, EPI_ISL_437810, EPI_ISL_437811, EPI_ISL_437812, EPI_ISL_437813, EPI_ISL_437814, EPI_ISL_437815, EPI_ISL_437816, EPI_ISL_437817, EPI_ISL_437818, EPI_ISL_437819, EPI_ISL_437820, EPI_ISL_437821, EPI_ISL_437822, EPI_ISL_437823, EPI_ISL_437824, EPI_ISL_437825, EPI_ISL_437826, EPI_ISL_437827, EPI_ISL_437828, EPI_ISL_437829, EPI_ISL_437830, EPI_ISL_437831, EPI_ISL_437832, EPI_ISL_437833, EPI_ISL_437834, EPI_ISL_437835, EPI_ISL_437836, EPI_ISL_437837, EPI_ISL_437838, EPI_ISL_437839, EPI_ISL_437840, EPI_ISL_437841, EPI_ISL_437842, EPI_ISL_437843, EPI_ISL_437844, EPI_ISL_437845, EPI_ISL_437846, EPI_ISL_437847, EPI_ISL_437848, EPI_ISL_437849, EPI_ISL_437850, EPI_ISL_437851, EPI_ISL_437852, EPI_ISL_437853, EPI_ISL_437854, EPI_ISL_437855, EPI_ISL_437856, EPI_ISL_437857, EPI_ISL_437858, EPI_ISL_437859, EPI_ISL_437860, EPI_ISL_437861, EPI_ISL_437862, EPI_ISL_437863, EPI_ISL_437864, EPI_ISL_437865, EPI_ISL_437866, EPI_ISL_437867, EPI_ISL_437868, EPI_ISL_437869, EPI_ISL_437870, EPI_ISL_437871, EPI_ISL_437872 | | | |
| see above | UW Virology Lab | UW Virology Lab | Pavitra Roychoudhury, Hong Xie, Keith Jerome, Alexander Greninger |
| EPI_ISL_437873 | Alaska State Virology Laboratory | Alaska State Virology Laboratory | Jack Chen, Ph.D. |
| EPI_ISL_437874, EPI_ISL_437875, EPI_ISL_437876, EPI_ISL_437877, EPI_ISL_437878, EPI_ISL_437879, EPI_ISL_437880, EPI_ISL_437881, EPI_ISL_437882, EPI_ISL_437883, EPI_ISL_437884, EPI_ISL_437885, EPI_ISL_437886, EPI_ISL_437887, EPI_ISL_437888, EPI_ISL_437889, EPI_ISL_437890, EPI_ISL_437891, EPI_ISL_437892, EPI_ISL_437893, EPI_ISL_437894, EPI_ISL_437895, EPI_ISL_437896, EPI_ISL_437897, EPI_ISL_437898, EPI_ISL_437899, EPI_ISL_437900, EPI_ISL_437901, EPI_ISL_437902, EPI_ISL_437903, EPI_ISL_437904, EPI_ISL_437905, EPI_ISL_437906, EPI_ISL_437907, EPI_ISL_437908, EPI_ISL_437909, EPI_ISL_437910, EPI_ISL_437911 | | | |
| see above | Laboratory of Microbiology, Medical School, National and Kapodistrian University of Athens | Laboratory of Biology, Department of Medicine, Democritus University of Thrace | Kassela K., Dovrolis,N., Bampali,M., Gatzidou,E., Froukala,E., Stavropoulou,A., Veletza,S., Tsakris,A., Spanakis,N. and Karakasiliotis,I. |
| EPI_ISL_437912 | Child Health Research Foundation | Child Health Research Lab | Senjuti Saha, Roly Malaker, Md Saiful Islam Sajib, Md Hasanuzzaman, Md Hafizur Rahman, Md Shahidul Islam, Zabed B Ahmed, Maksuda Islam, Samir K Saha |
| EPI_ISL_444273 | State Key Laboratory of Respiratory Disease, National Clinical Research Center for Respiratory Disease, Guangzhou Institute of Respiratory Health, the First Affiliated Hospital of Guangzhou Medical University | State Key Laboratory of Respiratory Disease, National Clinical Research Center for Respiratory Disease, Guangzhou Institute of Respiratory Health, the First Affiliated Hospital of Guangzhou Medical University | Sun,J., Shi,Y., Zheng,K., Huang,J. and Zhao,J. |
| EPI_ISL_444518 | unknown | Molecular Infectious Disease | Anderson,B.P., Rosenthal,S.H., Gerasimova,A., Kagan,R.M. and Owen,R. |
| EPI_ISL_445086 | Laboratory Diagnostic, Veterinary Specialized Institute Kraljevo | Laboratory Diagnostic, Veterinary Specialized Institute Kraljevo | Vidanovic,D., Tesovic,B., Sekler,M., Dmitric,M., Debeljak,Z., Matovic,K., Vaskovic,N., Petrovic,T., Volkening,J. and Alfonso,C.L. |
| EPI_ISL_445183 | Takayuki Hishiki Kanagawa Prefectural Institute of Public Health | Takayuki Hishiki Kanagawa Prefectural Institute of Public Health | Hishiki,T., Suzuki,R., Sakuragi,J., Usui,K., Tanaka,Y., Kawai,J., Kogo,Y., Matsuki,Y., An,T., Hayashizaki,Y. and Takasaki,T. |
| EPI_ISL_447898 | Tumor Immunology Unit, Department of Health Sciences, University of Palermo School of Medicine and National, Research Council of Italy - High Performance Computing and Networking Institute (CNR-ICAR) | Tumor Immunology Unit, Department of Health Sciences, University of Palermo School of Medicine and National, Research Council of Italy - High Performance Computing and Networking Institute (CNR-ICAR) | Vacca,D., Fiannaca,A., Tramuto,F., Cancila,V., La Paglia,L., Mazzucco,W., Gulino,A., La Rosa,M., Maida,C.M., Morello,G., Belmonte,B., Casuccio,A., Urso,A., Vitale,F. and Tripodo,C. |
| EPI_ISL_450212, EPI_ISL_450213, EPI_ISL_450214, EPI_ISL_450215, EPI_ISL_450216 | unknown | Microbiological Diagnostic Unit Public Health Laboratory (MDU-PHL) and Victorian Infectious Disease Reference Laboratory (VIDRL) | Seemann,T., Lane,C.R., Sherry,N.L., Duchene,S., Goncalves da Silva,A., Caly,L., Sait,M., Ballard,S.A., Horan,K., Schultz,M.B., Hoang,T., Easton,M., Dougal,S., Stinear,T.P., Druce,J., Catton,M., Sutton,B., van Diemen,A., Alpren,C., Williamson,D.A., Howden,B.P. |
| EPI_ISL_450217, EPI_ISL_450218, EPI_ISL_450219, EPI_ISL_450220, EPI_ISL_450221, EPI_ISL_450222, EPI_ISL_450223, EPI_ISL_450224, EPI_ISL_450225, EPI_ISL_450226, EPI_ISL_450227, EPI_ISL_450228, EPI_ISL_450229, EPI_ISL_450230 | | | |
| see above | unknown | Hamadan University of Medical Sciences | Teimoori,A., Azizi Jallilian,F., Ansari,N., Jamehdor,S., Nazari,A., Saadat,N., Mazaheri,Z., Zanjani,M. |
| EPI_ISL_450406 | Molecular Diagnostics, Antech Diagnostics | Molecular Diagnostics, Antech Diagnostics | Leutenegger,C.M., Lozoya,C.E., Tereski,J.L., Wyler,D. and Moroff,S. |
| EPI_ISL_450407 | Molecular Diagnostics, Antech Diagnostics | Molecular Diagnostics, Antech Diagnostics | Leutenegger,C.M., Lozoya,C.E., Tereski,J.L. and Moroff,S. |
| EPI_ISL_450408, EPI_ISL_450409, EPI_ISL_450410, EPI_ISL_450411, EPI_ISL_450412 | unknown | Microbiology | To,K.K.W., Yuen,K.-Y. |
| EPI_ISL_450414 | Microbiology, Regional Medical Research Centre (ICMR) | Microbiology, Regional Medical Research Centre (ICMR) | Borkakoty,B., Bali,N.K., Barua,P., Hazarika,R., Sharma,M.D. and Phukon,P. |
| EPI_ISL_450415 | Laboratory Diagnostic, Veterinary Specialized Institute Kraljevo | Laboratory Diagnostic, Veterinary Specialized Institute Kraljevo | Vidanovic,D., Skadric,I., Tesovic,B., Tolic,A., Sekler,M., Petrovic,T., Matovic,K., Dmitric,M., Debeljak,Z. and Vaskovic,N. |
| EPI_ISL_450416 | unknown | Anhui Provincial Center for Disease Control and Prevention | Yuan,Y., He,J., Gong,L., Li,W., Jiang,L., Liu,J., Chen,Q., Yu,J., Hou,S., Shi,Y., Lu,S., Zhang,Z., Ge,Y., Sa,N., He,L., Wu,J., Sun,Y., Liu,Z. |
| EPI_ISL_450417, EPI_ISL_450418, EPI_ISL_450419, EPI_ISL_450420, EPI_ISL_450421, EPI_ISL_450422, EPI_ISL_450423, EPI_ISL_450424, EPI_ISL_450425, EPI_ISL_450426 | unknown | Anhui Provincial Center for Disease Control | Yuan,Y., He,J., Gong,L., Li,W., Jiang,L., Liu,J., Chen,Q., Yu,J., Hou,S., Shi,Y., Lu,S., Zhang,Z., Ge,Y., Sa,N., He,L., Wu,J., Sun,Y., Liu,Z. |
| EPI_ISL_450427 | Anhui Provincial Center for Disease Control | Anhui Provincial CDC, Acute Infectious Disease Prevention & Ctrl | Yuan,Y., He,J., Gong,L., Li,W., Jiang,L., Liu,J., Chen,Q., Yu,J., Hou,S., Shi,Y., Lu,S., Zhang,Z., Ge,Y., Sa,N., He,L., Wu,J., Sun,Y., Liu,Z. |
| EPI_ISL_450428, EPI_ISL_450429, EPI_ISL_450430, EPI_ISL_450431, EPI_ISL_450432, EPI_ISL_450433, EPI_ISL_450434, EPI_ISL_450435, EPI_ISL_450436 | unknown | Central laboratory | Yan,Y. |
| EPI_ISL_450437 | Infectious Disease Hospital, Central Laboratory | Infectious Disease Hospital, Central Laboratory | Yan,Y. |
| EPI_ISL_450438, EPI_ISL_450439, EPI_ISL_450440, EPI_ISL_450441 | unknown | Central laboratory | Yan,Y. |
| EPI_ISL_450442 | The Department of Infectious Disease Prevention and Control, Henan Provincial Center for Disease Control and Prevention | The Department of Infectious Disease Prevention and Control, Henan Provincial Center for Disease Control and Prevention | Li,X., Lu,S., Wu,B., Hu,X., Li,D., Huang,X. and Guo,W. |
| EPI_ISL_450443 | Institute for Forensic Medicine, Faculty of Medicine, University of Belgrade | Institute for Forensic Medicine, Faculty of Medicine, University of Belgrade | Vidanovic,D., Skadric,I.R., Dordevic,N., Tolic,A., Tesovic,B., Sekler,M., Dmitric,M., Debeljak,Z., Zarkovic,A., Kolarevic,M., Petrovic,T. and Baskic,D. |
| EPI_ISL_450484, EPI_ISL_450485, EPI_ISL_450486, EPI_ISL_450487 | unknown | Data Science | Carroll,T.D., Tran,N.K., Cohen,S.H., Miller,C.J. |
| EPI_ISL_450499 | Molecular Pathology, Mehr Pathobiology Lab | Molecular Pathology, Mehr Pathobiology Lab | Shabadori,R., Soleimani Dodaran,M., Mirzapour,Z., Kamali,M. and Hamedi,D. |
| EPI_ISL_450505 | Molecular Pathology, Mehr Pathobiology Lab | Molecular Pathology, Mehr Pathobiology Lab | Soleimani Dodaran,M., Soleimani Dodaran,M., Mirzapour,Z., Shabadori,R., Kamali,M. and Hamedi,D. |
| EPI_ISL_454692, EPI_ISL_454693 | Quest Diagnostics | Quest Diagnostics | Anderson,B.P., Rosenthal,S.H., Gerasimova,A., Kagan,R.M. and Owen, R. |

| | | | |
|---|---|---|---|
| EPI_ISL_468063, EPI_ISL_468064, EPI_ISL_468065 | unknown | Computer Science and Engineering | Rouchka,E.C., Chariker,J.H., Chung,D., Ramirez,J., Palmer,K.E., Lasnik,A.B., Carrico,R., Arnold,F.W., Adcock,R.S., Zhang,M., Alejandro,B., Wolf,L.A., Hwang,J.Y., Park,J.W., Waigel,S., Zacharias,W. |
| EPI_ISL_524433, EPI_ISL_524434 | Environmental and Global Health, University of Florida - Gainesville | University of Florida | Elbadry,M.A., Subramaniam,K., Waltzek,T.B., Gibson,J.C., Stephenson,C.J., Alam,M.M., Morris,J.G. Jr., Lednicky,J.A. |
| EPI_ISL_529147, EPI_ISL_529148 | Democritus University of Thrace, Department of Medicine | Democritus University of Thrace, Department of Medicine | Kassela,K., Dovrolis,N., Bampali,M., Gatzidou,E., Froukala,E., Stavropoulou,A., Veletza,S., Tsakris,A., Spanakis,N., Karakasiliotis,I. |
| EPI_ISL_529202, EPI_ISL_529203, EPI_ISL_529205 | Utah Public Health Laboratory | Utah Public Health Laboratory | Erin Young, Kelly Oakeson |

We gratefully acknowledge the following Authors from the Originating laboratories responsible for obtaining the specimens, as well as the Submitting laboratories where the genome data were generated and shared via GISAID, on which this research is based.

All Submitters of data may be contacted directly via www.gisaid.org

| Accession ID | Originating Laboratory | Submitting Laboratory | Authors |
|---|---|---|---|
| EPI_ISL_437913, EPI_ISL_437914, EPI_ISL_437915, EPI_ISL_437916, EPI_ISL_437917, EPI_ISL_437918, EPI_ISL_437919, EPI_ISL_437920, EPI_ISL_437921, EPI_ISL_437922, EPI_ISL_437923, EPI_ISL_437924, EPI_ISL_437925, EPI_ISL_437926, EPI_ISL_437927, EPI_ISL_437928, EPI_ISL_437929, EPI_ISL_437930, EPI_ISL_437931, EPI_ISL_437932 | | | |
| see above | Institut für Virologie am Department für Hygiene, Mikrobiologie und Public Health | Bergthaler laboratory, CeMM Research Center for Molecular Medicine of the Austrian Academy of Sciences | Alexandra Popa, Benedikt Agerer, Henrique Colaco, Lukas Endler, Jakob-Wendelin Genger, Alexander Lercher, Mark Smyth, Thomas Penz, Michael Schuster, Jan Laine, Martin Senekowitsch, Judith Aberle, Stephan Aberle, Elisabeth Puchhammer-Stoeckl, Manfred Nairz, Guenter Weiss, Wegene Borena, Dorothee von Laer, Christoph Bock, Andreas Bergthaler |
| EPI_ISL_437933, EPI_ISL_437934, EPI_ISL_437935, EPI_ISL_437936, EPI_ISL_437937, EPI_ISL_437938, EPI_ISL_437939, EPI_ISL_437940, EPI_ISL_437941, EPI_ISL_437942, EPI_ISL_437943, EPI_ISL_437944, EPI_ISL_437945, EPI_ISL_437946, EPI_ISL_437947, EPI_ISL_437948, EPI_ISL_437949, EPI_ISL_437950, EPI_ISL_437951, EPI_ISL_437952, EPI_ISL_437953, EPI_ISL_437954, EPI_ISL_437955, EPI_ISL_437956, EPI_ISL_437957, EPI_ISL_437958, EPI_ISL_437959, EPI_ISL_437960, EPI_ISL_437961, EPI_ISL_437962, EPI_ISL_437963, EPI_ISL_437964, EPI_ISL_437965, EPI_ISL_437966, EPI_ISL_437967, EPI_ISL_437968, EPI_ISL_437969, EPI_ISL_437970, EPI_ISL_437971, EPI_ISL_437972, EPI_ISL_437973 | | | |
| see above | Universitaetsklinik für Innere Medizin II Innsbruck | Bergthaler laboratory, CeMM Research Center for Molecular Medicine of the Austrian Academy of Sciences | Alexandra Popa, Benedikt Agerer, Henrique Colaco, Lukas Endler, Jakob-Wendelin Genger, Alexander Lercher, Mark Smyth, Thomas Penz, Michael Schuster, Jan Laine, Martin Senekowitsch, Judith Aberle, Stephan Aberle, Elisabeth Puchhammer-Stoeckl, Manfred Nairz, Guenter Weiss, Wegene Borena, Dorothee von Laer, Christoph Bock, Andreas Bergthaler |
| EPI_ISL_437974, EPI_ISL_437975, EPI_ISL_437976, EPI_ISL_437977, EPI_ISL_437978, EPI_ISL_437979, EPI_ISL_437980, EPI_ISL_437981, EPI_ISL_437982, EPI_ISL_437983, EPI_ISL_437984, EPI_ISL_437985, EPI_ISL_437986, EPI_ISL_437987, EPI_ISL_437988, EPI_ISL_437989, EPI_ISL_437990, EPI_ISL_437991, EPI_ISL_437992 | | | |
| see above | Institut für Virologie am Department für Hygiene, Mikrobiologie und Public Health | Bergthaler laboratory, CeMM Research Center for Molecular Medicine of the Austrian Academy of Sciences | Alexandra Popa, Benedikt Agerer, Henrique Colaco, Lukas Endler, Jakob-Wendelin Genger, Alexander Lercher, Mark Smyth, Thomas Penz, Michael Schuster, Jan Laine, Martin Senekowitsch, Judith Aberle, Stephan Aberle, Elisabeth Puchhammer-Stoeckl, Manfred Nairz, Guenter Weiss, Wegene Borena, Dorothee von Laer, Christoph Bock, Andreas Bergthaler |
| EPI_ISL_437993, EPI_ISL_437994, EPI_ISL_437995, EPI_ISL_437996, EPI_ISL_437997, EPI_ISL_437998, EPI_ISL_437999, EPI_ISL_438000, EPI_ISL_438001, EPI_ISL_438002, EPI_ISL_438003, EPI_ISL_438004, EPI_ISL_438005, EPI_ISL_438006, EPI_ISL_438007, EPI_ISL_438008, EPI_ISL_438009, EPI_ISL_438010, EPI_ISL_438011, EPI_ISL_438012, EPI_ISL_438013, EPI_ISL_438014, EPI_ISL_438015, EPI_ISL_438016, EPI_ISL_438017, EPI_ISL_438018, EPI_ISL_438019, EPI_ISL_438020, EPI_ISL_438021, EPI_ISL_438022, EPI_ISL_438023, EPI_ISL_438024, EPI_ISL_438025, EPI_ISL_438026, EPI_ISL_438027, EPI_ISL_438028, EPI_ISL_438029, EPI_ISL_438030, EPI_ISL_438031, EPI_ISL_438032, EPI_ISL_438033, EPI_ISL_438034, EPI_ISL_438035, EPI_ISL_438036, EPI_ISL_438037, EPI_ISL_438038, EPI_ISL_438039, EPI_ISL_438040, EPI_ISL_438041, EPI_ISL_438042, EPI_ISL_438043, EPI_ISL_438044, EPI_ISL_438045, EPI_ISL_438046, EPI_ISL_438047, EPI_ISL_438048, EPI_ISL_438049, EPI_ISL_438050, EPI_ISL_438051, EPI_ISL_438052, EPI_ISL_438053, EPI_ISL_438054, EPI_ISL_438055, EPI_ISL_438056, EPI_ISL_438057, EPI_ISL_438058, EPI_ISL_438059, EPI_ISL_438060, EPI_ISL_438061, EPI_ISL_438062, EPI_ISL_438063, EPI_ISL_438064, EPI_ISL_438065, EPI_ISL_438066, EPI_ISL_438067, EPI_ISL_438068, EPI_ISL_438069, EPI_ISL_438070, EPI_ISL_438071, EPI_ISL_438072, EPI_ISL_438073, EPI_ISL_438074, EPI_ISL_438075, EPI_ISL_438076, EPI_ISL_438077, EPI_ISL_438078, EPI_ISL_438079, EPI_ISL_438080, EPI_ISL_438081, EPI_ISL_438082, EPI_ISL_438083, EPI_ISL_438084, EPI_ISL_438085, EPI_ISL_438086, EPI_ISL_438087, EPI_ISL_438088, EPI_ISL_438089, EPI_ISL_438090, EPI_ISL_438091, EPI_ISL_438092, EPI_ISL_438093, EPI_ISL_438094, EPI_ISL_438095, EPI_ISL_438096, EPI_ISL_438097, EPI_ISL_438098, EPI_ISL_438099, EPI_ISL_438100, EPI_ISL_438101, EPI_ISL_438102, EPI_ISL_438103, EPI_ISL_438104, EPI_ISL_438105, EPI_ISL_438106, EPI_ISL_438107, EPI_ISL_438108, EPI_ISL_438109, EPI_ISL_438110, EPI_ISL_438111, EPI_ISL_438112, EPI_ISL_438113, EPI_ISL_438114, EPI_ISL_438115, EPI_ISL_438116, EPI_ISL_438117, EPI_ISL_438118, EPI_ISL_438119, EPI_ISL_438120, EPI_ISL_438121, EPI_ISL_438122, EPI_ISL_438123, EPI_ISL_438124, EPI_ISL_438125, EPI_ISL_438126, EPI_ISL_438127, EPI_ISL_438128 | | | |
| see above | Center for Virology, Medical University of Vienna | Bergthaler laboratory, CeMM Research Center for Molecular Medicine of the Austrian Academy of Sciences | Alexandra Popa, Benedikt Agerer, Henrique Colaco, Lukas Endler, Jakob-Wendelin Genger, Alexander Lercher, Mark Smyth, Thomas Penz, Michael Schuster, Jan Laine, Martin Senekowitsch, Judith Aberle, Stephan Aberle, Elisabeth Puchhammer-Stoeckl, Manfred Nairz, Guenter Weiss, Wegene Borena, Dorothee von Laer, Christoph Bock, Andreas Bergthaler |
| EPI_ISL_438138 | Department of Microbiology,Gandhi Medical College and Hospital | Department of Microbiology, Gandhi Medical College and Hospital Secendrabad, Hyderbad, India | Raja Rao Mesipogu, Muttineni Radhakrishna, Nagamani K, Thrilok Chander B, Kalyani Putty, Ravikumar P, Sunitha P, Pankaj Singh D, Anand Kumar K, Amit A. Upadhyay, Steven Bosinger, Rama Amara |
| EPI_ISL_438139 | Department of Microbiology,Gandhi Medical College and Hospital,Hyderabad | Virus Research Laboratory, Department of Zoology, Osmania University, Hyderabad, India | Muttineni Radhakrishna, Nagamani K, Thrilok Chander B, Raja Rao M, Kalyani Putty, Ravikumar P, Sunitha P, Pankaj Singh D, Anand Kumar K, Amit A. Upadhyay, Steven Bosinger, Rama Amara |
| EPI_ISL_438140, EPI_ISL_438141, EPI_ISL_438142, EPI_ISL_438143, EPI_ISL_438144, EPI_ISL_438145, EPI_ISL_438146, EPI_ISL_438147, EPI_ISL_438148, EPI_ISL_438149, EPI_ISL_438150, EPI_ISL_438151, EPI_ISL_438152, EPI_ISL_438153, EPI_ISL_438154, EPI_ISL_438155, EPI_ISL_438156, EPI_ISL_438157, EPI_ISL_438158, EPI_ISL_438159, EPI_ISL_438160, EPI_ISL_438161, EPI_ISL_438162, EPI_ISL_438163, EPI_ISL_438164, EPI_ISL_438165, EPI_ISL_438166, EPI_ISL_438167, EPI_ISL_438168, EPI_ISL_438169, EPI_ISL_438170, EPI_ISL_438171, EPI_ISL_438172, EPI_ISL_438173, EPI_ISL_438174, EPI_ISL_438175 | | | |
| see above | Seattle Flu Study | Seattle Flu Study | Chu et al |
| EPI_ISL_438176, EPI_ISL_438177, EPI_ISL_438178, EPI_ISL_438179, EPI_ISL_438180, EPI_ISL_438181, EPI_ISL_438182, EPI_ISL_438183, EPI_ISL_438184, EPI_ISL_438185, EPI_ISL_438186, EPI_ISL_438187, EPI_ISL_438188, EPI_ISL_438189, EPI_ISL_438190, EPI_ISL_438191, EPI_ISL_438192, EPI_ISL_438193, EPI_ISL_438194, EPI_ISL_438195, EPI_ISL_438196, EPI_ISL_438197, EPI_ISL_438198, EPI_ISL_438199, EPI_ISL_438200, EPI_ISL_438201, EPI_ISL_438202, EPI_ISL_438203, EPI_ISL_438204, EPI_ISL_438205, EPI_ISL_438206, EPI_ISL_438207, EPI_ISL_438208, EPI_ISL_438209, EPI_ISL_438210, EPI_ISL_438211, EPI_ISL_438212, EPI_ISL_438213, EPI_ISL_438214, EPI_ISL_438215, EPI_ISL_438216, EPI_ISL_438217, EPI_ISL_438218, EPI_ISL_438219, EPI_ISL_438220, EPI_ISL_438221 | | | |
| see above | Washington State Department of Health | Seattle Flu Study | Chu et al |
| EPI_ISL_438222, EPI_ISL_438223, EPI_ISL_438224, EPI_ISL_438225, EPI_ISL_438226, EPI_ISL_438227, EPI_ISL_438228, EPI_ISL_438229, EPI_ISL_438230, EPI_ISL_438231, EPI_ISL_438232, EPI_ISL_438233, EPI_ISL_438234 | | | |
| see above | Johns Hopkins Hospital Department of Pathology | Johns Hopkins Hospital Department of Pathology | Peter M. Thielen, Thomas Mehoke, Shirlee Wohl, Srividya Ramakrishnan, Melanie Kirsche,_Amanda Ernlund,_Oluwaseun Falade-Nwulia, Timothy Gilpatrick, Paul Morris, Norah Sadowski, N_di_ Trovao, Victoria Gniazdowski, Michael Schatz, Stuart C. Ray, Winston Timp, Heba Mostafa |
| EPI_ISL_438235, EPI_ISL_438236, EPI_ISL_438237, EPI_ISL_438238, EPI_ISL_438239, EPI_ISL_438240, EPI_ISL_438241, EPI_ISL_438242, EPI_ISL_438243, EPI_ISL_438244, EPI_ISL_438245, EPI_ISL_438246, EPI_ISL_438247 | | | |
| see above | Johns Hopkins Hospital Department of Pathology | Johns Hopkins Hospital Department of Pathology | Peter M. Thielen, Thomas Mehoke, Shirlee Wohl, Srividya Ramakrishnan, Melanie Kirsche, Amanda Ernlund, Oluwaseun Falade-Nwulia, Timothy Gilpatrick, Paul Morris, Norah Sadowski, Nidia Trovao, Victoria Gniazdowski, Michael Schatz, Stuart C. Ray, Winston Timp, Heba Mostafa |
| EPI_ISL_438248, EPI_ISL_438249, EPI_ISL_438250, EPI_ISL_438251, EPI_ISL_438252, EPI_ISL_438253, EPI_ISL_438254, EPI_ISL_438255, EPI_ISL_438256, EPI_ISL_438257, EPI_ISL_438258, EPI_ISL_438259, EPI_ISL_438260, EPI_ISL_438261, EPI_ISL_438262, EPI_ISL_438263, EPI_ISL_438264, EPI_ISL_438265, EPI_ISL_438266, EPI_ISL_438267, EPI_ISL_438268, EPI_ISL_438269, EPI_ISL_438270, EPI_ISL_438271, EPI_ISL_438272, EPI_ISL_438273, EPI_ISL_438274, EPI_ISL_438275, EPI_ISL_438276, EPI_ISL_438277, EPI_ISL_438278, EPI_ISL_438279, EPI_ISL_438280, EPI_ISL_438281, EPI_ISL_438282, EPI_ISL_438283, EPI_ISL_438284, EPI_ISL_438285, EPI_ISL_438286, EPI_ISL_438287, EPI_ISL_438288, EPI_ISL_438289, EPI_ISL_438290, EPI_ISL_438291, EPI_ISL_438292, EPI_ISL_438293, EPI_ISL_438294, EPI_ISL_438295, EPI_ISL_438296, EPI_ISL_438297, EPI_ISL_438298, EPI_ISL_438299, EPI_ISL_438300, EPI_ISL_438301, EPI_ISL_438302, EPI_ISL_438303, EPI_ISL_438304, EPI_ISL_438305, EPI_ISL_438306, EPI_ISL_438307, EPI_ISL_438308, EPI_ISL_438309, EPI_ISL_438310, EPI_ISL_438311, EPI_ISL_438312, EPI_ISL_438313, EPI_ISL_438314, EPI_ISL_438315, EPI_ISL_438316, EPI_ISL_438317, EPI_ISL_438318, EPI_ISL_438319, EPI_ISL_438320, EPI_ISL_438321, EPI_ISL_438322, EPI_ISL_438323, EPI_ISL_438324, EPI_ISL_438325, EPI_ISL_438326, EPI_ISL_438327, EPI_ISL_438328, EPI_ISL_438329, EPI_ISL_438330, EPI_ISL_438331, EPI_ISL_438332, EPI_ISL_438333, EPI_ISL_438334, EPI_ISL_438335, EPI_ISL_438336, EPI_ISL_438337, EPI_ISL_438338, EPI_ISL_438339, EPI_ISL_438340, EPI_ISL_438341, EPI_ISL_438342, EPI_ISL_438343, EPI_ISL_438344, EPI_ISL_438345, EPI_ISL_438346, EPI_ISL_438347, EPI_ISL_438348, EPI_ISL_438349, EPI_ISL_438350, EPI_ISL_438351, EPI_ISL_438352, EPI_ISL_438353, EPI_ISL_438354, EPI_ISL_438355, EPI_ISL_438356, EPI_ISL_438357, EPI_ISL_438358, EPI_ISL_438359, EPI_ISL_438360, EPI_ISL_438361, EPI_ISL_438362, EPI_ISL_438363, EPI_ISL_438364, EPI_ISL_438365, EPI_ISL_438366, EPI_ISL_438367, EPI_ISL_438368, EPI_ISL_438369, EPI_ISL_438370, EPI_ISL_438371, EPI_ISL_438372, EPI_ISL_438373, EPI_ISL_438374, EPI_ISL_438375, EPI_ISL_438376, EPI_ISL_438377, EPI_ISL_438378, EPI_ISL_438379, EPI_ISL_438380, EPI_ISL_438381, EPI_ISL_438382, EPI_ISL_438383, EPI_ISL_438384, EPI_ISL_438385, EPI_ISL_438386, EPI_ISL_438387, EPI_ISL_438388, EPI_ISL_438389, EPI_ISL_438390, EPI_ISL_438391, EPI_ISL_438392, EPI_ISL_438393, EPI_ISL_438394, EPI_ISL_438395, EPI_ISL_438396, EPI_ISL_438397, EPI_ISL_438398, EPI_ISL_438399, EPI_ISL_438400, EPI_ISL_438401, EPI_ISL_438402, EPI_ISL_438403, EPI_ISL_438404, EPI_ISL_438405, EPI_ISL_438406, EPI_ISL_438407, EPI_ISL_438408, EPI_ISL_438409, EPI_ISL_438410, EPI_ISL_438411, EPI_ISL_438412, EPI_ISL_438413, EPI_ISL_438414, EPI_ISL_438415, EPI_ISL_438416, EPI_ISL_438417, EPI_ISL_438418, EPI_ISL_438419, EPI_ISL_438420, EPI_ISL_438421, EPI_ISL_438422, EPI_ISL_438423, EPI_ISL_438424, EPI_ISL_438425, EPI_ISL_438426, EPI_ISL_438427, EPI_ISL_438428, EPI_ISL_438429, EPI_ISL_438430, EPI_ISL_438431, EPI_ISL_438432, EPI_ISL_438433, EPI_ISL_438434, EPI_ISL_438435, EPI_ISL_438436, EPI_ISL_438437, EPI_ISL_438438, EPI_ISL_438439, EPI_ISL_438440, EPI_ISL_438441, EPI_ISL_438442, EPI_ISL_438443, EPI_ISL_438444, EPI_ISL_438445, EPI_ISL_438446, EPI_ISL_438447, EPI_ISL_438448, EPI_ISL_438449, EPI_ISL_438450, EPI_ISL_438451, EPI_ISL_438452, EPI_ISL_438453, EPI_ISL_438454, EPI_ISL_438455, EPI_ISL_438456, EPI_ISL_438457, EPI_ISL_438458, EPI_ISL_438459, EPI_ISL_438460, EPI_ISL_438461, EPI_ISL_438462, EPI_ISL_438463, EPI_ISL_438464, EPI_ISL_438465, EPI_ISL_438466, EPI_ISL_438467, EPI_ISL_438468, EPI_ISL_438469, EPI_ISL_438470, EPI_ISL_438471, EPI_ISL_438472, EPI_ISL_438473, EPI_ISL_438474, EPI_ISL_438475, EPI_ISL_438476, EPI_ISL_438477, EPI_ISL_438478, EPI_ISL_438479, EPI_ISL_438480, EPI_ISL_438481, EPI_ISL_438482, EPI_ISL_438483, EPI_ISL_438484, EPI_ISL_438485, EPI_ISL_438486, EPI_ISL_438487, EPI_ISL_438488, EPI_ISL_438489, EPI_ISL_438490, EPI_ISL_438491, EPI_ISL_438492, EPI_ISL_438493, EPI_ISL_438494, EPI_ISL_438495, EPI_ISL_438496, EPI_ISL_438497, EPI_ISL_438498, EPI_ISL_438499, EPI_ISL_438500, EPI_ISL_438501, EPI_ISL_438502, EPI_ISL_438503, EPI_ISL_438504, EPI_ISL_438505, EPI_ISL_438506, EPI_ISL_438507, EPI_ISL_438508, EPI_ISL_438509, EPI_ISL_438510, EPI_ISL_438511, EPI_ISL_438512, EPI_ISL_438513, EPI_ISL_438514, EPI_ISL_438515, EPI_ISL_438516, EPI_ISL_438517, EPI_ISL_438518, EPI_ISL_438519, EPI_ISL_438520, EPI_ISL_438521, EPI_ISL_438522, EPI_ISL_438523, EPI_ISL_438524, EPI_ISL_438525, EPI_ISL_438526, EPI_ISL_438527, EPI_ISL_438528, EPI_ISL_438529, EPI_ISL_438530, EPI_ISL_438531, EPI_ISL_438532, EPI_ISL_438533, EPI_ISL_438534, EPI_ISL_438535, EPI_ISL_438536, EPI_ISL_438537, EPI_ISL_438538, EPI_ISL_438539, EPI_ISL_438540, EPI_ISL_438541, EPI_ISL_438542, EPI_ISL_438543, EPI_ISL_438544, EPI_ISL_438545 | | | |
| see above | Department of Pathology, University of Cambridge | Wellcome Sanger Institute for the COVID-19 Genomics UK Consortium | Luke W Meredith, M. Estée Török , Myra Hosmillo, William L. Hamilton, Martin D. Curran, Theresa Feltwell, Grant Hall, Anna Yakovleva, Fahad A Khokhar, Charlotte J. Houldcroft, Laura G Caller, Aminu S. Jahun, Sarah L. Caddy, Ian Goodfellow, Alex Alderton, Roberto Amato, Sonia Goncalves, Ewan Harrison, David K. Jackson, Ian Johnston, Dominic Kwiatkowski, Cordelia Langford, John Sillitoe on behalf of the Wellcome Sanger Institute COVID-19 Surveillance Team (http://www.sanger.ac.uk/covid-team) |
| EPI_ISL_438546, EPI_ISL_438547, EPI_ISL_438548, EPI_ISL_438549 | Siloam Hospital Lippo Village | Mochtar Riady Institute for Nanotechnology, Universitas Pelita Harapan | Aksar C Lages, David Rustandi, Febi Andriani, Ivet M Suriapranata, Riska N Taufik, Tri Shinta Kurniasih, Irawan Yusuf |
| EPI_ISL_438550, EPI_ISL_438551, EPI_ISL_438552, EPI_ISL_438553, EPI_ISL_438554, EPI_ISL_438555, EPI_ISL_438556, EPI_ISL_438557, EPI_ISL_438558, EPI_ISL_438559, EPI_ISL_438560, EPI_ISL_438561, EPI_ISL_438562, EPI_ISL_438563, EPI_ISL_438564, EPI_ISL_438565, EPI_ISL_438566, EPI_ISL_438567, EPI_ISL_438568, EPI_ISL_438569, EPI_ISL_438570, EPI_ISL_438571, EPI_ISL_438572, EPI_ISL_438573, EPI_ISL_438574, EPI_ISL_438575, EPI_ISL_438576, EPI_ISL_438577, EPI_ISL_438578, EPI_ISL_438579, EPI_ISL_438580, EPI_ISL_438581, EPI_ISL_438582, EPI_ISL_438583, EPI_ISL_438584, EPI_ISL_438585, EPI_ISL_438586, EPI_ISL_438587, EPI_ISL_438588, EPI_ISL_438589, EPI_ISL_438590, EPI_ISL_438591, EPI_ISL_438592, EPI_ISL_438593, EPI_ISL_438594, EPI_ISL_438595, EPI_ISL_438596, EPI_ISL_438597, EPI_ISL_438598, EPI_ISL_438599, EPI_ISL_438600, EPI_ISL_438601, EPI_ISL_438602, EPI_ISL_438603, EPI_ISL_438604, EPI_ISL_438605, EPI_ISL_438606, EPI_ISL_438607, EPI_ISL_438608, EPI_ISL_438609, EPI_ISL_438610, EPI_ISL_438611, EPI_ISL_438612, EPI_ISL_438613, EPI_ISL_438614, EPI_ISL_438615, EPI_ISL_438616, EPI_ISL_438617, EPI_ISL_438618, EPI_ISL_438619, EPI_ISL_438620, EPI_ISL_438621, EPI_ISL_438622, EPI_ISL_438623, EPI_ISL_438624, EPI_ISL_438625, EPI_ISL_438626, EPI_ISL_438627, EPI_ISL_438628, EPI_ISL_438629, EPI_ISL_438630, EPI_ISL_438631, EPI_ISL_438632, EPI_ISL_438633, EPI_ISL_438634, EPI_ISL_438635, EPI_ISL_438636, EPI_ISL_438637, EPI_ISL_438638, EPI_ISL_438639, EPI_ISL_438640, EPI_ISL_438641, EPI_ISL_438642, EPI_ISL_438643, EPI_ISL_438644, EPI_ISL_438645, EPI_ISL_438646, EPI_ISL_438647, EPI_ISL_438648, EPI_ISL_438649, EPI_ISL_438650, EPI_ISL_438651, EPI_ISL_438652, EPI_ISL_438653, EPI_ISL_438654, EPI_ISL_438655, EPI_ISL_438656, EPI_ISL_438657, EPI_ISL_438658, EPI_ISL_438659, EPI_ISL_438660, EPI_ISL_438661, EPI_ISL_438662, EPI_ISL_438663, EPI_ISL_438664, EPI_ISL_438665, EPI_ISL_438666, EPI_ISL_438667, EPI_ISL_438668, EPI_ISL_438669, EPI_ISL_438670, EPI_ISL_438671, EPI_ISL_438672, EPI_ISL_438673, EPI_ISL_438674, EPI_ISL_438675, EPI_ISL_438676, EPI_ISL_438677, EPI_ISL_438678, EPI_ISL_438679, EPI_ISL_438680, EPI_ISL_438681, EPI_ISL_438682, EPI_ISL_438683, EPI_ISL_438684, EPI_ISL_438685, EPI_ISL_438686, EPI_ISL_438687, EPI_ISL_438688, EPI_ISL_438689, EPI_ISL_438690, EPI_ISL_438691, EPI_ISL_438692, EPI_ISL_438693, EPI_ISL_438694, EPI_ISL_438695, EPI_ISL_438696, EPI_ISL_438697, EPI_ISL_438698, EPI_ISL_438699, EPI_ISL_438700, EPI_ISL_438701, EPI_ISL_438702, EPI_ISL_438703, EPI_ISL_438704, EPI_ISL_438705, EPI_ISL_438706, EPI_ISL_438707, EPI_ISL_438708, EPI_ISL_438709, EPI_ISL_438710, EPI_ISL_438711, EPI_ISL_438712, EPI_ISL_438713, EPI_ISL_438714, EPI_ISL_438715, EPI_ISL_438716, EPI_ISL_438717, EPI_ISL_438718, EPI_ISL_438719, EPI_ISL_438720, EPI_ISL_438721, EPI_ISL_438722, EPI_ISL_438723, EPI_ISL_438724, EPI_ISL_438725, EPI_ISL_438726, EPI_ISL_438727, EPI_ISL_438728, EPI_ISL_438729, EPI_ISL_438730, EPI_ISL_438731, EPI_ISL_438732, EPI_ISL_438733, EPI_ISL_438734, EPI_ISL_438735, EPI_ISL_438736, EPI_ISL_438737, EPI_ISL_438738, EPI_ISL_438739, EPI_ISL_438740, EPI_ISL_438741, EPI_ISL_438742, EPI_ISL_438743, EPI_ISL_438744, EPI_ISL_438745, EPI_ISL_438746, EPI_ISL_438747, EPI_ISL_438748 | | | |
| see above | Department of Pathology, University of Cambridge | COVID-19 Genomics UK (COG-UK) Consortium | Luke W Meredith, M. Est√©e T√∂r√∂k , Myra Hosmillo, William L. Hamilton, Martin D. Curran, Theresa Feltwell, Grant Hall, Anna Yakovleva, Fahad A Khokhar, Charlotte J. Houldcroft, Laura G Caller, Aminu S. Jahun, Sarah L. Caddy, Ian Goodfellow |
| EPI_ISL_438749, EPI_ISL_438750, EPI_ISL_438751, EPI_ISL_438752, EPI_ISL_438753, EPI_ISL_438754, EPI_ISL_438755, EPI_ISL_438756, EPI_ISL_438757, EPI_ISL_438758, EPI_ISL_438759, EPI_ISL_438760, EPI_ISL_438761, EPI_ISL_438762, EPI_ISL_438763, EPI_ISL_438764, EPI_ISL_438765, EPI_ISL_438766, EPI_ISL_438767, EPI_ISL_438768, EPI_ISL_438769, EPI_ISL_438770, EPI_ISL_438771, EPI_ISL_438772, EPI_ISL_438773, EPI_ISL_438774, EPI_ISL_438775, EPI_ISL_438776, EPI_ISL_438777, EPI_ISL_438778, EPI_ISL_438779, EPI_ISL_438780, EPI_ISL_438781, EPI_ISL_438782, EPI_ISL_438783, EPI_ISL_438784, EPI_ISL_438785, EPI_ISL_438786, EPI_ISL_438787, EPI_ISL_438788, EPI_ISL_438789, EPI_ISL_438790, EPI_ISL_438791, EPI_ISL_438792, EPI_ISL_438793, EPI_ISL_438794, EPI_ISL_438795, EPI_ISL_438796, EPI_ISL_438797, EPI_ISL_438798, EPI_ISL_438799, EPI_ISL_438800, EPI_ISL_438801, EPI_ISL_438802, EPI_ISL_438803, EPI_ISL_438804, EPI_ISL_438805, EPI_ISL_438806, EPI_ISL_438807, EPI_ISL_438808, EPI_ISL_438809, EPI_ISL_438810, EPI_ISL_438811, EPI_ISL_438812, EPI_ISL_438813, EPI_ISL_438814, EPI_ISL_438815, EPI_ISL_438816, EPI_ISL_438817, EPI_ISL_438818, EPI_ISL_438819, EPI_ISL_438820, EPI_ISL_438821, EPI_ISL_438822, EPI_ISL_438823, EPI_ISL_438824, EPI_ISL_438825, EPI_ISL_438826, EPI_ISL_438827, EPI_ISL_438828, EPI_ISL_438829, EPI_ISL_438830, EPI_ISL_438831, EPI_ISL_438832, EPI_ISL_438833, EPI_ISL_438834, EPI_ISL_438835, EPI_ISL_438836, EPI_ISL_438837, EPI_ISL_438838, EPI_ISL_438839, EPI_ISL_438840, EPI_ISL_438841, EPI_ISL_438842, EPI_ISL_438843, EPI_ISL_438844, EPI_ISL_438845, EPI_ISL_438846, EPI_ISL_438847, EPI_ISL_438848, EPI_ISL_438849, EPI_ISL_438850, EPI_ISL_438851, EPI_ISL_438852, EPI_ISL_438853, EPI_ISL_438854, EPI_ISL_438855, EPI_ISL_438856, EPI_ISL_438857, EPI_ISL_438858, EPI_ISL_438859, EPI_ISL_438860, EPI_ISL_438861, EPI_ISL_438862, EPI_ISL_438863, EPI_ISL_438864, EPI_ISL_438865, EPI_ISL_438866, EPI_ISL_438867, EPI_ISL_438868, EPI_ISL_438869, EPI_ISL_438870, EPI_ISL_438871, EPI_ISL_438872, EPI_ISL_438873, EPI_ISL_438874, EPI_ISL_438875, EPI_ISL_438876, EPI_ISL_438877, EPI_ISL_438878, EPI_ISL_438879, EPI_ISL_438880, EPI_ISL_438881, EPI_ISL_438882, EPI_ISL_438883, EPI_ISL_438884, EPI_ISL_438885, EPI_ISL_438886, EPI_ISL_438887, EPI_ISL_438888, EPI_ISL_438889, EPI_ISL_438890, EPI_ISL_438891, EPI_ISL_438892, EPI_ISL_438893, EPI_ISL_438894, EPI_ISL_438895, EPI_ISL_438896, EPI_ISL_438897, EPI_ISL_438898, EPI_ISL_438899, EPI_ISL_438900, EPI_ISL_438901, EPI_ISL_438902, EPI_ISL_438903, EPI_ISL_438904, EPI_ISL_438905, EPI_ISL_438906, EPI_ISL_438907, EPI_ISL_438908, EPI_ISL_438909, EPI_ISL_438910, EPI_ISL_438911, EPI_ISL_438912, EPI_ISL_438913, EPI_ISL_438914, EPI_ISL_438915, EPI_ISL_438916, EPI_ISL_438917, EPI_ISL_438918, EPI_ISL_438919, EPI_ISL_438920, EPI_ISL_438921, EPI_ISL_438922, EPI_ISL_438923, EPI_ISL_438924, EPI_ISL_438925, EPI_ISL_438926, EPI_ISL_438927, EPI_ISL_438928, EPI_ISL_438929, EPI_ISL_438930, EPI_ISL_438931, EPI_ISL_438932, EPI_ISL_438933, EPI_ISL_438934, EPI_ISL_438935, EPI_ISL_438936, EPI_ISL_438937, EPI_ISL_438938, EPI_ISL_438939, EPI_ISL_438940, EPI_ISL_438941, EPI_ISL_438942, EPI_ISL_438943, EPI_ISL_438944, EPI_ISL_438945, EPI_ISL_438946 | | | |
| see above | West of Scotland Specialist Virology Centre, NHSGGC / MRC-University of Glasgow Centre for Virus Research | COVID-19 Genomics UK (COG-UK) Consortium | Ana da Silva Filipe, Natasha Johnson, Kathy Smollett, Daniel Mair, Stephen Carmichael, Lily Tong, Jenna Nichols, Elihu Aranday-Cortes, Kirstyn Brunker, Yasmin Parr, Kyriaki Nomikou; Sarah McDonald, Marc Niebel, Patawee Asamaphan; Richard Orton, Joseph Hughes, Sreenu Vattipally, David L Robertson; Alasdair MacLean, Rory Gunson; Kathy Li, Natasha Jesudason, Rajiv Shah, James Shepherd, Antonia Ho, Emma Thomson |
| EPI_ISL_438947, EPI_ISL_438948, EPI_ISL_438949, EPI_ISL_438951, EPI_ISL_438952, EPI_ISL_438953, EPI_ISL_438954, EPI_ISL_438955, EPI_ISL_438956, EPI_ISL_438958, EPI_ISL_438959, EPI_ISL_438961, EPI_ISL_438962, EPI_ISL_438963, EPI_ISL_438964, EPI_ISL_438965, EPI_ISL_438966, EPI_ISL_438967, EPI_ISL_438968, EPI_ISL_438969, EPI_ISL_438970, EPI_ISL_438971, EPI_ISL_438972 | | | |
| see above | Keio University School of Medicine | Keio University School of Medicine | Kenjiro Kosaki |
| EPI_ISL_438973, EPI_ISL_438974, EPI_ISL_438975, EPI_ISL_438976, EPI_ISL_438977, EPI_ISL_438978, EPI_ISL_438979, EPI_ISL_438980, EPI_ISL_438981, EPI_ISL_438982, EPI_ISL_438983, EPI_ISL_438984, EPI_ISL_438985, EPI_ISL_438986, EPI_ISL_438987, EPI_ISL_438988, EPI_ISL_438989, EPI_ISL_438990, EPI_ISL_438991, EPI_ISL_438992, EPI_ISL_438993, EPI_ISL_438994, EPI_ISL_438995, EPI_ISL_438996, EPI_ISL_438997, EPI_ISL_438998, EPI_ISL_438999, EPI_ISL_439000, EPI_ISL_439001, EPI_ISL_439002, EPI_ISL_439003, EPI_ISL_439004, EPI_ISL_439005, EPI_ISL_439006, EPI_ISL_439007, EPI_ISL_439008, EPI_ISL_439009, EPI_ISL_439010, EPI_ISL_439011, EPI_ISL_439012, EPI_ISL_439013, EPI_ISL_439014, EPI_ISL_439015, EPI_ISL_439016, EPI_ISL_439017, EPI_ISL_439018, EPI_ISL_439019, EPI_ISL_439020, EPI_ISL_439021, EPI_ISL_439022, EPI_ISL_439023, EPI_ISL_439024, EPI_ISL_439025, EPI_ISL_439026, EPI_ISL_439027, EPI_ISL_439028, EPI_ISL_439029, EPI_ISL_439030, EPI_ISL_439031, EPI_ISL_439032, EPI_ISL_439033, EPI_ISL_439034, EPI_ISL_439035, EPI_ISL_439036, EPI_ISL_439037, EPI_ISL_439038, EPI_ISL_439039, EPI_ISL_439040, EPI_ISL_439041, EPI_ISL_439042, EPI_ISL_439043, EPI_ISL_439044, EPI_ISL_439045, EPI_ISL_439046, EPI_ISL_439047, EPI_ISL_439048, EPI_ISL_439049, EPI_ISL_439050, EPI_ISL_439051, EPI_ISL_439052, EPI_ISL_439053, EPI_ISL_439054, EPI_ISL_439055, EPI_ISL_439056, EPI_ISL_439057, EPI_ISL_439058, EPI_ISL_439059, EPI_ISL_439060, EPI_ISL_439061, EPI_ISL_439062, EPI_ISL_439063, EPI_ISL_439064, EPI_ISL_439065, EPI_ISL_439066, EPI_ISL_439067, EPI_ISL_439068, EPI_ISL_439069, EPI_ISL_439070, EPI_ISL_439071, EPI_ISL_439072, EPI_ISL_439073, EPI_ISL_439074, EPI_ISL_439075, EPI_ISL_439076, EPI_ISL_439077, EPI_ISL_439078, EPI_ISL_439079, EPI_ISL_439080, EPI_ISL_439081, EPI_ISL_439082, EPI_ISL_439083, EPI_ISL_439084, EPI_ISL_439085, EPI_ISL_439086, EPI_ISL_439087, EPI_ISL_439088, EPI_ISL_439089, EPI_ISL_439090, EPI_ISL_439091, EPI_ISL_439092, EPI_ISL_439093, EPI_ISL_439094, EPI_ISL_439095, EPI_ISL_439096, EPI_ISL_439097, EPI_ISL_439098, EPI_ISL_439099, EPI_ISL_439100, EPI_ISL_439101, EPI_ISL_439102, EPI_ISL_439103, EPI_ISL_439104, EPI_ISL_439105, EPI_ISL_439106, EPI_ISL_439107, EPI_ISL_439108, EPI_ISL_439109, EPI_ISL_439110, EPI_ISL_439111, EPI_ISL_439112, EPI_ISL_439113, EPI_ISL_439114, EPI_ISL_439115, EPI_ISL_439116, EPI_ISL_439117, EPI_ISL_439118, EPI_ISL_439119, EPI_ISL_439120, EPI_ISL_439121, EPI_ISL_439122, EPI_ISL_439123, EPI_ISL_439124, EPI_ISL_439125, EPI_ISL_439126, EPI_ISL_439127, EPI_ISL_439128, EPI_ISL_439129, EPI_ISL_439130, EPI_ISL_439131, EPI_ISL_439132, EPI_ISL_439133, EPI_ISL_439134, EPI_ISL_439135, EPI_ISL_439136, EPI_ISL_439137, EPI_ISL_439138, EPI_ISL_439139, EPI_ISL_439140, EPI_ISL_439141, EPI_ISL_439142, EPI_ISL_439143 | | | |

| | | | |
|---|---|---|---|
| see above | West of Scotland Specialist Virology Centre, NHSGGC / MRC-University of Glasgow Centre for Virus Research | COVID-19 Genomics UK (COG-UK) Consortium | Ana da Silva Filipe, Natasha Johnson, Kathy Smollett, Daniel Mair, Stephen Carmichael, Lily Tong, Jenna Nichols, Elihu Aranday-Cortes, Kirstyn Brunker, Yasmin Parr, Kyriaki Nomikou; Sarah McDonald, Marc Niebel, Patawee Asamaphan; Richard Orton, Joseph Hughes, Sreenu Vattipally, David L Robertson; Alasdair MacLean, Rory Gunson; Kathy Li, Natasha Jesudason, Rajiv Shah, James Shepherd, Antonia Ho, Emma Thomson |
| EPI_ISL_439144, EPI_ISL_439145, EPI_ISL_439146, EPI_ISL_439147, EPI_ISL_439148, EPI_ISL_439149, EPI_ISL_439150, EPI_ISL_439151, EPI_ISL_439152, EPI_ISL_439153, EPI_ISL_439154, EPI_ISL_439155, EPI_ISL_439156, EPI_ISL_439157, EPI_ISL_439158, EPI_ISL_439159, EPI_ISL_439160, EPI_ISL_439161, EPI_ISL_439162, EPI_ISL_439163, EPI_ISL_439164, EPI_ISL_439165, EPI_ISL_439166, EPI_ISL_439167, EPI_ISL_439168, EPI_ISL_439169, EPI_ISL_439170, EPI_ISL_439171, EPI_ISL_439172, EPI_ISL_439173, EPI_ISL_439174, EPI_ISL_439175, EPI_ISL_439176, EPI_ISL_439177, EPI_ISL_439178, EPI_ISL_439179, EPI_ISL_439180, EPI_ISL_439181, EPI_ISL_439182, EPI_ISL_439183, EPI_ISL_439184, EPI_ISL_439185, EPI_ISL_439186, EPI_ISL_439187, EPI_ISL_439188, EPI_ISL_439189, EPI_ISL_439190, EPI_ISL_439191, EPI_ISL_439192, EPI_ISL_439193, EPI_ISL_439194, EPI_ISL_439195, EPI_ISL_439196, EPI_ISL_439197, EPI_ISL_439198, EPI_ISL_439199, EPI_ISL_439200, EPI_ISL_439201, EPI_ISL_439202, EPI_ISL_439203, EPI_ISL_439204, EPI_ISL_439205, EPI_ISL_439206, EPI_ISL_439207, EPI_ISL_439208, EPI_ISL_439209, EPI_ISL_439210, EPI_ISL_439211, EPI_ISL_439212, EPI_ISL_439213, EPI_ISL_439214, EPI_ISL_439215, EPI_ISL_439216, EPI_ISL_439217, EPI_ISL_439218, EPI_ISL_439219, EPI_ISL_439220, EPI_ISL_439221, EPI_ISL_439222, EPI_ISL_439223, EPI_ISL_439224, EPI_ISL_439225, EPI_ISL_439226, EPI_ISL_439227, EPI_ISL_439228, EPI_ISL_439229, EPI_ISL_439230, EPI_ISL_439231, EPI_ISL_439232, EPI_ISL_439233, EPI_ISL_439234, EPI_ISL_439235, EPI_ISL_439236, EPI_ISL_439237, EPI_ISL_439238, EPI_ISL_439239, EPI_ISL_439240, EPI_ISL_439241, EPI_ISL_439242, EPI_ISL_439243, EPI_ISL_439244, EPI_ISL_439245, EPI_ISL_439246, EPI_ISL_439247, EPI_ISL_439248, EPI_ISL_439249, EPI_ISL_439250, EPI_ISL_439251, EPI_ISL_439252, EPI_ISL_439253, EPI_ISL_439254, EPI_ISL_439255, EPI_ISL_439256, EPI_ISL_439257, EPI_ISL_439258, EPI_ISL_439259, EPI_ISL_439260, EPI_ISL_439261, EPI_ISL_439262, EPI_ISL_439263, EPI_ISL_439264, EPI_ISL_439265, EPI_ISL_439266, EPI_ISL_439267, EPI_ISL_439268, EPI_ISL_439269, EPI_ISL_439270, EPI_ISL_439271, EPI_ISL_439272, EPI_ISL_439273, EPI_ISL_439274, EPI_ISL_439275, EPI_ISL_439276, EPI_ISL_439277, EPI_ISL_439278, EPI_ISL_439279, EPI_ISL_439280, EPI_ISL_439281, EPI_ISL_439282, EPI_ISL_439283, EPI_ISL_439284, EPI_ISL_439285, EPI_ISL_439286, EPI_ISL_439287, EPI_ISL_439288, EPI_ISL_439289, EPI_ISL_439290, EPI_ISL_439291, EPI_ISL_439292, EPI_ISL_439293, EPI_ISL_439294, EPI_ISL_439295, EPI_ISL_439296, EPI_ISL_439297, EPI_ISL_439298, EPI_ISL_439299, EPI_ISL_439300, EPI_ISL_439301, EPI_ISL_439302, EPI_ISL_439303, EPI_ISL_439304, EPI_ISL_439305, EPI_ISL_439306, EPI_ISL_439307, EPI_ISL_439308, EPI_ISL_439309, EPI_ISL_439310, EPI_ISL_439311, EPI_ISL_439312, EPI_ISL_439313, EPI_ISL_439314, EPI_ISL_439315, EPI_ISL_439316, EPI_ISL_439317, EPI_ISL_439318, EPI_ISL_439319, EPI_ISL_439320, EPI_ISL_439321, EPI_ISL_439322, EPI_ISL_439323, EPI_ISL_439324, EPI_ISL_439325, EPI_ISL_439326, EPI_ISL_439327, EPI_ISL_439328, EPI_ISL_439329, EPI_ISL_439330, EPI_ISL_439331, EPI_ISL_439332, EPI_ISL_439333, EPI_ISL_439334, EPI_ISL_439335, EPI_ISL_439336, EPI_ISL_439337, EPI_ISL_439338, EPI_ISL_439339, EPI_ISL_439340, EPI_ISL_439341, EPI_ISL_439342, EPI_ISL_439343, EPI_ISL_439344, EPI_ISL_439345, EPI_ISL_439346, EPI_ISL_439347, EPI_ISL_439348, EPI_ISL_439349, EPI_ISL_439350, EPI_ISL_439351, EPI_ISL_439352, EPI_ISL_439353, EPI_ISL_439354, EPI_ISL_439355, EPI_ISL_439356, EPI_ISL_439357, EPI_ISL_439358, EPI_ISL_439359, EPI_ISL_439360, EPI_ISL_439361, EPI_ISL_439362, EPI_ISL_439363, EPI_ISL_439364, EPI_ISL_439365, EPI_ISL_439366, EPI_ISL_439367 | | | |
| see above | Virology Department, Royal Infirmary of Edinburgh, NHS Lothian / School of Biological Sciences, University of Edinburgh / Institute of Genetics and Molecular Medicine, University of Edinburgh | COVID-19 Genomics UK (COG-UK) Consortium | McHugh M, Dewar R, Rooke S, Gallagher M, Balcaza C, O‚ÄôToole √Å, Scher E, Hill V, McCrone JT, Colquhoun R, Yu X, Jackson B, Rambaut A, Williams TC, Templeton K |
| EPI_ISL_439368, EPI_ISL_439369, EPI_ISL_439370, EPI_ISL_439371, EPI_ISL_439372, EPI_ISL_439373, EPI_ISL_439374, EPI_ISL_439375, EPI_ISL_439376, EPI_ISL_439377, EPI_ISL_439378, EPI_ISL_439379, EPI_ISL_439380, EPI_ISL_439381, EPI_ISL_439382, EPI_ISL_439383, EPI_ISL_439384, EPI_ISL_439385, EPI_ISL_439386, EPI_ISL_439387, EPI_ISL_439388, EPI_ISL_439389, EPI_ISL_439390, EPI_ISL_439391, EPI_ISL_439392, EPI_ISL_439393, EPI_ISL_439394, EPI_ISL_439395, EPI_ISL_439396, EPI_ISL_439397, EPI_ISL_439398, EPI_ISL_439399, EPI_ISL_439400, EPI_ISL_439401, EPI_ISL_439402, EPI_ISL_439403, EPI_ISL_439404, EPI_ISL_439405, EPI_ISL_439406, EPI_ISL_439407, EPI_ISL_439408, EPI_ISL_439409, EPI_ISL_439410, EPI_ISL_439411, EPI_ISL_439412, EPI_ISL_439413, EPI_ISL_439414, EPI_ISL_439415, EPI_ISL_439416, EPI_ISL_439417, EPI_ISL_439418, EPI_ISL_439419, EPI_ISL_439420, EPI_ISL_439421, EPI_ISL_439422, EPI_ISL_439423, EPI_ISL_439424, EPI_ISL_439425, EPI_ISL_439426, EPI_ISL_439427, EPI_ISL_439428, EPI_ISL_439429, EPI_ISL_439430, EPI_ISL_439431, EPI_ISL_439432, EPI_ISL_439433, EPI_ISL_439434, EPI_ISL_439435, EPI_ISL_439436, EPI_ISL_439437, EPI_ISL_439438, EPI_ISL_439439, EPI_ISL_439440, EPI_ISL_439441, EPI_ISL_439442, EPI_ISL_439443, EPI_ISL_439444, EPI_ISL_439445, EPI_ISL_439446, EPI_ISL_439447, EPI_ISL_439448, EPI_ISL_439449, EPI_ISL_439450, EPI_ISL_439451, EPI_ISL_439452, EPI_ISL_439453, EPI_ISL_439454, EPI_ISL_439455, EPI_ISL_439456, EPI_ISL_439457, EPI_ISL_439458, EPI_ISL_439459, EPI_ISL_439460, EPI_ISL_439461, EPI_ISL_439462, EPI_ISL_439463, EPI_ISL_439464, EPI_ISL_439465, EPI_ISL_439466, EPI_ISL_439467, EPI_ISL_439468, EPI_ISL_439469, EPI_ISL_439470, EPI_ISL_439471, EPI_ISL_439472, EPI_ISL_439473, EPI_ISL_439474, EPI_ISL_439475, EPI_ISL_439476, EPI_ISL_439477, EPI_ISL_439478, EPI_ISL_439479, EPI_ISL_439480, EPI_ISL_439481, EPI_ISL_439482, EPI_ISL_439483, EPI_ISL_439484, EPI_ISL_439485, EPI_ISL_439486, EPI_ISL_439487, EPI_ISL_439488, EPI_ISL_439489, EPI_ISL_439490, EPI_ISL_439491, EPI_ISL_439492, EPI_ISL_439493, EPI_ISL_439494, EPI_ISL_439495, EPI_ISL_439496, EPI_ISL_439497, EPI_ISL_439498, EPI_ISL_439499, EPI_ISL_439500, EPI_ISL_439501, EPI_ISL_439502, EPI_ISL_439503, EPI_ISL_439504, EPI_ISL_439505, EPI_ISL_439506, EPI_ISL_439507, EPI_ISL_439508, EPI_ISL_439509, EPI_ISL_439510, EPI_ISL_439511, EPI_ISL_439512, EPI_ISL_439513, EPI_ISL_439514, EPI_ISL_439515, EPI_ISL_439516, EPI_ISL_439517, EPI_ISL_439518, EPI_ISL_439519, EPI_ISL_439520, EPI_ISL_439521, EPI_ISL_439522, EPI_ISL_439523, EPI_ISL_439524, EPI_ISL_439525, EPI_ISL_439526, EPI_ISL_439527, EPI_ISL_439528, EPI_ISL_439529, EPI_ISL_439530, EPI_ISL_439531, EPI_ISL_439532, EPI_ISL_439533, EPI_ISL_439534, EPI_ISL_439535, EPI_ISL_439536, EPI_ISL_439537, EPI_ISL_439538, EPI_ISL_439539, EPI_ISL_439540, EPI_ISL_439541, EPI_ISL_439542, EPI_ISL_439543, EPI_ISL_439544, EPI_ISL_439545, EPI_ISL_439546, EPI_ISL_439547, EPI_ISL_439548, EPI_ISL_439549, EPI_ISL_439550, EPI_ISL_439551, EPI_ISL_439552, EPI_ISL_439553, EPI_ISL_439554, EPI_ISL_439555, EPI_ISL_439556, EPI_ISL_439557, EPI_ISL_439558, EPI_ISL_439559, EPI_ISL_439560, EPI_ISL_439561, EPI_ISL_439562, EPI_ISL_439563, EPI_ISL_439564, EPI_ISL_439565, EPI_ISL_439566, EPI_ISL_439567, EPI_ISL_439568, EPI_ISL_439569, EPI_ISL_439570, EPI_ISL_439571, EPI_ISL_439572, EPI_ISL_439573, EPI_ISL_439574, EPI_ISL_439575, EPI_ISL_439576, EPI_ISL_439577, EPI_ISL_439578, EPI_ISL_439579, EPI_ISL_439580, EPI_ISL_439581, EPI_ISL_439582, EPI_ISL_439583, EPI_ISL_439584, EPI_ISL_439585, EPI_ISL_439586, EPI_ISL_439587, EPI_ISL_439588, EPI_ISL_439589, EPI_ISL_439590, EPI_ISL_439591, EPI_ISL_439592, EPI_ISL_439593, EPI_ISL_439594, EPI_ISL_439595, EPI_ISL_439596, EPI_ISL_439597, EPI_ISL_439598, EPI_ISL_439599, EPI_ISL_439600, EPI_ISL_439601, EPI_ISL_439602, EPI_ISL_439603, EPI_ISL_439604, EPI_ISL_439605, EPI_ISL_439606, EPI_ISL_439607, EPI_ISL_439608, EPI_ISL_439609, EPI_ISL_439610, EPI_ISL_439611, EPI_ISL_439612, EPI_ISL_439613, EPI_ISL_439614, EPI_ISL_439615, EPI_ISL_439616, EPI_ISL_439617, EPI_ISL_439618, EPI_ISL_439619, EPI_ISL_439620, EPI_ISL_439621, EPI_ISL_439622, EPI_ISL_439623, EPI_ISL_439624, EPI_ISL_439625, EPI_ISL_439626, EPI_ISL_439627, EPI_ISL_439628, EPI_ISL_439629, EPI_ISL_439630, EPI_ISL_439631, EPI_ISL_439632, EPI_ISL_439633, EPI_ISL_439634, EPI_ISL_439635, EPI_ISL_439636, EPI_ISL_439637, EPI_ISL_439638, EPI_ISL_439639, EPI_ISL_439640, EPI_ISL_439641, EPI_ISL_439642, EPI_ISL_439643, EPI_ISL_439644, EPI_ISL_439645, EPI_ISL_439646, EPI_ISL_439647, EPI_ISL_439648, EPI_ISL_439649, EPI_ISL_439650, EPI_ISL_439651, EPI_ISL_439652, EPI_ISL_439653, EPI_ISL_439654, EPI_ISL_439655, EPI_ISL_439656, EPI_ISL_439657, EPI_ISL_439658, EPI_ISL_439659, EPI_ISL_439660, EPI_ISL_439661, EPI_ISL_439662, EPI_ISL_439663, EPI_ISL_439664 | | | |
| see above | Department of Pathology, University of Cambridge | Wellcome Sanger Institute for the COVID-19 Genomics UK Consortium | Luke W Meredith, M. Estée Török , Myra Hosmillo, William L. Hamilton, Martin D. Curran, Theresa Feltwell, Grant Hall, Anna Yakovleva, Fahad A Khokhar, Charlotte J. Houldcroft, Laura G Caller, Aminu S. Jahun, Sarah L. Caddy, Ian Goodfellow, Alex Alderton, Roberto Amato, Sonia Goncalves, Ewan Harrison, David K. Jackson, Ian Johnston, Dominic Kwiatkowski, Cordelia Langford, John Sillitoe on behalf of the Wellcome Sanger Institute COVID-19 Surveillance Team (http://www.sanger.ac.uk/covid-team) |
| EPI_ISL_439665, EPI_ISL_439666, EPI_ISL_439667, EPI_ISL_439668, EPI_ISL_439669, EPI_ISL_439670, EPI_ISL_439671 | Virology Department, Royal Infirmary of Edinburgh, NHS Lothian / School of Biological Sciences, University of Edinburgh / Institute of Genetics and Molecular Medicine, University of Edinburgh | COVID-19 Genomics UK (COG-UK) Consortium | McHugh M, Dewar R, Rooke S, Gallagher M, Balcaza C, O‚ÄôToole √Å, Scher E, Hill V, McCrone JT, Colquhoun R, Yu X, Jackson B, Rambaut A, Williams TC, Templeton K |
| EPI_ISL_439672, EPI_ISL_439673, EPI_ISL_439674, EPI_ISL_439675, EPI_ISL_439676, EPI_ISL_439677, EPI_ISL_439678, EPI_ISL_439679, EPI_ISL_439680, EPI_ISL_439681, EPI_ISL_439682, EPI_ISL_439683, EPI_ISL_439684, EPI_ISL_439685, EPI_ISL_439686, EPI_ISL_439687, EPI_ISL_439688, EPI_ISL_439689, EPI_ISL_439690, EPI_ISL_439691, EPI_ISL_439692, EPI_ISL_439693, EPI_ISL_439694, EPI_ISL_439695, EPI_ISL_439696, EPI_ISL_439697, EPI_ISL_439698, EPI_ISL_439699, EPI_ISL_439700, EPI_ISL_439701, EPI_ISL_439702, EPI_ISL_439703, EPI_ISL_439704, EPI_ISL_439705, EPI_ISL_439706, EPI_ISL_439707, EPI_ISL_439708, EPI_ISL_439709, EPI_ISL_439710, EPI_ISL_439711, EPI_ISL_439712, EPI_ISL_439713, EPI_ISL_439714, EPI_ISL_439715, EPI_ISL_439716, EPI_ISL_439717, EPI_ISL_439718, EPI_ISL_439719, EPI_ISL_439720, EPI_ISL_439721, EPI_ISL_439722, EPI_ISL_439723, EPI_ISL_439724, EPI_ISL_439725, EPI_ISL_439726, EPI_ISL_439727, EPI_ISL_439728, EPI_ISL_439729, EPI_ISL_439730, EPI_ISL_439731, EPI_ISL_439732, EPI_ISL_439733, EPI_ISL_439734, EPI_ISL_439735, EPI_ISL_439736, EPI_ISL_439737, EPI_ISL_439738, EPI_ISL_439739, EPI_ISL_439740, EPI_ISL_439741, EPI_ISL_439742, EPI_ISL_439743, EPI_ISL_439744, EPI_ISL_439745, EPI_ISL_439746, EPI_ISL_439747, EPI_ISL_439748, EPI_ISL_439749, EPI_ISL_439750, EPI_ISL_439751, EPI_ISL_439752, EPI_ISL_439753, EPI_ISL_439754, EPI_ISL_439755, EPI_ISL_439756, EPI_ISL_439757, EPI_ISL_439758, EPI_ISL_439759, EPI_ISL_439760, EPI_ISL_439761, EPI_ISL_439762, EPI_ISL_439763, EPI_ISL_439764, EPI_ISL_439765, EPI_ISL_439766, EPI_ISL_439767, EPI_ISL_439768, EPI_ISL_439769, EPI_ISL_439770, EPI_ISL_439771, EPI_ISL_439772, EPI_ISL_439773, EPI_ISL_439774, EPI_ISL_439775, EPI_ISL_439776, EPI_ISL_439777, EPI_ISL_439778, EPI_ISL_439779, EPI_ISL_439780, EPI_ISL_439781, EPI_ISL_439782, EPI_ISL_439783, EPI_ISL_439784, EPI_ISL_439785, EPI_ISL_439786, EPI_ISL_439787, EPI_ISL_439788, EPI_ISL_439789, EPI_ISL_439790, EPI_ISL_439791, EPI_ISL_439792, EPI_ISL_439793, EPI_ISL_439794, EPI_ISL_439795, EPI_ISL_439796, EPI_ISL_439797, EPI_ISL_439798, EPI_ISL_439799, EPI_ISL_439800, EPI_ISL_439801, EPI_ISL_439802, EPI_ISL_439803, EPI_ISL_439804, EPI_ISL_439805, EPI_ISL_439806, EPI_ISL_439807, EPI_ISL_439808, EPI_ISL_439809, EPI_ISL_439810, EPI_ISL_439811, EPI_ISL_439812, EPI_ISL_439813, EPI_ISL_439814, EPI_ISL_439815, EPI_ISL_439816, EPI_ISL_439817, EPI_ISL_439818, EPI_ISL_439819, EPI_ISL_439820, EPI_ISL_439821, EPI_ISL_439822, EPI_ISL_439823, EPI_ISL_439824, EPI_ISL_439825, EPI_ISL_439826, EPI_ISL_439827, EPI_ISL_439828, EPI_ISL_439829, EPI_ISL_439830, EPI_ISL_439831, EPI_ISL_439832, EPI_ISL_439833, EPI_ISL_439834, EPI_ISL_439835, EPI_ISL_439836, EPI_ISL_439837, EPI_ISL_439838, EPI_ISL_439839, EPI_ISL_439840, EPI_ISL_439841, EPI_ISL_439842, EPI_ISL_439843, EPI_ISL_439844, EPI_ISL_439845, EPI_ISL_439846, EPI_ISL_439847, EPI_ISL_439848, EPI_ISL_439849, EPI_ISL_439850, EPI_ISL_439851, EPI_ISL_439852, EPI_ISL_439853, EPI_ISL_439854, EPI_ISL_439855, EPI_ISL_439856, EPI_ISL_439857, EPI_ISL_439858, EPI_ISL_439859, EPI_ISL_439860, EPI_ISL_439861, EPI_ISL_439862 | | | |
| see above | Liverpool Clinical Laboratories | COVID-19 Genomics UK (COG-UK) Consortium | Sam Haldenby, Anita Lucaci, Steve Paterson, Julian Hiscox, Alistair Darby, M Almsaud, A Alrezaihi, Muhannad Alruwaili, Stuart D Armstrong, Jones Benjamin , Eleanor G Bentley, Anu Chawla, Jordan J Clark, Angela Cowell, Richard Eccles, Isabel Garc√≠a-Dorival, Matthew Gemmell, Alessandro Gerada, PKF Gilmore, Richard Gregory, Ximeng Han, Catherine Hartley, Margaret Hughes, Miren Iturriza-Gomara, James Johnson, L Luu, Jenifer Manson , Charlotte Nelson, Elaine O‚ÄôToole, Cassie Olateju, Rebekah Penrice-Randal¬†, Lucille Rainbow, N.P Randle, Trevor Ian Robinson, Parul Sharma, Ghada T Shawli, James P Stewart , Neil Swainston, Ecaterina Vamos, Joanne Watts, Mark Whitehead |
| EPI_ISL_439863, EPI_ISL_439864, EPI_ISL_439865, EPI_ISL_439866, EPI_ISL_439867, EPI_ISL_439868, EPI_ISL_439869, EPI_ISL_439870, EPI_ISL_439871, EPI_ISL_439872, EPI_ISL_439873, EPI_ISL_439874, EPI_ISL_439875, EPI_ISL_439876, EPI_ISL_439877, EPI_ISL_439878, EPI_ISL_439879, EPI_ISL_439880, EPI_ISL_439881, EPI_ISL_439882, EPI_ISL_439883, EPI_ISL_439884, EPI_ISL_439885, EPI_ISL_439886, EPI_ISL_439887, EPI_ISL_439888, EPI_ISL_439889, EPI_ISL_439890, EPI_ISL_439891, EPI_ISL_439892, EPI_ISL_439893, EPI_ISL_439894, EPI_ISL_439895, EPI_ISL_439896, EPI_ISL_439897, EPI_ISL_439898, EPI_ISL_439899, EPI_ISL_439900, EPI_ISL_439901, EPI_ISL_439902, EPI_ISL_439903, EPI_ISL_439904, EPI_ISL_439905, EPI_ISL_439906, EPI_ISL_439907, EPI_ISL_439908, EPI_ISL_439909, EPI_ISL_439910, EPI_ISL_439911, EPI_ISL_439912, EPI_ISL_439913, EPI_ISL_439914, EPI_ISL_439915, EPI_ISL_439916, EPI_ISL_439917, EPI_ISL_439918, EPI_ISL_439919, EPI_ISL_439920, EPI_ISL_439921, EPI_ISL_439922, EPI_ISL_439923, EPI_ISL_439924, EPI_ISL_439925, EPI_ISL_439926, EPI_ISL_439927, EPI_ISL_439928, EPI_ISL_439929, EPI_ISL_439930, EPI_ISL_439931, EPI_ISL_439932, EPI_ISL_439933, EPI_ISL_439934, EPI_ISL_439935, EPI_ISL_439936, EPI_ISL_439937, EPI_ISL_439938, EPI_ISL_439939, EPI_ISL_439940, EPI_ISL_439941, EPI_ISL_439942, EPI_ISL_439943, EPI_ISL_439944, EPI_ISL_439945, EPI_ISL_439946, EPI_ISL_439947, EPI_ISL_439948, EPI_ISL_439949, EPI_ISL_439950, EPI_ISL_439951, EPI_ISL_439952 | | | |
| see above | Department of Pathology, University of Cambridge | Wellcome Sanger Institute for the COVID-19 Genomics UK Consortium | Luke W Meredith, M. Estée Török , Myra Hosmillo, William L. Hamilton, Martin D. Curran, Theresa Feltwell, Grant Hall, Anna Yakovleva, Fahad A Khokhar, Charlotte J. Houldcroft, Laura G Caller, Aminu S. Jahun, Sarah L. Caddy, Ian Goodfellow, Alex Alderton, Roberto Amato, Sonia Goncalves, Ewan Harrison, David K. Jackson, Ian Johnston, Dominic Kwiatkowski, Cordelia Langford, John Sillitoe on behalf of the Wellcome Sanger Institute COVID-19 Surveillance Team (http://www.sanger.ac.uk/covid-team) |
| EPI_ISL_439953 | PHE South West Regional Laboratory, National Infection Service | Wellcome Sanger Institute for the COVID-19 Genomics UK Consortium | Stephanie Hutchings, Hannah Pymont, Dr Peter Muir, Barry Vipond, Rich Hopes, Alex Alderton, Roberto Amato, Sonia Goncalves, Ewan Harrison, David K. Jackson, Ian Johnston, Dominic Kwiatkowski, Cordelia Langford, John Sillitoe on behalf of the Wellcome Sanger Institute COVID-19 Surveillance Team (http://www.sanger.ac.uk/covid-team) |
| EPI_ISL_439954, EPI_ISL_439955, EPI_ISL_439956 | Department of Pathology, University of Cambridge | Wellcome Sanger Institute for the COVID-19 Genomics UK Consortium | Luke W Meredith, M. Estée Török , Myra Hosmillo, William L. Hamilton, Martin D. Curran, Theresa Feltwell, Grant Hall, Anna Yakovleva, Fahad A Khokhar, Charlotte J. Houldcroft, Laura G Caller, Aminu S. Jahun, Sarah L. Caddy, Ian Goodfellow, Alex Alderton, Roberto Amato, Sonia Goncalves, Ewan Harrison, David K. Jackson, Ian Johnston, Dominic Kwiatkowski, Cordelia Langford, John Sillitoe on behalf of the Wellcome Sanger Institute COVID-19 Surveillance Team (http://www.sanger.ac.uk/covid-team) |
| EPI_ISL_439957 | PHE South West Regional Laboratory, National Infection Service | Wellcome Sanger Institute for the COVID-19 Genomics UK Consortium | Stephanie Hutchings, Hannah Pymont, Dr Peter Muir, Barry Vipond, Rich Hopes, Alex Alderton, Roberto Amato, Sonia Goncalves, Ewan Harrison, David K. Jackson, Ian Johnston, Dominic Kwiatkowski, Cordelia Langford, John Sillitoe on behalf of the Wellcome Sanger Institute COVID-19 Surveillance Team (http://www.sanger.ac.uk/covid-team) |
| EPI_ISL_439958 | Department of Pathology, University of Cambridge | Wellcome Sanger Institute for the COVID-19 Genomics UK Consortium | Luke W Meredith, M. Estée Török , Myra Hosmillo, William L. Hamilton, Martin D. Curran, Theresa Feltwell, Grant Hall, Anna Yakovleva, Fahad A Khokhar, Charlotte J. Houldcroft, Laura G Caller, Aminu S. Jahun, Sarah L. Caddy, Ian Goodfellow, Alex Alderton, Roberto Amato, Sonia Goncalves, Ewan Harrison, David K. Jackson, Ian Johnston, Dominic Kwiatkowski, Cordelia Langford, John Sillitoe on behalf of the Wellcome Sanger Institute COVID-19 Surveillance Team (http://www.sanger.ac.uk/covid-team) |
| EPI_ISL_439959, EPI_ISL_439960 | PHE South West Regional Laboratory, National Infection Service | Wellcome Sanger Institute for the COVID-19 Genomics UK Consortium | Stephanie Hutchings, Hannah Pymont, Dr Peter Muir, Barry Vipond, Rich Hopes, Alex Alderton, Roberto Amato, Sonia Goncalves, Ewan Harrison, David K. Jackson, Ian Johnston, Dominic Kwiatkowski, Cordelia Langford, John Sillitoe on behalf of the Wellcome Sanger Institute COVID-19 Surveillance Team (http://www.sanger.ac.uk/covid-team) |
| EPI_ISL_439961 | Department of Pathology, University of Cambridge | Wellcome Sanger Institute for the COVID-19 Genomics UK Consortium | Luke W Meredith, M. Estée Török , Myra Hosmillo, William L. Hamilton, Martin D. Curran, Theresa Feltwell, Grant Hall, Anna Yakovleva, Fahad A Khokhar, Charlotte J. Houldcroft, Laura G Caller, Aminu S. Jahun, Sarah L. Caddy, Ian Goodfellow, Alex Alderton, Roberto Amato, Sonia Goncalves, Ewan Harrison, David K. Jackson, Ian Johnston, Dominic Kwiatkowski, Cordelia Langford, John Sillitoe on behalf of the Wellcome Sanger Institute COVID-19 Surveillance Team (http://www.sanger.ac.uk/covid-team) |
| EPI_ISL_439962 | PHE South West Regional Laboratory, National Infection Service | Wellcome Sanger Institute for the COVID-19 Genomics UK Consortium | Stephanie Hutchings, Hannah Pymont, Dr Peter Muir, Barry Vipond, Rich Hopes, Alex Alderton, Roberto Amato, Sonia Goncalves, Ewan Harrison, David K. Jackson, Ian Johnston, Dominic Kwiatkowski, Cordelia Langford, John Sillitoe on behalf of the Wellcome Sanger Institute COVID-19 Surveillance Team (http://www.sanger.ac.uk/covid-team) |
| EPI_ISL_439963 | Department of Pathology, University of Cambridge | Wellcome Sanger Institute for the COVID-19 Genomics UK Consortium | Luke W Meredith, M. Estée Török , Myra Hosmillo, William L. Hamilton, Martin D. Curran, Theresa Feltwell, Grant Hall, Anna Yakovleva, Fahad A Khokhar, Charlotte J. Houldcroft, Laura G Caller, Aminu S. Jahun, Sarah L. Caddy, Ian Goodfellow, Alex Alderton, Roberto Amato, Sonia Goncalves, Ewan Harrison, David K. Jackson, Ian Johnston, Dominic Kwiatkowski, Cordelia Langford, John Sillitoe on behalf of the Wellcome Sanger Institute COVID-19 Surveillance Team (http://www.sanger.ac.uk/covid-team) |
| EPI_ISL_439964, EPI_ISL_439965, EPI_ISL_439966, EPI_ISL_439967 | PHE South West Regional Laboratory, National Infection Service | Wellcome Sanger Institute for the COVID-19 Genomics UK Consortium | Stephanie Hutchings, Hannah Pymont, Dr Peter Muir, Barry Vipond, Rich Hopes, Alex Alderton, Roberto Amato, Sonia Goncalves, Ewan Harrison, David K. Jackson, Ian Johnston, Dominic Kwiatkowski, Cordelia Langford, John Sillitoe on behalf of the Wellcome Sanger Institute COVID-19 Surveillance Team (http://www.sanger.ac.uk/covid-team) |
| EPI_ISL_439968 | Department of Pathology, University of Cambridge | Wellcome Sanger Institute for the COVID-19 Genomics UK Consortium | Luke W Meredith, M. Estée Török , Myra Hosmillo, William L. Hamilton, Martin D. Curran, Theresa Feltwell, Grant Hall, Anna Yakovleva, Fahad A Khokhar, Charlotte J. Houldcroft, Laura G Caller, Aminu S. Jahun, Sarah L. Caddy, Ian Goodfellow, Alex Alderton, Roberto Amato, Sonia Goncalves, Ewan Harrison, David K. Jackson, Ian Johnston, Dominic Kwiatkowski, Cordelia Langford, John Sillitoe on behalf of the Wellcome Sanger Institute COVID-19 Surveillance Team (http://www.sanger.ac.uk/covid-team) |
| EPI_ISL_439969 | PHE South West Regional Laboratory, National Infection Service | Wellcome Sanger Institute for the COVID-19 Genomics UK Consortium | Stephanie Hutchings, Hannah Pymont, Dr Peter Muir, Barry Vipond, Rich Hopes, Alex Alderton, Roberto Amato, Sonia Goncalves, Ewan Harrison, David K. Jackson, Ian Johnston, Dominic Kwiatkowski, Cordelia Langford, John Sillitoe on behalf of the Wellcome Sanger Institute COVID-19 Surveillance Team (http://www.sanger.ac.uk/covid-team) |
| EPI_ISL_439970 | Department of Pathology, University of Cambridge | Wellcome Sanger Institute for the COVID-19 Genomics UK Consortium | Luke W Meredith, M. Estée Török , Myra Hosmillo, William L. Hamilton, Martin D. Curran, Theresa Feltwell, Grant Hall, Anna Yakovleva, Fahad A Khokhar, Charlotte J. Houldcroft, Laura G Caller, Aminu S. Jahun, Sarah L. Caddy, Ian Goodfellow, Alex Alderton, Roberto Amato, Sonia Goncalves, Ewan Harrison, David K. Jackson, Ian Johnston, Dominic Kwiatkowski, Cordelia Langford, John Sillitoe on behalf of the Wellcome Sanger Institute COVID-19 Surveillance Team (http://www.sanger.ac.uk/covid-team) |
| EPI_ISL_439971, EPI_ISL_439972, EPI_ISL_439973, EPI_ISL_439974 | PHE South West Regional Laboratory, National Infection Service | Wellcome Sanger Institute for the COVID-19 Genomics UK Consortium | Stephanie Hutchings, Hannah Pymont, Dr Peter Muir, Barry Vipond, Rich Hopes, Alex Alderton, Roberto Amato, Sonia Goncalves, Ewan Harrison, David K. Jackson, Ian Johnston, Dominic Kwiatkowski, Cordelia Langford, John Sillitoe on behalf of the Wellcome Sanger Institute COVID-19 Surveillance Team (http://www.sanger.ac.uk/covid-team) |
| EPI_ISL_439975 | Department of Pathology, University of Cambridge | Wellcome Sanger Institute for the COVID-19 Genomics UK Consortium | Luke W Meredith, M. Estée Török , Myra Hosmillo, William L. Hamilton, Martin D. Curran, Theresa Feltwell, Grant Hall, Anna Yakovleva, Fahad A Khokhar, Charlotte J. Houldcroft, Laura G Caller, Aminu S. Jahun, Sarah L. Caddy, Ian Goodfellow, Alex Alderton, Roberto Amato, Sonia Goncalves, Ewan Harrison, David K. Jackson, Ian Johnston, Dominic Kwiatkowski, Cordelia Langford, John Sillitoe on behalf of the Wellcome Sanger Institute COVID-19 Surveillance Team (http://www.sanger.ac.uk/covid-team) |
| EPI_ISL_439976 | PHE South West Regional Laboratory, National Infection Service | Wellcome Sanger Institute for the COVID-19 Genomics UK Consortium | Stephanie Hutchings, Hannah Pymont, Dr Peter Muir, Barry Vipond, Rich Hopes, Alex Alderton, Roberto Amato, Sonia Goncalves, Ewan Harrison, David K. Jackson, Ian Johnston, Dominic Kwiatkowski, Cordelia Langford, John Sillitoe on behalf of the Wellcome Sanger Institute COVID-19 Surveillance Team (http://www.sanger.ac.uk/covid-team) |
| EPI_ISL_439977 | Department of Pathology, University of Cambridge | Wellcome Sanger Institute for the COVID-19 Genomics UK Consortium | Luke W Meredith, M. Estée Török , Myra Hosmillo, William L. Hamilton, Martin D. Curran, Theresa Feltwell, Grant Hall, Anna Yakovleva, Fahad A Khokhar, Charlotte J. Houldcroft, Laura G Caller, Aminu S. Jahun, Sarah L. Caddy, Ian Goodfellow, Alex Alderton, Roberto Amato, Sonia Goncalves, Ewan Harrison, David K. Jackson, Ian Johnston, Dominic Kwiatkowski, Cordelia Langford, John Sillitoe on behalf of the Wellcome Sanger Institute COVID-19 Surveillance Team (http://www.sanger.ac.uk/covid-team) |
| EPI_ISL_439978, EPI_ISL_439979 | PHE South West Regional Laboratory, National Infection | Wellcome Sanger Institute for the COVID-19 Genomics UK Consortium | Stephanie Hutchings, Hannah Pymont, Dr Peter Muir, Barry Vipond, Rich Hopes, Alex Alderton, Roberto Amato, Sonia Goncalves, Ewan Harrison, David K. Jackson, Ian Johnston, Dominic Kwiatkowski, Cordelia |

| | | | |
|---|---|---|---|
| | Service | | Langford, John Sillitoe on behalf of the Wellcome Sanger Institute COVID-19 Surveillance Team (http://www.sanger.ac.uk/covid-team) |
| EPI_ISL_439980 | Department of Pathology, University of Cambridge | Wellcome Sanger Institute for the COVID-19 Genomics UK Consortium | Luke W Meredith, M. Estée Török , Myra Hosmillo, William L. Hamilton, Martin D. Curran, Theresa Feltwell, Grant Hall, Anna Yakovleva, Fahad A Khokhar, Charlotte J. Houldcroft, Laura G Caller, Aminu S. Jahun, Sarah L. Caddy, Ian Goodfellow, Alex Alderton, Roberto Amato, Sonia Goncalves, Ewan Harrison, David K. Jackson, Ian Johnston, Dominic Kwiatkowski, Cordelia Langford, John Sillitoe on behalf of the Wellcome Sanger Institute COVID-19 Surveillance Team (http://www.sanger.ac.uk/covid-team) |
| EPI_ISL_439981, EPI_ISL_439982, EPI_ISL_439983, EPI_ISL_439984, EPI_ISL_439985, EPI_ISL_439986 | PHE South West Regional Laboratory, National Infection Service | Wellcome Sanger Institute for the COVID-19 Genomics UK Consortium | Stephanie Hutchings, Hannah Pymont, Dr Peter Muir, Barry Vipond, Rich Hopes, Alex Alderton, Roberto Amato, Sonia Goncalves, Ewan Harrison, David K. Jackson, Ian Johnston, Dominic Kwiatkowski, Cordelia Langford, John Sillitoe on behalf of the Wellcome Sanger Institute COVID-19 Surveillance Team (http://www.sanger.ac.uk/covid-team) |
| EPI_ISL_439987 | Department of Pathology, University of Cambridge | Wellcome Sanger Institute for the COVID-19 Genomics UK Consortium | Luke W Meredith, M. Estée Török , Myra Hosmillo, William L. Hamilton, Martin D. Curran, Theresa Feltwell, Grant Hall, Anna Yakovleva, Fahad A Khokhar, Charlotte J. Houldcroft, Laura G Caller, Aminu S. Jahun, Sarah L. Caddy, Ian Goodfellow, Alex Alderton, Roberto Amato, Sonia Goncalves, Ewan Harrison, David K. Jackson, Ian Johnston, Dominic Kwiatkowski, Cordelia Langford, John Sillitoe on behalf of the Wellcome Sanger Institute COVID-19 Surveillance Team (http://www.sanger.ac.uk/covid-team) |
| EPI_ISL_439988 | PHE South West Regional Laboratory, National Infection Service | Wellcome Sanger Institute for the COVID-19 Genomics UK Consortium | Stephanie Hutchings, Hannah Pymont, Dr Peter Muir, Barry Vipond, Rich Hopes, Alex Alderton, Roberto Amato, Sonia Goncalves, Ewan Harrison, David K. Jackson, Ian Johnston, Dominic Kwiatkowski, Cordelia Langford, John Sillitoe on behalf of the Wellcome Sanger Institute COVID-19 Surveillance Team (http://www.sanger.ac.uk/covid-team) |
| EPI_ISL_439989, EPI_ISL_439990, EPI_ISL_439991 | Department of Pathology, University of Cambridge | Wellcome Sanger Institute for the COVID-19 Genomics UK Consortium | Luke W Meredith, M. Estée Török , Myra Hosmillo, William L. Hamilton, Martin D. Curran, Theresa Feltwell, Grant Hall, Anna Yakovleva, Fahad A Khokhar, Charlotte J. Houldcroft, Laura G Caller, Aminu S. Jahun, Sarah L. Caddy, Ian Goodfellow, Alex Alderton, Roberto Amato, Sonia Goncalves, Ewan Harrison, David K. Jackson, Ian Johnston, Dominic Kwiatkowski, Cordelia Langford, John Sillitoe on behalf of the Wellcome Sanger Institute COVID-19 Surveillance Team (http://www.sanger.ac.uk/covid-team) |
| EPI_ISL_439992 | PHE South West Regional Laboratory, National Infection Service | Wellcome Sanger Institute for the COVID-19 Genomics UK Consortium | Stephanie Hutchings, Hannah Pymont, Dr Peter Muir, Barry Vipond, Rich Hopes, Alex Alderton, Roberto Amato, Sonia Goncalves, Ewan Harrison, David K. Jackson, Ian Johnston, Dominic Kwiatkowski, Cordelia Langford, John Sillitoe on behalf of the Wellcome Sanger Institute COVID-19 Surveillance Team (http://www.sanger.ac.uk/covid-team) |
| EPI_ISL_439993, EPI_ISL_439994, EPI_ISL_439995, EPI_ISL_439996, EPI_ISL_439997 | Department of Pathology, University of Cambridge | Wellcome Sanger Institute for the COVID-19 Genomics UK Consortium | Luke W Meredith, M. Estée Török , Myra Hosmillo, William L. Hamilton, Martin D. Curran, Theresa Feltwell, Grant Hall, Anna Yakovleva, Fahad A Khokhar, Charlotte J. Houldcroft, Laura G Caller, Aminu S. Jahun, Sarah L. Caddy, Ian Goodfellow, Alex Alderton, Roberto Amato, Sonia Goncalves, Ewan Harrison, David K. Jackson, Ian Johnston, Dominic Kwiatkowski, Cordelia Langford, John Sillitoe on behalf of the Wellcome Sanger Institute COVID-19 Surveillance Team (http://www.sanger.ac.uk/covid-team) |
| EPI_ISL_439998 | PHE South West Regional Laboratory, National Infection Service | Wellcome Sanger Institute for the COVID-19 Genomics UK Consortium | Stephanie Hutchings, Hannah Pymont, Dr Peter Muir, Barry Vipond, Rich Hopes, Alex Alderton, Roberto Amato, Sonia Goncalves, Ewan Harrison, David K. Jackson, Ian Johnston, Dominic Kwiatkowski, Cordelia Langford, John Sillitoe on behalf of the Wellcome Sanger Institute COVID-19 Surveillance Team (http://www.sanger.ac.uk/covid-team) |
| EPI_ISL_439999 | Department of Pathology, University of Cambridge | Wellcome Sanger Institute for the COVID-19 Genomics UK Consortium | Luke W Meredith, M. Estée Török , Myra Hosmillo, William L. Hamilton, Martin D. Curran, Theresa Feltwell, Grant Hall, Anna Yakovleva, Fahad A Khokhar, Charlotte J. Houldcroft, Laura G Caller, Aminu S. Jahun, Sarah L. Caddy, Ian Goodfellow, Alex Alderton, Roberto Amato, Sonia Goncalves, Ewan Harrison, David K. Jackson, Ian Johnston, Dominic Kwiatkowski, Cordelia Langford, John Sillitoe on behalf of the Wellcome Sanger Institute COVID-19 Surveillance Team (http://www.sanger.ac.uk/covid-team) |
| EPI_ISL_440000, EPI_ISL_440001, EPI_ISL_440002 | PHE South West Regional Laboratory, National Infection Service | Wellcome Sanger Institute for the COVID-19 Genomics UK Consortium | Stephanie Hutchings, Hannah Pymont, Dr Peter Muir, Barry Vipond, Rich Hopes, Alex Alderton, Roberto Amato, Sonia Goncalves, Ewan Harrison, David K. Jackson, Ian Johnston, Dominic Kwiatkowski, Cordelia Langford, John Sillitoe on behalf of the Wellcome Sanger Institute COVID-19 Surveillance Team (http://www.sanger.ac.uk/covid-team) |
| EPI_ISL_440003, EPI_ISL_440004 | Department of Pathology, University of Cambridge | Wellcome Sanger Institute for the COVID-19 Genomics UK Consortium | Luke W Meredith, M. Estée Török , Myra Hosmillo, William L. Hamilton, Martin D. Curran, Theresa Feltwell, Grant Hall, Anna Yakovleva, Fahad A Khokhar, Charlotte J. Houldcroft, Laura G Caller, Aminu S. Jahun, Sarah L. Caddy, Ian Goodfellow, Alex Alderton, Roberto Amato, Sonia Goncalves, Ewan Harrison, David K. Jackson, Ian Johnston, Dominic Kwiatkowski, Cordelia Langford, John Sillitoe on behalf of the Wellcome Sanger Institute COVID-19 Surveillance Team (http://www.sanger.ac.uk/covid-team) |
| EPI_ISL_440005, EPI_ISL_440006 | PHE South West Regional Laboratory, National Infection Service | Wellcome Sanger Institute for the COVID-19 Genomics UK Consortium | Stephanie Hutchings, Hannah Pymont, Dr Peter Muir, Barry Vipond, Rich Hopes, Alex Alderton, Roberto Amato, Sonia Goncalves, Ewan Harrison, David K. Jackson, Ian Johnston, Dominic Kwiatkowski, Cordelia Langford, John Sillitoe on behalf of the Wellcome Sanger Institute COVID-19 Surveillance Team (http://www.sanger.ac.uk/covid-team) |
| EPI_ISL_440007 | Department of Pathology, University of Cambridge | Wellcome Sanger Institute for the COVID-19 Genomics UK Consortium | Luke W Meredith, M. Estée Török , Myra Hosmillo, William L. Hamilton, Martin D. Curran, Theresa Feltwell, Grant Hall, Anna Yakovleva, Fahad A Khokhar, Charlotte J. Houldcroft, Laura G Caller, Aminu S. Jahun, Sarah L. Caddy, Ian Goodfellow, Alex Alderton, Roberto Amato, Sonia Goncalves, Ewan Harrison, David K. Jackson, Ian Johnston, Dominic Kwiatkowski, Cordelia Langford, John Sillitoe on behalf of the Wellcome Sanger Institute COVID-19 Surveillance Team (http://www.sanger.ac.uk/covid-team) |
| EPI_ISL_440008 | PHE South West Regional Laboratory, National Infection Service | Wellcome Sanger Institute for the COVID-19 Genomics UK Consortium | Stephanie Hutchings, Hannah Pymont, Dr Peter Muir, Barry Vipond, Rich Hopes, Alex Alderton, Roberto Amato, Sonia Goncalves, Ewan Harrison, David K. Jackson, Ian Johnston, Dominic Kwiatkowski, Cordelia Langford, John Sillitoe on behalf of the Wellcome Sanger Institute COVID-19 Surveillance Team (http://www.sanger.ac.uk/covid-team) |
| EPI_ISL_440009, EPI_ISL_440010, EPI_ISL_440011, EPI_ISL_440012 | Department of Pathology, University of Cambridge | Wellcome Sanger Institute for the COVID-19 Genomics UK Consortium | Luke W Meredith, M. Estée Török , Myra Hosmillo, William L. Hamilton, Martin D. Curran, Theresa Feltwell, Grant Hall, Anna Yakovleva, Fahad A Khokhar, Charlotte J. Houldcroft, Laura G Caller, Aminu S. Jahun, Sarah L. Caddy, Ian Goodfellow, Alex Alderton, Roberto Amato, Sonia Goncalves, Ewan Harrison, David K. Jackson, Ian Johnston, Dominic Kwiatkowski, Cordelia Langford, John Sillitoe on behalf of the Wellcome Sanger Institute COVID-19 Surveillance Team (http://www.sanger.ac.uk/covid-team) |
| EPI_ISL_440013 | PHE South West Regional Laboratory, National Infection Service | Wellcome Sanger Institute for the COVID-19 Genomics UK Consortium | Stephanie Hutchings, Hannah Pymont, Dr Peter Muir, Barry Vipond, Rich Hopes, Alex Alderton, Roberto Amato, Sonia Goncalves, Ewan Harrison, David K. Jackson, Ian Johnston, Dominic Kwiatkowski, Cordelia Langford, John Sillitoe on behalf of the Wellcome Sanger Institute COVID-19 Surveillance Team (http://www.sanger.ac.uk/covid-team) |
| EPI_ISL_440014, EPI_ISL_440015 | Department of Pathology, University of Cambridge | Wellcome Sanger Institute for the COVID-19 Genomics UK Consortium | Luke W Meredith, M. Estée Török , Myra Hosmillo, William L. Hamilton, Martin D. Curran, Theresa Feltwell, Grant Hall, Anna Yakovleva, Fahad A Khokhar, Charlotte J. Houldcroft, Laura G Caller, Aminu S. Jahun, Sarah L. Caddy, Ian Goodfellow, Alex Alderton, Roberto Amato, Sonia Goncalves, Ewan Harrison, David K. Jackson, Ian Johnston, Dominic Kwiatkowski, Cordelia Langford, John Sillitoe on behalf of the Wellcome Sanger Institute COVID-19 Surveillance Team (http://www.sanger.ac.uk/covid-team) |
| EPI_ISL_440016 | PHE South West Regional Laboratory, National Infection Service | Wellcome Sanger Institute for the COVID-19 Genomics UK Consortium | Stephanie Hutchings, Hannah Pymont, Dr Peter Muir, Barry Vipond, Rich Hopes, Alex Alderton, Roberto Amato, Sonia Goncalves, Ewan Harrison, David K. Jackson, Ian Johnston, Dominic Kwiatkowski, Cordelia Langford, John Sillitoe on behalf of the Wellcome Sanger Institute COVID-19 Surveillance Team (http://www.sanger.ac.uk/covid-team) |
| EPI_ISL_440017 | Department of Pathology, University of Cambridge | Wellcome Sanger Institute for the COVID-19 Genomics UK Consortium | Luke W Meredith, M. Estée Török , Myra Hosmillo, William L. Hamilton, Martin D. Curran, Theresa Feltwell, Grant Hall, Anna Yakovleva, Fahad A Khokhar, Charlotte J. Houldcroft, Laura G Caller, Aminu S. Jahun, Sarah L. Caddy, Ian Goodfellow, Alex Alderton, Roberto Amato, Sonia Goncalves, Ewan Harrison, David K. Jackson, Ian Johnston, Dominic Kwiatkowski, Cordelia Langford, John Sillitoe on behalf of the Wellcome Sanger Institute COVID-19 Surveillance Team (http://www.sanger.ac.uk/covid-team) |
| EPI_ISL_440018 | PHE South West Regional Laboratory, National Infection Service | Wellcome Sanger Institute for the COVID-19 Genomics UK Consortium | Stephanie Hutchings, Hannah Pymont, Dr Peter Muir, Barry Vipond, Rich Hopes, Alex Alderton, Roberto Amato, Sonia Goncalves, Ewan Harrison, David K. Jackson, Ian Johnston, Dominic Kwiatkowski, Cordelia Langford, John Sillitoe on behalf of the Wellcome Sanger Institute COVID-19 Surveillance Team (http://www.sanger.ac.uk/covid-team) |
| EPI_ISL_440019 | Department of Pathology, University of Cambridge | Wellcome Sanger Institute for the COVID-19 Genomics UK Consortium | Luke W Meredith, M. Estée Török , Myra Hosmillo, William L. Hamilton, Martin D. Curran, Theresa Feltwell, Grant Hall, Anna Yakovleva, Fahad A Khokhar, Charlotte J. Houldcroft, Laura G Caller, Aminu S. Jahun, Sarah L. Caddy, Ian Goodfellow, Alex Alderton, Roberto Amato, Sonia Goncalves, Ewan Harrison, David K. Jackson, Ian Johnston, Dominic Kwiatkowski, Cordelia Langford, John Sillitoe on behalf of the Wellcome Sanger Institute COVID-19 Surveillance Team (http://www.sanger.ac.uk/covid-team) |
| EPI_ISL_440020, EPI_ISL_440021, EPI_ISL_440022, EPI_ISL_440023 | PHE South West Regional Laboratory, National Infection Service | Wellcome Sanger Institute for the COVID-19 Genomics UK Consortium | Stephanie Hutchings, Hannah Pymont, Dr Peter Muir, Barry Vipond, Rich Hopes, Alex Alderton, Roberto Amato, Sonia Goncalves, Ewan Harrison, David K. Jackson, Ian Johnston, Dominic Kwiatkowski, Cordelia Langford, John Sillitoe on behalf of the Wellcome Sanger Institute COVID-19 Surveillance Team (http://www.sanger.ac.uk/covid-team) |
| EPI_ISL_440024, EPI_ISL_440025 | Department of Pathology, University of Cambridge | Wellcome Sanger Institute for the COVID-19 Genomics UK Consortium | Luke W Meredith, M. Estée Török , Myra Hosmillo, William L. Hamilton, Martin D. Curran, Theresa Feltwell, Grant Hall, Anna Yakovleva, Fahad A Khokhar, Charlotte J. Houldcroft, Laura G Caller, Aminu S. Jahun, Sarah L. Caddy, Ian Goodfellow, Alex Alderton, Roberto Amato, Sonia Goncalves, Ewan Harrison, David K. Jackson, Ian Johnston, Dominic Kwiatkowski, Cordelia Langford, John Sillitoe on behalf of the Wellcome Sanger Institute COVID-19 Surveillance Team (http://www.sanger.ac.uk/covid-team) |
| EPI_ISL_440026, EPI_ISL_440027, EPI_ISL_440028, EPI_ISL_440029 | PHE South West Regional Laboratory, National Infection Service | Wellcome Sanger Institute for the COVID-19 Genomics UK Consortium | Stephanie Hutchings, Hannah Pymont, Dr Peter Muir, Barry Vipond, Rich Hopes, Alex Alderton, Roberto Amato, Sonia Goncalves, Ewan Harrison, David K. Jackson, Ian Johnston, Dominic Kwiatkowski, Cordelia Langford, John Sillitoe on behalf of the Wellcome Sanger Institute COVID-19 Surveillance Team (http://www.sanger.ac.uk/covid-team) |
| EPI_ISL_440030 | Department of Pathology, University of Cambridge | Wellcome Sanger Institute for the COVID-19 Genomics UK Consortium | Luke W Meredith, M. Estée Török , Myra Hosmillo, William L. Hamilton, Martin D. Curran, Theresa Feltwell, Grant Hall, Anna Yakovleva, Fahad A Khokhar, Charlotte J. Houldcroft, Laura G Caller, Aminu S. Jahun, Sarah L. Caddy, Ian Goodfellow, Alex Alderton, Roberto Amato, Sonia Goncalves, Ewan Harrison, David K. Jackson, Ian Johnston, Dominic Kwiatkowski, Cordelia Langford, John Sillitoe on behalf of the Wellcome Sanger Institute COVID-19 Surveillance Team (http://www.sanger.ac.uk/covid-team) |
| EPI_ISL_440031, EPI_ISL_440032, EPI_ISL_440033 | PHE South West Regional Laboratory, National Infection Service | Wellcome Sanger Institute for the COVID-19 Genomics UK Consortium | Stephanie Hutchings, Hannah Pymont, Dr Peter Muir, Barry Vipond, Rich Hopes, Alex Alderton, Roberto Amato, Sonia Goncalves, Ewan Harrison, David K. Jackson, Ian Johnston, Dominic Kwiatkowski, Cordelia Langford, John Sillitoe on behalf of the Wellcome Sanger Institute COVID-19 Surveillance Team (http://www.sanger.ac.uk/covid-team) |
| EPI_ISL_440034 | Department of Pathology, University of Cambridge | Wellcome Sanger Institute for the COVID-19 Genomics UK Consortium | Luke W Meredith, M. Estée Török , Myra Hosmillo, William L. Hamilton, Martin D. Curran, Theresa Feltwell, Grant Hall, Anna Yakovleva, Fahad A Khokhar, Charlotte J. Houldcroft, Laura G Caller, Aminu S. Jahun, Sarah L. Caddy, Ian Goodfellow, Alex Alderton, Roberto Amato, Sonia Goncalves, Ewan Harrison, David K. Jackson, Ian Johnston, Dominic Kwiatkowski, Cordelia Langford, John Sillitoe on behalf of the Wellcome Sanger Institute COVID-19 Surveillance Team (http://www.sanger.ac.uk/covid-team) |
| EPI_ISL_440035, EPI_ISL_440036, EPI_ISL_440037, EPI_ISL_440038, EPI_ISL_440039 | PHE South West Regional Laboratory, National Infection Service | Wellcome Sanger Institute for the COVID-19 Genomics UK Consortium | Stephanie Hutchings, Hannah Pymont, Dr Peter Muir, Barry Vipond, Rich Hopes, Alex Alderton, Roberto Amato, Sonia Goncalves, Ewan Harrison, David K. Jackson, Ian Johnston, Dominic Kwiatkowski, Cordelia Langford, John Sillitoe on behalf of the Wellcome Sanger Institute COVID-19 Surveillance Team (http://www.sanger.ac.uk/covid-team) |
| EPI_ISL_440040, EPI_ISL_440041 | Department of Pathology, University of Cambridge | Wellcome Sanger Institute for the COVID-19 Genomics UK Consortium | Luke W Meredith, M. Estée Török , Myra Hosmillo, William L. Hamilton, Martin D. Curran, Theresa Feltwell, Grant Hall, Anna Yakovleva, Fahad A Khokhar, Charlotte J. Houldcroft, Laura G Caller, Aminu S. Jahun, Sarah L. Caddy, Ian Goodfellow, Alex Alderton, Roberto Amato, Sonia Goncalves, Ewan Harrison, David K. Jackson, Ian Johnston, Dominic Kwiatkowski, Cordelia Langford, John Sillitoe on behalf of the Wellcome Sanger Institute COVID-19 Surveillance Team (http://www.sanger.ac.uk/covid-team) |
| EPI_ISL_440042, EPI_ISL_440043 | PHE South West Regional Laboratory, National Infection Service | Wellcome Sanger Institute for the COVID-19 Genomics UK Consortium | Stephanie Hutchings, Hannah Pymont, Dr Peter Muir, Barry Vipond, Rich Hopes, Alex Alderton, Roberto Amato, Sonia Goncalves, Ewan Harrison, David K. Jackson, Ian Johnston, Dominic Kwiatkowski, Cordelia Langford, John Sillitoe on behalf of the Wellcome Sanger Institute COVID-19 Surveillance Team (http://www.sanger.ac.uk/covid-team) |
| EPI_ISL_440044, EPI_ISL_440045, EPI_ISL_440046, EPI_ISL_440047, EPI_ISL_440048, EPI_ISL_440049 | Department of Pathology, University of Cambridge | Wellcome Sanger Institute for the COVID-19 Genomics UK Consortium | Luke W Meredith, M. Estée Török , Myra Hosmillo, William L. Hamilton, Martin D. Curran, Theresa Feltwell, Grant Hall, Anna Yakovleva, Fahad A Khokhar, Charlotte J. Houldcroft, Laura G Caller, Aminu S. Jahun, Sarah L. Caddy, Ian Goodfellow, Alex Alderton, Roberto Amato, Sonia Goncalves, Ewan Harrison, David K. Jackson, Ian Johnston, Dominic Kwiatkowski, Cordelia Langford, John Sillitoe on behalf of the Wellcome Sanger Institute COVID-19 Surveillance Team (http://www.sanger.ac.uk/covid-team) |
| EPI_ISL_440050, EPI_ISL_440051, EPI_ISL_440052 | PHE South West Regional Laboratory, National Infection Service | Wellcome Sanger Institute for the COVID-19 Genomics UK Consortium | Stephanie Hutchings, Hannah Pymont, Dr Peter Muir, Barry Vipond, Rich Hopes, Alex Alderton, Roberto Amato, Sonia Goncalves, Ewan Harrison, David K. Jackson, Ian Johnston, Dominic Kwiatkowski, Cordelia Langford, John Sillitoe on behalf of the Wellcome Sanger Institute COVID-19 Surveillance Team (http://www.sanger.ac.uk/covid-team) |
| EPI_ISL_440053 | Department of Pathology, University of Cambridge | Wellcome Sanger Institute for the COVID-19 Genomics UK Consortium | Luke W Meredith, M. Estée Török , Myra Hosmillo, William L. Hamilton, Martin D. Curran, Theresa Feltwell, Grant Hall, Anna Yakovleva, Fahad A Khokhar, Charlotte J. Houldcroft, Laura G Caller, Aminu S. Jahun, Sarah L. Caddy, Ian Goodfellow, Alex Alderton, Roberto Amato, Sonia Goncalves, Ewan Harrison, David K. Jackson, Ian Johnston, Dominic Kwiatkowski, Cordelia Langford, John Sillitoe on behalf of the Wellcome Sanger Institute COVID-19 Surveillance Team (http://www.sanger.ac.uk/covid-team) |
| EPI_ISL_440054 | PHE South West Regional Laboratory, National Infection Service | Wellcome Sanger Institute for the COVID-19 Genomics UK Consortium | Stephanie Hutchings, Hannah Pymont, Dr Peter Muir, Barry Vipond, Rich Hopes, Alex Alderton, Roberto Amato, Sonia Goncalves, Ewan Harrison, David K. Jackson, Ian Johnston, Dominic Kwiatkowski, Cordelia Langford, John Sillitoe on behalf of the Wellcome Sanger Institute COVID-19 Surveillance Team (http://www.sanger.ac.uk/covid-team) |
| EPI_ISL_440055 | Department of Pathology, University of Cambridge | Wellcome Sanger Institute for the COVID-19 Genomics UK Consortium | Luke W Meredith, M. Estée Török , Myra Hosmillo, William L. Hamilton, Martin D. Curran, Theresa Feltwell, Grant Hall, Anna Yakovleva, Fahad A Khokhar, Charlotte J. Houldcroft, Laura G Caller, Aminu S. Jahun, Sarah L. Caddy, Ian Goodfellow, Alex Alderton, Roberto Amato, Sonia Goncalves, Ewan Harrison, David K. Jackson, Ian Johnston, Dominic Kwiatkowski, Cordelia Langford, John Sillitoe on behalf of the Wellcome Sanger Institute COVID-19 Surveillance Team (http://www.sanger.ac.uk/covid-team) |
| EPI_ISL_440056 | PHE South West Regional Laboratory, National Infection Service | Wellcome Sanger Institute for the COVID-19 Genomics UK Consortium | Stephanie Hutchings, Hannah Pymont, Dr Peter Muir, Barry Vipond, Rich Hopes, Alex Alderton, Roberto Amato, Sonia Goncalves, Ewan Harrison, David K. Jackson, Ian Johnston, Dominic Kwiatkowski, Cordelia Langford, John Sillitoe on behalf of the Wellcome Sanger Institute COVID-19 Surveillance Team (http://www.sanger.ac.uk/covid-team) |
| EPI_ISL_440057 | Department of Pathology, University of Cambridge | Wellcome Sanger Institute for the COVID-19 Genomics UK Consortium | Luke W Meredith, M. Estée Török , Myra Hosmillo, William L. Hamilton, Martin D. Curran, Theresa Feltwell, Grant Hall, Anna Yakovleva, Fahad A Khokhar, Charlotte J. Houldcroft, Laura G Caller, Aminu S. Jahun, Sarah L. Caddy, Ian Goodfellow, Alex Alderton, Roberto Amato, Sonia Goncalves, Ewan Harrison, David K. Jackson, Ian Johnston, Dominic Kwiatkowski, Cordelia Langford, John Sillitoe on behalf of the Wellcome Sanger Institute COVID-19 Surveillance Team (http://www.sanger.ac.uk/covid-team) |

| | | | |
|---|---|---|---|
| EPI_ISL_440058 | PHE South West Regional Laboratory, National Infection Service | Wellcome Sanger Institute for the COVID-19 Genomics UK Consortium | Stephanie Hutchings, Hannah Pymont, Dr Peter Muir, Barry Vipond, Rich Hopes, Alex Alderton, Roberto Amato, Sonia Goncalves, Ewan Harrison, David K. Jackson, Ian Johnston, Dominic Kwiatkowski, Cordelia Langford, John Sillitoe on behalf of the Wellcome Sanger Institute COVID-19 Surveillance Team (http://www.sanger.ac.uk/covid-team) |
| EPI_ISL_440059, EPI_ISL_440060, EPI_ISL_440061, EPI_ISL_440062 | Department of Pathology, University of Cambridge | Wellcome Sanger Institute for the COVID-19 Genomics UK Consortium | Luke W Meredith, M. Estée Török , Myra Hosmillo, William L. Hamilton, Martin D. Curran, Theresa Feltwell, Grant Hall, Anna Yakovleva, Fahad A Khokhar, Charlotte J. Houldcroft, Laura G Caller, Aminu S. Jahun, Sarah L. Caddy, Ian Goodfellow, Alex Alderton, Roberto Amato, Sonia Goncalves, Ewan Harrison, David K. Jackson, Ian Johnston, Dominic Kwiatkowski, Cordelia Langford, John Sillitoe on behalf of the Wellcome Sanger Institute COVID-19 Surveillance Team (http://www.sanger.ac.uk/covid-team) |
| EPI_ISL_440063 | PHE South West Regional Laboratory, National Infection Service | Wellcome Sanger Institute for the COVID-19 Genomics UK Consortium | Stephanie Hutchings, Hannah Pymont, Dr Peter Muir, Barry Vipond, Rich Hopes, Alex Alderton, Roberto Amato, Sonia Goncalves, Ewan Harrison, David K. Jackson, Ian Johnston, Dominic Kwiatkowski, Cordelia Langford, John Sillitoe on behalf of the Wellcome Sanger Institute COVID-19 Surveillance Team (http://www.sanger.ac.uk/covid-team) |
| EPI_ISL_440064 | Department of Pathology, University of Cambridge | Wellcome Sanger Institute for the COVID-19 Genomics UK Consortium | Luke W Meredith, M. Estée Török , Myra Hosmillo, William L. Hamilton, Martin D. Curran, Theresa Feltwell, Grant Hall, Anna Yakovleva, Fahad A Khokhar, Charlotte J. Houldcroft, Laura G Caller, Aminu S. Jahun, Sarah L. Caddy, Ian Goodfellow, Alex Alderton, Roberto Amato, Sonia Goncalves, Ewan Harrison, David K. Jackson, Ian Johnston, Dominic Kwiatkowski, Cordelia Langford, John Sillitoe on behalf of the Wellcome Sanger Institute COVID-19 Surveillance Team (http://www.sanger.ac.uk/covid-team) |
| EPI_ISL_440065 | PHE South West Regional Laboratory, National Infection Service | Wellcome Sanger Institute for the COVID-19 Genomics UK Consortium | Stephanie Hutchings, Hannah Pymont, Dr Peter Muir, Barry Vipond, Rich Hopes, Alex Alderton, Roberto Amato, Sonia Goncalves, Ewan Harrison, David K. Jackson, Ian Johnston, Dominic Kwiatkowski, Cordelia Langford, John Sillitoe on behalf of the Wellcome Sanger Institute COVID-19 Surveillance Team (http://www.sanger.ac.uk/covid-team) |
| EPI_ISL_440066, EPI_ISL_440067 | Department of Pathology, University of Cambridge | Wellcome Sanger Institute for the COVID-19 Genomics UK Consortium | Luke W Meredith, M. Estée Török , Myra Hosmillo, William L. Hamilton, Martin D. Curran, Theresa Feltwell, Grant Hall, Anna Yakovleva, Fahad A Khokhar, Charlotte J. Houldcroft, Laura G Caller, Aminu S. Jahun, Sarah L. Caddy, Ian Goodfellow, Alex Alderton, Roberto Amato, Sonia Goncalves, Ewan Harrison, David K. Jackson, Ian Johnston, Dominic Kwiatkowski, Cordelia Langford, John Sillitoe on behalf of the Wellcome Sanger Institute COVID-19 Surveillance Team (http://www.sanger.ac.uk/covid-team) |
| EPI_ISL_440068 | PHE South West Regional Laboratory, National Infection Service | Wellcome Sanger Institute for the COVID-19 Genomics UK Consortium | Stephanie Hutchings, Hannah Pymont, Dr Peter Muir, Barry Vipond, Rich Hopes, Alex Alderton, Roberto Amato, Sonia Goncalves, Ewan Harrison, David K. Jackson, Ian Johnston, Dominic Kwiatkowski, Cordelia Langford, John Sillitoe on behalf of the Wellcome Sanger Institute COVID-19 Surveillance Team (http://www.sanger.ac.uk/covid-team) |
| EPI_ISL_440069 | Department of Pathology, University of Cambridge | Wellcome Sanger Institute for the COVID-19 Genomics UK Consortium | Luke W Meredith, M. Estée Török , Myra Hosmillo, William L. Hamilton, Martin D. Curran, Theresa Feltwell, Grant Hall, Anna Yakovleva, Fahad A Khokhar, Charlotte J. Houldcroft, Laura G Caller, Aminu S. Jahun, Sarah L. Caddy, Ian Goodfellow, Alex Alderton, Roberto Amato, Sonia Goncalves, Ewan Harrison, David K. Jackson, Ian Johnston, Dominic Kwiatkowski, Cordelia Langford, John Sillitoe on behalf of the Wellcome Sanger Institute COVID-19 Surveillance Team (http://www.sanger.ac.uk/covid-team) |
| EPI_ISL_440070, EPI_ISL_440071 | PHE South West Regional Laboratory, National Infection Service | Wellcome Sanger Institute for the COVID-19 Genomics UK Consortium | Stephanie Hutchings, Hannah Pymont, Dr Peter Muir, Barry Vipond, Rich Hopes, Alex Alderton, Roberto Amato, Sonia Goncalves, Ewan Harrison, David K. Jackson, Ian Johnston, Dominic Kwiatkowski, Cordelia Langford, John Sillitoe on behalf of the Wellcome Sanger Institute COVID-19 Surveillance Team (http://www.sanger.ac.uk/covid-team) |
| EPI_ISL_440072, EPI_ISL_440073, EPI_ISL_440074, EPI_ISL_440075, EPI_ISL_440076, EPI_ISL_440077, EPI_ISL_440078, EPI_ISL_440079, EPI_ISL_440080 | Department of Pathology, University of Cambridge | Wellcome Sanger Institute for the COVID-19 Genomics UK Consortium | Luke W Meredith, M. Estée Török , Myra Hosmillo, William L. Hamilton, Martin D. Curran, Theresa Feltwell, Grant Hall, Anna Yakovleva, Fahad A Khokhar, Charlotte J. Houldcroft, Laura G Caller, Aminu S. Jahun, Sarah L. Caddy, Ian Goodfellow, Alex Alderton, Roberto Amato, Sonia Goncalves, Ewan Harrison, David K. Jackson, Ian Johnston, Dominic Kwiatkowski, Cordelia Langford, John Sillitoe on behalf of the Wellcome Sanger Institute COVID-19 Surveillance Team (http://www.sanger.ac.uk/covid-team) |
| EPI_ISL_440081 | PHE South West Regional Laboratory, National Infection Service | Wellcome Sanger Institute for the COVID-19 Genomics UK Consortium | Stephanie Hutchings, Hannah Pymont, Dr Peter Muir, Barry Vipond, Rich Hopes, Alex Alderton, Roberto Amato, Sonia Goncalves, Ewan Harrison, David K. Jackson, Ian Johnston, Dominic Kwiatkowski, Cordelia Langford, John Sillitoe on behalf of the Wellcome Sanger Institute COVID-19 Surveillance Team (http://www.sanger.ac.uk/covid-team) |
| EPI_ISL_440082, EPI_ISL_440083 | Department of Pathology, University of Cambridge | Wellcome Sanger Institute for the COVID-19 Genomics UK Consortium | Luke W Meredith, M. Estée Török , Myra Hosmillo, William L. Hamilton, Martin D. Curran, Theresa Feltwell, Grant Hall, Anna Yakovleva, Fahad A Khokhar, Charlotte J. Houldcroft, Laura G Caller, Aminu S. Jahun, Sarah L. Caddy, Ian Goodfellow, Alex Alderton, Roberto Amato, Sonia Goncalves, Ewan Harrison, David K. Jackson, Ian Johnston, Dominic Kwiatkowski, Cordelia Langford, John Sillitoe on behalf of the Wellcome Sanger Institute COVID-19 Surveillance Team (http://www.sanger.ac.uk/covid-team) |
| EPI_ISL_440084 | PHE South West Regional Laboratory, National Infection Service | Wellcome Sanger Institute for the COVID-19 Genomics UK Consortium | Stephanie Hutchings, Hannah Pymont, Dr Peter Muir, Barry Vipond, Rich Hopes, Alex Alderton, Roberto Amato, Sonia Goncalves, Ewan Harrison, David K. Jackson, Ian Johnston, Dominic Kwiatkowski, Cordelia Langford, John Sillitoe on behalf of the Wellcome Sanger Institute COVID-19 Surveillance Team (http://www.sanger.ac.uk/covid-team) |
| EPI_ISL_440085, EPI_ISL_440086 | Department of Pathology, University of Cambridge | Wellcome Sanger Institute for the COVID-19 Genomics UK Consortium | Luke W Meredith, M. Estée Török , Myra Hosmillo, William L. Hamilton, Martin D. Curran, Theresa Feltwell, Grant Hall, Anna Yakovleva, Fahad A Khokhar, Charlotte J. Houldcroft, Laura G Caller, Aminu S. Jahun, Sarah L. Caddy, Ian Goodfellow, Alex Alderton, Roberto Amato, Sonia Goncalves, Ewan Harrison, David K. Jackson, Ian Johnston, Dominic Kwiatkowski, Cordelia Langford, John Sillitoe on behalf of the Wellcome Sanger Institute COVID-19 Surveillance Team (http://www.sanger.ac.uk/covid-team) |
| EPI_ISL_440087 | PHE South West Regional Laboratory, National Infection Service | Wellcome Sanger Institute for the COVID-19 Genomics UK Consortium | Stephanie Hutchings, Hannah Pymont, Dr Peter Muir, Barry Vipond, Rich Hopes, Alex Alderton, Roberto Amato, Sonia Goncalves, Ewan Harrison, David K. Jackson, Ian Johnston, Dominic Kwiatkowski, Cordelia Langford, John Sillitoe on behalf of the Wellcome Sanger Institute COVID-19 Surveillance Team (http://www.sanger.ac.uk/covid-team) |
| EPI_ISL_440088, EPI_ISL_440089, EPI_ISL_440090 | Department of Pathology, University of Cambridge | Wellcome Sanger Institute for the COVID-19 Genomics UK Consortium | Luke W Meredith, M. Estée Török , Myra Hosmillo, William L. Hamilton, Martin D. Curran, Theresa Feltwell, Grant Hall, Anna Yakovleva, Fahad A Khokhar, Charlotte J. Houldcroft, Laura G Caller, Aminu S. Jahun, Sarah L. Caddy, Ian Goodfellow, Alex Alderton, Roberto Amato, Sonia Goncalves, Ewan Harrison, David K. Jackson, Ian Johnston, Dominic Kwiatkowski, Cordelia Langford, John Sillitoe on behalf of the Wellcome Sanger Institute COVID-19 Surveillance Team (http://www.sanger.ac.uk/covid-team) |
| EPI_ISL_440091, EPI_ISL_440092, EPI_ISL_440093, EPI_ISL_440094, EPI_ISL_440095 | PHE South West Regional Laboratory, National Infection Service | Wellcome Sanger Institute for the COVID-19 Genomics UK Consortium | Stephanie Hutchings, Hannah Pymont, Dr Peter Muir, Barry Vipond, Rich Hopes, Alex Alderton, Roberto Amato, Sonia Goncalves, Ewan Harrison, David K. Jackson, Ian Johnston, Dominic Kwiatkowski, Cordelia Langford, John Sillitoe on behalf of the Wellcome Sanger Institute COVID-19 Surveillance Team (http://www.sanger.ac.uk/covid-team) |
| EPI_ISL_440096, EPI_ISL_440097 | Department of Pathology, University of Cambridge | Wellcome Sanger Institute for the COVID-19 Genomics UK Consortium | Luke W Meredith, M. Estée Török , Myra Hosmillo, William L. Hamilton, Martin D. Curran, Theresa Feltwell, Grant Hall, Anna Yakovleva, Fahad A Khokhar, Charlotte J. Houldcroft, Laura G Caller, Aminu S. Jahun, Sarah L. Caddy, Ian Goodfellow, Alex Alderton, Roberto Amato, Sonia Goncalves, Ewan Harrison, David K. Jackson, Ian Johnston, Dominic Kwiatkowski, Cordelia Langford, John Sillitoe on behalf of the Wellcome Sanger Institute COVID-19 Surveillance Team (http://www.sanger.ac.uk/covid-team) |
| EPI_ISL_440098 | PHE South West Regional Laboratory, National Infection Service | Wellcome Sanger Institute for the COVID-19 Genomics UK Consortium | Stephanie Hutchings, Hannah Pymont, Dr Peter Muir, Barry Vipond, Rich Hopes, Alex Alderton, Roberto Amato, Sonia Goncalves, Ewan Harrison, David K. Jackson, Ian Johnston, Dominic Kwiatkowski, Cordelia Langford, John Sillitoe on behalf of the Wellcome Sanger Institute COVID-19 Surveillance Team (http://www.sanger.ac.uk/covid-team) |
| EPI_ISL_440099 | Department of Pathology, University of Cambridge | Wellcome Sanger Institute for the COVID-19 Genomics UK Consortium | Luke W Meredith, M. Estée Török , Myra Hosmillo, William L. Hamilton, Martin D. Curran, Theresa Feltwell, Grant Hall, Anna Yakovleva, Fahad A Khokhar, Charlotte J. Houldcroft, Laura G Caller, Aminu S. Jahun, Sarah L. Caddy, Ian Goodfellow, Alex Alderton, Roberto Amato, Sonia Goncalves, Ewan Harrison, David K. Jackson, Ian Johnston, Dominic Kwiatkowski, Cordelia Langford, John Sillitoe on behalf of the Wellcome Sanger Institute COVID-19 Surveillance Team (http://www.sanger.ac.uk/covid-team) |
| EPI_ISL_440100, EPI_ISL_440101, EPI_ISL_440102, EPI_ISL_440103 | PHE South West Regional Laboratory, National Infection Service | Wellcome Sanger Institute for the COVID-19 Genomics UK Consortium | Stephanie Hutchings, Hannah Pymont, Dr Peter Muir, Barry Vipond, Rich Hopes, Alex Alderton, Roberto Amato, Sonia Goncalves, Ewan Harrison, David K. Jackson, Ian Johnston, Dominic Kwiatkowski, Cordelia Langford, John Sillitoe on behalf of the Wellcome Sanger Institute COVID-19 Surveillance Team (http://www.sanger.ac.uk/covid-team) |
| EPI_ISL_440104 | Department of Pathology, University of Cambridge | Wellcome Sanger Institute for the COVID-19 Genomics UK Consortium | Luke W Meredith, M. Estée Török , Myra Hosmillo, William L. Hamilton, Martin D. Curran, Theresa Feltwell, Grant Hall, Anna Yakovleva, Fahad A Khokhar, Charlotte J. Houldcroft, Laura G Caller, Aminu S. Jahun, Sarah L. Caddy, Ian Goodfellow, Alex Alderton, Roberto Amato, Sonia Goncalves, Ewan Harrison, David K. Jackson, Ian Johnston, Dominic Kwiatkowski, Cordelia Langford, John Sillitoe on behalf of the Wellcome Sanger Institute COVID-19 Surveillance Team (http://www.sanger.ac.uk/covid-team) |
| EPI_ISL_440105, EPI_ISL_440106 | PHE South West Regional Laboratory, National Infection Service | Wellcome Sanger Institute for the COVID-19 Genomics UK Consortium | Stephanie Hutchings, Hannah Pymont, Dr Peter Muir, Barry Vipond, Rich Hopes, Alex Alderton, Roberto Amato, Sonia Goncalves, Ewan Harrison, David K. Jackson, Ian Johnston, Dominic Kwiatkowski, Cordelia Langford, John Sillitoe on behalf of the Wellcome Sanger Institute COVID-19 Surveillance Team (http://www.sanger.ac.uk/covid-team) |
| EPI_ISL_440107, EPI_ISL_440108, EPI_ISL_440109, EPI_ISL_440110, EPI_ISL_440111, EPI_ISL_440112 | Department of Pathology, University of Cambridge | Wellcome Sanger Institute for the COVID-19 Genomics UK Consortium | Luke W Meredith, M. Estée Török , Myra Hosmillo, William L. Hamilton, Martin D. Curran, Theresa Feltwell, Grant Hall, Anna Yakovleva, Fahad A Khokhar, Charlotte J. Houldcroft, Laura G Caller, Aminu S. Jahun, Sarah L. Caddy, Ian Goodfellow, Alex Alderton, Roberto Amato, Sonia Goncalves, Ewan Harrison, David K. Jackson, Ian Johnston, Dominic Kwiatkowski, Cordelia Langford, John Sillitoe on behalf of the Wellcome Sanger Institute COVID-19 Surveillance Team (http://www.sanger.ac.uk/covid-team) |
| EPI_ISL_440113, EPI_ISL_440114, EPI_ISL_440115, EPI_ISL_440116 | PHE South West Regional Laboratory, National Infection Service | Wellcome Sanger Institute for the COVID-19 Genomics UK Consortium | Stephanie Hutchings, Hannah Pymont, Dr Peter Muir, Barry Vipond, Rich Hopes, Alex Alderton, Roberto Amato, Sonia Goncalves, Ewan Harrison, David K. Jackson, Ian Johnston, Dominic Kwiatkowski, Cordelia Langford, John Sillitoe on behalf of the Wellcome Sanger Institute COVID-19 Surveillance Team (http://www.sanger.ac.uk/covid-team) |
| EPI_ISL_440117 | Department of Pathology, University of Cambridge | Wellcome Sanger Institute for the COVID-19 Genomics UK Consortium | Luke W Meredith, M. Estée Török , Myra Hosmillo, William L. Hamilton, Martin D. Curran, Theresa Feltwell, Grant Hall, Anna Yakovleva, Fahad A Khokhar, Charlotte J. Houldcroft, Laura G Caller, Aminu S. Jahun, Sarah L. Caddy, Ian Goodfellow, Alex Alderton, Roberto Amato, Sonia Goncalves, Ewan Harrison, David K. Jackson, Ian Johnston, Dominic Kwiatkowski, Cordelia Langford, John Sillitoe on behalf of the Wellcome Sanger Institute COVID-19 Surveillance Team (http://www.sanger.ac.uk/covid-team) |
| EPI_ISL_440118, EPI_ISL_440119 | PHE South West Regional Laboratory, National Infection Service | Wellcome Sanger Institute for the COVID-19 Genomics UK Consortium | Stephanie Hutchings, Hannah Pymont, Dr Peter Muir, Barry Vipond, Rich Hopes, Alex Alderton, Roberto Amato, Sonia Goncalves, Ewan Harrison, David K. Jackson, Ian Johnston, Dominic Kwiatkowski, Cordelia Langford, John Sillitoe on behalf of the Wellcome Sanger Institute COVID-19 Surveillance Team (http://www.sanger.ac.uk/covid-team) |
| EPI_ISL_440120, EPI_ISL_440121, EPI_ISL_440122 | Department of Pathology, University of Cambridge | Wellcome Sanger Institute for the COVID-19 Genomics UK Consortium | Luke W Meredith, M. Estée Török , Myra Hosmillo, William L. Hamilton, Martin D. Curran, Theresa Feltwell, Grant Hall, Anna Yakovleva, Fahad A Khokhar, Charlotte J. Houldcroft, Laura G Caller, Aminu S. Jahun, Sarah L. Caddy, Ian Goodfellow, Alex Alderton, Roberto Amato, Sonia Goncalves, Ewan Harrison, David K. Jackson, Ian Johnston, Dominic Kwiatkowski, Cordelia Langford, John Sillitoe on behalf of the Wellcome Sanger Institute COVID-19 Surveillance Team (http://www.sanger.ac.uk/covid-team) |
| EPI_ISL_440123, EPI_ISL_440124 | PHE South West Regional Laboratory, National Infection Service | Wellcome Sanger Institute for the COVID-19 Genomics UK Consortium | Stephanie Hutchings, Hannah Pymont, Dr Peter Muir, Barry Vipond, Rich Hopes, Alex Alderton, Roberto Amato, Sonia Goncalves, Ewan Harrison, David K. Jackson, Ian Johnston, Dominic Kwiatkowski, Cordelia Langford, John Sillitoe on behalf of the Wellcome Sanger Institute COVID-19 Surveillance Team (http://www.sanger.ac.uk/covid-team) |
| EPI_ISL_440125 | Department of Pathology, University of Cambridge | Wellcome Sanger Institute for the COVID-19 Genomics UK Consortium | Luke W Meredith, M. Estée Török , Myra Hosmillo, William L. Hamilton, Martin D. Curran, Theresa Feltwell, Grant Hall, Anna Yakovleva, Fahad A Khokhar, Charlotte J. Houldcroft, Laura G Caller, Aminu S. Jahun, Sarah L. Caddy, Ian Goodfellow, Alex Alderton, Roberto Amato, Sonia Goncalves, Ewan Harrison, David K. Jackson, Ian Johnston, Dominic Kwiatkowski, Cordelia Langford, John Sillitoe on behalf of the Wellcome Sanger Institute COVID-19 Surveillance Team (http://www.sanger.ac.uk/covid-team) |
| EPI_ISL_440126, EPI_ISL_440127, EPI_ISL_440128 | PHE South West Regional Laboratory, National Infection Service | Wellcome Sanger Institute for the COVID-19 Genomics UK Consortium | Stephanie Hutchings, Hannah Pymont, Dr Peter Muir, Barry Vipond, Rich Hopes, Alex Alderton, Roberto Amato, Sonia Goncalves, Ewan Harrison, David K. Jackson, Ian Johnston, Dominic Kwiatkowski, Cordelia Langford, John Sillitoe on behalf of the Wellcome Sanger Institute COVID-19 Surveillance Team (http://www.sanger.ac.uk/covid-team) |
| EPI_ISL_440129 | Department of Pathology, University of Cambridge | Wellcome Sanger Institute for the COVID-19 Genomics UK Consortium | Luke W Meredith, M. Estée Török , Myra Hosmillo, William L. Hamilton, Martin D. Curran, Theresa Feltwell, Grant Hall, Anna Yakovleva, Fahad A Khokhar, Charlotte J. Houldcroft, Laura G Caller, Aminu S. Jahun, Sarah L. Caddy, Ian Goodfellow, Alex Alderton, Roberto Amato, Sonia Goncalves, Ewan Harrison, David K. Jackson, Ian Johnston, Dominic Kwiatkowski, Cordelia Langford, John Sillitoe on behalf of the Wellcome Sanger Institute COVID-19 Surveillance Team (http://www.sanger.ac.uk/covid-team) |
| EPI_ISL_440130, EPI_ISL_440131 | PHE South West Regional Laboratory, National Infection Service | Wellcome Sanger Institute for the COVID-19 Genomics UK Consortium | Stephanie Hutchings, Hannah Pymont, Dr Peter Muir, Barry Vipond, Rich Hopes, Alex Alderton, Roberto Amato, Sonia Goncalves, Ewan Harrison, David K. Jackson, Ian Johnston, Dominic Kwiatkowski, Cordelia Langford, John Sillitoe on behalf of the Wellcome Sanger Institute COVID-19 Surveillance Team (http://www.sanger.ac.uk/covid-team) |
| EPI_ISL_440132, EPI_ISL_440133 | Department of Pathology, University of Cambridge | Wellcome Sanger Institute for the COVID-19 Genomics UK Consortium | Luke W Meredith, M. Estée Török , Myra Hosmillo, William L. Hamilton, Martin D. Curran, Theresa Feltwell, Grant Hall, Anna Yakovleva, Fahad A Khokhar, Charlotte J. Houldcroft, Laura G Caller, Aminu S. Jahun, Sarah L. Caddy, Ian Goodfellow, Alex Alderton, Roberto Amato, Sonia Goncalves, Ewan Harrison, David K. Jackson, Ian Johnston, Dominic Kwiatkowski, Cordelia Langford, John Sillitoe on behalf of the Wellcome Sanger Institute COVID-19 Surveillance Team (http://www.sanger.ac.uk/covid-team) |
| EPI_ISL_440134, EPI_ISL_440135, EPI_ISL_440136, EPI_ISL_440137 | PHE South West Regional Laboratory, National Infection Service | Wellcome Sanger Institute for the COVID-19 Genomics UK Consortium | Stephanie Hutchings, Hannah Pymont, Dr Peter Muir, Barry Vipond, Rich Hopes, Alex Alderton, Roberto Amato, Sonia Goncalves, Ewan Harrison, David K. Jackson, Ian Johnston, Dominic Kwiatkowski, Cordelia Langford, John Sillitoe on behalf of the Wellcome Sanger Institute COVID-19 Surveillance Team (http://www.sanger.ac.uk/covid-team) |
| EPI_ISL_440138 | Department of Pathology, University of Cambridge | Wellcome Sanger Institute for the COVID-19 Genomics UK Consortium | Luke W Meredith, M. Estée Török , Myra Hosmillo, William L. Hamilton, Martin D. Curran, Theresa Feltwell, Grant Hall, Anna Yakovleva, Fahad A Khokhar, Charlotte J. Houldcroft, Laura G Caller, Aminu S. Jahun, Sarah L. Caddy, Ian Goodfellow, Alex Alderton, Roberto Amato, Sonia Goncalves, Ewan Harrison, David K. Jackson, Ian Johnston, Dominic Kwiatkowski, Cordelia Langford, John Sillitoe on behalf of the Wellcome Sanger Institute COVID-19 Surveillance Team (http://www.sanger.ac.uk/covid-team) |
| EPI_ISL_440139, EPI_ISL_440140, EPI_ISL_440141, EPI_ISL_440142 | PHE South West Regional Laboratory, National Infection Service | Wellcome Sanger Institute for the COVID-19 Genomics UK Consortium | Stephanie Hutchings, Hannah Pymont, Dr Peter Muir, Barry Vipond, Rich Hopes, Alex Alderton, Roberto Amato, Sonia Goncalves, Ewan Harrison, David K. Jackson, Ian Johnston, Dominic Kwiatkowski, Cordelia Langford, John Sillitoe on behalf of the Wellcome Sanger Institute COVID-19 Surveillance Team (http://www.sanger.ac.uk/covid-team) |
| EPI_ISL_440143, EPI_ISL_440144 | Department of Pathology, University of Cambridge | Wellcome Sanger Institute for the COVID-19 Genomics UK Consortium | Luke W Meredith, M. Estée Török , Myra Hosmillo, William L. Hamilton, Martin D. Curran, Theresa Feltwell, Grant Hall, Anna Yakovleva, Fahad A Khokhar, Charlotte J. Houldcroft, Laura G Caller, Aminu S. Jahun, |

| | | | |
|---|---|---|---|
| | | | Sarah L. Caddy, Ian Goodfellow, Alex Alderton, Roberto Amato, Sonia Goncalves, Ewan Harrison, David K. Jackson, Ian Johnston, Dominic Kwiatkowski, Cordelia Langford, John Sillitoe on behalf of the Wellcome Sanger Institute COVID-19 Surveillance Team (http://www.sanger.ac.uk/covid-team) |
| EPI_ISL_440145 | PHE South West Regional Laboratory, National Infection Service | Wellcome Sanger Institute for the COVID-19 Genomics UK Consortium | Stephanie Hutchings, Hannah Pymont, Dr Peter Muir, Barry Vipond, Rich Hopes, Alex Alderton, Roberto Amato, Sonia Goncalves, Ewan Harrison, David K. Jackson, Ian Johnston, Dominic Kwiatkowski, Cordelia Langford, John Sillitoe on behalf of the Wellcome Sanger Institute COVID-19 Surveillance Team (http://www.sanger.ac.uk/covid-team) |
| EPI_ISL_440146, EPI_ISL_440147 | Department of Pathology, University of Cambridge | Wellcome Sanger Institute for the COVID-19 Genomics UK Consortium | Luke W Meredith, M. Estée Török , Myra Hosmillo, William L. Hamilton, Martin D. Curran, Theresa Feltwell, Grant Hall, Anna Yakovleva, Fahad A Khokhar, Charlotte J. Houldcroft, Laura G Caller, Aminu S. Jahun, Sarah L. Caddy, Ian Goodfellow, Alex Alderton, Roberto Amato, Sonia Goncalves, Ewan Harrison, David K. Jackson, Ian Johnston, Dominic Kwiatkowski, Cordelia Langford, John Sillitoe on behalf of the Wellcome Sanger Institute COVID-19 Surveillance Team (http://www.sanger.ac.uk/covid-team) |
| EPI_ISL_440148, EPI_ISL_440149 | PHE South West Regional Laboratory, National Infection Service | Wellcome Sanger Institute for the COVID-19 Genomics UK Consortium | Stephanie Hutchings, Hannah Pymont, Dr Peter Muir, Barry Vipond, Rich Hopes, Alex Alderton, Roberto Amato, Sonia Goncalves, Ewan Harrison, David K. Jackson, Ian Johnston, Dominic Kwiatkowski, Cordelia Langford, John Sillitoe on behalf of the Wellcome Sanger Institute COVID-19 Surveillance Team (http://www.sanger.ac.uk/covid-team) |
| EPI_ISL_440150 | Department of Pathology, University of Cambridge | Wellcome Sanger Institute for the COVID-19 Genomics UK Consortium | Luke W Meredith, M. Estée Török , Myra Hosmillo, William L. Hamilton, Martin D. Curran, Theresa Feltwell, Grant Hall, Anna Yakovleva, Fahad A Khokhar, Charlotte J. Houldcroft, Laura G Caller, Aminu S. Jahun, Sarah L. Caddy, Ian Goodfellow, Alex Alderton, Roberto Amato, Sonia Goncalves, Ewan Harrison, David K. Jackson, Ian Johnston, Dominic Kwiatkowski, Cordelia Langford, John Sillitoe on behalf of the Wellcome Sanger Institute COVID-19 Surveillance Team (http://www.sanger.ac.uk/covid-team) |
| EPI_ISL_440151 | PHE South West Regional Laboratory, National Infection Service | Wellcome Sanger Institute for the COVID-19 Genomics UK Consortium | Stephanie Hutchings, Hannah Pymont, Dr Peter Muir, Barry Vipond, Rich Hopes, Alex Alderton, Roberto Amato, Sonia Goncalves, Ewan Harrison, David K. Jackson, Ian Johnston, Dominic Kwiatkowski, Cordelia Langford, John Sillitoe on behalf of the Wellcome Sanger Institute COVID-19 Surveillance Team (http://www.sanger.ac.uk/covid-team) |
| EPI_ISL_440152 | Department of Pathology, University of Cambridge | Wellcome Sanger Institute for the COVID-19 Genomics UK Consortium | Luke W Meredith, M. Estée Török , Myra Hosmillo, William L. Hamilton, Martin D. Curran, Theresa Feltwell, Grant Hall, Anna Yakovleva, Fahad A Khokhar, Charlotte J. Houldcroft, Laura G Caller, Aminu S. Jahun, Sarah L. Caddy, Ian Goodfellow, Alex Alderton, Roberto Amato, Sonia Goncalves, Ewan Harrison, David K. Jackson, Ian Johnston, Dominic Kwiatkowski, Cordelia Langford, John Sillitoe on behalf of the Wellcome Sanger Institute COVID-19 Surveillance Team (http://www.sanger.ac.uk/covid-team) |
| EPI_ISL_440153, EPI_ISL_440154 | PHE South West Regional Laboratory, National Infection Service | Wellcome Sanger Institute for the COVID-19 Genomics UK Consortium | Stephanie Hutchings, Hannah Pymont, Dr Peter Muir, Barry Vipond, Rich Hopes, Alex Alderton, Roberto Amato, Sonia Goncalves, Ewan Harrison, David K. Jackson, Ian Johnston, Dominic Kwiatkowski, Cordelia Langford, John Sillitoe on behalf of the Wellcome Sanger Institute COVID-19 Surveillance Team (http://www.sanger.ac.uk/covid-team) |
| EPI_ISL_440155 | Department of Pathology, University of Cambridge | Wellcome Sanger Institute for the COVID-19 Genomics UK Consortium | Luke W Meredith, M. Estée Török , Myra Hosmillo, William L. Hamilton, Martin D. Curran, Theresa Feltwell, Grant Hall, Anna Yakovleva, Fahad A Khokhar, Charlotte J. Houldcroft, Laura G Caller, Aminu S. Jahun, Sarah L. Caddy, Ian Goodfellow, Alex Alderton, Roberto Amato, Sonia Goncalves, Ewan Harrison, David K. Jackson, Ian Johnston, Dominic Kwiatkowski, Cordelia Langford, John Sillitoe on behalf of the Wellcome Sanger Institute COVID-19 Surveillance Team (http://www.sanger.ac.uk/covid-team) |
| EPI_ISL_440156, EPI_ISL_440157, EPI_ISL_440158 | PHE South West Regional Laboratory, National Infection Service | Wellcome Sanger Institute for the COVID-19 Genomics UK Consortium | Stephanie Hutchings, Hannah Pymont, Dr Peter Muir, Barry Vipond, Rich Hopes, Alex Alderton, Roberto Amato, Sonia Goncalves, Ewan Harrison, David K. Jackson, Ian Johnston, Dominic Kwiatkowski, Cordelia Langford, John Sillitoe on behalf of the Wellcome Sanger Institute COVID-19 Surveillance Team (http://www.sanger.ac.uk/covid-team) |
| EPI_ISL_440159 | Department of Pathology, University of Cambridge | Wellcome Sanger Institute for the COVID-19 Genomics UK Consortium | Luke W Meredith, M. Estée Török , Myra Hosmillo, William L. Hamilton, Martin D. Curran, Theresa Feltwell, Grant Hall, Anna Yakovleva, Fahad A Khokhar, Charlotte J. Houldcroft, Laura G Caller, Aminu S. Jahun, Sarah L. Caddy, Ian Goodfellow, Alex Alderton, Roberto Amato, Sonia Goncalves, Ewan Harrison, David K. Jackson, Ian Johnston, Dominic Kwiatkowski, Cordelia Langford, John Sillitoe on behalf of the Wellcome Sanger Institute COVID-19 Surveillance Team (http://www.sanger.ac.uk/covid-team) |
| EPI_ISL_440160 | PHE South West Regional Laboratory, National Infection Service | Wellcome Sanger Institute for the COVID-19 Genomics UK Consortium | Stephanie Hutchings, Hannah Pymont, Dr Peter Muir, Barry Vipond, Rich Hopes, Alex Alderton, Roberto Amato, Sonia Goncalves, Ewan Harrison, David K. Jackson, Ian Johnston, Dominic Kwiatkowski, Cordelia Langford, John Sillitoe on behalf of the Wellcome Sanger Institute COVID-19 Surveillance Team (http://www.sanger.ac.uk/covid-team) |
| EPI_ISL_440161 | Department of Pathology, University of Cambridge | Wellcome Sanger Institute for the COVID-19 Genomics UK Consortium | Luke W Meredith, M. Estée Török , Myra Hosmillo, William L. Hamilton, Martin D. Curran, Theresa Feltwell, Grant Hall, Anna Yakovleva, Fahad A Khokhar, Charlotte J. Houldcroft, Laura G Caller, Aminu S. Jahun, Sarah L. Caddy, Ian Goodfellow, Alex Alderton, Roberto Amato, Sonia Goncalves, Ewan Harrison, David K. Jackson, Ian Johnston, Dominic Kwiatkowski, Cordelia Langford, John Sillitoe on behalf of the Wellcome Sanger Institute COVID-19 Surveillance Team (http://www.sanger.ac.uk/covid-team) |
| EPI_ISL_440162 | PHE South West Regional Laboratory, National Infection Service | Wellcome Sanger Institute for the COVID-19 Genomics UK Consortium | Stephanie Hutchings, Hannah Pymont, Dr Peter Muir, Barry Vipond, Rich Hopes, Alex Alderton, Roberto Amato, Sonia Goncalves, Ewan Harrison, David K. Jackson, Ian Johnston, Dominic Kwiatkowski, Cordelia Langford, John Sillitoe on behalf of the Wellcome Sanger Institute COVID-19 Surveillance Team (http://www.sanger.ac.uk/covid-team) |
| EPI_ISL_440163, EPI_ISL_440164 | Department of Pathology, University of Cambridge | Wellcome Sanger Institute for the COVID-19 Genomics UK Consortium | Luke W Meredith, M. Estée Török , Myra Hosmillo, William L. Hamilton, Martin D. Curran, Theresa Feltwell, Grant Hall, Anna Yakovleva, Fahad A Khokhar, Charlotte J. Houldcroft, Laura G Caller, Aminu S. Jahun, Sarah L. Caddy, Ian Goodfellow, Alex Alderton, Roberto Amato, Sonia Goncalves, Ewan Harrison, David K. Jackson, Ian Johnston, Dominic Kwiatkowski, Cordelia Langford, John Sillitoe on behalf of the Wellcome Sanger Institute COVID-19 Surveillance Team (http://www.sanger.ac.uk/covid-team) |
| EPI_ISL_440165 | PHE South West Regional Laboratory, National Infection Service | Wellcome Sanger Institute for the COVID-19 Genomics UK Consortium | Stephanie Hutchings, Hannah Pymont, Dr Peter Muir, Barry Vipond, Rich Hopes, Alex Alderton, Roberto Amato, Sonia Goncalves, Ewan Harrison, David K. Jackson, Ian Johnston, Dominic Kwiatkowski, Cordelia Langford, John Sillitoe on behalf of the Wellcome Sanger Institute COVID-19 Surveillance Team (http://www.sanger.ac.uk/covid-team) |
| EPI_ISL_440166 | Department of Pathology, University of Cambridge | Wellcome Sanger Institute for the COVID-19 Genomics UK Consortium | Luke W Meredith, M. Estée Török , Myra Hosmillo, William L. Hamilton, Martin D. Curran, Theresa Feltwell, Grant Hall, Anna Yakovleva, Fahad A Khokhar, Charlotte J. Houldcroft, Laura G Caller, Aminu S. Jahun, Sarah L. Caddy, Ian Goodfellow, Alex Alderton, Roberto Amato, Sonia Goncalves, Ewan Harrison, David K. Jackson, Ian Johnston, Dominic Kwiatkowski, Cordelia Langford, John Sillitoe on behalf of the Wellcome Sanger Institute COVID-19 Surveillance Team (http://www.sanger.ac.uk/covid-team) |
| EPI_ISL_440167, EPI_ISL_440168 | PHE South West Regional Laboratory, National Infection Service | Wellcome Sanger Institute for the COVID-19 Genomics UK Consortium | Stephanie Hutchings, Hannah Pymont, Dr Peter Muir, Barry Vipond, Rich Hopes, Alex Alderton, Roberto Amato, Sonia Goncalves, Ewan Harrison, David K. Jackson, Ian Johnston, Dominic Kwiatkowski, Cordelia Langford, John Sillitoe on behalf of the Wellcome Sanger Institute COVID-19 Surveillance Team (http://www.sanger.ac.uk/covid-team) |
| EPI_ISL_440169, EPI_ISL_440170, EPI_ISL_440171 | Department of Pathology, University of Cambridge | Wellcome Sanger Institute for the COVID-19 Genomics UK Consortium | Luke W Meredith, M. Estée Török , Myra Hosmillo, William L. Hamilton, Martin D. Curran, Theresa Feltwell, Grant Hall, Anna Yakovleva, Fahad A Khokhar, Charlotte J. Houldcroft, Laura G Caller, Aminu S. Jahun, Sarah L. Caddy, Ian Goodfellow, Alex Alderton, Roberto Amato, Sonia Goncalves, Ewan Harrison, David K. Jackson, Ian Johnston, Dominic Kwiatkowski, Cordelia Langford, John Sillitoe on behalf of the Wellcome Sanger Institute COVID-19 Surveillance Team (http://www.sanger.ac.uk/covid-team) |
| EPI_ISL_440172 | PHE South West Regional Laboratory, National Infection Service | Wellcome Sanger Institute for the COVID-19 Genomics UK Consortium | Stephanie Hutchings, Hannah Pymont, Dr Peter Muir, Barry Vipond, Rich Hopes, Alex Alderton, Roberto Amato, Sonia Goncalves, Ewan Harrison, David K. Jackson, Ian Johnston, Dominic Kwiatkowski, Cordelia Langford, John Sillitoe on behalf of the Wellcome Sanger Institute COVID-19 Surveillance Team (http://www.sanger.ac.uk/covid-team) |
| EPI_ISL_440173 | Department of Pathology, University of Cambridge | Wellcome Sanger Institute for the COVID-19 Genomics UK Consortium | Luke W Meredith, M. Estée Török , Myra Hosmillo, William L. Hamilton, Martin D. Curran, Theresa Feltwell, Grant Hall, Anna Yakovleva, Fahad A Khokhar, Charlotte J. Houldcroft, Laura G Caller, Aminu S. Jahun, Sarah L. Caddy, Ian Goodfellow, Alex Alderton, Roberto Amato, Sonia Goncalves, Ewan Harrison, David K. Jackson, Ian Johnston, Dominic Kwiatkowski, Cordelia Langford, John Sillitoe on behalf of the Wellcome Sanger Institute COVID-19 Surveillance Team (http://www.sanger.ac.uk/covid-team) |
| EPI_ISL_440174, EPI_ISL_440175 | PHE South West Regional Laboratory, National Infection Service | Wellcome Sanger Institute for the COVID-19 Genomics UK Consortium | Stephanie Hutchings, Hannah Pymont, Dr Peter Muir, Barry Vipond, Rich Hopes, Alex Alderton, Roberto Amato, Sonia Goncalves, Ewan Harrison, David K. Jackson, Ian Johnston, Dominic Kwiatkowski, Cordelia Langford, John Sillitoe on behalf of the Wellcome Sanger Institute COVID-19 Surveillance Team (http://www.sanger.ac.uk/covid-team) |
| EPI_ISL_440176, EPI_ISL_440177 | Department of Pathology, University of Cambridge | Wellcome Sanger Institute for the COVID-19 Genomics UK Consortium | Luke W Meredith, M. Estée Török , Myra Hosmillo, William L. Hamilton, Martin D. Curran, Theresa Feltwell, Grant Hall, Anna Yakovleva, Fahad A Khokhar, Charlotte J. Houldcroft, Laura G Caller, Aminu S. Jahun, Sarah L. Caddy, Ian Goodfellow, Alex Alderton, Roberto Amato, Sonia Goncalves, Ewan Harrison, David K. Jackson, Ian Johnston, Dominic Kwiatkowski, Cordelia Langford, John Sillitoe on behalf of the Wellcome Sanger Institute COVID-19 Surveillance Team (http://www.sanger.ac.uk/covid-team) |
| EPI_ISL_440178 | PHE South West Regional Laboratory, National Infection Service | Wellcome Sanger Institute for the COVID-19 Genomics UK Consortium | Stephanie Hutchings, Hannah Pymont, Dr Peter Muir, Barry Vipond, Rich Hopes, Alex Alderton, Roberto Amato, Sonia Goncalves, Ewan Harrison, David K. Jackson, Ian Johnston, Dominic Kwiatkowski, Cordelia Langford, John Sillitoe on behalf of the Wellcome Sanger Institute COVID-19 Surveillance Team (http://www.sanger.ac.uk/covid-team) |
| EPI_ISL_440179, EPI_ISL_440180, EPI_ISL_440181, EPI_ISL_440182, EPI_ISL_440183, EPI_ISL_440184, EPI_ISL_440185, EPI_ISL_440186 | Department of Pathology, University of Cambridge | Wellcome Sanger Institute for the COVID-19 Genomics UK Consortium | Luke W Meredith, M. Estée Török , Myra Hosmillo, William L. Hamilton, Martin D. Curran, Theresa Feltwell, Grant Hall, Anna Yakovleva, Fahad A Khokhar, Charlotte J. Houldcroft, Laura G Caller, Aminu S. Jahun, Sarah L. Caddy, Ian Goodfellow, Alex Alderton, Roberto Amato, Sonia Goncalves, Ewan Harrison, David K. Jackson, Ian Johnston, Dominic Kwiatkowski, Cordelia Langford, John Sillitoe on behalf of the Wellcome Sanger Institute COVID-19 Surveillance Team (http://www.sanger.ac.uk/covid-team) |
| EPI_ISL_440187 | PHE South West Regional Laboratory, National Infection Service | Wellcome Sanger Institute for the COVID-19 Genomics UK Consortium | Stephanie Hutchings, Hannah Pymont, Dr Peter Muir, Barry Vipond, Rich Hopes, Alex Alderton, Roberto Amato, Sonia Goncalves, Ewan Harrison, David K. Jackson, Ian Johnston, Dominic Kwiatkowski, Cordelia Langford, John Sillitoe on behalf of the Wellcome Sanger Institute COVID-19 Surveillance Team (http://www.sanger.ac.uk/covid-team) |
| EPI_ISL_440188 | Department of Pathology, University of Cambridge | Wellcome Sanger Institute for the COVID-19 Genomics UK Consortium | Luke W Meredith, M. Estée Török , Myra Hosmillo, William L. Hamilton, Martin D. Curran, Theresa Feltwell, Grant Hall, Anna Yakovleva, Fahad A Khokhar, Charlotte J. Houldcroft, Laura G Caller, Aminu S. Jahun, Sarah L. Caddy, Ian Goodfellow, Alex Alderton, Roberto Amato, Sonia Goncalves, Ewan Harrison, David K. Jackson, Ian Johnston, Dominic Kwiatkowski, Cordelia Langford, John Sillitoe on behalf of the Wellcome Sanger Institute COVID-19 Surveillance Team (http://www.sanger.ac.uk/covid-team) |
| EPI_ISL_440189, EPI_ISL_440190, EPI_ISL_440191, EPI_ISL_440192 | PHE South West Regional Laboratory, National Infection Service | Wellcome Sanger Institute for the COVID-19 Genomics UK Consortium | Stephanie Hutchings, Hannah Pymont, Dr Peter Muir, Barry Vipond, Rich Hopes, Alex Alderton, Roberto Amato, Sonia Goncalves, Ewan Harrison, David K. Jackson, Ian Johnston, Dominic Kwiatkowski, Cordelia Langford, John Sillitoe on behalf of the Wellcome Sanger Institute COVID-19 Surveillance Team (http://www.sanger.ac.uk/covid-team) |
| EPI_ISL_440193 | Department of Pathology, University of Cambridge | Wellcome Sanger Institute for the COVID-19 Genomics UK Consortium | Luke W Meredith, M. Estée Török , Myra Hosmillo, William L. Hamilton, Martin D. Curran, Theresa Feltwell, Grant Hall, Anna Yakovleva, Fahad A Khokhar, Charlotte J. Houldcroft, Laura G Caller, Aminu S. Jahun, Sarah L. Caddy, Ian Goodfellow, Alex Alderton, Roberto Amato, Sonia Goncalves, Ewan Harrison, David K. Jackson, Ian Johnston, Dominic Kwiatkowski, Cordelia Langford, John Sillitoe on behalf of the Wellcome Sanger Institute COVID-19 Surveillance Team (http://www.sanger.ac.uk/covid-team) |
| EPI_ISL_440194 | PHE South West Regional Laboratory, National Infection Service | Wellcome Sanger Institute for the COVID-19 Genomics UK Consortium | Stephanie Hutchings, Hannah Pymont, Dr Peter Muir, Barry Vipond, Rich Hopes, Alex Alderton, Roberto Amato, Sonia Goncalves, Ewan Harrison, David K. Jackson, Ian Johnston, Dominic Kwiatkowski, Cordelia Langford, John Sillitoe on behalf of the Wellcome Sanger Institute COVID-19 Surveillance Team (http://www.sanger.ac.uk/covid-team) |
| EPI_ISL_440195 | Department of Pathology, University of Cambridge | Wellcome Sanger Institute for the COVID-19 Genomics UK Consortium | Luke W Meredith, M. Estée Török , Myra Hosmillo, William L. Hamilton, Martin D. Curran, Theresa Feltwell, Grant Hall, Anna Yakovleva, Fahad A Khokhar, Charlotte J. Houldcroft, Laura G Caller, Aminu S. Jahun, Sarah L. Caddy, Ian Goodfellow, Alex Alderton, Roberto Amato, Sonia Goncalves, Ewan Harrison, David K. Jackson, Ian Johnston, Dominic Kwiatkowski, Cordelia Langford, John Sillitoe on behalf of the Wellcome Sanger Institute COVID-19 Surveillance Team (http://www.sanger.ac.uk/covid-team) |
| EPI_ISL_440196 | PHE South West Regional Laboratory, National Infection Service | Wellcome Sanger Institute for the COVID-19 Genomics UK Consortium | Stephanie Hutchings, Hannah Pymont, Dr Peter Muir, Barry Vipond, Rich Hopes, Alex Alderton, Roberto Amato, Sonia Goncalves, Ewan Harrison, David K. Jackson, Ian Johnston, Dominic Kwiatkowski, Cordelia Langford, John Sillitoe on behalf of the Wellcome Sanger Institute COVID-19 Surveillance Team (http://www.sanger.ac.uk/covid-team) |
| EPI_ISL_440197 | Department of Pathology, University of Cambridge | Wellcome Sanger Institute for the COVID-19 Genomics UK Consortium | Luke W Meredith, M. Estée Török , Myra Hosmillo, William L. Hamilton, Martin D. Curran, Theresa Feltwell, Grant Hall, Anna Yakovleva, Fahad A Khokhar, Charlotte J. Houldcroft, Laura G Caller, Aminu S. Jahun, Sarah L. Caddy, Ian Goodfellow, Alex Alderton, Roberto Amato, Sonia Goncalves, Ewan Harrison, David K. Jackson, Ian Johnston, Dominic Kwiatkowski, Cordelia Langford, John Sillitoe on behalf of the Wellcome Sanger Institute COVID-19 Surveillance Team (http://www.sanger.ac.uk/covid-team) |
| EPI_ISL_440198, EPI_ISL_440199, EPI_ISL_440200, EPI_ISL_440201 | PHE South West Regional Laboratory, National Infection Service | Wellcome Sanger Institute for the COVID-19 Genomics UK Consortium | Stephanie Hutchings, Hannah Pymont, Dr Peter Muir, Barry Vipond, Rich Hopes, Alex Alderton, Roberto Amato, Sonia Goncalves, Ewan Harrison, David K. Jackson, Ian Johnston, Dominic Kwiatkowski, Cordelia Langford, John Sillitoe on behalf of the Wellcome Sanger Institute COVID-19 Surveillance Team (http://www.sanger.ac.uk/covid-team) |
| EPI_ISL_440202, EPI_ISL_440203 | Department of Pathology, University of Cambridge | Wellcome Sanger Institute for the COVID-19 Genomics UK Consortium | Luke W Meredith, M. Estée Török , Myra Hosmillo, William L. Hamilton, Martin D. Curran, Theresa Feltwell, Grant Hall, Anna Yakovleva, Fahad A Khokhar, Charlotte J. Houldcroft, Laura G Caller, Aminu S. Jahun, Sarah L. Caddy, Ian Goodfellow, Alex Alderton, Roberto Amato, Sonia Goncalves, Ewan Harrison, David K. Jackson, Ian Johnston, Dominic Kwiatkowski, Cordelia Langford, John Sillitoe on behalf of the Wellcome Sanger Institute COVID-19 Surveillance Team (http://www.sanger.ac.uk/covid-team) |
| EPI_ISL_440204, EPI_ISL_440205, EPI_ISL_440206 | PHE South West Regional Laboratory, National Infection Service | Wellcome Sanger Institute for the COVID-19 Genomics UK Consortium | Stephanie Hutchings, Hannah Pymont, Dr Peter Muir, Barry Vipond, Rich Hopes, Alex Alderton, Roberto Amato, Sonia Goncalves, Ewan Harrison, David K. Jackson, Ian Johnston, Dominic Kwiatkowski, Cordelia Langford, John Sillitoe on behalf of the Wellcome Sanger Institute COVID-19 Surveillance Team (http://www.sanger.ac.uk/covid-team) |
| EPI_ISL_440207, EPI_ISL_440208, EPI_ISL_440209 | Department of Pathology, University of Cambridge | Wellcome Sanger Institute for the COVID-19 Genomics UK Consortium | Luke W Meredith, M. Estée Török , Myra Hosmillo, William L. Hamilton, Martin D. Curran, Theresa Feltwell, Grant Hall, Anna Yakovleva, Fahad A Khokhar, Charlotte J. Houldcroft, Laura G Caller, Aminu S. Jahun, Sarah L. Caddy, Ian Goodfellow, Alex Alderton, Roberto Amato, Sonia Goncalves, Ewan Harrison, David K. Jackson, Ian Johnston, Dominic Kwiatkowski, Cordelia Langford, John Sillitoe on behalf of the Wellcome Sanger Institute COVID-19 Surveillance Team (http://www.sanger.ac.uk/covid-team) |
| EPI_ISL_440210 | PHE South West Regional Laboratory, National Infection | Wellcome Sanger Institute for the COVID-19 Genomics UK Consortium | Stephanie Hutchings, Hannah Pymont, Dr Peter Muir, Barry Vipond, Rich Hopes, Alex Alderton, Roberto Amato, Sonia Goncalves, Ewan Harrison, David K. Jackson, Ian Johnston, Dominic Kwiatkowski, Cordelia |

| | | | |
|---|---|---|---|
| | Service | | Langford, John Sillitoe on behalf of the Wellcome Sanger Institute COVID-19 Surveillance Team (http://www.sanger.ac.uk/covid-team) |
| EPI_ISL_440211 | Department of Pathology, University of Cambridge | Wellcome Sanger Institute for the COVID-19 Genomics UK Consortium | Luke W Meredith, M. Estée Török , Myra Hosmillo, William L. Hamilton, Martin D. Curran, Theresa Feltwell, Grant Hall, Anna Yakovleva, Fahad A Khokhar, Charlotte J. Houldcroft, Laura G Caller, Aminu S. Jahun, Sarah L. Caddy, Ian Goodfellow, Alex Alderton, Roberto Amato, Sonia Goncalves, Ewan Harrison, David K. Jackson, Ian Johnston, Dominic Kwiatkowski, Cordelia Langford, John Sillitoe on behalf of the Wellcome Sanger Institute COVID-19 Surveillance Team (http://www.sanger.ac.uk/covid-team) |
| EPI_ISL_440212 | PHE South West Regional Laboratory, National Infection Service | Wellcome Sanger Institute for the COVID-19 Genomics UK Consortium | Stephanie Hutchings, Hannah Pymont, Dr Peter Muir, Barry Vipond, Rich Hopes, Alex Alderton, Roberto Amato, Sonia Goncalves, Ewan Harrison, David K. Jackson, Ian Johnston, Dominic Kwiatkowski, Cordelia Langford, John Sillitoe on behalf of the Wellcome Sanger Institute COVID-19 Surveillance Team (http://www.sanger.ac.uk/covid-team) |
| EPI_ISL_440213 | Department of Pathology, University of Cambridge | Wellcome Sanger Institute for the COVID-19 Genomics UK Consortium | Luke W Meredith, M. Estée Török , Myra Hosmillo, William L. Hamilton, Martin D. Curran, Theresa Feltwell, Grant Hall, Anna Yakovleva, Fahad A Khokhar, Charlotte J. Houldcroft, Laura G Caller, Aminu S. Jahun, Sarah L. Caddy, Ian Goodfellow, Alex Alderton, Roberto Amato, Sonia Goncalves, Ewan Harrison, David K. Jackson, Ian Johnston, Dominic Kwiatkowski, Cordelia Langford, John Sillitoe on behalf of the Wellcome Sanger Institute COVID-19 Surveillance Team (http://www.sanger.ac.uk/covid-team) |
| EPI_ISL_440214, EPI_ISL_440215, EPI_ISL_440216 | PHE South West Regional Laboratory, National Infection Service | Wellcome Sanger Institute for the COVID-19 Genomics UK Consortium | Stephanie Hutchings, Hannah Pymont, Dr Peter Muir, Barry Vipond, Rich Hopes, Alex Alderton, Roberto Amato, Sonia Goncalves, Ewan Harrison, David K. Jackson, Ian Johnston, Dominic Kwiatkowski, Cordelia Langford, John Sillitoe on behalf of the Wellcome Sanger Institute COVID-19 Surveillance Team (http://www.sanger.ac.uk/covid-team) |
| EPI_ISL_440217 | Department of Pathology, University of Cambridge | Wellcome Sanger Institute for the COVID-19 Genomics UK Consortium | Luke W Meredith, M. Estée Török , Myra Hosmillo, William L. Hamilton, Martin D. Curran, Theresa Feltwell, Grant Hall, Anna Yakovleva, Fahad A Khokhar, Charlotte J. Houldcroft, Laura G Caller, Aminu S. Jahun, Sarah L. Caddy, Ian Goodfellow, Alex Alderton, Roberto Amato, Sonia Goncalves, Ewan Harrison, David K. Jackson, Ian Johnston, Dominic Kwiatkowski, Cordelia Langford, John Sillitoe on behalf of the Wellcome Sanger Institute COVID-19 Surveillance Team (http://www.sanger.ac.uk/covid-team) |
| EPI_ISL_440218, EPI_ISL_440219, EPI_ISL_440220, EPI_ISL_440221, EPI_ISL_440222 | PHE South West Regional Laboratory, National Infection Service | Wellcome Sanger Institute for the COVID-19 Genomics UK Consortium | Stephanie Hutchings, Hannah Pymont, Dr Peter Muir, Barry Vipond, Rich Hopes, Alex Alderton, Roberto Amato, Sonia Goncalves, Ewan Harrison, David K. Jackson, Ian Johnston, Dominic Kwiatkowski, Cordelia Langford, John Sillitoe on behalf of the Wellcome Sanger Institute COVID-19 Surveillance Team (http://www.sanger.ac.uk/covid-team) |
| EPI_ISL_440223 | Department of Pathology, University of Cambridge | Wellcome Sanger Institute for the COVID-19 Genomics UK Consortium | Luke W Meredith, M. Estée Török , Myra Hosmillo, William L. Hamilton, Martin D. Curran, Theresa Feltwell, Grant Hall, Anna Yakovleva, Fahad A Khokhar, Charlotte J. Houldcroft, Laura G Caller, Aminu S. Jahun, Sarah L. Caddy, Ian Goodfellow, Alex Alderton, Roberto Amato, Sonia Goncalves, Ewan Harrison, David K. Jackson, Ian Johnston, Dominic Kwiatkowski, Cordelia Langford, John Sillitoe on behalf of the Wellcome Sanger Institute COVID-19 Surveillance Team (http://www.sanger.ac.uk/covid-team) |
| EPI_ISL_440224, EPI_ISL_440225 | PHE South West Regional Laboratory, National Infection Service | Wellcome Sanger Institute for the COVID-19 Genomics UK Consortium | Stephanie Hutchings, Hannah Pymont, Dr Peter Muir, Barry Vipond, Rich Hopes, Alex Alderton, Roberto Amato, Sonia Goncalves, Ewan Harrison, David K. Jackson, Ian Johnston, Dominic Kwiatkowski, Cordelia Langford, John Sillitoe on behalf of the Wellcome Sanger Institute COVID-19 Surveillance Team (http://www.sanger.ac.uk/covid-team) |
| EPI_ISL_440226 | Department of Pathology, University of Cambridge | Wellcome Sanger Institute for the COVID-19 Genomics UK Consortium | Luke W Meredith, M. Estée Török , Myra Hosmillo, William L. Hamilton, Martin D. Curran, Theresa Feltwell, Grant Hall, Anna Yakovleva, Fahad A Khokhar, Charlotte J. Houldcroft, Laura G Caller, Aminu S. Jahun, Sarah L. Caddy, Ian Goodfellow, Alex Alderton, Roberto Amato, Sonia Goncalves, Ewan Harrison, David K. Jackson, Ian Johnston, Dominic Kwiatkowski, Cordelia Langford, John Sillitoe on behalf of the Wellcome Sanger Institute COVID-19 Surveillance Team (http://www.sanger.ac.uk/covid-team) |
| EPI_ISL_440227, EPI_ISL_440228, EPI_ISL_440229 | PHE South West Regional Laboratory, National Infection Service | Wellcome Sanger Institute for the COVID-19 Genomics UK Consortium | Stephanie Hutchings, Hannah Pymont, Dr Peter Muir, Barry Vipond, Rich Hopes, Alex Alderton, Roberto Amato, Sonia Goncalves, Ewan Harrison, David K. Jackson, Ian Johnston, Dominic Kwiatkowski, Cordelia Langford, John Sillitoe on behalf of the Wellcome Sanger Institute COVID-19 Surveillance Team (http://www.sanger.ac.uk/covid-team) |
| EPI_ISL_440230, EPI_ISL_440231 | Department of Pathology, University of Cambridge | Wellcome Sanger Institute for the COVID-19 Genomics UK Consortium | Luke W Meredith, M. Estée Török , Myra Hosmillo, William L. Hamilton, Martin D. Curran, Theresa Feltwell, Grant Hall, Anna Yakovleva, Fahad A Khokhar, Charlotte J. Houldcroft, Laura G Caller, Aminu S. Jahun, Sarah L. Caddy, Ian Goodfellow, Alex Alderton, Roberto Amato, Sonia Goncalves, Ewan Harrison, David K. Jackson, Ian Johnston, Dominic Kwiatkowski, Cordelia Langford, John Sillitoe on behalf of the Wellcome Sanger Institute COVID-19 Surveillance Team (http://www.sanger.ac.uk/covid-team) |
| EPI_ISL_440232, EPI_ISL_440233 | PHE South West Regional Laboratory, National Infection Service | Wellcome Sanger Institute for the COVID-19 Genomics UK Consortium | Stephanie Hutchings, Hannah Pymont, Dr Peter Muir, Barry Vipond, Rich Hopes, Alex Alderton, Roberto Amato, Sonia Goncalves, Ewan Harrison, David K. Jackson, Ian Johnston, Dominic Kwiatkowski, Cordelia Langford, John Sillitoe on behalf of the Wellcome Sanger Institute COVID-19 Surveillance Team (http://www.sanger.ac.uk/covid-team) |
| EPI_ISL_440234 | Department of Pathology, University of Cambridge | Wellcome Sanger Institute for the COVID-19 Genomics UK Consortium | Luke W Meredith, M. Estée Török , Myra Hosmillo, William L. Hamilton, Martin D. Curran, Theresa Feltwell, Grant Hall, Anna Yakovleva, Fahad A Khokhar, Charlotte J. Houldcroft, Laura G Caller, Aminu S. Jahun, Sarah L. Caddy, Ian Goodfellow, Alex Alderton, Roberto Amato, Sonia Goncalves, Ewan Harrison, David K. Jackson, Ian Johnston, Dominic Kwiatkowski, Cordelia Langford, John Sillitoe on behalf of the Wellcome Sanger Institute COVID-19 Surveillance Team (http://www.sanger.ac.uk/covid-team) |
| EPI_ISL_440235, EPI_ISL_440236 | PHE South West Regional Laboratory, National Infection Service | Wellcome Sanger Institute for the COVID-19 Genomics UK Consortium | Stephanie Hutchings, Hannah Pymont, Dr Peter Muir, Barry Vipond, Rich Hopes, Alex Alderton, Roberto Amato, Sonia Goncalves, Ewan Harrison, David K. Jackson, Ian Johnston, Dominic Kwiatkowski, Cordelia Langford, John Sillitoe on behalf of the Wellcome Sanger Institute COVID-19 Surveillance Team (http://www.sanger.ac.uk/covid-team) |
| EPI_ISL_440237 | Department of Pathology, University of Cambridge | Wellcome Sanger Institute for the COVID-19 Genomics UK Consortium | Luke W Meredith, M. Estée Török , Myra Hosmillo, William L. Hamilton, Martin D. Curran, Theresa Feltwell, Grant Hall, Anna Yakovleva, Fahad A Khokhar, Charlotte J. Houldcroft, Laura G Caller, Aminu S. Jahun, Sarah L. Caddy, Ian Goodfellow, Alex Alderton, Roberto Amato, Sonia Goncalves, Ewan Harrison, David K. Jackson, Ian Johnston, Dominic Kwiatkowski, Cordelia Langford, John Sillitoe on behalf of the Wellcome Sanger Institute COVID-19 Surveillance Team (http://www.sanger.ac.uk/covid-team) |
| EPI_ISL_440238 | PHE South West Regional Laboratory, National Infection Service | Wellcome Sanger Institute for the COVID-19 Genomics UK Consortium | Stephanie Hutchings, Hannah Pymont, Dr Peter Muir, Barry Vipond, Rich Hopes, Alex Alderton, Roberto Amato, Sonia Goncalves, Ewan Harrison, David K. Jackson, Ian Johnston, Dominic Kwiatkowski, Cordelia Langford, John Sillitoe on behalf of the Wellcome Sanger Institute COVID-19 Surveillance Team (http://www.sanger.ac.uk/covid-team) |
| EPI_ISL_440239, EPI_ISL_440240 | Department of Pathology, University of Cambridge | Wellcome Sanger Institute for the COVID-19 Genomics UK Consortium | Luke W Meredith, M. Estée Török , Myra Hosmillo, William L. Hamilton, Martin D. Curran, Theresa Feltwell, Grant Hall, Anna Yakovleva, Fahad A Khokhar, Charlotte J. Houldcroft, Laura G Caller, Aminu S. Jahun, Sarah L. Caddy, Ian Goodfellow, Alex Alderton, Roberto Amato, Sonia Goncalves, Ewan Harrison, David K. Jackson, Ian Johnston, Dominic Kwiatkowski, Cordelia Langford, John Sillitoe on behalf of the Wellcome Sanger Institute COVID-19 Surveillance Team (http://www.sanger.ac.uk/covid-team) |
| EPI_ISL_440241, EPI_ISL_440242, EPI_ISL_440243 | PHE South West Regional Laboratory, National Infection Service | Wellcome Sanger Institute for the COVID-19 Genomics UK Consortium | Stephanie Hutchings, Hannah Pymont, Dr Peter Muir, Barry Vipond, Rich Hopes, Alex Alderton, Roberto Amato, Sonia Goncalves, Ewan Harrison, David K. Jackson, Ian Johnston, Dominic Kwiatkowski, Cordelia Langford, John Sillitoe on behalf of the Wellcome Sanger Institute COVID-19 Surveillance Team (http://www.sanger.ac.uk/covid-team) |
| EPI_ISL_440244 | Department of Pathology, University of Cambridge | Wellcome Sanger Institute for the COVID-19 Genomics UK Consortium | Luke W Meredith, M. Estée Török , Myra Hosmillo, William L. Hamilton, Martin D. Curran, Theresa Feltwell, Grant Hall, Anna Yakovleva, Fahad A Khokhar, Charlotte J. Houldcroft, Laura G Caller, Aminu S. Jahun, Sarah L. Caddy, Ian Goodfellow, Alex Alderton, Roberto Amato, Sonia Goncalves, Ewan Harrison, David K. Jackson, Ian Johnston, Dominic Kwiatkowski, Cordelia Langford, John Sillitoe on behalf of the Wellcome Sanger Institute COVID-19 Surveillance Team (http://www.sanger.ac.uk/covid-team) |
| EPI_ISL_440245 | PHE South West Regional Laboratory, National Infection Service | Wellcome Sanger Institute for the COVID-19 Genomics UK Consortium | Stephanie Hutchings, Hannah Pymont, Dr Peter Muir, Barry Vipond, Rich Hopes, Alex Alderton, Roberto Amato, Sonia Goncalves, Ewan Harrison, David K. Jackson, Ian Johnston, Dominic Kwiatkowski, Cordelia Langford, John Sillitoe on behalf of the Wellcome Sanger Institute COVID-19 Surveillance Team (http://www.sanger.ac.uk/covid-team) |
| EPI_ISL_440246, EPI_ISL_440247 | Department of Pathology, University of Cambridge | Wellcome Sanger Institute for the COVID-19 Genomics UK Consortium | Luke W Meredith, M. Estée Török , Myra Hosmillo, William L. Hamilton, Martin D. Curran, Theresa Feltwell, Grant Hall, Anna Yakovleva, Fahad A Khokhar, Charlotte J. Houldcroft, Laura G Caller, Aminu S. Jahun, Sarah L. Caddy, Ian Goodfellow, Alex Alderton, Roberto Amato, Sonia Goncalves, Ewan Harrison, David K. Jackson, Ian Johnston, Dominic Kwiatkowski, Cordelia Langford, John Sillitoe on behalf of the Wellcome Sanger Institute COVID-19 Surveillance Team (http://www.sanger.ac.uk/covid-team) |
| EPI_ISL_440248, EPI_ISL_440249, EPI_ISL_440250, EPI_ISL_440251, EPI_ISL_440252, EPI_ISL_440253 | PHE South West Regional Laboratory, National Infection Service | Wellcome Sanger Institute for the COVID-19 Genomics UK Consortium | Stephanie Hutchings, Hannah Pymont, Dr Peter Muir, Barry Vipond, Rich Hopes, Alex Alderton, Roberto Amato, Sonia Goncalves, Ewan Harrison, David K. Jackson, Ian Johnston, Dominic Kwiatkowski, Cordelia Langford, John Sillitoe on behalf of the Wellcome Sanger Institute COVID-19 Surveillance Team (http://www.sanger.ac.uk/covid-team) |
| EPI_ISL_440254 | Department of Pathology, University of Cambridge | Wellcome Sanger Institute for the COVID-19 Genomics UK Consortium | Luke W Meredith, M. Estée Török , Myra Hosmillo, William L. Hamilton, Martin D. Curran, Theresa Feltwell, Grant Hall, Anna Yakovleva, Fahad A Khokhar, Charlotte J. Houldcroft, Laura G Caller, Aminu S. Jahun, Sarah L. Caddy, Ian Goodfellow, Alex Alderton, Roberto Amato, Sonia Goncalves, Ewan Harrison, David K. Jackson, Ian Johnston, Dominic Kwiatkowski, Cordelia Langford, John Sillitoe on behalf of the Wellcome Sanger Institute COVID-19 Surveillance Team (http://www.sanger.ac.uk/covid-team) |
| EPI_ISL_440255, EPI_ISL_440256 | PHE South West Regional Laboratory, National Infection Service | Wellcome Sanger Institute for the COVID-19 Genomics UK Consortium | Stephanie Hutchings, Hannah Pymont, Dr Peter Muir, Barry Vipond, Rich Hopes, Alex Alderton, Roberto Amato, Sonia Goncalves, Ewan Harrison, David K. Jackson, Ian Johnston, Dominic Kwiatkowski, Cordelia Langford, John Sillitoe on behalf of the Wellcome Sanger Institute COVID-19 Surveillance Team (http://www.sanger.ac.uk/covid-team) |
| EPI_ISL_440257, EPI_ISL_440258, EPI_ISL_440259, EPI_ISL_440260, EPI_ISL_440261 | Department of Pathology, University of Cambridge | Wellcome Sanger Institute for the COVID-19 Genomics UK Consortium | Luke W Meredith, M. Estée Török , Myra Hosmillo, William L. Hamilton, Martin D. Curran, Theresa Feltwell, Grant Hall, Anna Yakovleva, Fahad A Khokhar, Charlotte J. Houldcroft, Laura G Caller, Aminu S. Jahun, Sarah L. Caddy, Ian Goodfellow, Alex Alderton, Roberto Amato, Sonia Goncalves, Ewan Harrison, David K. Jackson, Ian Johnston, Dominic Kwiatkowski, Cordelia Langford, John Sillitoe on behalf of the Wellcome Sanger Institute COVID-19 Surveillance Team (http://www.sanger.ac.uk/covid-team) |
| EPI_ISL_440262 | PHE South West Regional Laboratory, National Infection Service | Wellcome Sanger Institute for the COVID-19 Genomics UK Consortium | Stephanie Hutchings, Hannah Pymont, Dr Peter Muir, Barry Vipond, Rich Hopes, Alex Alderton, Roberto Amato, Sonia Goncalves, Ewan Harrison, David K. Jackson, Ian Johnston, Dominic Kwiatkowski, Cordelia Langford, John Sillitoe on behalf of the Wellcome Sanger Institute COVID-19 Surveillance Team (http://www.sanger.ac.uk/covid-team) |
| EPI_ISL_440263, EPI_ISL_440264, EPI_ISL_440265 | Department of Pathology, University of Cambridge | Wellcome Sanger Institute for the COVID-19 Genomics UK Consortium | Luke W Meredith, M. Estée Török , Myra Hosmillo, William L. Hamilton, Martin D. Curran, Theresa Feltwell, Grant Hall, Anna Yakovleva, Fahad A Khokhar, Charlotte J. Houldcroft, Laura G Caller, Aminu S. Jahun, Sarah L. Caddy, Ian Goodfellow, Alex Alderton, Roberto Amato, Sonia Goncalves, Ewan Harrison, David K. Jackson, Ian Johnston, Dominic Kwiatkowski, Cordelia Langford, John Sillitoe on behalf of the Wellcome Sanger Institute COVID-19 Surveillance Team (http://www.sanger.ac.uk/covid-team) |
| EPI_ISL_440266 | PHE South West Regional Laboratory, National Infection Service | Wellcome Sanger Institute for the COVID-19 Genomics UK Consortium | Stephanie Hutchings, Hannah Pymont, Dr Peter Muir, Barry Vipond, Rich Hopes, Alex Alderton, Roberto Amato, Sonia Goncalves, Ewan Harrison, David K. Jackson, Ian Johnston, Dominic Kwiatkowski, Cordelia Langford, John Sillitoe on behalf of the Wellcome Sanger Institute COVID-19 Surveillance Team (http://www.sanger.ac.uk/covid-team) |
| EPI_ISL_440267, EPI_ISL_440268 | Department of Pathology, University of Cambridge | Wellcome Sanger Institute for the COVID-19 Genomics UK Consortium | Luke W Meredith, M. Estée Török , Myra Hosmillo, William L. Hamilton, Martin D. Curran, Theresa Feltwell, Grant Hall, Anna Yakovleva, Fahad A Khokhar, Charlotte J. Houldcroft, Laura G Caller, Aminu S. Jahun, Sarah L. Caddy, Ian Goodfellow, Alex Alderton, Roberto Amato, Sonia Goncalves, Ewan Harrison, David K. Jackson, Ian Johnston, Dominic Kwiatkowski, Cordelia Langford, John Sillitoe on behalf of the Wellcome Sanger Institute COVID-19 Surveillance Team (http://www.sanger.ac.uk/covid-team) |
| EPI_ISL_440269 | PHE South West Regional Laboratory, National Infection Service | Wellcome Sanger Institute for the COVID-19 Genomics UK Consortium | Stephanie Hutchings, Hannah Pymont, Dr Peter Muir, Barry Vipond, Rich Hopes, Alex Alderton, Roberto Amato, Sonia Goncalves, Ewan Harrison, David K. Jackson, Ian Johnston, Dominic Kwiatkowski, Cordelia Langford, John Sillitoe on behalf of the Wellcome Sanger Institute COVID-19 Surveillance Team (http://www.sanger.ac.uk/covid-team) |
| EPI_ISL_440270, EPI_ISL_440271 | Department of Pathology, University of Cambridge | Wellcome Sanger Institute for the COVID-19 Genomics UK Consortium | Luke W Meredith, M. Estée Török , Myra Hosmillo, William L. Hamilton, Martin D. Curran, Theresa Feltwell, Grant Hall, Anna Yakovleva, Fahad A Khokhar, Charlotte J. Houldcroft, Laura G Caller, Aminu S. Jahun, Sarah L. Caddy, Ian Goodfellow, Alex Alderton, Roberto Amato, Sonia Goncalves, Ewan Harrison, David K. Jackson, Ian Johnston, Dominic Kwiatkowski, Cordelia Langford, John Sillitoe on behalf of the Wellcome Sanger Institute COVID-19 Surveillance Team (http://www.sanger.ac.uk/covid-team) |
| EPI_ISL_440272, EPI_ISL_440273, EPI_ISL_440274, EPI_ISL_440275, EPI_ISL_440276, EPI_ISL_440277 | PHE South West Regional Laboratory, National Infection Service | Wellcome Sanger Institute for the COVID-19 Genomics UK Consortium | Stephanie Hutchings, Hannah Pymont, Dr Peter Muir, Barry Vipond, Rich Hopes, Alex Alderton, Roberto Amato, Sonia Goncalves, Ewan Harrison, David K. Jackson, Ian Johnston, Dominic Kwiatkowski, Cordelia Langford, John Sillitoe on behalf of the Wellcome Sanger Institute COVID-19 Surveillance Team (http://www.sanger.ac.uk/covid-team) |
| EPI_ISL_440278, EPI_ISL_440279, EPI_ISL_440280, EPI_ISL_440281, EPI_ISL_440282, EPI_ISL_440283, EPI_ISL_440284, EPI_ISL_440285 | Department of Pathology, University of Cambridge | Wellcome Sanger Institute for the COVID-19 Genomics UK Consortium | Luke W Meredith, M. Estée Török , Myra Hosmillo, William L. Hamilton, Martin D. Curran, Theresa Feltwell, Grant Hall, Anna Yakovleva, Fahad A Khokhar, Charlotte J. Houldcroft, Laura G Caller, Aminu S. Jahun, Sarah L. Caddy, Ian Goodfellow, Alex Alderton, Roberto Amato, Sonia Goncalves, Ewan Harrison, David K. Jackson, Ian Johnston, Dominic Kwiatkowski, Cordelia Langford, John Sillitoe on behalf of the Wellcome Sanger Institute COVID-19 Surveillance Team (http://www.sanger.ac.uk/covid-team) |
| EPI_ISL_440286 | PHE South West Regional Laboratory, National Infection Service | Wellcome Sanger Institute for the COVID-19 Genomics UK Consortium | Stephanie Hutchings, Hannah Pymont, Dr Peter Muir, Barry Vipond, Rich Hopes, Alex Alderton, Roberto Amato, Sonia Goncalves, Ewan Harrison, David K. Jackson, Ian Johnston, Dominic Kwiatkowski, Cordelia Langford, John Sillitoe on behalf of the Wellcome Sanger Institute COVID-19 Surveillance Team (http://www.sanger.ac.uk/covid-team) |
| EPI_ISL_440287, EPI_ISL_440288, EPI_ISL_440289, EPI_ISL_440290 | Department of Pathology, University of Cambridge | Wellcome Sanger Institute for the COVID-19 Genomics UK Consortium | Luke W Meredith, M. Estée Török , Myra Hosmillo, William L. Hamilton, Martin D. Curran, Theresa Feltwell, Grant Hall, Anna Yakovleva, Fahad A Khokhar, Charlotte J. Houldcroft, Laura G Caller, Aminu S. Jahun, Sarah L. Caddy, Ian Goodfellow, Alex Alderton, Roberto Amato, Sonia Goncalves, Ewan Harrison, David K. Jackson, Ian Johnston, Dominic Kwiatkowski, Cordelia Langford, John Sillitoe on behalf of the Wellcome Sanger Institute COVID-19 Surveillance Team (http://www.sanger.ac.uk/covid-team) |
| EPI_ISL_440291, EPI_ISL_440292 | PHE South West Regional Laboratory, National Infection Service | Wellcome Sanger Institute for the COVID-19 Genomics UK Consortium | Stephanie Hutchings, Hannah Pymont, Dr Peter Muir, Barry Vipond, Rich Hopes, Alex Alderton, Roberto Amato, Sonia Goncalves, Ewan Harrison, David K. Jackson, Ian Johnston, Dominic Kwiatkowski, Cordelia Langford, John Sillitoe on behalf of the Wellcome Sanger Institute COVID-19 Surveillance Team (http://www.sanger.ac.uk/covid-team) |
| EPI_ISL_440293 | Department of Pathology, University of Cambridge | Wellcome Sanger Institute for the COVID-19 Genomics UK Consortium | Luke W Meredith, M. Estée Török , Myra Hosmillo, William L. Hamilton, Martin D. Curran, Theresa Feltwell, Grant Hall, Anna Yakovleva, Fahad A Khokhar, Charlotte J. Houldcroft, Laura G Caller, Aminu S. Jahun, Sarah L. Caddy, Ian Goodfellow, Alex Alderton, Roberto Amato, Sonia Goncalves, Ewan Harrison, David K. Jackson, Ian Johnston, Dominic Kwiatkowski, Cordelia Langford, John Sillitoe on behalf of the Wellcome Sanger Institute COVID-19 Surveillance Team (http://www.sanger.ac.uk/covid-team) |

| | | | |
|---|---|---|---|
| EPI_ISL_440294 | PHE South West Regional Laboratory, National Infection Service | Wellcome Sanger Institute for the COVID-19 Genomics UK Consortium | Stephanie Hutchings, Hannah Pymont, Dr Peter Muir, Barry Vipond, Rich Hopes, Alex Alderton, Roberto Amato, Sonia Goncalves, Ewan Harrison, David K. Jackson, Ian Johnston, Dominic Kwiatkowski, Cordelia Langford, John Sillitoe on behalf of the Wellcome Sanger Institute COVID-19 Surveillance Team (http://www.sanger.ac.uk/covid-team) |
| EPI_ISL_440295, EPI_ISL_440296, EPI_ISL_440297 | Department of Pathology, University of Cambridge | Wellcome Sanger Institute for the COVID-19 Genomics UK Consortium | Luke W Meredith, M. Estée Török , Myra Hosmillo, William L. Hamilton, Martin D. Curran, Theresa Feltwell, Grant Hall, Anna Yakovleva, Fahad A Khokhar, Charlotte J. Houldcroft, Laura G Caller, Aminu S. Jahun, Sarah L. Caddy, Ian Goodfellow, Alex Alderton, Roberto Amato, Sonia Goncalves, Ewan Harrison, David K. Jackson, Ian Johnston, Dominic Kwiatkowski, Cordelia Langford, John Sillitoe on behalf of the Wellcome Sanger Institute COVID-19 Surveillance Team (http://www.sanger.ac.uk/covid-team) |
| EPI_ISL_440298, EPI_ISL_440299 | PHE South West Regional Laboratory, National Infection Service | Wellcome Sanger Institute for the COVID-19 Genomics UK Consortium | Stephanie Hutchings, Hannah Pymont, Dr Peter Muir, Barry Vipond, Rich Hopes, Alex Alderton, Roberto Amato, Sonia Goncalves, Ewan Harrison, David K. Jackson, Ian Johnston, Dominic Kwiatkowski, Cordelia Langford, John Sillitoe on behalf of the Wellcome Sanger Institute COVID-19 Surveillance Team (http://www.sanger.ac.uk/covid-team) |
| EPI_ISL_440300, EPI_ISL_440301, EPI_ISL_440302, EPI_ISL_440303, EPI_ISL_440304, EPI_ISL_440305, EPI_ISL_440306, EPI_ISL_440307, EPI_ISL_440308, EPI_ISL_440309, EPI_ISL_440310, EPI_ISL_440311, EPI_ISL_440312, EPI_ISL_440313, EPI_ISL_440314, EPI_ISL_440315, EPI_ISL_440316, EPI_ISL_440317, EPI_ISL_440318, EPI_ISL_440319, EPI_ISL_440320, EPI_ISL_440321, EPI_ISL_440322, EPI_ISL_440323, EPI_ISL_440324, EPI_ISL_440325, EPI_ISL_440326, EPI_ISL_440327, EPI_ISL_440328, EPI_ISL_440329, EPI_ISL_440330, EPI_ISL_440331, EPI_ISL_440332, EPI_ISL_440333, EPI_ISL_440334, EPI_ISL_440335, EPI_ISL_440336, EPI_ISL_440337, EPI_ISL_440338, EPI_ISL_440339, EPI_ISL_440340, EPI_ISL_440341, EPI_ISL_440342, EPI_ISL_440343, EPI_ISL_440344, EPI_ISL_440345, EPI_ISL_440346, EPI_ISL_440347, EPI_ISL_440348, EPI_ISL_440349, EPI_ISL_440350, EPI_ISL_440351, EPI_ISL_440352, EPI_ISL_440353, EPI_ISL_440354, EPI_ISL_440355, EPI_ISL_440356, EPI_ISL_440357, EPI_ISL_440358, EPI_ISL_440359, EPI_ISL_440360, EPI_ISL_440361, EPI_ISL_440362, EPI_ISL_440363, EPI_ISL_440364, EPI_ISL_440365, EPI_ISL_440366, EPI_ISL_440367, EPI_ISL_440368, EPI_ISL_440369, EPI_ISL_440370, EPI_ISL_440371, EPI_ISL_440372, EPI_ISL_440373, EPI_ISL_440374, EPI_ISL_440375, EPI_ISL_440376, EPI_ISL_440377, EPI_ISL_440378, EPI_ISL_440379, EPI_ISL_440380, EPI_ISL_440381, EPI_ISL_440382, EPI_ISL_440383, EPI_ISL_440384, EPI_ISL_440385, EPI_ISL_440386, EPI_ISL_440387, EPI_ISL_440388, EPI_ISL_440389, EPI_ISL_440390, EPI_ISL_440391, EPI_ISL_440392, EPI_ISL_440393, EPI_ISL_440394, EPI_ISL_440395, EPI_ISL_440396, EPI_ISL_440397, EPI_ISL_440398, EPI_ISL_440399, EPI_ISL_440400, EPI_ISL_440401, EPI_ISL_440402, EPI_ISL_440403, EPI_ISL_440404, EPI_ISL_440405, EPI_ISL_440406, EPI_ISL_440407, EPI_ISL_440408, EPI_ISL_440409, EPI_ISL_440410, EPI_ISL_440411, EPI_ISL_440412, EPI_ISL_440413, EPI_ISL_440414, EPI_ISL_440415, EPI_ISL_440416, EPI_ISL_440417, EPI_ISL_440418, EPI_ISL_440419, EPI_ISL_440420, EPI_ISL_440421, EPI_ISL_440422, EPI_ISL_440423, EPI_ISL_440424, EPI_ISL_440425, EPI_ISL_440426, EPI_ISL_440427, EPI_ISL_440428, EPI_ISL_440429, EPI_ISL_440430, EPI_ISL_440431, EPI_ISL_440432, EPI_ISL_440433, EPI_ISL_440434, EPI_ISL_440435, EPI_ISL_440436, EPI_ISL_440437, EPI_ISL_440438, EPI_ISL_440439, EPI_ISL_440440, EPI_ISL_440441, EPI_ISL_440442, EPI_ISL_440443, EPI_ISL_440444, EPI_ISL_440445, EPI_ISL_440446, EPI_ISL_440447, EPI_ISL_440448, EPI_ISL_440449, EPI_ISL_440450, EPI_ISL_440451, EPI_ISL_440452, EPI_ISL_440453, EPI_ISL_440454, EPI_ISL_440455, EPI_ISL_440456, EPI_ISL_440457, EPI_ISL_440458, EPI_ISL_440459, EPI_ISL_440460, EPI_ISL_440461, EPI_ISL_440462, EPI_ISL_440463, EPI_ISL_440464, EPI_ISL_440465, EPI_ISL_440466, EPI_ISL_440467, EPI_ISL_440468, EPI_ISL_440469, EPI_ISL_440470, EPI_ISL_440471, EPI_ISL_440472, EPI_ISL_440473, EPI_ISL_440474, EPI_ISL_440475, EPI_ISL_440476, EPI_ISL_440477, EPI_ISL_440478, EPI_ISL_440479, EPI_ISL_440480, EPI_ISL_440481, EPI_ISL_440482, EPI_ISL_440483, EPI_ISL_440484, EPI_ISL_440485, EPI_ISL_440486, EPI_ISL_440487, EPI_ISL_440488, EPI_ISL_440489, EPI_ISL_440490, EPI_ISL_440491, EPI_ISL_440492, EPI_ISL_440493, EPI_ISL_440494, EPI_ISL_440495, EPI_ISL_440496, EPI_ISL_440497, EPI_ISL_440498, EPI_ISL_440499, EPI_ISL_440500, EPI_ISL_440501, EPI_ISL_440502, EPI_ISL_440503, EPI_ISL_440504, EPI_ISL_440505, EPI_ISL_440506, EPI_ISL_440507, EPI_ISL_440508, EPI_ISL_440509, EPI_ISL_440510, EPI_ISL_440511, EPI_ISL_440512, EPI_ISL_440513, EPI_ISL_440514, EPI_ISL_440515, EPI_ISL_440516, EPI_ISL_440517, EPI_ISL_440518, EPI_ISL_440519, EPI_ISL_440520, EPI_ISL_440521, EPI_ISL_440522, EPI_ISL_440523, EPI_ISL_440524, EPI_ISL_440525, EPI_ISL_440526, EPI_ISL_440527, EPI_ISL_440528, EPI_ISL_440529, EPI_ISL_440530, EPI_ISL_440531, EPI_ISL_440532, EPI_ISL_440533, EPI_ISL_440534, EPI_ISL_440535, EPI_ISL_440536, EPI_ISL_440537, EPI_ISL_440538, EPI_ISL_440539, EPI_ISL_440540, EPI_ISL_440541, EPI_ISL_440542, EPI_ISL_440543, EPI_ISL_440544, EPI_ISL_440545, EPI_ISL_440546, EPI_ISL_440547, EPI_ISL_440548, EPI_ISL_440549, EPI_ISL_440550, EPI_ISL_440551, EPI_ISL_440552, EPI_ISL_440553, EPI_ISL_440554, EPI_ISL_440555, EPI_ISL_440556, EPI_ISL_440557, EPI_ISL_440558, EPI_ISL_440559, EPI_ISL_440560, EPI_ISL_440561, EPI_ISL_440562, EPI_ISL_440563, EPI_ISL_440564, EPI_ISL_440565, EPI_ISL_440566, EPI_ISL_440567, EPI_ISL_440568, EPI_ISL_440569, EPI_ISL_440570, EPI_ISL_440571, EPI_ISL_440572, EPI_ISL_440573, EPI_ISL_440574, EPI_ISL_440575, EPI_ISL_440576, EPI_ISL_440577, EPI_ISL_440578, EPI_ISL_440579, EPI_ISL_440580, EPI_ISL_440581, EPI_ISL_440582, EPI_ISL_440583, EPI_ISL_440584, EPI_ISL_440585, EPI_ISL_440586, EPI_ISL_440587, EPI_ISL_440588, EPI_ISL_440589, EPI_ISL_440590, EPI_ISL_440591, EPI_ISL_440592, EPI_ISL_440593, EPI_ISL_440594, EPI_ISL_440595, EPI_ISL_440596, EPI_ISL_440597, EPI_ISL_440598, EPI_ISL_440599, EPI_ISL_440600, EPI_ISL_440601, EPI_ISL_440602, EPI_ISL_440603, EPI_ISL_440604, EPI_ISL_440605, EPI_ISL_440606, EPI_ISL_440607, EPI_ISL_440608, EPI_ISL_440609, EPI_ISL_440610, EPI_ISL_440611, EPI_ISL_440612, EPI_ISL_440613, EPI_ISL_440614, EPI_ISL_440615, EPI_ISL_440616, EPI_ISL_440617, EPI_ISL_440618, EPI_ISL_440619, EPI_ISL_440620, EPI_ISL_440621, EPI_ISL_440622 | | | |
| see above | Department of Pathology, University of Cambridge | Wellcome Sanger Institute for the COVID-19 Genomics UK Consortium | Luke W Meredith, M. Estée Török , Myra Hosmillo, William L. Hamilton, Martin D. Curran, Theresa Feltwell, Grant Hall, Anna Yakovleva, Fahad A Khokhar, Charlotte J. Houldcroft, Laura G Caller, Aminu S. Jahun, Sarah L. Caddy, Ian Goodfellow, Alex Alderton, Roberto Amato, Sonia Goncalves, Ewan Harrison, David K. Jackson, Ian Johnston, Dominic Kwiatkowski, Cordelia Langford, John Sillitoe on behalf of the Wellcome Sanger Institute COVID-19 Surveillance Team (http://www.sanger.ac.uk/covid-team) |
| EPI_ISL_440623, EPI_ISL_440624, EPI_ISL_440625, EPI_ISL_440626, EPI_ISL_440627, EPI_ISL_440628, EPI_ISL_440629, EPI_ISL_440630, EPI_ISL_440631, EPI_ISL_440632, EPI_ISL_440633, EPI_ISL_440634, EPI_ISL_440635, EPI_ISL_440636, EPI_ISL_440637, EPI_ISL_440638, EPI_ISL_440639, EPI_ISL_440640, EPI_ISL_440641, EPI_ISL_440642, EPI_ISL_440643, EPI_ISL_440644, EPI_ISL_440645, EPI_ISL_440646, EPI_ISL_440647, EPI_ISL_440648, EPI_ISL_440649, EPI_ISL_440650, EPI_ISL_440651, EPI_ISL_440652, EPI_ISL_440653, EPI_ISL_440654, EPI_ISL_440655, EPI_ISL_440656, EPI_ISL_440657, EPI_ISL_440658, EPI_ISL_440659, EPI_ISL_440660, EPI_ISL_440661, EPI_ISL_440662, EPI_ISL_440663, EPI_ISL_440664, EPI_ISL_440665, EPI_ISL_440666, EPI_ISL_440667, EPI_ISL_440668, EPI_ISL_440669, EPI_ISL_440670, EPI_ISL_440671, EPI_ISL_440672, EPI_ISL_440673, EPI_ISL_440674, EPI_ISL_440675, EPI_ISL_440676, EPI_ISL_440677, EPI_ISL_440678, EPI_ISL_440679, EPI_ISL_440680, EPI_ISL_440681, EPI_ISL_440682, EPI_ISL_440683, EPI_ISL_440684, EPI_ISL_440685, EPI_ISL_440686, EPI_ISL_440687, EPI_ISL_440688, EPI_ISL_440689, EPI_ISL_440690, EPI_ISL_440691, EPI_ISL_440692, EPI_ISL_440693, EPI_ISL_440694, EPI_ISL_440695, EPI_ISL_440696, EPI_ISL_440697, EPI_ISL_440698, EPI_ISL_440699, EPI_ISL_440700, EPI_ISL_440701, EPI_ISL_440702, EPI_ISL_440703, EPI_ISL_440704, EPI_ISL_440705, EPI_ISL_440706, EPI_ISL_440707, EPI_ISL_440708, EPI_ISL_440709, EPI_ISL_440710, EPI_ISL_440711, EPI_ISL_440712, EPI_ISL_440713, EPI_ISL_440714, EPI_ISL_440715, EPI_ISL_440716, EPI_ISL_440717, EPI_ISL_440718, EPI_ISL_440719, EPI_ISL_440720, EPI_ISL_440721, EPI_ISL_440722, EPI_ISL_440723, EPI_ISL_440724, EPI_ISL_440725, EPI_ISL_440726, EPI_ISL_440727, EPI_ISL_440728, EPI_ISL_440729, EPI_ISL_440730, EPI_ISL_440731, EPI_ISL_440732, EPI_ISL_440733, EPI_ISL_440734, EPI_ISL_440735, EPI_ISL_440736, EPI_ISL_440737, EPI_ISL_440738, EPI_ISL_440739, EPI_ISL_440740, EPI_ISL_440741, EPI_ISL_440742, EPI_ISL_440743, EPI_ISL_440744, EPI_ISL_440745, EPI_ISL_440746, EPI_ISL_440747, EPI_ISL_440748, EPI_ISL_440749, EPI_ISL_440750, EPI_ISL_440751, EPI_ISL_440752, EPI_ISL_440753, EPI_ISL_440754, EPI_ISL_440755, EPI_ISL_440756, EPI_ISL_440757, EPI_ISL_440758, EPI_ISL_440759, EPI_ISL_440760, EPI_ISL_440761, EPI_ISL_440762, EPI_ISL_440763, EPI_ISL_440764, EPI_ISL_440765, EPI_ISL_440766, EPI_ISL_440767, EPI_ISL_440768, EPI_ISL_440769, EPI_ISL_440770, EPI_ISL_440771, EPI_ISL_440772, EPI_ISL_440773, EPI_ISL_440774, EPI_ISL_440775, EPI_ISL_440776, EPI_ISL_440777, EPI_ISL_440778, EPI_ISL_440779, EPI_ISL_440780, EPI_ISL_440781, EPI_ISL_440782, EPI_ISL_440783, EPI_ISL_440784, EPI_ISL_440785, EPI_ISL_440786, EPI_ISL_440787, EPI_ISL_440788, EPI_ISL_440789, EPI_ISL_440790, EPI_ISL_440791, EPI_ISL_440792, EPI_ISL_440793, EPI_ISL_440794, EPI_ISL_440795, EPI_ISL_440796, EPI_ISL_440797, EPI_ISL_440798, EPI_ISL_440799, EPI_ISL_440800, EPI_ISL_440801, EPI_ISL_440802, EPI_ISL_440803, EPI_ISL_440804, EPI_ISL_440805, EPI_ISL_440806, EPI_ISL_440807, EPI_ISL_440808, EPI_ISL_440809 | | | |
| see above | PHE South West Regional Laboratory, National Infection Service | Wellcome Sanger Institute for the COVID-19 Genomics UK Consortium | Stephanie Hutchings, Hannah Pymont, Dr Peter Muir, Barry Vipond, Rich Hopes, Alex Alderton, Roberto Amato, Sonia Goncalves, Ewan Harrison, David K. Jackson, Ian Johnston, Dominic Kwiatkowski, Cordelia Langford, John Sillitoe on behalf of the Wellcome Sanger Institute COVID-19 Surveillance Team (http://www.sanger.ac.uk/covid-team) |
| EPI_ISL_440810, EPI_ISL_440811, EPI_ISL_440812, EPI_ISL_440813, EPI_ISL_440814, EPI_ISL_440815, EPI_ISL_440816, EPI_ISL_440817, EPI_ISL_440818, EPI_ISL_440819, EPI_ISL_440820, EPI_ISL_440821, EPI_ISL_440822, EPI_ISL_440823, EPI_ISL_440824, EPI_ISL_440825, EPI_ISL_440826, EPI_ISL_440827, EPI_ISL_440828, EPI_ISL_440829, EPI_ISL_440830, EPI_ISL_440831, EPI_ISL_440832, EPI_ISL_440833, EPI_ISL_440834, EPI_ISL_440835, EPI_ISL_440836, EPI_ISL_440837, EPI_ISL_440838, EPI_ISL_440839, EPI_ISL_440840, EPI_ISL_440841, EPI_ISL_440842, EPI_ISL_440843, EPI_ISL_440844, EPI_ISL_440845, EPI_ISL_440846, EPI_ISL_440847, EPI_ISL_440848, EPI_ISL_440849, EPI_ISL_440850, EPI_ISL_440851 | | | |
| see above | Department of Pathology, University of Cambridge | Wellcome Sanger Institute for the COVID-19 Genomics UK Consortium | Luke W Meredith, M. Estée Török , Myra Hosmillo, William L. Hamilton, Martin D. Curran, Theresa Feltwell, Grant Hall, Anna Yakovleva, Fahad A Khokhar, Charlotte J. Houldcroft, Laura G Caller, Aminu S. Jahun, Sarah L. Caddy, Ian Goodfellow, Alex Alderton, Roberto Amato, Sonia Goncalves, Ewan Harrison, David K. Jackson, Ian Johnston, Dominic Kwiatkowski, Cordelia Langford, John Sillitoe on behalf of the Wellcome Sanger Institute COVID-19 Surveillance Team (http://www.sanger.ac.uk/covid-team) |
| EPI_ISL_440852, EPI_ISL_440853, EPI_ISL_440854, EPI_ISL_440855, EPI_ISL_440856, EPI_ISL_440857, EPI_ISL_440858, EPI_ISL_440859, EPI_ISL_440860, EPI_ISL_440861, EPI_ISL_440862, EPI_ISL_440863, EPI_ISL_440864, EPI_ISL_440865, EPI_ISL_440866, EPI_ISL_440867, EPI_ISL_440868, EPI_ISL_440869, EPI_ISL_440870, EPI_ISL_440871, EPI_ISL_440872, EPI_ISL_440873, EPI_ISL_440874, EPI_ISL_440875, EPI_ISL_440876, EPI_ISL_440877, EPI_ISL_440878, EPI_ISL_440879, EPI_ISL_440880, EPI_ISL_440881, EPI_ISL_440882, EPI_ISL_440883, EPI_ISL_440884, EPI_ISL_440885, EPI_ISL_440886, EPI_ISL_440887, EPI_ISL_440888, EPI_ISL_440889, EPI_ISL_440890, EPI_ISL_440891, EPI_ISL_440892, EPI_ISL_440893, EPI_ISL_440894, EPI_ISL_440895, EPI_ISL_440896, EPI_ISL_440897, EPI_ISL_440898, EPI_ISL_440899, EPI_ISL_440900, EPI_ISL_440901, EPI_ISL_440902, EPI_ISL_440903, EPI_ISL_440904, EPI_ISL_440905, EPI_ISL_440906, EPI_ISL_440907, EPI_ISL_440908, EPI_ISL_440909, EPI_ISL_440910, EPI_ISL_440911, EPI_ISL_440912, EPI_ISL_440913, EPI_ISL_440914, EPI_ISL_440915, EPI_ISL_440916, EPI_ISL_440917, EPI_ISL_440918, EPI_ISL_440919, EPI_ISL_440920, EPI_ISL_440921, EPI_ISL_440922, EPI_ISL_440923, EPI_ISL_440924, EPI_ISL_440925, EPI_ISL_440926, EPI_ISL_440927, EPI_ISL_440928, EPI_ISL_440929, EPI_ISL_440930, EPI_ISL_440931, EPI_ISL_440932, EPI_ISL_440933, EPI_ISL_440934, EPI_ISL_440935, EPI_ISL_440936, EPI_ISL_440937, EPI_ISL_440938, EPI_ISL_440939, EPI_ISL_440940, EPI_ISL_440941, EPI_ISL_440942, EPI_ISL_440943, EPI_ISL_440944, EPI_ISL_440945, EPI_ISL_440946, EPI_ISL_440947, EPI_ISL_440948, EPI_ISL_440949, EPI_ISL_440950 | | | |
| see above | Liverpool Clinical Laboratories | COVID-19 Genomics UK (COG-UK) Consortium | Sam Haldenby, Anita Lucaci, Steve Paterson, Julian Hiscox, Alistair Darby, M Almsaud, A Alrezaihi, Muhannad Alruwaili, Stuart D Armstrong, Jones Benjamin , Eleanor G Bentley, Anu Chawla, Jordan J Clark, Angela Cowell, Richard Eccles, Isabel Garc√≠a-Dorival, Matthew Gemmell, Alessandro Gerada, PKF Gilmore, Richard Gregory, Ximeng Han, Catherine Hartley, Margaret Hughes, Miren Iturriza-Gomara, James Johnson, L Luu, Jenifer Manson , Charlotte Nelson, Elaine O‚ÄôToole, Cassie Olateju, Rebekah Penrice-Randal¬†, Lucille Rainbow, N.P Randle, Trevor Ian Robinson, Parul Sharma, Ghada T Shawli, James P Stewart , Neil Swainston, Ecaterina Vamos, Joanne Watts, Mark Whitehead |
| EPI_ISL_440951, EPI_ISL_440952, EPI_ISL_440953, EPI_ISL_440954, EPI_ISL_440955, EPI_ISL_440956, EPI_ISL_440957, EPI_ISL_440958, EPI_ISL_440959, EPI_ISL_440960, EPI_ISL_440961, EPI_ISL_440962, EPI_ISL_440963, EPI_ISL_440964, EPI_ISL_440965, EPI_ISL_440966, EPI_ISL_440967, EPI_ISL_440968, EPI_ISL_440969, EPI_ISL_440970, EPI_ISL_440971, EPI_ISL_440972, EPI_ISL_440973, EPI_ISL_440974, EPI_ISL_440975, EPI_ISL_440976, EPI_ISL_440977, EPI_ISL_440978, EPI_ISL_440979, EPI_ISL_440980, EPI_ISL_440981, EPI_ISL_440982, EPI_ISL_440983, EPI_ISL_440984, EPI_ISL_440985, EPI_ISL_440986, EPI_ISL_440987, EPI_ISL_440988, EPI_ISL_440989, EPI_ISL_440990, EPI_ISL_440991, EPI_ISL_440992, EPI_ISL_440993, EPI_ISL_440994, EPI_ISL_440995, EPI_ISL_440996, EPI_ISL_440997, EPI_ISL_440998, EPI_ISL_440999, EPI_ISL_441000, EPI_ISL_441001, EPI_ISL_441002, EPI_ISL_441003, EPI_ISL_441004, EPI_ISL_441005, EPI_ISL_441006, EPI_ISL_441007, EPI_ISL_441008, EPI_ISL_441009, EPI_ISL_441010, EPI_ISL_441011, EPI_ISL_441012, EPI_ISL_441013, EPI_ISL_441014, EPI_ISL_441015, EPI_ISL_441016, EPI_ISL_441017, EPI_ISL_441018, EPI_ISL_441019, EPI_ISL_441020, EPI_ISL_441021, EPI_ISL_441022, EPI_ISL_441023, EPI_ISL_441024, EPI_ISL_441025, EPI_ISL_441026, EPI_ISL_441027, EPI_ISL_441028, EPI_ISL_441029, EPI_ISL_441030, EPI_ISL_441031, EPI_ISL_441032, EPI_ISL_441033, EPI_ISL_441034, EPI_ISL_441035, EPI_ISL_441036, EPI_ISL_441037, EPI_ISL_441038, EPI_ISL_441039, EPI_ISL_441040, EPI_ISL_441041, EPI_ISL_441042, EPI_ISL_441043, EPI_ISL_441044, EPI_ISL_441045, EPI_ISL_441046, EPI_ISL_441047, EPI_ISL_441048, EPI_ISL_441049, EPI_ISL_441050, EPI_ISL_441051 | | | |
| see above | University College London, Great Ormond Street Hospital for Children NHS Foundation Trust, Imperial College Healthcare NHS Trust | COVID-19 Genomics UK (COG-UK) Consortium | Sergi Castellano, Rachel Williams, Mark Kristiansen, Paola Resende Silva, Sunando Roy, Tony Brooks, Helena Tutill, Paola Niola, Patricia Dyal, Charlotte Williams, Leysa Forrest, Yasmin Panchbhaya, Jacqueline Findlay, Sam Weeks, Julianne Brown, Kathryn Harris, Paul Randell, James Price, Alison Holmes, Judith Breuer |
| EPI_ISL_441052, EPI_ISL_441053, EPI_ISL_441054, EPI_ISL_441055, EPI_ISL_441056, EPI_ISL_441057, EPI_ISL_441058, EPI_ISL_441059, EPI_ISL_441060, EPI_ISL_441061, EPI_ISL_441062, EPI_ISL_441063, EPI_ISL_441064, EPI_ISL_441065, EPI_ISL_441066, EPI_ISL_441067, EPI_ISL_441068, EPI_ISL_441069, EPI_ISL_441070, EPI_ISL_441071, EPI_ISL_441072, EPI_ISL_441073, EPI_ISL_441074, EPI_ISL_441075, EPI_ISL_441076, EPI_ISL_441077, EPI_ISL_441078, EPI_ISL_441079, EPI_ISL_441080, EPI_ISL_441081, EPI_ISL_441082, EPI_ISL_441083, EPI_ISL_441084, EPI_ISL_441085, EPI_ISL_441086, EPI_ISL_441087, EPI_ISL_441088, EPI_ISL_441089, EPI_ISL_441090, EPI_ISL_441091, EPI_ISL_441092, EPI_ISL_441093, EPI_ISL_441094, EPI_ISL_441095, EPI_ISL_441096, EPI_ISL_441097, EPI_ISL_441098, EPI_ISL_441099, EPI_ISL_441100, EPI_ISL_441101, EPI_ISL_441102, EPI_ISL_441103, EPI_ISL_441104, EPI_ISL_441105, EPI_ISL_441106, EPI_ISL_441107, EPI_ISL_441108, EPI_ISL_441109, EPI_ISL_441110, EPI_ISL_441111, EPI_ISL_441112, EPI_ISL_441113, EPI_ISL_441114, EPI_ISL_441115, EPI_ISL_441116, EPI_ISL_441117, EPI_ISL_441118, EPI_ISL_441119, EPI_ISL_441120, EPI_ISL_441121, EPI_ISL_441122, EPI_ISL_441123, EPI_ISL_441124, EPI_ISL_441125, EPI_ISL_441126, EPI_ISL_441127, EPI_ISL_441128, EPI_ISL_441129, EPI_ISL_441130, EPI_ISL_441131, EPI_ISL_441132, EPI_ISL_441133, EPI_ISL_441134, EPI_ISL_441135, EPI_ISL_441136, EPI_ISL_441137, EPI_ISL_441138, EPI_ISL_441139, EPI_ISL_441140, EPI_ISL_441141, EPI_ISL_441142, EPI_ISL_441143, EPI_ISL_441144, EPI_ISL_441145, EPI_ISL_441146, EPI_ISL_441147, EPI_ISL_441148, EPI_ISL_441149, EPI_ISL_441150, EPI_ISL_441151, EPI_ISL_441152, EPI_ISL_441153, EPI_ISL_441154, EPI_ISL_441155, EPI_ISL_441156, EPI_ISL_441157, EPI_ISL_441158, EPI_ISL_441159, EPI_ISL_441160, EPI_ISL_441161, EPI_ISL_441162, EPI_ISL_441163, EPI_ISL_441164, EPI_ISL_441165, EPI_ISL_441166, EPI_ISL_441167, EPI_ISL_441168, EPI_ISL_441169, EPI_ISL_441170, EPI_ISL_441171, EPI_ISL_441172, EPI_ISL_441173, EPI_ISL_441174, EPI_ISL_441175, EPI_ISL_441176, EPI_ISL_441177, EPI_ISL_441178, EPI_ISL_441179, EPI_ISL_441180, EPI_ISL_441181, EPI_ISL_441182, EPI_ISL_441183, EPI_ISL_441184, EPI_ISL_441185, EPI_ISL_441186, EPI_ISL_441187, EPI_ISL_441188, EPI_ISL_441189, EPI_ISL_441190, EPI_ISL_441191, EPI_ISL_441192, EPI_ISL_441193, EPI_ISL_441194, EPI_ISL_441195, EPI_ISL_441196, EPI_ISL_441197, EPI_ISL_441198, EPI_ISL_441199, EPI_ISL_441200, EPI_ISL_441201, EPI_ISL_441202, EPI_ISL_441203, EPI_ISL_441204, EPI_ISL_441205, EPI_ISL_441206, EPI_ISL_441207, EPI_ISL_441208, EPI_ISL_441209, EPI_ISL_441210, EPI_ISL_441211, EPI_ISL_441212, EPI_ISL_441213, EPI_ISL_441214, EPI_ISL_441215, EPI_ISL_441216, EPI_ISL_441217, EPI_ISL_441218, EPI_ISL_441219, EPI_ISL_441220, EPI_ISL_441221, EPI_ISL_441222, EPI_ISL_441223, EPI_ISL_441224, EPI_ISL_441225, EPI_ISL_441226, EPI_ISL_441227, EPI_ISL_441228, EPI_ISL_441229, EPI_ISL_441230, EPI_ISL_441231, EPI_ISL_441232, EPI_ISL_441233, EPI_ISL_441234, EPI_ISL_441235, EPI_ISL_441236, EPI_ISL_441237, EPI_ISL_441238, EPI_ISL_441239, EPI_ISL_441240, EPI_ISL_441241, EPI_ISL_441242, EPI_ISL_441243, EPI_ISL_441244, EPI_ISL_441245, EPI_ISL_441246, EPI_ISL_441247, EPI_ISL_441248, EPI_ISL_441249, EPI_ISL_441250, EPI_ISL_441251, EPI_ISL_441252, EPI_ISL_441253, EPI_ISL_441254, EPI_ISL_441255, EPI_ISL_441256, EPI_ISL_441257, EPI_ISL_441258, EPI_ISL_441259, EPI_ISL_441260, EPI_ISL_441261, EPI_ISL_441262, EPI_ISL_441263, EPI_ISL_441264, EPI_ISL_441265, EPI_ISL_441266, EPI_ISL_441267, EPI_ISL_441268, EPI_ISL_441269, EPI_ISL_441270, EPI_ISL_441271, EPI_ISL_441272, EPI_ISL_441273, EPI_ISL_441274, EPI_ISL_441275, EPI_ISL_441276, EPI_ISL_441277, EPI_ISL_441278, EPI_ISL_441279, EPI_ISL_441280, EPI_ISL_441281, EPI_ISL_441282, EPI_ISL_441283, EPI_ISL_441284, EPI_ISL_441285, EPI_ISL_441286, EPI_ISL_441287, EPI_ISL_441288, EPI_ISL_441289, EPI_ISL_441290, EPI_ISL_441291, EPI_ISL_441292, EPI_ISL_441293, EPI_ISL_441294, EPI_ISL_441295, EPI_ISL_441296, EPI_ISL_441297, EPI_ISL_441298, EPI_ISL_441299, EPI_ISL_441300, EPI_ISL_441301, EPI_ISL_441302, EPI_ISL_441303, EPI_ISL_441304, EPI_ISL_441305, EPI_ISL_441306, EPI_ISL_441307, EPI_ISL_441308, EPI_ISL_441309, EPI_ISL_441310, EPI_ISL_441311, EPI_ISL_441312, EPI_ISL_441313, EPI_ISL_441314, EPI_ISL_441315, EPI_ISL_441316, EPI_ISL_441317, EPI_ISL_441318, EPI_ISL_441319, EPI_ISL_441320, EPI_ISL_441321, EPI_ISL_441322, EPI_ISL_441323, EPI_ISL_441324, EPI_ISL_441325, EPI_ISL_441326, EPI_ISL_441327, EPI_ISL_441328, EPI_ISL_441329, EPI_ISL_441330, EPI_ISL_441331, EPI_ISL_441332, EPI_ISL_441333, EPI_ISL_441334, EPI_ISL_441335, EPI_ISL_441336, EPI_ISL_441337, EPI_ISL_441338, EPI_ISL_441339, EPI_ISL_441340, EPI_ISL_441341, EPI_ISL_441342, EPI_ISL_441343, EPI_ISL_441344, EPI_ISL_441345, EPI_ISL_441346, EPI_ISL_441347, EPI_ISL_441348, EPI_ISL_441349 | | | |
| see above | Department of Pathology, University of Cambridge | Wellcome Sanger Institute for the COVID-19 Genomics UK Consortium | Luke W Meredith, M. Estée Török , Myra Hosmillo, William L. Hamilton, Martin D. Curran, Theresa Feltwell, Grant Hall, Anna Yakovleva, Fahad A Khokhar, Charlotte J. Houldcroft, Laura G Caller, Aminu S. Jahun, Sarah L. Caddy, Ian Goodfellow, Alex Alderton, Roberto Amato, Sonia Goncalves, Ewan Harrison, David K. Jackson, Ian Johnston, Dominic Kwiatkowski, Cordelia Langford, John Sillitoe on behalf of the Wellcome Sanger Institute COVID-19 Surveillance Team (http://www.sanger.ac.uk/covid-team) |
| EPI_ISL_441350, EPI_ISL_441351, EPI_ISL_441352, EPI_ISL_441353, EPI_ISL_441354 | University College London, Great Ormond Street Hospital for Children NHS Foundation Trust, Imperial College Healthcare NHS Trust | COVID-19 Genomics UK (COG-UK) Consortium | Sergi Castellano, Rachel Williams, Mark Kristiansen, Paola Resende Silva, Sunando Roy, Tony Brooks, Helena Tutill, Paola Niola, Patricia Dyal, Charlotte Williams, Leysa Forrest, Yasmin Panchbhaya, Jacqueline Findlay, Sam Weeks, Julianne Brown, Kathryn Harris, Paul Randell, James Price, Alison Holmes, Judith Breuer |
| EPI_ISL_441355, EPI_ISL_441356, EPI_ISL_441357, EPI_ISL_441358, EPI_ISL_441359, EPI_ISL_441360, EPI_ISL_441361, EPI_ISL_441362, EPI_ISL_441363, EPI_ISL_441364, EPI_ISL_441365, EPI_ISL_441366, EPI_ISL_441367, EPI_ISL_441368, EPI_ISL_441369, EPI_ISL_441370, EPI_ISL_441371, EPI_ISL_441372, EPI_ISL_441373, EPI_ISL_441374, EPI_ISL_441375, EPI_ISL_441376, EPI_ISL_441377, EPI_ISL_441378, EPI_ISL_441379, EPI_ISL_441380, EPI_ISL_441381, EPI_ISL_441382, EPI_ISL_441383, EPI_ISL_441384, EPI_ISL_441385, EPI_ISL_441386, EPI_ISL_441387, EPI_ISL_441388, EPI_ISL_441389, EPI_ISL_441390, EPI_ISL_441391, EPI_ISL_441392, EPI_ISL_441393, EPI_ISL_441394, EPI_ISL_441395, EPI_ISL_441396, EPI_ISL_441397, EPI_ISL_441398, EPI_ISL_441399, EPI_ISL_441400, EPI_ISL_441401, EPI_ISL_441402, EPI_ISL_441403, EPI_ISL_441404, EPI_ISL_441405, EPI_ISL_441406, EPI_ISL_441407, EPI_ISL_441408, EPI_ISL_441409, EPI_ISL_441410, EPI_ISL_441411, EPI_ISL_441412, EPI_ISL_441413, EPI_ISL_441414, EPI_ISL_441415, EPI_ISL_441416, EPI_ISL_441417, EPI_ISL_441418, EPI_ISL_441419, EPI_ISL_441420, EPI_ISL_441421, EPI_ISL_441422, EPI_ISL_441423, EPI_ISL_441424, EPI_ISL_441425, EPI_ISL_441426, EPI_ISL_441427, EPI_ISL_441428, EPI_ISL_441429, EPI_ISL_441430, EPI_ISL_441431, EPI_ISL_441432, EPI_ISL_441433, EPI_ISL_441434, EPI_ISL_441435, EPI_ISL_441436 | | | |
| see above | Regional Virus Laboratory, Belfast Health and Social Care Trust | COVID-19 Genomics UK (COG-UK) Consortium | Conall McCaughey, James McKenna, Tanya Curran, Susan Feeney, Alison Watt, Ciara Cox, Mairead Connor, Zoltan Molnar, David Simpson, Derek Fairley |
| EPI_ISL_441437, EPI_ISL_441438, EPI_ISL_441439, EPI_ISL_441440, EPI_ISL_441441, EPI_ISL_441442, EPI_ISL_441443, EPI_ISL_441444, EPI_ISL_441445, EPI_ISL_441446, EPI_ISL_441447, EPI_ISL_441448, EPI_ISL_441449, EPI_ISL_441450, EPI_ISL_441451, EPI_ISL_441452, EPI_ISL_441453, EPI_ISL_441454, EPI_ISL_441455, EPI_ISL_441456, EPI_ISL_441457, EPI_ISL_441458, EPI_ISL_441459, EPI_ISL_441460, EPI_ISL_441461, EPI_ISL_441462, EPI_ISL_441463, EPI_ISL_441464, EPI_ISL_441465, EPI_ISL_441466, EPI_ISL_441467, EPI_ISL_441468, EPI_ISL_441469, EPI_ISL_441470, EPI_ISL_441471, EPI_ISL_441472, EPI_ISL_441473, EPI_ISL_441474, EPI_ISL_441475, EPI_ISL_441476, EPI_ISL_441477, EPI_ISL_441478, EPI_ISL_441479, EPI_ISL_441480, EPI_ISL_441481, EPI_ISL_441482, EPI_ISL_441483, EPI_ISL_441484, EPI_ISL_441485, EPI_ISL_441486, EPI_ISL_441487, EPI_ISL_441488, EPI_ISL_441489, EPI_ISL_441490, EPI_ISL_441491, EPI_ISL_441492, EPI_ISL_441493, EPI_ISL_441494, EPI_ISL_441495, EPI_ISL_441496, EPI_ISL_441497, EPI_ISL_441498, EPI_ISL_441499, EPI_ISL_441500, EPI_ISL_441501, EPI_ISL_441502, EPI_ISL_441503, EPI_ISL_441504, EPI_ISL_441505, EPI_ISL_441506, EPI_ISL_441507, EPI_ISL_441508, EPI_ISL_441509, EPI_ISL_441510, EPI_ISL_441511, EPI_ISL_441512, EPI_ISL_441513, EPI_ISL_441514, EPI_ISL_441515, EPI_ISL_441516, EPI_ISL_441517, EPI_ISL_441518, EPI_ISL_441519, EPI_ISL_441520, EPI_ISL_441521, EPI_ISL_441522, EPI_ISL_441523, EPI_ISL_441524, EPI_ISL_441525, EPI_ISL_441526, EPI_ISL_441527, EPI_ISL_441528, EPI_ISL_441529, EPI_ISL_441530, EPI_ISL_441531, EPI_ISL_441532, EPI_ISL_441533, EPI_ISL_441534, EPI_ISL_441535, EPI_ISL_441536, EPI_ISL_441537, EPI_ISL_441538, EPI_ISL_441539, EPI_ISL_441540, EPI_ISL_441541, EPI_ISL_441542, EPI_ISL_441543, EPI_ISL_441544, EPI_ISL_441545, EPI_ISL_441546 | | | |
| see above | Queens Medical Centre, Clinical Microbiology Department / DeepSeq Nottingham | COVID-19 Genomics UK (COG-UK) Consortium | Gemma Clark, Wendy Smith, Manjinder Khakh, Hannah Howson-Wells, Jonathan Ball, Patrick McClure, Joseph Chappell, Theocharis Tsoleridis, Nadine Holmes, Matthew Carlisle, Christopher Moore, Fei Sang, Johnny Debebe, Victoria Wright, Matthew Loose |
| EPI_ISL_441547, EPI_ISL_441548, EPI_ISL_441549, EPI_ISL_441550, EPI_ISL_441551, EPI_ISL_441552, EPI_ISL_441553, EPI_ISL_441554, EPI_ISL_441555, EPI_ISL_441556, EPI_ISL_441557, EPI_ISL_441558, EPI_ISL_441559, EPI_ISL_441560, EPI_ISL_441561, EPI_ISL_441562, EPI_ISL_441563, EPI_ISL_441564, EPI_ISL_441565, EPI_ISL_441566, EPI_ISL_441567, EPI_ISL_441568, EPI_ISL_441569, EPI_ISL_441570, EPI_ISL_441571, EPI_ISL_441572, EPI_ISL_441573, EPI_ISL_441574, EPI_ISL_441575, EPI_ISL_441576, EPI_ISL_441577, EPI_ISL_441578, EPI_ISL_441579, EPI_ISL_441580, EPI_ISL_441581, EPI_ISL_441582, EPI_ISL_441583, EPI_ISL_441584, EPI_ISL_441585, EPI_ISL_441586, EPI_ISL_441587, EPI_ISL_441588, EPI_ISL_441589, EPI_ISL_441590, EPI_ISL_441591, EPI_ISL_441592, EPI_ISL_441593, EPI_ISL_441594, EPI_ISL_441595, EPI_ISL_441596, EPI_ISL_441597, EPI_ISL_441598, EPI_ISL_441599, EPI_ISL_441600, EPI_ISL_441601, EPI_ISL_441602, EPI_ISL_441603, EPI_ISL_441604, EPI_ISL_441605, EPI_ISL_441606, EPI_ISL_441607, EPI_ISL_441608, EPI_ISL_441609, EPI_ISL_441610, EPI_ISL_441611, EPI_ISL_441612, EPI_ISL_441613, EPI_ISL_441614, EPI_ISL_441615, EPI_ISL_441616, EPI_ISL_441617, EPI_ISL_441618, EPI_ISL_441619, EPI_ISL_441620, EPI_ISL_441621, EPI_ISL_441622, EPI_ISL_441623, EPI_ISL_441624, EPI_ISL_441625, EPI_ISL_441626, EPI_ISL_441627, EPI_ISL_441628, EPI_ISL_441629, EPI_ISL_441630, EPI_ISL_441631, EPI_ISL_441632, EPI_ISL_441633, EPI_ISL_441634, EPI_ISL_441635, EPI_ISL_441636, EPI_ISL_441637, EPI_ISL_441638, EPI_ISL_441639, EPI_ISL_441640, EPI_ISL_441641, EPI_ISL_441642, EPI_ISL_441643, EPI_ISL_441644, EPI_ISL_441645, EPI_ISL_441646, EPI_ISL_441647, EPI_ISL_441648, EPI_ISL_441649, EPI_ISL_441650, EPI_ISL_441651, EPI_ISL_441652, EPI_ISL_441653, EPI_ISL_441654, EPI_ISL_441655, EPI_ISL_441656, EPI_ISL_441657, EPI_ISL_441658 | | | |
| see above | Department of Pathology, University of Cambridge | Wellcome Sanger Institute for the COVID-19 Genomics UK Consortium | Luke W Meredith, M. Estée Török , Myra Hosmillo, William L. Hamilton, Martin D. Curran, Theresa Feltwell, Grant Hall, Anna Yakovleva, Fahad A Khokhar, Charlotte J. Houldcroft, Laura G Caller, Aminu S. Jahun, Sarah L. Caddy, Ian Goodfellow, Alex Alderton, Roberto Amato, Sonia Goncalves, Ewan Harrison, David K. Jackson, Ian Johnston, Dominic Kwiatkowski, Cordelia Langford, John Sillitoe on behalf of the Wellcome Sanger Institute COVID-19 Surveillance Team (http://www.sanger.ac.uk/covid-team) |
| EPI_ISL_441659 | Regional Virus Laboratory, Belfast Health and Social Care Trust | Wellcome Sanger Institute for the COVID-19 Genomics UK Consortium | Conall McCaughey, James McKenna, Tanya Curran, Susan Feeney, Alison Watt, Ciara Cox, Mairead Connor, Zoltan Molnar, David Simpson, Derek Fairley, Alex Alderton, Roberto Amato, Sonia Goncalves, Ewan Harrison, David K. Jackson, Ian Johnston, Dominic Kwiatkowski, Cordelia Langford, John Sillitoe on behalf of the Wellcome Sanger Institute COVID-19 Surveillance Team (http://www.sanger.ac.uk/covid-team) |
| EPI_ISL_441660, EPI_ISL_441661 | Department of Pathology, University of Cambridge | Wellcome Sanger Institute for the COVID-19 Genomics UK Consortium | Luke W Meredith, M. Estée Török , Myra Hosmillo, William L. Hamilton, Martin D. Curran, Theresa Feltwell, Grant Hall, Anna Yakovleva, Fahad A Khokhar, Charlotte J. Houldcroft, Laura G Caller, Aminu S. Jahun, |

| | | | |
|---|---|---|---|
| | | | Sarah L. Caddy, Ian Goodfellow, Alex Alderton, Roberto Amato, Sonia Goncalves, Ewan Harrison, David K. Jackson, Ian Johnston, Dominic Kwiatkowski, Cordelia Langford, John Sillitoe on behalf of the Wellcome Sanger Institute COVID-19 Surveillance Team (http://www.sanger.ac.uk/covid-team) |
| EPI_ISL_441662, EPI_ISL_441663 | Regional Virus Laboratory, Belfast Health and Social Care Trust | Wellcome Sanger Institute for the COVID-19 Genomics UK Consortium | Conall McCaughey, James McKenna, Tanya Curran, Susan Feeney, Alison Watt, Ciara Cox, Mairead Connor, Zoltan Molnar, David Simpson, Derek Fairley, Alex Alderton, Roberto Amato, Sonia Goncalves, Ewan Harrison, David K. Jackson, Ian Johnston, Dominic Kwiatkowski, Cordelia Langford, John Sillitoe on behalf of the Wellcome Sanger Institute COVID-19 Surveillance Team (http://www.sanger.ac.uk/covid-team) |
| EPI_ISL_441664, EPI_ISL_441665, EPI_ISL_441666, EPI_ISL_441667, EPI_ISL_441668, EPI_ISL_441669 | Department of Pathology, University of Cambridge | Wellcome Sanger Institute for the COVID-19 Genomics UK Consortium | Luke W Meredith, M. Estée Török , Myra Hosmillo, William L. Hamilton, Martin D. Curran, Theresa Feltwell, Grant Hall, Anna Yakovleva, Fahad A Khokhar, Charlotte J. Houldcroft, Laura G Caller, Aminu S. Jahun, Sarah L. Caddy, Ian Goodfellow, Alex Alderton, Roberto Amato, Sonia Goncalves, Ewan Harrison, David K. Jackson, Ian Johnston, Dominic Kwiatkowski, Cordelia Langford, John Sillitoe on behalf of the Wellcome Sanger Institute COVID-19 Surveillance Team (http://www.sanger.ac.uk/covid-team) |
| EPI_ISL_441670 | Regional Virus Laboratory, Belfast Health and Social Care Trust | Wellcome Sanger Institute for the COVID-19 Genomics UK Consortium | Conall McCaughey, James McKenna, Tanya Curran, Susan Feeney, Alison Watt, Ciara Cox, Mairead Connor, Zoltan Molnar, David Simpson, Derek Fairley, Alex Alderton, Roberto Amato, Sonia Goncalves, Ewan Harrison, David K. Jackson, Ian Johnston, Dominic Kwiatkowski, Cordelia Langford, John Sillitoe on behalf of the Wellcome Sanger Institute COVID-19 Surveillance Team (http://www.sanger.ac.uk/covid-team) |
| EPI_ISL_441671, EPI_ISL_441672, EPI_ISL_441673, EPI_ISL_441674 | Department of Pathology, University of Cambridge | Wellcome Sanger Institute for the COVID-19 Genomics UK Consortium | Luke W Meredith, M. Estée Török , Myra Hosmillo, William L. Hamilton, Martin D. Curran, Theresa Feltwell, Grant Hall, Anna Yakovleva, Fahad A Khokhar, Charlotte J. Houldcroft, Laura G Caller, Aminu S. Jahun, Sarah L. Caddy, Ian Goodfellow, Alex Alderton, Roberto Amato, Sonia Goncalves, Ewan Harrison, David K. Jackson, Ian Johnston, Dominic Kwiatkowski, Cordelia Langford, John Sillitoe on behalf of the Wellcome Sanger Institute COVID-19 Surveillance Team (http://www.sanger.ac.uk/covid-team) |
| EPI_ISL_441675, EPI_ISL_441676 | Regional Virus Laboratory, Belfast Health and Social Care Trust | Wellcome Sanger Institute for the COVID-19 Genomics UK Consortium | Conall McCaughey, James McKenna, Tanya Curran, Susan Feeney, Alison Watt, Ciara Cox, Mairead Connor, Zoltan Molnar, David Simpson, Derek Fairley, Alex Alderton, Roberto Amato, Sonia Goncalves, Ewan Harrison, David K. Jackson, Ian Johnston, Dominic Kwiatkowski, Cordelia Langford, John Sillitoe on behalf of the Wellcome Sanger Institute COVID-19 Surveillance Team (http://www.sanger.ac.uk/covid-team) |
| EPI_ISL_441677, EPI_ISL_441678, EPI_ISL_441679, EPI_ISL_441680, EPI_ISL_441681, EPI_ISL_441682, EPI_ISL_441683, EPI_ISL_441684 | Department of Pathology, University of Cambridge | Wellcome Sanger Institute for the COVID-19 Genomics UK Consortium | Luke W Meredith, M. Estée Török , Myra Hosmillo, William L. Hamilton, Martin D. Curran, Theresa Feltwell, Grant Hall, Anna Yakovleva, Fahad A Khokhar, Charlotte J. Houldcroft, Laura G Caller, Aminu S. Jahun, Sarah L. Caddy, Ian Goodfellow, Alex Alderton, Roberto Amato, Sonia Goncalves, Ewan Harrison, David K. Jackson, Ian Johnston, Dominic Kwiatkowski, Cordelia Langford, John Sillitoe on behalf of the Wellcome Sanger Institute COVID-19 Surveillance Team (http://www.sanger.ac.uk/covid-team) |
| EPI_ISL_441685 | Regional Virus Laboratory, Belfast Health and Social Care Trust | Wellcome Sanger Institute for the COVID-19 Genomics UK Consortium | Conall McCaughey, James McKenna, Tanya Curran, Susan Feeney, Alison Watt, Ciara Cox, Mairead Connor, Zoltan Molnar, David Simpson, Derek Fairley, Alex Alderton, Roberto Amato, Sonia Goncalves, Ewan Harrison, David K. Jackson, Ian Johnston, Dominic Kwiatkowski, Cordelia Langford, John Sillitoe on behalf of the Wellcome Sanger Institute COVID-19 Surveillance Team (http://www.sanger.ac.uk/covid-team) |
| EPI_ISL_441686, EPI_ISL_441687, EPI_ISL_441688, EPI_ISL_441689 | Department of Pathology, University of Cambridge | Wellcome Sanger Institute for the COVID-19 Genomics UK Consortium | Luke W Meredith, M. Estée Török , Myra Hosmillo, William L. Hamilton, Martin D. Curran, Theresa Feltwell, Grant Hall, Anna Yakovleva, Fahad A Khokhar, Charlotte J. Houldcroft, Laura G Caller, Aminu S. Jahun, Sarah L. Caddy, Ian Goodfellow, Alex Alderton, Roberto Amato, Sonia Goncalves, Ewan Harrison, David K. Jackson, Ian Johnston, Dominic Kwiatkowski, Cordelia Langford, John Sillitoe on behalf of the Wellcome Sanger Institute COVID-19 Surveillance Team (http://www.sanger.ac.uk/covid-team) |
| EPI_ISL_441690, EPI_ISL_441691 | Regional Virus Laboratory, Belfast Health and Social Care Trust | Wellcome Sanger Institute for the COVID-19 Genomics UK Consortium | Conall McCaughey, James McKenna, Tanya Curran, Susan Feeney, Alison Watt, Ciara Cox, Mairead Connor, Zoltan Molnar, David Simpson, Derek Fairley, Alex Alderton, Roberto Amato, Sonia Goncalves, Ewan Harrison, David K. Jackson, Ian Johnston, Dominic Kwiatkowski, Cordelia Langford, John Sillitoe on behalf of the Wellcome Sanger Institute COVID-19 Surveillance Team (http://www.sanger.ac.uk/covid-team) |
| EPI_ISL_441692 | Department of Pathology, University of Cambridge | Wellcome Sanger Institute for the COVID-19 Genomics UK Consortium | Luke W Meredith, M. Estée Török , Myra Hosmillo, William L. Hamilton, Martin D. Curran, Theresa Feltwell, Grant Hall, Anna Yakovleva, Fahad A Khokhar, Charlotte J. Houldcroft, Laura G Caller, Aminu S. Jahun, Sarah L. Caddy, Ian Goodfellow, Alex Alderton, Roberto Amato, Sonia Goncalves, Ewan Harrison, David K. Jackson, Ian Johnston, Dominic Kwiatkowski, Cordelia Langford, John Sillitoe on behalf of the Wellcome Sanger Institute COVID-19 Surveillance Team (http://www.sanger.ac.uk/covid-team) |
| EPI_ISL_441693, EPI_ISL_441694 | Regional Virus Laboratory, Belfast Health and Social Care Trust | Wellcome Sanger Institute for the COVID-19 Genomics UK Consortium | Conall McCaughey, James McKenna, Tanya Curran, Susan Feeney, Alison Watt, Ciara Cox, Mairead Connor, Zoltan Molnar, David Simpson, Derek Fairley, Alex Alderton, Roberto Amato, Sonia Goncalves, Ewan Harrison, David K. Jackson, Ian Johnston, Dominic Kwiatkowski, Cordelia Langford, John Sillitoe on behalf of the Wellcome Sanger Institute COVID-19 Surveillance Team (http://www.sanger.ac.uk/covid-team) |
| EPI_ISL_441695 | Department of Pathology, University of Cambridge | Wellcome Sanger Institute for the COVID-19 Genomics UK Consortium | Luke W Meredith, M. Estée Török , Myra Hosmillo, William L. Hamilton, Martin D. Curran, Theresa Feltwell, Grant Hall, Anna Yakovleva, Fahad A Khokhar, Charlotte J. Houldcroft, Laura G Caller, Aminu S. Jahun, Sarah L. Caddy, Ian Goodfellow, Alex Alderton, Roberto Amato, Sonia Goncalves, Ewan Harrison, David K. Jackson, Ian Johnston, Dominic Kwiatkowski, Cordelia Langford, John Sillitoe on behalf of the Wellcome Sanger Institute COVID-19 Surveillance Team (http://www.sanger.ac.uk/covid-team) |
| EPI_ISL_441696, EPI_ISL_441697, EPI_ISL_441698, EPI_ISL_441699 | Regional Virus Laboratory, Belfast Health and Social Care Trust | Wellcome Sanger Institute for the COVID-19 Genomics UK Consortium | Conall McCaughey, James McKenna, Tanya Curran, Susan Feeney, Alison Watt, Ciara Cox, Mairead Connor, Zoltan Molnar, David Simpson, Derek Fairley, Alex Alderton, Roberto Amato, Sonia Goncalves, Ewan Harrison, David K. Jackson, Ian Johnston, Dominic Kwiatkowski, Cordelia Langford, John Sillitoe on behalf of the Wellcome Sanger Institute COVID-19 Surveillance Team (http://www.sanger.ac.uk/covid-team) |
| EPI_ISL_441700, EPI_ISL_441701, EPI_ISL_441702, EPI_ISL_441703 | Department of Pathology, University of Cambridge | Wellcome Sanger Institute for the COVID-19 Genomics UK Consortium | Luke W Meredith, M. Estée Török , Myra Hosmillo, William L. Hamilton, Martin D. Curran, Theresa Feltwell, Grant Hall, Anna Yakovleva, Fahad A Khokhar, Charlotte J. Houldcroft, Laura G Caller, Aminu S. Jahun, Sarah L. Caddy, Ian Goodfellow, Alex Alderton, Roberto Amato, Sonia Goncalves, Ewan Harrison, David K. Jackson, Ian Johnston, Dominic Kwiatkowski, Cordelia Langford, John Sillitoe on behalf of the Wellcome Sanger Institute COVID-19 Surveillance Team (http://www.sanger.ac.uk/covid-team) |
| EPI_ISL_441704 | Regional Virus Laboratory, Belfast Health and Social Care Trust | Wellcome Sanger Institute for the COVID-19 Genomics UK Consortium | Conall McCaughey, James McKenna, Tanya Curran, Susan Feeney, Alison Watt, Ciara Cox, Mairead Connor, Zoltan Molnar, David Simpson, Derek Fairley, Alex Alderton, Roberto Amato, Sonia Goncalves, Ewan Harrison, David K. Jackson, Ian Johnston, Dominic Kwiatkowski, Cordelia Langford, John Sillitoe on behalf of the Wellcome Sanger Institute COVID-19 Surveillance Team (http://www.sanger.ac.uk/covid-team) |
| EPI_ISL_441705, EPI_ISL_441706, EPI_ISL_441707 | Department of Pathology, University of Cambridge | Wellcome Sanger Institute for the COVID-19 Genomics UK Consortium | Luke W Meredith, M. Estée Török , Myra Hosmillo, William L. Hamilton, Martin D. Curran, Theresa Feltwell, Grant Hall, Anna Yakovleva, Fahad A Khokhar, Charlotte J. Houldcroft, Laura G Caller, Aminu S. Jahun, Sarah L. Caddy, Ian Goodfellow, Alex Alderton, Roberto Amato, Sonia Goncalves, Ewan Harrison, David K. Jackson, Ian Johnston, Dominic Kwiatkowski, Cordelia Langford, John Sillitoe on behalf of the Wellcome Sanger Institute COVID-19 Surveillance Team (http://www.sanger.ac.uk/covid-team) |
| EPI_ISL_441708 | Regional Virus Laboratory, Belfast Health and Social Care Trust | Wellcome Sanger Institute for the COVID-19 Genomics UK Consortium | Conall McCaughey, James McKenna, Tanya Curran, Susan Feeney, Alison Watt, Ciara Cox, Mairead Connor, Zoltan Molnar, David Simpson, Derek Fairley, Alex Alderton, Roberto Amato, Sonia Goncalves, Ewan Harrison, David K. Jackson, Ian Johnston, Dominic Kwiatkowski, Cordelia Langford, John Sillitoe on behalf of the Wellcome Sanger Institute COVID-19 Surveillance Team (http://www.sanger.ac.uk/covid-team) |
| EPI_ISL_441709, EPI_ISL_441710, EPI_ISL_441711 | Department of Pathology, University of Cambridge | Wellcome Sanger Institute for the COVID-19 Genomics UK Consortium | Luke W Meredith, M. Estée Török , Myra Hosmillo, William L. Hamilton, Martin D. Curran, Theresa Feltwell, Grant Hall, Anna Yakovleva, Fahad A Khokhar, Charlotte J. Houldcroft, Laura G Caller, Aminu S. Jahun, Sarah L. Caddy, Ian Goodfellow, Alex Alderton, Roberto Amato, Sonia Goncalves, Ewan Harrison, David K. Jackson, Ian Johnston, Dominic Kwiatkowski, Cordelia Langford, John Sillitoe on behalf of the Wellcome Sanger Institute COVID-19 Surveillance Team (http://www.sanger.ac.uk/covid-team) |
| EPI_ISL_441712 | Regional Virus Laboratory, Belfast Health and Social Care Trust | Wellcome Sanger Institute for the COVID-19 Genomics UK Consortium | Conall McCaughey, James McKenna, Tanya Curran, Susan Feeney, Alison Watt, Ciara Cox, Mairead Connor, Zoltan Molnar, David Simpson, Derek Fairley, Alex Alderton, Roberto Amato, Sonia Goncalves, Ewan Harrison, David K. Jackson, Ian Johnston, Dominic Kwiatkowski, Cordelia Langford, John Sillitoe on behalf of the Wellcome Sanger Institute COVID-19 Surveillance Team (http://www.sanger.ac.uk/covid-team) |
| EPI_ISL_441713 | Department of Pathology, University of Cambridge | Wellcome Sanger Institute for the COVID-19 Genomics UK Consortium | Luke W Meredith, M. Estée Török , Myra Hosmillo, William L. Hamilton, Martin D. Curran, Theresa Feltwell, Grant Hall, Anna Yakovleva, Fahad A Khokhar, Charlotte J. Houldcroft, Laura G Caller, Aminu S. Jahun, Sarah L. Caddy, Ian Goodfellow, Alex Alderton, Roberto Amato, Sonia Goncalves, Ewan Harrison, David K. Jackson, Ian Johnston, Dominic Kwiatkowski, Cordelia Langford, John Sillitoe on behalf of the Wellcome Sanger Institute COVID-19 Surveillance Team (http://www.sanger.ac.uk/covid-team) |
| EPI_ISL_441714 | Regional Virus Laboratory, Belfast Health and Social Care Trust | Wellcome Sanger Institute for the COVID-19 Genomics UK Consortium | Conall McCaughey, James McKenna, Tanya Curran, Susan Feeney, Alison Watt, Ciara Cox, Mairead Connor, Zoltan Molnar, David Simpson, Derek Fairley, Alex Alderton, Roberto Amato, Sonia Goncalves, Ewan Harrison, David K. Jackson, Ian Johnston, Dominic Kwiatkowski, Cordelia Langford, John Sillitoe on behalf of the Wellcome Sanger Institute COVID-19 Surveillance Team (http://www.sanger.ac.uk/covid-team) |
| EPI_ISL_441715 | Department of Pathology, University of Cambridge | Wellcome Sanger Institute for the COVID-19 Genomics UK Consortium | Luke W Meredith, M. Estée Török , Myra Hosmillo, William L. Hamilton, Martin D. Curran, Theresa Feltwell, Grant Hall, Anna Yakovleva, Fahad A Khokhar, Charlotte J. Houldcroft, Laura G Caller, Aminu S. Jahun, Sarah L. Caddy, Ian Goodfellow, Alex Alderton, Roberto Amato, Sonia Goncalves, Ewan Harrison, David K. Jackson, Ian Johnston, Dominic Kwiatkowski, Cordelia Langford, John Sillitoe on behalf of the Wellcome Sanger Institute COVID-19 Surveillance Team (http://www.sanger.ac.uk/covid-team) |
| EPI_ISL_441716, EPI_ISL_441717 | Regional Virus Laboratory, Belfast Health and Social Care Trust | Wellcome Sanger Institute for the COVID-19 Genomics UK Consortium | Conall McCaughey, James McKenna, Tanya Curran, Susan Feeney, Alison Watt, Ciara Cox, Mairead Connor, Zoltan Molnar, David Simpson, Derek Fairley, Alex Alderton, Roberto Amato, Sonia Goncalves, Ewan Harrison, David K. Jackson, Ian Johnston, Dominic Kwiatkowski, Cordelia Langford, John Sillitoe on behalf of the Wellcome Sanger Institute COVID-19 Surveillance Team (http://www.sanger.ac.uk/covid-team) |
| EPI_ISL_441718 | Department of Pathology, University of Cambridge | Wellcome Sanger Institute for the COVID-19 Genomics UK Consortium | Luke W Meredith, M. Estée Török , Myra Hosmillo, William L. Hamilton, Martin D. Curran, Theresa Feltwell, Grant Hall, Anna Yakovleva, Fahad A Khokhar, Charlotte J. Houldcroft, Laura G Caller, Aminu S. Jahun, Sarah L. Caddy, Ian Goodfellow, Alex Alderton, Roberto Amato, Sonia Goncalves, Ewan Harrison, David K. Jackson, Ian Johnston, Dominic Kwiatkowski, Cordelia Langford, John Sillitoe on behalf of the Wellcome Sanger Institute COVID-19 Surveillance Team (http://www.sanger.ac.uk/covid-team) |
| EPI_ISL_441719 | Regional Virus Laboratory, Belfast Health and Social Care Trust | Wellcome Sanger Institute for the COVID-19 Genomics UK Consortium | Conall McCaughey, James McKenna, Tanya Curran, Susan Feeney, Alison Watt, Ciara Cox, Mairead Connor, Zoltan Molnar, David Simpson, Derek Fairley, Alex Alderton, Roberto Amato, Sonia Goncalves, Ewan Harrison, David K. Jackson, Ian Johnston, Dominic Kwiatkowski, Cordelia Langford, John Sillitoe on behalf of the Wellcome Sanger Institute COVID-19 Surveillance Team (http://www.sanger.ac.uk/covid-team) |
| EPI_ISL_441720, EPI_ISL_441721, EPI_ISL_441722, EPI_ISL_441723, EPI_ISL_441724, EPI_ISL_441725, EPI_ISL_441726, EPI_ISL_441727, EPI_ISL_441728, EPI_ISL_441729 | Department of Pathology, University of Cambridge | Wellcome Sanger Institute for the COVID-19 Genomics UK Consortium | Luke W Meredith, M. Estée Török , Myra Hosmillo, William L. Hamilton, Martin D. Curran, Theresa Feltwell, Grant Hall, Anna Yakovleva, Fahad A Khokhar, Charlotte J. Houldcroft, Laura G Caller, Aminu S. Jahun, Sarah L. Caddy, Ian Goodfellow, Alex Alderton, Roberto Amato, Sonia Goncalves, Ewan Harrison, David K. Jackson, Ian Johnston, Dominic Kwiatkowski, Cordelia Langford, John Sillitoe on behalf of the Wellcome Sanger Institute COVID-19 Surveillance Team (http://www.sanger.ac.uk/covid-team) |
| EPI_ISL_441730 | Regional Virus Laboratory, Belfast Health and Social Care Trust | Wellcome Sanger Institute for the COVID-19 Genomics UK Consortium | Conall McCaughey, James McKenna, Tanya Curran, Susan Feeney, Alison Watt, Ciara Cox, Mairead Connor, Zoltan Molnar, David Simpson, Derek Fairley, Alex Alderton, Roberto Amato, Sonia Goncalves, Ewan Harrison, David K. Jackson, Ian Johnston, Dominic Kwiatkowski, Cordelia Langford, John Sillitoe on behalf of the Wellcome Sanger Institute COVID-19 Surveillance Team (http://www.sanger.ac.uk/covid-team) |
| EPI_ISL_441731, EPI_ISL_441732 | Department of Pathology, University of Cambridge | Wellcome Sanger Institute for the COVID-19 Genomics UK Consortium | Luke W Meredith, M. Estée Török , Myra Hosmillo, William L. Hamilton, Martin D. Curran, Theresa Feltwell, Grant Hall, Anna Yakovleva, Fahad A Khokhar, Charlotte J. Houldcroft, Laura G Caller, Aminu S. Jahun, Sarah L. Caddy, Ian Goodfellow, Alex Alderton, Roberto Amato, Sonia Goncalves, Ewan Harrison, David K. Jackson, Ian Johnston, Dominic Kwiatkowski, Cordelia Langford, John Sillitoe on behalf of the Wellcome Sanger Institute COVID-19 Surveillance Team (http://www.sanger.ac.uk/covid-team) |
| EPI_ISL_441733 | Regional Virus Laboratory, Belfast Health and Social Care Trust | Wellcome Sanger Institute for the COVID-19 Genomics UK Consortium | Conall McCaughey, James McKenna, Tanya Curran, Susan Feeney, Alison Watt, Ciara Cox, Mairead Connor, Zoltan Molnar, David Simpson, Derek Fairley, Alex Alderton, Roberto Amato, Sonia Goncalves, Ewan Harrison, David K. Jackson, Ian Johnston, Dominic Kwiatkowski, Cordelia Langford, John Sillitoe on behalf of the Wellcome Sanger Institute COVID-19 Surveillance Team (http://www.sanger.ac.uk/covid-team) |
| EPI_ISL_441734, EPI_ISL_441735, EPI_ISL_441736 | Department of Pathology, University of Cambridge | Wellcome Sanger Institute for the COVID-19 Genomics UK Consortium | Luke W Meredith, M. Estée Török , Myra Hosmillo, William L. Hamilton, Martin D. Curran, Theresa Feltwell, Grant Hall, Anna Yakovleva, Fahad A Khokhar, Charlotte J. Houldcroft, Laura G Caller, Aminu S. Jahun, Sarah L. Caddy, Ian Goodfellow, Alex Alderton, Roberto Amato, Sonia Goncalves, Ewan Harrison, David K. Jackson, Ian Johnston, Dominic Kwiatkowski, Cordelia Langford, John Sillitoe on behalf of the Wellcome Sanger Institute COVID-19 Surveillance Team (http://www.sanger.ac.uk/covid-team) |
| EPI_ISL_441737 | Regional Virus Laboratory, Belfast Health and Social Care Trust | Wellcome Sanger Institute for the COVID-19 Genomics UK Consortium | Conall McCaughey, James McKenna, Tanya Curran, Susan Feeney, Alison Watt, Ciara Cox, Mairead Connor, Zoltan Molnar, David Simpson, Derek Fairley, Alex Alderton, Roberto Amato, Sonia Goncalves, Ewan Harrison, David K. Jackson, Ian Johnston, Dominic Kwiatkowski, Cordelia Langford, John Sillitoe on behalf of the Wellcome Sanger Institute COVID-19 Surveillance Team (http://www.sanger.ac.uk/covid-team) |
| EPI_ISL_441738, EPI_ISL_441739, EPI_ISL_441740, EPI_ISL_441741, EPI_ISL_441742, EPI_ISL_441743, EPI_ISL_441744, EPI_ISL_441745, EPI_ISL_441746 | Department of Pathology, University of Cambridge | Wellcome Sanger Institute for the COVID-19 Genomics UK Consortium | Luke W Meredith, M. Estée Török , Myra Hosmillo, William L. Hamilton, Martin D. Curran, Theresa Feltwell, Grant Hall, Anna Yakovleva, Fahad A Khokhar, Charlotte J. Houldcroft, Laura G Caller, Aminu S. Jahun, Sarah L. Caddy, Ian Goodfellow, Alex Alderton, Roberto Amato, Sonia Goncalves, Ewan Harrison, David K. Jackson, Ian Johnston, Dominic Kwiatkowski, Cordelia Langford, John Sillitoe on behalf of the Wellcome Sanger Institute COVID-19 Surveillance Team (http://www.sanger.ac.uk/covid-team) |
| EPI_ISL_441747 | Regional Virus Laboratory, Belfast Health and Social Care Trust | Wellcome Sanger Institute for the COVID-19 Genomics UK Consortium | Conall McCaughey, James McKenna, Tanya Curran, Susan Feeney, Alison Watt, Ciara Cox, Mairead Connor, Zoltan Molnar, David Simpson, Derek Fairley, Alex Alderton, Roberto Amato, Sonia Goncalves, Ewan Harrison, David K. Jackson, Ian Johnston, Dominic Kwiatkowski, Cordelia Langford, John Sillitoe on behalf of the Wellcome Sanger Institute COVID-19 Surveillance Team (http://www.sanger.ac.uk/covid-team) |
| EPI_ISL_441748, EPI_ISL_441749, EPI_ISL_441750 | Department of Pathology, University of Cambridge | Wellcome Sanger Institute for the COVID-19 Genomics UK Consortium | Luke W Meredith, M. Estée Török , Myra Hosmillo, William L. Hamilton, Martin D. Curran, Theresa Feltwell, Grant Hall, Anna Yakovleva, Fahad A Khokhar, Charlotte J. Houldcroft, Laura G Caller, Aminu S. Jahun, Sarah L. Caddy, Ian Goodfellow, Alex Alderton, Roberto Amato, Sonia Goncalves, Ewan Harrison, David K. Jackson, Ian Johnston, Dominic Kwiatkowski, Cordelia Langford, John Sillitoe on behalf of the Wellcome Sanger Institute COVID-19 Surveillance Team (http://www.sanger.ac.uk/covid-team) |
| EPI_ISL_441751 | Regional Virus Laboratory, Belfast Health and Social Care Trust | Wellcome Sanger Institute for the COVID-19 Genomics UK Consortium | Conall McCaughey, James McKenna, Tanya Curran, Susan Feeney, Alison Watt, Ciara Cox, Mairead Connor, Zoltan Molnar, David Simpson, Derek Fairley, Alex Alderton, Roberto Amato, Sonia Goncalves, Ewan Harrison, David K. Jackson, Ian Johnston, Dominic Kwiatkowski, Cordelia Langford, John Sillitoe on behalf of the Wellcome Sanger Institute COVID-19 Surveillance Team (http://www.sanger.ac.uk/covid-team) |
| EPI_ISL_441752 | Department of Pathology, University of Cambridge | Wellcome Sanger Institute for the COVID-19 Genomics UK Consortium | Luke W Meredith, M. Estée Török , Myra Hosmillo, William L. Hamilton, Martin D. Curran, Theresa Feltwell, Grant Hall, Anna Yakovleva, Fahad A Khokhar, Charlotte J. Houldcroft, Laura G Caller, Aminu S. Jahun, Sarah L. Caddy, Ian Goodfellow, Alex Alderton, Roberto Amato, Sonia Goncalves, Ewan Harrison, David K. Jackson, Ian Johnston, Dominic Kwiatkowski, Cordelia Langford, John Sillitoe on behalf of the Wellcome Sanger Institute COVID-19 Surveillance Team (http://www.sanger.ac.uk/covid-team) |
| EPI_ISL_441753 | Regional Virus Laboratory, Belfast Health and Social Care | Wellcome Sanger Institute for the COVID-19 Genomics UK Consortium | Conall McCaughey, James McKenna, Tanya Curran, Susan Feeney, Alison Watt, Ciara Cox, Mairead Connor, Zoltan Molnar, David Simpson, Derek Fairley, Alex Alderton, Roberto Amato, Sonia Goncalves, Ewan |

| | | | |
|---|---|---|---|
| | Trust | | Harrison, David K. Jackson, Ian Johnston, Dominic Kwiatkowski, Cordelia Langford, John Sillitoe on behalf of the Wellcome Sanger Institute COVID-19 Surveillance Team (http://www.sanger.ac.uk/covid-team) |
| EPI_ISL_441754, EPI_ISL_441755, EPI_ISL_441756, EPI_ISL_441757, EPI_ISL_441758, EPI_ISL_441759, EPI_ISL_441760 | Department of Pathology, University of Cambridge | Wellcome Sanger Institute for the COVID-19 Genomics UK Consortium | Luke W Meredith, M. Estée Török , Myra Hosmillo, William L. Hamilton, Martin D. Curran, Theresa Feltwell, Grant Hall, Anna Yakovleva, Fahad A Khokhar, Charlotte J. Houldcroft, Laura G Caller, Aminu S. Jahun, Sarah L. Caddy, Ian Goodfellow, Alex Alderton, Roberto Amato, Sonia Goncalves, Ewan Harrison, David K. Jackson, Ian Johnston, Dominic Kwiatkowski, Cordelia Langford, John Sillitoe on behalf of the Wellcome Sanger Institute COVID-19 Surveillance Team (http://www.sanger.ac.uk/covid-team) |
| EPI_ISL_441761 | Regional Virus Laboratory, Belfast Health and Social Care Trust | Wellcome Sanger Institute for the COVID-19 Genomics UK Consortium | Conall McCaughey, James McKenna, Tanya Curran, Susan Feeney, Alison Watt, Ciara Cox, Mairead Connor, Zoltan Molnar, David Simpson, Derek Fairley, Alex Alderton, Roberto Amato, Sonia Goncalves, Ewan Harrison, David K. Jackson, Ian Johnston, Dominic Kwiatkowski, Cordelia Langford, John Sillitoe on behalf of the Wellcome Sanger Institute COVID-19 Surveillance Team (http://www.sanger.ac.uk/covid-team) |
| EPI_ISL_441762, EPI_ISL_441763, EPI_ISL_441764 | Department of Pathology, University of Cambridge | Wellcome Sanger Institute for the COVID-19 Genomics UK Consortium | Luke W Meredith, M. Estée Török , Myra Hosmillo, William L. Hamilton, Martin D. Curran, Theresa Feltwell, Grant Hall, Anna Yakovleva, Fahad A Khokhar, Charlotte J. Houldcroft, Laura G Caller, Aminu S. Jahun, Sarah L. Caddy, Ian Goodfellow, Alex Alderton, Roberto Amato, Sonia Goncalves, Ewan Harrison, David K. Jackson, Ian Johnston, Dominic Kwiatkowski, Cordelia Langford, John Sillitoe on behalf of the Wellcome Sanger Institute COVID-19 Surveillance Team (http://www.sanger.ac.uk/covid-team) |
| EPI_ISL_441765, EPI_ISL_441766 | Regional Virus Laboratory, Belfast Health and Social Care Trust | Wellcome Sanger Institute for the COVID-19 Genomics UK Consortium | Conall McCaughey, James McKenna, Tanya Curran, Susan Feeney, Alison Watt, Ciara Cox, Mairead Connor, Zoltan Molnar, David Simpson, Derek Fairley, Alex Alderton, Roberto Amato, Sonia Goncalves, Ewan Harrison, David K. Jackson, Ian Johnston, Dominic Kwiatkowski, Cordelia Langford, John Sillitoe on behalf of the Wellcome Sanger Institute COVID-19 Surveillance Team (http://www.sanger.ac.uk/covid-team) |
| EPI_ISL_441767, EPI_ISL_441768, EPI_ISL_441769 | Department of Pathology, University of Cambridge | Wellcome Sanger Institute for the COVID-19 Genomics UK Consortium | Luke W Meredith, M. Estée Török , Myra Hosmillo, William L. Hamilton, Martin D. Curran, Theresa Feltwell, Grant Hall, Anna Yakovleva, Fahad A Khokhar, Charlotte J. Houldcroft, Laura G Caller, Aminu S. Jahun, Sarah L. Caddy, Ian Goodfellow, Alex Alderton, Roberto Amato, Sonia Goncalves, Ewan Harrison, David K. Jackson, Ian Johnston, Dominic Kwiatkowski, Cordelia Langford, John Sillitoe on behalf of the Wellcome Sanger Institute COVID-19 Surveillance Team (http://www.sanger.ac.uk/covid-team) |
| EPI_ISL_441770, EPI_ISL_441771 | Regional Virus Laboratory, Belfast Health and Social Care Trust | Wellcome Sanger Institute for the COVID-19 Genomics UK Consortium | Conall McCaughey, James McKenna, Tanya Curran, Susan Feeney, Alison Watt, Ciara Cox, Mairead Connor, Zoltan Molnar, David Simpson, Derek Fairley, Alex Alderton, Roberto Amato, Sonia Goncalves, Ewan Harrison, David K. Jackson, Ian Johnston, Dominic Kwiatkowski, Cordelia Langford, John Sillitoe on behalf of the Wellcome Sanger Institute COVID-19 Surveillance Team (http://www.sanger.ac.uk/covid-team) |
| EPI_ISL_441772, EPI_ISL_441773, EPI_ISL_441774 | Department of Pathology, University of Cambridge | Wellcome Sanger Institute for the COVID-19 Genomics UK Consortium | Luke W Meredith, M. Estée Török , Myra Hosmillo, William L. Hamilton, Martin D. Curran, Theresa Feltwell, Grant Hall, Anna Yakovleva, Fahad A Khokhar, Charlotte J. Houldcroft, Laura G Caller, Aminu S. Jahun, Sarah L. Caddy, Ian Goodfellow, Alex Alderton, Roberto Amato, Sonia Goncalves, Ewan Harrison, David K. Jackson, Ian Johnston, Dominic Kwiatkowski, Cordelia Langford, John Sillitoe on behalf of the Wellcome Sanger Institute COVID-19 Surveillance Team (http://www.sanger.ac.uk/covid-team) |
| EPI_ISL_441775, EPI_ISL_441776 | Regional Virus Laboratory, Belfast Health and Social Care Trust | Wellcome Sanger Institute for the COVID-19 Genomics UK Consortium | Conall McCaughey, James McKenna, Tanya Curran, Susan Feeney, Alison Watt, Ciara Cox, Mairead Connor, Zoltan Molnar, David Simpson, Derek Fairley, Alex Alderton, Roberto Amato, Sonia Goncalves, Ewan Harrison, David K. Jackson, Ian Johnston, Dominic Kwiatkowski, Cordelia Langford, John Sillitoe on behalf of the Wellcome Sanger Institute COVID-19 Surveillance Team (http://www.sanger.ac.uk/covid-team) |
| EPI_ISL_441777, EPI_ISL_441778, EPI_ISL_441779, EPI_ISL_441780 | Department of Pathology, University of Cambridge | Wellcome Sanger Institute for the COVID-19 Genomics UK Consortium | Luke W Meredith, M. Estée Török , Myra Hosmillo, William L. Hamilton, Martin D. Curran, Theresa Feltwell, Grant Hall, Anna Yakovleva, Fahad A Khokhar, Charlotte J. Houldcroft, Laura G Caller, Aminu S. Jahun, Sarah L. Caddy, Ian Goodfellow, Alex Alderton, Roberto Amato, Sonia Goncalves, Ewan Harrison, David K. Jackson, Ian Johnston, Dominic Kwiatkowski, Cordelia Langford, John Sillitoe on behalf of the Wellcome Sanger Institute COVID-19 Surveillance Team (http://www.sanger.ac.uk/covid-team) |
| EPI_ISL_441781 | Regional Virus Laboratory, Belfast Health and Social Care Trust | Wellcome Sanger Institute for the COVID-19 Genomics UK Consortium | Conall McCaughey, James McKenna, Tanya Curran, Susan Feeney, Alison Watt, Ciara Cox, Mairead Connor, Zoltan Molnar, David Simpson, Derek Fairley, Alex Alderton, Roberto Amato, Sonia Goncalves, Ewan Harrison, David K. Jackson, Ian Johnston, Dominic Kwiatkowski, Cordelia Langford, John Sillitoe on behalf of the Wellcome Sanger Institute COVID-19 Surveillance Team (http://www.sanger.ac.uk/covid-team) |
| EPI_ISL_441782, EPI_ISL_441783, EPI_ISL_441784, EPI_ISL_441785, EPI_ISL_441786, EPI_ISL_441787, EPI_ISL_441788, EPI_ISL_441789, EPI_ISL_441790, EPI_ISL_441791, EPI_ISL_441792, EPI_ISL_441793, EPI_ISL_441794, EPI_ISL_441795, EPI_ISL_441796, EPI_ISL_441797, EPI_ISL_441798, EPI_ISL_441799, EPI_ISL_441800, EPI_ISL_441801, EPI_ISL_441802, EPI_ISL_441803, EPI_ISL_441804, EPI_ISL_441805, EPI_ISL_441806, EPI_ISL_441807, EPI_ISL_441808, EPI_ISL_441809, EPI_ISL_441810, EPI_ISL_441811, EPI_ISL_441812, EPI_ISL_441813, EPI_ISL_441814, EPI_ISL_441815, EPI_ISL_441816, EPI_ISL_441817, EPI_ISL_441818, EPI_ISL_441819, EPI_ISL_441820, EPI_ISL_441821, EPI_ISL_441822, EPI_ISL_441823, EPI_ISL_441824, EPI_ISL_441825, EPI_ISL_441826, EPI_ISL_441827, EPI_ISL_441828, EPI_ISL_441829, EPI_ISL_441830, EPI_ISL_441831, EPI_ISL_441832, EPI_ISL_441833, EPI_ISL_441834, EPI_ISL_441835, EPI_ISL_441836, EPI_ISL_441837, EPI_ISL_441838, EPI_ISL_441839, EPI_ISL_441840, EPI_ISL_441841, EPI_ISL_441842, EPI_ISL_441843, EPI_ISL_441844, EPI_ISL_441845, EPI_ISL_441846 | | | |
| see above | Department of Pathology, University of Cambridge | Wellcome Sanger Institute for the COVID-19 Genomics UK Consortium | Luke W Meredith, M. Estée Török , Myra Hosmillo, William L. Hamilton, Martin D. Curran, Theresa Feltwell, Grant Hall, Anna Yakovleva, Fahad A Khokhar, Charlotte J. Houldcroft, Laura G Caller, Aminu S. Jahun, Sarah L. Caddy, Ian Goodfellow, Alex Alderton, Roberto Amato, Sonia Goncalves, Ewan Harrison, David K. Jackson, Ian Johnston, Dominic Kwiatkowski, Cordelia Langford, John Sillitoe on behalf of the Wellcome Sanger Institute COVID-19 Surveillance Team (http://www.sanger.ac.uk/covid-team) |
| EPI_ISL_441847, EPI_ISL_441848, EPI_ISL_441849, EPI_ISL_441850, EPI_ISL_441851, EPI_ISL_441852, EPI_ISL_441853, EPI_ISL_441854, EPI_ISL_441855, EPI_ISL_441856, EPI_ISL_441857, EPI_ISL_441858, EPI_ISL_441859, EPI_ISL_441860, EPI_ISL_441861, EPI_ISL_441862, EPI_ISL_441863, EPI_ISL_441864, EPI_ISL_441865, EPI_ISL_441866, EPI_ISL_441867, EPI_ISL_441868, EPI_ISL_441869, EPI_ISL_441870, EPI_ISL_441871, EPI_ISL_441872, EPI_ISL_441873, EPI_ISL_441874, EPI_ISL_441875, EPI_ISL_441876, EPI_ISL_441877, EPI_ISL_441878, EPI_ISL_441879, EPI_ISL_441880, EPI_ISL_441881, EPI_ISL_441882, EPI_ISL_441883, EPI_ISL_441884, EPI_ISL_441885, EPI_ISL_441886, EPI_ISL_441887, EPI_ISL_441888, EPI_ISL_441889, EPI_ISL_441890, EPI_ISL_441891, EPI_ISL_441892, EPI_ISL_441893, EPI_ISL_441894, EPI_ISL_441895, EPI_ISL_441896, EPI_ISL_441897 | | | |
| see above | Queens Medical Centre, Clinical Microbiology Department / DeepSeq Nottingham | COVID-19 Genomics UK (COG-UK) Consortium | Gemma Clark, Wendy Smith, Manjinder Khakh, Hannah Howson-Wells, Jonathan Ball, Patrick McClure, Joseph Chappell, Theocharis Tsoleridis, Nadine Holmes, Matthew Carlisle, Christopher Moore, Fei Sang, Johnny Debebe, Victoria Wright, Matthew Loose |
| EPI_ISL_441898, EPI_ISL_441899, EPI_ISL_441900, EPI_ISL_441901, EPI_ISL_441902, EPI_ISL_441903, EPI_ISL_441904, EPI_ISL_441905, EPI_ISL_441906, EPI_ISL_441907, EPI_ISL_441908, EPI_ISL_441909, EPI_ISL_441910, EPI_ISL_441911, EPI_ISL_441912, EPI_ISL_441913, EPI_ISL_441914, EPI_ISL_441915, EPI_ISL_441916, EPI_ISL_441917, EPI_ISL_441918, EPI_ISL_441919, EPI_ISL_441920, EPI_ISL_441921, EPI_ISL_441922, EPI_ISL_441923, EPI_ISL_441924, EPI_ISL_441925, EPI_ISL_441926, EPI_ISL_441927, EPI_ISL_441928, EPI_ISL_441929, EPI_ISL_441930, EPI_ISL_441931, EPI_ISL_441932, EPI_ISL_441933, EPI_ISL_441934, EPI_ISL_441935, EPI_ISL_441936, EPI_ISL_441937, EPI_ISL_441938, EPI_ISL_441939, EPI_ISL_441940, EPI_ISL_441941, EPI_ISL_441942, EPI_ISL_441943, EPI_ISL_441944, EPI_ISL_441945, EPI_ISL_441946, EPI_ISL_441947, EPI_ISL_441948, EPI_ISL_441949, EPI_ISL_441950, EPI_ISL_441951, EPI_ISL_441952, EPI_ISL_441953, EPI_ISL_441954, EPI_ISL_441955, EPI_ISL_441956, EPI_ISL_441957, EPI_ISL_441958, EPI_ISL_441959, EPI_ISL_441960, EPI_ISL_441961, EPI_ISL_441962, EPI_ISL_441963, EPI_ISL_441964, EPI_ISL_441965, EPI_ISL_441966, EPI_ISL_441967, EPI_ISL_441968, EPI_ISL_441969, EPI_ISL_441970, EPI_ISL_441971, EPI_ISL_441972, EPI_ISL_441973, EPI_ISL_441974, EPI_ISL_441975, EPI_ISL_441976, EPI_ISL_441977, EPI_ISL_441978, EPI_ISL_441979, EPI_ISL_441980, EPI_ISL_441981, EPI_ISL_441982, EPI_ISL_441983, EPI_ISL_441984, EPI_ISL_441985, EPI_ISL_441986, EPI_ISL_441987, EPI_ISL_441988, EPI_ISL_441989, EPI_ISL_441990, EPI_ISL_441991, EPI_ISL_441992, EPI_ISL_441993, EPI_ISL_441994, EPI_ISL_441995, EPI_ISL_441996, EPI_ISL_441997, EPI_ISL_441998, EPI_ISL_441999, EPI_ISL_442000, EPI_ISL_442001, EPI_ISL_442002, EPI_ISL_442003, EPI_ISL_442004, EPI_ISL_442005, EPI_ISL_442006, EPI_ISL_442007, EPI_ISL_442008, EPI_ISL_442009, EPI_ISL_442010, EPI_ISL_442011, EPI_ISL_442012, EPI_ISL_442013, EPI_ISL_442014, EPI_ISL_442015, EPI_ISL_442016, EPI_ISL_442017, EPI_ISL_442018, EPI_ISL_442019, EPI_ISL_442020, EPI_ISL_442021, EPI_ISL_442022, EPI_ISL_442023, EPI_ISL_442024, EPI_ISL_442025, EPI_ISL_442026, EPI_ISL_442027, EPI_ISL_442028, EPI_ISL_442029, EPI_ISL_442030, EPI_ISL_442031, EPI_ISL_442032, EPI_ISL_442033, EPI_ISL_442034, EPI_ISL_442035, EPI_ISL_442036, EPI_ISL_442037, EPI_ISL_442038, EPI_ISL_442039, EPI_ISL_442040, EPI_ISL_442041, EPI_ISL_442042, EPI_ISL_442043 | | | |
| see above | Virology Department, Sheffield Teaching Hospitals NHS Foundation Trust/Department of Infection, Immunity and Cardiovascular Disease, The Medical School, University of Sheffield | COVID-19 Genomics UK (COG-UK) Consortium | Thushan de Silva, Matthew Parker, Nikki Smith, Adri Angyal, Rebecca Brown, Luke Green, Rachel Tucker, Paul Parsons, Danielle Groves, Katie Johnson, Laura Carrilero, Alex Keeley, Dave Partridge, Matthew Wyles, Benjamin Lindsey, Mehmet Yavuz, Mohammad Raza, Cariad Evans |
| EPI_ISL_442044 | Kawsar Human Genetic Research Center | Kawsar Human Genetic Research Center | Mohammad Ali Khosravi, Maryam Abbasalipour Bashash, Sirous Zeinali, Solmaz Sabeghi, Yeganeh Keshvar, Fatemeh Hosseini, Yeganeh Haghdoost |
| EPI_ISL_442045, EPI_ISL_442046, EPI_ISL_442047, EPI_ISL_442048, EPI_ISL_442049, EPI_ISL_442050, EPI_ISL_442051, EPI_ISL_442052, EPI_ISL_442053, EPI_ISL_442054, EPI_ISL_442055, EPI_ISL_442056, EPI_ISL_442057, EPI_ISL_442058, EPI_ISL_442059, EPI_ISL_442060, EPI_ISL_442061, EPI_ISL_442062, EPI_ISL_442063, EPI_ISL_442064, EPI_ISL_442065, EPI_ISL_442066, EPI_ISL_442067, EPI_ISL_442068, EPI_ISL_442069, EPI_ISL_442070, EPI_ISL_442071, EPI_ISL_442072, EPI_ISL_442073, EPI_ISL_442074, EPI_ISL_442075, EPI_ISL_442076, EPI_ISL_442077, EPI_ISL_442078, EPI_ISL_442079, EPI_ISL_442080, EPI_ISL_442081, EPI_ISL_442082, EPI_ISL_442083, EPI_ISL_442084, EPI_ISL_442085, EPI_ISL_442086, EPI_ISL_442087, EPI_ISL_442088, EPI_ISL_442089, EPI_ISL_442090, EPI_ISL_442091, EPI_ISL_442092, EPI_ISL_442093, EPI_ISL_442094, EPI_ISL_442095, EPI_ISL_442096, EPI_ISL_442097, EPI_ISL_442098, EPI_ISL_442099, EPI_ISL_442100, EPI_ISL_442101, EPI_ISL_442102, EPI_ISL_442103, EPI_ISL_442104, EPI_ISL_442105, EPI_ISL_442106, EPI_ISL_442107, EPI_ISL_442108, EPI_ISL_442109, EPI_ISL_442110, EPI_ISL_442111, EPI_ISL_442112, EPI_ISL_442113, EPI_ISL_442114, EPI_ISL_442115, EPI_ISL_442116, EPI_ISL_442117, EPI_ISL_442118, EPI_ISL_442119, EPI_ISL_442120, EPI_ISL_442121, EPI_ISL_442122, EPI_ISL_442123, EPI_ISL_442124, EPI_ISL_442125, EPI_ISL_442126, EPI_ISL_442127, EPI_ISL_442128, EPI_ISL_442129, EPI_ISL_442130, EPI_ISL_442131, EPI_ISL_442132, EPI_ISL_442133, EPI_ISL_442134, EPI_ISL_442135, EPI_ISL_442136, EPI_ISL_442137, EPI_ISL_442138, EPI_ISL_442139, EPI_ISL_442140, EPI_ISL_442141, EPI_ISL_442142, EPI_ISL_442143, EPI_ISL_442144, EPI_ISL_442145, EPI_ISL_442146, EPI_ISL_442147, EPI_ISL_442148, EPI_ISL_442149, EPI_ISL_442150, EPI_ISL_442151, EPI_ISL_442152, EPI_ISL_442153, EPI_ISL_442154, EPI_ISL_442155, EPI_ISL_442156, EPI_ISL_442157, EPI_ISL_442158, EPI_ISL_442159, EPI_ISL_442160, EPI_ISL_442161, EPI_ISL_442162, EPI_ISL_442163, EPI_ISL_442164, EPI_ISL_442165, EPI_ISL_442166, EPI_ISL_442167, EPI_ISL_442168, EPI_ISL_442169, EPI_ISL_442170, EPI_ISL_442171, EPI_ISL_442172, EPI_ISL_442173, EPI_ISL_442174, EPI_ISL_442175, EPI_ISL_442176, EPI_ISL_442177, EPI_ISL_442178, EPI_ISL_442179, EPI_ISL_442180, EPI_ISL_442181, EPI_ISL_442182, EPI_ISL_442183, EPI_ISL_442184, EPI_ISL_442185, EPI_ISL_442186, EPI_ISL_442187, EPI_ISL_442188, EPI_ISL_442189, EPI_ISL_442190, EPI_ISL_442191, EPI_ISL_442192, EPI_ISL_442193, EPI_ISL_442194, EPI_ISL_442195, EPI_ISL_442196, EPI_ISL_442197, EPI_ISL_442198, EPI_ISL_442199, EPI_ISL_442200, EPI_ISL_442201, EPI_ISL_442202, EPI_ISL_442203, EPI_ISL_442204, EPI_ISL_442205, EPI_ISL_442206, EPI_ISL_442207, EPI_ISL_442208, EPI_ISL_442209, EPI_ISL_442210, EPI_ISL_442211, EPI_ISL_442212, EPI_ISL_442213, EPI_ISL_442214, EPI_ISL_442215, EPI_ISL_442216, EPI_ISL_442217, EPI_ISL_442218, EPI_ISL_442219, EPI_ISL_442220, EPI_ISL_442221, EPI_ISL_442222, EPI_ISL_442223, EPI_ISL_442224, EPI_ISL_442225, EPI_ISL_442226, EPI_ISL_442227, EPI_ISL_442228, EPI_ISL_442229, EPI_ISL_442230, EPI_ISL_442231, EPI_ISL_442232, EPI_ISL_442233, EPI_ISL_442234, EPI_ISL_442235, EPI_ISL_442236, EPI_ISL_442237, EPI_ISL_442238, EPI_ISL_442239, EPI_ISL_442240, EPI_ISL_442241, EPI_ISL_442242, EPI_ISL_442243, EPI_ISL_442244, EPI_ISL_442245, EPI_ISL_442246, EPI_ISL_442247, EPI_ISL_442248, EPI_ISL_442249, EPI_ISL_442250, EPI_ISL_442251, EPI_ISL_442252, EPI_ISL_442253, EPI_ISL_442254, EPI_ISL_442255, EPI_ISL_442256, EPI_ISL_442257, EPI_ISL_442258, EPI_ISL_442259, EPI_ISL_442260, EPI_ISL_442261, EPI_ISL_442262, EPI_ISL_442263, EPI_ISL_442264, EPI_ISL_442265, EPI_ISL_442266, EPI_ISL_442267, EPI_ISL_442268, EPI_ISL_442269, EPI_ISL_442270, EPI_ISL_442271, EPI_ISL_442272, EPI_ISL_442273, EPI_ISL_442274, EPI_ISL_442275, EPI_ISL_442276, EPI_ISL_442277, EPI_ISL_442278, EPI_ISL_442279, EPI_ISL_442280, EPI_ISL_442281, EPI_ISL_442282, EPI_ISL_442283, EPI_ISL_442284, EPI_ISL_442285, EPI_ISL_442286, EPI_ISL_442287, EPI_ISL_442288, EPI_ISL_442289, EPI_ISL_442290, EPI_ISL_442291, EPI_ISL_442292, EPI_ISL_442293, EPI_ISL_442294, EPI_ISL_442295, EPI_ISL_442296, EPI_ISL_442297, EPI_ISL_442298, EPI_ISL_442299, EPI_ISL_442300, EPI_ISL_442301, EPI_ISL_442302, EPI_ISL_442303, EPI_ISL_442304, EPI_ISL_442305, EPI_ISL_442306, EPI_ISL_442307, EPI_ISL_442308, EPI_ISL_442309, EPI_ISL_442310, EPI_ISL_442311, EPI_ISL_442312, EPI_ISL_442313, EPI_ISL_442314, EPI_ISL_442315, EPI_ISL_442316, EPI_ISL_442317, EPI_ISL_442318, EPI_ISL_442319, EPI_ISL_442320, EPI_ISL_442321, EPI_ISL_442322, EPI_ISL_442323, EPI_ISL_442324, EPI_ISL_442325, EPI_ISL_442326, EPI_ISL_442327, EPI_ISL_442328, EPI_ISL_442329, EPI_ISL_442330, EPI_ISL_442331, EPI_ISL_442332, EPI_ISL_442333, EPI_ISL_442334, EPI_ISL_442335, EPI_ISL_442336, EPI_ISL_442337, EPI_ISL_442338, EPI_ISL_442339, EPI_ISL_442340, EPI_ISL_442341, EPI_ISL_442342, EPI_ISL_442343 | | | |
| see above | Department of Pathology, University of Cambridge | Wellcome Sanger Institute for the COVID-19 Genomics UK Consortium | Luke W Meredith, M. Estée Török , Myra Hosmillo, William L. Hamilton, Martin D. Curran, Theresa Feltwell, Grant Hall, Anna Yakovleva, Fahad A Khokhar, Charlotte J. Houldcroft, Laura G Caller, Aminu S. Jahun, Sarah L. Caddy, Ian Goodfellow, Alex Alderton, Roberto Amato, Sonia Goncalves, Ewan Harrison, David K. Jackson, Ian Johnston, Dominic Kwiatkowski, Cordelia Langford, John Sillitoe on behalf of the Wellcome Sanger Institute COVID-19 Surveillance Team (http://www.sanger.ac.uk/covid-team) |
| EPI_ISL_442344, EPI_ISL_442345, EPI_ISL_442346, EPI_ISL_442347, EPI_ISL_442348, EPI_ISL_442349, EPI_ISL_442350, EPI_ISL_442351, EPI_ISL_442352, EPI_ISL_442353, EPI_ISL_442354, EPI_ISL_442355, EPI_ISL_442356, EPI_ISL_442357, EPI_ISL_442358, EPI_ISL_442359, EPI_ISL_442360, EPI_ISL_442361, EPI_ISL_442362, EPI_ISL_442363, EPI_ISL_442364, EPI_ISL_442365, EPI_ISL_442366, EPI_ISL_442367, EPI_ISL_442368, EPI_ISL_442369, EPI_ISL_442370, EPI_ISL_442371, EPI_ISL_442372, EPI_ISL_442373, EPI_ISL_442374, EPI_ISL_442375, EPI_ISL_442376, EPI_ISL_442377, EPI_ISL_442378, EPI_ISL_442379, EPI_ISL_442380, EPI_ISL_442381, EPI_ISL_442382, EPI_ISL_442383, EPI_ISL_442384, EPI_ISL_442385, EPI_ISL_442386, EPI_ISL_442387, EPI_ISL_442388, EPI_ISL_442389, EPI_ISL_442390, EPI_ISL_442391, EPI_ISL_442392, EPI_ISL_442393, EPI_ISL_442394, EPI_ISL_442395, EPI_ISL_442396, EPI_ISL_442397, EPI_ISL_442398, EPI_ISL_442399, EPI_ISL_442400, EPI_ISL_442401, EPI_ISL_442402, EPI_ISL_442403, EPI_ISL_442404, EPI_ISL_442405, EPI_ISL_442406, EPI_ISL_442407, EPI_ISL_442408, EPI_ISL_442409, EPI_ISL_442410, EPI_ISL_442411, EPI_ISL_442412, EPI_ISL_442413, EPI_ISL_442414, EPI_ISL_442415, EPI_ISL_442416, EPI_ISL_442417, EPI_ISL_442418, EPI_ISL_442419, EPI_ISL_442420, EPI_ISL_442421, EPI_ISL_442422, EPI_ISL_442423, EPI_ISL_442424, EPI_ISL_442425, EPI_ISL_442426, EPI_ISL_442427, EPI_ISL_442428, EPI_ISL_442429, EPI_ISL_442430, EPI_ISL_442431, EPI_ISL_442432, EPI_ISL_442433, EPI_ISL_442434, EPI_ISL_442435, EPI_ISL_442436, EPI_ISL_442437, EPI_ISL_442438, EPI_ISL_442439, EPI_ISL_442440, EPI_ISL_442441, EPI_ISL_442442, EPI_ISL_442443, EPI_ISL_442444, EPI_ISL_442445, EPI_ISL_442446, EPI_ISL_442447, EPI_ISL_442448, EPI_ISL_442449, EPI_ISL_442450, EPI_ISL_442451, EPI_ISL_442452, EPI_ISL_442453, EPI_ISL_442454, EPI_ISL_442455, EPI_ISL_442456, EPI_ISL_442457, EPI_ISL_442458, EPI_ISL_442459, EPI_ISL_442460, EPI_ISL_442461, EPI_ISL_442462, EPI_ISL_442463, EPI_ISL_442464, EPI_ISL_442465, EPI_ISL_442466, EPI_ISL_442467, EPI_ISL_442468, EPI_ISL_442469, EPI_ISL_442470, EPI_ISL_442471, EPI_ISL_442472, EPI_ISL_442473, EPI_ISL_442474, EPI_ISL_442475, EPI_ISL_442476, EPI_ISL_442477, EPI_ISL_442478, EPI_ISL_442479, EPI_ISL_442480, EPI_ISL_442481, EPI_ISL_442482, EPI_ISL_442483, EPI_ISL_442484, EPI_ISL_442485, EPI_ISL_442486, EPI_ISL_442487, EPI_ISL_442488, EPI_ISL_442489, EPI_ISL_442490, EPI_ISL_442491, EPI_ISL_442492, EPI_ISL_442493, EPI_ISL_442494, EPI_ISL_442495, EPI_ISL_442496, EPI_ISL_442497, EPI_ISL_442498, EPI_ISL_442499, EPI_ISL_442500, EPI_ISL_442501, EPI_ISL_442502, EPI_ISL_442503, EPI_ISL_442504, EPI_ISL_442505, EPI_ISL_442506, EPI_ISL_442507, EPI_ISL_442508, EPI_ISL_442509, EPI_ISL_442510, EPI_ISL_442511, EPI_ISL_442512, EPI_ISL_442513, EPI_ISL_442514, EPI_ISL_442515, EPI_ISL_442516, EPI_ISL_442517, EPI_ISL_442518, EPI_ISL_442519, EPI_ISL_442520, EPI_ISL_442521, EPI_ISL_442522 | | | |
| see above | Virology Department, Sheffield Teaching Hospitals NHS Foundation Trust/Department of Infection, Immunity and Cardiovascular Disease, The Medical School, University of Sheffield | COVID-19 Genomics UK (COG-UK) Consortium | Thushan de Silva, Matthew Parker, Nikki Smith, Adri Angyal, Rebecca Brown, Luke Green, Rachel Tucker, Paul Parsons, Danielle Groves, Katie Johnson, Laura Carrilero, Alex Keeley, Dave Partridge, Matthew Wyles, Benjamin Lindsey, Mehmet Yavuz, Mohammad Raza, Cariad Evans |
| EPI_ISL_442523 | Pasteur Institute of Iran | Kawsar Human Genetic Research Company | Sirous Zeinali, Mohammad Ali Khosravi,Maryam Abbasalipour Bashash, Sanaz Mostafavi Jabbari, Maraym Firoozi, Sormeh Pourtavakoli, Elmira Khateri, Razieh Zeinali and Fahimeh Hoseini |
| EPI_ISL_442524, EPI_ISL_442525, EPI_ISL_442526, EPI_ISL_442527, EPI_ISL_442528, EPI_ISL_442529, EPI_ISL_442530, EPI_ISL_442531, EPI_ISL_442532, EPI_ISL_442533, EPI_ISL_442534, EPI_ISL_442535, EPI_ISL_442536, EPI_ISL_442537, EPI_ISL_442538, EPI_ISL_442539, EPI_ISL_442540, EPI_ISL_442541, EPI_ISL_442542, EPI_ISL_442543, EPI_ISL_442544, EPI_ISL_442545, EPI_ISL_442546, EPI_ISL_442547, EPI_ISL_442548, EPI_ISL_442549, EPI_ISL_442550, EPI_ISL_442551, EPI_ISL_442552, EPI_ISL_442553, EPI_ISL_442554, EPI_ISL_442555, EPI_ISL_442556, EPI_ISL_442557, EPI_ISL_442558, EPI_ISL_442559, EPI_ISL_442560, EPI_ISL_442561, EPI_ISL_442562, EPI_ISL_442563, EPI_ISL_442564, EPI_ISL_442565, EPI_ISL_442566, EPI_ISL_442567, EPI_ISL_442568, EPI_ISL_442569, EPI_ISL_442570, EPI_ISL_442571, EPI_ISL_442572, EPI_ISL_442573, EPI_ISL_442574, EPI_ISL_442575, EPI_ISL_442576, EPI_ISL_442577, EPI_ISL_442578, EPI_ISL_442579, EPI_ISL_442580, EPI_ISL_442581, EPI_ISL_442582, EPI_ISL_442583, EPI_ISL_442584, EPI_ISL_442585, EPI_ISL_442586, EPI_ISL_442587, EPI_ISL_442588, EPI_ISL_442589, EPI_ISL_442590, EPI_ISL_442591, EPI_ISL_442592, EPI_ISL_442593, EPI_ISL_442594, EPI_ISL_442595, EPI_ISL_442596, EPI_ISL_442597, EPI_ISL_442598, EPI_ISL_442599, EPI_ISL_442600, EPI_ISL_442601, EPI_ISL_442602, EPI_ISL_442603, EPI_ISL_442604, EPI_ISL_442605, EPI_ISL_442606, EPI_ISL_442607, EPI_ISL_442608, EPI_ISL_442609, EPI_ISL_442610, EPI_ISL_442611, EPI_ISL_442612, EPI_ISL_442613, EPI_ISL_442614, EPI_ISL_442615, EPI_ISL_442616, EPI_ISL_442617, EPI_ISL_442618, EPI_ISL_442619, EPI_ISL_442620, EPI_ISL_442621, EPI_ISL_442622, EPI_ISL_442623 | | | |
| see above | Department of Pathology, University of Cambridge | Wellcome Sanger Institute for the COVID-19 Genomics UK Consortium | Luke W Meredith, M. Estée Török , Myra Hosmillo, William L. Hamilton, Martin D. Curran, Theresa Feltwell, Grant Hall, Anna Yakovleva, Fahad A Khokhar, Charlotte J. Houldcroft, Laura G Caller, Aminu S. Jahun, Sarah L. Caddy, Ian Goodfellow, Alex Alderton, Roberto Amato, Sonia Goncalves, Ewan Harrison, David K. Jackson, Ian Johnston, Dominic Kwiatkowski, Cordelia Langford, John Sillitoe on behalf of the Wellcome Sanger Institute COVID-19 Surveillance Team (http://www.sanger.ac.uk/covid-team) |
| EPI_ISL_442624, EPI_ISL_442625, EPI_ISL_442626, EPI_ISL_442627, EPI_ISL_442628, EPI_ISL_442629, EPI_ISL_442630, EPI_ISL_442631, EPI_ISL_442632, EPI_ISL_442633, EPI_ISL_442634, EPI_ISL_442635, EPI_ISL_442636, EPI_ISL_442637, EPI_ISL_442638, EPI_ISL_442639, EPI_ISL_442640, EPI_ISL_442641, EPI_ISL_442642, EPI_ISL_442643, EPI_ISL_442644, EPI_ISL_442645, EPI_ISL_442646, EPI_ISL_442647, EPI_ISL_442648, EPI_ISL_442649, EPI_ISL_442650, EPI_ISL_442651, EPI_ISL_442652, EPI_ISL_442653, EPI_ISL_442654, EPI_ISL_442655, EPI_ISL_442656, EPI_ISL_442657, EPI_ISL_442658, EPI_ISL_442659, EPI_ISL_442660, EPI_ISL_442661, EPI_ISL_442662, EPI_ISL_442663, EPI_ISL_442664, EPI_ISL_442665, EPI_ISL_442666, EPI_ISL_442667, EPI_ISL_442668, EPI_ISL_442669, EPI_ISL_442670, EPI_ISL_442671, EPI_ISL_442672, EPI_ISL_442673, EPI_ISL_442674, EPI_ISL_442675, EPI_ISL_442676, EPI_ISL_442677, EPI_ISL_442678, EPI_ISL_442679, EPI_ISL_442680, EPI_ISL_442681, EPI_ISL_442682, EPI_ISL_442683, EPI_ISL_442684, EPI_ISL_442685, EPI_ISL_442686, EPI_ISL_442687, EPI_ISL_442688, EPI_ISL_442689, EPI_ISL_442690, EPI_ISL_442691, EPI_ISL_442692, EPI_ISL_442693, EPI_ISL_442694, EPI_ISL_442695, EPI_ISL_442696, EPI_ISL_442697, EPI_ISL_442698, EPI_ISL_442699, EPI_ISL_442700, EPI_ISL_442701, EPI_ISL_442702, EPI_ISL_442703, EPI_ISL_442704, EPI_ISL_442705, EPI_ISL_442706, EPI_ISL_442707, EPI_ISL_442708, EPI_ISL_442709, EPI_ISL_442710, EPI_ISL_442711, EPI_ISL_442712, EPI_ISL_442713, EPI_ISL_442714, EPI_ISL_442715, EPI_ISL_442716, EPI_ISL_442717, EPI_ISL_442718, EPI_ISL_442719, EPI_ISL_442720, EPI_ISL_442721, EPI_ISL_442722, EPI_ISL_442723, EPI_ISL_442724, EPI_ISL_442725, EPI_ISL_442726, EPI_ISL_442727, EPI_ISL_442728, EPI_ISL_442729, EPI_ISL_442730, EPI_ISL_442731, EPI_ISL_442732, EPI_ISL_442733, EPI_ISL_442734, EPI_ISL_442735, EPI_ISL_442736, EPI_ISL_442737, EPI_ISL_442738, EPI_ISL_442739, EPI_ISL_442740, EPI_ISL_442741, EPI_ISL_442742, EPI_ISL_442743, EPI_ISL_442744, EPI_ISL_442745, EPI_ISL_442746, EPI_ISL_442747, EPI_ISL_442748, EPI_ISL_442749, EPI_ISL_442750, EPI_ISL_442751, EPI_ISL_442752, EPI_ISL_442753, EPI_ISL_442754, EPI_ISL_442755, EPI_ISL_442756, EPI_ISL_442757, EPI_ISL_442758, EPI_ISL_442759, EPI_ISL_442760, EPI_ISL_442761, EPI_ISL_442762, EPI_ISL_442763, EPI_ISL_442764, EPI_ISL_442765, EPI_ISL_442766, EPI_ISL_442767, EPI_ISL_442768, EPI_ISL_442769, EPI_ISL_442770, EPI_ISL_442771, EPI_ISL_442772, EPI_ISL_442773, EPI_ISL_442774, EPI_ISL_442775, EPI_ISL_442776, EPI_ISL_442777, EPI_ISL_442778, EPI_ISL_442779, EPI_ISL_442780, EPI_ISL_442781, EPI_ISL_442782, EPI_ISL_442783, EPI_ISL_442784, EPI_ISL_442785, EPI_ISL_442786, EPI_ISL_442787, EPI_ISL_442788, EPI_ISL_442789, EPI_ISL_442790, EPI_ISL_442791, EPI_ISL_442792, EPI_ISL_442793, EPI_ISL_442794, EPI_ISL_442795, EPI_ISL_442796, EPI_ISL_442797, EPI_ISL_442798, EPI_ISL_442799, EPI_ISL_442800, EPI_ISL_442801, EPI_ISL_442802, EPI_ISL_442803, EPI_ISL_442804, EPI_ISL_442805, EPI_ISL_442806, EPI_ISL_442807, EPI_ISL_442808, EPI_ISL_442809, EPI_ISL_442810 | | | |
| see above | PHE South West Regional Laboratory, National Infection Service | Wellcome Sanger Institute for the COVID-19 Genomics UK Consortium | Stephanie Hutchings, Hannah Pymont, Dr Peter Muir, Barry Vipond, Rich Hopes, Alex Alderton, Roberto Amato, Sonia Goncalves, Ewan Harrison, David K. Jackson, Ian Johnston, Dominic Kwiatkowski, Cordelia Langford, John Sillitoe on behalf of the Wellcome Sanger Institute COVID-19 Surveillance Team (http://www.sanger.ac.uk/covid-team) |
| EPI_ISL_442811, EPI_ISL_442812, EPI_ISL_442813, EPI_ISL_442814, EPI_ISL_442815, EPI_ISL_442816, EPI_ISL_442817, EPI_ISL_442818, EPI_ISL_442819, EPI_ISL_442820, EPI_ISL_442821, EPI_ISL_442822, EPI_ISL_442823, EPI_ISL_442824, EPI_ISL_442825, EPI_ISL_442826, EPI_ISL_442827, EPI_ISL_442828, EPI_ISL_442829, EPI_ISL_442830, EPI_ISL_442831, EPI_ISL_442832, EPI_ISL_442833, EPI_ISL_442834, EPI_ISL_442835, EPI_ISL_442836, EPI_ISL_442837, EPI_ISL_442838, EPI_ISL_442839, EPI_ISL_442840, EPI_ISL_442841, EPI_ISL_442842, EPI_ISL_442843, EPI_ISL_442844, EPI_ISL_442845, EPI_ISL_442846, EPI_ISL_442847, EPI_ISL_442848, EPI_ISL_442849, EPI_ISL_442850, EPI_ISL_442851, EPI_ISL_442852, EPI_ISL_442853, EPI_ISL_442854, EPI_ISL_442855, EPI_ISL_442856, EPI_ISL_442857, EPI_ISL_442858, | | | |

| | | | |
|---|---|---|---|
| EPI_ISL_442859, EPI_ISL_442860, EPI_ISL_442861, EPI_ISL_442862, EPI_ISL_442863, EPI_ISL_442864, EPI_ISL_442865, EPI_ISL_442866, EPI_ISL_442867, EPI_ISL_442868, EPI_ISL_442869, EPI_ISL_442870, EPI_ISL_442871, EPI_ISL_442872, EPI_ISL_442873, EPI_ISL_442874, EPI_ISL_442875, EPI_ISL_442876, EPI_ISL_442877, EPI_ISL_442878, EPI_ISL_442879, EPI_ISL_442880, EPI_ISL_442881, EPI_ISL_442882, EPI_ISL_442883, EPI_ISL_442884, EPI_ISL_442885, EPI_ISL_442886, EPI_ISL_442887, EPI_ISL_442888, EPI_ISL_442889, EPI_ISL_442890, EPI_ISL_442891, EPI_ISL_442892, EPI_ISL_442893, EPI_ISL_442894, EPI_ISL_442895, EPI_ISL_442896, EPI_ISL_442897, EPI_ISL_442898, EPI_ISL_442899, EPI_ISL_442900, EPI_ISL_442901, EPI_ISL_442902, EPI_ISL_442903, EPI_ISL_442904, EPI_ISL_442905, EPI_ISL_442906, EPI_ISL_442907, EPI_ISL_442908, EPI_ISL_442909, EPI_ISL_442910, EPI_ISL_442911, EPI_ISL_442912, EPI_ISL_442913, EPI_ISL_442914, EPI_ISL_442915, EPI_ISL_442916, EPI_ISL_442917, EPI_ISL_442918, EPI_ISL_442919, EPI_ISL_442920, EPI_ISL_442921, EPI_ISL_442922, EPI_ISL_442923, EPI_ISL_442924, EPI_ISL_442925, EPI_ISL_442926, EPI_ISL_442927, EPI_ISL_442928, EPI_ISL_442929, EPI_ISL_442930, EPI_ISL_442931, EPI_ISL_442932, EPI_ISL_442933, EPI_ISL_442934, EPI_ISL_442935, EPI_ISL_442936, EPI_ISL_442937, EPI_ISL_442938, EPI_ISL_442939, EPI_ISL_442940, EPI_ISL_442941, EPI_ISL_442942, EPI_ISL_442943, EPI_ISL_442944, EPI_ISL_442945, EPI_ISL_442946, EPI_ISL_442947, EPI_ISL_442948, EPI_ISL_442949, EPI_ISL_442950, EPI_ISL_442951, EPI_ISL_442952, EPI_ISL_442953, EPI_ISL_442954, EPI_ISL_442955, EPI_ISL_442956, EPI_ISL_442957, EPI_ISL_442958, EPI_ISL_442959, EPI_ISL_442960, EPI_ISL_442961, EPI_ISL_442962, EPI_ISL_442963, EPI_ISL_442964, EPI_ISL_442965, EPI_ISL_442966, EPI_ISL_442967, EPI_ISL_442968, EPI_ISL_442969, EPI_ISL_442970, EPI_ISL_442971, EPI_ISL_442972, EPI_ISL_442973, EPI_ISL_442974, EPI_ISL_442975, EPI_ISL_442976, EPI_ISL_442977, EPI_ISL_442978, EPI_ISL_442979, EPI_ISL_442980, EPI_ISL_442981, EPI_ISL_442982, EPI_ISL_442983, EPI_ISL_442984, EPI_ISL_442985, EPI_ISL_442986, EPI_ISL_442987, EPI_ISL_442988, EPI_ISL_442989, EPI_ISL_442990, EPI_ISL_442991, EPI_ISL_442992, EPI_ISL_442993, EPI_ISL_442994, EPI_ISL_442995, EPI_ISL_442996, EPI_ISL_442997, EPI_ISL_442998, EPI_ISL_442999, EPI_ISL_443000, EPI_ISL_443001, EPI_ISL_443002, EPI_ISL_443003, EPI_ISL_443004, EPI_ISL_443005, EPI_ISL_443006, EPI_ISL_443007, EPI_ISL_443008, EPI_ISL_443009, EPI_ISL_443010, EPI_ISL_443011, EPI_ISL_443012, EPI_ISL_443013, EPI_ISL_443014, EPI_ISL_443015, EPI_ISL_443016, EPI_ISL_443017, EPI_ISL_443018, EPI_ISL_443019, EPI_ISL_443020, EPI_ISL_443021, EPI_ISL_443022, EPI_ISL_443023, EPI_ISL_443024, EPI_ISL_443025, EPI_ISL_443026, EPI_ISL_443027, EPI_ISL_443028, EPI_ISL_443029, EPI_ISL_443030, EPI_ISL_443031, EPI_ISL_443032, EPI_ISL_443033, EPI_ISL_443034, EPI_ISL_443035, EPI_ISL_443036, EPI_ISL_443037, EPI_ISL_443038, EPI_ISL_443039, EPI_ISL_443040, EPI_ISL_443041, EPI_ISL_443042, EPI_ISL_443043, EPI_ISL_443044, EPI_ISL_443045, EPI_ISL_443046, EPI_ISL_443047, EPI_ISL_443048, EPI_ISL_443049, EPI_ISL_443050, EPI_ISL_443051, EPI_ISL_443052, EPI_ISL_443053, EPI_ISL_443054, EPI_ISL_443055, EPI_ISL_443056, EPI_ISL_443057, EPI_ISL_443058, EPI_ISL_443059, EPI_ISL_443060, EPI_ISL_443061, EPI_ISL_443062, EPI_ISL_443063, EPI_ISL_443064, EPI_ISL_443065, EPI_ISL_443066, EPI_ISL_443067, EPI_ISL_443068, EPI_ISL_443069, EPI_ISL_443070, EPI_ISL_443071, EPI_ISL_443072, EPI_ISL_443073, EPI_ISL_443074, EPI_ISL_443075, EPI_ISL_443076, EPI_ISL_443077, EPI_ISL_443078, EPI_ISL_443079, EPI_ISL_443080, EPI_ISL_443081, EPI_ISL_443082, EPI_ISL_443083, EPI_ISL_443084, EPI_ISL_443085, EPI_ISL_443086, EPI_ISL_443087, EPI_ISL_443088, EPI_ISL_443089, EPI_ISL_443090, EPI_ISL_443091, EPI_ISL_443092, EPI_ISL_443093, EPI_ISL_443094, EPI_ISL_443095, EPI_ISL_443096, EPI_ISL_443097, EPI_ISL_443098, EPI_ISL_443099, EPI_ISL_443100, EPI_ISL_443101, EPI_ISL_443102, EPI_ISL_443103, EPI_ISL_443104, EPI_ISL_443105, EPI_ISL_443106, EPI_ISL_443107, EPI_ISL_443108, EPI_ISL_443109, EPI_ISL_443110, EPI_ISL_443111, EPI_ISL_443112, EPI_ISL_443113, EPI_ISL_443114, EPI_ISL_443115, EPI_ISL_443116, EPI_ISL_443117, EPI_ISL_443118 | | | |
| see above | Department of Pathology, University of Cambridge | Wellcome Sanger Institute for the COVID-19 Genomics UK Consortium | Luke W Meredith, M. Estée Török , Myra Hosmillo, William L. Hamilton, Martin D. Curran, Theresa Feltwell, Grant Hall, Anna Yakovleva, Fahad A Khokhar, Charlotte J. Houldcroft, Laura G Caller, Aminu S. Jahun, Sarah L. Caddy, Ian Goodfellow, Alex Alderton, Roberto Amato, Sonia Goncalves, Ewan Harrison, David K. Jackson, Ian Johnston, Dominic Kwiatkowski, Cordelia Langford, John Sillitoe on behalf of the Wellcome Sanger Institute COVID-19 Surveillance Team (http://www.sanger.ac.uk/covid-team) |
| EPI_ISL_443183 | M Health Fairview | University of Minnesota Genomics Center | Daryl M. Gohl, John Garbe, Patrick Grady, Jerry Daniel, Ray Watson, Benjamin Auch, Andrew Nelson, Sophia Yohe, and Kenneth B. Beckman |
| EPI_ISL_443184, EPI_ISL_443185, EPI_ISL_443186 | UW Virology Lab | UW Virology Lab | Pavitra Roychoudhury, Hong Xie, Keith Jerome, Alexander Greninger |
| EPI_ISL_443187 | National Virology Reference Laboratory | National Public Health Laboratory, National Centre for Infectious Diseases | Mak Tze Minn, Octavia Sophie, Chavatte Jean-Marc, Zaini Zainun, Taib Surita, Cui Lin, Lin Raymond Tzer Pin |
| EPI_ISL_443188, EPI_ISL_443189, EPI_ISL_443190, EPI_ISL_443191, EPI_ISL_443192, EPI_ISL_443193, EPI_ISL_443194, EPI_ISL_443195, EPI_ISL_443196, EPI_ISL_443197, EPI_ISL_443198, EPI_ISL_443199, EPI_ISL_443200, EPI_ISL_443201, EPI_ISL_443202, EPI_ISL_443203, EPI_ISL_443204, EPI_ISL_443205, EPI_ISL_443206, EPI_ISL_443207, EPI_ISL_443208, EPI_ISL_443209, EPI_ISL_443210, EPI_ISL_443211, EPI_ISL_443212, EPI_ISL_443213, EPI_ISL_443214, EPI_ISL_443215, EPI_ISL_443216, EPI_ISL_443217, EPI_ISL_443218, EPI_ISL_443219, EPI_ISL_443220, EPI_ISL_443221, EPI_ISL_443222, EPI_ISL_443223, EPI_ISL_443224, EPI_ISL_443225, EPI_ISL_443226, EPI_ISL_443227, EPI_ISL_443228, EPI_ISL_443229, EPI_ISL_443230, EPI_ISL_443231, EPI_ISL_443232, EPI_ISL_443233, EPI_ISL_443234, EPI_ISL_443235, EPI_ISL_443236, EPI_ISL_443237, EPI_ISL_443238, EPI_ISL_443239, EPI_ISL_443240, EPI_ISL_443241, EPI_ISL_443242, EPI_ISL_443243, EPI_ISL_443244, EPI_ISL_443245, EPI_ISL_443246, EPI_ISL_443247, EPI_ISL_443248, EPI_ISL_443249 | | | |
| see above | National Public Health Laboratory, National Centre for Infectious Diseases | National Public Health Laboratory, National Centre for Infectious Diseases | Mak Tze Minn, Octavia Sophie, Chavatte Jean-Marc, Cui Lin, Lin Raymond Tzer Pin |
| EPI_ISL_443253, EPI_ISL_443254, EPI_ISL_443255, EPI_ISL_443256, EPI_ISL_443257 | M Health Fairview | University of Minnesota Genomics Center | Daryl M. Gohl, John Garbe, Patrick Grady, Jerry Daniel, Ray Watson, Benjamin Auch, Andrew Nelson, Sophia Yohe, and Kenneth B. Beckman |
| EPI_ISL_443258, EPI_ISL_443259 | Résidence Ornano | National Reference Center for Viruses of Respiratory Infections, Institut Pasteur, Paris | Mélanie Albert, Marion Barbet, Sylvie Behillil, Méline Bizard, Angela Brisebarre, Flora Donati, Etienne Simon-Lorière, Vincent Enouf, Maud Vanpeene, Sylvie van der Werf |
| EPI_ISL_443260 | LABM GH nord Essonne de Longjumeau - BP 125 | National Reference Center for Viruses of Respiratory Infections, Institut Pasteur, Paris | Mélanie Albert, Marion Barbet, Sylvie Behillil, Méline Bizard, Angela Brisebarre, Flora Donati, Etienne Simon-Lorière, Vincent Enouf, Maud Vanpeene, Sylvie van der Werf |
| EPI_ISL_443261, EPI_ISL_443262, EPI_ISL_443263, EPI_ISL_443264 | CHU de Dijon - Laboratoire de Virologie | National Reference Center for Viruses of Respiratory Infections, Institut Pasteur, Paris | Mélanie Albert, Marion Barbet, Sylvie Behillil, Méline Bizard, Angela Brisebarre, Flora Donati, Etienne Simon-Lorière, Vincent Enouf, Maud Vanpeene, Sylvie van der Werf, Jean-Baptiste Bour |
| EPI_ISL_443265, EPI_ISL_443266, EPI_ISL_443267, EPI_ISL_443268, EPI_ISL_443269, EPI_ISL_443270, EPI_ISL_443271, EPI_ISL_443272, EPI_ISL_443273, EPI_ISL_443274, EPI_ISL_443275, EPI_ISL_443276, EPI_ISL_443277, EPI_ISL_443278, EPI_ISL_443279, EPI_ISL_443280, EPI_ISL_443281, EPI_ISL_443282, EPI_ISL_443283 | | | |
| see above | CHU - Hôpital Cavale Blanche - Labo. de Virologie | National Reference Center for Viruses of Respiratory Infections, Institut Pasteur, Paris | Mélanie Albert, Marion Barbet, Sylvie Behillil, Méline Bizard, Angela Brisebarre, Flora Donati, Etienne Simon-Lorière, Vincent Enouf, Maud Vanpeene, Sylvie van der Werf, Léa Pilorge |
| EPI_ISL_443284, EPI_ISL_443285, EPI_ISL_443286, EPI_ISL_443287, EPI_ISL_443288 | Laboratoire de Microbiologie - Bât A - CH René Dubois | National Reference Center for Viruses of Respiratory Infections, Institut Pasteur, Paris | Mélanie Albert, Marion Barbet, Sylvie Behillil, Méline Bizard, Angela Brisebarre, Flora Donati, Etienne Simon-Lorière, Vincent Enouf, Maud Vanpeene, Sylvie van der Werf, Pascale Martres |
| EPI_ISL_443289, EPI_ISL_443290, EPI_ISL_443291, EPI_ISL_443292, EPI_ISL_443293, EPI_ISL_443294 | CHRU Pontchaillou - Laboratoire de Virologie | National Reference Center for Viruses of Respiratory Infections, Institut Pasteur, Paris | Mélanie Albert, Marion Barbet, Sylvie Behillil, Méline Bizard, Angela Brisebarre, Flora Donati, Etienne Simon-Lorière, Vincent Enouf, Maud Vanpeene, Sylvie van der Werf, Gisèle Lagathu |
| EPI_ISL_443295, EPI_ISL_443296, EPI_ISL_443297, EPI_ISL_443298, EPI_ISL_443299 | Hôpital Necker - Enfants - Malades Laboratoire de Virologie | National Reference Center for Viruses of Respiratory Infections, Institut Pasteur, Paris | Mélanie Albert, Marion Barbet, Sylvie Behillil, Méline Bizard, Angela Brisebarre, Flora Donati, Etienne Simon-Lorière, Vincent Enouf, Maud Vanpeene, Sylvie van der Werf, Marianne Leruez-Ville |
| EPI_ISL_443300, EPI_ISL_443301, EPI_ISL_443302 | Cabinet Médical | National Reference Center for Viruses of Respiratory Infections, Institut Pasteur, Paris | Mélanie Albert, Marion Barbet, Sylvie Behillil, Méline Bizard, Angela Brisebarre, Flora Donati, Etienne Simon-Lorière, Vincent Enouf, Maud Vanpeene, Sylvie van der Werf |
| EPI_ISL_443303 | Résidence Les Marines | National Reference Center for Viruses of Respiratory Infections, Institut Pasteur, Paris | Mélanie Albert, Marion Barbet, Sylvie Behillil, Méline Bizard, Angela Brisebarre, Flora Donati, Etienne Simon-Lorière, Vincent Enouf, Maud Vanpeene, Sylvie van der Werf |
| EPI_ISL_443304 | Résidence Esterel | National Reference Center for Viruses of Respiratory Infections, Institut Pasteur, Paris | Mélanie Albert, Marion Barbet, Sylvie Behillil, Méline Bizard, Angela Brisebarre, Flora Donati, Etienne Simon-Lorière, Vincent Enouf, Maud Vanpeene, Sylvie van der Werf |
| EPI_ISL_443305 | LABM GH nord Essonne de Longjumeau - BP 125 | National Reference Center for Viruses of Respiratory Infections, Institut Pasteur, Paris | Mélanie Albert, Marion Barbet, Sylvie Behillil, Méline Bizard, Angela Brisebarre, Flora Donati, Etienne Simon-Lorière, Vincent Enouf, Maud Vanpeene, Sylvie van der Werf |
| EPI_ISL_443306 | Cabinet Médical | National Reference Center for Viruses of Respiratory Infections, Institut Pasteur, Paris | Mélanie Albert, Marion Barbet, Sylvie Behillil, Méline Bizard, Angela Brisebarre, Flora Donati, Etienne Simon-Lorière, Vincent Enouf, Maud Vanpeene, Sylvie van der Werf |
| EPI_ISL_443307 | La Villa Papyri | National Reference Center for Viruses of Respiratory Infections, Institut Pasteur, Paris | Mélanie Albert, Marion Barbet, Sylvie Behillil, Méline Bizard, Angela Brisebarre, Flora Donati, Etienne Simon-Lorière, Vincent Enouf, Maud Vanpeene, Sylvie van der Werf |
| EPI_ISL_443308 | Plaisance | National Reference Center for Viruses of Respiratory Infections, Institut Pasteur, Paris | Mélanie Albert, Marion Barbet, Sylvie Behillil, Méline Bizard, Angela Brisebarre, Flora Donati, Etienne Simon-Lorière, Vincent Enouf, Maud Vanpeene, Sylvie van der Werf |
| EPI_ISL_443309 | CH Compiègne Laboratoire de Biologie | National Reference Center for Viruses of Respiratory Infections, Institut Pasteur, Paris | Mélanie Albert, Marion Barbet, Sylvie Behillil, Méline Bizard, Angela Brisebarre, Flora Donati, Etienne Simon-Lorière, Vincent Enouf, Maud Vanpeene, Sylvie van der Werf |
| EPI_ISL_443310 | Centre de santé Filieris | National Reference Center for Viruses of Respiratory Infections, Institut Pasteur, Paris | Mélanie Albert, Marion Barbet, Sylvie Behillil, Méline Bizard, Angela Brisebarre, Flora Donati, Etienne Simon-Lorière, Vincent Enouf, Maud Vanpeene, Sylvie van der Werf |
| EPI_ISL_443311, EPI_ISL_443312, EPI_ISL_443313 | Cabinet Médical | National Reference Center for Viruses of Respiratory Infections, Institut Pasteur, Paris | Mélanie Albert, Marion Barbet, Sylvie Behillil, Méline Bizard, Angela Brisebarre, Flora Donati, Etienne Simon-Lorière, Vincent Enouf, Maud Vanpeene, Sylvie van der Werf |
| EPI_ISL_443314 | LABM GH nord Essonne de Longjumeau - BP 125 | National Reference Center for Viruses of Respiratory Infections, Institut Pasteur, Paris | Mélanie Albert, Marion Barbet, Sylvie Behillil, Méline Bizard, Angela Brisebarre, Flora Donati, Etienne Simon-Lorière, Vincent Enouf, Maud Vanpeene, Sylvie van der Werf |
| EPI_ISL_443315 | Château de la Source | National Reference Center for Viruses of Respiratory Infections, Institut Pasteur, Paris | Mélanie Albert, Marion Barbet, Sylvie Behillil, Méline Bizard, Angela Brisebarre, Flora Donati, Etienne Simon-Lorière, Vincent Enouf, Maud Vanpeene, Sylvie van der Werf |
| EPI_ISL_443316 | CH Compiègne Laboratoire de Biologie | National Reference Center for Viruses of Respiratory Infections, Institut Pasteur, Paris | Mélanie Albert, Marion Barbet, Sylvie Behillil, Méline Bizard, Angela Brisebarre, Flora Donati, Etienne Simon-Lorière, Vincent Enouf, Maud Vanpeene, Sylvie van der Werf, Olivia Raulin |
| EPI_ISL_443317 | Cabinet Médical | National Reference Center for Viruses of Respiratory Infections, Institut Pasteur, Paris | Mélanie Albert, Marion Barbet, Sylvie Behillil, Méline Bizard, Angela Brisebarre, Flora Donati, Etienne Simon-Lorière, Vincent Enouf, Maud Vanpeene, Sylvie van der Werf |
| EPI_ISL_443318, EPI_ISL_443319, EPI_ISL_443320, EPI_ISL_443321, EPI_ISL_443322, EPI_ISL_443323, EPI_ISL_443324, EPI_ISL_443325, EPI_ISL_443326, EPI_ISL_443327, EPI_ISL_443328, EPI_ISL_443329, EPI_ISL_443330, EPI_ISL_443331, EPI_ISL_443332, EPI_ISL_443333, EPI_ISL_443334, EPI_ISL_443335, EPI_ISL_443336, EPI_ISL_443337, EPI_ISL_443338, EPI_ISL_443339, EPI_ISL_443340, EPI_ISL_443341, EPI_ISL_443342, EPI_ISL_443343, EPI_ISL_443344, EPI_ISL_443345, EPI_ISL_443346, EPI_ISL_443347, EPI_ISL_443348, EPI_ISL_443349, EPI_ISL_443350, EPI_ISL_443351, EPI_ISL_443352, EPI_ISL_443353, EPI_ISL_443354, EPI_ISL_443355, EPI_ISL_443356, EPI_ISL_443357, EPI_ISL_443358, EPI_ISL_443359, EPI_ISL_443360, EPI_ISL_443361, EPI_ISL_443362, EPI_ISL_443363, EPI_ISL_443364, EPI_ISL_443365, EPI_ISL_443366, EPI_ISL_443367, EPI_ISL_443368, EPI_ISL_443369, EPI_ISL_443370, EPI_ISL_443371, EPI_ISL_443372, EPI_ISL_443373, EPI_ISL_443374, EPI_ISL_443375, EPI_ISL_443376, EPI_ISL_443377, EPI_ISL_443378, EPI_ISL_443379, EPI_ISL_443380, EPI_ISL_443381, EPI_ISL_443382, EPI_ISL_443383, EPI_ISL_443384, EPI_ISL_443385, EPI_ISL_443386, EPI_ISL_443387, EPI_ISL_443388, EPI_ISL_443389, EPI_ISL_443390, EPI_ISL_443391, EPI_ISL_443392, EPI_ISL_443393, EPI_ISL_443394, EPI_ISL_443395, EPI_ISL_443396, EPI_ISL_443397, EPI_ISL_443398, EPI_ISL_443399, EPI_ISL_443400, EPI_ISL_443401, EPI_ISL_443402, EPI_ISL_443403, EPI_ISL_443404, EPI_ISL_443405, EPI_ISL_443406, EPI_ISL_443407, EPI_ISL_443408, EPI_ISL_443409, EPI_ISL_443410, EPI_ISL_443411, EPI_ISL_443412, EPI_ISL_443413, EPI_ISL_443414, EPI_ISL_443415, EPI_ISL_443416, EPI_ISL_443417, EPI_ISL_443418, EPI_ISL_443419, EPI_ISL_443420, EPI_ISL_443421, EPI_ISL_443422, EPI_ISL_443423, EPI_ISL_443424, EPI_ISL_443425, EPI_ISL_443426, EPI_ISL_443427, EPI_ISL_443428, EPI_ISL_443429, EPI_ISL_443430, EPI_ISL_443431, EPI_ISL_443432, EPI_ISL_443433, EPI_ISL_443434, EPI_ISL_443435, EPI_ISL_443436, EPI_ISL_443437, EPI_ISL_443438, EPI_ISL_443439, EPI_ISL_443440, EPI_ISL_443441, EPI_ISL_443442, EPI_ISL_443443, EPI_ISL_443444, EPI_ISL_443445, EPI_ISL_443446, EPI_ISL_443447, EPI_ISL_443448, EPI_ISL_443449, EPI_ISL_443450, EPI_ISL_443451, EPI_ISL_443452, EPI_ISL_443453, EPI_ISL_443454, EPI_ISL_443455, EPI_ISL_443456, EPI_ISL_443457, EPI_ISL_443458, EPI_ISL_443459, EPI_ISL_443460, EPI_ISL_443461, EPI_ISL_443462, EPI_ISL_443463, EPI_ISL_443464, EPI_ISL_443465, EPI_ISL_443466, EPI_ISL_443467, EPI_ISL_443468, EPI_ISL_443469, EPI_ISL_443470, EPI_ISL_443471, EPI_ISL_443472, EPI_ISL_443473, EPI_ISL_443474, EPI_ISL_443475, EPI_ISL_443476, EPI_ISL_443477, EPI_ISL_443478, EPI_ISL_443479, EPI_ISL_443480, EPI_ISL_443481, EPI_ISL_443482, EPI_ISL_443483, EPI_ISL_443484, EPI_ISL_443485, EPI_ISL_443486, EPI_ISL_443487, EPI_ISL_443488, EPI_ISL_443489, EPI_ISL_443490, EPI_ISL_443491, EPI_ISL_443492, EPI_ISL_443493, EPI_ISL_443494, EPI_ISL_443495, EPI_ISL_443496, EPI_ISL_443497, EPI_ISL_443498, EPI_ISL_443499, EPI_ISL_443500, EPI_ISL_443501, EPI_ISL_443502, EPI_ISL_443503, EPI_ISL_443504, EPI_ISL_443505, EPI_ISL_443506, EPI_ISL_443507, EPI_ISL_443508, EPI_ISL_443509, EPI_ISL_443510, EPI_ISL_443511, EPI_ISL_443512, EPI_ISL_443513, EPI_ISL_443514, EPI_ISL_443515, EPI_ISL_443516, EPI_ISL_443517, EPI_ISL_443518, EPI_ISL_443519, EPI_ISL_443520, EPI_ISL_443521, EPI_ISL_443522, EPI_ISL_443523, EPI_ISL_443524, EPI_ISL_443525, EPI_ISL_443526, EPI_ISL_443527, EPI_ISL_443528, EPI_ISL_443529, EPI_ISL_443530, EPI_ISL_443531, EPI_ISL_443532, EPI_ISL_443533, EPI_ISL_443534, EPI_ISL_443535, EPI_ISL_443536, EPI_ISL_443537, EPI_ISL_443538, EPI_ISL_443539, EPI_ISL_443540, EPI_ISL_443541, EPI_ISL_443542, EPI_ISL_443543, EPI_ISL_443544, EPI_ISL_443545, EPI_ISL_443546, EPI_ISL_443547, EPI_ISL_443548, EPI_ISL_443549, EPI_ISL_443550, EPI_ISL_443551, EPI_ISL_443552, EPI_ISL_443553, EPI_ISL_443554, EPI_ISL_443555, EPI_ISL_443556, EPI_ISL_443557, EPI_ISL_443558, EPI_ISL_443559, EPI_ISL_443560, EPI_ISL_443561, EPI_ISL_443562, EPI_ISL_443563, EPI_ISL_443564, EPI_ISL_443565, EPI_ISL_443566, EPI_ISL_443567, EPI_ISL_443568, EPI_ISL_443569, EPI_ISL_443570, EPI_ISL_443571, EPI_ISL_443572, EPI_ISL_443573, EPI_ISL_443574, EPI_ISL_443575, EPI_ISL_443576, EPI_ISL_443577, EPI_ISL_443578, EPI_ISL_443579, EPI_ISL_443580, EPI_ISL_443581, EPI_ISL_443582, EPI_ISL_443583, EPI_ISL_443584, EPI_ISL_443585, EPI_ISL_443586, EPI_ISL_443587, EPI_ISL_443588, EPI_ISL_443589, EPI_ISL_443590, EPI_ISL_443591, EPI_ISL_443592, EPI_ISL_443593, EPI_ISL_443594, EPI_ISL_443595, EPI_ISL_443596, EPI_ISL_443597, EPI_ISL_443598, EPI_ISL_443599, EPI_ISL_443600, EPI_ISL_443601, EPI_ISL_443602, EPI_ISL_443603, EPI_ISL_443604, EPI_ISL_443605, EPI_ISL_443606, EPI_ISL_443607, EPI_ISL_443608, EPI_ISL_443609, EPI_ISL_443610, EPI_ISL_443611, EPI_ISL_443612, EPI_ISL_443613, EPI_ISL_443614, EPI_ISL_443615, EPI_ISL_443616, EPI_ISL_443617, EPI_ISL_443618, EPI_ISL_443619, EPI_ISL_443620, EPI_ISL_443621, EPI_ISL_443622, EPI_ISL_443623, EPI_ISL_443624, EPI_ISL_443625, EPI_ISL_443626, EPI_ISL_443627, EPI_ISL_443628, EPI_ISL_443629, EPI_ISL_443630, EPI_ISL_443631, EPI_ISL_443632, EPI_ISL_443633, EPI_ISL_443634, EPI_ISL_443635, EPI_ISL_443636, EPI_ISL_443637, EPI_ISL_443638, EPI_ISL_443639, EPI_ISL_443640, EPI_ISL_443641, EPI_ISL_443642, EPI_ISL_443643, EPI_ISL_443644, EPI_ISL_443645, EPI_ISL_443646, EPI_ISL_443647, EPI_ISL_443648, EPI_ISL_443649, EPI_ISL_443650, EPI_ISL_443651, EPI_ISL_443652, EPI_ISL_443653, EPI_ISL_443654, EPI_ISL_443655, EPI_ISL_443656, EPI_ISL_443657, EPI_ISL_443658, EPI_ISL_443659, EPI_ISL_443660, EPI_ISL_443661, EPI_ISL_443662, EPI_ISL_443663, EPI_ISL_443664, EPI_ISL_443665, EPI_ISL_443666, EPI_ISL_443667, EPI_ISL_443668, EPI_ISL_443669, EPI_ISL_443670, EPI_ISL_443671, EPI_ISL_443672, EPI_ISL_443673, EPI_ISL_443674, EPI_ISL_443675, EPI_ISL_443676, EPI_ISL_443677, EPI_ISL_443678, EPI_ISL_443679, EPI_ISL_443680, EPI_ISL_443681, EPI_ISL_443682, EPI_ISL_443683, EPI_ISL_443684, EPI_ISL_443685 | | | |
| see above | Department of Pathology, University of Cambridge | Wellcome Sanger Institute for the COVID-19 Genomics UK Consortium | Luke W Meredith, M. Estée Török , Myra Hosmillo, William L. Hamilton, Martin D. Curran, Theresa Feltwell, Grant Hall, Anna Yakovleva, Fahad A Khokhar, Charlotte J. Houldcroft, Laura G Caller, Aminu S. Jahun, Sarah L. Caddy, Ian Goodfellow; and Alex Alderton, Roberto Amato, Sonia Goncalves, Ewan Harrison, David K. Jackson, Ian Johnston, Dominic Kwiatkowski, Cordelia Langford, John Sillitoe on behalf of the Wellcome Sanger Institute COVID-19 Surveillance Team (http://www.sanger.ac.uk/covid-team) |
| EPI_ISL_443686, EPI_ISL_443687, EPI_ISL_443688, EPI_ISL_443689, EPI_ISL_443690, EPI_ISL_443691, EPI_ISL_443692, EPI_ISL_443693, EPI_ISL_443694, EPI_ISL_443695, EPI_ISL_443696, EPI_ISL_443697, EPI_ISL_443698, EPI_ISL_443699, EPI_ISL_443700, EPI_ISL_443701, EPI_ISL_443702, EPI_ISL_443703, EPI_ISL_443704, EPI_ISL_443705, EPI_ISL_443706, EPI_ISL_443707, EPI_ISL_443708, EPI_ISL_443709, EPI_ISL_443710, EPI_ISL_443711, EPI_ISL_443712, EPI_ISL_443713, EPI_ISL_443714, EPI_ISL_443715, EPI_ISL_443716, EPI_ISL_443717, EPI_ISL_443718, EPI_ISL_443719, EPI_ISL_443720, EPI_ISL_443721, EPI_ISL_443722, EPI_ISL_443723, EPI_ISL_443724, EPI_ISL_443725, EPI_ISL_443726, EPI_ISL_443727, EPI_ISL_443728, EPI_ISL_443729, EPI_ISL_443730, EPI_ISL_443731, EPI_ISL_443732, EPI_ISL_443733, EPI_ISL_443734, EPI_ISL_443735, EPI_ISL_443736, EPI_ISL_443737, EPI_ISL_443738, EPI_ISL_443739, EPI_ISL_443740, EPI_ISL_443741, EPI_ISL_443742, EPI_ISL_443743, EPI_ISL_443744, EPI_ISL_443745, EPI_ISL_443746, EPI_ISL_443747, EPI_ISL_443748, EPI_ISL_443749, EPI_ISL_443750, EPI_ISL_443751, EPI_ISL_443752, EPI_ISL_443753, EPI_ISL_443754, EPI_ISL_443755, EPI_ISL_443756, EPI_ISL_443757, EPI_ISL_443758, EPI_ISL_443759, EPI_ISL_443760, EPI_ISL_443761, EPI_ISL_443762, EPI_ISL_443763, EPI_ISL_443764, EPI_ISL_443765, EPI_ISL_443766, EPI_ISL_443767, EPI_ISL_443768, EPI_ISL_443769, EPI_ISL_443770, EPI_ISL_443771, EPI_ISL_443772, EPI_ISL_443773, EPI_ISL_443774, EPI_ISL_443775, EPI_ISL_443776, EPI_ISL_443777, EPI_ISL_443778, EPI_ISL_443779, EPI_ISL_443780, EPI_ISL_443781, EPI_ISL_443782, EPI_ISL_443783, EPI_ISL_443784, EPI_ISL_443785, EPI_ISL_443786, EPI_ISL_443787, EPI_ISL_443788, EPI_ISL_443789, EPI_ISL_443790, EPI_ISL_443791, EPI_ISL_443792, EPI_ISL_443793, EPI_ISL_443794, EPI_ISL_443795, EPI_ISL_443796, EPI_ISL_443797, EPI_ISL_443798, EPI_ISL_443799, EPI_ISL_443800, EPI_ISL_443801, EPI_ISL_443802, EPI_ISL_443803, EPI_ISL_443804, EPI_ISL_443805, EPI_ISL_443806, EPI_ISL_443807, EPI_ISL_443808, EPI_ISL_443809, EPI_ISL_443810, EPI_ISL_443811, EPI_ISL_443812, EPI_ISL_443813, EPI_ISL_443814, EPI_ISL_443815, EPI_ISL_443816, EPI_ISL_443817, EPI_ISL_443818, EPI_ISL_443819, EPI_ISL_443820, EPI_ISL_443821, EPI_ISL_443822, EPI_ISL_443823, EPI_ISL_443824, EPI_ISL_443825, EPI_ISL_443826, EPI_ISL_443827, EPI_ISL_443828, EPI_ISL_443829, EPI_ISL_443830, EPI_ISL_443831, EPI_ISL_443832, EPI_ISL_443833, EPI_ISL_443834, EPI_ISL_443835, EPI_ISL_443836, EPI_ISL_443837, EPI_ISL_443838, EPI_ISL_443839, EPI_ISL_443840, EPI_ISL_443841, EPI_ISL_443842, EPI_ISL_443843, EPI_ISL_443844, EPI_ISL_443845, EPI_ISL_443846, EPI_ISL_443847, EPI_ISL_443848, EPI_ISL_443849, EPI_ISL_443850, EPI_ISL_443851, EPI_ISL_443852, EPI_ISL_443853, EPI_ISL_443854, EPI_ISL_443855, EPI_ISL_443856, EPI_ISL_443857, EPI_ISL_443858, EPI_ISL_443859, EPI_ISL_443860, EPI_ISL_443861, EPI_ISL_443862, EPI_ISL_443863, EPI_ISL_443864, EPI_ISL_443865, EPI_ISL_443866, EPI_ISL_443867, EPI_ISL_443868, EPI_ISL_443869, EPI_ISL_443870, EPI_ISL_443871, EPI_ISL_443872, EPI_ISL_443873, EPI_ISL_443874, EPI_ISL_443875, EPI_ISL_443876, EPI_ISL_443877, EPI_ISL_443878, EPI_ISL_443879, EPI_ISL_443880, EPI_ISL_443881, EPI_ISL_443882, EPI_ISL_443883, EPI_ISL_443884, EPI_ISL_443885, EPI_ISL_443886, EPI_ISL_443887, EPI_ISL_443888, EPI_ISL_443889, EPI_ISL_443890, EPI_ISL_443891, EPI_ISL_443892, EPI_ISL_443893, EPI_ISL_443894, EPI_ISL_443895, EPI_ISL_443896, EPI_ISL_443897, EPI_ISL_443898, EPI_ISL_443899, EPI_ISL_443900, EPI_ISL_443901, EPI_ISL_443902, EPI_ISL_443903, EPI_ISL_443904, EPI_ISL_443905, EPI_ISL_443906, EPI_ISL_443907, EPI_ISL_443908, EPI_ISL_443909, EPI_ISL_443910, EPI_ISL_443911, EPI_ISL_443912, EPI_ISL_443913, EPI_ISL_443914, EPI_ISL_443915, EPI_ISL_443916, EPI_ISL_443917, EPI_ISL_443918, EPI_ISL_443919, EPI_ISL_443920, EPI_ISL_443921, EPI_ISL_443922, EPI_ISL_443923, EPI_ISL_443924, EPI_ISL_443925, | | | |

| | | | |
|---|---|---|---|
| EPI_ISL_443926, EPI_ISL_443927, EPI_ISL_443928, EPI_ISL_443929, EPI_ISL_443930, EPI_ISL_443931, EPI_ISL_443932, EPI_ISL_443933, EPI_ISL_443934, EPI_ISL_443935, EPI_ISL_443936, EPI_ISL_443937, EPI_ISL_443938, EPI_ISL_443939, EPI_ISL_443940, EPI_ISL_443941, EPI_ISL_443942, EPI_ISL_443943, EPI_ISL_443944, EPI_ISL_443945, EPI_ISL_443946, EPI_ISL_443947, EPI_ISL_443948, EPI_ISL_443949, EPI_ISL_443950, EPI_ISL_443951, EPI_ISL_443952, EPI_ISL_443953, EPI_ISL_443954, EPI_ISL_443955, EPI_ISL_443956, EPI_ISL_443957, EPI_ISL_443958, EPI_ISL_443959, EPI_ISL_443960, EPI_ISL_443961, EPI_ISL_443962, EPI_ISL_443963, EPI_ISL_443964, EPI_ISL_443965, EPI_ISL_443966, EPI_ISL_443967, EPI_ISL_443968, EPI_ISL_443969, EPI_ISL_443970, EPI_ISL_443971, EPI_ISL_443972, EPI_ISL_443973, EPI_ISL_443974, EPI_ISL_443975, EPI_ISL_443976, EPI_ISL_443977, EPI_ISL_443978, EPI_ISL_443979, EPI_ISL_443980, EPI_ISL_443981, EPI_ISL_443982, EPI_ISL_443983, EPI_ISL_443984, EPI_ISL_443985, EPI_ISL_443986, EPI_ISL_443987, EPI_ISL_443988, EPI_ISL_443989, EPI_ISL_443990, EPI_ISL_443991, EPI_ISL_443992, EPI_ISL_443993, EPI_ISL_443994, EPI_ISL_443995, EPI_ISL_443996, EPI_ISL_443997, EPI_ISL_443998, EPI_ISL_443999, EPI_ISL_444000, EPI_ISL_444001, EPI_ISL_444002, EPI_ISL_444003, EPI_ISL_444004, EPI_ISL_444005, EPI_ISL_444006, EPI_ISL_444007, EPI_ISL_444008, EPI_ISL_444009, EPI_ISL_444010, EPI_ISL_444011, EPI_ISL_444012, EPI_ISL_444013, EPI_ISL_444014, EPI_ISL_444015, EPI_ISL_444016, EPI_ISL_444017, EPI_ISL_444018, EPI_ISL_444019, EPI_ISL_444020, EPI_ISL_444021 | | | |
| see above | PHE South West Regional Laboratory, National Infection Service | Wellcome Sanger Institute for the COVID-19 Genomics UK Consortium | Stephanie Hutchings, Hannah Pymont, Dr Peter Muir, Barry Vipond, Rich Hopes; and Alex Alderton, Roberto Amato, Sonia Goncalves, Ewan Harrison, David K. Jackson, Ian Johnston, Dominic Kwiatkowski, Cordelia Langford, John Sillitoe on behalf of the Wellcome Sanger Institute COVID-19 Surveillance Team (http://www.sanger.ac.uk/covid-team) |
| EPI_ISL_444022 | Baylor College of Medicine | Baylor College of Medicine: HGSC | Vasanthi Avadhanula, Erin Nicholson, David Henke, Pedro Piedra, Harsha Doddapaneni, Donna Muzny, Qingchang Meng, Hsu Chao, Zeineen Momin, Hua Shen, George Weissenberger, Kavya Kottapalli, Yimiti Meiheerguli, Sejal Salvi, Ginger Metcalf, Vipin Menon, Sara J.J. Cregeen, Matthew C. Ross, Tulin Ayvaz, Richard Sucgang, Kristi L. Hoffman, Matthew Wong, Joseph F. Petrosino |
| EPI_ISL_444023, EPI_ISL_444024, EPI_ISL_444025, EPI_ISL_444026 | County of Santa Clara Public Health | Chan-Zuckerberg Biohub | CZB Cliahub Consortium |
| EPI_ISL_444027 | Pamela Youde Nethersole Eastern Hospital | Hong Kong Department of Health | Mak Gannon C.K., Cheng Peter K.C., Lam Edman T.K., Chan Rickjason C.W., Tsang Dominic N.C. |
| EPI_ISL_444028 | Queen Elizabeth Hospital | Hong Kong Department of Health | Mak Gannon C.K., Cheng Peter K.C., Lam Edman T.K., Chan Rickjason C.W., Tsang Dominic N.C. |
| EPI_ISL_444029 | Prince of Wales Hospital | Hong Kong Department of Health | Mak Gannon C.K., Cheng Peter K.C., Lam Edman T.K., Chan Rickjason C.W., Tsang Dominic N.C. |
| EPI_ISL_444030 | United Christian Hospital | Hong Kong Department of Health | Mak Gannon C.K., Cheng Peter K.C., Lam Edman T.K., Chan Rickjason C.W., Tsang Dominic N.C. |
| EPI_ISL_444031 | Queen Mary Hospital | Hong Kong Department of Health | Mak Gannon C.K., Cheng Peter K.C., Lam Edman T.K., Chan Rickjason C.W., Tsang Dominic N.C. |
| EPI_ISL_444032 | North Lantau Hospital | Hong Kong Department of Health | Mak Gannon C.K., Cheng Peter K.C., Lam Edman T.K., Chan Rickjason C.W., Tsang Dominic N.C. |
| EPI_ISL_444033 | Queen Mary Hospital | Hong Kong Department of Health | Mak Gannon C.K., Cheng Peter K.C., Lam Edman T.K., Chan Rickjason C.W., Tsang Dominic N.C. |
| EPI_ISL_444034 | Prince of Wales Hospital | Hong Kong Department of Health | Mak Gannon C.K., Cheng Peter K.C., Lam Edman T.K., Chan Rickjason C.W., Tsang Dominic N.C. |
| EPI_ISL_444035 | Princess Margaret Hospital | Hong Kong Department of Health | Mak Gannon C.K., Cheng Peter K.C., Lam Edman T.K., Chan Rickjason C.W., Tsang Dominic N.C. |
| EPI_ISL_444036 | North Lantau Hospital | Hong Kong Department of Health | Mak Gannon C.K., Cheng Peter K.C., Lam Edman T.K., Chan Rickjason C.W., Tsang Dominic N.C. |
| EPI_ISL_444037 | Hong Kong Adventist Hospital | Hong Kong Department of Health | Mak Gannon C.K., Cheng Peter K.C., Lam Edman T.K., Chan Rickjason C.W., Tsang Dominic N.C. |
| EPI_ISL_444038 | Princess Margaret Hospital | Hong Kong Department of Health | Mak Gannon C.K., Cheng Peter K.C., Lam Edman T.K., Chan Rickjason C.W., Tsang Dominic N.C. |
| EPI_ISL_444039 | United Christian Hospital | Hong Kong Department of Health | Mak Gannon C.K., Cheng Peter K.C., Lam Edman T.K., Chan Rickjason C.W., Tsang Dominic N.C. |
| EPI_ISL_444040 | Queen Elizabeth Hospital | Hong Kong Department of Health | Mak Gannon C.K., Cheng Peter K.C., Lam Edman T.K., Chan Rickjason C.W., Tsang Dominic N.C. |
| EPI_ISL_444041 | Tuen Mun Hospital | Hong Kong Department of Health | Mak Gannon C.K., Cheng Peter K.C., Lam Edman T.K., Chan Rickjason C.W., Tsang Dominic N.C. |
| EPI_ISL_444042, EPI_ISL_444043 | Pamela Youde Nethersole Eastern Hospital | Hong Kong Department of Health | Mak Gannon C.K., Cheng Peter K.C., Lam Edman T.K., Chan Rickjason C.W., Tsang Dominic N.C. |
| EPI_ISL_444044 | Yan Chai Hospital | Hong Kong Department of Health | Mak Gannon C.K., Cheng Peter K.C., Lam Edman T.K., Chan Rickjason C.W., Tsang Dominic N.C. |
| EPI_ISL_444045, EPI_ISL_444046 | Queen Elizabeth Hospital | Hong Kong Department of Health | Mak Gannon C.K., Cheng Peter K.C., Lam Edman T.K., Chan Rickjason C.W., Tsang Dominic N.C. |
| EPI_ISL_444047 | United Christian Hospital | Hong Kong Department of Health | Mak Gannon C.K., Cheng Peter K.C., Lam Edman T.K., Chan Rickjason C.W., Tsang Dominic N.C. |
| EPI_ISL_444048 | Queen Elizabeth Hospital | Hong Kong Department of Health | Mak Gannon C.K., Cheng Peter K.C., Lam Edman T.K., Chan Rickjason C.W., Tsang Dominic N.C. |
| EPI_ISL_444049 | North Lantau Hospital | Hong Kong Department of Health | Mak Gannon C.K., Cheng Peter K.C., Lam Edman T.K., Chan Rickjason C.W., Tsang Dominic N.C. |
| EPI_ISL_444050 | Shek Wu Hui Jockey Club General Out-patient Clinic | Hong Kong Department of Health | Mak Gannon C.K., Cheng Peter K.C., Lam Edman T.K., Chan Rickjason C.W., Tsang Dominic N.C. |
| EPI_ISL_444051, EPI_ISL_444052, EPI_ISL_444053, EPI_ISL_444054, EPI_ISL_444055, EPI_ISL_444056, EPI_ISL_444057, EPI_ISL_444058, EPI_ISL_444059, EPI_ISL_444060, EPI_ISL_444061, EPI_ISL_444062, EPI_ISL_444063, EPI_ISL_444064, EPI_ISL_444065, EPI_ISL_444066, EPI_ISL_444067, EPI_ISL_444068, EPI_ISL_444069, EPI_ISL_444070, EPI_ISL_444071, EPI_ISL_444072, EPI_ISL_444073, EPI_ISL_444074, EPI_ISL_444075, EPI_ISL_444076, EPI_ISL_444077, EPI_ISL_444078 | | | |
| see above | UCSF Clinical Microbiology Laboratory | Chan-Zuckerberg Biohub | CZB Cliahub Consortium |
| EPI_ISL_444079, EPI_ISL_444080, EPI_ISL_444081, EPI_ISL_444082, EPI_ISL_444083, EPI_ISL_444084, EPI_ISL_444085, EPI_ISL_444086, EPI_ISL_444087, EPI_ISL_444088, EPI_ISL_444089, EPI_ISL_444090, EPI_ISL_444091, EPI_ISL_444092, EPI_ISL_444093, EPI_ISL_444094, EPI_ISL_444095, EPI_ISL_444096, EPI_ISL_444097, EPI_ISL_444098, EPI_ISL_444099, EPI_ISL_444100, EPI_ISL_444101, EPI_ISL_444102, EPI_ISL_444103, EPI_ISL_444104, EPI_ISL_444105, EPI_ISL_444106, EPI_ISL_444107, EPI_ISL_444108, EPI_ISL_444109, EPI_ISL_444110, EPI_ISL_444111, EPI_ISL_444112, EPI_ISL_444113, EPI_ISL_444114, EPI_ISL_444115, EPI_ISL_444116, EPI_ISL_444117, EPI_ISL_444118, EPI_ISL_444119, EPI_ISL_444120, EPI_ISL_444121, EPI_ISL_444122, EPI_ISL_444123, EPI_ISL_444124, EPI_ISL_444125, EPI_ISL_444126, EPI_ISL_444127, EPI_ISL_444128, EPI_ISL_444129, EPI_ISL_444130, EPI_ISL_444131, EPI_ISL_444132, EPI_ISL_444133, EPI_ISL_444134, EPI_ISL_444135, EPI_ISL_444136, EPI_ISL_444137, EPI_ISL_444138, EPI_ISL_444139, EPI_ISL_444140, EPI_ISL_444141, EPI_ISL_444142, EPI_ISL_444143, EPI_ISL_444144, EPI_ISL_444145, EPI_ISL_444146, EPI_ISL_444147, EPI_ISL_444148, EPI_ISL_444149, EPI_ISL_444150, EPI_ISL_444151, EPI_ISL_444152, EPI_ISL_444153, EPI_ISL_444154, EPI_ISL_444155, EPI_ISL_444156, EPI_ISL_444157, EPI_ISL_444158, EPI_ISL_444159, EPI_ISL_444160, EPI_ISL_444161, EPI_ISL_444162, EPI_ISL_444163, EPI_ISL_444164, EPI_ISL_444165, EPI_ISL_444166, EPI_ISL_444167, EPI_ISL_444168, EPI_ISL_444169, EPI_ISL_444170, EPI_ISL_444171, EPI_ISL_444172, EPI_ISL_444173, EPI_ISL_444174, EPI_ISL_444175, EPI_ISL_444176, EPI_ISL_444177, EPI_ISL_444178, EPI_ISL_444179, EPI_ISL_444180, EPI_ISL_444181, EPI_ISL_444182, EPI_ISL_444183, EPI_ISL_444184, EPI_ISL_444185, EPI_ISL_444186, EPI_ISL_444187, EPI_ISL_444188, EPI_ISL_444189, EPI_ISL_444190, EPI_ISL_444191, EPI_ISL_444192, EPI_ISL_444193, EPI_ISL_444194, EPI_ISL_444195, EPI_ISL_444196, EPI_ISL_444197, EPI_ISL_444198, EPI_ISL_444199, EPI_ISL_444200, EPI_ISL_444201, EPI_ISL_444202, EPI_ISL_444203, EPI_ISL_444204, EPI_ISL_444205, EPI_ISL_444206, EPI_ISL_444207, EPI_ISL_444208, EPI_ISL_444209, EPI_ISL_444210, EPI_ISL_444211, EPI_ISL_444212, EPI_ISL_444213, EPI_ISL_444214, EPI_ISL_444215, EPI_ISL_444216, EPI_ISL_444217, EPI_ISL_444218, EPI_ISL_444219, EPI_ISL_444220, EPI_ISL_444221, EPI_ISL_444222, EPI_ISL_444223, EPI_ISL_444224, EPI_ISL_444225, EPI_ISL_444226, EPI_ISL_444227, EPI_ISL_444228, EPI_ISL_444229, EPI_ISL_444230, EPI_ISL_444231, EPI_ISL_444232, EPI_ISL_444233, EPI_ISL_444234, EPI_ISL_444235, EPI_ISL_444236, EPI_ISL_444237, EPI_ISL_444238, EPI_ISL_444239, EPI_ISL_444240, EPI_ISL_444241, EPI_ISL_444242, EPI_ISL_444243, EPI_ISL_444244, EPI_ISL_444245, EPI_ISL_444246, EPI_ISL_444247, EPI_ISL_444248, EPI_ISL_444249, EPI_ISL_444250, EPI_ISL_444251, EPI_ISL_444252, EPI_ISL_444253, EPI_ISL_444254, EPI_ISL_444255, EPI_ISL_444256, EPI_ISL_444257, EPI_ISL_444258, EPI_ISL_444259, EPI_ISL_444260, EPI_ISL_444261, EPI_ISL_444262, EPI_ISL_444263, EPI_ISL_444264, EPI_ISL_444265, EPI_ISL_444266, EPI_ISL_444267, EPI_ISL_444268, EPI_ISL_444269, EPI_ISL_444270, EPI_ISL_444271, EPI_ISL_444272 | | | |
| see above | University College London, Great Ormond Street Hospital for Children NHS Foundation Trust, Imperial College Healthcare NHS Trust | COVID-19 Genomics UK (COG-UK) Consortium | Sergi Castellano, Rachel Williams, Mark Kristiansen, Paola Resende Silva, Sunando Roy, Tony Brooks, Helena Tutill, Paola Niola, Patricia Dyal, Charlotte Williams, Leysa Forrest, Yasmin Panchbhaya, Jacqueline Findlay, Sam Weeks, Julianne Brown, Kathryn Harris, Paul Randell, James Price, Alison Holmes, Judith Breuer |
| EPI_ISL_444274, EPI_ISL_444275, EPI_ISL_444276, EPI_ISL_444277, EPI_ISL_444278 | Laboratory Medicine | Department of Laboratory Medicine, Lin-Kou Chang Gung Memorial Hospital, Taoyuan, Taiwan | Kuo-Chien Tsao, Yu-Nong Gong, Shu-Li Yang, Yi-Chun Liu, Chung-Guei Huang, Mei-Jen Hsiao, Po-Wei Huang, Cheng-Ta Yang, Cheng-Hsun Chiu, Peng-Nien Huang, Kuo-Ming Lee, Guang-Wu Chen, Shin-Ru Shih |
| EPI_ISL_444279, EPI_ISL_444280, EPI_ISL_444281, EPI_ISL_444282, EPI_ISL_444283, EPI_ISL_444284, EPI_ISL_444285, EPI_ISL_444286, EPI_ISL_444287, EPI_ISL_444288, EPI_ISL_444289, EPI_ISL_444290, EPI_ISL_444291, EPI_ISL_444292, EPI_ISL_444293, EPI_ISL_444294, EPI_ISL_444295, EPI_ISL_444296, EPI_ISL_444297, EPI_ISL_444298, EPI_ISL_444299, EPI_ISL_444300, EPI_ISL_444301, EPI_ISL_444302, EPI_ISL_444303, EPI_ISL_444304, EPI_ISL_444305, EPI_ISL_444306, EPI_ISL_444307, EPI_ISL_444308, EPI_ISL_444309, EPI_ISL_444310, EPI_ISL_444311, EPI_ISL_444312 | | | |
| see above | University of Birmingham | COVID-19 Genomics UK (COG-UK) Consortium | Loman Lab: Claire McMurray, Joanne Stockton, Samuel Nicholls, Radoslaw Poplawski, Will Rowe, Josh Quick, Nicholas Loman // UHB Lab: Celina M Whalley, Andrew Bosworth, Charlotte Poxon, Kasun Wanigasooriya, Oliver Pickles, Mike Kidd, Alex Richter, Andrew D Beggs // PHE Heartlands Lab: Husam Osman, Andrew Bosworth |
| EPI_ISL_444313, EPI_ISL_444314, EPI_ISL_444315, EPI_ISL_444316, EPI_ISL_444317, EPI_ISL_444318, EPI_ISL_444319, EPI_ISL_444320, EPI_ISL_444321, EPI_ISL_444322, EPI_ISL_444323, EPI_ISL_444324, EPI_ISL_444325, EPI_ISL_444326, EPI_ISL_444327, EPI_ISL_444328, EPI_ISL_444329, EPI_ISL_444330, EPI_ISL_444331, EPI_ISL_444332, EPI_ISL_444333, EPI_ISL_444334, EPI_ISL_444335, EPI_ISL_444336, EPI_ISL_444337, EPI_ISL_444338, EPI_ISL_444339, EPI_ISL_444340, EPI_ISL_444341, EPI_ISL_444342, EPI_ISL_444343, EPI_ISL_444344, EPI_ISL_444345, EPI_ISL_444346, EPI_ISL_444347, EPI_ISL_444348, EPI_ISL_444349, EPI_ISL_444350, EPI_ISL_444351, EPI_ISL_444352, EPI_ISL_444353, EPI_ISL_444354, EPI_ISL_444355, EPI_ISL_444356, EPI_ISL_444357, EPI_ISL_444358, EPI_ISL_444359, EPI_ISL_444360, EPI_ISL_444361, EPI_ISL_444362, EPI_ISL_444363, EPI_ISL_444364, EPI_ISL_444365, EPI_ISL_444366, EPI_ISL_444367, EPI_ISL_444368, EPI_ISL_444369, EPI_ISL_444370, EPI_ISL_444371, EPI_ISL_444372, EPI_ISL_444373, EPI_ISL_444374, EPI_ISL_444375, EPI_ISL_444376, EPI_ISL_444377, EPI_ISL_444378, EPI_ISL_444379, EPI_ISL_444380, EPI_ISL_444381, EPI_ISL_444382, EPI_ISL_444383, EPI_ISL_444384, EPI_ISL_444385, EPI_ISL_444386, EPI_ISL_444387, EPI_ISL_444388, EPI_ISL_444389, EPI_ISL_444390, EPI_ISL_444391, EPI_ISL_444392, EPI_ISL_444393, EPI_ISL_444394, EPI_ISL_444395, EPI_ISL_444396, EPI_ISL_444397, EPI_ISL_444398, EPI_ISL_444399, EPI_ISL_444400, EPI_ISL_444401, EPI_ISL_444402, EPI_ISL_444403, EPI_ISL_444404, EPI_ISL_444405, EPI_ISL_444406, EPI_ISL_444407, EPI_ISL_444408, EPI_ISL_444409, EPI_ISL_444410, EPI_ISL_444411, EPI_ISL_444412, EPI_ISL_444413, EPI_ISL_444414, EPI_ISL_444415, EPI_ISL_444416, EPI_ISL_444417, EPI_ISL_444418, EPI_ISL_444419, EPI_ISL_444420, EPI_ISL_444421, EPI_ISL_444422, EPI_ISL_444423, EPI_ISL_444424, EPI_ISL_444425, EPI_ISL_444426, EPI_ISL_444427, EPI_ISL_444428, EPI_ISL_444429, EPI_ISL_444430, EPI_ISL_444431, EPI_ISL_444432, EPI_ISL_444433, EPI_ISL_444434, EPI_ISL_444435, EPI_ISL_444436, EPI_ISL_444437, EPI_ISL_444438, EPI_ISL_444439, EPI_ISL_444440, EPI_ISL_444441, EPI_ISL_444442, EPI_ISL_444443, EPI_ISL_444444, EPI_ISL_444445, EPI_ISL_444446, EPI_ISL_444447, EPI_ISL_444448, EPI_ISL_444449, EPI_ISL_444450, EPI_ISL_444451, EPI_ISL_444452, EPI_ISL_444453 | | | |
| see above | Department of Pathology, University of Cambridge | COVID-19 Genomics UK (COG-UK) Consortium | Luke W Meredith, M. Estée Török , Myra Hosmillo, William L. Hamilton, Martin D. Curran, Theresa Feltwell, Grant Hall, Anna Yakovleva, Fahad A Khokhar, Charlotte J. Houldcroft, Laura G Caller, Aminu S. Jahun, Sarah L. Caddy, Ian Goodfellow |
| EPI_ISL_444456 | B.J. Medical College and Civil hospital | Gujarat Biotechnology Research Centre | R D Dixit, Snehal Bagatharia, Kamlesh J Upadhyay, Ramesh Pandit, Tejas Shah, Ankit Hinsu, Pritesh Sabara, Apurvasinh Puvar, Janvi Raval, Monika Gandhi, Pinal Trivedi, Maharshi Pandya, Amit Kanani, Akanksha Verma, Nitin Savaliya, Raghawendra Kumar, Dinesh Kumar, Zuber Saiyed, Dipa Kinariwala, Disha Patel, Binita Aring, Neeta Khandelwal, Geeta Vaghela, Sonia Barve, Bhavesh Modi, Kairavi Joshi, Gaurishankar Shrimali, Nidhi Sood, Pranay Shah, Pooja P Doshi, Chaitanya Joshi, Madhvi Joshi |
| EPI_ISL_444457 | B.J. Medical College and Civil hospital | Gujarat Biotechnology Research Centre | Snehal Bagatharia, Kamlesh J Upadhyay, Ramesh Pandit, Tejas Shah, Ankit Hinsu, Pritesh Sabara, Apurvasinh Puvar, Janvi Raval, Monika Gandhi, Pinal Trivedi, Maharshi Pandya, Amit Kanani, Akanksha Verma, Nitin Savaliya, Raghawendra Kumar, Dinesh Kumar, Zuber Saiyed, Dipa Kinariwala, Disha Patel, Binita Aring, Neeta Khandelwal, Geeta Vaghela, Sonia Barve, Bhavesh Modi, Kairavi Joshi, Gaurishankar Shrimali, Nidhi Sood, Pranay Shah, R D Dixit, Nidhi Patel, Chaitanya Joshi, Madhvi Joshi |
| EPI_ISL_444458 | B.J. Medical College and Civil hospital | Gujarat Biotechnology Research Centre | Kamlesh J Upadhyay, Ramesh Pandit, Tejas Shah, Ankit Hinsu, Pritesh Sabara, Apurvasinh Puvar, Janvi Raval, Monika Gandhi, Pinal Trivedi, Maharshi Pandya, Amit Kanani, Akanksha Verma, Nitin Savaliya, Raghawendra Kumar, Dinesh Kumar, Zuber Saiyed, Dipa Kinariwala, Disha Patel, Binita Aring, Neeta Khandelwal, Geeta Vaghela, Sonia Barve, Bhavesh Modi, Kairavi Joshi, Gaurishankar Shrimali, Nidhi Sood, Pranay Shah, R D Dixit, Snehal Bagatharia, Priti Pandita, Chaitanya Joshi, Madhvi Joshi |
| EPI_ISL_444459 | B.J. Medical College and Civil hospital | Gujarat Biotechnology Research Centre | Ramesh Pandit, Tejas Shah, Ankit Hinsu, Pritesh Sabara, Apurvasinh Puvar, Janvi Raval, Monika Gandhi, Pinal Trivedi, Maharshi Pandya, Amit Kanani, Akanksha Verma, Nitin Savaliya, Raghawendra Kumar, Dinesh Kumar, Zuber Saiyed, Dipa Kinariwala, Disha Patel, Binita Aring, Neeta Khandelwal, Geeta Vaghela, Sonia Barve, Bhavesh Modi, Kairavi Joshi, Gaurishankar Shrimali, Nidhi Sood, Pranay Shah, R D Dixit, Snehal Bagatharia, Kamlesh J Upadhyay, Neha Rajpara, Chaitanya Joshi, Madhvi Joshi |
| EPI_ISL_444460 | B.J. Medical College and Civil hospital | Gujarat Biotechnology Research Centre | Tejas Shah, Ankit Hinsu, Pritesh Sabara, Apurvasinh Puvar, Janvi Raval, Monika Gandhi, Pinal Trivedi, Maharshi Pandya, Amit Kanani, Akanksha Verma, Nitin Savaliya, Raghawendra Kumar, Dinesh Kumar, Zuber Saiyed, Dipa Kinariwala, Disha Patel, Binita Aring, Neeta Khandelwal, Geeta Vaghela, Sonia Barve, Bhavesh Modi, Kairavi Joshi, Gaurishankar Shrimali, Nidhi Sood, Pranay Shah, R D Dixit, Snehal Bagatharia, Kamlesh J Upadhyay, Ramesh Pandit, Afzal Ansari, Chaitanya Joshi, Madhvi Joshi |
| EPI_ISL_444461 | B.J. Medical College and Civil hospital | Gujarat Biotechnology Research Centre | Ankit Hinsu, Pritesh Sabara, Apurvasinh Puvar, Janvi Raval, Monika Gandhi, Pinal Trivedi, Maharshi Pandya, Amit Kanani, Akanksha Verma, Nitin Savaliya, Raghawendra Kumar, Dinesh Kumar, Zuber Saiyed, Dipa Kinariwala, Disha Patel, Binita Aring, Neeta Khandelwal, Geeta Vaghela, Sonia Barve, Bhavesh Modi, Kairavi Joshi, Gaurishankar Shrimali, Nidhi Sood, Pranay Shah, R D Dixit, Snehal Bagatharia, Kamlesh J Upadhyay, Ramesh Pandit, Neelam Nathani, Chaitanya Joshi, Madhvi Joshi, Tejas Shah |
| EPI_ISL_444462 | B.J. Medical College and Civil hospital | Gujarat Biotechnology Research Centre | Pritesh Sabara, Apurvasinh Puvar, Janvi Raval, Monika Gandhi, Pinal Trivedi, Maharshi Pandya, Amit Kanani, Akanksha Verma, Nitin Savaliya, Raghawendra Kumar, Dinesh Kumar, Zuber Saiyed, Dipa Kinariwala, Disha Patel, Binita Aring, Neeta Khandelwal, Geeta Vaghela, Sonia Barve, Bhavesh Modi, Kairavi Joshi, Gaurishankar Shrimali, Nidhi Sood, Pranay Shah, R D Dixit, Snehal Bagatharia, Kamlesh J Upadhyay, Ramesh Pandit, Tejas Shah, Ankit Hinsu, Armi Chaudhari, Chaitanya Joshi, Madhvi Joshi |
| EPI_ISL_444463 | B.J. Medical College and Civil hospital | Gujarat Biotechnology Research Centre | Apurvasinh Puvar, Janvi Raval, Monika Gandhi, Pinal Trivedi, Maharshi Pandya, Amit Kanani, Akanksha Verma, Nitin Savaliya, Raghawendra Kumar, Dinesh Kumar, Zuber Saiyed, Dipa Kinariwala, Disha Patel, Binita Aring, Neeta Khandelwal, Geeta Vaghela, Sonia Barve, Bhavesh Modi, Kairavi Joshi, Gaurishankar Shrimali, Nidhi Sood, Pranay Shah, R D Dixit, Snehal Bagatharia, Kamlesh J Upadhyay, Ramesh Pandit, Tejas Shah, Ankit Hinsu, Pritesh Sabara, Bhavya Jindal, Chaitanya Joshi, Madhvi Joshi |
| EPI_ISL_444464 | B.J. Medical College and Civil hospital | Gujarat Biotechnology Research Centre | Janvi Raval, Monika Gandhi, Pinal Trivedi, Maharshi Pandya, Amit Kanani, Akanksha Verma, Nitin Savaliya, Raghawendra Kumar, Dinesh Kumar, Zuber Saiyed, Dipa Kinariwala, Disha Patel, Binita Aring, Neeta Khandelwal, Geeta Vaghela, Sonia Barve, Bhavesh Modi, Kairavi Joshi, Gaurishankar Shrimali, Nidhi Sood, Pranay Shah, R D Dixit, Snehal Bagatharia, Kamlesh J Upadhyay, Ramesh Pandit, Tejas Shah, Ankit Hinsu, Pritesh Sabara, Apurvasinh Puvar, Dipeshwari Shewale, Chaitanya Joshi, Madhvi Joshi |
| EPI_ISL_444465 | B.J. Medical College and Civil hospital | Gujarat Biotechnology Research Centre | Monika Gandhi, Pinal Trivedi, Maharshi Pandya, Amit Kanani, Akanksha Verma, Nitin Savaliya, Raghawendra Kumar, Dinesh Kumar, Zuber Saiyed, Dipa Kinariwala, Disha Patel, Binita Aring, Neeta Khandelwal, Geeta Vaghela, Sonia Barve, Bhavesh Modi, Kairavi Joshi, Gaurishankar Shrimali, Nidhi Sood, Pranay Shah, R D Dixit, Snehal Bagatharia, Kamlesh J Upadhyay, Ramesh Pandit, Tejas Shah, Ankit Hinsu, Pritesh |

| | | | |
|---|---|---|---|
| | | | Sabara, Apurvasinh Puvar, Janvi Raval, Anjali Rajwar, Chaitanya Joshi, Madhvi Joshi |
| EPI_ISL_444466 | B.J. Medical College and Civil hospital | Gujarat Biotechnology Research Centre | Pinal Trivedi, Maharshi Pandya, Amit Kanani, Akanksha Verma, Nitin Savaliya, Raghawendra Kumar, Dinesh Kumar, Zuber Saiyed, Dipa Kinariwala, Disha Patel, Binita Aring, Neeta Khandelwal, Geeta Vaghela, Sonia Barve, Bhavesh Modi, Kairavi Joshi, Gaurishankar Shrimali, Nidhi Sood, Pranay Shah, R D Dixit, Snehal Bagatharia, Kamlesh J Upadhyay, Ramesh Pandit, Tejas Shah, Ankit Hinsu, Pritesh Sabara, Apurvasinh Puvar, Janvi Raval, Monika Gandhi, Sharmistha Majumdar, Chaitanya Joshi, Madhvi Joshi |
| EPI_ISL_444467 | B.J. Medical College and Civil hospital | Gujarat Biotechnology Research Centre | Maharshi Pandya, Amit Kanani, Akanksha Verma, Nitin Savaliya, Raghawendra Kumar, Dinesh Kumar, Zuber Saiyed, Dipa Kinariwala, Disha Patel, Binita Aring, Neeta Khandelwal, Geeta Vaghela, Sonia Barve, Bhavesh Modi, Kairavi Joshi, Gaurishankar Shrimali, Nidhi Sood, Pranay Shah, R D Dixit, Snehal Bagatharia, Kamlesh J Upadhyay, Ramesh Pandit, Tejas Shah, Ankit Hinsu, Pritesh Sabara, Apurvasinh Puvar, Janvi Raval, Monika Gandhi, Pinal Trivedi, Pooja P Doshi, Chaitanya Joshi, Madhvi Joshi |
| EPI_ISL_444468 | B.J. Medical College and Civil hospital | Gujarat Biotechnology Research Centre | Amit Kanani, Akanksha Verma, Nitin Savaliya, Raghawendra Kumar, Dinesh Kumar, Zuber Saiyed, Dipa Kinariwala, Disha Patel, Binita Aring, Neeta Khandelwal, Geeta Vaghela, Sonia Barve, Bhavesh Modi, Kairavi Joshi, Gaurishankar Shrimali, Nidhi Sood, Pranay Shah, R D Dixit, Snehal Bagatharia, Kamlesh J Upadhyay, Ramesh Pandit, Tejas Shah, Ankit Hinsu, Pritesh Sabara, Apurvasinh Puvar, Janvi Raval, Monika Gandhi, Pinal Trivedi, Maharshi Pandya, Nidhi Patel, Chaitanya Joshi, Madhvi Joshi |
| EPI_ISL_444469 | B.J. Medical College and Civil hospital | Gujarat Biotechnology Research Centre | Akanksha Verma, Nitin Savaliya, Raghawendra Kumar, Dinesh Kumar, Zuber Saiyed, Dipa Kinariwala, Disha Patel, Binita Aring, Neeta Khandelwal, Geeta Vaghela, Sonia Barve, Bhavesh Modi, Kairavi Joshi, Gaurishankar Shrimali, Nidhi Sood, Pranay Shah, R D Dixit, Snehal Bagatharia, Kamlesh J Upadhyay, Ramesh Pandit, Tejas Shah, Ankit Hinsu, Pritesh Sabara, Apurvasinh Puvar, Janvi Raval, Monika Gandhi, Pinal Trivedi, Maharshi Pandya, Amit Kanani, Priti Pandita, Chaitanya Joshi, Madhvi Joshi |
| EPI_ISL_444470 | B.J. Medical College and Civil hospital | Gujarat Biotechnology Research Centre | Nitin Savaliya, Raghawendra Kumar, Dinesh Kumar, Zuber Saiyed, Dipa Kinariwala, Disha Patel, Binita Aring, Neeta Khandelwal, Geeta Vaghela, Sonia Barve, Bhavesh Modi, Kairavi Joshi, Gaurishankar Shrimali, Nidhi Sood, Pranay Shah, R D Dixit, Snehal Bagatharia, Kamlesh J Upadhyay, Ramesh Pandit, Tejas Shah, Ankit Hinsu, Pritesh Sabara, Apurvasinh Puvar, Janvi Raval, Monika Gandhi, Pinal Trivedi, Maharshi Pandya, Amit Kanani, Akanksha Verma, Neha Rajpara, Chaitanya Joshi, Madhvi Joshi |
| EPI_ISL_444471 | B.J. Medical College and Civil hospital | Gujarat Biotechnology Research Centre | Raghawendra Kumar, Dinesh Kumar, Zuber Saiyed, Dipa Kinariwala, Disha Patel, Binita Aring, Neeta Khandelwal, Geeta Vaghela, Sonia Barve, Bhavesh Modi, Kairavi Joshi, Gaurishankar Shrimali, Nidhi Sood, Pranay Shah, R D Dixit, Snehal Bagatharia, Kamlesh J Upadhyay, Ramesh Pandit, Tejas Shah, Ankit Hinsu, Pritesh Sabara, Apurvasinh Puvar, Janvi Raval, Monika Gandhi, Pinal Trivedi, Maharshi Pandya, Amit Kanani, Akanksha Verma, Nitin Savaliya, Afzal Ansari, Chaitanya Joshi, Madhvi Joshi |
| EPI_ISL_444472 | B.J. Medical College and Civil hospital | Gujarat Biotechnology Research Centre | Dinesh Kumar, Zuber Saiyed, Dipa Kinariwala, Disha Patel, Binita Aring, Neeta Khandelwal, Geeta Vaghela, Sonia Barve, Bhavesh Modi, Kairavi Joshi, Gaurishankar Shrimali, Nidhi Sood, Pranay Shah, R D Dixit, Snehal Bagatharia, Kamlesh J Upadhyay, Ramesh Pandit, Tejas Shah, Ankit Hinsu, Pritesh Sabara, Apurvasinh Puvar, Janvi Raval, Monika Gandhi, Pinal Trivedi, Maharshi Pandya, Amit Kanani, Akanksha Verma, Nitin Savaliya, Raghawendra Kumar, Neelam Nathani, Chaitanya Joshi, Madhvi Joshi |
| EPI_ISL_444473 | B.J. Medical College and Civil hospital | Gujarat Biotechnology Research Centre | Zuber Saiyed, Dipa Kinariwala, Disha Patel, Binita Aring, Neeta Khandelwal, Geeta Vaghela, Sonia Barve, Bhavesh Modi, Kairavi Joshi, Gaurishankar Shrimali, Nidhi Sood, Pranay Shah, R D Dixit, Snehal Bagatharia, Kamlesh J Upadhyay, Ramesh Pandit, Tejas Shah, Ankit Hinsu, Pritesh Sabara, Apurvasinh Puvar, Janvi Raval, Monika Gandhi, Pinal Trivedi, Maharshi Pandya, Amit Kanani, Akanksha Verma, Nitin Savaliya, Raghawendra Kumar, Dinesh Kumar, Armi Chaudhari, Chaitanya Joshi, Madhvi Joshi |
| EPI_ISL_444474 | B.J. Medical College and Civil hospital | Gujarat Biotechnology Research Centre | Dipa Kinariwala, Disha Patel, Binita Aring, Neeta Khandelwal, Geeta Vaghela, Sonia Barve, Bhavesh Modi, Kairavi Joshi, Gaurishankar Shrimali, Nidhi Sood, Pranay Shah, R D Dixit, Snehal Bagatharia, Kamlesh J Upadhyay, Ramesh Pandit, Tejas Shah, Ankit Hinsu, Pritesh Sabara, Apurvasinh Puvar, Janvi Raval, Monika Gandhi, Pinal Trivedi, Maharshi Pandya, Amit Kanani, Akanksha Verma, Nitin Savaliya, Raghawendra Kumar, Dinesh Kumar, Zuber Saiyed, Bhavya Jindal, Chaitanya Joshi, Madhvi Joshi |
| EPI_ISL_444475 | B.J. Medical College and Civil hospital | Gujarat Biotechnology Research Centre | Disha Patel, Binita Aring, Neeta Khandelwal, Geeta Vaghela, Sonia Barve, Bhavesh Modi, Kairavi Joshi, Gaurishankar Shrimali, Nidhi Sood, Pranay Shah, R D Dixit, Snehal Bagatharia, Kamlesh J Upadhyay, Ramesh Pandit, Tejas Shah, Ankit Hinsu, Pritesh Sabara, Apurvasinh Puvar, Janvi Raval, Monika Gandhi, Pinal Trivedi, Maharshi Pandya, Amit Kanani, Akanksha Verma, Nitin Savaliya, Raghawendra Kumar, Dinesh Kumar, Zuber Saiyed, Dipa Kinariwala, Dipeshwari Shewale, Chaitanya Joshi, Madhvi Joshi |
| EPI_ISL_444476 | B.J. Medical College and Civil hospital | Gujarat Biotechnology Research Centre | Binita Aring, Neeta Khandelwal, Geeta Vaghela, Sonia Barve, Bhavesh Modi, Kairavi Joshi, Gaurishankar Shrimali, Nidhi Sood, Pranay Shah, R D Dixit, Snehal Bagatharia, Kamlesh J Upadhyay, Ramesh Pandit, Tejas Shah, Ankit Hinsu, Pritesh Sabara, Apurvasinh Puvar, Janvi Raval, Monika Gandhi, Pinal Trivedi, Maharshi Pandya, Amit Kanani, Akanksha Verma, Nitin Savaliya, Raghawendra Kumar, Dinesh Kumar, Zuber Saiyed, Dipa Kinariwala, Disha Patel, Chaitanya Joshi, Madhvi Joshi, Dipeshwari Shewale |
| EPI_ISL_444477 | B.J. Medical College and Civil hospital | Gujarat Biotechnology Research Centre | Neeta Khandelwal, Geeta Vaghela, Sonia Barve, Bhavesh Modi, Kairavi Joshi, Gaurishankar Shrimali, Nidhi Sood, Pranay Shah, R D Dixit, Snehal Bagatharia, Kamlesh J Upadhyay, Ramesh Pandit, Tejas Shah, Ankit Hinsu, Pritesh Sabara, Apurvasinh Puvar, Janvi Raval, Monika Gandhi, Pinal Trivedi, Maharshi Pandya, Amit Kanani, Akanksha Verma, Nitin Savaliya, Raghawendra Kumar, Dinesh Kumar, Zuber Saiyed, Dipa Kinariwala, Disha Patel, Binita Aring, Sharmistha Majumdar, Chaitanya Joshi, Madhvi Joshi |
| EPI_ISL_444478 | B.J. Medical College and Civil hospital | Gujarat Biotechnology Research Centre | Geeta Vaghela, Sonia Barve, Bhavesh Modi, Kairavi Joshi, Gaurishankar Shrimali, Nidhi Sood, Pranay Shah, R D Dixit, Snehal Bagatharia, Kamlesh J Upadhyay, Ramesh Pandit, Tejas Shah, Ankit Hinsu, Pritesh Sabara, Apurvasinh Puvar, Janvi Raval, Monika Gandhi, Pinal Trivedi, Maharshi Pandya, Amit Kanani, Akanksha Verma, Nitin Savaliya, Raghawendra Kumar, Dinesh Kumar, Zuber Saiyed, Dipa Kinariwala, Disha Patel, Binita Aring, Neeta Khandelwal, Pooja P Doshi, Chaitanya Joshi, Madhvi Joshi |
| EPI_ISL_444479 | B.J. Medical College and Civil hospital | Gujarat Biotechnology Research Centre | Sonia Barve, Bhavesh Modi, Kairavi Joshi, Gaurishankar Shrimali, Nidhi Sood, Pranay Shah, R D Dixit, Snehal Bagatharia, Kamlesh J Upadhyay, Ramesh Pandit, Tejas Shah, Ankit Hinsu, Pritesh Sabara, Apurvasinh Puvar, Janvi Raval, Monika Gandhi, Pinal Trivedi, Maharshi Pandya, Amit Kanani, Akanksha Verma, Nitin Savaliya, Raghawendra Kumar, Dinesh Kumar, Zuber Saiyed, Dipa Kinariwala, Disha Patel, Binita Aring, Neeta Khandelwal, Geeta Vaghela, Nidhi Patel, Chaitanya Joshi, Madhvi Joshi |
| EPI_ISL_444480 | B.J. Medical College and Civil hospital | Gujarat Biotechnology Research Centre | Bhavesh Modi, Kairavi Joshi, Gaurishankar Shrimali, Nidhi Sood, Pranay Shah, R D Dixit, Snehal Bagatharia, Kamlesh J Upadhyay, Ramesh Pandit, Tejas Shah, Ankit Hinsu, Pritesh Sabara, Apurvasinh Puvar, Janvi Raval, Monika Gandhi, Pinal Trivedi, Maharshi Pandya, Amit Kanani, Akanksha Verma, Nitin Savaliya, Raghawendra Kumar, Dinesh Kumar, Zuber Saiyed, Dipa Kinariwala, Disha Patel, Binita Aring, Neeta Khandelwal, Geeta Vaghela, Sonia Barve, Priti Pandita, Chaitanya Joshi, Madhvi Joshi |
| EPI_ISL_444481 | B.J. Medical College and Civil hospital | Gujarat Biotechnology Research Centre | Kairavi Joshi, Gaurishankar Shrimali, Nidhi Sood, Pranay Shah, R D Dixit, Snehal Bagatharia, Kamlesh J Upadhyay, Ramesh Pandit, Tejas Shah, Ankit Hinsu, Pritesh Sabara, Apurvasinh Puvar, Janvi Raval, Monika Gandhi, Pinal Trivedi, Maharshi Pandya, Amit Kanani, Akanksha Verma, Nitin Savaliya, Raghawendra Kumar, Dinesh Kumar, Zuber Saiyed, Dipa Kinariwala, Disha Patel, Binita Aring, Neeta Khandelwal, Geeta Vaghela, Sonia Barve, Bhavesh Modi, Neha Rajpara, Chaitanya Joshi, Madhvi Joshi |
| EPI_ISL_444482 | Gujarat Biotechnology Research Centre | Gujarat Biotechnology Research Centre | Gaurishankar Shrimali, Nidhi Sood, Pranay Shah, R D Dixit, Snehal Bagatharia, Kamlesh J Upadhyay, Ramesh Pandit, Tejas Shah, Ankit Hinsu, Pritesh Sabara, Apurvasinh Puvar, Janvi Raval, Monika Gandhi, Pinal Trivedi, Maharshi Pandya, Amit Kanani, Akanksha Verma, Nitin Savaliya, Raghawendra Kumar, Dinesh Kumar, Zuber Saiyed, Dipa Kinariwala, Disha Patel, Binita Aring, Neeta Khandelwal, Geeta Vaghela, Sonia Barve, Bhavesh Modi, Kairavi Joshi, Afzal Ansari, Chaitanya Joshi, Madhvi Joshi |
| EPI_ISL_444483 | Gujarat Biotechnology Research Centre | Gujarat Biotechnology Research Centre | Nidhi Sood, Pranay Shah, R D Dixit, Snehal Bagatharia, Kamlesh J Upadhyay, Ramesh Pandit, Tejas Shah, Ankit Hinsu, Pritesh Sabara, Apurvasinh Puvar, Janvi Raval, Monika Gandhi, Pinal Trivedi, Maharshi Pandya, Amit Kanani, Akanksha Verma, Nitin Savaliya, Raghawendra Kumar, Dinesh Kumar, Zuber Saiyed, Dipa Kinariwala, Disha Patel, Binita Aring, Neeta Khandelwal, Geeta Vaghela, Sonia Barve, Bhavesh Modi, Kairavi Joshi, Gaurishankar Shrimali, Neelam Nathani, Chaitanya Joshi, Madhvi Joshi |
| EPI_ISL_444484 | Gujarat Biotechnology Research Centre | Gujarat Biotechnology Research Centre | Pranay Shah, R D Dixit, Snehal Bagatharia, Kamlesh J Upadhyay, Ramesh Pandit, Tejas Shah, Ankit Hinsu, Pritesh Sabara, Apurvasinh Puvar, Janvi Raval, Monika Gandhi, Pinal Trivedi, Maharshi Pandya, Amit Kanani, Akanksha Verma, Nitin Savaliya, Raghawendra Kumar, Dinesh Kumar, Zuber Saiyed, Dipa Kinariwala, Disha Patel, Binita Aring, Neeta Khandelwal, Geeta Vaghela, Sonia Barve, Bhavesh Modi, Kairavi Joshi, Gaurishankar Shrimali, Nidhi Sood, Armi Chaudhari, Chaitanya Joshi, Madhvi Joshi |
| EPI_ISL_444485 | Gujarat Biotechnology Research Centre | Gujarat Biotechnology Research Centre | R D Dixit, Snehal Bagatharia, Kamlesh J Upadhyay, Ramesh Pandit, Tejas Shah, Ankit Hinsu, Pritesh Sabara, Apurvasinh Puvar, Janvi Raval, Monika Gandhi, Pinal Trivedi, Maharshi Pandya, Amit Kanani, Akanksha Verma, Nitin Savaliya, Raghawendra Kumar, Dinesh Kumar, Zuber Saiyed, Dipa Kinariwala, Disha Patel, Binita Aring, Neeta Khandelwal, Geeta Vaghela, Sonia Barve, Bhavesh Modi, Kairavi Joshi, Gaurishankar Shrimali, Nidhi Sood, Pranay Shah, Bhavya Jindal, Chaitanya Joshi, Madhvi Joshi |
| EPI_ISL_444486 | Gujarat Biotechnology Research Centre | Gujarat Biotechnology Research Centre | Snehal Bagatharia, Kamlesh J Upadhyay, Ramesh Pandit, Tejas Shah, Ankit Hinsu, Pritesh Sabara, Apurvasinh Puvar, Janvi Raval, Monika Gandhi, Pinal Trivedi, Maharshi Pandya, Amit Kanani, Akanksha Verma, Nitin Savaliya, Raghawendra Kumar, Dinesh Kumar, Zuber Saiyed, Dipa Kinariwala, Disha Patel, Binita Aring, Neeta Khandelwal, Geeta Vaghela, Sonia Barve, Bhavesh Modi, Kairavi Joshi, Gaurishankar Shrimali, Nidhi Sood, Pranay Shah, R D Dixit, Dipeshwari Shewale, Chaitanya Joshi, Madhvi Joshi |
| EPI_ISL_444487, EPI_ISL_444488, EPI_ISL_444489, EPI_ISL_444490, EPI_ISL_444491, EPI_ISL_444492 | Karolinska Universitetslaboratoriet | CTMR, Karolinska Institutet, Stockholm, Sweden | Yue Hu, Stefanie Prast-Nielsen, Jingkai Ji, Fredrik Boulund, Jing Wang, Shuiqin Li, Yinghua Zha, Caroline Bjurnemark, Linnéa Påvénius, Marica Hamsten, Vivien Lan Yang Swartz, Lars Engstrand |
| EPI_ISL_444493 | Departamento de Laboratorios de Salud Publica (DLSP, Division Epidemiologia, Ministerio de Salud Publica) | Facultad de Ciencias (Sección Genética Evolutiva, Sección Virología). | Panzera,Y., Delfraro,A., Ramos,N., Frabasile,S., Calleros,L., Techera,C., Grecco,S., Fuques,E., Goni,N., Coppola,L., Ramos,V., Chiparelli,H., Arbiza,J. and Perez,R. |
| EPI_ISL_444494, EPI_ISL_444495, EPI_ISL_444496, EPI_ISL_444497, EPI_ISL_444498, EPI_ISL_444499, EPI_ISL_444500, EPI_ISL_444501, EPI_ISL_444502, EPI_ISL_444503, EPI_ISL_444504, EPI_ISL_444505, EPI_ISL_444506, EPI_ISL_444507, EPI_ISL_444508, EPI_ISL_444509, EPI_ISL_444510, EPI_ISL_444511, EPI_ISL_444512, EPI_ISL_444513, EPI_ISL_444514, EPI_ISL_444515, EPI_ISL_444516 | | | |
| see above | Laboratoire de microbiologie, Hopital de Verdun | Smith Laboratory, Centre de Recherche CHU Sainte-Justine | Martin Smith, Marieke Rozendaal, Ivan Pavlov |
| EPI_ISL_444520, EPI_ISL_444521, EPI_ISL_444522, EPI_ISL_444523, EPI_ISL_444524, EPI_ISL_444525, EPI_ISL_444526, EPI_ISL_444527, EPI_ISL_444528, EPI_ISL_444529, EPI_ISL_444530, EPI_ISL_444531, EPI_ISL_444532, EPI_ISL_444533, EPI_ISL_444534, EPI_ISL_444535, EPI_ISL_444536, EPI_ISL_444537, EPI_ISL_444538, EPI_ISL_444539, EPI_ISL_444540, EPI_ISL_444541, EPI_ISL_444542, EPI_ISL_444543, EPI_ISL_444544, EPI_ISL_444545, EPI_ISL_444546, EPI_ISL_444547, EPI_ISL_444548, EPI_ISL_444549, EPI_ISL_444550, EPI_ISL_444551, EPI_ISL_444552, EPI_ISL_444553, EPI_ISL_444554, EPI_ISL_444555, EPI_ISL_444556, EPI_ISL_444557, EPI_ISL_444558, EPI_ISL_444559, EPI_ISL_444560, EPI_ISL_444561, EPI_ISL_444562, EPI_ISL_444563, EPI_ISL_444564, EPI_ISL_444565, EPI_ISL_444566, EPI_ISL_444567, EPI_ISL_444568, EPI_ISL_444569, EPI_ISL_444570, EPI_ISL_444571, EPI_ISL_444572, EPI_ISL_444573, EPI_ISL_444574, EPI_ISL_444575, EPI_ISL_444576, EPI_ISL_444577, EPI_ISL_444578, EPI_ISL_444579, EPI_ISL_444580, EPI_ISL_444581, EPI_ISL_444582, EPI_ISL_444583, EPI_ISL_444584, EPI_ISL_444585, EPI_ISL_444586, EPI_ISL_444587, EPI_ISL_444588, EPI_ISL_444589, EPI_ISL_444590, EPI_ISL_444591, EPI_ISL_444592, EPI_ISL_444593, EPI_ISL_444594, EPI_ISL_444595, EPI_ISL_444596, EPI_ISL_444597, EPI_ISL_444598, EPI_ISL_444599, EPI_ISL_444600, EPI_ISL_444601, EPI_ISL_444602, EPI_ISL_444603, EPI_ISL_444604, EPI_ISL_444605, EPI_ISL_444606, EPI_ISL_444607, EPI_ISL_444608, EPI_ISL_444609 | | | |
| see above | Northwestern Memorial Hospital | Ozer Lab | Ramon Lorenzo-Redondo, Hannah H. Nam, Scott C. Roberts, Lacy M. Simons, Chad J. Achenbach, Lawrence J. Jennings, Chao Qi, Alan R. Hauser, Michael G. Ison, Judd F. Hultquist, Egon A. Ozer |
| EPI_ISL_444611 | Pathology Queensland | Public Health Virology Laboratory | Bixing Huang, Alyssa Pyke, Amanda De Jong, Andrew Van Den Hurk, Carmel Taylor, David Warrilow, Doris Genge, Elisabeth Gamez, Glen Hewitson, Ian Maxwell Mackay, Inga Sultana, Jamie McMahon, Jean Barcelon, Judy Northill, Mitchell Finger, Natalie Simpson, Neelima Nair, Peter Burtonclay, Peter Moore, Sarah Wheatley, Sean Moody, Sonja Hall-Mendelin, Timothy Gardam, and Frederick Moore |
| EPI_ISL_444612 | QML Pathology | Public Health Virology Laboratory | Bixing Huang, Alyssa Pyke, Amanda De Jong, Andrew Van Den Hurk, Carmel Taylor, David Warrilow, Doris Genge, Elisabeth Gamez, Glen Hewitson, Ian Maxwell Mackay, Inga Sultana, Jamie McMahon, Jean Barcelon, Judy Northill, Mitchell Finger, Natalie Simpson, Neelima Nair, Peter Burtonclay, Peter Moore, Sarah Wheatley, Sean Moody, Sonja Hall-Mendelin, Timothy Gardam, and Frederick Moore |
| EPI_ISL_444613, EPI_ISL_444614, EPI_ISL_444615, EPI_ISL_444616, EPI_ISL_444617, EPI_ISL_444618, EPI_ISL_444619, EPI_ISL_444620, EPI_ISL_444621, EPI_ISL_444622, EPI_ISL_444623, EPI_ISL_444624, EPI_ISL_444625, EPI_ISL_444626, EPI_ISL_444627, EPI_ISL_444628, EPI_ISL_444629, EPI_ISL_444630, EPI_ISL_444631, EPI_ISL_444632, EPI_ISL_444633, EPI_ISL_444634, EPI_ISL_444635, EPI_ISL_444636, EPI_ISL_444637, EPI_ISL_444638, EPI_ISL_444639, EPI_ISL_444640, EPI_ISL_444641, EPI_ISL_444642, EPI_ISL_444643, EPI_ISL_444644, EPI_ISL_444645, EPI_ISL_444646, EPI_ISL_444647, EPI_ISL_444648, EPI_ISL_444649, EPI_ISL_444650, EPI_ISL_444651, EPI_ISL_444652, EPI_ISL_444653, EPI_ISL_444654, EPI_ISL_444655, EPI_ISL_444656, EPI_ISL_444657, EPI_ISL_444658, EPI_ISL_444659, EPI_ISL_444660, EPI_ISL_444661, EPI_ISL_444662, EPI_ISL_444663, EPI_ISL_444664, EPI_ISL_444665, EPI_ISL_444666, EPI_ISL_444667, EPI_ISL_444668, EPI_ISL_444669, EPI_ISL_444670, EPI_ISL_444671, EPI_ISL_444672, EPI_ISL_444673, EPI_ISL_444674, EPI_ISL_444675, EPI_ISL_444676, EPI_ISL_444677, EPI_ISL_444678, EPI_ISL_444679, EPI_ISL_444680, EPI_ISL_444681, EPI_ISL_444682, EPI_ISL_444683, EPI_ISL_444684, EPI_ISL_444685, EPI_ISL_444686, EPI_ISL_444687, EPI_ISL_444688, EPI_ISL_444689, EPI_ISL_444690, EPI_ISL_444691, EPI_ISL_444692, EPI_ISL_444693, EPI_ISL_444694, EPI_ISL_444695, EPI_ISL_444696, EPI_ISL_444697, EPI_ISL_444698, EPI_ISL_444699, EPI_ISL_444700, EPI_ISL_444701, EPI_ISL_444702, EPI_ISL_444703, EPI_ISL_444704, EPI_ISL_444705, EPI_ISL_444706, EPI_ISL_444707, EPI_ISL_444708, EPI_ISL_444709, EPI_ISL_444710, EPI_ISL_444711, EPI_ISL_444712, EPI_ISL_444713, EPI_ISL_444714, EPI_ISL_444715, EPI_ISL_444716, EPI_ISL_444717, EPI_ISL_444718, EPI_ISL_444719, EPI_ISL_444720, EPI_ISL_444721, EPI_ISL_444722, EPI_ISL_444723, EPI_ISL_444724, EPI_ISL_444725, EPI_ISL_444726, EPI_ISL_444727, EPI_ISL_444728, EPI_ISL_444729, EPI_ISL_444730, EPI_ISL_444731, EPI_ISL_444732, EPI_ISL_444733, EPI_ISL_444734, EPI_ISL_444735, EPI_ISL_444736, EPI_ISL_444737, EPI_ISL_444738, EPI_ISL_444739, EPI_ISL_444740, EPI_ISL_444741, EPI_ISL_444742, EPI_ISL_444743, EPI_ISL_444744, EPI_ISL_444745, EPI_ISL_444746, EPI_ISL_444747, EPI_ISL_444748, EPI_ISL_444749, EPI_ISL_444750, EPI_ISL_444751, EPI_ISL_444752, EPI_ISL_444753, EPI_ISL_444754, EPI_ISL_444755, EPI_ISL_444756, EPI_ISL_444757, EPI_ISL_444758, EPI_ISL_444759, EPI_ISL_444760, EPI_ISL_444761, EPI_ISL_444762, EPI_ISL_444763, EPI_ISL_444764, EPI_ISL_444765, EPI_ISL_444766, EPI_ISL_444767, EPI_ISL_444768, EPI_ISL_444769, EPI_ISL_444770, EPI_ISL_444771, EPI_ISL_444772, EPI_ISL_444773, EPI_ISL_444774, EPI_ISL_444775, EPI_ISL_444776, EPI_ISL_444777, EPI_ISL_444778, EPI_ISL_444779, EPI_ISL_444780, EPI_ISL_444781, EPI_ISL_444782, EPI_ISL_444783, EPI_ISL_444784, EPI_ISL_444785, EPI_ISL_444786, EPI_ISL_444787, EPI_ISL_444788, EPI_ISL_444789, EPI_ISL_444790, EPI_ISL_444791, EPI_ISL_444792 | | | |
| see above | NYU Langone Health | Departments of Pathology and Medicine, New York University School of Medicine | Maria Aguero-Rosenfeld, Brendan Belovarac, Margaret Black, Ludovic Boytard, John Cadley, Paolo Cotzia, John Chen, Dacia Dimartino, Xiaojun Feng, Tatyana Gindin, Emily Guzman, Adriana Heguy, Megan Hogan, Emily Huang, George Jour, Alireza Khodadadi-Jamayran, Lawrence H. Lin, Raven Luther, Andrew Lytle, Christian Marier, Matthew T. Maurano, Mark J. Mulligan, Peter Meyn, Raquel Ordonez Ciriza, Iman Osman, Jared Pinnell, Vanessa Raabe, Sitharam Ramaswami, Amy Rapkiewicz, Andre M. Ribeiro-dos-Santos, Marie Samanovic-Golden, Antonio Serrano, Guomiao Shen, Matija Snuderl, Theodore Vougiouklakis, Nick Vulpescu, Gael Westby, Paul Zappile, Yutong Zhang |
| EPI_ISL_444793 | Pathology Queensland | Public Health Virology Laboratory | Bixing Huang, Alyssa Pyke, Amanda De Jong, Andrew Van Den Hurk, Carmel Taylor, David Warrilow, Doris Genge, Elisabeth Gamez, Glen Hewitson, Ian Maxwell Mackay, Inga Sultana, Jamie McMahon, Jean Barcelon, Judy Northill, Mitchell Finger, Natalie Simpson, Neelima Nair, Peter Burtonclay, Peter Moore, Sarah Wheatley, Sean Moody, Sonja Hall-Mendelin, Timothy Gardam, and Frederick Moore |
| EPI_ISL_444794 | Cairns Hospital | Public Health Virology Laboratory | Bixing Huang, Alyssa Pyke, Amanda De Jong, Andrew Van Den Hurk, Carmel Taylor, David Warrilow, Doris Genge, Elisabeth Gamez, Glen Hewitson, Ian Maxwell Mackay, Inga Sultana, Jamie McMahon, Jean Barcelon, Judy Northill, Mitchell Finger, Natalie Simpson, Neelima Nair, Peter Burtonclay, Peter Moore, Sarah Wheatley, Sean Moody, Sonja Hall-Mendelin, Timothy Gardam, and Frederick Moore |

| | | | |
|---|---|---|---|
| EPI_ISL_444817, EPI_ISL_444818, EPI_ISL_444819, EPI_ISL_444820, EPI_ISL_444821, EPI_ISL_444822, EPI_ISL_444823, EPI_ISL_444824, EPI_ISL_444825, EPI_ISL_444826, EPI_ISL_444827, EPI_ISL_444828, EPI_ISL_444829, EPI_ISL_444830, EPI_ISL_444831, EPI_ISL_444832, EPI_ISL_444833, EPI_ISL_444834, EPI_ISL_444835, EPI_ISL_444836, EPI_ISL_444837, EPI_ISL_444838, EPI_ISL_444839, EPI_ISL_444840, EPI_ISL_444841, EPI_ISL_444842, EPI_ISL_444843, EPI_ISL_444844, EPI_ISL_444845, EPI_ISL_444846, EPI_ISL_444847, EPI_ISL_444848, EPI_ISL_444849, EPI_ISL_444850, EPI_ISL_444851, EPI_ISL_444852, EPI_ISL_444853, EPI_ISL_444854, EPI_ISL_444855, EPI_ISL_444856, EPI_ISL_444857, EPI_ISL_444858, EPI_ISL_444859, EPI_ISL_444860, EPI_ISL_444861, EPI_ISL_444862, EPI_ISL_444863, EPI_ISL_444864, EPI_ISL_444865, EPI_ISL_444866, EPI_ISL_444867, EPI_ISL_444868, EPI_ISL_444869, EPI_ISL_444870, EPI_ISL_444871, EPI_ISL_444872, EPI_ISL_444873, EPI_ISL_444874, EPI_ISL_444875, EPI_ISL_444876, EPI_ISL_444877, EPI_ISL_444878, EPI_ISL_444879, EPI_ISL_444880, EPI_ISL_444881, EPI_ISL_444882, EPI_ISL_444883, EPI_ISL_444884, EPI_ISL_444885, EPI_ISL_444886, EPI_ISL_444887, EPI_ISL_444888, EPI_ISL_444889, EPI_ISL_444890, EPI_ISL_444891, EPI_ISL_444892, EPI_ISL_444893, EPI_ISL_444894, EPI_ISL_444895, EPI_ISL_444896, EPI_ISL_444897, EPI_ISL_444898, EPI_ISL_444899, EPI_ISL_444900, EPI_ISL_444901, EPI_ISL_444902, EPI_ISL_444903, EPI_ISL_444904, EPI_ISL_444905, EPI_ISL_444906, EPI_ISL_444907, EPI_ISL_444908, EPI_ISL_444909, EPI_ISL_444910, EPI_ISL_444911, EPI_ISL_444912, EPI_ISL_444913, EPI_ISL_444914, EPI_ISL_444915, EPI_ISL_444916, EPI_ISL_444917, EPI_ISL_444918, EPI_ISL_444919, EPI_ISL_444920, EPI_ISL_444921, EPI_ISL_444922, EPI_ISL_444923, EPI_ISL_444924, EPI_ISL_444925, EPI_ISL_444926, EPI_ISL_444927, EPI_ISL_444928, EPI_ISL_444929, EPI_ISL_444930, EPI_ISL_444931, EPI_ISL_444932, EPI_ISL_444933, EPI_ISL_444934, EPI_ISL_444935, EPI_ISL_444936, EPI_ISL_444937, EPI_ISL_444938, EPI_ISL_444939, EPI_ISL_444940, EPI_ISL_444941, EPI_ISL_444942, EPI_ISL_444943, EPI_ISL_444944, EPI_ISL_444945, EPI_ISL_444946, EPI_ISL_444947, EPI_ISL_444948, EPI_ISL_444949, EPI_ISL_444950, EPI_ISL_444951, EPI_ISL_444952, EPI_ISL_444953, EPI_ISL_444954, EPI_ISL_444955, EPI_ISL_444956, EPI_ISL_444957, EPI_ISL_444958, EPI_ISL_444959, EPI_ISL_444960, EPI_ISL_444961, EPI_ISL_444962, EPI_ISL_444963, EPI_ISL_444964, EPI_ISL_444965, EPI_ISL_444966, EPI_ISL_444967, EPI_ISL_444968 | | | |
| see above | Department of Virus and Microbiological Special Diagnostics, Statens Serum Institut, Copenhagen, Denmark, Artillerivej 5, 2300 Copenahgen S | Albertsen lab, Department of Chemistry and Bioscience, Aalborg University, Denmark | Rasmus Kirkegaard |
| EPI_ISL_444969 | Guangzhou Eighth People's Hospital (Jiahe Sector) | Institute of Human Virology, Zhongshan School of Medicine, Sun Yat-sen University | Junsong Zhang, Fei Yu, Jun Liu, Huimin Fan, Ruosu Ying, Feng Huang, Ting Pan, Bingfeng Liu,Yiwen Zhang, Xu Zhang, Mang Shi, Fengyu Hu, Fang Li, Kai Deng, Hui Zhang |
| EPI_ISL_444971 | Hospital Universitari Vall d'Hebron - Vall d'Hebron Institut de Recerca | Hospital Universitari Vall d'Hebron | Cristina Andrés, Maria Piñana, Damir Garcia-Cehic, Mercedes Guerrero-Murillo, Ariadna Rando, Juliana Esperalba, Maria Gema Codina, Tomàs Pumarola, Josep Quer, Andrés Antón |
| EPI_ISL_444972 | Hospital Universitari Vall d'Hebron - Vall d'Hebron Institut de Recerca | Hospital Universitari Vall d'Hebron | Cristina Andrés, Maria Piñana, DAmir Garcia-Cehic, Mercedes Guerrero-Murillo, Ariadna Rando, Juliana Esperalba, Maria Gema Codina, Tomàs Pumarola, Josep Quer, Andrés Antón |
| EPI_ISL_444973 | Hospital Universitari Vall d`Hebron - Vall d'Hebron Institut de Recerca | Hospital Universitari Vall d'Hebron | Cristina Andrés, Maria Piñana, DAmir Garcia-Cehic, Mercedes Guerrero-Murillo, Ariadna Rando, Juliana Esperalba, Maria Gema Codina, Tomàs Pumarola, Josep Quer, Andrés Antón |
| EPI_ISL_444974, EPI_ISL_444975, EPI_ISL_444976, EPI_ISL_444977, EPI_ISL_444978, EPI_ISL_444979 | Hospital Universitari Vall d'Hebron - Vall d'Hebron Institut de Recerca | Hospital Universitari Vall d'Hebron | Cristina Andrés, Maria Piñana, Damir Garcia-Cehic, Mercedes Guerrero-Murillo, Ariadna Rando, Juliana Esperalba, Maria Gema Codina, Tomàs Pumarola, Josep Quer, Andrés Antón |
| EPI_ISL_444980, EPI_ISL_444981, EPI_ISL_444982, EPI_ISL_444983 | Hospital Universitari Vall d'Hebron - Vall d"hebron Institut de Recerca | Hospital Universitari Vall d'Hebron | Cristina Andrés, Maria Piñana, Damir Garcia-Cehic, Mercedes Guerrero-Murillo, Ariadna Rando, Juliana Esperalba, Maria Gema Codina, Tomàs Pumarola, Josep Quer, Andrés Antón |
| EPI_ISL_444984, EPI_ISL_444985, EPI_ISL_444986, EPI_ISL_444987, EPI_ISL_444988, EPI_ISL_444989, EPI_ISL_444990 | Hospital Universitari Vall d'Hebron - Vall d'Hebron Institut de Recerca | Hospital Universitari Vall d'Hebron | Cristina Andrés, Maria Piñana, Damir Garcia-Cehic, Mercedes Guerrero-Murillo, Ariadna Rando, Juliana Esperalba, Maria Gema Codina, Tomàs Pumarola, Josep Quer, Andrés Antón |
| EPI_ISL_444994, EPI_ISL_444995, EPI_ISL_444996, EPI_ISL_444997, EPI_ISL_444998, EPI_ISL_444999, EPI_ISL_445000 | Naval Health Research Center | Naval Medical Research Center Biological Defense Research Directorate | Logan Voegtly, Regina Cer, Dessiree Pena-Gomez, Adrian Paskey,Kyle Long, Roger Pan, Melinda Balansay-Ames, Chris Myers, Ewell Hollis, Nathaniel Christy, Kimberly Bishop-Lilly |
| EPI_ISL_445003, EPI_ISL_445004, EPI_ISL_445005, EPI_ISL_445006, EPI_ISL_445007, EPI_ISL_445008, EPI_ISL_445009, EPI_ISL_445010, EPI_ISL_445011, EPI_ISL_445012, EPI_ISL_445013, EPI_ISL_445014, EPI_ISL_445015, EPI_ISL_445016, EPI_ISL_445017, EPI_ISL_445018, EPI_ISL_445019, EPI_ISL_445020, EPI_ISL_445021, EPI_ISL_445022, EPI_ISL_445023, EPI_ISL_445024, EPI_ISL_445025, EPI_ISL_445026, EPI_ISL_445027, EPI_ISL_445028, EPI_ISL_445029, EPI_ISL_445030, EPI_ISL_445031, EPI_ISL_445032, EPI_ISL_445033, EPI_ISL_445034, EPI_ISL_445035, EPI_ISL_445036, EPI_ISL_445037, EPI_ISL_445038, EPI_ISL_445039, EPI_ISL_445040, EPI_ISL_445041, EPI_ISL_445042, EPI_ISL_445043, EPI_ISL_445044, EPI_ISL_445045, EPI_ISL_445046, EPI_ISL_445047, EPI_ISL_445048, EPI_ISL_445049, EPI_ISL_445050 | | | |
| see above | Florida Bureau of Public Health Laboratories | Florida Bureau of Public Health Laboratories | Sarah Schmedes, Jason Blanton |
| EPI_ISL_445053, EPI_ISL_445054, EPI_ISL_445055, EPI_ISL_445056, EPI_ISL_445057, EPI_ISL_445058, EPI_ISL_445059, EPI_ISL_445060, EPI_ISL_445061, EPI_ISL_445062, EPI_ISL_445063, EPI_ISL_445064, EPI_ISL_445065, EPI_ISL_445066, EPI_ISL_445067, EPI_ISL_445068, EPI_ISL_445069, EPI_ISL_445070, EPI_ISL_445071, EPI_ISL_445072, EPI_ISL_445073, EPI_ISL_445074, EPI_ISL_445075, EPI_ISL_445076 | | | |
| see above | Laboratoire National de Sante, Microbiology, Virology | Laboratoire National de Sante, Microbiology, Epidemiology and Microbial Genomics | Anke Wienecke-Baldacchino, Ardashel Latsuzbaia, Jessica Tapp, Catherine Ragimbeau, Guillaume Fournier, Tamir Abdelrahman, Trung Nguyen Nguyen, Joel Mossong |
| EPI_ISL_445077 | M Health Fairview | University of Minnesota Genomics Center | Daryl M. Gohl, John Garbe, Patrick Grady, Jerry Daniel, Ray Watson, Benjamin Auch, Andrew Nelson, Sophia Yohe, and Kenneth B. Beckman |
| EPI_ISL_445078, EPI_ISL_445079, EPI_ISL_445080, EPI_ISL_445081, EPI_ISL_445082, EPI_ISL_445083, EPI_ISL_445084 | Baylor College of Medicine | Baylor College of Medicine: HGSC | Vasanthi Avadhanula, Erin Nicholson, David Henke, Pedro Piedra, Harsha Doddapaneni, Donna Muzny, Qingchang Meng, Hsu Chao, Zeineen Momin, Hua Shen, George Weissenberger, Kavya Kottapalli, Yimiti Meiheerguli, Sejal Salvi, Ginger Metcalf, Vipin Menon, Sara J.J. Cregeen, Matthew C. Ross, Tulin Ayvaz, Richard Sucgang, Kristi L. Hoffman, Matthew Wong, Joseph F. Petrosino |
| EPI_ISL_445085 | Virology Unit, Agrobiodiversity and Biotechnology Project, CIAT - International Center for Tropical Agriculture | Virology Unit, Agrobiodiversity and Biotechnology Project, CIAT - International Center for Tropical Agriculture | Lopez,D., Parra,B. and Cuellar,W.J. |
| EPI_ISL_445087 | Laboratory Diagnostic, Veterinary Specialized Institute Kraljevo | Laboratory Diagnostic, Veterinary Specialized Institute Kraljevo | Vidanovic,D., Tesovic,B., Sekler,M., Dmitric,M., Debeljak,Z., Matovic,K., Vaskovic,N., Petrovic,T., Volkening,J. and Alfonso,C. |
| EPI_ISL_445088 | Human Genetic Research Center, Kawsar Biotech Company | Human Genetic Research Center, Kawsar Biotech Company | Abbasalipour Bashash,M., Khosravi,M.A., Zeinali,S., Keshvar,Y., Sabeghi,S., Jadaliha,M. and Yazdani,R. |
| EPI_ISL_445094, EPI_ISL_445095, EPI_ISL_445096, EPI_ISL_445097, EPI_ISL_445098, EPI_ISL_445099, EPI_ISL_445100, EPI_ISL_445101, EPI_ISL_445102, EPI_ISL_445103, EPI_ISL_445104, EPI_ISL_445105, EPI_ISL_445106, EPI_ISL_445107, EPI_ISL_445108, EPI_ISL_445109, EPI_ISL_445110, EPI_ISL_445111, EPI_ISL_445112, EPI_ISL_445113, EPI_ISL_445114, EPI_ISL_445115, EPI_ISL_445116, EPI_ISL_445117 | | | |
| see above | UC San Diego Center for Advanced Laboratory Medicine | Andersen lab at Scripps Research | SEARCH Alliance San Diego with David Pride, Ji H Shin |
| EPI_ISL_445118 | Rady's Childrens Hospital | Andersen lab at Scripps Research | SEARCH Alliance San Diego |
| EPI_ISL_445119, EPI_ISL_445120, EPI_ISL_445121, EPI_ISL_445122, EPI_ISL_445123, EPI_ISL_445124, EPI_ISL_445125, EPI_ISL_445126, EPI_ISL_445127, EPI_ISL_445128, EPI_ISL_445129, EPI_ISL_445130, EPI_ISL_445131, EPI_ISL_445132, EPI_ISL_445133, EPI_ISL_445134, EPI_ISL_445135, EPI_ISL_445136, EPI_ISL_445137, EPI_ISL_445138, EPI_ISL_445139, EPI_ISL_445140, EPI_ISL_445141, EPI_ISL_445142, EPI_ISL_445143, EPI_ISL_445144, EPI_ISL_445145, EPI_ISL_445146, EPI_ISL_445147, EPI_ISL_445148, EPI_ISL_445149, EPI_ISL_445150, EPI_ISL_445151, EPI_ISL_445152, EPI_ISL_445153, EPI_ISL_445154, EPI_ISL_445155, EPI_ISL_445156, EPI_ISL_445157, EPI_ISL_445158, EPI_ISL_445159, EPI_ISL_445160, EPI_ISL_445161, EPI_ISL_445162, EPI_ISL_445163 | | | |
| see above | Robert Garry lab | Andersen lab at Scripps Research | Allison Smither, Gilberto Sabino-Santos, Patricia Snarski, Lilia Melnik, Antoinette Bell, Kaylynn Genemaras, Arnaud Drouin, Dahlene Fusco, Robert Garry with SEARCH Alliance San Diego |
| EPI_ISL_445164, EPI_ISL_445165, EPI_ISL_445166, EPI_ISL_445167, EPI_ISL_445168 | Scripps Medical Laboratory | Andersen lab at Scripps Research | SEARCH Alliance San Diego with Michael Quigley, Ellen Stefanski, Ian Mchardy |
| EPI_ISL_445169, EPI_ISL_445170, EPI_ISL_445171, EPI_ISL_445172, EPI_ISL_445173, EPI_ISL_445174, EPI_ISL_445175, EPI_ISL_445176, EPI_ISL_445177, EPI_ISL_445178, EPI_ISL_445179, EPI_ISL_445180, EPI_ISL_445181, EPI_ISL_445182 | | | |
| see above | UCSF Clinical Microbiology Laboratory | Chan-Zuckerberg Biohub | CZB Cliahub Consortium |
| EPI_ISL_445213 | DNA Solution Ltd | DNA Solution Ltd | Md. Imran Khan, Kazi Nadim Hasan, Abu Sufian, Mohammed Nafiz Imtiaz Polol, Abdul Khaleque, Mizanur Rahman, MSM Chowdhury, Hasan Ul Haider, Mamudul Hasan Razu, Mala Khan, Mohammad Fazle Alam Rabbi |
| EPI_ISL_445214, EPI_ISL_445215, EPI_ISL_445216, EPI_ISL_445217 | DNA Solution Ltd. | DNA Solution Ltd. | Md. Imran Khan, Kazi Nadim Hasan, Abu Sufian, Mohammed Nafiz Imtiaz Polol, Abdul Khaleque, Mizanur Rahman, MSM Chowdhury, Hasan Ul Haider, Mamudul Hasan Razu, Mala Khan, Mohammad Fazle Alam Rabbi |
| EPI_ISL_445219 | Universidad del Valle, Laboratorio de Microbiologia, VIREM | Universidad del Valle, Universidad Nacional de Colombia-Sede Palmira, International Center for Tropical Agriculture | Beatriz Parra, Diana López-Alvarez, Wilmer J. Cuellar |
| EPI_ISL_445220 | Laborarory for Respiratory Viruses, "Cantacuzino" National Military-Medical Institute for Resararch and Development | Cantacuzino Institute | M.Lazar, L.Ustea, A.Cretu |
| EPI_ISL_445221 | Wasterlakarna | The Public Health Agency of Sweden | Frida Ahlfors, Oskar Karlsson Lindsjo, Maria Lind Karlberg, Anna-Malin Linde, Olov Svartstrom, Anna Risberg, Theresa Enkirch, Mia Brytting, Karin Tegmark-Wisell |
| EPI_ISL_445222 | Saroledens Familjelakare | The Public Health Agency of Sweden | Katarina Jarbur, Oskar Karlsson Lindsjo, Maria Lind Karlberg, Anna-Malin Linde, Olov Svartstrom, Anna Risberg, Theresa Enkirch, Mia Brytting, Karin Tegmark-Wisell |
| EPI_ISL_445223 | Victoria Vard och Halsa | The Public Health Agency of Sweden | Sarah Henriksson, Oskar Karlsson Lindsjo, Maria Lind Karlberg, Anna-Malin Linde, Olov Svartstrom, Anna Risberg, Theresa Enkirch, Mia Brytting, Karin Tegmark-Wisell |
| EPI_ISL_445224 | Narhalsan Olskroken VC | The Public Health Agency of Sweden | Mahin Ghoroghi, Oskar Karlsson Lindsjo, Maria Lind Karlberg, Anna-Malin Linde, Olov Svartstrom, Anna Risberg, Theresa Enkirch, Mia Brytting, Karin Tegmark-Wisell |
| EPI_ISL_445225 | Surbrunns VC | The Public Health Agency of Sweden | Erik Embring, Oskar Karlsson Lindsjo, Maria Lind Karlberg, Anna-Malin Linde, Olov Svartstrom, Anna Risberg, Theresa Enkirch, Mia Brytting, Karin Tegmark-Wisell |
| EPI_ISL_445226 | Saroledens Familjelakare | The Public Health Agency of Sweden | Katarina Jarbur, Oskar Karlsson Lindsjo, Maria Lind Karlberg, Anna-Malin Linde, Olov Svartstrom, Anna Risberg, Theresa Enkirch, Mia Brytting, Karin Tegmark-Wisell |
| EPI_ISL_445227 | Uppsala Narakut Aleris | The Public Health Agency of Sweden | Annika Nilsson, Oskar Karlsson Lindsjo, Maria Lind Karlberg, Anna-Malin Linde, Olov Svartstrom, Anna Risberg, Theresa Enkirch, Mia Brytting, Karin Tegmark-Wisell |
| EPI_ISL_445228 | Ulltuna Vardcentral | The Public Health Agency of Sweden | Heidi Lindback, Oskar Karlsson Lindsjo, Maria Lind Karlberg, Anna-Malin Linde, Olov Svartstrom, Anna Risberg, Theresa Enkirch, Mia Brytting, Karin Tegmark-Wisell |
| EPI_ISL_445229 | Narhalsan Backa vardcentral | The Public Health Agency of Sweden | Mats Olsson, Oskar Karlsson Lindsjo, Maria Lind Karlberg, Anna-Malin Linde, Olov Svartstrom, Anna Risberg, Theresa Enkirch, Mia Brytting, Karin Tegmark-Wisell |
| EPI_ISL_445230, EPI_ISL_445231 | Uppsala Narakut Aleris | The Public Health Agency of Sweden | Annika Nilsson, Oskar Karlsson Lindsjo, Maria Lind Karlberg, Anna-Malin Linde, Olov Svartstrom, Anna Risberg, Theresa Enkirch, Mia Brytting, Karin Tegmark-Wisell |
| EPI_ISL_445232 | Kungsors VC | The Public Health Agency of Sweden | Jessica Karlsson, Oskar Karlsson Lindsjo, Maria Lind Karlberg, Anna-Malin Linde, Olov Svartstrom, Anna Risberg, Theresa Enkirch, Mia Brytting, Karin Tegmark-Wisell |
| EPI_ISL_445233 | Vardcentralen Brinken | The Public Health Agency of Sweden | Agnes Wigh, Oskar Karlsson Lindsjo, Maria Lind Karlberg, Anna-Malin Linde, Olov Svartstrom, Anna Risberg, Theresa Enkirch, Mia Brytting, Karin Tegmark-Wisell |
| EPI_ISL_445234, EPI_ISL_445235 | Wasterlakarna | The Public Health Agency of Sweden | Frida Ahlfors, Oskar Karlsson Lindsjo, Maria Lind Karlberg, Anna-Malin Linde, Olov Svartstrom, Anna Risberg, Theresa Enkirch, Mia Brytting, Karin Tegmark-Wisell |
| EPI_ISL_445236 | Narhalsan Backa vardcentral | The Public Health Agency of Sweden | Mats Olsson, Oskar Karlsson Lindsjo, Maria Lind Karlberg, Anna-Malin Linde, Olov Svartstrom, Anna Risberg, Theresa Enkirch, Mia Brytting, Karin Tegmark-Wisell |
| EPI_ISL_445237 | Narhalsan Molnlycke, Barn och ungdomsmedicin | The Public Health Agency of Sweden | Mats Reimer, Oskar Karlsson Lindsjo, Maria Lind Karlberg, Anna-Malin Linde, Olov Svartstrom, Anna Risberg, Theresa Enkirch, Mia Brytting, Karin Tegmark-Wisell |
| EPI_ISL_445238 | Ã–resundslakarna | The Public Health Agency of Sweden | Del Akrawi, Oskar Karlsson Lindsjo, Maria Lind Karlberg, Anna-Malin Linde, Olov Svartstrom, Anna Risberg, Theresa Enkirch, Mia Brytting, Karin Tegmark-Wisell |
| EPI_ISL_445239 | Uppsala Narakut Aleris | The Public Health Agency of Sweden | Annika Nilsson, Oskar Karlsson Lindsjo, Maria Lind Karlberg, Anna-Malin Linde, Olov Svartstrom, Anna Risberg, Theresa Enkirch, Mia Brytting, Karin Tegmark-Wisell |
| EPI_ISL_445240 | Ulltuna Vardcentral | The Public Health Agency of Sweden | Heidi Lindback, Oskar Karlsson Lindsjo, Maria Lind Karlberg, Anna-Malin Linde, Olov Svartstrom, Anna Risberg, Theresa Enkirch, Mia Brytting, Karin Tegmark-Wisell |
| EPI_ISL_445241 | Ã–restadsklinikens VC | The Public Health Agency of Sweden | Lisa Kjellberg / Laura Plavitu, Oskar Karlsson Lindsjo, Maria Lind Karlberg, Anna-Malin Linde, Olov Svartstrom, Anna Risberg, Theresa Enkirch, Mia Brytting, Karin Tegmark-Wisell |
| EPI_ISL_445242 | Jokkmokks Halsocentral | The Public Health Agency of Sweden | Markus Beland, Oskar Karlsson Lindsjo, Maria Lind Karlberg, Anna-Malin Linde, Olov Svartstrom, Anna Risberg, Theresa Enkirch, Mia Brytting, Karin Tegmark-Wisell |
| EPI_ISL_445243 | Laboratory for Respiratory Viruses, Cantacuzino National Military-Medical Institute for Research and Development | Cantacuzino Institute | M.Lazar, L.Ustea, A.Cretu |
| EPI_ISL_445244 | Akbiomed lab | Tejgaon College bmb lab | Md.Abdul kaium,Md.Easin Arafat |
| EPI_ISL_445245 | CLINICA ALEMANA DE SANTIAGO S.A. | Instituto de Salud Publica de Chile | Andrés E Castillo, Bárbara Parra,Paz Tapia, Jaime Lagos, Loredana Arata, Alejandra Acevedo, Winston Andrade, Gabriel Leal, Carolina Tambley, Patricia Bustos, Rodrigo Fasce, Jorge Fernandez |

| | | | |
|---|---|---|---|
| EPI_ISL_445246 | HOSPITAL PUERTO MONTT | Instituto de Salud Publica de Chile | Andrés E Castillo, Bárbara Parra,Paz Tapia, Jaime Lagos, Loredana Arata, Alejandra Acevedo, Winston Andrade, Gabriel Leal, Carolina Tambley, Patricia Bustos, Rodrigo Fasce, Jorge Fernandez |
| EPI_ISL_445247 | UNIVERSIDAD DE LOS ANDES | Instituto de Salud Publica de Chile | Andrés E Castillo, Bárbara Parra,Paz Tapia, Jaime Lagos, Loredana Arata, Alejandra Acevedo, Winston Andrade, Gabriel Leal, Carolina Tambley, Patricia Bustos, Rodrigo Fasce, Jorge Fernandez |
| EPI_ISL_445248 | CLINICA ALEMANA DE SANTIAGO S.A. | Instituto de Salud Publica de Chile | Andrés E Castillo, Bárbara Parra,Paz Tapia, Jaime Lagos, Loredana Arata, Alejandra Acevedo, Winston Andrade, Gabriel Leal, Carolina Tambley, Patricia Bustos, Rodrigo Fasce, Jorge Fernandez |
| EPI_ISL_445249 | CLINICA SANTA MARIA S.A. | Instituto de Salud Publica de Chile | Andrés E Castillo, Bárbara Parra,Paz Tapia, Jaime Lagos, Loredana Arata, Alejandra Acevedo, Winston Andrade, Gabriel Leal, Carolina Tambley, Patricia Bustos, Rodrigo Fasce, Jorge Fernandez |
| EPI_ISL_445250 | CLINICA ALEMANA DE SANTIAGO S.A. | Instituto de Salud Publica de Chile | Andrés E Castillo, Bárbara Parra,Paz Tapia, Jaime Lagos, Loredana Arata, Alejandra Acevedo, Winston Andrade, Gabriel Leal, Carolina Tambley, Patricia Bustos, Rodrigo Fasce, Jorge Fernandez |
| EPI_ISL_445251 | HOSPITAL DE CARABINEROS | Instituto de Salud Publica de Chile | Andrés E Castillo, Bárbara Parra,Paz Tapia, Jaime Lagos, Loredana Arata, Alejandra Acevedo, Winston Andrade, Gabriel Leal, Carolina Tambley, Patricia Bustos, Rodrigo Fasce, Jorge Fernandez |
| EPI_ISL_445252 | PONTIFICIA U. CATOLICA FAC. MEDICINA | Instituto de Salud Publica de Chile | Andrés E Castillo, Bárbara Parra,Paz Tapia, Jaime Lagos, Loredana Arata, Alejandra Acevedo, Winston Andrade, Gabriel Leal, Carolina Tambley, Patricia Bustos, Rodrigo Fasce, Jorge Fernandez |
| EPI_ISL_445253, EPI_ISL_445254, EPI_ISL_445255 | CLINICA ALEMANA DE SANTIAGO S.A. | Instituto de Salud Publica de Chile | Andrés E Castillo, Bárbara Parra,Paz Tapia, Jaime Lagos, Loredana Arata, Alejandra Acevedo, Winston Andrade, Gabriel Leal, Carolina Tambley, Patricia Bustos, Rodrigo Fasce, Jorge Fernandez |
| EPI_ISL_445256 | CLINICA LAS CONDES S.A. | Instituto de Salud Publica de Chile | Andrés E Castillo, Bárbara Parra,Paz Tapia, Jaime Lagos, Loredana Arata, Alejandra Acevedo, Winston Andrade, Gabriel Leal, Carolina Tambley, Patricia Bustos, Rodrigo Fasce, Jorge Fernandez |
| EPI_ISL_445257 | CLINICA TABANCURA | Instituto de Salud Publica de Chile | Andrés E Castillo, Bárbara Parra,Paz Tapia, Jaime Lagos, Loredana Arata, Alejandra Acevedo, Winston Andrade, Gabriel Leal, Carolina Tambley, Patricia Bustos, Rodrigo Fasce, Jorge Fernandez |
| EPI_ISL_445258 | CLINICA ALEMANA DE SANTIAGO S.A. | Instituto de Salud Publica de Chile | Andrés E Castillo, Bárbara Parra,Paz Tapia, Jaime Lagos, Loredana Arata, Alejandra Acevedo, Winston Andrade, Gabriel Leal, Carolina Tambley, Patricia Bustos, Rodrigo Fasce, Jorge Fernandez |
| EPI_ISL_445259 | CLINICA LAS CONDES S.A. | Instituto de Salud Publica de Chile | Andrés E Castillo, Bárbara Parra,Paz Tapia, Jaime Lagos, Loredana Arata, Alejandra Acevedo, Winston Andrade, Gabriel Leal, Carolina Tambley, Patricia Bustos, Rodrigo Fasce, Jorge Fernandez |
| EPI_ISL_445260 | CLINICA ALEMANA DE SANTIAGO S.A. | Instituto de Salud Publica de Chile | Andrés E Castillo, Bárbara Parra,Paz Tapia, Jaime Lagos, Loredana Arata, Alejandra Acevedo, Winston Andrade, Gabriel Leal, Carolina Tambley, Patricia Bustos, Rodrigo Fasce, Jorge Fernandez |
| EPI_ISL_445261 | INTEGRAMEDICA LAB. CLINICO LTDA. | Instituto de Salud Publica de Chile | Andrés E Castillo, Bárbara Parra,Paz Tapia, Jaime Lagos, Loredana Arata, Alejandra Acevedo, Winston Andrade, Gabriel Leal, Carolina Tambley, Patricia Bustos, Rodrigo Fasce, Jorge Fernandez |
| EPI_ISL_445262 | CLINICA TABANCURA | Instituto de Salud Publica de Chile | Andrés E Castillo, Bárbara Parra,Paz Tapia, Jaime Lagos, Loredana Arata, Alejandra Acevedo, Winston Andrade, Gabriel Leal, Carolina Tambley, Patricia Bustos, Rodrigo Fasce, Jorge Fernandez |
| EPI_ISL_445263 | MEGASALUD SPA. | Instituto de Salud Publica de Chile | Andrés E Castillo, Bárbara Parra,Paz Tapia, Jaime Lagos, Loredana Arata, Alejandra Acevedo, Winston Andrade, Gabriel Leal, Carolina Tambley, Patricia Bustos, Rodrigo Fasce, Jorge Fernandez |
| EPI_ISL_445264 | CLINICA REDSALUD VITACURA. | Instituto de Salud Publica de Chile | Andrés E Castillo, Bárbara Parra,Paz Tapia, Jaime Lagos, Loredana Arata, Alejandra Acevedo, Winston Andrade, Gabriel Leal, Carolina Tambley, Patricia Bustos, Rodrigo Fasce, Jorge Fernandez |
| EPI_ISL_445265 | PONTIFICIA UNIVERSIDAD CATOLICA DE CHILE | Instituto de Salud Publica de Chile | Andrés E Castillo, Bárbara Parra,Paz Tapia, Jaime Lagos, Loredana Arata, Alejandra Acevedo, Winston Andrade, Gabriel Leal, Carolina Tambley, Patricia Bustos, Rodrigo Fasce, Jorge Fernandez |
| EPI_ISL_445266, EPI_ISL_445267 | CENTRO ONCOLOGICO DEL NORTE | Instituto de Salud Publica de Chile | Andrés E Castillo, Bárbara Parra,Paz Tapia, Jaime Lagos, Loredana Arata, Alejandra Acevedo, Winston Andrade, Gabriel Leal, Carolina Tambley, Patricia Bustos, Rodrigo Fasce, Jorge Fernandez |
| EPI_ISL_445268, EPI_ISL_445269 | HOSPITAL REG.LAUTARO NAVARRO AVARIA | Instituto de Salud Publica de Chile | Andrés E Castillo, Bárbara Parra,Paz Tapia, Jaime Lagos, Loredana Arata, Alejandra Acevedo, Winston Andrade, Gabriel Leal, Carolina Tambley, Patricia Bustos, Rodrigo Fasce, Jorge Fernandez |
| EPI_ISL_445270 | HOSPITAL DR.HERNAN HENRIQUEZ ARAVENA | Instituto de Salud Publica de Chile | Andrés E Castillo, Bárbara Parra,Paz Tapia, Jaime Lagos, Loredana Arata, Alejandra Acevedo, Winston Andrade, Gabriel Leal, Carolina Tambley, Patricia Bustos, Rodrigo Fasce, Jorge Fernandez |
| EPI_ISL_445271 | LABORATORIO CLINICA CHILLAN | Instituto de Salud Publica de Chile | Andrés E Castillo, Bárbara Parra,Paz Tapia, Jaime Lagos, Loredana Arata, Alejandra Acevedo, Winston Andrade, Gabriel Leal, Carolina Tambley, Patricia Bustos, Rodrigo Fasce, Jorge Fernandez |
| EPI_ISL_445272 | CLINICA CIUDAD DEL MAR | Instituto de Salud Publica de Chile | Andrés E Castillo, Bárbara Parra,Paz Tapia, Jaime Lagos, Loredana Arata, Alejandra Acevedo, Winston Andrade, Gabriel Leal, Carolina Tambley, Patricia Bustos, Rodrigo Fasce, Jorge Fernandez |
| EPI_ISL_445273, EPI_ISL_445274 | LABORATORIO TORRE MEDICA LTDA. | Instituto de Salud Publica de Chile | Andrés E Castillo, Bárbara Parra,Paz Tapia, Jaime Lagos, Loredana Arata, Alejandra Acevedo, Winston Andrade, Gabriel Leal, Carolina Tambley, Patricia Bustos, Rodrigo Fasce, Jorge Fernandez |
| EPI_ISL_445275, EPI_ISL_445276 | HOSPITAL CLINICO FUSAT | Instituto de Salud Publica de Chile | Andrés E Castillo, Bárbara Parra,Paz Tapia, Jaime Lagos, Loredana Arata, Alejandra Acevedo, Winston Andrade, Gabriel Leal, Carolina Tambley, Patricia Bustos, Rodrigo Fasce, Jorge Fernandez |
| EPI_ISL_445277 | FUNDACION DE SALUD EL TENIENTE | Instituto de Salud Publica de Chile | Andrés E Castillo, Bárbara Parra,Paz Tapia, Jaime Lagos, Loredana Arata, Alejandra Acevedo, Winston Andrade, Gabriel Leal, Carolina Tambley, Patricia Bustos, Rodrigo Fasce, Jorge Fernandez |
| EPI_ISL_445278 | LABORATORIO TORRE MEDICA LTDA. | Instituto de Salud Publica de Chile | Andrés E Castillo, Bárbara Parra,Paz Tapia, Jaime Lagos, Loredana Arata, Alejandra Acevedo, Winston Andrade, Gabriel Leal, Carolina Tambley, Patricia Bustos, Rodrigo Fasce, Jorge Fernandez |
| EPI_ISL_445279 | LABORATORIO INMUNOLAB SPA | Instituto de Salud Publica de Chile | Andrés E Castillo, Bárbara Parra,Paz Tapia, Jaime Lagos, Loredana Arata, Alejandra Acevedo, Winston Andrade, Gabriel Leal, Carolina Tambley, Patricia Bustos, Rodrigo Fasce, Jorge Fernandez |
| EPI_ISL_445280 | HOSPITAL REG.LAUTARO NAVARRO AVARIA | Instituto de Salud Publica de Chile | Andrés E Castillo, Bárbara Parra,Paz Tapia, Jaime Lagos, Loredana Arata, Alejandra Acevedo, Winston Andrade, Gabriel Leal, Carolina Tambley, Patricia Bustos, Rodrigo Fasce, Jorge Fernandez |
| EPI_ISL_445281 | HOSPITAL CLINICO DEL SUR | Instituto de Salud Publica de Chile | Andrés E Castillo, Bárbara Parra,Paz Tapia, Jaime Lagos, Loredana Arata, Alejandra Acevedo, Winston Andrade, Gabriel Leal, Carolina Tambley, Patricia Bustos, Rodrigo Fasce, Jorge Fernandez |
| EPI_ISL_445282, EPI_ISL_445283 | CLINICA MAGALLANES S.A. | Instituto de Salud Publica de Chile | Andrés E Castillo, Bárbara Parra,Paz Tapia, Jaime Lagos, Loredana Arata, Alejandra Acevedo, Winston Andrade, Gabriel Leal, Carolina Tambley, Patricia Bustos, Rodrigo Fasce, Jorge Fernandez |
| EPI_ISL_445284 | HOSPITAL REG.LAUTARO NAVARRO AVARIA | Instituto de Salud Publica de Chile | Andrés E Castillo, Bárbara Parra,Paz Tapia, Jaime Lagos, Loredana Arata, Alejandra Acevedo, Winston Andrade, Gabriel Leal, Carolina Tambley, Patricia Bustos, Rodrigo Fasce, Jorge Fernandez |
| EPI_ISL_445285 | CLINICA CIUDAD DEL MAR | Instituto de Salud Publica de Chile | Andrés E Castillo, Bárbara Parra,Paz Tapia, Jaime Lagos, Loredana Arata, Alejandra Acevedo, Winston Andrade, Gabriel Leal, Carolina Tambley, Patricia Bustos, Rodrigo Fasce, Jorge Fernandez |
| EPI_ISL_445286 | HOSPITAL HANGA ROA | Instituto de Salud Publica de Chile | Andrés E Castillo, Bárbara Parra,Paz Tapia, Jaime Lagos, Loredana Arata, Alejandra Acevedo, Winston Andrade, Gabriel Leal, Carolina Tambley, Patricia Bustos, Rodrigo Fasce, Jorge Fernandez |
| EPI_ISL_445287 | CLINICA CIUDAD DEL MAR | Instituto de Salud Publica de Chile | Andrés E Castillo, Bárbara Parra,Paz Tapia, Jaime Lagos, Loredana Arata, Alejandra Acevedo, Winston Andrade, Gabriel Leal, Carolina Tambley, Patricia Bustos, Rodrigo Fasce, Jorge Fernandez |
| EPI_ISL_445288 | HOSPITAL REG.LAUTARO NAVARRO AVARIA | Instituto de Salud Publica de Chile | Andrés E Castillo, Bárbara Parra,Paz Tapia, Jaime Lagos, Loredana Arata, Alejandra Acevedo, Winston Andrade, Gabriel Leal, Carolina Tambley, Patricia Bustos, Rodrigo Fasce, Jorge Fernandez |
| EPI_ISL_445289 | HOSPITAL NAVAL PUERTO WILLIAMS | Instituto de Salud Publica de Chile | Andrés E Castillo, Bárbara Parra,Paz Tapia, Jaime Lagos, Loredana Arata, Alejandra Acevedo, Winston Andrade, Gabriel Leal, Carolina Tambley, Patricia Bustos, Rodrigo Fasce, Jorge Fernandez |
| EPI_ISL_445290, EPI_ISL_445291, EPI_ISL_445292 | CLINICA MAGALLANES S.A. | Instituto de Salud Publica de Chile | Andrés E Castillo, Bárbara Parra,Paz Tapia, Jaime Lagos, Loredana Arata, Alejandra Acevedo, Winston Andrade, Gabriel Leal, Carolina Tambley, Patricia Bustos, Rodrigo Fasce, Jorge Fernandez |
| EPI_ISL_445293, EPI_ISL_445294, EPI_ISL_445295 | HOSPITAL REG.LAUTARO NAVARRO AVARIA | Instituto de Salud Publica de Chile | Andrés E Castillo, Bárbara Parra,Paz Tapia, Jaime Lagos, Loredana Arata, Alejandra Acevedo, Winston Andrade, Gabriel Leal, Carolina Tambley, Patricia Bustos, Rodrigo Fasce, Jorge Fernandez |
| EPI_ISL_445296 | HOSPITAL REGIONAL DE COYHAIQUE | Instituto de Salud Publica de Chile | Andrés E Castillo, Bárbara Parra,Paz Tapia, Jaime Lagos, Loredana Arata, Alejandra Acevedo, Winston Andrade, Gabriel Leal, Carolina Tambley, Patricia Bustos, Rodrigo Fasce, Jorge Fernandez |
| EPI_ISL_445297 | CLINICA INTEGRAL S.A. | Instituto de Salud Publica de Chile | Andrés E Castillo, Bárbara Parra,Paz Tapia, Jaime Lagos, Loredana Arata, Alejandra Acevedo, Winston Andrade, Gabriel Leal, Carolina Tambley, Patricia Bustos, Rodrigo Fasce, Jorge Fernandez |
| EPI_ISL_445298 | HOSPITAL DE SAN FERNANDO | Instituto de Salud Publica de Chile | Andrés E Castillo, Bárbara Parra,Paz Tapia, Jaime Lagos, Loredana Arata, Alejandra Acevedo, Winston Andrade, Gabriel Leal, Carolina Tambley, Patricia Bustos, Rodrigo Fasce, Jorge Fernandez |
| EPI_ISL_445299 | CLINICA MAGALLANES S.A. | Instituto de Salud Publica de Chile | Andrés E Castillo, Bárbara Parra,Paz Tapia, Jaime Lagos, Loredana Arata, Alejandra Acevedo, Winston Andrade, Gabriel Leal, Carolina Tambley, Patricia Bustos, Rodrigo Fasce, Jorge Fernandez |
| EPI_ISL_445300 | HOSPITAL DE RANCAGUA | Instituto de Salud Publica de Chile | Andrés E Castillo, Bárbara Parra,Paz Tapia, Jaime Lagos, Loredana Arata, Alejandra Acevedo, Winston Andrade, Gabriel Leal, Carolina Tambley, Patricia Bustos, Rodrigo Fasce, Jorge Fernandez |
| EPI_ISL_445301 | HOSPITAL NAVAL PUERTO WILLIAMS | Instituto de Salud Publica de Chile | Andrés E Castillo, Bárbara Parra,Paz Tapia, Jaime Lagos, Loredana Arata, Alejandra Acevedo, Winston Andrade, Gabriel Leal, Carolina Tambley, Patricia Bustos, Rodrigo Fasce, Jorge Fernandez |
| EPI_ISL_445302 | INSTITUTO MEDICO LEGAL | Instituto de Salud Publica de Chile | Andrés E Castillo, Bárbara Parra,Paz Tapia, Jaime Lagos, Loredana Arata, Alejandra Acevedo, Winston Andrade, Gabriel Leal, Carolina Tambley, Patricia Bustos, Rodrigo Fasce, Jorge Fernandez |
| EPI_ISL_445303 | CTRO.DE SALUD FAMILIAR DR. RAUL YAZIGI | Instituto de Salud Publica de Chile | Andrés E Castillo, Bárbara Parra,Paz Tapia, Jaime Lagos, Loredana Arata, Alejandra Acevedo, Winston Andrade, Gabriel Leal, Carolina Tambley, Patricia Bustos, Rodrigo Fasce, Jorge Fernandez |
| EPI_ISL_445304 | HOSPITAL SAN JUAN DE DIOS | Instituto de Salud Publica de Chile | Andrés E Castillo, Bárbara Parra,Paz Tapia, Jaime Lagos, Loredana Arata, Alejandra Acevedo, Winston Andrade, Gabriel Leal, Carolina Tambley, Patricia Bustos, Rodrigo Fasce, Jorge Fernandez |
| EPI_ISL_445305 | HOSPITAL DE CARABINEROS | Instituto de Salud Publica de Chile | Andrés E Castillo, Bárbara Parra,Paz Tapia, Jaime Lagos, Loredana Arata, Alejandra Acevedo, Winston Andrade, Gabriel Leal, Carolina Tambley, Patricia Bustos, Rodrigo Fasce, Jorge Fernandez |
| EPI_ISL_445306 | CLINICA UC SAN CARLOS DE APOQUINDO | Instituto de Salud Publica de Chile | Andrés E Castillo, Bárbara Parra,Paz Tapia, Jaime Lagos, Loredana Arata, Alejandra Acevedo, Winston Andrade, Gabriel Leal, Carolina Tambley, Patricia Bustos, Rodrigo Fasce, Jorge Fernandez |
| EPI_ISL_445307 | HOSPITAL SAN JOSE DE MAIPO | Instituto de Salud Publica de Chile | Andrés E Castillo, Bárbara Parra,Paz Tapia, Jaime Lagos, Loredana Arata, Alejandra Acevedo, Winston Andrade, Gabriel Leal, Carolina Tambley, Patricia Bustos, Rodrigo Fasce, Jorge Fernandez |
| EPI_ISL_445308 | HOSPITAL EL CARMEN DR.LUIS VALENTIN F. | Instituto de Salud Publica de Chile | Andrés E Castillo, Bárbara Parra,Paz Tapia, Jaime Lagos, Loredana Arata, Alejandra Acevedo, Winston Andrade, Gabriel Leal, Carolina Tambley, Patricia Bustos, Rodrigo Fasce, Jorge Fernandez |
| EPI_ISL_445309 | MUTUAL DE SEGURIDAD C.CH.C. | Instituto de Salud Publica de Chile | Andrés E Castillo, Bárbara Parra,Paz Tapia, Jaime Lagos, Loredana Arata, Alejandra Acevedo, Winston Andrade, Gabriel Leal, Carolina Tambley, Patricia Bustos, Rodrigo Fasce, Jorge Fernandez |
| EPI_ISL_445310 | HOSPITAL DR.SOTERO DEL RIO | Instituto de Salud Publica de Chile | Andrés E Castillo, Bárbara Parra,Paz Tapia, Jaime Lagos, Loredana Arata, Alejandra Acevedo, Winston Andrade, Gabriel Leal, Carolina Tambley, Patricia Bustos, Rodrigo Fasce, Jorge Fernandez |
| EPI_ISL_445311 | MEGASALUD S.A. | Instituto de Salud Publica de Chile | Andrés E Castillo, Bárbara Parra,Paz Tapia, Jaime Lagos, Loredana Arata, Alejandra Acevedo, Winston Andrade, Gabriel Leal, Carolina Tambley, Patricia Bustos, Rodrigo Fasce, Jorge Fernandez |
| EPI_ISL_445312 | CLINICA UC SAN CARLOS DE APOQUINDO | Instituto de Salud Publica de Chile | Andrés E Castillo, Bárbara Parra,Paz Tapia, Jaime Lagos, Loredana Arata, Alejandra Acevedo, Winston Andrade, Gabriel Leal, Carolina Tambley, Patricia Bustos, Rodrigo Fasce, Jorge Fernandez |
| EPI_ISL_445313, EPI_ISL_445314 | HOSPITAL EL CARMEN DR.LUIS VALENTIN F. | Instituto de Salud Publica de Chile | Andrés E Castillo, Bárbara Parra,Paz Tapia, Jaime Lagos, Loredana Arata, Alejandra Acevedo, Winston Andrade, Gabriel Leal, Carolina Tambley, Patricia Bustos, Rodrigo Fasce, Jorge Fernandez |
| EPI_ISL_445315 | CLINICA UC SAN CARLOS DE APOQUINDO | Instituto de Salud Publica de Chile | Andrés E Castillo, Bárbara Parra,Paz Tapia, Jaime Lagos, Loredana Arata, Alejandra Acevedo, Winston Andrade, Gabriel Leal, Carolina Tambley, Patricia Bustos, Rodrigo Fasce, Jorge Fernandez |
| EPI_ISL_445316 | CESFAM BALMACEDA DE RENCA | Instituto de Salud Publica de Chile | Andrés E Castillo, Bárbara Parra,Paz Tapia, Jaime Lagos, Loredana Arata, Alejandra Acevedo, Winston Andrade, Gabriel Leal, Carolina Tambley, Patricia Bustos, Rodrigo Fasce, Jorge Fernandez |
| EPI_ISL_445317 | UNIV.DE CHILE HOSP.CLINICO | Instituto de Salud Publica de Chile | Andrés E Castillo, Bárbara Parra,Paz Tapia, Jaime Lagos, Loredana Arata, Alejandra Acevedo, Winston Andrade, Gabriel Leal, Carolina Tambley, Patricia Bustos, Rodrigo Fasce, Jorge Fernandez |
| EPI_ISL_445318 | C.DE SALUD FAMILIAR PABLO NERUDA | Instituto de Salud Publica de Chile | Andrés E Castillo, Bárbara Parra,Paz Tapia, Jaime Lagos, Loredana Arata, Alejandra Acevedo, Winston Andrade, Gabriel Leal, Carolina Tambley, Patricia Bustos, Rodrigo Fasce, Jorge Fernandez |
| EPI_ISL_445319 | HOSPITAL FELIX BULNES | Instituto de Salud Publica de Chile | Andrés E Castillo, Bárbara Parra,Paz Tapia, Jaime Lagos, Loredana Arata, Alejandra Acevedo, Winston Andrade, Gabriel Leal, Carolina Tambley, Patricia Bustos, Rodrigo Fasce, Jorge Fernandez |
| EPI_ISL_445320 | C.C.SALUD FAMILIAR PADRE FELIX DONOSO G. | Instituto de Salud Publica de Chile | Andrés E Castillo, Bárbara Parra,Paz Tapia, Jaime Lagos, Loredana Arata, Alejandra Acevedo, Winston Andrade, Gabriel Leal, Carolina Tambley, Patricia Bustos, Rodrigo Fasce, Jorge Fernandez |
| EPI_ISL_445321 | HOSPITAL FELIX BULNES | Instituto de Salud Publica de Chile | Andrés E Castillo, Bárbara Parra,Paz Tapia, Jaime Lagos, Loredana Arata, Alejandra Acevedo, Winston Andrade, Gabriel Leal, Carolina Tambley, Patricia Bustos, Rodrigo Fasce, Jorge Fernandez |
| EPI_ISL_445322 | UNIV.DE CHILE HOSP.CLINICO | Instituto de Salud Publica de Chile | Andrés E Castillo, Bárbara Parra,Paz Tapia, Jaime Lagos, Loredana Arata, Alejandra Acevedo, Winston Andrade, Gabriel Leal, Carolina Tambley, Patricia Bustos, Rodrigo Fasce, Jorge Fernandez |
| EPI_ISL_445323 | HOSP.ENFERMEDADES INFECCIOSAS | Instituto de Salud Publica de Chile | Andrés E Castillo, Bárbara Parra,Paz Tapia, Jaime Lagos, Loredana Arata, Alejandra Acevedo, Winston Andrade, Gabriel Leal, Carolina Tambley, Patricia Bustos, Rodrigo Fasce, Jorge Fernandez |
| EPI_ISL_445324 | INTEGRAMEDICA S.A | Instituto de Salud Publica de Chile | Andrés E Castillo, Bárbara Parra,Paz Tapia, Jaime Lagos, Loredana Arata, Alejandra Acevedo, Winston Andrade, Gabriel Leal, Carolina Tambley, Patricia Bustos, Rodrigo Fasce, Jorge Fernandez |
| EPI_ISL_445325 | HOSPITAL DR.SOTERO DEL RIO | Instituto de Salud Publica de Chile | Andrés E Castillo, Bárbara Parra,Paz Tapia, Jaime Lagos, Loredana Arata, Alejandra Acevedo, Winston Andrade, Gabriel Leal, Carolina Tambley, Patricia Bustos, Rodrigo Fasce, Jorge Fernandez |
| EPI_ISL_445326 | ASISTENCIA PUBLICA DR.ALEJANDRO DEL RIO | Instituto de Salud Publica de Chile | Andrés E Castillo, Bárbara Parra,Paz Tapia, Jaime Lagos, Loredana Arata, Alejandra Acevedo, Winston Andrade, Gabriel Leal, Carolina Tambley, Patricia Bustos, Rodrigo Fasce, Jorge Fernandez |
| EPI_ISL_445327 | HOSPITAL PADRE HURTADO | Instituto de Salud Publica de Chile | Andrés E Castillo, Bárbara Parra,Paz Tapia, Jaime Lagos, Loredana Arata, Alejandra Acevedo, Winston Andrade, Gabriel Leal, Carolina Tambley, Patricia Bustos, Rodrigo Fasce, Jorge Fernandez |
| EPI_ISL_445328 | MEGASALUD S.A. | Instituto de Salud Publica de Chile | Andrés E Castillo, Bárbara Parra,Paz Tapia, Jaime Lagos, Loredana Arata, Alejandra Acevedo, Winston Andrade, Gabriel Leal, Carolina Tambley, Patricia Bustos, Rodrigo Fasce, Jorge Fernandez |
| EPI_ISL_445329 | HOSPITAL DR.SOTERO DEL RIO | Instituto de Salud Publica de Chile | Andrés E Castillo, Bárbara Parra,Paz Tapia, Jaime Lagos, Loredana Arata, Alejandra Acevedo, Winston Andrade, Gabriel Leal, Carolina Tambley, Patricia Bustos, Rodrigo Fasce, Jorge Fernandez |
| EPI_ISL_445330 | CLINICA VESPUCIO S. A. | Instituto de Salud Publica de Chile | Andrés E Castillo, Bárbara Parra,Paz Tapia, Jaime Lagos, Loredana Arata, Alejandra Acevedo, Winston Andrade, Gabriel Leal, Carolina Tambley, Patricia Bustos, Rodrigo Fasce, Jorge Fernandez |
| EPI_ISL_445331, EPI_ISL_445332 | HOSPITAL HERMINDA MARTIN CHILLAN | Instituto de Salud Publica de Chile | Andrés E Castillo, Bárbara Parra,Paz Tapia, Jaime Lagos, Loredana Arata, Alejandra Acevedo, Winston Andrade, Gabriel Leal, Carolina Tambley, Patricia Bustos, Rodrigo Fasce, Jorge Fernandez |
| EPI_ISL_445333 | LABORATORIO CLINICA UNIVERSITARIA DE CONCEPCION | Instituto de Salud Publica de Chile | Andrés E Castillo, Bárbara Parra,Paz Tapia, Jaime Lagos, Loredana Arata, Alejandra Acevedo, Winston Andrade, Gabriel Leal, Carolina Tambley, Patricia Bustos, Rodrigo Fasce, Jorge Fernandez |
| EPI_ISL_445334 | CLINICA UNIVERSITARIA DE PUERTO MONTT S.A. | Instituto de Salud Publica de Chile | Andrés E Castillo, Bárbara Parra,Paz Tapia, Jaime Lagos, Loredana Arata, Alejandra Acevedo, Winston Andrade, Gabriel Leal, Carolina Tambley, Patricia Bustos, Rodrigo Fasce, Jorge Fernandez |

| | | | |
|---|---|---|---|
| EPI_ISL_445335 | HOSPITAL DE CALBUCO | Instituto de Salud Publica de Chile | Andrés E Castillo, Bárbara Parra,Paz Tapia, Jaime Lagos, Loredana Arata, Alejandra Acevedo, Winston Andrade, Gabriel Leal, Carolina Tambley, Patricia Bustos, Rodrigo Fasce, Jorge Fernandez |
| EPI_ISL_445336 | HOSPITAL LAS HIGUERAS DE TALCAHUANO | Instituto de Salud Publica de Chile | Andrés E Castillo, Bárbara Parra,Paz Tapia, Jaime Lagos, Loredana Arata, Alejandra Acevedo, Winston Andrade, Gabriel Leal, Carolina Tambley, Patricia Bustos, Rodrigo Fasce, Jorge Fernandez |
| EPI_ISL_445337 | HOSPITAL HANGA ROA | Instituto de Salud Publica de Chile | Andrés E Castillo, Bárbara Parra,Paz Tapia, Jaime Lagos, Loredana Arata, Alejandra Acevedo, Winston Andrade, Gabriel Leal, Carolina Tambley, Patricia Bustos, Rodrigo Fasce, Jorge Fernandez |
| EPI_ISL_445338, EPI_ISL_445339, EPI_ISL_445340, EPI_ISL_445341, EPI_ISL_445342, EPI_ISL_445343, EPI_ISL_445344, EPI_ISL_445345, EPI_ISL_445346, EPI_ISL_445347, EPI_ISL_445348 | | | |
| see above | HOSPITAL DR.HERNAN HENRIQUEZ ARAVENA | Instituto de Salud Publica de Chile | Andrés E Castillo, Bárbara Parra,Paz Tapia, Jaime Lagos, Loredana Arata, Alejandra Acevedo, Winston Andrade, Gabriel Leal, Carolina Tambley, Patricia Bustos, Rodrigo Fasce, Jorge Fernandez |
| EPI_ISL_445349, EPI_ISL_445350, EPI_ISL_445351 | HOSPITAL SAN JUAN DE DIOS | Instituto de Salud Publica de Chile | Andrés E Castillo, Bárbara Parra,Paz Tapia, Jaime Lagos, Loredana Arata, Alejandra Acevedo, Winston Andrade, Gabriel Leal, Carolina Tambley, Patricia Bustos, Rodrigo Fasce, Jorge Fernandez |
| EPI_ISL_445352 | HOSPITAL DEL PROFESOR | Instituto de Salud Publica de Chile | Andrés E Castillo, Bárbara Parra,Paz Tapia, Jaime Lagos, Loredana Arata, Alejandra Acevedo, Winston Andrade, Gabriel Leal, Carolina Tambley, Patricia Bustos, Rodrigo Fasce, Jorge Fernandez |
| EPI_ISL_445353 | HOSPITAL PADRE HURTADO | Instituto de Salud Publica de Chile | Andrés E Castillo, Bárbara Parra,Paz Tapia, Jaime Lagos, Loredana Arata, Alejandra Acevedo, Winston Andrade, Gabriel Leal, Carolina Tambley, Patricia Bustos, Rodrigo Fasce, Jorge Fernandez |
| EPI_ISL_445354 | HOSPITAL DE CARABINEROS | Instituto de Salud Publica de Chile | Andrés E Castillo, Bárbara Parra,Paz Tapia, Jaime Lagos, Loredana Arata, Alejandra Acevedo, Winston Andrade, Gabriel Leal, Carolina Tambley, Patricia Bustos, Rodrigo Fasce, Jorge Fernandez |
| EPI_ISL_445355 | MUTUAL DE SEGURIDAD C.CH.C. | Instituto de Salud Publica de Chile | Andrés E Castillo, Bárbara Parra,Paz Tapia, Jaime Lagos, Loredana Arata, Alejandra Acevedo, Winston Andrade, Gabriel Leal, Carolina Tambley, Patricia Bustos, Rodrigo Fasce, Jorge Fernandez |
| EPI_ISL_445356 | PONTIFICIA U. CATOLICA SERV. LABORATORIO | Instituto de Salud Publica de Chile | Andrés E Castillo, Bárbara Parra,Paz Tapia, Jaime Lagos, Loredana Arata, Alejandra Acevedo, Winston Andrade, Gabriel Leal, Carolina Tambley, Patricia Bustos, Rodrigo Fasce, Jorge Fernandez |
| EPI_ISL_445357 | INTEGRAMEDICA CENTROS MEDICOS S.A. | Instituto de Salud Publica de Chile | Andrés E Castillo, Bárbara Parra,Paz Tapia, Jaime Lagos, Loredana Arata, Alejandra Acevedo, Winston Andrade, Gabriel Leal, Carolina Tambley, Patricia Bustos, Rodrigo Fasce, Jorge Fernandez |
| EPI_ISL_445358 | MEGASALUD SPA. | Instituto de Salud Publica de Chile | Andrés E Castillo, Bárbara Parra,Paz Tapia, Jaime Lagos, Loredana Arata, Alejandra Acevedo, Winston Andrade, Gabriel Leal, Carolina Tambley, Patricia Bustos, Rodrigo Fasce, Jorge Fernandez |
| EPI_ISL_445359 | HOSP. SANTIAGO ORIENTE DR. LUIS TISNE B. | Instituto de Salud Publica de Chile | Andrés E Castillo, Bárbara Parra,Paz Tapia, Jaime Lagos, Loredana Arata, Alejandra Acevedo, Winston Andrade, Gabriel Leal, Carolina Tambley, Patricia Bustos, Rodrigo Fasce, Jorge Fernandez |
| EPI_ISL_445360 | HOSPITAL DEL PROFESOR | Instituto de Salud Publica de Chile | Andrés E Castillo, Bárbara Parra,Paz Tapia, Jaime Lagos, Loredana Arata, Alejandra Acevedo, Winston Andrade, Gabriel Leal, Carolina Tambley, Patricia Bustos, Rodrigo Fasce, Jorge Fernandez |
| EPI_ISL_445361 | CLINICA UC SAN CARLOS DE APOQUINDO | Instituto de Salud Publica de Chile | Andrés E Castillo, Bárbara Parra,Paz Tapia, Jaime Lagos, Loredana Arata, Alejandra Acevedo, Winston Andrade, Gabriel Leal, Carolina Tambley, Patricia Bustos, Rodrigo Fasce, Jorge Fernandez |
| EPI_ISL_445362 | BUPA SERVICIOS CLINICOS S.A | Instituto de Salud Publica de Chile | Andrés E Castillo, Bárbara Parra,Paz Tapia, Jaime Lagos, Loredana Arata, Alejandra Acevedo, Winston Andrade, Gabriel Leal, Carolina Tambley, Patricia Bustos, Rodrigo Fasce, Jorge Fernandez |
| EPI_ISL_445363 | ASISTENCIA PUBLICA DR.ALEJANDRO DEL RIO | Instituto de Salud Publica de Chile | Andrés E Castillo, Bárbara Parra,Paz Tapia, Jaime Lagos, Loredana Arata, Alejandra Acevedo, Winston Andrade, Gabriel Leal, Carolina Tambley, Patricia Bustos, Rodrigo Fasce, Jorge Fernandez |
| EPI_ISL_445364 | HOSPITAL EL CARMEN DR.LUIS VALENTIN F. | Instituto de Salud Publica de Chile | Andrés E Castillo, Bárbara Parra,Paz Tapia, Jaime Lagos, Loredana Arata, Alejandra Acevedo, Winston Andrade, Gabriel Leal, Carolina Tambley, Patricia Bustos, Rodrigo Fasce, Jorge Fernandez |
| EPI_ISL_445365, EPI_ISL_445366 | HOSPITAL DR.SOTERO DEL RIO | Instituto de Salud Publica de Chile | Andrés E Castillo, Bárbara Parra,Paz Tapia, Jaime Lagos, Loredana Arata, Alejandra Acevedo, Winston Andrade, Gabriel Leal, Carolina Tambley, Patricia Bustos, Rodrigo Fasce, Jorge Fernandez |
| EPI_ISL_445367 | ASISTENCIA PUBLICA DR.ALEJANDRO DEL RIO | Instituto de Salud Publica de Chile | Andrés E Castillo, Bárbara Parra,Paz Tapia, Jaime Lagos, Loredana Arata, Alejandra Acevedo, Winston Andrade, Gabriel Leal, Carolina Tambley, Patricia Bustos, Rodrigo Fasce, Jorge Fernandez |
| EPI_ISL_445368 | HOSPITAL DEL PROFESOR | Instituto de Salud Publica de Chile | Andrés E Castillo, Bárbara Parra,Paz Tapia, Jaime Lagos, Loredana Arata, Alejandra Acevedo, Winston Andrade, Gabriel Leal, Carolina Tambley, Patricia Bustos, Rodrigo Fasce, Jorge Fernandez |
| EPI_ISL_445369, EPI_ISL_445370 | HOSPITAL DE CARABINEROS | Instituto de Salud Publica de Chile | Andrés E Castillo, Bárbara Parra,Paz Tapia, Jaime Lagos, Loredana Arata, Alejandra Acevedo, Winston Andrade, Gabriel Leal, Carolina Tambley, Patricia Bustos, Rodrigo Fasce, Jorge Fernandez |
| EPI_ISL_445371 | HOSPITAL DR.SOTERO DEL RIO | Instituto de Salud Publica de Chile | Andrés E Castillo, Bárbara Parra,Paz Tapia, Jaime Lagos, Loredana Arata, Alejandra Acevedo, Winston Andrade, Gabriel Leal, Carolina Tambley, Patricia Bustos, Rodrigo Fasce, Jorge Fernandez |
| EPI_ISL_445372 | HOSPITAL FF.AA. "CIRUJANO C. GUZMAN | Instituto de Salud Publica de Chile | Andrés E Castillo, Bárbara Parra,Paz Tapia, Jaime Lagos, Loredana Arata, Alejandra Acevedo, Winston Andrade, Gabriel Leal, Carolina Tambley, Patricia Bustos, Rodrigo Fasce, Jorge Fernandez |
| EPI_ISL_445373, EPI_ISL_445374, EPI_ISL_445375, EPI_ISL_445376, EPI_ISL_445377 | HOSPITAL SAN JUAN DE DIOS | Instituto de Salud Publica de Chile | Andrés E Castillo, Bárbara Parra,Paz Tapia, Jaime Lagos, Loredana Arata, Alejandra Acevedo, Winston Andrade, Gabriel Leal, Carolina Tambley, Patricia Bustos, Rodrigo Fasce, Jorge Fernandez |
| EPI_ISL_445378 | HOSPITAL DE BULNES | Instituto de Salud Publica de Chile | Andrés E Castillo, Bárbara Parra,Paz Tapia, Jaime Lagos, Loredana Arata, Alejandra Acevedo, Winston Andrade, Gabriel Leal, Carolina Tambley, Patricia Bustos, Rodrigo Fasce, Jorge Fernandez |
| EPI_ISL_445379 | IMALAB- HOSPITAL FACH | Instituto de Salud Publica de Chile | Andrés E Castillo, Bárbara Parra,Paz Tapia, Jaime Lagos, Loredana Arata, Alejandra Acevedo, Winston Andrade, Gabriel Leal, Carolina Tambley, Patricia Bustos, Rodrigo Fasce, Jorge Fernandez |
| EPI_ISL_445380 | Ramathibodi Hospital | COVID-19 Network Investigations (CONI) Alliance | Elizabeth Batty, Wasun Chantratita, Thanat Chookajorn, Stefan Fernandez, Angkana Huang, Anthony R. Jones, Khajohn Joonsalak, Chonticha Klungtong, Theerarat Kochakarn, Namfon Kotanan, Krittikorn Kumpornsin, Wudtichai Manasatienkij, Bhakbhoom Panthan, Ekawat Pasomsub, Kingkan Rakmanee, Insee Sensorn, Janjira Thaipadungpanit, Arporn Wangwiwatsin, Treewat Watthanachockchai |

EPI_ISL_445381, EPI_ISL_445382, EPI_ISL_445383, EPI_ISL_445384, EPI_ISL_445385, EPI_ISL_445386, EPI_ISL_445387, EPI_ISL_445388, EPI_ISL_445389, EPI_ISL_445390, EPI_ISL_445391, EPI_ISL_445392, EPI_ISL_445393, EPI_ISL_445394, EPI_ISL_445395, EPI_ISL_445396, EPI_ISL_445397, EPI_ISL_445398, EPI_ISL_445399, EPI_ISL_445400, EPI_ISL_445401, EPI_ISL_445402, EPI_ISL_445403, EPI_ISL_445404,
EPI_ISL_445405, EPI_ISL_445406, EPI_ISL_445407, EPI_ISL_445408, EPI_ISL_445409, EPI_ISL_445410, EPI_ISL_445411, EPI_ISL_445412, EPI_ISL_445413, EPI_ISL_445414, EPI_ISL_445415, EPI_ISL_445416, EPI_ISL_445417, EPI_ISL_445418, EPI_ISL_445419, EPI_ISL_445420, EPI_ISL_445421, EPI_ISL_445422, EPI_ISL_445423, EPI_ISL_445424, EPI_ISL_445425, EPI_ISL_445426, EPI_ISL_445427, EPI_ISL_445428,
EPI_ISL_445429, EPI_ISL_445430, EPI_ISL_445431, EPI_ISL_445432, EPI_ISL_445433, EPI_ISL_445434, EPI_ISL_445435, EPI_ISL_445436, EPI_ISL_445437, EPI_ISL_445438, EPI_ISL_445439, EPI_ISL_445440, EPI_ISL_445441, EPI_ISL_445442, EPI_ISL_445443, EPI_ISL_445444, EPI_ISL_445445, EPI_ISL_445446, EPI_ISL_445447, EPI_ISL_445448, EPI_ISL_445449, EPI_ISL_445450, EPI_ISL_445451, EPI_ISL_445452,
EPI_ISL_445453, EPI_ISL_445454, EPI_ISL_445455, EPI_ISL_445456, EPI_ISL_445457, EPI_ISL_445458, EPI_ISL_445459, EPI_ISL_445460, EPI_ISL_445461, EPI_ISL_445462, EPI_ISL_445463, EPI_ISL_445464, EPI_ISL_445465, EPI_ISL_445466, EPI_ISL_445467, EPI_ISL_445468, EPI_ISL_445469, EPI_ISL_445470, EPI_ISL_445471, EPI_ISL_445472, EPI_ISL_445473, EPI_ISL_445474, EPI_ISL_445475, EPI_ISL_445476,
EPI_ISL_445477, EPI_ISL_445478, EPI_ISL_445479, EPI_ISL_445480, EPI_ISL_445481, EPI_ISL_445482, EPI_ISL_445483, EPI_ISL_445484, EPI_ISL_445485, EPI_ISL_445486, EPI_ISL_445487, EPI_ISL_445488, EPI_ISL_445489, EPI_ISL_445490, EPI_ISL_445491, EPI_ISL_445492, EPI_ISL_445493, EPI_ISL_445494, EPI_ISL_445495, EPI_ISL_445496, EPI_ISL_445497, EPI_ISL_445498, EPI_ISL_445499, EPI_ISL_445500,
EPI_ISL_445501, EPI_ISL_445502, EPI_ISL_445503, EPI_ISL_445504, EPI_ISL_445505, EPI_ISL_445506, EPI_ISL_445507, EPI_ISL_445508, EPI_ISL_445509, EPI_ISL_445510, EPI_ISL_445511, EPI_ISL_445512, EPI_ISL_445513, EPI_ISL_445514, EPI_ISL_445515, EPI_ISL_445516, EPI_ISL_445517, EPI_ISL_445518, EPI_ISL_445519, EPI_ISL_445520, EPI_ISL_445521, EPI_ISL_445522, EPI_ISL_445523, EPI_ISL_445524,
EPI_ISL_445525, EPI_ISL_445526, EPI_ISL_445527, EPI_ISL_445528, EPI_ISL_445529, EPI_ISL_445530, EPI_ISL_445531, EPI_ISL_445532, EPI_ISL_445533, EPI_ISL_445534, EPI_ISL_445535, EPI_ISL_445536, EPI_ISL_445537, EPI_ISL_445538, EPI_ISL_445539, EPI_ISL_445540, EPI_ISL_445541, EPI_ISL_445542, EPI_ISL_445543, EPI_ISL_445544, EPI_ISL_445545, EPI_ISL_445546, EPI_ISL_445547, EPI_ISL_445548,
EPI_ISL_445549, EPI_ISL_445550, EPI_ISL_445551, EPI_ISL_445552, EPI_ISL_445553, EPI_ISL_445554, EPI_ISL_445555, EPI_ISL_445556, EPI_ISL_445557, EPI_ISL_445558, EPI_ISL_445559, EPI_ISL_445560, EPI_ISL_445561, EPI_ISL_445562, EPI_ISL_445563, EPI_ISL_445564, EPI_ISL_445565, EPI_ISL_445566, EPI_ISL_445567, EPI_ISL_445568, EPI_ISL_445569, EPI_ISL_445570, EPI_ISL_445571, EPI_ISL_445572,
EPI_ISL_445573, EPI_ISL_445574, EPI_ISL_445575, EPI_ISL_445576, EPI_ISL_445577, EPI_ISL_445578, EPI_ISL_445579, EPI_ISL_445580, EPI_ISL_445581, EPI_ISL_445582, EPI_ISL_445583, EPI_ISL_445584, EPI_ISL_445585, EPI_ISL_445586, EPI_ISL_445587, EPI_ISL_445588, EPI_ISL_445589, EPI_ISL_445590, EPI_ISL_445591, EPI_ISL_445592, EPI_ISL_445593, EPI_ISL_445594, EPI_ISL_445595, EPI_ISL_445596,
EPI_ISL_445597, EPI_ISL_445598, EPI_ISL_445599, EPI_ISL_445600, EPI_ISL_445601, EPI_ISL_445602, EPI_ISL_445603, EPI_ISL_445604, EPI_ISL_445605, EPI_ISL_445606, EPI_ISL_445607, EPI_ISL_445608, EPI_ISL_445609, EPI_ISL_445610, EPI_ISL_445611, EPI_ISL_445612, EPI_ISL_445613, EPI_ISL_445614, EPI_ISL_445615, EPI_ISL_445616, EPI_ISL_445617, EPI_ISL_445618, EPI_ISL_445619, EPI_ISL_445620,
EPI_ISL_445621, EPI_ISL_445622, EPI_ISL_445623, EPI_ISL_445624, EPI_ISL_445625, EPI_ISL_445626, EPI_ISL_445627, EPI_ISL_445628, EPI_ISL_445629, EPI_ISL_445630, EPI_ISL_445631, EPI_ISL_445632, EPI_ISL_445633, EPI_ISL_445634, EPI_ISL_445635, EPI_ISL_445636, EPI_ISL_445637, EPI_ISL_445638, EPI_ISL_445639, EPI_ISL_445640, EPI_ISL_445641, EPI_ISL_445642, EPI_ISL_445643, EPI_ISL_445644,
EPI_ISL_445645, EPI_ISL_445646, EPI_ISL_445647, EPI_ISL_445648, EPI_ISL_445649, EPI_ISL_445650, EPI_ISL_445651, EPI_ISL_445652, EPI_ISL_445653, EPI_ISL_445654, EPI_ISL_445655, EPI_ISL_445656, EPI_ISL_445657, EPI_ISL_445658, EPI_ISL_445659, EPI_ISL_445660, EPI_ISL_445661, EPI_ISL_445662, EPI_ISL_445663, EPI_ISL_445664, EPI_ISL_445665, EPI_ISL_445666, EPI_ISL_445667, EPI_ISL_445668,
EPI_ISL_445669, EPI_ISL_445670, EPI_ISL_445671, EPI_ISL_445672, EPI_ISL_445673, EPI_ISL_445674, EPI_ISL_445675, EPI_ISL_445676, EPI_ISL_445677, EPI_ISL_445678, EPI_ISL_445679, EPI_ISL_445680, EPI_ISL_445681, EPI_ISL_445682, EPI_ISL_445683, EPI_ISL_445684, EPI_ISL_445685, EPI_ISL_445686, EPI_ISL_445687, EPI_ISL_445688, EPI_ISL_445689, EPI_ISL_445690, EPI_ISL_445691, EPI_ISL_445692,
EPI_ISL_445693, EPI_ISL_445694, EPI_ISL_445695, EPI_ISL_445696, EPI_ISL_445697, EPI_ISL_445698, EPI_ISL_445699, EPI_ISL_445700, EPI_ISL_445701, EPI_ISL_445702, EPI_ISL_445703, EPI_ISL_445704, EPI_ISL_445705, EPI_ISL_445706, EPI_ISL_445707, EPI_ISL_445708, EPI_ISL_445709, EPI_ISL_445710, EPI_ISL_445711, EPI_ISL_445712, EPI_ISL_445713, EPI_ISL_445714, EPI_ISL_445715, EPI_ISL_445716,
EPI_ISL_445717, EPI_ISL_445718, EPI_ISL_445719, EPI_ISL_445720, EPI_ISL_445721, EPI_ISL_445722, EPI_ISL_445723, EPI_ISL_445724, EPI_ISL_445725, EPI_ISL_445726, EPI_ISL_445727, EPI_ISL_445728, EPI_ISL_445729, EPI_ISL_445730, EPI_ISL_445731, EPI_ISL_445732, EPI_ISL_445733, EPI_ISL_445734, EPI_ISL_445735, EPI_ISL_445736, EPI_ISL_445737, EPI_ISL_445738, EPI_ISL_445739, EPI_ISL_445740,
EPI_ISL_445741, EPI_ISL_445742, EPI_ISL_445743, EPI_ISL_445744, EPI_ISL_445745, EPI_ISL_445746, EPI_ISL_445747, EPI_ISL_445748, EPI_ISL_445749, EPI_ISL_445750, EPI_ISL_445751, EPI_ISL_445752, EPI_ISL_445753, EPI_ISL_445754, EPI_ISL_445755, EPI_ISL_445756, EPI_ISL_445757, EPI_ISL_445758, EPI_ISL_445759, EPI_ISL_445760, EPI_ISL_445761, EPI_ISL_445762, EPI_ISL_445763, EPI_ISL_445764,
EPI_ISL_445765, EPI_ISL_445766, EPI_ISL_445767, EPI_ISL_445768, EPI_ISL_445769, EPI_ISL_445770, EPI_ISL_445771, EPI_ISL_445772, EPI_ISL_445773, EPI_ISL_445774, EPI_ISL_445775, EPI_ISL_445776, EPI_ISL_445777, EPI_ISL_445778, EPI_ISL_445779, EPI_ISL_445780, EPI_ISL_445781, EPI_ISL_445782, EPI_ISL_445783, EPI_ISL_445784, EPI_ISL_445785, EPI_ISL_445786, EPI_ISL_445787, EPI_ISL_445788,
EPI_ISL_445789, EPI_ISL_445790, EPI_ISL_445791, EPI_ISL_445792, EPI_ISL_445793, EPI_ISL_445794, EPI_ISL_445795, EPI_ISL_445796, EPI_ISL_445797, EPI_ISL_445798, EPI_ISL_445799, EPI_ISL_445800, EPI_ISL_445801, EPI_ISL_445802, EPI_ISL_445803, EPI_ISL_445804, EPI_ISL_445805, EPI_ISL_445806, EPI_ISL_445807, EPI_ISL_445808, EPI_ISL_445809, EPI_ISL_445810, EPI_ISL_445811, EPI_ISL_445812,
EPI_ISL_445813, EPI_ISL_445814, EPI_ISL_445815, EPI_ISL_445816, EPI_ISL_445817, EPI_ISL_445818, EPI_ISL_445819, EPI_ISL_445820, EPI_ISL_445821, EPI_ISL_445822, EPI_ISL_445823, EPI_ISL_445824, EPI_ISL_445825, EPI_ISL_445826, EPI_ISL_445827, EPI_ISL_445828, EPI_ISL_445829, EPI_ISL_445830, EPI_ISL_445831, EPI_ISL_445832, EPI_ISL_445833, EPI_ISL_445834, EPI_ISL_445835, EPI_ISL_445836,
EPI_ISL_445837, EPI_ISL_445838, EPI_ISL_445839, EPI_ISL_445840, EPI_ISL_445841, EPI_ISL_445842, EPI_ISL_445843, EPI_ISL_445844, EPI_ISL_445845, EPI_ISL_445846, EPI_ISL_445847, EPI_ISL_445848, EPI_ISL_445849, EPI_ISL_445850, EPI_ISL_445851, EPI_ISL_445852, EPI_ISL_445853, EPI_ISL_445854, EPI_ISL_445855, EPI_ISL_445856, EPI_ISL_445857, EPI_ISL_445858, EPI_ISL_445859, EPI_ISL_445860,
EPI_ISL_445861, EPI_ISL_445862, EPI_ISL_445863, EPI_ISL_445864, EPI_ISL_445865, EPI_ISL_445866, EPI_ISL_445867, EPI_ISL_445868, EPI_ISL_445869, EPI_ISL_445870, EPI_ISL_445871, EPI_ISL_445872, EPI_ISL_445873, EPI_ISL_445874, EPI_ISL_445875, EPI_ISL_445876, EPI_ISL_445877, EPI_ISL_445878, EPI_ISL_445879, EPI_ISL_445880, EPI_ISL_445881, EPI_ISL_445882, EPI_ISL_445883, EPI_ISL_445884,
EPI_ISL_445885, EPI_ISL_445886, EPI_ISL_445887, EPI_ISL_445888, EPI_ISL_445889, EPI_ISL_445890, EPI_ISL_445891, EPI_ISL_445892, EPI_ISL_445893, EPI_ISL_445894, EPI_ISL_445895, EPI_ISL_445896, EPI_ISL_445897, EPI_ISL_445898, EPI_ISL_445899, EPI_ISL_445900, EPI_ISL_445901, EPI_ISL_445902, EPI_ISL_445903, EPI_ISL_445904, EPI_ISL_445905, EPI_ISL_445906, EPI_ISL_445907, EPI_ISL_445908,
EPI_ISL_445909, EPI_ISL_445910, EPI_ISL_445911, EPI_ISL_445912, EPI_ISL_445913, EPI_ISL_445914, EPI_ISL_445915, EPI_ISL_445916, EPI_ISL_445917, EPI_ISL_445918, EPI_ISL_445919, EPI_ISL_445920, EPI_ISL_445921, EPI_ISL_445922, EPI_ISL_445923, EPI_ISL_445924, EPI_ISL_445925, EPI_ISL_445926, EPI_ISL_445927, EPI_ISL_445928, EPI_ISL_445929, EPI_ISL_445930, EPI_ISL_445931, EPI_ISL_445932,
EPI_ISL_445933, EPI_ISL_445934, EPI_ISL_445935, EPI_ISL_445936, EPI_ISL_445937, EPI_ISL_445938, EPI_ISL_445939, EPI_ISL_445940, EPI_ISL_445941, EPI_ISL_445942, EPI_ISL_445943, EPI_ISL_445944, EPI_ISL_445945, EPI_ISL_445946, EPI_ISL_445947, EPI_ISL_445948, EPI_ISL_445949, EPI_ISL_445950, EPI_ISL_445951, EPI_ISL_445952, EPI_ISL_445953, EPI_ISL_445954, EPI_ISL_445955, EPI_ISL_445956,
EPI_ISL_445957, EPI_ISL_445958, EPI_ISL_445959, EPI_ISL_445960, EPI_ISL_445961, EPI_ISL_445962, EPI_ISL_445963, EPI_ISL_445964, EPI_ISL_445965, EPI_ISL_445966, EPI_ISL_445967, EPI_ISL_445968, EPI_ISL_445969, EPI_ISL_445970, EPI_ISL_445971, EPI_ISL_445972, EPI_ISL_445973, EPI_ISL_445974, EPI_ISL_445975, EPI_ISL_445976, EPI_ISL_445977, EPI_ISL_445978, EPI_ISL_445979, EPI_ISL_445980,
EPI_ISL_445981, EPI_ISL_445982, EPI_ISL_445983, EPI_ISL_445984, EPI_ISL_445985, EPI_ISL_445986, EPI_ISL_445987, EPI_ISL_445988, EPI_ISL_445989, EPI_ISL_445990, EPI_ISL_445991, EPI_ISL_445992, EPI_ISL_445993, EPI_ISL_445994, EPI_ISL_445995, EPI_ISL_445996, EPI_ISL_445997, EPI_ISL_445998, EPI_ISL_445999, EPI_ISL_446000, EPI_ISL_446001, EPI_ISL_446002, EPI_ISL_446003, EPI_ISL_446004,
EPI_ISL_446005, EPI_ISL_446006, EPI_ISL_446007, EPI_ISL_446008, EPI_ISL_446009, EPI_ISL_446010, EPI_ISL_446011, EPI_ISL_446012, EPI_ISL_446013, EPI_ISL_446014, EPI_ISL_446015, EPI_ISL_446016, EPI_ISL_446017, EPI_ISL_446018, EPI_ISL_446019, EPI_ISL_446020, EPI_ISL_446021, EPI_ISL_446022, EPI_ISL_446023, EPI_ISL_446024, EPI_ISL_446025, EPI_ISL_446026, EPI_ISL_446027, EPI_ISL_446028,
EPI_ISL_446029, EPI_ISL_446030, EPI_ISL_446031, EPI_ISL_446032, EPI_ISL_446033, EPI_ISL_446034, EPI_ISL_446035, EPI_ISL_446036, EPI_ISL_446037, EPI_ISL_446038, EPI_ISL_446039, EPI_ISL_446040, EPI_ISL_446041, EPI_ISL_446042, EPI_ISL_446043, EPI_ISL_446044, EPI_ISL_446045, EPI_ISL_446046, EPI_ISL_446047, EPI_ISL_446048, EPI_ISL_446049, EPI_ISL_446050, EPI_ISL_446051, EPI_ISL_446052,
EPI_ISL_446053, EPI_ISL_446054, EPI_ISL_446055, EPI_ISL_446056, EPI_ISL_446057, EPI_ISL_446058, EPI_ISL_446059, EPI_ISL_446060, EPI_ISL_446061, EPI_ISL_446062, EPI_ISL_446063, EPI_ISL_446064, EPI_ISL_446065, EPI_ISL_446066, EPI_ISL_446067, EPI_ISL_446068, EPI_ISL_446069, EPI_ISL_446070, EPI_ISL_446071, EPI_ISL_446072, EPI_ISL_446073, EPI_ISL_446074, EPI_ISL_446075, EPI_ISL_446076,
EPI_ISL_446077, EPI_ISL_446078, EPI_ISL_446079, EPI_ISL_446080, EPI_ISL_446081, EPI_ISL_446082, EPI_ISL_446083, EPI_ISL_446084, EPI_ISL_446085, EPI_ISL_446086, EPI_ISL_446087, EPI_ISL_446088, EPI_ISL_446089, EPI_ISL_446090, EPI_ISL_446091, EPI_ISL_446092, EPI_ISL_446093, EPI_ISL_446094, EPI_ISL_446095, EPI_ISL_446096, EPI_ISL_446097, EPI_ISL_446098, EPI_ISL_446099, EPI_ISL_446100,
EPI_ISL_446101, EPI_ISL_446102, EPI_ISL_446103, EPI_ISL_446104, EPI_ISL_446105, EPI_ISL_446106, EPI_ISL_446107, EPI_ISL_446108, EPI_ISL_446109, EPI_ISL_446110, EPI_ISL_446111, EPI_ISL_446112, EPI_ISL_446113, EPI_ISL_446114, EPI_ISL_446115, EPI_ISL_446116, EPI_ISL_446117, EPI_ISL_446118, EPI_ISL_446119, EPI_ISL_446120, EPI_ISL_446121, EPI_ISL_446122, EPI_ISL_446123, EPI_ISL_446124,
EPI_ISL_446125, EPI_ISL_446126, EPI_ISL_446127, EPI_ISL_446128, EPI_ISL_446129, EPI_ISL_446130, EPI_ISL_446131, EPI_ISL_446132, EPI_ISL_446133, EPI_ISL_446134, EPI_ISL_446135, EPI_ISL_446136, EPI_ISL_446137, EPI_ISL_446138, EPI_ISL_446139, EPI_ISL_446140, EPI_ISL_446141, EPI_ISL_446142, EPI_ISL_446143, EPI_ISL_446144, EPI_ISL_446145, EPI_ISL_446146, EPI_ISL_446147, EPI_ISL_446148,
EPI_ISL_446149, EPI_ISL_446150, EPI_ISL_446151, EPI_ISL_446152, EPI_ISL_446153, EPI_ISL_446154, EPI_ISL_446155, EPI_ISL_446156, EPI_ISL_446157, EPI_ISL_446158, EPI_ISL_446159, EPI_ISL_446160, EPI_ISL_446161, EPI_ISL_446162, EPI_ISL_446163, EPI_ISL_446164, EPI_ISL_446165, EPI_ISL_446166, EPI_ISL_446167, EPI_ISL_446168, EPI_ISL_446169, EPI_ISL_446170, EPI_ISL_446171, EPI_ISL_446172,
EPI_ISL_446173, EPI_ISL_446174, EPI_ISL_446175, EPI_ISL_446176, EPI_ISL_446177, EPI_ISL_446178, EPI_ISL_446179, EPI_ISL_446180, EPI_ISL_446181, EPI_ISL_446182, EPI_ISL_446183, EPI_ISL_446184, EPI_ISL_446185, EPI_ISL_446186, EPI_ISL_446187, EPI_ISL_446188, EPI_ISL_446189, EPI_ISL_446190, EPI_ISL_446191, EPI_ISL_446192, EPI_ISL_446193, EPI_ISL_446194, EPI_ISL_446195, EPI_ISL_446196,
EPI_ISL_446197, EPI_ISL_446198, EPI_ISL_446199, EPI_ISL_446200, EPI_ISL_446201, EPI_ISL_446202, EPI_ISL_446203, EPI_ISL_446204, EPI_ISL_446205, EPI_ISL_446206, EPI_ISL_446207, EPI_ISL_446208, EPI_ISL_446209, EPI_ISL_446210, EPI_ISL_446211, EPI_ISL_446212, EPI_ISL_446213, EPI_ISL_446214, EPI_ISL_446215, EPI_ISL_446216, EPI_ISL_446217, EPI_ISL_446218, EPI_ISL_446219, EPI_ISL_446220,
EPI_ISL_446221, EPI_ISL_446222, EPI_ISL_446223, EPI_ISL_446224, EPI_ISL_446225, EPI_ISL_446226, EPI_ISL_446227, EPI_ISL_446228, EPI_ISL_446229, EPI_ISL_446230, EPI_ISL_446231, EPI_ISL_446232, EPI_ISL_446233, EPI_ISL_446234, EPI_ISL_446235, EPI_ISL_446236, EPI_ISL_446237, EPI_ISL_446238, EPI_ISL_446239, EPI_ISL_446240, EPI_ISL_446241, EPI_ISL_446242, EPI_ISL_446243, EPI_ISL_446244,
EPI_ISL_446245, EPI_ISL_446246, EPI_ISL_446247, EPI_ISL_446248, EPI_ISL_446249, EPI_ISL_446250, EPI_ISL_446251, EPI_ISL_446252, EPI_ISL_446253, EPI_ISL_446254, EPI_ISL_446255, EPI_ISL_446256, EPI_ISL_446257, EPI_ISL_446258, EPI_ISL_446259, EPI_ISL_446260, EPI_ISL_446261, EPI_ISL_446262, EPI_ISL_446263, EPI_ISL_446264, EPI_ISL_446265, EPI_ISL_446266, EPI_ISL_446267, EPI_ISL_446268,
EPI_ISL_446269, EPI_ISL_446270, EPI_ISL_446271, EPI_ISL_446272, EPI_ISL_446273, EPI_ISL_446274, EPI_ISL_446275, EPI_ISL_446276, EPI_ISL_446277, EPI_ISL_446278, EPI_ISL_446279, EPI_ISL_446280, EPI_ISL_446281, EPI_ISL_446282, EPI_ISL_446283, EPI_ISL_446284, EPI_ISL_446285, EPI_ISL_446286, EPI_ISL_446287, EPI_ISL_446288, EPI_ISL_446289, EPI_ISL_446290, EPI_ISL_446291, EPI_ISL_446292,
EPI_ISL_446293, EPI_ISL_446294, EPI_ISL_446295, EPI_ISL_446296, EPI_ISL_446297, EPI_ISL_446298, EPI_ISL_446299, EPI_ISL_446300, EPI_ISL_446301, EPI_ISL_446302, EPI_ISL_446303, EPI_ISL_446304, EPI_ISL_446305, EPI_ISL_446306, EPI_ISL_446307, EPI_ISL_446308, EPI_ISL_446309, EPI_ISL_446310, EPI_ISL_446311, EPI_ISL_446312, EPI_ISL_446313, EPI_ISL_446314, EPI_ISL_446315, EPI_ISL_446316,
EPI_ISL_446317, EPI_ISL_446318, EPI_ISL_446319, EPI_ISL_446320, EPI_ISL_446321, EPI_ISL_446322, EPI_ISL_446323, EPI_ISL_446324, EPI_ISL_446325, EPI_ISL_446326, EPI_ISL_446327, EPI_ISL_446328, EPI_ISL_446329, EPI_ISL_446330, EPI_ISL_446331, EPI_ISL_446332, EPI_ISL_446333, EPI_ISL_446334, EPI_ISL_446335, EPI_ISL_446336, EPI_ISL_446337, EPI_ISL_446338, EPI_ISL_446339, EPI_ISL_446340,
EPI_ISL_446341, EPI_ISL_446342, EPI_ISL_446343, EPI_ISL_446344, EPI_ISL_446345, EPI_ISL_446346, EPI_ISL_446347, EPI_ISL_446348, EPI_ISL_446349, EPI_ISL_446350, EPI_ISL_446351, EPI_ISL_446352, EPI_ISL_446353, EPI_ISL_446354, EPI_ISL_446355, EPI_ISL_446356, EPI_ISL_446357, EPI_ISL_446358, EPI_ISL_446359, EPI_ISL_446360, EPI_ISL_446361, EPI_ISL_446362, EPI_ISL_446363, EPI_ISL_446364,
EPI_ISL_446365, EPI_ISL_446366, EPI_ISL_446367, EPI_ISL_446368, EPI_ISL_446369, EPI_ISL_446370, EPI_ISL_446371, EPI_ISL_446372, EPI_ISL_446373, EPI_ISL_446374, EPI_ISL_446375, EPI_ISL_446376, EPI_ISL_446377, EPI_ISL_446378, EPI_ISL_446379, EPI_ISL_446380, EPI_ISL_446381, EPI_ISL_446382, EPI_ISL_446383, EPI_ISL_446384, EPI_ISL_446385, EPI_ISL_446386, EPI_ISL_446387, EPI_ISL_446388,
EPI_ISL_446389, EPI_ISL_446390, EPI_ISL_446391, EPI_ISL_446392, EPI_ISL_446393, EPI_ISL_446394, EPI_ISL_446395, EPI_ISL_446396, EPI_ISL_446397, EPI_ISL_446398, EPI_ISL_446399, EPI_ISL_446400, EPI_ISL_446401, EPI_ISL_446402, EPI_ISL_446403, EPI_ISL_446404, EPI_ISL_446405, EPI_ISL_446406, EPI_ISL_446407, EPI_ISL_446408, EPI_ISL_446409, EPI_ISL_446410, EPI_ISL_446411, EPI_ISL_446412,
EPI_ISL_446413, EPI_ISL_446414, EPI_ISL_446415, EPI_ISL_446416, EPI_ISL_446417, EPI_ISL_446418, EPI_ISL_446419, EPI_ISL_446420, EPI_ISL_446421, EPI_ISL_446422, EPI_ISL_446423, EPI_ISL_446424, EPI_ISL_446425, EPI_ISL_446426, EPI_ISL_446427, EPI_ISL_446428, EPI_ISL_446429, EPI_ISL_446430, EPI_ISL_446431, EPI_ISL_446432, EPI_ISL_446433, EPI_ISL_446434, EPI_ISL_446435, EPI_ISL_446436,
EPI_ISL_446437, EPI_ISL_446438, EPI_ISL_446439, EPI_ISL_446440, EPI_ISL_446441, EPI_ISL_446442, EPI_ISL_446443, EPI_ISL_446444, EPI_ISL_446445, EPI_ISL_446446, EPI_ISL_446447, EPI_ISL_446448, EPI_ISL_446449, EPI_ISL_446450, EPI_ISL_446451, EPI_ISL_446452, EPI_ISL_446453, EPI_ISL_446454, EPI_ISL_446455, EPI_ISL_446456, EPI_ISL_446457, EPI_ISL_446458, EPI_ISL_446459, EPI_ISL_446460,
EPI_ISL_446461, EPI_ISL_446462, EPI_ISL_446463, EPI_ISL_446464, EPI_ISL_446465, EPI_ISL_446466, EPI_ISL_446467, EPI_ISL_446468, EPI_ISL_446469, EPI_ISL_446470, EPI_ISL_446471, EPI_ISL_446472, EPI_ISL_446473, EPI_ISL_446474, EPI_ISL_446475, EPI_ISL_446476, EPI_ISL_446477, EPI_ISL_446478, EPI_ISL_446479, EPI_ISL_446480, EPI_ISL_446481, EPI_ISL_446482, EPI_ISL_446483, EPI_ISL_446484,
EPI_ISL_446485, EPI_ISL_446486, EPI_ISL_446487, EPI_ISL_446488, EPI_ISL_446489, EPI_ISL_446490, EPI_ISL_446491, EPI_ISL_446492, EPI_ISL_446493, EPI_ISL_446494, EPI_ISL_446495, EPI_ISL_446496, EPI_ISL_446497, EPI_ISL_446498, EPI_ISL_446499, EPI_ISL_446500, EPI_ISL_446501, EPI_ISL_446502, EPI_ISL_446503, EPI_ISL_446504, EPI_ISL_446505, EPI_ISL_446506, EPI_ISL_446507, EPI_ISL_446508,
EPI_ISL_446509, EPI_ISL_446510, EPI_ISL_446511, EPI_ISL_446512, EPI_ISL_446513, EPI_ISL_446514, EPI_ISL_446515, EPI_ISL_446516, EPI_ISL_446517, EPI_ISL_446518, EPI_ISL_446519, EPI_ISL_446520, EPI_ISL_446521, EPI_ISL_446522, EPI_ISL_446523, EPI_ISL_446524, EPI_ISL_446525, EPI_ISL_446526, EPI_ISL_446527, EPI_ISL_446528, EPI_ISL_446529, EPI_ISL_446530, EPI_ISL_446531, EPI_ISL_446532,
EPI_ISL_446533, EPI_ISL_446534, EPI_ISL_446535, EPI_ISL_446536, EPI_ISL_446537, EPI_ISL_446538, EPI_ISL_446539, EPI_ISL_446540, EPI_ISL_446541, EPI_ISL_446542, EPI_ISL_446543, EPI_ISL_446544, EPI_ISL_446545, EPI_ISL_446546, EPI_ISL_446547, EPI_ISL_446548, EPI_ISL_446549, EPI_ISL_446550, EPI_ISL_446551, EPI_ISL_446552, EPI_ISL_446553, EPI_ISL_446554, EPI_ISL_446555, EPI_ISL_446556,
EPI_ISL_446557, EPI_ISL_446558, EPI_ISL_446559, EPI_ISL_446560, EPI_ISL_446561, EPI_ISL_446562, EPI_ISL_446563, EPI_ISL_446564, EPI_ISL_446565, EPI_ISL_446566, EPI_ISL_446567, EPI_ISL_446568, EPI_ISL_446569, EPI_ISL_446570, EPI_ISL_446571, EPI_ISL_446572, EPI_ISL_446573, EPI_ISL_446574, EPI_ISL_446575, EPI_ISL_446576, EPI_ISL_446577, EPI_ISL_446578, EPI_ISL_446579, EPI_ISL_446580,
EPI_ISL_446581, EPI_ISL_446582, EPI_ISL_446583, EPI_ISL_446584, EPI_ISL_446585, EPI_ISL_446586, EPI_ISL_446587, EPI_ISL_446588, EPI_ISL_446589, EPI_ISL_446590, EPI_ISL_446591, EPI_ISL_446592, EPI_ISL_446593, EPI_ISL_446594, EPI_ISL_446595, EPI_ISL_446596, EPI_ISL_446597, EPI_ISL_446598, EPI_ISL_446599, EPI_ISL_446600, EPI_ISL_446601, EPI_ISL_446602, EPI_ISL_446603, EPI_ISL_446604,
EPI_ISL_446605, EPI_ISL_446606, EPI_ISL_446607, EPI_ISL_446608, EPI_ISL_446609, EPI_ISL_446610, EPI_ISL_446611, EPI_ISL_446612, EPI_ISL_446613, EPI_ISL_446614, EPI_ISL_446615, EPI_ISL_446616, EPI_ISL_446617, EPI_ISL_446618, EPI_ISL_446619, EPI_ISL_446620, EPI_ISL_446621, EPI_ISL_446622, EPI_ISL_446623, EPI_ISL_446624, EPI_ISL_446625, EPI_ISL_446626, EPI_ISL_446627, EPI_ISL_446628,
EPI_ISL_446629, EPI_ISL_446630, EPI_ISL_446631, EPI_ISL_446632, EPI_ISL_446633, EPI_ISL_446634, EPI_ISL_446635, EPI_ISL_446636, EPI_ISL_446637, EPI_ISL_446638, EPI_ISL_446639, EPI_ISL_446640, EPI_ISL_446641, EPI_ISL_446642, EPI_ISL_446643, EPI_ISL_446644, EPI_ISL_446645, EPI_ISL_446646, EPI_ISL_446647, EPI_ISL_446648, EPI_ISL_446649, EPI_ISL_446650, EPI_ISL_446651, EPI_ISL_446652,
EPI_ISL_446653, EPI_ISL_446654, EPI_ISL_446655, EPI_ISL_446656, EPI_ISL_446657, EPI_ISL_446658, EPI_ISL_446659, EPI_ISL_446660, EPI_ISL_446661, EPI_ISL_446662, EPI_ISL_446663, EPI_ISL_446664, EPI_ISL_446665, EPI_ISL_446666, EPI_ISL_446667, EPI_ISL_446668, EPI_ISL_446669, EPI_ISL_446670, EPI_ISL_446671, EPI_ISL_446672, EPI_ISL_446673, EPI_ISL_446674, EPI_ISL_446675, EPI_ISL_446676,
EPI_ISL_446677, EPI_ISL_446678, EPI_ISL_446679, EPI_ISL_446680, EPI_ISL_446681, EPI_ISL_446682, EPI_ISL_446683, EPI_ISL_446684, EPI_ISL_446685, EPI_ISL_446686, EPI_ISL_446687, EPI_ISL_446688, EPI_ISL_446689, EPI_ISL_446690, EPI_ISL_446691, EPI_ISL_446692, EPI_ISL_446693, EPI_ISL_446694, EPI_ISL_446695, EPI_ISL_446696, EPI_ISL_446697, EPI_ISL_446698, EPI_ISL_446699, EPI_ISL_446700,
EPI_ISL_446701, EPI_ISL_446702, EPI_ISL_446703, EPI_ISL_446704, EPI_ISL_446705, EPI_ISL_446706, EPI_ISL_446707, EPI_ISL_446708, EPI_ISL_446709, EPI_ISL_446710, EPI_ISL_446711, EPI_ISL_446712, EPI_ISL_446713, EPI_ISL_446714, EPI_ISL_446715, EPI_ISL_446716, EPI_ISL_446717, EPI_ISL_446718, EPI_ISL_446719, EPI_ISL_446720, EPI_ISL_446721, EPI_ISL_446722, EPI_ISL_446723, EPI_ISL_446724,
EPI_ISL_446725, EPI_ISL_446726, EPI_ISL_446727, EPI_ISL_446728, EPI_ISL_446729, EPI_ISL_446730, EPI_ISL_446731, EPI_ISL_446732, EPI_ISL_446733, EPI_ISL_446734, EPI_ISL_446735, EPI_ISL_446736, EPI_ISL_446737, EPI_ISL_446738, EPI_ISL_446739, EPI_ISL_446740, EPI_ISL_446741, EPI_ISL_446742, EPI_ISL_446743, EPI_ISL_446744, EPI_ISL_446745, EPI_ISL_446746, EPI_ISL_446747, EPI_ISL_446748,
EPI_ISL_446749, EPI_ISL_446750, EPI_ISL_446751, EPI_ISL_446752, EPI_ISL_446753, EPI_ISL_446754, EPI_ISL_446755, EPI_ISL_446756, EPI_ISL_446757, EPI_ISL_446758, EPI_ISL_446759, EPI_ISL_446760, EPI_ISL_446761, EPI_ISL_446762, EPI_ISL_446763, EPI_ISL_446764, EPI_ISL_446765, EPI_ISL_446766, EPI_ISL_446767, EPI_ISL_446768, EPI_ISL_446769, EPI_ISL_446770, EPI_ISL_446771, EPI_ISL_446772,
EPI_ISL_446773, EPI_ISL_446774, EPI_ISL_446775, EPI_ISL_446776, EPI_ISL_446777, EPI_ISL_446778, EPI_ISL_446779, EPI_ISL_446780, EPI_ISL_446781, EPI_ISL_446782, EPI_ISL_446783, EPI_ISL_446784, EPI_ISL_446785, EPI_ISL_446786, EPI_ISL_446787, EPI_ISL_446788, EPI_ISL_446789, EPI_ISL_446790, EPI_ISL_446791, EPI_ISL_446792, EPI_ISL_446793, EPI_ISL_446794, EPI_ISL_446795, EPI_ISL_446796,
EPI_ISL_446797, EPI_ISL_446798, EPI_ISL_446799, EPI_ISL_446800, EPI_ISL_446801, EPI_ISL_446802, EPI_ISL_446803, EPI_ISL_446804, EPI_ISL_446805, EPI_ISL_446806, EPI_ISL_446807, EPI_ISL_446808, EPI_ISL_446809, EPI_ISL_446810, EPI_ISL_446811, EPI_ISL_446812, EPI_ISL_446813, EPI_ISL_446814, EPI_ISL_446815, EPI_ISL_446816, EPI_ISL_446817, EPI_ISL_446818, EPI_ISL_446819, EPI_ISL_446820,
EPI_ISL_446821, EPI_ISL_446822, EPI_ISL_446823, EPI_ISL_446824, EPI_ISL_446825, EPI_ISL_446826, EPI_ISL_446827, EPI_ISL_446828, EPI_ISL_446829, EPI_ISL_446830, EPI_ISL_446831, EPI_ISL_446832, EPI_ISL_446833, EPI_ISL_446834, EPI_ISL_446835, EPI_ISL_446836, EPI_ISL_446837, EPI_ISL_446838, EPI_ISL_446839, EPI_ISL_446840, EPI_ISL_446841, EPI_ISL_446842, EPI_ISL_446843, EPI_ISL_446844,
EPI_ISL_446845, EPI_ISL_446846, EPI_ISL_446847, EPI_ISL_446848, EPI_ISL_446849, EPI_ISL_446850, EPI_ISL_446851, EPI_ISL_446852, EPI_ISL_446853, EPI_ISL_446854, EPI_ISL_446855, EPI_ISL_446856, EPI_ISL_446857, EPI_ISL_446858, EPI_ISL_446859, EPI_ISL_446860, EPI_ISL_446861, EPI_ISL_446862, EPI_ISL_446863, EPI_ISL_446864, EPI_ISL_446865, EPI_ISL_446866, EPI_ISL_446867, EPI_ISL_446868,
EPI_ISL_446869, EPI_ISL_446870, EPI_ISL_446871, EPI_ISL_446872, EPI_ISL_446873, EPI_ISL_446874, EPI_ISL_446875, EPI_ISL_446876, EPI_ISL_446877, EPI_ISL_446878, EPI_ISL_446879, EPI_ISL_446880, EPI_ISL_446881, EPI_ISL_446882, EPI_ISL_446883, EPI_ISL_446884, EPI_ISL_446885, EPI_ISL_446886, EPI_ISL_446887, EPI_ISL_446888, EPI_ISL_446889, EPI_ISL_446890, EPI_ISL_446891, EPI_ISL_446892,
EPI_ISL_446893, EPI_ISL_446894, EPI_ISL_446895, EPI_ISL_446896, EPI_ISL_446897, EPI_ISL_446898, EPI_ISL_446899, EPI_ISL_446900, EPI_ISL_446901, EPI_ISL_446902, EPI_ISL_446903, EPI_ISL_446904, EPI_ISL_446905, EPI_ISL_446906, EPI_ISL_446907, EPI_ISL_446908, EPI_ISL_446909, EPI_ISL_446910, EPI_ISL_446911, EPI_ISL_446912, EPI_ISL_446913, EPI_ISL_446914, EPI_ISL_446915, EPI_ISL_446916,
EPI_ISL_446917, EPI_ISL_446918, EPI_ISL_446919, EPI_ISL_446920, EPI_ISL_446921, EPI_ISL_446922, EPI_ISL_446923, EPI_ISL_446924, EPI_ISL_446925, EPI_ISL_446926, EPI_ISL_446927, EPI_ISL_446928, EPI_ISL_446929, EPI_ISL_446930, EPI_ISL_446931, EPI_ISL_446932, EPI_ISL_446933, EPI_ISL_446934, EPI_ISL_446935, EPI_ISL_446936, EPI_ISL_446937, EPI_ISL_446938, EPI_ISL_446939, EPI_ISL_446940,
EPI_ISL_446941, EPI_ISL_446942, EPI_ISL_446943, EPI_ISL_446944, EPI_ISL_446945, EPI_ISL_446946, EPI_ISL_446947, EPI_ISL_446948, EPI_ISL_446949, EPI_ISL_446950, EPI_ISL_446951, EPI_ISL_446952, EPI_ISL_446953, EPI_ISL_446954, EPI_ISL_446955, EPI_ISL_446956, EPI_ISL_446957, EPI_ISL_446958, EPI_ISL_446959, EPI_ISL_446960, EPI_ISL_446961, EPI_ISL_446962, EPI_ISL_446963, EPI_ISL_446964,

| | | | |
|---|---|---|---|
| EPI_ISL_446965, EPI_ISL_446966, EPI_ISL_446967, EPI_ISL_446968, EPI_ISL_446969, EPI_ISL_446970, EPI_ISL_446971, EPI_ISL_446972, EPI_ISL_446973, EPI_ISL_446974, EPI_ISL_446975, EPI_ISL_446976, EPI_ISL_446977, EPI_ISL_446978, EPI_ISL_446979, EPI_ISL_446980, EPI_ISL_446981, EPI_ISL_446982, EPI_ISL_446983, EPI_ISL_446984, EPI_ISL_446985, EPI_ISL_446986, EPI_ISL_446987, EPI_ISL_446988, EPI_ISL_446989, EPI_ISL_446990, EPI_ISL_446991, EPI_ISL_446992, EPI_ISL_446993, EPI_ISL_446994, EPI_ISL_446995 | | | |
| see above | Wales Specialist Virology Centre | Public Health Wales Microbiology Cardiff | Catherine Moore, Johnathan Evans, Laura Gifford, Malorie Perry, Simon Cottrell, Alec Birchley, Alexander Adams, Amy Gaskin, Bree Gatica-Wilcox, Jason Coombes, Lauren Gilbert, Lee Graham, Nicole Pacchiarini, Sara Kumziene-Summerhayes, Sarah Taylor, Sophie Jones, Sara Rey, Matthew Bull, Joanne Watkins, Sally Corden, Tom Connor |
| EPI_ISL_446996, EPI_ISL_446997, EPI_ISL_446998, EPI_ISL_446999, EPI_ISL_447000, EPI_ISL_447001, EPI_ISL_447002, EPI_ISL_447003, EPI_ISL_447004, EPI_ISL_447005, EPI_ISL_447006, EPI_ISL_447007, EPI_ISL_447008, EPI_ISL_447009, EPI_ISL_447010, EPI_ISL_447011, EPI_ISL_447012, EPI_ISL_447013, EPI_ISL_447014, EPI_ISL_447015, EPI_ISL_447016, EPI_ISL_447017, EPI_ISL_447018, EPI_ISL_447019, EPI_ISL_447020, EPI_ISL_447021, EPI_ISL_447022, EPI_ISL_447023, EPI_ISL_447024, EPI_ISL_447025, EPI_ISL_447026, EPI_ISL_447027, EPI_ISL_447028, EPI_ISL_447029 | | | |
| see above | Ramathibodi Hospital | COVID-19 Network Investigations (CONI) Alliance | Elizabeth Batty, Wasun Chantratita, Thanat Chookajorn, Stefan Fernandez, Angkana Huang, Anthony R. Jones, Khajohn Joonsalak, Chonticha Klungtong, Theerarat Kochakarn, Namfon Kotanan, Krittikorn Kumpornsin, Wudtichai Manasatienkij, Bhakbhoom Panthan, Ekawat Pasomsub, Kingkan Rakmanee, Insee Sensorn, Janjira Thaipadungpanit, Arporn Wangwiwatsin,Treewat Watthanachockchai |
| EPI_ISL_447030 | B.J. Medical College and Civil hospital | Gujarat Biotechnology Research Centre | Kamlesh J Upadhyay, Ramesh Pandit, Tejas Shah, Ankit Hinsu, Pritesh Sabara, Apurvasinh Puvar, Janvi Raval, Monika Gandhi, Pinal Trivedi, Maharshi Pandya, Amit Kanani, Akanksha Verma, Nitin Savaliya, Raghawendra Kumar, Dinesh Kumar, Zuber Saiyed, Dipa Kinariwala, Disha Patel, Binita Aring, Neeta Khandelwal, Geeta Vaghela, Sonia Barve, Bhavesh Modi, Kairavi Joshi, Gaurishankar Shrimali, Nidhi Sood, Pranay Shah, R D Dixit, Snehal Bagatharia, Anjali Rajwar, Chaitanya Joshi, Madhvi Joshi |
| EPI_ISL_447031 | B.J. Medical College and Civil hospital | Gujarat Biotechnology Research Centre | Ramesh Pandit, Tejas Shah, Ankit Hinsu, Pritesh Sabara, Apurvasinh Puvar, Janvi Raval, Monika Gandhi, Pinal Trivedi, Maharshi Pandya, Amit Kanani, Akanksha Verma, Nitin Savaliya, Raghawendra Kumar, Dinesh Kumar, Zuber Saiyed, Dipa Kinariwala, Disha Patel, Binita Aring, Neeta Khandelwal, Geeta Vaghela, Sonia Barve, Bhavesh Modi, Kairavi Joshi, Gaurishankar Shrimali, Nidhi Sood, Pranay Shah, R D Dixit, Snehal Bagatharia, Kamlesh J Upadhyay, Sharmistha Majumdar, Chaitanya Joshi, Madhvi Joshi |
| EPI_ISL_447032 | B.J. Medical College and Civil hospital | Gujarat Biotechnology Research Centre | Tejas Shah, Ankit Hinsu, Pritesh Sabara, Apurvasinh Puvar, Janvi Raval, Monika Gandhi, Pinal Trivedi, Maharshi Pandya, Amit Kanani, Akanksha Verma, Nitin Savaliya, Raghawendra Kumar, Dinesh Kumar, Zuber Saiyed, Dipa Kinariwala, Disha Patel, Binita Aring, Neeta Khandelwal, Geeta Vaghela, Sonia Barve, Bhavesh Modi, Kairavi Joshi, Gaurishankar Shrimali, Nidhi Sood, Pranay Shah, R D Dixit, Snehal Bagatharia, Ramesh Pandit, Kamlesh J Upadhyay, Pooja P Doshi, Chaitanya Joshi, Madhvi Joshi |
| EPI_ISL_447033 | B.J. Medical College and Civil hospital | Gujarat Biotechnology Research Centre | Ankit Hinsu, Pritesh Sabara, Apurvasinh Puvar, Janvi Raval, Monika Gandhi, Pinal Trivedi, Maharshi Pandya, Amit Kanani, Akanksha Verma, Nitin Savaliya, Raghawendra Kumar, Dinesh Kumar, Zuber Saiyed, Dipa Kinariwala, Disha Patel, Binita Aring, Neeta Khandelwal, Geeta Vaghela, Sonia Barve, Bhavesh Modi, Kairavi Joshi, Gaurishankar Shrimali, Nidhi Sood, Pranay Shah, R D Dixit, Snehal Bagatharia, Kamlesh J Upadhyay, Ramesh Pandit, Tejas Shah, Nidhi Patel, Chaitanya Joshi, Madhvi Joshi |
| EPI_ISL_447034 | B.J. Medical College and Civil hospital | Gujarat Biotechnology Research Centre | Pritesh Sabara, Apurvasinh Puvar, Janvi Raval, Monika Gandhi, Pinal Trivedi, Maharshi Pandya, Amit Kanani, Akanksha Verma, Nitin Savaliya, Raghawendra Kumar, Dinesh Kumar, Zuber Saiyed, Dipa Kinariwala, Disha Patel, Binita Aring, Neeta Khandelwal, Geeta Vaghela, Sonia Barve, Bhavesh Modi, Kairavi Joshi, Gaurishankar Shrimali, Nidhi Sood, Pranay Shah, R D Dixit, Snehal Bagatharia, Kamlesh J Upadhyay, Ramesh Pandit, Tejas Shah, Ankit Hinsu, Priti Pandita, Chaitanya Joshi, Madhvi Joshi |
| EPI_ISL_447035 | B.J. Medical College and Civil hospital | Gujarat Biotechnology Research Centre | Apurvasinh Puvar, Janvi Raval, Monika Gandhi, Pinal Trivedi, Maharshi Pandya, Amit Kanani, Akanksha Verma, Nitin Savaliya, Raghawendra Kumar, Dinesh Kumar, Zuber Saiyed, Dipa Kinariwala, Disha Patel, Binita Aring, Neeta Khandelwal, Geeta Vaghela, Sonia Barve, Bhavesh Modi, Kairavi Joshi, Gaurishankar Shrimali, Nidhi Sood, Pranay Shah, R D Dixit, Snehal Bagatharia, Kamlesh J Upadhyay, Ramesh Pandit, Tejas Shah, Ankit Hinsu, Pritesh Sabara, Neha Rajpara, Chaitanya Joshi, Madhvi Joshi |
| EPI_ISL_447036 | B.J. Medical College and Civil hospital | Gujarat Biotechnology Research Centre | Janvi Raval, Monika Gandhi, Pinal Trivedi, Maharshi Pandya, Amit Kanani, Akanksha Verma, Nitin Savaliya, Raghawendra Kumar, Dinesh Kumar, Zuber Saiyed, Dipa Kinariwala, Disha Patel, Binita Aring, Neeta Khandelwal, Geeta Vaghela, Sonia Barve, Bhavesh Modi, Kairavi Joshi, Gaurishankar Shrimali, Nidhi Sood, Pranay Shah, R D Dixit, Snehal Bagatharia, Kamlesh J Upadhyay, Ramesh Pandit, Tejas Shah, Ankit Hinsu, Pritesh Sabara, Apurvasinh Puvar, Afzal Ansari, Chaitanya Joshi, Madhvi Joshi |
| EPI_ISL_447037 | B.J. Medical College and Civil hospital | Gujarat Biotechnology Research Centre | Monika Gandhi, Pinal Trivedi, Maharshi Pandya, Amit Kanani, Akanksha Verma, Nitin Savaliya, Raghawendra Kumar, Dinesh Kumar, Zuber Saiyed, Dipa Kinariwala, Disha Patel, Binita Aring, Neeta Khandelwal, Geeta Vaghela, Sonia Barve, Bhavesh Modi, Kairavi Joshi, Gaurishankar Shrimali, Nidhi Sood, Pranay Shah, R D Dixit, Snehal Bagatharia, Kamlesh J Upadhyay, Ramesh Pandit, Tejas Shah, Ankit Hinsu, Pritesh Sabara, Apurvasinh Puvar, Janvi Raval, Neelam Nathani, Chaitanya Joshi, Madhvi Joshi |
| EPI_ISL_447038 | B.J. Medical College and Civil hospital | Gujarat Biotechnology Research Centre | Pinal Trivedi, Maharshi Pandya, Amit Kanani, Akanksha Verma, Nitin Savaliya, Raghawendra Kumar, Dinesh Kumar, Zuber Saiyed, Dipa Kinariwala, Disha Patel, Binita Aring, Neeta Khandelwal, Geeta Vaghela, Sonia Barve, Bhavesh Modi, Kairavi Joshi, Gaurishankar Shrimali, Nidhi Sood, Pranay Shah, R D Dixit, Snehal Bagatharia, Kamlesh J Upadhyay, Ramesh Pandit, Tejas Shah, Ankit Hinsu, Pritesh Sabara, Apurvasinh Puvar, Janvi Raval, Monika Gandhi, Armi Chaudhari, Chaitanya Joshi, Madhvi Joshi |
| EPI_ISL_447039 | B.J. Medical College and Civil hospital | Gujarat Biotechnology Research Centre | Maharshi Pandya, Amit Kanani, Akanksha Verma, Nitin Savaliya, Raghawendra Kumar, Dinesh Kumar, Zuber Saiyed, Dipa Kinariwala, Disha Patel, Binita Aring, Neeta Khandelwal, Geeta Vaghela, Sonia Barve, Bhavesh Modi, Kairavi Joshi, Gaurishankar Shrimali, Nidhi Sood, Pranay Shah, R D Dixit, Snehal Bagatharia, Kamlesh J Upadhyay, Ramesh Pandit, Tejas Shah, Ankit Hinsu, Pritesh Sabara, Apurvasinh Puvar, Janvi Raval, Monika Gandhi, Pinal Trivedi, Bhavya Jindal, Chaitanya Joshi, Madhvi Joshi |
| EPI_ISL_447040 | B.J. Medical College and Civil hospital | Gujarat Biotechnology Research Centre | Amit Kanani, Akanksha Verma, Nitin Savaliya, Raghawendra Kumar, Dinesh Kumar, Zuber Saiyed, Dipa Kinariwala, Disha Patel, Binita Aring, Neeta Khandelwal, Geeta Vaghela, Sonia Barve, Bhavesh Modi, Kairavi Joshi, Gaurishankar Shrimali, Nidhi Sood, Pranay Shah, R D Dixit, Snehal Bagatharia, Kamlesh J Upadhyay, Ramesh Pandit, Tejas Shah, Ankit Hinsu, Pritesh Sabara, Apurvasinh Puvar, Janvi Raval, Monika Gandhi, Pinal Trivedi, Maharshi Pandya, Anjali Rajwar, Chaitanya Joshi, Madhvi Joshi |
| EPI_ISL_447041 | B.J. Medical College and Civil hospital | Gujarat Biotechnology Research Centre | Akanksha Verma, Nitin Savaliya, Raghawendra Kumar, Dinesh Kumar, Zuber Saiyed, Dipa Kinariwala, Disha Patel, Binita Aring, Neeta Khandelwal, Geeta Vaghela, Sonia Barve, Bhavesh Modi, Kairavi Joshi, Gaurishankar Shrimali, Nidhi Sood, Pranay Shah, R D Dixit, Snehal Bagatharia, Kamlesh J Upadhyay, Ramesh Pandit, Tejas Shah, Ankit Hinsu, Pritesh Sabara, Apurvasinh Puvar, Janvi Raval, Monika Gandhi, Pinal Trivedi, Maharshi Pandya, Amit Kanani, Sharmistha Majumdar, Chaitanya Joshi, Madhvi Joshi |
| EPI_ISL_447042 | B.J. Medical College and Civil hospital | Gujarat Biotechnology Research Centre | Nitin Savaliya, Raghawendra Kumar, Dinesh Kumar, Zuber Saiyed, Dipa Kinariwala, Disha Patel, Binita Aring, Neeta Khandelwal, Geeta Vaghela, Sonia Barve, Bhavesh Modi, Kairavi Joshi, Gaurishankar Shrimali, Nidhi Sood, Pranay Shah, R D Dixit, Snehal Bagatharia, Kamlesh J Upadhyay, Ramesh Pandit, Tejas Shah, Ankit Hinsu, Pritesh Sabara, Apurvasinh Puvar, Janvi Raval, Monika Gandhi, Pinal Trivedi, Maharshi Pandya, Amit Kanani, Akanksha Verma, Pooja P Doshi, Chaitanya Joshi, Madhvi Joshi |
| EPI_ISL_447043 | B.J. Medical College and Civil hospital | Gujarat Biotechnology Research Centre | Raghawendra Kumar, Dinesh Kumar, Zuber Saiyed, Dipa Kinariwala, Disha Patel, Binita Aring, Neeta Khandelwal, Geeta Vaghela, Sonia Barve, Bhavesh Modi, Kairavi Joshi, Gaurishankar Shrimali, Nidhi Sood, Pranay Shah, R D Dixit, Snehal Bagatharia, Kamlesh J Upadhyay, Ramesh Pandit, Tejas Shah, Ankit Hinsu, Pritesh Sabara, Apurvasinh Puvar, Janvi Raval, Monika Gandhi, Pinal Trivedi, Maharshi Pandya, Amit Kanani, Akanksha Verma, Nitin Savaliya, Nidhi Patel, Chaitanya Joshi, Madhvi Joshi |
| EPI_ISL_447044 | B.J. Medical College and Civil hospital | Gujarat Biotechnology Research Centre | Dinesh Kumar, Zuber Saiyed, Dipa Kinariwala, Disha Patel, Binita Aring, Neeta Khandelwal, Geeta Vaghela, Sonia Barve, Bhavesh Modi, Kairavi Joshi, Gaurishankar Shrimali, Nidhi Sood, Pranay Shah, R D Dixit, Snehal Bagatharia, Kamlesh J Upadhyay, Ramesh Pandit, Tejas Shah, Ankit Hinsu, Pritesh Sabara, Apurvasinh Puvar, Janvi Raval, Monika Gandhi, Pinal Trivedi, Maharshi Pandya, Amit Kanani, Akanksha Verma, Nitin Savaliya, Raghawendra Kumar, Priti Pandita, Chaitanya Joshi, Madhvi Joshi |
| EPI_ISL_447045 | B.J. Medical College and Civil hospital | Gujarat Biotechnology Research Centre | Zuber Saiyed, Dipa Kinariwala, Disha Patel, Binita Aring, Neeta Khandelwal, Geeta Vaghela, Sonia Barve, Bhavesh Modi, Kairavi Joshi, Gaurishankar Shrimali, Nidhi Sood, Pranay Shah, R D Dixit, Snehal Bagatharia, Kamlesh J Upadhyay, Ramesh Pandit, Tejas Shah, Ankit Hinsu, Pritesh Sabara, Apurvasinh Puvar, Janvi Raval, Monika Gandhi, Pinal Trivedi, Maharshi Pandya, Amit Kanani, Akanksha Verma, Nitin Savaliya, Raghawendra Kumar, Dinesh Kumar, Neha Rajpara, Chaitanya Joshi, Madhvi Joshi |
| EPI_ISL_447046 | B.J. Medical College and Civil hospital | Gujarat Biotechnology Research Centre | Dipa Kinariwala, Disha Patel, Binita Aring, Neeta Khandelwal, Geeta Vaghela, Sonia Barve, Bhavesh Modi, Kairavi Joshi, Gaurishankar Shrimali, Nidhi Sood, Pranay Shah, R D Dixit, Snehal Bagatharia, Kamlesh J Upadhyay, Ramesh Pandit, Tejas Shah, Ankit Hinsu, Pritesh Sabara, Apurvasinh Puvar, Janvi Raval, Monika Gandhi, Pinal Trivedi, Maharshi Pandya, Amit Kanani, Akanksha Verma, Nitin Savaliya, Raghawendra Kumar, Dinesh Kumar, Zuber Saiyed, Afzal Ansari, Chaitanya Joshi, Madhvi Joshi |
| EPI_ISL_447047 | GMERS Medical College and Hospital, Gandhinagar | Gujarat Biotechnology Research Centre | Disha Patel, Binita Aring, Neeta Khandelwal, Geeta Vaghela, Sonia Barve, Bhavesh Modi, Kairavi Joshi, Gaurishankar Shrimali, Nidhi Sood, Pranay Shah, R D Dixit, Snehal Bagatharia, Kamlesh J Upadhyay, Ramesh Pandit, Tejas Shah, Ankit Hinsu, Pritesh Sabara, Apurvasinh Puvar, Janvi Raval, Monika Gandhi, Pinal Trivedi, Maharshi Pandya, Amit Kanani, Akanksha Verma, Nitin Savaliya, Raghawendra Kumar, Dinesh Kumar, Zuber Saiyed, Dipa Kinariwala, Neelam Nathani, Chaitanya Joshi, Madhvi Joshi |
| EPI_ISL_447048 | GMERS Medical College and Hospital, Gandhinagar | Gujarat Biotechnology Research Centre | Binita Aring, Neeta Khandelwal, Geeta Vaghela, Sonia Barve, Bhavesh Modi, Kairavi Joshi, Gaurishankar Shrimali, Nidhi Sood, Pranay Shah, R D Dixit, Snehal Bagatharia, Kamlesh J Upadhyay, Ramesh Pandit, Tejas Shah, Ankit Hinsu, Pritesh Sabara, Apurvasinh Puvar, Janvi Raval, Monika Gandhi, Pinal Trivedi, Maharshi Pandya, Amit Kanani, Akanksha Verma, Nitin Savaliya, Raghawendra Kumar, Dinesh Kumar, Zuber Saiyed, Dipa Kinariwala, Disha Patel, Armi Chaudhari, Chaitanya Joshi, Madhvi Joshi |
| EPI_ISL_447049 | GMERS Medical College and Hospital, Gandhinagar | Gujarat Biotechnology Research Centre | Neeta Khandelwal, Geeta Vaghela, Sonia Barve, Bhavesh Modi, Kairavi Joshi, Gaurishankar Shrimali, Nidhi Sood, Pranay Shah, R D Dixit, Snehal Bagatharia, Kamlesh J Upadhyay, Ramesh Pandit, Tejas Shah, Ankit Hinsu, Pritesh Sabara, Apurvasinh Puvar, Janvi Raval, Monika Gandhi, Pinal Trivedi, Maharshi Pandya, Amit Kanani, Akanksha Verma, Nitin Savaliya, Raghawendra Kumar, Dinesh Kumar, Zuber Saiyed, Dipa Kinariwala, Disha Patel, Binita Aring, Bhavya Jindal, Chaitanya Joshi, Madhvi Joshi |
| EPI_ISL_447050 | GMERS Medical College and Hospital, Gandhinagar | Gujarat Biotechnology Research Centre | Geeta Vaghela, Sonia Barve, Bhavesh Modi, Kairavi Joshi, Gaurishankar Shrimali, Nidhi Sood, Pranay Shah, R D Dixit, Snehal Bagatharia, Kamlesh J Upadhyay, Ramesh Pandit, Tejas Shah, Ankit Hinsu, Pritesh Sabara, Apurvasinh Puvar, Janvi Raval, Monika Gandhi, Pinal Trivedi, Maharshi Pandya, Amit Kanani, Akanksha Verma, Nitin Savaliya, Raghawendra Kumar, Dinesh Kumar, Zuber Saiyed, Dipa Kinariwala, Disha Patel, Binita Aring, Neeta Khandelwal, Dipeshwari Shewale, Chaitanya Joshi, Madhvi Joshi |
| EPI_ISL_447051 | GMERS Medical College and Hospital, Gandhinagar | Gujarat Biotechnology Research Centre | Sonia Barve, Bhavesh Modi, Kairavi Joshi, Gaurishankar Shrimali, Nidhi Sood, Pranay Shah, R D Dixit, Snehal Bagatharia, Kamlesh J Upadhyay, Ramesh Pandit, Tejas Shah, Ankit Hinsu, Pritesh Sabara, Apurvasinh Puvar, Janvi Raval, Monika Gandhi, Pinal Trivedi, Maharshi Pandya, Amit Kanani, Akanksha Verma, Nitin Savaliya, Raghawendra Kumar, Dinesh Kumar, Zuber Saiyed, Dipa Kinariwala, Disha Patel, Binita Aring, Neeta Khandelwal, Geeta Vaghela, Anjali Rajwar, Chaitanya Joshi, Madhvi Joshi |
| EPI_ISL_447052 | GMERS Medical College and Hospital, Gandhinagar | Gujarat Biotechnology Research Centre | Bhavesh Modi, Kairavi Joshi, Gaurishankar Shrimali, Nidhi Sood, Pranay Shah, R D Dixit, Snehal Bagatharia, Kamlesh J Upadhyay, Ramesh Pandit, Tejas Shah, Ankit Hinsu, Pritesh Sabara, Apurvasinh Puvar, Janvi Raval, Monika Gandhi, Pinal Trivedi, Maharshi Pandya, Amit Kanani, Akanksha Verma, Nitin Savaliya, Raghawendra Kumar, Dinesh Kumar, Zuber Saiyed, Dipa Kinariwala, Disha Patel, Binita Aring, Neeta Khandelwal, Geeta Vaghela, Sonia Barve, Sharmistha Majumdar, Chaitanya Joshi, Madhvi Joshi |
| EPI_ISL_447053 | GMERS Medical College and Hospital, Gandhinagar | Gujarat Biotechnology Research Centre | Kairavi Joshi, Gaurishankar Shrimali, Nidhi Sood, Pranay Shah, R D Dixit, Snehal Bagatharia, Kamlesh J Upadhyay, Ramesh Pandit, Tejas Shah, Ankit Hinsu, Pritesh Sabara, Apurvasinh Puvar, Janvi Raval, Monika Gandhi, Pinal Trivedi, Maharshi Pandya, Amit Kanani, Akanksha Verma, Nitin Savaliya, Raghawendra Kumar, Dinesh Kumar, Zuber Saiyed, Dipa Kinariwala, Disha Patel, Binita Aring, Neeta Khandelwal, Geeta Vaghela, Sonia Barve, Bhavesh Modi, Pooja P Doshi, Chaitanya Joshi, Madhvi Joshi |
| EPI_ISL_447054 | Cantacuzino National Military-Medical Institute for Research and Development | Cantacuzino Institute | M.Lazar, L.Ustea, A.Cretu |
| EPI_ISL_447055 | Department for Virology, Molecular Biology and Genome Research, R. G. Lugar Center for Public Health Research, National Center for Disease Control and Public Health (NCDC) of Georgia. | Department for Virology, Molecular Biology and Genome Research, R. G. Lugar Center for Public Health Research, National Center for Disease Control and Public Health (NCDC) of Georgia. | Meri Pantsulaia, Gvantsa Brachveli, Giorgi Tomashvili, Gvantsa Chanturia, Ann Machablishvili, Nato Kotaria, Marine Murtskhvaladze, Lela Sabadze, Mari Gavashelidze, Ana Papkiauri, Tata Imnadze, Tamar Jashiashvili, Tea Tevdoradze, Ketevan Sidamonidze, Ekaterine Khmaladze, Ekaterine Zhghenti, Roena Sukhiashvili, Mariam Zakalashvili, Lela Urushadze, Magda Dgebuadze, Davit Tsaguria, Ekaterine Zangaladze, Nino Berishvili, Adam Kotorashvili, Maia Alkhazashvili, Irma Burjanadze, Anna Kasradze, Khatuna Zakhashvili, Paata Imnadze, Amiran Gamkrelidze. |
| EPI_ISL_447056 | Department for Virology, Molecular Biology and Genome Research, R. G. Lugar Center for Public Health Research, National Center for Disease Control and Public Health (NCDC) of Georgia. | Department for Virology, Molecular Biology and Genome Research, R. G. Lugar Center for Public Health Research, National Center for Disease Control and Public Health (NCDC) of Georgia. | Gvantsa Brachveli, Meri Pantsulaia, Giorgi Tomashvili, Gvantsa Chanturia, Ann Machablishvili, Nato Kotaria, Marine Murtskhvaladze, Lela Sabadze, Mari Gavashelidze, Ana Papkiauri, Gvantsa Brachveli, Tata Imnadze, Tamar Jashiashvili, Tea Tevdoradze, Ketevan Sidamonidze, Ekaterine Khmaladze, Ekaterine Zhghenti, Roena Sukhiashvili, Mariam Zakalashvili, Lela Urushadze, Magda Dgebuadze, Davit Tsaguria, Ekaterine Zangaladze, Nino Berishvili, Adam Kotorashvili, Maia Alkhazashvili, Irma Burjanadze, Anna Kasradze, Khatuna Zakhashvili, Paata Imnadze, Amiran Gamkrelidze. |
| EPI_ISL_447057, EPI_ISL_447058, EPI_ISL_447059, EPI_ISL_447060, EPI_ISL_447061, EPI_ISL_447062, EPI_ISL_447063, EPI_ISL_447064, EPI_ISL_447065, EPI_ISL_447066, EPI_ISL_447067, EPI_ISL_447068, EPI_ISL_447069, EPI_ISL_447070, EPI_ISL_447071, EPI_ISL_447072, EPI_ISL_447073, EPI_ISL_447074, EPI_ISL_447075, EPI_ISL_447076, EPI_ISL_447077, EPI_ISL_447078, EPI_ISL_447079, EPI_ISL_447080, EPI_ISL_447081, EPI_ISL_447082, EPI_ISL_447083, EPI_ISL_447084, EPI_ISL_447085, EPI_ISL_447086, EPI_ISL_447087, EPI_ISL_447088, EPI_ISL_447089, EPI_ISL_447090, EPI_ISL_447091, EPI_ISL_447092, EPI_ISL_447093, EPI_ISL_447094, EPI_ISL_447095, EPI_ISL_447096, EPI_ISL_447097, EPI_ISL_447098, EPI_ISL_447099, EPI_ISL_447100, EPI_ISL_447101, EPI_ISL_447102, EPI_ISL_447103, EPI_ISL_447104, EPI_ISL_447105, EPI_ISL_447106, EPI_ISL_447107, EPI_ISL_447108, EPI_ISL_447109, EPI_ISL_447110, EPI_ISL_447111, EPI_ISL_447112, EPI_ISL_447113, EPI_ISL_447114, EPI_ISL_447115, EPI_ISL_447116, EPI_ISL_447117, EPI_ISL_447118 | | | |
| see above | Michigan Department of Health and Human Services, Bureau of Laboratories | Michigan Department of Health and Human Services, Bureau of Laboratories | Blankenship HM, Riner D, Soehnlen MK |

| | | | |
|---|---|---|---|
| EPI_ISL_447119 | HOSPITAL DR.HERNAN HENRIQUEZ ARAVENA | Instituto de Salud Publica de Chile | Andrés E Castillo, Bárbara Parra,Paz Tapia, Jaime Lagos, Loredana Arata, Alejandra Acevedo, Winston Andrade, Gabriel Leal, Carolina Tambley, Patricia Bustos, Rodrigo Fasce, Jorge Fernandez |
| EPI_ISL_447120, EPI_ISL_447121, EPI_ISL_447122, EPI_ISL_447123, EPI_ISL_447124, EPI_ISL_447125, EPI_ISL_447126, EPI_ISL_447127, EPI_ISL_447128, EPI_ISL_447129, EPI_ISL_447130, EPI_ISL_447131, EPI_ISL_447132, EPI_ISL_447133, EPI_ISL_447134, EPI_ISL_447135, EPI_ISL_447136, EPI_ISL_447137, EPI_ISL_447138, EPI_ISL_447139, EPI_ISL_447140, EPI_ISL_447141, EPI_ISL_447142, EPI_ISL_447143, EPI_ISL_447144, EPI_ISL_447145, EPI_ISL_447146, EPI_ISL_447147, EPI_ISL_447148, EPI_ISL_447149, EPI_ISL_447150, EPI_ISL_447151, EPI_ISL_447152, EPI_ISL_447153, EPI_ISL_447154, EPI_ISL_447155, EPI_ISL_447156, EPI_ISL_447157, EPI_ISL_447158, EPI_ISL_447159, EPI_ISL_447160, EPI_ISL_447161, EPI_ISL_447162 | | | |
| see above | Department of Clinical Microbiology | GIGA Medical Genomics | Keith Durkin, Maria Artesi, Sébastien Bontems, Raphaël Boreux, Cécile Meex, Pierrette Melin, Marie-Pierre Hayette, Vincent Bours. |
| EPI_ISL_447163, EPI_ISL_447164, EPI_ISL_447165, EPI_ISL_447166, EPI_ISL_447167, EPI_ISL_447168, EPI_ISL_447169, EPI_ISL_447170, EPI_ISL_447171, EPI_ISL_447172, EPI_ISL_447173, EPI_ISL_447174, EPI_ISL_447175, EPI_ISL_447176, EPI_ISL_447177, EPI_ISL_447178, EPI_ISL_447179, EPI_ISL_447180, EPI_ISL_447181, EPI_ISL_447182, EPI_ISL_447183, EPI_ISL_447184, EPI_ISL_447185, EPI_ISL_447186, EPI_ISL_447187, EPI_ISL_447188, EPI_ISL_447189, EPI_ISL_447190, EPI_ISL_447191, EPI_ISL_447192, EPI_ISL_447193, EPI_ISL_447194, EPI_ISL_447195, EPI_ISL_447196, EPI_ISL_447197, EPI_ISL_447198, EPI_ISL_447199, EPI_ISL_447200, EPI_ISL_447201, EPI_ISL_447202, EPI_ISL_447203, EPI_ISL_447204, EPI_ISL_447205, EPI_ISL_447206, EPI_ISL_447207, EPI_ISL_447208, EPI_ISL_447209, EPI_ISL_447210, EPI_ISL_447211, EPI_ISL_447212, EPI_ISL_447213, EPI_ISL_447214, EPI_ISL_447215, EPI_ISL_447216, EPI_ISL_447217, EPI_ISL_447218, EPI_ISL_447219, EPI_ISL_447220, EPI_ISL_447221, EPI_ISL_447222, EPI_ISL_447223, EPI_ISL_447224, EPI_ISL_447225, EPI_ISL_447226, EPI_ISL_447227, EPI_ISL_447228, EPI_ISL_447229 | | | |
| see above | Michigan Department of Health and Human Services, Bureau of Laboratories | Michigan Department of Health and Human Services, Bureau of Laboratories | Blankenship HM, Riner D, Soehnlen MK |
| EPI_ISL_447230, EPI_ISL_447231, EPI_ISL_447232, EPI_ISL_447233, EPI_ISL_447234, EPI_ISL_447235, EPI_ISL_447236, EPI_ISL_447237, EPI_ISL_447238, EPI_ISL_447239, EPI_ISL_447240, EPI_ISL_447241, EPI_ISL_447242, EPI_ISL_447243, EPI_ISL_447244, EPI_ISL_447245, EPI_ISL_447246, EPI_ISL_447247, EPI_ISL_447248, EPI_ISL_447249 | | | |
| see above | Viral Respiratory Lab, National Institute for Biomedical Research (INRB) | Pathogen Sequencing Lab, National Institute for Biomedical Research (INRB) | Placide Mbala-Kingebeni, Edith Nkwembe, Eddy Kinganda-Lusamaki, Amuri Aziza, Francisca Muyembe Mawete, Catherine Pratt, Matthias Pauthner, Josh Quick, Allison Black, James Hadfield, Trevor Bedford, Ian Goodfellow, Andrew Rambaut, Nick Loman, Kristian Andersen, Michael Wiley, Steve Ahuka-Mundeke, Jean-Jacques Muyembe Tamfum |
| EPI_ISL_447250 | Central Virology Laboratory | Central Virology Laboratory | Neta Zuckerman, Efrat Bucris, Oran Erster, Danit Sofer, Orna Mor, Ella Mendelson, Michal Mandelboim |
| EPI_ISL_447251 | Central Virology Laboratory | Central Virology Laboratory | Neta Zuckerman, Efrat Bucris, Oran Erster, Danit Sofer, Orna Mor, Ella Mendelson, Michal Mandelboim |
| EPI_ISL_447252, EPI_ISL_447253, EPI_ISL_447254, EPI_ISL_447255, EPI_ISL_447256, EPI_ISL_447257 | TSGH-CP molecular lab | TSGH-CP molecular lab | Cherng-Lih Perng, Ming-Jr JIAN, Chih-Kai Chang, Jung-Chung Lin, Kuo-Ming Yeh, Chien-Wen Chen, Sheng-Kang Chiu, Hsing-Yi Chung, Shih-Hung Tsai, Kuo-Sheng Hung, Tien-Yao Chang, Feng-Yee Chang, Hung-Sheng Shang |
| EPI_ISL_447258, EPI_ISL_447259, EPI_ISL_447260, EPI_ISL_447261, EPI_ISL_447262, EPI_ISL_447263, EPI_ISL_447264, EPI_ISL_447265, EPI_ISL_447266, EPI_ISL_447267, EPI_ISL_447268, EPI_ISL_447269, EPI_ISL_447270, EPI_ISL_447271, EPI_ISL_447272, EPI_ISL_447273, EPI_ISL_447274, EPI_ISL_447275, EPI_ISL_447276, EPI_ISL_447277, EPI_ISL_447278, EPI_ISL_447279, EPI_ISL_447280 | | | |
| see above | Microbiology laboratory, Assuta Ashdod University-Affiliated Hospital | Stern Lab | Stern Lab |
| EPI_ISL_447281, EPI_ISL_447282, EPI_ISL_447283, EPI_ISL_447284, EPI_ISL_447285, EPI_ISL_447286, EPI_ISL_447287, EPI_ISL_447288, EPI_ISL_447289, EPI_ISL_447290, EPI_ISL_447291, EPI_ISL_447292, EPI_ISL_447293, EPI_ISL_447294, EPI_ISL_447295, EPI_ISL_447296, EPI_ISL_447297, EPI_ISL_447298, EPI_ISL_447299, EPI_ISL_447300, EPI_ISL_447301, EPI_ISL_447302, EPI_ISL_447303, EPI_ISL_447304, EPI_ISL_447305, EPI_ISL_447306, EPI_ISL_447307, EPI_ISL_447308, EPI_ISL_447309, EPI_ISL_447310 | | | |
| see above | Microbiology Division, Barzilai University Medical Center | Stern Lab | Stern Lab |
| EPI_ISL_447311, EPI_ISL_447312, EPI_ISL_447313, EPI_ISL_447314, EPI_ISL_447315, EPI_ISL_447316, EPI_ISL_447317, EPI_ISL_447318, EPI_ISL_447319, EPI_ISL_447320, EPI_ISL_447321, EPI_ISL_447322, EPI_ISL_447323, EPI_ISL_447324, EPI_ISL_447325, EPI_ISL_447326, EPI_ISL_447327, EPI_ISL_447328, EPI_ISL_447329, EPI_ISL_447330 | | | |
| see above | Clinical Virology Laboratory, Soroka Medical Center and the Faculty of Health Sciences, Ben-Gurion University of the Negev | Stern Lab | Stern Lab |
| EPI_ISL_447331, EPI_ISL_447332, EPI_ISL_447333, EPI_ISL_447334, EPI_ISL_447335, EPI_ISL_447336, EPI_ISL_447337, EPI_ISL_447338, EPI_ISL_447339, EPI_ISL_447340, EPI_ISL_447341, EPI_ISL_447342, EPI_ISL_447343, EPI_ISL_447344, EPI_ISL_447345, EPI_ISL_447346, EPI_ISL_447347, EPI_ISL_447348, EPI_ISL_447349, EPI_ISL_447350, EPI_ISL_447351, EPI_ISL_447352, EPI_ISL_447353, EPI_ISL_447354, EPI_ISL_447355, EPI_ISL_447356, EPI_ISL_447357, EPI_ISL_447358, EPI_ISL_447359, EPI_ISL_447360, EPI_ISL_447361, EPI_ISL_447362, EPI_ISL_447363, EPI_ISL_447364, EPI_ISL_447365, EPI_ISL_447366, EPI_ISL_447367, EPI_ISL_447368, EPI_ISL_447369, EPI_ISL_447370, EPI_ISL_447371, EPI_ISL_447372, EPI_ISL_447373, EPI_ISL_447374, EPI_ISL_447375, EPI_ISL_447376, EPI_ISL_447377, EPI_ISL_447378, EPI_ISL_447379, EPI_ISL_447380, EPI_ISL_447381, EPI_ISL_447382 | | | |
| see above | Clinical Virology Unit, Hadassah Hebrew University Medical Center | Stern Lab | Stern Lab |
| EPI_ISL_447383, EPI_ISL_447384, EPI_ISL_447385, EPI_ISL_447386, EPI_ISL_447387, EPI_ISL_447388, EPI_ISL_447389, EPI_ISL_447390, EPI_ISL_447391, EPI_ISL_447392, EPI_ISL_447393, EPI_ISL_447394, EPI_ISL_447395, EPI_ISL_447396, EPI_ISL_447397, EPI_ISL_447398, EPI_ISL_447399, EPI_ISL_447400, EPI_ISL_447401, EPI_ISL_447402, EPI_ISL_447403, EPI_ISL_447404, EPI_ISL_447405, EPI_ISL_447406 | | | |
| see above | Clinical Microbiology Laboratory, The Baruch Padeh Medical Center, Poriya | Stern Lab | Stern Lab |
| EPI_ISL_447407, EPI_ISL_447408, EPI_ISL_447409, EPI_ISL_447410, EPI_ISL_447411, EPI_ISL_447412, EPI_ISL_447413, EPI_ISL_447414, EPI_ISL_447415, EPI_ISL_447416 | Clinical Virology Unit, Hadassah Hebrew University Medical Center | Stern Lab | Stern Lab |
| EPI_ISL_447417, EPI_ISL_447418 | Clinical Microbiology Laboratory, The Baruch Padeh Medical Center, Poriya | Stern Lab | Stern Lab |
| EPI_ISL_447419, EPI_ISL_447420, EPI_ISL_447421, EPI_ISL_447422, EPI_ISL_447423, EPI_ISL_447424, EPI_ISL_447425, EPI_ISL_447426, EPI_ISL_447427, EPI_ISL_447428, EPI_ISL_447429, EPI_ISL_447430, EPI_ISL_447431, EPI_ISL_447432, EPI_ISL_447433, EPI_ISL_447434, EPI_ISL_447435, EPI_ISL_447436, EPI_ISL_447437, EPI_ISL_447438, EPI_ISL_447439, EPI_ISL_447440, EPI_ISL_447441, EPI_ISL_447442, EPI_ISL_447443, EPI_ISL_447444, EPI_ISL_447445, EPI_ISL_447446, EPI_ISL_447447, EPI_ISL_447448, EPI_ISL_447449, EPI_ISL_447450, EPI_ISL_447451, EPI_ISL_447452, EPI_ISL_447453, EPI_ISL_447454, EPI_ISL_447455, EPI_ISL_447456, EPI_ISL_447457, EPI_ISL_447458, EPI_ISL_447459, EPI_ISL_447460, EPI_ISL_447461, EPI_ISL_447462, EPI_ISL_447463, EPI_ISL_447464, EPI_ISL_447465, EPI_ISL_447466, EPI_ISL_447467, EPI_ISL_447468, EPI_ISL_447469 | | | |
| see above | Clinical Microbiology Laboratory, Sheba Medical Center | Stern Lab | Stern Lab |
| EPI_ISL_447470 | Servicio de Microbiologia. Hospital Clinico Universitario de Valencia | Sequencing and Bioinformatics Service and Molecular Epidemiology Research Group. FISABIO-Public Health | David Navarro, Eliseo Albert, Maria Alma Bracho, Griselda De Marco, Lidia Ruiz Roldan, Neris Garcia-Gonzalez, Inma Galán Vendrell, Sandra Carbo, Loreto Ferrús Abad, Paula Ruiz-Hueso, Mariana Reyes-Prieto, Vicente Soriano Chirona, Ivan Ansari, Llúcia Martínez-Priego, Giuseppe 'Auria, Fernando Gonzalez-Candelas |
| EPI_ISL_447471 | Servicio de Microbiologia. Hospital Clinico Universitario de Valencia | Sequencing and Bioinformatics Service and Molecular Epidemiology Research Group. FISABIO-Public Health | Eliseo Albert, Maria Alma Bracho, Griselda De Marco, Lidia Ruiz Roldan, Neris Garcia-Gonzalez, Inma Galán Vendrell, Sandra Carbo, Loreto Ferrús Abad, Paula Ruiz-Hueso, Mariana Reyes-Prieto, Vicente Soriano Chirona, Ivan Ansari, Llúcia Martínez-Priego, Giuseppe 'Auria, David Navarro, Fernando Gonzalez-Candelas |
| EPI_ISL_447472 | Servicio de Microbiologia. Hospital Clinico Universitario de Valencia | Sequencing and Bioinformatics Service and Molecular Epidemiology Research Group. FISABIO-Public Health | Maria Alma Bracho, Griselda De Marco, Lidia Ruiz Roldan, Neris Garcia-Gonzalez, Inma Galán Vendrell, Sandra Carbo, Loreto Ferrús Abad, Paula Ruiz-Hueso, Mariana Reyes-Prieto, Vicente Soriano Chirona, Ivan Ansari, Llúcia Martínez-Priego, Giuseppe 'Auria, David Navarro, Eliseo Albert, Fernando Gonzalez-Candelas |
| EPI_ISL_447473 | Servicio de Microbiologia. Hospital Clinico Universitario de Valencia | Sequencing and Bioinformatics Service and Molecular Epidemiology Research Group. FISABIO-Public Health | Griselda De Marco, Lidia Ruiz Roldan, Neris Garcia-Gonzalez, Inma Galán Vendrell, Sandra Carbo, Loreto Ferrús Abad, Paula Ruiz-Hueso, Mariana Reyes-Prieto, Vicente Soriano Chirona, Ivan Ansari, Llúcia Martínez-Priego, Giuseppe 'Auria, David Navarro, Eliseo Albert, Maria Alma Bracho, Fernando Gonzalez-Candelas |
| EPI_ISL_447474 | Servicio de Microbiologia. Hospital Clinico Universitario de Valencia | Sequencing and Bioinformatics Service and Molecular Epidemiology Research Group. FISABIO-Public Health | Lidia Ruiz Roldan, Neris Garcia-Gonzalez, Inma Galán Vendrell, Sandra Carbo, Loreto Ferrús Abad, Paula Ruiz-Hueso, Mariana Reyes-Prieto, Vicente Soriano Chirona, Ivan Ansari, Llúcia Martínez-Priego, Giuseppe 'Auria, David Navarro, Eliseo Albert, Maria Alma Bracho, Fernando Gonzalez-Candelas |
| EPI_ISL_447475 | Servicio de Microbiologia. Hospital Clinico Universitario de Valencia | Sequencing and Bioinformatics Service and Molecular Epidemiology Research Group. FISABIO-Public Health | Neris Garcia-Gonzalez, Inma Galán Vendrell, Sandra Carbo, Loreto Ferrús Abad, Paula Ruiz-Hueso, Mariana Reyes-Prieto, Vicente Soriano Chirona, Ivan Ansari, Llúcia Martínez-Priego, Giuseppe 'Auria, David Navarro, Eliseo Albert, Maria Alma Bracho, Lidia Ruiz Roldan, Fernando Gonzalez-Candelas |
| EPI_ISL_447476 | Servicio de Microbiologia. Hospital Clinico Universitario de Valencia | Sequencing and Bioinformatics Service and Molecular Epidemiology Research Group. FISABIO-Public Health | Inma Galán Vendrell, Sandra Carbo, Loreto Ferrús Abad, Paula Ruiz-Hueso, Mariana Reyes-Prieto, Vicente Soriano Chirona, Ivan Ansari, Llúcia Martínez-Priego, Giuseppe 'Auria, David Navarro, Eliseo Albert, Maria Alma Bracho, Lidia Ruiz Roldan, Neris Garcia-Gonzalez, Fernando Gonzalez-Candelas |
| EPI_ISL_447477 | Servicio de Microbiologia. Hospital Clinico Universitario de Valencia | Sequencing and Bioinformatics Service and Molecular Epidemiology Research Group. FISABIO-Public Health | Sandra Carbo, Loreto Ferrús Abad, Paula Ruiz-Hueso, Mariana Reyes-Prieto, Vicente Soriano Chirona, Ivan Ansari, Llúcia Martínez-Priego, Giuseppe 'Auria, David Navarro, Eliseo Albert, Maria Alma Bracho, Lidia Ruiz Roldan, Neris Garcia-Gonzalez, Inma Galán Vendrell, Fernando Gonzalez-Candelas |
| EPI_ISL_447478 | Servicio de Microbiologia. Hospital Clinico Universitario de Valencia | Sequencing and Bioinformatics Service and Molecular Epidemiology Research Group. FISABIO-Public Health | Loreto Ferrús Abad, Paula Ruiz-Hueso, Mariana Reyes-Prieto, Vicente Soriano Chirona, Ivan Ansari, Llúcia Martínez-Priego, Giuseppe 'Auria, David Navarro, Eliseo Albert, Maria Alma Bracho, Lidia Ruiz Roldan, Neris Garcia-Gonzalez, Inma Galán Vendrell, Sandra Carbo, Fernando Gonzalez-Candelas |
| EPI_ISL_447479 | Servicio de Microbiologia. Hospital Clinico Universitario de Valencia | Sequencing and Bioinformatics Service and Molecular Epidemiology Research Group. FISABIO-Public Health | Paula Ruiz-Hueso, Mariana Reyes-Prieto, Vicente Soriano Chirona, Ivan Ansari, Llúcia Martínez-Priego, Giuseppe 'Auria, David Navarro, Eliseo Albert, Maria Alma Bracho, Lidia Ruiz Roldan, Neris Garcia-Gonzalez, Inma Galán Vendrell, Sandra Carbo, Loreto Ferrús Abad, Fernando Gonzalez-Candelas |
| EPI_ISL_447480 | Servicio de Microbiologia. Hospital Clinico Universitario de Valencia | Sequencing and Bioinformatics Service and Molecular Epidemiology Research Group. FISABIO-Public Health | Mariana Reyes-Prieto, Vicente Soriano Chirona, Ivan Ansari, Llúcia Martínez-Priego, Giuseppe 'Auria, David Navarro, Eliseo Albert, Maria Alma Bracho, Lidia Ruiz Roldan, Neris Garcia-Gonzalez, Inma Galán Vendrell, Sandra Carbo, Loreto Ferrús Abad, Paula Ruiz-Hueso, Fernando Gonzalez-Candelas |
| EPI_ISL_447481 | Servicio de Microbiologia. Hospital Clinico Universitario de Valencia | Sequencing and Bioinformatics Service and Molecular Epidemiology Research Group. FISABIO-Public Health | Vicente Soriano Chirona, Ivan Ansari, Llúcia Martínez-Priego, Giuseppe 'Auria, David Navarro, Eliseo Albert, Maria Alma Bracho, Lidia Ruiz Roldan, Neris Garcia-Gonzalez, Inma Galán Vendrell, Sandra Carbo, Loreto Ferrús Abad, Paula Ruiz-Hueso, Mariana Reyes-Prieto, Fernando Gonzalez-Candelas |
| EPI_ISL_447482 | Servicio de Microbiologia. Hospital Clinico Universitario de Valencia | Sequencing and Bioinformatics Service and Molecular Epidemiology Research Group. FISABIO-Public Health | Giuseppe 'Auria, David Navarro, Eliseo Albert, Maria Alma Bracho, Lidia Ruiz Roldan, Neris Garcia-Gonzalez, Inma Galán Vendrell, Sandra Carbo, Loreto Ferrús Abad, Paula Ruiz-Hueso, Mariana Reyes-Prieto, Vicente Soriano Chirona, Ivan Ansari, Llúcia Martínez-Priego, Fernando Gonzalez-Candelas |
| EPI_ISL_447483 | Servicio de Microbiologia. Hospital Clinico Universitario de Valencia | Sequencing and Bioinformatics Service and Molecular Epidemiology Research Group. FISABIO-Public Health | Llúcia Martínez-Priego, Giuseppe 'Auria, David Navarro, Eliseo Albert, Maria Alma Bracho, Lidia Ruiz Roldan, Neris Garcia-Gonzalez, Inma Galán Vendrell, Sandra Carbo, Loreto Ferrús Abad, Paula Ruiz-Hueso, Mariana Reyes-Prieto, Vicente Soriano Chirona, Ivan Ansari, Fernando Gonzalez-Candelas |
| EPI_ISL_447484 | Servicio de Microbiologia. Hospital Clinico Universitario de Valencia | Sequencing and Bioinformatics Service and Molecular Epidemiology Research Group. FISABIO-Public Health | David Navarro, Eliseo Albert, Maria Alma Bracho, Griselda De Marco, Lidia Ruiz Roldan, Neris Garcia-Gonzalez, Inma Galán Vendrell, Sandra Carbo, Loreto Ferrús Abad, Paula Ruiz-Hueso, Mariana Reyes-Prieto, Vicente Soriano Chirona, Ivan Ansari, Llúcia Martínez-Priego, Giuseppe 'Auria, Fernando Gonzalez-Candelas |
| EPI_ISL_447485 | Servicio de Microbiologia. Hospital Clinico Universitario de Valencia | Sequencing and Bioinformatics Service and Molecular Epidemiology Research Group. FISABIO-Public Health | Eliseo Albert, Maria Alma Bracho, Griselda De Marco, Lidia Ruiz Roldan, Neris Garcia-Gonzalez, Inma Galán Vendrell, Sandra Carbo, Loreto Ferrús Abad, Paula Ruiz-Hueso, Mariana Reyes-Prieto, Vicente Soriano Chirona, Ivan Ansari, Llúcia Martínez-Priego, Giuseppe 'Auria, David Navarro, Fernando Gonzalez-Candelas |
| EPI_ISL_447486 | Servicio de Microbiologia. Hospital Clinico Universitario de Valencia | Sequencing and Bioinformatics Service and Molecular Epidemiology Research Group. FISABIO-Public Health | Maria Alma Bracho, Griselda De Marco, Lidia Ruiz Roldan, Neris Garcia-Gonzalez, Inma Galán Vendrell, Sandra Carbo, Loreto Ferrús Abad, Paula Ruiz-Hueso, Mariana Reyes-Prieto, Vicente Soriano Chirona, Ivan Ansari, Llúcia Martínez-Priego, Giuseppe 'Auria, David Navarro, Eliseo Albert, Fernando Gonzalez-Candelas |
| EPI_ISL_447487 | Servicio de Microbiologia. Hospital Clinico Universitario de Valencia | Sequencing and Bioinformatics Service and Molecular Epidemiology Research Group. FISABIO-Public Health | Griselda De Marco, Lidia Ruiz Roldan, Neris Garcia-Gonzalez, Inma Galán Vendrell, Sandra Carbo, Loreto Ferrús Abad, Paula Ruiz-Hueso, Mariana Reyes-Prieto, Vicente Soriano Chirona, Ivan Ansari, Llúcia Martínez-Priego, Giuseppe 'Auria, David Navarro, Eliseo Albert, Maria Alma Bracho, Fernando Gonzalez-Candelas |
| EPI_ISL_447488 | Servicio de Microbiologia. Hospital Clinico Universitario de Valencia | Sequencing and Bioinformatics Service and Molecular Epidemiology Research Group. FISABIO-Public Health | Lidia Ruiz Roldan, Neris Garcia-Gonzalez, Inma Galán Vendrell, Sandra Carbo, Loreto Ferrús Abad, Paula Ruiz-Hueso, Mariana Reyes-Prieto, Vicente Soriano Chirona, Ivan Ansari, Llúcia Martínez-Priego, Giuseppe 'Auria, David Navarro, Eliseo Albert, Maria Alma Bracho, Fernando Gonzalez-Candelas |
| EPI_ISL_447489 | Servicio de Microbiologia. Hospital Clinico Universitario de Valencia | Sequencing and Bioinformatics Service and Molecular Epidemiology Research Group. FISABIO-Public Health | Neris Garcia-Gonzalez, Inma Galán Vendrell, Sandra Carbo, Loreto Ferrús Abad, Paula Ruiz-Hueso, Mariana Reyes-Prieto, Vicente Soriano Chirona, Ivan Ansari, Llúcia Martínez-Priego, Giuseppe 'Auria, David Navarro, Eliseo Albert, Maria Alma Bracho, Lidia Ruiz Roldan, Fernando Gonzalez-Candelas |
| EPI_ISL_447490 | Servicio de Microbiologia. Hospital Clinico Universitario de Valencia | Sequencing and Bioinformatics Service and Molecular Epidemiology Research Group. FISABIO-Public Health | Inma Galán Vendrell, Sandra Carbo, Loreto Ferrús Abad, Paula Ruiz-Hueso, Mariana Reyes-Prieto, Vicente Soriano Chirona, Ivan Ansari, Llúcia Martínez-Priego, Giuseppe 'Auria, David Navarro, Eliseo Albert, Maria Alma Bracho, Lidia Ruiz Roldan, Neris Garcia-Gonzalez, Fernando Gonzalez-Candelas |
| EPI_ISL_447491 | Servicio de Microbiologia. Hospital Clinico Universitario de Valencia | Sequencing and Bioinformatics Service and Molecular Epidemiology Research Group. FISABIO-Public Health | Sandra Carbo, Loreto Ferrús Abad, Paula Ruiz-Hueso, Mariana Reyes-Prieto, Vicente Soriano Chirona, Ivan Ansari, Llúcia Martínez-Priego, Giuseppe 'Auria, David Navarro, Eliseo Albert, Maria Alma Bracho, Lidia Ruiz Roldan, Neris Garcia-Gonzalez, Inma Galán Vendrell, Fernando Gonzalez-Candelas |
| EPI_ISL_447492 | Servicio de Microbiologia. Hospital Clinico Universitario de Valencia | Sequencing and Bioinformatics Service and Molecular Epidemiology Research Group. FISABIO-Public Health | Loreto Ferrús Abad, Paula Ruiz-Hueso, Mariana Reyes-Prieto, Vicente Soriano Chirona, Ivan Ansari, Llúcia Martínez-Priego, Giuseppe 'Auria, David Navarro, Eliseo Albert, Maria Alma Bracho, Lidia Ruiz Roldan, Neris Garcia-Gonzalez, Inma Galán Vendrell, Sandra Carbo, Fernando Gonzalez-Candelas |

| | | | |
|---|---|---|---|
| EPI_ISL_447493 | Servicio de Microbiologia. Hospital Clinico Universitario de Valencia | Sequencing and Bioinformatics Service and Molecular Epidemiology Research Group. FISABIO-Public Health | Paula Ruiz-Hueso, Mariana Reyes-Prieto, Vicente Soriano Chirona, Ivan Ansari, Llúcia Martínez-Priego, Giuseppe 'Auria, David Navarro, Eliseo Albert, Maria Alma Bracho, Lidia Ruiz Roldan, Neris Garcia-Gonzalez, Inma Galán Vendrell, Sandra Carbo, Loreto Ferrús Abad, Fernando Gonzalez-Candelas |
| EPI_ISL_447494 | Servicio de Microbiologia. Hospital Clinico Universitario de Valencia | Sequencing and Bioinformatics Service and Molecular Epidemiology Research Group. FISABIO-Public Health | Mariana Reyes-Prieto, Vicente Soriano Chirona, Ivan Ansari, Llúcia Martínez-Priego, Giuseppe 'Auria, David Navarro, Eliseo Albert, Maria Alma Bracho, Lidia Ruiz Roldan, Neris Garcia-Gonzalez, Inma Galán Vendrell, Sandra Carbo, Loreto Ferrús Abad, Paula Ruiz-Hueso, Fernando Gonzalez-Candelas |
| EPI_ISL_447495 | Servicio de Microbiologia. Hospital Clinico Universitario de Valencia | Sequencing and Bioinformatics Service and Molecular Epidemiology Research Group. FISABIO-Public Health | Vicente Soriano Chirona, Ivan Ansari, Llúcia Martínez-Priego, Giuseppe 'Auria, David Navarro, Eliseo Albert, Maria Alma Bracho, Lidia Ruiz Roldan, Neris Garcia-Gonzalez, Inma Galán Vendrell, Sandra Carbo, Loreto Ferrús Abad, Paula Ruiz-Hueso, Mariana Reyes-Prieto, Fernando Gonzalez-Candelas |
| EPI_ISL_447496 | Servicio de Microbiologia. Hospital Clinico Universitario de Valencia | Sequencing and Bioinformatics Service and Molecular Epidemiology Research Group. FISABIO-Public Health | Giuseppe 'Auria, David Navarro, Eliseo Albert, Maria Alma Bracho, Lidia Ruiz Roldan, Neris Garcia-Gonzalez, Inma Galán Vendrell, Sandra Carbo, Loreto Ferrús Abad, Paula Ruiz-Hueso, Mariana Reyes-Prieto, Vicente Soriano Chirona, Ivan Ansari, Llúcia Martínez-Priego, Fernando Gonzalez-Candelas |
| EPI_ISL_447497 | Servicio de Microbiologia. Hospital Clinico Universitario de Valencia | Sequencing and Bioinformatics Service and Molecular Epidemiology Research Group. FISABIO-Public Health | Llúcia Martínez-Priego, Giuseppe 'Auria, David Navarro, Eliseo Albert, Maria Alma Bracho, Lidia Ruiz Roldan, Neris Garcia-Gonzalez, Inma Galán Vendrell, Sandra Carbo, Loreto Ferrús Abad, Paula Ruiz-Hueso, Mariana Reyes-Prieto, Vicente Soriano Chirona, Ivan Ansari, Fernando Gonzalez-Candelas |
| EPI_ISL_447498 | Servicio de Microbiologia. Hospital Clinico Universitario de Valencia | Sequencing and Bioinformatics Service and Molecular Epidemiology Research Group. FISABIO-Public Health | David Navarro, Eliseo Albert, Maria Alma Bracho, Griselda De Marco, Lidia Ruiz Roldan, Neris Garcia-Gonzalez, Inma Galán Vendrell, Sandra Carbo, Loreto Ferrús Abad, Paula Ruiz-Hueso, Mariana Reyes-Prieto, Vicente Soriano Chirona, Ivan Ansari, Llúcia Martínez-Priego, Giuseppe 'Auria, Fernando Gonzalez-Candelas |
| EPI_ISL_447499 | Servicio de Microbiologia. Hospital Clinico Universitario de Valencia | Sequencing and Bioinformatics Service and Molecular Epidemiology Research Group. FISABIO-Public Health | Eliseo Albert, Maria Alma Bracho, Griselda De Marco, Lidia Ruiz Roldan, Neris Garcia-Gonzalez, Inma Galán Vendrell, Sandra Carbo, Loreto Ferrús Abad, Paula Ruiz-Hueso, Mariana Reyes-Prieto, Vicente Soriano Chirona, Ivan Ansari, Llúcia Martínez-Priego, Giuseppe 'Auria, David Navarro, Fernando Gonzalez-Candelas |
| EPI_ISL_447500 | Servicio de Microbiologia. Hospital Clinico Universitario de Valencia | Sequencing and Bioinformatics Service and Molecular Epidemiology Research Group. FISABIO-Public Health | Maria Alma Bracho, Griselda De Marco, Lidia Ruiz Roldan, Neris Garcia-Gonzalez, Inma Galán Vendrell, Sandra Carbo, Loreto Ferrús Abad, Paula Ruiz-Hueso, Mariana Reyes-Prieto, Vicente Soriano Chirona, Ivan Ansari, Llúcia Martínez-Priego, Giuseppe 'Auria, David Navarro, Eliseo Albert, Fernando Gonzalez-Candelas |
| EPI_ISL_447501 | Servicio de Microbiologia. Hospital Clinico Universitario de Valencia | Sequencing and Bioinformatics Service and Molecular Epidemiology Research Group. FISABIO-Public Health | Griselda De Marco, Lidia Ruiz Roldan, Neris Garcia-Gonzalez, Inma Galán Vendrell, Sandra Carbo, Loreto Ferrús Abad, Paula Ruiz-Hueso, Mariana Reyes-Prieto, Vicente Soriano Chirona, Ivan Ansari, Llúcia Martínez-Priego, Giuseppe 'Auria, David Navarro, Eliseo Albert, Maria Alma Bracho, Fernando Gonzalez-Candelas |
| EPI_ISL_447502 | Servicio de Microbiologia. Hospital Clinico Universitario de Valencia | Sequencing and Bioinformatics Service and Molecular Epidemiology Research Group. FISABIO-Public Health | Lidia Ruiz Roldan, Neris Garcia-Gonzalez, Inma Galán Vendrell, Sandra Carbo, Loreto Ferrús Abad, Paula Ruiz-Hueso, Mariana Reyes-Prieto, Vicente Soriano Chirona, Ivan Ansari, Llúcia Martínez-Priego, Giuseppe 'Auria, David Navarro, Eliseo Albert, Maria Alma Bracho, Fernando Gonzalez-Candelas |
| EPI_ISL_447503 | Servicio de Microbiologia. Hospital Clinico Universitario de Valencia | Sequencing and Bioinformatics Service and Molecular Epidemiology Research Group. FISABIO-Public Health | Neris Garcia-Gonzalez, Inma Galán Vendrell, Sandra Carbo, Loreto Ferrús Abad, Paula Ruiz-Hueso, Mariana Reyes-Prieto, Vicente Soriano Chirona, Ivan Ansari, Llúcia Martínez-Priego, Giuseppe 'Auria, David Navarro, Eliseo Albert, Maria Alma Bracho, Lidia Ruiz Roldan, Fernando Gonzalez-Candelas |
| EPI_ISL_447504 | Servicio de Microbiologia. Hospital Clinico Universitario de Valencia | Sequencing and Bioinformatics Service and Molecular Epidemiology Research Group. FISABIO-Public Health | Inma Galán Vendrell, Sandra Carbo, Loreto Ferrús Abad, Paula Ruiz-Hueso, Mariana Reyes-Prieto, Vicente Soriano Chirona, Ivan Ansari, Llúcia Martínez-Priego, Giuseppe 'Auria, David Navarro, Eliseo Albert, Maria Alma Bracho, Lidia Ruiz Roldan, Neris Garcia-Gonzalez, Fernando Gonzalez-Candelas |
| EPI_ISL_447505 | Servicio de Microbiologia. Hospital Clinico Universitario de Valencia | Sequencing and Bioinformatics Service and Molecular Epidemiology Research Group. FISABIO-Public Health | Sandra Carbo, Loreto Ferrús Abad, Paula Ruiz-Hueso, Mariana Reyes-Prieto, Vicente Soriano Chirona, Ivan Ansari, Llúcia Martínez-Priego, Giuseppe 'Auria, David Navarro, Eliseo Albert, Maria Alma Bracho, Lidia Ruiz Roldan, Neris Garcia-Gonzalez, Inma Galán Vendrell, Fernando Gonzalez-Candelas |
| EPI_ISL_447506 | Servicio de Microbiologia. Hospital Clinico Universitario de Valencia | Sequencing and Bioinformatics Service and Molecular Epidemiology Research Group. FISABIO-Public Health | Loreto Ferrús Abad, Paula Ruiz-Hueso, Mariana Reyes-Prieto, Vicente Soriano Chirona, Ivan Ansari, Llúcia Martínez-Priego, Giuseppe 'Auria, David Navarro, Eliseo Albert, Maria Alma Bracho, Lidia Ruiz Roldan, Neris Garcia-Gonzalez, Inma Galán Vendrell, Sandra Carbo, Fernando Gonzalez-Candelas |
| EPI_ISL_447507 | Servicio de Microbiologia. Hospital Clinico Universitario de Valencia | Sequencing and Bioinformatics Service and Molecular Epidemiology Research Group. FISABIO-Public Health | Paula Ruiz-Hueso, Mariana Reyes-Prieto, Vicente Soriano Chirona, Ivan Ansari, Llúcia Martínez-Priego, Giuseppe 'Auria, David Navarro, Eliseo Albert, Maria Alma Bracho, Lidia Ruiz Roldan, Neris Garcia-Gonzalez, Inma Galán Vendrell, Sandra Carbo, Loreto Ferrús Abad, Fernando Gonzalez-Candelas |
| EPI_ISL_447508 | Servicio de Microbiologia. Hospital Clinico Universitario de Valencia | Sequencing and Bioinformatics Service and Molecular Epidemiology Research Group. FISABIO-Public Health | Mariana Reyes-Prieto, Vicente Soriano Chirona, Ivan Ansari, Llúcia Martínez-Priego, Giuseppe 'Auria, David Navarro, Eliseo Albert, Maria Alma Bracho, Lidia Ruiz Roldan, Neris Garcia-Gonzalez, Inma Galán Vendrell, Sandra Carbo, Loreto Ferrús Abad, Paula Ruiz-Hueso, Fernando Gonzalez-Candelas |
| EPI_ISL_447509 | Servicio de Microbiologia. Hospital Clinico Universitario de Valencia | Sequencing and Bioinformatics Service and Molecular Epidemiology Research Group. FISABIO-Public Health | Vicente Soriano Chirona, Ivan Ansari, Llúcia Martínez-Priego, Giuseppe 'Auria, David Navarro, Eliseo Albert, Maria Alma Bracho, Lidia Ruiz Roldan, Neris Garcia-Gonzalez, Inma Galán Vendrell, Sandra Carbo, Loreto Ferrús Abad, Paula Ruiz-Hueso, Mariana Reyes-Prieto, Fernando Gonzalez-Candelas |
| EPI_ISL_447510 | Servicio de Microbiologia. Hospital Clinico Universitario de Valencia | Sequencing and Bioinformatics Service and Molecular Epidemiology Research Group. FISABIO-Public Health | Giuseppe 'Auria, David Navarro, Eliseo Albert, Maria Alma Bracho, Lidia Ruiz Roldan, Neris Garcia-Gonzalez, Inma Galán Vendrell, Sandra Carbo, Loreto Ferrús Abad, Paula Ruiz-Hueso, Mariana Reyes-Prieto, Vicente Soriano Chirona, Ivan Ansari, Llúcia Martínez-Priego, Fernando Gonzalez-Candelas |
| EPI_ISL_447511 | Servicio de Microbiologia. Hospital Clinico Universitario de Valencia | Sequencing and Bioinformatics Service and Molecular Epidemiology Research Group. FISABIO-Public Health | Llúcia Martínez-Priego, Giuseppe 'Auria, David Navarro, Eliseo Albert, Maria Alma Bracho, Lidia Ruiz Roldan, Neris Garcia-Gonzalez, Inma Galán Vendrell, Sandra Carbo, Loreto Ferrús Abad, Paula Ruiz-Hueso, Mariana Reyes-Prieto, Vicente Soriano Chirona, Ivan Ansari, Fernando Gonzalez-Candelas |
| EPI_ISL_447512 | Servicio de Microbiologia. Hospital Clinico Universitario de Valencia | Sequencing and Bioinformatics Service and Molecular Epidemiology Research Group. FISABIO-Public Health | David Navarro, Eliseo Albert, Maria Alma Bracho, Griselda De Marco, Lidia Ruiz Roldan, Neris Garcia-Gonzalez, Inma Galán Vendrell, Sandra Carbo, Loreto Ferrús Abad, Paula Ruiz-Hueso, Mariana Reyes-Prieto, Vicente Soriano Chirona, Ivan Ansari, Llúcia Martínez-Priego, Giuseppe 'Auria, Fernando Gonzalez-Candelas |
| EPI_ISL_447513 | Servicio de Microbiologia. Hospital Clinico Universitario de Valencia | Sequencing and Bioinformatics Service and Molecular Epidemiology Research Group. FISABIO-Public Health | Eliseo Albert, Maria Alma Bracho, Griselda De Marco, Lidia Ruiz Roldan, Neris Garcia-Gonzalez, Inma Galán Vendrell, Sandra Carbo, Loreto Ferrús Abad, Paula Ruiz-Hueso, Mariana Reyes-Prieto, Vicente Soriano Chirona, Ivan Ansari, Llúcia Martínez-Priego, Giuseppe 'Auria, David Navarro, Fernando Gonzalez-Candelas |
| EPI_ISL_447514 | Servicio de Microbiologia. Hospital Clinico Universitario de Valencia | Sequencing and Bioinformatics Service and Molecular Epidemiology Research Group. FISABIO-Public Health | Maria Alma Bracho, Griselda De Marco, Lidia Ruiz Roldan, Neris Garcia-Gonzalez, Inma Galán Vendrell, Sandra Carbo, Loreto Ferrús Abad, Paula Ruiz-Hueso, Mariana Reyes-Prieto, Vicente Soriano Chirona, Ivan Ansari, Llúcia Martínez-Priego, Giuseppe 'Auria, David Navarro, Eliseo Albert, Fernando Gonzalez-Candelas |
| EPI_ISL_447515 | Servicio de Microbiologia. Hospital Clinico Universitario de Valencia | Sequencing and Bioinformatics Service and Molecular Epidemiology Research Group. FISABIO-Public Health | Griselda De Marco, Lidia Ruiz Roldan, Neris Garcia-Gonzalez, Inma Galán Vendrell, Sandra Carbo, Loreto Ferrús Abad, Paula Ruiz-Hueso, Mariana Reyes-Prieto, Vicente Soriano Chirona, Ivan Ansari, Llúcia Martínez-Priego, Giuseppe 'Auria, David Navarro, Eliseo Albert, Maria Alma Bracho, Fernando Gonzalez-Candelas |
| EPI_ISL_447516 | Servicio de Microbiologia. Hospital Clinico Universitario de Valencia | Sequencing and Bioinformatics Service and Molecular Epidemiology Research Group. FISABIO-Public Health | Lidia Ruiz Roldan, Neris Garcia-Gonzalez, Inma Galán Vendrell, Sandra Carbo, Loreto Ferrús Abad, Paula Ruiz-Hueso, Mariana Reyes-Prieto, Vicente Soriano Chirona, Ivan Ansari, Llúcia Martínez-Priego, Giuseppe 'Auria, David Navarro, Eliseo Albert, Maria Alma Bracho, Fernando Gonzalez-Candelas |
| EPI_ISL_447517 | Servicio de Microbiologia. Hospital Clinico Universitario de Valencia | Sequencing and Bioinformatics Service and Molecular Epidemiology Research Group. FISABIO-Public Health | Neris Garcia-Gonzalez, Inma Galán Vendrell, Sandra Carbo, Loreto Ferrús Abad, Paula Ruiz-Hueso, Mariana Reyes-Prieto, Vicente Soriano Chirona, Ivan Ansari, Llúcia Martínez-Priego, Giuseppe 'Auria, David Navarro, Eliseo Albert, Maria Alma Bracho, Lidia Ruiz Roldan, Fernando Gonzalez-Candelas |
| EPI_ISL_447518 | Servicio de Microbiologia. Hospital Clinico Universitario de Valencia | Sequencing and Bioinformatics Service and Molecular Epidemiology Research Group. FISABIO-Public Health | Inma Galán Vendrell, Sandra Carbo, Loreto Ferrús Abad, Paula Ruiz-Hueso, Mariana Reyes-Prieto, Vicente Soriano Chirona, Ivan Ansari, Llúcia Martínez-Priego, Giuseppe 'Auria, David Navarro, Eliseo Albert, Maria Alma Bracho, Lidia Ruiz Roldan, Neris Garcia-Gonzalez, Fernando Gonzalez-Candelas |
| EPI_ISL_447519 | Servicio de Microbiologia. Hospital Clinico Universitario de Valencia | Sequencing and Bioinformatics Service and Molecular Epidemiology Research Group. FISABIO-Public Health | Sandra Carbo, Loreto Ferrús Abad, Paula Ruiz-Hueso, Mariana Reyes-Prieto, Vicente Soriano Chirona, Ivan Ansari, Llúcia Martínez-Priego, Giuseppe 'Auria, David Navarro, Eliseo Albert, Maria Alma Bracho, Lidia Ruiz Roldan, Neris Garcia-Gonzalez, Inma Galán Vendrell, Fernando Gonzalez-Candelas |
| EPI_ISL_447520 | Servicio de Microbiologia. Hospital Clinico Universitario de Valencia | Sequencing and Bioinformatics Service and Molecular Epidemiology Research Group. FISABIO-Public Health | Loreto Ferrús Abad, Paula Ruiz-Hueso, Mariana Reyes-Prieto, Vicente Soriano Chirona, Ivan Ansari, Llúcia Martínez-Priego, Giuseppe 'Auria, David Navarro, Eliseo Albert, Maria Alma Bracho, Lidia Ruiz Roldan, Neris Garcia-Gonzalez, Inma Galán Vendrell, Sandra Carbo, Fernando Gonzalez-Candelas |
| EPI_ISL_447521 | Servicio de Microbiologia. Hospital Clinico Universitario de Valencia | Sequencing and Bioinformatics Service and Molecular Epidemiology Research Group. FISABIO-Public Health | Paula Ruiz-Hueso, Mariana Reyes-Prieto, Vicente Soriano Chirona, Ivan Ansari, Llúcia Martínez-Priego, Giuseppe 'Auria, David Navarro, Eliseo Albert, Maria Alma Bracho, Lidia Ruiz Roldan, Neris Garcia-Gonzalez, Inma Galán Vendrell, Sandra Carbo, Loreto Ferrús Abad, Fernando Gonzalez-Candelas |
| EPI_ISL_447522 | Servicio de Microbiologia. Hospital Clinico Universitario de Valencia | Sequencing and Bioinformatics Service and Molecular Epidemiology Research Group. FISABIO-Public Health | Vicente Soriano Chirona, Ivan Ansari, Llúcia Martínez-Priego, Giuseppe 'Auria, David Navarro, Eliseo Albert, Maria Alma Bracho, Lidia Ruiz Roldan, Neris Garcia-Gonzalez, Inma Galán Vendrell, Sandra Carbo, Loreto Ferrús Abad, Paula Ruiz-Hueso, Mariana Reyes-Prieto, Fernando Gonzalez-Candelas |
| EPI_ISL_447523 | Servicio de Microbiologia. Hospital Clinico Universitario de Valencia | Sequencing and Bioinformatics Service and Molecular Epidemiology Research Group. FISABIO-Public Health | Giuseppe 'Auria, David Navarro, Eliseo Albert, Maria Alma Bracho, Lidia Ruiz Roldan, Neris Garcia-Gonzalez, Inma Galán Vendrell, Sandra Carbo, Loreto Ferrús Abad, Paula Ruiz-Hueso, Mariana Reyes-Prieto, Vicente Soriano Chirona, Ivan Ansari, Llúcia Martínez-Priego, Fernando Gonzalez-Candelas |
| EPI_ISL_447524 | Servicio de Microbiologia. Hospital Clinico Universitario de Valencia | Sequencing and Bioinformatics Service and Molecular Epidemiology Research Group. FISABIO-Public Health | Llúcia Martínez-Priego, Giuseppe 'Auria, David Navarro, Eliseo Albert, Maria Alma Bracho, Lidia Ruiz Roldan, Neris Garcia-Gonzalez, Inma Galán Vendrell, Sandra Carbo, Loreto Ferrús Abad, Paula Ruiz-Hueso, Mariana Reyes-Prieto, Vicente Soriano Chirona, Ivan Ansari, Fernando Gonzalez-Candelas |
| EPI_ISL_447525 | Servicio de Microbiologia. Hospital Clinico Universitario de Valencia | Sequencing and Bioinformatics Service and Molecular Epidemiology Research Group. FISABIO-Public Health | David Navarro, Eliseo Albert, Maria Alma Bracho, Griselda De Marco, Lidia Ruiz Roldan, Neris Garcia-Gonzalez, Inma Galán Vendrell, Sandra Carbo, Loreto Ferrús Abad, Paula Ruiz-Hueso, Mariana Reyes-Prieto, Vicente Soriano Chirona, Ivan Ansari, Llúcia Martínez-Priego, Giuseppe 'Auria, Fernando Gonzalez-Candelas |
| EPI_ISL_447526 | Servicio de Microbiologia. Hospital Clinico Universitario de Valencia | Sequencing and Bioinformatics Service and Molecular Epidemiology Research Group. FISABIO-Public Health | Eliseo Albert, Maria Alma Bracho, Griselda De Marco, Lidia Ruiz Roldan, Neris Garcia-Gonzalez, Inma Galán Vendrell, Sandra Carbo, Loreto Ferrús Abad, Paula Ruiz-Hueso, Mariana Reyes-Prieto, Vicente Soriano Chirona, Ivan Ansari, Llúcia Martínez-Priego, Giuseppe 'Auria, David Navarro, Fernando Gonzalez-Candelas |
| EPI_ISL_447527 | Servicio de Microbiologia. Hospital Clinico Universitario de Valencia | Sequencing and Bioinformatics Service and Molecular Epidemiology Research Group. FISABIO-Public Health | Maria Alma Bracho, Griselda De Marco, Lidia Ruiz Roldan, Neris Garcia-Gonzalez, Inma Galán Vendrell, Sandra Carbo, Loreto Ferrús Abad, Paula Ruiz-Hueso, Mariana Reyes-Prieto, Vicente Soriano Chirona, Ivan Ansari, Llúcia Martínez-Priego, Giuseppe 'Auria, David Navarro, Eliseo Albert, Fernando Gonzalez-Candelas |
| EPI_ISL_447528 | Servicio de Microbiologia. Hospital Clinico Universitario de Valencia | Sequencing and Bioinformatics Service and Molecular Epidemiology Research Group. FISABIO-Public Health | Griselda De Marco, Lidia Ruiz Roldan, Neris Garcia-Gonzalez, Inma Galán Vendrell, Sandra Carbo, Loreto Ferrús Abad, Paula Ruiz-Hueso, Mariana Reyes-Prieto, Vicente Soriano Chirona, Ivan Ansari, Llúcia Martínez-Priego, Giuseppe 'Auria, David Navarro, Eliseo Albert, Maria Alma Bracho, Fernando Gonzalez-Candelas |
| EPI_ISL_447529 | Servicio de Microbiologia. Hospital Clinico Universitario de Valencia | Sequencing and Bioinformatics Service and Molecular Epidemiology Research Group. FISABIO-Public Health | Lidia Ruiz Roldan, Neris Garcia-Gonzalez, Inma Galán Vendrell, Sandra Carbo, Loreto Ferrús Abad, Paula Ruiz-Hueso, Mariana Reyes-Prieto, Vicente Soriano Chirona, Ivan Ansari, Llúcia Martínez-Priego, Giuseppe 'Auria, David Navarro, Eliseo Albert, Maria Alma Bracho, Fernando Gonzalez-Candelas |
| EPI_ISL_447530 | Servicio de Microbiologia. Hospital Clinico Universitario de Valencia | Sequencing and Bioinformatics Service and Molecular Epidemiology Research Group. FISABIO-Public Health | Neris Garcia-Gonzalez, Inma Galán Vendrell, Sandra Carbo, Loreto Ferrús Abad, Paula Ruiz-Hueso, Mariana Reyes-Prieto, Vicente Soriano Chirona, Ivan Ansari, Llúcia Martínez-Priego, Giuseppe 'Auria, David Navarro, Eliseo Albert, Maria Alma Bracho, Lidia Ruiz Roldan, Fernando Gonzalez-Candelas |
| EPI_ISL_447531 | Servicio de Microbiologia. Hospital Clinico Universitario de Valencia | Sequencing and Bioinformatics Service and Molecular Epidemiology Research Group. FISABIO-Public Health | Inma Galán Vendrell, Sandra Carbo, Loreto Ferrús Abad, Paula Ruiz-Hueso, Mariana Reyes-Prieto, Vicente Soriano Chirona, Ivan Ansari, Llúcia Martínez-Priego, Giuseppe 'Auria, David Navarro, Eliseo Albert, Maria Alma Bracho, Lidia Ruiz Roldan, Neris Garcia-Gonzalez, Fernando Gonzalez-Candelas |
| EPI_ISL_447532, EPI_ISL_447533 | Hospital Universitari Vall d'Hebron - Vall d'Hebron Institut de Recerca | Hospital Universitari Vall d'Hebron | Cristina Andrés, Maria Piñana, Damir Garcia-Cehic, Mercedes Guerrero-Murillo, Ariadna Rando, Juliana Esperalba, Maria Gema Codina, Tomàs Pumarola, Josep Quer, Andrés Antón |
| EPI_ISL_447534 | Gujarat Biotechnology Research Centre | Gujarat Biotechnology Research Centre | Gaurishankar Shrimali, Nidhi Sood, Pranay Shah, R D Dixit, Snehal Bagatharia, Kamlesh J Upadhyay, Ramesh Pandit, Tejas Shah, Ankit Hinsu, Pritesh Sabara, Apurvasinh Puvar, Janvi Raval, Monika Gandhi, Pinal Trivedi, Maharshi Pandya, Amit Kanani, Akanksha Verma, Nitin Savaliya, Raghawendra Kumar, Dinesh Kumar, Zuber Saiyed, Dipa Kinariwala, Disha Patel, Binita Aring, Neeta Khandelwal, Geeta Vaghela, Sonia Barve, Bhavesh Modi, Kairavi Joshi, Nidhi Patel, Chaitanya Joshi, Madhvi Joshi |
| EPI_ISL_447535 | Gujarat Biotechnology Research Centre | Gujarat Biotechnology Research Centre | Nidhi Sood, Pranay Shah, R D Dixit, Snehal Bagatharia, Kamlesh J Upadhyay, Ramesh Pandit, Tejas Shah, Ankit Hinsu, Pritesh Sabara, Apurvasinh Puvar, Janvi Raval, Monika Gandhi, Pinal Trivedi, Maharshi Pandya, Amit Kanani, Akanksha Verma, Nitin Savaliya, Raghawendra Kumar, Dinesh Kumar, Zuber Saiyed, Dipa Kinariwala, Disha Patel, Binita Aring, Neeta Khandelwal, Geeta Vaghela, Sonia Barve, Bhavesh Modi, Kairavi Joshi, Gaurishankar Shrimali, Priti Pandita, Chaitanya Joshi, Madhvi Joshi |
| EPI_ISL_447536 | Gujarat Biotechnology Research Centre | Gujarat Biotechnology Research Centre | Pranay Shah, R D Dixit, Snehal Bagatharia, Kamlesh J Upadhyay, Ramesh Pandit, Tejas Shah, Ankit Hinsu, Pritesh Sabara, Apurvasinh Puvar, Janvi Raval, Monika Gandhi, Pinal Trivedi, Maharshi Pandya, Amit Kanani, Akanksha Verma, Nitin Savaliya, Raghawendra Kumar, Dinesh Kumar, Zuber Saiyed, Dipa Kinariwala, Disha Patel, Binita Aring, Neeta Khandelwal, Geeta Vaghela, Sonia Barve, Bhavesh Modi, Kairavi Joshi, Gaurishankar Shrimali, Nidhi Sood, Neha Rajpara, Chaitanya Joshi, Madhvi Joshi |
| EPI_ISL_447537 | Gujarat Biotechnology Research Centre | Gujarat Biotechnology Research Centre | R D Dixit, Snehal Bagatharia, Kamlesh J Upadhyay, Ramesh Pandit, Tejas Shah, Ankit Hinsu, Pritesh Sabara, Apurvasinh Puvar, Janvi Raval, Monika Gandhi, Pinal Trivedi, Maharshi Pandya, Amit Kanani, Akanksha |

| | | | |
|---|---|---|---|
| | | | Verma, Nitin Savaliya, Raghawendra Kumar, Dinesh Kumar, Zuber Saiyed, Dipa Kinariwala, Disha Patel, Binita Aring, Neeta Khandelwal, Geeta Vaghela, Sonia Barve, Bhavesh Modi, Kairavi Joshi, Gaurishankar Shrimali, Nidhi Sood, Pranay Shah, Afzal Ansari, Chaitanya Joshi, Madhvi Joshi |
| EPI_ISL_447538 | Gujarat Biotechnology Research Centre | Gujarat Biotechnology Research Centre | Snehal Bagatharia, Kamlesh J Upadhyay, Ramesh Pandit, Tejas Shah, Ankit Hinsu, Pritesh Sabara, Apurvasinh Puvar, Janvi Raval, Monika Gandhi, Pinal Trivedi, Maharshi Pandya, Amit Kanani, Akanksha Verma, Nitin Savaliya, Raghawendra Kumar, Dinesh Kumar, Zuber Saiyed, Dipa Kinariwala, Disha Patel, Binita Aring, Neeta Khandelwal, Geeta Vaghela, Sonia Barve, Bhavesh Modi, Kairavi Joshi, Gaurishankar Shrimali, Nidhi Sood, Pranay Shah, R D Dixit, Neelam Nathani, Chaitanya Joshi, Madhvi Joshi |
| EPI_ISL_447539 | Gujarat Biotechnology Research Centre | Gujarat Biotechnology Research Centre | Kamlesh J Upadhyay, Ramesh Pandit, Tejas Shah, Ankit Hinsu, Pritesh Sabara, Apurvasinh Puvar, Janvi Raval, Monika Gandhi, Pinal Trivedi, Maharshi Pandya, Amit Kanani, Akanksha Verma, Nitin Savaliya, Raghawendra Kumar, Dinesh Kumar, Zuber Saiyed, Dipa Kinariwala, Disha Patel, Binita Aring, Neeta Khandelwal, Geeta Vaghela, Sonia Barve, Bhavesh Modi, Kairavi Joshi, Gaurishankar Shrimali, Nidhi Sood, Pranay Shah, R D Dixit, Snehal Bagatharia, Armi Chaudhari, Chaitanya Joshi, Madhvi Joshi |
| EPI_ISL_447540 | Gujarat Biotechnology Research Centre | Gujarat Biotechnology Research Centre | Ramesh Pandit, Tejas Shah, Ankit Hinsu, Pritesh Sabara, Apurvasinh Puvar, Janvi Raval, Monika Gandhi, Pinal Trivedi, Maharshi Pandya, Amit Kanani, Akanksha Verma, Nitin Savaliya, Raghawendra Kumar, Dinesh Kumar, Zuber Saiyed, Dipa Kinariwala, Disha Patel, Binita Aring, Neeta Khandelwal, Geeta Vaghela, Sonia Barve, Bhavesh Modi, Kairavi Joshi, Gaurishankar Shrimali, Nidhi Sood, Pranay Shah, R D Dixit, Snehal Bagatharia, Kamlesh J Upadhyay, Bhavya Jindal, Chaitanya Joshi, Madhvi Joshi |
| EPI_ISL_447541 | Gujarat Biotechnology Research Centre | Gujarat Biotechnology Research Centre | Tejas Shah, Ankit Hinsu, Pritesh Sabara, Apurvasinh Puvar, Janvi Raval, Monika Gandhi, Pinal Trivedi, Maharshi Pandya, Amit Kanani, Akanksha Verma, Nitin Savaliya, Raghawendra Kumar, Dinesh Kumar, Zuber Saiyed, Dipa Kinariwala, Disha Patel, Binita Aring, Neeta Khandelwal, Geeta Vaghela, Sonia Barve, Bhavesh Modi, Kairavi Joshi, Gaurishankar Shrimali, Nidhi Sood, Pranay Shah, R D Dixit, Snehal Bagatharia, Kamlesh J Upadhyay, Ramesh Pandit, Anjali Rajwar, Chaitanya Joshi, Madhvi Joshi |
| EPI_ISL_447542 | Gujarat Biotechnology Research Centre | Gujarat Biotechnology Research Centre | Ankit Hinsu, Pritesh Sabara, Apurvasinh Puvar, Janvi Raval, Monika Gandhi, Pinal Trivedi, Maharshi Pandya, Amit Kanani, Akanksha Verma, Nitin Savaliya, Raghawendra Kumar, Dinesh Kumar, Zuber Saiyed, Dipa Kinariwala, Disha Patel, Binita Aring, Neeta Khandelwal, Geeta Vaghela, Sonia Barve, Bhavesh Modi, Kairavi Joshi, Gaurishankar Shrimali, Nidhi Sood, Pranay Shah, R D Dixit, Snehal Bagatharia, Kamlesh J Upadhyay, Ramesh Pandit, Tejas Shah, Dipeshwari Shewale, Chaitanya Joshi, Madhvi Joshi |
| EPI_ISL_447543 | Gujarat Biotechnology Research Centre | Gujarat Biotechnology Research Centre | Pritesh Sabara, Apurvasinh Puvar, Janvi Raval, Monika Gandhi, Pinal Trivedi, Maharshi Pandya, Amit Kanani, Akanksha Verma, Nitin Savaliya, Raghawendra Kumar, Dinesh Kumar, Zuber Saiyed, Dipa Kinariwala, Disha Patel, Binita Aring, Neeta Khandelwal, Geeta Vaghela, Sonia Barve, Bhavesh Modi, Kairavi Joshi, Gaurishankar Shrimali, Nidhi Sood, Pranay Shah, R D Dixit, Snehal Bagatharia, Kamlesh J Upadhyay, Ramesh Pandit, Tejas Shah, Ankit Hinsu, Sharmistha Majumdar, Chaitanya Joshi, Madhvi Joshi |
| EPI_ISL_447544 | Gujarat Biotechnology Research Centre | Gujarat Biotechnology Research Centre | Apurvasinh Puvar, Janvi Raval, Monika Gandhi, Pinal Trivedi, Maharshi Pandya, Amit Kanani, Akanksha Verma, Nitin Savaliya, Raghawendra Kumar, Dinesh Kumar, Zuber Saiyed, Dipa Kinariwala, Disha Patel, Binita Aring, Neeta Khandelwal, Geeta Vaghela, Sonia Barve, Bhavesh Modi, Kairavi Joshi, Gaurishankar Shrimali, Nidhi Sood, Pranay Shah, R D Dixit, Snehal Bagatharia, Kamlesh J Upadhyay, Ramesh Pandit, Tejas Shah, Ankit Hinsu, Pritesh Sabara, Pooja P Doshi, Chaitanya Joshi, Madhvi Joshi |
| EPI_ISL_447545 | Gujarat Biotechnology Research Centre | Gujarat Biotechnology Research Centre | Janvi Raval, Monika Gandhi, Pinal Trivedi, Maharshi Pandya, Amit Kanani, Akanksha Verma, Nitin Savaliya, Raghawendra Kumar, Dinesh Kumar, Zuber Saiyed, Dipa Kinariwala, Disha Patel, Binita Aring, Neeta Khandelwal, Geeta Vaghela, Sonia Barve, Bhavesh Modi, Kairavi Joshi, Gaurishankar Shrimali, Nidhi Sood, Pranay Shah, R D Dixit, Snehal Bagatharia, Kamlesh J Upadhyay, Ramesh Pandit, Tejas Shah, Ankit Hinsu, Pritesh Sabara, Apurvasinh Puvar, Nidhi Patel, Chaitanya Joshi, Madhvi Joshi |
| EPI_ISL_447546 | Gujarat Biotechnology Research Centre | Gujarat Biotechnology Research Centre | Monika Gandhi, Pinal Trivedi, Maharshi Pandya, Amit Kanani, Akanksha Verma, Nitin Savaliya, Raghawendra Kumar, Dinesh Kumar, Zuber Saiyed, Dipa Kinariwala, Disha Patel, Binita Aring, Neeta Khandelwal, Geeta Vaghela, Sonia Barve, Bhavesh Modi, Kairavi Joshi, Gaurishankar Shrimali, Nidhi Sood, Pranay Shah, R D Dixit, Snehal Bagatharia, Kamlesh J Upadhyay, Ramesh Pandit, Tejas Shah, Ankit Hinsu, Pritesh Sabara, Apurvasinh Puvar, Janvi Raval, Priti Pandita, Chaitanya Joshi, Madhvi Joshi |
| EPI_ISL_447547 | GMERS Medical College and Hospital, Gandhinagar | Gujarat Biotechnology Research Centre | Pinal Trivedi, Maharshi Pandya, Amit Kanani, Akanksha Verma, Nitin Savaliya, Raghawendra Kumar, Dinesh Kumar, Zuber Saiyed, Dipa Kinariwala, Disha Patel, Binita Aring, Neeta Khandelwal, Geeta Vaghela, Sonia Barve, Bhavesh Modi, Kairavi Joshi, Gaurishankar Shrimali, Nidhi Sood, Pranay Shah, R D Dixit, Snehal Bagatharia, Kamlesh J Upadhyay, Ramesh Pandit, Tejas Shah, Ankit Hinsu, Pritesh Sabara, Apurvasinh Puvar, Janvi Raval, Monika Gandhi, Neha Rajpara, Chaitanya Joshi, Madhvi Joshi |
| EPI_ISL_447548 | GMERS Medical College and Hospital, Gandhinagar | Gujarat Biotechnology Research Centre | Maharshi Pandya, Amit Kanani, Akanksha Verma, Nitin Savaliya, Raghawendra Kumar, Dinesh Kumar, Zuber Saiyed, Dipa Kinariwala, Disha Patel, Binita Aring, Neeta Khandelwal, Geeta Vaghela, Sonia Barve, Bhavesh Modi, Kairavi Joshi, Gaurishankar Shrimali, Nidhi Sood, Pranay Shah, R D Dixit, Snehal Bagatharia, Kamlesh J Upadhyay, Ramesh Pandit, Tejas Shah, Ankit Hinsu, Pritesh Sabara, Apurvasinh Puvar, Janvi Raval, Monika Gandhi, Pinal Trivedi, Afzal Ansari, Chaitanya Joshi, Madhvi Joshi |
| EPI_ISL_447549 | GMERS Medical College and Hospital, Gandhinagar | Gujarat Biotechnology Research Centre | Amit Kanani, Akanksha Verma, Nitin Savaliya, Raghawendra Kumar, Dinesh Kumar, Zuber Saiyed, Dipa Kinariwala, Disha Patel, Binita Aring, Neeta Khandelwal, Geeta Vaghela, Sonia Barve, Bhavesh Modi, Kairavi Joshi, Gaurishankar Shrimali, Nidhi Sood, Pranay Shah, R D Dixit, Snehal Bagatharia, Kamlesh J Upadhyay, Ramesh Pandit, Tejas Shah, Ankit Hinsu, Pritesh Sabara, Apurvasinh Puvar, Janvi Raval, Monika Gandhi, Pinal Trivedi, Maharshi Pandya, Neelam Nathani, Chaitanya Joshi, Madhvi Joshi |
| EPI_ISL_447550 | GMERS Medical College and Hospital, Gandhinagar | Gujarat Biotechnology Research Centre | Akanksha Verma, Nitin Savaliya, Raghawendra Kumar, Dinesh Kumar, Zuber Saiyed, Dipa Kinariwala, Disha Patel, Binita Aring, Neeta Khandelwal, Geeta Vaghela, Sonia Barve, Bhavesh Modi, Kairavi Joshi, Gaurishankar Shrimali, Nidhi Sood, Pranay Shah, R D Dixit, Snehal Bagatharia, Kamlesh J Upadhyay, Ramesh Pandit, Tejas Shah, Ankit Hinsu, Pritesh Sabara, Apurvasinh Puvar, Janvi Raval, Monika Gandhi, Pinal Trivedi, Maharshi Pandya, Amit Kanani, Armi Chaudhari, Chaitanya Joshi, Madhvi Joshi |
| EPI_ISL_447551 | GMERS Medical College and Hospital, Gandhinagar | Gujarat Biotechnology Research Centre | Nitin Savaliya, Raghawendra Kumar, Dinesh Kumar, Zuber Saiyed, Dipa Kinariwala, Disha Patel, Binita Aring, Neeta Khandelwal, Geeta Vaghela, Sonia Barve, Bhavesh Modi, Kairavi Joshi, Gaurishankar Shrimali, Nidhi Sood, Pranay Shah, R D Dixit, Snehal Bagatharia, Kamlesh J Upadhyay, Ramesh Pandit, Tejas Shah, Ankit Hinsu, Pritesh Sabara, Apurvasinh Puvar, Janvi Raval, Monika Gandhi, Pinal Trivedi, Maharshi Pandya, Amit Kanani, Akanksha Verma, Bhavya Jindal, Chaitanya Joshi, Madhvi Joshi |
| EPI_ISL_447552 | GMERS Medical College and Hospital, Gandhinagar | Gujarat Biotechnology Research Centre | Raghawendra Kumar, Dinesh Kumar, Zuber Saiyed, Dipa Kinariwala, Disha Patel, Binita Aring, Neeta Khandelwal, Geeta Vaghela, Sonia Barve, Bhavesh Modi, Kairavi Joshi, Gaurishankar Shrimali, Nidhi Sood, Pranay Shah, R D Dixit, Snehal Bagatharia, Kamlesh J Upadhyay, Ramesh Pandit, Tejas Shah, Ankit Hinsu, Pritesh Sabara, Apurvasinh Puvar, Janvi Raval, Monika Gandhi, Pinal Trivedi, Maharshi Pandya, Amit Kanani, Akanksha Verma, Nitin Savaliya, Anjali Rajwar, Chaitanya Joshi, Madhvi Joshi |
| EPI_ISL_447553 | GMERS Medical College and Hospital, Gandhinagar | Gujarat Biotechnology Research Centre | Dinesh Kumar, Zuber Saiyed, Dipa Kinariwala, Disha Patel, Binita Aring, Neeta Khandelwal, Geeta Vaghela, Sonia Barve, Bhavesh Modi, Kairavi Joshi, Gaurishankar Shrimali, Nidhi Sood, Pranay Shah, R D Dixit, Snehal Bagatharia, Kamlesh J Upadhyay, Ramesh Pandit, Tejas Shah, Ankit Hinsu, Pritesh Sabara, Apurvasinh Puvar, Janvi Raval, Monika Gandhi, Pinal Trivedi, Maharshi Pandya, Amit Kanani, Akanksha Verma, Nitin Savaliya, Raghawendra Kumar, Dipeshwari Shewale, Chaitanya Joshi, Madhvi Joshi |
| EPI_ISL_447554 | GMERS Medical College and Hospital, Gandhinagar | Gujarat Biotechnology Research Centre | Zuber Saiyed, Dipa Kinariwala, Disha Patel, Binita Aring, Neeta Khandelwal, Geeta Vaghela, Sonia Barve, Bhavesh Modi, Kairavi Joshi, Gaurishankar Shrimali, Nidhi Sood, Pranay Shah, R D Dixit, Snehal Bagatharia, Kamlesh J Upadhyay, Ramesh Pandit, Tejas Shah, Ankit Hinsu, Pritesh Sabara, Apurvasinh Puvar, Janvi Raval, Monika Gandhi, Pinal Trivedi, Maharshi Pandya, Amit Kanani, Akanksha Verma, Nitin Savaliya, Raghawendra Kumar, Dinesh Kumar, Sharmistha Majumdar, Chaitanya Joshi, Madhvi Joshi |
| EPI_ISL_447555 | GMERS Medical College and Hospital, Gandhinagar | Gujarat Biotechnology Research Centre | Dipa Kinariwala, Disha Patel, Binita Aring, Neeta Khandelwal, Geeta Vaghela, Sonia Barve, Bhavesh Modi, Kairavi Joshi, Gaurishankar Shrimali, Nidhi Sood, Pranay Shah, R D Dixit, Snehal Bagatharia, Kamlesh J Upadhyay, Ramesh Pandit, Tejas Shah, Ankit Hinsu, Pritesh Sabara, Apurvasinh Puvar, Janvi Raval, Monika Gandhi, Pinal Trivedi, Maharshi Pandya, Amit Kanani, Akanksha Verma, Nitin Savaliya, Raghawendra Kumar, Dinesh Kumar, Zuber Saiyed, Pooja P Doshi, Chaitanya Joshi, Madhvi Joshi |
| EPI_ISL_447556 | CSIR-Centre for Cellular and Molecular Biology | CSIR-Centre for Cellular and Molecular Biology | Sofia Banu, Payel Mukherjee, Priya Singh, Dhiviya Vedagiri, Divya Gupta, Vishal Sah, Santosh Kumar Kuncha, Krishnan Harinivas Harshan, Archana Bharadwaj Siva, Karthik Bharadwaj Tallapaka, Shagufta Khan, Lamuk Zaveri, Namami Gaur, Sakshi Shambhavi, Tulasi Nagabandi, Purushotham Vodnala, Rakesh K Mishra, Divya Tej Sowpati |
| EPI_ISL_447557, EPI_ISL_447558 | CSIR-Centre for Cellular and Molecular Biology | CSIR-Centre for Cellular and Molecular Biology | Payel Mukherjee, Sofia Banu, Priya Singh, Dhiviya Vedagiri, Divya Gupta, Vishal Sah, Santosh Kumar Kuncha, Krishnan Harinivas Harshan, Archana Bharadwaj Siva, Karthik Bharadwaj Tallapaka, Shagufta Khan, Lamuk Zaveri, Namami Gaur, Sakshi Shambhavi, Tulasi Nagabandi, Purushotham Vodnala, Rakesh K Mishra, Divya Tej Sowpati |
| EPI_ISL_447559 | CSIR-Centre for Cellular and Molecular Biology | CSIR-Centre for Cellular and Molecular Biology | Sofia Banu, Payel Mukherjee, Priya Singh, Dhiviya Vedagiri, Divya Gupta, Vishal Sah, Santosh Kumar Kuncha, Krishnan Harinivas Harshan, Archana Bharadwaj Siva, Karthik Bharadwaj Tallapaka, Shagufta Khan, Lamuk Zaveri, Namami Gaur, Sakshi Shambhavi, Tulasi Nagabandi, Purushotham Vodnala, Rakesh K Mishra, Divya Tej Sowpati |
| EPI_ISL_447560 | CSIR-Centre for Cellular and Molecular Biology | CSIR-Centre for Cellular and Molecular Biology | Payel Mukherjee, Sofia Banu, Priya Singh, Dhiviya Vedagiri, Divya Gupta, Vishal Sah, Santosh Kumar Kuncha, Krishnan Harinivas Harshan, Archana Bharadwaj Siva, Karthik Bharadwaj Tallapaka, Shagufta Khan, Lamuk Zaveri, Namami Gaur, Sakshi Shambhavi, Tulasi Nagabandi, Purushotham Vodnala, Rakesh K Mishra, Divya Tej Sowpati |
| EPI_ISL_447561, EPI_ISL_447562 | CSIR-Centre for Cellular and Molecular Biology | CSIR-Centre for Cellular and Molecular Biology | Shagufta Khan, Lamuk Zaveri, Namami Gaur, Sakshi Shambhavi, Tulasi Nagabandi, Purushotham Vodnala, Payel Mukherjee, Sofia Banu, Priya Singh, Dhiviya Vedagiri, Divya Gupta, Vishal Sah, Santosh Kumar Kuncha, Krishnan Harinivas Harshan, Archana Bharadwaj Siva, Karthik Bharadwaj Tallapaka, Rakesh K Mishra, Divya Tej Sowpati |
| EPI_ISL_447563 | CSIR-Centre for Cellular and Molecular Biology | CSIR-Centre for Cellular and Molecular Biology | Sakshi Shambhavi, Lamuk Zaveri, Shagufta Khan, Namami Gaur, Tulasi Nagabandi, Purushotham Vodnala, Payel Mukherjee, Sofia Banu, Priya Singh, Dhiviya Vedagiri, Divya Gupta, Vishal Sah, Santosh Kumar Kuncha, Krishnan Harinivas Harshan, Archana Bharadwaj Siva, Karthik Bharadwaj Tallapaka, Rakesh K Mishra, Divya Tej Sowpati |
| EPI_ISL_447564, EPI_ISL_447565, EPI_ISL_447566 | CSIR-Centre for Cellular and Molecular Biology | CSIR-Centre for Cellular and Molecular Biology | Sofia Banu, Payel Mukherjee, Priya Singh, Dhiviya Vedagiri, Divya Gupta, Vishal Sah, Santosh Kumar Kuncha, Krishnan Harinivas Harshan, Archana Bharadwaj Siva, Karthik Bharadwaj Tallapaka, Shagufta Khan, Lamuk Zaveri, Namami Gaur, Sakshi Shambhavi, Tulasi Nagabandi, Purushotham Vodnala, Rakesh K Mishra, Divya Tej Sowpati |
| EPI_ISL_447567 | CSIR-Centre for Cellular and Molecular Biology | CSIR-Centre for Cellular and Molecular Biology | Payel Mukherjee, Sofia Banu, Priya Singh, Dhiviya Vedagiri, Divya Gupta, Vishal Sah, Santosh Kumar Kuncha, Krishnan Harinivas Harshan, Archana Bharadwaj Siva, Karthik Bharadwaj Tallapaka, Shagufta Khan, Lamuk Zaveri, Namami Gaur, Sakshi Shambhavi, Tulasi Nagabandi, Purushotham Vodnala, Rakesh K Mishra, Divya Tej Sowpati |
| EPI_ISL_447568, EPI_ISL_447569 | CSIR-Centre for Cellular and Molecular Biology | CSIR-Centre for Cellular and Molecular Biology | Shagufta Khan, Lamuk Zaveri, Namami Gaur, Sakshi Shambhavi, Tulasi Nagabandi, Purushotham Vodnala, Payel Mukherjee, Sofia Banu, Priya Singh, Dhiviya Vedagiri, Divya Gupta, Vishal Sah, Santosh Kumar Kuncha, Krishnan Harinivas Harshan, Archana Bharadwaj Siva, Karthik Bharadwaj Tallapaka, Rakesh K Mishra, Divya Tej Sowpati |
| EPI_ISL_447570, EPI_ISL_447571, EPI_ISL_447572 | CSIR-Centre for Cellular and Molecular Biology | CSIR-Centre for Cellular and Molecular Biology | Lamuk Zaveri, Shagufta Khan, Namami Gaur, Sakshi Shambhavi, Tulasi Nagabandi, Purushotham Vodnala, Payel Mukherjee, Sofia Banu, Priya Singh, Dhiviya Vedagiri, Divya Gupta, Vishal Sah, Santosh Kumar Kuncha, Krishnan Harinivas Harshan, Archana Bharadwaj Siva, Karthik Bharadwaj Tallapaka, Rakesh K Mishra, Divya Tej Sowpati |
| EPI_ISL_447573 | CSIR-Centre for Cellular and Molecular Biology | CSIR-Centre for Cellular and Molecular Biology | Sakshi Shambhavi, Lamuk Zaveri, Shagufta Khan, Namami Gaur, Tulasi Nagabandi, Purushotham Vodnala, Payel Mukherjee, Sofia Banu, Priya Singh, Dhiviya Vedagiri, Divya Gupta, Vishal Sah, Santosh Kumar Kuncha, Krishnan Harinivas Harshan, Archana Bharadwaj Siva, Karthik Bharadwaj Tallapaka, Rakesh K Mishra, Divya Tej Sowpati |
| EPI_ISL_447574 | CSIR-Centre for Cellular and Molecular Biology | CSIR-Centre for Cellular and Molecular Biology | Namami Gaur, Sakshi Shambhavi, Lamuk Zaveri, Shagufta Khan, Tulasi Nagabandi, Purushotham Vodnala, Payel Mukherjee, Sofia Banu, Priya Singh, Dhiviya Vedagiri, Divya Gupta, Vishal Sah, Santosh Kumar Kuncha, Krishnan Harinivas Harshan, Archana Bharadwaj Siva, Karthik Bharadwaj Tallapaka, Rakesh K Mishra, Divya Tej Sowpati |
| EPI_ISL_447575 | CSIR-Centre for Cellular and Molecular Biology | CSIR-Centre for Cellular and Molecular Biology | Sofia Banu, Payel Mukherjee, Priya Singh, Dhiviya Vedagiri, Divya Gupta, Vishal Sah, Santosh Kumar Kuncha, Krishnan Harinivas Harshan, Archana Bharadwaj Siva, Karthik Bharadwaj Tallapaka, Shagufta Khan, Lamuk Zaveri, Namami Gaur, Sakshi Shambhavi, Tulasi Nagabandi, Purushotham Vodnala, Rakesh K Mishra, Divya Tej Sowpati |
| EPI_ISL_447576, EPI_ISL_447577, EPI_ISL_447578 | CSIR-Centre for Cellular and Molecular Biology | CSIR-Centre for Cellular and Molecular Biology | Namami Gaur, Sakshi Shambhavi, Lamuk Zaveri, Shagufta Khan, Tulasi Nagabandi, Purushotham Vodnala, Payel Mukherjee, Sofia Banu, Priya Singh, Dhiviya Vedagiri, Divya Gupta, Vishal Sah, Santosh Kumar Kuncha, Krishnan Harinivas Harshan, Archana Bharadwaj Siva, Karthik Bharadwaj Tallapaka, Rakesh K Mishra, Divya Tej Sowpati |
| EPI_ISL_447579, EPI_ISL_447580 | CSIR-Centre for Cellular and Molecular Biology | CSIR-Centre for Cellular and Molecular Biology | Tulasi Nagabandi, Namami Gaur, Sakshi Shambhavi, Lamuk Zaveri, Shagufta Khan, Purushotham Vodnala, Payel Mukherjee, Sofia Banu, Priya Singh, Dhiviya Vedagiri, Divya Gupta, Vishal Sah, Santosh Kumar Kuncha, Krishnan Harinivas Harshan, Archana Bharadwaj Siva, Karthik Bharadwaj Tallapaka, Rakesh K Mishra, Divya Tej Sowpati |
| EPI_ISL_447581 | CSIR-Centre for Cellular and Molecular Biology | CSIR-Centre for Cellular and Molecular Biology | Sakshi Shambhavi, Lamuk Zaveri, Shagufta Khan, Namami Gaur, Tulasi Nagabandi, Purushotham Vodnala, Payel Mukherjee, Sofia Banu, Priya Singh, Dhiviya Vedagiri, Divya Gupta, Vishal Sah, Santosh Kumar Kuncha, Krishnan Harinivas Harshan, Archana Bharadwaj Siva, Karthik Bharadwaj Tallapaka, Rakesh K Mishra, Divya Tej Sowpati |
| EPI_ISL_447582, EPI_ISL_447583 | CSIR-Centre for Cellular and Molecular Biology | CSIR-Centre for Cellular and Molecular Biology | Tulasi Nagabandi, Namami Gaur, Sakshi Shambhavi, Lamuk Zaveri, Shagufta Khan, Purushotham Vodnala, Payel Mukherjee, Sofia Banu, Priya Singh, Dhiviya Vedagiri, Divya Gupta, Vishal Sah, Santosh Kumar Kuncha, Krishnan Harinivas Harshan, Archana Bharadwaj Siva, Karthik Bharadwaj Tallapaka, Rakesh K Mishra, Divya Tej Sowpati |
| EPI_ISL_447584, EPI_ISL_447585, EPI_ISL_447586, EPI_ISL_447587 | Tamil Nadu Veterinary and Animal Sciences University | CSIR-Centre for Cellular and Molecular Biology | K Kaveri, S Sivasubramanian, S Vennila, P Padmapriya, R Kiruba, S Magesh, G Dhinakar Raj, G Ravi Kumar, Payel Mukherjee, Tulasi Nagabandi, Namami Gaur, Sakshi Shambhavi, Lamuk Zaveri, Shagufta Khan, Purushotham Vodnala, Sofia Banu, Priya Singh, Dhiviya Vedagiri, Divya Gupta, Vishal Sah, Santosh Kumar Kuncha, Krishnan Harinivas Harshan, Archana Bharadwaj Siva, Karthik Bharadwaj Tallapaka, Kumarasamy Thangaraj, Rakesh K Mishra, Divya Tej Sowpati |

| | | | |
|---|---|---|---|
| EPI_ISL_447588 | Lednicky Lab | Lednicky lab | Elbadry,M.A., Subramaniam,K., Waltzek,T.B., Gibson,J.C., Stephenson,C.J., Alam,M.M., Morris,J.G. Jr. and Lednicky,J.A. |
| EPI_ISL_447589 | University of Florida, Lednicky Lab | University of Florida, Lednicky Lab | Elbadry,M.A., Subramaniam,K., Waltzek,T.B., Gibson,J.C., Stephenson,C.J., Alam,M.M., Morris,J.G. Jr. and Lednicky,J.A. |
| EPI_ISL_447590 | Genome Centre | Genome Centre | A. S. M. Rubayet Ul Alam, M. Rafiul Islam, M. Shaminur Rahman, Md. Tanvir Islam, Md. Shazid Hasan, Pravas Chandra Roy, Habiba Ibnat, MD. Ali Ahasan Setu, Tanay Chakrovarty, Sourav Dutta Dip, Ruhul Amin, Md Nur Kabidul Azam, Ovinu Kibria Islam, Hassan M. Al-Emran, Shireen Nigar, Selina Akter, Md. Nazmul Hasan, Iqbal Kabir Jahid, M. Anwar Hossain |
| EPI_ISL_447591 | TSGH-CP molecular lab | TSGH-CP molecular lab | Cherng-Lih Perng, Ming-Jr JIAN, Chih-Kai Chang, Jung-Chung Lin, Kuo-Ming Yeh, Chien-Wen Chen, Sheng-Kang Chiu, Hsing-Yi Chung, Shih-Hung Tsai, Kuo-Sheng Hung, Tien-Yao Chang, Feng-Yee Chang, Hung-Sheng Shang |
| EPI_ISL_447592, EPI_ISL_447593 | TSGH-CP molecular lab | TSGH-CP molecular lab | Cherng-Lih Perng, Ming-Jr JIAN, Chih-Kai Chang, Jung-Chung Lin, Kuo-Ming Yeh, Chien-Wen Chen, Sheng-Kang Chiu, Hsing-Yi Chung, Shih-Hung Tsai, Kuo-Sheng Hung, Tien-Yao Chang, Feng-Yee Chang, Hung-Sheng Shang |
| EPI_ISL_447594 | Caloundra Hospital | Public Health Virology Laboratory | Bixing Huang, Alyssa Pyke, Amanda De Jong, Andrew Van Den Hurk, Carmel Taylor, David Warrilow, Doris Genge, Elisabeth Gamez, Glen Hewitson, Ian Maxwell Mackay, Inga Sultana, Jamie McMahon, Jean Barcelon, Judy Northill, Mitchell Finger, Natalie Simpson, Neelima Nair, Peter Burtonclay, Peter Moore, Sarah Wheatley, Sean Moody, Sonja Hall-Mendelin, Timothy Gardam, and Frederick Moore |
| EPI_ISL_447595 | Pathology Queensland, Sunshine Coast University Hospital | Public Health Virology Laboratory | Bixing Huang, Alyssa Pyke, Amanda De Jong, Andrew Van Den Hurk, Carmel Taylor, David Warrilow, Doris Genge, Elisabeth Gamez, Glen Hewitson, Ian Maxwell Mackay, Inga Sultana, Jamie McMahon, Jean Barcelon, Judy Northill, Mitchell Finger, Natalie Simpson, Neelima Nair, Peter Burtonclay, Peter Moore, Sarah Wheatley, Sean Moody, Sonja Hall-Mendelin, Timothy Gardam, and Frederick Moore |
| EPI_ISL_447596, EPI_ISL_447597, EPI_ISL_447598, EPI_ISL_447599, EPI_ISL_447600, EPI_ISL_447601, EPI_ISL_447602, EPI_ISL_447603, EPI_ISL_447604, EPI_ISL_447605, EPI_ISL_447606, EPI_ISL_447607 | | | |
| see above | Viral Respiratory Lab, National Institute for Biomedical Research (INRB) | Pathogen Sequencing Lab, National Institute for Biomedical Research (INRB) | Placide Mbala-Kingebeni, Edith Nkwembe, Eddy Kinganda-Lusamaki, Amuri Aziza, Francisca Muyembe Mawete, Catherine Pratt, Matthias Pauthner, Josh Quick, Allison Black, James Hadfield, Trevor Bedford, Ian Goodfellow, Andrew Rambaut, Nick Loman, Kristian Andersen, Michael Wiley, Steve Ahuka-Mundeke, Jean-Jacques Muyembe Tamfum |
| EPI_ISL_447608, EPI_ISL_447609, EPI_ISL_447610, EPI_ISL_447611, EPI_ISL_447612, EPI_ISL_447613 | Goethe University Hospital Frankfurt | Institute for Medical Virology, Goethe University Hospital Frankfurt | Tuna Toptan, Sebastian Hoehl, Sandra Westhaus, Denisa Bojkova, Annemarie Berger, Björn Rotter, Klaus Hoffmeier, Jindrich Cinatl, Sandra Ciesek, and Marek Widera |
| EPI_ISL_447614, EPI_ISL_447615, EPI_ISL_447616, EPI_ISL_447617, EPI_ISL_447618, EPI_ISL_447619, EPI_ISL_447620, EPI_ISL_447621, EPI_ISL_447622 | Department of Laboratory Medicine, National Taiwan University Hospital | Microbial Genomics Core Lab, National Taiwan University Centers of Genomic and Precision Medicine | Shiou-Hwei Yeh, You-Yu Lin, Ya-Yun Lai, Chiao-Ling Li, Shan-Chwen Chang, Pei-Jer Chen, Sui-Yuan Chang |
| EPI_ISL_447623, EPI_ISL_447624, EPI_ISL_447625, EPI_ISL_447626, EPI_ISL_447627, EPI_ISL_447628, EPI_ISL_447629, EPI_ISL_447630 | unknown | Virology | van der Poel,W.H.M., Hakze van der Honing,R.W., Harders,F. |
| EPI_ISL_447631 | unknown | Virology | Oreshkova,N., Vreman,S., Molenaar,R.J., Harders,F., Hakze van der Honing,R.W., Gerhards,N., Bouwstra,R., Hissink,H., Smit,L., Tacken,M., Weesendorp,E., Stegeman,A., van der Poel,W.H.M., Engelsma,M.Y. |
| EPI_ISL_447632, EPI_ISL_447633, EPI_ISL_447634 | unknown | Virology | Oreshkova,N., Vreman,S., Molenaar,R.J., Harders,F., Hakze van der Honing,R.W., Gerhards,N., Bouwstra,R., Hissink,H., Smit,L., Tacken,M., Weesendorp,E., Stegeman,A., van der Poel,W., Engelsma,M.Y. |
| EPI_ISL_447635, EPI_ISL_447636, EPI_ISL_447637, EPI_ISL_447638, EPI_ISL_447639, EPI_ISL_447640, EPI_ISL_447641, EPI_ISL_447642, EPI_ISL_447643, EPI_ISL_447644, EPI_ISL_447645, EPI_ISL_447646, EPI_ISL_447647, EPI_ISL_447648, EPI_ISL_447649, EPI_ISL_447650, EPI_ISL_447651, EPI_ISL_447652, EPI_ISL_447653 | | | |
| see above | unknown | Department of Medicine | Kassela,K., Dovrolis,N., Bampali,M., Gatzidou,E., Froukala,E., Stavropoulou,A., Veletza,S., Tsakris,A., Spanakis,N. and Karakasiliotis,I. |
| EPI_ISL_447654, EPI_ISL_447655 | Hôpital Henri-Mondor Ap-Hp | Hôpital Henri-Mondor Ap-Hp | Rodriguez,C., De Prost,N., Fourati,S., Lamoureux,C., Schmitz,D., Deveaux,I., Picard,O., Lepeule,R., Surgers,L., Mekontso-Dessap,A., Woerther,P.-L., Canoui-Poitrine,F., Pawlotsky,J.-M., Clinical Study Group,C., Gricourt,G., N'debi,M., Demontant,V., Trawinski,E. |
| EPI_ISL_447656 | unknown | Genomic platform | De Prost,N., Fourati,S., Lamoureux,C., Schmitz,D., Deveaux,I., Picard,O., Lepeule,R., Surgers,L., Mekontso-Dessap,A., Woerther,P.-L., Canoui-Poitrine,F., Pawlotsky,J.-M., Clinical Study Group,C., Rodrigue,C., Gricourt,G., N'debi,M., Demontant,V., Trawinski,E. |
| EPI_ISL_447657, EPI_ISL_447658, EPI_ISL_447659, EPI_ISL_447660, EPI_ISL_447661, EPI_ISL_447662, EPI_ISL_447663, EPI_ISL_447664, EPI_ISL_447665, EPI_ISL_447666, EPI_ISL_447667, EPI_ISL_447668, EPI_ISL_447669, EPI_ISL_447670, EPI_ISL_447671, EPI_ISL_447672, EPI_ISL_447673, EPI_ISL_447674, EPI_ISL_447675, EPI_ISL_447676, EPI_ISL_447677, EPI_ISL_447678, EPI_ISL_447679, EPI_ISL_447680, EPI_ISL_447681, EPI_ISL_447682, EPI_ISL_447683, EPI_ISL_447684, EPI_ISL_447685, EPI_ISL_447686, EPI_ISL_447687, EPI_ISL_447688, EPI_ISL_447689, EPI_ISL_447690, EPI_ISL_447691, EPI_ISL_447692, EPI_ISL_447693, EPI_ISL_447694, EPI_ISL_447695, EPI_ISL_447696, EPI_ISL_447697, EPI_ISL_447698, EPI_ISL_447699, EPI_ISL_447700, EPI_ISL_447701, EPI_ISL_447702, EPI_ISL_447703, EPI_ISL_447704, EPI_ISL_447705, EPI_ISL_447706, EPI_ISL_447707, EPI_ISL_447708, EPI_ISL_447709, EPI_ISL_447710, EPI_ISL_447711, EPI_ISL_447712, EPI_ISL_447713, EPI_ISL_447714, EPI_ISL_447715, EPI_ISL_447716, EPI_ISL_447717, EPI_ISL_447718, EPI_ISL_447719, EPI_ISL_447720, EPI_ISL_447721, EPI_ISL_447722, EPI_ISL_447723, EPI_ISL_447724, EPI_ISL_447725, EPI_ISL_447726, EPI_ISL_447727, EPI_ISL_447728, EPI_ISL_447729, EPI_ISL_447730, EPI_ISL_447731, EPI_ISL_447732, EPI_ISL_447733 | | | |
| see above | Hôpital Henri-Mondor Ap-Hp | Hôpital Henri-Mondor Ap-Hp | Rodriguez,C., De Prost,N., Fourati,S., Lamoureux,C., Schmitz,D., Deveaux,I., Picard,O., Lepeule,R., Surgers,L., Mekontso-Dessap,A., Woerther,P.-L., Canoui-Poitrine,F., Pawlotsky,J.-M., Clinical Study Group,C., Gricourt,G., N'debi,M., Demontant,V., Trawinski,E. |
| EPI_ISL_447734, EPI_ISL_447735, EPI_ISL_447736, EPI_ISL_447737, EPI_ISL_447738, EPI_ISL_447739, EPI_ISL_447740, EPI_ISL_447741, EPI_ISL_447742, EPI_ISL_447743, EPI_ISL_447744, EPI_ISL_447745, EPI_ISL_447746, EPI_ISL_447747, EPI_ISL_447748, EPI_ISL_447749, EPI_ISL_447750, EPI_ISL_447751, EPI_ISL_447752, EPI_ISL_447753, EPI_ISL_447754 | | | |
| see above | Grupo de Investigaciones Microbiológicas-UR (GIMUR), Departamento de Biología, Facultad de Ciencias Naturales, Universidad del Rosario, Bogotá, Colombia | Grupo de Investigaciones Microbiológicas-UR (GIMUR), Departamento de Biología, Facultad de Ciencias Naturales, Universidad del Rosario, Bogotá, Colombia Instituto Nacional de Salud, Bogotá, Colombia Icahn School of Medicine at Mount Sinai, New York, USA | Juan David Ramírez, Carolina Florez, Marina Muñoz, Carolina Hernandez, Adriana Castillo, Sergio Castañeda, Nathalia Ballesteros, David Martínez, Laura Vega, Jesús E. Jaimes, Sergio Gomez, Angelica Rico, Lisseth Pardo, Esther C. Barros, Martha L. Ospina, Aníbal A. Teherán, Ana S. Gonzalez-Reiche, Matthew M. Hernandez, Emilia Mia Sordillo, Viviana Simon, Harm van Bakel, Alberto Paniz-Mondolfi |
| EPI_ISL_447755, EPI_ISL_447756, EPI_ISL_447757, EPI_ISL_447758, EPI_ISL_447759, EPI_ISL_447760, EPI_ISL_447761, EPI_ISL_447762, EPI_ISL_447763, EPI_ISL_447764, EPI_ISL_447765, EPI_ISL_447766, EPI_ISL_447767, EPI_ISL_447768, EPI_ISL_447769, EPI_ISL_447771, EPI_ISL_447772, EPI_ISL_447774, EPI_ISL_447775, EPI_ISL_447776, EPI_ISL_447777, EPI_ISL_447778, EPI_ISL_447779, EPI_ISL_447780, EPI_ISL_447781, EPI_ISL_447782, EPI_ISL_447783, EPI_ISL_447784, EPI_ISL_447785, EPI_ISL_447786, EPI_ISL_447787, EPI_ISL_447789, EPI_ISL_447790, EPI_ISL_447791, EPI_ISL_447792, EPI_ISL_447793, EPI_ISL_447794, EPI_ISL_447795, EPI_ISL_447796, EPI_ISL_447797, EPI_ISL_447798, EPI_ISL_447799, EPI_ISL_447800, EPI_ISL_447801, EPI_ISL_447802, EPI_ISL_447803, EPI_ISL_447804, EPI_ISL_447805, EPI_ISL_447806, EPI_ISL_447807, EPI_ISL_447808, EPI_ISL_447809, EPI_ISL_447810, EPI_ISL_447811, EPI_ISL_447812, EPI_ISL_447813, EPI_ISL_447814, EPI_ISL_447815, EPI_ISL_447816, EPI_ISL_447817 | | | |
| see above | Instituto Nacional de Salud, Bogotá, Colombia | Grupo de Investigaciones Microbiológicas-UR (GIMUR), Departamento de Biología, Facultad de Ciencias Naturales, Universidad del Rosario, Bogotá, Colombia Instituto Nacional de Salud, Bogotá, Colombia Icahn School of Medicine at Mount Sinai, New York, USA | Juan David Ramírez, Carolina Florez, Marina Muñoz, Carolina Hernandez, Adriana Castillo, Sergio Castañeda, Nathalia Ballesteros, David Martínez, Laura Vega, Jesús E. Jaimes, Sergio Gomez, Angelica Rico, Lisseth Pardo, Esther C. Barros, Martha L. Ospina, Aníbal A. Teherán, Ana S. Gonzalez-Reiche, Matthew M. Hernandez, Emilia Mia Sordillo, Viviana Simon, Harm van Bakel, Alberto Paniz-Mondolfi |
| EPI_ISL_447832, EPI_ISL_447833, EPI_ISL_447834, EPI_ISL_447835, EPI_ISL_447836 | unknown | Department of Medicine | Kassela,K., Dovrolis,N., Bampali,M., Gatzidou,E., Froukala,E., Stavropoulou,A., Veletza,S., Tsakris,A., Spanakis,N. and Karakasiliotis,I. |
| EPI_ISL_447837 | Dept. of Medical Microbiology, Stavanger University Hospital, Helse Stavanger HF, | Norwegian Institute of Public Health, Department of Virology | Kathrine Stene-Johansen, Kamilla Heddeland Instefjord, Hilde Elshaug, Rasmus Riis Kopperud, Karoline Bragstad, Olav Hungnes |
| EPI_ISL_447838, EPI_ISL_447839 | Medical Microbiology Unit, Department for Laboratory Medicine, Drammen Hospital, Vestre Viken Health Trust, | Norwegian Institute of Public Health, Department of Virology | Kathrine Stene-Johansen, Kamilla Heddeland Instefjord, Hilde Elshaug, Rasmus Riis Kopperud, Karoline Bragstad, Olav Hungnes |
| EPI_ISL_447840 | DC Public Health Lab/ Dept. of Forensic Sciences | Pathogen Discovery, Respiratory Viruses Branch, Division of Viral Diseases, Centers for Disease Control and Prevention | Krista Queen, Yan Li, Anna Uehara, Jing Zhang, Ying Tao, Clinton R. Paden, Haibin Wang, Jasmine Padilla, Mary S. Keckler, Alison S. Laufer Halpin, Justin Lee, Christopher A. Elkins, Suxiang Tong |
| EPI_ISL_447841 | FL Bureau of Public Health Laboratories-Tampa | Pathogen Discovery, Respiratory Viruses Branch, Division of Viral Diseases, Centers for Disease Control and Prevention | Krista Queen, Yan Li, Anna Uehara, Jing Zhang, Ying Tao, Clinton R. Paden, Haibin Wang, Jasmine Padilla, Mary S. Keckler, Alison S. Laufer Halpin, Justin Lee, Christopher A. Elkins, Suxiang Tong |
| EPI_ISL_447842 | IA State Hygienic Laboratory | Pathogen Discovery, Respiratory Viruses Branch, Division of Viral Diseases, Centers for Disease Control and Prevention | Krista Queen, Yan Li, Anna Uehara, Jing Zhang, Ying Tao, Clinton R. Paden, Haibin Wang, Jasmine Padilla, Mary S. Keckler, Alison S. Laufer Halpin, Justin Lee, Christopher A. Elkins, Suxiang Tong |
| EPI_ISL_447843 | MD DOH Laboratories Administration | Pathogen Discovery, Respiratory Viruses Branch, Division of Viral Diseases, Centers for Disease Control and Prevention | Krista Queen, Yan Li, Anna Uehara, Jing Zhang, Ying Tao, Clinton R. Paden, Haibin Wang, Jasmine Padilla, Mary S. Keckler, Alison S. Laufer Halpin, Justin Lee, Christopher A. Elkins, Suxiang Tong |
| EPI_ISL_447844 | PA Department of Health, Bureau of Laboratories | Pathogen Discovery, Respiratory Viruses Branch, Division of Viral Diseases, Centers for Disease Control and Prevention | Krista Queen, Yan Li, Anna Uehara, Jing Zhang, Ying Tao, Clinton R. Paden, Haibin Wang, Jasmine Padilla, Mary S. Keckler, Alison S. Laufer Halpin, Justin Lee, Christopher A. Elkins, Suxiang Tong |
| EPI_ISL_447845 | PR - Biological and Chemical Emergencies Lab Office of Public Health Preparedness and Response | Pathogen Discovery, Respiratory Viruses Branch, Division of Viral Diseases, Centers for Disease Control and Prevention | Krista Queen, Yan Li, Anna Uehara, Jing Zhang, Ying Tao, Clinton R. Paden, Haibin Wang, Jasmine Padilla, Mary S. Keckler, Alison S. Laufer Halpin, Justin Lee, Christopher A. Elkins, Suxiang Tong |
| EPI_ISL_447846 | VT Dept. of Health Laboratory | Pathogen Discovery, Respiratory Viruses Branch, Division of Viral Diseases, Centers for Disease Control and Prevention | Krista Queen, Yan Li, Anna Uehara, Jing Zhang, Ying Tao, Clinton R. Paden, Haibin Wang, Jasmine Padilla, Mary S. Keckler, Alison S. Laufer Halpin, Justin Lee, Christopher A. Elkins, Suxiang Tong |
| EPI_ISL_447847 | CSIR-Centre for Cellular and Molecular Biology | CSIR-Centre for Cellular and Molecular Biology | Payel Mukherjee, Sofia Banu, Priya Singh, Dhiviya Vedagiri, Divya Gupta, Vishal Sah, Santosh Kumar Kuncha, Krishnan Harinivas Harshan, Archana Bharadwaj Siva, Karthik Bharadwaj Tallapaka, Shagufta Khan, Lamuk Zaveri, Namami Gaur, Sakshi Shambhavi, Tulasi Nagabandi, Purushotham Vodnala, Rakesh K Mishra, Divya Tej Sowpati |
| EPI_ISL_447848 | CSIR-Centre for Cellular and Molecular Biology | CSIR-Centre for Cellular and Molecular Biology | Sofia Banu, Payel Mukherjee, Priya Singh, Dhiviya Vedagiri, Divya Gupta, Vishal Sah, Santosh Kumar Kuncha, Krishnan Harinivas Harshan, Archana Bharadwaj Siva, Karthik Bharadwaj Tallapaka, Shagufta Khan, Lamuk Zaveri, Namami Gaur, Sakshi Shambhavi, Tulasi Nagabandi, Purushotham Vodnala, Rakesh K Mishra, Divya Tej Sowpati |
| EPI_ISL_447849, EPI_ISL_447850 | CSIR-Centre for Cellular and Molecular Biology | CSIR-Centre for Cellular and Molecular Biology | Shagufta Khan, Lamuk Zaveri, Namami Gaur, Sakshi Shambhavi, Tulasi Nagabandi, Purushotham Vodnala, Payel Mukherjee, Sofia Banu, Priya Singh, Dhiviya Vedagiri, Divya Gupta, Vishal Sah, Santosh Kumar Kuncha, Krishnan Harinivas Harshan, Archana Bharadwaj Siva, Karthik Bharadwaj Tallapaka, Rakesh K Mishra, Divya Tej Sowpati |
| EPI_ISL_447851, EPI_ISL_447852 | CSIR-Centre for Cellular and Molecular Biology | CSIR-Centre for Cellular and Molecular Biology | Lamuk Zaveri, Shagufta Khan, Namami Gaur, Sakshi Shambhavi, Tulasi Nagabandi, Purushotham Vodnala, Payel Mukherjee, Sofia Banu, Priya Singh, Dhiviya Vedagiri, Divya Gupta, Vishal Sah, Santosh Kumar Kuncha, Krishnan Harinivas Harshan, Archana Bharadwaj Siva, Karthik Bharadwaj Tallapaka, Rakesh K Mishra, Divya Tej Sowpati |
| EPI_ISL_447853 | CSIR-Centre for Cellular and Molecular Biology | CSIR-Centre for Cellular and Molecular Biology | Namami Gaur, Sakshi Shambhavi, Lamuk Zaveri, Shagufta Khan, Tulasi Nagabandi, Purushotham Vodnala, Payel Mukherjee, Sofia Banu, Priya Singh, Dhiviya Vedagiri, Divya Gupta, Vishal Sah, Santosh Kumar Kuncha, Krishnan Harinivas Harshan, Archana Bharadwaj Siva, Karthik Bharadwaj Tallapaka, Rakesh K Mishra, Divya Tej Sowpati |
| EPI_ISL_447854 | CSIR-Centre for Cellular and Molecular Biology | CSIR-Centre for Cellular and Molecular Biology | Payel Mukherjee, Sofia Banu, Priya Singh, Dhiviya Vedagiri, Divya Gupta, Vishal Sah, Santosh Kumar Kuncha, Krishnan Harinivas Harshan, Archana Bharadwaj Siva, Karthik Bharadwaj Tallapaka, Shagufta Khan, Lamuk Zaveri, Namami Gaur, Sakshi Shambhavi, Tulasi Nagabandi, Purushotham Vodnala, Rakesh K Mishra, Divya Tej Sowpati |
| EPI_ISL_447855 | CSIR-Centre for Cellular and Molecular Biology | CSIR-Centre for Cellular and Molecular Biology | Lamuk Zaveri, Shagufta Khan, Namami Gaur, Sakshi Shambhavi, Tulasi Nagabandi, Purushotham Vodnala, Payel Mukherjee, Sofia Banu, Priya Singh, Dhiviya Vedagiri, Divya Gupta, Vishal Sah, Santosh Kumar Kuncha, Krishnan Harinivas Harshan, Archana Bharadwaj Siva, Karthik Bharadwaj Tallapaka, Rakesh K Mishra, Divya Tej Sowpati |
| EPI_ISL_447856, EPI_ISL_447857, EPI_ISL_447858 | CSIR-Centre for Cellular and Molecular Biology | CSIR-Centre for Cellular and Molecular Biology | Sakshi Shambhavi, Lamuk Zaveri, Shagufta Khan, Namami Gaur, Tulasi Nagabandi, Purushotham Vodnala, Payel Mukherjee, Sofia Banu, Priya Singh, Dhiviya Vedagiri, Divya Gupta, Vishal Sah, Santosh Kumar Kuncha, Krishnan Harinivas Harshan, Archana Bharadwaj Siva, Karthik Bharadwaj Tallapaka, Rakesh K Mishra, Divya Tej Sowpati |
| EPI_ISL_447859 | CSIR-Centre for Cellular and Molecular Biology | CSIR-Centre for Cellular and Molecular Biology | Payel Mukherjee, Sofia Banu, Priya Singh, Dhiviya Vedagiri, Divya Gupta, Vishal Sah, Santosh Kumar Kuncha, Krishnan Harinivas Harshan, Archana Bharadwaj Siva, Karthik Bharadwaj Tallapaka, Shagufta Khan, Lamuk Zaveri, Namami Gaur, Sakshi Shambhavi, Tulasi Nagabandi, Purushotham Vodnala, Rakesh K Mishra, Divya Tej Sowpati |
| EPI_ISL_447860, EPI_ISL_447861 | CSIR-Centre for Cellular and Molecular Biology | CSIR-Centre for Cellular and Molecular Biology | Tulasi Nagabandi, Namami Gaur, Sakshi Shambhavi, Lamuk Zaveri, Shagufta Khan, Purushotham Vodnala, Payel Mukherjee, Sofia Banu, Priya Singh, Dhiviya Vedagiri, Divya Gupta, Vishal Sah, Santosh Kumar Kuncha, Krishnan Harinivas Harshan, Archana Bharadwaj Siva, Karthik Bharadwaj Tallapaka, Rakesh K Mishra, Divya Tej Sowpati |
| EPI_ISL_447862, EPI_ISL_447863, EPI_ISL_447864 | CSIR-Centre for Cellular and Molecular Biology | CSIR-Centre for Cellular and Molecular Biology | Payel Mukherjee, Sofia Banu, Priya Singh, Dhiviya Vedagiri, Divya Gupta, Vishal Sah, Santosh Kumar Kuncha, Krishnan Harinivas Harshan, Archana Bharadwaj Siva, Karthik Bharadwaj Tallapaka, Shagufta Khan, |

| | | | |
|---|---|---|---|
| | | | Lamuk Zaveri, Namami Gaur, Sakshi Shambhavi, Tulasi Nagabandi, Purushotham Vodnala, Rakesh K Mishra, Divya Tej Sowpati |
| EPI_ISL_447865, EPI_ISL_447866 | CSIR-Centre for Cellular and Molecular Biology | CSIR-Centre for Cellular and Molecular Biology | Sofia Banu, Payel Mukherjee, Priya Singh, Dhiviya Vedagiri, Divya Gupta, Vishal Sah, Santosh Kumar Kuncha, Krishnan Harinivas Harshan, Archana Bharadwaj Siva, Karthik Bharadwaj Tallapaka, Shagufta Khan, Lamuk Zaveri, Namami Gaur, Sakshi Shambhavi, Tulasi Nagabandi, Purushotham Vodnala, Rakesh K Mishra, Divya Tej Sowpati |
| EPI_ISL_447886 | unknown | Pathogen Discovery | Ying Tao, Yan Li, Jing Zhang, Clinton R. Paden, Krista Queen, Anna Uehara, Haibin Wang, Julu Bhatnagar, Suxiang Tong |
| EPI_ISL_447887, EPI_ISL_447888, EPI_ISL_447889, EPI_ISL_447890, EPI_ISL_447891, EPI_ISL_447892, EPI_ISL_447893, EPI_ISL_447894, EPI_ISL_447895, EPI_ISL_447896 | University of California, Davis | Chan-Zuckerberg Biohub | CZB Cliahub Consortium |
| EPI_ISL_447897 | Genome Centre | Genome Centre | A. S. M. Rubayet Ul Alam, M. Rafiul Islam, M. Shaminur Rahman, Md. Tanvir Islam, Md. Shazid Hasan, Pravas Chandra Roy, Habiba Ibnat, MD. Ali Ahasan Setu, Tanay Chakrovarty, Sourav Dutta Dip, Ruhul Amin, Md. Nur Kabidul Islam, Ovinu Kibria Islam, Hassan Md. Al-Emran, Shireen Nigar, Selina Akter, Md. Nazmul Hasan, Iqbal Kabir Jahid, Md. Anwar Hossain |
| EPI_ISL_447900 | Lednicky Laboratory at Emerging Pathogens Institute | University of Florida | Lednicky,J.A., Gibson,J.C., Alam,M.M., Stephenson,C.J., Elbadry,M.A. and Morris,J.G. |
| EPI_ISL_447903 | University of Florida | University of Florida | Elbadry,M.A., Subramaniam,K., Waltzek,T.B., Lauzardo,M., Gibson,J.C., Stephenson,C.J., Alam,M.M., Morris,J.G. Jr. and Lednicky,J.A. |
| EPI_ISL_447905 | University of Florida | University of Florida | Elbadry,M.A., Subramaniam,K., Waltzek,T.B., Gibson,J.C., Stephenson,C.J., Alam,M.M., Morris,J.G. Jr. and Lednicky,J.A. |
| EPI_ISL_449476, EPI_ISL_449477, EPI_ISL_449478, EPI_ISL_449479, EPI_ISL_449480, EPI_ISL_449481, EPI_ISL_449482, EPI_ISL_449483, EPI_ISL_449484, EPI_ISL_449485, EPI_ISL_449486, EPI_ISL_449487 | | | |
| see above | unknown | Department of Respiratory and Critical Care | Wang,X., Zhou,Q., He,Y., Liu,L., Ma,X., Wei,X., Jiang,N., Liang,L., Zheng,Y., Ma,L., Xu,Y., Yang,D., Zhang,J., Yang,B., Jiang,N., Zheng,Y., Ma,L., Xu,Y., Yang,D., Zhang,J., Yang,B., Jiang,N., Deng,T., Zhai,B., Gao,Y., Liu,W., Bai,X., Pan,T., Wang,G., Chang,Y., Zhang,Z., Shi,H., Ma,W.L. and Gao,Z. |
| EPI_ISL_467667 | Florida Bureau of Public Health Laboratories | Florida Bureau of Public Health Laboratories | Schmedes,S. and Blanton,J. |

We gratefully acknowledge the following Authors from the Originating laboratories responsible for obtaining the specimens, as well as the Submitting laboratories where the genome data were generated and shared via GISAID, on which this research is based.

All Submitters of data may be contacted directly via www.gisaid.org

| Accession ID | Originating Laboratory | Submitting Laboratory | Authors |
|---|---|---|---|
| EPI_ISL_447904 | National Institute of Biotechnology | National Institute of Biotechnology | Md. Moniruzzaman, Mohammad Uzzal Hossain, Md. Nazrul Islam, Md. Hadisur Rahman, Irfan Ahmed, Tahia Anan Rahman, Arittra Bhattacharjee, Md. Ruhul Amin, Asif Rashid, Chaman Ara Keya, Keshob Chandra Das, Md. Salimullah |
| EPI_ISL_447906, EPI_ISL_447907, EPI_ISL_447908 | Siriraj hospital | National Institute of Health. Department of medical Sciences, Ministry of Public Health, Thailand | Pilailuk,Okada; Navin Horthongkham, Siripaporn,Phuygun; Thanutsapa,Thanadachakul; Sittiporn,Parnmen;Warawan,Wongboot; Sunthareeya,Waicharoen; Malinee,Chittaganpitch |
| EPI_ISL_447909, EPI_ISL_447910, EPI_ISL_447911, EPI_ISL_447912, EPI_ISL_447913, EPI_ISL_447914, EPI_ISL_447915, EPI_ISL_447916, EPI_ISL_447917, EPI_ISL_447918, EPI_ISL_447919, EPI_ISL_447920, EPI_ISL_447921 | | | |
| see above | n/a | National Institute of Health. Department of medical Sciences, Ministry of Public Health, Thailand | Pilailuk,Okada; Siripaporn,Phuygun; Thanutsapa,Thanadachakul; Sittiporn,Parnmen;Warawan,Wongboot; Sunthareeya,Waicharoen; Malinee,Chittaganpitch |
| EPI_ISL_447922, EPI_ISL_447923, EPI_ISL_447924, EPI_ISL_447925, EPI_ISL_447926, EPI_ISL_447927, EPI_ISL_447928, EPI_ISL_447929, EPI_ISL_447930, EPI_ISL_447931, EPI_ISL_447932, EPI_ISL_447933, EPI_ISL_447934, EPI_ISL_447935, EPI_ISL_447936, EPI_ISL_447937, EPI_ISL_447938, EPI_ISL_447939, EPI_ISL_447940, EPI_ISL_447941, EPI_ISL_447942, EPI_ISL_447943, EPI_ISL_447944 | | | |
| see above | University of Birmingham | COVID-19 Genomics UK (COG-UK) Consortium | Claire McMurray, Joanne Stockton, Samuel Nicholls, Radoslaw Poplawski, Will Rowe, Josh Quick, Nicholas Loman, Celina M Whalley, Andrew Bosworth, Charlotte Poxon, Kasun Wanigasooriya, Oliver Pickles, Mike Kidd, Alex Richter, Andrew D Beggs, Husam Osman, Andrew Bosworth |
| EPI_ISL_447945, EPI_ISL_447946, EPI_ISL_447947, EPI_ISL_447948, EPI_ISL_447949, EPI_ISL_447950, EPI_ISL_447951, EPI_ISL_447952, EPI_ISL_447953, EPI_ISL_447954, EPI_ISL_447955, EPI_ISL_447956, EPI_ISL_447957, EPI_ISL_447958, EPI_ISL_447959, EPI_ISL_447960, EPI_ISL_447961, EPI_ISL_447962, EPI_ISL_447963, EPI_ISL_447964, EPI_ISL_447965, EPI_ISL_447966, EPI_ISL_447967, EPI_ISL_447968, EPI_ISL_447969, EPI_ISL_447970, EPI_ISL_447971, EPI_ISL_447972, EPI_ISL_447973, EPI_ISL_447974, EPI_ISL_447975, EPI_ISL_447976, EPI_ISL_447977, EPI_ISL_447978, EPI_ISL_447979, EPI_ISL_447980, EPI_ISL_447981, EPI_ISL_447982, EPI_ISL_447983, EPI_ISL_447984, EPI_ISL_447985, EPI_ISL_447986, EPI_ISL_447987, EPI_ISL_447988, EPI_ISL_447989, EPI_ISL_447990, EPI_ISL_447991, EPI_ISL_447992, EPI_ISL_447993, EPI_ISL_447994, EPI_ISL_447995, EPI_ISL_447996, EPI_ISL_447997, EPI_ISL_447998, EPI_ISL_447999, EPI_ISL_448000, EPI_ISL_448001, EPI_ISL_448002, EPI_ISL_448003, EPI_ISL_448004, EPI_ISL_448005, EPI_ISL_448006, EPI_ISL_448007, EPI_ISL_448008, EPI_ISL_448009, EPI_ISL_448010, EPI_ISL_448011, EPI_ISL_448012, EPI_ISL_448013, EPI_ISL_448014, EPI_ISL_448015, EPI_ISL_448016, EPI_ISL_448017, EPI_ISL_448018, EPI_ISL_448019, EPI_ISL_448020, EPI_ISL_448021, EPI_ISL_448022, EPI_ISL_448023, EPI_ISL_448024, EPI_ISL_448025, EPI_ISL_448026, EPI_ISL_448027, EPI_ISL_448028, EPI_ISL_448029, EPI_ISL_448030, EPI_ISL_448031, EPI_ISL_448032, EPI_ISL_448033, EPI_ISL_448034, EPI_ISL_448035, EPI_ISL_448036, EPI_ISL_448037, EPI_ISL_448038, EPI_ISL_448039, EPI_ISL_448040, EPI_ISL_448041, EPI_ISL_448042, EPI_ISL_448043, EPI_ISL_448044, EPI_ISL_448045, EPI_ISL_448046, EPI_ISL_448047, EPI_ISL_448048, EPI_ISL_448049, EPI_ISL_448050, EPI_ISL_448051, EPI_ISL_448052, EPI_ISL_448053, EPI_ISL_448054, EPI_ISL_448055, EPI_ISL_448056, EPI_ISL_448057, EPI_ISL_448058, EPI_ISL_448059, EPI_ISL_448060, EPI_ISL_448061, EPI_ISL_448062, EPI_ISL_448063, EPI_ISL_448064, EPI_ISL_448065, EPI_ISL_448066, EPI_ISL_448067, EPI_ISL_448068, EPI_ISL_448069, EPI_ISL_448070, EPI_ISL_448071, EPI_ISL_448072, EPI_ISL_448073, EPI_ISL_448074, EPI_ISL_448075, EPI_ISL_448076, EPI_ISL_448077, EPI_ISL_448078, EPI_ISL_448079, EPI_ISL_448080, EPI_ISL_448081, EPI_ISL_448082, EPI_ISL_448083, EPI_ISL_448084, EPI_ISL_448085, EPI_ISL_448086, EPI_ISL_448087, EPI_ISL_448088, EPI_ISL_448089, EPI_ISL_448090, EPI_ISL_448091, EPI_ISL_448092, EPI_ISL_448093, EPI_ISL_448094, EPI_ISL_448095, EPI_ISL_448096, EPI_ISL_448097, EPI_ISL_448098, EPI_ISL_448099, EPI_ISL_448100, EPI_ISL_448101, EPI_ISL_448102, EPI_ISL_448103, EPI_ISL_448104, EPI_ISL_448105, EPI_ISL_448106, EPI_ISL_448107, EPI_ISL_448108, EPI_ISL_448109, EPI_ISL_448110, EPI_ISL_448111, EPI_ISL_448112, EPI_ISL_448113, EPI_ISL_448114, EPI_ISL_448115 | | | |
| see above | Department of Pathology, University of Cambridge | COVID-19 Genomics UK (COG-UK) Consortium | Luke W Meredith, M. Estée Török , Myra Hosmillo, William L. Hamilton, Martin D. Curran, Theresa Feltwell, Grant Hall, Anna Yakovleva, Fahad A Khokhar, Charlotte J. Houldcroft, Laura G Caller, Aminu S. Jahun, Sarah L. Caddy, Ian Goodfellow |
| EPI_ISL_448116, EPI_ISL_448117, EPI_ISL_448118, EPI_ISL_448119, EPI_ISL_448120, EPI_ISL_448121, EPI_ISL_448122, EPI_ISL_448123, EPI_ISL_448124, EPI_ISL_448125, EPI_ISL_448126, EPI_ISL_448127, EPI_ISL_448128, EPI_ISL_448129, EPI_ISL_448130, EPI_ISL_448131, EPI_ISL_448132, EPI_ISL_448133, EPI_ISL_448134, EPI_ISL_448135, EPI_ISL_448136, EPI_ISL_448137, EPI_ISL_448138, EPI_ISL_448139, EPI_ISL_448140, EPI_ISL_448141, EPI_ISL_448142, EPI_ISL_448143, EPI_ISL_448144, EPI_ISL_448145, EPI_ISL_448146, EPI_ISL_448147, EPI_ISL_448148, EPI_ISL_448149, EPI_ISL_448150, EPI_ISL_448151, EPI_ISL_448152, EPI_ISL_448153, EPI_ISL_448154, EPI_ISL_448155, EPI_ISL_448156, EPI_ISL_448157, EPI_ISL_448158, EPI_ISL_448159, EPI_ISL_448160, EPI_ISL_448161, EPI_ISL_448162, EPI_ISL_448163, EPI_ISL_448164, EPI_ISL_448165, EPI_ISL_448166, EPI_ISL_448167, EPI_ISL_448168, EPI_ISL_448169, EPI_ISL_448170, EPI_ISL_448171, EPI_ISL_448172, EPI_ISL_448173, EPI_ISL_448174, EPI_ISL_448175, EPI_ISL_448176, EPI_ISL_448177, EPI_ISL_448178, EPI_ISL_448179, EPI_ISL_448180, EPI_ISL_448181, EPI_ISL_448182, EPI_ISL_448183, EPI_ISL_448184, EPI_ISL_448185, EPI_ISL_448186, EPI_ISL_448187, EPI_ISL_448188, EPI_ISL_448189, EPI_ISL_448190, EPI_ISL_448191, EPI_ISL_448192, EPI_ISL_448193, EPI_ISL_448194, EPI_ISL_448195, EPI_ISL_448196, EPI_ISL_448197, EPI_ISL_448198, EPI_ISL_448199, EPI_ISL_448200, EPI_ISL_448201, EPI_ISL_448202, EPI_ISL_448203, EPI_ISL_448204, EPI_ISL_448205, EPI_ISL_448206, EPI_ISL_448207, EPI_ISL_448208, EPI_ISL_448209, EPI_ISL_448210, EPI_ISL_448211, EPI_ISL_448212, EPI_ISL_448213, EPI_ISL_448214, EPI_ISL_448215, EPI_ISL_448216, EPI_ISL_448217, EPI_ISL_448218, EPI_ISL_448219, EPI_ISL_448220, EPI_ISL_448221 | | | |
| see above | West of Scotland Specialist Virology Centre, NHSGGC / MRC-University of Glasgow Centre for Virus Research | COVID-19 Genomics UK (COG-UK) Consortium | Ana da Silva Filipe, Natasha Johnson, Kathy Smollett, Daniel Mair, Stephen Carmichael, Lily Tong, Jenna Nichols, Elihu Aranday-Cortes, Kirstyn Brunker, Yasmin Parr, Kyriaki Nomikou, Sarah McDonald, Marc Niebel, Patawee Asamaphan, Richard Orton, Joseph Hughes, Sreenu Vattipally, David L Robertson, Alasdair MacLean, Rory Gunson, Kathy Li, Natasha Jesudason, Rajiv Shah, James Shepherd, Antonia Ho, Emma Thomson |
| EPI_ISL_448222 | Pasteur Insitute Ho Chi Minh City | National Key Laboratory of Gene Technology, Institute of Biotechnology, Vietnam Academy of Science and Technology | Le Tung Lam, Nguyen Trung Hieu, Nguyen Hong Trang, Ho Thi Thuong, Nguyen Thi Ngoc Thao, Huynh Thi Kim Loan, Luu Thuy Tien, Tran Huyen Linh, Pham Duy Quang, Luong Chan Quang, Cao Minh Thang, Nguyen Vu Thuong, Hoang Ha, Chu Hoang Ha, Phan Trong Lan, Truong Nam Hai |
| EPI_ISL_448223, EPI_ISL_448224, EPI_ISL_448225, EPI_ISL_448226, EPI_ISL_448227, EPI_ISL_448228, EPI_ISL_448229, EPI_ISL_448230, EPI_ISL_448231, EPI_ISL_448232, EPI_ISL_448233, EPI_ISL_448234, EPI_ISL_448235, EPI_ISL_448236, EPI_ISL_448237, EPI_ISL_448238, EPI_ISL_448239, EPI_ISL_448240, EPI_ISL_448241, EPI_ISL_448242, EPI_ISL_448243, EPI_ISL_448244, EPI_ISL_448245, EPI_ISL_448246, EPI_ISL_448247, EPI_ISL_448248, EPI_ISL_448249, EPI_ISL_448250, EPI_ISL_448251, EPI_ISL_448252, EPI_ISL_448253, EPI_ISL_448254, EPI_ISL_448255, EPI_ISL_448256, EPI_ISL_448257, EPI_ISL_448258, EPI_ISL_448259, EPI_ISL_448260, EPI_ISL_448261, EPI_ISL_448262, EPI_ISL_448263, EPI_ISL_448264, EPI_ISL_448265, EPI_ISL_448266, EPI_ISL_448267, EPI_ISL_448268, EPI_ISL_448269, EPI_ISL_448270, EPI_ISL_448271, EPI_ISL_448272, EPI_ISL_448273, EPI_ISL_448274, EPI_ISL_448275, EPI_ISL_448276, EPI_ISL_448277, EPI_ISL_448278, EPI_ISL_448279, EPI_ISL_448280, EPI_ISL_448281, EPI_ISL_448282, EPI_ISL_448283, EPI_ISL_448284, EPI_ISL_448285, EPI_ISL_448286, EPI_ISL_448287, EPI_ISL_448288, EPI_ISL_448289, EPI_ISL_448290, EPI_ISL_448291, EPI_ISL_448292, EPI_ISL_448293, EPI_ISL_448294, EPI_ISL_448295, EPI_ISL_448296, EPI_ISL_448297, EPI_ISL_448298, EPI_ISL_448299, EPI_ISL_448300, EPI_ISL_448301, EPI_ISL_448302, EPI_ISL_448303, EPI_ISL_448304, EPI_ISL_448305, EPI_ISL_448306, EPI_ISL_448307, EPI_ISL_448308, EPI_ISL_448309, EPI_ISL_448310, EPI_ISL_448311, EPI_ISL_448312, EPI_ISL_448313, EPI_ISL_448314, EPI_ISL_448315, EPI_ISL_448316, EPI_ISL_448317, EPI_ISL_448318, EPI_ISL_448319, EPI_ISL_448320, EPI_ISL_448321, EPI_ISL_448322, EPI_ISL_448323, EPI_ISL_448324, EPI_ISL_448325, EPI_ISL_448326, EPI_ISL_448327, EPI_ISL_448328, EPI_ISL_448329, EPI_ISL_448330, EPI_ISL_448331, EPI_ISL_448332, EPI_ISL_448333, EPI_ISL_448334, EPI_ISL_448335, EPI_ISL_448336, EPI_ISL_448337, EPI_ISL_448338, EPI_ISL_448339, EPI_ISL_448340, EPI_ISL_448341, EPI_ISL_448342, EPI_ISL_448343, EPI_ISL_448344, EPI_ISL_448345, EPI_ISL_448346, EPI_ISL_448347, EPI_ISL_448348, EPI_ISL_448349, EPI_ISL_448350, EPI_ISL_448351, EPI_ISL_448352, EPI_ISL_448353, EPI_ISL_448354, EPI_ISL_448355, EPI_ISL_448356, EPI_ISL_448357, EPI_ISL_448358, EPI_ISL_448359, EPI_ISL_448360, EPI_ISL_448361, EPI_ISL_448362, EPI_ISL_448363, EPI_ISL_448364, EPI_ISL_448365, EPI_ISL_448366, EPI_ISL_448367, EPI_ISL_448368, EPI_ISL_448369, EPI_ISL_448370, EPI_ISL_448371, EPI_ISL_448372, EPI_ISL_448373, EPI_ISL_448374, EPI_ISL_448375, EPI_ISL_448376, EPI_ISL_448377, EPI_ISL_448378, EPI_ISL_448379, EPI_ISL_448380, EPI_ISL_448381, EPI_ISL_448382, EPI_ISL_448383, EPI_ISL_448384, EPI_ISL_448385, EPI_ISL_448386, EPI_ISL_448387, EPI_ISL_448388 | | | |
| see above | Quadram Institute Bioscience | COVID-19 Genomics UK (COG-UK) Consortium | Dave J. Baker, Gemma L. Kay, Alp Aydin, Thanh Le-Viet, Steven Rudder, Ana P. Tedim, Anastasia Kolyva, Maria Diaz, Leonardo de Oliveira Martins, Nabil-Fareed Alikhan, Lizzie Meadows, Rachael Stanley, Ngozi Elumogo, Muhammed Yasir, Nicholas M. Thomson, Alexander J Trotter, Rachel Gilroy, Samuel Bloomfield, Claire Stuart, Andrew Bell, Reenesh Prakash, Samir Dervisevic, Alison E. Mather, John Wain, Mark Webber, Andrew J. Page, Justin O'Grady |
| EPI_ISL_448389, EPI_ISL_448390, EPI_ISL_448391, EPI_ISL_448392, EPI_ISL_448393, EPI_ISL_448394, EPI_ISL_448395, EPI_ISL_448396, EPI_ISL_448397, EPI_ISL_448398, EPI_ISL_448399, EPI_ISL_448400, EPI_ISL_448401, EPI_ISL_448402, EPI_ISL_448403, EPI_ISL_448404, EPI_ISL_448405, EPI_ISL_448406, EPI_ISL_448407, EPI_ISL_448408, EPI_ISL_448409, EPI_ISL_448410, EPI_ISL_448411, EPI_ISL_448412, EPI_ISL_448413, EPI_ISL_448414, EPI_ISL_448415, EPI_ISL_448416, EPI_ISL_448417, EPI_ISL_448418, EPI_ISL_448419, EPI_ISL_448420, EPI_ISL_448421, EPI_ISL_448422, EPI_ISL_448423, EPI_ISL_448424, EPI_ISL_448425, EPI_ISL_448426, EPI_ISL_448427, EPI_ISL_448428, EPI_ISL_448429, EPI_ISL_448430, EPI_ISL_448431, EPI_ISL_448432, EPI_ISL_448433, EPI_ISL_448434, EPI_ISL_448435, EPI_ISL_448436, EPI_ISL_448437, EPI_ISL_448438, EPI_ISL_448439, EPI_ISL_448440, EPI_ISL_448441, EPI_ISL_448442, EPI_ISL_448443, EPI_ISL_448444, EPI_ISL_448445, EPI_ISL_448446, EPI_ISL_448447, EPI_ISL_448448 | | | |
| see above | Queens Medical Centre, Clinical Microbiology Department / DeepSeq Nottingham | COVID-19 Genomics UK (COG-UK) Consortium | Gemma Clark, Wendy Smith, Manjinder Khakh, Hannah Howson-Wells, Jonathan Ball, Patrick McClure, Joseph Chappell, Theocharis Tsoleridis, Nadine Holmes, Matthew Carlisle, Christopher Moore, Fei Sang, Johnny Debebe, Victoria Wright, Matthew Loose |
| EPI_ISL_448449, EPI_ISL_448450, EPI_ISL_448451, EPI_ISL_448452, EPI_ISL_448453, EPI_ISL_448454, EPI_ISL_448455, EPI_ISL_448456, EPI_ISL_448457, EPI_ISL_448458, EPI_ISL_448459, EPI_ISL_448460, EPI_ISL_448461, EPI_ISL_448462, EPI_ISL_448463, EPI_ISL_448464, EPI_ISL_448465, EPI_ISL_448466, EPI_ISL_448467, EPI_ISL_448468, EPI_ISL_448469, EPI_ISL_448470, EPI_ISL_448471, EPI_ISL_448472, EPI_ISL_448473, EPI_ISL_448474, EPI_ISL_448475, EPI_ISL_448476, EPI_ISL_448477, EPI_ISL_448478, EPI_ISL_448479, EPI_ISL_448480, EPI_ISL_448481, EPI_ISL_448482, EPI_ISL_448483, EPI_ISL_448484, EPI_ISL_448485, EPI_ISL_448486, EPI_ISL_448487, EPI_ISL_448488, EPI_ISL_448489, EPI_ISL_448490, EPI_ISL_448491, EPI_ISL_448492, EPI_ISL_448493, EPI_ISL_448494, EPI_ISL_448495, EPI_ISL_448496, EPI_ISL_448497, EPI_ISL_448498, EPI_ISL_448499, EPI_ISL_448500, EPI_ISL_448501, EPI_ISL_448502, EPI_ISL_448503, EPI_ISL_448504, EPI_ISL_448505, EPI_ISL_448506, EPI_ISL_448507, EPI_ISL_448508, EPI_ISL_448509, EPI_ISL_448510, EPI_ISL_448511, EPI_ISL_448512, EPI_ISL_448513, EPI_ISL_448514, EPI_ISL_448515, EPI_ISL_448516, EPI_ISL_448517, EPI_ISL_448518, EPI_ISL_448519, EPI_ISL_448520, EPI_ISL_448521, EPI_ISL_448522, EPI_ISL_448523, EPI_ISL_448524, EPI_ISL_448525, EPI_ISL_448526, EPI_ISL_448527, EPI_ISL_448528, EPI_ISL_448529, EPI_ISL_448530, EPI_ISL_448531, EPI_ISL_448532, EPI_ISL_448533, EPI_ISL_448534, EPI_ISL_448535, EPI_ISL_448536, EPI_ISL_448537, EPI_ISL_448538, EPI_ISL_448539, EPI_ISL_448540, EPI_ISL_448541, EPI_ISL_448542, EPI_ISL_448543, EPI_ISL_448544, EPI_ISL_448545, EPI_ISL_448546, EPI_ISL_448547, EPI_ISL_448548, EPI_ISL_448549, EPI_ISL_448550, EPI_ISL_448551, EPI_ISL_448552, EPI_ISL_448553, EPI_ISL_448554, EPI_ISL_448555, EPI_ISL_448556, EPI_ISL_448557, EPI_ISL_448558, EPI_ISL_448559, EPI_ISL_448560, EPI_ISL_448561, EPI_ISL_448562, EPI_ISL_448563, EPI_ISL_448564, EPI_ISL_448565, EPI_ISL_448566, EPI_ISL_448567, EPI_ISL_448568, EPI_ISL_448569, EPI_ISL_448570, EPI_ISL_448571, EPI_ISL_448572, EPI_ISL_448573, EPI_ISL_448574, EPI_ISL_448575, EPI_ISL_448576, EPI_ISL_448577, EPI_ISL_448578, EPI_ISL_448579, EPI_ISL_448580, EPI_ISL_448581, EPI_ISL_448582, EPI_ISL_448583, EPI_ISL_448584, EPI_ISL_448585, EPI_ISL_448586, EPI_ISL_448587, EPI_ISL_448588, EPI_ISL_448589, EPI_ISL_448590, EPI_ISL_448591, EPI_ISL_448592, EPI_ISL_448593, EPI_ISL_448594, EPI_ISL_448595, EPI_ISL_448596, EPI_ISL_448597, EPI_ISL_448598, EPI_ISL_448599, EPI_ISL_448600, EPI_ISL_448601, EPI_ISL_448602, EPI_ISL_448603, EPI_ISL_448604, EPI_ISL_448605, EPI_ISL_448606, EPI_ISL_448607, EPI_ISL_448608, EPI_ISL_448609, EPI_ISL_448610, EPI_ISL_448611, EPI_ISL_448612, EPI_ISL_448613, EPI_ISL_448614, EPI_ISL_448615, EPI_ISL_448616, EPI_ISL_448617, EPI_ISL_448618, EPI_ISL_448619, EPI_ISL_448620, EPI_ISL_448621, EPI_ISL_448622, EPI_ISL_448623, EPI_ISL_448624, EPI_ISL_448625, EPI_ISL_448626, EPI_ISL_448627, EPI_ISL_448628, EPI_ISL_448629, EPI_ISL_448630, EPI_ISL_448631, EPI_ISL_448632, EPI_ISL_448633, EPI_ISL_448634, EPI_ISL_448635, EPI_ISL_448636, EPI_ISL_448637, EPI_ISL_448638, EPI_ISL_448639, EPI_ISL_448640, EPI_ISL_448641, EPI_ISL_448642, EPI_ISL_448643, EPI_ISL_448644, EPI_ISL_448645, EPI_ISL_448646, EPI_ISL_448647, EPI_ISL_448648, EPI_ISL_448649, EPI_ISL_448650, EPI_ISL_448651, EPI_ISL_448652, EPI_ISL_448653, EPI_ISL_448654, EPI_ISL_448655, EPI_ISL_448656, EPI_ISL_448657, EPI_ISL_448658, EPI_ISL_448659, EPI_ISL_448660, EPI_ISL_448661, EPI_ISL_448662, EPI_ISL_448663, EPI_ISL_448664, EPI_ISL_448665, EPI_ISL_448666, EPI_ISL_448667, EPI_ISL_448668, EPI_ISL_448669, EPI_ISL_448670, EPI_ISL_448671, EPI_ISL_448672, EPI_ISL_448673, EPI_ISL_448674, EPI_ISL_448675, EPI_ISL_448676, EPI_ISL_448677, EPI_ISL_448678, EPI_ISL_448679, EPI_ISL_448680, EPI_ISL_448681, EPI_ISL_448682, EPI_ISL_448683, EPI_ISL_448684, EPI_ISL_448685, EPI_ISL_448686, EPI_ISL_448687, EPI_ISL_448688, EPI_ISL_448689, EPI_ISL_448690, EPI_ISL_448691, EPI_ISL_448692, EPI_ISL_448693, EPI_ISL_448694, EPI_ISL_448695, EPI_ISL_448696, EPI_ISL_448697, EPI_ISL_448698, EPI_ISL_448699, EPI_ISL_448700, EPI_ISL_448701, EPI_ISL_448702, EPI_ISL_448703, EPI_ISL_448704, EPI_ISL_448705, EPI_ISL_448706, EPI_ISL_448707, EPI_ISL_448708, EPI_ISL_448709, EPI_ISL_448710, EPI_ISL_448711, EPI_ISL_448712, EPI_ISL_448713, EPI_ISL_448714, EPI_ISL_448715, EPI_ISL_448716, EPI_ISL_448717, EPI_ISL_448718, EPI_ISL_448719, EPI_ISL_448720, EPI_ISL_448721, EPI_ISL_448722, EPI_ISL_448723, EPI_ISL_448724, EPI_ISL_448725, EPI_ISL_448726, EPI_ISL_448727, EPI_ISL_448728, EPI_ISL_448729, EPI_ISL_448730, EPI_ISL_448731, EPI_ISL_448732, EPI_ISL_448733, EPI_ISL_448734, EPI_ISL_448735, EPI_ISL_448736, EPI_ISL_448737, EPI_ISL_448738, EPI_ISL_448739, EPI_ISL_448740, EPI_ISL_448741, EPI_ISL_448742, EPI_ISL_448743, EPI_ISL_448744, EPI_ISL_448745, EPI_ISL_448746, EPI_ISL_448747, EPI_ISL_448748, EPI_ISL_448749, EPI_ISL_448750, EPI_ISL_448751, EPI_ISL_448752, EPI_ISL_448753, EPI_ISL_448754, EPI_ISL_448755, EPI_ISL_448756, EPI_ISL_448757, EPI_ISL_448758, EPI_ISL_448759, EPI_ISL_448760, EPI_ISL_448761, EPI_ISL_448762, EPI_ISL_448763, EPI_ISL_448764, EPI_ISL_448765, EPI_ISL_448766, EPI_ISL_448767, EPI_ISL_448768, EPI_ISL_448769, EPI_ISL_448770, EPI_ISL_448771, EPI_ISL_448772, EPI_ISL_448773, EPI_ISL_448774, EPI_ISL_448775, EPI_ISL_448776, EPI_ISL_448777, EPI_ISL_448778, EPI_ISL_448779, EPI_ISL_448780, EPI_ISL_448781, EPI_ISL_448782, EPI_ISL_448783, EPI_ISL_448784, EPI_ISL_448785, EPI_ISL_448786, EPI_ISL_448787, EPI_ISL_448788, EPI_ISL_448789, EPI_ISL_448790, EPI_ISL_448791, EPI_ISL_448792, EPI_ISL_448793, EPI_ISL_448794, EPI_ISL_448795, EPI_ISL_448796, EPI_ISL_448797, EPI_ISL_448798, EPI_ISL_448799, EPI_ISL_448800, EPI_ISL_448801, EPI_ISL_448802, EPI_ISL_448803, EPI_ISL_448804, EPI_ISL_448805, EPI_ISL_448806, EPI_ISL_448807, EPI_ISL_448808, EPI_ISL_448809, EPI_ISL_448810, EPI_ISL_448811, EPI_ISL_448812, EPI_ISL_448813, EPI_ISL_448814, EPI_ISL_448815, EPI_ISL_448816, EPI_ISL_448817, EPI_ISL_448818, EPI_ISL_448819, EPI_ISL_448820, EPI_ISL_448821, EPI_ISL_448822, EPI_ISL_448823, EPI_ISL_448824, EPI_ISL_448825 | | | |
| see above | Oxford Viromics, NDM, University of Oxford; Oxford University Hospitals; Basingstoke and North Hampshire Hospital | COVID-19 Genomics UK (COG-UK) Consortium | Tanya Golubchik, David Bonsall, George Macintyre, Amy Trebes, Mariateresa de Cesare, Catrin Moore, Alex Mobbs, Anita Justice, Robert Shaw, Monique Andersson, Emma Wise, Nathan Moore, Jessica Lynch, Nick Cortes, Stephen Kidd, David Buck, John Todd, Christophe Fraser |
| EPI_ISL_448826, EPI_ISL_448827, EPI_ISL_448828, EPI_ISL_448829, EPI_ISL_448830, EPI_ISL_448831, EPI_ISL_448832, EPI_ISL_448833, EPI_ISL_448834, EPI_ISL_448835, EPI_ISL_448836, EPI_ISL_448837, EPI_ISL_448838, EPI_ISL_448839, EPI_ISL_448840, EPI_ISL_448841, EPI_ISL_448842, EPI_ISL_448843 | | | |
| see above | Virology Department, Sheffield Teaching Hospitals NHS Foundation Trust/Department of Infection, Immunity and Cardiovascular Disease, The Medical School, University of Sheffield | COVID-19 Genomics UK (COG-UK) Consortium | Thushan de Silva, Matthew Parker, Nikki Smith, Adri Angyal, Rebecca Brown, Luke Green, Rachel Tucker, Paul Parsons, Danielle Groves, Katie Johnson, Laura Carrilero, Alex Keeley, Dave Partridge, Matthew Wyles, Benjamin Lindsey, Mehmet Yavuz, Mohammad Raza, Cariad Evans |
| EPI_ISL_448844 | Virology Laboratory, Castle Hill Hospital, Hull University Teaching Hospitals NHS Trust/Department of Infection, Immunity and Cardiovascular Disease, The Medical School, University of Sheffield | COVID-19 Genomics UK (COG-UK) Consortium | Thushan de Silva, Matthew Parker, Nikki Smith, Adri Angyal, Rebecca Brown, Luke Green, Rachel Tucker, Paul Parsons, Danielle Groves, Katie Johnson, Laura Carrilero, Alex Keeley, Dave Partridge, Matthew Wyles, Benjamin Lindsey, Mehmet Yavuz, Mohammad Raza, Cariad Evans |
| EPI_ISL_448845, EPI_ISL_448846, EPI_ISL_448847, EPI_ISL_448848, EPI_ISL_448849 | Virology Department, Sheffield Teaching Hospitals NHS Foundation Trust/Department of Infection, Immunity and Cardiovascular Disease, The Medical School, University of Sheffield | COVID-19 Genomics UK (COG-UK) Consortium | Thushan de Silva, Matthew Parker, Nikki Smith, Adri Angyal, Rebecca Brown, Luke Green, Rachel Tucker, Paul Parsons, Danielle Groves, Katie Johnson, Laura Carrilero, Alex Keeley, Dave Partridge, Matthew Wyles, Benjamin Lindsey, Mehmet Yavuz, Mohammad Raza, Cariad Evans |
| EPI_ISL_448850 | Virology Laboratory, Castle Hill Hospital, Hull University Teaching Hospitals NHS Trust/Department of Infection, Immunity and Cardiovascular Disease, The Medical School, University of Sheffield | COVID-19 Genomics UK (COG-UK) Consortium | Thushan de Silva, Matthew Parker, Nikki Smith, Adri Angyal, Rebecca Brown, Luke Green, Rachel Tucker, Paul Parsons, Danielle Groves, Katie Johnson, Laura Carrilero, Alex Keeley, Dave Partridge, Matthew Wyles, Benjamin Lindsey, Mehmet Yavuz, Mohammad Raza, Cariad Evans |
| EPI_ISL_448851, EPI_ISL_448852, EPI_ISL_448853, EPI_ISL_448854, EPI_ISL_448855, EPI_ISL_448856, EPI_ISL_448857, EPI_ISL_448858, EPI_ISL_448859, EPI_ISL_448860, EPI_ISL_448861, EPI_ISL_448862, EPI_ISL_448863 | | | |
| see above | Virology Department, Sheffield Teaching Hospitals NHS Foundation Trust/Department of Infection, Immunity and Cardiovascular Disease, The Medical School, University of Sheffield | COVID-19 Genomics UK (COG-UK) Consortium | Thushan de Silva, Matthew Parker, Nikki Smith, Adri Angyal, Rebecca Brown, Luke Green, Rachel Tucker, Paul Parsons, Danielle Groves, Katie Johnson, Laura Carrilero, Alex Keeley, Dave Partridge, Matthew Wyles, Benjamin Lindsey, Mehmet Yavuz, Mohammad Raza, Cariad Evans |
| EPI_ISL_448864 | Virology Laboratory, Castle Hill Hospital, Hull University Teaching Hospitals NHS Trust/Department of Infection, Immunity and Cardiovascular Disease, The Medical School, University of Sheffield | COVID-19 Genomics UK (COG-UK) Consortium | Thushan de Silva, Matthew Parker, Nikki Smith, Adri Angyal, Rebecca Brown, Luke Green, Rachel Tucker, Paul Parsons, Danielle Groves, Katie Johnson, Laura Carrilero, Alex Keeley, Dave Partridge, Matthew Wyles, Benjamin Lindsey, Mehmet Yavuz, Mohammad Raza, Cariad Evans |
| EPI_ISL_448865 | Virology Department, Sheffield Teaching Hospitals NHS Foundation Trust/Department of Infection, Immunity and Cardiovascular Disease, The Medical School, University of Sheffield | COVID-19 Genomics UK (COG-UK) Consortium | Thushan de Silva, Matthew Parker, Nikki Smith, Adri Angyal, Rebecca Brown, Luke Green, Rachel Tucker, Paul Parsons, Danielle Groves, Katie Johnson, Laura Carrilero, Alex Keeley, Dave Partridge, Matthew Wyles, Benjamin Lindsey, Mehmet Yavuz, Mohammad Raza, Cariad Evans |

| | | | |
|---|---|---|---|
| EPI_ISL_448866 | Virology Laboratory, Castle Hill Hospital, Hull University Teaching Hospitals NHS Trust/Department of Infection, Immunity and Cardiovascular Disease, The Medical School, University of Sheffield | COVID-19 Genomics UK (COG-UK) Consortium | Thushan de Silva, Matthew Parker, Nikki Smith, Adri Angyal, Rebecca Brown, Luke Green, Rachel Tucker, Paul Parsons, Danielle Groves, Katie Johnson, Laura Carrilero, Alex Keeley, Dave Partridge, Matthew Wyles, Benjamin Lindsey, Mehmet Yavuz, Mohammad Raza, Cariad Evans |
| EPI_ISL_448867, EPI_ISL_448868, EPI_ISL_448869, EPI_ISL_448870, EPI_ISL_448871, EPI_ISL_448872, EPI_ISL_448873, EPI_ISL_448874, EPI_ISL_448875, EPI_ISL_448876, EPI_ISL_448877, EPI_ISL_448878, EPI_ISL_448879, EPI_ISL_448880, EPI_ISL_448881, EPI_ISL_448882, EPI_ISL_448883, EPI_ISL_448884, EPI_ISL_448885, EPI_ISL_448886, EPI_ISL_448887, EPI_ISL_448888, EPI_ISL_448889, EPI_ISL_448890, EPI_ISL_448891, EPI_ISL_448892, EPI_ISL_448893 | | | |
| see above | Virology Department, Sheffield Teaching Hospitals NHS Foundation Trust/Department of Infection, Immunity and Cardiovascular Disease, The Medical School, University of Sheffield | COVID-19 Genomics UK (COG-UK) Consortium | Thushan de Silva, Matthew Parker, Nikki Smith, Adri Angyal, Rebecca Brown, Luke Green, Rachel Tucker, Paul Parsons, Danielle Groves, Katie Johnson, Laura Carrilero, Alex Keeley, Dave Partridge, Matthew Wyles, Benjamin Lindsey, Mehmet Yavuz, Mohammad Raza, Cariad Evans |
| EPI_ISL_448894 | Virology Laboratory, Castle Hill Hospital, Hull University Teaching Hospitals NHS Trust/Department of Infection, Immunity and Cardiovascular Disease, The Medical School, University of Sheffield | COVID-19 Genomics UK (COG-UK) Consortium | Thushan de Silva, Matthew Parker, Nikki Smith, Adri Angyal, Rebecca Brown, Luke Green, Rachel Tucker, Paul Parsons, Danielle Groves, Katie Johnson, Laura Carrilero, Alex Keeley, Dave Partridge, Matthew Wyles, Benjamin Lindsey, Mehmet Yavuz, Mohammad Raza, Cariad Evans |
| EPI_ISL_448895 | Virology Department, Sheffield Teaching Hospitals NHS Foundation Trust/Department of Infection, Immunity and Cardiovascular Disease, The Medical School, University of Sheffield | COVID-19 Genomics UK (COG-UK) Consortium | Thushan de Silva, Matthew Parker, Nikki Smith, Adri Angyal, Rebecca Brown, Luke Green, Rachel Tucker, Paul Parsons, Danielle Groves, Katie Johnson, Laura Carrilero, Alex Keeley, Dave Partridge, Matthew Wyles, Benjamin Lindsey, Mehmet Yavuz, Mohammad Raza, Cariad Evans |
| EPI_ISL_448896 | Virology Laboratory, Castle Hill Hospital, Hull University Teaching Hospitals NHS Trust/Department of Infection, Immunity and Cardiovascular Disease, The Medical School, University of Sheffield | COVID-19 Genomics UK (COG-UK) Consortium | Thushan de Silva, Matthew Parker, Nikki Smith, Adri Angyal, Rebecca Brown, Luke Green, Rachel Tucker, Paul Parsons, Danielle Groves, Katie Johnson, Laura Carrilero, Alex Keeley, Dave Partridge, Matthew Wyles, Benjamin Lindsey, Mehmet Yavuz, Mohammad Raza, Cariad Evans |
| EPI_ISL_448897, EPI_ISL_448898, EPI_ISL_448899, EPI_ISL_448900, EPI_ISL_448901 | Virology Department, Sheffield Teaching Hospitals NHS Foundation Trust/Department of Infection, Immunity and Cardiovascular Disease, The Medical School, University of Sheffield | COVID-19 Genomics UK (COG-UK) Consortium | Thushan de Silva, Matthew Parker, Nikki Smith, Adri Angyal, Rebecca Brown, Luke Green, Rachel Tucker, Paul Parsons, Danielle Groves, Katie Johnson, Laura Carrilero, Alex Keeley, Dave Partridge, Matthew Wyles, Benjamin Lindsey, Mehmet Yavuz, Mohammad Raza, Cariad Evans |
| EPI_ISL_448902 | Virology Laboratory, Castle Hill Hospital, Hull University Teaching Hospitals NHS Trust/Department of Infection, Immunity and Cardiovascular Disease, The Medical School, University of Sheffield | COVID-19 Genomics UK (COG-UK) Consortium | Thushan de Silva, Matthew Parker, Nikki Smith, Adri Angyal, Rebecca Brown, Luke Green, Rachel Tucker, Paul Parsons, Danielle Groves, Katie Johnson, Laura Carrilero, Alex Keeley, Dave Partridge, Matthew Wyles, Benjamin Lindsey, Mehmet Yavuz, Mohammad Raza, Cariad Evans |
| EPI_ISL_448903, EPI_ISL_448904, EPI_ISL_448905 | Virology Department, Sheffield Teaching Hospitals NHS Foundation Trust/Department of Infection, Immunity and Cardiovascular Disease, The Medical School, University of Sheffield | COVID-19 Genomics UK (COG-UK) Consortium | Thushan de Silva, Matthew Parker, Nikki Smith, Adri Angyal, Rebecca Brown, Luke Green, Rachel Tucker, Paul Parsons, Danielle Groves, Katie Johnson, Laura Carrilero, Alex Keeley, Dave Partridge, Matthew Wyles, Benjamin Lindsey, Mehmet Yavuz, Mohammad Raza, Cariad Evans |
| EPI_ISL_448906 | Virology Laboratory, Castle Hill Hospital, Hull University Teaching Hospitals NHS Trust/Department of Infection, Immunity and Cardiovascular Disease, The Medical School, University of Sheffield | COVID-19 Genomics UK (COG-UK) Consortium | Thushan de Silva, Matthew Parker, Nikki Smith, Adri Angyal, Rebecca Brown, Luke Green, Rachel Tucker, Paul Parsons, Danielle Groves, Katie Johnson, Laura Carrilero, Alex Keeley, Dave Partridge, Matthew Wyles, Benjamin Lindsey, Mehmet Yavuz, Mohammad Raza, Cariad Evans |
| EPI_ISL_448907, EPI_ISL_448908, EPI_ISL_448909, EPI_ISL_448910, EPI_ISL_448911, EPI_ISL_448912, EPI_ISL_448913, EPI_ISL_448914, EPI_ISL_448915, EPI_ISL_448916, EPI_ISL_448917 | | | |
| see above | Virology Department, Sheffield Teaching Hospitals NHS Foundation Trust/Department of Infection, Immunity and Cardiovascular Disease, The Medical School, University of Sheffield | COVID-19 Genomics UK (COG-UK) Consortium | Thushan de Silva, Matthew Parker, Nikki Smith, Adri Angyal, Rebecca Brown, Luke Green, Rachel Tucker, Paul Parsons, Danielle Groves, Katie Johnson, Laura Carrilero, Alex Keeley, Dave Partridge, Matthew Wyles, Benjamin Lindsey, Mehmet Yavuz, Mohammad Raza, Cariad Evans |
| EPI_ISL_448918, EPI_ISL_448919, EPI_ISL_448920, EPI_ISL_448921, EPI_ISL_448922, EPI_ISL_448923, EPI_ISL_448924, EPI_ISL_448925, EPI_ISL_448926, EPI_ISL_448927, EPI_ISL_448928, EPI_ISL_448929, EPI_ISL_448930, EPI_ISL_448931, EPI_ISL_448932, EPI_ISL_448933, EPI_ISL_448934, EPI_ISL_448935, EPI_ISL_448936, EPI_ISL_448937, EPI_ISL_448938, EPI_ISL_448939, EPI_ISL_448940, EPI_ISL_448941, EPI_ISL_448942, EPI_ISL_448943, EPI_ISL_448944, EPI_ISL_448945, EPI_ISL_448946, EPI_ISL_448947, EPI_ISL_448948, EPI_ISL_448949, EPI_ISL_448950, EPI_ISL_448951, EPI_ISL_448952, EPI_ISL_448953, EPI_ISL_448954, EPI_ISL_448955, EPI_ISL_448956, EPI_ISL_448957, EPI_ISL_448958, EPI_ISL_448959, EPI_ISL_448960, EPI_ISL_448961, EPI_ISL_448962, EPI_ISL_448963, EPI_ISL_448964, EPI_ISL_448965, EPI_ISL_448966, EPI_ISL_448967, EPI_ISL_448968, EPI_ISL_448969, EPI_ISL_448970, EPI_ISL_448971, EPI_ISL_448972, EPI_ISL_448973, EPI_ISL_448974, EPI_ISL_448975, EPI_ISL_448976, EPI_ISL_448977 | | | |
| see above | Regional Virus Laboratory, Belfast Health and Social Care Trust | COVID-19 Genomics UK (COG-UK) Consortium | Conall McCaughey, James McKenna, Tanya Curran, Susan Feeney, Alison Watt, Ciara Cox, Mairead Connor, Zoltan Molnar, David Simpson, Derek Fairley |
| EPI_ISL_448978, EPI_ISL_448979, EPI_ISL_448980, EPI_ISL_448981, EPI_ISL_448982, EPI_ISL_448983, EPI_ISL_448984, EPI_ISL_448985, EPI_ISL_448986, EPI_ISL_448987, EPI_ISL_448988, EPI_ISL_448989, EPI_ISL_448990, EPI_ISL_448991, EPI_ISL_448992, EPI_ISL_448993, EPI_ISL_448994, EPI_ISL_448995, EPI_ISL_448996, EPI_ISL_448997, EPI_ISL_448998, EPI_ISL_448999, EPI_ISL_449000, EPI_ISL_449001, EPI_ISL_449002, EPI_ISL_449003, EPI_ISL_449004, EPI_ISL_449005, EPI_ISL_449006, EPI_ISL_449007, EPI_ISL_449008, EPI_ISL_449009, EPI_ISL_449010, EPI_ISL_449011, EPI_ISL_449012, EPI_ISL_449013, EPI_ISL_449014, EPI_ISL_449015, EPI_ISL_449016, EPI_ISL_449017, EPI_ISL_449018, EPI_ISL_449019, EPI_ISL_449020, EPI_ISL_449021, EPI_ISL_449022, EPI_ISL_449023, EPI_ISL_449024, EPI_ISL_449025, EPI_ISL_449026, EPI_ISL_449027, EPI_ISL_449028, EPI_ISL_449029, EPI_ISL_449030, EPI_ISL_449031, EPI_ISL_449032, EPI_ISL_449033, EPI_ISL_449034, EPI_ISL_449035, EPI_ISL_449036, EPI_ISL_449037, EPI_ISL_449038, EPI_ISL_449039, EPI_ISL_449040, EPI_ISL_449041, EPI_ISL_449042, EPI_ISL_449043, EPI_ISL_449044, EPI_ISL_449045, EPI_ISL_449046, EPI_ISL_449047, EPI_ISL_449048, EPI_ISL_449049, EPI_ISL_449050, EPI_ISL_449051, EPI_ISL_449052, EPI_ISL_449053, EPI_ISL_449054, EPI_ISL_449055, EPI_ISL_449056, EPI_ISL_449057, EPI_ISL_449058, EPI_ISL_449059, EPI_ISL_449060, EPI_ISL_449061, EPI_ISL_449062, EPI_ISL_449063, EPI_ISL_449064, EPI_ISL_449065, EPI_ISL_449066, EPI_ISL_449067, EPI_ISL_449068, EPI_ISL_449069, EPI_ISL_449070, EPI_ISL_449071, EPI_ISL_449072, EPI_ISL_449073, EPI_ISL_449074, EPI_ISL_449075, EPI_ISL_449076, EPI_ISL_449077, EPI_ISL_449078, EPI_ISL_449079, EPI_ISL_449080, EPI_ISL_449081, EPI_ISL_449082, EPI_ISL_449083, EPI_ISL_449084, EPI_ISL_449085, EPI_ISL_449086, EPI_ISL_449087, EPI_ISL_449088, EPI_ISL_449089, EPI_ISL_449090, EPI_ISL_449091, EPI_ISL_449092, EPI_ISL_449093, EPI_ISL_449094, EPI_ISL_449095, EPI_ISL_449096, EPI_ISL_449097, EPI_ISL_449098, EPI_ISL_449099, EPI_ISL_449100, EPI_ISL_449101, EPI_ISL_449102, EPI_ISL_449103, EPI_ISL_449104, EPI_ISL_449105, EPI_ISL_449106, EPI_ISL_449107, EPI_ISL_449108, EPI_ISL_449109, EPI_ISL_449110, EPI_ISL_449111, EPI_ISL_449112, EPI_ISL_449113, EPI_ISL_449114, EPI_ISL_449115, EPI_ISL_449116, EPI_ISL_449117, EPI_ISL_449118, EPI_ISL_449119, EPI_ISL_449120, EPI_ISL_449121, EPI_ISL_449122, EPI_ISL_449123, EPI_ISL_449124, EPI_ISL_449125, EPI_ISL_449126, EPI_ISL_449127, EPI_ISL_449128, EPI_ISL_449129, EPI_ISL_449130, EPI_ISL_449131, EPI_ISL_449132, EPI_ISL_449133, EPI_ISL_449134, EPI_ISL_449135, EPI_ISL_449136, EPI_ISL_449137, EPI_ISL_449138, EPI_ISL_449139, EPI_ISL_449140, EPI_ISL_449141, EPI_ISL_449142, EPI_ISL_449143, EPI_ISL_449144, EPI_ISL_449145, EPI_ISL_449146, EPI_ISL_449147, EPI_ISL_449148, EPI_ISL_449149, EPI_ISL_449150, EPI_ISL_449151, EPI_ISL_449152, EPI_ISL_449153, EPI_ISL_449154, EPI_ISL_449155, EPI_ISL_449156, EPI_ISL_449157, EPI_ISL_449158, EPI_ISL_449159, EPI_ISL_449160, EPI_ISL_449161, EPI_ISL_449162, EPI_ISL_449163, EPI_ISL_449164, EPI_ISL_449165, EPI_ISL_449166, EPI_ISL_449167, EPI_ISL_449168, EPI_ISL_449169, EPI_ISL_449170, EPI_ISL_449171, EPI_ISL_449172, EPI_ISL_449173, EPI_ISL_449174, EPI_ISL_449175 | | | |
| see above | Quadram Institute Bioscience | COVID-19 Genomics UK (COG-UK) Consortium | Dave J. Baker, Gemma L. Kay, Alp Aydin, Thanh Le-Viet, Steven Rudder, Ana P. Tedim, Anastasia Kolyva, Maria Diaz, Leonardo de Oliveira Martins, Nabil-Fareed Alikhan, Lizzie Meadows, Rachael Stanley, Ngozi Elumogo, Muhammed Yasir, Nicholas M. Thomson, Alexander J Trotter, Rachel Gilroy, Samuel Bloomfield, Claire Stuart, Andrew Bell, Reenesh Prakash, Samir Dervisevic, Alison E. Mather, John Wain, Mark Webber, Andrew J. Page, Justin O'Grady |
| EPI_ISL_449176, EPI_ISL_449177, EPI_ISL_449178, EPI_ISL_449179, EPI_ISL_449180, EPI_ISL_449181, EPI_ISL_449182, EPI_ISL_449183, EPI_ISL_449184, EPI_ISL_449185, EPI_ISL_449186, EPI_ISL_449187, EPI_ISL_449188, EPI_ISL_449189, EPI_ISL_449190, EPI_ISL_449191, EPI_ISL_449192, EPI_ISL_449193, EPI_ISL_449194, EPI_ISL_449195, EPI_ISL_449196, EPI_ISL_449197, EPI_ISL_449198, EPI_ISL_449199, EPI_ISL_449200, EPI_ISL_449201, EPI_ISL_449202, EPI_ISL_449203, EPI_ISL_449204, EPI_ISL_449205, EPI_ISL_449206, EPI_ISL_449207, EPI_ISL_449208, EPI_ISL_449209, EPI_ISL_449210, EPI_ISL_449211, EPI_ISL_449212, EPI_ISL_449213, EPI_ISL_449214, EPI_ISL_449215, EPI_ISL_449216, EPI_ISL_449217, EPI_ISL_449218, EPI_ISL_449219, EPI_ISL_449220, EPI_ISL_449221, EPI_ISL_449222, EPI_ISL_449223, EPI_ISL_449224, EPI_ISL_449225, EPI_ISL_449226, EPI_ISL_449227, EPI_ISL_449228, EPI_ISL_449229, EPI_ISL_449230, EPI_ISL_449231, EPI_ISL_449232, EPI_ISL_449233, EPI_ISL_449234, EPI_ISL_449235, EPI_ISL_449236, EPI_ISL_449237, EPI_ISL_449238, EPI_ISL_449239, EPI_ISL_449240, EPI_ISL_449241, EPI_ISL_449242, EPI_ISL_449243, EPI_ISL_449244, EPI_ISL_449245, EPI_ISL_449246, EPI_ISL_449247, EPI_ISL_449248, EPI_ISL_449249, EPI_ISL_449250, EPI_ISL_449251, EPI_ISL_449252, EPI_ISL_449253, EPI_ISL_449254, EPI_ISL_449255, EPI_ISL_449256, EPI_ISL_449257, EPI_ISL_449258 | | | |
| see above | West of Scotland Specialist Virology Centre, NHSGGC / MRC-University of Glasgow Centre for Virus Research | COVID-19 Genomics UK (COG-UK) Consortium | Ana da Silva Filipe, Natasha Johnson, Kathy Smollett, Daniel Mair, Stephen Carmichael, Lily Tong, Jenna Nichols, Elihu Aranday-Cortes, Kirstyn Brunker, Yasmin Parr, Kyriaki Nomikou, Sarah McDonald, Marc Niebel, Patawee Asamaphan, Richard Orton, Joseph Hughes, Sreenu Vattipally, David L Robertson, Alasdair MacLean, Rory Gunson, Kathy Li, Natasha Jesudason, Rajiv Shah, James Shepherd, Antonia Ho, Emma Thomson |
| EPI_ISL_449259, EPI_ISL_449260, EPI_ISL_449261, EPI_ISL_449262, EPI_ISL_449263, EPI_ISL_449264, EPI_ISL_449265, EPI_ISL_449266, EPI_ISL_449267, EPI_ISL_449268, EPI_ISL_449269, EPI_ISL_449270, EPI_ISL_449271, EPI_ISL_449272, EPI_ISL_449273, EPI_ISL_449274, EPI_ISL_449275, EPI_ISL_449276, EPI_ISL_449277, EPI_ISL_449278, EPI_ISL_449279, EPI_ISL_449280, EPI_ISL_449281, EPI_ISL_449282, EPI_ISL_449283, EPI_ISL_449284, EPI_ISL_449285, EPI_ISL_449286, EPI_ISL_449287, EPI_ISL_449288, EPI_ISL_449289, EPI_ISL_449290, EPI_ISL_449291, EPI_ISL_449292, EPI_ISL_449293, EPI_ISL_449294, EPI_ISL_449295, EPI_ISL_449296, EPI_ISL_449297, EPI_ISL_449298, EPI_ISL_449299, EPI_ISL_449300, EPI_ISL_449301, EPI_ISL_449302, EPI_ISL_449303, EPI_ISL_449304, EPI_ISL_449305, EPI_ISL_449306, EPI_ISL_449307, EPI_ISL_449308, EPI_ISL_449309, EPI_ISL_449310, EPI_ISL_449311, EPI_ISL_449312, EPI_ISL_449313, EPI_ISL_449314, EPI_ISL_449315, EPI_ISL_449316, EPI_ISL_449317, EPI_ISL_449318, EPI_ISL_449319, EPI_ISL_449320, EPI_ISL_449321, EPI_ISL_449322, EPI_ISL_449323, EPI_ISL_449324, EPI_ISL_449325, EPI_ISL_449326, EPI_ISL_449327, EPI_ISL_449328, EPI_ISL_449329 | | | |
| see above | Virology Department, Royal Infirmary of Edinburgh, NHS Lothian / School of Biological Sciences, University of Edinburgh / Institute of Genetics and Molecular Medicine, University of Edinburgh | COVID-19 Genomics UK (COG-UK) Consortium | McHugh M, Dewar R, Rooke S, Gallagher M, Balcaza C, O'Toole Á, Scher E, Hill V, McCrone JT, Colquhoun R, Yu X, Jackson B, Rambaut A, Williams TC, Templeton K |
| EPI_ISL_449330, EPI_ISL_449331, EPI_ISL_449332, EPI_ISL_449333, EPI_ISL_449334, EPI_ISL_449335, EPI_ISL_449336, EPI_ISL_449337, EPI_ISL_449338, EPI_ISL_449339, EPI_ISL_449340, EPI_ISL_449341, EPI_ISL_449342, EPI_ISL_449343, EPI_ISL_449344, EPI_ISL_449345, EPI_ISL_449346, EPI_ISL_449347, EPI_ISL_449348, EPI_ISL_449349, EPI_ISL_449350, EPI_ISL_449351, EPI_ISL_449352, EPI_ISL_449353, EPI_ISL_449354, EPI_ISL_449355, EPI_ISL_449356, EPI_ISL_449357, EPI_ISL_449358, EPI_ISL_449359, EPI_ISL_449360, EPI_ISL_449361, EPI_ISL_449362, EPI_ISL_449363, EPI_ISL_449364, EPI_ISL_449365, EPI_ISL_449366, EPI_ISL_449367, EPI_ISL_449368, EPI_ISL_449369, EPI_ISL_449370, EPI_ISL_449371, EPI_ISL_449372, EPI_ISL_449373, EPI_ISL_449374, EPI_ISL_449375, EPI_ISL_449376, EPI_ISL_449377, EPI_ISL_449378, EPI_ISL_449379, EPI_ISL_449380, EPI_ISL_449381, EPI_ISL_449382, EPI_ISL_449383, EPI_ISL_449384, EPI_ISL_449385, EPI_ISL_449386, EPI_ISL_449387, EPI_ISL_449388, EPI_ISL_449389, EPI_ISL_449390, EPI_ISL_449391, EPI_ISL_449392, EPI_ISL_449393, EPI_ISL_449394, EPI_ISL_449395, EPI_ISL_449396, EPI_ISL_449397, EPI_ISL_449398, EPI_ISL_449399, EPI_ISL_449400, EPI_ISL_449401, EPI_ISL_449402, EPI_ISL_449403, EPI_ISL_449404, EPI_ISL_449405, EPI_ISL_449406, EPI_ISL_449407, EPI_ISL_449408, EPI_ISL_449409, EPI_ISL_449410, EPI_ISL_449411, EPI_ISL_449412, EPI_ISL_449413, EPI_ISL_449414, EPI_ISL_449415, EPI_ISL_449416, EPI_ISL_449417, EPI_ISL_449418, EPI_ISL_449419, EPI_ISL_449420, EPI_ISL_449421, EPI_ISL_449422, EPI_ISL_449423, EPI_ISL_449424, EPI_ISL_449425, EPI_ISL_449426, EPI_ISL_449427, EPI_ISL_449428, EPI_ISL_449429, EPI_ISL_449430, EPI_ISL_449431, EPI_ISL_449432, EPI_ISL_449433, EPI_ISL_449434, EPI_ISL_449435, EPI_ISL_449436, EPI_ISL_449437, EPI_ISL_449438, EPI_ISL_449439, EPI_ISL_449440, EPI_ISL_449441, EPI_ISL_449442, EPI_ISL_449443, EPI_ISL_449444, EPI_ISL_449445, EPI_ISL_449446, EPI_ISL_449447, EPI_ISL_449448, EPI_ISL_449449, EPI_ISL_449450, EPI_ISL_449451, EPI_ISL_449452, EPI_ISL_449453, EPI_ISL_449454, EPI_ISL_449455, EPI_ISL_449456, EPI_ISL_449457, EPI_ISL_449458, EPI_ISL_449459, EPI_ISL_449460, EPI_ISL_449461, EPI_ISL_449462, EPI_ISL_449463, EPI_ISL_449464, EPI_ISL_449465, EPI_ISL_449466, EPI_ISL_449467, EPI_ISL_449468, EPI_ISL_449469, EPI_ISL_449470, EPI_ISL_449471, EPI_ISL_449472, EPI_ISL_449473, EPI_ISL_449474, EPI_ISL_449475, EPI_ISL_449488, EPI_ISL_449489, EPI_ISL_449490, EPI_ISL_449491, EPI_ISL_449492, EPI_ISL_449493, EPI_ISL_449494, EPI_ISL_449495, EPI_ISL_449496, EPI_ISL_449497, EPI_ISL_449498, EPI_ISL_449499, EPI_ISL_449500, EPI_ISL_449501, EPI_ISL_449502, EPI_ISL_449503, EPI_ISL_449504, EPI_ISL_449505, EPI_ISL_449506, EPI_ISL_449507, EPI_ISL_449508, EPI_ISL_449509, EPI_ISL_449510, EPI_ISL_449511, EPI_ISL_449512, EPI_ISL_449513, EPI_ISL_449514, EPI_ISL_449515, EPI_ISL_449516, EPI_ISL_449517, EPI_ISL_449518, EPI_ISL_449519, EPI_ISL_449520, EPI_ISL_449521, EPI_ISL_449522, EPI_ISL_449523, EPI_ISL_449524, EPI_ISL_449525, EPI_ISL_449526, EPI_ISL_449527, EPI_ISL_449528, EPI_ISL_449529, EPI_ISL_449530, EPI_ISL_449531, EPI_ISL_449532, EPI_ISL_449533, EPI_ISL_449534, EPI_ISL_449535, EPI_ISL_449536, EPI_ISL_449537, EPI_ISL_449538, EPI_ISL_449539, EPI_ISL_449540, EPI_ISL_449541, EPI_ISL_449542, EPI_ISL_449543, EPI_ISL_449544, EPI_ISL_449545, EPI_ISL_449546, EPI_ISL_449547, EPI_ISL_449548, EPI_ISL_449549, EPI_ISL_449550, EPI_ISL_449551, EPI_ISL_449552, EPI_ISL_449553, EPI_ISL_449554, EPI_ISL_449555, EPI_ISL_449556, EPI_ISL_449557, EPI_ISL_449558, EPI_ISL_449559, EPI_ISL_449560, EPI_ISL_449561, EPI_ISL_449562, EPI_ISL_449563, EPI_ISL_449564, EPI_ISL_449565, EPI_ISL_449566, EPI_ISL_449567, EPI_ISL_449568, EPI_ISL_449569, EPI_ISL_449570, EPI_ISL_449571, EPI_ISL_449572, EPI_ISL_449573, EPI_ISL_449574, EPI_ISL_449575, EPI_ISL_449576, EPI_ISL_449577, EPI_ISL_449578, EPI_ISL_449579, EPI_ISL_449580, EPI_ISL_449581, EPI_ISL_449582, EPI_ISL_449583, EPI_ISL_449584, EPI_ISL_449585, EPI_ISL_449586, EPI_ISL_449587, EPI_ISL_449588, EPI_ISL_449589, EPI_ISL_449590, EPI_ISL_449591, EPI_ISL_449592, EPI_ISL_449593, EPI_ISL_449594, EPI_ISL_449595, EPI_ISL_449596, EPI_ISL_449597, EPI_ISL_449598, EPI_ISL_449599, EPI_ISL_449600, EPI_ISL_449601, EPI_ISL_449602, EPI_ISL_449603, EPI_ISL_449604, EPI_ISL_449605, EPI_ISL_449606, EPI_ISL_449607, EPI_ISL_449608, EPI_ISL_449609, EPI_ISL_449610, EPI_ISL_449611, EPI_ISL_449612, EPI_ISL_449613, EPI_ISL_449614, EPI_ISL_449615, EPI_ISL_449616, EPI_ISL_449617, EPI_ISL_449618, EPI_ISL_449619, EPI_ISL_449620, EPI_ISL_449621, EPI_ISL_449622, EPI_ISL_449623, EPI_ISL_449624, EPI_ISL_449625 | | | |
| see above | Liverpool Clinical Laboratories | COVID-19 Genomics UK (COG-UK) Consortium | Sam Haldenby, Anita Lucaci, Steve Paterson, Julian Hiscox, Alistair Darby, M Almsaud, A Alrezaihi, Muhannad Alruwaili, Stuart D Armstrong, Jones Benjamin , Eleanor G Bentley, Anu Chawla, Jordan J Clark, Angela Cowell, Richard Eccles, Isabel García-Dorival, Matthew Gemmell, Alessandro Gerada, PKF Gilmore, Richard Gregory, Ximeng Han, Catherine Hartley, Margaret Hughes, Miren Iturriza-Gomara, James Johnson, L Luu, Jenifer Manson , Charlotte Nelson, Elaine O'Toole, Cassie Olateju, Rebekah Penrice-Randal , Lucille Rainbow, N.P Randle, Trevor Ian Robinson, Parul Sharma, Ghada T Shawli, James P Stewart , Neil Swainston, Ecaterina Vamos, Joanne Watts, Mark Whitehead |
| EPI_ISL_449626, EPI_ISL_449627, EPI_ISL_449628, EPI_ISL_449629, EPI_ISL_449630, EPI_ISL_449631, EPI_ISL_449632, EPI_ISL_449633, EPI_ISL_449634, EPI_ISL_449635, EPI_ISL_449636, EPI_ISL_449637, EPI_ISL_449638, EPI_ISL_449639, EPI_ISL_449640, EPI_ISL_449641, EPI_ISL_449642, EPI_ISL_449643, EPI_ISL_449644, EPI_ISL_449645, EPI_ISL_449646, EPI_ISL_449647, EPI_ISL_449648, EPI_ISL_449649, EPI_ISL_449650, EPI_ISL_449651, EPI_ISL_449652, EPI_ISL_449653, EPI_ISL_449654, EPI_ISL_449655, EPI_ISL_449656, EPI_ISL_449657, EPI_ISL_449658, EPI_ISL_449659, EPI_ISL_449660, EPI_ISL_449661, EPI_ISL_449662, EPI_ISL_449663, EPI_ISL_449664, EPI_ISL_449665, EPI_ISL_449666, EPI_ISL_449667, EPI_ISL_449668, EPI_ISL_449669, EPI_ISL_449670, EPI_ISL_449671, EPI_ISL_449672, EPI_ISL_449673, EPI_ISL_449674, EPI_ISL_449675, EPI_ISL_449676, EPI_ISL_449677, EPI_ISL_449678, EPI_ISL_449679, EPI_ISL_449680, EPI_ISL_449681, EPI_ISL_449682, EPI_ISL_449683, EPI_ISL_449684, EPI_ISL_449685, EPI_ISL_449686, EPI_ISL_449687, EPI_ISL_449688, EPI_ISL_449689, EPI_ISL_449690, EPI_ISL_449691, EPI_ISL_449692, EPI_ISL_449693, EPI_ISL_449694, EPI_ISL_449695, EPI_ISL_449696, EPI_ISL_449697, EPI_ISL_449698, EPI_ISL_449699, EPI_ISL_449700, EPI_ISL_449701, EPI_ISL_449702, EPI_ISL_449703, EPI_ISL_449704, EPI_ISL_449705, EPI_ISL_449706, EPI_ISL_449707, EPI_ISL_449708, EPI_ISL_449709, EPI_ISL_449710, EPI_ISL_449711, EPI_ISL_449712, EPI_ISL_449713, EPI_ISL_449714, EPI_ISL_449715, EPI_ISL_449716, EPI_ISL_449717, EPI_ISL_449718, EPI_ISL_449719, EPI_ISL_449720, EPI_ISL_449721, EPI_ISL_449722, EPI_ISL_449723, EPI_ISL_449724, EPI_ISL_449725, EPI_ISL_449726, EPI_ISL_449727, EPI_ISL_449728, EPI_ISL_449729 | | | |
| see above | University College London, Great Ormond Street Hospital for Children NHS Foundation Trust, Imperial College Healthcare NHS Trust | COVID-19 Genomics UK (COG-UK) Consortium | Sergi Castellano, Rachel Williams, Mark Kristiansen, Paola Resende Silva, Sunando Roy, Tony Brooks, Helena Tutill, Paola Niola, Patricia Dyal, Charlotte Williams, Leysa Forrest, Yasmin Panchbhaya, Jacqueline Findlay, Sam Weeks, Julianne Brown, Kathryn Harris, Paul Randell, James Price, Alison Holmes, Judith Breuer |
| EPI_ISL_449786 | Ostfold Hospital Trust - Kalnes, Centre for Laboratory Medicine, Section for gene technology and infection serology | Norwegian Institute of Public Health, Department of Virology | Kathrine Stene-Johansen, Kamilla Heddeland Instefjord, Hilde Elshaug, Rasmus Riis Kopperud, Karoline Bragstad, Olav Hungnes |
| EPI_ISL_449787, EPI_ISL_449788 | Furst Medical Laboratory | Norwegian Institute of Public Health, Department of Virology | Kathrine Stene-Johansen, Kamilla Heddeland Instefjord, Hilde Elshaug, Rasmus Riis Kopperud, Karoline Bragstad, Olav Hungnes |
| EPI_ISL_449789, EPI_ISL_449790, EPI_ISL_449791, EPI_ISL_449792, EPI_ISL_449793, EPI_ISL_449794 | Dept. of Medical Microbiology, Stavanger University Hospital, Helse Stavanger HF | Norwegian Institute of Public Health, Department of Virology | Kathrine Stene-Johansen, Kamilla Heddeland Instefjord, Hilde Elshaug, Rasmus Riis Kopperud, Karoline Bragstad, Olav Hungnes |
| EPI_ISL_449799 | National Laboratory for Health, Environment and Food | National Laboratory for Health, Environment and Food | Mahnic A., Hedzet S., Janezic S., Duh D., Zavrsnik J., Blazun Vosner H., Rupnik M. |
| EPI_ISL_449800 | HOSPITAL SAN JUAN DE DIOS | Instituto de Salud Publica de Chile | Andrés E Castillo, Bárbara Parra,Paz Tapia, Jaime Lagos, Loredana Arata, Alejandra Acevedo, Winston Andrade, Gabriel Leal, Carolina Tambley, Patricia Bustos, Rodrigo Fasce, Jorge Fernandez |
| EPI_ISL_449801, EPI_ISL_449802, EPI_ISL_449803, EPI_ISL_449804, EPI_ISL_449805, EPI_ISL_449806, EPI_ISL_449807, EPI_ISL_449808, EPI_ISL_449809, EPI_ISL_449810, EPI_ISL_449811, EPI_ISL_449812, EPI_ISL_449813, EPI_ISL_449814, EPI_ISL_449815, EPI_ISL_449816, EPI_ISL_449817, EPI_ISL_449818, EPI_ISL_449819, EPI_ISL_449820, EPI_ISL_449821, EPI_ISL_449822, EPI_ISL_449823, EPI_ISL_449824, EPI_ISL_449825, EPI_ISL_449826, EPI_ISL_449827, EPI_ISL_449828, EPI_ISL_449829, EPI_ISL_449830, EPI_ISL_449831, EPI_ISL_449832, EPI_ISL_449833, EPI_ISL_449834, EPI_ISL_449835, EPI_ISL_449836, EPI_ISL_449837, EPI_ISL_449838 | | | |
| see above | Utah Public Health Laboratory | Utah Public Health Laboratory | Erin Young, Kelly Oakeson |
| EPI_ISL_449839, EPI_ISL_449840, EPI_ISL_449841, EPI_ISL_449842, EPI_ISL_449843, EPI_ISL_449844, EPI_ISL_449845, EPI_ISL_449846, EPI_ISL_449847, EPI_ISL_449848, EPI_ISL_449849, EPI_ISL_449850, EPI_ISL_449851, EPI_ISL_449852, EPI_ISL_449853, EPI_ISL_449854, EPI_ISL_449855, EPI_ISL_449856, EPI_ISL_449857, EPI_ISL_449858, EPI_ISL_449859, EPI_ISL_449860, EPI_ISL_449861, EPI_ISL_449862, | | | |

| | | | |
|---|---|---|---|
| EPI_ISL_449863, EPI_ISL_449864, EPI_ISL_449865, EPI_ISL_449866, EPI_ISL_449867, EPI_ISL_449868, EPI_ISL_449869, EPI_ISL_449870, EPI_ISL_449871, EPI_ISL_449872, EPI_ISL_449873, EPI_ISL_449874, EPI_ISL_449875, EPI_ISL_449876, EPI_ISL_449877, EPI_ISL_449878, EPI_ISL_449879, EPI_ISL_449880, EPI_ISL_449881, EPI_ISL_449882, EPI_ISL_449883, EPI_ISL_449884, EPI_ISL_449885, EPI_ISL_449886, EPI_ISL_449887, EPI_ISL_449888, EPI_ISL_449889, EPI_ISL_449890, EPI_ISL_449891, EPI_ISL_449892, EPI_ISL_449893, EPI_ISL_449894, EPI_ISL_449895, EPI_ISL_449896, EPI_ISL_449897, EPI_ISL_449898, EPI_ISL_449899, EPI_ISL_449900, EPI_ISL_449901, EPI_ISL_449902, EPI_ISL_449903, EPI_ISL_449904, EPI_ISL_449905, EPI_ISL_449906, EPI_ISL_449907, EPI_ISL_449908, EPI_ISL_449909, EPI_ISL_449910, EPI_ISL_449911, EPI_ISL_449912, EPI_ISL_449913, EPI_ISL_449914, EPI_ISL_449915, EPI_ISL_449916, EPI_ISL_449917, EPI_ISL_449918, EPI_ISL_449919, EPI_ISL_449920, EPI_ISL_449921, EPI_ISL_449922, EPI_ISL_449923, EPI_ISL_449924, EPI_ISL_449925, EPI_ISL_449926, EPI_ISL_449927, EPI_ISL_449928, EPI_ISL_449929, EPI_ISL_449930, EPI_ISL_449931, EPI_ISL_449932, EPI_ISL_449933, EPI_ISL_449934, EPI_ISL_449935, EPI_ISL_449936, EPI_ISL_449937, EPI_ISL_449938, EPI_ISL_449939, EPI_ISL_449940, EPI_ISL_449941, EPI_ISL_449942, EPI_ISL_449943, EPI_ISL_449944, EPI_ISL_449945, EPI_ISL_449946, EPI_ISL_449947, EPI_ISL_449948, EPI_ISL_449949, EPI_ISL_449950, EPI_ISL_449951, EPI_ISL_449952, EPI_ISL_449953, EPI_ISL_449954, EPI_ISL_449955, EPI_ISL_449956, EPI_ISL_449957, EPI_ISL_449958, EPI_ISL_449959, EPI_ISL_449960, EPI_ISL_449961, EPI_ISL_449962, EPI_ISL_449963, EPI_ISL_449964, EPI_ISL_449965, EPI_ISL_449966, EPI_ISL_449967, EPI_ISL_449968, EPI_ISL_449969, EPI_ISL_449970, EPI_ISL_449971, EPI_ISL_449972, EPI_ISL_449973, EPI_ISL_449974, EPI_ISL_449975, EPI_ISL_449976, EPI_ISL_449977, EPI_ISL_449978, EPI_ISL_449979, EPI_ISL_449980, EPI_ISL_449981, EPI_ISL_449982, EPI_ISL_449983, EPI_ISL_449984, EPI_ISL_449985, EPI_ISL_449986, EPI_ISL_449987, EPI_ISL_449988, EPI_ISL_449989, EPI_ISL_449990, EPI_ISL_449991, EPI_ISL_449992, EPI_ISL_449993, EPI_ISL_449994, EPI_ISL_449995, EPI_ISL_449996, EPI_ISL_449997, EPI_ISL_449998, EPI_ISL_449999, EPI_ISL_450000, EPI_ISL_450001, EPI_ISL_450002, EPI_ISL_450003 | | | |
| see above | Washington State Department of Health | Seattle Flu Study | Chu et al |
| EPI_ISL_450004, EPI_ISL_450005, EPI_ISL_450006, EPI_ISL_450007, EPI_ISL_450008, EPI_ISL_450009, EPI_ISL_450010, EPI_ISL_450011, EPI_ISL_450012, EPI_ISL_450013, EPI_ISL_450014, EPI_ISL_450015, EPI_ISL_450016, EPI_ISL_450017, EPI_ISL_450018, EPI_ISL_450019, EPI_ISL_450020, EPI_ISL_450021, EPI_ISL_450022, EPI_ISL_450023, EPI_ISL_450024, EPI_ISL_450025, EPI_ISL_450026, EPI_ISL_450027, EPI_ISL_450028, EPI_ISL_450029, EPI_ISL_450030, EPI_ISL_450031, EPI_ISL_450032, EPI_ISL_450033, EPI_ISL_450034, EPI_ISL_450035, EPI_ISL_450036, EPI_ISL_450037, EPI_ISL_450038, EPI_ISL_450039, EPI_ISL_450040, EPI_ISL_450041, EPI_ISL_450042, EPI_ISL_450043, EPI_ISL_450044, EPI_ISL_450045, EPI_ISL_450046, EPI_ISL_450047, EPI_ISL_450048, EPI_ISL_450049, EPI_ISL_450050, EPI_ISL_450051, EPI_ISL_450052, EPI_ISL_450053, EPI_ISL_450054, EPI_ISL_450055, EPI_ISL_450056, EPI_ISL_450057, EPI_ISL_450058, EPI_ISL_450059, EPI_ISL_450060, EPI_ISL_450061, EPI_ISL_450062, EPI_ISL_450063, EPI_ISL_450064, EPI_ISL_450065, EPI_ISL_450066, EPI_ISL_450067, EPI_ISL_450068, EPI_ISL_450069, EPI_ISL_450070, EPI_ISL_450071, EPI_ISL_450072, EPI_ISL_450073, EPI_ISL_450074, EPI_ISL_450075, EPI_ISL_450076, EPI_ISL_450077, EPI_ISL_450078, EPI_ISL_450079, EPI_ISL_450080, EPI_ISL_450081, EPI_ISL_450082, EPI_ISL_450083, EPI_ISL_450084, EPI_ISL_450085, EPI_ISL_450086, EPI_ISL_450087, EPI_ISL_450088, EPI_ISL_450089, EPI_ISL_450090, EPI_ISL_450091, EPI_ISL_450092, EPI_ISL_450093, EPI_ISL_450094, EPI_ISL_450095, EPI_ISL_450096, EPI_ISL_450097, EPI_ISL_450098, EPI_ISL_450099, EPI_ISL_450100, EPI_ISL_450101, EPI_ISL_450102, EPI_ISL_450103, EPI_ISL_450104, EPI_ISL_450105, EPI_ISL_450106, EPI_ISL_450107, EPI_ISL_450108, EPI_ISL_450109, EPI_ISL_450110, EPI_ISL_450111, EPI_ISL_450112, EPI_ISL_450113, EPI_ISL_450114, EPI_ISL_450115, EPI_ISL_450116, EPI_ISL_450117, EPI_ISL_450118, EPI_ISL_450119, EPI_ISL_450120, EPI_ISL_450121, EPI_ISL_450122, EPI_ISL_450123, EPI_ISL_450124, EPI_ISL_450125, EPI_ISL_450126, EPI_ISL_450127, EPI_ISL_450128, EPI_ISL_450129, EPI_ISL_450130, EPI_ISL_450131, EPI_ISL_450132, EPI_ISL_450133, EPI_ISL_450134, EPI_ISL_450135, EPI_ISL_450136, EPI_ISL_450137, EPI_ISL_450138, EPI_ISL_450139, EPI_ISL_450140, EPI_ISL_450141, EPI_ISL_450142, EPI_ISL_450143, EPI_ISL_450144, EPI_ISL_450145, EPI_ISL_450146, EPI_ISL_450147, EPI_ISL_450148, EPI_ISL_450149, EPI_ISL_450150, EPI_ISL_450151, EPI_ISL_450152, EPI_ISL_450153, EPI_ISL_450154, EPI_ISL_450155, EPI_ISL_450156, EPI_ISL_450157, EPI_ISL_450158, EPI_ISL_450159, EPI_ISL_450160, EPI_ISL_450161, EPI_ISL_450162, EPI_ISL_450163, EPI_ISL_450164, EPI_ISL_450165, EPI_ISL_450166, EPI_ISL_450167, EPI_ISL_450168, EPI_ISL_450169, EPI_ISL_450170 | | | |
| see above | MSHS Clinical Microbiology Laboratories | MSHS Pathogen Surveillance Program | Ana S. Gonzalez-Reiche, Mitchell Sullivan, Ajay Obla, Gopi Patel, Emilia Sordillo, Melissa Gitman, Alberto Paniz-mondolfi, Matthew Hernandez, Shelcie Fabre, Jose Polanco, Zenab Khan, Bremy Alburquerque, Jayeeta Dutta, Juan Soto, Shwetha Sridhar Hara, Ying-Chih Wang, Melissa Smith, Robert Sebra, Lisa Miorin, Wen-chun Liu, Randy Albrecht, Judith Aberg, Florian Krammer, Adolfo Garcia-Sastre, Viviana Simon, Harm van Bakel |
| EPI_ISL_450171, EPI_ISL_450172 | National Influenza centre, Sri Lanka | The University of Hong Kong | Jude Jayamaha, Daniel KW Chu, Malik Peiris |
| EPI_ISL_450173, EPI_ISL_450174, EPI_ISL_450175, EPI_ISL_450176, EPI_ISL_450177, EPI_ISL_450178, EPI_ISL_450179, EPI_ISL_450180, EPI_ISL_450181, EPI_ISL_450182, EPI_ISL_450183, EPI_ISL_450184, EPI_ISL_450185 | | | |
| see above | Robert Garry lab | Andersen lab at Scripps Research | Allison Smither, Gilberto Sabino-Santos, Patricia Snarski, Lilia Melnik, Antoinette Bell, Kaylynn Genemaras, Arnaud Drouin, Dahlene Fusco, Robert Garry with SEARCH Alliance San Diego |
| EPI_ISL_450186, EPI_ISL_450187, EPI_ISL_450188, EPI_ISL_450189 | Biolab Diagnostic Laboratories | Andersen lab at Scripps Research | Issa Abu-Dayyeh, Ahmad Tibi, Lama Hussein, Lina Mohammad, Zein Naber, Amid Abdelnour with SEARCH Alliance San Diego |
| EPI_ISL_450190 | Rady's Childrens Hospital | Andersen lab at Scripps Research | SEARCH Alliance San Diego |
| EPI_ISL_450191, EPI_ISL_450192 | Scripps Medical Laboratory | Andersen lab at Scripps Research | SEARCH Alliance San Diego with Michael Quigley, Ellen Stefanski, Ian Mchardy |
| EPI_ISL_450193, EPI_ISL_450194, EPI_ISL_450195 | SA Pathology | SA Pathology | Lex Leong |
| EPI_ISL_450196 | bumrungrad international hospital | National Institute of Health. Department of medical Sciences, Ministry of Public Health, Thailand | Pilailuk,Okada; Siripaporn,Phuygun; Thanutsapa,Thanadachakul; Sittiporn,Parnmen;Warawan,Wongboot; Sunthareeya,Waicharoen; Malinee,Chittaganpitch |
| EPI_ISL_450197 | National Institute of Health. Department of medical Sciences, Ministry of Public Health, Thailand | National Institute of Health. Department of medical Sciences, Ministry of Public Health, Thailand | Pilailuk,Okada; Siripaporn,Phuygun; Thanutsapa,Thanadachakul; Sittiporn,Parnmen;Warawan,Wongboot; Sunthareeya,Waicharoen; Malinee,Chittaganpitch |
| EPI_ISL_450231 | Robert Garry lab | Andersen lab at Scripps Research | Allison Smither, Gilberto Sabino-Santos, Patricia Snarski, Lilia Melnik, Antoinette Bell, Kaylynn Genemaras, Arnaud Drouin, Dahlene Fusco, Robert Garry with SEARCH Alliance San Diego |
| EPI_ISL_450232, EPI_ISL_450233, EPI_ISL_450234, EPI_ISL_450235, EPI_ISL_450236, EPI_ISL_450237, EPI_ISL_450238, EPI_ISL_450239, EPI_ISL_450240 | UCSF Clinical Microbiology Laboratory | Chiu Laboratory, University of California, San Francisco | Xianding Deng, Scot Federman, Wei Gu, and Charles Y. Chiu |
| EPI_ISL_450241, EPI_ISL_450242, EPI_ISL_450243, EPI_ISL_450244, EPI_ISL_450245, EPI_ISL_450246, EPI_ISL_450247, EPI_ISL_450248, EPI_ISL_450249, EPI_ISL_450250, EPI_ISL_450251, EPI_ISL_450252, EPI_ISL_450253, EPI_ISL_450254, EPI_ISL_450255, EPI_ISL_450256, EPI_ISL_450257, EPI_ISL_450258, EPI_ISL_450259, EPI_ISL_450260, EPI_ISL_450261, EPI_ISL_450262, EPI_ISL_450263, EPI_ISL_450264, EPI_ISL_450265, EPI_ISL_450266, EPI_ISL_450267, EPI_ISL_450268, EPI_ISL_450269, EPI_ISL_450270, EPI_ISL_450271, EPI_ISL_450272, EPI_ISL_450273, EPI_ISL_450274, EPI_ISL_450275, EPI_ISL_450276, EPI_ISL_450277, EPI_ISL_450278, EPI_ISL_450279, EPI_ISL_450280, EPI_ISL_450281, EPI_ISL_450282, EPI_ISL_450283, EPI_ISL_450284, EPI_ISL_450285, EPI_ISL_450286, EPI_ISL_450287, EPI_ISL_450288, EPI_ISL_450289, EPI_ISL_450290, EPI_ISL_450291, EPI_ISL_450292, EPI_ISL_450293 | | | |
| see above | WHO National Influenza Centre Russian Federation | WHO National Influenza Centre Russian Federation | Andrey Komissarov, Artem Fadeev, Mariia Sergeeva, Anna Ivanova, Tamila Musaeva, Ksenia Komissarova, Mariia Timofeeva, Veronica Eder, Mariia Pisareva, Daria Danilenko |
| EPI_ISL_450294 | Institute of Human Genetics, Polish Academy of Sciences Sanitary and Epidemiological Station in Poznań | Institute of Human Genetics, Polish Academy of Sciences | Szymon Hryhorowicz, Adam Ustaszewski, Emilia Lis, Marta Kaczmarek-Ryś, Michał Witt, Andrzej Pławski |
| EPI_ISL_450295 | Institute of Human Genetics, Polish Academy of Sciences; Sanitary and Epidemiological Station in Poznań | Institute of Human Genetics, Polish Academy of Sciences | Szymon Hryhorowicz, Adam Ustaszewski, Emilia Lis, Marta Kaczmarek-Ryś, Michał Witt, Andrzej Pławski |
| EPI_ISL_450296, EPI_ISL_450297, EPI_ISL_450298, EPI_ISL_450299, EPI_ISL_450300, EPI_ISL_450301 | National Institute for Communicable Diseases of the National Health Laboratory Service | National Institute for Communicable Diseases of the National Health Laboratory Service | Allam M, Ismail A, Khumalo Z, Kwenda S, van Heusden P, Mtshali P, Mnyameni F, Mohale T, Subramoney K, Bhiman JN |
| EPI_ISL_450302, EPI_ISL_450303 | Centre hospitalier Anna-Laberge | Laboratoire de santé publique du Québec | Sandrine Moreira, Ioannis Ragoussis, Guillaume Bourque, Jesse Shapiro, Mark Lathrop and Michel Roger on behalf of the CoVSeQ research group (http://covseq.ca/researchgroup) |
| EPI_ISL_450304, EPI_ISL_450305 | Hôpital Charles-LeMoyne | Laboratoire de santé publique du Québec | Sandrine Moreira, Ioannis Ragoussis, Guillaume Bourque, Jesse Shapiro, Mark Lathrop and Michel Roger on behalf of the CoVSeQ research group (http://covseq.ca/researchgroup) |
| EPI_ISL_450306 | CSSS Haut-Richelieu/Rouville (Hôpital) | Laboratoire de santé publique du Québec | Sandrine Moreira, Ioannis Ragoussis, Guillaume Bourque, Jesse Shapiro, Mark Lathrop and Michel Roger on behalf of the CoVSeQ research group (http://covseq.ca/researchgroup) |
| EPI_ISL_450307 | Hôpital du Suroît | Laboratoire de santé publique du Québec | Sandrine Moreira, Ioannis Ragoussis, Guillaume Bourque, Jesse Shapiro, Mark Lathrop and Michel Roger on behalf of the CoVSeQ research group (http://covseq.ca/researchgroup) |
| EPI_ISL_450308 | Hôpital Charles-LeMoyne | Laboratoire de santé publique du Québec | Sandrine Moreira, Ioannis Ragoussis, Guillaume Bourque, Jesse Shapiro, Mark Lathrop and Michel Roger on behalf of the CoVSeQ research group (http://covseq.ca/researchgroup) |
| EPI_ISL_450309 | Hôpital Pierre-Boucher | Laboratoire de santé publique du Québec | Sandrine Moreira, Ioannis Ragoussis, Guillaume Bourque, Jesse Shapiro, Mark Lathrop and Michel Roger on behalf of the CoVSeQ research group (http://covseq.ca/researchgroup) |
| EPI_ISL_450310 | Hôpital Charles-LeMoyne | Laboratoire de santé publique du Québec | Sandrine Moreira, Ioannis Ragoussis, Guillaume Bourque, Jesse Shapiro, Mark Lathrop and Michel Roger on behalf of the CoVSeQ research group (http://covseq.ca/researchgroup) |
| EPI_ISL_450311, EPI_ISL_450312, EPI_ISL_450313 | Hôpital du Suroît | Laboratoire de santé publique du Québec | Sandrine Moreira, Ioannis Ragoussis, Guillaume Bourque, Jesse Shapiro, Mark Lathrop and Michel Roger on behalf of the CoVSeQ research group (http://covseq.ca/researchgroup) |
| EPI_ISL_450314, EPI_ISL_450315 | Hôpital Pierre-Boucher | Laboratoire de santé publique du Québec | Sandrine Moreira, Ioannis Ragoussis, Guillaume Bourque, Jesse Shapiro, Mark Lathrop and Michel Roger on behalf of the CoVSeQ research group (http://covseq.ca/researchgroup) |
| EPI_ISL_450316 | Hôpital Charles-LeMoyne | Laboratoire de santé publique du Québec | Sandrine Moreira, Ioannis Ragoussis, Guillaume Bourque, Jesse Shapiro, Mark Lathrop and Michel Roger on behalf of the CoVSeQ research group (http://covseq.ca/researchgroup) |
| EPI_ISL_450317, EPI_ISL_450318 | Hôpital Pierre-Boucher | Laboratoire de santé publique du Québec | Sandrine Moreira, Ioannis Ragoussis, Guillaume Bourque, Jesse Shapiro, Mark Lathrop and Michel Roger on behalf of the CoVSeQ research group (http://covseq.ca/researchgroup) |
| EPI_ISL_450321 | NIV Pune | CSIR-Centre for Cellular and Molecular Biology | Dr V A Potdar, Dr ML Choudhary,Dr Priya Abraham,V. Vipat, S. Jadhav, U. Saha, H. Kengle, A. Awhale, A. Jagtap, A. Gondhalikar, V Malik, N Srivastava, S. Digraskar, P. Malsane, S. Hundekar, K. Patel, Yogesh Balakartik, M. Kakade, S. Jadhav, R. Gunjikar, V. Awtade, S. Bhorekar, P Shinde, S. Salve, B. Minhas S. Bharadwaj, H Kaushal Y. Gurav, S. Tomar,Payel Mukherjee, Sofia Banu, Priya Singh, Dhiviya Vedagiri, Divya Gupta, Vishal Sah, Santosh Kumar Kuncha, Krishnan Harinivas Harshan, Archana Bharadwaj Siva, Karthik Bharadwaj Tallapaka, Shagufta Khan, Lamuk Zaveri, Namami Gaur, Sakshi Shambhavi, Tulasi Nagabandi, Purushotham Vodnala,G. Aditya Kumar, Koushick Sivakumar, Pooja Ramesh Gupta, Rajan Kumar Jha, Shraddha Vijay Lahoti, Deepak Kumar, Devi Prasad Vijayashankara, Disha Nanda, Divya Das, Jotin Gogoi, Manish |
| EPI_ISL_450322 | NIV Pune | CSIR-Centre for Cellular and Molecular Biology | Dr V A Potdar, Dr ML Choudhary,Dr Priya Abraham,V. Vipat, S. Jadhav, U. Saha, H. Kengle, A. Awhale, A. Jagtap, A. Gondhalikar, V Malik, N Srivastava, S. Digraskar, P. Malsane, S. Hundekar, K. Patel, Yogesh Balakartik, M. Kakade, S. Jadhav, R. Gunjikar, V. Awtade, S. Bhorekar, P Shinde, S. Salve, B. Minhas S. Bharadwaj, H Kaushal Y. Gurav, S. Tomar,Sofia Banu, Payel Mukherjee, Priya Singh, Dhiviya Vedagiri, Divya Gupta, Vishal Sah, Santosh Kumar Kuncha, Krishnan Harinivas Harshan, Archana Bharadwaj Siva, Karthik Bharadwaj Tallapaka, Shagufta Khan, Lamuk Zaveri, Namami Gaur, Sakshi Shambhavi, Tulasi Nagabandi, Purushotham Vodnala, Disha Nanda, Divya Das, Jotin Gogoi, Manish Bhattacharjee, Ravi Prasad Mukku, Renu Sudhakar, Somesh Gorde, Gangumala Srinivas Reddy, Sujoy Deb, Swati Bayyana, Zeba Rizvi, Rakesh K Mishra |
| EPI_ISL_450323 | NIV Pune | CSIR-Centre for Cellular and Molecular Biology | Dr V A Potdar, Dr ML Choudhary,Dr Priya Abraham,V. Vipat, S. Jadhav, U. Saha, H. Kengle, A. Awhale, A. Jagtap, A. Gondhalikar, V Malik, N Srivastava, S. Digraskar, P. Malsane, S. Hundekar, K. Patel, Yogesh Balakartik, M. Kakade, S. Jadhav, R. Gunjikar, V. Awtade, S. Bhorekar, P Shinde, S. Salve, B. Minhas S. Bharadwaj, H Kaushal Y. Gurav, S. Tomar,Payel Mukherjee, Sofia Banu, Priya Singh, Dhiviya Vedagiri, Divya Gupta, Vishal Sah, Santosh Kumar Kuncha, Krishnan Harinivas Harshan, Archana Bharadwaj Siva, Karthik Bharadwaj Tallapaka, Shagufta Khan, Lamuk Zaveri, Namami Gaur, Sakshi Shambhavi, Tulasi Nagabandi, Purushotham Vodnala,G. Aditya Kumar, Koushick Sivakumar, Pooja Ramesh Gupta, Rajan Kumar Jha, Shraddha Vijay Lahoti, Deepak Kumar, Devi Prasad Vijayashankara, Disha Nanda, Divya Das, Jotin Gogoi, Manish |
| EPI_ISL_450324 | NIV Pune | CSIR-Centre for Cellular and Molecular Biology | Dr V A Potdar, Dr ML Choudhary,Dr Priya Abraham,V. Vipat, S. Jadhav, U. Saha, H. Kengle, A. Awhale, A. Jagtap, A. Gondhalikar, V Malik, N Srivastava, S. Digraskar, P. Malsane, S. Hundekar, K. Patel, Yogesh Balakartik, M. Kakade, S. Jadhav, R. Gunjikar, V. Awtade, S. Bhorekar, P Shinde, S. Salve, B. Minhas S. Bharadwaj, H Kaushal Y. Gurav, S. Tomar,Sofia Banu, Payel Mukherjee, Priya Singh, Dhiviya Vedagiri, Divya Gupta, Vishal Sah, Santosh Kumar Kuncha, Krishnan Harinivas Harshan, Archana Bharadwaj Siva, Karthik Bharadwaj Tallapaka, Shagufta Khan, Lamuk Zaveri, Namami Gaur, Sakshi Shambhavi, Tulasi Nagabandi, Purushotham Vodnala, Disha Nanda, Divya Das, Jotin Gogoi, Manish Bhattacharjee, Ravi Prasad Mukku, Renu Sudhakar, Somesh Gorde, Gangumala Srinivas Reddy, Sujoy Deb, Swati Bayyana, Zeba Rizvi, Rakesh K Mishra |
| EPI_ISL_450325 | NIV Pune | CSIR-Centre for Cellular and Molecular Biology | Dr V A Potdar, Dr ML Choudhary,Dr Priya Abraham,V. Vipat, S. Jadhav, U. Saha, H. Kengle, A. Awhale, A. Jagtap, A. Gondhalikar, V Malik, N Srivastava, S. Digraskar, P. Malsane, S. Hundekar, K. Patel, Yogesh Balakartik, M. Kakade, S. Jadhav, R. Gunjikar, V. Awtade, S. Bhorekar, P Shinde, S. Salve, B. Minhas S. Bharadwaj, H Kaushal Y. Gurav, S. Tomar,Payel Mukherjee, Sofia Banu, Priya Singh, Dhiviya Vedagiri, Divya Gupta, Vishal Sah, Santosh Kumar Kuncha, Krishnan Harinivas Harshan, Archana Bharadwaj Siva, Karthik Bharadwaj Tallapaka, Shagufta Khan, Lamuk Zaveri, Namami Gaur, Sakshi Shambhavi, Tulasi Nagabandi, Purushotham Vodnala,G. Aditya Kumar, Koushick Sivakumar, Pooja Ramesh Gupta, Rajan Kumar Jha, Shraddha Vijay Lahoti, Deepak Kumar, Devi Prasad Vijayashankara, Disha Nanda, Divya Das, Jotin Gogoi, Manish |
| EPI_ISL_450326 | CSIR-Centre for Cellular and Molecular Biology | CSIR-Centre for Cellular and Molecular Biology | Payel Mukherjee, Sofia Banu, Priya Singh, Dhiviya Vedagiri, Divya Gupta, Vishal Sah, Santosh Kumar Kuncha, Krishnan Harinivas Harshan, Archana Bharadwaj Siva, Karthik Bharadwaj Tallapaka, Shagufta Khan, Lamuk Zaveri, Namami Gaur, Sakshi Shambhavi, Tulasi Nagabandi, Purushotham Vodnala,G. Aditya Kumar, Koushick Sivakumar, Pooja Ramesh Gupta, Rajan Kumar Jha, Shraddha Vijay Lahoti, Deepak Kumar, Devi Prasad Vijayashankara, Disha Nanda, Divya Das, Jotin Gogoi, Manish Bhattacharjee, Rakesh K Mishra, Divya Tej Sowpati |
| EPI_ISL_450327 | CSIR-Centre for Cellular and Molecular Biology | CSIR-Centre for Cellular and Molecular Biology | Sofia Banu, Payel Mukherjee, Priya Singh, Dhiviya Vedagiri, Divya Gupta, Vishal Sah, Santosh Kumar Kuncha, Krishnan Harinivas Harshan, Archana Bharadwaj Siva, Karthik Bharadwaj Tallapaka, Shagufta Khan, Lamuk Zaveri, Namami Gaur, Sakshi Shambhavi, Tulasi Nagabandi, Purushotham Vodnala, Disha Nanda, Divya Das, Jotin Gogoi, Manish Bhattacharjee, Ravi Prasad Mukku, Renu Sudhakar, Somesh Gorde, Gangumala Srinivas Reddy, Sujoy Deb, Swati Bayyana, Zeba Rizvi, Rakesh K Mishra, Divya Tej Sowpati |
| EPI_ISL_450328 | CSIR-Centre for Cellular and Molecular Biology | CSIR-Centre for Cellular and Molecular Biology | Shagufta Khan, Lamuk Zaveri, Namami Gaur, Sakshi Shambhavi, Tulasi Nagabandi, Purushotham Vodnala, Payel Mukherjee, Sofia Banu, Priya Singh, Dhiviya Vedagiri, Divya Gupta, Vishal Sah, Santosh Kumar Kuncha, Krishnan Harinivas Harshan, Archana Bharadwaj Siva, Karthik Bharadwaj Tallapaka, Zeba Rizvi, Zuberwasim Sayyad, Kakade Aishwarya Arun, Amrutha H C, Ananga Ghosh, Kezia J Ann, Radhika Khandelwal, Roshan Maku Venkata, Shemin Mansuri, Sonu Uday, Sudipta Mondal, Rakesh K Mishra, Divya Tej Sowpati |
| EPI_ISL_450329 | CSIR-Centre for Cellular and Molecular Biology | CSIR-Centre for Cellular and Molecular Biology | Namami Gaur, Sakshi Shambhavi, Lamuk Zaveri, Shagufta Khan, Tulasi Nagabandi, Purushotham Vodnala, Payel Mukherjee, Sofia Banu, Priya Singh, Dhiviya Vedagiri, Divya Gupta, Vishal Sah, Santosh Kumar Kuncha, Krishnan Harinivas Harshan, Archana Bharadwaj Siva, Karthik Bharadwaj Tallapaka, Sonu Uday, Sudipta Mondal, Annapoorna P Karthyayani, Debabrata Jana, Debrya Saha, Gokulan C G, Gunjan Purohit, |

| | | | |
|---|---|---|---|
| | | | Hanuman Tulashiram Kale, Pankaj Kumar, Prachand Issarapu, Preethi Jampala Rakesh K Mishra, Divya Tej Sowpati |
| EPI_ISL_450330 | CSIR-Centre for Cellular and Molecular Biology | CSIR-Centre for Cellular and Molecular Biology | Sakshi Shambhavi, Lamuk Zaveri, Shagufta Khan, Namami Gaur, Tulasi Nagabandi, Purushotham Vodnala, Payel Mukherjee, Sofia Banu, Priya Singh, Dhiviya Vedagiri, Divya Gupta, Vishal Sah, Santosh Kumar Kuncha, Krishnan Harinivas Harshan, Archana Bharadwaj Siva, Karthik Bharadwaj Tallapaka,Preethi Jampala, Sharada Ravi Iyer, Sulagana Mukherjee, Swetha Sundar, Peddapuvala Sai Uday Kiran, Umesh Kumar, Unis Ahmad Bhat, Ajay Sarawagi, Priyanka Pant, Rajkanwar Nathawat, Nikhil Hajirnis, Pratheusa Maccha, M Soujanya Reddy Rakesh K Mishra, Divya Tej Sowpati |
| EPI_ISL_450331 | CSIR-Centre for Cellular and Molecular Biology | CSIR-Centre for Cellular and Molecular Biology | Tulasi Nagabandi, Namami Gaur, Sakshi Shambhavi, Lamuk Zaveri, Shagufta Khan, Purushotham Vodnala, Payel Mukherjee, Sofia Banu, Priya Singh, Dhiviya Vedagiri, Divya Gupta, Vishal Sah, Santosh Kumar Kuncha, Krishnan Harinivas Harshan, Archana Bharadwaj Siva, Karthik Bharadwaj Tallapaka,G. Aditya Kumar, Koushick Sivakumar, Pooja Ramesh Gupta, Rajan Kumar Jha, Shraddha Vijay Lahoti, Deepak Kumar, Devi Prasad Vijayashankara, Disha Nanda, Divya Das, Jotin Gogoi, Manish Bhattacharjee, Rakesh K Mishra, Divya Tej Sowpati |
| EPI_ISL_450332 | CSIR-Centre for Cellular and Molecular Biology | CSIR-Centre for Cellular and Molecular Biology | Payel Mukherjee, Sofia Banu, Priya Singh, Dhiviya Vedagiri, Divya Gupta, Vishal Sah, Santosh Kumar Kuncha, Krishnan Harinivas Harshan, Archana Bharadwaj Siva, Karthik Bharadwaj Tallapaka, Shagufta Khan, Lamuk Zaveri, Namami Gaur, Sakshi Shambhavi, Tulasi Nagabandi, Purushotham Vodnala,G. Aditya Kumar, Koushick Sivakumar, Pooja Ramesh Gupta, Rajan Kumar Jha, Shraddha Vijay Lahoti, Deepak Kumar, Devi Prasad Vijayashankara, Disha Nanda, Divya Das, Jotin Gogoi, Manish Bhattacharjee, Rakesh K Mishra, Divya Tej Sowpati |
| EPI_ISL_450337 | Hospital Universitari Vall d'Hebron - Vall d'Hebron Institut de Recerca | Hospital Universitari Vall d'Hebron | Cristina Andrés, Maria Piñana, Damir Garcia-Cehic, Mercedes Guerrero-Murillo, Ariadna Rando, Juliana Esperalba, Maria Gema Codina, Maria Carmen Martín, Tomàs Pumarola, Josep Quer, Andrés Antón |
| EPI_ISL_450338 | Institute of Human Genetics, Polish Academy of Sciences; Sanitary and Epidemiological Station in Poznań | Institute of Human Genetics, Polish Academy of Sciences | Szymon Hryhorowicz, Adam Ustaszewski, Emilia Lis, Marta Kaczmarek-Ryś, Michał Witt, Andrzej Pławski |
| EPI_ISL_450339, EPI_ISL_450340 | Bangladesh Institute of Tropical & Infectious Diseases, COVID-19 Testing Laboratory | Basic and Applied Research on Jute Project | Rasel Ahmed, Md. Sabbir Hossain, Shah Md Tamim Kabir, Emdadul Mannan Emdad, Md. Nazmul Haq Rony, Eaftekhar Ahmed Rana, Paritous Kumar Biswas, M A Hassan Chowdhury, Md. Shakeel Ahmed, Md. Samiul Haque, Md. Monjurul Alam, Md. Sharifur Rahman, A S M Anwarul Huq, Md. Shahidul Islam, Goutam Buddha Das, AMAM Zonaed Siddiki |
| EPI_ISL_450341 | Bangladesh Institute of Tropical & Infectious Diseases, COVID-19 Testing Laboratory | Basic and Applied Research on Jute Project | Md. Sabbir Hossain, Rasel Ahmed, Shah Md Tamim Kabir, Emdadul Mannan Emdad, Md. Nazmul Haq Rony, Eaftekhar Ahmed Rana, Paritous Kumar Biswas, M A Hassan Chowdhury, Md. Shakeel Ahmed, Md. Samiul Haque, Md. Monjurul Alam, Md. Sharifur Rahman, A S M Anwarul Huq, Md. Shahidul Islam, Goutam Buddha Das, AMAM Zonaed Siddiki |
| EPI_ISL_450342 | Bangladesh Institute of Tropical & Infectious Diseases, COVID-19 Testing Laboratory | Basic and Applied Research on Jute Project | Rasel Ahmed, Md. Sabbir Hossain, Shah Md Tamim Kabir, Emdadul Mannan Emdad, Md. Nazmul Haq Rony, Eaftekhar Ahmed Rana, Paritous Kumar Biswas, M A Hassan Chowdhury, Md. Shakeel Ahmed, Md. Samiul Haque, Md. Monjurul Alam, Md. Sharifur Rahman, A S M Anwarul Huq, Md. Shahidul Islam, Goutam Buddha Das, AMAM Zonaed Siddiki |
| EPI_ISL_450343 | Bangladesh Institute of Tropical & Infectious Diseases, COVID-19 Testing Laboratory | Basic and Applied Research on Jute Project | Md. Sabbir Hossain, Rasel Ahmed, Shah Md Tamim Kabir, Emdadul Mannan Emdad, Md. Nazmul Haq Rony, Eaftekhar Ahmed Rana, Paritous Kumar Biswas, M A Hassan Chowdhury, Md. Shakeel Ahmed, Md. Samiul Haque, Md. Monjurul Alam, Md. Sharifur Rahman, A S M Anwarul Huq, Md. Shahidul Islam, Goutam Buddha Das, AMAM Zonaed Siddiki |
| EPI_ISL_450344 | Bangladesh Institute of Tropical & Infectious Diseases, COVID-19 Testing Laboratory | Basic and Applied Research on Jute Project | Rasel Ahmed, Md. Sabbir Hossain, Shah Md Tamim Kabir, Emdadul Mannan Emdad, Md. Nazmul Haq Rony, Eaftekhar Ahmed Rana, Paritous Kumar Biswas, M A Hassan Chowdhury, Md. Shakeel Ahmed, Md. Samiul Haque, Md. Monjurul Alam, Md. Sharifur Rahman, A S M Anwarul Huq, Md. Shahidul Islam, Goutam Buddha Das, AMAM Zonaed Siddiki |
| EPI_ISL_450345 | Bangladesh Institute of Tropical & Infectious Diseases, COVID-19 Testing Laboratory | Basic and Applied Research on Jute Project | Md. Sabbir Hossain, Rasel Ahmed, Shah Md Tamim Kabir, Emdadul Mannan Emdad, Md. Nazmul Haq Rony, Eaftekhar Ahmed Rana, Paritous Kumar Biswas, M A Hassan Chowdhury, Md. Shakeel Ahmed, Md. Samiul Haque, Md. Monjurul Alam, Md. Sharifur Rahman, A S M Anwarul Huq, Md. Shahidul Islam, Goutam Buddha Das, AMAM Zonaed Siddiki |
| EPI_ISL_450346, EPI_ISL_450347, EPI_ISL_450348, EPI_ISL_450349, EPI_ISL_450350, EPI_ISL_450351, EPI_ISL_450352 | St.Olavs hospital/NTNU | Institute of Genomics Core Facility, University of Tartu | Aleksandr Ianevski, Tuuli Reisberg, Janne-Fossum Malmring, Svein Arne Nordbø, Denis Kainov |
| EPI_ISL_450393, EPI_ISL_450394, EPI_ISL_450395, EPI_ISL_450396, EPI_ISL_450397, EPI_ISL_450398, EPI_ISL_450399, EPI_ISL_450400, EPI_ISL_450401, EPI_ISL_450402 | NYU Langone Health | Departments of Pathology and Medicine, New York University School of Medicine | Maria Aguero-Rosenfeld, Brendan Belovarac, Margaret Black, Ludovic Boytard, John Cadley, Paolo Cotzia, John Chen, Dacia Dimartino, Xiaojun Feng, Tatyana Gindin, Emily Guzman, Adriana Heguy, Megan Hogan, Emily Huang, George Jour, Alireza Khodadadi-Jamayran, Lawrence H. Lin, Raven Luther, Andrew Lytle, Christian Marier, Matthew T. Maurano, Mark J. Mulligan, Peter Meyn, Raquel Ordonez Ciriza, Iman Osman, Jared Pinnell, Vanessa Raabe, Sitharam Ramaswami, Amy Rapkiewicz, Andre M. Ribeiro-dos-Santos, Marie Samanovic-Golden, Antonio Serrano, Guomiao Shen, Matija Snuderl, Theodore Vougiouklakis, Nick Vulpescu, Gael Westby, Paul Zappile, Yutong Zhang |
| EPI_ISL_450445, EPI_ISL_450446, EPI_ISL_450447, EPI_ISL_450448, EPI_ISL_450449, EPI_ISL_450450, EPI_ISL_450451, EPI_ISL_450452, EPI_ISL_450453, EPI_ISL_450454, EPI_ISL_450455, EPI_ISL_450456, EPI_ISL_450457, EPI_ISL_450458, EPI_ISL_450459, EPI_ISL_450460, EPI_ISL_450461, EPI_ISL_450462, EPI_ISL_450463, EPI_ISL_450464, EPI_ISL_450465, EPI_ISL_450466, EPI_ISL_450467, EPI_ISL_450468, EPI_ISL_450469, EPI_ISL_450470, EPI_ISL_450471, EPI_ISL_450472, EPI_ISL_450473, EPI_ISL_450474, EPI_ISL_450475, EPI_ISL_450476, EPI_ISL_450477, EPI_ISL_450478, EPI_ISL_450479, EPI_ISL_450480, EPI_ISL_450481 | | | |
| see above | Stanford clinical virology lab | Chan-Zuckerberg Biohub | Benjamin Pinksy, Katharine Walter, Victoria N. Parikh, John Gorzynski, Hannah N. DeJong, Matthew T. Wheeler, Jason Andrews, Manuel Rivas, Carlos Bustamante, Euan Ashley, with CZB Cliahub Consortium |
| EPI_ISL_450495 | National Institute for Communicable Diseases of the National Health Laboratory Service | National Institute for Communicable Diseases of the National Health Laboratory Service | Allam M, Ismail A, Khumalo Z, Kwenda S, van Heusden P, Mtshali P, Mnyameni F, Mohale T, Subramoney K, Bhiman JN |
| EPI_ISL_450496, EPI_ISL_450497 | National Public Health Surveillance Laboratory, Vilnius, Lithuania | Charite Universitaetsmedizin Berlin, Institute of Virology | Victor M Corman, Jorn Beheim-Schwarzbach, Talitha Veith, Barbara Muehlemann, Julia Schneider, Terry Jones, Ana Steponkiene, Christian Drosten |
| EPI_ISL_450498 | Health Board Laboratory of Communicable Diseases | Charite Universitatsmedizin Berlin, Institute of Virology | Victor M Corman, Jörn Beheim-Schwarzbach, Barbara Mühlemann, Talitha Veith, Julia Schneider, Liidia Dotsenko, Natalja Kuznetsova, Terry Jones, Christian Drosten |
| EPI_ISL_450508, EPI_ISL_450509, EPI_ISL_450510, EPI_ISL_450511, EPI_ISL_450512, EPI_ISL_450513, EPI_ISL_450514, EPI_ISL_450515, EPI_ISL_450516, EPI_ISL_450517 | Rafik Hariri University Hospital | Rafik Hariri University Hospital | Rita Feghali |
| EPI_ISL_450518, EPI_ISL_450519 | E. Gulbja Laboratorija | Latvian Biomedical Research and Study Centre | Ivars Silamiķelis, Kaspars Megnis, Monta Ustinova, Ņikita Zrelovs, Vita Rovīte, Mikus Gavars, Dmitrijs Perminovs, Uga Dumpis, Jānis Kloviņš |
| EPI_ISL_450520, EPI_ISL_450521, EPI_ISL_450522, EPI_ISL_450523, EPI_ISL_450524 | Centrālā Laboratorija | Latvian Biomedical Research and Study Centre | Ivars Silamiķelis, Kaspars Megnis, Monta Ustinova, Ņikita Zrelovs, Vita Rovīte, Stella Lapiņa, Jana Osīte, Marta Priedīte, Uga Dumpis, Jānis Kloviņš |
| EPI_ISL_450525, EPI_ISL_450526, EPI_ISL_450527, EPI_ISL_450528, EPI_ISL_450529, EPI_ISL_450530 | Hematology Laboratory, Section of Molecular Diagnostics, University Clinical Centre, Medical University of Gdansk | Department of Virology, Faculty of Medicine, University of Helsinki, Helsinki, Finland | Maciej Grzybek, Marlena Robakowska, Aneta Szulc, Olli Vapalahti, Teemu Smura |
| EPI_ISL_450538, EPI_ISL_450539, EPI_ISL_450540, EPI_ISL_450541, EPI_ISL_450542, EPI_ISL_450543, EPI_ISL_450544, EPI_ISL_450545, EPI_ISL_450546, EPI_ISL_450547, EPI_ISL_450548, EPI_ISL_450549, EPI_ISL_450550, EPI_ISL_450551, EPI_ISL_450552, EPI_ISL_450553, EPI_ISL_450554, EPI_ISL_450555, EPI_ISL_450556, EPI_ISL_450557, EPI_ISL_450559, EPI_ISL_450560, EPI_ISL_450561, EPI_ISL_450562, EPI_ISL_450563, EPI_ISL_450564, EPI_ISL_450565, EPI_ISL_450566, EPI_ISL_450567, EPI_ISL_450568 | | | |
| see above | Utah Public Health Laboratory | Utah Public Health Laboratory | Erin Young, Kelly Oakeson |
| EPI_ISL_450570, EPI_ISL_450571, EPI_ISL_450572, EPI_ISL_450573, EPI_ISL_450574, EPI_ISL_450575, EPI_ISL_450576, EPI_ISL_450577, EPI_ISL_450578, EPI_ISL_450579, EPI_ISL_450580, EPI_ISL_450581, EPI_ISL_450582, EPI_ISL_450583, EPI_ISL_450584, EPI_ISL_450585, EPI_ISL_450586, EPI_ISL_450587, EPI_ISL_450588, EPI_ISL_450589, EPI_ISL_450590, EPI_ISL_450591, EPI_ISL_450592, EPI_ISL_450593, EPI_ISL_450594, EPI_ISL_450595, EPI_ISL_450596, EPI_ISL_450597, EPI_ISL_450598, EPI_ISL_450599, EPI_ISL_450600 | | | |
| see above | Michigan Department of Health and Human Services, Bureau of Laboratories | Michigan Department of Health and Human Services, Bureau of Laboratories | Blankenship HM; Riner D; Soehnlen MK |
| EPI_ISL_450601, EPI_ISL_450602, EPI_ISL_450603, EPI_ISL_450604, EPI_ISL_450605, EPI_ISL_450606, EPI_ISL_450607, EPI_ISL_450608, EPI_ISL_450609, EPI_ISL_450610, EPI_ISL_450611, EPI_ISL_450612, EPI_ISL_450613, EPI_ISL_450614, EPI_ISL_450615, EPI_ISL_450616, EPI_ISL_450617, EPI_ISL_450618, EPI_ISL_450619, EPI_ISL_450620, EPI_ISL_450621, EPI_ISL_450622, EPI_ISL_450623, EPI_ISL_450624, EPI_ISL_450625, EPI_ISL_450626, EPI_ISL_450627, EPI_ISL_450628, EPI_ISL_450629, EPI_ISL_450630, EPI_ISL_450631, EPI_ISL_450632, EPI_ISL_450633, EPI_ISL_450634, EPI_ISL_450635, EPI_ISL_450636, EPI_ISL_450637 | | | |
| see above | Michigan Department of Health and Human Services, Bureau of Laboratories | Michigan Department of Health and Human Services, Bureau of Laboratories | Blankenship HM, Riner D, Soehnlen MK |
| EPI_ISL_450638, EPI_ISL_450639, EPI_ISL_450640, EPI_ISL_450641, EPI_ISL_450642, EPI_ISL_450643, EPI_ISL_450644, EPI_ISL_450645, EPI_ISL_450646, EPI_ISL_450647, EPI_ISL_450648, EPI_ISL_450649, EPI_ISL_450650, EPI_ISL_450651, EPI_ISL_450652, EPI_ISL_450653 | | | |
| see above | Laboratoire de microbiologie, Hopital de Verdun | Smith Laboratory, Centre de Recherche CHU Sainte-Justine | Martin Smith, Marieke Rozendaal, Ivan Pavlov |
| EPI_ISL_450700, EPI_ISL_450701, EPI_ISL_450702, EPI_ISL_450703, EPI_ISL_450704, EPI_ISL_450705, EPI_ISL_450706, EPI_ISL_450707, EPI_ISL_450708, EPI_ISL_450709, EPI_ISL_450710, EPI_ISL_450711, EPI_ISL_450712, EPI_ISL_450713, EPI_ISL_450714, EPI_ISL_450715, EPI_ISL_450716, EPI_ISL_450717, EPI_ISL_450718, EPI_ISL_450719, EPI_ISL_450720, EPI_ISL_450721, EPI_ISL_450722 | | | |
| see above | University of Wisconsin-Madison AIDS Vaccine Research Laboratories | University of Wisconsin-Madison AIDS Vaccine Research Laboratories | Gage Moreno, Katarina Braun, et al. AIDS Vaccine Research Laboratories |
| EPI_ISL_450723 | Ramathibodi Hospital | COVID-19 Network Investigations (CONI) Alliance | Elizabeth Batty, Wasun Chantratita, Thanat Chookajorn, Stefan Fernandez, Angkana Huang, Anthony R. Jones, Khajohn Joonsalak, Chonticha Klungtong, Theerarat Kochakarn, Namfon Kotanan, Krittikorn Kumpornsin, Wudtichai Manasatienkij, Bhakbhoom Panthan, Ekawat Pasomsub, Insee Sensorn, Arporn Wangwiwatsin |
| EPI_ISL_450724, EPI_ISL_450725, EPI_ISL_450726, EPI_ISL_450727, EPI_ISL_450728, EPI_ISL_450729, EPI_ISL_450730, EPI_ISL_450731, EPI_ISL_450732, EPI_ISL_450733, EPI_ISL_450734, EPI_ISL_450735, EPI_ISL_450736, EPI_ISL_450737 | | | |
| see above | Hospital AZ Rivierenland | Institute of Tropical Medicine | Philippe Selhorst, Colin Anthony |
| EPI_ISL_450738, EPI_ISL_450739, EPI_ISL_450740, EPI_ISL_450741, EPI_ISL_450742, EPI_ISL_450743, EPI_ISL_450744, EPI_ISL_450745 | OUCRU/HTD | OUCRU/HTD | Nguyen Van Vinh Chau, Nguyen Thi Thu Hong, Nguyen Thi Han Ny, Le Nguyen Truc Nhu, Nghiem My Ngoc, Vo Thanh Lam, Nguyen Thanh Dung, Lam Minh Yen, Ngo Ngoc Quang Minh, Le Manh Hung, Nguyen Tri Dung, Dinh Nguyen Huy Man, Lam Anh Nguyet, Tran Chanh Xuan, Tran Tinh Hien, Nguyen Thanh Phong, Tran Nguyen Hoang Tu, Tran Tan Thanh, Nguyen Thanh Truong, Nguyen Tan Binh, Tang Chi Thuong, Guy Thwaites, and Le Van Tan, for OUCRU COVID-19 research group* |
| EPI_ISL_450746 | Laboratory of Molecular Biology, Diagnostyka sp. z o.o. | Laboratory of Recombinant Vaccines | Lukasz Rabalski, Anna Piotrowska-Mietelska, Maciej Kosinski, Boguslaw Szewczyk, Krystyna Bienkowska-Szewczyk |
| EPI_ISL_450747 | Sunnybrook Health Sciences Centre | Department of Laboratory Medicine and Molecular Diagnostics, Sunnybrook Health Sciences Centre | Jalees A. Nasir, Robert A. Kozak, Patryk Aftanas, Amogelang R. Raphenya, Kendrick M. Smith, Finlay Maguire, Hassaan Maan, Muhannad Alruwaili, Arinjay Banerjee, Hamza Mbareche, Brian P. Alcock, Natalie C. Knox, Karen Mossman, Bo Wang, Julian A. Hiscox, Andrew G. McArthur, Samira Mubareka |
| EPI_ISL_450748, EPI_ISL_450749, EPI_ISL_450750, EPI_ISL_450751, EPI_ISL_450752, EPI_ISL_450753, EPI_ISL_450754, EPI_ISL_450755, EPI_ISL_450756, EPI_ISL_450757, EPI_ISL_450758, EPI_ISL_450759, EPI_ISL_450760, EPI_ISL_450761, EPI_ISL_450762, EPI_ISL_450763, EPI_ISL_450764, EPI_ISL_450765, EPI_ISL_450766, EPI_ISL_450767, EPI_ISL_450768, EPI_ISL_450769, EPI_ISL_450770, EPI_ISL_450771, EPI_ISL_450772, EPI_ISL_450773, EPI_ISL_450774, EPI_ISL_450775, EPI_ISL_450776, EPI_ISL_450777, EPI_ISL_450778, EPI_ISL_450779, EPI_ISL_450780 | | | |
| see above | Minnesota Department of Health, Public Health Laboratory | Minnesota Department of Health, Public Health Laboratory | Matt Plumb, Jacob Garfin, and Xiong Wang |
| EPI_ISL_450781 | Government Medical College-Bhavnagar | Gujarat Biotechnology Research Centre | Kairavi Desai, Saklain Malek, Shirish Patel, Ramesh Pandit, Tejas Shah, Ankit Hinsu, Pritesh Sabara, Apurvasinh Puvar, Janvi Raval, Zarna Patel, Monika Gandhi, Pinal Trivedi, Maharshi Pandya, Amit Kanani, Nidhi Patel, Nitin Savaliya, Raghawendra Kumar, Dinesh Kumar, Zuber Saiyed, Komal Patel, Labdhi Pandya, Snehal Bagatharia, Bhavesh Modi, Gaurishankar Shrimali, R D Dixit, A M Kadri, Akanksha Verma, Chaitanya Joshi, Madhvi Joshi |
| EPI_ISL_450782 | Government Medical College-Bhavnagar | Gujarat Biotechnology Research Centre | Saklain Malek, Shirish Patel, Kairavi Desai, Tejas Shah, Ankit Hinsu, Pritesh Sabara, Apurvasinh Puvar, Janvi Raval, Zarna Patel, Monika Gandhi, Pinal Trivedi, Maharshi Pandya, Amit Kanani, Nidhi Patel, Nitin Savaliya, Raghawendra Kumar, Dinesh Kumar, Zuber Saiyed, Komal Patel, Labdhi Pandya, Snehal Bagatharia, Ramesh Pandit, Bhavesh Modi, Gaurishankar Shrimali, R D Dixit, A M Kadri, Priti Pandita, Chaitanya Joshi, Madhvi Joshi |
| EPI_ISL_450783 | Government Medical College-Bhavnagar | Gujarat Biotechnology Research Centre | Shirish Patel, Kairavi Desai, Saklain Malek, Ankit Hinsu, Pritesh Sabara, Apurvasinh Puvar, Janvi Raval, Zarna Patel, Monika Gandhi, Pinal Trivedi, Maharshi Pandya, Amit Kanani, Nidhi Patel, Nitin Savaliya, Raghawendra Kumar, Dinesh Kumar, Zuber Saiyed, Komal Patel, Labdhi Pandya, Snehal Bagatharia, Ramesh Pandit, Tejas Shah, Bhavesh Modi, Gaurishankar Shrimali, R D Dixit, A M Kadri, Neha Rajpara, Chaitanya Joshi, Madhvi Joshi |

| | | | |
|---|---|---|---|
| EPI_ISL_450784 | Government Medical College-Bhavnagar | Gujarat Biotechnology Research Centre | Zarna Patel, Ramesh Pandit, Tejas Shah, Ankit Hinsu, Pritesh Sabara, Apurvasinh Puvar, Janvi Raval, Monika Gandhi, Pinal Trivedi, Maharshi Pandya, Amit Kanani, Nidhi Patel, Nitin Savaliya, Raghawendra Kumar, Dinesh Kumar, Zuber Saiyed, Komal Patel, Labdhi Pandya, Snehal Bagatharia, Kairavi Desai, Saklain Malek, Shirish Patel, Bhavesh Modi, Gaurishankar Shrimali, R D Dixit, A M Kadri, Afzal Ansari, Chaitanya Joshi, Madhvi Joshi |
| EPI_ISL_450785 | Pandit Deendayal Upadhyay Government Medical College, Rajkot | Gujarat Biotechnology Research Centre | Prakash Modi, Sejul Antala, Manish Pattani, Apurvasinh Puvar, Janvi Raval, Zarna Patel, Monika Gandhi, Pinal Trivedi, Maharshi Pandya, Amit Kanani, Nidhi Patel, Nitin Savaliya, Raghawendra Kumar, Dinesh Kumar, Zuber Saiyed, Komal Patel, Labdhi Pandya, Snehal Bagatharia, Ramesh Pandit, Tejas Shah, Ankit Hinsu, Pritesh Sabara, Bhavesh Modi, Gaurishankar Shrimali, R D Dixit, A M Kadri, Neelam Nathani, Chaitanya Joshi, Madhvi Joshi |
| EPI_ISL_450786 | Pandit Deendayal Upadhyay Government Medical College, Rajkot | Gujarat Biotechnology Research Centre | Sejul Antala, Manish Pattani, Prakash Modi, Janvi Raval, Zarna Patel, Monika Gandhi, Pinal Trivedi, Maharshi Pandya, Amit Kanani, Nidhi Patel, Nitin Savaliya, Raghawendra Kumar, Dinesh Kumar, Zuber Saiyed, Komal Patel, Labdhi Pandya, Snehal Bagatharia, Ramesh Pandit, Tejas Shah, Ankit Hinsu, Pritesh Sabara, Apurvasinh Puvar, Bhavesh Modi, Gaurishankar Shrimali, R D Dixit, A M Kadri, Armi Chaudhari, Chaitanya Joshi, Madhvi Joshi |
| EPI_ISL_450787 | Pandit Deendayal Upadhyay Government Medical College, Rajkot | Gujarat Biotechnology Research Centre | Manish Pattani, Prakash Modi, Sejul Antala, Zarna Patel, Monika Gandhi, Pinal Trivedi, Maharshi Pandya, Amit Kanani, Nidhi Patel, Nitin Savaliya, Raghawendra Kumar, Dinesh Kumar, Zuber Saiyed, Komal Patel, Labdhi Pandya, Snehal Bagatharia, Ramesh Pandit, Tejas Shah, Ankit Hinsu, Pritesh Sabara, Apurvasinh Puvar, Janvi Raval, Bhavesh Modi, Gaurishankar Shrimali, R D Dixit, A M Kadri, Bhavya Jindal, Chaitanya Joshi, Madhvi Joshi |
| EPI_ISL_450788 | Pandit Deendayal Upadhyay Government Medical College, Rajkot | Gujarat Biotechnology Research Centre | Zarna Patel, Tejas Shah, Ankit Hinsu, Pritesh Sabara, Apurvasinh Puvar, Janvi Raval, Monika Gandhi, Pinal Trivedi, Maharshi Pandya, Amit Kanani, Nidhi Patel, Nitin Savaliya, Raghawendra Kumar, Dinesh Kumar, Zuber Saiyed, Komal Patel, Labdhi Pandya, Snehal Bagatharia, Prakash Modi, Sejul Antala, Manish Pattani, Ramesh Pandit, Bhavesh Modi, Gaurishankar Shrimali, R D Dixit, A M Kadri, Camellia Chakraborty, Chaitanya Joshi, Madhvi Joshi |
| EPI_ISL_450789 | Pandit Deendayal Upadhyay Government Medical College, Rajkot | Gujarat Biotechnology Research Centre | Ankit Hinsu, Pritesh Sabara, Apurvasinh Puvar, Janvi Raval, Zarna Patel, Monika Gandhi, Pinal Trivedi, Maharshi Pandya, Amit Kanani, Nidhi Patel, Nitin Savaliya, Raghawendra Kumar, Dinesh Kumar, Zuber Saiyed, Komal Patel, Labdhi Pandya, Snehal Bagatharia, Prakash Modi, Sejul Antala, Manish Pattani, Ramesh Pandit, Tejas Shah, Bhavesh Modi, Gaurishankar Shrimali, R D Dixit, A M Kadri, Siddhant Kumar, Chaitanya Joshi, Madhvi Joshi |
| EPI_ISL_450790 | Pandit Deendayal Upadhyay Government Medical College, Rajkot | Gujarat Biotechnology Research Centre | Zarna Patel, Pritesh Sabara, Apurvasinh Puvar, Janvi Raval, Monika Gandhi, Pinal Trivedi, Maharshi Pandya, Amit Kanani, Nidhi Patel, Nitin Savaliya, Raghawendra Kumar, Dinesh Kumar, Zuber Saiyed, Komal Patel, Labdhi Pandya, Snehal Bagatharia, Prakash Modi, Sejul Antala, Manish Pattani, Ramesh Pandit, Tejas Shah, Ankit Hinsu, Bhavesh Modi, Gaurishankar Shrimali, R D Dixit, A M Kadri, Sharmistha Majumdar, Chaitanya Joshi, Madhvi Joshi |
| EPI_ISL_450791 | Pandit Deendayal Upadhyay Government Medical College, Rajkot | Gujarat Biotechnology Research Centre | Zarna Patel, Apurvasinh Puvar, Janvi Raval, Monika Gandhi, Pinal Trivedi, Maharshi Pandya, Amit Kanani, Nidhi Patel, Nitin Savaliya, Raghawendra Kumar, Dinesh Kumar, Zuber Saiyed, Komal Patel, Labdhi Pandya, Snehal Bagatharia, Prakash Modi, Sejul Antala, Manish Pattani, Ramesh Pandit, Tejas Shah, Ankit Hinsu, Pritesh Sabara, Bhavesh Modi, Gaurishankar Shrimali, R D Dixit, A M Kadri, Pooja P Doshi, Chaitanya Joshi, Madhvi Joshi |
| EPI_ISL_450792 | Jamaica Ministry of Health and Wellness | Pathogen Discovery, Respiratory Viruses Branch, Division of Viral Diseases, Centers for Disease Control and Prevention | Krista Queen, Yan Li, Anna Montmayeur, Ying Tao, Jing Zhang, Anna Uehara, Clinton R. Paden, Rachel Marine, Haibin Wang, Jasmine Padilla, Justin Lee, Zachary Weiner, Bettina Bankamp, Suxiang Tong |
| EPI_ISL_450793, EPI_ISL_450794, EPI_ISL_450795, EPI_ISL_450796, EPI_ISL_450797 | Jamaica Ministry of Health and Wellness | Pathogen Discovery, Respiratory Viruses Branch, Division of Viral Diseases, Centers for Disease Control and Prevention | Yan Li, Anna Montmayeur, Ying Tao, Krista Queen, Jing Zhang, Anna Uehara, Clinton R. Paden, Rachel Marine, Haibin Wang, Zachary Weiner, Bettina Bankamp, Suxiang Tong |
| EPI_ISL_450798 | Jamaica Ministry of Health and Wellness | Pathogen Discovery, Respiratory Viruses Branch, Division of Viral Diseases, Centers for Disease Control and Prevention | Krista Queen, Yan Li, Anna Montmayeur, Ying Tao, Jing Zhang, Anna Uehara, Clinton R. Paden, Rachel Marine, Haibin Wang, Jasmine Padilla, Justin Lee, Zachary Weiner, Bettina Bankamp, Suxiang Tong |
| EPI_ISL_450799 | Jamaica Ministry of Health and Wellness | Pathogen Discovery, Respiratory Viruses Branch, Division of Viral Diseases, Centers for Disease Control and Prevention | Yan Li, Anna Montmayeur, Ying Tao, Krista Queen, Jing Zhang, Anna Uehara, Clinton R. Paden, Rachel Marine, Haibin Wang, Zachary Weiner, Bettina Bankamp, Suxiang Tong |
| EPI_ISL_450800 | AR Dept. of Health-Public Health Lab | Pathogen Discovery, Respiratory Viruses Branch, Division of Viral Diseases, Centers for Disease Control and Prevention | Yan Li, Anna Montmayeur, Ying Tao, Krista Queen, Jing Zhang, Anna Uehara, Clinton R. Paden, Rachel Marine, Haibin Wang, Zachary Weiner, Bettina Bankamp, Suxiang Tong |
| EPI_ISL_450801 | Georgia Department of Health | Pathogen Discovery, Respiratory Viruses Branch, Division of Viral Diseases, Centers for Disease Control and Prevention | Yan Li, Anna Montmayeur, Ying Tao, Krista Queen, Jing Zhang, Anna Uehara, Clinton R. Paden, Rachel Marine, Haibin Wang, Zachary Weiner, Bettina Bankamp, Suxiang Tong |
| EPI_ISL_450802, EPI_ISL_450803 | PA Department of Health, Bureau of Laboratories | Pathogen Discovery, Respiratory Viruses Branch, Division of Viral Diseases, Centers for Disease Control and Prevention | Yan Li, Anna Montmayeur, Ying Tao, Krista Queen, Jing Zhang, Anna Uehara, Clinton R. Paden, Rachel Marine, Haibin Wang, Zachary Weiner, Bettina Bankamp, Suxiang Tong |
| EPI_ISL_450804 | VI-US Virgin Islands Department of Health | Pathogen Discovery, Respiratory Viruses Branch, Division of Viral Diseases, Centers for Disease Control and Prevention | Yan Li, Anna Montmayeur, Ying Tao, Krista Queen, Jing Zhang, Anna Uehara, Clinton R. Paden, Rachel Marine, Haibin Wang, Zachary Weiner, Bettina Bankamp, Suxiang Tong |
| EPI_ISL_450805 | VI-US Virgin Islands Department of Health | Pathogen Discovery, Respiratory Viruses Branch, Division of Viral Diseases, Centers for Disease Control and Prevention | Krista Queen, Yan Li, Anna Montmayeur, Ying Tao, Jing Zhang, Anna Uehara, Clinton R. Paden, Rachel Marine, Haibin Wang, Jasmine Padilla, Justin Lee, Zachary Weiner, Bettina Bankamp, Suxiang Tong |
| EPI_ISL_450806 | VI-US Virgin Islands Department of Health | Pathogen Discovery, Respiratory Viruses Branch, Division of Viral Diseases, Centers for Disease Control and Prevention | Yan Li, Anna Montmayeur, Ying Tao, Krista Queen, Jing Zhang, Anna Uehara, Clinton R. Paden, Rachel Marine, Haibin Wang, Zachary Weiner, Bettina Bankamp, Suxiang Tong |
| EPI_ISL_450807 | Victoria Vard och Halsa | The Public Health Agency of Sweden | Sarah Henriksson, Anna-Malin Linde, Maria Lind Karlberg, Oskar Karlsson Lindsjo, Olov Svartstrom, Anna Risberg, Theresa Enkirch, Mia Brytting, Karin Tegmark-Wisell |
| EPI_ISL_450808 | Saroledens Familjelakare | The Public Health Agency of Sweden | Katarina Jarbur, Anna-Malin Linde, Maria Lind Karlberg, Oskar Karlsson Lindsjo, Olov Svartstrom, Anna Risberg, Theresa Enkirch, Mia Brytting, Karin Tegmark-Wisell |
| EPI_ISL_450809 | Kungsholmsdoktorn | The Public Health Agency of Sweden | Linus Hammar, Anna-Malin Linde, Maria Lind Karlberg, Oskar Karlsson Lindsjo, Olov Svartstrom, Anna Risberg, Theresa Enkirch, Mia Brytting, Karin Tegmark-Wisell |
| EPI_ISL_450810 | Saroledens Familjelakare | The Public Health Agency of Sweden | Katarina Jarbur, Anna-Malin Linde, Maria Lind Karlberg, Oskar Karlsson Lindsjo, Olov Svartstrom, Anna Risberg, Theresa Enkirch, Mia Brytting, Karin Tegmark-Wisell |
| EPI_ISL_450811 | Knivsta VC | The Public Health Agency of Sweden | Johanna Carlson, Anna-Malin Linde, Maria Lind Karlberg, Oskar Karlsson Lindsjo, Olov Svartstrom, Anna Risberg, Theresa Enkirch, Mia Brytting, Karin Tegmark-Wisell |
| EPI_ISL_450812 | Uppsala Narakut Aleris | The Public Health Agency of Sweden | Annika Nilsson, Anna-Malin Linde, Maria Lind Karlberg, Oskar Karlsson Lindsjo, Olov Svartstrom, Anna Risberg, Theresa Enkirch, Mia Brytting, Karin Tegmark-Wisell |
| EPI_ISL_450813 | Bla Kustens halsocentral | The Public Health Agency of Sweden | Olof Norrby, Anna-Malin Linde, Maria Lind Karlberg, Oskar Karlsson Lindsjo, Olov Svartstrom, Anna Risberg, Theresa Enkirch, Mia Brytting, Karin Tegmark-Wisell |
| EPI_ISL_450814 | Huslakarna Varmbadhuset Varberg | The Public Health Agency of Sweden | Johanna Hilmersson, Anna-Malin Linde, Maria Lind Karlberg, Oskar Karlsson Lindsjo, Olov Svartstrom, Anna Risberg, Theresa Enkirch, Mia Brytting, Karin Tegmark-Wisell |
| EPI_ISL_450815 | Narhalsan Molnlycke, Barn och ungdomsmedicin | The Public Health Agency of Sweden | Mats Reimer, Anna-Malin Linde, Maria Lind Karlberg, Oskar Karlsson Lindsjo, Olov Svartstrom, Anna Risberg, Theresa Enkirch, Mia Brytting, Karin Tegmark-Wisell |
| EPI_ISL_450816 | orestadsklinikens VC | The Public Health Agency of Sweden | Lisa Kjellberg / Laura Plavitu, Anna-Malin Linde, Maria Lind Karlberg, Oskar Karlsson Lindsjo, Olov Svartstrom, Anna Risberg, Theresa Enkirch, Mia Brytting, Karin Tegmark-Wisell |
| EPI_ISL_450817 | Narhalsan Backa vardcentral | The Public Health Agency of Sweden | Mats Olsson, Anna-Malin Linde, Maria Lind Karlberg, Oskar Karlsson Lindsjo, Olov Svartstrom, Anna Risberg, Theresa Enkirch, Mia Brytting, Karin Tegmark-Wisell |
| EPI_ISL_450818 | Aneby VC | The Public Health Agency of Sweden | Ken Granath, Anna-Malin Linde, Maria Lind Karlberg, Oskar Karlsson Lindsjo, Olov Svartstrom, Anna Risberg, Theresa Enkirch, Mia Brytting, Karin Tegmark-Wisell |
| EPI_ISL_450819 | Hovas Askim Familjelakare och BVC | The Public Health Agency of Sweden | Anna Wendel, Anna-Malin Linde, Maria Lind Karlberg, Oskar Karlsson Lindsjo, Olov Svartstrom, Anna Risberg, Theresa Enkirch, Mia Brytting, Karin Tegmark-Wisell |
| EPI_ISL_450820 | Narhalsan Molnlycke, Barn och ungdomsmedicin | The Public Health Agency of Sweden | Mats Reimer, Anna-Malin Linde, Maria Lind Karlberg, Oskar Karlsson Lindsjo, Olov Svartstrom, Anna Risberg, Theresa Enkirch, Mia Brytting, Karin Tegmark-Wisell |
| EPI_ISL_450821, EPI_ISL_450822 | Jarpens HC | The Public Health Agency of Sweden | Gunilla Johansson, Anna-Malin Linde, Maria Lind Karlberg, Oskar Karlsson Lindsjo, Olov Svartstrom, Anna Risberg, Theresa Enkirch, Mia Brytting, Karin Tegmark-Wisell |
| EPI_ISL_450823 | Narhalsan Backa vardcentral | The Public Health Agency of Sweden | Mats Olsson, Anna-Malin Linde, Maria Lind Karlberg, Oskar Karlsson Lindsjo, Olov Svartstrom, Anna Risberg, Theresa Enkirch, Mia Brytting, Karin Tegmark-Wisell |
| EPI_ISL_450824 | Ulltuna Vardcentral | The Public Health Agency of Sweden | Heidi Lindback, Anna-Malin Linde, Maria Lind Karlberg, Oskar Karlsson Lindsjo, Olov Svartstrom, Anna Risberg, Theresa Enkirch, Mia Brytting, Karin Tegmark-Wisell |
| EPI_ISL_450825 | Narhalsan Backa vardcentral | The Public Health Agency of Sweden | Mats Olsson, Anna-Malin Linde, Maria Lind Karlberg, Oskar Karlsson Lindsjo, Olov Svartstrom, Anna Risberg, Theresa Enkirch, Mia Brytting, Karin Tegmark-Wisell |
| EPI_ISL_450826, EPI_ISL_450827, EPI_ISL_450828 | Uppsala Narakut Aleris | The Public Health Agency of Sweden | Annika Nilsson, Anna-Malin Linde, Maria Lind Karlberg, Oskar Karlsson Lindsjo, Olov Svartstrom, Anna Risberg, Theresa Enkirch, Mia Brytting, Karin Tegmark-Wisell |
| EPI_ISL_450829 | Narhalsan Sjobo vardcentral | The Public Health Agency of Sweden | Lovisa Hjerten, Anna-Malin Linde, Maria Lind Karlberg, Oskar Karlsson Lindsjo, Olov Svartstrom, Anna Risberg, Theresa Enkirch, Mia Brytting, Karin Tegmark-Wisell |
| EPI_ISL_450830 | Narhalsan Olskroken VC | The Public Health Agency of Sweden | Mahin Ghoroghi, Anna-Malin Linde, Maria Lind Karlberg, Oskar Karlsson Lindsjo, Olov Svartstrom, Anna Risberg, Theresa Enkirch, Mia Brytting, Karin Tegmark-Wisell |
| EPI_ISL_450831 | Wetterhalsan | The Public Health Agency of Sweden | Anders Tengblad, Anna-Malin Linde, Maria Lind Karlberg, Oskar Karlsson Lindsjo, Olov Svartstrom, Anna Risberg, Theresa Enkirch, Mia Brytting, Karin Tegmark-Wisell |
| EPI_ISL_450832 | Byjorden vardcentral | The Public Health Agency of Sweden | Pernilla Brunman, Anna-Malin Linde, Maria Lind Karlberg, Oskar Karlsson Lindsjo, Olov Svartstrom, Anna Risberg, Theresa Enkirch, Mia Brytting, Karin Tegmark-Wisell |
| EPI_ISL_450833 | Wetterhalsan | The Public Health Agency of Sweden | Anders Tengblad, Anna-Malin Linde, Maria Lind Karlberg, Oskar Karlsson Lindsjo, Olov Svartstrom, Anna Risberg, Theresa Enkirch, Mia Brytting, Karin Tegmark-Wisell |
| EPI_ISL_450834 | Klinisk mikrobiologi och vardhygien Halmstad | The Public Health Agency of Sweden | Arne Kotz, Anna-Malin Linde, Maria Lind Karlberg, Oskar Karlsson Lindsjo, Olov Svartstrom, Anna Risberg, Theresa Enkirch, Mia Brytting, Karin Tegmark-Wisell |
| EPI_ISL_450835 | Unilabs Skovde | The Public Health Agency of Sweden | Tobias Kollberg, Helena Enroth, Anna-Malin Linde, Maria Lind Karlberg, Oskar Karlsson Lindsjo, Olov Svartstrom, Anna Risberg, Theresa Enkirch, Mia Brytting, Karin Tegmark-Wisell |
| EPI_ISL_450836, EPI_ISL_450837 | Laboratoriemedicin | The Public Health Agency of Sweden | Anna-Malin Linde, Maria Lind Karlberg, Oskar Karlsson Lindsjo, Olov Svartstrom, Anna Risberg, Theresa Enkirch, Mia Brytting, Karin Tegmark-Wisell |
| EPI_ISL_450838 | Utah Public Health Laboratory | Utah Public Health Laboratory | Erin Young, Kelly Oakeson |
| EPI_ISL_450839 | COVID-19 Laboratory Centre for Advanced Research in Sciences (CARS), University of Dhaka, Dhaka-1000, Bangladesh | DNA Solution Ltd | Sharif Akhteruzzaman, Zeba Islam Seraj, Nazmul Ahsan, Md Imdadul Hoque, MA Malek, Shahryar Nabi, Sabrina Moriom Elius, ABM Khademul Islam, Richard Malo, Imran Khan, Abu Sufian, Sabita Rezwana Rahman, Habibul Bari Shozib, Mamun Ahmed, AHM Nurun Nabi, Mohammad Riazul Islam, Md Mizanur Rahman, Md Ismail Hosen, Latiful Bari, Gazi Nurun Nahar, Haseena Khan, M Anwar Hossain. |
| EPI_ISL_450840 | COVID-19 Laboratory | DNA Solution Ltd. L-5 | Sharif Akhteruzzaman, Zeba Islam Seraj, Nazmul Ahsan, Md Imdadul Hoque, MA Malek, Shahryar Nabi, Sabrina Moriom Elius, ABM Khademul Islam, Richard Malo, Imran Khan, Abu Sufian, Sabita Rezwana Rahman, Habibul Bari Shozib, Mamun Ahmed, AHM Nurun Nabi, Mohammad Riazul Islam, Md Mizanur Rahman, Md Ismail Hosen, Latiful Bari, Gazi Nurun Nahar, Haseena Khan, M Anwar Hossain. |
| EPI_ISL_450841 | COVID-19 Laboratory | DNA Solution Ltd | Sharif Akhteruzzaman, Zeba Islam Seraj, Nazmul Ahsan, Md Imdadul Hoque, MA Malek, Shahryar Nabi, Sabrina Moriom Elius, ABM Khademul Islam, Richard Malo, Imran Khan, Abu Sufian, Sabita Rezwana Rahman, Habibul Bari Shozib, Mamun Ahmed, AHM Nurun Nabi, Mohammad Riazul Islam, Md Mizanur Rahman, Md Ismail Hosen, Latiful Bari, Gazi Nurun Nahar, Haseena Khan, M Anwar Hossain. |
| EPI_ISL_450842 | COVID-19 Laboratory | DNA Solution Ltd. | Sharif Akhteruzzaman, Zeba Islam Seraj, Nazmul Ahsan, Md Imdadul Hoque, MA Malek, Shahryar Nabi, Sabrina Moriom Elius, ABM Khademul Islam, Richard Malo, Imran Khan, Abu Sufian, Sabita Rezwana Rahman, Habibul Bari Shozib, Mamun Ahmed, AHM Nurun Nabi, Mohammad Riazul Islam, Md Mizanur Rahman, Md Ismail Hosen, Latiful Bari, Gazi Nurun Nahar, Haseena Khan, M Anwar Hossain. |
| EPI_ISL_450843 | COVID-19 Laboratory | DNA Solution Ltd. | Sharif Akhteruzzaman, Zeba Islam Seraj, Nazmul Ahsan, Md Imdadul Hoque, MA Malek, Shahryar Nabi, Sabrina Moriom Elius, ABM Khademul Islam, Richard Malo, Imran Khan, Abu Sufian, Sabita Rezwana Rahman, Habibul Bari Shozib, Mamun Ahmed, AHM Nurun Nabi, Mohammad Riazul Islam, Md Mizanur Rahman, Md Ismail Hosen, Latiful Bari, Gazi Nurun Nahar, Haseena Khan, M Anwar Hossain. |
| EPI_ISL_450844, EPI_ISL_450845, EPI_ISL_450846, EPI_ISL_450847, EPI_ISL_450848, EPI_ISL_450849, EPI_ISL_450850, EPI_ISL_450851, EPI_ISL_450852, EPI_ISL_450853, EPI_ISL_450854, EPI_ISL_450855, EPI_ISL_450856, EPI_ISL_450857, EPI_ISL_450858, EPI_ISL_450859, EPI_ISL_450860, EPI_ISL_450861, EPI_ISL_450862, EPI_ISL_450863, EPI_ISL_450864, EPI_ISL_450865, EPI_ISL_450866, EPI_ISL_450867, EPI_ISL_450868, EPI_ISL_450869, EPI_ISL_450870, EPI_ISL_450871, EPI_ISL_450872 | | | |
| see above | Florida Bureau of Public Health Laboratories | Florida Bureau of Public Health Laboratories | Sarah Schmedes, Jason Blanton |

| | | | |
|---|---|---|---|
| EPI_ISL_450873 | Evandro Chagas Institute | Evandro Chagas Institute | Santos, M.C.; Silva, A.M.; Junior, W.D.C.; Barbagelata, L.S.; Ferreira, J.A.; Sousa, E.M.A.; da Silva, P.S.; Martins, L.C.; Sousa Junior, E.C.; Viana, G.M.R |
| EPI_ISL_450874 | Evandro Chagas Institute | Evandro Chagas Institute | Santos, M.C.; Silva, A.M.; Junior, W.D.C.; Barbagelata, L.S.; Ferreira, J.A.; Sousa, E.M.A.; da Silva, P.S.; Martins, L.C.;Sousa Junior, E.C.;Viana, G.M.R |
| EPI_ISL_450875, EPI_ISL_450876, EPI_ISL_450877, EPI_ISL_450878, EPI_ISL_450879, EPI_ISL_450880, EPI_ISL_450881, EPI_ISL_450882, EPI_ISL_450883, EPI_ISL_450884, EPI_ISL_450885, EPI_ISL_450886, EPI_ISL_450887, EPI_ISL_450888, EPI_ISL_450889, EPI_ISL_450890, EPI_ISL_450891, EPI_ISL_450892, EPI_ISL_450893, EPI_ISL_450894, EPI_ISL_450895, EPI_ISL_450896, EPI_ISL_450897, EPI_ISL_450898, EPI_ISL_450899, EPI_ISL_450900, EPI_ISL_450901, EPI_ISL_450902, EPI_ISL_450903, EPI_ISL_450904, EPI_ISL_450905, EPI_ISL_450906, EPI_ISL_450907, EPI_ISL_450908, EPI_ISL_450909, EPI_ISL_450910, EPI_ISL_450911, EPI_ISL_450912, EPI_ISL_450913, EPI_ISL_450914, EPI_ISL_450915, EPI_ISL_450916, EPI_ISL_450917, EPI_ISL_450918, EPI_ISL_450919, EPI_ISL_450920, EPI_ISL_450921, EPI_ISL_450922, EPI_ISL_450923, EPI_ISL_450924, EPI_ISL_450925, EPI_ISL_450926, EPI_ISL_450927, EPI_ISL_450928, EPI_ISL_450929, EPI_ISL_450930, EPI_ISL_450931, EPI_ISL_450932, EPI_ISL_450933, EPI_ISL_450934, EPI_ISL_450935, EPI_ISL_450936, EPI_ISL_450937, EPI_ISL_450938, EPI_ISL_450939, EPI_ISL_450940, EPI_ISL_450941, EPI_ISL_450942, EPI_ISL_450943, EPI_ISL_450944, EPI_ISL_450945, EPI_ISL_450946, EPI_ISL_450947, EPI_ISL_450948, EPI_ISL_450949, EPI_ISL_450950, EPI_ISL_450951, EPI_ISL_450952, EPI_ISL_450953, EPI_ISL_450954, EPI_ISL_450955, EPI_ISL_450956, EPI_ISL_450957, EPI_ISL_450958, EPI_ISL_450959, EPI_ISL_450960, EPI_ISL_450961, EPI_ISL_450962, EPI_ISL_450963, EPI_ISL_450964, EPI_ISL_450965, EPI_ISL_450966, EPI_ISL_450967, EPI_ISL_450968, EPI_ISL_450969, EPI_ISL_450970, EPI_ISL_450971, EPI_ISL_450972, EPI_ISL_450973, EPI_ISL_450974, EPI_ISL_450975, EPI_ISL_450976, EPI_ISL_450977, EPI_ISL_450978, EPI_ISL_450979, EPI_ISL_450980, EPI_ISL_450981, EPI_ISL_450982, EPI_ISL_450983, EPI_ISL_450984, EPI_ISL_450985, EPI_ISL_450986, EPI_ISL_450987, EPI_ISL_450988, EPI_ISL_450989, EPI_ISL_450990, EPI_ISL_450991, EPI_ISL_450992, EPI_ISL_450993, EPI_ISL_450994, EPI_ISL_450995, EPI_ISL_450996, EPI_ISL_450997, EPI_ISL_450998, EPI_ISL_450999, EPI_ISL_451000, EPI_ISL_451001, EPI_ISL_451002, EPI_ISL_451003, EPI_ISL_451004, EPI_ISL_451005, EPI_ISL_451006, EPI_ISL_451007, EPI_ISL_451008, EPI_ISL_451009, EPI_ISL_451010, EPI_ISL_451011, EPI_ISL_451012, EPI_ISL_451013, EPI_ISL_451014, EPI_ISL_451015, EPI_ISL_451016, EPI_ISL_451017, EPI_ISL_451018, EPI_ISL_451019, EPI_ISL_451020, EPI_ISL_451021, EPI_ISL_451022, EPI_ISL_451023, EPI_ISL_451024, EPI_ISL_451025, EPI_ISL_451026, EPI_ISL_451027, EPI_ISL_451028, EPI_ISL_451029, EPI_ISL_451030, EPI_ISL_451031, EPI_ISL_451032, EPI_ISL_451033, EPI_ISL_451034, EPI_ISL_451035, EPI_ISL_451036, EPI_ISL_451037, EPI_ISL_451038, EPI_ISL_451039, EPI_ISL_451040, EPI_ISL_451041, EPI_ISL_451042, EPI_ISL_451043, EPI_ISL_451044, EPI_ISL_451045, EPI_ISL_451046, EPI_ISL_451047, EPI_ISL_451048, EPI_ISL_451049, EPI_ISL_451050, EPI_ISL_451051, EPI_ISL_451052, EPI_ISL_451053, EPI_ISL_451054, EPI_ISL_451055, EPI_ISL_451056, EPI_ISL_451057, EPI_ISL_451058, EPI_ISL_451059, EPI_ISL_451060, EPI_ISL_451061, EPI_ISL_451062, EPI_ISL_451063, EPI_ISL_451064, EPI_ISL_451065, EPI_ISL_451066, EPI_ISL_451067, EPI_ISL_451068, EPI_ISL_451069, EPI_ISL_451070, EPI_ISL_451071, EPI_ISL_451072, EPI_ISL_451073, EPI_ISL_451074 | | | |
| see above | Center of Excellence in Clinical Virology | Center of Excellence in Clinical Virology | Puenpa,J., Chansaenroj,J., Nilyanimit,P., Auphimai,C., Yorsaeng,R., Suwannakarn,K., Poovorawan,Y. |
| EPI_ISL_451076 | West China Hospital of Sichuan University | State Key Laboratory of Biotherapy of Sichuan University | Baowen Du, Minjin Wang, Chao Tanga, Chuan Chena, Yongzhao Zhou, Mingxia Yu, Han-Cheng Wei, Weimin Li, Jing-wen Lin, Jia Geng, Binwu Ying, Lu Chen |
| EPI_ISL_451077, EPI_ISL_451078, EPI_ISL_451079, EPI_ISL_451080, EPI_ISL_451081, EPI_ISL_451082, EPI_ISL_451083, EPI_ISL_451084, EPI_ISL_451085, EPI_ISL_451086, EPI_ISL_451087, EPI_ISL_451088, EPI_ISL_451089, EPI_ISL_451090, EPI_ISL_451091, EPI_ISL_451092, EPI_ISL_451093, EPI_ISL_451094, EPI_ISL_451095, EPI_ISL_451096, EPI_ISL_451097, EPI_ISL_451098, EPI_ISL_451099, EPI_ISL_451100, EPI_ISL_451101, EPI_ISL_451102, EPI_ISL_451103, EPI_ISL_451104, EPI_ISL_451105, EPI_ISL_451106, EPI_ISL_451107, EPI_ISL_451108, EPI_ISL_451109, EPI_ISL_451110, EPI_ISL_451111, EPI_ISL_451112, EPI_ISL_451113, EPI_ISL_451114, EPI_ISL_451115, EPI_ISL_451116, EPI_ISL_451117, EPI_ISL_451118, EPI_ISL_451119, EPI_ISL_451120, EPI_ISL_451121, EPI_ISL_451122, EPI_ISL_451123, EPI_ISL_451124, EPI_ISL_451125, EPI_ISL_451126, EPI_ISL_451127, EPI_ISL_451128, EPI_ISL_451129, EPI_ISL_451130, EPI_ISL_451131, EPI_ISL_451132, EPI_ISL_451133, EPI_ISL_451134, EPI_ISL_451135, EPI_ISL_451136, EPI_ISL_451137, EPI_ISL_451138, EPI_ISL_451139, EPI_ISL_451140, EPI_ISL_451141, EPI_ISL_451142, EPI_ISL_451143, EPI_ISL_451144, EPI_ISL_451145, EPI_ISL_451146, EPI_ISL_451147, EPI_ISL_451148 | | | |
| see above | SA Pathology | SA Pathology | Lex Leong, Chuan Kok Lim, Mark Turra, Ivan Bastian, Geoff Higgins |
| EPI_ISL_451149 | M.P Shah Government Medocal college Jamnagar | Gujarat Biotechnology Research Centre | Janvi Raval, Zarna Patel, Monika Gandhi, Pinal Trivedi, Maharshi Pandya, Amit Kanani, Nidhi Patel, Nitin Savaliya, Raghawendra Kumar, Dinesh Kumar, Zuber Saiyed, Komal Patel, Labdhi Pandya, Snehal Bagatharia, Ramesh Pandit, Tejas Shah, Ankit Hinsu, Pritesh Sabara, Apurvasinh Puvar, Binita Aring, Bhavesh Modi, Gaurishankar Shrimali, R D Dixit, A M Kadri, Priti Pandita, Chaitanya Joshi, Madhvi Joshi, |
| EPI_ISL_451150 | M.P Shah Government Medocal college Jamnagar | Gujarat Biotechnology Research Centre | Zarna Patel, Monika Gandhi, Pinal Trivedi, Maharshi Pandya, Amit Kanani, Nidhi Patel, Nitin Savaliya, Raghawendra Kumar, Dinesh Kumar, Zuber Saiyed, Komal Patel, Labdhi Pandya, Snehal Bagatharia, Ramesh Pandit, Tejas Shah, Ankit Hinsu, Pritesh Sabara, Apurvasinh Puvar, Binita Aring, Janvi Raval, Bhavesh Modi, Gaurishankar Shrimali, R D Dixit, A M Kadri, Pragya Sharma, Chaitanya Joshi, Madhvi Joshi, |
| EPI_ISL_451151 | M.P Shah Government Medocal college Jamnagar | Gujarat Biotechnology Research Centre | Monika Gandhi, Pinal Trivedi, Maharshi Pandya, Amit Kanani, Nidhi Patel, Nitin Savaliya, Raghawendra Kumar, Dinesh Kumar, Zuber Saiyed, Komal Patel, Labdhi Pandya, Snehal Bagatharia, Ramesh Pandit, Tejas Shah, Ankit Hinsu, Pritesh Sabara, Apurvasinh Puvar, Binita Aring, Janvi Raval, Zarna Patel, Bhavesh Modi, Gaurishankar Shrimali, R D Dixit, A M Kadri, Neha Rajpara, Chaitanya Joshi, Madhvi Joshi, |
| EPI_ISL_451152 | M.P Shah Government Medocal college Jamnagar | Gujarat Biotechnology Research Centre | Pinal Trivedi, Maharshi Pandya, Amit Kanani, Nidhi Patel, Nitin Savaliya, Raghawendra Kumar, Dinesh Kumar, Zuber Saiyed, Komal Patel, Labdhi Pandya, Snehal Bagatharia, Ramesh Pandit, Tejas Shah, Ankit Hinsu, Pritesh Sabara, Apurvasinh Puvar, Binita Aring, Janvi Raval, Zarna Patel, Monika Gandhi, Bhavesh Modi, Gaurishankar Shrimali, R D Dixit, A M Kadri, Afzal Ansari, Chaitanya Joshi, Madhvi Joshi, |
| EPI_ISL_451153 | M.P Shah Government Medocal college Jamnagar | Gujarat Biotechnology Research Centre | Maharshi Pandya, Amit Kanani, Nidhi Patel, Nitin Savaliya, Raghawendra Kumar, Dinesh Kumar, Zuber Saiyed, Komal Patel, Labdhi Pandya, Snehal Bagatharia, Ramesh Pandit, Tejas Shah, Ankit Hinsu, Pritesh Sabara, Apurvasinh Puvar, Binita Aring, Janvi Raval, Zarna Patel, Monika Gandhi, Pinal Trivedi, Bhavesh Modi, Gaurishankar Shrimali, R D Dixit, A M Kadri, Fenil Patel, Chaitanya Joshi, Madhvi Joshi, |
| EPI_ISL_451154 | Government Medical College, Vadodara | Gujarat Biotechnology Research Centre | Manish Pattani, Tanuja Javadekar , Maharshi Pandya, Amit Kanani, Nidhi Patel, Nitin Savaliya, Raghawendra Kumar, Dinesh Kumar, Zuber Saiyed, Komal Patel, Labdhi Pandya, Snehal Bagatharia, Ramesh Pandit, Tejas Shah, Ankit Hinsu, Pritesh Sabara, Apurvasinh Puvar, Janvi Raval, Zarna Patel, Monika Gandhi, Pinal Trivedi, Bhavesh Modi, Gaurishankar Shrimali, R D Dixit, A M Kadri, Neelam Nathani, Chaitanya Joshi, Madhvi Joshi |
| EPI_ISL_451155 | Government Medical College, Vadodara | Gujarat Biotechnology Research Centre | Tanuja Javadekar , Manish Pattani, Amit Kanani, Nidhi Patel, Nitin Savaliya, Raghawendra Kumar, Dinesh Kumar, Zuber Saiyed, Komal Patel, Labdhi Pandya, Snehal Bagatharia, Ramesh Pandit, Tejas Shah, Ankit Hinsu, Pritesh Sabara, Apurvasinh Puvar, Janvi Raval, Zarna Patel, Monika Gandhi, Pinal Trivedi, Maharshi Pandya, Bhavesh Modi, Gaurishankar Shrimali, R D Dixit, A M Kadri, Armi Chaudhari, Chaitanya Joshi, Madhvi Joshi |
| EPI_ISL_451156 | Government Medical College, Vadodara | Gujarat Biotechnology Research Centre | Amit Kanani, Nidhi Patel, Nitin Savaliya, Raghawendra Kumar, Dinesh Kumar, Zuber Saiyed, Komal Patel, Labdhi Pandya, Snehal Bagatharia, Ramesh Pandit, Tejas Shah, Ankit Hinsu, Pritesh Sabara, Apurvasinh Puvar, Janvi Raval, Zarna Patel, Monika Gandhi, Pinal Trivedi, Maharshi Pandya, Manish Pattani, Tanuja Javadekar , Bhavesh Modi, Gaurishankar Shrimali, R D Dixit, A M Kadri, Bhavya Jindal, Chaitanya Joshi, Madhvi Joshi |
| EPI_ISL_451157 | Government Medical College, Vadodara | Gujarat Biotechnology Research Centre | Nidhi Patel, Nitin Savaliya, Raghawendra Kumar, Dinesh Kumar, Zuber Saiyed, Komal Patel, Labdhi Pandya, Snehal Bagatharia, Ramesh Pandit, Tejas Shah, Ankit Hinsu, Pritesh Sabara, Apurvasinh Puvar, Janvi Raval, Zarna Patel, Monika Gandhi, Pinal Trivedi, Maharshi Pandya, Manish Pattani, Tanuja Javadekar , Amit Kanani, Bhavesh Modi, Gaurishankar Shrimali, R D Dixit, A M Kadri, Camellia Chakraborty, Chaitanya Joshi, Madhvi Joshi |
| EPI_ISL_451158 | Government Medical College, Vadodara | Gujarat Biotechnology Research Centre | Nitin Savaliya, Raghawendra Kumar, Dinesh Kumar, Zuber Saiyed, Komal Patel, Labdhi Pandya, Snehal Bagatharia, Ramesh Pandit, Tejas Shah, Ankit Hinsu, Pritesh Sabara, Apurvasinh Puvar, Janvi Raval, Zarna Patel, Monika Gandhi, Pinal Trivedi, Maharshi Pandya, Manish Pattani, Tanuja Javadekar , Amit Kanani, Nidhi Patel, Bhavesh Modi, Gaurishankar Shrimali, R D Dixit, A M Kadri, Siddhant Kumar, Chaitanya Joshi, Madhvi Joshi |
| EPI_ISL_451159 | Government Medical College, Vadodara | Gujarat Biotechnology Research Centre | Raghawendra Kumar, Dinesh Kumar, Zuber Saiyed, Komal Patel, Labdhi Pandya, Snehal Bagatharia, Ramesh Pandit, Tejas Shah, Ankit Hinsu, Pritesh Sabara, Apurvasinh Puvar, Janvi Raval, Zarna Patel, Monika Gandhi, Pinal Trivedi, Maharshi Pandya, Manish Pattani, Tanuja Javadekar , Amit Kanani, Nidhi Patel, Nitin Savaliya, Bhavesh Modi, Gaurishankar Shrimali, R D Dixit, A M Kadri, Sharmistha Majumdar, Chaitanya Joshi, Madhvi Joshi |
| EPI_ISL_451160 | Government Medical College, Vadodara | Gujarat Biotechnology Research Centre | Dinesh Kumar, Zuber Saiyed, Komal Patel, Labdhi Pandya, Snehal Bagatharia, Ramesh Pandit, Tejas Shah, Ankit Hinsu, Pritesh Sabara, Apurvasinh Puvar, Janvi Raval, Zarna Patel, Monika Gandhi, Pinal Trivedi, Maharshi Pandya, Manish Pattani, Tanuja Javadekar , Amit Kanani, Nidhi Patel, Nitin Savaliya, Raghawendra Kumar, Bhavesh Modi, Gaurishankar Shrimali, R D Dixit, A M Kadri, Pooja P Doshi, Chaitanya Joshi, Madhvi Joshi |
| EPI_ISL_451161 | Government Medical College, Vadodara | Gujarat Biotechnology Research Centre | Zuber Saiyed, Komal Patel, Labdhi Pandya, Snehal Bagatharia, Ramesh Pandit, Tejas Shah, Ankit Hinsu, Pritesh Sabara, Apurvasinh Puvar, Janvi Raval, Zarna Patel, Monika Gandhi, Pinal Trivedi, Maharshi Pandya, Manish Pattani, Tanuja Javadekar , Amit Kanani, Nidhi Patel, Nitin Savaliya, Raghawendra Kumar, Dinesh Kumar, Bhavesh Modi, Gaurishankar Shrimali, R D Dixit, A M Kadri, Akanksha Verma, Chaitanya Joshi, Madhvi Joshi |
| EPI_ISL_451162 | Government Medical College, Vadodara | Gujarat Biotechnology Research Centre | Komal Patel, Labdhi Pandya, Snehal Bagatharia, Ramesh Pandit, Tejas Shah, Ankit Hinsu, Pritesh Sabara, Apurvasinh Puvar, Janvi Raval, Zarna Patel, Monika Gandhi, Pinal Trivedi, Maharshi Pandya, Manish Pattani, Tanuja Javadekar , Amit Kanani, Nidhi Patel, Nitin Savaliya, Raghawendra Kumar, Dinesh Kumar, Zuber Saiyed, Bhavesh Modi, Gaurishankar Shrimali, R D Dixit, A M Kadri, Priti Pandita, Chaitanya Joshi, Madhvi Joshi |
| EPI_ISL_451163 | Government Medical College, Vadodara | Gujarat Biotechnology Research Centre | Labdhi Pandya, Snehal Bagatharia, Ramesh Pandit, Tejas Shah, Ankit Hinsu, Pritesh Sabara, Apurvasinh Puvar, Janvi Raval, Zarna Patel, Monika Gandhi, Pinal Trivedi, Maharshi Pandya, Manish Pattani, Tanuja Javadekar , Amit Kanani, Nidhi Patel, Nitin Savaliya, Raghawendra Kumar, Dinesh Kumar, Zuber Saiyed, Komal Patel, Bhavesh Modi, Gaurishankar Shrimali, R D Dixit, A M Kadri, Pragya Sharma, Chaitanya Joshi, Madhvi Joshi |
| EPI_ISL_451164, EPI_ISL_451165, EPI_ISL_451166, EPI_ISL_451167, EPI_ISL_451168, EPI_ISL_451169, EPI_ISL_451170, EPI_ISL_451171, EPI_ISL_451172, EPI_ISL_451173 | Lab voor klinische biologie | Onderzoeksgroep Virologie | Laurens Lambrechts, Nick Vereecke, Marthe Pauwels, Jozefien De Clercq, Bruno Verhasselt, Linos Vandekerckhove, Hans Nauwynck, Sebastiaan Theuns |
| EPI_ISL_451174, EPI_ISL_451175, EPI_ISL_451176, EPI_ISL_451177, EPI_ISL_451178, EPI_ISL_451179, EPI_ISL_451180, EPI_ISL_451181, EPI_ISL_451182 | Lab voor klinische biologie | Onderzoeksgroep Virologie | Nick Vereecke, Laurens Lambrechts, Marthe Pauwels, Jozefien De Clercq, Bruno Verhasselt, Linos Vandekerckhove, Hans Nauwynck, Sebastiaan Theuns |
| EPI_ISL_451183, EPI_ISL_451184, EPI_ISL_451185, EPI_ISL_451186, EPI_ISL_451187, EPI_ISL_451188, EPI_ISL_451189, EPI_ISL_451190, EPI_ISL_451191, EPI_ISL_451192, EPI_ISL_451193, EPI_ISL_451194, EPI_ISL_451195, EPI_ISL_451196, EPI_ISL_451197, EPI_ISL_451198, EPI_ISL_451199, EPI_ISL_451200, EPI_ISL_451201, EPI_ISL_451202 | | | |
| see above | Uganda Virus Research Institute | MRC/UVRI & LSHTM Uganda Research Unit | Dan Lule Bugembe, John Kayiwa, My V.T Phan, Phionah Tushabe, Stephen Balinandi, Beatrice Dhaala, Deogratius Ssemwanga, Jonas Lexow, Henry Mwebesa, Jane Aceng, Henry Kyobe, Julius Lutwama, Pontiano Kaleebu, Matthew Cotten |
| EPI_ISL_451203, EPI_ISL_451204, EPI_ISL_451205, EPI_ISL_451206, EPI_ISL_451207, EPI_ISL_451208, EPI_ISL_451209, EPI_ISL_451210, EPI_ISL_451211, EPI_ISL_451212, EPI_ISL_451213, EPI_ISL_451214, EPI_ISL_451215, EPI_ISL_451216, EPI_ISL_451217, EPI_ISL_451218 | | | |
| see above | LSUHS Emerging Viral Threat Laboratory | Microbial Genome Sequencing Center | Jeremy P. Kamil, John A. Vanchiere, Rona S. Scott, Camille F. Abshire, Abida Siddiqa, Byeong-Jae Lee, Chan-ki Min, Md Maksudul Alam, Monica Gestal-Cartele, Edna Ondari, Adam Greer, Malgorzata Bienkowska-Haba, Katarzyna Zwolinska, Jason M. Bodily, Andrew D. Yurochko, Paul M. Weinberger, Christopher G. Kevil, Martin J. Sapp, Daniel J. Snyder, Vaughn S. Cooper |
| EPI_ISL_451219, EPI_ISL_451220, EPI_ISL_451221, EPI_ISL_451222, EPI_ISL_451223, EPI_ISL_451224, EPI_ISL_451225, EPI_ISL_451226, EPI_ISL_451227, EPI_ISL_451228, EPI_ISL_451229 | | | |
| see above | LSUHS Emerging Viral Threat Laboratory | Microbial Genome Sequencing Center | Rona S. Scott, Jeremy P. Kamil, John A. Vanchiere, Camille F. Abshire, Abida Siddiqa, Byeong-Jae Lee, Chan-ki Min, Md Maksudul Alam, Monica Gestal-Cartele, Edna Ondari, Adam Greer, Malgorzata Bienkowska-Haba, Katarzyna Zwolinska, Jason M. Bodily, Andrew D. Yurochko, Paul M. Weinberger, Christopher G. Kevil, Martin J. Sapp, Daniel J. Snyder, Vaughn S. Cooper |
| EPI_ISL_451230, EPI_ISL_451231, EPI_ISL_451232, EPI_ISL_451233, EPI_ISL_451234, EPI_ISL_451235, EPI_ISL_451236, EPI_ISL_451237, EPI_ISL_451238, EPI_ISL_451239, EPI_ISL_451240, EPI_ISL_451241, EPI_ISL_451242, EPI_ISL_451243, EPI_ISL_451244, EPI_ISL_451245, EPI_ISL_451246, EPI_ISL_451247 | | | |
| see above | LSUHS Emerging Viral Threat Laboratory | Microbial Genome Sequencing Center | John A. Vanchiere, Jeremy P. Kamil, Rona S. Scott, Camille F. Abshire, Abida Siddiqa, Byeong-Jae Lee, Chan-ki Min, Md Maksudul Alam, Monica Gestal-Cartele, Edna Ondari, Adam Greer, Malgorzata Bienkowska-Haba, Katarzyna Zwolinska, Jason M. Bodily, Andrew D. Yurochko, Paul M. Weinberger, Christopher G. Kevil, Martin J. Sapp, Daniel J. Snyder, Vaughn S. Cooper |
| EPI_ISL_451248, EPI_ISL_451249, EPI_ISL_451250, EPI_ISL_451251, EPI_ISL_451252, EPI_ISL_451253, EPI_ISL_451254, EPI_ISL_451255, EPI_ISL_451256, EPI_ISL_451257, EPI_ISL_451258, EPI_ISL_451259, EPI_ISL_451260, EPI_ISL_451261, EPI_ISL_451262, EPI_ISL_451263, EPI_ISL_451264, EPI_ISL_451265, EPI_ISL_451266, EPI_ISL_451267, EPI_ISL_451268, EPI_ISL_451269, EPI_ISL_451270, EPI_ISL_451271, EPI_ISL_451272, EPI_ISL_451273, EPI_ISL_451274, EPI_ISL_451275, EPI_ISL_451276, EPI_ISL_451277, EPI_ISL_451278, EPI_ISL_451279, EPI_ISL_451280, EPI_ISL_451281, EPI_ISL_451282, EPI_ISL_451283, EPI_ISL_451284, EPI_ISL_451285, EPI_ISL_451286, EPI_ISL_451287, EPI_ISL_451288, EPI_ISL_451289, EPI_ISL_451290, EPI_ISL_451291, EPI_ISL_451292, EPI_ISL_451293, EPI_ISL_451294, EPI_ISL_451295, EPI_ISL_451296, EPI_ISL_451297 | | | |
| see above | LSUHS Emerging Viral Threat Laboratory | Microbial Genome Sequencing Center | Jeremy P. Kamil, John A. Vanchiere, Rona S. Scott, Camille F. Abshire, Abida Siddiqa, Byeong-Jae Lee, Chan-ki Min, Md Maksudul Alam, Monica Gestal-Cartele, Edna Ondari, Adam Greer, Malgorzata Bienkowska-Haba, Katarzyna Zwolinska, Jason M. Bodily, Andrew D. Yurochko, Paul M. Weinberger, Christopher G. Kevil, Martin J. Sapp, Daniel J. Snyder, Vaughn S. Cooper |
| EPI_ISL_451298 | Laboratory of Virology, INMI Lazzaro Spallanzani IRCCS | Laboratory of Virology, INMI Lazzaro Spallanzani IRCCS | Cesare E.M. Gruber, Martina Rueca, Barbara Bartolini, Francesco Messina, Antonino Di Caro, Maria R. Capobianchi, Giuseppe Ippolito |
| EPI_ISL_451299 | Laboratory of Virology, INMI Lazzaro Spallanzani IRCCS | Laboratory of Virology, INMI Lazzaro Spallanzani IRCCS | Martina Rueca, Cesare E.M. Gruber, Barbara Bartolini, Francesco Messina, Antonino Di Caro, Maria R. Capobianchi, Giuseppe Ippolito |
| EPI_ISL_451300 | Laboratory of Virology, INMI Lazzaro Spallanzani IRCCS | Laboratory of Virology, INMI Lazzaro Spallanzani IRCCS | Cesare E.M. Gruber, Martina Rueca, Barbara Bartolini, Francesco Messina, Antonino Di Caro, Maria R. Capobianchi, Giuseppe Ippolito |
| EPI_ISL_451301 | Laboratory of Virology, INMI Lazzaro Spallanzani IRCCS | Laboratory of Virology, INMI Lazzaro Spallanzani IRCCS | Martina Rueca, Cesare E.M. Gruber, Barbara Bartolini, Francesco Messina, Antonino Di Caro, Maria R. Capobianchi, Giuseppe Ippolito |
| EPI_ISL_451302 | Laboratory of Virology, INMI Lazzaro Spallanzani IRCCS | Laboratory of Virology, INMI Lazzaro Spallanzani IRCCS | Cesare E.M. Gruber, Martina Rueca, Barbara Bartolini, Francesco Messina, Antonino Di Caro, Maria R. Capobianchi, Giuseppe Ippolito |
| EPI_ISL_451303 | Laboratory of Virology, INMI Lazzaro Spallanzani IRCCS | Laboratory of Virology, INMI Lazzaro Spallanzani IRCCS | Martina Rueca, Cesare E.M. Gruber, Barbara Bartolini, Francesco Messina, Antonino Di Caro, Maria R. Capobianchi, Giuseppe Ippolito |
| EPI_ISL_451304 | Laboratory of Virology, INMI Lazzaro Spallanzani IRCCS | Laboratory of Virology, INMI Lazzaro Spallanzani IRCCS | Cesare E.M. Gruber, Martina Rueca, Barbara Bartolini, Francesco Messina, Antonino Di Caro, Maria R. Capobianchi, Giuseppe Ippolito |
| EPI_ISL_451305 | Laboratory of Virology, INMI Lazzaro Spallanzani IRCCS | Laboratory of Virology, INMI Lazzaro Spallanzani IRCCS | Martina Rueca, Cesare E.M. Gruber, Barbara Bartolini, Francesco Messina, Antonino Di Caro, Maria R. Capobianchi, Giuseppe Ippolito |

| | | | |
|---|---|---|---|
| EPI_ISL_451306 | Molecular Virology Unit, Fondazione IRCCS Policlinico San Matteo , Pavia | Laboratory of Virology, INMI Lazzaro Spallanzani IRCCS | Antonio Piralla, Fausto Baldanti, Martina Rueca, Antonino Di Caro, Maria R. Capobianchi, Cesare E.M. Gruber, Barbara Bartolini |
| EPI_ISL_451307 | Molecular Virology Unit, Fondazione IRCCS Policlinico San Matteo , Pavia | Laboratory of Virology, INMI Lazzaro Spallanzani IRCCS | Fausto Baldanti, Antonio Piralla, Antonino Di Caro, Cesare E.M. Gruber, Martina Rueca, Barbara Bartolini, Maria R. Capobianchi |
| EPI_ISL_451308 | Molecular Virology Unit, Fondazione IRCCS Policlinico San Matteo , Pavia | Laboratory of Virology, INMI Lazzaro Spallanzani IRCCS | Antonio Piralla, Fausto Baldanti, Maria R. Capobianchi, Cesare E.M. Gruber, Martina Rueca, Barbara Bartolini, Antonino Di Caro |
| EPI_ISL_451309 | Molecular Virology Unit, Fondazione IRCCS Policlinico San Matteo , Pavia | Laboratory of Virology, INMI Lazzaro Spallanzani IRCCS | Fausto Baldanti, Antonio Piralla, Cesare E.M. Gruber, Maria R. Capobianchi, Antonino Di Caro, Martina Rueca, Barbara Bartolini |
| EPI_ISL_451310, EPI_ISL_451311 | Hellenic Pasteur Institute, National Influenza Reference laboratory of Southern Greece & Unit of Bioinformatics and Applied Genomics | Hellenic Pasteur Institute, National Influenza Reference laboratory of Southern Greece & Unit of Bioinformatics and Applied Genomics | Vasiliki Pogka, Timokratis Karamitros, Athanasios Kossyvakis, Antonios Kalliaropoulos, Horefti Elina, Evangelidou Maria, Androniki Voulgari-Kokota, Aspasia Kontou, Andreas Mentis |
| EPI_ISL_451312, EPI_ISL_451313, EPI_ISL_451314, EPI_ISL_451315, EPI_ISL_451316, EPI_ISL_451317, EPI_ISL_451318, EPI_ISL_451319, EPI_ISL_451320, EPI_ISL_451321, EPI_ISL_451322, EPI_ISL_451323, EPI_ISL_451324, EPI_ISL_451325, EPI_ISL_451326, EPI_ISL_451327, EPI_ISL_451328, EPI_ISL_451329, EPI_ISL_451330, EPI_ISL_451331, EPI_ISL_451332, EPI_ISL_451333, EPI_ISL_451334, EPI_ISL_451335, EPI_ISL_451336, EPI_ISL_451337, EPI_ISL_451338, EPI_ISL_451339, EPI_ISL_451340, EPI_ISL_451341, EPI_ISL_451342, EPI_ISL_451343, EPI_ISL_451344, EPI_ISL_451345, EPI_ISL_451346, EPI_ISL_451347, EPI_ISL_451348, EPI_ISL_451349, EPI_ISL_451350, EPI_ISL_451351, EPI_ISL_451352, EPI_ISL_451353, EPI_ISL_451354, EPI_ISL_451355, EPI_ISL_451356, EPI_ISL_451357, EPI_ISL_451358, EPI_ISL_451359, EPI_ISL_451360, EPI_ISL_451361, EPI_ISL_451362, EPI_ISL_451363, EPI_ISL_451364, EPI_ISL_451365, EPI_ISL_451366, EPI_ISL_451367, EPI_ISL_451368, EPI_ISL_451369, EPI_ISL_451370, EPI_ISL_451371, EPI_ISL_451372, EPI_ISL_451373, EPI_ISL_451374, EPI_ISL_451375, EPI_ISL_451376, EPI_ISL_451377, EPI_ISL_451378, EPI_ISL_451379, EPI_ISL_451380, EPI_ISL_451381, EPI_ISL_451382, EPI_ISL_451383, EPI_ISL_451384, EPI_ISL_451385, EPI_ISL_451386, EPI_ISL_451387, EPI_ISL_451388, EPI_ISL_451389, EPI_ISL_451390, EPI_ISL_451391, EPI_ISL_451392, EPI_ISL_451393, EPI_ISL_451394, EPI_ISL_451395, EPI_ISL_451396, EPI_ISL_451397, EPI_ISL_451398, EPI_ISL_451399 | | | |
| see above | West China Hospital of Sichuan University | State Key Laboratory of Biotherapy of Sichuan University | Baowen Du, Minjin Wang, Chao Tang, Chuan Chen, Yongzhao Zhou, Mingxia Yu, Hancheng Wei, Weimin Li, Jing-wen Lin, Jia Geng, Binwu Ying, Lu Chen |
| EPI_ISL_451400 | Laboratoire de Recherche et d'Analyse Médicale de la Gendarmerie Royale | Laboratoire de Recherche et d'Analyse Médicale de la Gendarmerie Royale | Sanaâ LEMRISS, Amal SOUIRI, Saâd EL KABBAJ |
| EPI_ISL_451401, EPI_ISL_451402, EPI_ISL_451403, EPI_ISL_451404, EPI_ISL_451405, EPI_ISL_451406, EPI_ISL_451407, EPI_ISL_451408, EPI_ISL_451409, EPI_ISL_451410, EPI_ISL_451411, EPI_ISL_451412, EPI_ISL_451413, EPI_ISL_451414, EPI_ISL_451415, EPI_ISL_451416, EPI_ISL_451417, EPI_ISL_451418, EPI_ISL_451419, EPI_ISL_451420, EPI_ISL_451421, EPI_ISL_451422, EPI_ISL_451423, EPI_ISL_451424, EPI_ISL_451425, EPI_ISL_451426, EPI_ISL_451427, EPI_ISL_451428, EPI_ISL_451429, EPI_ISL_451430, EPI_ISL_451431, EPI_ISL_451432, EPI_ISL_451433, EPI_ISL_451434, EPI_ISL_451435, EPI_ISL_451436, EPI_ISL_451437, EPI_ISL_451438, EPI_ISL_451439, EPI_ISL_451440, EPI_ISL_451441, EPI_ISL_451442, EPI_ISL_451443, EPI_ISL_451444, EPI_ISL_451445, EPI_ISL_451446, EPI_ISL_451447, EPI_ISL_451448, EPI_ISL_451449, EPI_ISL_451450, EPI_ISL_451451, EPI_ISL_451452, EPI_ISL_451453, EPI_ISL_451454, EPI_ISL_451455, EPI_ISL_451456, EPI_ISL_451457, EPI_ISL_451458, EPI_ISL_451459, EPI_ISL_451460, EPI_ISL_451461, EPI_ISL_451462, EPI_ISL_451463, EPI_ISL_451464, EPI_ISL_451465, EPI_ISL_451466, EPI_ISL_451467, EPI_ISL_451468, EPI_ISL_451469, EPI_ISL_451470, EPI_ISL_451471, EPI_ISL_451472, EPI_ISL_451473, EPI_ISL_451474, EPI_ISL_451475, EPI_ISL_451476, EPI_ISL_451477, EPI_ISL_451478, EPI_ISL_451479 | | | |
| see above | NYU Langone Health | Departments of Pathology and Medicine, New York University School of Medicine | Maria Aguero-Rosenfeld, Brendan Belovarac, Margaret Black, Ludovic Boytard, John Cadley, Paolo Cotzia, John Chen, Dacia Dimartino, Xiaojun Feng, Tatyana Gindin, Emily Guzman, Adriana Heguy, Megan Hogan, Emily Huang, George Jour, Alireza Khodadadi-Jamayran, Lawrence H. Lin, Raven Luther, Andrew Lytle, Christian Marier, Matthew T. Maurano, Mark J. Mulligan, Peter Meyn, Raquel Ordonez Ciriza, Iman Osman, Jared Pinnell, Vanessa Raabe, Sitharam Ramaswami, Amy Rapkiewicz, Andre M. Ribeiro-dos-Santos, Marie Samanovic-Golden, Antonio Serrano, Guomiao Shen, Matija Snuderl, Theodore Vougiouklakis, Nick Vulpescu, Gael Westby, Paul Zappile, Yutong Zhang |
| EPI_ISL_451486 | Australian Clinical Labs | NSW Health Pathology - Institute of Clinical Pathology and Medical Research; Westmead Hospital; University of Sydney | CIDM-PH et al. |
| EPI_ISL_451487, EPI_ISL_451488 | Pathology North - NSW Health Pathology | NSW Health Pathology - Institute of Clinical Pathology and Medical Research; Westmead Hospital; University of Sydney | CIDM-PH et al. |
| EPI_ISL_451489 | Laverty Pathology | NSW Health Pathology - Institute of Clinical Pathology and Medical Research; Westmead Hospital; University of Sydney | CIDM-PH et al. |
| EPI_ISL_451490, EPI_ISL_451491, EPI_ISL_451492, EPI_ISL_451493, EPI_ISL_451494, EPI_ISL_451495, EPI_ISL_451496, EPI_ISL_451497, EPI_ISL_451498, EPI_ISL_451499, EPI_ISL_451500, EPI_ISL_451501, EPI_ISL_451502, EPI_ISL_451503, EPI_ISL_451504, EPI_ISL_451505, EPI_ISL_451506, EPI_ISL_451507, EPI_ISL_451508, EPI_ISL_451509, EPI_ISL_451510, EPI_ISL_451511, EPI_ISL_451512, EPI_ISL_451513, EPI_ISL_451514, EPI_ISL_451515, EPI_ISL_451516 | | | |
| see above | Pathology West - NSW Health Pathology | NSW Health Pathology - Institute of Clinical Pathology and Medical Research; Westmead Hospital; University of Sydney | CIDM-PH et al. |
| EPI_ISL_451517, EPI_ISL_451518, EPI_ISL_451519, EPI_ISL_451520, EPI_ISL_451521 | South Eastern Area Laboratory Services | NSW Health Pathology - Institute of Clinical Pathology and Medical Research; Westmead Hospital; University of Sydney | CIDM-PH et al. |
| EPI_ISL_451522, EPI_ISL_451523, EPI_ISL_451524, EPI_ISL_451525, EPI_ISL_451526, EPI_ISL_451527, EPI_ISL_451528, EPI_ISL_451529 | Pathology West - NSW Health Pathology | NSW Health Pathology - Institute of Clinical Pathology and Medical Research; Westmead Hospital; University of Sydney | CIDM-PH et al. |
| EPI_ISL_451530, EPI_ISL_451531 | Pathology Sydney South West - NSW Health Pathology | NSW Health Pathology - Institute of Clinical Pathology and Medical Research; Westmead Hospital; University of Sydney | CIDM-PH et al. |
| EPI_ISL_451532 | Pathology West - NSW Health Pathology | NSW Health Pathology - Institute of Clinical Pathology and Medical Research; Westmead Hospital; University of Sydney | CIDM-PH et al. |
| EPI_ISL_451533 | Pathology Sydney South West - NSW Health Pathology | NSW Health Pathology - Institute of Clinical Pathology and Medical Research; Westmead Hospital; University of Sydney | CIDM-PH et al. |
| EPI_ISL_451534 | Medlab Pathology | NSW Health Pathology - Institute of Clinical Pathology and Medical Research; Westmead Hospital; University of Sydney | CIDM-PH et al. |
| EPI_ISL_451535 | Pathology West - NSW Health Pathology | NSW Health Pathology - Institute of Clinical Pathology and Medical Research; Westmead Hospital; University of Sydney | CIDM-PH et al. |
| EPI_ISL_451536 | Pathology Sydney South West - NSW Health Pathology | NSW Health Pathology - Institute of Clinical Pathology and Medical Research; Westmead Hospital; University of Sydney | CIDM-PH et al. |
| EPI_ISL_451537, EPI_ISL_451538, EPI_ISL_451539 | Pathology West - NSW Health Pathology | NSW Health Pathology - Institute of Clinical Pathology and Medical Research; Westmead Hospital; University of Sydney | CIDM-PH et al. |
| EPI_ISL_451540 | ACT pathology | NSW Health Pathology - Institute of Clinical Pathology and Medical Research; Westmead Hospital; University of Sydney | CIDM-PH et al. |
| EPI_ISL_451541 | Pathology West - NSW Health Pathology | NSW Health Pathology - Institute of Clinical Pathology and Medical Research; Westmead Hospital; University of Sydney | CIDM-PH et al. |
| EPI_ISL_451542, EPI_ISL_451543, EPI_ISL_451544 | Pathology Sydney South West - NSW Health Pathology | NSW Health Pathology - Institute of Clinical Pathology and Medical Research; Westmead Hospital; University of Sydney | CIDM-PH et al. |
| EPI_ISL_451545 | Laverty Pathology | NSW Health Pathology - Institute of Clinical Pathology and Medical Research; Westmead Hospital; University of Sydney | CIDM-PH et al. |
| EPI_ISL_451546, EPI_ISL_451547 | Pathology West - NSW Health Pathology | NSW Health Pathology - Institute of Clinical Pathology and Medical Research; Westmead Hospital; University of Sydney | CIDM-PH et al. |
| EPI_ISL_451548 | Childrens Hospital Westmead | NSW Health Pathology - Institute of Clinical Pathology and Medical Research; Westmead Hospital; University of Sydney | CIDM-PH et al. |
| EPI_ISL_451549 | Pathology Sydney South West - NSW Health Pathology | NSW Health Pathology - Institute of Clinical Pathology and Medical Research; Westmead Hospital; University of Sydney | CIDM-PH et al. |
| EPI_ISL_451550, EPI_ISL_451551 | Pathology West - NSW Health Pathology | NSW Health Pathology - Institute of Clinical Pathology and Medical Research; Westmead Hospital; University of Sydney | CIDM-PH et al. |
| EPI_ISL_451552 | Pathology North - NSW Health Pathology | NSW Health Pathology - Institute of Clinical Pathology and Medical Research; Westmead Hospital; University of Sydney | CIDM-PH et al. |
| EPI_ISL_451553, EPI_ISL_451554, EPI_ISL_451555 | Medlab Pathology | NSW Health Pathology - Institute of Clinical Pathology and Medical Research; Westmead Hospital; University of Sydney | CIDM-PH et al. |
| EPI_ISL_451556, EPI_ISL_451557 | Pathology West - NSW Health Pathology | NSW Health Pathology - Institute of Clinical Pathology and Medical Research; Westmead Hospital; University of Sydney | CIDM-PH et al. |
| EPI_ISL_451558 | Medlab Pathology | NSW Health Pathology - Institute of Clinical Pathology and Medical Research; Westmead Hospital; University of Sydney | CIDM-PH et al. |
| EPI_ISL_451559, EPI_ISL_451560, EPI_ISL_451561, EPI_ISL_451562, EPI_ISL_451563, EPI_ISL_451564, EPI_ISL_451565 | Pathology Sydney South West - NSW Health Pathology | NSW Health Pathology - Institute of Clinical Pathology and Medical Research; Westmead Hospital; University of Sydney | CIDM-PH et al. |
| EPI_ISL_451566 | Pathology West - NSW Health Pathology | NSW Health Pathology - Institute of Clinical Pathology and Medical Research; Westmead Hospital; University of Sydney | CIDM-PH et al. |
| EPI_ISL_451567 | Medlab Pathology | NSW Health Pathology - Institute of Clinical Pathology and Medical Research; Westmead Hospital; University of Sydney | CIDM-PH et al. |
| EPI_ISL_451568, EPI_ISL_451569, EPI_ISL_451570, EPI_ISL_451571 | Pathology West - NSW Health Pathology | NSW Health Pathology - Institute of Clinical Pathology and Medical Research; Westmead Hospital; University of Sydney | CIDM-PH et al. |
| EPI_ISL_451572 | Laverty Pathology | NSW Health Pathology - Institute of Clinical Pathology and Medical Research; Westmead Hospital; University of Sydney | CIDM-PH et al. |
| EPI_ISL_451573 | Pathology West - NSW Health Pathology | NSW Health Pathology - Institute of Clinical Pathology and Medical Research; Westmead Hospital; University of Sydney | CIDM-PH et al. |
| EPI_ISL_451574 | Australian Clinical Labs | NSW Health Pathology - Institute of Clinical Pathology and Medical Research; Westmead Hospital; University of Sydney | CIDM-PH et al. |

| | | | |
|---|---|---|---|
| EPI_ISL_451575, EPI_ISL_451576, EPI_ISL_451577 | Pathology West - NSW Health Pathology | NSW Health Pathology - Institute of Clinical Pathology and Medical Research; Westmead Hospital; University of Sydney | CIDM-PH et al. |
| EPI_ISL_451578 | Pathology Sydney South West - NSW Health Pathology | NSW Health Pathology - Institute of Clinical Pathology and Medical Research; Westmead Hospital; University of Sydney | CIDM-PH et al. |
| EPI_ISL_451579, EPI_ISL_451580 | Pathology West - NSW Health Pathology | NSW Health Pathology - Institute of Clinical Pathology and Medical Research; Westmead Hospital; University of Sydney | CIDM-PH et al. |
| EPI_ISL_451581 | Pathology Sydney South West - NSW Health Pathology | NSW Health Pathology - Institute of Clinical Pathology and Medical Research; Westmead Hospital; University of Sydney | CIDM-PH et al. |
| EPI_ISL_451582 | Pathology West - NSW Health Pathology | NSW Health Pathology - Institute of Clinical Pathology and Medical Research; Westmead Hospital; University of Sydney | CIDM-PH et al. |
| EPI_ISL_451583 | Medlab Pathology | NSW Health Pathology - Institute of Clinical Pathology and Medical Research; Westmead Hospital; University of Sydney | CIDM-PH et al. |
| EPI_ISL_451584, EPI_ISL_451585, EPI_ISL_451586 | Pathology West - NSW Health Pathology | NSW Health Pathology - Institute of Clinical Pathology and Medical Research; Westmead Hospital; University of Sydney | CIDM-PH et al. |
| EPI_ISL_451587 | Pathology North Hunter- NSW Health Pathology | NSW Health Pathology - Institute of Clinical Pathology and Medical Research; Westmead Hospital; University of Sydney | CIDM-PH et al. |
| EPI_ISL_451588 | ACT pathology | NSW Health Pathology - Institute of Clinical Pathology and Medical Research; Westmead Hospital; University of Sydney | CIDM-PH et al. |
| EPI_ISL_451589 | Pathology North Hunter- NSW Health Pathology | NSW Health Pathology - Institute of Clinical Pathology and Medical Research; Westmead Hospital; University of Sydney | CIDM-PH et al. |
| EPI_ISL_451590, EPI_ISL_451591, EPI_ISL_451592, EPI_ISL_451593 | ACT pathology | NSW Health Pathology - Institute of Clinical Pathology and Medical Research; Westmead Hospital; University of Sydney | CIDM-PH et al. |
| EPI_ISL_451594 | Childrens Hospital Westmead | NSW Health Pathology - Institute of Clinical Pathology and Medical Research; Westmead Hospital; University of Sydney | CIDM-PH et al. |
| EPI_ISL_451595, EPI_ISL_451596 | ACT pathology | NSW Health Pathology - Institute of Clinical Pathology and Medical Research; Westmead Hospital; University of Sydney | CIDM-PH et al. |
| EPI_ISL_451597 | Medlab Pathology | NSW Health Pathology - Institute of Clinical Pathology and Medical Research; Westmead Hospital; University of Sydney | CIDM-PH et al. |
| EPI_ISL_451598 | ACT pathology | NSW Health Pathology - Institute of Clinical Pathology and Medical Research; Westmead Hospital; University of Sydney | CIDM-PH et al. |
| EPI_ISL_451599 | Australian Clinical Labs | NSW Health Pathology - Institute of Clinical Pathology and Medical Research; Westmead Hospital; University of Sydney | CIDM-PH et al. |
| EPI_ISL_451600 | Pathology North Hunter- NSW Health Pathology | NSW Health Pathology - Institute of Clinical Pathology and Medical Research; Westmead Hospital; University of Sydney | CIDM-PH et al. |
| EPI_ISL_451601 | Douglas Hanly Moir Pathology | NSW Health Pathology - Institute of Clinical Pathology and Medical Research; Westmead Hospital; University of Sydney | CIDM-PH et al. |
| EPI_ISL_451602 | Laverty Pathology | NSW Health Pathology - Institute of Clinical Pathology and Medical Research; Westmead Hospital; University of Sydney | CIDM-PH et al. |
| EPI_ISL_451603 | Pathology Sydney South West - NSW Health Pathology | NSW Health Pathology - Institute of Clinical Pathology and Medical Research; Westmead Hospital; University of Sydney | CIDM-PH et al. |
| EPI_ISL_451604 | Pathology North - NSW Health Pathology | NSW Health Pathology - Institute of Clinical Pathology and Medical Research; Westmead Hospital; University of Sydney | CIDM-PH et al. |
| EPI_ISL_451605 | Childrens Hospital Westmead | NSW Health Pathology - Institute of Clinical Pathology and Medical Research; Westmead Hospital; University of Sydney | CIDM-PH et al. |
| EPI_ISL_451606, EPI_ISL_451607 | Pathology West - NSW Health Pathology | NSW Health Pathology - Institute of Clinical Pathology and Medical Research; Westmead Hospital; University of Sydney | CIDM-PH et al. |
| EPI_ISL_451608 | Pathology Sydney South West - NSW Health Pathology | NSW Health Pathology - Institute of Clinical Pathology and Medical Research; Westmead Hospital; University of Sydney | CIDM-PH et al. |
| EPI_ISL_451609 | Laverty Pathology | NSW Health Pathology - Institute of Clinical Pathology and Medical Research; Westmead Hospital; University of Sydney | CIDM-PH et al. |
| EPI_ISL_451610 | Medlab Pathology | NSW Health Pathology - Institute of Clinical Pathology and Medical Research; Westmead Hospital; University of Sydney | CIDM-PH et al. |
| EPI_ISL_451611 | Laverty Pathology | NSW Health Pathology - Institute of Clinical Pathology and Medical Research; Westmead Hospital; University of Sydney | CIDM-PH et al. |
| EPI_ISL_451612 | ACT pathology | NSW Health Pathology - Institute of Clinical Pathology and Medical Research; Westmead Hospital; University of Sydney | CIDM-PH et al. |
| EPI_ISL_451613, EPI_ISL_451614, EPI_ISL_451615, EPI_ISL_451616, EPI_ISL_451617, EPI_ISL_451618, EPI_ISL_451619, EPI_ISL_451620, EPI_ISL_451621, EPI_ISL_451622, EPI_ISL_451623, EPI_ISL_451624, EPI_ISL_451625, EPI_ISL_451626, EPI_ISL_451627, EPI_ISL_451628, EPI_ISL_451629, EPI_ISL_451630 | | | |
| see above | Pathology West - NSW Health Pathology | NSW Health Pathology - Institute of Clinical Pathology and Medical Research; Westmead Hospital; University of Sydney | CIDM-PH et al. |
| EPI_ISL_451631 | Laverty Pathology | NSW Health Pathology - Institute of Clinical Pathology and Medical Research; Westmead Hospital; University of Sydney | CIDM-PH et al. |
| EPI_ISL_451632, EPI_ISL_451633, EPI_ISL_451634, EPI_ISL_451635, EPI_ISL_451636 | South Eastern Area Laboratory Services | NSW Health Pathology - Institute of Clinical Pathology and Medical Research; Westmead Hospital; University of Sydney | CIDM-PH et al. |
| EPI_ISL_451637, EPI_ISL_451638, EPI_ISL_451639, EPI_ISL_451640 | Pathology West - NSW Health Pathology | NSW Health Pathology - Institute of Clinical Pathology and Medical Research; Westmead Hospital; University of Sydney | CIDM-PH et al. |
| EPI_ISL_451641 | Laverty Pathology | NSW Health Pathology - Institute of Clinical Pathology and Medical Research; Westmead Hospital; University of Sydney | CIDM-PH et al. |
| EPI_ISL_451642, EPI_ISL_451643 | Pathology Sydney South West - NSW Health Pathology | NSW Health Pathology - Institute of Clinical Pathology and Medical Research; Westmead Hospital; University of Sydney | CIDM-PH et al. |
| EPI_ISL_451644, EPI_ISL_451645, EPI_ISL_451646, EPI_ISL_451647 | Laboratory of Molecular Biology, Diagnostyka sp. z o.o. | Laboratory of Recombinant Vaccines | Lukasz Rabalski, Anna Piotrowska-Mietelska, Maciej Kosinski, Boguslaw Szewczyk, Krystyna Bienkowska-Szewczyk |
| EPI_ISL_451648, EPI_ISL_451649, EPI_ISL_451650, EPI_ISL_451651, EPI_ISL_451652, EPI_ISL_451653, EPI_ISL_451654 | Hematology Laboratory, Section of Molecular Diagnostics, University Clinical Centre, Medical University of Gdansk | Laboratory of Recombinant Vaccines | Lukasz Rabalski, Adam Sodol, Aneta Szulc, Krzysztof Lewandowski, Ewa Milosz, Marlena Robakowska, Boguslaw Szewczyk, Krystyna Bienkowska-Szewczyk |
| EPI_ISL_451655, EPI_ISL_451656, EPI_ISL_451657, EPI_ISL_451658, EPI_ISL_451659, EPI_ISL_451660, EPI_ISL_451661, EPI_ISL_451662, EPI_ISL_451663, EPI_ISL_451664 | State Sanitary Inspectorate | Laboratory of Recombinant Vaccines | Lukasz Rabalski, Boguslaw Szewczyk, Krystyna Bienkowska-Szewczyk, Jaroslaw Pinkas |
| EPI_ISL_451665 | NYU Langone Health | Departments of Pathology and Medicine, New York University School of Medicine | Maria Aguero-Rosenfeld, Brendan Belovarac, Margaret Black, Ludovic Boytard, John Cadley, Paolo Cotzia, John Chen, Dacia Dimartino, Xiaojun Feng, Tatyana Gindin, Emily Guzman, Adriana Heguy, Megan Hogan, Emily Huang, George Jour, Alireza Khodadadi-Jamayran, Lawrence H. Lin, Raven Luther, Andrew Lytle, Christian Marier, Matthew T. Maurano, Mark J. Mulligan, Peter Meyn, Raquel Ordonez Ciriza, Iman Osman, Jared Pinnell, Vanessa Raabe, Sitharam Ramaswami, Amy Rapkiewicz, Andre M. Ribeiro-dos-Santos, Marie Samanovic-Golden, Antonio Serrano, Guomiao Shen, Matija Snuderl, Theodore Vougiouklakis, Nick Vulpescu, Gael Westby, Paul Zappile, Yutong Zhang |
| EPI_ISL_451666 | M.P Shah Government Medocal college Jamnagar | Gujarat Biotechnology Research Centre | Binita Aring, Janvi Raval, Zarna Patel, Monika Gandhi, Pinal Trivedi, Maharshi Pandya, Amit Kanani, Nidhi Patel, Nitin Savaliya, Raghawendra Kumar, Dinesh Kumar, Zuber Saiyed, Komal Patel, Labdhi Pandya, Snehal Bagatharia, Ramesh Pandit, Tejas Shah, Ankit Hinsu, Pritesh Sabara, Apurvasinh Puvar, Bhavesh Modi, Gaurishankar Shrimali, R D Dixit, A M Kadri, Akanksha Verma, Chaitanya Joshi, Madhvi Joshi, |
| EPI_ISL_451668, EPI_ISL_451669, EPI_ISL_451670, EPI_ISL_451671, EPI_ISL_451672, EPI_ISL_451673, EPI_ISL_451674, EPI_ISL_451675, EPI_ISL_451676, EPI_ISL_451677, EPI_ISL_451678, EPI_ISL_451679, EPI_ISL_451680, EPI_ISL_451681, EPI_ISL_451682, EPI_ISL_451683, EPI_ISL_451684, EPI_ISL_451685, EPI_ISL_451686, EPI_ISL_451687, EPI_ISL_451688, EPI_ISL_451689, EPI_ISL_451690, EPI_ISL_451691, EPI_ISL_451692, EPI_ISL_451693, EPI_ISL_451694, EPI_ISL_451695, EPI_ISL_451696, EPI_ISL_451697, EPI_ISL_451698, EPI_ISL_451699, EPI_ISL_451700, EPI_ISL_451701, EPI_ISL_451702, EPI_ISL_451703, EPI_ISL_451704, EPI_ISL_451705, EPI_ISL_451706, EPI_ISL_451707, EPI_ISL_451708, EPI_ISL_451709, EPI_ISL_451710, EPI_ISL_451711, EPI_ISL_451712, EPI_ISL_451713, EPI_ISL_451714, EPI_ISL_451715, EPI_ISL_451716, EPI_ISL_451717, EPI_ISL_451718, EPI_ISL_451719, EPI_ISL_451720, EPI_ISL_451721, EPI_ISL_451722, EPI_ISL_451723, EPI_ISL_451724, EPI_ISL_451725, EPI_ISL_451726, EPI_ISL_451727, EPI_ISL_451728, EPI_ISL_451729, EPI_ISL_451730, EPI_ISL_451731, EPI_ISL_451732, EPI_ISL_451733, EPI_ISL_451734, EPI_ISL_451735, EPI_ISL_451736, EPI_ISL_451737, EPI_ISL_451738, EPI_ISL_451739, EPI_ISL_451740, EPI_ISL_451741, EPI_ISL_451742, EPI_ISL_451743, EPI_ISL_451744, EPI_ISL_451745, EPI_ISL_451746, EPI_ISL_451747, EPI_ISL_451748, EPI_ISL_451749, EPI_ISL_451750, EPI_ISL_451751, EPI_ISL_451752, EPI_ISL_451753, EPI_ISL_451754, EPI_ISL_451755, EPI_ISL_451756, EPI_ISL_451757, EPI_ISL_451758, EPI_ISL_451759, EPI_ISL_451760, EPI_ISL_451761, EPI_ISL_451762, EPI_ISL_451763, EPI_ISL_451764, EPI_ISL_451765, EPI_ISL_451766, EPI_ISL_451767, EPI_ISL_451768, EPI_ISL_451769, EPI_ISL_451770, EPI_ISL_451771, EPI_ISL_451772, EPI_ISL_451773, EPI_ISL_451774, EPI_ISL_451775, EPI_ISL_451776, EPI_ISL_451777, EPI_ISL_451778, EPI_ISL_451779, EPI_ISL_451780, EPI_ISL_451781, EPI_ISL_451782, EPI_ISL_451783, EPI_ISL_451784, EPI_ISL_451785, EPI_ISL_451786, EPI_ISL_451787, | | | |

| | | | |
|---|---|---|---|
| EPI_ISL_451788, EPI_ISL_451789, EPI_ISL_451790, EPI_ISL_451791, EPI_ISL_451792, EPI_ISL_451793, EPI_ISL_451794, EPI_ISL_451795, EPI_ISL_451796, EPI_ISL_451797, EPI_ISL_451798, EPI_ISL_451799, EPI_ISL_451800, EPI_ISL_451801, EPI_ISL_451802, EPI_ISL_451803, EPI_ISL_451804, EPI_ISL_451805, EPI_ISL_451806, EPI_ISL_451807, EPI_ISL_451808, EPI_ISL_451809, EPI_ISL_451810, EPI_ISL_451811, EPI_ISL_451812, EPI_ISL_451813, EPI_ISL_451814, EPI_ISL_451815, EPI_ISL_451816, EPI_ISL_451817, EPI_ISL_451818, EPI_ISL_451819, EPI_ISL_451820, EPI_ISL_451821, EPI_ISL_451822, EPI_ISL_451823, EPI_ISL_451824, EPI_ISL_451825, EPI_ISL_451826, EPI_ISL_451827, EPI_ISL_451828, EPI_ISL_451829, EPI_ISL_451830, EPI_ISL_451831, EPI_ISL_451832, EPI_ISL_451833, EPI_ISL_451834, EPI_ISL_451835, EPI_ISL_451836, EPI_ISL_451837, EPI_ISL_451838, EPI_ISL_451839, EPI_ISL_451840, EPI_ISL_451841, EPI_ISL_451842, EPI_ISL_451843, EPI_ISL_451844, EPI_ISL_451845, EPI_ISL_451846, EPI_ISL_451847, EPI_ISL_451848, EPI_ISL_451849, EPI_ISL_451850, EPI_ISL_451851, EPI_ISL_451852, EPI_ISL_451853, EPI_ISL_451854, EPI_ISL_451855, EPI_ISL_451856, EPI_ISL_451857, EPI_ISL_451858, EPI_ISL_451859, EPI_ISL_451860, EPI_ISL_451861, EPI_ISL_451862, EPI_ISL_451863, EPI_ISL_451864, EPI_ISL_451865, EPI_ISL_451866, EPI_ISL_451867, EPI_ISL_451868, EPI_ISL_451869, EPI_ISL_451870, EPI_ISL_451871, EPI_ISL_451872, EPI_ISL_451873, EPI_ISL_451874, EPI_ISL_451875, EPI_ISL_451876, EPI_ISL_451877, EPI_ISL_451878, EPI_ISL_451879, EPI_ISL_451880, EPI_ISL_451881, EPI_ISL_451882, EPI_ISL_451883, EPI_ISL_451884, EPI_ISL_451885, EPI_ISL_451886, EPI_ISL_451887, EPI_ISL_451888, EPI_ISL_451889, EPI_ISL_451890, EPI_ISL_451891, EPI_ISL_451892, EPI_ISL_451893, EPI_ISL_451894, EPI_ISL_451895, EPI_ISL_451896, EPI_ISL_451897, EPI_ISL_451898, EPI_ISL_451899, EPI_ISL_451900, EPI_ISL_451901, EPI_ISL_451902, EPI_ISL_451903, EPI_ISL_451904, EPI_ISL_451905, EPI_ISL_451906, EPI_ISL_451907, EPI_ISL_451908, EPI_ISL_451909, EPI_ISL_451910, EPI_ISL_451911, EPI_ISL_451912, EPI_ISL_451913, EPI_ISL_451914, EPI_ISL_451915, EPI_ISL_451916 | | | |
| see above | Viollier AG | Department of Biosystems Science and Engineering, ETH Zürich | Christian Beisel, Sarah Nadeau, Ivan Topolsky, Pedro Ferreira, Philipp Jablonski, Susana Posada-Céspedes, Tobias Schär, Ina Nissen, Natascha Santacroce, Elodie Burcklen, Christiane Beckmann, Maurice Redondo, Olivier Kobel, Christoph Noppen, Sophie Seidel, Noemie Santamaria de Souza, Niko Beerenwinkel, Tanja Stadler |
| EPI_ISL_451934 | Research Unit, University Hospital for Infectious Diseases "Dr. Fran Mihaljević" | Cicin Sain lab, Helmholtz Centre for Infection Research | Zeeshan Chaudhry, Kathrin Eschke, Željka Mačak Šafranko, Ivan-Christian Kurolt |
| EPI_ISL_451935 | CUB Hopital Erasme Laboratoire d'Anatomie Pathologique | CUB Hopital Erasme Laboratoire d'Anatomie Pathologique | Prof. Isabelle Salmon, Dr. Nicky D'Haene |
| EPI_ISL_451936, EPI_ISL_451937, EPI_ISL_451938, EPI_ISL_451939, EPI_ISL_451940, EPI_ISL_451941, EPI_ISL_451942, EPI_ISL_451943, EPI_ISL_451944, EPI_ISL_451945, EPI_ISL_451946, EPI_ISL_451947 | | | |
| see above | Max von Pettenkofer Institute, Virology, National Reference Center for Retroviruses, LMU München | Laboratory for Functional Genome Analysis, Dept. Genomics, Gene Center of the LMU Munich | Max Muenchhoff, Stefan Krebs, Alexander Graf, Oliver Keppler, Helmut Blum |
| EPI_ISL_451948 | The Republican Research and Practical Center for Epidemiology and Microbiology | Charite Universitatsmedizin Berlin, Institute of Virology | Victor M Corman, Barbara Mühlemann, Talitha Veith, Jorn Beheim-Schwarzbach, Julia Schneider, Terry Jones, Natallia Shmialiova, Natallia Sivets, Christian Drosten |
| EPI_ISL_451957 | Molecular Pathology Division, Department of Pathology, Hong Kong Sanatorium & Hospital | Molecular Pathology Division, Department of Pathology, Hong Kong Sanatorium & Hospital | Chun Hang AU, Wai Sing CHAN, Ho Yin LAM, Dona N. HO, Simon Y.M. LAM, Jonpaul S.T. ZEE, Tsun Leung CHAN, Edmond S.K. MA |
| EPI_ISL_451958 | Jamil-ur-Rahman Center for Genome Research, Dr. Panjwani Center for Molecular Medicine and Drug Research, International Center for Chemical and Biological Sciences, University of Karachi | Jamil-ur-Rahman Center for Genome Research, Dr. Panjwani Center for Molecular Medicine and Drug Research, International Center for Chemical and Biological Sciences, University of Karachi | Shakeel,M., Raza,S.A., Khan,S., Khan,B.A., Zahid,M., Qureshi,M.A.and Khan,I.A |
| EPI_ISL_451960 | VI-US Virgin Islands Department of Health | Centers for Disease Control and Prevention | Queen,K., Li,Y., Montmayeur,A., Tao,Y., Zhang,J., Uehara,A., Paden,C.R., Marine,R., Wang,H., Padilla,J., Lee,J., Weiner,Z., Bankamp,B. and Tong,S. |
| EPI_ISL_451961 | Istituto Zooprofilattico Sperimentale Puglia e Basilicata; Dipartimento di Bioscienze, Biotecnologie e Biofarmaceutica dell'Università degli Studi di Bari "A.Moro"; Istituto di Biomembrane. Bioenergetica e Biotecnologie Molecolari del Consiglio Nazionale delle Ricerche di Bari | Beaconlab (Bioinformatics Evolution and Comparative Genomics lab), Dept of Biosciences, University of Milan | Parisi A.,Pesole G., Manzari C., Chiara M. |
| EPI_ISL_451962 | Istituto Zooprofilattico Sperimentale Puglia e Basilicata; Dipartimento di Bioscienze, Biotecnologie e Biofarmaceutica dell'Università degli Studi di Bari "A.Moro"; Istituto di Biomembrane. Bioenergetica e Biotecnologie Molecolari del Consiglio Nazionale delle Ricerche di Bari | Beaconlab (Bioinformatics, Evolution and Comparative Genomics lab), Dept of Biosciences, University on Milan | Parisi A.,Pesole G., Manzari C., Chiara M. |
| EPI_ISL_451963 | Federal Budget Institution of Science, State Research Center for Applied Microbiology & Biotechnology | Federal Budget Institution of Science, State Research Center for Applied Microbiology & Biotechnology | Dyatlov I, Shemyakin I, Khramov M, Bogun A, Kislichkina A, Frolov V, Shishkina L, Sizova A, Chekan L, Blagodatskikh S, Podkopaev Y, Kosilova I, Koroleva-Ushakova A, Tyurin E, Galkina E, Slukina N, Shaikhutdinova R, Kalmantayev T, Kalmantayeva O, Fursova N, Silkina M, Gorbatov A, Titareva G, Firstova V, Makarova M, Gapelchenkova T, Solovieva A, Slukin P, Dentovskaya S, Detushev K, Vagayskaya A, Kartsev N, Detusheva E, Zeninskaya N, Ivanov S, Kartseva A, Platonov M, Hlyntseva A, Khomyakov A, Chernysh S, Krasilnikova E, Ryabko A, Solomentsev V, Teymurazov M, Bakhteeva I, Borzilov A, Skryabin Y, Kanashenko M, Abaimova A, Kolchanova A, Novikova T, Goncharova J, Timofeev V, Kuzina E, Fursov M, Zhumakaev R, Marin M, Denisenko E, Trunyakova A, Kuzin V |
| EPI_ISL_451964, EPI_ISL_451965, EPI_ISL_451966, EPI_ISL_451967, EPI_ISL_451968, EPI_ISL_451969, EPI_ISL_451970 | Federal Budget Institution of Science, State Research Center for Applied Microbiology & Biotechnology | Federal Budget Institution of Science, State Research Center for Applied Microbiology & Biotechnology | Dyatlov I, Shemyakin I, Khramov M, Bogun A, Kislichkina A, Frolov V, Shishkina L, Sizova A, Chekan L, Blagodatskikh S, Podkopaev Y, Kosilova I, Koroleva-Ushakova A, Tyurin E, Galkina E, Slukina N, Shaikhutdinova R, Kalmantayev T, Kalmantayeva O, Fursova N, Silkina M, Gorbatov A, Titareva G, Firstova V, Makarova M, Gapelchenkova T, Solovieva A, Slukin P, Dentovskaya S, Detushev K, Vagayskaya A, Kartsev N, Detusheva E, Zeninskaya N, Ivanov S, Kartseva A, Platonov M, Hlyntseva A, Khomyakov A, Chernysh S, Krasilnikova E, Ryabko A, Solomentsev V, Teymurazov M, Bakhteeva I, Borzilov A, Skryabin Y, Kanashenko M, Abaimova A, Kolchanova A, Novikova T, Goncharova J, Timofeev V, Kuzina E, Fursov M, Zhumakaev R, Marin M, Denisenko E, Trunyakova A, Kuzin V |
| EPI_ISL_451971 | 1. ViroGenetics - BSL3 Laboratory of Virology, Małopolska Centre of Biotechnology, Jagiellonian University; 2. II Department of Internal Medicine, Faculty of Medicine, Jagiellonian University Medical College; 3. DIAGNOSTYKA Ltd. | 1. ViroGenetics - BSL3 Laboratory of Virology, Małopolska Centre of Biotechnology, Jagiellonian University; 2. II Department of Internal Medicine, Faculty of Medicine, Jagiellonian University Medical College. | Marek Sanak, Marcin Surmiak, Monika Gąsecka-Czapla, Wojciech Branicki, Paweł P Łabaj, Marta Rogalska-Kupiec, Jakub Swadźba, Krzysztof Pyrć |
| EPI_ISL_451972 | 1. ViroGenetics - BSL3 Laboratory of Virology, Małopolska Centre of Biotechnology, Jagiellonian University; 2. II Department of Internal Medicine, Faculty of Medicine, Jagiellonian University Medical College; 3. DIAGNOSTYKA Ltd. | 1. ViroGenetics - BSL3 Laboratory of Virology, Małopolska Centre of Biotechnology, Jagiellonian University; 2. II Department of Internal Medicine, Faculty of Medicine, Jagiellonian University Medical College. | Marek Sanak, Marcin Surmiak, Monika Gąsecka-Czapla, Wojciech Branicki, Paweł P Łabaj, Marta Rogalska-Kupiec, Jakub Swadźba, Krzysztof Pyrć |
| EPI_ISL_451973, EPI_ISL_451974 | 1. ViroGenetics - BSL3 Laboratory of Virology, Małopolska Centre of Biotechnology, Jagiellonian University; 2. II Department of Internal Medicine, Faculty of Medicine, Jagiellonian University Medical College; 3. DIAGNOSTYKA Ltd. | 1. ViroGenetics - BSL3 Laboratory of Virology, Małopolska Centre of Biotechnology, Jagiellonian University; 2. II Department of Internal Medicine, Faculty of Medicine, Jagiellonian University Medical College. | Marek Sanak, Marcin Surmiak, Monika Gąsecka-Czapla, Wojciech Branicki, Paweł P Łabaj, Marta Rogalska-Kupiec, Jakub Swadźba, Krzysztof Pyrć |
| EPI_ISL_451975, EPI_ISL_451976, EPI_ISL_451977, EPI_ISL_451978, EPI_ISL_451979, EPI_ISL_451980, EPI_ISL_451981, EPI_ISL_451982, EPI_ISL_451983, EPI_ISL_451984, EPI_ISL_451985, EPI_ISL_451986, EPI_ISL_451987 | | | |
| see above | 1. ViroGenetics - BSL3 Laboratory of Virology, Małopolska Centre of Biotechnology, Jagiellonian University; 2. II Department of Internal Medicine, Faculty of Medicine, Jagiellonian University Medical College; 3. DIAGNOSTYKA Ltd. | 1. ViroGenetics - BSL3 Laboratory of Virology, Małopolska Centre of Biotechnology, Jagiellonian University; 2. II Department of Internal Medicine, Faculty of Medicine, Jagiellonian University Medical College. | Marek Sanak, Marcin Surmiak, Monika Gąsecka-Czapla, Wojciech Branicki, Paweł P Łabaj, Marta Rogalska-Kupiec, Jakub Swadźba, Krzysztof Pyrć |
| EPI_ISL_451988, EPI_ISL_451989, EPI_ISL_451990, EPI_ISL_451991, EPI_ISL_451992, EPI_ISL_451993, EPI_ISL_451994, EPI_ISL_451995, EPI_ISL_451996, EPI_ISL_451997, EPI_ISL_451998, EPI_ISL_451999, EPI_ISL_452000, EPI_ISL_452001, EPI_ISL_452002, EPI_ISL_452003, EPI_ISL_452004, EPI_ISL_452005, EPI_ISL_452006, EPI_ISL_452007, EPI_ISL_452008, EPI_ISL_452009, EPI_ISL_452010, EPI_ISL_452011, EPI_ISL_452012, EPI_ISL_452013, EPI_ISL_452014, EPI_ISL_452015, EPI_ISL_452016, EPI_ISL_452017, EPI_ISL_452018, EPI_ISL_452019, EPI_ISL_452020, EPI_ISL_452021, EPI_ISL_452022, EPI_ISL_452023, EPI_ISL_452024, EPI_ISL_452025, EPI_ISL_452026, EPI_ISL_452027, EPI_ISL_452028, EPI_ISL_452029, EPI_ISL_452030, EPI_ISL_452031, EPI_ISL_452032, EPI_ISL_452033, EPI_ISL_452034, EPI_ISL_452035, EPI_ISL_452036, EPI_ISL_452037, EPI_ISL_452038, EPI_ISL_452039, EPI_ISL_452040, EPI_ISL_452041, EPI_ISL_452042, EPI_ISL_452043, EPI_ISL_452044, EPI_ISL_452045, EPI_ISL_452046, EPI_ISL_452047, EPI_ISL_452048, EPI_ISL_452049, EPI_ISL_452050, EPI_ISL_452051, EPI_ISL_452052, EPI_ISL_452053, EPI_ISL_452054, EPI_ISL_452055, EPI_ISL_452056, EPI_ISL_452057, EPI_ISL_452058, EPI_ISL_452059, EPI_ISL_452060, EPI_ISL_452061, EPI_ISL_452062, EPI_ISL_452063, EPI_ISL_452064, EPI_ISL_452065, EPI_ISL_452066, EPI_ISL_452067, EPI_ISL_452068, EPI_ISL_452069, EPI_ISL_452070, EPI_ISL_452071, EPI_ISL_452072, EPI_ISL_452073, EPI_ISL_452074, EPI_ISL_452075, EPI_ISL_452076, EPI_ISL_452077, EPI_ISL_452078, EPI_ISL_452079, EPI_ISL_452080, EPI_ISL_452081, EPI_ISL_452082, EPI_ISL_452083, EPI_ISL_452084, EPI_ISL_452085, EPI_ISL_452086, EPI_ISL_452087, EPI_ISL_452088, EPI_ISL_452089, EPI_ISL_452090, EPI_ISL_452091, EPI_ISL_452092, EPI_ISL_452093, EPI_ISL_452094, EPI_ISL_452095, EPI_ISL_452096, EPI_ISL_452097, EPI_ISL_452098, EPI_ISL_452099, EPI_ISL_452100 | | | |
| see above | Department of Clinical Microbiology, Copenhagen University Hospital, Hvidovre, Kettegaard Alle 30, 2650 Hvidovre. | Albertsen lab, Department of Chemistry and Bioscience, Aalborg University, Denmark | Rasmus Kirkegaard |
| EPI_ISL_452101, EPI_ISL_452102 | Department of Virus and Microbiological Special Diagnostics, Statens Serum Institut, Copenhagen, Denmark, Artillerivej 5, 2300 Copenahgen S | Albertsen lab, Department of Chemistry and Bioscience, Aalborg University, Denmark | Rasmus Kirkegaard |
| EPI_ISL_452103, EPI_ISL_452104 | Max von Pettenkofer Institute, Virology, National Reference Center for Retroviruses, LMU München | Laboratory for Functional Genome Analysis, Dept. Genomics, Gene Center of the LMU Munich | Max Muenchhoff, Stefan Krebs, Alexander Graf, Oliver Keppler, Helmut Blum |
| EPI_ISL_452105 | Texas DSHS Lab Services | Pathogen Discovery, Respiratory Viruses Branch, Division of Viral Diseases, Centers for Disease Control and Prevention | Yan Li, Anna Montmayeur, Ying Tao, Krista Queen, Jing Zhang, Anna Uehara, Clinton R. Paden, Rachel Marine, Mary S. Keckler, Alison S. Laufer Halpin, Haibin Wang, Christopher A. Elkins, Zachary Weiner, Suxiang Tong |
| EPI_ISL_452106, EPI_ISL_452107 | LA Office of Public Health Laboratories | Pathogen Discovery, Respiratory Viruses Branch, Division of Viral Diseases, Centers for Disease Control and Prevention | Yan Li, Anna Montmayeur, Ying Tao, Krista Queen, Jing Zhang, Anna Uehara, Clinton R. Paden, Rachel Marine, Mary S. Keckler, Alison S. Laufer Halpin, Haibin Wang, Christopher A. Elkins, Zachary Weiner, Suxiang Tong |
| EPI_ISL_452108 | MN Department of Health | Pathogen Discovery, Respiratory Viruses Branch, Division of Viral Diseases, Centers for Disease Control and Prevention | Yan Li, Anna Montmayeur, Ying Tao, Krista Queen, Jing Zhang, Anna Uehara, Clinton R. Paden, Rachel Marine, Mary S. Keckler, Alison S. Laufer Halpin, Haibin Wang, Christopher A. Elkins, Zachary Weiner, Suxiang Tong |
| EPI_ISL_452109 | NM Department of Health | Pathogen Discovery, Respiratory Viruses Branch, Division of Viral Diseases, Centers for Disease Control and Prevention | Yan Li, Anna Montmayeur, Ying Tao, Krista Queen, Jing Zhang, Anna Uehara, Clinton R. Paden, Rachel Marine, Mary S. Keckler, Alison S. Laufer Halpin, Haibin Wang, Christopher A. Elkins, Zachary Weiner, Suxiang Tong |
| EPI_ISL_452110, EPI_ISL_452111 | FL Bureau of Public Health Laboratories | Pathogen Discovery, Respiratory Viruses Branch, Division of Viral Diseases, Centers for Disease Control and Prevention | Yan Li, Anna Montmayeur, Ying Tao, Krista Queen, Jing Zhang, Anna Uehara, Clinton R. Paden, Rachel Marine, Mary S. Keckler, Alison S. Laufer Halpin, Haibin Wang, Christopher A. Elkins, Zachary Weiner, Suxiang Tong |
| EPI_ISL_452112 | Texas DSHS Lab Services | Pathogen Discovery, Respiratory Viruses Branch, Division of Viral Diseases, Centers for Disease Control and Prevention | Yan Li, Anna Montmayeur, Ying Tao, Krista Queen, Jing Zhang, Anna Uehara, Clinton R. Paden, Rachel Marine, Mary S. Keckler, Alison S. Laufer Halpin, Haibin Wang, Christopher A. Elkins, Zachary Weiner, Suxiang Tong |
| EPI_ISL_452113 | MN Department of Health | Pathogen Discovery, Respiratory Viruses Branch, Division of Viral Diseases, Centers for Disease Control and Prevention | Yan Li, Anna Montmayeur, Ying Tao, Krista Queen, Jing Zhang, Anna Uehara, Clinton R. Paden, Rachel Marine, Mary S. Keckler, Alison S. Laufer Halpin, Haibin Wang, Christopher A. Elkins, Zachary Weiner, Suxiang Tong |
| EPI_ISL_452114 | KS Health and Environmental Laboratories | Pathogen Discovery, Respiratory Viruses Branch, Division of Viral Diseases, Centers for Disease Control and Prevention | Yan Li, Anna Montmayeur, Ying Tao, Krista Queen, Jing Zhang, Anna Uehara, Clinton R. Paden, Rachel Marine, Mary S. Keckler, Alison S. Laufer Halpin, Haibin Wang, Christopher A. Elkins, Zachary Weiner, Suxiang Tong |
| EPI_ISL_452115 | Texas DSHS Lab Services | Pathogen Discovery, Respiratory Viruses Branch, Division of Viral Diseases, Centers for Disease Control and Prevention | Yan Li, Anna Montmayeur, Ying Tao, Krista Queen, Jing Zhang, Anna Uehara, Clinton R. Paden, Rachel Marine, Mary S. Keckler, Alison S. Laufer Halpin, Haibin Wang, Christopher A. Elkins, Zachary Weiner, Suxiang Tong |
| EPI_ISL_452116, EPI_ISL_452117 | NJ Public Health and Environmental Laboratories | Pathogen Discovery, Respiratory Viruses Branch, Division of Viral Diseases, Centers for Disease Control and Prevention | Yan Li, Anna Montmayeur, Ying Tao, Krista Queen, Jing Zhang, Anna Uehara, Clinton R. Paden, Rachel Marine, Mary S. Keckler, Alison S. Laufer Halpin, Haibin Wang, Christopher A. Elkins, Zachary Weiner, Suxiang Tong |
| EPI_ISL_452118 | AZ SPHL, Arizona Department of Health Services | Pathogen Discovery, Respiratory Viruses Branch, Division of Viral Diseases, Centers for Disease Control and Prevention | Yan Li, Anna Montmayeur, Ying Tao, Krista Queen, Jing Zhang, Anna Uehara, Clinton R. Paden, Rachel Marine, Mary S. Keckler, Alison S. Laufer Halpin, Haibin Wang, Christopher A. Elkins, Zachary Weiner, Suxiang Tong |
| EPI_ISL_452119 | VI-US Virgin Islands Department of Health | Pathogen Discovery, Respiratory Viruses Branch, Division of Viral Diseases, Centers for Disease Control and Prevention | Yan Li, Anna Montmayeur, Ying Tao, Krista Queen, Jing Zhang, Anna Uehara, Clinton R. Paden, Rachel Marine, Mary S. Keckler, Alison S. Laufer Halpin, Haibin Wang, Christopher A. Elkins, Zachary Weiner, Suxiang Tong |
| EPI_ISL_452120 | Georgia Department of Health | Pathogen Discovery, Respiratory Viruses Branch, Division of Viral Diseases, Centers for Disease Control and Prevention | Jing Zhang, Anna Montmayeur, Yan Li, Ying Tao, Krista Queen, Anna Uehara, Clinton R. Paden, Rachel Marine, Mary S. Keckler, Alison S. Laufer Halpin, Haibin Wang, Christopher A. Elkins, Zachary Weiner, Suxiang Tong |
| EPI_ISL_452121, EPI_ISL_452122, | VI-US Virgin Islands Department of Health | Pathogen Discovery, Respiratory Viruses Branch, Division of Viral | Jing Zhang, Anna Montmayeur, Yan Li, Ying Tao, Krista Queen, Anna Uehara, Clinton R. Paden, Rachel Marine, Mary S. Keckler, Alison S. Laufer Halpin, Haibin Wang, Christopher A. Elkins, Zachary Weiner, Suxiang |

| | | | |
|---|---|---|---|
| EPI_ISL_452123 | | Diseases, Centers for Disease Control and Prevention | Tong |
| EPI_ISL_452124 | NC State Laboratory of Public Health | Pathogen Discovery, Respiratory Viruses Branch, Division of Viral Diseases, Centers for Disease Control and Prevention | Jing Zhang, Anna Montmayeur, Yan Li, Ying Tao, Krista Queen, Anna Uehara, Clinton R. Paden, Rachel Marine, Mary S. Keckler, Alison S. Laufer Halpin, Haibin Wang, Christopher A. Elkins, Zachary Weiner, Suxiang Tong |
| EPI_ISL_452125 | Georgia Department of Health | Pathogen Discovery, Respiratory Viruses Branch, Division of Viral Diseases, Centers for Disease Control and Prevention | Jing Zhang, Anna Montmayeur, Yan Li, Ying Tao, Krista Queen, Anna Uehara, Clinton R. Paden, Rachel Marine, Mary S. Keckler, Alison S. Laufer Halpin, Haibin Wang, Christopher A. Elkins, Zachary Weiner, Suxiang Tong |
| EPI_ISL_452126 | NC State Laboratory of Public Health | Pathogen Discovery, Respiratory Viruses Branch, Division of Viral Diseases, Centers for Disease Control and Prevention | Jing Zhang, Anna Montmayeur, Yan Li, Ying Tao, Krista Queen, Anna Uehara, Clinton R. Paden, Rachel Marine, Mary S. Keckler, Alison S. Laufer Halpin, Haibin Wang, Christopher A. Elkins, Zachary Weiner, Suxiang Tong |
| EPI_ISL_452127, EPI_ISL_452128, EPI_ISL_452129, EPI_ISL_452130 | CO Department of Public Health and Environment | Pathogen Discovery, Respiratory Viruses Branch, Division of Viral Diseases, Centers for Disease Control and Prevention | Jing Zhang, Anna Montmayeur, Yan Li, Ying Tao, Krista Queen, Anna Uehara, Clinton R. Paden, Rachel Marine, Mary S. Keckler, Alison S. Laufer Halpin, Haibin Wang, Christopher A. Elkins, Zachary Weiner, Suxiang Tong |
| EPI_ISL_452131 | IN State Department of Health Laboratory Services | Pathogen Discovery, Respiratory Viruses Branch, Division of Viral Diseases, Centers for Disease Control and Prevention | Jing Zhang, Anna Montmayeur, Yan Li, Ying Tao, Krista Queen, Anna Uehara, Clinton R. Paden, Rachel Marine, Mary S. Keckler, Alison S. Laufer Halpin, Haibin Wang, Christopher A. Elkins, Zachary Weiner, Suxiang Tong |
| EPI_ISL_452132 | FL Bureau of Public Health Laboratories | Pathogen Discovery, Respiratory Viruses Branch, Division of Viral Diseases, Centers for Disease Control and Prevention | Krista Queen, Yan Li, Anna Montmayeur, Ying Tao, Jing Zhang, Anna Uehara, Clinton R. Paden, Rachel Marine, Mary S. Keckler, Alison S. Laufer Halpin, Haibin Wang, Christopher A. Elkins, Zachary Weiner, Suxiang Tong |
| EPI_ISL_452133 | MN Department of Health | Pathogen Discovery, Respiratory Viruses Branch, Division of Viral Diseases, Centers for Disease Control and Prevention | Krista Queen, Yan Li, Anna Montmayeur, Ying Tao, Jing Zhang, Anna Uehara, Clinton R. Paden, Rachel Marine, Mary S. Keckler, Alison S. Laufer Halpin, Haibin Wang, Christopher A. Elkins, Zachary Weiner, Suxiang Tong |
| EPI_ISL_452134 | IL Department of Public Health Chicago Laboratory | Pathogen Discovery, Respiratory Viruses Branch, Division of Viral Diseases, Centers for Disease Control and Prevention | Krista Queen, Yan Li, Anna Montmayeur, Ying Tao, Jing Zhang, Anna Uehara, Clinton R. Paden, Rachel Marine, Mary S. Keckler, Alison S. Laufer Halpin, Haibin Wang, Christopher A. Elkins, Zachary Weiner, Suxiang Tong |
| EPI_ISL_452135, EPI_ISL_452136 | IN State Department of Health Laboratory Services | Pathogen Discovery, Respiratory Viruses Branch, Division of Viral Diseases, Centers for Disease Control and Prevention | Krista Queen, Yan Li, Anna Montmayeur, Ying Tao, Jing Zhang, Anna Uehara, Clinton R. Paden, Rachel Marine, Mary S. Keckler, Alison S. Laufer Halpin, Haibin Wang, Christopher A. Elkins, Zachary Weiner, Suxiang Tong |
| EPI_ISL_452137, EPI_ISL_452138 | VI-US Virgin Islands Department of Health | Pathogen Discovery, Respiratory Viruses Branch, Division of Viral Diseases, Centers for Disease Control and Prevention | Anna Uehara, Yan Li, Anna Montmayeur, Ying Tao, Krista Queen, Jing Zhang, Clinton R. Paden, Rachel Marine, Haibin Wang, Bettina Bankamp, Zachary Weiner, Suxiang Tong |
| EPI_ISL_452139 | Instituto de Diagnostico y Referencia Epidemiologicos (INDRE) | Instituto de diagnóstico y Referencia Epidemiologicos (INDRE) | Ramirez-Gonzalez Ernesto, Garces-Ayala Fabiola, Araiza-Rodriguez Adnan, Mendieta-Condado Edgar, Rodriguez-Maldonado Abril, Wong-Arambula Claudia, Barrera-Badillo Gisela, Hernandez-Rivas Lucia, Lopez-Martinez Irma |
| EPI_ISL_452140 | CUB Hopital Erasme Laboratoire d'Anatomie Pathologique | CUB Hopital Erasme Laboratoire d'Anatomie Pathologique | Isabelle Salmon, Nicky D'Haene |
| EPI_ISL_452141 | Instituto de Diagnostico y Referencia Epidemiologicos (INDRE) | Instituto de diagnóstico y Referencia Epidemiologicos (INDRE) | Ramirez-Gonzalez Ernesto, Garces-Ayala Fabiola, Araiza-Rodriguez Adnan, Mendieta-Condado Edgar, Rodriguez-Maldonado Abril, Wong-Arambula Claudia, Barrera-Badillo Gisela, Hernandez-Rivas Lucia, Lopez-Martinez Irma. |
| EPI_ISL_452142 | CUB Hopital Erasme Laboratoire d'Anatomie Pathologique | CUB Hopital Erasme Laboratoire d'Anatomie Pathologique | Isabelle Salmon, Nikcy D'Haene |
| EPI_ISL_452143, EPI_ISL_452144, EPI_ISL_452145, EPI_ISL_452146, EPI_ISL_452147 | Yale COVID-19 Biorepository | Grubaugh Lab - Yale School of Public Health | Joseph Fauver, Tara Alpert, Anderson Brito, Anne Wyllie, Chantal Vogels, Mary Petrone, Cole Jensen, Chaney Kalinich, Isabel Ott, Arnau Casanovas, Catherine Muenker, Adam Moore, Alice Lu, Maria Tokuyama, Patrick Wong, Peiwen Lu, Saad Omer, Richard Martinello, Allison Nelson, Shelli Farhadian, Akiko Iwasaki, Charlese Dela Cruz, Albert Ko, Nathan Grubaugh |
| EPI_ISL_452148, EPI_ISL_452149 | CUB Hopital Erasme Laboratoire d'Anatomie Pathologique | CUB Hopital Erasme Laboratoire d'Anatomie Pathologique | Isabelle Salmon, Nicky D'Haene |
| EPI_ISL_452150 | CUB Hopital Erasme Laboratoire d'Anatomie Pathologique | CUB Hopital Erasme Laboratoire d'Anatomie Pathologique | Prof. Isabelle Salmon, Dr Nicky D'Haene |
| EPI_ISL_452151 | CUB Hopital Erasme Laboratoire d'Anatomie Pathologique | CUB Hopital Erasme Laboratoire d'Anatomie Pathologique | Prof. Isabelle Salmon, Dr.Nikcy D'Haene |
| EPI_ISL_452152 | CUB Hopital Erasme Laboratoire d'Anatomie Pathologique | CUB Hopital Erasme Laboratoire d'Anatomie Pathologique | Prof. Isabelle Salmon, Dr Nicky D'Haene |
| EPI_ISL_452153, EPI_ISL_452154, EPI_ISL_452155, EPI_ISL_452156, EPI_ISL_452157, EPI_ISL_452158, EPI_ISL_452159, EPI_ISL_452160, EPI_ISL_452161, EPI_ISL_452162, EPI_ISL_452163, EPI_ISL_452164, EPI_ISL_452165, EPI_ISL_452166, EPI_ISL_452167, EPI_ISL_452168, EPI_ISL_452169, EPI_ISL_452170, EPI_ISL_452171, EPI_ISL_452172, EPI_ISL_452173, EPI_ISL_452174, EPI_ISL_452175, EPI_ISL_452176, EPI_ISL_452177 | | | |
| see above | Utah Public Health Laboratory | Utah Public Health Laboratory | Erin Young, Kelly Oakeson |
| EPI_ISL_452178, EPI_ISL_452179 | Laboratory Medicine | Department of Laboratory Medicine, Lin-Kou Chang Gung Memorial Hospital, Taoyuan, Taiwan | Kuo-Chien Tsao, Yu-Nong Gong, Shu-Li Yang, Yi-Chun Liu, Chung-Guei Huang, Mei-Jen Hsiao, Po-Wei Huang, Cheng-Ta Yang, Cheng-Hsun Chiu, Peng-Nien Huang, Kuo-Ming Lee, Guang-Wu Chen, Shin-Ru Shih |
| EPI_ISL_452180 | Center of Medical Microbiology, Virology, and Hospital Hygiene, University of Duesseldorf | Center of Medical Microbiology, Virology, and Hospital Hygiene, University of Duesseldorf | Ortwin Adams, Marcel Andree, Alexander Dilthey, Torsten Feldt, Sandra Hauka, Torsten Houwaart, Björn-ErikJensen, Detlef Kindgen-Milles, Malte Kohns Vasconcelos, Klaus Pfeffer, Tina Senff, Daniel Strelow, Jörg Timm, Andreas Walker, Tobias Wienemann |
| EPI_ISL_452181, EPI_ISL_452182, EPI_ISL_452183, EPI_ISL_452184, EPI_ISL_452185, EPI_ISL_452186, EPI_ISL_452187, EPI_ISL_452188, EPI_ISL_452189 | ULSS9 Distretto di Bussolengo | Istituto Zooprofilattico Sperimentale delle Venezie | Adelaide Milani, Alessia Schivo, Annalisa Salviato, Erika Giorgia Quaranta, Gianpiero Zamperin, Ambra Pastori, Bianca Zecchin, Alice Fusaro, Calogero Terregino, Antonia Ricci |
| EPI_ISL_452190, EPI_ISL_452191 | ULSS9 Distretto di San Bonifacio | Istituto Zooprofilattico Sperimentale delle Venezie | Adelaide Milani, Alessia Schivo, Annalisa Salviato, Erika Giorgia Quaranta, Gianpiero Zamperin, Ambra Pastori, Bianca Zecchin, Alice Fusaro, Calogero Terregino, Antonia Ricci |
| EPI_ISL_452192, EPI_ISL_452193, EPI_ISL_452194, EPI_ISL_452195, EPI_ISL_452196, EPI_ISL_452197, EPI_ISL_452198, EPI_ISL_452199, EPI_ISL_452200, EPI_ISL_452201, EPI_ISL_452202, EPI_ISL_452203, EPI_ISL_452204, EPI_ISL_452205, EPI_ISL_452206, EPI_ISL_452207, EPI_ISL_452208, EPI_ISL_452209, EPI_ISL_452210, EPI_ISL_452211, EPI_ISL_452212, EPI_ISL_452213, EPI_ISL_452214, EPI_ISL_452215, EPI_ISL_452216, EPI_ISL_452217 | | | |
| see above | NIV Influenza | NIV Influenza | Potdar V |
| EPI_ISL_452218, EPI_ISL_452219, EPI_ISL_452220, EPI_ISL_452221, EPI_ISL_452222, EPI_ISL_452223 | Goethe University Hospital Frankfurt | Institute for Medical Virology, Goethe University Hospital Frankfurt | Tuna Toptan, Sebastian Hoehl, Sandra Westhaus, Denisa Bojkova, Annemarie Berger, Björn Rotter, Klaus Hoffmeier, Jindrich Cinatl, Sandra Ciesek, and Marek Widera |
| EPI_ISL_452229, EPI_ISL_452230 | Narhalsan Backa vardcentral | The Public Health Agency of Sweden | Mats Olsson, Anna-Malin Linde, Maria Lind Karlberg, Oskar Karlsson Lindsjo, Olov Svartstrom, Anna Risberg, Theresa Enkirch, Mia Brytting, Karin Tegmark-Wisell |
| EPI_ISL_452231, EPI_ISL_452232 | Saroledens Familjelakare | The Public Health Agency of Sweden | Katarina Jarbur, Anna-Malin Linde, Maria Lind Karlberg, Oskar Karlsson Lindsjo, Olov Svartstrom, Anna Risberg, Theresa Enkirch, Mia Brytting, Karin Tegmark-Wisell |
| EPI_ISL_452233 | Victoria Vard och Halsa | The Public Health Agency of Sweden | Sarah Henriksson, Anna-Malin Linde, Maria Lind Karlberg, Oskar Karlsson Lindsjo, Olov Svartstrom, Anna Risberg, Theresa Enkirch, Mia Brytting, Karin Tegmark-Wisell |
| EPI_ISL_452234 | Din Klinik | The Public Health Agency of Sweden | Helene Warnborg, Anna-Malin Linde, Maria Lind Karlberg, Oskar Karlsson Lindsjo, Olov Svartstrom, Anna Risberg, Theresa Enkirch, Mia Brytting, Karin Tegmark-Wisell |
| EPI_ISL_452235 | Narhalsan Backa vardcentral | The Public Health Agency of Sweden | Mats Olsson, Anna-Malin Linde, Maria Lind Karlberg, Oskar Karlsson Lindsjo, Olov Svartstrom, Anna Risberg, Theresa Enkirch, Mia Brytting, Karin Tegmark-Wisell |
| EPI_ISL_452236 | VC Sorgenfrimottagningen | The Public Health Agency of Sweden | Lisa Esbjornsson Klemendz, Anna-Malin Linde, Maria Lind Karlberg, Oskar Karlsson Lindsjo, Olov Svartstrom, Anna Risberg, Theresa Enkirch, Mia Brytting, Karin Tegmark-Wisell |
| EPI_ISL_452237 | Huddinge VC | The Public Health Agency of Sweden | Anders Johansson, Anna-Malin Linde, Maria Lind Karlberg, Oskar Karlsson Lindsjo, Olov Svartstrom, Anna Risberg, Theresa Enkirch, Mia Brytting, Karin Tegmark-Wisell |
| EPI_ISL_452238 | Narhalsan Sjobo vardcentral | The Public Health Agency of Sweden | Lovisa Hjerten, Anna-Malin Linde, Maria Lind Karlberg, Oskar Karlsson Lindsjo, Olov Svartstrom, Anna Risberg, Theresa Enkirch, Mia Brytting, Karin Tegmark-Wisell |
| EPI_ISL_452239 | Narhalsan Backa vardcentral | The Public Health Agency of Sweden | Mats Olsson, Anna-Malin Linde, Maria Lind Karlberg, Oskar Karlsson Lindsjo, Olov Svartstrom, Anna Risberg, Theresa Enkirch, Mia Brytting, Karin Tegmark-Wisell |
| EPI_ISL_452240 | Huddinge VC | The Public Health Agency of Sweden | Anders Johansson, Anna-Malin Linde, Maria Lind Karlberg, Oskar Karlsson Lindsjo, Olov Svartstrom, Anna Risberg, Theresa Enkirch, Mia Brytting, Karin Tegmark-Wisell |
| EPI_ISL_452241 | Huslakarna Varmbadhuset Varberg | The Public Health Agency of Sweden | Johanna Hilmersson, Anna-Malin Linde, Maria Lind Karlberg, Oskar Karlsson Lindsjo, Olov Svartstrom, Anna Risberg, Theresa Enkirch, Mia Brytting, Karin Tegmark-Wisell |
| EPI_ISL_452242 | Wernstedt Medical AB | The Public Health Agency of Sweden | Eva Sandberg, Anna-Malin Linde, Maria Lind Karlberg, Oskar Karlsson Lindsjo, Olov Svartstrom, Anna Risberg, Theresa Enkirch, Mia Brytting, Karin Tegmark-Wisell |
| EPI_ISL_452249, EPI_ISL_452250, EPI_ISL_452251, EPI_ISL_452252, EPI_ISL_452253, EPI_ISL_452254, EPI_ISL_452255, EPI_ISL_452256, EPI_ISL_452257, EPI_ISL_452258 | Respiratory Virus Unit, Microbiology Services Colindale, Public Health England | Respiratory Virus Unit, Microbiology Services Colindale, Public Health England | Steven Platt, Shahjahan Miah, Angie Lackenby, Omolola Akinbami, Tina Talts, Leena Bhaw, Richard Myers, Monica Galiano, Kirstin Edwards, Jonathan Hubb, Joanna Ellis, Maria Zambon |
| EPI_ISL_452262, EPI_ISL_452263, EPI_ISL_452264, EPI_ISL_452265, EPI_ISL_452266, EPI_ISL_452267, EPI_ISL_452268, EPI_ISL_452269, EPI_ISL_452270, EPI_ISL_452271, EPI_ISL_452272, EPI_ISL_452273, EPI_ISL_452274, EPI_ISL_452275, EPI_ISL_452276, EPI_ISL_452277, EPI_ISL_452278, EPI_ISL_452279, EPI_ISL_452280, EPI_ISL_452281, EPI_ISL_452282, EPI_ISL_452283, EPI_ISL_452284, EPI_ISL_452285, EPI_ISL_452286, EPI_ISL_452287, EPI_ISL_452288, EPI_ISL_452289, EPI_ISL_452290, EPI_ISL_452291, EPI_ISL_452292, EPI_ISL_452293, EPI_ISL_452294, EPI_ISL_452295, EPI_ISL_452296, EPI_ISL_452297, EPI_ISL_452298, EPI_ISL_452299, EPI_ISL_452300, EPI_ISL_452301, EPI_ISL_452302, EPI_ISL_452303, EPI_ISL_452304, EPI_ISL_452305, EPI_ISL_452306, EPI_ISL_452307, EPI_ISL_452308, EPI_ISL_452309, EPI_ISL_452310, EPI_ISL_452311, EPI_ISL_452312, EPI_ISL_452313, EPI_ISL_452314, EPI_ISL_452315, EPI_ISL_452316, EPI_ISL_452317, EPI_ISL_452318, EPI_ISL_452319, EPI_ISL_452320, EPI_ISL_452321, EPI_ISL_452322, EPI_ISL_452323, EPI_ISL_452324, EPI_ISL_452325, EPI_ISL_452326 | | | |
| see above | Michigan Department of Health and Human Services, Bureau of Laboratories | Michigan Department of Health and Human Services, Bureau of Laboratories | Blankenship HM, Riner D, Soehnlen MK |
| EPI_ISL_452327, EPI_ISL_452328, EPI_ISL_452329, EPI_ISL_452330, EPI_ISL_452331, EPI_ISL_452332, EPI_ISL_452333, EPI_ISL_452334, EPI_ISL_452335, EPI_ISL_452336, EPI_ISL_452337, EPI_ISL_452338, EPI_ISL_452339, EPI_ISL_452340, EPI_ISL_452341, EPI_ISL_452342, EPI_ISL_452343, EPI_ISL_452344, EPI_ISL_452345, EPI_ISL_452346, EPI_ISL_452347, EPI_ISL_452348, EPI_ISL_452349, EPI_ISL_452350, EPI_ISL_452351, EPI_ISL_452352, EPI_ISL_452353, EPI_ISL_452354, EPI_ISL_452355, EPI_ISL_452356, EPI_ISL_452357, EPI_ISL_452358, EPI_ISL_452359, EPI_ISL_452360, EPI_ISL_452361, EPI_ISL_452362, EPI_ISL_452363, EPI_ISL_452364 | | | |
| see above | Laboratory of Infectious Diseases Center of Beijing Ditan Hospital | Laboratory of Infectious Diseases Center of Beijing Ditan Hospital | Siyuan Yang, Chengjie Jie, Fengting Yu, Yunxia Tang, Liting Yan, Linghang Wang |
| EPI_ISL_452365, EPI_ISL_452366, EPI_ISL_452367, EPI_ISL_452368, EPI_ISL_452369, EPI_ISL_452370, EPI_ISL_452371, EPI_ISL_452372, EPI_ISL_452373, EPI_ISL_452374, EPI_ISL_452375, EPI_ISL_452376, EPI_ISL_452377, EPI_ISL_452378, EPI_ISL_452379, EPI_ISL_452380, EPI_ISL_452381, EPI_ISL_452382, EPI_ISL_452383, EPI_ISL_452384, EPI_ISL_452385, EPI_ISL_452386, EPI_ISL_452387, EPI_ISL_452388, EPI_ISL_452389, EPI_ISL_452390, EPI_ISL_452391, EPI_ISL_452392, EPI_ISL_452393, EPI_ISL_452394, EPI_ISL_452395, EPI_ISL_452396, EPI_ISL_452397, EPI_ISL_452398, EPI_ISL_452399, EPI_ISL_452400, EPI_ISL_452401, EPI_ISL_452402, EPI_ISL_452403, EPI_ISL_452404, EPI_ISL_452405, EPI_ISL_452406, EPI_ISL_452407, EPI_ISL_452408, EPI_ISL_452409, EPI_ISL_452410, EPI_ISL_452411, EPI_ISL_452412, EPI_ISL_452413, EPI_ISL_452414, EPI_ISL_452415, EPI_ISL_452416, EPI_ISL_452417, EPI_ISL_452418, EPI_ISL_452419, EPI_ISL_452420, EPI_ISL_452421, EPI_ISL_452422, EPI_ISL_452423, EPI_ISL_452424, EPI_ISL_452425, EPI_ISL_452426, EPI_ISL_452427, EPI_ISL_452428, EPI_ISL_452429, EPI_ISL_452430, EPI_ISL_452431, EPI_ISL_452432, EPI_ISL_452433, EPI_ISL_452434, EPI_ISL_452435, EPI_ISL_452436, EPI_ISL_452437, EPI_ISL_452438, EPI_ISL_452439, EPI_ISL_452440, EPI_ISL_452441, EPI_ISL_452442, EPI_ISL_452443, EPI_ISL_452444, EPI_ISL_452445, EPI_ISL_452446, EPI_ISL_452447, EPI_ISL_452448, EPI_ISL_452449, EPI_ISL_452450, EPI_ISL_452451, EPI_ISL_452452 | | | |
| see above | Servicio de Microbiología. HRU de Málaga. Servicio Andaluz de Salud | SeqCOVID-SPAIN consortium/IBV(CSIC) | Inmaculada de Toro Peinado, Maria Concepción Mediavilla Gradolph, Begoña Palop Borrás and SeqCOVID-SPAIN consortium |
| EPI_ISL_452453, EPI_ISL_452454, EPI_ISL_452455, EPI_ISL_452456, EPI_ISL_452457, EPI_ISL_452458, EPI_ISL_452459, EPI_ISL_452460, EPI_ISL_452461, EPI_ISL_452462, EPI_ISL_452463, EPI_ISL_452464, EPI_ISL_452465, EPI_ISL_452466, EPI_ISL_452467, EPI_ISL_452468, EPI_ISL_452469, EPI_ISL_452470, EPI_ISL_452471 | | | |
| see above | Hospital Universitario Puerta del Mar de Cádiz - INIBICA | SeqCOVID-SPAIN consortium/IBV(CSIC) | Salud Rodríguez-Pallares, Fátima-Galán-Sánchez, Manuel Rodríguez-Iglesias and SeqCOVID-SPAIN consortium |
| EPI_ISL_452472, EPI_ISL_452473, EPI_ISL_452474, EPI_ISL_452475, EPI_ISL_452476, EPI_ISL_452477, EPI_ISL_452478, EPI_ISL_452479, EPI_ISL_452480, EPI_ISL_452481, EPI_ISL_452482, EPI_ISL_452483, EPI_ISL_452484, EPI_ISL_452485, EPI_ISL_452486, EPI_ISL_452487, EPI_ISL_452488, EPI_ISL_452489, EPI_ISL_452490, EPI_ISL_452491, EPI_ISL_452492, EPI_ISL_452493, EPI_ISL_452494, EPI_ISL_452495, EPI_ISL_452496, EPI_ISL_452497, EPI_ISL_452498, EPI_ISL_452499, EPI_ISL_452500, EPI_ISL_452501, EPI_ISL_452502, EPI_ISL_452503, EPI_ISL_452504, EPI_ISL_452505, EPI_ISL_452506, EPI_ISL_452507, EPI_ISL_452508, EPI_ISL_452509, EPI_ISL_452510, EPI_ISL_452511, EPI_ISL_452512, EPI_ISL_452513, EPI_ISL_452514, EPI_ISL_452515, EPI_ISL_452516, EPI_ISL_452517, EPI_ISL_452518, EPI_ISL_452519, EPI_ISL_452520, EPI_ISL_452521, EPI_ISL_452522, EPI_ISL_452523, EPI_ISL_452524, EPI_ISL_452525, EPI_ISL_452526, EPI_ISL_452527, EPI_ISL_452528, EPI_ISL_452529, EPI_ISL_452530, EPI_ISL_452531, EPI_ISL_452532, EPI_ISL_452533, EPI_ISL_452534, EPI_ISL_452535, EPI_ISL_452536, EPI_ISL_452537, EPI_ISL_452538, EPI_ISL_452539, EPI_ISL_452540, EPI_ISL_452541, EPI_ISL_452542, EPI_ISL_452543 | | | |

| | | | |
|---|---|---|---|
| see above | Clínica Universidad de Navarra. Servicio de Enfermedades Infecciosas y Microbiología clínica | SeqCOVID-SPAIN consortium/IBV(CSIC) | Mirian Fernández-Alonso, Jose Luis del Pozo and SeqCOVID-SPAIN consortium |
| EPI_ISL_452544, EPI_ISL_452545, EPI_ISL_452546, EPI_ISL_452547, EPI_ISL_452548, EPI_ISL_452549, EPI_ISL_452550, EPI_ISL_452551, EPI_ISL_452552, EPI_ISL_452553, EPI_ISL_452554, EPI_ISL_452555, EPI_ISL_452556, EPI_ISL_452557, EPI_ISL_452558, EPI_ISL_452559, EPI_ISL_452560, EPI_ISL_452561, EPI_ISL_452562, EPI_ISL_452563, EPI_ISL_452564, EPI_ISL_452565, EPI_ISL_452566, EPI_ISL_452567, EPI_ISL_452568, EPI_ISL_452569, EPI_ISL_452570, EPI_ISL_452571, EPI_ISL_452572, EPI_ISL_452573, EPI_ISL_452574, EPI_ISL_452575, EPI_ISL_452576, EPI_ISL_452577, EPI_ISL_452578, EPI_ISL_452579, EPI_ISL_452580, EPI_ISL_452581, EPI_ISL_452582, EPI_ISL_452583, EPI_ISL_452584, EPI_ISL_452585, EPI_ISL_452586, EPI_ISL_452587, EPI_ISL_452588, EPI_ISL_452589, EPI_ISL_452590, EPI_ISL_452591, EPI_ISL_452592, EPI_ISL_452593, EPI_ISL_452594, EPI_ISL_452595, EPI_ISL_452596, EPI_ISL_452597, EPI_ISL_452598, EPI_ISL_452599, EPI_ISL_452600, EPI_ISL_452601, EPI_ISL_452602, EPI_ISL_452603, EPI_ISL_452604, EPI_ISL_452605, EPI_ISL_452606, EPI_ISL_452607, EPI_ISL_452608, EPI_ISL_452609, EPI_ISL_452610, EPI_ISL_452611, EPI_ISL_452612, EPI_ISL_452613, EPI_ISL_452614, EPI_ISL_452615, EPI_ISL_452616 | | | |
| see above | Servicio de Microbiología y Parasitología clínica. UCEIMP. Hospital Universitario Virgen del Rocío/IBIS/CSIC/US. | SeqCOVID-SPAIN consortium/IBV(CSIC) | Guillermo Martín Gutiérrez, Ángel Rodríguez Villodres, Lidia Gálvez Benitez, Verónica González Galán, Javier Aznar Martín and SeqCOVID-SPAIN consortium |
| EPI_ISL_452617, EPI_ISL_452618, EPI_ISL_452619, EPI_ISL_452620, EPI_ISL_452621, EPI_ISL_452622, EPI_ISL_452623, EPI_ISL_452624, EPI_ISL_452625, EPI_ISL_452626, EPI_ISL_452627, EPI_ISL_452628, EPI_ISL_452629, EPI_ISL_452630, EPI_ISL_452631, EPI_ISL_452632, EPI_ISL_452633, EPI_ISL_452634, EPI_ISL_452635, EPI_ISL_452636, EPI_ISL_452637, EPI_ISL_452638, EPI_ISL_452639, EPI_ISL_452640, EPI_ISL_452641, EPI_ISL_452642, EPI_ISL_452643, EPI_ISL_452644, EPI_ISL_452645, EPI_ISL_452646, EPI_ISL_452647, EPI_ISL_452648, EPI_ISL_452649, EPI_ISL_452650, EPI_ISL_452651, EPI_ISL_452652, EPI_ISL_452653, EPI_ISL_452654, EPI_ISL_452655, EPI_ISL_452656, EPI_ISL_452657, EPI_ISL_452658, EPI_ISL_452659, EPI_ISL_452660, EPI_ISL_452661, EPI_ISL_452662, EPI_ISL_452663, EPI_ISL_452664, EPI_ISL_452665, EPI_ISL_452666, EPI_ISL_452667, EPI_ISL_452668, EPI_ISL_452669, EPI_ISL_452670, EPI_ISL_452671, EPI_ISL_452672, EPI_ISL_452673, EPI_ISL_452674, EPI_ISL_452675, EPI_ISL_452676, EPI_ISL_452677, EPI_ISL_452678, EPI_ISL_452679, EPI_ISL_452680, EPI_ISL_452681, EPI_ISL_452682, EPI_ISL_452683, EPI_ISL_452684, EPI_ISL_452685, EPI_ISL_452686, EPI_ISL_452687, EPI_ISL_452688, EPI_ISL_452689, EPI_ISL_452690, EPI_ISL_452691 | | | |
| see above | Servicio de Microbiología. Hospital Universitario Donostia. OSI Donostialdea. Área de Enfermedades Infecciosas, Grupo de Infección Respiratoria y Resistencia Antimicrobiana. Instituto de Investigación Sanitaria Biodonostia. | SeqCOVID-SPAIN consortium/IBV(CSIC) | Gustavo Cilla, Milagrosa Montes, Luis Piñeiro, Jose Maria Marimón and SeqCOVID-SPAIN consortium |
| EPI_ISL_452692, EPI_ISL_452693, EPI_ISL_452694, EPI_ISL_452695, EPI_ISL_452696, EPI_ISL_452697, EPI_ISL_452698, EPI_ISL_452699, EPI_ISL_452700, EPI_ISL_452701, EPI_ISL_452702, EPI_ISL_452703, EPI_ISL_452704, EPI_ISL_452705, EPI_ISL_452706, EPI_ISL_452707, EPI_ISL_452708, EPI_ISL_452709, EPI_ISL_452710, EPI_ISL_452711, EPI_ISL_452712, EPI_ISL_452713, EPI_ISL_452714, EPI_ISL_452715, EPI_ISL_452716, EPI_ISL_452717, EPI_ISL_452718, EPI_ISL_452719, EPI_ISL_452720, EPI_ISL_452721, EPI_ISL_452722, EPI_ISL_452723, EPI_ISL_452724, EPI_ISL_452725, EPI_ISL_452726, EPI_ISL_452727, EPI_ISL_452728, EPI_ISL_452729, EPI_ISL_452730, EPI_ISL_452731, EPI_ISL_452732, EPI_ISL_452733, EPI_ISL_452734, EPI_ISL_452735, EPI_ISL_452736, EPI_ISL_452737, EPI_ISL_452738, EPI_ISL_452739, EPI_ISL_452740, EPI_ISL_452741, EPI_ISL_452742, EPI_ISL_452743, EPI_ISL_452744, EPI_ISL_452745, EPI_ISL_452746, EPI_ISL_452747, EPI_ISL_452748, EPI_ISL_452749, EPI_ISL_452750, EPI_ISL_452751, EPI_ISL_452752, EPI_ISL_452753, EPI_ISL_452754, EPI_ISL_452755, EPI_ISL_452756, EPI_ISL_452757, EPI_ISL_452758, EPI_ISL_452759, EPI_ISL_452760, EPI_ISL_452761, EPI_ISL_452762, EPI_ISL_452763, EPI_ISL_452764, EPI_ISL_452765, EPI_ISL_452766, EPI_ISL_452767, EPI_ISL_452768, EPI_ISL_452769, EPI_ISL_452770, EPI_ISL_452771, EPI_ISL_452772, EPI_ISL_452773, EPI_ISL_452774, EPI_ISL_452775, EPI_ISL_452776, EPI_ISL_452777, EPI_ISL_452778, EPI_ISL_452779, EPI_ISL_452780, EPI_ISL_452781, EPI_ISL_452782, EPI_ISL_452783, EPI_ISL_452784, EPI_ISL_452785, EPI_ISL_452786 | | | |
| see above | Hospital Universitario Araba. Vitoria-Gasteiz, | SeqCOVID-SPAIN consortium/IBV(CSIC) | Silvia Hernáez Crespo, Carmen Gómez González, Amaia Aguirre Quiñonero, Marina Fernández Torres, Maria Rosario Almela Ferrer, Maria Concepción Lecaroz Agara, Andrés Canut Blasco and SeqCOVID-SPAIN consortium |
| EPI_ISL_452787, EPI_ISL_452788, EPI_ISL_452789, EPI_ISL_452790, EPI_ISL_452791, EPI_ISL_452792, EPI_ISL_452793, EPI_ISL_452794, EPI_ISL_452795 | ICAR-National Institute of High Security Animal Diseases | ICAR-National Institute of High Security Animal Diseases | Anamika Mishra, Ashutosh Aasdev, Sandeep Bhatia, Harshad Murugkar, Chakradhar Tosh, Niranjan Mishra, Shanmugasundaram Nagarajan, Katherukamem Rajukumar, Richa Sood, G Venkatesh, Atul Kumar Pateriya, Manoj Kumar, Shashi Bhushan Sudhakar, Fateh Singh, Sethil Kumar D, Senmannan Kalaiyarasu, Pradeep Gandhale, Naveen Kumar, Chandan Kumar Dubey, Sushil Tripathi, Sandeep Kumar Jhade, Meghna Tripathi, Suman Kumari Shah, Pushpendra Singh, Pushpendra Namdeo, Suman Mishra, Rupal Singh, Vishnupriya Patil, Dipesh Kumar Nayak, Vijendra Pal Singh, Ashwin Ashok Raut |
| EPI_ISL_452796, EPI_ISL_452797, EPI_ISL_452798, EPI_ISL_452799, EPI_ISL_452800, EPI_ISL_452801, EPI_ISL_452802, EPI_ISL_452803, EPI_ISL_452804, EPI_ISL_452805, EPI_ISL_452806, EPI_ISL_452807, EPI_ISL_452808, EPI_ISL_452809, EPI_ISL_452810, EPI_ISL_452811, EPI_ISL_452812, EPI_ISL_452813, EPI_ISL_452814, EPI_ISL_452815, EPI_ISL_452816, EPI_ISL_452817, EPI_ISL_452818, EPI_ISL_452819, EPI_ISL_452820, EPI_ISL_452821, EPI_ISL_452822, EPI_ISL_452823, EPI_ISL_452824, EPI_ISL_452825, EPI_ISL_452826, EPI_ISL_452827, EPI_ISL_452828, EPI_ISL_452829, EPI_ISL_452830, EPI_ISL_452831, EPI_ISL_452832, EPI_ISL_452833, EPI_ISL_452834, EPI_ISL_452835, EPI_ISL_452836, EPI_ISL_452837, EPI_ISL_452838, EPI_ISL_452839, EPI_ISL_452840, EPI_ISL_452841, EPI_ISL_452842, EPI_ISL_452843, EPI_ISL_452844, EPI_ISL_452845, EPI_ISL_452846, EPI_ISL_452847, EPI_ISL_452848, EPI_ISL_452849, EPI_ISL_452850, EPI_ISL_452851, EPI_ISL_452852, EPI_ISL_452853, EPI_ISL_452854, EPI_ISL_452855, EPI_ISL_452856 | | | |
| see above | Virginia DCLS | Virginia DCLS | Virginia DCLS |
| EPI_ISL_452857, EPI_ISL_452858, EPI_ISL_452859, EPI_ISL_452860, EPI_ISL_452861, EPI_ISL_452862, EPI_ISL_452863, EPI_ISL_452864, EPI_ISL_452865, EPI_ISL_452866, EPI_ISL_452867, EPI_ISL_452868, EPI_ISL_452869, EPI_ISL_452870, EPI_ISL_452871, EPI_ISL_452872, EPI_ISL_452873, EPI_ISL_452874, EPI_ISL_452875, EPI_ISL_452876, EPI_ISL_452877, EPI_ISL_452878, EPI_ISL_452879, EPI_ISL_452880, EPI_ISL_452881, EPI_ISL_452882, EPI_ISL_452883, EPI_ISL_452884, EPI_ISL_452885, EPI_ISL_452886, EPI_ISL_452887, EPI_ISL_452888, EPI_ISL_452889, EPI_ISL_452890, EPI_ISL_452891, EPI_ISL_452892, EPI_ISL_452893, EPI_ISL_452894, EPI_ISL_452895, EPI_ISL_452896, EPI_ISL_452897, EPI_ISL_452898, EPI_ISL_452899, EPI_ISL_452900, EPI_ISL_452901, EPI_ISL_452902, EPI_ISL_452903, EPI_ISL_452904, EPI_ISL_452905, EPI_ISL_452906, EPI_ISL_452907, EPI_ISL_452908, EPI_ISL_452909, EPI_ISL_452910, EPI_ISL_452911, EPI_ISL_452912, EPI_ISL_452913, EPI_ISL_452914, EPI_ISL_452915, EPI_ISL_452916, EPI_ISL_452917, EPI_ISL_452918, EPI_ISL_452919, EPI_ISL_452920, EPI_ISL_452921, EPI_ISL_452922, EPI_ISL_452923, EPI_ISL_452924, EPI_ISL_452925, EPI_ISL_452926, EPI_ISL_452927, EPI_ISL_452928, EPI_ISL_452929, EPI_ISL_452930, EPI_ISL_452931, EPI_ISL_452932, EPI_ISL_452933, EPI_ISL_452934, EPI_ISL_452935, EPI_ISL_452936, EPI_ISL_452937, EPI_ISL_452938, EPI_ISL_452939, EPI_ISL_452940, EPI_ISL_452941, EPI_ISL_452942, EPI_ISL_452943, EPI_ISL_452944, EPI_ISL_452945, EPI_ISL_452946, EPI_ISL_452947, EPI_ISL_452948, EPI_ISL_452949, EPI_ISL_452950, EPI_ISL_452951, EPI_ISL_452952, EPI_ISL_452953, EPI_ISL_452954, EPI_ISL_452955, EPI_ISL_452956, EPI_ISL_452957, EPI_ISL_452958, EPI_ISL_452959, EPI_ISL_452960, EPI_ISL_452961, EPI_ISL_452962, EPI_ISL_452963, EPI_ISL_452964, EPI_ISL_452965, EPI_ISL_452966, EPI_ISL_452967, EPI_ISL_452968, EPI_ISL_452969, EPI_ISL_452970, EPI_ISL_452971, EPI_ISL_452972, EPI_ISL_452973, EPI_ISL_452974, EPI_ISL_452975, EPI_ISL_452976, EPI_ISL_452977, EPI_ISL_452978, EPI_ISL_452979, EPI_ISL_452980, EPI_ISL_452981, EPI_ISL_452982, EPI_ISL_452983, EPI_ISL_452984, EPI_ISL_452985, EPI_ISL_452986, EPI_ISL_452987, EPI_ISL_452988, EPI_ISL_452989, EPI_ISL_452990, EPI_ISL_452991, EPI_ISL_452992, EPI_ISL_452993, EPI_ISL_452994, EPI_ISL_452995, EPI_ISL_452996, EPI_ISL_452997, EPI_ISL_452998, EPI_ISL_452999, EPI_ISL_453000, EPI_ISL_453001, EPI_ISL_453002, EPI_ISL_453003, EPI_ISL_453004, EPI_ISL_453005 | | | |
| see above | Department of Pathology, University of Cambridge | COVID-19 Genomics UK (COG-UK) Consortium | Luke W Meredith, M. Estée Török , Myra Hosmillo, William L. Hamilton, Martin D. Curran, Theresa Feltwell, Grant Hall, Anna Yakovleva, Fahad A Khokhar, Charlotte J. Houldcroft, Laura G Caller, Aminu S. Jahun, Sarah L. Caddy, Ian Goodfellow |
| EPI_ISL_453006, EPI_ISL_453007, EPI_ISL_453008, EPI_ISL_453009, EPI_ISL_453010, EPI_ISL_453011, EPI_ISL_453012, EPI_ISL_453013, EPI_ISL_453014, EPI_ISL_453015, EPI_ISL_453016, EPI_ISL_453017, EPI_ISL_453018, EPI_ISL_453019, EPI_ISL_453020, EPI_ISL_453021, EPI_ISL_453022, EPI_ISL_453023, EPI_ISL_453024, EPI_ISL_453025, EPI_ISL_453026, EPI_ISL_453027, EPI_ISL_453028, EPI_ISL_453029, EPI_ISL_453030, EPI_ISL_453031, EPI_ISL_453032, EPI_ISL_453033, EPI_ISL_453034, EPI_ISL_453035, EPI_ISL_453036, EPI_ISL_453037, EPI_ISL_453038, EPI_ISL_453039, EPI_ISL_453040, EPI_ISL_453041, EPI_ISL_453042, EPI_ISL_453043, EPI_ISL_453044, EPI_ISL_453045, EPI_ISL_453046, EPI_ISL_453047, EPI_ISL_453048, EPI_ISL_453049, EPI_ISL_453050, EPI_ISL_453051, EPI_ISL_453052, EPI_ISL_453053, EPI_ISL_453054, EPI_ISL_453055, EPI_ISL_453056, EPI_ISL_453057, EPI_ISL_453058, EPI_ISL_453059, EPI_ISL_453060, EPI_ISL_453061, EPI_ISL_453062, EPI_ISL_453063, EPI_ISL_453064, EPI_ISL_453065, EPI_ISL_453066, EPI_ISL_453067, EPI_ISL_453068, EPI_ISL_453069, EPI_ISL_453070, EPI_ISL_453071, EPI_ISL_453072, EPI_ISL_453073, EPI_ISL_453074, EPI_ISL_453075, EPI_ISL_453076, EPI_ISL_453077, EPI_ISL_453078, EPI_ISL_453079, EPI_ISL_453080, EPI_ISL_453081, EPI_ISL_453082, EPI_ISL_453083, EPI_ISL_453084, EPI_ISL_453085, EPI_ISL_453086, EPI_ISL_453087, EPI_ISL_453088, EPI_ISL_453089, EPI_ISL_453090, EPI_ISL_453091, EPI_ISL_453092, EPI_ISL_453093, EPI_ISL_453094, EPI_ISL_453095, EPI_ISL_453096, EPI_ISL_453097, EPI_ISL_453098 | | | |
| see above | West of Scotland Specialist Virology Centre, NHSGGC / MRC-University of Glasgow Centre for Virus Research | COVID-19 Genomics UK (COG-UK) Consortium | Ana da Silva Filipe, Natasha Johnson, Kathy Smollett, Daniel Mair, Stephen Carmichael, Lily Tong, Jenna Nichols, Elihu Aranday-Cortes, Kirstyn Brunker, Yasmin Parr, Kyriaki Nomikou; Sarah McDonald, Marc Niebel, Patawee Asamaphan; Richard Orton, Joseph Hughes, Sreenu Vattipally, David L Robertson; Alasdair MacLean, Rory Gunson; Kathy Li, Natasha Jesudason, Rajiv Shah, James Shepherd, Antonia Ho, Emma Thomson |
| EPI_ISL_453099, EPI_ISL_453100, EPI_ISL_453101, EPI_ISL_453102, EPI_ISL_453103, EPI_ISL_453104, EPI_ISL_453105, EPI_ISL_453106, EPI_ISL_453107, EPI_ISL_453108, EPI_ISL_453109, EPI_ISL_453110, EPI_ISL_453111, EPI_ISL_453112, EPI_ISL_453113, EPI_ISL_453114, EPI_ISL_453115, EPI_ISL_453116, EPI_ISL_453117, EPI_ISL_453118, EPI_ISL_453119, EPI_ISL_453120, EPI_ISL_453121, EPI_ISL_453122, EPI_ISL_453123, EPI_ISL_453124, EPI_ISL_453125, EPI_ISL_453126, EPI_ISL_453127, EPI_ISL_453128, EPI_ISL_453129, EPI_ISL_453130, EPI_ISL_453131, EPI_ISL_453132, EPI_ISL_453133, EPI_ISL_453134, EPI_ISL_453135, EPI_ISL_453136, EPI_ISL_453137, EPI_ISL_453138, EPI_ISL_453139, EPI_ISL_453140, EPI_ISL_453141, EPI_ISL_453142, EPI_ISL_453143, EPI_ISL_453144, EPI_ISL_453145, EPI_ISL_453146, EPI_ISL_453147, EPI_ISL_453148, EPI_ISL_453149, EPI_ISL_453150, EPI_ISL_453151, EPI_ISL_453152, EPI_ISL_453153, EPI_ISL_453154, EPI_ISL_453155, EPI_ISL_453156, EPI_ISL_453157, EPI_ISL_453158, EPI_ISL_453159, EPI_ISL_453160, EPI_ISL_453161, EPI_ISL_453162, EPI_ISL_453163, EPI_ISL_453164, EPI_ISL_453165, EPI_ISL_453166, EPI_ISL_453167, EPI_ISL_453168, EPI_ISL_453169, EPI_ISL_453170, EPI_ISL_453171, EPI_ISL_453172, EPI_ISL_453173, EPI_ISL_453174, EPI_ISL_453175, EPI_ISL_453176, EPI_ISL_453177, EPI_ISL_453178, EPI_ISL_453179, EPI_ISL_453180, EPI_ISL_453181, EPI_ISL_453182, EPI_ISL_453183, EPI_ISL_453184, EPI_ISL_453185, EPI_ISL_453186, EPI_ISL_453187, EPI_ISL_453188, EPI_ISL_453189, EPI_ISL_453190, EPI_ISL_453191, EPI_ISL_453192, EPI_ISL_453193, EPI_ISL_453194 | | | |
| see above | Virology Department, Royal Infirmary of Edinburgh, NHS Lothian / School of Biological Sciences, University of Edinburgh / Institute of Genetics and Molecular Medicine, University of Edinburgh | COVID-19 Genomics UK (COG-UK) Consortium | McHugh M, Dewar R, Rooke S, Gallagher M, Balcaza C, O'Toole Á, Scher E, Hill V, McCrone JT, Colquhoun R, Yu X, Jackson B, Rambaut A, Williams TC, Templeton K |
| EPI_ISL_453195, EPI_ISL_453196, EPI_ISL_453197, EPI_ISL_453198, EPI_ISL_453199, EPI_ISL_453200, EPI_ISL_453201, EPI_ISL_453202, EPI_ISL_453203, EPI_ISL_453204, EPI_ISL_453205, EPI_ISL_453206, EPI_ISL_453207, EPI_ISL_453208, EPI_ISL_453209, EPI_ISL_453210, EPI_ISL_453211, EPI_ISL_453212, EPI_ISL_453213, EPI_ISL_453214, EPI_ISL_453215, EPI_ISL_453216, EPI_ISL_453217, EPI_ISL_453218, EPI_ISL_453219, EPI_ISL_453220, EPI_ISL_453221, EPI_ISL_453222, EPI_ISL_453223, EPI_ISL_453224, EPI_ISL_453225, EPI_ISL_453226, EPI_ISL_453227, EPI_ISL_453228, EPI_ISL_453229, EPI_ISL_453230, EPI_ISL_453231, EPI_ISL_453232, EPI_ISL_453233, EPI_ISL_453234, EPI_ISL_453235, EPI_ISL_453236, EPI_ISL_453237, EPI_ISL_453238, EPI_ISL_453239, EPI_ISL_453240, EPI_ISL_453241, EPI_ISL_453242, EPI_ISL_453243, EPI_ISL_453244, EPI_ISL_453245, EPI_ISL_453246, EPI_ISL_453247, EPI_ISL_453248, EPI_ISL_453249, EPI_ISL_453250, EPI_ISL_453251, EPI_ISL_453252, EPI_ISL_453253, EPI_ISL_453254, EPI_ISL_453255, EPI_ISL_453256, EPI_ISL_453257, EPI_ISL_453258, EPI_ISL_453259, EPI_ISL_453260, EPI_ISL_453261, EPI_ISL_453262, EPI_ISL_453263, EPI_ISL_453264, EPI_ISL_453265, EPI_ISL_453266, EPI_ISL_453267, EPI_ISL_453268, EPI_ISL_453269, EPI_ISL_453270, EPI_ISL_453271, EPI_ISL_453272, EPI_ISL_453273, EPI_ISL_453274, EPI_ISL_453275, EPI_ISL_453276, EPI_ISL_453277, EPI_ISL_453278, EPI_ISL_453279, EPI_ISL_453280, EPI_ISL_453281, EPI_ISL_453282, EPI_ISL_453283, EPI_ISL_453284, EPI_ISL_453285, EPI_ISL_453286, EPI_ISL_453287, EPI_ISL_453288, EPI_ISL_453289, EPI_ISL_453290, EPI_ISL_453291, EPI_ISL_453292, EPI_ISL_453293, EPI_ISL_453294, EPI_ISL_453295, EPI_ISL_453296, EPI_ISL_453297, EPI_ISL_453298, EPI_ISL_453299, EPI_ISL_453300, EPI_ISL_453301, EPI_ISL_453302, EPI_ISL_453303, EPI_ISL_453304, EPI_ISL_453305, EPI_ISL_453306, EPI_ISL_453307, EPI_ISL_453308, EPI_ISL_453309, EPI_ISL_453310, EPI_ISL_453311, EPI_ISL_453312, EPI_ISL_453313, EPI_ISL_453314, EPI_ISL_453315, EPI_ISL_453316, EPI_ISL_453317, EPI_ISL_453318, EPI_ISL_453319, EPI_ISL_453320, EPI_ISL_453321, EPI_ISL_453322, EPI_ISL_453323, EPI_ISL_453324, EPI_ISL_453325, EPI_ISL_453326, EPI_ISL_453327, EPI_ISL_453328, EPI_ISL_453329, EPI_ISL_453330, EPI_ISL_453331, EPI_ISL_453332, EPI_ISL_453333, EPI_ISL_453334, EPI_ISL_453335, EPI_ISL_453336, EPI_ISL_453337, EPI_ISL_453338, EPI_ISL_453339, EPI_ISL_453340, EPI_ISL_453341, EPI_ISL_453342, EPI_ISL_453343, EPI_ISL_453344, EPI_ISL_453345, EPI_ISL_453346, EPI_ISL_453347, EPI_ISL_453348, EPI_ISL_453349, EPI_ISL_453350, EPI_ISL_453351, EPI_ISL_453352, EPI_ISL_453353, EPI_ISL_453354, EPI_ISL_453355, EPI_ISL_453356, EPI_ISL_453357, EPI_ISL_453358, EPI_ISL_453359, EPI_ISL_453360, EPI_ISL_453361, EPI_ISL_453362, EPI_ISL_453363, EPI_ISL_453364, EPI_ISL_453365, EPI_ISL_453366, EPI_ISL_453367, EPI_ISL_453368, EPI_ISL_453369, EPI_ISL_453370, EPI_ISL_453371, EPI_ISL_453372, EPI_ISL_453373, EPI_ISL_453374, EPI_ISL_453375, EPI_ISL_453376, EPI_ISL_453377, EPI_ISL_453378, EPI_ISL_453379, EPI_ISL_453380, EPI_ISL_453381, EPI_ISL_453382, EPI_ISL_453383, EPI_ISL_453384, EPI_ISL_453385, EPI_ISL_453386, EPI_ISL_453387, EPI_ISL_453388, EPI_ISL_453389, EPI_ISL_453390, EPI_ISL_453391, EPI_ISL_453392, EPI_ISL_453393, EPI_ISL_453394, EPI_ISL_453395, EPI_ISL_453396, EPI_ISL_453397, EPI_ISL_453398, EPI_ISL_453399, EPI_ISL_453400, EPI_ISL_453401, EPI_ISL_453402, EPI_ISL_453403, EPI_ISL_453404, EPI_ISL_453405, EPI_ISL_453406, EPI_ISL_453407, EPI_ISL_453408, EPI_ISL_453409, EPI_ISL_453410, EPI_ISL_453411, EPI_ISL_453412, EPI_ISL_453413, EPI_ISL_453414, EPI_ISL_453415, EPI_ISL_453416, EPI_ISL_453417, EPI_ISL_453418, EPI_ISL_453419, EPI_ISL_453420, EPI_ISL_453421, EPI_ISL_453422, EPI_ISL_453423, EPI_ISL_453424, EPI_ISL_453425, EPI_ISL_453426, EPI_ISL_453427, EPI_ISL_453428, EPI_ISL_453429, EPI_ISL_453430, EPI_ISL_453431, EPI_ISL_453432, EPI_ISL_453433, EPI_ISL_453434, EPI_ISL_453435, EPI_ISL_453436, EPI_ISL_453437, EPI_ISL_453438, EPI_ISL_453439, EPI_ISL_453440, EPI_ISL_453441, EPI_ISL_453442, EPI_ISL_453443, EPI_ISL_453444, EPI_ISL_453445, EPI_ISL_453446, EPI_ISL_453447, EPI_ISL_453448, EPI_ISL_453449, EPI_ISL_453450, EPI_ISL_453451, EPI_ISL_453452, EPI_ISL_453453, EPI_ISL_453454, EPI_ISL_453455, EPI_ISL_453456, EPI_ISL_453457, EPI_ISL_453458, EPI_ISL_453459 | | | |
| see above | Liverpool Clinical Laboratories | COVID-19 Genomics UK (COG-UK) Consortium | Sam Haldenby, Anita Lucaci, Steve Paterson, Julian Hiscox, Alistair Darby, M Almsaud, A Alrezaihi, Muhannad Alruwaili, Stuart D Armstrong, Jones Benjamin , Eleanor G Bentley, Anu Chawla, Jordan J Clark, Angela Cowell, Richard Eccles, Isabel García-Dorival, Matthew Gemmell, Alessandro Gerada, PKF Gilmore, Richard Gregory, Ximeng Han, Catherine Hartley, Margaret Hughes, Miren Iturriza-Gomara, James Johnson, L Luu, Jenifer Manson , Charlotte Nelson, Elaine O'Toole, Cassie Olateju, Rebekah Penrice-Randal , Lucille Rainbow, N.P Randle, Trevor Ian Robinson, Parul Sharma, Ghada T Shawli, James P Stewart , Neil Swainston, Ecaterina Vamos, Joanne Watts, Mark Whitehead |
| EPI_ISL_453460, EPI_ISL_453461, EPI_ISL_453462, EPI_ISL_453463, EPI_ISL_453464 | University College London, Great Ormond Street Hospital for Children NHS Foundation Trust, Imperial College Healthcare NHS Trust | COVID-19 Genomics UK (COG-UK) Consortium | Sergi Castellano, Rachel Williams, Mark Kristiansen, Paola Resende Silva, Sunando Roy, Tony Brooks, Helena Tutill, Paola Niola, Patricia Dyal, Charlotte Williams, Leysa Forrest, Yasmin Panchbhaya, Jacqueline Findlay, Sam Weeks, Julianne Brown, Kathryn Harris, Paul Randell, James Price, Alison Holmes, Judith Breuer |
| EPI_ISL_453465, EPI_ISL_453466, EPI_ISL_453467, EPI_ISL_453468, EPI_ISL_453469, EPI_ISL_453470, EPI_ISL_453471, EPI_ISL_453472, EPI_ISL_453473, EPI_ISL_453474, EPI_ISL_453475, EPI_ISL_453476, EPI_ISL_453477, EPI_ISL_453478, EPI_ISL_453479, EPI_ISL_453480, EPI_ISL_453481, EPI_ISL_453482, EPI_ISL_453483, EPI_ISL_453484, EPI_ISL_453485 | | | |
| see above | Regional Virus Laboratory, Belfast Health and Social Care Trust | COVID-19 Genomics UK (COG-UK) Consortium | Conall McCaughey, James McKenna, Tanya Curran, Susan Feeney, Alison Watt, Ciara Cox, Mairead Connor, Zoltan Molnar, David Simpson, Derek Fairley |
| EPI_ISL_453486, EPI_ISL_453487, EPI_ISL_453488, EPI_ISL_453489, EPI_ISL_453490, EPI_ISL_453491, EPI_ISL_453492, EPI_ISL_453493, EPI_ISL_453494, EPI_ISL_453495, EPI_ISL_453496, EPI_ISL_453497, EPI_ISL_453498, EPI_ISL_453499, EPI_ISL_453500, EPI_ISL_453501, EPI_ISL_453502, EPI_ISL_453503, EPI_ISL_453504, EPI_ISL_453505, EPI_ISL_453506, EPI_ISL_453507, EPI_ISL_453508, EPI_ISL_453509, EPI_ISL_453510, EPI_ISL_453511, EPI_ISL_453512, EPI_ISL_453513, EPI_ISL_453514, EPI_ISL_453515, EPI_ISL_453516, EPI_ISL_453517, EPI_ISL_453518, EPI_ISL_453519, EPI_ISL_453520, EPI_ISL_453521, EPI_ISL_453522, EPI_ISL_453523, EPI_ISL_453524, EPI_ISL_453525, EPI_ISL_453526, EPI_ISL_453527, EPI_ISL_453528, EPI_ISL_453529, EPI_ISL_453530, EPI_ISL_453531, EPI_ISL_453532, EPI_ISL_453533, EPI_ISL_453534, EPI_ISL_453535, EPI_ISL_453536, EPI_ISL_453537, EPI_ISL_453538, EPI_ISL_453539, EPI_ISL_453540, EPI_ISL_453541, EPI_ISL_453542 | | | |
| see above | Northumbria University / South Tees Hospitals NHS Foundation Trust / North Cumbria Integrated Care NHS Foundation Trust / North Tees and Hartlepool NHS Foundation Trust / Newcastle Hospitals NHS Foundation Trust | COVID-19 Genomics UK (COG-UK) Consortium | Darren L Smith,Andrew Nelson,Matthew Bashton,Greg R Young,Joshua Loh,John Allan,Mohammad A Tariq,Giles S Holt,Gary Black,Wen C Yew,Lynn Dover ,Paul Baker,Steve Liggett,Sarah Essex,Jane Greenaway ,Debra Padgett,Clive Graham,Garren Scott,Edward Barton ,Emma Swindells ,Brendan Payne,Jennifer Collins,Yusri Taha,Gary Eltringham |
| EPI_ISL_453543 | Northumbria University / South Tees Hospitals NHS Foundation Trust / North Cumbria Integrated Care NHS Foundation Trust / North Tees and Hartlepool NHS Foundation Trust / Newcastle Hospitals NHS Foundation Trust | Northumbria University | Darren L Smith,Andrew Nelson,Matthew Bashton,Greg R Young,Joshua Loh,John Allan,Mohammad A Tariq,Giles S Holt,Gary Black,Wen C Yew,Lynn Dover ,Paul Baker,Steve Liggett,Sarah Essex,Jane Greenaway ,Debra Padgett,Clive Graham,Garren Scott,Edward Barton ,Emma Swindells ,Brendan Payne,Jennifer Collins,Yusri Taha,Gary Eltringham |
| EPI_ISL_453544 | Northumbria University / South Tees Hospitals NHS Foundation Trust / North Cumbria Integrated Care NHS Foundation Trust / North Tees and Hartlepool NHS Foundation Trust / Newcastle Hospitals NHS Foundation Trust | COVID-19 Genomics UK (COG-UK) Consortium | Darren L Smith,Andrew Nelson,Matthew Bashton,Greg R Young,Joshua Loh,John Allan,Mohammad A Tariq,Giles S Holt,Gary Black,Wen C Yew,Lynn Dover ,Paul Baker,Steve Liggett,Sarah Essex,Jane Greenaway ,Debra Padgett,Clive Graham,Garren Scott,Edward Barton ,Emma Swindells ,Brendan Payne,Jennifer Collins,Yusri Taha,Gary Eltringham |
| EPI_ISL_453545, EPI_ISL_453546, EPI_ISL_453547, EPI_ISL_453548, EPI_ISL_453549, EPI_ISL_453550, EPI_ISL_453551, EPI_ISL_453552, EPI_ISL_453553, EPI_ISL_453554, EPI_ISL_453555, EPI_ISL_453556, EPI_ISL_453557, EPI_ISL_453558, EPI_ISL_453559, EPI_ISL_453560, EPI_ISL_453561, EPI_ISL_453562, EPI_ISL_453563, EPI_ISL_453564, EPI_ISL_453565, EPI_ISL_453566, EPI_ISL_453567, EPI_ISL_453568, EPI_ISL_453569, EPI_ISL_453570, EPI_ISL_453571, EPI_ISL_453572, EPI_ISL_453573, EPI_ISL_453574, EPI_ISL_453575, EPI_ISL_453576, EPI_ISL_453577, EPI_ISL_453578, EPI_ISL_453579, EPI_ISL_453580, EPI_ISL_453581, EPI_ISL_453582, EPI_ISL_453583, EPI_ISL_453584, EPI_ISL_453585, EPI_ISL_453586, EPI_ISL_453587, EPI_ISL_453588, EPI_ISL_453589, EPI_ISL_453590, EPI_ISL_453591, EPI_ISL_453592, EPI_ISL_453593, EPI_ISL_453594, EPI_ISL_453595, EPI_ISL_453596, EPI_ISL_453597, EPI_ISL_453598, EPI_ISL_453599, EPI_ISL_453600, EPI_ISL_453601, EPI_ISL_453602, EPI_ISL_453603, EPI_ISL_453604, EPI_ISL_453605, EPI_ISL_453606, EPI_ISL_453607, EPI_ISL_453608, EPI_ISL_453609, EPI_ISL_453610, EPI_ISL_453611, EPI_ISL_453612, EPI_ISL_453613 | | | |
| see above | Quadram Institute Bioscience | COVID-19 Genomics UK (COG-UK) Consortium | Dave J. Baker, Gemma L. Kay, Alp Aydin, Thanh Le-Viet, Steven Rudder, Ana P. Tedim, Anastasia Kolyva, Maria Diaz, Leonardo de Oliveira Martins, Nabil-Fareed Alikhan, Lizzie Meadows, Rachael Stanley, Ngozi Elumogo, Muhammed Yasir, Nicholas M. Thomson, Alexander J Trotter, Rachel Gilroy, Samuel Bloomfield, Claire Stuart, Andrew Bell, Reenesh Prakash, Samir Dervisevic, Alison E. Mather, John Wain, Mark Webber, Andrew J. Page, Justin O'Grady |
| EPI_ISL_453614, EPI_ISL_453615, EPI_ISL_453616, EPI_ISL_453617, EPI_ISL_453618, EPI_ISL_453619, EPI_ISL_453620, EPI_ISL_453621, EPI_ISL_453622, EPI_ISL_453623, EPI_ISL_453624, EPI_ISL_453625, EPI_ISL_453626, EPI_ISL_453627, EPI_ISL_453628, EPI_ISL_453629, EPI_ISL_453630, EPI_ISL_453631, EPI_ISL_453632, EPI_ISL_453633, EPI_ISL_453634, EPI_ISL_453635, EPI_ISL_453636, EPI_ISL_453637, EPI_ISL_453638, EPI_ISL_453639, EPI_ISL_453640, EPI_ISL_453641, EPI_ISL_453642, EPI_ISL_453643, EPI_ISL_453644, EPI_ISL_453645, EPI_ISL_453646, EPI_ISL_453647, EPI_ISL_453648, EPI_ISL_453649, EPI_ISL_453650, EPI_ISL_453651, EPI_ISL_453652, EPI_ISL_453653, EPI_ISL_453654, EPI_ISL_453655, EPI_ISL_453656, EPI_ISL_453657, EPI_ISL_453658, EPI_ISL_453659, EPI_ISL_453660, EPI_ISL_453661, EPI_ISL_453662, EPI_ISL_453663, EPI_ISL_453664 | | | |

| | | | |
|---|---|---|---|
| see above | Queens Medical Centre, Clinical Microbiology Department / DeepSeq Nottingham | COVID-19 Genomics UK (COG-UK) Consortium | Gemma Clark, Wendy Smith, Manjinder Khakh, Hannah Howson-Wells, Jonathan Ball, Patrick McClure, Joseph Chappell, Theocharis Tsoleridis, Nadine Holmes, Matthew Carlisle, Christopher Moore, Fei Sang, Johnny Debebe, Victoria Wright, Matthew Loose |
| EPI_ISL_453665 | Centre for Enzyme Innovation, University of Portsmouth / Translational Research Laboratory, Portsmouth Hospitals NHS Trust | University of Portsmouth | Angela Beckett,,Yann Bourgeois,,Garry Scarlett,,Sharon Glaysher,,Scott Elliott,,Kelly Bicknell,,Robert Impey,,Allyson Lloyd,,Sarah Wyllie,,Ethan Butcher,,Anoop Chauhan,,Samuel Robson |
| EPI_ISL_453666, EPI_ISL_453667, EPI_ISL_453668, EPI_ISL_453669, EPI_ISL_453670, EPI_ISL_453671, EPI_ISL_453672, EPI_ISL_453673, EPI_ISL_453674, EPI_ISL_453675, EPI_ISL_453676, EPI_ISL_453677, EPI_ISL_453678, EPI_ISL_453679, EPI_ISL_453680, EPI_ISL_453681, EPI_ISL_453682, EPI_ISL_453683, EPI_ISL_453684, EPI_ISL_453685, EPI_ISL_453686, EPI_ISL_453687, EPI_ISL_453688, EPI_ISL_453689, EPI_ISL_453690, EPI_ISL_453691, EPI_ISL_453692, EPI_ISL_453693, EPI_ISL_453694, EPI_ISL_453695 | | | |
| see above | Centre for Enzyme Innovation, University of Portsmouth / Translational Research Laboratory, Portsmouth Hospitals NHS Trust | COVID-19 Genomics UK (COG-UK) Consortium | Angela Beckett,,Yann Bourgeois,,Garry Scarlett,,Sharon Glaysher,,Scott Elliott,,Kelly Bicknell,,Robert Impey,,Allyson Lloyd,,Sarah Wyllie,,Ethan Butcher,,Anoop Chauhan,,Samuel Robson |
| EPI_ISL_453696, EPI_ISL_453697, EPI_ISL_453698, EPI_ISL_453699, EPI_ISL_453700, EPI_ISL_453701, EPI_ISL_453702, EPI_ISL_453703, EPI_ISL_453704, EPI_ISL_453705, EPI_ISL_453706, EPI_ISL_453707, EPI_ISL_453708, EPI_ISL_453709, EPI_ISL_453710, EPI_ISL_453711, EPI_ISL_453712, EPI_ISL_453713, EPI_ISL_453714, EPI_ISL_453715, EPI_ISL_453716, EPI_ISL_453717, EPI_ISL_453718, EPI_ISL_453719, EPI_ISL_453720, EPI_ISL_453721, EPI_ISL_453722, EPI_ISL_453723, EPI_ISL_453724, EPI_ISL_453725, EPI_ISL_453726, EPI_ISL_453727, EPI_ISL_453728, EPI_ISL_453729, EPI_ISL_453730, EPI_ISL_453731, EPI_ISL_453732, EPI_ISL_453733, EPI_ISL_453734, EPI_ISL_453735, EPI_ISL_453736, EPI_ISL_453737, EPI_ISL_453738, EPI_ISL_453739, EPI_ISL_453740, EPI_ISL_453741, EPI_ISL_453742, EPI_ISL_453743, EPI_ISL_453744, EPI_ISL_453745, EPI_ISL_453746, EPI_ISL_453747, EPI_ISL_453748, EPI_ISL_453749, EPI_ISL_453750, EPI_ISL_453751, EPI_ISL_453752, EPI_ISL_453753, EPI_ISL_453754, EPI_ISL_453755, EPI_ISL_453756, EPI_ISL_453757, EPI_ISL_453758, EPI_ISL_453759, EPI_ISL_453760, EPI_ISL_453761, EPI_ISL_453762, EPI_ISL_453763, EPI_ISL_453764, EPI_ISL_453765, EPI_ISL_453766, EPI_ISL_453767, EPI_ISL_453768, EPI_ISL_453769, EPI_ISL_453770, EPI_ISL_453771, EPI_ISL_453772, EPI_ISL_453773, EPI_ISL_453774, EPI_ISL_453775, EPI_ISL_453776, EPI_ISL_453777, EPI_ISL_453778 | | | |
| see above | Virology Department, Sheffield Teaching Hospitals NHS Foundation Trust/Department of Infection, Immunity and Cardiovascular Disease, The Medical School, University of Sheffield | COVID-19 Genomics UK (COG-UK) Consortium | Thushan de Silva, Matthew Parker, Nikki Smith, Adri Angyal, Rebecca Brown, Luke Green, Rachel Tucker, Paul Parsons, Danielle Groves, Katie Johnson, Laura Carrilero, Alex Keeley, Dave Partridge, Matthew Wyles, Benjamin Lindsey, Mehmet Yavuz, Mohammad Raza, Cariad Evans |
| EPI_ISL_453813, EPI_ISL_453814, EPI_ISL_453815, EPI_ISL_453816, EPI_ISL_453817, EPI_ISL_453818, EPI_ISL_453819, EPI_ISL_453820, EPI_ISL_453821, EPI_ISL_453822, EPI_ISL_453823, EPI_ISL_453824, EPI_ISL_453825, EPI_ISL_453826, EPI_ISL_453827, EPI_ISL_453828, EPI_ISL_453829, EPI_ISL_453830, EPI_ISL_453831, EPI_ISL_453832, EPI_ISL_453833, EPI_ISL_453834, EPI_ISL_453835, EPI_ISL_453836, EPI_ISL_453837, EPI_ISL_453838, EPI_ISL_453839, EPI_ISL_453840, EPI_ISL_453841, EPI_ISL_453842, EPI_ISL_453843, EPI_ISL_453844, EPI_ISL_453845, EPI_ISL_453846, EPI_ISL_453847, EPI_ISL_453848, EPI_ISL_453849, EPI_ISL_453850, EPI_ISL_453851, EPI_ISL_453852, EPI_ISL_453853, EPI_ISL_453854, EPI_ISL_453855, EPI_ISL_453856, EPI_ISL_453857, EPI_ISL_453858, EPI_ISL_453859, EPI_ISL_453860, EPI_ISL_453861, EPI_ISL_453862, EPI_ISL_453863, EPI_ISL_453864, EPI_ISL_453865, EPI_ISL_453866, EPI_ISL_453867, EPI_ISL_453868, EPI_ISL_453869, EPI_ISL_453870, EPI_ISL_453871, EPI_ISL_453872, EPI_ISL_453873, EPI_ISL_453874, EPI_ISL_453875, EPI_ISL_453876, EPI_ISL_453877, EPI_ISL_453878, EPI_ISL_453879, EPI_ISL_453880, EPI_ISL_453881, EPI_ISL_453882, EPI_ISL_453883, EPI_ISL_453884, EPI_ISL_453885, EPI_ISL_453886, EPI_ISL_453887, EPI_ISL_453888, EPI_ISL_453889, EPI_ISL_453890, EPI_ISL_453891, EPI_ISL_453892, EPI_ISL_453893, EPI_ISL_453894, EPI_ISL_453895, EPI_ISL_453896, EPI_ISL_453897, EPI_ISL_453898, EPI_ISL_453899, EPI_ISL_453900, EPI_ISL_453901, EPI_ISL_453902, EPI_ISL_453903, EPI_ISL_453904, EPI_ISL_453905, EPI_ISL_453906, EPI_ISL_453907, EPI_ISL_453908, EPI_ISL_453909, EPI_ISL_453910, EPI_ISL_453911, EPI_ISL_453912, EPI_ISL_453913, EPI_ISL_453914, EPI_ISL_453915, EPI_ISL_453916, EPI_ISL_453917, EPI_ISL_453918, EPI_ISL_453919, EPI_ISL_453920, EPI_ISL_453921, EPI_ISL_453922, EPI_ISL_453923, EPI_ISL_453924, EPI_ISL_453925, EPI_ISL_453926, EPI_ISL_453927, EPI_ISL_453928, EPI_ISL_453929, EPI_ISL_453930, EPI_ISL_453931, EPI_ISL_453932, EPI_ISL_453933, EPI_ISL_453934, EPI_ISL_453935, EPI_ISL_453936, EPI_ISL_453937, EPI_ISL_453938, EPI_ISL_453939, EPI_ISL_453940, EPI_ISL_453941, EPI_ISL_453942, EPI_ISL_453943, EPI_ISL_453944, EPI_ISL_453945, EPI_ISL_453946, EPI_ISL_453947, EPI_ISL_453948, EPI_ISL_453949, EPI_ISL_453950, EPI_ISL_453951, EPI_ISL_453952, EPI_ISL_453953, EPI_ISL_453954, EPI_ISL_453955, EPI_ISL_453956, EPI_ISL_453957, EPI_ISL_453958, EPI_ISL_453959, EPI_ISL_453960, EPI_ISL_453961, EPI_ISL_453962, EPI_ISL_453963, EPI_ISL_453964, EPI_ISL_453965, EPI_ISL_453966, EPI_ISL_453967, EPI_ISL_453968, EPI_ISL_453969, EPI_ISL_453970, EPI_ISL_453971, EPI_ISL_453972, EPI_ISL_453973, EPI_ISL_453974, EPI_ISL_453975, EPI_ISL_453976, EPI_ISL_453977, EPI_ISL_453978, EPI_ISL_453979, EPI_ISL_453980, EPI_ISL_453981, EPI_ISL_453982, EPI_ISL_453983, EPI_ISL_453984, EPI_ISL_453985, EPI_ISL_453986, EPI_ISL_453987, EPI_ISL_453988, EPI_ISL_453989, EPI_ISL_453990, EPI_ISL_453991, EPI_ISL_453992, EPI_ISL_453993, EPI_ISL_453994, EPI_ISL_453995, EPI_ISL_453996, EPI_ISL_453997, EPI_ISL_453998, EPI_ISL_453999, EPI_ISL_454000, EPI_ISL_454001, EPI_ISL_454002, EPI_ISL_454003, EPI_ISL_454004, EPI_ISL_454005, EPI_ISL_454006, EPI_ISL_454007, EPI_ISL_454008, EPI_ISL_454009, EPI_ISL_454010, EPI_ISL_454011, EPI_ISL_454012, EPI_ISL_454013, EPI_ISL_454014, EPI_ISL_454015, EPI_ISL_454016, EPI_ISL_454017, EPI_ISL_454018, EPI_ISL_454019, EPI_ISL_454020, EPI_ISL_454021, EPI_ISL_454022, EPI_ISL_454023, EPI_ISL_454024, EPI_ISL_454025, EPI_ISL_454026, EPI_ISL_454027, EPI_ISL_454028, EPI_ISL_454029, EPI_ISL_454030, EPI_ISL_454031, EPI_ISL_454032, EPI_ISL_454033, EPI_ISL_454034, EPI_ISL_454035, EPI_ISL_454036, EPI_ISL_454037, EPI_ISL_454038, EPI_ISL_454039, EPI_ISL_454040, EPI_ISL_454041, EPI_ISL_454042, EPI_ISL_454043, EPI_ISL_454044, EPI_ISL_454045, EPI_ISL_454046, EPI_ISL_454047, EPI_ISL_454048, EPI_ISL_454049, EPI_ISL_454050, EPI_ISL_454051, EPI_ISL_454052, EPI_ISL_454053, EPI_ISL_454054, EPI_ISL_454055, EPI_ISL_454056, EPI_ISL_454057, EPI_ISL_454058, EPI_ISL_454059, EPI_ISL_454060, EPI_ISL_454061, EPI_ISL_454062, EPI_ISL_454063, EPI_ISL_454064, EPI_ISL_454065, EPI_ISL_454066, EPI_ISL_454067, EPI_ISL_454068, EPI_ISL_454069, EPI_ISL_454070, EPI_ISL_454071, EPI_ISL_454072, EPI_ISL_454073, EPI_ISL_454074, EPI_ISL_454075, EPI_ISL_454076, EPI_ISL_454077, EPI_ISL_454078, EPI_ISL_454079, EPI_ISL_454080, EPI_ISL_454081, EPI_ISL_454082, EPI_ISL_454083, EPI_ISL_454084, EPI_ISL_454085, EPI_ISL_454086, EPI_ISL_454087, EPI_ISL_454088, EPI_ISL_454089, EPI_ISL_454090, EPI_ISL_454091, EPI_ISL_454092, EPI_ISL_454093, EPI_ISL_454094, EPI_ISL_454095, EPI_ISL_454096, EPI_ISL_454097, EPI_ISL_454098, EPI_ISL_454099, EPI_ISL_454100, EPI_ISL_454101, EPI_ISL_454102, EPI_ISL_454103, EPI_ISL_454104, EPI_ISL_454105, EPI_ISL_454106, EPI_ISL_454107, EPI_ISL_454108, EPI_ISL_454109, EPI_ISL_454110, EPI_ISL_454111, EPI_ISL_454112, EPI_ISL_454113, EPI_ISL_454114, EPI_ISL_454115, EPI_ISL_454116, EPI_ISL_454117, EPI_ISL_454118, EPI_ISL_454119, EPI_ISL_454120, EPI_ISL_454121, EPI_ISL_454122, EPI_ISL_454123, EPI_ISL_454124, EPI_ISL_454125, EPI_ISL_454126, EPI_ISL_454127, EPI_ISL_454128, EPI_ISL_454129, EPI_ISL_454130, EPI_ISL_454131, EPI_ISL_454132, EPI_ISL_454133, EPI_ISL_454134, EPI_ISL_454135, EPI_ISL_454136, EPI_ISL_454137, EPI_ISL_454138, EPI_ISL_454139, EPI_ISL_454140, EPI_ISL_454141, EPI_ISL_454142, EPI_ISL_454143, EPI_ISL_454144, EPI_ISL_454145, EPI_ISL_454146, EPI_ISL_454147, EPI_ISL_454148, EPI_ISL_454149, EPI_ISL_454150, EPI_ISL_454151, EPI_ISL_454152, EPI_ISL_454153, EPI_ISL_454154, EPI_ISL_454155, EPI_ISL_454156, EPI_ISL_454157, EPI_ISL_454158, EPI_ISL_454159, EPI_ISL_454160, EPI_ISL_454161, EPI_ISL_454162, EPI_ISL_454163, EPI_ISL_454164, EPI_ISL_454165, EPI_ISL_454166, EPI_ISL_454167, EPI_ISL_454168, EPI_ISL_454169, EPI_ISL_454170, EPI_ISL_454171, EPI_ISL_454172, EPI_ISL_454173, EPI_ISL_454174, EPI_ISL_454175, EPI_ISL_454176, EPI_ISL_454177, EPI_ISL_454178, EPI_ISL_454179, EPI_ISL_454180, EPI_ISL_454181, EPI_ISL_454182, EPI_ISL_454183, EPI_ISL_454184, EPI_ISL_454185, EPI_ISL_454186, EPI_ISL_454187, EPI_ISL_454188, EPI_ISL_454189, EPI_ISL_454190, EPI_ISL_454191, EPI_ISL_454192, EPI_ISL_454193, EPI_ISL_454194, EPI_ISL_454195, EPI_ISL_454196, EPI_ISL_454197, EPI_ISL_454198, EPI_ISL_454199, EPI_ISL_454200, EPI_ISL_454201, EPI_ISL_454202, EPI_ISL_454203, EPI_ISL_454204, EPI_ISL_454205, EPI_ISL_454206, EPI_ISL_454207, EPI_ISL_454208, EPI_ISL_454209, EPI_ISL_454210, EPI_ISL_454211, EPI_ISL_454212, EPI_ISL_454213, EPI_ISL_454214, EPI_ISL_454215, EPI_ISL_454216, EPI_ISL_454217, EPI_ISL_454218, EPI_ISL_454219, EPI_ISL_454220, EPI_ISL_454221, EPI_ISL_454222, EPI_ISL_454223, EPI_ISL_454224, EPI_ISL_454225, EPI_ISL_454226, EPI_ISL_454227, EPI_ISL_454228, EPI_ISL_454229, EPI_ISL_454230, EPI_ISL_454231, EPI_ISL_454232, EPI_ISL_454233, EPI_ISL_454234, EPI_ISL_454235, EPI_ISL_454236, EPI_ISL_454237, EPI_ISL_454238, EPI_ISL_454239, EPI_ISL_454240, EPI_ISL_454241, EPI_ISL_454242, EPI_ISL_454243, EPI_ISL_454244, EPI_ISL_454245, EPI_ISL_454246, EPI_ISL_454247, EPI_ISL_454248, EPI_ISL_454249, EPI_ISL_454250, EPI_ISL_454251, EPI_ISL_454252, EPI_ISL_454253, EPI_ISL_454254, EPI_ISL_454255, EPI_ISL_454256, EPI_ISL_454257, EPI_ISL_454258, EPI_ISL_454259, EPI_ISL_454260, EPI_ISL_454261, EPI_ISL_454262, EPI_ISL_454263, EPI_ISL_454264, EPI_ISL_454265, EPI_ISL_454266, EPI_ISL_454267, EPI_ISL_454268, EPI_ISL_454269, EPI_ISL_454270, EPI_ISL_454271, EPI_ISL_454272, EPI_ISL_454273, EPI_ISL_454274, EPI_ISL_454275, EPI_ISL_454276, EPI_ISL_454277, EPI_ISL_454278, EPI_ISL_454279, EPI_ISL_454280, EPI_ISL_454281, EPI_ISL_454282, EPI_ISL_454283, EPI_ISL_454284, EPI_ISL_454285, EPI_ISL_454286, EPI_ISL_454287, EPI_ISL_454288, EPI_ISL_454289, EPI_ISL_454290, EPI_ISL_454291, EPI_ISL_454292, EPI_ISL_454293, EPI_ISL_454294, EPI_ISL_454295, EPI_ISL_454296, EPI_ISL_454297, EPI_ISL_454298, EPI_ISL_454299, EPI_ISL_454300, EPI_ISL_454301, EPI_ISL_454302, EPI_ISL_454303, EPI_ISL_454304, EPI_ISL_454305, EPI_ISL_454306, EPI_ISL_454307, EPI_ISL_454308, EPI_ISL_454309, EPI_ISL_454310, EPI_ISL_454311, EPI_ISL_454312, EPI_ISL_454313, EPI_ISL_454314, EPI_ISL_454315, EPI_ISL_454316, EPI_ISL_454317, EPI_ISL_454318, EPI_ISL_454319, EPI_ISL_454320, EPI_ISL_454321, EPI_ISL_454322, EPI_ISL_454323, EPI_ISL_454324, EPI_ISL_454325, EPI_ISL_454326, EPI_ISL_454327, EPI_ISL_454328, EPI_ISL_454329, EPI_ISL_454330, EPI_ISL_454331, EPI_ISL_454332, EPI_ISL_454333, EPI_ISL_454334, EPI_ISL_454335, EPI_ISL_454336, EPI_ISL_454337, EPI_ISL_454338, EPI_ISL_454339, EPI_ISL_454340, EPI_ISL_454341, EPI_ISL_454342, EPI_ISL_454343, EPI_ISL_454344, EPI_ISL_454345, EPI_ISL_454346, EPI_ISL_454347, EPI_ISL_454348, EPI_ISL_454349, EPI_ISL_454350, EPI_ISL_454351, EPI_ISL_454352 | | | |
| see above | unknown | Instituto Nacional de Saude (INSA) | Borges et al |
| EPI_ISL_454353 | Microbial Genome Sequencing Center; Microbial Genomic Epidemiology Laboratory | Microbial Genomic Epidemiology Laboratory, University of Pittsburgh | Mustapha M. Mustapha, Jane W. Marsh, Dan Snyder, Marissa P. Griffith, Stephanie L. Mitchell, Vatsala R. Srinivasa, Kady D. Waggle, Chinelo Ezeonwuku, Vaughn S. Cooper, Lee H. Harrison |
| EPI_ISL_454354 | UPMC Clinical Microbiology Laboratory | Microbial Genome Sequencing Center; Microbial Genomic Epidemiology Laboratory, University of Pittsburgh | Mustapha M. Mustapha, Jane W. Marsh, Dan Snyder, Marissa P. Griffith, Stephanie L. Mitchell, Vatsala R. Srinivasa, Kady D. Waggle, Chinelo Ezeonwuku, Vaughn S. Cooper, Lee H. Harrison |
| EPI_ISL_454355, EPI_ISL_454356, EPI_ISL_454357, EPI_ISL_454358, EPI_ISL_454359, EPI_ISL_454360, EPI_ISL_454361, EPI_ISL_454362, EPI_ISL_454363, EPI_ISL_454364, EPI_ISL_454365, EPI_ISL_454366, EPI_ISL_454367, EPI_ISL_454368, EPI_ISL_454369, EPI_ISL_454370, EPI_ISL_454371, EPI_ISL_454372, EPI_ISL_454373, EPI_ISL_454374, EPI_ISL_454375, EPI_ISL_454376, EPI_ISL_454377, EPI_ISL_454378, EPI_ISL_454379, EPI_ISL_454380, EPI_ISL_454381, EPI_ISL_454382, EPI_ISL_454383, EPI_ISL_454384, EPI_ISL_454385, EPI_ISL_454386, EPI_ISL_454387, EPI_ISL_454388, EPI_ISL_454389, EPI_ISL_454390, EPI_ISL_454391, EPI_ISL_454392, EPI_ISL_454393, EPI_ISL_454394, EPI_ISL_454395, EPI_ISL_454396, EPI_ISL_454397, EPI_ISL_454398, EPI_ISL_454399, EPI_ISL_454400, EPI_ISL_454401, EPI_ISL_454402, EPI_ISL_454403, EPI_ISL_454404, EPI_ISL_454405, EPI_ISL_454406, EPI_ISL_454407, EPI_ISL_454408, EPI_ISL_454409, EPI_ISL_454410, EPI_ISL_454411 | | | |
| see above | UPMC Clinical Microbiology Laboratory | Microbial Genome Sequencing Center, Microbial Genomic Epidemiological Laboratory | Mustapha M. Mustapha, Jane W. Marsh, Dan Snyder, Marissa P. Griffith, Stephanie L. Mitchell, Vatsala R. Srinivasa, Kady D. Waggle, Chinelo Ezeonwuku, Vaughn S. Cooper, Lee H. Harrison |
| EPI_ISL_454412, EPI_ISL_454413, EPI_ISL_454414, EPI_ISL_454415 | Dirk Dittmer | Dirk Dittmer | Bailey,A.G., Caro-Vegas,C.P., Dittmer,D., Eason,A.B., Juarez,A., Landis,J.T., McNamara,R.P., Miller,M.B., Moorad,R., Pluta,L.J., Seltzer,T.A., Thompson,C., Vahrson,W., Villamor,F. |
| EPI_ISL_454416 | Department of Medical Microbiology, Leiden University Medical Center | Department of Medical Microbiology, Leiden University Medical Center | Snijder,E.J., Ogando,N.S., Zevenhoven,J.C., Dalebout,T.J., de Vries,J.J. and Sidorov,I.I. |
| EPI_ISL_454417, EPI_ISL_454418 | Research and Experiment Center, Meizhou People Hospital | Research and Experiment Center, Meizhou People Hospital | Guo,X., Zeng,L. and Yu,Z. |
| EPI_ISL_454419 | Institute for Forensic Medicine, Faculty of Medicine, University of Belgrade | Institute for Forensic Medicine, Faculty of Medicine, University of Belgrade | Skadric,I., Stojkovic,O., Petrovic,T. and Tolic,A. |
| EPI_ISL_454420 | Rafik Hariri University Hospital | Rafik Hariri University Hospital | Rita Feghali |
| EPI_ISL_454421, EPI_ISL_454422, EPI_ISL_454423, EPI_ISL_454424, EPI_ISL_454425, EPI_ISL_454426, EPI_ISL_454427, EPI_ISL_454428, EPI_ISL_454429, EPI_ISL_454430, EPI_ISL_454431, EPI_ISL_454432, EPI_ISL_454433, EPI_ISL_454434, EPI_ISL_454435 | | | |
| see above | Maryland Department of Health Laboratories Administration | Maryland Department of Health Laboratories Administration | MDH Laboratories Administration |
| EPI_ISL_454436, EPI_ISL_454437, EPI_ISL_454438, EPI_ISL_454439, EPI_ISL_454440, EPI_ISL_454441, EPI_ISL_454442, EPI_ISL_454443, EPI_ISL_454444, EPI_ISL_454445, EPI_ISL_454446 | | | |
| see above | Halmstad klinisk mikrobiologi | The Public Health Agency of Sweden | Anna-Malin Linde, Maria Lind Karlberg, Mattias Haukland, Reza Advani, Olov Svartstrom, Oskar Karlsson Lindsjo, Petra Edquist, Shamam Muradrasoli, Anna Risberg, Karin Tegmark-Wisell |
| EPI_ISL_454447, EPI_ISL_454448, EPI_ISL_454449, EPI_ISL_454450, EPI_ISL_454451, EPI_ISL_454452, EPI_ISL_454453, EPI_ISL_454454, EPI_ISL_454455, EPI_ISL_454456, EPI_ISL_454457, EPI_ISL_454458, EPI_ISL_454459, EPI_ISL_454460, EPI_ISL_454461, EPI_ISL_454462, EPI_ISL_454463, EPI_ISL_454464, EPI_ISL_454465, EPI_ISL_454466, EPI_ISL_454467, EPI_ISL_454468, EPI_ISL_454469, EPI_ISL_454470, EPI_ISL_454471, EPI_ISL_454472, EPI_ISL_454473, EPI_ISL_454474, EPI_ISL_454475, EPI_ISL_454476, EPI_ISL_454477, EPI_ISL_454478, EPI_ISL_454479, EPI_ISL_454480, EPI_ISL_454481, EPI_ISL_454482, EPI_ISL_454483, EPI_ISL_454484, EPI_ISL_454485, EPI_ISL_454486, EPI_ISL_454487, EPI_ISL_454488, EPI_ISL_454489, EPI_ISL_454490, EPI_ISL_454491, EPI_ISL_454492, EPI_ISL_454493 | | | |
| see above | Karolinska Universitetslaboratoriet | The Public Health Agency of Sweden | Anna-Malin Linde, Maria Lind Karlberg, Mattias Haukland, Reza Advani, Olov Svartstrom, Oskar Karlsson Lindsjo, Petra Edquist, Shamam Muradrasoli, Anna Risberg, Karin Tegmark-Wisell |
| EPI_ISL_454494, EPI_ISL_454495, EPI_ISL_454496 | Skovde/Unilabs | The Public Health Agency of Sweden | Anna-Malin Linde, Maria Lind Karlberg, Mattias Haukland, Reza Advani, Olov Svartstrom, Oskar Karlsson Lindsjo, Petra Edquist, Shamam Muradrasoli, Anna Risberg, Karin Tegmark-Wisell |
| EPI_ISL_454497, EPI_ISL_454498 | RSE "National Center for Biotechnology" | RSE "National Center for Biotechnology" | Alexandr Shevtsov, Ilyas Akhmetollayev, Viktoriya Lutsay, Asylulan Amirgazin, Askar Abdaliyev, Akbota Rakhmetova, Zabira Aushakhmetova, Ruslan Kalendar, Yerlan Ramankulov |
| EPI_ISL_454499, EPI_ISL_454500, EPI_ISL_454501, EPI_ISL_454502, EPI_ISL_454503, EPI_ISL_454504, EPI_ISL_454505, EPI_ISL_454506, EPI_ISL_454507, EPI_ISL_454508, EPI_ISL_454509, EPI_ISL_454510, EPI_ISL_454511, EPI_ISL_454512, EPI_ISL_454513, EPI_ISL_454514, EPI_ISL_454515, EPI_ISL_454516, EPI_ISL_454517, EPI_ISL_454518, EPI_ISL_454519, EPI_ISL_454520 | | | |
| see above | RSE "National Center for Biotechnology" | RSE "National Center for Biotechnology" | Alexandr Shevtsov, Ilyas Akhmetollayev, Viktoriya Lutsay, Asylulan Amirgazin, Askar Abdaliyev, Akbota Rakhmetova, Zabira Aushakhmetova, Ruslan Kalendar, Yerlan Ramankulov |
| EPI_ISL_454521, EPI_ISL_454522, EPI_ISL_454523, EPI_ISL_454524, EPI_ISL_454525, EPI_ISL_454526, EPI_ISL_454527, EPI_ISL_454528, EPI_ISL_454529, EPI_ISL_454530, EPI_ISL_454531, EPI_ISL_454532, EPI_ISL_454533, EPI_ISL_454534, EPI_ISL_454535, EPI_ISL_454536, EPI_ISL_454537, EPI_ISL_454538, EPI_ISL_454539, EPI_ISL_454540, EPI_ISL_454541, EPI_ISL_454542, EPI_ISL_454543, EPI_ISL_454544, EPI_ISL_454545, EPI_ISL_454546, EPI_ISL_454547, EPI_ISL_454548, EPI_ISL_454549, EPI_ISL_454550, EPI_ISL_454551, EPI_ISL_454552, EPI_ISL_454553, EPI_ISL_454554, EPI_ISL_454555, EPI_ISL_454556, EPI_ISL_454557, EPI_ISL_454558, EPI_ISL_454559, EPI_ISL_454560, EPI_ISL_454561, EPI_ISL_454562, EPI_ISL_454563, EPI_ISL_454564, EPI_ISL_454565, EPI_ISL_454566, EPI_ISL_454567, EPI_ISL_454568, EPI_ISL_454569, EPI_ISL_454570 | | | |
| see above | NIV Influenza | NIV Influenza | Potdar V |
| EPI_ISL_454571 | National Center of Expertise | National Center for Expertise, National Center for Biotechnology, Kazakhstan | Abdaliyev Askar, Shevtsov Alexandr, Akhmetollayev Ilyas, Kalendar Ruslan, Rakhmetova Akbota, , Lutsay Viktoriya, Amirgazin Asylulan, Aushakhmetova Zabira, Ramankulov Yerlan |
| EPI_ISL_454572 | National Center of Expertise | National Center for Expertise, Kazakhstan National Center for Biotechnology, Kazakhstan | Abdaliyev Askar, Shevtsov Alexandr, Akhmetollayev Ilyas, Kalendar Ruslan, Rakhmetova Akbota, , Lutsay Viktoriya, Amirgazin Asylulan, Aushakhmetova Zabira, Ramankulov Yerlan |
| EPI_ISL_454574 | nstitute for Public Health | Laboratory for advanced genomics | Filip Rokić, Lovro Trgovec-Greif, Neven Sučić, Tomislav Rukavina, Igor Jurak, Oliver Vugrek |
| EPI_ISL_454575 | Laboratory of virology, National Center of Expertise | Laboratory of molecular-genetic research, National Center for Expertise, Kazakhstan National Center for Biotechnology, Kazakhstan | Abdaliyev Askar, Shevtsov Alexandr, Akhmetollayev Ilyas, Kalendar Ruslan, Rakhmetova Akbota, , Lutsay Viktoriya, Amirgazin Asylulan, Aushakhmetova Zabira, Ramankulov Yerlan |
| EPI_ISL_454576, EPI_ISL_454577 | Laboratory of virology, National Center of Expertise | Laboratory of molecular-genetic research, National Center for Expertise, Kazakhstan National Center for Biotechnology, Kazakhstan | Abdaliyev Askar, Shevtsov Alexandr, Akhmetollayev Ilyas, Kalendar Ruslan, Rakhmetova Akbota, , Lutsay Viktoriya, Amirgazin Asylulan, Aushakhmetova Zabira, Ramankulov Yerlan |
| EPI_ISL_454578 | University Hospital for Infectious Diseases "Dr. Fran Mihaljević", Research Unit | University of Zagreb, Centre for research and knowledge transfer in biotechnology | Ivan-Christian Kurolt, Jelena Ivancic Jelecki, Anamarija Slovic |
| EPI_ISL_454579, EPI_ISL_454580 | Laboratory of virology, National Center of Expertise | Laboratory of molecular-genetic research, National Center of Expertise, Kazakhstan National Center for Biotechnology, Kazakhstan | Abdaliyev Askar, Shevtsov Alexandr, Akhmetollayev Ilyas, Kalendar Ruslan, Rakhmetova Akbota, , Lutsay Viktoriya, Amirgazin Asylulan, Aushakhmetova Zabira, Ramankulov Yerlan |
| EPI_ISL_454581 | University Hospital for Infectious Diseases "Dr. Fran Mihaljević", Research Unit | University of Zagreb, Centre for research and knowledge transfer in biotechnology | Ivan-Christian Kurolt, Jelena Ivancic Jelecki, Anamarija Slovic |
| EPI_ISL_454582 | Laboratory of virology, National Center of Expertise | Laboratory of molecular-genetic research, National Center of Expertise, Kazakhstan National Center for Biotechnology, Kazakhstan | Abdaliyev Askar, Shevtsov Alexandr, Akhmetollayev Ilyas, Kalendar Ruslan, Rakhmetova Akbota, , Lutsay Viktoriya, Amirgazin Asylulan, Aushakhmetova Zabira, Ramankulov Yerlan |
| EPI_ISL_454583 | University Hospital for Infectious Diseases "Dr. Fran Mihaljević", Research Unit | University of Zagreb, Centre for research and knowledge transfer in biotechnology | Ivan-Christian Kurolt, Jelena Ivancic Jelecki, Anamarija Slovic |
| EPI_ISL_454584, EPI_ISL_454585, EPI_ISL_454586 | Laboratory of virology, National Center of Expertise | Laboratory of molecular-genetic research, National Center for Expertise, Kazakhstan National Center for Biotechnology, Kazakhstan | Abdaliyev Askar, Shevtsov Alexandr, Akhmetollayev Ilyas, Kalendar Ruslan, Rakhmetova Akbota, , Lutsay Viktoriya, Amirgazin Asylulan, Aushakhmetova Zabira, Ramankulov Yerlan |
| EPI_ISL_454587 | Laboratory of virology, National Center of Expertise | Laboratory of molecular-genetic research, National Center of Expertise, | Abdaliyev Askar, Shevtsov Alexandr, Akhmetollayev Ilyas, Kalendar Ruslan, Rakhmetova Akbota, , Lutsay Viktoriya, Amirgazin Asylulan, Aushakhmetova Zabira, Ramankulov Yerlan |

| | | | |
|---|---|---|---|
| | | Kazakhstan National Center for Biotechnology, Kazakhstan | |
| EPI_ISL_454588 | University Hospital for Infectious Diseases "Dr. Fran Mihaljević", Research Unit | University of Zagreb, Centre for research and knowledge transfer in biotechnology | Ivan-Christian Kurolt, Jelena Ivancic Jelecki, Anamarija Slovic |
| EPI_ISL_454589 | Laboratory of virology, National Center of Expertise | Laboratory of molecular-genetic research, National Center for Expertise, Kazakhstan National Center for Biotechnology, Kazakhstan | Abdaliyev Askar, Shevtsov Alexandr, Akhmetollayev Ilyas, Kalendar Ruslan, Rakhmetova Akbota, , Lutsay Viktoriya, Amirgazin Asylulan, Aushakhmetova Zabira, Ramankulov Yerlan |
| EPI_ISL_454590, EPI_ISL_454591 | Laboratory of virology, National Center of Expertise | Laboratory of molecular-genetic research, National Center of Expertise, Kazakhstan National Center for Biotechnology, Kazakhstan | Abdaliyev Askar, Shevtsov Alexandr, Akhmetollayev Ilyas, Kalendar Ruslan, Rakhmetova Akbota, , Lutsay Viktoriya, Amirgazin Asylulan, Aushakhmetova Zabira, Ramankulov Yerlan |
| EPI_ISL_454592 | University Hospital for Infectious Diseases "Dr. Fran Mihaljević", Research Unit | University of Zagreb, Centre for research and knowledge transfer in biotechnology | Ivan-Christian Kurolt, Jelena Ivancic Jelecki, Anamarija Slovic |
| EPI_ISL_454593, EPI_ISL_454594 | Laboratory of virology, National Center of Expertise | Laboratory of molecular-genetic research, National Center of Expertise, Kazakhstan National Center for Biotechnology, Kazakhstan | Abdaliyev Askar, Shevtsov Alexandr, Akhmetollayev Ilyas, Kalendar Ruslan, Rakhmetova Akbota, , Lutsay Viktoriya, Amirgazin Asylulan, Aushakhmetova Zabira, Ramankulov Yerlan |
| EPI_ISL_454595 | University Hospital for Infectious Diseases "Dr. Fran Mihaljević", Research Unit | University of Zagreb, Centre for research and knowledge transfer in biotechnology | Ivan-Christian Kurolt, Jelena Ivancic Jelecki, Anamarija Slovic |
| EPI_ISL_454596, EPI_ISL_454597 | Laboratory of virology, National Center of Expertise | Laboratory of molecular-genetic research, National Center for Expertise, Kazakhstan National Center for Biotechnology, Kazakhstan | Abdaliyev Askar, Shevtsov Alexandr, Akhmetollayev Ilyas, Kalendar Ruslan, Rakhmetova Akbota, , Lutsay Viktoriya, Amirgazin Asylulan, Aushakhmetova Zabira, Ramankulov Yerlan |
| EPI_ISL_454598 | Laboratory of virology, National Center of Expertise | Laboratory of molecular-genetic research, National Center of Expertise, Kazakhstan National Center for Biotechnology, Kazakhstan | Abdaliyev Askar, Shevtsov Alexandr, Akhmetollayev Ilyas, Kalendar Ruslan, Rakhmetova Akbota, , Lutsay Viktoriya, Amirgazin Asylulan, Aushakhmetova Zabira, Ramankulov Yerlan |
| EPI_ISL_454599, EPI_ISL_454600 | Laboratory of virology, National Center of Expertise | Laboratory of molecular-genetic research, National Center of Expertise, Kazakhstan National Center for Biotechnology, Kazakhstan | Abdaliyev Askar, Shevtsov Alexandr, Akhmetollayev Ilyas, Kalendar Ruslan, Rakhmetova Akbota, , Lutsay Viktoriya, Amirgazin Asylulan, Aushakhmetova Zabira, Ramankulov Yerlan |
| EPI_ISL_454601 | Laboratory of virology, National Center of Expertise | Laboratory of molecular-genetic research, National Center of Expertise, Kazakhstan National Center for Biotechnology, Kazakhstan | Abdaliyev Askar, Shevtsov Alexandr, Akhmetollayev Ilyas, Kalendar Ruslan, Rakhmetova Akbota, , Lutsay Viktoriya, Amirgazin Asylulan, Aushakhmetova Zabira, Ramankulov Yerlan |
| EPI_ISL_454602 | Croatian Institute of Public Health | University of Zagreb, Centre for research and knowledge transfer in biotechnology | Irena Tabain, Tatjana Vilibic-Cavlek, Jelena Ivancic Jelecki, Anamarija Slovic |
| EPI_ISL_454603 | Laboratory of virology, National Center of Expertise | Laboratory of molecular-genetic research, National Center of Expertise, Kazakhstan National Center for Biotechnology, Kazakhstan | Abdaliyev Askar, Shevtsov Alexandr, Akhmetollayev Ilyas, Kalendar Ruslan, Rakhmetova Akbota, , Lutsay Viktoriya, Amirgazin Asylulan, Aushakhmetova Zabira, Ramankulov Yerlan |
| EPI_ISL_454604 | Laboratory of virology, National Center of Expertise | Laboratory of molecular-genetic research, National Center for Expertise, Kazakhstan National Center for Biotechnology, Kazakhstan | Abdaliyev Askar, Shevtsov Alexandr, Akhmetollayev Ilyas, Kalendar Ruslan, Rakhmetova Akbota, , Lutsay Viktoriya, Amirgazin Asylulan, Aushakhmetova Zabira, Ramankulov Yerlan |
| EPI_ISL_454605, EPI_ISL_454606 | Institute for Public Health | Laboratory for advanced genomics | Filip Rokić, Lovro Trgovec-Greif, Neven Sučić, Tomislav Rukavina, Igor Jurak, Oliver Vugrek |
| EPI_ISL_454607, EPI_ISL_454608, EPI_ISL_454609, EPI_ISL_454610, EPI_ISL_454611, EPI_ISL_454612, EPI_ISL_454613 | Alameda County Public Health Lab | Chan-Zuckerberg Biohub | CZB Cliahub Consortium |
| EPI_ISL_454614, EPI_ISL_454615, EPI_ISL_454616, EPI_ISL_454617, EPI_ISL_454618, EPI_ISL_454619, EPI_ISL_454620, EPI_ISL_454621, EPI_ISL_454622, EPI_ISL_454623, EPI_ISL_454624, EPI_ISL_454625, EPI_ISL_454626, EPI_ISL_454627, EPI_ISL_454628, EPI_ISL_454629, EPI_ISL_454630, EPI_ISL_454631, EPI_ISL_454632, EPI_ISL_454633, EPI_ISL_454634 | | | |
| see above | UCSF Clinical Microbiology Laboratory | Chan-Zuckerberg Biohub | CZB Cliahub Consortium |
| EPI_ISL_454635 | County Of San Luis Obispo Public Health Laboratory | Chan-Zuckerberg Biohub | CZB Cliahub Consortium |
| EPI_ISL_454636, EPI_ISL_454637, EPI_ISL_454638, EPI_ISL_454639, EPI_ISL_454640, EPI_ISL_454641 | Humboldt County Public Health Laboratory | Chan-Zuckerberg Biohub | CZB Cliahub Consortium |
| EPI_ISL_454642 | CT-Dr. Katherine A. Kelley State Public Health Lab | Pathogen Discovery, Respiratory Viruses Branch, Division of Viral Diseases, Centers for Disease Control and Prevention | Jing Zhang, Ying Tao, Clinton R. Paden, Anna Uehara, Krista Queen, Yan Li, Haibin Wang, Zachary Weiner, Bettina Bankamp, Suxiang Tong |
| EPI_ISL_454643, EPI_ISL_454644 | VI-US Virgin Islands Department of Health | Pathogen Discovery, Respiratory Viruses Branch, Division of Viral Diseases, Centers for Disease Control and Prevention | Jing Zhang, Ying Tao, Clinton R. Paden, Anna Uehara, Krista Queen, Yan Li, Haibin Wang, Zachary Weiner, Bettina Bankamp, Suxiang Tong |
| EPI_ISL_454645 | CT-Dr. Katherine A. Kelley State Public Health Lab | Pathogen Discovery, Respiratory Viruses Branch, Division of Viral Diseases, Centers for Disease Control and Prevention | Jing Zhang, Ying Tao, Clinton R. Paden, Anna Uehara, Krista Queen, Yan Li, Haibin Wang, Zachary Weiner, Bettina Bankamp, Suxiang Tong |
| EPI_ISL_454646 | City of El Paso Department of Public Health Laboratory | Pathogen Discovery, Respiratory Viruses Branch, Division of Viral Diseases, Centers for Disease Control and Prevention | Ying Tao, Clinton R. Paden, Jing Zhang, Anna Uehara, Krista Queen, Yan Li, Haibin Wang, Zachary Weiner, Michael Bowen, Suxiang Tong |
| EPI_ISL_454647, EPI_ISL_454648, EPI_ISL_454649, EPI_ISL_454650, EPI_ISL_454651, EPI_ISL_454652 | VI-US Virgin Islands Department of Health | Pathogen Discovery, Respiratory Viruses Branch, Division of Viral Diseases, Centers for Disease Control and Prevention | Ying Tao, Clinton R. Paden, Jing Zhang, Anna Uehara, Krista Queen, Yan Li, Haibin Wang, Zachary Weiner, Bettina Bankamp, Suxiang Tong |
| EPI_ISL_454653, EPI_ISL_454654, EPI_ISL_454655, EPI_ISL_454656, EPI_ISL_454657, EPI_ISL_454658, EPI_ISL_454659, EPI_ISL_454660, EPI_ISL_454661, EPI_ISL_454662, EPI_ISL_454663, EPI_ISL_454664, EPI_ISL_454665, EPI_ISL_454666, EPI_ISL_454667, EPI_ISL_454668, EPI_ISL_454669, EPI_ISL_454670, EPI_ISL_454671, EPI_ISL_454672, EPI_ISL_454673, EPI_ISL_454674, EPI_ISL_454675, EPI_ISL_454676, EPI_ISL_454677, EPI_ISL_454678, EPI_ISL_454679, EPI_ISL_454680, EPI_ISL_454681, EPI_ISL_454682, EPI_ISL_454683, EPI_ISL_454684, EPI_ISL_454685, EPI_ISL_454686, EPI_ISL_454687, EPI_ISL_454688, EPI_ISL_454689 | | | |
| see above | County of Santa Clara Public Health Department | Chan-Zuckerberg Biohub | CZB Cliahub Consortium |
| EPI_ISL_454690 | Emory Molecular Diagnostics Laboratory, Emory Healthcare | Piantadosi Lab, Emory Department of Pathology | Ahmed Babiker, Anne Piantadosi |
| EPI_ISL_454691 | Quest Diagnostics | Quest Diagnostics | Anderson,B.P., Rosenthal,S.H., Gerasimova,A., Kagan,R.M. and Owen, R. |
| EPI_ISL_454707, EPI_ISL_454708, EPI_ISL_454709, EPI_ISL_454710, EPI_ISL_454711, EPI_ISL_454712, EPI_ISL_454713, EPI_ISL_454714, EPI_ISL_454715, EPI_ISL_454716, EPI_ISL_454717, EPI_ISL_454718, EPI_ISL_454719, EPI_ISL_454720, EPI_ISL_454721, EPI_ISL_454722, EPI_ISL_454723, EPI_ISL_454724, EPI_ISL_454725, EPI_ISL_454726, EPI_ISL_454727, EPI_ISL_454728, EPI_ISL_454729, EPI_ISL_454730, EPI_ISL_454731 | | | |
| see above | Utah Public Health Laboratory | Utah Public Health Laboratory | Erin Young, Kelly Oakeson |
| EPI_ISL_454732 | Russian State Collection of Viruses | Pathogenic Microorganisms Variability Laboratory | Denis Protsenko, Alexey Shchetinin, Maria Nikiforova, Elena Shidlovskaya, Nadezhda Kuznetsova, Vladimir Gushchin, Inna Dolzhikova, Daria Grousova, Andrey Botikov, Denis Logunov, Alexander Gintsburg, Alexey Mazus |
| EPI_ISL_454733 | Department of Medical, Biotechnologies University of Siena | Department of Medical, Biotechnologies University of Siena | Cusi,M.G., Pinzauti,D., Gandolfo,C., Anichini,G., Pozzi,G. and Santoro,F. |
| EPI_ISL_454749 | Japanese Quarantine Stations | Pathogen Genomics Center, National Institute of Infectious Diseases | Tsuyoshi Sekizuka, Kentaro Itokawa, Rina Tanaka, Masanori Hashino, Tsutomu Kageyama, Shinji Saito, Ikuyo Takayama, Hideki Hasegawa, Takuri Takahashi, Hajime Kamiya, Takuya Yamagishi, Motoi Suzuki, Takaji Wakita, Makoto Kuroda |
| EPI_ISL_454750, EPI_ISL_454751, EPI_ISL_454752, EPI_ISL_454753, EPI_ISL_454754, EPI_ISL_454755, EPI_ISL_454756, EPI_ISL_454757, EPI_ISL_454758, EPI_ISL_454759, EPI_ISL_454760, EPI_ISL_454761, EPI_ISL_454762, EPI_ISL_454763, EPI_ISL_454764, EPI_ISL_454765, EPI_ISL_454766, EPI_ISL_454767, EPI_ISL_454768, EPI_ISL_454769, EPI_ISL_454770, EPI_ISL_454771, EPI_ISL_454772, EPI_ISL_454773, EPI_ISL_454774, EPI_ISL_454775, EPI_ISL_454776, EPI_ISL_454777, EPI_ISL_454778, EPI_ISL_454779, EPI_ISL_454780, EPI_ISL_454781, EPI_ISL_454782, EPI_ISL_454783, EPI_ISL_454784, EPI_ISL_454785, EPI_ISL_454786, EPI_ISL_454787, EPI_ISL_454788, EPI_ISL_454789, EPI_ISL_454790, EPI_ISL_454791, EPI_ISL_454792, EPI_ISL_454793, EPI_ISL_454794 | | | |
| see above | Dutch COVID-19 response team | National Institute for Public Health and the Environment (RIVM) | Adam Meijer, Harry Vennema, Jeroen Cremer, Sharon van den Brink, Pieter Overduin, Florian Zwagemaker, Dennis Schmitz, Chantal Reusken, on behalf of the national COVID-19 response team |
| EPI_ISL_454795 | Veterinary Specialized Institute Kraljevo | Veterinary Specialized Institute Kraljevo | Vidanovic,D., Tesovic,B., Sekler,M., Dmitric,M., Debeljak,Z., Matovic,K., Vaskovic,N., Petrovic,T., Volkening,J. and Alfonso,C.L. |
| EPI_ISL_454796, EPI_ISL_454797, EPI_ISL_454798, EPI_ISL_454799, EPI_ISL_454800, EPI_ISL_454801, EPI_ISL_454802, EPI_ISL_454803, EPI_ISL_454804, EPI_ISL_454805, EPI_ISL_454806, EPI_ISL_454807, EPI_ISL_454808, EPI_ISL_454809, EPI_ISL_454810, EPI_ISL_454811, EPI_ISL_454812, EPI_ISL_454813, EPI_ISL_454814, EPI_ISL_454815, EPI_ISL_454816, EPI_ISL_454817, EPI_ISL_454818, EPI_ISL_454819, EPI_ISL_454820, EPI_ISL_454821, EPI_ISL_454822, EPI_ISL_454823, EPI_ISL_454824, EPI_ISL_454825, EPI_ISL_454826, EPI_ISL_454827, EPI_ISL_454828, EPI_ISL_454829 | | | |
| see above | Dirk Dittmer | Dirk Dittmer | Bailey,A.G., Caro-Vegas,C.P., Dittmer,D., Eason,A.B., Juarez,A., Landis,J.T., McNamara,R.P., Miller,M.B., Moorad,R., Pluta,L.J., Seltzer,T.A., Thompson,C., Vahrson,W., Villamor,F. |
| EPI_ISL_454830, EPI_ISL_454831, EPI_ISL_454832, EPI_ISL_454833 | SMS Medical College, Jaipur | CSIR Institute of Genomics and Integrative Biology | Sudhir Bhandari, Rahul Bhoyar, Mohammed Imran, Mohit Divakar, Disha Sharma, Anshul Kumar, Bani Jolly, Rahul Sahlot, Abhinav Jain, Paras Sehgal, Gyan Ranjan, Vinod Scaria, Sridhar Sivasubbu, Sandeep K Mathur |
| EPI_ISL_454858, EPI_ISL_454859, EPI_ISL_454860, EPI_ISL_454861, EPI_ISL_454862, EPI_ISL_454863, EPI_ISL_454864, EPI_ISL_454865, EPI_ISL_454866, EPI_ISL_454867 | Translational Health Science and Technology Institute -ESIC medical college and hospital, Faridabad | THSTI Bioassay laboratory | Saurabh Kumar, Jigme Wangchuk, Anil Kumar Pandey, Asim Das, Guruprasad R. Medigeshi |
| EPI_ISL_454868, EPI_ISL_454869, EPI_ISL_454870, EPI_ISL_454871, EPI_ISL_454872, EPI_ISL_454873, EPI_ISL_454874, EPI_ISL_454875, EPI_ISL_454876, EPI_ISL_454877, EPI_ISL_454878, EPI_ISL_454879, EPI_ISL_454880, EPI_ISL_454881, EPI_ISL_454882, EPI_ISL_454883, EPI_ISL_454884, EPI_ISL_454885, EPI_ISL_454886, EPI_ISL_454887, EPI_ISL_454888, EPI_ISL_454889, EPI_ISL_454890, EPI_ISL_454891, EPI_ISL_454892, EPI_ISL_454893, EPI_ISL_454894, EPI_ISL_454895, EPI_ISL_454896, EPI_ISL_454897, EPI_ISL_454898, EPI_ISL_454899, EPI_ISL_454900, EPI_ISL_454901, EPI_ISL_454902 | | | |
| see above | Karolinska Universitetslaboratoriet | The Public Health Agency of Sweden | Anna-Malin Linde, Maria Lind Karlberg, Mattias Haukland, Reza Advani, Olov Svartstrom, Oskar Karlsson Lindsjo, Petra Edquist, Shamam Muradrasoli, Anna Risberg, Karin Tegmark-Wisell |
| EPI_ISL_454903 | Klinisk mikrobiologi, UAS | The Public Health Agency of Sweden | Anna-Malin Linde, Maria Lind Karlberg, Mattias Haukland, Reza Advani, Olov Svartstrom, Oskar Karlsson Lindsjo, Petra Edquist, Shamam Muradrasoli, Anna Risberg, Karin Tegmark-Wisell |
| EPI_ISL_454904, EPI_ISL_454905, EPI_ISL_454906, EPI_ISL_454907, EPI_ISL_454908, EPI_ISL_454909, EPI_ISL_454910, EPI_ISL_454911, EPI_ISL_454912, EPI_ISL_454913, EPI_ISL_454914, EPI_ISL_454915, EPI_ISL_454916, EPI_ISL_454917, EPI_ISL_454918, EPI_ISL_454919, EPI_ISL_454920, EPI_ISL_454921, EPI_ISL_454922, EPI_ISL_454923, EPI_ISL_454924, EPI_ISL_454925, EPI_ISL_454926, EPI_ISL_454927, EPI_ISL_454928, EPI_ISL_454929, EPI_ISL_454930, EPI_ISL_454931, EPI_ISL_454932, EPI_ISL_454933, EPI_ISL_454934, EPI_ISL_454935, EPI_ISL_454936, EPI_ISL_454937, EPI_ISL_454938, EPI_ISL_454939, EPI_ISL_454940, EPI_ISL_454941, EPI_ISL_454942, EPI_ISL_454943, EPI_ISL_454944, EPI_ISL_454945, EPI_ISL_454946, EPI_ISL_454947, EPI_ISL_454948, EPI_ISL_454949, EPI_ISL_454950, EPI_ISL_454951, EPI_ISL_454952, EPI_ISL_454953, EPI_ISL_454954, EPI_ISL_454955, EPI_ISL_454956, EPI_ISL_454957, EPI_ISL_454958, EPI_ISL_454959, EPI_ISL_454960, EPI_ISL_454961, EPI_ISL_454962, EPI_ISL_454963, EPI_ISL_454964, EPI_ISL_454965, EPI_ISL_454966, EPI_ISL_454967, EPI_ISL_454968, EPI_ISL_454969, EPI_ISL_454970, EPI_ISL_454971, EPI_ISL_454972, EPI_ISL_454973, EPI_ISL_454974, EPI_ISL_454975, EPI_ISL_454976, EPI_ISL_454977, EPI_ISL_454978, EPI_ISL_454979, EPI_ISL_454980, EPI_ISL_454981, EPI_ISL_454982, EPI_ISL_454983, EPI_ISL_454984, EPI_ISL_454985, EPI_ISL_454986, EPI_ISL_454987, EPI_ISL_454988, EPI_ISL_454989, EPI_ISL_454990, EPI_ISL_454991, EPI_ISL_454992, EPI_ISL_454993, EPI_ISL_454994, EPI_ISL_454995, EPI_ISL_454996, EPI_ISL_454997, EPI_ISL_454998, EPI_ISL_454999, EPI_ISL_455000, EPI_ISL_455001, EPI_ISL_455002, EPI_ISL_455003, EPI_ISL_455004, EPI_ISL_455005, EPI_ISL_455006, EPI_ISL_455007, EPI_ISL_455008, EPI_ISL_455009, EPI_ISL_455010, EPI_ISL_455011, EPI_ISL_455012, EPI_ISL_455013, EPI_ISL_455014 | | | |
| see above | Wuhan Chain Medical Labs (CMLabs) | State Key Laboratory of Biotherapy of Sichuan University | Baowen Du, Minjin Wang, Chao Tang, Chuan Chen, Yongzhao Zhou, Mingxia Yu, Hancheng Wei, Weimin Li, Jing-wen Lin, Jia Geng, Binwu Ying, Lu Chen |
| EPI_ISL_455015 | Pandit Deendayal Upadhyay Government Medical College, Rajkot | Gujarat Biotechnology Research Centre | Snehal Bagatharia, Prakash Modi, Sejul Antala, Manish Pattani, Ramesh Pandit, Tejas Shah, Ankit Hinsu, Pritesh Sabara, Apurvasinh Puvar, Janvi Raval, Zarna Patel, Monika Gandhi, Pinal Trivedi, Maharshi Pandya, Amit Kanani, Nidhi Patel, Nitin Savaliya, Raghawendra Kumar, Dinesh Kumar, Zuber Saiyed, Komal Patel, Labdhi Pandya, Neha Rajpara, Bhavesh Modi, Gaurishankar Shrimali, R D Dixit, A M Kadri, Umang Mishra, Chaitanya Joshi, Madhvi Joshi |

| | | | |
|---|---|---|---|
| EPI_ISL_455016 | Pandit Deendayal Upadhyay Government Medical College, Rajkot | Gujarat Biotechnology Research Centre | Prakash Modi, Sejul Antala, Manish Pattani, Ramesh Pandit, Tejas Shah, Ankit Hinsu, Pritesh Sabara, Apurvasinh Puvar, Janvi Raval, Zarna Patel, Monika Gandhi, Pinal Trivedi, Maharshi Pandya, Amit Kanani, Nidhi Patel, Nitin Savaliya, Raghawendra Kumar, Dinesh Kumar, Zuber Saiyed, Komal Patel, Labdhi Pandya, Snehal Bagatharia, Afzal Ansari, Bhavesh Modi, Gaurishankar Shrimali, R D Dixit, A M Kadri, Umang Mishra, Chaitanya Joshi, Madhvi Joshi |
| EPI_ISL_455017 | Government Medical College, Vadodara | Gujarat Biotechnology Research Centre | Tanuja Javadekar , R N Daveshwar, Ramesh Pandit, Tejas Shah, Ankit Hinsu, Pritesh Sabara, Apurvasinh Puvar, Janvi Raval, Zarna Patel, Monika Gandhi, Pinal Trivedi, Maharshi Pandya, Amit Kanani, Nidhi Patel, Nitin Savaliya, Raghawendra Kumar, Dinesh Kumar, Zuber Saiyed, Komal Patel, Labdhi Pandya, Snehal Bagatharia, Fenil Patel, Bhavesh Modi, Gaurishankar Shrimali, R D Dixit, A M Kadri, Umang Mishra, Chaitanya Joshi, Madhvi Joshi, |
| EPI_ISL_455018 | Government Medical College, Vadodara | Gujarat Biotechnology Research Centre | R N Daveshwar, Ramesh Pandit, Tejas Shah, Ankit Hinsu, Pritesh Sabara, Apurvasinh Puvar, Janvi Raval, Zarna Patel, Monika Gandhi, Pinal Trivedi, Maharshi Pandya, Amit Kanani, Nidhi Patel, Nitin Savaliya, Raghawendra Kumar, Dinesh Kumar, Zuber Saiyed, Komal Patel, Labdhi Pandya, Snehal Bagatharia, Tanuja Javadekar , Neelam Nathani, Bhavesh Modi, Gaurishankar Shrimali, R D Dixit, A M Kadri, Umang Mishra, Chaitanya Joshi, Madhvi Joshi, |
| EPI_ISL_455019 | Government Medical College, Vadodara | Gujarat Biotechnology Research Centre | Ramesh Pandit, Tejas Shah, Ankit Hinsu, Pritesh Sabara, Apurvasinh Puvar, Janvi Raval, Zarna Patel, Monika Gandhi, Pinal Trivedi, Maharshi Pandya, Amit Kanani, Nidhi Patel, Nitin Savaliya, Raghawendra Kumar, Dinesh Kumar, Zuber Saiyed, Komal Patel, Labdhi Pandya, Snehal Bagatharia, Tanuja Javadekar , R N Daveshwar, Armi Chaudhari, Bhavesh Modi, Gaurishankar Shrimali, R D Dixit, A M Kadri, Umang Mishra, Chaitanya Joshi, Madhvi Joshi, |
| EPI_ISL_455020 | Government Medical College, Vadodara | Gujarat Biotechnology Research Centre | Tejas Shah, Ankit Hinsu, Pritesh Sabara, Apurvasinh Puvar, Janvi Raval, Zarna Patel, Monika Gandhi, Pinal Trivedi, Maharshi Pandya, Amit Kanani, Nidhi Patel, Nitin Savaliya, Raghawendra Kumar, Dinesh Kumar, Zuber Saiyed, Komal Patel, Labdhi Pandya, Snehal Bagatharia, Tanuja Javadekar , R N Daveshwar, Ramesh Pandit, Bhavya Jindal, Bhavesh Modi, Gaurishankar Shrimali, R D Dixit, A M Kadri, Umang Mishra, Chaitanya Joshi, Madhvi Joshi, |
| EPI_ISL_455021 | Government Medical College, Vadodara | Gujarat Biotechnology Research Centre | Ankit Hinsu, Pritesh Sabara, Apurvasinh Puvar, Janvi Raval, Zarna Patel, Monika Gandhi, Pinal Trivedi, Maharshi Pandya, Amit Kanani, Nidhi Patel, Nitin Savaliya, Raghawendra Kumar, Dinesh Kumar, Zuber Saiyed, Komal Patel, Labdhi Pandya, Snehal Bagatharia, Tanuja Javadekar , R N Daveshwar, Ramesh Pandit, Tejas Shah, Camellia Chakraborty, Bhavesh Modi, Gaurishankar Shrimali, R D Dixit, A M Kadri, Umang Mishra, Chaitanya Joshi, Madhvi Joshi, |
| EPI_ISL_455022 | Government Medical College, Vadodara | Gujarat Biotechnology Research Centre | Pritesh Sabara, Apurvasinh Puvar, Janvi Raval, Zarna Patel, Monika Gandhi, Pinal Trivedi, Maharshi Pandya, Amit Kanani, Nidhi Patel, Nitin Savaliya, Raghawendra Kumar, Dinesh Kumar, Zuber Saiyed, Komal Patel, Labdhi Pandya, Snehal Bagatharia, Tanuja Javadekar , R N Daveshwar, Ramesh Pandit, Tejas Shah, Ankit Hinsu, Siddhant Kumar, Bhavesh Modi, Gaurishankar Shrimali, R D Dixit, A M Kadri, Umang Mishra, Chaitanya Joshi, Madhvi Joshi, |
| EPI_ISL_455023 | Government Medical College, Vadodara | Gujarat Biotechnology Research Centre | Apurvasinh Puvar, Janvi Raval, Zarna Patel, Monika Gandhi, Pinal Trivedi, Maharshi Pandya, Amit Kanani, Nidhi Patel, Nitin Savaliya, Raghawendra Kumar, Dinesh Kumar, Zuber Saiyed, Komal Patel, Labdhi Pandya, Snehal Bagatharia, Tanuja Javadekar , R N Daveshwar, Ramesh Pandit, Tejas Shah, Ankit Hinsu, Pritesh Sabara, Priyanka P Vatsa, Bhavesh Modi, Gaurishankar Shrimali, R D Dixit, A M Kadri, Umang Mishra, Chaitanya Joshi, Madhvi Joshi, |
| EPI_ISL_455024 | Government Medical College, Vadodara | Gujarat Biotechnology Research Centre | Janvi Raval, Zarna Patel, Monika Gandhi, Pinal Trivedi, Maharshi Pandya, Amit Kanani, Nidhi Patel, Nitin Savaliya, Raghawendra Kumar, Dinesh Kumar, Zuber Saiyed, Komal Patel, Labdhi Pandya, Snehal Bagatharia, Tanuja Javadekar , R N Daveshwar, Ramesh Pandit, Tejas Shah, Ankit Hinsu, Pritesh Sabara, Apurvasinh Puvar, Pooja P Doshi, Bhavesh Modi, Gaurishankar Shrimali, R D Dixit, A M Kadri, Umang Mishra, Chaitanya Joshi, Madhvi Joshi, |
| EPI_ISL_455025 | Government Medical College, Vadodara | Gujarat Biotechnology Research Centre | Zarna Patel, Monika Gandhi, Pinal Trivedi, Maharshi Pandya, Amit Kanani, Nidhi Patel, Nitin Savaliya, Raghawendra Kumar, Dinesh Kumar, Zuber Saiyed, Komal Patel, Labdhi Pandya, Snehal Bagatharia, Tanuja Javadekar , R N Daveshwar, Ramesh Pandit, Tejas Shah, Ankit Hinsu, Pritesh Sabara, Apurvasinh Puvar, Janvi Raval, Akanksha Verma, Bhavesh Modi, Gaurishankar Shrimali, R D Dixit, A M Kadri, Umang Mishra, Chaitanya Joshi, Madhvi Joshi, |
| EPI_ISL_455026 | Government Medical College, Vadodara | Gujarat Biotechnology Research Centre | Monika Gandhi, Pinal Trivedi, Maharshi Pandya, Amit Kanani, Nidhi Patel, Nitin Savaliya, Raghawendra Kumar, Dinesh Kumar, Zuber Saiyed, Komal Patel, Labdhi Pandya, Snehal Bagatharia, Tanuja Javadekar , R N Daveshwar, Ramesh Pandit, Tejas Shah, Ankit Hinsu, Pritesh Sabara, Apurvasinh Puvar, Janvi Raval, Zarna Patel, Priti Pandita, Bhavesh Modi, Gaurishankar Shrimali, R D Dixit, A M Kadri, Umang Mishra, Chaitanya Joshi, Madhvi Joshi, |
| EPI_ISL_455027 | Government Medical College, Vadodara | Gujarat Biotechnology Research Centre | Pinal Trivedi, Maharshi Pandya, Amit Kanani, Nidhi Patel, Nitin Savaliya, Raghawendra Kumar, Dinesh Kumar, Zuber Saiyed, Komal Patel, Labdhi Pandya, Snehal Bagatharia, Tanuja Javadekar , R N Daveshwar, Ramesh Pandit, Tejas Shah, Ankit Hinsu, Pritesh Sabara, Apurvasinh Puvar, Janvi Raval, Zarna Patel, Monika Gandhi, Pragya Sharma, Bhavesh Modi, Gaurishankar Shrimali, R D Dixit, A M Kadri, Umang Mishra, Chaitanya Joshi, Madhvi Joshi, |
| EPI_ISL_455028, EPI_ISL_455029, EPI_ISL_455030, EPI_ISL_455031, EPI_ISL_455032, EPI_ISL_455033, EPI_ISL_455034, EPI_ISL_455035 | Pathology West - NSW Health Pathology | NSW Health Pathology - Institute of Clinical Pathology and Medical Research; Westmead Hospital; University of Sydney | CIDM-PH et al. |
| EPI_ISL_455036 | South Eastern Area Laboratory Services | NSW Health Pathology - Institute of Clinical Pathology and Medical Research; Westmead Hospital; University of Sydney | CIDM-PH et al. |
| EPI_ISL_455037 | Pathology Sydney South West - NSW Health Pathology | NSW Health Pathology - Institute of Clinical Pathology and Medical Research; Westmead Hospital; University of Sydney | CIDM-PH et al. |
| EPI_ISL_455038, EPI_ISL_455039, EPI_ISL_455040 | Pathology West - NSW Health Pathology | NSW Health Pathology - Institute of Clinical Pathology and Medical Research; Westmead Hospital; University of Sydney | CIDM-PH et al. |
| EPI_ISL_455041 | Laverty Pathology | NSW Health Pathology - Institute of Clinical Pathology and Medical Research; Westmead Hospital; University of Sydney | CIDM-PH et al. |
| EPI_ISL_455042, EPI_ISL_455043 | ACT Pathology | NSW Health Pathology - Institute of Clinical Pathology and Medical Research; Westmead Hospital; University of Sydney | CIDM-PH et al. |
| EPI_ISL_455044, EPI_ISL_455045, EPI_ISL_455046, EPI_ISL_455047 | Pathology West - NSW Health Pathology | NSW Health Pathology - Institute of Clinical Pathology and Medical Research; Westmead Hospital; University of Sydney | CIDM-PH et al. |
| EPI_ISL_455048 | Laverty Pathology | NSW Health Pathology - Institute of Clinical Pathology and Medical Research; Westmead Hospital; University of Sydney | CIDM-PH et al. |
| EPI_ISL_455049 | Pathology West - NSW Health Pathology | NSW Health Pathology - Institute of Clinical Pathology and Medical Research; Westmead Hospital; University of Sydney | CIDM-PH et al. |
| EPI_ISL_455050 | ACT Pathology | NSW Health Pathology - Institute of Clinical Pathology and Medical Research; Westmead Hospital; University of Sydney | CIDM-PH et al. |
| EPI_ISL_455051, EPI_ISL_455052, EPI_ISL_455053 | Douglas Hanly Moir Pathology | NSW Health Pathology - Institute of Clinical Pathology and Medical Research; Westmead Hospital; University of Sydney | CIDM-PH et al. |
| EPI_ISL_455054 | South Eastern Area Laboratory Services | NSW Health Pathology - Institute of Clinical Pathology and Medical Research; Westmead Hospital; University of Sydney | CIDM-PH et al. |
| EPI_ISL_455055, EPI_ISL_455056, EPI_ISL_455057, EPI_ISL_455058, EPI_ISL_455059, EPI_ISL_455060, EPI_ISL_455061, EPI_ISL_455062 | Pathology West - NSW Health Pathology | NSW Health Pathology - Institute of Clinical Pathology and Medical Research; Westmead Hospital; University of Sydney | CIDM-PH et al. |
| EPI_ISL_455063 | Laverty Pathology | NSW Health Pathology - Institute of Clinical Pathology and Medical Research; Westmead Hospital; University of Sydney | CIDM-PH et al. |
| EPI_ISL_455064 | Pathology West - NSW Health Pathology | NSW Health Pathology - Institute of Clinical Pathology and Medical Research; Westmead Hospital; University of Sydney | CIDM-PH et al. |
| EPI_ISL_455065 | Douglas Hanly Moir Pathology | NSW Health Pathology - Institute of Clinical Pathology and Medical Research; Westmead Hospital; University of Sydney | CIDM-PH et al. |
| EPI_ISL_455066, EPI_ISL_455067 | Pathology West - NSW Health Pathology | NSW Health Pathology - Institute of Clinical Pathology and Medical Research; Westmead Hospital; University of Sydney | CIDM-PH et al. |
| EPI_ISL_455068 | Childrens Hospital Westmead | NSW Health Pathology - Institute of Clinical Pathology and Medical Research; Westmead Hospital; University of Sydney | CIDM-PH et al. |
| EPI_ISL_455069, EPI_ISL_455070 | Pathology West - NSW Health Pathology | NSW Health Pathology - Institute of Clinical Pathology and Medical Research; Westmead Hospital; University of Sydney | CIDM-PH et al. |
| EPI_ISL_455071 | Pathology Sydney South West - NSW Health Pathology | NSW Health Pathology - Institute of Clinical Pathology and Medical Research; Westmead Hospital; University of Sydney | CIDM-PH et al. |
| EPI_ISL_455072, EPI_ISL_455073 | Sullivan Nicolaides Pathology | NSW Health Pathology - Institute of Clinical Pathology and Medical Research; Westmead Hospital; University of Sydney | CIDM-PH et al. |
| EPI_ISL_455074 | ACT Pathology | NSW Health Pathology - Institute of Clinical Pathology and Medical Research; Westmead Hospital; University of Sydney | CIDM-PH et al. |
| EPI_ISL_455075, EPI_ISL_455076, EPI_ISL_455077, EPI_ISL_455078, EPI_ISL_455079, EPI_ISL_455080, EPI_ISL_455081, EPI_ISL_455082, EPI_ISL_455083, EPI_ISL_455084, EPI_ISL_455085, EPI_ISL_455086, EPI_ISL_455087, EPI_ISL_455088, EPI_ISL_455089, EPI_ISL_455090, EPI_ISL_455091, EPI_ISL_455092, EPI_ISL_455093 | | | |
| see above | South Eastern Area Laboratory Services | NSW Health Pathology - Institute of Clinical Pathology and Medical Research; Westmead Hospital; University of Sydney | CIDM-PH et al. |
| EPI_ISL_455094 | Pathology Sydney South West - NSW Health Pathology | NSW Health Pathology - Institute of Clinical Pathology and Medical Research; Westmead Hospital; University of Sydney | CIDM-PH et al. |
| EPI_ISL_455095 | Pathology West - NSW Health Pathology | NSW Health Pathology - Institute of Clinical Pathology and Medical Research; Westmead Hospital; University of Sydney | CIDM-PH et al. |
| EPI_ISL_455096, EPI_ISL_455097, EPI_ISL_455098, EPI_ISL_455099 | South Eastern Area Laboratory Services | NSW Health Pathology - Institute of Clinical Pathology and Medical Research; Westmead Hospital; University of Sydney | CIDM-PH et al. |

| | | | |
|---|---|---|---|
| EPI_ISL_455100 | Orsa VC | The Public Health Agency of Sweden | Anna-Karin Lundqvist, Anna-Malin Linde, Maria Lind Karlberg, Oskar Karlsson Lindsjo, Olov Svartstrom, Anna Risberg, Theresa Enkirch, Mia Brytting, Karin Tegmark-Wisell |
| EPI_ISL_455101 | Jourcentralen | The Public Health Agency of Sweden | Salvatore Ascione, Anna-Malin Linde, Maria Lind Karlberg, Oskar Karlsson Lindsjo, Olov Svartstrom, Anna Risberg, Theresa Enkirch, Mia Brytting, Karin Tegmark-Wisell |
| EPI_ISL_455102 | Kungsors VC | The Public Health Agency of Sweden | Jessica Karlsson, Anna-Malin Linde, Maria Lind Karlberg, Oskar Karlsson Lindsjo, Olov Svartstrom, Anna Risberg, Theresa Enkirch, Mia Brytting, Karin Tegmark-Wisell |
| EPI_ISL_455103 | Huslakarna Varmbadhuset Varberg | The Public Health Agency of Sweden | Johanna Hilmersson, Anna-Malin Linde, Maria Lind Karlberg, Oskar Karlsson Lindsjo, Olov Svartstrom, Anna Risberg, Theresa Enkirch, Mia Brytting, Karin Tegmark-Wisell |
| EPI_ISL_455104 | Ulltuna Vardcentral | The Public Health Agency of Sweden | Heidi Lindback, Anna-Malin Linde, Maria Lind Karlberg, Oskar Karlsson Lindsjo, Olov Svartstrom, Anna Risberg, Theresa Enkirch, Mia Brytting, Karin Tegmark-Wisell |
| EPI_ISL_455105 | Wetterhalsan | The Public Health Agency of Sweden | Anders Tengblad, Anna-Malin Linde, Maria Lind Karlberg, Oskar Karlsson Lindsjo, Olov Svartstrom, Anna Risberg, Theresa Enkirch, Mia Brytting, Karin Tegmark-Wisell |
| EPI_ISL_455106 | Surbrunns VC | The Public Health Agency of Sweden | Erik Embring, Anna-Malin Linde, Maria Lind Karlberg, Oskar Karlsson Lindsjo, Olov Svartstrom, Anna Risberg, Theresa Enkirch, Mia Brytting, Karin Tegmark-Wisell |
| EPI_ISL_455107 | Narhalsan Mellerud | The Public Health Agency of Sweden | Maria Nykvist, Anna-Malin Linde, Maria Lind Karlberg, Oskar Karlsson Lindsjo, Olov Svartstrom, Anna Risberg, Theresa Enkirch, Mia Brytting, Karin Tegmark-Wisell |
| EPI_ISL_455108 | Smedby HC | The Public Health Agency of Sweden | Susanne Brunby, Anna-Malin Linde, Maria Lind Karlberg, Oskar Karlsson Lindsjo, Olov Svartstrom, Anna Risberg, Theresa Enkirch, Mia Brytting, Karin Tegmark-Wisell |
| EPI_ISL_455109 | Ulltuna Vardcentral | The Public Health Agency of Sweden | Heidi Lindback, Anna-Malin Linde, Maria Lind Karlberg, Oskar Karlsson Lindsjo, Olov Svartstrom, Anna Risberg, Theresa Enkirch, Mia Brytting, Karin Tegmark-Wisell |
| EPI_ISL_455110 | Scania Halsocenter, B288 | The Public Health Agency of Sweden | Christina Lergin, Anna-Malin Linde, Maria Lind Karlberg, Oskar Karlsson Lindsjo, Olov Svartstrom, Anna Risberg, Theresa Enkirch, Mia Brytting, Karin Tegmark-Wisell |
| EPI_ISL_455111 | Olof Norrby | The Public Health Agency of Sweden | Bla Kustens halsocentral, Anna-Malin Linde, Maria Lind Karlberg, Oskar Karlsson Lindsjo, Olov Svartstrom, Anna Risberg, Theresa Enkirch, Mia Brytting, Karin Tegmark-Wisell |
| EPI_ISL_455112, EPI_ISL_455113, EPI_ISL_455114, EPI_ISL_455115, EPI_ISL_455116, EPI_ISL_455117, EPI_ISL_455118, EPI_ISL_455119, EPI_ISL_455120, EPI_ISL_455121, EPI_ISL_455122, EPI_ISL_455123, EPI_ISL_455124, EPI_ISL_455125, EPI_ISL_455126, EPI_ISL_455127, EPI_ISL_455128, EPI_ISL_455129, EPI_ISL_455130, EPI_ISL_455131, EPI_ISL_455132, EPI_ISL_455133, EPI_ISL_455134, EPI_ISL_455135, EPI_ISL_455136, EPI_ISL_455137, EPI_ISL_455138, EPI_ISL_455139, EPI_ISL_455140, EPI_ISL_455141, EPI_ISL_455142, EPI_ISL_455143, EPI_ISL_455144, EPI_ISL_455145, EPI_ISL_455146, EPI_ISL_455147, EPI_ISL_455148, EPI_ISL_455149, EPI_ISL_455150, EPI_ISL_455151, EPI_ISL_455152, EPI_ISL_455153, EPI_ISL_455154, EPI_ISL_455155, EPI_ISL_455156, EPI_ISL_455157, EPI_ISL_455158, EPI_ISL_455159, EPI_ISL_455160, EPI_ISL_455161, EPI_ISL_455162, EPI_ISL_455163, EPI_ISL_455164, EPI_ISL_455165, EPI_ISL_455166, EPI_ISL_455167, EPI_ISL_455168, EPI_ISL_455169, EPI_ISL_455170, EPI_ISL_455171, EPI_ISL_455172, EPI_ISL_455173, EPI_ISL_455174, EPI_ISL_455175, EPI_ISL_455176, EPI_ISL_455177, EPI_ISL_455178, EPI_ISL_455179, EPI_ISL_455180, EPI_ISL_455181, EPI_ISL_455182, EPI_ISL_455183, EPI_ISL_455184, EPI_ISL_455185, EPI_ISL_455186, EPI_ISL_455187, EPI_ISL_455188, EPI_ISL_455189, EPI_ISL_455190, EPI_ISL_455191, EPI_ISL_455192, EPI_ISL_455193, EPI_ISL_455194, EPI_ISL_455195, EPI_ISL_455196, EPI_ISL_455197, EPI_ISL_455198, EPI_ISL_455199, EPI_ISL_455200, EPI_ISL_455201, EPI_ISL_455202, EPI_ISL_455203, EPI_ISL_455204, EPI_ISL_455205, EPI_ISL_455206, EPI_ISL_455207, EPI_ISL_455208, EPI_ISL_455209, EPI_ISL_455210, EPI_ISL_455211, EPI_ISL_455212, EPI_ISL_455213, EPI_ISL_455214, EPI_ISL_455215, EPI_ISL_455216, EPI_ISL_455217, EPI_ISL_455218, EPI_ISL_455219, EPI_ISL_455220, EPI_ISL_455221, EPI_ISL_455222, EPI_ISL_455223, EPI_ISL_455224, EPI_ISL_455225, EPI_ISL_455226, EPI_ISL_455227, EPI_ISL_455228, EPI_ISL_455229, EPI_ISL_455230, EPI_ISL_455231, EPI_ISL_455232, EPI_ISL_455233, EPI_ISL_455234, EPI_ISL_455235, EPI_ISL_455236, EPI_ISL_455237, EPI_ISL_455238, EPI_ISL_455239, EPI_ISL_455240, EPI_ISL_455241, EPI_ISL_455242, EPI_ISL_455243, EPI_ISL_455244, EPI_ISL_455245, EPI_ISL_455246, EPI_ISL_455247, EPI_ISL_455248, EPI_ISL_455249, EPI_ISL_455250, EPI_ISL_455251, EPI_ISL_455252, EPI_ISL_455253, EPI_ISL_455254, EPI_ISL_455255, EPI_ISL_455256, EPI_ISL_455257, EPI_ISL_455258, EPI_ISL_455259, EPI_ISL_455260, EPI_ISL_455261, EPI_ISL_455262, EPI_ISL_455263, EPI_ISL_455264, EPI_ISL_455265, EPI_ISL_455266, EPI_ISL_455267, EPI_ISL_455268, EPI_ISL_455269, EPI_ISL_455270, EPI_ISL_455271, EPI_ISL_455272, EPI_ISL_455273, EPI_ISL_455274, EPI_ISL_455275, EPI_ISL_455276, EPI_ISL_455277, EPI_ISL_455278, EPI_ISL_455279, EPI_ISL_455280, EPI_ISL_455281, EPI_ISL_455282, EPI_ISL_455283, EPI_ISL_455284, EPI_ISL_455285, EPI_ISL_455286, EPI_ISL_455287, EPI_ISL_455288, EPI_ISL_455289, EPI_ISL_455290, EPI_ISL_455291, EPI_ISL_455292, EPI_ISL_455293, EPI_ISL_455294, EPI_ISL_455295, EPI_ISL_455296, EPI_ISL_455297, EPI_ISL_455298, EPI_ISL_455299, EPI_ISL_455300, EPI_ISL_455301, EPI_ISL_455302, EPI_ISL_455303, EPI_ISL_455304, EPI_ISL_455305, EPI_ISL_455306, EPI_ISL_455307 | | | |
| see above | Dutch COVID-19 response team | Erasmus Medical Center | Bas Oude Munnink, David Nieuwenhuijse, Reina Sikkema, Claudia Schapendonk, Irina Chestakova, Anne van der Linden, Theo Bestebroer, Stefan van Nieuwkoop, Mark Pronk, Pascal Lexmond, Corien Swaan, Manon Haverkate, Madelief Mollers, Mart Stein, Sandra Kengne Kamga Mobou, Jeroen van Kampen, Jolanda Voermans, Aura Timen, Corine GeurtsvanKessel, Annemiek van der Eijk, Richard Molenkamp, Marion Koopmans, on behalf of the Dutch national COVID-19 response team. |
| EPI_ISL_455308, EPI_ISL_455309, EPI_ISL_455310, EPI_ISL_455311 | REGIONAL VRDL,ICMR-RMRC BBSR | Immunogenomics group, Institute of Life Sciences, Bhubaneswar | Sunil Raghav, Jyotirmayee Turuk, Arup Ghosh, Atimukta Jha, Viplov K. Biswas, Swati Madhulika, Manasi Priyadarshini, Shuchi Smita, Jaya Singh Khastri, Rupesh Dash, Soma Chattopadhyay, Ghulam Hussain Syed, Shanti Senapati, Tushar K. Beuria, Debdutta Bhattacharya, Rajeeb Swain, Punit Prasad, COVID-19 team of ILS & RMRC, Orissa COVID-19 study group, DBT's PAN-INDIA 1000 SARS-CoV2 RNA genome sequencing consortium, Sanghamitra Pati, Ajay Parida |
| EPI_ISL_455312 | Microbiology Unit, Department of Pathology & Laboratory Medicine, IIUM Medical Centre | SEA Microbiome Unit, Faculty of Industrial Sciences & Technology, Universiti Malaysia Pahang | Norhidayah Binti Kamarudin, Ahmad Hafiz Bin Zulkifly, Hajar Fauzan Ahmad, Muhammad Adam Lee Abdullah, Mohd Fazli Farida Asras, Ahmad Mahfuz Gazali, Mohd Nazli Bin Kamarulzaman, IIUM Medical Centre Covid19 Taskforce, UMP Covid19 Team |
| EPI_ISL_455313 | Microbiology Unit, Department of Pathology & Laboratory Medicine, IIUM Medical Centre | SEA Microbiome Unit, Faculty of Industrial Sciences & Technology, Universiti Malaysia Pahang | Hajar Fauzan Ahmad, Norhidayah Kamarudin, Ahmad Hafiz Zulkifly, IIUM Medical Centre Covid19 Taskforce, UMP Covid19 Team |
| EPI_ISL_455314 | Hospital Virgen del Rocio | Instituto de Salud Carlos III | Iglesias-Caballero, M. Molinero Calamita, M. González-Esguevillas, M. Camarero, S. Pozo, F. Casas, I. Jiménez, P. Jiménez, M. Zaballos, A. Monzón, S. Varona, S. Juliá, M. Cuesta, I, J. Lepe |
| EPI_ISL_455315, EPI_ISL_455316, EPI_ISL_455317, EPI_ISL_455318, EPI_ISL_455319, EPI_ISL_455320, EPI_ISL_455321, EPI_ISL_455322 | Hospital Virgen de las Nieves | Instituto de Salud Carlos III | Iglesias-Caballero, M. Molinero Calamita, M. González-Esguevillas, M. Camarero, S. Pozo, F. Casas, I. Jiménez, P. Jiménez, M. Zaballos, A. Monzón, S. Varona, S. Juliá, M. Cuesta, I, S. Sanbonmatsu |
| EPI_ISL_455323 | Hospital Virgen del Rocio | Instituto de Salud Carlos III | Iglesias-Caballero, M. Molinero Calamita, M. González-Esguevillas, M. Camarero, S. Pozo, F. Casas, I. Jiménez, P. Jiménez, M. Zaballos, A. Monzón, S. Varona, S. Juliá, M. Cuesta, I, J. Lepe |
| EPI_ISL_455324 | Hospital Virgen de las Nieves | Instituto de Salud Carlos III | Iglesias-Caballero, M. Molinero Calamita, M. González-Esguevillas, M. Camarero, S. Pozo, F. Casas, I. Jiménez, P. Jiménez, M. Zaballos, A. Monzón, S. Varona, S. Juliá, M. Cuesta, I, S. Sanbonmatsu |
| EPI_ISL_455325 | Hospital Universitario de Canarias | Instituto de Salud Carlos III | Iglesias-Caballero, M. Molinero Calamita, M. González-Esguevillas, M. Camarero, S. Pozo, F. Casas, I. Jiménez, P. Jiménez, M. Zaballos, A. Monzón, S. Varona, S. Juliá, M. Cuesta, I, B. Castro |
| EPI_ISL_455326 | Hospital Universitario Insular de Gran Canaria | Instituto de Salud Carlos III | Iglesias-Caballero, M. Molinero Calamita, M. González-Esguevillas, M. Camarero, S. Pozo, F. Casas, I. Jiménez, P. Jiménez, M. Zaballos, A. Monzón, S. Varona, S. Juliá, M. Cuesta, I, A. Hernández |
| EPI_ISL_455327 | Consejeria de Sanidad y Asuntos Sociales | Instituto de Salud Carlos III | Iglesias-Caballero, M. Molinero Calamita, M. González-Esguevillas, M. Camarero, S. Pozo, F. Casas, I. Jiménez, P. Jiménez, M. Zaballos, A. Monzón, S. Varona, S. Juliá, M. Cuesta, I, G. Gutiérrez |
| EPI_ISL_455328, EPI_ISL_455329, EPI_ISL_455330, EPI_ISL_455331 | Complejo Hospitalario Universitario La Coruna | Instituto de Salud Carlos III | Iglesias-Caballero, M. Molinero Calamita, M. González-Esguevillas, M. Camarero, S. Pozo, F. Casas, I. Jiménez, P. Jiménez, M. Zaballos, A. Monzón, S. Varona, S. Juliá, M. Cuesta, I, M.A Canizares |
| EPI_ISL_455332 | Xerencia de Xestión Integrada de Pontevedra e o Salnés | Instituto de Salud Carlos III | Iglesias-Caballero, M. Molinero Calamita, M. González-Esguevillas, M. Camarero, S. Pozo, F. Casas, I. Jiménez, P. Jiménez, M. Zaballos, A. Monzón, S. Varona, S. Juliá, M. Cuesta, I, M. Garcia |
| EPI_ISL_455333 | Complejo Hospitalario Universitario de Santiago | Instituto de Salud Carlos III | Iglesias-Caballero, M. Molinero Calamita, M. González-Esguevillas, M. Camarero, S. Pozo, F. Casas, I. Jiménez, P. Jiménez, M. Zaballos, A. Monzón, S. Varona, S. Juliá, M. Cuesta, I, J. Llovo |
| EPI_ISL_455334, EPI_ISL_455335 | Complejo Hospitalario de Orense | Instituto de Salud Carlos III | Iglesias-Caballero, M. Molinero Calamita, M. González-Esguevillas, M. Camarero, S. Pozo, F. Casas, I. Jiménez, P. Jiménez, M. Zaballos, A. Monzón, S. Varona, S. Juliá, M. Cuesta, I, M. Paz |
| EPI_ISL_455336, EPI_ISL_455337, EPI_ISL_455338, EPI_ISL_455339, EPI_ISL_455340, EPI_ISL_455341, EPI_ISL_455342, EPI_ISL_455343 | Hospital San Pedro | Instituto de Salud Carlos III | Iglesias-Caballero, M. Molinero Calamita, M. González-Esguevillas, M. Camarero, S. Pozo, F. Casas, I. Jiménez, P. Jiménez, M. Zaballos, A. Monzón, S. Varona, S. Juliá, M. Cuesta, I, C. Alonso |
| EPI_ISL_455344, EPI_ISL_455345, EPI_ISL_455346, EPI_ISL_455347, EPI_ISL_455348, EPI_ISL_455349 | Hospital Comarcal de Melilla | Instituto de Salud Carlos III | Iglesias-Caballero, M. Molinero Calamita, M. González-Esguevillas, M. Camarero, S. Pozo, F. Casas, I. Jiménez, P. Jiménez, M. Zaballos, A. Monzón, S. Varona, S. Juliá, M. Cuesta, I, I. Pérez |
| EPI_ISL_455350, EPI_ISL_455351 | Hospital Txagorritxu | Instituto de Salud Carlos III | Iglesias-Caballero, M. Molinero Calamita, M. González-Esguevillas, M. Camarero, S. Pozo, F. Casas, I. Jiménez, P. Jiménez, M. Zaballos, A. Monzón, S. Varona, S. Juliá, M. Cuesta, I, C. Gómez |
| EPI_ISL_455352, EPI_ISL_455353, EPI_ISL_455354 | Hospital de Cruces | Instituto de Salud Carlos III | Iglesias-Caballero, M. Molinero Calamita, M. González-Esguevillas, M. Camarero, S. Pozo, F. Casas, I. Jiménez, P. Jiménez, M. Zaballos, A. Monzón, S. Varona, S. Juliá, M. Cuesta, I, M. Aranzamendi |
| EPI_ISL_455355, EPI_ISL_455356, EPI_ISL_455357, EPI_ISL_455358, EPI_ISL_455359, EPI_ISL_455360, EPI_ISL_455361 | Emory Molecular Diagnostics Laboratory, Emory Healthcare | Piantadosi Lab, Emory Department of Pathology | Ahmed Babiker, Anne Piantadosi |
| EPI_ISL_455362 | Nigeria Centre for Disease Control (NCDC) | African Centre of Excellence for Genomics of Infectious Diseases (ACEGID), Redeemer's University, Ede, Osun State, Nigeria | Oluniyi P.E., Ajogbasile F.V., Kayode A., Olawoye I., Uwanibe J., Oguzie J., Olumade T., Folarin O.A., Ihekweazu C., Happi C.T. |
| EPI_ISL_455363, EPI_ISL_455364, EPI_ISL_455365, EPI_ISL_455366, EPI_ISL_455367, EPI_ISL_455368, EPI_ISL_455369, EPI_ISL_455370, EPI_ISL_455371, EPI_ISL_455372, EPI_ISL_455373, EPI_ISL_455374, EPI_ISL_455375, EPI_ISL_455376, EPI_ISL_455377, EPI_ISL_455378, EPI_ISL_455379, EPI_ISL_455380, EPI_ISL_455381, EPI_ISL_455382, EPI_ISL_455383, EPI_ISL_455384, EPI_ISL_455385, EPI_ISL_455386, EPI_ISL_455387, EPI_ISL_455388, EPI_ISL_455389, EPI_ISL_455390, EPI_ISL_455391, EPI_ISL_455392, EPI_ISL_455393, EPI_ISL_455394, EPI_ISL_455395, EPI_ISL_455396, EPI_ISL_455397, EPI_ISL_455398, EPI_ISL_455399, EPI_ISL_455400, EPI_ISL_455401, EPI_ISL_455402, EPI_ISL_455403, EPI_ISL_455404, EPI_ISL_455405, EPI_ISL_455406, EPI_ISL_455407, EPI_ISL_455408, EPI_ISL_455409, EPI_ISL_455410, EPI_ISL_455411 | | | |
| see above | Wuhan Chain Medical Labs (CMLabs) | State Key Laboratory of Biotherapy of Sichuan University | Baowen Du, Minjin Wang, Chao Tang, Chuan Chen, Yongzhao Zhou, Mingxia Yu, Hancheng Wei, Weimin Li, Jing-wen Lin, Jia Geng, Binwu Ying, Lu Chen |
| EPI_ISL_455412, EPI_ISL_455413, EPI_ISL_455414 | Nigeria Centre for Disease Control (NCDC) | African Centre of Excellence for Genomics of Infectious Diseases (ACEGID), Redeemer's University, Ede, Osun State, Nigeria | Oluniyi P.E., Ajogbasile F.V., Kayode A., Oguzie J., Olawoye I., Uwanibe J., Olumade T., Folarin O.A., Ihekweazu C., Happi C.T. |
| EPI_ISL_455415 | Nigeria Centre for Disease Control (NCDC) | African Centre of Excellence for Genomics of Infectious Diseases (ACEGID), Redeemer's University, Ede, Osun State, Nigeria | Oluniyi P.E., Ajogbasile F.V., Kayode A., Oguzie J., Olawoye I., Uwanibe J., Olumade T., Folarin O.A., Ihekweazu C., Happi C.T. |
| EPI_ISL_455418, EPI_ISL_455419 | Nigeria Centre for Disease Control (NCDC) | African Centre of Excellence for Genomics of Infectious Diseases (ACEGID), Redeemer's University, Ede, Osun State, Nigeria | Oluniyi P.E., Ajogbasile F.V., Kayode A., Oguzie J., Olawoye I., Uwanibe J., Olumade T., Folarin O.A., Ihekweazu C., Happi C.T. |
| EPI_ISL_455420 | National Institute of Laboratory Medicine and Referral Center | Genomic Research Lab, BCSIR | Abu Sayeed Mohammad Mahmud, Mohammad Samir Uzzaman, Eshrar Osman, Md. Ahasan Habib, Shahina Akhter, Tanjina Akhter Banu, Barna Goswami, Iffat Jahan, Tasnim Nafisa, Md. Maruf Ahmed Molla, MahmudaYeasmin, Sheikh Md. Selim Al Din, Utpal Chandra Ray, Md. Salim Khan |
| EPI_ISL_455422 | Nigeria Centre for Disease Control | African Centre of Excellence for Genomics of Infectious Diseases (ACEGID), Redeemer's University, Ede, Osun State, Nigeria | Oluniyi P.E., Ajogbasile F.V., Kayode A., Oguzie J., Olawoye I., Uwanibe J., Olumade T., Folarin O.A., Ihekweazu C., Happi C.T. |
| EPI_ISL_455423, EPI_ISL_455424, EPI_ISL_455425 | Nigeria Centre for Disease Control (NCDC) | African Centre of Excellence for Genomics of Infectious Diseases (ACEGID), Redeemer's University, Ede, Osun State, Nigeria | Oluniyi P.E., Ajogbasile F.V., Kayode A., Oguzie J., Olawoye I., Uwanibe J., Olumade T., Folarin O.A., Ihekweazu C., Happi C.T. |
| EPI_ISL_455426 | Nigeria Centre for Disease Control | African Centre of Excellence for Genomics of Infectious Diseases (ACEGID), Redeemer's University, Ede, Osun State, Nigeria | Oluniyi P.E., Ajogbasile F.V., Kayode A., Oguzie J., Olawoye I., Uwanibe J., Olumade T., Folarin O.A., Ihekweazu C., Happi C.T. |
| EPI_ISL_455427, EPI_ISL_455429, EPI_ISL_455430, EPI_ISL_455431 | Nigeria Centre for Disease Control (NCDC) | African Centre of Excellence for Genomics of Infectious Diseases (ACEGID), Redeemer's University, Ede, Osun State, Nigeria | Oluniyi P.E., Ajogbasile F.V., Kayode A., Oguzie J., Olawoye I., Uwanibe J., Olumade T., Folarin O.A., Ihekweazu C., Happi C.T. |
| EPI_ISL_455432 | Instituto de Diagnostico y Referencia Epidemiologicos (INDRE) | Instituto de Diagnostico y Referencia Epidemiologicos (INDRE) | Taboada Ramírez Blanca. Ramirez-Gonzalez Ernesto, Garces-Ayala Fabiola, Araiza-Rodriguez Adnan, Mendieta-Condado Edgar, Rodriguez-Maldonado Abril, Wong-Arambula Claudia, Barrera-Badillo Gisela, Hernandez-Rivas Lucia, Lopez-Martinez Irma. |
| EPI_ISL_455434 | Instituto de Diagnostico y Referencia Epidemiologicos (INDRE) | Instituto de Diagnostico y Referencia Epidemiologicos (INDRE) | Taboada Ramírez Blanca. Ramirez-Gonzalez Ernesto, Garces-Ayala Fabiola, Araiza-Rodriguez Adnan, Mendieta-Condado Edgar, Rodriguez-Maldonado Abril, Wong-Arambula Claudia, Barrera-Badillo Gisela, Hernandez-Rivas Lucia, Lopez-Martinez Irma. |
| EPI_ISL_455435 | Instituto de Diagnostico y Referencia Epidemiologicos (INDRE) | Instituto de Diagnostico y Referencia Epidemiologicos (INDRE) | Garces-Ayala Fabiola. Taboada Ramírez Blanca. Ramirez-Gonzalez Ernesto, Araiza-Rodriguez Adnan , Mendieta-Condado Edgar, Rodriguez-Maldonado Abril, Wong-Arambula Claudia, Barrera-Badillo Gisela, Hernandez-Rivas Lucia, Lopez-Martinez Irma |
| EPI_ISL_455436 | Instituto de Diagnostico y Referencia Epidemiologicos (INDRE) | Instituto de Diagnostico y Referencia Epidemiologicos (INDRE) | Garces-Ayala Fabiola. Taboada Ramírez Blanca. Ramirez-Gonzalez Ernesto, Araiza-Rodriguez Adnan, Mendieta-Condado Edgar, Rodriguez-Maldonado Abril, Wong-Arambula Claudia, Barrera-Badillo Gisela, |

| | | | |
|---|---|---|---|
| | | | Hernandez-Rivas Lucia, Lopez-Martinez Irma. |
| EPI_ISL_455437, EPI_ISL_455438 | Instituto de Diagnostico y Referencia Epidemiologicos (INDRE) | Instituto de Diagnostico y Referencia Epidemiologicos (INDRE) | Araiza-Rodriguez Adnan, Garces-Ayala Fabiola. Ramirez-Gonzalez Ernesto, Mendieta-Condado Edgar, Rodriguez-Maldonado Abril, Wong-Arambula Claudia, Barrera-Badillo Gisela, Hernandez-Rivas Lucia, Lopez-Martinez Irma, Taboada Ramírez Blanca. |
| EPI_ISL_455439 | Instituto de Diagnostico y Referencia Epidemiologicos (INDRE) | Instituto de Diagnostico y Referencia Epidemiologicos (INDRE) | Mendieta-Condado Edgar, Araiza-Rodriguez Adnan, Garces-Ayala Fabiola, Rodriguez-Maldonado Abril, Wong-Arambula Claudia, Barrera-Badillo Gisela, Hernandez-Rivas Lucia, Lopez-Martinez Irma, Taboada Ramirez Blanca, Ramirez-Gonzalez Ernesto. |
| EPI_ISL_455440, EPI_ISL_455441, EPI_ISL_455442, EPI_ISL_455443, EPI_ISL_455444, EPI_ISL_455445, EPI_ISL_455446, EPI_ISL_455447, EPI_ISL_455448, EPI_ISL_455449, EPI_ISL_455450, EPI_ISL_455451, EPI_ISL_455452, EPI_ISL_455453 | | | |
| see above | 1. ViroGenetics - BSL3 Laboratory of Virology, Małopolska Centre of Biotechnology, Jagiellonian University; 2. II Department of Internal Medicine, Faculty of Medicine, Jagiellonian University Medical College; 3. Narodowy Instytut Zdrowia Publicznego – Państwowy Zakład Higieny (NIZP-PZH) | 1. ViroGenetics - BSL3 Laboratory of Virology, Małopolska Centre of Biotechnology, Jagiellonian University; 2. II Department of Internal Medicine, Faculty of Medicine, Jagiellonian University Medical College; 3. Narodowy Instytut Zdrowia Publicznego – Państwowy Zakład Higieny (NIZP-PZH). | Katarzyna Pancer, Marek Sanak, Aleksandra A. Zasada, Magdalena Rzeczkowska, Tomasz Wołkowicz, Katarzyna Zacharczuk, Agnieszka Kołakowska-Kulesza, Katarzyna Owczarek, Aleksandra Milewska, Natalia Wolaniuk, Ewelina Hallman-Szelińska, Paweł P Łabaj, Wojciech Branicki, Krzysztof Pyrć |
| EPI_ISL_455454 | Instituto de Diagnostico y Referencia Epidemiologicos (INDRE) | Instituto de Diagnostico y Referencia Epidemiologicos (INDRE) | Mendieta-Condado Edgar, Araiza-Rodriguez Adnan, Garces-Ayala Fabiola, , Rodriguez-Maldonado Abril, Wong-Arambula Claudia, Barrera-Badillo Gisela, Taboada Ramírez Blanca, Ramirez-Gonzalez Ernesto, Hernandez-Rivas Lucia, Lopez-Martinez Irma. |
| EPI_ISL_455455 | Instituto de Diagnostico y Referencia Epidemiologicos (INDRE) | Instituto de Diagnostico y Referencia Epidemiologicos (INDRE) | Rodriguez-Maldonado Abril, Mendieta-Condado Edgar, Araiza-Rodriguez Adnan, Garces-Ayala Fabiola. Taboada Ramirez Blanca. Ramirez-Gonzalez Ernesto, Barrera-Badillo Gisela, Hernandez-Rivas Lucia, Lopez-Martinez Irma, Wong-Arambula Claudia. |
| EPI_ISL_455456 | Instituto de Diagnostico y Referencia Epidemiologicos (INDRE) | Instituto de Diagnostico y Referencia Epidemiologicos (INDRE) | Rodriguez-Maldonado Abril, Mendieta-Condado Edgar, Araiza-Rodriguez Adnan, Garces-Ayala Fabiola. Taboada Ramírez Blanca. Ramirez-Gonzalez Ernesto, , Barrera-Badillo Gisela, Hernandez-Rivas Lucia, Lopez-Martinez Irma, Wong-Arambula Claudia. |
| EPI_ISL_455458 | National Institute of Laboratory Medicine and Referral Center | Genomic Research Lab, BCSIR | Abu Sayeed Mohammad Mahmud, Mohammad Samir Uzzaman, Eshrar Osman, Md. Ahasan Habib, Shahina Akhter, Tanjina Akhter Banu, Barna Goswami, Iffat Jahan, Tasnim Nafisa, Md. Maruf Ahmed Molla, MahmudaYeasmin, Sheikh Md. Selim Al Din, Utpal Chandra Ray, Md. Salim Khan |
| EPI_ISL_455459 | National Institute of Laboratory Medicine and Referral Center | Genomic Research Lab, BCSIR | Abu Sayeed Mohammad Mahmud, Mohammad Samir Uzzaman, Eshrar Osman, Md. Ahasan Habib, Shahina Akhter, Tanjina Akhter Banu, Barna Goswami, Iffat Jahan, Tasnim Nafisa, Md. Maruf Ahmed Molla, MahmudaYeasmin, Sheikh Md. Selim Al Din, Utpal Chandra Ray, Md. Salim Khan |
| EPI_ISL_455460, EPI_ISL_455461, EPI_ISL_455462, EPI_ISL_455463, EPI_ISL_455464, EPI_ISL_455465, EPI_ISL_455466, EPI_ISL_455467 | Jiangxi Province Center for Disease Control and Prevention | Jiangxi Province Center for Disease Control and Prevention | JianXiong Li,Ying Xiong,Tian Gong,Yong Shi,Jun Zhou,Fang Xiao,ShiWen Liu,XiaoQing Liu,Gang Xu,DaJin Xiao,Xin Ran,YanNi Zhang |
| EPI_ISL_455468, EPI_ISL_455469, EPI_ISL_455470, EPI_ISL_455471, EPI_ISL_455472, EPI_ISL_455473, EPI_ISL_455474 | Laboratory for Respiratory Viruses, Cantacuzino National Military-Medical Institute for Research and Development | Cantacuzino Institute | M.Lazar, L.Ustea, A.Cretu, Tim Durfee |
| EPI_ISL_455475, EPI_ISL_455476, EPI_ISL_455477 | Laboratory for Respiratory Viruses, Cantacuzino National Military-Medical Institute for Research and Development | Cantacuzino Institute | M.Lazar, L.Ustea, A.Cretu, T.Durfee |
| EPI_ISL_455478 | REGIONAL VRDL,ICMR-RMRC BBSR | Immunogenomics group, Institute of Life Sciences, Bhubaneswar | Sunil Raghav, Jyotirmayee Turuk, Arup Ghosh, Atimukta Jha, Viplov K. Biswas, Swati Madhulika, Manasi Priyadarshini, Shuchi Smita, Jaya Singh Khastri, Rupesh Dash, Soma Chattopadhyay, Ghulam Hussain Syed, Shanti Senapati, Tushar K. Beuria, Debdutta Bhattacharya, Rajeeb Swain, Punit Prasad, COVID-19 team of ILS & RMRC, Orissa COVID-19 study group, DBT's PAN-INDIA 1000 SARS-CoV2 RNA genome sequencing consortium, Sanghamitra Pati, Ajay Parida |
| EPI_ISL_455479 | Laboratory for Respiratory Viruses, Cantacuzino National Military-Medical Institute for Research and Development | Cantacuzino Institute | M.Lazar, L.Ustea, A.Cretu, T.Durfee |
| EPI_ISL_455480 | Veterinary Specialized Institute Kraljevo | Veterinary Specialized Institute Kraljevo | Vidanovic,D., Tesovic,B., Banovic Djeri,B., Sekler,M., Dmitric,M., Debeljak,Z., Matovic,K., Vaskovic,N., Petrovic,T., Volkening,J. and Alfonso,C.L. |
| EPI_ISL_455566, EPI_ISL_455567 | Institute for Public Health | Laboratory for advanced genomics | Filip Rokić, Lovro Trgovec-Greif, Neven Sučić, Tomislav Rukavina, Igor Jurak, Oliver Vugrek |
| EPI_ISL_455568, EPI_ISL_455569, EPI_ISL_455570, EPI_ISL_455571, EPI_ISL_455572 | Gundersen Molecular Diagnostics Laboratory | Kabara Cancer Research Institute | Craig S. Richmond, Paraic A. Kenny |
| EPI_ISL_455573, EPI_ISL_455574 | Gundersen Clinical Microbiology Laboratory | Kabara Cancer Research Institute | Craig S. Richmond, Paraic A. Kenny |
| EPI_ISL_455575, EPI_ISL_455576, EPI_ISL_455577, EPI_ISL_455578, EPI_ISL_455579 | Gundersen Molecular Diagnostics Laboratory | Kabara Cancer Research Institute | Craig S. Richmond, Paraic A. Kenny |
| EPI_ISL_455580, EPI_ISL_455581, EPI_ISL_455582 | Gundersen Clinical Microbiology Laboratory | Kabara Cancer Research Institute | Craig S. Richmond, Paraic A. Kenny |
| EPI_ISL_455583 | Central Chest Institute of Thailand | National Institute of Health. Department of medical Sciences, Ministry of Public Health, Thailand | Pilailuk,Okada; Siripaporn,Phuygun; Thanutsapa,Thanadachakul; Sittiporn,Parnmen;Warawan,Wongboot; Sunthareeya,Waicharoen; Malinee,Chittaganpitch |
| EPI_ISL_455584 | National Institute of Health. Department of medical Sciences, Ministry of Public Health, Thailand | National Institute of Health. Department of medical Sciences, Ministry of Public Health, Thailand | Pilailuk,Okada; Siripaporn,Phuygun; Thanutsapa,Thanadachakul; Sittiporn,Parnmen;Warawan,Wongboot; Sunthareeya,Waicharoen; Malinee,Chittaganpitch |
| EPI_ISL_455585 | Phramongkutklao Hospital | National Institute of Health. Department of medical Sciences, Ministry of Public Health, Thailand | Pilailuk,Okada; Siripaporn,Phuygun; Thanutsapa,Thanadachakul; Sittiporn,Parnmen;Warawan,Wongboot; Sunthareeya,Waicharoen; Malinee,Chittaganpitch |
| EPI_ISL_455586 | Siriraj hospital | National Institute of Health. Department of medical Sciences, Ministry of Public Health, Thailand | Pilailuk,Okada; Siripaporn,Phuygun; Thanutsapa,Thanadachakul; Sittiporn,Parnmen;Warawan,Wongboot; Sunthareeya,Waicharoen; Malinee,Chittaganpitch |
| EPI_ISL_455587 | H.R.H. Maha Chakri Sirindhorn Medical Center | National Institute of Health. Department of medical Sciences, Ministry of Public Health, Thailand | Pilailuk,Okada; Siripaporn,Phuygun; Thanutsapa,Thanadachakul; Sittiporn,Parnmen;Warawan,Wongboot; Sunthareeya,Waicharoen; Malinee,Chittaganpitch |
| EPI_ISL_455588 | Trang Hospital | National Institute of Health. Department of medical Sciences, Ministry of Public Health, Thailand | Pilailuk,Okada; Siripaporn,Phuygun; Thanutsapa,Thanadachakul; Sittiporn,Parnmen;Warawan,Wongboot; Sunthareeya,Waicharoen; Malinee,Chittaganpitch |
| EPI_ISL_455589, EPI_ISL_455590, EPI_ISL_455591, EPI_ISL_455592, EPI_ISL_455593 | National Institute of Health. Department of medical Sciences, Ministry of Public Health, Thailand | National Institute of Health. Department of medical Sciences, Ministry of Public Health, Thailand | Pilailuk,Okada; Siripaporn,Phuygun; Thanutsapa,Thanadachakul; Sittiporn,Parnmen;Warawan,Wongboot; Sunthareeya,Waicharoen; Malinee,Chittaganpitch |
| EPI_ISL_455594 | Central Chest Institute of Thailand | National Institute of Health. Department of medical Sciences, Ministry of Public Health, Thailand | Pilailuk,Okada; Siripaporn,Phuygun; Thanutsapa,Thanadachakul; Sittiporn,Parnmen;Warawan,Wongboot; Sunthareeya,Waicharoen; Malinee,Chittaganpitch |
| EPI_ISL_455595, EPI_ISL_455596, EPI_ISL_455597, EPI_ISL_455598, EPI_ISL_455599, EPI_ISL_455600, EPI_ISL_455601, EPI_ISL_455602, EPI_ISL_455603 | SA Pathology | VPRL | Beard, MR., Van Der Hoek, K., Lim, C.K., Leong, L.E.X., Coldbeck-Shackley, R., Shue, B., Kirby, E., Merrett, J., Llamas, B. |
| EPI_ISL_455604 | Ramkhamhaeng Hospital | National Institute of Health. Department of medical Sciences, Ministry of Public Health, Thailand | Pilailuk,Okada; Siripaporn,Phuygun; Thanutsapa,Thanadachakul; Sittiporn,Parnmen;Warawan,Wongboot; Sunthareeya,Waicharoen; Malinee,Chittaganpitch |
| EPI_ISL_455605 | Panyananthaphikkhu Chonprathan Medical Center | National Institute of Health. Department of medical Sciences, Ministry of Public Health, Thailand | Pilailuk,Okada; Siripaporn,Phuygun; Thanutsapa,Thanadachakul; Sittiporn,Parnmen;Warawan,Wongboot; Sunthareeya,Waicharoen; Malinee,Chittaganpitch |
| EPI_ISL_455606, EPI_ISL_455607 | Praram 9 Hospital | National Institute of Health. Department of medical Sciences, Ministry of Public Health, Thailand | Pilailuk,Okada; Siripaporn,Phuygun; Thanutsapa,Thanadachakul; Sittiporn,Parnmen;Warawan,Wongboot; Sunthareeya,Waicharoen; Malinee,Chittaganpitch |
| EPI_ISL_455608 | Phramongkutklao Hospital | National Institute of Health. Department of medical Sciences, Ministry of Public Health, Thailand | Pilailuk,Okada; Siripaporn,Phuygun; Thanutsapa,Thanadachakul; Sittiporn,Parnmen;Warawan,Wongboot; Sunthareeya,Waicharoen; Malinee,Chittaganpitch |
| EPI_ISL_455609, EPI_ISL_455610, EPI_ISL_455611, EPI_ISL_455612, EPI_ISL_455613, EPI_ISL_455614, EPI_ISL_455615, EPI_ISL_455616, EPI_ISL_455617, EPI_ISL_455618, EPI_ISL_455619, EPI_ISL_455620, EPI_ISL_455621, EPI_ISL_455622, EPI_ISL_455623 | | | |
| see above | Ochsner Health | Bioinfoexperts, LLC | Susanna L. Lamers, David J. Nolan, Rebecca Rose, Sissy Cross, David Moraga Amador, Tong Yang, Luke Caruso, Wayra Navia, Lydia Von Borstel, Xiao Hui Zhou, Amy Feehan, Julia-Garcia-Diaz |
| EPI_ISL_455624 | National Institute of Health. Department of medical Sciences, Ministry of Public Health, Thailand | National Institute of Health. Department of medical Sciences, Ministry of Public Health, Thailand | Pilailuk,Okada; Siripaporn,Phuygun; Thanutsapa,Thanadachakul; Sittiporn,Parnmen;Warawan,Wongboot; Sunthareeya,Waicharoen; Malinee,Chittaganpitch |
| EPI_ISL_455625, EPI_ISL_455626, EPI_ISL_455627 | unknown | Instituto Nacional de Saude (INSA) | Borges et al |
| EPI_ISL_455629, EPI_ISL_455630, EPI_ISL_455631, EPI_ISL_455632, EPI_ISL_455633, EPI_ISL_455634, EPI_ISL_455635, EPI_ISL_455636, EPI_ISL_455637, EPI_ISL_455638, EPI_ISL_455639 | | | |
| see above | KRISP, KZN Research Innovation and Sequencing Platform | KRISP, KZN Research Innovation and Sequencing Platform | Giandhari J, Pillay S, Lessells R, Chimukangara B, Deforche K, Tegally H, Wilkinson E, de Oliveira T |
| EPI_ISL_455640, EPI_ISL_455641, EPI_ISL_455642, EPI_ISL_455643, EPI_ISL_455644, EPI_ISL_455645, EPI_ISL_455646, EPI_ISL_455647, EPI_ISL_455648, EPI_ISL_455649, EPI_ISL_455650, EPI_ISL_455651, EPI_ISL_455652, EPI_ISL_455653, EPI_ISL_455654, EPI_ISL_455655, EPI_ISL_455656, EPI_ISL_455657, EPI_ISL_455658, EPI_ISL_455659, EPI_ISL_455660, EPI_ISL_455661, EPI_ISL_455662, EPI_ISL_455663, EPI_ISL_455664, EPI_ISL_455665, EPI_ISL_455666, EPI_ISL_455667, EPI_ISL_455668, EPI_ISL_455669, EPI_ISL_455670, EPI_ISL_455671, EPI_ISL_455672, EPI_ISL_455673, EPI_ISL_455674, EPI_ISL_455675, EPI_ISL_455676, EPI_ISL_455677, EPI_ISL_455678, EPI_ISL_455679 | | | |
| see above | ICMR-National Institute of Cholera and Enteric Diseases | National Institute of Biomedical Genomics | Arindam Maitra, Mamta Chawla Sarkar, Sreedhar Chinnaswamy, Hasina Banu, Ananya Chatterjee, Shanta Dutta, Saumitra Das |
| EPI_ISL_455680 | Institute of pathogenic microbiology, Jiangsu Provincial Center for Disease Control and Prevention | Institute of pathogenic microbiology, Jiangsu Provincial Center for Disease Control and Prevention | Cui,L. |
| EPI_ISL_455681 | National Institute of Health, WHO Regional Reference Laboratory for | National Institute of Health, WHO Regional Reference Laboratory for | Sharif,S., Khurshid,A., Mahmood,N., Arshad,Y., Salman,M., Ikram,A., Badar,N., Umair,M., Tamim,S., Angez,M., Alam,M. and Ahad,A. |

| | | | |
|---|---|---|---|
| | Polio Eradication, Virology Department | Polio Eradication, Virology Department | |
| EPI_ISL_455682 | University of Florida | University of Florida | Lednicky,J.A., Wu,C.-Y., Lauzardo,M. and Morris,J.G. |
| EPI_ISL_455683, EPI_ISL_455684, EPI_ISL_455685, EPI_ISL_455686, EPI_ISL_455687, EPI_ISL_455688, EPI_ISL_455689, EPI_ISL_455690, EPI_ISL_455691, EPI_ISL_455692, EPI_ISL_455693 | | | |
| see above | unknown | Department of Microbiology | Gao,Q., Bao,L., Mao,H., Wang,L., Xu,K., Yang,M., Li,Y., Zhu,L., Wang,N., Lv,Z., Gao,H., Ge,X., Kan,B., Hu,Y., Liu,J., Cai,F., Jiang,D., Yin,Y., Qin,C., Li,J., Gong,X., Lou,X., Shi,W., Wu,D., Zhang,H., Deng,W., Lu,J., Li,C., Wang,X., Yin,W., Zhang,Y., Sun,Y. |
| EPI_ISL_455694, EPI_ISL_455695, EPI_ISL_455696, EPI_ISL_455697, EPI_ISL_455698, EPI_ISL_455699, EPI_ISL_455700, EPI_ISL_455701, EPI_ISL_455702, EPI_ISL_455703, EPI_ISL_455704, EPI_ISL_455705, EPI_ISL_455706, EPI_ISL_455707, EPI_ISL_455708, EPI_ISL_455709, EPI_ISL_455710, EPI_ISL_455711, EPI_ISL_455712 | | | |
| see above | National Hospital of Tropical Diseases | Oxford University Clinical Research Unit, Hanoi, Vietnam | Nguyen Thi Tam, Van Dinh Trang, Nguyen Thu Trang, Nguyen Thi Ngoc Diep, Le Nguyen Minh Hoa, Pham Ngoc Thach, H. Rogier van Doorn, on behalf of the OUCRU COVID-19 research group |
| EPI_ISL_455713, EPI_ISL_455714, EPI_ISL_455715, EPI_ISL_455716, EPI_ISL_455717, EPI_ISL_455718 | National Hospital of Tropical Diseases | Oxford University Clinical Research Unit, Hanoi, Vietnam | Nguyen Thi Tam, Van Dinh Trang, Nguyen Thi Hong Thuong, Vu Thi Ngoc Bich, Nguyen Thu Trang, Nguyen Thi Ngoc Diep, Le Nguyen Minh Hoa, Pham Ngoc Thach, H. Rogier van Doorn, on behalf of the OUCRU COVID-19 research group |
| EPI_ISL_455719 | T.C. Sağlık Bakanlığı Adıyaman İl Sağlık Müdürlüğü Adıyaman Eğitim Ve Araştırma Hastanesi | VETAL Animal Health Products Company, BSL3+ Production Laboratuary /Turkey | Mehmet Turgut, Muhittin Önderci, Fatma Nilay Tutak, Abidin Ercan Yonucu, Murat Dönen, Haluk Uluca, Fethiye Sevimli, O. Ugur Sezerman |
| EPI_ISL_455721 | Servicio de Microbiologia. Hospital Clinico Universitario de Valencia | Sequencing and Bioinformatics Service and Molecular Epidemiology Research Group. FISABIO-Public Health, and SeqCOVID-Spain Consortium | Sandra Carbo, Loreto Ferrús Abad, Paula Ruiz-Hueso, Mariana Reyes-Prieto, Vicente Soriano Chirona, Ivan Ansari, Llúcia Martínez-Priego, Giuseppe 'Auria, David Navarro, Eliseo Albert, Maria Alma Bracho, Lidia Ruiz Roldan, Neris Garcia-Gonzalez, Inma Galán Vendrell, Fernando Gonzalez-Candelas |
| EPI_ISL_455722 | Servicio de Microbiologia. Hospital Clinico Universitario de Valencia | Sequencing and Bioinformatics Service and Molecular Epidemiology Research Group. FISABIO-Public Health, and SeqCOVID-Spain Consortium | Giuseppe 'Auria, David Navarro, Eliseo Albert, Maria Alma Bracho, Lidia Ruiz Roldan, Neris Garcia-Gonzalez, Inma Galán Vendrell, Sandra Carbo, Loreto Ferrús Abad, Paula Ruiz-Hueso, Mariana Reyes-Prieto, Vicente Soriano Chirona, Ivan Ansari, Llúcia Martínez-Priego, Fernando Gonzalez-Candelas |
| EPI_ISL_455723 | Servicio de Microbiologia. Hospital Clinico Universitario de Valencia | Sequencing and Bioinformatics Service and Molecular Epidemiology Research Group. FISABIO-Public Health, and SeqCOVID-Spain Consortium | Vicente Soriano Chirona, Ivan Ansari, Llúcia Martínez-Priego, Giuseppe 'Auria, David Navarro, Eliseo Albert, Maria Alma Bracho, Lidia Ruiz Roldan, Neris Garcia-Gonzalez, Inma Galán Vendrell, Sandra Carbo, Loreto Ferrús Abad, Paula Ruiz-Hueso, Mariana Reyes-Prieto, Fernando Gonzalez-Candelas |
| EPI_ISL_455724 | Servicio de Microbiologia. Hospital Clinico Universitario de Valencia | Sequencing and Bioinformatics Service and Molecular Epidemiology Research Group. FISABIO-Public Health, and SeqCOVID-Spain Consortium | Loreto Ferrús Abad, Paula Ruiz-Hueso, Mariana Reyes-Prieto, Vicente Soriano Chirona, Ivan Ansari, Llúcia Martínez-Priego, Giuseppe 'Auria, David Navarro, Eliseo Albert, Maria Alma Bracho, Lidia Ruiz Roldan, Neris Garcia-Gonzalez, Inma Galán Vendrell, Sandra Carbo, Fernando Gonzalez-Candelas |
| EPI_ISL_455725 | Servicio de Microbiologia. Hospital Clinico Universitario de Valencia | Sequencing and Bioinformatics Service and Molecular Epidemiology Research Group. FISABIO-Public Health, and SeqCOVID-Spain Consortium | Eliseo Albert, Maria Alma Bracho, Griselda De Marco, Lidia Ruiz Roldan, Neris Garcia-Gonzalez, Inma Galán Vendrell, Sandra Carbo, Loreto Ferrús Abad, Paula Ruiz-Hueso, Mariana Reyes-Prieto, Vicente Soriano Chirona, Ivan Ansari, Llúcia Martínez-Priego, Giuseppe 'Auria, David Navarro, Fernando Gonzalez-Candelas |
| EPI_ISL_455726 | Servicio de Microbiologia. Hospital Clinico Universitario de Valencia | Sequencing and Bioinformatics Service and Molecular Epidemiology Research Group. FISABIO-Public Health, and SeqCOVID-Spain Consortium | Inma Galán Vendrell, Sandra Carbo, Loreto Ferrús Abad, Paula Ruiz-Hueso, Mariana Reyes-Prieto, Vicente Soriano Chirona, Ivan Ansari, Llúcia Martínez-Priego, Giuseppe 'Auria, David Navarro, Eliseo Albert, Maria Alma Bracho, Lidia Ruiz Roldan, Neris Garcia-Gonzalez, Fernando Gonzalez-Candelas |
| EPI_ISL_455727 | Servicio de Microbiologia. Hospital Clinico Universitario de Valencia | Sequencing and Bioinformatics Service and Molecular Epidemiology Research Group. FISABIO-Public Health, and SeqCOVID-Spain Consortium | David Navarro, Eliseo Albert, Maria Alma Bracho, Griselda De Marco, Lidia Ruiz Roldan, Neris Garcia-Gonzalez, Inma Galán Vendrell, Sandra Carbo, Loreto Ferrús Abad, Paula Ruiz-Hueso, Mariana Reyes-Prieto, Vicente Soriano Chirona, Ivan Ansari, Llúcia Martínez-Priego, Giuseppe 'Auria, Fernando Gonzalez-Candelas |
| EPI_ISL_455728 | Servicio de Microbiologia. Hospital Clinico Universitario de Valencia | Sequencing and Bioinformatics Service and Molecular Epidemiology Research Group. FISABIO-Public Health, and SeqCOVID-Spain Consortium | Maria Alma Bracho, Griselda De Marco, Lidia Ruiz Roldan, Neris Garcia-Gonzalez, Inma Galán Vendrell, Sandra Carbo, Loreto Ferrús Abad, Paula Ruiz-Hueso, Mariana Reyes-Prieto, Vicente Soriano Chirona, Ivan Ansari, Llúcia Martínez-Priego, Giuseppe 'Auria, David Navarro, Eliseo Albert, Fernando Gonzalez-Candelas |
| EPI_ISL_455729 | Servicio de Microbiologia. Hospital Clinico Universitario de Valencia | Sequencing and Bioinformatics Service and Molecular Epidemiology Research Group. FISABIO-Public Health, and SeqCOVID-Spain Consortium | Sandra Carbo, Loreto Ferrús Abad, Paula Ruiz-Hueso, Mariana Reyes-Prieto, Vicente Soriano Chirona, Ivan Ansari, Llúcia Martínez-Priego, Giuseppe 'Auria, David Navarro, Eliseo Albert, Maria Alma Bracho, Lidia Ruiz Roldan, Neris Garcia-Gonzalez, Inma Galán Vendrell, Fernando Gonzalez-Candelas |
| EPI_ISL_455730, EPI_ISL_455731 | Servicio de Microbiologia. Hospital Clinico Universitario de Valencia | Sequencing and Bioinformatics Service and Molecular Epidemiology Research Group. FISABIO-Public Health, and SeqCOVID-Spain Consortium | Mariana Reyes-Prieto, Vicente Soriano Chirona, Ivan Ansari, Llúcia Martínez-Priego, Giuseppe 'Auria, David Navarro, Eliseo Albert, Maria Alma Bracho, Lidia Ruiz Roldan, Neris Garcia-Gonzalez, Inma Galán Vendrell, Sandra Carbo, Loreto Ferrús Abad, Paula Ruiz-Hueso, Fernando Gonzalez-Candelas |
| EPI_ISL_455732 | Servicio de Microbiologia. Hospital Clinico Universitario de Valencia | Sequencing and Bioinformatics Service and Molecular Epidemiology Research Group. FISABIO-Public Health, and SeqCOVID-Spain Consortium | Lidia Ruiz Roldan, Neris Garcia-Gonzalez, Inma Galán Vendrell, Sandra Carbo, Loreto Ferrús Abad, Paula Ruiz-Hueso, Mariana Reyes-Prieto, Vicente Soriano Chirona, Ivan Ansari, Llúcia Martínez-Priego, Giuseppe 'Auria, David Navarro, Eliseo Albert, Maria Alma Bracho, Fernando Gonzalez-Candelas |
| EPI_ISL_455733 | Servicio de Microbiologia. Hospital Clinico Universitario de Valencia | Sequencing and Bioinformatics Service and Molecular Epidemiology Research Group. FISABIO-Public Health, and SeqCOVID-Spain Consortium | Loreto Ferrús Abad, Paula Ruiz-Hueso, Mariana Reyes-Prieto, Vicente Soriano Chirona, Ivan Ansari, Llúcia Martínez-Priego, Giuseppe 'Auria, David Navarro, Eliseo Albert, Maria Alma Bracho, Lidia Ruiz Roldan, Neris Garcia-Gonzalez, Inma Galán Vendrell, Sandra Carbo, Fernando Gonzalez-Candelas |
| EPI_ISL_455734 | Servicio de Microbiologia. Hospital Clinico Universitario de Valencia | Sequencing and Bioinformatics Service and Molecular Epidemiology Research Group. FISABIO-Public Health, and SeqCOVID-Spain Consortium | Griselda De Marco, Lidia Ruiz Roldan, Neris Garcia-Gonzalez, Inma Galán Vendrell, Sandra Carbo, Loreto Ferrús Abad, Paula Ruiz-Hueso, Mariana Reyes-Prieto, Vicente Soriano Chirona, Ivan Ansari, Llúcia Martínez-Priego, Giuseppe 'Auria, David Navarro, Eliseo Albert, Maria Alma Bracho, Fernando Gonzalez-Candelas |
| EPI_ISL_455735, EPI_ISL_455736 | Servicio de Microbiologia. Hospital_Arnau_de_Vilanova | Sequencing and Bioinformatics Service and Molecular Epidemiology Research Group. FISABIO-Public Health, and SeqCOVID-Spain Consortium | Victoria Dominguez, Maria Alma Bracho, Griselda De Marco, Lidia Ruiz Roldan, Neris Garcia-Gonzalez, Inma Galán Vendrell, Sandra Carbo, Loreto Ferrús Abad, Paula Ruiz-Hueso, Mariana Reyes-Prieto, Vicente Soriano Chirona, Ivan Ansari, Llúcia Martínez-Priego, Giuseppe 'Auria, Fernando Gonzalez-Candelas |
| EPI_ISL_455737 | Servicio de Microbiologia. Hospital Clinico Universitario de Valencia | Sequencing and Bioinformatics Service and Molecular Epidemiology Research Group. FISABIO-Public Health, and SeqCOVID-Spain Consortium | Griselda De Marco, Lidia Ruiz Roldan, Neris Garcia-Gonzalez, Inma Galán Vendrell, Sandra Carbo, Loreto Ferrús Abad, Paula Ruiz-Hueso, Mariana Reyes-Prieto, Vicente Soriano Chirona, Ivan Ansari, Llúcia Martínez-Priego, Giuseppe 'Auria, David Navarro, Eliseo Albert, Maria Alma Bracho, Fernando Gonzalez-Candelas |
| EPI_ISL_455738 | Servicio de Microbiologia. Hospital Clinico Universitario de Valencia | Sequencing and Bioinformatics Service and Molecular Epidemiology Research Group. FISABIO-Public Health, and SeqCOVID-Spain Consortium | Maria Alma Bracho, Griselda De Marco, Lidia Ruiz Roldan, Neris Garcia-Gonzalez, Inma Galán Vendrell, Sandra Carbo, Loreto Ferrús Abad, Paula Ruiz-Hueso, Mariana Reyes-Prieto, Vicente Soriano Chirona, Ivan Ansari, Llúcia Martínez-Priego, Giuseppe 'Auria, David Navarro, Eliseo Albert, Fernando Gonzalez-Candelas |
| EPI_ISL_455739 | Servicio de Microbiologia. Hospital Clinico Universitario de Valencia | Sequencing and Bioinformatics Service and Molecular Epidemiology Research Group. FISABIO-Public Health, and SeqCOVID-Spain Consortium | Lidia Ruiz Roldan, Neris Garcia-Gonzalez, Inma Galán Vendrell, Sandra Carbo, Loreto Ferrús Abad, Paula Ruiz-Hueso, Mariana Reyes-Prieto, Vicente Soriano Chirona, Ivan Ansari, Llúcia Martínez-Priego, Giuseppe 'Auria, David Navarro, Eliseo Albert, Maria Alma Bracho, Fernando Gonzalez-Candelas |
| EPI_ISL_455740 | Servicio de Microbiologia. Hospital Clinico Universitario de Valencia | Sequencing and Bioinformatics Service and Molecular Epidemiology Research Group. FISABIO-Public Health, and SeqCOVID-Spain Consortium | Paula Ruiz-Hueso, Mariana Reyes-Prieto, Vicente Soriano Chirona, Ivan Ansari, Llúcia Martínez-Priego, Giuseppe 'Auria, David Navarro, Eliseo Albert, Maria Alma Bracho, Lidia Ruiz Roldan, Neris Garcia-Gonzalez, Inma Galán Vendrell, Sandra Carbo, Loreto Ferrús Abad, Fernando Gonzalez-Candelas |
| EPI_ISL_455741 | Servicio de Microbiologia. Hospital Clinico Universitario de Valencia | Sequencing and Bioinformatics Service and Molecular Epidemiology Research Group. FISABIO-Public Health, and SeqCOVID-Spain Consortium | David Navarro, Eliseo Albert, Maria Alma Bracho, Griselda De Marco, Lidia Ruiz Roldan, Neris Garcia-Gonzalez, Inma Galán Vendrell, Sandra Carbo, Loreto Ferrús Abad, Paula Ruiz-Hueso, Mariana Reyes-Prieto, Vicente Soriano Chirona, Ivan Ansari, Llúcia Martínez-Priego, Giuseppe 'Auria, Fernando Gonzalez-Candelas |
| EPI_ISL_455742 | Servicio de Microbiologia. Hospital Clinico Universitario de Valencia | Sequencing and Bioinformatics Service and Molecular Epidemiology Research Group. FISABIO-Public Health, and SeqCOVID-Spain Consortium | Neris Garcia-Gonzalez, Inma Galán Vendrell, Sandra Carbo, Loreto Ferrús Abad, Paula Ruiz-Hueso, Mariana Reyes-Prieto, Vicente Soriano Chirona, Ivan Ansari, Llúcia Martínez-Priego, Giuseppe 'Auria, David Navarro, Eliseo Albert, Maria Alma Bracho, Lidia Ruiz Roldan, Fernando Gonzalez-Candelas |
| EPI_ISL_455743 | Servicio de Microbiologia. Hospital Clinico Universitario de Valencia | Sequencing and Bioinformatics Service and Molecular Epidemiology Research Group. FISABIO-Public Health, and SeqCOVID-Spain Consortium | Eliseo Albert, Maria Alma Bracho, Griselda De Marco, Lidia Ruiz Roldan, Neris Garcia-Gonzalez, Inma Galán Vendrell, Sandra Carbo, Loreto Ferrús Abad, Paula Ruiz-Hueso, Mariana Reyes-Prieto, Vicente Soriano Chirona, Ivan Ansari, Llúcia Martínez-Priego, Giuseppe 'Auria, David Navarro, Fernando Gonzalez-Candelas |
| EPI_ISL_455744 | Servicio de Microbiologia. Hospital Clinico Universitario de Valencia | Sequencing and Bioinformatics Service and Molecular Epidemiology Research Group. FISABIO-Public Health, and SeqCOVID-Spain Consortium | Llúcia Martínez-Priego, Giuseppe 'Auria, David Navarro, Eliseo Albert, Maria Alma Bracho, Lidia Ruiz Roldan, Neris Garcia-Gonzalez, Inma Galán Vendrell, Sandra Carbo, Loreto Ferrús Abad, Paula Ruiz-Hueso, Mariana Reyes-Prieto, Vicente Soriano Chirona, Ivan Ansari, Fernando Gonzalez-Candelas |
| EPI_ISL_455749, EPI_ISL_455750, EPI_ISL_455751, EPI_ISL_455752, EPI_ISL_455753, EPI_ISL_455754, EPI_ISL_455755, EPI_ISL_455756, EPI_ISL_455757, EPI_ISL_455758, EPI_ISL_455759, EPI_ISL_455760, EPI_ISL_455761, EPI_ISL_455762, EPI_ISL_455763, EPI_ISL_455764, EPI_ISL_455765, EPI_ISL_455766, EPI_ISL_455767, EPI_ISL_455768, EPI_ISL_455769, EPI_ISL_455770, EPI_ISL_455771, EPI_ISL_455772, EPI_ISL_455773, EPI_ISL_455774, EPI_ISL_455775, EPI_ISL_455776, EPI_ISL_455777, EPI_ISL_455778, EPI_ISL_455779, EPI_ISL_455780, EPI_ISL_455781, EPI_ISL_455782, EPI_ISL_455783, EPI_ISL_455784, EPI_ISL_455785, EPI_ISL_455786, EPI_ISL_455787, EPI_ISL_455788 | | | |
| see above | REGIONAL VRDL,ICMR-RMRC BBSR | Immunogenomics lab, Institute of Life Sciences, Bhubaneswar | Sunil Raghav, Jyotirmayee Turuk, Arup Ghosh, Atimukta Jha, Viplov K. Biswas, Swati Madhulika, Manasi Priyadarshini, Shuchi Smita, Jaya Singh Khastri, Rupesh Dash, Soma Chattopadhyay, Ghulam Hussain Syed, Shanti Senapati, Tushar K. Beuria, Debdutta Bhattacharya, Rajeeb Swain, Punit Prasad, COVID-19 team of ILS & RMRC, Orissa COVID-19 study group, DBT's PAN-INDIA 1000 SARS-CoV2 RNA genome sequencing consortium, Sanghamitra Pati, Ajay Parida |
| EPI_ISL_455790 | Institute for Medical Research, Infectious Disease Research Centre, National Institutes of Health, Ministry of Health Malaysia | Malaysia Genome Institute | Mohd Noor Mat Isa, Irni Suhayu Sapian, Yusuf Muhammad Noor, Jeyanthi Suppiah, Nurhezreen Md Iqbal, Enizza Kasim, Zarina Mohd Zawawi, Siti Noraini Othman, Mohd Faizal Abu Bakar, Shamsidar Sopie, Azrin Ahmad, Ravindran Thayan, Norazah Ahmad, Tahir Aris, Shahrul Hisham Zainal Ariffin |
| EPI_ISL_455791 | Institute for Medical Research, Infectious Disease Research Centre, National Institutes of Health, Ministry of Health Malaysia | Malaysia Genome Institute | Mohd Noor Mat Isa, Irni Suhayu Sapian, Yusuf Muhammad Noor, Jeyanthi Suppiah, Nurhezreen Md Iqbal, Enizza Kasim, Zarina Mohd Zawawi, Siti Noraini Othman, Mohd Faizal Abu Bakar, Shamsidar Sopie, Azrin Ahmad, Ravindran Thayan, Norazah Ahmad, Tahir Aris, Shahrul Hisham Zainal Ariffin |
| EPI_ISL_455792 | Institute for Medical Research, Infectious Disease Research Centre, National Institutes of Health, Ministry of Health Malaysia | Malaysia Genome Institute | Mohd Noor Mat Isa, Irni Suhayu Sapian, Yusuf Muhammad Noor, Jeyanthi Suppiah, Nurhezreen Md Iqbal, Enizza Kasim, Zarina Mohd Zawawi, Siti Noraini Othman, Mohd Faizal Abu Bakar, Shamsidar Sopie, Azrin Ahmad, Ravindran Thayan, Norazah Ahmad, Tahir Aris, Shahrul Hisham Zainal Ariffin |
| EPI_ISL_455793 | Institute for Medical Research, Infectious Disease Research Centre, National Institutes of Health, Ministry of Health Malaysia | Malaysia Genome Institute | Mohd Noor Mat Isa, Irni Suhayu Sapian, Yusuf Muhammad Noor, Jeyanthi Suppiah, Nurhezreen Md Iqbal, Enizza Kasim, Zarina Mohd Zawawi, Siti Noraini Othman, Mohd Faizal Abu Bakar, Shamsidar Sopie, Azrin Ahmad, Ravindran Thayan, Norazah Ahmad, Tahir Aris, Shahrul Hisham Zainal Ariffin |
| EPI_ISL_455840, EPI_ISL_455841 | Orebro klinisk mikrobiologi | The Public Health Agency of Sweden | Anna-Malin Linde, Maria Lind Karlberg, Mattias Haukland, Reza Advani, Olov Svartstrom, Oskar Karlsson Lindsjo, Petra Edquist, Shamam Muradrasoli, Anna Risberg, Karin Tegmark-Wisell |

| | | | |
|---|---|---|---|
| EPI_ISL_455842, EPI_ISL_455843, EPI_ISL_455844, EPI_ISL_455845 | Skovde/Unilabs | The Public Health Agency of Sweden | Anna-Malin Linde, Maria Lind Karlberg, Mattias Haukland, Reza Advani, Olov Svartstrom, Oskar Karlsson Lindsjo, Petra Edquist, Shamam Muradrasoli, Anna Risberg, Karin Tegmark-Wisell |
| EPI_ISL_455846, EPI_ISL_455847, EPI_ISL_455848, EPI_ISL_455849, EPI_ISL_455850, EPI_ISL_455851, EPI_ISL_455852, EPI_ISL_455853, EPI_ISL_455854, EPI_ISL_455855, EPI_ISL_455856, EPI_ISL_455857, EPI_ISL_455858, EPI_ISL_455859, EPI_ISL_455860, EPI_ISL_455861, EPI_ISL_455862, EPI_ISL_455863, EPI_ISL_455864, EPI_ISL_455865, EPI_ISL_455866, EPI_ISL_455867, EPI_ISL_455868, EPI_ISL_455869, EPI_ISL_455870, EPI_ISL_455871, EPI_ISL_455872, EPI_ISL_455873, EPI_ISL_455874, EPI_ISL_455875, EPI_ISL_455876, EPI_ISL_455877, EPI_ISL_455878, EPI_ISL_455879, EPI_ISL_455880, EPI_ISL_455881, EPI_ISL_455882, EPI_ISL_455883, EPI_ISL_455884, EPI_ISL_455885, EPI_ISL_455886, EPI_ISL_455887, EPI_ISL_455888, EPI_ISL_455889, EPI_ISL_455890, EPI_ISL_455891, EPI_ISL_455892, EPI_ISL_455893, EPI_ISL_455894, EPI_ISL_455895, EPI_ISL_455896, EPI_ISL_455897, EPI_ISL_455898, EPI_ISL_455899, EPI_ISL_455900, EPI_ISL_455901 | | | |
| see above | Karolinska Universitetslaboratoriet | The Public Health Agency of Sweden | Anna-Malin Linde, Maria Lind Karlberg, Mattias Haukland, Reza Advani, Olov Svartstrom, Oskar Karlsson Lindsjo, Petra Edquist, Shamam Muradrasoli, Anna Risberg, Karin Tegmark-Wisell |
| EPI_ISL_455902 | Halmstad klinisk mikrobiologi | The Public Health Agency of Sweden | Anna-Malin Linde, Maria Lind Karlberg, Mattias Haukland, Reza Advani, Olov Svartstrom, Oskar Karlsson Lindsjo, Petra Edquist, Shamam Muradrasoli, Anna Risberg, Karin Tegmark-Wisell |
| EPI_ISL_455903 | Folkhalsomyndigheten | The Public Health Agency of Sweden | Anna-Malin Linde, Maria Lind Karlberg, Mattias Haukland, Reza Advani, Olov Svartstrom, Oskar Karlsson Lindsjo, Petra Edquist, Shamam Muradrasoli, Anna Risberg, Karin Tegmark-Wisell |
| EPI_ISL_455904, EPI_ISL_455905, EPI_ISL_455906, EPI_ISL_455907 | Klinisk mikrobiologi, UAS | The Public Health Agency of Sweden | Anna-Malin Linde, Maria Lind Karlberg, Mattias Haukland, Reza Advani, Olov Svartstrom, Oskar Karlsson Lindsjo, Petra Edquist, Shamam Muradrasoli, Anna Risberg, Karin Tegmark-Wisell |
| EPI_ISL_455908, EPI_ISL_455909, EPI_ISL_455910, EPI_ISL_455911, EPI_ISL_455912, EPI_ISL_455913, EPI_ISL_455914, EPI_ISL_455915, EPI_ISL_455916, EPI_ISL_455917, EPI_ISL_455918, EPI_ISL_455919, EPI_ISL_455920, EPI_ISL_455921, EPI_ISL_455922, EPI_ISL_455923, EPI_ISL_455924, EPI_ISL_455925, EPI_ISL_455926, EPI_ISL_455927, EPI_ISL_455928, EPI_ISL_455929, EPI_ISL_455930, EPI_ISL_455931, EPI_ISL_455932, EPI_ISL_455933, EPI_ISL_455934, EPI_ISL_455935, EPI_ISL_455936, EPI_ISL_455937, EPI_ISL_455938, EPI_ISL_455939, EPI_ISL_455940, EPI_ISL_455941, EPI_ISL_455942, EPI_ISL_455943, EPI_ISL_455944, EPI_ISL_455945, EPI_ISL_455946, EPI_ISL_455947, EPI_ISL_455948 | | | |
| see above | Ramathibodi Hospital | COVID-19 Network Investigations (CONI) Alliance | Elizabeth Batty, Wasun Chantratita, Thanat Chookajorn, Stefan Fernandez, Angkana Huang, Anthony R. Jones, Khajohn Joonsalak, Chonticha Klungtong, Theerarat Kochakarn, Namfon Kotanan, Krittikorn Kumpornsin, Wudtichai Manasatienkij, Bhakbhoom Panthan, Ekawat Pasomsub, Kingkan Rakmanee, Insee Sensorn, Janjira Thaipadungpanit, Arporn Wangwiwatsin,Treewat Watthanachockchai |
| EPI_ISL_455958, EPI_ISL_455959, EPI_ISL_455960, EPI_ISL_455961, EPI_ISL_455962, EPI_ISL_455963, EPI_ISL_455964, EPI_ISL_455965, EPI_ISL_455966, EPI_ISL_455967, EPI_ISL_455968, EPI_ISL_455969, EPI_ISL_455970, EPI_ISL_455971, EPI_ISL_455972, EPI_ISL_455973, EPI_ISL_455974, EPI_ISL_455975, EPI_ISL_455976, EPI_ISL_455977, EPI_ISL_455978, EPI_ISL_455979 | | | |
| see above | Department of Clinical Microbiology | GIGA Medical Genomics | Keith Durkin, Maria Artesi, Sébastien Bontems, Raphaël Boreux, Cécile Meex, Pierrette Melin, Marie-Pierre Hayette, Vincent Bours. |
| EPI_ISL_455980, EPI_ISL_455981, EPI_ISL_455982, EPI_ISL_455983, EPI_ISL_455984, EPI_ISL_455985, EPI_ISL_455986, EPI_ISL_455987, EPI_ISL_455988, EPI_ISL_455989 | LSUHS Emerging Viral Threat Laboratory | Microbial Genome Sequencing Center | Jeremy P. Kamil, John A. Vanchiere, Rona S. Scott, Camille F. Abshire, Abida Siddiqa, Byeong-Jae Lee, Chan-ki Min, Md Maksudul Alam, Monica Gestal-Cartele, Edna Ondari, Adam Greer, Malgorzata Bienkowska-Haba, Katarzyna Zwolinska, Michelle M. Arnold, Jason M. Bodily, Andrew D. Yurochko, Paul M. Weinberger, Christopher G. Kevil, Martin J. Sapp, Daniel J. Snyder, Vaughn S. Cooper |
| EPI_ISL_455990, EPI_ISL_455991, EPI_ISL_455992, EPI_ISL_455993, EPI_ISL_455994, EPI_ISL_455995, EPI_ISL_455996, EPI_ISL_455997, EPI_ISL_455998, EPI_ISL_455999 | LSUHS Emerging Viral Threat Laboratory | Microbial Genome Sequencing Center | John A. Vanchiere, Jeremy P. Kamil, Rona S. Scott, Camille F. Abshire, Abida Siddiqa, Byeong-Jae Lee, Chan-ki Min, Md Maksudul Alam, Monica Gestal-Cartele, Edna Ondari, Adam Greer, Malgorzata Bienkowska-Haba, Katarzyna Zwolinska, Michelle M. Arnold, Jason M. Bodily, Andrew D. Yurochko, Paul M. Weinberger, Christopher G. Kevil, Martin J. Sapp, Daniel J. Snyder, Vaughn S. Cooper |
| EPI_ISL_456000, EPI_ISL_456001, EPI_ISL_456002 | LSUHS Emerging Viral Threat Laboratory | Microbial Genome Sequencing Center | Rona S. Scott, Jeremy P. Kamil, John A. Vanchiere, Camille F. Abshire, Abida Siddiqa, Byeong-Jae Lee, Chan-ki Min, Md Maksudul Alam, Monica Gestal-Cartele, Edna Ondari, Adam Greer, Malgorzata Bienkowska-Haba, Katarzyna Zwolinska, Michelle M. Arnold, Jason M. Bodily, Andrew D. Yurochko, Paul M. Weinberger, Christopher G. Kevil, Martin J. Sapp, Daniel J. Snyder, Vaughn S. Cooper |
| EPI_ISL_456003, EPI_ISL_456004, EPI_ISL_456005, EPI_ISL_456006, EPI_ISL_456007, EPI_ISL_456008, EPI_ISL_456009, EPI_ISL_456010, EPI_ISL_456011, EPI_ISL_456012, EPI_ISL_456013, EPI_ISL_456014, EPI_ISL_456015, EPI_ISL_456016, EPI_ISL_456017, EPI_ISL_456018, EPI_ISL_456019, EPI_ISL_456020, EPI_ISL_456021, EPI_ISL_456022, EPI_ISL_456023, EPI_ISL_456024, EPI_ISL_456025, EPI_ISL_456026, EPI_ISL_456027, EPI_ISL_456028, EPI_ISL_456029, EPI_ISL_456030, EPI_ISL_456031, EPI_ISL_456032, EPI_ISL_456033, EPI_ISL_456034, EPI_ISL_456035, EPI_ISL_456036, EPI_ISL_456037, EPI_ISL_456038, EPI_ISL_456039, EPI_ISL_456040, EPI_ISL_456041, EPI_ISL_456042, EPI_ISL_456043, EPI_ISL_456044, EPI_ISL_456045, EPI_ISL_456046, EPI_ISL_456047, EPI_ISL_456048, EPI_ISL_456049, EPI_ISL_456050, EPI_ISL_456051, EPI_ISL_456052, EPI_ISL_456053, EPI_ISL_456054, EPI_ISL_456055, EPI_ISL_456056, EPI_ISL_456057, EPI_ISL_456058, EPI_ISL_456059, EPI_ISL_456060, EPI_ISL_456061, EPI_ISL_456062, EPI_ISL_456063, EPI_ISL_456064, EPI_ISL_456065, EPI_ISL_456066, EPI_ISL_456067, EPI_ISL_456068, EPI_ISL_456069, EPI_ISL_456070 | | | |
| see above | NYU Langone Health | Departments of Pathology and Medicine, New York University School of Medicine | Maria Aguero-Rosenfeld, Brendan Belovarac, Margaret Black, Ludovic Boytard, John Cadley, Paolo Cotzia, John Chen, Dacia Dimartino, Xiaojun Feng, Tatyana Gindin, Emily Guzman, Adriana Heguy, Megan Hogan, Emily Huang, George Jour, Alireza Khodadadi-Jamayran, Lawrence H. Lin, Raven Luther, Andrew Lytle, Christian Marier, Matthew T. Maurano, Mark J. Mulligan, Peter Meyn, Raquel Ordonez Ciriza, Iman Osman, Jared Pinnell, Vanessa Raabe, Sitharam Ramaswami, Amy Rapkiewicz, Andre M. Ribeiro-dos-Santos, Marie Samanovic-Golden, Antonio Serrano, Guomiao Shen, Matija Snuderl, Theodore Vougiouklakis, Nick Vulpescu, Gael Westby, Paul Zappile, Yutong Zhang |
| EPI_ISL_456071, EPI_ISL_456072, EPI_ISL_456073, EPI_ISL_456074, EPI_ISL_456075 | Laboratory of Respiratory Viruses and Measles, Oswaldo Cruz Institute, FIOCRUZ | Laboratory of Respiratory Viruses and Measles, Oswaldo Cruz Institute, FIOCRUZ | Paola Resende, Luciana Appolinario, Fernando Motta, Aline Mattos, Milene Miranda, Cristiana Garcia, Braulia Caetano, Maria Ogrzewalska, Jonathan Lopes, Marilda Siqueira |
| EPI_ISL_456076, EPI_ISL_456077 | LACEN RJ - Laboratório Central de Saúde Pública Noel Nutels | Laboratory of Respiratory Viruses and Measles, Oswaldo Cruz Institute, FIOCRUZ | Paola Resende, Luciana Appolinario, Fernando Motta, Aline Mattos, Milene Miranda, Cristiana Garcia, Braulia Caetano, Maria Ogrzewalska, Jonathan Lopes, Marilda Siqueira |
| EPI_ISL_456079, EPI_ISL_456080, EPI_ISL_456081 | Laboratory of Respiratory Viruses and Measles, Oswaldo Cruz Institute, FIOCRUZ | Laboratory of Respiratory Viruses and Measles, Oswaldo Cruz Institute, FIOCRUZ | Paola Resende, Luciana Appolinario, Fernando Motta, Aline Mattos, Milene Miranda, Cristiana Garcia, Braulia Caetano, Maria Ogrzewalska, Jonathan Lopes, Marilda Siqueira |
| EPI_ISL_456082, EPI_ISL_456083 | LACEN RJ - Laboratório Central de Saúde Pública Noel Nutels | Laboratory of Respiratory Viruses and Measles, Oswaldo Cruz Institute, FIOCRUZ | Paola Resende, Luciana Appolinario, Fernando Motta, Aline Mattos, Milene Miranda, Cristiana Garcia, Braulia Caetano, Maria Ogrzewalska, Jonathan Lopes, Marilda Siqueira |
| EPI_ISL_456084, EPI_ISL_456085, EPI_ISL_456086, EPI_ISL_456087 | Laboratory of Respiratory Viruses and Measles, Oswaldo Cruz Institute, FIOCRUZ | Laboratory of Respiratory Viruses and Measles, Oswaldo Cruz Institute, FIOCRUZ | Paola Resende, Luciana Appolinario, Fernando Motta, Aline Mattos, Milene Miranda, Cristiana Garcia, Braulia Caetano, Maria Ogrzewalska, Jonathan Lopes, Marilda Siqueira |
| EPI_ISL_456088 | LACEN RJ - Laboratório Central de Saúde Pública Noel Nutels | Laboratory of Respiratory Viruses and Measles, Oswaldo Cruz Institute, FIOCRUZ | Paola Resende, Luciana Appolinario, Fernando Motta, Aline Mattos, Milene Miranda, Cristiana Garcia, Braulia Caetano, Maria Ogrzewalska, Jonathan Lopes, Marilda Siqueira |
| EPI_ISL_456089, EPI_ISL_456090, EPI_ISL_456091, EPI_ISL_456092, EPI_ISL_456093, EPI_ISL_456094, EPI_ISL_456095, EPI_ISL_456096, EPI_ISL_456097, EPI_ISL_456098, EPI_ISL_456099, EPI_ISL_456100, EPI_ISL_456101, EPI_ISL_456102, EPI_ISL_456103, EPI_ISL_456104, EPI_ISL_456105, EPI_ISL_456106 | | | |
| see above | Laboratory of Respiratory Viruses and Measles, Oswaldo Cruz Institute, FIOCRUZ | Laboratory of Respiratory Viruses and Measles, Oswaldo Cruz Institute, FIOCRUZ | Paola Resende, Luciana Appolinario, Fernando Motta, Aline Mattos, Milene Miranda, Cristiana Garcia, Braulia Caetano, Maria Ogrzewalska, Jonathan Lopes, Marilda Siqueira |
| EPI_ISL_456107, EPI_ISL_456108, EPI_ISL_456109, EPI_ISL_456110, EPI_ISL_456111, EPI_ISL_456112, EPI_ISL_456113, EPI_ISL_456114, EPI_ISL_456115 | NYU Langone Health | Departments of Pathology and Medicine, New York University School of Medicine | Maria Aguero-Rosenfeld, Brendan Belovarac, Margaret Black, Ludovic Boytard, John Cadley, Paolo Cotzia, John Chen, Dacia Dimartino, Xiaojun Feng, Tatyana Gindin, Emily Guzman, Adriana Heguy, Megan Hogan, Emily Huang, George Jour, Alireza Khodadadi-Jamayran, Lawrence H. Lin, Raven Luther, Andrew Lytle, Christian Marier, Matthew T. Maurano, Mark J. Mulligan, Peter Meyn, Raquel Ordonez Ciriza, Iman Osman, Jared Pinnell, Vanessa Raabe, Sitharam Ramaswami, Amy Rapkiewicz, Andre M. Ribeiro-dos-Santos, Marie Samanovic-Golden, Antonio Serrano, Guomiao Shen, Matija Snuderl, Theodore Vougiouklakis, Nick Vulpescu, Gael Westby, Paul Zappile, Yutong Zhang |
| EPI_ISL_456116, EPI_ISL_456117, EPI_ISL_456118, EPI_ISL_456119, EPI_ISL_456120, EPI_ISL_456121, EPI_ISL_456122, EPI_ISL_456123, EPI_ISL_456124, EPI_ISL_456125, EPI_ISL_456126, EPI_ISL_456127, EPI_ISL_456128, EPI_ISL_456129, EPI_ISL_456130, EPI_ISL_456131, EPI_ISL_456132, EPI_ISL_456133, EPI_ISL_456134, EPI_ISL_456135, EPI_ISL_456136, EPI_ISL_456137, EPI_ISL_456138, EPI_ISL_456139, EPI_ISL_456140, EPI_ISL_456141, EPI_ISL_456142, EPI_ISL_456143, EPI_ISL_456144, EPI_ISL_456145, EPI_ISL_456146, EPI_ISL_456147, EPI_ISL_456148, EPI_ISL_456149, EPI_ISL_456150, EPI_ISL_456151, EPI_ISL_456152, EPI_ISL_456153, EPI_ISL_456154, EPI_ISL_456155, EPI_ISL_456156 | | | |
| see above | Instituto Nacional de Salud - Unidad de Secuenciación y Análisis Genómico | Instituto Nacional de Salud, Universidad Cooperativa de Colombia, Instituto Alexander von Humboldt, Imperial College-London, London School of Hygiene & Tropical Medicine | Katherine Laiton-Donato, Diego A. Álvarez-Díaz, Carlos Franco-Muñoz, Jose A. Usme-Ciro, Gloria Puerto, Nicolas D. Franco-Sierra, Mailyn A.Gonzalez, Zulma M. Cucunubá, Christian Julian Villabona-Arenas, Liz Villabona-Arenas, Sussy Echeverria, Astrid C. Flórez, Sergio Gomez-Rangel, Luz Dary Rodriguez, Juliana Barbosa, Erika Ospitia, Diana Marcela Walteros-Acero, Martha Lucia Ospina Martinez, Marcela Mercado-Reyes. |
| EPI_ISL_456157 | LabPLUS | Institute of Environmental Science and Research (ESR) | Matt Storey, Xiaoyun Ren, Anja Werno, Antje van der Linden, Arlo Upton, Chris Mansell, David Hammer, Dragana Drinkovic, Erasmus Smit, Gary McAuliffe, Hana Sofia Andersson, James Ussher, Jill Sherwood, Josh Freeman, Julia Howard, Juliet Elvy, Mary DeAlmeida, Matt Blakiston, Matthew Rogers, Max Bloomfield, Michael Addidle, Michelle Balm, Sally Roberts, Sarah Jefferies, Sharmini Muttaiyah, Susan Morpeth, Susan Taylor, Timothy Blackmore, Vani Sathyendran, Veronica Playle, Virginia Hope, Erasmus Smit, Lauren Jelly, Joep de Ligt |
| EPI_ISL_456158, EPI_ISL_456159 | Southern Community Labs Dunedin | Institute of Environmental Science and Research (ESR) | Matt Storey, Xiaoyun Ren, Anja Werno, Antje van der Linden, Arlo Upton, Chris Mansell, David Hammer, Dragana Drinkovic, Erasmus Smit, Gary McAuliffe, Hana Sofia Andersson, James Ussher, Jill Sherwood, Josh Freeman, Julia Howard, Juliet Elvy, Mary DeAlmeida, Matt Blakiston, Matthew Rogers, Max Bloomfield, Michael Addidle, Michelle Balm, Sally Roberts, Sarah Jefferies, Sharmini Muttaiyah, Susan Morpeth, Susan Taylor, Timothy Blackmore, Vani Sathyendran, Veronica Playle, Virginia Hope, Erasmus Smit, Lauren Jelly, Joep de Ligt |
| EPI_ISL_456160 | Waikato Hospital | Institute of Environmental Science and Research (ESR) | Matt Storey, Xiaoyun Ren, Anja Werno, Antje van der Linden, Arlo Upton, Chris Mansell, David Hammer, Dragana Drinkovic, Erasmus Smit, Gary McAuliffe, Hana Sofia Andersson, James Ussher, Jill Sherwood, Josh Freeman, Julia Howard, Juliet Elvy, Mary DeAlmeida, Matt Blakiston, Matthew Rogers, Max Bloomfield, Michael Addidle, Michelle Balm, Sally Roberts, Sarah Jefferies, Sharmini Muttaiyah, Susan Morpeth, Susan Taylor, Timothy Blackmore, Vani Sathyendran, Veronica Playle, Virginia Hope, Erasmus Smit, Lauren Jelly, Joep de Ligt |
| EPI_ISL_456161 | Wellington SCL | Institute of Environmental Science and Research (ESR) | Matt Storey, Xiaoyun Ren, Anja Werno, Antje van der Linden, Arlo Upton, Chris Mansell, David Hammer, Dragana Drinkovic, Erasmus Smit, Gary McAuliffe, Hana Sofia Andersson, James Ussher, Jill Sherwood, Josh Freeman, Julia Howard, Juliet Elvy, Mary DeAlmeida, Matt Blakiston, Matthew Rogers, Max Bloomfield, Michael Addidle, Michelle Balm, Sally Roberts, Sarah Jefferies, Sharmini Muttaiyah, Susan Morpeth, Susan Taylor, Timothy Blackmore, Vani Sathyendran, Veronica Playle, Virginia Hope, Erasmus Smit, Lauren Jelly, Joep de Ligt |
| EPI_ISL_456162 | Southern Community Labs Dunedin | Institute of Environmental Science and Research (ESR) | Matt Storey, Xiaoyun Ren, Anja Werno, Antje van der Linden, Arlo Upton, Chris Mansell, David Hammer, Dragana Drinkovic, Erasmus Smit, Gary McAuliffe, Hana Sofia Andersson, James Ussher, Jill Sherwood, Josh Freeman, Julia Howard, Juliet Elvy, Mary DeAlmeida, Matt Blakiston, Matthew Rogers, Max Bloomfield, Michael Addidle, Michelle Balm, Sally Roberts, Sarah Jefferies, Sharmini Muttaiyah, Susan Morpeth, Susan Taylor, Timothy Blackmore, Vani Sathyendran, Veronica Playle, Virginia Hope, Erasmus Smit, Lauren Jelly, Joep de Ligt |
| EPI_ISL_456163 | PathLab Bay of Plenty | Institute of Environmental Science and Research (ESR) | Matt Storey, Xiaoyun Ren, Anja Werno, Antje van der Linden, Arlo Upton, Chris Mansell, David Hammer, Dragana Drinkovic, Erasmus Smit, Gary McAuliffe, Hana Sofia Andersson, James Ussher, Jill Sherwood, Josh Freeman, Julia Howard, Juliet Elvy, Mary DeAlmeida, Matt Blakiston, Matthew Rogers, Max Bloomfield, Michael Addidle, Michelle Balm, Sally Roberts, Sarah Jefferies, Sharmini Muttaiyah, Susan Morpeth, Susan Taylor, Timothy Blackmore, Vani Sathyendran, Veronica Playle, Virginia Hope, Erasmus Smit, Lauren Jelly, Joep de Ligt |
| EPI_ISL_456164, EPI_ISL_456165, EPI_ISL_456166 | Wellington SCL | Institute of Environmental Science and Research (ESR) | Matt Storey, Xiaoyun Ren, Anja Werno, Antje van der Linden, Arlo Upton, Chris Mansell, David Hammer, Dragana Drinkovic, Erasmus Smit, Gary McAuliffe, Hana Sofia Andersson, James Ussher, Jill Sherwood, Josh Freeman, Julia Howard, Juliet Elvy, Mary DeAlmeida, Matt Blakiston, Matthew Rogers, Max Bloomfield, Michael Addidle, Michelle Balm, Sally Roberts, Sarah Jefferies, Sharmini Muttaiyah, Susan Morpeth, Susan Taylor, Timothy Blackmore, Vani Sathyendran, Veronica Playle, Virginia Hope, Erasmus Smit, Lauren Jelly, Joep de Ligt |
| EPI_ISL_456167 | Southern Community Labs Dunedin | Institute of Environmental Science and Research (ESR) | Matt Storey, Xiaoyun Ren, Anja Werno, Antje van der Linden, Arlo Upton, Chris Mansell, David Hammer, Dragana Drinkovic, Erasmus Smit, Gary McAuliffe, Hana Sofia Andersson, James Ussher, Jill Sherwood, Josh Freeman, Julia Howard, Juliet Elvy, Mary DeAlmeida, Matt Blakiston, Matthew Rogers, Max Bloomfield, Michael Addidle, Michelle Balm, Sally Roberts, Sarah Jefferies, Sharmini Muttaiyah, Susan Morpeth, Susan Taylor, Timothy Blackmore, Vani Sathyendran, Veronica Playle, Virginia Hope, Erasmus Smit, Lauren Jelly, Joep de Ligt |
| EPI_ISL_456168 | Waikato Hospital | Institute of Environmental Science and Research (ESR) | Matt Storey, Xiaoyun Ren, Anja Werno, Antje van der Linden, Arlo Upton, Chris Mansell, David Hammer, Dragana Drinkovic, Erasmus Smit, Gary McAuliffe, Hana Sofia Andersson, James Ussher, Jill Sherwood, Josh Freeman, Julia Howard, Juliet Elvy, Mary DeAlmeida, Matt Blakiston, Matthew Rogers, Max Bloomfield, Michael Addidle, Michelle Balm, Sally Roberts, Sarah Jefferies, Sharmini Muttaiyah, Susan Morpeth, Susan Taylor, Timothy Blackmore, Vani Sathyendran, Veronica Playle, Virginia Hope, Erasmus Smit, Lauren Jelly, Joep de Ligt |
| EPI_ISL_456169, EPI_ISL_456170 | Wellington SCL | Institute of Environmental Science and Research (ESR) | Matt Storey, Xiaoyun Ren, Anja Werno, Antje van der Linden, Arlo Upton, Chris Mansell, David Hammer, Dragana Drinkovic, Erasmus Smit, Gary McAuliffe, Hana Sofia Andersson, James Ussher, Jill Sherwood, Josh Freeman, Julia Howard, Juliet Elvy, Mary DeAlmeida, Matt Blakiston, Matthew Rogers, Max Bloomfield, Michael Addidle, Michelle Balm, Sally Roberts, Sarah Jefferies, Sharmini Muttaiyah, Susan Morpeth, Susan Taylor, Timothy Blackmore, Vani Sathyendran, Veronica Playle, Virginia Hope, Erasmus Smit, Lauren Jelly, Joep de Ligt |
| EPI_ISL_456171 | PathLab Bay of Plenty | Institute of Environmental Science and Research (ESR) | Matt Storey, Xiaoyun Ren, Anja Werno, Antje van der Linden, Arlo Upton, Chris Mansell, David Hammer, Dragana Drinkovic, Erasmus Smit, Gary McAuliffe, Hana Sofia Andersson, James Ussher, Jill Sherwood, Josh Freeman, Julia Howard, Juliet Elvy, Mary DeAlmeida, Matt Blakiston, Matthew Rogers, Max Bloomfield, Michael Addidle, Michelle Balm, Sally Roberts, Sarah Jefferies, Sharmini Muttaiyah, Susan Morpeth, Susan Taylor, Timothy Blackmore, Vani Sathyendran, Veronica Playle, Virginia Hope, Erasmus Smit, Lauren Jelly, Joep de Ligt |

| | | | |
|---|---|---|---|
| EPI_ISL_456172, EPI_ISL_456173, EPI_ISL_456174, EPI_ISL_456175, EPI_ISL_456176, EPI_ISL_456177, EPI_ISL_456178, EPI_ISL_456179, EPI_ISL_456180 | LabPLUS | Institute of Environmental Science and Research (ESR) | Matt Storey, Xiaoyun Ren, Anja Werno, Antje van der Linden, Arlo Upton, Chris Mansell, David Hammer, Dragana Drinkovic, Erasmus Smit, Gary McAuliffe, Hana Sofia Andersson, James Ussher, Jill Sherwood, Josh Freeman, Julia Howard, Juliet Elvy, Mary DeAlmeida, Matt Blakiston, Matthew Rogers, Max Bloomfield, Michael Addidle, Michelle Balm, Sally Roberts, Sarah Jefferies, Sharmini Muttaiyah, Susan Morpeth, Susan Taylor, Timothy Blackmore, Vani Sathyendran, Veronica Playle, Virginia Hope, Erasmus Smit, Lauren Jelly, Joep de Ligt |
| EPI_ISL_456181 | Southern Community Labs Dunedin | Institute of Environmental Science and Research (ESR) | Matt Storey, Xiaoyun Ren, Anja Werno, Antje van der Linden, Arlo Upton, Chris Mansell, David Hammer, Dragana Drinkovic, Erasmus Smit, Gary McAuliffe, Hana Sofia Andersson, James Ussher, Jill Sherwood, Josh Freeman, Julia Howard, Juliet Elvy, Mary DeAlmeida, Matt Blakiston, Matthew Rogers, Max Bloomfield, Michael Addidle, Michelle Balm, Sally Roberts, Sarah Jefferies, Sharmini Muttaiyah, Susan Morpeth, Susan Taylor, Timothy Blackmore, Vani Sathyendran, Veronica Playle, Virginia Hope, Erasmus Smit, Lauren Jelly, Joep de Ligt |
| EPI_ISL_456182, EPI_ISL_456183, EPI_ISL_456184, EPI_ISL_456185, EPI_ISL_456186, EPI_ISL_456187, EPI_ISL_456188 | Wellington SCL | Institute of Environmental Science and Research (ESR) | Matt Storey, Xiaoyun Ren, Anja Werno, Antje van der Linden, Arlo Upton, Chris Mansell, David Hammer, Dragana Drinkovic, Erasmus Smit, Gary McAuliffe, Hana Sofia Andersson, James Ussher, Jill Sherwood, Josh Freeman, Julia Howard, Juliet Elvy, Mary DeAlmeida, Matt Blakiston, Matthew Rogers, Max Bloomfield, Michael Addidle, Michelle Balm, Sally Roberts, Sarah Jefferies, Sharmini Muttaiyah, Susan Morpeth, Susan Taylor, Timothy Blackmore, Vani Sathyendran, Veronica Playle, Virginia Hope, Erasmus Smit, Lauren Jelly, Joep de Ligt |
| EPI_ISL_456189, EPI_ISL_456190, EPI_ISL_456191, EPI_ISL_456192, EPI_ISL_456193 | Waikato Hospital | Institute of Environmental Science and Research (ESR) | Matt Storey, Xiaoyun Ren, Anja Werno, Antje van der Linden, Arlo Upton, Chris Mansell, David Hammer, Dragana Drinkovic, Erasmus Smit, Gary McAuliffe, Hana Sofia Andersson, James Ussher, Jill Sherwood, Josh Freeman, Julia Howard, Juliet Elvy, Mary DeAlmeida, Matt Blakiston, Matthew Rogers, Max Bloomfield, Michael Addidle, Michelle Balm, Sally Roberts, Sarah Jefferies, Sharmini Muttaiyah, Susan Morpeth, Susan Taylor, Timothy Blackmore, Vani Sathyendran, Veronica Playle, Virginia Hope, Erasmus Smit, Lauren Jelly, Joep de Ligt |
| EPI_ISL_456194, EPI_ISL_456195 | PathLab Bay of Plenty | Institute of Environmental Science and Research (ESR) | Matt Storey, Xiaoyun Ren, Anja Werno, Antje van der Linden, Arlo Upton, Chris Mansell, David Hammer, Dragana Drinkovic, Erasmus Smit, Gary McAuliffe, Hana Sofia Andersson, James Ussher, Jill Sherwood, Josh Freeman, Julia Howard, Juliet Elvy, Mary DeAlmeida, Matt Blakiston, Matthew Rogers, Max Bloomfield, Michael Addidle, Michelle Balm, Sally Roberts, Sarah Jefferies, Sharmini Muttaiyah, Susan Morpeth, Susan Taylor, Timothy Blackmore, Vani Sathyendran, Veronica Playle, Virginia Hope, Erasmus Smit, Lauren Jelly, Joep de Ligt |
| EPI_ISL_456196, EPI_ISL_456197, EPI_ISL_456198, EPI_ISL_456199, EPI_ISL_456200, EPI_ISL_456201, EPI_ISL_456202, EPI_ISL_456203, EPI_ISL_456204, EPI_ISL_456205, EPI_ISL_456206, EPI_ISL_456207, EPI_ISL_456208, EPI_ISL_456209, EPI_ISL_456210, EPI_ISL_456211, EPI_ISL_456212 | | | |
| see above | LabPLUS | Institute of Environmental Science and Research (ESR) | Matt Storey, Xiaoyun Ren, Anja Werno, Antje van der Linden, Arlo Upton, Chris Mansell, David Hammer, Dragana Drinkovic, Erasmus Smit, Gary McAuliffe, Hana Sofia Andersson, James Ussher, Jill Sherwood, Josh Freeman, Julia Howard, Juliet Elvy, Mary DeAlmeida, Matt Blakiston, Matthew Rogers, Max Bloomfield, Michael Addidle, Michelle Balm, Sally Roberts, Sarah Jefferies, Sharmini Muttaiyah, Susan Morpeth, Susan Taylor, Timothy Blackmore, Vani Sathyendran, Veronica Playle, Virginia Hope, Erasmus Smit, Lauren Jelly, Joep de Ligt |
| EPI_ISL_456213, EPI_ISL_456214 | Middlemore Hospital | Institute of Environmental Science and Research (ESR) | Matt Storey, Xiaoyun Ren, Anja Werno, Antje van der Linden, Arlo Upton, Chris Mansell, David Hammer, Dragana Drinkovic, Erasmus Smit, Gary McAuliffe, Hana Sofia Andersson, James Ussher, Jill Sherwood, Josh Freeman, Julia Howard, Juliet Elvy, Mary DeAlmeida, Matt Blakiston, Matthew Rogers, Max Bloomfield, Michael Addidle, Michelle Balm, Sally Roberts, Sarah Jefferies, Sharmini Muttaiyah, Susan Morpeth, Susan Taylor, Timothy Blackmore, Vani Sathyendran, Veronica Playle, Virginia Hope, Erasmus Smit, Lauren Jelly, Joep de Ligt |
| EPI_ISL_456215 | PathLab Bay of Plenty | Institute of Environmental Science and Research (ESR) | Matt Storey, Xiaoyun Ren, Anja Werno, Antje van der Linden, Arlo Upton, Chris Mansell, David Hammer, Dragana Drinkovic, Erasmus Smit, Gary McAuliffe, Hana Sofia Andersson, James Ussher, Jill Sherwood, Josh Freeman, Julia Howard, Juliet Elvy, Mary DeAlmeida, Matt Blakiston, Matthew Rogers, Max Bloomfield, Michael Addidle, Michelle Balm, Sally Roberts, Sarah Jefferies, Sharmini Muttaiyah, Susan Morpeth, Susan Taylor, Timothy Blackmore, Vani Sathyendran, Veronica Playle, Virginia Hope, Erasmus Smit, Lauren Jelly, Joep de Ligt |
| EPI_ISL_456216, EPI_ISL_456217, EPI_ISL_456218, EPI_ISL_456219, EPI_ISL_456220 | Southern Community Labs Dunedin | Institute of Environmental Science and Research (ESR) | Matt Storey, Xiaoyun Ren, Anja Werno, Antje van der Linden, Arlo Upton, Chris Mansell, David Hammer, Dragana Drinkovic, Erasmus Smit, Gary McAuliffe, Hana Sofia Andersson, James Ussher, Jill Sherwood, Josh Freeman, Julia Howard, Juliet Elvy, Mary DeAlmeida, Matt Blakiston, Matthew Rogers, Max Bloomfield, Michael Addidle, Michelle Balm, Sally Roberts, Sarah Jefferies, Sharmini Muttaiyah, Susan Morpeth, Susan Taylor, Timothy Blackmore, Vani Sathyendran, Veronica Playle, Virginia Hope, Erasmus Smit, Lauren Jelly, Joep de Ligt |
| EPI_ISL_456221, EPI_ISL_456222, EPI_ISL_456223, EPI_ISL_456224 | PathLab Bay of Plenty | Institute of Environmental Science and Research (ESR) | Matt Storey, Xiaoyun Ren, Anja Werno, Antje van der Linden, Arlo Upton, Chris Mansell, David Hammer, Dragana Drinkovic, Erasmus Smit, Gary McAuliffe, Hana Sofia Andersson, James Ussher, Jill Sherwood, Josh Freeman, Julia Howard, Juliet Elvy, Mary DeAlmeida, Matt Blakiston, Matthew Rogers, Max Bloomfield, Michael Addidle, Michelle Balm, Sally Roberts, Sarah Jefferies, Sharmini Muttaiyah, Susan Morpeth, Susan Taylor, Timothy Blackmore, Vani Sathyendran, Veronica Playle, Virginia Hope, Erasmus Smit, Lauren Jelly, Joep de Ligt |
| EPI_ISL_456225, EPI_ISL_456226, EPI_ISL_456227, EPI_ISL_456228, EPI_ISL_456229, EPI_ISL_456230, EPI_ISL_456231, EPI_ISL_456232, EPI_ISL_456233, EPI_ISL_456234, EPI_ISL_456235, EPI_ISL_456236, EPI_ISL_456237, EPI_ISL_456238, EPI_ISL_456239, EPI_ISL_456240 | | | |
| see above | Southern Community Labs Dunedin | Institute of Environmental Science and Research (ESR) | Matt Storey, Xiaoyun Ren, Anja Werno, Antje van der Linden, Arlo Upton, Chris Mansell, David Hammer, Dragana Drinkovic, Erasmus Smit, Gary McAuliffe, Hana Sofia Andersson, James Ussher, Jill Sherwood, Josh Freeman, Julia Howard, Juliet Elvy, Mary DeAlmeida, Matt Blakiston, Matthew Rogers, Max Bloomfield, Michael Addidle, Michelle Balm, Sally Roberts, Sarah Jefferies, Sharmini Muttaiyah, Susan Morpeth, Susan Taylor, Timothy Blackmore, Vani Sathyendran, Veronica Playle, Virginia Hope, Erasmus Smit, Lauren Jelly, Joep de Ligt |
| EPI_ISL_456241 | Waikato Hospital | Institute of Environmental Science and Research (ESR) | Matt Storey, Xiaoyun Ren, Anja Werno, Antje van der Linden, Arlo Upton, Chris Mansell, David Hammer, Dragana Drinkovic, Erasmus Smit, Gary McAuliffe, Hana Sofia Andersson, James Ussher, Jill Sherwood, Josh Freeman, Julia Howard, Juliet Elvy, Mary DeAlmeida, Matt Blakiston, Matthew Rogers, Max Bloomfield, Michael Addidle, Michelle Balm, Sally Roberts, Sarah Jefferies, Sharmini Muttaiyah, Susan Morpeth, Susan Taylor, Timothy Blackmore, Vani Sathyendran, Veronica Playle, Virginia Hope, Erasmus Smit, Lauren Jelly, Joep de Ligt |
| EPI_ISL_456242, EPI_ISL_456243, EPI_ISL_456244, EPI_ISL_456245, EPI_ISL_456246, EPI_ISL_456247, EPI_ISL_456248, EPI_ISL_456249, EPI_ISL_456250, EPI_ISL_456251, EPI_ISL_456252, EPI_ISL_456253, EPI_ISL_456254 | | | |
| see above | Southern Community Labs Dunedin | Institute of Environmental Science and Research (ESR) | Matt Storey, Xiaoyun Ren, Anja Werno, Antje van der Linden, Arlo Upton, Chris Mansell, David Hammer, Dragana Drinkovic, Erasmus Smit, Gary McAuliffe, Hana Sofia Andersson, James Ussher, Jill Sherwood, Josh Freeman, Julia Howard, Juliet Elvy, Mary DeAlmeida, Matt Blakiston, Matthew Rogers, Max Bloomfield, Michael Addidle, Michelle Balm, Sally Roberts, Sarah Jefferies, Sharmini Muttaiyah, Susan Morpeth, Susan Taylor, Timothy Blackmore, Vani Sathyendran, Veronica Playle, Virginia Hope, Erasmus Smit, Lauren Jelly, Joep de Ligt |
| EPI_ISL_456255, EPI_ISL_456256, EPI_ISL_456257, EPI_ISL_456258 | Middlemore Hospital | Institute of Environmental Science and Research (ESR) | Matt Storey, Xiaoyun Ren, Anja Werno, Antje van der Linden, Arlo Upton, Chris Mansell, David Hammer, Dragana Drinkovic, Erasmus Smit, Gary McAuliffe, Hana Sofia Andersson, James Ussher, Jill Sherwood, Josh Freeman, Julia Howard, Juliet Elvy, Mary DeAlmeida, Matt Blakiston, Matthew Rogers, Max Bloomfield, Michael Addidle, Michelle Balm, Sally Roberts, Sarah Jefferies, Sharmini Muttaiyah, Susan Morpeth, Susan Taylor, Timothy Blackmore, Vani Sathyendran, Veronica Playle, Virginia Hope, Erasmus Smit, Lauren Jelly, Joep de Ligt |
| EPI_ISL_456259, EPI_ISL_456260 | Waikato Hospital | Institute of Environmental Science and Research (ESR) | Matt Storey, Xiaoyun Ren, Anja Werno, Antje van der Linden, Arlo Upton, Chris Mansell, David Hammer, Dragana Drinkovic, Erasmus Smit, Gary McAuliffe, Hana Sofia Andersson, James Ussher, Jill Sherwood, Josh Freeman, Julia Howard, Juliet Elvy, Mary DeAlmeida, Matt Blakiston, Matthew Rogers, Max Bloomfield, Michael Addidle, Michelle Balm, Sally Roberts, Sarah Jefferies, Sharmini Muttaiyah, Susan Morpeth, Susan Taylor, Timothy Blackmore, Vani Sathyendran, Veronica Playle, Virginia Hope, Erasmus Smit, Lauren Jelly, Joep de Ligt |
| EPI_ISL_456261, EPI_ISL_456262, EPI_ISL_456263, EPI_ISL_456264, EPI_ISL_456265, EPI_ISL_456266, EPI_ISL_456267, EPI_ISL_456268, EPI_ISL_456269, EPI_ISL_456270, EPI_ISL_456271, EPI_ISL_456272, EPI_ISL_456273, EPI_ISL_456274, EPI_ISL_456275, EPI_ISL_456276, EPI_ISL_456277, EPI_ISL_456278, EPI_ISL_456279, EPI_ISL_456280, EPI_ISL_456281, EPI_ISL_456282, EPI_ISL_456283, EPI_ISL_456284, EPI_ISL_456285, EPI_ISL_456286, EPI_ISL_456287, EPI_ISL_456288, EPI_ISL_456289, EPI_ISL_456290, EPI_ISL_456291, EPI_ISL_456292, EPI_ISL_456293 | | | |
| see above | Southern Community Labs Dunedin | Institute of Environmental Science and Research (ESR) | Matt Storey, Xiaoyun Ren, Anja Werno, Antje van der Linden, Arlo Upton, Chris Mansell, David Hammer, Dragana Drinkovic, Erasmus Smit, Gary McAuliffe, Hana Sofia Andersson, James Ussher, Jill Sherwood, Josh Freeman, Julia Howard, Juliet Elvy, Mary DeAlmeida, Matt Blakiston, Matthew Rogers, Max Bloomfield, Michael Addidle, Michelle Balm, Sally Roberts, Sarah Jefferies, Sharmini Muttaiyah, Susan Morpeth, Susan Taylor, Timothy Blackmore, Vani Sathyendran, Veronica Playle, Virginia Hope, Erasmus Smit, Lauren Jelly, Joep de Ligt |
| EPI_ISL_456294 | North Shore Hospital | Institute of Environmental Science and Research (ESR) | Matt Storey, Xiaoyun Ren, Anja Werno, Antje van der Linden, Arlo Upton, Chris Mansell, David Hammer, Dragana Drinkovic, Erasmus Smit, Gary McAuliffe, Hana Sofia Andersson, James Ussher, Jill Sherwood, Josh Freeman, Julia Howard, Juliet Elvy, Mary DeAlmeida, Matt Blakiston, Matthew Rogers, Max Bloomfield, Michael Addidle, Michelle Balm, Sally Roberts, Sarah Jefferies, Sharmini Muttaiyah, Susan Morpeth, Susan Taylor, Timothy Blackmore, Vani Sathyendran, Veronica Playle, Virginia Hope, Erasmus Smit, Lauren Jelly, Joep de Ligt |
| EPI_ISL_456295, EPI_ISL_456296, EPI_ISL_456297, EPI_ISL_456298, EPI_ISL_456299 | Southern Community Labs Dunedin | Institute of Environmental Science and Research (ESR) | Matt Storey, Xiaoyun Ren, Anja Werno, Antje van der Linden, Arlo Upton, Chris Mansell, David Hammer, Dragana Drinkovic, Erasmus Smit, Gary McAuliffe, Hana Sofia Andersson, James Ussher, Jill Sherwood, Josh Freeman, Julia Howard, Juliet Elvy, Mary DeAlmeida, Matt Blakiston, Matthew Rogers, Max Bloomfield, Michael Addidle, Michelle Balm, Sally Roberts, Sarah Jefferies, Sharmini Muttaiyah, Susan Morpeth, Susan Taylor, Timothy Blackmore, Vani Sathyendran, Veronica Playle, Virginia Hope, Erasmus Smit, Lauren Jelly, Joep de Ligt |
| EPI_ISL_456300 | LabPLUS | Institute of Environmental Science and Research (ESR) | Matt Storey, Xiaoyun Ren, Anja Werno, Antje van der Linden, Arlo Upton, Chris Mansell, David Hammer, Dragana Drinkovic, Erasmus Smit, Gary McAuliffe, Hana Sofia Andersson, James Ussher, Jill Sherwood, Josh Freeman, Julia Howard, Juliet Elvy, Mary DeAlmeida, Matt Blakiston, Matthew Rogers, Max Bloomfield, Michael Addidle, Michelle Balm, Sally Roberts, Sarah Jefferies, Sharmini Muttaiyah, Susan Morpeth, Susan Taylor, Timothy Blackmore, Vani Sathyendran, Veronica Playle, Virginia Hope, Erasmus Smit, Lauren Jelly, Joep de Ligt |
| EPI_ISL_456301 | PathLab Bay of Plenty | Institute of Environmental Science and Research (ESR) | Matt Storey, Xiaoyun Ren, Anja Werno, Antje van der Linden, Arlo Upton, Chris Mansell, David Hammer, Dragana Drinkovic, Erasmus Smit, Gary McAuliffe, Hana Sofia Andersson, James Ussher, Jill Sherwood, Josh Freeman, Julia Howard, Juliet Elvy, Mary DeAlmeida, Matt Blakiston, Matthew Rogers, Max Bloomfield, Michael Addidle, Michelle Balm, Sally Roberts, Sarah Jefferies, Sharmini Muttaiyah, Susan Morpeth, Susan Taylor, Timothy Blackmore, Vani Sathyendran, Veronica Playle, Virginia Hope, Erasmus Smit, Lauren Jelly, Joep de Ligt |
| EPI_ISL_456302, EPI_ISL_456303, EPI_ISL_456304, EPI_ISL_456305, EPI_ISL_456306, EPI_ISL_456307, EPI_ISL_456308, EPI_ISL_456309, EPI_ISL_456310, EPI_ISL_456311, EPI_ISL_456312, EPI_ISL_456313, EPI_ISL_456314, EPI_ISL_456315, EPI_ISL_456316, EPI_ISL_456317, EPI_ISL_456318 | | | |
| see above | Southern Community Labs Dunedin | Institute of Environmental Science and Research (ESR) | Matt Storey, Xiaoyun Ren, Anja Werno, Antje van der Linden, Arlo Upton, Chris Mansell, David Hammer, Dragana Drinkovic, Erasmus Smit, Gary McAuliffe, Hana Sofia Andersson, James Ussher, Jill Sherwood, Josh Freeman, Julia Howard, Juliet Elvy, Mary DeAlmeida, Matt Blakiston, Matthew Rogers, Max Bloomfield, Michael Addidle, Michelle Balm, Sally Roberts, Sarah Jefferies, Sharmini Muttaiyah, Susan Morpeth, Susan Taylor, Timothy Blackmore, Vani Sathyendran, Veronica Playle, Virginia Hope, Erasmus Smit, Lauren Jelly, Joep de Ligt |
| EPI_ISL_456319, EPI_ISL_456320 | Wellington SCL | Institute of Environmental Science and Research (ESR) | Matt Storey, Xiaoyun Ren, Anja Werno, Antje van der Linden, Arlo Upton, Chris Mansell, David Hammer, Dragana Drinkovic, Erasmus Smit, Gary McAuliffe, Hana Sofia Andersson, James Ussher, Jill Sherwood, Josh Freeman, Julia Howard, Juliet Elvy, Mary DeAlmeida, Matt Blakiston, Matthew Rogers, Max Bloomfield, Michael Addidle, Michelle Balm, Sally Roberts, Sarah Jefferies, Sharmini Muttaiyah, Susan Morpeth, Susan Taylor, Timothy Blackmore, Vani Sathyendran, Veronica Playle, Virginia Hope, Erasmus Smit, Lauren Jelly, Joep de Ligt |
| EPI_ISL_456321, EPI_ISL_456322 | Canterbury Health Laboratories | Institute of Environmental Science and Research (ESR) | Matt Storey, Xiaoyun Ren, Anja Werno, Antje van der Linden, Arlo Upton, Chris Mansell, David Hammer, Dragana Drinkovic, Erasmus Smit, Gary McAuliffe, Hana Sofia Andersson, James Ussher, Jill Sherwood, Josh Freeman, Julia Howard, Juliet Elvy, Mary DeAlmeida, Matt Blakiston, Matthew Rogers, Max Bloomfield, Michael Addidle, Michelle Balm, Sally Roberts, Sarah Jefferies, Sharmini Muttaiyah, Susan Morpeth, Susan Taylor, Timothy Blackmore, Vani Sathyendran, Veronica Playle, Virginia Hope, Erasmus Smit, Lauren Jelly, Joep de Ligt |
| EPI_ISL_456323 | North Shore Hospital | Institute of Environmental Science and Research (ESR) | Matt Storey, Xiaoyun Ren, Anja Werno, Antje van der Linden, Arlo Upton, Chris Mansell, David Hammer, Dragana Drinkovic, Erasmus Smit, Gary McAuliffe, Hana Sofia Andersson, James Ussher, Jill Sherwood, Josh Freeman, Julia Howard, Juliet Elvy, Mary DeAlmeida, Matt Blakiston, Matthew Rogers, Max Bloomfield, Michael Addidle, Michelle Balm, Sally Roberts, Sarah Jefferies, Sharmini Muttaiyah, Susan Morpeth, Susan Taylor, Timothy Blackmore, Vani Sathyendran, Veronica Playle, Virginia Hope, Erasmus Smit, Lauren Jelly, Joep de Ligt |
| EPI_ISL_456324, EPI_ISL_456325, EPI_ISL_456326, EPI_ISL_456327, EPI_ISL_456328, EPI_ISL_456329, EPI_ISL_456330, EPI_ISL_456331, EPI_ISL_456332, EPI_ISL_456333, EPI_ISL_456334, EPI_ISL_456335, EPI_ISL_456336, EPI_ISL_456337, EPI_ISL_456338, EPI_ISL_456339, EPI_ISL_456340, EPI_ISL_456341, EPI_ISL_456342, EPI_ISL_456343 | | | |
| see above | Wellington SCL | Institute of Environmental Science and Research (ESR) | Matt Storey, Xiaoyun Ren, Anja Werno, Antje van der Linden, Arlo Upton, Chris Mansell, David Hammer, Dragana Drinkovic, Erasmus Smit, Gary McAuliffe, Hana Sofia Andersson, James Ussher, Jill Sherwood, Josh Freeman, Julia Howard, Juliet Elvy, Mary DeAlmeida, Matt Blakiston, Matthew Rogers, Max Bloomfield, Michael Addidle, Michelle Balm, Sally Roberts, Sarah Jefferies, Sharmini Muttaiyah, Susan Morpeth, Susan Taylor, Timothy Blackmore, Vani Sathyendran, Veronica Playle, Virginia Hope, Erasmus Smit, Lauren Jelly, Joep de Ligt |
| EPI_ISL_456344, EPI_ISL_456345, EPI_ISL_456346, EPI_ISL_456347 | Southern Community Labs Dunedin | Institute of Environmental Science and Research (ESR) | Matt Storey, Xiaoyun Ren, Anja Werno, Antje van der Linden, Arlo Upton, Chris Mansell, David Hammer, Dragana Drinkovic, Erasmus Smit, Gary McAuliffe, Hana Sofia Andersson, James Ussher, Jill Sherwood, Josh Freeman, Julia Howard, Juliet Elvy, Mary DeAlmeida, Matt Blakiston, Matthew Rogers, Max Bloomfield, Michael Addidle, Michelle Balm, Sally Roberts, Sarah Jefferies, Sharmini Muttaiyah, Susan Morpeth, Susan Taylor, Timothy Blackmore, Vani Sathyendran, Veronica Playle, Virginia Hope, Erasmus Smit, Lauren Jelly, Joep de Ligt |
| EPI_ISL_456348, EPI_ISL_456349, EPI_ISL_456350, EPI_ISL_456351, EPI_ISL_456352, EPI_ISL_456353, EPI_ISL_456354, EPI_ISL_456355, EPI_ISL_456356, EPI_ISL_456357, EPI_ISL_456358, EPI_ISL_456359, EPI_ISL_456360, EPI_ISL_456361, EPI_ISL_456362, EPI_ISL_456363, EPI_ISL_456364, EPI_ISL_456365, EPI_ISL_456366, EPI_ISL_456367, EPI_ISL_456368, EPI_ISL_456369, EPI_ISL_456370, EPI_ISL_456371, EPI_ISL_456372, EPI_ISL_456373, EPI_ISL_456374, EPI_ISL_456375 | | | |
| see above | Canterbury Health Laboratories | Institute of Environmental Science and Research (ESR) | Matt Storey, Xiaoyun Ren, Anja Werno, Antje van der Linden, Arlo Upton, Chris Mansell, David Hammer, Dragana Drinkovic, Erasmus Smit, Gary McAuliffe, Hana Sofia Andersson, James Ussher, Jill Sherwood, Josh Freeman, Julia Howard, Juliet Elvy, Mary DeAlmeida, Matt Blakiston, Matthew Rogers, Max Bloomfield, Michael Addidle, Michelle Balm, Sally Roberts, Sarah Jefferies, Sharmini Muttaiyah, Susan Morpeth, Susan Taylor, Timothy Blackmore, Vani Sathyendran, Veronica Playle, Virginia Hope, Erasmus Smit, Lauren Jelly, Joep de Ligt |
| EPI_ISL_456376 | Middlemore Hospital | Institute of Environmental Science and Research (ESR) | Matt Storey, Xiaoyun Ren, Anja Werno, Antje van der Linden, Arlo Upton, Chris Mansell, David Hammer, Dragana Drinkovic, Erasmus Smit, Gary McAuliffe, Hana Sofia Andersson, James Ussher, Jill Sherwood, Josh Freeman, Julia Howard, Juliet Elvy, Mary DeAlmeida, Matt Blakiston, Matthew Rogers, Max Bloomfield, Michael Addidle, Michelle Balm, Sally Roberts, Sarah Jefferies, Sharmini Muttaiyah, Susan Morpeth, Susan |

| | | | |
|---|---|---|---|
| | | | Taylor, Timothy Blackmore, Vani Sathyendran, Veronica Playle, Virginia Hope, Erasmus Smit, Lauren Jelly, Joep de Ligt |
| EPI_ISL_456377, EPI_ISL_456378 | MedLab Central Ltd | Institute of Environmental Science and Research (ESR) | Matt Storey, Xiaoyun Ren, Anja Werno, Antje van der Linden, Arlo Upton, Chris Mansell, David Hammer, Dragana Drinkovic, Erasmus Smit, Gary McAuliffe, Hana Sofia Andersson, James Ussher, Jill Sherwood, Josh Freeman, Julia Howard, Juliet Elvy, Mary DeAlmeida, Matt Blakiston, Matthew Rogers, Max Bloomfield, Michael Addidle, Michelle Balm, Sally Roberts, Sarah Jefferies, Sharmini Muttaiyah, Susan Morpeth, Susan Taylor, Timothy Blackmore, Vani Sathyendran, Veronica Playle, Virginia Hope, Erasmus Smit, Lauren Jelly, Joep de Ligt |
| EPI_ISL_456379, EPI_ISL_456380, EPI_ISL_456381, EPI_ISL_456382, EPI_ISL_456383, EPI_ISL_456384 | LabPLUS | Institute of Environmental Science and Research (ESR) | Matt Storey, Xiaoyun Ren, Anja Werno, Antje van der Linden, Arlo Upton, Chris Mansell, David Hammer, Dragana Drinkovic, Erasmus Smit, Gary McAuliffe, Hana Sofia Andersson, James Ussher, Jill Sherwood, Josh Freeman, Julia Howard, Juliet Elvy, Mary DeAlmeida, Matt Blakiston, Matthew Rogers, Max Bloomfield, Michael Addidle, Michelle Balm, Sally Roberts, Sarah Jefferies, Sharmini Muttaiyah, Susan Morpeth, Susan Taylor, Timothy Blackmore, Vani Sathyendran, Veronica Playle, Virginia Hope, Erasmus Smit, Lauren Jelly, Joep de Ligt |
| EPI_ISL_456385, EPI_ISL_456386, EPI_ISL_456387 | Middlemore Hospital | Institute of Environmental Science and Research (ESR) | Matt Storey, Xiaoyun Ren, Anja Werno, Antje van der Linden, Arlo Upton, Chris Mansell, David Hammer, Dragana Drinkovic, Erasmus Smit, Gary McAuliffe, Hana Sofia Andersson, James Ussher, Jill Sherwood, Josh Freeman, Julia Howard, Juliet Elvy, Mary DeAlmeida, Matt Blakiston, Matthew Rogers, Max Bloomfield, Michael Addidle, Michelle Balm, Sally Roberts, Sarah Jefferies, Sharmini Muttaiyah, Susan Morpeth, Susan Taylor, Timothy Blackmore, Vani Sathyendran, Veronica Playle, Virginia Hope, Erasmus Smit, Lauren Jelly, Joep de Ligt |
| EPI_ISL_456388, EPI_ISL_456389, EPI_ISL_456390, EPI_ISL_456391, EPI_ISL_456392, EPI_ISL_456393, EPI_ISL_456394 | LabPLUS | Institute of Environmental Science and Research (ESR) | Matt Storey, Xiaoyun Ren, Anja Werno, Antje van der Linden, Arlo Upton, Chris Mansell, David Hammer, Dragana Drinkovic, Erasmus Smit, Gary McAuliffe, Hana Sofia Andersson, James Ussher, Jill Sherwood, Josh Freeman, Julia Howard, Juliet Elvy, Mary DeAlmeida, Matt Blakiston, Matthew Rogers, Max Bloomfield, Michael Addidle, Michelle Balm, Sally Roberts, Sarah Jefferies, Sharmini Muttaiyah, Susan Morpeth, Susan Taylor, Timothy Blackmore, Vani Sathyendran, Veronica Playle, Virginia Hope, Erasmus Smit, Lauren Jelly, Joep de Ligt |
| EPI_ISL_456395, EPI_ISL_456396, EPI_ISL_456397 | North Shore Hospital | Institute of Environmental Science and Research (ESR) | Matt Storey, Xiaoyun Ren, Anja Werno, Antje van der Linden, Arlo Upton, Chris Mansell, David Hammer, Dragana Drinkovic, Erasmus Smit, Gary McAuliffe, Hana Sofia Andersson, James Ussher, Jill Sherwood, Josh Freeman, Julia Howard, Juliet Elvy, Mary DeAlmeida, Matt Blakiston, Matthew Rogers, Max Bloomfield, Michael Addidle, Michelle Balm, Sally Roberts, Sarah Jefferies, Sharmini Muttaiyah, Susan Morpeth, Susan Taylor, Timothy Blackmore, Vani Sathyendran, Veronica Playle, Virginia Hope, Erasmus Smit, Lauren Jelly, Joep de Ligt |
| EPI_ISL_456398, EPI_ISL_456399, EPI_ISL_456400, EPI_ISL_456401, EPI_ISL_456402 | Wellington SCL | Institute of Environmental Science and Research (ESR) | Matt Storey, Xiaoyun Ren, Anja Werno, Antje van der Linden, Arlo Upton, Chris Mansell, David Hammer, Dragana Drinkovic, Erasmus Smit, Gary McAuliffe, Hana Sofia Andersson, James Ussher, Jill Sherwood, Josh Freeman, Julia Howard, Juliet Elvy, Mary DeAlmeida, Matt Blakiston, Matthew Rogers, Max Bloomfield, Michael Addidle, Michelle Balm, Sally Roberts, Sarah Jefferies, Sharmini Muttaiyah, Susan Morpeth, Susan Taylor, Timothy Blackmore, Vani Sathyendran, Veronica Playle, Virginia Hope, Erasmus Smit, Lauren Jelly, Joep de Ligt |
| EPI_ISL_456403 | Middlemore Hospital | Institute of Environmental Science and Research (ESR) | Matt Storey, Xiaoyun Ren, Anja Werno, Antje van der Linden, Arlo Upton, Chris Mansell, David Hammer, Dragana Drinkovic, Erasmus Smit, Gary McAuliffe, Hana Sofia Andersson, James Ussher, Jill Sherwood, Josh Freeman, Julia Howard, Juliet Elvy, Mary DeAlmeida, Matt Blakiston, Matthew Rogers, Max Bloomfield, Michael Addidle, Michelle Balm, Sally Roberts, Sarah Jefferies, Sharmini Muttaiyah, Susan Morpeth, Susan Taylor, Timothy Blackmore, Vani Sathyendran, Veronica Playle, Virginia Hope, Erasmus Smit, Lauren Jelly, Joep de Ligt |
| EPI_ISL_456404 | unknown | Research Center Of Tropical and Infectious Of Medical Sciences | Mollaei,H.R., Aghaei-Afshar,A., Kalantar-Neyestanaki,D., Tabatabaeifar,F., Morones Ramirez,J.R. |
| EPI_ISL_456405 | unknown | Research Center Of Tropical and Infectious Of Medical Sciences | Mollaei,H.R., Kalantar-Neyestanaki,D., Aghaei-Afshar,A. |
| EPI_ISL_456406 | unknown | Research Center Of Tropical and Infectious Of Medical Sciences | Mollaei,H.R., Aghaei-Afshar,A., Kalantar-Neyestanaki,D., Morones Ramirez,J.R. |
| EPI_ISL_456407, EPI_ISL_456408, EPI_ISL_456409 | unknown | Research Center Of Tropical and Infectious Of Medical Sciences | Mollaei,H.R., Aghaei-Afshar,A., Kalantar-Neyestanaki,D. |
| EPI_ISL_456410, EPI_ISL_456411, EPI_ISL_456412, EPI_ISL_456413, EPI_ISL_456414, EPI_ISL_456415, EPI_ISL_456416, EPI_ISL_456417, EPI_ISL_456418, EPI_ISL_456419, EPI_ISL_456420, EPI_ISL_456421, EPI_ISL_456422, EPI_ISL_456423, EPI_ISL_456424, EPI_ISL_456425, EPI_ISL_456426, EPI_ISL_456427, EPI_ISL_456428, EPI_ISL_456429, EPI_ISL_456430, EPI_ISL_456431, EPI_ISL_456432, EPI_ISL_456433, EPI_ISL_456434, EPI_ISL_456435, EPI_ISL_456436, EPI_ISL_456437, EPI_ISL_456438, EPI_ISL_456439, EPI_ISL_456440, EPI_ISL_456441, EPI_ISL_456442, EPI_ISL_456443, EPI_ISL_456444, EPI_ISL_456445, EPI_ISL_456446, EPI_ISL_456447, EPI_ISL_456448, EPI_ISL_456449, EPI_ISL_456450, EPI_ISL_456451, EPI_ISL_456452, EPI_ISL_456453, EPI_ISL_456454, EPI_ISL_456455, EPI_ISL_456456, EPI_ISL_456457, EPI_ISL_456458, EPI_ISL_456459, EPI_ISL_456460, EPI_ISL_456461, EPI_ISL_456462, EPI_ISL_456463, EPI_ISL_456464, EPI_ISL_456465, EPI_ISL_456466, EPI_ISL_456467, EPI_ISL_456468, EPI_ISL_456469, EPI_ISL_456470, EPI_ISL_456471, EPI_ISL_456472, EPI_ISL_456473, EPI_ISL_456474, EPI_ISL_456475, EPI_ISL_456476, EPI_ISL_456477 | | | |
| see above | Victorian Infectious Diseases Reference Laboratory (VIDRL) | Microbiological Diagnostic Unit Public Health Laboratory and Victorian Infectious Diseases Reference Laboratory, Doherty Institute | Caly L., Seemann T., Sait, M., Schultz M., Druce J., Sherry, N. |
| EPI_ISL_456478, EPI_ISL_456479 | Microbiological Diagnostic Unit Public Health Laboratory | Microbiological Diagnostic Unit Public Health Laboratory, The Peter Doherty Institute for Infection and Immunity | Seemann T., Schultz M., Sait, M., Sherry, N. |
| EPI_ISL_456480, EPI_ISL_456481, EPI_ISL_456482, EPI_ISL_456483, EPI_ISL_456484, EPI_ISL_456485, EPI_ISL_456486, EPI_ISL_456487, EPI_ISL_456488, EPI_ISL_456489, EPI_ISL_456490, EPI_ISL_456491, EPI_ISL_456492, EPI_ISL_456493, EPI_ISL_456494, EPI_ISL_456495, EPI_ISL_456496 | | | |
| see above | Victorian Infectious Diseases Reference Laboratory (VIDRL) | Microbiological Diagnostic Unit Public Health Laboratory and Victorian Infectious Diseases Reference Laboratory, Doherty Institute | Caly L., Seemann T., Sait, M., Schultz M., Druce J., Sherry, N. |
| EPI_ISL_456497, EPI_ISL_456498 | Royal Darwin Hospital Pathology | Microbiological Diagnostic Unit Public Health Laboratory, The Peter Doherty Institute for Infection and Immunity | Meumann, E., Caly L., Seemann T., Sait, M., Schultz M., Druce J., Sherry, N. |
| EPI_ISL_456499, EPI_ISL_456500, EPI_ISL_456501, EPI_ISL_456502, EPI_ISL_456503, EPI_ISL_456504, EPI_ISL_456505, EPI_ISL_456506, EPI_ISL_456507, EPI_ISL_456508, EPI_ISL_456509, EPI_ISL_456510, EPI_ISL_456511, EPI_ISL_456512, EPI_ISL_456513, EPI_ISL_456514, EPI_ISL_456515, EPI_ISL_456516, EPI_ISL_456517, EPI_ISL_456518, EPI_ISL_456519, EPI_ISL_456520, EPI_ISL_456521, EPI_ISL_456522, EPI_ISL_456523, EPI_ISL_456524, EPI_ISL_456525, EPI_ISL_456526, EPI_ISL_456527, EPI_ISL_456528, EPI_ISL_456529, EPI_ISL_456530, EPI_ISL_456531, EPI_ISL_456532, EPI_ISL_456533, EPI_ISL_456534, EPI_ISL_456535, EPI_ISL_456536, EPI_ISL_456537, EPI_ISL_456538, EPI_ISL_456539, EPI_ISL_456540, EPI_ISL_456541, EPI_ISL_456542, EPI_ISL_456543, EPI_ISL_456544, EPI_ISL_456545, EPI_ISL_456546, EPI_ISL_456547, EPI_ISL_456548, EPI_ISL_456549, EPI_ISL_456550, EPI_ISL_456551, EPI_ISL_456552, EPI_ISL_456553, EPI_ISL_456554, EPI_ISL_456555, EPI_ISL_456556, EPI_ISL_456557, EPI_ISL_456558, EPI_ISL_456559, EPI_ISL_456560, EPI_ISL_456561, EPI_ISL_456562, EPI_ISL_456563, EPI_ISL_456564, EPI_ISL_456565, EPI_ISL_456566, EPI_ISL_456567, EPI_ISL_456568, EPI_ISL_456569, EPI_ISL_456570, EPI_ISL_456571, EPI_ISL_456572, EPI_ISL_456573, EPI_ISL_456574, EPI_ISL_456575, EPI_ISL_456576, EPI_ISL_456577, EPI_ISL_456578, EPI_ISL_456579, EPI_ISL_456580, EPI_ISL_456581, EPI_ISL_456582, EPI_ISL_456583, EPI_ISL_456584, EPI_ISL_456585, EPI_ISL_456586, EPI_ISL_456587, EPI_ISL_456588, EPI_ISL_456589, EPI_ISL_456590, EPI_ISL_456591, EPI_ISL_456592, EPI_ISL_456593 | | | |
| see above | Victorian Infectious Diseases Reference Laboratory (VIDRL) | Microbiological Diagnostic Unit Public Health Laboratory and Victorian Infectious Diseases Reference Laboratory, Doherty Institute | Caly L., Seemann T., Sait, M., Schultz M., Druce J., Sherry, N. |
| EPI_ISL_456594, EPI_ISL_456595 | Microbiological Diagnostic Unit Public Health Laboratory | Microbiological Diagnostic Unit Public Health Laboratory, The Peter Doherty Institute for Infection and Immunity | Seemann T., Schultz M., Sait, M., Sherry, N. |
| EPI_ISL_456596, EPI_ISL_456597, EPI_ISL_456598, EPI_ISL_456599, EPI_ISL_456600, EPI_ISL_456601, EPI_ISL_456602, EPI_ISL_456603, EPI_ISL_456604, EPI_ISL_456605, EPI_ISL_456606, EPI_ISL_456607, EPI_ISL_456608, EPI_ISL_456609, EPI_ISL_456610, EPI_ISL_456611, EPI_ISL_456612 | | | |
| see above | National Health Laboratory, Timor-Leste | Microbiological Diagnostic Unit Public Health Laboratory, The Peter Doherty Institute for Infection and Immunity | Soares da Silva, E., Dolores de Jesus da Costa, M., Salles de Sousa, A., Jayanti Pereira Tilman, A., Antonia da Costa, E., Barreto, I., Marr, I., Wapling, J., Francis, J., Ximenes, J., Canisia, D., Freeman, K., Dakh, F., Douglas, N., Baird, R., Caly, L., Seemann, T., Sait, M., Schultz, M., Sherry, N. |
| EPI_ISL_456613, EPI_ISL_456614, EPI_ISL_456615, EPI_ISL_456616, EPI_ISL_456617, EPI_ISL_456618, EPI_ISL_456619, EPI_ISL_456620, EPI_ISL_456621, EPI_ISL_456622, EPI_ISL_456623, EPI_ISL_456624, EPI_ISL_456625, EPI_ISL_456626, EPI_ISL_456627, EPI_ISL_456628, EPI_ISL_456629, EPI_ISL_456630, EPI_ISL_456631, EPI_ISL_456632, EPI_ISL_456633, EPI_ISL_456634, EPI_ISL_456635, EPI_ISL_456636, EPI_ISL_456637, EPI_ISL_456638, EPI_ISL_456639, EPI_ISL_456640, EPI_ISL_456641, EPI_ISL_456642, EPI_ISL_456643, EPI_ISL_456644, EPI_ISL_456645, EPI_ISL_456646, EPI_ISL_456647, EPI_ISL_456648, EPI_ISL_456649, EPI_ISL_456650, EPI_ISL_456651, EPI_ISL_456652, EPI_ISL_456653, EPI_ISL_456654, EPI_ISL_456655 | | | |
| see above | Victorian Infectious Diseases Reference Laboratory (VIDRL) | Microbiological Diagnostic Unit Public Health Laboratory and Victorian Infectious Diseases Reference Laboratory, Doherty Institute | Caly L., Seemann T., Sait, M., Schultz M., Druce J., Sherry, N. |
| EPI_ISL_456656 | American Type Culture Collection Inc. (ATCC) | American Type Culture Collection Inc. (ATCC) | Riojas, M.A., Frank,A.M., Puthuveetil, N.P., Benton,B., Peiris, J.S.M., Chu, D.K.W., King,S.P., Flores, B., Parker, M. and Rashid, S. |
| EPI_ISL_456657, EPI_ISL_456658, EPI_ISL_456659, EPI_ISL_456660, EPI_ISL_456661, EPI_ISL_456662, EPI_ISL_456663, EPI_ISL_456664, EPI_ISL_456665, EPI_ISL_456666, EPI_ISL_456667, EPI_ISL_456668, EPI_ISL_456669, EPI_ISL_456670, EPI_ISL_456671, EPI_ISL_456672, EPI_ISL_456673, EPI_ISL_456674, EPI_ISL_456675, EPI_ISL_456676 | | | |
| see above | University of Birmingham | COVID-19 Genomics UK (COG-UK) Consortium | Loman Lab: Claire McMurray, Joanne Stockton, Samuel Nicholls, Radoslaw Poplawski, Will Rowe, Josh Quick, Nicholas Loman // UHB Lab: Celina M Whalley, Andrew Bosworth, Charlotte Poxon, Kasun Wanigasooriya, Oliver Pickles, Mike Kidd, Alex Richter, Andrew D Beggs // PHE Heartlands Lab: Husam Osman, Andrew Bosworth |
| EPI_ISL_456677, EPI_ISL_456678, EPI_ISL_456679, EPI_ISL_456680, EPI_ISL_456681, EPI_ISL_456682, EPI_ISL_456683, EPI_ISL_456684, EPI_ISL_456685, EPI_ISL_456686, EPI_ISL_456687, EPI_ISL_456688, EPI_ISL_456689, EPI_ISL_456690, EPI_ISL_456691, EPI_ISL_456692, EPI_ISL_456693, EPI_ISL_456694, EPI_ISL_456695, EPI_ISL_456696, EPI_ISL_456697, EPI_ISL_456698, EPI_ISL_456699, EPI_ISL_456700, EPI_ISL_456701, EPI_ISL_456702, EPI_ISL_456703, EPI_ISL_456704, EPI_ISL_456705, EPI_ISL_456706, EPI_ISL_456707, EPI_ISL_456708, EPI_ISL_456709, EPI_ISL_456710, EPI_ISL_456711, EPI_ISL_456712, EPI_ISL_456713, EPI_ISL_456714, EPI_ISL_456715, EPI_ISL_456716, EPI_ISL_456717, EPI_ISL_456718, EPI_ISL_456719, EPI_ISL_456720, EPI_ISL_456721, EPI_ISL_456722, EPI_ISL_456723, EPI_ISL_456724, EPI_ISL_456725, EPI_ISL_456726, EPI_ISL_456727, EPI_ISL_456728, EPI_ISL_456729, EPI_ISL_456730, EPI_ISL_456731, EPI_ISL_456732, EPI_ISL_456733, EPI_ISL_456734, EPI_ISL_456735, EPI_ISL_456736, EPI_ISL_456737, EPI_ISL_456738, EPI_ISL_456739, EPI_ISL_456740, EPI_ISL_456741, EPI_ISL_456742, EPI_ISL_456743, EPI_ISL_456744, EPI_ISL_456745, EPI_ISL_456746, EPI_ISL_456747, EPI_ISL_456748, EPI_ISL_456749, EPI_ISL_456750, EPI_ISL_456751, EPI_ISL_456752, EPI_ISL_456753, EPI_ISL_456754, EPI_ISL_456755 | | | |
| see above | Department of Pathology, University of Cambridge | COVID-19 Genomics UK (COG-UK) Consortium | Luke W Meredith, M. Estée Török, Myra Hosmillo, William L. Hamilton, Martin D. Curran, Theresa Feltwell, Grant Hall, Anna Yakovleva, Fahad A Khokhar, Charlotte J. Houldcroft, Laura G Caller, Aminu S. Jahun, Sarah L. Caddy, Ian Goodfellow |
| EPI_ISL_456756, EPI_ISL_456757, EPI_ISL_456758, EPI_ISL_456759, EPI_ISL_456760, EPI_ISL_456761, EPI_ISL_456762, EPI_ISL_456763, EPI_ISL_456764, EPI_ISL_456765, EPI_ISL_456766, EPI_ISL_456767, EPI_ISL_456768, EPI_ISL_456769, EPI_ISL_456770, EPI_ISL_456771, EPI_ISL_456772, EPI_ISL_456773, EPI_ISL_456774, EPI_ISL_456775, EPI_ISL_456776, EPI_ISL_456777, EPI_ISL_456778, EPI_ISL_456779, EPI_ISL_456780, EPI_ISL_456781, EPI_ISL_456782, EPI_ISL_456783, EPI_ISL_456784, EPI_ISL_456785, EPI_ISL_456786, EPI_ISL_456787, EPI_ISL_456788, EPI_ISL_456789, EPI_ISL_456790, EPI_ISL_456791, EPI_ISL_456792, EPI_ISL_456793, EPI_ISL_456794, EPI_ISL_456795, EPI_ISL_456796, EPI_ISL_456797, EPI_ISL_456798, EPI_ISL_456799, EPI_ISL_456800, EPI_ISL_456801, EPI_ISL_456802, EPI_ISL_456803, EPI_ISL_456804, EPI_ISL_456805, EPI_ISL_456806, EPI_ISL_456807, EPI_ISL_456808, EPI_ISL_456809, EPI_ISL_456810, EPI_ISL_456811, EPI_ISL_456812, EPI_ISL_456813, EPI_ISL_456814, EPI_ISL_456815, EPI_ISL_456816, EPI_ISL_456817, EPI_ISL_456818, EPI_ISL_456819, EPI_ISL_456820, EPI_ISL_456821, EPI_ISL_456822, EPI_ISL_456823, EPI_ISL_456824, EPI_ISL_456825, EPI_ISL_456826, EPI_ISL_456827, EPI_ISL_456828, EPI_ISL_456829, EPI_ISL_456830, EPI_ISL_456831, EPI_ISL_456832, EPI_ISL_456833, EPI_ISL_456834, EPI_ISL_456835, EPI_ISL_456836, EPI_ISL_456837, EPI_ISL_456838, EPI_ISL_456839, EPI_ISL_456840, EPI_ISL_456841, EPI_ISL_456842, EPI_ISL_456843, EPI_ISL_456844, EPI_ISL_456845, EPI_ISL_456846, EPI_ISL_456847, EPI_ISL_456848, EPI_ISL_456849, EPI_ISL_456850, EPI_ISL_456851, EPI_ISL_456852, EPI_ISL_456853, EPI_ISL_456854, EPI_ISL_456855, EPI_ISL_456856, EPI_ISL_456857, EPI_ISL_456858, EPI_ISL_456859, EPI_ISL_456860, EPI_ISL_456861, EPI_ISL_456862, EPI_ISL_456863, EPI_ISL_456864, EPI_ISL_456865, EPI_ISL_456866, EPI_ISL_456867, EPI_ISL_456868, EPI_ISL_456869, EPI_ISL_456870, EPI_ISL_456871, EPI_ISL_456872, EPI_ISL_456873, EPI_ISL_456874, EPI_ISL_456875, EPI_ISL_456876, EPI_ISL_456877, EPI_ISL_456878, EPI_ISL_456879, EPI_ISL_456880, EPI_ISL_456881, EPI_ISL_456882, EPI_ISL_456883, EPI_ISL_456884, EPI_ISL_456885, EPI_ISL_456886, EPI_ISL_456887, EPI_ISL_456888, EPI_ISL_456889 | | | |
| see above | West of Scotland Specialist Virology Centre, NHSGGC / MRC-University of Glasgow Centre for Virus Research | COVID-19 Genomics UK (COG-UK) Consortium | Ana da Silva Filipe, Natasha Johnson, Kathy Smollett, Daniel Mair, Stephen Carmichael, Lily Tong, Jenna Nichols, Elihu Aranday-Cortes, Kirstyn Brunker, Yasmin Parr, Kyriaki Nomikou; Sarah McDonald, Marc Niebel, Patawee Asamaphan; Richard Orton, Joseph Hughes, Sreenu Vattipally, David L Robertson; Alasdair MacLean, Rory Gunson; Kathy Li, Natasha Jesudason, Rajiv Shah, James Shepherd, Antonia Ho, Emma Thomson |
| EPI_ISL_456890, EPI_ISL_456891, EPI_ISL_456892, EPI_ISL_456893, EPI_ISL_456894, EPI_ISL_456895, EPI_ISL_456896, EPI_ISL_456897, EPI_ISL_456898, EPI_ISL_456899, EPI_ISL_456900, EPI_ISL_456901, EPI_ISL_456902, EPI_ISL_456903, EPI_ISL_456904, EPI_ISL_456905, EPI_ISL_456906, EPI_ISL_456907, EPI_ISL_456908, EPI_ISL_456909, EPI_ISL_456910, EPI_ISL_456911, EPI_ISL_456912, EPI_ISL_456913, EPI_ISL_456914, EPI_ISL_456915, EPI_ISL_456916, EPI_ISL_456917, EPI_ISL_456918, EPI_ISL_456919, EPI_ISL_456920, EPI_ISL_456921, EPI_ISL_456922, EPI_ISL_456923, EPI_ISL_456924, EPI_ISL_456925, EPI_ISL_456926, EPI_ISL_456927, EPI_ISL_456928, EPI_ISL_456929, EPI_ISL_456930, EPI_ISL_456931, EPI_ISL_456932, EPI_ISL_456933, EPI_ISL_456934, EPI_ISL_456935, EPI_ISL_456936, EPI_ISL_456937, EPI_ISL_456938, EPI_ISL_456939, EPI_ISL_456940, EPI_ISL_456941, EPI_ISL_456942, EPI_ISL_456943, EPI_ISL_456944, EPI_ISL_456945, EPI_ISL_456946, EPI_ISL_456947, EPI_ISL_456948, EPI_ISL_456949, EPI_ISL_456950, EPI_ISL_456951, EPI_ISL_456952, EPI_ISL_456953, EPI_ISL_456954, EPI_ISL_456955, EPI_ISL_456956, EPI_ISL_456957, EPI_ISL_456958, EPI_ISL_456959, EPI_ISL_456960, EPI_ISL_456961, EPI_ISL_456962, EPI_ISL_456963, EPI_ISL_456964, EPI_ISL_456965, EPI_ISL_456966, EPI_ISL_456967, EPI_ISL_456968, EPI_ISL_456969, EPI_ISL_456970, EPI_ISL_456971, EPI_ISL_456972, EPI_ISL_456973, EPI_ISL_456974, EPI_ISL_456975, EPI_ISL_456976, EPI_ISL_456977, EPI_ISL_456978, EPI_ISL_456979, EPI_ISL_456980, EPI_ISL_456981, EPI_ISL_456982, EPI_ISL_456983, EPI_ISL_456984, EPI_ISL_456985, EPI_ISL_456986, EPI_ISL_456987, EPI_ISL_456988, EPI_ISL_456989, EPI_ISL_456990, EPI_ISL_456991, EPI_ISL_456992, EPI_ISL_456993, EPI_ISL_456994, EPI_ISL_456995, EPI_ISL_456996, EPI_ISL_456997, EPI_ISL_456998, EPI_ISL_456999, EPI_ISL_457000, EPI_ISL_457001, EPI_ISL_457002, EPI_ISL_457003, EPI_ISL_457004, EPI_ISL_457005, EPI_ISL_457006, EPI_ISL_457007, EPI_ISL_457008, EPI_ISL_457009, EPI_ISL_457010, EPI_ISL_457011, EPI_ISL_457012, EPI_ISL_457013, EPI_ISL_457014, EPI_ISL_457015, EPI_ISL_457016, EPI_ISL_457017, EPI_ISL_457018, EPI_ISL_457019, EPI_ISL_457020, EPI_ISL_457021, EPI_ISL_457022, EPI_ISL_457023 | | | |
| see above | Virology Department, Royal Infirmary of Edinburgh, NHS Lothian / School of Biological Sciences, University of Edinburgh / Institute of Genetics and Molecular Medicine, University of Edinburgh | COVID-19 Genomics UK (COG-UK) Consortium | McHugh M, Dewar R, Rooke S, Gallagher M, Balcaza C, O'Toole Á, Scher E, Hill V, McCrone JT, Colquhoun R, Yu X, Jackson B, Rambaut A, Williams TC, Templeton K |
| EPI_ISL_457024, EPI_ISL_457025, EPI_ISL_457026, EPI_ISL_457027, EPI_ISL_457028, EPI_ISL_457029, EPI_ISL_457030, EPI_ISL_457031, EPI_ISL_457032, EPI_ISL_457033, EPI_ISL_457034, EPI_ISL_457035, EPI_ISL_457036, EPI_ISL_457037, EPI_ISL_457038, EPI_ISL_457039, EPI_ISL_457040, EPI_ISL_457041, EPI_ISL_457042, EPI_ISL_457043, EPI_ISL_457044, EPI_ISL_457045, EPI_ISL_457046, EPI_ISL_457047, EPI_ISL_457048, EPI_ISL_457049, EPI_ISL_457050, EPI_ISL_457051, EPI_ISL_457052, EPI_ISL_457053, EPI_ISL_457054, EPI_ISL_457055, EPI_ISL_457056, EPI_ISL_457057, EPI_ISL_457058, EPI_ISL_457059, EPI_ISL_457060, EPI_ISL_457061, EPI_ISL_457062, EPI_ISL_457063, EPI_ISL_457064, EPI_ISL_457065, EPI_ISL_457066, EPI_ISL_457067, EPI_ISL_457068, EPI_ISL_457069, EPI_ISL_457070, EPI_ISL_457071, EPI_ISL_457072, EPI_ISL_457073, EPI_ISL_457074, EPI_ISL_457075, EPI_ISL_457076, EPI_ISL_457077, EPI_ISL_457078, EPI_ISL_457079, EPI_ISL_457080, EPI_ISL_457081, EPI_ISL_457082, EPI_ISL_457083, EPI_ISL_457084, EPI_ISL_457085, EPI_ISL_457086, EPI_ISL_457087, EPI_ISL_457088, EPI_ISL_457089, EPI_ISL_457090, EPI_ISL_457091, EPI_ISL_457092, EPI_ISL_457093, EPI_ISL_457094, EPI_ISL_457095, EPI_ISL_457096, EPI_ISL_457097, EPI_ISL_457098, EPI_ISL_457099, EPI_ISL_457100, EPI_ISL_457101, EPI_ISL_457102, EPI_ISL_457103, EPI_ISL_457104, EPI_ISL_457105, EPI_ISL_457106, EPI_ISL_457107, EPI_ISL_457108, EPI_ISL_457109, EPI_ISL_457110, EPI_ISL_457111, EPI_ISL_457112, EPI_ISL_457113, EPI_ISL_457114, EPI_ISL_457115, EPI_ISL_457116, EPI_ISL_457117, EPI_ISL_457118, EPI_ISL_457119, EPI_ISL_457120, EPI_ISL_457121, EPI_ISL_457122, EPI_ISL_457123, EPI_ISL_457124, EPI_ISL_457125, EPI_ISL_457126, EPI_ISL_457127, EPI_ISL_457128, EPI_ISL_457129, EPI_ISL_457130, EPI_ISL_457131, EPI_ISL_457132, EPI_ISL_457133, EPI_ISL_457134, EPI_ISL_457135, EPI_ISL_457136, EPI_ISL_457137, EPI_ISL_457138, EPI_ISL_457139, EPI_ISL_457140, EPI_ISL_457141, EPI_ISL_457142, EPI_ISL_457143, EPI_ISL_457144, EPI_ISL_457145, EPI_ISL_457146, EPI_ISL_457147, EPI_ISL_457148, EPI_ISL_457149, EPI_ISL_457150, EPI_ISL_457151, EPI_ISL_457152, EPI_ISL_457153, EPI_ISL_457154, EPI_ISL_457155, EPI_ISL_457156, EPI_ISL_457157, EPI_ISL_457158, EPI_ISL_457159, EPI_ISL_457160, EPI_ISL_457161, EPI_ISL_457162, EPI_ISL_457163, EPI_ISL_457164, EPI_ISL_457165, EPI_ISL_457166, EPI_ISL_457167, EPI_ISL_457168, EPI_ISL_457169, EPI_ISL_457170, EPI_ISL_457171, EPI_ISL_457172, EPI_ISL_457173, EPI_ISL_457174, EPI_ISL_457175, EPI_ISL_457176, EPI_ISL_457177, EPI_ISL_457178, EPI_ISL_457179, EPI_ISL_457180, EPI_ISL_457181, EPI_ISL_457182, EPI_ISL_457183, EPI_ISL_457184, EPI_ISL_457185, EPI_ISL_457186, EPI_ISL_457187, EPI_ISL_457188, EPI_ISL_457189, EPI_ISL_457190, EPI_ISL_457191, EPI_ISL_457192, EPI_ISL_457193, EPI_ISL_457194, EPI_ISL_457195, EPI_ISL_457196, EPI_ISL_457197, EPI_ISL_457198, EPI_ISL_457199, EPI_ISL_457200, EPI_ISL_457201, EPI_ISL_457202, EPI_ISL_457203, EPI_ISL_457204, EPI_ISL_457205, EPI_ISL_457206, EPI_ISL_457207, EPI_ISL_457208, EPI_ISL_457209, EPI_ISL_457210, EPI_ISL_457211, EPI_ISL_457212, EPI_ISL_457213, EPI_ISL_457214, EPI_ISL_457215, EPI_ISL_457216, EPI_ISL_457217, EPI_ISL_457218, EPI_ISL_457219, EPI_ISL_457220, EPI_ISL_457221, EPI_ISL_457222, EPI_ISL_457223, EPI_ISL_457224, EPI_ISL_457225, EPI_ISL_457226, EPI_ISL_457227, EPI_ISL_457228, EPI_ISL_457229, EPI_ISL_457230, EPI_ISL_457231, EPI_ISL_457232, EPI_ISL_457233, EPI_ISL_457234, EPI_ISL_457235, EPI_ISL_457236, EPI_ISL_457237, EPI_ISL_457238, EPI_ISL_457239, EPI_ISL_457240, EPI_ISL_457241, EPI_ISL_457242, EPI_ISL_457243, EPI_ISL_457244, EPI_ISL_457245, EPI_ISL_457246, EPI_ISL_457247, EPI_ISL_457248, EPI_ISL_457249, EPI_ISL_457250, EPI_ISL_457251, EPI_ISL_457252, EPI_ISL_457253, EPI_ISL_457254, EPI_ISL_457255, EPI_ISL_457256, EPI_ISL_457257, EPI_ISL_457258 | | | |
| see above | University of Exeter | COVID-19 Genomics UK (COG-UK) Consortium | Ben Temperton,Aaron Jeffries,Michelle Michelsen,Joanna Warwick-Dugdale,Audrey Farbos,Robyn Manley,Stephen Michell,Jane Masoli |
| EPI_ISL_457259, EPI_ISL_457260, EPI_ISL_457261, EPI_ISL_457262, EPI_ISL_457263, EPI_ISL_457264, EPI_ISL_457265, EPI_ISL_457266, EPI_ISL_457267, EPI_ISL_457268, EPI_ISL_457269 | | | |

| | | | |
|---|---|---|---|
| see above | Liverpool Clinical Laboratories | COVID-19 Genomics UK (COG-UK) Consortium | Sam Haldenby, Anita Lucaci, Steve Paterson, Julian Hiscox, Alistair Darby, M Almsaud, A Alrezaihi, Muhannad Alruwaili, Stuart D Armstrong, Jones Benjamin, Eleanor G Bentley, Anu Chawla, Jordan J Clark, Angela Cowell, Richard Eccles, Isabel García-Dorival, Matthew Gemmell, Alessandro Gerada, PKF Gilmore, Richard Gregory, Ximeng Han, Catherine Hartley, Margaret Hughes, Miren Iturriza-Gomara, James Johnson, L Luu, Jenifer Manson, Charlotte Nelson, Elaine O'Toole, Cassie Olateju, Rebekah Penrice-Randal , Lucille Rainbow, N.P Randle, Trevor Ian Robinson, Parul Sharma, Ghada T Shawli, James P Stewart, Neil Swainston, Ecaterina Vamos, Joanne Watts, Mark Whitehead |
| EPI_ISL_457270, EPI_ISL_457271, EPI_ISL_457272, EPI_ISL_457273, EPI_ISL_457274, EPI_ISL_457275, EPI_ISL_457276, EPI_ISL_457277, EPI_ISL_457278, EPI_ISL_457279, EPI_ISL_457280, EPI_ISL_457281, EPI_ISL_457282, EPI_ISL_457283, EPI_ISL_457284, EPI_ISL_457285, EPI_ISL_457286, EPI_ISL_457287, EPI_ISL_457288, EPI_ISL_457289, EPI_ISL_457290, EPI_ISL_457291, EPI_ISL_457292, EPI_ISL_457293, EPI_ISL_457294, EPI_ISL_457295, EPI_ISL_457296, EPI_ISL_457297, EPI_ISL_457298, EPI_ISL_457299, EPI_ISL_457300, EPI_ISL_457301, EPI_ISL_457302 | | | |
| see above | University College London, Great Ormond Street Hospital for Children NHS Foundation Trust, Imperial College Healthcare NHS Trust | COVID-19 Genomics UK (COG-UK) Consortium | Sergi Castellano, Rachel Williams, Mark Kristiansen, Paola Resende Silva, Sunando Roy, Tony Brooks, Helena Tutill, Paola Niola, Patricia Dyal, Charlotte Williams, Leysa Forrest, Yasmin Panchbhaya, Jacqueline Findlay, Sam Weeks, Julianne Brown, Kathryn Harris, Paul Randell, James Price, Alison Holmes, Judith Breuer |
| EPI_ISL_457303, EPI_ISL_457304, EPI_ISL_457305, EPI_ISL_457306, EPI_ISL_457307, EPI_ISL_457308, EPI_ISL_457309, EPI_ISL_457310, EPI_ISL_457311, EPI_ISL_457312, EPI_ISL_457313, EPI_ISL_457314, EPI_ISL_457315, EPI_ISL_457316, EPI_ISL_457317, EPI_ISL_457318, EPI_ISL_457319, EPI_ISL_457320, EPI_ISL_457321, EPI_ISL_457322, EPI_ISL_457323, EPI_ISL_457324, EPI_ISL_457325 | | | |
| see above | Northumbria University / South Tees Hospitals NHS Foundation Trust / North Cumbria Integrated Care NHS Foundation Trust / North Tees and Hartlepool NHS Foundation Trust / Newcastle Hospitals NHS Foundation Trust | COVID-19 Genomics UK (COG-UK) Consortium | Darren L Smith,Andrew Nelson,Matthew Bashton,Greg R Young,Joshua Loh,John Allan,Mohammad A Tariq,Giles S Holt,Gary Black,Wen C Yew,Lynn Dover,Paul Baker,Steve Liggett,Sarah Essex,Jane Greenaway,Debra Padgett,Clive Graham,Garren Scott,Edward Barton,Emma Swindells,Brendan Payne,Jennifer Collins,Yusri Taha,Gary Eltringham |
| EPI_ISL_457326, EPI_ISL_457327, EPI_ISL_457328, EPI_ISL_457329, EPI_ISL_457330, EPI_ISL_457331, EPI_ISL_457332, EPI_ISL_457333, EPI_ISL_457334, EPI_ISL_457335, EPI_ISL_457336, EPI_ISL_457337, EPI_ISL_457338, EPI_ISL_457339, EPI_ISL_457340, EPI_ISL_457341, EPI_ISL_457342, EPI_ISL_457343, EPI_ISL_457344, EPI_ISL_457345, EPI_ISL_457346, EPI_ISL_457347, EPI_ISL_457348, EPI_ISL_457349, EPI_ISL_457350, EPI_ISL_457351, EPI_ISL_457352, EPI_ISL_457353, EPI_ISL_457354, EPI_ISL_457355, EPI_ISL_457356, EPI_ISL_457357, EPI_ISL_457358, EPI_ISL_457359, EPI_ISL_457360, EPI_ISL_457361, EPI_ISL_457362, EPI_ISL_457363, EPI_ISL_457364, EPI_ISL_457365, EPI_ISL_457366, EPI_ISL_457367, EPI_ISL_457368, EPI_ISL_457369, EPI_ISL_457370, EPI_ISL_457371, EPI_ISL_457372, EPI_ISL_457373, EPI_ISL_457374, EPI_ISL_457375, EPI_ISL_457376, EPI_ISL_457377, EPI_ISL_457378, EPI_ISL_457379, EPI_ISL_457380, EPI_ISL_457381, EPI_ISL_457382, EPI_ISL_457383, EPI_ISL_457384, EPI_ISL_457385, EPI_ISL_457386, EPI_ISL_457387, EPI_ISL_457388, EPI_ISL_457389, EPI_ISL_457390, EPI_ISL_457391, EPI_ISL_457392, EPI_ISL_457393, EPI_ISL_457394, EPI_ISL_457395, EPI_ISL_457396, EPI_ISL_457397, EPI_ISL_457398, EPI_ISL_457399, EPI_ISL_457400, EPI_ISL_457401, EPI_ISL_457402, EPI_ISL_457403, EPI_ISL_457404, EPI_ISL_457405, EPI_ISL_457406, EPI_ISL_457407, EPI_ISL_457408, EPI_ISL_457409, EPI_ISL_457410, EPI_ISL_457411, EPI_ISL_457412, EPI_ISL_457413, EPI_ISL_457414, EPI_ISL_457415, EPI_ISL_457416, EPI_ISL_457417, EPI_ISL_457418, EPI_ISL_457419, EPI_ISL_457420, EPI_ISL_457421, EPI_ISL_457422, EPI_ISL_457423, EPI_ISL_457424, EPI_ISL_457425, EPI_ISL_457426, EPI_ISL_457427, EPI_ISL_457428, EPI_ISL_457429, EPI_ISL_457430, EPI_ISL_457431, EPI_ISL_457432, EPI_ISL_457433, EPI_ISL_457434, EPI_ISL_457435, EPI_ISL_457436, EPI_ISL_457437, EPI_ISL_457438, EPI_ISL_457439, EPI_ISL_457440, EPI_ISL_457441, EPI_ISL_457442, EPI_ISL_457443, EPI_ISL_457444, EPI_ISL_457445, EPI_ISL_457446, EPI_ISL_457447, EPI_ISL_457448, EPI_ISL_457449, EPI_ISL_457450, EPI_ISL_457451, EPI_ISL_457452, EPI_ISL_457453, EPI_ISL_457454, EPI_ISL_457455, EPI_ISL_457456, EPI_ISL_457457, EPI_ISL_457458, EPI_ISL_457459, EPI_ISL_457460, EPI_ISL_457461, EPI_ISL_457462, EPI_ISL_457463, EPI_ISL_457464, EPI_ISL_457465, EPI_ISL_457466, EPI_ISL_457467, EPI_ISL_457468, EPI_ISL_457469, EPI_ISL_457470, EPI_ISL_457471, EPI_ISL_457472, EPI_ISL_457473, EPI_ISL_457474, EPI_ISL_457475, EPI_ISL_457476, EPI_ISL_457477, EPI_ISL_457478, EPI_ISL_457479, EPI_ISL_457480, EPI_ISL_457481, EPI_ISL_457482, EPI_ISL_457483, EPI_ISL_457484, EPI_ISL_457485, EPI_ISL_457486, EPI_ISL_457487, EPI_ISL_457488, EPI_ISL_457489, EPI_ISL_457490, EPI_ISL_457491, EPI_ISL_457492, EPI_ISL_457493, EPI_ISL_457494, EPI_ISL_457495, EPI_ISL_457496, EPI_ISL_457497, EPI_ISL_457498, EPI_ISL_457499, EPI_ISL_457500, EPI_ISL_457501, EPI_ISL_457502, EPI_ISL_457503, EPI_ISL_457504, EPI_ISL_457505, EPI_ISL_457506, EPI_ISL_457507, EPI_ISL_457508, EPI_ISL_457509, EPI_ISL_457510, EPI_ISL_457511, EPI_ISL_457512, EPI_ISL_457513, EPI_ISL_457514, EPI_ISL_457515, EPI_ISL_457516, EPI_ISL_457517, EPI_ISL_457518, EPI_ISL_457519, EPI_ISL_457520, EPI_ISL_457521, EPI_ISL_457522, EPI_ISL_457523, EPI_ISL_457524, EPI_ISL_457525, EPI_ISL_457526, EPI_ISL_457527, EPI_ISL_457528, EPI_ISL_457529, EPI_ISL_457530, EPI_ISL_457531, EPI_ISL_457532, EPI_ISL_457533, EPI_ISL_457534, EPI_ISL_457535, EPI_ISL_457536, EPI_ISL_457537, EPI_ISL_457538, EPI_ISL_457539, EPI_ISL_457540, EPI_ISL_457541, EPI_ISL_457542, EPI_ISL_457543, EPI_ISL_457544, EPI_ISL_457545, EPI_ISL_457546, EPI_ISL_457547, EPI_ISL_457548, EPI_ISL_457549, EPI_ISL_457550, EPI_ISL_457551, EPI_ISL_457552, EPI_ISL_457553, EPI_ISL_457554, EPI_ISL_457555, EPI_ISL_457556, EPI_ISL_457557, EPI_ISL_457558, EPI_ISL_457559, EPI_ISL_457560, EPI_ISL_457561, EPI_ISL_457562, EPI_ISL_457563, EPI_ISL_457564, EPI_ISL_457565, EPI_ISL_457566, EPI_ISL_457567, EPI_ISL_457568, EPI_ISL_457569, EPI_ISL_457570, EPI_ISL_457571, EPI_ISL_457572 | | | |
| see above | Quadram Institute Bioscience | COVID-19 Genomics UK (COG-UK) Consortium | Dave J. Baker, Gemma L. Kay, Alp Aydin, Thanh Le-Viet, Steven Rudder, Ana P. Tedim, Anastasia Kolyva, Maria Diaz, Leonardo de Oliveira Martins, Nabil-Fareed Alikhan, Lizzie Meadows, Rachael Stanley, Ngozi Elumogo, Muhammed Yasir, Nicholas M. Thomson, Alexander J Trotter, Rachel Gilroy, Samuel Bloomfield, Claire Stuart, Andrew Bell, Reenesh Prakash, Samir Dervisevic, Alison E. Mather, John Wain, Mark Webber, Andrew J. Page, Justin O'Grady |
| EPI_ISL_457573, EPI_ISL_457574, EPI_ISL_457575, EPI_ISL_457576, EPI_ISL_457577, EPI_ISL_457578, EPI_ISL_457579, EPI_ISL_457580 | Queens Medical Centre, Clinical Microbiology Department / DeepSeq Nottingham | COVID-19 Genomics UK (COG-UK) Consortium | Gemma Clark, Wendy Smith, Manjinder Khakh, Hannah Howson-Wells, Jonathan Ball, Patrick McClure, Joseph Chappell, Theocharis Tsoleridis, Nadine Holmes, Matthew Carlisle, Christopher Moore, Fei Sang, Johnny Debebe, Victoria Wright, Matthew Loose |
| EPI_ISL_457581, EPI_ISL_457582, EPI_ISL_457583, EPI_ISL_457584, EPI_ISL_457585, EPI_ISL_457586, EPI_ISL_457587, EPI_ISL_457588, EPI_ISL_457589, EPI_ISL_457590, EPI_ISL_457591, EPI_ISL_457592, EPI_ISL_457593, EPI_ISL_457594, EPI_ISL_457595, EPI_ISL_457596, EPI_ISL_457597, EPI_ISL_457598, EPI_ISL_457599, EPI_ISL_457600, EPI_ISL_457601, EPI_ISL_457602, EPI_ISL_457603, EPI_ISL_457604, EPI_ISL_457605, EPI_ISL_457606, EPI_ISL_457607, EPI_ISL_457608, EPI_ISL_457609, EPI_ISL_457610, EPI_ISL_457611, EPI_ISL_457612, EPI_ISL_457613, EPI_ISL_457614, EPI_ISL_457615, EPI_ISL_457616, EPI_ISL_457617, EPI_ISL_457618, EPI_ISL_457619, EPI_ISL_457620, EPI_ISL_457621, EPI_ISL_457622, EPI_ISL_457623, EPI_ISL_457624, EPI_ISL_457625, EPI_ISL_457626, EPI_ISL_457627, EPI_ISL_457628, EPI_ISL_457629, EPI_ISL_457630, EPI_ISL_457631, EPI_ISL_457632, EPI_ISL_457633, EPI_ISL_457634, EPI_ISL_457635, EPI_ISL_457636, EPI_ISL_457637, EPI_ISL_457638, EPI_ISL_457639, EPI_ISL_457640, EPI_ISL_457641, EPI_ISL_457642, EPI_ISL_457643, EPI_ISL_457644, EPI_ISL_457645, EPI_ISL_457646, EPI_ISL_457647, EPI_ISL_457648, EPI_ISL_457649, EPI_ISL_457650, EPI_ISL_457651, EPI_ISL_457652, EPI_ISL_457653, EPI_ISL_457654, EPI_ISL_457655, EPI_ISL_457656, EPI_ISL_457657, EPI_ISL_457658, EPI_ISL_457659, EPI_ISL_457660, EPI_ISL_457661, EPI_ISL_457662, EPI_ISL_457663, EPI_ISL_457664, EPI_ISL_457665, EPI_ISL_457666, EPI_ISL_457667, EPI_ISL_457668, EPI_ISL_457669, EPI_ISL_457670, EPI_ISL_457671, EPI_ISL_457672, EPI_ISL_457673, EPI_ISL_457674, EPI_ISL_457675, EPI_ISL_457676, EPI_ISL_457677, EPI_ISL_457678, EPI_ISL_457679, EPI_ISL_457680, EPI_ISL_457681, EPI_ISL_457682, EPI_ISL_457683, EPI_ISL_457684, EPI_ISL_457685, EPI_ISL_457686 | | | |
| see above | Virology Department, Sheffield Teaching Hospitals NHS Foundation Trust/Department of Infection, Immunity and Cardiovascular Disease, The Medical School, University of Sheffield | COVID-19 Genomics UK (COG-UK) Consortium | Thushan de Silva, Matthew Parker, Nikki Smith, Adri Angyal, Rebecca Brown, Luke Green, Rachel Tucker, Paul Parsons, Danielle Groves, Katie Johnson, Laura Carrilero, Alex Keeley, Dave Partridge, Matthew Wyles, Benjamin Lindsey, Mehmet Yavuz, Mohammad Raza, Cariad Evans |
| EPI_ISL_457687, EPI_ISL_457688, EPI_ISL_457689, EPI_ISL_457690, EPI_ISL_457691, EPI_ISL_457692, EPI_ISL_457693, EPI_ISL_457694, EPI_ISL_457695, EPI_ISL_457696, EPI_ISL_457697, EPI_ISL_457698 | | | |
| see above | The First Affiliated Hospital of Guangzhou Medical University, Guangzhou, China | BGI-shenzhen & The First Affiliated Hospital of Guangzhou Medical University | Yanqun Wang, Daxi Wang, Lu Zhang, Wanying Sun, Zhaoyong Zhang et al. |
| EPI_ISL_457699, EPI_ISL_457700 | Department of Infectious Diseases, Istituto Superiore di Sanità, Roma , Italy | Army Medical and Veterinary Research Center | Paola Stefanelli, Alessandra Lo Presti, Stefano Fiore, Antonella Marchi, Eleonora Benedetti, Concetta Fabiani Silvia Fillo, Giovanni Faggioni, Riccardo De Sanctis, Antonella Fortunato, Anna Anselmo, Francesco Giordani, Vanessa Vera Fain, Nino D'Amore, Florigio Lista |
| EPI_ISL_457701 | Oman-NIC | Oman-NIC | Samira Al-Maruqi, Fahad Zadjali, Amina Al Jardani, Khulood Al-Mammary, Hanan Al-kindi, Fatma BaAlawi, Hamida AL Barwani, Zeyana AL-Dahmani, Intisar Al-Shukri, Aisha Al-Busaidi, Aisha Al-Amri, Ahlam Al-Amri, Mohammed Al-Tobi, Samiha Al Kharusi, Abdulla Balkhair |
| EPI_ISL_457702 | Oman-NIC | Microbiology laboratory- Sultan Qaboos University Hospital | Fahad Zadjali, Samira Al-Maruqi, Amina Al Jardani, Khulood Al-Mammary, Hanan Al-kindi, Fatma BaAlawi, Hamida AL Barwani, Zeyana AL-Dahmani, Intisar Al-Shukri, Aisha Al-Busaidi, Aisha Al-Amri, Ahlam Al-Amri, Mohammed Al-Tobi, Samiha Al Kharusi, Abdulla Balkhair |
| EPI_ISL_457703 | Oman-NIC | Department of Microbiology and Immunology- SQUH | Fahad Zadjali, Samira Al-Maruqi, Amina Al Jardani, Khulood Al-Mammary, Hanan Al-kindi, Fatma BaAlawi, Hamida AL Barwani, Zeyana AL-Dahmani, Intisar Al-Shukri, Aisha Al-Busaidi, Aisha Al-Amri, Ahlam Al-Amri, Mohammed Al-Tobi, Samiha Al Kharusi, Abdulla Balkhair |
| EPI_ISL_457704 | Oman-NIC | Oman-NIC | Samira Al-Maruqi, Fahad Zadjali, Amina Al Jardani, Khulood Al-Mammary, Hanan Al-kindi, Fatma BaAlawi, Hamida AL Barwani, Zeyana AL-Dahmani, Intisar Al-Shukri, Aisha Al-Busaidi, Aisha Al-Amri, Ahlam Al-Amri, Mohammed Al-Tobi, Samiha Al Kharusi, Abdulla Balkhair |
| EPI_ISL_457705 | OMAN-NIC | Department of Microbiology and Immunology- SQUH | Fahad Zadjali, Samira Al-Maruqi, Amina Al Jardani, Khulood Al-Mammary, Hanan Al-kindi, Fatma BaAlawi, Hamida AL Barwani, Zeyana AL-Dahmani, Intisar Al-Shukri, Aisha Al-Busaidi, Aisha Al-Amri, Ahlam Al-Amri, Mohammed Al-Tobi, Samiha Al Kharusi, Abdulla Balkhair |
| EPI_ISL_457706 | Oman-NIC | Oman-NIC | Samira Al-Maruqi, Fahad Zadjali, Amina Al Jardani, Khulood Al-Mammary, Hanan Al-kindi, Fatma BaAlawi, Hamida AL Barwani, Zeyana AL-Dahmani, Intisar Al-Shukri, Aisha Al-Busaidi, Aisha Al-Amri, Ahlam Al-Amri, Mohammed Al-Tobi, Samiha Al Kharusi, Abdulla Balkhair |
| EPI_ISL_457707 | Oman-NIC | Department of Microbiology and Immunology- SQUH | Fahad Zadjali, Samira Al-Maruqi, Amina Al Jardani, Khulood Al-Mammary, Hanan Al-kindi, Fatma BaAlawi, Hamida AL Barwani, Zeyana AL-Dahmani, Intisar Al-Shukri, Aisha Al-Busaidi, Aisha Al-Amri, Ahlam Al-Amri, Mohammed Al-Tobi, Samiha Al Kharusi, Abdulla Balkhair |
| EPI_ISL_457719, EPI_ISL_457720 | SYNLAB Eesti OÜ | Charite Universitatsmedizin Berlin, Institute of Virology | Victor M Corman, Jorn Beheim-Schwarzbach, Barbara Mühlemann, Talitha Veith, Julia Schneider, Paul Naaber, Terry Jones, Christian Drosten |
| EPI_ISL_457721 | Department of Infectious Diseases, Istituto Superiore di Sanità, Roma , Italy | Army Medical and Veterinary Research Center | Paola Stefanelli, Alessandra Lo Presti, Stefano Fiore, Antonella Marchi, Eleonora Benedetti, Concetta Fabiani Silvia Fillo, Giovanni Faggioni, Riccardo De Sanctis, Antonella Fortunato, Anna Anselmo, Francesco Giordani, Vanessa Vera Fain, Nino D'Amore, Florigio Lista |
| EPI_ISL_457722, EPI_ISL_457723 | SYNLAB Eesti OÜ | Charite Universitatsmedizin Berlin, Institute of Virology | Victor M Corman, Jorn Beheim-Schwarzbach, Barbara Mühlemann, Talitha Veith, Julia Schneider, Paul Naaber, Terry Jones, Christian Drosten |
| EPI_ISL_457724 | Department of Infectious Diseases, Istituto Superiore di Sanità, Roma , Italy | Army Medical and Veterinary Research Center | Paola Stefanelli, Alessandra Lo Presti, Stefano Fiore, Antonella Marchi, Eleonora Benedetti, Concetta Fabiani Silvia Fillo, Giovanni Faggioni, Riccardo De Sanctis, Antonella Fortunato, Anna Anselmo, Francesco Giordani, Vanessa Vera Fain, Nino D'Amore, Florigio Lista |
| EPI_ISL_457725 | SYNLAB Eesti OÜ | Charite Universitatsmedizin Berlin, Institute of Virology | Victor M Corman, Jorn Beheim-Schwarzbach, Barbara Mühlemann, Talitha Veith, Julia Schneider, Paul Naaber, Terry Jones, Christian Drosten |
| EPI_ISL_457726 | TSGH-CP molecular lab | TSGH-CP molecular lab | Cherng-Lih Perng, Ming-Jr JIAN, Chih-Kai Chang, Jung-Chung Lin, Kuo-Ming Yeh, Chien-Wen Chen, Sheng-Kang Chiu, Hsing-Yi Chung, Shih-Hung Tsai, Kuo-Sheng Hung, Tien-Yao Chang, Feng-Yee Chang, Hung-Sheng Shang |
| EPI_ISL_457727 | SYNLAB Eesti OÜ | Charite Universitatsmedizin Berlin, Institute of Virology | Victor M Corman, Jorn Beheim-Schwarzbach, Barbara Mühlemann, Talitha Veith, Julia Schneider, Paul Naaber, Terry Jones, Christian Drosten |
| EPI_ISL_457728 | Department of Infectious Diseases, Istituto Superiore di Sanità, Roma , Italy | Army Medical and Veterinary Research Center | Paola Stefanelli, Alessandra Lo Presti, Stefano Fiore, Antonella Marchi, Eleonora Benedetti, Concetta Fabiani Silvia Fillo, Giovanni Faggioni, Riccardo De Sanctis, Antonella Fortunato, Anna Anselmo, Francesco Giordani, Vanessa Vera Fain, Nino D'Amore, Florigio Lista |
| EPI_ISL_457729 | SYNLAB Eesti OÜ | Charite Universitatsmedizin Berlin, Institute of Virology | Victor M Corman, Jorn Beheim-Schwarzbach, Barbara Mühlemann, Talitha Veith, Julia Schneider, Paul Naaber, Terry Jones, Christian Drosten |
| EPI_ISL_457730 | TSGH-CP molecular lab | TSGH-CP molecular lab | Cherng-Lih Perng, Ming-Jr JIAN, Chih-Kai Chang, Jung-Chung Lin, Kuo-Ming Yeh, Chien-Wen Chen, Sheng-Kang Chiu, Hsing-Yi Chung, Shih-Hung Tsai, Kuo-Sheng Hung, Tien-Yao Chang, Feng-Yee Chang, Hung-Sheng Shang |
| EPI_ISL_457731 | SYNLAB Eesti OÜ | Charite Universitatsmedizin Berlin, Institute of Virology | Victor M Corman, Jorn Beheim-Schwarzbach, Barbara Mühlemann, Talitha Veith, Julia Schneider, Paul Naaber, Terry Jones, Christian Drosten |
| EPI_ISL_457732 | Department of Infectious Diseases, Istituto Superiore di Sanità, Roma , Italy | Army Medical and Veterinary Research Center | Paola Stefanelli, Alessandra Lo Presti, Stefano Fiore, Antonella Marchi, Eleonora Benedetti, Concetta Fabiani Silvia Fillo, Giovanni Faggioni, Riccardo De Sanctis, Antonella Fortunato, Anna Anselmo, Francesco Giordani, Vanessa Vera Fain, Nino D'Amore, Florigio Lista |
| EPI_ISL_457733 | TSGH-CP molecular lab | TSGH-CP molecular lab | Cherng-Lih Perng, Ming-Jr JIAN, Chih-Kai Chang, Jung-Chung Lin, Kuo-Ming Yeh, Chien-Wen Chen, Sheng-Kang Chiu, Hsing-Yi Chung, Shih-Hung Tsai, Kuo-Sheng Hung, Tien-Yao Chang, Feng-Yee Chang, Hung-Sheng Shang |
| EPI_ISL_457734, EPI_ISL_457735 | SYNLAB Eesti OÜ | Charite Universitatsmedizin Berlin, Institute of Virology | Victor M Corman, Jorn Beheim-Schwarzbach, Barbara Mühlemann, Talitha Veith, Julia Schneider, Paul Naaber, Terry Jones, Christian Drosten |
| EPI_ISL_457736 | Department of Infectious Diseases, Istituto Superiore di Sanità, Roma , Italy | Army Medical and Veterinary Research Center | Paola Stefanelli, Alessandra Lo Presti, Stefano Fiore, Antonella Marchi, Eleonora Benedetti, Concetta Fabiani Silvia Fillo, Giovanni Faggioni, Riccardo De Sanctis, Antonella Fortunato, Anna Anselmo, Francesco Giordani, Vanessa Vera Fain, Nino D'Amore, Florigio Lista |
| EPI_ISL_457737, EPI_ISL_457738, EPI_ISL_457739, EPI_ISL_457740 | SYNLAB Eesti OÜ | Charite Universitatsmedizin Berlin, Institute of Virology | Victor M Corman, Jorn Beheim-Schwarzbach, Barbara Mühlemann, Talitha Veith, Julia Schneider, Paul Naaber, Terry Jones, Christian Drosten |
| EPI_ISL_457749 | Department of Infectious Diseases, Istituto Superiore di Sanità, Roma , Italy | Army Medical and Veterinary Research Center | Paola Stefanelli, Alessandra Lo Presti, Stefano Fiore, Antonella Marchi, Eleonora Benedetti, Concetta Fabiani Silvia Fillo, Giovanni Faggioni, Riccardo De Sanctis, Antonella Fortunato, Anna Anselmo, Francesco Giordani, Vanessa Vera Fain, Nino D'Amore, Florigio Lista |
| EPI_ISL_457750 | Centogene AG | Centogene AG | Prof. Dr. Peter Bauer, Dr. Krishna Kumar Kandaswamy |
| EPI_ISL_457751, EPI_ISL_457752, EPI_ISL_457753, EPI_ISL_457754, EPI_ISL_457755, EPI_ISL_457756, EPI_ISL_457757, EPI_ISL_457758, EPI_ISL_457759, EPI_ISL_457760, EPI_ISL_457761, EPI_ISL_457762, EPI_ISL_457763, EPI_ISL_457764, EPI_ISL_457765, EPI_ISL_457766, EPI_ISL_457767, EPI_ISL_457768, EPI_ISL_457769, EPI_ISL_457770, EPI_ISL_457771, EPI_ISL_457772, EPI_ISL_457773, EPI_ISL_457774, EPI_ISL_457775, EPI_ISL_457776, EPI_ISL_457777, EPI_ISL_457778, EPI_ISL_457779, EPI_ISL_457780, EPI_ISL_457781, EPI_ISL_457782, EPI_ISL_457783, EPI_ISL_457784, EPI_ISL_457785, EPI_ISL_457786, EPI_ISL_457787, EPI_ISL_457788, EPI_ISL_457789, EPI_ISL_457790, EPI_ISL_457791, EPI_ISL_457792, EPI_ISL_457793, EPI_ISL_457794, EPI_ISL_457795, EPI_ISL_457796, EPI_ISL_457797, EPI_ISL_457798, EPI_ISL_457799, EPI_ISL_457800, EPI_ISL_457801, EPI_ISL_457802, EPI_ISL_457803, EPI_ISL_457804, EPI_ISL_457805, EPI_ISL_457806, EPI_ISL_457807, EPI_ISL_457808, EPI_ISL_457809, EPI_ISL_457810, EPI_ISL_457811, EPI_ISL_457812, EPI_ISL_457813, EPI_ISL_457814, EPI_ISL_457815, EPI_ISL_457816, EPI_ISL_457817, EPI_ISL_457818, EPI_ISL_457819, EPI_ISL_457820, EPI_ISL_457821, EPI_ISL_457822, EPI_ISL_457823 | | | |
| see above | Johns Hopkins Hospital Department of Pathology | Johns Hopkins Hospital Department of Pathology | Peter M. Thielen, Thomas Mehoke, Shirlee Wohl, Srividya Ramakrishnan, Melanie Kirsche, Amanda Ernlund, Craig Howser, Kristina Zudock, Oluwaseun Falade-Nwulia, Norah Sadowski, Paul Morris, Mark Hopkins, Yunfan Fan, Nidia Trovao, Victoria Gniazdowski, Michael C. Schatz, Stuart C. Ray, Winston Timp, Heba H. Mostafa |

| | | | |
|---|---|---|---|
| EPI_ISL_457824 | Bezmialem Vakif University, Dept Microbiology, Medical School, Fatih, Istanbul, Turkey | Bezmialem Vakif University, Medical School & Beykoz Institute of Life Sciences & Biotechnology | Mehmet Z. Doymaz, Merve Kalkan, Nesibe Cetin, Elif Karaaslan, Bilge Sumbul, Filiz Guney |
| EPI_ISL_457825 | Army Medical Research Center - Scientific Department | Army Medical and Veterinary Research Center | Silvia Fillo, Giovanni Faggioni, Riccardo De Sanctis, Antonella Fortunato, Anna Anselmo, Francesco Giordani, Vanessa Vera Fain, Nino D'Amore, Florigio Lista |
| EPI_ISL_457826 | Army Medical Center - Scientific Department | Army Medical and Veterinary Research Center | Silvia Fillo, Giovanni Faggioni, Riccardo De Sanctis, Antonella Fortunato, Anna Anselmo, Francesco Giordani, Vanessa Vera Fain, Nino D'Amore, Florigio Lista |
| EPI_ISL_457827, EPI_ISL_457828, EPI_ISL_457829, EPI_ISL_457830, EPI_ISL_457831, EPI_ISL_457832, EPI_ISL_457833, EPI_ISL_457834, EPI_ISL_457835, EPI_ISL_457836, EPI_ISL_457837, EPI_ISL_457838, EPI_ISL_457839, EPI_ISL_457840, EPI_ISL_457841, EPI_ISL_457842, EPI_ISL_457843, EPI_ISL_457844 | | | |
| see above | National Public Health Laboratory | KEMRI-Wellcome Trust Research Programme/KEMRI-CGMR-C Kilifi | Githinji G. et al 2020 |
| EPI_ISL_457845, EPI_ISL_457846, EPI_ISL_457847, EPI_ISL_457848, EPI_ISL_457849, EPI_ISL_457850, EPI_ISL_457851, EPI_ISL_457852, EPI_ISL_457853, EPI_ISL_457854, EPI_ISL_457855, EPI_ISL_457856, EPI_ISL_457857, EPI_ISL_457858, EPI_ISL_457859, EPI_ISL_457860, EPI_ISL_457861, EPI_ISL_457862, EPI_ISL_457863, EPI_ISL_457864, EPI_ISL_457865, EPI_ISL_457866, EPI_ISL_457867, EPI_ISL_457868, EPI_ISL_457869, EPI_ISL_457870, EPI_ISL_457871, EPI_ISL_457872, EPI_ISL_457873, EPI_ISL_457874, EPI_ISL_457875, EPI_ISL_457876, EPI_ISL_457877, EPI_ISL_457878, EPI_ISL_457879, EPI_ISL_457880, EPI_ISL_457881, EPI_ISL_457882, EPI_ISL_457883, EPI_ISL_457884, EPI_ISL_457885, EPI_ISL_457886, EPI_ISL_457887, EPI_ISL_457888, EPI_ISL_457889, EPI_ISL_457890, EPI_ISL_457891, EPI_ISL_457892, EPI_ISL_457893, EPI_ISL_457894, EPI_ISL_457895, EPI_ISL_457896, EPI_ISL_457897, EPI_ISL_457898, EPI_ISL_457899, EPI_ISL_457900, EPI_ISL_457901, EPI_ISL_457902, EPI_ISL_457903, EPI_ISL_457904, EPI_ISL_457905, EPI_ISL_457906, EPI_ISL_457907, EPI_ISL_457908, EPI_ISL_457909, EPI_ISL_457910, EPI_ISL_457911, EPI_ISL_457912, EPI_ISL_457913, EPI_ISL_457914, EPI_ISL_457915, EPI_ISL_457916, EPI_ISL_457917, EPI_ISL_457918, EPI_ISL_457919, EPI_ISL_457920, EPI_ISL_457921, EPI_ISL_457922, EPI_ISL_457923, EPI_ISL_457924, EPI_ISL_457925, EPI_ISL_457926, EPI_ISL_457927, EPI_ISL_457928, EPI_ISL_457929, EPI_ISL_457930, EPI_ISL_457931 | | | |
| see above | KEMRI-CGMR-C | KEMRI-Wellcome Trust Research Programme/KEMRI-CGMR-C Kilifi | Githinji G. et al 2020 |
| EPI_ISL_457932, EPI_ISL_457933, EPI_ISL_457934, EPI_ISL_457935, EPI_ISL_457936 | KEMRI-Centre for Virus Research | KEMRI-Wellcome Trust Research Programme/KEMRI-CGMR-C Kilifi | Githinji G. et al 2020 |
| EPI_ISL_457937, EPI_ISL_457938, EPI_ISL_457939 | Oman-NIC | Oman-NIC | Samira Al-Maruqi, Fahad Zadjali, Amina Al Jardani, Khulood Al-Mammary, Hanan Al-kindi, Fatma BaAlawi, Hamida AL Barwani, Zeyana AL-Dahmani, Intisar Al-Shukri, Aisha Al-Busaidi, Aisha Al-Amri, Ahlam Al-Amri, Mohammed Al-Tobi, Samiha Al Kharusi, Abdulla Balkhair |
| EPI_ISL_457940, EPI_ISL_457941, EPI_ISL_457942, EPI_ISL_457943, EPI_ISL_457944, EPI_ISL_457945, EPI_ISL_457946, EPI_ISL_457947, EPI_ISL_457948, EPI_ISL_457949, EPI_ISL_457950, EPI_ISL_457951, EPI_ISL_457952, EPI_ISL_457953, EPI_ISL_457954, EPI_ISL_457955, EPI_ISL_457956, EPI_ISL_457957, EPI_ISL_457958, EPI_ISL_457959, EPI_ISL_457960, EPI_ISL_457961, EPI_ISL_457962, EPI_ISL_457963, EPI_ISL_457964, EPI_ISL_457965, EPI_ISL_457966, EPI_ISL_457967, EPI_ISL_457968, EPI_ISL_457969, EPI_ISL_457970, EPI_ISL_457971, EPI_ISL_457972, EPI_ISL_457973 | | | |
| see above | Laboratorio de Biología Molecular Asociación Española Primera en Salud | Departments of Pathology and Medicine, New York University School of Medicine | Maria Victoria Elizondo, Maria Noel Zubillaga, Gonzalo Manrique, Paul Zappile, Gael Westby, Matthew T Maurano, Christian Marier, Adriana Heguy |
| EPI_ISL_457974, EPI_ISL_457975, EPI_ISL_457976, EPI_ISL_457977, EPI_ISL_457978, EPI_ISL_457979, EPI_ISL_457980 | Oman-NIC | Oman-NIC | Samira Al-Maruqi, Fahad Zadjali, Amina Al Jardani, Khulood Al-Mammary, Hanan Al-kindi, Fatma BaAlawi, Hamida AL Barwani, Zeyana AL-Dahmani, Intisar Al-Shukri, Aisha Al-Busaidi, Aisha Al-Amri, Ahlam Al-Amri, Mohammed Al-Tobi, Samiha Al Kharusi, Abdulla Balkhair |
| EPI_ISL_457981, EPI_ISL_457982, EPI_ISL_457983, EPI_ISL_457984 | Oman-NIC | Department of Microbiology and Immunology-SQUH | Fahad Zadjali, Samira Al-Maruqi, Amina Al Jardani, Khulood Al-Mammary, Hanan Al-kindi, Fatma BaAlawi, Hamida AL Barwani, Zeyana AL-Dahmani, Intisar Al-Shukri, Aisha Al-Busaidi, Aisha Al-Amri, Ahlam Al-Amri, Mohammed Al-Tobi, Samiha Al Kharusi, Abdulla Balkhair |
| EPI_ISL_457985, EPI_ISL_457986, EPI_ISL_457987, EPI_ISL_457988, EPI_ISL_457989, EPI_ISL_457990, EPI_ISL_457991, EPI_ISL_457992, EPI_ISL_457993, EPI_ISL_457994, EPI_ISL_457995, EPI_ISL_457996, EPI_ISL_457997, EPI_ISL_457998 | | | |
| see above | Oman-NIC | Oman-NIC | Samira Al-Maruqi, Fahad Zadjali, Amina Al Jardani, Khulood Al-Mammary, Hanan Al-kindi, Fatma BaAlawi, Hamida AL Barwani, Zeyana AL-Dahmani, Intisar Al-Shukri, Aisha Al-Busaidi, Aisha Al-Amri, Ahlam Al-Amri, Mohammed Al-Tobi, Samiha Al Kharusi, Abdulla Balkhair |
| EPI_ISL_457999 | unknown | Centre For Biotechnology Research and Development | Matoke-Muhia,D., Symeker,S.L., Muuo,S.N., Ochwoto,M., Zablon,J.O., Kimotho,J., Waruhiu,C.N. and Michuki,G.N. |
| EPI_ISL_458000 | unknown | Centre For Biotechnology Research and Development | Matoke-Muhia,D.K., Symeker,S.L., Muuo,S.N., Ochwoto,M., Zablon,J.O., Kimotho,J., Waruhiu,C.N. and Michuki,G.N. |
| EPI_ISL_458001, EPI_ISL_458002, EPI_ISL_458003, EPI_ISL_458004, EPI_ISL_458005, EPI_ISL_458006 | Dirk Dittmer | Dirk Dittmer | Bailey,A.G., Caro-Vegas,C.P., Dittmer,D., Eason,A.B., Juarez,A., Landis,J.T., McNamara,R.P., Miller,M.B., Moorad,R., Pluta,L.J., Seltzer,T.A., Thompson,C., Vahrson,W., Villamor,F. |
| EPI_ISL_458007 | Dirk Dittmer | Dirk Dittmer | Aubrey,B.G., Caro-Vegas,C.P., Dittmer,D., Eason,A.B., Juarez,A., Landis,J.T., McNamara,R.P., Miller,M.B., Moorad,R., Pluta,L.J., Seltzer,T.A., Thompson,C., Vahrson,W., Villamor,F. |
| EPI_ISL_458008, EPI_ISL_458009, EPI_ISL_458010, EPI_ISL_458011, EPI_ISL_458012, EPI_ISL_458013, EPI_ISL_458014, EPI_ISL_458015 | Department of Food Safety, Nutrition and Veterinary public health, Istituto Superiore di Sanita' | Department of Food Safety, Nutrition and Veterinary public health, Istituto Superiore di Sanita' | La Rosa,G., Iaconelli,M., Mancini,P., Bonanno Ferraro,G., Veneri,C., Bonadonna,L., Lucentini,L., Suffredini,E. |
| EPI_ISL_460097, EPI_ISL_460098, EPI_ISL_460099, EPI_ISL_460100, EPI_ISL_460101, EPI_ISL_460102, EPI_ISL_460103, EPI_ISL_460104, EPI_ISL_460105, EPI_ISL_460106, EPI_ISL_460107, EPI_ISL_460108, EPI_ISL_460109, EPI_ISL_460110, EPI_ISL_460111, EPI_ISL_460112, EPI_ISL_460113, EPI_ISL_460114, EPI_ISL_460115, EPI_ISL_460116, EPI_ISL_460117, EPI_ISL_460118, EPI_ISL_460119, EPI_ISL_460120, EPI_ISL_460121, EPI_ISL_460122, EPI_ISL_460123, EPI_ISL_460124, EPI_ISL_460125, EPI_ISL_460126, EPI_ISL_460127, EPI_ISL_460128, EPI_ISL_460129, EPI_ISL_460130, EPI_ISL_460131, EPI_ISL_460132, EPI_ISL_460133, EPI_ISL_460134, EPI_ISL_460135, EPI_ISL_460136, EPI_ISL_460137, EPI_ISL_460138, EPI_ISL_460139, EPI_ISL_460140, EPI_ISL_460141, EPI_ISL_460142, EPI_ISL_460143, EPI_ISL_460144, EPI_ISL_460145, EPI_ISL_460146, EPI_ISL_460147, EPI_ISL_460148, EPI_ISL_460149, EPI_ISL_460150, EPI_ISL_460151, EPI_ISL_460152, EPI_ISL_460153, EPI_ISL_460154, EPI_ISL_460155, EPI_ISL_460156, EPI_ISL_460157, EPI_ISL_460158, EPI_ISL_460159, EPI_ISL_460160, EPI_ISL_460161, EPI_ISL_460162, EPI_ISL_460163, EPI_ISL_460164, EPI_ISL_460165, EPI_ISL_460166, EPI_ISL_460167, EPI_ISL_460168, EPI_ISL_460169, EPI_ISL_460170, EPI_ISL_460171, EPI_ISL_460172, EPI_ISL_460173, EPI_ISL_460174, EPI_ISL_460175, EPI_ISL_460176, EPI_ISL_460177, EPI_ISL_460178, EPI_ISL_460179, EPI_ISL_460180, EPI_ISL_460181, EPI_ISL_460182, EPI_ISL_460183, EPI_ISL_460184, EPI_ISL_460185, EPI_ISL_460186, EPI_ISL_460187, EPI_ISL_460188, EPI_ISL_460189, EPI_ISL_460190, EPI_ISL_460191, EPI_ISL_460192, EPI_ISL_460193, EPI_ISL_460194, EPI_ISL_460195, EPI_ISL_460196, EPI_ISL_460197, EPI_ISL_460198, EPI_ISL_460199, EPI_ISL_460200, EPI_ISL_460201, EPI_ISL_460202, EPI_ISL_460203, EPI_ISL_460204, EPI_ISL_460205, EPI_ISL_460206, EPI_ISL_460207, EPI_ISL_460208, EPI_ISL_460209, EPI_ISL_460210, EPI_ISL_460211, EPI_ISL_460212, EPI_ISL_460213, EPI_ISL_460214, EPI_ISL_460215, EPI_ISL_460216, EPI_ISL_460217, EPI_ISL_460218, EPI_ISL_460219, EPI_ISL_460220, EPI_ISL_460221, EPI_ISL_460222, EPI_ISL_460223, EPI_ISL_460224, EPI_ISL_460225, EPI_ISL_460226, EPI_ISL_460227, EPI_ISL_460228, EPI_ISL_460229, EPI_ISL_460230, EPI_ISL_460231, EPI_ISL_460232, EPI_ISL_460233, EPI_ISL_460234, EPI_ISL_460235, EPI_ISL_460236, EPI_ISL_460237, EPI_ISL_460238, EPI_ISL_460239, EPI_ISL_460240, EPI_ISL_460241, EPI_ISL_460242, EPI_ISL_460243, EPI_ISL_460244, EPI_ISL_460245, EPI_ISL_460246, EPI_ISL_460247, EPI_ISL_460248, EPI_ISL_460249, EPI_ISL_460250, EPI_ISL_460251, EPI_ISL_460252, EPI_ISL_460253, EPI_ISL_460254, EPI_ISL_460255, EPI_ISL_460256, EPI_ISL_460257, EPI_ISL_460258, EPI_ISL_460259, EPI_ISL_460260, EPI_ISL_460261, EPI_ISL_460262, EPI_ISL_460263, EPI_ISL_460264, EPI_ISL_460265, EPI_ISL_460266, EPI_ISL_460267, EPI_ISL_460268, EPI_ISL_460269, EPI_ISL_460270, EPI_ISL_460271, EPI_ISL_460272, EPI_ISL_460273, EPI_ISL_460274, EPI_ISL_460275, EPI_ISL_460276, EPI_ISL_460277, EPI_ISL_460278, EPI_ISL_460279, EPI_ISL_460280, EPI_ISL_460281, EPI_ISL_460282, EPI_ISL_460283, EPI_ISL_460284, EPI_ISL_460285, EPI_ISL_460286, EPI_ISL_460287, EPI_ISL_460288, EPI_ISL_460289, EPI_ISL_460290, EPI_ISL_460291, EPI_ISL_460292, EPI_ISL_460293, EPI_ISL_460294, EPI_ISL_460295, EPI_ISL_460296, EPI_ISL_460297, EPI_ISL_460298, EPI_ISL_460299, EPI_ISL_460300, EPI_ISL_460301, EPI_ISL_460302, EPI_ISL_460303, EPI_ISL_460304, EPI_ISL_460305, EPI_ISL_460306, EPI_ISL_460307, EPI_ISL_460308, EPI_ISL_460309, EPI_ISL_460310, EPI_ISL_460311, EPI_ISL_460312, EPI_ISL_460313, EPI_ISL_460314, EPI_ISL_460315, EPI_ISL_460316, EPI_ISL_460317, EPI_ISL_460318, EPI_ISL_460319, EPI_ISL_460320, EPI_ISL_460321, EPI_ISL_460322, EPI_ISL_460323, EPI_ISL_460324, EPI_ISL_460325, EPI_ISL_460326, EPI_ISL_460327, EPI_ISL_460328, EPI_ISL_460329, EPI_ISL_460330, EPI_ISL_460331, EPI_ISL_460332, EPI_ISL_460333, EPI_ISL_460334, EPI_ISL_460335, EPI_ISL_460336, EPI_ISL_460337, EPI_ISL_460338, EPI_ISL_460339, EPI_ISL_460340, EPI_ISL_460341, EPI_ISL_460342, EPI_ISL_460343, EPI_ISL_460344, EPI_ISL_460345, EPI_ISL_460346, EPI_ISL_460347, EPI_ISL_460348, EPI_ISL_460349, EPI_ISL_460350, EPI_ISL_460351, EPI_ISL_460352, EPI_ISL_460353, EPI_ISL_460354, EPI_ISL_460355, EPI_ISL_460356, EPI_ISL_460357, EPI_ISL_460358, EPI_ISL_460359, EPI_ISL_460360, EPI_ISL_460361, EPI_ISL_460362, EPI_ISL_460363, EPI_ISL_460364, EPI_ISL_460365, EPI_ISL_460366, EPI_ISL_460367, EPI_ISL_460368, EPI_ISL_460369, EPI_ISL_460370, EPI_ISL_460371, EPI_ISL_460372, EPI_ISL_460373, EPI_ISL_460374, EPI_ISL_460375, EPI_ISL_460376, EPI_ISL_460377, EPI_ISL_460378, EPI_ISL_460379, EPI_ISL_460380, EPI_ISL_460381, EPI_ISL_460382, EPI_ISL_460383, EPI_ISL_460384, EPI_ISL_460385, EPI_ISL_460386, EPI_ISL_460387, EPI_ISL_460388, EPI_ISL_460389, EPI_ISL_460390, EPI_ISL_460391, EPI_ISL_460392, EPI_ISL_460393, EPI_ISL_460394, EPI_ISL_460395, EPI_ISL_460396, EPI_ISL_460397, EPI_ISL_460398, EPI_ISL_460399, EPI_ISL_460400, EPI_ISL_460401, EPI_ISL_460402, EPI_ISL_460403, EPI_ISL_460404, EPI_ISL_460405, EPI_ISL_460406, EPI_ISL_460407, EPI_ISL_460408, EPI_ISL_460409, EPI_ISL_460410, EPI_ISL_460411, EPI_ISL_460412, EPI_ISL_460413, EPI_ISL_460414, EPI_ISL_460415, EPI_ISL_460416, EPI_ISL_460417, EPI_ISL_460418, EPI_ISL_460419, EPI_ISL_460420, EPI_ISL_460421, EPI_ISL_460422, EPI_ISL_460423, EPI_ISL_460424, EPI_ISL_460425, EPI_ISL_460426, EPI_ISL_460427, EPI_ISL_460428, EPI_ISL_460429, EPI_ISL_460430, EPI_ISL_460431, EPI_ISL_460432, EPI_ISL_460433, EPI_ISL_460434, EPI_ISL_460435, EPI_ISL_460436, EPI_ISL_460437, EPI_ISL_460438, EPI_ISL_460439, EPI_ISL_460440, EPI_ISL_460441, EPI_ISL_460442, EPI_ISL_460443, EPI_ISL_460444, EPI_ISL_460445, EPI_ISL_460446, EPI_ISL_460447, EPI_ISL_460448, EPI_ISL_460449, EPI_ISL_460450, EPI_ISL_460451, EPI_ISL_460452, EPI_ISL_460453, EPI_ISL_460454, EPI_ISL_460455, EPI_ISL_460456, EPI_ISL_460457, EPI_ISL_460458, EPI_ISL_460459, EPI_ISL_460460, EPI_ISL_460461, EPI_ISL_460462, EPI_ISL_460463, EPI_ISL_460464, EPI_ISL_460465, EPI_ISL_460466, EPI_ISL_460467, EPI_ISL_460468, EPI_ISL_460469, EPI_ISL_460470, EPI_ISL_460471, EPI_ISL_460472 | | | |
| see above | Massachusetts General Hospital | Infectious Disease Program, Broad Institute of Harvard and MIT | Lemieux,J.E., Siddle,K.J., Shaw,B., Adams,G., Pierce,V., Turbett,S., Anahtar,M., Branda,J., Slater,D., Harris,J., Lin,A.E., Gladden-Young,A., Lagerborg,K., Rudy,M., DeRuff,K., Carter,A., Normandin,E., Bauer,M., Reilly,S., Tomkins-Tinch,C., Loreth,C., Chaluvadi,S., Neumann,A., Cusick,C., Chapman,S.B., Gnirke,A., Flowers,K., Cerrato,F., Birren,B.W., Gallagher,G., Smole,S., Park,D.J., MacInnis,B.L., Ryan,E., LaRocque,R., Rosenberg,E., Sabeti,P.C. |
| EPI_ISL_462446 | unknown | Ryota Kumagai Tokyo Metropolitan Institute of Public Health | Kumagai,R., Yoshida,I., Asakura,H., Nagashima,M., Chiba,T., Sadamasu,K. |
| EPI_ISL_463001, EPI_ISL_463002, EPI_ISL_463003, EPI_ISL_463004, EPI_ISL_463005, EPI_ISL_463006 | unknown | Clinical virology | Fares,W., Triki,H. |
| EPI_ISL_468044, EPI_ISL_468045, EPI_ISL_468046, EPI_ISL_468047, EPI_ISL_468048, EPI_ISL_468049, EPI_ISL_468050, EPI_ISL_468051, EPI_ISL_468052, EPI_ISL_468053, EPI_ISL_468054, EPI_ISL_468055 | | | |
| see above | Egyptian National Cancer Institute (ENCI) | Egyptian National Cancer Institute (ENCI) | Zekri, Abdel Rahman N, Amer,K.E., Ahmed,O.S., Soliman,H.K., Hafez,M.M., Bahnassy,A.A., Abdelhamid,W., Gad,A., Ali,M., Hassan,W., Samir,M., Raouf,A., Hamdy,M.S., Soliman,M.S., Elsissy,M.H., Elkhateeb,S.M., Ezzelarab,M.H., Abouelhoda, Mohamed |
| EPI_ISL_468056 | Egyptian National Cancer Institute (ENCI) | Egyptian National Cancer Institute (ENCI) | Zekri, Abdel Rahman N., Amer,K.E., Ahmed,O.S., Soliman,H.K., Ali,M.A., Hassan,W.A., Mahmoud,A.A., Khattab,A.A., Hafez,M.M., Abouelhoda, Mohamed |
| EPI_ISL_468057, EPI_ISL_468058, EPI_ISL_468059 | Egyptian National Cancer Institute (ENCI) | Egyptian National Cancer Institute (ENCI) | Zekri, Abdel Rahman N, Amer,K.E., Ahmed,O.S., Soliman,H.K., Hafez,M.M., Bahnassy,A.A., Abdelhamid,W., Gad,A., Ali,M., Hassan,W., Samir,M., Raouf,A., Hamdy,M.S., Soliman,M.S., Elsissy,M.H., Elkhateeb,S.M., Ezzelarab,M.H., Abouelhoda, Mohamed |
| EPI_ISL_468060, EPI_ISL_468061, EPI_ISL_468062 | Egyptian National Cancer Institute (ENCI) | Egyptian National Cancer Institute (ENCI) | Zekri, Abdel Rahman N., Amer,K.E., Ahmed,O.S., Soliman,H.K., Ali,M.A., Hassan,W.A., Mahmoud,A.A., Khattab,A.A., Hafez,M.M., Abouelhoda, Mohamed |
| EPI_ISL_475722 | Egyptian National Cancer Institute (ENCI) | Egyptian National Cancer Institute (ENCI) | Zekri, Abdel Rahman N, Amer,K.E., Ahmed,O.S., Soliman,H.K., Hafez,M.M., Bahnassy,A.A., Abdelhamid,W., Gad,A., Ali,M., Hassan,W., Samir,M., Raouf,A., Hamdy,M.S., Soliman,M.S., Elsissy,M.H., Elkhateeb,S.M., Ezzelarab,M.H., Abouelhoda, Mohamed |
| EPI_ISL_529146 | Goethe University Hospital Frankfurt, Institute for Medical Virology | Goethe University Hospital Frankfurt, Institute for Medical Virology | Grabmair,T.T., Hoehl,S., Westhaus,S., Bojkova,D., Berger,A., Rotter,B., Hoffmeier,K., Cinatl,J., Ciesek,S., Widera,M. |
| EPI_ISL_529206, EPI_ISL_529207 | Utah Public Health Laboratory | Utah Public Health Laboratory | Erin Young, Kelly Oakeson |

We gratefully acknowledge the following Authors from the Originating laboratories responsible for obtaining the specimens, as well as the Submitting laboratories where the genome data were generated and shared via GISAID, on which this research is based.

All Submitters of data may be contacted directly via www.gisaid.org

| Accession ID | Originating Laboratory | Submitting Laboratory | Authors |
|---|---|---|---|
| EPI_ISL_458017, EPI_ISL_458018, EPI_ISL_458019, EPI_ISL_458020, EPI_ISL_458021, EPI_ISL_458022 | NYU Langone Health | Departments of Pathology and Medicine, New York University School of Medicine | Maria Aguero-Rosenfeld, Brendan Belovarac, Margaret Black, Ludovic Boytard, John Cadley, Paolo Cotzia, John Chen, Dacia Dimartino, Xiaojun Feng, Tatyana Gindin, Emily Guzman, Adriana Heguy, Megan Hogan, Emily Huang, George Jour, Alireza Khodadadi-Jamayran, Lawrence H. Lin, Raven Luther, Andrew Lytle, Christian Marier, Matthew T. Maurano, Mark J. Mulligan, Peter Meyn, Raquel Ordonez Ciriza, Iman Osman, Jared Pinnell, Vanessa Raabe, Sitharam Ramaswami, Amy Rapkiewicz, Andre M. Ribeiro-dos-Santos, Marie Samanovic-Golden, Antonio Serrano, Guomiao Shen, Matija Snuderl, Theodore Vougiouklakis, Nick Vulpescu, Gael Westby, Paul Zappile, Yutong Zhang |
| EPI_ISL_458023, EPI_ISL_458024, EPI_ISL_458025, EPI_ISL_458026, EPI_ISL_458027, EPI_ISL_458028 | Hospital for Tropical Diseases | COVID-19 Network Investigations (CONI) Alliance | Elizabeth Batty, Nantarat Chantawat, Wasun Chantratita, Thanat Chookajorn, Stefan Fernandez, Angkana Huang, Weena Janwitthayanan, Akanitt Jittmittraphap, Anthony R. Jones, Khajohn Joonsalak, Chonticha Klungtong, Theerarat Kochakarn, Namfon Kotanan, Krittikorn Kumpornsin, Pornsawan Leaungwutiwong, Wudtichai Manasatienkij, Bhakbhoom Panthan, Ekawat Pasomsub, Kingkan Rakmanee, Insee Sensorn, Janjira Thaipadungpanit, Arporn Wangwiwatsin,Treewat Watthanachockchai |
| EPI_ISL_458029 | TSGH-CP molecular lab | TSGH-CP molecular lab | Cherng-Lih Perng, Ming-Jr JIAN, Chih-Kai Chang, Jung-Chung Lin, Kuo-Ming Yeh, Chien-Wen Chen, Sheng-Kang Chiu, Hsing-Yi Chung, Shih-Hung Tsai, Kuo-Sheng Hung, Tien-Yao Chang, Feng-Yee Chang, Hung-Sheng Shang |
| EPI_ISL_458030 | King Institute of Preventive Medicine & Research | CSIR-Centre for Cellular and Molecular Biology | K.Kaveri,S.Sivasubramanian,S.Vennila,P.Padmapriya,R.Kiruba,S.Magesh,G. Dhinakar Raj, G. Ravikumar, P. Azhahianambi,K Thangaraj,Payel Mukherjee, Sofia Banu, Priya Singh, Dhiviya Vedagiri, Divya Gupta, Vishal Sah, Santosh Kumar Kuncha, Krishnan Harinivas Harshan, Archana Bharadwaj Siva, Karthik Bharadwaj Tallapaka, Shagufta Khan, Lamuk Zaveri, Namami Gaur, Sakshi Shambhavi, Tulasi Nagabandi, Purushotham Vodnala, Rakesh K Mishra, Divya Tej Sowpati |
| EPI_ISL_458031 | King Institute of Preventive Medicine & Research | CSIR-Centre for Cellular and Molecular Biology | K.Kaveri,S.Sivasubramanian,S.Vennila,P.Padmapriya,R.Kiruba,S.Magesh,G. Dhinakar Raj, G. Ravikumar, P. Azhahianambi, K Thangaraj,Sofia Banu, Payel Mukherjee, Priya Singh, Dhiviya Vedagiri, Divya Gupta, Vishal Sah, Santosh Kumar Kuncha, Krishnan Harinivas Harshan, Archana Bharadwaj Siva, Karthik Bharadwaj Tallapaka, Shagufta Khan, Lamuk Zaveri, Namami Gaur, Sakshi Shambhavi, Tulasi Nagabandi, Purushotham Vodnala, Rakesh K Mishra, Divya Tej Sowpati |
| EPI_ISL_458032 | King Institute of Preventive Medicine & Research | CSIR-Centre for Cellular and Molecular Biology | K.Kaveri,S.Sivasubramanian,S.Vennila,P.Padmapriya,R.Kiruba,S.Magesh,G. Dhinakar Raj, G. Ravikumar, P. Azhahianambi, K Thangaraj,Shagufta Khan, Lamuk Zaveri, Namami Gaur, Sakshi Shambhavi, Tulasi Nagabandi, Purushotham Vodnala, Payel Mukherjee, Sofia Banu, Priya Singh, Dhiviya Vedagiri, Divya Gupta, Vishal Sah, Santosh Kumar Kuncha, Krishnan Harinivas Harshan, Archana Bharadwaj Siva, Karthik Bharadwaj Tallapaka, Rakesh K Mishra, Divya Tej Sowpati |
| EPI_ISL_458033 | King Institute of Preventive Medicine & Research | CSIR-Centre for Cellular and Molecular Biology | K.Kaveri,S.Sivasubramanian,S.Vennila,P.Padmapriya,R.Kiruba,S.Magesh,G. Dhinakar Raj, G. Ravikumar, P. Azhahianambi, K Thangaraj,Lamuk Zaveri, Shagufta Khan, Namami Gaur, Sakshi Shambhavi, Tulasi Nagabandi, Purushotham Vodnala, Payel Mukherjee, Sofia Banu, Priya Singh, Dhiviya Vedagiri, Divya Gupta, Vishal Sah, Santosh Kumar Kuncha, Krishnan Harinivas Harshan, Archana Bharadwaj Siva, Karthik Bharadwaj Tallapaka, Rakesh K Mishra, Divya Tej Sowpati |
| EPI_ISL_458034 | King Institute of Preventive Medicine & Research | CSIR-Centre for Cellular and Molecular Biology | K.Kaveri,S.Sivasubramanian,S.Vennila,P.Padmapriya,R.Kiruba,S.Magesh,G. Dhinakar Raj, G. Ravikumar, P. Azhahianambi, K Thangaraj, Namami Gaur, Sakshi Shambhavi, Lamuk Zaveri, Shagufta Khan, Tulasi Nagabandi, Purushotham Vodnala, Payel Mukherjee, Sofia Banu, Priya Singh, Dhiviya Vedagiri, Divya Gupta, Vishal Sah, Santosh Kumar Kuncha, Krishnan Harinivas Harshan, Archana Bharadwaj Siva, Karthik Bharadwaj Tallapaka, Rakesh K Mishra, Divya Tej Sowpati |
| EPI_ISL_458035 | King Institute of Preventive Medicine & Research | CSIR-Centre for Cellular and Molecular Biology | K.Kaveri,S.Sivasubramanian,S.Vennila,P.Padmapriya,R.Kiruba,S.Magesh,G. Dhinakar Raj, G. Ravikumar, P. Azhahianambi, K Thangaraj, Tulasi Nagabandi, Namami Gaur, Sakshi Shambhavi, Lamuk Zaveri, Shagufta Khan, Purushotham Vodnala, Payel Mukherjee, Sofia Banu, Priya Singh, Dhiviya Vedagiri, Divya Gupta, Vishal Sah, Santosh Kumar Kuncha, Krishnan Harinivas Harshan, Archana Bharadwaj Siva, Karthik Bharadwaj Tallapaka, Rakesh K Mishra, Divya Tej Sowpati |
| EPI_ISL_458036 | King Institute of Preventive Medicine & Research | CSIR-Centre for Cellular and Molecular Biology | K.Kaveri,S.Sivasubramanian,S.Vennila,P.Padmapriya,R.Kiruba,S.Magesh,G. Dhinakar Raj, G. Ravikumar, R. P. Aravindh Babu, K Thangaraj, Payel Mukherjee, Sofia Banu, Priya Singh, Dhiviya Vedagiri, Divya Gupta, Vishal Sah, Santosh Kumar Kuncha, Krishnan Harinivas Harshan, Archana Bharadwaj Siva, Karthik Bharadwaj Tallapaka, Shagufta Khan, Lamuk Zaveri, Namami Gaur, Sakshi Shambhavi, Tulasi Nagabandi, Purushotham Vodnala, Rakesh K Mishra, Divya Tej Sowpati |
| EPI_ISL_458037 | King Institute of Preventive Medicine & Research | CSIR-Centre for Cellular and Molecular Biology | K.Kaveri,S.Sivasubramanian,S.Vennila,P.Padmapriya,R.Kiruba,S.Magesh,G. Dhinakar Raj, G. Ravikumar, R. P. Aravindh Babu, K Thangaraj, Sofia Banu, Payel Mukherjee, Priya Singh, Dhiviya Vedagiri, Divya Gupta, Vishal Sah, Santosh Kumar Kuncha, Krishnan Harinivas Harshan, Archana Bharadwaj Siva, Karthik Bharadwaj Tallapaka, Shagufta Khan, Lamuk Zaveri, Namami Gaur, Sakshi Shambhavi, Tulasi Nagabandi, Purushotham Vodnala, Rakesh K Mishra, Divya Tej Sowpati |
| EPI_ISL_458038 | King Institute of Preventive Medicine & Research | CSIR-Centre for Cellular and Molecular Biology | K.Kaveri,S.Sivasubramanian,S.Vennila,P.Padmapriya,R.Kiruba,S.Magesh,G. Dhinakar Raj, G. Ravikumar, R. P. Aravindh Babu, K Thangaraj, Shagufta Khan, Lamuk Zaveri, Namami Gaur, Sakshi Shambhavi, Tulasi Nagabandi, Purushotham Vodnala, Payel Mukherjee, Sofia Banu, Priya Singh, Dhiviya Vedagiri, Divya Gupta, Vishal Sah, Santosh Kumar Kuncha, Krishnan Harinivas Harshan, Archana Bharadwaj Siva, Karthik Bharadwaj Tallapaka, Rakesh K Mishra, Divya Tej Sowpati |
| EPI_ISL_458039 | King Institute of Preventive Medicine & Research | CSIR-Centre for Cellular and Molecular Biology | K.Kaveri,S.Sivasubramanian,S.Vennila,P.Padmapriya,R.Kiruba,S.Magesh,G. Dhinakar Raj, G. Ravikumar, R. P. Aravindh Babu, K Thangaraj, Lamuk Zaveri, Shagufta Khan, Namami Gaur, Sakshi Shambhavi, Tulasi Nagabandi, Purushotham Vodnala, Payel Mukherjee, Sofia Banu, Priya Singh, Dhiviya Vedagiri, Divya Gupta, Vishal Sah, Santosh Kumar Kuncha, Krishnan Harinivas Harshan, Archana Bharadwaj Siva, Karthik Bharadwaj Tallapaka, Rakesh K Mishra, Divya Tej Sowpati |
| EPI_ISL_458040 | King Institute of Preventive Medicine & Research | CSIR-Centre for Cellular and Molecular Biology | K.Kaveri,S.Sivasubramanian,S.Vennila,P.Padmapriya,R.Kiruba,S.Magesh,G. Dhinakar Raj, G. Ravikumar, R. P. Aravindh Babu, K Thangaraj, Namami Gaur, Sakshi Shambhavi, Lamuk Zaveri, Shagufta Khan, Tulasi Nagabandi, Purushotham Vodnala, Payel Mukherjee, Sofia Banu, Priya Singh, Dhiviya Vedagiri, Divya Gupta, Vishal Sah, Santosh Kumar Kuncha, Krishnan Harinivas Harshan, Archana Bharadwaj Siva, Karthik Bharadwaj Tallapaka, Rakesh K Mishra, Divya Tej Sowpati |
| EPI_ISL_458041 | King Institute of Preventive Medicine & Research | CSIR-Centre for Cellular and Molecular Biology | K.Kaveri,S.Sivasubramanian,S.Vennila,P.Padmapriya,R.Kiruba,S.Magesh,G. Dhinakar Raj, G. Ravikumar, R. P. Aravindh Babu, K Thangaraj, Tulasi Nagabandi, Namami Gaur, Sakshi Shambhavi, Lamuk Zaveri, Shagufta Khan, Purushotham Vodnala, Payel Mukherjee, Sofia Banu, Priya Singh, Dhiviya Vedagiri, Divya Gupta, Vishal Sah, Santosh Kumar Kuncha, Krishnan Harinivas Harshan, Archana Bharadwaj Siva, Karthik Bharadwaj Tallapaka, Rakesh K Mishra, Divya Tej Sowpati |
| EPI_ISL_458042 | King Institute of Preventive Medicine & Research | CSIR-Centre for Cellular and Molecular Biology | K.Kaveri,S.Sivasubramanian,S.Vennila,P.Padmapriya,R.Kiruba,S.Magesh,G. Dhinakar Raj, G. Ravikumar, M. Sekar, K Thangaraj, Payel Mukherjee, Sofia Banu, Priya Singh, Dhiviya Vedagiri, Divya Gupta, Vishal Sah, Santosh Kumar Kuncha, Krishnan Harinivas Harshan, Archana Bharadwaj Siva, Karthik Bharadwaj Tallapaka, Shagufta Khan, Lamuk Zaveri, Namami Gaur, Sakshi Shambhavi, Tulasi Nagabandi, Purushotham Vodnala, Rakesh K Mishra, Divya Tej Sowpati |
| EPI_ISL_458043 | King Institute of Preventive Medicine & Research | CSIR-Centre for Cellular and Molecular Biology | K.Kaveri,S.Sivasubramanian,S.Vennila,P.Padmapriya,R.Kiruba,S.Magesh,G. Dhinakar Raj, G. Ravikumar, M. Sekar, K Thangaraj,Sofia Banu, Payel Mukherjee, Priya Singh, Dhiviya Vedagiri, Divya Gupta, Vishal Sah, Santosh Kumar Kuncha, Krishnan Harinivas Harshan, Archana Bharadwaj Siva, Karthik Bharadwaj Tallapaka, Shagufta Khan, Lamuk Zaveri, Namami Gaur, Sakshi Shambhavi, Tulasi Nagabandi, Purushotham Vodnala, Rakesh K Mishra, Divya Tej Sowpati |
| EPI_ISL_458044 | King Institute of Preventive Medicine & Research | CSIR-Centre for Cellular and Molecular Biology | K.Kaveri,S.Sivasubramanian,S.Vennila,P.Padmapriya,R.Kiruba,S.Magesh,G. Dhinakar Raj, G. Ravikumar, M. Sekar, K Thangaraj,Shagufta Khan, Lamuk Zaveri, Namami Gaur, Sakshi Shambhavi, Tulasi Nagabandi, Purushotham Vodnala, Payel Mukherjee, Sofia Banu, Priya Singh, Dhiviya Vedagiri, Divya Gupta, Vishal Sah, Santosh Kumar Kuncha, Krishnan Harinivas Harshan, Archana Bharadwaj Siva, Karthik Bharadwaj Tallapaka, Rakesh K Mishra, Divya Tej Sowpati |
| EPI_ISL_458045 | CSIR-Centre for Cellular and Molecular Biology | CSIR-Centre for Cellular and Molecular Biology | Payel Mukherjee, Sofia Banu, Priya Singh, Dhiviya Vedagiri, Divya Gupta, Vishal Sah, Santosh Kumar Kuncha, Krishnan Harinivas Harshan, Archana Bharadwaj Siva, Karthik Bharadwaj Tallapaka, Shagufta Khan, Lamuk Zaveri, Namami Gaur, Sakshi Shambhavi, Tulasi Nagabandi, Purushotham Vodnala, G. Aditya Kumar, Koushick Sivakumar, Pooja Ramesh Gupta, Rajan Kumar Jha, Shraddha Vijay Lahoti, Rakesh K Mishra, Divya Tej Sowpati |
| EPI_ISL_458046 | CSIR-Centre for Cellular and Molecular Biology | CSIR-Centre for Cellular and Molecular Biology | Sofia Banu, Payel Mukherjee, Priya Singh, Dhiviya Vedagiri, Divya Gupta, Vishal Sah, Santosh Kumar Kuncha, Krishnan Harinivas Harshan, Archana Bharadwaj Siva, Karthik Bharadwaj Tallapaka, Shagufta Khan, Lamuk Zaveri, Namami Gaur, Sakshi Shambhavi, Tulasi Nagabandi, Purushotham Vodnala, Deepak Kumar, Devi Prasad Vijayashankar, Disha Nanda, Divya Das, Jotin Gogoi, Manish Bhattacharjee, Rakesh K Mishra, Divya Tej Sowpati |
| EPI_ISL_458047 | CSIR-Centre for Cellular and Molecular Biology | CSIR-Centre for Cellular and Molecular Biology | Shagufta Khan, Lamuk Zaveri, Namami Gaur, Sakshi Shambhavi, Tulasi Nagabandi, Purushotham Vodnala, Payel Mukherjee, Sofia Banu, Priya Singh, Dhiviya Vedagiri, Divya Gupta, Vishal Sah, Santosh Kumar Kuncha, Krishnan Harinivas Harshan, Archana Bharadwaj Siva, Karthik Bharadwaj Tallapaka, Disha Nanda, Divya Das, Jotin Gogoi, Manish Bhattacharjee, Ravi Prasad Mukku, Rakesh K Mishra, Divya Tej Sowpati |
| EPI_ISL_458048 | CSIR-Centre for Cellular and Molecular Biology | CSIR-Centre for Cellular and Molecular Biology | Lamuk Zaveri, Shagufta Khan, Namami Gaur, Sakshi Shambhavi, Tulasi Nagabandi, Purushotham Vodnala, Payel Mukherjee, Sofia Banu, Priya Singh, Dhiviya Vedagiri, Divya Gupta, Vishal Sah, Santosh Kumar Kuncha, Krishnan Harinivas Harshan, Archana Bharadwaj Siva, Karthik Bharadwaj Tallapaka, Renu Sudhakar, Somesh Gorde, Gangumala Srinivas Reddy, Sujoy Deb, Swati Bayyana, Rakesh K Mishra, Divya Tej Sowpati |
| EPI_ISL_458049 | CSIR-Centre for Cellular and Molecular Biology | CSIR-Centre for Cellular and Molecular Biology | Namami Gaur, Sakshi Shambhavi, Lamuk Zaveri, Shagufta Khan, Tulasi Nagabandi, Purushotham Vodnala, Payel Mukherjee, Sofia Banu, Priya Singh, Dhiviya Vedagiri, Divya Gupta, Vishal Sah, Santosh Kumar Kuncha, Krishnan Harinivas Harshan, Archana Bharadwaj Siva, Karthik Bharadwaj Tallapaka, Zeba Rizvi, Zuberwasim Sayyad, Kakade Aishwarya Arun, Amrutha H C, Ananga Ghosh, Rakesh K Mishra, Divya Tej Sowpati |
| EPI_ISL_458050 | CSIR-Centre for Cellular and Molecular Biology | CSIR-Centre for Cellular and Molecular Biology | Tulasi Nagabandi, Namami Gaur, Sakshi Shambhavi, Lamuk Zaveri, Shagufta Khan, Purushotham Vodnala, Payel Mukherjee, Sofia Banu, Priya Singh, Dhiviya Vedagiri, Divya Gupta, Vishal Sah, Santosh Kumar Kuncha, Krishnan Harinivas Harshan, Archana Bharadwaj Siva, Karthik Bharadwaj Tallapaka,Kezia J Ann, Radhika Khandelwal, Roshan Maku Venkata, Shemin Mansuri, Sonu Uday, Rakesh K Mishra, Divya Tej Sowpati |
| EPI_ISL_458051 | CSIR-Centre for Cellular and Molecular Biology | CSIR-Centre for Cellular and Molecular Biology | Payel Mukherjee, Sofia Banu, Priya Singh, Dhiviya Vedagiri, Divya Gupta, Vishal Sah, Santosh Kumar Kuncha, Krishnan Harinivas Harshan, Archana Bharadwaj Siva, Karthik Bharadwaj Tallapaka, Shagufta Khan, Lamuk Zaveri, Namami Gaur, Sakshi Shambhavi, Tulasi Nagabandi, Purushotham Vodnala, Gokulan C G, Gunjan Purohit, Hanuman Tulashiram Kale, Pankaj Kumar, Prachand Issarapu, Rakesh K Mishra, Divya Tej Sowpati |
| EPI_ISL_458052 | CSIR-Centre for Cellular and Molecular Biology | CSIR-Centre for Cellular and Molecular Biology | Sofia Banu, Payel Mukherjee, Priya Singh, Dhiviya Vedagiri, Divya Gupta, Vishal Sah, Santosh Kumar Kuncha, Krishnan Harinivas Harshan, Archana Bharadwaj Siva, Karthik Bharadwaj Tallapaka, Shagufta Khan, Lamuk Zaveri, Namami Gaur, Sakshi Shambhavi, Tulasi Nagabandi, Purushotham Vodnala,Preethi Jampala, Sharada Ravi Iyer, Sulagana Mukherjee, Swetha Sundar, Peddapuvala Sai Uday Kiran, Rakesh K Mishra, Divya Tej Sowpati |
| EPI_ISL_458053 | CSIR-Centre for Cellular and Molecular Biology | CSIR-Centre for Cellular and Molecular Biology | Shagufta Khan, Lamuk Zaveri, Namami Gaur, Sakshi Shambhavi, Tulasi Nagabandi, Purushotham Vodnala, Payel Mukherjee, Sofia Banu, Priya Singh, Dhiviya Vedagiri, Divya Gupta, Vishal Sah, Santosh Kumar Kuncha, Krishnan Harinivas Harshan, Archana Bharadwaj Siva, Karthik Bharadwaj Tallapaka,Umesh Kumar, Unis Ahmad Bhat, Ajay Sarawagi, Priyanka Pant, Rajkanwar Nathawat, Rakesh K Mishra, Divya Tej Sowpati |
| EPI_ISL_458054 | CSIR-Centre for Cellular and Molecular Biology | CSIR-Centre for Cellular and Molecular Biology | Lamuk Zaveri, Shagufta Khan, Namami Gaur, Sakshi Shambhavi, Tulasi Nagabandi, Purushotham Vodnala, Payel Mukherjee, Sofia Banu, Priya Singh, Dhiviya Vedagiri, Divya Gupta, Vishal Sah, Santosh Kumar Kuncha, Krishnan Harinivas Harshan, Archana Bharadwaj Siva, Karthik Bharadwaj Tallapaka,Umesh Kumar, Unis Ahmad Bhat, Ajay Sarawagi, Priyanka Pant, Rajkanwar Nathawat, Rakesh K Mishra, Divya Tej Sowpati |
| EPI_ISL_458055 | CSIR-Centre for Cellular and Molecular Biology | CSIR-Centre for Cellular and Molecular Biology | Namami Gaur, Sakshi Shambhavi, Lamuk Zaveri, Shagufta Khan, Tulasi Nagabandi, Purushotham Vodnala, Payel Mukherjee, Sofia Banu, Priya Singh, Dhiviya Vedagiri, Divya Gupta, Vishal Sah, Santosh Kumar Kuncha, Krishnan Harinivas Harshan, Archana Bharadwaj Siva, Karthik Bharadwaj Tallapaka, Nikhil Hajirnis, Pratheusa Maccha, M Soujanya Reddy,G. Aditya Kumar, Koushick Sivakumar, Rakesh K Mishra, Divya Tej Sowpati |
| EPI_ISL_458056 | CSIR-Centre for Cellular and Molecular Biology | CSIR-Centre for Cellular and Molecular Biology | Tulasi Nagabandi, Namami Gaur, Sakshi Shambhavi, Lamuk Zaveri, Shagufta Khan, Purushotham Vodnala, Payel Mukherjee, Sofia Banu, Priya Singh, Dhiviya Vedagiri, Divya Gupta, Vishal Sah, Santosh Kumar Kuncha, Krishnan Harinivas Harshan, Archana Bharadwaj Siva, Karthik Bharadwaj Tallapaka,G. Aditya Kumar, Koushick Sivakumar, Pooja Ramesh Gupta, Rajan Kumar Jha, Shraddha Vijay Lahoti, Rakesh K Mishra, Divya Tej Sowpati |
| EPI_ISL_458057 | CSIR-Centre for Cellular and Molecular Biology | CSIR-Centre for Cellular and Molecular Biology | Payel Mukherjee, Sofia Banu, Priya Singh, Dhiviya Vedagiri, Divya Gupta, Vishal Sah, Santosh Kumar Kuncha, Krishnan Harinivas Harshan, Archana Bharadwaj Siva, Karthik Bharadwaj Tallapaka, Shagufta Khan, Lamuk Zaveri, Namami Gaur, Sakshi Shambhavi, Tulasi Nagabandi, Purushotham Vodnala,Deepak Kumar, Devi Prasad Vijayashankar, Disha Nanda, Divya Das, Jotin Gogoi, Manish Bhattacharjee, Rakesh K Mishra, Divya Tej Sowpati |
| EPI_ISL_458058 | CSIR-Centre for Cellular and Molecular Biology | CSIR-Centre for Cellular and Molecular Biology | Sofia Banu, Payel Mukherjee, Priya Singh, Dhiviya Vedagiri, Divya Gupta, Vishal Sah, Santosh Kumar Kuncha, Krishnan Harinivas Harshan, Archana Bharadwaj Siva, Karthik Bharadwaj Tallapaka, Shagufta Khan, Lamuk Zaveri, Namami Gaur, Sakshi Shambhavi, Tulasi Nagabandi, Purushotham Vodnala, Disha Nanda, Divya Das, Jotin Gogoi, Manish Bhattacharjee, Ravi Prasad Mukku, Rakesh K Mishra, Divya Tej Sowpati |
| EPI_ISL_458059 | CSIR-Centre for Cellular and Molecular Biology | CSIR-Centre for Cellular and Molecular Biology | Shagufta Khan, Lamuk Zaveri, Namami Gaur, Sakshi Shambhavi, Tulasi Nagabandi, Purushotham Vodnala, Payel Mukherjee, Sofia Banu, Priya Singh, Dhiviya Vedagiri, Divya Gupta, Vishal Sah, Santosh Kumar Kuncha, Krishnan Harinivas Harshan, Archana Bharadwaj Siva, Karthik Bharadwaj Tallapaka, Renu Sudhakar, Somesh Gorde, Gangumala Srinivas Reddy, Sujoy Deb, Swati Bayyana, Rakesh K Mishra, Divya Tej Sowpati |
| EPI_ISL_458060 | CSIR-Centre for Cellular and Molecular Biology | CSIR-Centre for Cellular and Molecular Biology | Lamuk Zaveri, Shagufta Khan, Namami Gaur, Sakshi Shambhavi, Tulasi Nagabandi, Purushotham Vodnala, Payel Mukherjee, Sofia Banu, Priya Singh, Dhiviya Vedagiri, Divya Gupta, Vishal Sah, Santosh Kumar Kuncha, Krishnan Harinivas Harshan, Archana Bharadwaj Siva, Karthik Bharadwaj Tallapaka,Zeba Rizvi, Zuberwasim Sayyad, Kakade Aishwarya Arun, Amrutha H C, Ananga Ghosh, Rakesh K Mishra, Divya Tej Sowpati |
| EPI_ISL_458061 | CSIR-Centre for Cellular and Molecular Biology | CSIR-Centre for Cellular and Molecular Biology | Namami Gaur, Sakshi Shambhavi, Lamuk Zaveri, Shagufta Khan, Tulasi Nagabandi, Purushotham Vodnala, Payel Mukherjee, Sofia Banu, Priya Singh, Dhiviya Vedagiri, Divya Gupta, Vishal Sah, Santosh Kumar Kuncha, Krishnan Harinivas Harshan, Archana Bharadwaj Siva, Karthik Bharadwaj Tallapaka,Kezia J Ann, Radhika Khandelwal, Roshan Maku Venkata, Shemin Mansuri, Sonu Uday, Rakesh K Mishra, Divya Tej Sowpati |
| EPI_ISL_458062 | CSIR-Centre for Cellular and Molecular Biology | CSIR-Centre for Cellular and Molecular Biology | Payel Mukherjee, Sofia Banu, Priya Singh, Dhiviya Vedagiri, Divya Gupta, Vishal Sah, Santosh Kumar Kuncha, Krishnan Harinivas Harshan, Archana Bharadwaj Siva, Karthik Bharadwaj Tallapaka, Shagufta Khan, Lamuk Zaveri, Namami Gaur, Sakshi Shambhavi, Tulasi Nagabandi, Purushotham Vodnala, Rakesh K Mishra, Sonu Uday, Sudipta Mondal, Annapoorna P Karthyayani, Debabrata Jana, Debrya Saha, Divya Tej Sowpati |
| EPI_ISL_458063 | CSIR-Centre for Cellular and Molecular Biology | CSIR-Centre for Cellular and Molecular Biology | Sofia Banu, Payel Mukherjee, Priya Singh, Dhiviya Vedagiri, Divya Gupta, Vishal Sah, Santosh Kumar Kuncha, Krishnan Harinivas Harshan, Archana Bharadwaj Siva, Karthik Bharadwaj Tallapaka, Shagufta Khan, Lamuk Zaveri, Namami Gaur, Sakshi Shambhavi, Tulasi Nagabandi, Purushotham Vodnala, Gokulan C G, Gunjan Purohit, Hanuman Tulashiram Kale, Pankaj Kumar, Prachand Issarapu, Rakesh K Mishra, Divya Tej Sowpati |
| EPI_ISL_458064 | CSIR-Centre for Cellular and Molecular Biology | CSIR-Centre for Cellular and Molecular Biology | Shagufta Khan, Lamuk Zaveri, Namami Gaur, Sakshi Shambhavi, Tulasi Nagabandi, Purushotham Vodnala, Payel Mukherjee, Sofia Banu, Priya Singh, Dhiviya Vedagiri, Divya Gupta, Vishal Sah, Santosh Kumar Kuncha, Krishnan Harinivas Harshan, Archana Bharadwaj Siva, Karthik Bharadwaj Tallapaka,Preethi Jampala, Sharada Ravi Iyer, Sulagana Mukherjee, Swetha Sundar, Peddapuvala Sai Uday Kiran Rakesh K Mishra, Divya Tej Sowpati |
| EPI_ISL_458065 | CSIR-Centre for Cellular and Molecular Biology | CSIR-Centre for Cellular and Molecular Biology | Lamuk Zaveri, Shagufta Khan, Namami Gaur, Sakshi Shambhavi, Tulasi Nagabandi, Purushotham Vodnala, Payel Mukherjee, Sofia Banu, Priya Singh, Dhiviya Vedagiri, Divya Gupta, Vishal Sah, Santosh Kumar Kuncha, Krishnan Harinivas Harshan, Archana Bharadwaj Siva, Karthik Bharadwaj Tallapaka,Umesh Kumar, Unis Ahmad Bhat, Ajay Sarawagi, Priyanka Pant, Rajkanwar Nathawat, Rakesh K Mishra, Divya Tej Sowpati |

| | | | |
|---|---|---|---|
| EPI_ISL_458066, EPI_ISL_458067, EPI_ISL_458068, EPI_ISL_458069 | Osmania Medical College | CSIR-Centre for Cellular and Molecular Biology | Shashikala Reddy, Mahboob Khan,Payel Mukherjee, Sofia Banu, Priya Singh, Dhiviya Vedagiri, Divya Gupta, Vishal Sah, Santosh Kumar Kuncha, Krishnan Harinivas Harshan, Archana Bharadwaj Siva, Karthik Bharadwaj Tallapaka, Shagufta Khan, Lamuk Zaveri, Namami Gaur, Sakshi Shambhavi, Tulasi Nagabandi, Purushotham Vodnala, Rakesh K Mishra, Divya Tej Sowpati |
| EPI_ISL_458070 | CSIR-Centre for Cellular and Molecular Biology | CSIR-Centre for Cellular and Molecular Biology | Sakshi Shambhavi, Lamuk Zaveri, Shagufta Khan, Namami Gaur, Tulasi Nagabandi, Purushotham Vodnala, Payel Mukherjee, Sofia Banu, Priya Singh, Dhiviya Vedagiri, Divya Gupta, Vishal Sah, Santosh Kumar Kuncha, Krishnan Harinivas Harshan, Archana Bharadwaj Siva, Karthik Bharadwaj Tallapaka,Nikhil Hajirnis, Pratheusa Maccha, M Soujanya Reddy,G. Aditya Kumar, Koushick Sivakumar,Disha Nanda, Divya Das, Jotin Gogoi, Manish Bhattacharjee, Ravi Prasad Mukku, Rakesh K Mishra, Divya Tej Sowpati |
| EPI_ISL_458071 | CSIR-Centre for Cellular and Molecular Biology | CSIR-Centre for Cellular and Molecular Biology | Sakshi Shambhavi, Lamuk Zaveri, Shagufta Khan, Namami Gaur, Tulasi Nagabandi, Purushotham Vodnala, Payel Mukherjee, Sofia Banu, Priya Singh, Dhiviya Vedagiri, Divya Gupta, Vishal Sah, Santosh Kumar Kuncha, Krishnan Harinivas Harshan, Archana Bharadwaj Siva, Karthik Bharadwaj Tallapaka,Nikhil Hajirnis, Pratheusa Maccha, M Soujanya Reddy,G. Aditya Kumar, Koushick Sivakumar, Rakesh K Mishra, Divya Tej Sowpati |
| EPI_ISL_458072, EPI_ISL_458073, EPI_ISL_458074, EPI_ISL_458075, EPI_ISL_458076, EPI_ISL_458077 | CSIR-Centre for Cellular and Molecular Biology | CSIR-Centre for Cellular and Molecular Biology | Dhiviya Vedagiri, Divya Gupta, Vishal Sah, Payel Mukherjee, Sofia Banu, Priya Singh, Santosh Kumar Kuncha, Archana Bharadwaj Siva, Karthik Bharadwaj Tallapaka, Shagufta Khan, Lamuk Zaveri, Namami Gaur, Sakshi Shambhavi, Tulasi Nagabandi, Purushotham Vodnala, Rakesh K Mishra, Divya Tej Sowpati, Krishnan Harinivas Harshan |
| EPI_ISL_458079 | Mitra Keluarga Hospital Kenjeran | Institute of Tropical Disease, Universitas Airlangga | Aldise M Nastri, Jezzy R Dewantari, Rima R Prasetya, Krisnoadi Rahardjo, Anastasia W Jefuna, Gatot Soegiarto, Laksmi Wulandari, Retno A Setyoningrum, Resti Yudhawati, Yohko K Shimizu, Mitsuhiro Nishimura, Yasuko Mori, Soetjipto, Kazufumi Shimizu, Maria I Lusida |
| EPI_ISL_458080 | CSIR-Centre for Cellular and Molecular Biology | CSIR-Centre for Cellular and Molecular Biology | Sakshi Shambhavi, Lamuk Zaveri, Shagufta Khan, Namami Gaur, Tulasi Nagabandi, Purushotham Vodnala, Payel Mukherjee, Sofia Banu, Priya Singh, Dhiviya Vedagiri, Divya Gupta, Vishal Sah, Santosh Kumar Kuncha, Krishnan Harinivas Harshan, Archana Bharadwaj Siva, Karthik Bharadwaj Tallapaka, G. Aditya Kumar, Koushick Sivakumar, Pooja Ramesh Gupta, Rajan Kumar Jha, Shraddha Vijay Lahoti, Rakesh K Mishra, Divya Tej Sowpati |
| EPI_ISL_458081 | RSUD Bangil Pasuruan | Institute of Tropical Disease, Universitas Airlangga | Jezzy R Dewantari, Rima R Prasetya, Krisnoadi Rahardjo, Aldise M Nastri, Arma Roosalina, Gatot Soegiarto, Laksmi Wulandari, Retno A Setyoningrum, Resti Yudhawati, Yohko K Shimizu, Mitsuhiro Nishimura, Yasuko Mori, Soetjipto, Kazufumi Shimizu, Maria I Lusida |
| EPI_ISL_458082 | Universitas Airlangga Hospital | Institute of Tropical Disease, Universitas Airlangga | Kazufumi Shimizu, Krisnoadi Rahardjo, Aldise M Nastri, Jezzy R Dewantari, Rima R Prasetya, Nasronudin, Gatot Soegiarto, Laksmi Wulandari, Retno A Setyoningrum, Resti Yudhawati, Yohko K Shimizu, Mitsuhiro Nishimura, Yasuko Mori, Soetjipto, Maria I Lusida |
| EPI_ISL_458083 | Adi Husada Undaan Hospital | Institute of Tropical Disease, Universitas Airlangga | Rima R Prasetya, Krisnoadi Rahardjo, Aldise M Nastri, Jezzy R Dewantari, Irawati Marga, Gatot Soegiarto, Laksmi Wulandari, Retno A Setyoningrum, Resti Yudhawati, Yohko K Shimizu, Mitsuhiro Nishimura, Yasuko Mori, Soetjipto, Kazufumi Shimizu, Maria I Lusida |
| EPI_ISL_458084 | Laboratorio Biologia Molecolare Sars Cov2 - UOC Laboratorio Analisi - Servizio Medicina di Laboratorio, Ospedale "San Francesco" - ATS-ASSL Nuoro | Laboratorio specialistico UOC Ematologia - Ospedale "San Francesco" - ATS-ASSL Nuoro | Piras Giovanna, Fancello Tatiana, Asproni Rosanna, Fiamma Maura, Monne Maria Itria, Toja Alessandro, Sanna Filomena, Floris Anna Rita, Sulis Vincenzo, Palmas Angelo Domenico, Casu Gavino, Lo Maglio Iana, Mameli Giuseppe. |
| EPI_ISL_458085 | Laboratorio Biologia Molecolare Sars Cov2 - UOC Laboratorio Analisi - Servizio Medicina di Laboratorio , Ospedale "San Francesco" - ATS- ASSL Nuoro | Laboratorio specialistico UOC Ematologia - Ospedale "San Francesco" - ATS-ASSL Nuoro | Piras Giovanna, Fancello Tatiana, Asproni Rosanna, Fiamma Maura, Monne Maria Itria, Toja Alessandro, Sanna Filomena, Floris Anna Rita, Sulis Vincenzo, Palmas Angelo Domenico, Casu Gavino, Lo Maglio Iana, Mameli Giuseppe. |
| EPI_ISL_458086 | B.J. Medical College and Civil hospital | Gujarat Biotechnology Research Centre | Dhaval Vaghela, Ramesh Patel, Pranay Shah, Kamlesh J Upadhyay, Ramesh Pandit, Tejas Shah, Ankit Hinsu, Pritesh Sabara, Apurvasinh Puvar, Janvi Raval, Zarna Patel, Monika Gandhi, Pinal Trivedi, Maharshi Pandya, Amit Kanani, Nidhi Patel, Nitin Savaliya, Raghawendra Kumar, Dinesh Kumar, Zuber Saiyed, Komal Patel, Labdhi Pandya, Snehal Bagatharia, Neha Rajpara, Bhavesh Modi, Gaurishankar Shrimali, R D Dixit, A M Kadri, Umang Mishra, Chaitanya Joshi, Madhvi Joshi |
| EPI_ISL_458087 | B.J. Medical College and Civil hospital | Gujarat Biotechnology Research Centre | Ramesh Patel, Pranay Shah, Kamlesh J Upadhyay, Ramesh Pandit, Tejas Shah, Ankit Hinsu, Pritesh Sabara, Apurvasinh Puvar, Janvi Raval, Zarna Patel, Monika Gandhi, Pinal Trivedi, Maharshi Pandya, Amit Kanani, Nidhi Patel, Nitin Savaliya, Raghawendra Kumar, Dinesh Kumar, Zuber Saiyed, Komal Patel, Labdhi Pandya, Snehal Bagatharia, Dhaval Vaghela, Afzal Ansari, Bhavesh Modi, Gaurishankar Shrimali, R D Dixit, A M Kadri, Umang Mishra, Chaitanya Joshi, Madhvi Joshi |
| EPI_ISL_458088 | B.J. Medical College and Civil hospital | Gujarat Biotechnology Research Centre | Pranay Shah, Kamlesh J Upadhyay, Ramesh Pandit, Tejas Shah, Ankit Hinsu, Pritesh Sabara, Apurvasinh Puvar, Janvi Raval, Zarna Patel, Monika Gandhi, Pinal Trivedi, Maharshi Pandya, Amit Kanani, Nidhi Patel, Nitin Savaliya, Raghawendra Kumar, Dinesh Kumar, Zuber Saiyed, Komal Patel, Labdhi Pandya, Snehal Bagatharia, Dhaval Vaghela, Ramesh Patel, Fenil Patel, Bhavesh Modi, Gaurishankar Shrimali, R D Dixit, A M Kadri, Umang Mishra, Chaitanya Joshi, Madhvi Joshi |
| EPI_ISL_458089 | B.J. Medical College and Civil hospital | Gujarat Biotechnology Research Centre | Pranay Shah, Kamlesh J Upadhyay, Ramesh Pandit, Tejas Shah, Ankit Hinsu, Pritesh Sabara, Apurvasinh Puvar, Janvi Raval, Zarna Patel, Monika Gandhi, Pinal Trivedi, Maharshi Pandya, Amit Kanani, Nidhi Patel, Nitin Savaliya, Raghawendra Kumar, Dinesh Kumar, Zuber Saiyed, Komal Patel, Labdhi Pandya, Snehal Bagatharia, Dhaval Vaghela, Ramesh Patel, Neelam Nathani, Bhavesh Modi, Gaurishankar Shrimali, R D Dixit, A M Kadri, Umang Mishra, Chaitanya Joshi, Madhvi Joshi |
| EPI_ISL_458090 | B.J. Medical College and Civil hospital | Gujarat Biotechnology Research Centre | Kamlesh J Upadhyay, Ramesh Pandit, Tejas Shah, Ankit Hinsu, Pritesh Sabara, Apurvasinh Puvar, Janvi Raval, Zarna Patel, Monika Gandhi, Pinal Trivedi, Maharshi Pandya, Amit Kanani, Nidhi Patel, Nitin Savaliya, Raghawendra Kumar, Dinesh Kumar, Zuber Saiyed, Komal Patel, Labdhi Pandya, Snehal Bagatharia, Dhaval Vaghela, Ramesh Patel, Pranay Shah, Armi Chaudhari, Bhavesh Modi, Gaurishankar Shrimali, R D Dixit, A M Kadri, Umang Mishra, Chaitanya Joshi, Madhvi Joshi |
| EPI_ISL_458091 | B.J. Medical College and Civil hospital | Gujarat Biotechnology Research Centre | Maharshi Pandya, Amit Kanani, Nidhi Patel, Nitin Savaliya, Raghawendra Kumar, Dinesh Kumar, Zuber Saiyed, Komal Patel, Labdhi Pandya, Snehal Bagatharia, Dhaval Vaghela, Ramesh Patel, Pranay Shah, Kamlesh J Upadhyay, Ramesh Pandit, Tejas Shah, Ankit Hinsu, Pritesh Sabara, Apurvasinh Puvar, Janvi Raval, Zarna Patel, Monika Gandhi, Pinal Trivedi, Bhavya Jindal, Bhavesh Modi, Gaurishankar Shrimali, R D Dixit, A M Kadri, Umang Mishra, Chaitanya Joshi, Madhvi Joshi |
| EPI_ISL_458092 | B.J. Medical College and Civil hospital | Gujarat Biotechnology Research Centre | Amit Kanani, Nidhi Patel, Nitin Savaliya, Raghawendra Kumar, Dinesh Kumar, Zuber Saiyed, Komal Patel, Labdhi Pandya, Snehal Bagatharia, Dhaval Vaghela, Ramesh Patel, Pranay Shah, Kamlesh J Upadhyay, Ramesh Pandit, Tejas Shah, Ankit Hinsu, Pritesh Sabara, Apurvasinh Puvar, Janvi Raval, Zarna Patel, Monika Gandhi, Pinal Trivedi, Maharshi Pandya, Camellia Chakraborty, Bhavesh Modi, Gaurishankar Shrimali, R D Dixit, A M Kadri, Umang Mishra, Chaitanya Joshi, Madhvi Joshi |
| EPI_ISL_458093 | B.J. Medical College and Civil hospital | Gujarat Biotechnology Research Centre | Nidhi Patel, Nitin Savaliya, Raghawendra Kumar, Dinesh Kumar, Zuber Saiyed, Komal Patel, Labdhi Pandya, Snehal Bagatharia, Dhaval Vaghela, Ramesh Patel, Pranay Shah, Kamlesh J Upadhyay, Ramesh Pandit, Tejas Shah, Ankit Hinsu, Pritesh Sabara, Apurvasinh Puvar, Janvi Raval, Zarna Patel, Monika Gandhi, Pinal Trivedi, Maharshi Pandya, Amit Kanani, Siddhant Kumar, Bhavesh Modi, Gaurishankar Shrimali, R D Dixit, A M Kadri, Umang Mishra, Chaitanya Joshi, Madhvi Joshi |
| EPI_ISL_458094 | B.J. Medical College and Civil hospital | Gujarat Biotechnology Research Centre | Nitin Savaliya, Raghawendra Kumar, Dinesh Kumar, Zuber Saiyed, Komal Patel, Labdhi Pandya, Snehal Bagatharia, Dhaval Vaghela, Ramesh Patel, Pranay Shah, Kamlesh J Upadhyay, Ramesh Pandit, Tejas Shah, Ankit Hinsu, Pritesh Sabara, Apurvasinh Puvar, Janvi Raval, Zarna Patel, Monika Gandhi, Pinal Trivedi, Maharshi Pandya, Amit Kanani, Nidhi Patel, Priyanka P Vatsa, Bhavesh Modi, Gaurishankar Shrimali, R D Dixit, A M Kadri, Umang Mishra, Chaitanya Joshi, Madhvi Joshi |
| EPI_ISL_458095 | B.J. Medical College and Civil hospital | Gujarat Biotechnology Research Centre | Raghawendra Kumar, Dinesh Kumar, Zuber Saiyed, Komal Patel, Labdhi Pandya, Snehal Bagatharia, Dhaval Vaghela, Ramesh Patel, Pranay Shah, Kamlesh J Upadhyay, Ramesh Pandit, Tejas Shah, Ankit Hinsu, Pritesh Sabara, Apurvasinh Puvar, Janvi Raval, Zarna Patel, Monika Gandhi, Pinal Trivedi, Maharshi Pandya, Amit Kanani, Nidhi Patel, Nitin Savaliya, Pooja P Doshi, Bhavesh Modi, Gaurishankar Shrimali, R D Dixit, A M Kadri, Umang Mishra, Chaitanya Joshi, Madhvi Joshi |
| EPI_ISL_458096 | B.J. Medical College and Civil hospital | Gujarat Biotechnology Research Centre | Dinesh Kumar, Zuber Saiyed, Komal Patel, Labdhi Pandya, Snehal Bagatharia, Dhaval Vaghela, Ramesh Patel, Pranay Shah, Kamlesh J Upadhyay, Ramesh Pandit, Tejas Shah, Ankit Hinsu, Pritesh Sabara, Apurvasinh Puvar, Janvi Raval, Zarna Patel, Monika Gandhi, Pinal Trivedi, Maharshi Pandya, Amit Kanani, Nidhi Patel, Nitin Savaliya, Raghawendra Kumar, Akanksha Verma, Bhavesh Modi, Gaurishankar Shrimali, R D Dixit, A M Kadri, Umang Mishra, Chaitanya Joshi, Madhvi Joshi |
| EPI_ISL_458097 | B.J. Medical College and Civil hospital | Gujarat Biotechnology Research Centre | Zuber Saiyed, Komal Patel, Labdhi Pandya, Snehal Bagatharia, Dhaval Vaghela, Ramesh Patel, Pranay Shah, Kamlesh J Upadhyay, Ramesh Pandit, Tejas Shah, Ankit Hinsu, Pritesh Sabara, Apurvasinh Puvar, Janvi Raval, Zarna Patel, Monika Gandhi, Pinal Trivedi, Maharshi Pandya, Amit Kanani, Nidhi Patel, Nitin Savaliya, Raghawendra Kumar, Dinesh Kumar, Priti Pandita, Bhavesh Modi, Gaurishankar Shrimali, R D Dixit, A M Kadri, Umang Mishra, Chaitanya Joshi, Madhvi Joshi |
| EPI_ISL_458098 | B.J. Medical College and Civil hospital | Gujarat Biotechnology Research Centre | Komal Patel, Labdhi Pandya, Snehal Bagatharia, Dhaval Vaghela, Ramesh Patel, Pranay Shah, Kamlesh J Upadhyay, Ramesh Pandit, Tejas Shah, Ankit Hinsu, Pritesh Sabara, Apurvasinh Puvar, Janvi Raval, Zarna Patel, Monika Gandhi, Pinal Trivedi, Maharshi Pandya, Amit Kanani, Nidhi Patel, Nitin Savaliya, Raghawendra Kumar, Dinesh Kumar, Zuber Saiyed, Pragya Sharma, Bhavesh Modi, Gaurishankar Shrimali, R D Dixit, A M Kadri, Umang Mishra, Chaitanya Joshi, Madhvi Joshi |
| EPI_ISL_458099 | B.J. Medical College and Civil hospital | Gujarat Biotechnology Research Centre | Labdhi Pandya, Snehal Bagatharia, Dhaval Vaghela, Ramesh Patel, Pranay Shah, Kamlesh J Upadhyay, Ramesh Pandit, Tejas Shah, Ankit Hinsu, Pritesh Sabara, Apurvasinh Puvar, Janvi Raval, Zarna Patel, Monika Gandhi, Pinal Trivedi, Maharshi Pandya, Amit Kanani, Nidhi Patel, Nitin Savaliya, Raghawendra Kumar, Dinesh Kumar, Zuber Saiyed, Komal Patel, Neha Rajpara, Bhavesh Modi, Gaurishankar Shrimali, R D Dixit, A M Kadri, Umang Mishra, Chaitanya Joshi, Madhvi Joshi |
| EPI_ISL_458100 | B.J. Medical College and Civil hospital | Gujarat Biotechnology Research Centre | Snehal Bagatharia, Dhaval Vaghela, Ramesh Patel, Pranay Shah, Kamlesh J Upadhyay, Ramesh Pandit, Tejas Shah, Ankit Hinsu, Pritesh Sabara, Apurvasinh Puvar, Janvi Raval, Zarna Patel, Monika Gandhi, Pinal Trivedi, Maharshi Pandya, Amit Kanani, Nidhi Patel, Nitin Savaliya, Raghawendra Kumar, Dinesh Kumar, Zuber Saiyed, Komal Patel, Labdhi Pandya, Afzal Ansari, Bhavesh Modi, Gaurishankar Shrimali, R D Dixit, A M Kadri, Umang Mishra, Chaitanya Joshi, Madhvi Joshi |
| EPI_ISL_458101 | B.J. Medical College and Civil hospital | Gujarat Biotechnology Research Centre | Dhaval Vaghela, Ramesh Patel, Pranay Shah, Kamlesh J Upadhyay, Ramesh Pandit, Tejas Shah, Ankit Hinsu, Pritesh Sabara, Apurvasinh Puvar, Janvi Raval, Zarna Patel, Monika Gandhi, Pinal Trivedi, Maharshi Pandya, Amit Kanani, Nidhi Patel, Nitin Savaliya, Raghawendra Kumar, Dinesh Kumar, Zuber Saiyed, Komal Patel, Labdhi Pandya, Snehal Bagatharia, Fenil Patel, Bhavesh Modi, Gaurishankar Shrimali, R D Dixit, A M Kadri, Umang Mishra, Chaitanya Joshi, Madhvi Joshi |
| EPI_ISL_458102 | B.J. Medical College and Civil hospital | Gujarat Biotechnology Research Centre | Ramesh Patel, Pranay Shah, Kamlesh J Upadhyay, Ramesh Pandit, Tejas Shah, Ankit Hinsu, Pritesh Sabara, Apurvasinh Puvar, Janvi Raval, Zarna Patel, Monika Gandhi, Pinal Trivedi, Maharshi Pandya, Amit Kanani, Nidhi Patel, Nitin Savaliya, Raghawendra Kumar, Dinesh Kumar, Zuber Saiyed, Komal Patel, Labdhi Pandya, Snehal Bagatharia, Dhaval Vaghela, Neelam Nathani, Bhavesh Modi, Gaurishankar Shrimali, R D Dixit, A M Kadri, Umang Mishra, Chaitanya Joshi, Madhvi Joshi |
| EPI_ISL_458103 | Gujarat Biotechnology Research Centre | Gujarat Biotechnology Research Centre | Ramesh Pandit, Tejas Shah, Ankit Hinsu, Pritesh Sabara, Apurvasinh Puvar, Janvi Raval, Zarna Patel, Monika Gandhi, Pinal Trivedi, Maharshi Pandya, Amit Kanani, Nidhi Patel, Nitin Savaliya, Raghawendra Kumar, Dinesh Kumar, Zuber Saiyed, Komal Patel, Labdhi Pandya, Snehal Bagatharia, Armi Chaudhari, Bhavesh Modi, Gaurishankar Shrimali, R D Dixit, A M Kadri, Umang Mishra, Chaitanya Joshi, Madhvi Joshi, , , , , |
| EPI_ISL_458104 | Gujarat Biotechnology Research Centre | Gujarat Biotechnology Research Centre | Tejas Shah, Ankit Hinsu, Pritesh Sabara, Apurvasinh Puvar, Janvi Raval, Zarna Patel, Monika Gandhi, Pinal Trivedi, Maharshi Pandya, Amit Kanani, Nidhi Patel, Nitin Savaliya, Raghawendra Kumar, Dinesh Kumar, Zuber Saiyed, Komal Patel, Labdhi Pandya, Snehal Bagatharia, Ramesh Patel, Bhavya Jindal, Bhavesh Modi, Gaurishankar Shrimali, R D Dixit, A M Kadri, Umang Mishra, Chaitanya Joshi, Madhvi Joshi, , , , , |
| EPI_ISL_458105 | Gujarat Biotechnology Research Centre | Gujarat Biotechnology Research Centre | Ankit Hinsu, Pritesh Sabara, Apurvasinh Puvar, Janvi Raval, Zarna Patel, Monika Gandhi, Pinal Trivedi, Maharshi Pandya, Amit Kanani, Nidhi Patel, Nitin Savaliya, Raghawendra Kumar, Dinesh Kumar, Zuber Saiyed, Komal Patel, Labdhi Pandya, Snehal Bagatharia, Ramesh Pandit, Tejas Shah, Camellia Chakraborty, Bhavesh Modi, Gaurishankar Shrimali, R D Dixit, A M Kadri, Umang Mishra, Chaitanya Joshi, Madhvi Joshi, , , , , |
| EPI_ISL_458106 | Gujarat Biotechnology Research Centre | Gujarat Biotechnology Research Centre | Pritesh Sabara, Apurvasinh Puvar, Janvi Raval, Zarna Patel, Monika Gandhi, Pinal Trivedi, Maharshi Pandya, Amit Kanani, Nidhi Patel, Nitin Savaliya, Raghawendra Kumar, Dinesh Kumar, Zuber Saiyed, Komal Patel, Labdhi Pandya, Snehal Bagatharia, Ramesh Pandit, Tejas Shah, Ankit Hinsu, Siddhant Kumar, Bhavesh Modi, Gaurishankar Shrimali, R D Dixit, A M Kadri, Umang Mishra, Chaitanya Joshi, Madhvi Joshi, , , , , |
| EPI_ISL_458107 | Gujarat Biotechnology Research Centre | Gujarat Biotechnology Research Centre | Apurvasinh Puvar, Janvi Raval, Zarna Patel, Monika Gandhi, Pinal Trivedi, Maharshi Pandya, Amit Kanani, Nidhi Patel, Nitin Savaliya, Raghawendra Kumar, Dinesh Kumar, Zuber Saiyed, Komal Patel, Labdhi Pandya, Snehal Bagatharia, Ramesh Pandit, Tejas Shah, Ankit Hinsu, Pritesh Sabara, Priyanka P Vatsa, Bhavesh Modi, Gaurishankar Shrimali, R D Dixit, A M Kadri, Umang Mishra, Chaitanya Joshi, Madhvi Joshi, , , , , |
| EPI_ISL_458108 | Gujarat Biotechnology Research Centre | Gujarat Biotechnology Research Centre | Janvi Raval, Zarna Patel, Monika Gandhi, Pinal Trivedi, Maharshi Pandya, Amit Kanani, Nidhi Patel, Nitin Savaliya, Raghawendra Kumar, Dinesh Kumar, Zuber Saiyed, Komal Patel, Labdhi Pandya, Snehal Bagatharia, Ramesh Pandit, Tejas Shah, Ankit Hinsu, Pritesh Sabara, Apurvasinh Puvar, Pooja P Doshi, Bhavesh Modi, Gaurishankar Shrimali, R D Dixit, A M Kadri, Umang Mishra, Chaitanya Joshi, Madhvi Joshi, , , , , |
| EPI_ISL_458109 | Gujarat Biotechnology Research Centre | Gujarat Biotechnology Research Centre | Zarna Patel, Monika Gandhi, Pinal Trivedi, Maharshi Pandya, Amit Kanani, Nidhi Patel, Nitin Savaliya, Raghawendra Kumar, Dinesh Kumar, Zuber Saiyed, Komal Patel, Labdhi Pandya, Snehal Bagatharia, Ramesh Pandit, Tejas Shah, Ankit Hinsu, Pritesh Sabara, Apurvasinh Puvar, Janvi Raval, Akanksha Verma, Bhavesh Modi, Gaurishankar Shrimali, R D Dixit, A M Kadri, Umang Mishra, Chaitanya Joshi, Madhvi Joshi, , , , , |
| EPI_ISL_458110 | Gujarat Biotechnology Research Centre | Gujarat Biotechnology Research Centre | Monika Gandhi, Pinal Trivedi, Maharshi Pandya, Amit Kanani, Nidhi Patel, Nitin Savaliya, Raghawendra Kumar, Dinesh Kumar, Zuber Saiyed, Komal Patel, Labdhi Pandya, Snehal Bagatharia, Ramesh Pandit, Tejas Shah, Ankit Hinsu, Pritesh Sabara, Apurvasinh Puvar, Janvi Raval, Zarna Patel, Priti Pandita, Bhavesh Modi, Gaurishankar Shrimali, R D Dixit, A M Kadri, Umang Mishra, Chaitanya Joshi, Madhvi Joshi, , , , , |
| EPI_ISL_458111 | Gujarat Biotechnology Research Centre | Gujarat Biotechnology Research Centre | Pinal Trivedi, Maharshi Pandya, Amit Kanani, Nidhi Patel, Nitin Savaliya, Raghawendra Kumar, Dinesh Kumar, Zuber Saiyed, Komal Patel, Labdhi Pandya, Snehal Bagatharia, Ramesh Pandit, Tejas Shah, Ankit Hinsu, Pritesh Sabara, Apurvasinh Puvar, Janvi Raval, Zarna Patel, Monika Gandhi, Pragya Sharma, Bhavesh Modi, Gaurishankar Shrimali, R D Dixit, A M Kadri, Umang Mishra, Chaitanya Joshi, Madhvi Joshi, , , , , |
| EPI_ISL_458112 | Gujarat Biotechnology Research Centre | Gujarat Biotechnology Research Centre | Maharshi Pandya, Amit Kanani, Nidhi Patel, Nitin Savaliya, Raghawendra Kumar, Dinesh Kumar, Zuber Saiyed, Komal Patel, Labdhi Pandya, Snehal Bagatharia, Ramesh Pandit, Tejas Shah, Ankit Hinsu, Pritesh Sabara, Apurvasinh Puvar, Janvi Raval, Zarna Patel, Monika Gandhi, Pinal Trivedi, Neha Rajpara, Bhavesh Modi, Gaurishankar Shrimali, R D Dixit, A M Kadri, Umang Mishra, Chaitanya Joshi, Madhvi Joshi, , , , , |
| EPI_ISL_458113 | Gujarat Biotechnology Research Centre | Gujarat Biotechnology Research Centre | Amit Kanani, Nidhi Patel, Nitin Savaliya, Raghawendra Kumar, Dinesh Kumar, Zuber Saiyed, Komal Patel, Labdhi Pandya, Snehal Bagatharia, Ramesh Pandit, Tejas Shah, Ankit Hinsu, Pritesh Sabara, Apurvasinh Puvar, Janvi Raval, Zarna Patel, Monika Gandhi, Pinal Trivedi, Maharshi Pandya, Afzal Ansari, Bhavesh Modi, Gaurishankar Shrimali, R D Dixit, A M Kadri, Umang Mishra, Chaitanya Joshi, Madhvi Joshi, , , , , |
| EPI_ISL_458116, EPI_ISL_458117, EPI_ISL_458118, EPI_ISL_458119, EPI_ISL_458120, EPI_ISL_458121, EPI_ISL_458122, EPI_ISL_458123, EPI_ISL_458124, EPI_ISL_458125, EPI_ISL_458126, EPI_ISL_458127, EPI_ISL_458128 | | | |
| see above | Oman National Influenza Centre | Department of Microbiology and Immunology-SQUH | Fahad Zadjali, Samira Al-Maruqi, Amina Al Jardani, Khulood Al-Mammary, Hanan Al-kindi, Fatma BaAlawi, Hamida AL Barwani, Zeyana AL-Dahmani, Intisar Al-Shukri, Aisha Al-Busaidi, Aisha Al-Amri, Ahlam Al-Amri, Mohammed Al-Tobi, Samiha Al Kharusi, Abdulla Balkhair |
| EPI_ISL_458130, EPI_ISL_458131 | Hospital Universitari Vall d'Hebron - Vall d'Hebron Institut de Recerca | Hospital Universitari Vall d'Hebron | Cristina Andrés, Maria Piñana, Damir Garcia-Cehic, Mercedes Guerrero-Murillo, Ariadna Rando, Josep Gregori, Juliana Esperalba, Maria Gema Codina, Maria Carmen Martin, Tomàs Pumarola, Josep Quer, Andrés Antón |
| EPI_ISL_458132 | Hospital Universitari Vall d'Hebron - Vall d'Hebron Institut de Rercerca | Hospital Universitari Vall d'Hebron | Cristina Andrés, Maria Piñana, Damir Garcia-Cehic, Mercedes Guerrero-Murillo, Ariadna Rando, Josep Gregori, Juliana Esperalba, Maria Gema Codina, Maria Carmen Martin, Tomàs Pumarola, Josep Quer, Andrés Antón |
| EPI_ISL_458133 | National Institute of Biotechnology | Bioinformatics Division, National Institute of Biotechnology | Mohammad Uzzal Hossain, Md. Moniruzzaman, Md. Salim Khan, Md. Nazrul Islam, Md. Hadisur Rahman, Arittra Bhattacharjee, Md. Ruhul Amin, Asif Rashid, Chaman Ara Keya, Keshob Chandra Das, Md. Salimullah |
| EPI_ISL_458137 | Oman National Influenza Centre | Department of Microbiology and | Fahad Zadjali, Samira Al-Maruqi, Amina Al Jardani, Khulood Al-Mammary, Hanan Al-kindi, Fatma BaAlawi, Hamida AL Barwani, Zeyana AL-Dahmani, Intisar Al-Shukri, Aisha Al-Busaidi, Aisha Al-Amri, Ahlam Al-Amri, Mohammed Al-Tobi, Samiha Al Kharusi, Abdulla |

| | | | |
|---|---|---|---|
| | | Immunology | Balkhair |
| EPI_ISL_458138, EPI_ISL_458139, EPI_ISL_458140, EPI_ISL_458141, EPI_ISL_458142, EPI_ISL_458143, EPI_ISL_458144, EPI_ISL_458145, EPI_ISL_458146, EPI_ISL_458147, EPI_ISL_458148, EPI_ISL_458149 | | | |
| see above | Evandro Chagas Institute | Evandro Chagas Institute | Santos, M.C.; Silva, A.M.; Junior, W.D.C.; Barbagelata, L.S.; Ferreira, J.A.; Sousa, E.M.A.; da Silva, P.S.; Resque, H.R; Martins, L.C.; Sousa Junior, E.C.;Viana, G.M.R |
| EPI_ISL_458150 | ANOUAL | ANOUAL | Jouali Farah, El Ansari Fatima Zahra, Marchoudi Nabila, Kasmi Yassine, Chenaoui Mohamed, El Aliani Aissam, Benhida Rachid, Azami Nawfel, Kitane Driss Lahlou, Loukman Salma, Fekkak Jamal |
| EPI_ISL_458156, EPI_ISL_458157, EPI_ISL_458158, EPI_ISL_458159, EPI_ISL_458160, EPI_ISL_458161, EPI_ISL_458162, EPI_ISL_458163, EPI_ISL_458164, EPI_ISL_458165, EPI_ISL_458166, EPI_ISL_458167, EPI_ISL_458168, EPI_ISL_458169, EPI_ISL_458170, EPI_ISL_458171, EPI_ISL_458172, EPI_ISL_458173, EPI_ISL_458174, EPI_ISL_458175, EPI_ISL_458176, EPI_ISL_458177, EPI_ISL_458178, EPI_ISL_458179, EPI_ISL_458180, EPI_ISL_458181, EPI_ISL_458182, EPI_ISL_458183, EPI_ISL_458184, EPI_ISL_458185, EPI_ISL_458186, EPI_ISL_458187, EPI_ISL_458188, EPI_ISL_458189, EPI_ISL_458190, EPI_ISL_458191, EPI_ISL_458192, EPI_ISL_458193, EPI_ISL_458194, EPI_ISL_458195, EPI_ISL_458196, EPI_ISL_458197, EPI_ISL_458198, EPI_ISL_458199, EPI_ISL_458200, EPI_ISL_458201, EPI_ISL_458202, EPI_ISL_458203, EPI_ISL_458204, EPI_ISL_458205, EPI_ISL_458206, EPI_ISL_458207, EPI_ISL_458208, EPI_ISL_458209, EPI_ISL_458210, EPI_ISL_458211, EPI_ISL_458212, EPI_ISL_458213, EPI_ISL_458214, EPI_ISL_458215, EPI_ISL_458216, EPI_ISL_458217, EPI_ISL_458218, EPI_ISL_458219, EPI_ISL_458220, EPI_ISL_458221, EPI_ISL_458222, EPI_ISL_458223, EPI_ISL_458224, EPI_ISL_458225, EPI_ISL_458226, EPI_ISL_458227, EPI_ISL_458228, EPI_ISL_458229, EPI_ISL_458230, EPI_ISL_458231, EPI_ISL_458232, EPI_ISL_458233, EPI_ISL_458234, EPI_ISL_458235 | | | |
| see above | KU Leuven, Rega Institute, Clinical and Epidemiological Virology | KU Leuven, Rega Institute, Clinical and Epidemiological Virology | Tony Wawina-Bokalanga, Bert Vanmechelen, Joan Marti-Carerras, Piet Maes |
| EPI_ISL_458236 | Hospital Mexico | Charité Virology-University of Costa Rica | Andres Moreira-Soto, Eugenia Corrales-Aguilar, Teresita Somogyi, Jan Felix Drexler |
| EPI_ISL_458237, EPI_ISL_458238, EPI_ISL_458239, EPI_ISL_458240, EPI_ISL_458241, EPI_ISL_458242, EPI_ISL_458243, EPI_ISL_458244, EPI_ISL_458245, EPI_ISL_458246, EPI_ISL_458247, EPI_ISL_458248, EPI_ISL_458249, EPI_ISL_458250, EPI_ISL_458251, EPI_ISL_458252, EPI_ISL_458253, EPI_ISL_458254, EPI_ISL_458255, EPI_ISL_458256, EPI_ISL_458257, EPI_ISL_458258, EPI_ISL_458259, EPI_ISL_458260, EPI_ISL_458261, EPI_ISL_458262, EPI_ISL_458263, EPI_ISL_458264, EPI_ISL_458265, EPI_ISL_458266, EPI_ISL_458267, EPI_ISL_458268, EPI_ISL_458269, EPI_ISL_458270, EPI_ISL_458271, EPI_ISL_458272, EPI_ISL_458273, EPI_ISL_458274, EPI_ISL_458275, EPI_ISL_458276, EPI_ISL_458277, EPI_ISL_458278, EPI_ISL_458279, EPI_ISL_458280, EPI_ISL_458281, EPI_ISL_458282, EPI_ISL_458283, EPI_ISL_458284 | | | |
| see above | Scripps Medical Laboratory | Andersen lab at Scripps Research | SEARCH Alliance San Diego with Michael Quigley, Ellen Stefanski, Ian Mchardy |
| EPI_ISL_458285, EPI_ISL_458286 | unknown | Bundeswehr Institute of Microbiology | Handrick,S., Bestehorn-Willmann,M.S., Eckstein,S., Walter,M.C., Antwerpen,M.H., Rehn,A., Naija,H., Stoecker,K., Woelfel,R. and Ben Moussa,M. |
| EPI_ISL_458287 | Biosafety Department PCL3 | Biosafety Department PCL3 | Lemriss,S., Souiri,A. and El Kabbaj,S. |
| EPI_ISL_458291 | Dirk Dittmer | Dirk Dittmer | Bailey,A.G., Caro-Vegas,C.P., Dittmer,D., Eason,A.B., Juarez,A., Landis,J.T., McNamara,R.P., Miller,M.B., Moorad,R., Pluta,L.J., Seltzer,T.A., Thompson,C., Vahrson,W., Villamor,F. |
| EPI_ISL_458292, EPI_ISL_458293 | Dirk Dittmer | Dirk Dittmer | Aubrey,B.G., Caro-Vegas,C.P., Dittmer,D., Eason,A.B., Juarez,A., Landis,J.T., Mcnamara,R.P., Miller,M.B., Moorad,R., Pluta,L.J., Seltzer,T.A., Thompson,C., Vahrson,W., Villamor,F. |
| EPI_ISL_458294, EPI_ISL_458295, EPI_ISL_458296, EPI_ISL_458297 | Dirk Dittmer | Dirk Dittmer | Bailey,A.G., Caro-Vegas,C.P., Dittmer,D., Eason,A.B., Juarez,A., Landis,J.T., McNamara,R.P., Miller,M.B., Moorad,R., Pluta,L.J., Seltzer,T.A., Thompson,C., Vahrson,W., Villamor,F. |
| EPI_ISL_458298 | CSIR-Centre for Cellular and Molecular Biology | CSIR-Centre for Cellular and Molecular Biology | Sakshi Shambhavi, Lamuk Zaveri, Shagufta Khan, Namami Gaur, Tulasi Nagabandi, Purushotham Vodnala, Payel Mukherjee, Sofia Banu, Priya Singh, Dhiviya Vedagiri, Divya Gupta, Vishal Sah, Santosh Kumar Kuncha, Krishnan Harinivas Harshan, Archana Bharadwaj Siva, Karthik Bharadwaj Tallapaka, Deepak Kumar, Devi Prasad Vijayashankar, Disha Nanda, Divya Das, Jotin Gogoi, Manish Bhattacharjee, Rakesh K Mishra, Divya Tej Sowpati |
| EPI_ISL_458299, EPI_ISL_458300, EPI_ISL_458301, EPI_ISL_458302, EPI_ISL_458303, EPI_ISL_458304, EPI_ISL_458305, EPI_ISL_458306, EPI_ISL_458307, EPI_ISL_458308, EPI_ISL_458309, EPI_ISL_458310, EPI_ISL_458311, EPI_ISL_458312, EPI_ISL_458313, EPI_ISL_458314, EPI_ISL_458315, EPI_ISL_458316, EPI_ISL_458317, EPI_ISL_458318, EPI_ISL_458319, EPI_ISL_458320, EPI_ISL_458321, EPI_ISL_458322, EPI_ISL_458323, EPI_ISL_458324, EPI_ISL_458325, EPI_ISL_458326, EPI_ISL_458327, EPI_ISL_458328, EPI_ISL_458329, EPI_ISL_458330, EPI_ISL_458331, EPI_ISL_458332, EPI_ISL_458333, EPI_ISL_458334, EPI_ISL_458335, EPI_ISL_458336, EPI_ISL_458337, EPI_ISL_458338, EPI_ISL_458339, EPI_ISL_458340, EPI_ISL_458341, EPI_ISL_458342, EPI_ISL_458343, EPI_ISL_458344, EPI_ISL_458345, EPI_ISL_458346, EPI_ISL_458347, EPI_ISL_458348, EPI_ISL_458349, EPI_ISL_458350, EPI_ISL_458351, EPI_ISL_458352, EPI_ISL_458353, EPI_ISL_458354, EPI_ISL_458355, EPI_ISL_458356, EPI_ISL_458357, EPI_ISL_458358, EPI_ISL_458359, EPI_ISL_458360, EPI_ISL_458361, EPI_ISL_458362, EPI_ISL_458363, EPI_ISL_458364, EPI_ISL_458365, EPI_ISL_458366, EPI_ISL_458367, EPI_ISL_458368, EPI_ISL_458369, EPI_ISL_458370, EPI_ISL_458371, EPI_ISL_458372, EPI_ISL_458373, EPI_ISL_458374, EPI_ISL_458375, EPI_ISL_458376, EPI_ISL_458377, EPI_ISL_458378, EPI_ISL_458379, EPI_ISL_458380, EPI_ISL_458381, EPI_ISL_458382, EPI_ISL_458383, EPI_ISL_458384, EPI_ISL_458385, EPI_ISL_458386, EPI_ISL_458387, EPI_ISL_458388, EPI_ISL_458389, EPI_ISL_458390, EPI_ISL_458391, EPI_ISL_458392, EPI_ISL_458393, EPI_ISL_458394, EPI_ISL_458395, EPI_ISL_458396, EPI_ISL_458397, EPI_ISL_458398, EPI_ISL_458399, EPI_ISL_458400, EPI_ISL_458401, EPI_ISL_458402, EPI_ISL_458403, EPI_ISL_458404, EPI_ISL_458405, EPI_ISL_458406, EPI_ISL_458407, EPI_ISL_458408, EPI_ISL_458409, EPI_ISL_458410, EPI_ISL_458411, EPI_ISL_458412, EPI_ISL_458413, EPI_ISL_458414, EPI_ISL_458415, EPI_ISL_458416, EPI_ISL_458417, EPI_ISL_458418, EPI_ISL_458419, EPI_ISL_458420, EPI_ISL_458421, EPI_ISL_458422, EPI_ISL_458423, EPI_ISL_458424, EPI_ISL_458425, EPI_ISL_458426, EPI_ISL_458427, EPI_ISL_458428, EPI_ISL_458429, EPI_ISL_458430, EPI_ISL_458431, EPI_ISL_458432, EPI_ISL_458433, EPI_ISL_458434, EPI_ISL_458435, EPI_ISL_458436, EPI_ISL_458437, EPI_ISL_458438, EPI_ISL_458439, EPI_ISL_458440, EPI_ISL_458441, EPI_ISL_458442, EPI_ISL_458443, EPI_ISL_458444, EPI_ISL_458445, EPI_ISL_458446, EPI_ISL_458447, EPI_ISL_458448, EPI_ISL_458449, EPI_ISL_458450, EPI_ISL_458451, EPI_ISL_458452, EPI_ISL_458453, EPI_ISL_458454, EPI_ISL_458455, EPI_ISL_458456, EPI_ISL_458457, EPI_ISL_458458, EPI_ISL_458459, EPI_ISL_458460, EPI_ISL_458461, EPI_ISL_458462, EPI_ISL_458463, EPI_ISL_458464, EPI_ISL_458465, EPI_ISL_458466, EPI_ISL_458467, EPI_ISL_458468, EPI_ISL_458469, EPI_ISL_458470, EPI_ISL_458471, EPI_ISL_458472, EPI_ISL_458473, EPI_ISL_458474, EPI_ISL_458475, EPI_ISL_458476, EPI_ISL_458477, EPI_ISL_458478, EPI_ISL_458479, EPI_ISL_458480, EPI_ISL_458481, EPI_ISL_458482, EPI_ISL_458483, EPI_ISL_458484, EPI_ISL_458485, EPI_ISL_458486, EPI_ISL_458487, EPI_ISL_458488, EPI_ISL_458489, EPI_ISL_458490, EPI_ISL_458491, EPI_ISL_458492, EPI_ISL_458493, EPI_ISL_458494, EPI_ISL_458495, EPI_ISL_458496, EPI_ISL_458497, EPI_ISL_458498, EPI_ISL_458499, EPI_ISL_458500, EPI_ISL_458501, EPI_ISL_458502, EPI_ISL_458503, EPI_ISL_458504, EPI_ISL_458505, EPI_ISL_458506, EPI_ISL_458507, EPI_ISL_458508, EPI_ISL_458509, EPI_ISL_458510, EPI_ISL_458511, EPI_ISL_458512, EPI_ISL_458513, EPI_ISL_458514, EPI_ISL_458515 | | | |
| see above | PHE South West Regional Laboratory, National Infection Service | Wellcome Sanger Institute for the COVID-19 Genomics UK Consortium | Stephanie Hutchings, Hannah Pymont, Dr Peter Muir, Barry Vipond, Rich Hopes; and Alex Alderton, Roberto Amato, Sonia Goncalves, Ewan Harrison, David K. Jackson, Ian Johnston, Dominic Kwiatkowski, Cordelia Langford, John Sillitoe on behalf of the Wellcome Sanger Institute COVID-19 Surveillance Team (http://www.sanger.ac.uk/covid-team) |
| EPI_ISL_458516, EPI_ISL_458517, EPI_ISL_458518, EPI_ISL_458519, EPI_ISL_458520, EPI_ISL_458521, EPI_ISL_458522, EPI_ISL_458523, EPI_ISL_458524, EPI_ISL_458525, EPI_ISL_458526, EPI_ISL_458527, EPI_ISL_458528, EPI_ISL_458529, EPI_ISL_458530, EPI_ISL_458531, EPI_ISL_458532, EPI_ISL_458533, EPI_ISL_458534, EPI_ISL_458535, EPI_ISL_458536, EPI_ISL_458537, EPI_ISL_458538, EPI_ISL_458539, EPI_ISL_458540, EPI_ISL_458541, EPI_ISL_458542, EPI_ISL_458543, EPI_ISL_458544, EPI_ISL_458545, EPI_ISL_458546, EPI_ISL_458547, EPI_ISL_458548, EPI_ISL_458549, EPI_ISL_458550, EPI_ISL_458551, EPI_ISL_458552, EPI_ISL_458553, EPI_ISL_458554, EPI_ISL_458555, EPI_ISL_458556, EPI_ISL_458557, EPI_ISL_458558, EPI_ISL_458559, EPI_ISL_458560, EPI_ISL_458561, EPI_ISL_458562, EPI_ISL_458563, EPI_ISL_458564, EPI_ISL_458565, EPI_ISL_458566, EPI_ISL_458567, EPI_ISL_458568, EPI_ISL_458569, EPI_ISL_458570, EPI_ISL_458571, EPI_ISL_458572, EPI_ISL_458573, EPI_ISL_458574, EPI_ISL_458575, EPI_ISL_458576, EPI_ISL_458577 | | | |
| see above | Department of Pathology, University of Cambridge | Wellcome Sanger Institute for the COVID-19 Genomics UK Consortium | Luke W Meredith, M. Estée Török , Myra Hosmillo, William L. Hamilton, Martin D. Curran, Theresa Feltwell, Grant Hall, Anna Yakovleva, Fahad A Khokhar, Charlotte J. Houldcroft, Laura G Caller, Aminu S. Jahun, Sarah L. Caddy, Ian Goodfellow; and Alex Alderton, Roberto Amato, Sonia Goncalves, Ewan Harrison, David K. Jackson, Ian Johnston, Dominic Kwiatkowski, Cordelia Langford, John Sillitoe on behalf of the Wellcome Sanger Institute COVID-19 Surveillance Team (http://www.sanger.ac.uk/covid-team) |
| EPI_ISL_458578, EPI_ISL_458579, EPI_ISL_458580, EPI_ISL_458581, EPI_ISL_458582, EPI_ISL_458583, EPI_ISL_458584, EPI_ISL_458585, EPI_ISL_458586, EPI_ISL_458587, EPI_ISL_458588, EPI_ISL_458589, EPI_ISL_458590, EPI_ISL_458591, EPI_ISL_458592, EPI_ISL_458593, EPI_ISL_458594, EPI_ISL_458595, EPI_ISL_458596, EPI_ISL_458597, EPI_ISL_458598, EPI_ISL_458599, EPI_ISL_458600, EPI_ISL_458601, EPI_ISL_458602, EPI_ISL_458603, EPI_ISL_458604, EPI_ISL_458605, EPI_ISL_458606, EPI_ISL_458607, EPI_ISL_458608, EPI_ISL_458609, EPI_ISL_458610, EPI_ISL_458611, EPI_ISL_458612, EPI_ISL_458613, EPI_ISL_458614, EPI_ISL_458615, EPI_ISL_458616, EPI_ISL_458617, EPI_ISL_458618, EPI_ISL_458619, EPI_ISL_458620, EPI_ISL_458621, EPI_ISL_458622, EPI_ISL_458623, EPI_ISL_458624, EPI_ISL_458625, EPI_ISL_458626, EPI_ISL_458627, EPI_ISL_458628, EPI_ISL_458629, EPI_ISL_458630, EPI_ISL_458631, EPI_ISL_458632, EPI_ISL_458633, EPI_ISL_458634, EPI_ISL_458635, EPI_ISL_458636, EPI_ISL_458637, EPI_ISL_458638, EPI_ISL_458639, EPI_ISL_458640, EPI_ISL_458641, EPI_ISL_458642, EPI_ISL_458643, EPI_ISL_458644, EPI_ISL_458645, EPI_ISL_458646, EPI_ISL_458647, EPI_ISL_458648, EPI_ISL_458649, EPI_ISL_458650, EPI_ISL_458651, EPI_ISL_458652, EPI_ISL_458653, EPI_ISL_458654, EPI_ISL_458655, EPI_ISL_458656, EPI_ISL_458657, EPI_ISL_458658, EPI_ISL_458659, EPI_ISL_458660, EPI_ISL_458661, EPI_ISL_458662, EPI_ISL_458663, EPI_ISL_458664, EPI_ISL_458665, EPI_ISL_458666, EPI_ISL_458667, EPI_ISL_458668, EPI_ISL_458669, EPI_ISL_458670, EPI_ISL_458671, EPI_ISL_458672, EPI_ISL_458673, EPI_ISL_458674, EPI_ISL_458675, EPI_ISL_458676, EPI_ISL_458677, EPI_ISL_458678, EPI_ISL_458679, EPI_ISL_458680, EPI_ISL_458681, EPI_ISL_458682, EPI_ISL_458683, EPI_ISL_458684, EPI_ISL_458685, EPI_ISL_458686, EPI_ISL_458687, EPI_ISL_458688, EPI_ISL_458689, EPI_ISL_458690, EPI_ISL_458691, EPI_ISL_458692, EPI_ISL_458693, EPI_ISL_458694, EPI_ISL_458695, EPI_ISL_458696, EPI_ISL_458697, EPI_ISL_458698, EPI_ISL_458699, EPI_ISL_458700, EPI_ISL_458701, EPI_ISL_458702, EPI_ISL_458703, EPI_ISL_458704, EPI_ISL_458705, EPI_ISL_458706, EPI_ISL_458707, EPI_ISL_458708, EPI_ISL_458709, EPI_ISL_458710, EPI_ISL_458711, EPI_ISL_458712, EPI_ISL_458713, EPI_ISL_458714, EPI_ISL_458715, EPI_ISL_458716, EPI_ISL_458717, EPI_ISL_458718 | | | |
| see above | NU-OMICS DNA Sequencing research facility, Northumbria University | Wellcome Sanger Institute for the COVID-19 Genomics UK Consortium | Chris Duncan, Sheia Waugh, Shirelle Burton-Fanning, Gary Eltringham, Jennifer Collins, Brendan Payne, Yusri Taha, Emma Swindells, Jane Greenaway, Edward Barton, Garren Scott, Debra Padgett, Clive Graham, Sarah Essex, Steve Ligget, Paul Baker, Lynn Dover, Wen Yew, Gary Black, John Allan, Joshua Loh, Greg Young, Matthew Bashton, Andrew Nelson, Darren Smith and Alex Alderton, Roberto Amato, Sonia Goncalves, Ewan Harrison, David K. Jackson, Ian Johnston, Dominic Kwiatkowski, Cordelia Langford, John Sillitoe on behalf of the Wellcome Sanger Institute COVID-19 Surveillance Team (http://www.sanger.ac.uk/covid-team) |
| EPI_ISL_458719, EPI_ISL_458720, EPI_ISL_458721, EPI_ISL_458722, EPI_ISL_458723, EPI_ISL_458724, EPI_ISL_458725, EPI_ISL_458726, EPI_ISL_458727, EPI_ISL_458728, EPI_ISL_458729, EPI_ISL_458730, EPI_ISL_458731, EPI_ISL_458732, EPI_ISL_458733, EPI_ISL_458734, EPI_ISL_458735, EPI_ISL_458736, EPI_ISL_458737, EPI_ISL_458738, EPI_ISL_458739, EPI_ISL_458740, EPI_ISL_458741, EPI_ISL_458742, EPI_ISL_458743, EPI_ISL_458744, EPI_ISL_458745, EPI_ISL_458746, EPI_ISL_458747, EPI_ISL_458748, EPI_ISL_458749, EPI_ISL_458750, EPI_ISL_458751, EPI_ISL_458752, EPI_ISL_458753, EPI_ISL_458754, EPI_ISL_458755, EPI_ISL_458756, EPI_ISL_458757, EPI_ISL_458758, EPI_ISL_458759, EPI_ISL_458760, EPI_ISL_458761, EPI_ISL_458762, EPI_ISL_458763, EPI_ISL_458764, EPI_ISL_458765, EPI_ISL_458766, EPI_ISL_458767, EPI_ISL_458768, EPI_ISL_458769, EPI_ISL_458770, EPI_ISL_458771, EPI_ISL_458772, EPI_ISL_458773, EPI_ISL_458774, EPI_ISL_458775, EPI_ISL_458776, EPI_ISL_458777, EPI_ISL_458778, EPI_ISL_458779, EPI_ISL_458780, EPI_ISL_458781, EPI_ISL_458782, EPI_ISL_458783, EPI_ISL_458784, EPI_ISL_458785, EPI_ISL_458786, EPI_ISL_458787, EPI_ISL_458788, EPI_ISL_458789, EPI_ISL_458790, EPI_ISL_458791, EPI_ISL_458792, EPI_ISL_458793, EPI_ISL_458794, EPI_ISL_458795, EPI_ISL_458796, EPI_ISL_458797, EPI_ISL_458798, EPI_ISL_458799, EPI_ISL_458800, EPI_ISL_458801, EPI_ISL_458802, EPI_ISL_458803, EPI_ISL_458804, EPI_ISL_458805, EPI_ISL_458806, EPI_ISL_458807, EPI_ISL_458808, EPI_ISL_458809, EPI_ISL_458810, EPI_ISL_458811, EPI_ISL_458812, EPI_ISL_458813, EPI_ISL_458814, EPI_ISL_458815, EPI_ISL_458816, EPI_ISL_458817, EPI_ISL_458818, EPI_ISL_458819, EPI_ISL_458820, EPI_ISL_458821, EPI_ISL_458822, EPI_ISL_458823, EPI_ISL_458824, EPI_ISL_458825, EPI_ISL_458826, EPI_ISL_458827, EPI_ISL_458828, EPI_ISL_458829, EPI_ISL_458830, EPI_ISL_458831, EPI_ISL_458832, EPI_ISL_458833, EPI_ISL_458834, EPI_ISL_458835, EPI_ISL_458836, EPI_ISL_458837, EPI_ISL_458838, EPI_ISL_458839, EPI_ISL_458840, EPI_ISL_458841, EPI_ISL_458842, EPI_ISL_458843, EPI_ISL_458844, EPI_ISL_458845, EPI_ISL_458846, EPI_ISL_458847, EPI_ISL_458848, EPI_ISL_458849, EPI_ISL_458850, EPI_ISL_458851, EPI_ISL_458852, EPI_ISL_458853, EPI_ISL_458854, EPI_ISL_458855, EPI_ISL_458856, EPI_ISL_458857, EPI_ISL_458858, EPI_ISL_458859, EPI_ISL_458860, EPI_ISL_458861, EPI_ISL_458862, EPI_ISL_458863, EPI_ISL_458864, EPI_ISL_458865, EPI_ISL_458866, EPI_ISL_458867, EPI_ISL_458868, EPI_ISL_458869, EPI_ISL_458870, EPI_ISL_458871, EPI_ISL_458872, EPI_ISL_458873, EPI_ISL_458874, EPI_ISL_458875, EPI_ISL_458876, EPI_ISL_458877, EPI_ISL_458878, EPI_ISL_458879, EPI_ISL_458880, EPI_ISL_458881, EPI_ISL_458882, EPI_ISL_458883, EPI_ISL_458884, EPI_ISL_458885, EPI_ISL_458886, EPI_ISL_458887, EPI_ISL_458888, EPI_ISL_458889, EPI_ISL_458890, EPI_ISL_458891, EPI_ISL_458892, EPI_ISL_458893, EPI_ISL_458894, EPI_ISL_458895, EPI_ISL_458896, EPI_ISL_458897, EPI_ISL_458898, EPI_ISL_458899, EPI_ISL_458900, EPI_ISL_458901, EPI_ISL_458902, EPI_ISL_458903, EPI_ISL_458904, EPI_ISL_458905, EPI_ISL_458906, EPI_ISL_458907, EPI_ISL_458908, EPI_ISL_458909, EPI_ISL_458910, EPI_ISL_458911, EPI_ISL_458912, EPI_ISL_458913, EPI_ISL_458914, EPI_ISL_458915, EPI_ISL_458916, EPI_ISL_458917, EPI_ISL_458918, EPI_ISL_458919, EPI_ISL_458920, EPI_ISL_458921, EPI_ISL_458922, EPI_ISL_458923, EPI_ISL_458924, EPI_ISL_458925, EPI_ISL_458926, EPI_ISL_458927, EPI_ISL_458928, EPI_ISL_458929, EPI_ISL_458930, EPI_ISL_458931, EPI_ISL_458932, EPI_ISL_458933, EPI_ISL_458934, EPI_ISL_458935, EPI_ISL_458936, EPI_ISL_458937, EPI_ISL_458938, EPI_ISL_458939, EPI_ISL_458940, EPI_ISL_458941, EPI_ISL_458942, EPI_ISL_458943, EPI_ISL_458944, EPI_ISL_458945, EPI_ISL_458946, EPI_ISL_458947, EPI_ISL_458948, EPI_ISL_458949, EPI_ISL_458950, EPI_ISL_458951, EPI_ISL_458952, EPI_ISL_458953, EPI_ISL_458954, EPI_ISL_458955, EPI_ISL_458956, EPI_ISL_458957, EPI_ISL_458958, EPI_ISL_458959, EPI_ISL_458960, EPI_ISL_458961, EPI_ISL_458962, EPI_ISL_458963, EPI_ISL_458964, EPI_ISL_458965, EPI_ISL_458966, EPI_ISL_458967, EPI_ISL_458968, EPI_ISL_458969, EPI_ISL_458970, EPI_ISL_458971, EPI_ISL_458972, EPI_ISL_458973, EPI_ISL_458974, EPI_ISL_458975, EPI_ISL_458976, EPI_ISL_458977, EPI_ISL_458978, EPI_ISL_458979, EPI_ISL_458980, EPI_ISL_458981, EPI_ISL_458982, EPI_ISL_458983, EPI_ISL_458984, EPI_ISL_458985, EPI_ISL_458986, EPI_ISL_458987, EPI_ISL_458988, EPI_ISL_458989, EPI_ISL_458990, EPI_ISL_458991, EPI_ISL_458992, EPI_ISL_458993, EPI_ISL_458994, EPI_ISL_458995, EPI_ISL_458996, EPI_ISL_458997, EPI_ISL_458998, EPI_ISL_458999, EPI_ISL_459000, EPI_ISL_459001, EPI_ISL_459002, EPI_ISL_459003, EPI_ISL_459004, EPI_ISL_459005, EPI_ISL_459006, EPI_ISL_459007, EPI_ISL_459008, EPI_ISL_459009, EPI_ISL_459010, EPI_ISL_459011, EPI_ISL_459012, EPI_ISL_459013, EPI_ISL_459014, EPI_ISL_459015, EPI_ISL_459016, EPI_ISL_459017, EPI_ISL_459018, EPI_ISL_459019, EPI_ISL_459020, EPI_ISL_459021, EPI_ISL_459022, EPI_ISL_459023, EPI_ISL_459024, EPI_ISL_459025, EPI_ISL_459026, EPI_ISL_459027, EPI_ISL_459028, EPI_ISL_459029, EPI_ISL_459030, EPI_ISL_459031, EPI_ISL_459032, EPI_ISL_459033, EPI_ISL_459034, EPI_ISL_459035, EPI_ISL_459036, EPI_ISL_459037, EPI_ISL_459038, EPI_ISL_459039, EPI_ISL_459040, EPI_ISL_459041, EPI_ISL_459042, EPI_ISL_459043, EPI_ISL_459044, EPI_ISL_459045, EPI_ISL_459046, EPI_ISL_459047, EPI_ISL_459048, EPI_ISL_459049, EPI_ISL_459050, EPI_ISL_459051, EPI_ISL_459052, EPI_ISL_459053, EPI_ISL_459054, EPI_ISL_459055, EPI_ISL_459056, EPI_ISL_459057, EPI_ISL_459058, EPI_ISL_459059, EPI_ISL_459060, EPI_ISL_459061, EPI_ISL_459062, EPI_ISL_459063, EPI_ISL_459064, EPI_ISL_459065, EPI_ISL_459066, EPI_ISL_459067, EPI_ISL_459068, EPI_ISL_459069, EPI_ISL_459070, EPI_ISL_459071, EPI_ISL_459072, EPI_ISL_459073, EPI_ISL_459074, EPI_ISL_459075, EPI_ISL_459076, EPI_ISL_459077, EPI_ISL_459078, EPI_ISL_459079, EPI_ISL_459080, EPI_ISL_459081, EPI_ISL_459082, EPI_ISL_459083, EPI_ISL_459084, EPI_ISL_459085, EPI_ISL_459086, EPI_ISL_459087, EPI_ISL_459088, EPI_ISL_459089, EPI_ISL_459090, EPI_ISL_459091, EPI_ISL_459092, EPI_ISL_459093, EPI_ISL_459094 | | | |
| see above | PHE South West Regional Laboratory, National Infection Service | Wellcome Sanger Institute for the COVID-19 Genomics UK Consortium | Stephanie Hutchings, Hannah Pymont, Dr Peter Muir, Barry Vipond, Rich Hopes; and Alex Alderton, Roberto Amato, Sonia Goncalves, Ewan Harrison, David K. Jackson, Ian Johnston, Dominic Kwiatkowski, Cordelia Langford, John Sillitoe on behalf of the Wellcome Sanger Institute COVID-19 Surveillance Team (http://www.sanger.ac.uk/covid-team) |
| EPI_ISL_459095, EPI_ISL_459096, EPI_ISL_459097, EPI_ISL_459098, EPI_ISL_459099, EPI_ISL_459100, EPI_ISL_459101, EPI_ISL_459102, EPI_ISL_459103, EPI_ISL_459104, EPI_ISL_459105, EPI_ISL_459106, EPI_ISL_459107, EPI_ISL_459108, EPI_ISL_459109, EPI_ISL_459110, EPI_ISL_459111, EPI_ISL_459112, EPI_ISL_459113, EPI_ISL_459114, EPI_ISL_459115, EPI_ISL_459116, EPI_ISL_459117, EPI_ISL_459118, EPI_ISL_459119, EPI_ISL_459120, EPI_ISL_459121, EPI_ISL_459122, EPI_ISL_459123, EPI_ISL_459124, EPI_ISL_459125, EPI_ISL_459126, EPI_ISL_459127, EPI_ISL_459128, EPI_ISL_459129, EPI_ISL_459130, EPI_ISL_459131, EPI_ISL_459132, EPI_ISL_459133, EPI_ISL_459134, EPI_ISL_459135, EPI_ISL_459136, EPI_ISL_459137, EPI_ISL_459138, EPI_ISL_459139, EPI_ISL_459140, EPI_ISL_459141, EPI_ISL_459142, EPI_ISL_459143, EPI_ISL_459144, EPI_ISL_459145, EPI_ISL_459146, EPI_ISL_459147, EPI_ISL_459148, EPI_ISL_459149, EPI_ISL_459150, EPI_ISL_459151, EPI_ISL_459152, EPI_ISL_459153, EPI_ISL_459154, EPI_ISL_459155, EPI_ISL_459156, EPI_ISL_459157, EPI_ISL_459158, EPI_ISL_459159, EPI_ISL_459160, EPI_ISL_459161, EPI_ISL_459162, EPI_ISL_459163, EPI_ISL_459164, EPI_ISL_459165 | | | |
| see above | NHSGGC West of Scotland Specialist Virology Centre / MRC-University of Glasgow Centre for Virus Research | Wellcome Sanger Institute for the COVID-19 Genomics UK Consortium | Ana da Silva Filipe, Natasha Johnson, Kathy Smollett, Daniel Mair, Stephen Carmichael, Lily Tong, Jenna Nichols, Elihu Aranday-Cortes, Kirstyn Brunker, Yasmin Parr, Kyriaki Nomikou; Sarah McDonald, Marc Niebel, Patawee Asamaphan; Richard Orton, Joseph Hughes, Sreenu Vattipally, David L Robertson; Alasdair MacLean, Rory Gunson; Kathy Li, Natasha Jesudason, Rajiv Shah, James Shepherd, Antonia Ho, Alice Broos, Emma Thomson and Alex Alderton, Roberto Amato, Sonia Goncalves, Ewan Harrison, David K. Jackson, Ian Johnston, Dominic Kwiatkowski, Cordelia Langford, John Sillitoe on behalf of the Wellcome Sanger Institute COVID-19 Surveillance Team (http://www.sanger.ac.uk/covid-team) |
| EPI_ISL_459166, EPI_ISL_459167, EPI_ISL_459168, EPI_ISL_459169, EPI_ISL_459170, EPI_ISL_459171, EPI_ISL_459172, EPI_ISL_459173, EPI_ISL_459174, EPI_ISL_459175, EPI_ISL_459176, EPI_ISL_459177, EPI_ISL_459178, EPI_ISL_459179, EPI_ISL_459180, EPI_ISL_459181, EPI_ISL_459182, EPI_ISL_459183, EPI_ISL_459184, EPI_ISL_459185, EPI_ISL_459186, EPI_ISL_459187, EPI_ISL_459188, EPI_ISL_459189, EPI_ISL_459190, EPI_ISL_459191, EPI_ISL_459192, EPI_ISL_459193, EPI_ISL_459194, EPI_ISL_459195, EPI_ISL_459196, EPI_ISL_459197, EPI_ISL_459198, EPI_ISL_459199, EPI_ISL_459200, EPI_ISL_459201, EPI_ISL_459202, EPI_ISL_459203, EPI_ISL_459204, EPI_ISL_459205, EPI_ISL_459206, EPI_ISL_459207, EPI_ISL_459208, EPI_ISL_459209, EPI_ISL_459210, EPI_ISL_459211, EPI_ISL_459212, EPI_ISL_459213, EPI_ISL_459214, EPI_ISL_459215, EPI_ISL_459216, EPI_ISL_459217, EPI_ISL_459218, EPI_ISL_459219, EPI_ISL_459220, EPI_ISL_459221, EPI_ISL_459222, EPI_ISL_459223, EPI_ISL_459224, EPI_ISL_459225, EPI_ISL_459226, EPI_ISL_459227, EPI_ISL_459228, EPI_ISL_459229, EPI_ISL_459230, EPI_ISL_459231, EPI_ISL_459232, EPI_ISL_459233, EPI_ISL_459234, EPI_ISL_459235, EPI_ISL_459236, EPI_ISL_459237, EPI_ISL_459238, EPI_ISL_459239, EPI_ISL_459240, EPI_ISL_459241, EPI_ISL_459242, EPI_ISL_459243, EPI_ISL_459244, EPI_ISL_459245, EPI_ISL_459246, EPI_ISL_459247, EPI_ISL_459248, EPI_ISL_459249, EPI_ISL_459250, EPI_ISL_459251, EPI_ISL_459252, EPI_ISL_459253, EPI_ISL_459254, EPI_ISL_459255, EPI_ISL_459256, EPI_ISL_459257, EPI_ISL_459258, EPI_ISL_459259, EPI_ISL_459260, EPI_ISL_459261, EPI_ISL_459262, EPI_ISL_459263, EPI_ISL_459264, EPI_ISL_459265, EPI_ISL_459266, EPI_ISL_459267, EPI_ISL_459268, EPI_ISL_459269, EPI_ISL_459270, EPI_ISL_459271, EPI_ISL_459272, EPI_ISL_459273, EPI_ISL_459274, EPI_ISL_459275, EPI_ISL_459276, EPI_ISL_459277, EPI_ISL_459278, EPI_ISL_459279, EPI_ISL_459280, EPI_ISL_459281, EPI_ISL_459282, EPI_ISL_459283, EPI_ISL_459284, EPI_ISL_459285, EPI_ISL_459286, EPI_ISL_459287, EPI_ISL_459288, EPI_ISL_459289, EPI_ISL_459290, EPI_ISL_459291, EPI_ISL_459292, EPI_ISL_459293, EPI_ISL_459294, EPI_ISL_459295, EPI_ISL_459296, EPI_ISL_459297, EPI_ISL_459304, EPI_ISL_459305, EPI_ISL_459306, EPI_ISL_459307, EPI_ISL_459308, EPI_ISL_459309, EPI_ISL_459310, EPI_ISL_459311, EPI_ISL_459312, EPI_ISL_459313, EPI_ISL_459314, EPI_ISL_459315, EPI_ISL_459316, EPI_ISL_459317, EPI_ISL_459318, EPI_ISL_459319, EPI_ISL_459320, EPI_ISL_459321, EPI_ISL_459322, EPI_ISL_459323 | | | |
| see above | Department of Pathology, University of Cambridge | Wellcome Sanger Institute for the COVID-19 Genomics UK Consortium | Luke W Meredith, M. Estée Török , Myra Hosmillo, William L. Hamilton, Martin D. Curran, Theresa Feltwell, Grant Hall, Anna Yakovleva, Fahad A Khokhar, Charlotte J. Houldcroft, Laura G Caller, Aminu S. Jahun, Sarah L. Caddy, Ian Goodfellow; and Alex Alderton, Roberto Amato, Sonia Goncalves, Ewan Harrison, David K. Jackson, Ian Johnston, Dominic Kwiatkowski, Cordelia Langford, John Sillitoe on behalf of the Wellcome Sanger Institute COVID-19 Surveillance Team (http://www.sanger.ac.uk/covid-team) |
| EPI_ISL_459324, EPI_ISL_459325, EPI_ISL_459326, EPI_ISL_459327, EPI_ISL_459328, EPI_ISL_459329, EPI_ISL_459330, EPI_ISL_459331, EPI_ISL_459332, EPI_ISL_459333, EPI_ISL_459334, EPI_ISL_459335, EPI_ISL_459336, EPI_ISL_459337, EPI_ISL_459338, EPI_ISL_459339, EPI_ISL_459340, EPI_ISL_459341, EPI_ISL_459342, EPI_ISL_459343, EPI_ISL_459344, EPI_ISL_459345, EPI_ISL_459346, EPI_ISL_459347, EPI_ISL_459348, EPI_ISL_459349, EPI_ISL_459350, EPI_ISL_459351, EPI_ISL_459352, EPI_ISL_459353, EPI_ISL_459354, EPI_ISL_459355, EPI_ISL_459356, EPI_ISL_459357, EPI_ISL_459358, EPI_ISL_459359, EPI_ISL_459360, EPI_ISL_459361, EPI_ISL_459362, EPI_ISL_459363, EPI_ISL_459364, EPI_ISL_459365, EPI_ISL_459366, EPI_ISL_459367, EPI_ISL_459368, EPI_ISL_459369, EPI_ISL_459370, EPI_ISL_459371, EPI_ISL_459372, EPI_ISL_459373, EPI_ISL_459374, EPI_ISL_459375, EPI_ISL_459376, EPI_ISL_459377, EPI_ISL_459378, EPI_ISL_459379, EPI_ISL_459380, EPI_ISL_459381, EPI_ISL_459382, EPI_ISL_459383, EPI_ISL_459384, EPI_ISL_459385, EPI_ISL_459386, EPI_ISL_459387, EPI_ISL_459388, EPI_ISL_459389, EPI_ISL_459390, EPI_ISL_459391, EPI_ISL_459392, EPI_ISL_459393, EPI_ISL_459394, EPI_ISL_459395, EPI_ISL_459396, EPI_ISL_459397, EPI_ISL_459398, EPI_ISL_459399, EPI_ISL_459400, EPI_ISL_459401, EPI_ISL_459402, EPI_ISL_459403, EPI_ISL_459404, EPI_ISL_459405, EPI_ISL_459406, EPI_ISL_459407, EPI_ISL_459408, EPI_ISL_459409 | | | |
| see above | Regional Virus Laboratory, Belfast Health and Social Care Trust | Wellcome Sanger Institute for the COVID-19 Genomics UK Consortium | Conall McCaughey, James McKenna, Tanya Curran, Susan Feeney, Alison Watt, Ciara Cox, Mairead Connor, Zoltan Molnar, David Simpson, Derek Fairley; and Alex Alderton, Roberto Amato, Sonia Goncalves, Ewan Harrison, David K. Jackson, Ian Johnston, Dominic Kwiatkowski, Cordelia Langford, John Sillitoe on behalf of the Wellcome Sanger Institute COVID-19 Surveillance Team (http://www.sanger.ac.uk/covid-team) |
| EPI_ISL_459410, EPI_ISL_459411, EPI_ISL_459412, EPI_ISL_459413, EPI_ISL_459414, EPI_ISL_459415, EPI_ISL_459416, EPI_ISL_459417, EPI_ISL_459418, EPI_ISL_459419, EPI_ISL_459420, EPI_ISL_459421, EPI_ISL_459422, EPI_ISL_459423, EPI_ISL_459424, EPI_ISL_459425, EPI_ISL_459426, EPI_ISL_459427, EPI_ISL_459428, EPI_ISL_459429, EPI_ISL_459430, EPI_ISL_459431, EPI_ISL_459432, EPI_ISL_459433, EPI_ISL_459434, EPI_ISL_459435, EPI_ISL_459436, EPI_ISL_459437, EPI_ISL_459438, EPI_ISL_459439, EPI_ISL_459440, EPI_ISL_459441, EPI_ISL_459442, EPI_ISL_459443, EPI_ISL_459444, EPI_ISL_459445, EPI_ISL_459446, EPI_ISL_459447, EPI_ISL_459448, EPI_ISL_459449, EPI_ISL_459450, EPI_ISL_459451, EPI_ISL_459452, EPI_ISL_459453, EPI_ISL_459454, EPI_ISL_459455, EPI_ISL_459456, EPI_ISL_459457, EPI_ISL_459458, EPI_ISL_459459, EPI_ISL_459460, EPI_ISL_459461, EPI_ISL_459462, EPI_ISL_459463, EPI_ISL_459464, EPI_ISL_459465, EPI_ISL_459466, EPI_ISL_459467, EPI_ISL_459468, EPI_ISL_459469, EPI_ISL_459470, EPI_ISL_459471, EPI_ISL_459472, EPI_ISL_459473, EPI_ISL_459474, EPI_ISL_459475, EPI_ISL_459476, EPI_ISL_459477, EPI_ISL_459478, EPI_ISL_459479, EPI_ISL_459480, EPI_ISL_459481, | | | |

| | | | |
|---|---|---|---|
| EPI_ISL_459482, EPI_ISL_459483, EPI_ISL_459484, EPI_ISL_459485, EPI_ISL_459486, EPI_ISL_459487, EPI_ISL_459488, EPI_ISL_459489, EPI_ISL_459490, EPI_ISL_459491, EPI_ISL_459492, EPI_ISL_459493, EPI_ISL_459494, EPI_ISL_459495, EPI_ISL_459496, EPI_ISL_459497, EPI_ISL_459498, EPI_ISL_459499, EPI_ISL_459500, EPI_ISL_459501, EPI_ISL_459502, EPI_ISL_459503, EPI_ISL_459504 | | | |
| see above | Department of Pathology, University of Cambridge | Wellcome Sanger Institute for the COVID-19 Genomics UK Consortium | Luke W Meredith, M. Estée Török , Myra Hosmillo, William L. Hamilton, Martin D. Curran, Theresa Feltwell, Grant Hall, Anna Yakovleva, Fahad A Khokhar, Charlotte J. Houldcroft, Laura G Caller, Aminu S. Jahun, Sarah L. Caddy, Ian Goodfellow; and Alex Alderton, Roberto Amato, Sonia Goncalves, Ewan Harrison, David K. Jackson, Ian Johnston, Dominic Kwiatkowski, Cordelia Langford, John Sillitoe on behalf of the Wellcome Sanger Institute COVID-19 Surveillance Team (http://www.sanger.ac.uk/covid-team) |
| EPI_ISL_459505, EPI_ISL_459506, EPI_ISL_459507, EPI_ISL_459508, EPI_ISL_459509, EPI_ISL_459510, EPI_ISL_459511, EPI_ISL_459512, EPI_ISL_459513, EPI_ISL_459514, EPI_ISL_459515, EPI_ISL_459516, EPI_ISL_459517, EPI_ISL_459518, EPI_ISL_459519, EPI_ISL_459520, EPI_ISL_459521, EPI_ISL_459522, EPI_ISL_459523, EPI_ISL_459524, EPI_ISL_459525, EPI_ISL_459526, EPI_ISL_459527, EPI_ISL_459528, EPI_ISL_459529, EPI_ISL_459530, EPI_ISL_459531, EPI_ISL_459532, EPI_ISL_459533, EPI_ISL_459534, EPI_ISL_459535, EPI_ISL_459536, EPI_ISL_459537, EPI_ISL_459538, EPI_ISL_459539, EPI_ISL_459540, EPI_ISL_459541, EPI_ISL_459542, EPI_ISL_459543, EPI_ISL_459544, EPI_ISL_459545, EPI_ISL_459546, EPI_ISL_459547, EPI_ISL_459548, EPI_ISL_459549, EPI_ISL_459550, EPI_ISL_459551, EPI_ISL_459552, EPI_ISL_459553, EPI_ISL_459554, EPI_ISL_459555, EPI_ISL_459556, EPI_ISL_459557, EPI_ISL_459558, EPI_ISL_459559, EPI_ISL_459560, EPI_ISL_459561, EPI_ISL_459562, EPI_ISL_459563, EPI_ISL_459564, EPI_ISL_459565, EPI_ISL_459566, EPI_ISL_459567, EPI_ISL_459568, EPI_ISL_459569, EPI_ISL_459570, EPI_ISL_459571, EPI_ISL_459572, EPI_ISL_459573, EPI_ISL_459574, EPI_ISL_459575, EPI_ISL_459576, EPI_ISL_459577, EPI_ISL_459578, EPI_ISL_459579, EPI_ISL_459580, EPI_ISL_459581, EPI_ISL_459582, EPI_ISL_459583, EPI_ISL_459584, EPI_ISL_459585, EPI_ISL_459586, EPI_ISL_459587, EPI_ISL_459588, EPI_ISL_459589, EPI_ISL_459590, EPI_ISL_459591, EPI_ISL_459592, EPI_ISL_459593, EPI_ISL_459594, EPI_ISL_459595, EPI_ISL_459596, EPI_ISL_459597, EPI_ISL_459598, EPI_ISL_459599, EPI_ISL_459600, EPI_ISL_459601, EPI_ISL_459602, EPI_ISL_459603, EPI_ISL_459604, EPI_ISL_459605, EPI_ISL_459606, EPI_ISL_459607, EPI_ISL_459608, EPI_ISL_459609, EPI_ISL_459610, EPI_ISL_459611, EPI_ISL_459612, EPI_ISL_459613, EPI_ISL_459614, EPI_ISL_459615, EPI_ISL_459616, EPI_ISL_459617, EPI_ISL_459618, EPI_ISL_459619, EPI_ISL_459620, EPI_ISL_459621, EPI_ISL_459622, EPI_ISL_459623, EPI_ISL_459624, EPI_ISL_459625, EPI_ISL_459626, EPI_ISL_459627, EPI_ISL_459628, EPI_ISL_459629, EPI_ISL_459630, EPI_ISL_459631, EPI_ISL_459632, EPI_ISL_459633, EPI_ISL_459634, EPI_ISL_459635, EPI_ISL_459636, EPI_ISL_459637, EPI_ISL_459638, EPI_ISL_459639, EPI_ISL_459640, EPI_ISL_459641, EPI_ISL_459642, EPI_ISL_459643, EPI_ISL_459644, EPI_ISL_459645, EPI_ISL_459646, EPI_ISL_459647, EPI_ISL_459648, EPI_ISL_459649, EPI_ISL_459650, EPI_ISL_459651, EPI_ISL_459652, EPI_ISL_459653, EPI_ISL_459654, EPI_ISL_459655, EPI_ISL_459656, EPI_ISL_459657, EPI_ISL_459658, EPI_ISL_459659, EPI_ISL_459660, EPI_ISL_459661, EPI_ISL_459662, EPI_ISL_459663, EPI_ISL_459664, EPI_ISL_459665, EPI_ISL_459666, EPI_ISL_459667, EPI_ISL_459668, EPI_ISL_459669, EPI_ISL_459670, EPI_ISL_459671, EPI_ISL_459672, EPI_ISL_459673, EPI_ISL_459674, EPI_ISL_459675, EPI_ISL_459676, EPI_ISL_459677, EPI_ISL_459678, EPI_ISL_459679, EPI_ISL_459680, EPI_ISL_459681, EPI_ISL_459682, EPI_ISL_459683, EPI_ISL_459684, EPI_ISL_459685, EPI_ISL_459686, EPI_ISL_459687, EPI_ISL_459688, EPI_ISL_459689, EPI_ISL_459690, EPI_ISL_459691, EPI_ISL_459692, EPI_ISL_459693, EPI_ISL_459694, EPI_ISL_459695, EPI_ISL_459696, EPI_ISL_459697, EPI_ISL_459698, EPI_ISL_459699, EPI_ISL_459700, EPI_ISL_459701, EPI_ISL_459702, EPI_ISL_459703, EPI_ISL_459704, EPI_ISL_459705, EPI_ISL_459706, EPI_ISL_459707, EPI_ISL_459708, EPI_ISL_459709, EPI_ISL_459710, EPI_ISL_459711, EPI_ISL_459712, EPI_ISL_459713, EPI_ISL_459714, EPI_ISL_459715, EPI_ISL_459716, EPI_ISL_459717, EPI_ISL_459718, EPI_ISL_459719, EPI_ISL_459720, EPI_ISL_459721, EPI_ISL_459722, EPI_ISL_459723, EPI_ISL_459724 | | | |
| see above | NHSGGC West of Scotland Specialist Virology Centre / MRC-University of Glasgow Centre for Virus Research | Wellcome Sanger Institute for the COVID-19 Genomics UK Consortium | Ana da Silva Filipe, Natasha Johnson, Kathy Smollett, Daniel Mair, Stephen Carmichael, Lily Tong, Jenna Nichols, Elihu Aranday-Cortes, Kirstyn Brunker, Yasmin Parr, Kyriaki Nomikou; Sarah McDonald, Marc Niebel, Patawee Asamaphan; Richard Orton, Joseph Hughes, Sreenu Vattipally, David L Robertson; Alasdair MacLean, Rory Gunson; Kathy Li, Natasha Jesudason, Rajiv Shah, James Shepherd, Antonia Ho, Alice Broos, Emma Thomson and Alex Alderton, Roberto Amato, Sonia Goncalves, Ewan Harrison, David K. Jackson, Ian Johnston, Dominic Kwiatkowski, Cordelia Langford, John Sillitoe on behalf of the Wellcome Sanger Institute COVID-19 Surveillance Team (http://www.sanger.ac.uk/covid-team) |
| EPI_ISL_459725, EPI_ISL_459726, EPI_ISL_459727 | PHE South West Regional Laboratory, National Infection Service | Wellcome Sanger Institute for the COVID-19 Genomics UK Consortium | Stephanie Hutchings, Hannah Pymont, Dr Peter Muir, Barry Vipond, Rich Hopes; and Alex Alderton, Roberto Amato, Sonia Goncalves, Ewan Harrison, David K. Jackson, Ian Johnston, Dominic Kwiatkowski, Cordelia Langford, John Sillitoe on behalf of the Wellcome Sanger Institute COVID-19 Surveillance Team (http://www.sanger.ac.uk/covid-team) |
| EPI_ISL_459856, EPI_ISL_459857, EPI_ISL_459858, EPI_ISL_459859, EPI_ISL_459860, EPI_ISL_459861, EPI_ISL_459862, EPI_ISL_459863, EPI_ISL_459864 | Center for Genome Regulation (CRG) | Center for Mathematical Modeling and Center for Genome Regulation. Santiago, Chile | Gaete A, Travisany D, Palma R, Urra C, Varas M, Allende ML, Maass A, González M. |
| EPI_ISL_459866 | Kingston Health Sciences Center | Queen's Genomics Lab at Ongwanada (Q-GLO) | Sjaarda CP, Rustom N, Huang D, Perez-Patrigeon S, Hudson ML, Wong H,Guan H, Ayub M, Soares CN, Colautti R, Evans GA, Sheth P |
| EPI_ISL_459867 | Kingston Health Sciences Center | Queen's Genomics Lab at Ongwanada (Q-GLO) | Sjaarda CP, Rustom N, Huang D, Perez-Patrigeon S, Hudson ML, Wong H,Guan H, Ayub M, Soares CN, Colautti R, Evans GA, Sheth P |
| EPI_ISL_459868, EPI_ISL_459869, EPI_ISL_459871, EPI_ISL_459872, EPI_ISL_459873, EPI_ISL_459874, EPI_ISL_459875, EPI_ISL_459877, EPI_ISL_459878, EPI_ISL_459879, EPI_ISL_459880, EPI_ISL_459881, EPI_ISL_459882, EPI_ISL_459883, EPI_ISL_459884, EPI_ISL_459885, EPI_ISL_459886, EPI_ISL_459887, EPI_ISL_459888, EPI_ISL_459889, EPI_ISL_459890, EPI_ISL_459891, EPI_ISL_459892 | | | |
| see above | Kingston Health Sciences Center | Queen's Genomics Lab at Ongwanada (Q-GLO) | Sjaarda CP, Rustom N, Huang D, Perez-Patrigeon S, Hudson ML, Wong H,Guan H, Ayub M, Soares CN, Colautti R, Evans GA, Sheth P |
| EPI_ISL_459893, EPI_ISL_459894, EPI_ISL_459895, EPI_ISL_459896, EPI_ISL_459897, EPI_ISL_459898, EPI_ISL_459899, EPI_ISL_459900, EPI_ISL_459901, EPI_ISL_459902, EPI_ISL_459903, EPI_ISL_459904, EPI_ISL_459905, EPI_ISL_459906 | | | |
| see above | Laboratoire National de Sante, Microbiology, Virology | Laboratoire National de Sante, Microbiology, Epidemiology and Microbial Genomics | Anke Wienecke-Baldacchino, Jessica Tapp, Guillaume Fournier, Tamir Abdelrahman, Trung Nguyen Nguyen, Catherine Ragimbeau |
| EPI_ISL_459909 | Zoonotic and Exotic infection Diseases Division, Harbin Veterinary Research Institute, CAAS | Zoonotic and Exotic infection Diseases Division, Harbin Veterinary Research Institute, CAAS | Zhigao Bu, Jinliang Wang |
| EPI_ISL_459910 | Zoonotic and Exotic infection Diseases Division, Harbin Veterinary Research Institute, CAAS | Zoonotic and Exotic infection Diseases Division, Harbin Veterinary Resrarch Institute, CAAS | Jinliang Wang, Lei Shuai, Chong Wang, Renqiang Liu, Xijun He, Xianfeng Zhang, Ziruo Sun, Dan Shan, Jinying Ge, Xijun Wang, Gongxun Zhong, Zhiyuan Wen, Zhigao Bu |
| EPI_ISL_459911 | Devki Devi Foundation, a unit of Max Healthcare | CSIR-IGIB/Max | Rajesh Pandey#, Samreen Siddiqui, Pooja Sharma, Bansidhar Tarai, Vivekanand A, Bharathram Uppili, Saruchi Wadhwa, Nishu Tyagi, Mitali Mukerji, Poonam Das, Sujeet Jha, Mohammed Faruq, Vinita Jha, Anurag Agrawal |
| EPI_ISL_459912, EPI_ISL_459913, EPI_ISL_459914, EPI_ISL_459915, EPI_ISL_459916, EPI_ISL_459917, EPI_ISL_459918, EPI_ISL_459919, EPI_ISL_459920, EPI_ISL_459921, EPI_ISL_459922, EPI_ISL_459923, EPI_ISL_459924, EPI_ISL_459925, EPI_ISL_459926, EPI_ISL_459927, EPI_ISL_459928, EPI_ISL_459929, EPI_ISL_459930, EPI_ISL_459931, EPI_ISL_459932, EPI_ISL_459933, EPI_ISL_459934, EPI_ISL_459935, EPI_ISL_459936, EPI_ISL_459937, EPI_ISL_459938, EPI_ISL_459939, EPI_ISL_459940, EPI_ISL_459941, EPI_ISL_459942, EPI_ISL_459943, EPI_ISL_459944, EPI_ISL_459945, EPI_ISL_459946, EPI_ISL_459947, EPI_ISL_459948, EPI_ISL_459949, EPI_ISL_459950, EPI_ISL_459951, EPI_ISL_459952 | | | |
| see above | Devki Devi Foundation, a unit of Max Healthcare | CSIR-IGIB/Max | Rajesh Pandey#, Samreen Siddiqui, Pooja Sharma, Bansidhar Tarai, Vivekanand A, Bharathram Uppili, Saruchi Wadhwa, Nishu Tyagi, Mitali Mukerji, Bansidhar Tarai, Poonam Das, Sujeet Jha, Mohammed Faruq, Vinita Jha, Anurag Agrawal |
| EPI_ISL_459953 | Institute for Medical Research, Infectious Disease Research Centre, National Institutes of Health, Ministry of Health Malaysia | Institute for Medical Research Infectious Disease Research Centre, National Institutes of Health, Ministry of Health Malaysia | Suppiah J, Mohd-Zawawi Z, Kamel KA, Ellan K, Kalyanasundram J, Mohd-Zain R, Thayan R |
| EPI_ISL_459954 | Institute for Medical Research, Infectious Disease Research Centre, National Institutes of Health, Ministry of Health Malaysia | Institute for Medical Research, Infectious Disease Research Centre, National Institutes of Health, Ministry of Health Malaysia | Suppiah J, Mohd-Zawawi Z, Kamel KA, Ellan K, Kalyanasundram J, Mohd-Zain R, Thayan R |
| EPI_ISL_459955, EPI_ISL_459956 | Institute for Medical Research, Infectious Disease Research Centre, National Institutes of Health, Ministry of Health Malaysia | Institute for Medical Research, Infectious Disease Research Centre, National Institutes of Health, Ministry of Health Malaysia | Suppiah J, Mohd-Zawawi Z, Kamel KA, Ellan K, Kalyanasundram J, Mohd-Zain R, Thayan R |
| EPI_ISL_459957 | Institute for Medical Research, Infectious Disease Research Centre, National Institutes of Health, Minis | Institute for Medical Research, Infectious Disease Research Centre, National Institutes of Health, Minis | Suppiah J, Mohd-Zawawi Z, Kamel KA, Ellan K, Kalyanasundram J, Mohd-Zain R, Thayan R |
| EPI_ISL_459958, EPI_ISL_459959, EPI_ISL_459960, EPI_ISL_459961 | Respiratory Virus Unit, Microbiology Services Colindale, Public Health England | Respiratory Virus Unit, Microbiology Services Colindale, Public Health England | Steven Platt, Shahjahan Miah, Angie Lackenby, Omolola Akinbami, Tina Talts, Leena Bhaw, Richard Myers, Monica Galiano, Kirstin Edwards, Jonathan Hubb, Joanna Ellis, Maria Zambon |
| EPI_ISL_459962, EPI_ISL_459963 | Centogene AG | Centogene AG | Prof. Dr. Peter Bauer, Dr. Krishna Kumar Kandaswamy |
| EPI_ISL_459964 | Centogene AG | Centogene AG | Prof. Dr. Peter Bauer, Dr. Krishna Kumar Kandaswamy |
| EPI_ISL_459965, EPI_ISL_459966, EPI_ISL_459967, EPI_ISL_459968, EPI_ISL_459969, EPI_ISL_459970, EPI_ISL_459971, EPI_ISL_459972, EPI_ISL_459973, EPI_ISL_459974, EPI_ISL_459975, EPI_ISL_459976, EPI_ISL_459977, EPI_ISL_459978, EPI_ISL_459979, EPI_ISL_459980, EPI_ISL_459981, EPI_ISL_459982, EPI_ISL_459983, EPI_ISL_459984 | | | |
| see above | Institut Pasteur du Maroc | Institut Pasteur du Maroc | Marion Barbet, Sylvie Behillil, Méline Bizard, Angela Brisebarre, Camille Capel, Etienne Simon-Lorière, Vincent Enouf, Maud Vanpeene, Sylvie van der Werf, Latifa Anga, Abdellah Faouzi, Anass Abbad, Mjid Eloualid, Jalal Nourlil, Anderrahmane Maaroufi |
| EPI_ISL_459992, EPI_ISL_459993, EPI_ISL_459994, EPI_ISL_459995, EPI_ISL_459996, EPI_ISL_459997, EPI_ISL_459998, EPI_ISL_459999, EPI_ISL_460000, EPI_ISL_460001, EPI_ISL_460002, EPI_ISL_460003, EPI_ISL_460004, EPI_ISL_460005, EPI_ISL_460006, EPI_ISL_460007, EPI_ISL_460008, EPI_ISL_460009, EPI_ISL_460010, EPI_ISL_460011, EPI_ISL_460012, EPI_ISL_460013, EPI_ISL_460014, EPI_ISL_460015, EPI_ISL_460016, EPI_ISL_460017, EPI_ISL_460018, EPI_ISL_460019, EPI_ISL_460020, EPI_ISL_460021, EPI_ISL_460022, EPI_ISL_460023, EPI_ISL_460024, EPI_ISL_460025, EPI_ISL_460026, EPI_ISL_460027, EPI_ISL_460028, EPI_ISL_460029, EPI_ISL_460030, EPI_ISL_460031, EPI_ISL_460032, EPI_ISL_460033, EPI_ISL_460034, EPI_ISL_460035, EPI_ISL_460036, EPI_ISL_460037, EPI_ISL_460038, EPI_ISL_460039, EPI_ISL_460040, EPI_ISL_460041, EPI_ISL_460042, EPI_ISL_460043 | | | |
| see above | Michigan Department of Health and Human Services, Bureau of Laboratories | Michigan Department of Health and Human Services, Bureau of Laboratories | Blankenship HM, Riner D, Soehnlen MK |
| EPI_ISL_460045, EPI_ISL_460046, EPI_ISL_460047, EPI_ISL_460048, EPI_ISL_460049, EPI_ISL_460050, EPI_ISL_460051, EPI_ISL_460052, EPI_ISL_460053, EPI_ISL_460054, EPI_ISL_460055, EPI_ISL_460056, EPI_ISL_460057, EPI_ISL_460058, EPI_ISL_460059, EPI_ISL_460060, EPI_ISL_460061, EPI_ISL_460062, EPI_ISL_460063, EPI_ISL_460064, EPI_ISL_460065, EPI_ISL_460066, EPI_ISL_460067, EPI_ISL_460068, EPI_ISL_460069, EPI_ISL_460070, EPI_ISL_460071, EPI_ISL_460072, EPI_ISL_460073, EPI_ISL_460074, EPI_ISL_460075, EPI_ISL_460076, EPI_ISL_460077, EPI_ISL_460078 | | | |
| see above | Minnesota Department of Health, Public Health Laboratory | Minnesota Department of Health, Public Health Laboratory | Matt Plumb, Jacob Garfin, and Xiong Wang |
| EPI_ISL_460079 | Molecular Virology Unit, Fondazione IRCCS Policlinico San Matteo , Pavia | Laboratory of Virology, INMI Lazzaro Spallanzani IRCCS | Barbara Bartolini, Cesare E.M. Gruber, Maria R. Capobianchi, Martina Rueca, Antonio Piralla, Fausto Baldanti, Antonino Di Caro |
| EPI_ISL_460080 | Molecular Virology Unit, Fondazione IRCCS Policlinico San Matteo , Pavia | Laboratory of Virology, INMI Lazzaro Spallanzani IRCCS | Antonio Piralla, Barbara Bartolini, Fausto Baldanti, Martina Rueca, Antonino Di Caro, Cesare E.M. Gruber, Maria R. Capobianchi |
| EPI_ISL_460081 | Molecular Virology Unit, Fondazione IRCCS Policlinico San Matteo , Pavia | Laboratory of Virology, INMI Lazzaro Spallanzani IRCCS | Fausto Baldanti, Martina Rueca, Antonio Piralla, Antonino Di Caro, Maria R. Capobianchi, Cesare E.M. Gruber, Barbara Bartolini |
| EPI_ISL_460082 | Molecular Virology Unit, Fondazione IRCCS Policlinico San Matteo , Pavia | Laboratory of Virology, INMI Lazzaro Spallanzani IRCCS | Martina Rueca, Cesare E.M. Gruber, Antonio Piralla, Antonino Di Caro, Barbara Bartolini, Maria R. Capobianchi, Fausto Baldanti |
| EPI_ISL_460083 | Molecular Virology Unit, Fondazione IRCCS Policlinico San Matteo , Pavia | Laboratory of Virology, INMI Lazzaro Spallanzani IRCCS | Martina Rueca, Antonino Di Caro, Cesare E.M. Gruber, Barbara Bartolini, Fausto Baldanti, Antonio Piralla, Maria R. Capobianchi |
| EPI_ISL_460084 | Molecular Virology Unit, Fondazione IRCCS Policlinico San Matteo , Pavia | Laboratory of Virology, INMI Lazzaro Spallanzani IRCCS | Fausto Baldanti, Antonio Piralla, Martina Rueca, Barbara Bartolini, Maria R. Capobianchi, Cesare E.M. Gruber, Antonino Di Caro |
| EPI_ISL_460085 | Molecular Virology Unit, Fondazione IRCCS Policlinico San | Laboratory of Virology, INMI | Cesare E.M. Gruber, Maria R. Capobianchi, Barbara Bartolini, Fausto Baldanti, Martina Rueca, Antonio Piralla, Antonino Di Caro |

| | | | |
|---|---|---|---|
| | Matteo , Pavia | Lazzaro Spallanzani IRCCS | |
| EPI_ISL_460086 | Molecular Virology Unit, Fondazione IRCCS Policlinico San Matteo , Pavia | Laboratory of Virology, INMI Lazzaro Spallanzani IRCCS | Maria R. Capobianchi, Fausto Baldanti, Antonio Piralla, Antonino Di Caro, Barbara Bartolini, Cesare E.M. Gruber, Martina Rueca |
| EPI_ISL_460087 | Molecular Virology Unit, Fondazione IRCCS Policlinico San Matteo , Pavia | Laboratory of Virology, INMI Lazzaro Spallanzani IRCCS | Cesare E.M. Gruber, Maria R. Capobianchi, Martina Rueca, Barbara Bartolini, Antonino Di Caro, Antonio Piralla, Fausto Baldanti |
| EPI_ISL_460088 | Molecular Virology Unit, Fondazione IRCCS Policlinico San Matteo , Pavia | Laboratory of Virology, INMI Lazzaro Spallanzani IRCCS | Martina Rueca, Barbara Bartolini, Fausto Baldanti, Maria R. Capobianchi, Cesare E.M. Gruber, Antonino Di Caro, Antonio Piralla |
| EPI_ISL_460089 | Molecular Virology Unit, Fondazione IRCCS Policlinico San Matteo , Pavia | Laboratory of Virology, INMI Lazzaro Spallanzani IRCCS | Antonino Di Caro, Barbara Bartolini, Martina Rueca, Cesare E.M. Gruber, Antonio Piralla, Fausto Baldanti, Maria R. Capobianchi |
| EPI_ISL_460090 | Molecular Virology Unit, Fondazione IRCCS Policlinico San Matteo , Pavia | Laboratory of Virology, INMI Lazzaro Spallanzani IRCCS | Antonio Piralla, Cesare E.M. Gruber, Antonino Di Caro, Maria R. Capobianchi, Martina Rueca, Barbara Bartolini, Fausto Baldanti |
| EPI_ISL_460091 | Molecular Virology Unit, Fondazione IRCCS Policlinico San Matteo , Pavia | Laboratory of Virology, INMI Lazzaro Spallanzani IRCCS | Antonino Di Caro, Antonio Piralla, Martina Rueca, Fausto Baldanti, Barbara Bartolini, Maria R. Capobianchi, Cesare E.M. Gruber |
| EPI_ISL_460092 | Molecular Virology Unit, Fondazione IRCCS Policlinico San Matteo , Pavia | Laboratory of Virology, INMI Lazzaro Spallanzani IRCCS | Cesare E.M. Gruber, Martina Rueca, Maria R. Capobianchi, Antonino Di Caro, Antonio Piralla, Barbara Bartolini, Fausto Baldanti |
| EPI_ISL_460093 | Molecular Virology Unit, Fondazione IRCCS Policlinico San Matteo , Pavia | Laboratory of Virology, INMI Lazzaro Spallanzani IRCCS | Maria R. Capobianchi, Antonio Piralla, Antonino Di Caro, Fausto Baldanti, Martina Rueca, Cesare E.M. Gruber, Barbara Bartolini |
| EPI_ISL_460094 | Molecular Virology Unit, Fondazione IRCCS Policlinico San Matteo , Pavia | Laboratory of Virology, INMI Lazzaro Spallanzani IRCCS | Barbara Bartolini, Maria R. Capobianchi, Antonino Di Caro, Antonio Piralla, Cesare E.M. Gruber, Martina Rueca, Fausto Baldanti |
| EPI_ISL_460095 | Molecular Virology Unit, Fondazione IRCCS Policlinico San Matteo , Pavia | Laboratory of Virology, INMI Lazzaro Spallanzani IRCCS | Barbara Bartolini, Antonino Di Caro, Fausto Baldanti, Cesare E.M. Gruber, Maria R. Capobianchi, Martina Rueca, Antonio Piralla |
| EPI_ISL_460096 | Molecular diagnostic laboratory of Federal Budget Institution of Science "Central Research Institute of Epidemiology" of The Federal Service on Customers' Rights Protection and Human Well-being Surveillance | Group of Genomics and Postgenomic Technologies of Central Research Institute of Epidemilology | Speranskaya AS, Kaptelova VV, Samoilov AE, Korneenko EV, Tivanova EV, Shipulina OY, Akimkin VG |
| EPI_ISL_460554, EPI_ISL_460555, EPI_ISL_460556, EPI_ISL_460557, EPI_ISL_460558, EPI_ISL_460559, EPI_ISL_460560, EPI_ISL_460561, EPI_ISL_460562, EPI_ISL_460563, EPI_ISL_460564, EPI_ISL_460565, EPI_ISL_460566, EPI_ISL_460567, EPI_ISL_460568, EPI_ISL_460569, EPI_ISL_460570, EPI_ISL_460571, EPI_ISL_460572, EPI_ISL_460573, EPI_ISL_460574, EPI_ISL_460575, EPI_ISL_460576, EPI_ISL_460577, EPI_ISL_460578, EPI_ISL_460579, EPI_ISL_460580, EPI_ISL_460581, EPI_ISL_460582, EPI_ISL_460583, EPI_ISL_460584, EPI_ISL_460585, EPI_ISL_460586, EPI_ISL_460587, EPI_ISL_460588, EPI_ISL_460589, EPI_ISL_460590, EPI_ISL_460591, EPI_ISL_460592, EPI_ISL_460593, EPI_ISL_460594, EPI_ISL_460595, EPI_ISL_460596, EPI_ISL_460597, EPI_ISL_460598, EPI_ISL_460599, EPI_ISL_460600, EPI_ISL_460601, EPI_ISL_460602 | | | |
| see above | Michigan Department of Health and Human Services, Bureau of Laboratories | Michigan Department of Health and Human Services, Bureau of Laboratories | Blankenship HM, Riner D, Soehnlen MK |
| EPI_ISL_460603 | NYU Langone Health | Departments of Pathology and Medicine, New York University School of Medicine | Maria Aguero-Rosenfeld, Brendan Belovarac, Margaret Black, Ludovic Boytard, John Cadley, Paolo Cotzia, John Chen, Dacia Dimartino, Xiaojun Feng, Tatyana Gindin, Emily Guzman, Adriana Heguy, Megan Hogan, Emily Huang, George Jour, Alireza Khodadadi-Jamayran, Lawrence H. Lin, Raven Luther, Andrew Lytle, Christian Marier, Matthew T. Maurano, Mark J. Mulligan, Peter Meyn, Raquel Ordonez Ciriza, Iman Osman, Jared Pinnell, Vanessa Raabe, Sitharam Ramaswami, Amy Rapkiewicz, Andre M. Ribeiro-dos-Santos, Marie Samanovic-Golden, Antonio Serrano, Guomiao Shen, Matija Snuderl, Theodore Vougiouklakis, Nick Vulpescu, Gael Westby, Paul Zappile, Yutong Zhang |
| EPI_ISL_460604 | Molecular diagnostic laboratory of Federal Budget Institution of Science "Central Research Institute of Epidemiology" of The Federal Service on Customers' Rights Protection and Human Well-being Surveillance | Group of Genomics and Postgenomic Technologies of Central Research Institute of Epidemilology | Speranskaya AS, Kaptelova VV, Samoilov AE, Korneenko EV, Tivanova EV, Shipulina OY, Akimkin VG |
| EPI_ISL_460605 | Molecular diagnostic laboratory of Federal Budget Institution of Science "Central Research Institute of Epidemiology" of The Federal Service on Customers' Rights Protection and Human Well-being Surveillance | Group of Genomics and Postgenomic Technologies of Central Research Institute of Epidemilology | Speranskaya AS, Kaptelova VV, Samoilov AE, Korneenko EV, Tivanova EV, Shipulina OY, Akimkin VG |
| EPI_ISL_460606, EPI_ISL_460607, EPI_ISL_460608, EPI_ISL_460609, EPI_ISL_460610, EPI_ISL_460611, EPI_ISL_460612, EPI_ISL_460613, EPI_ISL_460614, EPI_ISL_460615, EPI_ISL_460616 | | | |
| see above | BCCDC Public Health Laboratory | BCCDC Public Health Laboratory | Harrigan, Prystajecky, Krajden, Lee, Kamelian, Lapointe, Choi, Hoang, Sekirov, Levett, Tyson, Li, Gilmour |
| EPI_ISL_460617, EPI_ISL_460618, EPI_ISL_460619 | unknown | Physiology | Pence,S., Caykara,B., Pence,H.H., Tekin,S., Yiyit,N., Cevher Keskin,B., Kara,A. |
| EPI_ISL_460621, EPI_ISL_460622, EPI_ISL_460623, EPI_ISL_460624, EPI_ISL_460625, EPI_ISL_460626, EPI_ISL_460627, EPI_ISL_460628, EPI_ISL_460629, EPI_ISL_460630, EPI_ISL_460631, EPI_ISL_460632, EPI_ISL_460633, EPI_ISL_460634 | | | |
| see above | UW Virology Lab | UW Virology Lab | Pavitra Roychoudhury, Amin Addetia, Hong Xie, Lasata Shrestha, Truong Nguyen, Meei-Li Huang, Keith Jerome, Alexander Greninger |
| EPI_ISL_460635 | UHCW Pathology / University of Warwick | University of Warwick, for the COVID-19 Genomics (COG) UK Consortium | Richard Stark, Chrystala Constantinidou, Meera Unnikrishnan, Laura Baxter, Jeff Cheng, Grace Taylor-Joyce, Hannah Elizabeth Bridgewater, Lucy Frost, Sarojini Pandey, Paul Brown, Tauqeer Alam, Sascha Ott, Dimitris Grammatopoulos |
| EPI_ISL_460636, EPI_ISL_460637, EPI_ISL_460638, EPI_ISL_460639, EPI_ISL_460640, EPI_ISL_460641, EPI_ISL_460642, EPI_ISL_460643, EPI_ISL_460644, EPI_ISL_460645, EPI_ISL_460646, EPI_ISL_460647, EPI_ISL_460648, EPI_ISL_460649, EPI_ISL_460650, EPI_ISL_460651, EPI_ISL_460652, EPI_ISL_460653, EPI_ISL_460654, EPI_ISL_460655, EPI_ISL_460656, EPI_ISL_460657, EPI_ISL_460658, EPI_ISL_460659, EPI_ISL_460660, EPI_ISL_460661, EPI_ISL_460662, EPI_ISL_460663, EPI_ISL_460664, EPI_ISL_460665, EPI_ISL_460666, EPI_ISL_460667, EPI_ISL_460668, EPI_ISL_460669, EPI_ISL_460670, EPI_ISL_460671, EPI_ISL_460672, EPI_ISL_460673, EPI_ISL_460674, EPI_ISL_460675, EPI_ISL_460676, EPI_ISL_460677, EPI_ISL_460678, EPI_ISL_460679, EPI_ISL_460680, EPI_ISL_460681, EPI_ISL_460682, EPI_ISL_460683, EPI_ISL_460684, EPI_ISL_460685, EPI_ISL_460686, EPI_ISL_460687, EPI_ISL_460688, EPI_ISL_460689, EPI_ISL_460690, EPI_ISL_460691, EPI_ISL_460692, EPI_ISL_460693, EPI_ISL_460694, EPI_ISL_460695, EPI_ISL_460696, EPI_ISL_460697, EPI_ISL_460698, EPI_ISL_460699, EPI_ISL_460700, EPI_ISL_460701, EPI_ISL_460702, EPI_ISL_460703, EPI_ISL_460704, EPI_ISL_460705, EPI_ISL_460706, EPI_ISL_460707, EPI_ISL_460708, EPI_ISL_460709, EPI_ISL_460710, EPI_ISL_460711, EPI_ISL_460712, EPI_ISL_460713, EPI_ISL_460714, EPI_ISL_460715, EPI_ISL_460716, EPI_ISL_460717, EPI_ISL_460718, EPI_ISL_460719, EPI_ISL_460720, EPI_ISL_460721, EPI_ISL_460722, EPI_ISL_460723, EPI_ISL_460724, EPI_ISL_460725, EPI_ISL_460726, EPI_ISL_460727, EPI_ISL_460728, EPI_ISL_460729, EPI_ISL_460730, EPI_ISL_460731, EPI_ISL_460732, EPI_ISL_460733, EPI_ISL_460734, EPI_ISL_460735, EPI_ISL_460736, EPI_ISL_460737, EPI_ISL_460738, EPI_ISL_460739, EPI_ISL_460740, EPI_ISL_460741, EPI_ISL_460742, EPI_ISL_460743, EPI_ISL_460744, EPI_ISL_460745, EPI_ISL_460746, EPI_ISL_460747, EPI_ISL_460748, EPI_ISL_460749, EPI_ISL_460750, EPI_ISL_460751, EPI_ISL_460752, EPI_ISL_460753, EPI_ISL_460754, EPI_ISL_460755, EPI_ISL_460756, EPI_ISL_460757, EPI_ISL_460758, EPI_ISL_460759, EPI_ISL_460760, EPI_ISL_460761, EPI_ISL_460762, EPI_ISL_460763, EPI_ISL_460764, EPI_ISL_460765, EPI_ISL_460766, EPI_ISL_460767, EPI_ISL_460768, EPI_ISL_460769, EPI_ISL_460770, EPI_ISL_460771, EPI_ISL_460772, EPI_ISL_460773, EPI_ISL_460774, EPI_ISL_460775, EPI_ISL_460776, EPI_ISL_460777, EPI_ISL_460778, EPI_ISL_460779, EPI_ISL_460780, EPI_ISL_460781, EPI_ISL_460782, EPI_ISL_460783, EPI_ISL_460784, EPI_ISL_460785, EPI_ISL_460786, EPI_ISL_460787, EPI_ISL_460788, EPI_ISL_460789, EPI_ISL_460790, EPI_ISL_460791, EPI_ISL_460792, EPI_ISL_460793, EPI_ISL_460794, EPI_ISL_460795, EPI_ISL_460796, EPI_ISL_460797, EPI_ISL_460798, EPI_ISL_460799, EPI_ISL_460800, EPI_ISL_460801, EPI_ISL_460802, EPI_ISL_460803, EPI_ISL_460804, EPI_ISL_460805, EPI_ISL_460806, EPI_ISL_460807, EPI_ISL_460808, EPI_ISL_460809, EPI_ISL_460810, EPI_ISL_460811, EPI_ISL_460812, EPI_ISL_460813, EPI_ISL_460814, EPI_ISL_460815, EPI_ISL_460816, EPI_ISL_460817, EPI_ISL_460818, EPI_ISL_460819, EPI_ISL_460820, EPI_ISL_460821, EPI_ISL_460822, EPI_ISL_460823, EPI_ISL_460824, EPI_ISL_460825, EPI_ISL_460826, EPI_ISL_460827, EPI_ISL_460828, EPI_ISL_460829, EPI_ISL_460830, EPI_ISL_460831, EPI_ISL_460832, EPI_ISL_460833, EPI_ISL_460834, EPI_ISL_460835, EPI_ISL_460836, EPI_ISL_460837, EPI_ISL_460838, EPI_ISL_460839, EPI_ISL_460840, EPI_ISL_460841, EPI_ISL_460842, EPI_ISL_460843, EPI_ISL_460844, EPI_ISL_460845, EPI_ISL_460846, EPI_ISL_460847, EPI_ISL_460848, EPI_ISL_460849, EPI_ISL_460850, EPI_ISL_460851, EPI_ISL_460852, EPI_ISL_460853, EPI_ISL_460854, EPI_ISL_460855, EPI_ISL_460856, EPI_ISL_460857, EPI_ISL_460858, EPI_ISL_460859, EPI_ISL_460860, EPI_ISL_460861, EPI_ISL_460862, EPI_ISL_460863, EPI_ISL_460864, EPI_ISL_460865, EPI_ISL_460866, EPI_ISL_460867, EPI_ISL_460868, EPI_ISL_460869, EPI_ISL_460870, EPI_ISL_460871, EPI_ISL_460872, EPI_ISL_460873, EPI_ISL_460874, EPI_ISL_460875, EPI_ISL_460876, EPI_ISL_460877, EPI_ISL_460878, EPI_ISL_460879, EPI_ISL_460880, EPI_ISL_460881, EPI_ISL_460882, EPI_ISL_460883, EPI_ISL_460884, EPI_ISL_460885, EPI_ISL_460886, EPI_ISL_460887, EPI_ISL_460888, EPI_ISL_460889, EPI_ISL_460890, EPI_ISL_460891, EPI_ISL_460892, EPI_ISL_460893, EPI_ISL_460894, EPI_ISL_460895, EPI_ISL_460896, EPI_ISL_460897, EPI_ISL_460898, EPI_ISL_460899, EPI_ISL_460900, EPI_ISL_460901, EPI_ISL_460902, EPI_ISL_460903, EPI_ISL_460904, EPI_ISL_460905, EPI_ISL_460906, EPI_ISL_460907, EPI_ISL_460908, EPI_ISL_460909, EPI_ISL_460910, EPI_ISL_460911, EPI_ISL_460912, EPI_ISL_460913, EPI_ISL_460914, EPI_ISL_460915, EPI_ISL_460916, EPI_ISL_460917, EPI_ISL_460918, EPI_ISL_460919, EPI_ISL_460920, EPI_ISL_460921, EPI_ISL_460922, EPI_ISL_460923, EPI_ISL_460924, EPI_ISL_460925, EPI_ISL_460926, EPI_ISL_460927, EPI_ISL_460928, EPI_ISL_460929, EPI_ISL_460930, EPI_ISL_460931, EPI_ISL_460932, EPI_ISL_460933, EPI_ISL_460934, EPI_ISL_460935, EPI_ISL_460936, EPI_ISL_460937, EPI_ISL_460938, EPI_ISL_460939, EPI_ISL_460940, EPI_ISL_460941, EPI_ISL_460942, EPI_ISL_460943, EPI_ISL_460944, EPI_ISL_460945, EPI_ISL_460946, EPI_ISL_460947, EPI_ISL_460948, EPI_ISL_460949, EPI_ISL_460950, EPI_ISL_460951, EPI_ISL_460952, EPI_ISL_460953, EPI_ISL_460954, EPI_ISL_460955, EPI_ISL_460956, EPI_ISL_460957, EPI_ISL_460958, EPI_ISL_460959, EPI_ISL_460960, EPI_ISL_460961, EPI_ISL_460962, EPI_ISL_460963, EPI_ISL_460964, EPI_ISL_460965, EPI_ISL_460966, EPI_ISL_460967, EPI_ISL_460968, EPI_ISL_460969, EPI_ISL_460970, EPI_ISL_460971, EPI_ISL_460972, EPI_ISL_460973, EPI_ISL_460974, EPI_ISL_460975, EPI_ISL_460976, EPI_ISL_460977, EPI_ISL_460978, EPI_ISL_460979, EPI_ISL_460980, EPI_ISL_460981, EPI_ISL_460982, EPI_ISL_460983, EPI_ISL_460984, EPI_ISL_460985, EPI_ISL_460986, EPI_ISL_460987, EPI_ISL_460988, EPI_ISL_460989, EPI_ISL_460990, EPI_ISL_460991, EPI_ISL_460992, EPI_ISL_460993, EPI_ISL_460994, EPI_ISL_460995, EPI_ISL_460996, EPI_ISL_460997, EPI_ISL_460998, EPI_ISL_460999, EPI_ISL_461000, EPI_ISL_461001, EPI_ISL_461002, EPI_ISL_461003, EPI_ISL_461004, EPI_ISL_461005, EPI_ISL_461006, EPI_ISL_461007, EPI_ISL_461008, EPI_ISL_461009, EPI_ISL_461010, EPI_ISL_461011, EPI_ISL_461012, EPI_ISL_461013, EPI_ISL_461014, EPI_ISL_461015, EPI_ISL_461016, EPI_ISL_461017, EPI_ISL_461018, EPI_ISL_461019, EPI_ISL_461020, EPI_ISL_461021, EPI_ISL_461022, EPI_ISL_461023, EPI_ISL_461024, EPI_ISL_461025, EPI_ISL_461026, EPI_ISL_461027, EPI_ISL_461028, EPI_ISL_461029, EPI_ISL_461030, EPI_ISL_461031, EPI_ISL_461032, EPI_ISL_461033, EPI_ISL_461034, EPI_ISL_461035, EPI_ISL_461036, EPI_ISL_461037, EPI_ISL_461038, EPI_ISL_461039, EPI_ISL_461040, EPI_ISL_461041, EPI_ISL_461042, EPI_ISL_461043, EPI_ISL_461044, EPI_ISL_461045, EPI_ISL_461046, EPI_ISL_461047, EPI_ISL_461048, EPI_ISL_461049, EPI_ISL_461050, EPI_ISL_461051, EPI_ISL_461052, EPI_ISL_461053, EPI_ISL_461054, EPI_ISL_461055, EPI_ISL_461056, EPI_ISL_461057, EPI_ISL_461058, EPI_ISL_461059, EPI_ISL_461060, EPI_ISL_461061, EPI_ISL_461062, EPI_ISL_461063, EPI_ISL_461064, EPI_ISL_461065, EPI_ISL_461066, EPI_ISL_461067, EPI_ISL_461068, EPI_ISL_461069, EPI_ISL_461070, EPI_ISL_461071, EPI_ISL_461072, EPI_ISL_461073, EPI_ISL_461074, EPI_ISL_461075, EPI_ISL_461076, EPI_ISL_461077, EPI_ISL_461078, EPI_ISL_461079, EPI_ISL_461080, EPI_ISL_461081, EPI_ISL_461082, EPI_ISL_461083, EPI_ISL_461084, EPI_ISL_461085, EPI_ISL_461086, EPI_ISL_461087, EPI_ISL_461088, EPI_ISL_461089, EPI_ISL_461090, EPI_ISL_461091, EPI_ISL_461092, EPI_ISL_461093, EPI_ISL_461094, EPI_ISL_461095, EPI_ISL_461096, EPI_ISL_461097, EPI_ISL_461098, EPI_ISL_461099, EPI_ISL_461100, EPI_ISL_461101, EPI_ISL_461102, EPI_ISL_461103, EPI_ISL_461104, EPI_ISL_461105, EPI_ISL_461106, EPI_ISL_461107, EPI_ISL_461108, EPI_ISL_461109, EPI_ISL_461110, EPI_ISL_461111, EPI_ISL_461112, EPI_ISL_461113, EPI_ISL_461114, EPI_ISL_461115, EPI_ISL_461116, EPI_ISL_461117, EPI_ISL_461118, EPI_ISL_461119, EPI_ISL_461120, EPI_ISL_461121, EPI_ISL_461122, EPI_ISL_461123, EPI_ISL_461124, EPI_ISL_461125, EPI_ISL_461126, EPI_ISL_461127, EPI_ISL_461128, EPI_ISL_461129, EPI_ISL_461130, EPI_ISL_461131, EPI_ISL_461132, EPI_ISL_461133, EPI_ISL_461134, EPI_ISL_461135, EPI_ISL_461136, EPI_ISL_461137, EPI_ISL_461138, EPI_ISL_461139, EPI_ISL_461140, EPI_ISL_461141, EPI_ISL_461142, EPI_ISL_461143, EPI_ISL_461144, EPI_ISL_461145, EPI_ISL_461146, EPI_ISL_461147, EPI_ISL_461148, EPI_ISL_461149, EPI_ISL_461150, EPI_ISL_461151, EPI_ISL_461152, EPI_ISL_461153, EPI_ISL_461154, EPI_ISL_461155, EPI_ISL_461156, EPI_ISL_461157, EPI_ISL_461158, EPI_ISL_461159, EPI_ISL_461160, EPI_ISL_461161, EPI_ISL_461162, EPI_ISL_461163, EPI_ISL_461164, EPI_ISL_461165, EPI_ISL_461166, EPI_ISL_461167, EPI_ISL_461168, EPI_ISL_461169, EPI_ISL_461170, EPI_ISL_461171, EPI_ISL_461172, EPI_ISL_461173, EPI_ISL_461174, EPI_ISL_461175, EPI_ISL_461176, EPI_ISL_461177, EPI_ISL_461178, EPI_ISL_461179, EPI_ISL_461180, EPI_ISL_461181, EPI_ISL_461182, EPI_ISL_461183, EPI_ISL_461184, EPI_ISL_461185, EPI_ISL_461186, EPI_ISL_461187, EPI_ISL_461188, EPI_ISL_461189, EPI_ISL_461190, EPI_ISL_461191, EPI_ISL_461192, EPI_ISL_461193, EPI_ISL_461194, EPI_ISL_461195, EPI_ISL_461196, EPI_ISL_461197, EPI_ISL_461198, EPI_ISL_461199, EPI_ISL_461200, EPI_ISL_461201, EPI_ISL_461202, EPI_ISL_461203, EPI_ISL_461204, EPI_ISL_461205, EPI_ISL_461206, EPI_ISL_461207, EPI_ISL_461208, EPI_ISL_461209, EPI_ISL_461210, EPI_ISL_461211, EPI_ISL_461212, EPI_ISL_461213, EPI_ISL_461214, EPI_ISL_461215, EPI_ISL_461216, EPI_ISL_461217, EPI_ISL_461218, EPI_ISL_461219, EPI_ISL_461220, EPI_ISL_461221, EPI_ISL_461222, EPI_ISL_461223, EPI_ISL_461224, EPI_ISL_461225, EPI_ISL_461226, EPI_ISL_461227, EPI_ISL_461228, EPI_ISL_461229, EPI_ISL_461230, EPI_ISL_461231, EPI_ISL_461232, EPI_ISL_461233, EPI_ISL_461234, EPI_ISL_461235, EPI_ISL_461236, EPI_ISL_461237, EPI_ISL_461238, EPI_ISL_461239, EPI_ISL_461240, EPI_ISL_461241, EPI_ISL_461242, EPI_ISL_461243, EPI_ISL_461244, EPI_ISL_461245, EPI_ISL_461246, EPI_ISL_461247, EPI_ISL_461248, EPI_ISL_461249, EPI_ISL_461250, EPI_ISL_461251, EPI_ISL_461252, EPI_ISL_461253, EPI_ISL_461254, EPI_ISL_461255, EPI_ISL_461256, EPI_ISL_461257, EPI_ISL_461258, EPI_ISL_461259, EPI_ISL_461260, EPI_ISL_461261, EPI_ISL_461262, EPI_ISL_461263, EPI_ISL_461264, EPI_ISL_461265, EPI_ISL_461266, EPI_ISL_461267, EPI_ISL_461268, EPI_ISL_461269, EPI_ISL_461270, EPI_ISL_461271, EPI_ISL_461272, EPI_ISL_461273, EPI_ISL_461274, EPI_ISL_461275, EPI_ISL_461276, EPI_ISL_461277, EPI_ISL_461278, EPI_ISL_461279, EPI_ISL_461280, EPI_ISL_461281, EPI_ISL_461282, EPI_ISL_461283, EPI_ISL_461284, EPI_ISL_461285, EPI_ISL_461286, EPI_ISL_461287, EPI_ISL_461288, EPI_ISL_461289, EPI_ISL_461290, EPI_ISL_461291, EPI_ISL_461292, EPI_ISL_461293, EPI_ISL_461294, EPI_ISL_461295, EPI_ISL_461296, EPI_ISL_461297, EPI_ISL_461298, EPI_ISL_461299, EPI_ISL_461300, EPI_ISL_461301, EPI_ISL_461302, EPI_ISL_461303, EPI_ISL_461304, EPI_ISL_461305, EPI_ISL_461306, EPI_ISL_461307, EPI_ISL_461308, EPI_ISL_461309, EPI_ISL_461310, EPI_ISL_461311, EPI_ISL_461312, EPI_ISL_461313, EPI_ISL_461314, EPI_ISL_461315, EPI_ISL_461316, EPI_ISL_461317, EPI_ISL_461318, EPI_ISL_461319, EPI_ISL_461320, EPI_ISL_461321, EPI_ISL_461322, EPI_ISL_461323, EPI_ISL_461324, EPI_ISL_461325, EPI_ISL_461326, EPI_ISL_461327, EPI_ISL_461328, EPI_ISL_461329, EPI_ISL_461330, EPI_ISL_461331, EPI_ISL_461332, EPI_ISL_461333, EPI_ISL_461334, EPI_ISL_461335, EPI_ISL_461336, EPI_ISL_461337, EPI_ISL_461338, EPI_ISL_461339, EPI_ISL_461340, EPI_ISL_461341, EPI_ISL_461342, EPI_ISL_461343, EPI_ISL_461344, EPI_ISL_461345, EPI_ISL_461346, EPI_ISL_461347, EPI_ISL_461348, EPI_ISL_461349, EPI_ISL_461350, EPI_ISL_461351, EPI_ISL_461352, EPI_ISL_461353, EPI_ISL_461354, EPI_ISL_461355, EPI_ISL_461356, EPI_ISL_461357, EPI_ISL_461358, EPI_ISL_461359, EPI_ISL_461360, EPI_ISL_461361, EPI_ISL_461362, EPI_ISL_461363, EPI_ISL_461364, EPI_ISL_461365, EPI_ISL_461366, EPI_ISL_461367, EPI_ISL_461368, EPI_ISL_461369, EPI_ISL_461370, EPI_ISL_461371, EPI_ISL_461372, EPI_ISL_461373, EPI_ISL_461374, EPI_ISL_461375, EPI_ISL_461376, EPI_ISL_461377, EPI_ISL_461378, EPI_ISL_461379, EPI_ISL_461380, EPI_ISL_461381, EPI_ISL_461382, EPI_ISL_461383, EPI_ISL_461384, EPI_ISL_461385, EPI_ISL_461386, EPI_ISL_461387, EPI_ISL_461388, EPI_ISL_461389, EPI_ISL_461390, EPI_ISL_461391, EPI_ISL_461392, EPI_ISL_461393, EPI_ISL_461394, EPI_ISL_461395, EPI_ISL_461396, EPI_ISL_461397, EPI_ISL_461398 | | | |
| see above | Dutch COVID-19 response team | Erasmus Medical Center | Bas Oude Munnink, David Nieuwenhuijse, Reina Sikkema, Claudia Schapendonk, Irina Chestakova, Anne van der Linden, Theo Bestebroer, Stefan van Nieuwkoop, Mark Pronk, Pascal Lexmond, Corien Swaan, Manon Haverkate, Madelief Mollers, Mart Stein, Sandra Kengne Kamga Mobou, Jeroen van Kampen, Jolanda Voermans, Aura Timen, Corine GeurtsvanKessel, Annemiek van der Eijk, Richard Molenkamp, Marion Koopmans, on behalf of the Dutch national COVID-19 response team. |
| EPI_ISL_461399, EPI_ISL_461400, EPI_ISL_461401, EPI_ISL_461402, EPI_ISL_461403, EPI_ISL_461404, EPI_ISL_461405, EPI_ISL_461406, EPI_ISL_461407, EPI_ISL_461408, EPI_ISL_461409, EPI_ISL_461410, EPI_ISL_461411, EPI_ISL_461412, EPI_ISL_461413, EPI_ISL_461414, EPI_ISL_461415, EPI_ISL_461416, EPI_ISL_461417, EPI_ISL_461418, EPI_ISL_461419, EPI_ISL_461420, EPI_ISL_461421, EPI_ISL_461422, EPI_ISL_461423, EPI_ISL_461424, EPI_ISL_461425, EPI_ISL_461426, EPI_ISL_461427, EPI_ISL_461428, EPI_ISL_461429, EPI_ISL_461430, EPI_ISL_461431, EPI_ISL_461432, EPI_ISL_461433, EPI_ISL_461434, EPI_ISL_461435, EPI_ISL_461436, EPI_ISL_461437, EPI_ISL_461438, EPI_ISL_461439, EPI_ISL_461440, EPI_ISL_461441, EPI_ISL_461442, EPI_ISL_461443, EPI_ISL_461444, EPI_ISL_461445, EPI_ISL_461446, EPI_ISL_461447, EPI_ISL_461448, EPI_ISL_461449, EPI_ISL_461450, EPI_ISL_461451, EPI_ISL_461452, EPI_ISL_461453, EPI_ISL_461454, EPI_ISL_461455, EPI_ISL_461456, EPI_ISL_461457, EPI_ISL_461458, EPI_ISL_461459, EPI_ISL_461460, EPI_ISL_461461, EPI_ISL_461462, EPI_ISL_461463, EPI_ISL_461464, EPI_ISL_461465, EPI_ISL_461466, EPI_ISL_461467, EPI_ISL_461468, EPI_ISL_461469, EPI_ISL_461470, EPI_ISL_461471, EPI_ISL_461472, EPI_ISL_461473, EPI_ISL_461474, EPI_ISL_461475, EPI_ISL_461476, EPI_ISL_461477 | | | |
| see above | UW Virology Lab | UW Virology Lab | Pavitra Roychoudhury, Amin Addetia, Hong Xie, Lasata Shrestha, Truong Nguyen, Meei-Li Huang, Keith Jerome, Alexander Greninger |
| EPI_ISL_461478 | Government Medical College, Vadodara | Gujarat Biotechnology Research Centre | Fenil Patel, Nidhi Patel, Nitin Savaliya, Raghawendra Kumar, Dinesh Kumar, Zuber Saiyed, Komal Patel, Labdhi Pandya, Snehal Bagatharia, Tanuja Javadekar , R N Daveshwar, Tejas Shah, Ankit Hinsu, Pritesh Sabara, Apurvasinh Puvar, Janvi Raval, Zarna Patel, Monika Gandhi, Pinal Trivedi, Maharshi Pandya, R D Dixit, A M Kadri, Harsh Bakshi, Chaitanya Joshi, Madhvi Joshi, |
| EPI_ISL_461479 | Government Medical College, Vadodara | Gujarat Biotechnology Research Centre | Neelam Nathani, Nitin Savaliya, Raghawendra Kumar, Dinesh Kumar, Zuber Saiyed, Komal Patel, Labdhi Pandya, Snehal Bagatharia, Tanuja Javadekar , R N Daveshwar, Tejas Shah, Ankit Hinsu, Pritesh Sabara, Apurvasinh Puvar, Janvi Raval, Zarna Patel, Monika Gandhi, Pinal Trivedi, Maharshi Pandya, Nidhi Patel, R D Dixit, A M Kadri, Harsh Bakshi, Chaitanya Joshi, Madhvi Joshi, |
| EPI_ISL_461480 | Government Medical College, Vadodara | Gujarat Biotechnology Research Centre | Armi Chaudhari, Raghawendra Kumar, Dinesh Kumar, Zuber Saiyed, Komal Patel, Labdhi Pandya, Snehal Bagatharia, Tanuja Javadekar , R N Daveshwar, Tejas Shah, Ankit Hinsu, Pritesh Sabara, Apurvasinh Puvar, Janvi Raval, Zarna Patel, Monika Gandhi, Pinal Trivedi, Maharshi Pandya, Nidhi Patel, Nitin Savaliya, R D Dixit, A M Kadri, Harsh Bakshi, Chaitanya Joshi, Madhvi Joshi, |
| EPI_ISL_461481 | Pandit Deendayal Upadhyay Government Medical College, | Gujarat Biotechnology Research | Bhavya Jindal, Dinesh Kumar, Zuber Saiyed, Komal Patel, Labdhi Pandya, Snehal Bagatharia, Prakash Modi, Sejul Antala, Manish Pattani, Tejas Shah, Ankit Hinsu, Pritesh Sabara, Apurvasinh Puvar, Janvi Raval, Zarna Patel, Monika Gandhi, Pinal Trivedi, Maharshi |

| | | | |
|---|---|---|---|
| | Rajkot | Centre | Pandya, Nidhi Patel, Nitin Savaliya, Raghawendra Kumar, R D Dixit, A M Kadri, Harsh Bakshi, Chaitanya Joshi, Madhvi Joshi |
| EPI_ISL_461482 | Pandit Deendayal Upadhyay Government Medical College, Rajkot | Gujarat Biotechnology Research Centre | Anjali Rajwar, Zuber Saiyed, Komal Patel, Labdhi Pandya, Snehal Bagatharia, Prakash Modi, Sejul Antala, Manish Pattani, Tejas Shah, Ankit Hinsu, Pritesh Sabara, Apurvasinh Puvar, Janvi Raval, Zarna Patel, Monika Gandhi, Pinal Trivedi, Maharshi Pandya, Nidhi Patel, Nitin Savaliya, Raghawendra Kumar, Dinesh Kumar, R D Dixit, A M Kadri, Harsh Bakshi, Chaitanya Joshi, Madhvi Joshi |
| EPI_ISL_461483 | B.J. Medical College and Civil hospital | Gujarat Biotechnology Research Centre | Dipeshwari Shewale, Komal Patel, Labdhi Pandya, Snehal Bagatharia, Pranay Shah, Kamlesh J Upadhyay, Tejas Shah, Ankit Hinsu, Pritesh Sabara, Apurvasinh Puvar, Janvi Raval, Zarna Patel, Monika Gandhi, Pinal Trivedi, Maharshi Pandya, Nidhi Patel, Nitin Savaliya, Raghawendra Kumar, Dinesh Kumar, Zuber Saiyed, R D Dixit, A M Kadri, Harsh Bakshi, Chaitanya Joshi, Madhvi Joshi, |
| EPI_ISL_461484 | B.J. Medical College and Civil hospital | Gujarat Biotechnology Research Centre | Priyanka P Vatsa, Labdhi Pandya, Snehal Bagatharia, Pranay Shah, Kamlesh J Upadhyay, Tejas Shah, Ankit Hinsu, Pritesh Sabara, Apurvasinh Puvar, Janvi Raval, Zarna Patel, Monika Gandhi, Pinal Trivedi, Maharshi Pandya, Nidhi Patel, Nitin Savaliya, Raghawendra Kumar, Dinesh Kumar, Zuber Saiyed, Komal Patel, R D Dixit, A M Kadri, Harsh Bakshi, Chaitanya Joshi, Madhvi Joshi, |
| EPI_ISL_461485 | B.J. Medical College and Civil hospital | Gujarat Biotechnology Research Centre | Pooja P Doshi, Snehal Bagatharia, Pranay Shah, Kamlesh J Upadhyay, Tejas Shah, Ankit Hinsu, Pritesh Sabara, Apurvasinh Puvar, Janvi Raval, Zarna Patel, Monika Gandhi, Pinal Trivedi, Maharshi Pandya, Nidhi Patel, Nitin Savaliya, Raghawendra Kumar, Dinesh Kumar, Zuber Saiyed, Komal Patel, Labdhi Pandya, R D Dixit, A M Kadri, Harsh Bakshi, Chaitanya Joshi, Madhvi Joshi, |
| EPI_ISL_461486 | B.J. Medical College and Civil hospital | Gujarat Biotechnology Research Centre | Akanksha Verma, Pranay Shah, Kamlesh J Upadhyay, Tejas Shah, Ankit Hinsu, Pritesh Sabara, Apurvasinh Puvar, Janvi Raval, Zarna Patel, Monika Gandhi, Pinal Trivedi, Maharshi Pandya, Nidhi Patel, Nitin Savaliya, Raghawendra Kumar, Dinesh Kumar, Zuber Saiyed, Komal Patel, Labdhi Pandya, Snehal Bagatharia, R D Dixit, A M Kadri, Harsh Bakshi, Chaitanya Joshi, Madhvi Joshi, |
| EPI_ISL_461487 | B.J. Medical College and Civil hospital | Gujarat Biotechnology Research Centre | Priti Pandita, Kamlesh J Upadhyay, Tejas Shah, Ankit Hinsu, Pritesh Sabara, Apurvasinh Puvar, Janvi Raval, Zarna Patel, Monika Gandhi, Pinal Trivedi, Maharshi Pandya, Nidhi Patel, Nitin Savaliya, Raghawendra Kumar, Dinesh Kumar, Zuber Saiyed, Komal Patel, Labdhi Pandya, Snehal Bagatharia, Pranay Shah, R D Dixit, A M Kadri, Harsh Bakshi, Chaitanya Joshi, Madhvi Joshi, |
| EPI_ISL_461488 | B.J. Medical College and Civil hospital | Gujarat Biotechnology Research Centre | Pragya Sharma, Tejas Shah, Ankit Hinsu, Pritesh Sabara, Apurvasinh Puvar, Janvi Raval, Zarna Patel, Monika Gandhi, Pinal Trivedi, Maharshi Pandya, Nidhi Patel, Nitin Savaliya, Raghawendra Kumar, Dinesh Kumar, Zuber Saiyed, Komal Patel, Labdhi Pandya, Snehal Bagatharia, Pranay Shah, Kamlesh J Upadhyay, R D Dixit, A M Kadri, Harsh Bakshi, Chaitanya Joshi, Madhvi Joshi, |
| EPI_ISL_461489 | B.J. Medical College and Civil hospital | Gujarat Biotechnology Research Centre | Neha Rajpara, Ankit Hinsu, Pritesh Sabara, Apurvasinh Puvar, Janvi Raval, Zarna Patel, Monika Gandhi, Pinal Trivedi, Maharshi Pandya, Nidhi Patel, Nitin Savaliya, Raghawendra Kumar, Dinesh Kumar, Zuber Saiyed, Komal Patel, Labdhi Pandya, Snehal Bagatharia, Pranay Shah, Kamlesh J Upadhyay, Tejas Shah, R D Dixit, A M Kadri, Harsh Bakshi, Chaitanya Joshi, Madhvi Joshi, |
| EPI_ISL_461490 | B.J. Medical College and Civil hospital | Gujarat Biotechnology Research Centre | Afzal Ansari, Pritesh Sabara, Apurvasinh Puvar, Janvi Raval, Zarna Patel, Monika Gandhi, Pinal Trivedi, Maharshi Pandya, Nidhi Patel, Nitin Savaliya, Raghawendra Kumar, Dinesh Kumar, Zuber Saiyed, Komal Patel, Labdhi Pandya, Snehal Bagatharia, Pranay Shah, Kamlesh J Upadhyay, Tejas Shah, Ankit Hinsu, R D Dixit, A M Kadri, Harsh Bakshi, Chaitanya Joshi, Madhvi Joshi, |
| EPI_ISL_461491 | B.J. Medical College and Civil hospital | Gujarat Biotechnology Research Centre | Fenil Patel, Apurvasinh Puvar, Janvi Raval, Zarna Patel, Monika Gandhi, Pinal Trivedi, Maharshi Pandya, Nidhi Patel, Nitin Savaliya, Raghawendra Kumar, Dinesh Kumar, Zuber Saiyed, Komal Patel, Labdhi Pandya, Snehal Bagatharia, Pranay Shah, Kamlesh J Upadhyay, Tejas Shah, Ankit Hinsu, Pritesh Sabara, R D Dixit, A M Kadri, Harsh Bakshi, Chaitanya Joshi, Madhvi Joshi, |
| EPI_ISL_461492 | B.J. Medical College and Civil hospital | Gujarat Biotechnology Research Centre | Neelam Nathani, Janvi Raval, Zarna Patel, Monika Gandhi, Pinal Trivedi, Maharshi Pandya, Nidhi Patel, Nitin Savaliya, Raghawendra Kumar, Dinesh Kumar, Zuber Saiyed, Komal Patel, Labdhi Pandya, Snehal Bagatharia, Pranay Shah, Kamlesh J Upadhyay, Tejas Shah, Ankit Hinsu, Pritesh Sabara, Apurvasinh Puvar, R D Dixit, A M Kadri, Harsh Bakshi, Chaitanya Joshi, Madhvi Joshi, |
| EPI_ISL_461493 | B.J. Medical College and Civil hospital | Gujarat Biotechnology Research Centre | Armi Chaudhari, Zarna Patel, Monika Gandhi, Pinal Trivedi, Maharshi Pandya, Nidhi Patel, Nitin Savaliya, Raghawendra Kumar, Dinesh Kumar, Zuber Saiyed, Komal Patel, Labdhi Pandya, Snehal Bagatharia, Pranay Shah, Kamlesh J Upadhyay, Tejas Shah, Ankit Hinsu, Pritesh Sabara, Apurvasinh Puvar, Janvi Raval, R D Dixit, A M Kadri, Harsh Bakshi, Chaitanya Joshi, Madhvi Joshi, |
| EPI_ISL_461494 | B.J. Medical College and Civil hospital | Gujarat Biotechnology Research Centre | Bhavya Jindal, Monika Gandhi, Pinal Trivedi, Maharshi Pandya, Nidhi Patel, Nitin Savaliya, Raghawendra Kumar, Dinesh Kumar, Zuber Saiyed, Komal Patel, Labdhi Pandya, Snehal Bagatharia, Pranay Shah, Kamlesh J Upadhyay, Tejas Shah, Ankit Hinsu, Pritesh Sabara, Apurvasinh Puvar, Janvi Raval, Zarna Patel, R D Dixit, A M Kadri, Harsh Bakshi, Chaitanya Joshi, Madhvi Joshi, |
| EPI_ISL_461495 | B.J. Medical College and Civil hospital | Gujarat Biotechnology Research Centre | Anjali Rajwar, Pinal Trivedi, Maharshi Pandya, Nidhi Patel, Nitin Savaliya, Raghawendra Kumar, Dinesh Kumar, Zuber Saiyed, Komal Patel, Labdhi Pandya, Snehal Bagatharia, Pranay Shah, Kamlesh J Upadhyay, Tejas Shah, Ankit Hinsu, Pritesh Sabara, Apurvasinh Puvar, Janvi Raval, Zarna Patel, Monika Gandhi, R D Dixit, A M Kadri, Harsh Bakshi, Chaitanya Joshi, Madhvi Joshi, |
| EPI_ISL_461496 | B.J. Medical College and Civil hospital | Gujarat Biotechnology Research Centre | Dipeshwari Shewale, Maharshi Pandya, Nidhi Patel, Nitin Savaliya, Raghawendra Kumar, Dinesh Kumar, Zuber Saiyed, Komal Patel, Labdhi Pandya, Snehal Bagatharia, Pranay Shah, Kamlesh J Upadhyay, Tejas Shah, Ankit Hinsu, Pritesh Sabara, Apurvasinh Puvar, Janvi Raval, Zarna Patel, Monika Gandhi, Pinal Trivedi, R D Dixit, A M Kadri, Harsh Bakshi, Chaitanya Joshi, Madhvi Joshi, |
| EPI_ISL_461497 | B.J. Medical College and Civil hospital | Gujarat Biotechnology Research Centre | Priyanka P Vatsa, Nidhi Patel, Nitin Savaliya, Raghawendra Kumar, Dinesh Kumar, Zuber Saiyed, Komal Patel, Labdhi Pandya, Snehal Bagatharia, Pranay Shah, Kamlesh J Upadhyay, Tejas Shah, Ankit Hinsu, Pritesh Sabara, Apurvasinh Puvar, Janvi Raval, Zarna Patel, Monika Gandhi, Pinal Trivedi, Maharshi Pandya, R D Dixit, A M Kadri, Harsh Bakshi, Chaitanya Joshi, Madhvi Joshi, |
| EPI_ISL_461498 | B.J. Medical College and Civil hospital | Gujarat Biotechnology Research Centre | Pooja P Doshi, Nitin Savaliya, Raghawendra Kumar, Dinesh Kumar, Zuber Saiyed, Komal Patel, Labdhi Pandya, Snehal Bagatharia, Pranay Shah, Kamlesh J Upadhyay, Tejas Shah, Ankit Hinsu, Pritesh Sabara, Apurvasinh Puvar, Janvi Raval, Zarna Patel, Monika Gandhi, Pinal Trivedi, Maharshi Pandya, Nidhi Patel, R D Dixit, A M Kadri, Harsh Bakshi, Chaitanya Joshi, Madhvi Joshi, |
| EPI_ISL_461499 | B.J. Medical College and Civil hospital | Gujarat Biotechnology Research Centre | Akanksha Verma, Raghawendra Kumar, Dinesh Kumar, Zuber Saiyed, Komal Patel, Labdhi Pandya, Snehal Bagatharia, Pranay Shah, Kamlesh J Upadhyay, Tejas Shah, Ankit Hinsu, Pritesh Sabara, Apurvasinh Puvar, Janvi Raval, Zarna Patel, Monika Gandhi, Pinal Trivedi, Maharshi Pandya, Nidhi Patel, Nitin Savaliya, R D Dixit, A M Kadri, Harsh Bakshi, Chaitanya Joshi, Madhvi Joshi, |
| EPI_ISL_461500 | B.J. Medical College and Civil hospital | Gujarat Biotechnology Research Centre | Priti Pandita, Dinesh Kumar, Zuber Saiyed, Komal Patel, Labdhi Pandya, Snehal Bagatharia, Pranay Shah, Kamlesh J Upadhyay, Tejas Shah, Ankit Hinsu, Pritesh Sabara, Apurvasinh Puvar, Janvi Raval, Zarna Patel, Monika Gandhi, Pinal Trivedi, Maharshi Pandya, Nidhi Patel, Nitin Savaliya, Raghawendra Kumar, R D Dixit, A M Kadri, Harsh Bakshi, Chaitanya Joshi, Madhvi Joshi, |
| EPI_ISL_461501 | B.J. Medical College and Civil hospital | Gujarat Biotechnology Research Centre | Pragya Sharma, Zuber Saiyed, Komal Patel, Labdhi Pandya, Snehal Bagatharia, Pranay Shah, Kamlesh J Upadhyay, Tejas Shah, Ankit Hinsu, Pritesh Sabara, Apurvasinh Puvar, Janvi Raval, Zarna Patel, Monika Gandhi, Pinal Trivedi, Maharshi Pandya, Nidhi Patel, Nitin Savaliya, Raghawendra Kumar, Dinesh Kumar, R D Dixit, A M Kadri, Harsh Bakshi, Chaitanya Joshi, Madhvi Joshi, |
| EPI_ISL_461502 | B.J. Medical College and Civil hospital | Gujarat Biotechnology Research Centre | Neha Rajpara, Komal Patel, Labdhi Pandya, Snehal Bagatharia, Pranay Shah, Kamlesh J Upadhyay, Tejas Shah, Ankit Hinsu, Pritesh Sabara, Apurvasinh Puvar, Janvi Raval, Zarna Patel, Monika Gandhi, Pinal Trivedi, Maharshi Pandya, Nidhi Patel, Nitin Savaliya, Raghawendra Kumar, Dinesh Kumar, Zuber Saiyed, R D Dixit, A M Kadri, Harsh Bakshi, Chaitanya Joshi, Madhvi Joshi, |
| EPI_ISL_461503 | B.J. Medical College and Civil hospital | Gujarat Biotechnology Research Centre | Afzal Ansari, Labdhi Pandya, Snehal Bagatharia, Pranay Shah, Kamlesh J Upadhyay, Tejas Shah, Ankit Hinsu, Pritesh Sabara, Apurvasinh Puvar, Janvi Raval, Zarna Patel, Monika Gandhi, Pinal Trivedi, Maharshi Pandya, Nidhi Patel, Nitin Savaliya, Raghawendra Kumar, Dinesh Kumar, Zuber Saiyed, Komal Patel, R D Dixit, A M Kadri, Harsh Bakshi, Chaitanya Joshi, Madhvi Joshi, |
| EPI_ISL_461504 | B.J. Medical College and Civil hospital | Gujarat Biotechnology Research Centre | Snehal Bagatharia, Pranay Shah, Kamlesh J Upadhyay, Tejas Shah, Ankit Hinsu, Pritesh Sabara, Apurvasinh Puvar, Janvi Raval, Zarna Patel, Monika Gandhi, Pinal Trivedi, Maharshi Pandya, Nidhi Patel, Nitin Savaliya, Raghawendra Kumar, Dinesh Kumar, Zuber Saiyed, Komal Patel, Labdhi Pandya, Fenil Patel, R D Dixit, A M Kadri, Harsh Bakshi, Chaitanya Joshi, Madhvi Joshi, |
| EPI_ISL_461505 | B.J. Medical College and Civil hospital | Gujarat Biotechnology Research Centre | Pranay Shah, Kamlesh J Upadhyay, Tejas Shah, Ankit Hinsu, Pritesh Sabara, Apurvasinh Puvar, Janvi Raval, Zarna Patel, Monika Gandhi, Pinal Trivedi, Maharshi Pandya, Nidhi Patel, Nitin Savaliya, Raghawendra Kumar, Dinesh Kumar, Zuber Saiyed, Komal Patel, Labdhi Pandya, Snehal Bagatharia, Neelam Nathani, R D Dixit, A M Kadri, Harsh Bakshi, Chaitanya Joshi, Madhvi Joshi, |
| EPI_ISL_461506 | B.J. Medical College and Civil hospital | Gujarat Biotechnology Research Centre | Kamlesh J Upadhyay, Tejas Shah, Ankit Hinsu, Pritesh Sabara, Apurvasinh Puvar, Janvi Raval, Zarna Patel, Monika Gandhi, Pinal Trivedi, Maharshi Pandya, Nidhi Patel, Nitin Savaliya, Raghawendra Kumar, Dinesh Kumar, Zuber Saiyed, Komal Patel, Labdhi Pandya, Snehal Bagatharia, Pranay Shah, Armi Chaudhari, R D Dixit, A M Kadri, Harsh Bakshi, Chaitanya Joshi, Madhvi Joshi, |
| EPI_ISL_461507, EPI_ISL_461508, EPI_ISL_461509, EPI_ISL_461510, EPI_ISL_461511, EPI_ISL_461512, EPI_ISL_461513, EPI_ISL_461514, EPI_ISL_461515, EPI_ISL_461516, EPI_ISL_461517, EPI_ISL_461518, EPI_ISL_461519, EPI_ISL_461520, EPI_ISL_461521, EPI_ISL_461522, EPI_ISL_461523, EPI_ISL_461524, EPI_ISL_461525, EPI_ISL_461526, EPI_ISL_461527, EPI_ISL_461528, EPI_ISL_461529, EPI_ISL_461530, EPI_ISL_461531, EPI_ISL_461532, EPI_ISL_461533, EPI_ISL_461534, EPI_ISL_461535, EPI_ISL_461536, EPI_ISL_461537, EPI_ISL_461538, EPI_ISL_461539, EPI_ISL_461540, EPI_ISL_461541, EPI_ISL_461542, EPI_ISL_461543, EPI_ISL_461544, EPI_ISL_461545 | | | |
| see above | University of Birmingham | COVID-19 Genomics UK (COG-UK) Consortium | Loman Lab: Claire McMurray, Joanne Stockton, Samuel Nicholls, Radoslaw Poplawski, Will Rowe, Josh Quick, Nicholas Loman // UHB Lab: Celina M Whalley, Andrew Bosworth, Charlotte Poxon, Kasun Wanigasooriya, Oliver Pickles, Mike Kidd, Alex Richter, Andrew D Beggs // PHE Heartlands Lab: Husam Osman, Andrew Bosworth |
| EPI_ISL_461546, EPI_ISL_461547, EPI_ISL_461548, EPI_ISL_461549, EPI_ISL_461550, EPI_ISL_461551, EPI_ISL_461552, EPI_ISL_461553, EPI_ISL_461554, EPI_ISL_461555, EPI_ISL_461556, EPI_ISL_461557, EPI_ISL_461558, EPI_ISL_461559, EPI_ISL_461560, EPI_ISL_461561, EPI_ISL_461562, EPI_ISL_461563, EPI_ISL_461564, EPI_ISL_461565, EPI_ISL_461566, EPI_ISL_461567, EPI_ISL_461568, EPI_ISL_461569, EPI_ISL_461570, EPI_ISL_461571, EPI_ISL_461572, EPI_ISL_461573, EPI_ISL_461574, EPI_ISL_461575, EPI_ISL_461576, EPI_ISL_461577, EPI_ISL_461578, EPI_ISL_461579, EPI_ISL_461580, EPI_ISL_461581, EPI_ISL_461582, EPI_ISL_461583, EPI_ISL_461584, EPI_ISL_461585, EPI_ISL_461586, EPI_ISL_461587, EPI_ISL_461588 | | | |
| see above | Department of Pathology, University of Cambridge | COVID-19 Genomics UK (COG-UK) Consortium | Luke W Meredith, M. Estée Török, Myra Hosmillo, William L. Hamilton, Martin D. Curran, Theresa Feltwell, Grant Hall, Anna Yakovleva, Fahad A Khokhar, Charlotte J. Houldcroft, Laura G Caller, Aminu S. Jahun, Sarah L. Caddy, Ian Goodfellow |
| EPI_ISL_461589, EPI_ISL_461590, EPI_ISL_461591, EPI_ISL_461592, EPI_ISL_461593, EPI_ISL_461594, EPI_ISL_461595, EPI_ISL_461596, EPI_ISL_461597, EPI_ISL_461598, EPI_ISL_461599, EPI_ISL_461600, EPI_ISL_461601, EPI_ISL_461602, EPI_ISL_461603, EPI_ISL_461604, EPI_ISL_461605, EPI_ISL_461606, EPI_ISL_461607, EPI_ISL_461608, EPI_ISL_461609, EPI_ISL_461610, EPI_ISL_461611, EPI_ISL_461612, EPI_ISL_461613, EPI_ISL_461614, EPI_ISL_461615, EPI_ISL_461616, EPI_ISL_461617, EPI_ISL_461618, EPI_ISL_461619, EPI_ISL_461620, EPI_ISL_461621, EPI_ISL_461622, EPI_ISL_461623, EPI_ISL_461624, EPI_ISL_461625, EPI_ISL_461626, EPI_ISL_461627, EPI_ISL_461628, EPI_ISL_461629, EPI_ISL_461630, EPI_ISL_461631, EPI_ISL_461632, EPI_ISL_461633, EPI_ISL_461634, EPI_ISL_461635, EPI_ISL_461636, EPI_ISL_461637, EPI_ISL_461638, EPI_ISL_461639, EPI_ISL_461640, EPI_ISL_461641, EPI_ISL_461642, EPI_ISL_461643, EPI_ISL_461644, EPI_ISL_461645, EPI_ISL_461646, EPI_ISL_461647, EPI_ISL_461648, EPI_ISL_461649, EPI_ISL_461650, EPI_ISL_461651, EPI_ISL_461652, EPI_ISL_461653, EPI_ISL_461654, EPI_ISL_461655, EPI_ISL_461656, EPI_ISL_461657, EPI_ISL_461658, EPI_ISL_461659, EPI_ISL_461660, EPI_ISL_461661, EPI_ISL_461662, EPI_ISL_461663, EPI_ISL_461664, EPI_ISL_461665, EPI_ISL_461666, EPI_ISL_461667, EPI_ISL_461668, EPI_ISL_461669, EPI_ISL_461670, EPI_ISL_461671, EPI_ISL_461672, EPI_ISL_461673, EPI_ISL_461674, EPI_ISL_461675, EPI_ISL_461676, EPI_ISL_461677, EPI_ISL_461678, EPI_ISL_461679, EPI_ISL_461680, EPI_ISL_461681, EPI_ISL_461682, EPI_ISL_461683, EPI_ISL_461684, EPI_ISL_461685, EPI_ISL_461686, EPI_ISL_461687, EPI_ISL_461688, EPI_ISL_461689, EPI_ISL_461690, EPI_ISL_461691, EPI_ISL_461692, EPI_ISL_461693, EPI_ISL_461694, EPI_ISL_461695, EPI_ISL_461696, EPI_ISL_461697, EPI_ISL_461698, EPI_ISL_461699, EPI_ISL_461700, EPI_ISL_461701, EPI_ISL_461702, EPI_ISL_461703, EPI_ISL_461704, EPI_ISL_461705 | | | |
| see above | West of Scotland Specialist Virology Centre, NHSGGC / MRC-University of Glasgow Centre for Virus Research | COVID-19 Genomics UK (COG-UK) Consortium | Ana da Silva Filipe, Natasha Johnson, Kathy Smollett, Daniel Mair, Stephen Carmichael, Lily Tong, Jenna Nichols, Elihu Aranday-Cortes, Kirstyn Brunker, Yasmin Parr, Kyriaki Nomikou; Sarah McDonald, Marc Niebel, Patawee Asamaphan; Richard Orton, Joseph Hughes, Sreenu Vattipally, David L Robertson; Alasdair MacLean, Rory Gunson; Kathy Li, Natasha Jesudason, Rajiv Shah, James Shepherd, Antonia Ho, Emma Thomson |
| EPI_ISL_461706, EPI_ISL_461707, EPI_ISL_461708, EPI_ISL_461709, EPI_ISL_461710, EPI_ISL_461711, EPI_ISL_461712, EPI_ISL_461713, EPI_ISL_461714, EPI_ISL_461715, EPI_ISL_461716, EPI_ISL_461717, EPI_ISL_461718, EPI_ISL_461719, EPI_ISL_461720, EPI_ISL_461721, EPI_ISL_461722, EPI_ISL_461723, EPI_ISL_461724, EPI_ISL_461725, EPI_ISL_461726, EPI_ISL_461727, EPI_ISL_461728, EPI_ISL_461729, EPI_ISL_461730, EPI_ISL_461731, EPI_ISL_461732, EPI_ISL_461733, EPI_ISL_461734, EPI_ISL_461735, EPI_ISL_461736, EPI_ISL_461737, EPI_ISL_461738, EPI_ISL_461739, EPI_ISL_461740, EPI_ISL_461741, EPI_ISL_461742, EPI_ISL_461743, EPI_ISL_461744, EPI_ISL_461745, EPI_ISL_461746, EPI_ISL_461747, EPI_ISL_461748, EPI_ISL_461749, EPI_ISL_461750, EPI_ISL_461751, EPI_ISL_461752, EPI_ISL_461753, EPI_ISL_461754, EPI_ISL_461755, EPI_ISL_461756, EPI_ISL_461757, EPI_ISL_461758, EPI_ISL_461759, EPI_ISL_461760, EPI_ISL_461761, EPI_ISL_461762 | | | |
| see above | Virology Department, Royal Infirmary of Edinburgh, NHS Lothian / School of Biological Sciences, University of Edinburgh / Institute of Genetics and Molecular Medicine, University of Edinburgh | COVID-19 Genomics UK (COG-UK) Consortium | McHugh M, Dewar R, Rooke S, Gallagher M, Balcaza C, O'Toole Á, Scher E, Hill V, McCrone JT, Colquhoun R, Yu X, Jackson B, Rambaut A, Williams TC, Templeton K |
| EPI_ISL_461763, EPI_ISL_461764, EPI_ISL_461765, EPI_ISL_461766, EPI_ISL_461767, EPI_ISL_461768, EPI_ISL_461769, EPI_ISL_461770, EPI_ISL_461771, EPI_ISL_461772 | University College London, Great Ormond Street Hospital for Children NHS Foundation Trust, Imperial College Healthcare NHS Trust | COVID-19 Genomics UK (COG-UK) Consortium | Sergi Castellano, Rachel Williams, Mark Kristiansen, Paola Resende Silva, Sunando Roy, Tony Brooks, Helena Tutill, Paola Niola, Patricia Dyal, Charlotte Williams, Leysa Forrest, Yasmin Panchbhaya, Jacqueline Findlay, Sam Weeks, Julianne Brown, Kathryn Harris, Paul Randell, James Price, Alison Holmes, Judith Breuer |
| EPI_ISL_461773, EPI_ISL_461774, EPI_ISL_461775, EPI_ISL_461776, EPI_ISL_461777, EPI_ISL_461778, EPI_ISL_461779, EPI_ISL_461780, EPI_ISL_461781, EPI_ISL_461782, EPI_ISL_461783, EPI_ISL_461784, EPI_ISL_461785, EPI_ISL_461786, EPI_ISL_461787, EPI_ISL_461788, EPI_ISL_461789, EPI_ISL_461790 | | | |
| see above | Regional Virus Laboratory, Belfast Health and Social Care Trust | COVID-19 Genomics UK (COG-UK) Consortium | Conall McCaughey, James McKenna, Tanya Curran, Susan Feeney, Alison Watt, Ciara Cox, Mairead Connor, Zoltan Molnar, David Simpson, Derek Fairley |
| EPI_ISL_461791, EPI_ISL_461792, EPI_ISL_461793, EPI_ISL_461794, EPI_ISL_461795, EPI_ISL_461796, EPI_ISL_461797, EPI_ISL_461798, EPI_ISL_461799, EPI_ISL_461800, EPI_ISL_461801, EPI_ISL_461802, EPI_ISL_461803, EPI_ISL_461804 | | | |
| see above | Northumbria University / South Tees Hospitals NHS Foundation Trust / North Cumbria Integrated Care NHS Foundation Trust / North Tees and Hartlepool NHS Foundation Trust / Newcastle Hospitals NHS Foundation Trust | COVID-19 Genomics UK (COG-UK) Consortium | Darren L Smith,Andrew Nelson,Matthew Bashton,Greg R Young,Joshua Loh,John Allan,Mohammad A Tariq,Giles S Holt,Gary Black,Wen C Yew,Lynn Dover,Paul Baker,Steve Liggett,Sarah Essex,Jane Greenaway,Debra Padgett,Clive Graham,Garren Scott,Edward Barton,Emma Swindells,Brendan Payne,Jennifer Collins,Yusri Taha,Gary Eltringham |
| EPI_ISL_461805, EPI_ISL_461806, EPI_ISL_461807, EPI_ISL_461808, EPI_ISL_461809, EPI_ISL_461810, EPI_ISL_461811, EPI_ISL_461812, EPI_ISL_461813, EPI_ISL_461814, EPI_ISL_461815, EPI_ISL_461816, EPI_ISL_461817, EPI_ISL_461818, EPI_ISL_461819, EPI_ISL_461820, EPI_ISL_461821, EPI_ISL_461822, EPI_ISL_461823, EPI_ISL_461824, EPI_ISL_461825, EPI_ISL_461826, EPI_ISL_461827, EPI_ISL_461828, EPI_ISL_461829, EPI_ISL_461830, EPI_ISL_461831, EPI_ISL_461832, EPI_ISL_461833, EPI_ISL_461834, EPI_ISL_461835, EPI_ISL_461836, EPI_ISL_461837, EPI_ISL_461838, EPI_ISL_461839, EPI_ISL_461840, EPI_ISL_461841, EPI_ISL_461842, EPI_ISL_461843, EPI_ISL_461844, EPI_ISL_461845, EPI_ISL_461846, EPI_ISL_461847, EPI_ISL_461848, EPI_ISL_461849, EPI_ISL_461850, EPI_ISL_461851, EPI_ISL_461852, EPI_ISL_461853, EPI_ISL_461854, EPI_ISL_461855, EPI_ISL_461856, EPI_ISL_461857, EPI_ISL_461858, EPI_ISL_461859, EPI_ISL_461860, EPI_ISL_461861, EPI_ISL_461862, EPI_ISL_461863, EPI_ISL_461864, EPI_ISL_461865, EPI_ISL_461866, EPI_ISL_461867, EPI_ISL_461868, EPI_ISL_461869, EPI_ISL_461870, EPI_ISL_461871, EPI_ISL_461872, EPI_ISL_461873, EPI_ISL_461874, EPI_ISL_461875, EPI_ISL_461876, EPI_ISL_461877, EPI_ISL_461878, EPI_ISL_461879, EPI_ISL_461880, EPI_ISL_461881, EPI_ISL_461882, EPI_ISL_461883, EPI_ISL_461884, EPI_ISL_461885, EPI_ISL_461886, EPI_ISL_461887, EPI_ISL_461888, EPI_ISL_461889, EPI_ISL_461890, EPI_ISL_461891, EPI_ISL_461892, EPI_ISL_461893 | | | |
| see above | Quadram Institute Bioscience | COVID-19 Genomics UK (COG-UK) | Dave J. Baker, Gemma L. Kay, Alp Aydin, Thanh Le-Viet, Steven Rudder, Ana P. Tedim, Anastasia Kolyva, Maria Diaz, Leonardo de Oliveira Martins, Nabil-Fareed Alikhan, Lizzie Meadows, Rachael Stanley, Ngozi Elumogo, Muhammed Yasir, Nicholas M. Thomson, |

| | | | |
|---|---|---|---|
| | | Consortium | Alexander J Trotter, Rachel Gilroy, Samuel Bloomfield, Claire Stuart, Andrew Bell, Reenesh Prakash, Samir Dervisevic, Alison E. Mather, John Wain, Mark Webber, Andrew J. Page, Justin O'Grady |
| EPI_ISL_461894, EPI_ISL_461895, EPI_ISL_461896, EPI_ISL_461897, EPI_ISL_461898, EPI_ISL_461899, EPI_ISL_461900, EPI_ISL_461901, EPI_ISL_461902, EPI_ISL_461903, EPI_ISL_461904, EPI_ISL_461905, EPI_ISL_461906, EPI_ISL_461907, EPI_ISL_461908, EPI_ISL_461909, EPI_ISL_461910, EPI_ISL_461911, EPI_ISL_461912, EPI_ISL_461913, EPI_ISL_461914, EPI_ISL_461915, EPI_ISL_461916, EPI_ISL_461917, EPI_ISL_461918, EPI_ISL_461919, EPI_ISL_461920, EPI_ISL_461921, EPI_ISL_461922, EPI_ISL_461923, EPI_ISL_461924, EPI_ISL_461925, EPI_ISL_461926, EPI_ISL_461927, EPI_ISL_461928, EPI_ISL_461929, EPI_ISL_461930, EPI_ISL_461931, EPI_ISL_461932, EPI_ISL_461933, EPI_ISL_461934, EPI_ISL_461935, EPI_ISL_461936, EPI_ISL_461937, EPI_ISL_461938, EPI_ISL_461939, EPI_ISL_461940, EPI_ISL_461941, EPI_ISL_461942, EPI_ISL_461943, EPI_ISL_461944, EPI_ISL_461945, EPI_ISL_461946, EPI_ISL_461947, EPI_ISL_461948, EPI_ISL_461949, EPI_ISL_461950, EPI_ISL_461951, EPI_ISL_461952, EPI_ISL_461953, EPI_ISL_461954, EPI_ISL_461955, EPI_ISL_461956, EPI_ISL_461957, EPI_ISL_461958, EPI_ISL_461959, EPI_ISL_461960, EPI_ISL_461961, EPI_ISL_461962, EPI_ISL_461963, EPI_ISL_461964, EPI_ISL_461965, EPI_ISL_461966, EPI_ISL_461967, EPI_ISL_461968, EPI_ISL_461969 | | | |
| see above | Queens Medical Centre, Clinical Microbiology Department / DeepSeq Nottingham | COVID-19 Genomics UK (COG-UK) Consortium | Gemma Clark, Wendy Smith, Manjinder Khakh, Hannah Howson-Wells, Jonathan Ball, Patrick McClure, Joseph Chappell, Theocharis Tsoleridis, Nadine Holmes, Matthew Carlisle, Christopher Moore, Fei Sang, Johnny Debebe, Victoria Wright, Matthew Loose |
| EPI_ISL_461970, EPI_ISL_461971, EPI_ISL_461972, EPI_ISL_461973, EPI_ISL_461974, EPI_ISL_461975, EPI_ISL_461976, EPI_ISL_461977, EPI_ISL_461978, EPI_ISL_461979, EPI_ISL_461980, EPI_ISL_461981, EPI_ISL_461982, EPI_ISL_461983, EPI_ISL_461984, EPI_ISL_461985, EPI_ISL_461986, EPI_ISL_461987, EPI_ISL_461988, EPI_ISL_461989, EPI_ISL_461990, EPI_ISL_461991, EPI_ISL_461992, EPI_ISL_461993, EPI_ISL_461994, EPI_ISL_461995, EPI_ISL_461996, EPI_ISL_461997, EPI_ISL_461998 | | | |
| see above | Centre for Enzyme Innovation, University of Portsmouth / Translational Research Laboratory, Portsmouth Hospitals NHS Trust | COVID-19 Genomics UK (COG-UK) Consortium | Angela Beckett,Yann Bourgeois,Garry Scarlett,Sharon Glaysher,Scott Elliott,Kelly Bicknell,Robert Impey,Allyson Lloyd,Sarah Wyllie,Ethan Butcher,Anoop Chauhan,Samuel Robson |
| EPI_ISL_461999, EPI_ISL_462000, EPI_ISL_462001, EPI_ISL_462002, EPI_ISL_462003, EPI_ISL_462004, EPI_ISL_462005, EPI_ISL_462006, EPI_ISL_462007, EPI_ISL_462008, EPI_ISL_462009, EPI_ISL_462010, EPI_ISL_462011, EPI_ISL_462012, EPI_ISL_462013, EPI_ISL_462014, EPI_ISL_462015, EPI_ISL_462016, EPI_ISL_462017, EPI_ISL_462018, EPI_ISL_462019, EPI_ISL_462020, EPI_ISL_462021, EPI_ISL_462022, EPI_ISL_462023, EPI_ISL_462024, EPI_ISL_462025, EPI_ISL_462026, EPI_ISL_462027, EPI_ISL_462028, EPI_ISL_462029, EPI_ISL_462030, EPI_ISL_462031, EPI_ISL_462032, EPI_ISL_462033, EPI_ISL_462034, EPI_ISL_462035, EPI_ISL_462036, EPI_ISL_462037, EPI_ISL_462038, EPI_ISL_462039, EPI_ISL_462040, EPI_ISL_462041, EPI_ISL_462042, EPI_ISL_462043, EPI_ISL_462044, EPI_ISL_462045, EPI_ISL_462046, EPI_ISL_462047, EPI_ISL_462048, EPI_ISL_462049, EPI_ISL_462050, EPI_ISL_462051, EPI_ISL_462052, EPI_ISL_462053, EPI_ISL_462054, EPI_ISL_462055, EPI_ISL_462056, EPI_ISL_462057, EPI_ISL_462058, EPI_ISL_462059, EPI_ISL_462060, EPI_ISL_462061, EPI_ISL_462062, EPI_ISL_462063, EPI_ISL_462064, EPI_ISL_462065, EPI_ISL_462066, EPI_ISL_462067, EPI_ISL_462068, EPI_ISL_462069, EPI_ISL_462070, EPI_ISL_462071, EPI_ISL_462072, EPI_ISL_462073, EPI_ISL_462074, EPI_ISL_462075, EPI_ISL_462076, EPI_ISL_462077, EPI_ISL_462078, EPI_ISL_462079, EPI_ISL_462080, EPI_ISL_462081, EPI_ISL_462082, EPI_ISL_462083, EPI_ISL_462084 | | | |
| see above | Virology Department, Sheffield Teaching Hospitals NHS Foundation Trust/Department of Infection, Immunity and Cardiovascular Disease, The Medical School, University of Sheffield | COVID-19 Genomics UK (COG-UK) Consortium | Thushan de Silva, Matthew Parker, Nikki Smith, Adri Angyal, Rebecca Brown, Luke Green, Rachel Tucker, Paul Parsons, Danielle Groves, Katie Johnson, Laura Carrilero, Alex Keeley, Dave Partridge, Matthew Wyles, Benjamin Lindsey, Mehmet Yavuz, Mohammad Raza, Cariad Evans |
| EPI_ISL_462085, EPI_ISL_462086, EPI_ISL_462087, EPI_ISL_462088, EPI_ISL_462089 | Singapore General Hospital | Department of Microbiology | Nurdyana Abdul Rahman, Kun Lee Lim, Chenhao Li, Kian Sing Chan, Lynette Oon, Kern Rei Chng, Niranjan Nagarajan, Karrie Ko |
| EPI_ISL_462090 | National Institute of Laboratory Medicine and Referral Center | Genomic Research Lab, BCSIR | Barna Goswami, Abu Sayeed Mohammad Mahmud, Mohammad Samir Uzzaman, Eshrar Osman, Md. Ahasan Habib, Shahina Akter, Tanjina Akhter Banu, Iffat Jahan, Md. Saddam Hossain, Tasnim Nafisa, Md. Maruf Ahmed Molla, Mahmuda Yeasmin, Asish Kumar Ghos, Bayzid Bin Monir, Arifa Akram, Sheikh Md. Selim Al Din, Salek Ahmed Sajib, Utpal Chandra Ray, Md. Salim Khan |
| EPI_ISL_462091 | National Institute of Laboratory Medicine and Referral Center | Genomic Research Lab, BCSIR | Iffat Jahan, Abu Sayeed Mohammad Mahmud, Mohammad Samir Uzzaman, Eshrar Osman, Md. Ahasan Habib, Shahina Akter, Tanjina Akhter Banu, Barna Goswami, Md. Saddam Hossain, Tasnim Nafisa, Md. Maruf Ahmed Molla, Mahmuda Yeasmin, Asish Kumar Ghos, Bayzid Bin Monir, Arifa Akram, Sheikh Md. Selim Al Din, Salek Ahmed Sajib, Utpal Chandra Ray, Md. Salim Khan |
| EPI_ISL_462092 | National Institute of Laboratory Medicine and Referral Center | Genomic Research Lab, BCSIR | Shahina Akter, Abu Sayeed Mohammad Mahmud, Mohammad Samir Uzzaman, Eshrar Osman, Md. Ahasan Habib, Tanjina Akhter Banu, Barna Goswami, Iffat Jahan, Md. Saddam Hossain, Tasnim Nafisa, Md. Maruf Ahmed Molla, Mahmuda Yeasmin, Asish Kumar Ghos, Bayzid Bin Monir, Arifa Akram, Sheikh Md. Selim Al Din, Salek Ahmed Sajib, Utpal Chandra Ray, Md. Salim Khan |
| EPI_ISL_462093 | National Institute of Laboratory Medicine and Referral Center | Genomic Research Lab, BCSIR | Abu Sayeed Mohammad Mahmud, Mohammad Samir Uzzaman, Eshrar Osman, Md. Ahasan Habib, Tanjina Akhter Banu, Shahina Akter, Barna Goswami, Iffat Jahan, Md. Saddam Hossain, Tasnim Nafisa, Md. Maruf Ahmed Molla, Mahmuda Yeasmin, Asish Kumar Ghosh, Bayzid Bin Monir, Arifa Akram, Sheikh Md. Selim Al Din, Salek Ahmed Sajib, Utpal Chandra Ray, Md. Salim Khan |
| EPI_ISL_462094, EPI_ISL_462095 | National Institute of Laboratory Medicine and Referral Center | Genomic Research Lab, BCSIR | Abu Sayeed Mohammad Mahmud, Mohammad Samir Uzzaman, Eshrar Osman, Md. Ahasan Habib, Tanjina Akhter Banu, Shahina Akter, Barna Goswami, Iffat Jahan, Md. Saddam Hossain, Tasnim Nafisa, Md. Maruf Ahmed Molla, Mahmuda Yeasmin, Asish Kumar Ghosh, Bayzid Bin Monir, Arifa Akram, Sheikh Md. Selim Al Din, Salek Ahmed Sajib, Utpal Chandra Ray, Md. Salim Khan |
| EPI_ISL_462096 | National Institute of Laboratory Medicine and Referral Center | Genomic Research Lab, BCSIR | Abu Sayeed Mohammad Mahmud, Mohammad Samir Uzzaman, Eshrar Osman, Md. Ahasan Habib, Tanjina Akhter Banu, Shahina Akter, Barna Goswami, Iffat Jahan, Md. Saddam Hossain, Tasnim Nafisa, Md. Maruf Ahmed Molla, Mahmuda Yeasmin, Asish Kumar Ghosh, Bayzid Bin Monir, Arifa Akram, Sheikh Md. Selim Al Din, Salek Ahmed Sajib, Utpal Chandra Ray, Md. Salim Khan |
| EPI_ISL_462097 | National Institute of Laboratory Medicine and Referral Center | Genomic Research Lab, BCSIR | Abu Sayeed Mohammad Mahmud, Mohammad Samir Uzzaman, Eshrar Osman, Md. Ahasan Habib, Tanjina Akhter Banu, Shahina Akter, Barna Goswami, Iffat Jahan, Md. Saddam Hossain, Tasnim Nafisa, Md. Maruf Ahmed Molla, Mahmuda Yeasmin, Asish Kumar Ghosh, Bayzid Bin Monir, Arifa Akram, Sheikh Md. Selim Al Din, Salek Ahmed Sajib, Utpal Chandra Ray, Md. Salim Khan |
| EPI_ISL_462098 | National Institute of Laboratory Medicine and Referral Center | Genomic Research Lab, BCSIR | Abu Sayeed Mohammad Mahmud, Mohammad Samir Uzzaman, Eshrar Osman, Md. Ahasan Habib, Tanjina Akhter Banu, Shahina Akter, Barna Goswami, Iffat Jahan, Md. Saddam Hossain, Tasnim Nafisa, Md. Maruf Ahmed Molla, Mahmuda Yeasmin, Asish Kumar Ghosh, Bayzid Bin Monir, Arifa Akram, Sheikh Md. Selim Al Din, Salek Ahmed Sajib, Utpal Chandra Ray, Md. Salim Khan |
| EPI_ISL_462149, EPI_ISL_462150 | Molecular diagnostic laboratory of Federal Budget Institution of Science "Central Research Institute of Epidemiology" of The Federal Service on Customers' Rights Protection and Human Well-being Surveillance | Group of Genomics and Postgenomic Technologies of Central Research Institute of Epidemiology | Speranskaya AS, Kaptelova VV, Samoilov AE, Korneenko EV, Sizova TV, Tivanova EV, Shipulina OY, Akimkin VG |
| EPI_ISL_462151, EPI_ISL_462152, EPI_ISL_462153, EPI_ISL_462154, EPI_ISL_462155, EPI_ISL_462156, EPI_ISL_462157, EPI_ISL_462158, EPI_ISL_462159, EPI_ISL_462160, EPI_ISL_462161, EPI_ISL_462162, EPI_ISL_462163, EPI_ISL_462164, EPI_ISL_462165, EPI_ISL_462166, EPI_ISL_462167, EPI_ISL_462168, EPI_ISL_462169, EPI_ISL_462170, EPI_ISL_462171, EPI_ISL_462172, EPI_ISL_462173, EPI_ISL_462174, EPI_ISL_462175, EPI_ISL_462176, EPI_ISL_462177, EPI_ISL_462178, EPI_ISL_462179, EPI_ISL_462180, EPI_ISL_462181, EPI_ISL_462182, EPI_ISL_462183, EPI_ISL_462184, EPI_ISL_462185, EPI_ISL_462186, EPI_ISL_462187, EPI_ISL_462188, EPI_ISL_462189, EPI_ISL_462190, EPI_ISL_462191, EPI_ISL_462192, EPI_ISL_462193, EPI_ISL_462194, EPI_ISL_462195, EPI_ISL_462196, EPI_ISL_462197, EPI_ISL_462198, EPI_ISL_462199, EPI_ISL_462200, EPI_ISL_462201, EPI_ISL_462202, EPI_ISL_462203, EPI_ISL_462204, EPI_ISL_462205, EPI_ISL_462206, EPI_ISL_462207, EPI_ISL_462208, EPI_ISL_462209, EPI_ISL_462210, EPI_ISL_462211, EPI_ISL_462212, EPI_ISL_462213, EPI_ISL_462214, EPI_ISL_462215, EPI_ISL_462216, EPI_ISL_462217, EPI_ISL_462218, EPI_ISL_462219, EPI_ISL_462220, EPI_ISL_462221, EPI_ISL_462222, EPI_ISL_462223, EPI_ISL_462224, EPI_ISL_462225, EPI_ISL_462226, EPI_ISL_462227, EPI_ISL_462228, EPI_ISL_462229, EPI_ISL_462230, EPI_ISL_462231, EPI_ISL_462232, EPI_ISL_462233, EPI_ISL_462234, EPI_ISL_462235, EPI_ISL_462236, EPI_ISL_462237, EPI_ISL_462238, EPI_ISL_462239, EPI_ISL_462240, EPI_ISL_462241, EPI_ISL_462242, EPI_ISL_462243, EPI_ISL_462244, EPI_ISL_462245, EPI_ISL_462246, EPI_ISL_462247, EPI_ISL_462248, EPI_ISL_462249, EPI_ISL_462250, EPI_ISL_462251, EPI_ISL_462252, EPI_ISL_462253, EPI_ISL_462254, EPI_ISL_462255, EPI_ISL_462256, EPI_ISL_462257, EPI_ISL_462258, EPI_ISL_462259, EPI_ISL_462260, EPI_ISL_462261, EPI_ISL_462262, EPI_ISL_462263, EPI_ISL_462264, EPI_ISL_462265, EPI_ISL_462266, EPI_ISL_462267, EPI_ISL_462268, EPI_ISL_462269, EPI_ISL_462270, EPI_ISL_462271, EPI_ISL_462272, EPI_ISL_462273, EPI_ISL_462274, EPI_ISL_462275 | | | |
| see above | KU Leuven, Rega Institute, Clinical and Epidemiological Virology | KU Leuven, Rega Institute, Clinical and Epidemiological Virology | Tony Wawina-Bokalanga, Bert Vanmechelen, Joan Marti-Carerras, Piet Maes |
| EPI_ISL_462276, EPI_ISL_462277, EPI_ISL_462278, EPI_ISL_462279, EPI_ISL_462280, EPI_ISL_462281, EPI_ISL_462282, EPI_ISL_462283, EPI_ISL_462284, EPI_ISL_462285, EPI_ISL_462286, EPI_ISL_462287, EPI_ISL_462288, EPI_ISL_462289, EPI_ISL_462290, EPI_ISL_462291, EPI_ISL_462292, EPI_ISL_462293, EPI_ISL_462294, EPI_ISL_462295, EPI_ISL_462296, EPI_ISL_462297, EPI_ISL_462298, EPI_ISL_462299, EPI_ISL_462300, EPI_ISL_462301, EPI_ISL_462302, EPI_ISL_462303, EPI_ISL_462304, EPI_ISL_462305, EPI_ISL_462306, EPI_ISL_462307, EPI_ISL_462308, EPI_ISL_462309, EPI_ISL_462310, EPI_ISL_462311, EPI_ISL_462312, EPI_ISL_462313, EPI_ISL_462314, EPI_ISL_462315, EPI_ISL_462316, EPI_ISL_462317, EPI_ISL_462318, EPI_ISL_462319, EPI_ISL_462320, EPI_ISL_462321, EPI_ISL_462322, EPI_ISL_462323, EPI_ISL_462324, EPI_ISL_462325, EPI_ISL_462326, EPI_ISL_462327, EPI_ISL_462328, EPI_ISL_462329, EPI_ISL_462330, EPI_ISL_462331, EPI_ISL_462332, EPI_ISL_462333, EPI_ISL_462334, EPI_ISL_462335, EPI_ISL_462336, EPI_ISL_462337, EPI_ISL_462338, EPI_ISL_462339, EPI_ISL_462340, EPI_ISL_462341, EPI_ISL_462342, EPI_ISL_462343, EPI_ISL_462344, EPI_ISL_462345, EPI_ISL_462346, EPI_ISL_462347, EPI_ISL_462348, EPI_ISL_462349, EPI_ISL_462350, EPI_ISL_462351, EPI_ISL_462352, EPI_ISL_462353, EPI_ISL_462354, EPI_ISL_462355, EPI_ISL_462356, EPI_ISL_462357, EPI_ISL_462358, EPI_ISL_462359, EPI_ISL_462360, EPI_ISL_462361, EPI_ISL_462362, EPI_ISL_462363, EPI_ISL_462364, EPI_ISL_462365, EPI_ISL_462366, EPI_ISL_462367, EPI_ISL_462368, EPI_ISL_462369, EPI_ISL_462370, EPI_ISL_462371, EPI_ISL_462372, EPI_ISL_462373, EPI_ISL_462374, EPI_ISL_462375, EPI_ISL_462376, EPI_ISL_462377, EPI_ISL_462378, EPI_ISL_462379, EPI_ISL_462380, EPI_ISL_462381, EPI_ISL_462382, EPI_ISL_462383, EPI_ISL_462384, EPI_ISL_462385, EPI_ISL_462386, EPI_ISL_462387, EPI_ISL_462388, EPI_ISL_462389, EPI_ISL_462390, EPI_ISL_462391, EPI_ISL_462392, EPI_ISL_462393, EPI_ISL_462394, EPI_ISL_462395, EPI_ISL_462396, EPI_ISL_462397, EPI_ISL_462398, EPI_ISL_462399, EPI_ISL_462400, EPI_ISL_462401, EPI_ISL_462402, EPI_ISL_462403, EPI_ISL_462404, EPI_ISL_462405, EPI_ISL_462406, EPI_ISL_462407, EPI_ISL_462408, EPI_ISL_462409, EPI_ISL_462410, EPI_ISL_462411, EPI_ISL_462412, EPI_ISL_462413, EPI_ISL_462414, EPI_ISL_462415, EPI_ISL_462416, EPI_ISL_462417, EPI_ISL_462418, EPI_ISL_462419, EPI_ISL_462420, EPI_ISL_462421, EPI_ISL_462422, EPI_ISL_462423, EPI_ISL_462424, EPI_ISL_462425, EPI_ISL_462426, EPI_ISL_462427, EPI_ISL_462428, EPI_ISL_462429, EPI_ISL_462430, EPI_ISL_462431, EPI_ISL_462432, EPI_ISL_462433 | | | |
| see above | National Public Health Laboratory, National Centre for Infectious Diseases | National Public Health Laboratory, National Centre for Infectious Diseases | Mak TM, Octavia S, Chavatte JM, Cui L, Lin RTP |
| EPI_ISL_462434, EPI_ISL_462435, EPI_ISL_462436, EPI_ISL_462437, EPI_ISL_462438 | unknown | Laboratory Diagnostic | Vidanovic,D., Tesovic,B., Banovic Djeri,B., Knezevic,A., Jankovic,M., Sekler,M., Dmitric,M., Petrovic,T., Volkening,J., Afonso,C.L. |
| EPI_ISL_462439, EPI_ISL_462440, EPI_ISL_462441, EPI_ISL_462442, EPI_ISL_462443, EPI_ISL_462444, EPI_ISL_462445 | unknown | Ryota Kumagai Tokyo Metropolitan Institute of Public Health | Asakura,H., Kumagai,R., Yoshida,I., Nagashima,M., Chiba,T., Sadamasu,K. |
| EPI_ISL_462447, EPI_ISL_462448 | Fundació Lluita contra la SIDA (FLSida)/Hospital Universitari Germans Trias i Pujol | IrsiCaixa AIDS Research Lab | Marc Noguera-Julian, Mariona Parera, Maria Pilar Armengol, Marc Corbacho, Maria Ubals, Oriol Mitjà, Lidia Ruiz, Nuria Izquierdo, Jorge Carrillo, Roger Paredes, Julia Blanco, Joaquim Segalés, Bonaventura Clotet |
| EPI_ISL_462449 | Fundació Lluita contra la SIDA (FLSida)/Hospital Universitari Germans Trias i Pujol | IrsiCaixa AIDS Research Lab | Marc Noguera-Julian, Mariona Parera, Maria Pilar Armengol, Marc Corbacho, Maria Ubals, Oriol Mitjà, Lidia Ruiz, Nuria Izquierdo, Jorge Carrillo, Roger Paredes, Julia Blanco, Bonaventura Clotet |
| EPI_ISL_462450, EPI_ISL_462451, EPI_ISL_462452, EPI_ISL_462453, EPI_ISL_462454, EPI_ISL_462455, EPI_ISL_462456, EPI_ISL_462457, EPI_ISL_462458, EPI_ISL_462459, EPI_ISL_462460, EPI_ISL_462461, EPI_ISL_462462, EPI_ISL_462463, EPI_ISL_462464, EPI_ISL_462465, EPI_ISL_462466, EPI_ISL_462467, EPI_ISL_462468, EPI_ISL_462469, EPI_ISL_462470, EPI_ISL_462471, EPI_ISL_462472, EPI_ISL_462473, EPI_ISL_462474, EPI_ISL_462475, EPI_ISL_462476 | | | |
| see above | Clinical Center, University of Sarajevo | Charite Universitatsmedizin Berlin, Institute of Virology | Victor M Corman, Jorn Beheim-Schwarzbach, Barbara Muehlemann, Talitha Veith, Julia Schneider, Terry Jones, Amela Dedeic-Ljubovic, Irma Salimovic-Besic, Suzana Arapcic, Almedina Hadzihasanovic-Moro, Selma Mutevelic, Christian Drosten |
| EPI_ISL_462477 | Hospital Costa del Sol | Instituto de Salud Carlos III | Iglesias-Caballero, M. Molinero Calamita, M. González-Esguevillas, M. Camarero, S. Pozo, F. Casas, I. Jiménez, P. Jiménez, M. Zaballos, A. Monzón, S. Varona, S. Juliá, M. Cuesta, I, F. Fernández |
| EPI_ISL_462480 | Institute of Human Genetics, Polish Academy of Sciences | Institute of Human Genetics, Polish Academy of Sciences | Szymon Hryhorowicz, Adam Ustaszewski, Emilia Lis, Marta Kaczmarek-Ryś, Michał Witt, Andrzej Pławski |
| EPI_ISL_462636, EPI_ISL_462637, EPI_ISL_462638, EPI_ISL_462639, EPI_ISL_462640, EPI_ISL_462641, EPI_ISL_462642, EPI_ISL_462643, EPI_ISL_462644, EPI_ISL_462645, EPI_ISL_462646, EPI_ISL_462647, EPI_ISL_462648, EPI_ISL_462649, EPI_ISL_462650, EPI_ISL_462651, EPI_ISL_462652, EPI_ISL_462653, EPI_ISL_462654, EPI_ISL_462655, EPI_ISL_462656, EPI_ISL_462657, EPI_ISL_462658, EPI_ISL_462659, EPI_ISL_462660, EPI_ISL_462661, EPI_ISL_462662, EPI_ISL_462663, EPI_ISL_462664, EPI_ISL_462665, EPI_ISL_462666, EPI_ISL_462667, EPI_ISL_462668, EPI_ISL_462669, EPI_ISL_462670, EPI_ISL_462671, EPI_ISL_462672, EPI_ISL_462673, EPI_ISL_462674, EPI_ISL_462675, EPI_ISL_462676, EPI_ISL_462677, EPI_ISL_462678, EPI_ISL_462679, EPI_ISL_462680, EPI_ISL_462681, EPI_ISL_462682, EPI_ISL_462683, EPI_ISL_462684, EPI_ISL_462685, EPI_ISL_462686, EPI_ISL_462687, EPI_ISL_462688, EPI_ISL_462689, EPI_ISL_462690, EPI_ISL_462691, EPI_ISL_462692, EPI_ISL_462693, EPI_ISL_462694, EPI_ISL_462695, EPI_ISL_462696, EPI_ISL_462697, EPI_ISL_462698, EPI_ISL_462699, EPI_ISL_462700, EPI_ISL_462701, EPI_ISL_462702, EPI_ISL_462703, EPI_ISL_462704, EPI_ISL_462705, EPI_ISL_462706, EPI_ISL_462707, EPI_ISL_462708, EPI_ISL_462709, EPI_ISL_462710, EPI_ISL_462711, EPI_ISL_462712, EPI_ISL_462713, EPI_ISL_462714, EPI_ISL_462715, EPI_ISL_462716, EPI_ISL_462717, EPI_ISL_462718, EPI_ISL_462719, EPI_ISL_462720, EPI_ISL_462721, EPI_ISL_462722, EPI_ISL_462723, EPI_ISL_462724, EPI_ISL_462725, EPI_ISL_462726, EPI_ISL_462727, EPI_ISL_462728, EPI_ISL_462729, EPI_ISL_462730, EPI_ISL_462731, EPI_ISL_462732, EPI_ISL_462733, EPI_ISL_462734, EPI_ISL_462735, EPI_ISL_462736 | | | |
| see above | Michigan Department of Health and Human Services, Bureau of Laboratories | Michigan Department of Health and Human Services, Bureau of Laboratories | Blankenship HM, Riner D, Soehnlen MK |
| EPI_ISL_462753 | University Clinical Hospital of Mostar | University of Sarajevo Veterinary Faculty | Goletic, T., Softic, A., Goletic, S., Ostojic, M., Hukic, M., Eterovic, T., Seho-Alic, A. |
| EPI_ISL_462754, EPI_ISL_462755, EPI_ISL_462756, EPI_ISL_462757, EPI_ISL_462758, EPI_ISL_462759, EPI_ISL_462760, EPI_ISL_462761, EPI_ISL_462762, EPI_ISL_462763, EPI_ISL_462764, EPI_ISL_462765, EPI_ISL_462766, EPI_ISL_462767, EPI_ISL_462768, EPI_ISL_462769, EPI_ISL_462770, EPI_ISL_462771, EPI_ISL_462772, EPI_ISL_462773, EPI_ISL_462774, EPI_ISL_462775, EPI_ISL_462776, EPI_ISL_462777, EPI_ISL_462778, EPI_ISL_462779, EPI_ISL_462780, EPI_ISL_462781, EPI_ISL_462782, EPI_ISL_462783, EPI_ISL_462784, EPI_ISL_462785, EPI_ISL_462786, EPI_ISL_462787, EPI_ISL_462788, EPI_ISL_462789, EPI_ISL_462790, EPI_ISL_462791, EPI_ISL_462792, EPI_ISL_462793, EPI_ISL_462794, EPI_ISL_462795, EPI_ISL_462796, EPI_ISL_462797, EPI_ISL_462798, EPI_ISL_462799, EPI_ISL_462800, EPI_ISL_462801, EPI_ISL_462802, EPI_ISL_462803, EPI_ISL_462804, EPI_ISL_462805, EPI_ISL_462806, EPI_ISL_462807, EPI_ISL_462808, EPI_ISL_462809, EPI_ISL_462810, EPI_ISL_462811, EPI_ISL_462812, EPI_ISL_462813, EPI_ISL_462814, EPI_ISL_462815, EPI_ISL_462816, EPI_ISL_462817, EPI_ISL_462818, EPI_ISL_462819, EPI_ISL_462820, EPI_ISL_462821, EPI_ISL_462822, EPI_ISL_462823, EPI_ISL_462824, EPI_ISL_462825, EPI_ISL_462826, EPI_ISL_462827, EPI_ISL_462828, EPI_ISL_462829, EPI_ISL_462830, EPI_ISL_462831, EPI_ISL_462832, EPI_ISL_462833, EPI_ISL_462834, EPI_ISL_462835, EPI_ISL_462836, EPI_ISL_462837, EPI_ISL_462838, EPI_ISL_462839, EPI_ISL_462840, EPI_ISL_462841, EPI_ISL_462842, EPI_ISL_462843, EPI_ISL_462844 | | | |
| see above | BCCDC Public Health Laboratory | BCCDC Public Health Laboratory | Harrigan, Prystajecky, Krajden, Lee, Kamelian, Lapointe, Choi, Hoang, Sekirov, Levett, Tyson, Li, Gilmour |

| | | | |
|---|---|---|---|
| EPI_ISL_462845, EPI_ISL_462846, EPI_ISL_462847, EPI_ISL_462848, EPI_ISL_462849, EPI_ISL_462850, EPI_ISL_462851, EPI_ISL_462852, EPI_ISL_462853, EPI_ISL_462854, EPI_ISL_462855, EPI_ISL_462856, EPI_ISL_462857, EPI_ISL_462858, EPI_ISL_462859, EPI_ISL_462860, EPI_ISL_462861, EPI_ISL_462862, EPI_ISL_462863, EPI_ISL_462864, EPI_ISL_462865, EPI_ISL_462866, EPI_ISL_462867, EPI_ISL_462868, EPI_ISL_462869, EPI_ISL_462870, EPI_ISL_462871, EPI_ISL_462872, EPI_ISL_462873, EPI_ISL_462874, EPI_ISL_462875, EPI_ISL_462876, EPI_ISL_462877, EPI_ISL_462878, EPI_ISL_462879, EPI_ISL_462880, EPI_ISL_462881, EPI_ISL_462882, EPI_ISL_462883, EPI_ISL_462884, EPI_ISL_462885, EPI_ISL_462886, EPI_ISL_462887, EPI_ISL_462888, EPI_ISL_462889, EPI_ISL_462890, EPI_ISL_462891, EPI_ISL_462892, EPI_ISL_462893, EPI_ISL_462894, EPI_ISL_462895, EPI_ISL_462896, EPI_ISL_462897, EPI_ISL_462898, EPI_ISL_462899, EPI_ISL_462900, EPI_ISL_462901, EPI_ISL_462902, EPI_ISL_462903, EPI_ISL_462904, EPI_ISL_462905, EPI_ISL_462906, EPI_ISL_462907, EPI_ISL_462908, EPI_ISL_462909, EPI_ISL_462910, EPI_ISL_462911 | | | |
| see above | Minnesota Department of Health, Public Health Laboratory | Minnesota Department of Health, Public Health Laboratory | Matt Plumb, Jacob Garfin, and Xiong Wang |
| EPI_ISL_462912, EPI_ISL_462913, EPI_ISL_462914, EPI_ISL_462915, EPI_ISL_462916, EPI_ISL_462917, EPI_ISL_462918, EPI_ISL_462919, EPI_ISL_462920, EPI_ISL_462921, EPI_ISL_462922, EPI_ISL_462923, EPI_ISL_462924, EPI_ISL_462925, EPI_ISL_462926, EPI_ISL_462927, EPI_ISL_462928, EPI_ISL_462929, EPI_ISL_462930, EPI_ISL_462931, EPI_ISL_462932, EPI_ISL_462933, EPI_ISL_462934, EPI_ISL_462935, EPI_ISL_462936, EPI_ISL_462937, EPI_ISL_462938, EPI_ISL_462939, EPI_ISL_462940, EPI_ISL_462941, EPI_ISL_462942, EPI_ISL_462943, EPI_ISL_462944, EPI_ISL_462945, EPI_ISL_462946, EPI_ISL_462947, EPI_ISL_462948, EPI_ISL_462949, EPI_ISL_462950, EPI_ISL_462951, EPI_ISL_462952, EPI_ISL_462953, EPI_ISL_462954, EPI_ISL_462955, EPI_ISL_462956, EPI_ISL_462957, EPI_ISL_462958, EPI_ISL_462959, EPI_ISL_462960, EPI_ISL_462961, EPI_ISL_462962, EPI_ISL_462963, EPI_ISL_462964, EPI_ISL_462965, EPI_ISL_462966, EPI_ISL_462967, EPI_ISL_462968, EPI_ISL_462969, EPI_ISL_462970, EPI_ISL_462971, EPI_ISL_462972, EPI_ISL_462973, EPI_ISL_462974, EPI_ISL_462975, EPI_ISL_462976, EPI_ISL_462977, EPI_ISL_462978, EPI_ISL_462979, EPI_ISL_462980, EPI_ISL_462981, EPI_ISL_462982, EPI_ISL_462983, EPI_ISL_462984, EPI_ISL_462985, EPI_ISL_462986, EPI_ISL_462987, EPI_ISL_462988, EPI_ISL_462989 | | | |
| see above | Wyoming Public Health Laboratory | Center for Global Health, University of New Mexico Health Sciences Center | Daryl Domman, Kurt Schwalm, Rob Christensen, Wanda Manley, Cari Sloma, Noah Hull, Darrell Dinwiddie |
| EPI_ISL_462990 | University Clinical Centre of the Republic of Srpska | University of Sarajevo, Veterinary Faculty | Teufik, G., Šejla, G., Toni, E., Maja, T., Mirsada, H., Aida, K., Alma, Š.A. |
| EPI_ISL_462991 | Microbiology Division | Microbiology Division | Flores,H. |
| EPI_ISL_462992 | Nigerian Institute of Medical Research | Nigerian Institute of Medical Research | Saibu,J.O., Onwuamah,C.K., Okwuraiwe,A.P., Amoo,O.S., Salu,O.B., Ige,F.A., Liboro,G., Odewale,E., Adesegun,A., Abosede,O., Ahmed,R., Sokei,J., Oyefolu,A., Adegbola,R., Salako,B., Omilabu,S. and Audu,R. |
| EPI_ISL_462995, EPI_ISL_462996, EPI_ISL_462997, EPI_ISL_462998, EPI_ISL_462999, EPI_ISL_463000 | Molecular Genetics | Molecular Genetics | Gomez,J., Coto,E. |
| EPI_ISL_463007 | Department of Laboratory Medicine, National Taiwan University Hospital | Microbial Genomics Core Lab, National Taiwan University Centers of Genomic and Precision Medicine | Shiou-Hwei Yeh, You-Yu Lin, Ya-Yun Lai, Chiao-Ling Li, Shan-Chwen Chang, Pei-Jer Chen, Sui-Yuan Chang |
| EPI_ISL_463008 | Institute of Molecular Virology, University Münster | Institute of Molecular Virology, University Münster | Angeles Mecate Zambrano, Linda Brunotte, Stephan Ludwig, Joachim Kühn, Alexander Mellmann |
| EPI_ISL_463010, EPI_ISL_463011, EPI_ISL_463012, EPI_ISL_463013, EPI_ISL_463014, EPI_ISL_463015, EPI_ISL_463016, EPI_ISL_463017, EPI_ISL_463018, EPI_ISL_463019, EPI_ISL_463020, EPI_ISL_463021, EPI_ISL_463022, EPI_ISL_463023, EPI_ISL_463024, EPI_ISL_463025, EPI_ISL_463026, EPI_ISL_463027, EPI_ISL_463028, EPI_ISL_463029, EPI_ISL_463030 | | | |
| see above | Institute of Life Sciences, Bhubaneswar | Immunogenomics lab, Institute of Life Sciences, Bhubaneswar | Sunil Raghav, Arup Ghosh, Atimukta Jha, Viplov K. Biswas, Swati Madhulika, Manasi Priyadarshini, Shuchi Smita, Kaushik Sen, Hiren G. Dodia, Deepak Singh, Jeky Chawla, Shamima Ansari, Rupesh Dash, Soma Chattopadhyay, Ghulam Hussain Syed, Shanti Senapati, Tushar K. Beuria, Rajeeb Swain, Punit Prasad, ILS COVID-19 TEAM, Orissa COVID-19 Study Group, DBT's PAN-INDIA 1000 SARS-CoV2 RNA genome sequencing consortium, Ajay Parida |
| EPI_ISL_463031, EPI_ISL_463032, EPI_ISL_463033, EPI_ISL_463034, EPI_ISL_463035, EPI_ISL_463036, EPI_ISL_463037, EPI_ISL_463038, EPI_ISL_463039, EPI_ISL_463040, EPI_ISL_463041, EPI_ISL_463042, EPI_ISL_463043, EPI_ISL_463044, EPI_ISL_463045, EPI_ISL_463046, EPI_ISL_463047, EPI_ISL_463048, EPI_ISL_463049, EPI_ISL_463050, EPI_ISL_463051 | | | |
| see above | Institute of Life Sciences, Bhubaneswar | Immunogenomics lab, Institute of Life Sciences, Bhubaneswar | Sunil Raghav, Arup Ghosh, Atimukta Jha, Viplov K. Biswas, Swati Madhulika, Manasi Priyadarshini, Shuchi Smita, O. P. Shriwas, Priyanka Mohapatra, Satya Ranjan Sahu, Aliva Minz, Debyashrita Barik, Rupesh Dash, Soma Chattopadhyay, Ghulam Hussain Syed, Shanti Senapati, Tushar K. Beuria, Rajeeb Swain, Punit Prasad, ILS COVID-19 TEAM, Orissa COVID-19 Study Group, DBT's PAN-INDIA 1000 SARS-CoV2 RNA genome sequencing consortium, Ajay Parida |
| EPI_ISL_463052, EPI_ISL_463053, EPI_ISL_463054, EPI_ISL_463055, EPI_ISL_463056, EPI_ISL_463057, EPI_ISL_463058, EPI_ISL_463059, EPI_ISL_463060, EPI_ISL_463061, EPI_ISL_463062, EPI_ISL_463063, EPI_ISL_463064, EPI_ISL_463065, EPI_ISL_463066, EPI_ISL_463067, EPI_ISL_463068, EPI_ISL_463069, EPI_ISL_463070, EPI_ISL_463071, EPI_ISL_463072 | | | |
| see above | Institute of Life Sciences, Bhubaneswar | Immunogenomics lab, Institute of Life Sciences, Bhubaneswar | Sunil Raghav, Arup Ghosh, Atimukta Jha, Viplov K. Biswas, Swati Madhulika, Manasi Priyadarshini, Shuchi Smita, Sifu Agarwal, Sanchari Chatterjee, Avula Kiran, Parej Nath, Supriya Suman, Rina Yadav, Rupesh Dash, Soma Chattopadhyay, Ghulam Hussain Syed, Shanti Senapati, Tushar K. Beuria, Rajeeb Swain, Punit Prasad, ILS COVID-19 TEAM, Orissa COVID-19 Study Group, DBT's PAN-INDIA 1000 SARS-CoV2 RNA genome sequencing consortium, Ajay Parida |
| EPI_ISL_463073, EPI_ISL_463074, EPI_ISL_463075, EPI_ISL_463076, EPI_ISL_463077, EPI_ISL_463078, EPI_ISL_463079, EPI_ISL_463080, EPI_ISL_463081, EPI_ISL_463082, EPI_ISL_463083, EPI_ISL_463084, EPI_ISL_463085, EPI_ISL_463086, EPI_ISL_463087, EPI_ISL_463088, EPI_ISL_463089, EPI_ISL_463090, EPI_ISL_463091, EPI_ISL_463092, EPI_ISL_463093 | | | |
| see above | Institute of Life Sciences, Bhubaneswar | Immunogenomics lab, Institute of Life Sciences, Bhubaneswar | Sunil Raghav, Arup Ghosh, Atimukta Jha, Viplov K. Biswas, Swati Madhulika, Manasi Priyadarshini, Shuchi Smita, Kautilya Kumar Jena, Sandhya Suranjika, Neha Singh, Eshna Laha, Saiket De, Rupesh Dash, Soma Chattopadhyay, Ghulam Hussain Syed, Shanti Senapati, Tushar K. Beuria, Rajeeb Swain, Punit Prasad, ILS COVID-19 TEAM, Orissa COVID-19 Study Group, DBT's PAN-INDIA 1000 SARS-CoV2 RNA genome sequencing consortium, Ajay Parida |
| EPI_ISL_463094, EPI_ISL_463095, EPI_ISL_463096, EPI_ISL_463097, EPI_ISL_463098, EPI_ISL_463099, EPI_ISL_463100, EPI_ISL_463101, EPI_ISL_463102, EPI_ISL_463103, EPI_ISL_463104, EPI_ISL_463105, EPI_ISL_463106, EPI_ISL_463107, EPI_ISL_463108, EPI_ISL_463109, EPI_ISL_463110, EPI_ISL_463111, EPI_ISL_463112, EPI_ISL_463113, EPI_ISL_463114, EPI_ISL_463115, EPI_ISL_463116, EPI_ISL_463117, EPI_ISL_463118, EPI_ISL_463119, EPI_ISL_463120, EPI_ISL_463121, EPI_ISL_463122, EPI_ISL_463123, EPI_ISL_463124, EPI_ISL_463125, EPI_ISL_463126, EPI_ISL_463127, EPI_ISL_463128, EPI_ISL_463129, EPI_ISL_463130, EPI_ISL_463131, EPI_ISL_463132, EPI_ISL_463133, EPI_ISL_463134, EPI_ISL_463135, EPI_ISL_463136, EPI_ISL_463137 | | | |
| see above | Virginia DCLS | Virginia DCLS | Virginia DCLS |
| EPI_ISL_463138, EPI_ISL_463139, EPI_ISL_463140, EPI_ISL_463141, EPI_ISL_463142, EPI_ISL_463143, EPI_ISL_463144, EPI_ISL_463145, EPI_ISL_463146, EPI_ISL_463147, EPI_ISL_463148, EPI_ISL_463149, EPI_ISL_463150, EPI_ISL_463151, EPI_ISL_463152, EPI_ISL_463153, EPI_ISL_463154, EPI_ISL_463155, EPI_ISL_463156, EPI_ISL_463157, EPI_ISL_463158, EPI_ISL_463159, EPI_ISL_463160, EPI_ISL_463161, EPI_ISL_463162, EPI_ISL_463163, EPI_ISL_463164, EPI_ISL_463165, EPI_ISL_463166, EPI_ISL_463167, EPI_ISL_463168, EPI_ISL_463169, EPI_ISL_463170, EPI_ISL_463171, EPI_ISL_463172, EPI_ISL_463173, EPI_ISL_463174 | | | |
| see above | Yale Clinical Virology Laboratory | Grubaugh Lab - Yale School of Public Health | Joseph Fauver, Tara Alpert, Anderson Brito, Anne Wyllie, Chantal Vogels, Mary Petrone, Cole Jensen, Chaney Kalinich, Isabel Ott, Arnau Casanovas, Catherine Muenker, Adam Moore, Alice Lu, Maria Tokuyama, Patrick Wong, Peiwen Lu, Saad Omer, Richard Martinello, Allison Nelson, Shelli Farhadian, Akiko Iwasaki, Charlese Dela Cruz, Albert Ko, Nathan Grubaugh |
| EPI_ISL_463186, EPI_ISL_463187, EPI_ISL_463188, EPI_ISL_463189, EPI_ISL_463190, EPI_ISL_463191, EPI_ISL_463192, EPI_ISL_463193, EPI_ISL_463194, EPI_ISL_463195, EPI_ISL_463196, EPI_ISL_463197, EPI_ISL_463198, EPI_ISL_463199, EPI_ISL_463200, EPI_ISL_463201, EPI_ISL_463202, EPI_ISL_463203, EPI_ISL_463204, EPI_ISL_463205, EPI_ISL_463206, EPI_ISL_463207, EPI_ISL_463208, EPI_ISL_463209, EPI_ISL_463210, EPI_ISL_463211, EPI_ISL_463212, EPI_ISL_463213, EPI_ISL_463214, EPI_ISL_463215, EPI_ISL_463216, EPI_ISL_463217, EPI_ISL_463218, EPI_ISL_463219, EPI_ISL_463220, EPI_ISL_463221, EPI_ISL_463222, EPI_ISL_463223, EPI_ISL_463224, EPI_ISL_463225, EPI_ISL_463226, EPI_ISL_463227, EPI_ISL_463228, EPI_ISL_463229, EPI_ISL_463230, EPI_ISL_463231, EPI_ISL_463232, EPI_ISL_463233, EPI_ISL_463234, EPI_ISL_463235, EPI_ISL_463236, EPI_ISL_463237, EPI_ISL_463238, EPI_ISL_463239, EPI_ISL_463240, EPI_ISL_463241, EPI_ISL_463242, EPI_ISL_463243, EPI_ISL_463244, EPI_ISL_463245, EPI_ISL_463246, EPI_ISL_463247, EPI_ISL_463248, EPI_ISL_463249, EPI_ISL_463250, EPI_ISL_463251, EPI_ISL_463252, EPI_ISL_463253, EPI_ISL_463254, EPI_ISL_463255, EPI_ISL_463256, EPI_ISL_463257, EPI_ISL_463258, EPI_ISL_463259, EPI_ISL_463260, EPI_ISL_463261, EPI_ISL_463262, EPI_ISL_463263, EPI_ISL_463264, EPI_ISL_463265, EPI_ISL_463266, EPI_ISL_463267, EPI_ISL_463268, EPI_ISL_463269, EPI_ISL_463270, EPI_ISL_463271, EPI_ISL_463272, EPI_ISL_463273, EPI_ISL_463274, EPI_ISL_463275, EPI_ISL_463276 | | | |
| see above | BCCDC Public Health Laboratory | BCCDC Public Health Laboratory | Richard Harrigan, Hope Lapointe, Jinny Choi, Kimia Kamelian, John Tyson,Terry Snutch, Linda Hoang, Inna Sekirov, Paul Levett, Mel Krajden, Natalie Prystajecky |
| EPI_ISL_463277, EPI_ISL_463278, EPI_ISL_463279, EPI_ISL_463280, EPI_ISL_463281, EPI_ISL_463282, EPI_ISL_463283, EPI_ISL_463284, EPI_ISL_463285, EPI_ISL_463286, EPI_ISL_463287, EPI_ISL_463288, EPI_ISL_463289, EPI_ISL_463290, EPI_ISL_463291, EPI_ISL_463292, EPI_ISL_463293, EPI_ISL_463294, EPI_ISL_463295, EPI_ISL_463296, EPI_ISL_463297, EPI_ISL_463298, EPI_ISL_463299, EPI_ISL_463300 | | | |
| see above | Ochsner Health | Bioinfoexperts, LLC | Susanna L. Lamers, David J. Nolan, Rebecca Rose, Sissy Cross, David Moraga Amador, Tong Yang, Luke Caruso, Wayra Navia, Lydia Von Borstel, Xiao Hui Zhou, Amy Feehan, Julia-Garcia-Diaz |
| EPI_ISL_463301 | Tuen Mun Hospital | Hong Kong Department of Health | Mak Gannon C.K., Cheng Peter K.C., Lam Edman T.K., Chan Rickjason C.W., Tsang Dominic N.C. |
| EPI_ISL_463302, EPI_ISL_463303 | United Christian Hospital | Hong Kong Department of Health | Mak Gannon C.K., Cheng Peter K.C., Lam Edman T.K., Chan Rickjason C.W., Tsang Dominic N.C. |
| EPI_ISL_463304 | Mrs Wu York Yu GOPC | Hong Kong Department of Health | Mak Gannon C.K., Cheng Peter K.C., Lam Edman T.K., Chan Rickjason C.W., Tsang Dominic N.C. |
| EPI_ISL_463305 | Queen Elizabeth Hospital | Hong Kong Department of Health | Mak Gannon C.K., Cheng Peter K.C., Lam Edman T.K., Chan Rickjason C.W., Tsang Dominic N.C. |
| EPI_ISL_463306 | Prince of Wales Hospital | Hong Kong Department of Health | Mak Gannon C.K., Cheng Peter K.C., Lam Edman T.K., Chan Rickjason C.W., Tsang Dominic N.C. |
| EPI_ISL_463307 | HK Molecular Pathology Disgnostic Centre | Hong Kong Department of Health | Mak Gannon C.K., Cheng Peter K.C., Lam Edman T.K., Chan Rickjason C.W., Tsang Dominic N.C. |
| EPI_ISL_463308 | Princess Margaret Hospital | Hong Kong Department of Health | Mak Gannon C.K., Cheng Peter K.C., Lam Edman T.K., Chan Rickjason C.W., Tsang Dominic N.C. |
| EPI_ISL_463309 | Prince of Wales Hospital | Hong Kong Department of Health | Mak Gannon C.K., Cheng Peter K.C., Lam Edman T.K., Chan Rickjason C.W., Tsang Dominic N.C. |
| EPI_ISL_463310 | Tseung Kwan O Hospital | Hong Kong Department of Health | Mak Gannon C.K., Cheng Peter K.C., Lam Edman T.K., Chan Rickjason C.W., Tsang Dominic N.C. |
| EPI_ISL_463311, EPI_ISL_463312 | Queen Elizabeth Hospital | Hong Kong Department of Health | Mak Gannon C.K., Cheng Peter K.C., Lam Edman T.K., Chan Rickjason C.W., Tsang Dominic N.C. |
| EPI_ISL_463313 | Central Health Medical Practice | Hong Kong Department of Health | Mak Gannon C.K., Cheng Peter K.C., Lam Edman T.K., Chan Rickjason C.W., Tsang Dominic N.C. |
| EPI_ISL_463314 | Queen Elizabeth Hospital | Hong Kong Department of Health | Mak Gannon C.K., Cheng Peter K.C., Lam Edman T.K., Chan Rickjason C.W., Tsang Dominic N.C. |
| EPI_ISL_463315 | Princess Margaret Hospital | Hong Kong Department of Health | Mak Gannon C.K., Cheng Peter K.C., Lam Edman T.K., Chan Rickjason C.W., Tsang Dominic N.C. |
| EPI_ISL_463316, EPI_ISL_463317 | Yan Chai Hospital | Hong Kong Department of Health | Mak Gannon C.K., Cheng Peter K.C., Lam Edman T.K., Chan Rickjason C.W., Tsang Dominic N.C. |
| EPI_ISL_463318 | Pamela Youde Nethersole Eastern Hospital | Hong Kong Department of Health | Mak Gannon C.K., Cheng Peter K.C., Lam Edman T.K., Chan Rickjason C.W., Tsang Dominic N.C. |
| EPI_ISL_463319 | Prince of Wales Hospital | Hong Kong Department of Health | Mak Gannon C.K., Cheng Peter K.C., Lam Edman T.K., Chan Rickjason C.W., Tsang Dominic N.C. |
| EPI_ISL_463320 | Tseung Kwan O Hospital | Hong Kong Department of Health | Mak Gannon C.K., Cheng Peter K.C., Lam Edman T.K., Chan Rickjason C.W., Tsang Dominic N.C. |
| EPI_ISL_463321 | Queen Mary Hospital | Hong Kong Department of Health | Mak Gannon C.K., Cheng Peter K.C., Lam Edman T.K., Chan Rickjason C.W., Tsang Dominic N.C. |
| EPI_ISL_463322 | North Lantau Hospital | Hong Kong Department of Health | Mak Gannon C.K., Cheng Peter K.C., Lam Edman T.K., Chan Rickjason C.W., Tsang Dominic N.C. |
| EPI_ISL_463323 | Yuen Long JC GOPC | Hong Kong Department of Health | Mak Gannon C.K., Cheng Peter K.C., Lam Edman T.K., Chan Rickjason C.W., Tsang Dominic N.C. |
| EPI_ISL_463324, EPI_ISL_463325, EPI_ISL_463326, EPI_ISL_463327, EPI_ISL_463328, EPI_ISL_463329, EPI_ISL_463330, EPI_ISL_463331, EPI_ISL_463332, EPI_ISL_463333, EPI_ISL_463334, EPI_ISL_463335, EPI_ISL_463336, EPI_ISL_463337, EPI_ISL_463338, EPI_ISL_463339, EPI_ISL_463340, EPI_ISL_463341, EPI_ISL_463342, EPI_ISL_463343, EPI_ISL_463344, EPI_ISL_463345, EPI_ISL_463346, EPI_ISL_463347, EPI_ISL_463348, EPI_ISL_463349, EPI_ISL_463350, EPI_ISL_463351, EPI_ISL_463352, EPI_ISL_463353, EPI_ISL_463354, EPI_ISL_463355, EPI_ISL_463356, EPI_ISL_463357, EPI_ISL_463358, EPI_ISL_463359, EPI_ISL_463360, EPI_ISL_463361, EPI_ISL_463362, EPI_ISL_463363, EPI_ISL_463364, EPI_ISL_463365, EPI_ISL_463366, EPI_ISL_463367, EPI_ISL_463368, EPI_ISL_463369, EPI_ISL_463370, EPI_ISL_463371, EPI_ISL_463372, EPI_ISL_463373, EPI_ISL_463374, EPI_ISL_463375, EPI_ISL_463376, EPI_ISL_463377, EPI_ISL_463378, EPI_ISL_463379, EPI_ISL_463380, EPI_ISL_463381, EPI_ISL_463382, EPI_ISL_463383, EPI_ISL_463384, EPI_ISL_463385, EPI_ISL_463386, EPI_ISL_463387, EPI_ISL_463388, EPI_ISL_463389, EPI_ISL_463390, EPI_ISL_463391, EPI_ISL_463392, EPI_ISL_463393, EPI_ISL_463394, EPI_ISL_463395, EPI_ISL_463396, EPI_ISL_463397, EPI_ISL_463398, EPI_ISL_463399, EPI_ISL_463400, EPI_ISL_463401, EPI_ISL_463402, EPI_ISL_463403, EPI_ISL_463404, EPI_ISL_463405, EPI_ISL_463406, EPI_ISL_463407, EPI_ISL_463408, EPI_ISL_463409, EPI_ISL_463410, EPI_ISL_463411, EPI_ISL_463412, EPI_ISL_463413, EPI_ISL_463414, EPI_ISL_463415, EPI_ISL_463416, EPI_ISL_463417, EPI_ISL_463418, EPI_ISL_463419, EPI_ISL_463420, EPI_ISL_463421, EPI_ISL_463422, EPI_ISL_463423, EPI_ISL_463424, EPI_ISL_463425, EPI_ISL_463426, EPI_ISL_463427, EPI_ISL_463428, EPI_ISL_463429, EPI_ISL_463430, EPI_ISL_463431, EPI_ISL_463432, EPI_ISL_463433, EPI_ISL_463434, EPI_ISL_463435, EPI_ISL_463436, EPI_ISL_463437, EPI_ISL_463438, EPI_ISL_463439, EPI_ISL_463440, EPI_ISL_463441, EPI_ISL_463442, EPI_ISL_463443, EPI_ISL_463444, EPI_ISL_463445, EPI_ISL_463446, EPI_ISL_463447, EPI_ISL_463448, EPI_ISL_463449, EPI_ISL_463450, EPI_ISL_463451, EPI_ISL_463452, EPI_ISL_463453, EPI_ISL_463454, EPI_ISL_463455, EPI_ISL_463456, EPI_ISL_463457, EPI_ISL_463458, EPI_ISL_463459, EPI_ISL_463460, EPI_ISL_463461, EPI_ISL_463462, EPI_ISL_463463, EPI_ISL_463464, EPI_ISL_463465, EPI_ISL_463466, EPI_ISL_463467, EPI_ISL_463468, EPI_ISL_463469, EPI_ISL_463470, EPI_ISL_463471, EPI_ISL_463472, EPI_ISL_463473, EPI_ISL_463474, EPI_ISL_463475, EPI_ISL_463476, EPI_ISL_463477, EPI_ISL_463478, EPI_ISL_463479, EPI_ISL_463480, EPI_ISL_463481, EPI_ISL_463482, EPI_ISL_463483, EPI_ISL_463484, EPI_ISL_463485, EPI_ISL_463486, EPI_ISL_463487, EPI_ISL_463488, EPI_ISL_463489, EPI_ISL_463490, EPI_ISL_463491, EPI_ISL_463492, EPI_ISL_463493, EPI_ISL_463494, EPI_ISL_463495, EPI_ISL_463496, EPI_ISL_463497, EPI_ISL_463498, EPI_ISL_463499, EPI_ISL_463500, EPI_ISL_463501, EPI_ISL_463502, EPI_ISL_463503, EPI_ISL_463504, EPI_ISL_463505, EPI_ISL_463506, EPI_ISL_463507, EPI_ISL_463508, EPI_ISL_463509, EPI_ISL_463510, EPI_ISL_463511, EPI_ISL_463512, EPI_ISL_463513, EPI_ISL_463514, EPI_ISL_463515, EPI_ISL_463516, EPI_ISL_463517, EPI_ISL_463518, EPI_ISL_463519, EPI_ISL_463520, EPI_ISL_463521, EPI_ISL_463522, EPI_ISL_463523, EPI_ISL_463524, EPI_ISL_463525, EPI_ISL_463526, EPI_ISL_463527, EPI_ISL_463528, EPI_ISL_463529, EPI_ISL_463530, EPI_ISL_463531, EPI_ISL_463532, EPI_ISL_463533, EPI_ISL_463534, EPI_ISL_463535, EPI_ISL_463536, EPI_ISL_463537, EPI_ISL_463538, EPI_ISL_463539, EPI_ISL_463540, EPI_ISL_463541, EPI_ISL_463542, EPI_ISL_463543, EPI_ISL_463544, EPI_ISL_463545, EPI_ISL_463546, EPI_ISL_463547, EPI_ISL_463548, EPI_ISL_463549, EPI_ISL_463550, EPI_ISL_463551, EPI_ISL_463552, EPI_ISL_463553, EPI_ISL_463554, EPI_ISL_463555, EPI_ISL_463556, EPI_ISL_463557, EPI_ISL_463558, EPI_ISL_463559, EPI_ISL_463560, EPI_ISL_463561, EPI_ISL_463562, EPI_ISL_463563, EPI_ISL_463564, EPI_ISL_463565, EPI_ISL_463566, EPI_ISL_463567, EPI_ISL_463568, EPI_ISL_463569, EPI_ISL_463570, EPI_ISL_463571, EPI_ISL_463572, EPI_ISL_463573, EPI_ISL_463574, EPI_ISL_463575, EPI_ISL_463576, EPI_ISL_463577, EPI_ISL_463578, EPI_ISL_463579, EPI_ISL_463580, EPI_ISL_463581, EPI_ISL_463582, EPI_ISL_463583, EPI_ISL_463584, EPI_ISL_463585, EPI_ISL_463586, EPI_ISL_463587, EPI_ISL_463588, EPI_ISL_463589, EPI_ISL_463590, EPI_ISL_463591, EPI_ISL_463592, EPI_ISL_463593, EPI_ISL_463594, EPI_ISL_463595, EPI_ISL_463596, EPI_ISL_463597, EPI_ISL_463598, EPI_ISL_463599, EPI_ISL_463600, EPI_ISL_463601, EPI_ISL_463602, EPI_ISL_463603, EPI_ISL_463604, EPI_ISL_463605, EPI_ISL_463606, EPI_ISL_463607, EPI_ISL_463608, EPI_ISL_463609, EPI_ISL_463610, EPI_ISL_463611, EPI_ISL_463612, EPI_ISL_463613, EPI_ISL_463614, EPI_ISL_463615, EPI_ISL_463616, EPI_ISL_463617, EPI_ISL_463618, EPI_ISL_463619, EPI_ISL_463620, EPI_ISL_463621, EPI_ISL_463622, EPI_ISL_463623, EPI_ISL_463624, EPI_ISL_463625, EPI_ISL_463626, EPI_ISL_463627, EPI_ISL_463628, EPI_ISL_463629, EPI_ISL_463630, EPI_ISL_463631, EPI_ISL_463632, EPI_ISL_463633, EPI_ISL_463634, EPI_ISL_463635, EPI_ISL_463636, EPI_ISL_463637, EPI_ISL_463638, EPI_ISL_463639, EPI_ISL_463640, EPI_ISL_463641, EPI_ISL_463642, EPI_ISL_463643, EPI_ISL_463644, EPI_ISL_463645, EPI_ISL_463646, EPI_ISL_463647, EPI_ISL_463648, EPI_ISL_463649, EPI_ISL_463650, EPI_ISL_463651, EPI_ISL_463652, EPI_ISL_463653, EPI_ISL_463654, EPI_ISL_463655, EPI_ISL_463656, EPI_ISL_463657, EPI_ISL_463658, EPI_ISL_463659, EPI_ISL_463660, EPI_ISL_463661, EPI_ISL_463662, EPI_ISL_463663, EPI_ISL_463664, EPI_ISL_463665, EPI_ISL_463666, EPI_ISL_463667, EPI_ISL_463668, EPI_ISL_463669, EPI_ISL_463670, EPI_ISL_463671, EPI_ISL_463672, EPI_ISL_463673, EPI_ISL_463674, EPI_ISL_463675, EPI_ISL_463676, EPI_ISL_463677, EPI_ISL_463678, EPI_ISL_463679, EPI_ISL_463680, EPI_ISL_463681, EPI_ISL_463682, EPI_ISL_463683, EPI_ISL_463684, EPI_ISL_463685, EPI_ISL_463686, EPI_ISL_463687, EPI_ISL_463688, EPI_ISL_463689, EPI_ISL_463690, EPI_ISL_463691, EPI_ISL_463692, EPI_ISL_463693, EPI_ISL_463694, EPI_ISL_463695, EPI_ISL_463696, EPI_ISL_463697, EPI_ISL_463698, EPI_ISL_463699, EPI_ISL_463700, EPI_ISL_463701, EPI_ISL_463702, EPI_ISL_463703, EPI_ISL_463704, EPI_ISL_463705 | | | |
| see above | Washington State Department of Health | Seattle Flu Study | Chu et al |
| EPI_ISL_463740 | Mohammed Bin Rashid University of Medicine and Health | Al Jalila Genomics Center | Ahmad Abou Tayoun, Tom Loney, Hamda Khansaheb, Sathishkumar Ramaswamy, Divinlal Harilal, Zulfa Omar Deesi, Rupa Murthy Varghese, Hanan Al Suwaidi, Abdulmajeed Alkhaja, Mohammed Uddin, Rifat Hamoudi, Rabih Halwani, Abiola Catherine Senok, |

| | | | |
|---|---|---|---|
| | Sciences | | Qutayba Hamid, Norbert Nowotny, Alawi Alsheikh-Ali |
| EPI_ISL_463741, EPI_ISL_463742, EPI_ISL_463743, EPI_ISL_463744, EPI_ISL_463745, EPI_ISL_463746, EPI_ISL_463747, EPI_ISL_463748 | Department of Molecular Virology, Cyprus Institute of Neurology and Genetics | Department of Molecular Virology, Cyprus Institute of Neurology and Genetics | Jan Richter, George Krashias, Christina Tryfonos, Stavros Bashiardes, Dana Koptides, Christina Christodoulou |
| EPI_ISL_463749 | Pasteur Institute of Iran | Rapid Response Team | Mahboobeh Rafigh, Kayhan Azadmanesh, Tahmineh Jalali, Fatemeh Fotouhi-Chahooki, Mohammad Hassan Pouriayevali, Arash Arashkia, Zahra Ahmadi, Mohammad Sadegh Shams Nosrati, Ali Maleki, Zabihollah Shoja, Sanam Azad-Mazjiri, Mehdi Rohani, Saber Esmaeili, Ahmad Ghasemi, Amir Hesam Nemati, Ahmad Mahmoudi, Zahra Fereydouni, Mahsa Tavakolirad, Tahereh Mohammadi, Sahar Khakifirouz, Mehdi Fazlalipour, Hesam Karimi, Kazem Baesi, Seyed Dawood Mousavi Nasab, Mahmood Barati, Mohammad Reza Asadi Karam, Mehri Habibi, Neda Afzali, Ali Torabi, Azita Eshratkhah mohammadnejad, Seyedeh Sahar Bathaeian, Mohamad Mahdi Mortazavipour, Seyedeh Atefe Hosseini, Farideh niknam oskouei, Zahra Nejatipour, Parastoo Yekta Sanati, Hadiseh Shokouhi Targhi, Mahsa Ghalejoogh, Azam Amirian, Afsaneh Zokaei, Hajarossadat Ghaderi, Elmira Vadaye kheiri, Mina Agharezaei, Akram Abouie Mehrizi, Seyedeh Zahra Moravej, Mostafa Salehi-Vaziri |
| EPI_ISL_463889 | Shaoxing Center for Disease Control and Prevention | Department of Pathology and Laboratory Medicine, University of California Los Angeles | Jinkun Chen, Evann E. Hilt, Huan Wu, Zhuojing Jiang, QinChao Zhang, JiLing Wang, Yifang Wang, Fan Li, Ziqin Li, Jialiang Tang, Shangxin Yang |
| EPI_ISL_463893 | University Clinical Center Tuzla | Alea Genetički Centar | Konjhodžić,R;Salihefendić,L;Goletić,T;Pećar,D;Tihić,N;Marjanović,D;Hukić,M. |
| EPI_ISL_463894, EPI_ISL_463895, EPI_ISL_463896, EPI_ISL_463897, EPI_ISL_463898, EPI_ISL_463899, EPI_ISL_463900, EPI_ISL_463901, EPI_ISL_463902 | Shaoxing Center for Disease Control and Prevention | Department of Pathology and Laboratory Medicine, University of California Los Angeles | Jinkun Chen, Evann E. Hilt, Huan Wu, Zhuojing Jiang, QinChao Zhang, JiLing Wang, Yifang Wang, Fan Li, Ziqin Li, Jialiang Tang, Shangxin Yang |
| EPI_ISL_463903, EPI_ISL_463904, EPI_ISL_463905, EPI_ISL_463906, EPI_ISL_463907, EPI_ISL_463908, EPI_ISL_463909, EPI_ISL_463910, EPI_ISL_463911, EPI_ISL_463912, EPI_ISL_463913, EPI_ISL_463914, EPI_ISL_463915, EPI_ISL_463916, EPI_ISL_463917, EPI_ISL_463918, EPI_ISL_463919, EPI_ISL_463920, EPI_ISL_463921, EPI_ISL_463922, EPI_ISL_463923, EPI_ISL_463924, EPI_ISL_463925, EPI_ISL_463926, EPI_ISL_463927, EPI_ISL_463928, EPI_ISL_463929, EPI_ISL_463930, EPI_ISL_463931, EPI_ISL_463932, EPI_ISL_463933, EPI_ISL_463934, EPI_ISL_463935, EPI_ISL_463936, EPI_ISL_463937, EPI_ISL_463938, EPI_ISL_463939, EPI_ISL_463940, EPI_ISL_463941, EPI_ISL_463942, EPI_ISL_463943, EPI_ISL_463944, EPI_ISL_463945, EPI_ISL_463946, EPI_ISL_463947, EPI_ISL_463948, EPI_ISL_463949, EPI_ISL_463950, EPI_ISL_463951, EPI_ISL_463952, EPI_ISL_463953, EPI_ISL_463954, EPI_ISL_463955, EPI_ISL_463956, EPI_ISL_463957, EPI_ISL_463958, EPI_ISL_463959, EPI_ISL_463960, EPI_ISL_463961, EPI_ISL_463962, EPI_ISL_463963, EPI_ISL_463964, EPI_ISL_463965, EPI_ISL_463966, EPI_ISL_463967, EPI_ISL_463968, EPI_ISL_463969 | | | |
| see above | Laboratoire de microbiologie, Hopital de Verdun | Smith Laboratory, Centre de Recherche CHU Sainte-Justine | Martin Smith, Marieke Rozendaal, Ivan Pavlov |
| EPI_ISL_463970, EPI_ISL_463971, EPI_ISL_463972, EPI_ISL_463973, EPI_ISL_463974, EPI_ISL_463975, EPI_ISL_463976, EPI_ISL_463977, EPI_ISL_463978, EPI_ISL_463979, EPI_ISL_463980, EPI_ISL_463981, EPI_ISL_463982, EPI_ISL_463983, EPI_ISL_463984, EPI_ISL_463985, EPI_ISL_463986, EPI_ISL_463987, EPI_ISL_463988, EPI_ISL_463989, EPI_ISL_463990, EPI_ISL_463991, EPI_ISL_463992, EPI_ISL_463993, EPI_ISL_463994 | | | |
| see above | Toronto Invasive Bacterial Diseases Network | McMaster University | Allison McGeer, Patryk Aftanas, Angel Li, Kuganya Nirmalarajah, Samira Mubareka, Andrew G. McArthur |
| EPI_ISL_463995, EPI_ISL_463996, EPI_ISL_463997, EPI_ISL_463998, EPI_ISL_463999, EPI_ISL_464000, EPI_ISL_464001, EPI_ISL_464002, EPI_ISL_464003, EPI_ISL_464004, EPI_ISL_464005, EPI_ISL_464006, EPI_ISL_464007, EPI_ISL_464008, EPI_ISL_464009, EPI_ISL_464010, EPI_ISL_464011, EPI_ISL_464012, EPI_ISL_464013, EPI_ISL_464014, EPI_ISL_464015, EPI_ISL_464016, EPI_ISL_464017, EPI_ISL_464018, EPI_ISL_464019, EPI_ISL_464020, EPI_ISL_464021, EPI_ISL_464022, EPI_ISL_464023, EPI_ISL_464024, EPI_ISL_464025, EPI_ISL_464026, EPI_ISL_464027, EPI_ISL_464028, EPI_ISL_464029, EPI_ISL_464030, EPI_ISL_464031, EPI_ISL_464032, EPI_ISL_464033, EPI_ISL_464034, EPI_ISL_464035, EPI_ISL_464036, EPI_ISL_464037, EPI_ISL_464038, EPI_ISL_464039, EPI_ISL_464040, EPI_ISL_464041, EPI_ISL_464042, EPI_ISL_464043, EPI_ISL_464044, EPI_ISL_464045, EPI_ISL_464046, EPI_ISL_464047, EPI_ISL_464048, EPI_ISL_464049, EPI_ISL_464050, EPI_ISL_464051, EPI_ISL_464052, EPI_ISL_464053, EPI_ISL_464054, EPI_ISL_464055, EPI_ISL_464056, EPI_ISL_464057, EPI_ISL_464058, EPI_ISL_464059, EPI_ISL_464060, EPI_ISL_464061, EPI_ISL_464062, EPI_ISL_464063, EPI_ISL_464064 | | | |
| see above | Unity Health Toronto | Ontario Institute for Cancer Research | Ramzi Fattouh,Larissa M. Matukas,Mark Downing,Annette Gower,Karel Boissinot,Samira Mubareka,TIBDN,Ilinca Lungu,Bernard Lam,Jeremy Johns,Paul Krzyzanowski,Richard de Borja,Philip Zuzarte,Jared Simpson |
| EPI_ISL_464065, EPI_ISL_464066, EPI_ISL_464067, EPI_ISL_464068, EPI_ISL_464069, EPI_ISL_464070, EPI_ISL_464071, EPI_ISL_464072, EPI_ISL_464073, EPI_ISL_464074, EPI_ISL_464075, EPI_ISL_464076, EPI_ISL_464077, EPI_ISL_464078, EPI_ISL_464079, EPI_ISL_464080, EPI_ISL_464081, EPI_ISL_464082, EPI_ISL_464083, EPI_ISL_464084, EPI_ISL_464085, EPI_ISL_464086, EPI_ISL_464087, EPI_ISL_464088, EPI_ISL_464089, EPI_ISL_464090 | | | |
| see above | KU Leuven, Rega Institute, Clinical and Epidemiological Virology | KU Leuven, Rega Institute, Clinical and Epidemiological Virology | Tony Wawina-Bokalanga, Bert Vanmechelen, Joan Marti-Carerras, Piet Maes |
| EPI_ISL_464092, EPI_ISL_464093, EPI_ISL_464094 | Laboratory Medicine | Department of Laboratory Medicine, Lin-Kou Chang Gung Memorial Hospital, Taoyuan, Taiwan | Kuo-Chien Tsao, Yu-Nong Gong, Shu-Li Yang, Yi-Chun Liu, Chung-Guei Huang, Mei-Jen Hsiao, Po-Wei Huang, Cheng-Ta Yang, Cheng-Hsun Chiu, Peng-Nien Huang, Kuo-Ming Lee, Guang-Wu Chen, Shin-Ru Shih |
| EPI_ISL_464112, EPI_ISL_464113, EPI_ISL_464114, EPI_ISL_464115, EPI_ISL_464116, EPI_ISL_464117, EPI_ISL_464118, EPI_ISL_464119, EPI_ISL_464120, EPI_ISL_464121, EPI_ISL_464122, EPI_ISL_464123, EPI_ISL_464124, EPI_ISL_464125, EPI_ISL_464126, EPI_ISL_464127, EPI_ISL_464128, EPI_ISL_464129, EPI_ISL_464130, EPI_ISL_464131, EPI_ISL_464132, EPI_ISL_464133 | | | |
| see above | National Health Laboratory Service (NHLS), Tygerberg | Division of Medical Virology, Stellenbosch University and National Health Laboratory Service (NHLS) | Susan Engelbrecht, Kayla Delaney, Bronwyn Kleinhans, Houriiyah Tegally, Eduan Wilkindon, Gert van Zyl, Wolfgang Preiser, Tulio de Oliveira |
| EPI_ISL_464134 | National Health Laboratory Service (NHLS), Tygerberg | Stellenbosch University and NHLS | Susan Engelbrecht, Kayla Delaney, Bronwyn Kleinhans, Houriiyah Tegally, Eduan Wilkindon, Gert van Zyl, Wolfgang Preiser, Tulio de Oliveira |
| EPI_ISL_464135, EPI_ISL_464136, EPI_ISL_464137, EPI_ISL_464138 | National Health Laboratory Service (NHLS), Tygerberg | Division of Medical Virology, Stellenbosch University and National Health Laboratory Service (NHLS) | Susan Engelbrecht, Kayla Delaney, Bronwyn Kleinhans, Houriiyah Tegally, Eduan Wilkindon, Gert van Zyl, Wolfgang Preiser, Tulio de Oliveira |
| EPI_ISL_464139 | National Health Laboratory Service (NHLS), Tygerberg | Division of Medical Virology, Stellenbosch University and National Health Laboratory Service (NHLS) | Susan Engelbrecht, Kayla Delaney, Bronwyn Kleinhans, Houriiyah Tegally, Eduan Wilkindon, Gert van Zyl, Wolfgang Preiser, Tulio de Oliveira |
| EPI_ISL_464140 | National Health Laboratory Service (NHLS), Tygerberg | Division of Medical Virology, Stellenbosch University and National Health Laboratory Service (NHLS) | Susan Engelbrecht, Kayla Delaney, Bronwyn Kleinhans, Houriiyah Tegally, Eduan Wilkindon, Gert van Zyl, Wolfgang Preiser, Tulio de Oliveira |
| EPI_ISL_464141, EPI_ISL_464142, EPI_ISL_464143 | National Health Laboratory Service (NHLS), Tygerberg | Division of Medical Virology, Stellenbosch University and National Health Laboratory Service (NHLS) | Susan Engelbrecht, Kayla Delaney, Bronwyn Kleinhans, Houriiyah Tegally, Eduan Wilkindon, Gert van Zyl, Wolfgang Preiser, Tulio de Oliveira |
| EPI_ISL_464144, EPI_ISL_464145, EPI_ISL_464146, EPI_ISL_464147, EPI_ISL_464148, EPI_ISL_464149, EPI_ISL_464150, EPI_ISL_464151, EPI_ISL_464152, EPI_ISL_464153 | National Health Laboratory Service (NHLS), Tygerberg | Division of Medical Virology, Stellenbosch University and National Health Laboratory Service (NHLS) | Susan Engelbrecht, Kayla Delaney, Bronwyn Kleinhans, Houriiyah Tegally, Eduan Wilkindon, Gert van Zyl, Wolfgang Preiser, Tulio de Oliveira |
| EPI_ISL_464154 | National Health Laboratory Service (NHLS), Tygerberg | Division of Medical Virology, Stellenbosch University and National Health Laboratory Service (NHLS) | Susan Engelbrecht, Kayla Delaney, Bronwyn Kleinhans, Houriiyah Tegally, Eduan Wilkindon, Gert van Zyl, Wolfgang Preiser, Tulio de Oliveira |
| EPI_ISL_464155, EPI_ISL_464156, EPI_ISL_464157, EPI_ISL_464158 | National Health Laboratory Service (NHLS), Tygerberg | Division of Medical Virology, Stellenbosch University and National Health Laboratory Service (NHLS) | Susan Engelbrecht, Kayla Delaney, Bronwyn Kleinhans, Houriiyah Tegally, Eduan Wilkindon, Gert van Zyl, Wolfgang Preiser, Tulio de Oliveira |
| EPI_ISL_464159, EPI_ISL_464160 | National Institute of Laboratory Medicine and Referral Center | Genomic Research Lab, BCSIR | Shahina Akter, Abu Sayeed Mohammad Mahmud, Mohammad Samir Uzzaman, Eshrar Osman, Md. Ahasan Habib, Tanjina Akhter Banu, Md. Murshed Hasan Sarker, Barna Goswami, Iffat Jahan, Md. Saddam Hossain, Tasnim Nafisa, Md. Maruf Ahmed Molla, Mahmuda Yeasmin, Asish Kumar Ghosh, Arifa Akram, A. K. M. Shamsuzzaman, Sheikh Md. Selim Al Din, Utpal Chandra Ray, Salek Ahmed Sajib, Md. Salim Khan |
| EPI_ISL_464161, EPI_ISL_464162 | National Institute of Laboratory Medicine and Referral Center | Genomic Research Lab, BCSIR | Md. Ahasan Habib, Abu Sayeed Mohammad Mahmud, Mohammad Samir Uzzaman, Eshrar Osman, Shahina Akter, Tanjina Akhter Banu, Md. Murshed Hasan Sarker, Barna Goswami, Iffat Jahan, Md. Saddam Hossain, Tasnim Nafisa, Md. Maruf Ahmed Molla, Mahmuda Yeasmin, Asish Kumar Ghosh, Arifa Akram, A. K. M. Shamsuzzaman, Sheikh Md. Selim Al Din, Utpal Chandra Ray, Salek Ahmed Sajib, Md. Salim Khan |
| EPI_ISL_464163, EPI_ISL_464164 | National Institute of Laboratory Medicine and Referral Center | Genomic Research Lab, BCSIR | Tanjina Akhter Banu, Abu Sayeed Mohammad Mahmud, Mohammad Samir Uzzaman, Eshrar Osman, Md. Ahasan Habib, Shahina Akter, Md. Murshed Hasan Sarker, Barna Goswami, Iffat Jahan, Md. Saddam Hossain, Tasnim Nafisa, Md. Maruf Ahmed Molla, Mahmuda Yeasmin, Asish Kumar Ghosh, Arifa Akram, A. K. M. Shamsuzzaman, Sheikh Md. Selim Al Din, Utpal Chandra Ray, Salek Ahmed Sajib, Md. Salim Khan |
| EPI_ISL_464165, EPI_ISL_464166 | National Institute of Laboratory Medicine and Referral Center | Genomic Research Lab, BCSIR | Barna Goswami, Abu Sayeed Mohammad Mahmud, Mohammad Samir Uzzaman, Eshrar Osman, Md. Ahasan Habib, Shahina Akter, Tanjina Akhter Banu, Md. Murshed Hasan Sarker, Iffat Jahan, Md. Saddam Hossain, Tasnim Nafisa, Md. Maruf Ahmed Molla, Mahmuda Yeasmin, Asish Kumar Ghosh, Arifa Akram, A. K. M. Shamsuzzaman, Sheikh Md. Selim Al Din, Utpal Chandra Ray, Salek Ahmed Sajib, Md. Salim Khan |
| EPI_ISL_464167 | VI-US Virgin Islands Department of Health | Pathogen Discovery, Respiratory Viruses Branch, Division of Viral Diseases, Centers for Disease Control and Prevention | Krista Queen, Ying Tao, Jing Zhang, Yan Li, Anna Uehara, Clinton R. Paden, Mary S. Keckler, Alison S. Laufer Halpin, Haibin Wang, Jasmine Padilla, Justin Lee, Christopher A. Elkins, Suxiang Tong |
| EPI_ISL_464168, EPI_ISL_464169, EPI_ISL_464170, EPI_ISL_464171, EPI_ISL_464172, EPI_ISL_464173, EPI_ISL_464174, EPI_ISL_464175, EPI_ISL_464176, EPI_ISL_464177, EPI_ISL_464178, EPI_ISL_464179, EPI_ISL_464180, EPI_ISL_464181, EPI_ISL_464182, EPI_ISL_464183, EPI_ISL_464184, EPI_ISL_464185, EPI_ISL_464186, EPI_ISL_464187, EPI_ISL_464188, EPI_ISL_464189, EPI_ISL_464190, EPI_ISL_464191, EPI_ISL_464192, EPI_ISL_464193, EPI_ISL_464194, EPI_ISL_464195, EPI_ISL_464196, EPI_ISL_464197, EPI_ISL_464198, EPI_ISL_464199, EPI_ISL_464200, EPI_ISL_464201, EPI_ISL_464202, EPI_ISL_464203, EPI_ISL_464204, EPI_ISL_464205, EPI_ISL_464206, EPI_ISL_464207, EPI_ISL_464208, EPI_ISL_464209, EPI_ISL_464210, EPI_ISL_464211, EPI_ISL_464212, EPI_ISL_464213, EPI_ISL_464214, EPI_ISL_464215, EPI_ISL_464216, EPI_ISL_464217, EPI_ISL_464218, EPI_ISL_464219, EPI_ISL_464220, EPI_ISL_464221, EPI_ISL_464222, EPI_ISL_464223, EPI_ISL_464224, EPI_ISL_464225, EPI_ISL_464226, EPI_ISL_464227, EPI_ISL_464228, EPI_ISL_464229, EPI_ISL_464230, EPI_ISL_464231, EPI_ISL_464232, EPI_ISL_464233, EPI_ISL_464234, EPI_ISL_464235, EPI_ISL_464236, EPI_ISL_464237, EPI_ISL_464238, EPI_ISL_464239, EPI_ISL_464240, EPI_ISL_464241, EPI_ISL_464242, EPI_ISL_464243, EPI_ISL_464244, EPI_ISL_464245, EPI_ISL_464246, EPI_ISL_464247, EPI_ISL_464248, EPI_ISL_464249, EPI_ISL_464250, EPI_ISL_464251, EPI_ISL_464252, EPI_ISL_464253, EPI_ISL_464254, EPI_ISL_464255, EPI_ISL_464256, EPI_ISL_464257, EPI_ISL_464258, EPI_ISL_464259, EPI_ISL_464260, EPI_ISL_464261, EPI_ISL_464262, EPI_ISL_464263, EPI_ISL_464264, EPI_ISL_464265, EPI_ISL_464266, EPI_ISL_464267, EPI_ISL_464268, EPI_ISL_464269, EPI_ISL_464270, EPI_ISL_464271, EPI_ISL_464272, EPI_ISL_464273, EPI_ISL_464274, EPI_ISL_464275, EPI_ISL_464276, EPI_ISL_464277, EPI_ISL_464278, EPI_ISL_464279, EPI_ISL_464280, EPI_ISL_464281, EPI_ISL_464282, EPI_ISL_464283, EPI_ISL_464284, EPI_ISL_464285, EPI_ISL_464286, EPI_ISL_464287, EPI_ISL_464288, EPI_ISL_464289, EPI_ISL_464290, EPI_ISL_464291, EPI_ISL_464292, EPI_ISL_464293, EPI_ISL_464294, EPI_ISL_464295, EPI_ISL_464296, EPI_ISL_464297, EPI_ISL_464298, EPI_ISL_464299, EPI_ISL_464300, EPI_ISL_464301, EPI_ISL_464302, EPI_ISL_464303, EPI_ISL_464304, EPI_ISL_464305, EPI_ISL_464306, EPI_ISL_464307, EPI_ISL_464308, EPI_ISL_464309, EPI_ISL_464310, EPI_ISL_464311, EPI_ISL_464312, EPI_ISL_464313, EPI_ISL_464314, EPI_ISL_464315, EPI_ISL_464316, EPI_ISL_464317, EPI_ISL_464318, EPI_ISL_464319, EPI_ISL_464320, EPI_ISL_464321, EPI_ISL_464322, EPI_ISL_464323, EPI_ISL_464324, EPI_ISL_464325, EPI_ISL_464326, EPI_ISL_464327, EPI_ISL_464328, EPI_ISL_464329, EPI_ISL_464330, EPI_ISL_464331, EPI_ISL_464332, EPI_ISL_464333, EPI_ISL_464334, EPI_ISL_464335, EPI_ISL_464336, EPI_ISL_464337, EPI_ISL_464338, EPI_ISL_464339, EPI_ISL_464340, EPI_ISL_464341, EPI_ISL_464342, EPI_ISL_464343, EPI_ISL_464344, EPI_ISL_464345, EPI_ISL_464346, EPI_ISL_464347, EPI_ISL_464348, EPI_ISL_464349, EPI_ISL_464350, EPI_ISL_464351, EPI_ISL_464352, EPI_ISL_464353, EPI_ISL_464354, EPI_ISL_464355, EPI_ISL_464356, EPI_ISL_464357, EPI_ISL_464358, EPI_ISL_464359, EPI_ISL_464360, EPI_ISL_464361, EPI_ISL_464362, EPI_ISL_464363, EPI_ISL_464364, EPI_ISL_464365, EPI_ISL_464366, EPI_ISL_464367, EPI_ISL_464368, EPI_ISL_464369, EPI_ISL_464370, EPI_ISL_464371, EPI_ISL_464372, EPI_ISL_464373, EPI_ISL_464374, EPI_ISL_464375, EPI_ISL_464376, EPI_ISL_464377, EPI_ISL_464378, EPI_ISL_464379, EPI_ISL_464380, EPI_ISL_464381, EPI_ISL_464382, EPI_ISL_464383, EPI_ISL_464384, EPI_ISL_464385, EPI_ISL_464386, EPI_ISL_464387, EPI_ISL_464388, EPI_ISL_464389, EPI_ISL_464390, EPI_ISL_464391, EPI_ISL_464392, EPI_ISL_464393, EPI_ISL_464394, EPI_ISL_464395, EPI_ISL_464396, EPI_ISL_464397, EPI_ISL_464398, EPI_ISL_464399, EPI_ISL_464400, EPI_ISL_464401, EPI_ISL_464402, EPI_ISL_464403, EPI_ISL_464404, EPI_ISL_464405, EPI_ISL_464406, EPI_ISL_464407, EPI_ISL_464408, EPI_ISL_464409, EPI_ISL_464410, EPI_ISL_464411, EPI_ISL_464412, EPI_ISL_464413, EPI_ISL_464414, EPI_ISL_464415, EPI_ISL_464416, EPI_ISL_464417, EPI_ISL_464418, EPI_ISL_464419, EPI_ISL_464420, EPI_ISL_464421, EPI_ISL_464422, EPI_ISL_464423, EPI_ISL_464424, EPI_ISL_464425, EPI_ISL_464426, EPI_ISL_464427, EPI_ISL_464428, EPI_ISL_464429, EPI_ISL_464430, EPI_ISL_464431, | | | |

| | | | |
|---|---|---|---|
| EPI_ISL_464432, EPI_ISL_464433, EPI_ISL_464434, EPI_ISL_464435, EPI_ISL_464436, EPI_ISL_464437, EPI_ISL_464438, EPI_ISL_464439, EPI_ISL_464440, EPI_ISL_464441, EPI_ISL_464442, EPI_ISL_464443, EPI_ISL_464444, EPI_ISL_464445, EPI_ISL_464446, EPI_ISL_464447, EPI_ISL_464448, EPI_ISL_464449, EPI_ISL_464450, EPI_ISL_464451, EPI_ISL_464452, EPI_ISL_464453, EPI_ISL_464454, EPI_ISL_464455, EPI_ISL_464456, EPI_ISL_464457, EPI_ISL_464458, EPI_ISL_464459, EPI_ISL_464460, EPI_ISL_464461, EPI_ISL_464462, EPI_ISL_464463, EPI_ISL_464464, EPI_ISL_464465, EPI_ISL_464466, EPI_ISL_464467, EPI_ISL_464468, EPI_ISL_464469, EPI_ISL_464470, EPI_ISL_464471, EPI_ISL_464472, EPI_ISL_464473, EPI_ISL_464474, EPI_ISL_464475, EPI_ISL_464476, EPI_ISL_464477, EPI_ISL_464478, EPI_ISL_464479, EPI_ISL_464480, EPI_ISL_464481, EPI_ISL_464482, EPI_ISL_464483, EPI_ISL_464484, EPI_ISL_464485, EPI_ISL_464486, EPI_ISL_464487, EPI_ISL_464488, EPI_ISL_464489, EPI_ISL_464490, EPI_ISL_464491, EPI_ISL_464492, EPI_ISL_464493, EPI_ISL_464494, EPI_ISL_464495, EPI_ISL_464496, EPI_ISL_464497, EPI_ISL_464498, EPI_ISL_464499, EPI_ISL_464500, EPI_ISL_464501, EPI_ISL_464502, EPI_ISL_464503, EPI_ISL_464504, EPI_ISL_464505, EPI_ISL_464506, EPI_ISL_464507, EPI_ISL_464508, EPI_ISL_464509, EPI_ISL_464510, EPI_ISL_464511, EPI_ISL_464512, EPI_ISL_464513, EPI_ISL_464514, EPI_ISL_464515, EPI_ISL_464516, EPI_ISL_464517, EPI_ISL_464518, EPI_ISL_464519, EPI_ISL_464520, EPI_ISL_464521, EPI_ISL_464522, EPI_ISL_464523, EPI_ISL_464524, EPI_ISL_464525, EPI_ISL_464526, EPI_ISL_464527, EPI_ISL_464528, EPI_ISL_464529, EPI_ISL_464530, EPI_ISL_464531, EPI_ISL_464532, EPI_ISL_464533, EPI_ISL_464534, EPI_ISL_464535, EPI_ISL_464536, EPI_ISL_464537, EPI_ISL_464538, EPI_ISL_464539, EPI_ISL_464540, EPI_ISL_464541, EPI_ISL_464542, EPI_ISL_464543, EPI_ISL_464544, EPI_ISL_464545, EPI_ISL_464546, EPI_ISL_464547, EPI_ISL_464548, EPI_ISL_464549, EPI_ISL_464550, EPI_ISL_464551, EPI_ISL_464552, EPI_ISL_464553, EPI_ISL_464554, EPI_ISL_464555, EPI_ISL_464556, EPI_ISL_464557, EPI_ISL_464558, EPI_ISL_464559, EPI_ISL_464560, EPI_ISL_464561, EPI_ISL_464562, EPI_ISL_464563, EPI_ISL_464564, EPI_ISL_464565, EPI_ISL_464566, EPI_ISL_464567, EPI_ISL_464568, EPI_ISL_464569, EPI_ISL_464570, EPI_ISL_464571, EPI_ISL_464572, EPI_ISL_464573, EPI_ISL_464574, EPI_ISL_464575, EPI_ISL_464576, EPI_ISL_464577, EPI_ISL_464578, EPI_ISL_464579, EPI_ISL_464580, EPI_ISL_464581, EPI_ISL_464582, EPI_ISL_464583, EPI_ISL_464584, EPI_ISL_464585, EPI_ISL_464586, EPI_ISL_464587, EPI_ISL_464588, EPI_ISL_464589, EPI_ISL_464590, EPI_ISL_464591, EPI_ISL_464592, EPI_ISL_464593, EPI_ISL_464594, EPI_ISL_464595, EPI_ISL_464596, EPI_ISL_464597, EPI_ISL_464598, EPI_ISL_464599, EPI_ISL_464600, EPI_ISL_464601, EPI_ISL_464602, EPI_ISL_464603, EPI_ISL_464604, EPI_ISL_464605, EPI_ISL_464606, EPI_ISL_464607, EPI_ISL_464608, EPI_ISL_464609, EPI_ISL_464610, EPI_ISL_464611, EPI_ISL_464612, EPI_ISL_464613, EPI_ISL_464614, EPI_ISL_464615, EPI_ISL_464616, EPI_ISL_464617, EPI_ISL_464618, EPI_ISL_464619, EPI_ISL_464620, EPI_ISL_464621, EPI_ISL_464622, EPI_ISL_464623, EPI_ISL_464624, EPI_ISL_464625, EPI_ISL_464626, EPI_ISL_464627, EPI_ISL_464628, EPI_ISL_464629, EPI_ISL_464630, EPI_ISL_464631, EPI_ISL_464632, EPI_ISL_464633, EPI_ISL_464634, EPI_ISL_464635, EPI_ISL_464636, EPI_ISL_464637, EPI_ISL_464638, EPI_ISL_464639, EPI_ISL_464640, EPI_ISL_464641, EPI_ISL_464642, EPI_ISL_464643, EPI_ISL_464644, EPI_ISL_464645, EPI_ISL_464646, EPI_ISL_464647, EPI_ISL_464648, EPI_ISL_464649, EPI_ISL_464650, EPI_ISL_464651, EPI_ISL_464652, EPI_ISL_464653, EPI_ISL_464654, EPI_ISL_464655, EPI_ISL_464656, EPI_ISL_464657, EPI_ISL_464658, EPI_ISL_464659, EPI_ISL_464660, EPI_ISL_464661, EPI_ISL_464662, EPI_ISL_464663, EPI_ISL_464664, EPI_ISL_464665, EPI_ISL_464666, EPI_ISL_464667, EPI_ISL_464668, EPI_ISL_464669, EPI_ISL_464670, EPI_ISL_464671, EPI_ISL_464672, EPI_ISL_464673, EPI_ISL_464674, EPI_ISL_464675, EPI_ISL_464676, EPI_ISL_464677, EPI_ISL_464678, EPI_ISL_464679, EPI_ISL_464680, EPI_ISL_464681, EPI_ISL_464682, EPI_ISL_464683, EPI_ISL_464684, EPI_ISL_464685, EPI_ISL_464686, EPI_ISL_464687, EPI_ISL_464688, EPI_ISL_464689, EPI_ISL_464690, EPI_ISL_464691, EPI_ISL_464692, EPI_ISL_464693, EPI_ISL_464694, EPI_ISL_464695, EPI_ISL_464696, EPI_ISL_464697, EPI_ISL_464698, EPI_ISL_464699, EPI_ISL_464700, EPI_ISL_464701, EPI_ISL_464702, EPI_ISL_464703, EPI_ISL_464704, EPI_ISL_464705, EPI_ISL_464706, EPI_ISL_464707, EPI_ISL_464708, EPI_ISL_464709, EPI_ISL_464710, EPI_ISL_464711, EPI_ISL_464712, EPI_ISL_464713, EPI_ISL_464714, EPI_ISL_464715, EPI_ISL_464716, EPI_ISL_464717, EPI_ISL_464718, EPI_ISL_464719, EPI_ISL_464720, EPI_ISL_464721, EPI_ISL_464722, EPI_ISL_464723, EPI_ISL_464724, EPI_ISL_464725, EPI_ISL_464726, EPI_ISL_464727, EPI_ISL_464728, EPI_ISL_464729, EPI_ISL_464730, EPI_ISL_464731, EPI_ISL_464732, EPI_ISL_464733, EPI_ISL_464734, EPI_ISL_464735, EPI_ISL_464736, EPI_ISL_464737, EPI_ISL_464738, EPI_ISL_464739, EPI_ISL_464740, EPI_ISL_464741, EPI_ISL_464742, EPI_ISL_464743, EPI_ISL_464744, EPI_ISL_464745, EPI_ISL_464746, EPI_ISL_464747, EPI_ISL_464748, EPI_ISL_464749, EPI_ISL_464750, EPI_ISL_464751, EPI_ISL_464752, EPI_ISL_464753, EPI_ISL_464754, EPI_ISL_464755, EPI_ISL_464756, EPI_ISL_464757, EPI_ISL_464758, EPI_ISL_464759, EPI_ISL_464760, EPI_ISL_464761, EPI_ISL_464762, EPI_ISL_464763, EPI_ISL_464764, EPI_ISL_464765, EPI_ISL_464766, EPI_ISL_464767, EPI_ISL_464768, EPI_ISL_464769, EPI_ISL_464770, EPI_ISL_464771, EPI_ISL_464772, EPI_ISL_464773, EPI_ISL_464774, EPI_ISL_464775, EPI_ISL_464776, EPI_ISL_464777, EPI_ISL_464778, EPI_ISL_464779, EPI_ISL_464780, EPI_ISL_464781, EPI_ISL_464782, EPI_ISL_464783, EPI_ISL_464784, EPI_ISL_464785, EPI_ISL_464786, EPI_ISL_464787, EPI_ISL_464788, EPI_ISL_464789, EPI_ISL_464790, EPI_ISL_464791, EPI_ISL_464792, EPI_ISL_464793, EPI_ISL_464794, EPI_ISL_464795, EPI_ISL_464796, EPI_ISL_464797, EPI_ISL_464798, EPI_ISL_464799, EPI_ISL_464800, EPI_ISL_464801, EPI_ISL_464802, EPI_ISL_464803, EPI_ISL_464804, EPI_ISL_464805, EPI_ISL_464806, EPI_ISL_464807, EPI_ISL_464808, EPI_ISL_464809, EPI_ISL_464810, EPI_ISL_464811, EPI_ISL_464812, EPI_ISL_464813, EPI_ISL_464814, EPI_ISL_464815, EPI_ISL_464816, EPI_ISL_464817, EPI_ISL_464818, EPI_ISL_464819, EPI_ISL_464820, EPI_ISL_464821, EPI_ISL_464822, EPI_ISL_464823, EPI_ISL_464824, EPI_ISL_464825, EPI_ISL_464826, EPI_ISL_464827, EPI_ISL_464828, EPI_ISL_464829, EPI_ISL_464830, EPI_ISL_464831, EPI_ISL_464832, EPI_ISL_464833, EPI_ISL_464834, EPI_ISL_464835, EPI_ISL_464836, EPI_ISL_464837, EPI_ISL_464838, EPI_ISL_464839, EPI_ISL_464840, EPI_ISL_464841, EPI_ISL_464842, EPI_ISL_464843, EPI_ISL_464844, EPI_ISL_464845, EPI_ISL_464846, EPI_ISL_464847, EPI_ISL_464848, EPI_ISL_464849, EPI_ISL_464850, EPI_ISL_464851, EPI_ISL_464852, EPI_ISL_464853, EPI_ISL_464854, EPI_ISL_464855, EPI_ISL_464856, EPI_ISL_464857, EPI_ISL_464858, EPI_ISL_464859, EPI_ISL_464860, EPI_ISL_464861, EPI_ISL_464862, EPI_ISL_464863, EPI_ISL_464864, EPI_ISL_464865, EPI_ISL_464866, EPI_ISL_464867, EPI_ISL_464868, EPI_ISL_464869, EPI_ISL_464870, EPI_ISL_464871, EPI_ISL_464872, EPI_ISL_464873, EPI_ISL_464874, EPI_ISL_464875, EPI_ISL_464876, EPI_ISL_464877, EPI_ISL_464878, EPI_ISL_464879, EPI_ISL_464880, EPI_ISL_464881, EPI_ISL_464882, EPI_ISL_464883, EPI_ISL_464884, EPI_ISL_464885, EPI_ISL_464886, EPI_ISL_464887, EPI_ISL_464888, EPI_ISL_464889, EPI_ISL_464890, EPI_ISL_464891, EPI_ISL_464892, EPI_ISL_464893, EPI_ISL_464894, EPI_ISL_464895, EPI_ISL_464896, EPI_ISL_464897, EPI_ISL_464898, EPI_ISL_464899, EPI_ISL_464900, EPI_ISL_464901, EPI_ISL_464902, EPI_ISL_464903, EPI_ISL_464904, EPI_ISL_464905, EPI_ISL_464906, EPI_ISL_464907, EPI_ISL_464908, EPI_ISL_464909, EPI_ISL_464910, EPI_ISL_464911, EPI_ISL_464912, EPI_ISL_464913, EPI_ISL_464914, EPI_ISL_464915, EPI_ISL_464916, EPI_ISL_464917, EPI_ISL_464918, EPI_ISL_464919, EPI_ISL_464920, EPI_ISL_464921, EPI_ISL_464922, EPI_ISL_464923, EPI_ISL_464924, EPI_ISL_464925, EPI_ISL_464926, EPI_ISL_464927, EPI_ISL_464928, EPI_ISL_464929, EPI_ISL_464930, EPI_ISL_464931, EPI_ISL_464932, EPI_ISL_464933, EPI_ISL_464934, EPI_ISL_464935, EPI_ISL_464936, EPI_ISL_464937, EPI_ISL_464938, EPI_ISL_464939, EPI_ISL_464940, EPI_ISL_464941, EPI_ISL_464942, EPI_ISL_464943, EPI_ISL_464944, EPI_ISL_464945, EPI_ISL_464946, EPI_ISL_464947, EPI_ISL_464948, EPI_ISL_464949, EPI_ISL_464950, EPI_ISL_464951, EPI_ISL_464952, EPI_ISL_464953, EPI_ISL_464954, EPI_ISL_464955, EPI_ISL_464956, EPI_ISL_464957, EPI_ISL_464958, EPI_ISL_464959, EPI_ISL_464960, EPI_ISL_464961, EPI_ISL_464962, EPI_ISL_464963, EPI_ISL_464964, EPI_ISL_464965, EPI_ISL_464966, EPI_ISL_464967, EPI_ISL_464968, EPI_ISL_464969, EPI_ISL_464970, EPI_ISL_464971, EPI_ISL_464972, EPI_ISL_464973, EPI_ISL_464974, EPI_ISL_464975, EPI_ISL_464976, EPI_ISL_464977, EPI_ISL_464978, EPI_ISL_464979, EPI_ISL_464980, EPI_ISL_464981, EPI_ISL_464982, EPI_ISL_464983, EPI_ISL_464984, EPI_ISL_464985, EPI_ISL_464986, EPI_ISL_464987, EPI_ISL_464988, EPI_ISL_464989, EPI_ISL_464990, EPI_ISL_464991, EPI_ISL_464992, EPI_ISL_464993, EPI_ISL_464994, EPI_ISL_464995, EPI_ISL_464996, EPI_ISL_464997, EPI_ISL_464998, EPI_ISL_464999, EPI_ISL_465000, EPI_ISL_465001, EPI_ISL_465002, EPI_ISL_465003, EPI_ISL_465004, EPI_ISL_465005, EPI_ISL_465006, EPI_ISL_465007, EPI_ISL_465008, EPI_ISL_465009, EPI_ISL_465010, EPI_ISL_465011, EPI_ISL_465012, EPI_ISL_465013, EPI_ISL_465014, EPI_ISL_465015, EPI_ISL_465016, EPI_ISL_465017, EPI_ISL_465018, EPI_ISL_465019, EPI_ISL_465020, EPI_ISL_465021, EPI_ISL_465022, EPI_ISL_465023, EPI_ISL_465024, EPI_ISL_465025, EPI_ISL_465026, EPI_ISL_465027, EPI_ISL_465028, EPI_ISL_465029, EPI_ISL_465030, EPI_ISL_465031, EPI_ISL_465032, EPI_ISL_465033, EPI_ISL_465034, EPI_ISL_465035, EPI_ISL_465036, EPI_ISL_465037, EPI_ISL_465038, EPI_ISL_465039, EPI_ISL_465040, EPI_ISL_465041, EPI_ISL_465042, EPI_ISL_465043, EPI_ISL_465044, EPI_ISL_465045, EPI_ISL_465046, EPI_ISL_465047, EPI_ISL_465048, EPI_ISL_465049, EPI_ISL_465050, EPI_ISL_465051, EPI_ISL_465052, EPI_ISL_465053, EPI_ISL_465054, EPI_ISL_465055, EPI_ISL_465056, EPI_ISL_465057, EPI_ISL_465058, EPI_ISL_465059, EPI_ISL_465060, EPI_ISL_465061, EPI_ISL_465062, EPI_ISL_465063, EPI_ISL_465064, EPI_ISL_465065, EPI_ISL_465066, EPI_ISL_465067, EPI_ISL_465068, EPI_ISL_465069, EPI_ISL_465070, EPI_ISL_465071, EPI_ISL_465072, EPI_ISL_465073, EPI_ISL_465074, EPI_ISL_465075, EPI_ISL_465076, EPI_ISL_465077, EPI_ISL_465078, EPI_ISL_465079, EPI_ISL_465080, EPI_ISL_465081, EPI_ISL_465082, EPI_ISL_465083, EPI_ISL_465084, EPI_ISL_465085, EPI_ISL_465086, EPI_ISL_465087, EPI_ISL_465088, EPI_ISL_465089, EPI_ISL_465090, EPI_ISL_465091, EPI_ISL_465092, EPI_ISL_465093, EPI_ISL_465094, EPI_ISL_465095, EPI_ISL_465096, EPI_ISL_465097, EPI_ISL_465098, EPI_ISL_465099, EPI_ISL_465100, EPI_ISL_465101, EPI_ISL_465102, EPI_ISL_465103, EPI_ISL_465104, EPI_ISL_465105, EPI_ISL_465106, EPI_ISL_465107, EPI_ISL_465108, EPI_ISL_465109, EPI_ISL_465110, EPI_ISL_465111, EPI_ISL_465112, EPI_ISL_465113, EPI_ISL_465114, EPI_ISL_465115, EPI_ISL_465116, EPI_ISL_465117, EPI_ISL_465118, EPI_ISL_465119, EPI_ISL_465120, EPI_ISL_465121, EPI_ISL_465122, EPI_ISL_465123, EPI_ISL_465124, EPI_ISL_465125, EPI_ISL_465126, EPI_ISL_465127, EPI_ISL_465128, EPI_ISL_465129, EPI_ISL_465130, EPI_ISL_465131, EPI_ISL_465132, EPI_ISL_465133, EPI_ISL_465134, EPI_ISL_465135, EPI_ISL_465136, EPI_ISL_465137, EPI_ISL_465138, EPI_ISL_465139, EPI_ISL_465140, EPI_ISL_465141, EPI_ISL_465142, EPI_ISL_465143, EPI_ISL_465144, EPI_ISL_465145, EPI_ISL_465146, EPI_ISL_465147, EPI_ISL_465148, EPI_ISL_465149, EPI_ISL_465150, EPI_ISL_465151, EPI_ISL_465152, EPI_ISL_465153, EPI_ISL_465154, EPI_ISL_465155, EPI_ISL_465156, EPI_ISL_465157, EPI_ISL_465158, EPI_ISL_465159, EPI_ISL_465160, EPI_ISL_465161, EPI_ISL_465162 | | | |
| see above | Respiratory Virus Unit, Microbiology Services Colindale, Public Health England | Respiratory Virus Unit, Microbiology Services Colindale, Public Health England | PHE Covid Sequencing Team |
| EPI_ISL_465163, EPI_ISL_465164 | National Institute of Laboratory Medicine and Referral Center | Genomic Research Lab, BCSIR | Iffat Jahan, Abu Sayeed Mohammad Mahmud, Mohammad Samir Uzzaman, Eshrar Osman, Md. Ahasan Habib, Shahina Akter, Tanjina Akhter Banu, Md. Murshed Hasan Sarker, Barna Goswami, Md. Saddam Hossain, Tasnim Nafisa, Md. Maruf Ahmed Molla, Mahmuda Yeasmin, Asish Kumar Ghosh, Arifa Akram, A. K. M. Shamsuzzaman, Sheikh Md. Selim Al Din, Utpal Chandra Ray, Salek Ahmed Sajib, Md. Salim Khan |
| EPI_ISL_465165, EPI_ISL_465166, EPI_ISL_465167, EPI_ISL_465168, EPI_ISL_465169, EPI_ISL_465170, EPI_ISL_465171, EPI_ISL_465172, EPI_ISL_465173, EPI_ISL_465174, EPI_ISL_465175, EPI_ISL_465176, EPI_ISL_465177, EPI_ISL_465178, EPI_ISL_465179, EPI_ISL_465180, EPI_ISL_465181, EPI_ISL_465182, EPI_ISL_465183, EPI_ISL_465184, EPI_ISL_465185, EPI_ISL_465186, EPI_ISL_465187, EPI_ISL_465188, EPI_ISL_465189, EPI_ISL_465190, EPI_ISL_465191, EPI_ISL_465192, EPI_ISL_465193, EPI_ISL_465194, EPI_ISL_465195, EPI_ISL_465196, EPI_ISL_465197, EPI_ISL_465198, EPI_ISL_465199, EPI_ISL_465200, EPI_ISL_465201, EPI_ISL_465202, EPI_ISL_465203, EPI_ISL_465204, EPI_ISL_465205, EPI_ISL_465206, EPI_ISL_465207, EPI_ISL_465208, EPI_ISL_465209, EPI_ISL_465210, EPI_ISL_465211, EPI_ISL_465212, EPI_ISL_465213, EPI_ISL_465214, EPI_ISL_465215, EPI_ISL_465216, EPI_ISL_465217, EPI_ISL_465218, EPI_ISL_465219, EPI_ISL_465220, EPI_ISL_465221, EPI_ISL_465222, EPI_ISL_465223, EPI_ISL_465224, EPI_ISL_465225, EPI_ISL_465226, EPI_ISL_465227, EPI_ISL_465228, EPI_ISL_465229, EPI_ISL_465230, EPI_ISL_465231, EPI_ISL_465232, EPI_ISL_465233, EPI_ISL_465234, EPI_ISL_465235, EPI_ISL_465236, EPI_ISL_465237, EPI_ISL_465238, EPI_ISL_465239, EPI_ISL_465240, EPI_ISL_465241, EPI_ISL_465242, EPI_ISL_465243, EPI_ISL_465244, EPI_ISL_465245, EPI_ISL_465246, EPI_ISL_465247, EPI_ISL_465248, EPI_ISL_465249, EPI_ISL_465250, EPI_ISL_465251, EPI_ISL_465252, EPI_ISL_465253, EPI_ISL_465254, EPI_ISL_465255, EPI_ISL_465256, EPI_ISL_465257, EPI_ISL_465258, EPI_ISL_465259, EPI_ISL_465260, EPI_ISL_465261, EPI_ISL_465262, EPI_ISL_465263, EPI_ISL_465264, EPI_ISL_465265, EPI_ISL_465266, EPI_ISL_465267, EPI_ISL_465268, EPI_ISL_465269, EPI_ISL_465270, EPI_ISL_465271, EPI_ISL_465272, EPI_ISL_465273, EPI_ISL_465274, EPI_ISL_465275, EPI_ISL_465276, EPI_ISL_465277, EPI_ISL_465278, EPI_ISL_465279, EPI_ISL_465280, EPI_ISL_465281, EPI_ISL_465282, EPI_ISL_465283, EPI_ISL_465284, EPI_ISL_465285, EPI_ISL_465286, EPI_ISL_465287, EPI_ISL_465288, EPI_ISL_465289, EPI_ISL_465290, EPI_ISL_465291, EPI_ISL_465292, EPI_ISL_465293, EPI_ISL_465294, EPI_ISL_465295, EPI_ISL_465296, EPI_ISL_465297, EPI_ISL_465298, EPI_ISL_465299, EPI_ISL_465300, EPI_ISL_465301, EPI_ISL_465302, EPI_ISL_465303, EPI_ISL_465304, EPI_ISL_465305, EPI_ISL_465306, EPI_ISL_465307, EPI_ISL_465308, EPI_ISL_465309, EPI_ISL_465310, EPI_ISL_465311, EPI_ISL_465312, EPI_ISL_465313, EPI_ISL_465314, EPI_ISL_465315, EPI_ISL_465316, EPI_ISL_465317, EPI_ISL_465318, EPI_ISL_465319, EPI_ISL_465320, EPI_ISL_465321, EPI_ISL_465322, EPI_ISL_465323, EPI_ISL_465324, EPI_ISL_465325, EPI_ISL_465326, EPI_ISL_465327, EPI_ISL_465328, EPI_ISL_465329, EPI_ISL_465330, EPI_ISL_465331, EPI_ISL_465332, EPI_ISL_465333, EPI_ISL_465334, EPI_ISL_465335, EPI_ISL_465336, EPI_ISL_465337, EPI_ISL_465338, EPI_ISL_465339, EPI_ISL_465340, EPI_ISL_465341, EPI_ISL_465342, EPI_ISL_465343, EPI_ISL_465344, EPI_ISL_465345, EPI_ISL_465346, EPI_ISL_465347, EPI_ISL_465348, EPI_ISL_465349, EPI_ISL_465350, EPI_ISL_465351, EPI_ISL_465352, EPI_ISL_465353, EPI_ISL_465354, EPI_ISL_465355, EPI_ISL_465356, EPI_ISL_465357, EPI_ISL_465358, EPI_ISL_465359, EPI_ISL_465360, EPI_ISL_465361, EPI_ISL_465362, EPI_ISL_465363, EPI_ISL_465364, EPI_ISL_465365, EPI_ISL_465366, EPI_ISL_465367, EPI_ISL_465368, EPI_ISL_465369, EPI_ISL_465370, EPI_ISL_465371, EPI_ISL_465372, EPI_ISL_465373, EPI_ISL_465374, EPI_ISL_465375, EPI_ISL_465376, EPI_ISL_465377, EPI_ISL_465378, EPI_ISL_465379, EPI_ISL_465380, EPI_ISL_465381, EPI_ISL_465382, EPI_ISL_465383, EPI_ISL_465384, EPI_ISL_465385, EPI_ISL_465386, EPI_ISL_465387, EPI_ISL_465388, EPI_ISL_465389, EPI_ISL_465390, EPI_ISL_465391, EPI_ISL_465392, EPI_ISL_465393, EPI_ISL_465394, EPI_ISL_465395, EPI_ISL_465396, EPI_ISL_465397, EPI_ISL_465398, EPI_ISL_465399, EPI_ISL_465400, EPI_ISL_465401, EPI_ISL_465402, EPI_ISL_465403, EPI_ISL_465404, EPI_ISL_465405, EPI_ISL_465406, EPI_ISL_465407, EPI_ISL_465408, EPI_ISL_465409, EPI_ISL_465410, EPI_ISL_465411, EPI_ISL_465412, EPI_ISL_465413, EPI_ISL_465414, EPI_ISL_465415, EPI_ISL_465416, EPI_ISL_465417, EPI_ISL_465418, EPI_ISL_465419, EPI_ISL_465420, EPI_ISL_465421, EPI_ISL_465422, EPI_ISL_465423, EPI_ISL_465424, EPI_ISL_465425, EPI_ISL_465426, EPI_ISL_465427, EPI_ISL_465428, EPI_ISL_465429, EPI_ISL_465430, EPI_ISL_465431, EPI_ISL_465432, EPI_ISL_465433, EPI_ISL_465434, EPI_ISL_465435, EPI_ISL_465436, EPI_ISL_465437, EPI_ISL_465438, EPI_ISL_465439, EPI_ISL_465440, EPI_ISL_465441, EPI_ISL_465442, EPI_ISL_465443, EPI_ISL_465444, EPI_ISL_465445, EPI_ISL_465446, EPI_ISL_465447, EPI_ISL_465448, EPI_ISL_465449, EPI_ISL_465450, EPI_ISL_465451, EPI_ISL_465452, EPI_ISL_465453, EPI_ISL_465454, EPI_ISL_465455, EPI_ISL_465456, EPI_ISL_465457, EPI_ISL_465458, EPI_ISL_465459, EPI_ISL_465460, EPI_ISL_465461, EPI_ISL_465462, EPI_ISL_465463, EPI_ISL_465464, EPI_ISL_465465, EPI_ISL_465466, EPI_ISL_465467, EPI_ISL_465468, EPI_ISL_465469, EPI_ISL_465470, EPI_ISL_465471, EPI_ISL_465472, EPI_ISL_465473, EPI_ISL_465474, EPI_ISL_465475, EPI_ISL_465476, EPI_ISL_465477, EPI_ISL_465478, EPI_ISL_465479, EPI_ISL_465480, EPI_ISL_465481, EPI_ISL_465482, EPI_ISL_465483, EPI_ISL_465484, EPI_ISL_465485, EPI_ISL_465486, EPI_ISL_465487, EPI_ISL_465488, EPI_ISL_465489, EPI_ISL_465490, EPI_ISL_465491, EPI_ISL_465492, EPI_ISL_465493, EPI_ISL_465494, EPI_ISL_465495, EPI_ISL_465496, EPI_ISL_465497, EPI_ISL_465498, EPI_ISL_465499, EPI_ISL_465500, EPI_ISL_465501, EPI_ISL_465502, EPI_ISL_465503, EPI_ISL_465504, EPI_ISL_465505, EPI_ISL_465506, EPI_ISL_465507, EPI_ISL_465508, EPI_ISL_465509, EPI_ISL_465510, EPI_ISL_465511, EPI_ISL_465512, EPI_ISL_465513, EPI_ISL_465514, EPI_ISL_465515, EPI_ISL_465516, EPI_ISL_465517, EPI_ISL_465518, EPI_ISL_465519, EPI_ISL_465520, EPI_ISL_465521, EPI_ISL_465522, EPI_ISL_465523, EPI_ISL_465524, EPI_ISL_465525, EPI_ISL_465526, EPI_ISL_465527, EPI_ISL_465528, EPI_ISL_465529, EPI_ISL_465530, EPI_ISL_465531, EPI_ISL_465532, EPI_ISL_465533, EPI_ISL_465534, EPI_ISL_465535, EPI_ISL_465536, EPI_ISL_465537, EPI_ISL_465538, EPI_ISL_465539, EPI_ISL_465540, EPI_ISL_465541, EPI_ISL_465542, EPI_ISL_465543, EPI_ISL_465544, EPI_ISL_465545, EPI_ISL_465546, EPI_ISL_465547, EPI_ISL_465548, EPI_ISL_465549, EPI_ISL_465550, EPI_ISL_465551, EPI_ISL_465552, EPI_ISL_465553, EPI_ISL_465554, EPI_ISL_465555, EPI_ISL_465556, EPI_ISL_465557, EPI_ISL_465558, EPI_ISL_465559, EPI_ISL_465560, EPI_ISL_465561, EPI_ISL_465562, EPI_ISL_465563, EPI_ISL_465564, EPI_ISL_465565, EPI_ISL_465566, EPI_ISL_465567, EPI_ISL_465568, EPI_ISL_465569, EPI_ISL_465570, EPI_ISL_465571, EPI_ISL_465572, EPI_ISL_465573, EPI_ISL_465574, EPI_ISL_465575, EPI_ISL_465576, EPI_ISL_465577, EPI_ISL_465578, EPI_ISL_465579, EPI_ISL_465580, EPI_ISL_465581, EPI_ISL_465582, EPI_ISL_465583, EPI_ISL_465584, EPI_ISL_465585, EPI_ISL_465586, EPI_ISL_465587, EPI_ISL_465588, EPI_ISL_465589, EPI_ISL_465590, EPI_ISL_465591, EPI_ISL_465592, EPI_ISL_465593, EPI_ISL_465594, EPI_ISL_465595, EPI_ISL_465596, EPI_ISL_465597, EPI_ISL_465598, EPI_ISL_465599, EPI_ISL_465600, EPI_ISL_465601, EPI_ISL_465602, EPI_ISL_465603, EPI_ISL_465604, EPI_ISL_465605, EPI_ISL_465606, EPI_ISL_465607, EPI_ISL_465608, EPI_ISL_465609, EPI_ISL_465610, EPI_ISL_465611, EPI_ISL_465612, EPI_ISL_465613, EPI_ISL_465614, EPI_ISL_465615, EPI_ISL_465616, EPI_ISL_465617, EPI_ISL_465618, EPI_ISL_465619, EPI_ISL_465620, EPI_ISL_465621, EPI_ISL_465622, EPI_ISL_465623, EPI_ISL_465624, EPI_ISL_465625, EPI_ISL_465626, EPI_ISL_465627, EPI_ISL_465628, EPI_ISL_465629, EPI_ISL_465630, EPI_ISL_465631, EPI_ISL_465632, EPI_ISL_465633, EPI_ISL_465634, EPI_ISL_465635, EPI_ISL_465636, EPI_ISL_465637, EPI_ISL_465638, EPI_ISL_465639, EPI_ISL_465640, EPI_ISL_465641, EPI_ISL_465642, EPI_ISL_465643, EPI_ISL_465644, EPI_ISL_465645, EPI_ISL_465646, EPI_ISL_465647, EPI_ISL_465648, EPI_ISL_465649, EPI_ISL_465650, EPI_ISL_465651, EPI_ISL_465652, EPI_ISL_465653, EPI_ISL_465654, EPI_ISL_465655, EPI_ISL_465656, EPI_ISL_465657, EPI_ISL_465658, EPI_ISL_465659, EPI_ISL_465660, EPI_ISL_465661, EPI_ISL_465662, EPI_ISL_465663, EPI_ISL_465664, EPI_ISL_465665, EPI_ISL_465666, EPI_ISL_465667, EPI_ISL_465668, EPI_ISL_465669, EPI_ISL_465670, EPI_ISL_465671, EPI_ISL_465672, EPI_ISL_465673, EPI_ISL_465674, EPI_ISL_465675, EPI_ISL_465676, EPI_ISL_465677, EPI_ISL_465678 | | | |
| see above | Respiratory Virus Unit, Microbiology Services Colindale, Public Health England | Respiratory Virus Unit, Microbiology Services Colindale, Public Health England | PHE Covid Sequencing Team |
| EPI_ISL_465679, EPI_ISL_465680 | Hôpital du Suroît | Laboratoire de santé publique du Québec | Sandrine Moreira, Ioannis Ragoussis, Guillaume Bourque, Jesse Shapiro, Mark Lathrop and Michel Roger on behalf of the CoVSeQ research group (http://covseq.ca/researchgroup) |
| EPI_ISL_465681, EPI_ISL_465682 | Hôpital Charles-LeMoyne | Laboratoire de santé publique du Québec | Sandrine Moreira, Ioannis Ragoussis, Guillaume Bourque, Jesse Shapiro, Mark Lathrop and Michel Roger on behalf of the CoVSeQ research group (http://covseq.ca/researchgroup) |
| EPI_ISL_465683, EPI_ISL_465684 | Hôpital de Maria | Laboratoire de santé publique du Québec | Sandrine Moreira, Ioannis Ragoussis, Guillaume Bourque, Jesse Shapiro, Mark Lathrop and Michel Roger on behalf of the CoVSeQ research group (http://covseq.ca/researchgroup) |
| EPI_ISL_465685 | CSSS Haut-Richelieu/Rouville (Hôpital) | Laboratoire de santé publique du Québec | Sandrine Moreira, Ioannis Ragoussis, Guillaume Bourque, Jesse Shapiro, Mark Lathrop and Michel Roger on behalf of the CoVSeQ research group (http://covseq.ca/researchgroup) |
| EPI_ISL_465686 | Hôpital de Hull | Laboratoire de santé publique du Québec | Sandrine Moreira, Ioannis Ragoussis, Guillaume Bourque, Jesse Shapiro, Mark Lathrop and Michel Roger on behalf of the CoVSeQ research group (http://covseq.ca/researchgroup) |
| EPI_ISL_465687 | CSSS Haut-Richelieu/Rouville (Hôpital) | Laboratoire de santé publique du Québec | Sandrine Moreira, Ioannis Ragoussis, Guillaume Bourque, Jesse Shapiro, Mark Lathrop and Michel Roger on behalf of the CoVSeQ research group (http://covseq.ca/researchgroup) |
| EPI_ISL_465688, EPI_ISL_465689, EPI_ISL_465690 | Centre hospitalier Anna-Laberge | Laboratoire de santé publique du Québec | Sandrine Moreira, Ioannis Ragoussis, Guillaume Bourque, Jesse Shapiro, Mark Lathrop and Michel Roger on behalf of the CoVSeQ research group (http://covseq.ca/researchgroup) |
| EPI_ISL_465691 | Hôpital de Gatineau | Laboratoire de santé publique du Québec | Sandrine Moreira, Ioannis Ragoussis, Guillaume Bourque, Jesse Shapiro, Mark Lathrop and Michel Roger on behalf of the CoVSeQ research group (http://covseq.ca/researchgroup) |
| EPI_ISL_465692 | CSSS Haut-Richelieu/Rouville (Hôpital) | Laboratoire de santé publique du Québec | Sandrine Moreira, Ioannis Ragoussis, Guillaume Bourque, Jesse Shapiro, Mark Lathrop and Michel Roger on behalf of the CoVSeQ research group (http://covseq.ca/researchgroup) |
| EPI_ISL_465693 | Hôpital du Suroît | Laboratoire de santé publique du Québec | Sandrine Moreira, Ioannis Ragoussis, Guillaume Bourque, Jesse Shapiro, Mark Lathrop and Michel Roger on behalf of the CoVSeQ research group (http://covseq.ca/researchgroup) |
| EPI_ISL_465694 | Hôpital Charles-LeMoyne | Laboratoire de santé publique du Québec | Sandrine Moreira, Ioannis Ragoussis, Guillaume Bourque, Jesse Shapiro, Mark Lathrop and Michel Roger on behalf of the CoVSeQ research group (http://covseq.ca/researchgroup) |
| EPI_ISL_465695, EPI_ISL_465696 | Hôpital Pierre-Boucher | Laboratoire de santé publique du Québec | Sandrine Moreira, Ioannis Ragoussis, Guillaume Bourque, Jesse Shapiro, Mark Lathrop and Michel Roger on behalf of the CoVSeQ research group (http://covseq.ca/researchgroup) |
| EPI_ISL_465697 | Centre de santé Inuulitsivik | Laboratoire de santé publique du Québec | Sandrine Moreira, Ioannis Ragoussis, Guillaume Bourque, Jesse Shapiro, Mark Lathrop and Michel Roger on behalf of the CoVSeQ research group (http://covseq.ca/researchgroup) |
| EPI_ISL_465698, EPI_ISL_465699 | Hôpital Charles-LeMoyne | Laboratoire de santé publique du Québec | Sandrine Moreira, Ioannis Ragoussis, Guillaume Bourque, Jesse Shapiro, Mark Lathrop and Michel Roger on behalf of the CoVSeQ research group (http://covseq.ca/researchgroup) |
| EPI_ISL_465700 | Hôpital Pierre-Boucher | Laboratoire de santé publique du Québec | Sandrine Moreira, Ioannis Ragoussis, Guillaume Bourque, Jesse Shapiro, Mark Lathrop and Michel Roger on behalf of the CoVSeQ research group (http://covseq.ca/researchgroup) |
| EPI_ISL_465701 | Hôpital du Suroît | Laboratoire de santé publique du Québec | Sandrine Moreira, Ioannis Ragoussis, Guillaume Bourque, Jesse Shapiro, Mark Lathrop and Michel Roger on behalf of the CoVSeQ research group (http://covseq.ca/researchgroup) |
| EPI_ISL_465702, EPI_ISL_465703 | Hôpital Pierre-Boucher | Laboratoire de santé publique du Québec | Sandrine Moreira, Ioannis Ragoussis, Guillaume Bourque, Jesse Shapiro, Mark Lathrop and Michel Roger on behalf of the CoVSeQ research group (http://covseq.ca/researchgroup) |
| EPI_ISL_465704, EPI_ISL_465705, EPI_ISL_465706, EPI_ISL_465707, EPI_ISL_465708, EPI_ISL_465709, EPI_ISL_465710, EPI_ISL_465711, EPI_ISL_465712, EPI_ISL_465713, EPI_ISL_465714, EPI_ISL_465715, EPI_ISL_465716, EPI_ISL_465717, EPI_ISL_465718, EPI_ISL_465719, EPI_ISL_465720, EPI_ISL_465721, EPI_ISL_465722, EPI_ISL_465723, EPI_ISL_465724, EPI_ISL_465725, EPI_ISL_465726, EPI_ISL_465727, EPI_ISL_465728, EPI_ISL_465729, EPI_ISL_465730, EPI_ISL_465731, EPI_ISL_465732, EPI_ISL_465733, EPI_ISL_465734, EPI_ISL_465735, EPI_ISL_465736, EPI_ISL_465737, EPI_ISL_465738, EPI_ISL_465739, EPI_ISL_465740, EPI_ISL_465741, EPI_ISL_465742, EPI_ISL_465743, EPI_ISL_465744, EPI_ISL_465745, EPI_ISL_465746, EPI_ISL_465747, EPI_ISL_465748, EPI_ISL_465749, EPI_ISL_465750, EPI_ISL_465751, EPI_ISL_465752, EPI_ISL_465753, EPI_ISL_465754, EPI_ISL_465755, EPI_ISL_465756, EPI_ISL_465757, EPI_ISL_465758, EPI_ISL_465759, EPI_ISL_465760, EPI_ISL_465761, EPI_ISL_465762, EPI_ISL_465763, EPI_ISL_465764, EPI_ISL_465765, EPI_ISL_465766, EPI_ISL_465767, EPI_ISL_465768, EPI_ISL_465769, EPI_ISL_465770, EPI_ISL_465771, EPI_ISL_465772, EPI_ISL_465773, EPI_ISL_465774, EPI_ISL_465775, EPI_ISL_465776, EPI_ISL_465777, EPI_ISL_465778, EPI_ISL_465779, EPI_ISL_465780, EPI_ISL_465781, EPI_ISL_465782, EPI_ISL_465783, EPI_ISL_465784, EPI_ISL_465785, EPI_ISL_465786, EPI_ISL_465787, EPI_ISL_465788, EPI_ISL_465789, EPI_ISL_465790, EPI_ISL_465791, EPI_ISL_465792, EPI_ISL_465793, EPI_ISL_465794, EPI_ISL_465795, EPI_ISL_465796, EPI_ISL_465797, EPI_ISL_465798, EPI_ISL_465799, EPI_ISL_465800, EPI_ISL_465801, EPI_ISL_465802, EPI_ISL_465803, EPI_ISL_465804, EPI_ISL_465805, EPI_ISL_465806, EPI_ISL_465807, EPI_ISL_465808, EPI_ISL_465809, EPI_ISL_465810, EPI_ISL_465811, EPI_ISL_465812, EPI_ISL_465813, EPI_ISL_465814, EPI_ISL_465815, EPI_ISL_465816, EPI_ISL_465817, EPI_ISL_465818, EPI_ISL_465819, EPI_ISL_465820, EPI_ISL_465821, EPI_ISL_465822, EPI_ISL_465823, | | | |

| | | | |
|---|---|---|---|
| EPI_ISL_465824, EPI_ISL_465825, EPI_ISL_465826, EPI_ISL_465827, EPI_ISL_465828, EPI_ISL_465829, EPI_ISL_465830, EPI_ISL_465831, EPI_ISL_465832, EPI_ISL_465833, EPI_ISL_465834, EPI_ISL_465835, EPI_ISL_465836, EPI_ISL_465837, EPI_ISL_465838, EPI_ISL_465839, EPI_ISL_465840, EPI_ISL_465841, EPI_ISL_465842, EPI_ISL_465843, EPI_ISL_465844, EPI_ISL_465845, EPI_ISL_465846, EPI_ISL_465847, EPI_ISL_465848, EPI_ISL_465849, EPI_ISL_465850, EPI_ISL_465851, EPI_ISL_465852, EPI_ISL_465853, EPI_ISL_465854, EPI_ISL_465855, EPI_ISL_465856, EPI_ISL_465857, EPI_ISL_465858, EPI_ISL_465859, EPI_ISL_465860, EPI_ISL_465861, EPI_ISL_465862, EPI_ISL_465863, EPI_ISL_465864, EPI_ISL_465865, EPI_ISL_465866, EPI_ISL_465867, EPI_ISL_465868, EPI_ISL_465869, EPI_ISL_465870, EPI_ISL_465871, EPI_ISL_465872, EPI_ISL_465873, EPI_ISL_465874, EPI_ISL_465875, EPI_ISL_465876, EPI_ISL_465877, EPI_ISL_465878, EPI_ISL_465879, EPI_ISL_465880, EPI_ISL_465881, EPI_ISL_465882, EPI_ISL_465883, EPI_ISL_465884, EPI_ISL_465885, EPI_ISL_465886, EPI_ISL_465887, EPI_ISL_465888, EPI_ISL_465889, EPI_ISL_465890, EPI_ISL_465891, EPI_ISL_465892, EPI_ISL_465893, EPI_ISL_465894, EPI_ISL_465895, EPI_ISL_465896, EPI_ISL_465897, EPI_ISL_465898, EPI_ISL_465899, EPI_ISL_465900, EPI_ISL_465901, EPI_ISL_465902, EPI_ISL_465903, EPI_ISL_465904, EPI_ISL_465905, EPI_ISL_465906, EPI_ISL_465907, EPI_ISL_465908, EPI_ISL_465909, EPI_ISL_465910, EPI_ISL_465911, EPI_ISL_465912, EPI_ISL_465913, EPI_ISL_465914, EPI_ISL_465915, EPI_ISL_465916, EPI_ISL_465917, EPI_ISL_465918, EPI_ISL_465919, EPI_ISL_465920, EPI_ISL_465921, EPI_ISL_465922, EPI_ISL_465923, EPI_ISL_465924, EPI_ISL_465925, EPI_ISL_465926, EPI_ISL_465927, EPI_ISL_465928, EPI_ISL_465929, EPI_ISL_465930, EPI_ISL_465931, EPI_ISL_465932, EPI_ISL_465933, EPI_ISL_465934, EPI_ISL_465935, EPI_ISL_465936, EPI_ISL_465937, EPI_ISL_465938, EPI_ISL_465939, EPI_ISL_465940, EPI_ISL_465941, EPI_ISL_465942, EPI_ISL_465943, EPI_ISL_465944, EPI_ISL_465945, EPI_ISL_465946, EPI_ISL_465947, EPI_ISL_465948, EPI_ISL_465949, EPI_ISL_465950, EPI_ISL_465951, EPI_ISL_465952, EPI_ISL_465953, EPI_ISL_465954, EPI_ISL_465955, EPI_ISL_465956, EPI_ISL_465957, EPI_ISL_465958, EPI_ISL_465959, EPI_ISL_465960, EPI_ISL_465961, EPI_ISL_465962, EPI_ISL_465963, EPI_ISL_465964, EPI_ISL_465965, EPI_ISL_465966, EPI_ISL_465967, EPI_ISL_465968, EPI_ISL_465969, EPI_ISL_465970, EPI_ISL_465971, EPI_ISL_465972, EPI_ISL_465973, EPI_ISL_465974, EPI_ISL_465975, EPI_ISL_465976, EPI_ISL_465977, EPI_ISL_465978, EPI_ISL_465979, EPI_ISL_465980, EPI_ISL_465981, EPI_ISL_465982, EPI_ISL_465983, EPI_ISL_465984, EPI_ISL_465985, EPI_ISL_465986, EPI_ISL_465987, EPI_ISL_465988, EPI_ISL_465989, EPI_ISL_465990, EPI_ISL_465991, EPI_ISL_465992, EPI_ISL_465993, EPI_ISL_465994, EPI_ISL_465995, EPI_ISL_465996, EPI_ISL_465997, EPI_ISL_465998, EPI_ISL_465999, EPI_ISL_466000, EPI_ISL_466001, EPI_ISL_466002, EPI_ISL_466003, EPI_ISL_466004, EPI_ISL_466005, EPI_ISL_466006, EPI_ISL_466007, EPI_ISL_466008, EPI_ISL_466009, EPI_ISL_466010, EPI_ISL_466011, EPI_ISL_466012, EPI_ISL_466013, EPI_ISL_466014, EPI_ISL_466015, EPI_ISL_466016, EPI_ISL_466017, EPI_ISL_466018, EPI_ISL_466019, EPI_ISL_466020, EPI_ISL_466021, EPI_ISL_466022, EPI_ISL_466023, EPI_ISL_466024, EPI_ISL_466025, EPI_ISL_466026, EPI_ISL_466027, EPI_ISL_466028, EPI_ISL_466029, EPI_ISL_466030, EPI_ISL_466031, EPI_ISL_466032, EPI_ISL_466033, EPI_ISL_466034, EPI_ISL_466035, EPI_ISL_466036, EPI_ISL_466037, EPI_ISL_466038, EPI_ISL_466039, EPI_ISL_466040, EPI_ISL_466041, EPI_ISL_466042, EPI_ISL_466043, EPI_ISL_466044, EPI_ISL_466045, EPI_ISL_466046, EPI_ISL_466047, EPI_ISL_466048, EPI_ISL_466049, EPI_ISL_466050, EPI_ISL_466051, EPI_ISL_466052, EPI_ISL_466053, EPI_ISL_466054, EPI_ISL_466055, EPI_ISL_466056, EPI_ISL_466057, EPI_ISL_466058, EPI_ISL_466059, EPI_ISL_466060, EPI_ISL_466061, EPI_ISL_466062, EPI_ISL_466063, EPI_ISL_466064, EPI_ISL_466065, EPI_ISL_466066, EPI_ISL_466067, EPI_ISL_466068, EPI_ISL_466069, EPI_ISL_466070, EPI_ISL_466071, EPI_ISL_466072, EPI_ISL_466073, EPI_ISL_466074, EPI_ISL_466075, EPI_ISL_466076, EPI_ISL_466077, EPI_ISL_466078, EPI_ISL_466079, EPI_ISL_466080, EPI_ISL_466081, EPI_ISL_466082, EPI_ISL_466083, EPI_ISL_466084, EPI_ISL_466085, EPI_ISL_466086, EPI_ISL_466087, EPI_ISL_466088, EPI_ISL_466089, EPI_ISL_466090, EPI_ISL_466091, EPI_ISL_466092, EPI_ISL_466093, EPI_ISL_466094, EPI_ISL_466095, EPI_ISL_466096, EPI_ISL_466097, EPI_ISL_466098, EPI_ISL_466099, EPI_ISL_466100, EPI_ISL_466101, EPI_ISL_466102, EPI_ISL_466103, EPI_ISL_466104, EPI_ISL_466105, EPI_ISL_466106, EPI_ISL_466107, EPI_ISL_466108, EPI_ISL_466109, EPI_ISL_466110, EPI_ISL_466111, EPI_ISL_466112, EPI_ISL_466113, EPI_ISL_466114, EPI_ISL_466115, EPI_ISL_466116, EPI_ISL_466117, EPI_ISL_466118, EPI_ISL_466119, EPI_ISL_466120, EPI_ISL_466121, EPI_ISL_466122, EPI_ISL_466123, EPI_ISL_466124, EPI_ISL_466125, EPI_ISL_466126, EPI_ISL_466127, EPI_ISL_466128, EPI_ISL_466129, EPI_ISL_466130, EPI_ISL_466131, EPI_ISL_466132, EPI_ISL_466133, EPI_ISL_466134, EPI_ISL_466135, EPI_ISL_466136, EPI_ISL_466137, EPI_ISL_466138, EPI_ISL_466139, EPI_ISL_466140, EPI_ISL_466141, EPI_ISL_466142, EPI_ISL_466143, EPI_ISL_466144, EPI_ISL_466145, EPI_ISL_466146, EPI_ISL_466147, EPI_ISL_466148, EPI_ISL_466149, EPI_ISL_466150, EPI_ISL_466151, EPI_ISL_466152, EPI_ISL_466153, EPI_ISL_466154, EPI_ISL_466155, EPI_ISL_466156, EPI_ISL_466157, EPI_ISL_466158, EPI_ISL_466159, EPI_ISL_466160, EPI_ISL_466161, EPI_ISL_466162, EPI_ISL_466163, EPI_ISL_466164, EPI_ISL_466165, EPI_ISL_466166, EPI_ISL_466167, EPI_ISL_466168, EPI_ISL_466169, EPI_ISL_466170, EPI_ISL_466171, EPI_ISL_466172, EPI_ISL_466173, EPI_ISL_466174, EPI_ISL_466175, EPI_ISL_466176, EPI_ISL_466177, EPI_ISL_466178, EPI_ISL_466179, EPI_ISL_466180, EPI_ISL_466181, EPI_ISL_466182, EPI_ISL_466183, EPI_ISL_466184, EPI_ISL_466185, EPI_ISL_466186, EPI_ISL_466187, EPI_ISL_466188, EPI_ISL_466189, EPI_ISL_466190, EPI_ISL_466191, EPI_ISL_466192, EPI_ISL_466193, EPI_ISL_466194, EPI_ISL_466195, EPI_ISL_466196, EPI_ISL_466197, EPI_ISL_466198, EPI_ISL_466199, EPI_ISL_466200, EPI_ISL_466201, EPI_ISL_466202, EPI_ISL_466203, EPI_ISL_466204, EPI_ISL_466205, EPI_ISL_466206, EPI_ISL_466207, EPI_ISL_466208, EPI_ISL_466209, EPI_ISL_466210, EPI_ISL_466211, EPI_ISL_466212, EPI_ISL_466213, EPI_ISL_466214, EPI_ISL_466215, EPI_ISL_466216, EPI_ISL_466217, EPI_ISL_466218, EPI_ISL_466219, EPI_ISL_466220, EPI_ISL_466221, EPI_ISL_466222, EPI_ISL_466223, EPI_ISL_466224, EPI_ISL_466225, EPI_ISL_466226, EPI_ISL_466227, EPI_ISL_466228, EPI_ISL_466229, EPI_ISL_466230, EPI_ISL_466231, EPI_ISL_466232, EPI_ISL_466233, EPI_ISL_466234, EPI_ISL_466235, EPI_ISL_466236, EPI_ISL_466237, EPI_ISL_466238, EPI_ISL_466239, EPI_ISL_466240, EPI_ISL_466241, EPI_ISL_466242, EPI_ISL_466243, EPI_ISL_466244, EPI_ISL_466245, EPI_ISL_466246, EPI_ISL_466247, EPI_ISL_466248, EPI_ISL_466249, EPI_ISL_466250, EPI_ISL_466251, EPI_ISL_466252, EPI_ISL_466253, EPI_ISL_466254, EPI_ISL_466255, EPI_ISL_466256, EPI_ISL_466257, EPI_ISL_466258, EPI_ISL_466259, EPI_ISL_466260, EPI_ISL_466261, EPI_ISL_466262, EPI_ISL_466263, EPI_ISL_466264, EPI_ISL_466265, EPI_ISL_466266, EPI_ISL_466267, EPI_ISL_466268, EPI_ISL_466269, EPI_ISL_466270, EPI_ISL_466271, EPI_ISL_466272, EPI_ISL_466273, EPI_ISL_466274, EPI_ISL_466275, EPI_ISL_466276, EPI_ISL_466277, EPI_ISL_466278, EPI_ISL_466279, EPI_ISL_466280, EPI_ISL_466281, EPI_ISL_466282, EPI_ISL_466283, EPI_ISL_466284, EPI_ISL_466285, EPI_ISL_466286, EPI_ISL_466287, EPI_ISL_466288, EPI_ISL_466289, EPI_ISL_466290, EPI_ISL_466291, EPI_ISL_466292, EPI_ISL_466293, EPI_ISL_466294, EPI_ISL_466295, EPI_ISL_466296, EPI_ISL_466297, EPI_ISL_466298, EPI_ISL_466299, EPI_ISL_466300, EPI_ISL_466301, EPI_ISL_466302, EPI_ISL_466303, EPI_ISL_466304, EPI_ISL_466305, EPI_ISL_466306, EPI_ISL_466307, EPI_ISL_466308, EPI_ISL_466309, EPI_ISL_466310, EPI_ISL_466311, EPI_ISL_466312, EPI_ISL_466313, EPI_ISL_466314, EPI_ISL_466315, EPI_ISL_466316, EPI_ISL_466317, EPI_ISL_466318, EPI_ISL_466319, EPI_ISL_466320, EPI_ISL_466321, EPI_ISL_466322, EPI_ISL_466323, EPI_ISL_466324, EPI_ISL_466325, EPI_ISL_466326, EPI_ISL_466327, EPI_ISL_466328, EPI_ISL_466329, EPI_ISL_466330, EPI_ISL_466331, EPI_ISL_466332, EPI_ISL_466333, EPI_ISL_466334, EPI_ISL_466335, EPI_ISL_466336, EPI_ISL_466337, EPI_ISL_466338, EPI_ISL_466339, EPI_ISL_466340, EPI_ISL_466341, EPI_ISL_466342, EPI_ISL_466343, EPI_ISL_466344, EPI_ISL_466345, EPI_ISL_466346, EPI_ISL_466347, EPI_ISL_466348, EPI_ISL_466349, EPI_ISL_466350, EPI_ISL_466351, EPI_ISL_466352, EPI_ISL_466353, EPI_ISL_466354, EPI_ISL_466355, EPI_ISL_466356, EPI_ISL_466357, EPI_ISL_466358, EPI_ISL_466359, EPI_ISL_466360, EPI_ISL_466361, EPI_ISL_466362, EPI_ISL_466363, EPI_ISL_466364, EPI_ISL_466365, EPI_ISL_466366, EPI_ISL_466367, EPI_ISL_466368, EPI_ISL_466369, EPI_ISL_466370, EPI_ISL_466371, EPI_ISL_466372, EPI_ISL_466373, EPI_ISL_466374, EPI_ISL_466375, EPI_ISL_466376, EPI_ISL_466377, EPI_ISL_466378, EPI_ISL_466379, EPI_ISL_466380, EPI_ISL_466381, EPI_ISL_466382, EPI_ISL_466383, EPI_ISL_466384, EPI_ISL_466385, EPI_ISL_466386, EPI_ISL_466387, EPI_ISL_466388, EPI_ISL_466389, EPI_ISL_466390, EPI_ISL_466391, EPI_ISL_466392, EPI_ISL_466393, EPI_ISL_466394, EPI_ISL_466395, EPI_ISL_466396, EPI_ISL_466397, EPI_ISL_466398, EPI_ISL_466399, EPI_ISL_466400, EPI_ISL_466401, EPI_ISL_466402, EPI_ISL_466403, EPI_ISL_466404, EPI_ISL_466405, EPI_ISL_466406, EPI_ISL_466407, EPI_ISL_466408, EPI_ISL_466409, EPI_ISL_466410, EPI_ISL_466411, EPI_ISL_466412, EPI_ISL_466413, EPI_ISL_466414, EPI_ISL_466415, EPI_ISL_466416, EPI_ISL_466417, EPI_ISL_466418, EPI_ISL_466419, EPI_ISL_466420, EPI_ISL_466421, EPI_ISL_466422, EPI_ISL_466423, EPI_ISL_466424, EPI_ISL_466425, EPI_ISL_466426, EPI_ISL_466427, EPI_ISL_466428, EPI_ISL_466429, EPI_ISL_466430, EPI_ISL_466431, EPI_ISL_466432, EPI_ISL_466433, EPI_ISL_466434, EPI_ISL_466435, EPI_ISL_466436, EPI_ISL_466437, EPI_ISL_466438, EPI_ISL_466439, EPI_ISL_466440, EPI_ISL_466441, EPI_ISL_466442, EPI_ISL_466443, EPI_ISL_466444, EPI_ISL_466445, EPI_ISL_466446, EPI_ISL_466447, EPI_ISL_466448, EPI_ISL_466449, EPI_ISL_466450, EPI_ISL_466451, EPI_ISL_466452, EPI_ISL_466453, EPI_ISL_466454, EPI_ISL_466455, EPI_ISL_466456, EPI_ISL_466457, EPI_ISL_466458, EPI_ISL_466459, EPI_ISL_466460, EPI_ISL_466461, EPI_ISL_466462, EPI_ISL_466463, EPI_ISL_466464, EPI_ISL_466465, EPI_ISL_466466, EPI_ISL_466467, EPI_ISL_466468, EPI_ISL_466469, EPI_ISL_466470, EPI_ISL_466471, EPI_ISL_466472, EPI_ISL_466473, EPI_ISL_466474, EPI_ISL_466475, EPI_ISL_466476, EPI_ISL_466477, EPI_ISL_466478, EPI_ISL_466479, EPI_ISL_466480, EPI_ISL_466481, EPI_ISL_466482, EPI_ISL_466483, EPI_ISL_466484, EPI_ISL_466485, EPI_ISL_466486, EPI_ISL_466487, EPI_ISL_466488, EPI_ISL_466489, EPI_ISL_466490, EPI_ISL_466491, EPI_ISL_466492, EPI_ISL_466493, EPI_ISL_466494, EPI_ISL_466495, EPI_ISL_466496, EPI_ISL_466497, EPI_ISL_466498, EPI_ISL_466499, EPI_ISL_466500, EPI_ISL_466501, EPI_ISL_466502, EPI_ISL_466503, EPI_ISL_466504, EPI_ISL_466505, EPI_ISL_466506, EPI_ISL_466507, EPI_ISL_466508, EPI_ISL_466509, EPI_ISL_466510, EPI_ISL_466511, EPI_ISL_466512, EPI_ISL_466513, EPI_ISL_466514, EPI_ISL_466515, EPI_ISL_466516, EPI_ISL_466517, EPI_ISL_466518, EPI_ISL_466519, EPI_ISL_466520, EPI_ISL_466521, EPI_ISL_466522, EPI_ISL_466523, EPI_ISL_466524, EPI_ISL_466525, EPI_ISL_466526, EPI_ISL_466527, EPI_ISL_466528, EPI_ISL_466529, EPI_ISL_466530, EPI_ISL_466531, EPI_ISL_466532, EPI_ISL_466533, EPI_ISL_466534, EPI_ISL_466535, EPI_ISL_466536, EPI_ISL_466537, EPI_ISL_466538, EPI_ISL_466539, EPI_ISL_466540, EPI_ISL_466541, EPI_ISL_466542, EPI_ISL_466543, EPI_ISL_466544, EPI_ISL_466545, EPI_ISL_466546, EPI_ISL_466547, EPI_ISL_466548, EPI_ISL_466549, EPI_ISL_466550, EPI_ISL_466551, EPI_ISL_466552, EPI_ISL_466553, EPI_ISL_466554, EPI_ISL_466555, EPI_ISL_466556, EPI_ISL_466557, EPI_ISL_466558, EPI_ISL_466559, EPI_ISL_466560, EPI_ISL_466561, EPI_ISL_466562, EPI_ISL_466563, EPI_ISL_466564, EPI_ISL_466565, EPI_ISL_466566, EPI_ISL_466567, EPI_ISL_466568, EPI_ISL_466569, EPI_ISL_466570, EPI_ISL_466571, EPI_ISL_466572, EPI_ISL_466573, EPI_ISL_466574, EPI_ISL_466575, EPI_ISL_466576, EPI_ISL_466577, EPI_ISL_466578, EPI_ISL_466579, EPI_ISL_466580, EPI_ISL_466581, EPI_ISL_466582, EPI_ISL_466583, EPI_ISL_466584, EPI_ISL_466585, EPI_ISL_466586, EPI_ISL_466587, EPI_ISL_466588, EPI_ISL_466589, EPI_ISL_466590, EPI_ISL_466591, EPI_ISL_466592, EPI_ISL_466593, EPI_ISL_466594, EPI_ISL_466595, EPI_ISL_466596, EPI_ISL_466597, EPI_ISL_466598, EPI_ISL_466599, EPI_ISL_466600, EPI_ISL_466601, EPI_ISL_466602, EPI_ISL_466603, EPI_ISL_466604, EPI_ISL_466605, EPI_ISL_466606, EPI_ISL_466607, EPI_ISL_466608, EPI_ISL_466609, EPI_ISL_466610, EPI_ISL_466611, EPI_ISL_466612, EPI_ISL_466613, EPI_ISL_466614, EPI_ISL_466615, EPI_ISL_466616, EPI_ISL_466617, EPI_ISL_466618, EPI_ISL_466619, EPI_ISL_466620, EPI_ISL_466621, EPI_ISL_466622, EPI_ISL_466623, EPI_ISL_466624, EPI_ISL_466625 | | | |
| see above | Respiratory Virus Unit, Microbiology Services Colindale, Public Health England | Respiratory Virus Unit, Microbiology Services Colindale, Public Health England | PHE Covid Sequencing Team |
| EPI_ISL_466626, EPI_ISL_466627, EPI_ISL_466628, EPI_ISL_466629, EPI_ISL_466630, EPI_ISL_466636, EPI_ISL_466637, EPI_ISL_466638, EPI_ISL_466639, EPI_ISL_466644, EPI_ISL_466645 | | | |
| see above | National Institute of Laboratory Medicine and Referral Center | Genomic Research Lab, BCSIR | Abu Sayeed Mohammad Mahmud, Mohammad Samir Uzzaman, Eshrar Osman, Md. Ahasan Habib, Shahina Akter, Tanjina Akhter Banu, Md. Murshed Hasan Sarker, Iffat Jahan, Barna Goswami, Md. Saddam Hossain, Tasnim Nafisa, Md. Maruf Ahmed Molla, Mahmuda Yeasmin, Asish Kumar Ghosh, Arifa Akram, A. K. M. Shamsuzzaman, Sheikh Md. Selim Al Din, Utpal Chandra Ray, Salek Ahmed Sajib, Md. Salim Khan |
| EPI_ISL_466648 | Innovative Genomics Institute, UCB | Innovative Genomics Institute, UCB | Stacia Wyman, Haridha Shivram, Liana Lareau, Shana McDevitt, Justin Choi |
| EPI_ISL_466649, EPI_ISL_466650 | National Institute of Laboratory Medicine and Referral Center | Genomic Research Lab, BCSIR | Abu Sayeed Mohammad Mahmud, Mohammad Samir Uzzaman, Eshrar Osman, Md. Ahasan Habib, Shahina Akter, Tanjina Akhter Banu, Md. Murshed Hasan Sarker, Iffat Jahan, Barna Goswami, Md. Saddam Hossain, Tasnim Nafisa, Md. Maruf Ahmed Molla, Mahmuda Yeasmin, Asish Kumar Ghosh, Arifa Akram, A. K. M. Shamsuzzaman, Sheikh Md. Selim Al Din, Utpal Chandra Ray, Salek Ahmed Sajib, Md. Salim Khan |
| EPI_ISL_466652, EPI_ISL_466653, EPI_ISL_466654, EPI_ISL_466655, EPI_ISL_466656, EPI_ISL_466657, EPI_ISL_466658, EPI_ISL_466659, EPI_ISL_466660, EPI_ISL_466661, EPI_ISL_466662, EPI_ISL_466663, EPI_ISL_466664, EPI_ISL_466665, EPI_ISL_466666, EPI_ISL_466667, EPI_ISL_466668, EPI_ISL_466669, EPI_ISL_466670, EPI_ISL_466671, EPI_ISL_466672, EPI_ISL_466673, EPI_ISL_466674, EPI_ISL_466675, EPI_ISL_466676, EPI_ISL_466677, EPI_ISL_466678, EPI_ISL_466679, EPI_ISL_466680, EPI_ISL_466681, EPI_ISL_466682, EPI_ISL_466683 | | | |
| see above | Nebraska Public Health Laboratory | UNMC COVID-19 Response Team | UNMC COVID-19 Response Team |
| EPI_ISL_466686, EPI_ISL_466687, EPI_ISL_466688, EPI_ISL_466689, EPI_ISL_466690, EPI_ISL_466691, EPI_ISL_466692, EPI_ISL_466693, EPI_ISL_466694 | National Institute of Laboratory Medicine and Referral Center | Genomic Research Lab, BCSIR | Abu Sayeed Mohammad Mahmud, Mohammad Samir Uzzaman, Eshrar Osman, Md. Ahasan Habib, Shahina Akter, Tanjina Akhter Banu, Md. Murshed Hasan Sarker, Iffat Jahan, Barna Goswami, Md. Saddam Hossain, Tasnim Nafisa, Md. Maruf Ahmed Molla, Mahmuda Yeasmin, Asish Kumar Ghosh, Arifa Akram, A. K. M. Shamsuzzaman, Sheikh Md. Selim Al Din, Utpal Chandra Ray, Salek Ahmed Sajib, Md. Salim Khan |
| EPI_ISL_466696, EPI_ISL_466697, EPI_ISL_466698, EPI_ISL_466699, EPI_ISL_466700, EPI_ISL_466701, EPI_ISL_466702, EPI_ISL_466703, EPI_ISL_466704, EPI_ISL_466705, EPI_ISL_466706, EPI_ISL_466707, EPI_ISL_466708, EPI_ISL_466709, EPI_ISL_466710, EPI_ISL_466711, EPI_ISL_466712, EPI_ISL_466713, EPI_ISL_466714, EPI_ISL_466715, EPI_ISL_466716, EPI_ISL_466717, EPI_ISL_466718, EPI_ISL_466719, EPI_ISL_466720, EPI_ISL_466721, EPI_ISL_466722, EPI_ISL_466723, EPI_ISL_466724, EPI_ISL_466725, EPI_ISL_466726, EPI_ISL_466727, EPI_ISL_466728, EPI_ISL_466729, EPI_ISL_466730, EPI_ISL_466731, EPI_ISL_466732, EPI_ISL_466733, EPI_ISL_466734, EPI_ISL_466735, EPI_ISL_466736, EPI_ISL_466737, EPI_ISL_466738, EPI_ISL_466739, EPI_ISL_466740, EPI_ISL_466741, EPI_ISL_466742, EPI_ISL_466743, EPI_ISL_466744, EPI_ISL_466745, EPI_ISL_466746, EPI_ISL_466747, EPI_ISL_466748, EPI_ISL_466749, EPI_ISL_466750, EPI_ISL_466751, EPI_ISL_466752, EPI_ISL_466753, EPI_ISL_466754, EPI_ISL_466755, EPI_ISL_466756, EPI_ISL_466757, EPI_ISL_466758, EPI_ISL_466759, EPI_ISL_466760, EPI_ISL_466761, EPI_ISL_466762, EPI_ISL_466763, EPI_ISL_466764, EPI_ISL_466765, EPI_ISL_466766, EPI_ISL_466767, EPI_ISL_466768, EPI_ISL_466769, EPI_ISL_466770, EPI_ISL_466771, EPI_ISL_466772, EPI_ISL_466773, EPI_ISL_466774, EPI_ISL_466775, EPI_ISL_466776, EPI_ISL_466777, EPI_ISL_466778, EPI_ISL_466779, EPI_ISL_466780, EPI_ISL_466781, EPI_ISL_466782, EPI_ISL_466783, EPI_ISL_466784, EPI_ISL_466785, EPI_ISL_466786, EPI_ISL_466787, EPI_ISL_466788, EPI_ISL_466789, EPI_ISL_466790, EPI_ISL_466791, EPI_ISL_466792, EPI_ISL_466793, EPI_ISL_466794, EPI_ISL_466795, EPI_ISL_466796, EPI_ISL_466797, EPI_ISL_466798, EPI_ISL_466799, EPI_ISL_466800, EPI_ISL_466801, EPI_ISL_466802, EPI_ISL_466803, EPI_ISL_466804, EPI_ISL_466805, EPI_ISL_466806, EPI_ISL_466807, EPI_ISL_466808, EPI_ISL_466809, EPI_ISL_466810, EPI_ISL_466811, EPI_ISL_466812, EPI_ISL_466813, EPI_ISL_466814, EPI_ISL_466815, EPI_ISL_466816, EPI_ISL_466817, EPI_ISL_466818, EPI_ISL_466819, EPI_ISL_466820, EPI_ISL_466821, EPI_ISL_466822, EPI_ISL_466823, EPI_ISL_466824, EPI_ISL_466825, EPI_ISL_466826, EPI_ISL_466827, EPI_ISL_466828, EPI_ISL_466829, EPI_ISL_466830, EPI_ISL_466831, EPI_ISL_466832, EPI_ISL_466833, EPI_ISL_466834, EPI_ISL_466835, EPI_ISL_466836, EPI_ISL_466837, EPI_ISL_466838 | | | |
| see above | BCCDC Public Health Laboratory | BCCDC Public Health Laboratory | Richard Harrigan, Hope Lapointe, Jinny Choi, Kimia Kamelian, John Tyson,Terry Snutch, Linda Hoang, Inna Sekirov, Paul Levett, Mel Krajden, Natalie Prystajecky |
| EPI_ISL_466839 | National Genomics Core-Center for DNA Fingerprinting and Diagnostics | National Genomics Core- Center for DNA Fingerprinting and Diagnostics (NGC-CDFD)- DBT's PAN-INDIA-1000 Genome consortium | Bala Pratyusha, Vinay Donipadi, G Shashikanth, Amrita Bhattacherjee, Rajeshree Sanyal, Raju Kumar, Ajay Kumar Chaudhary, Akash Chinchole, Brahmaji Sontyana, C. Arun Kumar, R HARINARAYANAN, RASHNA BHANDARI, MURALI DHARAN BASHYAM, DEBASHIS MITRA, DIVYA VASHISHT, ASHWIN DALAL |
| EPI_ISL_466840, EPI_ISL_466841, EPI_ISL_466842, EPI_ISL_466843, EPI_ISL_466844 | National Genomics Core-Center for DNA Fingerprinting and Diagnostics | National Genomics Core- Center for DNA Fingerprinting and Diagnostics (NGC-CDFD)- DBT's PAN-INDIA-1000 Genome consortium | Bala Pratyusha, Vinay Donipadi, G Shashikanth, Amrita Bhattacherjee, Nalini Raghunathan, Rajeshree Sanyal, Raju Kumar, Ajay Kumar Chaudhary, Akash Chinchole, Brahmaji Sontyana, C. Arun Kumar, R Harinarayanan, Rashna Bhandari, Murali Dharan Bashyam, Debashis Mitra, Divya Vashisht, Ashwin Dalal |
| EPI_ISL_466845, EPI_ISL_466846, EPI_ISL_466847 | National Genomics Core-Center for DNA Fingerprinting and Diagnostics | National Genomics Core- Center for DNA Fingerprinting and Diagnostics (NGC-CDFD)- DBT's PAN-INDIA-1000 Genome consortium | Bala Pratyusha, Vinay Donipadi, G Shashikanth, Amrita Bhattacherjee, Chandra Shekhar V, Chilakala Gangi Reddy, Chinthakindi KrishnaPrasad, Edurugatla Dinesh, Guru Raja, Hilal Ahmad Reshi, R HARINARAYANAN, RASHNA BHANDARI, MURALI DHARAN BASHYAM, DEBASHIS MITRA, DIVYA VASHISHT, ASHWIN DALAL |
| EPI_ISL_466848, EPI_ISL_466849, EPI_ISL_466850, EPI_ISL_466851, EPI_ISL_466852 | National Genomics Core-Center for DNA Fingerprinting and Diagnostics | National Genomics Core- Center for DNA Fingerprinting and Diagnostics (NGC-CDFD)- DBT's PAN-INDIA-1000 Genome consortium | Bala Pratyusha, Vinay Donipadi, G Shashikanth, Amrita Bhattacherjee, J. Mallikarjun, K. Viswakalyan, Kaisar Ahmad Lone, Kausika Kumar Malik, N. Sudheer, Neeraj Kumar, R HARINARAYANAN, RASHNA BHANDARI, MURALI DHARAN BASHYAM, DEBASHIS MITRA, DIVYA VASHISHT, ASHWIN DALAL |
| EPI_ISL_466853, EPI_ISL_466854, EPI_ISL_466855, EPI_ISL_466856, EPI_ISL_466857 | National Genomics Core-Center for DNA Fingerprinting and Diagnostics | National Genomics Core- Center for DNA Fingerprinting and Diagnostics (NGC-CDFD)- DBT's PAN-INDIA-1000 Genome consortium | Bala Pratyusha, Vinay Donipadi, G Shashikanth, Amrita Bhattacherjee, Niteen Pathak, Pradipta Hore, Rahul Baroi, Sayantan Goswami, Shaffiqu T S, Shalini Aricthota, R HARINARAYANAN, RASHNA BHANDARI, MURALI DHARAN BASHYAM, DEBASHIS MITRA, DIVYA VASHISHT, ASHWIN DALAL |
| EPI_ISL_466858, EPI_ISL_466859, EPI_ISL_466860, EPI_ISL_466861, EPI_ISL_466862 | National Genomics Core-Center for DNA Fingerprinting and Diagnostics | National Genomics Core- Center for DNA Fingerprinting and Diagnostics (NGC-CDFD)- DBT's PAN-INDIA-1000 Genome consortium | Bala Pratyusha, Vinay Donipadi, G Shashikanth, Amrita Bhattacherjee, Sobhan Babu, SPR Prasad, Yogesh Patidar, Arjita Jaiswal, Arpita Singh, Devanshi Gupta, R HARINARAYANAN, RASHNA BHANDARI, MURALI DHARAN BASHYAM, DEBASHIS MITRA, DIVYA VASHISHT, ASHWIN DALAL |
| EPI_ISL_466863, EPI_ISL_466864, EPI_ISL_466865, EPI_ISL_466866, EPI_ISL_466867 | National Genomics Core-Center for DNA Fingerprinting and Diagnostics | National Genomics Core- Center for DNA Fingerprinting and Diagnostics (NGC-CDFD)- DBT's PAN-INDIA-1000 Genome consortium | Bala Pratyusha, Vinay Donipadi, G Shashikanth, Amrita Bhattacherjee, Romila Moirangthem, Sanjana Sarkar, Shivani Yadav, Shubhra Ganguli, Suchitra Upreti, Swathi Chodisetty , R HARINARAYANAN, RASHNA BHANDARI, MURALI DHARAN BASHYAM, DEBASHIS MITRA, DIVYA VASHISHT, ASHWIN DALAL |
| EPI_ISL_466868, EPI_ISL_466869, EPI_ISL_466870, EPI_ISL_466871, EPI_ISL_466872 | National Genomics Core-Center for DNA Fingerprinting and Diagnostics | National Genomics Core- Center for DNA Fingerprinting and Diagnostics (NGC-CDFD)- DBT's PAN-INDIA-1000 Genome consortium | Bala Pratyusha, Vinay Donipadi, G Shashikanth, Amrita Bhattacherjee, Vani Singh, Shubhra Ganguli, Suchitra Upreti, Swathi Chodisetty , Vani Singh , R HARINARAYANAN, RASHNA BHANDARI, MURALI DHARAN BASHYAM, DEBASHIS MITRA, DIVYA VASHISHT, ASHWIN DALAL |
| EPI_ISL_466873 | Centre for Clinical Infection and Diagnostics Research and Genomics Innovation Unit | Respiratory Virus Unit, Microbiology Services Colindale, Public Health England | PHE Covid Sequencing Team, Chloe Fisher, Luke Snell, Gaia Nebbia, Ali Awan |

| | | | |
|---|---|---|---|
| EPI_ISL_466874, EPI_ISL_466875, EPI_ISL_466876, EPI_ISL_466877, EPI_ISL_466878, EPI_ISL_466879, EPI_ISL_466880, EPI_ISL_466881, EPI_ISL_466882, EPI_ISL_466883, EPI_ISL_466884, EPI_ISL_466885, EPI_ISL_466886, EPI_ISL_466887, EPI_ISL_466888, EPI_ISL_466889, EPI_ISL_466890, EPI_ISL_466891, EPI_ISL_466892, EPI_ISL_466893, EPI_ISL_466894, EPI_ISL_466895, EPI_ISL_466896, EPI_ISL_466897, EPI_ISL_466898, EPI_ISL_466899, EPI_ISL_466900, EPI_ISL_466901, EPI_ISL_466902, EPI_ISL_466903, EPI_ISL_466904, EPI_ISL_466905, EPI_ISL_466906, EPI_ISL_466907, EPI_ISL_466908, EPI_ISL_466909, EPI_ISL_466910, EPI_ISL_466911, EPI_ISL_466912, EPI_ISL_466913, EPI_ISL_466914, EPI_ISL_466915, EPI_ISL_466916, EPI_ISL_466917, EPI_ISL_466918, EPI_ISL_466919, EPI_ISL_466920, EPI_ISL_466921, EPI_ISL_466922, EPI_ISL_466923, EPI_ISL_466924, EPI_ISL_466925 | | | |
| see above | Max von Pettenkofer Institute, Virology, National Reference Center for Retroviruses, LMU München | Laboratory for Functional Genome Analysis, Dept. Genomics, Gene Center of the LMU Munich | Max Muenchhoff, Stefan Krebs, Alexander Graf, Oliver Keppler, Helmut Blum |
| EPI_ISL_466927, EPI_ISL_466928, EPI_ISL_466929, EPI_ISL_466930, EPI_ISL_466931, EPI_ISL_466932, EPI_ISL_466933, EPI_ISL_466934, EPI_ISL_466935, EPI_ISL_466936, EPI_ISL_466937, EPI_ISL_466938, EPI_ISL_466939, EPI_ISL_466940, EPI_ISL_466941, EPI_ISL_466942, EPI_ISL_466943, EPI_ISL_466944, EPI_ISL_466945, EPI_ISL_466946, EPI_ISL_466947, EPI_ISL_466948, EPI_ISL_466949, EPI_ISL_466950, EPI_ISL_466951, EPI_ISL_466952, EPI_ISL_466953, EPI_ISL_466954, EPI_ISL_466955, EPI_ISL_466956, EPI_ISL_466957, EPI_ISL_466958, EPI_ISL_466959, EPI_ISL_466960, EPI_ISL_466961, EPI_ISL_466962, EPI_ISL_466963, EPI_ISL_466964, EPI_ISL_466965, EPI_ISL_466966, EPI_ISL_466967, EPI_ISL_466968, EPI_ISL_466969, EPI_ISL_466970, EPI_ISL_466971, EPI_ISL_466972, EPI_ISL_466973, EPI_ISL_466974, EPI_ISL_466975, EPI_ISL_466976, EPI_ISL_466977, EPI_ISL_466978, EPI_ISL_466979, EPI_ISL_466980, EPI_ISL_466981, EPI_ISL_466982, EPI_ISL_466983, EPI_ISL_466984, EPI_ISL_466985, EPI_ISL_466986, EPI_ISL_466987, EPI_ISL_466988, EPI_ISL_466989, EPI_ISL_466990, EPI_ISL_466991, EPI_ISL_466992, EPI_ISL_466993, EPI_ISL_466994, EPI_ISL_466995, EPI_ISL_466996, EPI_ISL_466997, EPI_ISL_466998, EPI_ISL_466999, EPI_ISL_467000, EPI_ISL_467001, EPI_ISL_467002, EPI_ISL_467003, EPI_ISL_467004, EPI_ISL_467005, EPI_ISL_467006, EPI_ISL_467007, EPI_ISL_467008, EPI_ISL_467009, EPI_ISL_467010, EPI_ISL_467011, EPI_ISL_467012, EPI_ISL_467013, EPI_ISL_467014, EPI_ISL_467015, EPI_ISL_467016 | | | |
| see above | Viollier AG | Department of Biosystems Science and Engineering, ETH Zürich | Christian Beisel, Sarah Nadeau, Ivan Topolsky, Pedro Ferreira, Philipp Jablonski, Susana Posada-Céspedes, Tobias Schär, Ina Nissen, Natascha Santacroce, Elodie Burcklen, Christiane Beckmann, Maurice Redondo, Olivier Kobel, Christoph Noppen, Sophie Seidel, Noemie Santamaria de Souza, Niko Beerenwinkel, Tanja Stadler |
| EPI_ISL_467029 | GMERS Medical College and Hospital, Gandhinagar | Gujarat Biotechnology Research Centre | Seema Bhatt, Gaurishankar Shrimali, Bhavesh Modi, Bharti Rajani, Tejas Shah, Ankit Hinsu, Pritesh Sabara, Apurvasinh Puvar, Janvi Raval, Zarna Patel, Monika Gandhi, Pinal Trivedi, Maharshi Pandya, Nidhi Patel, Nitin Savaliya, Raghawendra Kumar, Dinesh Kumar, Zuber Saiyed, Komal Patel, Labdhi Pandya, Snehal Bagatharia, Bhavya Jindal, R D Dixit, A M Kadri, Harsh Bakshi, Chaitanya Joshi, Madhvi Joshi |
| EPI_ISL_467030 | GMERS Medical College and Hospital, Gandhinagar | Gujarat Biotechnology Research Centre | Gaurishankar Shrimali, Bhavesh Modi, Bharti Rajani, Tejas Shah, Ankit Hinsu, Pritesh Sabara, Apurvasinh Puvar, Janvi Raval, Zarna Patel, Monika Gandhi, Pinal Trivedi, Maharshi Pandya, Nidhi Patel, Nitin Savaliya, Raghawendra Kumar, Dinesh Kumar, Zuber Saiyed, Komal Patel, Labdhi Pandya, Snehal Bagatharia, Seema Bhatt, Priyanka P Vatsa, R D Dixit, A M Kadri, Harsh Bakshi, Chaitanya Joshi, Madhvi Joshi |
| EPI_ISL_467031 | GMERS Medical College and Hospital, Gandhinagar | Gujarat Biotechnology Research Centre | Bhavesh Modi, Bharti Rajani, Tejas Shah, Ankit Hinsu, Pritesh Sabara, Apurvasinh Puvar, Janvi Raval, Zarna Patel, Monika Gandhi, Pinal Trivedi, Maharshi Pandya, Nidhi Patel, Nitin Savaliya, Raghawendra Kumar, Dinesh Kumar, Zuber Saiyed, Komal Patel, Labdhi Pandya, Snehal Bagatharia, Seema Bhatt, Gaurishankar Shrimali, Pooja P Doshi, R D Dixit, A M Kadri, Harsh Bakshi, Chaitanya Joshi, Madhvi Joshi |
| EPI_ISL_467032 | GMERS Medical College and Hospital, Gandhinagar | Gujarat Biotechnology Research Centre | Bharti Rajani, Tejas Shah, Ankit Hinsu, Pritesh Sabara, Apurvasinh Puvar, Janvi Raval, Zarna Patel, Monika Gandhi, Pinal Trivedi, Maharshi Pandya, Nidhi Patel, Nitin Savaliya, Raghawendra Kumar, Dinesh Kumar, Zuber Saiyed, Komal Patel, Labdhi Pandya, Snehal Bagatharia, Seema Bhatt, Gaurishankar Shrimali, Bhavesh Modi, Akanksha Verma, R D Dixit, A M Kadri, Harsh Bakshi, Chaitanya Joshi, Madhvi Joshi |
| EPI_ISL_467033 | GMERS Medical College and Hospital, Gandhinagar | Gujarat Biotechnology Research Centre | Tejas Shah, Ankit Hinsu, Pritesh Sabara, Apurvasinh Puvar, Janvi Raval, Zarna Patel, Monika Gandhi, Pinal Trivedi, Maharshi Pandya, Nidhi Patel, Nitin Savaliya, Raghawendra Kumar, Dinesh Kumar, Zuber Saiyed, Komal Patel, Labdhi Pandya, Snehal Bagatharia, Seema Bhatt, Gaurishankar Shrimali, Bhavesh Modi, Bharti Rajani, Priti Pandita, R D Dixit, A M Kadri, Harsh Bakshi, Chaitanya Joshi, Madhvi Joshi |
| EPI_ISL_467034 | GMERS Medical College and Hospital, Gandhinagar | Gujarat Biotechnology Research Centre | Ankit Hinsu, Pritesh Sabara, Apurvasinh Puvar, Janvi Raval, Zarna Patel, Monika Gandhi, Pinal Trivedi, Maharshi Pandya, Nidhi Patel, Nitin Savaliya, Raghawendra Kumar, Dinesh Kumar, Zuber Saiyed, Komal Patel, Labdhi Pandya, Snehal Bagatharia, Seema Bhatt, Gaurishankar Shrimali, Bhavesh Modi, Bharti Rajani, Tejas Shah, Pragya Sharma, R D Dixit, A M Kadri, Harsh Bakshi, Chaitanya Joshi, Madhvi Joshi |
| EPI_ISL_467035 | GMERS Medical College and Hospital, Gandhinagar | Gujarat Biotechnology Research Centre | Pritesh Sabara, Apurvasinh Puvar, Janvi Raval, Zarna Patel, Monika Gandhi, Pinal Trivedi, Maharshi Pandya, Nidhi Patel, Nitin Savaliya, Raghawendra Kumar, Dinesh Kumar, Zuber Saiyed, Komal Patel, Labdhi Pandya, Snehal Bagatharia, Seema Bhatt, Gaurishankar Shrimali, Bhavesh Modi, Bharti Rajani, Tejas Shah, Ankit Hinsu, Neha Rajpara, R D Dixit, A M Kadri, Harsh Bakshi, Chaitanya Joshi, Madhvi Joshi |
| EPI_ISL_467036 | GMERS Medical College and Hospital, Gandhinagar | Gujarat Biotechnology Research Centre | Apurvasinh Puvar, Janvi Raval, Zarna Patel, Monika Gandhi, Pinal Trivedi, Maharshi Pandya, Nidhi Patel, Nitin Savaliya, Raghawendra Kumar, Dinesh Kumar, Zuber Saiyed, Komal Patel, Labdhi Pandya, Snehal Bagatharia, Seema Bhatt, Gaurishankar Shrimali, Bhavesh Modi, Bharti Rajani, Tejas Shah, Ankit Hinsu, Pritesh Sabara, Afzal Ansari, R D Dixit, A M Kadri, Harsh Bakshi, Chaitanya Joshi, Madhvi Joshi |
| EPI_ISL_467037 | GMERS Medical College and Hospital, Gandhinagar | Gujarat Biotechnology Research Centre | Janvi Raval, Zarna Patel, Monika Gandhi, Pinal Trivedi, Maharshi Pandya, Nidhi Patel, Nitin Savaliya, Raghawendra Kumar, Dinesh Kumar, Zuber Saiyed, Komal Patel, Labdhi Pandya, Snehal Bagatharia, Seema Bhatt, Gaurishankar Shrimali, Bhavesh Modi, Bharti Rajani, Tejas Shah, Ankit Hinsu, Pritesh Sabara, Apurvasinh Puvar, Fenil Patel, R D Dixit, A M Kadri, Harsh Bakshi, Chaitanya Joshi, Madhvi Joshi |
| EPI_ISL_467038 | GMERS Medical College and Hospital, Gandhinagar | Gujarat Biotechnology Research Centre | Zarna Patel, Monika Gandhi, Pinal Trivedi, Maharshi Pandya, Nidhi Patel, Nitin Savaliya, Raghawendra Kumar, Dinesh Kumar, Zuber Saiyed, Komal Patel, Labdhi Pandya, Snehal Bagatharia, Seema Bhatt, Gaurishankar Shrimali, Bhavesh Modi, Bharti Rajani, Tejas Shah, Ankit Hinsu, Pritesh Sabara, Apurvasinh Puvar, Janvi Raval, Neelam Nathani, R D Dixit, A M Kadri, Harsh Bakshi, Chaitanya Joshi, Madhvi Joshi |
| EPI_ISL_467039 | Government Medical College, Vadodara | Gujarat Biotechnology Research Centre | Meenakshi Shah, Neena Doshi, Varsha Godbole, Tejas Shah, Ankit Hinsu, Pritesh Sabara, Apurvasinh Puvar, Janvi Raval, Zarna Patel, Monika Gandhi, Pinal Trivedi, Maharshi Pandya, Nidhi Patel, Nitin Savaliya, Raghawendra Kumar, Dinesh Kumar, Zuber Saiyed, Komal Patel, Labdhi Pandya, Snehal Bagatharia, Armi Chaudhari, R D Dixit, A M Kadri, Harsh Bakshi, Chaitanya Joshi, Madhvi Joshi, |
| EPI_ISL_467040 | Government Medical College, Vadodara | Gujarat Biotechnology Research Centre | Neena Doshi, Varsha Godbole, Tejas Shah, Ankit Hinsu, Pritesh Sabara, Apurvasinh Puvar, Janvi Raval, Zarna Patel, Monika Gandhi, Pinal Trivedi, Maharshi Pandya, Nidhi Patel, Nitin Savaliya, Raghawendra Kumar, Dinesh Kumar, Zuber Saiyed, Komal Patel, Labdhi Pandya, Snehal Bagatharia, Meenakshi Shah, Bhavya Jindal, R D Dixit, A M Kadri, Harsh Bakshi, Chaitanya Joshi, Madhvi Joshi, |
| EPI_ISL_467041 | B.J. Medical College and Civil hospital | Gujarat Biotechnology Research Centre | Monika Gandhi, Pinal Trivedi, Maharshi Pandya, Nidhi Patel, Nitin Savaliya, Raghawendra Kumar, Dinesh Kumar, Zuber Saiyed, Komal Patel, Labdhi Pandya, Snehal Bagatharia, Pranay Shah, Kamlesh J Upadhyay, Nirav Mungalpara, Tejas Shah, Ankit Hinsu, Pritesh Sabara, Apurvasinh Puvar, Janvi Raval, Zarna Patel, Priyanka P Vatsa, R D Dixit, A M Kadri, Harsh Bakshi, Chaitanya Joshi, Madhvi Joshi, |
| EPI_ISL_467042 | B.J. Medical College and Civil hospital | Gujarat Biotechnology Research Centre | Pinal Trivedi, Maharshi Pandya, Nidhi Patel, Nitin Savaliya, Raghawendra Kumar, Dinesh Kumar, Zuber Saiyed, Komal Patel, Labdhi Pandya, Snehal Bagatharia, Pranay Shah, Kamlesh J Upadhyay, Nirav Mungalpara, Tejas Shah, Ankit Hinsu, Pritesh Sabara, Apurvasinh Puvar, Janvi Raval, Zarna Patel, Monika Gandhi, Pooja P Doshi, R D Dixit, A M Kadri, Harsh Bakshi, Chaitanya Joshi, Madhvi Joshi, |
| EPI_ISL_467043 | B.J. Medical College and Civil hospital | Gujarat Biotechnology Research Centre | Maharshi Pandya, Nidhi Patel, Nitin Savaliya, Raghawendra Kumar, Dinesh Kumar, Zuber Saiyed, Komal Patel, Labdhi Pandya, Snehal Bagatharia, Pranay Shah, Kamlesh J Upadhyay, Nirav Mungalpara, Tejas Shah, Ankit Hinsu, Pritesh Sabara, Apurvasinh Puvar, Janvi Raval, Zarna Patel, Monika Gandhi, Pinal Trivedi, Akanksha Verma, R D Dixit, A M Kadri, Harsh Bakshi, Chaitanya Joshi, Madhvi Joshi, |
| EPI_ISL_467044 | B.J. Medical College and Civil hospital | Gujarat Biotechnology Research Centre | Nidhi Patel, Nitin Savaliya, Raghawendra Kumar, Dinesh Kumar, Zuber Saiyed, Komal Patel, Labdhi Pandya, Snehal Bagatharia, Pranay Shah, Kamlesh J Upadhyay, Nirav Mungalpara, Tejas Shah, Ankit Hinsu, Pritesh Sabara, Apurvasinh Puvar, Janvi Raval, Zarna Patel, Monika Gandhi, Pinal Trivedi, Maharshi Pandya, Priti Pandita, R D Dixit, A M Kadri, Harsh Bakshi, Chaitanya Joshi, Madhvi Joshi, |
| EPI_ISL_467045 | B.J. Medical College and Civil hospital | Gujarat Biotechnology Research Centre | Nitin Savaliya, Raghawendra Kumar, Dinesh Kumar, Zuber Saiyed, Komal Patel, Labdhi Pandya, Snehal Bagatharia, Pranay Shah, Kamlesh J Upadhyay, Nirav Mungalpara, Tejas Shah, Ankit Hinsu, Pritesh Sabara, Apurvasinh Puvar, Janvi Raval, Zarna Patel, Monika Gandhi, Pinal Trivedi, Maharshi Pandya, Nidhi Patel, Pragya Sharma, R D Dixit, A M Kadri, Harsh Bakshi, Chaitanya Joshi, Madhvi Joshi, |
| EPI_ISL_467046 | B.J. Medical College and Civil hospital | Gujarat Biotechnology Research Centre | Raghawendra Kumar, Dinesh Kumar, Zuber Saiyed, Komal Patel, Labdhi Pandya, Snehal Bagatharia, Pranay Shah, Kamlesh J Upadhyay, Nirav Mungalpara, Tejas Shah, Ankit Hinsu, Pritesh Sabara, Apurvasinh Puvar, Janvi Raval, Zarna Patel, Monika Gandhi, Pinal Trivedi, Maharshi Pandya, Nidhi Patel, Nitin Savaliya, Neha Rajpara, R D Dixit, A M Kadri, Harsh Bakshi, Chaitanya Joshi, Madhvi Joshi, |
| EPI_ISL_467047 | B.J. Medical College and Civil hospital | Gujarat Biotechnology Research Centre | Dinesh Kumar, Zuber Saiyed, Komal Patel, Labdhi Pandya, Snehal Bagatharia, Pranay Shah, Kamlesh J Upadhyay, Nirav Mungalpara, Tejas Shah, Ankit Hinsu, Pritesh Sabara, Apurvasinh Puvar, Janvi Raval, Zarna Patel, Monika Gandhi, Pinal Trivedi, Maharshi Pandya, Nidhi Patel, Nitin Savaliya, Raghawendra Kumar, Afzal Ansari, R D Dixit, A M Kadri, Harsh Bakshi, Chaitanya Joshi, Madhvi Joshi, |
| EPI_ISL_467048 | B.J. Medical College and Civil hospital | Gujarat Biotechnology Research Centre | Zuber Saiyed, Komal Patel, Labdhi Pandya, Snehal Bagatharia, Pranay Shah, Kamlesh J Upadhyay, Nirav Mungalpara, Tejas Shah, Ankit Hinsu, Pritesh Sabara, Apurvasinh Puvar, Janvi Raval, Zarna Patel, Monika Gandhi, Pinal Trivedi, Maharshi Pandya, Nidhi Patel, Nitin Savaliya, Raghawendra Kumar, Dinesh Kumar, Fenil Patel, R D Dixit, A M Kadri, Harsh Bakshi, Chaitanya Joshi, Madhvi Joshi, |
| EPI_ISL_467049 | B.J. Medical College and Civil hospital | Gujarat Biotechnology Research Centre | Komal Patel, Labdhi Pandya, Snehal Bagatharia, Pranay Shah, Kamlesh J Upadhyay, Nirav Mungalpara, Tejas Shah, Ankit Hinsu, Pritesh Sabara, Apurvasinh Puvar, Janvi Raval, Zarna Patel, Monika Gandhi, Pinal Trivedi, Maharshi Pandya, Nidhi Patel, Nitin Savaliya, Raghawendra Kumar, Dinesh Kumar, Zuber Saiyed, Neelam Nathani, R D Dixit, A M Kadri, Harsh Bakshi, Chaitanya Joshi, Madhvi Joshi, |
| EPI_ISL_467050 | B.J. Medical College and Civil hospital | Gujarat Biotechnology Research Centre | Labdhi Pandya, Snehal Bagatharia, Pranay Shah, Kamlesh J Upadhyay, Nirav Mungalpara, Tejas Shah, Ankit Hinsu, Pritesh Sabara, Apurvasinh Puvar, Janvi Raval, Zarna Patel, Monika Gandhi, Pinal Trivedi, Maharshi Pandya, Nidhi Patel, Nitin Savaliya, Raghawendra Kumar, Dinesh Kumar, Zuber Saiyed, Komal Patel, Armi Chaudhari, R D Dixit, A M Kadri, Harsh Bakshi, Chaitanya Joshi, Madhvi Joshi, |
| EPI_ISL_467051 | B.J. Medical College and Civil hospital | Gujarat Biotechnology Research Centre | Snehal Bagatharia, Pranay Shah, Kamlesh J Upadhyay, Nirav Mungalpara, Tejas Shah, Ankit Hinsu, Pritesh Sabara, Apurvasinh Puvar, Janvi Raval, Zarna Patel, Monika Gandhi, Pinal Trivedi, Maharshi Pandya, Nidhi Patel, Nitin Savaliya, Raghawendra Kumar, Dinesh Kumar, Zuber Saiyed, Komal Patel, Labdhi Pandya, Bhavya Jindal, R D Dixit, A M Kadri, Harsh Bakshi, Chaitanya Joshi, Madhvi Joshi, |
| EPI_ISL_467052 | B.J. Medical College and Civil hospital | Gujarat Biotechnology Research Centre | Pranay Shah, Kamlesh J Upadhyay, Nirav Mungalpara, Tejas Shah, Ankit Hinsu, Pritesh Sabara, Apurvasinh Puvar, Janvi Raval, Zarna Patel, Monika Gandhi, Pinal Trivedi, Maharshi Pandya, Nidhi Patel, Nitin Savaliya, Raghawendra Kumar, Dinesh Kumar, Zuber Saiyed, Komal Patel, Labdhi Pandya, Snehal Bagatharia, Priyanka P Vatsa, R D Dixit, A M Kadri, Harsh Bakshi, Chaitanya Joshi, Madhvi Joshi, |
| EPI_ISL_467053 | B.J. Medical College and Civil hospital | Gujarat Biotechnology Research Centre | Kamlesh J Upadhyay, Nirav Mungalpara, Tejas Shah, Ankit Hinsu, Pritesh Sabara, Apurvasinh Puvar, Janvi Raval, Zarna Patel, Monika Gandhi, Pinal Trivedi, Maharshi Pandya, Nidhi Patel, Nitin Savaliya, Raghawendra Kumar, Dinesh Kumar, Zuber Saiyed, Komal Patel, Labdhi Pandya, Snehal Bagatharia, Pranay Shah, Pooja P Doshi, R D Dixit, A M Kadri, Harsh Bakshi, Chaitanya Joshi, Madhvi Joshi, |
| EPI_ISL_467054 | B.J. Medical College and Civil hospital | Gujarat Biotechnology Research Centre | Nirav Mungalpara, Tejas Shah, Ankit Hinsu, Pritesh Sabara, Apurvasinh Puvar, Janvi Raval, Zarna Patel, Monika Gandhi, Pinal Trivedi, Maharshi Pandya, Nidhi Patel, Nitin Savaliya, Raghawendra Kumar, Dinesh Kumar, Zuber Saiyed, Komal Patel, Labdhi Pandya, Snehal Bagatharia, Pranay Shah, Kamlesh J Upadhyay, Akanksha Verma, R D Dixit, A M Kadri, Harsh Bakshi, Chaitanya Joshi, Madhvi Joshi, |
| EPI_ISL_467055, EPI_ISL_467056, EPI_ISL_467057, EPI_ISL_467058 | Servicio de Microbiología, Hospital Universitario Son Espases | SeqCOVID-SPAIN consortium/IBV(CSIC) | Carla López-Causapé, Jordi Reina y Antonio Oliver and SeqCOVID-SPAIN consortium |
| EPI_ISL_467059, EPI_ISL_467060, EPI_ISL_467061, EPI_ISL_467062, EPI_ISL_467063 | Hospital Universitario Virgen de las Nieves de Granada-SAS | SeqCOVID-SPAIN consortium/IBV(CSIC) | Mercedes Pérez Ruiz, Sara Sanbonmatsu Gámez, Irene Pedrosa Corral, José M. Navarro-Marí and SeqCOVID-SPAIN consortium |
| EPI_ISL_467064, EPI_ISL_467065, EPI_ISL_467066, EPI_ISL_467067, EPI_ISL_467068, EPI_ISL_467069, EPI_ISL_467070, EPI_ISL_467071, EPI_ISL_467072, EPI_ISL_467073, EPI_ISL_467074, EPI_ISL_467075, EPI_ISL_467076, EPI_ISL_467077, EPI_ISL_467078, EPI_ISL_467079, EPI_ISL_467080, EPI_ISL_467081, EPI_ISL_467082, EPI_ISL_467083, EPI_ISL_467084, EPI_ISL_467085 | | | |
| see above | Hospital Universitario Puerta del Mar de Cádiz - INIBICA | SeqCOVID-SPAIN consortium/IBV(CSIC) | Salud Rodríguez-Pallares, Fátima-Galán-Sánchez, Manuel Rodríguez-Iglesias and SeqCOVID-SPAIN consortium |
| EPI_ISL_467086, EPI_ISL_467087, EPI_ISL_467088, EPI_ISL_467089, EPI_ISL_467090, EPI_ISL_467091 | Hospital Universitario de Gran Canaria Dr. Negrín | SeqCOVID-SPAIN consortium/IBV(CSIC) | M. Carmen Pérez González, Francisco J. Chamizo López, Ana Bordes Benítez and SeqCOVID-SPAIN consortium |
| EPI_ISL_467092, EPI_ISL_467093, EPI_ISL_467094, EPI_ISL_467095, EPI_ISL_467096, EPI_ISL_467097, EPI_ISL_467098, EPI_ISL_467099, EPI_ISL_467100, EPI_ISL_467101, EPI_ISL_467102, EPI_ISL_467103, EPI_ISL_467104, EPI_ISL_467105, EPI_ISL_467106, EPI_ISL_467107, EPI_ISL_467108, EPI_ISL_467109, EPI_ISL_467110, EPI_ISL_467111, EPI_ISL_467112, EPI_ISL_467113, EPI_ISL_467114, EPI_ISL_467115, EPI_ISL_467116, EPI_ISL_467117, EPI_ISL_467118, EPI_ISL_467119, EPI_ISL_467120, EPI_ISL_467121, EPI_ISL_467122, EPI_ISL_467123, EPI_ISL_467124, EPI_ISL_467125, EPI_ISL_467126, EPI_ISL_467127, EPI_ISL_467128, EPI_ISL_467129, EPI_ISL_467130, EPI_ISL_467131, EPI_ISL_467132, EPI_ISL_467133, EPI_ISL_467134, EPI_ISL_467135, EPI_ISL_467136, EPI_ISL_467137, EPI_ISL_467138, EPI_ISL_467139, EPI_ISL_467140, EPI_ISL_467141, EPI_ISL_467142, EPI_ISL_467143, EPI_ISL_467144, EPI_ISL_467145, EPI_ISL_467146, EPI_ISL_467147, EPI_ISL_467148, EPI_ISL_467149, EPI_ISL_467150, EPI_ISL_467151, EPI_ISL_467152, EPI_ISL_467153, EPI_ISL_467154, EPI_ISL_467155, EPI_ISL_467156, EPI_ISL_467157, EPI_ISL_467158, EPI_ISL_467159, EPI_ISL_467160, EPI_ISL_467161, EPI_ISL_467162, EPI_ISL_467163, EPI_ISL_467164, EPI_ISL_467165, EPI_ISL_467166, EPI_ISL_467167, EPI_ISL_467168, EPI_ISL_467169, EPI_ISL_467170, EPI_ISL_467171, EPI_ISL_467172, EPI_ISL_467173, EPI_ISL_467174, EPI_ISL_467175, EPI_ISL_467176, EPI_ISL_467177, EPI_ISL_467178, EPI_ISL_467179, EPI_ISL_467180, EPI_ISL_467181, EPI_ISL_467182, EPI_ISL_467183 | | | |
| see above | Hospital Universitario Araba. Vitoria-Gasteiz | SeqCOVID-SPAIN consortium/IBV(CSIC) | Silvia Hernáez Crespo, Carmen Gómez González, Amaia Aguirre Quiñonero, Marina Fernández Torres, Mª Rosario Almela Ferrer, Mª Concepción Lecaroz Agara, Andrés Canut Blasco. and SeqCOVID-SPAIN consortium |
| EPI_ISL_467184, EPI_ISL_467185, EPI_ISL_467186, EPI_ISL_467187, EPI_ISL_467188, EPI_ISL_467189, EPI_ISL_467190, EPI_ISL_467191, EPI_ISL_467192, EPI_ISL_467193, EPI_ISL_467194, EPI_ISL_467195, EPI_ISL_467196, EPI_ISL_467197, EPI_ISL_467198, EPI_ISL_467199, EPI_ISL_467200, EPI_ISL_467201, EPI_ISL_467202, EPI_ISL_467203, EPI_ISL_467204, EPI_ISL_467205, EPI_ISL_467206, EPI_ISL_467207, EPI_ISL_467208, EPI_ISL_467209, EPI_ISL_467210, EPI_ISL_467211, EPI_ISL_467212, EPI_ISL_467213, EPI_ISL_467214, EPI_ISL_467215, EPI_ISL_467216, EPI_ISL_467217, EPI_ISL_467218, EPI_ISL_467219, EPI_ISL_467220, EPI_ISL_467221, EPI_ISL_467222, EPI_ISL_467223, EPI_ISL_467224, EPI_ISL_467225, EPI_ISL_467226, EPI_ISL_467227, EPI_ISL_467228, EPI_ISL_467229, EPI_ISL_467230, EPI_ISL_467231, EPI_ISL_467232, EPI_ISL_467233, EPI_ISL_467234, EPI_ISL_467235, EPI_ISL_467236, EPI_ISL_467237, EPI_ISL_467238, EPI_ISL_467239, EPI_ISL_467240, EPI_ISL_467241, EPI_ISL_467242, EPI_ISL_467243, EPI_ISL_467244, EPI_ISL_467245, EPI_ISL_467246, EPI_ISL_467247, EPI_ISL_467248, EPI_ISL_467249, EPI_ISL_467250, EPI_ISL_467251, EPI_ISL_467252, EPI_ISL_467253, EPI_ISL_467254, EPI_ISL_467255, EPI_ISL_467256, EPI_ISL_467257, EPI_ISL_467258, EPI_ISL_467259, EPI_ISL_467260, EPI_ISL_467261 | | | |
| see above | Hospital General Universitario Gregorio Marañón | SeqCOVID-SPAIN consortium/IBV(CSIC) | Laura Pérez-Lago, Marta Herranz, Jon Sicilia, Julia Suárez, Pilar Catalán, Patricia Muñoz, Darío García de Viedma and SeqCOVID-SPAIN consortium |
| EPI_ISL_467262, EPI_ISL_467263, EPI_ISL_467264, EPI_ISL_467265, EPI_ISL_467266, EPI_ISL_467267, EPI_ISL_467268, EPI_ISL_467269, EPI_ISL_467270, EPI_ISL_467271, EPI_ISL_467272, EPI_ISL_467273, EPI_ISL_467274, EPI_ISL_467275, EPI_ISL_467276, EPI_ISL_467277, EPI_ISL_467278, EPI_ISL_467279, EPI_ISL_467280, EPI_ISL_467281, EPI_ISL_467282, EPI_ISL_467283, EPI_ISL_467284, EPI_ISL_467285, EPI_ISL_467286, EPI_ISL_467287, EPI_ISL_467288, EPI_ISL_467289, EPI_ISL_467290, EPI_ISL_467291, EPI_ISL_467292, EPI_ISL_467293, EPI_ISL_467294, EPI_ISL_467295, EPI_ISL_467296, EPI_ISL_467297 | | | |
| see above | Hospital Clínico Universitario de Santiago de Compostela | SeqCOVID-SPAIN consortium/IBV(CSIC) | José Javier Costa Alcalde, Antonio Aguilera Guirao, Mª Luisa Pérez del Molino Bernal, Amparo Coira Nieto, Gema Barbeito Castiñeiras, Rocio Trastoy Pena and SeqCOVID-SPAIN consortium |
| EPI_ISL_467298 | Nebraska Public Health Laboratory | UNMC COVID-19 Response Team | UNMC COVID-19 Response Team |

| | | | |
|---|---|---|---|
| EPI_ISL_467299 | Research and Medical Analysis Laboratory of Gendarmerie Royale | Research and Medical Analysis Laboratory of Gendarmerie Royale | Sanaâ LEMRISS Amal SOUIRI Hicham EL OSSMANI Saâd EL Kabbaj |
| EPI_ISL_467300 | General Hospital "Abdulah Nakas" | Alea Genetic Center | Rijad Konjhodzic; Lana Salihefendic; Teufik Goletic; Sead Jazic; Dino Pecar; Nihad Fejzic; Damir Marjanovic; Enis Kandic |
| EPI_ISL_467301 | Washington University in St. Louis | Washington University in St. Louis | David Wang, Carey-Ann Burnham, Scott Handley, Lindsay Droit, Stephen Tahan |
| EPI_ISL_467302 | Washington University in St. Louis | Washington University in St. Louis | David Wang, Carey-Ann Burnham, Scott Handley, Lindsay Droit, Stephen Tahan |
| EPI_ISL_467303 | BCCDC Public Health Laboratory | BCCDC Public Health Laboratory | Richard Harrigan, Hope Lapointe, Jinny Choi, Kimia Kamelian, John Tyson,Terry Snutch, Linda Hoang, Inna Sekirov, Paul Levett, Mel Krajden, Natalie Prystajecky |
| EPI_ISL_467304 | Washington University in St. Louis | Washington University in St. Louis | David Wang, Carey-Ann Burnham, Scott Handley, Lindsay Droit, Stephen Tahan |
| EPI_ISL_467305, EPI_ISL_467306, EPI_ISL_467307, EPI_ISL_467308, EPI_ISL_467309, EPI_ISL_467310, EPI_ISL_467311, EPI_ISL_467312, EPI_ISL_467313, EPI_ISL_467314, EPI_ISL_467315, EPI_ISL_467316, EPI_ISL_467317, EPI_ISL_467318, EPI_ISL_467319, EPI_ISL_467320, EPI_ISL_467321, EPI_ISL_467322, EPI_ISL_467323, EPI_ISL_467324, EPI_ISL_467325, EPI_ISL_467326, EPI_ISL_467327, EPI_ISL_467328, EPI_ISL_467329, EPI_ISL_467330, EPI_ISL_467331, EPI_ISL_467332, EPI_ISL_467333, EPI_ISL_467334, EPI_ISL_467335, EPI_ISL_467336, EPI_ISL_467337, EPI_ISL_467338, EPI_ISL_467339, EPI_ISL_467340, EPI_ISL_467341, EPI_ISL_467342, EPI_ISL_467343 | | | |
| see above | BCCDC Public Health Laboratory | BCCDC Public Health Laboratory | Richard Harrigan, Hope Lapointe, Jinny Choi, Kimia Kamelian, John Tyson,Terry Snutch, Linda Hoang, Inna Sekirov, Paul Levett, Mel Krajden, Natalie Prystajecky |
| EPI_ISL_467344, EPI_ISL_467345, EPI_ISL_467346, EPI_ISL_467347, EPI_ISL_467348, EPI_ISL_467349, EPI_ISL_467350, EPI_ISL_467351, EPI_ISL_467352, EPI_ISL_467353, EPI_ISL_467354, EPI_ISL_467355, EPI_ISL_467356, EPI_ISL_467357, EPI_ISL_467358, EPI_ISL_467359, EPI_ISL_467360, EPI_ISL_467361, EPI_ISL_467362, EPI_ISL_467363, EPI_ISL_467364, EPI_ISL_467365, EPI_ISL_467366, EPI_ISL_467367, EPI_ISL_467368, EPI_ISL_467369, EPI_ISL_467370, EPI_ISL_467371 | | | |
| see above | Laboratory of Respiratory Viruses and Measles, Oswaldo Cruz Institute, FIOCRUZ | Laboratory of Respiratory Viruses and Measles, Oswaldo Cruz Institute, FIOCRUZ | Paola Resende, Luciana Appolinario, Fernando Motta, Anna Carolina Paixão, Ana Carolina Mendonça, Aline Mattos, Milene Miranda, Cristiana Garcia, Braulia Caetano, Maria Ogrzewalska, Jonathan Lopes, Marilda Siqueira |
| EPI_ISL_467372 | Arizona State University Health Services | Arizona State University | Peter T. Skidmore, Rabia Maqsood, LaRinda A. Holland, Emily A. Kaelin, Lily I. Wu, Arvind Varsani, Rolf U. Halden, Brenda G. Hogue, Matthew Scotch, Efrem S. Lim |
| EPI_ISL_467373 | Arizona State University Health Services | Arizona State University | Peter T. Skidmore, Rabia Maqsood, LaRinda A. Holland, Emily A. Kaelin, Lily I. Wu, Arvind Varsani, Rolf U. Halden, Brenda G. Hogue, Matthew Scotch, Efrem S. Lim |
| EPI_ISL_467374 | Dinkes Samarinda | Eijkman Institute for Molecular Biology, Ministry of Research and Technology/National Agency for Research and Innovation | Edison Johar, Frilasita A Yudhaputri, Hidayat Trimarsanto, David H Muljono, Safarina G Malik, Khin Saw Myint, Amin Soebandrio |
| EPI_ISL_467375 | RSUP Prof. Dr. R. Kandou Manado | Eijkman Institute for Molecular Biology, Ministry of Research and Technology/National Agency for Research and Innovation | Edison Johar, Frilasita A Yudhaputri, Hidayat Trimarsanto, David H Muljono, Safarina G Malik, Khin Saw Myint, Amin Soebandrio |
| EPI_ISL_467376 | RSUP Fatmawati | Eijkman Institute for Molecular Biology, Ministry of Research and Technology/National Agency for Research and Innovation | Edison Johar, Frilasita A Yudhaputri, Hidayat Trimarsanto, David H Muljono, Safarina G Malik, Khin Saw Myint, Amin Soebandrio |
| EPI_ISL_467377, EPI_ISL_467378, EPI_ISL_467379, EPI_ISL_467380, EPI_ISL_467381, EPI_ISL_467382, EPI_ISL_467383, EPI_ISL_467384, EPI_ISL_467385, EPI_ISL_467386, EPI_ISL_467387, EPI_ISL_467388, EPI_ISL_467389, EPI_ISL_467390, EPI_ISL_467391, EPI_ISL_467392, EPI_ISL_467393, EPI_ISL_467394, EPI_ISL_467395, EPI_ISL_467396, EPI_ISL_467397, EPI_ISL_467398, EPI_ISL_467399, EPI_ISL_467400, EPI_ISL_467401, EPI_ISL_467402, EPI_ISL_467403, EPI_ISL_467404, EPI_ISL_467405, EPI_ISL_467406, EPI_ISL_467407, EPI_ISL_467408, EPI_ISL_467409, EPI_ISL_467410, EPI_ISL_467411, EPI_ISL_467412, EPI_ISL_467413, EPI_ISL_467414, EPI_ISL_467415, EPI_ISL_467416, EPI_ISL_467417, EPI_ISL_467418, EPI_ISL_467419, EPI_ISL_467420, EPI_ISL_467421, EPI_ISL_467422 | | | |
| see above | NYU Langone Health | Departments of Pathology and Medicine, New York University School of Medicine | Maria Aguero-Rosenfeld, Brendan Belovarac, Margaret Black, Ludovic Boytard, John Cadley, Paolo Cotzia, John Chen, Dacia Dimartino, Xiaojun Feng, Tatyana Gindin, Emily Guzman, Adriana Heguy, Megan Hogan, Emily Huang, George Jour, Alireza Khodadadi-Jamayran, Lawrence H. Lin, Raven Luther, Andrew Lytle, Christian Marier, Matthew T. Maurano, Mark J. Mulligan, Peter Meyn, Raquel Ordonez Ciriza, Iman Osman, Jared Pinnell, Vanessa Raabe, Sitharam Ramaswami, Amy Rapkiewicz, Andre M. Ribeiro-dos-Santos, Marie Samanovic-Golden, Antonio Serrano, Guomiao Shen, Matija Snuderl, Theodore Vougiouklakis, Nick Vulpescu, Gael Westby, Paul Zappile, Yutong Zhang |
| EPI_ISL_467423, EPI_ISL_467424, EPI_ISL_467425, EPI_ISL_467426, EPI_ISL_467427, EPI_ISL_467428, EPI_ISL_467429 | BCCDC Public Health Laboratory | BCCDC Public Health Laboratory | Richard Harrigan, Hope Lapointe, Jinny Choi, Kimia Kamelian, John Tyson,Terry Snutch, Linda Hoang, Inna Sekirov, Paul Levett, Mel Krajden, Natalie Prystajecky |
| EPI_ISL_467430 | Zoonotic and Exotic infection Diseases Division | Zoontic and Exotic infection Diseases Division | Jinlang Wang, Zhigao Bu |
| EPI_ISL_467431 | Molecular Diagnostics Services (MDS) | KRISP, KZN Research Innovation and Sequencing Platform | Giandhari J, Pillay S, Lessells R, Chimukangara B, Mdlalose K, York D, Khan S, Tegally H, Wilkinson E, de Oliveira T |
| EPI_ISL_467432, EPI_ISL_467433, EPI_ISL_467434, EPI_ISL_467435 | AMPATH-DBN | KRISP, KZN Research Innovation and Sequencing Platform | Giandhari J, Pillay S, Lessells R, Chimukangara B, Mdlalose K, York D, Khan S, Tegally H, Wilkinson E, de Oliveira T |
| EPI_ISL_467436, EPI_ISL_467437, EPI_ISL_467438, EPI_ISL_467439, EPI_ISL_467440, EPI_ISL_467441, EPI_ISL_467442, EPI_ISL_467443 | NHLS-IALCH | KRISP, KZN Research Innovation and Sequencing Platform | Giandhari J, Pillay S, Lessells R, Chimukangara B, Mdlalose K, York D, Khan S, Tegally H, Wilkinson E, de Oliveira T |
| EPI_ISL_467444, EPI_ISL_467445, EPI_ISL_467446, EPI_ISL_467447, EPI_ISL_467448 | Molecular Diagnostics Services (MDS) | KRISP, KZN Research Innovation and Sequencing Platform | Giandhari J, Pillay S, Lessells R, Chimukangara B, Mdlalose K, York D, Khan S, Tegally H, Wilkinson E, de Oliveira T |
| EPI_ISL_467449, EPI_ISL_467450, EPI_ISL_467451, EPI_ISL_467452, EPI_ISL_467453, EPI_ISL_467454, EPI_ISL_467455, EPI_ISL_467456, EPI_ISL_467457, EPI_ISL_467458, EPI_ISL_467459, EPI_ISL_467460, EPI_ISL_467461, EPI_ISL_467462, EPI_ISL_467463, EPI_ISL_467464, EPI_ISL_467465, EPI_ISL_467466, EPI_ISL_467467, EPI_ISL_467468, EPI_ISL_467469, EPI_ISL_467470, EPI_ISL_467471, EPI_ISL_467472, EPI_ISL_467473, EPI_ISL_467474 | | | |
| see above | AMPATH-DBN | KRISP, KZN Research Innovation and Sequencing Platform | Giandhari J, Pillay S, Lessells R, Chimukangara B, Mdlalose K, York D, Khan S, Tegally H, Wilkinson E, de Oliveira T |
| EPI_ISL_467475, EPI_ISL_467476, EPI_ISL_467477, EPI_ISL_467478, EPI_ISL_467479, EPI_ISL_467480, EPI_ISL_467481, EPI_ISL_467482, EPI_ISL_467483, EPI_ISL_467484, EPI_ISL_467485, EPI_ISL_467486, EPI_ISL_467487, EPI_ISL_467488, EPI_ISL_467489, EPI_ISL_467490, EPI_ISL_467491 | | | |
| see above | Molecular Diagnostics Services (MDS) | KRISP, KZN Research Innovation and Sequencing Platform | Giandhari J, Pillay S, Lessells R, Chimukangara B, Mdlalose K, York D, Khan S, Tegally H, Wilkinson E, de Oliveira T |
| EPI_ISL_467492, EPI_ISL_467493 | NHLS-IALCH | KRISP, KZN Research Innovation and Sequencing Platform | Giandhari J, Pillay S, Lessells R, Chimukangara B, Mdlalose K, York D, Khan S, Tegally H, Wilkinson E, de Oliveira T |
| EPI_ISL_467494, EPI_ISL_467495, EPI_ISL_467496, EPI_ISL_467497, EPI_ISL_467498, EPI_ISL_467499, EPI_ISL_467500, EPI_ISL_467501, EPI_ISL_467502, EPI_ISL_467503, EPI_ISL_467504, EPI_ISL_467505, EPI_ISL_467506 | | | |
| see above | Molecular Diagnostics Services (MDS) | KRISP, KZN Research Innovation and Sequencing Platform | Giandhari J, Pillay S, Lessells R, Chimukangara B, Mdlalose K, York D, Khan S, Tegally H, Wilkinson E, de Oliveira T |
| EPI_ISL_467507, EPI_ISL_467508, EPI_ISL_467509, EPI_ISL_467510, EPI_ISL_467511, EPI_ISL_467512, EPI_ISL_467513, EPI_ISL_467514, EPI_ISL_467515 | NHLS-IALCH | KRISP, KZN Research Innovation and Sequencing Platform | Giandhari J, Pillay S, Lessells R, Chimukangara B, Mdlalose K, York D, Khan S, Tegally H, Wilkinson E, de Oliveira T |
| EPI_ISL_467516 | CAPRISA | KRISP, KZN Research Innovation and Sequencing Platform | Giandhari J, Pillay S, Lessells R, Chimukangara B, Mdlalose K, York D, Khan S, Tegally H, Wilkinson E, de Oliveira T |
| EPI_ISL_467517, EPI_ISL_467518, EPI_ISL_467519, EPI_ISL_467520, EPI_ISL_467521, EPI_ISL_467522, EPI_ISL_467523, EPI_ISL_467524 | NHLS-IALCH | KRISP, KZN Research Innovation and Sequencing Platform | Giandhari J, Pillay S, Lessells R, Chimukangara B, Mdlalose K, York D, Khan S, Tegally H, Wilkinson E, de Oliveira T |
| EPI_ISL_467525, EPI_ISL_467526, EPI_ISL_467527, EPI_ISL_467528, EPI_ISL_467529, EPI_ISL_467530, EPI_ISL_467531, EPI_ISL_467532, EPI_ISL_467533, EPI_ISL_467534, EPI_ISL_467535, EPI_ISL_467536, EPI_ISL_467537, EPI_ISL_467538, EPI_ISL_467539, EPI_ISL_467540, EPI_ISL_467541, EPI_ISL_467542, EPI_ISL_467543, EPI_ISL_467544, EPI_ISL_467545, EPI_ISL_467546, EPI_ISL_467547, EPI_ISL_467548, EPI_ISL_467549, EPI_ISL_467550, EPI_ISL_467551, EPI_ISL_467552, EPI_ISL_467553, EPI_ISL_467554, EPI_ISL_467555, EPI_ISL_467556, EPI_ISL_467557, EPI_ISL_467558, EPI_ISL_467559, EPI_ISL_467560, EPI_ISL_467561, EPI_ISL_467562, EPI_ISL_467563, EPI_ISL_467564, EPI_ISL_467565, EPI_ISL_467566, EPI_ISL_467567, EPI_ISL_467568, EPI_ISL_467569, EPI_ISL_467570, EPI_ISL_467571, EPI_ISL_467572, EPI_ISL_467573, EPI_ISL_467574, EPI_ISL_467575, EPI_ISL_467576, EPI_ISL_467577, EPI_ISL_467578, EPI_ISL_467579, EPI_ISL_467580, EPI_ISL_467581, EPI_ISL_467582, EPI_ISL_467583, EPI_ISL_467584, EPI_ISL_467585, EPI_ISL_467586, EPI_ISL_467587, EPI_ISL_467588, EPI_ISL_467589, EPI_ISL_467590, EPI_ISL_467591, EPI_ISL_467592, EPI_ISL_467593, EPI_ISL_467594, EPI_ISL_467595, EPI_ISL_467596, EPI_ISL_467597, EPI_ISL_467598, EPI_ISL_467599, EPI_ISL_467600, EPI_ISL_467601, EPI_ISL_467602, EPI_ISL_467603, EPI_ISL_467604, EPI_ISL_467605, EPI_ISL_467606, EPI_ISL_467607, EPI_ISL_467608, EPI_ISL_467609, EPI_ISL_467610, EPI_ISL_467611, EPI_ISL_467612, EPI_ISL_467613, EPI_ISL_467614, EPI_ISL_467615, EPI_ISL_467616, EPI_ISL_467617, EPI_ISL_467618, EPI_ISL_467619, EPI_ISL_467620, EPI_ISL_467621, EPI_ISL_467622, EPI_ISL_467623, EPI_ISL_467624, EPI_ISL_467625, EPI_ISL_467626, EPI_ISL_467627, EPI_ISL_467628, EPI_ISL_467629, EPI_ISL_467630, EPI_ISL_467631, EPI_ISL_467632, EPI_ISL_467633, EPI_ISL_467634, EPI_ISL_467635, EPI_ISL_467636, EPI_ISL_467637, EPI_ISL_467638, EPI_ISL_467639, EPI_ISL_467640, EPI_ISL_467641, EPI_ISL_467642, EPI_ISL_467643, EPI_ISL_467644, EPI_ISL_467645, EPI_ISL_467646, EPI_ISL_467647, EPI_ISL_467648, EPI_ISL_467649, EPI_ISL_467650, EPI_ISL_467651, EPI_ISL_467652, EPI_ISL_467653, EPI_ISL_467654, EPI_ISL_467655, EPI_ISL_467656, EPI_ISL_467657, EPI_ISL_467658, EPI_ISL_467659, EPI_ISL_467660, EPI_ISL_467661, EPI_ISL_467662, EPI_ISL_467663, EPI_ISL_467664 | | | |
| see above | New Mexico Department of Health Scientific Laboratory Division | Center for Global Health, University of New Mexico Health Sciences Center | Daryl Domman, Kurt Schwalm, Twila Kunde, Joseph Hicks, Michael Edwards, Darrell Dinwiddie |
| EPI_ISL_467666 | Virology lab, NIC, NCCD, Ulaanbaatar, Mongolia | National Centre for Communicable Deseases (NCCD) | Naranzul Ts,Darmaa B,Bayasgalan N,Ankhbayar S,Tsogtbaatar B, Erdene-Ochir Ts,Nymadawa P |
| EPI_ISL_467668, EPI_ISL_467669, EPI_ISL_467670, EPI_ISL_467671, EPI_ISL_467672, EPI_ISL_467673, EPI_ISL_467674, EPI_ISL_467675, EPI_ISL_467676, EPI_ISL_467677, EPI_ISL_467678, EPI_ISL_467679, EPI_ISL_467680, EPI_ISL_467681, EPI_ISL_467682, EPI_ISL_467683, EPI_ISL_467684, EPI_ISL_467685, EPI_ISL_467686, EPI_ISL_467687, EPI_ISL_467688, EPI_ISL_467689, EPI_ISL_467690, EPI_ISL_467691, EPI_ISL_467692 | | | |
| see above | unknown | Rapid Response Team | Shoja,Z., Fazlalipour,M., Salehi Vaziri,M., Azadmanesh,K., Jalali,T., Arashkia,A., Rohani,M., Esmaeili,S., Fotouhi-Chahooki,F., Maleki,A., Baesi,K., Pouriayevali,M.H., Ghasemi,A., Mahmoudi,A., Mostafavi,E., Fereydouni,Z., Tavakolirad,M., Khakifirouz,S., Mohammadi,T., Asadi Karam,M.R.A.K., Habibi,M., Mousavi Nasab,S.D., Ahmadi,Z., Azad-Mazjiri,S., Rafigh,M., Nemati,A.H., Shams Nosrati,M.S., Parikhani,A., Bathaeian,S.S., Nejatipour,Z., Yekta Sanati,P., Ghalejoogh,M. |
| EPI_ISL_467693, EPI_ISL_467694, EPI_ISL_467695, EPI_ISL_467696, EPI_ISL_467697, EPI_ISL_467698, EPI_ISL_467699, EPI_ISL_467700, EPI_ISL_467701, EPI_ISL_467702, EPI_ISL_467703, EPI_ISL_467704, EPI_ISL_467705, EPI_ISL_467706, EPI_ISL_467707, EPI_ISL_467708, EPI_ISL_467709, EPI_ISL_467710, EPI_ISL_467711, EPI_ISL_467712, EPI_ISL_467713, EPI_ISL_467714, EPI_ISL_467715, EPI_ISL_467716, EPI_ISL_467717, EPI_ISL_467718, EPI_ISL_467719 | | | |
| see above | PHE South West Regional Laboratory, National Infection Service | Wellcome Sanger Institute for the COVID-19 Genomics UK | Stephanie Hutchings, Hannah Pymont, Dr Peter Muir, Barry Vipond, Rich Hopes; and Alex Alderton, Roberto Amato, Sonia Goncalves, Ewan Harrison, David K. Jackson, Ian Johnston, Dominic Kwiatkowski, Cordelia Langford, John Sillitoe on behalf of the Wellcome Sanger Institute COVID-19 Surveillance Team (http://www.sanger.ac.uk/covid-team) |

| | | Consortium | |
|---|---|---|---|
| EPI_ISL_467774, EPI_ISL_467775 | Molecular diagnostic laboratory of Federal Budget Institution of Science "Central Research Institute of Epidemiology" of The Federal Service on Customers' Rights Protection and Human Well-being Surveillance | Group of Genomics and Postgenomic Technologies of Central Research Institute of Epidemilology | Speranskaya AS, Kaptelova VV, Samoilov AE, Korneenko EV, Sizova TV, Tivanova EV, Shipulina OY, Akimkin VG |
| EPI_ISL_467778, EPI_ISL_467779, EPI_ISL_467780, EPI_ISL_467781 | National Influenza Centre Romania | Charite Universitatsmedizin Berlin, Institute of Virology | Victor M Corman, Jorn Beheim-Schwarzbach, Barbara Muehlemann, Talitha Veith, Julia Schneider, Terry Jones, L. Ustea, N. Paraschiv, M. Lazar, Christian Drosten |
| EPI_ISL_467782, EPI_ISL_467783, EPI_ISL_467784, EPI_ISL_467785, EPI_ISL_467786, EPI_ISL_467787, EPI_ISL_467788, EPI_ISL_467789, EPI_ISL_467790, EPI_ISL_467791, EPI_ISL_467792, EPI_ISL_467793, EPI_ISL_467794, EPI_ISL_467795, EPI_ISL_467796, EPI_ISL_467797, EPI_ISL_467798, EPI_ISL_467799, EPI_ISL_467800, EPI_ISL_467801, EPI_ISL_467802, EPI_ISL_467803, EPI_ISL_467804, EPI_ISL_467805, EPI_ISL_467806, EPI_ISL_467807, EPI_ISL_467808 | | | |
| see above | Virginia DCLS | Virginia DCLS | Virginia DCLS |
| EPI_ISL_467809 | Cedars-Sinai Medical Center, Department of Pathology & Laboratory Medicine, Molecular Pathology Laboratory | Cedars-Sinai Medical Center, Molecular Pathology Laboratory of Department of Pathology & Laboratory Medicine and Genomic Core | Wenjuan Zhang, John Paul Govindavari, Brian Davis, Stephanie Chen, Jong Taek Kim, Jianbo Song, Jean Lopategui, Jasmine T Plummer, Eric Vail |
| EPI_ISL_467811, EPI_ISL_467812, EPI_ISL_467813, EPI_ISL_467814, EPI_ISL_467815, EPI_ISL_467816, EPI_ISL_467817, EPI_ISL_467818, EPI_ISL_467819, EPI_ISL_467820, EPI_ISL_467821, EPI_ISL_467822, EPI_ISL_467823, EPI_ISL_467824, EPI_ISL_467825, EPI_ISL_467826, EPI_ISL_467827, EPI_ISL_467828, EPI_ISL_467829, EPI_ISL_467830, EPI_ISL_467831, EPI_ISL_467832, EPI_ISL_467833, EPI_ISL_467834, EPI_ISL_467835, EPI_ISL_467836, EPI_ISL_467837, EPI_ISL_467838, EPI_ISL_467839, EPI_ISL_467840, EPI_ISL_467841, EPI_ISL_467842, EPI_ISL_467843, EPI_ISL_467844, EPI_ISL_467845, EPI_ISL_467846, EPI_ISL_467847, EPI_ISL_467848, EPI_ISL_467849, EPI_ISL_467850, EPI_ISL_467851, EPI_ISL_467852, EPI_ISL_467853, EPI_ISL_467854, EPI_ISL_467855, EPI_ISL_467856, EPI_ISL_467857, EPI_ISL_467858, EPI_ISL_467859, EPI_ISL_467860, EPI_ISL_467861, EPI_ISL_467862, EPI_ISL_467863, EPI_ISL_467864, EPI_ISL_467865, EPI_ISL_467866, EPI_ISL_467867, EPI_ISL_467868, EPI_ISL_467869, EPI_ISL_467870, EPI_ISL_467871, EPI_ISL_467872, EPI_ISL_467873, EPI_ISL_467874, EPI_ISL_467875, EPI_ISL_467876, EPI_ISL_467877, EPI_ISL_467878, EPI_ISL_467879, EPI_ISL_467880, EPI_ISL_467881, EPI_ISL_467882, EPI_ISL_467883, EPI_ISL_467884, EPI_ISL_467885, EPI_ISL_467886, EPI_ISL_467887, EPI_ISL_467888, EPI_ISL_467889, EPI_ISL_467890, EPI_ISL_467891, EPI_ISL_467892, EPI_ISL_467893, EPI_ISL_467894, EPI_ISL_467895, EPI_ISL_467896, EPI_ISL_467897, EPI_ISL_467898, EPI_ISL_467899, EPI_ISL_467900, EPI_ISL_467901, EPI_ISL_467902, EPI_ISL_467903, EPI_ISL_467904, EPI_ISL_467905, EPI_ISL_467906, EPI_ISL_467907, EPI_ISL_467908, EPI_ISL_467909, EPI_ISL_467910, EPI_ISL_467911, EPI_ISL_467912, EPI_ISL_467913, EPI_ISL_467914, EPI_ISL_467915, EPI_ISL_467916, EPI_ISL_467917, EPI_ISL_467918, EPI_ISL_467919, EPI_ISL_467920, EPI_ISL_467921, EPI_ISL_467922, EPI_ISL_467923, EPI_ISL_467924, EPI_ISL_467925, EPI_ISL_467926, EPI_ISL_467927 | | | |
| see above | Quest Diagnostics | Quest Diagnostics | Anderson,B.P., Rosenthal,S.H., Gerasimova,A., Kagan,R.M. and Owen, R. |
| EPI_ISL_467928, EPI_ISL_467929, EPI_ISL_467930, EPI_ISL_467931, EPI_ISL_467932, EPI_ISL_467933, EPI_ISL_467934, EPI_ISL_467935, EPI_ISL_467936, EPI_ISL_467937, EPI_ISL_467938, EPI_ISL_467939, EPI_ISL_467940, EPI_ISL_467941, EPI_ISL_467942, EPI_ISL_467943, EPI_ISL_467944 | | | |
| see above | Virginia DCLS | Virginia DCLS | Virginia DCLS |
| EPI_ISL_467945 | Montefiore Medical Center, Dept. of Pathology, Clinical Virology | Albert Einstein College of Medicine, Dept. of Microbiology & Immunology, Chandran lab | J. Maximilian Fels, Saad Khan, Ryan Forster, Karin A. Skalina, Ariel S. Wirchnianski, Denise Haslwanter, Catalina Florez, Robert H. Bortz III, M. Eugenia Dieterle, Ethan Laudermilch, Rohit K. Jangra, Amanda Mengotto, Duncan Kimmel, Shahina B. Maqbool, John M. Greally, Wendy A. Szymczak, Amy S. Fox, Michael B. Prystowsky, D. Yitzchak Goldstein, Johanna P. Daily, Libusha Kelly, Kartik Chandran |
| EPI_ISL_467946, EPI_ISL_467947, EPI_ISL_467948, EPI_ISL_467949 | Innovative Genomics Institute, UC Berkeley | Innovative Genomics Institute, UC Berkeley | Stacia Wyman, Haridha Shivram, Liana Lareau, Shana McDevitt, Justin Choi |
| EPI_ISL_467950, EPI_ISL_467951, EPI_ISL_467952, EPI_ISL_467953, EPI_ISL_467954, EPI_ISL_467955, EPI_ISL_467956, EPI_ISL_467957, EPI_ISL_467958, EPI_ISL_467959, EPI_ISL_467960, EPI_ISL_467961, EPI_ISL_467962 | | | |
| see above | San Diego County Public Health Laboratory | Andersen lab at Scripps Research | SEARCH Alliance San Diego with Tracy Basler, Jovan Shephard, Brett Austin |
| EPI_ISL_467963 | San Diego County Public Health Laboratory | Andersen lab at Scripps Research | SEARCH Alliance San Diego with Michael Quigley, Ellen Stefanski, Ian Mchardy |
| EPI_ISL_467964, EPI_ISL_467965, EPI_ISL_467966 | San Diego County Public Health Laboratory | Andersen lab at Scripps Research | SEARCH Alliance San Diego with Tracy Basler, Jovan Shephard, Brett Austin |
| EPI_ISL_467967 | Scripps Medical Laboratory | Andersen lab at Scripps Research | SEARCH Alliance San Diego with Tracy Basler, Jovan Shephard, Brett Austin |
| EPI_ISL_467968, EPI_ISL_467969, EPI_ISL_467970, EPI_ISL_467971 | San Diego County Public Health Laboratory | Andersen lab at Scripps Research | SEARCH Alliance San Diego with Tracy Basler, Jovan Shephard, Brett Austin |
| EPI_ISL_467972 | Scripps Medical Laboratory | Andersen lab at Scripps Research | SEARCH Alliance San Diego with Michael Quigley, Ellen Stefanski, Ian Mchardy |
| EPI_ISL_467973, EPI_ISL_467974, EPI_ISL_467975 | San Diego County Public Health Laboratory | Andersen lab at Scripps Research | SEARCH Alliance San Diego with Tracy Basler, Jovan Shephard, Brett Austin |
| EPI_ISL_467976 | Rady's Childrens Hospital | Andersen lab at Scripps Research | SEARCH Alliance San Diego |
| EPI_ISL_467977, EPI_ISL_467978, EPI_ISL_467979, EPI_ISL_467980 | San Diego County Public Health Laboratory | Andersen lab at Scripps Research | SEARCH Alliance San Diego with Tracy Basler, Jovan Shephard, Brett Austin |
| EPI_ISL_467981 | Rady's Childrens Hospital | Andersen lab at Scripps Research | SEARCH Alliance San Diego |
| EPI_ISL_467982, EPI_ISL_467983 | San Diego County Public Health Laboratory | Andersen lab at Scripps Research | SEARCH Alliance San Diego with Tracy Basler, Jovan Shephard, Brett Austin |
| EPI_ISL_467984 | Rady's Childrens Hospital | Andersen lab at Scripps Research | SEARCH Alliance San Diego |
| EPI_ISL_467985, EPI_ISL_467986, EPI_ISL_467987, EPI_ISL_467988, EPI_ISL_467989 | SA Pathology | SA Pathology | Lex Leong, Chuan Kok Lim, Mark Turra, Ivan Bastian, Geoff Higgins |
| EPI_ISL_467990 | SA Pathology | SA Pathology | Leong, LEX, Lim, CK, Turra, M, Bastian, I, Higgins, G |
| EPI_ISL_467991, EPI_ISL_467992, EPI_ISL_467993, EPI_ISL_467994, EPI_ISL_467995, EPI_ISL_467996, EPI_ISL_467997, EPI_ISL_467998, EPI_ISL_467999, EPI_ISL_468000, EPI_ISL_468001, EPI_ISL_468002, EPI_ISL_468003, EPI_ISL_468004, EPI_ISL_468005, EPI_ISL_468006, EPI_ISL_468007, EPI_ISL_468008, EPI_ISL_468009, EPI_ISL_468010, EPI_ISL_468011, EPI_ISL_468012, EPI_ISL_468013, EPI_ISL_468014, EPI_ISL_468015, EPI_ISL_468016, EPI_ISL_468017, EPI_ISL_468018, EPI_ISL_468019, EPI_ISL_468020, EPI_ISL_468021, EPI_ISL_468022, EPI_ISL_468023, EPI_ISL_468024, EPI_ISL_468025, EPI_ISL_468026, EPI_ISL_468027, EPI_ISL_468028, EPI_ISL_468029, EPI_ISL_468030, EPI_ISL_468031, EPI_ISL_468032, EPI_ISL_468033, EPI_ISL_468034, EPI_ISL_468035, EPI_ISL_468036, EPI_ISL_468037, EPI_ISL_468038, EPI_ISL_468039, EPI_ISL_468040, EPI_ISL_468041, EPI_ISL_468042, EPI_ISL_468043 | | | |
| see above | SA Pathology | SA Pathology | Lex Leong, Chuan Kok Lim, Mark Turra, Ivan Bastian, Geoff Higgins |
| EPI_ISL_468066 | Physiology, Istanbul Medeniyet University | Physiology, Istanbul Medeniyet University | Pence,S., Caykara,B., Pence,H.H., Tekin,S., Yiyit,N., Cevher Keskin,B. and Kara,A. |
| EPI_ISL_468067 | Microbiology and Immunology, The Peter Doherty Institute for Infection and Immunity | Microbiology and Immunology, The Peter Doherty Institute for Infection and Immunity | Caly,L., Seemann,T., Sait,M., Schultz,M.B., Druce,J., Sherry,N., Meumann,E., Soares da Silva,E., Dolores de Jesus da Costa,M., Salles de Sousa,A., Jayanti Pereira Tilman,A., Antonia da Costa,E., Barreto,I., Marr,I., Wapling,J., Francis,J., Ximenes,J., Canisia,D., Freeman,K., Dakh,F., Douglas,N. and Baird,R. |
| EPI_ISL_468070, EPI_ISL_468071, EPI_ISL_468072, EPI_ISL_468073 | Child Health Research Foundation | Child Health Research Foundation | Senjuti Saha, Roly Malaker, Md Saiful Islam Sajib, Hafizur Rahman, Maksuda Islam, Samir K Saha |
| EPI_ISL_468074, EPI_ISL_468075, EPI_ISL_468076, EPI_ISL_468077, EPI_ISL_468078 | Child Health Research Foundation | Child Health Research Foundation | Senjuti Saha, Roly Malaker, Md Saiful Islam Sajib, Hafizur Rahman, Afroza Akter Tanni, Syed Muktadir Al Sium, Maksuda Islam, Samir K Saha |
| EPI_ISL_468080, EPI_ISL_468081, EPI_ISL_468082, EPI_ISL_468083, EPI_ISL_468084, EPI_ISL_468085, EPI_ISL_468086, EPI_ISL_468087, EPI_ISL_468088, EPI_ISL_468089, EPI_ISL_468090, EPI_ISL_468091, EPI_ISL_468092, EPI_ISL_468093, EPI_ISL_468094, EPI_ISL_468095, EPI_ISL_468096, EPI_ISL_468097, EPI_ISL_468098, EPI_ISL_468099, EPI_ISL_468100, EPI_ISL_468101, EPI_ISL_468102, EPI_ISL_468103, EPI_ISL_468104, EPI_ISL_468105, EPI_ISL_468106, EPI_ISL_468107, EPI_ISL_468108, EPI_ISL_468109, EPI_ISL_468110, EPI_ISL_468111, EPI_ISL_468112, EPI_ISL_468113, EPI_ISL_468114, EPI_ISL_468115, EPI_ISL_468116, EPI_ISL_468117, EPI_ISL_468118, EPI_ISL_468119, EPI_ISL_468120, EPI_ISL_468121, EPI_ISL_468122, EPI_ISL_468123, EPI_ISL_468124, EPI_ISL_468125, EPI_ISL_468126, EPI_ISL_468127, EPI_ISL_468128, EPI_ISL_468129 | | | |
| see above | OHSU Lab Services Molecular Microbiology Lab | Oregon SARS-CoV-2 Genome Sequencing Center | Brendan L. O'Connell, Ruth V. Nichols, Alec J. Hirsch, Guang Fan, Daniel N. Streblow, William B. Messer, Andrew C. Adey, Benjamin N. Bimber, Brian J. O'Roak |
| EPI_ISL_468134, EPI_ISL_468135, EPI_ISL_468136, EPI_ISL_468137, EPI_ISL_468138, EPI_ISL_468139, EPI_ISL_468140, EPI_ISL_468141, EPI_ISL_468142, EPI_ISL_468143, EPI_ISL_468144, EPI_ISL_468145, EPI_ISL_468146, EPI_ISL_468147, EPI_ISL_468148, EPI_ISL_468149, EPI_ISL_468150, EPI_ISL_468151, EPI_ISL_468152, EPI_ISL_468153, EPI_ISL_468154, EPI_ISL_468155, EPI_ISL_468156, EPI_ISL_468157, EPI_ISL_468158 | | | |
| see above | [Romania, Bucharest] National Institute for Infectious Diseases "Prof. Dr. Matei Balș" | [Romania, Bucharest] National Institute for Infectious Diseases "Prof. Dr. Matei Balș" | Leontina Banica, Marius Cotic, Corina Casangiu, Marius Surleac, Simona Paraschiv |
| EPI_ISL_468159 | unknown | Department of Virology, Public Health Laboratories Division, National Institute of Health | Massab Umair, Aamer Ikram, Muhammad Salman, Adnan Khurshid, Nazish Badar, Shannon Whitmer, John Klena |
| EPI_ISL_468160 | unknown | Department of Virology, Public Health Laboratories Division, National Institute of Health | Massab Umair, Aamer Ikram, Muhammad Salman, Adnan Khurshid, Nazish Badar, Shannon Whitmer, John Klena |
| EPI_ISL_468161 | Department of Virology, Public Health Laboratories Division, National Institute of Health | Department of Virology, Public Health Laboratories Division, National Institute of Health | Massab Umair, Aamer Ikram, Muhammad Salman, Adnan Khurshid, Nazish Badar, Shannon Whitmer, John Klena |
| EPI_ISL_468162 | unknown | Department of Virology, Public Health Laboratories Division, National Institute of Health | Massab Umair, Aamer Ikram, Muhammad Salman, Adnan Khurshid, Nazish Badar, Shannon Whitmer, John Klena |
| EPI_ISL_468163 | Department of Virology, Public Health Laboratories Division, National Institute of Health | Department of Virology, Public Health Laboratories Division, National Institute of Health | Massab Umair, Aamer Ikram, Muhammad Salman, Adnan Khurshid, Nazish Badar, Shannon Whitmer, John Klena |
| EPI_ISL_468202, EPI_ISL_468203, EPI_ISL_468204, EPI_ISL_468205, EPI_ISL_468206, EPI_ISL_468207, EPI_ISL_468208, EPI_ISL_468209, EPI_ISL_468210, EPI_ISL_468211, EPI_ISL_468212, EPI_ISL_468213, EPI_ISL_468214, EPI_ISL_468215, EPI_ISL_468216, EPI_ISL_468217, EPI_ISL_468218, EPI_ISL_468219, EPI_ISL_468220, EPI_ISL_468221, EPI_ISL_468222, EPI_ISL_468223, EPI_ISL_468224, EPI_ISL_468225, EPI_ISL_468226, EPI_ISL_468227, EPI_ISL_468228, EPI_ISL_468229, EPI_ISL_468230, EPI_ISL_468231, EPI_ISL_468232, EPI_ISL_468233, EPI_ISL_468234, EPI_ISL_468235, EPI_ISL_468236, EPI_ISL_468237, EPI_ISL_468238, EPI_ISL_468239, EPI_ISL_468240, EPI_ISL_468241, EPI_ISL_468242, EPI_ISL_468243, EPI_ISL_468244, EPI_ISL_468245, EPI_ISL_468246, EPI_ISL_468247, EPI_ISL_468248, EPI_ISL_468249, EPI_ISL_468250, EPI_ISL_468251, EPI_ISL_468252, EPI_ISL_468253, EPI_ISL_468254, EPI_ISL_468255, EPI_ISL_468256, EPI_ISL_468257, EPI_ISL_468258, EPI_ISL_468259, EPI_ISL_468260, EPI_ISL_468261, EPI_ISL_468262, EPI_ISL_468263, EPI_ISL_468264, EPI_ISL_468265, EPI_ISL_468266, EPI_ISL_468267, EPI_ISL_468268, EPI_ISL_468269, EPI_ISL_468270, EPI_ISL_468271, EPI_ISL_468272, EPI_ISL_468273, | | | |

| | | | |
|---|---|---|---|
| EPI_ISL_468274, EPI_ISL_468275, EPI_ISL_468276, EPI_ISL_468277, EPI_ISL_468278, EPI_ISL_468279, EPI_ISL_468280, EPI_ISL_468281, EPI_ISL_468282, EPI_ISL_468283, EPI_ISL_468284, EPI_ISL_468285, EPI_ISL_468286, EPI_ISL_468287, EPI_ISL_468288, EPI_ISL_468289, EPI_ISL_468290, EPI_ISL_468291, EPI_ISL_468292, EPI_ISL_468293, EPI_ISL_468294, EPI_ISL_468295, EPI_ISL_468296, EPI_ISL_468297, EPI_ISL_468298, EPI_ISL_468299, EPI_ISL_468300, EPI_ISL_468301, EPI_ISL_468302, EPI_ISL_468303, EPI_ISL_468304 | | | |
| see above | Viollier AG | Department of Biosystems Science and Engineering, ETH Zürich | Christian Beisel, Sarah Nadeau, Ivan Topolsky, Pedro Ferreira, Philipp Jablonski, Susana Posada-Céspedes, Tobias Schär, Ina Nissen, Natascha Santacroce, Elodie Burcklen, Christiane Beckmann, Maurice Redondo, Olivier Kobel, Christoph Noppen, Sophie Seidel, Noemie Santamaria de Souza, Niko Beerenwinkel, Tanja Stadler |
| EPI_ISL_468305, EPI_ISL_468307 | Centro de Vigilancia a Saude de Diadema | Instituto Adolfo Lutz, Interdiciplinary Procedures Center, Strategic Laboratory | Claudio Tavares Sacchi, Claudia Regina Gonçalves, Erica Valessa Ramos Gomes |
| EPI_ISL_468308 | Hospital Municipal do Tatuape Carmino Caricchio | Instituto Adolfo Lutz, Interdiciplinary Procedures Center, Strategic Laboratory | Claudio Tavares Sacchi, Claudia Regina Gonçalves, Erica Valessa Ramos Gomes |
| EPI_ISL_468310 | Hospital Sao Paulo de Ensino da UNIFESP | Instituto Adolfo Lutz, Interdiciplinary Procedures Center, Strategic Laboratory | Claudio Tavares Sacchi, Claudia Regina Gonçalves, Erica Valessa Ramos Gomes |
| EPI_ISL_468311, EPI_ISL_468312 | Hospital Municipal Dr Ignacio Proenca de Gouvea | Instituto Adolfo Lutz, Interdiciplinary Procedures Center, Strategic Laboratory | Claudio Tavares Sacchi, Claudia Regina Gonçalves, Erica Valessa Ramos Gomes |
| EPI_ISL_468313 | Vigilancia Epidemiologica de São Bernardo do Campo | Instituto Adolfo Lutz, Interdiciplinary Procedures Center, Strategic Laboratory | Claudio Tavares Sacchi, Claudia Regina Gonçalves, Erica Valessa Ramos Gomes |
| EPI_ISL_468314 | CTA Centro de Testagem e Aconselhamento | Instituto Adolfo Lutz, Interdiciplinary Procedures Center, Strategic Laboratory | Claudio Tavares Sacchi, Claudia Regina Gonçalves, Erica Valessa Ramos Gomes |
| EPI_ISL_468315 | Hospital Municipal do Tatuape Carmino Caricchio | Instituto Adolfo Lutz, Interdiciplinary Procedures Center, Strategic Laboratory | Claudio Tavares Sacchi, Claudia Regina Gonçalves, Erica Valessa Ramos Gomes |
| EPI_ISL_468316 | UPA Vila Assis | Instituto Adolfo Lutz, Interdiciplinary Procedures Center, Strategic Laboratory | Claudio Tavares Sacchi, Claudia Regina Gonçalves, Erica Valessa Ramos Gomes |
| EPI_ISL_468318 | Hospital Universitario da USP | Instituto Adolfo Lutz, Interdiciplinary Procedures Center, Strategic Laboratory | Claudio Tavares Sacchi, Claudia Regina Gonçalves, Erica Valessa Ramos Gomes |
| EPI_ISL_468319 | Vigilancia Epidemiologica de São Bernardo do Campo | Instituto Adolfo Lutz, Interdiciplinary Procedures Center, Strategic Laboratory | Claudio Tavares Sacchi, Claudia Regina Gonçalves, Erica Valessa Ramos Gomes |
| EPI_ISL_468320 | Secretaria Municipal de Saude de Hortolandia | Instituto Adolfo Lutz, Interdiciplinary Procedures Center, Strategic Laboratory | Claudio Tavares Sacchi, Claudia Regina Gonçalves, Erica Valessa Ramos Gomes |
| EPI_ISL_468321 | Hospital Universitario da USP | Instituto Adolfo Lutz, Interdiciplinary Procedures Center, Strategic Laboratory | Claudio Tavares Sacchi, Claudia Regina Gonçalves, Erica Valessa Ramos Gomes |
| EPI_ISL_468331, EPI_ISL_468332, EPI_ISL_468333, EPI_ISL_468334, EPI_ISL_468335, EPI_ISL_468336, EPI_ISL_468337, EPI_ISL_468338, EPI_ISL_468339, EPI_ISL_468340, EPI_ISL_468341, EPI_ISL_468342, EPI_ISL_468343, EPI_ISL_468344 | | | |
| see above | Microbiology Service, University Hospital of A Coruna-Biomedical Research Institute | Genomes & Disease, Center for Research in Molecular Medicine and Chronic Diseases, University of Santiago de Compostela | Kelly Conde, Jorge Arca, Soraya Rumbo, Juan A. Vallejo, M Poza, G Bou, Ana Pequeno-Valtierra, Jorge Rodriguez-Castro, Javier Temes, Daniel Garcia-Souto, Martin Santamarina, Cristina Gomez, Jose M. C. Tubio |
| EPI_ISL_468345, EPI_ISL_468346, EPI_ISL_468347, EPI_ISL_468348, EPI_ISL_468349, EPI_ISL_468350, EPI_ISL_468351, EPI_ISL_468352, EPI_ISL_468353, EPI_ISL_468354, EPI_ISL_468355, EPI_ISL_468356 | | | |
| see above | County of Santa Clara Public Health Department | Chan-Zuckerberg Biohub | CZB Cliahub Consortium |
| EPI_ISL_468357, EPI_ISL_468358, EPI_ISL_468359, EPI_ISL_468360, EPI_ISL_468361, EPI_ISL_468362, EPI_ISL_468363, EPI_ISL_468364, EPI_ISL_468365, EPI_ISL_468366, EPI_ISL_468367, EPI_ISL_468368, EPI_ISL_468369, EPI_ISL_468370, EPI_ISL_468371, EPI_ISL_468372, EPI_ISL_468373, EPI_ISL_468374, EPI_ISL_468375, EPI_ISL_468376, EPI_ISL_468377, EPI_ISL_468378, EPI_ISL_468379, EPI_ISL_468380, EPI_ISL_468381, EPI_ISL_468382, EPI_ISL_468383, EPI_ISL_468384, EPI_ISL_468385, EPI_ISL_468386, EPI_ISL_468387 | | | |
| see above | Alameda County Public Health Lab | Chan-Zuckerberg Biohub | CZB Cliahub Consortium |
| EPI_ISL_468388, EPI_ISL_468389, EPI_ISL_468390, EPI_ISL_468391, EPI_ISL_468392, EPI_ISL_468393, EPI_ISL_468394, EPI_ISL_468395, EPI_ISL_468396, EPI_ISL_468397, EPI_ISL_468398, EPI_ISL_468399, EPI_ISL_468400, EPI_ISL_468401, EPI_ISL_468402, EPI_ISL_468403, EPI_ISL_468404, EPI_ISL_468405, EPI_ISL_468406, EPI_ISL_468407, EPI_ISL_468408, EPI_ISL_468409, EPI_ISL_468410, EPI_ISL_468411, EPI_ISL_468412, EPI_ISL_468413, EPI_ISL_468414, EPI_ISL_468415, EPI_ISL_468416, EPI_ISL_468417, EPI_ISL_468418, EPI_ISL_468419, EPI_ISL_468420, EPI_ISL_468421, EPI_ISL_468422, EPI_ISL_468423, EPI_ISL_468424, EPI_ISL_468425, EPI_ISL_468426, EPI_ISL_468427, EPI_ISL_468428, EPI_ISL_468429, EPI_ISL_468430, EPI_ISL_468431, EPI_ISL_468432, EPI_ISL_468433, EPI_ISL_468434, EPI_ISL_468435, EPI_ISL_468436, EPI_ISL_468437 | | | |
| see above | County of San Luis Obispo Public Health Laboratory | Chan-Zuckerberg Biohub | CZB Cliahub Consortium |
| EPI_ISL_468438, EPI_ISL_468439, EPI_ISL_468440, EPI_ISL_468441, EPI_ISL_468442, EPI_ISL_468443, EPI_ISL_468444, EPI_ISL_468445, EPI_ISL_468446, EPI_ISL_468447, EPI_ISL_468448, EPI_ISL_468449, EPI_ISL_468450, EPI_ISL_468451, EPI_ISL_468452, EPI_ISL_468453, EPI_ISL_468454, EPI_ISL_468455, EPI_ISL_468456, EPI_ISL_468457, EPI_ISL_468458, EPI_ISL_468459, EPI_ISL_468460, EPI_ISL_468461 | | | |
| see above | Humboldt County Public Health Laboratory | Chan-Zuckerberg Biohub | CZB Cliahub Consortium |
| EPI_ISL_468462, EPI_ISL_468463, EPI_ISL_468464, EPI_ISL_468465, EPI_ISL_468466, EPI_ISL_468467, EPI_ISL_468468, EPI_ISL_468469, EPI_ISL_468470, EPI_ISL_468471, EPI_ISL_468472, EPI_ISL_468473, EPI_ISL_468474, EPI_ISL_468475, EPI_ISL_468476, EPI_ISL_468477, EPI_ISL_468478, EPI_ISL_468479, EPI_ISL_468480, EPI_ISL_468481, EPI_ISL_468482, EPI_ISL_468483, EPI_ISL_468484, EPI_ISL_468485, EPI_ISL_468486, EPI_ISL_468487, EPI_ISL_468488, EPI_ISL_468489, EPI_ISL_468490, EPI_ISL_468491, EPI_ISL_468492, EPI_ISL_468493, EPI_ISL_468494, EPI_ISL_468495, EPI_ISL_468496, EPI_ISL_468497, EPI_ISL_468498, EPI_ISL_468499, EPI_ISL_468500, EPI_ISL_468501, EPI_ISL_468502, EPI_ISL_468503, EPI_ISL_468504, EPI_ISL_468505 | | | |
| see above | Ventura County Public Health Lab | Chan-Zuckerberg Biohub | CZB Cliahub Consortium |
| EPI_ISL_468506, EPI_ISL_468507, EPI_ISL_468508, EPI_ISL_468509, EPI_ISL_468510, EPI_ISL_468511, EPI_ISL_468512, EPI_ISL_468513, EPI_ISL_468514, EPI_ISL_468515, EPI_ISL_468516, EPI_ISL_468517, EPI_ISL_468518, EPI_ISL_468519, EPI_ISL_468520, EPI_ISL_468521, EPI_ISL_468522, EPI_ISL_468523, EPI_ISL_468524, EPI_ISL_468525, EPI_ISL_468526, EPI_ISL_468527, EPI_ISL_468528, EPI_ISL_468529, EPI_ISL_468530, EPI_ISL_468531, EPI_ISL_468532, EPI_ISL_468533, EPI_ISL_468534, EPI_ISL_468535, EPI_ISL_468536, EPI_ISL_468537, EPI_ISL_468538, EPI_ISL_468539, EPI_ISL_468540, EPI_ISL_468541, EPI_ISL_468542, EPI_ISL_468543, EPI_ISL_468544, EPI_ISL_468545, EPI_ISL_468546, EPI_ISL_468547, EPI_ISL_468548, EPI_ISL_468549, EPI_ISL_468550, EPI_ISL_468551, EPI_ISL_468552, EPI_ISL_468553, EPI_ISL_468554, EPI_ISL_468555, EPI_ISL_468556, EPI_ISL_468557, EPI_ISL_468558, EPI_ISL_468559 | | | |
| see above | San Joaquin County Public Health Lab | Chan-Zuckerberg Biohub | CZB Cliahub Consortium |
| EPI_ISL_468560, EPI_ISL_468561, EPI_ISL_468562, EPI_ISL_468563, EPI_ISL_468564, EPI_ISL_468565, EPI_ISL_468566, EPI_ISL_468567, EPI_ISL_468568, EPI_ISL_468569, EPI_ISL_468570, EPI_ISL_468571, EPI_ISL_468572, EPI_ISL_468573, EPI_ISL_468574, EPI_ISL_468575, EPI_ISL_468576, EPI_ISL_468577, EPI_ISL_468578, EPI_ISL_468579, EPI_ISL_468580, EPI_ISL_468581, EPI_ISL_468582, EPI_ISL_468583, EPI_ISL_468584, EPI_ISL_468585, EPI_ISL_468586, EPI_ISL_468587, EPI_ISL_468588, EPI_ISL_468589, EPI_ISL_468590 | | | |
| see above | Quest Diagnostics | Quest Diagnostics | Anderson,B.P., Rosenthal,S.H., Gerasimova,A., Kagan,R.M. and Owen, R. |
| EPI_ISL_468591 | Institute for Public Health | Laboratory for advanced genomics | Filip Rokić, Lovro Trgovec-Greif, Neven Sučić, Tomislav Rukavina, Igor Jurak, Oliver Vugrek |
| EPI_ISL_468592, EPI_ISL_468593, EPI_ISL_468594, EPI_ISL_468595, EPI_ISL_468596, EPI_ISL_468597, EPI_ISL_468598, EPI_ISL_468599, EPI_ISL_468600, EPI_ISL_468601, EPI_ISL_468602, EPI_ISL_468603, EPI_ISL_468604, EPI_ISL_468605, EPI_ISL_468606, EPI_ISL_468607, EPI_ISL_468608, EPI_ISL_468609, EPI_ISL_468610, EPI_ISL_468611, EPI_ISL_468612, EPI_ISL_468613, EPI_ISL_468614 | | | |
| see above | Orange County Public Health Lab | Chan-Zuckerberg Biohub | CZB Cliahub Consortium |
| EPI_ISL_468615, EPI_ISL_468616, EPI_ISL_468617, EPI_ISL_468618, EPI_ISL_468619, EPI_ISL_468620, EPI_ISL_468621, EPI_ISL_468622, EPI_ISL_468623, EPI_ISL_468624, EPI_ISL_468625, EPI_ISL_468626, EPI_ISL_468627, EPI_ISL_468628, EPI_ISL_468629, EPI_ISL_468630, EPI_ISL_468631, EPI_ISL_468632, EPI_ISL_468633, EPI_ISL_468634, EPI_ISL_468635, EPI_ISL_468636, EPI_ISL_468637, EPI_ISL_468638, EPI_ISL_468639, EPI_ISL_468640, EPI_ISL_468641, EPI_ISL_468642, EPI_ISL_468643, EPI_ISL_468644, EPI_ISL_468645, EPI_ISL_468646, EPI_ISL_468647, EPI_ISL_468648, EPI_ISL_468649, EPI_ISL_468650, EPI_ISL_468651, EPI_ISL_468652, EPI_ISL_468653, EPI_ISL_468654, EPI_ISL_468655 | | | |
| see above | Contra Costa Public Health Lab | Chan-Zuckerberg Biohub | CZB Cliahub Consortium |
| EPI_ISL_468656 | Institute for Public Health | Laboratory for advanced genomics | Filip Rokić, Lovro Trgovec-Greif, Neven Sučić, Tomislav Rukavina, Igor Jurak, Oliver Vugrek |
| EPI_ISL_468718 | unknown | Environmental and Global Health | Stephenson,C.J., Subramaniam,K., Waltzek,T.B., Merck,L.H., Gibson,J.C., Morris,J.G. |
| EPI_ISL_468719 | University of Florida | University of Florida | Elbadry,M.A., Subramaniam,K., Waltzek,T.B., Stephenson,C.J., Gibson,J.C., Alam,M.M., Lauzardo,M., Morris,J.G., Lednicky,J.A. |
| EPI_ISL_468720 | University of Florida | University of Florida | Stephenson,C.J., Subramaniam,K., Waltzek,T.B., Lauzardo,M., Morris,J.G., Lednicky,J.A. |
| EPI_ISL_468721 | University of Florida | University of Florida | Stephenson,C.J., Subramaniam,K., Waltzek,T.B., Lauzardo,M., Gibson,J.C., Morris,J.G., Lednicky,J.A. |
| EPI_ISL_468722 | University of Florida | University of Florida | Elbadry,M.A., Subramaniam,K., Waltzek,T.B., Lauzardo,M., Morris,J.G., Lednicky,J.A. |
| EPI_ISL_468723 | University of Florida | University of Florida | Stephenson,C.J., Subramaniam,K., Waltzek,T.B., Lauzardo,M., Morris,J.G., Lednicky,J.A. |
| EPI_ISL_468724, EPI_ISL_468725 | unknown | Contact:Ryota Kumagai Tokyo Metropolitan Institute of Public Health | Kumagai,R., Yoshida,I., Asakura,H., Nagashima,M., Chiba,T., Sadamasu,K. |
| EPI_ISL_468726 | unknown | Department of Microbiology | Peng,H., Tang,H., Jiang,L., Qi,Z., Zhao,P. |
| EPI_ISL_469241, EPI_ISL_469242, EPI_ISL_469243, EPI_ISL_469244, EPI_ISL_469245, EPI_ISL_469246, EPI_ISL_469247, EPI_ISL_469248, EPI_ISL_469249, EPI_ISL_469250, EPI_ISL_469251, EPI_ISL_469252 | | | |
| see above | Special Infectious Agents Unit | Special Infectious Agents Unit | Azhar,E.I., Hassan,A.M., Tolah,A.M., Uthman,N.A., Al-Sobahy,T.L., Farraj,S.A., El-Kafrawy,S.A. |
| EPI_ISL_469253 | Second Military Medical University, Department of Microbiology | Second Military Medical University, Department of Microbiology | Peng,H., Tang,H., Jiang,L., Qi,Z. and Zhao,P. |
| EPI_ISL_478672 | Egyptian National Cancer Institute (ENCI) | Egyptian National Cancer Institute (ENCI) | Zekri, Abdel Rahman N, Amer,K.E., Ahmed,O.S., Soliman,H.K., Hafez,M.M., Bahnassy,A.A., Abdelhamid,W., Gad,A., Ali,M., Hassan,W., Samir,M., Raouf,A., Hamdy,M.S., Soliman,M.S., Elsissy,M.H., Elkhateeb,S.M., Ezzelarab,M.H., Abouelhoda, Mohamed |

| | | | |
|---|---|---|---|
| EPI_ISL_479676, EPI_ISL_479677, EPI_ISL_479678, EPI_ISL_479679, EPI_ISL_479680, EPI_ISL_479681, EPI_ISL_479682, EPI_ISL_479683, EPI_ISL_479684, EPI_ISL_479685 | unknown | Contact:Hiroyuki Asakura Tokyo Metropolitan Institute of Public Health, Department of Microbiology | Asakura,H., Yoshida,I., Kumagai,R., Nagashima,M., Chiba,T., Sadamasu,K. |
| EPI_ISL_479686, EPI_ISL_479687, EPI_ISL_479688, EPI_ISL_479689, EPI_ISL_479690, EPI_ISL_479691, EPI_ISL_479692, EPI_ISL_479693, EPI_ISL_479694, EPI_ISL_479695, EPI_ISL_479696, EPI_ISL_479697, EPI_ISL_479698, EPI_ISL_479699, EPI_ISL_479700, EPI_ISL_479701, EPI_ISL_479702, EPI_ISL_479703, EPI_ISL_479704, EPI_ISL_479705, EPI_ISL_479706, EPI_ISL_479707, EPI_ISL_479708, EPI_ISL_479709, EPI_ISL_479710, EPI_ISL_479711, EPI_ISL_479712, EPI_ISL_479713, EPI_ISL_479714, EPI_ISL_479715, EPI_ISL_479716, EPI_ISL_479717, EPI_ISL_479718, EPI_ISL_479719, EPI_ISL_479720, EPI_ISL_479721, EPI_ISL_479722, EPI_ISL_479723, EPI_ISL_479724, EPI_ISL_479725, EPI_ISL_479726, EPI_ISL_479727 | | | |
| see above | Egyptian National Cancer Institute (ENCI) | Egyptian National Cancer Institute (ENCI) | Zekri, Abdel Rahman N, Amer,K.E., Ahmed,O.S., Soliman,H.K., Hafez,M.M., Bahnassy,A.A., Abdelhamid,W., Gad,A., Ali,M., Hassan,W., Samir,M., Raouf,A., Hamdy,M.S., Soliman,M.S., Elsissy,M.H., Elkhateeb,S.M., Ezzelarab,M.H., Abouelhoda, Mohamed |
| EPI_ISL_524450, EPI_ISL_524451, EPI_ISL_524452, EPI_ISL_524453, EPI_ISL_524454 | Microbiology & Immunology, University of North Carolina | Microbiology & Immunology, University of North Carolina | Bailey,A.G., Caro-Vegas,C.P., Dittmer,D., Eason,A.B., Juarez,A., Landis,J.T., McNamara,R.P., Miller,M.B., Moorad,R., Pluta,L.J., Seltzer,T.A., Thompson,C., Vahrson,W., Villamor,F. |

We gratefully acknowledge the following Authors from the Originating laboratories responsible for obtaining the specimens, as well as the Submitting laboratories where the genome data were generated and shared via GISAID, on which this research is based.

All Submitters of data may be contacted directly via www.gisaid.org

| Accession ID | Originating Laboratory | Submitting Laboratory | Authors |
|---|---|---|---|
| EPI_ISL_468657, EPI_ISL_468658, EPI_ISL_468659, EPI_ISL_468660, EPI_ISL_468661, EPI_ISL_468662, EPI_ISL_468663, EPI_ISL_468664, EPI_ISL_468665, EPI_ISL_468666, EPI_ISL_468667, EPI_ISL_468668, EPI_ISL_468669, EPI_ISL_468670, EPI_ISL_468671, EPI_ISL_468672, EPI_ISL_468673, EPI_ISL_468674, EPI_ISL_468675, EPI_ISL_468676, EPI_ISL_468677, EPI_ISL_468678, EPI_ISL_468679, EPI_ISL_468680, EPI_ISL_468681, EPI_ISL_468682, EPI_ISL_468683, EPI_ISL_468684, EPI_ISL_468685, EPI_ISL_468686, EPI_ISL_468687, EPI_ISL_468688, EPI_ISL_468689, EPI_ISL_468690, EPI_ISL_468691, EPI_ISL_468692, EPI_ISL_468693, EPI_ISL_468694, EPI_ISL_468695, EPI_ISL_468696, EPI_ISL_468697, EPI_ISL_468698, EPI_ISL_468699, EPI_ISL_468700 | | | |
| see above | BCCDC Public Health Laboratory | BCCDC Public Health Laboratory | Richard Harrigan, Hope Lapointe, Jinny Choi, Kimia Kamelian, John Tyson,Terry Snutch, Linda Hoang, Inna Sekirov, Paul Levett, Mel Krajden, Natalie Prystajecky |
| EPI_ISL_468701, EPI_ISL_468702, EPI_ISL_468703, EPI_ISL_468704, EPI_ISL_468705, EPI_ISL_468706, EPI_ISL_468707, EPI_ISL_468708, EPI_ISL_468709, EPI_ISL_468710, EPI_ISL_468711, EPI_ISL_468712, EPI_ISL_468713, EPI_ISL_468714, EPI_ISL_468715, EPI_ISL_468716, EPI_ISL_468717 | | | |
| see above | Ochsner Health | Bioinfoexperts, LLC | Rebecca Rose, Amy Feehan, David J. Nolan, Sissy Cross, David Moraga Amador, Tong Yang, Luke Caruso, Wayra Navia, Lydia Von Borstel, Xiao Hui Zhou, Julia-Garcia-Diaz, Susanna L. Lamers |
| EPI_ISL_468727, EPI_ISL_468728, EPI_ISL_468729, EPI_ISL_468730, EPI_ISL_468731, EPI_ISL_468732, EPI_ISL_468733, EPI_ISL_468734, EPI_ISL_468735 | Lab voor klinische biologie | Onderzoeksgroep Virologie | Laurens Lambrechts, Nick Vereecke, Marthe Pauwels, Bruno Verhasselt, Linos Vandekerckhove, Hans Nauwynck, Sebastiaan Theuns |
| EPI_ISL_468736, EPI_ISL_468737, EPI_ISL_468738, EPI_ISL_468739, EPI_ISL_468740, EPI_ISL_468741, EPI_ISL_468742, EPI_ISL_468743, EPI_ISL_468744, EPI_ISL_468745, EPI_ISL_468746 | | | |
| see above | Lab voor klinische biologie | Onderzoeksgroep Virologie | Nick Vereecke, Laurens Lambrechts, Marthe Pauwels, Bruno Verhasselt, Linos Vandekerckhove, Hans Nauwynck, Sebastiaan Theuns |
| EPI_ISL_468747, EPI_ISL_468748, EPI_ISL_468749, EPI_ISL_468750, EPI_ISL_468751 | Facultad de Medicina UC | Center for Mathematical Modeling and Center for Genome Regulation. Santiago, Chile | Gaete A, Travisany D, Palma R, Urra C, Varas M, Allende ML, Maass A, González M, Ferres M. |
| EPI_ISL_468752 | Center for Genome Regulation (CRG) | Center for Mathematical Modeling and Center for Genome Regulation. Santiago, Chile | Gaete A, Travisany D, Palma R, Urra C, Varas M, Allende ML, Maass A, González M. |
| EPI_ISL_468753, EPI_ISL_468754, EPI_ISL_468755, EPI_ISL_468756, EPI_ISL_468757, EPI_ISL_468758, EPI_ISL_468759 | Laboratorio de Biologia Molecular, Facultad de Medicina, Universidad de Atacama | Center for Mathematical Modeling and Center for Genome Regulation. Santiago, Chile | Gaete A, Travisany D, Palma R, Urra C, Varas M, Allende ML, Maass A, González M, C Echeverria |
| EPI_ISL_468760 | Center for Genome Regulation (CRG) | Center for Mathematical Modeling and Center for Genome Regulation. Santiago, Chile | Gaete A, Travisany D, Palma R, Urra C, Varas M, Allende ML, Maass A, González M. |
| EPI_ISL_468761, EPI_ISL_468762, EPI_ISL_468763, EPI_ISL_468764 | Centro de Investigación Biomédica de La Rioja - Hospital San Pedro Logroño | SeqCOVID-SPAIN consortium/IBV(CSIC) | María de Toro, José Manuel Azcona Gutiérrez, María Pilar Bea Escudero, Miriam Blasco Alberdi and SeqCOVID-SPAIN consortium |
| EPI_ISL_468765, EPI_ISL_468766, EPI_ISL_468767, EPI_ISL_468768, EPI_ISL_468769, EPI_ISL_468770, EPI_ISL_468771, EPI_ISL_468772, EPI_ISL_468773, EPI_ISL_468774, EPI_ISL_468775, EPI_ISL_468776, EPI_ISL_468777, EPI_ISL_468778, EPI_ISL_468779, EPI_ISL_468780, EPI_ISL_468781, EPI_ISL_468782, EPI_ISL_468783, EPI_ISL_468784, EPI_ISL_468785, EPI_ISL_468786, EPI_ISL_468787, EPI_ISL_468788, EPI_ISL_468789, EPI_ISL_468790, EPI_ISL_468791, EPI_ISL_468792, EPI_ISL_468793, EPI_ISL_468794, EPI_ISL_468795, EPI_ISL_468796, EPI_ISL_468797, EPI_ISL_468798, EPI_ISL_468799, EPI_ISL_468800, EPI_ISL_468801, EPI_ISL_468802, EPI_ISL_468803, EPI_ISL_468804, EPI_ISL_468805, EPI_ISL_468806, EPI_ISL_468807, EPI_ISL_468808, EPI_ISL_468809, EPI_ISL_468810, EPI_ISL_468811, EPI_ISL_468812, EPI_ISL_468813, EPI_ISL_468814, EPI_ISL_468815, EPI_ISL_468816, EPI_ISL_468817, EPI_ISL_468818, EPI_ISL_468819, EPI_ISL_468820, EPI_ISL_468821, EPI_ISL_468822, EPI_ISL_468823, EPI_ISL_468824, EPI_ISL_468825, EPI_ISL_468826, EPI_ISL_468827, EPI_ISL_468828, EPI_ISL_468829, EPI_ISL_468830, EPI_ISL_468831, EPI_ISL_468832, EPI_ISL_468833, EPI_ISL_468834, EPI_ISL_468835, EPI_ISL_468836, EPI_ISL_468837, EPI_ISL_468838, EPI_ISL_468839, EPI_ISL_468840, EPI_ISL_468841, EPI_ISL_468842, EPI_ISL_468843, EPI_ISL_468844, EPI_ISL_468845, EPI_ISL_468846, EPI_ISL_468847, EPI_ISL_468848, EPI_ISL_468849, EPI_ISL_468850, EPI_ISL_468851, EPI_ISL_468852, EPI_ISL_468853, EPI_ISL_468854, EPI_ISL_468855, EPI_ISL_468856, EPI_ISL_468857, EPI_ISL_468858, EPI_ISL_468859 | | | |
| see above | Servicio de Microbiología, Hospital Miguel Servet, Zaragoza | SeqCOVID-SPAIN consortium/IBV(CSIC) | Antonio Rezusta López, Alexander Tristancho Baró, Ana Milagro, Yolanda Gracia Grataloup, Nieves Martínez Cameo and SeqCOVID-SPAIN consortium |
| EPI_ISL_468860, EPI_ISL_468861, EPI_ISL_468862, EPI_ISL_468863, EPI_ISL_468864, EPI_ISL_468865, EPI_ISL_468866, EPI_ISL_468867, EPI_ISL_468868, EPI_ISL_468869, EPI_ISL_468870, EPI_ISL_468871, EPI_ISL_468872, EPI_ISL_468873, EPI_ISL_468874, EPI_ISL_468875, EPI_ISL_468876, EPI_ISL_468877, EPI_ISL_468878, EPI_ISL_468879, EPI_ISL_468880, EPI_ISL_468881, EPI_ISL_468882, EPI_ISL_468883, EPI_ISL_468884, EPI_ISL_468885, EPI_ISL_468886, EPI_ISL_468887, EPI_ISL_468888, EPI_ISL_468889, EPI_ISL_468890, EPI_ISL_468891, EPI_ISL_468892, EPI_ISL_468893, EPI_ISL_468894, EPI_ISL_468895, EPI_ISL_468896, EPI_ISL_468897, EPI_ISL_468898, EPI_ISL_468899, EPI_ISL_468900, EPI_ISL_468901, EPI_ISL_468902, EPI_ISL_468903, EPI_ISL_468904, EPI_ISL_468905, EPI_ISL_468906, EPI_ISL_468907, EPI_ISL_468908, EPI_ISL_468909, EPI_ISL_468910, EPI_ISL_468911, EPI_ISL_468912, EPI_ISL_468913 | | | |
| see above | Servicio de Microbiología. Hospital Universitario Donostia. OSI Donostialdea. Área de Enfermedades Infecciosas, Grupo de Infección Respiratoria y Resistencia Antimicrobiana. Instituto de Investigación Sanitaria Biodonostia. | SeqCOVID-SPAIN consortium/IBV(CSIC) | Gustavo Cilla, Milagrosa Montes, Luis Piñeiro, Jose Maria Marimón and SeqCOVID-SPAIN consortium |
| EPI_ISL_468914 | Istituto Zooprofilattico Sperimentale Puglia e Basilicata; Dipartimento di Bioscienze, Biotecnologie e Biofarmaceutica dell'Università degli Studi di Bari "A.Moro"; Istituto di Biomembrane. Bioenergetica e Biotecnologie Molecolari del Consiglio Nazionale delle Ricerche di Bari | Beaconlab (Bioinformatics, Evolution and Comparative Genomics lab), Dept of Biosciences, University on Milan | Parisi A.,Pesole G., Manzari C., Chiara M. |
| EPI_ISL_468915, EPI_ISL_468916, EPI_ISL_468917, EPI_ISL_468918, EPI_ISL_468919, EPI_ISL_468920, EPI_ISL_468921, EPI_ISL_468922, EPI_ISL_468923, EPI_ISL_468924, EPI_ISL_468925, EPI_ISL_468926, EPI_ISL_468927, EPI_ISL_468928, EPI_ISL_468929, EPI_ISL_468930, EPI_ISL_468931, EPI_ISL_468932, EPI_ISL_468933, EPI_ISL_468934, EPI_ISL_468935, EPI_ISL_468936, EPI_ISL_468937, EPI_ISL_468938, EPI_ISL_468939, EPI_ISL_468940, EPI_ISL_468941, EPI_ISL_468942, EPI_ISL_468943, EPI_ISL_468944, EPI_ISL_468945, EPI_ISL_468946, EPI_ISL_468947, EPI_ISL_468948, EPI_ISL_468949, EPI_ISL_468950, EPI_ISL_468951 | | | |
| see above | Servicio de Microbiología. Hospital Universitario Donostia. OSI Donostialdea. Área de Enfermedades Infecciosas, Grupo de Infección Respiratoria y Resistencia Antimicrobiana. Instituto de Investigación Sanitaria Biodonostia. | SeqCOVID-SPAIN consortium/IBV(CSIC) | Gustavo Cilla, Milagrosa Montes, Luis Piñeiro, Jose Maria Marimón and SeqCOVID-SPAIN consortium |
| EPI_ISL_468952, EPI_ISL_468953, EPI_ISL_468954, EPI_ISL_468955, EPI_ISL_468956, EPI_ISL_468957, EPI_ISL_468958, EPI_ISL_468959, EPI_ISL_468960, EPI_ISL_468961, EPI_ISL_468962, EPI_ISL_468963, EPI_ISL_468964, EPI_ISL_468965, EPI_ISL_468966, EPI_ISL_468967, EPI_ISL_468968, EPI_ISL_468969, EPI_ISL_468970, EPI_ISL_468971, EPI_ISL_468972, EPI_ISL_468973, EPI_ISL_468974, EPI_ISL_468975, EPI_ISL_468976, EPI_ISL_468977, EPI_ISL_468978, EPI_ISL_468979, EPI_ISL_468980, EPI_ISL_468981, EPI_ISL_468982, EPI_ISL_468983, EPI_ISL_468984, EPI_ISL_468985, EPI_ISL_468986, EPI_ISL_468987, EPI_ISL_468988, EPI_ISL_468989, EPI_ISL_468990, EPI_ISL_468991, EPI_ISL_468992, EPI_ISL_468993, EPI_ISL_468994, EPI_ISL_468995, EPI_ISL_468996, EPI_ISL_468997, EPI_ISL_468998, EPI_ISL_468999, EPI_ISL_469000, EPI_ISL_469001, EPI_ISL_469002, EPI_ISL_469003, EPI_ISL_469004, EPI_ISL_469005, EPI_ISL_469006, EPI_ISL_469007, EPI_ISL_469008, EPI_ISL_469009, EPI_ISL_469010, EPI_ISL_469011, EPI_ISL_469012, EPI_ISL_469013, EPI_ISL_469014, EPI_ISL_469015 | | | |
| see above | Servicio de Microbiología, Hospital Universitario Son Espases | SeqCOVID-SPAIN consortium/IBV(CSIC) | Carla López-Causapé, Jordi Reina, Antonio Oliver and SeqCOVID-SPAIN consortium |
| EPI_ISL_469016 | Istituto Zooprofilattico Sperimentale Puglia e Basilicata; Dipartimento di Bioscienze, Biotecnologie e Biofarmaceutica dell'Università degli Studi di Bari "A.Moro"; Istituto di Biomembrane. Bioenergetica e Biotecnologie Molecolari del Consiglio Nazionale delle Ricerche di Bari | Beaconlab (Bioinformatics, Evolution and Comparative Genomics lab), Dept of Biosciences, University on Milan | Parisi A.,Pesole G., Manzari C., Chiara M. |
| EPI_ISL_469017 | LNR National Reference Laboratory, Mohammed VI University of Health Sciences | Medical Biotechnology Laboratory, Rabat Medical and Pharmacy School, Mohammed The Vth University in Rabat | Meriem LAAMARTI, Souad KARTTI, Rokaia LAAMRTI , M.W. CHEMAO-ELFIHRI, Loubna ALLAM, Mouna OUADGHIRI, Imane SMYEJ, Jalila RAHOUI, Houda BENRAHMA, Jalil El Atar, Idrissa Diawara, Rachid EL JAOUDI, Laila SBABOU, Chakib NEJJARI, Saaid AMZAZI, Rachid MENTAG, Lahcen BELYAMANI and Azeddine IBRAHIMI |
| EPI_ISL_469018 | Istituto Zooprofilattico Sperimentale Puglia e Basilicata; Dipartimento di Bioscienze, Biotecnologie e Biofarmaceutica dell'Università degli Studi di Bari "A.Moro"; Istituto di Biomembrane. Bioenergetica e Biotecnologie Molecolari del Consiglio Nazionale delle Ricerche di Bari | Beaconlab (Bioinformatics, Evolution and Comparative Genomics lab), Dept of Biosciences, University on Milan | Parisi A.,Pesole G., Manzari C., Chiara M. |
| EPI_ISL_469019, EPI_ISL_469020, EPI_ISL_469021, EPI_ISL_469022 | Istituto Zooprofilattico Sperimentale Puglia e Basilicata; Dipartimento di Bioscienze, Biotecnologie e Biofarmaceutica dell'Università degli Studi di Bari "A.Moro"; Istituto di Biomembrane. Bioenergetica e Biotecnologie Molecolari del Consiglio Nazionale delle Ricerche di Bari | Beaconlab (Bioinformatics, Evolution and Comparative Genomics lab), Dept of Biosciences, University on Milan | Parisi A.,Pesole G., Manzari C., Chiara M. |
| EPI_ISL_469023 | Istituto Zooprofilattico Sperimentale Puglia e Basilicata; Dipartimento di Bioscienze, Biotecnologie e Biofarmaceutica dell'Università degli Studi di Bari "A.Moro"; Istituto di Biomembrane. Bioenergetica e Biotecnologie Molecolari del Consiglio Nazionale delle Ricerche di Bari | Beaconlab (Bioinformatics, Evolution and Comparative Genomics lab), Dept of Biosciences, University on Milan | Parisi A.,Pesole G., Manzari C., Chiara M |
| EPI_ISL_469024 | B.J. Medical College and Civil hospital | Gujarat Biotechnology Research Centre | Tejas Shah, Ankit Hinsu, Pritesh Sabara, Apurvasinh Puvar, Janvi Raval, Zarna Patel, Monika Gandhi, Pinal Trivedi, Maharshi Pandya, Nidhi Patel, Nitin Savaliya, Raghawendra Kumar, Dinesh Kumar, Zuber Saiyed, Komal Patel, Labdhi Pandya, Snehal Bagatharia, Pranay Shah, Kamlesh J Upadhyay, Nirav Mungalpara, Priti Pandita, R D Dixit, A M Kadri, Harsh Bakshi, Chaitanya Joshi, Madhvi Joshi |
| EPI_ISL_469025 | B.J. Medical College and Civil hospital | Gujarat Biotechnology Research Centre | Ankit Hinsu, Pritesh Sabara, Apurvasinh Puvar, Janvi Raval, Zarna Patel, Monika Gandhi, Pinal Trivedi, Maharshi Pandya, Nidhi Patel, Nitin Savaliya, Raghawendra Kumar, Dinesh Kumar, Zuber Saiyed, Komal Patel, Labdhi Pandya, Snehal Bagatharia, Pranay Shah, Kamlesh J Upadhyay, Nirav Mungalpara, Tejas Shah, Pragya Sharma, R D Dixit, A M Kadri, Harsh Bakshi, Chaitanya Joshi, Madhvi Joshi |
| EPI_ISL_469026 | B.J. Medical College and Civil hospital | Gujarat Biotechnology Research Centre | Pritesh Sabara, Apurvasinh Puvar, Janvi Raval, Zarna Patel, Monika Gandhi, Pinal Trivedi, Maharshi Pandya, Nidhi Patel, Nitin Savaliya, Raghawendra Kumar, Dinesh Kumar, Zuber Saiyed, Komal Patel, Labdhi Pandya, Snehal Bagatharia, Pranay Shah, Kamlesh J Upadhyay, Nirav Mungalpara, Tejas Shah, Ankit Hinsu, Neha Rajpara, R D Dixit, A M Kadri, Harsh Bakshi, Chaitanya Joshi, Madhvi Joshi |
| EPI_ISL_469027 | B.J. Medical College and Civil hospital | Gujarat Biotechnology Research Centre | Apurvasinh Puvar, Janvi Raval, Zarna Patel, Monika Gandhi, Pinal Trivedi, Maharshi Pandya, Nidhi Patel, Nitin Savaliya, Raghawendra Kumar, Dinesh Kumar, Zuber Saiyed, Komal Patel, Labdhi Pandya, Snehal Bagatharia, Pranay Shah, Kamlesh J Upadhyay, Nirav Mungalpara, Tejas Shah, Ankit Hinsu, Pritesh Sabara, Afzal Ansari, R D Dixit, A M Kadri, Harsh Bakshi, Chaitanya Joshi, Madhvi Joshi |
| EPI_ISL_469028 | B.J. Medical College and Civil hospital | Gujarat Biotechnology Research Centre | Janvi Raval, Zarna Patel, Monika Gandhi, Pinal Trivedi, Maharshi Pandya, Nidhi Patel, Nitin Savaliya, Raghawendra Kumar, Dinesh Kumar, Zuber Saiyed, Komal Patel, Labdhi Pandya, Snehal Bagatharia, Pranay Shah, Kamlesh J |

| | | | |
|---|---|---|---|
| | | | Upadhyay, Nirav Mungalpara, Tejas Shah, Ankit Hinsu, Pritesh Sabara, Apurvasinh Puvar, Fenil Patel, R D Dixit, A M Kadri, Harsh Bakshi, Chaitanya Joshi, Madhvi Joshi |
| EPI_ISL_469029 | Government Medical College, Vadodara | Gujarat Biotechnology Research Centre | Zarna Patel, Monika Gandhi, Pinal Trivedi, Maharshi Pandya, Nidhi Patel, Nitin Savaliya, Raghawendra Kumar, Dinesh Kumar, Zuber Saiyed, Komal Patel, Labdhi Pandya, Snehal Bagatharia, Meenakshi Shah, Neena Doshi, Varsha Godbole, Tejas Shah, Ankit Hinsu, Pritesh Sabara, Apurvasinh Puvar, Janvi Raval, Neelam Nathani, R D Dixit, A M Kadri, Harsh Bakshi, Chaitanya Joshi, Madhvi Joshi |
| EPI_ISL_469030 | Government Medical College, Vadodara | Gujarat Biotechnology Research Centre | Monika Gandhi, Pinal Trivedi, Maharshi Pandya, Nidhi Patel, Nitin Savaliya, Raghawendra Kumar, Dinesh Kumar, Zuber Saiyed, Komal Patel, Labdhi Pandya, Snehal Bagatharia, Meenakshi Shah, Neena Doshi, Varsha Godbole, Tejas Shah, Ankit Hinsu, Pritesh Sabara, Apurvasinh Puvar, Janvi Raval, Zarna Patel, Armi Chaudhari, R D Dixit, A M Kadri, Harsh Bakshi, Chaitanya Joshi, Madhvi Joshi |
| EPI_ISL_469031 | Government Medical College, Vadodara | Gujarat Biotechnology Research Centre | Pinal Trivedi, Maharshi Pandya, Nidhi Patel, Nitin Savaliya, Raghawendra Kumar, Dinesh Kumar, Zuber Saiyed, Komal Patel, Labdhi Pandya, Snehal Bagatharia, Meenakshi Shah, Neena Doshi, Varsha Godbole, Tejas Shah, Ankit Hinsu, Pritesh Sabara, Apurvasinh Puvar, Janvi Raval, Zarna Patel, Monika Gandhi, Bhavya Jindal, R D Dixit, A M Kadri, Harsh Bakshi, Chaitanya Joshi, Madhvi Joshi |
| EPI_ISL_469032 | Government Medical College, Vadodara | Gujarat Biotechnology Research Centre | Maharshi Pandya, Nidhi Patel, Nitin Savaliya, Raghawendra Kumar, Dinesh Kumar, Zuber Saiyed, Komal Patel, Labdhi Pandya, Snehal Bagatharia, Meenakshi Shah, Neena Doshi, Varsha Godbole, Tejas Shah, Ankit Hinsu, Pritesh Sabara, Apurvasinh Puvar, Janvi Raval, Zarna Patel, Monika Gandhi, Pinal Trivedi, Pragya Sharma, R D Dixit, A M Kadri, Harsh Bakshi, Chaitanya Joshi, Madhvi Joshi |
| EPI_ISL_469033 | Government Medical College, Vadodara | Gujarat Biotechnology Research Centre | Nidhi Patel, Nitin Savaliya, Raghawendra Kumar, Dinesh Kumar, Zuber Saiyed, Komal Patel, Labdhi Pandya, Snehal Bagatharia, Meenakshi Shah, Neena Doshi, Varsha Godbole, Tejas Shah, Ankit Hinsu, Pritesh Sabara, Apurvasinh Puvar, Janvi Raval, Zarna Patel, Monika Gandhi, Pinal Trivedi, Maharshi Pandya, Priyanka P Vatsa, R D Dixit, A M Kadri, Harsh Bakshi, Chaitanya Joshi, Madhvi Joshi |
| EPI_ISL_469034 | Government Medical College, Vadodara | Gujarat Biotechnology Research Centre | Nitin Savaliya, Raghawendra Kumar, Dinesh Kumar, Zuber Saiyed, Komal Patel, Labdhi Pandya, Snehal Bagatharia, Meenakshi Shah, Neena Doshi, Varsha Godbole, Tejas Shah, Ankit Hinsu, Pritesh Sabara, Apurvasinh Puvar, Janvi Raval, Zarna Patel, Monika Gandhi, Pinal Trivedi, Maharshi Pandya, Nidhi Patel, Pooja P Doshi, R D Dixit, A M Kadri, Harsh Bakshi, Chaitanya Joshi, Madhvi Joshi |
| EPI_ISL_469035 | Government Medical College, Vadodara | Gujarat Biotechnology Research Centre | Raghawendra Kumar, Dinesh Kumar, Zuber Saiyed, Komal Patel, Labdhi Pandya, Snehal Bagatharia, Meenakshi Shah, Neena Doshi, Varsha Godbole, Tejas Shah, Ankit Hinsu, Pritesh Sabara, Apurvasinh Puvar, Janvi Raval, Zarna Patel, Monika Gandhi, Pinal Trivedi, Maharshi Pandya, Nidhi Patel, Nitin Savaliya, Akanksha Verma, R D Dixit, A M Kadri, Harsh Bakshi, Chaitanya Joshi, Madhvi Joshi |
| EPI_ISL_469036 | Government Medical College, Vadodara | Gujarat Biotechnology Research Centre | Dinesh Kumar, Zuber Saiyed, Komal Patel, Labdhi Pandya, Snehal Bagatharia, Meenakshi Shah, Neena Doshi, Varsha Godbole, Tejas Shah, Ankit Hinsu, Pritesh Sabara, Apurvasinh Puvar, Janvi Raval, Zarna Patel, Monika Gandhi, Pinal Trivedi, Maharshi Pandya, Nidhi Patel, Nitin Savaliya, Raghawendra Kumar, Priti Pandita, R D Dixit, A M Kadri, Harsh Bakshi, Chaitanya Joshi, Madhvi Joshi |
| EPI_ISL_469037 | GMERS Medical College & Hospital | Gujarat Biotechnology Research Centre | Zuber Saiyed, Komal Patel, Labdhi Pandya, Snehal Bagatharia, Meenakshi Shah, Neena Doshi, Varsha Godbole, Tejas Shah, Ankit Hinsu, Pritesh Sabara, Apurvasinh Puvar, Janvi Raval, Zarna Patel, Monika Gandhi, Pinal Trivedi, Maharshi Pandya, Nidhi Patel, Nitin Savaliya, Raghawendra Kumar, Dinesh Kumar, Pragya Sharma, R D Dixit, A M Kadri, Harsh Bakshi, Chaitanya Joshi, Madhvi Joshi |
| EPI_ISL_469038 | GMERS Medical College & Hospital | Gujarat Biotechnology Research Centre | Komal Patel, Labdhi Pandya, Snehal Bagatharia, Meenakshi Shah, Neena Doshi, Varsha Godbole, Tejas Shah, Ankit Hinsu, Pritesh Sabara, Apurvasinh Puvar, Janvi Raval, Zarna Patel, Monika Gandhi, Pinal Trivedi, Maharshi Pandya, Nidhi Patel, Nitin Savaliya, Raghawendra Kumar, Dinesh Kumar, Zuber Saiyed, Neha Rajpara, R D Dixit, A M Kadri, Harsh Bakshi, Chaitanya Joshi, Madhvi Joshi |
| EPI_ISL_469039 | GMERS Medical College & Hospital | Gujarat Biotechnology Research Centre | Labdhi Pandya, Snehal Bagatharia, Meenakshi Shah, Neena Doshi, Varsha Godbole, Tejas Shah, Ankit Hinsu, Pritesh Sabara, Apurvasinh Puvar, Janvi Raval, Zarna Patel, Monika Gandhi, Pinal Trivedi, Maharshi Pandya, Nidhi Patel, Nitin Savaliya, Raghawendra Kumar, Dinesh Kumar, Zuber Saiyed, Komal Patel, Afzal Ansari, R D Dixit, A M Kadri, Harsh Bakshi, Chaitanya Joshi, Madhvi Joshi |
| EPI_ISL_469040 | GMERS Medical College & Hospital | Gujarat Biotechnology Research Centre | Snehal Bagatharia, Meenakshi Shah, Neena Doshi, Varsha Godbole, Tejas Shah, Ankit Hinsu, Pritesh Sabara, Apurvasinh Puvar, Janvi Raval, Zarna Patel, Monika Gandhi, Pinal Trivedi, Maharshi Pandya, Nidhi Patel, Nitin Savaliya, Raghawendra Kumar, Dinesh Kumar, Zuber Saiyed, Komal Patel, Labdhi Pandya, Fenil Patel, R D Dixit, A M Kadri, Harsh Bakshi, Chaitanya Joshi, Madhvi Joshi |
| EPI_ISL_469041 | GMERS Medical College & Hospital | Gujarat Biotechnology Research Centre | Meenakshi Shah, Neena Doshi, Varsha Godbole, Tejas Shah, Ankit Hinsu, Pritesh Sabara, Apurvasinh Puvar, Janvi Raval, Zarna Patel, Monika Gandhi, Pinal Trivedi, Maharshi Pandya, Nidhi Patel, Nitin Savaliya, Raghawendra Kumar, Dinesh Kumar, Zuber Saiyed, Komal Patel, Labdhi Pandya, Snehal Bagatharia, Neelam Nathani, R D Dixit, A M Kadri, Harsh Bakshi, Chaitanya Joshi, Madhvi Joshi |
| EPI_ISL_469042 | GMERS Medical College & Hospital | Gujarat Biotechnology Research Centre | Neena Doshi, Varsha Godbole, Tejas Shah, Ankit Hinsu, Pritesh Sabara, Apurvasinh Puvar, Janvi Raval, Zarna Patel, Monika Gandhi, Pinal Trivedi, Maharshi Pandya, Nidhi Patel, Nitin Savaliya, Raghawendra Kumar, Dinesh Kumar, Zuber Saiyed, Komal Patel, Labdhi Pandya, Snehal Bagatharia, Meenakshi Shah, Armi Chaudhari, R D Dixit, A M Kadri, Harsh Bakshi, Chaitanya Joshi, Madhvi Joshi |
| EPI_ISL_469043 | Dr. N. D. Desai Medical College & Hospital | Gujarat Biotechnology Research Centre | J G Buch, Jigar Gusani, Supreet Prabhu, Tejas Shah, Ankit Hinsu, Pritesh Sabara, Apurvasinh Puvar, Janvi Raval, Zarna Patel, Monika Gandhi, Pinal Trivedi, Maharshi Pandya, Nidhi Patel, Nitin Savaliya, Raghawendra Kumar, Dinesh Kumar, Zuber Saiyed, Komal Patel, Labdhi Pandya, Snehal Bagatharia, Bhavya Jindal, R D Dixit, A M Kadri, Harsh Bakshi, Chaitanya Joshi, Madhvi Joshi |
| EPI_ISL_469044 | Dr. N. D. Desai Medical College & Hospital | Gujarat Biotechnology Research Centre | Jigar Gusani, Supreet Prabhu, Tejas Shah, Ankit Hinsu, Pritesh Sabara, Apurvasinh Puvar, Janvi Raval, Zarna Patel, Monika Gandhi, Pinal Trivedi, Maharshi Pandya, Nidhi Patel, Nitin Savaliya, Raghawendra Kumar, Dinesh Kumar, Zuber Saiyed, Komal Patel, Labdhi Pandya, Snehal Bagatharia, J G Buch, Neha Rajpara, R D Dixit, A M Kadri, Harsh Bakshi, Chaitanya Joshi, Madhvi Joshi |
| EPI_ISL_469045 | Dr. N. D. Desai Medical College & Hospital | Gujarat Biotechnology Research Centre | Supreet Prabhu, Tejas Shah, Ankit Hinsu, Pritesh Sabara, Apurvasinh Puvar, Janvi Raval, Zarna Patel, Monika Gandhi, Pinal Trivedi, Maharshi Pandya, Nidhi Patel, Nitin Savaliya, Raghawendra Kumar, Dinesh Kumar, Zuber Saiyed, Komal Patel, Labdhi Pandya, Snehal Bagatharia, J G Buch, Jigar Gusani, Priyanka P Vatsa, R D Dixit, A M Kadri, Harsh Bakshi, Chaitanya Joshi, Madhvi Joshi |
| EPI_ISL_469046 | Dr. N. D. Desai Medical College & Hospital | Gujarat Biotechnology Research Centre | Tejas Shah, Ankit Hinsu, Pritesh Sabara, Apurvasinh Puvar, Janvi Raval, Zarna Patel, Monika Gandhi, Pinal Trivedi, Maharshi Pandya, Nidhi Patel, Nitin Savaliya, Raghawendra Kumar, Dinesh Kumar, Zuber Saiyed, Komal Patel, Labdhi Pandya, Snehal Bagatharia, J G Buch, Jigar Gusani, Supreet Prabhu, Pooja P Doshi, R D Dixit, A M Kadri, Harsh Bakshi, Chaitanya Joshi, Madhvi Joshi |
| EPI_ISL_469047 | Dr. N. D. Desai Medical College & Hospital | Gujarat Biotechnology Research Centre | Ankit Hinsu, Pritesh Sabara, Apurvasinh Puvar, Janvi Raval, Zarna Patel, Monika Gandhi, Pinal Trivedi, Maharshi Pandya, Nidhi Patel, Nitin Savaliya, Raghawendra Kumar, Dinesh Kumar, Zuber Saiyed, Komal Patel, Labdhi Pandya, Snehal Bagatharia, J G Buch, Jigar Gusani, Supreet Prabhu, Tejas Shah, Akanksha Verma, R D Dixit, A M Kadri, Harsh Bakshi, Chaitanya Joshi, Madhvi Joshi |
| EPI_ISL_469048 | Banas Medical College and Research Institute | Gujarat Biotechnology Research Centre | Radhika Khara, Sunil R Joshi, Viren s Doshi, Zarna Patel, Monika Gandhi, Pinal Trivedi, Maharshi Pandya, Nidhi Patel, Nitin Savaliya, Raghawendra Kumar, Dinesh Kumar, Zuber Saiyed, Komal Patel, Labdhi Pandya, Snehal Bagatharia, Tejas Shah, Ankit Hinsu, Pritesh Sabara, Apurvasinh Puvar, Janvi Raval, Priti Pandita, R D Dixit, A M Kadri, Harsh Bakshi, Chaitanya Joshi, Madhvi Joshi |
| EPI_ISL_469049 | LNR National Reference Laboratory, Mohammed VI University of Health Sciences | Medical Biotechnology Laboratory, Rabat Medical and Pharmacy School, Mohammed The Vth University in Rabat | Meriem LAAMARTI, Souad KARTTI, Rokaia LAAMRTI , M.W. CHEMAO-ELFIHRI, Loubna ALLAM, Mouna OUADGHIRI, Imane SMYEJ, Jalila RAHOUI, Houda BENRAHMA, Jalil El Atar, Idrissa Diawara, Rachid EL JAOUDI, Laila SBABOU, Chakib NEJJARI, Saaid AMZAZI, Rachid MENTAG, Lahcen BELYAMANI and Azeddine IBRAHIMI |
| EPI_ISL_469050 | Istituto Zooprofilattico Sperimentale Puglia e Basilicata; Dipartimento di Bioscienze, Biotecnologie e Biofarmaceutica dell'Università degli Studi di Bari "A.Moro"; Istituto di Biomembrane. Bioenergetica e Biotecnologie Molecolari del Consiglio Nazionale delle Ricerche di Bari | Beaconlab (Bioinformatics, Evolution and Comparative Genomics lab), Dept of Biosciences, University on Milan | Parisi A.,Pesole G., Manzari C., Chiara M. |
| EPI_ISL_469051, EPI_ISL_469052, EPI_ISL_469053, EPI_ISL_469054 | LNR National Reference Laboratory, Mohammed VI University of Health Sciences | Medical Biotechnology Laboratory, Rabat Medical and Pharmacy School, Mohammed The Vth University in Rabat | Meriem LAAMARTI, Souad KARTTI, Rokaia LAAMRTI , M.W. CHEMAO-ELFIHRI, Loubna ALLAM, Mouna OUADGHIRI, Imane SMYEJ, Jalila RAHOUI, Houda BENRAHMA, Jalil El Atar, Idrissa Diawara, Rachid EL JAOUDI, Laila SBABOU, Chakib NEJJARI, Saaid AMZAZI, Rachid MENTAG, Lahcen BELYAMANI and Azeddine IBRAHIMI |
| EPI_ISL_469055, EPI_ISL_469056 | Jourcentralen | The Public Health Agency of Sweden | Oskar Karlsson Lindsjo, Maria Lind Karlberg, Mattias Haukland, Reza Advani, Olov Svartstrom, Anna-Malin Linde, Sandra Broddesson, Petra Edquist, Shamam Muradrasoli, Anna Risberg, Karin Tegmark-Wisell |
| EPI_ISL_469057 | Inger Landgren | The Public Health Agency of Sweden | Oskar Karlsson Lindsjo, Maria Lind Karlberg, Mattias Haukland, Reza Advani, Olov Svartstrom, Anna-Malin Linde, Sandra Broddesson, Petra Edquist, Shamam Muradrasoli, Anna Risberg, Karin Tegmark-Wisell |
| EPI_ISL_469058 | Narhalsan Sjobo vardcentral | The Public Health Agency of Sweden | Oskar Karlsson Lindsjo, Maria Lind Karlberg, Mattias Haukland, Reza Advani, Olov Svartstrom, Anna-Malin Linde, Sandra Broddesson, Petra Edquist, Shamam Muradrasoli, Anna Risberg, Karin Tegmark-Wisell |
| EPI_ISL_469059 | Hovas Askim Familjelakare och BVC | The Public Health Agency of Sweden | Oskar Karlsson Lindsjo, Maria Lind Karlberg, Mattias Haukland, Reza Advani, Olov Svartstrom, Anna-Malin Linde, Sandra Broddesson, Petra Edquist, Shamam Muradrasoli, Anna Risberg, Karin Tegmark-Wisell |
| EPI_ISL_469060, EPI_ISL_469061 | Narhalsan Sjobo vardcentral | The Public Health Agency of Sweden | Oskar Karlsson Lindsjo, Maria Lind Karlberg, Mattias Haukland, Reza Advani, Olov Svartstrom, Anna-Malin Linde, Sandra Broddesson, Petra Edquist, Shamam Muradrasoli, Anna Risberg, Karin Tegmark-Wisell |
| EPI_ISL_469062 | Huddinge VC | The Public Health Agency of Sweden | Oskar Karlsson Lindsjo, Maria Lind Karlberg, Mattias Haukland, Reza Advani, Olov Svartstrom, Anna-Malin Linde, Sandra Broddesson, Petra Edquist, Shamam Muradrasoli, Anna Risberg, Karin Tegmark-Wisell |
| EPI_ISL_469063 | Ulltuna Vardcentral | The Public Health Agency of Sweden | Oskar Karlsson Lindsjo, Maria Lind Karlberg, Mattias Haukland, Reza Advani, Olov Svartstrom, Anna-Malin Linde, Sandra Broddesson, Petra Edquist, Shamam Muradrasoli, Anna Risberg, Karin Tegmark-Wisell |
| EPI_ISL_469064, EPI_ISL_469065 | Huddinge VC | The Public Health Agency of Sweden | Oskar Karlsson Lindsjo, Maria Lind Karlberg, Mattias Haukland, Reza Advani, Olov Svartstrom, Anna-Malin Linde, Sandra Broddesson, Petra Edquist, Shamam Muradrasoli, Anna Risberg, Karin Tegmark-Wisell |
| EPI_ISL_469066 | Surbrunns VC | The Public Health Agency of Sweden | Oskar Karlsson Lindsjo, Maria Lind Karlberg, Mattias Haukland, Reza Advani, Olov Svartstrom, Anna-Malin Linde, Sandra Broddesson, Petra Edquist, Shamam Muradrasoli, Anna Risberg, Karin Tegmark-Wisell |
| EPI_ISL_469067 | Kungsholmsdoktorn | The Public Health Agency of Sweden | Oskar Karlsson Lindsjo, Maria Lind Karlberg, Mattias Haukland, Reza Advani, Olov Svartstrom, Anna-Malin Linde, Sandra Broddesson, Petra Edquist, Shamam Muradrasoli, Anna Risberg, Karin Tegmark-Wisell |
| EPI_ISL_469068 | Hovas Askim Familjelakare och BVC | The Public Health Agency of Sweden | Oskar Karlsson Lindsjo, Maria Lind Karlberg, Mattias Haukland, Reza Advani, Olov Svartstrom, Anna-Malin Linde, Sandra Broddesson, Petra Edquist, Shamam Muradrasoli, Anna Risberg, Karin Tegmark-Wisell |
| EPI_ISL_469069 | Narhalsan Olskroken VC | The Public Health Agency of Sweden | Oskar Karlsson Lindsjo, Maria Lind Karlberg, Mattias Haukland, Reza Advani, Olov Svartstrom, Anna-Malin Linde, Sandra Broddesson, Petra Edquist, Shamam Muradrasoli, Anna Risberg, Karin Tegmark-Wisell |
| EPI_ISL_469070 | Surbrunns VC | The Public Health Agency of Sweden | Oskar Karlsson Lindsjo, Maria Lind Karlberg, Mattias Haukland, Reza Advani, Olov Svartstrom, Anna-Malin Linde, Sandra Broddesson, Petra Edquist, Shamam Muradrasoli, Anna Risberg, Karin Tegmark-Wisell |
| EPI_ISL_469071 | Wasterlakarna | The Public Health Agency of Sweden | Oskar Karlsson Lindsjo, Maria Lind Karlberg, Mattias Haukland, Reza Advani, Olov Svartstrom, Anna-Malin Linde, Sandra Broddesson, Petra Edquist, Shamam Muradrasoli, Anna Risberg, Karin Tegmark-Wisell |
| EPI_ISL_469072 | Ulltuna Vardcentral | The Public Health Agency of Sweden | Oskar Karlsson Lindsjo, Maria Lind Karlberg, Mattias Haukland, Reza Advani, Olov Svartstrom, Anna-Malin Linde, Sandra Broddesson, Petra Edquist, Shamam Muradrasoli, Anna Risberg, Karin Tegmark-Wisell |
| EPI_ISL_469073, EPI_ISL_469074 | Halmstad klinisk mikrobiologi | The Public Health Agency of Sweden | Oskar Karlsson Lindsjo, Maria Lind Karlberg, Mattias Haukland, Reza Advani, Olov Svartstrom, Anna-Malin Linde, Sandra Broddesson, Petra Edquist, Shamam Muradrasoli, Anna Risberg, Karin Tegmark-Wisell |
| EPI_ISL_469075 | Karolinska Universitetslaboratoriet | The Public Health Agency of Sweden | Oskar Karlsson Lindsjo, Maria Lind Karlberg, Mattias Haukland, Reza Advani, Olov Svartstrom, Anna-Malin Linde, Sandra Broddesson, Petra Edquist, Shamam Muradrasoli, Anna Risberg, Karin Tegmark-Wisell |
| EPI_ISL_469076 | Uppsala klinisk mikrobiologi | The Public Health Agency of Sweden | Oskar Karlsson Lindsjo, Maria Lind Karlberg, Mattias Haukland, Reza Advani, Olov Svartstrom, Anna-Malin Linde, Sandra Broddesson, Petra Edquist, Shamam Muradrasoli, Anna Risberg, Karin Tegmark-Wisell |
| EPI_ISL_469077, EPI_ISL_469078, EPI_ISL_469079 | Karolinska Universitetslaboratoriet | The Public Health Agency of Sweden | Oskar Karlsson Lindsjo, Maria Lind Karlberg, Mattias Haukland, Reza Advani, Olov Svartstrom, Anna-Malin Linde, Sandra Broddesson, Petra Edquist, Shamam Muradrasoli, Anna Risberg, Karin Tegmark-Wisell |
| EPI_ISL_469080, EPI_ISL_469081, EPI_ISL_469082, EPI_ISL_469083, EPI_ISL_469084, EPI_ISL_469085, EPI_ISL_469086, EPI_ISL_469087, EPI_ISL_469088, EPI_ISL_469089, EPI_ISL_469090, EPI_ISL_469091, EPI_ISL_469092, EPI_ISL_469093, EPI_ISL_469094, EPI_ISL_469095, EPI_ISL_469096, EPI_ISL_469097, EPI_ISL_469098, EPI_ISL_469099, EPI_ISL_469100, EPI_ISL_469101, EPI_ISL_469102, EPI_ISL_469103, EPI_ISL_469104, EPI_ISL_469105, EPI_ISL_469106, EPI_ISL_469107, EPI_ISL_469108, EPI_ISL_469109, EPI_ISL_469110, EPI_ISL_469111, EPI_ISL_469112, EPI_ISL_469113, EPI_ISL_469114, EPI_ISL_469115, EPI_ISL_469116, EPI_ISL_469117, EPI_ISL_469118, EPI_ISL_469119, EPI_ISL_469120, EPI_ISL_469121, EPI_ISL_469122, EPI_ISL_469123, EPI_ISL_469124, EPI_ISL_469125, EPI_ISL_469126, EPI_ISL_469127, EPI_ISL_469128, EPI_ISL_469129, EPI_ISL_469130, EPI_ISL_469131, EPI_ISL_469132, EPI_ISL_469133, EPI_ISL_469134, EPI_ISL_469135, EPI_ISL_469136, EPI_ISL_469137, EPI_ISL_469138, EPI_ISL_469139, EPI_ISL_469140, EPI_ISL_469141, EPI_ISL_469142, EPI_ISL_469143, EPI_ISL_469144, EPI_ISL_469145, EPI_ISL_469146, EPI_ISL_469147, EPI_ISL_469148, EPI_ISL_469149, EPI_ISL_469150, EPI_ISL_469151, EPI_ISL_469152, EPI_ISL_469153, EPI_ISL_469154, EPI_ISL_469155 | | | |
| see above | National Public Health Laboratory, National Centre for Infectious Diseases | National Public Health Laboratory, National Centre for Infectious Diseases | Mak TM, Octavia S, Chavatte JM, Cui L, Lin RTP |
| EPI_ISL_469171, EPI_ISL_469172, EPI_ISL_469173, EPI_ISL_469174, EPI_ISL_469175, EPI_ISL_469176, EPI_ISL_469177, EPI_ISL_469178, EPI_ISL_469179, EPI_ISL_469180, EPI_ISL_469181, EPI_ISL_469182, EPI_ISL_469183, EPI_ISL_469184, EPI_ISL_469185, EPI_ISL_469186, EPI_ISL_469187, EPI_ISL_469188, EPI_ISL_469189, EPI_ISL_469190, EPI_ISL_469191, EPI_ISL_469192, EPI_ISL_469193, EPI_ISL_469194, EPI_ISL_469195, EPI_ISL_469196, EPI_ISL_469197, EPI_ISL_469198, EPI_ISL_469199, EPI_ISL_469200, EPI_ISL_469201, EPI_ISL_469202, EPI_ISL_469203, EPI_ISL_469204, EPI_ISL_469205, EPI_ISL_469206, EPI_ISL_469207, EPI_ISL_469208 | | | |
| see above | Yale Clinical Virology Laboratory | Grubaugh Lab - Yale School of Public Health | Joseph Fauver, Tara Alpert, Anderson Brito, Anne Wyllie, Chantal Vogels, Mary Petrone, Cole Jensen, Chaney Kalinich, Isabel Ott, Arnau Casanovas, Catherine Muenker, Adam Moore, Alice Lu, Maria Tokuyama, Patrick Wong, Peiwen Lu, Saad Omer, Richard Martinello, Allison Nelson, Shelli Farhadian, Akiko Iwasaki, Charlese Dela Cruz, Albert Ko, Nathan Grubaugh |
| EPI_ISL_469209, EPI_ISL_469210, EPI_ISL_469211, EPI_ISL_469212, EPI_ISL_469213, EPI_ISL_469214, EPI_ISL_469215, EPI_ISL_469216, EPI_ISL_469217, EPI_ISL_469218, EPI_ISL_469219, EPI_ISL_469220, EPI_ISL_469221, EPI_ISL_469222, EPI_ISL_469223 | | | |
| see above | BCCDC Public Health Laboratory | BCCDC Public Health Laboratory | Richard Harrigan, Hope Lapointe, Jinny Choi, Kimia Kamelian, John Tyson,Terry Snutch, Linda Hoang, Inna Sekirov, Paul Levett, Mel Krajden, Natalie Prystajecky |
| EPI_ISL_469224, EPI_ISL_469225, EPI_ISL_469226, EPI_ISL_469227, EPI_ISL_469228, EPI_ISL_469229, EPI_ISL_469230, EPI_ISL_469231, EPI_ISL_469232, EPI_ISL_469233, EPI_ISL_469234, EPI_ISL_469235, EPI_ISL_469236, EPI_ISL_469237, EPI_ISL_469238, EPI_ISL_469239, EPI_ISL_469240 | | | |
| see above | Public Health Laboratory | National Microbiology Laboratory | Anna Majer, Shari Tyson, Grace Seo, Kristyn Burak, Philip Mabon, Elsie Grudeski, Rhiannon Huzarewich, Russell Mandes, Jennifer Tanner, Natalie Knox, Morag Graham, Gary Van Domselaar, Robert Needle, Yang Yu, Adel Malek, Laura Gilbert, George Zahariadis, Nathalie Bastien, Yan Li, Timothy Booth, Matthew Gilmour |
| EPI_ISL_469254 | National Institute for Viral Disease Control and Prevention, China CDC | Institute of Viral Disease Control and Prevention, China CDC | Wenjie Tan, Lijuan Chen, Peihua NiuBaoying Huang, Li Zhao, Yubai Bi, Wenling Wang, Roujian Lu, Dayan Wang, Wenbo Xu, George Fu Gao, Chun Huang, Guizhen Wu |

| | | | |
|---|---|---|---|
| EPI_ISL_469255 | National Institute for Viral Disease Control and Prevention, China CDC | Institute of Viral Disease Control and Prevention, China CDC | Xiang ZhaoLijuan Chen, Dayan Wang, Yong Zhang, Yao MengZhixiao ChenYuchao Wu, Jun Han, Weifeng Shi, Yanhai Wang, William J. Liu, Shiwen Wang, George F. Gao, Wenbo Xu, Chun Huang, Guizhen Wu |
| EPI_ISL_469256 | National Institute for Viral Disease Control and Prevention, China CDC | National Institute for Viral Disease Control and Prevention, China CDC | Xiang ZhaoLijuan Chen, Dayan Wang, Yong Zhang, Yao MengZhixiao ChenYuchao Wu, Jun Han, Weifeng Shi, Yanhai Wang, William J. Liu, Shiwen Wang, George F. Gao, Wenbo Xu, Chun Huang, Guizhen Wu |
| EPI_ISL_469274 | National Public Health Laboratory, National Centre for Infectious Diseases | National Public Health Laboratory, National Centre for Infectious Diseases | Mak TM, Octavia S, Chavatte JM, Cui L, Lin RTP |
| EPI_ISL_469275 | Egyptian National Cancer Institute (ENCI) | Human Genome Center | Zekri, Abdel Rahman N, Amer,K.E., Ahmed,O.S., Soliman,H.K., Hafez,M.M., Bahnassy,A.A., Abdelhamid,W., Gad,A., Ali,M., Hassan,W., Samir,M., Raouf,A., Hamdy,M.S., Soliman,M.S., Elsissy,M.H., Elkhateeb,S.M., Ezzelarab,M.H., Abouelhoda, Mohamed |
| EPI_ISL_469276, EPI_ISL_469277, EPI_ISL_469278, EPI_ISL_469279, EPI_ISL_469280, EPI_ISL_469281 | Mohammed Bin Rashid University of Medicine and Health Sciences | Al Jalila Genomics Center | Ahmad Abou Tayoun, Tom Loney, Hamda Khansaheb, Sathishkumar Ramaswamy, Divinlal Harilal, Zulfa Omar Deesi, Rupa Murthy Varghese, Hanan Al Suwaidi, Abdulmajeed Alkhaja, Mohammed Uddin, Rifat Hamoudi, Rabih Halwani, Abiola Catherine Senok, Qutayba Hamid, Norbert Nowotny, Alawi Alsheikh-Ali |
| EPI_ISL_469282 | Service de Virologie Hôpital Saint-Louis | Laboratory of Cell Biology of viral infection, Unit INSERM-U944 | Laurent Meertens, Lucie Bonnet-Madin, Constance Delaugerre, Ali Amara |
| EPI_ISL_469283, EPI_ISL_469284 | Service de Virologie Hôpital Saint-Louis | Laboratory Cell Biology of Viral Infection-INSERM unit 944 | Laurent Meertens, Lucie Bonnet-Madin, Séverine Mercier-Delarue, Maud SALMONA, Constance Delaugerre, Ali Amara |
| EPI_ISL_469285 | National Institute of Laboratory Medicine and Referral Center | Genomic Research Lab, BCSIR | Shahina Akter, Abu Sayeed Mohammad Mahmud, Mohammad Samir Uzzaman, Eshrar Osman, Md. Ahasan Habib, Tanjina Akhter Banu, Md. Murshed Hasan Sarkar, Iffat Jahan, Barna Goswami, Md. Saddam Hossain, Tasnim Nafisa, Md. Maruf Ahmed Molla, Mahmuda Yeasmin, Asish Kumar Ghosh, Bayzid Bin Monir, A. K. M. Shamsuzzaman, Sheikh Md. Selim Al Din, Utpal Chandra Ray, Salek Ahmed Sajib, Md. Salim Khan |
| EPI_ISL_469286 | National Institute of Laboratory Medicine and Referral Center | Genomic Research Lab, BCSIR | Tanjina Akhter Banu, Abu Sayeed Mohammad Mahmud, Mohammad Samir Uzzaman, Eshrar Osman, Md. Ahasan Habib, Shahina Akter, Md. Murshed Hasan Sarkar, Iffat Jahan, Barna Goswami, Md. Saddam Hossain, Tasnim Nafisa, Md. Maruf Ahmed Molla, Mahmuda Yeasmin, Asish Kumar Ghosh, Bayzid Bin Monir, A. K. M. Shamsuzzaman, Sheikh Md. Selim Al Din, Utpal Chandra Ray, Salek Ahmed Sajib, Md. Salim Khan |
| EPI_ISL_469287, EPI_ISL_469288, EPI_ISL_469289, EPI_ISL_469290, EPI_ISL_469291, EPI_ISL_469292, EPI_ISL_469293, EPI_ISL_469294, EPI_ISL_469295, EPI_ISL_469296 | Keio University Hospital | Keio University Hospital | Kenjiro Kosaki |
| EPI_ISL_469297 | National Institute of Laboratory Medicine and Referral Center | Genomic Research Lab, BCSIR | Barna Goswami, Abu Sayeed Mohammad Mahmud, Mohammad Samir Uzzaman, Eshrar Osman, Md. Ahasan Habib, Shahina Akter, Tanjina Akhter Banu, Md. Murshed Hasan Sarkar, Iffat Jahan, Md. Saddam Hossain, Tasnim Nafisa, Md. Maruf Ahmed Molla, Mahmuda Yeasmin, Asish Kumar Ghosh, Bayzid Bin Monir, A. K. M. Shamsuzzaman, Sheikh Md. Selim Al Din, Utpal Chandra Ray, Salek Ahmed Sajib, Md. Salim Khan |
| EPI_ISL_469298 | National Institute of Laboratory Medicine and Referral Center | Genomic Research Lab, BCSIR | Md. Murshed Hasan Sarkar, Abu Sayeed Mohammad Mahmud, Mohammad Samir Uzzaman, Eshrar Osman, Md. Ahasan Habib, Shahina Akter, Tanjina Akhter Banu, Barna Goswami, Iffat Jahan, Md. Saddam Hossain, Tasnim Nafisa, Md. Maruf Ahmed Molla, Mahmuda Yeasmin, Asish Kumar Ghosh, Bayzid Bin Monir, A. K. M. Shamsuzzaman, Sheikh Md. Selim Al Din, Utpal Chandra Ray, Salek Ahmed Sajib, Md. Salim Khan |
| EPI_ISL_469299 | National Institute of Laboratory Medicine and Referral Center | Genomic Research Lab, BCSIR | Iffat Jahan, Abu Sayeed Mohammad Mahmud, Mohammad Samir Uzzaman, Eshrar Osman, Md. Ahasan Habib, Shahina Akter, Tanjina Akhter Banu, Md. Murshed Hasan Sarkar, Barna Goswami, Md. Saddam Hossain, Tasnim Nafisa, Md. Maruf Ahmed Molla, Mahmuda Yeasmin, Asish Kumar Ghosh, Bayzid Bin Monir, A. K. M. Shamsuzzaman, Sheikh Md. Selim Al Din, Utpal Chandra Ray, Salek Ahmed Sajib, Md. Salim Khan |
| EPI_ISL_469300 | National Institute of Laboratory Medicine and Referral Center | Genomic Research Lab, BCSIR | Abu Sayeed Mohammad Mahmud, Mohammad Samir Uzzaman, Eshrar Osman, Md. Ahasan Habib, Shahina Akter, Tanjina Akhter Banu, Md. Murshed Hasan Sarkar, Barna Goswami, Iffat Jahan, Md. Saddam Hossain, Tasnim Nafisa, Md. Maruf Ahmed Molla, Mahmuda Yeasmin, Asish Kumar Ghosh, Bayzid Bin Monir, A. K. M. Shamsuzzaman, Sheikh Md. Selim Al Din, Utpal Chandra Ray, Salek Ahmed Sajib, Md. Salim Khan |
| EPI_ISL_469302, EPI_ISL_469303, EPI_ISL_469304, EPI_ISL_469305, EPI_ISL_469306, EPI_ISL_469307, EPI_ISL_469308, EPI_ISL_469309, EPI_ISL_469310, EPI_ISL_469311, EPI_ISL_469312, EPI_ISL_469313, EPI_ISL_469314, EPI_ISL_469315, EPI_ISL_469316, EPI_ISL_469317, EPI_ISL_469318, EPI_ISL_469319, EPI_ISL_469320, EPI_ISL_469321, EPI_ISL_469322, EPI_ISL_469323, EPI_ISL_469324, EPI_ISL_469325, EPI_ISL_469326, EPI_ISL_469327, EPI_ISL_469328, EPI_ISL_469329, EPI_ISL_469330, EPI_ISL_469331, EPI_ISL_469332, EPI_ISL_469333, EPI_ISL_469334, EPI_ISL_469335, EPI_ISL_469336, EPI_ISL_469337, EPI_ISL_469338, EPI_ISL_469339, EPI_ISL_469340, EPI_ISL_469341, EPI_ISL_469342, EPI_ISL_469343, EPI_ISL_469344 | | | |
| see above | NU-OMICS DNA Sequencing research facility, Northumbria University | Wellcome Sanger Institute for the COVID-19 Genomics UK Consortium | Chris Duncan, Sheia Waugh, Shirelle Burton-Fanning, Gary Eltringham, Jennifer Collins, Brendan Payne, Yusri Taha, Emma Swindells, Jane Greenaway, Edward Barton, Garren Scott, Debra Padgett, Clive Graham, Sarah Essex, Steve Ligget, Paul Baker, Lynn Dover, Wen Yew, Gary Black, John Allan, Joshua Loh, Greg Young, Matthew Bashton, Andrew Nelson, Darren Smith and Alex Alderton, Roberto Amato, Sonia Goncalves, Ewan Harrison, David K. Jackson, Ian Johnston, Dominic Kwiatkowski, Cordelia Langford, John Sillitoe on behalf of the Wellcome Sanger Institute COVID-19 Surveillance Team (http://www.sanger.ac.uk/covid-team) |
| EPI_ISL_469345, EPI_ISL_469346, EPI_ISL_469347, EPI_ISL_469348, EPI_ISL_469349, EPI_ISL_469350, EPI_ISL_469351, EPI_ISL_469352, EPI_ISL_469353, EPI_ISL_469354, EPI_ISL_469355, EPI_ISL_469356, EPI_ISL_469357, EPI_ISL_469358, EPI_ISL_469359, EPI_ISL_469360, EPI_ISL_469361, EPI_ISL_469362, EPI_ISL_469363, EPI_ISL_469364, EPI_ISL_469365, EPI_ISL_469366, EPI_ISL_469367, EPI_ISL_469368, EPI_ISL_469369, EPI_ISL_469370, EPI_ISL_469371, EPI_ISL_469372, EPI_ISL_469373, EPI_ISL_469374, EPI_ISL_469375, EPI_ISL_469376, EPI_ISL_469377, EPI_ISL_469378, EPI_ISL_469379, EPI_ISL_469380, EPI_ISL_469381, EPI_ISL_469382, EPI_ISL_469383, EPI_ISL_469384, EPI_ISL_469385, EPI_ISL_469386, EPI_ISL_469387, EPI_ISL_469388, EPI_ISL_469389, EPI_ISL_469390, EPI_ISL_469391, EPI_ISL_469392, EPI_ISL_469393, EPI_ISL_469394, EPI_ISL_469395, EPI_ISL_469396, EPI_ISL_469397, EPI_ISL_469398, EPI_ISL_469399, EPI_ISL_469400, EPI_ISL_469401, EPI_ISL_469402, EPI_ISL_469403, EPI_ISL_469404, EPI_ISL_469405, EPI_ISL_469406, EPI_ISL_469407, EPI_ISL_469408, EPI_ISL_469409, EPI_ISL_469410, EPI_ISL_469411, EPI_ISL_469412, EPI_ISL_469413, EPI_ISL_469414, EPI_ISL_469415, EPI_ISL_469416, EPI_ISL_469417, EPI_ISL_469418, EPI_ISL_469419, EPI_ISL_469420, EPI_ISL_469421, EPI_ISL_469422, EPI_ISL_469423, EPI_ISL_469424, EPI_ISL_469425, EPI_ISL_469426, EPI_ISL_469427, EPI_ISL_469428, EPI_ISL_469429, EPI_ISL_469430, EPI_ISL_469431, EPI_ISL_469432, EPI_ISL_469433, EPI_ISL_469434, EPI_ISL_469435, EPI_ISL_469436, EPI_ISL_469437, EPI_ISL_469438, EPI_ISL_469439, EPI_ISL_469440, EPI_ISL_469441, EPI_ISL_469442, EPI_ISL_469443, EPI_ISL_469444, EPI_ISL_469445, EPI_ISL_469446, EPI_ISL_469447, EPI_ISL_469448, EPI_ISL_469449, EPI_ISL_469450, EPI_ISL_469451, EPI_ISL_469452, EPI_ISL_469453, EPI_ISL_469454, EPI_ISL_469455, EPI_ISL_469456, EPI_ISL_469457, EPI_ISL_469458, EPI_ISL_469459, EPI_ISL_469460, EPI_ISL_469461, EPI_ISL_469462, EPI_ISL_469463, EPI_ISL_469464, EPI_ISL_469465, EPI_ISL_469466, EPI_ISL_469467, EPI_ISL_469468, EPI_ISL_469469, EPI_ISL_469470, EPI_ISL_469471, EPI_ISL_469472, EPI_ISL_469473, EPI_ISL_469474, EPI_ISL_469475, EPI_ISL_469476, EPI_ISL_469477, EPI_ISL_469478, EPI_ISL_469479, EPI_ISL_469480, EPI_ISL_469481, EPI_ISL_469482, EPI_ISL_469483, EPI_ISL_469484, EPI_ISL_469485, EPI_ISL_469486, EPI_ISL_469487, EPI_ISL_469488, EPI_ISL_469489, EPI_ISL_469490, EPI_ISL_469491, EPI_ISL_469492, EPI_ISL_469493, EPI_ISL_469494, EPI_ISL_469495, EPI_ISL_469496, EPI_ISL_469497, EPI_ISL_469498, EPI_ISL_469499, EPI_ISL_469500, EPI_ISL_469501, EPI_ISL_469502, EPI_ISL_469503, EPI_ISL_469504, EPI_ISL_469505, EPI_ISL_469506, EPI_ISL_469507, EPI_ISL_469508, EPI_ISL_469509, EPI_ISL_469510, EPI_ISL_469511, EPI_ISL_469512, EPI_ISL_469513, EPI_ISL_469514, EPI_ISL_469515, EPI_ISL_469516, EPI_ISL_469517, EPI_ISL_469518, EPI_ISL_469519, EPI_ISL_469520, EPI_ISL_469521, EPI_ISL_469522, EPI_ISL_469523, EPI_ISL_469524, EPI_ISL_469525, EPI_ISL_469526, EPI_ISL_469527, EPI_ISL_469528, EPI_ISL_469529, EPI_ISL_469530, EPI_ISL_469531, EPI_ISL_469532, EPI_ISL_469533, EPI_ISL_469534, EPI_ISL_469535, EPI_ISL_469536, EPI_ISL_469537, EPI_ISL_469538, EPI_ISL_469539, EPI_ISL_469540, EPI_ISL_469541, EPI_ISL_469542, EPI_ISL_469543, EPI_ISL_469544, EPI_ISL_469545, EPI_ISL_469546, EPI_ISL_469547, EPI_ISL_469548, EPI_ISL_469549, EPI_ISL_469550, EPI_ISL_469551, EPI_ISL_469552, EPI_ISL_469553, EPI_ISL_469554, EPI_ISL_469555, EPI_ISL_469556, EPI_ISL_469557, EPI_ISL_469558, EPI_ISL_469559, EPI_ISL_469560, EPI_ISL_469561, EPI_ISL_469562, EPI_ISL_469563, EPI_ISL_469564, EPI_ISL_469565, EPI_ISL_469566, EPI_ISL_469567, EPI_ISL_469568, EPI_ISL_469569, EPI_ISL_469570, EPI_ISL_469571, EPI_ISL_469572, EPI_ISL_469573, EPI_ISL_469574, EPI_ISL_469575, EPI_ISL_469576, EPI_ISL_469577, EPI_ISL_469578, EPI_ISL_469579, EPI_ISL_469580, EPI_ISL_469581, EPI_ISL_469582, EPI_ISL_469583, EPI_ISL_469584, EPI_ISL_469585, EPI_ISL_469586, EPI_ISL_469587, EPI_ISL_469588, EPI_ISL_469589, EPI_ISL_469590, EPI_ISL_469591, EPI_ISL_469592, EPI_ISL_469593, EPI_ISL_469594, EPI_ISL_469595, EPI_ISL_469596, EPI_ISL_469597, EPI_ISL_469598, EPI_ISL_469599, EPI_ISL_469600, EPI_ISL_469601, EPI_ISL_469602, EPI_ISL_469603, EPI_ISL_469604, EPI_ISL_469605, EPI_ISL_469606, EPI_ISL_469607, EPI_ISL_469608, EPI_ISL_469609, EPI_ISL_469610, EPI_ISL_469611, EPI_ISL_469612, EPI_ISL_469613, EPI_ISL_469614, EPI_ISL_469615, EPI_ISL_469616, EPI_ISL_469617, EPI_ISL_469618, EPI_ISL_469619, EPI_ISL_469620, EPI_ISL_469621, EPI_ISL_469622, EPI_ISL_469623, EPI_ISL_469624, EPI_ISL_469625, EPI_ISL_469626, EPI_ISL_469627, EPI_ISL_469628, EPI_ISL_469629, EPI_ISL_469630, EPI_ISL_469631, EPI_ISL_469632, EPI_ISL_469633, EPI_ISL_469634, EPI_ISL_469635, EPI_ISL_469636, EPI_ISL_469637, EPI_ISL_469638, EPI_ISL_469639, EPI_ISL_469640, EPI_ISL_469641, EPI_ISL_469642, EPI_ISL_469643, EPI_ISL_469644, EPI_ISL_469645, EPI_ISL_469646, EPI_ISL_469647, EPI_ISL_469648, EPI_ISL_469649, EPI_ISL_469650, EPI_ISL_469651, EPI_ISL_469652, EPI_ISL_469653, EPI_ISL_469654, EPI_ISL_469655, EPI_ISL_469656, EPI_ISL_469657, EPI_ISL_469658, EPI_ISL_469659, EPI_ISL_469660, EPI_ISL_469661, EPI_ISL_469662, EPI_ISL_469663, EPI_ISL_469664, EPI_ISL_469665, EPI_ISL_469666, EPI_ISL_469667, EPI_ISL_469668, EPI_ISL_469669, EPI_ISL_469670, EPI_ISL_469671, EPI_ISL_469672, EPI_ISL_469673, EPI_ISL_469674, EPI_ISL_469675, EPI_ISL_469676, EPI_ISL_469677, EPI_ISL_469678, EPI_ISL_469679, EPI_ISL_469680, EPI_ISL_469681, EPI_ISL_469682, EPI_ISL_469683, EPI_ISL_469684, EPI_ISL_469685, EPI_ISL_469686, EPI_ISL_469687, EPI_ISL_469688, EPI_ISL_469689, EPI_ISL_469690, EPI_ISL_469691, EPI_ISL_469692, EPI_ISL_469693, EPI_ISL_469694, EPI_ISL_469695, EPI_ISL_469696, EPI_ISL_469697, EPI_ISL_469698, EPI_ISL_469699, EPI_ISL_469700, EPI_ISL_469701, EPI_ISL_469702, EPI_ISL_469703, EPI_ISL_469704, EPI_ISL_469705, EPI_ISL_469706, EPI_ISL_469707, EPI_ISL_469708, EPI_ISL_469709, EPI_ISL_469710, EPI_ISL_469711, EPI_ISL_469712, EPI_ISL_469713, EPI_ISL_469714, EPI_ISL_469715, EPI_ISL_469716, EPI_ISL_469717, EPI_ISL_469718, EPI_ISL_469719, EPI_ISL_469720, EPI_ISL_469721, EPI_ISL_469722, EPI_ISL_469723, EPI_ISL_469724, EPI_ISL_469725, EPI_ISL_469726, EPI_ISL_469727, EPI_ISL_469728, EPI_ISL_469729, EPI_ISL_469730, EPI_ISL_469731, EPI_ISL_469732, EPI_ISL_469733, EPI_ISL_469734, EPI_ISL_469735, EPI_ISL_469736, EPI_ISL_469737, EPI_ISL_469738, EPI_ISL_469739, EPI_ISL_469740, EPI_ISL_469741, EPI_ISL_469742, EPI_ISL_469743, EPI_ISL_469744, EPI_ISL_469745, EPI_ISL_469746, EPI_ISL_469747, EPI_ISL_469748, EPI_ISL_469749, EPI_ISL_469750, EPI_ISL_469751, EPI_ISL_469752, EPI_ISL_469753, EPI_ISL_469754, EPI_ISL_469755, EPI_ISL_469756, EPI_ISL_469757, EPI_ISL_469758, EPI_ISL_469759, EPI_ISL_469760, EPI_ISL_469761, EPI_ISL_469762, EPI_ISL_469763, EPI_ISL_469764, EPI_ISL_469765, EPI_ISL_469766, EPI_ISL_469767, EPI_ISL_469768, EPI_ISL_469769, EPI_ISL_469770, EPI_ISL_469771, EPI_ISL_469772, EPI_ISL_469773, EPI_ISL_469774, EPI_ISL_469775, EPI_ISL_469776, EPI_ISL_469777, EPI_ISL_469778, EPI_ISL_469779, EPI_ISL_469780, EPI_ISL_469781, EPI_ISL_469782, EPI_ISL_469783, EPI_ISL_469784, EPI_ISL_469785, EPI_ISL_469786, EPI_ISL_469787, EPI_ISL_469788, EPI_ISL_469789, EPI_ISL_469790, EPI_ISL_469791, EPI_ISL_469792, EPI_ISL_469793, EPI_ISL_469794, EPI_ISL_469795, EPI_ISL_469796, EPI_ISL_469797, EPI_ISL_469798, EPI_ISL_469799, EPI_ISL_469800, EPI_ISL_469801, EPI_ISL_469802, EPI_ISL_469803, EPI_ISL_469804 | | | |
| see above | PHE South West Regional Laboratory, National Infection Service | Wellcome Sanger Institute for the COVID-19 Genomics UK Consortium | Stephanie Hutchings, Hannah Pymont, Dr Peter Muir, Barry Vipond, Rich Hopes; and Alex Alderton, Roberto Amato, Sonia Goncalves, Ewan Harrison, David K. Jackson, Ian Johnston, Dominic Kwiatkowski, Cordelia Langford, John Sillitoe on behalf of the Wellcome Sanger Institute COVID-19 Surveillance Team (http://www.sanger.ac.uk/covid-team) |
| EPI_ISL_469805 | NU-OMICS DNA Sequencing research facility, Northumbria University | Wellcome Sanger Institute for the COVID-19 Genomics UK Consortium | Chris Duncan, Sheia Waugh, Shirelle Burton-Fanning, Gary Eltringham, Jennifer Collins, Brendan Payne, Yusri Taha, Emma Swindells, Jane Greenaway, Edward Barton, Garren Scott, Debra Padgett, Clive Graham, Sarah Essex, Steve Ligget, Paul Baker, Lynn Dover, Wen Yew, Gary Black, John Allan, Joshua Loh, Greg Young, Matthew Bashton, Andrew Nelson, Darren Smith and Alex Alderton, Roberto Amato, Sonia Goncalves, Ewan Harrison, David K. Jackson, Ian Johnston, Dominic Kwiatkowski, Cordelia Langford, John Sillitoe on behalf of the Wellcome Sanger Institute COVID-19 Surveillance Team (http://www.sanger.ac.uk/covid-team) |
| EPI_ISL_469806, EPI_ISL_469807, EPI_ISL_469808, EPI_ISL_469809, EPI_ISL_469810, EPI_ISL_469811, EPI_ISL_469812, EPI_ISL_469813, EPI_ISL_469814, EPI_ISL_469815, EPI_ISL_469816, EPI_ISL_469817, EPI_ISL_469818, EPI_ISL_469819, EPI_ISL_469820, EPI_ISL_469821, EPI_ISL_469822, EPI_ISL_469823, EPI_ISL_469824, EPI_ISL_469825, EPI_ISL_469826, EPI_ISL_469827, EPI_ISL_469828, EPI_ISL_469829, EPI_ISL_469830, EPI_ISL_469831, EPI_ISL_469832, EPI_ISL_469833, EPI_ISL_469834, EPI_ISL_469835, EPI_ISL_469836, EPI_ISL_469837, EPI_ISL_469838, EPI_ISL_469839, EPI_ISL_469840, EPI_ISL_469841, EPI_ISL_469842, EPI_ISL_469843 | | | |
| see above | PHE South West Regional Laboratory, National Infection Service | Wellcome Sanger Institute for the COVID-19 Genomics UK Consortium | Stephanie Hutchings, Hannah Pymont, Dr Peter Muir, Barry Vipond, Rich Hopes; and Alex Alderton, Roberto Amato, Sonia Goncalves, Ewan Harrison, David K. Jackson, Ian Johnston, Dominic Kwiatkowski, Cordelia Langford, John Sillitoe on behalf of the Wellcome Sanger Institute COVID-19 Surveillance Team (http://www.sanger.ac.uk/covid-team) |
| EPI_ISL_469844, EPI_ISL_469845, EPI_ISL_469846, EPI_ISL_469847, EPI_ISL_469848, EPI_ISL_469849, EPI_ISL_469850, EPI_ISL_469851, EPI_ISL_469852, EPI_ISL_469853, EPI_ISL_469854, EPI_ISL_469855, EPI_ISL_469856, EPI_ISL_469857, EPI_ISL_469858, EPI_ISL_469859, EPI_ISL_469860, EPI_ISL_469861, EPI_ISL_469862, EPI_ISL_469863, EPI_ISL_469864, EPI_ISL_469865, EPI_ISL_469866, EPI_ISL_469867, EPI_ISL_469868, EPI_ISL_469869, EPI_ISL_469870, EPI_ISL_469871, EPI_ISL_469872, EPI_ISL_469873, EPI_ISL_469874, EPI_ISL_469875, EPI_ISL_469876, EPI_ISL_469877, EPI_ISL_469878, EPI_ISL_469879, EPI_ISL_469880, EPI_ISL_469881, EPI_ISL_469882, EPI_ISL_469883, EPI_ISL_469884, EPI_ISL_469885, EPI_ISL_469886, EPI_ISL_469887, EPI_ISL_469888, EPI_ISL_469889, EPI_ISL_469890 | | | |
| see above | Regional Virus Laboratory, Belfast Health and Social Care Trust | Wellcome Sanger Institute for the COVID-19 Genomics UK Consortium | Conall McCaughey, James McKenna, Tanya Curran, Susan Feeney, Alison Watt, Ciara Cox, Mairead Connor, Zoltan Molnar, David Simpson, Derek Fairley; and Alex Alderton, Roberto Amato, Sonia Goncalves, Ewan Harrison, David K. Jackson, Ian Johnston, Dominic Kwiatkowski, Cordelia Langford, John Sillitoe on behalf of the Wellcome Sanger Institute COVID-19 Surveillance Team (http://www.sanger.ac.uk/covid-team) |
| EPI_ISL_469891, EPI_ISL_469892, EPI_ISL_469893, EPI_ISL_469894, EPI_ISL_469895, EPI_ISL_469896, EPI_ISL_469897, EPI_ISL_469898, EPI_ISL_469899, EPI_ISL_469900, EPI_ISL_469901, EPI_ISL_469902, EPI_ISL_469903, EPI_ISL_469904, EPI_ISL_469905, EPI_ISL_469906, EPI_ISL_469907, EPI_ISL_469908, EPI_ISL_469909, EPI_ISL_469910, EPI_ISL_469911, EPI_ISL_469912, EPI_ISL_469913 | | | |
| see above | Department of Pathology, University of Cambridge | Wellcome Sanger Institute for the COVID-19 Genomics UK Consortium | Luke W Meredith, M. Estée Török , Myra Hosmillo, William L. Hamilton, Martin D. Curran, Theresa Feltwell, Grant Hall, Anna Yakovleva, Fahad A Khokhar, Charlotte J. Houldcroft, Laura G Caller, Aminu S. Jahun, Sarah L. Caddy, Ian Goodfellow; and Alex Alderton, Roberto Amato, Sonia Goncalves, Ewan Harrison, David K. Jackson, Ian Johnston, Dominic Kwiatkowski, Cordelia Langford, John Sillitoe on behalf of the Wellcome Sanger Institute COVID-19 Surveillance Team (http://www.sanger.ac.uk/covid-team) |
| EPI_ISL_469914 | PHE South West Regional Laboratory, National Infection Service | Wellcome Sanger Institute for the COVID-19 Genomics UK Consortium | Stephanie Hutchings, Hannah Pymont, Dr Peter Muir, Barry Vipond, Rich Hopes; and Alex Alderton, Roberto Amato, Sonia Goncalves, Ewan Harrison, David K. Jackson, Ian Johnston, Dominic Kwiatkowski, Cordelia Langford, John Sillitoe on behalf of the Wellcome Sanger Institute COVID-19 Surveillance Team (http://www.sanger.ac.uk/covid-team) |
| EPI_ISL_469915 | NU-OMICS DNA Sequencing research facility, Northumbria University | Wellcome Sanger Institute for the COVID-19 Genomics UK Consortium | Chris Duncan, Sheia Waugh, Shirelle Burton-Fanning, Gary Eltringham, Jennifer Collins, Brendan Payne, Yusri Taha, Emma Swindells, Jane Greenaway, Edward Barton, Garren Scott, Debra Padgett, Clive Graham, Sarah Essex, Steve Ligget, Paul Baker, Lynn Dover, Wen Yew, Gary Black, John Allan, Joshua Loh, Greg Young, Matthew Bashton, Andrew Nelson, Darren Smith and Alex Alderton, Roberto Amato, Sonia Goncalves, Ewan Harrison, David K. Jackson, Ian Johnston, Dominic Kwiatkowski, Cordelia Langford, John Sillitoe on behalf of the Wellcome Sanger Institute COVID-19 Surveillance Team (http://www.sanger.ac.uk/covid-team) |
| EPI_ISL_469916, EPI_ISL_469917 | PHE South West Regional Laboratory, National Infection Service | Wellcome Sanger Institute for the COVID-19 Genomics UK Consortium | Stephanie Hutchings, Hannah Pymont, Dr Peter Muir, Barry Vipond, Rich Hopes; and Alex Alderton, Roberto Amato, Sonia Goncalves, Ewan Harrison, David K. Jackson, Ian Johnston, Dominic Kwiatkowski, Cordelia Langford, John Sillitoe on behalf of the Wellcome Sanger Institute COVID-19 Surveillance Team (http://www.sanger.ac.uk/covid-team) |
| EPI_ISL_469918, EPI_ISL_469919, EPI_ISL_469920, EPI_ISL_469921, EPI_ISL_469922, EPI_ISL_469923, EPI_ISL_469924, EPI_ISL_469925, EPI_ISL_469926, EPI_ISL_469927, EPI_ISL_469928, EPI_ISL_469929 | | | |
| see above | Virology Department, Sheffield Teaching Hospitals NHS Foundation Trust / Department of Infection, Immunity and Cardiovascular Disease, The Medical School, University of Sheffield | Wellcome Sanger Institute for the COVID-19 Genomics UK Consortium | Thushan de Silva, Matthew Parker,Adri Angyal, Rebecca Brown, Luke Green, Rachel Tucker, Paul Parsons, Danielle Groves, Alex Keeley, Dave Partridge, Matthew Wyles, Benjamin Lindsey, Mehmet Yavuz, Mohammad Raza, Cariad Evans and Alex Alderton, Roberto Amato, Sonia Goncalves, Ewan Harrison, David K. Jackson, Ian Johnston, Dominic Kwiatkowski, Cordelia Langford, John Sillitoe on behalf of the Wellcome Sanger Institute COVID-19 Surveillance Team (http://www.sanger.ac.uk/covid-team) |
| EPI_ISL_469930, EPI_ISL_469931, EPI_ISL_469932, EPI_ISL_469933, EPI_ISL_469934, EPI_ISL_469935, EPI_ISL_469936, EPI_ISL_469937, EPI_ISL_469938, EPI_ISL_469939, EPI_ISL_469940, EPI_ISL_469941, EPI_ISL_469942, EPI_ISL_469943, EPI_ISL_469944, EPI_ISL_469945, EPI_ISL_469946, EPI_ISL_469947, EPI_ISL_469948, EPI_ISL_469949, EPI_ISL_469950, EPI_ISL_469951, EPI_ISL_469952, EPI_ISL_469953, EPI_ISL_469954, EPI_ISL_469955, EPI_ISL_469956, EPI_ISL_469957, EPI_ISL_469958, EPI_ISL_469959, EPI_ISL_469960, EPI_ISL_469961, EPI_ISL_469962, EPI_ISL_469963, EPI_ISL_469964, EPI_ISL_469965, EPI_ISL_469966, EPI_ISL_469967, EPI_ISL_469968, EPI_ISL_469969, EPI_ISL_469970, EPI_ISL_469971, EPI_ISL_469972, EPI_ISL_469973, EPI_ISL_469974, EPI_ISL_469975, EPI_ISL_469976, EPI_ISL_469977, EPI_ISL_469978, EPI_ISL_469979, EPI_ISL_469980, EPI_ISL_469981, EPI_ISL_469982, EPI_ISL_469983, EPI_ISL_469984, EPI_ISL_469985, EPI_ISL_469986, EPI_ISL_469987, EPI_ISL_469988, EPI_ISL_469989, EPI_ISL_469990, EPI_ISL_469991, EPI_ISL_469992, EPI_ISL_469993, EPI_ISL_469994, EPI_ISL_469995, EPI_ISL_469996, EPI_ISL_469997, EPI_ISL_469998, EPI_ISL_469999, EPI_ISL_470000, EPI_ISL_470001, EPI_ISL_470002, EPI_ISL_470003, EPI_ISL_470004, EPI_ISL_470005, EPI_ISL_470006, EPI_ISL_470007, EPI_ISL_470008, EPI_ISL_470009, EPI_ISL_470010, EPI_ISL_470011, EPI_ISL_470012 | | | |
| see above | NHSGGC West of Scotland Specialist Virology Centre / MRC-University of Glasgow Centre for Virus Research | Wellcome Sanger Institute for the COVID-19 Genomics UK Consortium | Ana da Silva Filipe, Natasha Johnson, Kathy Smollett, Daniel Mair, Stephen Carmichael, Lily Tong, Jenna Nichols, Elihu Aranday-Cortes, Kirstyn Brunker, Yasmin Parr, Kyriaki Nomikou; Sarah McDonald, Marc Niebel, Patawee Asamaphan; Richard Orton, Joseph Hughes, Sreenu Vattipally, David L Robertson; Alasdair MacLean, Rory Gunson; Kathy Li, Natasha Jesudason, Rajiv Shah, James Shepherd, Antonia Ho, Alice Broos, Emma Thomson and Alex Alderton, Roberto Amato, Sonia Goncalves, Ewan Harrison, David K. Jackson, Ian Johnston, Dominic Kwiatkowski, Cordelia Langford, John Sillitoe on behalf of the Wellcome Sanger Institute COVID-19 Surveillance Team (http://www.sanger.ac.uk/covid-team) |
| EPI_ISL_470013, EPI_ISL_470014, EPI_ISL_470015, EPI_ISL_470016, EPI_ISL_470017, EPI_ISL_470018, EPI_ISL_470019, EPI_ISL_470020, EPI_ISL_470021, EPI_ISL_470022, EPI_ISL_470023, EPI_ISL_470024, EPI_ISL_470025, EPI_ISL_470026, EPI_ISL_470027, EPI_ISL_470028, EPI_ISL_470029, EPI_ISL_470030, EPI_ISL_470031, EPI_ISL_470032, EPI_ISL_470033, EPI_ISL_470034, EPI_ISL_470035, EPI_ISL_470036, EPI_ISL_470037, EPI_ISL_470038, EPI_ISL_470039, EPI_ISL_470040, EPI_ISL_470041, EPI_ISL_470042, EPI_ISL_470043, EPI_ISL_470044, EPI_ISL_470045, EPI_ISL_470046, EPI_ISL_470047, EPI_ISL_470048, EPI_ISL_470049, EPI_ISL_470050, EPI_ISL_470051, EPI_ISL_470052, EPI_ISL_470053, EPI_ISL_470054, EPI_ISL_470055, EPI_ISL_470056, EPI_ISL_470057, EPI_ISL_470058, EPI_ISL_470059, EPI_ISL_470060, EPI_ISL_470061, EPI_ISL_470062, EPI_ISL_470063, EPI_ISL_470064, EPI_ISL_470065, EPI_ISL_470066, EPI_ISL_470067, EPI_ISL_470068, EPI_ISL_470069, EPI_ISL_470070, EPI_ISL_470071, EPI_ISL_470072, EPI_ISL_470073, EPI_ISL_470074, EPI_ISL_470075, EPI_ISL_470076, EPI_ISL_470077, EPI_ISL_470078, EPI_ISL_470079, EPI_ISL_470080, EPI_ISL_470081, EPI_ISL_470082, EPI_ISL_470083, EPI_ISL_470084, EPI_ISL_470085, EPI_ISL_470086, EPI_ISL_470087, EPI_ISL_470088, EPI_ISL_470089 | | | |

| | | | |
|---|---|---|---|
| see above | Regional Virus Laboratory, Belfast Health and Social Care Trust | Wellcome Sanger Institute for the COVID-19 Genomics UK Consortium | Conall McCaughey, James McKenna, Tanya Curran, Susan Feeney, Alison Watt, Ciara Cox, Mairead Connor, Zoltan Molnar, David Simpson, Derek Fairley; and Alex Alderton, Roberto Amato, Sonia Goncalves, Ewan Harrison, David K. Jackson, Ian Johnston, Dominic Kwiatkowski, Cordelia Langford, John Sillitoe on behalf of the Wellcome Sanger Institute COVID-19 Surveillance Team (http://www.sanger.ac.uk/covid-team) |
| EPI_ISL_470090, EPI_ISL_470091, EPI_ISL_470092, EPI_ISL_470093, EPI_ISL_470094, EPI_ISL_470095, EPI_ISL_470096, EPI_ISL_470097, EPI_ISL_470098, EPI_ISL_470099, EPI_ISL_470100, EPI_ISL_470101, EPI_ISL_470102, EPI_ISL_470103, EPI_ISL_470104, EPI_ISL_470105, EPI_ISL_470106, EPI_ISL_470107, EPI_ISL_470108, EPI_ISL_470109, EPI_ISL_470110, EPI_ISL_470111, EPI_ISL_470112, EPI_ISL_470113, EPI_ISL_470114, EPI_ISL_470115, EPI_ISL_470116, EPI_ISL_470117, EPI_ISL_470118, EPI_ISL_470119, EPI_ISL_470120, EPI_ISL_470121, EPI_ISL_470122, EPI_ISL_470123, EPI_ISL_470124, EPI_ISL_470125, EPI_ISL_470126, EPI_ISL_470127, EPI_ISL_470128, EPI_ISL_470129, EPI_ISL_470130, EPI_ISL_470131, EPI_ISL_470132, EPI_ISL_470133, EPI_ISL_470134, EPI_ISL_470135, EPI_ISL_470136, EPI_ISL_470137, EPI_ISL_470138, EPI_ISL_470139, EPI_ISL_470140, EPI_ISL_470141, EPI_ISL_470142, EPI_ISL_470143, EPI_ISL_470144, EPI_ISL_470145, EPI_ISL_470146, EPI_ISL_470147, EPI_ISL_470148, EPI_ISL_470149, EPI_ISL_470150, EPI_ISL_470151, EPI_ISL_470152, EPI_ISL_470153, EPI_ISL_470154, EPI_ISL_470155, EPI_ISL_470156, EPI_ISL_470157, EPI_ISL_470158, EPI_ISL_470159, EPI_ISL_470160, EPI_ISL_470161, EPI_ISL_470162, EPI_ISL_470163, EPI_ISL_470164, EPI_ISL_470165, EPI_ISL_470166, EPI_ISL_470167, EPI_ISL_470168, EPI_ISL_470169, EPI_ISL_470170, EPI_ISL_470171, EPI_ISL_470172, EPI_ISL_470173, EPI_ISL_470174, EPI_ISL_470175, EPI_ISL_470176, EPI_ISL_470177, EPI_ISL_470178, EPI_ISL_470179, EPI_ISL_470180, EPI_ISL_470181, EPI_ISL_470182, EPI_ISL_470183, EPI_ISL_470184, EPI_ISL_470185, EPI_ISL_470186, EPI_ISL_470187, EPI_ISL_470188, EPI_ISL_470189, EPI_ISL_470190, EPI_ISL_470191, EPI_ISL_470192, EPI_ISL_470193, EPI_ISL_470194, EPI_ISL_470195, EPI_ISL_470196, EPI_ISL_470197, EPI_ISL_470198, EPI_ISL_470199, EPI_ISL_470200, EPI_ISL_470201, EPI_ISL_470202, EPI_ISL_470203, EPI_ISL_470204, EPI_ISL_470205, EPI_ISL_470206, EPI_ISL_470207, EPI_ISL_470208, EPI_ISL_470209, EPI_ISL_470210, EPI_ISL_470211, EPI_ISL_470212, EPI_ISL_470213, EPI_ISL_470214, EPI_ISL_470215, EPI_ISL_470216, EPI_ISL_470217, EPI_ISL_470218, EPI_ISL_470219, EPI_ISL_470220, EPI_ISL_470221, EPI_ISL_470222, EPI_ISL_470223, EPI_ISL_470224, EPI_ISL_470225, EPI_ISL_470226, EPI_ISL_470227, EPI_ISL_470228, EPI_ISL_470229, EPI_ISL_470230, EPI_ISL_470231, EPI_ISL_470232, EPI_ISL_470233, EPI_ISL_470234, EPI_ISL_470235, EPI_ISL_470236, EPI_ISL_470237, EPI_ISL_470238, EPI_ISL_470239, EPI_ISL_470240, EPI_ISL_470241, EPI_ISL_470242, EPI_ISL_470243, EPI_ISL_470244, EPI_ISL_470245, EPI_ISL_470246, EPI_ISL_470247, EPI_ISL_470248, EPI_ISL_470249, EPI_ISL_470250, EPI_ISL_470251, EPI_ISL_470252, EPI_ISL_470253, EPI_ISL_470254, EPI_ISL_470255, EPI_ISL_470256, EPI_ISL_470257, EPI_ISL_470258, EPI_ISL_470259, EPI_ISL_470260, EPI_ISL_470261, EPI_ISL_470262, EPI_ISL_470263, EPI_ISL_470264, EPI_ISL_470265, EPI_ISL_470266, EPI_ISL_470267, EPI_ISL_470268, EPI_ISL_470269, EPI_ISL_470270, EPI_ISL_470271, EPI_ISL_470272, EPI_ISL_470273, EPI_ISL_470274, EPI_ISL_470275, EPI_ISL_470276, EPI_ISL_470277, EPI_ISL_470278, EPI_ISL_470279, EPI_ISL_470280, EPI_ISL_470281, EPI_ISL_470282, EPI_ISL_470283, EPI_ISL_470284, EPI_ISL_470285, EPI_ISL_470286, EPI_ISL_470287, EPI_ISL_470288, EPI_ISL_470289, EPI_ISL_470290, EPI_ISL_470291, EPI_ISL_470292, EPI_ISL_470293, EPI_ISL_470294, EPI_ISL_470295, EPI_ISL_470296, EPI_ISL_470297, EPI_ISL_470298, EPI_ISL_470299, EPI_ISL_470300, EPI_ISL_470301, EPI_ISL_470302, EPI_ISL_470303, EPI_ISL_470304, EPI_ISL_470305, EPI_ISL_470306, EPI_ISL_470307, EPI_ISL_470308, EPI_ISL_470309, EPI_ISL_470310, EPI_ISL_470311, EPI_ISL_470312, EPI_ISL_470313, EPI_ISL_470314, EPI_ISL_470315, EPI_ISL_470316, EPI_ISL_470317, EPI_ISL_470318, EPI_ISL_470319, EPI_ISL_470320, EPI_ISL_470321, EPI_ISL_470322, EPI_ISL_470323, EPI_ISL_470324, EPI_ISL_470325, EPI_ISL_470326, EPI_ISL_470327, EPI_ISL_470328, EPI_ISL_470329, EPI_ISL_470330, EPI_ISL_470331, EPI_ISL_470332, EPI_ISL_470333, EPI_ISL_470334, EPI_ISL_470335, EPI_ISL_470336, EPI_ISL_470337, EPI_ISL_470338, EPI_ISL_470339, EPI_ISL_470340, EPI_ISL_470341, EPI_ISL_470342, EPI_ISL_470343, EPI_ISL_470344, EPI_ISL_470345, EPI_ISL_470346, EPI_ISL_470347, EPI_ISL_470348, EPI_ISL_470349, EPI_ISL_470350, EPI_ISL_470351, EPI_ISL_470352, EPI_ISL_470353, EPI_ISL_470354, EPI_ISL_470355 | | | |
| see above | Department of Pathology, University of Cambridge | Wellcome Sanger Institute for the COVID-19 Genomics UK Consortium | Luke W Meredith, M. Estée Török , Myra Hosmillo, William L. Hamilton, Martin D. Curran, Theresa Feltwell, Grant Hall, Anna Yakovleva, Fahad A Khokhar, Charlotte J. Houldcroft, Laura G Caller, Aminu S. Jahun, Sarah L. Caddy, Ian Goodfellow; and Alex Alderton, Roberto Amato, Sonia Goncalves, Ewan Harrison, David K. Jackson, Ian Johnston, Dominic Kwiatkowski, Cordelia Langford, John Sillitoe on behalf of the Wellcome Sanger Institute COVID-19 Surveillance Team (http://www.sanger.ac.uk/covid-team) |
| EPI_ISL_470356, EPI_ISL_470357 | PHE South West Regional Laboratory, National Infection Service | Wellcome Sanger Institute for the COVID-19 Genomics UK Consortium | Stephanie Hutchings, Hannah Pymont, Dr Peter Muir, Barry Vipond, Rich Hopes; and Alex Alderton, Roberto Amato, Sonia Goncalves, Ewan Harrison, David K. Jackson, Ian Johnston, Dominic Kwiatkowski, Cordelia Langford, John Sillitoe on behalf of the Wellcome Sanger Institute COVID-19 Surveillance Team (http://www.sanger.ac.uk/covid-team) |
| EPI_ISL_470358, EPI_ISL_470359, EPI_ISL_470360, EPI_ISL_470361, EPI_ISL_470362, EPI_ISL_470363, EPI_ISL_470364, EPI_ISL_470365, EPI_ISL_470366, EPI_ISL_470367, EPI_ISL_470368, EPI_ISL_470369, EPI_ISL_470370, EPI_ISL_470371, EPI_ISL_470372, EPI_ISL_470373, EPI_ISL_470374, EPI_ISL_470375, EPI_ISL_470376, EPI_ISL_470377, EPI_ISL_470378, EPI_ISL_470379, EPI_ISL_470380, EPI_ISL_470381, EPI_ISL_470382, EPI_ISL_470383, EPI_ISL_470384, EPI_ISL_470385, EPI_ISL_470386, EPI_ISL_470387, EPI_ISL_470388, EPI_ISL_470389, EPI_ISL_470390, EPI_ISL_470391, EPI_ISL_470392, EPI_ISL_470393, EPI_ISL_470394, EPI_ISL_470395, EPI_ISL_470396, EPI_ISL_470397, EPI_ISL_470398, EPI_ISL_470399, EPI_ISL_470400, EPI_ISL_470401, EPI_ISL_470402, EPI_ISL_470403, EPI_ISL_470404, EPI_ISL_470405, EPI_ISL_470406, EPI_ISL_470407, EPI_ISL_470408, EPI_ISL_470409, EPI_ISL_470410, EPI_ISL_470411, EPI_ISL_470412, EPI_ISL_470413, EPI_ISL_470414, EPI_ISL_470415, EPI_ISL_470416, EPI_ISL_470417, EPI_ISL_470418, EPI_ISL_470419, EPI_ISL_470420, EPI_ISL_470421, EPI_ISL_470422, EPI_ISL_470423, EPI_ISL_470424, EPI_ISL_470425, EPI_ISL_470426, EPI_ISL_470427, EPI_ISL_470428, EPI_ISL_470429, EPI_ISL_470430, EPI_ISL_470431, EPI_ISL_470432, EPI_ISL_470433, EPI_ISL_470434, EPI_ISL_470435, EPI_ISL_470436, EPI_ISL_470437, EPI_ISL_470438, EPI_ISL_470439, EPI_ISL_470440, EPI_ISL_470441, EPI_ISL_470442, EPI_ISL_470443, EPI_ISL_470444, EPI_ISL_470445, EPI_ISL_470446, EPI_ISL_470447, EPI_ISL_470448, EPI_ISL_470449, EPI_ISL_470450, EPI_ISL_470451, EPI_ISL_470452, EPI_ISL_470453, EPI_ISL_470454, EPI_ISL_470455, EPI_ISL_470456, EPI_ISL_470457, EPI_ISL_470458, EPI_ISL_470459, EPI_ISL_470460, EPI_ISL_470461, EPI_ISL_470462, EPI_ISL_470463, EPI_ISL_470464, EPI_ISL_470465, EPI_ISL_470466, EPI_ISL_470467, EPI_ISL_470468, EPI_ISL_470469, EPI_ISL_470470, EPI_ISL_470471, EPI_ISL_470472, EPI_ISL_470473, EPI_ISL_470474, EPI_ISL_470475, EPI_ISL_470476, EPI_ISL_470477, EPI_ISL_470478, EPI_ISL_470479, EPI_ISL_470480, EPI_ISL_470481, EPI_ISL_470482, EPI_ISL_470483, EPI_ISL_470484, EPI_ISL_470485, EPI_ISL_470486, EPI_ISL_470487, EPI_ISL_470488, EPI_ISL_470489, EPI_ISL_470490, EPI_ISL_470491, EPI_ISL_470492, EPI_ISL_470493, EPI_ISL_470494, EPI_ISL_470495, EPI_ISL_470496, EPI_ISL_470497, EPI_ISL_470498, EPI_ISL_470499, EPI_ISL_470500, EPI_ISL_470501, EPI_ISL_470502, EPI_ISL_470503, EPI_ISL_470504, EPI_ISL_470505, EPI_ISL_470506, EPI_ISL_470507, EPI_ISL_470508, EPI_ISL_470509, EPI_ISL_470510, EPI_ISL_470511, EPI_ISL_470512, EPI_ISL_470513, EPI_ISL_470514, EPI_ISL_470515, EPI_ISL_470516, EPI_ISL_470517, EPI_ISL_470518, EPI_ISL_470519, EPI_ISL_470520, EPI_ISL_470521, EPI_ISL_470522, EPI_ISL_470523, EPI_ISL_470524, EPI_ISL_470525, EPI_ISL_470526, EPI_ISL_470527, EPI_ISL_470528 | | | |
| see above | Department of Pathology, University of Cambridge | Wellcome Sanger Institute for the COVID-19 Genomics UK Consortium | Luke W Meredith, M. Estée Török , Myra Hosmillo, William L. Hamilton, Martin D. Curran, Theresa Feltwell, Grant Hall, Anna Yakovleva, Fahad A Khokhar, Charlotte J. Houldcroft, Laura G Caller, Aminu S. Jahun, Sarah L. Caddy, Ian Goodfellow; and Alex Alderton, Roberto Amato, Sonia Goncalves, Ewan Harrison, David K. Jackson, Ian Johnston, Dominic Kwiatkowski, Cordelia Langford, John Sillitoe on behalf of the Wellcome Sanger Institute COVID-19 Surveillance Team (http://www.sanger.ac.uk/covid-team) |
| EPI_ISL_470529, EPI_ISL_470530 | PHE South West Regional Laboratory, National Infection Service | Wellcome Sanger Institute for the COVID-19 Genomics UK Consortium | Stephanie Hutchings, Hannah Pymont, Dr Peter Muir, Barry Vipond, Rich Hopes; and Alex Alderton, Roberto Amato, Sonia Goncalves, Ewan Harrison, David K. Jackson, Ian Johnston, Dominic Kwiatkowski, Cordelia Langford, John Sillitoe on behalf of the Wellcome Sanger Institute COVID-19 Surveillance Team (http://www.sanger.ac.uk/covid-team) |
| EPI_ISL_470531, EPI_ISL_470532, EPI_ISL_470533 | Department of Pathology, University of Cambridge | Wellcome Sanger Institute for the COVID-19 Genomics UK Consortium | Luke W Meredith, M. Estée Török , Myra Hosmillo, William L. Hamilton, Martin D. Curran, Theresa Feltwell, Grant Hall, Anna Yakovleva, Fahad A Khokhar, Charlotte J. Houldcroft, Laura G Caller, Aminu S. Jahun, Sarah L. Caddy, Ian Goodfellow; and Alex Alderton, Roberto Amato, Sonia Goncalves, Ewan Harrison, David K. Jackson, Ian Johnston, Dominic Kwiatkowski, Cordelia Langford, John Sillitoe on behalf of the Wellcome Sanger Institute COVID-19 Surveillance Team (http://www.sanger.ac.uk/covid-team) |
| EPI_ISL_470534 | Regional Virus Laboratory, Belfast Health and Social Care Trust | Wellcome Sanger Institute for the COVID-19 Genomics UK Consortium | Conall McCaughey, James McKenna, Tanya Curran, Susan Feeney, Alison Watt, Ciara Cox, Mairead Connor, Zoltan Molnar, David Simpson, Derek Fairley; and Alex Alderton, Roberto Amato, Sonia Goncalves, Ewan Harrison, David K. Jackson, Ian Johnston, Dominic Kwiatkowski, Cordelia Langford, John Sillitoe on behalf of the Wellcome Sanger Institute COVID-19 Surveillance Team (http://www.sanger.ac.uk/covid-team) |
| EPI_ISL_470535, EPI_ISL_470536, EPI_ISL_470537, EPI_ISL_470538 | Department of Pathology, University of Cambridge | Wellcome Sanger Institute for the COVID-19 Genomics UK Consortium | Luke W Meredith, M. Estée Török , Myra Hosmillo, William L. Hamilton, Martin D. Curran, Theresa Feltwell, Grant Hall, Anna Yakovleva, Fahad A Khokhar, Charlotte J. Houldcroft, Laura G Caller, Aminu S. Jahun, Sarah L. Caddy, Ian Goodfellow; and Alex Alderton, Roberto Amato, Sonia Goncalves, Ewan Harrison, David K. Jackson, Ian Johnston, Dominic Kwiatkowski, Cordelia Langford, John Sillitoe on behalf of the Wellcome Sanger Institute COVID-19 Surveillance Team (http://www.sanger.ac.uk/covid-team) |
| EPI_ISL_470539 | Molecular diagnostic laboratory of Federal Budget Institution of Science "Central Research Institute of Epidemiology" of The Federal Service on Customers' Rights Protection and Human Well-being Surveillance | Group of Genomics and Postgenomic Technologies of Central Research Institute of Epidemilology | Speranskaya AS, Kaptelova VV, Samoilov AE, Korneenko EV, Sizova TV, Tivanova EV, Shipulina OY, Akimkin VG |
| EPI_ISL_470540 | Wisconsin State Laboratory of Hygiene Communicable Disease Division | Wisconsin State Laboratory of Hygiene Communicable Disease Division | Kelsey R Florek |
| EPI_ISL_470541, EPI_ISL_470542, EPI_ISL_470543, EPI_ISL_470544, EPI_ISL_470545, EPI_ISL_470546, EPI_ISL_470547, EPI_ISL_470548, EPI_ISL_470549, EPI_ISL_470550, EPI_ISL_470551, EPI_ISL_470552, EPI_ISL_470553, EPI_ISL_470554, EPI_ISL_470555, EPI_ISL_470556, EPI_ISL_470557, EPI_ISL_470558, EPI_ISL_470559, EPI_ISL_470560, EPI_ISL_470561, EPI_ISL_470562, EPI_ISL_470563, EPI_ISL_470564, EPI_ISL_470565, EPI_ISL_470566, EPI_ISL_470567 | | | |
| see above | Utah Public Health Laboratory | Utah Public Health Laboratory | Erin Young, Kelly Oakeson |
| EPI_ISL_470568, EPI_ISL_470569, EPI_ISL_470570, EPI_ISL_470571, EPI_ISL_470572, EPI_ISL_470573, EPI_ISL_470574, EPI_ISL_470575, EPI_ISL_470576, EPI_ISL_470577, EPI_ISL_470578, EPI_ISL_470579, EPI_ISL_470580, EPI_ISL_470581, EPI_ISL_470582, EPI_ISL_470583, EPI_ISL_470584, EPI_ISL_470585, EPI_ISL_470586, EPI_ISL_470587, EPI_ISL_470588 | | | |
| see above | Hermes Pardini | Bioinformatics Laboratory / LNCC | Alexandra Gerber, Ana Paula Guimarães, Luiz Gonzaga Paula de Almeida, Ronaldo da Silva Francisco Junior, Mariane Talon, Filipe Romero, Átila Duque Rossi, Terezinha Marta Pereira, working group UFRJ, Jaqueline Goes de Jesus, Ingra Morales Claro, Ester Cerdeira Sabino, Nuno Rodrigues Faria, CADDE-group, Laboratorio Hermes Pardini, Laboratorio Simile, working group UFMG, Amilcar Tanuri, Carolina Voloch, Renato Santana Aguiar e Ana Tereza Vasconcelos |
| EPI_ISL_470589, EPI_ISL_470590, EPI_ISL_470591, EPI_ISL_470592, EPI_ISL_470593, EPI_ISL_470594, EPI_ISL_470595, EPI_ISL_470596, EPI_ISL_470597 | Simile | Bioinformatics Laboratory / LNCC | Alexandra Gerber, Ana Paula Guimarães, Luiz Gonzaga Paula de Almeida, Ronaldo da Silva Francisco Junior, Mariane Talon, Filipe Romero, Átila Duque Rossi, Terezinha Marta Pereira, working group UFRJ, Jaqueline Goes de Jesus, Ingra Morales Claro, Ester Cerdeira Sabino, Nuno Rodrigues Faria, CADDE-group, Laboratorio Hermes Pardini, Laboratorio Simile, working group UFMG, Amilcar Tanuri, Carolina Voloch, Renato Santana Aguiar e Ana Tereza Vasconcelos |
| EPI_ISL_470598, EPI_ISL_470599, EPI_ISL_470600, EPI_ISL_470601, EPI_ISL_470602, EPI_ISL_470603, EPI_ISL_470604, EPI_ISL_470605, EPI_ISL_470606, EPI_ISL_470607, EPI_ISL_470608, EPI_ISL_470609, EPI_ISL_470610, EPI_ISL_470611, EPI_ISL_470612, EPI_ISL_470613, EPI_ISL_470614 | | | |
| see above | Hermes Pardini | Bioinformatics Laboratory / LNCC | Alexandra Gerber, Ana Paula Guimarães, Luiz Gonzaga Paula de Almeida, Ronaldo da Silva Francisco Junior, Mariane Talon, Filipe Romero, Átila Duque Rossi, Terezinha Marta Pereira, working group UFRJ, Jaqueline Goes de Jesus, Ingra Morales Claro, Ester Cerdeira Sabino, Nuno Rodrigues Faria, CADDE-group, Laboratorio Hermes Pardini, Laboratorio Simile, working group UFMG, Amilcar Tanuri, Carolina Voloch, Renato Santana Aguiar e Ana Tereza Vasconcelos |
| EPI_ISL_470615, EPI_ISL_470616, EPI_ISL_470617, EPI_ISL_470618, EPI_ISL_470619, EPI_ISL_470620, EPI_ISL_470621, EPI_ISL_470622, EPI_ISL_470623, EPI_ISL_470624, EPI_ISL_470625, EPI_ISL_470626, EPI_ISL_470627, EPI_ISL_470628, EPI_ISL_470629, EPI_ISL_470630, EPI_ISL_470631, EPI_ISL_470632, EPI_ISL_470633, EPI_ISL_470634, EPI_ISL_470635, EPI_ISL_470636, EPI_ISL_470637, EPI_ISL_470638, EPI_ISL_470639, EPI_ISL_470640, EPI_ISL_470641, EPI_ISL_470642, EPI_ISL_470643, EPI_ISL_470644, EPI_ISL_470645, EPI_ISL_470646, EPI_ISL_470647, EPI_ISL_470648, EPI_ISL_470649, EPI_ISL_470650 | | | |
| see above | Laboratorio de Virologia Molecular / UFRJ | Bioinformatics Laboratory / LNCC | Alexandra Gerber, Ana Paula Guimarães, Luiz Gonzaga Paula de Almeida, Ronaldo da Silva Francisco Junior, Mariane Talon, Filipe Romero, Átila Duque Rossi, Terezinha Marta Pereira, working group UFRJ, Jaqueline Goes de Jesus, Ingra Morales Claro, Ester Cerdeira Sabino, Nuno Rodrigues Faria, CADDE-group, Laboratorio Hermes Pardini, Laboratorio Simile, working group UFMG, Amilcar Tanuri, Carolina Voloch, Renato Santana Aguiar e Ana Tereza Vasconcelos |
| EPI_ISL_470651, EPI_ISL_470652, EPI_ISL_470653, EPI_ISL_470654, EPI_ISL_470655 | Hermes Pardini | Bioinformatics Laboratory / LNCC | Alexandra Gerber, Ana Paula Guimarães, Luiz Gonzaga Paula de Almeida, Ronaldo da Silva Francisco Junior, Mariane Talon, Filipe Romero, Átila Duque Rossi, Terezinha Marta Pereira, working group UFRJ, Jaqueline Goes de Jesus, Ingra Morales Claro, Ester Cerdeira Sabino, Nuno Rodrigues Faria, CADDE-group, Laboratorio Hermes Pardini, Laboratorio Simile, working group UFMG, Amilcar Tanuri, Carolina Voloch, Renato Santana Aguiar e Ana Tereza Vasconcelos |
| EPI_ISL_470656, EPI_ISL_470657, EPI_ISL_470658, EPI_ISL_470659, EPI_ISL_470660, EPI_ISL_470661, EPI_ISL_470662, EPI_ISL_470663, EPI_ISL_470664, EPI_ISL_470665, EPI_ISL_470666, EPI_ISL_470667, EPI_ISL_470668, EPI_ISL_470669, EPI_ISL_470670, EPI_ISL_470671, EPI_ISL_470672, EPI_ISL_470673, EPI_ISL_470674, EPI_ISL_470675, EPI_ISL_470676, EPI_ISL_470677, EPI_ISL_470678, EPI_ISL_470679, EPI_ISL_470680, EPI_ISL_470681, EPI_ISL_470682, EPI_ISL_470683, EPI_ISL_470684, EPI_ISL_470685, EPI_ISL_470686, EPI_ISL_470687, EPI_ISL_470688, EPI_ISL_470689, EPI_ISL_470690, EPI_ISL_470691, EPI_ISL_470692, EPI_ISL_470693, EPI_ISL_470694, EPI_ISL_470695, EPI_ISL_470696, EPI_ISL_470697, EPI_ISL_470698, EPI_ISL_470699, EPI_ISL_470700, EPI_ISL_470701, EPI_ISL_470702, EPI_ISL_470703, EPI_ISL_470704, EPI_ISL_470705, EPI_ISL_470706, EPI_ISL_470707, EPI_ISL_470708, EPI_ISL_470709, EPI_ISL_470710, EPI_ISL_470711, EPI_ISL_470712, EPI_ISL_470713, EPI_ISL_470714, EPI_ISL_470715, EPI_ISL_470716, EPI_ISL_470717, EPI_ISL_470718 | | | |
| see above | Utah Public Health Laboratory | Utah Public Health Laboratory | Erin Young, Kelly Oakeson |
| EPI_ISL_470719, EPI_ISL_470720, EPI_ISL_470721, EPI_ISL_470722, EPI_ISL_470723, EPI_ISL_470724, EPI_ISL_470725, EPI_ISL_470726, EPI_ISL_470727, EPI_ISL_470728, EPI_ISL_470729, EPI_ISL_470730, EPI_ISL_470731, EPI_ISL_470732, EPI_ISL_470733, EPI_ISL_470734, EPI_ISL_470735, EPI_ISL_470736, EPI_ISL_470737, EPI_ISL_470738, EPI_ISL_470739, EPI_ISL_470740, EPI_ISL_470741, EPI_ISL_470742, EPI_ISL_470743, EPI_ISL_470744, EPI_ISL_470745, EPI_ISL_470746 | | | |
| see above | Utah Public Health Laboratory | Utah Public Health Laboratory | Heidi Butz, Erin Young, Kelly Oakeson |
| EPI_ISL_470747, EPI_ISL_470748, EPI_ISL_470749, EPI_ISL_470750, EPI_ISL_470751, EPI_ISL_470752, EPI_ISL_470753, EPI_ISL_470754, EPI_ISL_470755, EPI_ISL_470756, EPI_ISL_470757, EPI_ISL_470758, EPI_ISL_470759, EPI_ISL_470760, EPI_ISL_470761, EPI_ISL_470762, EPI_ISL_470763, EPI_ISL_470764, EPI_ISL_470765, EPI_ISL_470766, EPI_ISL_470767, EPI_ISL_470768, EPI_ISL_470769, EPI_ISL_470770, EPI_ISL_470771, EPI_ISL_470772, EPI_ISL_470773, EPI_ISL_470774, EPI_ISL_470775, EPI_ISL_470776, EPI_ISL_470777, EPI_ISL_470778, EPI_ISL_470779, EPI_ISL_470780, EPI_ISL_470781, EPI_ISL_470782, EPI_ISL_470783, EPI_ISL_470784, EPI_ISL_470785, EPI_ISL_470786, EPI_ISL_470787, EPI_ISL_470788, EPI_ISL_470789 | | | |
| see above | Minnesota Department of Health, Public Health Laboratory | Minnesota Department of Health, Public Health Laboratory | Matt Plumb, Jacob Garfin, and Xiong Wang |
| EPI_ISL_470790, EPI_ISL_470791, EPI_ISL_470792, EPI_ISL_470793, EPI_ISL_470794, EPI_ISL_470795, EPI_ISL_470796, EPI_ISL_470797, EPI_ISL_470798, EPI_ISL_470799, EPI_ISL_470800 | | | |
| see above | M Health Fairview | Minnesota Department of Health, Public Health Laboratory | Matt Plumb, Jacob Garfin, and Xiong Wang |
| EPI_ISL_470801 | Virology | Virology | Hossain,M.E., Hasan,R., Miah,M., Hasan,M.M., Sumaiya,M.K., Rahman,M.M., Alam,M.S., Clemens,J.D., Ahmed,T., Rahman,M.Z. and Rahman,M. |
| EPI_ISL_470802 | State Key Laboratory of Agriculture Microbiology | State Key Laboratory of Agriculture Microbiology, Huazhong Agric | Zhong Zou |
| EPI_ISL_470830, EPI_ISL_470831, EPI_ISL_470832, EPI_ISL_470833, EPI_ISL_470834, EPI_ISL_470835, EPI_ISL_470836, EPI_ISL_470837, EPI_ISL_470838, EPI_ISL_470839, EPI_ISL_470840, EPI_ISL_470841, EPI_ISL_470842, EPI_ISL_470843, EPI_ISL_470844, EPI_ISL_470845, EPI_ISL_470846, EPI_ISL_470847, EPI_ISL_470848, EPI_ISL_470849, EPI_ISL_470850, EPI_ISL_470851, EPI_ISL_470852, EPI_ISL_470853, EPI_ISL_470854, EPI_ISL_470855, EPI_ISL_470856, EPI_ISL_470857, EPI_ISL_470858, EPI_ISL_470859, EPI_ISL_470860, EPI_ISL_470861, EPI_ISL_470862, EPI_ISL_470863, EPI_ISL_470864, EPI_ISL_470865, EPI_ISL_470866, EPI_ISL_470867, EPI_ISL_470868, EPI_ISL_470869, EPI_ISL_470870, EPI_ISL_470871, EPI_ISL_470872, EPI_ISL_470873, EPI_ISL_470874, EPI_ISL_470875 | | | |
| see above | PathWest Laboratory Medicine WA | PathWest Laboratory Medicine WA | Chisha Sikazwe, Jurissa Lang, Avram Levy, David Smith and David Speers |
| EPI_ISL_470876 | Department for Virology, Molecular Biology and Genome Research, R. G. Lugar Center for Public Health Research, National Center for Disease Control and Public Health (NCDC) of Georgia. | Department for Virology, Molecular Biology and Genome Research, R. G. Lugar Center for Public Health Research, National Center for Disease Control and Public Health (NCDC) of Georgia. | Giorgi Tomashvili, Meri Pantsulaia, Gvantsa Brachveli, Gvantsa Chanturia, Ann Machablishvili, Nato Kotaria, Marine Murtskhvaladze, Lela Sabadze, Mari Gavashelidze, Ana Papkiauri, Tata Imnadze, Tamar Jashiashvili, Tea Tevdoradze, Ketevan Sidamonidze, Ekaterine Khmaladze, Ekaterine Zhghenti, Roena Sukhiashvili, Mariam Zakalashvili, Lela Urushadze, Magda Dgebuadze, Davit Tsaguria, Ekaterine Zangaladze, Nino Berishvili, Adam Kotorashvili, Maia Alkhazashvili, Irma Burjanadze, Anna Kasradze, Khatuna Zakhashvili, Paata Imnadze, Amiran Gamkrelidze. |
| EPI_ISL_470877 | Department for Virology, Molecular Biology and Genome Research, R. G. Lugar Center for Public Health Research, National Center for Disease Control and Public Health (NCDC) of Georgia. | Department for Virology, Molecular Biology and Genome Research, R. G. Lugar Center for Public Health Research, National Center for Disease Control and Public Health (NCDC) of Georgia. | Gvantsa Brachveli, Meri Pantsulaia, Giorgi Tomashvili, Gvantsa Chanturia, Ann Machablishvili, Nato Kotaria, Marine Murtskhvaladze, Lela Sabadze, Mari Gavashelidze, Ana Papkiauri, Tata Imnadze, Tamar Jashiashvili, Tea Tevdoradze, Ketevan Sidamonidze, Ekaterine Khmaladze, Ekaterine Zhghenti, Roena Sukhiashvili, Mariam Zakalashvili, Lela Urushadze, Magda Dgebuadze, Davit Tsaguria, Ekaterine Zangaladze, Nino Berishvili, Adam Kotorashvili, Maia Alkhazashvili, Irma Burjanadze, Anna Kasradze, Khatuna Zakhashvili, Paata Imnadze, Amiran Gamkrelidze. |

| | | | |
|---|---|---|---|
| EPI_ISL_470878, EPI_ISL_470879, EPI_ISL_470880 | National Institute for Communicable Diseases of the National Health Laboratory Service | National Institute for Communicable Diseases of the National Health Laboratory Service | Allam M, Ismail A, Khumalo Z, Kwenda S, van Heusden P, Mtshali P, Mnyameni F, Mohale T, Subramoney K, Bhiman JN |
| EPI_ISL_470882 | Foerde Hospital, Department of Microbiology | Norwegian Institute of Public Health, Department of Virology | Kathrine Stene-Johansen, Kamilla Heddeland Instefjord, Hilde Elshaug, Rasmus Riis Kopperud, Karoline Bragstad, Olav Hungnes |
| EPI_ISL_470896 | Russian State Collection of Viruses | Pathogenic Microorganisms Variability Laboratory | Alexey Shchetinin, Maria Nikiforova, Elena Shidlovskaya, Nadezhda Kuznetsova, Inna Dolzhikova, Daria Grousova, Andrey Botikov, Denis Logunov, Alexander Gintsburg, Vladimir Gushchin |
| EPI_ISL_470897, EPI_ISL_470898, EPI_ISL_470899 | Pathogenic Microorganisms Variability Laboratory | Pathogenic Microorganisms Variability Laboratory | Alexey Shchetinin, Maria Nikiforova, Elena Shidlovskaya, Nadezhda Kuznetsova, Andrey Botikov, Alexander Gintsburg, Vladimir Gushchin |
| EPI_ISL_470900, EPI_ISL_470901, EPI_ISL_470902 | Influenza etiology and epidemiology laboratory | Pathogenic Microorganisms Variability Laboratory | Alexey Shchetinin, Maria Nikiforova, Elena Shidlovskaya, Nadezhda Kuznetsova, Vladimir Gushchin, Inna Dolzhikova, Daria Grousova, Andrey Botikov, Denis Logunov, Kirill Krasnoslobotsev, Svetlana Trushakova, Elena Burtseva, Ludmila Kolobukhina, Svetlana Smetanina, Alexander Gintsburg |
| EPI_ISL_470903, EPI_ISL_470904 | Influenza etiology and epidemiology laboratory | Pathogenic Microorganisms Variability Laboratory | Alexey Shchetinin, Maria Nikiforova, Elena Shidlovskaya, Nadezhda Kuznetsova, Vladimir Gushchin, Inna Dolzhikova, Daria Grousova, Andrey Botikov, Denis Logunov, Anna Ignatjeva, Evgeniya Mukasheva, Elena Burtseva, Ludmila Kolobukhina, Svetlana Smetanina, Alexander Gintsburg |
| EPI_ISL_471144, EPI_ISL_471145, EPI_ISL_471146, EPI_ISL_471147, EPI_ISL_471148, EPI_ISL_471149, EPI_ISL_471150, EPI_ISL_471151, EPI_ISL_471152, EPI_ISL_471153, EPI_ISL_471154, EPI_ISL_471155, EPI_ISL_471156 | | | |
| see above | Gundersen Molecular Diagnostics Laboratory | Kabara Cancer Research Institute | Craig S. Richmond, Paraic A. Kenny |
| EPI_ISL_471157 | Gundersen Clinical Microbiology Laboratory | Kabara Cancer Research Institute | Craig S. Richmond, Paraic A. Kenny |
| EPI_ISL_471158, EPI_ISL_471159, EPI_ISL_471160, EPI_ISL_471161, EPI_ISL_471162, EPI_ISL_471163, EPI_ISL_471164, EPI_ISL_471165, EPI_ISL_471166, EPI_ISL_471167, EPI_ISL_471168, EPI_ISL_471169, EPI_ISL_471170, EPI_ISL_471171 | | | |
| see above | MRCG at LSHTM Genomics lab | MRCG at LSHTM Genomics lab | Sesay et al |
| EPI_ISL_471172 | Unilabs Laboratory Medicine | Norwegian Institute of Public Health, Department of Virology | Kathrine Stene-Johansen, Kamilla Heddeland Instefjord, Hilde Elshaug, Rasmus Riis Kopperud, Karoline Bragstad, Olav Hungnes |
| EPI_ISL_471173 | Hospital of Southern Norway - Kristiansand, Department of Medical Microbiology | Norwegian Institute of Public Health, Department of Virology | Kathrine Stene-Johansen, Kamilla Heddeland Instefjord, Hilde Elshaug, Rasmus Riis Kopperud, Karoline Bragstad, Olav Hungnes |
| EPI_ISL_471174 | Ostfold Hospital Trust - Kalnes, Centre for Laboratory Medicine, Section for gene technology and infection serology | Norwegian Institute of Public Health, Department of Virology | Kathrine Stene-Johansen, Kamilla Heddeland Instefjord, Hilde Elshaug, Rasmus Riis Kopperud, Karoline Bragstad, Olav Hungnes |
| EPI_ISL_471175 | Oslo University Hospital, Department of Medical Microbiology | Norwegian Institute of Public Health, Department of Virology | Kathrine Stene-Johansen, Kamilla Heddeland Instefjord, Hilde Elshaug, Rasmus Riis Kopperud, Karoline Bragstad, Olav Hungnes |
| EPI_ISL_471176 | Hospital of Southern Norway - Kristiansand, Department of Medical Microbiology | Norwegian Institute of Public Health, Department of Virology | Kathrine Stene-Johansen, Kamilla Heddeland Instefjord, Hilde Elshaug, Rasmus Riis Kopperud, Karoline Bragstad, Olav Hungnes |
| EPI_ISL_471177 | Oslo University Hospital, Department of Medical Microbiology | Norwegian Institute of Public Health, Department of Virology | Kathrine Stene-Johansen, Kamilla Heddeland Instefjord, Hilde Elshaug, Rasmus Riis Kopperud, Karoline Bragstad, Olav Hungnes |
| EPI_ISL_471178, EPI_ISL_471179, EPI_ISL_471180, EPI_ISL_471181, EPI_ISL_471182, EPI_ISL_471183, EPI_ISL_471184, EPI_ISL_471185, EPI_ISL_471186, EPI_ISL_471187, EPI_ISL_471188, EPI_ISL_471189, EPI_ISL_471190, EPI_ISL_471191, EPI_ISL_471192, EPI_ISL_471193, EPI_ISL_471194, EPI_ISL_471195, EPI_ISL_471196, EPI_ISL_471197, EPI_ISL_471198, EPI_ISL_471199, EPI_ISL_471200, EPI_ISL_471201, EPI_ISL_471202, EPI_ISL_471203, EPI_ISL_471204, EPI_ISL_471205, EPI_ISL_471206, EPI_ISL_471207, EPI_ISL_471208, EPI_ISL_471209, EPI_ISL_471210, EPI_ISL_471211, EPI_ISL_471212, EPI_ISL_471213, EPI_ISL_471214, EPI_ISL_471215, EPI_ISL_471216, EPI_ISL_471217, EPI_ISL_471218, EPI_ISL_471219, EPI_ISL_471220, EPI_ISL_471221, EPI_ISL_471222, EPI_ISL_471223, EPI_ISL_471224, EPI_ISL_471225, EPI_ISL_471226, EPI_ISL_471227, EPI_ISL_471228, EPI_ISL_471229, EPI_ISL_471230, EPI_ISL_471231, EPI_ISL_471232, EPI_ISL_471233, EPI_ISL_471234, EPI_ISL_471235, EPI_ISL_471236, EPI_ISL_471237, EPI_ISL_471238, EPI_ISL_471239, EPI_ISL_471240, EPI_ISL_471241, EPI_ISL_471242, EPI_ISL_471243, EPI_ISL_471244, EPI_ISL_471245, EPI_ISL_471246, EPI_ISL_471247, EPI_ISL_471248, EPI_ISL_471249, EPI_ISL_471250, EPI_ISL_471251, EPI_ISL_471252, EPI_ISL_471253, EPI_ISL_471254, EPI_ISL_471255, EPI_ISL_471256, EPI_ISL_471257, EPI_ISL_471258, EPI_ISL_471259, EPI_ISL_471260, EPI_ISL_471261, EPI_ISL_471262, EPI_ISL_471263, EPI_ISL_471264, EPI_ISL_471265, EPI_ISL_471266 | | | |
| see above | Wisconsin State Laboratory of Hygiene Communicable Disease Division | Wisconsin State Laboratory of Hygiene Communicable Disease Division | Kelsey R. Florek, Abigail C. Shockey |
| EPI_ISL_471267 | Hospital IESS Babahoyo | Institute of Microbiology, Universidad San Francisco de Quito | Sully Márquez, Belén Prado-Vivar, Juan José Guadalupe, Bernardo Gutiérrez, Francisco Cordova, Ninfa Henriquez, Killen Briones-Zamora, Killen Briones-Claudette, Verónica Barragán, Patricio Rojas-Silva, Gabriel Trueba, Michelle Grunauer, Paúl Cárdenas |
| EPI_ISL_471268 | Hospital IESS Babahoyo | Institute of Microbiology, Universidad San Francisco de Quito | Belén Prado-Vivar, Sully Márquez, Juan José Guadalupe, Bernardo Gutiérrez, Francisco Cordova, Ninfa Henriquez, Killen Briones-Zamora, Killen Briones-Claudette, Verónica Barragán, Patricio Rojas-Silva, Gabriel Trueba, Michelle Grunauer, Paúl Cárdenas |
| EPI_ISL_471269 | Hospital Oncológico Solca Núcleo de Quito | Institute of Microbiology, Universidad San Francisco de Quito | Sully Márquez, Belén Prado-Vivar, Juan José Guadalupe, Bernardo Gutiérrez, Marcos Di Stefano, Grace Salazar, Verónica Barragán, Patricio Rojas-Silva, Gabriel Trueba, Michelle Grunauer, Paúl Cárdenas |
| EPI_ISL_471270 | Hospital Oncológico Solca Núcleo de Quito | Institute of Microbiology, Universidad San Francisco de Quito | Sully Márquez, Belén Prado-Vivar, Juan José Guadalupe, Bernardo Gutiérrez, Marcos Di Stefano, Grace Salazar, Verónica Barragán, Patricio Rojas-Silva, Gabriel Trueba, Michelle Grunauer, Paúl Cárdenas |
| EPI_ISL_471271 | Hospital Oncológico Solca Núcleo de Quito | Institute of Microbiology, Universidad San Francisco de Quito | Sully Márquez, Belén Prado-Vivar, Juan José Guadalupe, Bernardo Gutiérrez, Marcos Di Stefano, Grace Salazar, Verónica Barragán, Patricio Rojas-Silva, Gabriel Trueba, Michelle Grunauer, Paúl Cárdenas |
| EPI_ISL_471396, EPI_ISL_471397, EPI_ISL_471398, EPI_ISL_471399, EPI_ISL_471400, EPI_ISL_471401, EPI_ISL_471402, EPI_ISL_471403, EPI_ISL_471404, EPI_ISL_471405, EPI_ISL_471406, EPI_ISL_471407, EPI_ISL_471408, EPI_ISL_471409, EPI_ISL_471410, EPI_ISL_471411, EPI_ISL_471412, EPI_ISL_471413, EPI_ISL_471414, EPI_ISL_471415 | | | |
| see above | Viral Respiratory Lab, National Institute for Biomedical Research (INRB) | Pathogen Sequencing Lab, National Institute for Biomedical Research (INRB) | Placide Mbala-Kingebeni, Edith Nkwembe, Eddy Kinganda-Lusamaki, Amuri Aziza, Francisca Muyembe Mawete, Catherine Pratt, Matthias Pauthner, Josh Quick, Allison Black, James Hadfield, Trevor Bedford, Ian Goodfellow, Andrew Rambaut, Nick Loman, Kristian Andersen, Michael Wiley, Steve Ahuka-Mundeke, Jean-Jacques Muyembe Tamfum |
| EPI_ISL_471416, EPI_ISL_471417, EPI_ISL_471418, EPI_ISL_471419, EPI_ISL_471420, EPI_ISL_471421, EPI_ISL_471422, EPI_ISL_471423, EPI_ISL_471424 | Laboratory for Respiratory Viruses, National Influenza Centre, Cantacuzino National Military-Medical Institute for Research and Development | Cantacuzino Institute | Luiza Ustea, Nicoleta Paraschiv, Tim Durfee, Mihaela Lazar |
| EPI_ISL_471425 | Division of Viral Diseases, Center for Laboratory Control of Infectious Diseases, Korea Centers for Diseases Control and Prevention | Division of Viral Diseases, Center for Laboratory Control of Infectious Diseases, Korea Centers for Diseases Control and Prevention | Jeong-Min Kim, Yoon-Seok Chung, Namjoo Lee, Mi-Seon Kim, Sang Hee Woo, Hye-Jun Jo, Sehee Park, Heui Man Kim, Jun-Sub Kim, Junhyeong Jang, Dong Hyun Song, Daesang Lee, Seong Tae Jeong, Myung Guk Han |
| EPI_ISL_471426 | Division of Viral Diseases, Center for Laboratory Control of Infectious Diseases, Korea Centers for Diseases Control and Prevention | Division of Viral Diseases, Center for Laboratory Control of Infectious Diseases, Korea Centers for Diseases Control and Prevention | Jeong-Min Kim, Yoon-Seok Chung, Namjoo Lee, Mi-Seon Kim, Sang Hee Woo, Hye-Jun Jo, Sehee Park, Heui Man Kim, Jun-Sub Kim, Junhyeong Jang, Dong Hyun Song, Daesang Lee, Seong Tae Jeong, Myung Guk Han |
| EPI_ISL_471427, EPI_ISL_471428, EPI_ISL_471429, EPI_ISL_471430, EPI_ISL_471431, EPI_ISL_471432, EPI_ISL_471433, EPI_ISL_471434, EPI_ISL_471435, EPI_ISL_471436, EPI_ISL_471437 | | | |
| see above | Department of Clinical Microbiology | GIGA Medical Genomics | Keith Durkin, Maria Artesi, Sébastien Bontems, Raphaël Boreux, Cécile Meex, Axelle Chaslain, Céline Fombellida-Lopez, Pierrette Melin, Marie-Pierre Hayette, Vincent Bours. |
| EPI_ISL_471438, EPI_ISL_471439, EPI_ISL_471440, EPI_ISL_471441, EPI_ISL_471442, EPI_ISL_471443, EPI_ISL_471444 | Division of Viral Diseases, Center for Laboratory Control of Infectious Diseases, Korea Centers for Diseases Control and Prevention | Division of Viral Diseases, Center for Laboratory Control of Infectious Diseases, Korea Centers for Diseases Control and Prevention | Jeong-Min Kim, Yoon-Seok Chung, Namjoo Lee, Sang Hee Woo, Hye-Jun Jo, Heui Man Kim, Jun-Sub Kim, Dong Hyun Song, Daesang Lee, Seong Tae Jeong, Myung Guk Han |
| EPI_ISL_471445 | Division of Viral Diseases, Center for Laboratory Control of Infectious Diseases, Korea Centers for Diseases Control and Prevention | Division of Viral Diseases, Center for Laboratory Control of Infectious Diseases, Korea Centers for Diseases Control and Prevention | Jeong-Min Kim, Yoon-Seok Chung, Namjoo Lee, Sang Hee Woo, Hye-Jun Jo, Heui Man Kim, Jun-Sub Kim, Myung Guk Han |
| EPI_ISL_471446, EPI_ISL_471447, EPI_ISL_471448, EPI_ISL_471449, EPI_ISL_471450, EPI_ISL_471451, EPI_ISL_471452 | Division of Viral Diseases, Center for Laboratory Control of Infectious Diseases, Korea Centers for Diseases Control and Prevention | Division of Viral Diseases, Center for Laboratory Control of Infectious Diseases, Korea Centers for Diseases Control and Prevention | Jeong-Min Kim, Yoon-Seok Chung, Namjoo Lee, Sang Hee Woo, Hye-Jun Jo, Heui Man Kim, Jun-Sub Kim, Dong Hyun Song, Daesang Lee, Seong Tae Jeong, Myung Guk Han |
| EPI_ISL_471453 | Division of Viral Diseases, Center for Laboratory Control of Infectious Diseases, Korea Centers for Diseases Control and Prevention | Division of Viral Diseases, Center for Laboratory Control of Infectious Diseases, Korea Centers for Diseases Control and Prevention | Jeong-Min Kim, Yoon-Seok Chung, Namjoo Lee, Sang Hee Woo, Hye-Jun Jo, Heui Man Kim, Jun-Sub Kim, Myung Guk Han |
| EPI_ISL_471454, EPI_ISL_471455 | Division of Viral Diseases, Center for Laboratory Control of Infectious Diseases, Korea Centers for Diseases Control and Prevention | Division of Viral Diseases, Center for Laboratory Control of Infectious Diseases, Korea Centers for Diseases Control and Prevention | Jeong-Min Kim, Yoon-Seok Chung, Namjoo Lee, Sang Hee Woo, Hye-Jun Jo, Heui Man Kim, Jun-Sub Kim, Dong Hyun Song, Daesang Lee, Seong Tae Jeong, Myung Guk Han |
| EPI_ISL_471456, EPI_ISL_471457, EPI_ISL_471458, EPI_ISL_471459, EPI_ISL_471460 | Centre de Virologie des Maladies Tropicales | Functional Genomic Platform/Service Analyses Biologique/UATRS/ Centre National Pour la Recherche Scientifique Et Technique (CNRST) | Hicham ANNAZ, Elmostafa EL FAHIME, Marouane MELLOUL, Yassine AKHOUAD, Mly Abdelaziz ELALAOUI, Ahmed REGGAD, Sanaa ALAOUI-Amine , Rachid ABI, Rida TAGAJDID, Zhor KASMY, Safaa ELKORCHI, Nadia TOUIL, Farida HILALI, Abdelkader LAATIRIS , Abdelillah LARAQUI, Tahra BAJJOU , Yassine SEKHSOKH , Idriss-Amine LAHLOU, Mostafa ELOUENNASS, Khalid ENNIBI |
| EPI_ISL_471461 | South China Agricultural University | South China Agricultural University | Yongyi Shen, Lihua Xiao, Wu Chen |
| EPI_ISL_471462, EPI_ISL_471463 | South China Agricultural University | South China Agricultural University | Yongyi Shen, Lihua Xiao, Wu Chen |
| EPI_ISL_471464 | South China Agricultural University | South China Agricultural University | Yongyi Shen, Lihua Xiao, Wu Chen |
| EPI_ISL_471465, EPI_ISL_471466 | South China Agricultural University | South China Agricultural University | Yongyi Shen, Lihua Xiao, Wu Chen |
| EPI_ISL_471467, EPI_ISL_471468, EPI_ISL_471469, EPI_ISL_471470 | South China Agricultural University | South China Agricultural University | Yongyi Shen, Wu Chen |
| EPI_ISL_471471 | Victorian Infectious Diseases Reference Laboratory | Victorian Infectious Diseases Reference Laboratory | Caly,L., Seemann,T., Sait,M., Schultz,M.B., Druce,J., Sherry,N. Meumann,E., Soares da Silva,E., Dolores de Jesus da Costa,M., Salles de Sousa,A., Jayanti Pereira Tilman,A., Antonia da Costa,E., Barreto,I., Marr,I., Wapling,J., Francis,J., Ximenes,J., Canisia,D., Freeman,K., Dakh,F., Douglas,N. and Baird,R. |
| EPI_ISL_471472 | Hospital Universitari Germans Trias i Pujol(HUGTiP)/Fundació Lluita contra la SIDA (FLSida)/IRTA-CReSA | IrsiCaixa AIDS Research Lab | Marc Noguera-Julian, Pilar Armengol, Jordi Rodón, Julia Vergara, Lidia Ruiz, Nuria Izquierdo, Jorge Carrillo, Roger Paredes, Albert Bensaid, Julia Blanco, Joaquim Segalés, Bonaventura Clotet |
| EPI_ISL_471510, EPI_ISL_471511, EPI_ISL_471512, EPI_ISL_471513, EPI_ISL_471514, EPI_ISL_471515, EPI_ISL_471516, EPI_ISL_471517, EPI_ISL_471518, EPI_ISL_471519, EPI_ISL_471520, EPI_ISL_471521, EPI_ISL_471522, EPI_ISL_471523, EPI_ISL_471524, EPI_ISL_471525, EPI_ISL_471526, EPI_ISL_471527 | | | |
| see above | Respiratory Virus Unit, Microbiology Services Colindale, Public Health England | Respiratory Virus Unit, Microbiology Services Colindale, Public Health England | PHE Covid Sequencing Team |
| EPI_ISL_471528 | The National Institute of Public Health | State Veterinary Institute Prague and The National Institute of Public Health | Nagy,A;Jirincova,H;Novakova,L;Trnka,D;Vecerova,J |
| EPI_ISL_471529 | Department for Virology, Molecular Biology and Genome | Department for Virology, Molecular Biology and Genome Research, R. | Meri Pantsulaia, Gvantsa Brachveli, Giorgi Tomashvili, Gvantsa Chanturia, Ann Machablishvili, Nato Kotaria, Marine Murtskhvaladze, Lela Sabadze, Mari Gavashelidze, Ana Papkiauri, Gvantsa Brachveli, Tata Imnadze, Tamar |

| | | | |
|---|---|---|---|
| | Research, R. G. Lugar Center for Public Health Research, National Center for Disease Control and Public Health (NCDC) of Georgia. | G. Lugar Center for Public Health Research, National Center for Disease Control and Public Health (NCDC) of Georgia. | Jashiashvili, Tea Tevdoradze, Ketevan Sidamonidze, Ekaterine Khmaladze, Ekaterine Zhghenti, Roena Sukhiashvili, Mariam Zakalashvili, Lela Urushadze, Magda Dgebuadze, Davit Tsaguria, Ekaterine Zangaladze, Nino Berishvili, Adam Kotorashvili, Maia Alkhazashvili, Irma Burjanadze, Anna Kasradze, Khatuna Zakhashvili, Paata Imnadze, Amiran Gamkrelidze. |
| EPI_ISL_471530 | The National Institute of Public Health | State Veterinary Institute Prague and The National Institute of Public Health | Nagy,A;Jirincova,H;Novakova,L;Trnka,D;Vecerova,J |
| EPI_ISL_471539 | Hospital Universitario da USP Sao Paulo | Instituto Adolfo Lutz, Interdiciplinary Procedures Center, Strategic Laboratory | Claudio Tavares Sacchi, Claudia Regina Gonçalves, Erica Valessa Ramos Gomes |
| EPI_ISL_471540 | The National Institute of Public Health | State Veterinary Institute Prague and The National Institute of Public Health | Nagy,A;Jirincova,H;Novakova,L;Trnka,D;Vecerova,J |
| EPI_ISL_471541 | Hospital Geral Santa Marcelina | Instituto Adolfo Lutz, Interdiciplinary Procedures Center, Strategic Laboratory | Claudio Tavares Sacchi, Claudia Regina Gonçalves, Erica Valessa Ramos Gomes |
| EPI_ISL_471542 | Secretaria de Saude de Mogi das Cruzes | Instituto Adolfo Lutz, Interdiciplinary Procedures Center, Strategic Laboratory | Claudio Tavares Sacchi, Claudia Regina Gonçalves, Erica Valessa Ramos Gomes |
| EPI_ISL_471543 | Centro de Saude I Tacito Leite de Carvalho e Silva | Instituto Adolfo Lutz, Interdiciplinary Procedures Center, Strategic Laboratory | Claudio Tavares Sacchi, Claudia Regina Gonçalves, Erica Valessa Ramos Gomes |
| EPI_ISL_471544 | The National Institute of Public Health | State Veterinary Institute Prague and The National Institute of Public Health | Nagy,A;Jirincova,H;Novakova,L;Trnka,D;Vecerova,J |
| EPI_ISL_471545 | Hospital Sao Paulo de Ensino da Unifesp | Instituto Adolfo Lutz, Interdiciplinary Procedures Center, Strategic Laboratory | Claudio Tavares Sacchi, Claudia Regina Gonçalves, Erica Valessa Ramos Gomes |
| EPI_ISL_471546 | AMA DR Jose Soares Hungria | Instituto Adolfo Lutz, Interdiciplinary Procedures Center, Strategic Laboratory | Claudio Tavares Sacchi, Claudia Regina Gonçalves, Erica Valessa Ramos Gomes |
| EPI_ISL_471547 | The National Institute of Public Health | State Veterinary Institute Prague and The National Institute of Public Health | Nagy,A;Jirincova,H;Novakova,L;Trnka,D;Vecerova,J |
| EPI_ISL_471548 | Hospital do Servidor Público Estadual Francisco Morato de Oliveira | Instituto Adolfo Lutz, Interdiciplinary Procedures Center, Strategic Laboratory | Claudio Tavares Sacchi, Claudia Regina Gonçalves, Erica Valessa Ramos Gomes |
| EPI_ISL_471549 | Hospital Municipal Carmen Prudente | Instituto Adolfo Lutz, Interdiciplinary Procedures Center, Strategic Laboratory | Claudio Tavares Sacchi, Claudia Regina Gonçalves, Erica Valessa Ramos Gomes |
| EPI_ISL_471550 | The National Institute of Public Health | State Veterinary Institute Prague and The National Institute of Public Health | Nagy,A;Jirincova,H;Novakova,L;Trnka,D;Vecerova,J |
| EPI_ISL_471551 | Hospital Sao Paulo de Ensino da Unifesp | Instituto Adolfo Lutz, Interdiciplinary Procedures Center, Strategic Laboratory | Claudio Tavares Sacchi, Claudia Regina Gonçalves, Erica Valessa Ramos Gomes |
| EPI_ISL_471552 | Hospital Sancta Maggiore | Instituto Adolfo Lutz, Interdiciplinary Procedures Center, Strategic Laboratory | Claudio Tavares Sacchi, Claudia Regina Gonçalves, Erica Valessa Ramos Gomes |
| EPI_ISL_471553 | The National Institute of Public Health | State Veterinary Institute Prague and The National Institute of Public Health | Nagy,A;Jirincova,H;Novakova,L;Trnka,D;Vecerova,J |
| EPI_ISL_471554 | Hospital Bosque da Saúde | Instituto Adolfo Lutz, Interdiciplinary Procedures Center, Strategic Laboratory | Claudio Tavares Sacchi, Claudia Regina Gonçalves, Erica Valessa Ramos Gomes |
| EPI_ISL_471555 | The National Institute of Public Health | State Veterinary Institute Prague and The National Institute of Public Health | Nagy,A;Jirincova,H;Novakova,L;Trnka,D;Vecerova,J |
| EPI_ISL_471556 | Pronto Socorro Jose Ibrahin | Instituto Adolfo Lutz, Interdiciplinary Procedures Center, Strategic Laboratory | Claudio Tavares Sacchi, Claudia Regina Gonçalves, Erica Valessa Ramos Gomes |
| EPI_ISL_471562, EPI_ISL_471581, EPI_ISL_471582 | Hosp. Municipal Prof. Dr. Alípio Corrêa Netto | Instituto Adolfo Lutz, Interdiciplinary Procedures Center, Strategic Laboratory | Claudio Tavares Sacchi, Claudia Regina Gonçalves, Erica Valessa Ramos Gomes |
| EPI_ISL_471583, EPI_ISL_471584 | King Institute of Preventive Medicine & Research | CSIR-Centre for Cellular and Molecular Biology | K.Kaveri,S.Sivasubramanian,S.Vennila,P.Padmapriya,R.Kiruba,S.Magesh,G. Dhinakar Raj, G. Ravikumar, P. Azhahianambi,K Thangaraj,Payel Mukherjee, Sofia Banu, Priya Singh, Dhiviya Vedagiri, Divya Gupta, Vishal Sah, Santosh Kumar Kuncha, Krishnan Harinivas Harshan, Archana Bharadwaj Siva, Karthik Bharadwaj Tallapaka, Shagufta Khan, Lamuk Zaveri, Namami Gaur, Sakshi Shambhavi, Tulasi Nagabandi, Purushotham Vodnala, Rakesh K Mishra, Divya Tej Sowpati |
| EPI_ISL_471585 | CSIR-Centre for Cellular and Molecular Biology | CSIR-Centre for Cellular and Molecular Biology | Dhiviya Vedagiri, Divya Gupta, Vishal Sah, Payel Mukherjee, Sofia Banu, Priya Singh, Santosh Kumar Kuncha, Archana Bharadwaj Siva, Karthik Bharadwaj Tallapaka, Shagufta Khan, Lamuk Zaveri, Namami Gaur, Sakshi Shambhavi, Tulasi Nagabandi, Purushotham Vodnala, Rakesh K Mishra, Divya Tej Sowpati, Krishnan Harinivas Harshan |
| EPI_ISL_471586 | CSIR-Centre for Cellular and Molecular Biology | CSIR-Centre for Cellular and Molecular Biology | Lamuk Zaveri, Shagufta Khan, Namami Gaur, Sakshi Shambhavi, Tulasi Nagabandi, Purushotham Vodnala, Payel Mukherjee, Sofia Banu, Priya Singh, Dhiviya Vedagiri, Divya Gupta, Vishal Sah, Santosh Kumar Kuncha, Krishnan Harinivas Harshan, Archana Bharadwaj Siva, Karthik Bharadwaj Tallapaka,Zeba Rizvi, Zuberwasim Sayyad, Kakade Aishwarya Arun, Amrutha H C, Ananga Ghosh, Rakesh K Mishra, Divya Tej Sowpati |
| EPI_ISL_471587 | CSIR-Centre for Cellular and Molecular Biology | CSIR-Centre for Cellular and Molecular Biology | Dhiviya Vedagiri, Divya Gupta, Vishal Sah, Payel Mukherjee, Sofia Banu, Priya Singh, Santosh Kumar Kuncha, Archana Bharadwaj Siva, Karthik Bharadwaj Tallapaka, Shagufta Khan, Lamuk Zaveri, Namami Gaur, Sakshi Shambhavi, Tulasi Nagabandi, Purushotham Vodnala, Rakesh K Mishra, Divya Tej Sowpati, Krishnan Harinivas Harshan |
| EPI_ISL_471588 | CSIR-Centre for Cellular and Molecular Biology | CSIR-Centre for Cellular and Molecular Biology | Lamuk Zaveri, Shagufta Khan, Namami Gaur, Sakshi Shambhavi, Tulasi Nagabandi, Purushotham Vodnala, Payel Mukherjee, Sofia Banu, Priya Singh, Dhiviya Vedagiri, Divya Gupta, Vishal Sah, Santosh Kumar Kuncha, Krishnan Harinivas Harshan, Archana Bharadwaj Siva, Karthik Bharadwaj Tallapaka, Renu Sudhakar, Somesh Gorde, Gangumala Srinivas Reddy, Sujoy Deb, Swati Bayyana, Rakesh K Mishra, Divya Tej Sowpati |
| EPI_ISL_471589 | CSIR-Centre for Cellular and Molecular Biology | CSIR-Centre for Cellular and Molecular Biology | Lamuk Zaveri, Shagufta Khan, Namami Gaur, Sakshi Shambhavi, Tulasi Nagabandi, Purushotham Vodnala, Payel Mukherjee, Sofia Banu, Priya Singh, Dhiviya Vedagiri, Divya Gupta, Vishal Sah, Santosh Kumar Kuncha, Krishnan Harinivas Harshan, Archana Bharadwaj Siva, Karthik Bharadwaj Tallapaka,Umesh Kumar, Unis Ahmad Bhat, Ajay Sarawagi, Priyanka Pant, Rajkanwar Nathawat, Rakesh K Mishra, Divya Tej Sowpati |
| EPI_ISL_471590 | CSIR-Centre for Cellular and Molecular Biology | CSIR-Centre for Cellular and Molecular Biology | Lamuk Zaveri, Shagufta Khan, Namami Gaur, Sakshi Shambhavi, Tulasi Nagabandi, Purushotham Vodnala, Payel Mukherjee, Sofia Banu, Priya Singh, Dhiviya Vedagiri, Divya Gupta, Vishal Sah, Santosh Kumar Kuncha, Krishnan Harinivas Harshan, Archana Bharadwaj Siva, Karthik Bharadwaj Tallapaka,Zeba Rizvi, Zuberwasim Sayyad, Kakade Aishwarya Arun, Amrutha H C, Ananga Ghosh, Rakesh K Mishra, Divya Tej Sowpati |
| EPI_ISL_471591 | CSIR-Centre for Cellular and Molecular Biology | CSIR-Centre for Cellular and Molecular Biology | Namami Gaur, Sakshi Shambhavi, Lamuk Zaveri, Shagufta Khan, Tulasi Nagabandi, Purushotham Vodnala, Payel Mukherjee, Sofia Banu, Priya Singh, Dhiviya Vedagiri, Divya Gupta, Vishal Sah, Santosh Kumar Kuncha, Krishnan Harinivas Harshan, Archana Bharadwaj Siva, Karthik Bharadwaj Tallapaka, Zeba Rizvi, Zuberwasim Sayyad, Kakade Aishwarya Arun, Amrutha H C, Ananga Ghosh, Rakesh K Mishra, Divya Tej Sowpati |
| EPI_ISL_471592 | CSIR-Centre for Cellular and Molecular Biology | CSIR-Centre for Cellular and Molecular Biology | Namami Gaur, Sakshi Shambhavi, Lamuk Zaveri, Shagufta Khan, Tulasi Nagabandi, Purushotham Vodnala, Payel Mukherjee, Sofia Banu, Priya Singh, Dhiviya Vedagiri, Divya Gupta, Vishal Sah, Santosh Kumar Kuncha, Krishnan Harinivas Harshan, Archana Bharadwaj Siva, Karthik Bharadwaj Tallapaka, Nikhil Hajirnis, Pratheusa Maccha, M Soujanya Reddy,G. Aditya Kumar, Koushick Sivakumar, Rakesh K Mishra, Divya Tej Sowpati |
| EPI_ISL_471593 | CSIR-Centre for Cellular and Molecular Biology | CSIR-Centre for Cellular and Molecular Biology | Namami Gaur, Sakshi Shambhavi, Lamuk Zaveri, Shagufta Khan, Tulasi Nagabandi, Purushotham Vodnala, Payel Mukherjee, Sofia Banu, Priya Singh, Dhiviya Vedagiri, Divya Gupta, Vishal Sah, Santosh Kumar Kuncha, Krishnan Harinivas Harshan, Archana Bharadwaj Siva, Karthik Bharadwaj Tallapaka, Zeba Rizvi, Zuberwasim Sayyad, Kakade Aishwarya Arun, Amrutha H C, Ananga Ghosh, Rakesh K Mishra, Divya Tej Sowpati |
| EPI_ISL_471594 | CSIR-Centre for Cellular and Molecular Biology | CSIR-Centre for Cellular and Molecular Biology | Namami Gaur, Sakshi Shambhavi, Lamuk Zaveri, Shagufta Khan, Tulasi Nagabandi, Purushotham Vodnala, Payel Mukherjee, Sofia Banu, Priya Singh, Dhiviya Vedagiri, Divya Gupta, Vishal Sah, Santosh Kumar Kuncha, Krishnan Harinivas Harshan, Archana Bharadwaj Siva, Karthik Bharadwaj Tallapaka,Kezia J Ann, Radhika Khandelwal, Roshan Maku Venkata, Shemin Mansuri, Sonu Uday, Rakesh K Mishra, Divya Tej Sowpati |
| EPI_ISL_471595 | CSIR-Centre for Cellular and Molecular Biology | CSIR-Centre for Cellular and Molecular Biology | Payel Mukherjee, Sofia Banu, Priya Singh, Dhiviya Vedagiri, Divya Gupta, Vishal Sah, Santosh Kumar Kuncha, Krishnan Harinivas Harshan, Archana Bharadwaj Siva, Karthik Bharadwaj Tallapaka, Shagufta Khan, Lamuk Zaveri, Namami Gaur, Sakshi Shambhavi, Tulasi Nagabandi, Purushotham Vodnala, G. Aditya Kumar, Koushick Sivakumar, Pooja Ramesh Gupta, Rajan Kumar Jha, Shraddha Vijay Lahoti, Rakesh K Mishra, Divya Tej Sowpati |
| EPI_ISL_471596 | CSIR-Centre for Cellular and Molecular Biology | CSIR-Centre for Cellular and Molecular Biology | Payel Mukherjee, Sofia Banu, Priya Singh, Dhiviya Vedagiri, Divya Gupta, Vishal Sah, Santosh Kumar Kuncha, Krishnan Harinivas Harshan, Archana Bharadwaj Siva, Karthik Bharadwaj Tallapaka, Shagufta Khan, Lamuk Zaveri, Namami Gaur, Sakshi Shambhavi, Tulasi Nagabandi, Purushotham Vodnala, Gokulan C G, Gunjan Purohit, Hanuman Tulashiram Kale, Pankaj Kumar, Prachand Issarapu, Rakesh K Mishra, Divya Tej Sowpati |
| EPI_ISL_471597 | CSIR-Centre for Cellular and Molecular Biology | CSIR-Centre for Cellular and Molecular Biology | Payel Mukherjee, Sofia Banu, Priya Singh, Dhiviya Vedagiri, Divya Gupta, Vishal Sah, Santosh Kumar Kuncha, Krishnan Harinivas Harshan, Archana Bharadwaj Siva, Karthik Bharadwaj Tallapaka, Shagufta Khan, Lamuk Zaveri, Namami Gaur, Sakshi Shambhavi, Tulasi Nagabandi, Purushotham Vodnala, Rakesh K Mishra, Sonu Uday, Sudipta Mondal, Annapoorna P Karthyayani, Debabrata Jana, Debrya Saha, Divya Tej Sowpati |
| EPI_ISL_471598 | CSIR-Centre for Cellular and Molecular Biology | CSIR-Centre for Cellular and Molecular Biology | Payel Mukherjee, Sofia Banu, Priya Singh, Dhiviya Vedagiri, Divya Gupta, Vishal Sah, Santosh Kumar Kuncha, Krishnan Harinivas Harshan, Archana Bharadwaj Siva, Karthik Bharadwaj Tallapaka, Shagufta Khan, Lamuk Zaveri, Namami Gaur, Sakshi Shambhavi, Tulasi Nagabandi, Purushotham Vodnala,Deepak Kumar, Devi Prasad Vijayashankar, Disha Nanda, Divya Das, Jotin Gogoi, Manish Bhattacharjee, Rakesh K Mishra, Divya Tej Sowpati |
| EPI_ISL_471599 | CSIR-Centre for Cellular and Molecular Biology | CSIR-Centre for Cellular and Molecular Biology | Sakshi Shambhavi, Lamuk Zaveri, Shagufta Khan, Namami Gaur, Tulasi Nagabandi, Purushotham Vodnala, Payel Mukherjee, Sofia Banu, Priya Singh, Dhiviya Vedagiri, Divya Gupta, Vishal Sah, Santosh Kumar Kuncha, Krishnan Harinivas Harshan, Archana Bharadwaj Siva, Karthik Bharadwaj Tallapaka, Deepak Kumar, Devi Prasad Vijayashankar, Disha Nanda, Divya Das, Jotin Gogoi, Manish Bhattacharjee, Rakesh K Mishra, Divya Tej Sowpati |
| EPI_ISL_471600 | CSIR-Centre for Cellular and Molecular Biology | CSIR-Centre for Cellular and Molecular Biology | Sakshi Shambhavi, Lamuk Zaveri, Shagufta Khan, Namami Gaur, Tulasi Nagabandi, Purushotham Vodnala, Payel Mukherjee, Sofia Banu, Priya Singh, Dhiviya Vedagiri, Divya Gupta, Vishal Sah, Santosh Kumar Kuncha, Krishnan Harinivas Harshan, Archana Bharadwaj Siva, Karthik Bharadwaj Tallapaka, G. Aditya Kumar, Koushick Sivakumar, Pooja Ramesh Gupta, Rajan Kumar Jha, Shraddha Vijay Lahoti, Rakesh K Mishra, Divya Tej Sowpati |
| EPI_ISL_471601 | CSIR-Centre for Cellular and Molecular Biology | CSIR-Centre for Cellular and Molecular Biology | Sakshi Shambhavi, Lamuk Zaveri, Shagufta Khan, Namami Gaur, Tulasi Nagabandi, Purushotham Vodnala, Payel Mukherjee, Sofia Banu, Priya Singh, Dhiviya Vedagiri, Divya Gupta, Vishal Sah, Santosh Kumar Kuncha, Krishnan Harinivas Harshan, Archana Bharadwaj Siva, Karthik Bharadwaj Tallapaka,Nikhil Hajirnis, Pratheusa Maccha, M Soujanya Reddy,G. Aditya Kumar, Koushick Sivakumar, Rakesh K Mishra, Divya Tej Sowpati |
| EPI_ISL_471602 | CSIR-Centre for Cellular and Molecular Biology | CSIR-Centre for Cellular and Molecular Biology | Sakshi Shambhavi, Lamuk Zaveri, Shagufta Khan, Namami Gaur, Tulasi Nagabandi, Purushotham Vodnala, Payel Mukherjee, Sofia Banu, Priya Singh, Dhiviya Vedagiri, Divya Gupta, Vishal Sah, Santosh Kumar Kuncha, Krishnan Harinivas Harshan, Archana Bharadwaj Siva, Karthik Bharadwaj Tallapaka,Nikhil Hajirnis, Pratheusa Maccha, M Soujanya Reddy,G. Aditya Kumar, Koushick Sivakumar,Disha Nanda, Divya Das, Jotin Gogoi, Manish Bhattacharjee, Ravi Prasad Mukku, Rakesh K Mishra, Divya Tej Sowpati |
| EPI_ISL_471603 | CSIR-Centre for Cellular and Molecular Biology | CSIR-Centre for Cellular and Molecular Biology | Shagufta Khan, Lamuk Zaveri, Namami Gaur, Sakshi Shambhavi, Tulasi Nagabandi, Purushotham Vodnala, Payel Mukherjee, Sofia Banu, Priya Singh, Dhiviya Vedagiri, Divya Gupta, Vishal Sah, Santosh Kumar Kuncha, Krishnan Harinivas Harshan, Archana Bharadwaj Siva, Karthik Bharadwaj Tallapaka, Disha Nanda, Divya Das, Jotin Gogoi, Manish Bhattacharjee, Ravi Prasad Mukku, Rakesh K Mishra, Divya Tej Sowpati |
| EPI_ISL_471604 | CSIR-Centre for Cellular and Molecular Biology | CSIR-Centre for Cellular and Molecular Biology | Shagufta Khan, Lamuk Zaveri, Namami Gaur, Sakshi Shambhavi, Tulasi Nagabandi, Purushotham Vodnala, Payel Mukherjee, Sofia Banu, Priya Singh, Dhiviya Vedagiri, Divya Gupta, Vishal Sah, Santosh Kumar Kuncha, Krishnan Harinivas Harshan, Archana Bharadwaj Siva, Karthik Bharadwaj Tallapaka, Renu Sudhakar, Somesh Gorde, Gangumala Srinivas Reddy, Sujoy Deb, Swati Bayyana, Rakesh K Mishra, Divya Tej Sowpati |
| EPI_ISL_471605 | CSIR-Centre for Cellular and Molecular Biology | CSIR-Centre for Cellular and Molecular Biology | Shagufta Khan, Lamuk Zaveri, Namami Gaur, Sakshi Shambhavi, Tulasi Nagabandi, Purushotham Vodnala, Payel Mukherjee, Sofia Banu, Priya Singh, Dhiviya Vedagiri, Divya Gupta, Vishal Sah, Santosh Kumar Kuncha, Krishnan Harinivas Harshan, Archana Bharadwaj Siva, Karthik Bharadwaj Tallapaka,Preethi Jampala, Sharada Ravi Iyer, Sulagana Mukherjee, Swetha Sundar, Peddapuvala Sai Uday Kiran Rakesh K Mishra, Divya Tej Sowpati |
| EPI_ISL_471606 | CSIR-Centre for Cellular and Molecular Biology | CSIR-Centre for Cellular and Molecular Biology | Shagufta Khan, Lamuk Zaveri, Namami Gaur, Sakshi Shambhavi, Tulasi Nagabandi, Purushotham Vodnala, Payel Mukherjee, Sofia Banu, Priya Singh, Dhiviya Vedagiri, Divya Gupta, Vishal Sah, Santosh Kumar Kuncha, Krishnan Harinivas Harshan, Archana Bharadwaj Siva, Karthik Bharadwaj Tallapaka,Umesh Kumar, Unis Ahmad Bhat, Ajay Sarawagi, Priyanka Pant, Rajkanwar Nathawat, Rakesh K Mishra, Divya Tej Sowpati |

| | | | |
|---|---|---|---|
| EPI_ISL_471607 | CSIR-Centre for Cellular and Molecular Biology | CSIR-Centre for Cellular and Molecular Biology | Sofia Banu, Payel Mukherjee, Priya Singh, Dhiviya Vedagiri, Divya Gupta, Vishal Sah, Santosh Kumar Kuncha, Krishnan Harinivas Harshan, Archana Bharadwaj Siva, Karthik Bharadwaj Tallapaka, Shagufta Khan, Lamuk Zaveri, Namami Gaur, Sakshi Shambhavi, Tulasi Nagabandi, Purushotham Vodnala, Deepak Kumar, Devi Prasad Vijayashankar, Disha Nanda, Divya Das, Jotin Gogoi, Manish Bhattacharjee, Rakesh K Mishra, Divya Tej Sowpati |
| EPI_ISL_471608 | CSIR-Centre for Cellular and Molecular Biology | CSIR-Centre for Cellular and Molecular Biology | Sofia Banu, Payel Mukherjee, Priya Singh, Dhiviya Vedagiri, Divya Gupta, Vishal Sah, Santosh Kumar Kuncha, Krishnan Harinivas Harshan, Archana Bharadwaj Siva, Karthik Bharadwaj Tallapaka, Shagufta Khan, Lamuk Zaveri, Namami Gaur, Sakshi Shambhavi, Tulasi Nagabandi, Purushotham Vodnala, Disha Nanda, Divya Das, Jotin Gogoi, Manish Bhattacharjee, Ravi Prasad Mukku, Rakesh K Mishra, Divya Tej Sowpati |
| EPI_ISL_471609 | CSIR-Centre for Cellular and Molecular Biology | CSIR-Centre for Cellular and Molecular Biology | Sofia Banu, Payel Mukherjee, Priya Singh, Dhiviya Vedagiri, Divya Gupta, Vishal Sah, Santosh Kumar Kuncha, Krishnan Harinivas Harshan, Archana Bharadwaj Siva, Karthik Bharadwaj Tallapaka, Shagufta Khan, Lamuk Zaveri, Namami Gaur, Sakshi Shambhavi, Tulasi Nagabandi, Purushotham Vodnala, Gokulan C G, Gunjan Purohit, Hanuman Tulashiram Kale, Pankaj Kumar, Prachand Issarapu, Rakesh K Mishra, Divya Tej Sowpati |
| EPI_ISL_471610 | CSIR-Centre for Cellular and Molecular Biology | CSIR-Centre for Cellular and Molecular Biology | Sofia Banu, Payel Mukherjee, Priya Singh, Dhiviya Vedagiri, Divya Gupta, Vishal Sah, Santosh Kumar Kuncha, Krishnan Harinivas Harshan, Archana Bharadwaj Siva, Karthik Bharadwaj Tallapaka, Shagufta Khan, Lamuk Zaveri, Namami Gaur, Sakshi Shambhavi, Tulasi Nagabandi, Purushotham Vodnala,Preethi Jampala, Sharada Ravi Iyer, Sulagana Mukherjee, Swetha Sundar, Peddapuvala Sai Uday Kiran, Rakesh K Mishra, Divya Tej Sowpati |
| EPI_ISL_471611 | CSIR-Centre for Cellular and Molecular Biology | CSIR-Centre for Cellular and Molecular Biology | Tulasi Nagabandi, Namami Gaur, Sakshi Shambhavi, Lamuk Zaveri, Shagufta Khan, Purushotham Vodnala, Payel Mukherjee, Sofia Banu, Priya Singh, Dhiviya Vedagiri, Divya Gupta, Vishal Sah, Santosh Kumar Kuncha, Krishnan Harinivas Harshan, Archana Bharadwaj Siva, Karthik Bharadwaj Tallapaka,G. Aditya Kumar, Koushick Sivakumar, Pooja Ramesh Gupta, Rajan Kumar Jha, Shraddha Vijay Lahoti, Rakesh K Mishra, Divya Tej Sowpati |
| EPI_ISL_471612 | CSIR-Centre for Cellular and Molecular Biology | CSIR-Centre for Cellular and Molecular Biology | Tulasi Nagabandi, Namami Gaur, Sakshi Shambhavi, Lamuk Zaveri, Shagufta Khan, Purushotham Vodnala, Payel Mukherjee, Sofia Banu, Priya Singh, Dhiviya Vedagiri, Divya Gupta, Vishal Sah, Santosh Kumar Kuncha, Krishnan Harinivas Harshan, Archana Bharadwaj Siva, Karthik Bharadwaj Tallapaka,Kezia J Ann, Radhika Khandelwal, Roshan Maku Venkata, Shemin Mansuri, Sonu Uday, Rakesh K Mishra, Divya Tej Sowpati |
| EPI_ISL_471613 | CSIR-Centre for Cellular and Molecular Biology | CSIR-Centre for Cellular and Molecular Biology | Tulasi Nagabandi, Namami Gaur, Sakshi Shambhavi, Lamuk Zaveri, Shagufta Khan, Purushotham Vodnala, Payel Mukherjee, Sofia Banu, Priya Singh, Dhiviya Vedagiri, Divya Gupta, Vishal Sah, Santosh Kumar Kuncha, Krishnan Harinivas Harshan, Archana Bharadwaj Siva, Karthik Bharadwaj Tallapaka,G. Aditya Kumar, Koushick Sivakumar, Pooja Ramesh Gupta, Rajan Kumar Jha, Shraddha Vijay Lahoti, Rakesh K Mishra, Divya Tej Sowpati |
| EPI_ISL_471614 | CSIR-Centre for Cellular and Molecular Biology | CSIR-Centre for Cellular and Molecular Biology | Tulasi Nagabandi, Namami Gaur, Sakshi Shambhavi, Lamuk Zaveri, Shagufta Khan, Purushotham Vodnala, Payel Mukherjee, Sofia Banu, Priya Singh, Dhiviya Vedagiri, Divya Gupta, Vishal Sah, Santosh Kumar Kuncha, Krishnan Harinivas Harshan, Archana Bharadwaj Siva, Karthik Bharadwaj Tallapaka,Kezia J Ann, Radhika Khandelwal, Roshan Maku Venkata, Shemin Mansuri, Sonu Uday, Rakesh K Mishra, Divya Tej Sowpati |
| EPI_ISL_471615 | CSIR-Centre for Cellular and Molecular Biology | CSIR-Centre for Cellular and Molecular Biology | Lamuk Zaveri, Shagufta Khan, Namami Gaur, Sakshi Shambhavi, Tulasi Nagabandi, Purushotham Vodnala, Payel Mukherjee, Sofia Banu, Priya Singh, Dhiviya Vedagiri, Divya Gupta, Vishal Sah, Santosh Kumar Kuncha, Krishnan Harinivas Harshan, Archana Bharadwaj Siva, Karthik Bharadwaj Tallapaka,Zeba Rizvi, Zuberwasim Sayyad, Kakade Aishwarya Arun, Amrutha H C, Ananga Ghosh, Rakesh K Mishra, Divya Tej Sowpati |
| EPI_ISL_471616 | CSIR-Centre for Cellular and Molecular Biology | CSIR-Centre for Cellular and Molecular Biology | Lamuk Zaveri, Shagufta Khan, Namami Gaur, Sakshi Shambhavi, Tulasi Nagabandi, Purushotham Vodnala, Payel Mukherjee, Sofia Banu, Priya Singh, Dhiviya Vedagiri, Divya Gupta, Vishal Sah, Santosh Kumar Kuncha, Krishnan Harinivas Harshan, Archana Bharadwaj Siva, Karthik Bharadwaj Tallapaka, Renu Sudhakar, Somesh Gorde, Gangumala Srinivas Reddy, Sujoy Deb, Swati Bayyana, Rakesh K Mishra, Divya Tej Sowpati |
| EPI_ISL_471617 | CSIR-Centre for Cellular and Molecular Biology | CSIR-Centre for Cellular and Molecular Biology | Lamuk Zaveri, Shagufta Khan, Namami Gaur, Sakshi Shambhavi, Tulasi Nagabandi, Purushotham Vodnala, Payel Mukherjee, Sofia Banu, Priya Singh, Dhiviya Vedagiri, Divya Gupta, Vishal Sah, Santosh Kumar Kuncha, Krishnan Harinivas Harshan, Archana Bharadwaj Siva, Karthik Bharadwaj Tallapaka,Umesh Kumar, Unis Ahmad Bhat, Ajay Sarawagi, Priyanka Pant, Rajkanwar Nathawat, Rakesh K Mishra, Divya Tej Sowpati |
| EPI_ISL_471618 | CSIR-Centre for Cellular and Molecular Biology | CSIR-Centre for Cellular and Molecular Biology | Lamuk Zaveri, Shagufta Khan, Namami Gaur, Sakshi Shambhavi, Tulasi Nagabandi, Purushotham Vodnala, Payel Mukherjee, Sofia Banu, Priya Singh, Dhiviya Vedagiri, Divya Gupta, Vishal Sah, Santosh Kumar Kuncha, Krishnan Harinivas Harshan, Archana Bharadwaj Siva, Karthik Bharadwaj Tallapaka,Zeba Rizvi, Zuberwasim Sayyad, Kakade Aishwarya Arun, Amrutha H C, Ananga Ghosh, Rakesh K Mishra, Divya Tej Sowpati |
| EPI_ISL_471619 | CSIR-Centre for Cellular and Molecular Biology | CSIR-Centre for Cellular and Molecular Biology | Namami Gaur, Sakshi Shambhavi, Lamuk Zaveri, Shagufta Khan, Tulasi Nagabandi, Purushotham Vodnala, Payel Mukherjee, Sofia Banu, Priya Singh, Dhiviya Vedagiri, Divya Gupta, Vishal Sah, Santosh Kumar Kuncha, Krishnan Harinivas Harshan, Archana Bharadwaj Siva, Karthik Bharadwaj Tallapaka, Zeba Rizvi, Zuberwasim Sayyad, Kakade Aishwarya Arun, Amrutha H C, Ananga Ghosh, Rakesh K Mishra, Divya Tej Sowpati |
| EPI_ISL_471620 | CSIR-Centre for Cellular and Molecular Biology | CSIR-Centre for Cellular and Molecular Biology | Namami Gaur, Sakshi Shambhavi, Lamuk Zaveri, Shagufta Khan, Tulasi Nagabandi, Purushotham Vodnala, Payel Mukherjee, Sofia Banu, Priya Singh, Dhiviya Vedagiri, Divya Gupta, Vishal Sah, Santosh Kumar Kuncha, Krishnan Harinivas Harshan, Archana Bharadwaj Siva, Karthik Bharadwaj Tallapaka, Nikhil Hajirnis, Pratheusa Maccha, M Soujanya Reddy,G. Aditya Kumar, Koushick Sivakumar, Rakesh K Mishra, Divya Tej Sowpati |
| EPI_ISL_471621 | CSIR-Centre for Cellular and Molecular Biology | CSIR-Centre for Cellular and Molecular Biology | Namami Gaur, Sakshi Shambhavi, Lamuk Zaveri, Shagufta Khan, Tulasi Nagabandi, Purushotham Vodnala, Payel Mukherjee, Sofia Banu, Priya Singh, Dhiviya Vedagiri, Divya Gupta, Vishal Sah, Santosh Kumar Kuncha, Krishnan Harinivas Harshan, Archana Bharadwaj Siva, Karthik Bharadwaj Tallapaka, Zeba Rizvi, Zuberwasim Sayyad, Kakade Aishwarya Arun, Amrutha H C, Ananga Ghosh, Rakesh K Mishra, Divya Tej Sowpati |
| EPI_ISL_471622 | CSIR-Centre for Cellular and Molecular Biology | CSIR-Centre for Cellular and Molecular Biology | Namami Gaur, Sakshi Shambhavi, Lamuk Zaveri, Shagufta Khan, Tulasi Nagabandi, Purushotham Vodnala, Payel Mukherjee, Sofia Banu, Priya Singh, Dhiviya Vedagiri, Divya Gupta, Vishal Sah, Santosh Kumar Kuncha, Krishnan Harinivas Harshan, Archana Bharadwaj Siva, Karthik Bharadwaj Tallapaka,Kezia J Ann, Radhika Khandelwal, Roshan Maku Venkata, Shemin Mansuri, Sonu Uday, Rakesh K Mishra, Divya Tej Sowpati |
| EPI_ISL_471623 | CSIR-Centre for Cellular and Molecular Biology | CSIR-Centre for Cellular and Molecular Biology | Payel Mukherjee, Sofia Banu, Priya Singh, Dhiviya Vedagiri, Divya Gupta, Vishal Sah, Santosh Kumar Kuncha, Krishnan Harinivas Harshan, Archana Bharadwaj Siva, Karthik Bharadwaj Tallapaka, Shagufta Khan, Lamuk Zaveri, Namami Gaur, Sakshi Shambhavi, Tulasi Nagabandi, Purushotham Vodnala, G. Aditya Kumar, Koushick Sivakumar, Pooja Ramesh Gupta, Rajan Kumar Jha, Shraddha Vijay Lahoti, Rakesh K Mishra, Divya Tej Sowpati |
| EPI_ISL_471624 | CSIR-Centre for Cellular and Molecular Biology | CSIR-Centre for Cellular and Molecular Biology | Payel Mukherjee, Sofia Banu, Priya Singh, Dhiviya Vedagiri, Divya Gupta, Vishal Sah, Santosh Kumar Kuncha, Krishnan Harinivas Harshan, Archana Bharadwaj Siva, Karthik Bharadwaj Tallapaka, Shagufta Khan, Lamuk Zaveri, Namami Gaur, Sakshi Shambhavi, Tulasi Nagabandi, Purushotham Vodnala, Gokulan C G, Gunjan Purohit, Hanuman Tulashiram Kale, Pankaj Kumar, Prachand Issarapu, Rakesh K Mishra, Divya Tej Sowpati |
| EPI_ISL_471625 | CSIR-Centre for Cellular and Molecular Biology | CSIR-Centre for Cellular and Molecular Biology | Payel Mukherjee, Sofia Banu, Priya Singh, Dhiviya Vedagiri, Divya Gupta, Vishal Sah, Santosh Kumar Kuncha, Krishnan Harinivas Harshan, Archana Bharadwaj Siva, Karthik Bharadwaj Tallapaka, Shagufta Khan, Lamuk Zaveri, Namami Gaur, Sakshi Shambhavi, Tulasi Nagabandi, Purushotham Vodnala, Rakesh K Mishra, Sonu Uday, Sudipta Mondal, Annapoorna P Karthyayani, Debabrata Jana, Debrya Saha, Divya Tej Sowpati |
| EPI_ISL_471626 | CSIR-Centre for Cellular and Molecular Biology | CSIR-Centre for Cellular and Molecular Biology | Payel Mukherjee, Sofia Banu, Priya Singh, Dhiviya Vedagiri, Divya Gupta, Vishal Sah, Santosh Kumar Kuncha, Krishnan Harinivas Harshan, Archana Bharadwaj Siva, Karthik Bharadwaj Tallapaka, Shagufta Khan, Lamuk Zaveri, Namami Gaur, Sakshi Shambhavi, Tulasi Nagabandi, Purushotham Vodnala,Deepak Kumar, Devi Prasad Vijayashankar, Disha Nanda, Divya Das, Jotin Gogoi, Manish Bhattacharjee, Rakesh K Mishra, Divya Tej Sowpati |
| EPI_ISL_471627 | CSIR-Centre for Cellular and Molecular Biology | CSIR-Centre for Cellular and Molecular Biology | Sakshi Shambhavi, Lamuk Zaveri, Shagufta Khan, Namami Gaur, Tulasi Nagabandi, Purushotham Vodnala, Payel Mukherjee, Sofia Banu, Priya Singh, Dhiviya Vedagiri, Divya Gupta, Vishal Sah, Santosh Kumar Kuncha, Krishnan Harinivas Harshan, Archana Bharadwaj Siva, Karthik Bharadwaj Tallapaka, Deepak Kumar, Devi Prasad Vijayashankar, Disha Nanda, Divya Das, Jotin Gogoi, Manish Bhattacharjee, Rakesh K Mishra, Divya Tej Sowpati |
| EPI_ISL_471628 | CSIR-Centre for Cellular and Molecular Biology | CSIR-Centre for Cellular and Molecular Biology | Sakshi Shambhavi, Lamuk Zaveri, Shagufta Khan, Namami Gaur, Tulasi Nagabandi, Purushotham Vodnala, Payel Mukherjee, Sofia Banu, Priya Singh, Dhiviya Vedagiri, Divya Gupta, Vishal Sah, Santosh Kumar Kuncha, Krishnan Harinivas Harshan, Archana Bharadwaj Siva, Karthik Bharadwaj Tallapaka, G. Aditya Kumar, Koushick Sivakumar, Pooja Ramesh Gupta, Rajan Kumar Jha, Shraddha Vijay Lahoti, Rakesh K Mishra, Divya Tej Sowpati |
| EPI_ISL_471629 | CSIR-Centre for Cellular and Molecular Biology | CSIR-Centre for Cellular and Molecular Biology | Sakshi Shambhavi, Lamuk Zaveri, Shagufta Khan, Namami Gaur, Tulasi Nagabandi, Purushotham Vodnala, Payel Mukherjee, Sofia Banu, Priya Singh, Dhiviya Vedagiri, Divya Gupta, Vishal Sah, Santosh Kumar Kuncha, Krishnan Harinivas Harshan, Archana Bharadwaj Siva, Karthik Bharadwaj Tallapaka,Nikhil Hajirnis, Pratheusa Maccha, M Soujanya Reddy,G. Aditya Kumar, Koushick Sivakumar, Rakesh K Mishra, Divya Tej Sowpati |
| EPI_ISL_471630 | CSIR-Centre for Cellular and Molecular Biology | CSIR-Centre for Cellular and Molecular Biology | Sakshi Shambhavi, Lamuk Zaveri, Shagufta Khan, Namami Gaur, Tulasi Nagabandi, Purushotham Vodnala, Payel Mukherjee, Sofia Banu, Priya Singh, Dhiviya Vedagiri, Divya Gupta, Vishal Sah, Santosh Kumar Kuncha, Krishnan Harinivas Harshan, Archana Bharadwaj Siva, Karthik Bharadwaj Tallapaka,Nikhil Hajirnis, Pratheusa Maccha, M Soujanya Reddy,G. Aditya Kumar, Koushick Sivakumar,Disha Nanda, Divya Das, Jotin Gogoi, Manish Bhattacharjee, Ravi Prasad Mukku, Rakesh K Mishra, Divya Tej Sowpati |
| EPI_ISL_471631 | CSIR-Centre for Cellular and Molecular Biology | CSIR-Centre for Cellular and Molecular Biology | Shagufta Khan, Lamuk Zaveri, Namami Gaur, Sakshi Shambhavi, Tulasi Nagabandi, Purushotham Vodnala, Payel Mukherjee, Sofia Banu, Priya Singh, Dhiviya Vedagiri, Divya Gupta, Vishal Sah, Santosh Kumar Kuncha, Krishnan Harinivas Harshan, Archana Bharadwaj Siva, Karthik Bharadwaj Tallapaka, Disha Nanda, Divya Das, Jotin Gogoi, Manish Bhattacharjee, Ravi Prasad Mukku, Rakesh K Mishra, Divya Tej Sowpati |
| EPI_ISL_471632 | CSIR-Centre for Cellular and Molecular Biology | CSIR-Centre for Cellular and Molecular Biology | Shagufta Khan, Lamuk Zaveri, Namami Gaur, Sakshi Shambhavi, Tulasi Nagabandi, Purushotham Vodnala, Payel Mukherjee, Sofia Banu, Priya Singh, Dhiviya Vedagiri, Divya Gupta, Vishal Sah, Santosh Kumar Kuncha, Krishnan Harinivas Harshan, Archana Bharadwaj Siva, Karthik Bharadwaj Tallapaka, Renu Sudhakar, Somesh Gorde, Gangumala Srinivas Reddy, Sujoy Deb, Swati Bayyana, Rakesh K Mishra, Divya Tej Sowpati |
| EPI_ISL_471633 | CSIR-Centre for Cellular and Molecular Biology | CSIR-Centre for Cellular and Molecular Biology | Shagufta Khan, Lamuk Zaveri, Namami Gaur, Sakshi Shambhavi, Tulasi Nagabandi, Purushotham Vodnala, Payel Mukherjee, Sofia Banu, Priya Singh, Dhiviya Vedagiri, Divya Gupta, Vishal Sah, Santosh Kumar Kuncha, Krishnan Harinivas Harshan, Archana Bharadwaj Siva, Karthik Bharadwaj Tallapaka,Preethi Jampala, Sharada Ravi Iyer, Sulagana Mukherjee, Swetha Sundar, Peddapuvala Sai Uday Kiran Rakesh K Mishra, Divya Tej Sowpati |
| EPI_ISL_471634 | CSIR-Centre for Cellular and Molecular Biology | CSIR-Centre for Cellular and Molecular Biology | Shagufta Khan, Lamuk Zaveri, Namami Gaur, Sakshi Shambhavi, Tulasi Nagabandi, Purushotham Vodnala, Payel Mukherjee, Sofia Banu, Priya Singh, Dhiviya Vedagiri, Divya Gupta, Vishal Sah, Santosh Kumar Kuncha, Krishnan Harinivas Harshan, Archana Bharadwaj Siva, Karthik Bharadwaj Tallapaka,Umesh Kumar, Unis Ahmad Bhat, Ajay Sarawagi, Priyanka Pant, Rajkanwar Nathawat, Rakesh K Mishra, Divya Tej Sowpati |
| EPI_ISL_471635 | CSIR-Centre for Cellular and Molecular Biology | CSIR-Centre for Cellular and Molecular Biology | Sofia Banu, Payel Mukherjee, Priya Singh, Dhiviya Vedagiri, Divya Gupta, Vishal Sah, Santosh Kumar Kuncha, Krishnan Harinivas Harshan, Archana Bharadwaj Siva, Karthik Bharadwaj Tallapaka, Shagufta Khan, Lamuk Zaveri, Namami Gaur, Sakshi Shambhavi, Tulasi Nagabandi, Purushotham Vodnala, Deepak Kumar, Devi Prasad Vijayashankar, Disha Nanda, Divya Das, Jotin Gogoi, Manish Bhattacharjee, Rakesh K Mishra, Divya Tej Sowpati |
| EPI_ISL_471636 | CSIR-Centre for Cellular and Molecular Biology | CSIR-Centre for Cellular and Molecular Biology | Sofia Banu, Payel Mukherjee, Priya Singh, Dhiviya Vedagiri, Divya Gupta, Vishal Sah, Santosh Kumar Kuncha, Krishnan Harinivas Harshan, Archana Bharadwaj Siva, Karthik Bharadwaj Tallapaka, Shagufta Khan, Lamuk Zaveri, Namami Gaur, Sakshi Shambhavi, Tulasi Nagabandi, Purushotham Vodnala, Disha Nanda, Divya Das, Jotin Gogoi, Manish Bhattacharjee, Ravi Prasad Mukku, Rakesh K Mishra, Divya Tej Sowpati |
| EPI_ISL_471637 | CSIR-Centre for Cellular and Molecular Biology | CSIR-Centre for Cellular and Molecular Biology | Sofia Banu, Payel Mukherjee, Priya Singh, Dhiviya Vedagiri, Divya Gupta, Vishal Sah, Santosh Kumar Kuncha, Krishnan Harinivas Harshan, Archana Bharadwaj Siva, Karthik Bharadwaj Tallapaka, Shagufta Khan, Lamuk Zaveri, Namami Gaur, Sakshi Shambhavi, Tulasi Nagabandi, Purushotham Vodnala, Gokulan C G, Gunjan Purohit, Hanuman Tulashiram Kale, Pankaj Kumar, Prachand Issarapu, Rakesh K Mishra, Divya Tej Sowpati |
| EPI_ISL_471638 | CSIR-Centre for Cellular and Molecular Biology | CSIR-Centre for Cellular and Molecular Biology | Sofia Banu, Payel Mukherjee, Priya Singh, Dhiviya Vedagiri, Divya Gupta, Vishal Sah, Santosh Kumar Kuncha, Krishnan Harinivas Harshan, Archana Bharadwaj Siva, Karthik Bharadwaj Tallapaka, Shagufta Khan, Lamuk Zaveri, Namami Gaur, Sakshi Shambhavi, Tulasi Nagabandi, Purushotham Vodnala,Preethi Jampala, Sharada Ravi Iyer, Sulagana Mukherjee, Swetha Sundar, Peddapuvala Sai Uday Kiran, Rakesh K Mishra, Divya Tej Sowpati |
| EPI_ISL_471639 | CSIR-Centre for Cellular and Molecular Biology | CSIR-Centre for Cellular and Molecular Biology | Tulasi Nagabandi, Namami Gaur, Sakshi Shambhavi, Lamuk Zaveri, Shagufta Khan, Purushotham Vodnala, Payel Mukherjee, Sofia Banu, Priya Singh, Dhiviya Vedagiri, Divya Gupta, Vishal Sah, Santosh Kumar Kuncha, Krishnan Harinivas Harshan, Archana Bharadwaj Siva, Karthik Bharadwaj Tallapaka,G. Aditya Kumar, Koushick Sivakumar, Pooja Ramesh Gupta, Rajan Kumar Jha, Shraddha Vijay Lahoti, Rakesh K Mishra, Divya Tej Sowpati |
| EPI_ISL_471640 | CSIR-Centre for Cellular and Molecular Biology | CSIR-Centre for Cellular and Molecular Biology | Tulasi Nagabandi, Namami Gaur, Sakshi Shambhavi, Lamuk Zaveri, Shagufta Khan, Purushotham Vodnala, Payel Mukherjee, Sofia Banu, Priya Singh, Dhiviya Vedagiri, Divya Gupta, Vishal Sah, Santosh Kumar Kuncha, Krishnan Harinivas Harshan, Archana Bharadwaj Siva, Karthik Bharadwaj Tallapaka,Kezia J Ann, Radhika Khandelwal, Roshan Maku Venkata, Shemin Mansuri, Sonu Uday, Rakesh K Mishra, Divya Tej Sowpati |
| EPI_ISL_471641, EPI_ISL_471642 | CSIR-Centre for Cellular and Molecular Biology | CSIR-Centre for Cellular and Molecular Biology | Dhiviya Vedagiri, Divya Gupta, Vishal Sah, Payel Mukherjee, Sofia Banu, Priya Singh, Santosh Kumar Kuncha, Archana Bharadwaj Siva, Karthik Bharadwaj Tallapaka, Shagufta Khan, Lamuk Zaveri, Namami Gaur, Sakshi Shambhavi, Tulasi Nagabandi, Purushotham Vodnala, Rakesh K Mishra, Divya Tej Sowpati, Krishnan Harinivas Harshan |
| EPI_ISL_471643 | CSIR-Centre for Cellular and Molecular Biology | CSIR-Centre for Cellular and Molecular Biology | Tulasi Nagabandi, Namami Gaur, Sakshi Shambhavi, Lamuk Zaveri, Shagufta Khan, Purushotham Vodnala, Payel Mukherjee, Sofia Banu, Priya Singh, Dhiviya Vedagiri, Divya Gupta, Vishal Sah, Santosh Kumar Kuncha, Krishnan Harinivas Harshan, Archana Bharadwaj Siva, Karthik Bharadwaj Tallapaka,G. Aditya Kumar, Koushick Sivakumar, Pooja Ramesh Gupta, Rajan Kumar Jha, Shraddha Vijay Lahoti, Rakesh K Mishra, Divya Tej Sowpati |
| EPI_ISL_471644 | CSIR-Centre for Cellular and Molecular Biology | CSIR-Centre for Cellular and Molecular Biology | Tulasi Nagabandi, Namami Gaur, Sakshi Shambhavi, Lamuk Zaveri, Shagufta Khan, Purushotham Vodnala, Payel Mukherjee, Sofia Banu, Priya Singh, Dhiviya Vedagiri, Divya Gupta, Vishal Sah, Santosh Kumar Kuncha, Krishnan Harinivas Harshan, Archana Bharadwaj Siva, Karthik Bharadwaj Tallapaka,Kezia J Ann, Radhika Khandelwal, Roshan Maku Venkata, Shemin Mansuri, Sonu Uday, Rakesh K Mishra, Divya Tej Sowpati |
| EPI_ISL_471645, EPI_ISL_471646 | CSIR-Centre for Cellular and Molecular Biology | CSIR-Centre for Cellular and Molecular Biology | Dhiviya Vedagiri, Divya Gupta, Vishal Sah, Payel Mukherjee, Sofia Banu, Priya Singh, Santosh Kumar Kuncha, Archana Bharadwaj Siva, Karthik Bharadwaj Tallapaka, Shagufta Khan, Lamuk Zaveri, Namami Gaur, Sakshi Shambhavi, Tulasi Nagabandi, Purushotham Vodnala, Rakesh K Mishra, Divya Tej Sowpati, Krishnan Harinivas Harshan |
| EPI_ISL_471647 | Hospital Municipal de Barueri Dr. Francisco Moran | Instituto Adolfo Lutz, Interdiciplinary Procedures Center, Strategic Laboratory | Claudio Tavares Sacchi, Claudia Regina Gonçalves, Erica Valessa Ramos Gomes |
| EPI_ISL_471648 | UBS e Pronto Socorro Jd. Jacira | Instituto Adolfo Lutz, Interdiciplinary Procedures Center, Strategic Laboratory | Claudio Tavares Sacchi, Claudia Regina Gonçalves, Erica Valessa Ramos Gomes |
| EPI_ISL_471676, EPI_ISL_471677, EPI_ISL_471678, EPI_ISL_471679, EPI_ISL_471680, EPI_ISL_471681, EPI_ISL_471682, EPI_ISL_471683, EPI_ISL_471684, EPI_ISL_471685, EPI_ISL_471686, EPI_ISL_471687, EPI_ISL_471688, EPI_ISL_471689, EPI_ISL_471690, EPI_ISL_471691, EPI_ISL_471692, EPI_ISL_471693, EPI_ISL_471694, EPI_ISL_471695, EPI_ISL_471696, EPI_ISL_471697, EPI_ISL_471698, EPI_ISL_471699, EPI_ISL_471700, EPI_ISL_471701, EPI_ISL_471702, EPI_ISL_471703, EPI_ISL_471704, EPI_ISL_471705, EPI_ISL_471706, EPI_ISL_471707, EPI_ISL_471708, EPI_ISL_471709, EPI_ISL_471710, EPI_ISL_471711, EPI_ISL_471712, EPI_ISL_471713, EPI_ISL_471714, EPI_ISL_471715, EPI_ISL_471716, EPI_ISL_471717, EPI_ISL_471718, EPI_ISL_471719, EPI_ISL_471720, EPI_ISL_471721, EPI_ISL_471722, EPI_ISL_471723, EPI_ISL_471724, EPI_ISL_471725, EPI_ISL_471726, EPI_ISL_471727, EPI_ISL_471728, EPI_ISL_471729, EPI_ISL_471730, EPI_ISL_471731, EPI_ISL_471732, EPI_ISL_471733, EPI_ISL_471734, EPI_ISL_471735, EPI_ISL_471736, EPI_ISL_471737, EPI_ISL_471738, EPI_ISL_471739, EPI_ISL_471740, EPI_ISL_471741, EPI_ISL_471742, EPI_ISL_471743, EPI_ISL_471744, EPI_ISL_471745, EPI_ISL_471746, EPI_ISL_471747, EPI_ISL_471748, EPI_ISL_471749, EPI_ISL_471750, EPI_ISL_471751, EPI_ISL_471752, EPI_ISL_471753, EPI_ISL_471754, EPI_ISL_471755, EPI_ISL_471756, EPI_ISL_471757, EPI_ISL_471758, EPI_ISL_471759, EPI_ISL_471760, EPI_ISL_471761, EPI_ISL_471762, EPI_ISL_471763, EPI_ISL_471764, EPI_ISL_471765, EPI_ISL_471766, EPI_ISL_471767, EPI_ISL_471768, EPI_ISL_471769, EPI_ISL_471770, EPI_ISL_471771, EPI_ISL_471772, EPI_ISL_471773, EPI_ISL_471774, EPI_ISL_471775, EPI_ISL_471776, EPI_ISL_471777, EPI_ISL_471778, EPI_ISL_471779, EPI_ISL_471780, EPI_ISL_471781, EPI_ISL_471782, EPI_ISL_471783, EPI_ISL_471784, EPI_ISL_471785, EPI_ISL_471786, EPI_ISL_471787, EPI_ISL_471788, EPI_ISL_471789, EPI_ISL_471790, EPI_ISL_471791, EPI_ISL_471792, EPI_ISL_471793, EPI_ISL_471794, EPI_ISL_471795, EPI_ISL_471796, EPI_ISL_471797, EPI_ISL_471798, EPI_ISL_471799, EPI_ISL_471800, EPI_ISL_471801, EPI_ISL_471802, EPI_ISL_471803, EPI_ISL_471804, EPI_ISL_471805, EPI_ISL_471806, EPI_ISL_471807, EPI_ISL_471808, EPI_ISL_471809, EPI_ISL_471810, EPI_ISL_471811, EPI_ISL_471812, EPI_ISL_471813, EPI_ISL_471814, EPI_ISL_471815, EPI_ISL_471816, EPI_ISL_471817, EPI_ISL_471818, EPI_ISL_471819, EPI_ISL_471820, EPI_ISL_471821, EPI_ISL_471822, EPI_ISL_471823, EPI_ISL_471824, EPI_ISL_471825, EPI_ISL_471826, EPI_ISL_471827, EPI_ISL_471828, EPI_ISL_471829, EPI_ISL_471830, EPI_ISL_471831, EPI_ISL_471832, EPI_ISL_471833, EPI_ISL_471834, EPI_ISL_471835, EPI_ISL_471836, EPI_ISL_471837, EPI_ISL_471838, EPI_ISL_471839, EPI_ISL_471840, EPI_ISL_471841, EPI_ISL_471842, EPI_ISL_471843, EPI_ISL_471844, EPI_ISL_471845, EPI_ISL_471846, EPI_ISL_471847, EPI_ISL_471848, EPI_ISL_471849, EPI_ISL_471850, EPI_ISL_471851, EPI_ISL_471852, EPI_ISL_471853, EPI_ISL_471854, EPI_ISL_471855, EPI_ISL_471856, EPI_ISL_471857, EPI_ISL_471858, EPI_ISL_471859, EPI_ISL_471860, EPI_ISL_471861, EPI_ISL_471862, EPI_ISL_471863, EPI_ISL_471864, EPI_ISL_471865, EPI_ISL_471866, EPI_ISL_471867, EPI_ISL_471868, EPI_ISL_471869, EPI_ISL_471870, EPI_ISL_471871, EPI_ISL_471872, EPI_ISL_471873, EPI_ISL_471874, EPI_ISL_471875, EPI_ISL_471876, EPI_ISL_471877, EPI_ISL_471878, EPI_ISL_471879, EPI_ISL_471880, EPI_ISL_471881, EPI_ISL_471882, EPI_ISL_471883, EPI_ISL_471884, EPI_ISL_471885, EPI_ISL_471886, EPI_ISL_471887, EPI_ISL_471888, EPI_ISL_471889, EPI_ISL_471890, EPI_ISL_471891, EPI_ISL_471892, EPI_ISL_471893, EPI_ISL_471894, EPI_ISL_471895, EPI_ISL_471896, EPI_ISL_471897, EPI_ISL_471898, EPI_ISL_471899, EPI_ISL_471900, EPI_ISL_471901, EPI_ISL_471902, EPI_ISL_471903, EPI_ISL_471904, EPI_ISL_471905, EPI_ISL_471906, EPI_ISL_471907, EPI_ISL_471908, EPI_ISL_471909, EPI_ISL_471910 | | | |

| | | | |
|---|---|---|---|
| see above | Michigan Department of Health and Human Services, Bureau of Laboratories | Michigan Department of Health and Human Services, Bureau of Laboratories | Blankenship HM, Riner D, Soehnlen MK |
| EPI_ISL_471911, EPI_ISL_471912, EPI_ISL_471913, EPI_ISL_471914, EPI_ISL_471915, EPI_ISL_471916, EPI_ISL_471917, EPI_ISL_471918, EPI_ISL_471919, EPI_ISL_471920, EPI_ISL_471921, EPI_ISL_471922, EPI_ISL_471923, EPI_ISL_471924, EPI_ISL_471925, EPI_ISL_471926, EPI_ISL_471927, EPI_ISL_471928, EPI_ISL_471929, EPI_ISL_471930, EPI_ISL_471931, EPI_ISL_471932, EPI_ISL_471933, EPI_ISL_471934, EPI_ISL_471935, EPI_ISL_471936, EPI_ISL_471937, EPI_ISL_471938, EPI_ISL_471939, EPI_ISL_471940, EPI_ISL_471941, EPI_ISL_471942, EPI_ISL_471943, EPI_ISL_471944, EPI_ISL_471945, EPI_ISL_471946, EPI_ISL_471947, EPI_ISL_471948, EPI_ISL_471949, EPI_ISL_471950, EPI_ISL_471951, EPI_ISL_471952, EPI_ISL_471953, EPI_ISL_471954, EPI_ISL_471955, EPI_ISL_471956, EPI_ISL_471957, EPI_ISL_471958, EPI_ISL_471959, EPI_ISL_471960, EPI_ISL_471961, EPI_ISL_471962, EPI_ISL_471963, EPI_ISL_471964, EPI_ISL_471965, EPI_ISL_471966, EPI_ISL_471967, EPI_ISL_471968, EPI_ISL_471969, EPI_ISL_471970, EPI_ISL_471971, EPI_ISL_471972, EPI_ISL_471973, EPI_ISL_471974, EPI_ISL_471975, EPI_ISL_471976, EPI_ISL_471977, EPI_ISL_471978, EPI_ISL_471979, EPI_ISL_471980, EPI_ISL_471981, EPI_ISL_471982, EPI_ISL_471983, EPI_ISL_471984, EPI_ISL_471985, EPI_ISL_471986, EPI_ISL_471987, EPI_ISL_471988, EPI_ISL_471989, EPI_ISL_471990, EPI_ISL_471991, EPI_ISL_471992 | | | |
| see above | University of Exeter | COVID-19 Genomics UK (COG-UK) Consortium | Ben Temperton,Aaron Jeffries,Michelle Michelsen,Joanna Warwick-Dugdale,Audrey Farbos,Robyn Manley,Stephen Michell,Jane Masoli |
| EPI_ISL_471993, EPI_ISL_471994, EPI_ISL_471995, EPI_ISL_471996, EPI_ISL_471997, EPI_ISL_471998, EPI_ISL_471999, EPI_ISL_472000, EPI_ISL_472001, EPI_ISL_472002, EPI_ISL_472003, EPI_ISL_472004, EPI_ISL_472005, EPI_ISL_472006, EPI_ISL_472007, EPI_ISL_472008, EPI_ISL_472009, EPI_ISL_472010, EPI_ISL_472011, EPI_ISL_472012, EPI_ISL_472013, EPI_ISL_472014, EPI_ISL_472015, EPI_ISL_472016, EPI_ISL_472017, EPI_ISL_472018, EPI_ISL_472019, EPI_ISL_472020, EPI_ISL_472021, EPI_ISL_472022, EPI_ISL_472023, EPI_ISL_472024, EPI_ISL_472025, EPI_ISL_472026, EPI_ISL_472027, EPI_ISL_472028, EPI_ISL_472029, EPI_ISL_472030, EPI_ISL_472031, EPI_ISL_472032, EPI_ISL_472033, EPI_ISL_472034, EPI_ISL_472035, EPI_ISL_472036, EPI_ISL_472037, EPI_ISL_472038, EPI_ISL_472039, EPI_ISL_472040, EPI_ISL_472041, EPI_ISL_472042, EPI_ISL_472043, EPI_ISL_472044, EPI_ISL_472045, EPI_ISL_472046, EPI_ISL_472047, EPI_ISL_472048, EPI_ISL_472049, EPI_ISL_472050, EPI_ISL_472051, EPI_ISL_472052, EPI_ISL_472053, EPI_ISL_472054, EPI_ISL_472055, EPI_ISL_472056, EPI_ISL_472057, EPI_ISL_472058, EPI_ISL_472059, EPI_ISL_472060, EPI_ISL_472061, EPI_ISL_472062, EPI_ISL_472063, EPI_ISL_472064, EPI_ISL_472065, EPI_ISL_472066, EPI_ISL_472067, EPI_ISL_472068, EPI_ISL_472069, EPI_ISL_472070, EPI_ISL_472071, EPI_ISL_472072, EPI_ISL_472073, EPI_ISL_472074, EPI_ISL_472075, EPI_ISL_472076, EPI_ISL_472077, EPI_ISL_472078, EPI_ISL_472079, EPI_ISL_472080, EPI_ISL_472081, EPI_ISL_472082, EPI_ISL_472083, EPI_ISL_472084, EPI_ISL_472085, EPI_ISL_472086, EPI_ISL_472087, EPI_ISL_472088, EPI_ISL_472089, EPI_ISL_472090, EPI_ISL_472091, EPI_ISL_472092, EPI_ISL_472093, EPI_ISL_472094, EPI_ISL_472095, EPI_ISL_472096, EPI_ISL_472097, EPI_ISL_472098, EPI_ISL_472099, EPI_ISL_472100, EPI_ISL_472101, EPI_ISL_472102, EPI_ISL_472103, EPI_ISL_472104, EPI_ISL_472105, EPI_ISL_472106, EPI_ISL_472107, EPI_ISL_472108, EPI_ISL_472109, EPI_ISL_472110, EPI_ISL_472111, EPI_ISL_472112, EPI_ISL_472113, EPI_ISL_472114, EPI_ISL_472115, EPI_ISL_472116, EPI_ISL_472117, EPI_ISL_472118, EPI_ISL_472119, EPI_ISL_472120, EPI_ISL_472121, EPI_ISL_472122, EPI_ISL_472123, EPI_ISL_472124, EPI_ISL_472125, EPI_ISL_472126, EPI_ISL_472127, EPI_ISL_472128, EPI_ISL_472129, EPI_ISL_472130, EPI_ISL_472131, EPI_ISL_472132, EPI_ISL_472133, EPI_ISL_472134, EPI_ISL_472135 | | | |
| see above | Liverpool Clinical Laboratories | COVID-19 Genomics UK (COG-UK) Consortium | Sam Haldenby, Anita Lucaci, Steve Paterson, Julian Hiscox, Alistair Darby, M Almsaud, A Alrezaihi, Muhannad Alruwaili, Stuart D Armstrong, Jones Benjamin, Eleanor G Bentley, Anu Chawla, Jordan J Clark, Angela Cowell, Richard Eccles, Isabel García-Dorival, Matthew Gemmell, Alessandro Gerada, PKF Gilmore, Richard Gregory, Ximeng Han, Catherine Hartley, Margaret Hughes, Miren Iturriza-Gomara, James Johnson, L Luu, Jenifer Manson, Charlotte Nelson, Elaine O'Toole, Cassie Olateju, Rebekah Penrice-Randal , Lucille Rainbow, N.P Randle, Trevor Ian Robinson, Parul Sharma, Ghada T Shawli, James P Stewart, Neil Swainston, Ecaterina Vamos, Joanne Watts, Mark Whitehead |
| EPI_ISL_472136, EPI_ISL_472137, EPI_ISL_472138, EPI_ISL_472139, EPI_ISL_472140, EPI_ISL_472141, EPI_ISL_472142, EPI_ISL_472143, EPI_ISL_472144, EPI_ISL_472145, EPI_ISL_472146, EPI_ISL_472147, EPI_ISL_472148, EPI_ISL_472149, EPI_ISL_472150, EPI_ISL_472151, EPI_ISL_472152, EPI_ISL_472153, EPI_ISL_472154 | | | |
| see above | Regional Virus Laboratory, Belfast Health and Social Care Trust | COVID-19 Genomics UK (COG-UK) Consortium | Conall McCaughey, James McKenna, Tanya Curran, Susan Feeney, Alison Watt, Ciara Cox, Mairead Connor, Zoltan Molnar, David Simpson, Derek Fairley |
| EPI_ISL_472155, EPI_ISL_472156, EPI_ISL_472157, EPI_ISL_472158, EPI_ISL_472159, EPI_ISL_472160, EPI_ISL_472161, EPI_ISL_472162, EPI_ISL_472163, EPI_ISL_472164, EPI_ISL_472165, EPI_ISL_472166, EPI_ISL_472167, EPI_ISL_472168, EPI_ISL_472169, EPI_ISL_472170, EPI_ISL_472171, EPI_ISL_472172, EPI_ISL_472173, EPI_ISL_472174, EPI_ISL_472175, EPI_ISL_472176, EPI_ISL_472177, EPI_ISL_472178, EPI_ISL_472179, EPI_ISL_472180, EPI_ISL_472181, EPI_ISL_472182, EPI_ISL_472183, EPI_ISL_472184, EPI_ISL_472185, EPI_ISL_472186, EPI_ISL_472187, EPI_ISL_472188, EPI_ISL_472189, EPI_ISL_472190, EPI_ISL_472191, EPI_ISL_472192, EPI_ISL_472193, EPI_ISL_472194, EPI_ISL_472195, EPI_ISL_472196, EPI_ISL_472197, EPI_ISL_472198, EPI_ISL_472199, EPI_ISL_472200, EPI_ISL_472201, EPI_ISL_472202, EPI_ISL_472203, EPI_ISL_472204, EPI_ISL_472205, EPI_ISL_472206, EPI_ISL_472207, EPI_ISL_472208, EPI_ISL_472209, EPI_ISL_472210, EPI_ISL_472211, EPI_ISL_472212, EPI_ISL_472213, EPI_ISL_472214, EPI_ISL_472215, EPI_ISL_472216, EPI_ISL_472217, EPI_ISL_472218, EPI_ISL_472219, EPI_ISL_472220, EPI_ISL_472221, EPI_ISL_472222, EPI_ISL_472223, EPI_ISL_472224, EPI_ISL_472225, EPI_ISL_472226, EPI_ISL_472227, EPI_ISL_472228, EPI_ISL_472229, EPI_ISL_472230, EPI_ISL_472231, EPI_ISL_472232, EPI_ISL_472233, EPI_ISL_472234, EPI_ISL_472235, EPI_ISL_472236, EPI_ISL_472237, EPI_ISL_472238, EPI_ISL_472239, EPI_ISL_472240, EPI_ISL_472241, EPI_ISL_472242, EPI_ISL_472243, EPI_ISL_472244, EPI_ISL_472245, EPI_ISL_472246, EPI_ISL_472247, EPI_ISL_472248, EPI_ISL_472249, EPI_ISL_472250, EPI_ISL_472251, EPI_ISL_472252, EPI_ISL_472253, EPI_ISL_472254, EPI_ISL_472255, EPI_ISL_472256, EPI_ISL_472257, EPI_ISL_472258, EPI_ISL_472259, EPI_ISL_472260, EPI_ISL_472261, EPI_ISL_472262, EPI_ISL_472263, EPI_ISL_472264, EPI_ISL_472265, EPI_ISL_472266, EPI_ISL_472267, EPI_ISL_472268, EPI_ISL_472269, EPI_ISL_472270, EPI_ISL_472271, EPI_ISL_472272, EPI_ISL_472273, EPI_ISL_472274, EPI_ISL_472275, EPI_ISL_472276, EPI_ISL_472277, EPI_ISL_472278, EPI_ISL_472279, EPI_ISL_472280, EPI_ISL_472281, EPI_ISL_472282, EPI_ISL_472283, EPI_ISL_472284, EPI_ISL_472285, EPI_ISL_472286, EPI_ISL_472287, EPI_ISL_472288, EPI_ISL_472289, EPI_ISL_472290 | | | |
| see above | Northumbria University / South Tees Hospitals NHS Foundation Trust / North Cumbria Integrated Care NHS Foundation Trust / North Tees and Hartlepool NHS Foundation Trust / Newcastle Hospitals NHS Foundation Trust | COVID-19 Genomics UK (COG-UK) Consortium | Darren L Smith,Andrew Nelson,Matthew Bashton,Greg R Young,Joshua Loh,John Allan,Mohammad A Tariq,Giles S Holt,Gary Black,Wen C Yew,Lynn Dover,Paul Baker,Steve Liggett,Sarah Essex,Jane Greenaway,Debra Padgett,Clive Graham,Garren Scott,Edward Barton,Emma Swindells,Brendan Payne,Jennifer Collins,Yusri Taha,Gary Eltringham |
| EPI_ISL_472291, EPI_ISL_472292, EPI_ISL_472293, EPI_ISL_472294, EPI_ISL_472295, EPI_ISL_472296, EPI_ISL_472297, EPI_ISL_472298, EPI_ISL_472299, EPI_ISL_472300, EPI_ISL_472301, EPI_ISL_472302, EPI_ISL_472303, EPI_ISL_472304, EPI_ISL_472305, EPI_ISL_472306, EPI_ISL_472307, EPI_ISL_472308, EPI_ISL_472309, EPI_ISL_472310, EPI_ISL_472311, EPI_ISL_472312, EPI_ISL_472313, EPI_ISL_472314, EPI_ISL_472315, EPI_ISL_472316, EPI_ISL_472317, EPI_ISL_472318, EPI_ISL_472319, EPI_ISL_472320, EPI_ISL_472321, EPI_ISL_472322, EPI_ISL_472323, EPI_ISL_472324, EPI_ISL_472325, EPI_ISL_472326, EPI_ISL_472327, EPI_ISL_472328, EPI_ISL_472329, EPI_ISL_472330, EPI_ISL_472331, EPI_ISL_472332, EPI_ISL_472333, EPI_ISL_472334, EPI_ISL_472335, EPI_ISL_472336, EPI_ISL_472337, EPI_ISL_472338, EPI_ISL_472339, EPI_ISL_472340, EPI_ISL_472341, EPI_ISL_472342, EPI_ISL_472343, EPI_ISL_472344, EPI_ISL_472345, EPI_ISL_472346, EPI_ISL_472347, EPI_ISL_472348, EPI_ISL_472349, EPI_ISL_472350, EPI_ISL_472351, EPI_ISL_472352, EPI_ISL_472353, EPI_ISL_472354, EPI_ISL_472355, EPI_ISL_472356, EPI_ISL_472357, EPI_ISL_472358, EPI_ISL_472359, EPI_ISL_472360, EPI_ISL_472361, EPI_ISL_472362, EPI_ISL_472363, EPI_ISL_472364, EPI_ISL_472365, EPI_ISL_472366, EPI_ISL_472367, EPI_ISL_472368, EPI_ISL_472369, EPI_ISL_472370, EPI_ISL_472371, EPI_ISL_472372, EPI_ISL_472373, EPI_ISL_472374, EPI_ISL_472375, EPI_ISL_472376, EPI_ISL_472377, EPI_ISL_472378, EPI_ISL_472379, EPI_ISL_472380, EPI_ISL_472381, EPI_ISL_472382, EPI_ISL_472383 | | | |
| see above | Quadram Institute Bioscience | COVID-19 Genomics UK (COG-UK) Consortium | Dave J. Baker, Gemma L. Kay, Alp Aydin, Thanh Le-Viet, Steven Rudder, Ana P. Tedim, Anastasia Kolyva, Maria Diaz, Leonardo de Oliveira Martins, Nabil-Fareed Alikhan, Lizzie Meadows, Rachael Stanley, Ngozi Elumogo, Muhammed Yasir, Nicholas M. Thomson, Alexander J Trotter, Rachel Gilroy, Samuel Bloomfield, Claire Stuart, Andrew Bell, Reenesh Prakash, Samir Dervisevic, Alison E. Mather, John Wain, Mark Webber, Andrew J. Page, Justin O'Grady |
| EPI_ISL_472384, EPI_ISL_472385, EPI_ISL_472386, EPI_ISL_472387, EPI_ISL_472388, EPI_ISL_472389, EPI_ISL_472390, EPI_ISL_472391, EPI_ISL_472392, EPI_ISL_472393, EPI_ISL_472394, EPI_ISL_472395, EPI_ISL_472396, EPI_ISL_472397, EPI_ISL_472398, EPI_ISL_472399, EPI_ISL_472400, EPI_ISL_472401, EPI_ISL_472402, EPI_ISL_472403, EPI_ISL_472404, EPI_ISL_472405, EPI_ISL_472406, EPI_ISL_472407, EPI_ISL_472408, EPI_ISL_472409, EPI_ISL_472410, EPI_ISL_472411, EPI_ISL_472412, EPI_ISL_472413, EPI_ISL_472414, EPI_ISL_472415, EPI_ISL_472416, EPI_ISL_472417, EPI_ISL_472418, EPI_ISL_472419, EPI_ISL_472420, EPI_ISL_472421, EPI_ISL_472422, EPI_ISL_472423, EPI_ISL_472424, EPI_ISL_472425, EPI_ISL_472426, EPI_ISL_472427, EPI_ISL_472428, EPI_ISL_472429 | | | |
| see above | Queens Medical Centre, Clinical Microbiology Department / DeepSeq Nottingham | COVID-19 Genomics UK (COG-UK) Consortium | Gemma Clark, Wendy Smith, Manjinder Khakh, Vicki M Fleming, Michelle M Lister, Hannah Howson-Wells, Jonathan Ball, Patrick McClure, Joseph Chappell, Theocharis Tsoleridis, Nadine Holmes, Matthew Carlisle, Christopher Moore, Fei Sang, Johnny Debebe, Victoria Wright, Matthew Loose |
| EPI_ISL_472430, EPI_ISL_472431 | Queens Medical Centre, Clinical Microbiology Department / DeepSeq Nottingham | COVID-19 Genomics UK (COG-UK) Consortium | Nichola Duckworth, Tim Sloan, Sarah Walsh, Jonathan Ball, Patrick McClure, Joeseph Chappell, Nadine Holmes, Matthew Carlisle, Christopher Moore, Fei Sang, Johnny Debebe, Victoria Wright, Matthew Loose |
| EPI_ISL_472432, EPI_ISL_472433, EPI_ISL_472434, EPI_ISL_472435, EPI_ISL_472436, EPI_ISL_472437, EPI_ISL_472438, EPI_ISL_472439, EPI_ISL_472440, EPI_ISL_472441, EPI_ISL_472442, EPI_ISL_472443, EPI_ISL_472444, EPI_ISL_472445, EPI_ISL_472446, EPI_ISL_472447, EPI_ISL_472448, EPI_ISL_472449, EPI_ISL_472450, EPI_ISL_472451, EPI_ISL_472452, EPI_ISL_472453, EPI_ISL_472454, EPI_ISL_472455, EPI_ISL_472456, EPI_ISL_472457, EPI_ISL_472458, EPI_ISL_472459, EPI_ISL_472460, EPI_ISL_472461, EPI_ISL_472462, EPI_ISL_472463, EPI_ISL_472464, EPI_ISL_472465, EPI_ISL_472466, EPI_ISL_472467, EPI_ISL_472468, EPI_ISL_472469, EPI_ISL_472470, EPI_ISL_472471, EPI_ISL_472472, EPI_ISL_472473, EPI_ISL_472474, EPI_ISL_472475, EPI_ISL_472476, EPI_ISL_472477, EPI_ISL_472478, EPI_ISL_472479, EPI_ISL_472480, EPI_ISL_472481, EPI_ISL_472482, EPI_ISL_472483, EPI_ISL_472484, EPI_ISL_472485, EPI_ISL_472486, EPI_ISL_472487, EPI_ISL_472488, EPI_ISL_472489, EPI_ISL_472490, EPI_ISL_472491, EPI_ISL_472492, EPI_ISL_472493, EPI_ISL_472494, EPI_ISL_472495, EPI_ISL_472496, EPI_ISL_472497, EPI_ISL_472498, EPI_ISL_472499, EPI_ISL_472500, EPI_ISL_472501, EPI_ISL_472502, EPI_ISL_472503, EPI_ISL_472504, EPI_ISL_472505, EPI_ISL_472506, EPI_ISL_472507, EPI_ISL_472508, EPI_ISL_472509, EPI_ISL_472510, EPI_ISL_472511, EPI_ISL_472512, EPI_ISL_472513, EPI_ISL_472514, EPI_ISL_472515, EPI_ISL_472516, EPI_ISL_472517, EPI_ISL_472518, EPI_ISL_472519, EPI_ISL_472520, EPI_ISL_472521, EPI_ISL_472522, EPI_ISL_472523, EPI_ISL_472524, EPI_ISL_472525, EPI_ISL_472526, EPI_ISL_472527, EPI_ISL_472528, EPI_ISL_472529, EPI_ISL_472530, EPI_ISL_472531, EPI_ISL_472532, EPI_ISL_472533, EPI_ISL_472534, EPI_ISL_472535, EPI_ISL_472536, EPI_ISL_472537, EPI_ISL_472538, EPI_ISL_472539, EPI_ISL_472540, EPI_ISL_472541, EPI_ISL_472542, EPI_ISL_472543, EPI_ISL_472544, EPI_ISL_472545, EPI_ISL_472546, EPI_ISL_472547, EPI_ISL_472548, EPI_ISL_472549, EPI_ISL_472550, EPI_ISL_472551, EPI_ISL_472552, EPI_ISL_472553, EPI_ISL_472554, EPI_ISL_472555, EPI_ISL_472556, EPI_ISL_472557, EPI_ISL_472558, EPI_ISL_472559, EPI_ISL_472560, EPI_ISL_472561, EPI_ISL_472562, EPI_ISL_472563, EPI_ISL_472564, EPI_ISL_472565, EPI_ISL_472566, EPI_ISL_472567, EPI_ISL_472568, EPI_ISL_472569, EPI_ISL_472570, EPI_ISL_472571, EPI_ISL_472572, EPI_ISL_472573, EPI_ISL_472574, EPI_ISL_472575, EPI_ISL_472576, EPI_ISL_472577, EPI_ISL_472578, EPI_ISL_472579, EPI_ISL_472580, EPI_ISL_472581, EPI_ISL_472582, EPI_ISL_472583, EPI_ISL_472584, EPI_ISL_472585, EPI_ISL_472586, EPI_ISL_472587, EPI_ISL_472588, EPI_ISL_472589, EPI_ISL_472590, EPI_ISL_472591, EPI_ISL_472592, EPI_ISL_472593, EPI_ISL_472594, EPI_ISL_472595, EPI_ISL_472596, EPI_ISL_472597, EPI_ISL_472598, EPI_ISL_472599, EPI_ISL_472600, EPI_ISL_472601, EPI_ISL_472602, EPI_ISL_472603, EPI_ISL_472604, EPI_ISL_472605, EPI_ISL_472606, EPI_ISL_472607, EPI_ISL_472608, EPI_ISL_472609, EPI_ISL_472610, EPI_ISL_472611, EPI_ISL_472612, EPI_ISL_472613, EPI_ISL_472614, EPI_ISL_472615, EPI_ISL_472616, EPI_ISL_472617, EPI_ISL_472618, EPI_ISL_472619, EPI_ISL_472620, EPI_ISL_472621, EPI_ISL_472622, EPI_ISL_472623, EPI_ISL_472624, EPI_ISL_472625, EPI_ISL_472626, EPI_ISL_472627, EPI_ISL_472628, EPI_ISL_472629, EPI_ISL_472630, EPI_ISL_472631, EPI_ISL_472632, EPI_ISL_472633, EPI_ISL_472634, EPI_ISL_472635, EPI_ISL_472636, EPI_ISL_472637, EPI_ISL_472638, EPI_ISL_472639, EPI_ISL_472640, EPI_ISL_472641, EPI_ISL_472642, EPI_ISL_472643, EPI_ISL_472644, EPI_ISL_472645, EPI_ISL_472646, EPI_ISL_472647, EPI_ISL_472648, EPI_ISL_472649, EPI_ISL_472650, EPI_ISL_472651, EPI_ISL_472652, EPI_ISL_472653, EPI_ISL_472654, EPI_ISL_472655, EPI_ISL_472656, EPI_ISL_472657, EPI_ISL_472658, EPI_ISL_472659, EPI_ISL_472660, EPI_ISL_472661, EPI_ISL_472662, EPI_ISL_472663, EPI_ISL_472664, EPI_ISL_472665, EPI_ISL_472666, EPI_ISL_472667, EPI_ISL_472668, EPI_ISL_472669, EPI_ISL_472670, EPI_ISL_472671, EPI_ISL_472672, EPI_ISL_472673, EPI_ISL_472674, EPI_ISL_472675, EPI_ISL_472676, EPI_ISL_472677, EPI_ISL_472678, EPI_ISL_472679, EPI_ISL_472680, EPI_ISL_472681, EPI_ISL_472682, EPI_ISL_472683, EPI_ISL_472684, EPI_ISL_472685, EPI_ISL_472686, EPI_ISL_472687, EPI_ISL_472688, EPI_ISL_472689, EPI_ISL_472690, EPI_ISL_472691, EPI_ISL_472692, EPI_ISL_472693, EPI_ISL_472694, EPI_ISL_472695, EPI_ISL_472696, EPI_ISL_472697, EPI_ISL_472698, EPI_ISL_472699, EPI_ISL_472700, EPI_ISL_472701, EPI_ISL_472702, EPI_ISL_472703, EPI_ISL_472704, EPI_ISL_472705, EPI_ISL_472706, EPI_ISL_472707, EPI_ISL_472708, EPI_ISL_472709, EPI_ISL_472710, EPI_ISL_472711, EPI_ISL_472712, EPI_ISL_472713, EPI_ISL_472714, EPI_ISL_472715, EPI_ISL_472716, EPI_ISL_472717, EPI_ISL_472718, EPI_ISL_472719, EPI_ISL_472720, EPI_ISL_472721, EPI_ISL_472722, EPI_ISL_472723, EPI_ISL_472724, EPI_ISL_472725, EPI_ISL_472726, EPI_ISL_472727, EPI_ISL_472728, EPI_ISL_472729, EPI_ISL_472730, EPI_ISL_472731, EPI_ISL_472732, EPI_ISL_472733, EPI_ISL_472734, EPI_ISL_472735, EPI_ISL_472736, EPI_ISL_472737, EPI_ISL_472738, EPI_ISL_472739, EPI_ISL_472740, EPI_ISL_472741, EPI_ISL_472742, EPI_ISL_472743, EPI_ISL_472744, EPI_ISL_472745, EPI_ISL_472746, EPI_ISL_472747, EPI_ISL_472748, EPI_ISL_472749, EPI_ISL_472750, EPI_ISL_472751, EPI_ISL_472752, EPI_ISL_472753, EPI_ISL_472754, EPI_ISL_472755, EPI_ISL_472756, EPI_ISL_472757, EPI_ISL_472758, EPI_ISL_472759, EPI_ISL_472760, EPI_ISL_472761, EPI_ISL_472762, EPI_ISL_472763, EPI_ISL_472764, EPI_ISL_472765, EPI_ISL_472766, EPI_ISL_472767, EPI_ISL_472768, EPI_ISL_472769, EPI_ISL_472770, EPI_ISL_472771, EPI_ISL_472772, EPI_ISL_472773, EPI_ISL_472774, EPI_ISL_472775, EPI_ISL_472776, EPI_ISL_472777, EPI_ISL_472778, EPI_ISL_472779, EPI_ISL_472780, EPI_ISL_472781, EPI_ISL_472782, EPI_ISL_472783, EPI_ISL_472784, EPI_ISL_472785, EPI_ISL_472786, EPI_ISL_472787, EPI_ISL_472788, EPI_ISL_472789, EPI_ISL_472790, EPI_ISL_472791, EPI_ISL_472792, EPI_ISL_472793, EPI_ISL_472794, EPI_ISL_472795, EPI_ISL_472796, EPI_ISL_472797, EPI_ISL_472798, EPI_ISL_472799, EPI_ISL_472800, EPI_ISL_472801, EPI_ISL_472802, EPI_ISL_472803, EPI_ISL_472804, EPI_ISL_472805, EPI_ISL_472806, EPI_ISL_472807, EPI_ISL_472808, EPI_ISL_472809, EPI_ISL_472810, EPI_ISL_472811, EPI_ISL_472812, EPI_ISL_472813, EPI_ISL_472814, EPI_ISL_472815, EPI_ISL_472816, EPI_ISL_472817, EPI_ISL_472818, EPI_ISL_472819, EPI_ISL_472820, EPI_ISL_472821, EPI_ISL_472822, EPI_ISL_472823, EPI_ISL_472824, EPI_ISL_472825, EPI_ISL_472826, EPI_ISL_472827, EPI_ISL_472828, EPI_ISL_472829, EPI_ISL_472830, EPI_ISL_472831, EPI_ISL_472832, EPI_ISL_472833, EPI_ISL_472834, EPI_ISL_472835, EPI_ISL_472836, EPI_ISL_472837, EPI_ISL_472838, EPI_ISL_472839, EPI_ISL_472840, EPI_ISL_472841, EPI_ISL_472842, EPI_ISL_472843, EPI_ISL_472844, EPI_ISL_472845, EPI_ISL_472846, EPI_ISL_472847, EPI_ISL_472848, EPI_ISL_472849, EPI_ISL_472850, EPI_ISL_472851, EPI_ISL_472852, EPI_ISL_472853, EPI_ISL_472854, EPI_ISL_472855, EPI_ISL_472856, EPI_ISL_472857, EPI_ISL_472858, EPI_ISL_472859, EPI_ISL_472860, EPI_ISL_472861, EPI_ISL_472862, EPI_ISL_472863, EPI_ISL_472864, EPI_ISL_472865, EPI_ISL_472866, EPI_ISL_472867, EPI_ISL_472868, EPI_ISL_472869, EPI_ISL_472870, EPI_ISL_472871, EPI_ISL_472872, EPI_ISL_472873, EPI_ISL_472874, EPI_ISL_472875, EPI_ISL_472876, EPI_ISL_472877, EPI_ISL_472878, EPI_ISL_472879, EPI_ISL_472880, EPI_ISL_472881, EPI_ISL_472882, EPI_ISL_472883, EPI_ISL_472884, EPI_ISL_472885, EPI_ISL_472886, EPI_ISL_472887, EPI_ISL_472888, EPI_ISL_472889, EPI_ISL_472890, EPI_ISL_472891, EPI_ISL_472892, EPI_ISL_472893, EPI_ISL_472894, EPI_ISL_472895, EPI_ISL_472896, EPI_ISL_472897, EPI_ISL_472898, EPI_ISL_472899, EPI_ISL_472900, EPI_ISL_472901, EPI_ISL_472902, EPI_ISL_472903, EPI_ISL_472904, EPI_ISL_472905, EPI_ISL_472906, EPI_ISL_472907, EPI_ISL_472908, EPI_ISL_472909, EPI_ISL_472910, EPI_ISL_472911, EPI_ISL_472912, EPI_ISL_472913, EPI_ISL_472914, EPI_ISL_472915, EPI_ISL_472916, EPI_ISL_472917, EPI_ISL_472918, EPI_ISL_472919, EPI_ISL_472920, EPI_ISL_472921, EPI_ISL_472922, EPI_ISL_472923, EPI_ISL_472924, EPI_ISL_472925, EPI_ISL_472926, EPI_ISL_472927, EPI_ISL_472928, EPI_ISL_472929, EPI_ISL_472930, EPI_ISL_472931, EPI_ISL_472932, EPI_ISL_472933, EPI_ISL_472934, EPI_ISL_472935, EPI_ISL_472936, EPI_ISL_472937, EPI_ISL_472938, EPI_ISL_472939, EPI_ISL_472940, EPI_ISL_472941, EPI_ISL_472942, EPI_ISL_472943, EPI_ISL_472944, EPI_ISL_472945, EPI_ISL_472946, EPI_ISL_472947, EPI_ISL_472948, EPI_ISL_472949, EPI_ISL_472950, EPI_ISL_472951, EPI_ISL_472952, EPI_ISL_472953, EPI_ISL_472954, EPI_ISL_472955, EPI_ISL_472956, EPI_ISL_472957, EPI_ISL_472958, EPI_ISL_472959, EPI_ISL_472960, EPI_ISL_472961, EPI_ISL_472962, EPI_ISL_472963, EPI_ISL_472964, EPI_ISL_472965, EPI_ISL_472966, EPI_ISL_472967, EPI_ISL_472968, EPI_ISL_472969, EPI_ISL_472970, EPI_ISL_472971, EPI_ISL_472972, EPI_ISL_472973, EPI_ISL_472974, EPI_ISL_472975, EPI_ISL_472976, EPI_ISL_472977, EPI_ISL_472978, EPI_ISL_472979, EPI_ISL_472980, EPI_ISL_472981, EPI_ISL_472982, EPI_ISL_472983, EPI_ISL_472984, EPI_ISL_472985, EPI_ISL_472986, EPI_ISL_472987, EPI_ISL_472988, EPI_ISL_472989, EPI_ISL_472990, EPI_ISL_472991, EPI_ISL_472992, EPI_ISL_472993, EPI_ISL_472994, EPI_ISL_472995, EPI_ISL_472996, EPI_ISL_472997, EPI_ISL_472998, EPI_ISL_472999, EPI_ISL_473000, EPI_ISL_473001, EPI_ISL_473002, EPI_ISL_473003, EPI_ISL_473004, EPI_ISL_473005, EPI_ISL_473006, EPI_ISL_473007, EPI_ISL_473008, EPI_ISL_473009, EPI_ISL_473010, EPI_ISL_473011, EPI_ISL_473012, EPI_ISL_473013, EPI_ISL_473014, EPI_ISL_473015, EPI_ISL_473016, EPI_ISL_473017, EPI_ISL_473018, EPI_ISL_473019, EPI_ISL_473020, EPI_ISL_473021, EPI_ISL_473022, EPI_ISL_473023, EPI_ISL_473024, EPI_ISL_473025, EPI_ISL_473026, EPI_ISL_473027, EPI_ISL_473028, EPI_ISL_473029, EPI_ISL_473030, EPI_ISL_473031, EPI_ISL_473032, EPI_ISL_473033, EPI_ISL_473034, EPI_ISL_473035, EPI_ISL_473036, EPI_ISL_473037, EPI_ISL_473038, EPI_ISL_473039, EPI_ISL_473040, EPI_ISL_473041, EPI_ISL_473042, EPI_ISL_473043, EPI_ISL_473044, EPI_ISL_473045, EPI_ISL_473046, EPI_ISL_473047, EPI_ISL_473048, EPI_ISL_473049, EPI_ISL_473050, EPI_ISL_473051, EPI_ISL_473052, EPI_ISL_473053, EPI_ISL_473054, EPI_ISL_473055, EPI_ISL_473056, EPI_ISL_473057, EPI_ISL_473058, EPI_ISL_473059, EPI_ISL_473060, EPI_ISL_473061, EPI_ISL_473062, EPI_ISL_473063, EPI_ISL_473064, EPI_ISL_473065, EPI_ISL_473066, EPI_ISL_473067, EPI_ISL_473068, EPI_ISL_473069, EPI_ISL_473070, EPI_ISL_473071, EPI_ISL_473072, EPI_ISL_473073, EPI_ISL_473074, EPI_ISL_473075, EPI_ISL_473076, EPI_ISL_473077, EPI_ISL_473078, EPI_ISL_473079, EPI_ISL_473080, EPI_ISL_473081, EPI_ISL_473082, EPI_ISL_473083, EPI_ISL_473084, EPI_ISL_473085, EPI_ISL_473086, EPI_ISL_473087, EPI_ISL_473088, EPI_ISL_473089, EPI_ISL_473090, EPI_ISL_473091, EPI_ISL_473092, EPI_ISL_473093, EPI_ISL_473094, EPI_ISL_473095, EPI_ISL_473096, EPI_ISL_473097, EPI_ISL_473098, EPI_ISL_473099, EPI_ISL_473100, EPI_ISL_473101, EPI_ISL_473102, EPI_ISL_473103, EPI_ISL_473104, EPI_ISL_473105, EPI_ISL_473106, EPI_ISL_473107, EPI_ISL_473108, EPI_ISL_473109, EPI_ISL_473110, EPI_ISL_473111, EPI_ISL_473112, EPI_ISL_473113, EPI_ISL_473114, EPI_ISL_473115, EPI_ISL_473116, EPI_ISL_473117, EPI_ISL_473118, EPI_ISL_473119, EPI_ISL_473120, EPI_ISL_473121, EPI_ISL_473122, EPI_ISL_473123, EPI_ISL_473124, EPI_ISL_473125, EPI_ISL_473126, EPI_ISL_473127, EPI_ISL_473128, EPI_ISL_473129, EPI_ISL_473130, EPI_ISL_473131, EPI_ISL_473132, EPI_ISL_473133, EPI_ISL_473134, EPI_ISL_473135, EPI_ISL_473136, EPI_ISL_473137, EPI_ISL_473138, EPI_ISL_473139, EPI_ISL_473140, EPI_ISL_473141, EPI_ISL_473142, EPI_ISL_473143, EPI_ISL_473144, EPI_ISL_473145, EPI_ISL_473146, EPI_ISL_473147, EPI_ISL_473148, EPI_ISL_473149, EPI_ISL_473150, EPI_ISL_473151, EPI_ISL_473152, EPI_ISL_473153, EPI_ISL_473154, EPI_ISL_473155, EPI_ISL_473156, EPI_ISL_473157, EPI_ISL_473158, EPI_ISL_473159, EPI_ISL_473160, EPI_ISL_473161, EPI_ISL_473162, EPI_ISL_473163, EPI_ISL_473164, EPI_ISL_473165, EPI_ISL_473166, EPI_ISL_473167, EPI_ISL_473168, EPI_ISL_473169, EPI_ISL_473170, EPI_ISL_473171, EPI_ISL_473172, EPI_ISL_473173, EPI_ISL_473174, EPI_ISL_473175, EPI_ISL_473176, EPI_ISL_473177, EPI_ISL_473178, EPI_ISL_473179, EPI_ISL_473180, EPI_ISL_473181, EPI_ISL_473182, EPI_ISL_473183, EPI_ISL_473184, EPI_ISL_473185, EPI_ISL_473186, EPI_ISL_473187, EPI_ISL_473188, EPI_ISL_473189, EPI_ISL_473190, EPI_ISL_473191, EPI_ISL_473192, EPI_ISL_473193, EPI_ISL_473194, EPI_ISL_473195, EPI_ISL_473196, EPI_ISL_473197, EPI_ISL_473198, EPI_ISL_473199, EPI_ISL_473200, EPI_ISL_473201, EPI_ISL_473202, EPI_ISL_473203, EPI_ISL_473204, EPI_ISL_473205, EPI_ISL_473206, EPI_ISL_473207, EPI_ISL_473208, EPI_ISL_473209, EPI_ISL_473210, EPI_ISL_473211, EPI_ISL_473212, EPI_ISL_473213, EPI_ISL_473214, EPI_ISL_473215, EPI_ISL_473216, EPI_ISL_473217, EPI_ISL_473218, EPI_ISL_473219, EPI_ISL_473220, EPI_ISL_473221, EPI_ISL_473222, EPI_ISL_473223, EPI_ISL_473224, EPI_ISL_473225, EPI_ISL_473226, EPI_ISL_473227, EPI_ISL_473228, EPI_ISL_473229, EPI_ISL_473230, EPI_ISL_473231, EPI_ISL_473232, EPI_ISL_473233, EPI_ISL_473234, EPI_ISL_473235, EPI_ISL_473236, EPI_ISL_473237, EPI_ISL_473238, EPI_ISL_473239, EPI_ISL_473240, EPI_ISL_473241, EPI_ISL_473242, EPI_ISL_473243, EPI_ISL_473244, EPI_ISL_473245, EPI_ISL_473246, EPI_ISL_473247, EPI_ISL_473248, EPI_ISL_473249, EPI_ISL_473250, EPI_ISL_473251, EPI_ISL_473252, EPI_ISL_473253, EPI_ISL_473254, EPI_ISL_473255, EPI_ISL_473256, EPI_ISL_473257, EPI_ISL_473258, EPI_ISL_473259, EPI_ISL_473260, EPI_ISL_473261, EPI_ISL_473262, EPI_ISL_473263, EPI_ISL_473264, EPI_ISL_473265, EPI_ISL_473266, EPI_ISL_473267, EPI_ISL_473268, EPI_ISL_473269, EPI_ISL_473270, EPI_ISL_473271, EPI_ISL_473272, EPI_ISL_473273, EPI_ISL_473274, EPI_ISL_473275, EPI_ISL_473276, EPI_ISL_473277, EPI_ISL_473278, EPI_ISL_473279, EPI_ISL_473280, EPI_ISL_473281, EPI_ISL_473282, EPI_ISL_473283, EPI_ISL_473284, EPI_ISL_473285, EPI_ISL_473286, EPI_ISL_473287, EPI_ISL_473288, EPI_ISL_473289, EPI_ISL_473290, EPI_ISL_473291, EPI_ISL_473292, EPI_ISL_473293, EPI_ISL_473294, EPI_ISL_473295, EPI_ISL_473296, EPI_ISL_473297, EPI_ISL_473298, EPI_ISL_473299, EPI_ISL_473300, EPI_ISL_473301, EPI_ISL_473302, EPI_ISL_473303, EPI_ISL_473304, EPI_ISL_473305 | | | |
| see above | Wales Specialist Virology Centre Sequencing lab: Pathogen Genomics Unit | COVID-19 Genomics UK (COG-UK) Consortium | Catherine Moore, Johnathan Evans, Laura Gifford, Malorie Perry, Simon Cottrell, Angela Marchbank, Alec Birchley, Alexander Adams, Amy Gaskin, Bree Gatica-Wilcox, Jason Coombes, Joel Southgate, Lauren Gilbert, Lee Graham, Nicole Pacchiarini, Sara Kumziene-Summerhayes, Sarah Taylor, Sophie Jones, Sara Rey, Matthew Bull, Joanne Watkins, Sally Corden, Tom Connor |
| EPI_ISL_473306, EPI_ISL_473307, EPI_ISL_473308, EPI_ISL_473309, EPI_ISL_473310, EPI_ISL_473311, EPI_ISL_473312, EPI_ISL_473313, EPI_ISL_473314, EPI_ISL_473315, EPI_ISL_473316, EPI_ISL_473317, EPI_ISL_473318, EPI_ISL_473319, EPI_ISL_473320, EPI_ISL_473321, EPI_ISL_473322, EPI_ISL_473323, EPI_ISL_473324, EPI_ISL_473325, EPI_ISL_473326, EPI_ISL_473327, EPI_ISL_473328, EPI_ISL_473329, EPI_ISL_473330, EPI_ISL_473331, EPI_ISL_473332, EPI_ISL_473333, EPI_ISL_473334, EPI_ISL_473335, EPI_ISL_473336, EPI_ISL_473337, EPI_ISL_473338, EPI_ISL_473339, EPI_ISL_473340, EPI_ISL_473341, EPI_ISL_473342, EPI_ISL_473343, EPI_ISL_473344, EPI_ISL_473345, EPI_ISL_473346, EPI_ISL_473347, EPI_ISL_473348, EPI_ISL_473349, EPI_ISL_473350, EPI_ISL_473351, EPI_ISL_473352, EPI_ISL_473353, EPI_ISL_473354, EPI_ISL_473355, EPI_ISL_473356, EPI_ISL_473357, EPI_ISL_473358, EPI_ISL_473359, EPI_ISL_473360, EPI_ISL_473361, EPI_ISL_473362, EPI_ISL_473363, EPI_ISL_473364, EPI_ISL_473365, EPI_ISL_473366, EPI_ISL_473367, EPI_ISL_473368, EPI_ISL_473369, EPI_ISL_473370, EPI_ISL_473371, EPI_ISL_473372, EPI_ISL_473373, EPI_ISL_473374, EPI_ISL_473375, EPI_ISL_473376, EPI_ISL_473377, EPI_ISL_473378, EPI_ISL_473379, EPI_ISL_473380, EPI_ISL_473381, EPI_ISL_473382, EPI_ISL_473383, EPI_ISL_473384, EPI_ISL_473385, EPI_ISL_473386, EPI_ISL_473387, EPI_ISL_473388, EPI_ISL_473389, EPI_ISL_473390, EPI_ISL_473391, EPI_ISL_473392, EPI_ISL_473393, EPI_ISL_473394, EPI_ISL_473395, EPI_ISL_473396, EPI_ISL_473397, EPI_ISL_473398, EPI_ISL_473399, EPI_ISL_473400, EPI_ISL_473401, EPI_ISL_473402, EPI_ISL_473403, EPI_ISL_473404, EPI_ISL_473405, EPI_ISL_473406, EPI_ISL_473407, EPI_ISL_473408, EPI_ISL_473409, EPI_ISL_473410, EPI_ISL_473411, EPI_ISL_473412, EPI_ISL_473413, EPI_ISL_473414, EPI_ISL_473415, EPI_ISL_473416, EPI_ISL_473417, EPI_ISL_473418, EPI_ISL_473419, EPI_ISL_473420, EPI_ISL_473421, EPI_ISL_473422, EPI_ISL_473423, EPI_ISL_473424, EPI_ISL_473425, EPI_ISL_473426, EPI_ISL_473427, EPI_ISL_473428, EPI_ISL_473429, EPI_ISL_473430, EPI_ISL_473431, EPI_ISL_473432, EPI_ISL_473433, EPI_ISL_473434, EPI_ISL_473435, EPI_ISL_473436, EPI_ISL_473437, EPI_ISL_473438, EPI_ISL_473439, EPI_ISL_473440, EPI_ISL_473441, EPI_ISL_473442, EPI_ISL_473443, EPI_ISL_473444, EPI_ISL_473445, EPI_ISL_473446, EPI_ISL_473447, EPI_ISL_473448, EPI_ISL_473449, EPI_ISL_473450 | | | |
| see above | University of Birmingham | COVID-19 Genomics UK (COG-UK) Consortium | Institute of Microbiology, University of Birmingham: Claire McMurray, Joanne Stockton, Samuel Nicholls, Radoslaw Poplawski, Will Rowe, Josh Quick, Nicholas Loman, University of Birmingham Testing Laboratory: Celina M Whalley, Andrew Bosworth, Charlotte Poxon, Kasun Wanigasooriya, Oliver Pickles, Mike Kidd, Alex Richter, Andrew D Beggs PHE Heartlands Lab: Husam Osman, Andrew Bosworth. Queen Elizabeth Hospital: Anna Casey |
| EPI_ISL_473451, EPI_ISL_473452, EPI_ISL_473453, EPI_ISL_473454, EPI_ISL_473455, EPI_ISL_473456, EPI_ISL_473457, EPI_ISL_473458, EPI_ISL_473459, EPI_ISL_473460, EPI_ISL_473461, EPI_ISL_473462, EPI_ISL_473463, EPI_ISL_473464, EPI_ISL_473465, EPI_ISL_473466, EPI_ISL_473467, EPI_ISL_473468, EPI_ISL_473469, EPI_ISL_473470, EPI_ISL_473471, EPI_ISL_473472, EPI_ISL_473473, EPI_ISL_473474, EPI_ISL_473475, EPI_ISL_473476, EPI_ISL_473477, EPI_ISL_473478, EPI_ISL_473479, EPI_ISL_473480, EPI_ISL_473481, EPI_ISL_473482, EPI_ISL_473483, EPI_ISL_473484, EPI_ISL_473485, EPI_ISL_473486, EPI_ISL_473487, EPI_ISL_473488, EPI_ISL_473489, EPI_ISL_473490, EPI_ISL_473491, EPI_ISL_473492, EPI_ISL_473493, EPI_ISL_473494, EPI_ISL_473495, EPI_ISL_473496, EPI_ISL_473497, EPI_ISL_473498, EPI_ISL_473499, EPI_ISL_473500, EPI_ISL_473501, EPI_ISL_473502, EPI_ISL_473503, EPI_ISL_473504, EPI_ISL_473505, EPI_ISL_473506, EPI_ISL_473507 | | | |
| see above | Department of Pathology, University of Cambridge | COVID-19 Genomics UK (COG-UK) Consortium | Luke W Meredith, M. Estée Török, Myra Hosmillo, William L. Hamilton, Martin D. Curran, Theresa Feltwell, Grant Hall, Anna Yakovleva, Fahad A Khokhar, Charlotte J. Houldcroft, Laura G Caller, Aminu S. Jahun, Sarah L. Caddy, Yasmin Chaudhry, Malte Pinckert, Ian Goodfellow |
| EPI_ISL_473508, EPI_ISL_473509, EPI_ISL_473510, EPI_ISL_473511, EPI_ISL_473512, EPI_ISL_473513, EPI_ISL_473514, EPI_ISL_473515, EPI_ISL_473516, EPI_ISL_473517, EPI_ISL_473518, EPI_ISL_473519, EPI_ISL_473520, EPI_ISL_473521, EPI_ISL_473522, EPI_ISL_473523, EPI_ISL_473524, EPI_ISL_473525, EPI_ISL_473526, EPI_ISL_473527, EPI_ISL_473528, EPI_ISL_473529, EPI_ISL_473530, EPI_ISL_473531, EPI_ISL_473532, EPI_ISL_473533, EPI_ISL_473534, EPI_ISL_473535, EPI_ISL_473536, EPI_ISL_473537, EPI_ISL_473538, EPI_ISL_473539, EPI_ISL_473540, EPI_ISL_473541, EPI_ISL_473542, EPI_ISL_473543, EPI_ISL_473544, EPI_ISL_473545, EPI_ISL_473546, EPI_ISL_473547, EPI_ISL_473548, EPI_ISL_473549, EPI_ISL_473550, EPI_ISL_473551, EPI_ISL_473552, EPI_ISL_473553, EPI_ISL_473554, EPI_ISL_473555, EPI_ISL_473556, EPI_ISL_473557, EPI_ISL_473558, EPI_ISL_473559, EPI_ISL_473560, EPI_ISL_473561, EPI_ISL_473562, EPI_ISL_473563, EPI_ISL_473564, EPI_ISL_473565, EPI_ISL_473566, EPI_ISL_473567, EPI_ISL_473568, EPI_ISL_473569, EPI_ISL_473570, EPI_ISL_473571, EPI_ISL_473572, EPI_ISL_473573, EPI_ISL_473574, EPI_ISL_473575, EPI_ISL_473576, EPI_ISL_473577, EPI_ISL_473578, EPI_ISL_473579, EPI_ISL_473580, EPI_ISL_473581, EPI_ISL_473582, EPI_ISL_473583, EPI_ISL_473584, EPI_ISL_473585, EPI_ISL_473586, EPI_ISL_473587, EPI_ISL_473588, EPI_ISL_473589, EPI_ISL_473590, EPI_ISL_473591, EPI_ISL_473592, EPI_ISL_473593, EPI_ISL_473594, EPI_ISL_473595, EPI_ISL_473596, EPI_ISL_473597, EPI_ISL_473598, EPI_ISL_473599, EPI_ISL_473600, EPI_ISL_473601, EPI_ISL_473602, EPI_ISL_473603, EPI_ISL_473604, EPI_ISL_473605, EPI_ISL_473606, EPI_ISL_473607, EPI_ISL_473608, EPI_ISL_473609, EPI_ISL_473610, EPI_ISL_473611, EPI_ISL_473612, EPI_ISL_473613, EPI_ISL_473614, EPI_ISL_473615, EPI_ISL_473616, EPI_ISL_473617, EPI_ISL_473618, EPI_ISL_473619, EPI_ISL_473620, EPI_ISL_473621, EPI_ISL_473622, EPI_ISL_473623, EPI_ISL_473624, EPI_ISL_473625, EPI_ISL_473626, EPI_ISL_473627, EPI_ISL_473628, EPI_ISL_473629, EPI_ISL_473630, EPI_ISL_473631, EPI_ISL_473632, EPI_ISL_473633, EPI_ISL_473634, EPI_ISL_473635, EPI_ISL_473636, EPI_ISL_473637, EPI_ISL_473638, EPI_ISL_473639, EPI_ISL_473640, EPI_ISL_473641, EPI_ISL_473642, EPI_ISL_473643, EPI_ISL_473644, EPI_ISL_473645, EPI_ISL_473646, EPI_ISL_473647, EPI_ISL_473648, EPI_ISL_473649, EPI_ISL_473650, EPI_ISL_473651, EPI_ISL_473652, EPI_ISL_473653, EPI_ISL_473654, EPI_ISL_473655, EPI_ISL_473656, EPI_ISL_473657, EPI_ISL_473658, EPI_ISL_473659, EPI_ISL_473660, EPI_ISL_473661, EPI_ISL_473662, EPI_ISL_473663, EPI_ISL_473664, EPI_ISL_473665, EPI_ISL_473666, EPI_ISL_473667, EPI_ISL_473668, EPI_ISL_473669, EPI_ISL_473670, EPI_ISL_473671, EPI_ISL_473672, EPI_ISL_473673, EPI_ISL_473674, EPI_ISL_473675, | | | |

| | | | |
|---|---|---|---|
| EPI_ISL_473676, EPI_ISL_473677, EPI_ISL_473678, EPI_ISL_473679, EPI_ISL_473680, EPI_ISL_473681, EPI_ISL_473682, EPI_ISL_473683, EPI_ISL_473684, EPI_ISL_473685, EPI_ISL_473686, EPI_ISL_473687, EPI_ISL_473688, EPI_ISL_473689, EPI_ISL_473690, EPI_ISL_473691, EPI_ISL_473692, EPI_ISL_473693, EPI_ISL_473694, EPI_ISL_473695, EPI_ISL_473696, EPI_ISL_473697, EPI_ISL_473698, EPI_ISL_473699, EPI_ISL_473700, EPI_ISL_473701, EPI_ISL_473702, EPI_ISL_473703, EPI_ISL_473704, EPI_ISL_473705, EPI_ISL_473706, EPI_ISL_473707, EPI_ISL_473708, EPI_ISL_473709, EPI_ISL_473710, EPI_ISL_473711, EPI_ISL_473712, EPI_ISL_473713, EPI_ISL_473714, EPI_ISL_473715, EPI_ISL_473716, EPI_ISL_473717, EPI_ISL_473718, EPI_ISL_473719, EPI_ISL_473720, EPI_ISL_473721, EPI_ISL_473722, EPI_ISL_473723, EPI_ISL_473724, EPI_ISL_473725, EPI_ISL_473726, EPI_ISL_473727, EPI_ISL_473728, EPI_ISL_473729, EPI_ISL_473730, EPI_ISL_473731, EPI_ISL_473732, EPI_ISL_473733, EPI_ISL_473734, EPI_ISL_473735, EPI_ISL_473736, EPI_ISL_473737, EPI_ISL_473738, EPI_ISL_473739, EPI_ISL_473740, EPI_ISL_473741, EPI_ISL_473742, EPI_ISL_473743, EPI_ISL_473744, EPI_ISL_473745, EPI_ISL_473746, EPI_ISL_473747, EPI_ISL_473748, EPI_ISL_473749, EPI_ISL_473750, EPI_ISL_473751, EPI_ISL_473752, EPI_ISL_473753, EPI_ISL_473754, EPI_ISL_473755, EPI_ISL_473756, EPI_ISL_473757, EPI_ISL_473758, EPI_ISL_473759, EPI_ISL_473760, EPI_ISL_473761, EPI_ISL_473762, EPI_ISL_473763, EPI_ISL_473764, EPI_ISL_473765, EPI_ISL_473766, EPI_ISL_473767, EPI_ISL_473768, EPI_ISL_473769, EPI_ISL_473770, EPI_ISL_473771, EPI_ISL_473772, EPI_ISL_473773, EPI_ISL_473774, EPI_ISL_473775, EPI_ISL_473776, EPI_ISL_473777, EPI_ISL_473778, EPI_ISL_473779, EPI_ISL_473780, EPI_ISL_473781, EPI_ISL_473782 | | | |
| see above | West of Scotland Specialist Virology Centre, NHSGGC / MRC-University of Glasgow Centre for Virus Research | COVID-19 Genomics UK (COG-UK) Consortium | Ana da Silva Filipe, Natasha Johnson, Kathy Smollett, Daniel Mair, Stephen Carmichael, Lily Tong, Jenna Nichols, Elihu Aranday-Cortes, Kirstyn Brunker, Yasmin Parr, Alice Broos, Kyriaki Nomikou; Sarah McDonald, Marc Niebel, Patawee Asamaphan; Richard Orton, Joseph Hughes, Sreenu Vattipally, David L Robertson; Alasdair MacLean, Rory Gunson; Kathy Li, Natasha Jesudason, Rajiv Shah, James Shepherd, Antonia Ho, Emma Thomson |
| EPI_ISL_473783, EPI_ISL_473784, EPI_ISL_473785, EPI_ISL_473786, EPI_ISL_473787, EPI_ISL_473788, EPI_ISL_473789, EPI_ISL_473790, EPI_ISL_473791, EPI_ISL_473792, EPI_ISL_473793, EPI_ISL_473794, EPI_ISL_473795, EPI_ISL_473796, EPI_ISL_473797, EPI_ISL_473798, EPI_ISL_473799, EPI_ISL_473800, EPI_ISL_473801, EPI_ISL_473802, EPI_ISL_473803, EPI_ISL_473804, EPI_ISL_473805, EPI_ISL_473806, EPI_ISL_473807, EPI_ISL_473808, EPI_ISL_473809, EPI_ISL_473810, EPI_ISL_473811, EPI_ISL_473812, EPI_ISL_473813, EPI_ISL_473814, EPI_ISL_473815, EPI_ISL_473816, EPI_ISL_473817, EPI_ISL_473818, EPI_ISL_473819, EPI_ISL_473820, EPI_ISL_473821, EPI_ISL_473822, EPI_ISL_473823, EPI_ISL_473824, EPI_ISL_473825, EPI_ISL_473826, EPI_ISL_473827, EPI_ISL_473828, EPI_ISL_473829, EPI_ISL_473830, EPI_ISL_473831, EPI_ISL_473832, EPI_ISL_473833, EPI_ISL_473834, EPI_ISL_473835, EPI_ISL_473836, EPI_ISL_473837, EPI_ISL_473838, EPI_ISL_473839, EPI_ISL_473840, EPI_ISL_473841, EPI_ISL_473842, EPI_ISL_473843, EPI_ISL_473844, EPI_ISL_473845, EPI_ISL_473846, EPI_ISL_473847, EPI_ISL_473848, EPI_ISL_473849, EPI_ISL_473850, EPI_ISL_473851, EPI_ISL_473852, EPI_ISL_473853, EPI_ISL_473854, EPI_ISL_473855, EPI_ISL_473856, EPI_ISL_473857, EPI_ISL_473858, EPI_ISL_473859, EPI_ISL_473860, EPI_ISL_473861, EPI_ISL_473862, EPI_ISL_473863, EPI_ISL_473864, EPI_ISL_473865, EPI_ISL_473866, EPI_ISL_473867, EPI_ISL_473868, EPI_ISL_473869, EPI_ISL_473870, EPI_ISL_473871, EPI_ISL_473872, EPI_ISL_473873, EPI_ISL_473874, EPI_ISL_473875, EPI_ISL_473876, EPI_ISL_473877, EPI_ISL_473878, EPI_ISL_473879, EPI_ISL_473880, EPI_ISL_473881, EPI_ISL_473882, EPI_ISL_473883, EPI_ISL_473884, EPI_ISL_473885, EPI_ISL_473886, EPI_ISL_473887, EPI_ISL_473888, EPI_ISL_473889, EPI_ISL_473890, EPI_ISL_473891, EPI_ISL_473892, EPI_ISL_473893, EPI_ISL_473894, EPI_ISL_473895, EPI_ISL_473896, EPI_ISL_473897, EPI_ISL_473898, EPI_ISL_473899, EPI_ISL_473900, EPI_ISL_473901, EPI_ISL_473902, EPI_ISL_473903, EPI_ISL_473904, EPI_ISL_473905, EPI_ISL_473906, EPI_ISL_473907, EPI_ISL_473908, EPI_ISL_473909, EPI_ISL_473910, EPI_ISL_473911, EPI_ISL_473912, EPI_ISL_473913, EPI_ISL_473914, EPI_ISL_473915, EPI_ISL_473916, EPI_ISL_473917, EPI_ISL_473918, EPI_ISL_473919, EPI_ISL_473920, EPI_ISL_473921, EPI_ISL_473922, EPI_ISL_473923, EPI_ISL_473924, EPI_ISL_473925, EPI_ISL_473926, EPI_ISL_473927, EPI_ISL_473928, EPI_ISL_473929, EPI_ISL_473930, EPI_ISL_473931, EPI_ISL_473932, EPI_ISL_473933, EPI_ISL_473934, EPI_ISL_473935, EPI_ISL_473936, EPI_ISL_473937, EPI_ISL_473938, EPI_ISL_473939, EPI_ISL_473940, EPI_ISL_473941, EPI_ISL_473942, EPI_ISL_473943, EPI_ISL_473944, EPI_ISL_473945, EPI_ISL_473946, EPI_ISL_473947, EPI_ISL_473948, EPI_ISL_473949, EPI_ISL_473950, EPI_ISL_473951, EPI_ISL_473952, EPI_ISL_473953, EPI_ISL_473954, EPI_ISL_473955 | | | |
| see above | Virology Department, Royal Infirmary of Edinburgh, NHS Lothian / School of Biological Sciences, University of Edinburgh / Institute of Genetics and Molecular Medicine, University of Edinburgh | COVID-19 Genomics UK (COG-UK) Consortium | McHugh M, Dewar R, Rooke S, Gallagher M, Balcaza C, O'Toole Á, Scher E, Hill V, McCrone JT, Colquhoun R, Yu X, Jackson B, Rambaut A, Williams TC, Templeton K |
| EPI_ISL_473956, EPI_ISL_473957, EPI_ISL_473958, EPI_ISL_473959, EPI_ISL_473960, EPI_ISL_473961, EPI_ISL_473962, EPI_ISL_473963, EPI_ISL_473964, EPI_ISL_473965, EPI_ISL_473966, EPI_ISL_473967, EPI_ISL_473968, EPI_ISL_473969, EPI_ISL_473970, EPI_ISL_473971, EPI_ISL_473972, EPI_ISL_473973, EPI_ISL_473974, EPI_ISL_473975, EPI_ISL_473976, EPI_ISL_473977, EPI_ISL_473978, EPI_ISL_473979, EPI_ISL_473980, EPI_ISL_473981, EPI_ISL_473982, EPI_ISL_473983, EPI_ISL_473984, EPI_ISL_473985, EPI_ISL_473986, EPI_ISL_473987, EPI_ISL_473988, EPI_ISL_473989, EPI_ISL_473990, EPI_ISL_473991, EPI_ISL_473992, EPI_ISL_473993, EPI_ISL_473994, EPI_ISL_473995, EPI_ISL_473996, EPI_ISL_473997, EPI_ISL_473998, EPI_ISL_473999, EPI_ISL_474000, EPI_ISL_474001, EPI_ISL_474002, EPI_ISL_474003, EPI_ISL_474004, EPI_ISL_474005, EPI_ISL_474006, EPI_ISL_474007, EPI_ISL_474008, EPI_ISL_474009, EPI_ISL_474010, EPI_ISL_474011, EPI_ISL_474012, EPI_ISL_474013, EPI_ISL_474014, EPI_ISL_474015, EPI_ISL_474016, EPI_ISL_474017, EPI_ISL_474018, EPI_ISL_474019, EPI_ISL_474020, EPI_ISL_474021, EPI_ISL_474022, EPI_ISL_474023, EPI_ISL_474024, EPI_ISL_474025, EPI_ISL_474026, EPI_ISL_474027, EPI_ISL_474028, EPI_ISL_474029, EPI_ISL_474030, EPI_ISL_474031, EPI_ISL_474032, EPI_ISL_474033, EPI_ISL_474034, EPI_ISL_474035, EPI_ISL_474036, EPI_ISL_474037, EPI_ISL_474038, EPI_ISL_474039, EPI_ISL_474040, EPI_ISL_474041, EPI_ISL_474042, EPI_ISL_474043, EPI_ISL_474044, EPI_ISL_474045, EPI_ISL_474046, EPI_ISL_474047, EPI_ISL_474048, EPI_ISL_474049, EPI_ISL_474050, EPI_ISL_474051, EPI_ISL_474052, EPI_ISL_474053, EPI_ISL_474054, EPI_ISL_474055, EPI_ISL_474056, EPI_ISL_474057, EPI_ISL_474058, EPI_ISL_474059, EPI_ISL_474060, EPI_ISL_474061, EPI_ISL_474062, EPI_ISL_474063, EPI_ISL_474064, EPI_ISL_474065, EPI_ISL_474066, EPI_ISL_474067, EPI_ISL_474068, EPI_ISL_474069, EPI_ISL_474070, EPI_ISL_474071, EPI_ISL_474072, EPI_ISL_474073, EPI_ISL_474074, EPI_ISL_474075, EPI_ISL_474076, EPI_ISL_474077, EPI_ISL_474078, EPI_ISL_474079, EPI_ISL_474080, EPI_ISL_474081, EPI_ISL_474082, EPI_ISL_474083, EPI_ISL_474084, EPI_ISL_474085, EPI_ISL_474086, EPI_ISL_474087, EPI_ISL_474088, EPI_ISL_474089, EPI_ISL_474090, EPI_ISL_474091, EPI_ISL_474092, EPI_ISL_474093, EPI_ISL_474094, EPI_ISL_474095, EPI_ISL_474096, EPI_ISL_474097, EPI_ISL_474098, EPI_ISL_474099, EPI_ISL_474100, EPI_ISL_474101, EPI_ISL_474102, EPI_ISL_474103, EPI_ISL_474104, EPI_ISL_474105, EPI_ISL_474106, EPI_ISL_474107, EPI_ISL_474108, EPI_ISL_474109, EPI_ISL_474110, EPI_ISL_474111, EPI_ISL_474112, EPI_ISL_474113, EPI_ISL_474114, EPI_ISL_474115, EPI_ISL_474116, EPI_ISL_474117, EPI_ISL_474118, EPI_ISL_474119, EPI_ISL_474120, EPI_ISL_474121, EPI_ISL_474122, EPI_ISL_474123, EPI_ISL_474124, EPI_ISL_474125, EPI_ISL_474126, EPI_ISL_474127, EPI_ISL_474128, EPI_ISL_474129, EPI_ISL_474130, EPI_ISL_474131, EPI_ISL_474132, EPI_ISL_474133, EPI_ISL_474134, EPI_ISL_474135, EPI_ISL_474136, EPI_ISL_474137, EPI_ISL_474138, EPI_ISL_474139, EPI_ISL_474140, EPI_ISL_474141, EPI_ISL_474142, EPI_ISL_474143, EPI_ISL_474144, EPI_ISL_474145, EPI_ISL_474146, EPI_ISL_474147, EPI_ISL_474148, EPI_ISL_474149, EPI_ISL_474150, EPI_ISL_474151, EPI_ISL_474152, EPI_ISL_474153, EPI_ISL_474154, EPI_ISL_474155, EPI_ISL_474156, EPI_ISL_474157, EPI_ISL_474158, EPI_ISL_474159, EPI_ISL_474160, EPI_ISL_474161, EPI_ISL_474162, EPI_ISL_474163, EPI_ISL_474164, EPI_ISL_474165, EPI_ISL_474166, EPI_ISL_474167, EPI_ISL_474168, EPI_ISL_474169, EPI_ISL_474170, EPI_ISL_474171, EPI_ISL_474172, EPI_ISL_474173, EPI_ISL_474174, EPI_ISL_474175, EPI_ISL_474176, EPI_ISL_474177, EPI_ISL_474178, EPI_ISL_474179, EPI_ISL_474180, EPI_ISL_474181, EPI_ISL_474182, EPI_ISL_474183, EPI_ISL_474184, EPI_ISL_474185, EPI_ISL_474186, EPI_ISL_474187, EPI_ISL_474188, EPI_ISL_474189, EPI_ISL_474190, EPI_ISL_474191, EPI_ISL_474192, EPI_ISL_474193, EPI_ISL_474194, EPI_ISL_474195, EPI_ISL_474196, EPI_ISL_474197, EPI_ISL_474198, EPI_ISL_474199, EPI_ISL_474200, EPI_ISL_474201, EPI_ISL_474202, EPI_ISL_474203, EPI_ISL_474204, EPI_ISL_474205, EPI_ISL_474206, EPI_ISL_474207, EPI_ISL_474208, EPI_ISL_474209, EPI_ISL_474210, EPI_ISL_474211, EPI_ISL_474212, EPI_ISL_474213, EPI_ISL_474214, EPI_ISL_474215, EPI_ISL_474216, EPI_ISL_474217, EPI_ISL_474218, EPI_ISL_474219, EPI_ISL_474220, EPI_ISL_474221, EPI_ISL_474222, EPI_ISL_474223, EPI_ISL_474224, EPI_ISL_474225, EPI_ISL_474226, EPI_ISL_474227, EPI_ISL_474228, EPI_ISL_474229, EPI_ISL_474230, EPI_ISL_474231, EPI_ISL_474232, EPI_ISL_474233, EPI_ISL_474234, EPI_ISL_474235, EPI_ISL_474236, EPI_ISL_474237, EPI_ISL_474238, EPI_ISL_474239, EPI_ISL_474240, EPI_ISL_474241, EPI_ISL_474242, EPI_ISL_474243, EPI_ISL_474244, EPI_ISL_474245, EPI_ISL_474246, EPI_ISL_474247, EPI_ISL_474248, EPI_ISL_474249, EPI_ISL_474250, EPI_ISL_474251, EPI_ISL_474252, EPI_ISL_474253, EPI_ISL_474254, EPI_ISL_474255, EPI_ISL_474256, EPI_ISL_474257, EPI_ISL_474258, EPI_ISL_474259, EPI_ISL_474260, EPI_ISL_474261, EPI_ISL_474262, EPI_ISL_474263, EPI_ISL_474264, EPI_ISL_474265, EPI_ISL_474266, EPI_ISL_474267, EPI_ISL_474268, EPI_ISL_474269, EPI_ISL_474270, EPI_ISL_474271, EPI_ISL_474272, EPI_ISL_474273, EPI_ISL_474274, EPI_ISL_474275, EPI_ISL_474276, EPI_ISL_474277, EPI_ISL_474278, EPI_ISL_474279, EPI_ISL_474280, EPI_ISL_474281, EPI_ISL_474282, EPI_ISL_474283, EPI_ISL_474284, EPI_ISL_474285, EPI_ISL_474286, EPI_ISL_474287, EPI_ISL_474288, EPI_ISL_474289, EPI_ISL_474290, EPI_ISL_474291, EPI_ISL_474292, EPI_ISL_474293, EPI_ISL_474294, EPI_ISL_474295, EPI_ISL_474296, EPI_ISL_474297, EPI_ISL_474298, EPI_ISL_474299, EPI_ISL_474300, EPI_ISL_474301, EPI_ISL_474302, EPI_ISL_474303, EPI_ISL_474304, EPI_ISL_474305, EPI_ISL_474306, EPI_ISL_474307, EPI_ISL_474308, EPI_ISL_474309, EPI_ISL_474310, EPI_ISL_474311, EPI_ISL_474312, EPI_ISL_474313, EPI_ISL_474314, EPI_ISL_474315, EPI_ISL_474316, EPI_ISL_474317, EPI_ISL_474318, EPI_ISL_474319, EPI_ISL_474320, EPI_ISL_474321, EPI_ISL_474322, EPI_ISL_474323, EPI_ISL_474324, EPI_ISL_474325, EPI_ISL_474326, EPI_ISL_474327, EPI_ISL_474328, EPI_ISL_474329, EPI_ISL_474330, EPI_ISL_474331, EPI_ISL_474332, EPI_ISL_474333, EPI_ISL_474334, EPI_ISL_474335, EPI_ISL_474336, EPI_ISL_474337, EPI_ISL_474338, EPI_ISL_474339, EPI_ISL_474340, EPI_ISL_474341, EPI_ISL_474342, EPI_ISL_474343, EPI_ISL_474344, EPI_ISL_474345, EPI_ISL_474346, EPI_ISL_474347, EPI_ISL_474348, EPI_ISL_474349, EPI_ISL_474350, EPI_ISL_474351, EPI_ISL_474352, EPI_ISL_474353, EPI_ISL_474354, EPI_ISL_474355, EPI_ISL_474356, EPI_ISL_474357, EPI_ISL_474358, EPI_ISL_474359, EPI_ISL_474360, EPI_ISL_474361, EPI_ISL_474362, EPI_ISL_474363, EPI_ISL_474364, EPI_ISL_474365, EPI_ISL_474366, EPI_ISL_474367, EPI_ISL_474368, EPI_ISL_474369, EPI_ISL_474370, EPI_ISL_474371, EPI_ISL_474372, EPI_ISL_474373, EPI_ISL_474374, EPI_ISL_474375, EPI_ISL_474376, EPI_ISL_474377, EPI_ISL_474378, EPI_ISL_474379, EPI_ISL_474380, EPI_ISL_474381, EPI_ISL_474382, EPI_ISL_474383, EPI_ISL_474384, EPI_ISL_474385, EPI_ISL_474386, EPI_ISL_474387, EPI_ISL_474388, EPI_ISL_474389, EPI_ISL_474390, EPI_ISL_474391, EPI_ISL_474392, EPI_ISL_474393, EPI_ISL_474394, EPI_ISL_474395, EPI_ISL_474396, EPI_ISL_474397, EPI_ISL_474398, EPI_ISL_474399, EPI_ISL_474400, EPI_ISL_474401, EPI_ISL_474402, EPI_ISL_474403, EPI_ISL_474404, EPI_ISL_474405, EPI_ISL_474406, EPI_ISL_474407, EPI_ISL_474408, EPI_ISL_474409, EPI_ISL_474410, EPI_ISL_474411, EPI_ISL_474412, EPI_ISL_474413, EPI_ISL_474414, EPI_ISL_474415, EPI_ISL_474416, EPI_ISL_474417, EPI_ISL_474418, EPI_ISL_474419, EPI_ISL_474420, EPI_ISL_474421, EPI_ISL_474422, EPI_ISL_474423, EPI_ISL_474424, EPI_ISL_474425, EPI_ISL_474426, EPI_ISL_474427, EPI_ISL_474428, EPI_ISL_474429, EPI_ISL_474430, EPI_ISL_474431, EPI_ISL_474432, EPI_ISL_474433, EPI_ISL_474434, EPI_ISL_474435, EPI_ISL_474436, EPI_ISL_474437, EPI_ISL_474438, EPI_ISL_474439, EPI_ISL_474440, EPI_ISL_474441, EPI_ISL_474442, EPI_ISL_474443, EPI_ISL_474444, EPI_ISL_474445, EPI_ISL_474446, EPI_ISL_474447, EPI_ISL_474448, EPI_ISL_474449, EPI_ISL_474450, EPI_ISL_474451, EPI_ISL_474452, EPI_ISL_474453, EPI_ISL_474454, EPI_ISL_474455, EPI_ISL_474456, EPI_ISL_474457, EPI_ISL_474458, EPI_ISL_474459, EPI_ISL_474460, EPI_ISL_474461, EPI_ISL_474462, EPI_ISL_474463, EPI_ISL_474464, EPI_ISL_474465, EPI_ISL_474466, EPI_ISL_474467, EPI_ISL_474468, EPI_ISL_474469, EPI_ISL_474470, EPI_ISL_474471, EPI_ISL_474472, EPI_ISL_474473, EPI_ISL_474474, EPI_ISL_474475, EPI_ISL_474476, EPI_ISL_474477, EPI_ISL_474478, EPI_ISL_474479, EPI_ISL_474480, EPI_ISL_474481, EPI_ISL_474482, EPI_ISL_474483, EPI_ISL_474484, EPI_ISL_474485, EPI_ISL_474486, EPI_ISL_474487, EPI_ISL_474488, EPI_ISL_474489, EPI_ISL_474490, EPI_ISL_474491, EPI_ISL_474492, EPI_ISL_474493, EPI_ISL_474494, EPI_ISL_474495, EPI_ISL_474496, EPI_ISL_474497, EPI_ISL_474498, EPI_ISL_474499, EPI_ISL_474500, EPI_ISL_474501, EPI_ISL_474502, EPI_ISL_474503, EPI_ISL_474504, EPI_ISL_474505, EPI_ISL_474506, EPI_ISL_474507, EPI_ISL_474508, EPI_ISL_474509, EPI_ISL_474510, EPI_ISL_474511, EPI_ISL_474512, EPI_ISL_474513, EPI_ISL_474514, EPI_ISL_474515, EPI_ISL_474516, EPI_ISL_474517, EPI_ISL_474518, EPI_ISL_474519, EPI_ISL_474520, EPI_ISL_474521, EPI_ISL_474522, EPI_ISL_474523, EPI_ISL_474524, EPI_ISL_474525, EPI_ISL_474526, EPI_ISL_474527, EPI_ISL_474528, EPI_ISL_474529, EPI_ISL_474530, EPI_ISL_474531, EPI_ISL_474532, EPI_ISL_474533, EPI_ISL_474534, EPI_ISL_474535, EPI_ISL_474536, EPI_ISL_474537, EPI_ISL_474538, EPI_ISL_474539, EPI_ISL_474540, EPI_ISL_474541, EPI_ISL_474542, EPI_ISL_474543, EPI_ISL_474544, EPI_ISL_474545, EPI_ISL_474546, EPI_ISL_474547, EPI_ISL_474548, EPI_ISL_474549, EPI_ISL_474550, EPI_ISL_474551, EPI_ISL_474552, EPI_ISL_474553, EPI_ISL_474554, EPI_ISL_474555, EPI_ISL_474556, EPI_ISL_474557, EPI_ISL_474558, EPI_ISL_474559, EPI_ISL_474560, EPI_ISL_474561, EPI_ISL_474562, EPI_ISL_474563, EPI_ISL_474564, EPI_ISL_474565, EPI_ISL_474566, EPI_ISL_474567, EPI_ISL_474568, EPI_ISL_474569, EPI_ISL_474570, EPI_ISL_474571, EPI_ISL_474572, EPI_ISL_474573, EPI_ISL_474574, EPI_ISL_474575, EPI_ISL_474576, EPI_ISL_474577, EPI_ISL_474578, EPI_ISL_474579, EPI_ISL_474580, EPI_ISL_474581, EPI_ISL_474582, EPI_ISL_474583, EPI_ISL_474584, EPI_ISL_474585, EPI_ISL_474586, EPI_ISL_474587, EPI_ISL_474588, EPI_ISL_474589, EPI_ISL_474590, EPI_ISL_474591, EPI_ISL_474592, EPI_ISL_474593, EPI_ISL_474594, EPI_ISL_474595, EPI_ISL_474596, EPI_ISL_474597, EPI_ISL_474598, EPI_ISL_474599, EPI_ISL_474600, EPI_ISL_474601, EPI_ISL_474602, EPI_ISL_474603, EPI_ISL_474604, EPI_ISL_474605, EPI_ISL_474606, EPI_ISL_474607, EPI_ISL_474608, EPI_ISL_474609, EPI_ISL_474610, EPI_ISL_474611, EPI_ISL_474612, EPI_ISL_474613, EPI_ISL_474614, EPI_ISL_474615, EPI_ISL_474616, EPI_ISL_474617, EPI_ISL_474618, EPI_ISL_474619, EPI_ISL_474620, EPI_ISL_474621, EPI_ISL_474622, EPI_ISL_474623, EPI_ISL_474624, EPI_ISL_474625, EPI_ISL_474626, EPI_ISL_474627, EPI_ISL_474628, EPI_ISL_474629, EPI_ISL_474630, EPI_ISL_474631, EPI_ISL_474632, EPI_ISL_474633, EPI_ISL_474634, EPI_ISL_474635, EPI_ISL_474636, EPI_ISL_474637, EPI_ISL_474638, EPI_ISL_474639, EPI_ISL_474640, EPI_ISL_474641, EPI_ISL_474642, EPI_ISL_474643, EPI_ISL_474644, EPI_ISL_474645, EPI_ISL_474646, EPI_ISL_474647, EPI_ISL_474648, EPI_ISL_474649, EPI_ISL_474650, EPI_ISL_474651, EPI_ISL_474652, EPI_ISL_474653, EPI_ISL_474654, EPI_ISL_474655, EPI_ISL_474656, EPI_ISL_474657, EPI_ISL_474658, EPI_ISL_474659, EPI_ISL_474660, EPI_ISL_474661, EPI_ISL_474662, EPI_ISL_474663, EPI_ISL_474664, EPI_ISL_474665, EPI_ISL_474666, EPI_ISL_474667, EPI_ISL_474668, EPI_ISL_474669, EPI_ISL_474670, EPI_ISL_474671, EPI_ISL_474672, EPI_ISL_474673, EPI_ISL_474674, EPI_ISL_474675, EPI_ISL_474676, EPI_ISL_474677, EPI_ISL_474678, EPI_ISL_474679, EPI_ISL_474680, EPI_ISL_474681, EPI_ISL_474682, EPI_ISL_474683, EPI_ISL_474684, EPI_ISL_474685, EPI_ISL_474686, EPI_ISL_474687, EPI_ISL_474688, EPI_ISL_474689, EPI_ISL_474690, EPI_ISL_474691, EPI_ISL_474692, EPI_ISL_474693, EPI_ISL_474694, EPI_ISL_474695, EPI_ISL_474696, EPI_ISL_474697, EPI_ISL_474698, EPI_ISL_474699, EPI_ISL_474700, EPI_ISL_474701, EPI_ISL_474702, EPI_ISL_474703, EPI_ISL_474704, EPI_ISL_474705, EPI_ISL_474706, EPI_ISL_474707, EPI_ISL_474708, EPI_ISL_474709, EPI_ISL_474710, EPI_ISL_474711, EPI_ISL_474712, EPI_ISL_474713, EPI_ISL_474714, EPI_ISL_474715, EPI_ISL_474716, EPI_ISL_474717, EPI_ISL_474718, EPI_ISL_474719, EPI_ISL_474720, EPI_ISL_474721, EPI_ISL_474722, EPI_ISL_474723, EPI_ISL_474724, EPI_ISL_474725, EPI_ISL_474726, EPI_ISL_474727, EPI_ISL_474728, EPI_ISL_474729, EPI_ISL_474730, EPI_ISL_474731, EPI_ISL_474732, EPI_ISL_474733, EPI_ISL_474734, EPI_ISL_474735, EPI_ISL_474736, EPI_ISL_474737, EPI_ISL_474738, EPI_ISL_474739, EPI_ISL_474740, EPI_ISL_474741, EPI_ISL_474742, EPI_ISL_474743, EPI_ISL_474744, EPI_ISL_474745, EPI_ISL_474746, EPI_ISL_474747, EPI_ISL_474748, EPI_ISL_474749, EPI_ISL_474750, EPI_ISL_474751, EPI_ISL_474752, EPI_ISL_474753, EPI_ISL_474754, EPI_ISL_474755, EPI_ISL_474756, EPI_ISL_474757, EPI_ISL_474758, EPI_ISL_474759, EPI_ISL_474760, EPI_ISL_474761, EPI_ISL_474762, EPI_ISL_474763, EPI_ISL_474764, EPI_ISL_474765, EPI_ISL_474766, EPI_ISL_474767, EPI_ISL_474768, EPI_ISL_474769, EPI_ISL_474770, EPI_ISL_474771, EPI_ISL_474772, EPI_ISL_474773, EPI_ISL_474774, EPI_ISL_474775, EPI_ISL_474776, EPI_ISL_474777, EPI_ISL_474778, EPI_ISL_474779, EPI_ISL_474780, EPI_ISL_474781, EPI_ISL_474782, EPI_ISL_474783, EPI_ISL_474784, EPI_ISL_474785, EPI_ISL_474786, EPI_ISL_474787, EPI_ISL_474788, EPI_ISL_474789, EPI_ISL_474790, EPI_ISL_474791, EPI_ISL_474792, EPI_ISL_474793, EPI_ISL_474794, EPI_ISL_474795, EPI_ISL_474796 | | | |
| see above | Originating lab: Wales Specialist Virology Centre Sequencing lab: Pathogen Genomics Unit | COVID-19 Genomics UK (COG-UK) Consortium | Catherine Moore, Johnathan Evans, Laura Gifford, Malorie Perry, Simon Cottrell, Angela Marchbank, Alec Birchley, Alexander Adams, Amy Gaskin, Bree Gatica-Wilcox, Jason Coombes, Joel Southgate, Lauren Gilbert, Lee Graham, Nicole Pacchiarini, Sara Kumziene-Summerhayes, Sarah Taylor, Sophie Jones, Sara Rey, Matthew Bull, Joanne Watkins, Sally Corden, Tom Connor |
| EPI_ISL_474797 | Hospital Universitario Virgen de las Nieves de Granada-SAS | SeqCOVID-SPAIN consortium/IBV(CSIC) | Mercedes Pérez Ruiz, Sara Sanbonmatsu Gámez, Irene Pedrosa Corral, José M. Navarro-Marí and SeqCOVID-SPAIN consortium |
| EPI_ISL_474798, EPI_ISL_474799, EPI_ISL_474800, EPI_ISL_474801, EPI_ISL_474802, EPI_ISL_474803, EPI_ISL_474804, EPI_ISL_474805, EPI_ISL_474806, EPI_ISL_474807, EPI_ISL_474808, EPI_ISL_474809, EPI_ISL_474810, EPI_ISL_474811, EPI_ISL_474812, EPI_ISL_474813, EPI_ISL_474814, EPI_ISL_474815, EPI_ISL_474816, EPI_ISL_474817, EPI_ISL_474818, EPI_ISL_474819, EPI_ISL_474820, EPI_ISL_474821, EPI_ISL_474822, EPI_ISL_474823, EPI_ISL_474824, EPI_ISL_474825, EPI_ISL_474826, EPI_ISL_474827, EPI_ISL_474828, EPI_ISL_474829, EPI_ISL_474830, EPI_ISL_474831 | | | |
| see above | Complejo Hospitalario Universitario de Albacete | SeqCOVID-SPAIN consortium/IBV(CSIC) | Encarnacion Simarro Córdoba, Julia Lozano Serra, Lorena Robles Fonseca , Monica Parra Grandes, Caridad Sainz de Baranda Camino and SeqCOVID-SPAIN consortium |
| EPI_ISL_474832, EPI_ISL_474833, EPI_ISL_474834, EPI_ISL_474835, EPI_ISL_474836 | Hospital Universitario Virgen de las Nieves de Granada-SAS | SeqCOVID-SPAIN consortium/IBV(CSIC) | Mercedes Pérez Ruiz, Sara Sanbonmatsu Gámez, Irene Pedrosa Corral, José M. Navarro-Marí and SeqCOVID-SPAIN consortium |
| EPI_ISL_474837, EPI_ISL_474838, EPI_ISL_474839, EPI_ISL_474840 | Complejo Hospitalario Universitario de Albacete | SeqCOVID-SPAIN consortium/IBV(CSIC) | Encarnacion Simarro Córdoba, Julia Lozano Serra, Lorena Robles Fonseca , Monica Parra Grandes, Caridad Sainz de Baranda Camino and SeqCOVID-SPAIN consortium |
| EPI_ISL_474841, EPI_ISL_474842, EPI_ISL_474843, EPI_ISL_474844, EPI_ISL_474845, EPI_ISL_474846 | Hospital Universitario Virgen de las Nieves de Granada-SAS | SeqCOVID-SPAIN consortium/IBV(CSIC) | Mercedes Pérez Ruiz, Sara Sanbonmatsu Gámez, Irene Pedrosa Corral, José M. Navarro-Marí and SeqCOVID-SPAIN consortium |
| EPI_ISL_474847, EPI_ISL_474848, EPI_ISL_474849 | Complejo Hospitalario Universitario de Albacete | SeqCOVID-SPAIN consortium/IBV(CSIC) | Encarnacion Simarro Córdoba, Julia Lozano Serra, Lorena Robles Fonseca , Monica Parra Grandes, Caridad Sainz de Baranda Camino and SeqCOVID-SPAIN consortium |
| EPI_ISL_474850, EPI_ISL_474851, EPI_ISL_474852 | Hospital Universitario Virgen de las Nieves de Granada-SAS | SeqCOVID-SPAIN consortium/IBV(CSIC) | Mercedes Pérez Ruiz, Sara Sanbonmatsu Gámez, Irene Pedrosa Corral, José M. Navarro-Marí and SeqCOVID-SPAIN consortium |
| EPI_ISL_474853 | Complejo Hospitalario Universitario de Albacete | SeqCOVID-SPAIN consortium/IBV(CSIC) | Encarnacion Simarro Córdoba, Julia Lozano Serra, Lorena Robles Fonseca , Monica Parra Grandes, Caridad Sainz de Baranda Camino and SeqCOVID-SPAIN consortium |
| EPI_ISL_474854, EPI_ISL_474855, EPI_ISL_474856, EPI_ISL_474857, EPI_ISL_474858, EPI_ISL_474859, EPI_ISL_474860, EPI_ISL_474861, EPI_ISL_474862, EPI_ISL_474863, EPI_ISL_474864, EPI_ISL_474865, EPI_ISL_474866, EPI_ISL_474867, EPI_ISL_474868, EPI_ISL_474869, EPI_ISL_474870, EPI_ISL_474871, EPI_ISL_474872, EPI_ISL_474873, EPI_ISL_474874, EPI_ISL_474875, EPI_ISL_474876, EPI_ISL_474877, EPI_ISL_474878, EPI_ISL_474879, EPI_ISL_474880, EPI_ISL_474881, EPI_ISL_474882, EPI_ISL_474883, EPI_ISL_474884, EPI_ISL_474885, EPI_ISL_474886, EPI_ISL_474887, EPI_ISL_474888, EPI_ISL_474889, EPI_ISL_474890, EPI_ISL_474891, EPI_ISL_474892, EPI_ISL_474893, EPI_ISL_474894, EPI_ISL_474895, EPI_ISL_474896, EPI_ISL_474897, EPI_ISL_474898, EPI_ISL_474899, EPI_ISL_474900 | | | |
| see above | Hospital Universitario Virgen de las Nieves de Granada-SAS | SeqCOVID-SPAIN consortium/IBV(CSIC) | Mercedes Pérez Ruiz, Sara Sanbonmatsu Gámez, Irene Pedrosa Corral, José M. Navarro-Marí and SeqCOVID-SPAIN consortium |
| EPI_ISL_474901, EPI_ISL_474902, EPI_ISL_474903, EPI_ISL_474904, EPI_ISL_474905, EPI_ISL_474906 | Complejo Hospitalario Universitario de Albacete | SeqCOVID-SPAIN consortium/IBV(CSIC) | Encarnacion Simarro Córdoba, Julia Lozano Serra, Lorena Robles Fonseca , Monica Parra Grandes, Caridad Sainz de Baranda Camino and SeqCOVID-SPAIN consortium |
| EPI_ISL_474907, EPI_ISL_474908, EPI_ISL_474909 | Hospital Universitario Virgen de las Nieves de Granada-SAS | SeqCOVID-SPAIN consortium/IBV(CSIC) | Mercedes Pérez Ruiz, Sara Sanbonmatsu Gámez, Irene Pedrosa Corral, José M. Navarro-Marí and SeqCOVID-SPAIN consortium |
| EPI_ISL_474910, EPI_ISL_474911, EPI_ISL_474912, EPI_ISL_474913, EPI_ISL_474914, EPI_ISL_474915, EPI_ISL_474916, EPI_ISL_474917, EPI_ISL_474918 | Hospital Universitario de Gran Canaria Dr. Negrín | SeqCOVID-SPAIN consortium/IBV(CSIC) | M. Carmen Pérez González, Francisco J. Chamizo López, Ana Bordes Benítez and SeqCOVID-SPAIN consortium |
| EPI_ISL_474919 | Complejo Hospitalario Universitario de Albacete | SeqCOVID-SPAIN consortium/IBV(CSIC) | Encarnacion Simarro Córdoba, Julia Lozano Serra, Lorena Robles Fonseca , Monica Parra Grandes, Caridad Sainz de Baranda Camino and SeqCOVID-SPAIN consortium |
| EPI_ISL_474920 | Hospital Universitario Virgen de las Nieves de Granada-SAS | SeqCOVID-SPAIN consortium/IBV(CSIC) | Mercedes Pérez Ruiz, Sara Sanbonmatsu Gámez, Irene Pedrosa Corral, José M. Navarro-Marí and SeqCOVID-SPAIN consortium |
| EPI_ISL_474921 | Complejo Hospitalario Universitario de Albacete | SeqCOVID-SPAIN consortium/IBV(CSIC) | Encarnacion Simarro Córdoba, Julia Lozano Serra, Lorena Robles Fonseca , Monica Parra Grandes, Caridad Sainz de Baranda Camino and SeqCOVID-SPAIN consortium |
| EPI_ISL_474922, EPI_ISL_474923, EPI_ISL_474924, EPI_ISL_474925, EPI_ISL_474926, EPI_ISL_474927, EPI_ISL_474928, EPI_ISL_474929, EPI_ISL_474930, EPI_ISL_474931, EPI_ISL_474932 | | | |
| see above | Hospital Universitario Virgen de las Nieves de Granada-SAS | SeqCOVID-SPAIN consortium/IBV(CSIC) | Mercedes Pérez Ruiz, Sara Sanbonmatsu Gámez, Irene Pedrosa Corral, José M. Navarro-Marí and SeqCOVID-SPAIN consortium |
| EPI_ISL_474933 | Complejo Hospitalario Universitario de Albacete | SeqCOVID-SPAIN consortium/IBV(CSIC) | Encarnacion Simarro Córdoba, Julia Lozano Serra, Lorena Robles Fonseca , Monica Parra Grandes, Caridad Sainz de Baranda Camino and SeqCOVID-SPAIN consortium |
| EPI_ISL_474934, EPI_ISL_474935, EPI_ISL_474936, EPI_ISL_474937, EPI_ISL_474938, EPI_ISL_474939 | Hospital Universitario Virgen de las Nieves de Granada-SAS | SeqCOVID-SPAIN consortium/IBV(CSIC) | Mercedes Pérez Ruiz, Sara Sanbonmatsu Gámez, Irene Pedrosa Corral, José M. Navarro-Marí and SeqCOVID-SPAIN consortium |
| EPI_ISL_474940, EPI_ISL_474941 | Complejo Hospitalario Universitario de Albacete | SeqCOVID-SPAIN consortium/IBV(CSIC) | Encarnacion Simarro Córdoba, Julia Lozano Serra, Lorena Robles Fonseca , Monica Parra Grandes, Caridad Sainz de Baranda Camino and SeqCOVID-SPAIN consortium |

| | | | |
|---|---|---|---|
| EPI_ISL_474942, EPI_ISL_474943, EPI_ISL_474944 | Hospital Universitario Virgen de las Nieves de Granada-SAS | SeqCOVID-SPAIN consortium/IBV(CSIC) | Mercedes Pérez Ruiz, Sara Sanbonmatsu Gámez, Irene Pedrosa Corral, José M. Navarro-Marí and SeqCOVID-SPAIN consortium |
| EPI_ISL_474945, EPI_ISL_474946, EPI_ISL_474947 | Complejo Hospitalario Universitario de Albacete | SeqCOVID-SPAIN consortium/IBV(CSIC) | Encarnacion Simarro Córdoba, Julia Lozano Serra, Lorena Robles Fonseca , Monica Parra Grandes, Caridad Sainz de Baranda Camino and SeqCOVID-SPAIN consortium |
| EPI_ISL_474948, EPI_ISL_474949, EPI_ISL_474950 | Hospital Universitario Virgen de las Nieves de Granada-SAS | SeqCOVID-SPAIN consortium/IBV(CSIC) | Mercedes Pérez Ruiz, Sara Sanbonmatsu Gámez, Irene Pedrosa Corral, José M. Navarro-Marí and SeqCOVID-SPAIN consortium |
| EPI_ISL_474951, EPI_ISL_474952, EPI_ISL_474953, EPI_ISL_474954, EPI_ISL_474955, EPI_ISL_474956 | Complejo Hospitalario Universitario de Albacete | SeqCOVID-SPAIN consortium/IBV(CSIC) | Encarnacion Simarro Córdoba, Julia Lozano Serra, Lorena Robles Fonseca , Monica Parra Grandes, Caridad Sainz de Baranda Camino and SeqCOVID-SPAIN consortium |
| EPI_ISL_474957 | Hospital Universitario Virgen de las Nieves de Granada-SAS | SeqCOVID-SPAIN consortium/IBV(CSIC) | Mercedes Pérez Ruiz, Sara Sanbonmatsu Gámez, Irene Pedrosa Corral, José M. Navarro-Marí and SeqCOVID-SPAIN consortium |
| EPI_ISL_474958 | Israeli Central Virology laboratory | Israel Central Virology laboratory | Neta Zuckerman, Efrat Dahan Bucris, Oran Erster, Ella Mendelson, Michal Mandelboim |
| EPI_ISL_474959, EPI_ISL_474960, EPI_ISL_474961, EPI_ISL_474962, EPI_ISL_474963, EPI_ISL_474964, EPI_ISL_474965, EPI_ISL_474966, EPI_ISL_474967, EPI_ISL_474968, EPI_ISL_474969, EPI_ISL_474970, EPI_ISL_474971, EPI_ISL_474972, EPI_ISL_474973, EPI_ISL_474974, EPI_ISL_474975, EPI_ISL_474976, EPI_ISL_474977, EPI_ISL_474978, EPI_ISL_474979, EPI_ISL_474980, EPI_ISL_474981, EPI_ISL_474982, EPI_ISL_474983, EPI_ISL_474984, EPI_ISL_474985, EPI_ISL_474986, EPI_ISL_474987, EPI_ISL_474988, EPI_ISL_474989, EPI_ISL_474990, EPI_ISL_474991, EPI_ISL_474992, EPI_ISL_474993, EPI_ISL_474994, EPI_ISL_474995, EPI_ISL_474996, EPI_ISL_474997, EPI_ISL_474998, EPI_ISL_474999, EPI_ISL_475000, EPI_ISL_475001, EPI_ISL_475002, EPI_ISL_475003, EPI_ISL_475004, EPI_ISL_475005, EPI_ISL_475006, EPI_ISL_475007, EPI_ISL_475008, EPI_ISL_475009, EPI_ISL_475010, EPI_ISL_475011, EPI_ISL_475012, EPI_ISL_475013, EPI_ISL_475014, EPI_ISL_475015, EPI_ISL_475016, EPI_ISL_475017, EPI_ISL_475018, EPI_ISL_475019, EPI_ISL_475020, EPI_ISL_475021, EPI_ISL_475022, EPI_ISL_475023, EPI_ISL_475024, EPI_ISL_475025 | | | |
| see above | Israel Central Virology laboratory | Israel Central Virology laboratory | Neta Zuckerman, Efrat Dahan Bucris, Oran Erster, Ella Mendelson, Michal Mandelboim |
| EPI_ISL_475026 | Banas Medical College and Research Institute | Gujarat Biotechnology Research Centre | Sunil R Joshi, Viren s Doshi, Pritesh Sabara, Apurvasinh Puvar, Janvi Raval, Zarna Patel, Monika Gandhi, Pinal Trivedi, Maharshi Pandya, Nidhi Patel, Nitin Savaliya, Raghawendra Kumar, Dinesh Kumar, Zuber Saiyed, Komal Patel, Labdhi Pandya, Snehal Bagatharia, Radhika Khara, Neha Rajpara, R D Dixit, A M Kadri, Harsh Bakshi, Chaitanya Joshi, Madhvi Joshi |
| EPI_ISL_475027 | Banas Medical College and Research Institute | Gujarat Biotechnology Research Centre | Viren s Doshi, Pritesh Sabara, Apurvasinh Puvar, Janvi Raval, Zarna Patel, Monika Gandhi, Pinal Trivedi, Maharshi Pandya, Nidhi Patel, Nitin Savaliya, Raghawendra Kumar, Dinesh Kumar, Zuber Saiyed, Komal Patel, Labdhi Pandya, Snehal Bagatharia, Radhika Khara, Sunil R Joshi, Afzal Ansari, R D Dixit, A M Kadri, Harsh Bakshi, Chaitanya Joshi, Madhvi Joshi |
| EPI_ISL_475028 | Banas Medical College and Research Institute | Gujarat Biotechnology Research Centre | Pritesh Sabara, Apurvasinh Puvar, Janvi Raval, Zarna Patel, Monika Gandhi, Pinal Trivedi, Maharshi Pandya, Nidhi Patel, Nitin Savaliya, Raghawendra Kumar, Dinesh Kumar, Zuber Saiyed, Komal Patel, Labdhi Pandya, Snehal Bagatharia, Radhika Khara, Sunil R Joshi, Viren s Doshi, Fenil Patel, R D Dixit, A M Kadri, Harsh Bakshi, Chaitanya Joshi, Madhvi Joshi |
| EPI_ISL_475029 | Banas Medical College and Research Institute | Gujarat Biotechnology Research Centre | Apurvasinh Puvar, Janvi Raval, Zarna Patel, Monika Gandhi, Pinal Trivedi, Maharshi Pandya, Nidhi Patel, Nitin Savaliya, Raghawendra Kumar, Dinesh Kumar, Zuber Saiyed, Komal Patel, Labdhi Pandya, Snehal Bagatharia, Radhika Khara, Sunil R Joshi, Viren s Doshi, Pritesh Sabara, Neelam Nathani, R D Dixit, A M Kadri, Harsh Bakshi, Chaitanya Joshi, Madhvi Joshi |
| EPI_ISL_475030 | Department of MicroBiology, Government Medical College, Surat | Gujarat Biotechnology Research Centre | Janvi Raval, Zarna Patel, Monika Gandhi, Pinal Trivedi, Maharshi Pandya, Nidhi Patel, Nitin Savaliya, Raghawendra Kumar, Dinesh Kumar, Zuber Saiyed, Komal Patel, Labdhi Pandya, Snehal Bagatharia, Naresh Chauhan, Summaiya Mullan, Amit gamit, Pritesh Sabara, Apurvasinh Puvar, Armi Chaudhari, R D Dixit, A M Kadri, Harsh Bakshi, Chaitanya Joshi, Madhvi Joshi |
| EPI_ISL_475031 | Department of MicroBiology, Government Medical College, Surat | Gujarat Biotechnology Research Centre | Zarna Patel, Monika Gandhi, Pinal Trivedi, Maharshi Pandya, Nidhi Patel, Nitin Savaliya, Raghawendra Kumar, Dinesh Kumar, Zuber Saiyed, Komal Patel, Labdhi Pandya, Snehal Bagatharia, Naresh Chauhan, Summaiya Mullan, Amit gamit, Pritesh Sabara, Apurvasinh Puvar, Janvi Raval, Bhavya Jindal, R D Dixit, A M Kadri, Harsh Bakshi, Chaitanya Joshi, Madhvi Joshi |
| EPI_ISL_475032 | Department of MicroBiology, Government Medical College, Surat | Gujarat Biotechnology Research Centre | Monika Gandhi, Pinal Trivedi, Maharshi Pandya, Nidhi Patel, Nitin Savaliya, Raghawendra Kumar, Dinesh Kumar, Zuber Saiyed, Komal Patel, Labdhi Pandya, Snehal Bagatharia, Naresh Chauhan, Summaiya Mullan, Amit gamit, Pritesh Sabara, Apurvasinh Puvar, Janvi Raval, Zarna Patel, Priyanka P Vatsa, R D Dixit, A M Kadri, Harsh Bakshi, Chaitanya Joshi, Madhvi Joshi |
| EPI_ISL_475033 | Department of MicroBiology, Government Medical College, Surat | Gujarat Biotechnology Research Centre | Pinal Trivedi, Maharshi Pandya, Nidhi Patel, Nitin Savaliya, Raghawendra Kumar, Dinesh Kumar, Zuber Saiyed, Komal Patel, Labdhi Pandya, Snehal Bagatharia, Naresh Chauhan, Summaiya Mullan, Amit gamit, Pritesh Sabara, Apurvasinh Puvar, Janvi Raval, Zarna Patel, Monika Gandhi, Pooja P Doshi, R D Dixit, A M Kadri, Harsh Bakshi, Chaitanya Joshi, Madhvi Joshi |
| EPI_ISL_475034 | Department of MicroBiology, Government Medical College, Surat | Gujarat Biotechnology Research Centre | Maharshi Pandya, Nidhi Patel, Nitin Savaliya, Raghawendra Kumar, Dinesh Kumar, Zuber Saiyed, Komal Patel, Labdhi Pandya, Snehal Bagatharia, Naresh Chauhan, Summaiya Mullan, Amit gamit, Pritesh Sabara, Apurvasinh Puvar, Janvi Raval, Zarna Patel, Monika Gandhi, Pinal Trivedi, Akanksha Verma, R D Dixit, A M Kadri, Harsh Bakshi, Chaitanya Joshi, Madhvi Joshi |
| EPI_ISL_475035 | Department of MicroBiology, Government Medical College, Surat | Gujarat Biotechnology Research Centre | Nidhi Patel, Nitin Savaliya, Raghawendra Kumar, Dinesh Kumar, Zuber Saiyed, Komal Patel, Labdhi Pandya, Snehal Bagatharia, Naresh Chauhan, Summaiya Mullan, Amit gamit, Pritesh Sabara, Apurvasinh Puvar, Janvi Raval, Zarna Patel, Monika Gandhi, Pinal Trivedi, Maharshi Pandya, Priti Pandita, R D Dixit, A M Kadri, Harsh Bakshi, Chaitanya Joshi, Madhvi Joshi |
| EPI_ISL_475036 | Department of MicroBiology, Government Medical College, Surat | Gujarat Biotechnology Research Centre | Nitin Savaliya, Raghawendra Kumar, Dinesh Kumar, Zuber Saiyed, Komal Patel, Labdhi Pandya, Snehal Bagatharia, Naresh Chauhan, Summaiya Mullan, Amit gamit, Pritesh Sabara, Apurvasinh Puvar, Janvi Raval, Zarna Patel, Monika Gandhi, Pinal Trivedi, Maharshi Pandya, Nidhi Patel, Pragya Sharma, R D Dixit, A M Kadri, Harsh Bakshi, Chaitanya Joshi, Madhvi Joshi |
| EPI_ISL_475037 | Department of MicroBiology, Government Medical College, Surat | Gujarat Biotechnology Research Centre | Raghawendra Kumar, Dinesh Kumar, Zuber Saiyed, Komal Patel, Labdhi Pandya, Snehal Bagatharia, Naresh Chauhan, Summaiya Mullan, Amit gamit, Pritesh Sabara, Apurvasinh Puvar, Janvi Raval, Zarna Patel, Monika Gandhi, Pinal Trivedi, Maharshi Pandya, Nidhi Patel, Nitin Savaliya, Neha Rajpara, R D Dixit, A M Kadri, Harsh Bakshi, Chaitanya Joshi, Madhvi Joshi |
| EPI_ISL_475038 | Department of MicroBiology, Government Medical College, Surat | Gujarat Biotechnology Research Centre | Dinesh Kumar, Zuber Saiyed, Komal Patel, Labdhi Pandya, Snehal Bagatharia, Naresh Chauhan, Summaiya Mullan, Amit gamit, Pritesh Sabara, Apurvasinh Puvar, Janvi Raval, Zarna Patel, Monika Gandhi, Pinal Trivedi, Maharshi Pandya, Nidhi Patel, Nitin Savaliya, Raghawendra Kumar, Afzal Ansari, R D Dixit, A M Kadri, Harsh Bakshi, Chaitanya Joshi, Madhvi Joshi |
| EPI_ISL_475039 | Department of MicroBiology, Government Medical College, Surat | Gujarat Biotechnology Research Centre | Zuber Saiyed, Komal Patel, Labdhi Pandya, Snehal Bagatharia, Naresh Chauhan, Summaiya Mullan, Amit gamit, Pritesh Sabara, Apurvasinh Puvar, Janvi Raval, Zarna Patel, Monika Gandhi, Pinal Trivedi, Maharshi Pandya, Nidhi Patel, Nitin Savaliya, Raghawendra Kumar, Dinesh Kumar, Fenil Patel, R D Dixit, A M Kadri, Harsh Bakshi, Chaitanya Joshi, Madhvi Joshi |
| EPI_ISL_475040 | Department of MicroBiology, Government Medical College, Surat | Gujarat Biotechnology Research Centre | Komal Patel, Labdhi Pandya, Snehal Bagatharia, Naresh Chauhan, Summaiya Mullan, Amit gamit, Pritesh Sabara, Apurvasinh Puvar, Janvi Raval, Zarna Patel, Monika Gandhi, Pinal Trivedi, Maharshi Pandya, Nidhi Patel, Nitin Savaliya, Raghawendra Kumar, Dinesh Kumar, Zuber Saiyed, Neelam Nathani, R D Dixit, A M Kadri, Harsh Bakshi, Chaitanya Joshi, Madhvi Joshi |
| EPI_ISL_475041 | Department of MicroBiology, Government Medical College, Surat | Gujarat Biotechnology Research Centre | Labdhi Pandya, Snehal Bagatharia, Naresh Chauhan, Summaiya Mullan, Amit gamit, Pritesh Sabara, Apurvasinh Puvar, Janvi Raval, Zarna Patel, Monika Gandhi, Pinal Trivedi, Maharshi Pandya, Nidhi Patel, Nitin Savaliya, Raghawendra Kumar, Dinesh Kumar, Zuber Saiyed, Komal Patel, Armi Chaudhari, R D Dixit, A M Kadri, Harsh Bakshi, Chaitanya Joshi, Madhvi Joshi |
| EPI_ISL_475042 | Department of MicroBiology, Government Medical College, Surat | Gujarat Biotechnology Research Centre | Snehal Bagatharia, Naresh Chauhan, Summaiya Mullan, Amit gamit, Pritesh Sabara, Apurvasinh Puvar, Janvi Raval, Zarna Patel, Monika Gandhi, Pinal Trivedi, Maharshi Pandya, Nidhi Patel, Nitin Savaliya, Raghawendra Kumar, Dinesh Kumar, Zuber Saiyed, Komal Patel, Labdhi Pandya, Bhavya Jindal, R D Dixit, A M Kadri, Harsh Bakshi, Chaitanya Joshi, Madhvi Joshi |
| EPI_ISL_475043 | Department of MicroBiology, Government Medical College, Surat | Gujarat Biotechnology Research Centre | Naresh Chauhan, Summaiya Mullan, Amit gamit, Pritesh Sabara, Apurvasinh Puvar, Janvi Raval, Zarna Patel, Monika Gandhi, Pinal Trivedi, Maharshi Pandya, Nidhi Patel, Nitin Savaliya, Raghawendra Kumar, Dinesh Kumar, Zuber Saiyed, Komal Patel, Labdhi Pandya, Snehal Bagatharia, Priyanka P Vatsa, R D Dixit, A M Kadri, Harsh Bakshi, Chaitanya Joshi, Madhvi Joshi |
| EPI_ISL_475044 | Department of MicroBiology, Government Medical College, Surat | Gujarat Biotechnology Research Centre | Summaiya Mullan, Amit gamit, Pritesh Sabara, Apurvasinh Puvar, Janvi Raval, Zarna Patel, Monika Gandhi, Pinal Trivedi, Maharshi Pandya, Nidhi Patel, Nitin Savaliya, Raghawendra Kumar, Dinesh Kumar, Zuber Saiyed, Komal Patel, Labdhi Pandya, Snehal Bagatharia, Naresh Chauhan, Pooja P Doshi, R D Dixit, A M Kadri, Harsh Bakshi, Chaitanya Joshi, Madhvi Joshi |
| EPI_ISL_475045 | Department of MicroBiology, Government Medical College, Surat | Gujarat Biotechnology Research Centre | Amit gamit, Pritesh Sabara, Apurvasinh Puvar, Janvi Raval, Zarna Patel, Monika Gandhi, Pinal Trivedi, Maharshi Pandya, Nidhi Patel, Nitin Savaliya, Raghawendra Kumar, Dinesh Kumar, Zuber Saiyed, Komal Patel, Labdhi Pandya, Snehal Bagatharia, Naresh Chauhan, Summaiya Mullan, Akanksha Verma, R D Dixit, A M Kadri, Harsh Bakshi, Chaitanya Joshi, Madhvi Joshi |
| EPI_ISL_475046 | Department of MicroBiology, Government Medical College, Surat | Gujarat Biotechnology Research Centre | Pritesh Sabara, Apurvasinh Puvar, Janvi Raval, Zarna Patel, Monika Gandhi, Pinal Trivedi, Maharshi Pandya, Nidhi Patel, Nitin Savaliya, Raghawendra Kumar, Dinesh Kumar, Zuber Saiyed, Komal Patel, Labdhi Pandya, Snehal Bagatharia, Naresh Chauhan, Summaiya Mullan, Amit gamit, Priti Pandita, R D Dixit, A M Kadri, Harsh Bakshi, Chaitanya Joshi, Madhvi Joshi |
| EPI_ISL_475047 | GMERS Medical College & Hospital | Gujarat Biotechnology Research Centre | Apurvasinh Puvar, Janvi Raval, Zarna Patel, Monika Gandhi, Pinal Trivedi, Maharshi Pandya, Nidhi Patel, Nitin Savaliya, Raghawendra Kumar, Dinesh Kumar, Zuber Saiyed, Komal Patel, Labdhi Pandya, Snehal Bagatharia, Meenakshi Shah, Neena Doshi, Varsha Godbole, Pritesh Sabara, Pragya Sharma, R D Dixit, A M Kadri, Harsh Bakshi, Chaitanya Joshi, Madhvi Joshi |
| EPI_ISL_475048 | GMERS Medical College & Hospital | Gujarat Biotechnology Research Centre | Janvi Raval, Zarna Patel, Monika Gandhi, Pinal Trivedi, Maharshi Pandya, Nidhi Patel, Nitin Savaliya, Raghawendra Kumar, Dinesh Kumar, Zuber Saiyed, Komal Patel, Labdhi Pandya, Snehal Bagatharia, Meenakshi Shah, Neena Doshi, Varsha Godbole, Pritesh Sabara, Apurvasinh Puvar, Neha Rajpara, R D Dixit, A M Kadri, Harsh Bakshi, Chaitanya Joshi, Madhvi Joshi |
| EPI_ISL_475049 | GMERS Medical College & Hospital | Gujarat Biotechnology Research Centre | Zarna Patel, Monika Gandhi, Pinal Trivedi, Maharshi Pandya, Nidhi Patel, Nitin Savaliya, Raghawendra Kumar, Dinesh Kumar, Zuber Saiyed, Komal Patel, Labdhi Pandya, Snehal Bagatharia, Meenakshi Shah, Neena Doshi, Varsha Godbole, Pritesh Sabara, Apurvasinh Puvar, Janvi Raval, Afzal Ansari, R D Dixit, A M Kadri, Harsh Bakshi, Chaitanya Joshi, Madhvi Joshi |
| EPI_ISL_475050 | GMERS Medical College & Hospital | Gujarat Biotechnology Research Centre | Monika Gandhi, Pinal Trivedi, Maharshi Pandya, Nidhi Patel, Nitin Savaliya, Raghawendra Kumar, Dinesh Kumar, Zuber Saiyed, Komal Patel, Labdhi Pandya, Snehal Bagatharia, Meenakshi Shah, Neena Doshi, Varsha Godbole, Pritesh Sabara, Apurvasinh Puvar, Janvi Raval, Zarna Patel, Fenil Patel, R D Dixit, A M Kadri, Harsh Bakshi, Chaitanya Joshi, Madhvi Joshi |
| EPI_ISL_475051 | GMERS Medical College & Hospital | Gujarat Biotechnology Research Centre | Pinal Trivedi, Maharshi Pandya, Nidhi Patel, Nitin Savaliya, Raghawendra Kumar, Dinesh Kumar, Zuber Saiyed, Komal Patel, Labdhi Pandya, Snehal Bagatharia, Meenakshi Shah, Neena Doshi, Varsha Godbole, Pritesh Sabara, Apurvasinh Puvar, Janvi Raval, Zarna Patel, Monika Gandhi, Neelam Nathani, R D Dixit, A M Kadri, Harsh Bakshi, Chaitanya Joshi, Madhvi Joshi |
| EPI_ISL_475052 | GMERS Medical College & Hospital | Gujarat Biotechnology Research Centre | Maharshi Pandya, Nidhi Patel, Nitin Savaliya, Raghawendra Kumar, Dinesh Kumar, Zuber Saiyed, Komal Patel, Labdhi Pandya, Snehal Bagatharia, Meenakshi Shah, Neena Doshi, Varsha Godbole, Pritesh Sabara, Apurvasinh Puvar, Janvi Raval, Zarna Patel, Monika Gandhi, Pinal Trivedi, Armi Chaudhari, R D Dixit, A M Kadri, Harsh Bakshi, Chaitanya Joshi, Madhvi Joshi |
| EPI_ISL_475053 | GMERS Medical College & Hospital | Gujarat Biotechnology Research Centre | Nidhi Patel, Nitin Savaliya, Raghawendra Kumar, Dinesh Kumar, Zuber Saiyed, Komal Patel, Labdhi Pandya, Snehal Bagatharia, Meenakshi Shah, Neena Doshi, Varsha Godbole, Pritesh Sabara, Apurvasinh Puvar, Janvi Raval, Zarna Patel, Monika Gandhi, Pinal Trivedi, Maharshi Pandya, Bhavya Jindal, R D Dixit, A M Kadri, Harsh Bakshi, Chaitanya Joshi, Madhvi Joshi |
| EPI_ISL_475054 | GMERS Medical College & Hospital | Gujarat Biotechnology Research Centre | Nitin Savaliya, Raghawendra Kumar, Dinesh Kumar, Zuber Saiyed, Komal Patel, Labdhi Pandya, Snehal Bagatharia, Meenakshi Shah, Neena Doshi, Varsha Godbole, Pritesh Sabara, Apurvasinh Puvar, Janvi Raval, Zarna Patel, Monika Gandhi, Pinal Trivedi, Maharshi Pandya, Nidhi Patel, Priyanka P Vatsa, R D Dixit, A M Kadri, Harsh Bakshi, Chaitanya Joshi, Madhvi Joshi |
| EPI_ISL_475055 | GMERS Medical College & Hospital | Gujarat Biotechnology Research Centre | Raghawendra Kumar, Dinesh Kumar, Zuber Saiyed, Komal Patel, Labdhi Pandya, Snehal Bagatharia, Meenakshi Shah, Neena Doshi, Varsha Godbole, Pritesh Sabara, Apurvasinh Puvar, Janvi Raval, Zarna Patel, Monika Gandhi, Pinal Trivedi, Maharshi Pandya, Nidhi Patel, Nitin Savaliya, Pooja P Doshi, R D Dixit, A M Kadri, Harsh Bakshi, Chaitanya Joshi, Madhvi Joshi |
| EPI_ISL_475056 | Dr. N. D. Desai Medical College & Hospital | Gujarat Biotechnology Research Centre | Dinesh Kumar, Zuber Saiyed, Komal Patel, Labdhi Pandya, Snehal Bagatharia, J G Buch, Jigar Gusani, Supreet Prabhu, Pritesh Sabara, Apurvasinh Puvar, Janvi Raval, Zarna Patel, Monika Gandhi, Pinal Trivedi, Maharshi Pandya, Nidhi Patel, Nitin Savaliya, Raghawendra Kumar, Akanksha Verma, R D Dixit, A M Kadri, Harsh Bakshi, Chaitanya Joshi, Madhvi Joshi |
| EPI_ISL_475057 | Dr. N. D. Desai Medical College & Hospital | Gujarat Biotechnology Research Centre | Zuber Saiyed, Komal Patel, Labdhi Pandya, Supreet Prabhu, Snehal Bagatharia, Jigar Gusani, J G Buch, Pritesh Sabara, Apurvasinh Puvar, Janvi Raval, Zarna Patel, Monika Gandhi, Pinal Trivedi, Maharshi Pandya, Nidhi Patel, Nitin Savaliya, Raghawendra Kumar, Dinesh Kumar, Priti Pandita, R D Dixit, A M Kadri, Harsh Bakshi, Chaitanya Joshi, Madhvi Joshi |
| EPI_ISL_475058 | GAIMS & G K General Hospital | Gujarat Biotechnology Research Centre | Babulal Babhoria, Hitesh Assudani, Komal Patel, Labdhi Pandya, Snehal Bagatharia, Pritesh Sabara, Apurvasinh Puvar, Janvi Raval, Zarna Patel, Monika Gandhi, Pinal Trivedi, Maharshi Pandya, Nidhi Patel, Nitin Savaliya, Raghawendra Kumar, Dinesh Kumar, Zuber Saiyed, Pragya Sharma, R D Dixit, A M Kadri, Harsh Bakshi, Chaitanya Joshi, Madhvi Joshi |
| EPI_ISL_475059 | GAIMS & G K General Hospital | Gujarat Biotechnology Research Centre | Hitesh Assudani, Babulal Babhoria, Labdhi Pandya, Snehal Bagatharia, Pritesh Sabara, Apurvasinh Puvar, Janvi Raval, Zarna Patel, Monika Gandhi, Pinal Trivedi, Maharshi Pandya, Nidhi Patel, Nitin Savaliya, Raghawendra Kumar, Dinesh Kumar, Zuber Saiyed, Komal Patel, Neha Rajpara, R D Dixit, A M Kadri, Harsh Bakshi, Chaitanya Joshi, Madhvi Joshi |
| EPI_ISL_475060, EPI_ISL_475061, EPI_ISL_475062, EPI_ISL_475063, EPI_ISL_475064, EPI_ISL_475065, EPI_ISL_475066, EPI_ISL_475067, EPI_ISL_475068, EPI_ISL_475069, EPI_ISL_475070 | | | |
| see above | Lab voor klinische biologie | Onderzoeksgroep Virologie | Laurens Lambrechts, Nick Vereecke, Marthe Pauwels, Bruno Verhasselt, Linos Vandekerckhove, Hans Nauwynck, Sebastiaan Theuns |
| EPI_ISL_475071, EPI_ISL_475072, EPI_ISL_475073, EPI_ISL_475074, EPI_ISL_475075, EPI_ISL_475076, EPI_ISL_475077, EPI_ISL_475078, EPI_ISL_475079, EPI_ISL_475080, EPI_ISL_475081, EPI_ISL_475082 | | | |
| see above | Lab voor klinische biologie | Onderzoeksgroep Virologie | Nick Vereecke, Laurens Lambrechts, Marthe Pauwels, Bruno Verhasselt, Linos Vandekerckhove, Hans Nauwynck, Sebastiaan Theuns |
| EPI_ISL_475083, EPI_ISL_475084 | National Institute of Laboratory Medicine and Referral Center | Genomic Research Lab, BCSIR | Md. Murshed Hasan Sarkar, Abu Sayeed Mohammad Mahmud, Mohammad Samir Uzzaman, Eshrar Osman, Md. Ahasan Habib, Shahina Akter, Tanjina Akhter Banu, Barna Goswami, Iffat Jahan, Md. Saddam Hossain, Tasnim Nafisa, Md. Maruf Ahmed Molla, Mahmuda Yeasmin, Asish Kumar Ghosh, Bayzid Bin Monir, A. K. M. Shamsuzzaman, Sheikh Md. Selim Al Din, Utpal Chandra Ray, Salek Ahmed Sajib, Md. Salim Khan |

| | | | |
|---|---|---|---|
| EPI_ISL_475085 | Skovde/Unilabs | The Public Health Agency of Sweden | Oskar Karlsson Lindsjo, Maria Lind Karlberg, Mattias Haukland, Reza Advani, Olov Svartstrom, Anna-Malin Linde, Sandra Broddesson, Petra Edquist, Shamam Muradrasoli, Anna Risberg, Karin Tegmark-Wisell |
| EPI_ISL_475086, EPI_ISL_475087, EPI_ISL_475088, EPI_ISL_475089, EPI_ISL_475090, EPI_ISL_475091 | Karolinska Universitetslaboratoriet | The Public Health Agency of Sweden | Oskar Karlsson Lindsjo, Maria Lind Karlberg, Mattias Haukland, Reza Advani, Olov Svartstrom, Anna-Malin Linde, Sandra Broddesson, Petra Edquist, Shamam Muradrasoli, Anna Risberg, Karin Tegmark-Wisell |
| EPI_ISL_475092 | Skovde/Unilabs | The Public Health Agency of Sweden | Oskar Karlsson Lindsjo, Maria Lind Karlberg, Mattias Haukland, Reza Advani, Olov Svartstrom, Anna-Malin Linde, Sandra Broddesson, Petra Edquist, Shamam Muradrasoli, Anna Risberg, Karin Tegmark-Wisell |
| EPI_ISL_475093, EPI_ISL_475094 | Halmstad klinisk mikrobiologi | The Public Health Agency of Sweden | Oskar Karlsson Lindsjo, Maria Lind Karlberg, Mattias Haukland, Reza Advani, Olov Svartstrom, Anna-Malin Linde, Sandra Broddesson, Petra Edquist, Shamam Muradrasoli, Anna Risberg, Karin Tegmark-Wisell |
| EPI_ISL_475095 | Karolinska Universitetslaboratoriet | The Public Health Agency of Sweden | Oskar Karlsson Lindsjo, Maria Lind Karlberg, Mattias Haukland, Reza Advani, Olov Svartstrom, Anna-Malin Linde, Sandra Broddesson, Petra Edquist, Shamam Muradrasoli, Anna Risberg, Karin Tegmark-Wisell |
| EPI_ISL_475096, EPI_ISL_475097, EPI_ISL_475098 | Halmstad klinisk mikrobiologi | The Public Health Agency of Sweden | Oskar Karlsson Lindsjo, Maria Lind Karlberg, Mattias Haukland, Reza Advani, Olov Svartstrom, Anna-Malin Linde, Sandra Broddesson, Petra Edquist, Shamam Muradrasoli, Anna Risberg, Karin Tegmark-Wisell |
| EPI_ISL_475099 | Skovde/Unilabs | The Public Health Agency of Sweden | Oskar Karlsson Lindsjo, Maria Lind Karlberg, Mattias Haukland, Reza Advani, Olov Svartstrom, Anna-Malin Linde, Sandra Broddesson, Petra Edquist, Shamam Muradrasoli, Anna Risberg, Karin Tegmark-Wisell |
| EPI_ISL_475100 | Halmstad klinisk mikrobiologi | The Public Health Agency of Sweden | Oskar Karlsson Lindsjo, Maria Lind Karlberg, Mattias Haukland, Reza Advani, Olov Svartstrom, Anna-Malin Linde, Sandra Broddesson, Petra Edquist, Shamam Muradrasoli, Anna Risberg, Karin Tegmark-Wisell |
| EPI_ISL_475101, EPI_ISL_475102, EPI_ISL_475103, EPI_ISL_475104 | Klinisk Mikrobiologi | The Public Health Agency of Sweden | Oskar Karlsson Lindsjo, Maria Lind Karlberg, Mattias Haukland, Reza Advani, Olov Svartstrom, Anna-Malin Linde, Sandra Broddesson, Petra Edquist, Shamam Muradrasoli, Anna Risberg, Karin Tegmark-Wisell |
| EPI_ISL_475105, EPI_ISL_475106, EPI_ISL_475107, EPI_ISL_475108, EPI_ISL_475109 | Skovde/Unilabs | The Public Health Agency of Sweden | Oskar Karlsson Lindsjo, Maria Lind Karlberg, Mattias Haukland, Reza Advani, Olov Svartstrom, Anna-Malin Linde, Sandra Broddesson, Petra Edquist, Shamam Muradrasoli, Anna Risberg, Karin Tegmark-Wisell |
| EPI_ISL_475110 | Gavle klinisk mikrobiologi | The Public Health Agency of Sweden | Oskar Karlsson Lindsjo, Maria Lind Karlberg, Mattias Haukland, Reza Advani, Olov Svartstrom, Anna-Malin Linde, Sandra Broddesson, Petra Edquist, Shamam Muradrasoli, Anna Risberg, Karin Tegmark-Wisell |
| EPI_ISL_475111, EPI_ISL_475112, EPI_ISL_475113 | Skovde/Unilabs | The Public Health Agency of Sweden | Oskar Karlsson Lindsjo, Maria Lind Karlberg, Mattias Haukland, Reza Advani, Olov Svartstrom, Anna-Malin Linde, Sandra Broddesson, Petra Edquist, Shamam Muradrasoli, Anna Risberg, Karin Tegmark-Wisell |
| EPI_ISL_475114 | Halmstad klinisk mikrobiologi | The Public Health Agency of Sweden | Oskar Karlsson Lindsjo, Maria Lind Karlberg, Mattias Haukland, Reza Advani, Olov Svartstrom, Anna-Malin Linde, Sandra Broddesson, Petra Edquist, Shamam Muradrasoli, Anna Risberg, Karin Tegmark-Wisell |
| EPI_ISL_475115 | Gavle klinisk mikrobiologi | The Public Health Agency of Sweden | Oskar Karlsson Lindsjo, Maria Lind Karlberg, Mattias Haukland, Reza Advani, Olov Svartstrom, Anna-Malin Linde, Sandra Broddesson, Petra Edquist, Shamam Muradrasoli, Anna Risberg, Karin Tegmark-Wisell |
| EPI_ISL_475116, EPI_ISL_475117 | Halmstad klinisk mikrobiologi | The Public Health Agency of Sweden | Oskar Karlsson Lindsjo, Maria Lind Karlberg, Mattias Haukland, Reza Advani, Olov Svartstrom, Anna-Malin Linde, Sandra Broddesson, Petra Edquist, Shamam Muradrasoli, Anna Risberg, Karin Tegmark-Wisell |
| EPI_ISL_475118, EPI_ISL_475119 | Uppsala klinisk mikrobiologi | The Public Health Agency of Sweden | Oskar Karlsson Lindsjo, Maria Lind Karlberg, Mattias Haukland, Reza Advani, Olov Svartstrom, Anna-Malin Linde, Sandra Broddesson, Petra Edquist, Shamam Muradrasoli, Anna Risberg, Karin Tegmark-Wisell |
| EPI_ISL_475120, EPI_ISL_475121 | Halmstad klinisk mikrobiologi | The Public Health Agency of Sweden | Oskar Karlsson Lindsjo, Maria Lind Karlberg, Mattias Haukland, Reza Advani, Olov Svartstrom, Anna-Malin Linde, Sandra Broddesson, Petra Edquist, Shamam Muradrasoli, Anna Risberg, Karin Tegmark-Wisell |
| EPI_ISL_475122 | Umea klinisk mikrobiologi | The Public Health Agency of Sweden | Oskar Karlsson Lindsjo, Maria Lind Karlberg, Mattias Haukland, Reza Advani, Olov Svartstrom, Anna-Malin Linde, Sandra Broddesson, Petra Edquist, Shamam Muradrasoli, Anna Risberg, Karin Tegmark-Wisell |
| EPI_ISL_475123 | Halmstad klinisk mikrobiologi | The Public Health Agency of Sweden | Oskar Karlsson Lindsjo, Maria Lind Karlberg, Mattias Haukland, Reza Advani, Olov Svartstrom, Anna-Malin Linde, Sandra Broddesson, Petra Edquist, Shamam Muradrasoli, Anna Risberg, Karin Tegmark-Wisell |
| EPI_ISL_475124 | Ostersund klinisk mikrobiologi | The Public Health Agency of Sweden | Oskar Karlsson Lindsjo, Maria Lind Karlberg, Mattias Haukland, Reza Advani, Olov Svartstrom, Anna-Malin Linde, Sandra Broddesson, Petra Edquist, Shamam Muradrasoli, Anna Risberg, Karin Tegmark-Wisell |
| EPI_ISL_475125 | Halmstad klinisk mikrobiologi | The Public Health Agency of Sweden | Oskar Karlsson Lindsjo, Maria Lind Karlberg, Mattias Haukland, Reza Advani, Olov Svartstrom, Anna-Malin Linde, Sandra Broddesson, Petra Edquist, Shamam Muradrasoli, Anna Risberg, Karin Tegmark-Wisell |
| EPI_ISL_475126, EPI_ISL_475127 | Umea klinisk mikrobiologi | The Public Health Agency of Sweden | Oskar Karlsson Lindsjo, Maria Lind Karlberg, Mattias Haukland, Reza Advani, Olov Svartstrom, Anna-Malin Linde, Sandra Broddesson, Petra Edquist, Shamam Muradrasoli, Anna Risberg, Karin Tegmark-Wisell |
| EPI_ISL_475128, EPI_ISL_475129, EPI_ISL_475130, EPI_ISL_475131, EPI_ISL_475132, EPI_ISL_475133, EPI_ISL_475134, EPI_ISL_475135 | Orebro klinisk mikrobiologi | The Public Health Agency of Sweden | Oskar Karlsson Lindsjo, Maria Lind Karlberg, Mattias Haukland, Reza Advani, Olov Svartstrom, Anna-Malin Linde, Sandra Broddesson, Petra Edquist, Shamam Muradrasoli, Anna Risberg, Karin Tegmark-Wisell |
| EPI_ISL_475136 | Karolinska Universitetslaboratoriet | The Public Health Agency of Sweden | Oskar Karlsson Lindsjo, Maria Lind Karlberg, Mattias Haukland, Reza Advani, Olov Svartstrom, Anna-Malin Linde, Sandra Broddesson, Petra Edquist, Shamam Muradrasoli, Anna Risberg, Karin Tegmark-Wisell |
| EPI_ISL_475137, EPI_ISL_475138 | Kalmar klinisk mikrobiologi | The Public Health Agency of Sweden | Oskar Karlsson Lindsjo, Maria Lind Karlberg, Mattias Haukland, Reza Advani, Olov Svartstrom, Anna-Malin Linde, Sandra Broddesson, Petra Edquist, Shamam Muradrasoli, Anna Risberg, Karin Tegmark-Wisell |
| EPI_ISL_475139 | Orebro klinisk mikrobiologi | The Public Health Agency of Sweden | Oskar Karlsson Lindsjo, Maria Lind Karlberg, Mattias Haukland, Reza Advani, Olov Svartstrom, Anna-Malin Linde, Sandra Broddesson, Petra Edquist, Shamam Muradrasoli, Anna Risberg, Karin Tegmark-Wisell |
| EPI_ISL_475140 | Skovde/Unilabs | The Public Health Agency of Sweden | Oskar Karlsson Lindsjo, Maria Lind Karlberg, Mattias Haukland, Reza Advani, Olov Svartstrom, Anna-Malin Linde, Sandra Broddesson, Petra Edquist, Shamam Muradrasoli, Anna Risberg, Karin Tegmark-Wisell |
| EPI_ISL_475141, EPI_ISL_475142 | Karolinska Universitetslaboratoriet | The Public Health Agency of Sweden | Oskar Karlsson Lindsjo, Maria Lind Karlberg, Mattias Haukland, Reza Advani, Olov Svartstrom, Anna-Malin Linde, Sandra Broddesson, Petra Edquist, Shamam Muradrasoli, Anna Risberg, Karin Tegmark-Wisell |
| EPI_ISL_475143 | Umea klinisk mikrobiologi | The Public Health Agency of Sweden | Oskar Karlsson Lindsjo, Maria Lind Karlberg, Mattias Haukland, Reza Advani, Olov Svartstrom, Anna-Malin Linde, Sandra Broddesson, Petra Edquist, Shamam Muradrasoli, Anna Risberg, Karin Tegmark-Wisell |
| EPI_ISL_475144, EPI_ISL_475145, EPI_ISL_475146, EPI_ISL_475147, EPI_ISL_475148 | Klinisk mikrobiologi Vasternorrland | The Public Health Agency of Sweden | Oskar Karlsson Lindsjo, Maria Lind Karlberg, Mattias Haukland, Reza Advani, Olov Svartstrom, Anna-Malin Linde, Sandra Broddesson, Petra Edquist, Shamam Muradrasoli, Anna Risberg, Karin Tegmark-Wisell |
| EPI_ISL_475149, EPI_ISL_475150 | Karolinska Universitetslaboratoriet | The Public Health Agency of Sweden | Oskar Karlsson Lindsjo, Maria Lind Karlberg, Mattias Haukland, Reza Advani, Olov Svartstrom, Anna-Malin Linde, Sandra Broddesson, Petra Edquist, Shamam Muradrasoli, Anna Risberg, Karin Tegmark-Wisell |
| EPI_ISL_475151 | Uppsala klinisk mikrobiologi | The Public Health Agency of Sweden | Oskar Karlsson Lindsjo, Maria Lind Karlberg, Mattias Haukland, Reza Advani, Olov Svartstrom, Anna-Malin Linde, Sandra Broddesson, Petra Edquist, Shamam Muradrasoli, Anna Risberg, Karin Tegmark-Wisell |
| EPI_ISL_475152 | Folkhalsomyndigheten | The Public Health Agency of Sweden | Oskar Karlsson Lindsjo, Maria Lind Karlberg, Mattias Haukland, Reza Advani, Olov Svartstrom, Anna-Malin Linde, Sandra Broddesson, Petra Edquist, Shamam Muradrasoli, Anna Risberg, Karin Tegmark-Wisell |
| EPI_ISL_475153, EPI_ISL_475154, EPI_ISL_475155 | Klinisk mikrobiologi Vasternorrland | The Public Health Agency of Sweden | Oskar Karlsson Lindsjo, Maria Lind Karlberg, Mattias Haukland, Reza Advani, Olov Svartstrom, Anna-Malin Linde, Sandra Broddesson, Petra Edquist, Shamam Muradrasoli, Anna Risberg, Karin Tegmark-Wisell |
| EPI_ISL_475156, EPI_ISL_475157, EPI_ISL_475158, EPI_ISL_475159 | Halmstad klinisk mikrobiologi | The Public Health Agency of Sweden | Oskar Karlsson Lindsjo, Maria Lind Karlberg, Mattias Haukland, Reza Advani, Olov Svartstrom, Anna-Malin Linde, Sandra Broddesson, Petra Edquist, Shamam Muradrasoli, Anna Risberg, Karin Tegmark-Wisell |
| EPI_ISL_475160, EPI_ISL_475161, EPI_ISL_475162, EPI_ISL_475163 | Klinisk mikrobiologi Vasternorrland | The Public Health Agency of Sweden | Oskar Karlsson Lindsjo, Maria Lind Karlberg, Mattias Haukland, Reza Advani, Olov Svartstrom, Anna-Malin Linde, Sandra Broddesson, Petra Edquist, Shamam Muradrasoli, Anna Risberg, Karin Tegmark-Wisell |
| EPI_ISL_475164 | Halmstad klinisk mikrobiologi | The Public Health Agency of Sweden | Oskar Karlsson Lindsjo, Maria Lind Karlberg, Mattias Haukland, Reza Advani, Olov Svartstrom, Anna-Malin Linde, Sandra Broddesson, Petra Edquist, Shamam Muradrasoli, Anna Risberg, Karin Tegmark-Wisell |
| EPI_ISL_475165 | National Institute of Laboratory Medicine and Referral Center | Genomic Research Lab, BCSIR | Shahina Akter, Abu Sayeed Mohammad Mahmud, Mohammad Samir Uzzaman, Eshrar Osman, Md. Ahasan Habib, Tanjina Akhter Banu, Md. Murshed Hasan Sarkar, Barna Goswami, Iffat Jahan, Md. Saddam Hossain, Tasnim Nafisa, Md. Maruf Ahmed Molla, Mahmuda Yeasmin, Asish Kumar Ghosh, Bayzid Bin Monir, A. K. M. Shamsuzzaman, Sheikh Md. Selim Al Din, Utpal Chandra Ray, Salek Ahmed Sajib, Md. Salim Khan |
| EPI_ISL_475166 | National Institute of Laboratory Medicine and Referral Center | Genomic Research Lab, BCSIR | Tanjina Akhter Banu, Abu Sayeed Mohammad Mahmud, Mohammad Samir Uzzaman, Eshrar Osman, Md. Ahasan Habib, Shahina Akter, Md. Murshed Hasan Sarkar, Barna Goswami, Iffat Jahan, Md. Saddam Hossain, Tasnim Nafisa, Md. Maruf Ahmed Molla, Mahmuda Yeasmin, Asish Kumar Ghosh, Bayzid Bin Monir, A. K. M. Shamsuzzaman, Sheikh Md. Selim Al Din, Utpal Chandra Ray, Salek Ahmed Sajib, Md. Salim Khan |
| EPI_ISL_475167 | National Institute of Laboratory Medicine and Referral Center | Genomic Research Lab, BCSIR | Barna Goswami, Abu Sayeed Mohammad Mahmud, Mohammad Samir Uzzaman, Eshrar Osman, Md. Ahasan Habib, Shahina Akter, Tanjina Akhter Banu, Md. Murshed Hasan Sarkar, Iffat Jahan, Md. Saddam Hossain, Tasnim Nafisa, Md. Maruf Ahmed Molla, Mahmuda Yeasmin, Asish Kumar Ghosh, Bayzid Bin Monir, A. K. M. Shamsuzzaman, Sheikh Md. Selim Al Din, Utpal Chandra Ray, Salek Ahmed Sajib, Md. Salim Khan |
| EPI_ISL_475168 | National Institute of Laboratory Medicine and Referral Center | Genomic Research Lab, BCSIR | Iffat Jahan, Abu Sayeed Mohammad Mahmud, Mohammad Samir Uzzaman, Eshrar Osman, Md. Ahasan Habib, Shahina Akter, Tanjina Akhter Banu, Md. Murshed Hasan Sarkar, Barna Goswami, Md. Saddam Hossain, Tasnim Nafisa, Md. Maruf Ahmed Molla, Mahmuda Yeasmin, Asish Kumar Ghosh, Bayzid Bin Monir, A. K. M. Shamsuzzaman, Sheikh Md. Selim Al Din, Utpal Chandra Ray, Salek Ahmed Sajib, Md. Salim Khan |
| EPI_ISL_475169 | National Institute of Laboratory Medicine and Referral Center | Genomic Research Lab, BCSIR | Md. Saddam Hossain, Abu Sayeed Mohammad Mahmud, Mohammad Samir Uzzaman, Eshrar Osman, Md. Ahasan Habib, Shahina Akter, Tanjina Akhter Banu, Md. Murshed Hasan Sarkar, Barna Goswami, Iffat Jahan, Tasnim Nafisa, Md. Maruf Ahmed Molla, Mahmuda Yeasmin, Asish Kumar Ghosh, Bayzid Bin Monir, A. K. M. Shamsuzzaman, Sheikh Md. Selim Al Din, Utpal Chandra Ray, Salek Ahmed Sajib, Md. Salim Khan |
| EPI_ISL_475170, EPI_ISL_475171, EPI_ISL_475172, EPI_ISL_475173 | National Institute of Laboratory Medicine and Referral Center | Genomic Research Lab, BCSIR | Abu Sayeed Mohammad Mahmud, Mohammad Samir Uzzaman, Eshrar Osman, Md. Ahasan Habib, Shahina Akter, Tanjina Akhter Banu, Md. Murshed Hasan Sarkar, Barna Goswami, Iffat Jahan, Md. Saddam Hossain, Tasnim Nafisa, Md. Maruf Ahmed Molla, Mahmuda Yeasmin, Asish Kumar Ghosh, Bayzid Bin Monir, A. K. M. Shamsuzzaman, Sheikh Md. Selim Al Din, Utpal Chandra Ray, Salek Ahmed Sajib, Md. Salim Khan |
| EPI_ISL_475174, EPI_ISL_475175, EPI_ISL_475176, EPI_ISL_475177, EPI_ISL_475178, EPI_ISL_475179, EPI_ISL_475180, EPI_ISL_475181, EPI_ISL_475182, EPI_ISL_475183, EPI_ISL_475184, EPI_ISL_475185, EPI_ISL_475186, EPI_ISL_475187, EPI_ISL_475188, EPI_ISL_475189, EPI_ISL_475190, EPI_ISL_475191, EPI_ISL_475192, EPI_ISL_475193, EPI_ISL_475194, EPI_ISL_475195, EPI_ISL_475196, EPI_ISL_475197, EPI_ISL_475198, EPI_ISL_475199, EPI_ISL_475200, EPI_ISL_475201, EPI_ISL_475202, EPI_ISL_475203, EPI_ISL_475204, EPI_ISL_475205, EPI_ISL_475206, EPI_ISL_475207, EPI_ISL_475208, EPI_ISL_475209, EPI_ISL_475210, EPI_ISL_475211, EPI_ISL_475212, EPI_ISL_475213, EPI_ISL_475214, EPI_ISL_475215, EPI_ISL_475216, EPI_ISL_475217, EPI_ISL_475218, EPI_ISL_475219, EPI_ISL_475220, EPI_ISL_475221, EPI_ISL_475222, EPI_ISL_475223, EPI_ISL_475224, EPI_ISL_475225, EPI_ISL_475226, EPI_ISL_475227, EPI_ISL_475228, EPI_ISL_475229, EPI_ISL_475230, EPI_ISL_475231, EPI_ISL_475232, EPI_ISL_475233, EPI_ISL_475234, EPI_ISL_475235, EPI_ISL_475236, EPI_ISL_475237 | | | |
| see above | Nebraska Public Health Laboratory | UNMC COVID-19 Response Team | UNMC COVID-19 Response Team |
| EPI_ISL_475238 | National Institute of Laboratory Medicine and Referral Center | Genomic Research Lab, BCSIR | Abu Sayeed Mohammad Mahmud, Mohammad Samir Uzzaman, Eshrar Osman, Md. Ahasan Habib, Shahina Akter, Tanjina Akhter Banu, Md. Murshed Hasan Sarkar, Barna Goswami, Iffat Jahan, Md. Saddam Hossain, Tasnim Nafisa, Md. Maruf Ahmed Molla, Mahmuda Yeasmin, Asish Kumar Ghosh, Bayzid Bin Monir, A. K. M. Shamsuzzaman, Sheikh Md. Selim Al Din, Utpal Chandra Ray, Salek Ahmed Sajib, Md. Salim Khan |
| EPI_ISL_475239, EPI_ISL_475240, EPI_ISL_475241, EPI_ISL_475242, EPI_ISL_475243, EPI_ISL_475244, EPI_ISL_475245, EPI_ISL_475246, EPI_ISL_475247, EPI_ISL_475248, EPI_ISL_475249, EPI_ISL_475250, EPI_ISL_475251, EPI_ISL_475252, EPI_ISL_475253, EPI_ISL_475254, EPI_ISL_475255, EPI_ISL_475256, EPI_ISL_475257, EPI_ISL_475258, EPI_ISL_475259, EPI_ISL_475260, EPI_ISL_475261, EPI_ISL_475262, EPI_ISL_475263, EPI_ISL_475264, EPI_ISL_475265, EPI_ISL_475266, EPI_ISL_475267, EPI_ISL_475268, EPI_ISL_475269, EPI_ISL_475270, EPI_ISL_475271, EPI_ISL_475272, EPI_ISL_475273, EPI_ISL_475274, EPI_ISL_475275, EPI_ISL_475276, EPI_ISL_475277, EPI_ISL_475278, EPI_ISL_475279, EPI_ISL_475280, EPI_ISL_475281, EPI_ISL_475282, EPI_ISL_475283, EPI_ISL_475284, EPI_ISL_475285, EPI_ISL_475286, EPI_ISL_475287, EPI_ISL_475288, EPI_ISL_475289, EPI_ISL_475290, EPI_ISL_475291, EPI_ISL_475292, EPI_ISL_475293, EPI_ISL_475294, EPI_ISL_475295, EPI_ISL_475296, EPI_ISL_475297, EPI_ISL_475298, EPI_ISL_475299, EPI_ISL_475300, EPI_ISL_475301, EPI_ISL_475302, EPI_ISL_475303, EPI_ISL_475304, EPI_ISL_475305, EPI_ISL_475306, EPI_ISL_475307, EPI_ISL_475308, EPI_ISL_475309, EPI_ISL_475310, EPI_ISL_475311, EPI_ISL_475312, EPI_ISL_475313, EPI_ISL_475314, EPI_ISL_475315, EPI_ISL_475316, EPI_ISL_475317, EPI_ISL_475318, EPI_ISL_475319, EPI_ISL_475320, EPI_ISL_475321, EPI_ISL_475322, EPI_ISL_475323, EPI_ISL_475324, EPI_ISL_475325, EPI_ISL_475326, EPI_ISL_475327, EPI_ISL_475328, EPI_ISL_475329, EPI_ISL_475330, EPI_ISL_475331, EPI_ISL_475332, EPI_ISL_475333, EPI_ISL_475334, EPI_ISL_475335, EPI_ISL_475336, EPI_ISL_475337, EPI_ISL_475338, EPI_ISL_475339, EPI_ISL_475340, EPI_ISL_475341 | | | |
| see above | Centre for Enzyme Innovation, University of Portsmouth / Translational Research Laboratory, Portsmouth Hospitals NHS Trust | COVID-19 Genomics UK (COG-UK) Consortium | Angela Beckett,Yann Bourgeois,Garry Scarlett,Sharon Glaysher,Scott Elliott,Kelly Bicknell,Robert Impey,Allyson Lloyd,Sarah Wyllie,Ethan Butcher,Anoop Chauhan,Samuel Robson |
| EPI_ISL_475342, EPI_ISL_475343, EPI_ISL_475344, EPI_ISL_475345, EPI_ISL_475346, EPI_ISL_475347, EPI_ISL_475348, EPI_ISL_475349, EPI_ISL_475350, EPI_ISL_475351, EPI_ISL_475352, EPI_ISL_475353, EPI_ISL_475354, EPI_ISL_475355, EPI_ISL_475356, EPI_ISL_475357, EPI_ISL_475358, EPI_ISL_475359, EPI_ISL_475360, EPI_ISL_475361, EPI_ISL_475362, EPI_ISL_475363, EPI_ISL_475364, EPI_ISL_475365, EPI_ISL_475366, EPI_ISL_475367, EPI_ISL_475368, EPI_ISL_475369, EPI_ISL_475370, EPI_ISL_475371, EPI_ISL_475372, EPI_ISL_475373, EPI_ISL_475374, EPI_ISL_475375, EPI_ISL_475376, EPI_ISL_475377, EPI_ISL_475378, EPI_ISL_475379, EPI_ISL_475380, EPI_ISL_475381, EPI_ISL_475382, EPI_ISL_475383, EPI_ISL_475384, EPI_ISL_475385, EPI_ISL_475386, EPI_ISL_475387, EPI_ISL_475388, EPI_ISL_475389, EPI_ISL_475390, EPI_ISL_475391, EPI_ISL_475392, EPI_ISL_475393, EPI_ISL_475394, EPI_ISL_475395, EPI_ISL_475396, EPI_ISL_475397, EPI_ISL_475398, EPI_ISL_475399, EPI_ISL_475400, EPI_ISL_475401, EPI_ISL_475402, EPI_ISL_475403, EPI_ISL_475404, EPI_ISL_475405, EPI_ISL_475406, EPI_ISL_475407, EPI_ISL_475408, EPI_ISL_475409, EPI_ISL_475410, EPI_ISL_475411, EPI_ISL_475412, EPI_ISL_475413, EPI_ISL_475414, EPI_ISL_475415, EPI_ISL_475416, EPI_ISL_475417, EPI_ISL_475418, EPI_ISL_475419, EPI_ISL_475420, EPI_ISL_475421, EPI_ISL_475422, EPI_ISL_475423, EPI_ISL_475424, EPI_ISL_475425, EPI_ISL_475426, EPI_ISL_475427, EPI_ISL_475428, EPI_ISL_475429, EPI_ISL_475430, EPI_ISL_475431, EPI_ISL_475432, EPI_ISL_475433, EPI_ISL_475434, EPI_ISL_475435, EPI_ISL_475436, EPI_ISL_475437, EPI_ISL_475438, EPI_ISL_475439, EPI_ISL_475440, EPI_ISL_475441, EPI_ISL_475442, EPI_ISL_475443, EPI_ISL_475444, EPI_ISL_475445, EPI_ISL_475446, EPI_ISL_475447, EPI_ISL_475448, EPI_ISL_475449, EPI_ISL_475450, EPI_ISL_475451, EPI_ISL_475452, EPI_ISL_475453, EPI_ISL_475454, EPI_ISL_475455, EPI_ISL_475456, EPI_ISL_475457, EPI_ISL_475458, EPI_ISL_475459, EPI_ISL_475460, EPI_ISL_475461, EPI_ISL_475462, EPI_ISL_475463, EPI_ISL_475464, EPI_ISL_475465, EPI_ISL_475466, EPI_ISL_475467, EPI_ISL_475468, EPI_ISL_475469, EPI_ISL_475470, EPI_ISL_475471, EPI_ISL_475472, EPI_ISL_475473, EPI_ISL_475474, EPI_ISL_475475, EPI_ISL_475476, EPI_ISL_475477, EPI_ISL_475478, EPI_ISL_475479, EPI_ISL_475480, EPI_ISL_475481, EPI_ISL_475482, EPI_ISL_475483, EPI_ISL_475484, EPI_ISL_475485, EPI_ISL_475486, EPI_ISL_475487, EPI_ISL_475488, EPI_ISL_475489, EPI_ISL_475490, EPI_ISL_475491, EPI_ISL_475492, EPI_ISL_475493, EPI_ISL_475494, EPI_ISL_475495, EPI_ISL_475496, EPI_ISL_475497, EPI_ISL_475498, EPI_ISL_475499, EPI_ISL_475500, EPI_ISL_475501, EPI_ISL_475502, EPI_ISL_475503, EPI_ISL_475504, EPI_ISL_475505, EPI_ISL_475506, EPI_ISL_475507, EPI_ISL_475508, EPI_ISL_475509, EPI_ISL_475510 | | | |
| see above | Virology Department, Sheffield Teaching Hospitals NHS Foundation Trust/Department of Infection, Immunity and Cardiovascular Disease, The Medical School, University of | COVID-19 Genomics UK (COG-UK) Consortium | Thushan de Silva, Matthew Parker, Nikki Smith, Adri Angyal, Rebecca Brown, Luke Green, Rachel Tucker, Paul Parsons, Danielle Groves, Katie Johnson, Laura Carrilero, Alex Keeley, Dave Partridge, Matthew Wyles, Benjamin Lindsey, Mehmet Yavuz, Mohammad Raza, Cariad Evans |

| | | | |
|---|---|---|---|
| | Sheffield | | |
| EPI_ISL_475511 | Orestadsklinikens VC | The Public Health Agency of Sweden | Oskar Karlsson Lindsjo, Maria Lind Karlberg, Mattias Haukland, Reza Advani, Olov Svartstrom, Anna-Malin Linde, Sandra Broddesson, Mia Brytting, Anna Risberg, Karin Tegmark-Wisell |
| EPI_ISL_475512 | Din Klinik | The Public Health Agency of Sweden | Oskar Karlsson Lindsjo, Maria Lind Karlberg, Mattias Haukland, Reza Advani, Olov Svartstrom, Anna-Malin Linde, Sandra Broddesson, Mia Brytting, Anna Risberg, Karin Tegmark-Wisell |
| EPI_ISL_475513 | Huddinge VC | The Public Health Agency of Sweden | Oskar Karlsson Lindsjo, Maria Lind Karlberg, Mattias Haukland, Reza Advani, Olov Svartstrom, Anna-Malin Linde, Sandra Broddesson, Mia Brytting, Anna Risberg, Karin Tegmark-Wisell |
| EPI_ISL_475514 | Uppsala Narakut Aleris | The Public Health Agency of Sweden | Oskar Karlsson Lindsjo, Maria Lind Karlberg, Mattias Haukland, Reza Advani, Olov Svartstrom, Anna-Malin Linde, Sandra Broddesson, Mia Brytting, Anna Risberg, Karin Tegmark-Wisell |
| EPI_ISL_475515 | Lakargruppen | The Public Health Agency of Sweden | Oskar Karlsson Lindsjo, Maria Lind Karlberg, Mattias Haukland, Reza Advani, Olov Svartstrom, Anna-Malin Linde, Sandra Broddesson, Mia Brytting, Anna Risberg, Karin Tegmark-Wisell |
| EPI_ISL_475516, EPI_ISL_475517 | Uppsala Narakut Aleris | The Public Health Agency of Sweden | Oskar Karlsson Lindsjo, Maria Lind Karlberg, Mattias Haukland, Reza Advani, Olov Svartstrom, Anna-Malin Linde, Sandra Broddesson, Mia Brytting, Anna Risberg, Karin Tegmark-Wisell |
| EPI_ISL_475518 | Trollbackens VC | The Public Health Agency of Sweden | Oskar Karlsson Lindsjo, Maria Lind Karlberg, Mattias Haukland, Reza Advani, Olov Svartstrom, Anna-Malin Linde, Sandra Broddesson, Mia Brytting, Anna Risberg, Karin Tegmark-Wisell |
| EPI_ISL_475519 | Orsa VC | The Public Health Agency of Sweden | Oskar Karlsson Lindsjo, Maria Lind Karlberg, Mattias Haukland, Reza Advani, Olov Svartstrom, Anna-Malin Linde, Sandra Broddesson, Mia Brytting, Anna Risberg, Karin Tegmark-Wisell |
| EPI_ISL_475520 | Vardcentralen Brinken | The Public Health Agency of Sweden | Oskar Karlsson Lindsjo, Maria Lind Karlberg, Mattias Haukland, Reza Advani, Olov Svartstrom, Anna-Malin Linde, Sandra Broddesson, Mia Brytting, Anna Risberg, Karin Tegmark-Wisell |
| EPI_ISL_475521 | Ulltuna Vardcentral | The Public Health Agency of Sweden | Oskar Karlsson Lindsjo, Maria Lind Karlberg, Mattias Haukland, Reza Advani, Olov Svartstrom, Anna-Malin Linde, Sandra Broddesson, Mia Brytting, Anna Risberg, Karin Tegmark-Wisell |
| EPI_ISL_475522, EPI_ISL_475523 | Huddinge VC | The Public Health Agency of Sweden | Oskar Karlsson Lindsjo, Maria Lind Karlberg, Mattias Haukland, Reza Advani, Olov Svartstrom, Anna-Malin Linde, Sandra Broddesson, Mia Brytting, Anna Risberg, Karin Tegmark-Wisell |
| EPI_ISL_475524 | Narhalsan Sjobo vardcentral | The Public Health Agency of Sweden | Oskar Karlsson Lindsjo, Maria Lind Karlberg, Mattias Haukland, Reza Advani, Olov Svartstrom, Anna-Malin Linde, Sandra Broddesson, Mia Brytting, Anna Risberg, Karin Tegmark-Wisell |
| EPI_ISL_475525 | Huddinge VC | The Public Health Agency of Sweden | Oskar Karlsson Lindsjo, Maria Lind Karlberg, Mattias Haukland, Reza Advani, Olov Svartstrom, Anna-Malin Linde, Sandra Broddesson, Mia Brytting, Anna Risberg, Karin Tegmark-Wisell |
| EPI_ISL_475526, EPI_ISL_475527 | Uppsala Narakut Aleris | The Public Health Agency of Sweden | Oskar Karlsson Lindsjo, Maria Lind Karlberg, Mattias Haukland, Reza Advani, Olov Svartstrom, Anna-Malin Linde, Sandra Broddesson, Mia Brytting, Anna Risberg, Karin Tegmark-Wisell |
| EPI_ISL_475528 | Omtanken Grimmered | The Public Health Agency of Sweden | Oskar Karlsson Lindsjo, Maria Lind Karlberg, Mattias Haukland, Reza Advani, Olov Svartstrom, Anna-Malin Linde, Sandra Broddesson, Mia Brytting, Anna Risberg, Karin Tegmark-Wisell |
| EPI_ISL_475529, EPI_ISL_475530, EPI_ISL_475531, EPI_ISL_475532 | Kungsors VC | The Public Health Agency of Sweden | Oskar Karlsson Lindsjo, Maria Lind Karlberg, Mattias Haukland, Reza Advani, Olov Svartstrom, Anna-Malin Linde, Sandra Broddesson, Mia Brytting, Anna Risberg, Karin Tegmark-Wisell |
| EPI_ISL_475533, EPI_ISL_475534, EPI_ISL_475535 | Omtanken Grimmered | The Public Health Agency of Sweden | Oskar Karlsson Lindsjo, Maria Lind Karlberg, Mattias Haukland, Reza Advani, Olov Svartstrom, Anna-Malin Linde, Sandra Broddesson, Mia Brytting, Anna Risberg, Karin Tegmark-Wisell |
| EPI_ISL_475536 | Follinge Halsocentral | The Public Health Agency of Sweden | Oskar Karlsson Lindsjo, Maria Lind Karlberg, Mattias Haukland, Reza Advani, Olov Svartstrom, Anna-Malin Linde, Sandra Broddesson, Mia Brytting, Anna Risberg, Karin Tegmark-Wisell |
| EPI_ISL_475537 | Narhalsan Oden VC | The Public Health Agency of Sweden | Oskar Karlsson Lindsjo, Maria Lind Karlberg, Mattias Haukland, Reza Advani, Olov Svartstrom, Anna-Malin Linde, Sandra Broddesson, Mia Brytting, Anna Risberg, Karin Tegmark-Wisell |
| EPI_ISL_475538 | Omtanken Grimmered | The Public Health Agency of Sweden | Oskar Karlsson Lindsjo, Maria Lind Karlberg, Mattias Haukland, Reza Advani, Olov Svartstrom, Anna-Malin Linde, Sandra Broddesson, Mia Brytting, Anna Risberg, Karin Tegmark-Wisell |
| EPI_ISL_475539 | Narhalsan Sjobo vardcentral | The Public Health Agency of Sweden | Oskar Karlsson Lindsjo, Maria Lind Karlberg, Mattias Haukland, Reza Advani, Olov Svartstrom, Anna-Malin Linde, Sandra Broddesson, Mia Brytting, Anna Risberg, Karin Tegmark-Wisell |
| EPI_ISL_475540 | Bla Kustens halsocentral | The Public Health Agency of Sweden | Oskar Karlsson Lindsjo, Maria Lind Karlberg, Mattias Haukland, Reza Advani, Olov Svartstrom, Anna-Malin Linde, Sandra Broddesson, Mia Brytting, Anna Risberg, Karin Tegmark-Wisell |
| EPI_ISL_475541 | Follinge Halsocentral | The Public Health Agency of Sweden | Oskar Karlsson Lindsjo, Maria Lind Karlberg, Mattias Haukland, Reza Advani, Olov Svartstrom, Anna-Malin Linde, Sandra Broddesson, Mia Brytting, Anna Risberg, Karin Tegmark-Wisell |
| EPI_ISL_475542 | Kungsholmsdoktorn | The Public Health Agency of Sweden | Oskar Karlsson Lindsjo, Maria Lind Karlberg, Mattias Haukland, Reza Advani, Olov Svartstrom, Anna-Malin Linde, Sandra Broddesson, Mia Brytting, Anna Risberg, Karin Tegmark-Wisell |
| EPI_ISL_475543 | Surbrunns VC | The Public Health Agency of Sweden | Oskar Karlsson Lindsjo, Maria Lind Karlberg, Mattias Haukland, Reza Advani, Olov Svartstrom, Anna-Malin Linde, Sandra Broddesson, Mia Brytting, Anna Risberg, Karin Tegmark-Wisell |
| EPI_ISL_475544, EPI_ISL_475545, EPI_ISL_475546, EPI_ISL_475547 | Karolinska Universitetslaboratoriet | The Public Health Agency of Sweden | Oskar Karlsson Lindsjo, Maria Lind Karlberg, Mattias Haukland, Reza Advani, Olov Svartstrom, Anna-Malin Linde, Sandra Broddesson, Shaman Muradrasoli, Anna Risberg, Karin Tegmark-Wisell |
| EPI_ISL_475548 | Halmstad klinisk mikrobiologi | The Public Health Agency of Sweden | Oskar Karlsson Lindsjo, Maria Lind Karlberg, Mattias Haukland, Reza Advani, Olov Svartstrom, Anna-Malin Linde, Sandra Broddesson, Shaman Muradrasoli, Anna Risberg, Karin Tegmark-Wisell |
| EPI_ISL_475549, EPI_ISL_475550 | Skovde/Unilabs | The Public Health Agency of Sweden | Oskar Karlsson Lindsjo, Maria Lind Karlberg, Mattias Haukland, Reza Advani, Olov Svartstrom, Anna-Malin Linde, Sandra Broddesson, Shaman Muradrasoli, Anna Risberg, Karin Tegmark-Wisell |
| EPI_ISL_475551, EPI_ISL_475552 | Karolinska Universitetslaboratoriet | The Public Health Agency of Sweden | Oskar Karlsson Lindsjo, Maria Lind Karlberg, Mattias Haukland, Reza Advani, Olov Svartstrom, Anna-Malin Linde, Sandra Broddesson, Shaman Muradrasoli, Anna Risberg, Karin Tegmark-Wisell |
| EPI_ISL_475553 | Halmstad klinisk mikrobiologi | The Public Health Agency of Sweden | Oskar Karlsson Lindsjo, Maria Lind Karlberg, Mattias Haukland, Reza Advani, Olov Svartstrom, Anna-Malin Linde, Sandra Broddesson, Shaman Muradrasoli, Anna Risberg, Karin Tegmark-Wisell |
| EPI_ISL_475554, EPI_ISL_475555 | Skovde/Unilabs | The Public Health Agency of Sweden | Oskar Karlsson Lindsjo, Maria Lind Karlberg, Mattias Haukland, Reza Advani, Olov Svartstrom, Anna-Malin Linde, Sandra Broddesson, Shaman Muradrasoli, Anna Risberg, Karin Tegmark-Wisell |
| EPI_ISL_475556, EPI_ISL_475557 | Halmstad klinisk mikrobiologi | The Public Health Agency of Sweden | Oskar Karlsson Lindsjo, Maria Lind Karlberg, Mattias Haukland, Reza Advani, Olov Svartstrom, Anna-Malin Linde, Sandra Broddesson, Shaman Muradrasoli, Anna Risberg, Karin Tegmark-Wisell |
| EPI_ISL_475558, EPI_ISL_475559, EPI_ISL_475560, EPI_ISL_475561 | Karolinska Universitetslaboratoriet | The Public Health Agency of Sweden | Oskar Karlsson Lindsjo, Maria Lind Karlberg, Mattias Haukland, Reza Advani, Olov Svartstrom, Anna-Malin Linde, Sandra Broddesson, Shaman Muradrasoli, Anna Risberg, Karin Tegmark-Wisell |
| EPI_ISL_475562, EPI_ISL_475563 | Din Klinik | The Public Health Agency of Sweden | Oskar Karlsson Lindsjo, Maria Lind Karlberg, Mattias Haukland, Reza Advani, Olov Svartstrom, Anna-Malin Linde, Sandra Broddesson, Mia Brytting, Anna Risberg, Karin Tegmark-Wisell |
| EPI_ISL_475564 | Surbrunns VC | The Public Health Agency of Sweden | Oskar Karlsson Lindsjo, Maria Lind Karlberg, Mattias Haukland, Reza Advani, Olov Svartstrom, Anna-Malin Linde, Sandra Broddesson, Mia Brytting, Anna Risberg, Karin Tegmark-Wisell |
| EPI_ISL_475565 | Narhalsan Backa vardcentral | The Public Health Agency of Sweden | Oskar Karlsson Lindsjo, Maria Lind Karlberg, Mattias Haukland, Reza Advani, Olov Svartstrom, Anna-Malin Linde, Sandra Broddesson, Mia Brytting, Anna Risberg, Karin Tegmark-Wisell |
| EPI_ISL_475566 | Vardcentralen Brinken | The Public Health Agency of Sweden | Oskar Karlsson Lindsjo, Maria Lind Karlberg, Mattias Haukland, Reza Advani, Olov Svartstrom, Anna-Malin Linde, Sandra Broddesson, Mia Brytting, Anna Risberg, Karin Tegmark-Wisell |
| EPI_ISL_475567 | Huddinge VC | The Public Health Agency of Sweden | Oskar Karlsson Lindsjo, Maria Lind Karlberg, Mattias Haukland, Reza Advani, Olov Svartstrom, Anna-Malin Linde, Sandra Broddesson, Mia Brytting, Anna Risberg, Karin Tegmark-Wisell |
| EPI_ISL_475568 | Kungsors VC | The Public Health Agency of Sweden | Oskar Karlsson Lindsjo, Maria Lind Karlberg, Mattias Haukland, Reza Advani, Olov Svartstrom, Anna-Malin Linde, Sandra Broddesson, Mia Brytting, Anna Risberg, Karin Tegmark-Wisell |
| EPI_ISL_475569 | Kungsholmsdoktorn | The Public Health Agency of Sweden | Oskar Karlsson Lindsjo, Maria Lind Karlberg, Mattias Haukland, Reza Advani, Olov Svartstrom, Anna-Malin Linde, Sandra Broddesson, Mia Brytting, Anna Risberg, Karin Tegmark-Wisell |
| EPI_ISL_475570 | Genome Center | Genome Center | A. S. M. Rubayet- Ul- Alam, Ovinu Kibria Islam, Md. Shazid Hasan, Hassan M. Al-Emran, Shireen Nigar, Selina Akter, Pravas Chandra Roy, Md. Tanvir Islam, Shovon Lal Sarkar, M. Shaminur Rahman, M. Rafiul Islam, Habiba Ibnat, Md Nur Kabidul Azam, Chakrabortty Atonu, Proshanto Kumar Das, Md. Hasan al Pramanik, Md. Zannat Ali, Shohanur Rahaman, Md. Aminul Islam, Ashok Kumar, Md. Nazmul Hasan, Md. Iqbal Kabir Jahid, Md. Anwar Hossain |
| EPI_ISL_475571 | Genome Center | Genome Center | Hassan M. Al-Emran, Md. Shazid Hasan, Ovinu Kibria Islam, A. S. M. Rubayet- Ul- Alam, Pravas Chandra Roy, Selina Akter, Shireen Nigar, Shovon Lal Sarkar, Md. Tanvir Islam, Mithun Talukder Md. Tawyabur, Md. Taijul Islam, Provakar Mondol, Md. Muzahidul Islam, Md. Iqbal Kabir Jahid Md. Anwar Hossain |
| EPI_ISL_475572 | Imperial College London | Imperial College London | Jie Zhou, Wendy Barclay |
| EPI_ISL_475573 | Genome Center | Genome Center | Md. Shazid Hasan, Hassan M. Al-Emran, Ovinu Kibria Islam, A. S. M. Rubayet- Ul- Alam, Selina Akter, Shireen Nigar, Md. Tanvir Islam, Pravas Chandra Roy, Shovon Lal Sarkar, Md. Nazmul Hasan, Tanay Chakrovarty, Md. Ali Ahasan Setu, Sourav Dutta, Ruhul Amin, Md. Iqbal Kabir Jahid, Md. Anwar Hossain |
| EPI_ISL_475574, EPI_ISL_475575, EPI_ISL_475576, EPI_ISL_475577, EPI_ISL_475578, EPI_ISL_475579, EPI_ISL_475580, EPI_ISL_475581, EPI_ISL_475582, EPI_ISL_475583, EPI_ISL_475584, EPI_ISL_475585, EPI_ISL_475586, EPI_ISL_475587, EPI_ISL_475588, EPI_ISL_475589, EPI_ISL_475590, EPI_ISL_475591, EPI_ISL_475592, EPI_ISL_475593, EPI_ISL_475594, EPI_ISL_475595, EPI_ISL_475596, EPI_ISL_475597, EPI_ISL_475598, EPI_ISL_475599, EPI_ISL_475600, EPI_ISL_475601, EPI_ISL_475602, EPI_ISL_475603, EPI_ISL_475604, EPI_ISL_475605, EPI_ISL_475606, EPI_ISL_475607, EPI_ISL_475608, EPI_ISL_475609, EPI_ISL_475610, EPI_ISL_475611, EPI_ISL_475612, EPI_ISL_475613, EPI_ISL_475614, EPI_ISL_475615, EPI_ISL_475616, EPI_ISL_475617, EPI_ISL_475618, EPI_ISL_475619, EPI_ISL_475620, EPI_ISL_475621, EPI_ISL_475622, EPI_ISL_475623, EPI_ISL_475624, EPI_ISL_475625, EPI_ISL_475626, EPI_ISL_475627, EPI_ISL_475628, EPI_ISL_475629, EPI_ISL_475630, EPI_ISL_475631, EPI_ISL_475632, EPI_ISL_475633, EPI_ISL_475634, EPI_ISL_475635, EPI_ISL_475636, EPI_ISL_475637, EPI_ISL_475638, EPI_ISL_475639, EPI_ISL_475640, EPI_ISL_475641, EPI_ISL_475642, EPI_ISL_475643, EPI_ISL_475644, EPI_ISL_475645, EPI_ISL_475646, EPI_ISL_475647, EPI_ISL_475648, EPI_ISL_475649, EPI_ISL_475650, EPI_ISL_475651, EPI_ISL_475652, EPI_ISL_475653, EPI_ISL_475654, EPI_ISL_475655, EPI_ISL_475656, EPI_ISL_475657, EPI_ISL_475658, EPI_ISL_475659, EPI_ISL_475660, EPI_ISL_475661, EPI_ISL_475662, EPI_ISL_475663, EPI_ISL_475664, EPI_ISL_475665, EPI_ISL_475666, EPI_ISL_475667, EPI_ISL_475668, EPI_ISL_475669, EPI_ISL_475670, EPI_ISL_475671, EPI_ISL_475672, EPI_ISL_475673, EPI_ISL_475674, EPI_ISL_475675, EPI_ISL_475676, EPI_ISL_475677, EPI_ISL_475678, EPI_ISL_475679, EPI_ISL_475680, EPI_ISL_475681, EPI_ISL_475682, EPI_ISL_475683, EPI_ISL_475684, EPI_ISL_475685, EPI_ISL_475686, EPI_ISL_475687, EPI_ISL_475688, EPI_ISL_475689, EPI_ISL_475690, EPI_ISL_475691, EPI_ISL_475692, EPI_ISL_475693, EPI_ISL_475694, EPI_ISL_475695, EPI_ISL_475696, EPI_ISL_475697, EPI_ISL_475698, EPI_ISL_475699, EPI_ISL_475700, EPI_ISL_475701, EPI_ISL_475702, EPI_ISL_475703, EPI_ISL_475704, EPI_ISL_475705, EPI_ISL_475706, EPI_ISL_475707, EPI_ISL_475708, EPI_ISL_475709, EPI_ISL_475710, EPI_ISL_475711, EPI_ISL_475712, EPI_ISL_475713, EPI_ISL_475714, EPI_ISL_475715, EPI_ISL_475716 | | | |
| see above | Cedars-Sinai Medical Center, Department of Pathology & Laboratory Medicine, Molecular Pathology Laboratory | Cedars-Sinai Medical Center, Molecular Pathology Laboratory of Department of Pathology & Laboratory Medicine and Genomic Core | Wenjuan Zhang, John Paul Govindavari, Brian Davis, Stephanie Chen, Jong Taek Kim, Jianbo Song, Jean Lopategui, Jasmine T Plummer, Eric Vail |
| EPI_ISL_475717, EPI_ISL_475718, EPI_ISL_475719, EPI_ISL_475720, EPI_ISL_475721 | Microbiology, University Hospital Donostia | Microbiology, University Hospital Donostia | Cilla,G., Montes,M., Pineiro,L., Marimon,J.M. |
| EPI_ISL_475723, EPI_ISL_475724 | Egyptian National Cancer Institute (ENCI) | Egyptian National Cancer Institute (ENCI) | Zekri, Abdel Rahman N, Amer,K.E., Ahmed,O.S., Soliman,H.K., Hafez,M.M., Bahnassy,A.A., Abdelhamid,W., Gad,A., Ali,M., Hassan,W., Samir,M., Raouf,A., Hamdy,M.S., Soliman,M.S., Elsissy,M.H., Elkhateeb,S.M., Ezzelarab,M.H., Abouelhoda, Mohamed |
| EPI_ISL_475725, EPI_ISL_475726, EPI_ISL_475727, EPI_ISL_475728, EPI_ISL_475729, EPI_ISL_475730, EPI_ISL_475731, EPI_ISL_475732, EPI_ISL_475733, EPI_ISL_475734, EPI_ISL_475735, EPI_ISL_475736, EPI_ISL_475737, EPI_ISL_475738, EPI_ISL_475739, EPI_ISL_475740, EPI_ISL_475741, EPI_ISL_475742, EPI_ISL_475743, EPI_ISL_475744 | | | |
| see above | Utah Public Health Laboratory | Utah Public Health Laboratory | Erin Young, Kelly Oakeson |
| EPI_ISL_475745, EPI_ISL_475746, EPI_ISL_475747, EPI_ISL_475748, EPI_ISL_475749, EPI_ISL_475750, EPI_ISL_475751, EPI_ISL_475752, EPI_ISL_475753 | Medical Ain Shams Research Institute (MASRI), Ain Shams University | Medical Ain Shams Research Institute (MASRI), Ain Shams University | Hesham Elghazaly , Sara Hassan Agwa, Mahmoud Elmeteini , Ahmad Moustafa , Ashraf Omar, Osama Mansour, Samia Abdo, Hala Hafez, Ghada Ismael , Shaimaa Moustafa , Aya Mohamed, Reham Mamdouh , Hoda Abd Elsatar, Manal Hamdy Elsaid, Fatma Ebied |
| EPI_ISL_475754 | National Institute of Laboratory Medicine and Referral Center | Genomic Research Lab, BCSIR | Shahina Akter, Abu Sayeed Mohammad Mahmud, Mohammad Samir Uzzaman, Eshrar Osman, Md. Ahasan Habib, Tanjina Akhter Banu, Md. Murshed Hasan Sarkar, Barna Goswami, Iffat Jahan, Md. Saddam Hossain, Tasnim Nafisa, Md. Maruf Ahmed Molla, Mahmuda Yeasmin, Asish Kumar Ghosh, Arifa Akram, A. K. M. Shamsuzzaman, Sheikh Md. Selim Al Din, Utpal Chandra Ray, Salek Ahmed Sajib, Md. Salim Khan |
| EPI_ISL_475755 | National Institute of Laboratory Medicine and Referral Center | Genomic Research Lab, BCSIR | Md. Murshed Hasan Sarkar, Abu Sayeed Mohammad Mahmud, Mohammad Samir Uzzaman, Eshrar Osman, Md. Ahasan Habib, Shahina Akter, Tanjina Akhter Banu, Barna Goswami, Iffat Jahan, Md. Saddam Hossain, Tasnim Nafisa, Md. Maruf Ahmed Molla, Mahmuda Yeasmin, Asish Kumar Ghosh, Arifa Akram, A. K. M. Shamsuzzaman, Sheikh Md. Selim Al Din, Utpal Chandra Ray, Salek Ahmed Sajib, Md. Salim Khan |
| EPI_ISL_475756 | National Institute of Laboratory Medicine and Referral Center | Genomic Research Lab, BCSIR | Tanjina Akhter Banu, Abu Sayeed Mohammad Mahmud, Mohammad Samir Uzzaman, Eshrar Osman, Md. Ahasan Habib, Shahina Akter, Md. Murshed Hasan Sarkar, Barna Goswami, Iffat Jahan, Md. Saddam Hossain, Tasnim Nafisa, Md. Maruf Ahmed Molla, Mahmuda Yeasmin, Asish Kumar Ghosh, Arifa Akram, A. K. M. Shamsuzzaman, Sheikh Md. Selim Al Din, Utpal Chandra Ray, Salek Ahmed Sajib, Md. Salim Khan |
| EPI_ISL_475757 | National Institute of Laboratory Medicine and Referral Center | Genomic Research Lab, BCSIR | Barna Goswami, Abu Sayeed Mohammad Mahmud, Mohammad Samir Uzzaman, Eshrar Osman, Md. Ahasan Habib, Shahina Akter, Tanjina Akhter Banu, Md. Murshed Hasan Sarkar, Iffat Jahan, Md. Saddam Hossain, Tasnim Nafisa, Md. Maruf Ahmed Molla, Mahmuda Yeasmin, Asish Kumar Ghosh, Arifa Akram, A. K. M. Shamsuzzaman, Sheikh Md. Selim Al Din, Utpal Chandra Ray, Salek Ahmed Sajib, Md. Salim Khan |
| EPI_ISL_475758 | National Institute of Laboratory Medicine and Referral Center | Genomic Research Lab, BCSIR | Iffat Jahan, Abu Sayeed Mohammad Mahmud, Mohammad Samir Uzzaman, Eshrar Osman, Md. Ahasan Habib, Shahina Akter, Tanjina Akhter Banu, Md. Murshed Hasan Sarkar, Barna Goswami, Md. Saddam Hossain, Tasnim Nafisa, Md. Maruf Ahmed Molla, Mahmuda Yeasmin, Asish Kumar Ghosh, Arifa Akram, A. K. M. Shamsuzzaman, Sheikh Md. Selim Al Din, Utpal Chandra Ray, Salek Ahmed Sajib, Md. Salim Khan |
| EPI_ISL_475759 | National Institute of Laboratory Medicine and Referral Center | Genomic Research Lab, BCSIR | Md. Saddam Hossain, Abu Sayeed Mohammad Mahmud, Mohammad Samir Uzzaman, Eshrar Osman, Md. Ahasan Habib, Shahina Akter, Tanjina Akhter Banu, Md. Murshed Hasan Sarkar, Barna Goswami, Iffat Jahan, Tasnim Nafisa, Md. Maruf Ahmed Molla, Mahmuda Yeasmin, Asish Kumar Ghosh, Arifa Akram, A. K. M. Shamsuzzaman, Sheikh Md. Selim Al Din, Utpal Chandra Ray, Salek Ahmed Sajib, Md. Salim Khan |
| EPI_ISL_475760, EPI_ISL_475761 | National Institute of Laboratory Medicine and Referral Center | Genomic Research Lab, BCSIR | Abu Sayeed Mohammad Mahmud, Mohammad Samir Uzzaman, Eshrar Osman, Md. Ahasan Habib, Shahina Akter, Tanjina Akhter Banu, Md. Murshed Hasan Sarkar, Barna Goswami, Iffat Jahan, Md. Saddam Hossain, Tasnim Nafisa, |

| | | | |
|---|---|---|---|
| | | | Md. Maruf Ahmed Molla, Mahmuda Yeasmin, Asish Kumar Ghosh, Arifa Akram, A. K. M. Shamsuzzaman, Sheikh Md. Selim Al Din, Utpal Chandra Ray, Salek Ahmed Sajib, Md. Salim Khan |
| EPI_ISL_475762 | Oklahoma State Department of Health | França Lab | Caio Martinelle B. de França, Graham Wiley, Samuel T. Dunn, and Matthew J. Miller. |
| EPI_ISL_475763 | Institut für Virologie am Department für Hygiene, Mikrobiologie und Public Health | Bergthaler laboratory, CeMM Research Center for Molecular Medicine of the Austrian Academy of Sciences | Alexandra Popa, Benedikt Agerer, Henrique Colaco, Lukas Endler, Jakob-Wendelin Genger, Alexander Lercher, Mark Smyth, Thomas Penz, Michael Schuster, Jan Laine, Martin Senekowitsch, Judith Aberle, Stephan Aberle, Peter Hufnagl, Daniela Schmid, Franz Allerberger, Elisabeth Puchhammer-Stoeckl, Manfred Nairz, Guenter Weiss, Gregor Hörmann, Kinga Rigler-Hohenwarter, Rainer Gattringer, Wegene Borena, Dorothee von Laer, Christoph Bock, Andreas Bergthaler |
| EPI_ISL_475764, EPI_ISL_475765, EPI_ISL_475766, EPI_ISL_475767, EPI_ISL_475768 | Universitaetsklinik für Innere Medizin II Innsbruck | Bergthaler laboratory, CeMM Research Center for Molecular Medicine of the Austrian Academy of Sciences | Alexandra Popa, Benedikt Agerer, Henrique Colaco, Lukas Endler, Jakob-Wendelin Genger, Alexander Lercher, Mark Smyth, Thomas Penz, Michael Schuster, Jan Laine, Martin Senekowitsch, Judith Aberle, Stephan Aberle, Peter Hufnagl, Daniela Schmid, Franz Allerberger, Elisabeth Puchhammer-Stoeckl, Manfred Nairz, Guenter Weiss, Gregor Hörmann, Kinga Rigler-Hohenwarter, Rainer Gattringer, Wegene Borena, Dorothee von Laer, Christoph Bock, Andreas Bergthaler |
| EPI_ISL_475769 | Institut für Virologie am Department für Hygiene, Mikrobiologie und Public Health | Bergthaler laboratory, CeMM Research Center for Molecular Medicine of the Austrian Academy of Sciences | Alexandra Popa, Benedikt Agerer, Henrique Colaco, Lukas Endler, Jakob-Wendelin Genger, Alexander Lercher, Mark Smyth, Thomas Penz, Michael Schuster, Jan Laine, Martin Senekowitsch, Judith Aberle, Stephan Aberle, Peter Hufnagl, Daniela Schmid, Franz Allerberger, Elisabeth Puchhammer-Stoeckl, Manfred Nairz, Guenter Weiss, Gregor Hörmann, Kinga Rigler-Hohenwarter, Rainer Gattringer, Wegene Borena, Dorothee von Laer, Christoph Bock, Andreas Bergthaler |
| EPI_ISL_475770, EPI_ISL_475771, EPI_ISL_475772, EPI_ISL_475773, EPI_ISL_475774, EPI_ISL_475775, EPI_ISL_475776, EPI_ISL_475777, EPI_ISL_475778, EPI_ISL_475779, EPI_ISL_475780, EPI_ISL_475781, EPI_ISL_475782, EPI_ISL_475783, EPI_ISL_475784, EPI_ISL_475785, EPI_ISL_475786, EPI_ISL_475787, EPI_ISL_475788, EPI_ISL_475789, EPI_ISL_475790, EPI_ISL_475791, EPI_ISL_475792, EPI_ISL_475793, EPI_ISL_475794, EPI_ISL_475795, EPI_ISL_475796, EPI_ISL_475797, EPI_ISL_475798, EPI_ISL_475799, EPI_ISL_475800, EPI_ISL_475801, EPI_ISL_475802, EPI_ISL_475803, EPI_ISL_475804, EPI_ISL_475805, EPI_ISL_475806, EPI_ISL_475807, EPI_ISL_475808, EPI_ISL_475809, EPI_ISL_475810, EPI_ISL_475811, EPI_ISL_475812 | | | |
| see above | Center for Virology, Medical University of Vienna | Bergthaler laboratory, CeMM Research Center for Molecular Medicine of the Austrian Academy of Sciences | Alexandra Popa, Benedikt Agerer, Henrique Colaco, Lukas Endler, Jakob-Wendelin Genger, Alexander Lercher, Mark Smyth, Thomas Penz, Michael Schuster, Jan Laine, Martin Senekowitsch, Judith Aberle, Stephan Aberle, Peter Hufnagl, Daniela Schmid, Franz Allerberger, Elisabeth Puchhammer-Stoeckl, Manfred Nairz, Guenter Weiss, Gregor Hörmann, Kinga Rigler-Hohenwarter, Rainer Gattringer, Wegene Borena, Dorothee von Laer, Christoph Bock, Andreas Bergthaler |
| EPI_ISL_475813, EPI_ISL_475814, EPI_ISL_475815, EPI_ISL_475816, EPI_ISL_475817, EPI_ISL_475818, EPI_ISL_475819, EPI_ISL_475820, EPI_ISL_475821, EPI_ISL_475822, EPI_ISL_475823, EPI_ISL_475824, EPI_ISL_475825, EPI_ISL_475826, EPI_ISL_475827, EPI_ISL_475828, EPI_ISL_475829 | | | |
| see above | Institut für Virologie am Department für Hygiene, Mikrobiologie und Public Health | Bergthaler laboratory, CeMM Research Center for Molecular Medicine of the Austrian Academy of Sciences | Alexandra Popa, Benedikt Agerer, Henrique Colaco, Lukas Endler, Jakob-Wendelin Genger, Alexander Lercher, Mark Smyth, Thomas Penz, Michael Schuster, Jan Laine, Martin Senekowitsch, Judith Aberle, Stephan Aberle, Peter Hufnagl, Daniela Schmid, Franz Allerberger, Elisabeth Puchhammer-Stoeckl, Manfred Nairz, Guenter Weiss, Gregor Hörmann, Kinga Rigler-Hohenwarter, Rainer Gattringer, Wegene Borena, Dorothee von Laer, Christoph Bock, Andreas Bergthaler |
| EPI_ISL_475830, EPI_ISL_475831, EPI_ISL_475832, EPI_ISL_475833, EPI_ISL_475834, EPI_ISL_475835, EPI_ISL_475836, EPI_ISL_475837, EPI_ISL_475838, EPI_ISL_475839, EPI_ISL_475840, EPI_ISL_475841, EPI_ISL_475842, EPI_ISL_475843, EPI_ISL_475844, EPI_ISL_475845, EPI_ISL_475846, EPI_ISL_475847, EPI_ISL_475848, EPI_ISL_475849, EPI_ISL_475850, EPI_ISL_475851, EPI_ISL_475852, EPI_ISL_475853, EPI_ISL_475854, EPI_ISL_475855, EPI_ISL_475856, EPI_ISL_475857, EPI_ISL_475858, EPI_ISL_475859, EPI_ISL_475860, EPI_ISL_475861, EPI_ISL_475862, EPI_ISL_475863, EPI_ISL_475864, EPI_ISL_475865, EPI_ISL_475866, EPI_ISL_475867, EPI_ISL_475868, EPI_ISL_475869, EPI_ISL_475870, EPI_ISL_475871, EPI_ISL_475872, EPI_ISL_475873, EPI_ISL_475874, EPI_ISL_475875, EPI_ISL_475876, EPI_ISL_475877, EPI_ISL_475878, EPI_ISL_475879, EPI_ISL_475880, EPI_ISL_475881, EPI_ISL_475882, EPI_ISL_475883, EPI_ISL_475884, EPI_ISL_475885, EPI_ISL_475886 | | | |
| see above | Austrian Agency for Health and Food Safety (AGES) | Bergthaler laboratory, CeMM Research Center for Molecular Medicine of the Austrian Academy of Sciences | Alexandra Popa, Benedikt Agerer, Henrique Colaco, Lukas Endler, Jakob-Wendelin Genger, Alexander Lercher, Mark Smyth, Thomas Penz, Michael Schuster, Jan Laine, Martin Senekowitsch, Judith Aberle, Stephan Aberle, Peter Hufnagl, Daniela Schmid, Franz Allerberger, Elisabeth Puchhammer-Stoeckl, Manfred Nairz, Guenter Weiss, Gregor Hörmann, Kinga Rigler-Hohenwarter, Rainer Gattringer, Wegene Borena, Dorothee von Laer, Christoph Bock, Andreas Bergthaler |
| EPI_ISL_475887, EPI_ISL_475888, EPI_ISL_475889, EPI_ISL_475890, EPI_ISL_475891, EPI_ISL_475892, EPI_ISL_475893, EPI_ISL_475894, EPI_ISL_475895, EPI_ISL_475896, EPI_ISL_475897, EPI_ISL_475898, EPI_ISL_475899, EPI_ISL_475900, EPI_ISL_475901, EPI_ISL_475902, EPI_ISL_475903, EPI_ISL_475904, EPI_ISL_475905, EPI_ISL_475906, EPI_ISL_475907, EPI_ISL_475908, EPI_ISL_475909 | | | |
| see above | Zentralinstitut für medizinische und chemische Labordiagnostik, Universitätskliniken Insbruck | Bergthaler laboratory, CeMM Research Center for Molecular Medicine of the Austrian Academy of Sciences | Alexandra Popa, Benedikt Agerer, Henrique Colaco, Lukas Endler, Jakob-Wendelin Genger, Alexander Lercher, Mark Smyth, Thomas Penz, Michael Schuster, Jan Laine, Martin Senekowitsch, Judith Aberle, Stephan Aberle, Peter Hufnagl, Daniela Schmid, Franz Allerberger, Elisabeth Puchhammer-Stoeckl, Manfred Nairz, Guenter Weiss, Gregor Hörmann, Kinga Rigler-Hohenwarter, Rainer Gattringer, Wegene Borena, Dorothee von Laer, Christoph Bock, Andreas Bergthaler |
| EPI_ISL_475910, EPI_ISL_475911, EPI_ISL_475912, EPI_ISL_475913, EPI_ISL_475914, EPI_ISL_475915 | Klinikum Wels-Grieskirchen | Bergthaler laboratory, CeMM Research Center for Molecular Medicine of the Austrian Academy of Sciences | Alexandra Popa, Benedikt Agerer, Henrique Colaco, Lukas Endler, Jakob-Wendelin Genger, Alexander Lercher, Mark Smyth, Thomas Penz, Michael Schuster, Jan Laine, Martin Senekowitsch, Judith Aberle, Stephan Aberle, Peter Hufnagl, Daniela Schmid, Franz Allerberger, Elisabeth Puchhammer-Stoeckl, Manfred Nairz, Guenter Weiss, Gregor Hörmann, Kinga Rigler-Hohenwarter, Rainer Gattringer, Wegene Borena, Dorothee von Laer, Christoph Bock, Andreas Bergthaler |
| EPI_ISL_475916, EPI_ISL_475917, EPI_ISL_475918, EPI_ISL_475919, EPI_ISL_475920, EPI_ISL_475921, EPI_ISL_475922, EPI_ISL_475923, EPI_ISL_475924, EPI_ISL_475925, EPI_ISL_475926, EPI_ISL_475927, EPI_ISL_475928 | | | |
| see above | Institut für Virologie am Department für Hygiene, Mikrobiologie und Public Health | Bergthaler laboratory, CeMM Research Center for Molecular Medicine of the Austrian Academy of Sciences | Alexandra Popa, Benedikt Agerer, Henrique Colaco, Lukas Endler, Jakob-Wendelin Genger, Alexander Lercher, Mark Smyth, Thomas Penz, Michael Schuster, Jan Laine, Martin Senekowitsch, Judith Aberle, Stephan Aberle, Peter Hufnagl, Daniela Schmid, Franz Allerberger, Elisabeth Puchhammer-Stoeckl, Manfred Nairz, Guenter Weiss, Gregor Hörmann, Kinga Rigler-Hohenwarter, Rainer Gattringer, Wegene Borena, Dorothee von Laer, Christoph Bock, Andreas Bergthaler |
| EPI_ISL_475929, EPI_ISL_475930, EPI_ISL_475931, EPI_ISL_475932, EPI_ISL_475933, EPI_ISL_475934, EPI_ISL_475935, EPI_ISL_475936 | Universitaetsklinik für Innere Medizin II Innsbruck | Bergthaler laboratory, CeMM Research Center for Molecular Medicine of the Austrian Academy of Sciences | Alexandra Popa, Benedikt Agerer, Henrique Colaco, Lukas Endler, Jakob-Wendelin Genger, Alexander Lercher, Mark Smyth, Thomas Penz, Michael Schuster, Jan Laine, Martin Senekowitsch, Judith Aberle, Stephan Aberle, Peter Hufnagl, Daniela Schmid, Franz Allerberger, Elisabeth Puchhammer-Stoeckl, Manfred Nairz, Guenter Weiss, Gregor Hörmann, Kinga Rigler-Hohenwarter, Rainer Gattringer, Wegene Borena, Dorothee von Laer, Christoph Bock, Andreas Bergthaler |
| EPI_ISL_475937, EPI_ISL_475938, EPI_ISL_475939, EPI_ISL_475940, EPI_ISL_475941, EPI_ISL_475942, EPI_ISL_475943, EPI_ISL_475944, EPI_ISL_475945, EPI_ISL_475946, EPI_ISL_475947, EPI_ISL_475948, EPI_ISL_475949, EPI_ISL_475950, EPI_ISL_475951, EPI_ISL_475952, EPI_ISL_475953, EPI_ISL_475954, EPI_ISL_475955, EPI_ISL_475956, EPI_ISL_475957, EPI_ISL_475958, EPI_ISL_475959, EPI_ISL_475960, EPI_ISL_475961, EPI_ISL_475962, EPI_ISL_475963, EPI_ISL_475964, EPI_ISL_475965, EPI_ISL_475966, EPI_ISL_475967, EPI_ISL_475968, EPI_ISL_475969, EPI_ISL_475970, EPI_ISL_475971, EPI_ISL_475972, EPI_ISL_475973, EPI_ISL_475974, EPI_ISL_475975, EPI_ISL_475976, EPI_ISL_475977, EPI_ISL_475978, EPI_ISL_475979, EPI_ISL_475980, EPI_ISL_475981, EPI_ISL_475982, EPI_ISL_475983, EPI_ISL_475984, EPI_ISL_475985, EPI_ISL_475986, EPI_ISL_475987, EPI_ISL_475988, EPI_ISL_475989, EPI_ISL_475990, EPI_ISL_475991, EPI_ISL_475992, EPI_ISL_475993, EPI_ISL_475994, EPI_ISL_475995, EPI_ISL_475996, EPI_ISL_475997, EPI_ISL_475998, EPI_ISL_475999 | | | |
| see above | National Public Health Laboratory, National Centre for Infectious Diseases | National Public Health Laboratory, National Centre for Infectious Diseases | Mak TM, Octavia S, Chavatte JM, Cui L, Lin RTP |
| EPI_ISL_476018, EPI_ISL_476019, EPI_ISL_476020, EPI_ISL_476021 | Washington University in St. Louis | Washington University in St. Louis | David Wang, Carey-Ann Burnham, Scott Handley, Lindsay Droit, Stephen Tahan |
| EPI_ISL_476022 | Defence Research & Development Establishment | Defence Research & Development Establishment | Shashi Sharma, Paban Kumar Dash, Jyoti S Kumar, Sushil Kumar Sharma, Ambuj Shrivastava |
| EPI_ISL_476023 | Defence Research & Development Establishment (DRDE) | Defence Research & Development Establishment (DRDE) | Shashi Sharma, Paban Kumar Dash, Sushil Kumar Sharma, Ambuj Shrivastava, Jyoti S. Kumar |
| EPI_ISL_476024 | Laboratoire de Recherche et d'Analyses Médicales de la Gendarmerie Royale | Laboratoire de Recherche et d'Analyses Médicales de la Gendarmerie Royale | Sanaâ Lemriss, Amal SOUIRI, Nabil Lemzaoui, Omar Mestoui, Mohamed Labioui, Nabil Ouaariba, Ayoub Jibjibe, Mahmoud Yartaoui, Mohamed Chahmi, Marouane El Rhouila, Samiha Sellak, Nadia Kandoussi, Saâd El Kabbaj. |
| EPI_ISL_476025 | Laboratoire de Recherche et d'Analyses Médicales de la Gendarmerie Royale | Laboratoire de Recherche et d'Analyses Médicales de la Gendarmerie Royale | Sanaâ LEMRISS, Amal Souiri, Saâd EL KABBAJ |
| EPI_ISL_476026 | Laboratoire de Recherche et d'Analyses Médicales de la Gendarmerie Royale | Laboratoire de Recherche et d'Analyses Médicales de la Gendarmerie Royale | Sanaâ Lemriss, Amal SOUIRI, Saâd EL KABBAJ |
| EPI_ISL_476027, EPI_ISL_476028, EPI_ISL_476029, EPI_ISL_476030, EPI_ISL_476031, EPI_ISL_476032, EPI_ISL_476033, EPI_ISL_476034, EPI_ISL_476035, EPI_ISL_476036, EPI_ISL_476037, EPI_ISL_476038, EPI_ISL_476039, EPI_ISL_476040, EPI_ISL_476041, EPI_ISL_476042, EPI_ISL_476043, EPI_ISL_476044, EPI_ISL_476045, EPI_ISL_476046, EPI_ISL_476047, EPI_ISL_476048, EPI_ISL_476049, EPI_ISL_476050, EPI_ISL_476051, EPI_ISL_476052, EPI_ISL_476053, EPI_ISL_476054, EPI_ISL_476055, EPI_ISL_476056, EPI_ISL_476057, EPI_ISL_476058, EPI_ISL_476059, EPI_ISL_476060, EPI_ISL_476061, EPI_ISL_476062, EPI_ISL_476063, EPI_ISL_476064, EPI_ISL_476065, EPI_ISL_476066 | | | |
| see above | Michigan Department of Health and Human Services, Bureau of Laboratories | Michigan Department of Health and Human Services, Bureau of Laboratories | Blankenship HM, Riner D, Soehnlen MK |
| EPI_ISL_476067 | The National Institute of Public Health | State Veterinary Institute Prague and The National Institute of Public Health | Nagy,A;Jirincova,H;Novakova,L;Trnka,D;Vecerova,J |
| EPI_ISL_476068, EPI_ISL_476069, EPI_ISL_476070, EPI_ISL_476071, EPI_ISL_476072, EPI_ISL_476073, EPI_ISL_476074, EPI_ISL_476075, EPI_ISL_476076, EPI_ISL_476077 | University of Debrecen, Department of Medical Microbiology | National Laboratory of Virology, Szentágothai Research Centre | Endre Gábor Tóth, Balázs Somogyi, Brigitta Zana, Eszter Csoma, Ferenc Jakab, Gábor Kemenesi |
| EPI_ISL_476078 | University of Szeged, Institute of Clinical Microbiology | National Laboratory of Virology, Szentágothai Research Centre | Endre Gábor Tóth, Balázs Somogyi, Brigitta Zana, Terhes Gabriella, Ferenc Jakab, Gábor Kemenesi |
| EPI_ISL_476079, EPI_ISL_476080, EPI_ISL_476081, EPI_ISL_476082, EPI_ISL_476083, EPI_ISL_476084, EPI_ISL_476085, EPI_ISL_476086, EPI_ISL_476087, EPI_ISL_476088, EPI_ISL_476089, EPI_ISL_476090, EPI_ISL_476091, EPI_ISL_476092, EPI_ISL_476093, EPI_ISL_476094, EPI_ISL_476095, EPI_ISL_476096, EPI_ISL_476097, EPI_ISL_476098, EPI_ISL_476099, EPI_ISL_476100, EPI_ISL_476101, EPI_ISL_476102, EPI_ISL_476103, EPI_ISL_476104, EPI_ISL_476105, EPI_ISL_476106, EPI_ISL_476107, EPI_ISL_476108, EPI_ISL_476109, EPI_ISL_476110, EPI_ISL_476111, EPI_ISL_476112, EPI_ISL_476113, EPI_ISL_476114, EPI_ISL_476115, EPI_ISL_476116, EPI_ISL_476117, EPI_ISL_476118, EPI_ISL_476119, EPI_ISL_476120, EPI_ISL_476121, EPI_ISL_476122, EPI_ISL_476123, EPI_ISL_476124, EPI_ISL_476125, EPI_ISL_476126, EPI_ISL_476127, EPI_ISL_476128, EPI_ISL_476129, EPI_ISL_476130, EPI_ISL_476131, EPI_ISL_476132, EPI_ISL_476133, EPI_ISL_476134 | | | |
| see above | Viollier AG | Department of Biosystems Science and Engineering, ETH Zürich | Christian Beisel, Sarah Nadeau, Ivan Topolsky, Pedro Ferreira, Philipp Jablonski, Susana Posada-Céspedes, Tobias Schär, Ina Nissen, Natascha Santacroce, Elodie Burcklen, Christiane Beckmann, Maurice Redondo, Olivier Kobel, Christoph Noppen, Sophie Seidel, Noemie Santamaria de Souza, Niko Beerenwinkel, Tanja Stadler |
| EPI_ISL_476135 | Achima Care Fristadens VC | The Public Health Agency of Sweden | Oskar Karlsson Lindsjo, Maria Lind Karlberg, Mattias Haukland, Reza Advani, Olov Svartstrom, Anna-Malin Linde, Sandra Broddesson, Petra Edquist, Mia Brytting, Anna Risberg, Karin Tegmark-Wisell |
| EPI_ISL_476136 | Surbrunns VC | The Public Health Agency of Sweden | Oskar Karlsson Lindsjo, Maria Lind Karlberg, Mattias Haukland, Reza Advani, Olov Svartstrom, Anna-Malin Linde, Sandra Broddesson, Petra Edquist, Mia Brytting, Anna Risberg, Karin Tegmark-Wisell |
| EPI_ISL_476137 | Wasterlakarna | The Public Health Agency of Sweden | Oskar Karlsson Lindsjo, Maria Lind Karlberg, Mattias Haukland, Reza Advani, Olov Svartstrom, Anna-Malin Linde, Sandra Broddesson, Petra Edquist, Mia Brytting, Anna Risberg, Karin Tegmark-Wisell |
| EPI_ISL_476138 | Ulltuna Vardcentral | The Public Health Agency of Sweden | Oskar Karlsson Lindsjo, Maria Lind Karlberg, Mattias Haukland, Reza Advani, Olov Svartstrom, Anna-Malin Linde, Sandra Broddesson, Petra Edquist, Mia Brytting, Anna Risberg, Karin Tegmark-Wisell |
| EPI_ISL_476139 | Folkhalsomyndigheten | The Public Health Agency of Sweden | Oskar Karlsson Lindsjo, Maria Lind Karlberg, Mattias Haukland, Reza Advani, Olov Svartstrom, Anna-Malin Linde, Sandra Broddesson, Petra Edquist, Shamam Muradrasoli, Anna Risberg, Karin Tegmark-Wisell |
| EPI_ISL_476140, EPI_ISL_476141, EPI_ISL_476142 | Klinisk Mikrobiologi | The Public Health Agency of Sweden | Oskar Karlsson Lindsjo, Maria Lind Karlberg, Mattias Haukland, Reza Advani, Olov Svartstrom, Anna-Malin Linde, Sandra Broddesson, Petra Edquist, Shamam Muradrasoli, Anna Risberg, Karin Tegmark-Wisell |
| EPI_ISL_476143, EPI_ISL_476144 | Skovde/Unilabs | The Public Health Agency of Sweden | Oskar Karlsson Lindsjo, Maria Lind Karlberg, Mattias Haukland, Reza Advani, Olov Svartstrom, Anna-Malin Linde, Sandra Broddesson, Petra Edquist, Shamam Muradrasoli, Anna Risberg, Karin Tegmark-Wisell |
| EPI_ISL_476145, EPI_ISL_476146, EPI_ISL_476147 | Ostersund klinisk mikrobiologi | The Public Health Agency of Sweden | Oskar Karlsson Lindsjo, Maria Lind Karlberg, Mattias Haukland, Reza Advani, Olov Svartstrom, Anna-Malin Linde, Sandra Broddesson, Petra Edquist, Shamam Muradrasoli, Anna Risberg, Karin Tegmark-Wisell |
| EPI_ISL_476148, EPI_ISL_476149 | Institut Pasteur Dakar | Institut Pasteur de Dakar | Ndongo Dia, Moussa Moise Diagne, Mamadou Diop, Ousmane Faye, Amadou Alpha Sall |
| EPI_ISL_476150 | Institut Pasteur Dakar | Institut Pasteur de Dakar | Ndongo Dia, Moussa Moise Diagne, Mamadou diop, Ousmane Faye, Amadou Alpha Sall |
| EPI_ISL_476151 | Institut Pasteur Dakar | Institut Pasteur de Dakar | Ndongo Dia, Moussa Moise Diagne, Mamadou Diop, Ousmane faye, Amadou Alpha Sall |
| EPI_ISL_476152, EPI_ISL_476153, EPI_ISL_476154, EPI_ISL_476155, EPI_ISL_476156, EPI_ISL_476157, EPI_ISL_476158, EPI_ISL_476159, EPI_ISL_476160, EPI_ISL_476161, EPI_ISL_476162, EPI_ISL_476163, EPI_ISL_476164, EPI_ISL_476165, EPI_ISL_476166, EPI_ISL_476167, EPI_ISL_476168, EPI_ISL_476169, EPI_ISL_476170 | | | |
| see above | Laboratório de Patologia Clínica - UNICAMP | Laboratório de Estudos de Vírus Emergentes - UNICAMP | José Luiz Proença-Modena, Magnun Nueldo Nunes dos Santos, Angelica Schreiber, Julia Forato,Camila Simeoni, Marcilio Jorge Fumagalli, Mariene Ribeiro Amorim, Darlan da Silva Candido, Nuno Rodrigues Faria, Julien Theze, Luiz Gonzaga,Jaqueline Goes Jesus e William Marciel de Souza |
| EPI_ISL_476171, EPI_ISL_476172, EPI_ISL_476173, EPI_ISL_476174, EPI_ISL_476175, EPI_ISL_476176, EPI_ISL_476177, EPI_ISL_476178, EPI_ISL_476179, EPI_ISL_476180, EPI_ISL_476181, EPI_ISL_476182, EPI_ISL_476183, EPI_ISL_476184, EPI_ISL_476185, EPI_ISL_476186, EPI_ISL_476187, EPI_ISL_476188, EPI_ISL_476189, EPI_ISL_476190, EPI_ISL_476191, EPI_ISL_476192, EPI_ISL_476193, EPI_ISL_476194, | | | |

| | | | |
|---|---|---|---|
| EPI_ISL_476195, EPI_ISL_476196, EPI_ISL_476197, EPI_ISL_476198, EPI_ISL_476199, EPI_ISL_476200, EPI_ISL_476201, EPI_ISL_476202 | | | |
| see above | DB Diagnósticos do Brasil | Instituto de Medicina Tropical da Univesidade de São Paulo | Samples: Nelson Gaburo Jr; Sequencing: Ingra Morales Claro, Jaqueline Goes de Jesus, Erika Regina Manuli, Flavia Cristina da Silva Sales, Thais de Moura Coletti, Camila Alves Maia da Silva, Mariana Severo Ramundo, Giula Magalhaes Ferreira, Darlan da Silva Candido, Julien Theze, Nuno Faria, Ester Sabino |
| EPI_ISL_476203, EPI_ISL_476204, EPI_ISL_476205, EPI_ISL_476206, EPI_ISL_476207, EPI_ISL_476208 | Hospital da Clínicas da Faculdade de Medicina da Universidade de São Paulo | Instituto de Medicina Tropical da Univesidade de São Paulo | Samples: Ingra Morales Claro, Erika Regina Manuli, Cecilia Salete Alencar, Carolina S. Lazar, Silvia F. Costa; Sequencing: Ingra Morales Claro, Jaqueline Goes de Jesus, Erika Regina Manuli, Flavia Cristina da Silva Sales, Thais de Moura Coletti, Camila Alves Maia da Silva, Mariana Severo Ramundo, Giula Magalhaes Ferreira, Darlan da Silva Candido, Julien Theze, Nuno Faria, Ester Sabino |
| EPI_ISL_476209, EPI_ISL_476210, EPI_ISL_476211, EPI_ISL_476212, EPI_ISL_476213, EPI_ISL_476214, EPI_ISL_476215, EPI_ISL_476216 | DB Diagnósticos do Brasil | Instituto de Medicina Tropical da Univesidade de São Paulo | Samples: Nelson Gaburo Jr; Sequencing: Ingra Morales Claro, Jaqueline Goes de Jesus, Erika Regina Manuli, Flavia Cristina da Silva Sales, Thais de Moura Coletti, Camila Alves Maia da Silva, Mariana Severo Ramundo, Giula Magalhaes Ferreira, Darlan da Silva Candido, Julien Theze, Nuno Faria, Ester Sabino |
| EPI_ISL_476217 | Hospital da Clínicas da Faculdade de Medicina da Universidade de São Paulo | Instituto de Medicina Tropical da Univesidade de São Paulo | Samples: Ingra Morales Claro, Erika Regina Manuli, Cecilia Salete Alencar, Carolina S. Lazar, Silvia F. Costa; Sequencing: Ingra Morales Claro, Jaqueline Goes de Jesus, Erika Regina Manuli, Flavia Cristina da Silva Sales, Thais de Moura Coletti, Camila Alves Maia da Silva, Mariana Severo Ramundo, Giula Magalhaes Ferreira, Darlan da Silva Candido, Julien Theze, Nuno Faria, Ester Sabino |
| EPI_ISL_476218, EPI_ISL_476219 | DB Diagnósticos do Brasil | Instituto de Medicina Tropical da Univesidade de São Paulo | Samples: Nelson Gaburo Jr; Sequencing: Ingra Morales Claro, Jaqueline Goes de Jesus, Erika Regina Manuli, Flavia Cristina da Silva Sales, Thais de Moura Coletti, Camila Alves Maia da Silva, Mariana Severo Ramundo, Giula Magalhaes Ferreira, Darlan da Silva Candido, Julien Theze, Nuno Faria, Ester Sabino |
| EPI_ISL_476220, EPI_ISL_476221 | Laboratory Fleury | Instituto de Medicina Tropical da Univesidade de São Paulo | Samples: Celso Granato; Sequencing: Ingra Morales Claro, Jaqueline Goes de Jesus, Erika Regina Manuli, Flavia Cristina da Silva Sales, Thais de Moura Coletti, Camila Alves Maia da Silva, Mariana Severo Ramundo, Giula Magalhaes Ferreira, Darlan da Silva Candido, Julien Theze, Nuno Faria, Ester Sabino |
| EPI_ISL_476222, EPI_ISL_476223 | Hospital da Clínicas da Faculdade de Medicina da Universidade de São Paulo | Instituto de Medicina Tropical da Univesidade de São Paulo | Samples: Ingra Morales Claro, Erika Regina Manuli, Cecilia Salete Alencar, Carolina S. Lazar, Silvia F. Costa; Sequencing: Ingra Morales Claro, Jaqueline Goes de Jesus, Erika Regina Manuli, Flavia Cristina da Silva Sales, Thais de Moura Coletti, Camila Alves Maia da Silva, Mariana Severo Ramundo, Giula Magalhaes Ferreira, Darlan da Silva Candido, Julien Theze, Nuno Faria, Ester Sabino |
| EPI_ISL_476224, EPI_ISL_476225, EPI_ISL_476226, EPI_ISL_476227, EPI_ISL_476228, EPI_ISL_476229 | DB Diagnósticos do Brasil | Instituto de Medicina Tropical da Univesidade de São Paulo | Samples: Nelson Gaburo Jr; Sequencing: Ingra Morales Claro, Jaqueline Goes de Jesus, Erika Regina Manuli, Flavia Cristina da Silva Sales, Thais de Moura Coletti, Camila Alves Maia da Silva, Mariana Severo Ramundo, Giula Magalhaes Ferreira, Darlan da Silva Candido, Julien Theze, Nuno Faria, Ester Sabino |
| EPI_ISL_476230, EPI_ISL_476231 | Laboratory Fleury | Instituto de Medicina Tropical da Univesidade de São Paulo | Samples: Celso Granato; Sequencing: Ingra Morales Claro, Jaqueline Goes de Jesus, Erika Regina Manuli, Flavia Cristina da Silva Sales, Thais de Moura Coletti, Camila Alves Maia da Silva, Mariana Severo Ramundo, Giula Magalhaes Ferreira, Darlan da Silva Candido, Julien Theze, Nuno Faria, Ester Sabino |
| EPI_ISL_476232 | DB Diagnósticos do Brasil | Instituto de Medicina Tropical da Univesidade de São Paulo | Samples: Nelson Gaburo Jr; Sequencing: Ingra Morales Claro, Jaqueline Goes de Jesus, Erika Regina Manuli, Flavia Cristina da Silva Sales, Thais de Moura Coletti, Camila Alves Maia da Silva, Mariana Severo Ramundo, Giula Magalhaes Ferreira, Darlan da Silva Candido, Julien Theze, Nuno Faria, Ester Sabino |
| EPI_ISL_476233, EPI_ISL_476234, EPI_ISL_476235, EPI_ISL_476236 | Laboratory Fleury | Instituto de Medicina Tropical da Univesidade de São Paulo | Samples: Celso Granato; Sequencing: Ingra Morales Claro, Jaqueline Goes de Jesus, Erika Regina Manuli, Flavia Cristina da Silva Sales, Thais de Moura Coletti, Camila Alves Maia da Silva, Mariana Severo Ramundo, Giula Magalhaes Ferreira, Darlan da Silva Candido, Julien Theze, Nuno Faria, Ester Sabino |
| EPI_ISL_476237, EPI_ISL_476238, EPI_ISL_476239, EPI_ISL_476240, EPI_ISL_476241, EPI_ISL_476242, EPI_ISL_476243, EPI_ISL_476244, EPI_ISL_476245, EPI_ISL_476246, EPI_ISL_476247, EPI_ISL_476248, EPI_ISL_476249, EPI_ISL_476250, EPI_ISL_476251, EPI_ISL_476252, EPI_ISL_476253, EPI_ISL_476254, EPI_ISL_476255, EPI_ISL_476256, EPI_ISL_476257, EPI_ISL_476258, EPI_ISL_476259, EPI_ISL_476260, EPI_ISL_476261, EPI_ISL_476262, EPI_ISL_476263, EPI_ISL_476264, EPI_ISL_476265, EPI_ISL_476266, EPI_ISL_476267, EPI_ISL_476268, EPI_ISL_476269, EPI_ISL_476270, EPI_ISL_476271, EPI_ISL_476272, EPI_ISL_476273, EPI_ISL_476274, EPI_ISL_476275, EPI_ISL_476276, EPI_ISL_476277 | | | |
| see above | Hospital da Clínicas da Faculdade de Medicina da Universidade de São Paulo | Instituto de Medicina Tropical da Univesidade de São Paulo | Samples: Ingra Morales Claro, Erika Regina Manuli, Cecilia Salete Alencar, Carolina S. Lazar, Silvia F. Costa; Sequencing: Ingra Morales Claro, Jaqueline Goes de Jesus, Erika Regina Manuli, Flavia Cristina da Silva Sales, Thais de Moura Coletti, Camila Alves Maia da Silva, Mariana Severo Ramundo, Giula Magalhaes Ferreira, Darlan da Silva Candido, Julien Theze, Nuno Faria, Ester Sabino |
| EPI_ISL_476278, EPI_ISL_476279, EPI_ISL_476280, EPI_ISL_476281, EPI_ISL_476282, EPI_ISL_476283, EPI_ISL_476284, EPI_ISL_476285, EPI_ISL_476286, EPI_ISL_476287, EPI_ISL_476288, EPI_ISL_476289, EPI_ISL_476290, EPI_ISL_476291, EPI_ISL_476292, EPI_ISL_476293, EPI_ISL_476294, EPI_ISL_476295, EPI_ISL_476296, EPI_ISL_476297, EPI_ISL_476298, EPI_ISL_476299, EPI_ISL_476300, EPI_ISL_476301, EPI_ISL_476302, EPI_ISL_476303, EPI_ISL_476304, EPI_ISL_476305, EPI_ISL_476306, EPI_ISL_476307, EPI_ISL_476308, EPI_ISL_476309, EPI_ISL_476310, EPI_ISL_476311, EPI_ISL_476312, EPI_ISL_476313, EPI_ISL_476314, EPI_ISL_476315, EPI_ISL_476316, EPI_ISL_476317, EPI_ISL_476318, EPI_ISL_476319, EPI_ISL_476320, EPI_ISL_476321, EPI_ISL_476322, EPI_ISL_476323, EPI_ISL_476324, EPI_ISL_476325, EPI_ISL_476326, EPI_ISL_476327, EPI_ISL_476328, EPI_ISL_476329, EPI_ISL_476330, EPI_ISL_476331, EPI_ISL_476332, EPI_ISL_476333, EPI_ISL_476334, EPI_ISL_476335, EPI_ISL_476336 | | | |
| see above | DB Diagnósticos do Brasil | Instituto de Medicina Tropical da Univesidade de São Paulo | Samples: Nelson Gaburo Jr; Sequencing: Ingra Morales Claro, Jaqueline Goes de Jesus, Erika Regina Manuli, Flavia Cristina da Silva Sales, Thais de Moura Coletti, Camila Alves Maia da Silva, Mariana Severo Ramundo, Giula Magalhaes Ferreira, Darlan da Silva Candido, Julien Theze, Nuno Faria, Ester Sabino |
| EPI_ISL_476337, EPI_ISL_476338, EPI_ISL_476339, EPI_ISL_476340, EPI_ISL_476341, EPI_ISL_476342, EPI_ISL_476343, EPI_ISL_476344, EPI_ISL_476345, EPI_ISL_476346, EPI_ISL_476347, EPI_ISL_476348, EPI_ISL_476349 | | | |
| see above | Laboratório de Patologia Clínica - UNICAMP | Laboratório de Estudos de Vírus Emergentes - UNICAMP | José Luiz Proença-Modena, Magnun Nueldo Nunes dos Santos, Angelica Schreiber, Julia Forato,Camila Simeoni, Marcilio Jorge Fumagalli, Mariene Ribeiro Amorim, Darlan da Silva Candido, Nuno Rodrigues Faria, Julien Theze, Luiz Gonzaga,Jaqueline Goes Jesus e William Marciel de Souza |
| EPI_ISL_476350, EPI_ISL_476351, EPI_ISL_476352, EPI_ISL_476353, EPI_ISL_476354, EPI_ISL_476355, EPI_ISL_476356, EPI_ISL_476357, EPI_ISL_476358, EPI_ISL_476359, EPI_ISL_476360, EPI_ISL_476361, EPI_ISL_476362, EPI_ISL_476363, EPI_ISL_476364, EPI_ISL_476365, EPI_ISL_476366, EPI_ISL_476367, EPI_ISL_476368, EPI_ISL_476369, EPI_ISL_476370, EPI_ISL_476371 | | | |
| see above | DB Diagnósticos do Brasil | Instituto de Medicina Tropical da Univesidade de São Paulo | Samples: Nelson Gaburo Jr; Sequencing: Ingra Morales Claro, Jaqueline Goes de Jesus, Erika Regina Manuli, Flavia Cristina da Silva Sales, Thais de Moura Coletti, Camila Alves Maia da Silva, Mariana Severo Ramundo, Giula Magalhaes Ferreira, Darlan da Silva Candido, Julien Theze, Nuno Faria, Ester Sabino |
| EPI_ISL_476372, EPI_ISL_476373, EPI_ISL_476374, EPI_ISL_476375, EPI_ISL_476376, EPI_ISL_476377, EPI_ISL_476378, EPI_ISL_476379, EPI_ISL_476380, EPI_ISL_476381, EPI_ISL_476382, EPI_ISL_476383, EPI_ISL_476384, EPI_ISL_476385, EPI_ISL_476386 | | | |
| see above | Hospital da Clínicas da Faculdade de Medicina da Universidade de São Paulo | Instituto de Medicina Tropical da Univesidade de São Paulo | Samples: Ingra Morales Claro, Erika Regina Manuli, Cecilia Salete Alencar, Carolina S. Lazar, Silvia F. Costa; Sequencing: Ingra Morales Claro, Jaqueline Goes de Jesus, Erika Regina Manuli, Flavia Cristina da Silva Sales, Thais de Moura Coletti, Camila Alves Maia da Silva, Mariana Severo Ramundo, Giula Magalhaes Ferreira, Darlan da Silva Candido, Julien Theze, Nuno Faria, Ester Sabino |
| EPI_ISL_476387, EPI_ISL_476388, EPI_ISL_476389, EPI_ISL_476390, EPI_ISL_476391, EPI_ISL_476392, EPI_ISL_476393, EPI_ISL_476394, EPI_ISL_476395, EPI_ISL_476396, EPI_ISL_476397, EPI_ISL_476398, EPI_ISL_476399, EPI_ISL_476400, EPI_ISL_476401, EPI_ISL_476402 | | | |
| see above | Laboratório de Patologia Clínica - UNICAMP | Laboratório de Estudos de Vírus Emergentes - UNICAMP | José Luiz Proença-Modena, Magnun Nueldo Nunes dos Santos, Angelica Schreiber, Julia Forato,Camila Simeoni, Marcilio Jorge Fumagalli, Mariene Ribeiro Amorim, Darlan da Silva Candido, Nuno Rodrigues Faria, Julien Theze, Luiz Gonzaga,Jaqueline Goes Jesus e William Marciel de Souza |
| EPI_ISL_476403, EPI_ISL_476404, EPI_ISL_476405 | Laboratory DASA | Instituto de Medicina Tropical da Univesidade de São Paulo | Samples: José Eduardo Levi; Sequencing: Ingra Morales Claro, Jaqueline Goes de Jesus, Erika Regina Manuli, Flavia Cristina da Silva Sales, Thais de Moura Coletti, Camila Alves Maia da Silva, Mariana Severo Ramundo, Giula Magalhaes Ferreira, Darlan da Silva Candido, Julien Theze, Nuno Faria, Ester Sabino |
| EPI_ISL_476406, EPI_ISL_476407 | Laboratory Fleury | Instituto de Medicina Tropical da Univesidade de São Paulo | Samples: Celso Granato; Sequencing: Ingra Morales Claro, Jaqueline Goes de Jesus, Erika Regina Manuli, Flavia Cristina da Silva Sales, Thais de Moura Coletti, Camila Alves Maia da Silva, Mariana Severo Ramundo, Giula Magalhaes Ferreira, Darlan da Silva Candido, Julien Theze, Nuno Faria, Ester Sabino |
| EPI_ISL_476408, EPI_ISL_476409, EPI_ISL_476410, EPI_ISL_476411, EPI_ISL_476412, EPI_ISL_476413, EPI_ISL_476414, EPI_ISL_476415, EPI_ISL_476416, EPI_ISL_476417, EPI_ISL_476418, EPI_ISL_476419, EPI_ISL_476420, EPI_ISL_476421, EPI_ISL_476422, EPI_ISL_476423, EPI_ISL_476424, EPI_ISL_476425 | | | |
| see above | Laboratório de Patologia Clínica - UNICAMP | Laboratório de Estudos de Vírus Emergentes - UNICAMP | José Luiz Proença-Modena, Magnun Nueldo Nunes dos Santos, Angelica Schreiber, Julia Forato,Camila Simeoni, Marcilio Jorge Fumagalli, Mariene Ribeiro Amorim, Darlan da Silva Candido, Nuno Rodrigues Faria, Julien Theze, Luiz Gonzaga,Jaqueline Goes Jesus e William Marciel de Souza |
| EPI_ISL_476426, EPI_ISL_476427 | Laboratory Fleury | Instituto de Medicina Tropical da Univesidade de São Paulo | Samples: Celso Granato; Sequencing: Ingra Morales Claro, Jaqueline Goes de Jesus, Erika Regina Manuli, Flavia Cristina da Silva Sales, Thais de Moura Coletti, Camila Alves Maia da Silva, Mariana Severo Ramundo, Giula Magalhaes Ferreira, Darlan da Silva Candido, Julien Theze, Nuno Faria, Ester Sabino |
| EPI_ISL_476428, EPI_ISL_476429, EPI_ISL_476430, EPI_ISL_476431, EPI_ISL_476432, EPI_ISL_476433, EPI_ISL_476434, EPI_ISL_476435, EPI_ISL_476436, EPI_ISL_476437, EPI_ISL_476438, EPI_ISL_476439, EPI_ISL_476440, EPI_ISL_476441, EPI_ISL_476442, EPI_ISL_476443, EPI_ISL_476444, EPI_ISL_476445, EPI_ISL_476446, EPI_ISL_476447, EPI_ISL_476448, EPI_ISL_476449, EPI_ISL_476450, EPI_ISL_476451, EPI_ISL_476452, EPI_ISL_476453, EPI_ISL_476454, EPI_ISL_476455, EPI_ISL_476456, EPI_ISL_476457, EPI_ISL_476458, EPI_ISL_476459, EPI_ISL_476460, EPI_ISL_476461, EPI_ISL_476462, EPI_ISL_476463, EPI_ISL_476464, EPI_ISL_476465, EPI_ISL_476466, EPI_ISL_476467, EPI_ISL_476468, EPI_ISL_476469, EPI_ISL_476470, EPI_ISL_476471, EPI_ISL_476472, EPI_ISL_476473, EPI_ISL_476474, EPI_ISL_476475, EPI_ISL_476476, EPI_ISL_476477, EPI_ISL_476478, EPI_ISL_476479, EPI_ISL_476480, EPI_ISL_476481, EPI_ISL_476482, EPI_ISL_476483, EPI_ISL_476484, EPI_ISL_476485, EPI_ISL_476486, EPI_ISL_476487, EPI_ISL_476488, EPI_ISL_476489, EPI_ISL_476490 | | | |
| see above | Hospital da Clínicas da Faculdade de Medicina da Universidade de São Paulo | Instituto de Medicina Tropical da Univesidade de São Paulo | Samples: Ingra Morales Claro, Erika Regina Manuli, Cecilia Salete Alencar, Carolina S. Lazar, Silvia F. Costa; Sequencing: Ingra Morales Claro, Jaqueline Goes de Jesus, Erika Regina Manuli, Flavia Cristina da Silva Sales, Thais de Moura Coletti, Camila Alves Maia da Silva, Mariana Severo Ramundo, Giula Magalhaes Ferreira, Darlan da Silva Candido, Julien Theze, Nuno Faria, Ester Sabino |
| EPI_ISL_476491, EPI_ISL_476492 | Institut Pasteur Dakar | Institut Pasteur de Dakar | Ndongo Dia, Moussa Moise Diagne, Mamadou Diop, Ousmane Faye, Amadou Alpha Sall |
| EPI_ISL_476493 | Institut Pasteur Dakar | Institut Pasteur de Dakar | Ndongo Dia, Moussa Moise Diagne, Mamadou Diop, Ousmane Faye, Amadou alpha Sall |
| EPI_ISL_476494 | Institut Pasteur Dakar | Institut Pasteur de Dakar | Ndongo Dia, Moussa Moise Diagne, Mamadou Diop, Ousmane Faye, Amadou Alpha Sall |
| EPI_ISL_476495 | Institut Pasteur Dakar | Institut Pasteur de Dakar | Ndongo Dia, Moussa Moise Diagne, Mamadou Diop, Ousmane Faye, Amadou alpha Sall |
| EPI_ISL_476496 | Hospital Garrahan | Héritas | Dalmacio Pereyra, Roberta Crespo, Mauricio Grisolia, Cristian Rohr, Andrea Mangano, Maria Florencia Fernandez, Fabian Fay, Martin Vazquez |
| EPI_ISL_476497 | Institut Pasteur Dakar | Institut Pasteur de Dakar | Ndongo Dia, Moussa Moise Diagne, Mamadou Diop, Ousmane Faye, Amadou alpha Sall |
| EPI_ISL_476498, EPI_ISL_476499, EPI_ISL_476500, EPI_ISL_476501, EPI_ISL_476502, EPI_ISL_476503, EPI_ISL_476504, EPI_ISL_476505, EPI_ISL_476506, EPI_ISL_476507, EPI_ISL_476508, EPI_ISL_476509, EPI_ISL_476510, EPI_ISL_476511, EPI_ISL_476512, EPI_ISL_476513 | | | |
| see above | Laboratoire de microbiologie, Hopital de Verdun | Smith Laboratory, Centre de Recherche CHU Sainte-Justine | Martin Smith, Marieke Rozendaal, Ivan Pavlov |
| EPI_ISL_476514 | Institut Pasteur Dakar | Institut Pasteur de Dakar | Ndongo Dia, Moussa Moise Diagne, Mamadou Diop, Ousmane Faye, Amadou Alpha Sall |
| EPI_ISL_476515 | Institut Pasteur Dakar | Institut Pasteur de Dakar | Ndongo Dia, Moussa Moise Diagne, Mamadou diop, Ousmane Faye, Amadou alpha Sall |
| EPI_ISL_476516 | Institut Pasteur Dakar | Institut Pasteur de Dakar | Ndongo Dia, Moussa Moise Diagne, mamadou Diop, Ousmane Faye, Amadou Alpha Sall |
| EPI_ISL_476517, EPI_ISL_476518, EPI_ISL_476519, EPI_ISL_476520, EPI_ISL_476521, EPI_ISL_476522, EPI_ISL_476523, EPI_ISL_476524, EPI_ISL_476525, EPI_ISL_476526, EPI_ISL_476527, EPI_ISL_476528, EPI_ISL_476529, EPI_ISL_476530, EPI_ISL_476531, EPI_ISL_476532, EPI_ISL_476533, EPI_ISL_476534, EPI_ISL_476535, EPI_ISL_476536, EPI_ISL_476537, EPI_ISL_476538, EPI_ISL_476539, EPI_ISL_476540, EPI_ISL_476541, EPI_ISL_476542, EPI_ISL_476543, EPI_ISL_476544, EPI_ISL_476545, EPI_ISL_476546, EPI_ISL_476547, EPI_ISL_476548, EPI_ISL_476549, EPI_ISL_476550, EPI_ISL_476551, EPI_ISL_476552, EPI_ISL_476553, EPI_ISL_476554, EPI_ISL_476555, EPI_ISL_476556, EPI_ISL_476557 | | | |
| see above | Yale Clinical Virology Laboratory | Grubaugh Lab - Yale School of Public Health | Joseph Fauver, Tara Alpert, Anderson Brito, Anne Wyllie, Chantal Vogels, Mary Petrone, Cole Jensen, Chaney Kalinich, Isabel Ott, Arnau Casanovas, Catherine Muenker, Adam Moore, Alice Lu, Maria Tokuyama, Patrick Wong, Peiwen Lu, Saad Omer, Richard Martinello, Allison Nelson, Shelli Farhadian, Akiko Iwasaki, Charlese Dela Cruz, Albert Ko, Nathan Grubaugh |
| EPI_ISL_476558 | Institut Pasteur Dakar | Institut Pasteur de Dakar | Ndongo Dia, Moussa Moise Diagne, Mamadou Diop, Ousmane Faye, Amadou Alpha Sall |
| EPI_ISL_476559 | unknown | Laboratoire Sciences et Technologies de la Santé (STS) Institut Supérieur des Sciences de la Santé Université Hassan 1er, Settat, Morocco | Hajar Lemriss, Sanaâ Lemriss, Amal Souiri, Narjis Amar, Mustapha Mouallif, Touria Essayagh, Jawad Bouzid, Saâd EL Kabbaj, Abderraouf Hilali |
| EPI_ISL_476560 | Institut Pasteur Dakar | Institut Pasteur de Dakar | Ndongo Dia, Moussa Moise Diagne, Mamadou Diop, Ousmane Faye, Amadou Alpha Sall |
| EPI_ISL_476561 | Hospital Garrahan | Héritas | Roberta Crespo, Dalmacio Pereyra, Mauricio Grisolia, Cristian Rohr, Andrea Mangano, Maria Florencia Fernandez, Fabian Fay, Martin Vazquez |
| EPI_ISL_476562 | Institut Pasteur Dakar | Institut Pasteur de Dakar | Ndongo Dia, Moussa Moise Diagne, Mamadou Diop, Ousmane Faye, Amadou Alpha Sall |
| EPI_ISL_476563 | Hospital de Pediatría "Prof. Dr. Juan P Garrahan" | Héritas | Dalmacio Pereyra, Roberta Crespo, Mauricio Grisolia, Cristian Rohr, Andrea Mangano, Maria Florencia Fernandez, Fabian Fay, Martin Vazquez |
| EPI_ISL_476564 | Institut Pasteur Dakar | Institut Pasteur de Dakar | Ndongo Dia, Moussa Moise Diagne, Mamadou diop, Ousmane Faye, Amadou alpha Sall |
| EPI_ISL_476565 | Hospital de Pediatría "Prof. Dr. Juan P Garrahan" | Héritas | Andrea Mangano, Maria Florencia Fernandez, Dalmacio Pereyra, Roberta Crespo, Mauricio Grisolia, Cristian Rohr, Fabian Fay, Martin Vazquez |

| | | | |
|---|---|---|---|
| EPI_ISL_476566 | Institut pasteur Dakar | Institut Pasteur de Dakar | Ndongo Dia, Moussa Moise Diagne, Mamadou Diop, Ousmane Faye, Amadou Alpha Sall |
| EPI_ISL_476567 | Hospital de Pediatría "Prof. Dr. Juan P Garrahan" | Héritas | Dalmacio Pereyra, Roberta Crespo, Mauricio Grisolia, Cristian Rohr, Andrea Mangano, Maria Florencia Fernandez, Fabian Fay, Martin Vazquez |
| EPI_ISL_476568 | Hospital de Pediatría "Prof. Dr. Juan P Garrahan" | Héritas | Cristian Rohr, Andrea Mangano, Maria Florencia Fernandez, Dalmacio Pereyra, Roberta Crespo, Mauricio Grisolia, Fabian Fay, Martin Vazquez |
| EPI_ISL_476569 | Institut Pasteur Dakar | Institut Pasteur de Dakar | Ndongo Dia, Moussa Moise, Mamadou Diop, Ousmane Faye, Amadou Alpha Sall |
| EPI_ISL_476570 | Institut Pasteur Dakar | Institut Pasteur de Dakar | Ndongo Dia, Moussa Moise Diagne, Mamadou Diop, Ousmane Faye, Amadou Alpha Sall |
| EPI_ISL_476571 | Hospital de Pediatría "Prof. Dr. Juan P Garrahan" | Héritas | Dalmacio Pereyra, Roberta Crespo, Mauricio Grisolia, Cristian Rohr, Andrea Mangano, Maria Florencia Fernandez, Fabian Fay, Martin Vazquez |
| EPI_ISL_476572 | Institut Pasteur Dakar | Institut Pasteur de Dakar | Ndongo Dia, Moussa Moise Diagne, Mamadou Diop, Ousmane Faye, Amadou Alpha Sall |
| EPI_ISL_476573 | Hospital de Pediatría "Prof. Dr. Juan P Garrahan" | Héritas | Dalmacio Pereyra, Roberta Crespo, Mauricio Grisolia, Cristian Rohr, Andrea Mangano, Maria Florencia Fernandez, Fabian Fay, Martin Vazquez |
| EPI_ISL_476574 | Institut Pasteur Dakar | Institut Pasteur de Dakar | Ndongo Dia, Moussa Moise Diagne, Mamadou Diop, Ousmane Faye, Amadou Alpha Sall |
| EPI_ISL_476702 | Incubadora Venezolana de Ciencia, Venezuela | Incubadora Venezolana de Ciencia, Venezuela / Instituto Nacional de Salud, Bogotá, Colombia / Grupo de Investigaciones Microbiológicas-UR (GIMUR), Departamento de Biología, Facultad de Ciencias Naturales, Universidad del Rosario, Bogotá, Colombia / Icahn School of Medicine at Mount Sinai, New York, USA | Alberto Paniz-Mondolfi, Marina Muñoz, Luis Perez-Garcia, Lourdes Delgado, Carolina Florez, Sergio Gomez, Angelica Rico, Lisseth Pardo, Esther C. Barros, Carolina Hernández, Jesús E. Jaimes, Aníbal A. Teherán, Ana S. Gonzalez-Reiche, Matthew M. Hernandez, Emilia Mia Sordillo, Viviana Simon, Harm van Bakel, Juan David Ramírez |
| EPI_ISL_476703, EPI_ISL_476704 | Incubadora Venezolana de Ciencia, Venezuela | Incubadora Venezolana de Ciencia, Venezuela / Instituto Nacional de Salud, Bogotá, Colombia / Grupo de Investigaciones Microbiológicas-UR (GIMUR), Departamento de Biología, Facultad de Ciencias Naturales, Universidad del Rosario, Bogotá, Colombia / Icahn School of Medicine at Mount Sinai, New York, USA | Alberto Paniz-Mondolfi, Marina Muñoz, Luis Perez-Garcia, Lourdes Delgado, Carolina Florez, Sergio Gomez, Angelica Rico, Lisseth Pardo, Esther C. Barros, Carolina Hernández, Jesús E. Jaimes, Aníbal A. Teherán, Ana S. Gonzalez-Reiche, Matthew M. Hernandez, Emilia Mia Sordillo, Viviana Simon, Harm van Bakel, Juan David Ramírez |
| EPI_ISL_476705 | Labor Kneißler GmbH & Co. KG | Heinrich Pette Institute, Leibniz Institute for Experimental Virology | Thomas Günther, Adam Grundhoff, Manja Czech-Sioli, Nicole Fischer, Matthias Ottinger, Melanie M. Brinkmann |
| EPI_ISL_476706, EPI_ISL_476707, EPI_ISL_476708, EPI_ISL_476709, EPI_ISL_476710, EPI_ISL_476711, EPI_ISL_476712, EPI_ISL_476713, EPI_ISL_476714, EPI_ISL_476715, EPI_ISL_476716, EPI_ISL_476717, EPI_ISL_476718, EPI_ISL_476719, EPI_ISL_476720, EPI_ISL_476721, EPI_ISL_476722, EPI_ISL_476723, EPI_ISL_476724, EPI_ISL_476725, EPI_ISL_476726, EPI_ISL_476727, EPI_ISL_476728, EPI_ISL_476729, EPI_ISL_476730, EPI_ISL_476731, EPI_ISL_476732, EPI_ISL_476733, EPI_ISL_476734, EPI_ISL_476735, EPI_ISL_476736, EPI_ISL_476737, EPI_ISL_476738, EPI_ISL_476739, EPI_ISL_476740, EPI_ISL_476741, EPI_ISL_476742, EPI_ISL_476743, EPI_ISL_476744, EPI_ISL_476745, EPI_ISL_476746, EPI_ISL_476747, EPI_ISL_476748, EPI_ISL_476749, EPI_ISL_476750, EPI_ISL_476751, EPI_ISL_476752, EPI_ISL_476753, EPI_ISL_476754, EPI_ISL_476755, EPI_ISL_476756, EPI_ISL_476757, EPI_ISL_476758, EPI_ISL_476759, EPI_ISL_476760, EPI_ISL_476761, EPI_ISL_476762, EPI_ISL_476763, EPI_ISL_476764, EPI_ISL_476765, EPI_ISL_476766 | | | |
| see above | Minnesota Department of Health, Public Health Laboratory | Minnesota Department of Health, Public Health Laboratory | Matt Plumb, Jacob Garfin, and Xiong Wang |
| EPI_ISL_476767, EPI_ISL_476768, EPI_ISL_476769, EPI_ISL_476770, EPI_ISL_476771, EPI_ISL_476772, EPI_ISL_476773, EPI_ISL_476774, EPI_ISL_476775, EPI_ISL_476776, EPI_ISL_476777, EPI_ISL_476778, EPI_ISL_476779, EPI_ISL_476780, EPI_ISL_476781, EPI_ISL_476782, EPI_ISL_476783, EPI_ISL_476784, EPI_ISL_476785, EPI_ISL_476786, EPI_ISL_476787, EPI_ISL_476788, EPI_ISL_476789, EPI_ISL_476790, EPI_ISL_476791, EPI_ISL_476792, EPI_ISL_476793, EPI_ISL_476794 | | | |
| see above | Stanford clinical virology lab | Chan-Zuckerberg Biohub | Benjamin Pinksy, Katharine Walter, Victoria N. Parikh, John Gorzynski, Hannah N. DeJong, Matthew T. Wheeler, Jason Andrews, Manuel Rivas, Carlos Bustamante, Euan Ashley, with CZB Cliahub Consortium |
| EPI_ISL_476795 | Department of Laboratory Medicine, Tan Tock Seng Hospital | Department of Laboratory Medicine, Tan Tock Seng Hospital | Chen YYC, Zair X, Li C, Tang WY, Maurer-Stroh S, Barkham TMS, Nagarajan N, Sessions OM |
| EPI_ISL_476796, EPI_ISL_476797 | Department of Laboratory Medicine, Tan Tock Seng Hospital | Department of Laboratory Medicine, Tan Tock Seng Hospital | Chen YYC, Zair X, Li C, Tang WY, Maurer-Stroh S, Barkham TMS, Nagarajan N, Sessions OM |
| EPI_ISL_476801, EPI_ISL_476802, EPI_ISL_476803, EPI_ISL_476804 | Hong Kong Department of Health | School of Public Health, The University of Hong Kong | Dominic N.C. Tsang, Daniel K.W. Chu, Leo L.M. Poon, Malik Peiris |
| EPI_ISL_476805, EPI_ISL_476806, EPI_ISL_476807, EPI_ISL_476808, EPI_ISL_476809, EPI_ISL_476810, EPI_ISL_476811, EPI_ISL_476812 | Department of Laboratory Medicine Tan Tock Seng Hospital | Department of Laboratory Medicine Tan Tock Seng Hospital | Chen YYC, Zair X, Li C, Tang WY, Maurer-Stroh S, Barkham TMS, Nagarajan N, Sessions OM |
| EPI_ISL_476813, EPI_ISL_476814 | Department of Laboratory Medicine, Tan Tock Seng Hospital | Department of Laboratory Medicine, Tan Tock Seng Hospital | Chen YYC, Zair X, Li C, Tang WY, Maurer-Stroh S, Barkham TMS, Nagarajan N, Sessions OM |
| EPI_ISL_476815 | Department of Laboratory Medicine Tan Tock Seng Hospital | Department of Laboratory Medicine Tan Tock Seng Hospital | Chen YYC, Zair X, Li C, Tang WY, Maurer-Stroh S, Barkham TMS, Nagarajan N, Sessions OM |
| EPI_ISL_476816, EPI_ISL_476817, EPI_ISL_476818, EPI_ISL_476819 | Department of Laboratory Medicine, Tan Tock Seng Hospital | Department of Laboratory Medicine, Tan Tock Seng Hospital | Chen YYC, Zair X, Li C, Tang WY, Maurer-Stroh S, Barkham TMS, Nagarajan N, Sessions OM |
| EPI_ISL_476820 | Department of Laboratory Medicine, Tan Tock Seng Hospital | Department of Laboratory Medicine, Tan Tock Seng Hospital | Chen YYC, Zair X, Li C, Tang WY, Maurer-Stroh S, Barkham TMS, Nagarajan N, Sessions OM |
| EPI_ISL_476821 | Department of Laboratory Medicine, Tan Tock Seng Hospital | Department of Laboratory Medicine, Tan Tock Seng Hospital | Chen YYC, Zair X, Li C, Tang WY, Maurer-Stroh S, Barkham TMS, Nagarajan N, Sessions OM |
| EPI_ISL_476822, EPI_ISL_476823, EPI_ISL_476824, EPI_ISL_476825, EPI_ISL_476826, EPI_ISL_476827, EPI_ISL_476828, EPI_ISL_476829, EPI_ISL_476830, EPI_ISL_476831 | Laboratoire des Fièvres Hémorragiques Virales du Benin | Charité-Universitätsmedizin Berlin | Yadouleton, Anges; Sander Anna-Lena; Moreira-Soto Andres; Drexler, Jan Felix |
| EPI_ISL_476832 | Medical Biology Department, Kocaeli University | Medical Genetics Department, Kocaeli University | Savli H, Cine N, Sunnetci-Akkoyunlu D, Eren-Keskin S, Ilgazli A, Akhan S, Karadenizli A, Kasap M, Sayan M, Akpinar G, Canturk NZ. |
| EPI_ISL_476833, EPI_ISL_476834 | Laboratoire des Fièvres Hémorragiques Virales du Benin | Charité-Universitätsmedizin Berlin | Yadouleton, Anges; Sander Anna-Lena; Moreira-Soto Andres; Drexler, Jan Felix |
| EPI_ISL_476835 | National Influenza Centre for Northern Greece | National Influenza Centre for Northern Greece | Maria Christoforidi |
| EPI_ISL_476836 | National Influenza Centre for Nothern Greece | National Influenza Centre for Nothern Greece | Maria Christoforidi |
| EPI_ISL_476837, EPI_ISL_476838, EPI_ISL_476839 | National Influenza Centre for Northern Greece | National Influenza Centre for Northern Greece | Maria Christoforidi |
| EPI_ISL_476840 | Defence Research & Development Establishment (DRDE) | Defence Research & Development Establishment (DRDE) | Shashi Sharma, Paban Kumar Dash, Sushil Kumar Sharma, Ambuj Shrivastava, Jyoti S. Kumar |
| EPI_ISL_476841 | National Influenza Centre for Northern Greece | National Influenza Centre for Northern Greece | Maria Christoforidi |
| EPI_ISL_476842 | Defence Research & Development Establishment (DRDE) | Defence Research & Development Establishment (DRDE) | Shashi Sharma, Paban Kumar Dash, Sushil Kumar Sharma, Ambuj Shrivastava, Jyoti S. Kumar |
| EPI_ISL_476843 | National Influenza Centre for Northern Greece | National Influenza Centre for Northern Greece | Maria Christoforidi |
| EPI_ISL_476844 | Defence Research & Development Establishment (DRDE) | Defence Research & Development Establishment (DRDE) | Shashi Sharma, Paban Kumar Dash, Sushil Kumar Sharma, Ambuj Shrivastava, Jyoti S. Kumar |
| EPI_ISL_476845 | National Influenza Centre for Northern Greece | National Influenza Centre for Northern Greece | Maria Christoforidi |
| EPI_ISL_476846 | Defence Research & Development Establishment (DRDE) | Defence Research & Development Establishment (DRDE) | Shashi Sharma, Paban Kumar Dash, Sushil Kumar Sharma, Ambuj Shrivastava, Jyoti S. Kumar |
| EPI_ISL_476847 | National Influenza Centre for Northern Greece | National Influenza Centre for Northern Greece | Maria Christoforidi |
| EPI_ISL_476848, EPI_ISL_476849, EPI_ISL_476850 | Defence Research & Development Establishment (DRDE) | Defence Research & Development Establishment (DRDE) | Shashi Sharma, Paban Kumar Dash, Sushil Kumar Sharma, Ambuj Shrivastava, Jyoti S. Kumar |
| EPI_ISL_476851 | National Influenza Centre for Northern Greece | National Influenza Centre for Northern Greece | Maria Christoforidi |
| EPI_ISL_476852, EPI_ISL_476853, EPI_ISL_476854 | Defence Research & Development Establishment (DRDE) | Defence Research & Development Establishment (DRDE) | Shashi Sharma, Paban Kumar Dash, Sushil Kumar Sharma, Ambuj Shrivastava, Jyoti S. Kumar |
| EPI_ISL_476855 | GMERS Medical College & Hospital, Gotri, Vadodara | Gujarat Biotechnology Research Centre | Apurvasinh Puvar, Janvi Raval, Zarna Patel, Monika Gandhi, Pinal Trivedi, Maharshi Pandya, Nidhi Patel, Nitin Savaliya, Raghawendra Kumar, Dinesh Kumar, Zuber Saiyed, Komal Patel, Labdhi Pandya, Afzal Ansari, Nikha Trivedi, Meenakshi Shah, Neena Doshi, Varsha Godbole, R D Dixit, A M Kadri, Harsh Bakshi, Chaitanya Joshi, Madhvi Joshi |
| EPI_ISL_476856 | GMERS Medical College & Hospital, Gotri, Vadodara | Gujarat Biotechnology Research Centre | Janvi Raval, Zarna Patel, Monika Gandhi, Pinal Trivedi, Maharshi Pandya, Nidhi Patel, Nitin Savaliya, Raghawendra Kumar, Dinesh Kumar, Zuber Saiyed, Komal Patel, Labdhi Pandya, Afzal Ansari, Nikha Trivedi, Meenakshi Shah, Neena Doshi, Varsha Godbole, Apurvasinh Puvar, R D Dixit, A M Kadri, Harsh Bakshi, Chaitanya Joshi, Madhvi Joshi |
| EPI_ISL_476857 | GMERS Medical College & Hospital, Gotri, Vadodara | Gujarat Biotechnology Research Centre | Zarna Patel, Monika Gandhi, Pinal Trivedi, Maharshi Pandya, Nidhi Patel, Nitin Savaliya, Raghawendra Kumar, Dinesh Kumar, Zuber Saiyed, Komal Patel, Labdhi Pandya, Afzal Ansari, Nikha Trivedi, Meenakshi Shah, Neena Doshi, Varsha Godbole, Apurvasinh Puvar, Janvi Raval, R D Dixit, A M Kadri, Harsh Bakshi, Chaitanya Joshi, Madhvi Joshi |
| EPI_ISL_476858 | GMERS Medical College & Hospital, Gotri, Vadodara | Gujarat Biotechnology Research Centre | Monika Gandhi, Pinal Trivedi, Maharshi Pandya, Nidhi Patel, Nitin Savaliya, Raghawendra Kumar, Dinesh Kumar, Zuber Saiyed, Komal Patel, Labdhi Pandya, Afzal Ansari, Nikha Trivedi, Meenakshi Shah, Neena Doshi, Varsha Godbole, Apurvasinh Puvar, Janvi Raval, Zarna Patel, R D Dixit, A M Kadri, Harsh Bakshi, Chaitanya Joshi, Madhvi Joshi |
| EPI_ISL_476859 | GMERS Medical College & Hospital, Gotri, Vadodara | Gujarat Biotechnology Research Centre | Pinal Trivedi, Maharshi Pandya, Nidhi Patel, Nitin Savaliya, Raghawendra Kumar, Dinesh Kumar, Zuber Saiyed, Komal Patel, Labdhi Pandya, Afzal Ansari, Nikha Trivedi, Meenakshi Shah, Neena Doshi, Varsha Godbole, Apurvasinh Puvar, Janvi Raval, Zarna Patel, Monika Gandhi, R D Dixit, A M Kadri, Harsh Bakshi, Chaitanya Joshi, Madhvi Joshi |
| EPI_ISL_476860 | GMERS Medical College & Hospital, Gotri, Vadodara | Gujarat Biotechnology Research Centre | Maharshi Pandya, Nidhi Patel, Nitin Savaliya, Raghawendra Kumar, Dinesh Kumar, Zuber Saiyed, Komal Patel, Labdhi Pandya, Afzal Ansari, Nikha Trivedi, Meenakshi Shah, Neena Doshi, Varsha Godbole, Apurvasinh Puvar, Janvi Raval, Zarna Patel, Monika Gandhi, Pinal Trivedi, R D Dixit, A M Kadri, Harsh Bakshi, Chaitanya Joshi, Madhvi Joshi |
| EPI_ISL_476861 | GMERS Medical College & Hospital, Gotri, Vadodara | Gujarat Biotechnology Research Centre | Nidhi Patel, Nitin Savaliya, Raghawendra Kumar, Dinesh Kumar, Zuber Saiyed, Komal Patel, Labdhi Pandya, Afzal Ansari, Nikha Trivedi, Meenakshi Shah, Neena Doshi, Varsha Godbole, Apurvasinh Puvar, Janvi Raval, Zarna Patel, Monika Gandhi, Pinal Trivedi, Maharshi Pandya, R D Dixit, A M Kadri, Harsh Bakshi, Chaitanya Joshi, Madhvi Joshi |
| EPI_ISL_476862 | GMERS Medical College & Hospital, Gotri, Vadodara | Gujarat Biotechnology Research Centre | Nitin Savaliya, Raghawendra Kumar, Dinesh Kumar, Zuber Saiyed, Komal Patel, Labdhi Pandya, Afzal Ansari, Nikha Trivedi, Meenakshi Shah, Neena Doshi, Varsha Godbole, Apurvasinh Puvar, Janvi Raval, Zarna Patel, Monika Gandhi, Pinal Trivedi, Maharshi Pandya, Nidhi Patel, R D Dixit, A M Kadri, Harsh Bakshi, Chaitanya Joshi, Madhvi Joshi |
| EPI_ISL_476863 | GMERS Medical College and Hospital, Gandhinagar | Gujarat Biotechnology Research Centre | Raghawendra Kumar, Dinesh Kumar, Zuber Saiyed, Komal Patel, Labdhi Pandya, Afzal Ansari, Nikha Trivedi, Seema Bhatt, Gaurishankar Shrimali, Bhavesh Modi, Bharti Rajani, Apurvasinh Puvar, Janvi Raval, Zarna Patel, Monika Gandhi, Pinal Trivedi, Maharshi Pandya, Nidhi Patel, Nitin Savaliya, R D Dixit, A M Kadri, Harsh Bakshi, Chaitanya Joshi, Madhvi Joshi |
| EPI_ISL_476864 | GMERS Medical College and Hospital, Gandhinagar | Gujarat Biotechnology Research Centre | Dinesh Kumar, Zuber Saiyed, Komal Patel, Labdhi Pandya, Afzal Ansari, Nikha Trivedi, Seema Bhatt, Gaurishankar Shrimali, Bhavesh Modi, Bharti Rajani, Apurvasinh Puvar, Janvi Raval, Zarna Patel, Monika Gandhi, Pinal Trivedi, Maharshi Pandya, Nidhi Patel, Nitin Savaliya, Raghawendra Kumar, R D Dixit, A M Kadri, Harsh Bakshi, Chaitanya Joshi, Madhvi Joshi |
| EPI_ISL_476865 | GMERS Medical College and Hospital, Gandhinagar | Gujarat Biotechnology Research Centre | Zuber Saiyed, Komal Patel, Labdhi Pandya, Afzal Ansari, Nikha Trivedi, Seema Bhatt, Gaurishankar Shrimali, Bhavesh Modi, Bharti Rajani, Apurvasinh Puvar, Janvi Raval, Zarna Patel, Monika Gandhi, Pinal Trivedi, Maharshi Pandya, |

| | | | |
|---|---|---|---|
| | | | Nidhi Patel, Nitin Savaliya, Raghawendra Kumar, Dinesh Kumar, R D Dixit, A M Kadri, Harsh Bakshi, Chaitanya Joshi, Madhvi Joshi |
| EPI_ISL_476866 | GMERS Medical College and Hospital, Gandhinagar | Gujarat Biotechnology Research Centre | Komal Patel, Labdhi Pandya, Afzal Ansari, Nikha Trivedi, Seema Bhatt, Gaurishankar Shrimali, Bhavesh Modi, Bharti Rajani, Apurvasinh Puvar, Janvi Raval, Zarna Patel, Monika Gandhi, Pinal Trivedi, Maharshi Pandya, Nidhi Patel, Nitin Savaliya, Raghawendra Kumar, Dinesh Kumar, Zuber Saiyed, R D Dixit, A M Kadri, Harsh Bakshi, Chaitanya Joshi, Madhvi Joshi |
| EPI_ISL_476867 | Banas Medical College and Research Institute | Gujarat Biotechnology Research Centre | Labdhi Pandya, Afzal Ansari, Nikha Trivedi, Radhika Khara, Sunil R Joshi, Viren s Doshi, Apurvasinh Puvar, Janvi Raval, Zarna Patel, Monika Gandhi, Pinal Trivedi, Maharshi Pandya, Nidhi Patel, Nitin Savaliya, Raghawendra Kumar, Dinesh Kumar, Zuber Saiyed, Komal Patel, R D Dixit, A M Kadri, Harsh Bakshi, Chaitanya Joshi, Madhvi Joshi |
| EPI_ISL_476868 | Banas Medical College and Research Institute | Gujarat Biotechnology Research Centre | Afzal Ansari, Nikha Trivedi, Radhika Khara, Sunil R Joshi, Viren s Doshi, Apurvasinh Puvar, Janvi Raval, Zarna Patel, Monika Gandhi, Pinal Trivedi, Maharshi Pandya, Nidhi Patel, Nitin Savaliya, Raghawendra Kumar, Dinesh Kumar, Zuber Saiyed, Komal Patel, Labdhi Pandya, R D Dixit, A M Kadri, Harsh Bakshi, Chaitanya Joshi, Madhvi Joshi |
| EPI_ISL_476869 | Department of MicroBiology, Government Medical College, Surat | Gujarat Biotechnology Research Centre | Nikha Trivedi, Naresh Chauhan, Summaiya Mullan, Amit gamit, Apurvasinh Puvar, Janvi Raval, Zarna Patel, Monika Gandhi, Pinal Trivedi, Maharshi Pandya, Nidhi Patel, Nitin Savaliya, Raghawendra Kumar, Dinesh Kumar, Zuber Saiyed, Komal Patel, Labdhi Pandya, Afzal Ansari, R D Dixit, A M Kadri, Harsh Bakshi, Chaitanya Joshi, Madhvi Joshi |
| EPI_ISL_476870 | Department of MicroBiology, Government Medical College, Surat | Gujarat Biotechnology Research Centre | Naresh Chauhan, Summaiya Mullan, Amit gamit, Apurvasinh Puvar, Janvi Raval, Zarna Patel, Monika Gandhi, Pinal Trivedi, Maharshi Pandya, Nidhi Patel, Nitin Savaliya, Raghawendra Kumar, Dinesh Kumar, Zuber Saiyed, Komal Patel, Labdhi Pandya, Afzal Ansari, Nikha Trivedi, R D Dixit, A M Kadri, Harsh Bakshi, Chaitanya Joshi, Madhvi Joshi |
| EPI_ISL_476871 | Department of MicroBiology, Government Medical College, Surat | Gujarat Biotechnology Research Centre | Summaiya Mullan, Amit gamit, Apurvasinh Puvar, Janvi Raval, Zarna Patel, Monika Gandhi, Pinal Trivedi, Maharshi Pandya, Nidhi Patel, Nitin Savaliya, Raghawendra Kumar, Dinesh Kumar, Zuber Saiyed, Komal Patel, Labdhi Pandya, Afzal Ansari, Nikha Trivedi, Naresh Chauhan, R D Dixit, A M Kadri, Harsh Bakshi, Chaitanya Joshi, Madhvi Joshi |
| EPI_ISL_476872 | Department of MicroBiology, Government Medical College, Surat | Gujarat Biotechnology Research Centre | Amit gamit, Apurvasinh Puvar, Janvi Raval, Zarna Patel, Monika Gandhi, Pinal Trivedi, Maharshi Pandya, Nidhi Patel, Nitin Savaliya, Raghawendra Kumar, Dinesh Kumar, Zuber Saiyed, Komal Patel, Labdhi Pandya, Afzal Ansari, Nikha Trivedi, Naresh Chauhan, Summaiya Mullan, R D Dixit, A M Kadri, Harsh Bakshi, Chaitanya Joshi, Madhvi Joshi |
| EPI_ISL_476873 | Department of MicroBiology, Government Medical College, Surat | Gujarat Biotechnology Research Centre | Apurvasinh Puvar, Janvi Raval, Zarna Patel, Monika Gandhi, Pinal Trivedi, Maharshi Pandya, Nidhi Patel, Nitin Savaliya, Raghawendra Kumar, Dinesh Kumar, Zuber Saiyed, Komal Patel, Labdhi Pandya, Afzal Ansari, Nikha Trivedi, Naresh Chauhan, Summaiya Mullan, Amit gamit, R D Dixit, A M Kadri, Harsh Bakshi, Chaitanya Joshi, Madhvi Joshi |
| EPI_ISL_476874 | Department of MicroBiology, Government Medical College, Surat | Gujarat Biotechnology Research Centre | Janvi Raval, Zarna Patel, Monika Gandhi, Pinal Trivedi, Maharshi Pandya, Nidhi Patel, Nitin Savaliya, Raghawendra Kumar, Dinesh Kumar, Zuber Saiyed, Komal Patel, Labdhi Pandya, Afzal Ansari, Nikha Trivedi, Naresh Chauhan, Summaiya Mullan, Amit gamit, Apurvasinh Puvar, R D Dixit, A M Kadri, Harsh Bakshi, Chaitanya Joshi, Madhvi Joshi |
| EPI_ISL_476875 | Department of MicroBiology, Government Medical College, Surat | Gujarat Biotechnology Research Centre | Zarna Patel, Monika Gandhi, Pinal Trivedi, Maharshi Pandya, Nidhi Patel, Nitin Savaliya, Raghawendra Kumar, Dinesh Kumar, Zuber Saiyed, Komal Patel, Labdhi Pandya, Afzal Ansari, Nikha Trivedi, Naresh Chauhan, Summaiya Mullan, Amit gamit, Apurvasinh Puvar, Janvi Raval, R D Dixit, A M Kadri, Harsh Bakshi, Chaitanya Joshi, Madhvi Joshi |
| EPI_ISL_476876 | Department of MicroBiology, Government Medical College, Surat | Gujarat Biotechnology Research Centre | Pinal Trivedi, Maharshi Pandya, Nidhi Patel, Nitin Savaliya, Raghawendra Kumar, Dinesh Kumar, Zuber Saiyed, Komal Patel, Labdhi Pandya, Afzal Ansari, Nikha Trivedi, Naresh Chauhan, Summaiya Mullan, Amit gamit, Apurvasinh Puvar, Janvi Raval, Zarna Patel, Monika Gandhi, R D Dixit, A M Kadri, Harsh Bakshi, Chaitanya Joshi, Madhvi Joshi |
| EPI_ISL_476877 | Department of MicroBiology, Government Medical College, Surat | Gujarat Biotechnology Research Centre | Maharshi Pandya, Nidhi Patel, Nitin Savaliya, Raghawendra Kumar, Dinesh Kumar, Zuber Saiyed, Komal Patel, Labdhi Pandya, Afzal Ansari, Nikha Trivedi, Naresh Chauhan, Summaiya Mullan, Amit gamit, Apurvasinh Puvar, Janvi Raval, Zarna Patel, Monika Gandhi, Pinal Trivedi, R D Dixit, A M Kadri, Harsh Bakshi, Chaitanya Joshi, Madhvi Joshi |
| EPI_ISL_476878 | Department of MicroBiology, Government Medical College, Surat | Gujarat Biotechnology Research Centre | Nidhi Patel, Nitin Savaliya, Raghawendra Kumar, Dinesh Kumar, Zuber Saiyed, Komal Patel, Labdhi Pandya, Afzal Ansari, Nikha Trivedi, Naresh Chauhan, Summaiya Mullan, Amit gamit, Apurvasinh Puvar, Janvi Raval, Zarna Patel, Monika Gandhi, Pinal Trivedi, Maharshi Pandya, R D Dixit, A M Kadri, Harsh Bakshi, Chaitanya Joshi, Madhvi Joshi |
| EPI_ISL_476879 | Department of MicroBiology, Government Medical College, Surat | Gujarat Biotechnology Research Centre | Nitin Savaliya, Raghawendra Kumar, Dinesh Kumar, Zuber Saiyed, Komal Patel, Labdhi Pandya, Afzal Ansari, Nikha Trivedi, Naresh Chauhan, Summaiya Mullan, Amit gamit, Apurvasinh Puvar, Janvi Raval, Zarna Patel, Monika Gandhi, Pinal Trivedi, Maharshi Pandya, Nidhi Patel, R D Dixit, A M Kadri, Harsh Bakshi, Chaitanya Joshi, Madhvi Joshi |
| EPI_ISL_476880 | Department of MicroBiology, Government Medical College, Surat | Gujarat Biotechnology Research Centre | Raghawendra Kumar, Dinesh Kumar, Zuber Saiyed, Komal Patel, Labdhi Pandya, Afzal Ansari, Nikha Trivedi, Naresh Chauhan, Summaiya Mullan, Amit gamit, Apurvasinh Puvar, Janvi Raval, Zarna Patel, Monika Gandhi, Pinal Trivedi, Maharshi Pandya, Nidhi Patel, Nitin Savaliya, R D Dixit, A M Kadri, Harsh Bakshi, Chaitanya Joshi, Madhvi Joshi |
| EPI_ISL_476881 | Department of MicroBiology, Government Medical College, Surat | Gujarat Biotechnology Research Centre | Dinesh Kumar, Zuber Saiyed, Komal Patel, Labdhi Pandya, Afzal Ansari, Nikha Trivedi, Naresh Chauhan, Summaiya Mullan, Amit gamit, Apurvasinh Puvar, Janvi Raval, Zarna Patel, Monika Gandhi, Pinal Trivedi, Maharshi Pandya, Nidhi Patel, Nitin Savaliya, Raghawendra Kumar, R D Dixit, A M Kadri, Harsh Bakshi, Chaitanya Joshi, Madhvi Joshi |
| EPI_ISL_476882 | Department of MicroBiology, Government Medical College, Surat | Gujarat Biotechnology Research Centre | Zuber Saiyed, Komal Patel, Labdhi Pandya, Afzal Ansari, Nikha Trivedi, Naresh Chauhan, Summaiya Mullan, Amit gamit, Apurvasinh Puvar, Janvi Raval, Zarna Patel, Monika Gandhi, Pinal Trivedi, Maharshi Pandya, Nidhi Patel, Nitin Savaliya, Raghawendra Kumar, Dinesh Kumar, R D Dixit, A M Kadri, Harsh Bakshi, Chaitanya Joshi, Madhvi Joshi |
| EPI_ISL_476883, EPI_ISL_476884, EPI_ISL_476885, EPI_ISL_476886, EPI_ISL_476887, EPI_ISL_476888, EPI_ISL_476889, EPI_ISL_476890, EPI_ISL_476891, EPI_ISL_476892, EPI_ISL_476893, EPI_ISL_476894, EPI_ISL_476895, EPI_ISL_476896 | | | |
| see above | Defence Research & Development Establishment (DRDE) | Defence Research & Development Establishment (DRDE) | Shashi Sharma, Paban Kumar Dash, Sushil Kumar Sharma, Ambuj Shrivastava, Jyoti S. Kumar |
| EPI_ISL_476897 | University of South Carolina Functional Genomics Core | University of South Carolina Functional Genomics Core | Michael Shtutman |
| EPI_ISL_476898, EPI_ISL_476899 | Alaska State Virology Laboratory | Alaska State Virology Laboratory | Jack Chen, Ph.D. |
| EPI_ISL_476900, EPI_ISL_476901, EPI_ISL_476902, EPI_ISL_476903, EPI_ISL_476904, EPI_ISL_476905, EPI_ISL_476906, EPI_ISL_476907, EPI_ISL_476908, EPI_ISL_476909, EPI_ISL_476910, EPI_ISL_476911, EPI_ISL_476912, EPI_ISL_476913, EPI_ISL_476914, EPI_ISL_476915, EPI_ISL_476916, EPI_ISL_476917, EPI_ISL_476918, EPI_ISL_476919, EPI_ISL_476920, EPI_ISL_476921, EPI_ISL_476922, EPI_ISL_476923, EPI_ISL_476924, EPI_ISL_476925, EPI_ISL_476926, EPI_ISL_476927, EPI_ISL_476928, EPI_ISL_476929, EPI_ISL_476930, EPI_ISL_476931, EPI_ISL_476932, EPI_ISL_476933, EPI_ISL_476934, EPI_ISL_476935, EPI_ISL_476936, EPI_ISL_476937, EPI_ISL_476938, EPI_ISL_476939, EPI_ISL_476940 | | | |
| see above | UW Virology Lab | UW Virology Lab | Pavitra Roychoudhury, Hong Xie, Lasata Shrestha, Amin Addetia, Truong Nguyen, Victoria M Rachleff, Meei-Li Huang, Keith R Jerome, Alexander Greninger |
| EPI_ISL_476941, EPI_ISL_476942, EPI_ISL_476943, EPI_ISL_476944, EPI_ISL_476945, EPI_ISL_476946, EPI_ISL_476947, EPI_ISL_476948, EPI_ISL_476949, EPI_ISL_476950, EPI_ISL_476951, EPI_ISL_476952, EPI_ISL_476953, EPI_ISL_476954, EPI_ISL_476955, EPI_ISL_476956, EPI_ISL_476957, EPI_ISL_476958, EPI_ISL_476959, EPI_ISL_476960, EPI_ISL_476961, EPI_ISL_476962, EPI_ISL_476963, EPI_ISL_476964, EPI_ISL_476965, EPI_ISL_476966, EPI_ISL_476967, EPI_ISL_476968, EPI_ISL_476969, EPI_ISL_476970, EPI_ISL_476971, EPI_ISL_476972, EPI_ISL_476973, EPI_ISL_476974, EPI_ISL_476975, EPI_ISL_476976, EPI_ISL_476977, EPI_ISL_476978, EPI_ISL_476979, EPI_ISL_476980, EPI_ISL_476981, EPI_ISL_476982, EPI_ISL_476983, EPI_ISL_476984, EPI_ISL_476985, EPI_ISL_476986, EPI_ISL_476987, EPI_ISL_476988, EPI_ISL_476989, EPI_ISL_476990, EPI_ISL_476991, EPI_ISL_476992, EPI_ISL_476993, EPI_ISL_476994, EPI_ISL_476995, EPI_ISL_476996, EPI_ISL_476997, EPI_ISL_476998, EPI_ISL_476999, EPI_ISL_477000, EPI_ISL_477001, EPI_ISL_477002, EPI_ISL_477003, EPI_ISL_477004, EPI_ISL_477005, EPI_ISL_477006, EPI_ISL_477007 | | | |
| see above | KU Leuven, Rega Institute, Clinical and Epidemiological Virology | KU Leuven, Rega Institute, Clinical and Epidemiological Virology | Tony Wawina-Bokalanga, Joan Marti-Carerras, Bert Vanmechelen, Piet Maes |
| EPI_ISL_477008, EPI_ISL_477009, EPI_ISL_477010, EPI_ISL_477011, EPI_ISL_477012, EPI_ISL_477013 | University of Debrecen, Department of Medical Microbiology | National Laboratory of Virology, Szentágothai Research Centre | Endre Gábor Tóth, Balázs Somogyi, Brigitta Zana, Eszter Csoma, Ferenc Jakab, Gábor Kemenesi |
| EPI_ISL_477014 | Institute of Microbiology, Universidad San Francisco de Quito | Institute of Microbiology, Universidad San Francisco de Quito | Belen Prado-Vivar, Sully Marquez, Juan Jose Guadalupe, Monica Becerra-Wong, Carla Torres, Bernardo Gutierrez, Francisco Mora, Juan Gaviria, Alejandra Ramones, Franklin Espinoza, Edison Ligña, Jorge Reyes, Patricio Rojas-Silva, Veronica Barragan, Gabriel Trueba, Michelle Grunauer, Paul Cardenas |
| EPI_ISL_477015 | Institute of Microbiology, Universidad San Francisco de Quito | Institute of Microbiology, Universidad San Francisco de Quito | Sully Márquez, Belén Prado-Vivar, Juan José Guadalupe, Monica Becerra-Wong, Carla Torres, Bernardo Gutiérrez, Jorge Luis Velez, Verónica Barragán, Patricio Rojas-Silva, Gabriel Trueba, Michelle Grunauer, Paúl Cárdenas |
| EPI_ISL_477016 | Institute of Microbiology, Universidad San Francisco de Quito | Institute of Microbiology, Universidad San Francisco de Quito | Juan José Guadalupe, Sully Márquez, Belén Prado-Vivar, Monica Becerra-Wong, Carla Torres, Bernardo Gutiérrez, Jorge Luis Velez, Verónica Barragán, Patricio Rojas-Silva, Gabriel Trueba, Michelle Grunauer, Paúl Cárdenas |
| EPI_ISL_477017, EPI_ISL_477018, EPI_ISL_477019, EPI_ISL_477020, EPI_ISL_477021, EPI_ISL_477022, EPI_ISL_477023, EPI_ISL_477024, EPI_ISL_477025, EPI_ISL_477026, EPI_ISL_477027, EPI_ISL_477028, EPI_ISL_477029, EPI_ISL_477030, EPI_ISL_477031, EPI_ISL_477032, EPI_ISL_477033, EPI_ISL_477034, EPI_ISL_477035, EPI_ISL_477036, EPI_ISL_477037, EPI_ISL_477038, EPI_ISL_477039, EPI_ISL_477040, EPI_ISL_477041, EPI_ISL_477042, EPI_ISL_477043, EPI_ISL_477044, EPI_ISL_477045, EPI_ISL_477046, EPI_ISL_477047, EPI_ISL_477048, EPI_ISL_477049, EPI_ISL_477050, EPI_ISL_477051, EPI_ISL_477052, EPI_ISL_477053, EPI_ISL_477054, EPI_ISL_477055, EPI_ISL_477056, EPI_ISL_477057, EPI_ISL_477058, EPI_ISL_477059, EPI_ISL_477060, EPI_ISL_477061, EPI_ISL_477062, EPI_ISL_477063, EPI_ISL_477064, EPI_ISL_477065, EPI_ISL_477066, EPI_ISL_477067, EPI_ISL_477068, EPI_ISL_477069, EPI_ISL_477070, EPI_ISL_477071, EPI_ISL_477072, EPI_ISL_477073, EPI_ISL_477074, EPI_ISL_477075, EPI_ISL_477076, EPI_ISL_477077, EPI_ISL_477078, EPI_ISL_477079, EPI_ISL_477080, EPI_ISL_477081, EPI_ISL_477082, EPI_ISL_477083, EPI_ISL_477084, EPI_ISL_477085, EPI_ISL_477086, EPI_ISL_477087, EPI_ISL_477088, EPI_ISL_477089, EPI_ISL_477090, EPI_ISL_477091, EPI_ISL_477092, EPI_ISL_477093, EPI_ISL_477094, EPI_ISL_477095, EPI_ISL_477096, EPI_ISL_477097, EPI_ISL_477098, EPI_ISL_477099, EPI_ISL_477100, EPI_ISL_477101, EPI_ISL_477102, EPI_ISL_477103, EPI_ISL_477104, EPI_ISL_477105, EPI_ISL_477106, EPI_ISL_477107, EPI_ISL_477108, EPI_ISL_477109, EPI_ISL_477110, EPI_ISL_477111, EPI_ISL_477112, EPI_ISL_477113, EPI_ISL_477114, EPI_ISL_477115, EPI_ISL_477116, EPI_ISL_477117, EPI_ISL_477118, EPI_ISL_477119 | | | |
| see above | BCCDC Public Health Laboratory | BCCDC Public Health Laboratory | Richard Harrigan, Hope Lapointe, Jinny Choi, Kimia Kamelian, John Tyson,Terry Snutch, Linda Hoang, Inna Sekirov, Paul Levett, Mel Krajden, Natalie Prystajecky |
| EPI_ISL_477125, EPI_ISL_477126, EPI_ISL_477127, EPI_ISL_477128, EPI_ISL_477129, EPI_ISL_477130, EPI_ISL_477131, EPI_ISL_477132, EPI_ISL_477133, EPI_ISL_477134, EPI_ISL_477135, EPI_ISL_477136, EPI_ISL_477137, EPI_ISL_477138, EPI_ISL_477139, EPI_ISL_477140 | | | |
| see above | Child Health Research Foundation | Child Health Research Foundation | Senjuti Saha, Md Saiful Islam Sajib, Roly Malaker, Md Hafizur Rahman, Afroza Akter Tanni, Syed Muktadir Al Sium, Maksuda Islam, Samir K Saha |
| EPI_ISL_477141, EPI_ISL_477142, EPI_ISL_477143, EPI_ISL_477144, EPI_ISL_477145, EPI_ISL_477146, EPI_ISL_477147, EPI_ISL_477148, EPI_ISL_477149, EPI_ISL_477150, EPI_ISL_477151, EPI_ISL_477152, EPI_ISL_477153, EPI_ISL_477154, EPI_ISL_477155, EPI_ISL_477156, EPI_ISL_477157, EPI_ISL_477158, EPI_ISL_477159 | | | |
| see above | Institut Pasteur Dakar | Institut Pasteur de Dakar | Ndongo Dia, Moussa Moise Diagne, Mamadou Diop, Mamadou Malado Jallow, Marie Henriette Dior Ndione, Safietou Sankhe, Ousmane Faye, Amadou Alpha Sall. |
| EPI_ISL_477160 | Laboratory of Dr. John Lednicky | University of Florida | John A. Lednicky, Chang-Yu Wu, and John Glenn Morris, Jr. |
| EPI_ISL_477161 | Egyptian National Cancer Institute (ENCI) | Egyptian National Cancer Institute (ENCI) | Zekri, Abdel Rahman N, Amer,K.E., Ahmed,O.S., Soliman,H.K., Hafez,M.M., Bahnassy,A.A., Abdelhamid,W., Gad,A., Ali,M., Hassan,W., Samir,M., Raouf,A., Hamdy,M.S., Soliman,M.S., Elsissy,M.H., Elkhateeb,S.M., Ezzelarab,M.H., Abouelhoda, Mohamed |
| EPI_ISL_477163 | Laboratory of Dr. John Lednicky | University of Florida | John A. Lednicky, Maha A. Elbadry, Kuttichantran Subramaniam, Thomas B. Waltzek, John Glenn Morris, Jr. |
| EPI_ISL_477164 | Department of Virology, Public Health Laboratories Division, National Institute of Health | Department of Virology, Public Health Laboratories Division, National Institute of Health | Nazish Badar, Aamer Ikram, Muhammad Salman, Hamza Ahmed Mirza, Abdul Ahad, Yasir Arshad, Massab Umair |
| EPI_ISL_477165, EPI_ISL_477166, EPI_ISL_477167 | Department of Virology, Public Health Laboratories Division, National Institute of Health | Department of Virology, Public Health Laboratories Division, National Institute of Health | Nazish Badar,Aamer Ikram, Muhammad Salman, Massab Umair, Hamza Ahmed Mirza, Abdul Ahad, Yasir Arshad |
| EPI_ISL_477168 | Institute for Stem Cell Science and Regenerative Medicine | National Centre for Biological Sciences | Farhan Ali, Vanessa Molin Paynter, Srikar Krishna, Mohak Sharda, Shah-e-Jahan Gulzar, Awadhesh Pandit, Varadha Sundarmurthy, Uma Ramakrishnan, Dasaradhi Palakodeti, Aswin Seshasayee |
| EPI_ISL_477169 | Department for Virology, Molecular Biology and Genome Research, R. G. Lugar Center for Public Health Research, National Center for Disease Control and Public Health (NCDC) of Georgia. | Department for Virology, Molecular Biology and Genome Research, R. G. Lugar Center for Public Health Research, National Center for Disease Control and Public Health (NCDC) of Georgia. | Tata Imnadze, Giorgi Tomashvili, Meri Pantsulaia, Gvantsa Brachveli, Gvantsa Chanturia, Ann Machablishvili, Nato Kotaria, Marine Murtskhvaladze, Lela Sabadze, Mari Gavashelidze, Ana Papkiauri, Tamar Jashiashvili, Tea Tevdoradze, Ketevan Sidamonidze, Ekaterine Khmaladze, Ekaterine Zhghenti, Roena Sukhiashvili, Mariam Zakalashvili, Lela Urushadze, Magda Dgebuadze, Davit Tsaguria, Ekaterine Zangaladze, Nino Berishvili, Adam Kotorashvili, Maia Alkhazashvili, Irma Burjanadze, Anna Kasradze, Khatuna Zakhashvili, Paata Imnadze, Amiran Gamkrelidze. |
| EPI_ISL_477170 | Department of Laboratory, Medicine Tan Tock Seng Hospital | Department of Laboratory Medicine Tan Tock Seng Hospital | Chen YYC, Zair X, Li C, Tang WY, Maurer-Stroh S, Barkham TMS, Nagarajan N, Sessions OM |
| EPI_ISL_477171 | Department of Laboratory, Medicine Tan Tock Seng Hospital | Department of Laboratory, Medicine Tan Tock Seng Hospital | Chen YYC, Zair X, Li C, Tang WY, Maurer-Stroh S, Barkham TMS, Nagarajan N, Sessions OM |
| EPI_ISL_477172, EPI_ISL_477174, EPI_ISL_477175, EPI_ISL_477177, EPI_ISL_477178, EPI_ISL_477180, EPI_ISL_477182 | Department of Laboratory Medicine Tan Tock Seng Hospital | Department of Laboratory Medicine Tan Tock Seng Hospital | Chen YYC, Zair X, Li C, Tang WY, Maurer-Stroh S, Barkham TMS, Nagarajan N, Sessions OM |
| EPI_ISL_477183 | Department of MicroBiology, Government Medical College, Surat | Gujarat Biotechnology Research Centre | Monika Gandhi, Pinal Trivedi, Maharshi Pandya, Nidhi Patel, Nitin Savaliya, Raghawendra Kumar, Dinesh Kumar, Zuber Saiyed, Komal Patel, Labdhi Pandya, Afzal Ansari, Nikha Trivedi, Naresh Chauhan, Summaiya Mullan, Amit gamit, Apurvasinh Puvar, Janvi Raval, Zarna Patel, R D Dixit, A M Kadri, Harsh Bakshi, Chaitanya Joshi, Madhvi Joshi |
| EPI_ISL_477184, EPI_ISL_477187, EPI_ISL_477188, EPI_ISL_477189, EPI_ISL_477190, EPI_ISL_477191, EPI_ISL_477192 | Department of Laboratory Medicine Tan Tock Seng Hospital | Department of Laboratory Medicine Tan Tock Seng Hospital | Chen YYC, Zair X, Li C, Tang WY, Maurer-Stroh S, Barkham TMS, Nagarajan N, Sessions OM |

| | | | |
|---|---|---|---|
| EPI_ISL_477193, EPI_ISL_477194 | Istituto Zooprofilattico Sperimentale Puglia e Basilicata; | Beaconlab (Bioinformatics, Evolution and Comparative Genomics lab), Dept of Biosciences, University on Mila | Parisi A.,Pesole G., Manzari C., Chiara M. |
| EPI_ISL_477195, EPI_ISL_477196, EPI_ISL_477197, EPI_ISL_477198, EPI_ISL_477199, EPI_ISL_477200, EPI_ISL_477201 | Istituto Zooprofilattico Sperimentale Puglia e Basilicata; | Beaconlab (Bioinformatics, Evolution and Comparative Genomics lab), Dept of Biosciences, University on Milan | Parisi A.,Pesole G., Manzari C., Chiara M. |
| EPI_ISL_477202, EPI_ISL_477203 | Istituto Zooprofilattico Sperimentale Puglia e Basilicata; | Beaconlab (Bioinformatics, Evolution and Comparative Genomics lab), Dept of Biosciences, University on Mila | Parisi A.,Pesole G., Manzari C., Chiara M. |
| EPI_ISL_477204 | Prof. Massimo Zollo CEINGE TASK-FORCE COVID19 - Regione Campania | Prof. Massimo Zollo CEINGE TASK-FORCE COVID19 - Regione Campania | Veronica Ferrucci1,2, Dae young Kong8, Fatemeh asadzadeh1,2, Laura Marrone1,2, Roberto Siciliano1,2, Rino Cerino3, Giovanna Fusco3, Marika Comegna1,2, Angelo Boccia2, Maurizio Viscardi3, Giorgia Borriello3, Sergio Brandi3, Claudia Tiberio4, Luigi Atripaldi4, Giovanni Paolella1,2, Giuseppe Castaldo1,2, Stefano Pascarella4, Martina Bianchi4, Lorenzo Chiariotti1,2, Jae Myun Lee5, Jae Ho Jung6, Kyong Seop Yun7, Hong Yeoul Kim 7,8* and Massimo Zollo1,2* 1 CEINGE Biotecnologie Avanzate, Naples, Italia 2 Dipartimento di Medicina Molecolare e Biotecnologie Mediche DMMBM University of Naples Federico II, Italia 3 Istituto Zooprofilattico Sperimentale del Mezzogiorno, Naples, Italia 4 - U.O.C. di Patologia Clinica Ospedale D. Cotugno, Azienda Sanitaria Ospedali dei Colli, Naples, Italy. 5 Università La Sapienza di Roma, Italia 6 Department of Microbiology, Yonsei University College of Medicine, Seoul, Korea 7 Department of Surgery, Yonsei University College of Medicine, Seoul, Korea 8 Haim bio co., Ltd, , Indust |
| EPI_ISL_477205, EPI_ISL_477206, EPI_ISL_477207, EPI_ISL_477208, EPI_ISL_477209, EPI_ISL_477210, EPI_ISL_477211, EPI_ISL_477212, EPI_ISL_477213, EPI_ISL_477214, EPI_ISL_477215, EPI_ISL_477216, EPI_ISL_477217, EPI_ISL_477218, EPI_ISL_477219, EPI_ISL_477220, EPI_ISL_477221, EPI_ISL_477222, EPI_ISL_477223, EPI_ISL_477224, EPI_ISL_477225, EPI_ISL_477226, EPI_ISL_477227, EPI_ISL_477228, EPI_ISL_477229, EPI_ISL_477230, EPI_ISL_477231, EPI_ISL_477232, EPI_ISL_477233, EPI_ISL_477234, EPI_ISL_477235, EPI_ISL_477236, EPI_ISL_477237, EPI_ISL_477238, EPI_ISL_477239, EPI_ISL_477240, EPI_ISL_477241, EPI_ISL_477242, EPI_ISL_477243, EPI_ISL_477244, EPI_ISL_477245, EPI_ISL_477246, EPI_ISL_477247, EPI_ISL_477248, EPI_ISL_477249, EPI_ISL_477250, EPI_ISL_477251, EPI_ISL_477252, EPI_ISL_477253, EPI_ISL_477254, EPI_ISL_477255, EPI_ISL_477256, EPI_ISL_477257, EPI_ISL_477258, EPI_ISL_477259, EPI_ISL_477260, EPI_ISL_477261, EPI_ISL_477262, EPI_ISL_477263 | | | |
| see above | Institute for Stem Cell Science and Regenerative Medicine | National Centre for Biological Sciences | Farhan Ali, Vanessa Molin Paynter, Srikar Krishna, Mohak Sharda, Shah-e-Jahan Gulzar, Awadhesh Pandit, Varadha Sundarmurthy, Uma Ramakrishnan, Dasaradhi Palakodeti, Aswin Seshasayee |
| EPI_ISL_477272, EPI_ISL_477273, EPI_ISL_477274, EPI_ISL_477275, EPI_ISL_477276 | Mayo Clinic & Mayo Clinic Laboratories | Minnesota Department of Health, Public Health Laboratory | Matt Plumb, Jacob Garfin, Kelly Pung, and Xiong Wang |
| EPI_ISL_477277, EPI_ISL_477278, EPI_ISL_477279, EPI_ISL_477280, EPI_ISL_477281, EPI_ISL_477282, EPI_ISL_477283, EPI_ISL_477284, EPI_ISL_477285, EPI_ISL_477286, EPI_ISL_477287, EPI_ISL_477288, EPI_ISL_477289, EPI_ISL_477290 | | | |
| see above | M Health Fairview | Minnesota Department of Health, Public Health Laboratory | Matt Plumb, Jacob Garfin, Kelly Pung, and Xiong Wang |
| EPI_ISL_477291, EPI_ISL_477292, EPI_ISL_477293, EPI_ISL_477294, EPI_ISL_477295, EPI_ISL_477296, EPI_ISL_477297, EPI_ISL_477298, EPI_ISL_477299, EPI_ISL_477300, EPI_ISL_477301, EPI_ISL_477302, EPI_ISL_477303, EPI_ISL_477304, EPI_ISL_477305, EPI_ISL_477306, EPI_ISL_477307, EPI_ISL_477308, EPI_ISL_477309, EPI_ISL_477310, EPI_ISL_477311, EPI_ISL_477312 | | | |
| see above | Mayo Clinic & Mayo Clinic Laboratories | Minnesota Department of Health, Public Health Laboratory | Matt Plumb, Jacob Garfin, Kelly Pung, and Xiong Wang |
| EPI_ISL_477615, EPI_ISL_477616, EPI_ISL_477617, EPI_ISL_477618, EPI_ISL_477619, EPI_ISL_477620, EPI_ISL_477621, EPI_ISL_477622, EPI_ISL_477623, EPI_ISL_477624, EPI_ISL_477625 | | | |
| see above | University of Szeged, Institute of Clinical Microbiology | National Laboratory of Virology, Szentágothai Research Centre | Endre Gábor Tóth, Balázs Somogyi, Brigitta Zana, Terhes Gabriella, Ferenc Jakab, Gábor Kemenesi |
| EPI_ISL_477631, EPI_ISL_477632, EPI_ISL_477633, EPI_ISL_477634, EPI_ISL_477635, EPI_ISL_477636, EPI_ISL_477637, EPI_ISL_477638, EPI_ISL_477639, EPI_ISL_477640, EPI_ISL_477641, EPI_ISL_477642, EPI_ISL_477643, EPI_ISL_477644, EPI_ISL_477645, EPI_ISL_477646, EPI_ISL_477647, EPI_ISL_477648, EPI_ISL_477649, EPI_ISL_477650, EPI_ISL_477651, EPI_ISL_477652, EPI_ISL_477653, EPI_ISL_477654, EPI_ISL_477655, EPI_ISL_477656, EPI_ISL_477657, EPI_ISL_477658, EPI_ISL_477659, EPI_ISL_477660, EPI_ISL_477661, EPI_ISL_477662, EPI_ISL_477663, EPI_ISL_477664, EPI_ISL_477665, EPI_ISL_477666, EPI_ISL_477667, EPI_ISL_477668, EPI_ISL_477669, EPI_ISL_477670, EPI_ISL_477671, EPI_ISL_477672 | | | |
| see above | Virginia DCLS | Virginia DCLS | Virginia DCLS |
| EPI_ISL_477673, EPI_ISL_477674, EPI_ISL_477675, EPI_ISL_477676, EPI_ISL_477677, EPI_ISL_477678, EPI_ISL_477679, EPI_ISL_477680, EPI_ISL_477681, EPI_ISL_477682, EPI_ISL_477683, EPI_ISL_477684, EPI_ISL_477685, EPI_ISL_477686, EPI_ISL_477687, EPI_ISL_477688, EPI_ISL_477689, EPI_ISL_477690, EPI_ISL_477691, EPI_ISL_477692, EPI_ISL_477693, EPI_ISL_477694, EPI_ISL_477695, EPI_ISL_477696, EPI_ISL_477697, EPI_ISL_477698, EPI_ISL_477699, EPI_ISL_477700, EPI_ISL_477701, EPI_ISL_477702, EPI_ISL_477703, EPI_ISL_477704, EPI_ISL_477705, EPI_ISL_477706, EPI_ISL_477707, EPI_ISL_477708, EPI_ISL_477709, EPI_ISL_477710, EPI_ISL_477711, EPI_ISL_477712, EPI_ISL_477713, EPI_ISL_477714, EPI_ISL_477715, EPI_ISL_477716, EPI_ISL_477717, EPI_ISL_477718, EPI_ISL_477719, EPI_ISL_477720, EPI_ISL_477721, EPI_ISL_477722, EPI_ISL_477723, EPI_ISL_477724 | | | |
| see above | UW Virology Lab | UW Virology Lab | Pavitra Roychoudhury, Hong Xie, Lasata Shrestha, Amin Addetia, Truong Nguyen, Victoria M Rachleff, Meei-Li Huang, Keith R Jerome, Alexander Greninger |
| EPI_ISL_477727, EPI_ISL_477728, EPI_ISL_477729, EPI_ISL_477730, EPI_ISL_477731, EPI_ISL_477732, EPI_ISL_477733, EPI_ISL_477734, EPI_ISL_477735, EPI_ISL_477736, EPI_ISL_477737, EPI_ISL_477738, EPI_ISL_477739, EPI_ISL_477740, EPI_ISL_477741, EPI_ISL_477742, EPI_ISL_477743, EPI_ISL_477744, EPI_ISL_477745, EPI_ISL_477746, EPI_ISL_477747, EPI_ISL_477748, EPI_ISL_477749, EPI_ISL_477750, EPI_ISL_477751, EPI_ISL_477752, EPI_ISL_477753, EPI_ISL_477754, EPI_ISL_477755, EPI_ISL_477756, EPI_ISL_477757, EPI_ISL_477758, EPI_ISL_477759, EPI_ISL_477760, EPI_ISL_477761, EPI_ISL_477762, EPI_ISL_477763, EPI_ISL_477764, EPI_ISL_477765, EPI_ISL_477766, EPI_ISL_477767, EPI_ISL_477768, EPI_ISL_477769, EPI_ISL_477770, EPI_ISL_477771, EPI_ISL_477772, EPI_ISL_477773, EPI_ISL_477774, EPI_ISL_477775, EPI_ISL_477776, EPI_ISL_477777, EPI_ISL_477778, EPI_ISL_477779, EPI_ISL_477780, EPI_ISL_477781, EPI_ISL_477782 | | | |
| see above | University of Birmingham | COVID-19 Genomics UK (COG-UK) Consortium | Institute of Microbiology, University of Birmingham: Claire McMurray, Joanne Stockton, Samuel Nicholls, Radoslaw Poplawski, Will Rowe, Josh Quick, Nicholas Loman. University of Birmingham Testing Laboratory: Celina M Whalley, Andrew Bosworth, Charlotte Poxon, Kasun Wanigasooriya, Oliver Pickles, Mike Kidd, Alex Richter, Andrew D Beggs PHE Heartlands Lab: Husam Osman, Andrew Bosworth. Queen Elizabeth Hospital: Anna Casey |
| EPI_ISL_477783, EPI_ISL_477784, EPI_ISL_477785, EPI_ISL_477786, EPI_ISL_477787, EPI_ISL_477788, EPI_ISL_477789, EPI_ISL_477790, EPI_ISL_477791, EPI_ISL_477792, EPI_ISL_477793, EPI_ISL_477794, EPI_ISL_477795, EPI_ISL_477796, EPI_ISL_477797, EPI_ISL_477798, EPI_ISL_477799, EPI_ISL_477800, EPI_ISL_477801, EPI_ISL_477802, EPI_ISL_477803, EPI_ISL_477804, EPI_ISL_477805, EPI_ISL_477806, EPI_ISL_477807, EPI_ISL_477808, EPI_ISL_477809, EPI_ISL_477810, EPI_ISL_477811, EPI_ISL_477812, EPI_ISL_477813, EPI_ISL_477814, EPI_ISL_477815, EPI_ISL_477816, EPI_ISL_477817, EPI_ISL_477818, EPI_ISL_477819 | | | |
| see above | Department of Pathology, University of Cambridge | COVID-19 Genomics UK (COG-UK) Consortium | Luke W Meredith, M. Estée Török, Myra Hosmillo, William L. Hamilton, Martin D. Curran, Theresa Feltwell, Grant Hall, Anna Yakovleva, Fahad A Khokhar, Charlotte J. Houldcroft, Laura G Caller, Aminu S. Jahun, Sarah L. Caddy, Yasmin Chaudhry, Malte Pinckert, Ian Goodfellow |
| EPI_ISL_477820, EPI_ISL_477821, EPI_ISL_477822, EPI_ISL_477823, EPI_ISL_477824, EPI_ISL_477825, EPI_ISL_477826, EPI_ISL_477827, EPI_ISL_477828, EPI_ISL_477829, EPI_ISL_477830, EPI_ISL_477831, EPI_ISL_477832, EPI_ISL_477833, EPI_ISL_477834, EPI_ISL_477835, EPI_ISL_477836, EPI_ISL_477837, EPI_ISL_477838, EPI_ISL_477839, EPI_ISL_477840, EPI_ISL_477841, EPI_ISL_477842, EPI_ISL_477843, EPI_ISL_477844, EPI_ISL_477845, EPI_ISL_477846, EPI_ISL_477847, EPI_ISL_477848, EPI_ISL_477849, EPI_ISL_477850, EPI_ISL_477851, EPI_ISL_477852, EPI_ISL_477853, EPI_ISL_477854, EPI_ISL_477855, EPI_ISL_477856, EPI_ISL_477857, EPI_ISL_477858, EPI_ISL_477859, EPI_ISL_477860, EPI_ISL_477861, EPI_ISL_477862, EPI_ISL_477863, EPI_ISL_477864, EPI_ISL_477865, EPI_ISL_477866, EPI_ISL_477867, EPI_ISL_477868, EPI_ISL_477869, EPI_ISL_477870, EPI_ISL_477871, EPI_ISL_477872, EPI_ISL_477873, EPI_ISL_477874, EPI_ISL_477875, EPI_ISL_477876, EPI_ISL_477877, EPI_ISL_477878, EPI_ISL_477879, EPI_ISL_477880, EPI_ISL_477881, EPI_ISL_477882, EPI_ISL_477883, EPI_ISL_477884, EPI_ISL_477885, EPI_ISL_477886, EPI_ISL_477887, EPI_ISL_477888, EPI_ISL_477889, EPI_ISL_477890, EPI_ISL_477891, EPI_ISL_477892, EPI_ISL_477893, EPI_ISL_477894, EPI_ISL_477895, EPI_ISL_477896, EPI_ISL_477897, EPI_ISL_477898, EPI_ISL_477899, EPI_ISL_477900, EPI_ISL_477901, EPI_ISL_477902, EPI_ISL_477903, EPI_ISL_477904, EPI_ISL_477905, EPI_ISL_477906, EPI_ISL_477907, EPI_ISL_477908, EPI_ISL_477909, EPI_ISL_477910, EPI_ISL_477911, EPI_ISL_477912, EPI_ISL_477913, EPI_ISL_477914, EPI_ISL_477915, EPI_ISL_477916, EPI_ISL_477917, EPI_ISL_477918, EPI_ISL_477919, EPI_ISL_477920, EPI_ISL_477921, EPI_ISL_477922, EPI_ISL_477923, EPI_ISL_477924, EPI_ISL_477925, EPI_ISL_477926, EPI_ISL_477927, EPI_ISL_477928, EPI_ISL_477929, EPI_ISL_477930, EPI_ISL_477931, EPI_ISL_477932, EPI_ISL_477933, EPI_ISL_477934, EPI_ISL_477935, EPI_ISL_477936, EPI_ISL_477937, EPI_ISL_477938, EPI_ISL_477939, EPI_ISL_477940, EPI_ISL_477941, EPI_ISL_477942, EPI_ISL_477943, EPI_ISL_477944, EPI_ISL_477945, EPI_ISL_477946, EPI_ISL_477947, EPI_ISL_477948, EPI_ISL_477949, EPI_ISL_477950, EPI_ISL_477951, EPI_ISL_477952, EPI_ISL_477953, EPI_ISL_477954, EPI_ISL_477955, EPI_ISL_477956, EPI_ISL_477957, EPI_ISL_477958, EPI_ISL_477959, EPI_ISL_477960, EPI_ISL_477961, EPI_ISL_477962, EPI_ISL_477963, EPI_ISL_477964, EPI_ISL_477965, EPI_ISL_477966, EPI_ISL_477967, EPI_ISL_477968, EPI_ISL_477969, EPI_ISL_477970, EPI_ISL_477971, EPI_ISL_477972, EPI_ISL_477973, EPI_ISL_477974, EPI_ISL_477975, EPI_ISL_477976, EPI_ISL_477977, EPI_ISL_477978, EPI_ISL_477979, EPI_ISL_477980, EPI_ISL_477981, EPI_ISL_477982, EPI_ISL_477983, EPI_ISL_477984, EPI_ISL_477985, EPI_ISL_477986, EPI_ISL_477987, EPI_ISL_477988, EPI_ISL_477989, EPI_ISL_477990, EPI_ISL_477991, EPI_ISL_477992, EPI_ISL_477993, EPI_ISL_477994, EPI_ISL_477995, EPI_ISL_477996, EPI_ISL_477997, EPI_ISL_477998, EPI_ISL_477999, EPI_ISL_478000, EPI_ISL_478001, EPI_ISL_478002, EPI_ISL_478003, EPI_ISL_478004, EPI_ISL_478005, EPI_ISL_478006, EPI_ISL_478007, EPI_ISL_478008, EPI_ISL_478009, EPI_ISL_478010, EPI_ISL_478011, EPI_ISL_478012, EPI_ISL_478013, EPI_ISL_478014, EPI_ISL_478015, EPI_ISL_478016, EPI_ISL_478017, EPI_ISL_478018, EPI_ISL_478019, EPI_ISL_478020, EPI_ISL_478021, EPI_ISL_478022, EPI_ISL_478023, EPI_ISL_478024, EPI_ISL_478025, EPI_ISL_478026, EPI_ISL_478027, EPI_ISL_478028, EPI_ISL_478029, EPI_ISL_478030, EPI_ISL_478031, EPI_ISL_478032, EPI_ISL_478033, EPI_ISL_478034, EPI_ISL_478035, EPI_ISL_478036, EPI_ISL_478037, EPI_ISL_478038, EPI_ISL_478039, EPI_ISL_478040, EPI_ISL_478041, EPI_ISL_478042, EPI_ISL_478043, EPI_ISL_478044, EPI_ISL_478045, EPI_ISL_478046, EPI_ISL_478047, EPI_ISL_478048, EPI_ISL_478049, EPI_ISL_478050, EPI_ISL_478051, EPI_ISL_478052, EPI_ISL_478053, EPI_ISL_478054, EPI_ISL_478055, EPI_ISL_478056, EPI_ISL_478057, EPI_ISL_478058, EPI_ISL_478059, EPI_ISL_478060, EPI_ISL_478061, EPI_ISL_478062, EPI_ISL_478063, EPI_ISL_478064, EPI_ISL_478065, EPI_ISL_478066, EPI_ISL_478067, EPI_ISL_478068, EPI_ISL_478069, EPI_ISL_478070, EPI_ISL_478071, EPI_ISL_478072, EPI_ISL_478073, EPI_ISL_478074, EPI_ISL_478075, EPI_ISL_478076, EPI_ISL_478077, EPI_ISL_478078, EPI_ISL_478079, EPI_ISL_478080, EPI_ISL_478081, EPI_ISL_478082, EPI_ISL_478083, EPI_ISL_478084, EPI_ISL_478085, EPI_ISL_478086, EPI_ISL_478087, EPI_ISL_478088, EPI_ISL_478089, EPI_ISL_478090, EPI_ISL_478091, EPI_ISL_478092, EPI_ISL_478093, EPI_ISL_478094, EPI_ISL_478095, EPI_ISL_478096, EPI_ISL_478097, EPI_ISL_478098, EPI_ISL_478099, EPI_ISL_478100, EPI_ISL_478101, EPI_ISL_478102, EPI_ISL_478103, EPI_ISL_478104, EPI_ISL_478105, EPI_ISL_478106, EPI_ISL_478107, EPI_ISL_478108, EPI_ISL_478109, EPI_ISL_478110, EPI_ISL_478111, EPI_ISL_478112, EPI_ISL_478113, EPI_ISL_478114, EPI_ISL_478115, EPI_ISL_478116, EPI_ISL_478117, EPI_ISL_478118, EPI_ISL_478119, EPI_ISL_478120, EPI_ISL_478121, EPI_ISL_478122, EPI_ISL_478123, EPI_ISL_478124, EPI_ISL_478125, EPI_ISL_478126, EPI_ISL_478127, EPI_ISL_478128, EPI_ISL_478129, EPI_ISL_478130, EPI_ISL_478131, EPI_ISL_478132, EPI_ISL_478133, EPI_ISL_478134, EPI_ISL_478135, EPI_ISL_478136, EPI_ISL_478137, EPI_ISL_478138, EPI_ISL_478139, EPI_ISL_478140, EPI_ISL_478141, EPI_ISL_478142, EPI_ISL_478143, EPI_ISL_478144, EPI_ISL_478145, EPI_ISL_478146, EPI_ISL_478147, EPI_ISL_478148, EPI_ISL_478149, EPI_ISL_478150, EPI_ISL_478151, EPI_ISL_478152, EPI_ISL_478153, EPI_ISL_478154, EPI_ISL_478155, EPI_ISL_478156, EPI_ISL_478157, EPI_ISL_478158, EPI_ISL_478159, EPI_ISL_478160, EPI_ISL_478161, EPI_ISL_478162 | | | |
| see above | West of Scotland Specialist Virology Centre, NHSGGC / MRC-University of Glasgow Centre for Virus Research | COVID-19 Genomics UK (COG-UK) Consortium | Ana da Silva Filipe, Natasha Johnson, Kathy Smollett, Daniel Mair, Stephen Carmichael, Lily Tong, Jenna Nichols, Elihu Aranday-Cortes, Kirstyn Brunker, Yasmin Parr, Alice Broos, Kyriaki Nomikou; Sarah McDonald, Marc Niebel, Patawee Asamaphan; Richard Orton, Joseph Hughes, Sreenu Vattipally, David L Robertson; Alasdair MacLean, Rory Gunson; Kathy Li, Natasha Jesudason, Rajiv Shah, James Shepherd, Antonia Ho, Emma Thomson |
| EPI_ISL_478163, EPI_ISL_478164, EPI_ISL_478165, EPI_ISL_478166, EPI_ISL_478167, EPI_ISL_478168, EPI_ISL_478169, EPI_ISL_478170, EPI_ISL_478171, EPI_ISL_478172, EPI_ISL_478173, EPI_ISL_478174, EPI_ISL_478175, EPI_ISL_478176, EPI_ISL_478177, EPI_ISL_478178, EPI_ISL_478179, EPI_ISL_478180, EPI_ISL_478181, EPI_ISL_478182, EPI_ISL_478183, EPI_ISL_478184, EPI_ISL_478185, EPI_ISL_478186, EPI_ISL_478187, EPI_ISL_478188, EPI_ISL_478189, EPI_ISL_478190, EPI_ISL_478191, EPI_ISL_478192, EPI_ISL_478193, EPI_ISL_478194, EPI_ISL_478195, EPI_ISL_478196, EPI_ISL_478197, EPI_ISL_478198, EPI_ISL_478199, EPI_ISL_478200, EPI_ISL_478201, EPI_ISL_478202, EPI_ISL_478203, EPI_ISL_478204, EPI_ISL_478205, EPI_ISL_478206, EPI_ISL_478207, EPI_ISL_478208, EPI_ISL_478209, EPI_ISL_478210, EPI_ISL_478211, EPI_ISL_478212, EPI_ISL_478213, EPI_ISL_478214, EPI_ISL_478215, EPI_ISL_478216, EPI_ISL_478217, EPI_ISL_478218, EPI_ISL_478219, EPI_ISL_478220, EPI_ISL_478221, EPI_ISL_478222, EPI_ISL_478223, EPI_ISL_478224, EPI_ISL_478225, EPI_ISL_478226, EPI_ISL_478227, EPI_ISL_478228, EPI_ISL_478229, EPI_ISL_478230, EPI_ISL_478231, EPI_ISL_478232, EPI_ISL_478233, EPI_ISL_478234, EPI_ISL_478235, EPI_ISL_478236, EPI_ISL_478237, EPI_ISL_478238, EPI_ISL_478239, EPI_ISL_478240, EPI_ISL_478241, EPI_ISL_478242, EPI_ISL_478243, EPI_ISL_478244, EPI_ISL_478245, EPI_ISL_478246, EPI_ISL_478247, EPI_ISL_478248, EPI_ISL_478249, EPI_ISL_478250, EPI_ISL_478251, EPI_ISL_478252, EPI_ISL_478253, EPI_ISL_478254, EPI_ISL_478255, EPI_ISL_478256, EPI_ISL_478257, EPI_ISL_478258, EPI_ISL_478259, EPI_ISL_478260, EPI_ISL_478261, EPI_ISL_478262, EPI_ISL_478263, EPI_ISL_478264, EPI_ISL_478265, EPI_ISL_478266, EPI_ISL_478267, EPI_ISL_478268, EPI_ISL_478269, EPI_ISL_478270, EPI_ISL_478271, EPI_ISL_478272, EPI_ISL_478273 | | | |
| see above | Virology Department, Royal Infirmary of Edinburgh, NHS Lothian / School of Biological Sciences, University of Edinburgh / Institute of Genetics and Molecular Medicine, University of Edinburgh | COVID-19 Genomics UK (COG-UK) Consortium | McHugh M, Dewar R, Rooke S, Gallagher M, Balcaza C, O'Toole Á, Scher E, Hill V, McCrone JT, Colquhoun R, Yu X, Jackson B, Rambaut A, Williams TC, Templeton K |
| EPI_ISL_478274, EPI_ISL_478275, EPI_ISL_478276, EPI_ISL_478277, EPI_ISL_478278, EPI_ISL_478279, EPI_ISL_478280, EPI_ISL_478281, EPI_ISL_478282, EPI_ISL_478283, EPI_ISL_478284, EPI_ISL_478285, EPI_ISL_478286, EPI_ISL_478287, EPI_ISL_478288 | | | |
| see above | University of Exeter | COVID-19 Genomics UK (COG-UK) Consortium | Ben Temperton,Aaron Jeffries,Michelle Michelsen,Joanna Warwick-Dugdale,Audrey Farbos,Robyn Manley,Stephen Michell,Jane Masoli |
| EPI_ISL_478289, EPI_ISL_478290, EPI_ISL_478291, EPI_ISL_478292, EPI_ISL_478293, EPI_ISL_478294, EPI_ISL_478295, EPI_ISL_478296, EPI_ISL_478297, EPI_ISL_478298, EPI_ISL_478299, EPI_ISL_478300, EPI_ISL_478301, EPI_ISL_478302, EPI_ISL_478303, EPI_ISL_478304, EPI_ISL_478305, EPI_ISL_478306, EPI_ISL_478307, EPI_ISL_478308, EPI_ISL_478309, EPI_ISL_478310, EPI_ISL_478311, EPI_ISL_478312, EPI_ISL_478313, EPI_ISL_478314, EPI_ISL_478315, EPI_ISL_478316, EPI_ISL_478317, EPI_ISL_478318, EPI_ISL_478319, EPI_ISL_478320, EPI_ISL_478321, EPI_ISL_478322, EPI_ISL_478323, EPI_ISL_478324, EPI_ISL_478325, EPI_ISL_478326, EPI_ISL_478327, EPI_ISL_478328, EPI_ISL_478329, EPI_ISL_478330, EPI_ISL_478331, EPI_ISL_478332, EPI_ISL_478333, EPI_ISL_478334, EPI_ISL_478335, EPI_ISL_478336, EPI_ISL_478337, EPI_ISL_478338, EPI_ISL_478339, EPI_ISL_478340, EPI_ISL_478341, EPI_ISL_478342, EPI_ISL_478343, EPI_ISL_478344, EPI_ISL_478345, EPI_ISL_478346, EPI_ISL_478347, EPI_ISL_478348, EPI_ISL_478349, EPI_ISL_478350, EPI_ISL_478351, EPI_ISL_478352, EPI_ISL_478353, EPI_ISL_478354, EPI_ISL_478355, EPI_ISL_478356, EPI_ISL_478357, EPI_ISL_478358, EPI_ISL_478359, EPI_ISL_478360, EPI_ISL_478361, EPI_ISL_478362, EPI_ISL_478363, EPI_ISL_478364, EPI_ISL_478365, EPI_ISL_478366, EPI_ISL_478367, EPI_ISL_478368, EPI_ISL_478369, EPI_ISL_478370, EPI_ISL_478371 | | | |
| see above | University Hospitals Of Leicester NHS Trust and DeepSeq Nottingham | COVID-19 Genomics UK (COG-UK) Consortium | Christopher Holmes, Paul Bird, Thomas Helmer, Karlie Fallon, Julian Tang, Jonathan Ball, Patrick McClure, Joeseph Chappell, Nadine Holmes, Matthew Carlisle, Christopher Moore, Fei Sang, Johnny Debebe, Victoria Wright, Matthew Loose |
| EPI_ISL_478372, EPI_ISL_478373, EPI_ISL_478374, EPI_ISL_478375, EPI_ISL_478376, EPI_ISL_478377, EPI_ISL_478378, EPI_ISL_478379, EPI_ISL_478380, EPI_ISL_478381, EPI_ISL_478382, EPI_ISL_478383, EPI_ISL_478384, EPI_ISL_478385, EPI_ISL_478386, EPI_ISL_478387, EPI_ISL_478388, EPI_ISL_478389, EPI_ISL_478390, EPI_ISL_478391, EPI_ISL_478392, EPI_ISL_478393, EPI_ISL_478394, EPI_ISL_478395, EPI_ISL_478396, EPI_ISL_478397, EPI_ISL_478398, EPI_ISL_478399, EPI_ISL_478400, EPI_ISL_478401, EPI_ISL_478402, EPI_ISL_478403 | | | |
| see above | Liverpool Clinical Laboratories | COVID-19 Genomics UK (COG-UK) Consortium | Sam Haldenby, Anita Lucaci, Steve Paterson, Julian Hiscox, Alistair Darby, M Almsaud, A Alrezaihi, Muhannad Alruwaili, Stuart D Armstrong, Jones Benjamin, Eleanor G Bentley, Anu Chawla, Jordan J Clark, Angela Cowell, Richard Eccles, Isabel García-Dorival, Matthew Gemmell, Alessandro Gerada, PKF Gilmore, Richard Gregory, Ximeng Han, Catherine Hartley, Margaret Hughes, Miren Iturriza-Gomara, James Johnson, L Luu, Jenifer Manson, Charlotte Nelson, Elaine O'Toole, Cassie Olateju, Rebekah Penrice-Randal , Lucille Rainbow, N.P Randle, Trevor Ian Robinson, Parul Sharma, Ghada T Shawli, James P Stewart, Neil Swainston, Ecaterina Vamos, Joanne Watts, Mark Whitehead |
| EPI_ISL_478404, EPI_ISL_478405, EPI_ISL_478406, EPI_ISL_478407, EPI_ISL_478408, EPI_ISL_478409, EPI_ISL_478410, EPI_ISL_478411, EPI_ISL_478412, EPI_ISL_478413, EPI_ISL_478414, EPI_ISL_478415, EPI_ISL_478416, EPI_ISL_478417, EPI_ISL_478418, EPI_ISL_478419, EPI_ISL_478420, EPI_ISL_478421, EPI_ISL_478422, EPI_ISL_478423, EPI_ISL_478424, EPI_ISL_478425, EPI_ISL_478426, EPI_ISL_478427, EPI_ISL_478428, EPI_ISL_478429, EPI_ISL_478430, EPI_ISL_478431, EPI_ISL_478432, EPI_ISL_478433, EPI_ISL_478434, EPI_ISL_478435, EPI_ISL_478436, EPI_ISL_478437, EPI_ISL_478438, EPI_ISL_478439, EPI_ISL_478440, EPI_ISL_478441, EPI_ISL_478442, EPI_ISL_478443, EPI_ISL_478444, EPI_ISL_478445, EPI_ISL_478446, EPI_ISL_478447, EPI_ISL_478448, EPI_ISL_478449, EPI_ISL_478450, EPI_ISL_478451, EPI_ISL_478452, EPI_ISL_478453, EPI_ISL_478454, EPI_ISL_478455, EPI_ISL_478456, EPI_ISL_478457, EPI_ISL_478458, EPI_ISL_478459, EPI_ISL_478460, EPI_ISL_478461, EPI_ISL_478462, EPI_ISL_478463, EPI_ISL_478464, EPI_ISL_478465, EPI_ISL_478466, EPI_ISL_478467, EPI_ISL_478468, EPI_ISL_478469, EPI_ISL_478470, EPI_ISL_478471, EPI_ISL_478472 | | | |
| see above | University College London, Great Ormond Street Hospital for Children NHS Foundation Trust, Imperial College Healthcare NHS Trust | COVID-19 Genomics UK (COG-UK) Consortium | Sergi Castellano, Rachel Williams, Mark Kristiansen, Paola Resende Silva, Sunando Roy, Tony Brooks, Helena Tutill, Paola Niola, Patricia Dyal, Charlotte Williams, Leysa Forrest, Yasmin Panchbhaya, Jacqueline Findlay, Samuel Weeks, Julianne Brown, Kathryn Harris, Paul Randell, James Price, Alison Holmes, Judith Breuer |
| EPI_ISL_478473, EPI_ISL_478474, EPI_ISL_478475, EPI_ISL_478476, EPI_ISL_478477, EPI_ISL_478478, EPI_ISL_478479, EPI_ISL_478480, EPI_ISL_478481, EPI_ISL_478482, EPI_ISL_478483, EPI_ISL_478484, EPI_ISL_478485, EPI_ISL_478486, EPI_ISL_478487, EPI_ISL_478488, EPI_ISL_478489, EPI_ISL_478490, EPI_ISL_478491, EPI_ISL_478492, EPI_ISL_478493, EPI_ISL_478494, EPI_ISL_478495, EPI_ISL_478496, EPI_ISL_478497, EPI_ISL_478498, EPI_ISL_478499, EPI_ISL_478500, EPI_ISL_478501, EPI_ISL_478502, EPI_ISL_478503, EPI_ISL_478504, EPI_ISL_478505, EPI_ISL_478506, EPI_ISL_478507, EPI_ISL_478508, EPI_ISL_478509, EPI_ISL_478510, EPI_ISL_478511, EPI_ISL_478512, EPI_ISL_478513, EPI_ISL_478514, EPI_ISL_478515, EPI_ISL_478516, EPI_ISL_478517, EPI_ISL_478518, EPI_ISL_478519, EPI_ISL_478520, EPI_ISL_478521, EPI_ISL_478522, EPI_ISL_478523, EPI_ISL_478524, EPI_ISL_478525, EPI_ISL_478526, EPI_ISL_478527, EPI_ISL_478528, EPI_ISL_478529, EPI_ISL_478530, EPI_ISL_478531, EPI_ISL_478532, EPI_ISL_478533, EPI_ISL_478534, EPI_ISL_478535, EPI_ISL_478536, EPI_ISL_478537, EPI_ISL_478538, EPI_ISL_478539, EPI_ISL_478540, EPI_ISL_478541, EPI_ISL_478542, EPI_ISL_478543, EPI_ISL_478544, EPI_ISL_478545, EPI_ISL_478546, EPI_ISL_478547, EPI_ISL_478548, EPI_ISL_478549, EPI_ISL_478550, EPI_ISL_478551, EPI_ISL_478552, EPI_ISL_478553, EPI_ISL_478554, EPI_ISL_478555, EPI_ISL_478556, EPI_ISL_478557, EPI_ISL_478558, EPI_ISL_478559, EPI_ISL_478560, EPI_ISL_478561, EPI_ISL_478562, EPI_ISL_478563, EPI_ISL_478564, EPI_ISL_478565, EPI_ISL_478566, EPI_ISL_478567, EPI_ISL_478568, EPI_ISL_478569, EPI_ISL_478570, EPI_ISL_478571, EPI_ISL_478572, EPI_ISL_478573, EPI_ISL_478574, EPI_ISL_478575, EPI_ISL_478576, EPI_ISL_478577, EPI_ISL_478578, EPI_ISL_478579, EPI_ISL_478580, EPI_ISL_478581, EPI_ISL_478582, EPI_ISL_478583, EPI_ISL_478584, EPI_ISL_478585, EPI_ISL_478586, EPI_ISL_478587, EPI_ISL_478588, EPI_ISL_478589, EPI_ISL_478590, EPI_ISL_478591, EPI_ISL_478592, EPI_ISL_478593, EPI_ISL_478594, EPI_ISL_478595, EPI_ISL_478596, EPI_ISL_478597, EPI_ISL_478598, EPI_ISL_478599, EPI_ISL_478600, EPI_ISL_478601, EPI_ISL_478602, EPI_ISL_478603, EPI_ISL_478604, EPI_ISL_478605, EPI_ISL_478606, EPI_ISL_478607, EPI_ISL_478608, EPI_ISL_478609, EPI_ISL_478610, EPI_ISL_478611, EPI_ISL_478612, EPI_ISL_478613, EPI_ISL_478614, EPI_ISL_478615, EPI_ISL_478616, EPI_ISL_478617, EPI_ISL_478618, EPI_ISL_478619, EPI_ISL_478620, EPI_ISL_478621, EPI_ISL_478622, EPI_ISL_478623, EPI_ISL_478624, EPI_ISL_478625, EPI_ISL_478626, EPI_ISL_478627, EPI_ISL_478628, EPI_ISL_478629, EPI_ISL_478630, EPI_ISL_478631, EPI_ISL_478632, EPI_ISL_478633, EPI_ISL_478634, EPI_ISL_478635, EPI_ISL_478636, EPI_ISL_478637, EPI_ISL_478638, EPI_ISL_478639, EPI_ISL_478640, EPI_ISL_478641, EPI_ISL_478642, EPI_ISL_478643, EPI_ISL_478644, EPI_ISL_478645, EPI_ISL_478646, EPI_ISL_478647, EPI_ISL_478648, EPI_ISL_478649, EPI_ISL_478650, EPI_ISL_478651, EPI_ISL_478652, EPI_ISL_478653, EPI_ISL_478654, EPI_ISL_478655, EPI_ISL_478656, EPI_ISL_478657, EPI_ISL_478658, EPI_ISL_478659, EPI_ISL_478660, EPI_ISL_478661, EPI_ISL_478662, EPI_ISL_478663, EPI_ISL_478664, EPI_ISL_478665, EPI_ISL_478666, EPI_ISL_478667, EPI_ISL_478668 | | | |

| | | | |
|---|---|---|---|
| see above | Northumbria University / South Tees Hospitals NHS Foundation Trust / North Cumbria Integrated Care NHS Foundation Trust / North Tees and Hartlepool NHS Foundation Trust / Newcastle Hospitals NHS Foundation Trust | COVID-19 Genomics UK (COG-UK) Consortium | Darren L Smith,Andrew Nelson,Matthew Bashton,Greg R Young,Joshua Loh,John Allan,Mohammad A Tariq,Giles S Holt,Gary Black,Wen C Yew,Lynn Dover,Paul Baker,Steve Liggett,Sarah Essex,Jane Greenaway,Debra Padgett,Clive Graham,Garren Scott,Edward Barton,Emma Swindells,Brendan Payne,Jennifer Collins,Yusri Taha,Gary Eltringham |
| EPI_ISL_478669, EPI_ISL_478670 | unknown | Microbiology, Koc University | Can,F., Ozer,B., Nurtop,E., Dogan,O. |
| EPI_ISL_478671 | unknown | Molecular and Cell Biology | Baray,J.C., Mahmud,A., Khan,M.R., Nag,K., Sultana,N. |
| EPI_ISL_478673, EPI_ISL_478674 | Pathology North - Royal North Shore Hospital - NSW Health Pathology | NSW Health Pathology - Institute of Clinical Pathology and Medical Research; Westmead Hospital; University of Sydney | CIDM-PH et al. |
| EPI_ISL_478675, EPI_ISL_478676, EPI_ISL_478677, EPI_ISL_478678, EPI_ISL_478679, EPI_ISL_478680, EPI_ISL_478681, EPI_ISL_478682 | Sydney South West Pathology Service (SSWPS) - Liverpool Hospital - NSW Health Pathology | NSW Health Pathology - Institute of Clinical Pathology and Medical Research; Westmead Hospital; University of Sydney | CIDM-PH et al. |
| EPI_ISL_478683, EPI_ISL_478684, EPI_ISL_478685, EPI_ISL_478686, EPI_ISL_478687, EPI_ISL_478688, EPI_ISL_478689, EPI_ISL_478690, EPI_ISL_478691, EPI_ISL_478692, EPI_ISL_478693, EPI_ISL_478694, EPI_ISL_478695, EPI_ISL_478696, EPI_ISL_478697, EPI_ISL_478698, EPI_ISL_478699, EPI_ISL_478700, EPI_ISL_478701, EPI_ISL_478702, EPI_ISL_478703, EPI_ISL_478704, EPI_ISL_478705 | | | |
| see above | South Eastern Area Laboratory Services (SEALS) | NSW Health Pathology - Institute of Clinical Pathology and Medical Research; Westmead Hospital; University of Sydney | CIDM-PH et al. |
| EPI_ISL_478706, EPI_ISL_478707 | Sydney South West Pathology Service (SSWPS) - Liverpool Hospital - NSW Health Pathology | NSW Health Pathology - Institute of Clinical Pathology and Medical Research; Westmead Hospital; University of Sydney | CIDM-PH et al. |
| EPI_ISL_478708, EPI_ISL_478709, EPI_ISL_478710, EPI_ISL_478711 | South Eastern Area Laboratory Services (SEALS) | NSW Health Pathology - Institute of Clinical Pathology and Medical Research; Westmead Hospital; University of Sydney | CIDM-PH et al. |
| EPI_ISL_478712, EPI_ISL_478713, EPI_ISL_478714, EPI_ISL_478715, EPI_ISL_478716 | Sydney South West Pathology Service (SSWPS) - Liverpool Hospital - NSW Health Pathology | NSW Health Pathology - Institute of Clinical Pathology and Medical Research; Westmead Hospital; University of Sydney | CIDM-PH et al. |
| EPI_ISL_478717, EPI_ISL_478718 | Sydney South West Pathology Service (SSWPS) - Concord Repatriation General Hospital - NSW Health Pathology | NSW Health Pathology - Institute of Clinical Pathology and Medical Research; Westmead Hospital; University of Sydney | CIDM-PH et al. |
| EPI_ISL_478719, EPI_ISL_478720 | Quadram Institute Bioscience | COVID-19 Genomics UK (COG-UK) Consortium | Dave J. Baker, Gemma L. Kay, Alp Aydin, Thanh Le-Viet, Steven Rudder, Ana P. Tedim, Anastasia Kolyva, Maria Diaz, Leonardo de Oliveira Martins, Nabil-Fareed Alikhan, Lizzie Meadows, Rachael Stanley, Ngozi Elumogo, Muhammed Yasir, Nicholas M. Thomson, Alexander J Trotter, Rachel Gilroy, Samuel Bloomfield, Claire Stuart, Andrew Bell, Reenesh Prakash, Samir Dervisevic, Alison E. Mather, John Wain, Mark Webber, Andrew J. Page, Justin O'Grady |
| EPI_ISL_478721, EPI_ISL_478722, EPI_ISL_478723, EPI_ISL_478724, EPI_ISL_478725, EPI_ISL_478726, EPI_ISL_478727, EPI_ISL_478728 | Queens Medical Centre, Clinical Microbiology Department / DeepSeq Nottingham | COVID-19 Genomics UK (COG-UK) Consortium | Gemma Clark, Wendy Smith, Manjinder Khakh, Vicki M Fleming, Michelle M Lister, Hannah Howson-Wells, Jonathan Ball, Patrick McClure, Joseph Chappell, Theocharis Tsoleridis, Nadine Holmes, Matthew Carlisle, Christopher Moore, Fei Sang, Johnny Debebe, Victoria Wright, Matthew Loose |
| EPI_ISL_478729, EPI_ISL_478730, EPI_ISL_478731, EPI_ISL_478732, EPI_ISL_478733, EPI_ISL_478734, EPI_ISL_478735, EPI_ISL_478736, EPI_ISL_478737, EPI_ISL_478738, EPI_ISL_478739, EPI_ISL_478740, EPI_ISL_478741, EPI_ISL_478742, EPI_ISL_478743, EPI_ISL_478744, EPI_ISL_478745, EPI_ISL_478746, EPI_ISL_478747, EPI_ISL_478748, EPI_ISL_478749, EPI_ISL_478750, EPI_ISL_478751, EPI_ISL_478752, EPI_ISL_478753, EPI_ISL_478754, EPI_ISL_478755, EPI_ISL_478756, EPI_ISL_478757, EPI_ISL_478758, EPI_ISL_478759, EPI_ISL_478760, EPI_ISL_478761, EPI_ISL_478762, EPI_ISL_478763, EPI_ISL_478764, EPI_ISL_478765, EPI_ISL_478766, EPI_ISL_478767, EPI_ISL_478768, EPI_ISL_478769, EPI_ISL_478770, EPI_ISL_478771, EPI_ISL_478772, EPI_ISL_478773, EPI_ISL_478774, EPI_ISL_478775, EPI_ISL_478776, EPI_ISL_478777, EPI_ISL_478778, EPI_ISL_478779, EPI_ISL_478780, EPI_ISL_478781, EPI_ISL_478782, EPI_ISL_478783, EPI_ISL_478784, EPI_ISL_478785, EPI_ISL_478786, EPI_ISL_478787, EPI_ISL_478788, EPI_ISL_478789, EPI_ISL_478790, EPI_ISL_478791, EPI_ISL_478792, EPI_ISL_478793, EPI_ISL_478794, EPI_ISL_478795, EPI_ISL_478796, EPI_ISL_478797, EPI_ISL_478798, EPI_ISL_478799, EPI_ISL_478800, EPI_ISL_478801, EPI_ISL_478802, EPI_ISL_478803, EPI_ISL_478804, EPI_ISL_478805, EPI_ISL_478806, EPI_ISL_478807, EPI_ISL_478808, EPI_ISL_478809, EPI_ISL_478810, EPI_ISL_478811, EPI_ISL_478812, EPI_ISL_478813, EPI_ISL_478814, EPI_ISL_478815, EPI_ISL_478816, EPI_ISL_478817, EPI_ISL_478818, EPI_ISL_478819, EPI_ISL_478820, EPI_ISL_478821, EPI_ISL_478822, EPI_ISL_478823, EPI_ISL_478824, EPI_ISL_478825, EPI_ISL_478826, EPI_ISL_478827, EPI_ISL_478828, EPI_ISL_478829, EPI_ISL_478830, EPI_ISL_478831, EPI_ISL_478832, EPI_ISL_478833, EPI_ISL_478834, EPI_ISL_478835, EPI_ISL_478836, EPI_ISL_478837, EPI_ISL_478838, EPI_ISL_478839, EPI_ISL_478840, EPI_ISL_478841, EPI_ISL_478842, EPI_ISL_478843, EPI_ISL_478844, EPI_ISL_478845, EPI_ISL_478846, EPI_ISL_478847, EPI_ISL_478848, EPI_ISL_478849, EPI_ISL_478850, EPI_ISL_478851, EPI_ISL_478852, EPI_ISL_478853, EPI_ISL_478854, EPI_ISL_478855, EPI_ISL_478856, EPI_ISL_478857, EPI_ISL_478858, EPI_ISL_478859, EPI_ISL_478860, EPI_ISL_478861, EPI_ISL_478862, EPI_ISL_478863, EPI_ISL_478864, EPI_ISL_478865, EPI_ISL_478866, EPI_ISL_478867, EPI_ISL_478868, EPI_ISL_478869, EPI_ISL_478870, EPI_ISL_478871, EPI_ISL_478872, EPI_ISL_478873, EPI_ISL_478874, EPI_ISL_478875, EPI_ISL_478876, EPI_ISL_478877, EPI_ISL_478878, EPI_ISL_478879, EPI_ISL_478880, EPI_ISL_478881, EPI_ISL_478882, EPI_ISL_478883, EPI_ISL_478884, EPI_ISL_478885, EPI_ISL_478886, EPI_ISL_478887, EPI_ISL_478888, EPI_ISL_478889, EPI_ISL_478890, EPI_ISL_478891, EPI_ISL_478892, EPI_ISL_478893, EPI_ISL_478894, EPI_ISL_478895, EPI_ISL_478896, EPI_ISL_478897, EPI_ISL_478898, EPI_ISL_478899, EPI_ISL_478900, EPI_ISL_478901, EPI_ISL_478902, EPI_ISL_478903, EPI_ISL_478904, EPI_ISL_478905, EPI_ISL_478906, EPI_ISL_478907, EPI_ISL_478908, EPI_ISL_478909, EPI_ISL_478910, EPI_ISL_478911, EPI_ISL_478912, EPI_ISL_478913, EPI_ISL_478914, EPI_ISL_478915, EPI_ISL_478916, EPI_ISL_478917, EPI_ISL_478918, EPI_ISL_478919, EPI_ISL_478920, EPI_ISL_478921, EPI_ISL_478922, EPI_ISL_478923, EPI_ISL_478924, EPI_ISL_478925, EPI_ISL_478926, EPI_ISL_478927, EPI_ISL_478928, EPI_ISL_478929, EPI_ISL_478930, EPI_ISL_478931, EPI_ISL_478932, EPI_ISL_478933, EPI_ISL_478934, EPI_ISL_478935, EPI_ISL_478936, EPI_ISL_478937, EPI_ISL_478938, EPI_ISL_478939, EPI_ISL_478940, EPI_ISL_478941, EPI_ISL_478942, EPI_ISL_478943, EPI_ISL_478944, EPI_ISL_478945, EPI_ISL_478946, EPI_ISL_478947, EPI_ISL_478948, EPI_ISL_478949, EPI_ISL_478950, EPI_ISL_478951, EPI_ISL_478952, EPI_ISL_478953, EPI_ISL_478954, EPI_ISL_478955, EPI_ISL_478956, EPI_ISL_478957, EPI_ISL_478958, EPI_ISL_478959, EPI_ISL_478960, EPI_ISL_478961, EPI_ISL_478962, EPI_ISL_478963, EPI_ISL_478964, EPI_ISL_478965, EPI_ISL_478966, EPI_ISL_478967, EPI_ISL_478968, EPI_ISL_478969, EPI_ISL_478970, EPI_ISL_478971, EPI_ISL_478972, EPI_ISL_478973, EPI_ISL_478974, EPI_ISL_478975, EPI_ISL_478976, EPI_ISL_478977, EPI_ISL_478978, EPI_ISL_478979, EPI_ISL_478980, EPI_ISL_478981, EPI_ISL_478982, EPI_ISL_478983, EPI_ISL_478984, EPI_ISL_478985, EPI_ISL_478986, EPI_ISL_478987, EPI_ISL_478988, EPI_ISL_478989, EPI_ISL_478990, EPI_ISL_478991, EPI_ISL_478992, EPI_ISL_478993, EPI_ISL_478994, EPI_ISL_478995, EPI_ISL_478996, EPI_ISL_478997, EPI_ISL_478998, EPI_ISL_478999, EPI_ISL_479000, EPI_ISL_479001, EPI_ISL_479002, EPI_ISL_479003, EPI_ISL_479004, EPI_ISL_479005, EPI_ISL_479006, EPI_ISL_479007, EPI_ISL_479008, EPI_ISL_479009, EPI_ISL_479010, EPI_ISL_479011, EPI_ISL_479012, EPI_ISL_479013, EPI_ISL_479014, EPI_ISL_479015, EPI_ISL_479016, EPI_ISL_479017, EPI_ISL_479018, EPI_ISL_479019, EPI_ISL_479020, EPI_ISL_479021, EPI_ISL_479022, EPI_ISL_479023, EPI_ISL_479024, EPI_ISL_479025, EPI_ISL_479026, EPI_ISL_479027, EPI_ISL_479028, EPI_ISL_479029, EPI_ISL_479030, EPI_ISL_479031, EPI_ISL_479032, EPI_ISL_479033, EPI_ISL_479034, EPI_ISL_479035, EPI_ISL_479036, EPI_ISL_479037, EPI_ISL_479038, EPI_ISL_479039, EPI_ISL_479040, EPI_ISL_479041, EPI_ISL_479042, EPI_ISL_479043, EPI_ISL_479044, EPI_ISL_479045, EPI_ISL_479046, EPI_ISL_479047, EPI_ISL_479048, EPI_ISL_479049, EPI_ISL_479050, EPI_ISL_479051, EPI_ISL_479052, EPI_ISL_479053, EPI_ISL_479054, EPI_ISL_479055, EPI_ISL_479056, EPI_ISL_479057, EPI_ISL_479058, EPI_ISL_479059, EPI_ISL_479060, EPI_ISL_479061, EPI_ISL_479062, EPI_ISL_479063, EPI_ISL_479064, EPI_ISL_479065, EPI_ISL_479066, EPI_ISL_479067, EPI_ISL_479068, EPI_ISL_479069, EPI_ISL_479070, EPI_ISL_479071, EPI_ISL_479072, EPI_ISL_479073, EPI_ISL_479074, EPI_ISL_479075, EPI_ISL_479076, EPI_ISL_479077, EPI_ISL_479078, EPI_ISL_479079, EPI_ISL_479080, EPI_ISL_479081, EPI_ISL_479082, EPI_ISL_479083, EPI_ISL_479084, EPI_ISL_479085, EPI_ISL_479086, EPI_ISL_479087, EPI_ISL_479088, EPI_ISL_479089, EPI_ISL_479090, EPI_ISL_479091, EPI_ISL_479092, EPI_ISL_479093, EPI_ISL_479094, EPI_ISL_479095, EPI_ISL_479096, EPI_ISL_479097, EPI_ISL_479098, EPI_ISL_479099, EPI_ISL_479100, EPI_ISL_479101, EPI_ISL_479102, EPI_ISL_479103, EPI_ISL_479104, EPI_ISL_479105, EPI_ISL_479106, EPI_ISL_479107, EPI_ISL_479108, EPI_ISL_479109, EPI_ISL_479110, EPI_ISL_479111, EPI_ISL_479112, EPI_ISL_479113, EPI_ISL_479114, EPI_ISL_479115, EPI_ISL_479116, EPI_ISL_479117, EPI_ISL_479118, EPI_ISL_479119, EPI_ISL_479120, EPI_ISL_479121, EPI_ISL_479122, EPI_ISL_479123, EPI_ISL_479124, EPI_ISL_479125, EPI_ISL_479126, EPI_ISL_479127, EPI_ISL_479128, EPI_ISL_479129, EPI_ISL_479130, EPI_ISL_479131, EPI_ISL_479132, EPI_ISL_479133, EPI_ISL_479134, EPI_ISL_479135, EPI_ISL_479136, EPI_ISL_479137, EPI_ISL_479138, EPI_ISL_479139, EPI_ISL_479140, EPI_ISL_479141, EPI_ISL_479142, EPI_ISL_479143, EPI_ISL_479144, EPI_ISL_479145, EPI_ISL_479146, EPI_ISL_479147, EPI_ISL_479148, EPI_ISL_479149, EPI_ISL_479150, EPI_ISL_479151, EPI_ISL_479152, EPI_ISL_479153, EPI_ISL_479154, EPI_ISL_479155, EPI_ISL_479156, EPI_ISL_479157, EPI_ISL_479158, EPI_ISL_479159, EPI_ISL_479160, EPI_ISL_479161, EPI_ISL_479162, EPI_ISL_479163, EPI_ISL_479164, EPI_ISL_479165, EPI_ISL_479166, EPI_ISL_479167, EPI_ISL_479168, EPI_ISL_479169, EPI_ISL_479170, EPI_ISL_479171, EPI_ISL_479172, EPI_ISL_479173, EPI_ISL_479174 | | | |
| see above | Oxford Viromics, NDM, University of Oxford; Oxford University Hospitals; Basingstoke and North Hampshire Hospital | COVID-19 Genomics UK (COG-UK) Consortium | Tanya Golubchik, David Bonsall, George Macintyre, Amy Trebes, Mariateresa de Cesare, Catrin Moore, Alex Mobbs, Anita Justice, Robert Shaw, Monique Andersson, Timothy Peto, Emma Wise, Nathan Moore, Jessica Lynch, Nick Cortes, Matilde Mori, Stephen Kidd, David Buck, John Todd, Christophe Fraser |
| EPI_ISL_479175, EPI_ISL_479176, EPI_ISL_479177, EPI_ISL_479178, EPI_ISL_479179, EPI_ISL_479180, EPI_ISL_479181, EPI_ISL_479182, EPI_ISL_479183, EPI_ISL_479184, EPI_ISL_479185, EPI_ISL_479186, EPI_ISL_479187, EPI_ISL_479188, EPI_ISL_479189, EPI_ISL_479190, EPI_ISL_479191, EPI_ISL_479192, EPI_ISL_479193, EPI_ISL_479194 | | | |
| see above | Centre for Enzyme Innovation, University of Portsmouth / Translational Research Laboratory, Portsmouth Hospitals NHS Trust | COVID-19 Genomics UK (COG-UK) Consortium | Angela Beckett,Yann Bourgeois,Garry Scarlett,Sharon Glaysher,Scott Elliott,Kelly Bicknell,Robert Impey,Allyson Lloyd,Sarah Wyllie,Ethan Butcher,Anoop Chauhan,Samuel Robson |
| EPI_ISL_479195, EPI_ISL_479196, EPI_ISL_479197, EPI_ISL_479198, EPI_ISL_479199, EPI_ISL_479200, EPI_ISL_479201, EPI_ISL_479202, EPI_ISL_479203, EPI_ISL_479204, EPI_ISL_479205, EPI_ISL_479206, EPI_ISL_479207, EPI_ISL_479208, EPI_ISL_479209, EPI_ISL_479210, EPI_ISL_479211, EPI_ISL_479212, EPI_ISL_479213, EPI_ISL_479214, EPI_ISL_479215, EPI_ISL_479216, EPI_ISL_479217, EPI_ISL_479218, EPI_ISL_479219, EPI_ISL_479220, EPI_ISL_479221, EPI_ISL_479222, EPI_ISL_479223, EPI_ISL_479224, EPI_ISL_479225, EPI_ISL_479226, EPI_ISL_479227, EPI_ISL_479228, EPI_ISL_479229, EPI_ISL_479230, EPI_ISL_479231, EPI_ISL_479232, EPI_ISL_479233, EPI_ISL_479234, EPI_ISL_479235, EPI_ISL_479236, EPI_ISL_479237, EPI_ISL_479238, EPI_ISL_479239, EPI_ISL_479240, EPI_ISL_479241, EPI_ISL_479242, EPI_ISL_479243, EPI_ISL_479244, EPI_ISL_479245, EPI_ISL_479246, EPI_ISL_479247, EPI_ISL_479248, EPI_ISL_479249, EPI_ISL_479250, EPI_ISL_479251, EPI_ISL_479252, EPI_ISL_479253, EPI_ISL_479254, EPI_ISL_479255, EPI_ISL_479256, EPI_ISL_479257, EPI_ISL_479258, EPI_ISL_479259, EPI_ISL_479260, EPI_ISL_479261, EPI_ISL_479262, EPI_ISL_479263, EPI_ISL_479264, EPI_ISL_479265, EPI_ISL_479266, EPI_ISL_479267, EPI_ISL_479268, EPI_ISL_479269, EPI_ISL_479270, EPI_ISL_479271, EPI_ISL_479272, EPI_ISL_479273, EPI_ISL_479274, EPI_ISL_479275, EPI_ISL_479276, EPI_ISL_479277, EPI_ISL_479278, EPI_ISL_479279, EPI_ISL_479280, EPI_ISL_479281, EPI_ISL_479282, EPI_ISL_479283 | | | |
| see above | Virology Department, Sheffield Teaching Hospitals NHS Foundation Trust/Department of Infection, Immunity and Cardiovascular Disease, The Medical School, University of Sheffield | COVID-19 Genomics UK (COG-UK) Consortium | Thushan de Silva, Matthew Parker, Nikki Smith, Adri Angyal, Rebecca Brown, Luke Green, Rachel Tucker, Paul Parsons, Danielle Groves, Katie Johnson, Laura Carrilero, Alex Keeley, Dave Partridge, Matthew Wyles, Benjamin Lindsey, Mehmet Yavuz, Mohammad Raza, Cariad Evans |
| EPI_ISL_479284, EPI_ISL_479285, EPI_ISL_479286, EPI_ISL_479287, EPI_ISL_479288, EPI_ISL_479289, EPI_ISL_479290, EPI_ISL_479291, EPI_ISL_479292, EPI_ISL_479293, EPI_ISL_479294, EPI_ISL_479295, EPI_ISL_479296, EPI_ISL_479297, EPI_ISL_479298, EPI_ISL_479299, EPI_ISL_479300, EPI_ISL_479301, EPI_ISL_479302, EPI_ISL_479303, EPI_ISL_479304, EPI_ISL_479305, EPI_ISL_479306, EPI_ISL_479307, EPI_ISL_479308, EPI_ISL_479309, EPI_ISL_479310, EPI_ISL_479311, EPI_ISL_479312, EPI_ISL_479313, EPI_ISL_479314, EPI_ISL_479315, EPI_ISL_479316, EPI_ISL_479317, EPI_ISL_479318, EPI_ISL_479319, EPI_ISL_479320, EPI_ISL_479321, EPI_ISL_479322, EPI_ISL_479323, EPI_ISL_479324, EPI_ISL_479325, EPI_ISL_479326, EPI_ISL_479327, EPI_ISL_479328, EPI_ISL_479329, EPI_ISL_479330, EPI_ISL_479331, EPI_ISL_479332, EPI_ISL_479333, EPI_ISL_479334, EPI_ISL_479335, EPI_ISL_479336, EPI_ISL_479337, EPI_ISL_479338, EPI_ISL_479339, EPI_ISL_479340, EPI_ISL_479341, EPI_ISL_479342, EPI_ISL_479343, EPI_ISL_479344, EPI_ISL_479345, EPI_ISL_479346, EPI_ISL_479347, EPI_ISL_479348, EPI_ISL_479349, EPI_ISL_479350, EPI_ISL_479351, EPI_ISL_479352, EPI_ISL_479353, EPI_ISL_479354, EPI_ISL_479355, EPI_ISL_479356, EPI_ISL_479357, EPI_ISL_479358, EPI_ISL_479359, EPI_ISL_479360, EPI_ISL_479361, EPI_ISL_479362, EPI_ISL_479363, EPI_ISL_479364, EPI_ISL_479365, EPI_ISL_479366, EPI_ISL_479367, EPI_ISL_479368, EPI_ISL_479369, EPI_ISL_479370, EPI_ISL_479371, EPI_ISL_479372, EPI_ISL_479373, EPI_ISL_479374, EPI_ISL_479375, EPI_ISL_479376, EPI_ISL_479377, EPI_ISL_479378, EPI_ISL_479379, EPI_ISL_479380, EPI_ISL_479381, EPI_ISL_479382, EPI_ISL_479383, EPI_ISL_479384, EPI_ISL_479385, EPI_ISL_479386, EPI_ISL_479387, EPI_ISL_479388, EPI_ISL_479389, EPI_ISL_479390, EPI_ISL_479391, EPI_ISL_479392, EPI_ISL_479393, EPI_ISL_479394, EPI_ISL_479395, EPI_ISL_479396, EPI_ISL_479397, EPI_ISL_479398, EPI_ISL_479399, EPI_ISL_479400, EPI_ISL_479401, EPI_ISL_479402, EPI_ISL_479403, EPI_ISL_479404, EPI_ISL_479405, EPI_ISL_479406, EPI_ISL_479407, EPI_ISL_479408, EPI_ISL_479409, EPI_ISL_479410, EPI_ISL_479411, EPI_ISL_479412, EPI_ISL_479413, EPI_ISL_479414, EPI_ISL_479415, EPI_ISL_479416, EPI_ISL_479417, EPI_ISL_479418, EPI_ISL_479419, EPI_ISL_479420, EPI_ISL_479421, EPI_ISL_479422, EPI_ISL_479423, EPI_ISL_479424, EPI_ISL_479425, EPI_ISL_479426, EPI_ISL_479427, EPI_ISL_479428, EPI_ISL_479429, EPI_ISL_479430, EPI_ISL_479431, EPI_ISL_479432, EPI_ISL_479433, EPI_ISL_479434, EPI_ISL_479435, EPI_ISL_479436, EPI_ISL_479437, EPI_ISL_479438, EPI_ISL_479439, EPI_ISL_479440, EPI_ISL_479441, EPI_ISL_479442, EPI_ISL_479443, EPI_ISL_479444, EPI_ISL_479445, EPI_ISL_479446, EPI_ISL_479447, EPI_ISL_479448, EPI_ISL_479449, EPI_ISL_479450, EPI_ISL_479451, EPI_ISL_479452, EPI_ISL_479453, EPI_ISL_479454, EPI_ISL_479455, EPI_ISL_479456, EPI_ISL_479457, EPI_ISL_479458, EPI_ISL_479459, EPI_ISL_479460, EPI_ISL_479461, EPI_ISL_479462, EPI_ISL_479463, EPI_ISL_479464, EPI_ISL_479465, EPI_ISL_479466, EPI_ISL_479467, EPI_ISL_479468, EPI_ISL_479469, EPI_ISL_479470, EPI_ISL_479471, EPI_ISL_479472, EPI_ISL_479473, EPI_ISL_479474, EPI_ISL_479475, EPI_ISL_479476, EPI_ISL_479477, EPI_ISL_479478, EPI_ISL_479479, EPI_ISL_479480, EPI_ISL_479481 | | | |
| see above | Wales Specialist Virology Centre Sequencing lab: Pathogen Genomics Unit | COVID-19 Genomics UK (COG-UK) Consortium | Catherine Moore, Johnathan Evans, Laura Gifford, Malorie Perry, Simon Cottrell, Angela Marchbank, Alec Birchley, Alexander Adams, Amy Gaskin, Bree Gatica-Wilcox, Jason Coombes, Joel Southgate, Lauren Gilbert, Lee Graham, Nicole Pacchiarini, Sara Kumziene-Summerhayes, Sarah Taylor, Sophie Jones, Sara Rey, Matthew Bull, Joanne Watkins, Sally Corden, Tom Connor |
| EPI_ISL_479482, EPI_ISL_479483, EPI_ISL_479484, EPI_ISL_479485, EPI_ISL_479486, EPI_ISL_479487, EPI_ISL_479488, EPI_ISL_479489, EPI_ISL_479490, EPI_ISL_479491, EPI_ISL_479492 | | | |
| see above | Department of Laboratory Medicine Tan Tock Seng Hospital | Department of Laboratory Medicine Tan Tock Seng Hospital | Chen YYC, Zair X, Li C, Tang WY, Maurer-Stroh S, Barkham TMS, Nagarajan N, Sessions OM |
| EPI_ISL_479493, EPI_ISL_479494, EPI_ISL_479495, EPI_ISL_479496, EPI_ISL_479497, EPI_ISL_479498, EPI_ISL_479499, EPI_ISL_479500, EPI_ISL_479501, EPI_ISL_479502, EPI_ISL_479503, EPI_ISL_479504, EPI_ISL_479505, EPI_ISL_479506, EPI_ISL_479507, EPI_ISL_479508, EPI_ISL_479509, EPI_ISL_479510, EPI_ISL_479511, EPI_ISL_479512, EPI_ISL_479513, EPI_ISL_479514, EPI_ISL_479515, EPI_ISL_479516, EPI_ISL_479517, EPI_ISL_479518, EPI_ISL_479519, EPI_ISL_479520, EPI_ISL_479521, EPI_ISL_479522, EPI_ISL_479523, EPI_ISL_479524, EPI_ISL_479525, EPI_ISL_479526, EPI_ISL_479527, EPI_ISL_479528, EPI_ISL_479529, EPI_ISL_479530, EPI_ISL_479531, EPI_ISL_479532, EPI_ISL_479533, EPI_ISL_479534, EPI_ISL_479535, EPI_ISL_479536, EPI_ISL_479537, EPI_ISL_479538, EPI_ISL_479539, EPI_ISL_479540, EPI_ISL_479541, EPI_ISL_479542, EPI_ISL_479543, EPI_ISL_479544, EPI_ISL_479545, EPI_ISL_479546, EPI_ISL_479547, EPI_ISL_479548, EPI_ISL_479549, EPI_ISL_479550, EPI_ISL_479551, EPI_ISL_479552, EPI_ISL_479553, EPI_ISL_479554, EPI_ISL_479555, EPI_ISL_479556, EPI_ISL_479557, EPI_ISL_479558, EPI_ISL_479559, EPI_ISL_479560, EPI_ISL_479561, EPI_ISL_479562, EPI_ISL_479563, EPI_ISL_479564, EPI_ISL_479565, EPI_ISL_479566, EPI_ISL_479567, EPI_ISL_479568, EPI_ISL_479569, EPI_ISL_479570, EPI_ISL_479571, EPI_ISL_479572, EPI_ISL_479573 | | | |
| see above | NIV Influenza | NIV Influenza | Potdar V |
| EPI_ISL_479574, EPI_ISL_479575, EPI_ISL_479576, EPI_ISL_479577, EPI_ISL_479578, EPI_ISL_479579, EPI_ISL_479580, EPI_ISL_479581, EPI_ISL_479582, EPI_ISL_479583, EPI_ISL_479584, EPI_ISL_479585, EPI_ISL_479586, EPI_ISL_479587, EPI_ISL_479588, EPI_ISL_479589, EPI_ISL_479590, EPI_ISL_479591, EPI_ISL_479592, EPI_ISL_479593, EPI_ISL_479594, EPI_ISL_479595, EPI_ISL_479596, EPI_ISL_479597, EPI_ISL_479598, EPI_ISL_479599, EPI_ISL_479600, EPI_ISL_479601, EPI_ISL_479602, EPI_ISL_479603 | | | |
| see above | National Public Health Laboratory, National Centre for Infectious Diseases | National Public Health Laboratory, National Centre for Infectious Diseases | Mak TM, Octavia S, Zhou Z, Chavatte JM, Cui L, Lin RTP |
| EPI_ISL_479616, EPI_ISL_479617 | Laboratory of Molecular Virology of the International Centre for Genetic Engineering and Biotechnology (ICGEB) | ARGO Open Lab Platform for Genome Sequencing | Licastro, D, Rajasekharan S, Dal Monego S, Segat L, D'Agaro P, Salton F, Confalonieri P, Confalonieri M Marcello A |
| EPI_ISL_479618 | Laboratory of Molecular Virology of the International Centre for Genetic Engineering and Biotechnology (ICGEB) | ARGO Open Lab Platform for Genome Sequencing | Licastro, D, Rajasekharan S, Dal Monego S, Segat L, D'Agaro P, Salton F, Confalonieri P, Confalonieri M, Marcello A |
| EPI_ISL_479619 | Laboratory of Molecular Virology of the International Centre for Genetic Engineering and Biotechnology (ICGEB) | ARGO Open Lab Platform for Genome Sequencing | Licastro, D, Rajasekharan S, Dal Monego S, Segat L, D'Agaro P, Salton F, Confalonieri P, Confalonieri M, Marcello A |
| EPI_ISL_479620, EPI_ISL_479621, EPI_ISL_479622, EPI_ISL_479623, EPI_ISL_479624 | Molecular diagnostic laboratory of Federal Budget Institution of Science "Central Research Institute of Epidemiology" of The Federal Service on Customers' Rights Protection and Human Well-being Surveillance | Group of Genomics and Postgenomic Technologies of Central Research Institute of Epidemilology | Speranskaya AS, Kaptelova VV,Valdokhina AV, Bulanenko VP, Samoilov AE, Korneenko EV, Sizova TV, Tivanova EV, Shipulina OY, Akimkin VG |

| | | | |
|---|---|---|---|
| EPI_ISL_479662 | College of Veterinary Medicine, Chungnam National University | College of Veterinary Medicine, Chungnam National University | Seo,S. |
| EPI_ISL_479663, EPI_ISL_479664, EPI_ISL_479665, EPI_ISL_479666, EPI_ISL_479667, EPI_ISL_479668, EPI_ISL_479669, EPI_ISL_479670, EPI_ISL_479671, EPI_ISL_479672, EPI_ISL_479673, EPI_ISL_479674, EPI_ISL_479675 | | | |
| see above | Center for Genomics and System Biology, New York University | Center for Genomics and System Biology, New York University | Roder,A., Banakis,S., Johnson,K., Khalfan,M., Borenstein,E.S., Samanovic,M., Cornelius,A., Herati,R., Ulrich,R., Fleming,A., Kottkamp,A., Raabe,V., Mulligan,M.J., Gresham,D., Ghedin,E. |
| EPI_ISL_479728 | Egyptian National Cancer Institute (ENCI) | Egyptian National Cancer Institute (ENCI) | Zekri,A.N., Amer,K.E., Ahmed,O.S., Soliman,H.K., Bahnassy,A.A., Ali,M., Abdelhamid,W., Gad,A., Hassan,W., Samir,M., Raouf,A., Hamdy,M.S., Soliman,M.S., Elsissy,M.H., Elkhateeb,S.M., Ezzelarab,M.H., Abouelhoda,M. |
| EPI_ISL_479729, EPI_ISL_479730, EPI_ISL_479731, EPI_ISL_479732, EPI_ISL_479733, EPI_ISL_479734, EPI_ISL_479735 | Egyptian National Cancer Institute (ENCI) | Egyptian National Cancer Institute (ENCI) | Zekri, Abdel Rahman N, Amer,K.E., Ahmed,O.S., Soliman,H.K., Hafez,M.M., Bahnassy,A.A., Abdelhamid,W., Gad,A., Ali,M., Hassan,W., Samir,M., Raouf,A., Hamdy,M.S., Soliman,M.S., Elsissy,M.H., Elkhateeb,S.M., Ezzelarab,M.H., Abouelhoda, Mohamed |
| EPI_ISL_486890, EPI_ISL_486891, EPI_ISL_486892, EPI_ISL_486893, EPI_ISL_486894, EPI_ISL_486895, EPI_ISL_486896 | Tokyo Metoropolitan Institute of Public Health | Tokyo Metoropolitan Institute of Public Health | Asakura,H., Yoshida,I., Kumagai,R., Nagashima,M., Chiba,T., Sadamasu,K. |
| EPI_ISL_491111, EPI_ISL_491112 | Friedrich-Loeffler-Institut, Laboratory for NGS and Microarray Diagnostics | Friedrich-Loeffler-Institut, Laboratory for NGS and Microarray Diagnostics | Dirk Höper, Laboratory for NGS and Microarray Diagnostics |

We gratefully acknowledge the following Authors from the Originating laboratories responsible for obtaining the specimens, as well as the Submitting laboratories where the genome data were generated and shared via GISAID, on which this research is based.

All Submitters of data may be contacted directly via www.gisaid.org

| Accession ID | Originating Laboratory | Submitting Laboratory | Authors |
|---|---|---|---|
| EPI_ISL_491437 | Area de Salud Escazu (Coopesana) | Inciensa, Instituto Costarricense de Investigación y Enseñanza en Nutrición y Salud | Francisco Duarte, Hebleen Brenes, Claudio Soto-Garita, Estela Cordero, Adriana Godinez & Melany Calderon |
| EPI_ISL_491438 | Hospital San Rafael de Alajuela | Inciensa, Instituto Costarricense de Investigación y Enseñanza en Nutrición y Salud | Francisco Duarte, Hebleen Brenes, Claudio Soto-Garita, Estela Cordero, Adriana Godinez & Melany Calderon |
| EPI_ISL_491439 | Hospital Calderon Guardia | Inciensa, Instituto Costarricense de Investigación y Enseñanza en Nutrición y Salud | Francisco Duarte, Hebleen Brenes, Claudio Soto-Garita, Estela Cordero, Adriana Godinez & Melany Calderon |
| EPI_ISL_491440 | Hospital San Rafael de Alajuela | Inciensa, Instituto Costarricense de Investigación y Enseñanza en Nutrición y Salud | Francisco Duarte, Hebleen Brenes, Claudio Soto-Garita, Estela Cordero, Adriana Godinez & Melany Calderon |
| EPI_ISL_491441, EPI_ISL_491442 | Hospital Clinica Biblica | Inciensa, Instituto Costarricense de Investigación y Enseñanza en Nutrición y Salud | Francisco Duarte, Hebleen Brenes, Claudio Soto-Garita, Estela Cordero, Adriana Godinez & Melany Calderon |
| EPI_ISL_491443 | Hospital Fernando Escalante Pradilla | Inciensa, Instituto Costarricense de Investigación y Enseñanza en Nutrición y Salud | Francisco Duarte, Hebleen Brenes, Claudio Soto-Garita, Estela Cordero, Adriana Godinez & Melany Calderon |
| EPI_ISL_491444 | Area de Salud Escazu (Coopesana) | Inciensa, Instituto Costarricense de Investigación y Enseñanza en Nutrición y Salud | Francisco Duarte, Hebleen Brenes, Claudio Soto-Garita, Estela Cordero, Adriana Godinez & Melany Calderon |
| EPI_ISL_491445 | Area de Salud Mata Redonda | Inciensa, Instituto Costarricense de Investigación y Enseñanza en Nutrición y Salud | Francisco Duarte, Hebleen Brenes, Claudio Soto-Garita, Estela Cordero, Adriana Godinez & Melany Calderon |
| EPI_ISL_491446 | Area de Salud Alajuela Central | Inciensa, Instituto Costarricense de Investigación y Enseñanza en Nutrición y Salud | Francisco Duarte, Hebleen Brenes, Claudio Soto-Garita, Estela Cordero, Adriana Godinez & Melany Calderon |
| EPI_ISL_491447 | Hospital Fernando Escalante Pradilla | Inciensa, Instituto Costarricense de Investigación y Enseñanza en Nutrición y Salud | Francisco Duarte, Hebleen Brenes, Claudio Soto-Garita, Estela Cordero, Adriana Godinez & Melany Calderon |
| EPI_ISL_491448 | Hospital San Rafael de Alajuela | Inciensa, Instituto Costarricense de Investigación y Enseñanza en Nutrición y Salud | Francisco Duarte, Hebleen Brenes, Claudio Soto-Garita, Estela Cordero, Adriana Godinez & Melany Calderon |
| EPI_ISL_491449 | Area de Salud Alajuela Sur | Inciensa, Instituto Costarricense de Investigación y Enseñanza en Nutrición y Salud | Francisco Duarte, Hebleen Brenes, Claudio Soto-Garita, Estela Cordero, Adriana Godinez & Melany Calderon |
| EPI_ISL_491450 | Hospital San Juan de Dios | Inciensa, Instituto Costarricense de Investigación y Enseñanza en Nutrición y Salud | Francisco Duarte, Hebleen Brenes, Claudio Soto-Garita, Estela Cordero, Adriana Godinez & Melany Calderon |
| EPI_ISL_491451 | Hospital México | Inciensa, Instituto Costarricense de Investigación y Enseñanza en Nutrición y Salud | Francisco Duarte, Hebleen Brenes, Claudio Soto-Garita, Estela Cordero, Adriana Godinez & Melany Calderon |
| EPI_ISL_491452 | Hospital San Rafael de Alajuela | Inciensa, Instituto Costarricense de Investigación y Enseñanza en Nutrición y Salud | Francisco Duarte, Hebleen Brenes, Claudio Soto-Garita, Estela Cordero, Adriana Godinez & Melany Calderon |
| EPI_ISL_491453 | Hospital México | Inciensa, Instituto Costarricense de Investigación y Enseñanza en Nutrición y Salud | Francisco Duarte, Hebleen Brenes, Claudio Soto-Garita, Estela Cordero, Adriana Godinez & Melany Calderon |
| EPI_ISL_491454 | Hospital San Juan de Dios | Inciensa, Instituto Costarricense de Investigación y Enseñanza en Nutrición y Salud | Francisco Duarte, Hebleen Brenes, Claudio Soto-Garita, Estela Cordero, Adriana Godinez & Melany Calderon |
| EPI_ISL_491455 | Hospital Clinica Biblica | Inciensa, Instituto Costarricense de Investigación y Enseñanza en Nutrición y Salud | Francisco Duarte, Hebleen Brenes, Claudio Soto-Garita, Estela Cordero, Adriana Godinez & Melany Calderon |
| EPI_ISL_491456 | Hospital San Juan de Dios | Inciensa, Instituto Costarricense de Investigación y Enseñanza en Nutrición y Salud | Francisco Duarte, Hebleen Brenes, Claudio Soto-Garita, Estela Cordero, Adriana Godinez & Melany Calderon |
| EPI_ISL_491457 | Area de Salud Los Santos | Inciensa, Instituto Costarricense de Investigación y Enseñanza en Nutrición y Salud | Francisco Duarte, Hebleen Brenes, Claudio Soto-Garita, Estela Cordero, Adriana Godinez & Melany Calderon |
| EPI_ISL_491458, EPI_ISL_491459, EPI_ISL_491460, EPI_ISL_491461, EPI_ISL_491462 | Laboratorio de Referencia Nacional de Virus Respiratorio. Instituto Nacional de Salud Perú | Laboratorio de Referencia Nacional de Biotecnología y Biología Molecular. Instituto Nacional de Salud Perú | Carlos Padilla Rojas, Karolyn Vega Chozo, Priscila Lope Pari, Omar Caceres Rey, Marco Galarza Perez, Maribel Huaringa Nuñez, Johanna Balbuena Torrez, Henri Bailon Calderon, Nancy Rojas Serrano |
| EPI_ISL_491463 | Laboratorio de Referencia Nacional de Virus Respiratorio. Instituto Nacional de Salud Perú | Laboratorio de Referencia Nacional de Biotecnología y Biología Molecular. Instituto Nacional de Salud Perú | Carlos Padilla Rojas, Karolyn Vega Chozo, Priscila Lope Pari, Omar Caceres Rey, Marco Galarza Perez, Maribel Huaringa Nuñez, Johanna Balbuena Torres, Henri Bailon Calderon, Nancy Rojas Serrano |
| EPI_ISL_491464 | Laboratorio de Referencia Nacional de Virus Respiratorio. Instituto Nacional de Salud Perú | Laboratorio de Referencia Nacional de Biotecnología y Biología Molecular. Instituto Nacional de Salud Perú | Carlos Padilla Rojas, Karolyn Vega Chozo, Priscila Lope Pari, Omar Caceres Rey, Marco Galarza Perez, Maribel Huaringa Nuñez, Johanna Balbuena Torrez, Henri Bailon Calderon, Nancy Rojas Serrano |
| EPI_ISL_491465, EPI_ISL_491466 | San Lazaro Hospital | Research Institute for Tropical Medicine | Ma. Angelica Tujan, Othoniel Jan Onza, Francisco Gerardo Polotan, Inez Andrea Medado, Criselda Bautista, Kirstyn Brunker, Edelwisa Mercado, Daria Manalo, Catalino Demetria |
| EPI_ISL_491467, EPI_ISL_491468 | Research Institute for Tropical Medicine | Research Institute for Tropical Medicine | Ma. Angelica Tujan, Othoniel Jan Onza, Francisco Gerardo Polotan, Inez Andrea Medado, Criselda Bautista, Kirstyn Brunker, Edelwisa Mercado, Daria Manalo, Catalino Demetria |
| EPI_ISL_491469 | St. Luke's Medical Center - Quezon City | Research Institute for Tropical Medicine | Ma. Angelica Tujan, Othoniel Jan Onza, Francisco Gerardo Polotan, Inez Andrea Medado, Criselda Bautista, Kirstyn Brunker, Edelwisa Mercado, Daria Manalo, Catalino Demetria |
| EPI_ISL_491470 | Cardinal Santos Medical Center | Research Institute for Tropical Medicine | Ma. Angelica Tujan, Othoniel Jan Onza, Francisco Gerardo Polotan, Inez Andrea Medado, Criselda Bautista, Kirstyn Brunker, Edelwisa Mercado, Daria Manalo, Catalino Demetria |
| EPI_ISL_491471 | San Lazaro Hospital | Research Institute for Tropical Medicine | Ma. Angelica Tujan, Othoniel Jan Onza, Francisco Gerardo Polotan, Inez Andrea Medado, Criselda Bautista, Kirstyn Brunker, Edelwisa Mercado, Daria Manalo, Catalino Demetria |
| EPI_ISL_491472 | Cardinal Santos Medical Center | Research Institute for Tropical Medicine | Ma. Angelica Tujan, Othoniel Jan Onza, Francisco Gerardo Polotan, Inez Andrea Medado, Criselda Bautista, Kirstyn Brunker, Edelwisa Mercado, Daria Manalo, Catalino Demetria |
| EPI_ISL_491473 | The Medical City | Research Institute for Tropical Medicine | Ma. Angelica Tujan, Othoniel Jan Onza, Francisco Gerardo Polotan, Inez Andrea Medado, Criselda Bautista, Kirstyn Brunker, Edelwisa Mercado, Daria Manalo, Catalino Demetria |
| EPI_ISL_491474 | Research Institute for Tropical Medicine | Research Institute for Tropical Medicine | Ma. Angelica Tujan, Othoniel Jan Onza, Francisco Gerardo Polotan, Inez Andrea Medado, Criselda Bautista, Kirstyn Brunker, Edelwisa Mercado, Daria Manalo, Catalino Demetria |
| EPI_ISL_491475 | Protacio Hospital | Research Institute for Tropical Medicine | Ma. Angelica Tujan, Othoniel Jan Onza, Francisco Gerardo Polotan, Inez Andrea Medado, Criselda Bautista, Kirstyn Brunker, Edelwisa Mercado, Daria Manalo, Catalino Demetria |
| EPI_ISL_491476 | BSL3 Lab, Pendik Veterinary Control Enstitue | Genomic Laboratory (GLAB), Istanbul Technical University | Mustafa HASOKSUZ, Fahriye SARAC, Osman ERGANIS, Serdar UZAR, Hakan ENUL, Cumhur ADIAY, Ahmet SAIT, Orbay SAYI, Kadir YESILBAG, Oguz KARABEY |
| EPI_ISL_491477, EPI_ISL_491478, EPI_ISL_491479, EPI_ISL_491480 | CSIR-CDRI/SGPGI, Lucknow | CSIR-CDRI/SGPGI, Lucknow | Saumya Sarkar, Dharam Veer Singh, Rahul Vishvkarma, Ujjala Ghoshal, Uday Ghoshal, Ravishankar Ramachandran, Tapas Kumar Kundu, Rajender Singh |
| EPI_ISL_491481 | Functional Genomics Core University of South Carolina / Prisma Health–Midlands | Functional Genomics Core, Center For Targeted Therapeutics, | Hao Ji, Diego Altomare, B.Celia Cui, Mengqian Chen, Alyssa Clay-Glimour, Michael Wyatt, Phillip Buckhaults, Helmut Albrecht, Michael Shtutman |
| EPI_ISL_491494, EPI_ISL_491495, EPI_ISL_491496, EPI_ISL_491497, EPI_ISL_491498, EPI_ISL_491499, EPI_ISL_491500, EPI_ISL_491501, EPI_ISL_491502, EPI_ISL_491503, EPI_ISL_491504, EPI_ISL_491505, EPI_ISL_491506, EPI_ISL_491507, EPI_ISL_491508, EPI_ISL_491509, EPI_ISL_491510, EPI_ISL_491511, EPI_ISL_491512, EPI_ISL_491513, EPI_ISL_491514, EPI_ISL_491515, EPI_ISL_491516, EPI_ISL_491517, EPI_ISL_491518, EPI_ISL_491519, EPI_ISL_491520, EPI_ISL_491521, EPI_ISL_491522, EPI_ISL_491523, EPI_ISL_491524, EPI_ISL_491525, EPI_ISL_491526, EPI_ISL_491527, EPI_ISL_491528, EPI_ISL_491529, EPI_ISL_491530, EPI_ISL_491531, EPI_ISL_491532, EPI_ISL_491533, EPI_ISL_491534, EPI_ISL_491535, EPI_ISL_491536, EPI_ISL_491537, EPI_ISL_491538, EPI_ISL_491539, EPI_ISL_491540, EPI_ISL_491541, EPI_ISL_491542, EPI_ISL_491543, EPI_ISL_491544, EPI_ISL_491545, EPI_ISL_491546, EPI_ISL_491547, EPI_ISL_491548, EPI_ISL_491549, EPI_ISL_491550, EPI_ISL_491551, EPI_ISL_491552, EPI_ISL_491553, EPI_ISL_491554, EPI_ISL_491555, EPI_ISL_491556, EPI_ISL_491557, EPI_ISL_491558, EPI_ISL_491559, EPI_ISL_491560, EPI_ISL_491561, EPI_ISL_491562, EPI_ISL_491563, EPI_ISL_491564, EPI_ISL_491565, EPI_ISL_491566, EPI_ISL_491567, EPI_ISL_491568, EPI_ISL_491569, EPI_ISL_491570, EPI_ISL_491571, EPI_ISL_491573, EPI_ISL_491574, EPI_ISL_491575, EPI_ISL_491576, EPI_ISL_491577, EPI_ISL_491578, EPI_ISL_491579, EPI_ISL_491580, EPI_ISL_491581, EPI_ISL_491582, EPI_ISL_491583, EPI_ISL_491584, EPI_ISL_491585, EPI_ISL_491586, EPI_ISL_491587, EPI_ISL_491588, EPI_ISL_491589, EPI_ISL_491590, EPI_ISL_491591, EPI_ISL_491592, EPI_ISL_491593, EPI_ISL_491594, EPI_ISL_491595, EPI_ISL_491596, EPI_ISL_491597, EPI_ISL_491598, EPI_ISL_491599, EPI_ISL_491600, EPI_ISL_491601, EPI_ISL_491602, EPI_ISL_491603, EPI_ISL_491604, EPI_ISL_491605, EPI_ISL_491606, EPI_ISL_491607, EPI_ISL_491609, EPI_ISL_491610, EPI_ISL_491612, EPI_ISL_491613, EPI_ISL_491614, EPI_ISL_491615, EPI_ISL_491617, EPI_ISL_491618, EPI_ISL_491619, EPI_ISL_491620, EPI_ISL_491621, EPI_ISL_491622, EPI_ISL_491623, EPI_ISL_491624, EPI_ISL_491625, EPI_ISL_491626, EPI_ISL_491627, EPI_ISL_491628, EPI_ISL_491629, EPI_ISL_491630, EPI_ISL_491631, EPI_ISL_491632, EPI_ISL_491633, EPI_ISL_491634, EPI_ISL_491635, EPI_ISL_491636, EPI_ISL_491637, EPI_ISL_491638, EPI_ISL_491639, EPI_ISL_491640, EPI_ISL_491641, EPI_ISL_491642, EPI_ISL_491643, EPI_ISL_491644, EPI_ISL_491645, EPI_ISL_491646, EPI_ISL_491647, EPI_ISL_491648, EPI_ISL_491649, EPI_ISL_491650, EPI_ISL_491651, EPI_ISL_491652, EPI_ISL_491653, EPI_ISL_491654, EPI_ISL_491656, EPI_ISL_491657, EPI_ISL_491658, EPI_ISL_491659, EPI_ISL_491660, EPI_ISL_491661, EPI_ISL_491662, EPI_ISL_491663, EPI_ISL_491664, EPI_ISL_491665, EPI_ISL_491666, EPI_ISL_491667, EPI_ISL_491668, EPI_ISL_491669, EPI_ISL_491670, EPI_ISL_491671, EPI_ISL_491672, EPI_ISL_491673, EPI_ISL_491674, EPI_ISL_491675, EPI_ISL_491676, EPI_ISL_491677, EPI_ISL_491678, EPI_ISL_491679, EPI_ISL_491680, EPI_ISL_491681, EPI_ISL_491682, EPI_ISL_491683, EPI_ISL_491684, EPI_ISL_491685, EPI_ISL_491686, EPI_ISL_491687, EPI_ISL_491688, EPI_ISL_491689, EPI_ISL_491690, EPI_ISL_491691, EPI_ISL_491692, EPI_ISL_491693, EPI_ISL_491694, EPI_ISL_491695, EPI_ISL_491696, EPI_ISL_491697, EPI_ISL_491698, EPI_ISL_491699, EPI_ISL_491700 | | | |
| see above | Virology Department, Royal Infirmary of Edinburgh, NHS Lothian / School of Biological Sciences, University of Edinburgh | Wellcome Sanger Institute for the COVID-19 Genomics UK Consortium | McHugh M, Dewar R, Rooke S, O'Toole Á, Scher E, Hill V, McCrone JT, Colquhoun R, Yu X, Jackson B, Rambaut A, Templeton K and Alex Alderton, Roberto Amato, Sonia Goncalves, Ewan Harrison, David K. Jackson, Ian Johnston, Dominic Kwiatkowski, Cordelia Langford, John Sillitoe on behalf of the Wellcome Sanger Institute COVID-19 Surveillance Team (http://www.sanger.ac.uk/covid-team) |
| EPI_ISL_491709, EPI_ISL_491710, EPI_ISL_491711, EPI_ISL_491712, EPI_ISL_491713, EPI_ISL_491714, EPI_ISL_491715, EPI_ISL_491716, EPI_ISL_491717, EPI_ISL_491718, EPI_ISL_491719, EPI_ISL_491720, EPI_ISL_491721, EPI_ISL_491722, EPI_ISL_491723, EPI_ISL_491724, EPI_ISL_491725, EPI_ISL_491726, EPI_ISL_491727, EPI_ISL_491728, EPI_ISL_491729, EPI_ISL_491730, EPI_ISL_491731, EPI_ISL_491732, EPI_ISL_491733, EPI_ISL_491734, EPI_ISL_491735 | | | |
| see above | Respiratory Virus Unit, Microbiology Services Colindale, Public Health England | Respiratory Virus Unit, Microbiology Services Colindale, Public Health England | PHE Covid Sequencing Team |
| EPI_ISL_491907, EPI_ISL_491908, EPI_ISL_491909, EPI_ISL_491910, EPI_ISL_491911, EPI_ISL_491912, EPI_ISL_491913, EPI_ISL_491914, EPI_ISL_491915, EPI_ISL_491916, EPI_ISL_491917, EPI_ISL_491918, EPI_ISL_491919, EPI_ISL_491920, EPI_ISL_491921, EPI_ISL_491922, EPI_ISL_491923, EPI_ISL_491924, EPI_ISL_491925, EPI_ISL_491926, EPI_ISL_491927, EPI_ISL_491928, EPI_ISL_491929, EPI_ISL_491930, EPI_ISL_491931 | | | |
| see above | Naval Infectious Diseases Diagnostic Laboratory | Naval Medical Research Center Biological Defense Research Directorate | Logan Voegtly, Regina Cer, Lindsay Glang, Victor Sugiharto, Francisco Malgon Bautista, Hua Wei Chen, Dessiree Pena-Gomez, Megan Schilling, Adrian Paskey, Kyle Long, Mark Simons, Kimberly |

| | | | |
|---|---|---|---|
| | | | Bishop-Lilly |
| EPI_ISL_491932 | Institute of Microbiology, Universidad San Francisco de Quito | Institute of Microbiology, Universidad San Francisco de Quito | Belén Prado-Vivar, Sully Márquez, Juan José Guadalupe, Monica Becerra-Wong, Bernardo Gutiérrez, Carlos Mena, Nabih Dahik, Verónica Barragán, Patricio Rojas-Silva, Gabriel Trueba, Michelle Grunauer, Paúl Cárdenas |
| EPI_ISL_491933, EPI_ISL_491934, EPI_ISL_491935 | Centro de Investigaciones, Universidad de Especialidades Espíritu Santo | Institute of Microbiology, Universidad San Francisco de Quito | Derly Andrade, Juan Carlos Fernandez, Belén Prado-Vivar, Sully Márquez, Juan José Guadalupe, Monica Becerra-Wong, Bernardo Gutiérrez, Gabriel Morey, Ruben Armas, Jose Pedro Barberan, Fernando Espinoza, Edith Lopez, Verónica Barragán, Patricio Rojas-Silva, Gabriel Trueba, Michelle Grunauer, Paúl Cárdenas |
| EPI_ISL_491936 | Institute of Microbiology, Universidad San Francisco de Quito | Institute of Microbiology, Universidad San Francisco de Quito | Belén Prado-Vivar, Sully Márquez, Juan José Guadalupe, Monica Becerra-Wong, Bernardo Gutiérrez, Carlos Guerrero, Verónica Barragán, Patricio Rojas-Silva, Gabriel Trueba, Michelle Grunauer, Paúl Cárdenas |
| EPI_ISL_491937, EPI_ISL_491938 | Institute of Microbiology, Universidad San Francisco de Quito | Institute of Microbiology, Universidad San Francisco de Quito | Belén Prado-Vivar, Sully Márquez, Juan José Guadalupe, Monica Becerra-Wong, Bernardo Gutiérrez, Rosario Erazo, Verónica Barragán, Patricio Rojas-Silva, Gabriel Trueba, Michelle Grunauer, Paúl Cárdenas |
| EPI_ISL_491939, EPI_ISL_491940 | Institute of Microbiology, Universidad San Francisco de Quito | Institute of Microbiology, Universidad San Francisco de Quito | Belén Prado-Vivar, Sully Márquez, Juan José Guadalupe, Monica Becerra-Wong, Bernardo Gutiérrez, Carlos Mena, Nabih Dahik, Verónica Barragán, Patricio Rojas-Silva, Gabriel Trueba, Michelle Grunauer, Paúl Cárdenas |
| EPI_ISL_491941 | Centro de Investigaciones, Universidad de Especialidades Espíritu Santo | Institute of Microbiology, Universidad San Francisco de Quito | Derly Andrade, Juan Carlos Fernandez, Belén Prado-Vivar, Sully Márquez, Juan José Guadalupe, Monica Becerra-Wong, Bernardo Gutiérrez, Gabriel Morey, Ruben Armas, Jose Pedro Barberan, Fernando Espinoza, Edith Lopez, Verónica Barragán, Patricio Rojas-Silva, Gabriel Trueba, Michelle Grunauer, Paúl Cárdenas |
| EPI_ISL_491942, EPI_ISL_491943 | Naval Infectious Diseases Diagnostic Laboratory | Naval Medical Research Center Biological Defense Research Directorate | Logan Voegtly, Regina Cer, Lindsay Glang, Victor Sugiharto, Francisco Malgon Bautista, Hua Wei Chen, Dessiree Pena-Gomez, Megan Schilling, Adrian Paskey, Kyle Long, Mark Simons, Kimberly Bishop-Lilly |
| EPI_ISL_491944, EPI_ISL_491945, EPI_ISL_491946, EPI_ISL_491947, EPI_ISL_491948, EPI_ISL_491949, EPI_ISL_491950, EPI_ISL_491951, EPI_ISL_491952, EPI_ISL_491953, EPI_ISL_491954 | | | |
| see above | Instituto Nacional de Investigación en Salud Pública - INSPI | INSPI - Charité | Alfredo Bruno Caicedo, Domenica de Mora Coloma, Andres Moreira-Soto, Anna-Lena Sander, Nina Krause, Maritza Olmedo,Denisses Portugal, Manuel Gonzalez, Silvia Salgado, Alberto Orlando, Alexandra Usiña, Juan Carlos Zeballos, Jan Felix Drexler |
| EPI_ISL_491968, EPI_ISL_491969, EPI_ISL_491970, EPI_ISL_491971, EPI_ISL_491972, EPI_ISL_491973, EPI_ISL_491974, EPI_ISL_491975, EPI_ISL_491976, EPI_ISL_491977, EPI_ISL_491978, EPI_ISL_491979, EPI_ISL_491980, EPI_ISL_491981, EPI_ISL_491982, EPI_ISL_491983, EPI_ISL_491984, EPI_ISL_491985, EPI_ISL_491986, EPI_ISL_491987, EPI_ISL_491988, EPI_ISL_491989, EPI_ISL_491990, EPI_ISL_491991, EPI_ISL_491992, EPI_ISL_491993, EPI_ISL_491994, EPI_ISL_491995, EPI_ISL_491996, EPI_ISL_491997, EPI_ISL_491998, EPI_ISL_491999, EPI_ISL_492000, EPI_ISL_492001, EPI_ISL_492002, EPI_ISL_492003, EPI_ISL_492004, EPI_ISL_492005, EPI_ISL_492006, EPI_ISL_492007, EPI_ISL_492008, EPI_ISL_492009, EPI_ISL_492010, EPI_ISL_492011, EPI_ISL_492012, EPI_ISL_492013, EPI_ISL_492014, EPI_ISL_492015, EPI_ISL_492016, EPI_ISL_492017, EPI_ISL_492018, EPI_ISL_492019, EPI_ISL_492020, EPI_ISL_492021, EPI_ISL_492022, EPI_ISL_492023, EPI_ISL_492024, EPI_ISL_492025, EPI_ISL_492026 | | | |
| see above | Oman-NIC | Department of Microbiology and Immunology-SQUH | Fahad Zadjali, Samira Al-Maruqi, Amina Al Jardani, Khulood Al-Mammary, Hanan Al-kindi, Fatma BaAlawi, Hamida AL Barwani, Zeyana AL-Dahmani, Intisar Al-Shukri, Aisha Al-Busaidi, Aisha Al-Amri, Ahlam Al-Amri, Mohammed Al-Tobi, Samiha Al Kharusi, Abdulla Balkhair |
| EPI_ISL_492028, EPI_ISL_492029, EPI_ISL_492030, EPI_ISL_492031 | Child Health Research Foundation | Child Health Research Foundation | Senjuti Saha, Md Saiful Islam Sajib, Roly Malaker, Md Hafizur Rahman, Afroza Akter Tanni, Syed Muktadir Al Sium, Maksuda Islam, Samir K Saha |
| EPI_ISL_492032 | Instituto de Biologia do Exército | Laboratório Metabolismo Macromolecular FirminoTorres de Castro, Instituto de Biofísica Carlos Chagas Filho, Universidade Federal do Rio de Janeiro | Bianca Catarina Azevedo Cabral, Aline Rosa Vianna de Souza , Marcos Dornelas-Ribeiro, Tatiana LS Nogueira, Nádia Vaez Gonçalves da Cruz, Caleb GM Santos, Elizabeth Valentin, Marcio da Costa Cipitelli, Virginia Sara Grancieri do Amaral, Rodrigo Soares de Moura Neto, Clarissa Damaso, Rosane Silva |
| EPI_ISL_492033 | Instituto de Biologia do Exército | Laboratório Metabolismo Macromolecular FirminoTorres de Castro, Instituto de Biofísica Carlos Chagas Filho, Universidade Federal do Rio de Janeiro | Bianca Catarina Azevedo Cabral, Aline Rosa Vianna de Souza, Caleb GM Santos, Marcos Dornelas-Ribeiro, Tatiana LS Nogueira, Nádia Vaez Gonçalves da Cruz, Elizabeth Valentin, Marcio da Costa Cipitelli, Virginia Sara Grancieri do Amaral, Rodrigo Soares de Moura Neto, Clarissa Damaso, Rosane Silva |
| EPI_ISL_492034 | Instituto de Biologia do Exército | Laboratório Metabolismo Macromolecular FirminoTorres de Castro, Instituto de Biofísica Carlos Chagas Filho, Universidade Federal do Rio de Janeiro | Bianca Catarina Azevedo Cabral, Aline Rosa Vianna de Souza, Nádia Vaez Gonçalves da Cruz, Caleb GM Santos, Marcos Dornelas-Ribeiro, Tatiana LS Nogueira, Elizabeth Valentin, Marcio da Costa Cipitelli, Virginia Sara Grancieri do Amaral, Rodrigo Soares de Moura Neto, Clarissa Damaso, Rosane Silva |
| EPI_ISL_492035 | Instituto de Biologia do Exército | Laboratório Metabolismo Macromolecular FirminoTorres de Castro, Instituto de Biofísica Carlos Chagas Filho, Universidade Federal do Rio de Janeiro | Bianca Catarina Azevedo Cabral, Aline Rosa Vianna de Souza, Tatiana LS Nogueira, Nádia Vaez Gonçalves da Cruz, Caleb GM Santos, Marcos Dornelas-Ribeiro, Elizabeth Valentin, Marcio da Costa Cipitelli, Virginia Sara Grancieri do Amaral, Rodrigo Soares de Moura Neto, Clarissa Damaso, Rosane Silva |
| EPI_ISL_492036 | Instituto de Biologia do Exército | Laboratório Metabolismo Macromolecular FirminoTorres de Castro, Instituto de Biofísica Carlos Chagas Filho, Universidade Federal do Rio de Janeiro | Bianca Catarina Azevedo Cabral, Aline Rosa Vianna de Souza , Marcos Dornelas-Ribeiro, Tatiana LS Nogueira, Nádia Vaez Gonçalves da Cruz, Caleb GM Santos, Elizabeth Valentin, Marcio da Costa Cipitelli, Virginia Sara Grancieri do Amaral, Rodrigo Soares de Moura Neto, Clarissa Damaso, Rosane Silva |
| EPI_ISL_492037 | Instituto de Biologia do Exército | Laboratório Metabolismo Macromolecular FirminoTorres de Castro, Instituto de Biofísica Carlos Chagas Filho, Universidade Federal do Rio de Janeiro | Bianca Catarina Azevedo Cabral, Aline Rosa Vianna de Souza, Caleb GM Santos, Marcos Dornelas-Ribeiro, Tatiana LS Nogueira, Nádia Vaez Gonçalves da Cruz, Elizabeth Valentin, Marcio da Costa Cipitelli, Virginia Sara Grancieri do Amaral, Rodrigo Soares de Moura Neto, Clarissa Damaso, Rosane Silva |
| EPI_ISL_492038 | Instituto de Biologia do Exército | Laboratório Metabolismo Macromolecular FirminoTorres de Castro, Instituto de Biofísica Carlos Chagas Filho, Universidade Federal do Rio de Janeiro | Bianca Catarina Azevedo Cabral, Aline Rosa Vianna de Souza, Nádia Vaez Gonçalves da Cruz, Caleb GM Santos, Marcos Dornelas-Ribeiro, Tatiana LS Nogueira, Elizabeth Valentin, Marcio da Costa Cipitelli, Virginia Sara Grancieri do Amaral, Rodrigo Soares de Moura Neto, Clarissa Damaso, Rosane Silva |
| EPI_ISL_492039 | Instituto de Biologia do Exército | Laboratório Metabolismo Macromolecular FirminoTorres de Castro, Instituto de Biofísica Carlos Chagas Filho, Universidade Federal do Rio de Janeiro | Bianca Catarina Azevedo Cabral, Aline Rosa Vianna de Souza, Tatiana LS Nogueira, Nádia Vaez Gonçalves da Cruz, Caleb GM Santos, Marcos Dornelas-Ribeiro, Elizabeth Valentin, Marcio da Costa Cipitelli, Virginia Sara Grancieri do Amaral, Rodrigo Soares de Moura Neto, Clarissa Damaso, Rosane Silva |
| EPI_ISL_492040 | Instituto de Biologia do Exército | Laboratório Metabolismo Macromolecular FirminoTorres de Castro, Instituto de Biofísica Carlos Chagas Filho, Universidade Federal do Rio de Janeiro | Bianca Catarina Azevedo Cabral, Aline Rosa Vianna de Souza , Marcos Dornelas-Ribeiro, Tatiana LS Nogueira, Nádia Vaez Gonçalves da Cruz, Caleb GM Santos, Elizabeth Valentin, Marcio da Costa Cipitelli, Virginia Sara Grancieri do Amaral, Rodrigo Soares de Moura Neto, Clarissa Damaso, Rosane Silva |
| EPI_ISL_492041 | Instituto de Biologia do Exército | Laboratório Metabolismo Macromolecular FirminoTorres de Castro, Instituto de Biofísica Carlos Chagas Filho, Universidade Federal do Rio de Janeiro | Bianca Catarina Azevedo Cabral, Aline Rosa Vianna de Souza, Caleb GM Santos, Marcos Dornelas-Ribeiro, Tatiana LS Nogueira, Nádia Vaez Gonçalves da Cruz, Elizabeth Valentin, Marcio da Costa Cipitelli, Virginia Sara Grancieri do Amaral, Rodrigo Soares de Moura Neto, Clarissa Damaso, Rosane Silva |
| EPI_ISL_492042 | Instituto de Biologia do Exército | Laboratório Metabolismo Macromolecular FirminoTorres de Castro, Instituto de Biofísica Carlos Chagas Filho, Universidade Federal do Rio de Janeiro | Bianca Catarina Azevedo Cabral, Aline Rosa Vianna de Souza, Nádia Vaez Gonçalves da Cruz, Caleb GM Santos, Marcos Dornelas-Ribeiro, Tatiana LS Nogueira, Elizabeth Valentin, Marcio da Costa Cipitelli, Virginia Sara Grancieri do Amaral, Rodrigo Soares de Moura Neto, Clarissa Damaso, Rosane Silva |
| EPI_ISL_492043 | Instituto de Biologia do Exército | Laboratório Metabolismo Macromolecular FirminoTorres de Castro, Instituto de Biofísica Carlos Chagas Filho, Universidade Federal do Rio de Janeiro | Bianca Catarina Azevedo Cabral, Aline Rosa Vianna de Souza, Tatiana LS Nogueira, Nádia Vaez Gonçalves da Cruz, Caleb GM Santos, Marcos Dornelas-Ribeiro, Elizabeth Valentin, Marcio da Costa Cipitelli, Virginia Sara Grancieri do Amaral, Rodrigo Soares de Moura Neto, Clarissa Damaso, Rosane Silva |
| EPI_ISL_492044 | Instituto de Biologia do Exército | Laboratório Metabolismo Macromolecular FirminoTorres de Castro, Instituto de Biofísica Carlos Chagas Filho, Universidade Federal do Rio de Janeiro | Bianca Catarina Azevedo Cabral, Aline Rosa Vianna de Souza , Marcos Dornelas-Ribeiro, Tatiana LS Nogueira, Nádia Vaez Gonçalves da Cruz, Caleb GM Santos, Elizabeth Valentin, Marcio da Costa Cipitelli, Virginia Sara Grancieri do Amaral, Rodrigo Soares de Moura Neto, Clarissa Damaso, Rosane Silva |
| EPI_ISL_492045 | Instituto de Biologia do Exército | Laboratório Metabolismo Macromolecular FirminoTorres de Castro, Instituto de Biofísica Carlos Chagas Filho, Universidade Federal do Rio de Janeiro | Bianca Catarina Azevedo Cabral, Aline Rosa Vianna de Souza, Caleb GM Santos, Marcos Dornelas-Ribeiro, Tatiana LS Nogueira, Nádia Vaez Gonçalves da Cruz, Elizabeth Valentin, Marcio da Costa Cipitelli, Virginia Sara Grancieri do Amaral, Rodrigo Soares de Moura Neto, Clarissa Damaso, Rosane Silva |
| EPI_ISL_492046 | Instituto de Biologia do Exército | Laboratório Metabolismo Macromolecular FirminoTorres de Castro, Instituto de Biofísica Carlos Chagas Filho, Universidade Federal do Rio de Janeiro | Bianca Catarina Azevedo Cabral, Aline Rosa Vianna de Souza, Nádia Vaez Gonçalves da Cruz, Caleb GM Santos, Marcos Dornelas-Ribeiro, Tatiana LS Nogueira, Elizabeth Valentin, Marcio da Costa Cipitelli, Virginia Sara Grancieri do Amaral, Rodrigo Soares de Moura Neto, Clarissa Damaso, Rosane Silva |
| EPI_ISL_492047 | Instituto de Biologia do Exército | Laboratório Metabolismo Macromolecular FirminoTorres de Castro, Instituto de Biofísica Carlos Chagas Filho, Universidade Federal do Rio de Janeiro | Bianca Catarina Azevedo Cabral, Aline Rosa Vianna de Souza, Tatiana LS Nogueira, Nádia Vaez Gonçalves da Cruz, Caleb GM Santos, Marcos Dornelas-Ribeiro, Elizabeth Valentin, Marcio da Costa Cipitelli, Virginia Sara Grancieri do Amaral, Rodrigo Soares de Moura Neto, Clarissa Damaso, Rosane Silva |
| EPI_ISL_492048 | Instituto de Biologia do Exército | Laboratório Metabolismo Macromolecular FirminoTorres de Castro, Instituto de Biofísica Carlos Chagas Filho, Universidade Federal do Rio de Janeiro | Bianca Catarina Azevedo Cabral, Aline Rosa Vianna de Souza , Marcos Dornelas-Ribeiro, Tatiana LS Nogueira, Nádia Vaez Gonçalves da Cruz, Caleb GM Santos, Elizabeth Valentin, Marcio da Costa Cipitelli, Virginia Sara Grancieri do Amaral, Rodrigo Soares de Moura Neto, Clarissa Damaso, Rosane Silva |
| EPI_ISL_492049, EPI_ISL_492050, EPI_ISL_492051, EPI_ISL_492052, EPI_ISL_492053, EPI_ISL_492054, EPI_ISL_492055, EPI_ISL_492056, EPI_ISL_492057, EPI_ISL_492058, EPI_ISL_492059, EPI_ISL_492060, EPI_ISL_492061, EPI_ISL_492062, EPI_ISL_492063 | | | |
| see above | Alaska State Virology Laboratory | Alaska State Virology Laboratory | Chen J et al with Pathogenomics group Dagdag R, Redlinger M, Milton E, George W, Kovalenko A, Drown DM, Bortz E |
| EPI_ISL_492065 | Oman-National Influenza Center | Department of Microbiology and Immunology-SQUH Department of Microbiology and Immunology, Sultan Qaboos University Hopsital, P.O 35, Postal code 123 | Samira Al-Maruqi, Fahad Zadjali, Amina Al Jardani, Khulood Al-Mammary, Hanan Al-kindi, Fatma BaAlawi, Hamida AL Barwani, Zeyana AL-Dahmani, Intisar Al-Shukri, Azza Al-Rashdi, Samiha Al Kharusi, Abdulla Balkhair |
| EPI_ISL_492066, EPI_ISL_492067, EPI_ISL_492068, EPI_ISL_492069, EPI_ISL_492070, EPI_ISL_492071, EPI_ISL_492072, EPI_ISL_492073 | 1. ViroGenetics - BSL3 Laboratory of Virology, Małopolska Centre of Biotechnology, Jagiellonian University; 2. II Department of Internal Medicine, Faculty of Medicine, Jagiellonian University Medical College; 3. Narodowy Instytut Zdrowia Publicznego – Państwowy Zakład Higieny (NIZP-PZH) | 1. ViroGenetics - BSL3 Laboratory of Virology, Małopolska Centre of Biotechnology, Jagiellonian University; 2. II Department of Internal Medicine, Faculty of Medicine, Jagiellonian University Medical College; 3. Narodowy Instytut Zdrowia Publicznego – Państwowy Zakład Higieny (NIZP-PZH). | Katarzyna Pancer, Marek Sanak, Aleksandra A. Zasada, Magdalena Rzeczkowska, Tomasz Wołkowicz, Katarzyna Zacharczuk, Agnieszka Kołakowska-Kulesza, Katarzyna Owczarek, Aleksandra Milewska, Natalia Wolaniuk, Ewelina Hallman-Szelińska, Paweł P Łabaj, Wojciech Branicki, Krzysztof Pyrć |
| EPI_ISL_492074 | Functional Genomics Core University of South Carolina / Prisma Health–Midlands | Functional Genomics Core, University of South Carolina, | Hao Ji, Diego Altomare, B.Celia Cui, Mengqian Chen, Alyssa Clay-Glimour, Michael Wyatt, Phillip Buckhaults, Helmut Albrecht, Michael Shtutman |
| EPI_ISL_492075, EPI_ISL_492076, EPI_ISL_492077, EPI_ISL_492078, EPI_ISL_492079, EPI_ISL_492080, EPI_ISL_492081, EPI_ISL_492082, EPI_ISL_492083, EPI_ISL_492084, EPI_ISL_492085, EPI_ISL_492086 | | | |
| see above | Institute for Public Health of the Republic of North Macedonia | Charite Universitatsmedizin Berlin, Institute of Virology | Victor M Corman, Joern Beheim-Schwarzbach, Barbara Mühlemann, Talitha Veith, Julia Schneider, Elizabeta Jancheska, Maja Kuzmanovska, Golubinka Bosevska, Terry Jones, Christian Drosten |
| EPI_ISL_492087 | Alaska State Virology Laboratory | Alaska State Virology Laboratory | Chen J et al with Pathogenomics group Dagdag R, Redlinger M, Milton E, George W, Kovalenko A, Drown DM, Bortz E |
| EPI_ISL_492109, EPI_ISL_492110, EPI_ISL_492111, EPI_ISL_492112, EPI_ISL_492113, EPI_ISL_492114, EPI_ISL_492115, EPI_ISL_492116, EPI_ISL_492117, EPI_ISL_492118, EPI_ISL_492119, EPI_ISL_492120, EPI_ISL_492121, EPI_ISL_492122, EPI_ISL_492123, EPI_ISL_492124, EPI_ISL_492125, EPI_ISL_492126, EPI_ISL_492127, EPI_ISL_492128, EPI_ISL_492129, EPI_ISL_492130, EPI_ISL_492131, EPI_ISL_492132, EPI_ISL_492133, EPI_ISL_492134, EPI_ISL_492135, EPI_ISL_492136, EPI_ISL_492137, EPI_ISL_492138, EPI_ISL_492139, EPI_ISL_492140, EPI_ISL_492141, EPI_ISL_492142, EPI_ISL_492143, EPI_ISL_492144, EPI_ISL_492145, EPI_ISL_492146, EPI_ISL_492147, EPI_ISL_492148, EPI_ISL_492149, EPI_ISL_492150, EPI_ISL_492151, EPI_ISL_492152, EPI_ISL_492153, EPI_ISL_492154, EPI_ISL_492155, EPI_ISL_492156, EPI_ISL_492157, EPI_ISL_492158, EPI_ISL_492159, EPI_ISL_492160, EPI_ISL_492161, EPI_ISL_492162, EPI_ISL_492163, EPI_ISL_492164, EPI_ISL_492165, EPI_ISL_492166, EPI_ISL_492167, EPI_ISL_492168, EPI_ISL_492169, EPI_ISL_492170, EPI_ISL_492171, EPI_ISL_492172, EPI_ISL_492173, EPI_ISL_492174, EPI_ISL_492175, EPI_ISL_492176, EPI_ISL_492177, EPI_ISL_492178, EPI_ISL_492179, EPI_ISL_492180 | | | |
| see above | SA Pathology | SA Pathology | Lex Leong, Chuan Kok Lim, Mark Turra, Ivan Bastian, Geoff Higgins |
| EPI_ISL_492184 | INT Fondazione Pascale | INT Fondazione Pascale | Pascale |
| EPI_ISL_492185, EPI_ISL_492186, EPI_ISL_492187, EPI_ISL_492188 | PHE South West Regional Laboratory, National Infection Service | Wellcome Sanger Institute for the COVID-19 Genomics UK Consortium | Stephanie Hutchings, Hannah Pymont, Dr Peter Muir, Barry Vipond, Rich Hopes; and Alex Alderton, Roberto Amato, Sonia Goncalves, Ewan Harrison, David K. Jackson, Ian Johnston, Dominic Kwiatkowski, Cordelia Langford, John Sillitoe on behalf of the Wellcome Sanger Institute COVID-19 Surveillance Team (http://www.sanger.ac.uk/covid-team) |
| EPI_ISL_492189, EPI_ISL_492190, EPI_ISL_492191, EPI_ISL_492192, EPI_ISL_492193, EPI_ISL_492194 | Department of Pathology, University of Cambridge | Wellcome Sanger Institute for the COVID-19 Genomics UK Consortium | Luke W Meredith, M. Estée Török , Myra Hosmillo, William L. Hamilton, Martin D. Curran, Theresa Feltwell, Grant Hall, Anna Yakovleva, Fahad A Khokhar, Charlotte J. Houldcroft, Laura G Caller, Aminu S. Jahun, Sarah L. Caddy, Ian Goodfellow; and Alex Alderton, Roberto Amato, Sonia Goncalves, Ewan Harrison, David K. Jackson, Ian Johnston, Dominic Kwiatkowski, Cordelia Langford, John Sillitoe on behalf of the Wellcome Sanger Institute COVID-19 Surveillance Team (http://www.sanger.ac.uk/covid-team) |
| EPI_ISL_492195 | PHE South West Regional Laboratory, National Infection Service | Wellcome Sanger Institute for the COVID-19 Genomics UK Consortium | Stephanie Hutchings, Hannah Pymont, Dr Peter Muir, Barry Vipond, Rich Hopes; and Alex Alderton, Roberto Amato, Sonia Goncalves, Ewan Harrison, David K. Jackson, Ian Johnston, Dominic Kwiatkowski, Cordelia Langford, John Sillitoe on behalf of the Wellcome Sanger Institute COVID-19 Surveillance Team (http://www.sanger.ac.uk/covid-team) |
| EPI_ISL_492196 | Department of Pathology, University of Cambridge | Wellcome Sanger Institute for the COVID-19 Genomics UK Consortium | Luke W Meredith, M. Estée Török , Myra Hosmillo, William L. Hamilton, Martin D. Curran, Theresa Feltwell, Grant Hall, Anna Yakovleva, Fahad A Khokhar, Charlotte J. Houldcroft, Laura G Caller, Aminu S. Jahun, Sarah L. Caddy, Ian Goodfellow; and Alex Alderton, Roberto Amato, Sonia Goncalves, Ewan Harrison, David K. Jackson, Ian Johnston, Dominic Kwiatkowski, Cordelia Langford, John Sillitoe on behalf of the Wellcome Sanger Institute COVID-19 Surveillance Team (http://www.sanger.ac.uk/covid-team) |
| EPI_ISL_492197 | PHE South West Regional Laboratory, National Infection Service | Wellcome Sanger Institute for the COVID-19 Genomics UK Consortium | Stephanie Hutchings, Hannah Pymont, Dr Peter Muir, Barry Vipond, Rich Hopes; and Alex Alderton, Roberto Amato, Sonia Goncalves, Ewan Harrison, David K. Jackson, Ian Johnston, Dominic Kwiatkowski, Cordelia Langford, John Sillitoe on behalf of the Wellcome Sanger Institute COVID-19 Surveillance Team (http://www.sanger.ac.uk/covid-team) |

| | | | |
|---|---|---|---|
| EPI_ISL_492198, EPI_ISL_492199, EPI_ISL_492200, EPI_ISL_492201 | Department of Pathology, University of Cambridge | Wellcome Sanger Institute for the COVID-19 Genomics UK Consortium | Luke W Meredith, M. Estée Török , Myra Hosmillo, William L. Hamilton, Martin D. Curran, Theresa Feltwell, Grant Hall, Anna Yakovleva, Fahad A Khokhar, Charlotte J. Houldcroft, Laura G Caller, Aminu S. Jahun, Sarah L. Caddy, Ian Goodfellow; and Alex Alderton, Roberto Amato, Sonia Goncalves, Ewan Harrison, David K. Jackson, Ian Johnston, Dominic Kwiatkowski, Cordelia Langford, John Sillitoe on behalf of the Wellcome Sanger Institute COVID-19 Surveillance Team (http://www.sanger.ac.uk/covid-team) |
| EPI_ISL_492202, EPI_ISL_492203 | PHE South West Regional Laboratory, National Infection Service | Wellcome Sanger Institute for the COVID-19 Genomics UK Consortium | Stephanie Hutchings, Hannah Pymont, Dr Peter Muir, Barry Vipond, Rich Hopes; and Alex Alderton, Roberto Amato, Sonia Goncalves, Ewan Harrison, David K. Jackson, Ian Johnston, Dominic Kwiatkowski, Cordelia Langford, John Sillitoe on behalf of the Wellcome Sanger Institute COVID-19 Surveillance Team (http://www.sanger.ac.uk/covid-team) |
| EPI_ISL_492204, EPI_ISL_492205, EPI_ISL_492206 | Department of Pathology, University of Cambridge | Wellcome Sanger Institute for the COVID-19 Genomics UK Consortium | Luke W Meredith, M. Estée Török , Myra Hosmillo, William L. Hamilton, Martin D. Curran, Theresa Feltwell, Grant Hall, Anna Yakovleva, Fahad A Khokhar, Charlotte J. Houldcroft, Laura G Caller, Aminu S. Jahun, Sarah L. Caddy, Ian Goodfellow; and Alex Alderton, Roberto Amato, Sonia Goncalves, Ewan Harrison, David K. Jackson, Ian Johnston, Dominic Kwiatkowski, Cordelia Langford, John Sillitoe on behalf of the Wellcome Sanger Institute COVID-19 Surveillance Team (http://www.sanger.ac.uk/covid-team) |
| EPI_ISL_492207, EPI_ISL_492208, EPI_ISL_492209, EPI_ISL_492210 | NU-OMICS DNA Sequencing research facility, Northumbria University | Wellcome Sanger Institute for the COVID-19 Genomics UK Consortium | Chris Duncan, Sheia Waugh, Shirelle Burton-Fanning, Gary Eltringham, Jennifer Collins, Brendan Payne, Yusri Taha, Emma Swindells, Jane Greenaway, Edward Barton, Garren Scott, Debra Padgett, Clive Graham, Sarah Essex, Steve Ligget, Paul Baker, Lynn Dover, Wen Yew, Gary Black, John Allan, Joshua Loh, Greg Young, Matthew Bashton, Andrew Nelson, Darren Smith and Alex Alderton, Roberto Amato, Sonia Goncalves, Ewan Harrison, David K. Jackson, Ian Johnston, Dominic Kwiatkowski, Cordelia Langford, John Sillitoe on behalf of the Wellcome Sanger Institute COVID-19 Surveillance Team (http://www.sanger.ac.uk/covid-team) |
| EPI_ISL_492211, EPI_ISL_492212, EPI_ISL_492213, EPI_ISL_492214, EPI_ISL_492215, EPI_ISL_492216, EPI_ISL_492217, EPI_ISL_492218, EPI_ISL_492219, EPI_ISL_492220, EPI_ISL_492221, EPI_ISL_492222, EPI_ISL_492223, EPI_ISL_492224, EPI_ISL_492225, EPI_ISL_492226, EPI_ISL_492227, EPI_ISL_492228, EPI_ISL_492229, EPI_ISL_492230, EPI_ISL_492231, EPI_ISL_492232, EPI_ISL_492233, EPI_ISL_492234, EPI_ISL_492235, EPI_ISL_492236, EPI_ISL_492237, EPI_ISL_492238, EPI_ISL_492239, EPI_ISL_492240, EPI_ISL_492241, EPI_ISL_492242, EPI_ISL_492243, EPI_ISL_492244, EPI_ISL_492245, EPI_ISL_492246, EPI_ISL_492247, EPI_ISL_492248, EPI_ISL_492249, EPI_ISL_492250, EPI_ISL_492251, EPI_ISL_492252, EPI_ISL_492253, EPI_ISL_492254, EPI_ISL_492255, EPI_ISL_492256, EPI_ISL_492257, EPI_ISL_492258, EPI_ISL_492259, EPI_ISL_492260, EPI_ISL_492261, EPI_ISL_492262, EPI_ISL_492263, EPI_ISL_492264, EPI_ISL_492265, EPI_ISL_492266, EPI_ISL_492267, EPI_ISL_492268, EPI_ISL_492269, EPI_ISL_492270, EPI_ISL_492271, EPI_ISL_492272, EPI_ISL_492273, EPI_ISL_492274, EPI_ISL_492275, EPI_ISL_492276, EPI_ISL_492277, EPI_ISL_492278, EPI_ISL_492279, EPI_ISL_492280, EPI_ISL_492281, EPI_ISL_492282, EPI_ISL_492283, EPI_ISL_492284, EPI_ISL_492285, EPI_ISL_492286, EPI_ISL_492287, EPI_ISL_492288, EPI_ISL_492289, EPI_ISL_492290, EPI_ISL_492291, EPI_ISL_492292, EPI_ISL_492293, EPI_ISL_492294, EPI_ISL_492295, EPI_ISL_492296, EPI_ISL_492297, EPI_ISL_492298, EPI_ISL_492299, EPI_ISL_492300, EPI_ISL_492301, EPI_ISL_492302, EPI_ISL_492303, EPI_ISL_492304, EPI_ISL_492305, EPI_ISL_492306, EPI_ISL_492307, EPI_ISL_492308, EPI_ISL_492309, EPI_ISL_492310, EPI_ISL_492311, EPI_ISL_492312, EPI_ISL_492313, EPI_ISL_492314, EPI_ISL_492315, EPI_ISL_492316, EPI_ISL_492317, EPI_ISL_492318, EPI_ISL_492319, EPI_ISL_492320, EPI_ISL_492321, EPI_ISL_492322, EPI_ISL_492323, EPI_ISL_492324, EPI_ISL_492325, EPI_ISL_492326, EPI_ISL_492327, EPI_ISL_492328, EPI_ISL_492329, EPI_ISL_492330, EPI_ISL_492331, EPI_ISL_492332, EPI_ISL_492333, EPI_ISL_492334, EPI_ISL_492335, EPI_ISL_492336, EPI_ISL_492337, EPI_ISL_492338, EPI_ISL_492339, EPI_ISL_492340, EPI_ISL_492341, EPI_ISL_492342, EPI_ISL_492343, EPI_ISL_492344, EPI_ISL_492345, EPI_ISL_492346, EPI_ISL_492347, EPI_ISL_492348, EPI_ISL_492349, EPI_ISL_492350, EPI_ISL_492351, EPI_ISL_492352, EPI_ISL_492353, EPI_ISL_492354, EPI_ISL_492355, EPI_ISL_492356, EPI_ISL_492357, EPI_ISL_492358, EPI_ISL_492359, EPI_ISL_492360, EPI_ISL_492361, EPI_ISL_492362, EPI_ISL_492363, EPI_ISL_492364, EPI_ISL_492365, EPI_ISL_492366, EPI_ISL_492367, EPI_ISL_492368, EPI_ISL_492369, EPI_ISL_492370, EPI_ISL_492371, EPI_ISL_492372, EPI_ISL_492373, EPI_ISL_492374, EPI_ISL_492375, EPI_ISL_492376, EPI_ISL_492377, EPI_ISL_492378, EPI_ISL_492379, EPI_ISL_492380, EPI_ISL_492381, EPI_ISL_492382, EPI_ISL_492383, EPI_ISL_492384, EPI_ISL_492385, EPI_ISL_492386, EPI_ISL_492387, EPI_ISL_492388, EPI_ISL_492389, EPI_ISL_492390, EPI_ISL_492391, EPI_ISL_492392, EPI_ISL_492393, EPI_ISL_492394, EPI_ISL_492395, EPI_ISL_492396, EPI_ISL_492397, EPI_ISL_492398, EPI_ISL_492399, EPI_ISL_492400, EPI_ISL_492401, EPI_ISL_492402, EPI_ISL_492403, EPI_ISL_492404, EPI_ISL_492405, EPI_ISL_492406, EPI_ISL_492407, EPI_ISL_492408, EPI_ISL_492409, EPI_ISL_492410, EPI_ISL_492411, EPI_ISL_492412, EPI_ISL_492413, EPI_ISL_492414, EPI_ISL_492415, EPI_ISL_492416, EPI_ISL_492417, EPI_ISL_492418, EPI_ISL_492419, EPI_ISL_492420, EPI_ISL_492421, EPI_ISL_492422, EPI_ISL_492423, EPI_ISL_492424, EPI_ISL_492425, EPI_ISL_492426, EPI_ISL_492427, EPI_ISL_492428, EPI_ISL_492429, EPI_ISL_492430, EPI_ISL_492431, EPI_ISL_492432, EPI_ISL_492433, EPI_ISL_492434, EPI_ISL_492435, EPI_ISL_492436, EPI_ISL_492437, EPI_ISL_492438, EPI_ISL_492439, EPI_ISL_492440, EPI_ISL_492441, EPI_ISL_492442, EPI_ISL_492443, EPI_ISL_492444, EPI_ISL_492445, EPI_ISL_492446, EPI_ISL_492447, EPI_ISL_492448, EPI_ISL_492449 | | | |
| see above | PHE South West Regional Laboratory, National Infection Service | Wellcome Sanger Institute for the COVID-19 Genomics UK Consortium | Stephanie Hutchings, Hannah Pymont, Dr Peter Muir, Barry Vipond, Rich Hopes; and Alex Alderton, Roberto Amato, Sonia Goncalves, Ewan Harrison, David K. Jackson, Ian Johnston, Dominic Kwiatkowski, Cordelia Langford, John Sillitoe on behalf of the Wellcome Sanger Institute COVID-19 Surveillance Team (http://www.sanger.ac.uk/covid-team) |
| EPI_ISL_492450, EPI_ISL_492451, EPI_ISL_492452, EPI_ISL_492453, EPI_ISL_492454, EPI_ISL_492455, EPI_ISL_492456, EPI_ISL_492458, EPI_ISL_492459, EPI_ISL_492460, EPI_ISL_492461, EPI_ISL_492462, EPI_ISL_492463, EPI_ISL_492464, EPI_ISL_492465, EPI_ISL_492466, EPI_ISL_492468, EPI_ISL_492469, EPI_ISL_492470, EPI_ISL_492471, EPI_ISL_492472, EPI_ISL_492473, EPI_ISL_492474, EPI_ISL_492475, EPI_ISL_492476, EPI_ISL_492477, EPI_ISL_492478, EPI_ISL_492479, EPI_ISL_492480, EPI_ISL_492481, EPI_ISL_492482, EPI_ISL_492483, EPI_ISL_492484, EPI_ISL_492485, EPI_ISL_492486, EPI_ISL_492487, EPI_ISL_492488, EPI_ISL_492489, EPI_ISL_492490, EPI_ISL_492491, EPI_ISL_492492, EPI_ISL_492493, EPI_ISL_492494 | | | |
| see above | NU-OMICS DNA Sequencing research facility, Northumbria University | Wellcome Sanger Institute for the COVID-19 Genomics UK Consortium | Chris Duncan, Sheia Waugh, Shirelle Burton-Fanning, Gary Eltringham, Jennifer Collins, Brendan Payne, Yusri Taha, Emma Swindells, Jane Greenaway, Edward Barton, Garren Scott, Debra Padgett, Clive Graham, Sarah Essex, Steve Ligget, Paul Baker, Lynn Dover, Wen Yew, Gary Black, John Allan, Joshua Loh, Greg Young, Matthew Bashton, Andrew Nelson, Darren Smith and Alex Alderton, Roberto Amato, Sonia Goncalves, Ewan Harrison, David K. Jackson, Ian Johnston, Dominic Kwiatkowski, Cordelia Langford, John Sillitoe on behalf of the Wellcome Sanger Institute COVID-19 Surveillance Team (http://www.sanger.ac.uk/covid-team) |
| EPI_ISL_492496, EPI_ISL_492498, EPI_ISL_492499, EPI_ISL_492500, EPI_ISL_492501, EPI_ISL_492502, EPI_ISL_492503, EPI_ISL_492504 | University College London, Great Ormond Street Hospital for Children NHS Foundation Trust, Imperial College Healthcare NHS Trust | Wellcome Sanger Institute for the COVID-19 Genomics UK Consortium | Sergi Castellano, Rachel Williams, Mark Kristiansen, Paola Resende Silva, Sunando Roy, Tony Brooks, Helena Tutill, Paola Niola, Patricia Dyal, Charlotte Williams, Leysa Forrest, Yasmin Panchbhaya, Jacqueline Findlay, Sam Weeks, Julianne Brown, Kathryn Harris, Paul Randell, James Price, Alison Holmes, Judith Breuer and Alex Alderton, Roberto Amato, Sonia Goncalves, Ewan Harrison, David K. Jackson, Ian Johnston, Dominic Kwiatkowski, Cordelia Langford, John Sillitoe on behalf of the Wellcome Sanger Institute COVID-19 Surveillance Team (http://www.sanger.ac.uk/covid-team) |
| EPI_ISL_492505, EPI_ISL_492506, EPI_ISL_492507, EPI_ISL_492508, EPI_ISL_492509, EPI_ISL_492510, EPI_ISL_492511, EPI_ISL_492512, EPI_ISL_492513, EPI_ISL_492515, EPI_ISL_492516, EPI_ISL_492517, EPI_ISL_492518, EPI_ISL_492519, EPI_ISL_492520, EPI_ISL_492521, EPI_ISL_492522, EPI_ISL_492523, EPI_ISL_492524, EPI_ISL_492525, EPI_ISL_492526, EPI_ISL_492527, EPI_ISL_492528, EPI_ISL_492529, EPI_ISL_492530, EPI_ISL_492531, EPI_ISL_492532, EPI_ISL_492533, EPI_ISL_492534, EPI_ISL_492535, EPI_ISL_492536, EPI_ISL_492537, EPI_ISL_492538, EPI_ISL_492539, EPI_ISL_492540, EPI_ISL_492541, EPI_ISL_492542, EPI_ISL_492543, EPI_ISL_492544, EPI_ISL_492545, EPI_ISL_492546, EPI_ISL_492547, EPI_ISL_492548, EPI_ISL_492549, EPI_ISL_492550, EPI_ISL_492551, EPI_ISL_492552, EPI_ISL_492553, EPI_ISL_492554, EPI_ISL_492555, EPI_ISL_492556, EPI_ISL_492557, EPI_ISL_492558, EPI_ISL_492559, EPI_ISL_492560, EPI_ISL_492561, EPI_ISL_492562, EPI_ISL_492563, EPI_ISL_492564, EPI_ISL_492565, EPI_ISL_492566, EPI_ISL_492567, EPI_ISL_492568, EPI_ISL_492569, EPI_ISL_492570, EPI_ISL_492571, EPI_ISL_492572, EPI_ISL_492573, EPI_ISL_492574, EPI_ISL_492575, EPI_ISL_492576, EPI_ISL_492577, EPI_ISL_492578, EPI_ISL_492579, EPI_ISL_492580, EPI_ISL_492581, EPI_ISL_492582, EPI_ISL_492583, EPI_ISL_492584, EPI_ISL_492585, EPI_ISL_492586, EPI_ISL_492587, EPI_ISL_492588, EPI_ISL_492589, EPI_ISL_492590, EPI_ISL_492591, EPI_ISL_492592, EPI_ISL_492593, EPI_ISL_492595, EPI_ISL_492597, EPI_ISL_492598, EPI_ISL_492599, EPI_ISL_492600, EPI_ISL_492601, EPI_ISL_492602, EPI_ISL_492603, EPI_ISL_492604, EPI_ISL_492605, EPI_ISL_492606, EPI_ISL_492607, EPI_ISL_492608, EPI_ISL_492609, EPI_ISL_492610, EPI_ISL_492611, EPI_ISL_492612, EPI_ISL_492613, EPI_ISL_492614, EPI_ISL_492615, EPI_ISL_492616, EPI_ISL_492617, EPI_ISL_492618, EPI_ISL_492619, EPI_ISL_492620, EPI_ISL_492621, EPI_ISL_492622, EPI_ISL_492623, EPI_ISL_492624, EPI_ISL_492625, EPI_ISL_492626, EPI_ISL_492627, EPI_ISL_492628, EPI_ISL_492629, EPI_ISL_492630, EPI_ISL_492631, EPI_ISL_492632, EPI_ISL_492633, EPI_ISL_492634, EPI_ISL_492635, EPI_ISL_492636, EPI_ISL_492637, EPI_ISL_492638, EPI_ISL_492639, EPI_ISL_492640, EPI_ISL_492641, EPI_ISL_492642, EPI_ISL_492643, EPI_ISL_492644, EPI_ISL_492645, EPI_ISL_492646, EPI_ISL_492647, EPI_ISL_492648, EPI_ISL_492649, EPI_ISL_492650, EPI_ISL_492651, EPI_ISL_492652, EPI_ISL_492653, EPI_ISL_492654, EPI_ISL_492655, EPI_ISL_492656, EPI_ISL_492657, EPI_ISL_492658, EPI_ISL_492659, EPI_ISL_492660, EPI_ISL_492661, EPI_ISL_492662, EPI_ISL_492663, EPI_ISL_492664, EPI_ISL_492665, EPI_ISL_492666, EPI_ISL_492667, EPI_ISL_492668, EPI_ISL_492669, EPI_ISL_492670, EPI_ISL_492671, EPI_ISL_492672, EPI_ISL_492673, EPI_ISL_492674, EPI_ISL_492675, EPI_ISL_492676, EPI_ISL_492677, EPI_ISL_492678, EPI_ISL_492679, EPI_ISL_492680, EPI_ISL_492681, EPI_ISL_492682, EPI_ISL_492683, EPI_ISL_492684, EPI_ISL_492685, EPI_ISL_492686, EPI_ISL_492687, EPI_ISL_492688, EPI_ISL_492689, EPI_ISL_492690, EPI_ISL_492691, EPI_ISL_492692, EPI_ISL_492693, EPI_ISL_492694, EPI_ISL_492695, EPI_ISL_492696, EPI_ISL_492697, EPI_ISL_492698, EPI_ISL_492699, EPI_ISL_492700, EPI_ISL_492701, EPI_ISL_492702, EPI_ISL_492703, EPI_ISL_492704, EPI_ISL_492705, EPI_ISL_492706, EPI_ISL_492707, EPI_ISL_492708, EPI_ISL_492709, EPI_ISL_492710, EPI_ISL_492711, EPI_ISL_492712, EPI_ISL_492713, EPI_ISL_492714, EPI_ISL_492715, EPI_ISL_492716, EPI_ISL_492717, EPI_ISL_492718, EPI_ISL_492719, EPI_ISL_492720, EPI_ISL_492721, EPI_ISL_492722, EPI_ISL_492723, EPI_ISL_492724, EPI_ISL_492725, EPI_ISL_492726, EPI_ISL_492727, EPI_ISL_492728, EPI_ISL_492729, EPI_ISL_492730, EPI_ISL_492731, EPI_ISL_492732, EPI_ISL_492733, EPI_ISL_492734, EPI_ISL_492735, EPI_ISL_492736, EPI_ISL_492737, EPI_ISL_492738, EPI_ISL_492739, EPI_ISL_492740, EPI_ISL_492741, EPI_ISL_492742, EPI_ISL_492743, EPI_ISL_492744, EPI_ISL_492745 | | | |
| see above | PHE South West Regional Laboratory, National Infection Service | Wellcome Sanger Institute for the COVID-19 Genomics UK Consortium | Stephanie Hutchings, Hannah Pymont, Dr Peter Muir, Barry Vipond, Rich Hopes; and Alex Alderton, Roberto Amato, Sonia Goncalves, Ewan Harrison, David K. Jackson, Ian Johnston, Dominic Kwiatkowski, Cordelia Langford, John Sillitoe on behalf of the Wellcome Sanger Institute COVID-19 Surveillance Team (http://www.sanger.ac.uk/covid-team) |
| EPI_ISL_492747, EPI_ISL_492749, EPI_ISL_492751, EPI_ISL_492752, EPI_ISL_492753, EPI_ISL_492754, EPI_ISL_492755, EPI_ISL_492756, EPI_ISL_492757, EPI_ISL_492759, EPI_ISL_492760, EPI_ISL_492761, EPI_ISL_492762, EPI_ISL_492763 | | | |
| see above | University College London, Great Ormond Street Hospital for Children NHS Foundation Trust, Imperial College Healthcare NHS Trust | Wellcome Sanger Institute for the COVID-19 Genomics UK Consortium | Sergi Castellano, Rachel Williams, Mark Kristiansen, Paola Resende Silva, Sunando Roy, Tony Brooks, Helena Tutill, Paola Niola, Patricia Dyal, Charlotte Williams, Leysa Forrest, Yasmin Panchbhaya, Jacqueline Findlay, Sam Weeks, Julianne Brown, Kathryn Harris, Paul Randell, James Price, Alison Holmes, Judith Breuer and Alex Alderton, Roberto Amato, Sonia Goncalves, Ewan Harrison, David K. Jackson, Ian Johnston, Dominic Kwiatkowski, Cordelia Langford, John Sillitoe on behalf of the Wellcome Sanger Institute COVID-19 Surveillance Team (http://www.sanger.ac.uk/covid-team) |
| EPI_ISL_492764, EPI_ISL_492765, EPI_ISL_492766, EPI_ISL_492767, EPI_ISL_492768, EPI_ISL_492769, EPI_ISL_492770, EPI_ISL_492771, EPI_ISL_492772, EPI_ISL_492773, EPI_ISL_492774, EPI_ISL_492775, EPI_ISL_492776, EPI_ISL_492777, EPI_ISL_492778, EPI_ISL_492779, EPI_ISL_492780, EPI_ISL_492781, EPI_ISL_492782, EPI_ISL_492783, EPI_ISL_492784, EPI_ISL_492785, EPI_ISL_492786, EPI_ISL_492787, EPI_ISL_492788, EPI_ISL_492789, EPI_ISL_492790, EPI_ISL_492791, EPI_ISL_492792, EPI_ISL_492793, EPI_ISL_492794, EPI_ISL_492795, EPI_ISL_492796, EPI_ISL_492797, EPI_ISL_492798, EPI_ISL_492799, EPI_ISL_492800, EPI_ISL_492801, EPI_ISL_492802, EPI_ISL_492803, EPI_ISL_492804, EPI_ISL_492805, EPI_ISL_492806, EPI_ISL_492807, EPI_ISL_492808, EPI_ISL_492809, EPI_ISL_492810, EPI_ISL_492811, EPI_ISL_492812, EPI_ISL_492813, EPI_ISL_492814, EPI_ISL_492815, EPI_ISL_492816, EPI_ISL_492817, EPI_ISL_492818, EPI_ISL_492819, EPI_ISL_492820, EPI_ISL_492821, EPI_ISL_492822, EPI_ISL_492823, EPI_ISL_492824, EPI_ISL_492825, EPI_ISL_492826, EPI_ISL_492827, EPI_ISL_492828, EPI_ISL_492829, EPI_ISL_492830, EPI_ISL_492831, EPI_ISL_492832, EPI_ISL_492833, EPI_ISL_492834, EPI_ISL_492835 | | | |
| see above | PHE South West Regional Laboratory, National Infection Service | Wellcome Sanger Institute for the COVID-19 Genomics UK Consortium | Stephanie Hutchings, Hannah Pymont, Dr Peter Muir, Barry Vipond, Rich Hopes; and Alex Alderton, Roberto Amato, Sonia Goncalves, Ewan Harrison, David K. Jackson, Ian Johnston, Dominic Kwiatkowski, Cordelia Langford, John Sillitoe on behalf of the Wellcome Sanger Institute COVID-19 Surveillance Team (http://www.sanger.ac.uk/covid-team) |
| EPI_ISL_492836 | Department of Medical Microbiology, Western Sussex Hospitals NHS Foundation Trust, St Richard's Hospital | Wellcome Sanger Institute for the COVID-19 Genomics UK Consortium | Manasa Mutingwende, Sarah Lowdon, Olga Podplomyk, Michelle Erkiert, Jonathan Lewis, Paul Randell and Alex Alderton, Roberto Amato, Sonia Goncalves, Ewan Harrison, David K. Jackson, Ian Johnston, Dominic Kwiatkowski, Cordelia Langford, John Sillitoe on behalf of the Wellcome Sanger Institute COVID-19 Surveillance Team (http://www.sanger.ac.uk/covid-team) |
| EPI_ISL_492837 | Royal Free Hospital / Health Services Laboratories | Wellcome Sanger Institute for the COVID-19 Genomics UK Consortium | Tanzina Haque, Tabitha Mahungu, Dianne Irish, Cate Goodlad, Jenny Cross, Judith Heaney and Alex Alderton, Roberto Amato, Sonia Goncalves, Ewan Harrison, David K. Jackson, Ian Johnston, Dominic Kwiatkowski, Cordelia Langford, John Sillitoe on behalf of the Wellcome Sanger Institute COVID-19 Surveillance Team (http://www.sanger.ac.uk/covid-team) |
| EPI_ISL_492838, EPI_ISL_492839 | Department of Medical Microbiology, Western Sussex Hospitals NHS Foundation Trust, St Richard's Hospital | Wellcome Sanger Institute for the COVID-19 Genomics UK Consortium | Manasa Mutingwende, Sarah Lowdon, Olga Podplomyk, Michelle Erkiert, Jonathan Lewis, Paul Randell and Alex Alderton, Roberto Amato, Sonia Goncalves, Ewan Harrison, David K. Jackson, Ian Johnston, Dominic Kwiatkowski, Cordelia Langford, John Sillitoe on behalf of the Wellcome Sanger Institute COVID-19 Surveillance Team (http://www.sanger.ac.uk/covid-team) |
| EPI_ISL_492840 | Royal Free Hospital / Health Services Laboratories | Wellcome Sanger Institute for the COVID-19 Genomics UK Consortium | Tanzina Haque, Tabitha Mahungu, Dianne Irish, Cate Goodlad, Jenny Cross, Judith Heaney and Alex Alderton, Roberto Amato, Sonia Goncalves, Ewan Harrison, David K. Jackson, Ian Johnston, Dominic Kwiatkowski, Cordelia Langford, John Sillitoe on behalf of the Wellcome Sanger Institute COVID-19 Surveillance Team (http://www.sanger.ac.uk/covid-team) |
| EPI_ISL_492841, EPI_ISL_492842 | Department of Medical Microbiology, Western Sussex Hospitals NHS Foundation Trust, St Richard's Hospital | Wellcome Sanger Institute for the COVID-19 Genomics UK Consortium | Manasa Mutingwende, Sarah Lowdon, Olga Podplomyk, Michelle Erkiert, Jonathan Lewis, Paul Randell and Alex Alderton, Roberto Amato, Sonia Goncalves, Ewan Harrison, David K. Jackson, Ian Johnston, Dominic Kwiatkowski, Cordelia Langford, John Sillitoe on behalf of the Wellcome Sanger Institute COVID-19 Surveillance Team (http://www.sanger.ac.uk/covid-team) |
| EPI_ISL_492843, EPI_ISL_492844, EPI_ISL_492845, EPI_ISL_492846, EPI_ISL_492847 | Royal Free Hospital / Health Services Laboratories | Wellcome Sanger Institute for the COVID-19 Genomics UK Consortium | Tanzina Haque, Tabitha Mahungu, Dianne Irish, Cate Goodlad, Jenny Cross, Judith Heaney and Alex Alderton, Roberto Amato, Sonia Goncalves, Ewan Harrison, David K. Jackson, Ian Johnston, Dominic Kwiatkowski, Cordelia Langford, John Sillitoe on behalf of the Wellcome Sanger Institute COVID-19 Surveillance Team (http://www.sanger.ac.uk/covid-team) |
| EPI_ISL_492848, EPI_ISL_492849 | Department of Medical Microbiology, Western Sussex Hospitals NHS Foundation Trust, St Richard's Hospital | Wellcome Sanger Institute for the COVID-19 Genomics UK Consortium | Manasa Mutingwende, Sarah Lowdon, Olga Podplomyk, Michelle Erkiert, Jonathan Lewis, Paul Randell and Alex Alderton, Roberto Amato, Sonia Goncalves, Ewan Harrison, David K. Jackson, Ian Johnston, Dominic Kwiatkowski, Cordelia Langford, John Sillitoe on behalf of the Wellcome Sanger Institute COVID-19 Surveillance Team (http://www.sanger.ac.uk/covid-team) |
| EPI_ISL_492850, EPI_ISL_492851, EPI_ISL_492852 | Royal Free Hospital / Health Services Laboratories | Wellcome Sanger Institute for the COVID-19 Genomics UK Consortium | Tanzina Haque, Tabitha Mahungu, Dianne Irish, Cate Goodlad, Jenny Cross, Judith Heaney and Alex Alderton, Roberto Amato, Sonia Goncalves, Ewan Harrison, David K. Jackson, Ian Johnston, Dominic Kwiatkowski, Cordelia Langford, John Sillitoe on behalf of the Wellcome Sanger Institute COVID-19 Surveillance Team (http://www.sanger.ac.uk/covid-team) |
| EPI_ISL_492853, EPI_ISL_492854, EPI_ISL_492855 | Department of Medical Microbiology, Western Sussex Hospitals NHS Foundation Trust, St Richard's Hospital | Wellcome Sanger Institute for the COVID-19 Genomics UK Consortium | Manasa Mutingwende, Sarah Lowdon, Olga Podplomyk, Michelle Erkiert, Jonathan Lewis, Paul Randell and Alex Alderton, Roberto Amato, Sonia Goncalves, Ewan Harrison, David K. Jackson, Ian Johnston, Dominic Kwiatkowski, Cordelia Langford, John Sillitoe on behalf of the Wellcome Sanger Institute COVID-19 Surveillance Team (http://www.sanger.ac.uk/covid-team) |
| EPI_ISL_492856 | Royal Free Hospital / Health Services Laboratories | Wellcome Sanger Institute for the COVID-19 Genomics UK Consortium | Tanzina Haque, Tabitha Mahungu, Dianne Irish, Cate Goodlad, Jenny Cross, Judith Heaney and Alex Alderton, Roberto Amato, Sonia Goncalves, Ewan Harrison, David K. Jackson, Ian Johnston, Dominic Kwiatkowski, Cordelia Langford, John Sillitoe on behalf of the Wellcome Sanger Institute COVID-19 Surveillance Team (http://www.sanger.ac.uk/covid-team) |
| EPI_ISL_492857, EPI_ISL_492858, EPI_ISL_492859, EPI_ISL_492860, EPI_ISL_492861 | Department of Medical Microbiology, Western Sussex Hospitals NHS Foundation Trust, St Richard's Hospital | Wellcome Sanger Institute for the COVID-19 Genomics UK Consortium | Manasa Mutingwende, Sarah Lowdon, Olga Podplomyk, Michelle Erkiert, Jonathan Lewis, Paul Randell and Alex Alderton, Roberto Amato, Sonia Goncalves, Ewan Harrison, David K. Jackson, Ian Johnston, Dominic Kwiatkowski, Cordelia Langford, John Sillitoe on behalf of the Wellcome Sanger Institute COVID-19 Surveillance Team (http://www.sanger.ac.uk/covid-team) |
| EPI_ISL_492863, EPI_ISL_492864, EPI_ISL_492865 | Royal Free Hospital / Health Services Laboratories | Wellcome Sanger Institute for the COVID-19 Genomics UK Consortium | Tanzina Haque, Tabitha Mahungu, Dianne Irish, Cate Goodlad, Jenny Cross, Judith Heaney and Alex Alderton, Roberto Amato, Sonia Goncalves, Ewan Harrison, David K. Jackson, Ian Johnston, Dominic Kwiatkowski, Cordelia Langford, John Sillitoe on behalf of the Wellcome Sanger Institute COVID-19 Surveillance Team (http://www.sanger.ac.uk/covid-team) |
| EPI_ISL_492866, EPI_ISL_492867 | Department of Medical Microbiology, Western Sussex Hospitals NHS Foundation Trust, St Richard's Hospital | Wellcome Sanger Institute for the COVID-19 Genomics UK Consortium | Manasa Mutingwende, Sarah Lowdon, Olga Podplomyk, Michelle Erkiert, Jonathan Lewis, Paul Randell and Alex Alderton, Roberto Amato, Sonia Goncalves, Ewan Harrison, David K. Jackson, Ian Johnston, Dominic Kwiatkowski, Cordelia Langford, John Sillitoe on behalf of the Wellcome Sanger Institute COVID-19 Surveillance Team (http://www.sanger.ac.uk/covid-team) |
| EPI_ISL_492868, EPI_ISL_492869 | Royal Free Hospital / Health Services Laboratories | Wellcome Sanger Institute for the COVID-19 Genomics UK Consortium | Tanzina Haque, Tabitha Mahungu, Dianne Irish, Cate Goodlad, Jenny Cross, Judith Heaney and Alex Alderton, Roberto Amato, Sonia Goncalves, Ewan Harrison, David K. Jackson, Ian Johnston, Dominic Kwiatkowski, Cordelia Langford, John Sillitoe on behalf of the Wellcome Sanger Institute COVID-19 Surveillance Team (http://www.sanger.ac.uk/covid-team) |
| EPI_ISL_492870, EPI_ISL_492871 | Department of Medical Microbiology, Western Sussex Hospitals NHS Foundation Trust, St Richard's Hospital | Wellcome Sanger Institute for the COVID-19 Genomics UK Consortium | Manasa Mutingwende, Sarah Lowdon, Olga Podplomyk, Michelle Erkiert, Jonathan Lewis, Paul Randell and Alex Alderton, Roberto Amato, Sonia Goncalves, Ewan Harrison, David K. Jackson, Ian Johnston, Dominic Kwiatkowski, Cordelia Langford, John Sillitoe on behalf of the Wellcome Sanger Institute COVID-19 Surveillance Team (http://www.sanger.ac.uk/covid-team) |
| EPI_ISL_492872 | Royal Free Hospital / Health Services Laboratories | Wellcome Sanger Institute for the COVID-19 Genomics UK Consortium | Tanzina Haque, Tabitha Mahungu, Dianne Irish, Cate Goodlad, Jenny Cross, Judith Heaney and Alex Alderton, Roberto Amato, Sonia Goncalves, Ewan Harrison, David K. Jackson, Ian Johnston, Dominic Kwiatkowski, Cordelia Langford, John Sillitoe on behalf of the Wellcome Sanger Institute COVID-19 Surveillance Team (http://www.sanger.ac.uk/covid-team) |
| EPI_ISL_492873 | Department of Medical Microbiology, Western Sussex Hospitals NHS Foundation Trust, St Richard's Hospital | Wellcome Sanger Institute for the COVID-19 Genomics UK Consortium | Manasa Mutingwende, Sarah Lowdon, Olga Podplomyk, Michelle Erkiert, Jonathan Lewis, Paul Randell and Alex Alderton, Roberto Amato, Sonia Goncalves, Ewan Harrison, David K. Jackson, Ian Johnston, Dominic Kwiatkowski, Cordelia Langford, John Sillitoe on behalf of the Wellcome Sanger Institute COVID-19 Surveillance Team (http://www.sanger.ac.uk/covid-team) |
| EPI_ISL_492874 | Royal Free Hospital / Health Services Laboratories | Wellcome Sanger Institute for the COVID-19 Genomics UK Consortium | Tanzina Haque, Tabitha Mahungu, Dianne Irish, Cate Goodlad, Jenny Cross, Judith Heaney and Alex Alderton, Roberto Amato, Sonia Goncalves, Ewan Harrison, David K. Jackson, Ian Johnston, Dominic Kwiatkowski, Cordelia Langford, John Sillitoe on behalf of the Wellcome Sanger Institute COVID-19 Surveillance Team (http://www.sanger.ac.uk/covid-team) |
| EPI_ISL_492875 | Department of Medical Microbiology, Western Sussex Hospitals NHS Foundation Trust, St Richard's Hospital | Wellcome Sanger Institute for the COVID-19 Genomics UK Consortium | Manasa Mutingwende, Sarah Lowdon, Olga Podplomyk, Michelle Erkiert, Jonathan Lewis, Paul Randell and Alex Alderton, Roberto Amato, Sonia Goncalves, Ewan Harrison, David K. Jackson, Ian Johnston, Dominic Kwiatkowski, Cordelia Langford, John Sillitoe on behalf of the Wellcome Sanger Institute COVID-19 Surveillance Team (http://www.sanger.ac.uk/covid-team) |

| | | | |
|---|---|---|---|
| EPI_ISL_492876, EPI_ISL_492877 | Royal Free Hospital / Health Services Laboratories | Wellcome Sanger Institute for the COVID-19 Genomics UK Consortium | Tanzina Haque, Tabitha Mahungu, Dianne Irish, Cate Goodlad, Jenny Cross, Judith Heaney and Alex Alderton, Roberto Amato, Sonia Goncalves, Ewan Harrison, David K. Jackson, Ian Johnston, Dominic Kwiatkowski, Cordelia Langford, John Sillitoe on behalf of the Wellcome Sanger Institute COVID-19 Surveillance Team (http://www.sanger.ac.uk/covid-team) |
| EPI_ISL_492878 | Department of Medical Microbiology, Western Sussex Hospitals NHS Foundation Trust, St Richard's Hospital | Wellcome Sanger Institute for the COVID-19 Genomics UK Consortium | Manasa Mutingwende, Sarah Lowdon, Olga Podplomyk, Michelle Erkiert, Jonathan Lewis, Paul Randell and Alex Alderton, Roberto Amato, Sonia Goncalves, Ewan Harrison, David K. Jackson, Ian Johnston, Dominic Kwiatkowski, Cordelia Langford, John Sillitoe on behalf of the Wellcome Sanger Institute COVID-19 Surveillance Team (http://www.sanger.ac.uk/covid-team) |
| EPI_ISL_492879, EPI_ISL_492880 | Royal Free Hospital / Health Services Laboratories | Wellcome Sanger Institute for the COVID-19 Genomics UK Consortium | Tanzina Haque, Tabitha Mahungu, Dianne Irish, Cate Goodlad, Jenny Cross, Judith Heaney and Alex Alderton, Roberto Amato, Sonia Goncalves, Ewan Harrison, David K. Jackson, Ian Johnston, Dominic Kwiatkowski, Cordelia Langford, John Sillitoe on behalf of the Wellcome Sanger Institute COVID-19 Surveillance Team (http://www.sanger.ac.uk/covid-team) |
| EPI_ISL_492881 | Department of Medical Microbiology, Western Sussex Hospitals NHS Foundation Trust, St Richard's Hospital | Wellcome Sanger Institute for the COVID-19 Genomics UK Consortium | Manasa Mutingwende, Sarah Lowdon, Olga Podplomyk, Michelle Erkiert, Jonathan Lewis, Paul Randell and Alex Alderton, Roberto Amato, Sonia Goncalves, Ewan Harrison, David K. Jackson, Ian Johnston, Dominic Kwiatkowski, Cordelia Langford, John Sillitoe on behalf of the Wellcome Sanger Institute COVID-19 Surveillance Team (http://www.sanger.ac.uk/covid-team) |
| EPI_ISL_492882, EPI_ISL_492883 | Royal Free Hospital / Health Services Laboratories | Wellcome Sanger Institute for the COVID-19 Genomics UK Consortium | Tanzina Haque, Tabitha Mahungu, Dianne Irish, Cate Goodlad, Jenny Cross, Judith Heaney and Alex Alderton, Roberto Amato, Sonia Goncalves, Ewan Harrison, David K. Jackson, Ian Johnston, Dominic Kwiatkowski, Cordelia Langford, John Sillitoe on behalf of the Wellcome Sanger Institute COVID-19 Surveillance Team (http://www.sanger.ac.uk/covid-team) |
| EPI_ISL_492884 | Department of Medical Microbiology, Western Sussex Hospitals NHS Foundation Trust, St Richard's Hospital | Wellcome Sanger Institute for the COVID-19 Genomics UK Consortium | Manasa Mutingwende, Sarah Lowdon, Olga Podplomyk, Michelle Erkiert, Jonathan Lewis, Paul Randell and Alex Alderton, Roberto Amato, Sonia Goncalves, Ewan Harrison, David K. Jackson, Ian Johnston, Dominic Kwiatkowski, Cordelia Langford, John Sillitoe on behalf of the Wellcome Sanger Institute COVID-19 Surveillance Team (http://www.sanger.ac.uk/covid-team) |
| EPI_ISL_492885 | Royal Free Hospital / Health Services Laboratories | Wellcome Sanger Institute for the COVID-19 Genomics UK Consortium | Tanzina Haque, Tabitha Mahungu, Dianne Irish, Cate Goodlad, Jenny Cross, Judith Heaney and Alex Alderton, Roberto Amato, Sonia Goncalves, Ewan Harrison, David K. Jackson, Ian Johnston, Dominic Kwiatkowski, Cordelia Langford, John Sillitoe on behalf of the Wellcome Sanger Institute COVID-19 Surveillance Team (http://www.sanger.ac.uk/covid-team) |
| EPI_ISL_492886 | Department of Medical Microbiology, Western Sussex Hospitals NHS Foundation Trust, St Richard's Hospital | Wellcome Sanger Institute for the COVID-19 Genomics UK Consortium | Manasa Mutingwende, Sarah Lowdon, Olga Podplomyk, Michelle Erkiert, Jonathan Lewis, Paul Randell and Alex Alderton, Roberto Amato, Sonia Goncalves, Ewan Harrison, David K. Jackson, Ian Johnston, Dominic Kwiatkowski, Cordelia Langford, John Sillitoe on behalf of the Wellcome Sanger Institute COVID-19 Surveillance Team (http://www.sanger.ac.uk/covid-team) |
| EPI_ISL_492887, EPI_ISL_492888 | Royal Free Hospital / Health Services Laboratories | Wellcome Sanger Institute for the COVID-19 Genomics UK Consortium | Tanzina Haque, Tabitha Mahungu, Dianne Irish, Cate Goodlad, Jenny Cross, Judith Heaney and Alex Alderton, Roberto Amato, Sonia Goncalves, Ewan Harrison, David K. Jackson, Ian Johnston, Dominic Kwiatkowski, Cordelia Langford, John Sillitoe on behalf of the Wellcome Sanger Institute COVID-19 Surveillance Team (http://www.sanger.ac.uk/covid-team) |
| EPI_ISL_492889 | Department of Medical Microbiology, Western Sussex Hospitals NHS Foundation Trust, St Richard's Hospital | Wellcome Sanger Institute for the COVID-19 Genomics UK Consortium | Manasa Mutingwende, Sarah Lowdon, Olga Podplomyk, Michelle Erkiert, Jonathan Lewis, Paul Randell and Alex Alderton, Roberto Amato, Sonia Goncalves, Ewan Harrison, David K. Jackson, Ian Johnston, Dominic Kwiatkowski, Cordelia Langford, John Sillitoe on behalf of the Wellcome Sanger Institute COVID-19 Surveillance Team (http://www.sanger.ac.uk/covid-team) |
| EPI_ISL_492890 | Royal Free Hospital / Health Services Laboratories | Wellcome Sanger Institute for the COVID-19 Genomics UK Consortium | Tanzina Haque, Tabitha Mahungu, Dianne Irish, Cate Goodlad, Jenny Cross, Judith Heaney and Alex Alderton, Roberto Amato, Sonia Goncalves, Ewan Harrison, David K. Jackson, Ian Johnston, Dominic Kwiatkowski, Cordelia Langford, John Sillitoe on behalf of the Wellcome Sanger Institute COVID-19 Surveillance Team (http://www.sanger.ac.uk/covid-team) |
| EPI_ISL_492891, EPI_ISL_492892, EPI_ISL_492893 | Department of Medical Microbiology, Western Sussex Hospitals NHS Foundation Trust, St Richard's Hospital | Wellcome Sanger Institute for the COVID-19 Genomics UK Consortium | Manasa Mutingwende, Sarah Lowdon, Olga Podplomyk, Michelle Erkiert, Jonathan Lewis, Paul Randell and Alex Alderton, Roberto Amato, Sonia Goncalves, Ewan Harrison, David K. Jackson, Ian Johnston, Dominic Kwiatkowski, Cordelia Langford, John Sillitoe on behalf of the Wellcome Sanger Institute COVID-19 Surveillance Team (http://www.sanger.ac.uk/covid-team) |
| EPI_ISL_492894 | Royal Free Hospital / Health Services Laboratories | Wellcome Sanger Institute for the COVID-19 Genomics UK Consortium | Tanzina Haque, Tabitha Mahungu, Dianne Irish, Cate Goodlad, Jenny Cross, Judith Heaney and Alex Alderton, Roberto Amato, Sonia Goncalves, Ewan Harrison, David K. Jackson, Ian Johnston, Dominic Kwiatkowski, Cordelia Langford, John Sillitoe on behalf of the Wellcome Sanger Institute COVID-19 Surveillance Team (http://www.sanger.ac.uk/covid-team) |
| EPI_ISL_492895, EPI_ISL_492896, EPI_ISL_492897, EPI_ISL_492898, EPI_ISL_492899, EPI_ISL_492900 | Department of Medical Microbiology, Western Sussex Hospitals NHS Foundation Trust, St Richard's Hospital | Wellcome Sanger Institute for the COVID-19 Genomics UK Consortium | Manasa Mutingwende, Sarah Lowdon, Olga Podplomyk, Michelle Erkiert, Jonathan Lewis, Paul Randell and Alex Alderton, Roberto Amato, Sonia Goncalves, Ewan Harrison, David K. Jackson, Ian Johnston, Dominic Kwiatkowski, Cordelia Langford, John Sillitoe on behalf of the Wellcome Sanger Institute COVID-19 Surveillance Team (http://www.sanger.ac.uk/covid-team) |
| EPI_ISL_492901, EPI_ISL_492902 | Royal Free Hospital / Health Services Laboratories | Wellcome Sanger Institute for the COVID-19 Genomics UK Consortium | Tanzina Haque, Tabitha Mahungu, Dianne Irish, Cate Goodlad, Jenny Cross, Judith Heaney and Alex Alderton, Roberto Amato, Sonia Goncalves, Ewan Harrison, David K. Jackson, Ian Johnston, Dominic Kwiatkowski, Cordelia Langford, John Sillitoe on behalf of the Wellcome Sanger Institute COVID-19 Surveillance Team (http://www.sanger.ac.uk/covid-team) |
| EPI_ISL_492903, EPI_ISL_492904 | Department of Medical Microbiology, Western Sussex Hospitals NHS Foundation Trust, St Richard's Hospital | Wellcome Sanger Institute for the COVID-19 Genomics UK Consortium | Manasa Mutingwende, Sarah Lowdon, Olga Podplomyk, Michelle Erkiert, Jonathan Lewis, Paul Randell and Alex Alderton, Roberto Amato, Sonia Goncalves, Ewan Harrison, David K. Jackson, Ian Johnston, Dominic Kwiatkowski, Cordelia Langford, John Sillitoe on behalf of the Wellcome Sanger Institute COVID-19 Surveillance Team (http://www.sanger.ac.uk/covid-team) |
| EPI_ISL_492905, EPI_ISL_492906 | Royal Free Hospital / Health Services Laboratories | Wellcome Sanger Institute for the COVID-19 Genomics UK Consortium | Tanzina Haque, Tabitha Mahungu, Dianne Irish, Cate Goodlad, Jenny Cross, Judith Heaney and Alex Alderton, Roberto Amato, Sonia Goncalves, Ewan Harrison, David K. Jackson, Ian Johnston, Dominic Kwiatkowski, Cordelia Langford, John Sillitoe on behalf of the Wellcome Sanger Institute COVID-19 Surveillance Team (http://www.sanger.ac.uk/covid-team) |
| EPI_ISL_492907 | Department of Medical Microbiology, Western Sussex Hospitals NHS Foundation Trust, St Richard's Hospital | Wellcome Sanger Institute for the COVID-19 Genomics UK Consortium | Manasa Mutingwende, Sarah Lowdon, Olga Podplomyk, Michelle Erkiert, Jonathan Lewis, Paul Randell and Alex Alderton, Roberto Amato, Sonia Goncalves, Ewan Harrison, David K. Jackson, Ian Johnston, Dominic Kwiatkowski, Cordelia Langford, John Sillitoe on behalf of the Wellcome Sanger Institute COVID-19 Surveillance Team (http://www.sanger.ac.uk/covid-team) |
| EPI_ISL_492908 | Royal Free Hospital / Health Services Laboratories | Wellcome Sanger Institute for the COVID-19 Genomics UK Consortium | Tanzina Haque, Tabitha Mahungu, Dianne Irish, Cate Goodlad, Jenny Cross, Judith Heaney and Alex Alderton, Roberto Amato, Sonia Goncalves, Ewan Harrison, David K. Jackson, Ian Johnston, Dominic Kwiatkowski, Cordelia Langford, John Sillitoe on behalf of the Wellcome Sanger Institute COVID-19 Surveillance Team (http://www.sanger.ac.uk/covid-team) |
| EPI_ISL_492909, EPI_ISL_492910 | Department of Medical Microbiology, Western Sussex Hospitals NHS Foundation Trust, St Richard's Hospital | Wellcome Sanger Institute for the COVID-19 Genomics UK Consortium | Manasa Mutingwende, Sarah Lowdon, Olga Podplomyk, Michelle Erkiert, Jonathan Lewis, Paul Randell and Alex Alderton, Roberto Amato, Sonia Goncalves, Ewan Harrison, David K. Jackson, Ian Johnston, Dominic Kwiatkowski, Cordelia Langford, John Sillitoe on behalf of the Wellcome Sanger Institute COVID-19 Surveillance Team (http://www.sanger.ac.uk/covid-team) |
| EPI_ISL_492911 | Royal Free Hospital / Health Services Laboratories | Wellcome Sanger Institute for the COVID-19 Genomics UK Consortium | Tanzina Haque, Tabitha Mahungu, Dianne Irish, Cate Goodlad, Jenny Cross, Judith Heaney and Alex Alderton, Roberto Amato, Sonia Goncalves, Ewan Harrison, David K. Jackson, Ian Johnston, Dominic Kwiatkowski, Cordelia Langford, John Sillitoe on behalf of the Wellcome Sanger Institute COVID-19 Surveillance Team (http://www.sanger.ac.uk/covid-team) |
| EPI_ISL_492912, EPI_ISL_492913 | Department of Medical Microbiology, Western Sussex Hospitals NHS Foundation Trust, St Richard's Hospital | Wellcome Sanger Institute for the COVID-19 Genomics UK Consortium | Manasa Mutingwende, Sarah Lowdon, Olga Podplomyk, Michelle Erkiert, Jonathan Lewis, Paul Randell and Alex Alderton, Roberto Amato, Sonia Goncalves, Ewan Harrison, David K. Jackson, Ian Johnston, Dominic Kwiatkowski, Cordelia Langford, John Sillitoe on behalf of the Wellcome Sanger Institute COVID-19 Surveillance Team (http://www.sanger.ac.uk/covid-team) |
| EPI_ISL_492914 | Royal Free Hospital / Health Services Laboratories | Wellcome Sanger Institute for the COVID-19 Genomics UK Consortium | Tanzina Haque, Tabitha Mahungu, Dianne Irish, Cate Goodlad, Jenny Cross, Judith Heaney and Alex Alderton, Roberto Amato, Sonia Goncalves, Ewan Harrison, David K. Jackson, Ian Johnston, Dominic Kwiatkowski, Cordelia Langford, John Sillitoe on behalf of the Wellcome Sanger Institute COVID-19 Surveillance Team (http://www.sanger.ac.uk/covid-team) |
| EPI_ISL_492915 | Department of Medical Microbiology, Western Sussex Hospitals NHS Foundation Trust, St Richard's Hospital | Wellcome Sanger Institute for the COVID-19 Genomics UK Consortium | Manasa Mutingwende, Sarah Lowdon, Olga Podplomyk, Michelle Erkiert, Jonathan Lewis, Paul Randell and Alex Alderton, Roberto Amato, Sonia Goncalves, Ewan Harrison, David K. Jackson, Ian Johnston, Dominic Kwiatkowski, Cordelia Langford, John Sillitoe on behalf of the Wellcome Sanger Institute COVID-19 Surveillance Team (http://www.sanger.ac.uk/covid-team) |
| EPI_ISL_492916, EPI_ISL_492917, EPI_ISL_492918, EPI_ISL_492919, EPI_ISL_492920, EPI_ISL_492921, EPI_ISL_492922, EPI_ISL_492923, EPI_ISL_492924, EPI_ISL_492925, EPI_ISL_492926, EPI_ISL_492927, EPI_ISL_492928, EPI_ISL_492930, EPI_ISL_492932, EPI_ISL_492933, EPI_ISL_492934, EPI_ISL_492935, EPI_ISL_492936, EPI_ISL_492937, EPI_ISL_492938, EPI_ISL_492939, EPI_ISL_492940, EPI_ISL_492941, EPI_ISL_492942, EPI_ISL_492943, EPI_ISL_492944, EPI_ISL_492945, EPI_ISL_492946, EPI_ISL_492947, EPI_ISL_492948, EPI_ISL_492949, EPI_ISL_492950, EPI_ISL_492951, EPI_ISL_492952, EPI_ISL_492953, EPI_ISL_492954, EPI_ISL_492955, EPI_ISL_492956, EPI_ISL_492957, EPI_ISL_492958, EPI_ISL_492959, EPI_ISL_492960, EPI_ISL_492961, EPI_ISL_492962, EPI_ISL_492963, EPI_ISL_492964, EPI_ISL_492965, EPI_ISL_492966, EPI_ISL_492967, EPI_ISL_492968, EPI_ISL_492969, EPI_ISL_492970, EPI_ISL_492971, EPI_ISL_492972, EPI_ISL_492973, EPI_ISL_492974, EPI_ISL_492975, EPI_ISL_492976, EPI_ISL_492977 | | | |
| see above | NU-OMICS DNA Sequencing research facility, Northumbria University | Wellcome Sanger Institute for the COVID-19 Genomics UK Consortium | Chris Duncan, Sheia Waugh, Shirelle Burton-Fanning, Gary Eltringham, Jennifer Collins, Brendan Payne, Yusri Taha, Emma Swindells, Jane Greenaway, Edward Barton, Garren Scott, Debra Padgett, Clive Graham, Sarah Essex, Steve Ligget, Paul Baker, Lynn Dover, Wen Yew, Gary Black, John Allan, Joshua Loh, Greg Young, Matthew Bashton, Andrew Nelson, Darren Smith and Alex Alderton, Roberto Amato, Sonia Goncalves, Ewan Harrison, David K. Jackson, Ian Johnston, Dominic Kwiatkowski, Cordelia Langford, John Sillitoe on behalf of the Wellcome Sanger Institute COVID-19 Surveillance Team (http://www.sanger.ac.uk/covid-team) |
| EPI_ISL_492978, EPI_ISL_492979 | Department of Laboratory Medicine Tan Tock Seng Hospital | Department of Laboratory Medicine Tan Tock Seng Hospital | Chen YYC, Zair X, Li C, Tang WY, Maurer-Stroh S, Barkham TMS, Nagarajan N, Sessions OM |
| EPI_ISL_492980, EPI_ISL_492981 | IRCCS Sacro Cuore Don Calabria Hospital, Department of Infectious, Tropical Diseases & Microbiology | University of Verona, Department of Biotechnology | Antonio Mori, Michela Deiana, Elena Pomari, Chiara Piubelli; Giulia Lopatriello, Luca Marcolungo, Cristina Beltrami, Chiara Degli Esposti, Emanuela Cosentino, Massimo Delledonne |
| EPI_ISL_492982 | IRCCS Sacro Cuore Don Calabria Hospital, Department of Infectious, Tropical Diseases & Microbiology | University of Verona, Department of Biotechnology | Antonio Mori, Michela Deiana, Elena Pomari, Chiara Piubelli; Giulia Lopatriello, Luca Marcolungo, Cristina Beltrami, Chiara Degli Esposti, Emanuela Cosentino, Massimo Delledonne |
| EPI_ISL_492983, EPI_ISL_492984 | IRCCS Sacro Cuore Don Calabria Hospital, Department of Infectious, Tropical Diseases & Microbiology | University of Verona, Department of Biotechnology | Antonio Mori, Michela Deiana, Elena Pomari, Chiara Piubelli; Giulia Lopatriello, Luca Marcolungo, Cristina Beltrami, Chiara Degli Esposti, Emanuela Cosentino, Massimo Delledonne |
| EPI_ISL_492985, EPI_ISL_492986, EPI_ISL_492987 | IRCCS Sacro Cuore Don Calabria Hospital, Department of Infectious, Tropical Diseases & Microbiology | University of Verona, Department of Biotechnology | Antonio Mori, Michela Deiana, Elena Pomari, Chiara Piubelli; Giulia Lopatriello, Luca Marcolungo, Cristina Beltrami, Chiara Degli Esposti, Emanuela Cosentino, Massimo Delledonne |
| EPI_ISL_492988, EPI_ISL_492989 | Centrālā laboratorija | Latvian Biomedical Research and Study Centre | Ivars Silamiķelis, Kaspars Megnis, Monta Ustinova, Ņikita Zrelovs, Vita Rovīte, Stella Lapiņa, Jana Osīte, Marta Priedīte, Uga Dumpis, Jānis Kloviņš |
| EPI_ISL_492990 | Centrālā laboratorija | Latvian Biomedical Research and Study Centre | Ivars Silamiķelis, Kaspars Megnis, Monta Ustinova, Ņikita Zrelovs, Vita Rovīte, Stella Lapiņa, Jana Osīte, Marta Priedīte, Uga Dumpis, Jānis Kloviņš |
| EPI_ISL_492991, EPI_ISL_492992 | Centrālā laboratorija | Latvian Biomedical Research and Study Centre | Ivars Silamiķelis, Kaspars Megnis, Monta Ustinova, Ņikita Zrelovs, Vita Rovīte, Stella Lapiņa, Jana Osīte, Marta Priedīte, Uga Dumpis, Jānis Kloviņš |
| EPI_ISL_492993, EPI_ISL_492994, EPI_ISL_492995, EPI_ISL_492996, EPI_ISL_492997, EPI_ISL_492998, EPI_ISL_492999, EPI_ISL_493000 | E. Gulbja Laboratorija | Latvian Biomedical Research and Study Centre | Ivars Silamiķelis, Kaspars Megnis, Monta Ustinova, Ņikita Zrelovs, Vita Rovīte, Mikus Gavars, Dmitrijs Perminovs, Uga Dumpis, Jānis Kloviņš |
| EPI_ISL_493001 | Wyoming Public Health Laboratory | Wyoming Public Health Laboratory | Noah Hull, Rob Christensen, Jim Mildenberger, Joel Sevinsky, Cari Sloma, and Wanda Manley |
| EPI_ISL_493002, EPI_ISL_493003 | Respiratory Virus Unit, Microbiology Services Colindale, Public Health England | Respiratory Virus Unit, Microbiology Services Colindale, Public Health England | PHE Covid Sequencing Team |
| EPI_ISL_493004, EPI_ISL_493005, EPI_ISL_493006, EPI_ISL_493007, EPI_ISL_493008, EPI_ISL_493009, EPI_ISL_493010, EPI_ISL_493011, EPI_ISL_493012, EPI_ISL_493013, EPI_ISL_493014, EPI_ISL_493015, EPI_ISL_493016, EPI_ISL_493017, EPI_ISL_493018, EPI_ISL_493019, EPI_ISL_493020, EPI_ISL_493021, EPI_ISL_493022, EPI_ISL_493023, EPI_ISL_493024, EPI_ISL_493025, EPI_ISL_493026, EPI_ISL_493027, EPI_ISL_493028, EPI_ISL_493029, EPI_ISL_493030, EPI_ISL_493031, EPI_ISL_493032, EPI_ISL_493033, EPI_ISL_493034, EPI_ISL_493035, EPI_ISL_493036, EPI_ISL_493037, EPI_ISL_493038, EPI_ISL_493039, EPI_ISL_493040, EPI_ISL_493041, EPI_ISL_493042, EPI_ISL_493043, EPI_ISL_493044, EPI_ISL_493045, EPI_ISL_493046, EPI_ISL_493047, EPI_ISL_493048, EPI_ISL_493049, EPI_ISL_493050, EPI_ISL_493051, EPI_ISL_493052, EPI_ISL_493054, EPI_ISL_493055, EPI_ISL_493056, EPI_ISL_493057, EPI_ISL_493058, EPI_ISL_493059, EPI_ISL_493060 | | | |
| see above | Utah Public Health Laboratory | Utah Public Health Laboratory | Heidi Butz, Erin Young, Kelly Oakeson |
| EPI_ISL_493061, EPI_ISL_493062, EPI_ISL_493063, EPI_ISL_493064 | Wyoming Public Health Laboratory | Wyoming Public Health Laboratory | Noah Hull, Rob Christensen, Jim Mildenberger, Joel Sevinsky, Cari Sloma, and Wanda Manley |
| EPI_ISL_493069, EPI_ISL_493070, EPI_ISL_493071, EPI_ISL_493072, EPI_ISL_493073, EPI_ISL_493074, EPI_ISL_493075, EPI_ISL_493076, EPI_ISL_493077, EPI_ISL_493078, EPI_ISL_493079, EPI_ISL_493080, EPI_ISL_493081, EPI_ISL_493082, EPI_ISL_493083, EPI_ISL_493084, EPI_ISL_493085, EPI_ISL_493086 | | | |
| see above | Washington University in St. Louis | Washington University in St. Louis | David Wang, Carey-Ann Burnham, Scott Handley, Lindsay Droit, Stephen Tahan |
| EPI_ISL_493087, EPI_ISL_493088, EPI_ISL_493089, EPI_ISL_493090, EPI_ISL_493091, EPI_ISL_493092, EPI_ISL_493093, EPI_ISL_493094, EPI_ISL_493095, EPI_ISL_493096, EPI_ISL_493097, EPI_ISL_493098, EPI_ISL_493099, EPI_ISL_493100, EPI_ISL_493101, EPI_ISL_493102, EPI_ISL_493103, EPI_ISL_493104, EPI_ISL_493105, EPI_ISL_493106, EPI_ISL_493107, EPI_ISL_493108, EPI_ISL_493109, EPI_ISL_493110, EPI_ISL_493111, EPI_ISL_493112, EPI_ISL_493113, EPI_ISL_493114, EPI_ISL_493115, EPI_ISL_493116, EPI_ISL_493117, EPI_ISL_493118, EPI_ISL_493119, EPI_ISL_493120, EPI_ISL_493121, EPI_ISL_493122, EPI_ISL_493123, EPI_ISL_493124 | | | |
| see above | Utah Public Health Laboratory | Utah Public Health Laboratory | Heidi Butz, Erin Young, Kelly Oakeson |
| EPI_ISL_493125, EPI_ISL_493126, EPI_ISL_493127, EPI_ISL_493128 | Wyoming Public Health Laboratory | Wyoming Public Health Laboratory | Noah Hull, Rob Christensen, Jim Mildenberger, Joel Sevinsky, Cari Sloma, and Wanda Manley |
| EPI_ISL_493129, EPI_ISL_493130 | Functional Genomics Core University of South Carolina / Prisma Health–Midlands | Functional Genomics Core, University of South Carolina, | Hao Ji, Diego Altomare, B.Celia Cui, Mengqian Chen, Alyssa Clay-Glimour, Michael Wyatt, Phillip Buckhaults, Helmut Albrecht, Michael Shtutman |

| | | | |
|---|---|---|---|
| EPI_ISL_493137 | Center for Research and Innovation, Faculty of Medical Technology, Mahidol University | Center for Research and Innovation, Faculty of Medical Technology, Mahidol University | Kantima Sangsiriwut; Hatairat Lerdsamran; Jarunee Prasertsopon; Tipsuda Chanmanee; Anek Mungaomklang; Kamolthip Atsawawaranunt; Prabda Praphasiri; Somrak Sirikhetkon; Nattakan Thinpan; Pilaipan Puthavathana |
| EPI_ISL_493139 | Center for Research and Innovation, Faculty of Medical Technology, Mahidol University | Center for Research and Innovation, Faculty of Medical Technology, Mahidol University | Kantima Sangsiriwut; Hatairat Lerdsamran; Jarunee Prasertsopon; Tipsuda Chanmanee; Anek Mungaomklang; Kamolthip Atsawawaranunt; Prabda Praphasiri; Somrak Sirikhetkon; Nattakan Thinpan; Pilaipan Puthavathana |
| EPI_ISL_493149, EPI_ISL_493150, EPI_ISL_493151, EPI_ISL_493152, EPI_ISL_493153, EPI_ISL_493154, EPI_ISL_493155, EPI_ISL_493156, EPI_ISL_493157, EPI_ISL_493158, EPI_ISL_493159, EPI_ISL_493160, EPI_ISL_493161, EPI_ISL_493162, EPI_ISL_493163, EPI_ISL_493164, EPI_ISL_493165, EPI_ISL_493166, EPI_ISL_493167, EPI_ISL_493168, EPI_ISL_493169, EPI_ISL_493170, EPI_ISL_493171, EPI_ISL_493172, EPI_ISL_493173, EPI_ISL_493174, EPI_ISL_493175, EPI_ISL_493176, EPI_ISL_493177, EPI_ISL_493178, EPI_ISL_493179, EPI_ISL_493180, EPI_ISL_493181, EPI_ISL_493182, EPI_ISL_493183, EPI_ISL_493184, EPI_ISL_493185, EPI_ISL_493186, EPI_ISL_493187, EPI_ISL_493188, EPI_ISL_493189, EPI_ISL_493190 | | | |
| see above | National Virus Resource Center, Chinese Academy of Sciences, Wuhan 430071, China | Computational Virology Group, Center for Bacteria and Viruses Resources and Bioinformation, Wuhan Institute of Virology, Chinese Academy of Sciences Wuhan 430071, China | Jianjun Chen, Yi Yan, Yi Huang, Jin Xiong, Hongping Wei, Di Liu |
| EPI_ISL_493197 | INT Fondazione Pascale | INT Fondazione Pascale | Pascale |
| EPI_ISL_493198 | Virology Lab,Department of Pathology, National Cheng Kung University Hospital | Virology Lab,Department of Pathology, National Cheng Kung University Hospital | Huey-Pin Tsai, et al |
| EPI_ISL_493199 | Virology Lab,Department of Pathology, National Cheng Kung University Hospital | Virology Lab,Department of Pathology, National Cheng Kung University Hospital | Huey-Pin Tsai, et al |
| EPI_ISL_493200, EPI_ISL_493201, EPI_ISL_493202, EPI_ISL_493203, EPI_ISL_493204, EPI_ISL_493205, EPI_ISL_493206, EPI_ISL_493207 | Virology Lab,Department of Pathology, National Cheng Kung University Hospital | Virology Lab,Department of Pathology, National Cheng Kung University Hospital | Huey-Pin Tsai, et al |
| EPI_ISL_493208 | Virology Lab,Department of Pathology, National Cheng Kung University Hospital | Virology Lab,Department of Pathology, National Cheng Kung University Hospital | Huey-Pin Tsai et al |
| EPI_ISL_493209, EPI_ISL_493210, EPI_ISL_493211, EPI_ISL_493212 | Functional Genomics Core University of South Carolina / Prisma Health–Midlands | Functional Genomics Core, University of South Carolina | Hao Ji, Diego Altomare, B.Celia Cui, Mengqian Chen, Alyssa Clay-Glimour, Michael Wyatt, Phillip Buckhaults, Helmut Albrecht, Michael Shtutman |
| EPI_ISL_493213 | INT Fondazione Pascale | INT Fondazione Pascale | Pascale |
| EPI_ISL_493328 | INMI Lazzaro Spallanzani IRCCS | INMI Lazzaro Spallanzani IRCCS | Martina Rueca, Cesare E.M. Gruber, Barbara Bartolini, Francesco Messina, Maria R. Capobianchi, Antonino Di Caro |
| EPI_ISL_493329 | INMI Lazzaro Spallanzani IRCCS | INMI Lazzaro Spallanzani IRCCS | Barbara Bartolini, Martina Rueca, Cesare E.M. Gruber, Francesco Messina, Antonino Di Caro, Maria R. Capobianchi |
| EPI_ISL_493330 | INMI Lazzaro Spallanzani IRCCS | INMI Lazzaro Spallanzani IRCCS | Cesare E.M. Gruber, Martina Rueca, Barbara Bartolini, Francesco Messina, Maria R. Capobianchi, Antonino Di Caro |
| EPI_ISL_493331 | INMI Lazzaro Spallanzani IRCCS | INMI Lazzaro Spallanzani IRCCS | Martina Rueca, Cesare E.M. Gruber, Barbara Bartolini, Francesco Messina, Maria R. Capobianchi, Antonino Di Caro |
| EPI_ISL_493332 | Istituto Zooprofilattico Sperimentale del Mezzogiorno | INMI Lazzaro Spallanzani IRCCS | Cesare E.M. Gruber, Martina Rueca, Barbara Bartolini, Francesco Messina, Antonino Di Caro, Giovanna Fusco, Maurizio Viscardi, Giorgia Borriello, Maria R. Capobianchi |
| EPI_ISL_493333 | Istituto Zooprofilattico Sperimentale del Mezzogiorno | INMI Lazzaro Spallanzani IRCCS | Barbara Bartolini, Martina Rueca, Cesare E.M. Gruber, Francesco Messina, Antonino Di Caro, Giovanna Fusco, Maurizio Viscardi, Giorgia Borriello, Maria R. Capobianchi |
| EPI_ISL_493334, EPI_ISL_493335, EPI_ISL_493336, EPI_ISL_493337, EPI_ISL_493338 | Instituto de Diagnostico y Referencia Epidemiologicos (INDRE) | Instituto de Diagnostico y Referencia Epidemiologicos (INDRE) | Gisela Barrera-Badillo , Abril Rodriguez-Maldonado, Claudia Wong-Arambula , Natividad Cruz-Ortiz, Tatiana Nunez-Garcia, Dayanira Arellano-Suarez, Fabiola Garces-Ayala, Edgar Mendieta-Condado, Lucia Hernandez-Rivas, Irma Lopez-Martinez, Ernesto Ramirez-Gonzalez. |
| EPI_ISL_493339, EPI_ISL_493340, EPI_ISL_493341 | Instituto de Diagnostico y Referencia Epidemiologicos (INDRE) | Instituto de Diagnostico y Referencia Epidemiologicos (INDRE) | Gisela Barrera-Badillo , Abril Rodriguez-Maldonado, Claudia Wong-Arambula , Natividad Cruz-Ortiz, Tatiana Nunez-Garcia, Dayanira Arellano-Suarez, Adnan Araiza-Rodriguez, Edgar Mendieta-Condado, Lucia Hernandez-Rivas, Irma Lopez-Martinez, Ernesto Ramirez-Gonzalez. |
| EPI_ISL_493342, EPI_ISL_493343 | Instituto de Diagnostico y Referencia Epidemiologicos (INDRE) | Instituto de Diagnostico y Referencia Epidemiologicos (INDRE) | Ernesto Ramirez-Gonzalez, Abril Rodriguez-Maldonado, Claudia Wong-Arambula , Natividad Cruz-Ortiz, Tatiana Nunez-Garcia, Dayanira Arellano-Suarez, Adnan Araiza-Rodriguez, Edgar Mendieta-Condado, Lucia Hernandez-Rivas, Irma Lopez-Martinez, Gisela Barrera-Badillo. |
| EPI_ISL_493344, EPI_ISL_493345, EPI_ISL_493346, EPI_ISL_493347, EPI_ISL_493348, EPI_ISL_493349 | Instituto de Diagnostico y Referencia Epidemiologicos (INDRE) | Instituto de Diagnostico y Referencia Epidemiologicos (INDRE) | Ernesto Ramirez-Gonzalez, Abril Rodriguez-Maldonado, Claudia Wong-Arambula , Natividad Cruz-Ortiz, Tatiana Nunez-Garcia, Dayanira Arellano-Suarez, Adnan Araiza-Rodriguez, Fabiola Garces-Ayala, Lucia Hernandez-Rivas, Irma Lopez-Martinez, Gisela Barrera-Badillo. |
| EPI_ISL_493350 | Ostfold Hospital Trust - Kalnes, Centre for Laboratory Medicine, Section for gene technology and infection serology | Norwegian Institute of Public Health, Department of Virology | Kathrine Stene-Johansen, Kamilla Heddeland Instefjord, Hilde Elshaug, Rasmus Riis Kopperud, Karoline Bragstad, Olav Hungnes |
| EPI_ISL_493351, EPI_ISL_493352, EPI_ISL_493353, EPI_ISL_493354, EPI_ISL_493355 | Oslo University Hospital, Department of Medical Microbiology | Norwegian Institute of Public Health, Department of Virology | Kathrine Stene-Johansen, Kamilla Heddeland Instefjord, Hilde Elshaug, Rasmus Riis Kopperud, Karoline Bragstad, Olav Hungnes |
| EPI_ISL_493356, EPI_ISL_493357, EPI_ISL_493358, EPI_ISL_493359, EPI_ISL_493360, EPI_ISL_493361, EPI_ISL_493362, EPI_ISL_493363, EPI_ISL_493364, EPI_ISL_493365, EPI_ISL_493366, EPI_ISL_493367, EPI_ISL_493368, EPI_ISL_493369, EPI_ISL_493370, EPI_ISL_493371, EPI_ISL_493372, EPI_ISL_493373, EPI_ISL_493374, EPI_ISL_493375, EPI_ISL_493376, EPI_ISL_493377, EPI_ISL_493378, EPI_ISL_493379, EPI_ISL_493380 | | | |
| see above | Furst Medical Laboratory | Norwegian Institute of Public Health, Department of Virology | Kathrine Stene-Johansen, Kamilla Heddeland Instefjord, Hilde Elshaug, Rasmus Riis Kopperud, Karoline Bragstad, Olav Hungnes |
| EPI_ISL_493381, EPI_ISL_493382, EPI_ISL_493383 | Medical Microbiology Unit, Department for Laboratory Medicine, Drammen Hospital, Vestre Viken Health Trust, | Norwegian Institute of Public Health, Department of Virology | Kathrine Stene-Johansen, Kamilla Heddeland Instefjord, Hilde Elshaug, Rasmus Riis Kopperud, Karoline Bragstad, Olav Hungnes |
| EPI_ISL_493384 | Hospital of Southern Norway - Kristiansand, Department of Medical Microbiology | Norwegian Institute of Public Health, Department of Virology | Kathrine Stene-Johansen, Kamilla Heddeland Instefjord, Hilde Elshaug, Rasmus Riis Kopperud, Karoline Bragstad, Olav Hungnes |
| EPI_ISL_493385, EPI_ISL_493386, EPI_ISL_493387, EPI_ISL_493388, EPI_ISL_493389 | Oslo University Hospital, Department of Medical Microbiology | Norwegian Institute of Public Health, Department of Virology | Kathrine Stene-Johansen, Kamilla Heddeland Instefjord, Hilde Elshaug, Rasmus Riis Kopperud, Karoline Bragstad, Olav Hungnes |
| EPI_ISL_493390, EPI_ISL_493391, EPI_ISL_493392, EPI_ISL_493393, EPI_ISL_493394, EPI_ISL_493395, EPI_ISL_493396, EPI_ISL_493397, EPI_ISL_493398, EPI_ISL_493399, EPI_ISL_493400, EPI_ISL_493401, EPI_ISL_493402, EPI_ISL_493403, EPI_ISL_493404, EPI_ISL_493405, EPI_ISL_493406, EPI_ISL_493407, EPI_ISL_493408, EPI_ISL_493409, EPI_ISL_493410, EPI_ISL_493411, EPI_ISL_493412, EPI_ISL_493413, EPI_ISL_493414, EPI_ISL_493415, EPI_ISL_493416, EPI_ISL_493417, EPI_ISL_493418, EPI_ISL_493419, EPI_ISL_493420, EPI_ISL_493421, EPI_ISL_493422, EPI_ISL_493423, EPI_ISL_493424, EPI_ISL_493425 | | | |
| see above | National Public Health Laboratory, National Centre for Infectious Diseases | National Public Health Laboratory, National Centre for Infectious Diseases | Mak TM, Octavia S, Zhou Z, Chavatte JM, Cui L, Lin RTP |
| EPI_ISL_493426 | Functional Genomics Core University of South Carolina / Prisma Health–Midlands | Functional Genomics Core, University of South Carolina, | Hao Ji, Diego Altomare, B.Celia Cui, Mengqian Chen, Alyssa Clay-Glimour, Michael Wyatt, Phillip Buckhaults, Helmut Albrecht, Michael Shtutman |
| EPI_ISL_493427 | Functional Genomics Core University of South Carolina / Prisma Health–Midlands | Functional Genomics Core, University of South Carolina | Hao Ji, Diego Altomare, B.Celia Cui, Mengqian Chen, Alyssa Clay-Glimour, Michael Wyatt, Phillip Buckhaults, Helmut Albrecht, Michael Shtutman |
| EPI_ISL_493428, EPI_ISL_493429, EPI_ISL_493430, EPI_ISL_493431, EPI_ISL_493432, EPI_ISL_493433, EPI_ISL_493434, EPI_ISL_493435, EPI_ISL_493436, EPI_ISL_493437, EPI_ISL_493438, EPI_ISL_493439, EPI_ISL_493440, EPI_ISL_493441, EPI_ISL_493442, EPI_ISL_493443, EPI_ISL_493444, EPI_ISL_493445, EPI_ISL_493446, EPI_ISL_493447, EPI_ISL_493448, EPI_ISL_493449, EPI_ISL_493450, EPI_ISL_493451 | | | |
| see above | University of Birmingham | COVID-19 Genomics UK (COG-UK) Consortium | Institute of Microbiology, University of Birmingham: Claire McMurray, Joanne Stockton, Samuel Nicholls, Radoslaw Poplawski, Will Rowe, Josh Quick, Nicholas Loman. University of Birmingham Testing Laboratory: Celina M Whalley, Andrew Bosworth, Charlotte Poxon, Kasun Wanigasooriya, Oliver Pickles, Mike Kidd, Alex Richter, Andrew D Beggs PHE Heartlands Lab: Husam Osman, Andrew Bosworth. Queen Elizabeth Hospital: Anna Casey |
| EPI_ISL_493452, EPI_ISL_493453, EPI_ISL_493454, EPI_ISL_493455, EPI_ISL_493456, EPI_ISL_493457, EPI_ISL_493458, EPI_ISL_493459, EPI_ISL_493460, EPI_ISL_493461, EPI_ISL_493462, EPI_ISL_493463, EPI_ISL_493464, EPI_ISL_493465, EPI_ISL_493466, EPI_ISL_493467, EPI_ISL_493468, EPI_ISL_493469, EPI_ISL_493470, EPI_ISL_493471, EPI_ISL_493472, EPI_ISL_493473, EPI_ISL_493474, EPI_ISL_493475, EPI_ISL_493476, EPI_ISL_493477, EPI_ISL_493478, EPI_ISL_493479, EPI_ISL_493480, EPI_ISL_493481, EPI_ISL_493482, EPI_ISL_493483, EPI_ISL_493484, EPI_ISL_493485, EPI_ISL_493486, EPI_ISL_493487, EPI_ISL_493488, EPI_ISL_493489, EPI_ISL_493490, EPI_ISL_493491, EPI_ISL_493492, EPI_ISL_493493, EPI_ISL_493494, EPI_ISL_493495, EPI_ISL_493496, EPI_ISL_493497, EPI_ISL_493498, EPI_ISL_493499, EPI_ISL_493500, EPI_ISL_493501, EPI_ISL_493502, EPI_ISL_493503, EPI_ISL_493504, EPI_ISL_493505, EPI_ISL_493506, EPI_ISL_493507, EPI_ISL_493508, EPI_ISL_493509, EPI_ISL_493510, EPI_ISL_493511, EPI_ISL_493512, EPI_ISL_493513, EPI_ISL_493514, EPI_ISL_493515, EPI_ISL_493516, EPI_ISL_493517, EPI_ISL_493518, EPI_ISL_493519, EPI_ISL_493520, EPI_ISL_493521, EPI_ISL_493522, EPI_ISL_493523, EPI_ISL_493524, EPI_ISL_493525, EPI_ISL_493526, EPI_ISL_493527, EPI_ISL_493528, EPI_ISL_493529, EPI_ISL_493530, EPI_ISL_493531, EPI_ISL_493532, EPI_ISL_493533, EPI_ISL_493534, EPI_ISL_493535, EPI_ISL_493536, EPI_ISL_493537, EPI_ISL_493538, EPI_ISL_493539, EPI_ISL_493540 | | | |
| see above | Northumbria University / South Tees Hospitals NHS Foundation Trust / North Cumbria Integrated Care NHS Foundation Trust / North Tees and Hartlepool NHS Foundation Trust / Newcastle Hospitals NHS Foundation Trust | COVID-19 Genomics UK (COG-UK) Consortium | Darren L Smith,Andrew Nelson,Matthew Bashton,Greg R Young,Joshua Loh,John Allan,Mohammad A Tariq,Giles S Holt,Gary Black,Wen C Yew,Lynn Dover,Paul Baker,Steve Liggett,Sarah Essex,Jane Greenaway,Debra Padgett,Clive Graham,Garren Scott,Edward Barton,Emma Swindells,Brendan Payne,Jennifer Collins,Yusri Taha,Gary Eltringham |
| EPI_ISL_493541, EPI_ISL_493542, EPI_ISL_493543, EPI_ISL_493544, EPI_ISL_493545, EPI_ISL_493546 | Quadram Institute Bioscience | COVID-19 Genomics UK (COG-UK) Consortium | Dave J. Baker, Gemma L. Kay, Alp Aydin, Thanh Le-Viet, Steven Rudder, Ana P. Tedim, Anastasia Kolyva, Maria Diaz, Leonardo de Oliveira Martins, Nabil-Fareed Alikhan, Lizzie Meadows, Rachael Stanley, Ngozi Elumogo, Muhammed Yasir, Nicholas M. Thomson, Alexander J Trotter, Rachel Gilroy, Samuel Bloomfield, Claire Stuart, Andrew Bell, Reenesh Prakash, Samir Dervisevic, Alison E. Mather, John Wain, Mark Webber, Andrew J. Page, Justin O'Grady |
| EPI_ISL_493547, EPI_ISL_493548 | Queens Medical Centre, Clinical Microbiology Department / DeepSeq Nottingham | COVID-19 Genomics UK (COG-UK) Consortium | Gemma Clark, Wendy Smith, Manjinder Khakh, Vicki M Fleming, Michelle M Lister, Hannah Howson-Wells, Jonathan Ball, Patrick McClure, Joseph Chappell, Theocharis Tsoleridis, Nadine Holmes, Matthew Carlisle, Christopher Moore, Fei Sang, Johnny Debebe, Victoria Wright, Matthew Loose |
| EPI_ISL_493549, EPI_ISL_493550, EPI_ISL_493551, EPI_ISL_493552, EPI_ISL_493553, EPI_ISL_493554, EPI_ISL_493555, EPI_ISL_493556, EPI_ISL_493557, EPI_ISL_493558, EPI_ISL_493559, EPI_ISL_493560, EPI_ISL_493561, EPI_ISL_493562, EPI_ISL_493563, EPI_ISL_493564, EPI_ISL_493565, EPI_ISL_493566, EPI_ISL_493567, EPI_ISL_493568, EPI_ISL_493569, EPI_ISL_493570, EPI_ISL_493571, EPI_ISL_493572, EPI_ISL_493573, EPI_ISL_493574, EPI_ISL_493575, EPI_ISL_493576, EPI_ISL_493577, EPI_ISL_493578, EPI_ISL_493579, EPI_ISL_493580, EPI_ISL_493581, EPI_ISL_493582, EPI_ISL_493583, EPI_ISL_493584, EPI_ISL_493585, EPI_ISL_493586, EPI_ISL_493587, EPI_ISL_493588, EPI_ISL_493589, EPI_ISL_493590, EPI_ISL_493591, EPI_ISL_493592, EPI_ISL_493593, EPI_ISL_493594, EPI_ISL_493595, EPI_ISL_493596, EPI_ISL_493597, EPI_ISL_493598, EPI_ISL_493599, EPI_ISL_493600 | | | |
| see above | Lincolnshire Hospitals and DeepSeq Nottingham | COVID-19 Genomics UK (COG-UK) Consortium | Nichola Duckworth, Tim Sloan, Sarah Walsh, Jonathan Ball, Patrick McClure, Joeseph Chappell, Nadine Holmes, Matthew Carlisle, Christopher Moore, Fei Sang, Johnny Debebe, Victoria Wright, Matthew Loose |
| EPI_ISL_493601, EPI_ISL_493602 | Queens Medical Centre, Clinical Microbiology Department / DeepSeq Nottingham | COVID-19 Genomics UK (COG-UK) Consortium | Gemma Clark, Wendy Smith, Manjinder Khakh, Vicki M Fleming, Michelle M Lister, Hannah Howson-Wells, Jonathan Ball, Patrick McClure, Joseph Chappell, Theocharis Tsoleridis, Nadine Holmes, Matthew Carlisle, Christopher Moore, Fei Sang, Johnny Debebe, Victoria Wright, Matthew Loose |
| EPI_ISL_493603, EPI_ISL_493604, EPI_ISL_493605 | Lincolnshire Hospitals and DeepSeq Nottingham | COVID-19 Genomics UK (COG-UK) Consortium | Nichola Duckworth, Tim Sloan, Sarah Walsh, Jonathan Ball, Patrick McClure, Joeseph Chappell, Nadine Holmes, Matthew Carlisle, Christopher Moore, Fei Sang, Johnny Debebe, Victoria Wright, Matthew Loose |
| EPI_ISL_493606, EPI_ISL_493607, EPI_ISL_493608, EPI_ISL_493609 | Queens Medical Centre, Clinical Microbiology Department / DeepSeq Nottingham | COVID-19 Genomics UK (COG-UK) Consortium | Gemma Clark, Wendy Smith, Manjinder Khakh, Vicki M Fleming, Michelle M Lister, Hannah Howson-Wells, Jonathan Ball, Patrick McClure, Joseph Chappell, Theocharis Tsoleridis, Nadine Holmes, Matthew Carlisle, Christopher Moore, Fei Sang, Johnny Debebe, Victoria Wright, Matthew Loose |
| EPI_ISL_493610, EPI_ISL_493611, EPI_ISL_493612, EPI_ISL_493613 | Lincolnshire Hospitals and DeepSeq Nottingham | COVID-19 Genomics UK (COG-UK) Consortium | Nichola Duckworth, Tim Sloan, Sarah Walsh, Jonathan Ball, Patrick McClure, Joeseph Chappell, Nadine Holmes, Matthew Carlisle, Christopher Moore, Fei Sang, Johnny Debebe, Victoria Wright, Matthew Loose |
| EPI_ISL_493614, EPI_ISL_493615, EPI_ISL_493616, EPI_ISL_493617, EPI_ISL_493618, EPI_ISL_493619, EPI_ISL_493620, EPI_ISL_493621, EPI_ISL_493622, EPI_ISL_493623, EPI_ISL_493624, EPI_ISL_493625, EPI_ISL_493626, EPI_ISL_493627, EPI_ISL_493628, EPI_ISL_493629, EPI_ISL_493630, EPI_ISL_493631, EPI_ISL_493632, EPI_ISL_493633, EPI_ISL_493634, EPI_ISL_493635, EPI_ISL_493636, EPI_ISL_493637, EPI_ISL_493638, EPI_ISL_493639, EPI_ISL_493640, EPI_ISL_493641, EPI_ISL_493642, EPI_ISL_493643, EPI_ISL_493644, EPI_ISL_493645, EPI_ISL_493646, EPI_ISL_493647, EPI_ISL_493648, EPI_ISL_493649, EPI_ISL_493650, EPI_ISL_493651, EPI_ISL_493652, EPI_ISL_493653, EPI_ISL_493654, EPI_ISL_493655, EPI_ISL_493656, EPI_ISL_493657, EPI_ISL_493658, EPI_ISL_493659, EPI_ISL_493660, EPI_ISL_493661, EPI_ISL_493662, EPI_ISL_493663, EPI_ISL_493664, EPI_ISL_493665 | | | |

| | | | |
|---|---|---|---|
| see above | Centre for Enzyme Innovation, University of Portsmouth / Translational Research Laboratory, Portsmouth Hospitals NHS Trust | COVID-19 Genomics UK (COG-UK) Consortium | Angela Beckett,Yann Bourgeois,Garry Scarlett,Sharon Glaysher,Scott Elliott,Kelly Bicknell,Robert Impey,Allyson Lloyd,Sarah Wyllie,Ethan Butcher,Anoop Chauhan,Samuel Robson |
| EPI_ISL_493666, EPI_ISL_493667, EPI_ISL_493668, EPI_ISL_493669, EPI_ISL_493670, EPI_ISL_493671, EPI_ISL_493672, EPI_ISL_493673, EPI_ISL_493674, EPI_ISL_493675, EPI_ISL_493676, EPI_ISL_493677, EPI_ISL_493678, EPI_ISL_493679, EPI_ISL_493680, EPI_ISL_493681, EPI_ISL_493682, EPI_ISL_493683, EPI_ISL_493684, EPI_ISL_493685, EPI_ISL_493686, EPI_ISL_493687, EPI_ISL_493688, EPI_ISL_493689, EPI_ISL_493690, EPI_ISL_493691, EPI_ISL_493692, EPI_ISL_493693, EPI_ISL_493694, EPI_ISL_493695, EPI_ISL_493696, EPI_ISL_493697, EPI_ISL_493698, EPI_ISL_493699, EPI_ISL_493700, EPI_ISL_493701, EPI_ISL_493702, EPI_ISL_493703, EPI_ISL_493704, EPI_ISL_493705, EPI_ISL_493706, EPI_ISL_493707, EPI_ISL_493708, EPI_ISL_493709, EPI_ISL_493710, EPI_ISL_493711, EPI_ISL_493712, EPI_ISL_493713, EPI_ISL_493714, EPI_ISL_493715, EPI_ISL_493716, EPI_ISL_493717, EPI_ISL_493718, EPI_ISL_493719, EPI_ISL_493720, EPI_ISL_493721, EPI_ISL_493722, EPI_ISL_493723, EPI_ISL_493724, EPI_ISL_493725, EPI_ISL_493726, EPI_ISL_493727, EPI_ISL_493728, EPI_ISL_493729, EPI_ISL_493730, EPI_ISL_493731, EPI_ISL_493732, EPI_ISL_493733, EPI_ISL_493734, EPI_ISL_493735, EPI_ISL_493736, EPI_ISL_493737, EPI_ISL_493738, EPI_ISL_493739, EPI_ISL_493740 | | | |
| see above | Virology Department, Sheffield Teaching Hospitals NHS Foundation Trust/Department of Infection, Immunity and Cardiovascular Disease, The Medical School, University of Sheffield | COVID-19 Genomics UK (COG-UK) Consortium | Thushan de Silva, Matthew Parker, Nikki Smith, Adri Angyal, Rebecca Brown, Luke Green, Rachel Tucker, Paul Parsons, Danielle Groves, Katie Johnson, Laura Carrilero, Alex Keeley, Dave Partridge, Matthew Wyles, Benjamin Lindsey, Mehmet Yavuz, Mohammad Raza, Cariad Evans |
| EPI_ISL_493741, EPI_ISL_493742, EPI_ISL_493743, EPI_ISL_493744, EPI_ISL_493745, EPI_ISL_493746, EPI_ISL_493747, EPI_ISL_493748, EPI_ISL_493749, EPI_ISL_493750, EPI_ISL_493751, EPI_ISL_493752, EPI_ISL_493753, EPI_ISL_493754, EPI_ISL_493755, EPI_ISL_493756, EPI_ISL_493757, EPI_ISL_493758, EPI_ISL_493759, EPI_ISL_493760, EPI_ISL_493761, EPI_ISL_493762, EPI_ISL_493763, EPI_ISL_493764, EPI_ISL_493765, EPI_ISL_493766, EPI_ISL_493767, EPI_ISL_493768, EPI_ISL_493769, EPI_ISL_493770, EPI_ISL_493771, EPI_ISL_493772, EPI_ISL_493773, EPI_ISL_493774, EPI_ISL_493775, EPI_ISL_493776, EPI_ISL_493777, EPI_ISL_493778, EPI_ISL_493779, EPI_ISL_493780, EPI_ISL_493781, EPI_ISL_493782, EPI_ISL_493783, EPI_ISL_493784, EPI_ISL_493785, EPI_ISL_493786, EPI_ISL_493787, EPI_ISL_493788, EPI_ISL_493789, EPI_ISL_493790, EPI_ISL_493791, EPI_ISL_493792, EPI_ISL_493793, EPI_ISL_493794, EPI_ISL_493795, EPI_ISL_493796, EPI_ISL_493797, EPI_ISL_493798, EPI_ISL_493799, EPI_ISL_493800, EPI_ISL_493801, EPI_ISL_493802, EPI_ISL_493803, EPI_ISL_493804, EPI_ISL_493805, EPI_ISL_493806, EPI_ISL_493807, EPI_ISL_493808, EPI_ISL_493809, EPI_ISL_493810, EPI_ISL_493811, EPI_ISL_493812, EPI_ISL_493813, EPI_ISL_493814, EPI_ISL_493815, EPI_ISL_493816, EPI_ISL_493817, EPI_ISL_493818, EPI_ISL_493819, EPI_ISL_493820, EPI_ISL_493821, EPI_ISL_493822, EPI_ISL_493823, EPI_ISL_493824, EPI_ISL_493825, EPI_ISL_493826, EPI_ISL_493827, EPI_ISL_493828, EPI_ISL_493829, EPI_ISL_493830, EPI_ISL_493831, EPI_ISL_493832, EPI_ISL_493833, EPI_ISL_493834, EPI_ISL_493835, EPI_ISL_493836, EPI_ISL_493837, EPI_ISL_493838, EPI_ISL_493839, EPI_ISL_493840, EPI_ISL_493841, EPI_ISL_493842, EPI_ISL_493843, EPI_ISL_493844, EPI_ISL_493845, EPI_ISL_493846, EPI_ISL_493847, EPI_ISL_493848, EPI_ISL_493849, EPI_ISL_493850, EPI_ISL_493851, EPI_ISL_493852, EPI_ISL_493853, EPI_ISL_493854, EPI_ISL_493855, EPI_ISL_493856, EPI_ISL_493857, EPI_ISL_493858, EPI_ISL_493859, EPI_ISL_493860, EPI_ISL_493861, EPI_ISL_493862, EPI_ISL_493863, EPI_ISL_493864, EPI_ISL_493865, EPI_ISL_493866, EPI_ISL_493867, EPI_ISL_493868, EPI_ISL_493869, EPI_ISL_493870, EPI_ISL_493871, EPI_ISL_493872, EPI_ISL_493873, EPI_ISL_493874, EPI_ISL_493875, EPI_ISL_493876, EPI_ISL_493877, EPI_ISL_493878, EPI_ISL_493879, EPI_ISL_493880, EPI_ISL_493881, EPI_ISL_493882, EPI_ISL_493883, EPI_ISL_493884, EPI_ISL_493885, EPI_ISL_493886, EPI_ISL_493887, EPI_ISL_493888, EPI_ISL_493889, EPI_ISL_493890, EPI_ISL_493891, EPI_ISL_493892, EPI_ISL_493893, EPI_ISL_493894, EPI_ISL_493895, EPI_ISL_493896, EPI_ISL_493897 | | | |
| see above | West of Scotland Specialist Virology Centre, NHSGGC / MRC-University of Glasgow Centre for Virus Research | COVID-19 Genomics UK (COG-UK) Consortium | Ana da Silva Filipe, Natasha Johnson, Kathy Smollett, Daniel Mair, Stephen Carmichael, Lily Tong, Jenna Nichols, Elihu Aranday-Cortes, Kirstyn Brunker, Yasmin Parr, Alice Broos, Kyriaki Nomikou; Sarah McDonald, Marc Niebel, Patawee Asamaphan; Richard Orton, Joseph Hughes, Sreenu Vattipally, David L Robertson; Alasdair MacLean, Rory Gunson; Kathy Li, Natasha Jesudason, Rajiv Shah, James Shepherd, Antonia Ho, Emma Thomson |
| EPI_ISL_493898, EPI_ISL_493899, EPI_ISL_493900, EPI_ISL_493901, EPI_ISL_493902, EPI_ISL_493903, EPI_ISL_493904, EPI_ISL_493905, EPI_ISL_493906, EPI_ISL_493907, EPI_ISL_493908, EPI_ISL_493909, EPI_ISL_493910, EPI_ISL_493911, EPI_ISL_493912, EPI_ISL_493913, EPI_ISL_493914, EPI_ISL_493915, EPI_ISL_493916, EPI_ISL_493917, EPI_ISL_493918, EPI_ISL_493919, EPI_ISL_493920, EPI_ISL_493921, EPI_ISL_493922, EPI_ISL_493923, EPI_ISL_493924, EPI_ISL_493925, EPI_ISL_493926, EPI_ISL_493927, EPI_ISL_493928, EPI_ISL_493929, EPI_ISL_493930, EPI_ISL_493931, EPI_ISL_493932, EPI_ISL_493933, EPI_ISL_493934, EPI_ISL_493935, EPI_ISL_493936, EPI_ISL_493937, EPI_ISL_493938, EPI_ISL_493939, EPI_ISL_493940, EPI_ISL_493941, EPI_ISL_493942, EPI_ISL_493943, EPI_ISL_493944, EPI_ISL_493945, EPI_ISL_493946, EPI_ISL_493947, EPI_ISL_493948, EPI_ISL_493949, EPI_ISL_493950, EPI_ISL_493951, EPI_ISL_493952, EPI_ISL_493953, EPI_ISL_493954, EPI_ISL_493955, EPI_ISL_493956, EPI_ISL_493957, EPI_ISL_493958, EPI_ISL_493959, EPI_ISL_493960, EPI_ISL_493961, EPI_ISL_493962, EPI_ISL_493963, EPI_ISL_493964, EPI_ISL_493965, EPI_ISL_493966, EPI_ISL_493967, EPI_ISL_493968, EPI_ISL_493969, EPI_ISL_493970, EPI_ISL_493971, EPI_ISL_493972, EPI_ISL_493973, EPI_ISL_493974 | | | |
| see above | Virology Department, Royal Infirmary of Edinburgh, NHS Lothian / School of Biological Sciences, University of Edinburgh / Institute of Genetics and Molecular Medicine, University of Edinburgh | COVID-19 Genomics UK (COG-UK) Consortium | McHugh M, Dewar R, Rooke S, Gallagher M, Balcaza C, O'Toole Á, Scher E, Hill V, McCrone JT, Colquhoun R, Yu X, Jackson B, Rambaut A, Williams TC, Templeton K |
| EPI_ISL_493975, EPI_ISL_493976, EPI_ISL_493977, EPI_ISL_493978, EPI_ISL_493979, EPI_ISL_493980, EPI_ISL_493981, EPI_ISL_493982, EPI_ISL_493983, EPI_ISL_493984, EPI_ISL_493985, EPI_ISL_493986, EPI_ISL_493987, EPI_ISL_493988, EPI_ISL_493989, EPI_ISL_493990, EPI_ISL_493991, EPI_ISL_493992, EPI_ISL_493993, EPI_ISL_493994, EPI_ISL_493995, EPI_ISL_493996, EPI_ISL_493997, EPI_ISL_493998, EPI_ISL_493999, EPI_ISL_494000, EPI_ISL_494001, EPI_ISL_494002, EPI_ISL_494003, EPI_ISL_494004, EPI_ISL_494005, EPI_ISL_494006, EPI_ISL_494007, EPI_ISL_494008, EPI_ISL_494009, EPI_ISL_494010, EPI_ISL_494011, EPI_ISL_494012, EPI_ISL_494013, EPI_ISL_494014, EPI_ISL_494015, EPI_ISL_494016, EPI_ISL_494017, EPI_ISL_494018, EPI_ISL_494019, EPI_ISL_494020, EPI_ISL_494021, EPI_ISL_494022, EPI_ISL_494023, EPI_ISL_494024, EPI_ISL_494025, EPI_ISL_494026, EPI_ISL_494027, EPI_ISL_494028, EPI_ISL_494029, EPI_ISL_494030, EPI_ISL_494031, EPI_ISL_494032, EPI_ISL_494033, EPI_ISL_494034, EPI_ISL_494035, EPI_ISL_494036, EPI_ISL_494037, EPI_ISL_494038, EPI_ISL_494039, EPI_ISL_494040, EPI_ISL_494041, EPI_ISL_494042, EPI_ISL_494043, EPI_ISL_494044, EPI_ISL_494045, EPI_ISL_494046, EPI_ISL_494047, EPI_ISL_494048, EPI_ISL_494049, EPI_ISL_494050, EPI_ISL_494051, EPI_ISL_494052, EPI_ISL_494053, EPI_ISL_494054, EPI_ISL_494055, EPI_ISL_494056, EPI_ISL_494057, EPI_ISL_494058, EPI_ISL_494059, EPI_ISL_494060, EPI_ISL_494061, EPI_ISL_494062, EPI_ISL_494063, EPI_ISL_494064, EPI_ISL_494065, EPI_ISL_494066, EPI_ISL_494067, EPI_ISL_494068, EPI_ISL_494069, EPI_ISL_494070, EPI_ISL_494071, EPI_ISL_494072, EPI_ISL_494073, EPI_ISL_494074, EPI_ISL_494075, EPI_ISL_494076, EPI_ISL_494077, EPI_ISL_494078, EPI_ISL_494079, EPI_ISL_494080, EPI_ISL_494081, EPI_ISL_494082, EPI_ISL_494083, EPI_ISL_494084, EPI_ISL_494085, EPI_ISL_494086, EPI_ISL_494087, EPI_ISL_494088, EPI_ISL_494089, EPI_ISL_494090, EPI_ISL_494091, EPI_ISL_494092, EPI_ISL_494093, EPI_ISL_494094, EPI_ISL_494095, EPI_ISL_494096, EPI_ISL_494097, EPI_ISL_494098, EPI_ISL_494099, EPI_ISL_494100, EPI_ISL_494101, EPI_ISL_494102, EPI_ISL_494103, EPI_ISL_494104, EPI_ISL_494105, EPI_ISL_494106, EPI_ISL_494107, EPI_ISL_494108, EPI_ISL_494109, EPI_ISL_494110, EPI_ISL_494111, EPI_ISL_494112, EPI_ISL_494113, EPI_ISL_494114, EPI_ISL_494115, EPI_ISL_494116, EPI_ISL_494117, EPI_ISL_494118, EPI_ISL_494119, EPI_ISL_494120, EPI_ISL_494121, EPI_ISL_494122, EPI_ISL_494123, EPI_ISL_494124, EPI_ISL_494125, EPI_ISL_494126, EPI_ISL_494127, EPI_ISL_494128, EPI_ISL_494129, EPI_ISL_494130, EPI_ISL_494131, EPI_ISL_494132, EPI_ISL_494133, EPI_ISL_494134, EPI_ISL_494135, EPI_ISL_494136, EPI_ISL_494137, EPI_ISL_494138, EPI_ISL_494139, EPI_ISL_494140, EPI_ISL_494141, EPI_ISL_494142, EPI_ISL_494143, EPI_ISL_494144, EPI_ISL_494145, EPI_ISL_494146, EPI_ISL_494147, EPI_ISL_494148, EPI_ISL_494149, EPI_ISL_494150, EPI_ISL_494151, EPI_ISL_494152, EPI_ISL_494153, EPI_ISL_494154, EPI_ISL_494155, EPI_ISL_494156, EPI_ISL_494157, EPI_ISL_494158, EPI_ISL_494159, EPI_ISL_494160, EPI_ISL_494161, EPI_ISL_494162, EPI_ISL_494163, EPI_ISL_494164, EPI_ISL_494165, EPI_ISL_494166, EPI_ISL_494167, EPI_ISL_494168, EPI_ISL_494169, EPI_ISL_494170, EPI_ISL_494171, EPI_ISL_494172, EPI_ISL_494173, EPI_ISL_494174, EPI_ISL_494175, EPI_ISL_494176, EPI_ISL_494177, EPI_ISL_494178, EPI_ISL_494179, EPI_ISL_494180, EPI_ISL_494181, EPI_ISL_494182, EPI_ISL_494183, EPI_ISL_494184, EPI_ISL_494185, EPI_ISL_494186, EPI_ISL_494187, EPI_ISL_494188, EPI_ISL_494189, EPI_ISL_494190, EPI_ISL_494191, EPI_ISL_494192, EPI_ISL_494193, EPI_ISL_494194, EPI_ISL_494195, EPI_ISL_494196, EPI_ISL_494197, EPI_ISL_494198, EPI_ISL_494199, EPI_ISL_494200, EPI_ISL_494201, EPI_ISL_494202, EPI_ISL_494203, EPI_ISL_494204, EPI_ISL_494205, EPI_ISL_494206, EPI_ISL_494207, EPI_ISL_494208, EPI_ISL_494209, EPI_ISL_494210, EPI_ISL_494211, EPI_ISL_494212, EPI_ISL_494213, EPI_ISL_494214, EPI_ISL_494215, EPI_ISL_494216, EPI_ISL_494217, EPI_ISL_494218, EPI_ISL_494219, EPI_ISL_494220, EPI_ISL_494221, EPI_ISL_494222, EPI_ISL_494223, EPI_ISL_494224, EPI_ISL_494225, EPI_ISL_494226, EPI_ISL_494227, EPI_ISL_494228, EPI_ISL_494229, EPI_ISL_494230, EPI_ISL_494231, EPI_ISL_494232, EPI_ISL_494233, EPI_ISL_494234, EPI_ISL_494235, EPI_ISL_494236, EPI_ISL_494237, EPI_ISL_494238, EPI_ISL_494239, EPI_ISL_494240, EPI_ISL_494241, EPI_ISL_494242, EPI_ISL_494243, EPI_ISL_494244, EPI_ISL_494245, EPI_ISL_494246, EPI_ISL_494247, EPI_ISL_494248, EPI_ISL_494249, EPI_ISL_494250, EPI_ISL_494251, EPI_ISL_494252, EPI_ISL_494253, EPI_ISL_494254, EPI_ISL_494255, EPI_ISL_494256, EPI_ISL_494257, EPI_ISL_494258, EPI_ISL_494259, EPI_ISL_494260, EPI_ISL_494261, EPI_ISL_494262, EPI_ISL_494263, EPI_ISL_494264, EPI_ISL_494265, EPI_ISL_494266, EPI_ISL_494267, EPI_ISL_494268, EPI_ISL_494269, EPI_ISL_494270, EPI_ISL_494271, EPI_ISL_494272, EPI_ISL_494273, EPI_ISL_494274, EPI_ISL_494275, EPI_ISL_494276, EPI_ISL_494277, EPI_ISL_494278, EPI_ISL_494279, EPI_ISL_494280, EPI_ISL_494281, EPI_ISL_494282, EPI_ISL_494283, EPI_ISL_494284, EPI_ISL_494285, EPI_ISL_494286, EPI_ISL_494287, EPI_ISL_494288, EPI_ISL_494289, EPI_ISL_494290, EPI_ISL_494291, EPI_ISL_494292, EPI_ISL_494293, EPI_ISL_494294, EPI_ISL_494295, EPI_ISL_494296, EPI_ISL_494297, EPI_ISL_494298, EPI_ISL_494299, EPI_ISL_494300, EPI_ISL_494301, EPI_ISL_494302, EPI_ISL_494303, EPI_ISL_494304, EPI_ISL_494305, EPI_ISL_494306, EPI_ISL_494307, EPI_ISL_494308, EPI_ISL_494309, EPI_ISL_494310, EPI_ISL_494311, EPI_ISL_494312, EPI_ISL_494313, EPI_ISL_494314, EPI_ISL_494315, EPI_ISL_494316, EPI_ISL_494317, EPI_ISL_494318, EPI_ISL_494319, EPI_ISL_494320, EPI_ISL_494321, EPI_ISL_494322, EPI_ISL_494323, EPI_ISL_494324, EPI_ISL_494325, EPI_ISL_494326, EPI_ISL_494327, EPI_ISL_494328, EPI_ISL_494329, EPI_ISL_494330, EPI_ISL_494331, EPI_ISL_494332, EPI_ISL_494333, EPI_ISL_494334, EPI_ISL_494335, EPI_ISL_494336, EPI_ISL_494337, EPI_ISL_494338, EPI_ISL_494339, EPI_ISL_494340, EPI_ISL_494341, EPI_ISL_494342, EPI_ISL_494343, EPI_ISL_494344, EPI_ISL_494345, EPI_ISL_494346, EPI_ISL_494347, EPI_ISL_494348, EPI_ISL_494349, EPI_ISL_494350, EPI_ISL_494351, EPI_ISL_494352, EPI_ISL_494353, EPI_ISL_494354, EPI_ISL_494355, EPI_ISL_494356, EPI_ISL_494357, EPI_ISL_494358, EPI_ISL_494359, EPI_ISL_494360, EPI_ISL_494361, EPI_ISL_494362, EPI_ISL_494363, EPI_ISL_494364, EPI_ISL_494365, EPI_ISL_494366, EPI_ISL_494367, EPI_ISL_494368, EPI_ISL_494369, EPI_ISL_494370, EPI_ISL_494371 | | | |
| see above | Originating lab: Wales Specialist Virology Centre Sequencing lab: Pathogen Genomics Unit | COVID-19 Genomics UK (COG-UK) Consortium | Catherine Moore, Johnathan Evans, Laura Gifford, Malorie Perry, Simon Cottrell, Angela Marchbank, Alec Birchley, Alexander Adams, Amy Gaskin, Bree Gatica-Wilcox, Jason Coombes, Joel Southgate, Lauren Gilbert, Lee Graham, Nicole Pacchiarini, Sara Kumziene-Summerhayes, Sarah Taylor, Sophie Jones, Sara Rey, Matthew Bull, Joanne Watkins, Sally Corden, Tom Connor |
| EPI_ISL_494372, EPI_ISL_494373, EPI_ISL_494374, EPI_ISL_494375, EPI_ISL_494376, EPI_ISL_494377, EPI_ISL_494378, EPI_ISL_494379, EPI_ISL_494380, EPI_ISL_494381, EPI_ISL_494382, EPI_ISL_494383, EPI_ISL_494384, EPI_ISL_494385, EPI_ISL_494386, EPI_ISL_494387, EPI_ISL_494388, EPI_ISL_494389, EPI_ISL_494390, EPI_ISL_494391, EPI_ISL_494392, EPI_ISL_494393, EPI_ISL_494394, EPI_ISL_494395, EPI_ISL_494396, EPI_ISL_494397, EPI_ISL_494398, EPI_ISL_494399, EPI_ISL_494400, EPI_ISL_494401, EPI_ISL_494402, EPI_ISL_494403, EPI_ISL_494404, EPI_ISL_494405, EPI_ISL_494406, EPI_ISL_494407, EPI_ISL_494408, EPI_ISL_494409, EPI_ISL_494410, EPI_ISL_494411, EPI_ISL_494412, EPI_ISL_494413, EPI_ISL_494414, EPI_ISL_494415, EPI_ISL_494416, EPI_ISL_494417, EPI_ISL_494418, EPI_ISL_494419, EPI_ISL_494420, EPI_ISL_494421, EPI_ISL_494422, EPI_ISL_494423, EPI_ISL_494424, EPI_ISL_494425, EPI_ISL_494426, EPI_ISL_494427, EPI_ISL_494428, EPI_ISL_494429, EPI_ISL_494430, EPI_ISL_494431, EPI_ISL_494432, EPI_ISL_494433, EPI_ISL_494434, EPI_ISL_494435, EPI_ISL_494436, EPI_ISL_494437, EPI_ISL_494438, EPI_ISL_494439, EPI_ISL_494440, EPI_ISL_494441, EPI_ISL_494442, EPI_ISL_494443, EPI_ISL_494444, EPI_ISL_494445, EPI_ISL_494446, EPI_ISL_494447, EPI_ISL_494448, EPI_ISL_494449, EPI_ISL_494450, EPI_ISL_494451, EPI_ISL_494452, EPI_ISL_494453, EPI_ISL_494454, EPI_ISL_494455, EPI_ISL_494456, EPI_ISL_494457, EPI_ISL_494458, EPI_ISL_494459, EPI_ISL_494460, EPI_ISL_494461, EPI_ISL_494462, EPI_ISL_494463, EPI_ISL_494464, EPI_ISL_494465, EPI_ISL_494466, EPI_ISL_494467, EPI_ISL_494468, EPI_ISL_494469, EPI_ISL_494470, EPI_ISL_494471, EPI_ISL_494472, EPI_ISL_494473, EPI_ISL_494474, EPI_ISL_494475, EPI_ISL_494476, EPI_ISL_494477, EPI_ISL_494478, EPI_ISL_494479, EPI_ISL_494480, EPI_ISL_494481, EPI_ISL_494482, EPI_ISL_494483, EPI_ISL_494484, EPI_ISL_494485, EPI_ISL_494486, EPI_ISL_494487, EPI_ISL_494488, EPI_ISL_494489, EPI_ISL_494490, EPI_ISL_494491, EPI_ISL_494492, EPI_ISL_494493, EPI_ISL_494494, EPI_ISL_494495, EPI_ISL_494496, EPI_ISL_494497, EPI_ISL_494498, EPI_ISL_494499, EPI_ISL_494500, EPI_ISL_494501, EPI_ISL_494502, EPI_ISL_494503 | | | |
| see above | San Diego County Public Health Laboratory | Andersen lab at Scripps Research | SEARCH Alliance San Diego with Tracy Basler, Jovan Shephard, Brett Austin |
| EPI_ISL_494504, EPI_ISL_494505, EPI_ISL_494506, EPI_ISL_494507, EPI_ISL_494508, EPI_ISL_494509, EPI_ISL_494510, EPI_ISL_494511, EPI_ISL_494512, EPI_ISL_494513, EPI_ISL_494514, EPI_ISL_494515, EPI_ISL_494516, EPI_ISL_494517, EPI_ISL_494518, EPI_ISL_494519, EPI_ISL_494520, EPI_ISL_494521, EPI_ISL_494522, EPI_ISL_494523, EPI_ISL_494524, EPI_ISL_494525, EPI_ISL_494526, EPI_ISL_494527, EPI_ISL_494528, EPI_ISL_494529, EPI_ISL_494530, EPI_ISL_494531, EPI_ISL_494532, EPI_ISL_494533, EPI_ISL_494534, EPI_ISL_494535, EPI_ISL_494536, EPI_ISL_494537, EPI_ISL_494538, EPI_ISL_494539, EPI_ISL_494540, EPI_ISL_494541, EPI_ISL_494542, EPI_ISL_494543, EPI_ISL_494544, EPI_ISL_494545, EPI_ISL_494546, EPI_ISL_494547, EPI_ISL_494548, EPI_ISL_494549, EPI_ISL_494550, EPI_ISL_494551, EPI_ISL_494552 | | | |
| see above | Quest Diagnostics | Quest Diagnostics | Anderson,B.P., Rosenthal,S.H., Gerasimova,A., Kagan,R.M. and Owen, R. |
| EPI_ISL_494553, EPI_ISL_494554, EPI_ISL_494555, EPI_ISL_494558 | Functional Genomics Core University of South Carolina / Prisma Health–Midlands | Functional Genomics Core, University of South Carolina | Hao Ji, Diego Altomare, B.Celia Cui, Mengqian Chen, Alyssa Clay-Glimour, Michael Wyatt, Phillip Buckhaults, Helmut Albrecht, Michael Shtutman |
| EPI_ISL_494568, EPI_ISL_494569 | San Diego County Public Health Laboratory | Andersen lab at Scripps Research | SEARCH Alliance San Diego with Tracy Basler, Jovan Shephard, Brett Austin |
| EPI_ISL_494570 | Rady's Childrens Hospital | Andersen lab at Scripps Research | SEARCH Alliance San Diego |
| EPI_ISL_494571, EPI_ISL_494572, EPI_ISL_494573, EPI_ISL_494574, EPI_ISL_494575, EPI_ISL_494576, EPI_ISL_494577, EPI_ISL_494578, EPI_ISL_494579, EPI_ISL_494580, EPI_ISL_494581, EPI_ISL_494582, EPI_ISL_494583, EPI_ISL_494584, EPI_ISL_494585, EPI_ISL_494586, EPI_ISL_494587, EPI_ISL_494588, EPI_ISL_494589, EPI_ISL_494590, EPI_ISL_494591, EPI_ISL_494592, EPI_ISL_494593, EPI_ISL_494594, EPI_ISL_494595 | | | |
| see above | San Diego County Public Health Laboratory | Andersen lab at Scripps Research | SEARCH Alliance San Diego with Tracy Basler, Jovan Shephard, Brett Austin |
| EPI_ISL_494596, EPI_ISL_494597, EPI_ISL_494598, EPI_ISL_494599, EPI_ISL_494600, EPI_ISL_494601, EPI_ISL_494602, EPI_ISL_494603, EPI_ISL_494604, EPI_ISL_494605, EPI_ISL_494606, EPI_ISL_494607, EPI_ISL_494608, EPI_ISL_494609, EPI_ISL_494610, EPI_ISL_494611, EPI_ISL_494612, EPI_ISL_494613, EPI_ISL_494614, EPI_ISL_494615, EPI_ISL_494616, EPI_ISL_494617, EPI_ISL_494618, EPI_ISL_494619 | | | |
| see above | Scripps Medical Laboratory | Andersen lab at Scripps Research | SEARCH Alliance San Diego with Michael Quigley, Ellen Stefanski, Ian Mchardy |
| EPI_ISL_494620, EPI_ISL_494621, EPI_ISL_494622, EPI_ISL_494623, EPI_ISL_494624, EPI_ISL_494625, EPI_ISL_494626, EPI_ISL_494627, EPI_ISL_494628, EPI_ISL_494629, EPI_ISL_494630, EPI_ISL_494631, EPI_ISL_494632, EPI_ISL_494633, EPI_ISL_494634 | | | |
| see above | San Diego County Public Health Laboratory | Andersen lab at Scripps Research | SEARCH Alliance San Diego with Tracy Basler, Jovan Shephard, Brett Austin |
| EPI_ISL_494635, EPI_ISL_494636, EPI_ISL_494637, EPI_ISL_494638, EPI_ISL_494639, EPI_ISL_494640, EPI_ISL_494641, EPI_ISL_494642, EPI_ISL_494643, EPI_ISL_494644, EPI_ISL_494645, EPI_ISL_494646, EPI_ISL_494647, EPI_ISL_494648, EPI_ISL_494649, EPI_ISL_494650, EPI_ISL_494651, EPI_ISL_494652, EPI_ISL_494653, EPI_ISL_494654, EPI_ISL_494655, EPI_ISL_494656, EPI_ISL_494657, EPI_ISL_494658, EPI_ISL_494659, EPI_ISL_494660, EPI_ISL_494661, EPI_ISL_494662, EPI_ISL_494663, EPI_ISL_494664, EPI_ISL_494665, EPI_ISL_494666, EPI_ISL_494667, EPI_ISL_494668, EPI_ISL_494669, EPI_ISL_494670, EPI_ISL_494671, EPI_ISL_494672, EPI_ISL_494673, EPI_ISL_494674, EPI_ISL_494675, EPI_ISL_494676, EPI_ISL_494677, EPI_ISL_494678, EPI_ISL_494679, EPI_ISL_494680, EPI_ISL_494681, EPI_ISL_494682, EPI_ISL_494683, EPI_ISL_494684, EPI_ISL_494685, EPI_ISL_494686, EPI_ISL_494687, EPI_ISL_494688, EPI_ISL_494689, EPI_ISL_494690, EPI_ISL_494691, EPI_ISL_494692, EPI_ISL_494693, EPI_ISL_494694, EPI_ISL_494695, EPI_ISL_494696, EPI_ISL_494697, EPI_ISL_494698, EPI_ISL_494699, EPI_ISL_494700, EPI_ISL_494701, EPI_ISL_494702, EPI_ISL_494703, EPI_ISL_494704, EPI_ISL_494705, EPI_ISL_494706, EPI_ISL_494707, EPI_ISL_494708, EPI_ISL_494709, EPI_ISL_494710, EPI_ISL_494711, EPI_ISL_494712, EPI_ISL_494713 | | | |
| see above | Scripps Medical Laboratory | Andersen lab at Scripps Research | SEARCH Alliance San Diego with Michael Quigley, Ellen Stefanski, Ian Mchardy |
| EPI_ISL_494714, EPI_ISL_494715, EPI_ISL_494716, EPI_ISL_494717, EPI_ISL_494718, EPI_ISL_494719, EPI_ISL_494720, EPI_ISL_494721, EPI_ISL_494722, EPI_ISL_494723, EPI_ISL_494724, EPI_ISL_494725, EPI_ISL_494726, EPI_ISL_494727, EPI_ISL_494728, EPI_ISL_494729, EPI_ISL_494730, EPI_ISL_494731, EPI_ISL_494732, EPI_ISL_494733, EPI_ISL_494734, EPI_ISL_494735, EPI_ISL_494736, EPI_ISL_494737, EPI_ISL_494738, EPI_ISL_494739, EPI_ISL_494740, EPI_ISL_494741, EPI_ISL_494742, EPI_ISL_494743, EPI_ISL_494744, EPI_ISL_494745, EPI_ISL_494746 | | | |
| see above | San Diego County Public Health Laboratory | Andersen lab at Scripps Research | SEARCH Alliance San Diego with Tracy Basler, Jovan Shephard, Brett Austin |
| EPI_ISL_494747 | Virology Lab,Department of Pathology, National Cheng Kung University Hospital | Virology Lab,Department of Pathology, National Cheng Kung University Hospital | Huey-Pin Tsai, et al |
| EPI_ISL_494756, EPI_ISL_494757, EPI_ISL_494759, EPI_ISL_494761, EPI_ISL_494762, EPI_ISL_494763, EPI_ISL_494771, EPI_ISL_494774, EPI_ISL_494775, EPI_ISL_494776 | INT Fondazione Pascale | INT Fondazione Pascale | INT Fondazione Pascale |
| EPI_ISL_494970 | Dr. Tony Mazzulli Microbiologist-in-Chief | Dr. Jeff Wrana, Senior Investigator | Jeff Wrana, Jess Shen, Seda Barutcu, Kin Chan, Dan Trcka, Marie-Ming Aynaud, Javier Hernandez, Jessica Bourke, Christine Bruce, Bryn Hazlett, Laurence Pelletier, Sue Poutanen, Tony Mazzulli |
| EPI_ISL_495012 | Dr. Tony Mazzulli Microbiologist-in-Chief | Dr. Jeff Wrana, Senior Investigator | Jeff Wrana, Jess Shen, Seda Barutcu, Kin Chan, Dan Trcka, Marie-Ming Aynaud, Javier Hernandez, Jessica Bourke, Christine Bruce, Bryn Hazlett, Laurence Pelletier, Sue Poutanen, Tony Mazzulli |
| EPI_ISL_495013 | Dr. Tony Mazzulli Microbiologist-in-Chief | Dr. Jeff Wrana, Senior Investigator | Jeff Wrana, Jess Shen, Seda Barutcu, Kin Chan, Dan Trcka, Marie-Ming Aynaud, Javier Hernandez, Jessica Bourke, Christine Bruce, Bryn Hazlett, Laurence Pelletier, Sue Poutanen, Tony Mazzulli |
| EPI_ISL_495014 | B.J. Medical College and Civil hospital | Gujarat Biotechnology Research Centre | Janvi Raval, Zarna Patel, Monika Gandhi, Pinal Trivedi, Maharshi Pandya, Nidhi Patel, Nitin Savaliya, Raghawendra Kumar, Dinesh Kumar, Zuber Saiyed, Komal Patel, Labdhi Pandya, Afzal Ansari, Nikha Trivedi, Pranay Shah, Kamlesh J Upadhyay, Sanjay Kapadia, Apurvasinh Puvar, R D Dixit, A M Kadri, Harsh Bakshi, Chaitanya Joshi, Madhvi Joshi |
| EPI_ISL_495015 | B.J. Medical College and Civil hospital | Gujarat Biotechnology Research Centre | Zarna Patel, Monika Gandhi, Pinal Trivedi, Maharshi Pandya, Nidhi Patel, Nitin Savaliya, Raghawendra Kumar, Dinesh Kumar, Zuber Saiyed, Komal Patel, Labdhi Pandya, Afzal Ansari, Nikha Trivedi, Pranay Shah, Kamlesh J Upadhyay, Sanjay Kapadia, Apurvasinh Puvar, Janvi Raval, R D Dixit, A M Kadri, Harsh Bakshi, Chaitanya Joshi, Madhvi Joshi |
| EPI_ISL_495016 | B.J. Medical College and Civil hospital | Gujarat Biotechnology Research Centre | Monika Gandhi, Pinal Trivedi, Maharshi Pandya, Nidhi Patel, Nitin Savaliya, Raghawendra Kumar, Dinesh Kumar, Zuber Saiyed, Komal Patel, Labdhi Pandya, Afzal Ansari, Nikha Trivedi, Pranay Shah, Kamlesh J Upadhyay, Sanjay Kapadia, Apurvasinh Puvar, Janvi Raval, Zarna Patel, R D Dixit, A M Kadri, Harsh Bakshi, Chaitanya Joshi, Madhvi Joshi |
| EPI_ISL_495017 | B.J. Medical College and Civil hospital | Gujarat Biotechnology Research Centre | Pinal Trivedi, Maharshi Pandya, Nidhi Patel, Nitin Savaliya, Raghawendra Kumar, Dinesh Kumar, Zuber Saiyed, Komal Patel, Labdhi Pandya, Afzal Ansari, Nikha Trivedi, Pranay Shah, Kamlesh J Upadhyay, Sanjay Kapadia, Apurvasinh Puvar, Janvi Raval, Zarna Patel, Monika Gandhi, R D Dixit, A M Kadri, Harsh Bakshi, Chaitanya Joshi, Madhvi Joshi |
| EPI_ISL_495018 | B.J. Medical College and Civil hospital | Gujarat Biotechnology Research Centre | Maharshi Pandya, Nidhi Patel, Nitin Savaliya, Raghawendra Kumar, Dinesh Kumar, Zuber Saiyed, Komal Patel, Labdhi Pandya, Afzal Ansari, Nikha Trivedi, Pranay Shah, Kamlesh J Upadhyay, Sanjay Kapadia, Apurvasinh Puvar, Janvi Raval, Zarna Patel, Monika Gandhi, Pinal Trivedi, R D Dixit, A M Kadri, Harsh Bakshi, Chaitanya Joshi, Madhvi Joshi |

| | | | |
|---|---|---|---|
| EPI_ISL_495019 | B.J. Medical College and Civil hospital | Gujarat Biotechnology Research Centre | Nidhi Patel, Nitin Savaliya, Raghawendra Kumar, Dinesh Kumar, Zuber Saiyed, Komal Patel, Labdhi Pandya, Afzal Ansari, Nikha Trivedi, Pranay Shah, Kamlesh J Upadhyay, Sanjay Kapadia, Apurvasinh Puvar, Janvi Raval, Zarna Patel, Monika Gandhi, Pinal Trivedi, Maharshi Pandya, R D Dixit, A M Kadri, Harsh Bakshi, Chaitanya Joshi, Madhvi Joshi |
| EPI_ISL_495020 | Government Medical College, Bhavnagar | Gujarat Biotechnology Research Centre | Kairavi Desai, Saklin Malek, Shirish Patel, Nitin Savaliya, Raghawendra Kumar, Dinesh Kumar, Zuber Saiyed, Komal Patel, Labdhi Pandya, Afzal Ansari, Nikha Trivedi, Apurvasinh Puvar, Janvi Raval, Zarna Patel, Monika Gandhi, Pinal Trivedi, Maharshi Pandya, Nidhi Patel, R D Dixit, A M Kadri, Harsh Bakshi, Chaitanya Joshi, Madhvi Joshi |
| EPI_ISL_495021 | Government Medical College, Bhavnagar | Gujarat Biotechnology Research Centre | Saklin Malek, Shirish Patel, Kairavi Desai, Raghawendra Kumar, Dinesh Kumar, Zuber Saiyed, Komal Patel, Labdhi Pandya, Afzal Ansari, Nikha Trivedi, Apurvasinh Puvar, Janvi Raval, Zarna Patel, Monika Gandhi, Pinal Trivedi, Maharshi Pandya, Nidhi Patel, Nitin Savaliya, R D Dixit, A M Kadri, Harsh Bakshi, Chaitanya Joshi, Madhvi Joshi |
| EPI_ISL_495022 | Government Medical College, Bhavnagar | Gujarat Biotechnology Research Centre | Shirish Patel, Kairavi Desai, Saklin Malek, Dinesh Kumar, Zuber Saiyed, Komal Patel, Labdhi Pandya, Afzal Ansari, Nikha Trivedi, Apurvasinh Puvar, Janvi Raval, Zarna Patel, Monika Gandhi, Pinal Trivedi, Maharshi Pandya, Nidhi Patel, Nitin Savaliya, Raghawendra Kumar, R D Dixit, A M Kadri, Harsh Bakshi, Chaitanya Joshi, Madhvi Joshi |
| EPI_ISL_495023 | Government Medical College, Bhavnagar | Gujarat Biotechnology Research Centre | Nitin Savaliya, Raghawendra Kumar, Dinesh Kumar, Zuber Saiyed, Komal Patel, Labdhi Pandya, Afzal Ansari, Nikha Trivedi, Kairavi Desai, Saklin Malek, Shirish Patel, Apurvasinh Puvar, Janvi Raval, Zarna Patel, Monika Gandhi, Pinal Trivedi, Maharshi Pandya, Nidhi Patel, R D Dixit, A M Kadri, Harsh Bakshi, Chaitanya Joshi, Madhvi Joshi |
| EPI_ISL_495024 | Government Medical College, Bhavnagar | Gujarat Biotechnology Research Centre | Raghawendra Kumar, Dinesh Kumar, Zuber Saiyed, Komal Patel, Labdhi Pandya, Afzal Ansari, Nikha Trivedi, Kairavi Desai, Saklin Malek, Shirish Patel, Apurvasinh Puvar, Janvi Raval, Zarna Patel, Monika Gandhi, Pinal Trivedi, Maharshi Pandya, Nidhi Patel, Nitin Savaliya, R D Dixit, A M Kadri, Harsh Bakshi, Chaitanya Joshi, Madhvi Joshi |
| EPI_ISL_495025 | Government Medical College, Bhavnagar | Gujarat Biotechnology Research Centre | Dinesh Kumar, Zuber Saiyed, Komal Patel, Labdhi Pandya, Afzal Ansari, Nikha Trivedi, Kairavi Desai, Saklin Malek, Shirish Patel, Apurvasinh Puvar, Janvi Raval, Zarna Patel, Monika Gandhi, Pinal Trivedi, Maharshi Pandya, Nidhi Patel, Nitin Savaliya, Raghawendra Kumar, R D Dixit, A M Kadri, Harsh Bakshi, Chaitanya Joshi, Madhvi Joshi |
| EPI_ISL_495026 | GMERS Medical College & Hospital, Gotri, Vadodara | Gujarat Biotechnology Research Centre | Komal Patel, Labdhi Pandya, Afzal Ansari, Nikha Trivedi, Meenakshi Shah, Neena Doshi, Varsha Godbole, Apurvasinh Puvar, Janvi Raval, Zarna Patel, Monika Gandhi, Pinal Trivedi, Maharshi Pandya, Nidhi Patel, Nitin Savaliya, Raghawendra Kumar, Dinesh Kumar, Zuber Saiyed, R D Dixit, A M Kadri, Harsh Bakshi, Chaitanya Joshi, Madhvi Joshi |
| EPI_ISL_495027 | GMERS Medical College & Hospital, Gotri, Vadodara | Gujarat Biotechnology Research Centre | Labdhi Pandya, Afzal Ansari, Nikha Trivedi, Meenakshi Shah, Neena Doshi, Varsha Godbole, Apurvasinh Puvar, Janvi Raval, Zarna Patel, Monika Gandhi, Pinal Trivedi, Maharshi Pandya, Nidhi Patel, Nitin Savaliya, Raghawendra Kumar, Dinesh Kumar, Zuber Saiyed, Komal Patel, R D Dixit, A M Kadri, Harsh Bakshi, Chaitanya Joshi, Madhvi Joshi |
| EPI_ISL_495028 | GMERS Medical College & Hospital, Gotri, Vadodara | Gujarat Biotechnology Research Centre | Afzal Ansari, Nikha Trivedi, Meenakshi Shah, Neena Doshi, Varsha Godbole, Apurvasinh Puvar, Janvi Raval, Zarna Patel, Monika Gandhi, Pinal Trivedi, Maharshi Pandya, Nidhi Patel, Nitin Savaliya, Raghawendra Kumar, Dinesh Kumar, Zuber Saiyed, Komal Patel, Labdhi Pandya, R D Dixit, A M Kadri, Harsh Bakshi, Chaitanya Joshi, Madhvi Joshi |
| EPI_ISL_495029 | GMERS Medical College & Hospital, Gotri, Vadodara | Gujarat Biotechnology Research Centre | Nikha Trivedi, Meenakshi Shah, Neena Doshi, Varsha Godbole, Apurvasinh Puvar, Janvi Raval, Zarna Patel, Monika Gandhi, Pinal Trivedi, Maharshi Pandya, Nidhi Patel, Nitin Savaliya, Raghawendra Kumar, Dinesh Kumar, Zuber Saiyed, Komal Patel, Labdhi Pandya, Afzal Ansari, R D Dixit, A M Kadri, Harsh Bakshi, Chaitanya Joshi, Madhvi Joshi |
| EPI_ISL_495030 | GMERS Medical College & Hospital, Gotri, Vadodara | Gujarat Biotechnology Research Centre | Meenakshi Shah, Neena Doshi, Varsha Godbole, Apurvasinh Puvar, Janvi Raval, Zarna Patel, Monika Gandhi, Pinal Trivedi, Maharshi Pandya, Nidhi Patel, Nitin Savaliya, Raghawendra Kumar, Dinesh Kumar, Zuber Saiyed, Komal Patel, Labdhi Pandya, Afzal Ansari, Nikha Trivedi, R D Dixit, A M Kadri, Harsh Bakshi, Chaitanya Joshi, Madhvi Joshi |
| EPI_ISL_495031 | GMERS Medical College & Hospital, Gotri, Vadodara | Gujarat Biotechnology Research Centre | Neena Doshi, Varsha Godbole, Apurvasinh Puvar, Janvi Raval, Zarna Patel, Monika Gandhi, Pinal Trivedi, Maharshi Pandya, Nidhi Patel, Nitin Savaliya, Raghawendra Kumar, Dinesh Kumar, Zuber Saiyed, Komal Patel, Labdhi Pandya, Afzal Ansari, Nikha Trivedi, Meenakshi Shah, R D Dixit, A M Kadri, Harsh Bakshi, Chaitanya Joshi, Madhvi Joshi |
| EPI_ISL_495032 | GMERS Medical College & Hospital, Gotri, Vadodara | Gujarat Biotechnology Research Centre | Varsha Godbole, Apurvasinh Puvar, Janvi Raval, Zarna Patel, Monika Gandhi, Pinal Trivedi, Maharshi Pandya, Nidhi Patel, Nitin Savaliya, Raghawendra Kumar, Dinesh Kumar, Zuber Saiyed, Komal Patel, Labdhi Pandya, Afzal Ansari, Nikha Trivedi, Meenakshi Shah, Neena Doshi, R D Dixit, A M Kadri, Harsh Bakshi, Chaitanya Joshi, Madhvi Joshi |
| EPI_ISL_495033 | GMERS Medical College & Hospital, Gotri, Vadodara | Gujarat Biotechnology Research Centre | Apurvasinh Puvar, Janvi Raval, Zarna Patel, Monika Gandhi, Pinal Trivedi, Maharshi Pandya, Nidhi Patel, Nitin Savaliya, Raghawendra Kumar, Dinesh Kumar, Zuber Saiyed, Komal Patel, Labdhi Pandya, Afzal Ansari, Nikha Trivedi, Meenakshi Shah, Neena Doshi, Varsha Godbole, R D Dixit, A M Kadri, Harsh Bakshi, Chaitanya Joshi, Madhvi Joshi |
| EPI_ISL_495034 | GMERS Medical College & Hospital, Gotri, Vadodara | Gujarat Biotechnology Research Centre | Janvi Raval, Zarna Patel, Monika Gandhi, Pinal Trivedi, Maharshi Pandya, Nidhi Patel, Nitin Savaliya, Raghawendra Kumar, Dinesh Kumar, Zuber Saiyed, Komal Patel, Labdhi Pandya, Afzal Ansari, Nikha Trivedi, Meenakshi Shah, Neena Doshi, Varsha Godbole, Apurvasinh Puvar, R D Dixit, A M Kadri, Harsh Bakshi, Chaitanya Joshi, Madhvi Joshi |
| EPI_ISL_495035 | GMERS Medical College & Hospital, Gotri, Vadodara | Gujarat Biotechnology Research Centre | Zarna Patel, Monika Gandhi, Pinal Trivedi, Maharshi Pandya, Nidhi Patel, Nitin Savaliya, Raghawendra Kumar, Dinesh Kumar, Zuber Saiyed, Komal Patel, Labdhi Pandya, Afzal Ansari, Nikha Trivedi, Meenakshi Shah, Neena Doshi, Varsha Godbole, Apurvasinh Puvar, Janvi Raval, R D Dixit, A M Kadri, Harsh Bakshi, Chaitanya Joshi, Madhvi Joshi |
| EPI_ISL_495036 | GMERS Medical College & Hospital, Gotri, Vadodara | Gujarat Biotechnology Research Centre | Monika Gandhi, Pinal Trivedi, Maharshi Pandya, Nidhi Patel, Nitin Savaliya, Raghawendra Kumar, Dinesh Kumar, Zuber Saiyed, Komal Patel, Labdhi Pandya, Afzal Ansari, Nikha Trivedi, Meenakshi Shah, Neena Doshi, Varsha Godbole, Apurvasinh Puvar, Janvi Raval, Zarna Patel, R D Dixit, A M Kadri, Harsh Bakshi, Chaitanya Joshi, Madhvi Joshi |
| EPI_ISL_495037 | GMERS Medical College & Hospital, Gotri, Vadodara | Gujarat Biotechnology Research Centre | Pinal Trivedi, Maharshi Pandya, Nidhi Patel, Nitin Savaliya, Raghawendra Kumar, Dinesh Kumar, Zuber Saiyed, Komal Patel, Labdhi Pandya, Afzal Ansari, Nikha Trivedi, Meenakshi Shah, Neena Doshi, Varsha Godbole, Apurvasinh Puvar, Janvi Raval, Zarna Patel, Monika Gandhi, R D Dixit, A M Kadri, Harsh Bakshi, Chaitanya Joshi, Madhvi Joshi |
| EPI_ISL_495038 | GMERS Medical College & Hospital, Gotri, Vadodara | Gujarat Biotechnology Research Centre | Maharshi Pandya, Nidhi Patel, Nitin Savaliya, Raghawendra Kumar, Dinesh Kumar, Zuber Saiyed, Komal Patel, Labdhi Pandya, Afzal Ansari, Nikha Trivedi, Meenakshi Shah, Neena Doshi, Varsha Godbole, Apurvasinh Puvar, Janvi Raval, Zarna Patel, Monika Gandhi, Pinal Trivedi, R D Dixit, A M Kadri, Harsh Bakshi, Chaitanya Joshi, Madhvi Joshi |
| EPI_ISL_495039 | GMERS Medical College & Hospital, Gotri, Vadodara | Gujarat Biotechnology Research Centre | Nidhi Patel, Nitin Savaliya, Raghawendra Kumar, Dinesh Kumar, Zuber Saiyed, Komal Patel, Labdhi Pandya, Afzal Ansari, Nikha Trivedi, Meenakshi Shah, Neena Doshi, Varsha Godbole, Apurvasinh Puvar, Janvi Raval, Zarna Patel, Monika Gandhi, Pinal Trivedi, Maharshi Pandya, R D Dixit, A M Kadri, Harsh Bakshi, Chaitanya Joshi, Madhvi Joshi |
| EPI_ISL_495040 | GMERS Medical College & Hospital, Gotri, Vadodara | Gujarat Biotechnology Research Centre | Nitin Savaliya, Raghawendra Kumar, Dinesh Kumar, Zuber Saiyed, Komal Patel, Labdhi Pandya, Afzal Ansari, Nikha Trivedi, Meenakshi Shah, Neena Doshi, Varsha Godbole, Apurvasinh Puvar, Janvi Raval, Zarna Patel, Monika Gandhi, Pinal Trivedi, Maharshi Pandya, Nidhi Patel, R D Dixit, A M Kadri, Harsh Bakshi, Chaitanya Joshi, Madhvi Joshi |
| EPI_ISL_495041 | GMERS Medical College & Hospital, Gotri, Vadodara | Gujarat Biotechnology Research Centre | Raghawendra Kumar, Dinesh Kumar, Zuber Saiyed, Komal Patel, Labdhi Pandya, Afzal Ansari, Nikha Trivedi, Meenakshi Shah, Neena Doshi, Varsha Godbole, Apurvasinh Puvar, Janvi Raval, Zarna Patel, Monika Gandhi, Pinal Trivedi, Maharshi Pandya, Nidhi Patel, Nitin Savaliya, R D Dixit, A M Kadri, Harsh Bakshi, Chaitanya Joshi, Madhvi Joshi |
| EPI_ISL_495042 | GMERS Medical College & Hospital, Gotri, Vadodara | Gujarat Biotechnology Research Centre | Dinesh Kumar, Zuber Saiyed, Komal Patel, Labdhi Pandya, Afzal Ansari, Nikha Trivedi, Meenakshi Shah, Neena Doshi, Varsha Godbole, Apurvasinh Puvar, Janvi Raval, Zarna Patel, Monika Gandhi, Pinal Trivedi, Maharshi Pandya, Nidhi Patel, Nitin Savaliya, Raghawendra Kumar, R D Dixit, A M Kadri, Harsh Bakshi, Chaitanya Joshi, Madhvi Joshi |
| EPI_ISL_495043 | GMERS Medical College & Hospital, Gotri, Vadodara | Gujarat Biotechnology Research Centre | Zuber Saiyed, Komal Patel, Labdhi Pandya, Afzal Ansari, Nikha Trivedi, Meenakshi Shah, Neena Doshi, Varsha Godbole, Apurvasinh Puvar, Janvi Raval, Zarna Patel, Monika Gandhi, Pinal Trivedi, Maharshi Pandya, Nidhi Patel, Nitin Savaliya, Raghawendra Kumar, Dinesh Kumar, R D Dixit, A M Kadri, Harsh Bakshi, Chaitanya Joshi, Madhvi Joshi |
| EPI_ISL_495044 | GMERS Medical College & Hospital, Gotri, Vadodara | Gujarat Biotechnology Research Centre | Komal Patel, Labdhi Pandya, Afzal Ansari, Nikha Trivedi, Meenakshi Shah, Neena Doshi, Varsha Godbole, Apurvasinh Puvar, Janvi Raval, Zarna Patel, Monika Gandhi, Pinal Trivedi, Maharshi Pandya, Nidhi Patel, Nitin Savaliya, Raghawendra Kumar, Dinesh Kumar, Zuber Saiyed, R D Dixit, A M Kadri, Harsh Bakshi, Chaitanya Joshi, Madhvi Joshi |
| EPI_ISL_495045 | GMERS Medical College & Hospital, Gotri, Vadodara | Gujarat Biotechnology Research Centre | Labdhi Pandya, Afzal Ansari, Nikha Trivedi, Meenakshi Shah, Neena Doshi, Varsha Godbole, Apurvasinh Puvar, Janvi Raval, Zarna Patel, Monika Gandhi, Pinal Trivedi, Maharshi Pandya, Nidhi Patel, Nitin Savaliya, Raghawendra Kumar, Dinesh Kumar, Zuber Saiyed, Komal Patel, R D Dixit, A M Kadri, Harsh Bakshi, Chaitanya Joshi, Madhvi Joshi |
| EPI_ISL_495046 | GMERS Medical College & Hospital, Gotri, Vadodara | Gujarat Biotechnology Research Centre | Afzal Ansari, Nikha Trivedi, Meenakshi Shah, Neena Doshi, Varsha Godbole, Apurvasinh Puvar, Janvi Raval, Zarna Patel, Monika Gandhi, Pinal Trivedi, Maharshi Pandya, Nidhi Patel, Nitin Savaliya, Raghawendra Kumar, Dinesh Kumar, Zuber Saiyed, Komal Patel, Labdhi Pandya, R D Dixit, A M Kadri, Harsh Bakshi, Chaitanya Joshi, Madhvi Joshi |
| EPI_ISL_495047 | GMERS Medical College & Hospital, Gotri, Vadodara | Gujarat Biotechnology Research Centre | Nikha Trivedi, Meenakshi Shah, Neena Doshi, Varsha Godbole, Apurvasinh Puvar, Janvi Raval, Zarna Patel, Monika Gandhi, Pinal Trivedi, Maharshi Pandya, Nidhi Patel, Nitin Savaliya, Raghawendra Kumar, Dinesh Kumar, Zuber Saiyed, Komal Patel, Labdhi Pandya, Afzal Ansari, R D Dixit, A M Kadri, Harsh Bakshi, Chaitanya Joshi, Madhvi Joshi |
| EPI_ISL_495048 | GMERS Medical College & Hospital, Gotri, Vadodara | Gujarat Biotechnology Research Centre | Meenakshi Shah, Neena Doshi, Varsha Godbole, Apurvasinh Puvar, Janvi Raval, Zarna Patel, Monika Gandhi, Pinal Trivedi, Maharshi Pandya, Nidhi Patel, Nitin Savaliya, Raghawendra Kumar, Dinesh Kumar, Zuber Saiyed, Komal Patel, Labdhi Pandya, Afzal Ansari, Nikha Trivedi, R D Dixit, A M Kadri, Harsh Bakshi, Chaitanya Joshi, Madhvi Joshi |
| EPI_ISL_495049 | GMERS Medical College & Hospital, Gotri, Vadodara | Gujarat Biotechnology Research Centre | Neena Doshi, Varsha Godbole, Apurvasinh Puvar, Janvi Raval, Zarna Patel, Monika Gandhi, Pinal Trivedi, Maharshi Pandya, Nidhi Patel, Nitin Savaliya, Raghawendra Kumar, Dinesh Kumar, Zuber Saiyed, Komal Patel, Labdhi Pandya, Afzal Ansari, Nikha Trivedi, Meenakshi Shah, R D Dixit, A M Kadri, Harsh Bakshi, Chaitanya Joshi, Madhvi Joshi |
| EPI_ISL_495050 | GMERS Medical College & Hospital, Gotri, Vadodara | Gujarat Biotechnology Research Centre | Varsha Godbole, Apurvasinh Puvar, Janvi Raval, Zarna Patel, Monika Gandhi, Pinal Trivedi, Maharshi Pandya, Nidhi Patel, Nitin Savaliya, Raghawendra Kumar, Dinesh Kumar, Zuber Saiyed, Komal Patel, Labdhi Pandya, Afzal Ansari, Nikha Trivedi, Meenakshi Shah, Neena Doshi, R D Dixit, A M Kadri, Harsh Bakshi, Chaitanya Joshi, Madhvi Joshi |
| EPI_ISL_495051 | GMERS Medical College & Hospital, Gotri, Vadodara | Gujarat Biotechnology Research Centre | Apurvasinh Puvar, Janvi Raval, Zarna Patel, Monika Gandhi, Pinal Trivedi, Maharshi Pandya, Nidhi Patel, Nitin Savaliya, Raghawendra Kumar, Dinesh Kumar, Zuber Saiyed, Komal Patel, Labdhi Pandya, Afzal Ansari, Nikha Trivedi, Meenakshi Shah, Neena Doshi, Varsha Godbole, R D Dixit, A M Kadri, Harsh Bakshi, Chaitanya Joshi, Madhvi Joshi |
| EPI_ISL_495052 | GMERS Medical College & Hospital, Gotri, Vadodara | Gujarat Biotechnology Research Centre | Janvi Raval, Zarna Patel, Monika Gandhi, Pinal Trivedi, Maharshi Pandya, Nidhi Patel, Nitin Savaliya, Raghawendra Kumar, Dinesh Kumar, Zuber Saiyed, Komal Patel, Labdhi Pandya, Afzal Ansari, Nikha Trivedi, Meenakshi Shah, Neena Doshi, Varsha Godbole, Apurvasinh Puvar, R D Dixit, A M Kadri, Harsh Bakshi, Chaitanya Joshi, Madhvi Joshi |
| EPI_ISL_495053 | GMERS Medical College & Hospital, Gotri, Vadodara | Gujarat Biotechnology Research Centre | Zarna Patel, Monika Gandhi, Pinal Trivedi, Maharshi Pandya, Nidhi Patel, Nitin Savaliya, Raghawendra Kumar, Dinesh Kumar, Zuber Saiyed, Komal Patel, Labdhi Pandya, Afzal Ansari, Nikha Trivedi, Meenakshi Shah, Neena Doshi, Varsha Godbole, Apurvasinh Puvar, Janvi Raval, R D Dixit, A M Kadri, Harsh Bakshi, Chaitanya Joshi, Madhvi Joshi |
| EPI_ISL_495054 | GMERS Medical College & Hospital, Gotri, Vadodara | Gujarat Biotechnology Research Centre | Monika Gandhi, Pinal Trivedi, Maharshi Pandya, Nidhi Patel, Nitin Savaliya, Raghawendra Kumar, Dinesh Kumar, Zuber Saiyed, Komal Patel, Labdhi Pandya, Afzal Ansari, Nikha Trivedi, Meenakshi Shah, Neena Doshi, Varsha Godbole, Apurvasinh Puvar, Janvi Raval, Zarna Patel, R D Dixit, A M Kadri, Harsh Bakshi, Chaitanya Joshi, Madhvi Joshi |
| EPI_ISL_495055 | GMERS Medical College & Hospital, Gotri, Vadodara | Gujarat Biotechnology Research Centre | Pinal Trivedi, Maharshi Pandya, Nidhi Patel, Nitin Savaliya, Raghawendra Kumar, Dinesh Kumar, Zuber Saiyed, Komal Patel, Labdhi Pandya, Afzal Ansari, Nikha Trivedi, Meenakshi Shah, Neena Doshi, Varsha Godbole, Apurvasinh Puvar, Janvi Raval, Zarna Patel, Monika Gandhi, R D Dixit, A M Kadri, Harsh Bakshi, Chaitanya Joshi, Madhvi Joshi |
| EPI_ISL_495056 | GMERS Medical College & Hospital, Gotri, Vadodara | Gujarat Biotechnology Research Centre | Maharshi Pandya, Nidhi Patel, Nitin Savaliya, Raghawendra Kumar, Dinesh Kumar, Zuber Saiyed, Komal Patel, Labdhi Pandya, Afzal Ansari, Nikha Trivedi, Meenakshi Shah, Neena Doshi, Varsha Godbole, Apurvasinh Puvar, Janvi Raval, Zarna Patel, Monika Gandhi, Pinal Trivedi, R D Dixit, A M Kadri, Harsh Bakshi, Chaitanya Joshi, Madhvi Joshi |
| EPI_ISL_495057 | GMERS Medical College & Hospital, Gotri, Vadodara | Gujarat Biotechnology Research Centre | Nidhi Patel, Nitin Savaliya, Raghawendra Kumar, Dinesh Kumar, Zuber Saiyed, Komal Patel, Labdhi Pandya, Afzal Ansari, Nikha Trivedi, Meenakshi Shah, Neena Doshi, Varsha Godbole, Apurvasinh Puvar, Janvi Raval, Zarna Patel, Monika Gandhi, Pinal Trivedi, Maharshi Pandya, R D Dixit, A M Kadri, Harsh Bakshi, Chaitanya Joshi, Madhvi Joshi |
| EPI_ISL_495058 | GMERS Medical College & Hospital, Gotri, Vadodara | Gujarat Biotechnology Research Centre | Nitin Savaliya, Raghawendra Kumar, Dinesh Kumar, Zuber Saiyed, Komal Patel, Labdhi Pandya, Afzal Ansari, Nikha Trivedi, Meenakshi Shah, Neena Doshi, Varsha Godbole, Apurvasinh Puvar, Janvi Raval, Zarna Patel, Monika Gandhi, Pinal Trivedi, Maharshi Pandya, Nidhi Patel, R D Dixit, A M Kadri, Harsh Bakshi, Chaitanya Joshi, Madhvi Joshi |
| EPI_ISL_495059 | GMERS Medical College & Hospital, Gotri, Vadodara | Gujarat Biotechnology Research Centre | Raghawendra Kumar, Dinesh Kumar, Zuber Saiyed, Komal Patel, Labdhi Pandya, Afzal Ansari, Nikha Trivedi, Meenakshi Shah, Neena Doshi, Varsha Godbole, Apurvasinh Puvar, Janvi Raval, Zarna Patel, Monika Gandhi, Pinal Trivedi, Maharshi Pandya, Nidhi Patel, Nitin Savaliya, R D Dixit, A M Kadri, Harsh Bakshi, Chaitanya Joshi, Madhvi Joshi |
| EPI_ISL_495060 | GMERS Medical College & Hospital, Gotri, Vadodara | Gujarat Biotechnology Research Centre | Dinesh Kumar, Zuber Saiyed, Komal Patel, Labdhi Pandya, Afzal Ansari, Nikha Trivedi, Meenakshi Shah, Neena Doshi, Varsha Godbole, Apurvasinh Puvar, Janvi Raval, Zarna Patel, Monika Gandhi, Pinal Trivedi, Maharshi Pandya, Nidhi Patel, Nitin Savaliya, Raghawendra Kumar, R D Dixit, A M Kadri, Harsh Bakshi, Chaitanya Joshi, Madhvi Joshi |
| EPI_ISL_495061 | GMERS Medical College & Hospital, Gotri, Vadodara | Gujarat Biotechnology Research Centre | Zuber Saiyed, Komal Patel, Labdhi Pandya, Afzal Ansari, Nikha Trivedi, Meenakshi Shah, Neena Doshi, Varsha Godbole, Apurvasinh Puvar, Janvi Raval, Zarna Patel, Monika Gandhi, Pinal Trivedi, Maharshi Pandya, Nidhi Patel, Nitin Savaliya, Raghawendra Kumar, Dinesh Kumar, R D Dixit, A M Kadri, Harsh Bakshi, Chaitanya Joshi, Madhvi Joshi |
| EPI_ISL_495062 | GMERS Medical College & Hospital, Gotri, Vadodara | Gujarat Biotechnology Research Centre | Komal Patel, Labdhi Pandya, Afzal Ansari, Nikha Trivedi, Meenakshi Shah, Neena Doshi, Varsha Godbole, Apurvasinh Puvar, Janvi Raval, Zarna Patel, Monika Gandhi, Pinal Trivedi, Maharshi Pandya, Nidhi Patel, Nitin Savaliya, Raghawendra Kumar, Dinesh Kumar, Zuber Saiyed, R D Dixit, A M Kadri, Harsh Bakshi, Chaitanya Joshi, Madhvi Joshi |
| EPI_ISL_495063 | GMERS Medical College & Hospital, Himmatnagar | Gujarat Biotechnology Research Centre | Labdhi Pandya, Afzal Ansari, Nikha Trivedi, Himanshu Khatri, Mayur Gandhi, Apurvasinh Puvar, Janvi Raval, Zarna Patel, Monika Gandhi, Pinal Trivedi, Maharshi Pandya, Nidhi Patel, Nitin Savaliya, Raghawendra Kumar, Dinesh Kumar, Zuber Saiyed, Komal Patel, R D Dixit, A M Kadri, Harsh Bakshi, Chaitanya Joshi, Madhvi Joshi, |

| | | | |
|---|---|---|---|
| EPI_ISL_495064 | GMERS Medical College & Hospital, Himmatnagar | Gujarat Biotechnology Research Centre | Afzal Ansari, Nikha Trivedi, Himanshu Khatri, Mayur Gandhi, Apurvasinh Puvar, Janvi Raval, Zarna Patel, Monika Gandhi, Pinal Trivedi, Maharshi Pandya, Nidhi Patel, Nitin Savaliya, Raghawendra Kumar, Dinesh Kumar, Zuber Saiyed, Komal Patel, Labdhi Pandya, R D Dixit, A M Kadri, Harsh Bakshi, Chaitanya Joshi, Madhvi Joshi, |
| EPI_ISL_495065 | GAIMS & G K General Hospital | Gujarat Biotechnology Research Centre | Nikha Trivedi, Babulal Babhoria, Hitesh Assudani, Apurvasinh Puvar, Janvi Raval, Zarna Patel, Monika Gandhi, Pinal Trivedi, Maharshi Pandya, Nidhi Patel, Nitin Savaliya, Raghawendra Kumar, Dinesh Kumar, Zuber Saiyed, Komal Patel, Labdhi Pandya, Afzal Ansari, R D Dixit, A M Kadri, Harsh Bakshi, Chaitanya Joshi, Madhvi Joshi, |
| EPI_ISL_495066 | GAIMS & G K General Hospital | Gujarat Biotechnology Research Centre | Babulal Babhoria, Hitesh Assudani, Apurvasinh Puvar, Janvi Raval, Zarna Patel, Monika Gandhi, Pinal Trivedi, Maharshi Pandya, Nidhi Patel, Nitin Savaliya, Raghawendra Kumar, Dinesh Kumar, Zuber Saiyed, Komal Patel, Labdhi Pandya, Afzal Ansari, Nikha Trivedi, R D Dixit, A M Kadri, Harsh Bakshi, Chaitanya Joshi, Madhvi Joshi, |
| EPI_ISL_495067 | Dr. N. D. Desai Medical College & Hospital | Gujarat Biotechnology Research Centre | J G Buch, Jigar Gusani, Supreet Prabhu, Apurvasinh Puvar, Janvi Raval, Zarna Patel, Monika Gandhi, Pinal Trivedi, Maharshi Pandya, Nidhi Patel, Nitin Savaliya, Raghawendra Kumar, Dinesh Kumar, Zuber Saiyed, Komal Patel, Labdhi Pandya, Afzal Ansari, Nikha Trivedi, R D Dixit, A M Kadri, Harsh Bakshi, Chaitanya Joshi, Madhvi Joshi |
| EPI_ISL_495068 | Dr. N. D. Desai Medical College & Hospital | Gujarat Biotechnology Research Centre | Jigar Gusani, Supreet Prabhu, Apurvasinh Puvar, Janvi Raval, Zarna Patel, Monika Gandhi, Pinal Trivedi, Maharshi Pandya, Nidhi Patel, Nitin Savaliya, Raghawendra Kumar, Dinesh Kumar, Zuber Saiyed, Komal Patel, Labdhi Pandya, Afzal Ansari, Nikha Trivedi, J G Buch, R D Dixit, A M Kadri, Harsh Bakshi, Chaitanya Joshi, Madhvi Joshi |
| EPI_ISL_495069 | Dr. N. D. Desai Medical College & Hospital | Gujarat Biotechnology Research Centre | Supreet Prabhu, Apurvasinh Puvar, Janvi Raval, Zarna Patel, Monika Gandhi, Pinal Trivedi, Maharshi Pandya, Nidhi Patel, Nitin Savaliya, Raghawendra Kumar, Dinesh Kumar, Zuber Saiyed, Komal Patel, Labdhi Pandya, Afzal Ansari, Nikha Trivedi, J G Buch, Jigar Gusani, R D Dixit, A M Kadri, Harsh Bakshi, Chaitanya Joshi, Madhvi Joshi |
| EPI_ISL_495070 | Department of MicroBiology, Government Medical College, Surat | Gujarat Biotechnology Research Centre | Apurvasinh Puvar, Janvi Raval, Zarna Patel, Monika Gandhi, Pinal Trivedi, Maharshi Pandya, Nidhi Patel, Nitin Savaliya, Raghawendra Kumar, Dinesh Kumar, Zuber Saiyed, Komal Patel, Labdhi Pandya, Afzal Ansari, Nikha Trivedi, Naresh Chauhan, Summaiya Mullan, Amit gamit, R D Dixit, A M Kadri, Harsh Bakshi, Chaitanya Joshi, Madhvi Joshi |
| EPI_ISL_495071 | Department of MicroBiology, Government Medical College, Surat | Gujarat Biotechnology Research Centre | Janvi Raval, Zarna Patel, Monika Gandhi, Pinal Trivedi, Maharshi Pandya, Nidhi Patel, Nitin Savaliya, Raghawendra Kumar, Dinesh Kumar, Zuber Saiyed, Komal Patel, Labdhi Pandya, Afzal Ansari, Nikha Trivedi, Naresh Chauhan, Summaiya Mullan, Amit gamit, Apurvasinh Puvar, R D Dixit, A M Kadri, Harsh Bakshi, Chaitanya Joshi, Madhvi Joshi |
| EPI_ISL_495072 | Department of MicroBiology, Government Medical College, Surat | Gujarat Biotechnology Research Centre | Zarna Patel, Monika Gandhi, Pinal Trivedi, Maharshi Pandya, Nidhi Patel, Nitin Savaliya, Raghawendra Kumar, Dinesh Kumar, Zuber Saiyed, Komal Patel, Labdhi Pandya, Afzal Ansari, Nikha Trivedi, Naresh Chauhan, Summaiya Mullan, Amit gamit, Apurvasinh Puvar, Janvi Raval, R D Dixit, A M Kadri, Harsh Bakshi, Chaitanya Joshi, Madhvi Joshi |
| EPI_ISL_495073 | Department of MicroBiology, Government Medical College, Surat | Gujarat Biotechnology Research Centre | Monika Gandhi, Pinal Trivedi, Maharshi Pandya, Nidhi Patel, Nitin Savaliya, Raghawendra Kumar, Dinesh Kumar, Zuber Saiyed, Komal Patel, Labdhi Pandya, Afzal Ansari, Nikha Trivedi, Naresh Chauhan, Summaiya Mullan, Amit gamit, Apurvasinh Puvar, Janvi Raval, Zarna Patel, R D Dixit, A M Kadri, Harsh Bakshi, Chaitanya Joshi, Madhvi Joshi |
| EPI_ISL_495074 | Department of MicroBiology, Government Medical College, Surat | Gujarat Biotechnology Research Centre | Pinal Trivedi, Maharshi Pandya, Nidhi Patel, Nitin Savaliya, Raghawendra Kumar, Dinesh Kumar, Zuber Saiyed, Komal Patel, Labdhi Pandya, Afzal Ansari, Nikha Trivedi, Naresh Chauhan, Summaiya Mullan, Amit gamit, Apurvasinh Puvar, Janvi Raval, Zarna Patel, Monika Gandhi, R D Dixit, A M Kadri, Harsh Bakshi, Chaitanya Joshi, Madhvi Joshi |
| EPI_ISL_495075 | Department of MicroBiology, Government Medical College, Surat | Gujarat Biotechnology Research Centre | Nidhi Patel, Nitin Savaliya, Raghawendra Kumar, Dinesh Kumar, Zuber Saiyed, Komal Patel, Labdhi Pandya, Afzal Ansari, Nikha Trivedi, Naresh Chauhan, Summaiya Mullan, Amit gamit, Apurvasinh Puvar, Janvi Raval, Zarna Patel, Monika Gandhi, Pinal Trivedi, Maharshi Pandya, R D Dixit, A M Kadri, Harsh Bakshi, Chaitanya Joshi, Madhvi Joshi |
| EPI_ISL_495076 | Department of MicroBiology, Government Medical College, Surat | Gujarat Biotechnology Research Centre | Nitin Savaliya, Raghawendra Kumar, Dinesh Kumar, Zuber Saiyed, Komal Patel, Labdhi Pandya, Afzal Ansari, Nikha Trivedi, Naresh Chauhan, Summaiya Mullan, Amit gamit, Apurvasinh Puvar, Janvi Raval, Zarna Patel, Monika Gandhi, Pinal Trivedi, Maharshi Pandya, Nidhi Patel, R D Dixit, A M Kadri, Harsh Bakshi, Chaitanya Joshi, Madhvi Joshi |
| EPI_ISL_495077 | Department of MicroBiology, Government Medical College, Surat | Gujarat Biotechnology Research Centre | Raghawendra Kumar, Dinesh Kumar, Zuber Saiyed, Komal Patel, Labdhi Pandya, Afzal Ansari, Nikha Trivedi, Naresh Chauhan, Summaiya Mullan, Amit gamit, Apurvasinh Puvar, Janvi Raval, Zarna Patel, Monika Gandhi, Pinal Trivedi, Maharshi Pandya, Nidhi Patel, Nitin Savaliya, R D Dixit, A M Kadri, Harsh Bakshi, Chaitanya Joshi, Madhvi Joshi |
| EPI_ISL_495078 | Department of MicroBiology, Government Medical College, Surat | Gujarat Biotechnology Research Centre | Dinesh Kumar, Zuber Saiyed, Komal Patel, Labdhi Pandya, Afzal Ansari, Nikha Trivedi, Naresh Chauhan, Summaiya Mullan, Amit gamit, Apurvasinh Puvar, Janvi Raval, Zarna Patel, Monika Gandhi, Pinal Trivedi, Maharshi Pandya, Nidhi Patel, Nitin Savaliya, Raghawendra Kumar, R D Dixit, A M Kadri, Harsh Bakshi, Chaitanya Joshi, Madhvi Joshi |
| EPI_ISL_495079 | Department of MicroBiology, Government Medical College, Surat | Gujarat Biotechnology Research Centre | Afzal Ansari, Nikha Trivedi, Naresh Chauhan, Summaiya Mullan, Amit gamit, Apurvasinh Puvar, Janvi Raval, Zarna Patel, Monika Gandhi, Pinal Trivedi, Maharshi Pandya, Nidhi Patel, Nitin Savaliya, Raghawendra Kumar, Dinesh Kumar, Zuber Saiyed, Komal Patel, Labdhi Pandya, R D Dixit, A M Kadri, Harsh Bakshi, Chaitanya Joshi, Madhvi Joshi |
| EPI_ISL_495080 | Department of MicroBiology, Government Medical College, Surat | Gujarat Biotechnology Research Centre | Nikha Trivedi, Naresh Chauhan, Summaiya Mullan, Amit gamit, Apurvasinh Puvar, Janvi Raval, Zarna Patel, Monika Gandhi, Pinal Trivedi, Maharshi Pandya, Nidhi Patel, Nitin Savaliya, Raghawendra Kumar, Dinesh Kumar, Zuber Saiyed, Komal Patel, Labdhi Pandya, Afzal Ansari, R D Dixit, A M Kadri, Harsh Bakshi, Chaitanya Joshi, Madhvi Joshi |
| EPI_ISL_495081, EPI_ISL_495082, EPI_ISL_495083, EPI_ISL_495084, EPI_ISL_495085, EPI_ISL_495086, EPI_ISL_495087, EPI_ISL_495088, EPI_ISL_495089, EPI_ISL_495090, EPI_ISL_495091, EPI_ISL_495092, EPI_ISL_495093, EPI_ISL_495094, EPI_ISL_495095, EPI_ISL_495096 | | | |
| see above | Department of Medical Microbiology, Western Sussex Hospitals NHS Foundation Trust, St Richard's Hospital | Wellcome Sanger Institute for the COVID-19 Genomics UK Consortium | Manasa Mutingwende, Sarah Lowdon, Olga Podplomyk, Michelle Erkiert, Jonathan Lewis, Paul Randell and Alex Alderton, Roberto Amato, Sonia Goncalves, Ewan Harrison, David K. Jackson, Ian Johnston, Dominic Kwiatkowski, Cordelia Langford, John Sillitoe on behalf of the Wellcome Sanger Institute COVID-19 Surveillance Team (http://www.sanger.ac.uk/covid-team) |
| EPI_ISL_495097, EPI_ISL_495098, EPI_ISL_495099, EPI_ISL_495100, EPI_ISL_495101, EPI_ISL_495102, EPI_ISL_495103, EPI_ISL_495104, EPI_ISL_495105, EPI_ISL_495106, EPI_ISL_495107, EPI_ISL_495108, EPI_ISL_495109, EPI_ISL_495110, EPI_ISL_495111, EPI_ISL_495112, EPI_ISL_495113, EPI_ISL_495114, EPI_ISL_495115, EPI_ISL_495116, EPI_ISL_495117, EPI_ISL_495118, EPI_ISL_495119, EPI_ISL_495120, EPI_ISL_495121 | | | |
| see above | PHE South West Regional Laboratory, National Infection Service | Wellcome Sanger Institute for the COVID-19 Genomics UK Consortium | Stephanie Hutchings, Hannah Pymont, Dr Peter Muir, Barry Vipond, Rich Hopes; and Alex Alderton, Roberto Amato, Sonia Goncalves, Ewan Harrison, David K. Jackson, Ian Johnston, Dominic Kwiatkowski, Cordelia Langford, John Sillitoe on behalf of the Wellcome Sanger Institute COVID-19 Surveillance Team (http://www.sanger.ac.uk/covid-team) |
| EPI_ISL_495122, EPI_ISL_495123, EPI_ISL_495124, EPI_ISL_495125, EPI_ISL_495126, EPI_ISL_495127, EPI_ISL_495128, EPI_ISL_495129, EPI_ISL_495130, EPI_ISL_495131, EPI_ISL_495132, EPI_ISL_495133, EPI_ISL_495134, EPI_ISL_495135, EPI_ISL_495136, EPI_ISL_495137, EPI_ISL_495138, EPI_ISL_495139, EPI_ISL_495140, EPI_ISL_495141, EPI_ISL_495142, EPI_ISL_495143, EPI_ISL_495144, EPI_ISL_495145, EPI_ISL_495146, EPI_ISL_495147, EPI_ISL_495148, EPI_ISL_495149, EPI_ISL_495150, EPI_ISL_495151, EPI_ISL_495152, EPI_ISL_495153, EPI_ISL_495154, EPI_ISL_495155, EPI_ISL_495156, EPI_ISL_495157, EPI_ISL_495158, EPI_ISL_495159, EPI_ISL_495160 | | | |
| see above | Innovative Genomics Institute, UC Berkeley | Innovative Genomics Institute, UC Berkeley | Stacia Wyman, Haridha Shivram, Liana Lareau, Shana McDevitt, Justin Choi |
| EPI_ISL_495161 | CSIR-Centre for Cellular and Molecular Biology | CSIR-Centre for Cellular and Molecular Biology | Onkar Kulkarni,Sofia Banu, Payel Mukherjee, Priya Singh, Dhiviya Vedagiri, Divya Gupta, Vishal Sah, Santosh Kumar Kuncha, Krishnan Harinivas Harshan, Archana Bharadwaj Siva, Karthik Bharadwaj Tallapaka, Shagufta Khan, Lamuk Zaveri,Nikhil Hajirnis, M Soujanya Reddy, Pratheusa Maccha, Namami Gaur, Sakshi Shambhavi, Tulasi Nagabandi, Purushotham Vodnala, Deepak Kumar, Devi Prasad Vijayashankar, Disha Nanda, Divya Das, Jotin Gogoi, Manish Bhattacharjee, Rakesh K Mishra, Divya Tej Sowpati |
| EPI_ISL_495162 | CSIR-Centre for Cellular and Molecular Biology | CSIR-Centre for Cellular and Molecular Biology | Onkar Kulkarni, Payel Mukherjee, Sofia Banu, Priya Singh, Dhiviya Vedagiri, Divya Gupta, Vishal Sah, Santosh Kumar Kuncha, Krishnan Harinivas Harshan, Archana Bharadwaj Siva, Karthik Bharadwaj Tallapaka, Shagufta Khan, Lamuk Zaveri,Nikhil Hajirnis, M Soujanya Reddy, Pratheusa Maccha, Namami Gaur, Sakshi Shambhavi, Tulasi Nagabandi, Purushotham Vodnala, Deepak Kumar, Devi Prasad Vijayashankar, Disha Nanda, Divya Das, Jotin Gogoi, Manish Bhattacharjee, Rakesh K Mishra, Divya Tej Sowpati |
| EPI_ISL_495163 | CSIR-Centre for Cellular and Molecular Biology | CSIR-Centre for Cellular and Molecular Biology | Onkar Kulkarni, Payel Mukherjee, Sofia Banu, Priya Singh, Dhiviya Vedagiri, Divya Gupta, Vishal Sah, Santosh Kumar Kuncha, Krishnan Harinivas Harshan, Archana Bharadwaj Siva, Karthik Bharadwaj Tallapaka, Shagufta Khan, Lamuk Zaveri, Nikhil Hajirnis, M Soujanya Reddy, Pratheusa Maccha, Namami Gaur, Sakshi Shambhavi, Tulasi Nagabandi, Purushotham Vodnala, Deepak Kumar, Devi Prasad Vijayashankar, Disha Nanda, Divya Das, Jotin Gogoi, Manish Bhattacharjee, Rakesh K Mishra, Divya Tej Sowpati |
| EPI_ISL_495164 | CSIR-Centre for Cellular and Molecular Biology | CSIR-Centre for Cellular and Molecular Biology | Onkar Kulkarni, Payel Mukherjee, Sofia Banu, Priya Singh, Dhiviya Vedagiri, Divya Gupta, Vishal Sah, Santosh Kumar Kuncha, Krishnan Harinivas Harshan, Archana Bharadwaj Siva, Karthik Bharadwaj Tallapaka, Shagufta Khan, Lamuk Zaveri, Namami Gaur, Sakshi Shambhavi, Nikhil Hajirnis, M Soujanya Reddy, Pratheusa Maccha, Tulasi Nagabandi, Purushotham Vodnala,Preethi Jampala, Sharada Ravi Iyer, Sulagana Mukherjee, Swetha Sundar, Peddapuvala Sai Uday Kiran, Rakesh K Mishra, Divya Tej Sowpati |
| EPI_ISL_495165 | CSIR-Centre for Cellular and Molecular Biology | CSIR-Centre for Cellular and Molecular Biology | Onkar Kulkarni, Payel Mukherjee, Sofia Banu, Priya Singh, Dhiviya Vedagiri, Divya Gupta, Vishal Sah, Santosh Kumar Kuncha, Krishnan Harinivas Harshan, Archana Bharadwaj Siva, Karthik Bharadwaj Tallapaka, Shagufta Khan, Lamuk Zaveri, Namami Gaur, Sakshi Shambhavi,Nikhil Hajirnis, M Soujanya Reddy, Pratheusa Maccha, Tulasi Nagabandi, Purushotham Vodnala,Preethi Jampala, Sharada Ravi Iyer, Sulagana Mukherjee, Swetha Sundar, Peddapuvala Sai Uday Kiran, Rakesh K Mishra, Divya Tej Sowpati |
| EPI_ISL_495166 | CSIR-Centre for Cellular and Molecular Biology | CSIR-Centre for Cellular and Molecular Biology | Onkar Kulkarni,Sofia Banu, Payel Mukherjee, Priya Singh, Dhiviya Vedagiri, Divya Gupta, Vishal Sah, Santosh Kumar Kuncha, Krishnan Harinivas Harshan, Archana Bharadwaj Siva, Karthik Bharadwaj Tallapaka, Shagufta Khan, Lamuk Zaveri, Namami Gaur, Nikhil Hajirnis, M Soujanya Reddy, Pratheusa Maccha,Sakshi Shambhavi, Tulasi Nagabandi, Purushotham Vodnala, Deepak Kumar, Devi Prasad Vijayashankar, Disha Nanda, Divya Das, Jotin Gogoi, Manish Bhattacharjee, Rakesh K Mishra, Divya Tej Sowpati |
| EPI_ISL_495167 | CSIR-Centre for Cellular and Molecular Biology | CSIR-Centre for Cellular and Molecular Biology | Onkar Kulkarni, Payel Mukherjee, Sofia Banu, Priya Singh, Dhiviya Vedagiri, Divya Gupta, Vishal Sah, Santosh Kumar Kuncha, Krishnan Harinivas Harshan, Archana Bharadwaj Siva, Karthik Bharadwaj Tallapaka, Shagufta Khan, Lamuk Zaveri, Namami Gaur, Sakshi Shambhavi, Nikhil Hajirnis, M Soujanya Reddy, Pratheusa Maccha,Tulasi Nagabandi, Purushotham Vodnala, Deepak Kumar, Devi Prasad Vijayashankar, Disha Nanda, Divya Das, Jotin Gogoi, Manish Bhattacharjee, Rakesh K Mishra, Divya Tej Sowpati |
| EPI_ISL_495168 | CSIR-Centre for Cellular and Molecular Biology | CSIR-Centre for Cellular and Molecular Biology | Sofia Banu, Payel Mukherjee, Priya Singh,Onkar Kulkarni, Dhiviya Vedagiri, Divya Gupta, Vishal Sah, Santosh Kumar Kuncha, Krishnan Harinivas Harshan, Archana Bharadwaj Siva, Karthik Bharadwaj Tallapaka, Shagufta Khan, Lamuk Zaveri, Namami Gaur, Sakshi Shambhavi, Nikhil Hajirnis, M Soujanya Reddy, Pratheusa Maccha,Tulasi Nagabandi, Purushotham Vodnala, Deepak Kumar, Devi Prasad Vijayashankar, Disha Nanda, Divya Das, Jotin Gogoi, Manish Bhattacharjee, Rakesh K Mishra, Divya Tej Sowpati |
| EPI_ISL_495169 | CSIR-Centre for Cellular and Molecular Biology | CSIR-Centre for Cellular and Molecular Biology | Onkar Kulkarni, Payel Mukherjee, Sofia Banu, Priya Singh, Dhiviya Vedagiri, Divya Gupta, Vishal Sah, Santosh Kumar Kuncha, Krishnan Harinivas Harshan, Archana Bharadwaj Siva, Karthik Bharadwaj Tallapaka, Shagufta Khan, Lamuk Zaveri, Namami Gaur,Nikhil Hajirnis, M Soujanya Reddy, Pratheusa Maccha, Sakshi Shambhavi, Tulasi Nagabandi, Purushotham Vodnala, Deepak Kumar, Devi Prasad Vijayashankar, Disha Nanda, Divya Das, Jotin Gogoi, Manish Bhattacharjee, Rakesh K Mishra, Divya Tej Sowpati |
| EPI_ISL_495170 | CSIR-Centre for Cellular and Molecular Biology | CSIR-Centre for Cellular and Molecular Biology | Onkar Kulkarni, Tulasi Nagabandi, Namami Gaur, Sakshi Shambhavi, Lamuk Zaveri,Nikhil Hajirnis, M Soujanya Reddy, Pratheusa Maccha, Shagufta Khan, Purushotham Vodnala, Payel Mukherjee, Sofia Banu, Priya Singh, Onkar Kulkarni, Dhiviya Vedagiri, Divya Gupta, Vishal Sah, Santosh Kumar Kuncha, Krishnan Harinivas Harshan, Archana Bharadwaj Siva, Karthik Bharadwaj Tallapaka,G. Aditya Kumar, Koushick Sivakumar, Pooja Ramesh Gupta, Rajan Kumar Jha, Shraddha Vijay Lahoti, Rakesh K Mishra, Divya Tej Sowpati |
| EPI_ISL_495171 | CSIR-Centre for Cellular and Molecular Biology | CSIR-Centre for Cellular and Molecular Biology | M Soujanya Reddy, Nikhil Hajirnis, Pratheusa Maccha, Namami Gaur, Sakshi Shambhavi, Lamuk Zaveri, Shagufta Khan, Tulasi Nagabandi, Purushotham Vodnala, Payel Mukherjee, Sofia Banu, Priya Singh, Onkar Kulkarni, Dhiviya Vedagiri, Divya Gupta, Vishal Sah, Santosh Kumar Kuncha, Krishnan Harinivas Harshan, Archana Bharadwaj Siva, Karthik Bharadwaj Tallapaka, Zeba Rizvi, Zuberwasim Sayyad, Kakade Aishwarya Arun, Amrutha H C, Ananga Ghosh, Rakesh K Mishra, Divya Tej Sowpati |
| EPI_ISL_495172 | CSIR-Centre for Cellular and Molecular Biology | CSIR-Centre for Cellular and Molecular Biology | Onkar Kulkarni, Tulasi Nagabandi, Namami Gaur, Sakshi Shambhavi, Lamuk Zaveri, Nikhil Hajirnis, M Soujanya Reddy, Pratheusa Maccha, Shagufta Khan, Purushotham Vodnala, Payel Mukherjee, Sofia Banu, Priya Singh,Onkar Kulkarni, Dhiviya Vedagiri, Divya Gupta, Vishal Sah, Santosh Kumar Kuncha, Krishnan Harinivas Harshan, Archana Bharadwaj Siva, Karthik Bharadwaj Tallapaka,Kezia J Ann, Radhika Khandelwal, Roshan Maku Venkata, Shemin Mansuri, Sonu Uday, Rakesh K Mishra, Divya Tej Sowpati |
| EPI_ISL_495173 | CSIR-Centre for Cellular and Molecular Biology | CSIR-Centre for Cellular and Molecular Biology | Onkar Kulkarni, Lamuk Zaveri, Nikhil Hajirnis, M Soujanya Reddy, Pratheusa Maccha, Tulasi Nagabandi, Namami Gaur, Sakshi Shambhavi,Shagufta Khan, Purushotham Vodnala, Payel Mukherjee, Sofia Banu, Priya Singh, Dhiviya Vedagiri, Divya Gupta, Vishal Sah, Santosh Kumar Kuncha, Krishnan Harinivas Harshan, Archana Bharadwaj Siva, Karthik Bharadwaj Tallapaka,G. Aditya Kumar, Koushick Sivakumar, Pooja Ramesh Gupta, Rajan Kumar Jha, Shraddha Vijay Lahoti, Rakesh K Mishra, Divya Tej Sowpati |
| EPI_ISL_495174 | CSIR-Centre for Cellular and Molecular Biology | CSIR-Centre for Cellular and Molecular Biology | M Soujanya Reddy, Nikhil Hajirnis, Pratheusa Maccha, Payel Mukherjee, Sofia Banu, Priya Singh, Onkar Kulkarni,Tulasi Nagabandi, Namami Gaur, Sakshi Shambhavi, Lamuk Zaveri, Shagufta Khan, Purushotham Vodnala, Dhiviya Vedagiri, Divya Gupta, Vishal Sah, Santosh Kumar Kuncha, Krishnan Harinivas Harshan, Archana Bharadwaj Siva, Karthik Bharadwaj Tallapaka,Kezia J Ann, Radhika Khandelwal, Roshan Maku Venkata, Shemin Mansuri, Sonu Uday, Rakesh K Mishra, Divya Tej Sowpati |
| EPI_ISL_495175 | CSIR-Centre for Cellular and Molecular Biology | CSIR-Centre for Cellular and Molecular Biology | Onkar Kulkarni, Lamuk Zaveri, Tulasi Nagabandi, Namami Gaur, Sakshi Shambhavi,Shagufta Khan, Nikhil Hajirnis, M Soujanya Reddy, Pratheusa Maccha, Purushotham Vodnala, Payel Mukherjee, Sofia Banu, Priya Singh, Dhiviya Vedagiri, Divya Gupta, Vishal Sah, Santosh Kumar Kuncha, Krishnan Harinivas Harshan, Archana Bharadwaj Siva, Karthik Bharadwaj Tallapaka,G. Aditya Kumar, Koushick Sivakumar, Pooja Ramesh Gupta, Rajan Kumar Jha, Shraddha Vijay Lahoti, Rakesh K Mishra, Divya Tej Sowpati |
| EPI_ISL_495176 | CSIR-Centre for Cellular and Molecular Biology | CSIR-Centre for Cellular and Molecular Biology | Tulasi Nagabandi, Namami Gaur, Sakshi Shambhavi, Lamuk Zaveri, Shagufta Khan, Nikhil Hajirnis, M Soujanya Reddy, Pratheusa Maccha, Purushotham Vodnala, Payel Mukherjee, Sofia Banu, Priya Singh, Onkar Kulkarni, Dhiviya Vedagiri, Divya Gupta, Vishal Sah, Santosh Kumar Kuncha, Krishnan Harinivas Harshan, Archana Bharadwaj Siva, Karthik Bharadwaj Tallapaka,G. Aditya Kumar, Koushick Sivakumar, Pooja Ramesh Gupta, Rajan Kumar Jha, Shraddha Vijay Lahoti, Rakesh K Mishra, Divya Tej Sowpati |
| EPI_ISL_495177 | CSIR-Centre for Cellular and Molecular Biology | CSIR-Centre for Cellular and Molecular Biology | Sakshi Shambhavi, Lamuk Zaveri, Shagufta Khan, Namami Gaur, Nikhil Hajirnis, M Soujanya Reddy, Pratheusa Maccha,Tulasi Nagabandi, Purushotham Vodnala, Payel Mukherjee, Sofia Banu, |

| | | | |
|---|---|---|---|
| | | | Priya Singh,Onkar Kulkarni, Dhiviya Vedagiri, Divya Gupta, Vishal Sah, Santosh Kumar Kuncha, Krishnan Harinivas Harshan, Archana Bharadwaj Siva, Karthik Bharadwaj Tallapaka, G. Aditya Kumar, Koushick Sivakumar, Pooja Ramesh Gupta, Rajan Kumar Jha, Shraddha Vijay Lahoti, Rakesh K Mishra, Divya Tej Sowpati |
| EPI_ISL_495178 | CSIR-Centre for Cellular and Molecular Biology | CSIR-Centre for Cellular and Molecular Biology | Payel Mukherjee, Sofia Banu, Priya Singh, Onkar Kulkarni, Dhiviya Vedagiri, Divya Gupta, Vishal Sah, Santosh Kumar Kuncha, Krishnan Harinivas Harshan, Archana Bharadwaj Siva, Karthik Bharadwaj Tallapaka, Shagufta Khan, Lamuk Zaveri, Nikhil Hajirnis, M Soujanya Reddy, Pratheusa Maccha, Namami Gaur, Sakshi Shambhavi, Tulasi Nagabandi, Purushotham Vodnala, Rakesh K Mishra, Sonu Uday, Sudipta Mondal, Annapoorna P Karthyayani, Debabrata Jana, Debrya Saha, Divya Tej Sowpati |
| EPI_ISL_495179 | CSIR-Centre for Cellular and Molecular Biology | CSIR-Centre for Cellular and Molecular Biology | Pratheusa Maccha, Sakshi Shambhavi, Lamuk Zaveri, Shagufta Khan, Namami Gaur, Nikhil Hajirnis, M Soujanya Reddy, Tulasi Nagabandi, Purushotham Vodnala, Payel Mukherjee, Sofia Banu, Priya Singh,Onkar Kulkarni, Dhiviya Vedagiri, Divya Gupta, Vishal Sah, Santosh Kumar Kuncha, Krishnan Harinivas Harshan, Archana Bharadwaj Siva, Karthik Bharadwaj Tallapaka, G. Aditya Kumar, Koushick Sivakumar,Disha Nanda, Divya Das, Jotin Gogoi, Manish Bhattacharjee, Ravi Prasad Mukku, Rakesh K Mishra, Divya Tej Sowpati |
| EPI_ISL_495180 | CSIR-Centre for Cellular and Molecular Biology | CSIR-Centre for Cellular and Molecular Biology | M Soujanya Reddy, Nikhil Hajirnis, Pratheusa Maccha, Sakshi Shambhavi, Lamuk Zaveri, Shagufta Khan, Namami Gaur, Tulasi Nagabandi, Purushotham Vodnala, Payel Mukherjee, Sofia Banu, Priya Singh,Onkar Kulkarni, Dhiviya Vedagiri, Divya Gupta, Vishal Sah, Santosh Kumar Kuncha, Krishnan Harinivas Harshan, Archana Bharadwaj Siva, Karthik Bharadwaj Tallapaka, G. Aditya Kumar, Koushick Sivakumar, Pooja Ramesh Gupta, Rajan Kumar Jha, Shraddha Vijay Lahoti, Rakesh K Mishra, Divya Tej Sowpati |
| EPI_ISL_495181 | CSIR-Centre for Cellular and Molecular Biology | CSIR-Centre for Cellular and Molecular Biology | Pratheusa Maccha, Sofia Banu, Payel Mukherjee, Priya Singh, Onkar Kulkarni, Dhiviya Vedagiri, Divya Gupta, Vishal Sah, Santosh Kumar Kuncha, Krishnan Harinivas Harshan, Archana Bharadwaj Siva, Karthik Bharadwaj Tallapaka, Shagufta Khan, Lamuk Zaveri, Namami Gaur, Sakshi Shambhavi, Nikhil Hajirnis, M Soujanya Reddy, Tulasi Nagabandi, Purushotham Vodnala,Preethi Jampala, Sharada Ravi Iyer, Sulagana Mukherjee, Swetha Sundar, Peddapuvala Sai Uday Kiran, Rakesh K Mishra, Divya Tej Sowpati |
| EPI_ISL_495182 | CSIR-Centre for Cellular and Molecular Biology | CSIR-Centre for Cellular and Molecular Biology | Sakshi Shambhavi, Lamuk Zaveri, Shagufta Khan, Namami Gaur, Nikhil Hajirnis, M Soujanya Reddy, Pratheusa Maccha, Tulasi Nagabandi, Purushotham Vodnala, Payel Mukherjee, Sofia Banu, Priya Singh, Onkar Kulkarni, Dhiviya Vedagiri, Divya Gupta, Vishal Sah, Santosh Kumar Kuncha, Krishnan Harinivas Harshan, Archana Bharadwaj Siva, Karthik Bharadwaj Tallapaka, Deepak Kumar, Devi Prasad Vijayashankar, Disha Nanda, Divya Das, Jotin Gogoi, Manish Bhattacharjee, Rakesh K Mishra, Divya Tej Sowpati |
| EPI_ISL_495183 | CSIR-Centre for Cellular and Molecular Biology | CSIR-Centre for Cellular and Molecular Biology | Namami Gaur, Sakshi Shambhavi, Lamuk Zaveri, Shagufta Khan, Nikhil Hajirnis, M Soujanya Reddy, Pratheusa Maccha, Tulasi Nagabandi, Purushotham Vodnala, Payel Mukherjee, Sofia Banu, Priya Singh, Onkar Kulkarni, Dhiviya Vedagiri, Divya Gupta, Vishal Sah, Santosh Kumar Kuncha, Krishnan Harinivas Harshan, Archana Bharadwaj Siva, Karthik Bharadwaj Tallapaka, Zeba Rizvi, Zuberwasim Sayyad, Kakade Aishwarya Arun, Amrutha H C, Ananga Ghosh, Rakesh K Mishra, Divya Tej Sowpati |
| EPI_ISL_495184 | CSIR-Centre for Cellular and Molecular Biology | CSIR-Centre for Cellular and Molecular Biology | Namami Gaur, Sakshi Shambhavi, Lamuk Zaveri, Shagufta Khan, Nikhil Hajirnis, M Soujanya Reddy, Pratheusa Maccha, Tulasi Nagabandi, Purushotham Vodnala, Payel Mukherjee, Sofia Banu, Priya Singh,Onkar Kulkarni, Dhiviya Vedagiri, Divya Gupta, Vishal Sah, Santosh Kumar Kuncha, Krishnan Harinivas Harshan, Archana Bharadwaj Siva, Karthik Bharadwaj Tallapaka, Zeba Rizvi, Zuberwasim Sayyad, Kakade Aishwarya Arun, Amrutha H C, Ananga Ghosh, Rakesh K Mishra, Divya Tej Sowpati |
| EPI_ISL_495185, EPI_ISL_495186 | CSIR-Centre for Cellular and Molecular Biology | CSIR-Centre for Cellular and Molecular Biology | Shagufta Khan, Lamuk Zaveri, Namami Gaur, Sakshi Shambhavi, Nikhil Hajirnis, M Soujanya Reddy, Pratheusa Maccha, Tulasi Nagabandi, Purushotham Vodnala, Payel Mukherjee, Sofia Banu, Priya Singh, Onkar Kulkarni, Dhiviya Vedagiri, Divya Gupta, Vishal Sah, Santosh Kumar Kuncha, Krishnan Harinivas Harshan, Archana Bharadwaj Siva, Karthik Bharadwaj Tallapaka, Renu Sudhakar, Somesh Gorde, Gangumala Srinivas Reddy, Sujoy Deb, Swati Bayyana, Rakesh K Mishra, Divya Tej Sowpati |
| EPI_ISL_495187 | CSIR-Centre for Cellular and Molecular Biology | CSIR-Centre for Cellular and Molecular Biology | Shagufta Khan, Lamuk Zaveri, Namami Gaur, Sakshi Shambhavi, Nikhil Hajirnis, M Soujanya Reddy, Pratheusa Maccha,Tulasi Nagabandi, Purushotham Vodnala, Payel Mukherjee, Sofia Banu, Priya Singh, Onkar Kulkarni, Dhiviya Vedagiri, Divya Gupta, Vishal Sah, Santosh Kumar Kuncha, Krishnan Harinivas Harshan, Archana Bharadwaj Siva, Karthik Bharadwaj Tallapaka,Umesh Kumar, Unis Ahmad Bhat, Ajay Sarawagi, Priyanka Pant, Rajkanwar Nathawat, Rakesh K Mishra, Divya Tej Sowpati |
| EPI_ISL_495188 | CSIR-Centre for Cellular and Molecular Biology | CSIR-Centre for Cellular and Molecular Biology | Nikhil Hajirnis, M Soujanya Reddy, Pratheusa Maccha, Payel Mukherjee, Sofia Banu, Priya Singh,Onkar Kulkarni, Dhiviya Vedagiri, Divya Gupta, Vishal Sah, Santosh Kumar Kuncha, Krishnan Harinivas Harshan, Archana Bharadwaj Siva, Karthik Bharadwaj Tallapaka, Shagufta Khan, Lamuk Zaveri, Namami Gaur, Sakshi Shambhavi, Tulasi Nagabandi, Purushotham Vodnala,Deepak Kumar, Devi Prasad Vijayashankar, Disha Nanda, Divya Das, Jotin Gogoi, Manish Bhattacharjee, Rakesh K Mishra, Divya Tej Sowpati |
| EPI_ISL_495189 | CSIR-Centre for Cellular and Molecular Biology | CSIR-Centre for Cellular and Molecular Biology | Sofia Banu, Payel Mukherjee, Priya Singh,Onkar Kulkarni, Dhiviya Vedagiri, Divya Gupta, Vishal Sah, Santosh Kumar Kuncha, Krishnan Harinivas Harshan, Archana Bharadwaj Siva, Karthik Bharadwaj Tallapaka, Shagufta Khan, Lamuk Zaveri, Namami Gaur, Sakshi Shambhavi, Nikhil Hajirnis, M Soujanya Reddy, Pratheusa Maccha,Tulasi Nagabandi, Purushotham Vodnala, Deepak Kumar, Devi Prasad Vijayashankar, Disha Nanda, Divya Das, Jotin Gogoi, Manish Bhattacharjee, Rakesh K Mishra, Divya Tej Sowpati |
| EPI_ISL_495190 | CSIR-Centre for Cellular and Molecular Biology | CSIR-Centre for Cellular and Molecular Biology | Sakshi Shambhavi, Lamuk Zaveri, Shagufta Khan, Nikhil Hajirnis, M Soujanya Reddy, Pratheusa Maccha, Namami Gaur, Tulasi Nagabandi, Purushotham Vodnala, Payel Mukherjee, Sofia Banu, Priya Singh, Onkar Kulkarni, Dhiviya Vedagiri, Divya Gupta, Vishal Sah, Santosh Kumar Kuncha, Krishnan Harinivas Harshan, Archana Bharadwaj Siva, Karthik Bharadwaj Tallapaka,G. Aditya Kumar, Koushick Sivakumar, Rakesh K Mishra, Divya Tej Sowpati |
| EPI_ISL_495191 | CSIR-Centre for Cellular and Molecular Biology | CSIR-Centre for Cellular and Molecular Biology | Nikhil Hajirnis, M Soujanya Reddy, Pratheusa Maccha, Lamuk Zaveri, Shagufta Khan, Namami Gaur, Sakshi Shambhavi, Tulasi Nagabandi, Purushotham Vodnala, Payel Mukherjee, Sofia Banu, Priya Singh, Onkar Kulkarni, Dhiviya Vedagiri, Divya Gupta, Vishal Sah, Santosh Kumar Kuncha, Krishnan Harinivas Harshan, Archana Bharadwaj Siva, Karthik Bharadwaj Tallapaka,Zeba Rizvi, Zuberwasim Sayyad, Kakade Aishwarya Arun, Amrutha H C, Ananga Ghosh, Rakesh K Mishra, Divya Tej Sowpati |
| EPI_ISL_495192 | CSIR-Centre for Cellular and Molecular Biology | CSIR-Centre for Cellular and Molecular Biology | Sofia Banu, Payel Mukherjee, Priya Singh,Onkar Kulkarni, Dhiviya Vedagiri, Divya Gupta, Vishal Sah, Santosh Kumar Kuncha, Krishnan Harinivas Harshan, Archana Bharadwaj Siva, Karthik Bharadwaj Tallapaka, Shagufta Khan, Lamuk Zaveri, Namami Gaur, Sakshi Shambhavi, Tulasi Nagabandi, Nikhil Hajirnis, M Soujanya Reddy, Pratheusa Maccha, Purushotham Vodnala, Gokulan C G, Gunjan Purohit, Hanuman Tulashiram Kale, Pankaj Kumar, Prachand Issarapu, Rakesh K Mishra, Divya Tej Sowpati |
| EPI_ISL_495193 | CSIR-Centre for Cellular and Molecular Biology | CSIR-Centre for Cellular and Molecular Biology | Namami Gaur, Sakshi Shambhavi, Lamuk Zaveri, Shagufta Khan, Nikhil Hajirnis, M Soujanya Reddy, Pratheusa Maccha, Tulasi Nagabandi, Purushotham Vodnala, Payel Mukherjee, Sofia Banu, Priya Singh, Onkar Kulkarni, Dhiviya Vedagiri, Divya Gupta, Vishal Sah, Santosh Kumar Kuncha, Krishnan Harinivas Harshan, Archana Bharadwaj Siva, Karthik Bharadwaj Tallapaka, Zeba Rizvi, Zuberwasim Sayyad, Kakade Aishwarya Arun, Amrutha H C, Ananga Ghosh, Rakesh K Mishra, Divya Tej Sowpati |
| EPI_ISL_495194 | CSIR-Centre for Cellular and Molecular Biology | CSIR-Centre for Cellular and Molecular Biology | Sofia Banu, Payel Mukherjee, Priya Singh,Onkar Kulkarni, Dhiviya Vedagiri, Divya Gupta, Vishal Sah, Santosh Kumar Kuncha, Krishnan Harinivas Harshan, Archana Bharadwaj Siva, Karthik Bharadwaj Tallapaka, Shagufta Khan, Lamuk Zaveri, Namami Gaur, Sakshi Shambhavi, Nikhil Hajirnis, M Soujanya Reddy, Pratheusa Maccha,Tulasi Nagabandi, Purushotham Vodnala, Deepak Kumar, Devi Prasad Vijayashankar, Disha Nanda, Divya Das, Jotin Gogoi, Manish Bhattacharjee, Rakesh K Mishra, Divya Tej Sowpati |
| EPI_ISL_495195 | CSIR-Centre for Cellular and Molecular Biology | CSIR-Centre for Cellular and Molecular Biology | Nikhil Hajirnis, M Soujanya Reddy, Pratheusa Maccha, Namami Gaur, Sakshi Shambhavi, Lamuk Zaveri, Shagufta Khan, Tulasi Nagabandi, Purushotham Vodnala, Payel Mukherjee, Sofia Banu, Priya Singh,Onkar Kulkarni, Dhiviya Vedagiri, Divya Gupta, Vishal Sah, Santosh Kumar Kuncha, Krishnan Harinivas Harshan, Archana Bharadwaj Siva, Karthik Bharadwaj Tallapaka,Kezia J Ann, Radhika Khandelwal, Roshan Maku Venkata, Shemin Mansuri, Sonu Uday, Rakesh K Mishra, Divya Tej Sowpati |
| EPI_ISL_495196 | CSIR-Centre for Cellular and Molecular Biology | CSIR-Centre for Cellular and Molecular Biology | Payel Mukherjee, Sofia Banu, Priya Singh, Onkar Kulkarni, Dhiviya Vedagiri, Divya Gupta, Vishal Sah, Santosh Kumar Kuncha, Krishnan Harinivas Harshan, Archana Bharadwaj Siva, Karthik Bharadwaj Tallapaka, Shagufta Khan, Lamuk Zaveri, Nikhil Hajirnis, M Soujanya Reddy, Pratheusa Maccha, Namami Gaur, Sakshi Shambhavi, Tulasi Nagabandi, Purushotham Vodnala, G. Aditya Kumar, Koushick Sivakumar, Pooja Ramesh Gupta, Rajan Kumar Jha, Shraddha Vijay Lahoti, Rakesh K Mishra, Divya Tej Sowpati |
| EPI_ISL_495197 | CSIR-Centre for Cellular and Molecular Biology | CSIR-Centre for Cellular and Molecular Biology | Sofia Banu, Payel Mukherjee, Priya Singh,Onkar Kulkarni, Dhiviya Vedagiri, Divya Gupta, Vishal Sah, Santosh Kumar Kuncha, Krishnan Harinivas Harshan, Archana Bharadwaj Siva, Karthik Bharadwaj Tallapaka, Shagufta Khan, Lamuk Zaveri, Namami Gaur, Sakshi Shambhavi, Nikhil Hajirnis, M Soujanya Reddy, Pratheusa Maccha, Tulasi Nagabandi, Purushotham Vodnala, Disha Nanda, Divya Das, Jotin Gogoi, Manish Bhattacharjee, Ravi Prasad Mukku, Rakesh K Mishra, Divya Tej Sowpati |
| EPI_ISL_495198 | CSIR-Centre for Cellular and Molecular Biology | CSIR-Centre for Cellular and Molecular Biology | Tulasi Nagabandi, Namami Gaur, Sakshi Shambhavi, Lamuk Zaveri, Shagufta Khan, Nikhil Hajirnis, M Soujanya Reddy, Pratheusa Maccha, Purushotham Vodnala, Payel Mukherjee, Sofia Banu, Priya Singh,Onkar Kulkarni, Dhiviya Vedagiri, Divya Gupta, Vishal Sah, Santosh Kumar Kuncha, Krishnan Harinivas Harshan, Archana Bharadwaj Siva, Karthik Bharadwaj Tallapaka,Kezia J Ann, Radhika Khandelwal, Roshan Maku Venkata, Shemin Mansuri, Sonu Uday, Rakesh K Mishra, Divya Tej Sowpati |
| EPI_ISL_495199 | CSIR-Centre for Cellular and Molecular Biology | CSIR-Centre for Cellular and Molecular Biology | Pratheusa Maccha, Sakshi Shambhavi, Lamuk Zaveri, Shagufta Khan, Namami Gaur, Nikhil Hajirnis, M Soujanya Reddy, Tulasi Nagabandi, Purushotham Vodnala, Payel Mukherjee, Sofia Banu, Priya Singh,Onkar Kulkarni, Dhiviya Vedagiri, Divya Gupta, Vishal Sah, Santosh Kumar Kuncha, Krishnan Harinivas Harshan, Archana Bharadwaj Siva, Karthik Bharadwaj Tallapaka, G. Aditya Kumar, Koushick Sivakumar,Disha Nanda, Divya Das, Jotin Gogoi, Manish Bhattacharjee, Ravi Prasad Mukku, Rakesh K Mishra, Divya Tej Sowpati |
| EPI_ISL_495200 | CSIR-Centre for Cellular and Molecular Biology | CSIR-Centre for Cellular and Molecular Biology | Lamuk Zaveri, Shagufta Khan, Namami Gaur, Sakshi Shambhavi, Nikhil Hajirnis, M Soujanya Reddy, Pratheusa Maccha, Tulasi Nagabandi, Purushotham Vodnala, Payel Mukherjee, Sofia Banu, Priya Singh, Onkar Kulkarni, Dhiviya Vedagiri, Divya Gupta, Vishal Sah, Santosh Kumar Kuncha, Krishnan Harinivas Harshan, Archana Bharadwaj Siva, Karthik Bharadwaj Tallapaka, Renu Sudhakar, Somesh Gorde, Gangumala Srinivas Reddy, Sujoy Deb, Swati Bayyana, Rakesh K Mishra, Divya Tej Sowpati |
| EPI_ISL_495201 | CSIR-Centre for Cellular and Molecular Biology | CSIR-Centre for Cellular and Molecular Biology | Sakshi Shambhavi, Lamuk Zaveri, Shagufta Khan, Namami Gaur, Nikhil Hajirnis, M Soujanya Reddy, Pratheusa Maccha,Tulasi Nagabandi, Purushotham Vodnala, Payel Mukherjee, Sofia Banu, Priya Singh,Onkar Kulkarni, Dhiviya Vedagiri, Divya Gupta, Vishal Sah, Santosh Kumar Kuncha, Krishnan Harinivas Harshan, Archana Bharadwaj Siva, Karthik Bharadwaj Tallapaka, G. Aditya Kumar, Koushick Sivakumar, Pooja Ramesh Gupta, Rajan Kumar Jha, Shraddha Vijay Lahoti, Rakesh K Mishra, Divya Tej Sowpati |
| EPI_ISL_495202 | CSIR-Centre for Cellular and Molecular Biology | CSIR-Centre for Cellular and Molecular Biology | Shagufta Khan, Lamuk Zaveri, Namami Gaur, Sakshi Shambhavi, Nikhil Hajirnis, M Soujanya Reddy, Pratheusa Maccha, Tulasi Nagabandi, Purushotham Vodnala, Payel Mukherjee, Sofia Banu, Priya Singh, Onkar Kulkarni, Dhiviya Vedagiri, Divya Gupta, Vishal Sah, Santosh Kumar Kuncha, Krishnan Harinivas Harshan, Archana Bharadwaj Siva, Karthik Bharadwaj Tallapaka, Renu Sudhakar, Somesh Gorde, Gangumala Srinivas Reddy, Sujoy Deb, Swati Bayyana, Rakesh K Mishra, Divya Tej Sowpati |
| EPI_ISL_495203 | CSIR-Centre for Cellular and Molecular Biology | CSIR-Centre for Cellular and Molecular Biology | Tulasi Nagabandi, Namami Gaur, Sakshi Shambhavi, Lamuk Zaveri, Shagufta Khan, Nikhil Hajirnis, M Soujanya Reddy, Pratheusa Maccha, Purushotham Vodnala, Payel Mukherjee, Sofia Banu, Priya Singh,Onkar Kulkarni , Dhiviya Vedagiri, Divya Gupta, Vishal Sah, Santosh Kumar Kuncha, Krishnan Harinivas Harshan, Archana Bharadwaj Siva, Karthik Bharadwaj Tallapaka,G. Aditya Kumar, Koushick Sivakumar, Pooja Ramesh Gupta, Rajan Kumar Jha, Shraddha Vijay Lahoti, Rakesh K Mishra, Divya Tej Sowpati |
| EPI_ISL_495204 | CSIR-Centre for Cellular and Molecular Biology | CSIR-Centre for Cellular and Molecular Biology | Pratheusa Maccha, Shagufta Khan, Lamuk Zaveri, Namami Gaur, Sakshi Shambhavi, Tulasi Nagabandi, Nikhil Hajirnis, M Soujanya Reddy, Purushotham Vodnala, Payel Mukherjee, Sofia Banu, Priya Singh, Onkar Kulkarni, Dhiviya Vedagiri, Divya Gupta, Vishal Sah, Santosh Kumar Kuncha, Krishnan Harinivas Harshan, Archana Bharadwaj Siva, Karthik Bharadwaj Tallapaka, Disha Nanda, Divya Das, Jotin Gogoi, Manish Bhattacharjee, Ravi Prasad Mukku, Rakesh K Mishra, Divya Tej Sowpati |
| EPI_ISL_495205 | CSIR-Centre for Cellular and Molecular Biology | CSIR-Centre for Cellular and Molecular Biology | Shagufta Khan, Lamuk Zaveri, Namami Gaur, Sakshi Shambhavi, Nikhil Hajirnis, M Soujanya Reddy, Pratheusa Maccha, Tulasi Nagabandi, Purushotham Vodnala, Payel Mukherjee, Sofia Banu, Priya Singh, Onkar Kulkarni, Dhiviya Vedagiri, Divya Gupta, Vishal Sah, Santosh Kumar Kuncha, Krishnan Harinivas Harshan, Archana Bharadwaj Siva, Karthik Bharadwaj Tallapaka,Preethi Jampala, Sharada Ravi Iyer, Sulagana Mukherjee, Swetha Sundar, Peddapuvala Sai Uday Kiran Rakesh K Mishra, Divya Tej Sowpati |
| EPI_ISL_495206 | CSIR-Centre for Cellular and Molecular Biology | CSIR-Centre for Cellular and Molecular Biology | Shagufta Khan, Lamuk Zaveri, Namami Gaur, Sakshi Shambhavi, Nikhil Hajirnis, M Soujanya Reddy, Pratheusa Maccha,Tulasi Nagabandi, Purushotham Vodnala, Payel Mukherjee, Sofia Banu, Priya Singh, Onkar Kulkarni, Dhiviya Vedagiri, Divya Gupta, Vishal Sah, Santosh Kumar Kuncha, Krishnan Harinivas Harshan, Archana Bharadwaj Siva, Karthik Bharadwaj Tallapaka,Umesh Kumar, Unis Ahmad Bhat, Ajay Sarawagi, Priyanka Pant, Rajkanwar Nathawat, Rakesh K Mishra, Divya Tej Sowpati |
| EPI_ISL_495207 | CSIR-Centre for Cellular and Molecular Biology | CSIR-Centre for Cellular and Molecular Biology | Namami Gaur, Sakshi Shambhavi, Lamuk Zaveri, Shagufta Khan, Nikhil Hajirnis, M Soujanya Reddy, Pratheusa Maccha, Tulasi Nagabandi, Purushotham Vodnala, Payel Mukherjee, Sofia Banu, Priya Singh, Onkar Kulkarni, Dhiviya Vedagiri, Divya Gupta, Vishal Sah, Santosh Kumar Kuncha, Krishnan Harinivas Harshan, Archana Bharadwaj Siva, Karthik Bharadwaj Tallapaka,G. Aditya Kumar, Koushick Sivakumar, Rakesh K Mishra, Divya Tej Sowpati |
| EPI_ISL_495208 | CSIR-Centre for Cellular and Molecular Biology | CSIR-Centre for Cellular and Molecular Biology | Shagufta Khan, Lamuk Zaveri, Namami Gaur, Sakshi Shambhavi, Nikhil Hajirnis, M Soujanya Reddy, Pratheusa Maccha, Tulasi Nagabandi, Purushotham Vodnala, Payel Mukherjee, Sofia Banu, Priya Singh, Onkar Kulkarni, Dhiviya Vedagiri, Divya Gupta, Vishal Sah, Santosh Kumar Kuncha, Krishnan Harinivas Harshan, Archana Bharadwaj Siva, Karthik Bharadwaj Tallapaka, Renu Sudhakar, Somesh Gorde, Gangumala Srinivas Reddy, Sujoy Deb, Swati Bayyana, Rakesh K Mishra, Divya Tej Sowpati |
| EPI_ISL_495209 | CSIR-Centre for Cellular and Molecular Biology | CSIR-Centre for Cellular and Molecular Biology | Payel Mukherjee, Sofia Banu, Priya Singh, Onkar Kulkarni, Dhiviya Vedagiri, Divya Gupta, Vishal Sah, Santosh Kumar Kuncha, Krishnan Harinivas Harshan, Archana Bharadwaj Siva, Karthik Bharadwaj Tallapaka, Shagufta Khan, Lamuk Zaveri, Nikhil Hajirnis, M Soujanya Reddy, Pratheusa Maccha, Namami Gaur, Sakshi Shambhavi, Tulasi Nagabandi, Purushotham Vodnala, Gokulan C G, Gunjan Purohit, Hanuman Tulashiram Kale, Pankaj Kumar, Prachand Issarapu, Rakesh K Mishra, Divya Tej Sowpati |
| EPI_ISL_495210 | CSIR-Centre for Cellular and Molecular Biology | CSIR-Centre for Cellular and Molecular Biology | Sofia Banu, Payel Mukherjee, Priya Singh,Onkar Kulkarni, Dhiviya Vedagiri, Divya Gupta, Vishal Sah, Santosh Kumar Kuncha, Krishnan Harinivas Harshan, Archana Bharadwaj Siva, Karthik |

| | | | |
|---|---|---|---|
| | | | Bharadwaj Tallapaka, Shagufta Khan, Lamuk Zaveri, Namami Gaur, Sakshi Shambhavi, Nikhil Hajirnis, M Soujanya Reddy, Pratheusa Maccha,Tulasi Nagabandi, Purushotham Vodnala, Deepak Kumar, Devi Prasad Vijayashankar, Disha Nanda, Divya Das, Jotin Gogoi, Manish Bhattacharjee, Rakesh K Mishra, Divya Tej Sowpati |
| EPI_ISL_495211 | CSIR-Centre for Cellular and Molecular Biology | CSIR-Centre for Cellular and Molecular Biology | Shagufta Khan, Lamuk Zaveri, Namami Gaur, Sakshi Shambhavi, Nikhil Hajirnis, M Soujanya Reddy, Pratheusa Maccha, Tulasi Nagabandi, Purushotham Vodnala, Payel Mukherjee, Sofia Banu, Priya Singh, Onkar Kulkarni, Dhiviya Vedagiri, Divya Gupta, Vishal Sah, Santosh Kumar Kuncha, Krishnan Harinivas Harshan, Archana Bharadwaj Siva, Karthik Bharadwaj Tallapaka,Preethi Jampala, Sharada Ravi Iyer, Sulagana Mukherjee, Swetha Sundar, Peddapuvala Sai Uday Kiran Rakesh K Mishra, Divya Tej Sowpati |
| EPI_ISL_495212 | CSIR-Centre for Cellular and Molecular Biology | CSIR-Centre for Cellular and Molecular Biology | M Soujanya Reddy, Nikhil Hajirnis, Pratheusa Maccha, Sakshi Shambhavi, Lamuk Zaveri, Shagufta Khan, Namami Gaur, Tulasi Nagabandi, Purushotham Vodnala, Payel Mukherjee, Sofia Banu, Priya Singh,Onkar Kulkarni, Dhiviya Vedagiri, Divya Gupta, Vishal Sah, Santosh Kumar Kuncha, Krishnan Harinivas Harshan, Archana Bharadwaj Siva, Karthik Bharadwaj Tallapaka, G. Aditya Kumar, Koushick Sivakumar, Pooja Ramesh Gupta, Rajan Kumar Jha, Shraddha Vijay Lahoti, Rakesh K Mishra, Divya Tej Sowpati |
| EPI_ISL_495213 | CSIR-Centre for Cellular and Molecular Biology | CSIR-Centre for Cellular and Molecular Biology | Sakshi Shambhavi, Lamuk Zaveri, Shagufta Khan, Nikhil Hajirnis, M Soujanya Reddy, Pratheusa Maccha, Namami Gaur, Tulasi Nagabandi, Purushotham Vodnala, Payel Mukherjee, Sofia Banu, Priya Singh, Onkar Kulkarni, Dhiviya Vedagiri, Divya Gupta, Vishal Sah, Santosh Kumar Kuncha, Krishnan Harinivas Harshan, Archana Bharadwaj Siva, Karthik Bharadwaj Tallapaka,G. Aditya Kumar, Koushick Sivakumar, Rakesh K Mishra, Divya Tej Sowpati |
| EPI_ISL_495214 | CSIR-Centre for Cellular and Molecular Biology | CSIR-Centre for Cellular and Molecular Biology | Tulasi Nagabandi, Namami Gaur, Sakshi Shambhavi, Lamuk Zaveri, Shagufta Khan, Nikhil Hajirnis, M Soujanya Reddy, Pratheusa Maccha, Purushotham Vodnala, Payel Mukherjee, Sofia Banu, Priya Singh,Onkar Kulkarni , Dhiviya Vedagiri, Divya Gupta, Vishal Sah, Santosh Kumar Kuncha, Krishnan Harinivas Harshan, Archana Bharadwaj Siva, Karthik Bharadwaj Tallapaka,G. Aditya Kumar, Koushick Sivakumar, Pooja Ramesh Gupta, Rajan Kumar Jha, Shraddha Vijay Lahoti, Rakesh K Mishra, Divya Tej Sowpati |
| EPI_ISL_495215 | CSIR-Centre for Cellular and Molecular Biology | CSIR-Centre for Cellular and Molecular Biology | Lamuk Zaveri, Shagufta Khan,Nikhil Hajirnis, M Soujanya Reddy, Pratheusa Maccha, Namami Gaur, Sakshi Shambhavi, Tulasi Nagabandi, Purushotham Vodnala, Payel Mukherjee, Sofia Banu, Priya Singh, Onkar Kulkarni, Dhiviya Vedagiri, Divya Gupta, Vishal Sah, Santosh Kumar Kuncha, Krishnan Harinivas Harshan, Archana Bharadwaj Siva, Karthik Bharadwaj Tallapaka,Umesh Kumar, Unis Ahmad Bhat, Ajay Sarawagi, Priyanka Pant, Rajkanwar Nathawat, Rakesh K Mishra, Divya Tej Sowpati |
| EPI_ISL_495216 | CSIR-Centre for Cellular and Molecular Biology | CSIR-Centre for Cellular and Molecular Biology | Nikhil Hajirnis, M Soujanya Reddy, Pratheusa Maccha, Lamuk Zaveri, Shagufta Khan, Namami Gaur, Sakshi Shambhavi, Tulasi Nagabandi, Purushotham Vodnala, Payel Mukherjee, Sofia Banu, Priya Singh, Onkar Kulkarni, Dhiviya Vedagiri, Divya Gupta, Vishal Sah, Santosh Kumar Kuncha, Krishnan Harinivas Harshan, Archana Bharadwaj Siva, Karthik Bharadwaj Tallapaka,Zeba Rizvi, Zuberwasim Sayyad, Kakade Aishwarya Arun, Amrutha H C, Ananga Ghosh, Rakesh K Mishra, Divya Tej Sowpati |
| EPI_ISL_495217 | CSIR-Centre for Cellular and Molecular Biology | CSIR-Centre for Cellular and Molecular Biology | Nikhil Hajirnis, M Soujanya Reddy, Pratheusa Maccha, Namami Gaur, Sakshi Shambhavi, Lamuk Zaveri, Shagufta Khan, Tulasi Nagabandi, Purushotham Vodnala, Payel Mukherjee, Sofia Banu, Priya Singh,Onkar Kulkarni, Dhiviya Vedagiri, Divya Gupta, Vishal Sah, Santosh Kumar Kuncha, Krishnan Harinivas Harshan, Archana Bharadwaj Siva, Karthik Bharadwaj Tallapaka,Kezia J Ann, Radhika Khandelwal, Roshan Maku Venkata, Shemin Mansuri, Sonu Uday, Rakesh K Mishra, Divya Tej Sowpati |
| EPI_ISL_495218 | CSIR-Centre for Cellular and Molecular Biology | CSIR-Centre for Cellular and Molecular Biology | Sakshi Shambhavi, Lamuk Zaveri, Shagufta Khan, Namami Gaur, Nikhil Hajirnis, M Soujanya Reddy, Pratheusa Maccha, Tulasi Nagabandi, Purushotham Vodnala, Payel Mukherjee, Sofia Banu, Priya Singh, Onkar Kulkarni, Dhiviya Vedagiri, Divya Gupta, Vishal Sah, Santosh Kumar Kuncha, Krishnan Harinivas Harshan, Archana Bharadwaj Siva, Karthik Bharadwaj Tallapaka, Deepak Kumar, Devi Prasad Vijayashankar, Disha Nanda, Divya Das, Jotin Gogoi, Manish Bhattacharjee, Rakesh K Mishra, Divya Tej Sowpati |
| EPI_ISL_495219 | CSIR-Centre for Cellular and Molecular Biology | CSIR-Centre for Cellular and Molecular Biology | Sofia Banu, Payel Mukherjee, Priya Singh,Onkar Kulkarni, Dhiviya Vedagiri, Divya Gupta, Vishal Sah, Santosh Kumar Kuncha, Krishnan Harinivas Harshan, Archana Bharadwaj Siva, Karthik Bharadwaj Tallapaka, Shagufta Khan, Lamuk Zaveri, Namami Gaur, Sakshi Shambhavi, Nikhil Hajirnis, M Soujanya Reddy, Pratheusa Maccha, Tulasi Nagabandi, Purushotham Vodnala, Disha Nanda, Divya Das, Jotin Gogoi, Manish Bhattacharjee, Ravi Prasad Mukku, Rakesh K Mishra, Divya Tej Sowpati |
| EPI_ISL_495220 | CSIR-Centre for Cellular and Molecular Biology | CSIR-Centre for Cellular and Molecular Biology | Shagufta Khan, Lamuk Zaveri, Namami Gaur, Sakshi Shambhavi, Nikhil Hajirnis, M Soujanya Reddy, Pratheusa Maccha, Tulasi Nagabandi, Purushotham Vodnala, Payel Mukherjee, Sofia Banu, Priya Singh, Onkar Kulkarni, Dhiviya Vedagiri, Divya Gupta, Vishal Sah, Santosh Kumar Kuncha, Krishnan Harinivas Harshan, Archana Bharadwaj Siva, Karthik Bharadwaj Tallapaka,Preethi Jampala, Sharada Ravi Iyer, Sulagana Mukherjee, Swetha Sundar, Peddapuvala Sai Uday Kiran Rakesh K Mishra, Divya Tej Sowpati |
| EPI_ISL_495221 | CSIR-Centre for Cellular and Molecular Biology | CSIR-Centre for Cellular and Molecular Biology | Sofia Banu, Payel Mukherjee, Priya Singh,Onkar Kulkarni, Dhiviya Vedagiri, Divya Gupta, Vishal Sah, Santosh Kumar Kuncha, Krishnan Harinivas Harshan, Archana Bharadwaj Siva, Karthik Bharadwaj Tallapaka, Shagufta Khan, Lamuk Zaveri, Namami Gaur, Sakshi Shambhavi, Nikhil Hajirnis, M Soujanya Reddy, Pratheusa Maccha, Tulasi Nagabandi, Purushotham Vodnala, Disha Nanda, Divya Das, Jotin Gogoi, Manish Bhattacharjee, Ravi Prasad Mukku, Rakesh K Mishra, Divya Tej Sowpati |
| EPI_ISL_495222 | CSIR-Centre for Cellular and Molecular Biology | CSIR-Centre for Cellular and Molecular Biology | Pratheusa Maccha, Shagufta Khan, Lamuk Zaveri, Namami Gaur, Sakshi Shambhavi, Tulasi Nagabandi, Nikhil Hajirnis, M Soujanya Reddy, Purushotham Vodnala, Payel Mukherjee, Sofia Banu, Priya Singh, Onkar Kulkarni, Dhiviya Vedagiri, Divya Gupta, Vishal Sah, Santosh Kumar Kuncha, Krishnan Harinivas Harshan, Archana Bharadwaj Siva, Karthik Bharadwaj Tallapaka, Disha Nanda, Divya Das, Jotin Gogoi, Manish Bhattacharjee, Ravi Prasad Mukku, Rakesh K Mishra, Divya Tej Sowpati |
| EPI_ISL_495223 | CSIR-Centre for Cellular and Molecular Biology | CSIR-Centre for Cellular and Molecular Biology | Nikhil Hajirnis, M Soujanya Reddy, Pratheusa Maccha, Payel Mukherjee, Sofia Banu, Priya Singh,Onkar Kulkarni, Dhiviya Vedagiri, Divya Gupta, Vishal Sah, Santosh Kumar Kuncha, Krishnan Harinivas Harshan, Archana Bharadwaj Siva, Karthik Bharadwaj Tallapaka, Shagufta Khan, Lamuk Zaveri, Namami Gaur, Sakshi Shambhavi, Tulasi Nagabandi, Purushotham Vodnala,Deepak Kumar, Devi Prasad Vijayashankar, Disha Nanda, Divya Das, Jotin Gogoi, Manish Bhattacharjee, Rakesh K Mishra, Divya Tej Sowpati |
| EPI_ISL_495224 | CSIR-Centre for Cellular and Molecular Biology | CSIR-Centre for Cellular and Molecular Biology | Namami Gaur, Sakshi Shambhavi, Lamuk Zaveri, Shagufta Khan, Nikhil Hajirnis, M Soujanya Reddy, Pratheusa Maccha, Tulasi Nagabandi, Purushotham Vodnala, Payel Mukherjee, Sofia Banu, Priya Singh, Onkar Kulkarni, Dhiviya Vedagiri, Divya Gupta, Vishal Sah, Santosh Kumar Kuncha, Krishnan Harinivas Harshan, Archana Bharadwaj Siva, Karthik Bharadwaj Tallapaka,G. Aditya Kumar, Koushick Sivakumar, Rakesh K Mishra, Divya Tej Sowpati |
| EPI_ISL_495225 | CSIR-Centre for Cellular and Molecular Biology | CSIR-Centre for Cellular and Molecular Biology | Pratheusa Maccha, Sakshi Shambhavi, Lamuk Zaveri, Shagufta Khan, Namami Gaur, Nikhil Hajirnis, M Soujanya Reddy, Tulasi Nagabandi, Purushotham Vodnala, Payel Mukherjee, Sofia Banu, Priya Singh,Onkar Kulkarni, Dhiviya Vedagiri, Divya Gupta, Vishal Sah, Santosh Kumar Kuncha, Krishnan Harinivas Harshan, Archana Bharadwaj Siva, Karthik Bharadwaj Tallapaka, G. Aditya Kumar, Koushick Sivakumar,Disha Nanda, Divya Das, Jotin Gogoi, Manish Bhattacharjee, Ravi Prasad Mukku, Rakesh K Mishra, Divya Tej Sowpati |
| EPI_ISL_495226 | CSIR-Centre for Cellular and Molecular Biology | CSIR-Centre for Cellular and Molecular Biology | M Soujanya Reddy, Nikhil Hajirnis, Pratheusa Maccha, Namami Gaur, Sakshi Shambhavi, Lamuk Zaveri, Shagufta Khan, Tulasi Nagabandi, Purushotham Vodnala, Payel Mukherjee, Sofia Banu, Priya Singh, Onkar Kulkarni, Dhiviya Vedagiri, Divya Gupta, Vishal Sah, Santosh Kumar Kuncha, Krishnan Harinivas Harshan, Archana Bharadwaj Siva, Karthik Bharadwaj Tallapaka, Zeba Rizvi, Zuberwasim Sayyad, Kakade Aishwarya Arun, Amrutha H C, Ananga Ghosh, Rakesh K Mishra, Divya Tej Sowpati |
| EPI_ISL_495227 | CSIR-Centre for Cellular and Molecular Biology | CSIR-Centre for Cellular and Molecular Biology | Payel Mukherjee, Sofia Banu, Priya Singh, Onkar Kulkarni, Dhiviya Vedagiri, Divya Gupta, Vishal Sah, Santosh Kumar Kuncha, Krishnan Harinivas Harshan, Archana Bharadwaj Siva, Karthik Bharadwaj Tallapaka, Shagufta Khan, Lamuk Zaveri, Nikhil Hajirnis, M Soujanya Reddy, Pratheusa Maccha, Namami Gaur, Sakshi Shambhavi, Tulasi Nagabandi, Purushotham Vodnala, Gokulan C G, Gunjan Purohit, Hanuman Tulashiram Kale, Pankaj Kumar, Prachand Issarapu, Rakesh K Mishra, Divya Tej Sowpati |
| EPI_ISL_495228 | CSIR-Centre for Cellular and Molecular Biology | CSIR-Centre for Cellular and Molecular Biology | Payel Mukherjee, Sofia Banu, Priya Singh, Onkar Kulkarni, Dhiviya Vedagiri, Divya Gupta, Vishal Sah, Santosh Kumar Kuncha, Krishnan Harinivas Harshan, Archana Bharadwaj Siva, Karthik Bharadwaj Tallapaka, Shagufta Khan, Lamuk Zaveri, Nikhil Hajirnis, M Soujanya Reddy, Pratheusa Maccha, Namami Gaur, Sakshi Shambhavi, Tulasi Nagabandi, Purushotham Vodnala, Rakesh K Mishra, Sonu Uday, Sudipta Mondal, Annapoorna P Karthyayani, Debabrata Jana, Debrya Saha, Divya Tej Sowpati |
| EPI_ISL_495229 | CSIR-Centre for Cellular and Molecular Biology | CSIR-Centre for Cellular and Molecular Biology | Namami Gaur, Sakshi Shambhavi, Lamuk Zaveri, Shagufta Khan, Nikhil Hajirnis, M Soujanya Reddy, Pratheusa Maccha, Tulasi Nagabandi, Purushotham Vodnala, Payel Mukherjee, Sofia Banu, Priya Singh,Onkar Kulkarni, Dhiviya Vedagiri, Divya Gupta, Vishal Sah, Santosh Kumar Kuncha, Krishnan Harinivas Harshan, Archana Bharadwaj Siva, Karthik Bharadwaj Tallapaka, Zeba Rizvi, Zuberwasim Sayyad, Kakade Aishwarya Arun, Amrutha H C, Ananga Ghosh, Rakesh K Mishra, Divya Tej Sowpati |
| EPI_ISL_495230 | CSIR-Centre for Cellular and Molecular Biology | CSIR-Centre for Cellular and Molecular Biology | Sofia Banu, Payel Mukherjee, Priya Singh,Onkar Kulkarni, Dhiviya Vedagiri, Divya Gupta, Vishal Sah, Santosh Kumar Kuncha, Krishnan Harinivas Harshan, Archana Bharadwaj Siva, Karthik Bharadwaj Tallapaka, Shagufta Khan, Lamuk Zaveri, Namami Gaur, Sakshi Shambhavi, Tulasi Nagabandi, Nikhil Hajirnis, M Soujanya Reddy, Pratheusa Maccha, Purushotham Vodnala, Gokulan C G, Gunjan Purohit, Hanuman Tulashiram Kale, Pankaj Kumar, Prachand Issarapu, Rakesh K Mishra, Divya Tej Sowpati |
| EPI_ISL_495231 | CSIR-Centre for Cellular and Molecular Biology | CSIR-Centre for Cellular and Molecular Biology | Namami Gaur, Sakshi Shambhavi, Lamuk Zaveri, Shagufta Khan, Nikhil Hajirnis, M Soujanya Reddy, Pratheusa Maccha, Tulasi Nagabandi, Purushotham Vodnala, Payel Mukherjee, Sofia Banu, Priya Singh, Onkar Kulkarni, Dhiviya Vedagiri, Divya Gupta, Vishal Sah, Santosh Kumar Kuncha, Krishnan Harinivas Harshan, Archana Bharadwaj Siva, Karthik Bharadwaj Tallapaka, Zeba Rizvi, Zuberwasim Sayyad, Kakade Aishwarya Arun, Amrutha H C, Ananga Ghosh, Rakesh K Mishra, Divya Tej Sowpati |
| EPI_ISL_495232 | CSIR-Centre for Cellular and Molecular Biology | CSIR-Centre for Cellular and Molecular Biology | Sofia Banu, Payel Mukherjee, Priya Singh,Onkar Kulkarni, Dhiviya Vedagiri, Divya Gupta, Vishal Sah, Santosh Kumar Kuncha, Krishnan Harinivas Harshan, Archana Bharadwaj Siva, Karthik Bharadwaj Tallapaka, Shagufta Khan, Lamuk Zaveri, Namami Gaur, Sakshi Shambhavi, Nikhil Hajirnis, M Soujanya Reddy, Pratheusa Maccha, Tulasi Nagabandi, Purushotham Vodnala, Disha Nanda, Divya Das, Jotin Gogoi, Manish Bhattacharjee, Ravi Prasad Mukku, Rakesh K Mishra, Divya Tej Sowpati |
| EPI_ISL_495233 | CSIR-Centre for Cellular and Molecular Biology | CSIR-Centre for Cellular and Molecular Biology | Lamuk Zaveri, Shagufta Khan,Nikhil Hajirnis, M Soujanya Reddy, Pratheusa Maccha, Namami Gaur, Sakshi Shambhavi, Tulasi Nagabandi, Purushotham Vodnala, Payel Mukherjee, Sofia Banu, Priya Singh, Onkar Kulkarni, Dhiviya Vedagiri, Divya Gupta, Vishal Sah, Santosh Kumar Kuncha, Krishnan Harinivas Harshan, Archana Bharadwaj Siva, Karthik Bharadwaj Tallapaka,Umesh Kumar, Unis Ahmad Bhat, Ajay Sarawagi, Priyanka Pant, Rajkanwar Nathawat, Rakesh K Mishra, Divya Tej Sowpati |
| EPI_ISL_495234 | CSIR-Centre for Cellular and Molecular Biology | CSIR-Centre for Cellular and Molecular Biology | Lamuk Zaveri, Shagufta Khan,Nikhil Hajirnis, M Soujanya Reddy, Pratheusa Maccha, Namami Gaur, Sakshi Shambhavi, Tulasi Nagabandi, Purushotham Vodnala, Payel Mukherjee, Sofia Banu, Priya Singh, Onkar Kulkarni, Dhiviya Vedagiri, Divya Gupta, Vishal Sah, Santosh Kumar Kuncha, Krishnan Harinivas Harshan, Archana Bharadwaj Siva, Karthik Bharadwaj Tallapaka,Zeba Rizvi, Zuberwasim Sayyad, Kakade Aishwarya Arun, Amrutha H C, Ananga Ghosh, Rakesh K Mishra, Divya Tej Sowpati |
| EPI_ISL_495235 | CSIR-Centre for Cellular and Molecular Biology | CSIR-Centre for Cellular and Molecular Biology | Tulasi Nagabandi, Namami Gaur, Sakshi Shambhavi, Lamuk Zaveri, Shagufta Khan, Nikhil Hajirnis, M Soujanya Reddy, Pratheusa Maccha, Purushotham Vodnala, Payel Mukherjee, Sofia Banu, Priya Singh, Onkar Kulkarni, Dhiviya Vedagiri, Divya Gupta, Vishal Sah, Santosh Kumar Kuncha, Krishnan Harinivas Harshan, Archana Bharadwaj Siva, Karthik Bharadwaj Tallapaka,G. Aditya Kumar, Koushick Sivakumar, Pooja Ramesh Gupta, Rajan Kumar Jha, Shraddha Vijay Lahoti, Rakesh K Mishra, Divya Tej Sowpati |
| EPI_ISL_495236 | CSIR-Centre for Cellular and Molecular Biology | CSIR-Centre for Cellular and Molecular Biology | Nikhil Hajirnis, M Soujanya Reddy, Pratheusa Maccha, Lamuk Zaveri, Shagufta Khan, Namami Gaur, Sakshi Shambhavi, Tulasi Nagabandi, Purushotham Vodnala, Payel Mukherjee, Sofia Banu, Priya Singh, Onkar Kulkarni, Dhiviya Vedagiri, Divya Gupta, Vishal Sah, Santosh Kumar Kuncha, Krishnan Harinivas Harshan, Archana Bharadwaj Siva, Karthik Bharadwaj Tallapaka,Zeba Rizvi, Zuberwasim Sayyad, Kakade Aishwarya Arun, Amrutha H C, Ananga Ghosh, Rakesh K Mishra, Divya Tej Sowpati |
| EPI_ISL_495237 | CSIR-Centre for Cellular and Molecular Biology | CSIR-Centre for Cellular and Molecular Biology | Payel Mukherjee, Sofia Banu, Priya Singh, Onkar Kulkarni, Dhiviya Vedagiri, Divya Gupta, Vishal Sah, Santosh Kumar Kuncha, Krishnan Harinivas Harshan, Archana Bharadwaj Siva, Karthik Bharadwaj Tallapaka, Shagufta Khan, Lamuk Zaveri, Nikhil Hajirnis, M Soujanya Reddy, Pratheusa Maccha, Namami Gaur, Sakshi Shambhavi, Tulasi Nagabandi, Purushotham Vodnala, Rakesh K Mishra, Sonu Uday, Sudipta Mondal, Annapoorna P Karthyayani, Debabrata Jana, Debrya Saha, Divya Tej Sowpati |
| EPI_ISL_495238 | CSIR-Centre for Cellular and Molecular Biology | CSIR-Centre for Cellular and Molecular Biology | Pratheusa Maccha, Sakshi Shambhavi, Lamuk Zaveri, Shagufta Khan, Namami Gaur, Nikhil Hajirnis, M Soujanya Reddy, Tulasi Nagabandi, Purushotham Vodnala, Payel Mukherjee, Sofia Banu, Priya Singh,Onkar Kulkarni, Dhiviya Vedagiri, Divya Gupta, Vishal Sah, Santosh Kumar Kuncha, Krishnan Harinivas Harshan, Archana Bharadwaj Siva, Karthik Bharadwaj Tallapaka, G. Aditya Kumar, Koushick Sivakumar,Disha Nanda, Divya Das, Jotin Gogoi, Manish Bhattacharjee, Ravi Prasad Mukku, Rakesh K Mishra, Divya Tej Sowpati |
| EPI_ISL_495239 | CSIR-Centre for Cellular and Molecular Biology | CSIR-Centre for Cellular and Molecular Biology | Nikhil Hajirnis, M Soujanya Reddy, Pratheusa Maccha, Payel Mukherjee, Sofia Banu, Priya Singh,Onkar Kulkarni, Dhiviya Vedagiri, Divya Gupta, Vishal Sah, Santosh Kumar Kuncha, Krishnan Harinivas Harshan, Archana Bharadwaj Siva, Karthik Bharadwaj Tallapaka, Shagufta Khan, Lamuk Zaveri, Namami Gaur, Sakshi Shambhavi, Tulasi Nagabandi, Purushotham Vodnala,Deepak Kumar, Devi Prasad Vijayashankar, Disha Nanda, Divya Das, Jotin Gogoi, Manish Bhattacharjee, Rakesh K Mishra, Divya Tej Sowpati |
| EPI_ISL_495240 | CSIR-Centre for Cellular and Molecular Biology | CSIR-Centre for Cellular and Molecular Biology | M Soujanya Reddy, Nikhil Hajirnis, Pratheusa Maccha, Payel Mukherjee, Sofia Banu, Priya Singh, Onkar Kulkarni,Tulasi Nagabandi, Namami Gaur, Sakshi Shambhavi, Lamuk Zaveri, Shagufta Khan, Purushotham Vodnala, Dhiviya Vedagiri, Divya Gupta, Vishal Sah, Santosh Kumar Kuncha, Krishnan Harinivas Harshan, Archana Bharadwaj Siva, Karthik Bharadwaj Tallapaka,Kezia J Ann, Radhika Khandelwal, Roshan Maku Venkata, Shemin Mansuri, Sonu Uday, Rakesh K Mishra, Divya Tej Sowpati |
| EPI_ISL_495241 | CSIR-Centre for Cellular and Molecular Biology | CSIR-Centre for Cellular and Molecular Biology | Shagufta Khan, Lamuk Zaveri, Namami Gaur, Sakshi Shambhavi, Nikhil Hajirnis, M Soujanya Reddy, Pratheusa Maccha,Tulasi Nagabandi, Purushotham Vodnala, Payel Mukherjee, Sofia Banu, Priya Singh, Onkar Kulkarni, Dhiviya Vedagiri, Divya Gupta, Vishal Sah, Santosh Kumar Kuncha, Krishnan Harinivas Harshan, Archana Bharadwaj Siva, Karthik Bharadwaj Tallapaka,Umesh Kumar, Unis Ahmad Bhat, Ajay Sarawagi, Priyanka Pant, Rajkanwar Nathawat, Rakesh K Mishra, Divya Tej Sowpati |
| EPI_ISL_495242 | CSIR-Centre for Cellular and Molecular Biology | CSIR-Centre for Cellular and Molecular Biology | Tulasi Nagabandi, Namami Gaur, Sakshi Shambhavi, Lamuk Zaveri, Shagufta Khan, Nikhil Hajirnis, M Soujanya Reddy, Pratheusa Maccha, Purushotham Vodnala, Payel Mukherjee, Sofia Banu, |

| | | | |
|---|---|---|---|
| | | | Priya Singh,Onkar Kulkarni, Dhiviya Vedagiri, Divya Gupta, Vishal Sah, Santosh Kumar Kuncha, Krishnan Harinivas Harshan, Archana Bharadwaj Siva, Karthik Bharadwaj Tallapaka,Kezia J Ann, Radhika Khandelwal, Roshan Maku Venkata, Shemin Mansuri, Sonu Uday, Rakesh K Mishra, Divya Tej Sowpati |
| EPI_ISL_495243 | CSIR-Centre for Cellular and Molecular Biology | CSIR-Centre for Cellular and Molecular Biology | Lamuk Zaveri, Shagufta Khan,Nikhil Hajirnis, M Soujanya Reddy, Pratheusa Maccha, Namami Gaur, Sakshi Shambhavi, Tulasi Nagabandi, Purushotham Vodnala, Payel Mukherjee, Sofia Banu, Priya Singh, Onkar Kulkarni, Dhiviya Vedagiri, Divya Gupta, Vishal Sah, Santosh Kumar Kuncha, Krishnan Harinivas Harshan, Archana Bharadwaj Siva, Karthik Bharadwaj Tallapaka,Zeba Rizvi, Zuberwasim Sayyad, Kakade Aishwarya Arun, Amrutha H C, Ananga Ghosh, Rakesh K Mishra, Divya Tej Sowpati |
| EPI_ISL_495244 | CSIR-Centre for Cellular and Molecular Biology | CSIR-Centre for Cellular and Molecular Biology | Payel Mukherjee, Sofia Banu, Priya Singh, Onkar Kulkarni, Dhiviya Vedagiri, Divya Gupta, Vishal Sah, Santosh Kumar Kuncha, Krishnan Harinivas Harshan, Archana Bharadwaj Siva, Karthik Bharadwaj Tallapaka, Shagufta Khan, Lamuk Zaveri, Nikhil Hajirnis, M Soujanya Reddy, Pratheusa Maccha, Namami Gaur, Sakshi Shambhavi, Tulasi Nagabandi, Purushotham Vodnala, G. Aditya Kumar, Koushick Sivakumar, Pooja Ramesh Gupta, Rajan Kumar Jha, Shraddha Vijay Lahoti, Rakesh K Mishra, Divya Tej Sowpati |
| EPI_ISL_495245 | CSIR-Centre for Cellular and Molecular Biology | CSIR-Centre for Cellular and Molecular Biology | Payel Mukherjee, Sofia Banu, Priya Singh, Onkar Kulkarni, Dhiviya Vedagiri, Divya Gupta, Vishal Sah, Santosh Kumar Kuncha, Krishnan Harinivas Harshan, Archana Bharadwaj Siva, Karthik Bharadwaj Tallapaka, Shagufta Khan, Lamuk Zaveri, Nikhil Hajirnis, M Soujanya Reddy, Pratheusa Maccha, Namami Gaur, Sakshi Shambhavi, Tulasi Nagabandi, Purushotham Vodnala, Gokulan C G, Gunjan Purohit, Hanuman Tulashiram Kale, Pankaj Kumar, Prachand Issarapu, Rakesh K Mishra, Divya Tej Sowpati |
| EPI_ISL_495246 | CSIR-Centre for Cellular and Molecular Biology | CSIR-Centre for Cellular and Molecular Biology | Lamuk Zaveri, Shagufta Khan, Namami Gaur, Sakshi Shambhavi, Nikhil Hajirnis, M Soujanya Reddy, Pratheusa Maccha, Tulasi Nagabandi, Purushotham Vodnala, Payel Mukherjee, Sofia Banu, Priya Singh, Onkar Kulkarni, Dhiviya Vedagiri, Divya Gupta, Vishal Sah, Santosh Kumar Kuncha, Krishnan Harinivas Harshan, Archana Bharadwaj Siva, Karthik Bharadwaj Tallapaka, Renu Sudhakar, Somesh Gorde, Gangumala Srinivas Reddy, Sujoy Deb, Swati Bayyana, Rakesh K Mishra, Divya Tej Sowpati |
| EPI_ISL_495247 | CSIR-Centre for Cellular and Molecular Biology | CSIR-Centre for Cellular and Molecular Biology | Lamuk Zaveri, Shagufta Khan,Nikhil Hajirnis, M Soujanya Reddy, Pratheusa Maccha, Namami Gaur, Sakshi Shambhavi, Tulasi Nagabandi, Purushotham Vodnala, Payel Mukherjee, Sofia Banu, Priya Singh, Onkar Kulkarni, Dhiviya Vedagiri, Divya Gupta, Vishal Sah, Santosh Kumar Kuncha, Krishnan Harinivas Harshan, Archana Bharadwaj Siva, Karthik Bharadwaj Tallapaka,Zeba Rizvi, Zuberwasim Sayyad, Kakade Aishwarya Arun, Amrutha H C, Ananga Ghosh, Rakesh K Mishra, Divya Tej Sowpati |
| EPI_ISL_495248 | CSIR-Centre for Cellular and Molecular Biology | CSIR-Centre for Cellular and Molecular Biology | Sakshi Shambhavi, Lamuk Zaveri, Shagufta Khan, Nikhil Hajirnis, M Soujanya Reddy, Pratheusa Maccha, Namami Gaur, Tulasi Nagabandi, Purushotham Vodnala, Payel Mukherjee, Sofia Banu, Priya Singh, Onkar Kulkarni, Dhiviya Vedagiri, Divya Gupta, Vishal Sah, Santosh Kumar Kuncha, Krishnan Harinivas Harshan, Archana Bharadwaj Siva, Karthik Bharadwaj Tallapaka,G. Aditya Kumar, Koushick Sivakumar, Rakesh K Mishra, Divya Tej Sowpati |
| EPI_ISL_495249 | CSIR-Centre for Cellular and Molecular Biology | CSIR-Centre for Cellular and Molecular Biology | Payel Mukherjee, Sofia Banu, Priya Singh, Onkar Kulkarni, Dhiviya Vedagiri, Divya Gupta, Vishal Sah, Santosh Kumar Kuncha, Krishnan Harinivas Harshan, Archana Bharadwaj Siva, Karthik Bharadwaj Tallapaka, Shagufta Khan, Lamuk Zaveri, Nikhil Hajirnis, M Soujanya Reddy, Pratheusa Maccha, Namami Gaur, Sakshi Shambhavi, Tulasi Nagabandi, Purushotham Vodnala, Gokulan C G, Gunjan Purohit, Hanuman Tulashiram Kale, Pankaj Kumar, Prachand Issarapu, Rakesh K Mishra, Divya Tej Sowpati |
| EPI_ISL_495250 | CSIR-Centre for Cellular and Molecular Biology | CSIR-Centre for Cellular and Molecular Biology | Lamuk Zaveri, Shagufta Khan,Nikhil Hajirnis, M Soujanya Reddy, Pratheusa Maccha, Namami Gaur, Sakshi Shambhavi, Tulasi Nagabandi, Purushotham Vodnala, Payel Mukherjee, Sofia Banu, Priya Singh, Onkar Kulkarni, Dhiviya Vedagiri, Divya Gupta, Vishal Sah, Santosh Kumar Kuncha, Krishnan Harinivas Harshan, Archana Bharadwaj Siva, Karthik Bharadwaj Tallapaka,Umesh Kumar, Unis Ahmad Bhat, Ajay Sarawagi, Priyanka Pant, Rajkanwar Nathawat, Rakesh K Mishra, Divya Tej Sowpati |
| EPI_ISL_495251 | CSIR-Centre for Cellular and Molecular Biology | CSIR-Centre for Cellular and Molecular Biology | Pratheusa Maccha, Shagufta Khan, Lamuk Zaveri, Namami Gaur, Sakshi Shambhavi, Tulasi Nagabandi, Nikhil Hajirnis, M Soujanya Reddy, Purushotham Vodnala, Payel Mukherjee, Sofia Banu, Priya Singh, Onkar Kulkarni, Dhiviya Vedagiri, Divya Gupta, Vishal Sah, Santosh Kumar Kuncha, Krishnan Harinivas Harshan, Archana Bharadwaj Siva, Karthik Bharadwaj Tallapaka, Disha Nanda, Divya Das, Jotin Gogoi, Manish Bhattacharjee, Ravi Prasad Mukku, Rakesh K Mishra, Divya Tej Sowpati |
| EPI_ISL_495252 | CSIR-Centre for Cellular and Molecular Biology | CSIR-Centre for Cellular and Molecular Biology | Payel Mukherjee, Sofia Banu, Priya Singh, Onkar Kulkarni, Dhiviya Vedagiri, Divya Gupta, Vishal Sah, Santosh Kumar Kuncha, Krishnan Harinivas Harshan, Archana Bharadwaj Siva, Karthik Bharadwaj Tallapaka, Shagufta Khan, Lamuk Zaveri, Nikhil Hajirnis, M Soujanya Reddy, Pratheusa Maccha, Namami Gaur, Sakshi Shambhavi, Tulasi Nagabandi, Purushotham Vodnala, G. Aditya Kumar, Koushick Sivakumar, Pooja Ramesh Gupta, Rajan Kumar Jha, Shraddha Vijay Lahoti, Rakesh K Mishra, Divya Tej Sowpati |
| EPI_ISL_495253 | CSIR-Centre for Cellular and Molecular Biology | CSIR-Centre for Cellular and Molecular Biology | M Soujanya Reddy, Nikhil Hajirnis, Pratheusa Maccha, Namami Gaur, Sakshi Shambhavi, Lamuk Zaveri, Shagufta Khan, Tulasi Nagabandi, Purushotham Vodnala, Payel Mukherjee, Sofia Banu, Priya Singh, Onkar Kulkarni, Dhiviya Vedagiri, Divya Gupta, Vishal Sah, Santosh Kumar Kuncha, Krishnan Harinivas Harshan, Archana Bharadwaj Siva, Karthik Bharadwaj Tallapaka, Zeba Rizvi, Zuberwasim Sayyad, Kakade Aishwarya Arun, Amrutha H C, Ananga Ghosh, Rakesh K Mishra, Divya Tej Sowpati |
| EPI_ISL_495254 | CSIR-Centre for Cellular and Molecular Biology | CSIR-Centre for Cellular and Molecular Biology | Pratheusa Maccha, Sofia Banu, Payel Mukherjee, Priya Singh, Onkar Kulkarni, Dhiviya Vedagiri, Divya Gupta, Vishal Sah, Santosh Kumar Kuncha, Krishnan Harinivas Harshan, Archana Bharadwaj Siva, Karthik Bharadwaj Tallapaka, Shagufta Khan, Lamuk Zaveri, Namami Gaur, Sakshi Shambhavi, Nikhil Hajirnis, M Soujanya Reddy, Tulasi Nagabandi, Purushotham Vodnala,Preethi Jampala, Sharada Ravi Iyer, Sulagana Mukherjee, Swetha Sundar, Peddapuvala Sai Uday Kiran, Rakesh K Mishra, Divya Tej Sowpati |
| EPI_ISL_495255 | CSIR-Centre for Cellular and Molecular Biology | CSIR-Centre for Cellular and Molecular Biology | Tulasi Nagabandi, Namami Gaur, Sakshi Shambhavi, Lamuk Zaveri, Shagufta Khan, Nikhil Hajirnis, M Soujanya Reddy, Pratheusa Maccha, Purushotham Vodnala, Payel Mukherjee, Sofia Banu, Priya Singh, Onkar Kulkarni, Dhiviya Vedagiri, Divya Gupta, Vishal Sah, Santosh Kumar Kuncha, Krishnan Harinivas Harshan, Archana Bharadwaj Siva, Karthik Bharadwaj Tallapaka,G. Aditya Kumar, Koushick Sivakumar, Pooja Ramesh Gupta, Rajan Kumar Jha, Shraddha Vijay Lahoti, Rakesh K Mishra, Divya Tej Sowpati |
| EPI_ISL_495256 | CSIR-Centre for Cellular and Molecular Biology | CSIR-Centre for Cellular and Molecular Biology | Sakshi Shambhavi, Lamuk Zaveri, Shagufta Khan, Namami Gaur, Nikhil Hajirnis, M Soujanya Reddy, Pratheusa Maccha, Tulasi Nagabandi, Purushotham Vodnala, Payel Mukherjee, Sofia Banu, Priya Singh, Onkar Kulkarni, Dhiviya Vedagiri, Divya Gupta, Vishal Sah, Santosh Kumar Kuncha, Krishnan Harinivas Harshan, Archana Bharadwaj Siva, Karthik Bharadwaj Tallapaka, Deepak Kumar, Devi Prasad Vijayashankar, Disha Nanda, Divya Das, Jotin Gogoi, Manish Bhattacharjee, Rakesh K Mishra, Divya Tej Sowpati |
| EPI_ISL_495257 | CSIR-Centre for Cellular and Molecular Biology | CSIR-Centre for Cellular and Molecular Biology | M Soujanya Reddy, Nikhil Hajirnis, Pratheusa Maccha, Payel Mukherjee, Sofia Banu, Priya Singh, Onkar Kulkarni,Tulasi Nagabandi, Namami Gaur, Sakshi Shambhavi, Lamuk Zaveri, Shagufta Khan, Purushotham Vodnala, Dhiviya Vedagiri, Divya Gupta, Vishal Sah, Santosh Kumar Kuncha, Krishnan Harinivas Harshan, Archana Bharadwaj Siva, Karthik Bharadwaj Tallapaka,Kezia J Ann, Radhika Khandelwal, Roshan Maku Venkata, Shemin Mansuri, Sonu Uday, Rakesh K Mishra, Divya Tej Sowpati |
| EPI_ISL_495258 | CSIR-Centre for Cellular and Molecular Biology | CSIR-Centre for Cellular and Molecular Biology | Lamuk Zaveri, Shagufta Khan, Namami Gaur, Sakshi Shambhavi, Nikhil Hajirnis, M Soujanya Reddy, Pratheusa Maccha, Tulasi Nagabandi, Purushotham Vodnala, Payel Mukherjee, Sofia Banu, Priya Singh, Onkar Kulkarni, Dhiviya Vedagiri, Divya Gupta, Vishal Sah, Santosh Kumar Kuncha, Krishnan Harinivas Harshan, Archana Bharadwaj Siva, Karthik Bharadwaj Tallapaka, Renu Sudhakar, Somesh Gorde, Gangumala Srinivas Reddy, Sujoy Deb, Swati Bayyana, Rakesh K Mishra, Divya Tej Sowpati |
| EPI_ISL_495259 | CSIR-Centre for Cellular and Molecular Biology | CSIR-Centre for Cellular and Molecular Biology | Nikhil Hajirnis, M Soujanya Reddy, Pratheusa Maccha, Namami Gaur, Sakshi Shambhavi, Lamuk Zaveri, Shagufta Khan, Tulasi Nagabandi, Purushotham Vodnala, Payel Mukherjee, Sofia Banu, Priya Singh,Onkar Kulkarni, Dhiviya Vedagiri, Divya Gupta, Vishal Sah, Santosh Kumar Kuncha, Krishnan Harinivas Harshan, Archana Bharadwaj Siva, Karthik Bharadwaj Tallapaka,Kezia J Ann, Radhika Khandelwal, Roshan Maku Venkata, Shemin Mansuri, Sonu Uday, Rakesh K Mishra, Divya Tej Sowpati |
| EPI_ISL_495260 | CSIR-Centre for Cellular and Molecular Biology | CSIR-Centre for Cellular and Molecular Biology | Namami Gaur, Sakshi Shambhavi, Lamuk Zaveri, Shagufta Khan, Nikhil Hajirnis, M Soujanya Reddy, Pratheusa Maccha, Tulasi Nagabandi, Purushotham Vodnala, Payel Mukherjee, Sofia Banu, Priya Singh, Onkar Kulkarni, Dhiviya Vedagiri, Divya Gupta, Vishal Sah, Santosh Kumar Kuncha, Krishnan Harinivas Harshan, Archana Bharadwaj Siva, Karthik Bharadwaj Tallapaka,G. Aditya Kumar, Koushick Sivakumar, Rakesh K Mishra, Divya Tej Sowpati |
| EPI_ISL_495261 | CSIR-Centre for Cellular and Molecular Biology | CSIR-Centre for Cellular and Molecular Biology | Nikhil Hajirnis, M Soujanya Reddy, Pratheusa Maccha, Lamuk Zaveri, Shagufta Khan, Namami Gaur, Sakshi Shambhavi, Tulasi Nagabandi, Purushotham Vodnala, Payel Mukherjee, Sofia Banu, Priya Singh, Onkar Kulkarni, Dhiviya Vedagiri, Divya Gupta, Vishal Sah, Santosh Kumar Kuncha, Krishnan Harinivas Harshan, Archana Bharadwaj Siva, Karthik Bharadwaj Tallapaka,Zeba Rizvi, Zuberwasim Sayyad, Kakade Aishwarya Arun, Amrutha H C, Ananga Ghosh, Rakesh K Mishra, Divya Tej Sowpati |
| EPI_ISL_495262 | CSIR-Centre for Cellular and Molecular Biology | CSIR-Centre for Cellular and Molecular Biology | Sofia Banu, Payel Mukherjee, Priya Singh,Onkar Kulkarni, Dhiviya Vedagiri, Divya Gupta, Vishal Sah, Santosh Kumar Kuncha, Krishnan Harinivas Harshan, Archana Bharadwaj Siva, Karthik Bharadwaj Tallapaka, Shagufta Khan, Lamuk Zaveri, Namami Gaur, Sakshi Shambhavi, Tulasi Nagabandi, Nikhil Hajirnis, M Soujanya Reddy, Pratheusa Maccha, Purushotham Vodnala, Gokulan C G, Gunjan Purohit, Hanuman Tulashiram Kale, Pankaj Kumar, Prachand Issarapu, Rakesh K Mishra, Divya Tej Sowpati |
| EPI_ISL_495263 | CSIR-Centre for Cellular and Molecular Biology | CSIR-Centre for Cellular and Molecular Biology | Tulasi Nagabandi, Namami Gaur, Sakshi Shambhavi, Lamuk Zaveri, Shagufta Khan, Nikhil Hajirnis, M Soujanya Reddy, Pratheusa Maccha, Purushotham Vodnala, Payel Mukherjee, Sofia Banu, Priya Singh, Onkar Kulkarni, Dhiviya Vedagiri, Divya Gupta, Vishal Sah, Santosh Kumar Kuncha, Krishnan Harinivas Harshan, Archana Bharadwaj Siva, Karthik Bharadwaj Tallapaka,G. Aditya Kumar, Koushick Sivakumar, Pooja Ramesh Gupta, Rajan Kumar Jha, Shraddha Vijay Lahoti, Rakesh K Mishra, Divya Tej Sowpati |
| EPI_ISL_495264 | CSIR-Centre for Cellular and Molecular Biology | CSIR-Centre for Cellular and Molecular Biology | Sakshi Shambhavi, Lamuk Zaveri, Shagufta Khan, Namami Gaur, Nikhil Hajirnis, M Soujanya Reddy, Pratheusa Maccha, Tulasi Nagabandi, Purushotham Vodnala, Payel Mukherjee, Sofia Banu, Priya Singh, Onkar Kulkarni, Dhiviya Vedagiri, Divya Gupta, Vishal Sah, Santosh Kumar Kuncha, Krishnan Harinivas Harshan, Archana Bharadwaj Siva, Karthik Bharadwaj Tallapaka, Deepak Kumar, Devi Prasad Vijayashankar, Disha Nanda, Divya Das, Jotin Gogoi, Manish Bhattacharjee, Rakesh K Mishra, Divya Tej Sowpati |
| EPI_ISL_495265 | CSIR-Centre for Cellular and Molecular Biology | CSIR-Centre for Cellular and Molecular Biology | Lamuk Zaveri, Shagufta Khan, Namami Gaur, Sakshi Shambhavi, Nikhil Hajirnis, M Soujanya Reddy, Pratheusa Maccha, Tulasi Nagabandi, Purushotham Vodnala, Payel Mukherjee, Sofia Banu, Priya Singh, Onkar Kulkarni, Dhiviya Vedagiri, Divya Gupta, Vishal Sah, Santosh Kumar Kuncha, Krishnan Harinivas Harshan, Archana Bharadwaj Siva, Karthik Bharadwaj Tallapaka, Renu Sudhakar, Somesh Gorde, Gangumala Srinivas Reddy, Sujoy Deb, Swati Bayyana, Rakesh K Mishra, Divya Tej Sowpati |
| EPI_ISL_495266 | CSIR-Centre for Cellular and Molecular Biology | CSIR-Centre for Cellular and Molecular Biology | Namami Gaur, Sakshi Shambhavi, Lamuk Zaveri, Shagufta Khan, Nikhil Hajirnis, M Soujanya Reddy, Pratheusa Maccha, Tulasi Nagabandi, Purushotham Vodnala, Payel Mukherjee, Sofia Banu, Priya Singh,Onkar Kulkarni, Dhiviya Vedagiri, Divya Gupta, Vishal Sah, Santosh Kumar Kuncha, Krishnan Harinivas Harshan, Archana Bharadwaj Siva, Karthik Bharadwaj Tallapaka, Zeba Rizvi, Zuberwasim Sayyad, Kakade Aishwarya Arun, Amrutha H C, Ananga Ghosh, Rakesh K Mishra, Divya Tej Sowpati |
| EPI_ISL_495267 | CSIR-Centre for Cellular and Molecular Biology | CSIR-Centre for Cellular and Molecular Biology | Tulasi Nagabandi, Namami Gaur, Sakshi Shambhavi, Lamuk Zaveri, Shagufta Khan, Nikhil Hajirnis, M Soujanya Reddy, Pratheusa Maccha, Purushotham Vodnala, Payel Mukherjee, Sofia Banu, Priya Singh,Onkar Kulkarni, Dhiviya Vedagiri, Divya Gupta, Vishal Sah, Santosh Kumar Kuncha, Krishnan Harinivas Harshan, Archana Bharadwaj Siva, Karthik Bharadwaj Tallapaka,Kezia J Ann, Radhika Khandelwal, Roshan Maku Venkata, Shemin Mansuri, Sonu Uday, Rakesh K Mishra, Divya Tej Sowpati |
| EPI_ISL_495268 | CSIR-Centre for Cellular and Molecular Biology | CSIR-Centre for Cellular and Molecular Biology | Tulasi Nagabandi, Namami Gaur, Sakshi Shambhavi, Lamuk Zaveri, Shagufta Khan, Nikhil Hajirnis, M Soujanya Reddy, Pratheusa Maccha, Purushotham Vodnala, Payel Mukherjee, Sofia Banu, Priya Singh,Onkar Kulkarni , Dhiviya Vedagiri, Divya Gupta, Vishal Sah, Santosh Kumar Kuncha, Krishnan Harinivas Harshan, Archana Bharadwaj Siva, Karthik Bharadwaj Tallapaka,G. Aditya Kumar, Koushick Sivakumar, Pooja Ramesh Gupta, Rajan Kumar Jha, Shraddha Vijay Lahoti, Rakesh K Mishra, Divya Tej Sowpati |
| EPI_ISL_495269 | CSIR-Centre for Cellular and Molecular Biology | CSIR-Centre for Cellular and Molecular Biology | Sakshi Shambhavi, Lamuk Zaveri, Shagufta Khan, Namami Gaur, Nikhil Hajirnis, M Soujanya Reddy, Pratheusa Maccha,Tulasi Nagabandi, Purushotham Vodnala, Payel Mukherjee, Sofia Banu, Priya Singh,Onkar Kulkarni, Dhiviya Vedagiri, Divya Gupta, Vishal Sah, Santosh Kumar Kuncha, Krishnan Harinivas Harshan, Archana Bharadwaj Siva, Karthik Bharadwaj Tallapaka, G. Aditya Kumar, Koushick Sivakumar, Pooja Ramesh Gupta, Rajan Kumar Jha, Shraddha Vijay Lahoti, Rakesh K Mishra, Divya Tej Sowpati |
| EPI_ISL_495270 | CSIR-Centre for Cellular and Molecular Biology | CSIR-Centre for Cellular and Molecular Biology | Namami Gaur, Sakshi Shambhavi, Lamuk Zaveri, Shagufta Khan, Nikhil Hajirnis, M Soujanya Reddy, Pratheusa Maccha, Tulasi Nagabandi, Purushotham Vodnala, Payel Mukherjee, Sofia Banu, Priya Singh, Onkar Kulkarni, Dhiviya Vedagiri, Divya Gupta, Vishal Sah, Santosh Kumar Kuncha, Krishnan Harinivas Harshan, Archana Bharadwaj Siva, Karthik Bharadwaj Tallapaka, Zeba Rizvi, Zuberwasim Sayyad, Kakade Aishwarya Arun, Amrutha H C, Ananga Ghosh, Rakesh K Mishra, Divya Tej Sowpati |
| EPI_ISL_495271 | CSIR-Centre for Cellular and Molecular Biology | CSIR-Centre for Cellular and Molecular Biology | M Soujanya Reddy, Nikhil Hajirnis, Pratheusa Maccha, Sakshi Shambhavi, Lamuk Zaveri, Shagufta Khan, Namami Gaur, Tulasi Nagabandi, Purushotham Vodnala, Payel Mukherjee, Sofia Banu, Priya Singh,Onkar Kulkarni, Dhiviya Vedagiri, Divya Gupta, Vishal Sah, Santosh Kumar Kuncha, Krishnan Harinivas Harshan, Archana Bharadwaj Siva, Karthik Bharadwaj Tallapaka, G. Aditya Kumar, Koushick Sivakumar, Pooja Ramesh Gupta, Rajan Kumar Jha, Shraddha Vijay Lahoti, Rakesh K Mishra, Divya Tej Sowpati |
| EPI_ISL_495272 | CSIR-Centre for Cellular and Molecular Biology | CSIR-Centre for Cellular and Molecular Biology | M Soujanya Reddy, Nikhil Hajirnis, Pratheusa Maccha, Namami Gaur, Sakshi Shambhavi, Lamuk Zaveri, Shagufta Khan, Tulasi Nagabandi, Purushotham Vodnala, Payel Mukherjee, Sofia Banu, Priya Singh, Onkar Kulkarni, Dhiviya Vedagiri, Divya Gupta, Vishal Sah, Santosh Kumar Kuncha, Krishnan Harinivas Harshan, Archana Bharadwaj Siva, Karthik Bharadwaj Tallapaka, Zeba Rizvi, Zuberwasim Sayyad, Kakade Aishwarya Arun, Amrutha H C, Ananga Ghosh, Rakesh K Mishra, Divya Tej Sowpati |
| EPI_ISL_495273 | CSIR-Centre for Cellular and Molecular Biology | CSIR-Centre for Cellular and Molecular Biology | Pratheusa Maccha, Sofia Banu, Payel Mukherjee, Priya Singh, Onkar Kulkarni, Dhiviya Vedagiri, Divya Gupta, Vishal Sah, Santosh Kumar Kuncha, Krishnan Harinivas Harshan, Archana Bharadwaj Siva, Karthik Bharadwaj Tallapaka, Shagufta Khan, Lamuk Zaveri, Namami Gaur, Sakshi Shambhavi, Nikhil Hajirnis, M Soujanya Reddy, Tulasi Nagabandi, Purushotham Vodnala,Preethi Jampala, Sharada Ravi Iyer, Sulagana Mukherjee, Swetha Sundar, Peddapuvala Sai Uday Kiran, Rakesh K Mishra, Divya Tej Sowpati |
| EPI_ISL_495274 | Osmania Medical College | CSIR-Centre for Cellular and Molecular Biology | Shashikala Reddy, Mahboob Khan, Lamuk Zaveri, Sofia Banu,Payel Mukherjee, Shagufta Khan,Priya Singh, Onkar Kulkarni, Dhiviya Vedagiri, Divya Gupta, Vishal Sah, Santosh Kumar Kuncha, |

| | | | |
|---|---|---|---|
| | | | Krishnan Harinivas Harshan, Archana Bharadwaj Siva, Karthik Bharadwaj Tallapaka,Nikhil Hajirnis, M Soujanya Reddy, Pratheusa Maccha, Namami Gaur, Sakshi Shambhavi, Tulasi Nagabandi, Purushotham Vodnala, Rakesh K Mishra, Divya Tej Sowpati |
| EPI_ISL_495275 | Osmania Medical College | CSIR-Centre for Cellular and Molecular Biology | Shashikala Reddy, Mahboob Khan, Onkar Kulkarni, Payel Mukherjee, Sofia Banu, Priya Singh, Dhiviya Vedagiri, Divya Gupta, Vishal Sah, Santosh Kumar Kuncha, Krishnan Harinivas Harshan, Archana Bharadwaj Siva, Karthik Bharadwaj Tallapaka, Shagufta Khan, Lamuk Zaveri, Nikhil Hajirnis, M Soujanya Reddy, Pratheusa Maccha,Namami Gaur, Sakshi Shambhavi, Tulasi Nagabandi, Purushotham Vodnala, Rakesh K Mishra, Divya Tej Sowpati |
| EPI_ISL_495276 | Osmania Medical College | CSIR-Centre for Cellular and Molecular Biology | Shashikala Reddy, Mahboob Khan,Payel Mukherjee, Sofia Banu, Priya Singh, Onkar Kulkarni, Dhiviya Vedagiri, Divya Gupta, Vishal Sah, Santosh Kumar Kuncha, Krishnan Harinivas Harshan, Archana Bharadwaj Siva, Karthik Bharadwaj Tallapaka, Shagufta Khan, Lamuk Zaveri, Namami Gaur, Sakshi Shambhavi, Nikhil Hajirnis, M Soujanya Reddy, Pratheusa Maccha,Tulasi Nagabandi, Purushotham Vodnala, Rakesh K Mishra, Divya Tej Sowpati |
| EPI_ISL_495277 | Osmania Medical College | CSIR-Centre for Cellular and Molecular Biology | Shashikala Reddy, Mahboob Khan, Namami Gaur, Lamuk Zaveri,Shagufta Khan, Nikhil Hajirnis, M Soujanya Reddy, Pratheusa Maccha, Sofia Banu, Onkar Kulkarni, Payel Mukherjee, Priya Singh, Dhiviya Vedagiri, Divya Gupta, Vishal Sah, Santosh Kumar Kuncha, Krishnan Harinivas Harshan, Archana Bharadwaj Siva, Karthik Bharadwaj Tallapaka,Sakshi Shambhavi, Tulasi Nagabandi, Purushotham Vodnala, Rakesh K Mishra, Divya Tej Sowpati |
| EPI_ISL_495278 | Osmania Medical College | CSIR-Centre for Cellular and Molecular Biology | Shashikala Reddy, Mahboob Khan,Tulasi Nagabandi, Namami Gaur, Lamuk Zaveri,Shagufta Khan,Nikhil Hajirnis, M Soujanya Reddy, Pratheusa Maccha, Sofia Banu,Payel Mukherjee, Priya Singh, Onkar Kulkarni, Dhiviya Vedagiri, Divya Gupta, Vishal Sah, Santosh Kumar Kuncha, Krishnan Harinivas Harshan, Archana Bharadwaj Siva, Karthik Bharadwaj Tallapaka,Sakshi Shambhavi, Purushotham Vodnala, Rakesh K Mishra, Divya Tej Sowpati |
| EPI_ISL_495279 | Osmania Medical College | CSIR-Centre for Cellular and Molecular Biology | Shashikala Reddy, Mahboob Khan,Sakshi Shambhavi,Tulasi Nagabandi, Namami Gaur, Lamuk Zaveri,Shagufta Khan, Nikhil Hajirnis, M Soujanya Reddy, Pratheusa Maccha,Sofia Banu,Payel Mukherjee, Priya Singh,Onkar Kulkarni, Dhiviya Vedagiri, Divya Gupta, Vishal Sah, Santosh Kumar Kuncha, Krishnan Harinivas Harshan, Archana Bharadwaj Siva, Karthik Bharadwaj Tallapaka,Sakshi Shambhavi, Purushotham Vodnala, Rakesh K Mishra, Divya Tej Sowpati |
| EPI_ISL_495280 | Osmania Medical College | CSIR-Centre for Cellular and Molecular Biology | Shashikala Reddy, Mahboob Khan,Payel Mukherjee, Sofia Banu, Priya Singh, Onkar Kulkarni, Dhiviya Vedagiri, Divya Gupta, Vishal Sah, Santosh Kumar Kuncha, Krishnan Harinivas Harshan, Archana Bharadwaj Siva, Karthik Bharadwaj Tallapaka, Shagufta Khan, Lamuk Zaveri, Nikhil Hajirnis, M Soujanya Reddy, Pratheusa Maccha,Namami Gaur, Sakshi Shambhavi, Tulasi Nagabandi, Purushotham Vodnala, Rakesh K Mishra, Divya Tej Sowpati |
| EPI_ISL_495281 | Osmania Medical College | CSIR-Centre for Cellular and Molecular Biology | Shashikala Reddy, Mahboob Khan, Sofia Banu,Payel Mukherjee, Priya Singh,Onkar Kulkarni, Dhiviya Vedagiri, Divya Gupta, Vishal Sah, Santosh Kumar Kuncha, Krishnan Harinivas Harshan, Archana Bharadwaj Siva, Karthik Bharadwaj Tallapaka, Shagufta Khan, Lamuk Zaveri, Namami Gaur, Nikhil Hajirnis, M Soujanya Reddy, Pratheusa Maccha,Sakshi Shambhavi, Tulasi Nagabandi, Purushotham Vodnala, Rakesh K Mishra, Divya Tej Sowpati |
| EPI_ISL_495282 | Osmania Medical College | CSIR-Centre for Cellular and Molecular Biology | Shashikala Reddy, Mahboob Khan,Payel Mukherjee, Sofia Banu, Priya Singh, Onkar Kulkarni, Dhiviya Vedagiri, Divya Gupta, Vishal Sah, Santosh Kumar Kuncha, Krishnan Harinivas Harshan, Archana Bharadwaj Siva, Karthik Bharadwaj Tallapaka, Shagufta Khan, Lamuk Zaveri, Nikhil Hajirnis, M Soujanya Reddy, Pratheusa Maccha,Namami Gaur, Sakshi Shambhavi, Tulasi Nagabandi, Purushotham Vodnala, Rakesh K Mishra, Divya Tej Sowpati |
| EPI_ISL_495283 | Osmania Medical College | CSIR-Centre for Cellular and Molecular Biology | Shashikala Reddy, Mahboob Khan, Namami Gaur, Lamuk Zaveri,Shagufta Khan, Sofia Banu, Onkar Kulkarni, Payel Mukherjee, Priya Singh, Dhiviya Vedagiri, Divya Gupta, Vishal Sah, Santosh Kumar Kuncha, Krishnan Harinivas Harshan, Archana Bharadwaj Siva, Karthik Bharadwaj Tallapaka,Sakshi Shambhavi, Nikhil Hajirnis, M Soujanya Reddy, Pratheusa Maccha, Tulasi Nagabandi, Purushotham Vodnala, Rakesh K Mishra, Divya Tej Sowpati |
| EPI_ISL_495284 | Osmania Medical College | CSIR-Centre for Cellular and Molecular Biology | Shashikala Reddy, Mahboob Khan,Tulasi Nagabandi, Namami Gaur, Lamuk Zaveri,Shagufta Khan, Sofia Banu,Payel Mukherjee, Priya Singh, Onkar Kulkarni, Dhiviya Vedagiri, Divya Gupta, Vishal Sah, Santosh Kumar Kuncha, Krishnan Harinivas Harshan, Archana Bharadwaj Siva, Karthik Bharadwaj Tallapaka,Sakshi Shambhavi,Nikhil Hajirnis, M Soujanya Reddy, Pratheusa Maccha, Purushotham Vodnala, Rakesh K Mishra, Divya Tej Sowpati |
| EPI_ISL_495285 | Osmania Medical College | CSIR-Centre for Cellular and Molecular Biology | Shashikala Reddy, Mahboob Khan,Shagufta Khan, Sofia Banu,Payel Mukherjee, Priya Singh,Onkar Kulkarni, Dhiviya Vedagiri, Divya Gupta, Vishal Sah, Santosh Kumar Kuncha, Krishnan Harinivas Harshan, Archana Bharadwaj Siva, Karthik Bharadwaj Tallapaka, Lamuk Zaveri, Nikhil Hajirnis, M Soujanya Reddy, Pratheusa Maccha, Namami Gaur, Sakshi Shambhavi, Tulasi Nagabandi, Purushotham Vodnala, Rakesh K Mishra, Divya Tej Sowpati |
| EPI_ISL_495286 | Osmania Medical College | CSIR-Centre for Cellular and Molecular Biology | Shashikala Reddy, Mahboob Khan, Sofia Banu,Payel Mukherjee, Priya Singh,Onkar Kulkarni, Dhiviya Vedagiri, Divya Gupta, Vishal Sah, Santosh Kumar Kuncha, Krishnan Harinivas Harshan, Archana Bharadwaj Siva, Karthik Bharadwaj Tallapaka, Shagufta Khan, Lamuk Zaveri, Namami Gaur, Sakshi Shambhavi, Nikhil Hajirnis, M Soujanya Reddy, Pratheusa Maccha,Tulasi Nagabandi, Purushotham Vodnala, Rakesh K Mishra, Divya Tej Sowpati |
| EPI_ISL_495287 | Osmania Medical College | CSIR-Centre for Cellular and Molecular Biology | Shashikala Reddy, Mahboob Khan,Onkar Kulkarni, Sofia Banu,Payel Mukherjee, Priya Singh, Dhiviya Vedagiri, Divya Gupta, Vishal Sah, Santosh Kumar Kuncha, Krishnan Harinivas Harshan, Archana Bharadwaj Siva, Karthik Bharadwaj Tallapaka, Shagufta Khan, Nikhil Hajirnis, M Soujanya Reddy, Pratheusa Maccha, Lamuk Zaveri, Namami Gaur, Sakshi Shambhavi, Tulasi Nagabandi, Purushotham Vodnala, Rakesh K Mishra, Divya Tej Sowpati |
| EPI_ISL_495288 | Osmania Medical College | CSIR-Centre for Cellular and Molecular Biology | Shashikala Reddy, Mahboob Khan, Lamuk Zaveri, Sofia Banu,Payel Mukherjee, Shagufta Khan,Priya Singh, Onkar Kulkarni, Dhiviya Vedagiri, Divya Gupta, Vishal Sah, Santosh Kumar Kuncha, Krishnan Harinivas Harshan, Archana Bharadwaj Siva, Karthik Bharadwaj Tallapaka, Namami Gaur, Nikhil Hajirnis, M Soujanya Reddy, Pratheusa Maccha,Sakshi Shambhavi, Tulasi Nagabandi, Purushotham Vodnala, Rakesh K Mishra, Divya Tej Sowpati |
| EPI_ISL_495289 | Osmania Medical College | CSIR-Centre for Cellular and Molecular Biology | Shashikala Reddy, Mahboob Khan, Namami Gaur, Lamuk Zaveri,Shagufta Khan, Sofia Banu, Onkar Kulkarni, Payel Mukherjee, Priya Singh, Dhiviya Vedagiri, Divya Gupta, Vishal Sah, Santosh Kumar Kuncha, Krishnan Harinivas Harshan, Archana Bharadwaj Siva, Karthik Bharadwaj Tallapaka,Sakshi Shambhavi, Nikhil Hajirnis, M Soujanya Reddy, Pratheusa Maccha, Tulasi Nagabandi, Purushotham Vodnala, Rakesh K Mishra, Divya Tej Sowpati |
| EPI_ISL_495290 | Osmania Medical College | CSIR-Centre for Cellular and Molecular Biology | Shashikala Reddy, Mahboob Khan,Tulasi Nagabandi, Namami Gaur, Lamuk Zaveri,Shagufta Khan, Sofia Banu,Payel Mukherjee, Priya Singh, Onkar Kulkarni, Dhiviya Vedagiri, Divya Gupta, Vishal Sah, Santosh Kumar Kuncha, Krishnan Harinivas Harshan, Archana Bharadwaj Siva, Karthik Bharadwaj Tallapaka,Sakshi Shambhavi, Nikhil Hajirnis, M Soujanya Reddy, Pratheusa Maccha, Purushotham Vodnala, Rakesh K Mishra, Divya Tej Sowpati |
| EPI_ISL_495291 | Osmania Medical College | CSIR-Centre for Cellular and Molecular Biology | Shashikala Reddy, Mahboob Khan,Onkar Kulkarni,Shagufta Khan, Sofia Banu,Payel Mukherjee, Priya Singh, Dhiviya Vedagiri, Divya Gupta, Vishal Sah, Santosh Kumar Kuncha, Krishnan Harinivas Harshan, Archana Bharadwaj Siva, Karthik Bharadwaj Tallapaka, Lamuk Zaveri, Nikhil Hajirnis, M Soujanya Reddy, Pratheusa Maccha, Namami Gaur, Sakshi Shambhavi, Tulasi Nagabandi, Purushotham Vodnala, Rakesh K Mishra, Divya Tej Sowpati |
| EPI_ISL_495292 | Osmania Medical College | CSIR-Centre for Cellular and Molecular Biology | Shashikala Reddy, Mahboob Khan,Shagufta Khan, Sofia Banu,Payel Mukherjee, Priya Singh,Onkar Kulkarni, Dhiviya Vedagiri, Divya Gupta, Vishal Sah, Santosh Kumar Kuncha, Krishnan Harinivas Harshan, Archana Bharadwaj Siva, Karthik Bharadwaj Tallapaka, Lamuk Zaveri, Namami Gaur, Sakshi Shambhavi,Nikhil Hajirnis, M Soujanya Reddy, Pratheusa Maccha, Tulasi Nagabandi, Purushotham Vodnala, Rakesh K Mishra, Divya Tej Sowpati |
| EPI_ISL_495293 | Osmania Medical College | CSIR-Centre for Cellular and Molecular Biology | Shashikala Reddy, Mahboob Khan, Lamuk Zaveri, Sofia Banu,Payel Mukherjee, Shagufta Khan,Priya Singh, Onkar Kulkarni, Dhiviya Vedagiri, Divya Gupta, Vishal Sah, Santosh Kumar Kuncha, Krishnan Harinivas Harshan, Archana Bharadwaj Siva, Karthik Bharadwaj Tallapaka, Namami Gaur, Sakshi Shambhavi, Nikhil Hajirnis, M Soujanya Reddy, Pratheusa Maccha, Tulasi Nagabandi, Purushotham Vodnala, Rakesh K Mishra, Divya Tej Sowpati |
| EPI_ISL_495294 | Osmania Medical College | CSIR-Centre for Cellular and Molecular Biology | Shashikala Reddy, Mahboob Khan,Onkar Kulkarni, Lamuk Zaveri, Sofia Banu,Payel Mukherjee, Shagufta Khan,Priya Singh, Dhiviya Vedagiri, Divya Gupta, Vishal Sah, Santosh Kumar Kuncha, Krishnan Harinivas Harshan, Archana Bharadwaj Siva, Karthik Bharadwaj Tallapaka, Namami Gaur, Sakshi Shambhavi, Nikhil Hajirnis, M Soujanya Reddy, Pratheusa Maccha, Tulasi Nagabandi, Purushotham Vodnala, Rakesh K Mishra, Divya Tej Sowpati |
| EPI_ISL_495295 | Osmania Medical College | CSIR-Centre for Cellular and Molecular Biology | Shashikala Reddy, Mahboob Khan, Sofia Banu,Payel Mukherjee, Priya Singh,Onkar Kulkarni, Dhiviya Vedagiri, Divya Gupta, Vishal Sah, Santosh Kumar Kuncha, Krishnan Harinivas Harshan, Archana Bharadwaj Siva, Karthik Bharadwaj Tallapaka, Shagufta Khan, Lamuk Zaveri,Nikhil Hajirnis, M Soujanya Reddy, Pratheusa Maccha, Namami Gaur, Sakshi Shambhavi, Tulasi Nagabandi, Purushotham Vodnala, Rakesh K Mishra, Divya Tej Sowpati |
| EPI_ISL_495296 | Osmania Medical College | CSIR-Centre for Cellular and Molecular Biology | Shashikala Reddy, Mahboob Khan,Shagufta Khan, Sofia Banu,Payel Mukherjee, Priya Singh,Onkar Kulkarni, Dhiviya Vedagiri, Divya Gupta, Vishal Sah, Santosh Kumar Kuncha, Krishnan Harinivas Harshan, Archana Bharadwaj Siva, Karthik Bharadwaj Tallapaka, Lamuk Zaveri, Nikhil Hajirnis, M Soujanya Reddy, Pratheusa Maccha, Namami Gaur, Sakshi Shambhavi, Tulasi Nagabandi, Purushotham Vodnala, Rakesh K Mishra, Divya Tej Sowpati |
| EPI_ISL_495297 | Osmania Medical College | CSIR-Centre for Cellular and Molecular Biology | Shashikala Reddy, Mahboob Khan,Sakshi Shambhavi,Tulasi Nagabandi, Namami Gaur, Lamuk Zaveri,Shagufta Khan, Sofia Banu,Payel Mukherjee, Priya Singh,Onkar Kulkarni, Dhiviya Vedagiri, Divya Gupta, Vishal Sah, Santosh Kumar Kuncha, Krishnan Harinivas Harshan, Archana Bharadwaj Siva, Karthik Bharadwaj Tallapaka,Sakshi Shambhavi, Nikhil Hajirnis, M Soujanya Reddy, Pratheusa Maccha,Purushotham Vodnala, Rakesh K Mishra, Divya Tej Sowpati |
| EPI_ISL_495298, EPI_ISL_495299, EPI_ISL_495300, EPI_ISL_495301, EPI_ISL_495302, EPI_ISL_495303, EPI_ISL_495304, EPI_ISL_495305, EPI_ISL_495306, EPI_ISL_495307, EPI_ISL_495308, EPI_ISL_495309, EPI_ISL_495310, EPI_ISL_495311, EPI_ISL_495312, EPI_ISL_495313, EPI_ISL_495314, EPI_ISL_495315, EPI_ISL_495316, EPI_ISL_495317, EPI_ISL_495318, EPI_ISL_495319, EPI_ISL_495320, EPI_ISL_495321, EPI_ISL_495322, EPI_ISL_495323, EPI_ISL_495324, EPI_ISL_495325, EPI_ISL_495326, EPI_ISL_495327, EPI_ISL_495328, EPI_ISL_495329 | | | |
| see above | Virginia DCLS | Virginia DCLS | Virginia DCLS |
| EPI_ISL_495330, EPI_ISL_495331, EPI_ISL_495332, EPI_ISL_495333, EPI_ISL_495334, EPI_ISL_495335, EPI_ISL_495336, EPI_ISL_495337, EPI_ISL_495338, EPI_ISL_495339, EPI_ISL_495340, EPI_ISL_495341, EPI_ISL_495342, EPI_ISL_495343, EPI_ISL_495344, EPI_ISL_495345, EPI_ISL_495346, EPI_ISL_495347, EPI_ISL_495348, EPI_ISL_495349, EPI_ISL_495350, EPI_ISL_495351, EPI_ISL_495352, EPI_ISL_495353, EPI_ISL_495354, EPI_ISL_495355, EPI_ISL_495356, EPI_ISL_495357, EPI_ISL_495358, EPI_ISL_495359, EPI_ISL_495360, EPI_ISL_495361, EPI_ISL_495362, EPI_ISL_495363, EPI_ISL_495364, EPI_ISL_495365, EPI_ISL_495366, EPI_ISL_495367, EPI_ISL_495368, EPI_ISL_495369, EPI_ISL_495370, EPI_ISL_495371, EPI_ISL_495372, EPI_ISL_495373, EPI_ISL_495374, EPI_ISL_495375, EPI_ISL_495376, EPI_ISL_495377, EPI_ISL_495378, EPI_ISL_495379, EPI_ISL_495380, EPI_ISL_495381, EPI_ISL_495382, EPI_ISL_495383, EPI_ISL_495384, EPI_ISL_495385, EPI_ISL_495386, EPI_ISL_495387, EPI_ISL_495388, EPI_ISL_495389, EPI_ISL_495390, EPI_ISL_495391, EPI_ISL_495392, EPI_ISL_495393, EPI_ISL_495394, EPI_ISL_495395, EPI_ISL_495396, EPI_ISL_495397, EPI_ISL_495398, EPI_ISL_495399, EPI_ISL_495400, EPI_ISL_495401, EPI_ISL_495402, EPI_ISL_495403, EPI_ISL_495404, EPI_ISL_495405, EPI_ISL_495406, EPI_ISL_495407, EPI_ISL_495408, EPI_ISL_495409, EPI_ISL_495410 | | | |
| see above | Florida Bureau of Public Health Laboratories | Florida Bureau of Public Health Laboratories | Sarah Schmedes, Jason Blanton |
| EPI_ISL_495411, EPI_ISL_495412, EPI_ISL_495413, EPI_ISL_495414, EPI_ISL_495415, EPI_ISL_495416, EPI_ISL_495417, EPI_ISL_495418, EPI_ISL_495419, EPI_ISL_495420, EPI_ISL_495421, EPI_ISL_495422, EPI_ISL_495423, EPI_ISL_495424, EPI_ISL_495425, EPI_ISL_495426, EPI_ISL_495427, EPI_ISL_495428, EPI_ISL_495429, EPI_ISL_495430, EPI_ISL_495431, EPI_ISL_495432, EPI_ISL_495433, EPI_ISL_495434, EPI_ISL_495435, EPI_ISL_495436, EPI_ISL_495437, EPI_ISL_495438, EPI_ISL_495439, EPI_ISL_495440, EPI_ISL_495441, EPI_ISL_495442, EPI_ISL_495443, EPI_ISL_495444, EPI_ISL_495445, EPI_ISL_495446, EPI_ISL_495447, EPI_ISL_495448, EPI_ISL_495449, EPI_ISL_495450, EPI_ISL_495451, EPI_ISL_495452, EPI_ISL_495453, EPI_ISL_495454, EPI_ISL_495455, EPI_ISL_495456, EPI_ISL_495457 | | | |
| see above | Kafkas University, Faculty of Medicine, Department of Medical Microbiology | Kafkas University, Faculty of Medicine, Department of Medical Microbiology | Murat Karamese, Didem Ozgur, E. Ediz Tutuncu |
| EPI_ISL_495459 | Centers for Disease Control and Prevention of Lishui | Department of InspectionCenters for Disease Control and Prevention of Lishui | Wang Xiaoguang,Ji Qiaoying,Ji Jiansong,Ye Bifeng,Ye Ling |
| EPI_ISL_495460, EPI_ISL_495461, EPI_ISL_495462, EPI_ISL_495463, EPI_ISL_495464, EPI_ISL_495465, EPI_ISL_495466, EPI_ISL_495467, EPI_ISL_495468, EPI_ISL_495469, EPI_ISL_495470, EPI_ISL_495471, EPI_ISL_495472, EPI_ISL_495473, EPI_ISL_495474, EPI_ISL_495475, EPI_ISL_495476, EPI_ISL_495477, EPI_ISL_495478, EPI_ISL_495479, EPI_ISL_495480, EPI_ISL_495481, EPI_ISL_495482, EPI_ISL_495483, EPI_ISL_495484, EPI_ISL_495485, EPI_ISL_495486, EPI_ISL_495487, EPI_ISL_495488, EPI_ISL_495489, EPI_ISL_495490, EPI_ISL_495491, EPI_ISL_495492, EPI_ISL_495493, EPI_ISL_495494, EPI_ISL_495495, EPI_ISL_495496, EPI_ISL_495497, EPI_ISL_495498, EPI_ISL_495499, EPI_ISL_495500, EPI_ISL_495501, EPI_ISL_495502, EPI_ISL_495503, EPI_ISL_495504, EPI_ISL_495505, EPI_ISL_495506, EPI_ISL_495507, EPI_ISL_495508, EPI_ISL_495509 | | | |
| see above | University of Wisconsin-Madison AIDS Vaccine Research Laboratories | University of Wisconsin-Madison AIDS Vaccine Research Laboratories | Gage Moreno, Katarina Braun, et al. AIDS Vaccine Research Laboratories |
| EPI_ISL_495516, EPI_ISL_495517, EPI_ISL_495518, EPI_ISL_495519, EPI_ISL_495520, EPI_ISL_495521, EPI_ISL_495522, EPI_ISL_495523, EPI_ISL_495524, EPI_ISL_495525, EPI_ISL_495526, EPI_ISL_495527, EPI_ISL_495528, EPI_ISL_495529, EPI_ISL_495530, EPI_ISL_495531, EPI_ISL_495532, EPI_ISL_495533, EPI_ISL_495534 | | | |
| see above | NHLS-IALCH | KRISP, KZN Research Innovation and Sequencing Platform | Giandhari J, Pillay S, Lessells R, Chimukangara B, Mdlalose K, York D, Khan S, Tegally H, Wilkinson E, de Oliveira T |
| EPI_ISL_495535, EPI_ISL_495536, EPI_ISL_495537, EPI_ISL_495538, EPI_ISL_495539, EPI_ISL_495540, EPI_ISL_495541, EPI_ISL_495542 | Medical Disagnostics Services (MDS) | KRISP, KZN Research Innovation and Sequencing Platform | Giandhari J, Pillay S, Lessells R, Chimukangara B, Mdlalose K, York D, Khan S, Tegally H, Wilkinson E, de Oliveira T |
| EPI_ISL_495543, EPI_ISL_495544, EPI_ISL_495545, EPI_ISL_495546, EPI_ISL_495547, EPI_ISL_495548, EPI_ISL_495549, EPI_ISL_495550, EPI_ISL_495551, EPI_ISL_495552, EPI_ISL_495553, EPI_ISL_495554, EPI_ISL_495555, EPI_ISL_495556, EPI_ISL_495557, EPI_ISL_495558, EPI_ISL_495559, EPI_ISL_495560, EPI_ISL_495561, EPI_ISL_495562 | | | |

| | | | |
|---|---|---|---|
| see above | NHLS-IALCH | KRISP, KZN Research Innovation and Sequencing Platform | Giandhari J, Pillay S, Lessells R, Chimukangara B, Mdlalose K, York D, Khan S, Tegally H, Wilkinson E, de Oliveira T |
| EPI_ISL_495563, EPI_ISL_495564, EPI_ISL_495565, EPI_ISL_495566, EPI_ISL_495567, EPI_ISL_495568, EPI_ISL_495569, EPI_ISL_495570, EPI_ISL_495571, EPI_ISL_495572, EPI_ISL_495573, EPI_ISL_495574, EPI_ISL_495575, EPI_ISL_495576, EPI_ISL_495577, EPI_ISL_495578, EPI_ISL_495579, EPI_ISL_495580, EPI_ISL_495581, EPI_ISL_495582, EPI_ISL_495583, EPI_ISL_495584, EPI_ISL_495585, EPI_ISL_495586, EPI_ISL_495587, EPI_ISL_495588, EPI_ISL_495589, EPI_ISL_495590, EPI_ISL_495591, EPI_ISL_495592, EPI_ISL_495593, EPI_ISL_495594, EPI_ISL_495595, EPI_ISL_495596 | | | |
| see above | University of Michigan Clinical Microbiology Laboratory | Lauring Lab, University of Michigan, Department of Microbiology and Immunology | Valesano et al. |
| EPI_ISL_495597, EPI_ISL_495598, EPI_ISL_495599, EPI_ISL_495600, EPI_ISL_495601, EPI_ISL_495602, EPI_ISL_495603, EPI_ISL_495604, EPI_ISL_495605, EPI_ISL_495606, EPI_ISL_495607, EPI_ISL_495608, EPI_ISL_495609 | | | |
| see above | Mayo Clinic & Mayo Clinic Laboratories | Minnesota Department of Health, Public Health Laboratory | Matt Plumb, Jacob Garfin, and Xiong Wang |
| EPI_ISL_495610, EPI_ISL_495611, EPI_ISL_495612, EPI_ISL_495613, EPI_ISL_495614, EPI_ISL_495615, EPI_ISL_495616, EPI_ISL_495617, EPI_ISL_495618, EPI_ISL_495619, EPI_ISL_495620, EPI_ISL_495621, EPI_ISL_495622, EPI_ISL_495623, EPI_ISL_495624, EPI_ISL_495625, EPI_ISL_495626, EPI_ISL_495627, EPI_ISL_495628 | | | |
| see above | Minnesota Department of Health, Public Health Laboratory | Minnesota Department of Health, Public Health Laboratory | Matt Plumb, Jacob Garfin, and Xiong Wang |
| EPI_ISL_495629, EPI_ISL_495630, EPI_ISL_495631, EPI_ISL_495632, EPI_ISL_495633, EPI_ISL_495634, EPI_ISL_495635, EPI_ISL_495636, EPI_ISL_495637, EPI_ISL_495638, EPI_ISL_495639, EPI_ISL_495640, EPI_ISL_495641 | | | |
| see above | Viral Respiratory Lab, National Institute for Biomedical Research (INRB) | Pathogen Sequencing Lab, National Institute for Biomedical Research (INRB) | Placide Mbala-Kingebeni, Edith Nkwembe, Eddy Kinganda-Lusamaki, Amuri Aziza, Francisca Muyembe Mawete, Emmanuel Lokilo Lofiko, Catherine Pratt, Matthias Pauthner, Josh Quick, Allison Black, James Hadfield, Trevor Bedford, Ian Goodfellow, Andrew Rambaut, Nick Loman, Kristian Andersen, Michael Wiley, Steve Ahuka-Mundeke, Jean-Jacques Muyembe Tamfum |
| EPI_ISL_495642, EPI_ISL_495643, EPI_ISL_495644, EPI_ISL_495645, EPI_ISL_495646, EPI_ISL_495647, EPI_ISL_495648, EPI_ISL_495649, EPI_ISL_495650, EPI_ISL_495651, EPI_ISL_495652, EPI_ISL_495653 | | | |
| see above | Washington State Department of Health | Seattle Flu Study | Deborah A. Nickerson, Chris D. Frazar, Jover Lee, Benjamin Pelle, Matthew Richardson, Amanda Adler, Elisabeth Brandstetter, Peter D. Han, Kairsten Fay, Misja Ilcisin, Kirsten Lacombe, Thomas R. Sibley, Melissa Truong, Caitlin R. Wolf, Romesh Gautom, Geoff Melly, Brian Hiatt, Philip Dykema, Scott Lindquist, Michael Boeckh, Janet A. Englund, Michael Famulare, Barry R. Lutz, Mark J. Rieder, Lea M. Starita, Matthew Thompson, Helen Y. Chu, Jay Shendure, Trevor Bedford |
| EPI_ISL_495654, EPI_ISL_495655, EPI_ISL_495656, EPI_ISL_495657, EPI_ISL_495658, EPI_ISL_495659, EPI_ISL_495660, EPI_ISL_495661, EPI_ISL_495662 | Seattle Flu Study | Seattle Flu Study | Deborah A. Nickerson, Chris D. Frazar, Jover Lee, Benjamin Pelle, Matthew Richardson, Amanda Adler, Elisabeth Brandstetter, Peter D. Han, Kairsten Fay, Misja Ilcisin, Kirsten Lacombe, Thomas R. Sibley, Melissa Truong, Caitlin R. Wolf, Karen Cowgill, Stephanie Schrag, Jeff Duchin, Michael Boeckh, Janet A. Englund, Michael Famulare, Barry R. Lutz, Mark J. Rieder, Lea M. Starita, Matthew Thompson, Helen Y. Chu, Trevor Bedford, Jay Shendure |
| EPI_ISL_495663 | Seattle Flu Study | Seattle Flu Study | Deborah A. Nickerson, Chris D. Frazar, Jover Lee, Benjamin Pelle, Matthew Richardson, Amanda Adler, Elisabeth Brandstetter, Peter D. Han, Kairsten Fay, Misja Ilcisin, Kirsten Lacombe, Thomas R. Sibley, Melissa Truong, Caitlin R. Wolf, Michael Boeckh, Janet A. Englund, Michael Famulare, Barry R. Lutz, Mark J. Rieder, Lea M. Starita, Matthew Thompson, Jay Shendure, Trevor Bedford, Helen Y. Chu |
| EPI_ISL_495664, EPI_ISL_495665 | Seattle Flu Study | Seattle Flu Study | Deborah A. Nickerson, Chris D. Frazar, Jover Lee, Benjamin Pelle, Matthew Richardson, Amanda Adler, Elisabeth Brandstetter, Peter D. Han, Kairsten Fay, Misja Ilcisin, Kirsten Lacombe, Thomas R. Sibley, Melissa Truong, Caitlin R. Wolf, Karen Cowgill, Stephanie Schrag, Jeff Duchin, Michael Boeckh, Janet A. Englund, Michael Famulare, Barry R. Lutz, Mark J. Rieder, Lea M. Starita, Matthew Thompson, Helen Y. Chu, Trevor Bedford, Jay Shendure |
| EPI_ISL_495666, EPI_ISL_495667, EPI_ISL_495668, EPI_ISL_495669, EPI_ISL_495670, EPI_ISL_495671, EPI_ISL_495672, EPI_ISL_495673, EPI_ISL_495674, EPI_ISL_495675, EPI_ISL_495676, EPI_ISL_495677, EPI_ISL_495678, EPI_ISL_495679, EPI_ISL_495680, EPI_ISL_495681, EPI_ISL_495682, EPI_ISL_495683, EPI_ISL_495684, EPI_ISL_495685, EPI_ISL_495686, EPI_ISL_495687, EPI_ISL_495688, EPI_ISL_495689, EPI_ISL_495690, EPI_ISL_495691, EPI_ISL_495692, EPI_ISL_495693, EPI_ISL_495694, EPI_ISL_495695, EPI_ISL_495696, EPI_ISL_495697, EPI_ISL_495698, EPI_ISL_495699, EPI_ISL_495700, EPI_ISL_495701, EPI_ISL_495702, EPI_ISL_495703, EPI_ISL_495704, EPI_ISL_495705, EPI_ISL_495706, EPI_ISL_495707, EPI_ISL_495708, EPI_ISL_495709, EPI_ISL_495710, EPI_ISL_495711, EPI_ISL_495712, EPI_ISL_495713, EPI_ISL_495714, EPI_ISL_495715, EPI_ISL_495716, EPI_ISL_495717, EPI_ISL_495718, EPI_ISL_495719, EPI_ISL_495720, EPI_ISL_495721, EPI_ISL_495722, EPI_ISL_495723, EPI_ISL_495724, EPI_ISL_495725, EPI_ISL_495726, EPI_ISL_495727, EPI_ISL_495728, EPI_ISL_495729, EPI_ISL_495730, EPI_ISL_495731, EPI_ISL_495732, EPI_ISL_495733, EPI_ISL_495734, EPI_ISL_495735, EPI_ISL_495736, EPI_ISL_495737, EPI_ISL_495738, EPI_ISL_495739, EPI_ISL_495740, EPI_ISL_495741, EPI_ISL_495742, EPI_ISL_495743, EPI_ISL_495744, EPI_ISL_495745, EPI_ISL_495746, EPI_ISL_495747, EPI_ISL_495748, EPI_ISL_495749, EPI_ISL_495750, EPI_ISL_495751, EPI_ISL_495752, EPI_ISL_495753, EPI_ISL_495754, EPI_ISL_495755, EPI_ISL_495756, EPI_ISL_495757, EPI_ISL_495758, EPI_ISL_495759, EPI_ISL_495760, EPI_ISL_495761, EPI_ISL_495762, EPI_ISL_495763, EPI_ISL_495764, EPI_ISL_495765, EPI_ISL_495766, EPI_ISL_495767, EPI_ISL_495768, EPI_ISL_495769, EPI_ISL_495770, EPI_ISL_495771, EPI_ISL_495772, EPI_ISL_495773, EPI_ISL_495774, EPI_ISL_495775, EPI_ISL_495776, EPI_ISL_495777, EPI_ISL_495778, EPI_ISL_495779, EPI_ISL_495780, EPI_ISL_495781, EPI_ISL_495782, EPI_ISL_495783, EPI_ISL_495784, EPI_ISL_495785, EPI_ISL_495786, EPI_ISL_495787, EPI_ISL_495788, EPI_ISL_495789, EPI_ISL_495790, EPI_ISL_495791, EPI_ISL_495792, EPI_ISL_495793, EPI_ISL_495794, EPI_ISL_495795, EPI_ISL_495796, EPI_ISL_495797, EPI_ISL_495798, EPI_ISL_495799, EPI_ISL_495800, EPI_ISL_495801, EPI_ISL_495802, EPI_ISL_495803, EPI_ISL_495804, EPI_ISL_495805, EPI_ISL_495806, EPI_ISL_495807, EPI_ISL_495808, EPI_ISL_495809, EPI_ISL_495810, EPI_ISL_495811, EPI_ISL_495812, EPI_ISL_495813, EPI_ISL_495814, EPI_ISL_495815, EPI_ISL_495816, EPI_ISL_495817, EPI_ISL_495818, EPI_ISL_495819, EPI_ISL_495820, EPI_ISL_495821, EPI_ISL_495822, EPI_ISL_495823, EPI_ISL_495824, EPI_ISL_495825, EPI_ISL_495826, EPI_ISL_495827, EPI_ISL_495828, EPI_ISL_495829, EPI_ISL_495830, EPI_ISL_495831, EPI_ISL_495832, EPI_ISL_495833, EPI_ISL_495834, EPI_ISL_495835, EPI_ISL_495836, EPI_ISL_495837, EPI_ISL_495838, EPI_ISL_495839, EPI_ISL_495840, EPI_ISL_495841, EPI_ISL_495842, EPI_ISL_495843, EPI_ISL_495844, EPI_ISL_495845, EPI_ISL_495846, EPI_ISL_495847, EPI_ISL_495848, EPI_ISL_495849, EPI_ISL_495850, EPI_ISL_495851, EPI_ISL_495852, EPI_ISL_495853, EPI_ISL_495854, EPI_ISL_495855, EPI_ISL_495856, EPI_ISL_495857, EPI_ISL_495858, EPI_ISL_495859, EPI_ISL_495860, EPI_ISL_495861, EPI_ISL_495862, EPI_ISL_495863, EPI_ISL_495864, EPI_ISL_495865, EPI_ISL_495866, EPI_ISL_495867, EPI_ISL_495868, EPI_ISL_495869, EPI_ISL_495870, EPI_ISL_495871, EPI_ISL_495872, EPI_ISL_495873, EPI_ISL_495874, EPI_ISL_495875, EPI_ISL_495876, EPI_ISL_495877, EPI_ISL_495878, EPI_ISL_495879, EPI_ISL_495880, EPI_ISL_495881, EPI_ISL_495882, EPI_ISL_495883, EPI_ISL_495884, EPI_ISL_495885, EPI_ISL_495886, EPI_ISL_495887, EPI_ISL_495888, EPI_ISL_495889, EPI_ISL_495890, EPI_ISL_495891, EPI_ISL_495892, EPI_ISL_495893, EPI_ISL_495894, EPI_ISL_495895, EPI_ISL_495896, EPI_ISL_495897, EPI_ISL_495898, EPI_ISL_495899, EPI_ISL_495900, EPI_ISL_495901, EPI_ISL_495902, EPI_ISL_495903, EPI_ISL_495904, EPI_ISL_495905, EPI_ISL_495906, EPI_ISL_495907, EPI_ISL_495908, EPI_ISL_495909, EPI_ISL_495910, EPI_ISL_495911, EPI_ISL_495912, EPI_ISL_495913, EPI_ISL_495914, EPI_ISL_495915, EPI_ISL_495916 | | | |
| see above | Washington State Department of Health | Seattle Flu Study | Deborah A. Nickerson, Chris D. Frazar, Jover Lee, Benjamin Pelle, Matthew Richardson, Amanda Adler, Elisabeth Brandstetter, Peter D. Han, Kairsten Fay, Misja Ilcisin, Kirsten Lacombe, Thomas R. Sibley, Melissa Truong, Caitlin R. Wolf, Romesh Gautom, Geoff |
| EPI_ISL_495917, EPI_ISL_495918, EPI_ISL_495919, EPI_ISL_495920, EPI_ISL_495921, EPI_ISL_495922, EPI_ISL_495923, EPI_ISL_495924, EPI_ISL_495925, EPI_ISL_495926, EPI_ISL_495927, EPI_ISL_495928, EPI_ISL_495929, EPI_ISL_495930, EPI_ISL_495931, EPI_ISL_495932, EPI_ISL_495933, EPI_ISL_495934, EPI_ISL_495935, EPI_ISL_495936, EPI_ISL_495937, EPI_ISL_495938, EPI_ISL_495939, EPI_ISL_495940, EPI_ISL_495941, EPI_ISL_495942, EPI_ISL_495943, EPI_ISL_495944, EPI_ISL_495945, EPI_ISL_495946, EPI_ISL_495947, EPI_ISL_495948, EPI_ISL_495949, EPI_ISL_495950, EPI_ISL_495951, EPI_ISL_495952, EPI_ISL_495953, EPI_ISL_495954, EPI_ISL_495955, EPI_ISL_495956, EPI_ISL_495957, EPI_ISL_495958, EPI_ISL_495959, EPI_ISL_495960, EPI_ISL_495961, EPI_ISL_495962, EPI_ISL_495963, EPI_ISL_495964, EPI_ISL_495965, EPI_ISL_495966, EPI_ISL_495967, EPI_ISL_495968, EPI_ISL_495969, EPI_ISL_495970, EPI_ISL_495971, EPI_ISL_495972, EPI_ISL_495973, EPI_ISL_495974, EPI_ISL_495975, EPI_ISL_495976, EPI_ISL_495977, EPI_ISL_495978, EPI_ISL_495979, EPI_ISL_495980, EPI_ISL_495981, EPI_ISL_495982, EPI_ISL_495983, EPI_ISL_495984, EPI_ISL_495985, EPI_ISL_495986, EPI_ISL_495987, EPI_ISL_495988, EPI_ISL_495989, EPI_ISL_495990, EPI_ISL_495991, EPI_ISL_495992, EPI_ISL_495993, EPI_ISL_495994, EPI_ISL_495995, EPI_ISL_495996, EPI_ISL_495997, EPI_ISL_495998, EPI_ISL_495999, EPI_ISL_496000, EPI_ISL_496001, EPI_ISL_496002, EPI_ISL_496003, EPI_ISL_496004, EPI_ISL_496005, EPI_ISL_496006, EPI_ISL_496007, EPI_ISL_496008, EPI_ISL_496009, EPI_ISL_496010, EPI_ISL_496011, EPI_ISL_496012, EPI_ISL_496013, EPI_ISL_496014, EPI_ISL_496015, EPI_ISL_496016, EPI_ISL_496017, EPI_ISL_496018, EPI_ISL_496019, EPI_ISL_496020, EPI_ISL_496021, EPI_ISL_496022, EPI_ISL_496023, EPI_ISL_496024, EPI_ISL_496025, EPI_ISL_496026, EPI_ISL_496027, EPI_ISL_496028, EPI_ISL_496029, EPI_ISL_496030, EPI_ISL_496031, EPI_ISL_496032, EPI_ISL_496033, EPI_ISL_496034, EPI_ISL_496035, EPI_ISL_496036, EPI_ISL_496037, EPI_ISL_496038, EPI_ISL_496039, EPI_ISL_496040, EPI_ISL_496041, EPI_ISL_496042, EPI_ISL_496043, EPI_ISL_496044, EPI_ISL_496045, EPI_ISL_496046, EPI_ISL_496047, EPI_ISL_496048, EPI_ISL_496049, EPI_ISL_496050, EPI_ISL_496051, EPI_ISL_496052, EPI_ISL_496053, EPI_ISL_496054, EPI_ISL_496055, EPI_ISL_496056, EPI_ISL_496057, EPI_ISL_496058, EPI_ISL_496059, EPI_ISL_496060, EPI_ISL_496061, EPI_ISL_496062, EPI_ISL_496063, EPI_ISL_496064, EPI_ISL_496065, EPI_ISL_496066, EPI_ISL_496067, EPI_ISL_496068, EPI_ISL_496069, EPI_ISL_496070, EPI_ISL_496071, EPI_ISL_496072, EPI_ISL_496073, EPI_ISL_496074, EPI_ISL_496075, EPI_ISL_496076, EPI_ISL_496077, EPI_ISL_496078, EPI_ISL_496079, EPI_ISL_496080, EPI_ISL_496081, EPI_ISL_496082, EPI_ISL_496083, EPI_ISL_496084, EPI_ISL_496085, EPI_ISL_496086, EPI_ISL_496087, EPI_ISL_496088, EPI_ISL_496089, EPI_ISL_496090, EPI_ISL_496091, EPI_ISL_496092, EPI_ISL_496093, EPI_ISL_496094, EPI_ISL_496095, EPI_ISL_496096, EPI_ISL_496097, EPI_ISL_496098, EPI_ISL_496099, EPI_ISL_496100, EPI_ISL_496101, EPI_ISL_496102, EPI_ISL_496103, EPI_ISL_496104, EPI_ISL_496105, EPI_ISL_496106, EPI_ISL_496107, EPI_ISL_496108, EPI_ISL_496109, EPI_ISL_496110, EPI_ISL_496111, EPI_ISL_496112, EPI_ISL_496113, EPI_ISL_496114, EPI_ISL_496115, EPI_ISL_496116, EPI_ISL_496117, EPI_ISL_496118, EPI_ISL_496119, EPI_ISL_496120, EPI_ISL_496121, EPI_ISL_496122, EPI_ISL_496123, EPI_ISL_496124, EPI_ISL_496125, EPI_ISL_496126, EPI_ISL_496127, EPI_ISL_496128, EPI_ISL_496129, EPI_ISL_496130, EPI_ISL_496131, EPI_ISL_496132, EPI_ISL_496133, EPI_ISL_496134, EPI_ISL_496135, EPI_ISL_496136, EPI_ISL_496137, EPI_ISL_496138, EPI_ISL_496139, EPI_ISL_496140, EPI_ISL_496141, EPI_ISL_496142, EPI_ISL_496143, EPI_ISL_496144, EPI_ISL_496145, EPI_ISL_496146, EPI_ISL_496147, EPI_ISL_496148, EPI_ISL_496149, EPI_ISL_496150, EPI_ISL_496151, EPI_ISL_496152, EPI_ISL_496153, EPI_ISL_496154, EPI_ISL_496155, EPI_ISL_496156, EPI_ISL_496157, EPI_ISL_496158, EPI_ISL_496159, EPI_ISL_496160, EPI_ISL_496161, EPI_ISL_496162, EPI_ISL_496163, EPI_ISL_496164, EPI_ISL_496165, EPI_ISL_496166, EPI_ISL_496167, EPI_ISL_496168, EPI_ISL_496169, EPI_ISL_496170, EPI_ISL_496171, EPI_ISL_496172, EPI_ISL_496173, EPI_ISL_496174, EPI_ISL_496175, EPI_ISL_496176, EPI_ISL_496177, EPI_ISL_496178, EPI_ISL_496179, EPI_ISL_496180, EPI_ISL_496181, EPI_ISL_496182, EPI_ISL_496183, EPI_ISL_496184, EPI_ISL_496185, EPI_ISL_496186, EPI_ISL_496187, EPI_ISL_496188, EPI_ISL_496189, EPI_ISL_496190, EPI_ISL_496191, EPI_ISL_496192, EPI_ISL_496193, EPI_ISL_496194, EPI_ISL_496195, EPI_ISL_496196, EPI_ISL_496197, EPI_ISL_496198, EPI_ISL_496199, EPI_ISL_496200, EPI_ISL_496201, EPI_ISL_496202, EPI_ISL_496203, EPI_ISL_496204, EPI_ISL_496205, EPI_ISL_496206, EPI_ISL_496207, EPI_ISL_496208, EPI_ISL_496209, EPI_ISL_496210, EPI_ISL_496211, EPI_ISL_496212, EPI_ISL_496213, EPI_ISL_496214, EPI_ISL_496215, EPI_ISL_496216, EPI_ISL_496217, EPI_ISL_496218, EPI_ISL_496219, EPI_ISL_496220, EPI_ISL_496221, EPI_ISL_496222, EPI_ISL_496223, EPI_ISL_496224, EPI_ISL_496225, EPI_ISL_496226, EPI_ISL_496227, EPI_ISL_496228, EPI_ISL_496229, EPI_ISL_496230, EPI_ISL_496231, EPI_ISL_496232, EPI_ISL_496233, EPI_ISL_496234, EPI_ISL_496235, EPI_ISL_496236, EPI_ISL_496237, EPI_ISL_496238, EPI_ISL_496239, EPI_ISL_496240, EPI_ISL_496241, EPI_ISL_496242, EPI_ISL_496243, EPI_ISL_496244, EPI_ISL_496245, EPI_ISL_496246, EPI_ISL_496247, EPI_ISL_496248, EPI_ISL_496249, EPI_ISL_496250, EPI_ISL_496251, EPI_ISL_496252, EPI_ISL_496253, EPI_ISL_496254, EPI_ISL_496255, EPI_ISL_496256, EPI_ISL_496257, EPI_ISL_496258, EPI_ISL_496259, EPI_ISL_496260, EPI_ISL_496261, EPI_ISL_496262, EPI_ISL_496263, EPI_ISL_496264, EPI_ISL_496265, EPI_ISL_496266, EPI_ISL_496267, EPI_ISL_496268, EPI_ISL_496269, EPI_ISL_496270, EPI_ISL_496271, EPI_ISL_496272, EPI_ISL_496273, EPI_ISL_496274, EPI_ISL_496275, EPI_ISL_496276, EPI_ISL_496277, EPI_ISL_496278, EPI_ISL_496279, EPI_ISL_496280, EPI_ISL_496281, EPI_ISL_496282, EPI_ISL_496283, EPI_ISL_496284, EPI_ISL_496285, EPI_ISL_496286, EPI_ISL_496287, EPI_ISL_496288, EPI_ISL_496289, EPI_ISL_496290, EPI_ISL_496291, EPI_ISL_496292, EPI_ISL_496293, EPI_ISL_496294, EPI_ISL_496295, EPI_ISL_496296, EPI_ISL_496297, EPI_ISL_496298, EPI_ISL_496299, EPI_ISL_496300, EPI_ISL_496301, EPI_ISL_496302, EPI_ISL_496303, EPI_ISL_496304, EPI_ISL_496305, EPI_ISL_496306, EPI_ISL_496307, EPI_ISL_496308, EPI_ISL_496309, EPI_ISL_496310, EPI_ISL_496311, EPI_ISL_496312, EPI_ISL_496313, EPI_ISL_496314, EPI_ISL_496315, EPI_ISL_496316, EPI_ISL_496317, EPI_ISL_496318, EPI_ISL_496319, EPI_ISL_496320, EPI_ISL_496321, EPI_ISL_496322, EPI_ISL_496323, EPI_ISL_496324, EPI_ISL_496325, EPI_ISL_496326, EPI_ISL_496327, EPI_ISL_496328, EPI_ISL_496329, EPI_ISL_496330, EPI_ISL_496331, EPI_ISL_496332, EPI_ISL_496333, EPI_ISL_496334, EPI_ISL_496335, EPI_ISL_496336, EPI_ISL_496337, EPI_ISL_496338 | | | |
| see above | Washington State Department of Health | Seattle Flu Study | Deborah A. Nickerson, Chris D. Frazar, Jover Lee, Benjamin Pelle, Matthew Richardson, Amanda Adler, Elisabeth Brandstetter, Peter D. Han, Kairsten Fay, Misja Ilcisin, Kirsten Lacombe, Thomas R. Sibley, Melissa Truong, Caitlin R. Wolf, Romesh Gautom, Geoff Melly, Brian Hiatt, Philip Dykema, Scott Lindquist, Michael Boeckh, Janet A. Englund, Michael Famulare, Barry R. Lutz, Mark J. Rieder, Lea M. Starita, Matthew Thompson, Helen Y. Chu, Jay Shendure, Trevor Bedford |
| EPI_ISL_496339, EPI_ISL_496340, EPI_ISL_496341, EPI_ISL_496342, EPI_ISL_496343, EPI_ISL_496344, EPI_ISL_496345, EPI_ISL_496346, EPI_ISL_496347, EPI_ISL_496348, EPI_ISL_496349, EPI_ISL_496350, EPI_ISL_496351, EPI_ISL_496352, EPI_ISL_496353, EPI_ISL_496354, EPI_ISL_496355, EPI_ISL_496356, EPI_ISL_496357, EPI_ISL_496358, EPI_ISL_496359, EPI_ISL_496360, EPI_ISL_496361, EPI_ISL_496362, EPI_ISL_496363, EPI_ISL_496364, EPI_ISL_496365, EPI_ISL_496366, EPI_ISL_496367, EPI_ISL_496368, EPI_ISL_496369, EPI_ISL_496370, EPI_ISL_496371, EPI_ISL_496372, EPI_ISL_496373, EPI_ISL_496374, EPI_ISL_496375, EPI_ISL_496376 | | | |
| see above | Infectolab | Andersen lab at Scripps Research | SEARCH Alliance San Diego with Samuel Navarro Alvarez, Carlos A. Cota Haros, Octavio Renteria Pacheco |
| EPI_ISL_496482 | Dept. Infectious, Tropical Diseases & Microbiology, IRCCS Sacro Cuore Don Calabria Hospital | 1) Dept. Infectious, Tropical Diseases & Microbiology, IRCCS Sacro Cuore Don Calabria Hospital; 2) Centro Piattaforme Tecnologiche, University of Verona; 3) Dept. Neurosciences, Biomedicine and Movement Sciences, University of Verona. | 1) Antonio Mori, Michela Deiana, Elena Pomari, Chiara Piubelli; 2) Monica Castellucci and Francesca Griggio; 3) Giovanni Malerba |
| EPI_ISL_496491, EPI_ISL_496492, EPI_ISL_496493, EPI_ISL_496494, EPI_ISL_496495, EPI_ISL_496496, EPI_ISL_496497, EPI_ISL_496498, EPI_ISL_496499, EPI_ISL_496500, EPI_ISL_496501, EPI_ISL_496502, EPI_ISL_496503, EPI_ISL_496504, EPI_ISL_496505, EPI_ISL_496506, EPI_ISL_496507, EPI_ISL_496508, EPI_ISL_496509, EPI_ISL_496510, EPI_ISL_496511, EPI_ISL_496512, EPI_ISL_496513, EPI_ISL_496514, EPI_ISL_496515, EPI_ISL_496516, EPI_ISL_496517 | | | |
| see above | Viral Respiratory Lab, National Institute for Biomedical Research (INRB) | Pathogen Sequencing Lab, National Institute for Biomedical Research (INRB) | Placide Mbala-Kingebeni, Edith Nkwembe, Eddy Kinganda-Lusamaki, Amuri Aziza, Francisca Muyembe Mawete, Emmanuel Lokilo Lofiko, Catherine Pratt, Matthias Pauthner, Josh Quick, Allison Black, James Hadfield, Trevor Bedford, Ian Goodfellow, Andrew Rambaut, Nick Loman, Kristian Andersen, Michael Wiley, Steve Ahuka-Mundeke, Jean-Jacques Muyembe Tamfum |
| EPI_ISL_496518, EPI_ISL_496519, EPI_ISL_496520 | Armed Forces Medical College | National Centre For Cell Science | Dhiraj Paul, Kunal Jani, Radha Chauhan, Janesh Kumar, Vasudevan Seshadri, Girdhari Lal, Rajesh Karyakarte, Suvarna Joshi, Murlidhar Tambe, Sourav Sen, Santosh Karade, Kavita Bala Anand, Shelinder Pal Singh Shergill, Rajiv Mohan Gupta, Manoj Kumar Bhat, Arvind Sahu, Maharashtra COVID-19 Study Group, DBT's PAN-INDIA 1000 SARS-CoV2 RNA genome sequencing consortium, Yogesh S Shouche |
| EPI_ISL_496521, EPI_ISL_496522, EPI_ISL_496523 | B.J. Govt. Medical College | National Centre For Cell Science | Dhiraj Paul, Kunal Jani, Radha Chauhan, Janesh Kumar, Vasudevan Seshadri, Girdhari Lal, Rajesh Karyakarte, Suvarna Joshi, Murlidhar Tambe, Sourav Sen, Santosh Karade, Kavita Bala Anand, Shelinder Pal Singh Shergill, Rajiv Mohan Gupta, Manoj Kumar Bhat, Arvind Sahu, Maharashtra COVID-19 Study Group, DBT's PAN-INDIA 1000 SARS-CoV2 RNA genome sequencing consortium, Yogesh S Shouche |
| EPI_ISL_496524, EPI_ISL_496525, EPI_ISL_496526 | National Centre For Cell Science | National Centre For Cell Science | Dhiraj Paul, Kunal Jani, Radha Chauhan, Janesh Kumar, Vasudevan Seshadri, Girdhari Lal, Rajesh Karyakarte, Suvarna Joshi, Murlidhar Tambe, Sourav Sen, Santosh Karade, Kavita Bala Anand, Shelinder Pal Singh Shergill, Rajiv Mohan Gupta, Manoj Kumar Bhat, Arvind Sahu, Maharashtra COVID-19 Study Group, DBT's PAN-INDIA 1000 SARS-CoV2 RNA genome sequencing consortium, Yogesh S Shouche |
| EPI_ISL_496527, EPI_ISL_496528 | B.J. Govt. Medical College | National Centre For Cell Science | Dhiraj Paul, Kunal Jani, Radha Chauhan, Janesh Kumar, Vasudevan Seshadri, Girdhari Lal, Rajesh Karyakarte, Suvarna Joshi, Murlidhar Tambe, Sourav Sen, Santosh Karade, Kavita Bala Anand, Shelinder Pal Singh Shergill, Rajiv Mohan Gupta, Manoj Kumar Bhat, Arvind Sahu, Maharashtra COVID-19 Study Group, DBT's PAN-INDIA 1000 SARS-CoV2 RNA genome sequencing consortium, Yogesh S Shouche |
| EPI_ISL_496529 | Armed Forces Medical College | National Centre For Cell Science | Dhiraj Paul, Kunal Jani, Radha Chauhan, Janesh Kumar, Vasudevan Seshadri, Girdhari Lal, Rajesh Karyakarte, Suvarna Joshi, Murlidhar Tambe, Sourav Sen, Santosh Karade, Kavita Bala Anand, Shelinder Pal Singh Shergill, Rajiv Mohan Gupta, Manoj Kumar Bhat, Arvind Sahu, Maharashtra COVID-19 Study Group, DBT's PAN-INDIA 1000 SARS-CoV2 RNA genome sequencing consortium, Yogesh S Shouche |
| EPI_ISL_496530 | B.J. Govt. Medical College | National Centre For Cell Science | Dhiraj Paul, Kunal Jani, Radha Chauhan, Janesh Kumar, Vasudevan Seshadri, Girdhari Lal, Rajesh Karyakarte, Suvarna Joshi, Murlidhar Tambe, Sourav Sen, Santosh Karade, Kavita Bala Anand, Shelinder Pal Singh Shergill, Rajiv Mohan Gupta, Manoj Kumar Bhat, Arvind Sahu, Maharashtra COVID-19 Study Group, DBT's PAN-INDIA 1000 SARS-CoV2 RNA genome sequencing consortium, Yogesh S Shouche |
| EPI_ISL_496531, EPI_ISL_496532 | National Centre For Cell Science | National Centre For Cell Science | Dhiraj Paul, Kunal Jani, Radha Chauhan, Janesh Kumar, Vasudevan Seshadri, Girdhari Lal, Rajesh Karyakarte, Suvarna Joshi, Murlidhar Tambe, Sourav Sen, Santosh Karade, Kavita Bala Anand, Shelinder Pal Singh Shergill, Rajiv Mohan Gupta, Manoj Kumar Bhat, Arvind Sahu, Maharashtra COVID-19 Study Group, DBT's PAN-INDIA 1000 SARS-CoV2 RNA genome sequencing consortium, Yogesh S Shouche |
| EPI_ISL_496533 | Armed Forces Medical College | National Centre For Cell Science | Dhiraj Paul, Kunal Jani, Radha Chauhan, Janesh Kumar, Vasudevan Seshadri, Girdhari Lal, Rajesh Karyakarte, Suvarna Joshi, Murlidhar Tambe, Sourav Sen, Santosh Karade, Kavita Bala Anand, Shelinder Pal Singh Shergill, Rajiv Mohan Gupta, Manoj Kumar Bhat, Arvind Sahu, Maharashtra COVID-19 Study Group, DBT's PAN-INDIA 1000 SARS-CoV2 RNA genome sequencing consortium, Yogesh S Shouche |

| | | | |
|---|---|---|---|
| EPI_ISL_496534 | B.J. Govt. Medical College | National Centre For Cell Science | Dhiraj Paul, Kunal Jani, Radha Chauhan, Janesh Kumar, Vasudevan Seshadri, Girdhari Lal, Rajesh Karyakarte, Suvarna Joshi, Murlidhar Tambe, Sourav Sen, Santosh Karade, Kavita Bala Anand, Shelinder Pal Singh Shergill, Rajiv Mohan Gupta, Manoj Kumar Bhat, Arvind Sahu, Maharashtra COVID-19 Study Group, DBT's PAN-INDIA 1000 SARS-CoV2 RNA genome sequencing consortium, Yogesh S Shouche |
| EPI_ISL_496535, EPI_ISL_496536 | National Centre For Cell Science | National Centre For Cell Science | Dhiraj Paul, Kunal Jani, Radha Chauhan, Janesh Kumar, Vasudevan Seshadri, Girdhari Lal, Rajesh Karyakarte, Suvarna Joshi, Murlidhar Tambe, Sourav Sen, Santosh Karade, Kavita Bala Anand, Shelinder Pal Singh Shergill, Rajiv Mohan Gupta, Manoj Kumar Bhat, Arvind Sahu, Maharashtra COVID-19 Study Group, DBT's PAN-INDIA 1000 SARS-CoV2 RNA genome sequencing consortium, Yogesh S Shouche |
| EPI_ISL_496537, EPI_ISL_496538, EPI_ISL_496539, EPI_ISL_496540, EPI_ISL_496541, EPI_ISL_496542, EPI_ISL_496543, EPI_ISL_496544, EPI_ISL_496545 | Armed Forces Medical College | National Centre For Cell Science | Dhiraj Paul, Kunal Jani, Radha Chauhan, Janesh Kumar, Vasudevan Seshadri, Girdhari Lal, Rajesh Karyakarte, Suvarna Joshi, Murlidhar Tambe, Sourav Sen, Santosh Karade, Kavita Bala Anand, Shelinder Pal Singh Shergill, Rajiv Mohan Gupta, Manoj Kumar Bhat, Arvind Sahu, Maharashtra COVID-19 Study Group, DBT's PAN-INDIA 1000 SARS-CoV2 RNA genome sequencing consortium, Yogesh S Shouche |
| EPI_ISL_496546, EPI_ISL_496547, EPI_ISL_496548, EPI_ISL_496549, EPI_ISL_496550, EPI_ISL_496551, EPI_ISL_496552, EPI_ISL_496553, EPI_ISL_496554 | B.J. Govt. Medical College | National Centre For Cell Science | Dhiraj Paul, Kunal Jani, Radha Chauhan, Janesh Kumar, Vasudevan Seshadri, Girdhari Lal, Rajesh Karyakarte, Suvarna Joshi, Murlidhar Tambe, Sourav Sen, Santosh Karade, Kavita Bala Anand, Shelinder Pal Singh Shergill, Rajiv Mohan Gupta, Manoj Kumar Bhat, Arvind Sahu, Maharashtra COVID-19 Study Group, DBT's PAN-INDIA 1000 SARS-CoV2 RNA genome sequencing consortium, Yogesh S Shouche |
| EPI_ISL_496555, EPI_ISL_496556, EPI_ISL_496557, EPI_ISL_496558, EPI_ISL_496559, EPI_ISL_496560, EPI_ISL_496561, EPI_ISL_496562, EPI_ISL_496563, EPI_ISL_496564, EPI_ISL_496565, EPI_ISL_496566, EPI_ISL_496567, EPI_ISL_496568, EPI_ISL_496569, EPI_ISL_496570, EPI_ISL_496571, EPI_ISL_496572, EPI_ISL_496573, EPI_ISL_496574, EPI_ISL_496575, EPI_ISL_496576, EPI_ISL_496577, EPI_ISL_496578, EPI_ISL_496579, EPI_ISL_496580, EPI_ISL_496581, EPI_ISL_496582, EPI_ISL_496583, EPI_ISL_496584, EPI_ISL_496585, EPI_ISL_496586, EPI_ISL_496587 | | | |
| see above | National Centre For Cell Science | National Centre For Cell Science | Dhiraj Paul, Kunal Jani, Radha Chauhan, Janesh Kumar, Vasudevan Seshadri, Girdhari Lal, Rajesh Karyakarte, Suvarna Joshi, Murlidhar Tambe, Sourav Sen, Santosh Karade, Kavita Bala Anand, Shelinder Pal Singh Shergill, Rajiv Mohan Gupta, Manoj Kumar Bhat, Arvind Sahu, Maharashtra COVID-19 Study Group, DBT's PAN-INDIA 1000 SARS-CoV2 RNA genome sequencing consortium, Yogesh S Shouche |
| EPI_ISL_496602 | Armed Forces Medical College | National Centre For Cell Science | Dhiraj Paul, Kunal Jani, Radha Chauhan, Janesh Kumar, Vasudevan Seshadri, Girdhari Lal, Rajesh Karyakarte, Suvarna Joshi, Murlidhar Tambe, Sourav Sen, Santosh Karade, Kavita Bala Anand, Shelinder Pal Singh Shergill, Rajiv Mohan Gupta, Manoj Kumar Bhat, Arvind Sahu, Maharashtra COVID-19 Study Group, DBT's PAN-INDIA 1000 SARS-CoV2 RNA genome sequencing consortium, Yogesh S Shouche |
| EPI_ISL_496603, EPI_ISL_496604, EPI_ISL_496605, EPI_ISL_496606, EPI_ISL_496607, EPI_ISL_496608, EPI_ISL_496609, EPI_ISL_496610, EPI_ISL_496611, EPI_ISL_496612, EPI_ISL_496613, EPI_ISL_496614, EPI_ISL_496615, EPI_ISL_496616, EPI_ISL_496617, EPI_ISL_496618, EPI_ISL_496619, EPI_ISL_496620, EPI_ISL_496621, EPI_ISL_496622, EPI_ISL_496623, EPI_ISL_496624, EPI_ISL_496625, EPI_ISL_496626, EPI_ISL_496627, EPI_ISL_496628, EPI_ISL_496629, EPI_ISL_496630, EPI_ISL_496631, EPI_ISL_496632, EPI_ISL_496633, EPI_ISL_496634, EPI_ISL_496635, EPI_ISL_496636, EPI_ISL_496637, EPI_ISL_496638, EPI_ISL_496639, EPI_ISL_496640, EPI_ISL_496641, EPI_ISL_496642, EPI_ISL_496643, EPI_ISL_496644, EPI_ISL_496645, EPI_ISL_496646, EPI_ISL_496647, EPI_ISL_496648, EPI_ISL_496649, EPI_ISL_496650, EPI_ISL_496651, EPI_ISL_496652, EPI_ISL_496653, EPI_ISL_496654, EPI_ISL_496655, EPI_ISL_496656, EPI_ISL_496657, EPI_ISL_496658, EPI_ISL_496659, EPI_ISL_496660, EPI_ISL_496661, EPI_ISL_496662, EPI_ISL_496663, EPI_ISL_496664, EPI_ISL_496665, EPI_ISL_496666, EPI_ISL_496667, EPI_ISL_496668, EPI_ISL_496669, EPI_ISL_496670, EPI_ISL_496671, EPI_ISL_496672, EPI_ISL_496673, EPI_ISL_496674, EPI_ISL_496675, EPI_ISL_496676, EPI_ISL_496677, EPI_ISL_496678, EPI_ISL_496679, EPI_ISL_496680, EPI_ISL_496681, EPI_ISL_496682, EPI_ISL_496683, EPI_ISL_496684, EPI_ISL_496685, EPI_ISL_496686, EPI_ISL_496687, EPI_ISL_496688, EPI_ISL_496689, EPI_ISL_496690, EPI_ISL_496691, EPI_ISL_496692, EPI_ISL_496693, EPI_ISL_496694, EPI_ISL_496695, EPI_ISL_496696, EPI_ISL_496697, EPI_ISL_496698, EPI_ISL_496699, EPI_ISL_496700, EPI_ISL_496701, EPI_ISL_496702, EPI_ISL_496703, EPI_ISL_496704, EPI_ISL_496705, EPI_ISL_496706, EPI_ISL_496707, EPI_ISL_496708, EPI_ISL_496709, EPI_ISL_496710, EPI_ISL_496711, EPI_ISL_496712, EPI_ISL_496713, EPI_ISL_496714, EPI_ISL_496715, EPI_ISL_496716, EPI_ISL_496717, EPI_ISL_496718, EPI_ISL_496719, EPI_ISL_496720, EPI_ISL_496721, EPI_ISL_496722, EPI_ISL_496723, EPI_ISL_496724, EPI_ISL_496725, EPI_ISL_496726, EPI_ISL_496727, EPI_ISL_496728, EPI_ISL_496729, EPI_ISL_496730, EPI_ISL_496731, EPI_ISL_496732, EPI_ISL_496733, EPI_ISL_496734, EPI_ISL_496735, EPI_ISL_496736, EPI_ISL_496737, EPI_ISL_496738, EPI_ISL_496739, EPI_ISL_496740, EPI_ISL_496741, EPI_ISL_496742, EPI_ISL_496743, EPI_ISL_496744, EPI_ISL_496745, EPI_ISL_496746, EPI_ISL_496747, EPI_ISL_496748, EPI_ISL_496749, EPI_ISL_496750, EPI_ISL_496751, EPI_ISL_496752, EPI_ISL_496753, EPI_ISL_496754, EPI_ISL_496755, EPI_ISL_496756, EPI_ISL_496757, EPI_ISL_496758, EPI_ISL_496759, EPI_ISL_496760, EPI_ISL_496761, EPI_ISL_496762, EPI_ISL_496763, EPI_ISL_496764, EPI_ISL_496765, EPI_ISL_496766, EPI_ISL_496767, EPI_ISL_496768, EPI_ISL_496769, EPI_ISL_496770, EPI_ISL_496771, EPI_ISL_496772, EPI_ISL_496773, EPI_ISL_496774, EPI_ISL_496775, EPI_ISL_496776, EPI_ISL_496777, EPI_ISL_496778, EPI_ISL_496779, EPI_ISL_496780, EPI_ISL_496781, EPI_ISL_496782, EPI_ISL_496783, EPI_ISL_496784, EPI_ISL_496785, EPI_ISL_496786, EPI_ISL_496787, EPI_ISL_496788, EPI_ISL_496789, EPI_ISL_496790, EPI_ISL_496791, EPI_ISL_496792, EPI_ISL_496793, EPI_ISL_496794, EPI_ISL_496795, EPI_ISL_496796, EPI_ISL_496797, EPI_ISL_496798, EPI_ISL_496799, EPI_ISL_496800, EPI_ISL_496801, EPI_ISL_496802, EPI_ISL_496803, EPI_ISL_496804, EPI_ISL_496805, EPI_ISL_496806, EPI_ISL_496807, EPI_ISL_496808, EPI_ISL_496809, EPI_ISL_496810, EPI_ISL_496811, EPI_ISL_496812, EPI_ISL_496813, EPI_ISL_496814, EPI_ISL_496815, EPI_ISL_496816, EPI_ISL_496817, EPI_ISL_496818, EPI_ISL_496819, EPI_ISL_496820, EPI_ISL_496821, EPI_ISL_496822, EPI_ISL_496823, EPI_ISL_496824, EPI_ISL_496825, EPI_ISL_496826, EPI_ISL_496827, EPI_ISL_496828, EPI_ISL_496829, EPI_ISL_496830, EPI_ISL_496831, EPI_ISL_496832, EPI_ISL_496833, EPI_ISL_496834, EPI_ISL_496835, EPI_ISL_496836, EPI_ISL_496837, EPI_ISL_496838, EPI_ISL_496839, EPI_ISL_496840, EPI_ISL_496841, EPI_ISL_496842, EPI_ISL_496843, EPI_ISL_496844, EPI_ISL_496845, EPI_ISL_496846, EPI_ISL_496847, EPI_ISL_496848, EPI_ISL_496849, EPI_ISL_496850, EPI_ISL_496851, EPI_ISL_496852, EPI_ISL_496853, EPI_ISL_496854, EPI_ISL_496855, EPI_ISL_496856, EPI_ISL_496857, EPI_ISL_496858, EPI_ISL_496859, EPI_ISL_496860, EPI_ISL_496861, EPI_ISL_496862, EPI_ISL_496863, EPI_ISL_496864, EPI_ISL_496865, EPI_ISL_496866, EPI_ISL_496867, EPI_ISL_496868, EPI_ISL_496869, EPI_ISL_496870, EPI_ISL_496871, EPI_ISL_496872, EPI_ISL_496873, EPI_ISL_496874, EPI_ISL_496875, EPI_ISL_496876, EPI_ISL_496877, EPI_ISL_496878, EPI_ISL_496879, EPI_ISL_496880, EPI_ISL_496881, EPI_ISL_496882, EPI_ISL_496883, EPI_ISL_496884, EPI_ISL_496885, EPI_ISL_496886, EPI_ISL_496887, EPI_ISL_496888, EPI_ISL_496889, EPI_ISL_496890, EPI_ISL_496891, EPI_ISL_496892, EPI_ISL_496893, EPI_ISL_496894, EPI_ISL_496895, EPI_ISL_496896, EPI_ISL_496897, EPI_ISL_496898, EPI_ISL_496899, EPI_ISL_496900, EPI_ISL_496901, EPI_ISL_496902, EPI_ISL_496903, EPI_ISL_496904, EPI_ISL_496905, EPI_ISL_496906, EPI_ISL_496907, EPI_ISL_496908, EPI_ISL_496909, EPI_ISL_496910, EPI_ISL_496911, EPI_ISL_496912, EPI_ISL_496913, EPI_ISL_496914, EPI_ISL_496915 | | | |
| see above | Gorgas Memorial Laboratory of Health Studies | Gorgas Memorial Laboratory of Health Studies | Danilo Franco, Claudia Gonzalez Sandra Lopez-Verges, Alexander A Martinez |
| EPI_ISL_496916, EPI_ISL_496917 | Minnesota Department of Health, Public Health Laboratory | Minnesota Department of Health, Public Health Laboratory | Matt Plumb, Jacob Garfin, and Xiong Wang |
| EPI_ISL_496918 | Mayo Clinic & Mayo Clinic Laboratories | Minnesota Department of Health, Public Health Laboratory | Matt Plumb, Jacob Garfin, and Xiong Wang |
| EPI_ISL_496919, EPI_ISL_496920, EPI_ISL_496921, EPI_ISL_496922, EPI_ISL_496923, EPI_ISL_496924, EPI_ISL_496925, EPI_ISL_496926, EPI_ISL_496927, EPI_ISL_496928, EPI_ISL_496929, EPI_ISL_496930 | | | |
| see above | Minnesota Department of Health, Public Health Laboratory | Minnesota Department of Health, Public Health Laboratory | Matt Plumb, Jacob Garfin, and Xiong Wang |
| EPI_ISL_496931, EPI_ISL_496932, EPI_ISL_496933, EPI_ISL_496934, EPI_ISL_496935, EPI_ISL_496936, EPI_ISL_496937, EPI_ISL_496938, EPI_ISL_496939, EPI_ISL_496940, EPI_ISL_496941, EPI_ISL_496942, EPI_ISL_496943, EPI_ISL_496944, EPI_ISL_496945, EPI_ISL_496946, EPI_ISL_496947, EPI_ISL_496948, EPI_ISL_496949, EPI_ISL_496950, EPI_ISL_496951, EPI_ISL_496952, EPI_ISL_496953, EPI_ISL_496954, EPI_ISL_496955, EPI_ISL_496956, EPI_ISL_496957, EPI_ISL_496958, EPI_ISL_496959, EPI_ISL_496960, EPI_ISL_496961, EPI_ISL_496962, EPI_ISL_496963, EPI_ISL_496964, EPI_ISL_496965, EPI_ISL_496966, EPI_ISL_496967, EPI_ISL_496968, EPI_ISL_496969, EPI_ISL_496970, EPI_ISL_496971, EPI_ISL_496972, EPI_ISL_496973, EPI_ISL_496974, EPI_ISL_496975, EPI_ISL_496976, EPI_ISL_496977, EPI_ISL_496978, EPI_ISL_496979, EPI_ISL_496980, EPI_ISL_496981, EPI_ISL_496982, EPI_ISL_496983, EPI_ISL_496984, EPI_ISL_496985, EPI_ISL_496986, EPI_ISL_496987, EPI_ISL_496988, EPI_ISL_496989, EPI_ISL_496990, EPI_ISL_496991, EPI_ISL_496992, EPI_ISL_496993, EPI_ISL_496994, EPI_ISL_496995, EPI_ISL_496996, EPI_ISL_496997, EPI_ISL_496998, EPI_ISL_496999, EPI_ISL_497000, EPI_ISL_497001, EPI_ISL_497002, EPI_ISL_497003, EPI_ISL_497004, EPI_ISL_497005, EPI_ISL_497006, EPI_ISL_497007, EPI_ISL_497008, EPI_ISL_497009, EPI_ISL_497010, EPI_ISL_497011, EPI_ISL_497012, EPI_ISL_497013, EPI_ISL_497014, EPI_ISL_497015, EPI_ISL_497016, EPI_ISL_497017, EPI_ISL_497018, EPI_ISL_497019, EPI_ISL_497020, EPI_ISL_497021, EPI_ISL_497022, EPI_ISL_497023, EPI_ISL_497024, EPI_ISL_497025, EPI_ISL_497026, EPI_ISL_497027, EPI_ISL_497028, EPI_ISL_497029, EPI_ISL_497030, EPI_ISL_497031, EPI_ISL_497032, EPI_ISL_497033, EPI_ISL_497034, EPI_ISL_497035, EPI_ISL_497036, EPI_ISL_497037, EPI_ISL_497038, EPI_ISL_497039, EPI_ISL_497040, EPI_ISL_497041, EPI_ISL_497042, EPI_ISL_497043, EPI_ISL_497044, EPI_ISL_497045, EPI_ISL_497046, EPI_ISL_497047, EPI_ISL_497048, EPI_ISL_497049, EPI_ISL_497050, EPI_ISL_497051, EPI_ISL_497052, EPI_ISL_497053, EPI_ISL_497054, EPI_ISL_497055, EPI_ISL_497056, EPI_ISL_497057, EPI_ISL_497058, EPI_ISL_497059, EPI_ISL_497060, EPI_ISL_497061, EPI_ISL_497062, EPI_ISL_497063, EPI_ISL_497064, EPI_ISL_497065, EPI_ISL_497066, EPI_ISL_497067, EPI_ISL_497068, EPI_ISL_497069, EPI_ISL_497070, EPI_ISL_497071, EPI_ISL_497072, EPI_ISL_497073, EPI_ISL_497074, EPI_ISL_497075, EPI_ISL_497076, EPI_ISL_497077, EPI_ISL_497078, EPI_ISL_497079, EPI_ISL_497080, EPI_ISL_497081, EPI_ISL_497082, EPI_ISL_497083, EPI_ISL_497084, EPI_ISL_497085, EPI_ISL_497086, EPI_ISL_497087, EPI_ISL_497088, EPI_ISL_497089, EPI_ISL_497090, EPI_ISL_497091, EPI_ISL_497092, EPI_ISL_497093, EPI_ISL_497094, EPI_ISL_497095, EPI_ISL_497096, EPI_ISL_497097, EPI_ISL_497098, EPI_ISL_497099, EPI_ISL_497100, EPI_ISL_497101, EPI_ISL_497102, EPI_ISL_497103, EPI_ISL_497104, EPI_ISL_497105, EPI_ISL_497106, EPI_ISL_497107, EPI_ISL_497108, EPI_ISL_497109, EPI_ISL_497110, EPI_ISL_497111, EPI_ISL_497112, EPI_ISL_497113, EPI_ISL_497114, EPI_ISL_497115, EPI_ISL_497116, EPI_ISL_497117, EPI_ISL_497118, EPI_ISL_497119, EPI_ISL_497120, EPI_ISL_497121, EPI_ISL_497122, EPI_ISL_497123, EPI_ISL_497124, EPI_ISL_497125, EPI_ISL_497126, EPI_ISL_497127, EPI_ISL_497128, EPI_ISL_497129, EPI_ISL_497130, EPI_ISL_497131, EPI_ISL_497132, EPI_ISL_497133, EPI_ISL_497134, EPI_ISL_497135, EPI_ISL_497136, EPI_ISL_497137, EPI_ISL_497138, EPI_ISL_497139, EPI_ISL_497140, EPI_ISL_497141, EPI_ISL_497142, EPI_ISL_497143, EPI_ISL_497144, EPI_ISL_497145, EPI_ISL_497146, EPI_ISL_497147, EPI_ISL_497148, EPI_ISL_497149, EPI_ISL_497150, EPI_ISL_497151, EPI_ISL_497152, EPI_ISL_497153, EPI_ISL_497154, EPI_ISL_497155, EPI_ISL_497156, EPI_ISL_497157, EPI_ISL_497158, EPI_ISL_497159, EPI_ISL_497160, EPI_ISL_497161, EPI_ISL_497162, EPI_ISL_497163, EPI_ISL_497164, EPI_ISL_497165, EPI_ISL_497166, EPI_ISL_497167, EPI_ISL_497168, EPI_ISL_497169, EPI_ISL_497170, EPI_ISL_497171, EPI_ISL_497172, EPI_ISL_497173, EPI_ISL_497174, EPI_ISL_497175, EPI_ISL_497176, EPI_ISL_497177, EPI_ISL_497178, EPI_ISL_497179, EPI_ISL_497180, EPI_ISL_497181, EPI_ISL_497182, EPI_ISL_497183, EPI_ISL_497184, EPI_ISL_497185, EPI_ISL_497186, EPI_ISL_497187, EPI_ISL_497188, EPI_ISL_497189, EPI_ISL_497190, EPI_ISL_497191, EPI_ISL_497192, EPI_ISL_497193, EPI_ISL_497194, EPI_ISL_497195, EPI_ISL_497196, EPI_ISL_497197, EPI_ISL_497198, EPI_ISL_497199, EPI_ISL_497200, EPI_ISL_497201, EPI_ISL_497202, EPI_ISL_497203, EPI_ISL_497204, EPI_ISL_497205, EPI_ISL_497206, EPI_ISL_497207, EPI_ISL_497208, EPI_ISL_497209, EPI_ISL_497210, EPI_ISL_497211, EPI_ISL_497212, EPI_ISL_497213, EPI_ISL_497214, EPI_ISL_497215, EPI_ISL_497216, EPI_ISL_497217, EPI_ISL_497218, EPI_ISL_497219, EPI_ISL_497220, EPI_ISL_497221, EPI_ISL_497222, EPI_ISL_497223, EPI_ISL_497224, EPI_ISL_497225, EPI_ISL_497226, EPI_ISL_497227, EPI_ISL_497228, EPI_ISL_497229, EPI_ISL_497230, EPI_ISL_497231, EPI_ISL_497232, EPI_ISL_497233, EPI_ISL_497234, EPI_ISL_497235, EPI_ISL_497236, EPI_ISL_497237, EPI_ISL_497238, EPI_ISL_497239, EPI_ISL_497240, EPI_ISL_497241, EPI_ISL_497242, EPI_ISL_497243, EPI_ISL_497244, EPI_ISL_497245, EPI_ISL_497246, EPI_ISL_497247, EPI_ISL_497248, EPI_ISL_497249, EPI_ISL_497250, EPI_ISL_497251, EPI_ISL_497252, EPI_ISL_497253, EPI_ISL_497254, EPI_ISL_497255, EPI_ISL_497256, EPI_ISL_497257, EPI_ISL_497258, EPI_ISL_497259, EPI_ISL_497260, EPI_ISL_497261, EPI_ISL_497262, EPI_ISL_497263, EPI_ISL_497264, EPI_ISL_497265, EPI_ISL_497266, EPI_ISL_497267, EPI_ISL_497268, EPI_ISL_497269, EPI_ISL_497270, EPI_ISL_497271, EPI_ISL_497272, EPI_ISL_497273, EPI_ISL_497274, EPI_ISL_497275, EPI_ISL_497276, EPI_ISL_497277, EPI_ISL_497278, EPI_ISL_497279, EPI_ISL_497280, EPI_ISL_497281, EPI_ISL_497282, EPI_ISL_497283, EPI_ISL_497284, EPI_ISL_497285, EPI_ISL_497286, EPI_ISL_497287, EPI_ISL_497288, EPI_ISL_497289, EPI_ISL_497290, EPI_ISL_497291, EPI_ISL_497292, EPI_ISL_497293, EPI_ISL_497294, EPI_ISL_497295, EPI_ISL_497296, EPI_ISL_497297, EPI_ISL_497298, EPI_ISL_497299, EPI_ISL_497300, EPI_ISL_497301, EPI_ISL_497302, EPI_ISL_497303, EPI_ISL_497304, EPI_ISL_497305, EPI_ISL_497306, EPI_ISL_497307, EPI_ISL_497308, EPI_ISL_497309, EPI_ISL_497310, EPI_ISL_497311, EPI_ISL_497312, EPI_ISL_497313, EPI_ISL_497314, EPI_ISL_497315, EPI_ISL_497316, EPI_ISL_497317, EPI_ISL_497318, EPI_ISL_497319, EPI_ISL_497320, EPI_ISL_497321, EPI_ISL_497322, EPI_ISL_497323, EPI_ISL_497324, EPI_ISL_497325, EPI_ISL_497326, EPI_ISL_497327, EPI_ISL_497328, EPI_ISL_497329, EPI_ISL_497330, EPI_ISL_497331, EPI_ISL_497332, EPI_ISL_497333, EPI_ISL_497334, EPI_ISL_497335, EPI_ISL_497336, EPI_ISL_497337, EPI_ISL_497338, EPI_ISL_497339, EPI_ISL_497340, EPI_ISL_497341, EPI_ISL_497342, EPI_ISL_497343, EPI_ISL_497344, EPI_ISL_497345, EPI_ISL_497346, EPI_ISL_497347, EPI_ISL_497348, EPI_ISL_497349, EPI_ISL_497350, EPI_ISL_497351, EPI_ISL_497352, EPI_ISL_497353, EPI_ISL_497354, EPI_ISL_497355, EPI_ISL_497356, EPI_ISL_497357, EPI_ISL_497358, EPI_ISL_497359, EPI_ISL_497360, EPI_ISL_497361, EPI_ISL_497362, EPI_ISL_497363, EPI_ISL_497364, EPI_ISL_497365, EPI_ISL_497366, EPI_ISL_497367, EPI_ISL_497368, EPI_ISL_497369, EPI_ISL_497370, EPI_ISL_497371, EPI_ISL_497372, EPI_ISL_497373, EPI_ISL_497374, EPI_ISL_497375, EPI_ISL_497376, EPI_ISL_497377, EPI_ISL_497378, EPI_ISL_497379, EPI_ISL_497380, EPI_ISL_497381, EPI_ISL_497382, EPI_ISL_497383, EPI_ISL_497384, EPI_ISL_497385, EPI_ISL_497386, EPI_ISL_497387, EPI_ISL_497388, EPI_ISL_497389, EPI_ISL_497390, EPI_ISL_497391, EPI_ISL_497392, EPI_ISL_497393, EPI_ISL_497394, EPI_ISL_497395, EPI_ISL_497396, EPI_ISL_497397, EPI_ISL_497398, EPI_ISL_497399, EPI_ISL_497400, EPI_ISL_497401, EPI_ISL_497402, EPI_ISL_497403, EPI_ISL_497404, EPI_ISL_497405, EPI_ISL_497406, EPI_ISL_497407, EPI_ISL_497408, EPI_ISL_497409, EPI_ISL_497410, EPI_ISL_497411, EPI_ISL_497412, EPI_ISL_497413, EPI_ISL_497414, EPI_ISL_497415, EPI_ISL_497416, EPI_ISL_497417, EPI_ISL_497418, EPI_ISL_497419, EPI_ISL_497420, EPI_ISL_497421, EPI_ISL_497422, EPI_ISL_497423, EPI_ISL_497424, EPI_ISL_497425, EPI_ISL_497426, EPI_ISL_497427, EPI_ISL_497428, EPI_ISL_497429, EPI_ISL_497430, EPI_ISL_497431, EPI_ISL_497432, EPI_ISL_497433, EPI_ISL_497434, EPI_ISL_497435, EPI_ISL_497436, EPI_ISL_497437, EPI_ISL_497438, EPI_ISL_497439, EPI_ISL_497440, EPI_ISL_497441, EPI_ISL_497442, EPI_ISL_497443, EPI_ISL_497444, EPI_ISL_497445, EPI_ISL_497446, EPI_ISL_497447, EPI_ISL_497448, EPI_ISL_497449, EPI_ISL_497450, EPI_ISL_497451, EPI_ISL_497452, EPI_ISL_497453, EPI_ISL_497454, EPI_ISL_497455, EPI_ISL_497456, EPI_ISL_497457, EPI_ISL_497458, EPI_ISL_497459, EPI_ISL_497460, EPI_ISL_497461, EPI_ISL_497462, EPI_ISL_497463, EPI_ISL_497464, EPI_ISL_497465, EPI_ISL_497466, EPI_ISL_497467, EPI_ISL_497468, EPI_ISL_497469, EPI_ISL_497470, EPI_ISL_497471, EPI_ISL_497472, EPI_ISL_497473, EPI_ISL_497474, EPI_ISL_497475, EPI_ISL_497476, EPI_ISL_497477, EPI_ISL_497478, EPI_ISL_497479, EPI_ISL_497480, EPI_ISL_497481, EPI_ISL_497482, EPI_ISL_497483, EPI_ISL_497484, EPI_ISL_497485, EPI_ISL_497486, EPI_ISL_497487, EPI_ISL_497488, EPI_ISL_497489, EPI_ISL_497490, EPI_ISL_497491, EPI_ISL_497492, EPI_ISL_497493, EPI_ISL_497494, EPI_ISL_497495, EPI_ISL_497496, EPI_ISL_497497, EPI_ISL_497498, EPI_ISL_497499, EPI_ISL_497500, EPI_ISL_497501, EPI_ISL_497502, EPI_ISL_497503, EPI_ISL_497504, EPI_ISL_497505, EPI_ISL_497506, EPI_ISL_497507, EPI_ISL_497508, EPI_ISL_497509, EPI_ISL_497510, EPI_ISL_497511, EPI_ISL_497512, EPI_ISL_497513, EPI_ISL_497514, EPI_ISL_497515, EPI_ISL_497516, EPI_ISL_497517, EPI_ISL_497518, EPI_ISL_497519, EPI_ISL_497520, EPI_ISL_497521, EPI_ISL_497522, EPI_ISL_497523, EPI_ISL_497524, EPI_ISL_497525, EPI_ISL_497526, EPI_ISL_497527, EPI_ISL_497528, EPI_ISL_497529, EPI_ISL_497530, EPI_ISL_497531, EPI_ISL_497532, EPI_ISL_497533, EPI_ISL_497534, EPI_ISL_497535, EPI_ISL_497536, EPI_ISL_497537, EPI_ISL_497538, EPI_ISL_497539, EPI_ISL_497540, EPI_ISL_497541, EPI_ISL_497542, EPI_ISL_497543, EPI_ISL_497544, EPI_ISL_497545, EPI_ISL_497546, EPI_ISL_497547, EPI_ISL_497548, EPI_ISL_497549, EPI_ISL_497550, EPI_ISL_497551, EPI_ISL_497552, EPI_ISL_497553, EPI_ISL_497554, EPI_ISL_497555, EPI_ISL_497556, EPI_ISL_497557, EPI_ISL_497558, EPI_ISL_497559, EPI_ISL_497560, EPI_ISL_497561, EPI_ISL_497562, EPI_ISL_497563, EPI_ISL_497564, EPI_ISL_497565, EPI_ISL_497566, EPI_ISL_497567, EPI_ISL_497568, EPI_ISL_497569, EPI_ISL_497570, EPI_ISL_497571, EPI_ISL_497572, EPI_ISL_497573, EPI_ISL_497574, EPI_ISL_497575, EPI_ISL_497576, EPI_ISL_497577, EPI_ISL_497578, EPI_ISL_497579, EPI_ISL_497580, EPI_ISL_497581, EPI_ISL_497582, EPI_ISL_497583, EPI_ISL_497584, EPI_ISL_497585, EPI_ISL_497586, EPI_ISL_497587, EPI_ISL_497588, EPI_ISL_497589, EPI_ISL_497590, EPI_ISL_497591, EPI_ISL_497592, EPI_ISL_497593, EPI_ISL_497594, EPI_ISL_497595, EPI_ISL_497596, EPI_ISL_497597, EPI_ISL_497598, EPI_ISL_497599, EPI_ISL_497600, EPI_ISL_497601, EPI_ISL_497602, EPI_ISL_497603, EPI_ISL_497604, EPI_ISL_497605, EPI_ISL_497606, EPI_ISL_497607, EPI_ISL_497608, EPI_ISL_497609, EPI_ISL_497610, EPI_ISL_497611, EPI_ISL_497612, EPI_ISL_497613, EPI_ISL_497614, EPI_ISL_497615, EPI_ISL_497616, EPI_ISL_497617, EPI_ISL_497618, EPI_ISL_497619, EPI_ISL_497620, EPI_ISL_497621, EPI_ISL_497622, EPI_ISL_497623, EPI_ISL_497624, EPI_ISL_497625, EPI_ISL_497626, EPI_ISL_497627, EPI_ISL_497628, EPI_ISL_497629, EPI_ISL_497630, EPI_ISL_497631, EPI_ISL_497632, EPI_ISL_497633, EPI_ISL_497634, EPI_ISL_497635, EPI_ISL_497636, EPI_ISL_497637, EPI_ISL_497638, EPI_ISL_497639, EPI_ISL_497640, EPI_ISL_497641, EPI_ISL_497642, EPI_ISL_497643, EPI_ISL_497644, EPI_ISL_497645, EPI_ISL_497646, EPI_ISL_497647, EPI_ISL_497648, EPI_ISL_497649, EPI_ISL_497650, EPI_ISL_497651, EPI_ISL_497652, EPI_ISL_497653, EPI_ISL_497654, EPI_ISL_497655, EPI_ISL_497656, EPI_ISL_497657, EPI_ISL_497658, EPI_ISL_497659, EPI_ISL_497660, EPI_ISL_497661, EPI_ISL_497662, EPI_ISL_497663, EPI_ISL_497664, EPI_ISL_497665, EPI_ISL_497666, EPI_ISL_497667, EPI_ISL_497668, EPI_ISL_497669, EPI_ISL_497670, EPI_ISL_497671, EPI_ISL_497672, EPI_ISL_497673, EPI_ISL_497674, EPI_ISL_497675, EPI_ISL_497676, EPI_ISL_497677, EPI_ISL_497678, EPI_ISL_497679, EPI_ISL_497680, EPI_ISL_497681, EPI_ISL_497682, EPI_ISL_497683, EPI_ISL_497684, EPI_ISL_497685, EPI_ISL_497686, EPI_ISL_497687, EPI_ISL_497688, EPI_ISL_497689, EPI_ISL_497690, EPI_ISL_497691, EPI_ISL_497692, EPI_ISL_497693, EPI_ISL_497694, EPI_ISL_497695, EPI_ISL_497696, EPI_ISL_497697, EPI_ISL_497698, EPI_ISL_497699, EPI_ISL_497700, EPI_ISL_497701, EPI_ISL_497702, EPI_ISL_497703, EPI_ISL_497704, EPI_ISL_497705, EPI_ISL_497706, EPI_ISL_497707, EPI_ISL_497708, EPI_ISL_497709, EPI_ISL_497710, EPI_ISL_497711, EPI_ISL_497712, EPI_ISL_497713, EPI_ISL_497714, EPI_ISL_497715, EPI_ISL_497716, EPI_ISL_497717, EPI_ISL_497718, EPI_ISL_497719, EPI_ISL_497720, EPI_ISL_497721, EPI_ISL_497722, EPI_ISL_497723, EPI_ISL_497724, EPI_ISL_497725, EPI_ISL_497726 | | | |
| see above | Washington State Department of Health | Seattle Flu Study | Deborah A. Nickerson, Chris D. Frazar, Jover Lee, Benjamin Pelle, Matthew Richardson, Amanda Adler, Elisabeth Brandstetter, Peter D. Han, Kairsten Fay, Misja Ilcisin, Kirsten Lacombe, Thomas R. Sibley, Melissa Truong, Caitlin R. Wolf, Romesh Gautom, Geoff Melly, Brian Hiatt, Philip Dykema, Scott Lindquist, Michael Boeckh, Janet A. Englund, Michael Famulare, Barry R. Lutz, Mark J. Rieder, Lea M. Starita, Matthew Thompson, Helen Y. Chu, Jay Shendure, Trevor Bedford |
| EPI_ISL_497736, EPI_ISL_497738, EPI_ISL_497744, EPI_ISL_497745 | Instituto Nacional de Salud, Bogotá, Colombia | Instituto Nacional de Salud, Bogotá, Colombia | Katherine Laiton-Donato, Diego A. Álvarez-Díaz, Carlos Franco-Muñoz, Jonathan Reales, Diego Andrés Prada, Jose A. Usme-Ciro, Nicolas D. Franco-Sierra, Zulma M. Cucunubá, Christian Julian Villabona-Arenas, Liz Villabona-Arenas, Sussy Echeverria, Astrid C. Flórez, Carolina Ferro, Diana Marcela Walteros-Acero, Franklin Prieto, Carlos Andrés Durán, Martha Lucia Ospina Martinez, Marcela Mercado-Reyes |
| EPI_ISL_497758, EPI_ISL_497760 | CSIR-CDRI/SGPGI, Lucknow | CSIr-CDRI/SGPGI, Lucknow | Saumya Sarkar, Dharam Veer Singh, Rahul Vishvkarma, Ujjala Ghoshal, Uday Ghoshal, Ravishankar Ramachandran, Tapas Kumar Kundu, Rajender Singh |
| EPI_ISL_497762, EPI_ISL_497763 | CSIR-CDRI/SGPGI, Lucknow | CSIR-CDRI/SGPGI, Lucknow | Saumya Sarkar, Dharam Veer Singh, Rahul Vishvkarma, Ujjala Ghoshal, Uday Ghoshal, Ravishankar Ramachandran, Tapas Kumar Kundu, Rajender Singh |
| EPI_ISL_497764 | CSIR-CDRi/SGPGI, Lucknow | CSIR-CDRI/SGPGI, Lucknow | Saumya Sarkar, Dharam Veer Singh, Rahul Vishvkarma, Ujjala Ghoshal, Uday Ghoshal, Ravishankar Ramachandran, Tapas Kumar Kundu, Rajender Singh |
| EPI_ISL_497765 | CSIR-CDRI/SGPGI, Lucknow | CSIR-CDRi/SGPGI, Lucknow | Saumya Sarkar, Dharam Veer Singh, Rahul Vishvkarma, Ujjala Ghoshal, Uday Ghoshal, Ravishankar Ramachandran, Tapas Kumar Kundu, Rajender Singh |
| EPI_ISL_497766 | CSIR-CDRi/SGPGI, Lucknow | CSIR-CDRi/SGPGI, Lucknow | Saumya Sarkar, Dharam Veer Singh, Rahul Vishvkarma, Ujjala Ghoshal, Uday Ghoshal, Ravishankar Ramachandran, Tapas Kumar Kundu, Rajender Singh |
| EPI_ISL_497767 | CSIR-CDRI/SGPGI, Lucknow | CSIR-CDRI/SGPGI, Lucknow | Saumya Sarkar, Dharam Veer Singh, Rahul Vishvkarma, Ujjala Ghoshal, Uday Ghoshal, Ravishankar Ramachandran, Tapas Kumar Kundu, Rajender Singh |
| EPI_ISL_497768, EPI_ISL_497769, EPI_ISL_497770, EPI_ISL_497771, EPI_ISL_497772, EPI_ISL_497773, EPI_ISL_497774, EPI_ISL_497775, EPI_ISL_497776, EPI_ISL_497777, EPI_ISL_497778, EPI_ISL_497779, EPI_ISL_497780, EPI_ISL_497781, EPI_ISL_497782, EPI_ISL_497783, EPI_ISL_497784, EPI_ISL_497785, EPI_ISL_497786, EPI_ISL_497787, EPI_ISL_497788, EPI_ISL_497789, EPI_ISL_497790, EPI_ISL_497791, EPI_ISL_497792, EPI_ISL_497793, EPI_ISL_497794, EPI_ISL_497795, EPI_ISL_497796, EPI_ISL_497797, EPI_ISL_497798, EPI_ISL_497799, EPI_ISL_497800, EPI_ISL_497801, EPI_ISL_497802, EPI_ISL_497803, EPI_ISL_497804, EPI_ISL_497805, EPI_ISL_497806, EPI_ISL_497807, EPI_ISL_497808, EPI_ISL_497809, EPI_ISL_497810, EPI_ISL_497811, EPI_ISL_497812, EPI_ISL_497813, EPI_ISL_497814, EPI_ISL_497815, EPI_ISL_497816, EPI_ISL_497817, EPI_ISL_497818, EPI_ISL_497819, EPI_ISL_497820, EPI_ISL_497821, EPI_ISL_497822, EPI_ISL_497823, EPI_ISL_497824, EPI_ISL_497825, EPI_ISL_497826, EPI_ISL_497827, EPI_ISL_497828, EPI_ISL_497829, EPI_ISL_497830, EPI_ISL_497831, EPI_ISL_497832, EPI_ISL_497833, EPI_ISL_497834, EPI_ISL_497835, EPI_ISL_497836, EPI_ISL_497837, EPI_ISL_497838, EPI_ISL_497839, EPI_ISL_497840, EPI_ISL_497841, EPI_ISL_497842, EPI_ISL_497843, EPI_ISL_497844, EPI_ISL_497845, EPI_ISL_497846, EPI_ISL_497847, EPI_ISL_497848, EPI_ISL_497849, EPI_ISL_497850, EPI_ISL_497851, EPI_ISL_497852, EPI_ISL_497853, EPI_ISL_497854, EPI_ISL_497855, EPI_ISL_497856, EPI_ISL_497857, EPI_ISL_497858, EPI_ISL_497859, EPI_ISL_497860, EPI_ISL_497861, EPI_ISL_497862, EPI_ISL_497863, EPI_ISL_497864, EPI_ISL_497865, EPI_ISL_497866, EPI_ISL_497867, EPI_ISL_497868, EPI_ISL_497869, EPI_ISL_497870 | | | |
| see above | Department of Microbiology, The University of Hong Kong | Department of Microbiology, The University of Hong Kong | Kelvin K.W. To, Kwok-Yung Yuen |
| EPI_ISL_497872 | UW Virology Lab | UW Virology Lab | Pavitra Roychoudhury, Amin Addetia, Hong Xie, Lasata Shrestha, Truong Nguyen, Meei-Li Huang, Keith Jerome, Alexander Greninger |
| EPI_ISL_497873, EPI_ISL_497874, EPI_ISL_497875, EPI_ISL_497876, EPI_ISL_497877, EPI_ISL_497878, | National Centre For Cell Science | National Centre For Cell Science | Dhiraj Paul, Kunal Jani, Radha Chauhan, Janesh Kumar, Vasudevan Seshadri, Girdhari Lal, Rajesh Karyakarte, Suvarna Joshi, Murlidhar Tambe, Sourav Sen, Santosh Karade, Kavita Bala Anand, Shelinder Pal Singh Shergill, Rajiv Mohan Gupta, Manoj Kumar Bhat, Arvind Sahu, Maharashtra COVID-19 Study Group, DBT's PAN-INDIA 1000 SARS-CoV2 RNA genome sequencing consortium, Yogesh S Shouche |

| | | | |
|---|---|---|---|
| EPI_ISL_497879 | | | |
| EPI_ISL_497880, EPI_ISL_497881, EPI_ISL_497882, EPI_ISL_497883, EPI_ISL_497884, EPI_ISL_497885, EPI_ISL_497886, EPI_ISL_497887 | Armed Forces Medical College | National Centre For Cell Science | Dhiraj Paul, Kunal Jani, Radha Chauhan, Janesh Kumar, Vasudevan Seshadri, Girdhari Lal, Rajesh Karyakarte, Suvarna Joshi, Murlidhar Tambe, Sourav Sen, Santosh Karade, Kavita Bala Anand, Shelinder Pal Singh Shergill, Rajiv Mohan Gupta, Manoj Kumar Bhat, Arvind Sahu, Maharashtra COVID-19 Study Group, DBT's PAN-INDIA 1000 SARS-CoV2 RNA genome sequencing consortium, Yogesh S Shouche |
| EPI_ISL_497888, EPI_ISL_497889, EPI_ISL_497890, EPI_ISL_497891 | B.J. Govt. Medical College | National Centre For Cell Science | Dhiraj Paul, Kunal Jani, Radha Chauhan, Janesh Kumar, Vasudevan Seshadri, Girdhari Lal, Rajesh Karyakarte, Suvarna Joshi, Murlidhar Tambe, Sourav Sen, Santosh Karade, Kavita Bala Anand, Shelinder Pal Singh Shergill, Rajiv Mohan Gupta, Manoj Kumar Bhat, Arvind Sahu, Maharashtra COVID-19 Study Group, DBT's PAN-INDIA 1000 SARS-CoV2 RNA genome sequencing consortium, Yogesh S Shouche |
| EPI_ISL_497950 | Shaoxing CDC | Zhejiang Provincial Center for Disease Control and Prevention | Yin Chen, Yanjun Zhang, Haiyan Mao, Junhang Pan, Xiuyu Lou, Yi Sun, Hao Yan, Zhen Li, Wen Shi |
| EPI_ISL_497951 | Division of Viral Diseases, Center for Laboratory Control of Infectious Diseases, Korea Centers for Diseases Control and Prevention | Division of Viral Diseases, Center for Laboratory Control of Infectious Diseases, Korea Centers for Diseases Control and Prevention | Jeong-Min Kim, Yoon-Seok Chung, Namjoo Lee, Sang Hee Woo, Hye-Jun Jo, Heui Man Kim, Jun-Sub Kim, Myung Guk Han |
| EPI_ISL_497953, EPI_ISL_497954, EPI_ISL_497955, EPI_ISL_497956, EPI_ISL_497957, EPI_ISL_497958, EPI_ISL_497959, EPI_ISL_497960 | Division of Viral Diseases, Center for Laboratory Control of Infectious Diseases, Korea Centers for Diseases Control and Prevention | Division of Viral Diseases, Center for Laboratory Control of Infectious Diseases, Korea Centers for Diseases Control and Prevention | Jeong-Min Kim, Yoon-Seok Chung, Namjoo Lee, Sang Hee Woo, Hye-Jun Jo, Heui Man Kim, Jun-Sub Kim, Dong Hyun Song, Daesang Lee, Seong Tae Jeong, Myung Guk Han |
| EPI_ISL_497961, EPI_ISL_497962, EPI_ISL_497963, EPI_ISL_497964, EPI_ISL_497965, EPI_ISL_497966, EPI_ISL_497967 | Division of Viral Diseases, Center for Laboratory Control of Infectious Diseases, Korea Centers for Diseases Control and Prevention | Division of Viral Diseases, Center for Laboratory Control of Infectious Diseases, Korea Centers for Diseases Control and Prevention | Jeong-Min Kim, Yoon-Seok Chung, Namjoo Lee, Sang Hee Woo, Hye-Jun Jo, Heui Man Kim, Jun-Sub Kim, Myung Guk Han |
| EPI_ISL_497968, EPI_ISL_497969, EPI_ISL_497970, EPI_ISL_497971 | Division of Viral Diseases, Center for Laboratory Control of Infectious Diseases, Korea Centers for Diseases Control and Prevention | Division of Viral Diseases, Center for Laboratory Control of Infectious Diseases, Korea Centers for Diseases Control and Prevention | Jeong-Min Kim, Yoon-Seok Chung, Namjoo Lee, Sang Hee Woo, Hye-Jun Jo, Heui Man Kim, Jun-Sub Kim, Dong Hyun Song, Daesang Lee, Seong Tae Jeong, Myung Guk Han |
| EPI_ISL_497972, EPI_ISL_497973 | Division of Viral Diseases, Center for Laboratory Control of Infectious Diseases, Korea Centers for Diseases Control and Prevention | Division of Viral Diseases, Center for Laboratory Control of Infectious Diseases, Korea Centers for Diseases Control and Prevention | Jeong-Min Kim, Yoon-Seok Chung, Namjoo Lee, Sang Hee Woo, Hye-Jun Jo, Heui Man Kim, Jun-Sub Kim, Myung Guk Han |
| EPI_ISL_497974 | Division of Viral Diseases, Center for Laboratory Control of Infectious Diseases, Korea Centers for Diseases Control and Prevention | Division of Viral Diseases, Center for Laboratory Control of Infectious Diseases, Korea Centers for Diseases Control and Prevention | Jeong-Min Kim, Yoon-Seok Chung, Namjoo Lee, Sang Hee Woo, Hye-Jun Jo, Heui Man Kim, Jun-Sub Kim, Dong Hyun Song, Daesang Lee, Seong Tae Jeong, Myung Guk Han |
| EPI_ISL_497975, EPI_ISL_497976, EPI_ISL_497977, EPI_ISL_497978, EPI_ISL_497979, EPI_ISL_497980, EPI_ISL_497981, EPI_ISL_497982, EPI_ISL_497983, EPI_ISL_497984 | Division of Viral Diseases, Center for Laboratory Control of Infectious Diseases, Korea Centers for Diseases Control and Prevention | Division of Viral Diseases, Center for Laboratory Control of Infectious Diseases, Korea Centers for Diseases Control and Prevention | Jeong-Min Kim, Yoon-Seok Chung, Namjoo Lee, Sang Hee Woo, Hye-Jun Jo, Heui Man Kim, Jun-Sub Kim, Myung Guk Han |
| EPI_ISL_497985 | Division of Viral Diseases, Center for Laboratory Control of Infectious Diseases, Korea Centers for Diseases Control and Prevention | Division of Viral Diseases, Center for Laboratory Control of Infectious Diseases, Korea Centers for Diseases Control and Prevention | Jeong-Min Kim, Yoon-Seok Chung, Namjoo Lee, Sang Hee Woo, Hye-Jun Jo, Heui Man Kim, Jun-Sub Kim, Dong Hyun Song, Daesang Lee, Seong Tae Jeong, Myung Guk Han |
| EPI_ISL_497986 | Division of Viral Diseases, Center for Laboratory Control of Infectious Diseases, Korea Centers for Diseases Control and Prevention | Division of Viral Diseases, Center for Laboratory Control of Infectious Diseases, Korea Centers for Diseases Control and Prevention | Jeong-Min Kim, Yoon-Seok Chung, Namjoo Lee, Sang Hee Woo, Hye-Jun Jo, Heui Man Kim, Jun-Sub Kim, Myung Guk Han |
| EPI_ISL_497987, EPI_ISL_497988, EPI_ISL_497989, EPI_ISL_497990 | Division of Viral Diseases, Center for Laboratory Control of Infectious Diseases, Korea Centers for Diseases Control and Prevention | Division of Viral Diseases, Center for Laboratory Control of Infectious Diseases, Korea Centers for Diseases Control and Prevention | Jeong-Min Kim, Yoon-Seok Chung, Namjoo Lee, Sang Hee Woo, Hye-Jun Jo, Heui Man Kim, Jun-Sub Kim, Dong Hyun Song, Daesang Lee, Seong Tae Jeong, Myung Guk Han |
| EPI_ISL_497991, EPI_ISL_497992, EPI_ISL_497993 | Division of Viral Diseases, Center for Laboratory Control of Infectious Diseases, Korea Centers for Diseases Control and Prevention | Division of Viral Diseases, Center for Laboratory Control of Infectious Diseases, Korea Centers for Diseases Control and Prevention | Jeong-Min Kim, Yoon-Seok Chung, Namjoo Lee, Sang Hee Woo, Hye-Jun Jo, Heui Man Kim, Jun-Sub Kim, Myung Guk Han |
| EPI_ISL_497994, EPI_ISL_497995 | Division of Viral Diseases, Center for Laboratory Control of Infectious Diseases, Korea Centers for Diseases Control and Prevention | Division of Viral Diseases, Center for Laboratory Control of Infectious Diseases, Korea Centers for Diseases Control and Prevention | Jeong-Min Kim, Yoon-Seok Chung, Namjoo Lee, Sang Hee Woo, Hye-Jun Jo, Heui Man Kim, Jun-Sub Kim, Dong Hyun Song, Daesang Lee, Seong Tae Jeong, Myung Guk Han |
| EPI_ISL_497996 | Division of Viral Diseases, Center for Laboratory Control of Infectious Diseases, Korea Centers for Diseases Control and Prevention | Division of Viral Diseases, Center for Laboratory Control of Infectious Diseases, Korea Centers for Diseases Control and Prevention | Jeong-Min Kim, Yoon-Seok Chung, Namjoo Lee, Sang Hee Woo, Hye-Jun Jo, Heui Man Kim, Jun-Sub Kim, Myung Guk Han |
| EPI_ISL_497997 | Division of Viral Diseases, Center for Laboratory Control of Infectious Diseases, Korea Centers for Diseases Control and Prevention | Division of Viral Diseases, Center for Laboratory Control of Infectious Diseases, Korea Centers for Diseases Control and Prevention | Jeong-Min Kim, Yoon-Seok Chung, Namjoo Lee, Sang Hee Woo, Hye-Jun Jo, Heui Man Kim, Jun-Sub Kim, Dong Hyun Song, Daesang Lee, Seong Tae Jeong, Myung Guk Han |
| EPI_ISL_497998, EPI_ISL_497999, EPI_ISL_498000, EPI_ISL_498001, EPI_ISL_498002, EPI_ISL_498003 | Division of Viral Diseases, Center for Laboratory Control of Infectious Diseases, Korea Centers for Diseases Control and Prevention | Division of Viral Diseases, Center for Laboratory Control of Infectious Diseases, Korea Centers for Diseases Control and Prevention | Jeong-Min Kim, Yoon-Seok Chung, Namjoo Lee, Sang Hee Woo, Hye-Jun Jo, Heui Man Kim, Jun-Sub Kim, Myung Guk Han |
| EPI_ISL_498004 | Division of Viral Diseases, Center for Laboratory Control of Infectious Diseases, Korea Centers for Diseases Control and Prevention | Division of Viral Diseases, Center for Laboratory Control of Infectious Diseases, Korea Centers for Diseases Control and Prevention | Jeong-Min Kim, Yoon-Seok Chung, Namjoo Lee, Sang Hee Woo, Hye-Jun Jo, Heui Man Kim, Jun-Sub Kim, Dong Hyun Song, Daesang Lee, Seong Tae Jeong, Myung Guk Han |
| EPI_ISL_498005 | Division of Viral Diseases, Center for Laboratory Control of Infectious Diseases, Korea Centers for Diseases Control and Prevention | Division of Viral Diseases, Center for Laboratory Control of Infectious Diseases, Korea Centers for Diseases Control and Prevention | Jeong-Min Kim, Yoon-Seok Chung, Namjoo Lee, Sang Hee Woo, Hye-Jun Jo, Heui Man Kim, Jun-Sub Kim, Myung Guk Han |
| EPI_ISL_498006, EPI_ISL_498007, EPI_ISL_498008, EPI_ISL_498009, EPI_ISL_498010, EPI_ISL_498011, EPI_ISL_498012, EPI_ISL_498013, EPI_ISL_498014, EPI_ISL_498015, EPI_ISL_498016 | | | |
| see above | Division of Viral Diseases, Center for Laboratory Control of Infectious Diseases, Korea Centers for Diseases Control and Prevention | Division of Viral Diseases, Center for Laboratory Control of Infectious Diseases, Korea Centers for Diseases Control and Prevention | Jeong-Min Kim, Yoon-Seok Chung, Namjoo Lee, Sang Hee Woo, Hye-Jun Jo, Heui Man Kim, Jun-Sub Kim, Dong Hyun Song, Daesang Lee, Seong Tae Jeong, Myung Guk Han |
| EPI_ISL_498017 | Division of Viral Diseases, Center for Laboratory Control of Infectious Diseases, Korea Centers for Diseases Control and Prevention | Division of Viral Diseases, Center for Laboratory Control of Infectious Diseases, Korea Centers for Diseases Control and Prevention | Jeong-Min Kim, Yoon-Seok Chung, Namjoo Lee, Sang Hee Woo, Hye-Jun Jo, Heui Man Kim, Jun-Sub Kim, Myung Guk Han |
| EPI_ISL_498018, EPI_ISL_498019, EPI_ISL_498020, EPI_ISL_498021, EPI_ISL_498022, EPI_ISL_498023, EPI_ISL_498024, EPI_ISL_498025, EPI_ISL_498026, EPI_ISL_498027, EPI_ISL_498028, EPI_ISL_498029, EPI_ISL_498030, EPI_ISL_498031, EPI_ISL_498032, EPI_ISL_498033, EPI_ISL_498034, EPI_ISL_498035 | | | |
| see above | Division of Viral Diseases, Center for Laboratory Control of Infectious Diseases, Korea Centers for Diseases Control and Prevention | Division of Viral Diseases, Center for Laboratory Control of Infectious Diseases, Korea Centers for Diseases Control and Prevention | Jeong-Min Kim, Yoon-Seok Chung, Namjoo Lee, Sang Hee Woo, Hye-Jun Jo, Heui Man Kim, Jun-Sub Kim, Dong Hyun Song, Daesang Lee, Seong Tae Jeong, Myung Guk Han |
| EPI_ISL_498036 | Division of Viral Diseases, Center for Laboratory Control of Infectious Diseases, Korea Centers for Diseases Control and Prevention | Division of Viral Diseases, Center for Laboratory Control of Infectious Diseases, Korea Centers for Diseases Control and Prevention | Jeong-Min Kim, Yoon-Seok Chung, Namjoo Lee, Sang Hee Woo, Hye-Jun Jo, Heui Man Kim, Jun-Sub Kim, Myung Guk Han |
| EPI_ISL_498037, EPI_ISL_498038, EPI_ISL_498039, EPI_ISL_498040, EPI_ISL_498041, EPI_ISL_498042, EPI_ISL_498043, EPI_ISL_498044, EPI_ISL_498045, EPI_ISL_498046, EPI_ISL_498047, EPI_ISL_498048, EPI_ISL_498049, EPI_ISL_498050, EPI_ISL_498051, EPI_ISL_498052, EPI_ISL_498053 | | | |
| see above | Division of Viral Diseases, Center for Laboratory Control of Infectious Diseases, Korea Centers for Diseases Control and Prevention | Division of Viral Diseases, Center for Laboratory Control of Infectious Diseases, Korea Centers for Diseases Control and Prevention | Jeong-Min Kim, Yoon-Seok Chung, Namjoo Lee, Sang Hee Woo, Hye-Jun Jo, Heui Man Kim, Jun-Sub Kim, Dong Hyun Song, Daesang Lee, Seong Tae Jeong, Myung Guk Han |
| EPI_ISL_498054, EPI_ISL_498055, EPI_ISL_498056, EPI_ISL_498057, EPI_ISL_498058, EPI_ISL_498059, EPI_ISL_498060, EPI_ISL_498061, EPI_ISL_498062, EPI_ISL_498063, EPI_ISL_498064, EPI_ISL_498065, EPI_ISL_498066, EPI_ISL_498067, EPI_ISL_498068, EPI_ISL_498069, EPI_ISL_498070, EPI_ISL_498071, EPI_ISL_498072, EPI_ISL_498073, EPI_ISL_498074, EPI_ISL_498075, EPI_ISL_498076, EPI_ISL_498077, EPI_ISL_498078, EPI_ISL_498079, EPI_ISL_498080, EPI_ISL_498081, EPI_ISL_498082, EPI_ISL_498083, EPI_ISL_498084, EPI_ISL_498085, EPI_ISL_498086, EPI_ISL_498087, EPI_ISL_498088, EPI_ISL_498089, EPI_ISL_498090, EPI_ISL_498091, EPI_ISL_498092, EPI_ISL_498093, EPI_ISL_498094, EPI_ISL_498095, EPI_ISL_498096, EPI_ISL_498097, EPI_ISL_498098, EPI_ISL_498099, EPI_ISL_498100, EPI_ISL_498101, EPI_ISL_498102, EPI_ISL_498103, EPI_ISL_498104, EPI_ISL_498105, EPI_ISL_498106, EPI_ISL_498107, EPI_ISL_498108, EPI_ISL_498109, EPI_ISL_498110, EPI_ISL_498111, EPI_ISL_498112, EPI_ISL_498113, EPI_ISL_498114, EPI_ISL_498115, EPI_ISL_498116, EPI_ISL_498117, EPI_ISL_498118, EPI_ISL_498119, EPI_ISL_498120, EPI_ISL_498121, EPI_ISL_498122, EPI_ISL_498123, EPI_ISL_498124, EPI_ISL_498125, EPI_ISL_498126 | | | |
| see above | NHLS-IALCH | KRISP, KZN Research Innovation and Sequencing Platform | Giandhari J, Pillay S, Lessells R, Chimukangara B, Mdlalose K, York D, Khan S, Tegally H, Wilkinson E, de Oliveira T |
| EPI_ISL_498127, EPI_ISL_498128, EPI_ISL_498129, EPI_ISL_498130, EPI_ISL_498131, EPI_ISL_498132, EPI_ISL_498133, EPI_ISL_498134, EPI_ISL_498135, EPI_ISL_498136, EPI_ISL_498137, EPI_ISL_498138, EPI_ISL_498139, EPI_ISL_498140, EPI_ISL_498141, EPI_ISL_498142, EPI_ISL_498143, EPI_ISL_498144, EPI_ISL_498145, EPI_ISL_498146, EPI_ISL_498147, EPI_ISL_498148, EPI_ISL_498149, EPI_ISL_498150, EPI_ISL_498151 | | | |
| see above | Department of Clinical Microbiology | GIGA Medical Genomics | Keith Durkin, Maria Artesi, Sébastien Bontems, Raphaël Boreux, Cécile Meex, Axelle Chaslain, Céline Fombellida-Lopez, Pierrette Melin, Marie-Pierre Hayette, Vincent Bours. |
| EPI_ISL_498152, EPI_ISL_498153, EPI_ISL_498154, EPI_ISL_498155, EPI_ISL_498156, EPI_ISL_498157, EPI_ISL_498158, EPI_ISL_498159, EPI_ISL_498160, EPI_ISL_498161, EPI_ISL_498162, EPI_ISL_498163, EPI_ISL_498164, EPI_ISL_498165, EPI_ISL_498166, EPI_ISL_498167, EPI_ISL_498168, EPI_ISL_498169, EPI_ISL_498170 | | | |
| see above | Instituto Nacional de Salud, Bogotá, Colombia | Instituto Nacional de Salud, Bogotá, Colombia | Katherine Laiton-Donato, Diego A. Álvarez-Díaz, Carlos Franco-Muñoz, Jonathan Reales, Diego Andrés Prada, Jose A. Usme-Ciro, Nicolas D. Franco-Sierra, Zulma M. Cucunubá, Christian Julian Villabona-Arenas, Liz Villabona-Arenas, Sussy Echeverria, Astrid C. Flórez, Carolina Ferro, Diana Marcela Walteros-Acero, Franklin Prieto, Carlos Andrés Durán, Martha Lucia Ospina Martinez, Marcela Mercado-Reyes |
| EPI_ISL_498171, EPI_ISL_498172, EPI_ISL_498173, EPI_ISL_498174, EPI_ISL_498175, EPI_ISL_498176, EPI_ISL_498177, EPI_ISL_498178, EPI_ISL_498179, EPI_ISL_498180, EPI_ISL_498181, EPI_ISL_498182, EPI_ISL_498183, EPI_ISL_498184, EPI_ISL_498185, EPI_ISL_498186, EPI_ISL_498187, EPI_ISL_498188, EPI_ISL_498189, EPI_ISL_498190, EPI_ISL_498191, EPI_ISL_498192 | | | |
| see above | OUCRU | OUCRU | Nguyen Van Vinh Chau, Nguyen Thi Thu Hong, Nguyen Thi Han Ny, Le Nguyen Truc Nhu, Nghiem My Ngoc, Vo Thanh Lam, Nguyen Thanh Dung, Lam Minh Yen, Ngo Ngoc Quang Minh, Le Manh Hung, Nguyen Tri Dung, Dinh Nguyen Huy Man, Lam Anh Nguyet, Tran Chanh Xuan, Tran Tinh Hien, Nguyen Thanh Phong, Tran Nguyen Hoang Tu, Tran Tan Thanh, Nguyen Thanh Truong, Nguyen Tan Binh, Tang Chi Thuong, Guy Thwaites, and Le Van Tan, for OUCRU COVID-19 research group* |
| EPI_ISL_498193 | Molecular Pathology Division, Department of Pathology, Hong Kong Sanatorium & Hospital | Molecular Pathology Division, Department of Pathology, Hong Kong Sanatorium & Hospital | Chun Hang AU, Wai Sing CHAN, Ho Yin LAM, Dona N. HO, Simon Y.M. LAM, Jonpaul S.T. ZEE, Tsun Leung CHAN, Edmond S.K. MA |
| EPI_ISL_498226 | LIC | LIC | LIC |
| EPI_ISL_498227, EPI_ISL_498228 | National Institute of Laboratory Medicine and Referral Center | Genomic Research Lab, BCSIR | Shahina Akter, Abu Sayeed Mohammad Mahmud, Mohammad Samir Uzzaman, Eshrar Osman, Md. Ahasan Habib, Tanjina Akhter Banu, Md. Murshed Hasan Sarkar, Barna Goswami, Iffat Jahan, Md. Saddam Hossain, Tasnim Nafisa, Md. Maruf Ahmed Molla, Mahmuda Yeasmin, Asish Kumar Ghosh, A. K. M. Shamsuzzaman, Sheikh Md. Selim Al Din, Utpal Chandra Ray, Salek Ahmed Sajib, Md. Salim Khan |
| EPI_ISL_498229, EPI_ISL_498230, EPI_ISL_498231, EPI_ISL_498232, EPI_ISL_498233, EPI_ISL_498234, EPI_ISL_498235, EPI_ISL_498236, EPI_ISL_498237, EPI_ISL_498238, EPI_ISL_498239, EPI_ISL_498240, EPI_ISL_498241, EPI_ISL_498242, EPI_ISL_498243, EPI_ISL_498244, EPI_ISL_498245, EPI_ISL_498246, EPI_ISL_498247, EPI_ISL_498248, EPI_ISL_498249, EPI_ISL_498250, EPI_ISL_498251, EPI_ISL_498252 | | | |
| see above | Institut Pasteur de Dakar | Institut Pasteur de Dakar | Ndongo Dia, Moussa Moise Diagne, Mamadou Diop, Marie Henriette Dior Ndione, Mamadou Malado Jallow, Safietou Sankhe Mbengue, Ousmane Faye, Amadou Alpha Sall. |
| EPI_ISL_498253, EPI_ISL_498254 | National Institute of Laboratory Medicine and Referral Center | Genomic Research Lab, BCSIR | Tanjina Akhter Banu, Abu Sayeed Mohammad Mahmud, Mohammad Samir Uzzaman, Eshrar Osman, Md. Ahasan Habib, Shahina Akter, Md. Murshed Hasan Sarkar, Barna Goswami, Iffat Jahan, Md. Saddam Hossain, Tasnim Nafisa, Md. Maruf Ahmed Molla, Mahmuda Yeasmin, Asish Kumar Ghosh, A. K. M. Shamsuzzaman, Sheikh Md. Selim Al Din, Utpal Chandra Ray, Salek Ahmed Sajib, Md. Salim Khan |

| | | | |
|---|---|---|---|
| EPI_ISL_498255, EPI_ISL_498256, EPI_ISL_498257, EPI_ISL_498258, EPI_ISL_498259, EPI_ISL_498260, EPI_ISL_498261 | Hospital for Tropical Diseases | COVID-19 Network Investigations (CONI) Alliance | Elizabeth Batty, Nantarat Chantawat, Wasun Chantratita, Thanat Chookajorn, Stefan Fernandez, Angkana Huang, Weena Janwitthayanan, Akanitt Jittmittraphap, Anthony R. Jones, Khajohn Joonsalak, Chonticha Klungtong, Theerarat Kochakarn, Namfon Kotanan, Krittikorn Kumpornsin, Pornsawan Leaungwutiwong, Wudtichai Manasatienkij, Bhakbhoom Panthan, Ekawat Pasomsub, Kingkan Rakmanee, Insee Sensorn, Janjira Thaipadungpanit, Arporn Wangwiwatsin,Treewat Watthanachockchai |
| EPI_ISL_498262, EPI_ISL_498263, EPI_ISL_498264, EPI_ISL_498265, EPI_ISL_498266 | Ramathibodi Hospital | COVID-19 Network Investigations (CONI) Alliance | Elizabeth Batty, Wasun Chantratita, Thanat Chookajorn, Stefan Fernandez, Angkana Huang, Anthony R. Jones, Khajohn Joonsalak, Chonticha Klungtong, Theerarat Kochakarn, Namfon Kotanan, Krittikorn Kumpornsin, Wudtichai Manasatienkij, Bhakbhoom Panthan, Ekawat Pasomsub, Kingkan Rakmanee, Insee Sensorn, Janjira Thaipadungpanit, Arporn Wangwiwatsin,Treewat Watthanachockchai |
| EPI_ISL_498267, EPI_ISL_498268 | National Institute of Laboratory Medicine and Referral Center | Genomic Research Lab, BCSIR | Barna Goswami, Abu Sayeed Mohammad Mahmud, Mohammad Samir Uzzaman, Eshrar Osman, Md. Ahasan Habib, Shahina Akter, Tanjina Akhter Banu, Md. Murshed Hasan Sarkar, Iffat Jahan, Md. Saddam Hossain, Tasnim Nafisa, Md. Maruf Ahmed Molla, Mahmuda Yeasmin, Asish Kumar Ghosh, A. K. M. Shamsuzzaman, Sheikh Md. Selim Al Din, Utpal Chandra Ray, Salek Ahmed Sajib, Md. Salim Khan |
| EPI_ISL_498269, EPI_ISL_498270, EPI_ISL_498271 | Department of Microbiology, The University of Hong Kong | Department of Microbiology, The University of Hong Kong | Kelvin K.W. To, Kwok-Yung Yuen |
| EPI_ISL_498272, EPI_ISL_498273 | National Institute of Laboratory Medicine and Referral Center | Genomic Research Lab, BCSIR | Iffat Jahan, Abu Sayeed Mohammad Mahmud, Mohammad Samir Uzzaman, Eshrar Osman, Md. Ahasan Habib, Shahina Akter, Tanjina Akhter Banu, Md. Murshed Hasan Sarkar, Barna Goswami, Md. Saddam Hossain, Tasnim Nafisa, Md. Maruf Ahmed Molla, Mahmuda Yeasmin, Asish Kumar Ghosh, A. K. M. Shamsuzzaman, Sheikh Md. Selim Al Din, Utpal Chandra Ray, Salek Ahmed Sajib, Md. Salim Khan |
| EPI_ISL_498274, EPI_ISL_498417 | National Institute of Laboratory Medicine and Referral Center | Genomic Research Lab, BCSIR | Tasnim Nafisa, Abu Sayeed Mohammad Mahmud, Mohammad Samir Uzzaman, Eshrar Osman, Md. Ahasan Habib, Shahina Akter, Tanjina Akhter Banu, Md. Murshed Hasan Sarkar, Barna Goswami, Iffat Jahan, Md. Saddam Hossain, Md. Maruf Ahmed Molla, Mahmuda Yeasmin, Asish Kumar Ghosh, A. K. M. Shamsuzzaman, Sheikh Md. Selim Al Din, Utpal Chandra Ray, Salek Ahmed Sajib, Md. Salim Khan |
| EPI_ISL_498418, EPI_ISL_498419 | National Institute of Laboratory Medicine and Referral Center | Genomic Research Lab, BCSIR | Md. Maruf Ahmed Molla, Abu Sayeed Mohammad Mahmud, Mohammad Samir Uzzaman, Eshrar Osman, Md. Ahasan Habib, Shahina Akter, Tanjina Akhter Banu, Md. Murshed Hasan Sarkar, Barna Goswami, Iffat Jahan, Md. Saddam Hossain, Tasnim Nafisa, Mahmuda Yeasmin, Asish Kumar Ghosh, A. K. M. Shamsuzzaman, Sheikh Md. Selim Al Din, Utpal Chandra Ray, Salek Ahmed Sajib, Md. Salim Khan |
| EPI_ISL_498466, EPI_ISL_498467 | National Institute of Laboratory Medicine and Referral Center | Genomic Research Lab, BCSIR | Mahmuda Yeasmin, Abu Sayeed Mohammad Mahmud, Mohammad Samir Uzzaman, Eshrar Osman, Md. Ahasan Habib, Shahina Akter, Tanjina Akhter Banu, Md. Murshed Hasan Sarkar, Barna Goswami, Iffat Jahan, Md. Saddam Hossain, Tasnim Nafisa, Md. Maruf Ahmed Molla, Asish Kumar Ghosh, A. K. M. Shamsuzzaman, Sheikh Md. Selim Al Din, Utpal Chandra Ray, Salek Ahmed Sajib, Md. Salim Khan |
| EPI_ISL_498468, EPI_ISL_498469, EPI_ISL_498470, EPI_ISL_498471, EPI_ISL_498472, EPI_ISL_498473, EPI_ISL_498474, EPI_ISL_498475, EPI_ISL_498476, EPI_ISL_498477, EPI_ISL_498478, EPI_ISL_498479, EPI_ISL_498480, EPI_ISL_498481, EPI_ISL_498482, EPI_ISL_498483, EPI_ISL_498484, EPI_ISL_498485, EPI_ISL_498486, EPI_ISL_498487, EPI_ISL_498488, EPI_ISL_498489, EPI_ISL_498490, EPI_ISL_498491, EPI_ISL_498492, EPI_ISL_498493, EPI_ISL_498494, EPI_ISL_498495, EPI_ISL_498496, EPI_ISL_498497, EPI_ISL_498498, EPI_ISL_498499, EPI_ISL_498500, EPI_ISL_498501, EPI_ISL_498502, EPI_ISL_498503, EPI_ISL_498504, EPI_ISL_498505, EPI_ISL_498506, EPI_ISL_498507, EPI_ISL_498508, EPI_ISL_498509, EPI_ISL_498510, EPI_ISL_498511, EPI_ISL_498512, EPI_ISL_498513, EPI_ISL_498514, EPI_ISL_498515, EPI_ISL_498516, EPI_ISL_498517, EPI_ISL_498518, EPI_ISL_498519, EPI_ISL_498520, EPI_ISL_498521, EPI_ISL_498522, EPI_ISL_498523, EPI_ISL_498524, EPI_ISL_498525, EPI_ISL_498526, EPI_ISL_498527, EPI_ISL_498528, EPI_ISL_498529, EPI_ISL_498530, EPI_ISL_498531, EPI_ISL_498532, EPI_ISL_498533, EPI_ISL_498534, EPI_ISL_498535, EPI_ISL_498536, EPI_ISL_498537, EPI_ISL_498538, EPI_ISL_498539, EPI_ISL_498540, EPI_ISL_498541, EPI_ISL_498542, EPI_ISL_498543, EPI_ISL_498544, EPI_ISL_498545, EPI_ISL_498546, EPI_ISL_498547, EPI_ISL_498548 | | | |
| see above | ACT Pathology | Schwessinger Lab | Ashley Jones, Benjamin Schwessinger, Robert Lanfear, Robyn N Hall, Megan McDonald, Ming-Dao Chia, Kevin Murray, Craig Kennedy, Karina Kennedy |
| EPI_ISL_498549, EPI_ISL_498550 | National Institute of Laboratory Medicine and Referral Center | Genomic Research Lab, BCSIR | Asish Kumar Ghosh, Abu Sayeed Mohammad Mahmud, Mohammad Samir Uzzaman, Eshrar Osman, Md. Ahasan Habib, Shahina Akter, Tanjina Akhter Banu, Md. Murshed Hasan Sarkar, Barna Goswami, Iffat Jahan, Md. Saddam Hossain, Tasnim Nafisa, Md. Maruf Ahmed Molla, Mahmuda Yeasmin, A. K. M. Shamsuzzaman, Sheikh Md. Selim Al Din, Utpal Chandra Ray, Salek Ahmed Sajib, Md. Salim Khan |
| EPI_ISL_498551, EPI_ISL_498552, EPI_ISL_498554, EPI_ISL_498556 | Lebanese American University | Lebanese American University | Abi Habib,W., Abdallah,J., El Shesheny,R., Mokhbat,J., Webby,R.J., Goldstein,J. and Kayali,G. |
| EPI_ISL_498558, EPI_ISL_498559, EPI_ISL_498560, EPI_ISL_498561 | Laboratory of Molecular Virology International Center for Genetic Engineering and Biotechnology (ICGEB) | ARGO Open Lab Platform for Genome Sequencing | Licastro D, Rajasekharan S, Dal Monego S, Segat L, D'Agaro P, Marcello A |
| EPI_ISL_498562 | Laboratory of Molecular Virology International Center for Genetic Engineering and Biotechnology (ICGEB) | ARGO Open Lab Platform for Genome Sequencing | Licastro D, Rajasekharan S, Dal Monego S, Segat L, D'Agaro P, Marcello A |
| EPI_ISL_498563 | Laboratory of Molecular Virology International Center for Genetic Engineering and Biotechnology (ICGEB) | ARGO Open Lab Platform for Genome Sequencing | Licastro D, Rajasekharan S, Dal Monego S, Segat L, D'Agaro P, Marcello A |
| EPI_ISL_498564, EPI_ISL_498565, EPI_ISL_498566, EPI_ISL_498567, EPI_ISL_498568, EPI_ISL_498569, EPI_ISL_498570, EPI_ISL_498571, EPI_ISL_498572, EPI_ISL_498573, EPI_ISL_498574, EPI_ISL_498575, EPI_ISL_498576, EPI_ISL_498577, EPI_ISL_498578, EPI_ISL_498579, EPI_ISL_498580, EPI_ISL_498581, EPI_ISL_498582, EPI_ISL_498583, EPI_ISL_498584, EPI_ISL_498585, EPI_ISL_498586, EPI_ISL_498587, EPI_ISL_498588, EPI_ISL_498589, EPI_ISL_498590, EPI_ISL_498591, EPI_ISL_498592, EPI_ISL_498593, EPI_ISL_498594, EPI_ISL_498595, EPI_ISL_498596, EPI_ISL_498597, EPI_ISL_498598, EPI_ISL_498599, EPI_ISL_498600, EPI_ISL_498601, EPI_ISL_498602, EPI_ISL_498603, EPI_ISL_498604, EPI_ISL_498605, EPI_ISL_498606, EPI_ISL_498607, EPI_ISL_498608, EPI_ISL_498609, EPI_ISL_498610, EPI_ISL_498611, EPI_ISL_498612, EPI_ISL_498613, EPI_ISL_498614, EPI_ISL_498615, EPI_ISL_498616, EPI_ISL_498617, EPI_ISL_498618 | | | |
| see above | National Public Health Laboratory, National Centre for Infectious Diseases | National Public Health Laboratory, National Centre for Infectious Diseases | Mak TM, Octavia S, Zhou Z, Chavatte JM, Cui L, Lin RTP |
| EPI_ISL_498619, EPI_ISL_498620, EPI_ISL_498621, EPI_ISL_498622, EPI_ISL_498623, EPI_ISL_498624, EPI_ISL_498625, EPI_ISL_498626, EPI_ISL_498627 | Viollier AG | Department of Biosystems Science and Engineering, ETH Zürich | Christian Beisel, Sarah Nadeau, Ivan Topolsky, Pedro Ferreira, Philipp Jablonski, Susana Posada-Céspedes, Tobias Schär, Ina Nissen, Natascha Santacroce, Elodie Burcklen, Christiane Beckmann, Maurice Redondo, Olivier Kobel, Christoph Noppen, Sophie Seidel, Noemie Santamaria de Souza, Niko Beerenwinkel, Tanja Stadler |
| EPI_ISL_498628, EPI_ISL_498629 | Department of Clinical Microbiology | GIGA Medical Genomics | Keith Durkin, Maria Artesi, Sébastien Bontems, Raphaël Boreux, Cécile Meex, Axelle Chaslain, Céline Fombellida-Lopez, Pierrette Melin, Marie-Pierre Hayette, Vincent Bours. |
| EPI_ISL_498630, EPI_ISL_498631, EPI_ISL_498632, EPI_ISL_498633, EPI_ISL_498634, EPI_ISL_498635, EPI_ISL_498637, EPI_ISL_498638, EPI_ISL_498639, EPI_ISL_498640, EPI_ISL_498641, EPI_ISL_498642, EPI_ISL_498643, EPI_ISL_498644, EPI_ISL_498645, EPI_ISL_498646, EPI_ISL_498647, EPI_ISL_498648, EPI_ISL_498649, EPI_ISL_498650, EPI_ISL_498651, EPI_ISL_498652, EPI_ISL_498653, EPI_ISL_498654, EPI_ISL_498655, EPI_ISL_498656, EPI_ISL_498657, EPI_ISL_498658, EPI_ISL_498659, EPI_ISL_498660, EPI_ISL_498661, EPI_ISL_498662, EPI_ISL_498663, EPI_ISL_498664, EPI_ISL_498665, EPI_ISL_498666, EPI_ISL_498667, EPI_ISL_498668, EPI_ISL_498669, EPI_ISL_498670, EPI_ISL_498671, EPI_ISL_498672, EPI_ISL_498673, EPI_ISL_498674, EPI_ISL_498675, EPI_ISL_498676, EPI_ISL_498677, EPI_ISL_498678, EPI_ISL_498679, EPI_ISL_498680, EPI_ISL_498681, EPI_ISL_498682, EPI_ISL_498683, EPI_ISL_498684, EPI_ISL_498685, EPI_ISL_498686, EPI_ISL_498687, EPI_ISL_498688, EPI_ISL_498689, EPI_ISL_498690 | | | |
| see above | Utah Public Health Laboratory | Utah Public Health Laboratory | Heidi Butz, Erin Young, Kelly Oakeson |
| EPI_ISL_498691, EPI_ISL_498692, EPI_ISL_498693, EPI_ISL_498694 | National Institute for Viral Disease Control and Prevention, China CDC | National Institute for Viral Disease Control and Prevention, China CDC | Xiang Zhao,LingLing Mao,Yao Meng,Zhixiao Chen,Yuchao Wu,Yong ZhangBo ZhijianJianqun Zhang,Yang Song,Dayan Wang,WenQing YaoWenbo Xu |
| EPI_ISL_498695, EPI_ISL_498696, EPI_ISL_498697, EPI_ISL_498698, EPI_ISL_498699, EPI_ISL_498700, EPI_ISL_498701, EPI_ISL_498702, EPI_ISL_498703, EPI_ISL_498704, EPI_ISL_498705, EPI_ISL_498706, EPI_ISL_498707, EPI_ISL_498708, EPI_ISL_498709, EPI_ISL_498710, EPI_ISL_498711, EPI_ISL_498712, EPI_ISL_498713, EPI_ISL_498714, EPI_ISL_498715, EPI_ISL_498716, EPI_ISL_498717, EPI_ISL_498718, EPI_ISL_498719, EPI_ISL_498720, EPI_ISL_498721, EPI_ISL_498722, EPI_ISL_498723, EPI_ISL_498724, EPI_ISL_498725, EPI_ISL_498726, EPI_ISL_498727, EPI_ISL_498728, EPI_ISL_498729, EPI_ISL_498730, EPI_ISL_498731, EPI_ISL_498732, EPI_ISL_498733, EPI_ISL_498734, EPI_ISL_498735, EPI_ISL_498736, EPI_ISL_498737, EPI_ISL_498738, EPI_ISL_498739, EPI_ISL_498740, EPI_ISL_498741, EPI_ISL_498742, EPI_ISL_498743, EPI_ISL_498744, EPI_ISL_498745, EPI_ISL_498746, EPI_ISL_498747 | | | |
| see above | Quest Diagnostics | Quest Diagnostics | Rosenthal,S.H., Gerasimova,A., Kagan,R.M. and Owen, R. |
| EPI_ISL_498748, EPI_ISL_498749, EPI_ISL_498750 | Pathology West - NSW Health Pathology | NSW Health Pathology - Institute of Clinical Pathology and Medical Research; Westmead Hospital; University of Sydney | CIDM-PH et al. |
| EPI_ISL_498751 | South Eastern Area Laboratory Services (SEALS) | NSW Health Pathology - Institute of Clinical Pathology and Medical Research; Westmead Hospital; University of Sydney | CIDM-PH et al. |
| EPI_ISL_498761, EPI_ISL_498762 | Pathology West - NSW Health Pathology | NSW Health Pathology - Institute of Clinical Pathology and Medical Research; Westmead Hospital; University of Sydney | CIDM-PH et al. |
| EPI_ISL_498763, EPI_ISL_498764 | Sydney South West Pathology Service (SSWPS) - Liverpool Hospital - NSW Health Pathology | NSW Health Pathology - Institute of Clinical Pathology and Medical Research; Westmead Hospital; University of Sydney | CIDM-PH et al. |
| EPI_ISL_498765 | Sydney South West Pathology Service (SSWPS) - Concord Repatriation General Hospital - NSW Health Pathology | NSW Health Pathology - Institute of Clinical Pathology and Medical Research; Westmead Hospital; University of Sydney | CIDM-PH et al. |
| EPI_ISL_498766, EPI_ISL_498767 | Pathology West - NSW Health Pathology | NSW Health Pathology - Institute of Clinical Pathology and Medical Research; Westmead Hospital; University of Sydney | CIDM-PH et al. |
| EPI_ISL_498768 | South Eastern Area Laboratory Services (SEALS) | NSW Health Pathology - Institute of Clinical Pathology and Medical Research; Westmead Hospital; University of Sydney | CIDM-PH et al. |
| EPI_ISL_498769, EPI_ISL_498770, EPI_ISL_498771, EPI_ISL_498772 | Laverty Pathology | NSW Health Pathology - Institute of Clinical Pathology and Medical Research; Westmead Hospital; University of Sydney | CIDM-PH et al. |
| EPI_ISL_498773 | Douglass Hanly Moir Pathology | NSW Health Pathology - Institute of Clinical Pathology and Medical Research; Westmead Hospital; University of Sydney | CIDM-PH et al. |
| EPI_ISL_498774, EPI_ISL_498775 | Histopath | NSW Health Pathology - Institute of Clinical Pathology and Medical Research; Westmead Hospital; University of Sydney | CIDM-PH et al. |
| EPI_ISL_498776, EPI_ISL_498777 | Pathology West - NSW Health Pathology | NSW Health Pathology - Institute of Clinical Pathology and Medical Research; Westmead Hospital; University of Sydney | CIDM-PH et al. |
| EPI_ISL_498778 | Sydney South West Pathology Service (SSWPS) - Liverpool Hospital - NSW Health Pathology | NSW Health Pathology - Institute of Clinical Pathology and Medical Research; Westmead Hospital; University of Sydney | CIDM-PH et al. |
| EPI_ISL_498779, EPI_ISL_498780 | Pathology West - NSW Health Pathology | NSW Health Pathology - Institute of Clinical Pathology and Medical Research; Westmead Hospital; University of Sydney | CIDM-PH et al. |
| EPI_ISL_498781, EPI_ISL_498782 | Sydney South West Pathology Service (SSWPS) - Liverpool Hospital - NSW Health Pathology | NSW Health Pathology - Institute of Clinical Pathology and Medical Research; Westmead Hospital; University of Sydney | CIDM-PH et al. |
| EPI_ISL_498783, EPI_ISL_498784, EPI_ISL_498785, EPI_ISL_498786, EPI_ISL_498787, EPI_ISL_498788, EPI_ISL_498789, EPI_ISL_498790, EPI_ISL_498791, EPI_ISL_498792 | National Institute of Laboratory Medicine and Referral Center | Genomic Research Lab, BCSIR | Md. Saddam Hossain, Abu Sayeed Mohammad Mahmud, Mohammad Samir Uzzaman, Eshrar Osman, Md. Ahasan Habib, Shahina Akter, Tanjina Akhter Banu, Md. Murshed Hasan Sarkar, Barna Goswami, Iffat Jahan, Tasnim Nafisa, Md. Maruf Ahmed Molla, Mahmuda Yeasmin, Asish Kumar Ghosh, A. K. M. Shamsuzzaman, Sheikh Md. Selim Al Din, Utpal Chandra Ray, Salek Ahmed Sajib, Md. Salim Khan |

| | | | |
|---|---|---|---|
| EPI_ISL_498793, EPI_ISL_498794, EPI_ISL_498795, EPI_ISL_498796, EPI_ISL_498797, EPI_ISL_498800, EPI_ISL_498801, EPI_ISL_498802, EPI_ISL_498803, EPI_ISL_498804, EPI_ISL_498805, EPI_ISL_498806 | | | |
| see above | National Institute of Laboratory Medicine and Referral Center | Genomic Research Lab, BCSIR | Md. Murshed Hasan Sarkar, Abu Sayeed Mohammad Mahmud, Mohammad Samir Uzzaman, Eshrar Osman, Md. Ahasan Habib, Shahina Akter, Tanjina Akhter Banu, Barna Goswami, Iffat Jahan, Md. Saddam Hossain, Tasnim Nafisa, Md. Maruf Ahmed Molla, Mahmuda Yeasmin, Asish Kumar Ghosh, A. K. M. Shamsuzzaman, Sheikh Md. Selim Al Din, Utpal Chandra Ray, Salek Ahmed Sajib, Md. Salim Khan |
| EPI_ISL_498808, EPI_ISL_498809, EPI_ISL_498811, EPI_ISL_498814, EPI_ISL_498815, EPI_ISL_498816, EPI_ISL_498817, EPI_ISL_498818, EPI_ISL_498830, EPI_ISL_498892, EPI_ISL_498932, EPI_ISL_498933 | | | |
| see above | National Institute of Laboratory Medicine and Referral Center | Genomic Research Lab, BCSIR | Abu Sayeed Mohammad Mahmud, Mohammad Samir Uzzaman, Eshrar Osman, Md. Ahasan Habib, Shahina Akter, Tanjina Akhter Banu, Md. Murshed Hasan Sarkar, Barna Goswami, Iffat Jahan, Md. Saddam Hossain, Tasnim Nafisa, Md. Maruf Ahmed Molla, Mahmuda Yeasmin, Asish Kumar Ghosh, A. K. M. Shamsuzzaman, Sheikh Md. Selim Al Din, Utpal Chandra Ray, Salek Ahmed Sajib, Md. Salim Khan |
| EPI_ISL_499042 | National Institute for Biological Standards and Control (NIBSC) | National Institute for Biological Standards and Control (NIBSC) | Javier Martin, Dimitra Klapsa, Thomas Wilton |
| EPI_ISL_499083 | Instituto de Virologia "Dr. J. M. Vanella", Facultad de Ciencias Medicas, Universidad Nacional de Cordoba. Laboratorio Central de la Provincia de Cordoba, Argentina. Ministerio de Salud de la provincia de Cordoba, Argentina. | Laboratorio de Virología, Hospital de Niños Ricardo Gutiérrez, CABA, Argentina. | Sandra Gallego, Brenda Konigheim, Sebastian Blanco, Lorena Spinsanti, Javier Aguilar, Adrian Diaz, Gonzalo Castro, Gabriela Barbas, Mercedes Nabaes, Stephanie Goya, Monica Natale, Silvina Lusso, Mariana Viegas. |
| EPI_ISL_499266, EPI_ISL_499267, EPI_ISL_499268, EPI_ISL_499269 | Queens Medical Centre, Clinical Microbiology Department / DeepSeq Nottingham | COVID-19 Genomics UK (COG-UK) Consortium | Gemma Clark, Wendy Smith, Manjinder Khakh, Vicki M Fleming, Michelle M Lister, Hannah Howson-Wells, Jonathan Ball, Patrick McClure, Joseph Chappell, Theocharis Tsoleridis, Nadine Holmes, Matthew Carlisle, Christopher Moore, Fei Sang, Johnny Debebe, Victoria Wright, Matthew Loose |
| EPI_ISL_499270, EPI_ISL_499271, EPI_ISL_499272, EPI_ISL_499273, EPI_ISL_499274, EPI_ISL_499275, EPI_ISL_499276, EPI_ISL_499277, EPI_ISL_499278, EPI_ISL_499279, EPI_ISL_499280, EPI_ISL_499281, EPI_ISL_499282, EPI_ISL_499283, EPI_ISL_499284, EPI_ISL_499285, EPI_ISL_499286, EPI_ISL_499287, EPI_ISL_499288, EPI_ISL_499289, EPI_ISL_499290, EPI_ISL_499291, EPI_ISL_499292, EPI_ISL_499293, EPI_ISL_499294, EPI_ISL_499295, EPI_ISL_499296, EPI_ISL_499297, EPI_ISL_499298, EPI_ISL_499299, EPI_ISL_499300, EPI_ISL_499301, EPI_ISL_499302, EPI_ISL_499303, EPI_ISL_499304, EPI_ISL_499305, EPI_ISL_499306, EPI_ISL_499307, EPI_ISL_499308, EPI_ISL_499309, EPI_ISL_499310, EPI_ISL_499311, EPI_ISL_499312, EPI_ISL_499313, EPI_ISL_499314, EPI_ISL_499315, EPI_ISL_499316, EPI_ISL_499317, EPI_ISL_499318, EPI_ISL_499319, EPI_ISL_499320, EPI_ISL_499321, EPI_ISL_499322, EPI_ISL_499323, EPI_ISL_499324, EPI_ISL_499325, EPI_ISL_499326, EPI_ISL_499327, EPI_ISL_499328, EPI_ISL_499329 | | | |
| see above | Centre for Enzyme Innovation, University of Portsmouth / Translational Research Laboratory, Portsmouth Hospitals NHS Trust | COVID-19 Genomics UK (COG-UK) Consortium | Angela Beckett,Yann Bourgeois,Garry Scarlett,Sharon Glaysher,Scott Elliott,Kelly Bicknell,Robert Impey,Allyson Lloyd,Sarah Wyllie,Ethan Butcher,Anoop Chauhan,Samuel Robson |
| EPI_ISL_499330, EPI_ISL_499331, EPI_ISL_499332, EPI_ISL_499333, EPI_ISL_499334, EPI_ISL_499335, EPI_ISL_499336, EPI_ISL_499337, EPI_ISL_499338, EPI_ISL_499339, EPI_ISL_499340, EPI_ISL_499341, EPI_ISL_499342, EPI_ISL_499343, EPI_ISL_499344, EPI_ISL_499345, EPI_ISL_499346, EPI_ISL_499347, EPI_ISL_499348, EPI_ISL_499349, EPI_ISL_499350, EPI_ISL_499351 | | | |
| see above | Virology Department, Sheffield Teaching Hospitals NHS Foundation Trust/Department of Infection, Immunity and Cardiovascular Disease, The Medical School, University of Sheffield | COVID-19 Genomics UK (COG-UK) Consortium | Thushan de Silva, Matthew Parker, Nikki Smith, Adri Angyal, Rebecca Brown, Luke Green, Rachel Tucker, Paul Parsons, Danielle Groves, Katie Johnson, Laura Carrilero, Alex Keeley, Dave Partridge, Matthew Wyles, Benjamin Lindsey, Mehmet Yavuz, Mohammad Raza, Cariad Evans |
| EPI_ISL_499352, EPI_ISL_499353, EPI_ISL_499354 | West of Scotland Specialist Virology Centre, NHSGGC / MRC-University of Glasgow Centre for Virus Research | COVID-19 Genomics UK (COG-UK) Consortium | Ana da Silva Filipe, Natasha Johnson, Kathy Smollett, Daniel Mair, Stephen Carmichael, Lily Tong, Jenna Nichols, Elihu Aranday-Cortes, Kirstyn Brunker, Yasmin Parr, Alice Broos, Kyriaki Nomikou; Sarah McDonald, Marc Niebel, Patawee Asamaphan; Richard Orton, Joseph Hughes, Sreenu Vattipally, David L Robertson; Alasdair MacLean, Rory Gunson; Kathy Li, Natasha Jesudason, Rajiv Shah, James Shepherd, Antonia Ho, Emma Thomson |
| EPI_ISL_499355, EPI_ISL_499356, EPI_ISL_499357, EPI_ISL_499358, EPI_ISL_499359, EPI_ISL_499360, EPI_ISL_499361, EPI_ISL_499362, EPI_ISL_499363, EPI_ISL_499364, EPI_ISL_499365, EPI_ISL_499366, EPI_ISL_499367, EPI_ISL_499368, EPI_ISL_499369, EPI_ISL_499370, EPI_ISL_499371, EPI_ISL_499372, EPI_ISL_499373, EPI_ISL_499374, EPI_ISL_499375, EPI_ISL_499376, EPI_ISL_499377, EPI_ISL_499378, EPI_ISL_499379, EPI_ISL_499380, EPI_ISL_499381, EPI_ISL_499382, EPI_ISL_499383, EPI_ISL_499384, EPI_ISL_499385, EPI_ISL_499386, EPI_ISL_499387, EPI_ISL_499388, EPI_ISL_499389, EPI_ISL_499390, EPI_ISL_499391, EPI_ISL_499392, EPI_ISL_499393, EPI_ISL_499394, EPI_ISL_499395, EPI_ISL_499396, EPI_ISL_499397 | | | |
| see above | Originating lab: Wales Specialist Virology Centre Sequencing lab: Pathogen Genomics Unit | COVID-19 Genomics UK (COG-UK) Consortium | Catherine Moore, Johnathan Evans, Laura Gifford, Malorie Perry, Simon Cottrell, Angela Marchbank, Alec Birchley, Alexander Adams, Amy Gaskin, Bree Gatica-Wilcox, Jason Coombes, Joel Southgate, Lauren Gilbert, Lee Graham, Nicole Pacchiarini, Sara Kumziene-Summerhayes, Sarah Taylor, Sophie Jones, Sara Rey, Matthew Bull, Joanne Watkins, Sally Corden, Tom Connor |
| EPI_ISL_499398, EPI_ISL_499399, EPI_ISL_499400, EPI_ISL_499401, EPI_ISL_499402, EPI_ISL_499403, EPI_ISL_499404, EPI_ISL_499405, EPI_ISL_499406, EPI_ISL_499407, EPI_ISL_499408, EPI_ISL_499409, EPI_ISL_499410, EPI_ISL_499411, EPI_ISL_499412, EPI_ISL_499413, EPI_ISL_499414, EPI_ISL_499415, EPI_ISL_499416, EPI_ISL_499417, EPI_ISL_499418, EPI_ISL_499419, EPI_ISL_499420, EPI_ISL_499421, EPI_ISL_499422, EPI_ISL_499423, EPI_ISL_499424, EPI_ISL_499425, EPI_ISL_499426, EPI_ISL_499427, EPI_ISL_499428, EPI_ISL_499429, EPI_ISL_499430, EPI_ISL_499431, EPI_ISL_499432, EPI_ISL_499433, EPI_ISL_499434, EPI_ISL_499435, EPI_ISL_499436, EPI_ISL_499437, EPI_ISL_499438, EPI_ISL_499439, EPI_ISL_499440, EPI_ISL_499441, EPI_ISL_499442, EPI_ISL_499443, EPI_ISL_499444, EPI_ISL_499445, EPI_ISL_499446, EPI_ISL_499447, EPI_ISL_499448, EPI_ISL_499449, EPI_ISL_499450, EPI_ISL_499451, EPI_ISL_499452, EPI_ISL_499453, EPI_ISL_499454, EPI_ISL_499455, EPI_ISL_499456, EPI_ISL_499457, EPI_ISL_499458, EPI_ISL_499459 | | | |
| see above | Wales Specialist Virology Centre Sequencing lab: Pathogen Genomics Unit | COVID-19 Genomics UK (COG-UK) Consortium | Catherine Moore, Johnathan Evans, Laura Gifford, Malorie Perry, Simon Cottrell, Angela Marchbank, Alec Birchley, Alexander Adams, Amy Gaskin, Bree Gatica-Wilcox, Jason Coombes, Joel Southgate, Lauren Gilbert, Lee Graham, Nicole Pacchiarini, Sara Kumziene-Summerhayes, Sarah Taylor, Sophie Jones, Sara Rey, Matthew Bull, Joanne Watkins, Sally Corden, Tom Connor |
| EPI_ISL_499460, EPI_ISL_499461, EPI_ISL_499462, EPI_ISL_499463, EPI_ISL_499464, EPI_ISL_499465, EPI_ISL_499466, EPI_ISL_499467, EPI_ISL_499468, EPI_ISL_499469, EPI_ISL_499470, EPI_ISL_499471 | | | |
| see above | Liverpool Clinical Laboratories | COVID-19 Genomics UK (COG-UK) Consortium | Sam Haldenby, Anita Lucaci, Steve Paterson, Julian Hiscox, Alistair Darby, M Almsaud, A Alrezaihi, Muhannad Alruwaili, Stuart D Armstrong, Jones Benjamin, Eleanor G Bentley, Anu Chawla, Jordan J Clark, Angela Cowell, Richard Eccles, Isabel García-Dorival, Matthew Gemmell, Alessandro Gerada, PKF Gilmore, Richard Gregory, Ximeng Han, Catherine Hartley, Margaret Hughes, Miren Iturriza-Gomara, James Johnson, L Luu, Jenifer Manson, Charlotte Nelson, Elaine O'Toole, Cassie Olateju, Rebekah Penrice-Randal , Lucille Rainbow, N.P Randle, Trevor Ian Robinson, Parul Sharma, Ghada T Shawli, James P Stewart, Neil Swainston, Ecaterina Vamos, Joanne Watts, Mark Whitehead |
| EPI_ISL_499472 | Northumbria University / South Tees Hospitals NHS Foundation Trust / North Cumbria Integrated Care NHS Foundation Trust / North Tees and Hartlepool NHS Foundation Trust / Newcastle Hospitals NHS Foundation Trust | COVID-19 Genomics UK (COG-UK) Consortium | Darren L Smith,Andrew Nelson,Matthew Bashton,Greg R Young,Joshua Loh,John Allan,Mohammad A Tariq,Giles S Holt,Gary Black,Wen C Yew,Lynn Dover,Paul Baker,Steve Liggett,Sarah Essex,Jane Greenaway,Debra Padgett,Clive Graham,Garren Scott,Edward Barton,Emma Swindells,Brendan Payne,Jennifer Collins,Yusri Taha,Gary Eltringham |
| EPI_ISL_499473 | Liverpool Clinical Laboratories | COVID-19 Genomics UK (COG-UK) Consortium | Sam Haldenby, Anita Lucaci, Steve Paterson, Julian Hiscox, Alistair Darby, M Almsaud, A Alrezaihi, Muhannad Alruwaili, Stuart D Armstrong, Jones Benjamin, Eleanor G Bentley, Anu Chawla, Jordan J Clark, Angela Cowell, Richard Eccles, Isabel García-Dorival, Matthew Gemmell, Alessandro Gerada, PKF Gilmore, Richard Gregory, Ximeng Han, Catherine Hartley, Margaret Hughes, Miren Iturriza-Gomara, James Johnson, L Luu, Jenifer Manson, Charlotte Nelson, Elaine O'Toole, Cassie Olateju, Rebekah Penrice-Randal , Lucille Rainbow, N.P Randle, Trevor Ian Robinson, Parul Sharma, Ghada T Shawli, James P Stewart, Neil Swainston, Ecaterina Vamos, Joanne Watts, Mark Whitehead |
| EPI_ISL_499474 | Northumbria University / South Tees Hospitals NHS Foundation Trust / North Cumbria Integrated Care NHS Foundation Trust / North Tees and Hartlepool NHS Foundation Trust / Newcastle Hospitals NHS Foundation Trust | COVID-19 Genomics UK (COG-UK) Consortium | Darren L Smith,Andrew Nelson,Matthew Bashton,Greg R Young,Joshua Loh,John Allan,Mohammad A Tariq,Giles S Holt,Gary Black,Wen C Yew,Lynn Dover,Paul Baker,Steve Liggett,Sarah Essex,Jane Greenaway,Debra Padgett,Clive Graham,Garren Scott,Edward Barton,Emma Swindells,Brendan Payne,Jennifer Collins,Yusri Taha,Gary Eltringham |
| EPI_ISL_499475, EPI_ISL_499476, EPI_ISL_499477 | Liverpool Clinical Laboratories | COVID-19 Genomics UK (COG-UK) Consortium | Sam Haldenby, Anita Lucaci, Steve Paterson, Julian Hiscox, Alistair Darby, M Almsaud, A Alrezaihi, Muhannad Alruwaili, Stuart D Armstrong, Jones Benjamin, Eleanor G Bentley, Anu Chawla, Jordan J Clark, Angela Cowell, Richard Eccles, Isabel García-Dorival, Matthew Gemmell, Alessandro Gerada, PKF Gilmore, Richard Gregory, Ximeng Han, Catherine Hartley, Margaret Hughes, Miren Iturriza-Gomara, James Johnson, L Luu, Jenifer Manson, Charlotte Nelson, Elaine O'Toole, Cassie Olateju, Rebekah Penrice-Randal , Lucille Rainbow, N.P Randle, Trevor Ian Robinson, Parul Sharma, Ghada T Shawli, James P Stewart, Neil Swainston, Ecaterina Vamos, Joanne Watts, Mark Whitehead |
| EPI_ISL_499478 | Northumbria University / South Tees Hospitals NHS Foundation Trust / North Cumbria Integrated Care NHS Foundation Trust / North Tees and Hartlepool NHS Foundation Trust / Newcastle Hospitals NHS Foundation Trust | COVID-19 Genomics UK (COG-UK) Consortium | Darren L Smith,Andrew Nelson,Matthew Bashton,Greg R Young,Joshua Loh,John Allan,Mohammad A Tariq,Giles S Holt,Gary Black,Wen C Yew,Lynn Dover,Paul Baker,Steve Liggett,Sarah Essex,Jane Greenaway,Debra Padgett,Clive Graham,Garren Scott,Edward Barton,Emma Swindells,Brendan Payne,Jennifer Collins,Yusri Taha,Gary Eltringham |
| EPI_ISL_499479 | Quadram Institute Bioscience | COVID-19 Genomics UK (COG-UK) Consortium | Dave J. Baker, Gemma L. Kay, Alp Aydin, Thanh Le-Viet, Steven Rudder, Ana P. Tedim, Anastasia Kolyva, Maria Diaz, Leonardo de Oliveira Martins, Nabil-Fareed Alikhan, Lizzie Meadows, Rachael Stanley, Ngozi Elumogo, Muhammed Yasir, Nicholas M. Thomson, Alexander J Trotter, Rachel Gilroy, Samuel Bloomfield, Claire Stuart, Andrew Bell, Reenesh Prakash, Samir Dervisevic, Alison E. Mather, John Wain, Mark Webber, Andrew J. Page, Justin O'Grady |
| EPI_ISL_499480, EPI_ISL_499481 | Liverpool Clinical Laboratories | COVID-19 Genomics UK (COG-UK) Consortium | Sam Haldenby, Anita Lucaci, Steve Paterson, Julian Hiscox, Alistair Darby, M Almsaud, A Alrezaihi, Muhannad Alruwaili, Stuart D Armstrong, Jones Benjamin, Eleanor G Bentley, Anu Chawla, Jordan J Clark, Angela Cowell, Richard Eccles, Isabel García-Dorival, Matthew Gemmell, Alessandro Gerada, PKF Gilmore, Richard Gregory, Ximeng Han, Catherine Hartley, Margaret Hughes, Miren Iturriza-Gomara, James Johnson, L Luu, Jenifer Manson, Charlotte Nelson, Elaine O'Toole, Cassie Olateju, Rebekah Penrice-Randal , Lucille Rainbow, N.P Randle, Trevor Ian Robinson, Parul Sharma, Ghada T Shawli, James P Stewart, Neil Swainston, Ecaterina Vamos, Joanne Watts, Mark Whitehead |
| EPI_ISL_499482 | Northumbria University / South Tees Hospitals NHS Foundation Trust / North Cumbria Integrated Care NHS Foundation Trust / North Tees and Hartlepool NHS Foundation Trust / Newcastle Hospitals NHS Foundation Trust | COVID-19 Genomics UK (COG-UK) Consortium | Darren L Smith,Andrew Nelson,Matthew Bashton,Greg R Young,Joshua Loh,John Allan,Mohammad A Tariq,Giles S Holt,Gary Black,Wen C Yew,Lynn Dover,Paul Baker,Steve Liggett,Sarah Essex,Jane Greenaway,Debra Padgett,Clive Graham,Garren Scott,Edward Barton,Emma Swindells,Brendan Payne,Jennifer Collins,Yusri Taha,Gary Eltringham |
| EPI_ISL_499483, EPI_ISL_499484 | Liverpool Clinical Laboratories | COVID-19 Genomics UK (COG-UK) Consortium | Sam Haldenby, Anita Lucaci, Steve Paterson, Julian Hiscox, Alistair Darby, M Almsaud, A Alrezaihi, Muhannad Alruwaili, Stuart D Armstrong, Jones Benjamin, Eleanor G Bentley, Anu Chawla, Jordan J Clark, Angela Cowell, Richard Eccles, Isabel García-Dorival, Matthew Gemmell, Alessandro Gerada, PKF Gilmore, Richard Gregory, Ximeng Han, Catherine Hartley, Margaret Hughes, Miren Iturriza-Gomara, James Johnson, L Luu, Jenifer Manson, Charlotte Nelson, Elaine O'Toole, Cassie Olateju, Rebekah Penrice-Randal , Lucille Rainbow, N.P Randle, Trevor Ian Robinson, Parul Sharma, Ghada T Shawli, James P Stewart, Neil Swainston, Ecaterina Vamos, Joanne Watts, Mark Whitehead |
| EPI_ISL_499485 | Northumbria University / South Tees Hospitals NHS Foundation Trust / North Cumbria Integrated Care NHS Foundation Trust / North Tees and Hartlepool NHS Foundation Trust / Newcastle Hospitals NHS Foundation Trust | COVID-19 Genomics UK (COG-UK) Consortium | Darren L Smith,Andrew Nelson,Matthew Bashton,Greg R Young,Joshua Loh,John Allan,Mohammad A Tariq,Giles S Holt,Gary Black,Wen C Yew,Lynn Dover,Paul Baker,Steve Liggett,Sarah Essex,Jane Greenaway,Debra Padgett,Clive Graham,Garren Scott,Edward Barton,Emma Swindells,Brendan Payne,Jennifer Collins,Yusri Taha,Gary Eltringham |
| EPI_ISL_499486, EPI_ISL_499487, EPI_ISL_499488, EPI_ISL_499489 | Liverpool Clinical Laboratories | COVID-19 Genomics UK (COG-UK) Consortium | Sam Haldenby, Anita Lucaci, Steve Paterson, Julian Hiscox, Alistair Darby, M Almsaud, A Alrezaihi, Muhannad Alruwaili, Stuart D Armstrong, Jones Benjamin, Eleanor G Bentley, Anu Chawla, Jordan J Clark, Angela Cowell, Richard Eccles, Isabel García-Dorival, Matthew Gemmell, Alessandro Gerada, PKF Gilmore, Richard Gregory, Ximeng Han, Catherine Hartley, Margaret Hughes, Miren Iturriza-Gomara, James Johnson, L Luu, Jenifer Manson, Charlotte Nelson, Elaine O'Toole, Cassie Olateju, Rebekah Penrice-Randal , Lucille Rainbow, N.P Randle, Trevor Ian Robinson, Parul Sharma, Ghada T Shawli, James P Stewart, Neil Swainston, Ecaterina Vamos, Joanne Watts, Mark Whitehead |
| EPI_ISL_499490 | Quadram Institute Bioscience | COVID-19 Genomics UK (COG-UK) Consortium | Dave J. Baker, Gemma L. Kay, Alp Aydin, Thanh Le-Viet, Steven Rudder, Ana P. Tedim, Anastasia Kolyva, Maria Diaz, Leonardo de Oliveira Martins, Nabil-Fareed Alikhan, Lizzie Meadows, Rachael Stanley, Ngozi Elumogo, Muhammed Yasir, Nicholas M. Thomson, Alexander J Trotter, Rachel Gilroy, Samuel Bloomfield, Claire Stuart, Andrew Bell, Reenesh Prakash, Samir Dervisevic, Alison E. Mather, John Wain, Mark Webber, Andrew J. Page, Justin O'Grady |
| EPI_ISL_499491, EPI_ISL_499492 | Liverpool Clinical Laboratories | COVID-19 Genomics UK (COG-UK) Consortium | Sam Haldenby, Anita Lucaci, Steve Paterson, Julian Hiscox, Alistair Darby, M Almsaud, A Alrezaihi, Muhannad Alruwaili, Stuart D Armstrong, Jones Benjamin, Eleanor G Bentley, Anu Chawla, Jordan J Clark, Angela Cowell, Richard Eccles, Isabel García-Dorival, Matthew Gemmell, Alessandro Gerada, PKF Gilmore, Richard Gregory, Ximeng Han, Catherine Hartley, Margaret Hughes, Miren Iturriza-Gomara, James Johnson, L Luu, Jenifer Manson, Charlotte Nelson, Elaine O'Toole, Cassie Olateju, Rebekah Penrice-Randal , Lucille Rainbow, N.P Randle, Trevor Ian Robinson, Parul Sharma, Ghada T Shawli, James P Stewart, Neil Swainston, Ecaterina Vamos, Joanne Watts, Mark Whitehead |
| EPI_ISL_499493, EPI_ISL_499494 | Northumbria University / South Tees Hospitals NHS Foundation Trust / North Cumbria Integrated Care NHS Foundation Trust / North Tees and Hartlepool NHS Foundation Trust / Newcastle Hospitals NHS Foundation Trust | COVID-19 Genomics UK (COG-UK) Consortium | Darren L Smith,Andrew Nelson,Matthew Bashton,Greg R Young,Joshua Loh,John Allan,Mohammad A Tariq,Giles S Holt,Gary Black,Wen C Yew,Lynn Dover,Paul Baker,Steve Liggett,Sarah Essex,Jane Greenaway,Debra Padgett,Clive Graham,Garren Scott,Edward Barton,Emma Swindells,Brendan Payne,Jennifer Collins,Yusri Taha,Gary Eltringham |
| EPI_ISL_499495 | Quadram Institute Bioscience | COVID-19 Genomics UK (COG-UK) Consortium | Dave J. Baker, Gemma L. Kay, Alp Aydin, Thanh Le-Viet, Steven Rudder, Ana P. Tedim, Anastasia Kolyva, Maria Diaz, Leonardo de Oliveira Martins, Nabil-Fareed Alikhan, Lizzie Meadows, Rachael Stanley, Ngozi Elumogo, Muhammed Yasir, Nicholas M. Thomson, Alexander J Trotter, Rachel Gilroy, Samuel Bloomfield, Claire Stuart, Andrew Bell, Reenesh Prakash, Samir Dervisevic, Alison E. Mather, John Wain, Mark Webber, Andrew J. Page, Justin O'Grady |
| EPI_ISL_499496, EPI_ISL_499497, EPI_ISL_499498, EPI_ISL_499499, EPI_ISL_499500, EPI_ISL_499501 | Liverpool Clinical Laboratories | COVID-19 Genomics UK (COG-UK) Consortium | Sam Haldenby, Anita Lucaci, Steve Paterson, Julian Hiscox, Alistair Darby, M Almsaud, A Alrezaihi, Muhannad Alruwaili, Stuart D Armstrong, Jones Benjamin, Eleanor G Bentley, Anu Chawla, Jordan J Clark, Angela Cowell, Richard Eccles, Isabel García-Dorival, Matthew Gemmell, Alessandro Gerada, PKF Gilmore, Richard Gregory, Ximeng Han, Catherine Hartley, Margaret Hughes, Miren Iturriza-Gomara, James Johnson, L Luu, Jenifer Manson, Charlotte Nelson, Elaine O'Toole, Cassie Olateju, Rebekah Penrice-Randal , Lucille Rainbow, N.P Randle, Trevor Ian Robinson, Parul Sharma, Ghada T Shawli, James P Stewart, Neil Swainston, Ecaterina Vamos, Joanne Watts, Mark Whitehead |

| | | | |
|---|---|---|---|
| EPI_ISL_499502, EPI_ISL_499503, EPI_ISL_499504, EPI_ISL_499505, EPI_ISL_499506, EPI_ISL_499507, EPI_ISL_499508 | Northumbria University / South Tees Hospitals NHS Foundation Trust / North Cumbria Integrated Care NHS Foundation Trust / North Tees and Hartlepool NHS Foundation Trust / Newcastle Hospitals NHS Foundation Trust | COVID-19 Genomics UK (COG-UK) Consortium | Darren L Smith,Andrew Nelson,Matthew Bashton,Greg R Young,Joshua Loh,John Allan,Mohammad A Tariq,Giles S Holt,Gary Black,Wen C Yew,Lynn Dover,Paul Baker,Steve Liggett,Sarah Essex,Jane Greenaway,Debra Padgett,Clive Graham,Garren Scott,Edward Barton,Emma Swindells,Brendan Payne,Jennifer Collins,Yusri Taha,Gary Eltringham |
| EPI_ISL_499509 | Liverpool Clinical Laboratories | COVID-19 Genomics UK (COG-UK) Consortium | Sam Haldenby, Anita Lucaci, Steve Paterson, Julian Hiscox, Alistair Darby, M Almsaud, A Alrezaihi, Muhannad Alruwaili, Stuart D Armstrong, Jones Benjamin, Eleanor G Bentley, Anu Chawla, Jordan J Clark, Angela Cowell, Richard Eccles, Isabel García-Dorival, Matthew Gemmell, Alessandro Gerada, PKF Gilmore, Richard Gregory, Ximeng Han, Catherine Hartley, Margaret Hughes, Miren Iturriza-Gomara, James Johnson, L Luu, Jenifer Manson, Charlotte Nelson, Elaine O'Toole, Cassie Olateju, Rebekah Penrice-Randal , Lucille Rainbow, N.P Randle, Trevor Ian Robinson, Parul Sharma, Ghada T Shawli, James P Stewart, Neil Swainston, Ecaterina Vamos, Joanne Watts, Mark Whitehead |
| EPI_ISL_499510 | Northumbria University / South Tees Hospitals NHS Foundation Trust / North Cumbria Integrated Care NHS Foundation Trust / North Tees and Hartlepool NHS Foundation Trust / Newcastle Hospitals NHS Foundation Trust | COVID-19 Genomics UK (COG-UK) Consortium | Darren L Smith,Andrew Nelson,Matthew Bashton,Greg R Young,Joshua Loh,John Allan,Mohammad A Tariq,Giles S Holt,Gary Black,Wen C Yew,Lynn Dover,Paul Baker,Steve Liggett,Sarah Essex,Jane Greenaway,Debra Padgett,Clive Graham,Garren Scott,Edward Barton,Emma Swindells,Brendan Payne,Jennifer Collins,Yusri Taha,Gary Eltringham |
| EPI_ISL_499511, EPI_ISL_499512, EPI_ISL_499513, EPI_ISL_499514 | Liverpool Clinical Laboratories | COVID-19 Genomics UK (COG-UK) Consortium | Sam Haldenby, Anita Lucaci, Steve Paterson, Julian Hiscox, Alistair Darby, M Almsaud, A Alrezaihi, Muhannad Alruwaili, Stuart D Armstrong, Jones Benjamin, Eleanor G Bentley, Anu Chawla, Jordan J Clark, Angela Cowell, Richard Eccles, Isabel García-Dorival, Matthew Gemmell, Alessandro Gerada, PKF Gilmore, Richard Gregory, Ximeng Han, Catherine Hartley, Margaret Hughes, Miren Iturriza-Gomara, James Johnson, L Luu, Jenifer Manson, Charlotte Nelson, Elaine O'Toole, Cassie Olateju, Rebekah Penrice-Randal , Lucille Rainbow, N.P Randle, Trevor Ian Robinson, Parul Sharma, Ghada T Shawli, James P Stewart, Neil Swainston, Ecaterina Vamos, Joanne Watts, Mark Whitehead |
| EPI_ISL_499515 | Northumbria University / South Tees Hospitals NHS Foundation Trust / North Cumbria Integrated Care NHS Foundation Trust / North Tees and Hartlepool NHS Foundation Trust / Newcastle Hospitals NHS Foundation Trust | COVID-19 Genomics UK (COG-UK) Consortium | Darren L Smith,Andrew Nelson,Matthew Bashton,Greg R Young,Joshua Loh,John Allan,Mohammad A Tariq,Giles S Holt,Gary Black,Wen C Yew,Lynn Dover,Paul Baker,Steve Liggett,Sarah Essex,Jane Greenaway,Debra Padgett,Clive Graham,Garren Scott,Edward Barton,Emma Swindells,Brendan Payne,Jennifer Collins,Yusri Taha,Gary Eltringham |
| EPI_ISL_499516, EPI_ISL_499517, EPI_ISL_499518, EPI_ISL_499519, EPI_ISL_499520, EPI_ISL_499521, EPI_ISL_499522, EPI_ISL_499523, EPI_ISL_499524, EPI_ISL_499525, EPI_ISL_499526, EPI_ISL_499527, EPI_ISL_499528, EPI_ISL_499529, EPI_ISL_499530, EPI_ISL_499531, EPI_ISL_499532, EPI_ISL_499533, EPI_ISL_499534, EPI_ISL_499535, EPI_ISL_499536, EPI_ISL_499537, EPI_ISL_499538, EPI_ISL_499539, EPI_ISL_499540, EPI_ISL_499541, EPI_ISL_499542, EPI_ISL_499543, EPI_ISL_499544, EPI_ISL_499545 | | | |
| see above | Liverpool Clinical Laboratories | COVID-19 Genomics UK (COG-UK) Consortium | Sam Haldenby, Anita Lucaci, Steve Paterson, Julian Hiscox, Alistair Darby, M Almsaud, A Alrezaihi, Muhannad Alruwaili, Stuart D Armstrong, Jones Benjamin, Eleanor G Bentley, Anu Chawla, Jordan J Clark, Angela Cowell, Richard Eccles, Isabel García-Dorival, Matthew Gemmell, Alessandro Gerada, PKF Gilmore, Richard Gregory, Ximeng Han, Catherine Hartley, Margaret Hughes, Miren Iturriza-Gomara, James Johnson, L Luu, Jenifer Manson, Charlotte Nelson, Elaine O'Toole, Cassie Olateju, Rebekah Penrice-Randal , Lucille Rainbow, N.P Randle, Trevor Ian Robinson, Parul Sharma, Ghada T Shawli, James P Stewart, Neil Swainston, Ecaterina Vamos, Joanne Watts, Mark Whitehead |
| EPI_ISL_499546, EPI_ISL_499547, EPI_ISL_499548, EPI_ISL_499549, EPI_ISL_499550, EPI_ISL_499551, EPI_ISL_499552, EPI_ISL_499553, EPI_ISL_499554, EPI_ISL_499555, EPI_ISL_499556 | | | |
| see above | Northumbria University / South Tees Hospitals NHS Foundation Trust / North Cumbria Integrated Care NHS Foundation Trust / North Tees and Hartlepool NHS Foundation Trust / Newcastle Hospitals NHS Foundation Trust | COVID-19 Genomics UK (COG-UK) Consortium | Darren L Smith,Andrew Nelson,Matthew Bashton,Greg R Young,Joshua Loh,John Allan,Mohammad A Tariq,Giles S Holt,Gary Black,Wen C Yew,Lynn Dover,Paul Baker,Steve Liggett,Sarah Essex,Jane Greenaway,Debra Padgett,Clive Graham,Garren Scott,Edward Barton,Emma Swindells,Brendan Payne,Jennifer Collins,Yusri Taha,Gary Eltringham |
| EPI_ISL_499557 | Liverpool Clinical Laboratories | COVID-19 Genomics UK (COG-UK) Consortium | Sam Haldenby, Anita Lucaci, Steve Paterson, Julian Hiscox, Alistair Darby, M Almsaud, A Alrezaihi, Muhannad Alruwaili, Stuart D Armstrong, Jones Benjamin, Eleanor G Bentley, Anu Chawla, Jordan J Clark, Angela Cowell, Richard Eccles, Isabel García-Dorival, Matthew Gemmell, Alessandro Gerada, PKF Gilmore, Richard Gregory, Ximeng Han, Catherine Hartley, Margaret Hughes, Miren Iturriza-Gomara, James Johnson, L Luu, Jenifer Manson, Charlotte Nelson, Elaine O'Toole, Cassie Olateju, Rebekah Penrice-Randal , Lucille Rainbow, N.P Randle, Trevor Ian Robinson, Parul Sharma, Ghada T Shawli, James P Stewart, Neil Swainston, Ecaterina Vamos, Joanne Watts, Mark Whitehead |
| EPI_ISL_499558, EPI_ISL_499559, EPI_ISL_499560, EPI_ISL_499561 | Northumbria University / South Tees Hospitals NHS Foundation Trust / North Cumbria Integrated Care NHS Foundation Trust / North Tees and Hartlepool NHS Foundation Trust / Newcastle Hospitals NHS Foundation Trust | COVID-19 Genomics UK (COG-UK) Consortium | Darren L Smith,Andrew Nelson,Matthew Bashton,Greg R Young,Joshua Loh,John Allan,Mohammad A Tariq,Giles S Holt,Gary Black,Wen C Yew,Lynn Dover,Paul Baker,Steve Liggett,Sarah Essex,Jane Greenaway,Debra Padgett,Clive Graham,Garren Scott,Edward Barton,Emma Swindells,Brendan Payne,Jennifer Collins,Yusri Taha,Gary Eltringham |
| EPI_ISL_499562 | Quadram Institute Bioscience | COVID-19 Genomics UK (COG-UK) Consortium | Dave J. Baker, Gemma L. Kay, Alp Aydin, Thanh Le-Viet, Steven Rudder, Ana P. Tedim, Anastasia Kolyva, Maria Diaz, Leonardo de Oliveira Martins, Nabil-Fareed Alikhan, Lizzie Meadows, Rachael Stanley, Ngozi Elumogo, Muhammed Yasir, Nicholas M. Thomson, Alexander J Trotter, Rachel Gilroy, Samuel Bloomfield, Claire Stuart, Andrew Bell, Reenesh Prakash, Samir Dervisevic, Alison E. Mather, John Wain, Mark Webber, Andrew J. Page, Justin O'Grady |
| EPI_ISL_499563 | Northumbria University / South Tees Hospitals NHS Foundation Trust / North Cumbria Integrated Care NHS Foundation Trust / North Tees and Hartlepool NHS Foundation Trust / Newcastle Hospitals NHS Foundation Trust | COVID-19 Genomics UK (COG-UK) Consortium | Darren L Smith,Andrew Nelson,Matthew Bashton,Greg R Young,Joshua Loh,John Allan,Mohammad A Tariq,Giles S Holt,Gary Black,Wen C Yew,Lynn Dover,Paul Baker,Steve Liggett,Sarah Essex,Jane Greenaway,Debra Padgett,Clive Graham,Garren Scott,Edward Barton,Emma Swindells,Brendan Payne,Jennifer Collins,Yusri Taha,Gary Eltringham |
| EPI_ISL_499564, EPI_ISL_499565, EPI_ISL_499566, EPI_ISL_499567 | Liverpool Clinical Laboratories | COVID-19 Genomics UK (COG-UK) Consortium | Sam Haldenby, Anita Lucaci, Steve Paterson, Julian Hiscox, Alistair Darby, M Almsaud, A Alrezaihi, Muhannad Alruwaili, Stuart D Armstrong, Jones Benjamin, Eleanor G Bentley, Anu Chawla, Jordan J Clark, Angela Cowell, Richard Eccles, Isabel García-Dorival, Matthew Gemmell, Alessandro Gerada, PKF Gilmore, Richard Gregory, Ximeng Han, Catherine Hartley, Margaret Hughes, Miren Iturriza-Gomara, James Johnson, L Luu, Jenifer Manson, Charlotte Nelson, Elaine O'Toole, Cassie Olateju, Rebekah Penrice-Randal , Lucille Rainbow, N.P Randle, Trevor Ian Robinson, Parul Sharma, Ghada T Shawli, James P Stewart, Neil Swainston, Ecaterina Vamos, Joanne Watts, Mark Whitehead |
| EPI_ISL_499568 | Northumbria University / South Tees Hospitals NHS Foundation Trust / North Cumbria Integrated Care NHS Foundation Trust / North Tees and Hartlepool NHS Foundation Trust / Newcastle Hospitals NHS Foundation Trust | COVID-19 Genomics UK (COG-UK) Consortium | Darren L Smith,Andrew Nelson,Matthew Bashton,Greg R Young,Joshua Loh,John Allan,Mohammad A Tariq,Giles S Holt,Gary Black,Wen C Yew,Lynn Dover,Paul Baker,Steve Liggett,Sarah Essex,Jane Greenaway,Debra Padgett,Clive Graham,Garren Scott,Edward Barton,Emma Swindells,Brendan Payne,Jennifer Collins,Yusri Taha,Gary Eltringham |
| EPI_ISL_499569, EPI_ISL_499570, EPI_ISL_499571 | Liverpool Clinical Laboratories | COVID-19 Genomics UK (COG-UK) Consortium | Sam Haldenby, Anita Lucaci, Steve Paterson, Julian Hiscox, Alistair Darby, M Almsaud, A Alrezaihi, Muhannad Alruwaili, Stuart D Armstrong, Jones Benjamin, Eleanor G Bentley, Anu Chawla, Jordan J Clark, Angela Cowell, Richard Eccles, Isabel García-Dorival, Matthew Gemmell, Alessandro Gerada, PKF Gilmore, Richard Gregory, Ximeng Han, Catherine Hartley, Margaret Hughes, Miren Iturriza-Gomara, James Johnson, L Luu, Jenifer Manson, Charlotte Nelson, Elaine O'Toole, Cassie Olateju, Rebekah Penrice-Randal , Lucille Rainbow, N.P Randle, Trevor Ian Robinson, Parul Sharma, Ghada T Shawli, James P Stewart, Neil Swainston, Ecaterina Vamos, Joanne Watts, Mark Whitehead |
| EPI_ISL_499572 | Northumbria University / South Tees Hospitals NHS Foundation Trust / North Cumbria Integrated Care NHS Foundation Trust / North Tees and Hartlepool NHS Foundation Trust / Newcastle Hospitals NHS Foundation Trust | COVID-19 Genomics UK (COG-UK) Consortium | Darren L Smith,Andrew Nelson,Matthew Bashton,Greg R Young,Joshua Loh,John Allan,Mohammad A Tariq,Giles S Holt,Gary Black,Wen C Yew,Lynn Dover,Paul Baker,Steve Liggett,Sarah Essex,Jane Greenaway,Debra Padgett,Clive Graham,Garren Scott,Edward Barton,Emma Swindells,Brendan Payne,Jennifer Collins,Yusri Taha,Gary Eltringham |
| EPI_ISL_499573, EPI_ISL_499574, EPI_ISL_499575, EPI_ISL_499576, EPI_ISL_499577, EPI_ISL_499578, EPI_ISL_499579, EPI_ISL_499580, EPI_ISL_499581, EPI_ISL_499582, EPI_ISL_499583, EPI_ISL_499584, EPI_ISL_499585, EPI_ISL_499586 | | | |
| see above | Liverpool Clinical Laboratories | COVID-19 Genomics UK (COG-UK) Consortium | Sam Haldenby, Anita Lucaci, Steve Paterson, Julian Hiscox, Alistair Darby, M Almsaud, A Alrezaihi, Muhannad Alruwaili, Stuart D Armstrong, Jones Benjamin, Eleanor G Bentley, Anu Chawla, Jordan J Clark, Angela Cowell, Richard Eccles, Isabel García-Dorival, Matthew Gemmell, Alessandro Gerada, PKF Gilmore, Richard Gregory, Ximeng Han, Catherine Hartley, Margaret Hughes, Miren Iturriza-Gomara, James Johnson, L Luu, Jenifer Manson, Charlotte Nelson, Elaine O'Toole, Cassie Olateju, Rebekah Penrice-Randal , Lucille Rainbow, N.P Randle, Trevor Ian Robinson, Parul Sharma, Ghada T Shawli, James P Stewart, Neil Swainston, Ecaterina Vamos, Joanne Watts, Mark Whitehead |
| EPI_ISL_499587 | Northumbria University / South Tees Hospitals NHS Foundation Trust / North Cumbria Integrated Care NHS Foundation Trust / North Tees and Hartlepool NHS Foundation Trust / Newcastle Hospitals NHS Foundation Trust | COVID-19 Genomics UK (COG-UK) Consortium | Darren L Smith,Andrew Nelson,Matthew Bashton,Greg R Young,Joshua Loh,John Allan,Mohammad A Tariq,Giles S Holt,Gary Black,Wen C Yew,Lynn Dover,Paul Baker,Steve Liggett,Sarah Essex,Jane Greenaway,Debra Padgett,Clive Graham,Garren Scott,Edward Barton,Emma Swindells,Brendan Payne,Jennifer Collins,Yusri Taha,Gary Eltringham |
| EPI_ISL_499588 | Quadram Institute Bioscience | COVID-19 Genomics UK (COG-UK) Consortium | Dave J. Baker, Gemma L. Kay, Alp Aydin, Thanh Le-Viet, Steven Rudder, Ana P. Tedim, Anastasia Kolyva, Maria Diaz, Leonardo de Oliveira Martins, Nabil-Fareed Alikhan, Lizzie Meadows, Rachael Stanley, Ngozi Elumogo, Muhammed Yasir, Nicholas M. Thomson, Alexander J Trotter, Rachel Gilroy, Samuel Bloomfield, Claire Stuart, Andrew Bell, Reenesh Prakash, Samir Dervisevic, Alison E. Mather, John Wain, Mark Webber, Andrew J. Page, Justin O'Grady |
| EPI_ISL_499589, EPI_ISL_499590 | Liverpool Clinical Laboratories | COVID-19 Genomics UK (COG-UK) Consortium | Sam Haldenby, Anita Lucaci, Steve Paterson, Julian Hiscox, Alistair Darby, M Almsaud, A Alrezaihi, Muhannad Alruwaili, Stuart D Armstrong, Jones Benjamin, Eleanor G Bentley, Anu Chawla, Jordan J Clark, Angela Cowell, Richard Eccles, Isabel García-Dorival, Matthew Gemmell, Alessandro Gerada, PKF Gilmore, Richard Gregory, Ximeng Han, Catherine Hartley, Margaret Hughes, Miren Iturriza-Gomara, James Johnson, L Luu, Jenifer Manson, Charlotte Nelson, Elaine O'Toole, Cassie Olateju, Rebekah Penrice-Randal , Lucille Rainbow, N.P Randle, Trevor Ian Robinson, Parul Sharma, Ghada T Shawli, James P Stewart, Neil Swainston, Ecaterina Vamos, Joanne Watts, Mark Whitehead |
| EPI_ISL_499591, EPI_ISL_499592 | Quadram Institute Bioscience | COVID-19 Genomics UK (COG-UK) Consortium | Dave J. Baker, Gemma L. Kay, Alp Aydin, Thanh Le-Viet, Steven Rudder, Ana P. Tedim, Anastasia Kolyva, Maria Diaz, Leonardo de Oliveira Martins, Nabil-Fareed Alikhan, Lizzie Meadows, Rachael Stanley, Ngozi Elumogo, Muhammed Yasir, Nicholas M. Thomson, Alexander J Trotter, Rachel Gilroy, Samuel Bloomfield, Claire Stuart, Andrew Bell, Reenesh Prakash, Samir Dervisevic, Alison E. Mather, John Wain, Mark Webber, Andrew J. Page, Justin O'Grady |
| EPI_ISL_499593, EPI_ISL_499594 | Liverpool Clinical Laboratories | COVID-19 Genomics UK (COG-UK) Consortium | Sam Haldenby, Anita Lucaci, Steve Paterson, Julian Hiscox, Alistair Darby, M Almsaud, A Alrezaihi, Muhannad Alruwaili, Stuart D Armstrong, Jones Benjamin, Eleanor G Bentley, Anu Chawla, Jordan J Clark, Angela Cowell, Richard Eccles, Isabel García-Dorival, Matthew Gemmell, Alessandro Gerada, PKF Gilmore, Richard Gregory, Ximeng Han, Catherine Hartley, Margaret Hughes, Miren Iturriza-Gomara, James Johnson, L Luu, Jenifer Manson, Charlotte Nelson, Elaine O'Toole, Cassie Olateju, Rebekah Penrice-Randal , Lucille Rainbow, N.P Randle, Trevor Ian Robinson, Parul Sharma, Ghada T Shawli, James P Stewart, Neil Swainston, Ecaterina Vamos, Joanne Watts, Mark Whitehead |
| EPI_ISL_499595, EPI_ISL_499596, EPI_ISL_499597, EPI_ISL_499598 | Northumbria University / South Tees Hospitals NHS Foundation Trust / North Cumbria Integrated Care NHS Foundation Trust / North Tees and Hartlepool NHS Foundation Trust / Newcastle Hospitals NHS Foundation Trust | COVID-19 Genomics UK (COG-UK) Consortium | Darren L Smith,Andrew Nelson,Matthew Bashton,Greg R Young,Joshua Loh,John Allan,Mohammad A Tariq,Giles S Holt,Gary Black,Wen C Yew,Lynn Dover,Paul Baker,Steve Liggett,Sarah Essex,Jane Greenaway,Debra Padgett,Clive Graham,Garren Scott,Edward Barton,Emma Swindells,Brendan Payne,Jennifer Collins,Yusri Taha,Gary Eltringham |
| EPI_ISL_499599 | Quadram Institute Bioscience | COVID-19 Genomics UK (COG-UK) Consortium | Dave J. Baker, Gemma L. Kay, Alp Aydin, Thanh Le-Viet, Steven Rudder, Ana P. Tedim, Anastasia Kolyva, Maria Diaz, Leonardo de Oliveira Martins, Nabil-Fareed Alikhan, Lizzie Meadows, Rachael Stanley, Ngozi Elumogo, Muhammed Yasir, Nicholas M. Thomson, Alexander J Trotter, Rachel Gilroy, Samuel Bloomfield, Claire Stuart, Andrew Bell, Reenesh Prakash, Samir Dervisevic, Alison E. Mather, John Wain, Mark Webber, Andrew J. Page, Justin O'Grady |
| EPI_ISL_499600 | Liverpool Clinical Laboratories | COVID-19 Genomics UK (COG-UK) Consortium | Sam Haldenby, Anita Lucaci, Steve Paterson, Julian Hiscox, Alistair Darby, M Almsaud, A Alrezaihi, Muhannad Alruwaili, Stuart D Armstrong, Jones Benjamin, Eleanor G Bentley, Anu Chawla, Jordan J Clark, Angela Cowell, Richard Eccles, Isabel García-Dorival, Matthew Gemmell, Alessandro Gerada, PKF Gilmore, Richard Gregory, Ximeng Han, Catherine Hartley, Margaret Hughes, Miren Iturriza-Gomara, James Johnson, L Luu, Jenifer Manson, Charlotte Nelson, Elaine O'Toole, Cassie Olateju, Rebekah Penrice-Randal , Lucille Rainbow, N.P Randle, Trevor Ian Robinson, Parul Sharma, Ghada T Shawli, James P Stewart, Neil Swainston, Ecaterina Vamos, Joanne Watts, Mark Whitehead |
| EPI_ISL_499601 | Quadram Institute Bioscience | COVID-19 Genomics UK (COG-UK) Consortium | Dave J. Baker, Gemma L. Kay, Alp Aydin, Thanh Le-Viet, Steven Rudder, Ana P. Tedim, Anastasia Kolyva, Maria Diaz, Leonardo de Oliveira Martins, Nabil-Fareed Alikhan, Lizzie Meadows, Rachael Stanley, Ngozi Elumogo, Muhammed Yasir, Nicholas M. Thomson, Alexander J Trotter, Rachel Gilroy, Samuel Bloomfield, Claire Stuart, Andrew Bell, Reenesh Prakash, Samir Dervisevic, Alison E. Mather, John Wain, Mark Webber, Andrew J. Page, Justin O'Grady |
| EPI_ISL_499602 | Northumbria University / South Tees Hospitals NHS Foundation Trust / North Cumbria Integrated Care NHS Foundation Trust / North Tees and Hartlepool NHS Foundation Trust / Newcastle Hospitals NHS Foundation Trust | COVID-19 Genomics UK (COG-UK) Consortium | Darren L Smith,Andrew Nelson,Matthew Bashton,Greg R Young,Joshua Loh,John Allan,Mohammad A Tariq,Giles S Holt,Gary Black,Wen C Yew,Lynn Dover,Paul Baker,Steve Liggett,Sarah Essex,Jane Greenaway,Debra Padgett,Clive Graham,Garren Scott,Edward Barton,Emma Swindells,Brendan Payne,Jennifer Collins,Yusri Taha,Gary Eltringham |
| EPI_ISL_499603 | Quadram Institute Bioscience | COVID-19 Genomics UK (COG-UK) Consortium | Dave J. Baker, Gemma L. Kay, Alp Aydin, Thanh Le-Viet, Steven Rudder, Ana P. Tedim, Anastasia Kolyva, Maria Diaz, Leonardo de Oliveira Martins, Nabil-Fareed Alikhan, Lizzie Meadows, Rachael Stanley, Ngozi Elumogo, Muhammed Yasir, Nicholas M. Thomson, Alexander J Trotter, Rachel Gilroy, Samuel Bloomfield, Claire Stuart, Andrew Bell, Reenesh Prakash, Samir Dervisevic, Alison E. Mather, John Wain, Mark Webber, Andrew J. Page, Justin O'Grady |
| EPI_ISL_499604, EPI_ISL_499605 | Liverpool Clinical Laboratories | COVID-19 Genomics UK (COG-UK) Consortium | Sam Haldenby, Anita Lucaci, Steve Paterson, Julian Hiscox, Alistair Darby, M Almsaud, A Alrezaihi, Muhannad Alruwaili, Stuart D Armstrong, Jones Benjamin, Eleanor G Bentley, Anu Chawla, Jordan |

| | | | |
|---|---|---|---|
| | | | J Clark, Angela Cowell, Richard Eccles, Isabel García-Dorival, Matthew Gemmell, Alessandro Gerada, PKF Gilmore, Richard Gregory, Ximeng Han, Catherine Hartley, Margaret Hughes, Miren Iturriza-Gomara, James Johnson, L Luu, Jenifer Manson, Charlotte Nelson, Elaine O’Toole, Cassie Olateju, Rebekah Penrice-Randal , Lucille Rainbow, N.P Randle, Trevor Ian Robinson, Parul Sharma, Ghada T Shawli, James P Stewart, Neil Swainston, Ecaterina Vamos, Joanne Watts, Mark Whitehead |
| EPI_ISL_499606 | Northumbria University / South Tees Hospitals NHS Foundation Trust / North Cumbria Integrated Care NHS Foundation Trust / North Tees and Hartlepool NHS Foundation Trust / Newcastle Hospitals NHS Foundation Trust | COVID-19 Genomics UK (COG-UK) Consortium | Darren L Smith,Andrew Nelson,Matthew Bashton,Greg R Young,Joshua Loh,John Allan,Mohammad A Tariq,Giles S Holt,Gary Black,Wen C Yew,Lynn Dover,Paul Baker,Steve Liggett,Sarah Essex,Jane Greenaway,Debra Padgett,Clive Graham,Garren Scott,Edward Barton,Emma Swindells,Brendan Payne,Jennifer Collins,Yusri Taha,Gary Eltringham |
| EPI_ISL_499607, EPI_ISL_499608 | Liverpool Clinical Laboratories | COVID-19 Genomics UK (COG-UK) Consortium | Sam Haldenby, Anita Lucaci, Steve Paterson, Julian Hiscox, Alistair Darby, M Almsaud, A Alrezaihi, Muhannad Alruwaili, Stuart D Armstrong, Jones Benjamin, Eleanor G Bentley, Anu Chawla, Jordan J Clark, Angela Cowell, Richard Eccles, Isabel García-Dorival, Matthew Gemmell, Alessandro Gerada, PKF Gilmore, Richard Gregory, Ximeng Han, Catherine Hartley, Margaret Hughes, Miren Iturriza-Gomara, James Johnson, L Luu, Jenifer Manson, Charlotte Nelson, Elaine O’Toole, Cassie Olateju, Rebekah Penrice-Randal , Lucille Rainbow, N.P Randle, Trevor Ian Robinson, Parul Sharma, Ghada T Shawli, James P Stewart, Neil Swainston, Ecaterina Vamos, Joanne Watts, Mark Whitehead |
| EPI_ISL_499609, EPI_ISL_499610 | Northumbria University / South Tees Hospitals NHS Foundation Trust / North Cumbria Integrated Care NHS Foundation Trust / North Tees and Hartlepool NHS Foundation Trust / Newcastle Hospitals NHS Foundation Trust | COVID-19 Genomics UK (COG-UK) Consortium | Darren L Smith,Andrew Nelson,Matthew Bashton,Greg R Young,Joshua Loh,John Allan,Mohammad A Tariq,Giles S Holt,Gary Black,Wen C Yew,Lynn Dover,Paul Baker,Steve Liggett,Sarah Essex,Jane Greenaway,Debra Padgett,Clive Graham,Garren Scott,Edward Barton,Emma Swindells,Brendan Payne,Jennifer Collins,Yusri Taha,Gary Eltringham |
| EPI_ISL_499611, EPI_ISL_499612, EPI_ISL_499613, EPI_ISL_499614, EPI_ISL_499615, EPI_ISL_499616, EPI_ISL_499617, EPI_ISL_499618, EPI_ISL_499619, EPI_ISL_499620, EPI_ISL_499621, EPI_ISL_499622 | | | |
| see above | Liverpool Clinical Laboratories | COVID-19 Genomics UK (COG-UK) Consortium | Sam Haldenby, Anita Lucaci, Steve Paterson, Julian Hiscox, Alistair Darby, M Almsaud, A Alrezaihi, Muhannad Alruwaili, Stuart D Armstrong, Jones Benjamin, Eleanor G Bentley, Anu Chawla, Jordan J Clark, Angela Cowell, Richard Eccles, Isabel García-Dorival, Matthew Gemmell, Alessandro Gerada, PKF Gilmore, Richard Gregory, Ximeng Han, Catherine Hartley, Margaret Hughes, Miren Iturriza-Gomara, James Johnson, L Luu, Jenifer Manson, Charlotte Nelson, Elaine O’Toole, Cassie Olateju, Rebekah Penrice-Randal , Lucille Rainbow, N.P Randle, Trevor Ian Robinson, Parul Sharma, Ghada T Shawli, James P Stewart, Neil Swainston, Ecaterina Vamos, Joanne Watts, Mark Whitehead |
| EPI_ISL_499623 | Quadram Institute Bioscience | COVID-19 Genomics UK (COG-UK) Consortium | Dave J. Baker, Gemma L. Kay, Alp Aydin, Thanh Le-Viet, Steven Rudder, Ana P. Tedim, Anastasia Kolyva, Maria Diaz, Leonardo de Oliveira Martins, Nabil-Fareed Alikhan, Lizzie Meadows, Rachael Stanley, Ngozi Elumogo, Muhammed Yasir, Nicholas M. Thomson, Alexander J Trotter, Rachel Gilroy, Samuel Bloomfield, Claire Stuart, Andrew Bell, Reenesh Prakash, Samir Dervisevic, Alison E. Mather, John Wain, Mark Webber, Andrew J. Page, Justin O’Grady |
| EPI_ISL_499624, EPI_ISL_499625 | Liverpool Clinical Laboratories | COVID-19 Genomics UK (COG-UK) Consortium | Sam Haldenby, Anita Lucaci, Steve Paterson, Julian Hiscox, Alistair Darby, M Almsaud, A Alrezaihi, Muhannad Alruwaili, Stuart D Armstrong, Jones Benjamin, Eleanor G Bentley, Anu Chawla, Jordan J Clark, Angela Cowell, Richard Eccles, Isabel García-Dorival, Matthew Gemmell, Alessandro Gerada, PKF Gilmore, Richard Gregory, Ximeng Han, Catherine Hartley, Margaret Hughes, Miren Iturriza-Gomara, James Johnson, L Luu, Jenifer Manson, Charlotte Nelson, Elaine O’Toole, Cassie Olateju, Rebekah Penrice-Randal , Lucille Rainbow, N.P Randle, Trevor Ian Robinson, Parul Sharma, Ghada T Shawli, James P Stewart, Neil Swainston, Ecaterina Vamos, Joanne Watts, Mark Whitehead |
| EPI_ISL_499626 | Northumbria University / South Tees Hospitals NHS Foundation Trust / North Cumbria Integrated Care NHS Foundation Trust / North Tees and Hartlepool NHS Foundation Trust / Newcastle Hospitals NHS Foundation Trust | COVID-19 Genomics UK (COG-UK) Consortium | Darren L Smith,Andrew Nelson,Matthew Bashton,Greg R Young,Joshua Loh,John Allan,Mohammad A Tariq,Giles S Holt,Gary Black,Wen C Yew,Lynn Dover,Paul Baker,Steve Liggett,Sarah Essex,Jane Greenaway,Debra Padgett,Clive Graham,Garren Scott,Edward Barton,Emma Swindells,Brendan Payne,Jennifer Collins,Yusri Taha,Gary Eltringham |
| EPI_ISL_499627, EPI_ISL_499628, EPI_ISL_499629, EPI_ISL_499630 | Liverpool Clinical Laboratories | COVID-19 Genomics UK (COG-UK) Consortium | Sam Haldenby, Anita Lucaci, Steve Paterson, Julian Hiscox, Alistair Darby, M Almsaud, A Alrezaihi, Muhannad Alruwaili, Stuart D Armstrong, Jones Benjamin, Eleanor G Bentley, Anu Chawla, Jordan J Clark, Angela Cowell, Richard Eccles, Isabel García-Dorival, Matthew Gemmell, Alessandro Gerada, PKF Gilmore, Richard Gregory, Ximeng Han, Catherine Hartley, Margaret Hughes, Miren Iturriza-Gomara, James Johnson, L Luu, Jenifer Manson, Charlotte Nelson, Elaine O’Toole, Cassie Olateju, Rebekah Penrice-Randal , Lucille Rainbow, N.P Randle, Trevor Ian Robinson, Parul Sharma, Ghada T Shawli, James P Stewart, Neil Swainston, Ecaterina Vamos, Joanne Watts, Mark Whitehead |
| EPI_ISL_499631 | Quadram Institute Bioscience | COVID-19 Genomics UK (COG-UK) Consortium | Dave J. Baker, Gemma L. Kay, Alp Aydin, Thanh Le-Viet, Steven Rudder, Ana P. Tedim, Anastasia Kolyva, Maria Diaz, Leonardo de Oliveira Martins, Nabil-Fareed Alikhan, Lizzie Meadows, Rachael Stanley, Ngozi Elumogo, Muhammed Yasir, Nicholas M. Thomson, Alexander J Trotter, Rachel Gilroy, Samuel Bloomfield, Claire Stuart, Andrew Bell, Reenesh Prakash, Samir Dervisevic, Alison E. Mather, John Wain, Mark Webber, Andrew J. Page, Justin O’Grady |
| EPI_ISL_499632, EPI_ISL_499633, EPI_ISL_499634, EPI_ISL_499635, EPI_ISL_499636, EPI_ISL_499637, EPI_ISL_499638, EPI_ISL_499639, EPI_ISL_499640, EPI_ISL_499641, EPI_ISL_499642, EPI_ISL_499643, EPI_ISL_499644, EPI_ISL_499645, EPI_ISL_499646, EPI_ISL_499647, EPI_ISL_499648, EPI_ISL_499649, EPI_ISL_499650, EPI_ISL_499651, EPI_ISL_499652, EPI_ISL_499653, EPI_ISL_499654, EPI_ISL_499655, EPI_ISL_499656, EPI_ISL_499657, EPI_ISL_499658, EPI_ISL_499659, EPI_ISL_499660, EPI_ISL_499661, EPI_ISL_499662, EPI_ISL_499663, EPI_ISL_499664, EPI_ISL_499665, EPI_ISL_499666, EPI_ISL_499667, EPI_ISL_499668, EPI_ISL_499669, EPI_ISL_499670, EPI_ISL_499671, EPI_ISL_499672, EPI_ISL_499673, EPI_ISL_499674, EPI_ISL_499675, EPI_ISL_499676, EPI_ISL_499677, EPI_ISL_499678, EPI_ISL_499679, EPI_ISL_499680, EPI_ISL_499681, EPI_ISL_499682, EPI_ISL_499683, EPI_ISL_499684, EPI_ISL_499685, EPI_ISL_499686, EPI_ISL_499687, EPI_ISL_499688, EPI_ISL_499689, EPI_ISL_499690, EPI_ISL_499691, EPI_ISL_499692, EPI_ISL_499693, EPI_ISL_499694, EPI_ISL_499695, EPI_ISL_499696, EPI_ISL_499697, EPI_ISL_499698, EPI_ISL_499699, EPI_ISL_499700, EPI_ISL_499701, EPI_ISL_499702, EPI_ISL_499703, EPI_ISL_499704, EPI_ISL_499705, EPI_ISL_499706, EPI_ISL_499707, EPI_ISL_499708, EPI_ISL_499709, EPI_ISL_499710, EPI_ISL_499711, EPI_ISL_499712, EPI_ISL_499713, EPI_ISL_499714, EPI_ISL_499715, EPI_ISL_499716, EPI_ISL_499717, EPI_ISL_499718, EPI_ISL_499719, EPI_ISL_499720, EPI_ISL_499721, EPI_ISL_499722, EPI_ISL_499723, EPI_ISL_499724, EPI_ISL_499725, EPI_ISL_499726, EPI_ISL_499727, EPI_ISL_499728, EPI_ISL_499729, EPI_ISL_499730, EPI_ISL_499731, EPI_ISL_499732, EPI_ISL_499733, EPI_ISL_499734, EPI_ISL_499735, EPI_ISL_499736, EPI_ISL_499737, EPI_ISL_499738, EPI_ISL_499739, EPI_ISL_499740, EPI_ISL_499741, EPI_ISL_499742, EPI_ISL_499743, EPI_ISL_499744, EPI_ISL_499745, EPI_ISL_499746, EPI_ISL_499747, EPI_ISL_499748, EPI_ISL_499749, EPI_ISL_499750, EPI_ISL_499751, EPI_ISL_499752, EPI_ISL_499753, EPI_ISL_499754, EPI_ISL_499755, EPI_ISL_499756, EPI_ISL_499757, EPI_ISL_499758, EPI_ISL_499759, EPI_ISL_499760, EPI_ISL_499761, EPI_ISL_499762, EPI_ISL_499763, EPI_ISL_499764, EPI_ISL_499765, EPI_ISL_499766, EPI_ISL_499767, EPI_ISL_499768, EPI_ISL_499769, EPI_ISL_499770, EPI_ISL_499771, EPI_ISL_499772, EPI_ISL_499773, EPI_ISL_499774, EPI_ISL_499775, EPI_ISL_499776 | | | |
| see above | Liverpool Clinical Laboratories | COVID-19 Genomics UK (COG-UK) Consortium | Sam Haldenby, Anita Lucaci, Steve Paterson, Julian Hiscox, Alistair Darby, M Almsaud, A Alrezaihi, Muhannad Alruwaili, Stuart D Armstrong, Jones Benjamin, Eleanor G Bentley, Anu Chawla, Jordan J Clark, Angela Cowell, Richard Eccles, Isabel García-Dorival, Matthew Gemmell, Alessandro Gerada, PKF Gilmore, Richard Gregory, Ximeng Han, Catherine Hartley, Margaret Hughes, Miren Iturriza-Gomara, James Johnson, L Luu, Jenifer Manson, Charlotte Nelson, Elaine O’Toole, Cassie Olateju, Rebekah Penrice-Randal , Lucille Rainbow, N.P Randle, Trevor Ian Robinson, Parul Sharma, Ghada T Shawli, James P Stewart, Neil Swainston, Ecaterina Vamos, Joanne Watts, Mark Whitehead |
| EPI_ISL_499777, EPI_ISL_499778, EPI_ISL_499779, EPI_ISL_499780, EPI_ISL_499781, EPI_ISL_499782, EPI_ISL_499783, EPI_ISL_499784, EPI_ISL_499785, EPI_ISL_499786, EPI_ISL_499787, EPI_ISL_499788, EPI_ISL_499789, EPI_ISL_499790, EPI_ISL_499791, EPI_ISL_499792, EPI_ISL_499793, EPI_ISL_499794, EPI_ISL_499795, EPI_ISL_499796, EPI_ISL_499797, EPI_ISL_499798, EPI_ISL_499799, EPI_ISL_499800, EPI_ISL_499801, EPI_ISL_499802, EPI_ISL_499803, EPI_ISL_499804, EPI_ISL_499805, EPI_ISL_499806, EPI_ISL_499807 | | | |
| see above | Northumbria University / South Tees Hospitals NHS Foundation Trust / North Cumbria Integrated Care NHS Foundation Trust / North Tees and Hartlepool NHS Foundation Trust / Newcastle Hospitals NHS Foundation Trust | COVID-19 Genomics UK (COG-UK) Consortium | Darren L Smith,Andrew Nelson,Matthew Bashton,Greg R Young,Joshua Loh,John Allan,Mohammad A Tariq,Giles S Holt,Gary Black,Wen C Yew,Lynn Dover,Paul Baker,Steve Liggett,Sarah Essex,Jane Greenaway,Debra Padgett,Clive Graham,Garren Scott,Edward Barton,Emma Swindells,Brendan Payne,Jennifer Collins,Yusri Taha,Gary Eltringham |
| EPI_ISL_499808, EPI_ISL_499809 | Queens Medical Centre, Clinical Microbiology Department / DeepSeq Nottingham | COVID-19 Genomics UK (COG-UK) Consortium | Gemma Clark, Wendy Smith, Manjinder Khakh, Vicki M Fleming, Michelle M Lister, Hannah Howson-Wells, Jonathan Ball, Patrick McClure, Joseph Chappell, Theocharis Tsoleridis, Nadine Holmes, Matthew Carlisle, Christopher Moore, Fei Sang, Johnny Debebe, Victoria Wright, Matthew Loose |
| EPI_ISL_499810 | University Hospitals Of Leicester NHS Trust and DeepSeq Nottingham | COVID-19 Genomics UK (COG-UK) Consortium | Christopher Holmes, Paul Bird, Thomas Helmer, Karlie Fallon, Julian Tang, Jonathan Ball, Patrick McClure, Joeseph Chappell, Nadine Holmes, Matthew Carlisle, Christopher Moore, Fei Sang, Johnny Debebe, Victoria Wright, Matthew Loose |
| EPI_ISL_499811, EPI_ISL_499812, EPI_ISL_499813, EPI_ISL_499814, EPI_ISL_499815, EPI_ISL_499816, EPI_ISL_499817, EPI_ISL_499818, EPI_ISL_499819, EPI_ISL_499820, EPI_ISL_499821, EPI_ISL_499822, EPI_ISL_499823, EPI_ISL_499824, EPI_ISL_499825, EPI_ISL_499826, EPI_ISL_499827 | | | |
| see above | Liverpool Clinical Laboratories | COVID-19 Genomics UK (COG-UK) Consortium | Sam Haldenby, Anita Lucaci, Steve Paterson, Julian Hiscox, Alistair Darby, M Almsaud, A Alrezaihi, Muhannad Alruwaili, Stuart D Armstrong, Jones Benjamin, Eleanor G Bentley, Anu Chawla, Jordan J Clark, Angela Cowell, Richard Eccles, Isabel García-Dorival, Matthew Gemmell, Alessandro Gerada, PKF Gilmore, Richard Gregory, Ximeng Han, Catherine Hartley, Margaret Hughes, Miren Iturriza-Gomara, James Johnson, L Luu, Jenifer Manson, Charlotte Nelson, Elaine O’Toole, Cassie Olateju, Rebekah Penrice-Randal , Lucille Rainbow, N.P Randle, Trevor Ian Robinson, Parul Sharma, Ghada T Shawli, James P Stewart, Neil Swainston, Ecaterina Vamos, Joanne Watts, Mark Whitehead |
| EPI_ISL_499828, EPI_ISL_499829 | University Hospitals Of Leicester NHS Trust and DeepSeq Nottingham | COVID-19 Genomics UK (COG-UK) Consortium | Christopher Holmes, Paul Bird, Thomas Helmer, Karlie Fallon, Julian Tang, Jonathan Ball, Patrick McClure, Joeseph Chappell, Nadine Holmes, Matthew Carlisle, Christopher Moore, Fei Sang, Johnny Debebe, Victoria Wright, Matthew Loose |
| EPI_ISL_499830 | University of Birmingham | COVID-19 Genomics UK (COG-UK) Consortium | Institute of Microbiology, University of Birmingham: Claire McMurray, Joanne Stockton, Samuel Nicholls, Radoslaw Poplawski, Will Rowe, Josh Quick, Nicholas Loman. University of Birmingham Testing Laboratory: Celina M Whalley, Andrew Bosworth, Charlotte Poxon, Kasun Wanigasooriya, Oliver Pickles, Mike Kidd, Alex Richter, Andrew D Beggs PHE Heartlands Lab: Husam Osman, Andrew Bosworth. Queen Elizabeth Hospital: Anna Casey |
| EPI_ISL_499831, EPI_ISL_499832, EPI_ISL_499833, EPI_ISL_499834, EPI_ISL_499835, EPI_ISL_499836, EPI_ISL_499837, EPI_ISL_499838 | Liverpool Clinical Laboratories | COVID-19 Genomics UK (COG-UK) Consortium | Sam Haldenby, Anita Lucaci, Steve Paterson, Julian Hiscox, Alistair Darby, M Almsaud, A Alrezaihi, Muhannad Alruwaili, Stuart D Armstrong, Jones Benjamin, Eleanor G Bentley, Anu Chawla, Jordan J Clark, Angela Cowell, Richard Eccles, Isabel García-Dorival, Matthew Gemmell, Alessandro Gerada, PKF Gilmore, Richard Gregory, Ximeng Han, Catherine Hartley, Margaret Hughes, Miren Iturriza-Gomara, James Johnson, L Luu, Jenifer Manson, Charlotte Nelson, Elaine O’Toole, Cassie Olateju, Rebekah Penrice-Randal , Lucille Rainbow, N.P Randle, Trevor Ian Robinson, Parul Sharma, Ghada T Shawli, James P Stewart, Neil Swainston, Ecaterina Vamos, Joanne Watts, Mark Whitehead |
| EPI_ISL_499839 | University of Birmingham | COVID-19 Genomics UK (COG-UK) Consortium | Institute of Microbiology, University of Birmingham: Claire McMurray, Joanne Stockton, Samuel Nicholls, Radoslaw Poplawski, Will Rowe, Josh Quick, Nicholas Loman. University of Birmingham Testing Laboratory: Celina M Whalley, Andrew Bosworth, Charlotte Poxon, Kasun Wanigasooriya, Oliver Pickles, Mike Kidd, Alex Richter, Andrew D Beggs PHE Heartlands Lab: Husam Osman, Andrew Bosworth. Queen Elizabeth Hospital: Anna Casey |
| EPI_ISL_499840, EPI_ISL_499841, EPI_ISL_499842, EPI_ISL_499843, EPI_ISL_499844, EPI_ISL_499845, EPI_ISL_499846, EPI_ISL_499847, EPI_ISL_499848, EPI_ISL_499849, EPI_ISL_499850 | | | |
| see above | Liverpool Clinical Laboratories | COVID-19 Genomics UK (COG-UK) Consortium | Sam Haldenby, Anita Lucaci, Steve Paterson, Julian Hiscox, Alistair Darby, M Almsaud, A Alrezaihi, Muhannad Alruwaili, Stuart D Armstrong, Jones Benjamin, Eleanor G Bentley, Anu Chawla, Jordan J Clark, Angela Cowell, Richard Eccles, Isabel García-Dorival, Matthew Gemmell, Alessandro Gerada, PKF Gilmore, Richard Gregory, Ximeng Han, Catherine Hartley, Margaret Hughes, Miren Iturriza-Gomara, James Johnson, L Luu, Jenifer Manson, Charlotte Nelson, Elaine O’Toole, Cassie Olateju, Rebekah Penrice-Randal , Lucille Rainbow, N.P Randle, Trevor Ian Robinson, Parul Sharma, Ghada T Shawli, James P Stewart, Neil Swainston, Ecaterina Vamos, Joanne Watts, Mark Whitehead |
| EPI_ISL_499851, EPI_ISL_499852 | University of Birmingham | COVID-19 Genomics UK (COG-UK) Consortium | Institute of Microbiology, University of Birmingham: Claire McMurray, Joanne Stockton, Samuel Nicholls, Radoslaw Poplawski, Will Rowe, Josh Quick, Nicholas Loman. University of Birmingham Testing Laboratory: Celina M Whalley, Andrew Bosworth, Charlotte Poxon, Kasun Wanigasooriya, Oliver Pickles, Mike Kidd, Alex Richter, Andrew D Beggs PHE Heartlands Lab: Husam Osman, Andrew Bosworth. Queen Elizabeth Hospital: Anna Casey |
| EPI_ISL_499853 | Department of Pathology, University of Cambridge | COVID-19 Genomics UK (COG-UK) Consortium | Luke W Meredith, M. Estée Török, Myra Hosmillo, William L. Hamilton, Martin D. Curran, Theresa Feltwell, Grant Hall, Anna Yakovleva, Fahad A Khokhar, Charlotte J. Houldcroft, Laura G Caller, Aminu S. Jahun, Sarah L. Caddy, Yasmin Chaudhry, Malte Pinckert, Ian Goodfellow |
| EPI_ISL_499854 | Liverpool Clinical Laboratories | COVID-19 Genomics UK (COG-UK) Consortium | Sam Haldenby, Anita Lucaci, Steve Paterson, Julian Hiscox, Alistair Darby, M Almsaud, A Alrezaihi, Muhannad Alruwaili, Stuart D Armstrong, Jones Benjamin, Eleanor G Bentley, Anu Chawla, Jordan J Clark, Angela Cowell, Richard Eccles, Isabel García-Dorival, Matthew Gemmell, Alessandro Gerada, PKF Gilmore, Richard Gregory, Ximeng Han, Catherine Hartley, Margaret Hughes, Miren Iturriza-Gomara, James Johnson, L Luu, Jenifer Manson, Charlotte Nelson, Elaine O’Toole, Cassie Olateju, Rebekah Penrice-Randal , Lucille Rainbow, N.P Randle, Trevor Ian Robinson, Parul Sharma, Ghada T Shawli, James P Stewart, Neil Swainston, Ecaterina Vamos, Joanne Watts, Mark Whitehead |
| EPI_ISL_499855 | Department of Pathology, University of Cambridge | COVID-19 Genomics UK (COG-UK) Consortium | Luke W Meredith, M. Estée Török, Myra Hosmillo, William L. Hamilton, Martin D. Curran, Theresa Feltwell, Grant Hall, Anna Yakovleva, Fahad A Khokhar, Charlotte J. Houldcroft, Laura G Caller, Aminu S. Jahun, Sarah L. Caddy, Yasmin Chaudhry, Malte Pinckert, Ian Goodfellow |
| EPI_ISL_499856, EPI_ISL_499857 | Liverpool Clinical Laboratories | COVID-19 Genomics UK (COG-UK) Consortium | Sam Haldenby, Anita Lucaci, Steve Paterson, Julian Hiscox, Alistair Darby, M Almsaud, A Alrezaihi, Muhannad Alruwaili, Stuart D Armstrong, Jones Benjamin, Eleanor G Bentley, Anu Chawla, Jordan J Clark, Angela Cowell, Richard Eccles, Isabel García-Dorival, Matthew Gemmell, Alessandro Gerada, PKF Gilmore, Richard Gregory, Ximeng Han, Catherine Hartley, Margaret Hughes, Miren Iturriza-Gomara, James Johnson, L Luu, Jenifer Manson, Charlotte Nelson, Elaine O’Toole, Cassie Olateju, Rebekah Penrice-Randal , Lucille Rainbow, N.P Randle, Trevor Ian Robinson, Parul Sharma, Ghada T Shawli, James P Stewart, Neil Swainston, Ecaterina Vamos, Joanne Watts, Mark Whitehead |
| EPI_ISL_499858 | University Hospitals Of Leicester NHS Trust and DeepSeq Nottingham | COVID-19 Genomics UK (COG-UK) Consortium | Christopher Holmes, Paul Bird, Thomas Helmer, Karlie Fallon, Julian Tang, Jonathan Ball, Patrick McClure, Joeseph Chappell, Nadine Holmes, Matthew Carlisle, Christopher Moore, Fei Sang, Johnny Debebe, Victoria Wright, Matthew Loose |
| EPI_ISL_499859, EPI_ISL_499860 | Liverpool Clinical Laboratories | COVID-19 Genomics UK (COG-UK) Consortium | Sam Haldenby, Anita Lucaci, Steve Paterson, Julian Hiscox, Alistair Darby, M Almsaud, A Alrezaihi, Muhannad Alruwaili, Stuart D Armstrong, Jones Benjamin, Eleanor G Bentley, Anu Chawla, Jordan J Clark, Angela Cowell, Richard Eccles, Isabel García-Dorival, Matthew Gemmell, Alessandro Gerada, PKF Gilmore, Richard Gregory, Ximeng Han, Catherine Hartley, Margaret Hughes, Miren Iturriza-Gomara, James Johnson, L Luu, Jenifer Manson, Charlotte Nelson, Elaine O’Toole, Cassie Olateju, Rebekah Penrice-Randal , Lucille Rainbow, N.P Randle, Trevor Ian Robinson, Parul Sharma, |

| | | | |
|---|---|---|---|
| | | | Ghada T Shawli, James P Stewart, Neil Swainston, Ecaterina Vamos, Joanne Watts, Mark Whitehead |
| EPI_ISL_499861 | University Hospitals Of Leicester NHS Trust and DeepSeq Nottingham | COVID-19 Genomics UK (COG-UK) Consortium | Christopher Holmes, Paul Bird, Thomas Helmer, Karlie Fallon, Julian Tang, Jonathan Ball, Patrick McClure, Joeseph Chappell, Nadine Holmes, Matthew Carlisle, Christopher Moore, Fei Sang, Johnny Debebe, Victoria Wright, Matthew Loose |
| EPI_ISL_499862, EPI_ISL_499863, EPI_ISL_499864, EPI_ISL_499865, EPI_ISL_499866, EPI_ISL_499867, EPI_ISL_499868 | Liverpool Clinical Laboratories | COVID-19 Genomics UK (COG-UK) Consortium | Sam Haldenby, Anita Lucaci, Steve Paterson, Julian Hiscox, Alistair Darby, M Almsaud, A Alrezaihi, Muhannad Alruwaili, Stuart D Armstrong, Jones Benjamin, Eleanor G Bentley, Anu Chawla, Jordan J Clark, Angela Cowell, Richard Eccles, Isabel García-Dorival, Matthew Gemmell, Alessandro Gerada, PKF Gilmore, Richard Gregory, Ximeng Han, Catherine Hartley, Margaret Hughes, Miren Iturriza-Gomara, James Johnson, L Luu, Jenifer Manson, Charlotte Nelson, Elaine O'Toole, Cassie Olateju, Rebekah Penrice-Randal , Lucille Rainbow, N.P Randle, Trevor Ian Robinson, Parul Sharma, Ghada T Shawli, James P Stewart, Neil Swainston, Ecaterina Vamos, Joanne Watts, Mark Whitehead |
| EPI_ISL_499869 | University Hospitals Of Leicester NHS Trust and DeepSeq Nottingham | COVID-19 Genomics UK (COG-UK) Consortium | Christopher Holmes, Paul Bird, Thomas Helmer, Karlie Fallon, Julian Tang, Jonathan Ball, Patrick McClure, Joeseph Chappell, Nadine Holmes, Matthew Carlisle, Christopher Moore, Fei Sang, Johnny Debebe, Victoria Wright, Matthew Loose |
| EPI_ISL_499870, EPI_ISL_499871, EPI_ISL_499872, EPI_ISL_499873, EPI_ISL_499874, EPI_ISL_499875, EPI_ISL_499876, EPI_ISL_499877, EPI_ISL_499878, EPI_ISL_499879, EPI_ISL_499880, EPI_ISL_499881, EPI_ISL_499882, EPI_ISL_499883, EPI_ISL_499884, EPI_ISL_499885, EPI_ISL_499886, EPI_ISL_499887, EPI_ISL_499888, EPI_ISL_499889, EPI_ISL_499890, EPI_ISL_499891, EPI_ISL_499892, EPI_ISL_499893 | | | |
| see above | Liverpool Clinical Laboratories | COVID-19 Genomics UK (COG-UK) Consortium | Sam Haldenby, Anita Lucaci, Steve Paterson, Julian Hiscox, Alistair Darby, M Almsaud, A Alrezaihi, Muhannad Alruwaili, Stuart D Armstrong, Jones Benjamin, Eleanor G Bentley, Anu Chawla, Jordan J Clark, Angela Cowell, Richard Eccles, Isabel García-Dorival, Matthew Gemmell, Alessandro Gerada, PKF Gilmore, Richard Gregory, Ximeng Han, Catherine Hartley, Margaret Hughes, Miren Iturriza-Gomara, James Johnson, L Luu, Jenifer Manson, Charlotte Nelson, Elaine O'Toole, Cassie Olateju, Rebekah Penrice-Randal , Lucille Rainbow, N.P Randle, Trevor Ian Robinson, Parul Sharma, Ghada T Shawli, James P Stewart, Neil Swainston, Ecaterina Vamos, Joanne Watts, Mark Whitehead |
| EPI_ISL_499894, EPI_ISL_499895 | University Hospitals Of Leicester NHS Trust and DeepSeq Nottingham | COVID-19 Genomics UK (COG-UK) Consortium | Christopher Holmes, Paul Bird, Thomas Helmer, Karlie Fallon, Julian Tang, Jonathan Ball, Patrick McClure, Joeseph Chappell, Nadine Holmes, Matthew Carlisle, Christopher Moore, Fei Sang, Johnny Debebe, Victoria Wright, Matthew Loose |
| EPI_ISL_499896 | Liverpool Clinical Laboratories | COVID-19 Genomics UK (COG-UK) Consortium | Sam Haldenby, Anita Lucaci, Steve Paterson, Julian Hiscox, Alistair Darby, M Almsaud, A Alrezaihi, Muhannad Alruwaili, Stuart D Armstrong, Jones Benjamin, Eleanor G Bentley, Anu Chawla, Jordan J Clark, Angela Cowell, Richard Eccles, Isabel García-Dorival, Matthew Gemmell, Alessandro Gerada, PKF Gilmore, Richard Gregory, Ximeng Han, Catherine Hartley, Margaret Hughes, Miren Iturriza-Gomara, James Johnson, L Luu, Jenifer Manson, Charlotte Nelson, Elaine O'Toole, Cassie Olateju, Rebekah Penrice-Randal , Lucille Rainbow, N.P Randle, Trevor Ian Robinson, Parul Sharma, Ghada T Shawli, James P Stewart, Neil Swainston, Ecaterina Vamos, Joanne Watts, Mark Whitehead |
| EPI_ISL_499897 | University Hospitals Of Leicester NHS Trust and DeepSeq Nottingham | COVID-19 Genomics UK (COG-UK) Consortium | Christopher Holmes, Paul Bird, Thomas Helmer, Karlie Fallon, Julian Tang, Jonathan Ball, Patrick McClure, Joeseph Chappell, Nadine Holmes, Matthew Carlisle, Christopher Moore, Fei Sang, Johnny Debebe, Victoria Wright, Matthew Loose |
| EPI_ISL_499898, EPI_ISL_499899, EPI_ISL_499900, EPI_ISL_499901, EPI_ISL_499902, EPI_ISL_499903, EPI_ISL_499904 | Liverpool Clinical Laboratories | COVID-19 Genomics UK (COG-UK) Consortium | Sam Haldenby, Anita Lucaci, Steve Paterson, Julian Hiscox, Alistair Darby, M Almsaud, A Alrezaihi, Muhannad Alruwaili, Stuart D Armstrong, Jones Benjamin, Eleanor G Bentley, Anu Chawla, Jordan J Clark, Angela Cowell, Richard Eccles, Isabel García-Dorival, Matthew Gemmell, Alessandro Gerada, PKF Gilmore, Richard Gregory, Ximeng Han, Catherine Hartley, Margaret Hughes, Miren Iturriza-Gomara, James Johnson, L Luu, Jenifer Manson, Charlotte Nelson, Elaine O'Toole, Cassie Olateju, Rebekah Penrice-Randal , Lucille Rainbow, N.P Randle, Trevor Ian Robinson, Parul Sharma, Ghada T Shawli, James P Stewart, Neil Swainston, Ecaterina Vamos, Joanne Watts, Mark Whitehead |
| EPI_ISL_499905 | Department of Pathology, University of Cambridge | COVID-19 Genomics UK (COG-UK) Consortium | Luke W Meredith, M. Estée Török, Myra Hosmillo, William L. Hamilton, Martin D. Curran, Theresa Feltwell, Grant Hall, Anna Yakovleva, Fahad A Khokhar, Charlotte J. Houldcroft, Laura G Caller, Aminu S. Jahun, Sarah L. Caddy, Yasmin Chaudhry, Malte Pinckert, Ian Goodfellow |
| EPI_ISL_499906, EPI_ISL_499907 | Liverpool Clinical Laboratories | COVID-19 Genomics UK (COG-UK) Consortium | Sam Haldenby, Anita Lucaci, Steve Paterson, Julian Hiscox, Alistair Darby, M Almsaud, A Alrezaihi, Muhannad Alruwaili, Stuart D Armstrong, Jones Benjamin, Eleanor G Bentley, Anu Chawla, Jordan J Clark, Angela Cowell, Richard Eccles, Isabel García-Dorival, Matthew Gemmell, Alessandro Gerada, PKF Gilmore, Richard Gregory, Ximeng Han, Catherine Hartley, Margaret Hughes, Miren Iturriza-Gomara, James Johnson, L Luu, Jenifer Manson, Charlotte Nelson, Elaine O'Toole, Cassie Olateju, Rebekah Penrice-Randal , Lucille Rainbow, N.P Randle, Trevor Ian Robinson, Parul Sharma, Ghada T Shawli, James P Stewart, Neil Swainston, Ecaterina Vamos, Joanne Watts, Mark Whitehead |
| EPI_ISL_499908 | Department of Pathology, University of Cambridge | COVID-19 Genomics UK (COG-UK) Consortium | Luke W Meredith, M. Estée Török, Myra Hosmillo, William L. Hamilton, Martin D. Curran, Theresa Feltwell, Grant Hall, Anna Yakovleva, Fahad A Khokhar, Charlotte J. Houldcroft, Laura G Caller, Aminu S. Jahun, Sarah L. Caddy, Yasmin Chaudhry, Malte Pinckert, Ian Goodfellow |
| EPI_ISL_499909 | Liverpool Clinical Laboratories | COVID-19 Genomics UK (COG-UK) Consortium | Sam Haldenby, Anita Lucaci, Steve Paterson, Julian Hiscox, Alistair Darby, M Almsaud, A Alrezaihi, Muhannad Alruwaili, Stuart D Armstrong, Jones Benjamin, Eleanor G Bentley, Anu Chawla, Jordan J Clark, Angela Cowell, Richard Eccles, Isabel García-Dorival, Matthew Gemmell, Alessandro Gerada, PKF Gilmore, Richard Gregory, Ximeng Han, Catherine Hartley, Margaret Hughes, Miren Iturriza-Gomara, James Johnson, L Luu, Jenifer Manson, Charlotte Nelson, Elaine O'Toole, Cassie Olateju, Rebekah Penrice-Randal , Lucille Rainbow, N.P Randle, Trevor Ian Robinson, Parul Sharma, Ghada T Shawli, James P Stewart, Neil Swainston, Ecaterina Vamos, Joanne Watts, Mark Whitehead |
| EPI_ISL_499910 | University Hospitals Of Leicester NHS Trust and DeepSeq Nottingham | COVID-19 Genomics UK (COG-UK) Consortium | Christopher Holmes, Paul Bird, Thomas Helmer, Karlie Fallon, Julian Tang, Jonathan Ball, Patrick McClure, Joeseph Chappell, Nadine Holmes, Matthew Carlisle, Christopher Moore, Fei Sang, Johnny Debebe, Victoria Wright, Matthew Loose |
| EPI_ISL_499911 | Liverpool Clinical Laboratories | COVID-19 Genomics UK (COG-UK) Consortium | Sam Haldenby, Anita Lucaci, Steve Paterson, Julian Hiscox, Alistair Darby, M Almsaud, A Alrezaihi, Muhannad Alruwaili, Stuart D Armstrong, Jones Benjamin, Eleanor G Bentley, Anu Chawla, Jordan J Clark, Angela Cowell, Richard Eccles, Isabel García-Dorival, Matthew Gemmell, Alessandro Gerada, PKF Gilmore, Richard Gregory, Ximeng Han, Catherine Hartley, Margaret Hughes, Miren Iturriza-Gomara, James Johnson, L Luu, Jenifer Manson, Charlotte Nelson, Elaine O'Toole, Cassie Olateju, Rebekah Penrice-Randal , Lucille Rainbow, N.P Randle, Trevor Ian Robinson, Parul Sharma, Ghada T Shawli, James P Stewart, Neil Swainston, Ecaterina Vamos, Joanne Watts, Mark Whitehead |
| EPI_ISL_499912, EPI_ISL_499913 | University Hospitals Of Leicester NHS Trust and DeepSeq Nottingham | COVID-19 Genomics UK (COG-UK) Consortium | Christopher Holmes, Paul Bird, Thomas Helmer, Karlie Fallon, Julian Tang, Jonathan Ball, Patrick McClure, Joeseph Chappell, Nadine Holmes, Matthew Carlisle, Christopher Moore, Fei Sang, Johnny Debebe, Victoria Wright, Matthew Loose |
| EPI_ISL_499914, EPI_ISL_499915, EPI_ISL_499916, EPI_ISL_499917, EPI_ISL_499918, EPI_ISL_499919, EPI_ISL_499920, EPI_ISL_499921, EPI_ISL_499922, EPI_ISL_499923, EPI_ISL_499924, EPI_ISL_499925, EPI_ISL_499926, EPI_ISL_499927, EPI_ISL_499928, EPI_ISL_499929, EPI_ISL_499930, EPI_ISL_499931, EPI_ISL_499932 | | | |
| see above | Liverpool Clinical Laboratories | COVID-19 Genomics UK (COG-UK) Consortium | Sam Haldenby, Anita Lucaci, Steve Paterson, Julian Hiscox, Alistair Darby, M Almsaud, A Alrezaihi, Muhannad Alruwaili, Stuart D Armstrong, Jones Benjamin, Eleanor G Bentley, Anu Chawla, Jordan J Clark, Angela Cowell, Richard Eccles, Isabel García-Dorival, Matthew Gemmell, Alessandro Gerada, PKF Gilmore, Richard Gregory, Ximeng Han, Catherine Hartley, Margaret Hughes, Miren Iturriza-Gomara, James Johnson, L Luu, Jenifer Manson, Charlotte Nelson, Elaine O'Toole, Cassie Olateju, Rebekah Penrice-Randal , Lucille Rainbow, N.P Randle, Trevor Ian Robinson, Parul Sharma, Ghada T Shawli, James P Stewart, Neil Swainston, Ecaterina Vamos, Joanne Watts, Mark Whitehead |
| EPI_ISL_499933 | University Hospitals Of Leicester NHS Trust and DeepSeq Nottingham | COVID-19 Genomics UK (COG-UK) Consortium | Christopher Holmes, Paul Bird, Thomas Helmer, Karlie Fallon, Julian Tang, Jonathan Ball, Patrick McClure, Joeseph Chappell, Nadine Holmes, Matthew Carlisle, Christopher Moore, Fei Sang, Johnny Debebe, Victoria Wright, Matthew Loose |
| EPI_ISL_499934, EPI_ISL_499935, EPI_ISL_499936, EPI_ISL_499937, EPI_ISL_499938, EPI_ISL_499939, EPI_ISL_499940, EPI_ISL_499941, EPI_ISL_499942, EPI_ISL_499943, EPI_ISL_499944, EPI_ISL_499945, EPI_ISL_499946 | | | |
| see above | Liverpool Clinical Laboratories | COVID-19 Genomics UK (COG-UK) Consortium | Sam Haldenby, Anita Lucaci, Steve Paterson, Julian Hiscox, Alistair Darby, M Almsaud, A Alrezaihi, Muhannad Alruwaili, Stuart D Armstrong, Jones Benjamin, Eleanor G Bentley, Anu Chawla, Jordan J Clark, Angela Cowell, Richard Eccles, Isabel García-Dorival, Matthew Gemmell, Alessandro Gerada, PKF Gilmore, Richard Gregory, Ximeng Han, Catherine Hartley, Margaret Hughes, Miren Iturriza-Gomara, James Johnson, L Luu, Jenifer Manson, Charlotte Nelson, Elaine O'Toole, Cassie Olateju, Rebekah Penrice-Randal , Lucille Rainbow, N.P Randle, Trevor Ian Robinson, Parul Sharma, Ghada T Shawli, James P Stewart, Neil Swainston, Ecaterina Vamos, Joanne Watts, Mark Whitehead |
| EPI_ISL_499947 | University of Birmingham | COVID-19 Genomics UK (COG-UK) Consortium | Institute of Microbiology, University of Birmingham: Claire McMurray, Joanne Stockton, Samuel Nicholls, Radoslaw Poplawski, Will Rowe, Josh Quick, Nicholas Loman. University of Birmingham Testing Laboratory: Celina M Whalley, Andrew Bosworth, Charlotte Poxon, Kasun Wanigasooriya, Oliver Pickles, Mike Kidd, Alex Richter, Andrew D Beggs PHE Heartlands Lab: Husam Osman, Andrew Bosworth. Queen Elizabeth Hospital: Anna Casey |
| EPI_ISL_499948, EPI_ISL_499949 | Liverpool Clinical Laboratories | COVID-19 Genomics UK (COG-UK) Consortium | Sam Haldenby, Anita Lucaci, Steve Paterson, Julian Hiscox, Alistair Darby, M Almsaud, A Alrezaihi, Muhannad Alruwaili, Stuart D Armstrong, Jones Benjamin, Eleanor G Bentley, Anu Chawla, Jordan J Clark, Angela Cowell, Richard Eccles, Isabel García-Dorival, Matthew Gemmell, Alessandro Gerada, PKF Gilmore, Richard Gregory, Ximeng Han, Catherine Hartley, Margaret Hughes, Miren Iturriza-Gomara, James Johnson, L Luu, Jenifer Manson, Charlotte Nelson, Elaine O'Toole, Cassie Olateju, Rebekah Penrice-Randal , Lucille Rainbow, N.P Randle, Trevor Ian Robinson, Parul Sharma, Ghada T Shawli, James P Stewart, Neil Swainston, Ecaterina Vamos, Joanne Watts, Mark Whitehead |
| EPI_ISL_499950 | University Hospitals Of Leicester NHS Trust and DeepSeq Nottingham | COVID-19 Genomics UK (COG-UK) Consortium | Christopher Holmes, Paul Bird, Thomas Helmer, Karlie Fallon, Julian Tang, Jonathan Ball, Patrick McClure, Joeseph Chappell, Nadine Holmes, Matthew Carlisle, Christopher Moore, Fei Sang, Johnny Debebe, Victoria Wright, Matthew Loose |
| EPI_ISL_499951, EPI_ISL_499952, EPI_ISL_499953, EPI_ISL_499954, EPI_ISL_499955, EPI_ISL_499956, EPI_ISL_499957, EPI_ISL_499958, EPI_ISL_499959, EPI_ISL_499960, EPI_ISL_499961, EPI_ISL_499962, EPI_ISL_499963, EPI_ISL_499964, EPI_ISL_499965 | | | |
| see above | Liverpool Clinical Laboratories | COVID-19 Genomics UK (COG-UK) Consortium | Sam Haldenby, Anita Lucaci, Steve Paterson, Julian Hiscox, Alistair Darby, M Almsaud, A Alrezaihi, Muhannad Alruwaili, Stuart D Armstrong, Jones Benjamin, Eleanor G Bentley, Anu Chawla, Jordan J Clark, Angela Cowell, Richard Eccles, Isabel García-Dorival, Matthew Gemmell, Alessandro Gerada, PKF Gilmore, Richard Gregory, Ximeng Han, Catherine Hartley, Margaret Hughes, Miren Iturriza-Gomara, James Johnson, L Luu, Jenifer Manson, Charlotte Nelson, Elaine O'Toole, Cassie Olateju, Rebekah Penrice-Randal , Lucille Rainbow, N.P Randle, Trevor Ian Robinson, Parul Sharma, Ghada T Shawli, James P Stewart, Neil Swainston, Ecaterina Vamos, Joanne Watts, Mark Whitehead |
| EPI_ISL_499966 | Department of Pathology, University of Cambridge | COVID-19 Genomics UK (COG-UK) Consortium | Luke W Meredith, M. Estée Török, Myra Hosmillo, William L. Hamilton, Martin D. Curran, Theresa Feltwell, Grant Hall, Anna Yakovleva, Fahad A Khokhar, Charlotte J. Houldcroft, Laura G Caller, Aminu S. Jahun, Sarah L. Caddy, Yasmin Chaudhry, Malte Pinckert, Ian Goodfellow |
| EPI_ISL_499967 | Liverpool Clinical Laboratories | COVID-19 Genomics UK (COG-UK) Consortium | Sam Haldenby, Anita Lucaci, Steve Paterson, Julian Hiscox, Alistair Darby, M Almsaud, A Alrezaihi, Muhannad Alruwaili, Stuart D Armstrong, Jones Benjamin, Eleanor G Bentley, Anu Chawla, Jordan J Clark, Angela Cowell, Richard Eccles, Isabel García-Dorival, Matthew Gemmell, Alessandro Gerada, PKF Gilmore, Richard Gregory, Ximeng Han, Catherine Hartley, Margaret Hughes, Miren Iturriza-Gomara, James Johnson, L Luu, Jenifer Manson, Charlotte Nelson, Elaine O'Toole, Cassie Olateju, Rebekah Penrice-Randal , Lucille Rainbow, N.P Randle, Trevor Ian Robinson, Parul Sharma, Ghada T Shawli, James P Stewart, Neil Swainston, Ecaterina Vamos, Joanne Watts, Mark Whitehead |
| EPI_ISL_499968, EPI_ISL_499969, EPI_ISL_499970, EPI_ISL_499971, EPI_ISL_499972, EPI_ISL_499973 | Department of Pathology, University of Cambridge | COVID-19 Genomics UK (COG-UK) Consortium | Luke W Meredith, M. Estée Török, Myra Hosmillo, William L. Hamilton, Martin D. Curran, Theresa Feltwell, Grant Hall, Anna Yakovleva, Fahad A Khokhar, Charlotte J. Houldcroft, Laura G Caller, Aminu S. Jahun, Sarah L. Caddy, Yasmin Chaudhry, Malte Pinckert, Ian Goodfellow |
| EPI_ISL_499974, EPI_ISL_499975, EPI_ISL_499976, EPI_ISL_499977, EPI_ISL_499978, EPI_ISL_499979, EPI_ISL_499980, EPI_ISL_499981, EPI_ISL_499982, EPI_ISL_499983 | University Hospitals Of Leicester NHS Trust and DeepSeq Nottingham | COVID-19 Genomics UK (COG-UK) Consortium | Christopher Holmes, Paul Bird, Thomas Helmer, Karlie Fallon, Julian Tang, Jonathan Ball, Patrick McClure, Joeseph Chappell, Nadine Holmes, Matthew Carlisle, Christopher Moore, Fei Sang, Johnny Debebe, Victoria Wright, Matthew Loose |
| EPI_ISL_499984, EPI_ISL_499985, EPI_ISL_499986, EPI_ISL_499987, EPI_ISL_499988, EPI_ISL_499989, EPI_ISL_499990, EPI_ISL_499991, EPI_ISL_499992, EPI_ISL_499993, EPI_ISL_499994, EPI_ISL_499995, EPI_ISL_499996, EPI_ISL_499997, EPI_ISL_499998, EPI_ISL_499999, EPI_ISL_500000, EPI_ISL_500001, EPI_ISL_500002, EPI_ISL_500003, EPI_ISL_500004, EPI_ISL_500005, EPI_ISL_500006, EPI_ISL_500007, EPI_ISL_500008, EPI_ISL_500009, EPI_ISL_500010, EPI_ISL_500011, EPI_ISL_500012, EPI_ISL_500013, EPI_ISL_500014, EPI_ISL_500015, EPI_ISL_500016, EPI_ISL_500017, EPI_ISL_500018, EPI_ISL_500019, EPI_ISL_500020, EPI_ISL_500021, EPI_ISL_500022, EPI_ISL_500023, EPI_ISL_500024, EPI_ISL_500025, EPI_ISL_500026, EPI_ISL_500027, EPI_ISL_500028, EPI_ISL_500029, EPI_ISL_500030, EPI_ISL_500031, EPI_ISL_500032, EPI_ISL_500033, EPI_ISL_500034, EPI_ISL_500035, EPI_ISL_500036, EPI_ISL_500037, EPI_ISL_500038, EPI_ISL_500039, EPI_ISL_500040, EPI_ISL_500041, EPI_ISL_500042, EPI_ISL_500043, EPI_ISL_500044, EPI_ISL_500045, EPI_ISL_500046, EPI_ISL_500047, EPI_ISL_500048, EPI_ISL_500049, EPI_ISL_500050, EPI_ISL_500051, EPI_ISL_500052, EPI_ISL_500053, EPI_ISL_500054, EPI_ISL_500055, EPI_ISL_500056, EPI_ISL_500057, EPI_ISL_500058, EPI_ISL_500059, EPI_ISL_500060, EPI_ISL_500061, EPI_ISL_500062, EPI_ISL_500063, EPI_ISL_500064, EPI_ISL_500065, EPI_ISL_500066, EPI_ISL_500067, EPI_ISL_500068, EPI_ISL_500069, EPI_ISL_500070, EPI_ISL_500071, EPI_ISL_500072, EPI_ISL_500073, EPI_ISL_500074, EPI_ISL_500075, EPI_ISL_500076, EPI_ISL_500077, EPI_ISL_500078, EPI_ISL_500079, EPI_ISL_500080, EPI_ISL_500081, EPI_ISL_500082, EPI_ISL_500083, EPI_ISL_500084, EPI_ISL_500085, EPI_ISL_500086, EPI_ISL_500087, EPI_ISL_500088, EPI_ISL_500089, EPI_ISL_500090, EPI_ISL_500091, EPI_ISL_500092, EPI_ISL_500093, EPI_ISL_500094, EPI_ISL_500095, EPI_ISL_500096, EPI_ISL_500097, EPI_ISL_500098, EPI_ISL_500099, EPI_ISL_500100, EPI_ISL_500101, EPI_ISL_500102, EPI_ISL_500103, EPI_ISL_500104, EPI_ISL_500105, EPI_ISL_500106, EPI_ISL_500107, EPI_ISL_500108, EPI_ISL_500109, EPI_ISL_500110, EPI_ISL_500111, EPI_ISL_500112, EPI_ISL_500113, EPI_ISL_500114, EPI_ISL_500115, EPI_ISL_500116, EPI_ISL_500117, EPI_ISL_500118, EPI_ISL_500119, EPI_ISL_500120, EPI_ISL_500121, EPI_ISL_500122, EPI_ISL_500123, EPI_ISL_500124, EPI_ISL_500125, EPI_ISL_500126, EPI_ISL_500127, EPI_ISL_500128, EPI_ISL_500129, EPI_ISL_500130, EPI_ISL_500131, EPI_ISL_500132, EPI_ISL_500133, EPI_ISL_500134, EPI_ISL_500135, EPI_ISL_500136, EPI_ISL_500137, EPI_ISL_500138, EPI_ISL_500139, EPI_ISL_500140, EPI_ISL_500141, EPI_ISL_500142, EPI_ISL_500143, EPI_ISL_500144, EPI_ISL_500145, EPI_ISL_500146, EPI_ISL_500147, EPI_ISL_500148, EPI_ISL_500149, EPI_ISL_500150, EPI_ISL_500151, EPI_ISL_500152, EPI_ISL_500153, EPI_ISL_500154, EPI_ISL_500155, EPI_ISL_500156 | | | |
| see above | Liverpool Clinical Laboratories | COVID-19 Genomics UK (COG-UK) Consortium | Sam Haldenby, Anita Lucaci, Steve Paterson, Julian Hiscox, Alistair Darby, M Almsaud, A Alrezaihi, Muhannad Alruwaili, Stuart D Armstrong, Jones Benjamin, Eleanor G Bentley, Anu Chawla, Jordan |

| | | | |
|---|---|---|---|
| | | | J Clark, Angela Cowell, Richard Eccles, Isabel García-Dorival, Matthew Gemmell, Alessandro Gerada, PKF Gilmore, Richard Gregory, Ximeng Han, Catherine Hartley, Margaret Hughes, Miren Iturriza-Gomara, James Johnson, L Luu, Jenifer Manson, Charlotte Nelson, Elaine O'Toole, Cassie Olateju, Rebekah Penrice-Randal , Lucille Rainbow, N.P Randle, Trevor Ian Robinson, Parul Sharma, Ghada T Shawli, James P Stewart, Neil Swainston, Ecaterina Vamos, Joanne Watts, Mark Whitehead |
| EPI_ISL_500157, EPI_ISL_500158, EPI_ISL_500159 | Complejo Hospitalario Universitario de Albacete | SeqCOVID-SPAIN consortium/IBV(CSIC) | Encarnacion Simarro Córdoba, Julia Lozano Serra, Lorena Robles Fonseca , Monica Parra Grandes, Caridad Sainz de Baranda Camino and SeqCOVID-SPAIN consortium |
| EPI_ISL_500160, EPI_ISL_500161 | Hospital Universitario Virgen de las Nieves de Granada-SAS | SeqCOVID-SPAIN consortium/IBV(CSIC) | Mercedes Pérez Ruiz, Sara Sanbonmatsu Gámez, Irene Pedrosa Corral, José M. Navarro-Marí and SeqCOVID-SPAIN consortium |
| EPI_ISL_500162 | Hospital Clínico Universitario de Santiago de Compostela | SeqCOVID-SPAIN consortium/IBV(CSIC) | José Javier Costa Alcalde, Antonio Aguilera Guirao, Mª Luisa Pérez del Molino Bernal, Amparo Coira Nieto, Gema Barbeito Castiñeiras, Rocio Trastoy Pena and SeqCOVID-SPAIN consortium |
| EPI_ISL_500163, EPI_ISL_500164, EPI_ISL_500165 | Hospital Universitario Virgen de las Nieves de Granada-SAS | SeqCOVID-SPAIN consortium/IBV(CSIC) | Mercedes Pérez Ruiz, Sara Sanbonmatsu Gámez, Irene Pedrosa Corral, José M. Navarro-Marí and SeqCOVID-SPAIN consortium |
| EPI_ISL_500166 | Complejo Hospitalario Universitario de Albacete | SeqCOVID-SPAIN consortium/IBV(CSIC) | Encarnacion Simarro Córdoba, Julia Lozano Serra, Lorena Robles Fonseca , Monica Parra Grandes, Caridad Sainz de Baranda Camino and SeqCOVID-SPAIN consortium |
| EPI_ISL_500167 | Hospital Clínico Universitario de Santiago de Compostela | SeqCOVID-SPAIN consortium/IBV(CSIC) | José Javier Costa Alcalde, Antonio Aguilera Guirao, Mª Luisa Pérez del Molino Bernal, Amparo Coira Nieto, Gema Barbeito Castiñeiras, Rocio Trastoy Pena and SeqCOVID-SPAIN consortium |
| EPI_ISL_500168, EPI_ISL_500169, EPI_ISL_500170, EPI_ISL_500171, EPI_ISL_500172, EPI_ISL_500173, EPI_ISL_500174, EPI_ISL_500175, EPI_ISL_500176, EPI_ISL_500177, EPI_ISL_500178, EPI_ISL_500179, EPI_ISL_500180, EPI_ISL_500181, EPI_ISL_500182, EPI_ISL_500183, EPI_ISL_500184, EPI_ISL_500185, EPI_ISL_500186, EPI_ISL_500187, EPI_ISL_500188, EPI_ISL_500189, EPI_ISL_500190, EPI_ISL_500191, EPI_ISL_500192, EPI_ISL_500193, EPI_ISL_500194, EPI_ISL_500195, EPI_ISL_500196, EPI_ISL_500197, EPI_ISL_500198, EPI_ISL_500199, EPI_ISL_500200, EPI_ISL_500201, EPI_ISL_500202, EPI_ISL_500203, EPI_ISL_500204, EPI_ISL_500205, EPI_ISL_500206 | | | |
| see above | Hospital Universitario Virgen de las Nieves de Granada-SAS | SeqCOVID-SPAIN consortium/IBV(CSIC) | Mercedes Pérez Ruiz, Sara Sanbonmatsu Gámez, Irene Pedrosa Corral, José M. Navarro-Marí and SeqCOVID-SPAIN consortium |
| EPI_ISL_500207 | Complejo Hospitalario Universitario de Albacete | SeqCOVID-SPAIN consortium/IBV(CSIC) | Encarnacion Simarro Córdoba, Julia Lozano Serra, Lorena Robles Fonseca , Monica Parra Grandes, Caridad Sainz de Baranda Camino and SeqCOVID-SPAIN consortium |
| EPI_ISL_500208 | Hospital Universitario Virgen de las Nieves de Granada-SAS | SeqCOVID-SPAIN consortium/IBV(CSIC) | Mercedes Pérez Ruiz, Sara Sanbonmatsu Gámez, Irene Pedrosa Corral, José M. Navarro-Marí and SeqCOVID-SPAIN consortium |
| EPI_ISL_500209 | Hospital Clínico Universitario de Santiago de Compostela | SeqCOVID-SPAIN consortium/IBV(CSIC) | José Javier Costa Alcalde, Antonio Aguilera Guirao, Mª Luisa Pérez del Molino Bernal, Amparo Coira Nieto, Gema Barbeito Castiñeiras, Rocio Trastoy Pena and SeqCOVID-SPAIN consortium |
| EPI_ISL_500210, EPI_ISL_500211, EPI_ISL_500212, EPI_ISL_500213, EPI_ISL_500214, EPI_ISL_500215, EPI_ISL_500216, EPI_ISL_500217 | Hospital Universitario Virgen de las Nieves de Granada-SAS | SeqCOVID-SPAIN consortium/IBV(CSIC) | Mercedes Pérez Ruiz, Sara Sanbonmatsu Gámez, Irene Pedrosa Corral, José M. Navarro-Marí and SeqCOVID-SPAIN consortium |
| EPI_ISL_500218 | Complejo Hospitalario Universitario de Albacete | SeqCOVID-SPAIN consortium/IBV(CSIC) | Encarnacion Simarro Córdoba, Julia Lozano Serra, Lorena Robles Fonseca , Monica Parra Grandes, Caridad Sainz de Baranda Camino and SeqCOVID-SPAIN consortium |
| EPI_ISL_500219 | Hospital Clínico Universitario de Santiago de Compostela | SeqCOVID-SPAIN consortium/IBV(CSIC) | José Javier Costa Alcalde, Antonio Aguilera Guirao, Mª Luisa Pérez del Molino Bernal, Amparo Coira Nieto, Gema Barbeito Castiñeiras, Rocio Trastoy Pena and SeqCOVID-SPAIN consortium |
| EPI_ISL_500220 | Hospital Universitario Virgen de las Nieves de Granada-SAS | SeqCOVID-SPAIN consortium/IBV(CSIC) | Mercedes Pérez Ruiz, Sara Sanbonmatsu Gámez, Irene Pedrosa Corral, José M. Navarro-Marí and SeqCOVID-SPAIN consortium |
| EPI_ISL_500221 | Hospital Clínico Universitario de Santiago de Compostela | SeqCOVID-SPAIN consortium/IBV(CSIC) | José Javier Costa Alcalde, Antonio Aguilera Guirao, Mª Luisa Pérez del Molino Bernal, Amparo Coira Nieto, Gema Barbeito Castiñeiras, Rocio Trastoy Pena and SeqCOVID-SPAIN consortium |
| EPI_ISL_500222, EPI_ISL_500223, EPI_ISL_500224, EPI_ISL_500225, EPI_ISL_500226, EPI_ISL_500227 | Hospital Universitario Virgen de las Nieves de Granada-SAS | SeqCOVID-SPAIN consortium/IBV(CSIC) | Mercedes Pérez Ruiz, Sara Sanbonmatsu Gámez, Irene Pedrosa Corral, José M. Navarro-Marí and SeqCOVID-SPAIN consortium |
| EPI_ISL_500228, EPI_ISL_500229, EPI_ISL_500230, EPI_ISL_500231, EPI_ISL_500232, EPI_ISL_500233, EPI_ISL_500234, EPI_ISL_500235, EPI_ISL_500236, EPI_ISL_500237, EPI_ISL_500238, EPI_ISL_500239, EPI_ISL_500240, EPI_ISL_500241, EPI_ISL_500242, EPI_ISL_500243, EPI_ISL_500244, EPI_ISL_500245, EPI_ISL_500246, EPI_ISL_500247, EPI_ISL_500248, EPI_ISL_500249, EPI_ISL_500250, EPI_ISL_500251, EPI_ISL_500252, EPI_ISL_500253, EPI_ISL_500254, EPI_ISL_500255, EPI_ISL_500256, EPI_ISL_500257, EPI_ISL_500258, EPI_ISL_500259, EPI_ISL_500260, EPI_ISL_500261, EPI_ISL_500262, EPI_ISL_500263, EPI_ISL_500264, EPI_ISL_500265, EPI_ISL_500266, EPI_ISL_500267, EPI_ISL_500268, EPI_ISL_500269, EPI_ISL_500270, EPI_ISL_500271, EPI_ISL_500272, EPI_ISL_500273, EPI_ISL_500274, EPI_ISL_500275, EPI_ISL_500276, EPI_ISL_500277, EPI_ISL_500278, EPI_ISL_500279, EPI_ISL_500280, EPI_ISL_500281, EPI_ISL_500282, EPI_ISL_500283, EPI_ISL_500284, EPI_ISL_500285, EPI_ISL_500286 | | | |
| see above | Hospital Universitario Araba. Vitoria-Gasteiz | SeqCOVID-SPAIN consortium/IBV(CSIC) | Silvia Hernáez Crespo, Carmen Gómez González, Amaia Aguirre Quiñonero, Marina Fernández Torres, Mª Rosario Almela Ferrer, Mª Concepción Lecaroz Agara, Andrés Canut Blasco and SeqCOVID-SPAIN consortium |
| EPI_ISL_500287, EPI_ISL_500288, EPI_ISL_500290, EPI_ISL_500291 | Servicio de Microbiología, Hospital Miguel Servet, Zaragoza | SeqCOVID-SPAIN consortium/IBV(CSIC) | Antonio Rezusta López, Alexander Tristancho Baró, Ana Milagro, Yolanda Gracia Grataloup, Nieves Martínez Cameo and SeqCOVID-SPAIN consortium |
| EPI_ISL_500292, EPI_ISL_500293, EPI_ISL_500294, EPI_ISL_500295, EPI_ISL_500296, EPI_ISL_500297, EPI_ISL_500298, EPI_ISL_500299, EPI_ISL_500300, EPI_ISL_500301, EPI_ISL_500302, EPI_ISL_500303, EPI_ISL_500304, EPI_ISL_500305, EPI_ISL_500306, EPI_ISL_500307, EPI_ISL_500308, EPI_ISL_500309, EPI_ISL_500310, EPI_ISL_500311, EPI_ISL_500312, EPI_ISL_500313, EPI_ISL_500314, EPI_ISL_500315, EPI_ISL_500316, EPI_ISL_500317, EPI_ISL_500318, EPI_ISL_500319, EPI_ISL_500320, EPI_ISL_500321, EPI_ISL_500322, EPI_ISL_500323, EPI_ISL_500324, EPI_ISL_500325, EPI_ISL_500326, EPI_ISL_500327, EPI_ISL_500328, EPI_ISL_500329, EPI_ISL_500330, EPI_ISL_500331, EPI_ISL_500332, EPI_ISL_500333, EPI_ISL_500334, EPI_ISL_500335 | | | |
| see above | Servicio de Microbiología. Hospital Universitario Donostia. OSI Donostialdea. Área de Enfermedades Infecciosas, Grupo de Infección Respiratoria y Resistencia Antimicrobiana. Instituto de Investigación Sanitaria Biodonostia | SeqCOVID-SPAIN consortium/IBV(CSIC) | Gustavo Cilla, Milagrosa Montes, Luis Piñeiro, Jose Maria Marimón and SeqCOVID-SPAIN consortium |
| EPI_ISL_500336, EPI_ISL_500337, EPI_ISL_500338, EPI_ISL_500339, EPI_ISL_500340, EPI_ISL_500341, EPI_ISL_500343, EPI_ISL_500344, EPI_ISL_500345, EPI_ISL_500346, EPI_ISL_500347, EPI_ISL_500349, EPI_ISL_500350, EPI_ISL_500351, EPI_ISL_500352, EPI_ISL_500353, EPI_ISL_500354, EPI_ISL_500355, EPI_ISL_500356, EPI_ISL_500357, EPI_ISL_500359, EPI_ISL_500360, EPI_ISL_500361, EPI_ISL_500362, EPI_ISL_500363, EPI_ISL_500364, EPI_ISL_500365, EPI_ISL_500366, EPI_ISL_500367, EPI_ISL_500368 | | | |
| see above | Servicio de Microbiología, Hospital Miguel Servet, Zaragoza | SeqCOVID-SPAIN consortium/IBV(CSIC) | Antonio Rezusta López, Alexander Tristancho Baró, Ana Milagro, Yolanda Gracia Grataloup, Nieves Martínez Cameo and SeqCOVID-SPAIN consortium |
| EPI_ISL_500369, EPI_ISL_500370, EPI_ISL_500371, EPI_ISL_500372, EPI_ISL_500374, EPI_ISL_500375, EPI_ISL_500376, EPI_ISL_500377, EPI_ISL_500378, EPI_ISL_500379, EPI_ISL_500380, EPI_ISL_500381, EPI_ISL_500383, EPI_ISL_500385, EPI_ISL_500386, EPI_ISL_500387, EPI_ISL_500388, EPI_ISL_500389, EPI_ISL_500390, EPI_ISL_500391, EPI_ISL_500392, EPI_ISL_500393, EPI_ISL_500394, EPI_ISL_500395, EPI_ISL_500396, EPI_ISL_500397, EPI_ISL_500398, EPI_ISL_500399, EPI_ISL_500400, EPI_ISL_500401, EPI_ISL_500402, EPI_ISL_500403, EPI_ISL_500404, EPI_ISL_500405, EPI_ISL_500406, EPI_ISL_500407, EPI_ISL_500408, EPI_ISL_500409, EPI_ISL_500410, EPI_ISL_500411, EPI_ISL_500412, EPI_ISL_500413, EPI_ISL_500414, EPI_ISL_500415, EPI_ISL_500416, EPI_ISL_500417, EPI_ISL_500418, EPI_ISL_500419, EPI_ISL_500420, EPI_ISL_500421, EPI_ISL_500422, EPI_ISL_500423, EPI_ISL_500424, EPI_ISL_500425, EPI_ISL_500426, EPI_ISL_500427, EPI_ISL_500428, EPI_ISL_500429, EPI_ISL_500430, EPI_ISL_500431, EPI_ISL_500432, EPI_ISL_500433, EPI_ISL_500434, EPI_ISL_500435, EPI_ISL_500436, EPI_ISL_500437, EPI_ISL_500438, EPI_ISL_500439, EPI_ISL_500440, EPI_ISL_500441, EPI_ISL_500442, EPI_ISL_500443, EPI_ISL_500444, EPI_ISL_500445, EPI_ISL_500446, EPI_ISL_500447, EPI_ISL_500448, EPI_ISL_500449, EPI_ISL_500450, EPI_ISL_500451, EPI_ISL_500452, EPI_ISL_500453, EPI_ISL_500454, EPI_ISL_500455, EPI_ISL_500456, EPI_ISL_500457, EPI_ISL_500458 | | | |
| see above | Centro de Investigación Biomédica de La Rioja - Hospital San Pedro Logroño | SeqCOVID-SPAIN consortium/IBV(CSIC) | María de Toro, José Manuel Azcona Gutiérrez, María Pilar Bea Escudero, Miriam Blasco Alberdi and SeqCOVID-SPAIN consortium |
| EPI_ISL_500459 | Quest Diagnostics | Quest Diagnostics | Rosenthal,S.H., Gerasimova,A., Kagan,R.M., Owen, R. AND Lacbawan, F. |
| EPI_ISL_500460, EPI_ISL_500461, EPI_ISL_500462, EPI_ISL_500463, EPI_ISL_500464, EPI_ISL_500465, EPI_ISL_500466, EPI_ISL_500467, EPI_ISL_500468, EPI_ISL_500469, EPI_ISL_500470, EPI_ISL_500471, EPI_ISL_500472, EPI_ISL_500473, EPI_ISL_500474, EPI_ISL_500475, EPI_ISL_500476, EPI_ISL_500477, EPI_ISL_500478, EPI_ISL_500479, EPI_ISL_500480, EPI_ISL_500481, EPI_ISL_500482, EPI_ISL_500483, EPI_ISL_500484, EPI_ISL_500485, EPI_ISL_500486 | | | |
| see above | LACEN/PE | WallauLab, Aggeu Magalhaes Institute | Marcelo Henrique dos Santos Paiva, Duschinka Ribeiro Duarte Guedes, Cássia Docena, Matheus Filgueira Bezerra, Filipe Zimmer Dezordi, Laís Ceschini Machado, Larissa Krokovsky, Elisama Helvecio, Alexandre Freitas da Silva, Antonio Mauro Rezende5, Sinval Pinto Brandão Filho7, Constância Flávia Junqueira Ayres, Gabriel Luz Wallau |
| EPI_ISL_500500, EPI_ISL_500501, EPI_ISL_500502 | University of Washington Virology Lab | University of Washington Virology Lab | Pavitra Roychoudhury, Hong Xie, Lasata Shrestha, Amin Addetia, Truong Nguyen, Victoria M Rachleff, Meei-Li Huang, Keith R Jerome, Alexander Greninger |
| EPI_ISL_500503, EPI_ISL_500504, EPI_ISL_500505, EPI_ISL_500506, EPI_ISL_500507, EPI_ISL_500508, EPI_ISL_500509, EPI_ISL_500510, EPI_ISL_500511, EPI_ISL_500512, EPI_ISL_500513, EPI_ISL_500514, EPI_ISL_500515, EPI_ISL_500516, EPI_ISL_500517, EPI_ISL_500518, EPI_ISL_500519, EPI_ISL_500520, EPI_ISL_500521, EPI_ISL_500522, EPI_ISL_500523, EPI_ISL_500524, EPI_ISL_500525, EPI_ISL_500526, EPI_ISL_500527, EPI_ISL_500528, EPI_ISL_500529, EPI_ISL_500530, EPI_ISL_500531, EPI_ISL_500532, EPI_ISL_500533, EPI_ISL_500534, EPI_ISL_500535, EPI_ISL_500536, EPI_ISL_500537, EPI_ISL_500538 | | | |
| see above | Mayo Clinic Laboratories | University of Washington Virology Lab | Pavitra Roychoudhury, Hong Xie, Lasata Shrestha, Amin Addetia, Truong Nguyen, Victoria M Rachleff, Meei-Li Huang, Keith R Jerome, Alexander Greninger |
| EPI_ISL_500539, EPI_ISL_500540, EPI_ISL_500541, EPI_ISL_500542, EPI_ISL_500543, EPI_ISL_500544, EPI_ISL_500545, EPI_ISL_500546, EPI_ISL_500547, EPI_ISL_500548, EPI_ISL_500549, EPI_ISL_500550, EPI_ISL_500551, EPI_ISL_500552, EPI_ISL_500553, EPI_ISL_500554, EPI_ISL_500555, EPI_ISL_500556, EPI_ISL_500557, EPI_ISL_500558, EPI_ISL_500559, EPI_ISL_500560, EPI_ISL_500561, EPI_ISL_500562, EPI_ISL_500563, EPI_ISL_500564, EPI_ISL_500565, EPI_ISL_500566, EPI_ISL_500567, EPI_ISL_500568, EPI_ISL_500569, EPI_ISL_500570, EPI_ISL_500571, EPI_ISL_500572 | | | |
| see above | Singapore General Hospital | Department of Microbiology | Nurdyana Abdul Rahman, Kun Lee Lim, Chenhao Li, Kian Sing Chan, Lynette Oon, Kern Rei Chng, Niranjan Nagarajan, Karrie Ko |
| EPI_ISL_500573, EPI_ISL_500574, EPI_ISL_500575, EPI_ISL_500576, EPI_ISL_500577, EPI_ISL_500578, EPI_ISL_500579, EPI_ISL_500580, EPI_ISL_500581, EPI_ISL_500582, EPI_ISL_500583, EPI_ISL_500584, EPI_ISL_500585, EPI_ISL_500586, EPI_ISL_500587, EPI_ISL_500588, EPI_ISL_500589, EPI_ISL_500590, EPI_ISL_500591, EPI_ISL_500592, EPI_ISL_500593, EPI_ISL_500594, EPI_ISL_500595 | | | |
| see above | National Virus Reference Laboratory | National Virus Reference Laboratory | Michael Carr, Gabriel Gonzalez, Jonathan Dean, Suzie Coughlan, Cillian F De Gascun |
| EPI_ISL_500596, EPI_ISL_500597, EPI_ISL_500598, EPI_ISL_500599, EPI_ISL_500600, EPI_ISL_500601, EPI_ISL_500602, EPI_ISL_500603, EPI_ISL_500604, EPI_ISL_500605, EPI_ISL_500606, EPI_ISL_500607, EPI_ISL_500608, EPI_ISL_500609, EPI_ISL_500610, EPI_ISL_500611, EPI_ISL_500612, EPI_ISL_500613, EPI_ISL_500614, EPI_ISL_500615, EPI_ISL_500616, EPI_ISL_500617, EPI_ISL_500618, EPI_ISL_500619, EPI_ISL_500620, EPI_ISL_500621, EPI_ISL_500622, EPI_ISL_500623, EPI_ISL_500624, EPI_ISL_500625, EPI_ISL_500626, EPI_ISL_500627, EPI_ISL_500628, EPI_ISL_500629, EPI_ISL_500630, EPI_ISL_500631, EPI_ISL_500632, EPI_ISL_500633, EPI_ISL_500634, EPI_ISL_500635, EPI_ISL_500636, EPI_ISL_500637, EPI_ISL_500638, EPI_ISL_500639, EPI_ISL_500640, EPI_ISL_500641, EPI_ISL_500642, EPI_ISL_500643, EPI_ISL_500644, EPI_ISL_500645, EPI_ISL_500646, EPI_ISL_500647, EPI_ISL_500648, EPI_ISL_500649, EPI_ISL_500650, EPI_ISL_500651, EPI_ISL_500652, EPI_ISL_500653, EPI_ISL_500654, EPI_ISL_500655, EPI_ISL_500656, EPI_ISL_500657, EPI_ISL_500658, EPI_ISL_500659, EPI_ISL_500660, EPI_ISL_500661, EPI_ISL_500662, EPI_ISL_500663, EPI_ISL_500664, EPI_ISL_500665, EPI_ISL_500666, EPI_ISL_500667, EPI_ISL_500668, EPI_ISL_500669, EPI_ISL_500670, EPI_ISL_500671, EPI_ISL_500672, EPI_ISL_500673, EPI_ISL_500674, EPI_ISL_500675, EPI_ISL_500676, EPI_ISL_500677, EPI_ISL_500678, EPI_ISL_500679, EPI_ISL_500680, EPI_ISL_500681, EPI_ISL_500682, EPI_ISL_500683, EPI_ISL_500684, EPI_ISL_500685, EPI_ISL_500686, EPI_ISL_500687, EPI_ISL_500688, EPI_ISL_500689, EPI_ISL_500690, EPI_ISL_500691, EPI_ISL_500692, EPI_ISL_500693, EPI_ISL_500694, EPI_ISL_500695, EPI_ISL_500696, EPI_ISL_500697, EPI_ISL_500698, EPI_ISL_500699, EPI_ISL_500700, EPI_ISL_500701, EPI_ISL_500702, EPI_ISL_500703, EPI_ISL_500704, EPI_ISL_500705, EPI_ISL_500706 | | | |
| see above | Area of Virology, Serology and Virology Division (SAViD), New South Wales Health Pathology Randwick | Area of Virology, Serology and Virology Division (SAViD), New South Wales Health Pathology Randwick | Rawlinson, W. |
| EPI_ISL_500707, EPI_ISL_500708, EPI_ISL_500709, EPI_ISL_500710, EPI_ISL_500711, EPI_ISL_500712, EPI_ISL_500713, EPI_ISL_500714, EPI_ISL_500715 | Respiratory Virus Unit, Microbiology Services Colindale, Public Health England | Respiratory Virus Unit, Microbiology Services Colindale, Public Health England | PHE Covid Sequencing Team |
| EPI_ISL_500716 | BSL3 Lab Pendik Veterinary Control Institute | Department of Medicinal Genetics, Bursa Uludağ University, Faculty of medicine By Sehime Gülsün Temel, Adem Alemdar, Kadir Yeşilbağ | Mustafa HASOKSUZ, Fahriye SARAC, Osman ERGANIS, Serdar UZAR, Hakan ENUL, Cumhur ADIAY, Ahmet SAIT, Orbay SAYI, Kadir YESILBAG, Oguz KARABEY |
| EPI_ISL_500717 | Area of Virology, Serology and Virology Division (SAViD), New South Wales Health Pathology Randwick | Area of Virology, Serology and Virology Division (SAViD), New South Wales Health Pathology Randwick | Rawlinson, W |
| EPI_ISL_500719 | Respiratory Virus Unit, Microbiology Services Colindale, Public Health England | Respiratory Virus Unit, Microbiology Services Colindale, Public Health England | PHE Covid Sequencing Team |
| EPI_ISL_500768, EPI_ISL_500769, EPI_ISL_500770, EPI_ISL_500771, EPI_ISL_500772, EPI_ISL_500773, EPI_ISL_500774 | Furst Medical Laboratory | Norwegian Institute of Public Health, Department of Virology | Kathrine Stene-Johansen, Kamilla Heddeland Instefjord, Hilde Elshaug, Rasmus Riis Kopperud, Karoline Bragstad, Olav Hungnes |
| EPI_ISL_500775 | Hospital of Southern Norway - Kristiansand, Department of Medical Microbiology | Norwegian Institute of Public Health, Department of Virology | Kathrine Stene-Johansen, Kamilla Heddeland Instefjord, Hilde Elshaug, Rasmus Riis Kopperud, Karoline Bragstad, Olav Hungnes |
| EPI_ISL_500776, EPI_ISL_500777, EPI_ISL_500778 | Foerde Hospital, Department of Microbiology | Norwegian Institute of Public Health, Department of Virology | Kathrine Stene-Johansen, Kamilla Heddeland Instefjord, Hilde Elshaug, Rasmus Riis Kopperud, Karoline Bragstad, Olav Hungnes |
| EPI_ISL_500779, EPI_ISL_500780, EPI_ISL_500781, EPI_ISL_500782, | Akershus University Hospital, Department for Microbiology and Infectious Disease Control | Norwegian Institute of Public Health, Department of Virology | Kathrine Stene-Johansen, Kamilla Heddeland Instefjord, Hilde Elshaug, Rasmus Riis Kopperud, Karoline Bragstad, Olav Hungnes |

| | | | |
|---|---|---|---|
| EPI_ISL_500783 | | | |
| EPI_ISL_500784, EPI_ISL_500785, EPI_ISL_500786, EPI_ISL_500787, EPI_ISL_500788, EPI_ISL_500789, EPI_ISL_500790 | Furst Medical Laboratory | Norwegian Institute of Public Health, Department of Virology | Kathrine Stene-Johansen, Kamilla Heddeland Instefjord, Hilde Elshaug, Rasmus Riis Kopperud, Karoline Bragstad, Olav Hungnes |
| EPI_ISL_500791, EPI_ISL_500792 | Hospital of Southern Norway - Kristiansand, Department of Medical Microbiology | Norwegian Institute of Public Health, Department of Virology | Kathrine Stene-Johansen, Kamilla Heddeland Instefjord, Hilde Elshaug, Rasmus Riis Kopperud, Karoline Bragstad, Olav Hungnes |
| EPI_ISL_500793, EPI_ISL_500794, EPI_ISL_500795, EPI_ISL_500796 | Akershus University Hospital, Department for Microbiology and Infectious Disease Control | Norwegian Institute of Public Health, Department of Virology | Kathrine Stene-Johansen, Kamilla Heddeland Instefjord, Hilde Elshaug, Rasmus Riis Kopperud, Karoline Bragstad, Olav Hungnes |
| EPI_ISL_500797, EPI_ISL_500798, EPI_ISL_500799 | Hospital of Southern Norway - Kristiansand, Department of Medical Microbiology | Norwegian Institute of Public Health, Department of Virology | Kathrine Stene-Johansen, Kamilla Heddeland Instefjord, Hilde Elshaug, Rasmus Riis Kopperud, Karoline Bragstad, Olav Hungnes |
| EPI_ISL_500800 | Oslo University Hospital, Department of Medical Microbiology | Norwegian Institute of Public Health, Department of Virology | Kathrine Stene-Johansen, Kamilla Heddeland Instefjord, Hilde Elshaug, Rasmus Riis Kopperud, Karoline Bragstad, Olav Hungnes |
| EPI_ISL_500801, EPI_ISL_500802, EPI_ISL_500803, EPI_ISL_500804, EPI_ISL_500805, EPI_ISL_500806, EPI_ISL_500807, EPI_ISL_500808, EPI_ISL_500809, EPI_ISL_500810, EPI_ISL_500811, EPI_ISL_500812, EPI_ISL_500813, EPI_ISL_500814, EPI_ISL_500815, EPI_ISL_500816, EPI_ISL_500817, EPI_ISL_500818, EPI_ISL_500819, EPI_ISL_500820, EPI_ISL_500821, EPI_ISL_500822, EPI_ISL_500823, EPI_ISL_500824, EPI_ISL_500825, EPI_ISL_500826, EPI_ISL_500827, EPI_ISL_500828, EPI_ISL_500829, EPI_ISL_500830 | | | |
| see above | National Institute for Biological Standards and Control | National Institute for Biological Standards and Control | Javier Martin, Dimitra Klapsa, Thomas Wilton |
| EPI_ISL_500831, EPI_ISL_500832, EPI_ISL_500833, EPI_ISL_500834, EPI_ISL_500835, EPI_ISL_500836, EPI_ISL_500837, EPI_ISL_500838, EPI_ISL_500839, EPI_ISL_500840, EPI_ISL_500841, EPI_ISL_500842, EPI_ISL_500843, EPI_ISL_500844, EPI_ISL_500845, EPI_ISL_500846, EPI_ISL_500847, EPI_ISL_500848, EPI_ISL_500849, EPI_ISL_500850, EPI_ISL_500851, EPI_ISL_500852, EPI_ISL_500853, EPI_ISL_500854, EPI_ISL_500855, EPI_ISL_500856, EPI_ISL_500857, EPI_ISL_500858, EPI_ISL_500859, EPI_ISL_500860, EPI_ISL_500861, EPI_ISL_500862, EPI_ISL_500863 | | | |
| see above | Virginia DCLS | Virginia DCLS | Virginia DCLS |
| EPI_ISL_500864 | Instituto de Pesquisas Biomédicas, Hospital Naval Marcílio Dias | Bioinformatics Laboratory / LNCC | Andressa Rangel de Oliveira Lima, Luiz Gonzaga Paula de Almeida, Alexandra Lehmkuhl Gerber, Marlon Daniel Lima Tonin, Mônica Gomes Monteiro da R. Santos, Claudio Alberto Mule Monteiro, Átila Duque Rossi, Isabel de Medeiros Magalhaes de As, Luana Ferreira Martins de Toledo, Amilcar Tanuri, Albina Luciana da Silva Freitas, Fernanda Conceição Silva do Amorim, Shana Priscila Coutinho Barroso, Cynthia Chester Cardoso, Carolina Moreira Voloch, Ana Tereza Vasconcelos |
| EPI_ISL_500865, EPI_ISL_500866, EPI_ISL_500867, EPI_ISL_500868, EPI_ISL_500869, EPI_ISL_500870, EPI_ISL_500871, EPI_ISL_500872, EPI_ISL_500873, EPI_ISL_500874, EPI_ISL_500875 | | | |
| see above | LACEN/PE | WallauLab, Aggeu Magalhaes Institute | Marcelo Henrique dos Santos Paiva, Duschinka Ribeiro Duarte Guedes, Cássia Docena, Matheus Filgueira Bezerra, Filipe Zimmer Dezordi, Laís Ceschini Machado, Larissa Krokovsky, Elisama Helvecio, Alexandre Freitas da Silva, Antonio Mauro Rezende5, Sinval Pinto Brandão Filho7, Constância Flávia Junqueira Ayres, Gabriel Luz Wallau |
| EPI_ISL_500877, EPI_ISL_500878, EPI_ISL_500879, EPI_ISL_500880, EPI_ISL_500881, EPI_ISL_500882, EPI_ISL_500883, EPI_ISL_500884, EPI_ISL_500885, EPI_ISL_500886, EPI_ISL_500887, EPI_ISL_500888, EPI_ISL_500889, EPI_ISL_500890, EPI_ISL_500891, EPI_ISL_500892, EPI_ISL_500893, EPI_ISL_500894, EPI_ISL_500895, EPI_ISL_500896, EPI_ISL_500897, EPI_ISL_500898, EPI_ISL_500899, EPI_ISL_500900, EPI_ISL_500901, EPI_ISL_500902, EPI_ISL_500903, EPI_ISL_500904, EPI_ISL_500905, EPI_ISL_500906, EPI_ISL_500907, EPI_ISL_500908, EPI_ISL_500909, EPI_ISL_500910, EPI_ISL_500911, EPI_ISL_500912, EPI_ISL_500913, EPI_ISL_500914, EPI_ISL_500915, EPI_ISL_500916, EPI_ISL_500917, EPI_ISL_500918, EPI_ISL_500919, EPI_ISL_500920, EPI_ISL_500921, EPI_ISL_500922, EPI_ISL_500923, EPI_ISL_500924, EPI_ISL_500925, EPI_ISL_500926, EPI_ISL_500927, EPI_ISL_500928, EPI_ISL_500929, EPI_ISL_500930, EPI_ISL_500931, EPI_ISL_500932, EPI_ISL_500933, EPI_ISL_500934, EPI_ISL_500935, EPI_ISL_500936, EPI_ISL_500937, EPI_ISL_500938, EPI_ISL_500939, EPI_ISL_500940, EPI_ISL_500941, EPI_ISL_500942, EPI_ISL_500943, EPI_ISL_500944, EPI_ISL_500945 | | | |
| see above | Viollier AG | Department of Biosystems Science and Engineering, ETH Zürich | Christian Beisel, Sarah Nadeau, Ivan Topolsky, Pedro Ferreira, Philipp Jablonski, Susana Posada-Céspedes, Tobias Schär, Ina Nissen, Natascha Santacroce, Elodie Burcklen, Christiane Beckmann, Maurice Redondo, Olivier Kobel, Christoph Noppen, Sophie Seidel, Noemie Santamaria de Souza, Niko Beerenwinkel, Tanja Stadler |
| EPI_ISL_500946 | GMERS Medical College & Hospital, Gotri, Vadodara | Gujarat Biotechnology Research Centre | Zuber Saiyed, Komal Patel, Labdhi Pandya, Afzal Ansari, Nikha Trivedi, Meenakshi Shah, Neena Doshi, Varsha Godbole, Apurvasinh Puvar, Janvi Raval, Zarna Patel, Monika Gandhi, Pinal Trivedi, Maharshi Pandya, Nidhi Patel, Nitin Savaliya, Raghawendra Kumar, Dinesh Kumar, R D Dixit, A M Kadri, Harsh Bakshi, Chaitanya Joshi, Madhvi Joshi |
| EPI_ISL_500947 | Department of MicroBiology, Government Medical College, Surat | Gujarat Biotechnology Research Centre | Maharshi Pandya, Nidhi Patel, Nitin Savaliya, Raghawendra Kumar, Dinesh Kumar, Zuber Saiyed, Komal Patel, Labdhi Pandya, Afzal Ansari, Nikha Trivedi, Naresh Chauhan, Summaiya Mullan, Amit gamit, Apurvasinh Puvar, Janvi Raval, Zarna Patel, Monika Gandhi, Pinal Trivedi, R D Dixit, A M Kadri, Harsh Bakshi, Chaitanya Joshi, Madhvi Joshi |
| EPI_ISL_500948 | Department of MicroBiology, Government Medical College, Surat | Gujarat Biotechnology Research Centre | Zuber Saiyed, Komal Patel, Labdhi Pandya, Afzal Ansari, Nikha Trivedi, Naresh Chauhan, Summaiya Mullan, Amit gamit, Apurvasinh Puvar, Janvi Raval, Zarna Patel, Monika Gandhi, Pinal Trivedi, Maharshi Pandya, Nidhi Patel, Nitin Savaliya, Raghawendra Kumar, Dinesh Kumar, R D Dixit, A M Kadri, Harsh Bakshi, Chaitanya Joshi, Madhvi Joshi |
| EPI_ISL_500949 | Department of MicroBiology, Government Medical College, Surat | Gujarat Biotechnology Research Centre | Komal Patel, Labdhi Pandya, Afzal Ansari, Nikha Trivedi, Naresh Chauhan, Summaiya Mullan, Amit gamit, Apurvasinh Puvar, Janvi Raval, Zarna Patel, Monika Gandhi, Pinal Trivedi, Maharshi Pandya, Nidhi Patel, Nitin Savaliya, Raghawendra Kumar, Dinesh Kumar, Zuber Saiyed, R D Dixit, A M Kadri, Harsh Bakshi, Chaitanya Joshi, Madhvi Joshi |
| EPI_ISL_500950 | Department of MicroBiology, Government Medical College, Surat | Gujarat Biotechnology Research Centre | Labdhi Pandya, Afzal Ansari, Nikha Trivedi, Naresh Chauhan, Summaiya Mullan, Amit gamit, Apurvasinh Puvar, Janvi Raval, Zarna Patel, Monika Gandhi, Pinal Trivedi, Maharshi Pandya, Nidhi Patel, Nitin Savaliya, Raghawendra Kumar, Dinesh Kumar, Zuber Saiyed, Komal Patel, R D Dixit, A M Kadri, Harsh Bakshi, Chaitanya Joshi, Madhvi Joshi |
| EPI_ISL_500951, EPI_ISL_500952, EPI_ISL_500953 | Respiratory Virus Unit, Microbiology Services Colindale, Public Health England | Respiratory Virus Unit, Microbiology Services Colindale, Public Health England | PHE Covid Sequencing Team |
| EPI_ISL_500954, EPI_ISL_500955, EPI_ISL_500956, EPI_ISL_500957, EPI_ISL_500958, EPI_ISL_500959, EPI_ISL_500960, EPI_ISL_500961, EPI_ISL_500962, EPI_ISL_500963, EPI_ISL_500964, EPI_ISL_500965, EPI_ISL_500966, EPI_ISL_500967, EPI_ISL_500968, EPI_ISL_500969, EPI_ISL_500970, EPI_ISL_500971, EPI_ISL_500972, EPI_ISL_500973, EPI_ISL_500974, EPI_ISL_500975, EPI_ISL_500976, EPI_ISL_500977, EPI_ISL_500978, EPI_ISL_500979, EPI_ISL_500980, EPI_ISL_500981, EPI_ISL_500982, EPI_ISL_500983, EPI_ISL_500984, EPI_ISL_500985, EPI_ISL_500986, EPI_ISL_500987, EPI_ISL_500988, EPI_ISL_500989, EPI_ISL_500990, EPI_ISL_500991, EPI_ISL_500992, EPI_ISL_500993, EPI_ISL_500994, EPI_ISL_500995, EPI_ISL_500996, EPI_ISL_500997, EPI_ISL_500998, EPI_ISL_500999, EPI_ISL_501000, EPI_ISL_501001, EPI_ISL_501002, EPI_ISL_501003, EPI_ISL_501004, EPI_ISL_501005, EPI_ISL_501006, EPI_ISL_501007, EPI_ISL_501008, EPI_ISL_501009, EPI_ISL_501010, EPI_ISL_501011, EPI_ISL_501012, EPI_ISL_501013, EPI_ISL_501014, EPI_ISL_501015, EPI_ISL_501016, EPI_ISL_501017, EPI_ISL_501018, EPI_ISL_501019, EPI_ISL_501020, EPI_ISL_501021, EPI_ISL_501022, EPI_ISL_501023, EPI_ISL_501024, EPI_ISL_501025, EPI_ISL_501026, EPI_ISL_501027, EPI_ISL_501028, EPI_ISL_501029, EPI_ISL_501030, EPI_ISL_501031, EPI_ISL_501032, EPI_ISL_501033, EPI_ISL_501034, EPI_ISL_501035, EPI_ISL_501036, EPI_ISL_501037, EPI_ISL_501038, EPI_ISL_501039, EPI_ISL_501040, EPI_ISL_501041, EPI_ISL_501042, EPI_ISL_501043, EPI_ISL_501044, EPI_ISL_501045, EPI_ISL_501046, EPI_ISL_501047, EPI_ISL_501048, EPI_ISL_501049, EPI_ISL_501050, EPI_ISL_501051, EPI_ISL_501052, EPI_ISL_501053, EPI_ISL_501054, EPI_ISL_501055, EPI_ISL_501056, EPI_ISL_501057, EPI_ISL_501058, EPI_ISL_501059, EPI_ISL_501060, EPI_ISL_501061, EPI_ISL_501062, EPI_ISL_501063, EPI_ISL_501064, EPI_ISL_501065, EPI_ISL_501066, EPI_ISL_501067, EPI_ISL_501068, EPI_ISL_501069, EPI_ISL_501070, EPI_ISL_501071 | | | |
| see above | Regional Virus Laboratory, Belfast Health and Social Care Trust | Wellcome Sanger Institute for the COVID-19 Genomics UK Consortium | Conall McCaughey, James McKenna, Tanya Curran, Susan Feeney, Alison Watt, Ciara Cox, Mairead Connor, Zoltan Molnar, David Simpson, Derek Fairley; and Alex Alderton, Roberto Amato, Sonia Goncalves, Ewan Harrison, David K. Jackson, Ian Johnston, Dominic Kwiatkowski, Cordelia Langford, John Sillitoe on behalf of the Wellcome Sanger Institute COVID-19 Surveillance Team (http://www.sanger.ac.uk/covid-team) |
| EPI_ISL_501072, EPI_ISL_501073, EPI_ISL_501074, EPI_ISL_501075, EPI_ISL_501076, EPI_ISL_501077, EPI_ISL_501078, EPI_ISL_501079, EPI_ISL_501080, EPI_ISL_501081, EPI_ISL_501082 | | | |
| see above | Mayo Clinic Laboratories | University of Washington Virology Lab | Pavitra Roychoudhury, Hong Xie, Lasata Shrestha, Amin Addetia, Truong Nguyen, Victoria M Rachleff, Meei-Li Huang, Keith R Jerome, Alexander Greninger |
| EPI_ISL_501083, EPI_ISL_501084, EPI_ISL_501085, EPI_ISL_501086, EPI_ISL_501087, EPI_ISL_501088, EPI_ISL_501089, EPI_ISL_501090, EPI_ISL_501091, EPI_ISL_501092, EPI_ISL_501093, EPI_ISL_501094, EPI_ISL_501095, EPI_ISL_501096, EPI_ISL_501097, EPI_ISL_501098, EPI_ISL_501099, EPI_ISL_501100, EPI_ISL_501101, EPI_ISL_501102, EPI_ISL_501103, EPI_ISL_501104, EPI_ISL_501105, EPI_ISL_501106, EPI_ISL_501107, EPI_ISL_501108, EPI_ISL_501109, EPI_ISL_501110, EPI_ISL_501111, EPI_ISL_501112, EPI_ISL_501113, EPI_ISL_501114, EPI_ISL_501115, EPI_ISL_501116, EPI_ISL_501117, EPI_ISL_501118, EPI_ISL_501119, EPI_ISL_501120, EPI_ISL_501121, EPI_ISL_501122, EPI_ISL_501123, EPI_ISL_501124, EPI_ISL_501125, EPI_ISL_501126, EPI_ISL_501127, EPI_ISL_501128, EPI_ISL_501129, EPI_ISL_501130, EPI_ISL_501131, EPI_ISL_501132, EPI_ISL_501133, EPI_ISL_501134, EPI_ISL_501135, EPI_ISL_501136, EPI_ISL_501137, EPI_ISL_501138, EPI_ISL_501139, EPI_ISL_501140, EPI_ISL_501141, EPI_ISL_501142, EPI_ISL_501143, EPI_ISL_501144, EPI_ISL_501145, EPI_ISL_501146, EPI_ISL_501147, EPI_ISL_501148, EPI_ISL_501149, EPI_ISL_501150, EPI_ISL_501151, EPI_ISL_501152, EPI_ISL_501153, EPI_ISL_501154, EPI_ISL_501155, EPI_ISL_501156, EPI_ISL_501157, EPI_ISL_501158, EPI_ISL_501159, EPI_ISL_501160, EPI_ISL_501161, EPI_ISL_501162, EPI_ISL_501163, EPI_ISL_501164 | | | |
| see above | University of Washington Virology Lab | University of Washington Virology Lab | Pavitra Roychoudhury, Hong Xie, Lasata Shrestha, Amin Addetia, Truong Nguyen, Victoria M Rachleff, Meei-Li Huang, Keith R Jerome, Alexander Greninger |
| EPI_ISL_501165 | Mayo Clinic Laboratories | University of Washington Virology Lab | Pavitra Roychoudhury, Hong Xie, Lasata Shrestha, Amin Addetia, Truong Nguyen, Victoria M Rachleff, Meei-Li Huang, Keith R Jerome, Alexander Greninger |
| EPI_ISL_501167, EPI_ISL_501168, EPI_ISL_501169, EPI_ISL_501170, EPI_ISL_501171, EPI_ISL_501172, EPI_ISL_501173, EPI_ISL_501174 | Baylor College of Medicine | Baylor College of Medicine: HGSC | Vasanthi Avadhanula, Erin Nicholson, David Henke, Pedro Piedra, Harsha Doddapaneni, Donna Muzny, Qingchang Meng, Hsu Chao, Zeineen Momin, Hua Shen, George Weissenberger, Kavya Kottapalli, Yimiti Meiheerguli, Sejal Salvi, Ginger Metcalf, Vipin Menon, Sara J.J. Cregeen, Matthew C. Ross, Tulin Ayvaz, Richard Sucgang, Kristi L. Hoffman, Matthew Wong, Joseph F. Petrosino |
| EPI_ISL_501176, EPI_ISL_501177, EPI_ISL_501178, EPI_ISL_501179, EPI_ISL_501180, EPI_ISL_501181, EPI_ISL_501182, EPI_ISL_501183, EPI_ISL_501184, EPI_ISL_501185, EPI_ISL_501186, EPI_ISL_501187, EPI_ISL_501188, EPI_ISL_501189, EPI_ISL_501190, EPI_ISL_501191, EPI_ISL_501192, EPI_ISL_501193, EPI_ISL_501194, EPI_ISL_501195, EPI_ISL_501196, EPI_ISL_501197, EPI_ISL_501198, EPI_ISL_501199, EPI_ISL_501200, EPI_ISL_501201, EPI_ISL_501202, EPI_ISL_501203, EPI_ISL_501204, EPI_ISL_501205, EPI_ISL_501206, EPI_ISL_501207, EPI_ISL_501208, EPI_ISL_501209, EPI_ISL_501210, EPI_ISL_501211, EPI_ISL_501212, EPI_ISL_501213, EPI_ISL_501214, EPI_ISL_501215, EPI_ISL_501216, EPI_ISL_501217, EPI_ISL_501218, EPI_ISL_501219, EPI_ISL_501220, EPI_ISL_501221, EPI_ISL_501222, EPI_ISL_501223, EPI_ISL_501224, EPI_ISL_501225, EPI_ISL_501226, EPI_ISL_501227, EPI_ISL_501228 | | | |
| see above | Department of Medical Microbiology, University Malaya Medical Centre | Department of Medical Microbiology, Faculty of Medicine, University of Malaya | Yoong Min CHONG, Jennifer Chong, I-Ching SAM, Yoke Fun CHAN, University Malaya Medical Centre COVID Team |
| EPI_ISL_501230 | Hellenic Pasteur Institute, Public Health Laboratories | Hellenic Pasteur Institute, National Influenza Reference laboratory of Southern Greece & Unit of Bioinformatics and Applied Genomics | Vasiliki Pogka, Timokratis Karamitros, Athanasios Kossyvakis, Antonios Kalliaropoulos, Horefti Elina, Evangelidou Maria, Androniki Voulgari-Kokota, Aspasia Kontou, Andreas Mentis |
| EPI_ISL_501231, EPI_ISL_501232, EPI_ISL_501233, EPI_ISL_501234, EPI_ISL_501235, EPI_ISL_501236, EPI_ISL_501237, EPI_ISL_501238, EPI_ISL_501241, EPI_ISL_501242, EPI_ISL_501246, EPI_ISL_501248, EPI_ISL_501249, EPI_ISL_501250, EPI_ISL_501251 | | | |
| see above | Hellenic Pasteur Institute, National Influenza Reference laboratory of Southern Greece & Unit of Bioinformatics and Applied Genomics | Hellenic Pasteur Institute, National Influenza Reference laboratory of Southern Greece & Unit of Bioinformatics and Applied Genomics | Vasiliki Pogka, Timokratis Karamitros, Athanasios Kossyvakis, Antonios Kalliaropoulos, Horefti Elina, Evangelidou Maria, Androniki Voulgari-Kokota, Aspasia Kontou, Andreas Mentis |
| EPI_ISL_501252, EPI_ISL_501253, EPI_ISL_501254, EPI_ISL_501255, EPI_ISL_501256, EPI_ISL_501257, EPI_ISL_501258, EPI_ISL_501259, EPI_ISL_501260, EPI_ISL_501261, EPI_ISL_501262, EPI_ISL_501263, EPI_ISL_501264, EPI_ISL_501265, EPI_ISL_501266, EPI_ISL_501267, EPI_ISL_501268, EPI_ISL_501269, EPI_ISL_501270, EPI_ISL_501271, EPI_ISL_501272, EPI_ISL_501273, EPI_ISL_501274 | | | |
| see above | National Virus Reference Laboratory | National Virus Reference Laboratory | Michael Carr, Gabriel Gonzalez, Jonathan Dean, Suzie Coughlan, Cillian F De Gascun |
| EPI_ISL_501275 | E. Gulbja Laboratorija | Latvian Biomedical Research and Study Centre | Ivars Silamiķelis, Kaspars Megnis, Monta Ustinova, Ņikita Zrelovs, Vita Rovīte, Mikus Gavars, Dmitrijs Perminovs, Uga Dumpis, Jānis Kloviņš |
| EPI_ISL_501284, EPI_ISL_501285 | E. Gulbja Laboratorija | Latvian Biomedical Research and Study Centre | Ivars Silamiķelis, Kaspars Megnis, Monta Ustinova, Ņikita Zrelovs, Vita Rovīte, Mikus Gavars, Dmitrijs Perminovs, Uga Dumpis, Jānis Kloviņš |
| EPI_ISL_501286, EPI_ISL_501287, EPI_ISL_501288, EPI_ISL_501289 | Centrālā laboratorija | Latvian Biomedical Research and Study Centre | Ivars Silamiķelis, Kaspars Megnis, Monta Ustinova, Ņikita Zrelovs, Vita Rovīte, Stella Lapiņa, Jana Osīte, Marta Priedīte, Uga Dumpis, Jānis Kloviņš |
| EPI_ISL_501555, EPI_ISL_501556 | PHE South West Regional Laboratory, National Infection Service | Wellcome Sanger Institute for the COVID-19 Genomics UK Consortium | Stephanie Hutchings, Hannah Pymont, Dr Peter Muir, Barry Vipond, Rich Hopes; and Alex Alderton, Roberto Amato, Sonia Goncalves, Ewan Harrison, David K. Jackson, Ian Johnston, Dominic Kwiatkowski, Cordelia Langford, John Sillitoe on behalf of the Wellcome Sanger Institute COVID-19 Surveillance Team (http://www.sanger.ac.uk/covid-team) |
| EPI_ISL_501557 | Department of Medical Microbiology, Western Sussex Hospitals NHS Foundation Trust, St Richard's Hospital | Wellcome Sanger Institute for the COVID-19 Genomics UK Consortium | Manasa Mutingwende, Sarah Lowdon, Olga Podplomyk, Michelle Erkiert, Jonathan Lewis, Paul Randell and Alex Alderton, Roberto Amato, Sonia Goncalves, Ewan Harrison, David K. Jackson, Ian Johnston, Dominic Kwiatkowski, Cordelia Langford, John Sillitoe on behalf of the Wellcome Sanger Institute COVID-19 Surveillance Team (http://www.sanger.ac.uk/covid-team) |
| EPI_ISL_501558, EPI_ISL_501559, EPI_ISL_501560, EPI_ISL_501561 | PHE South West Regional Laboratory, National Infection Service | Wellcome Sanger Institute for the COVID-19 Genomics UK Consortium | Stephanie Hutchings, Hannah Pymont, Dr Peter Muir, Barry Vipond, Rich Hopes; and Alex Alderton, Roberto Amato, Sonia Goncalves, Ewan Harrison, David K. Jackson, Ian Johnston, Dominic Kwiatkowski, Cordelia Langford, John Sillitoe on behalf of the Wellcome Sanger Institute COVID-19 Surveillance Team (http://www.sanger.ac.uk/covid-team) |
| EPI_ISL_501562 | Department of Medical Microbiology, Western Sussex Hospitals NHS Foundation Trust, St Richard's Hospital | Wellcome Sanger Institute for the COVID-19 Genomics UK Consortium | Manasa Mutingwende, Sarah Lowdon, Olga Podplomyk, Michelle Erkiert, Jonathan Lewis, Paul Randell and Alex Alderton, Roberto Amato, Sonia Goncalves, Ewan Harrison, David K. Jackson, Ian Johnston, Dominic Kwiatkowski, Cordelia Langford, John Sillitoe on behalf of the Wellcome Sanger Institute COVID-19 Surveillance Team (http://www.sanger.ac.uk/covid-team) |
| EPI_ISL_501563, EPI_ISL_501564, EPI_ISL_501565, EPI_ISL_501566, EPI_ISL_501567, EPI_ISL_501568, EPI_ISL_501569, EPI_ISL_501570, EPI_ISL_501571, EPI_ISL_501572, EPI_ISL_501573, EPI_ISL_501574, EPI_ISL_501575, EPI_ISL_501576, EPI_ISL_501577, EPI_ISL_501578, EPI_ISL_501579, EPI_ISL_501580, EPI_ISL_501581, EPI_ISL_501582, EPI_ISL_501583, EPI_ISL_501584, EPI_ISL_501585, EPI_ISL_501586, EPI_ISL_501587, EPI_ISL_501588, EPI_ISL_501589, EPI_ISL_501590, EPI_ISL_501591, EPI_ISL_501592, EPI_ISL_501593, EPI_ISL_501594, EPI_ISL_501595, EPI_ISL_501596, EPI_ISL_501597, EPI_ISL_501598, EPI_ISL_501599, EPI_ISL_501601 | | | |
| see above | PHE South West Regional Laboratory, National Infection Service | Wellcome Sanger Institute for the COVID-19 Genomics UK Consortium | Stephanie Hutchings, Hannah Pymont, Dr Peter Muir, Barry Vipond, Rich Hopes; and Alex Alderton, Roberto Amato, Sonia Goncalves, Ewan Harrison, David K. Jackson, Ian Johnston, Dominic Kwiatkowski, Cordelia Langford, John Sillitoe on behalf of the Wellcome Sanger Institute COVID-19 Surveillance Team (http://www.sanger.ac.uk/covid-team) |
| EPI_ISL_501602 | Department of Medical Microbiology, Western Sussex Hospitals NHS Foundation Trust, St Richard's Hospital | Wellcome Sanger Institute for the COVID-19 Genomics UK Consortium | Manasa Mutingwende, Sarah Lowdon, Olga Podplomyk, Michelle Erkiert, Jonathan Lewis, Paul Randell and Alex Alderton, Roberto Amato, Sonia Goncalves, Ewan Harrison, David K. Jackson, Ian Johnston, Dominic Kwiatkowski, Cordelia Langford, John Sillitoe on behalf of the Wellcome Sanger Institute COVID-19 Surveillance Team (http://www.sanger.ac.uk/covid-team) |

| | | | |
|---|---|---|---|
| EPI_ISL_501603, EPI_ISL_501604, EPI_ISL_501605, EPI_ISL_501606, EPI_ISL_501607, EPI_ISL_501608, EPI_ISL_501609, EPI_ISL_501610, EPI_ISL_501611, EPI_ISL_501612 | PHE South West Regional Laboratory, National Infection Service | Wellcome Sanger Institute for the COVID-19 Genomics UK Consortium | Stephanie Hutchings, Hannah Pymont, Dr Peter Muir, Barry Vipond, Rich Hopes; and Alex Alderton, Roberto Amato, Sonia Goncalves, Ewan Harrison, David K. Jackson, Ian Johnston, Dominic Kwiatkowski, Cordelia Langford, John Sillitoe on behalf of the Wellcome Sanger Institute COVID-19 Surveillance Team (http://www.sanger.ac.uk/covid-team) |
| EPI_ISL_501613 | Department of Medical Microbiology, Western Sussex Hospitals NHS Foundation Trust, St Richard's Hospital | Wellcome Sanger Institute for the COVID-19 Genomics UK Consortium | Manasa Mutingwende, Sarah Lowdon, Olga Podplomyk, Michelle Erkiert, Jonathan Lewis, Paul Randell and Alex Alderton, Roberto Amato, Sonia Goncalves, Ewan Harrison, David K. Jackson, Ian Johnston, Dominic Kwiatkowski, Cordelia Langford, John Sillitoe on behalf of the Wellcome Sanger Institute COVID-19 Surveillance Team (http://www.sanger.ac.uk/covid-team) |
| EPI_ISL_501614, EPI_ISL_501615, EPI_ISL_501616, EPI_ISL_501617, EPI_ISL_501618, EPI_ISL_501619, EPI_ISL_501620, EPI_ISL_501621, EPI_ISL_501622, EPI_ISL_501623, EPI_ISL_501624, EPI_ISL_501625, EPI_ISL_501626, EPI_ISL_501627, EPI_ISL_501628, EPI_ISL_501629, EPI_ISL_501630, EPI_ISL_501631 | | | |
| see above | Lab Microbiology, Pathology Department, William Harvey Hospital | Wellcome Sanger Institute for the COVID-19 Genomics UK Consortium | Samuel Moses, Hannah Lowe, Felicity Ryan and Alex Alderton, Roberto Amato, Sonia Goncalves, Ewan Harrison, David K. Jackson, Ian Johnston, Dominic Kwiatkowski, Cordelia Langford, John Sillitoe on behalf of the Wellcome Sanger Institute COVID-19 Surveillance Team (http://www.sanger.ac.uk/covid-team) |
| EPI_ISL_501632 | Virology Department, Royal Infirmary of Edinburgh, NHS Lothian / School of Biological Sciences, University of Edinburgh | Wellcome Sanger Institute for the COVID-19 Genomics UK Consortium | McHugh M, Dewar R, Rooke S, O'Toole Á, Scher E, Hill V, McCrone JT, Colquhoun R, Yu X, Jackson B, Rambaut A, Templeton K and Alex Alderton, Roberto Amato, Sonia Goncalves, Ewan Harrison, David K. Jackson, Ian Johnston, Dominic Kwiatkowski, Cordelia Langford, John Sillitoe on behalf of the Wellcome Sanger Institute COVID-19 Surveillance Team (http://www.sanger.ac.uk/covid-team) |
| EPI_ISL_501633, EPI_ISL_501634, EPI_ISL_501635 | NHSGGC West of Scotland Specialist Virology Centre / MRC-University of Glasgow Centre for Virus Research | Wellcome Sanger Institute for the COVID-19 Genomics UK Consortium | Ana da Silva Filipe, Natasha Johnson, Kathy Smollett, Daniel Mair, Stephen Carmichael, Lily Tong, Jenna Nichols, Elihu Aranday-Cortes, Kirstyn Brunker, Yasmin Parr, Kyriaki Nomikou; Sarah McDonald, Marc Niebel, Patawee Asamaphan; Richard Orton, Joseph Hughes, Sreenu Vattipally, David L Robertson; Alasdair MacLean, Rory Gunson; Kathy Li, Natasha Jesudason, Rajiv Shah, James Shepherd, Antonia Ho, Alice Broos, Emma Thomson and Alex Alderton, Roberto Amato, Sonia Goncalves, Ewan Harrison, David K. Jackson, Ian Johnston, Dominic Kwiatkowski, Cordelia Langford, John Sillitoe on behalf of the Wellcome Sanger Institute COVID-19 Surveillance Team (http://www.sanger.ac.uk/covid-team) |
| EPI_ISL_501808, EPI_ISL_501817 | Centrālā laboratorija | Latvian Biomedical Research and Study Centre | Ivars Silamiķelis, Kaspars Megnis, Monta Ustinova, Ņikita Zrelovs, Vita Rovīte, Stella Lapiņa, Jana Osīte, Marta Priedīte, Uga Dumpis, Jānis Kloviņš |
| EPI_ISL_501823, EPI_ISL_501829, EPI_ISL_501833, EPI_ISL_501839, EPI_ISL_501849, EPI_ISL_501894 | E. Gulbja Laboratorija | Latvian Biomedical Research and Study Centre | Ivars Silamiķelis, Kaspars Megnis, Monta Ustinova, Ņikita Zrelovs, Vita Rovīte, Mikus Gavars, Dmitrijs Perminovs, Uga Dumpis, Jānis Kloviņš |
| EPI_ISL_501895, EPI_ISL_501896, EPI_ISL_501915, EPI_ISL_501922 | Centrālā laboratorija | Latvian Biomedical Research and Study Centre | Ivars Silamiķelis, Kaspars Megnis, Monta Ustinova, Ņikita Zrelovs, Vita Rovīte, Stella Lapiņa, Jana Osīte, Marta Priedīte, Uga Dumpis, Jānis Kloviņš |
| EPI_ISL_501929, EPI_ISL_501936 | E. Gulbja Laboratorija | Latvian Biomedical Research and Study Centre | Ivars Silamiķelis, Kaspars Megnis, Monta Ustinova, Ņikita Zrelovs, Vita Rovīte, Mikus Gavars, Dmitrijs Perminovs, Uga Dumpis, Jānis Kloviņš |
| EPI_ISL_502779 | LACEN/PE | LABBE, Federal University of Pernambuco | WILSON JOSE DA SILVA JUNIOR, HEIDI LACERDA ALVES DA CRUZ, MARCOS DA SILVEIRA REGUEIRA NETO, BRUNO SAMPAIO, SERGIO DE SA LEITAO PAIVA JUNIOR, ZILDENE DE SOUSA SILVEIRA, MAIRA GALDINO DA ROCHA PITTA, MICHELLY CRISTINY PEREIRA, MARCOS ANTONIO DE MORAIS JUNIOR, ANTONIO CARLOS DE FREITAS, VALDIR DE QUEIROZ BALBINO. |
| EPI_ISL_502875 | LACEN/PE | LABBE, Federal University of Pernambuco | WILSON JOSE DA SILVA JUNIOR, HEIDI LACERDA ALVES DA CRUZ, MARCOS DA SILVEIRA REGUEIRA NETO, BRUNO SAMPAIO, SERGIO DE SA LEITAO PAIVA JUNIOR, ZILDENE DE SOUSA SILVEIRA, MAIRA GALDINO DA ROCHA PITTA, MICHELLY CRISTINY PEREIRA, REGINALDO GONCALVES DE LIMA NETO, MARCOS ANTONIO DE MORAIS JUNIOR, ANTONIO CARLOS DE FREITAS, VALDIR DE QUEIROZ BALBINO. |
| EPI_ISL_503958, EPI_ISL_504040 | National Institute of Laboratory Medicine and Referral Center | Genomic Research Lab, BCSIR | Md. Ahasan Habib, Abu Sayeed Mohammad Mahmud, Mohammad Samir Uzzaman, Eshrar Osman, Shahina Akter, Tanjina Akhter Banu, Md. Murshed Hasan Sarkar, Barna Goswami, Iffat Jahan, Md. Saddam Hossain, Tarannum Taznin, Tasnim Nafisa, Md. Maruf Ahmed Molla, Mahmuda Yeasmin, Asish Kumar Ghosh, A. K. M. Shamsuzzaman, Sheikh Md. Selim Al Din, Utpal Chandra Ray, Salek Ahmed Sajib, Md. Salim Khan |
| EPI_ISL_504137, EPI_ISL_504176, EPI_ISL_504177, EPI_ISL_504178, EPI_ISL_504179, EPI_ISL_504180, EPI_ISL_504181, EPI_ISL_504182, EPI_ISL_504183, EPI_ISL_504184 | National Institute of Laboratory Medicine and Referral Center | Genomic Research Lab, BCSIR | Abu Sayeed Mohammad Mahmud, Mohammad Samir Uzzaman, Eshrar Osman, Md. Ahasan Habib, Shahina Akter, Tanjina Akhter Banu, Md. Murshed Hasan Sarkar, Barna Goswami, Iffat Jahan, Md. Saddam Hossain, Tarannum Taznin, Tasnim Nafisa, Md. Maruf Ahmed Molla, Mahmuda Yeasmin, Asish Kumar Ghosh, A. K. M. Shamsuzzaman, Sheikh Md. Selim Al Din, Utpal Chandra Ray, Salek Ahmed Sajib, Md. Salim Khan |
| EPI_ISL_504185 | Discovery DNA | Discovery DNA | Dustin Hittel, Marina Kerr, Leo Dimnik, Desmond Koo, Alice Li, Aneal Khan |
| EPI_ISL_504186, EPI_ISL_504187, EPI_ISL_504188, EPI_ISL_504189, EPI_ISL_504190, EPI_ISL_504191, EPI_ISL_504192, EPI_ISL_504193, EPI_ISL_504194, EPI_ISL_504195, EPI_ISL_504196, EPI_ISL_504197, EPI_ISL_504198, EPI_ISL_504199, EPI_ISL_504200, EPI_ISL_504201, EPI_ISL_504202, EPI_ISL_504203, EPI_ISL_504204, EPI_ISL_504205, EPI_ISL_504206, EPI_ISL_504207, EPI_ISL_504208, EPI_ISL_504209, EPI_ISL_504210, EPI_ISL_504211, EPI_ISL_504212, EPI_ISL_504213, EPI_ISL_504214, EPI_ISL_504215, EPI_ISL_504216, EPI_ISL_504217, EPI_ISL_504218, EPI_ISL_504219, EPI_ISL_504220, EPI_ISL_504221, EPI_ISL_504222, EPI_ISL_504223, EPI_ISL_504224, EPI_ISL_504225, EPI_ISL_504226, EPI_ISL_504227, EPI_ISL_504228, EPI_ISL_504229, EPI_ISL_504230, EPI_ISL_504231, EPI_ISL_504232, EPI_ISL_504233, EPI_ISL_504234, EPI_ISL_504235, EPI_ISL_504236, EPI_ISL_504237, EPI_ISL_504238, EPI_ISL_504239, EPI_ISL_504240, EPI_ISL_504241, EPI_ISL_504242, EPI_ISL_504243, EPI_ISL_504244 | | | |
| see above | National Institute for Communicable Diseases of the National Health Laboratory Service | National Institute for Communicable Diseases of the National Health Laboratory Service | Allam M, Ismail A, Khumalo Z, Kwenda S, Mtshali P, Mnyameni F, Mohale T, Bhiman JN |
| EPI_ISL_505003 | Biology Dpt | Microbiology and Infections Diseases | Emmanuelle Billon-Denis, Audrey Ferrier-Rembert, Annabelle Garnier, Laurence Cheutin, Clarisse Vigne, Emilie Tessier, Jessica Denis, Olivier Gorgé, Flora Nolent, Isabelle Drouet, Noémie Verguet, Olivier Ferraris, Jean-Nicolas Tournier |
| EPI_ISL_506041 | Biology Dpt, HIA Percy | Microbiology and Infectious Diseases Dpt | Emmanuelle Billon-Denis, Audrey Ferrier-Rembert, Annabelle Garnier, Laurence Cheutin, Clarisse Vigne, Emilie Tessier, Jessica Denis, Olivier Gorgé, Flora Nolent, Isabelle Drouet, Noémie Verguet, Olivier Ferraris, Jean-Nicolas Tournier |
| EPI_ISL_506947, EPI_ISL_506948, EPI_ISL_506949, EPI_ISL_506950, EPI_ISL_506951, EPI_ISL_506952, EPI_ISL_506953, EPI_ISL_506954, EPI_ISL_506955, EPI_ISL_506956 | Division of Viral Diseases, Center for Laboratory Control of Infectious Diseases, Korea Centers for Diseases Control and Prevention | Division of Viral Diseases, Center for Laboratory Control of Infectious Diseases, Korea Centers for Diseases Control and Prevention | Jeong-Min Kim, Yoon-Seok Chung, Namjoo Lee, Sang Hee Woo, Hye-Jun Jo, Heui Man Kim, Jun-Sub Kim, Dong Hyun Song, Daesang Lee, Seong Tae Jeong, Myung Guk Han |
| EPI_ISL_506957, EPI_ISL_506958, EPI_ISL_506959, EPI_ISL_506960, EPI_ISL_506961, EPI_ISL_506962, EPI_ISL_506963, EPI_ISL_506964 | Division of Viral Diseases, Center for Laboratory Control of Infectious Diseases, Korea Centers for Diseases Control and Prevention | Division of Viral Diseases, Center for Laboratory Control of Infectious Diseases, Korea Centers for Diseases Control and Prevention | Jeong-Min Kim, Yoon-Seok Chung, Namjoo Lee, Sang Hee Woo, Hye-Jun Jo, Heui Man Kim, Jun-Sub Kim, Myung Guk Han |
| EPI_ISL_506965, EPI_ISL_506966, EPI_ISL_506967, EPI_ISL_506968, EPI_ISL_506969, EPI_ISL_506970, EPI_ISL_506971, EPI_ISL_506972, EPI_ISL_506973, EPI_ISL_506974 | Division of Viral Diseases, Center for Laboratory Control of Infectious Diseases, Korea Centers for Diseases Control and Prevention | Division of Viral Diseases, Center for Laboratory Control of Infectious Diseases, Korea Centers for Diseases Control and Prevention | Jeong-Min Kim, Yoon-Seok Chung, Namjoo Lee, Sang Hee Woo, Hye-Jun Jo, Heui Man Kim, Jun-Sub Kim, Dong Hyun Song, Daesang Lee, Seong Tae Jeong, Myung Guk Han |
| EPI_ISL_506975 | Division of Viral Diseases, Center for Laboratory Control of Infectious Diseases, Korea Centers for Diseases Control and Prevention | Division of Viral Diseases, Center for Laboratory Control of Infectious Diseases, Korea Centers for Diseases Control and Prevention | Jeong-Min Kim, Yoon-Seok Chung, Namjoo Lee, Sang Hee Woo, Hye-Jun Jo, Heui Man Kim, Jun-Sub Kim, Myung Guk Han |
| EPI_ISL_506976, EPI_ISL_506977, EPI_ISL_506978, EPI_ISL_506979, EPI_ISL_506980, EPI_ISL_506981, EPI_ISL_506982, EPI_ISL_506983, EPI_ISL_506984, EPI_ISL_506985, EPI_ISL_506986, EPI_ISL_506987, EPI_ISL_506988, EPI_ISL_506989, EPI_ISL_506990, EPI_ISL_506991, EPI_ISL_506992, EPI_ISL_506993 | | | |
| see above | Division of Viral Diseases, Center for Laboratory Control of Infectious Diseases, Korea Centers for Diseases Control and Prevention | Division of Viral Diseases, Center for Laboratory Control of Infectious Diseases, Korea Centers for Diseases Control and Prevention | Jeong-Min Kim, Yoon-Seok Chung, Namjoo Lee, Sang Hee Woo, Hye-Jun Jo, Heui Man Kim, Jun-Sub Kim, Dong Hyun Song, Daesang Lee, Seong Tae Jeong, Myung Guk Han |
| EPI_ISL_506994, EPI_ISL_506995 | Division of Viral Diseases, Center for Laboratory Control of Infectious Diseases, Korea Centers for Diseases Control and Prevention | Division of Viral Diseases, Center for Laboratory Control of Infectious Diseases, Korea Centers for Diseases Control and Prevention | Jeong-Min Kim, Yoon-Seok Chung, Namjoo Lee, Sang Hee Woo, Hye-Jun Jo, Heui Man Kim, Jun-Sub Kim, Myung Guk Han |
| EPI_ISL_506996, EPI_ISL_506997, EPI_ISL_506998, EPI_ISL_506999, EPI_ISL_507000 | Department of Medical Microbiology, University Malaya Medical Centre | Department of Medical Microbiology, Faculty of Medicine, University of Malaya | Yoong Min CHONG, Jennifer Chong, I-Ching SAM, Yoke Fun CHAN, University Malaya Medical Centre COVID Team |
| EPI_ISL_507002, EPI_ISL_507003, EPI_ISL_507004, EPI_ISL_507005, EPI_ISL_507006 | Department of Laboratory Medicine, Tan Tock Seng Hospital | Department of Laboratory Medicine, Tan Tock Seng Hospital | Chen YYC, Zair X, Li C, Tang WY, Maurer-Stroh S, Barkham TMS, Nagarajan N, Sessions OM |
| EPI_ISL_507007 | Yaftabad Hospital, Covid Lab Center | Tehran University of Medical Sciences | Shahabzadeh,Z., Hosseinzadeh Gharajeh,N., Hashemian,S.M. and Barati,O. |
| EPI_ISL_507008 | National Veterinary Institute | National Veterinary Institute | Siamak Zohari |
| EPI_ISL_507010, EPI_ISL_507011, EPI_ISL_507012, EPI_ISL_507013, EPI_ISL_507014, EPI_ISL_507015, EPI_ISL_507016, EPI_ISL_507017, EPI_ISL_507018, EPI_ISL_507019, EPI_ISL_507020, EPI_ISL_507021, EPI_ISL_507022, EPI_ISL_507023, EPI_ISL_507024, EPI_ISL_507025, EPI_ISL_507026, EPI_ISL_507027, EPI_ISL_507028, EPI_ISL_507029, EPI_ISL_507030, EPI_ISL_507031, EPI_ISL_507032, EPI_ISL_507033, EPI_ISL_507034, EPI_ISL_507035, EPI_ISL_507036, EPI_ISL_507037, EPI_ISL_507038 | | | |
| see above | unknown | Infectious Diseases Research, King Abdullah International Medical Research Center (KAIMRC) | Alghoribi,M.F. |
| EPI_ISL_507039 | Department of Microbiology, College of Medicine and Medical Research Institute Chungbuk National University | Department of Microbiology, College of Medicine and Medical Research Institute Chungbuk National University | Young-Il Kim, Mark Anthony B. Casel, Se-Mi Kim, Seong-Gyu Kim, Su-Jin Park, Eun-Ha Kim, Hye Won Jeong, Young Ki Choi |
| EPI_ISL_507040, EPI_ISL_507041, EPI_ISL_507042, EPI_ISL_507043, EPI_ISL_507044, EPI_ISL_507045, EPI_ISL_507046, EPI_ISL_507047, EPI_ISL_507048, EPI_ISL_507049, EPI_ISL_507050, EPI_ISL_507051, EPI_ISL_507052, EPI_ISL_507053, EPI_ISL_507054, EPI_ISL_507055, EPI_ISL_507056, EPI_ISL_507057, EPI_ISL_507058, EPI_ISL_507059, EPI_ISL_507060, EPI_ISL_507061, EPI_ISL_507062, EPI_ISL_507063, EPI_ISL_507064, EPI_ISL_507065, EPI_ISL_507066, EPI_ISL_507067, EPI_ISL_507068, EPI_ISL_507069, EPI_ISL_507070, EPI_ISL_507071, EPI_ISL_507072, EPI_ISL_507073, EPI_ISL_507074, EPI_ISL_507075, EPI_ISL_507076, EPI_ISL_507077, EPI_ISL_507078, EPI_ISL_507079, EPI_ISL_507080, EPI_ISL_507081, EPI_ISL_507082, EPI_ISL_507083, EPI_ISL_507084, EPI_ISL_507085, EPI_ISL_507086, EPI_ISL_507087, EPI_ISL_507088, EPI_ISL_507089, EPI_ISL_507090, EPI_ISL_507091, EPI_ISL_507092, EPI_ISL_507093, EPI_ISL_507094, EPI_ISL_507095, EPI_ISL_507096, EPI_ISL_507097, EPI_ISL_507098, EPI_ISL_507099, EPI_ISL_507100, EPI_ISL_507101, EPI_ISL_507102, EPI_ISL_507103, EPI_ISL_507104, EPI_ISL_507105, EPI_ISL_507106, EPI_ISL_507107, EPI_ISL_507108, EPI_ISL_507109 | | | |
| see above | University College London Hospital | COVID-19 Genomics UK (COG-UK) Consortium | Judith Heaney, Matthew Byott, Catherine Houlihan, Dan Frampton, Stuart Kirk, Moira Spyer and Eleni Nastouli |
| EPI_ISL_507110, EPI_ISL_507111, EPI_ISL_507112, EPI_ISL_507113, EPI_ISL_507114, EPI_ISL_507115, EPI_ISL_507116, EPI_ISL_507117, EPI_ISL_507118, EPI_ISL_507119, EPI_ISL_507120, EPI_ISL_507121, EPI_ISL_507122, EPI_ISL_507123, EPI_ISL_507124, EPI_ISL_507125, EPI_ISL_507126, EPI_ISL_507127, EPI_ISL_507128, EPI_ISL_507129 | | | |
| see above | Northumbria University / South Tees Hospitals NHS Foundation Trust / North Cumbria Integrated Care NHS Foundation Trust / North Tees and Hartlepool NHS Foundation Trust / Newcastle Hospitals NHS Foundation Trust | COVID-19 Genomics UK (COG-UK) Consortium | Darren L Smith,Andrew Nelson,Matthew Bashton,Greg R Young,Joshua Loh,John Allan,Mohammad A Tariq,Giles S Holt,Gary Black,Wen C Yew,Lynn Dover,Paul Baker,Steve Liggett,Sarah Essex,Jane Greenaway,Debra Padgett,Clive Graham,Garren Scott,Edward Barton,Emma Swindells,Brendan Payne,Jennifer Collins,Yusri Taha,Gary Eltringham |
| EPI_ISL_507130 | Quadram Institute Bioscience | COVID-19 Genomics UK (COG-UK) Consortium | Dave J. Baker, Gemma L. Kay, Alp Aydin, Thanh Le-Viet, Steven Rudder, Ana P. Tedim, Anastasia Kolyva, Maria Diaz, Leonardo de Oliveira Martins, Nabil-Fareed Alikhan, Lizzie Meadows, Rachel Stanley, Ngozi Elumogo, Muhammed Yasir, Nicholas M. Thomson, Alexander J Trotter, Rachel Gilroy, Samuel Bloomfield, Claire Stuart, Andrew Bell, Reenesh Prakash, Samir Dervisevic, Alison E. Mather, John Wain, Mark Webber, Andrew J. Page, Justin O'Grady |

| | | | |
|---|---|---|---|
| EPI_ISL_507131, EPI_ISL_507132, EPI_ISL_507133, EPI_ISL_507134, EPI_ISL_507135, EPI_ISL_507136, EPI_ISL_507137, EPI_ISL_507138, EPI_ISL_507139, EPI_ISL_507140, EPI_ISL_507141, EPI_ISL_507142, EPI_ISL_507143, EPI_ISL_507144, EPI_ISL_507145, EPI_ISL_507146, EPI_ISL_507147, EPI_ISL_507148, EPI_ISL_507149, EPI_ISL_507150, EPI_ISL_507151, EPI_ISL_507152, EPI_ISL_507153, EPI_ISL_507154, EPI_ISL_507155, EPI_ISL_507156, EPI_ISL_507157, EPI_ISL_507158, EPI_ISL_507159 | | | |
| see above | Centre for Enzyme Innovation, University of Portsmouth / Translational Research Laboratory, Portsmouth Hospitals NHS Trust | COVID-19 Genomics UK (COG-UK) Consortium | Angela Beckett,Yann Bourgeois,Garry Scarlett,Sharon Glaysher,Scott Elliott,Kelly Bicknell,Robert Impey,Allyson Lloyd,Sarah Wyllie,Ethan Butcher,Anoop Chauhan,Samuel Robson |
| EPI_ISL_507160, EPI_ISL_507161, EPI_ISL_507162, EPI_ISL_507163, EPI_ISL_507164, EPI_ISL_507165, EPI_ISL_507166, EPI_ISL_507167, EPI_ISL_507168, EPI_ISL_507169, EPI_ISL_507170, EPI_ISL_507171, EPI_ISL_507172, EPI_ISL_507173, EPI_ISL_507174, EPI_ISL_507175, EPI_ISL_507176, EPI_ISL_507177, EPI_ISL_507178, EPI_ISL_507179 | | | |
| see above | Virology Department, Sheffield Teaching Hospitals NHS Foundation Trust/Department of Infection, Immunity and Cardiovascular Disease, The Medical School, University of Sheffield | COVID-19 Genomics UK (COG-UK) Consortium | Thushan de Silva, Matthew Parker, Nikki Smith, Adri Angyal, Rebecca Brown, Luke Green, Rachel Tucker, Paul Parsons, Danielle Groves, Katie Johnson, Laura Carrilero, Alex Keeley, Dave Partridge, Matthew Wyles, Benjamin Lindsey, Mehmet Yavuz, Mohammad Raza, Cariad Evans |
| EPI_ISL_507180, EPI_ISL_507181, EPI_ISL_507182, EPI_ISL_507183, EPI_ISL_507184, EPI_ISL_507185, EPI_ISL_507186, EPI_ISL_507187, EPI_ISL_507188, EPI_ISL_507189, EPI_ISL_507190, EPI_ISL_507191, EPI_ISL_507192, EPI_ISL_507193, EPI_ISL_507194, EPI_ISL_507195, EPI_ISL_507196, EPI_ISL_507197, EPI_ISL_507198, EPI_ISL_507199, EPI_ISL_507200, EPI_ISL_507201 | | | |
| see above | Virology Department, Royal Infirmary of Edinburgh, NHS Lothian / School of Biological Sciences, University of Edinburgh / Institute of Genetics and Molecular Medicine, University of Edinburgh | COVID-19 Genomics UK (COG-UK) Consortium | McHugh M, Dewar R, Rooke S, Gallagher M, Balcaza C, O'Toole Á, Scher E, Hill V, McCrone JT, Colquhoun R, Yu X, Jackson B, Rambaut A, Williams TC, Templeton K |
| EPI_ISL_507205 | WHO National Influenza Centre Russian Federation | WHO National Influenza Centre Russian Federation | Andrey Komissarov, Artem Fadeev, Mariia Sergeeva, Anna Ivanova, Daria Danilenko |
| EPI_ISL_507206, EPI_ISL_507207, EPI_ISL_507208, EPI_ISL_507209, EPI_ISL_507210, EPI_ISL_507211, EPI_ISL_507212, EPI_ISL_507213, EPI_ISL_507214, EPI_ISL_507215 | Department of Experimental Modeling and Pathogenesis of Infectious Diseases | WHO National Influenza Centre Russian Federation | Andrey Komissarov, Artem Fadeev, Mariia Sergeeva, Anna Ivanova, Daria Danilenko |
| EPI_ISL_507216, EPI_ISL_507217, EPI_ISL_507218, EPI_ISL_507219, EPI_ISL_507220, EPI_ISL_507221, EPI_ISL_507222, EPI_ISL_507223, EPI_ISL_507224, EPI_ISL_507225, EPI_ISL_507226, EPI_ISL_507227, EPI_ISL_507228, EPI_ISL_507229, EPI_ISL_507230, EPI_ISL_507231, EPI_ISL_507232, EPI_ISL_507233, EPI_ISL_507234, EPI_ISL_507235, EPI_ISL_507236, EPI_ISL_507237, EPI_ISL_507238, EPI_ISL_507239, EPI_ISL_507240, EPI_ISL_507241, EPI_ISL_507242, EPI_ISL_507243, EPI_ISL_507244, EPI_ISL_507245, EPI_ISL_507246, EPI_ISL_507247, EPI_ISL_507248, EPI_ISL_507249, EPI_ISL_507250, EPI_ISL_507251, EPI_ISL_507252, EPI_ISL_507253, EPI_ISL_507254, EPI_ISL_507255, EPI_ISL_507256, EPI_ISL_507257, EPI_ISL_507258, EPI_ISL_507259, EPI_ISL_507260, EPI_ISL_507261, EPI_ISL_507262, EPI_ISL_507263, EPI_ISL_507264, EPI_ISL_507265, EPI_ISL_507266, EPI_ISL_507267, EPI_ISL_507268, EPI_ISL_507269, EPI_ISL_507270, EPI_ISL_507271, EPI_ISL_507272, EPI_ISL_507273, EPI_ISL_507274, EPI_ISL_507275, EPI_ISL_507276, EPI_ISL_507277, EPI_ISL_507278, EPI_ISL_507279, EPI_ISL_507280, EPI_ISL_507281, EPI_ISL_507282, EPI_ISL_507283, EPI_ISL_507284, EPI_ISL_507285, EPI_ISL_507286, EPI_ISL_507287, EPI_ISL_507288, EPI_ISL_507289, EPI_ISL_507290, EPI_ISL_507291, EPI_ISL_507292, EPI_ISL_507293, EPI_ISL_507294, EPI_ISL_507295, EPI_ISL_507296 | | | |
| see above | WHO National Influenza Centre Russian Federation | WHO National Influenza Centre Russian Federation | Andrey Komissarov, Artem Fadeev, Mariia Sergeeva, Anna Ivanova, Daria Danilenko |
| EPI_ISL_507297, EPI_ISL_507298, EPI_ISL_507299 | Republican Medical Genetics Center | WHO National Influenza Centre Russian Federation | Andrey Komissarov, Artem Fadeev, Mariia Sergeeva, Anna Ivanova, Ildar Minniakhmetov, Rita Khusainova, Daria Danilenko |
| EPI_ISL_507300 | WHO National Influenza Centre Russian Federation | WHO National Influenza Centre Russian Federation | Andrey Komissarov, Artem Fadeev, Mariia Sergeeva, Anna Ivanova, Daria Danilenko |
| EPI_ISL_507415, EPI_ISL_507416, EPI_ISL_507417, EPI_ISL_507418, EPI_ISL_507419, EPI_ISL_507420, EPI_ISL_507421, EPI_ISL_507422, EPI_ISL_507423, EPI_ISL_507424, EPI_ISL_507425, EPI_ISL_507426, EPI_ISL_507427, EPI_ISL_507428, EPI_ISL_507429, EPI_ISL_507430, EPI_ISL_507431, EPI_ISL_507432, EPI_ISL_507433, EPI_ISL_507434, EPI_ISL_507435, EPI_ISL_507436, EPI_ISL_507437, EPI_ISL_507438, EPI_ISL_507439, EPI_ISL_507440, EPI_ISL_507441, EPI_ISL_507442, EPI_ISL_507443, EPI_ISL_507444, EPI_ISL_507445, EPI_ISL_507446, EPI_ISL_507447, EPI_ISL_507448, EPI_ISL_507449, EPI_ISL_507450, EPI_ISL_507451, EPI_ISL_507452, EPI_ISL_507453, EPI_ISL_507454, EPI_ISL_507455, EPI_ISL_507456, EPI_ISL_507457, EPI_ISL_507458, EPI_ISL_507459, EPI_ISL_507460, EPI_ISL_507461, EPI_ISL_507462, EPI_ISL_507463, EPI_ISL_507464, EPI_ISL_507465, EPI_ISL_507466, EPI_ISL_507467, EPI_ISL_507468, EPI_ISL_507469, EPI_ISL_507470, EPI_ISL_507471, EPI_ISL_507472, EPI_ISL_507473, EPI_ISL_507474, EPI_ISL_507475, EPI_ISL_507476, EPI_ISL_507477, EPI_ISL_507478, EPI_ISL_507479, EPI_ISL_507480, EPI_ISL_507481, EPI_ISL_507482, EPI_ISL_507483, EPI_ISL_507484, EPI_ISL_507485, EPI_ISL_507486, EPI_ISL_507487, EPI_ISL_507488, EPI_ISL_507489, EPI_ISL_507490, EPI_ISL_507491, EPI_ISL_507492, EPI_ISL_507493, EPI_ISL_507494, EPI_ISL_507495, EPI_ISL_507496, EPI_ISL_507497, EPI_ISL_507498, EPI_ISL_507499, EPI_ISL_507500, EPI_ISL_507501, EPI_ISL_507502, EPI_ISL_507503, EPI_ISL_507504, EPI_ISL_507505, EPI_ISL_507506, EPI_ISL_507507, EPI_ISL_507508, EPI_ISL_507509, EPI_ISL_507510, EPI_ISL_507511, EPI_ISL_507512, EPI_ISL_507513, EPI_ISL_507514, EPI_ISL_507515, EPI_ISL_507516, EPI_ISL_507517, EPI_ISL_507518, EPI_ISL_507519, EPI_ISL_507520, EPI_ISL_507521, EPI_ISL_507522, EPI_ISL_507523, EPI_ISL_507524, EPI_ISL_507525, EPI_ISL_507526, EPI_ISL_507527, EPI_ISL_507528, EPI_ISL_507529, EPI_ISL_507530, EPI_ISL_507531, EPI_ISL_507532, EPI_ISL_507533, EPI_ISL_507534, EPI_ISL_507535, EPI_ISL_507536, EPI_ISL_507537, EPI_ISL_507538, EPI_ISL_507539, EPI_ISL_507540, EPI_ISL_507541, EPI_ISL_507542, EPI_ISL_507543, EPI_ISL_507544, EPI_ISL_507545, EPI_ISL_507546, EPI_ISL_507547, EPI_ISL_507548, EPI_ISL_507549, EPI_ISL_507550, EPI_ISL_507551, EPI_ISL_507552, EPI_ISL_507553, EPI_ISL_507554, EPI_ISL_507555, EPI_ISL_507556, EPI_ISL_507557, EPI_ISL_507558, EPI_ISL_507559, EPI_ISL_507560, EPI_ISL_507561, EPI_ISL_507562, EPI_ISL_507563, EPI_ISL_507564, EPI_ISL_507565, EPI_ISL_507566, EPI_ISL_507567, EPI_ISL_507568, EPI_ISL_507569, EPI_ISL_507570, EPI_ISL_507571, EPI_ISL_507572, EPI_ISL_507573, EPI_ISL_507574, EPI_ISL_507575, EPI_ISL_507576, EPI_ISL_507577, EPI_ISL_507578, EPI_ISL_507579, EPI_ISL_507580, EPI_ISL_507581, EPI_ISL_507582, EPI_ISL_507583, EPI_ISL_507584, EPI_ISL_507585, EPI_ISL_507586, EPI_ISL_507587, EPI_ISL_507588, EPI_ISL_507589, EPI_ISL_507590, EPI_ISL_507591, EPI_ISL_507592, EPI_ISL_507593, EPI_ISL_507594, EPI_ISL_507595, EPI_ISL_507596, EPI_ISL_507597, EPI_ISL_507598, EPI_ISL_507599, EPI_ISL_507600, EPI_ISL_507601, EPI_ISL_507602, EPI_ISL_507603, EPI_ISL_507604, EPI_ISL_507605, EPI_ISL_507606, EPI_ISL_507607, EPI_ISL_507608, EPI_ISL_507609, EPI_ISL_507610, EPI_ISL_507611, EPI_ISL_507612, EPI_ISL_507613, EPI_ISL_507614, EPI_ISL_507615, EPI_ISL_507616, EPI_ISL_507617, EPI_ISL_507618, EPI_ISL_507619, EPI_ISL_507620, EPI_ISL_507621, EPI_ISL_507622, EPI_ISL_507623, EPI_ISL_507624, EPI_ISL_507625, EPI_ISL_507626, EPI_ISL_507627, EPI_ISL_507628, EPI_ISL_507629, EPI_ISL_507630, EPI_ISL_507631, EPI_ISL_507632, EPI_ISL_507633, EPI_ISL_507634, EPI_ISL_507635, EPI_ISL_507636, EPI_ISL_507637, EPI_ISL_507638, EPI_ISL_507639, EPI_ISL_507640, EPI_ISL_507641, EPI_ISL_507642, EPI_ISL_507643, EPI_ISL_507644, EPI_ISL_507645, EPI_ISL_507646, EPI_ISL_507647, EPI_ISL_507648, EPI_ISL_507649, EPI_ISL_507650, EPI_ISL_507651, EPI_ISL_507652, EPI_ISL_507653, EPI_ISL_507654, EPI_ISL_507655, EPI_ISL_507656, EPI_ISL_507657, EPI_ISL_507658, EPI_ISL_507659, EPI_ISL_507660, EPI_ISL_507661, EPI_ISL_507662, EPI_ISL_507663, EPI_ISL_507664, EPI_ISL_507665, EPI_ISL_507666, EPI_ISL_507667, EPI_ISL_507668, EPI_ISL_507669, EPI_ISL_507670, EPI_ISL_507671, EPI_ISL_507672, EPI_ISL_507673, EPI_ISL_507674, EPI_ISL_507675, EPI_ISL_507676, EPI_ISL_507677, EPI_ISL_507678, EPI_ISL_507679, EPI_ISL_507680, EPI_ISL_507681, EPI_ISL_507682, EPI_ISL_507683, EPI_ISL_507684, EPI_ISL_507685, EPI_ISL_507686, EPI_ISL_507687, EPI_ISL_507688, EPI_ISL_507689, EPI_ISL_507690, EPI_ISL_507691, EPI_ISL_507692, EPI_ISL_507693, EPI_ISL_507694, EPI_ISL_507695, EPI_ISL_507696, EPI_ISL_507697, EPI_ISL_507698, EPI_ISL_507699, EPI_ISL_507700, EPI_ISL_507701, EPI_ISL_507702, EPI_ISL_507703, EPI_ISL_507704, EPI_ISL_507705, EPI_ISL_507706, EPI_ISL_507707, EPI_ISL_507708, EPI_ISL_507709, EPI_ISL_507710, EPI_ISL_507711, EPI_ISL_507712, EPI_ISL_507713, EPI_ISL_507714, EPI_ISL_507715, EPI_ISL_507716, EPI_ISL_507717, EPI_ISL_507718, EPI_ISL_507719, EPI_ISL_507720, EPI_ISL_507721, EPI_ISL_507722, EPI_ISL_507723, EPI_ISL_507724, EPI_ISL_507725, EPI_ISL_507726, EPI_ISL_507727, EPI_ISL_507728, EPI_ISL_507729, EPI_ISL_507730, EPI_ISL_507731, EPI_ISL_507732, EPI_ISL_507733, EPI_ISL_507734, EPI_ISL_507735, EPI_ISL_507736, EPI_ISL_507737, EPI_ISL_507738, EPI_ISL_507739, EPI_ISL_507740, EPI_ISL_507741, EPI_ISL_507742, EPI_ISL_507743, EPI_ISL_507744, EPI_ISL_507745, EPI_ISL_507746, EPI_ISL_507747, EPI_ISL_507748, EPI_ISL_507749, EPI_ISL_507750, EPI_ISL_507751, EPI_ISL_507752, EPI_ISL_507753, EPI_ISL_507754, EPI_ISL_507755, EPI_ISL_507756, EPI_ISL_507757, EPI_ISL_507758, EPI_ISL_507759, EPI_ISL_507760, EPI_ISL_507761, EPI_ISL_507762, EPI_ISL_507763, EPI_ISL_507764, EPI_ISL_507765, EPI_ISL_507766, EPI_ISL_507767, EPI_ISL_507768, EPI_ISL_507769, EPI_ISL_507770, EPI_ISL_507771, EPI_ISL_507772, EPI_ISL_507773, EPI_ISL_507774, EPI_ISL_507775, EPI_ISL_507776, EPI_ISL_507777, EPI_ISL_507778, EPI_ISL_507779, EPI_ISL_507780, EPI_ISL_507781, EPI_ISL_507782, EPI_ISL_507783, EPI_ISL_507784, EPI_ISL_507785, EPI_ISL_507786, EPI_ISL_507787, EPI_ISL_507788, EPI_ISL_507789, EPI_ISL_507790, EPI_ISL_507791, EPI_ISL_507792, EPI_ISL_507793, EPI_ISL_507794, EPI_ISL_507795, EPI_ISL_507796, EPI_ISL_507797, EPI_ISL_507798, EPI_ISL_507799, EPI_ISL_507800, EPI_ISL_507801, EPI_ISL_507802, EPI_ISL_507803, EPI_ISL_507804, EPI_ISL_507805, EPI_ISL_507806, EPI_ISL_507807, EPI_ISL_507808, EPI_ISL_507809, EPI_ISL_507810, EPI_ISL_507811, EPI_ISL_507812, EPI_ISL_507813, EPI_ISL_507814, EPI_ISL_507815, EPI_ISL_507816, EPI_ISL_507817, EPI_ISL_507818, EPI_ISL_507819, EPI_ISL_507820, EPI_ISL_507821, EPI_ISL_507822, EPI_ISL_507823, EPI_ISL_507824, EPI_ISL_507825, EPI_ISL_507826, EPI_ISL_507827, EPI_ISL_507828, EPI_ISL_507829, EPI_ISL_507830, EPI_ISL_507831, EPI_ISL_507832, EPI_ISL_507833, EPI_ISL_507834, EPI_ISL_507835, EPI_ISL_507836, EPI_ISL_507837, EPI_ISL_507838, EPI_ISL_507839, EPI_ISL_507840, EPI_ISL_507841, EPI_ISL_507842, EPI_ISL_507843, EPI_ISL_507844, EPI_ISL_507845, EPI_ISL_507846, EPI_ISL_507847, EPI_ISL_507848, EPI_ISL_507849, EPI_ISL_507850, EPI_ISL_507851, EPI_ISL_507852, EPI_ISL_507853, EPI_ISL_507854, EPI_ISL_507855, EPI_ISL_507856, EPI_ISL_507857, EPI_ISL_507858, EPI_ISL_507859, EPI_ISL_507860, EPI_ISL_507861, EPI_ISL_507862, EPI_ISL_507863, EPI_ISL_507864, EPI_ISL_507865, EPI_ISL_507866, EPI_ISL_507867, EPI_ISL_507868, EPI_ISL_507869, EPI_ISL_507870, EPI_ISL_507871, EPI_ISL_507872, EPI_ISL_507873, EPI_ISL_507874, EPI_ISL_507875, EPI_ISL_507876, EPI_ISL_507877, EPI_ISL_507878, EPI_ISL_507879, EPI_ISL_507880, EPI_ISL_507881, EPI_ISL_507882, EPI_ISL_507883, EPI_ISL_507884, EPI_ISL_507885, EPI_ISL_507886, EPI_ISL_507887, EPI_ISL_507888, EPI_ISL_507889, EPI_ISL_507890, EPI_ISL_507891, EPI_ISL_507892, EPI_ISL_507893, EPI_ISL_507894, EPI_ISL_507895, EPI_ISL_507896, EPI_ISL_507897, EPI_ISL_507898, EPI_ISL_507899, EPI_ISL_507900, EPI_ISL_507901, EPI_ISL_507902, EPI_ISL_507903, EPI_ISL_507904, EPI_ISL_507905, EPI_ISL_507906, EPI_ISL_507907, EPI_ISL_507908, EPI_ISL_507909, EPI_ISL_507910, EPI_ISL_507911, EPI_ISL_507912, EPI_ISL_507913, EPI_ISL_507914, EPI_ISL_507915, EPI_ISL_507916, EPI_ISL_507917, EPI_ISL_507918, EPI_ISL_507919, EPI_ISL_507920, EPI_ISL_507921, EPI_ISL_507922, EPI_ISL_507923, EPI_ISL_507924, EPI_ISL_507925, EPI_ISL_507926, EPI_ISL_507927, EPI_ISL_507928, EPI_ISL_507929, EPI_ISL_507930, EPI_ISL_507931, EPI_ISL_507932, EPI_ISL_507933 | | | |
| see above | Michigan Department of Health and Human Services, Bureau of Laboratories | Michigan Department of Health and Human Services, Bureau of Laboratories | Blankenship HM, Riner D, Soehnlen MK |
| EPI_ISL_507934, EPI_ISL_507935, EPI_ISL_507936, EPI_ISL_507937, EPI_ISL_507938, EPI_ISL_507939, EPI_ISL_507940, EPI_ISL_507941, EPI_ISL_507942, EPI_ISL_507943, EPI_ISL_507944, EPI_ISL_507945, EPI_ISL_507946, EPI_ISL_507947, EPI_ISL_507948, EPI_ISL_507949, EPI_ISL_507950, EPI_ISL_507951, EPI_ISL_507952, EPI_ISL_507953, EPI_ISL_507954, EPI_ISL_507955, EPI_ISL_507956 | | | |
| see above | Minnesota Department of Health, Public Health Laboratory | Minnesota Department of Health, Public Health Laboratory | Matt Plumb, Jacob Garfin, and Xiong Wang |
| EPI_ISL_507957, EPI_ISL_507958, EPI_ISL_507959, EPI_ISL_507960, EPI_ISL_507961 | Mayo Clinic & Mayo Clinic Laboratories | Minnesota Department of Health, Public Health Laboratory | Matt Plumb, Jacob Garfin, and Xiong Wang |
| EPI_ISL_507962 | Children's Hospitals and Clinics of Minnesota | Minnesota Department of Health, Public Health Laboratory | Matt Plumb, Jacob Garfin, and Xiong Wang |
| EPI_ISL_507963, EPI_ISL_507964, EPI_ISL_507965, EPI_ISL_507966, EPI_ISL_507967, EPI_ISL_507968, EPI_ISL_507969, EPI_ISL_507970, EPI_ISL_507971, EPI_ISL_507972 | Avera Mckennan Laboratory | Minnesota Department of Health, Public Health Laboratory | Matt Plumb, Jacob Garfin, and Xiong Wang |
| EPI_ISL_507973, EPI_ISL_507974, EPI_ISL_507975, EPI_ISL_507976, EPI_ISL_507977, EPI_ISL_507978, EPI_ISL_507979, EPI_ISL_507980, EPI_ISL_507981, EPI_ISL_507982, EPI_ISL_507983 | | | |
| see above | Minnesota Department of Health, Public Health Laboratory | Minnesota Department of Health, Public Health Laboratory | Matt Plumb, Jacob Garfin, and Xiong Wang |
| EPI_ISL_507984, EPI_ISL_507985, EPI_ISL_507986, EPI_ISL_507987, EPI_ISL_507988, EPI_ISL_507989, EPI_ISL_507990, EPI_ISL_507991, EPI_ISL_507992, EPI_ISL_507993, EPI_ISL_507994, EPI_ISL_507995, EPI_ISL_507996, EPI_ISL_507997, EPI_ISL_507998, EPI_ISL_507999, EPI_ISL_508000, EPI_ISL_508001, EPI_ISL_508002, EPI_ISL_508003, EPI_ISL_508004, EPI_ISL_508005, EPI_ISL_508006, EPI_ISL_508007, EPI_ISL_508008, EPI_ISL_508009, EPI_ISL_508010, EPI_ISL_508011, EPI_ISL_508012, EPI_ISL_508013, EPI_ISL_508014, EPI_ISL_508015, EPI_ISL_508016, EPI_ISL_508017, EPI_ISL_508018, EPI_ISL_508019, EPI_ISL_508020, EPI_ISL_508021, EPI_ISL_508022, EPI_ISL_508023, EPI_ISL_508024, EPI_ISL_508025, EPI_ISL_508026, EPI_ISL_508027, EPI_ISL_508028, EPI_ISL_508029, EPI_ISL_508030, EPI_ISL_508031, EPI_ISL_508032, EPI_ISL_508033, EPI_ISL_508034, EPI_ISL_508035, EPI_ISL_508036, EPI_ISL_508037, EPI_ISL_508038, EPI_ISL_508039, EPI_ISL_508040, EPI_ISL_508041, EPI_ISL_508042, EPI_ISL_508043, EPI_ISL_508044, EPI_ISL_508045, EPI_ISL_508046, EPI_ISL_508047, EPI_ISL_508048, EPI_ISL_508049, EPI_ISL_508050, EPI_ISL_508051, EPI_ISL_508052, EPI_ISL_508053, EPI_ISL_508054, EPI_ISL_508055, EPI_ISL_508056, EPI_ISL_508057, EPI_ISL_508058, EPI_ISL_508059, EPI_ISL_508060, EPI_ISL_508061, EPI_ISL_508062, EPI_ISL_508063, EPI_ISL_508064, EPI_ISL_508065, EPI_ISL_508066, EPI_ISL_508067, EPI_ISL_508068, EPI_ISL_508069, EPI_ISL_508070, EPI_ISL_508071, EPI_ISL_508072, EPI_ISL_508073, EPI_ISL_508074, EPI_ISL_508075, EPI_ISL_508076, EPI_ISL_508077, EPI_ISL_508078, EPI_ISL_508079, EPI_ISL_508080, EPI_ISL_508081, EPI_ISL_508082, EPI_ISL_508083, EPI_ISL_508084, EPI_ISL_508085, EPI_ISL_508086, EPI_ISL_508087, EPI_ISL_508088, EPI_ISL_508089, EPI_ISL_508090, EPI_ISL_508091, EPI_ISL_508092, EPI_ISL_508093, EPI_ISL_508094, EPI_ISL_508095, EPI_ISL_508096, EPI_ISL_508097, EPI_ISL_508098, EPI_ISL_508099, EPI_ISL_508100, EPI_ISL_508101, EPI_ISL_508102, EPI_ISL_508103, EPI_ISL_508104, EPI_ISL_508105, EPI_ISL_508106, EPI_ISL_508107, EPI_ISL_508108, EPI_ISL_508109, EPI_ISL_508110, EPI_ISL_508111, EPI_ISL_508112, EPI_ISL_508113, EPI_ISL_508114, EPI_ISL_508115, EPI_ISL_508116, EPI_ISL_508117, EPI_ISL_508118, EPI_ISL_508119, EPI_ISL_508120, EPI_ISL_508121 | | | |
| see above | New Mexico Department of Health Scientific Laboratory Division | Center for Global Health, University of New Mexico Health Sciences Center | Daryl Domman, Kurt Schwalm, Twila Kunde, Joseph Hicks, Michael Edwards, Darrell Dinwiddie |
| EPI_ISL_510526 | Biological prevention, army | Biological prevention, army | Seadawy, M.G., Shamel,M.D., Harty,B.S., Elhoseny,M.M. and Gad,A.F. |
| EPI_ISL_510527 | Unite des virus emergents, UMR190 | Unite des virus emergents, UMR190 | Baronti, C., Piorkowski,G., Coutard,B., Charrel,R. and de Lamballerie,X. |
| EPI_ISL_510528 | Communicable Disease Laboratory, Public Health Directorate | Communicable Disease Laboratory, Public Health Directorate | Al Wasti,H. and AlTaif,Z. |
| EPI_ISL_510529 | School of Veterinary Medicine, Disease Control | School of Veterinary Medicine, Disease Control | Simulundu,E., Kapata,N., Mupeta,F., Kapata,P.C., Saasa,N., Changula,K., Muleya,W., Chitanga,S., Chambaro,H., Mubemba,B., Masahiro,K., Chanda,D., Mulenga,L., Fwoloshi,S., Shibemba,A.L., Kapaya,F., Zulu,P., Musonda,K., Monze,M., Sinyange,N., Liwewe,M.M., Kapin'a,M., Chipimo,P.J., Ngosa,W., Morales,A.N., Kayeyi,N., Malama,K., Tembo,J., Bates,M., Sawa,H., Takada,A., Nalubamba,K.S., Mukonka,V., Chilufya,C. and Zumla,A. |
| EPI_ISL_510530 | UMR 190 - Faculte de medecine, UMR 'Emergence des Pathologies Virales' (EPV: Aix-Marseille University - IRD 190 - Inserm 1207 - EHE | UMR 190 - Faculte de medecine, UMR 'Emergence des Pathologies Virales' (EPV: Aix-Marseille University - IRD 190 - Inserm 1207 - EHE | Durand, G.A., Piorkowski,G., Pommier De Santi,V., De Laval,F., Leparc Goffart,I., Gilles,M., Geulen,M., Peduzzi,F., Le Flem,F.-X. |
| EPI_ISL_510531 | Communicable Disease Laboratory, Public Health Directorate | Communicable Disease Laboratory, Public Health Directorate | Shehab,F., Altaif,Z., AlWasti,H., Zaed,A. |
| EPI_ISL_510532 | Biological prevention, army | Biological prevention, army | Seadawy,M.G., ELnabrawy,H.A., Shamel,M.D., Elhoseiny,M.F., Gad,A.F., Hassan,W.A., Raouf,A.A., Harty,B.E., ElGohary,A.A., Amer,k.E., Elnakeeb,M.A., Elnagdy,T.A., Ali,M.A., Kandeil,A.M. and Soliman,Y.A. |
| EPI_ISL_510533, EPI_ISL_510534 | Faculty of Medicine, Bursa Uludag University | Bursa Uludag University | Temel, S.G., Saglik, I., Özemri Sağ, Ş., Alemdar, A. |
| EPI_ISL_510535 | Molecular Virology, Instituto Carlos Chagas / Fiocruz Paraná | Universidade Federal do Parana (UFPR) | Suzukawa,A., Tscha,M., Zanluca,C., Raboni,S., Duarte dos Santos,C. |
| EPI_ISL_510545 | Takayuki Hishiki Kanagawa Prefectural Institute of Public Health | Takayuki Hishiki Kanagawa Prefectural Institute of Public Health | Hishiki,T., Suzuki,R., Sakuragi,J., Shimizu,H., Usui,K., Tanaka,Y., Kawai,J., Kogo,Y., Matsuki,Y., An,T., Hayashizaki,Y. and Takasaki,T. |
| EPI_ISL_524426 | Egyptian National Cancer Institute (ENCI) | Egyptian National Cancer Institute (ENCI) | Zekri, Abdel Rahman N., Amer,K.E., Ahmed,O.S., Soliman,H.K., Ali,M.A., Hassan,W.A., Mahmoud,A.A., Khattab,A.A., Hafez,M.M., Abouelhoda, Mohamed |
| EPI_ISL_524427 | Egyptian National Cancer Institute (ENCI) | Egyptian National Cancer Institute (ENCI) | Zekri, Abdel Rahman N, Amer,K.E., Ahmed,O.S., Soliman,H.K., Hafez,M.M., Bahnassy,A.A., Abdelhamid,W., Gad,A., Ali,M., Hassan,W., Samir,M., Raouf,A., Hamdy,M.S., Soliman,M.S., Elsissy,M.H., Elkhateeb,S.M., Ezzelarab,M.H., Abouelhoda, Mohamed |
| EPI_ISL_524435 | Department of Immunology, The Scripps Research Institute | Department of Immunology, The Scripps Research Institute | SEARCH Alliance,S.D., Pride,D., Shin,J.H. |
| EPI_ISL_524436 | Department of Immunology, The Scripps Research Institute | Department of Immunology, The Scripps Research Institute | Smither,A., Sabino-Santos,G., Snarski,P., Melnik,L., Bell,A., Genemaras,K., Drouin,A., Fusco,D., Garry,R., SEARCH Alliance,S.D. |
| EPI_ISL_529141, EPI_ISL_529142, EPI_ISL_529143, EPI_ISL_529144, EPI_ISL_529145 | Egyptian National Cancer Institute (ENCI) | Egyptian National Cancer Institute (ENCI) | Zekri, Abdel Rahman N., Amer,K.E., Ahmed,O.S., Soliman,H.K., Ali,M.A., Hassan,W.A., Mahmoud,A.A., Khattab,A.A., Hafez,M.M., Abouelhoda, Mohamed |
| EPI_ISL_529151, EPI_ISL_529152, EPI_ISL_529153 | Department of Immunology, The Scripps Research Institute | Andersen lab at Scripps Research | Quigley, M., Stefanski, E., Mchardy, I. with SEARCH Alliance San Diego |

| | | | |
|---|---|---|---|
| EPI_ISL_529154 | Department of Immunology, The Scripps Research Institute | Andersen lab at Scripps Research | Basler, T., Shephard, J., Austin, B. with SEARCH Alliance San Diego |
| EPI_ISL_529162, EPI_ISL_529163, EPI_ISL_529164, EPI_ISL_529165, EPI_ISL_529166 | Department of Immunology, The Scripps Research Institute | Andersen lab at Scripps Research | Allison Smither, Gilberto Sabino-Santos, Patricia Snarski, Lilia Melnik, Antoinette Bell, Kaylynn Genemaras, Arnaud Drouin, Dahlene Fusco, Robert Garry with SEARCH Alliance San Diego |
| EPI_ISL_529171, EPI_ISL_529172, EPI_ISL_529173, EPI_ISL_529174, EPI_ISL_529175, EPI_ISL_529176, EPI_ISL_529177, EPI_ISL_529178, EPI_ISL_529180, EPI_ISL_529181, EPI_ISL_529182, EPI_ISL_529183, EPI_ISL_529184, EPI_ISL_529185, EPI_ISL_529186, EPI_ISL_529187, EPI_ISL_529188, EPI_ISL_529189, EPI_ISL_529190, EPI_ISL_529191, EPI_ISL_529192, EPI_ISL_529193, EPI_ISL_529194, EPI_ISL_529195, EPI_ISL_529196, EPI_ISL_529197, EPI_ISL_529198, EPI_ISL_529199 | | | |
| see above | South Carolina Department of Health and Environmental Control | South Carolina Department of Health and Environmental Control | Haley V. Flores |

We gratefully acknowledge the following Authors from the Originating laboratories responsible for obtaining the specimens, as well as the Submitting laboratories where the genome data were generated and shared via GISAID, on which this research is based.

All Submitters of data may be contacted directly via www.gisaid.org

| Accession ID | Originating Laboratory | Submitting Laboratory | Authors |
|---|---|---|---|
| EPI_ISL_479625, EPI_ISL_479626, EPI_ISL_479627, EPI_ISL_479628, EPI_ISL_479629, EPI_ISL_479630, EPI_ISL_479631, EPI_ISL_479632, EPI_ISL_479633, EPI_ISL_479634, EPI_ISL_479635, EPI_ISL_479636, EPI_ISL_479637, EPI_ISL_479638, EPI_ISL_479639, EPI_ISL_479640, EPI_ISL_479641, EPI_ISL_479642, EPI_ISL_479643, EPI_ISL_479644, EPI_ISL_479645, EPI_ISL_479646, EPI_ISL_479647, EPI_ISL_479648, EPI_ISL_479649, EPI_ISL_479650, EPI_ISL_479651 | | | |
| see above | Dr. Georges-L.-Dumont University Hospital Centre | National Microbiology Laboratory | Anna Majer, Shari Tyson, Grace Seo, Kristyn Burak, Philip Mabon, Elsie Grudeski, Rhiannon Huzarewich, Russell Mandes, Jennifer Tanner, Natalie Knox, Morag Graham, Gary Van Domselaar, Richard Garceau, Guillaume Desnoyers, Nathalie Bastien, Yan Li, Timothy Booth |
| EPI_ISL_479657, EPI_ISL_479658, EPI_ISL_479659, EPI_ISL_479660, EPI_ISL_479661 | NIV Influenza | NIV Influenza | Potdar V |
| EPI_ISL_479736, EPI_ISL_479737, EPI_ISL_479738, EPI_ISL_479739, EPI_ISL_479740, EPI_ISL_479741, EPI_ISL_479742, EPI_ISL_479743, EPI_ISL_479744, EPI_ISL_479745, EPI_ISL_479746, EPI_ISL_479747, EPI_ISL_479748, EPI_ISL_479749, EPI_ISL_479750, EPI_ISL_479751, EPI_ISL_479752, EPI_ISL_479753, EPI_ISL_479754, EPI_ISL_479755 | | | |
| see above | Institute for Stem Cell Science and Regenerative Medicine | National Centre for Biological Sciences | Farhan Ali, Vanessa Molin Paynter, Srikar Krishna, Mohak Sharda, Shah-e-Jahan Gulzar, Awadhesh Pandit, Varadha Sundarmurthy, Uma Ramakrishnan, Dasaradhi Palakodeti, Aswin Seshasayee |
| EPI_ISL_479756, EPI_ISL_479757, EPI_ISL_479758 | National Institute of Hygiene and Epidemiology (NIHE) | National Key Laboratory of Gene Technology, Institute of Biotechnology (IBT) | Le Tung Lam, Nguyen Hong Trang, Ho Thi Thuong, Tran Huyen Linh, Ung Thi Hong Trang, Le Thi Thanh, Nguyen Vu Son, Vuong Duc Cuong, Tran Thu Huong, Pham Thi Hien, Nguyen Phuong Anh, Nguyen Le Khanh Hang, Hoang Vu Mai Phuong, Hoang Ha, Taichiro Takemura, Futoshi Hasebe, Chu Hoang Ha, Le Quynh Mai, Dang Duc Anh, Truong Nam Hai |
| EPI_ISL_479759, EPI_ISL_479760, EPI_ISL_479761, EPI_ISL_479762, EPI_ISL_479763, EPI_ISL_479764, EPI_ISL_479765, EPI_ISL_479766, EPI_ISL_479767, EPI_ISL_479768, EPI_ISL_479769, EPI_ISL_479770, EPI_ISL_479771, EPI_ISL_479772, EPI_ISL_479773, EPI_ISL_479774, EPI_ISL_479775 | | | |
| see above | University of Miami Immunology and Histocompatibility Laboratory | University of Miami Immunology and Histocompatibility Laboratory | Emilio Margolles-Clark, PhD and Phillip Ruiz, MD, PhD |
| EPI_ISL_479776 | NIV Influenza | NIV Influenza | Potdar V |
| EPI_ISL_479777, EPI_ISL_479778, EPI_ISL_479779, EPI_ISL_479780, EPI_ISL_479781, EPI_ISL_479782, EPI_ISL_479783, EPI_ISL_479784, EPI_ISL_479785, EPI_ISL_479786, EPI_ISL_479787, EPI_ISL_479788, EPI_ISL_479789 | | | |
| see above | Breuer Lab, UCL | Breuer Lab, UCL | Breuer Lab |
| EPI_ISL_479790 | Laboratory of Molecular Virology of the International Centre for Genetic Engineering and Biotechnology (ICGEB) | ARGO Open Lab Platform for Genome Sequencing | Licastro, D, Rajasekharan S, Dal Monego S, Segat L, D'Agaro P, Salton F, Confalonieri P, Confalonieri M, Marcello A |
| EPI_ISL_479791 | Laboratory of Molecular Virology of the International Centre for Genetic Engineering and Biotechnology (ICGEB) | ARGO Open Lab Platform for Genome Sequencing | Licastro, D, Rajasekharan S, Dal Monego S, Segat L, D'Agaro P, Salton F, Confalonieri P, Confalonieri M, Marcello A |
| EPI_ISL_479792, EPI_ISL_479793, EPI_ISL_479794, EPI_ISL_479795 | Hokkaido Institute of Public Health | Pathogen Genomics Center, National Institute of Infectious Diseases | Tsuyoshi Sekizuka, Rika Komagome, Kentaro Itokawa, Rina Tanaka, Masanori Hashino, Hajime Kamiya, Motoi Suzuki, Makoto Kuroda |
| EPI_ISL_479796 | Ishikawa Prefectural Institute of Public Health and Environmental Science | Pathogen Genomics Center, National Institute of Infectious Diseases | Tsuyoshi Sekizuka, Sanae Kuramoto, Eri Nariai, Kentaro Itokawa, Rina Tanaka, Masanori Hashino, Hajime Kamiya, Motoi Suzuki, Makoto Kuroda |
| EPI_ISL_479797, EPI_ISL_479798 | Sagamihara City Public Health Research Institute | Pathogen Genomics Center, National Institute of Infectious Diseases | Tsuyoshi Sekizuka, Hiroshi Nakamura, Kentaro Itokawa, Rina Tanaka, Masanori Hashino, Hajime Kamiya, Motoi Suzuki, Makoto Kuroda |
| EPI_ISL_479799, EPI_ISL_479800 | Sapporo City Institute of Public Health | Pathogen Genomics Center, National Institute of Infectious Diseases | Tsuyoshi Sekizuka, Asami Ohnishi, Kentaro Itokawa, Rina Tanaka, Masanori Hashino, Hajime Kamiya, Motoi Suzuki, Makoto Kuroda |
| EPI_ISL_479801 | Hokkaido Institute of Public Health | Pathogen Genomics Center, National Institute of Infectious Diseases | Tsuyoshi Sekizuka, Rika Komagome, Kentaro Itokawa, Rina Tanaka, Masanori Hashino, Hajime Kamiya, Motoi Suzuki, Makoto Kuroda |
| EPI_ISL_479802, EPI_ISL_479803, EPI_ISL_479804 | Sagamihara City Public Health Research Institute | Pathogen Genomics Center, National Institute of Infectious Diseases | Tsuyoshi Sekizuka, Hiroshi Nakamura, Kentaro Itokawa, Rina Tanaka, Masanori Hashino, Hajime Kamiya, Motoi Suzuki, Makoto Kuroda |
| EPI_ISL_479805, EPI_ISL_479806, EPI_ISL_479807, EPI_ISL_479808 | Saitama Prefectural Institute of Public Health | Pathogen Genomics Center, National Institute of Infectious Diseases | Tsuyoshi Sekizuka, Hayato Ehara, Kentaro Itokawa, Rina Tanaka, Masanori Hashino, Hajime Kamiya, Motoi Suzuki, Makoto Kuroda |
| EPI_ISL_479809, EPI_ISL_479810, EPI_ISL_479811 | Chiba Prefectural Institute of Public Health | Pathogen Genomics Center, National Institute of Infectious Diseases | Tsuyoshi Sekizuka, Masakatsu Taira, Kentaro Itokawa, Rina Tanaka, Masanori Hashino, Hajime Kamiya, Motoi Suzuki, Makoto Kuroda |
| EPI_ISL_479812, EPI_ISL_479813, EPI_ISL_479814, EPI_ISL_479815, EPI_ISL_479816, EPI_ISL_479817, EPI_ISL_479818, EPI_ISL_479819, EPI_ISL_479820 | Hokkaido Institute of Public Health | Pathogen Genomics Center, National Institute of Infectious Diseases | Tsuyoshi Sekizuka, Rika Komagome, Kentaro Itokawa, Rina Tanaka, Masanori Hashino, Hajime Kamiya, Motoi Suzuki, Makoto Kuroda |
| EPI_ISL_479821, EPI_ISL_479822 | Department of Infectious Diseases, Kobe Institute of Health | Pathogen Genomics Center, National Institute of Infectious Diseases | Tsuyoshi Sekizuka, Ryohei Nomoto, Kentaro Itokawa, Rina Tanaka, Masanori Hashino, Hajime Kamiya, Motoi Suzuki, Makoto Kuroda |
| EPI_ISL_479823 | Kochi Prefectural Institute of Public Health | Pathogen Genomics Center, National Institute of Infectious Diseases | Tsuyoshi Sekizuka, Akihiko Tokaji, Kentaro Itokawa, Rina Tanaka, Masanori Hashino, Hajime Kamiya, Motoi Suzuki, Makoto Kuroda |
| EPI_ISL_479824 | Kumamoto PrefecturalInstituteof Public-Health and Environmental Science | Pathogen Genomics Center, National Institute of Infectious Diseases | Tsuyoshi Sekizuka, Shunsuke Yahiro, Kentaro Itokawa, Rina Tanaka, Masanori Hashino, Hajime Kamiya, Motoi Suzuki, Makoto Kuroda |
| EPI_ISL_479825 | Tokyo Metropolitan Institute of Public Health | Pathogen Genomics Center, National Institute of Infectious Diseases | Tsuyoshi Sekizuka, Kenji Sadamasu, Takashi Chiba, Mami Nagashima, Kentaro Itokawa, Rina Tanaka, Masanori Hashino, Hajime Kamiya, Motoi Suzuki, Makoto Kuroda |
| EPI_ISL_479826, EPI_ISL_479827, EPI_ISL_479828, EPI_ISL_479829, EPI_ISL_479830, EPI_ISL_479831, EPI_ISL_479832, EPI_ISL_479833, EPI_ISL_479834, EPI_ISL_479835, EPI_ISL_479836, EPI_ISL_479837, EPI_ISL_479838, EPI_ISL_479839, EPI_ISL_479840, EPI_ISL_479841, EPI_ISL_479842, EPI_ISL_479843, EPI_ISL_479844, EPI_ISL_479845, EPI_ISL_479846, EPI_ISL_479847, EPI_ISL_479848, EPI_ISL_479849 | | | |
| see above | Sapporo City Institute of Public Health | Pathogen Genomics Center, National Institute of Infectious Diseases | Tsuyoshi Sekizuka, Asami Ohnishi, Kentaro Itokawa, Rina Tanaka, Masanori Hashino, Hajime Kamiya, Motoi Suzuki, Makoto Kuroda |
| EPI_ISL_479850, EPI_ISL_479851, EPI_ISL_479852, EPI_ISL_479853, EPI_ISL_479854 | Gunma Prefectural Institute of Public Health and Environmental Sciences | Pathogen Genomics Center, National Institute of Infectious Diseases | Tsuyoshi Sekizuka, Hiroyuki Tsukagoshi, Kentaro Itokawa, Rina Tanaka, Masanori Hashino, Hajime Kamiya, Motoi Suzuki, Makoto Kuroda |
| EPI_ISL_479855, EPI_ISL_479856, EPI_ISL_479857, EPI_ISL_479858, EPI_ISL_479859, EPI_ISL_479860, EPI_ISL_479861 | Department of Infectious Diseases, Kobe Institute of Health | Pathogen Genomics Center, National Institute of Infectious Diseases | Tsuyoshi Sekizuka, Ryohei Nomoto, Kentaro Itokawa, Rina Tanaka, Masanori Hashino, Hajime Kamiya, Motoi Suzuki, Makoto Kuroda |
| EPI_ISL_479862, EPI_ISL_479863, EPI_ISL_479864, EPI_ISL_479865, EPI_ISL_479866, EPI_ISL_479867 | Wakayama Prefectural Research Center of Environment and Public Health | Pathogen Genomics Center, National Institute of Infectious Diseases | Tsuyoshi Sekizuka, Fumio Terasoma, Yosuke Hamajima, Kentaro Itokawa, Rina Tanaka, Masanori Hashino, Hajime Kamiya, Motoi Suzuki, Makoto Kuroda |
| EPI_ISL_479868 | Department of Infectious Diseases, Kobe Institute of Health | Pathogen Genomics Center, National Institute of Infectious Diseases | Tsuyoshi Sekizuka, Ryohei Nomoto, Kentaro Itokawa, Rina Tanaka, Masanori Hashino, Hajime Kamiya, Motoi Suzuki, Makoto Kuroda |
| EPI_ISL_479869 | Niigata Prefectural Institute of Public Health and Environmental Sciences | Pathogen Genomics Center, National Institute of Infectious Diseases | Tsuyoshi Sekizuka, Reiko Arai, Kentaro Itokawa, Rina Tanaka, Masanori Hashino, Hajime Kamiya, Motoi Suzuki, Makoto Kuroda |
| EPI_ISL_479870, EPI_ISL_479871 | Sagamihara City Public Health Research Institute | Pathogen Genomics Center, National Institute of Infectious Diseases | Tsuyoshi Sekizuka, Hiroshi Nakamura, Kentaro Itokawa, Rina Tanaka, Masanori Hashino, Hajime Kamiya, Motoi Suzuki, Makoto Kuroda |
| EPI_ISL_479872, EPI_ISL_479873, EPI_ISL_479874, EPI_ISL_479875, EPI_ISL_479876, EPI_ISL_479877, EPI_ISL_479878, EPI_ISL_479879, EPI_ISL_479880, EPI_ISL_479881, EPI_ISL_479882, EPI_ISL_479883, EPI_ISL_479884, EPI_ISL_479885 | | | |
| see above | Sapporo City Institute of Public Health | Pathogen Genomics Center, National Institute of Infectious Diseases | Tsuyoshi Sekizuka, Asami Ohnishi, Kentaro Itokawa, Rina Tanaka, Masanori Hashino, Hajime Kamiya, Motoi Suzuki, Makoto Kuroda |
| EPI_ISL_479886, EPI_ISL_479887, EPI_ISL_479888, EPI_ISL_479889, EPI_ISL_479890, EPI_ISL_479891, EPI_ISL_479892, EPI_ISL_479893, EPI_ISL_479894, EPI_ISL_479895 | Tokyo Metropolitan Institute of Public Health | Pathogen Genomics Center, National Institute of Infectious Diseases | Tsuyoshi Sekizuka, Kenji Sadamasu, Takashi Chiba, Mami Nagashima, Kentaro Itokawa, Rina Tanaka, Masanori Hashino, Hajime Kamiya, Motoi Suzuki, Makoto Kuroda |

| | | | |
|---|---|---|---|
| EPI_ISL_479896, EPI_ISL_479897, EPI_ISL_479898, EPI_ISL_479899, EPI_ISL_479900, EPI_ISL_479901 | Gunma Prefectural Institute of Public Health and Environmental Sciences | Pathogen Genomics Center, National Institute of Infectious Diseases | Tsuyoshi Sekizuka, Hiroyuki Tsukagoshi, Kentaro Itokawa, Rina Tanaka, Masanori Hashino, Hajime Kamiya, Motoi Suzuki, Makoto Kuroda |
| EPI_ISL_479902 | Niigata Prefectural Institute of Public Health and Environmental Sciences | Pathogen Genomics Center, National Institute of Infectious Diseases | Tsuyoshi Sekizuka, Reiko Arai, Kentaro Itokawa, Rina Tanaka, Masanori Hashino, Hajime Kamiya, Motoi Suzuki, Makoto Kuroda |
| EPI_ISL_479903, EPI_ISL_479904, EPI_ISL_479905, EPI_ISL_479906, EPI_ISL_479907, EPI_ISL_479908, EPI_ISL_479909, EPI_ISL_479910, EPI_ISL_479911, EPI_ISL_479912 | Himeji City Institute of Environment and Health | Pathogen Genomics Center, National Institute of Infectious Diseases | Tsuyoshi Sekizuka, Kentaro Itokawa, Rina Tanaka, Masanori Hashino, Hajime Kamiya, Motoi Suzuki, Makoto Kuroda |
| EPI_ISL_479913, EPI_ISL_479914, EPI_ISL_479915, EPI_ISL_479916, EPI_ISL_479917, EPI_ISL_479918, EPI_ISL_479919, EPI_ISL_479920, EPI_ISL_479921, EPI_ISL_479922, EPI_ISL_479923, EPI_ISL_479924 | | | |
| see above | Niigata City Public Health Research Institute | Pathogen Genomics Center, National Institute of Infectious Diseases | Tsuyoshi Sekizuka, Yurie Takahashi, Kentaro Itokawa, Rina Tanaka, Masanori Hashino, Hajime Kamiya, Motoi Suzuki, Makoto Kuroda |
| EPI_ISL_479925, EPI_ISL_479926, EPI_ISL_479927 | Sakai City Institute of Public Health | Pathogen Genomics Center, National Institute of Infectious Diseases | Tsuyoshi Sekizuka, Tatsuya Miyoshi, Kentaro Itokawa, Rina Tanaka, Masanori Hashino, Hajime Kamiya, Motoi Suzuki, Makoto Kuroda |
| EPI_ISL_479928, EPI_ISL_479929, EPI_ISL_479930, EPI_ISL_479931, EPI_ISL_479932, EPI_ISL_479933, EPI_ISL_479934, EPI_ISL_479935 | Saitama Prefectural Institute of Public Health | Pathogen Genomics Center, National Institute of Infectious Diseases | Tsuyoshi Sekizuka, Hayato Ehara, Kentaro Itokawa, Rina Tanaka, Masanori Hashino, Hajime Kamiya, Motoi Suzuki, Makoto Kuroda |
| EPI_ISL_479936, EPI_ISL_479937, EPI_ISL_479938, EPI_ISL_479939, EPI_ISL_479940, EPI_ISL_479941, EPI_ISL_479942, EPI_ISL_479943 | Ibaraki Prefectural Institute of Public Health | Pathogen Genomics Center, National Institute of Infectious Diseases | Tsuyoshi Sekizuka, Keiko Goto, Kentaro Itokawa, Rina Tanaka, Masanori Hashino, Hajime Kamiya, Motoi Suzuki, Makoto Kuroda |
| EPI_ISL_479944, EPI_ISL_479945, EPI_ISL_479946, EPI_ISL_479947, EPI_ISL_479948, EPI_ISL_479949, EPI_ISL_479950, EPI_ISL_479951, EPI_ISL_479952, EPI_ISL_479953, EPI_ISL_479954, EPI_ISL_479955, EPI_ISL_479956, EPI_ISL_479957, EPI_ISL_479958 | | | |
| see above | Osaka Institute of Public Health | Pathogen Genomics Center, National Institute of Infectious Diseases | Tsuyoshi Sekizuka, Satoshi Hiroi, Saeko Morikawa, Kazushi Motomura, Kentaro Itokawa, Rina Tanaka, Masanori Hashino, Hajime Kamiya, Motoi Suzuki, Makoto Kuroda |
| EPI_ISL_479959, EPI_ISL_479960, EPI_ISL_479961, EPI_ISL_479962, EPI_ISL_479963, EPI_ISL_479964, EPI_ISL_479965 | Tokyo Metropolitan Institute of Public Health | Pathogen Genomics Center, National Institute of Infectious Diseases | Tsuyoshi Sekizuka, Kenji Sadamasu, Takashi Chiba, Mami Nagashima, Kentaro Itokawa, Rina Tanaka, Masanori Hashino, Hajime Kamiya, Motoi Suzuki, Makoto Kuroda |
| EPI_ISL_479966 | Osaka Institute of Public Health | Pathogen Genomics Center, National Institute of Infectious Diseases | Tsuyoshi Sekizuka, Satoshi Hiroi, Saeko Morikawa, Kazushi Motomura, Kentaro Itokawa, Rina Tanaka, Masanori Hashino, Hajime Kamiya, Motoi Suzuki, Makoto Kuroda |
| EPI_ISL_479967, EPI_ISL_479968, EPI_ISL_479969, EPI_ISL_479970, EPI_ISL_479971, EPI_ISL_479972, EPI_ISL_479973, EPI_ISL_479974, EPI_ISL_479975, EPI_ISL_479976, EPI_ISL_479977, EPI_ISL_479978 | | | |
| see above | Fukui Prefectural Institute of Public Health and Environmental Science | Pathogen Genomics Center, National Institute of Infectious Diseases | Tsuyoshi Sekizuka, Miho Toho, Kentaro Itokawa, Rina Tanaka, Masanori Hashino, Hajime Kamiya, Motoi Suzuki, Makoto Kuroda |
| EPI_ISL_479979, EPI_ISL_479980, EPI_ISL_479981, EPI_ISL_479982, EPI_ISL_479983, EPI_ISL_479984 | Oita Prefectural Institute of Public Health and Environmental Science | Pathogen Genomics Center, National Institute of Infectious Diseases | Tsuyoshi Sekizuka, Mari Sasaki, Kentaro Itokawa, Rina Tanaka, Masanori Hashino, Hajime Kamiya, Motoi Suzuki, Makoto Kuroda |
| EPI_ISL_479985 | Ibaraki Prefectural Institute of Public Health | Pathogen Genomics Center, National Institute of Infectious Diseases | Tsuyoshi Sekizuka, Keiko Goto, Kentaro Itokawa, Rina Tanaka, Masanori Hashino, Hajime Kamiya, Motoi Suzuki, Makoto Kuroda |
| EPI_ISL_479986, EPI_ISL_479987, EPI_ISL_479988, EPI_ISL_479989 | Department of Infectious Diseases, Kobe Institute of Health | Pathogen Genomics Center, National Institute of Infectious Diseases | Tsuyoshi Sekizuka, Ryohei Nomoto, Kentaro Itokawa, Rina Tanaka, Masanori Hashino, Hajime Kamiya, Motoi Suzuki, Makoto Kuroda |
| EPI_ISL_479990 | Kitakyushu City Institute of Health and Environmental Sciences | Pathogen Genomics Center, National Institute of Infectious Diseases | Tsuyoshi Sekizuka, Katsuya Obata, Asuka Kikuchi Kentaro Itokawa, Rina Tanaka, Masanori Hashino, Hajime Kamiya, Motoi Suzuki, Makoto Kuroda |
| EPI_ISL_479991, EPI_ISL_479992, EPI_ISL_479993, EPI_ISL_479994, EPI_ISL_479995, EPI_ISL_479996 | Kumamoto City Public Health Research Institute | Pathogen Genomics Center, National Institute of Infectious Diseases | Tsuyoshi Sekizuka, Kaori Tashiro, Kentaro Itokawa, Rina Tanaka, Masanori Hashino, Hajime Kamiya, Motoi Suzuki, Makoto Kuroda |
| EPI_ISL_479997, EPI_ISL_479998, EPI_ISL_479999, EPI_ISL_480000, EPI_ISL_480001 | Nagano Environmental Conservation Research Institute | Pathogen Genomics Center, National Institute of Infectious Diseases | Tsuyoshi Sekizuka, Naoko Shimodaira, Kentaro Itokawa, Rina Tanaka, Masanori Hashino, Hajime Kamiya, Motoi Suzuki, Makoto Kuroda |
| EPI_ISL_480002, EPI_ISL_480003 | Nagasaki Prefectural Institute for Environmental Research and Public Health | Pathogen Genomics Center, National Institute of Infectious Diseases | Tsuyoshi Sekizuka, Fumiaki Matsumoto, Kentaro Itokawa, Rina Tanaka, Masanori Hashino, Hajime Kamiya, Motoi Suzuki, Makoto Kuroda |
| EPI_ISL_480004, EPI_ISL_480005, EPI_ISL_480006, EPI_ISL_480007, EPI_ISL_480008, EPI_ISL_480009, EPI_ISL_480010, EPI_ISL_480011, EPI_ISL_480012, EPI_ISL_480013, EPI_ISL_480014 | | | |
| see above | Chiba Prefectural Institute of Public Health | Pathogen Genomics Center, National Institute of Infectious Diseases | Tsuyoshi Sekizuka, Masakatsu Taira, Kentaro Itokawa, Rina Tanaka, Masanori Hashino, Hajime Kamiya, Motoi Suzuki, Makoto Kuroda |
| EPI_ISL_480015, EPI_ISL_480016, EPI_ISL_480017, EPI_ISL_480018, EPI_ISL_480019, EPI_ISL_480020 | Gunma Prefectural Institute of Public Health and Environmental Sciences | Pathogen Genomics Center, National Institute of Infectious Diseases | Tsuyoshi Sekizuka, Hiroyuki Tsukagoshi, Kentaro Itokawa, Rina Tanaka, Masanori Hashino, Hajime Kamiya, Motoi Suzuki, Makoto Kuroda |
| EPI_ISL_480021, EPI_ISL_480022, EPI_ISL_480023, EPI_ISL_480024, EPI_ISL_480025, EPI_ISL_480026, EPI_ISL_480027, EPI_ISL_480028, EPI_ISL_480029 | Ibaraki Prefectural Institute of Public Health | Pathogen Genomics Center, National Institute of Infectious Diseases | Tsuyoshi Sekizuka, Keiko Goto, Kentaro Itokawa, Rina Tanaka, Masanori Hashino, Hajime Kamiya, Motoi Suzuki, Makoto Kuroda |
| EPI_ISL_480030, EPI_ISL_480031, EPI_ISL_480032, EPI_ISL_480033, EPI_ISL_480034, EPI_ISL_480035, EPI_ISL_480036, EPI_ISL_480037, EPI_ISL_480038, EPI_ISL_480039, EPI_ISL_480040, EPI_ISL_480041 | | | |
| see above | Tochigi Prefectural Institute of Public Health and Environmental Science | Pathogen Genomics Center, National Institute of Infectious Diseases | Tsuyoshi Sekizuka, Ako Nakajima, Kentaro Itokawa, Rina Tanaka, Masanori Hashino, Hajime Kamiya, Motoi Suzuki, Makoto Kuroda |
| EPI_ISL_480042, EPI_ISL_480043, EPI_ISL_480044, EPI_ISL_480045, EPI_ISL_480046, EPI_ISL_480047, EPI_ISL_480048, EPI_ISL_480049, EPI_ISL_480050, EPI_ISL_480051, EPI_ISL_480052, EPI_ISL_480053, EPI_ISL_480054, EPI_ISL_480055, EPI_ISL_480056, EPI_ISL_480057, EPI_ISL_480058, EPI_ISL_480059, EPI_ISL_480060, EPI_ISL_480061, EPI_ISL_480062, EPI_ISL_480063, EPI_ISL_480064 | | | |
| see above | Nagoya City Public Health Research Institute | Pathogen Genomics Center, National Institute of Infectious Diseases | Tsuyoshi Sekizuka, Takuya Miki, Shinichiro Shibata, Kentaro Itokawa, Rina Tanaka, Masanori Hashino, Hajime Kamiya, Motoi Suzuki, Makoto Kuroda |
| EPI_ISL_480065, EPI_ISL_480066, EPI_ISL_480067, EPI_ISL_480068, EPI_ISL_480069, EPI_ISL_480070, EPI_ISL_480071, EPI_ISL_480072 | Sakai City Institute of Public Health | Pathogen Genomics Center, National Institute of Infectious Diseases | Tsuyoshi Sekizuka, Tatsuya Miyoshi, Kentaro Itokawa, Rina Tanaka, Masanori Hashino, Hajime Kamiya, Motoi Suzuki, Makoto Kuroda |
| EPI_ISL_480073 | Tochigi Prefectural Institute of Public Health and Environmental Science | Pathogen Genomics Center, National Institute of Infectious Diseases | Tsuyoshi Sekizuka, Ako Nakajima, Kentaro Itokawa, Rina Tanaka, Masanori Hashino, Hajime Kamiya, Motoi Suzuki, Makoto Kuroda |
| EPI_ISL_480074, EPI_ISL_480075, EPI_ISL_480076, EPI_ISL_480077, EPI_ISL_480078, EPI_ISL_480079, EPI_ISL_480080, EPI_ISL_480081, EPI_ISL_480082 | Shizuoka City Institute of Environmental Sciences and Public Health | Pathogen Genomics Center, National Institute of Infectious Diseases | Tsuyoshi Sekizuka, Takaharu Maehata,Sou Okamura,Yuji Kanazawa,Kenji Yagi, Kentaro Itokawa, Rina Tanaka, Masanori Hashino, Hajime Kamiya, Motoi Suzuki, Makoto Kuroda |
| EPI_ISL_480083, EPI_ISL_480084, EPI_ISL_480085, EPI_ISL_480086, EPI_ISL_480087, EPI_ISL_480088, EPI_ISL_480089 | Gifu Prefectural Institute of Public Health and Environmental Sciences | Pathogen Genomics Center, National Institute of Infectious Diseases | Tsuyoshi Sekizuka, Yoshihiko Kameyama, Kentaro Itokawa, Rina Tanaka, Masanori Hashino, Hajime Kamiya, Motoi Suzuki, Makoto Kuroda |
| EPI_ISL_480090, EPI_ISL_480091, EPI_ISL_480092, EPI_ISL_480093, EPI_ISL_480094, EPI_ISL_480095, EPI_ISL_480096, EPI_ISL_480097, EPI_ISL_480098, EPI_ISL_480099, EPI_ISL_480100, EPI_ISL_480101, EPI_ISL_480102 | | | |
| see above | Department of Infectious Diseases, Kobe Institute of Health | Pathogen Genomics Center, National Institute of Infectious Diseases | Tsuyoshi Sekizuka, Ryohei Nomoto, Kentaro Itokawa, Rina Tanaka, Masanori Hashino, Hajime Kamiya, Motoi Suzuki, Makoto Kuroda |
| EPI_ISL_480103, EPI_ISL_480104, EPI_ISL_480105, EPI_ISL_480106, EPI_ISL_480107, EPI_ISL_480108 | Koshigaya City Public Health Center | Pathogen Genomics Center, National Institute of Infectious Diseases | Tsuyoshi Sekizuka, Yuka Furui, Aya Tamura, Kyohei Sakata, Takumi Daimon, Yoko Togawa, Yoshiko Hamada, Kentaro Itokawa, Rina Tanaka, Masanori Hashino, Hajime Kamiya, Motoi Suzuki, Makoto Kuroda |
| EPI_ISL_480109, EPI_ISL_480110, EPI_ISL_480111, EPI_ISL_480112, EPI_ISL_480113, EPI_ISL_480114, EPI_ISL_480115, EPI_ISL_480116, EPI_ISL_480117, EPI_ISL_480118, EPI_ISL_480119 | | | |
| see above | Oita Prefectural Institute of Public Health and Environmental Science | Pathogen Genomics Center, National Institute of Infectious Diseases | Tsuyoshi Sekizuka, Mari Sasaki, Kentaro Itokawa, Rina Tanaka, Masanori Hashino, Hajime Kamiya, Motoi Suzuki, Makoto Kuroda |

| | | | |
|---|---|---|---|
| EPI_ISL_480120, EPI_ISL_480121, EPI_ISL_480122, EPI_ISL_480123, EPI_ISL_480124, EPI_ISL_480125, EPI_ISL_480126, EPI_ISL_480127, EPI_ISL_480128, EPI_ISL_480129, EPI_ISL_480130, EPI_ISL_480131, EPI_ISL_480132, EPI_ISL_480133, EPI_ISL_480134, EPI_ISL_480135, EPI_ISL_480136, EPI_ISL_480137, EPI_ISL_480138, EPI_ISL_480139, EPI_ISL_480140, EPI_ISL_480141, EPI_ISL_480142, EPI_ISL_480143, EPI_ISL_480144, EPI_ISL_480145, EPI_ISL_480146, EPI_ISL_480147, EPI_ISL_480148, EPI_ISL_480149, EPI_ISL_480150, EPI_ISL_480151, EPI_ISL_480152, EPI_ISL_480153, EPI_ISL_480154, EPI_ISL_480155, EPI_ISL_480156, EPI_ISL_480157, EPI_ISL_480158, EPI_ISL_480159, EPI_ISL_480160, EPI_ISL_480161, EPI_ISL_480162, EPI_ISL_480163, EPI_ISL_480164, EPI_ISL_480165, EPI_ISL_480166, EPI_ISL_480167, EPI_ISL_480168 | | | |
| see above | Fukui Prefectural Institute of Public Health and Environmental Science | Pathogen Genomics Center, National Institute of Infectious Diseases | Tsuyoshi Sekizuka, Miho Toho, Kentaro Itokawa, Rina Tanaka, Masanori Hashino, Hajime Kamiya, Motoi Suzuki, Makoto Kuroda |
| EPI_ISL_480169, EPI_ISL_480170, EPI_ISL_480171, EPI_ISL_480172, EPI_ISL_480173, EPI_ISL_480174, EPI_ISL_480175, EPI_ISL_480176, EPI_ISL_480177 | Gunma Prefectural Institute of Public Health and Environmental Sciences | Pathogen Genomics Center, National Institute of Infectious Diseases | Tsuyoshi Sekizuka, Hiroyuki Tsukagoshi, Kentaro Itokawa, Rina Tanaka, Masanori Hashino, Hajime Kamiya, Motoi Suzuki, Makoto Kuroda |
| EPI_ISL_480178, EPI_ISL_480179 | Hiroshima City Institute of Public Health | Pathogen Genomics Center, National Institute of Infectious Diseases | Tsuyoshi Sekizuka, Kota Noritsune, Kentaro Itokawa, Rina Tanaka, Masanori Hashino, Hajime Kamiya, Motoi Suzuki, Makoto Kuroda |
| EPI_ISL_480180, EPI_ISL_480181, EPI_ISL_480182, EPI_ISL_480183, EPI_ISL_480184, EPI_ISL_480185, EPI_ISL_480186, EPI_ISL_480187, EPI_ISL_480188, EPI_ISL_480189 | Ibaraki Prefectural Institute of Public Health | Pathogen Genomics Center, National Institute of Infectious Diseases | Tsuyoshi Sekizuka, Keiko Goto, Kentaro Itokawa, Rina Tanaka, Masanori Hashino, Hajime Kamiya, Motoi Suzuki, Makoto Kuroda |
| EPI_ISL_480190, EPI_ISL_480191, EPI_ISL_480192, EPI_ISL_480193, EPI_ISL_480194, EPI_ISL_480195 | Ota Health Center Welfare Section | Pathogen Genomics Center, National Institute of Infectious Diseases | Tsuyoshi Sekizuka, Chika Takahashi, Kentaro Itokawa, Rina Tanaka, Masanori Hashino, Hajime Kamiya, Motoi Suzuki, Makoto Kuroda |
| EPI_ISL_480196, EPI_ISL_480197, EPI_ISL_480198, EPI_ISL_480199, EPI_ISL_480200, EPI_ISL_480201, EPI_ISL_480202, EPI_ISL_480203 | Toyama Institute of Health | Pathogen Genomics Center, National Institute of Infectious Diseases | Tsuyoshi Sekizuka, Masae Itamochi, Kazunori Oishi, Kentaro Itokawa, Rina Tanaka, Masanori Hashino, Hajime Kamiya, Motoi Suzuki, Makoto Kuroda |
| EPI_ISL_480204 | Akita City Public Health Center | Pathogen Genomics Center, National Institute of Infectious Diseases | Tsuyoshi Sekizuka, Koichi Ito, Kentaro Itokawa, Rina Tanaka, Masanori Hashino, Hajime Kamiya, Motoi Suzuki, Makoto Kuroda |
| EPI_ISL_480205, EPI_ISL_480206, EPI_ISL_480207, EPI_ISL_480208, EPI_ISL_480209, EPI_ISL_480210, EPI_ISL_480211, EPI_ISL_480212, EPI_ISL_480213, EPI_ISL_480214, EPI_ISL_480215, EPI_ISL_480216, EPI_ISL_480217, EPI_ISL_480218, EPI_ISL_480219, EPI_ISL_480220 | | | |
| see above | Department of Infectious Diseases, Kobe Institute of Health | Pathogen Genomics Center, National Institute of Infectious Diseases | Tsuyoshi Sekizuka, Ryohei Nomoto, Kentaro Itokawa, Rina Tanaka, Masanori Hashino, Hajime Kamiya, Motoi Suzuki, Makoto Kuroda |
| EPI_ISL_480221, EPI_ISL_480222, EPI_ISL_480223 | Koshigaya City Public Health Center | Pathogen Genomics Center, National Institute of Infectious Diseases | Tsuyoshi Sekizuka, Yuka Furui, Aya Tamura, Kyohei Sakata, Takumi Daimon, Yoko Togawa, Yoshiko Hamada, Kentaro Itokawa, Rina Tanaka, Masanori Hashino, Hajime Kamiya, Motoi Suzuki, Makoto Kuroda |
| EPI_ISL_480224 | National Reference Laboratory "Influenza and acute respiratory diseases" | NRL-HIV | Ivan Ivanov, Ivailo Alexiev, Ivva Philipova |
| EPI_ISL_480225 | Fukui Prefectural Institute of Public Health and Environmental Science | Pathogen Genomics Center, National Institute of Infectious Diseases | Tsuyoshi Sekizuka, Miho Toho, Kentaro Itokawa, Rina Tanaka, Masanori Hashino, Hajime Kamiya, Motoi Suzuki, Makoto Kuroda |
| EPI_ISL_480226 | Niigata Prefectural Institute of Public Health and Environmental Sciences | Pathogen Genomics Center, National Institute of Infectious Diseases | Tsuyoshi Sekizuka, Reiko Arai, Kentaro Itokawa, Rina Tanaka, Masanori Hashino, Hajime Kamiya, Motoi Suzuki, Makoto Kuroda |
| EPI_ISL_480227 | Tokyo Metropolitan Institute of Public Health | Pathogen Genomics Center, National Institute of Infectious Diseases | Tsuyoshi Sekizuka, Kenji Sadamasu, Takashi Chiba, Mami Nagashima, Kentaro Itokawa, Rina Tanaka, Masanori Hashino, Hajime Kamiya, Motoi Suzuki, Makoto Kuroda |
| EPI_ISL_480228, EPI_ISL_480229, EPI_ISL_480230, EPI_ISL_480231, EPI_ISL_480232, EPI_ISL_480233, EPI_ISL_480234, EPI_ISL_480235, EPI_ISL_480236, EPI_ISL_480237, EPI_ISL_480238, EPI_ISL_480239, EPI_ISL_480240, EPI_ISL_480241, EPI_ISL_480242, EPI_ISL_480243, EPI_ISL_480244, EPI_ISL_480245, EPI_ISL_480246, EPI_ISL_480247, EPI_ISL_480248, EPI_ISL_480249, EPI_ISL_480250, EPI_ISL_480251, EPI_ISL_480252, EPI_ISL_480253, EPI_ISL_480254, EPI_ISL_480255, EPI_ISL_480256, EPI_ISL_480257, EPI_ISL_480258, EPI_ISL_480259, EPI_ISL_480260, EPI_ISL_480261, EPI_ISL_480262, EPI_ISL_480263, EPI_ISL_480264, EPI_ISL_480265, EPI_ISL_480266, EPI_ISL_480267, EPI_ISL_480268, EPI_ISL_480269, EPI_ISL_480270, EPI_ISL_480271, EPI_ISL_480272, EPI_ISL_480273, EPI_ISL_480274, EPI_ISL_480275, EPI_ISL_480276, EPI_ISL_480277, EPI_ISL_480278, EPI_ISL_480279, EPI_ISL_480280, EPI_ISL_480281, EPI_ISL_480282, EPI_ISL_480283, EPI_ISL_480284, EPI_ISL_480285, EPI_ISL_480286, EPI_ISL_480287, EPI_ISL_480288, EPI_ISL_480289, EPI_ISL_480290, EPI_ISL_480291, EPI_ISL_480292 | | | |
| see above | Genomic Laboratory (GLAB) (Conjoint lab of Health Directorate of Istanbul and Istanbul Technical University) | Genomic Laboratory (GLAB), Istanbul Technical University | Ilker Karacan, Tugba Kizilboga Akgun, Bugra Agaoglu, Gizem Alkurt, Jale Yildiz, Betsi Köse, Elifnaz Çelik, Arzu Irvem, Yasemin Kendir Demirkol, Ozlem Akgun Dogan, Mehtap Aydın, Levent Doganay, Gizem Dinler Doganay |
| EPI_ISL_480293, EPI_ISL_480294, EPI_ISL_480295, EPI_ISL_480296 | Institute for Stem Cell Science and Regenerative Medicine | National Centre for Biological Sciences | Farhan Ali, Vanessa Molin Paynter, Srikar Krishna, Mohak Sharda, Shah-e-Jahan Gulzar, Awadhesh Pandit, Varadha Sundarmurthy, Uma Ramakrishnan, Dasaradhi Palakodeti, Aswin Seshasayee |
| EPI_ISL_480297, EPI_ISL_480298, EPI_ISL_480299, EPI_ISL_480300, EPI_ISL_480301, EPI_ISL_480302, EPI_ISL_480303, EPI_ISL_480304, EPI_ISL_480305, EPI_ISL_480306, EPI_ISL_480307, EPI_ISL_480308, EPI_ISL_480309, EPI_ISL_480310 | | | |
| see above | National Reference Laboratory "Influenza and acute respiratory diseases" | NRL-HIV | Ivan Ivanov, Ivailo Alexiev, Ivva Philipova |
| EPI_ISL_480312, EPI_ISL_480313, EPI_ISL_480314 | Hospital Mexico | Charité Virology-University of Costa Rica | Andres Moreira-Soto, Eugenia Corrales-Aguilar, Ignacio Postigo-Hidalgo, Teresita Somogyi, Jan Felix Drexler |
| EPI_ISL_480315, EPI_ISL_480316, EPI_ISL_480317, EPI_ISL_480318, EPI_ISL_480319, EPI_ISL_480320 | Hospital Clínica Bíblica | Charité Virology-University of Costa Rica | Andres Moreira-Soto, Eugenia Corrales-Aguilar, Ignacio Postigo-Hidalgo, Karla Sofía Gutiérrez, Jan Felix Drexler |
| EPI_ISL_480321 | Laboratorio Clínico San José | Charité Virology-University of Costa Rica | Andres Moreira-Soto, Eugenia Corrales-Aguilar, Ignacio Postigo-Hidalgo, Hugo Núñez Navas, Jan Felix Drexler |
| EPI_ISL_480322, EPI_ISL_480323, EPI_ISL_480324, EPI_ISL_480325, EPI_ISL_480326, EPI_ISL_480327 | Hospital Nacional de Niños | Charité Virology-University of Costa Rica | Andres Moreira-Soto, Eugenia Corrales-Aguilar, Ignacio Postigo-Hidalgo, Cristian Pérez Corrales, Andrei Montero Bonilla, Jan Felix Drexler |
| EPI_ISL_480328 | Laboratorio LABIN | Charité Virology-University of Costa Rica | Andres Moreira-Soto, Eugenia Corrales-Aguilar, Ignacio Postigo-Hidalgo, Ignacio Soto Pacheco, Jan Felix Drexler |
| EPI_ISL_480329, EPI_ISL_480330 | Quadram Institute Bioscience | COVID-19 Genomics UK (COG-UK) Consortium | Dave J. Baker, Gemma L. Kay, Alp Aydin, Thanh Le-Viet, Steven Rudder, Ana P. Tedim, Anastasia Kolyva, Maria Diaz, Leonardo de Oliveira Martins, Nabil-Fareed Alikhan, Lizzie Meadows, Rachael Stanley, Ngozi Elumogo, Muhammed Yasir, Nicholas M. Thomson, Alexander J Trotter, Rachel Gilroy, Samuel Bloomfield, Claire Stuart, Andrew Bell, Reenesh Prakash, Samir Dervisevic, Alison E. Mather, John Wain, Mark Webber, Andrew J. Page, Justin O'Grady |
| EPI_ISL_480331, EPI_ISL_480332, EPI_ISL_480333, EPI_ISL_480334, EPI_ISL_480335, EPI_ISL_480336, EPI_ISL_480337, EPI_ISL_480338, EPI_ISL_480339, EPI_ISL_480340, EPI_ISL_480341, EPI_ISL_480342, EPI_ISL_480343, EPI_ISL_480344, EPI_ISL_480345, EPI_ISL_480346, EPI_ISL_480347, EPI_ISL_480348 | | | |
| see above | Microbial Genomics Laboratory, Institut Pasteur de Montevideo | Microbial Genomics Laboratory, Institut Pasteur de Montevideo | Cecilia Salazar, Marianoel Pereira, Ignacio Ferrés, Gonzalo Moratorio, Pilar Moreno, Gregorio Iraola |
| EPI_ISL_480351, EPI_ISL_480352, EPI_ISL_480353, EPI_ISL_480354, EPI_ISL_480355, EPI_ISL_480356, EPI_ISL_480357, EPI_ISL_480358, EPI_ISL_480359, EPI_ISL_480360, EPI_ISL_480361, EPI_ISL_480362, EPI_ISL_480363, EPI_ISL_480364, EPI_ISL_480365, EPI_ISL_480366, EPI_ISL_480367, EPI_ISL_480368, EPI_ISL_480369, EPI_ISL_480370, EPI_ISL_480371, EPI_ISL_480372, EPI_ISL_480373, EPI_ISL_480374, EPI_ISL_480375, EPI_ISL_480376, EPI_ISL_480377, EPI_ISL_480378, EPI_ISL_480379, EPI_ISL_480380, EPI_ISL_480381, EPI_ISL_480382, EPI_ISL_480383, EPI_ISL_480384, EPI_ISL_480385, EPI_ISL_480386, EPI_ISL_480387, EPI_ISL_480388, EPI_ISL_480389, EPI_ISL_480390, EPI_ISL_480391, EPI_ISL_480392, EPI_ISL_480393, EPI_ISL_480394, EPI_ISL_480395, EPI_ISL_480396, EPI_ISL_480397, EPI_ISL_480398, EPI_ISL_480399, EPI_ISL_480400, EPI_ISL_480401, EPI_ISL_480402, EPI_ISL_480403, EPI_ISL_480404, EPI_ISL_480405, EPI_ISL_480406, EPI_ISL_480407, EPI_ISL_480408, EPI_ISL_480409, EPI_ISL_480410, EPI_ISL_480411, EPI_ISL_480412, EPI_ISL_480413 | | | |
| see above | University of Wisconsin-Madison AIDS Vaccine Research Laboratories | University of Wisconsin-Madison AIDS Vaccine Research Laboratories | Gage Moreno, Katarina Braun, et al. AIDS Vaccine Research Laboratories |
| EPI_ISL_480414, EPI_ISL_480415, EPI_ISL_480416, EPI_ISL_480417, EPI_ISL_480418 | National Institute of Laboratory Medicine and Referral Center | Bangladesh Council of Scientific and Industrial Research | Md. Saddam Hossain, Abu Sayeed Mohammad Mahmud, Mohammad Samir Uzzaman, Eshrar Osman, Md. Ahasan Habib, Shahina Akter, Tanjina Akhter Banu, Md. Murshed Hasan Sarkar, Barna Goswami, Iffat Jahan, Tasnim Nafisa, Md. Maruf Ahmed Molla, Mahmuda Yeasmin, Asish Kumar Ghosh, Shahjahan Siddike, A. K. M. Shamsuzzaman, Sheikh Md. Selim Al Din, Utpal Chandra Ray, Salek Ahmed Sajib, Md. Salim Khan |
| EPI_ISL_480419, EPI_ISL_480420, EPI_ISL_480421, EPI_ISL_480424, EPI_ISL_480425 | National Institute of Laboratory Medicine and Referral Center | Bangladesh Council of Scientific and Industrial Research | Md. Murshed Hasan Sarkar, Abu Sayeed Mohammad Mahmud, Mohammad Samir Uzzaman, Eshrar Osman, Md. Ahasan Habib, Shahina Akter, Tanjina Akhter Banu, Barna Goswami, Iffat Jahan, Md. Saddam Hossain, Tasnim Nafisa, Md. Maruf Ahmed Molla, Mahmuda Yeasmin, Asish Kumar Ghosh, Shahjahan Siddike, A. K. M. Shamsuzzaman, Sheikh Md. Selim Al Din, Utpal Chandra Ray, Salek Ahmed Sajib, Md. Salim Khan |
| EPI_ISL_480426, EPI_ISL_480427 | National Institute of Laboratory Medicine and Referral Center | Bangladesh Council of Scientific and Industrial Research | Shahina Akter, Abu Sayeed Mohammad Mahmud, Mohammad Samir Uzzaman, Eshrar Osman, Md. Ahasan Habib, Tanjina Akhter Banu, Md. Murshed Hasan Sarkar, Barna Goswami, Iffat Jahan, Md. Saddam Hossain, Tasnim Nafisa, Md. Maruf Ahmed Molla, Mahmuda Yeasmin, Asish Kumar Ghosh, Shahjahan Siddike, A. K. M. Shamsuzzaman, Sheikh Md. Selim Al Din, Utpal Chandra Ray, Salek Ahmed Sajib, Md. Salim Khan |
| EPI_ISL_480428, EPI_ISL_480429, EPI_ISL_480430, EPI_ISL_480431, EPI_ISL_480432, EPI_ISL_480433, EPI_ISL_480434, EPI_ISL_480435, EPI_ISL_480436, EPI_ISL_480437, EPI_ISL_480438 | | | |
| see above | Laboratorio de Biología Molecular Asociación Española Primera en Salud | Departments of Pathology and Medicine, New York University School of Medicine | Maria Victoria Elizondo, Maria Noel Zubillaga, Gonzalo Manrique, Paul Zappile, Gael Westby, Matthew T Maurano, Christian Marier, Adriana Heguy |
| EPI_ISL_480439, EPI_ISL_480440 | National Institute of Laboratory Medicine and Referral Center | Bangladesh Council of Scientific and Industrial Research | Tanjina Akhter Banu, Abu Sayeed Mohammad Mahmud, Mohammad Samir Uzzaman, Eshrar Osman, Md. Ahasan Habib, Shahina Akter, Md. Murshed Hasan Sarkar, Barna Goswami, Iffat Jahan, Md. Saddam Hossain, Tasnim Nafisa, Md. Maruf Ahmed Molla, Mahmuda Yeasmin, Asish Kumar Ghosh, Shahjahan Siddike, A. K. M. Shamsuzzaman, Sheikh Md. Selim Al Din, Utpal Chandra Ray, Salek Ahmed Sajib, Md. Salim Khan |
| EPI_ISL_480441, EPI_ISL_480442 | National Institute of Laboratory Medicine and Referral Center | Bangladesh Council of Scientific and Industrial Research | Barna Goswami, Abu Sayeed Mohammad Mahmud, Mohammad Samir Uzzaman, Eshrar Osman, Md. Ahasan Habib, Shahina Akter, Tanjina Akhter Banu, Md. Murshed Hasan Sarkar, Iffat Jahan, Md. Saddam Hossain, Tasnim Nafisa, Md. Maruf Ahmed Molla, Mahmuda Yeasmin, Asish Kumar Ghosh, Shahjahan Siddike, A. K. M. Shamsuzzaman, Sheikh Md. Selim Al Din, Utpal Chandra Ray, Salek Ahmed Sajib, Md. Salim Khan |
| EPI_ISL_480443, EPI_ISL_480444 | National Institute of Laboratory Medicine and Referral Center | Bangladesh Council of Scientific and Industrial Research | Iffat Jahan, Abu Sayeed Mohammad Mahmud, Mohammad Samir Uzzaman, Eshrar Osman, Md. Ahasan Habib, Shahina Akter, Tanjina Akhter Banu, Md. Murshed Hasan Sarkar, Barna Goswami, Md. Saddam Hossain, Tasnim Nafisa, Md. Maruf Ahmed Molla, Mahmuda Yeasmin, Asish Kumar Ghosh, Shahjahan Siddike, A. K. M. Shamsuzzaman, Sheikh Md. Selim Al Din, Utpal Chandra Ray, Salek Ahmed Sajib, Md. Salim Khan |
| EPI_ISL_480445 | National Institute of Laboratory Medicine and Referral Center | Genomic Research Lab, BCSIR | Md. Ahasan Habib, Abu Sayeed Mohammad Mahmud, Mohammad Samir Uzzaman, Eshrar Osman, , Shahina Akter, Tanjina Akhter Banu, Md. Murshed Hasan Sarkar, Barna Goswami, Iffat Jahan, Md. Saddam Hossain, Tasnim Nafisa, Md. Maruf Ahmed Molla, Mahmuda Yeasmin, Asish Kumar Ghosh, Shahjahan Siddike, A. K. M. Shamsuzzaman, Sheikh Md. Selim Al Din, Utpal Chandra Ray, Salek Ahmed Sajib, Md. Salim Khan |
| EPI_ISL_480446, EPI_ISL_480447, EPI_ISL_480448, EPI_ISL_480449, | National Institute of Laboratory Medicine and Referral Center | Genomic Research Lab, BCSIR | Abu Sayeed Mohammad Mahmud, Mohammad Samir Uzzaman, Eshrar Osman, Md. Ahasan Habib, Shahina Akter, Tanjina Akhter Banu, Md. Murshed Hasan Sarkar, Barna Goswami, Iffat Jahan, Md. Saddam Hossain, Tasnim Nafisa, Md. Maruf Ahmed Molla, Mahmuda Yeasmin, Asish Kumar Ghosh, Shahjahan Siddike, A. K. M. Shamsuzzaman, Sheikh Md. Selim Al Din, Utpal Chandra Ray, Salek Ahmed Sajib, Md. Salim Khan |

| | | | |
|---|---|---|---|
| EPI_ISL_480450 | | | |
| EPI_ISL_480554, EPI_ISL_480556 | Institut Pasteur Dakar | Institut Pasteur de Dakar | Ndongo Dia, Moussa Moise Diagne, Mamadou Diop, Marie Henriette Dior Ndione, Mamadou Malado Jallow, Safietou Sanke, Ousmane Faye, Amadou Alpha Sall. |
| EPI_ISL_480557 | Victorian Infectious Diseases Reference Laboratory (VIDRL) | VIDRL and MDU-PHL | Caly L., Seemann T., Sait, M., Schultz M., Druce J., Sherry, N. |
| EPI_ISL_480558, EPI_ISL_480559, EPI_ISL_480560, EPI_ISL_480561, EPI_ISL_480562 | Microbiological Diagnostic Unit - Public Health Laboratory (MDU-PHL) | MDU-PHL | Seemann T., Schultz M., Sait, M., Sherry, N. |
| EPI_ISL_480563, EPI_ISL_480564, EPI_ISL_480565, EPI_ISL_480566, EPI_ISL_480567, EPI_ISL_480568, EPI_ISL_480569, EPI_ISL_480570, EPI_ISL_480571, EPI_ISL_480572, EPI_ISL_480573, EPI_ISL_480574, EPI_ISL_480576, EPI_ISL_480577, EPI_ISL_480579, EPI_ISL_480581, EPI_ISL_480582, EPI_ISL_480583, EPI_ISL_480584, EPI_ISL_480585, EPI_ISL_480586 | | | |
| see above | Victorian Infectious Diseases Reference Laboratory (VIDRL) | VIDRL and MDU-PHL | Caly L., Seemann T., Sait, M., Schultz M., Druce J., Sherry, N. |
| EPI_ISL_480587 | Microbiological Diagnostic Unit - Public Health Laboratory (MDU-PHL) | MDU-PHL | Seemann T., Schultz M., Sait, M., Sherry, N. |
| EPI_ISL_480588, EPI_ISL_480589, EPI_ISL_480590, EPI_ISL_480591, EPI_ISL_480592, EPI_ISL_480593, EPI_ISL_480594, EPI_ISL_480595, EPI_ISL_480597, EPI_ISL_480598, EPI_ISL_480599, EPI_ISL_480600 | | | |
| see above | Victorian Infectious Diseases Reference Laboratory (VIDRL) | VIDRL and MDU-PHL | Caly L., Seemann T., Sait, M., Schultz M., Druce J., Sherry, N. |
| EPI_ISL_480601, EPI_ISL_480602 | National Health Laboratory, Timor-Leste | MDU-PHL | Soares da Silva, E., Dolores de Jesus da Costa, M., Salles de Sousa, A., Jayanti Pereira Tilman, A., Antonia da Costa, E., Barreto, I., Marr, I., Wapling, J., Francis, J., Ximenes, J., Canisia, D., Freeman, K., Dakh, F., Douglas, N., Baird, R., Caly, L., Seemann, T., Sait, M., Schultz, M., Sherry, N. |
| EPI_ISL_480603, EPI_ISL_480604, EPI_ISL_480605, EPI_ISL_480606, EPI_ISL_480607, EPI_ISL_480608, EPI_ISL_480609, EPI_ISL_480610, EPI_ISL_480611 | Victorian Infectious Diseases Reference Laboratory (VIDRL) | VIDRL and MDU-PHL | Caly L., Seemann T., Sait, M., Schultz M., Druce J., Sherry, N. |
| EPI_ISL_480612, EPI_ISL_480613, EPI_ISL_480614, EPI_ISL_480615, EPI_ISL_480616, EPI_ISL_480617, EPI_ISL_480618, EPI_ISL_480620 | Microbiological Diagnostic Unit - Public Health Laboratory (MDU-PHL) | MDU-PHL | Seemann T., Schultz M., Sait, M., Sherry, N. |
| EPI_ISL_480621, EPI_ISL_480622, EPI_ISL_480624, EPI_ISL_480625, EPI_ISL_480626, EPI_ISL_480627, EPI_ISL_480628, EPI_ISL_480629, EPI_ISL_480630, EPI_ISL_480631, EPI_ISL_480632, EPI_ISL_480633, EPI_ISL_480634, EPI_ISL_480635, EPI_ISL_480636, EPI_ISL_480637, EPI_ISL_480638, EPI_ISL_480639, EPI_ISL_480640, EPI_ISL_480641, EPI_ISL_480642, EPI_ISL_480643, EPI_ISL_480644, EPI_ISL_480645, EPI_ISL_480646, EPI_ISL_480647, EPI_ISL_480648, EPI_ISL_480649, EPI_ISL_480650, EPI_ISL_480651, EPI_ISL_480652, EPI_ISL_480653, EPI_ISL_480654, EPI_ISL_480655, EPI_ISL_480656, EPI_ISL_480657, EPI_ISL_480658, EPI_ISL_480659, EPI_ISL_480660, EPI_ISL_480661, EPI_ISL_480662, EPI_ISL_480663, EPI_ISL_480664, EPI_ISL_480665, EPI_ISL_480666, EPI_ISL_480667, EPI_ISL_480668, EPI_ISL_480669, EPI_ISL_480670, EPI_ISL_480671, EPI_ISL_480672, EPI_ISL_480673, EPI_ISL_480674, EPI_ISL_480675, EPI_ISL_480676, EPI_ISL_480677, EPI_ISL_480678, EPI_ISL_480679, EPI_ISL_480680, EPI_ISL_480681, EPI_ISL_480682, EPI_ISL_480683, EPI_ISL_480684, EPI_ISL_480685, EPI_ISL_480686 | | | |
| see above | Victorian Infectious Diseases Reference Laboratory (VIDRL) | VIDRL and MDU-PHL | Caly L., Seemann T., Sait, M., Schultz M., Druce J., Sherry, N. |
| EPI_ISL_480687 | Microbiological Diagnostic Unit - Public Health Laboratory (MDU-PHL) | MDU-PHL | Seemann T., Schultz M., Sait, M., Sherry, N. |
| EPI_ISL_480688, EPI_ISL_480689, EPI_ISL_480690 | Victorian Infectious Diseases Reference Laboratory (VIDRL) | VIDRL and MDU-PHL | Caly L., Seemann T., Sait, M., Schultz M., Druce J., Sherry, N. |
| EPI_ISL_480691, EPI_ISL_480692, EPI_ISL_480693, EPI_ISL_480694, EPI_ISL_480695, EPI_ISL_480696, EPI_ISL_480697 | Royal Darwin Hospital Pathology | MDU-PHL | Meumann, E., Caly L., Seemann T., Sait, M., Schultz M., Druce J., Sherry, N. |
| EPI_ISL_480698, EPI_ISL_480699, EPI_ISL_480700, EPI_ISL_480701, EPI_ISL_480702, EPI_ISL_480703, EPI_ISL_480704, EPI_ISL_480705, EPI_ISL_480706, EPI_ISL_480707, EPI_ISL_480708, EPI_ISL_480709, EPI_ISL_480710, EPI_ISL_480711, EPI_ISL_480712, EPI_ISL_480713, EPI_ISL_480714, EPI_ISL_480715, EPI_ISL_480716, EPI_ISL_480717, EPI_ISL_480718, EPI_ISL_480719, EPI_ISL_480720, EPI_ISL_480721, EPI_ISL_480722, EPI_ISL_480723, EPI_ISL_480724, EPI_ISL_480725, EPI_ISL_480726, EPI_ISL_480727, EPI_ISL_480728, EPI_ISL_480729, EPI_ISL_480730, EPI_ISL_480731, EPI_ISL_480732, EPI_ISL_480733, EPI_ISL_480734, EPI_ISL_480735, EPI_ISL_480736, EPI_ISL_480737, EPI_ISL_480738, EPI_ISL_480739, EPI_ISL_480740, EPI_ISL_480741, EPI_ISL_480742, EPI_ISL_480743, EPI_ISL_480744 | | | |
| see above | Victorian Infectious Diseases Reference Laboratory (VIDRL) | VIDRL and MDU-PHL | Caly L., Seemann T., Sait, M., Schultz M., Druce J., Sherry, N. |
| EPI_ISL_480745, EPI_ISL_480746, EPI_ISL_480747, EPI_ISL_480748, EPI_ISL_480749, EPI_ISL_480750, EPI_ISL_480751, EPI_ISL_480752, EPI_ISL_480753, EPI_ISL_480754, EPI_ISL_480755, EPI_ISL_480756, EPI_ISL_480757, EPI_ISL_480758, EPI_ISL_480759, EPI_ISL_480760, EPI_ISL_480761, EPI_ISL_480762, EPI_ISL_480763 | | | |
| see above | Microbiological Diagnostic Unit - Public Health Laboratory (MDU-PHL) | MDU-PHL | Seemann T., Schultz M., Sait, M., Sherry, N. |
| EPI_ISL_480764, EPI_ISL_480765, EPI_ISL_480766 | Victorian Infectious Diseases Reference Laboratory (VIDRL) | VIDRL and MDU-PHL | Caly L., Seemann T., Sait, M., Schultz M., Druce J., Sherry, N. |
| EPI_ISL_480767, EPI_ISL_480768, EPI_ISL_480769, EPI_ISL_480770, EPI_ISL_480771, EPI_ISL_480772, EPI_ISL_480773, EPI_ISL_480774, EPI_ISL_480775, EPI_ISL_480776, EPI_ISL_480777 | | | |
| see above | Microbiological Diagnostic Unit - Public Health Laboratory (MDU-PHL) | MDU-PHL | Seemann T., Schultz M., Sait, M., Sherry, N. |
| EPI_ISL_480778, EPI_ISL_480779, EPI_ISL_480780, EPI_ISL_480781 | Victorian Infectious Diseases Reference Laboratory (VIDRL) | VIDRL and MDU-PHL | Caly L., Seemann T., Sait, M., Schultz M., Druce J., Sherry, N. |
| EPI_ISL_480782, EPI_ISL_480783, EPI_ISL_480786, EPI_ISL_480787, EPI_ISL_480788, EPI_ISL_480789 | Institut Pasteur Dakar | Institut Pasteur de Dakar | Ndongo Dia, Moussa Moise Diagne, Mamadou Diop, Marie Henriette Dior Ndione, Mamadou Malado Jallow, Safietou Sanke, Ousmane Faye, Amadou Alpha Sall. |
| EPI_ISL_480790, EPI_ISL_480791, EPI_ISL_480792, EPI_ISL_480793, EPI_ISL_480794, EPI_ISL_480795, EPI_ISL_480796, EPI_ISL_480797, EPI_ISL_480798, EPI_ISL_480799, EPI_ISL_480800, EPI_ISL_480801, EPI_ISL_480802, EPI_ISL_480803, EPI_ISL_480804, EPI_ISL_480805, EPI_ISL_480806, EPI_ISL_480807, EPI_ISL_480808, EPI_ISL_480809, EPI_ISL_480810, EPI_ISL_480811, EPI_ISL_480812, EPI_ISL_480813, EPI_ISL_480814, EPI_ISL_480815, EPI_ISL_480816, EPI_ISL_480817, EPI_ISL_480818, EPI_ISL_480819, EPI_ISL_480820, EPI_ISL_480821, EPI_ISL_480822, EPI_ISL_480823, EPI_ISL_480824, EPI_ISL_480825, EPI_ISL_480826, EPI_ISL_480827, EPI_ISL_480828, EPI_ISL_480829, EPI_ISL_480830, EPI_ISL_480831, EPI_ISL_480832, EPI_ISL_480833, EPI_ISL_480834, EPI_ISL_480835, EPI_ISL_480836, EPI_ISL_480837, EPI_ISL_480838, EPI_ISL_480839, EPI_ISL_480840, EPI_ISL_480841, EPI_ISL_480842, EPI_ISL_480843, EPI_ISL_480844, EPI_ISL_480845, EPI_ISL_480846, EPI_ISL_480847, EPI_ISL_480848, EPI_ISL_480849, EPI_ISL_480850, EPI_ISL_480851, EPI_ISL_480852, EPI_ISL_480853, EPI_ISL_480854, EPI_ISL_480855, EPI_ISL_480856, EPI_ISL_480857, EPI_ISL_480858, EPI_ISL_480859, EPI_ISL_480860, EPI_ISL_480861, EPI_ISL_480862, EPI_ISL_480863, EPI_ISL_480864, EPI_ISL_480865, EPI_ISL_480866, EPI_ISL_480867, EPI_ISL_480868, EPI_ISL_480869, EPI_ISL_480870, EPI_ISL_480871, EPI_ISL_480872, EPI_ISL_480873, EPI_ISL_480874, EPI_ISL_480875, EPI_ISL_480876, EPI_ISL_480877, EPI_ISL_480878, EPI_ISL_480879, EPI_ISL_480880, EPI_ISL_480881, EPI_ISL_480882, EPI_ISL_480883, EPI_ISL_480884, EPI_ISL_480885, EPI_ISL_480886, EPI_ISL_480887, EPI_ISL_480888, EPI_ISL_480889, EPI_ISL_480890, EPI_ISL_480891, EPI_ISL_480892, EPI_ISL_480893, EPI_ISL_480894, EPI_ISL_480895, EPI_ISL_480896, EPI_ISL_480897, EPI_ISL_480898, EPI_ISL_480899, EPI_ISL_480900, EPI_ISL_480901, EPI_ISL_480902, EPI_ISL_480903, EPI_ISL_480904, EPI_ISL_480905, EPI_ISL_480906, EPI_ISL_480907, EPI_ISL_480908, EPI_ISL_480909, EPI_ISL_480910, EPI_ISL_480911, EPI_ISL_480912, EPI_ISL_480913, EPI_ISL_480914, EPI_ISL_480915, EPI_ISL_480916, EPI_ISL_480917, EPI_ISL_480918, EPI_ISL_480919, EPI_ISL_480920, EPI_ISL_480921, EPI_ISL_480922, EPI_ISL_480923, EPI_ISL_480924, EPI_ISL_480925, EPI_ISL_480926, EPI_ISL_480927, EPI_ISL_480928, EPI_ISL_480929, EPI_ISL_480930, EPI_ISL_480931, EPI_ISL_480932, EPI_ISL_480933, EPI_ISL_480934, EPI_ISL_480935, EPI_ISL_480936, EPI_ISL_480937, EPI_ISL_480938, EPI_ISL_480939, EPI_ISL_480940, EPI_ISL_480941, EPI_ISL_480942, EPI_ISL_480943, EPI_ISL_480944, EPI_ISL_480945, EPI_ISL_480946, EPI_ISL_480947, EPI_ISL_480948, EPI_ISL_480949, EPI_ISL_480950, EPI_ISL_480951 | | | |
| see above | Florida Bureau of Public Health Laboratories | Florida Bureau of Public Health Laboratories | Sarah Schmedes, Jason Blanton |
| EPI_ISL_480952, EPI_ISL_480953, EPI_ISL_480954, EPI_ISL_480955, EPI_ISL_480956, EPI_ISL_480957, EPI_ISL_480958, EPI_ISL_480959, EPI_ISL_480960 | Servicio de Microbiología. Hospital Universitario Donostia. OSI Donostialdea. Área de Enfermedades Infecciosas, Grupo de Infección Respiratoria y Resistencia Antimicrobiana. Instituto de Investigación Sanitaria Biodonostia | SeqCOVID-SPAIN consortium/IBV(CSIC) | Gustavo Cilla, Milagrosa Montes, Luis Piñeiro, Jose Maria Marimón and SeqCOVID-SPAIN consortium |
| EPI_ISL_480961 | ISGlobal, Institut de Salut Global de Barcelona | SeqCOVID-SPAIN consortium/IBV(CSIC) | Alfredo Mayor, Alberto L Garcia-Basteiro, Carlota Dobaño, Gemma Moncunill, Pau Cisteró and SeqCOVID-SPAIN consortium |
| EPI_ISL_480962, EPI_ISL_480963, EPI_ISL_480964, EPI_ISL_480965, EPI_ISL_480966, EPI_ISL_480967, EPI_ISL_480968, EPI_ISL_480969, EPI_ISL_480970, EPI_ISL_480971, EPI_ISL_480972, EPI_ISL_480973, EPI_ISL_480974 | | | |
| see above | Servicio de Microbiología. Hospital Universitario Donostia. OSI Donostialdea. Área de Enfermedades Infecciosas, Grupo de Infección Respiratoria y Resistencia Antimicrobiana. Instituto de Investigación Sanitaria Biodonostia | SeqCOVID-SPAIN consortium/IBV(CSIC) | Gustavo Cilla, Milagrosa Montes, Luis Piñeiro, Jose Maria Marimón and SeqCOVID-SPAIN consortium |
| EPI_ISL_480975 | ISGlobal, Institut de Salut Global de Barcelona | SeqCOVID-SPAIN consortium/IBV(CSIC) | Alfredo Mayor, Alberto L Garcia-Basteiro, Carlota Dobaño, Gemma Moncunill, Pau Cisteró and SeqCOVID-SPAIN consortium |
| EPI_ISL_480976, EPI_ISL_480977, EPI_ISL_480978, EPI_ISL_480979, EPI_ISL_480980 | Servicio de Microbiología. Hospital Universitario Donostia. OSI Donostialdea. Área de Enfermedades Infecciosas, Grupo de Infección Respiratoria y Resistencia Antimicrobiana. Instituto de Investigación Sanitaria Biodonostia | SeqCOVID-SPAIN consortium/IBV(CSIC) | Gustavo Cilla, Milagrosa Montes, Luis Piñeiro, Jose Maria Marimón and SeqCOVID-SPAIN consortium |
| EPI_ISL_480981 | ISGlobal, Institut de Salut Global de Barcelona | SeqCOVID-SPAIN consortium/IBV(CSIC) | Alfredo Mayor, Alberto L Garcia-Basteiro, Carlota Dobaño, Gemma Moncunill, Pau Cisteró and SeqCOVID-SPAIN consortium |
| EPI_ISL_480982, EPI_ISL_480983, EPI_ISL_480984, EPI_ISL_480985, EPI_ISL_480986, EPI_ISL_480987, EPI_ISL_480988 | Servicio de Microbiología. Hospital Universitario Donostia. OSI Donostialdea. Área de Enfermedades Infecciosas, Grupo de Infección Respiratoria y Resistencia Antimicrobiana. Instituto de Investigación Sanitaria Biodonostia | SeqCOVID-SPAIN consortium/IBV(CSIC) | Gustavo Cilla, Milagrosa Montes, Luis Piñeiro, Jose Maria Marimón and SeqCOVID-SPAIN consortium |
| EPI_ISL_480989 | ISGlobal, Institut de Salut Global de Barcelona | SeqCOVID-SPAIN consortium/IBV(CSIC) | Alfredo Mayor, Alberto L Garcia-Basteiro, Carlota Dobaño, Gemma Moncunill, Pau Cisteró and SeqCOVID-SPAIN consortium |
| EPI_ISL_480990, EPI_ISL_480991, EPI_ISL_480992, EPI_ISL_480993, EPI_ISL_480994 | Servicio de Microbiología. Hospital Universitario Donostia. OSI Donostialdea. Área de Enfermedades Infecciosas, Grupo de Infección Respiratoria y Resistencia Antimicrobiana. Instituto de Investigación Sanitaria Biodonostia | SeqCOVID-SPAIN consortium/IBV(CSIC) | Gustavo Cilla, Milagrosa Montes, Luis Piñeiro, Jose Maria Marimón and SeqCOVID-SPAIN consortium |

| | | | |
|---|---|---|---|
| EPI_ISL_480995 | ISGlobal, Institut de Salut Global de Barcelona | SeqCOVID-SPAIN consortium/IBV(CSIC) | Alfredo Mayor, Alberto L Garcia-Basteiro, Carlota Dobaño, Gemma Moncunill, Pau Cisteró and SeqCOVID-SPAIN consortium |
| EPI_ISL_480996, EPI_ISL_480997, EPI_ISL_480998, EPI_ISL_480999, EPI_ISL_481000, EPI_ISL_481001, EPI_ISL_481002 | Servicio de Microbiología. Hospital Universitario Donostia. OSI Donostialdea. Área de Enfermedades Infecciosas, Grupo de Infección Respiratoria y Resistencia Antimicrobiana. Instituto de Investigación Sanitaria Biodonostia | SeqCOVID-SPAIN consortium/IBV(CSIC) | Gustavo Cilla, Milagrosa Montes, Luis Piñeiro, Jose Maria Marimón and SeqCOVID-SPAIN consortium |
| EPI_ISL_481003 | ISGlobal, Institut de Salut Global de Barcelona | SeqCOVID-SPAIN consortium/IBV(CSIC) | Alfredo Mayor, Alberto L Garcia-Basteiro, Carlota Dobaño, Gemma Moncunill, Pau Cisteró and SeqCOVID-SPAIN consortium |
| EPI_ISL_481004, EPI_ISL_481005, EPI_ISL_481006, EPI_ISL_481007, EPI_ISL_481008, EPI_ISL_481009, EPI_ISL_481010, EPI_ISL_481011, EPI_ISL_481012, EPI_ISL_481013, EPI_ISL_481014, EPI_ISL_481015, EPI_ISL_481016 | | | |
| see above | Servicio de Microbiología. Hospital Universitario Donostia. OSI Donostialdea. Área de Enfermedades Infecciosas, Grupo de Infección Respiratoria y Resistencia Antimicrobiana. Instituto de Investigación Sanitaria Biodonostia | SeqCOVID-SPAIN consortium/IBV(CSIC) | Gustavo Cilla, Milagrosa Montes, Luis Piñeiro, Jose Maria Marimón and SeqCOVID-SPAIN consortium |
| EPI_ISL_481017 | ISGlobal, Institut de Salut Global de Barcelona | SeqCOVID-SPAIN consortium/IBV(CSIC) | Alfredo Mayor, Alberto L Garcia-Basteiro, Carlota Dobaño, Gemma Moncunill, Pau Cisteró and SeqCOVID-SPAIN consortium |
| EPI_ISL_481018, EPI_ISL_481019, EPI_ISL_481020, EPI_ISL_481021, EPI_ISL_481022, EPI_ISL_481023, EPI_ISL_481024 | Servicio de Microbiología. Hospital Universitario Donostia. OSI Donostialdea. Área de Enfermedades Infecciosas, Grupo de Infección Respiratoria y Resistencia Antimicrobiana. Instituto de Investigación Sanitaria Biodonostia | SeqCOVID-SPAIN consortium/IBV(CSIC) | Gustavo Cilla, Milagrosa Montes, Luis Piñeiro, Jose Maria Marimón and SeqCOVID-SPAIN consortium |
| EPI_ISL_481025 | ISGlobal, Institut de Salut Global de Barcelona | SeqCOVID-SPAIN consortium/IBV(CSIC) | Alfredo Mayor, Alberto L Garcia-Basteiro, Carlota Dobaño, Gemma Moncunill, Pau Cisteró and SeqCOVID-SPAIN consortium |
| EPI_ISL_481026, EPI_ISL_481027, EPI_ISL_481028 | Servicio de Microbiología. Hospital Universitario Donostia. OSI Donostialdea. Área de Enfermedades Infecciosas, Grupo de Infección Respiratoria y Resistencia Antimicrobiana. Instituto de Investigación Sanitaria Biodonostia | SeqCOVID-SPAIN consortium/IBV(CSIC) | Gustavo Cilla, Milagrosa Montes, Luis Piñeiro, Jose Maria Marimón and SeqCOVID-SPAIN consortium |
| EPI_ISL_481029 | ISGlobal, Institut de Salut Global de Barcelona | SeqCOVID-SPAIN consortium/IBV(CSIC) | Alfredo Mayor, Alberto L Garcia-Basteiro, Carlota Dobaño, Gemma Moncunill, Pau Cisteró and SeqCOVID-SPAIN consortium |
| EPI_ISL_481030, EPI_ISL_481031, EPI_ISL_481032, EPI_ISL_481033 | Servicio de Microbiología. Hospital Universitario Donostia. OSI Donostialdea. Área de Enfermedades Infecciosas, Grupo de Infección Respiratoria y Resistencia Antimicrobiana. Instituto de Investigación Sanitaria Biodonostia | SeqCOVID-SPAIN consortium/IBV(CSIC) | Gustavo Cilla, Milagrosa Montes, Luis Piñeiro, Jose Maria Marimón and SeqCOVID-SPAIN consortium |
| EPI_ISL_481034, EPI_ISL_481035 | ISGlobal, Institut de Salut Global de Barcelona | SeqCOVID-SPAIN consortium/IBV(CSIC) | Alfredo Mayor, Alberto L Garcia-Basteiro, Carlota Dobaño, Gemma Moncunill, Pau Cisteró and SeqCOVID-SPAIN consortium |
| EPI_ISL_481036, EPI_ISL_481037, EPI_ISL_481038, EPI_ISL_481039, EPI_ISL_481040 | Servicio de Microbiología. Hospital Universitario Donostia. OSI Donostialdea. Área de Enfermedades Infecciosas, Grupo de Infección Respiratoria y Resistencia Antimicrobiana. Instituto de Investigación Sanitaria Biodonostia | SeqCOVID-SPAIN consortium/IBV(CSIC) | Gustavo Cilla, Milagrosa Montes, Luis Piñeiro, Jose Maria Marimón and SeqCOVID-SPAIN consortium |
| EPI_ISL_481041, EPI_ISL_481042, EPI_ISL_481043, EPI_ISL_481044, EPI_ISL_481045, EPI_ISL_481046, EPI_ISL_481047, EPI_ISL_481048, EPI_ISL_481049, EPI_ISL_481050, EPI_ISL_481051, EPI_ISL_481052, EPI_ISL_481053, EPI_ISL_481054, EPI_ISL_481055, EPI_ISL_481056, EPI_ISL_481057, EPI_ISL_481058, EPI_ISL_481059, EPI_ISL_481060, EPI_ISL_481061, EPI_ISL_481062, EPI_ISL_481063, EPI_ISL_481064, EPI_ISL_481065, EPI_ISL_481066, EPI_ISL_481067, EPI_ISL_481068, EPI_ISL_481069, EPI_ISL_481070, EPI_ISL_481071, EPI_ISL_481072, EPI_ISL_481073, EPI_ISL_481074, EPI_ISL_481075, EPI_ISL_481076, EPI_ISL_481077, EPI_ISL_481078, EPI_ISL_481079, EPI_ISL_481080, EPI_ISL_481081, EPI_ISL_481082, EPI_ISL_481083, EPI_ISL_481084, EPI_ISL_481085, EPI_ISL_481086, EPI_ISL_481087, EPI_ISL_481088, EPI_ISL_481089, EPI_ISL_481090, EPI_ISL_481091, EPI_ISL_481092, EPI_ISL_481093, EPI_ISL_481094, EPI_ISL_481095, EPI_ISL_481096, EPI_ISL_481097, EPI_ISL_481098, EPI_ISL_481099, EPI_ISL_481100, EPI_ISL_481101, EPI_ISL_481102, EPI_ISL_481103, EPI_ISL_481104, EPI_ISL_481105, EPI_ISL_481106, EPI_ISL_481107, EPI_ISL_481108, EPI_ISL_481109 | | | |
| see above | Hospital General Universitario Gregorio Marañón | SeqCOVID-SPAIN consortium/IBV(CSIC) | Laura Pérez-Lago, Marta Herranz, Jon Sicilia, Julia Suárez, Pilar Catalán, Patricia Muñoz, Darío García de Viedma and SeqCOVID-SPAIN consortium |
| EPI_ISL_481110, EPI_ISL_481111, EPI_ISL_481112, EPI_ISL_481113, EPI_ISL_481114, EPI_ISL_481115, EPI_ISL_481116, EPI_ISL_481117, EPI_ISL_481118, EPI_ISL_481119, EPI_ISL_481120, EPI_ISL_481121, EPI_ISL_481122, EPI_ISL_481123, EPI_ISL_481124, EPI_ISL_481125, EPI_ISL_481126, EPI_ISL_481127, EPI_ISL_481128, EPI_ISL_481129, EPI_ISL_481130, EPI_ISL_481131, EPI_ISL_481132, EPI_ISL_481133 | | | |
| see above | Immunogenomics lab, Institute of Life Sciences, Bhubaneswar | Immunogenomics lab, Institute of Life Sciences, Bhubaneswar | Sunil Raghav, Arup Ghosh, Deepika Singh, Ankita Datey, P. Sushree Shyamli, Bharati Singh, Neha Singh, Atimukta Jha, Viplov K. Biswas, Swati Madhulika, Manasi Priyadarshini, Sneha Dutta, Auromira Khuntia, Rupesh Dash, Soma Chattopadhyay, Ghulam Hussain Syed, Shanti Senapati, Tushar K. Beuria, Rajeeb Swain, Punit Prasad, Orissa COVID-19 Study Group, DBT's PAN-INDIA 1000 SARS-CoV2 RNA genome sequencing consortium, Ajay Parida |
| EPI_ISL_481134, EPI_ISL_481135, EPI_ISL_481136, EPI_ISL_481137, EPI_ISL_481138, EPI_ISL_481139, EPI_ISL_481140, EPI_ISL_481141, EPI_ISL_481142, EPI_ISL_481143, EPI_ISL_481144, EPI_ISL_481145, EPI_ISL_481146, EPI_ISL_481147, EPI_ISL_481148, EPI_ISL_481149, EPI_ISL_481150, EPI_ISL_481151, EPI_ISL_481152, EPI_ISL_481153, EPI_ISL_481154, EPI_ISL_481155, EPI_ISL_481156, EPI_ISL_481157 | | | |
| see above | Immunogenomics lab, Institute of Life Sciences, Bhubaneswar | Immunogenomics lab, Institute of Life Sciences, Bhubaneswar | Sunil Raghav, Arup Ghosh, Ankita Datey, P. Sushree Shyamli, Bharati Singh, Neha Singh, Deepika Singh, Atimukta Jha, Viplov K. Biswas, Swati Madhulika, Manasi Priyadarshini, Aditi Chatterjee, Rahul Das, Soumyajit Ghosh, Rupesh Dash, Soma Chattopadhyay, Ghulam Hussain Syed, Shanti Senapati, Tushar K. Beuria, Rajeeb Swain, Punit Prasad, Amol Ratnakar Suryawanshi, Dileep Vasudeva, Orissa COVID-19 Study Group, DBT's PAN-INDIA 1000 SARS-CoV2 RNA genome sequencing consortium, Ajay Parida |
| EPI_ISL_481158, EPI_ISL_481159, EPI_ISL_481160, EPI_ISL_481161, EPI_ISL_481162, EPI_ISL_481163, EPI_ISL_481164, EPI_ISL_481165, EPI_ISL_481166, EPI_ISL_481167, EPI_ISL_481168, EPI_ISL_481169, EPI_ISL_481170, EPI_ISL_481171, EPI_ISL_481172, EPI_ISL_481173, EPI_ISL_481174, EPI_ISL_481175, EPI_ISL_481176, EPI_ISL_481177, EPI_ISL_481178, EPI_ISL_481179, EPI_ISL_481180, EPI_ISL_481181 | | | |
| see above | Immunogenomics lab, Institute of Life Sciences, Bhubaneswar | Immunogenomics lab, Institute of Life Sciences, Bhubaneswar | Sunil Raghav, Arup Ghosh, P. Sushree Shyamli, Bharati Singh, Neha Singh, Ankita Datey, Deepika Singh, Atimukta Jha, Viplov K. Biswas, Swati Madhulika, Manasi Priyadarshini, Tsheten Sheroa, Auromira Khuntia, Rupesh Dash, Soma Chattopadhyay, Ghulam Hussain Syed, Shanti Senapati, Tushar K. Beuria, Rajeeb Swain, Punit Prasad, Amol Ratnakar Suryawanshi, Dileep Vasudevan, Orissa COVID-19 Study Group, DBT's PAN-INDIA 1000 SARS-CoV2 RNA genome sequencing consortium, Ajay Parida |
| EPI_ISL_481182, EPI_ISL_481183, EPI_ISL_481184, EPI_ISL_481185, EPI_ISL_481186, EPI_ISL_481187, EPI_ISL_481188, EPI_ISL_481189, EPI_ISL_481190, EPI_ISL_481191, EPI_ISL_481192, EPI_ISL_481193, EPI_ISL_481194, EPI_ISL_481195, EPI_ISL_481196, EPI_ISL_481197, EPI_ISL_481198, EPI_ISL_481199, EPI_ISL_481200, EPI_ISL_481201, EPI_ISL_481202, EPI_ISL_481203, EPI_ISL_481204, EPI_ISL_481205 | | | |
| see above | Immunogenomics lab, Institute of Life Sciences, Bhubaneswar | Immunogenomics lab, Institute of Life Sciences, Bhubaneswar | Sunil Raghav, Arup Ghosh, Atimukta Jha, Viplov K. Biswas, Swati Madhulika, Manasi Priyadarshini, Ajit Singh, Sivaram Krishna, Naga Jogayya Kothakota, Rupesh Dash, Soma Chattopadhyay, Ghulam Hussain Syed, Shanti Senapati, Tushar K. Beuria, Rajeeb Swain, Punit Prasad, Amol Ratnakar Suryawanshi, Dileep Vasudevan, Orissa COVID-19 Study Group, DBT's PAN-INDIA 1000 SARS-CoV2 RNA genome sequencing consortium, Ajay Parida |
| EPI_ISL_481206 | Hospital of Southern Norway - Kristiansand, Department of Medical Microbiology | Norwegian Institute of Public Health, Department of Virology | Kathrine Stene-Johansen, Kamilla Heddeland Instefjord, Hilde Elshaug, Rasmus Riis Kopperud, Karoline Bragstad, Olav Hungnes |
| EPI_ISL_481207 | Ostfold Hospital Trust - Kalnes, Centre for Laboratory Medicine, Section for gene technology and infection serology | Norwegian Institute of Public Health, Department of Virology | Kathrine Stene-Johansen, Kamilla Heddeland Instefjord, Hilde Elshaug, Rasmus Riis Kopperud, Karoline Bragstad, Olav Hungnes |
| EPI_ISL_481208 | Furst Medical Laboratory | Norwegian Institute of Public Health, Department of Virology | Kathrine Stene-Johansen, Kamilla Heddeland Instefjord, Hilde Elshaug, Rasmus Riis Kopperud, Karoline Bragstad, Olav Hungnes |
| EPI_ISL_481209, EPI_ISL_481210, EPI_ISL_481211, EPI_ISL_481212, EPI_ISL_481213 | Ostfold Hospital Trust - Kalnes, Centre for Laboratory Medicine, Section for gene technology and infection serology | Norwegian Institute of Public Health, Department of Virology | Kathrine Stene-Johansen, Kamilla Heddeland Instefjord, Hilde Elshaug, Rasmus Riis Kopperud, Karoline Bragstad, Olav Hungnes |
| EPI_ISL_481214, EPI_ISL_481215 | Oslo University Hospital, Department of Medical Microbiology | Norwegian Institute of Public Health, Department of Virology | Kathrine Stene-Johansen, Kamilla Heddeland Instefjord, Hilde Elshaug, Rasmus Riis Kopperud, Karoline Bragstad, Olav Hungnes |
| EPI_ISL_481216 | Medical Microbiology Unit, Department for Laboratory Medicine, Drammen Hospital, Vestre Viken Health Trust, | Norwegian Institute of Public Health, Department of Virology | Kathrine Stene-Johansen, Kamilla Heddeland Instefjord, Hilde Elshaug, Rasmus Riis Kopperud, Karoline Bragstad, Olav Hungnes |
| EPI_ISL_481217, EPI_ISL_481218, EPI_ISL_481219 | Oslo University Hospital, Department of Medical Microbiology | Norwegian Institute of Public Health, Department of Virology | Kathrine Stene-Johansen, Kamilla Heddeland Instefjord, Hilde Elshaug, Rasmus Riis Kopperud, Karoline Bragstad, Olav Hungnes |
| EPI_ISL_481220 | Institut Pasteur Dakar | Institut Pasteur de Dakar | Ndongo Dia, Moussa Moise Diagne, Mamadou Diop, Marie Henriette Dior Ndione, Mamadou Malado Jallow, Safietou Sanke, Ousmane Faye, Amadou Alpha Sall. |
| EPI_ISL_481221, EPI_ISL_481222, EPI_ISL_481223, EPI_ISL_481224, EPI_ISL_481225, EPI_ISL_481226 | Lab voor klinische biologie | Onderzoeksgroep Virologie | Laurens Lambrechts, Nick Vereecke, Marthe Pauwels, Bruno Verhasselt, Linos Vandekerckhove, Hans Nauwynck, Sebastiaan Theuns |
| EPI_ISL_481227, EPI_ISL_481228, EPI_ISL_481229, EPI_ISL_481230, EPI_ISL_481231, EPI_ISL_481232, EPI_ISL_481233 | Lab voor klinische biologie | Onderzoeksgroep Virologie | Nick Vereecke, Laurens Lambrechts, Marthe Pauwels, Bruno Verhasselt, Linos Vandekerckhove, Hans Nauwynck, Sebastiaan Theuns |
| EPI_ISL_481234, EPI_ISL_481235, EPI_ISL_481236, EPI_ISL_481237, EPI_ISL_481238, EPI_ISL_481239, EPI_ISL_481240 | Institut Pasteur Dakar | Institut Pasteur de Dakar | Ndongo Dia, Moussa Moise Diagne, Mamadou Diop, Marie Henriette Dior Ndione, Mamadou Malado Jallow, Safietou Sanke, Ousmane Faye, Amadou Alpha Sall. |
| EPI_ISL_481241 | M Health Fairview | Minnesota Department of Health, Public Health Laboratory | Matt Plumb, Jacob Garfin, Kelly Pung, and Xiong Wang |
| EPI_ISL_481242 | Mayo Clinic & Mayo Clinic Laboratories | Minnesota Department of Health, Public Health Laboratory | Matt Plumb, Jacob Garfin, Kelly Pung, and Xiong Wang |
| EPI_ISL_481243 | Institut Pasteur Dakar | Institut Pasteur de Dakar | Ndongo Dia, Moussa Moise Diagne, Mamadou Diop, Marie Henriette Dior Ndione, Mamadou Malado Jallow, Safietou Sanke, Ousmane Faye, Amadou Alpha Sall. |
| EPI_ISL_481244, EPI_ISL_481245, EPI_ISL_481246, EPI_ISL_481247, EPI_ISL_481248 | Hospital IESS Babahoyo | Institute of Microbiology, Universidad San Francisco de Quito | Belén Prado-Vivar, Sully Márquez, Juan José Guadalupe, Monica Becerra-Wong, Carla Torres, Bernardo Gutiérrez, Francisco Cordova, Ninfa Henriquez, Killen Briones-Zamora, Killen Briones-Claudette, Verónica Barragán, Patricio Rojas-Silva, Gabriel Trueba, Michelle Grunauer, Paúl Cárdenas |
| EPI_ISL_481251, EPI_ISL_481252 | Department of Emerging Infectious Diseases, Institute of Tropical Medicine, Nagasaki University | Department of Emerging Infectious Diseases, Institute of Tropical Medicine, Nagasaki | Jiro Yasuda, Rokusuke Yoshikawa, Yuichiro Furusato, Haruka Abe |

| | | | |
|---|---|---|---|
| | | University | |
| EPI_ISL_481253 | Robert Koch Institute, National Reference center for Influenza, Berlin, Germany | Robert Koch Institute, Bioinformatics MF1, Berlin, Germany | Marianne Wedde, Oliver Drechsel, Andrea Thuermer, Rene Kmiecinski, Ralf Duerrwald, Thorsten Wolff, Stephan Fuchs, Max v. Kleist |
| EPI_ISL_481254, EPI_ISL_481255 | Department of Emerging Infectious Diseases, Institute of Tropical Medicine, Nagasaki University | Department of Emerging Infectious Diseases, Institute of Tropical Medicine, Nagasaki University | Jiro Yasuda, Rokusuke Yoshikawa, Yuichiro Furusato, Haruka Abe |
| EPI_ISL_481256 | Robert Koch Institute, National Reference center for Influenza, Berlin, Germany | Robert Koch Institute, Bioinformatics MF1, Berlin, Germany | Marianne Wedde, Oliver Drechsel, Andrea Thuermer, Rene Kmiecinski, Ralf Duerrwald, Thorsten Wolff, Stephan Fuchs, Max v. Kleist |
| EPI_ISL_481257, EPI_ISL_481258, EPI_ISL_481259, EPI_ISL_481260, EPI_ISL_481261 | Department of Emerging Infectious Diseases, Institute of Tropical Medicine, Nagasaki University | Department of Emerging Infectious Diseases, Institute of Tropical Medicine, Nagasaki University | Jiro Yasuda, Rokusuke Yoshikawa, Yuichiro Furusato, Haruka Abe |
| EPI_ISL_481262 | Robert Koch Institute, National Reference center for Influenza, Berlin, Germany | Robert Koch Institute, Bioinformatics MF1, Berlin, Germany | Marianne Wedde, Oliver Drechsel, Andrea Thuermer, Rene Kmiecinski, Ralf Duerrwald, Thorsten Wolff, Stephan Fuchs, Max v. Kleist |
| EPI_ISL_481263 | Department of Emerging Infectious Diseases, Institute of Tropical Medicine, Nagasaki University | Department of Emerging Infectious Diseases, Institute of Tropical Medicine, Nagasaki University | Jiro Yasuda, Rokusuke Yoshikawa, Yuichiro Furusato, Haruka Abe |
| EPI_ISL_481264 | Robert Koch Institute, National Reference center for Influenza, Berlin, Germany | Robert Koch Institute, Bioinformatics MF1, Berlin, Germany | Marianne Wedde, Oliver Drechsel, Andrea Thuermer, Rene Kmiecinski, Ralf Duerrwald, Thorsten Wolff, Stephan Fuchs, Max v. Kleist |
| EPI_ISL_481265, EPI_ISL_481266, EPI_ISL_481267, EPI_ISL_481268, EPI_ISL_481269, EPI_ISL_481270, EPI_ISL_481271, EPI_ISL_481272, EPI_ISL_481273, EPI_ISL_481274, EPI_ISL_481275, EPI_ISL_481276, EPI_ISL_481277, EPI_ISL_481278, EPI_ISL_481279, EPI_ISL_481280, EPI_ISL_481281, EPI_ISL_481282 | | | |
| see above | Maryland Department of Health | Maryland Department of Health | Keller,E. |
| EPI_ISL_481283 | Center for Genomics and System Biology, New York University | Center for Genomics and System Biology, New York University | Roder,A., Banakis,S., Johnson,K., Khalfan,M., Borenstein,E.S., Samanovic,M., Cornelius,A., Herati,R., Ulrich,R., Fleming,A., Kottkamp,A., Raabe,V., Mulligan,M.J., Gresham,D. and Ghedin,E. |
| EPI_ISL_481284 | Department of Experimental Modeling and Pathogenesis of Infectious Diseases, Federal Research Center of Fundamental and Translational Medicine | Department of Experimental Modeling and Pathogenesis of Infectious Diseases, Federal Research Center of Fundamental and Translational Medicine | Sobolev,I.A., Shanshin,D.V., Chepurnov,A.A., Kononova,J.V., Bondar,A.A., Alekseev,A.Y. and Shestopalov,A.M. |
| EPI_ISL_481370 | Division of Viral Diseases, Center for Laboratory Control of Infectious Diseases, Korea Centers for Diseases Control and Prevention | Division of Viral Diseases, Center for Laboratory Control of Infectious Diseases, Korea Centers for Diseases Control and Prevention | Jeong-Min Kim, Yoon-Seok Chung, Namjoo Lee, Sang Hee Woo, Hye-Jun Jo, Heui Man Kim, Jun-Sub Kim, Myung Guk Han |
| EPI_ISL_481371, EPI_ISL_481372, EPI_ISL_481373, EPI_ISL_481374, EPI_ISL_481375, EPI_ISL_481376, EPI_ISL_481377, EPI_ISL_481378, EPI_ISL_481379 | Division of Viral Diseases, Center for Laboratory Control of Infectious Diseases, Korea Centers for Diseases Control and Prevention | Division of Viral Diseases, Center for Laboratory Control of Infectious Diseases, Korea Centers for Diseases Control and Prevention | Jeong-Min Kim, Yoon-Seok Chung, Namjoo Lee, Sang Hee Woo, Hye-Jun Jo, Heui Man Kim, Jun-Sub Kim, Dong Hyun Song, Daesang Lee, Seong Tae Jeong, Myung Guk Han |
| EPI_ISL_481380 | Department for Virology, Molecular Biology and Genome Research, R. G. Lugar Center for Public Health Research, National Center for Disease Control and Public Health (NCDC) of Georgia. | Department for Virology, Molecular Biology and Genome Research, R. G. Lugar Center for Public Health Research, National Center for Disease Control and Public Health (NCDC) of Georgia. | Ana Papkiauri, Tata Imnadze, Giorgi Tomashvili, Meri Pantsulaia, Gvantsa Brachveli, Gvantsa Chanturia, Ann Machablishvili, Nato Kotaria, Marine Murtskhvaladze, Lela Sabadze, Mari Gavashelidze, Tamar Jashiashvili, Tea Tevdoradze, Ketevan Sidamonidze, Ekaterine Khmaladze, Ekaterine Zhghenti, Roena Sukhiashvili, Mariam Zakalashvili, Lela Urushadze, Magda Dgebuadze, Davit Tsaguria, Ekaterine Zangaladze, Nino Berishvili, Adam Kotorashvili, Maia Alkhazashvili, Irma Burjanadze, Anna Kasradze, Khatuna Zakhashvili, Paata Imnadze, Amiran Gamkrelidze. |
| EPI_ISL_481483 | Department for Virology, Molecular Biology and Genome Research, R. G. Lugar Center for Public Health Research, National Center for Disease Control and Public Health (NCDC) of Georgia. | Department for Virology, Molecular Biology and Genome Research, R. G. Lugar Center for Public Health Research, National Center for Disease Control and Public Health (NCDC) of Georgia. | Nino Berishvili, Tata Imnadze, Giorgi Tomashvili, Ana Papkiauri, Meri Pantsulaia, Gvantsa Brachveli, Gvantsa Chanturia, Ann Machablishvili, Nato Kotaria, Marine Murtskhvaladze, Lela Sabadze, Mari Gavashelidze, Tamar Jashiashvili, Tea Tevdoradze, Ketevan Sidamonidze, Ekaterine Khmaladze, Ekaterine Zhghenti, Roena Sukhiashvili, Mariam Zakalashvili, Lela Urushadze, Magda Dgebuadze, Davit Tsaguria, Ekaterine Zangaladze, Adam Kotorashvili, Maia Alkhazashvili, Irma Burjanadze, Anna Kasradze, Khatuna Zakhashvili, Paata Imnadze, Amiran Gamkrelidze. |
| EPI_ISL_481510, EPI_ISL_481511 | Prof. Massimo Zollo CEINGE TASK-FORCE COVID19 - Regione Campania | Prof. Massimo Zollo CEINGE TASK-FORCE COVID19 - Regione Campania | Veronica Ferrucci1,2, Dae young Kong8, Fatemeh asadzadeh1,2, Laura Marrone1,2, Roberto Siciliano1,2, Rino Cerino3, Giovanna Fusco3, Marika Comegna1,2, Angelo Boccia2, Maurizio Viscardi3, Giorgia Borriello3, Sergio Brandi3, Claudia Tiberio4, Luigi Atripaldi4, Giovanni Paolella1,2, Giuseppe Castaldo1,2, Stefano Pascarella4, Martina Bianchi4, Lorenzo Chiariotti1,2, Jae Myun Lee5, Jae Ho Jung6, Kyong Seop Yun7, Hong Yeoul Kim 7,8* and Massimo Zollo1,2* 1 CEINGE Biotecnologie Avanzate, Naples, Italia 2 Dipartimento di Medicina Molecolare e Biotecnologie Mediche DMMBM University of Naples Federico II, Italia 3 Istituto Zooprofilattico Sperimentale del Mezzogiorno, Naples, Italia 4 -U.O.C. di Patologia Clinica Ospedale D. Cotugno, Azienda Sanitaria Ospedali dei Colli, Naples, Italy. 5 Università La Sapienza di Roma, Italia 6 Department of Microbiology, Yonsei University College of Medicine, Seoul, Korea 7 Department of Surgery, Yonsei University College of Medicine, Seoul, Korea 8 Haim bio co., Ltd, , Indust |
| EPI_ISL_481512 | Prof. Massimo Zollo CEINGE TASK-FORCE COVID19 - Regione Campania | Prof. Massimo Zollo CEINGE TASK-FORCE COVID19 - Regione Campania | Veronica Ferrucci1,2, Dae young Kong8, Fatemeh asadzadeh1,2, Laura Marrone1,2, Roberto Siciliano1,2, Rino Cerino3, Giovanna Fusco3, Marika Comegna1,2, Angelo Boccia2, Maurizio Viscardi3, Giorgia Borriello3, Sergio Brandi3, Claudia Tiberio4, Luigi Atripaldi4, Giovanni Paolella1,2, Giuseppe Castaldo1,2, Stefano Pascarella4, Martina Bianchi4, Lorenzo Chiariotti1,2, Jae Myun Lee5, Jae Ho Jung6, Kyong Seop Yun7, Hong Yeoul Kim 7,8* and Massimo Zollo1,2* 1 CEINGE Biotecnologie Avanzate, Naples, Italia 2 Dipartimento di Medicina Molecolare e Biotecnologie Mediche DMMBM University of Naples Federico II, Italia 3 Istituto Zooprofilattico Sperimentale del Mezzogiorno, Naples, Italia 4 -U.O.C. di Patologia Clinica Ospedale D. Cotugno, Azienda Sanitaria Ospedali dei Colli, Naples, Italy. 5 Università La Sapienza di Roma, Italia 6 Department of Microbiology, Yonsei University College of Medicine, Seoul, Korea 7 Department of Surgery, Yonsei University College of Medicine, Seoul, Korea 8 Haim bio co., Ltd, , Indust |
| EPI_ISL_481513, EPI_ISL_481514, EPI_ISL_481515, EPI_ISL_481516, EPI_ISL_481517, EPI_ISL_481518, EPI_ISL_481519, EPI_ISL_481520, EPI_ISL_481521, EPI_ISL_481522, EPI_ISL_481523, EPI_ISL_481524, EPI_ISL_481525, EPI_ISL_481526, EPI_ISL_481527, EPI_ISL_481528, EPI_ISL_481529, EPI_ISL_481530, EPI_ISL_481531, EPI_ISL_481532, EPI_ISL_481533, EPI_ISL_481534, EPI_ISL_481535, EPI_ISL_481536, EPI_ISL_481537, EPI_ISL_481538, EPI_ISL_481539, EPI_ISL_481540, EPI_ISL_481541, EPI_ISL_481542, EPI_ISL_481543, EPI_ISL_481544, EPI_ISL_481545, EPI_ISL_481546, EPI_ISL_481547, EPI_ISL_481548, EPI_ISL_481549, EPI_ISL_481550, EPI_ISL_481551, EPI_ISL_481552, EPI_ISL_481553, EPI_ISL_481554, EPI_ISL_481555, EPI_ISL_481556, EPI_ISL_481557, EPI_ISL_481558, EPI_ISL_481559, EPI_ISL_481560, EPI_ISL_481561, EPI_ISL_481562, EPI_ISL_481563, EPI_ISL_481564, EPI_ISL_481565, EPI_ISL_481566, EPI_ISL_481567, EPI_ISL_481568, EPI_ISL_481569, EPI_ISL_481570, EPI_ISL_481571, EPI_ISL_481572, EPI_ISL_481573, EPI_ISL_481574, EPI_ISL_481575, EPI_ISL_481576, EPI_ISL_481577, EPI_ISL_481578, EPI_ISL_481579, EPI_ISL_481580, EPI_ISL_481581, EPI_ISL_481582, EPI_ISL_481583, EPI_ISL_481584, EPI_ISL_481585, EPI_ISL_481586, EPI_ISL_481587, EPI_ISL_481588, EPI_ISL_481589, EPI_ISL_481590, EPI_ISL_481591, EPI_ISL_481592, EPI_ISL_481593, EPI_ISL_481594, EPI_ISL_481595, EPI_ISL_481596, EPI_ISL_481597, EPI_ISL_481598, EPI_ISL_481599, EPI_ISL_481600, EPI_ISL_481601, EPI_ISL_481602, EPI_ISL_481603, EPI_ISL_481604, EPI_ISL_481605, EPI_ISL_481606, EPI_ISL_481607, EPI_ISL_481608, EPI_ISL_481609, EPI_ISL_481610, EPI_ISL_481611, EPI_ISL_481612, EPI_ISL_481613, EPI_ISL_481614, EPI_ISL_481615, EPI_ISL_481616, EPI_ISL_481617, EPI_ISL_481618, EPI_ISL_481619, EPI_ISL_481620, EPI_ISL_481621, EPI_ISL_481622, EPI_ISL_481623, EPI_ISL_481624, EPI_ISL_481625, EPI_ISL_481626, EPI_ISL_481627, EPI_ISL_481628, EPI_ISL_481629, EPI_ISL_481630, EPI_ISL_481631, EPI_ISL_481632, EPI_ISL_481633, EPI_ISL_481634, EPI_ISL_481635, EPI_ISL_481636, EPI_ISL_481637, EPI_ISL_481638, EPI_ISL_481639, EPI_ISL_481640, EPI_ISL_481641, EPI_ISL_481642, EPI_ISL_481643, EPI_ISL_481644, EPI_ISL_481645, EPI_ISL_481646, EPI_ISL_481647, EPI_ISL_481648, EPI_ISL_481649, EPI_ISL_481650, EPI_ISL_481651, EPI_ISL_481652, EPI_ISL_481653, EPI_ISL_481654, EPI_ISL_481655, EPI_ISL_481656, EPI_ISL_481657, EPI_ISL_481658, EPI_ISL_481659, EPI_ISL_481660, EPI_ISL_481661, EPI_ISL_481662, EPI_ISL_481663, EPI_ISL_481664, EPI_ISL_481665, EPI_ISL_481666, EPI_ISL_481667, EPI_ISL_481668, EPI_ISL_481669, EPI_ISL_481670, EPI_ISL_481671, EPI_ISL_481672, EPI_ISL_481673, EPI_ISL_481674, EPI_ISL_481675, EPI_ISL_481676, EPI_ISL_481677, EPI_ISL_481678, EPI_ISL_481679, EPI_ISL_481680, EPI_ISL_481681, EPI_ISL_481682, EPI_ISL_481683, EPI_ISL_481684, EPI_ISL_481685, EPI_ISL_481686, EPI_ISL_481687, EPI_ISL_481688, EPI_ISL_481689, EPI_ISL_481690, EPI_ISL_481691, EPI_ISL_481692, EPI_ISL_481693, EPI_ISL_481694, EPI_ISL_481695, EPI_ISL_481696, EPI_ISL_481697, EPI_ISL_481698, EPI_ISL_481699, EPI_ISL_481700, EPI_ISL_481701, EPI_ISL_481702, EPI_ISL_481703, EPI_ISL_481704, EPI_ISL_481705, EPI_ISL_481706, EPI_ISL_481707, EPI_ISL_481708, EPI_ISL_481709, EPI_ISL_481710, EPI_ISL_481711, EPI_ISL_481712, EPI_ISL_481713, EPI_ISL_481714, EPI_ISL_481715 | | | |
| see above | Department of Virology and Immunology, University of Helsinki and Helsinki University Hospital, Huslab Finland | Department of Virology, Faculty of Medicine, University of Helsinki, Helsinki, Finland | Teemu Smura, Hannimari Kallio-Kokko, Jenni Virtanen, Maija Suvanto, Sari Hannula, Harri Kangas, Pekka Ellonen, Olli Vapalahti |
| EPI_ISL_481716 | Prof. Massimo Zollo CEINGE TASK-FORCE COVID19 - Regione Campania | Prof. Massimo Zollo CEINGE TASK-FORCE COVID19 - Regione Campania | Veronica Ferrucci1,2, Dae young Kong8, Fatemeh asadzadeh1,2, Laura Marrone1,2, Roberto Siciliano1,2, Rino Cerino3, Giovanna Fusco3, Marika Comegna1,2, Angelo Boccia2, Maurizio Viscardi3, Giorgia Borriello3, Sergio Brandi3, Claudia Tiberio4, Luigi Atripaldi4, Giovanni Paolella1,2, Giuseppe Castaldo1,2, Stefano Pascarella4, Martina Bianchi4, Lorenzo Chiariotti1,2, Jae Myun Lee5, Jae Ho Jung6, Kyong Seop Yun7, Hong Yeoul Kim 7,8* and Massimo Zollo1,2* 1 CEINGE Biotecnologie Avanzate, Naples, Italia 2 Dipartimento di Medicina Molecolare e Biotecnologie Mediche DMMBM University of Naples Federico II, Italia 3 Istituto Zooprofilattico Sperimentale del Mezzogiorno, Naples, Italia 4 -U.O.C. di Patologia Clinica Ospedale D. Cotugno, Azienda Sanitaria Ospedali dei Colli, Naples, Italy. 5 Università La Sapienza di Roma, Italia 6 Department of Microbiology, Yonsei University College of Medicine, Seoul, Korea 7 Department of Surgery, Yonsei University College of Medicine, Seoul, Korea 8 Haim bio co., Ltd, , Indust |
| EPI_ISL_481717, EPI_ISL_481718, EPI_ISL_481719, EPI_ISL_481720, EPI_ISL_481721, EPI_ISL_481722, EPI_ISL_481723, EPI_ISL_481724, EPI_ISL_481725, EPI_ISL_481726, EPI_ISL_481727, EPI_ISL_481728, EPI_ISL_481729, EPI_ISL_481730, EPI_ISL_481731, EPI_ISL_481732, EPI_ISL_481733, EPI_ISL_481734, EPI_ISL_481735, EPI_ISL_481736, EPI_ISL_481737, EPI_ISL_481738, EPI_ISL_481739, EPI_ISL_481740 | | | |
| see above | Department of Virology and Immunology, University of Helsinki and Helsinki University Hospital, Huslab Finland | Department of Virology, Faculty of Medicine, University of Helsinki, Helsinki, Finland | Teemu Smura, Hannimari Kallio-Kokko, Jenni Virtanen, Maija Suvanto, Sari Hannula, Harri Kangas, Pekka Ellonen, Olli Vapalahti |
| EPI_ISL_481741 | Prof. Massimo Zollo CEINGE TASK-FORCE COVID19 - Regione Campania | Prof. Massimo Zollo CEINGE TASK-FORCE COVID19 - Regione Campania | Veronica Ferrucci1,2, Dae young Kong8, Fatemeh asadzadeh1,2, Laura Marrone1,2, Roberto Siciliano1,2, Rino Cerino3, Giovanna Fusco3, Marika Comegna1,2, Angelo Boccia2, Maurizio Viscardi3, Giorgia Borriello3, Sergio Brandi3, Claudia Tiberio4, Luigi Atripaldi4, Giovanni Paolella1,2, Giuseppe Castaldo1,2, Stefano Pascarella4, Martina Bianchi4, Lorenzo Chiariotti1,2, Jae Myun Lee5, Jae Ho Jung6, Kyong Seop Yun7, Hong Yeoul Kim 7,8* and Massimo Zollo1,2* 1 CEINGE Biotecnologie Avanzate, Naples, Italia 2 Dipartimento di Medicina Molecolare e Biotecnologie Mediche DMMBM University of Naples Federico II, Italia 3 Istituto Zooprofilattico Sperimentale del Mezzogiorno, Naples, Italia 4 -U.O.C. di Patologia Clinica Ospedale D. Cotugno, Azienda Sanitaria Ospedali dei Colli, Naples, Italy. 5 Università La Sapienza di Roma, Italia 6 Department of Microbiology, Yonsei University College of Medicine, Seoul, Korea 7 Department of Surgery, Yonsei University College of Medicine, Seoul, Korea 8 Haim bio co., Ltd, , Indust |
| EPI_ISL_481742, EPI_ISL_481743, EPI_ISL_481744, EPI_ISL_481745, EPI_ISL_481746, EPI_ISL_481747, EPI_ISL_481748, EPI_ISL_481749, EPI_ISL_481750, EPI_ISL_481751, EPI_ISL_481752, EPI_ISL_481753, EPI_ISL_481754, EPI_ISL_481755, EPI_ISL_481756, EPI_ISL_481757, EPI_ISL_481758 | | | |
| see above | Dr. Georges-L.-Dumont University Hospital Centre | National Microbiology Laboratory | Anna Majer, Shari Tyson, Grace Seo, Kristyn Burak, Philip Mabon, Elsie Grudeski, Rhiannon Huzarewich, Russell Mandes, Jennifer Tanner, Natalie Knox, Morag Graham, Gary Van Domselaar, Richard Garceau, Guillaume Desnoyers, Nathalie Bastien, Yan Li, Timothy Booth |
| EPI_ISL_481759 | Prof. Massimo Zollo CEINGE TASK-FORCE COVID19 - Regione Campania | Prof. Massimo Zollo CEINGE TASK-FORCE COVID19 - Regione Campania | Veronica Ferrucci1,2, Dae young Kong8, Fatemeh asadzadeh1,2, Laura Marrone1,2, Roberto Siciliano1,2, Rino Cerino3, Giovanna Fusco3, Marika Comegna1,2, Angelo Boccia2, Maurizio Viscardi3, Giorgia Borriello3, Sergio Brandi3, Claudia Tiberio4, Luigi Atripaldi4, Giovanni Paolella1,2, Giuseppe Castaldo1,2, Stefano Pascarella4, Martina Bianchi4, Lorenzo Chiariotti1,2, Jae Myun Lee5, Jae Ho Jung6, Kyong Seop Yun7, Hong Yeoul Kim 7,8* and Massimo Zollo1,2* 1 CEINGE Biotecnologie Avanzate, Naples, Italia 2 Dipartimento di Medicina Molecolare e Biotecnologie Mediche DMMBM University of Naples Federico II, Italia 3 Istituto Zooprofilattico Sperimentale del Mezzogiorno, Naples, Italia 4 -U.O.C. di Patologia Clinica Ospedale D. Cotugno, Azienda Sanitaria Ospedali dei Colli, Naples, Italy. 5 Università La Sapienza di Roma, Italia 6 Department of Microbiology, Yonsei University College of Medicine, Seoul, Korea 7 Department of Surgery, Yonsei University College of Medicine, Seoul, Korea 8 Haim bio co., Ltd, , Indust |
| EPI_ISL_481760, EPI_ISL_481761, EPI_ISL_481762 | Prof. Massimo Zollo CEINGE TASK-FORCE COVID19 - Regione Campania | Prof. Massimo Zollo CEINGE TASK-FORCE COVID19 - Regione Campania | Veronica Ferrucci1,2, Dae young Kong8, Fatemeh asadzadeh1,2, Laura Marrone1,2, Roberto Siciliano1,2, Rino Cerino3, Giovanna Fusco3, Marika Comegna1,2, Angelo Boccia2, Maurizio Viscardi3, Giorgia Borriello3, Sergio Brandi3, Claudia Tiberio4, Luigi Atripaldi4, Giovanni Paolella1,2, Giuseppe Castaldo1,2, Stefano Pascarella4, Martina Bianchi4, Lorenzo Chiariotti1,2, Jae Myun Lee5, Jae Ho Jung6, Kyong Seop Yun7, Hong Yeoul Kim 7,8* and Massimo Zollo1,2* 1 CEINGE Biotecnologie Avanzate, Naples, Italia 2 Dipartimento di Medicina Molecolare e Biotecnologie Mediche DMMBM University of Naples Federico II, Italia 3 Istituto Zooprofilattico Sperimentale del Mezzogiorno, Naples, Italia 4 -U.O.C. di Patologia Clinica Ospedale D. Cotugno, Azienda Sanitaria Ospedali dei Colli, Naples, Italy. 5 Università La Sapienza di Roma, Italia 6 Department of Microbiology, Yonsei University College of Medicine, Seoul, Korea 7 Department of Surgery, Yonsei University College of Medicine, Seoul, Korea 8 Haim bio co., Ltd, , Indust |
| EPI_ISL_481763, EPI_ISL_481764, EPI_ISL_481765, EPI_ISL_481766, EPI_ISL_481767, EPI_ISL_481768, EPI_ISL_481769, EPI_ISL_481770, EPI_ISL_481771, EPI_ISL_481772, EPI_ISL_481773, EPI_ISL_481774, EPI_ISL_481775, EPI_ISL_481776, EPI_ISL_481777, EPI_ISL_481778, EPI_ISL_481779, EPI_ISL_481780, EPI_ISL_481781, EPI_ISL_481782, EPI_ISL_481783, EPI_ISL_481784, EPI_ISL_481785, EPI_ISL_481786, EPI_ISL_481787, EPI_ISL_481788, EPI_ISL_481789, EPI_ISL_481790, EPI_ISL_481791, EPI_ISL_481792, EPI_ISL_481793, EPI_ISL_481794, EPI_ISL_481795, EPI_ISL_481796, EPI_ISL_481797, EPI_ISL_481798, EPI_ISL_481799, EPI_ISL_481800, EPI_ISL_481801, EPI_ISL_481802, EPI_ISL_481803, EPI_ISL_481804, EPI_ISL_481805, EPI_ISL_481806, EPI_ISL_481807, EPI_ISL_481808, EPI_ISL_481809, EPI_ISL_481810, EPI_ISL_481811, EPI_ISL_481812, EPI_ISL_481813, EPI_ISL_481814, EPI_ISL_481815, EPI_ISL_481816, EPI_ISL_481817, EPI_ISL_481818, EPI_ISL_481819, EPI_ISL_481820, EPI_ISL_481821, EPI_ISL_481822, EPI_ISL_481823, EPI_ISL_481824, EPI_ISL_481825, EPI_ISL_481826, EPI_ISL_481827, EPI_ISL_481828, EPI_ISL_481829, EPI_ISL_481830, EPI_ISL_481831, EPI_ISL_481832, EPI_ISL_481833, EPI_ISL_481834, | | | |

| | | | |
|---|---|---|---|
| EPI_ISL_481835, EPI_ISL_481836, EPI_ISL_481837, EPI_ISL_481838, EPI_ISL_481839, EPI_ISL_481840, EPI_ISL_481841, EPI_ISL_481842, EPI_ISL_481843, EPI_ISL_481844, EPI_ISL_481845, EPI_ISL_481846, EPI_ISL_481847, EPI_ISL_481848, EPI_ISL_481849, EPI_ISL_481850, EPI_ISL_481851, EPI_ISL_481852, EPI_ISL_481853, EPI_ISL_481854, EPI_ISL_481855, EPI_ISL_481856, EPI_ISL_481857, EPI_ISL_481858, EPI_ISL_481859, EPI_ISL_481860, EPI_ISL_481861, EPI_ISL_481862, EPI_ISL_481863, EPI_ISL_481864, EPI_ISL_481865, EPI_ISL_481866, EPI_ISL_481867, EPI_ISL_481868, EPI_ISL_481869, EPI_ISL_481870, EPI_ISL_481871, EPI_ISL_481872, EPI_ISL_481873, EPI_ISL_481874, EPI_ISL_481875, EPI_ISL_481876, EPI_ISL_481877, EPI_ISL_481878, EPI_ISL_481879, EPI_ISL_481880, EPI_ISL_481881, EPI_ISL_481882, EPI_ISL_481883, EPI_ISL_481884, EPI_ISL_481885, EPI_ISL_481886, EPI_ISL_481887, EPI_ISL_481888, EPI_ISL_481889, EPI_ISL_481890, EPI_ISL_481891, EPI_ISL_481892, EPI_ISL_481893, EPI_ISL_481894, EPI_ISL_481895, EPI_ISL_481896, EPI_ISL_481897, EPI_ISL_481898, EPI_ISL_481899, EPI_ISL_481900, EPI_ISL_481901, EPI_ISL_481902, EPI_ISL_481903, EPI_ISL_481904, EPI_ISL_481905, EPI_ISL_481906, EPI_ISL_481907, EPI_ISL_481908, EPI_ISL_481909, EPI_ISL_481910, EPI_ISL_481911, EPI_ISL_481912, EPI_ISL_481913, EPI_ISL_481914, EPI_ISL_481915, EPI_ISL_481916, EPI_ISL_481917, EPI_ISL_481918, EPI_ISL_481919, EPI_ISL_481920, EPI_ISL_481921, EPI_ISL_481922, EPI_ISL_481923, EPI_ISL_481924, EPI_ISL_481925, EPI_ISL_481926, EPI_ISL_481927, EPI_ISL_481928, EPI_ISL_481929, EPI_ISL_481930, EPI_ISL_481931, EPI_ISL_481932, EPI_ISL_481933, EPI_ISL_481934, EPI_ISL_481935, EPI_ISL_481936, EPI_ISL_481937, EPI_ISL_481938, EPI_ISL_481939, EPI_ISL_481940, EPI_ISL_481941, EPI_ISL_481942, EPI_ISL_481943, EPI_ISL_481944, EPI_ISL_481945, EPI_ISL_481946, EPI_ISL_481947, EPI_ISL_481948, EPI_ISL_481949, EPI_ISL_481950, EPI_ISL_481951, EPI_ISL_481952, EPI_ISL_481953, EPI_ISL_481954, EPI_ISL_481955, EPI_ISL_481956, EPI_ISL_481957, EPI_ISL_481958, EPI_ISL_481959, EPI_ISL_481960, EPI_ISL_481961, EPI_ISL_481962, EPI_ISL_481963, EPI_ISL_481964, EPI_ISL_481965, EPI_ISL_481966, EPI_ISL_481967, EPI_ISL_481968, EPI_ISL_481969, EPI_ISL_481970, EPI_ISL_481971, EPI_ISL_481972, EPI_ISL_481973, EPI_ISL_481974, EPI_ISL_481975, EPI_ISL_481976, EPI_ISL_481977, EPI_ISL_481978, EPI_ISL_481979, EPI_ISL_481980, EPI_ISL_481981, EPI_ISL_481982, EPI_ISL_481983, EPI_ISL_481984, EPI_ISL_481985, EPI_ISL_481986, EPI_ISL_481987, EPI_ISL_481988, EPI_ISL_481989, EPI_ISL_481990, EPI_ISL_481991, EPI_ISL_481992, EPI_ISL_481993, EPI_ISL_481994, EPI_ISL_481995, EPI_ISL_481996, EPI_ISL_481997, EPI_ISL_481998, EPI_ISL_481999, EPI_ISL_482000, EPI_ISL_482001, EPI_ISL_482002, EPI_ISL_482003, EPI_ISL_482004, EPI_ISL_482005, EPI_ISL_482006, EPI_ISL_482007, EPI_ISL_482008, EPI_ISL_482009, EPI_ISL_482010, EPI_ISL_482011, EPI_ISL_482012, EPI_ISL_482013, EPI_ISL_482014, EPI_ISL_482015, EPI_ISL_482016, EPI_ISL_482017, EPI_ISL_482018, EPI_ISL_482019, EPI_ISL_482020, EPI_ISL_482021, EPI_ISL_482022, EPI_ISL_482023, EPI_ISL_482024, EPI_ISL_482025, EPI_ISL_482026, EPI_ISL_482027, EPI_ISL_482029, EPI_ISL_482030, EPI_ISL_482031 | | | |
| see above | PHE South West Regional Laboratory, National Infection Service | Wellcome Sanger Institute for the COVID-19 Genomics UK Consortium | Stephanie Hutchings, Hannah Pymont, Dr Peter Muir, Barry Vipond, Rich Hopes; and Alex Alderton, Roberto Amato, Sonia Goncalves, Ewan Harrison, David K. Jackson, Ian Johnston, Dominic Kwiatkowski, Cordelia Langford, John Sillitoe on behalf of the Wellcome Sanger Institute COVID-19 Surveillance Team (http://www.sanger.ac.uk/covid-team) |
| EPI_ISL_482032, EPI_ISL_482033, EPI_ISL_482034, EPI_ISL_482035, EPI_ISL_482036, EPI_ISL_482037, EPI_ISL_482038, EPI_ISL_482039, EPI_ISL_482040, EPI_ISL_482041, EPI_ISL_482042, EPI_ISL_482043, EPI_ISL_482044, EPI_ISL_482045, EPI_ISL_482046, EPI_ISL_482047, EPI_ISL_482048, EPI_ISL_482049, EPI_ISL_482050, EPI_ISL_482051, EPI_ISL_482052, EPI_ISL_482053, EPI_ISL_482054, EPI_ISL_482055, EPI_ISL_482056 | | | |
| see above | Regional Virus Laboratory, Belfast Health and Social Care Trust | Wellcome Sanger Institute for the COVID-19 Genomics UK Consortium | Conall McCaughey, James McKenna, Tanya Curran, Susan Feeney, Alison Watt, Ciara Cox, Mairead Connor, Zoltan Molnar, David Simpson, Derek Fairley; and Alex Alderton, Roberto Amato, Sonia Goncalves, Ewan Harrison, David K. Jackson, Ian Johnston, Dominic Kwiatkowski, Cordelia Langford, John Sillitoe on behalf of the Wellcome Sanger Institute COVID-19 Surveillance Team (http://www.sanger.ac.uk/covid-team) |
| EPI_ISL_482057, EPI_ISL_482058, EPI_ISL_482059, EPI_ISL_482060, EPI_ISL_482061, EPI_ISL_482062, EPI_ISL_482063, EPI_ISL_482064, EPI_ISL_482065, EPI_ISL_482066, EPI_ISL_482067, EPI_ISL_482068 | | | |
| see above | The Department of Microbiology, Torbay and South Devon NHS Foundation Trust | Wellcome Sanger Institute for the COVID-19 Genomics UK Consortium | Amy Hurd, Sophie Lloyd, Anthony Mogridge, Jack Howe, Helen Brown, Gary Booth, Mel Brown, Cheryl Bailiss, Michelle Harrison and Alex Alderton, Roberto Amato, Sonia Goncalves, Ewan Harrison, David K. Jackson, Ian Johnston, Dominic Kwiatkowski, Cordelia Langford, John Sillitoe on behalf of the Wellcome Sanger Institute COVID-19 Surveillance Team (http://www.sanger.ac.uk/covid-team) |
| EPI_ISL_482069 | Microbiology Department, Hereford County Hospital | Wellcome Sanger Institute for the COVID-19 Genomics UK Consortium | Alison Johnson, Venkat Sivaprakasam, Fenella Halstead, Jane Thomas, Wendy Hogsden, Samantha Lamb and Alex Alderton, Roberto Amato, Sonia Goncalves, Ewan Harrison, David K. Jackson, Ian Johnston, Dominic Kwiatkowski, Cordelia Langford, John Sillitoe on behalf of the Wellcome Sanger Institute COVID-19 Surveillance Team (http://www.sanger.ac.uk/covid-team) |
| EPI_ISL_482070 | Regional Virus Laboratory, Belfast Health and Social Care Trust | Wellcome Sanger Institute for the COVID-19 Genomics UK Consortium | Conall McCaughey, James McKenna, Tanya Curran, Susan Feeney, Alison Watt, Ciara Cox, Mairead Connor, Zoltan Molnar, David Simpson, Derek Fairley; and Alex Alderton, Roberto Amato, Sonia Goncalves, Ewan Harrison, David K. Jackson, Ian Johnston, Dominic Kwiatkowski, Cordelia Langford, John Sillitoe on behalf of the Wellcome Sanger Institute COVID-19 Surveillance Team (http://www.sanger.ac.uk/covid-team) |
| EPI_ISL_482071, EPI_ISL_482072, EPI_ISL_482073, EPI_ISL_482074 | Microbiology Department, Hereford County Hospital | Wellcome Sanger Institute for the COVID-19 Genomics UK Consortium | Alison Johnson, Venkat Sivaprakasam, Fenella Halstead, Jane Thomas, Wendy Hogsden, Samantha Lamb and Alex Alderton, Roberto Amato, Sonia Goncalves, Ewan Harrison, David K. Jackson, Ian Johnston, Dominic Kwiatkowski, Cordelia Langford, John Sillitoe on behalf of the Wellcome Sanger Institute COVID-19 Surveillance Team (http://www.sanger.ac.uk/covid-team) |
| EPI_ISL_482075 | Regional Virus Laboratory, Belfast Health and Social Care Trust | Wellcome Sanger Institute for the COVID-19 Genomics UK Consortium | Conall McCaughey, James McKenna, Tanya Curran, Susan Feeney, Alison Watt, Ciara Cox, Mairead Connor, Zoltan Molnar, David Simpson, Derek Fairley; and Alex Alderton, Roberto Amato, Sonia Goncalves, Ewan Harrison, David K. Jackson, Ian Johnston, Dominic Kwiatkowski, Cordelia Langford, John Sillitoe on behalf of the Wellcome Sanger Institute COVID-19 Surveillance Team (http://www.sanger.ac.uk/covid-team) |
| EPI_ISL_482076 | Microbiology Department, Hereford County Hospital | Wellcome Sanger Institute for the COVID-19 Genomics UK Consortium | Alison Johnson, Venkat Sivaprakasam, Fenella Halstead, Jane Thomas, Wendy Hogsden, Samantha Lamb and Alex Alderton, Roberto Amato, Sonia Goncalves, Ewan Harrison, David K. Jackson, Ian Johnston, Dominic Kwiatkowski, Cordelia Langford, John Sillitoe on behalf of the Wellcome Sanger Institute COVID-19 Surveillance Team (http://www.sanger.ac.uk/covid-team) |
| EPI_ISL_482077 | Regional Virus Laboratory, Belfast Health and Social Care Trust | Wellcome Sanger Institute for the COVID-19 Genomics UK Consortium | Conall McCaughey, James McKenna, Tanya Curran, Susan Feeney, Alison Watt, Ciara Cox, Mairead Connor, Zoltan Molnar, David Simpson, Derek Fairley; and Alex Alderton, Roberto Amato, Sonia Goncalves, Ewan Harrison, David K. Jackson, Ian Johnston, Dominic Kwiatkowski, Cordelia Langford, John Sillitoe on behalf of the Wellcome Sanger Institute COVID-19 Surveillance Team (http://www.sanger.ac.uk/covid-team) |
| EPI_ISL_482078, EPI_ISL_482079, EPI_ISL_482080, EPI_ISL_482081 | Microbiology Department, Hereford County Hospital | Wellcome Sanger Institute for the COVID-19 Genomics UK Consortium | Alison Johnson, Venkat Sivaprakasam, Fenella Halstead, Jane Thomas, Wendy Hogsden, Samantha Lamb and Alex Alderton, Roberto Amato, Sonia Goncalves, Ewan Harrison, David K. Jackson, Ian Johnston, Dominic Kwiatkowski, Cordelia Langford, John Sillitoe on behalf of the Wellcome Sanger Institute COVID-19 Surveillance Team (http://www.sanger.ac.uk/covid-team) |
| EPI_ISL_482082 | Regional Virus Laboratory, Belfast Health and Social Care Trust | Wellcome Sanger Institute for the COVID-19 Genomics UK Consortium | Conall McCaughey, James McKenna, Tanya Curran, Susan Feeney, Alison Watt, Ciara Cox, Mairead Connor, Zoltan Molnar, David Simpson, Derek Fairley; and Alex Alderton, Roberto Amato, Sonia Goncalves, Ewan Harrison, David K. Jackson, Ian Johnston, Dominic Kwiatkowski, Cordelia Langford, John Sillitoe on behalf of the Wellcome Sanger Institute COVID-19 Surveillance Team (http://www.sanger.ac.uk/covid-team) |
| EPI_ISL_482083, EPI_ISL_482084, EPI_ISL_482085, EPI_ISL_482086, EPI_ISL_482087, EPI_ISL_482088, EPI_ISL_482089, EPI_ISL_482090 | Microbiology Department, Hereford County Hospital | Wellcome Sanger Institute for the COVID-19 Genomics UK Consortium | Alison Johnson, Venkat Sivaprakasam, Fenella Halstead, Jane Thomas, Wendy Hogsden, Samantha Lamb and Alex Alderton, Roberto Amato, Sonia Goncalves, Ewan Harrison, David K. Jackson, Ian Johnston, Dominic Kwiatkowski, Cordelia Langford, John Sillitoe on behalf of the Wellcome Sanger Institute COVID-19 Surveillance Team (http://www.sanger.ac.uk/covid-team) |
| EPI_ISL_482091 | Regional Virus Laboratory, Belfast Health and Social Care Trust | Wellcome Sanger Institute for the COVID-19 Genomics UK Consortium | Conall McCaughey, James McKenna, Tanya Curran, Susan Feeney, Alison Watt, Ciara Cox, Mairead Connor, Zoltan Molnar, David Simpson, Derek Fairley; and Alex Alderton, Roberto Amato, Sonia Goncalves, Ewan Harrison, David K. Jackson, Ian Johnston, Dominic Kwiatkowski, Cordelia Langford, John Sillitoe on behalf of the Wellcome Sanger Institute COVID-19 Surveillance Team (http://www.sanger.ac.uk/covid-team) |
| EPI_ISL_482092, EPI_ISL_482093, EPI_ISL_482094, EPI_ISL_482095 | Microbiology Department, Hereford County Hospital | Wellcome Sanger Institute for the COVID-19 Genomics UK Consortium | Alison Johnson, Venkat Sivaprakasam, Fenella Halstead, Jane Thomas, Wendy Hogsden, Samantha Lamb and Alex Alderton, Roberto Amato, Sonia Goncalves, Ewan Harrison, David K. Jackson, Ian Johnston, Dominic Kwiatkowski, Cordelia Langford, John Sillitoe on behalf of the Wellcome Sanger Institute COVID-19 Surveillance Team (http://www.sanger.ac.uk/covid-team) |
| EPI_ISL_482096, EPI_ISL_482097 | Regional Virus Laboratory, Belfast Health and Social Care Trust | Wellcome Sanger Institute for the COVID-19 Genomics UK Consortium | Conall McCaughey, James McKenna, Tanya Curran, Susan Feeney, Alison Watt, Ciara Cox, Mairead Connor, Zoltan Molnar, David Simpson, Derek Fairley; and Alex Alderton, Roberto Amato, Sonia Goncalves, Ewan Harrison, David K. Jackson, Ian Johnston, Dominic Kwiatkowski, Cordelia Langford, John Sillitoe on behalf of the Wellcome Sanger Institute COVID-19 Surveillance Team (http://www.sanger.ac.uk/covid-team) |
| EPI_ISL_482098, EPI_ISL_482099, EPI_ISL_482100 | Microbiology Department, Hereford County Hospital | Wellcome Sanger Institute for the COVID-19 Genomics UK Consortium | Alison Johnson, Venkat Sivaprakasam, Fenella Halstead, Jane Thomas, Wendy Hogsden, Samantha Lamb and Alex Alderton, Roberto Amato, Sonia Goncalves, Ewan Harrison, David K. Jackson, Ian Johnston, Dominic Kwiatkowski, Cordelia Langford, John Sillitoe on behalf of the Wellcome Sanger Institute COVID-19 Surveillance Team (http://www.sanger.ac.uk/covid-team) |
| EPI_ISL_482101, EPI_ISL_482102 | Regional Virus Laboratory, Belfast Health and Social Care Trust | Wellcome Sanger Institute for the COVID-19 Genomics UK Consortium | Conall McCaughey, James McKenna, Tanya Curran, Susan Feeney, Alison Watt, Ciara Cox, Mairead Connor, Zoltan Molnar, David Simpson, Derek Fairley; and Alex Alderton, Roberto Amato, Sonia Goncalves, Ewan Harrison, David K. Jackson, Ian Johnston, Dominic Kwiatkowski, Cordelia Langford, John Sillitoe on behalf of the Wellcome Sanger Institute COVID-19 Surveillance Team (http://www.sanger.ac.uk/covid-team) |
| EPI_ISL_482103, EPI_ISL_482104 | Microbiology Department, Hereford County Hospital | Wellcome Sanger Institute for the COVID-19 Genomics UK Consortium | Alison Johnson, Venkat Sivaprakasam, Fenella Halstead, Jane Thomas, Wendy Hogsden, Samantha Lamb and Alex Alderton, Roberto Amato, Sonia Goncalves, Ewan Harrison, David K. Jackson, Ian Johnston, Dominic Kwiatkowski, Cordelia Langford, John Sillitoe on behalf of the Wellcome Sanger Institute COVID-19 Surveillance Team (http://www.sanger.ac.uk/covid-team) |
| EPI_ISL_482105 | Regional Virus Laboratory, Belfast Health and Social Care Trust | Wellcome Sanger Institute for the COVID-19 Genomics UK Consortium | Conall McCaughey, James McKenna, Tanya Curran, Susan Feeney, Alison Watt, Ciara Cox, Mairead Connor, Zoltan Molnar, David Simpson, Derek Fairley; and Alex Alderton, Roberto Amato, Sonia Goncalves, Ewan Harrison, David K. Jackson, Ian Johnston, Dominic Kwiatkowski, Cordelia Langford, John Sillitoe on behalf of the Wellcome Sanger Institute COVID-19 Surveillance Team (http://www.sanger.ac.uk/covid-team) |
| EPI_ISL_482106, EPI_ISL_482107, EPI_ISL_482108, EPI_ISL_482109, EPI_ISL_482110, EPI_ISL_482111, EPI_ISL_482112, EPI_ISL_482113, EPI_ISL_482114 | Microbiology Department, Hereford County Hospital | Wellcome Sanger Institute for the COVID-19 Genomics UK Consortium | Alison Johnson, Venkat Sivaprakasam, Fenella Halstead, Jane Thomas, Wendy Hogsden, Samantha Lamb and Alex Alderton, Roberto Amato, Sonia Goncalves, Ewan Harrison, David K. Jackson, Ian Johnston, Dominic Kwiatkowski, Cordelia Langford, John Sillitoe on behalf of the Wellcome Sanger Institute COVID-19 Surveillance Team (http://www.sanger.ac.uk/covid-team) |
| EPI_ISL_482115 | Regional Virus Laboratory, Belfast Health and Social Care Trust | Wellcome Sanger Institute for the COVID-19 Genomics UK Consortium | Conall McCaughey, James McKenna, Tanya Curran, Susan Feeney, Alison Watt, Ciara Cox, Mairead Connor, Zoltan Molnar, David Simpson, Derek Fairley; and Alex Alderton, Roberto Amato, Sonia Goncalves, Ewan Harrison, David K. Jackson, Ian Johnston, Dominic Kwiatkowski, Cordelia Langford, John Sillitoe on behalf of the Wellcome Sanger Institute COVID-19 Surveillance Team (http://www.sanger.ac.uk/covid-team) |
| EPI_ISL_482116, EPI_ISL_482117, EPI_ISL_482118, EPI_ISL_482119 | Microbiology Department, Hereford County Hospital | Wellcome Sanger Institute for the COVID-19 Genomics UK Consortium | Alison Johnson, Venkat Sivaprakasam, Fenella Halstead, Jane Thomas, Wendy Hogsden, Samantha Lamb and Alex Alderton, Roberto Amato, Sonia Goncalves, Ewan Harrison, David K. Jackson, Ian Johnston, Dominic Kwiatkowski, Cordelia Langford, John Sillitoe on behalf of the Wellcome Sanger Institute COVID-19 Surveillance Team (http://www.sanger.ac.uk/covid-team) |
| EPI_ISL_482120, EPI_ISL_482121, EPI_ISL_482122, EPI_ISL_482123 | The Department of Microbiology, Torbay and South Devon NHS Foundation Trust | Wellcome Sanger Institute for the COVID-19 Genomics UK Consortium | Amy Hurd, Sophie Lloyd, Anthony Mogridge, Jack Howe, Helen Brown, Gary Booth, Mel Brown, Cheryl Bailiss, Michelle Harrison and Alex Alderton, Roberto Amato, Sonia Goncalves, Ewan Harrison, David K. Jackson, Ian Johnston, Dominic Kwiatkowski, Cordelia Langford, John Sillitoe on behalf of the Wellcome Sanger Institute COVID-19 Surveillance Team (http://www.sanger.ac.uk/covid-team) |
| EPI_ISL_482124 | North West London Pathology, Imperial College Healthcare NHS Trust | Wellcome Sanger Institute for the COVID-19 Genomics UK Consortium | Ling Li, Paul Randell, David Muir, Frankie Bolt, Alison Holmes, James Price, Aileen Rowan, Graham Taylor, Anjna Badhan, Carolina Herrera and Alex Alderton, Roberto Amato, Sonia Goncalves, Ewan Harrison, David K. Jackson, Ian Johnston, Dominic Kwiatkowski, Cordelia Langford, John Sillitoe on behalf of the Wellcome Sanger Institute COVID-19 Surveillance Team (http://www.sanger.ac.uk/covid-team) |
| EPI_ISL_482125, EPI_ISL_482126, EPI_ISL_482127, EPI_ISL_482128, EPI_ISL_482129, EPI_ISL_482130 | The Department of Microbiology, Torbay and South Devon NHS Foundation Trust | Wellcome Sanger Institute for the COVID-19 Genomics UK Consortium | Amy Hurd, Sophie Lloyd, Anthony Mogridge, Jack Howe, Helen Brown, Gary Booth, Mel Brown, Cheryl Bailiss, Michelle Harrison and Alex Alderton, Roberto Amato, Sonia Goncalves, Ewan Harrison, David K. Jackson, Ian Johnston, Dominic Kwiatkowski, Cordelia Langford, John Sillitoe on behalf of the Wellcome Sanger Institute COVID-19 Surveillance Team (http://www.sanger.ac.uk/covid-team) |
| EPI_ISL_482131 | North West London Pathology, Imperial College Healthcare NHS Trust | Wellcome Sanger Institute for the COVID-19 Genomics UK Consortium | Ling Li, Paul Randell, David Muir, Frankie Bolt, Alison Holmes, James Price, Aileen Rowan, Graham Taylor, Anjna Badhan, Carolina Herrera and Alex Alderton, Roberto Amato, Sonia Goncalves, Ewan Harrison, David K. Jackson, Ian Johnston, Dominic Kwiatkowski, Cordelia Langford, John Sillitoe on behalf of the Wellcome Sanger Institute COVID-19 Surveillance Team (http://www.sanger.ac.uk/covid-team) |
| EPI_ISL_482132, EPI_ISL_482133, EPI_ISL_482134, EPI_ISL_482135, EPI_ISL_482136, EPI_ISL_482137, EPI_ISL_482139, EPI_ISL_482140, EPI_ISL_482141, EPI_ISL_482142, EPI_ISL_482143, EPI_ISL_482144, EPI_ISL_482145, EPI_ISL_482146, EPI_ISL_482147, EPI_ISL_482148, EPI_ISL_482149, EPI_ISL_482150, EPI_ISL_482151, EPI_ISL_482152, EPI_ISL_482153, EPI_ISL_482154, EPI_ISL_482155, EPI_ISL_482156, EPI_ISL_482157, EPI_ISL_482158, EPI_ISL_482159 | | | |
| see above | Regional Virus Laboratory, Belfast Health and Social Care Trust | Wellcome Sanger Institute for the COVID-19 Genomics UK Consortium | Conall McCaughey, James McKenna, Tanya Curran, Susan Feeney, Alison Watt, Ciara Cox, Mairead Connor, Zoltan Molnar, David Simpson, Derek Fairley; and Alex Alderton, Roberto Amato, Sonia Goncalves, Ewan Harrison, David K. Jackson, Ian Johnston, Dominic Kwiatkowski, Cordelia Langford, John Sillitoe on behalf of the Wellcome Sanger Institute COVID-19 Surveillance Team (http://www.sanger.ac.uk/covid-team) |
| EPI_ISL_482160, EPI_ISL_482161, EPI_ISL_482162, EPI_ISL_482163, EPI_ISL_482164, EPI_ISL_482165, EPI_ISL_482166, EPI_ISL_482167, EPI_ISL_482168, EPI_ISL_482169, EPI_ISL_482170, EPI_ISL_482171, EPI_ISL_482172, EPI_ISL_482173, EPI_ISL_482174, EPI_ISL_482175, EPI_ISL_482176, EPI_ISL_482177, EPI_ISL_482178, EPI_ISL_482179, EPI_ISL_482180, EPI_ISL_482181, EPI_ISL_482182, EPI_ISL_482183, EPI_ISL_482184, EPI_ISL_482185, EPI_ISL_482186, EPI_ISL_482187, EPI_ISL_482188, EPI_ISL_482189, EPI_ISL_482190, EPI_ISL_482191, EPI_ISL_482192, EPI_ISL_482193, EPI_ISL_482194, EPI_ISL_482195, EPI_ISL_482196, EPI_ISL_482197, EPI_ISL_482198, EPI_ISL_482199, EPI_ISL_482200, EPI_ISL_482201, EPI_ISL_482202, EPI_ISL_482203, EPI_ISL_482204, EPI_ISL_482205, EPI_ISL_482206, EPI_ISL_482207, EPI_ISL_482208, EPI_ISL_482209, EPI_ISL_482210, EPI_ISL_482211, EPI_ISL_482212, EPI_ISL_482213, EPI_ISL_482214, EPI_ISL_482215, EPI_ISL_482216, EPI_ISL_482217, EPI_ISL_482218 | | | |
| see above | University College London, Great Ormond Street Hospital for Children NHS Foundation Trust, Imperial College Healthcare NHS Trust | Wellcome Sanger Institute for the COVID-19 Genomics UK Consortium | Sergi Castellano, Rachel Williams, Mark Kristiansen, Paola Resende Silva, Sunando Roy, Tony Brooks, Helena Tutill, Paola Niola, Patricia Dyal, Charlotte Williams, Leysa Forrest, Yasmin Panchbhaya, Jacqueline Findlay, Sam Weeks, Julianne Brown, Kathryn Harris, Paul Randell, James Price, Alison Holmes, Judith Breuer and Alex Alderton, Roberto Amato, Sonia Goncalves, Ewan Harrison, David K. Jackson, Ian Johnston, Dominic Kwiatkowski, Cordelia Langford, John Sillitoe on behalf of the Wellcome Sanger Institute COVID-19 Surveillance Team (http://www.sanger.ac.uk/covid-team) |
| EPI_ISL_482292, EPI_ISL_482293, EPI_ISL_482294, EPI_ISL_482295, EPI_ISL_482296, EPI_ISL_482297, EPI_ISL_482298, EPI_ISL_482299, EPI_ISL_482300, EPI_ISL_482301, EPI_ISL_482302, EPI_ISL_482303, EPI_ISL_482304, EPI_ISL_482305, EPI_ISL_482306, EPI_ISL_482307, EPI_ISL_482308, EPI_ISL_482309, EPI_ISL_482310, EPI_ISL_482311, EPI_ISL_482312, EPI_ISL_482313, EPI_ISL_482314, EPI_ISL_482315, EPI_ISL_482316, EPI_ISL_482317, EPI_ISL_482318, EPI_ISL_482319, EPI_ISL_482320, EPI_ISL_482321, EPI_ISL_482322, EPI_ISL_482323, EPI_ISL_482324, EPI_ISL_482325, EPI_ISL_482326, EPI_ISL_482327, EPI_ISL_482328, EPI_ISL_482329, EPI_ISL_482330, EPI_ISL_482331, EPI_ISL_482332, EPI_ISL_482333, EPI_ISL_482334, EPI_ISL_482335, EPI_ISL_482336, EPI_ISL_482337, EPI_ISL_482338, EPI_ISL_482339, EPI_ISL_482340, EPI_ISL_482341, EPI_ISL_482342, EPI_ISL_482343, EPI_ISL_482344, EPI_ISL_482345, EPI_ISL_482346, EPI_ISL_482347, EPI_ISL_482348, EPI_ISL_482349, EPI_ISL_482350, EPI_ISL_482351, EPI_ISL_482352, EPI_ISL_482353, EPI_ISL_482354, EPI_ISL_482355, EPI_ISL_482356, EPI_ISL_482357, EPI_ISL_482358, EPI_ISL_482359, EPI_ISL_482360, EPI_ISL_482361, EPI_ISL_482362, EPI_ISL_482363, EPI_ISL_482364, EPI_ISL_482365, EPI_ISL_482366, EPI_ISL_482367, EPI_ISL_482368, EPI_ISL_482369, EPI_ISL_482370, EPI_ISL_482371, EPI_ISL_482372, EPI_ISL_482373, EPI_ISL_482374, EPI_ISL_482375, EPI_ISL_482376, EPI_ISL_482377, EPI_ISL_482378, EPI_ISL_482379, EPI_ISL_482380, EPI_ISL_482381, EPI_ISL_482382, EPI_ISL_482383, EPI_ISL_482384, EPI_ISL_482385, EPI_ISL_482386, EPI_ISL_482387, EPI_ISL_482388, EPI_ISL_482389, EPI_ISL_482390, EPI_ISL_482391, EPI_ISL_482392, EPI_ISL_482393, EPI_ISL_482394, EPI_ISL_482395, EPI_ISL_482396, EPI_ISL_482397, EPI_ISL_482398, EPI_ISL_482399, EPI_ISL_482400, EPI_ISL_482401, EPI_ISL_482402, EPI_ISL_482403, EPI_ISL_482404, EPI_ISL_482405, EPI_ISL_482406, EPI_ISL_482407, EPI_ISL_482408, EPI_ISL_482409, EPI_ISL_482410, EPI_ISL_482411, EPI_ISL_482412, EPI_ISL_482413, EPI_ISL_482414, EPI_ISL_482415, EPI_ISL_482416, EPI_ISL_482417, EPI_ISL_482418, EPI_ISL_482419, EPI_ISL_482420, EPI_ISL_482421, EPI_ISL_482422, EPI_ISL_482423, EPI_ISL_482424, EPI_ISL_482425, EPI_ISL_482426, EPI_ISL_482427, EPI_ISL_482428, EPI_ISL_482429, EPI_ISL_482430, EPI_ISL_482431, EPI_ISL_482432, EPI_ISL_482433, EPI_ISL_482434, EPI_ISL_482435, EPI_ISL_482436, EPI_ISL_482437, EPI_ISL_482438, EPI_ISL_482439, EPI_ISL_482440, EPI_ISL_482441, EPI_ISL_482442, EPI_ISL_482443, EPI_ISL_482444, EPI_ISL_482445, EPI_ISL_482446, EPI_ISL_482447, EPI_ISL_482448, EPI_ISL_482449, EPI_ISL_482450, EPI_ISL_482451, EPI_ISL_482452, EPI_ISL_482453, EPI_ISL_482454, EPI_ISL_482455, EPI_ISL_482456, EPI_ISL_482457, EPI_ISL_482458, EPI_ISL_482459, EPI_ISL_482460, EPI_ISL_482461, EPI_ISL_482462, EPI_ISL_482463, EPI_ISL_482464, EPI_ISL_482465, EPI_ISL_482466, EPI_ISL_482467 | | | |
| see above | Providence St. Joseph Health Molecular Genomics Laboratory | Providence St. Joseph Health Molecular Genomics Laboratory | Alexa K Dowdell, Brian D Piening, Fred L Robinson, Carlo B Bifulco, Mary Campbell |
| EPI_ISL_482468 | Laboratorio de Referencia Nacional de Virus Respiratorios. Instituto Nacional de Salud Peru | Laboratorio de Referencia Nacional de Biotecnologia y Biologia Molecular. Instituto Nacional de Salud Peru | Carlos Padilla Rojas, Priscila Lope Pari, Karolyn Vega Chozo, Johanna Balbuena Torres, Omar Caceres Rey, Henri Bailon Calderon, Maribel Huaringa Nuñez, Nancy Rojas Serrano |
| EPI_ISL_482469, EPI_ISL_482470 | Queen Elizabeth II Health Science Centre | National Microbiology Laboratory | Anna Majer, Shari Tyson, Grace Seo, Kristyn Burak, Philip Mabon, Elsie Grudeski, Rhiannon Huzarewich, Russell Mandes, Jennifer Tanner, Natalie Knox, Morag Graham, Gary Van Domselaar, Todd Hatchette, Jason LeBlanc, Nathalie Bastien, Yan Li, Timothy |

| | | | |
|---|---|---|---|
| | | | Booth |
| EPI_ISL_482471, EPI_ISL_482472, EPI_ISL_482473 | Dr. Georges-L.-Dumont University Hospital Centre | National Microbiology Laboratory | Anna Majer, Shari Tyson, Grace Seo, Kristyn Burak, Philip Mabon, Elsie Grudeski, Rhiannon Huzarewich, Russell Mandes, Jennifer Tanner, Natalie Knox, Morag Graham, Gary Van Domselaar, Richard Garceau, Guillaume Desnoyers, Nathalie Bastien, Yan Li, Timothy Booth |
| EPI_ISL_482474, EPI_ISL_482475 | Cadham Provincial Laboratory | National Microbiology Laboratory | Anna Majer, Shari Tyson, Grace Seo, Kristyn Burak, Philip Mabon, Elsie Grudeski, Rhiannon Huzarewich, Russell Mandes, Jennifer Tanner, Natalie Knox, Morag Graham, Gary Van Domselaar, Paul Van Caeseele, Jared Bullard, David Alexander, Kerry Dust, Nathalie Bastien, Yan Li, Timothy Booth, |
| EPI_ISL_482476, EPI_ISL_482477 | Queen Elizabeth II Health Science Centre | National Microbiology Laboratory | Anna Majer, Shari Tyson, Grace Seo, Kristyn Burak, Philip Mabon, Elsie Grudeski, Rhiannon Huzarewich, Russell Mandes, Jennifer Tanner, Natalie Knox, Morag Graham, Gary Van Domselaar, Todd Hatchette, Jason LeBlanc, Nathalie Bastien, Yan Li, Timothy Booth |
| EPI_ISL_482478 | Cadham Provincial Laboratory | National Microbiology Laboratory | Anna Majer, Shari Tyson, Grace Seo, Kristyn Burak, Philip Mabon, Elsie Grudeski, Rhiannon Huzarewich, Russell Mandes, Jennifer Tanner, Natalie Knox, Morag Graham, Gary Van Domselaar, Paul Van Caeseele, Jared Bullard, David Alexander, Kerry Dust, Nathalie Bastien, Yan Li, Timothy Booth, |
| EPI_ISL_482479 | Public Health Laboratory | National Microbiology Laboratory | Anna Majer, Shari Tyson, Grace Seo, Kristyn Burak, Philip Mabon, Elsie Grudeski, Rhiannon Huzarewich, Russell Mandes, Jennifer Tanner, Natalie Knox, Morag Graham, Gary Van Domselaar, Robert Needle, Yang Yu, Adel Malek, Laura Gilbert, George Zahariadis, Nathalie Bastien, Yan Li, Timothy Booth |
| EPI_ISL_482480, EPI_ISL_482481, EPI_ISL_482482, EPI_ISL_482483, EPI_ISL_482484 | Cadham Provincial Laboratory | National Microbiology Laboratory | Anna Majer, Shari Tyson, Grace Seo, Kristyn Burak, Philip Mabon, Elsie Grudeski, Rhiannon Huzarewich, Russell Mandes, Jennifer Tanner, Natalie Knox, Morag Graham, Gary Van Domselaar, Paul Van Caeseele, Jared Bullard, David Alexander, Kerry Dust, Nathalie Bastien, Yan Li, Timothy Booth, |
| EPI_ISL_482485, EPI_ISL_482486, EPI_ISL_482487 | National Institute of Laboratory Medicine and Referral Center | Genomic Research Lab, BCSIR | Abu Sayeed Mohammad Mahmud, Mohammad Samir Uzzaman, Eshrar Osman, Md. Ahasan Habib, Shahina Akter, Tanjina Akhter Banu, Md. Murshed Hasan Sarkar, Barna Goswami, Iffat Jahan, Md. Saddam Hossain, Tasnim Nafisa, Md. Maruf Ahmed Molla, Mahmuda Yeasmin, Asish Kumar Ghosh, Shahjahan Siddike, A. K. M. Shamsuzzaman, Sheikh Md. Selim Al Din, Utpal Chandra Ray, Salek Ahmed Sajib, Md. Salim Khan |
| EPI_ISL_482488 | National Institute of Laboratory Medicine and Referral Center | Genomic Research Lab, BCSIR | Md. Murshed Hasan Sarkar, Abu Sayeed Mohammad Mahmud, Mohammad Samir Uzzaman, Eshrar Osman, Md. Ahasan Habib, Shahina Akter, Tanjina Akhter Banu, Barna Goswami, Iffat Jahan, Md. Saddam Hossain, Tasnim Nafisa, Md. Maruf Ahmed Molla, Mahmuda Yeasmin, Asish Kumar Ghosh, Shahjahan Siddike, A. K. M. Shamsuzzaman, Sheikh Md. Selim Al Din, Utpal Chandra Ray, Salek Ahmed Sajib, Md. Salim Khan |
| EPI_ISL_482489 | National Institute of Laboratory Medicine and Referral Center | Genomic Research Lab, BCSIR | Md. Ahasan Habib, Abu Sayeed Mohammad Mahmud, Mohammad Samir Uzzaman, Eshrar Osman, Shahina Akter, Tanjina Akhter Banu, Md. Murshed Hasan Sarkar, Barna Goswami, Iffat Jahan, Md. Saddam Hossain, Tasnim Nafisa, Md. Maruf Ahmed Molla, Mahmuda Yeasmin, Asish Kumar Ghosh, Shahjahan Siddike, A. K. M. Shamsuzzaman, Sheikh Md. Selim Al Din, Utpal Chandra Ray, Salek Ahmed Sajib, Md. Salim Khan |
| EPI_ISL_482491, EPI_ISL_482492, EPI_ISL_482493, EPI_ISL_482494, EPI_ISL_482495, EPI_ISL_482496, EPI_ISL_482497, EPI_ISL_482498, EPI_ISL_482499, EPI_ISL_482500, EPI_ISL_482501, EPI_ISL_482502, EPI_ISL_482503, EPI_ISL_482504, EPI_ISL_482505, EPI_ISL_482506, EPI_ISL_482507, EPI_ISL_482508, EPI_ISL_482509, EPI_ISL_482510, EPI_ISL_482511, EPI_ISL_482512, EPI_ISL_482513, EPI_ISL_482514, EPI_ISL_482515, EPI_ISL_482516, EPI_ISL_482517, EPI_ISL_482518, EPI_ISL_482519, EPI_ISL_482520, EPI_ISL_482521, EPI_ISL_482522, EPI_ISL_482523, EPI_ISL_482524, EPI_ISL_482525, EPI_ISL_482526, EPI_ISL_482527, EPI_ISL_482528, EPI_ISL_482529, EPI_ISL_482530, EPI_ISL_482531, EPI_ISL_482532, EPI_ISL_482533, EPI_ISL_482534, EPI_ISL_482535, EPI_ISL_482536, EPI_ISL_482537, EPI_ISL_482538, EPI_ISL_482539, EPI_ISL_482540, EPI_ISL_482541, EPI_ISL_482542, EPI_ISL_482543, EPI_ISL_482544, EPI_ISL_482545, EPI_ISL_482546, EPI_ISL_482547, EPI_ISL_482548, EPI_ISL_482549, EPI_ISL_482550, EPI_ISL_482551, EPI_ISL_482552, EPI_ISL_482553, EPI_ISL_482554, EPI_ISL_482555, EPI_ISL_482556, EPI_ISL_482557, EPI_ISL_482558, EPI_ISL_482559, EPI_ISL_482560, EPI_ISL_482561, EPI_ISL_482562, EPI_ISL_482563, EPI_ISL_482564, EPI_ISL_482565, EPI_ISL_482566, EPI_ISL_482567, EPI_ISL_482568, EPI_ISL_482569, EPI_ISL_482570, EPI_ISL_482571, EPI_ISL_482572, EPI_ISL_482573, EPI_ISL_482574 | | | |
| see above | National Centre for Disease control (NCDC) | NCDC/CSIR-IGIB | Pramod Kumar#, Rajesh Pandey#, Pooja Sharma, Mahesh S Dhar, Vivekanand A, Bharathram Uppili, Robin Marwal, Radhakrishanan VS, Saruchi Wadhwa, Nishu Tyagi, Uma Sharma, Priyanka Singh, Hemlata Lall, Meena Datta, Varun Jaiswal, Hema Gogia, Preeti Madan, Prateek Singh, Debasis Dash, Mitali Mukerji, Sandhya Kabra, Sujeet Singh, Mohammed Faruq, Anurag Agrawal*, Partha Rakshit* |
| EPI_ISL_482575, EPI_ISL_482576, EPI_ISL_482577, EPI_ISL_482578, EPI_ISL_482579, EPI_ISL_482580, EPI_ISL_482581, EPI_ISL_482582, EPI_ISL_482583, EPI_ISL_482584, EPI_ISL_482585, EPI_ISL_482586 | | | |
| see above | Hangzhou Center for Diseases Control and Prevention | Hangzhou Center for Diseases Control and Prevention | Jun Li, Haoqiu Wang, Lingfeng Mao, Hua Yu, Xinfen Yu, Zhou Sun, Xin Qian, Shuchang Chen, Junfang Chen, Xuchu Wang |
| EPI_ISL_482587, EPI_ISL_482588, EPI_ISL_482589, EPI_ISL_482590, EPI_ISL_482591, EPI_ISL_482592, EPI_ISL_482593, EPI_ISL_482594, EPI_ISL_482595, EPI_ISL_482596, EPI_ISL_482597, EPI_ISL_482598, EPI_ISL_482599, EPI_ISL_482600, EPI_ISL_482601, EPI_ISL_482602, EPI_ISL_482603, EPI_ISL_482604, EPI_ISL_482605, EPI_ISL_482606, EPI_ISL_482607, EPI_ISL_482608, EPI_ISL_482609, EPI_ISL_482610, EPI_ISL_482611, EPI_ISL_482612, EPI_ISL_482613, EPI_ISL_482614, EPI_ISL_482615, EPI_ISL_482616, EPI_ISL_482617, EPI_ISL_482618, EPI_ISL_482619, EPI_ISL_482620, EPI_ISL_482621, EPI_ISL_482622, EPI_ISL_482623, EPI_ISL_482624, EPI_ISL_482625, EPI_ISL_482626, EPI_ISL_482627, EPI_ISL_482628, EPI_ISL_482629, EPI_ISL_482630, EPI_ISL_482631, EPI_ISL_482632, EPI_ISL_482633, EPI_ISL_482634, EPI_ISL_482635, EPI_ISL_482636, EPI_ISL_482637, EPI_ISL_482638, EPI_ISL_482639, EPI_ISL_482640, EPI_ISL_482641, EPI_ISL_482642, EPI_ISL_482643, EPI_ISL_482644, EPI_ISL_482645, EPI_ISL_482646, EPI_ISL_482647, EPI_ISL_482648, EPI_ISL_482649, EPI_ISL_482650, EPI_ISL_482651, EPI_ISL_482652, EPI_ISL_482653, EPI_ISL_482654, EPI_ISL_482655, EPI_ISL_482656, EPI_ISL_482657, EPI_ISL_482658, EPI_ISL_482659, EPI_ISL_482660, EPI_ISL_482661, EPI_ISL_482662, EPI_ISL_482663, EPI_ISL_482664, EPI_ISL_482665, EPI_ISL_482666, EPI_ISL_482667, EPI_ISL_482668, EPI_ISL_482669, EPI_ISL_482670, EPI_ISL_482671 | | | |
| see above | National Centre for Disease control (NCDC) | NCDC/CSIR-IGIB | Pramod Kumar#, Rajesh Pandey#, Pooja Sharma, Mahesh S Dhar, Vivekanand A, Bharathram Uppili, Robin Marwal, Radhakrishanan VS, Saruchi Wadhwa, Nishu Tyagi, Uma Sharma, Priyanka Singh, Hemlata Lall, Meena Datta, Varun Jaiswal, Hema Gogia, Preeti Madan, Prateek Singh, Debasis Dash, Mitali Mukerji, Sandhya Kabra, Sujeet Singh, Mohammed Faruq, Anurag Agrawal*, Partha Rakshit* |
| EPI_ISL_482672, EPI_ISL_482673, EPI_ISL_482674, EPI_ISL_482675, EPI_ISL_482676, EPI_ISL_482677, EPI_ISL_482678, EPI_ISL_482679, EPI_ISL_482680, EPI_ISL_482681, EPI_ISL_482682, EPI_ISL_482683, EPI_ISL_482684, EPI_ISL_482685, EPI_ISL_482686, EPI_ISL_482687, EPI_ISL_482688, EPI_ISL_482689, EPI_ISL_482690, EPI_ISL_482691, EPI_ISL_482692, EPI_ISL_482693, EPI_ISL_482694, EPI_ISL_482695, EPI_ISL_482696, EPI_ISL_482697, EPI_ISL_482698, EPI_ISL_482699 | | | |
| see above | Singapore General Hospital | Department of Microbiology | Nurdyana Abdul Rahman, Kun Lee Lim, Chenhao Li, Kian Sing Chan, Lynette Oon, Kern Rei Chng, Niranjan Nagarajan, Karrie Ko |
| EPI_ISL_482700, EPI_ISL_482701 | National Institute of Laboratory Medicine and Referral Center | Genomic Research Lab, BCSIR | Abu Sayeed Mohammad Mahmud, Mohammad Samir Uzzaman, Eshrar Osman, Md. Ahasan Habib, Shahina Akter, Tanjina Akhter Banu, Md. Murshed Hasan Sarkar, Barna Goswami, Iffat Jahan, Md. Saddam Hossain, Tasnim Nafisa, Md. Maruf Ahmed Molla, Mahmuda Yeasmin, Asish Kumar Ghosh, Shahjahan Siddike, A. K. M. Shamsuzzaman, Sheikh Md. Selim Al Din, Utpal Chandra Ray, Salek Ahmed Sajib, Md. Salim Khan |
| EPI_ISL_482702, EPI_ISL_482703, EPI_ISL_482704, EPI_ISL_482705, EPI_ISL_482706, EPI_ISL_482707, EPI_ISL_482708, EPI_ISL_482709 | Molecular Diagnostics Services (MDS) | KRISP, KZN Research Innovation and Sequencing Platform | Giandhari J, Pillay S, Lessells R, Chimukangara B, Mdlalose K, York D, Khan S, Tegally H, Wilkinson E, de Oliveira T |
| EPI_ISL_482710, EPI_ISL_482711, EPI_ISL_482712, EPI_ISL_482713 | NHLS-IALCH | KRISP, KZN Research Innovation and Sequencing Platform | Giandhari J, Pillay S, Lessells R, Chimukangara B, Mdlalose K, York D, Khan S, Tegally H, Wilkinson E, de Oliveira T |
| EPI_ISL_482714, EPI_ISL_482715, EPI_ISL_482716, EPI_ISL_482717, EPI_ISL_482718, EPI_ISL_482719, EPI_ISL_482720, EPI_ISL_482721, EPI_ISL_482722, EPI_ISL_482723 | Molecular Diagnostics Services (MDS) | KRISP, KZN Research Innovation and Sequencing Platform | Giandhari J, Pillay S, Lessells R, Chimukangara B, Mdlalose K, York D, Khan S, Tegally H, Wilkinson E, de Oliveira T |
| EPI_ISL_482724, EPI_ISL_482725, EPI_ISL_482726, EPI_ISL_482727, EPI_ISL_482728, EPI_ISL_482729, EPI_ISL_482730, EPI_ISL_482731 | NHLS-IALCH | KRISP, KZN Research Innovation and Sequencing Platform | Giandhari J, Pillay S, Lessells R, Chimukangara B, Mdlalose K, York D, Khan S, Tegally H, Wilkinson E, de Oliveira T |
| EPI_ISL_482732, EPI_ISL_482733, EPI_ISL_482734, EPI_ISL_482735, EPI_ISL_482736, EPI_ISL_482737, EPI_ISL_482738, EPI_ISL_482739, EPI_ISL_482740 | LNR National Reference Laboratory, Mohammed VI University of Health Sciences | Medical Biotechnology Laboratory, Rabat Medical and Pharmacy School, Mohammed The Vth University in Rabat | Meriem LAAMARTI, Souad KARTTI, Rokia LAAMARTI , M.W. CHEMAO-ELFIHRI, Loubna ALLAM, Mouna OUADGHIRI, Imane SMYEJ, Jalila RAHOUI, Houda BENRAHMA, Jalil El ATAR, Idrissa DIAWARA, Rachid EL JAOUDI, Laila SBABOU, Chakib NEJJARI, Saaid AMZAZI, Rachid MENTAG, Lahcen BELYAMANI and Azeddine IBRAHIMI |
| EPI_ISL_482742 | Molecular and Cell Biology, Globe Biotech Limited | Molecular and Cell Biology, Globe Biotech Limited | Baray,J.C., Mahmud,A., Khan,M.R., Chowdhury,M.M.H., Roy,R., Islam,F., Nag,K. and Sultana,N. |
| EPI_ISL_482744 | Laboratory Diagnostic, Veterinary Specialized Institute Kraljevo | Laboratory Diagnostic, Veterinary Specialized Institute Kraljevo | Vidanovic,D., Tesovic,B., Banovic Djeri,B., Knezevic,A., Vidanovic,D., Tesovic,B., Banovic Djeri,B., Knezevic,A., Afonso,C. |
| EPI_ISL_482745 | Medical Microbiology, Leiden University Medical Center | Medical Microbiology, Leiden University Medical Center | Snijder,E.J., Ogando,N.S., Zevenhoven,J.C., Dalebout,T.J., de Vries,J.C. and Sidorov,I. |
| EPI_ISL_482746 | Medical Microbiology, Leiden University Medical Center | Medical Microbiology, Leiden University Medical Center | Snijder,E.J., Ogando,N.S., Zevenhoven,J.C., Dalebout,T.J., de Vries,J.J. and Sidorov,I. |
| EPI_ISL_482759, EPI_ISL_482760, EPI_ISL_482761, EPI_ISL_482762, EPI_ISL_482763, EPI_ISL_482764, EPI_ISL_482765, EPI_ISL_482766, EPI_ISL_482767, EPI_ISL_482768, EPI_ISL_482769, EPI_ISL_482770, EPI_ISL_482771, EPI_ISL_482772, EPI_ISL_482773, EPI_ISL_482774, EPI_ISL_482775 | | | |
| see above | Medical Ain Shams Research Institute (MASRI), Ain Shams University | Medical Ain Shams Research Institute (MASRI), Ain Shams University | Hesham Elghazaly, Sara Hassan Agwa, Ahmad Moustafa, Hala Hafez, Sara Elnakeep, Shaimaa Moustafa, Aya Mohamed, Reham Mamdouh, Ghada Ismael, Ashraf Omar, Osama Mansour, Mahmoud Elmeitini |
| EPI_ISL_482777 | Queen Elizabeth Hospital | Hong Kong Department of Health | Mak Gannon C.K., Cheng Peter K.C., Lam Edman T.K., Chan Rickjason C.W., Tsang Dominic N.C. |
| EPI_ISL_482778 | Tuen Mun Hospital | Hong Kong Department of Health | Mak Gannon C.K., Cheng Peter K.C., Lam Edman T.K., Chan Rickjason C.W., Tsang Dominic N.C. |
| EPI_ISL_482779, EPI_ISL_482780 | Prince of Wales Hospital | Hong Kong Department of Health | Mak Gannon C.K., Cheng Peter K.C., Lam Edman T.K., Chan Rickjason C.W., Tsang Dominic N.C. |
| EPI_ISL_482781 | Centre for Health Protection | Hong Kong Department of Health | Mak Gannon C.K., Cheng Peter K.C., Lam Edman T.K., Chan Rickjason C.W., Tsang Dominic N.C. |
| EPI_ISL_482782, EPI_ISL_482783 | Tuen Mun Hospital | Hong Kong Department of Health | Mak Gannon C.K., Cheng Peter K.C., Lam Edman T.K., Chan Rickjason C.W., Tsang Dominic N.C. |
| EPI_ISL_482784 | Prince of Wales Hospital | Hong Kong Department of Health | Mak Gannon C.K., Cheng Peter K.C., Lam Edman T.K., Chan Rickjason C.W., Tsang Dominic N.C. |
| EPI_ISL_482820 | Centre de Recerca en Sanitat Animal (IRTA-CReSA) | IrsiCaixa AIDS Research Lab | J. Segalés, M. Puig, J. Rodon, C. Avila-Nieto, J. Carrillo, G. Cantero, M.T. Terrón, S. Cruz, M. Parera ,M. Noguera-Julián, N. Izquierdo-Useros, V. Guallar, E. Vidal, A. Valencia, I. Blanco, J. Blanco, B. Clotet, J. Vergara-Alert |
| EPI_ISL_482848, EPI_ISL_482849, EPI_ISL_482850 | NHLS-IALCH | KRISP, KZN Research Innovation and Sequencing Platform | Giandhari J, Pillay S, Lessells R, Chimukangara B, Mdlalose K, York D, Khan S, Tegally H, Wilkinson E, de Oliveira T |
| EPI_ISL_482851, EPI_ISL_482852, EPI_ISL_482853, EPI_ISL_482854, EPI_ISL_482855, EPI_ISL_482856, EPI_ISL_482857, EPI_ISL_482858, EPI_ISL_482859, EPI_ISL_482860, EPI_ISL_482861, EPI_ISL_482862, EPI_ISL_482863, EPI_ISL_482864, EPI_ISL_482865, EPI_ISL_482866, EPI_ISL_482867, EPI_ISL_482868, EPI_ISL_482869, EPI_ISL_482870, EPI_ISL_482871, EPI_ISL_482872 | | | |
| see above | Molecular Diagnostics Services (MDS) | KRISP, KZN Research Innovation and Sequencing Platform | Giandhari J, Pillay S, Lessells R, Chimukangara B, Mdlalose K, York D, Khan S, Tegally H, Wilkinson E, de Oliveira T |
| EPI_ISL_482874, EPI_ISL_482875, EPI_ISL_482876, EPI_ISL_482877, EPI_ISL_482878 | Institut Pasteur Dakar | Institut Pasteur de Dakar | Ndongo Dia, Moussa Moise Diagne, Mamadou Diop, Marie Henriette Dior Ndione, Mamadou malado Jallow, Safietou Sankhe, Ousmane Faye, Amadou Alpha Sall. |
| EPI_ISL_482879, EPI_ISL_482880, EPI_ISL_482881, EPI_ISL_482882, EPI_ISL_482883, EPI_ISL_482884, EPI_ISL_482885, EPI_ISL_482886, EPI_ISL_482887, EPI_ISL_482888, EPI_ISL_482889 | | | |
| see above | CHU Purpan - Laboratoire de Virologie - Institut Fédératif de Biologie | Laboratoire de virologie - École Nationale Vétérinaire de Toulouse | Guillaume Croville, Jean-Luc Guérin, Jacques Izopet |
| EPI_ISL_482946, EPI_ISL_482947, EPI_ISL_482948, EPI_ISL_482949, EPI_ISL_482950, EPI_ISL_482951, EPI_ISL_482952, EPI_ISL_482953, EPI_ISL_482954, EPI_ISL_482955, EPI_ISL_482956, EPI_ISL_482957, EPI_ISL_482958, EPI_ISL_482959, EPI_ISL_482960, EPI_ISL_482961, EPI_ISL_482962, EPI_ISL_482963, EPI_ISL_482964, EPI_ISL_482965, EPI_ISL_482966 | | | |
| see above | Minnesota Department of Health, Public Health | Minnesota Department of Health, Public | Matt Plumb, Jacob Garfin, and Xiong Wang |

| | | | |
|---|---|---|---|
| | Laboratory | Health Laboratory | |
| EPI_ISL_482967, EPI_ISL_482968, EPI_ISL_482969, EPI_ISL_482970, EPI_ISL_482971, EPI_ISL_482972, EPI_ISL_482973, EPI_ISL_482974, EPI_ISL_482975, EPI_ISL_482976, EPI_ISL_482977, EPI_ISL_482978, EPI_ISL_482979, EPI_ISL_482980, EPI_ISL_482981, EPI_ISL_482982, EPI_ISL_482983, EPI_ISL_482984, EPI_ISL_482985, EPI_ISL_482986, EPI_ISL_482987 | | | |
| see above | Mayo Clinic & Mayo Clinic Laboratories | Minnesota Department of Health, Public Health Laboratory | Matt Plumb, Jacob Garfin, and Xiong Wang |
| EPI_ISL_482988, EPI_ISL_482989, EPI_ISL_482990, EPI_ISL_482991, EPI_ISL_482992, EPI_ISL_482993, EPI_ISL_482994, EPI_ISL_482995, EPI_ISL_482996, EPI_ISL_482997, EPI_ISL_482998, EPI_ISL_482999, EPI_ISL_483000, EPI_ISL_483001, EPI_ISL_483002, EPI_ISL_483003, EPI_ISL_483004, EPI_ISL_483005, EPI_ISL_483006, EPI_ISL_483007, EPI_ISL_483008, EPI_ISL_483009, EPI_ISL_483010, EPI_ISL_483011, EPI_ISL_483012, EPI_ISL_483013, EPI_ISL_483014, EPI_ISL_483015, EPI_ISL_483016, EPI_ISL_483017 | | | |
| see above | Minnesota Department of Health, Public Health Laboratory | Minnesota Department of Health, Public Health Laboratory | Matt Plumb, Jacob Garfin, and Xiong Wang |
| EPI_ISL_483018, EPI_ISL_483019, EPI_ISL_483020, EPI_ISL_483021, EPI_ISL_483022, EPI_ISL_483023, EPI_ISL_483024, EPI_ISL_483025, EPI_ISL_483026, EPI_ISL_483027, EPI_ISL_483028, EPI_ISL_483029, EPI_ISL_483030, EPI_ISL_483031, EPI_ISL_483032, EPI_ISL_483033 | | | |
| see above | Utah Public Health Laboratory | Utah Public Health Laboratory | Heidi Butz, Erin Young, Kelly Oakeson |
| EPI_ISL_483035, EPI_ISL_483036, EPI_ISL_483037, EPI_ISL_483038 | Medical Ain Shams Research Institute (MASRI), Ain Shams University | Medical Ain Shams Research Institute (MASRI), Ain Shams University | Hesham Elghazaly, Sara Hassan Agwa, Ahmad Moustafa, Hala Hafez, Sara Elnakeep, Shaimaa Moustafa, Aya Mohamed, Reham Mamdouh, Ghada Ismael, Ashraf Omar, Osama Mansour, Mahmoud Elmeitini |
| EPI_ISL_483059 | Hospital Universitari Germans Trias i Pujol | IrsiCaixa AIDS Research Lab | J. Segalés, M. Puig, J. Rodon, C. Avila-Nieto, J. Carrillo, G. Cantero, M.T. Terrón, S. Cruz, M. Parera ,M. Noguera-Julián, N. Izquierdo-Useros, V. Guallar, E. Vidal, A. Valencia, I. Blanco, J. Blanco, B. Clotet, J. Vergara-Alert |
| EPI_ISL_483060 | unknown | Microbiology, Canterbury Health Laboratories | Dilcher,M., Anderson,T. |
| EPI_ISL_483061, EPI_ISL_483062 | unknown | National Reference Center for transfusion infectious risks, INTS | Cappy,P., Candotti,D., Sauvage,V., Lucas,Q., Boizeau,L., Gomez,J., Laperche,S. |
| EPI_ISL_483063, EPI_ISL_483064 | unknown | Virology, Ecole Nationale Veterinaire de Toulouse | Bessiere,P., Cadiergues,M.-C., Croville,G., Walch,M., Dubois,M., Izopet,J., Guerin,J.-L. |
| EPI_ISL_483065 | Centro de Desenvolvimento Tecnologico em Saude, Fundacao Oswaldo Cruz | Centro de Desenvolvimento Tecnologico em Saude, Fundacao Oswaldo Cruz | Souza,T.M., Fintelman-Rodrigues,N., De Paula,A.D., Tschoeke,D., Barroso,S.P., Gregorio,M.L., Oliveira,J.S., Saraiva,F.B., Ferreira,M.A., Sacramento,C.Q. |
| EPI_ISL_483066, EPI_ISL_483067, EPI_ISL_483068, EPI_ISL_483069, EPI_ISL_483070, EPI_ISL_483071, EPI_ISL_483072, EPI_ISL_483073, EPI_ISL_483074, EPI_ISL_483075, EPI_ISL_483076, EPI_ISL_483077, EPI_ISL_483078, EPI_ISL_483079, EPI_ISL_483080, EPI_ISL_483081, EPI_ISL_483082, EPI_ISL_483083, EPI_ISL_483084, EPI_ISL_483085, EPI_ISL_483086, EPI_ISL_483087, EPI_ISL_483088, EPI_ISL_483089, EPI_ISL_483090, EPI_ISL_483091, EPI_ISL_483092, EPI_ISL_483093, EPI_ISL_483094, EPI_ISL_483095, EPI_ISL_483096, EPI_ISL_483097, EPI_ISL_483098, EPI_ISL_483099, EPI_ISL_483100, EPI_ISL_483101, EPI_ISL_483102, EPI_ISL_483103, EPI_ISL_483104, EPI_ISL_483105, EPI_ISL_483106, EPI_ISL_483107, EPI_ISL_483108, EPI_ISL_483109, EPI_ISL_483110, EPI_ISL_483111, EPI_ISL_483112, EPI_ISL_483113, EPI_ISL_483114, EPI_ISL_483115, EPI_ISL_483116, EPI_ISL_483117, EPI_ISL_483118, EPI_ISL_483119, EPI_ISL_483120, EPI_ISL_483121, EPI_ISL_483122, EPI_ISL_483123, EPI_ISL_483124, EPI_ISL_483125, EPI_ISL_483126, EPI_ISL_483127, EPI_ISL_483128, EPI_ISL_483129, EPI_ISL_483130, EPI_ISL_483131, EPI_ISL_483132, EPI_ISL_483133, EPI_ISL_483134, EPI_ISL_483135, EPI_ISL_483136, EPI_ISL_483137, EPI_ISL_483138 | | | |
| see above | SA Pathology | SA Pathology | Lex Leong, Chuan Kok Lim, Mark Turra, Ivan Bastian, Geoff Higgins |
| EPI_ISL_483139, EPI_ISL_483140, EPI_ISL_483141, EPI_ISL_483142, EPI_ISL_483143, EPI_ISL_483144, EPI_ISL_483145, EPI_ISL_483146, EPI_ISL_483147, EPI_ISL_483148, EPI_ISL_483149, EPI_ISL_483150, EPI_ISL_483151, EPI_ISL_483152, EPI_ISL_483153, EPI_ISL_483154, EPI_ISL_483155, EPI_ISL_483156, EPI_ISL_483157 | | | |
| see above | Robert Koch Institute, ZBS1 Highly Pathogenic Viruses, Berlin, Germany | Robert Koch Institute, Bioinformatics MF1, Berlin, Germany | Janine Michel, Andrea Thuermer, Oliver Drechsel, Rene Kmiecinski, Stephan Fuchs, Max v. Kleist, Andreas Nitsche |
| EPI_ISL_483158, EPI_ISL_483159, EPI_ISL_483160, EPI_ISL_483161, EPI_ISL_483162, EPI_ISL_483163, EPI_ISL_483164 | San Diego County Public Health Laboratory | Andersen lab at Scripps Research | SEARCH Alliance San Diego with Tracy Basler, Jovan Shephard, Brett Austin |
| EPI_ISL_483165, EPI_ISL_483166, EPI_ISL_483167, EPI_ISL_483168, EPI_ISL_483169, EPI_ISL_483170, EPI_ISL_483171, EPI_ISL_483172, EPI_ISL_483173, EPI_ISL_483174, EPI_ISL_483175, EPI_ISL_483176, EPI_ISL_483177, EPI_ISL_483178, EPI_ISL_483179, EPI_ISL_483180, EPI_ISL_483181, EPI_ISL_483182, EPI_ISL_483183, EPI_ISL_483184, EPI_ISL_483185, EPI_ISL_483186, EPI_ISL_483187, EPI_ISL_483188, EPI_ISL_483189, EPI_ISL_483190, EPI_ISL_483191, EPI_ISL_483192, EPI_ISL_483193, EPI_ISL_483194, EPI_ISL_483195, EPI_ISL_483196, EPI_ISL_483197, EPI_ISL_483198, EPI_ISL_483199, EPI_ISL_483200, EPI_ISL_483201, EPI_ISL_483202, EPI_ISL_483203, EPI_ISL_483204, EPI_ISL_483205, EPI_ISL_483206, EPI_ISL_483207, EPI_ISL_483208, EPI_ISL_483209, EPI_ISL_483210, EPI_ISL_483211, EPI_ISL_483212, EPI_ISL_483213, EPI_ISL_483214, EPI_ISL_483215, EPI_ISL_483216, EPI_ISL_483217, EPI_ISL_483218, EPI_ISL_483219, EPI_ISL_483220, EPI_ISL_483221, EPI_ISL_483222, EPI_ISL_483223, EPI_ISL_483224, EPI_ISL_483225, EPI_ISL_483226, EPI_ISL_483227, EPI_ISL_483228, EPI_ISL_483229, EPI_ISL_483230, EPI_ISL_483231, EPI_ISL_483232, EPI_ISL_483233, EPI_ISL_483234, EPI_ISL_483235, EPI_ISL_483236, EPI_ISL_483237, EPI_ISL_483238, EPI_ISL_483239, EPI_ISL_483240, EPI_ISL_483241, EPI_ISL_483242, EPI_ISL_483243, EPI_ISL_483244, EPI_ISL_483245, EPI_ISL_483246, EPI_ISL_483247, EPI_ISL_483248, EPI_ISL_483249, EPI_ISL_483250, EPI_ISL_483251, EPI_ISL_483252, EPI_ISL_483253, EPI_ISL_483254, EPI_ISL_483255, EPI_ISL_483256, EPI_ISL_483257, EPI_ISL_483258, EPI_ISL_483259, EPI_ISL_483260, EPI_ISL_483261, EPI_ISL_483262, EPI_ISL_483263, EPI_ISL_483264, EPI_ISL_483265, EPI_ISL_483266, EPI_ISL_483267, EPI_ISL_483268, EPI_ISL_483269, EPI_ISL_483270, EPI_ISL_483271, EPI_ISL_483272, EPI_ISL_483273, EPI_ISL_483274, EPI_ISL_483275, EPI_ISL_483276, EPI_ISL_483277, EPI_ISL_483278, EPI_ISL_483279, EPI_ISL_483280, EPI_ISL_483281, EPI_ISL_483282, EPI_ISL_483283, EPI_ISL_483284, EPI_ISL_483285, EPI_ISL_483286, EPI_ISL_483287, EPI_ISL_483288, EPI_ISL_483289, EPI_ISL_483290, EPI_ISL_483291, EPI_ISL_483292, EPI_ISL_483293, EPI_ISL_483294, EPI_ISL_483295, EPI_ISL_483296, EPI_ISL_483297, EPI_ISL_483298, EPI_ISL_483299, EPI_ISL_483300, EPI_ISL_483301, EPI_ISL_483302, EPI_ISL_483303, EPI_ISL_483304, EPI_ISL_483305, EPI_ISL_483306, EPI_ISL_483307, EPI_ISL_483308, EPI_ISL_483309, EPI_ISL_483310, EPI_ISL_483311, EPI_ISL_483312, EPI_ISL_483313, EPI_ISL_483314, EPI_ISL_483315, EPI_ISL_483316, EPI_ISL_483317, EPI_ISL_483318, EPI_ISL_483319, EPI_ISL_483320, EPI_ISL_483321, EPI_ISL_483322, EPI_ISL_483323, EPI_ISL_483324, EPI_ISL_483325, EPI_ISL_483326, EPI_ISL_483327, EPI_ISL_483328, EPI_ISL_483329, EPI_ISL_483330, EPI_ISL_483331, EPI_ISL_483332, EPI_ISL_483333, EPI_ISL_483334, EPI_ISL_483335, EPI_ISL_483336, EPI_ISL_483337, EPI_ISL_483338, EPI_ISL_483339, EPI_ISL_483340, EPI_ISL_483341, EPI_ISL_483342, EPI_ISL_483343, EPI_ISL_483344, EPI_ISL_483345, EPI_ISL_483346, EPI_ISL_483347, EPI_ISL_483348, EPI_ISL_483349, EPI_ISL_483350, EPI_ISL_483351, EPI_ISL_483352, EPI_ISL_483353, EPI_ISL_483354, EPI_ISL_483355, EPI_ISL_483356, EPI_ISL_483357, EPI_ISL_483358, EPI_ISL_483359, EPI_ISL_483360, EPI_ISL_483361, EPI_ISL_483362, EPI_ISL_483363, EPI_ISL_483364, EPI_ISL_483365, EPI_ISL_483366, EPI_ISL_483367, EPI_ISL_483368, EPI_ISL_483369, EPI_ISL_483370, EPI_ISL_483371, EPI_ISL_483372, EPI_ISL_483373, EPI_ISL_483374, EPI_ISL_483375, EPI_ISL_483376, EPI_ISL_483377, EPI_ISL_483378, EPI_ISL_483379, EPI_ISL_483380, EPI_ISL_483381, EPI_ISL_483382, EPI_ISL_483383, EPI_ISL_483384, EPI_ISL_483385, EPI_ISL_483386, EPI_ISL_483387, EPI_ISL_483388, EPI_ISL_483389, EPI_ISL_483390, EPI_ISL_483391, EPI_ISL_483392, EPI_ISL_483393, EPI_ISL_483394, EPI_ISL_483395, EPI_ISL_483396, EPI_ISL_483397 | | | |
| see above | UC San Diego Center for Advanced Laboratory Medicine | Andersen lab at Scripps Research | SEARCH Alliance San Diego with David Pride, Ji H Shin |
| EPI_ISL_483398, EPI_ISL_483399, EPI_ISL_483400, EPI_ISL_483401, EPI_ISL_483402 | UC San Diego Center for Advanced Laboratory Medicine | Andersen lab at Scripps Research | Allison Smither, Gilberto Sabino-Santos, Patricia Snarski, Lilia Melnik, Antoinette Bell, Kaylynn Genemaras, Arnaud Drouin, Dahlene Fusco, Robert Garry with SEARCH Alliance San Diego |
| EPI_ISL_483403, EPI_ISL_483404, EPI_ISL_483405, EPI_ISL_483406, EPI_ISL_483407, EPI_ISL_483408, EPI_ISL_483409, EPI_ISL_483410, EPI_ISL_483411, EPI_ISL_483412, EPI_ISL_483413, EPI_ISL_483414, EPI_ISL_483415, EPI_ISL_483416, EPI_ISL_483417, EPI_ISL_483418, EPI_ISL_483419, EPI_ISL_483420, EPI_ISL_483421, EPI_ISL_483422, EPI_ISL_483423, EPI_ISL_483424, EPI_ISL_483425, EPI_ISL_483426, EPI_ISL_483427, EPI_ISL_483428, EPI_ISL_483429, EPI_ISL_483430, EPI_ISL_483431, EPI_ISL_483432, EPI_ISL_483433, EPI_ISL_483434, EPI_ISL_483435, EPI_ISL_483436, EPI_ISL_483437, EPI_ISL_483438, EPI_ISL_483439, EPI_ISL_483440, EPI_ISL_483441, EPI_ISL_483442, EPI_ISL_483443, EPI_ISL_483444, EPI_ISL_483445, EPI_ISL_483446, EPI_ISL_483447, EPI_ISL_483448, EPI_ISL_483449, EPI_ISL_483450, EPI_ISL_483451, EPI_ISL_483452, EPI_ISL_483453, EPI_ISL_483454, EPI_ISL_483455, EPI_ISL_483456, EPI_ISL_483457, EPI_ISL_483458, EPI_ISL_483459, EPI_ISL_483460, EPI_ISL_483461, EPI_ISL_483462, EPI_ISL_483463, EPI_ISL_483464, EPI_ISL_483465, EPI_ISL_483466, EPI_ISL_483467, EPI_ISL_483468, EPI_ISL_483469, EPI_ISL_483470, EPI_ISL_483471, EPI_ISL_483472, EPI_ISL_483473, EPI_ISL_483474, EPI_ISL_483475, EPI_ISL_483476, EPI_ISL_483477, EPI_ISL_483478, EPI_ISL_483479, EPI_ISL_483480, EPI_ISL_483481, EPI_ISL_483482, EPI_ISL_483483, EPI_ISL_483484, EPI_ISL_483485, EPI_ISL_483486, EPI_ISL_483487, EPI_ISL_483488, EPI_ISL_483489, EPI_ISL_483490, EPI_ISL_483491, EPI_ISL_483492, EPI_ISL_483493, EPI_ISL_483494, EPI_ISL_483495, EPI_ISL_483496, EPI_ISL_483497, EPI_ISL_483498, EPI_ISL_483499, EPI_ISL_483500, EPI_ISL_483501, EPI_ISL_483502, EPI_ISL_483503, EPI_ISL_483504, EPI_ISL_483505, EPI_ISL_483506, EPI_ISL_483507, EPI_ISL_483508, EPI_ISL_483509, EPI_ISL_483510, EPI_ISL_483511, EPI_ISL_483512, EPI_ISL_483513, EPI_ISL_483514, EPI_ISL_483515, EPI_ISL_483516, EPI_ISL_483517, EPI_ISL_483518, EPI_ISL_483519, EPI_ISL_483520, EPI_ISL_483521, EPI_ISL_483522, EPI_ISL_483523, EPI_ISL_483524, EPI_ISL_483525, EPI_ISL_483526, EPI_ISL_483527, EPI_ISL_483528, EPI_ISL_483529, EPI_ISL_483530 | | | |
| see above | UC San Diego Center for Advanced Laboratory Medicine | Andersen lab at Scripps Research | SEARCH Alliance San Diego with David Pride, Ji H Shin |
| EPI_ISL_483531, EPI_ISL_483532, EPI_ISL_483533, EPI_ISL_483534, EPI_ISL_483535, EPI_ISL_483536, EPI_ISL_483537, EPI_ISL_483538, EPI_ISL_483539, EPI_ISL_483540, EPI_ISL_483541 | | | |
| see above | San Diego County Public Health Laboratory | Andersen lab at Scripps Research | SEARCH Alliance San Diego with Tracy Basler, Jovan Shephard, Brett Austin |
| EPI_ISL_483542, EPI_ISL_483543, EPI_ISL_483544, EPI_ISL_483545, EPI_ISL_483546, EPI_ISL_483547, EPI_ISL_483548, EPI_ISL_483549, EPI_ISL_483550, EPI_ISL_483551, EPI_ISL_483552, EPI_ISL_483553, EPI_ISL_483554, EPI_ISL_483555, EPI_ISL_483556, EPI_ISL_483557, EPI_ISL_483558, EPI_ISL_483559, EPI_ISL_483560, EPI_ISL_483561, EPI_ISL_483562, EPI_ISL_483563, EPI_ISL_483564, EPI_ISL_483565 | | | |
| see above | Kingdom of Bahrein Ministry of Health | Erasmus Medical Center | Bas Oude Munnink, David Nieuwenhuijse, Reina Sikkema, Fatema, Ebrahim Shehad, Amjad Ghanem Mohamed, Hashmeya Al Wasti, Claudia Schapendonk, Irina Chestakova, Anne van der Linden, Theo Bestebroer, Stefan van Nieuwkoop, Mark Pronk, Pascal Lexmond, Richard Molenkamp, Marion Koopmans, on behalf of the Dutch national COVID-19 response team. |
| EPI_ISL_483566 | Clinical Microbiology Laboratory- Basurto University Hospital | Biocruces-Bizkaia | Mikel J. Urrutikoetxea-Gutierrez, Ana Belén Belén de la Hoz, Matxalen Vidal-García, Mº Carmen Nieto Toboso, Estibaliz Ugalde-Zarraga, José Luis Díaz de Tuesta del Arco |
| EPI_ISL_483570 | Clinical Microbiology Laboratory- Basurto University Hospita | Biocruces-Bizkaia | Mikel J. Urrutikoetxea-Gutierrez, Ana Belén Belén de la Hoz, Matxalen Vidal-García, Mº Carmen Nieto Toboso, Estibaliz Ugalde-Zarraga, José Luis Díaz de Tuesta del Arco |
| EPI_ISL_483571 | Clinical Microbiology Laboratory- Basurto University Hospital | Biocruces-Bizkaia | Mikel J. Urrutikoetxea-Gutierrez, Ana Belén Belén de la Hoz, Matxalen Vidal-García, Mº Carmen Nieto Toboso, Estibaliz Ugalde-Zarraga, José Luis Díaz de Tuesta del Arco |
| EPI_ISL_483572 | Clinical Microbiology Laboratory- Basurto University Hospital | Biocruces-Bizkaia | Mikel J. Urrutikoetxea-Gutierrez, Ana Belén Belén de la Hoz, Matxalen Vidal-García, Mº Carmen Nieto Toboso, Estibaliz Ugalde-Zarraga, José Luis Díaz de Tuesta del Arco |
| EPI_ISL_483573 | Clinical Microbiology Laboratory- Basurto University Hospital | Biocruces-Bizkaia | Mikel J. Urrutikoetxea-Gutierrez, Ana Belén Belén de la Hoz, Matxalen Vidal-García, Mº Carmen Nieto Toboso, Estibaliz Ugalde-Zarraga, José Luis Díaz de Tuesta del Arco |
| EPI_ISL_483576, EPI_ISL_483577, EPI_ISL_483578, EPI_ISL_483579, EPI_ISL_483580, EPI_ISL_483581, EPI_ISL_483582, EPI_ISL_483583, EPI_ISL_483584, EPI_ISL_483585, EPI_ISL_483586, EPI_ISL_483587, EPI_ISL_483588, EPI_ISL_483589, EPI_ISL_483590, EPI_ISL_483591, EPI_ISL_483592, EPI_ISL_483593, EPI_ISL_483594, EPI_ISL_483595, EPI_ISL_483596, EPI_ISL_483597, EPI_ISL_483598, EPI_ISL_483599, EPI_ISL_483600, EPI_ISL_483601, EPI_ISL_483602, EPI_ISL_483603, EPI_ISL_483604, EPI_ISL_483605, EPI_ISL_483606, EPI_ISL_483607, EPI_ISL_483608, EPI_ISL_483609, EPI_ISL_483610, EPI_ISL_483611, EPI_ISL_483612, EPI_ISL_483613, EPI_ISL_483614, EPI_ISL_483615, EPI_ISL_483616, EPI_ISL_483617, EPI_ISL_483618, EPI_ISL_483619, EPI_ISL_483620, EPI_ISL_483621 | | | |
| see above | National Public Health Laboratory, National Centre for Infectious Diseases | National Public Health Laboratory, National Centre for Infectious Diseases | Mak TM, Octavia S, Zhou Z, Chavatte JM, Cui L, Lin RTP |
| EPI_ISL_483622, EPI_ISL_483623 | National Institute of Laboratory Medicine and Referral Center | Genomic Research Lab, BCSIR | Tasnim Nafisa, Abu Sayeed Mohammad Mahmud, Mohammad Samir Uzzaman, Eshrar Osman, Md. Ahasan Habib, Shahina Akter, Tanjina Akhter Banu, Md. Murshed Hasan Sarkar, Barna Goswami, Iffat Jahan, Md. Saddam Hossain, Md. Maruf Ahmed Molla, Mahmuda Yeasmin, Asish Kumar Ghosh, A. K. M. Shamsuzzaman, Sheikh Md. Selim Al Din, Utpal Chandra Ray, Salek Ahmed Sajib, Md. Salim Khan |
| EPI_ISL_483624 | National Institute of Laboratory Medicine and Referral Center | Genomic Research Lab, BCSIR | Md. Maruf Ahmed Molla, Abu Sayeed Mohammad Mahmud, Mohammad Samir Uzzaman, Eshrar Osman, Md. Ahasan Habib, Shahina Akter, Tanjina Akhter Banu, Md. Murshed Hasan Sarkar, Barna Goswami, Iffat Jahan, Md. Saddam Hossain, Tasnim Nafisa, Mahmuda Yeasmin, Asish Kumar Ghosh, A. K. M. Shamsuzzaman, Sheikh Md. Selim Al Din, Utpal Chandra Ray, Salek Ahmed Sajib, Md. Salim Khan |
| EPI_ISL_483625 | Molecular Microbiology of Laboratory, Institute of Biotechnology, Vietnam Academy of Science and Technology | Molecular Microbiology of Laboratory, Institute of Biotechnology, Vietnam Academy of Science and Technology | Bui Thi,D.T., Nguyen,H.T., Tran,T.X., Nguyen,D.D., Pham,L.T., Vu,H.T., Le Thi,Q.M., Nguyen Le,H.K., Hoang Vu,P.M., Dang,A.D., Dong,Q.V. and Dinh,K.D. |
| EPI_ISL_483626 | National Institute of Laboratory Medicine and Referral Center | Genomic Research Lab, BCSIR | Md. Maruf Ahmed Molla, Abu Sayeed Mohammad Mahmud, Mohammad Samir Uzzaman, Eshrar Osman, Md. Ahasan Habib, Shahina Akter, Tanjina Akhter Banu, Md. Murshed Hasan Sarkar, Barna Goswami, Iffat Jahan, Md. Saddam Hossain, Tasnim Nafisa, Mahmuda Yeasmin, Asish Kumar Ghosh, A. K. M. Shamsuzzaman, Sheikh Md. Selim Al Din, Utpal Chandra Ray, Salek Ahmed Sajib, Md. Salim Khan |
| EPI_ISL_483627, EPI_ISL_483628 | National Institute of Laboratory Medicine and Referral Center | Genomic Research Lab, BCSIR | Mahmuda Yeasmin, Abu Sayeed Mohammad Mahmud, Mohammad Samir Uzzaman, Eshrar Osman, Md. Ahasan Habib, Shahina Akter, Tanjina Akhter Banu, Md. Murshed Hasan Sarkar, Barna Goswami, Iffat Jahan, Md. Saddam Hossain, Tasnim Nafisa, Md. Maruf Ahmed Molla, Asish Kumar Ghosh, A. K. M. Shamsuzzaman, Sheikh Md. Selim Al Din, Utpal Chandra Ray, Salek Ahmed Sajib, Md. Salim Khan |
| EPI_ISL_483629, EPI_ISL_483630 | National Institute of Laboratory Medicine and Referral Center | Genomic Research Lab, BCSIR | Asish Kumar Ghosh, Abu Sayeed Mohammad Mahmud, Mohammad Samir Uzzaman, Eshrar Osman, Md. Ahasan Habib, Shahina Akter, Tanjina Akhter Banu, Md. Murshed Hasan Sarkar, Barna Goswami, Iffat Jahan, Md. Saddam Hossain, Tasnim Nafisa, Md. Maruf Ahmed Molla, Mahmuda Yeasmin, A. K. M. Shamsuzzaman, Sheikh Md. Selim Al Din, Utpal Chandra Ray, Salek Ahmed Sajib, Md. Salim Khan |
| EPI_ISL_483631, EPI_ISL_483632 | National Institute of Laboratory Medicine and Referral Center | Genomic Research Lab, BCSIR | Md. Ahasan Habib, Abu Sayeed Mohammad Mahmud, Mohammad Samir Uzzaman, Eshrar Osman, Shahina Akter, Tanjina Akhter Banu, Md. Murshed Hasan Sarkar, Barna Goswami, Iffat Jahan, Md. Saddam Hossain, Tasnim Nafisa, Md. Maruf Ahmed Molla, Mahmuda Yeasmin, Asish Kumar Ghosh, A. K. M. Shamsuzzaman, Sheikh Md. Selim Al Din, Utpal Chandra Ray, Salek Ahmed Sajib, Md. Salim Khan |
| EPI_ISL_483633, EPI_ISL_483634 | National Institute of Laboratory Medicine and Referral Center | Genomic Research Lab, BCSIR | Shahina Akter, Abu Sayeed Mohammad Mahmud, Mohammad Samir Uzzaman, Eshrar Osman, Md. Ahasan Habib, Tanjina Akhter Banu, Md. Murshed Hasan Sarkar, Barna Goswami, Iffat Jahan, Md. Saddam Hossain, Tasnim Nafisa, Md. Maruf Ahmed Molla, Mahmuda Yeasmin, Asish Kumar Ghosh, A. K. M. Shamsuzzaman, Sheikh Md. Selim Al Din, Utpal Chandra Ray, Salek Ahmed Sajib, Md. Salim Khan |
| EPI_ISL_483635, EPI_ISL_483636 | National Institute of Laboratory Medicine and Referral | Genomic Research Lab, BCSIR | Tanjina Akhter Banu, Abu Sayeed Mohammad Mahmud, Mohammad Samir Uzzaman, Eshrar Osman, Md. Ahasan Habib, Shahina Akter, Md. Murshed Hasan Sarkar, Barna Goswami, Iffat Jahan, Md. Saddam Hossain, Tasnim Nafisa, Md. Maruf Ahmed Molla, |

| | | | |
|---|---|---|---|
| | Center | | Mahmuda Yeasmin, Asish Kumar Ghosh, A. K. M. Shamsuzzaman, Sheikh Md. Selim Al Din, Utpal Chandra Ray, Salek Ahmed Sajib, Md. Salim Khan |
| EPI_ISL_483637 | National Laboratory of Virology, Szentágothai Research Centre | National Laboratory of Virology, Szentágothai Research Centre | Endre Gábor Tóth, Balázs Somogyi, Ferenc Jakab, Gábor Kemenesi |
| EPI_ISL_483638, EPI_ISL_483639, EPI_ISL_483640 | Kingdom of Bahrein Ministry of Health | Erasmus Medical Center | Bas Oude Munnink, David Nieuwenhuijse, Reina Sikkema, Fatema, Ebrahim Shehad, Amjad Ghanem Mohamed, Hashmeya Al Wasti, Claudia Schapendonk, Irina Chestakova, Anne van der Linden, Theo Bestebroer, Stefan van Nieuwkoop, Mark Pronk, Pascal Lexmond, Richard Molenkamp, Marion Koopmans, on behalf of the Dutch national COVID-19 response team. |
| EPI_ISL_483641, EPI_ISL_483642 | National Institute of Laboratory Medicine and Referral Center | Genomic Research Lab, BCSIR | Barna Goswami, Abu Sayeed Mohammad Mahmud, Mohammad Samir Uzzaman, Eshrar Osman, Md. Ahasan Habib, Shahina Akter, Tanjina Akhter Banu, Md. Murshed Hasan Sarkar, Iffat Jahan, Md. Saddam Hossain, Tasnim Nafisa, Md. Maruf Ahmed Molla, Mahmuda Yeasmin, Asish Kumar Ghosh, A. K. M. Shamsuzzaman, Sheikh Md. Selim Al Din, Utpal Chandra Ray, Salek Ahmed Sajib, Md. Salim Khan |
| EPI_ISL_483643, EPI_ISL_483644 | National Institute of Laboratory Medicine and Referral Center | Genomic Research Lab, BCSIR | Iffat Jahan, Abu Sayeed Mohammad Mahmud, Mohammad Samir Uzzaman, Eshrar Osman, Md. Ahasan Habib, Shahina Akter, Tanjina Akhter Banu, Md. Murshed Hasan Sarkar, Barna Goswami, Md. Saddam Hossain, Tasnim Nafisa, Md. Maruf Ahmed Molla, Mahmuda Yeasmin, Asish Kumar Ghosh, A. K. M. Shamsuzzaman, Sheikh Md. Selim Al Din, Utpal Chandra Ray, Salek Ahmed Sajib, Md. Salim Khan |
| EPI_ISL_483645, EPI_ISL_483646, EPI_ISL_483647 | National Institute of Laboratory Medicine and Referral Center | Genomic Research Lab, BCSIR | Md. Saddam Hossain, Abu Sayeed Mohammad Mahmud, Mohammad Samir Uzzaman, Eshrar Osman, Md. Ahasan Habib, Shahina Akter, Tanjina Akhter Banu, Md. Murshed Hasan Sarkar, Barna Goswami, Iffat Jahan, Tasnim Nafisa, Md. Maruf Ahmed Molla, Mahmuda Yeasmin, Asish Kumar Ghosh, A. K. M. Shamsuzzaman, Sheikh Md. Selim Al Din, Utpal Chandra Ray, Salek Ahmed Sajib, Md. Salim Khan |
| EPI_ISL_483648, EPI_ISL_483649, EPI_ISL_483650, EPI_ISL_483651, EPI_ISL_483652, EPI_ISL_483653, EPI_ISL_483654, EPI_ISL_483655, EPI_ISL_483656, EPI_ISL_483657, EPI_ISL_483658, EPI_ISL_483659, EPI_ISL_483660, EPI_ISL_483661, EPI_ISL_483662, EPI_ISL_483663, EPI_ISL_483664, EPI_ISL_483665, EPI_ISL_483666, EPI_ISL_483667 | | | |
| see above | Viollier AG | Department of Biosystems Science and Engineering, ETH Zürich | Christian Beisel, Sarah Nadeau, Ivan Topolsky, Pedro Ferreira, Philipp Jablonski, Susana Posada-Céspedes, Tobias Schär, Ina Nissen, Natascha Santacroce, Elodie Burcklen, Christiane Beckmann, Maurice Redondo, Olivier Kobel, Christoph Noppen, Sophie Seidel, Noemie Santamaria de Souza, Niko Beerenwinkel, Tanja Stadler |
| EPI_ISL_483668, EPI_ISL_483669, EPI_ISL_483670, EPI_ISL_483671, EPI_ISL_483672, EPI_ISL_483673, EPI_ISL_483674, EPI_ISL_483675, EPI_ISL_483676, EPI_ISL_483677, EPI_ISL_483678, EPI_ISL_483679, EPI_ISL_483680, EPI_ISL_483681, EPI_ISL_483682, EPI_ISL_483683, EPI_ISL_483684, EPI_ISL_483685 | | | |
| see above | University Hospital Zurich | Department of Biosystems Science and Engineering, ETH Zürich | Christian Beisel, Sarah Nadeau, Ivan Topolsky, Pedro Ferreira, Philipp Jablonski, Susana Posada-Céspedes, Tobias Schär, Ina Nissen, Natascha Santacroce, Elodie Burcklen, Julia Martinez-Gomez, Phil Cheng, Mitch Levesque, Philipp Bosshard, Niko Beerenwinkel, Tanja Stadler |
| EPI_ISL_483686, EPI_ISL_483687 | National Institute of Laboratory Medicine and Referral Center | Genomic Research Lab, BCSIR | Md. Murshed Hasan Sarkar, Abu Sayeed Mohammad Mahmud, Mohammad Samir Uzzaman, Eshrar Osman, Md. Ahasan Habib, Shahina Akter, Tanjina Akhter Banu, Barna Goswami, Iffat Jahan, Md. Saddam Hossain, Tasnim Nafisa, Md. Maruf Ahmed Molla, Mahmuda Yeasmin, Asish Kumar Ghosh, A. K. M. Shamsuzzaman, Sheikh Md. Selim Al Din, Utpal Chandra Ray, Salek Ahmed Sajib, Md. Salim Khan |
| EPI_ISL_483688 | Genomic Research Lab, BCSIR | Genomic Research Lab, BCSIR | Md. Murshed Hasan Sarkar, Abu Sayeed Mohammad Mahmud, Mohammad Samir Uzzaman, Eshrar Osman, Md. Ahasan Habib, Shahina Akter, Tanjina Akhter Banu, Barna Goswami, Iffat Jahan, Md. Saddam Hossain, Tasnim Nafisa, Md. Maruf Ahmed Molla, Mahmuda Yeasmin, Asish Kumar Ghosh, A. K. M. Shamsuzzaman, Sheikh Md. Selim Al Din, Utpal Chandra Ray, Salek Ahmed Sajib, Md. Salim Khan |
| EPI_ISL_483689, EPI_ISL_483690, EPI_ISL_483691, EPI_ISL_483692 | National Institute of Laboratory Medicine and Referral Center | Genomic Research Lab, BCSIR | Md. Murshed Hasan Sarkar, Abu Sayeed Mohammad Mahmud, Mohammad Samir Uzzaman, Eshrar Osman, Md. Ahasan Habib, Shahina Akter, Tanjina Akhter Banu, Barna Goswami, Iffat Jahan, Md. Saddam Hossain, Tasnim Nafisa, Md. Maruf Ahmed Molla, Mahmuda Yeasmin, Asish Kumar Ghosh, A. K. M. Shamsuzzaman, Sheikh Md. Selim Al Din, Utpal Chandra Ray, Salek Ahmed Sajib, Md. Salim Khan |
| EPI_ISL_483693, EPI_ISL_483694, EPI_ISL_483695, EPI_ISL_483699, EPI_ISL_483700 | National Institute of Laboratory Medicine and Referral Center | Genomic Research Lab, BCSIR | Abu Sayeed Mohammad Mahmud, Mohammad Samir Uzzaman, Eshrar Osman, Md. Ahasan Habib, Shahina Akter, Tanjina Akhter Banu, Md. Murshed Hasan Sarkar, Barna Goswami, Iffat Jahan, Md. Saddam Hossain, Tasnim Nafisa, Md. Maruf Ahmed Molla, Mahmuda Yeasmin, Asish Kumar Ghosh, A. K. M. Shamsuzzaman, Sheikh Md. Selim Al Din, Utpal Chandra Ray, Salek Ahmed Sajib, Md. Salim Khan |
| EPI_ISL_483701 | Israel Central Virology laboratory | Israel Central Virology laboratory | Neta Zuckerman, Efrat Dahan Bucris, Oran Erster, Ella Mendelson, Michal Mandelboim |
| EPI_ISL_483703 | National Institute of Laboratory Medicine and Referral Center | Genomic Research Lab, BCSIR | Abu Sayeed Mohammad Mahmud, Mohammad Samir Uzzaman, Eshrar Osman, Md. Ahasan Habib, Shahina Akter, Tanjina Akhter Banu, Md. Murshed Hasan Sarkar, Barna Goswami, Iffat Jahan, Md. Saddam Hossain, Tasnim Nafisa, Md. Maruf Ahmed Molla, Mahmuda Yeasmin, Asish Kumar Ghosh, A. K. M. Shamsuzzaman, Sheikh Md. Selim Al Din, Utpal Chandra Ray, Salek Ahmed Sajib, Md. Salim Khan |
| EPI_ISL_483704 | Israel Central Virology laboratory | Israel Central Virology laboratory | Neta Zuckerman, Efrat Dahan Bucris, Oran Erster, Ella Mendelson, Michal Mandelboim |
| EPI_ISL_483705 | National Institute of Laboratory Medicine and Referral Center | Genomic Research Lab, BCSIR | Abu Sayeed Mohammad Mahmud, Mohammad Samir Uzzaman, Eshrar Osman, Md. Ahasan Habib, Shahina Akter, Tanjina Akhter Banu, Md. Murshed Hasan Sarkar, Barna Goswami, Iffat Jahan, Md. Saddam Hossain, Tasnim Nafisa, Md. Maruf Ahmed Molla, Mahmuda Yeasmin, Asish Kumar Ghosh, A. K. M. Shamsuzzaman, Sheikh Md. Selim Al Din, Utpal Chandra Ray, Salek Ahmed Sajib, Md. Salim Khan |
| EPI_ISL_483706 | Israel Central Virology laboratory | Israel Central Virology laboratory | Neta Zuckerman, Efrat Dahan Bucris, Oran Erster, Ella Mendelson, Michal Mandelboim |
| EPI_ISL_483707 | National Institute of Laboratory Medicine and Referral Center | Genomic Research Lab, BCSIR | Abu Sayeed Mohammad Mahmud, Mohammad Samir Uzzaman, Eshrar Osman, Md. Ahasan Habib, Shahina Akter, Tanjina Akhter Banu, Md. Murshed Hasan Sarkar, Barna Goswami, Iffat Jahan, Md. Saddam Hossain, Tasnim Nafisa, Md. Maruf Ahmed Molla, Mahmuda Yeasmin, Asish Kumar Ghosh, A. K. M. Shamsuzzaman, Sheikh Md. Selim Al Din, Utpal Chandra Ray, Salek Ahmed Sajib, Md. Salim Khan |
| EPI_ISL_483708, EPI_ISL_483709 | Israel Central Virology laboratory | Israel Central Virology laboratory | Neta Zuckerman, Efrat Dahan Bucris, Oran Erster, Ella Mendelson, Michal Mandelboim |
| EPI_ISL_483710 | National Institute of Laboratory Medicine and Referral Center | Genomic Research Lab, BCSIR | Abu Sayeed Mohammad Mahmud, Mohammad Samir Uzzaman, Eshrar Osman, Md. Ahasan Habib, Shahina Akter, Tanjina Akhter Banu, Md. Murshed Hasan Sarkar, Barna Goswami, Iffat Jahan, Md. Saddam Hossain, Tasnim Nafisa, Md. Maruf Ahmed Molla, Mahmuda Yeasmin, Asish Kumar Ghosh, A. K. M. Shamsuzzaman, Sheikh Md. Selim Al Din, Utpal Chandra Ray, Salek Ahmed Sajib, Md. Salim Khan |
| EPI_ISL_483711, EPI_ISL_483712, EPI_ISL_483713, EPI_ISL_483714, EPI_ISL_483715, EPI_ISL_483716, EPI_ISL_483717, EPI_ISL_483718, EPI_ISL_483719, EPI_ISL_483720, EPI_ISL_483721, EPI_ISL_483722, EPI_ISL_483723, EPI_ISL_483724, EPI_ISL_483725 | | | |
| see above | Israel Central Virology laboratory | Israel Central Virology laboratory | Neta Zuckerman, Efrat Dahan Bucris, Oran Erster, Ella Mendelson, Michal Mandelboim |
| EPI_ISL_483820 | GMERS Medical College and Hospital, Gandhinagar | Gujarat Biotechnology Research Centre | Komal Patel, Labdhi Pandya, Afzal Ansari, Nikha Trivedi, Seema Bhatt, Gaurishankar Shrimali, Bhavesh Modi, Bharti Rajani, Apurvasinh Puvar, Janvi Raval, Zarna Patel, Monika Gandhi, Pinal Trivedi, Maharshi Pandya, Nidhi Patel, Nitin Savaliya, Raghawendra Kumar, Dinesh Kumar, Zuber Saiyed, R D Dixit, A M Kadri, Harsh Bakshi, Chaitanya Joshi, Madhvi Joshi |
| EPI_ISL_483821 | Government Medical College, Vadodara | Gujarat Biotechnology Research Centre | Labdhi Pandya, Afzal Ansari, Nikha Trivedi, Meenakshi Shah, Neena Doshi, Varsha Godbole, Apurvasinh Puvar, Janvi Raval, Zarna Patel, Monika Gandhi, Pinal Trivedi, Maharshi Pandya, Nidhi Patel, Nitin Savaliya, Raghawendra Kumar, Dinesh Kumar, Zuber Saiyed, Komal Patel, R D Dixit, A M Kadri, Harsh Bakshi, Chaitanya Joshi, Madhvi Joshi |
| EPI_ISL_483822 | Government Medical College, Vadodara | Gujarat Biotechnology Research Centre | Afzal Ansari, Nikha Trivedi, Meenakshi Shah, Neena Doshi, Varsha Godbole, Apurvasinh Puvar, Janvi Raval, Zarna Patel, Monika Gandhi, Pinal Trivedi, Maharshi Pandya, Nidhi Patel, Nitin Savaliya, Raghawendra Kumar, Dinesh Kumar, Zuber Saiyed, Komal Patel, Labdhi Pandya, R D Dixit, A M Kadri, Harsh Bakshi, Chaitanya Joshi, Madhvi Joshi |
| EPI_ISL_483823 | GMERS Medical College Himmatnagar | Gujarat Biotechnology Research Centre | Nikha Trivedi, Himanshu Khatri, Mayur Gandhi, Apurvasinh Puvar, Janvi Raval, Zarna Patel, Monika Gandhi, Pinal Trivedi, Maharshi Pandya, Nidhi Patel, Nitin Savaliya, Raghawendra Kumar, Dinesh Kumar, Zuber Saiyed, Komal Patel, Labdhi Pandya, Afzal Ansari, R D Dixit, A M Kadri, Harsh Bakshi, Chaitanya Joshi, Madhvi Joshi |
| EPI_ISL_483824 | GMERS Medical College Himmatnagar | Gujarat Biotechnology Research Centre | Himanshu Khatri, Mayur Gandhi, Apurvasinh Puvar, Janvi Raval, Zarna Patel, Monika Gandhi, Pinal Trivedi, Maharshi Pandya, Nidhi Patel, Nitin Savaliya, Raghawendra Kumar, Dinesh Kumar, Zuber Saiyed, Komal Patel, Labdhi Pandya, Afzal Ansari, Nikha Trivedi, R D Dixit, A M Kadri, Harsh Bakshi, Chaitanya Joshi, Madhvi Joshi |
| EPI_ISL_483825 | GMERS Medical College Himmatnagar | Gujarat Biotechnology Research Centre | Mayur Gandhi, Apurvasinh Puvar, Janvi Raval, Zarna Patel, Monika Gandhi, Pinal Trivedi, Maharshi Pandya, Nidhi Patel, Nitin Savaliya, Raghawendra Kumar, Dinesh Kumar, Zuber Saiyed, Komal Patel, Labdhi Pandya, Afzal Ansari, Nikha Trivedi, Himanshu Khatri, R D Dixit, A M Kadri, Harsh Bakshi, Chaitanya Joshi, Madhvi Joshi |
| EPI_ISL_483826 | GMERS Medical College Himmatnagar | Gujarat Biotechnology Research Centre | Apurvasinh Puvar, Janvi Raval, Zarna Patel, Monika Gandhi, Pinal Trivedi, Maharshi Pandya, Nidhi Patel, Nitin Savaliya, Raghawendra Kumar, Dinesh Kumar, Zuber Saiyed, Komal Patel, Labdhi Pandya, Afzal Ansari, Nikha Trivedi, Himanshu Khatri, Mayur Gandhi, R D Dixit, A M Kadri, Harsh Bakshi, Chaitanya Joshi, Madhvi Joshi |
| EPI_ISL_483827 | GMERS Medical College Himmatnagar | Gujarat Biotechnology Research Centre | Janvi Raval, Zarna Patel, Monika Gandhi, Pinal Trivedi, Maharshi Pandya, Nidhi Patel, Nitin Savaliya, Raghawendra Kumar, Dinesh Kumar, Zuber Saiyed, Komal Patel, Labdhi Pandya, Afzal Ansari, Nikha Trivedi, Himanshu Khatri, Mayur Gandhi, Apurvasinh Puvar, R D Dixit, A M Kadri, Harsh Bakshi, Chaitanya Joshi, Madhvi Joshi |
| EPI_ISL_483828 | GMERS Medical College Himmatnagar | Gujarat Biotechnology Research Centre | Zarna Patel, Monika Gandhi, Pinal Trivedi, Maharshi Pandya, Nidhi Patel, Nitin Savaliya, Raghawendra Kumar, Dinesh Kumar, Zuber Saiyed, Komal Patel, Labdhi Pandya, Afzal Ansari, Nikha Trivedi, Himanshu Khatri, Mayur Gandhi, Apurvasinh Puvar, Janvi Raval, R D Dixit, A M Kadri, Harsh Bakshi, Chaitanya Joshi, Madhvi Joshi |
| EPI_ISL_483829 | Pandit Deendayal Upadhyay Government Medical College, Rajkot | Gujarat Biotechnology Research Centre | Monika Gandhi, Pinal Trivedi, Maharshi Pandya, Nidhi Patel, Nitin Savaliya, Raghawendra Kumar, Dinesh Kumar, Zuber Saiyed, Komal Patel, Labdhi Pandya, Afzal Ansari, Nikha Trivedi, Gauravi Dhruv, Arti Trivedi, Apurvasinh Puvar, Janvi Raval, Zarna Patel, R D Dixit, A M Kadri, Harsh Bakshi, Chaitanya Joshi, Madhvi Joshi |
| EPI_ISL_483830 | Pandit Deendayal Upadhyay Government Medical College, Rajkot | Gujarat Biotechnology Research Centre | Pinal Trivedi, Maharshi Pandya, Nidhi Patel, Nitin Savaliya, Raghawendra Kumar, Dinesh Kumar, Zuber Saiyed, Komal Patel, Labdhi Pandya, Afzal Ansari, Nikha Trivedi, Gauravi Dhruv, Arti Trivedi, Apurvasinh Puvar, Janvi Raval, Zarna Patel, Monika Gandhi, R D Dixit, A M Kadri, Harsh Bakshi, Chaitanya Joshi, Madhvi Joshi |
| EPI_ISL_483831 | Pandit Deendayal Upadhyay Government Medical College, Rajkot | Gujarat Biotechnology Research Centre | Maharshi Pandya, Nidhi Patel, Nitin Savaliya, Raghawendra Kumar, Dinesh Kumar, Zuber Saiyed, Komal Patel, Labdhi Pandya, Afzal Ansari, Nikha Trivedi, Gauravi Dhruv, Arti Trivedi, Apurvasinh Puvar, Janvi Raval, Zarna Patel, Monika Gandhi, Pinal Trivedi, R D Dixit, A M Kadri, Harsh Bakshi, Chaitanya Joshi, Madhvi Joshi |
| EPI_ISL_483832 | Pandit Deendayal Upadhyay Government Medical College, Rajkot | Gujarat Biotechnology Research Centre | Nidhi Patel, Nitin Savaliya, Raghawendra Kumar, Dinesh Kumar, Zuber Saiyed, Komal Patel, Labdhi Pandya, Afzal Ansari, Nikha Trivedi, Gauravi Dhruv, Arti Trivedi, Apurvasinh Puvar, Janvi Raval, Zarna Patel, Monika Gandhi, Pinal Trivedi, Maharshi Pandya, R D Dixit, A M Kadri, Harsh Bakshi, Chaitanya Joshi, Madhvi Joshi |
| EPI_ISL_483833 | Pandit Deendayal Upadhyay Government Medical College, Rajkot | Gujarat Biotechnology Research Centre | Nitin Savaliya, Raghawendra Kumar, Dinesh Kumar, Zuber Saiyed, Komal Patel, Labdhi Pandya, Afzal Ansari, Nikha Trivedi, Gauravi Dhruv, Arti Trivedi, Apurvasinh Puvar, Janvi Raval, Zarna Patel, Monika Gandhi, Pinal Trivedi, Maharshi Pandya, Nidhi Patel, R D Dixit, A M Kadri, Harsh Bakshi, Chaitanya Joshi, Madhvi Joshi |
| EPI_ISL_483834 | Pandit Deendayal Upadhyay Government Medical College, Rajkot | Gujarat Biotechnology Research Centre | Raghawendra Kumar, Dinesh Kumar, Zuber Saiyed, Komal Patel, Labdhi Pandya, Afzal Ansari, Nikha Trivedi, Gauravi Dhruv, Arti Trivedi, Apurvasinh Puvar, Janvi Raval, Zarna Patel, Monika Gandhi, Pinal Trivedi, Maharshi Pandya, Nidhi Patel, Nitin Savaliya, R D Dixit, A M Kadri, Harsh Bakshi, Chaitanya Joshi, Madhvi Joshi |
| EPI_ISL_483835 | Pandit Deendayal Upadhyay Government Medical College, Rajkot | Gujarat Biotechnology Research Centre | Dinesh Kumar, Zuber Saiyed, Komal Patel, Labdhi Pandya, Afzal Ansari, Nikha Trivedi, Gauravi Dhruv, Arti Trivedi, Apurvasinh Puvar, Janvi Raval, Zarna Patel, Monika Gandhi, Pinal Trivedi, Maharshi Pandya, Nidhi Patel, Nitin Savaliya, Raghawendra Kumar, R D Dixit, A M Kadri, Harsh Bakshi, Chaitanya Joshi, Madhvi Joshi |
| EPI_ISL_483836 | Pandit Deendayal Upadhyay Government Medical College, Rajkot | Gujarat Biotechnology Research Centre | Zuber Saiyed, Komal Patel, Labdhi Pandya, Afzal Ansari, Nikha Trivedi, Gauravi Dhruv, Arti Trivedi, Apurvasinh Puvar, Janvi Raval, Zarna Patel, Monika Gandhi, Pinal Trivedi, Maharshi Pandya, Nidhi Patel, Nitin Savaliya, Raghawendra Kumar, Dinesh Kumar, R D Dixit, A M Kadri, Harsh Bakshi, Chaitanya Joshi, Madhvi Joshi |
| EPI_ISL_483837 | Pandit Deendayal Upadhyay Government Medical College, Rajkot | Gujarat Biotechnology Research Centre | Komal Patel, Labdhi Pandya, Afzal Ansari, Nikha Trivedi, Gauravi Dhruv, Arti Trivedi, Apurvasinh Puvar, Janvi Raval, Zarna Patel, Monika Gandhi, Pinal Trivedi, Maharshi Pandya, Nidhi Patel, Nitin Savaliya, Raghawendra Kumar, Dinesh Kumar, Zuber Saiyed, R D Dixit, A M Kadri, Harsh Bakshi, Chaitanya Joshi, Madhvi Joshi |
| EPI_ISL_483838 | Pandit Deendayal Upadhyay Government Medical College, Rajkot | Gujarat Biotechnology Research Centre | Labdhi Pandya, Afzal Ansari, Nikha Trivedi, Gauravi Dhruv, Arti Trivedi, Apurvasinh Puvar, Janvi Raval, Zarna Patel, Monika Gandhi, Pinal Trivedi, Maharshi Pandya, Nidhi Patel, Nitin Savaliya, Raghawendra Kumar, Dinesh Kumar, Zuber Saiyed, Komal Patel, R D Dixit, A M Kadri, Harsh Bakshi, Chaitanya Joshi, Madhvi Joshi |
| EPI_ISL_483839 | Pandit Deendayal Upadhyay Government Medical College, Rajkot | Gujarat Biotechnology Research Centre | Afzal Ansari, Nikha Trivedi, Gauravi Dhruv, Arti Trivedi, Apurvasinh Puvar, Janvi Raval, Zarna Patel, Monika Gandhi, Pinal Trivedi, Maharshi Pandya, Nidhi Patel, Nitin Savaliya, Raghawendra Kumar, Dinesh Kumar, Zuber Saiyed, Komal Patel, Labdhi Pandya, R D Dixit, A M Kadri, Harsh Bakshi, Chaitanya Joshi, Madhvi Joshi |
| EPI_ISL_483840 | Pandit Deendayal Upadhyay Government Medical College, Rajkot | Gujarat Biotechnology Research Centre | Nikha Trivedi, Gauravi Dhruv, Arti Trivedi, Apurvasinh Puvar, Janvi Raval, Zarna Patel, Monika Gandhi, Pinal Trivedi, Maharshi Pandya, Nidhi Patel, Nitin Savaliya, Raghawendra Kumar, Dinesh Kumar, Zuber Saiyed, Komal Patel, Labdhi Pandya, Afzal Ansari, R D Dixit, A M Kadri, Harsh Bakshi, Chaitanya Joshi, Madhvi Joshi |
| EPI_ISL_483841 | Pandit Deendayal Upadhyay Government Medical College, Rajkot | Gujarat Biotechnology Research Centre | Gauravi Dhruv, Arti Trivedi, Apurvasinh Puvar, Janvi Raval, Zarna Patel, Monika Gandhi, Pinal Trivedi, Maharshi Pandya, Nidhi Patel, Nitin Savaliya, Raghawendra Kumar, Dinesh Kumar, Zuber Saiyed, Komal Patel, Labdhi Pandya, Afzal Ansari, Nikha Trivedi, R D Dixit, A M Kadri, Harsh Bakshi, Chaitanya Joshi, Madhvi Joshi |
| EPI_ISL_483842 | Pandit Deendayal Upadhyay Government Medical College, Rajkot | Gujarat Biotechnology Research Centre | Arti Trivedi, Apurvasinh Puvar, Janvi Raval, Zarna Patel, Monika Gandhi, Pinal Trivedi, Maharshi Pandya, Nidhi Patel, Nitin Savaliya, Raghawendra Kumar, Dinesh Kumar, Zuber Saiyed, Komal Patel, Labdhi Pandya, Afzal Ansari, Nikha Trivedi, Gauravi Dhruv, R D Dixit, A M Kadri, Harsh Bakshi, Chaitanya Joshi, Madhvi Joshi |
| EPI_ISL_483843 | Pandit Deendayal Upadhyay Government Medical College, Rajkot | Gujarat Biotechnology Research Centre | Apurvasinh Puvar, Janvi Raval, Zarna Patel, Monika Gandhi, Pinal Trivedi, Maharshi Pandya, Nidhi Patel, Nitin Savaliya, Raghawendra Kumar, Dinesh Kumar, Zuber Saiyed, Komal Patel, Labdhi Pandya, Afzal Ansari, Nikha Trivedi, Gauravi Dhruv, Arti Trivedi, R D Dixit, A M Kadri, Harsh Bakshi, Chaitanya Joshi, Madhvi Joshi |

| | | | |
|---|---|---|---|
| EPI_ISL_483844 | Pandit Deendayal Upadhyay Government Medical College, Rajkot | Gujarat Biotechnology Research Centre | Janvi Raval, Zarna Patel, Monika Gandhi, Pinal Trivedi, Maharshi Pandya, Nidhi Patel, Nitin Savaliya, Raghawendra Kumar, Dinesh Kumar, Zuber Saiyed, Komal Patel, Labdhi Pandya, Afzal Ansari, Nikha Trivedi, Gauravi Dhruv, Arti Trivedi, Apurvasinh Puvar, R D Dixit, A M Kadri, Harsh Bakshi, Chaitanya Joshi, Madhvi Joshi |
| EPI_ISL_483845 | Pandit Deendayal Upadhyay Government Medical College, Rajkot | Gujarat Biotechnology Research Centre | Zarna Patel, Monika Gandhi, Pinal Trivedi, Maharshi Pandya, Nidhi Patel, Nitin Savaliya, Raghawendra Kumar, Dinesh Kumar, Zuber Saiyed, Komal Patel, Labdhi Pandya, Afzal Ansari, Nikha Trivedi, Gauravi Dhruv, Arti Trivedi, Apurvasinh Puvar, Janvi Raval, R D Dixit, A M Kadri, Harsh Bakshi, Chaitanya Joshi, Madhvi Joshi |
| EPI_ISL_483846 | Pandit Deendayal Upadhyay Government Medical College, Rajkot | Gujarat Biotechnology Research Centre | Monika Gandhi, Pinal Trivedi, Maharshi Pandya, Nidhi Patel, Nitin Savaliya, Raghawendra Kumar, Dinesh Kumar, Zuber Saiyed, Komal Patel, Labdhi Pandya, Afzal Ansari, Nikha Trivedi, Gauravi Dhruv, Arti Trivedi, Apurvasinh Puvar, Janvi Raval, Zarna Patel, R D Dixit, A M Kadri, Harsh Bakshi, Chaitanya Joshi, Madhvi Joshi |
| EPI_ISL_483847 | Pandit Deendayal Upadhyay Government Medical College, Rajkot | Gujarat Biotechnology Research Centre | Pinal Trivedi, Maharshi Pandya, Nidhi Patel, Nitin Savaliya, Raghawendra Kumar, Dinesh Kumar, Zuber Saiyed, Komal Patel, Labdhi Pandya, Afzal Ansari, Nikha Trivedi, Gauravi Dhruv, Arti Trivedi, Apurvasinh Puvar, Janvi Raval, Zarna Patel, Monika Gandhi, R D Dixit, A M Kadri, Harsh Bakshi, Chaitanya Joshi, Madhvi Joshi |
| EPI_ISL_483848 | Pandit Deendayal Upadhyay Government Medical College, Rajkot | Gujarat Biotechnology Research Centre | Maharshi Pandya, Nidhi Patel, Nitin Savaliya, Raghawendra Kumar, Dinesh Kumar, Zuber Saiyed, Komal Patel, Labdhi Pandya, Afzal Ansari, Nikha Trivedi, Gauravi Dhruv, Arti Trivedi, Apurvasinh Puvar, Janvi Raval, Zarna Patel, Monika Gandhi, Pinal Trivedi, R D Dixit, A M Kadri, Harsh Bakshi, Chaitanya Joshi, Madhvi Joshi |
| EPI_ISL_483849 | Pandit Deendayal Upadhyay Government Medical College, Rajkot | Gujarat Biotechnology Research Centre | Nidhi Patel, Nitin Savaliya, Raghawendra Kumar, Dinesh Kumar, Zuber Saiyed, Komal Patel, Labdhi Pandya, Afzal Ansari, Nikha Trivedi, Gauravi Dhruv, Arti Trivedi, Apurvasinh Puvar, Janvi Raval, Zarna Patel, Monika Gandhi, Pinal Trivedi, Maharshi Pandya, R D Dixit, A M Kadri, Harsh Bakshi, Chaitanya Joshi, Madhvi Joshi |
| EPI_ISL_483850 | Department of Microbiology, Government Medical College, Surat | Gujarat Biotechnology Research Centre | Nitin Savaliya, Raghawendra Kumar, Dinesh Kumar, Zuber Saiyed, Komal Patel, Labdhi Pandya, Afzal Ansari, Nikha Trivedi, Naresh Chauhan, Summaiya Mullan, Amit gamit, Apurvasinh Puvar, Janvi Raval, Zarna Patel, Monika Gandhi, Pinal Trivedi, Maharshi Pandya, Nidhi Patel, R D Dixit, A M Kadri, Harsh Bakshi, Chaitanya Joshi, Madhvi Joshi |
| EPI_ISL_483851 | Department of Microbiology, Government Medical College, Surat | Gujarat Biotechnology Research Centre | Raghawendra Kumar, Dinesh Kumar, Zuber Saiyed, Komal Patel, Labdhi Pandya, Afzal Ansari, Nikha Trivedi, Naresh Chauhan, Summaiya Mullan, Amit gamit, Apurvasinh Puvar, Janvi Raval, Zarna Patel, Monika Gandhi, Pinal Trivedi, Maharshi Pandya, Nidhi Patel, Nitin Savaliya, R D Dixit, A M Kadri, Harsh Bakshi, Chaitanya Joshi, Madhvi Joshi |
| EPI_ISL_483852 | Department of Microbiology, Government Medical College, Surat | Gujarat Biotechnology Research Centre | Dinesh Kumar, Zuber Saiyed, Komal Patel, Labdhi Pandya, Afzal Ansari, Nikha Trivedi, Naresh Chauhan, Summaiya Mullan, Amit gamit, Apurvasinh Puvar, Janvi Raval, Zarna Patel, Monika Gandhi, Pinal Trivedi, Maharshi Pandya, Nidhi Patel, Nitin Savaliya, Raghawendra Kumar, R D Dixit, A M Kadri, Harsh Bakshi, Chaitanya Joshi, Madhvi Joshi |
| EPI_ISL_483853 | Department of Microbiology, Government Medical College, Surat | Gujarat Biotechnology Research Centre | Zuber Saiyed, Komal Patel, Labdhi Pandya, Afzal Ansari, Nikha Trivedi, Naresh Chauhan, Summaiya Mullan, Amit gamit, Apurvasinh Puvar, Janvi Raval, Zarna Patel, Monika Gandhi, Pinal Trivedi, Maharshi Pandya, Nidhi Patel, Nitin Savaliya, Raghawendra Kumar, Dinesh Kumar, R D Dixit, A M Kadri, Harsh Bakshi, Chaitanya Joshi, Madhvi Joshi |
| EPI_ISL_483854 | Department of Microbiology, Government Medical College, Surat | Gujarat Biotechnology Research Centre | Komal Patel, Labdhi Pandya, Afzal Ansari, Nikha Trivedi, Naresh Chauhan, Summaiya Mullan, Amit gamit, Apurvasinh Puvar, Janvi Raval, Zarna Patel, Monika Gandhi, Pinal Trivedi, Maharshi Pandya, Nidhi Patel, Nitin Savaliya, Raghawendra Kumar, Dinesh Kumar, Zuber Saiyed, R D Dixit, A M Kadri, Harsh Bakshi, Chaitanya Joshi, Madhvi Joshi |
| EPI_ISL_483855 | Department of Microbiology, Government Medical College, Surat | Gujarat Biotechnology Research Centre | Labdhi Pandya, Afzal Ansari, Nikha Trivedi, Naresh Chauhan, Summaiya Mullan, Amit gamit, Apurvasinh Puvar, Janvi Raval, Zarna Patel, Monika Gandhi, Pinal Trivedi, Maharshi Pandya, Nidhi Patel, Nitin Savaliya, Raghawendra Kumar, Dinesh Kumar, Zuber Saiyed, Komal Patel, R D Dixit, A M Kadri, Harsh Bakshi, Chaitanya Joshi, Madhvi Joshi |
| EPI_ISL_483856 | Department of Microbiology, Government Medical College, Surat | Gujarat Biotechnology Research Centre | Afzal Ansari, Nikha Trivedi, Naresh Chauhan, Summaiya Mullan, Amit gamit, Apurvasinh Puvar, Janvi Raval, Zarna Patel, Monika Gandhi, Pinal Trivedi, Maharshi Pandya, Nidhi Patel, Nitin Savaliya, Raghawendra Kumar, Dinesh Kumar, Zuber Saiyed, Komal Patel, Labdhi Pandya, R D Dixit, A M Kadri, Harsh Bakshi, Chaitanya Joshi, Madhvi Joshi |
| EPI_ISL_483857 | Department of Microbiology, Government Medical College, Surat | Gujarat Biotechnology Research Centre | Nikha Trivedi, Naresh Chauhan, Summaiya Mullan, Amit gamit, Apurvasinh Puvar, Janvi Raval, Zarna Patel, Monika Gandhi, Pinal Trivedi, Maharshi Pandya, Nidhi Patel, Nitin Savaliya, Raghawendra Kumar, Dinesh Kumar, Zuber Saiyed, Komal Patel, Labdhi Pandya, Afzal Ansari, R D Dixit, A M Kadri, Harsh Bakshi, Chaitanya Joshi, Madhvi Joshi |
| EPI_ISL_483858 | Department of Microbiology, Government Medical College, Surat | Gujarat Biotechnology Research Centre | Naresh Chauhan, Summaiya Mullan, Amit gamit, Apurvasinh Puvar, Janvi Raval, Zarna Patel, Monika Gandhi, Pinal Trivedi, Maharshi Pandya, Nidhi Patel, Nitin Savaliya, Raghawendra Kumar, Dinesh Kumar, Zuber Saiyed, Komal Patel, Labdhi Pandya, Afzal Ansari, Nikha Trivedi, R D Dixit, A M Kadri, Harsh Bakshi, Chaitanya Joshi, Madhvi Joshi |
| EPI_ISL_483859 | Department of Microbiology, Government Medical College, Surat | Gujarat Biotechnology Research Centre | Summaiya Mullan, Amit gamit, Apurvasinh Puvar, Janvi Raval, Zarna Patel, Monika Gandhi, Pinal Trivedi, Maharshi Pandya, Nidhi Patel, Nitin Savaliya, Raghawendra Kumar, Dinesh Kumar, Zuber Saiyed, Komal Patel, Labdhi Pandya, Afzal Ansari, Nikha Trivedi, Naresh Chauhan, R D Dixit, A M Kadri, Harsh Bakshi, Chaitanya Joshi, Madhvi Joshi |
| EPI_ISL_483860 | Department of Microbiology, Government Medical College, Surat | Gujarat Biotechnology Research Centre | Amit gamit, Apurvasinh Puvar, Janvi Raval, Zarna Patel, Monika Gandhi, Pinal Trivedi, Maharshi Pandya, Nidhi Patel, Nitin Savaliya, Raghawendra Kumar, Dinesh Kumar, Zuber Saiyed, Komal Patel, Labdhi Pandya, Afzal Ansari, Nikha Trivedi, Naresh Chauhan, Summaiya Mullan, R D Dixit, A M Kadri, Harsh Bakshi, Chaitanya Joshi, Madhvi Joshi |
| EPI_ISL_483861 | Department of Microbiology, Government Medical College, Surat | Gujarat Biotechnology Research Centre | Apurvasinh Puvar, Janvi Raval, Zarna Patel, Monika Gandhi, Pinal Trivedi, Maharshi Pandya, Nidhi Patel, Nitin Savaliya, Raghawendra Kumar, Dinesh Kumar, Zuber Saiyed, Komal Patel, Labdhi Pandya, Afzal Ansari, Nikha Trivedi, Naresh Chauhan, Summaiya Mullan, Amit gamit, R D Dixit, A M Kadri, Harsh Bakshi, Chaitanya Joshi, Madhvi Joshi |
| EPI_ISL_483862 | Department of Microbiology, Government Medical College, Surat | Gujarat Biotechnology Research Centre | Janvi Raval, Zarna Patel, Monika Gandhi, Pinal Trivedi, Maharshi Pandya, Nidhi Patel, Nitin Savaliya, Raghawendra Kumar, Dinesh Kumar, Zuber Saiyed, Komal Patel, Labdhi Pandya, Afzal Ansari, Nikha Trivedi, Naresh Chauhan, Summaiya Mullan, Amit gamit, Apurvasinh Puvar, R D Dixit, A M Kadri, Harsh Bakshi, Chaitanya Joshi, Madhvi Joshi |
| EPI_ISL_483863 | Department of Microbiology, Government Medical College, Surat | Gujarat Biotechnology Research Centre | Zarna Patel, Monika Gandhi, Pinal Trivedi, Maharshi Pandya, Nidhi Patel, Nitin Savaliya, Raghawendra Kumar, Dinesh Kumar, Zuber Saiyed, Komal Patel, Labdhi Pandya, Afzal Ansari, Nikha Trivedi, Naresh Chauhan, Summaiya Mullan, Amit gamit, Apurvasinh Puvar, Janvi Raval, R D Dixit, A M Kadri, Harsh Bakshi, Chaitanya Joshi, Madhvi Joshi |
| EPI_ISL_483864 | Department of Microbiology, Government Medical College, Surat | Gujarat Biotechnology Research Centre | Monika Gandhi, Pinal Trivedi, Maharshi Pandya, Nidhi Patel, Nitin Savaliya, Raghawendra Kumar, Dinesh Kumar, Zuber Saiyed, Komal Patel, Labdhi Pandya, Afzal Ansari, Nikha Trivedi, Naresh Chauhan, Summaiya Mullan, Amit gamit, Apurvasinh Puvar, Janvi Raval, Zarna Patel, R D Dixit, A M Kadri, Harsh Bakshi, Chaitanya Joshi, Madhvi Joshi |
| EPI_ISL_483865 | Department of Microbiology, Government Medical College, Surat | Gujarat Biotechnology Research Centre | Pinal Trivedi, Maharshi Pandya, Nidhi Patel, Nitin Savaliya, Raghawendra Kumar, Dinesh Kumar, Zuber Saiyed, Komal Patel, Labdhi Pandya, Afzal Ansari, Nikha Trivedi, Naresh Chauhan, Summaiya Mullan, Amit gamit, Apurvasinh Puvar, Janvi Raval, Zarna Patel, Monika Gandhi, R D Dixit, A M Kadri, Harsh Bakshi, Chaitanya Joshi, Madhvi Joshi |
| EPI_ISL_483866 | Department of Microbiology, Government Medical College, Surat | Gujarat Biotechnology Research Centre | Maharshi Pandya, Nidhi Patel, Nitin Savaliya, Raghawendra Kumar, Dinesh Kumar, Zuber Saiyed, Komal Patel, Labdhi Pandya, Afzal Ansari, Nikha Trivedi, Naresh Chauhan, Summaiya Mullan, Amit gamit, Apurvasinh Puvar, Janvi Raval, Zarna Patel, Monika Gandhi, Pinal Trivedi, R D Dixit, A M Kadri, Harsh Bakshi, Chaitanya Joshi, Madhvi Joshi |
| EPI_ISL_483867 | Department of Microbiology, Government Medical College, Surat | Gujarat Biotechnology Research Centre | Nidhi Patel, Nitin Savaliya, Raghawendra Kumar, Dinesh Kumar, Zuber Saiyed, Komal Patel, Labdhi Pandya, Afzal Ansari, Nikha Trivedi, Naresh Chauhan, Summaiya Mullan, Amit gamit, Apurvasinh Puvar, Janvi Raval, Zarna Patel, Monika Gandhi, Pinal Trivedi, Maharshi Pandya, R D Dixit, A M Kadri, Harsh Bakshi, Chaitanya Joshi, Madhvi Joshi |
| EPI_ISL_483868 | Department of Microbiology, Government Medical College, Surat | Gujarat Biotechnology Research Centre | Nitin Savaliya, Raghawendra Kumar, Dinesh Kumar, Zuber Saiyed, Komal Patel, Labdhi Pandya, Afzal Ansari, Nikha Trivedi, Naresh Chauhan, Summaiya Mullan, Amit gamit, Apurvasinh Puvar, Janvi Raval, Zarna Patel, Monika Gandhi, Pinal Trivedi, Maharshi Pandya, Nidhi Patel, R D Dixit, A M Kadri, Harsh Bakshi, Chaitanya Joshi, Madhvi Joshi |
| EPI_ISL_483869 | Department of Microbiology, Government Medical College, Surat | Gujarat Biotechnology Research Centre | Raghawendra Kumar, Dinesh Kumar, Zuber Saiyed, Komal Patel, Labdhi Pandya, Afzal Ansari, Nikha Trivedi, Naresh Chauhan, Summaiya Mullan, Amit gamit, Apurvasinh Puvar, Janvi Raval, Zarna Patel, Monika Gandhi, Pinal Trivedi, Maharshi Pandya, Nidhi Patel, Nitin Savaliya, R D Dixit, A M Kadri, Harsh Bakshi, Chaitanya Joshi, Madhvi Joshi |
| EPI_ISL_483870 | Department of Microbiology, Government Medical College, Surat | Gujarat Biotechnology Research Centre | Dinesh Kumar, Zuber Saiyed, Komal Patel, Labdhi Pandya, Afzal Ansari, Nikha Trivedi, Naresh Chauhan, Summaiya Mullan, Amit gamit, Apurvasinh Puvar, Janvi Raval, Zarna Patel, Monika Gandhi, Pinal Trivedi, Maharshi Pandya, Nidhi Patel, Nitin Savaliya, Raghawendra Kumar, R D Dixit, A M Kadri, Harsh Bakshi, Chaitanya Joshi, Madhvi Joshi |
| EPI_ISL_483871 | Department of Microbiology, Government Medical College, Surat | Gujarat Biotechnology Research Centre | Zuber Saiyed, Komal Patel, Labdhi Pandya, Afzal Ansari, Nikha Trivedi, Naresh Chauhan, Summaiya Mullan, Amit gamit, Apurvasinh Puvar, Janvi Raval, Zarna Patel, Monika Gandhi, Pinal Trivedi, Maharshi Pandya, Nidhi Patel, Nitin Savaliya, Raghawendra Kumar, Dinesh Kumar, R D Dixit, A M Kadri, Harsh Bakshi, Chaitanya Joshi, Madhvi Joshi |
| EPI_ISL_483872 | Department of Microbiology, Government Medical College, Surat | Gujarat Biotechnology Research Centre | Komal Patel, Labdhi Pandya, Afzal Ansari, Nikha Trivedi, Naresh Chauhan, Summaiya Mullan, Amit gamit, Apurvasinh Puvar, Janvi Raval, Zarna Patel, Monika Gandhi, Pinal Trivedi, Maharshi Pandya, Nidhi Patel, Nitin Savaliya, Raghawendra Kumar, Dinesh Kumar, Zuber Saiyed, R D Dixit, A M Kadri, Harsh Bakshi, Chaitanya Joshi, Madhvi Joshi |
| EPI_ISL_483873 | Department of Microbiology, Government Medical College, Surat | Gujarat Biotechnology Research Centre | Labdhi Pandya, Afzal Ansari, Nikha Trivedi, Naresh Chauhan, Summaiya Mullan, Amit gamit, Apurvasinh Puvar, Janvi Raval, Zarna Patel, Monika Gandhi, Pinal Trivedi, Maharshi Pandya, Nidhi Patel, Nitin Savaliya, Raghawendra Kumar, Dinesh Kumar, Zuber Saiyed, Komal Patel, R D Dixit, A M Kadri, Harsh Bakshi, Chaitanya Joshi, Madhvi Joshi |
| EPI_ISL_483874 | Department of Microbiology, Government Medical College, Surat | Gujarat Biotechnology Research Centre | Afzal Ansari, Nikha Trivedi, Naresh Chauhan, Summaiya Mullan, Amit gamit, Apurvasinh Puvar, Janvi Raval, Zarna Patel, Monika Gandhi, Pinal Trivedi, Maharshi Pandya, Nidhi Patel, Nitin Savaliya, Raghawendra Kumar, Dinesh Kumar, Zuber Saiyed, Komal Patel, Labdhi Pandya, R D Dixit, A M Kadri, Harsh Bakshi, Chaitanya Joshi, Madhvi Joshi |
| EPI_ISL_483875 | Department of Microbiology, Government Medical College, Surat | Gujarat Biotechnology Research Centre | Nikha Trivedi, Naresh Chauhan, Summaiya Mullan, Amit gamit, Apurvasinh Puvar, Janvi Raval, Zarna Patel, Monika Gandhi, Pinal Trivedi, Maharshi Pandya, Nidhi Patel, Nitin Savaliya, Raghawendra Kumar, Dinesh Kumar, Zuber Saiyed, Komal Patel, Labdhi Pandya, Afzal Ansari, R D Dixit, A M Kadri, Harsh Bakshi, Chaitanya Joshi, Madhvi Joshi |
| EPI_ISL_483876 | Department of Microbiology, Government Medical College, Surat | Gujarat Biotechnology Research Centre | Naresh Chauhan, Summaiya Mullan, Amit gamit, Apurvasinh Puvar, Janvi Raval, Zarna Patel, Monika Gandhi, Pinal Trivedi, Maharshi Pandya, Nidhi Patel, Nitin Savaliya, Raghawendra Kumar, Dinesh Kumar, Zuber Saiyed, Komal Patel, Labdhi Pandya, Afzal Ansari, Nikha Trivedi, R D Dixit, A M Kadri, Harsh Bakshi, Chaitanya Joshi, Madhvi Joshi |
| EPI_ISL_483877 | Department of Microbiology, Government Medical College, Surat | Gujarat Biotechnology Research Centre | Summaiya Mullan, Amit gamit, Apurvasinh Puvar, Janvi Raval, Zarna Patel, Monika Gandhi, Pinal Trivedi, Maharshi Pandya, Nidhi Patel, Nitin Savaliya, Raghawendra Kumar, Dinesh Kumar, Zuber Saiyed, Komal Patel, Labdhi Pandya, Afzal Ansari, Nikha Trivedi, Naresh Chauhan, R D Dixit, A M Kadri, Harsh Bakshi, Chaitanya Joshi, Madhvi Joshi |
| EPI_ISL_483878 | Department of Microbiology, Government Medical College, Surat | Gujarat Biotechnology Research Centre | Amit gamit, Apurvasinh Puvar, Janvi Raval, Zarna Patel, Monika Gandhi, Pinal Trivedi, Maharshi Pandya, Nidhi Patel, Nitin Savaliya, Raghawendra Kumar, Dinesh Kumar, Zuber Saiyed, Komal Patel, Labdhi Pandya, Afzal Ansari, Nikha Trivedi, Naresh Chauhan, Summaiya Mullan, R D Dixit, A M Kadri, Harsh Bakshi, Chaitanya Joshi, Madhvi Joshi |
| EPI_ISL_483879 | Department of Microbiology, Government Medical College, Surat | Gujarat Biotechnology Research Centre | Apurvasinh Puvar, Janvi Raval, Zarna Patel, Monika Gandhi, Pinal Trivedi, Maharshi Pandya, Nidhi Patel, Nitin Savaliya, Raghawendra Kumar, Dinesh Kumar, Zuber Saiyed, Komal Patel, Labdhi Pandya, Afzal Ansari, Nikha Trivedi, Naresh Chauhan, Summaiya Mullan, Amit gamit, R D Dixit, A M Kadri, Harsh Bakshi, Chaitanya Joshi, Madhvi Joshi |
| EPI_ISL_483880, EPI_ISL_483881, EPI_ISL_483882, EPI_ISL_483883, EPI_ISL_483884, EPI_ISL_483885, EPI_ISL_483886, EPI_ISL_483887, EPI_ISL_483888, EPI_ISL_483889, EPI_ISL_483890, EPI_ISL_483891, EPI_ISL_483892, EPI_ISL_483893, EPI_ISL_483894, EPI_ISL_483895, EPI_ISL_483896, EPI_ISL_483897, EPI_ISL_483898, EPI_ISL_483899, EPI_ISL_483900, EPI_ISL_483901, EPI_ISL_483902, EPI_ISL_483903, EPI_ISL_483904, EPI_ISL_483905, EPI_ISL_483906, EPI_ISL_483907, EPI_ISL_483908, EPI_ISL_483909, EPI_ISL_483910, EPI_ISL_483911, EPI_ISL_483912 | | | |
| see above | University of Birmingham | COVID-19 Genomics UK (COG-UK) Consortium | Institute of Microbiology, University of Birmingham: Claire McMurray, Joanne Stockton, Samuel Nicholls, Radoslaw Poplawski, Will Rowe, Josh Quick, Nicholas Loman. University of Birmingham Testing Laboratory: Celina M Whalley, Andrew Bosworth, Charlotte Poxon, Kasun Wanigasooriya, Oliver Pickles, Mike Kidd, Alex Richter, Andrew D Beggs PHE Heartlands Lab: Husam Osman, Andrew Bosworth. Queen Elizabeth Hospital: Anna Casey |
| EPI_ISL_483913, EPI_ISL_483914, EPI_ISL_483915 | Department of Pathology, University of Cambridge | COVID-19 Genomics UK (COG-UK) Consortium | Luke W Meredith, M. Estée Török, Myra Hosmillo, William L. Hamilton, Martin D. Curran, Theresa Feltwell, Grant Hall, Anna Yakovleva, Fahad A Khokhar, Charlotte J. Houldcroft, Laura G Caller, Aminu S. Jahun, Sarah L. Caddy, Yasmin Chaudhry, Malte Pinckert, Ian Goodfellow |
| EPI_ISL_483916, EPI_ISL_483917, EPI_ISL_483918, EPI_ISL_483919, EPI_ISL_483920, EPI_ISL_483921, EPI_ISL_483922, EPI_ISL_483923, EPI_ISL_483924, EPI_ISL_483925, EPI_ISL_483926, EPI_ISL_483927, EPI_ISL_483928, EPI_ISL_483929, EPI_ISL_483930, EPI_ISL_483931, EPI_ISL_483932, EPI_ISL_483933, EPI_ISL_483934, EPI_ISL_483935, EPI_ISL_483936, EPI_ISL_483937, EPI_ISL_483938, EPI_ISL_483939, EPI_ISL_483940, EPI_ISL_483941, EPI_ISL_483942, EPI_ISL_483943, EPI_ISL_483944, EPI_ISL_483945, EPI_ISL_483946, EPI_ISL_483947, EPI_ISL_483948, EPI_ISL_483949, EPI_ISL_483950, EPI_ISL_483951, EPI_ISL_483952, EPI_ISL_483953, EPI_ISL_483954, EPI_ISL_483955, EPI_ISL_483956, EPI_ISL_483957, EPI_ISL_483958, EPI_ISL_483959, EPI_ISL_483960, EPI_ISL_483961, EPI_ISL_483962, EPI_ISL_483963, EPI_ISL_483964, EPI_ISL_483965, EPI_ISL_483966, EPI_ISL_483967, EPI_ISL_483968, EPI_ISL_483969, EPI_ISL_483970, EPI_ISL_483971, EPI_ISL_483972, EPI_ISL_483973, EPI_ISL_483974, EPI_ISL_483975, EPI_ISL_483976, EPI_ISL_483977, EPI_ISL_483978, EPI_ISL_483979, EPI_ISL_483980, EPI_ISL_483981, EPI_ISL_483982, EPI_ISL_483983, EPI_ISL_483984, EPI_ISL_483985, EPI_ISL_483986, EPI_ISL_483987, EPI_ISL_483988, EPI_ISL_483989, EPI_ISL_483990, EPI_ISL_483991, EPI_ISL_483992, EPI_ISL_483993, EPI_ISL_483994, EPI_ISL_483995, EPI_ISL_483996, EPI_ISL_483997, EPI_ISL_483998, EPI_ISL_483999, EPI_ISL_484000, EPI_ISL_484001, EPI_ISL_484002, EPI_ISL_484003, EPI_ISL_484004, EPI_ISL_484005, EPI_ISL_484006, EPI_ISL_484007, EPI_ISL_484008, EPI_ISL_484009, EPI_ISL_484010, EPI_ISL_484011, EPI_ISL_484012, EPI_ISL_484013, EPI_ISL_484014, EPI_ISL_484015, EPI_ISL_484016, EPI_ISL_484017, EPI_ISL_484018, EPI_ISL_484019, EPI_ISL_484020, EPI_ISL_484021, EPI_ISL_484022, EPI_ISL_484023, EPI_ISL_484024, EPI_ISL_484025, EPI_ISL_484026, EPI_ISL_484027, EPI_ISL_484028, EPI_ISL_484029, EPI_ISL_484030, EPI_ISL_484031, EPI_ISL_484032, EPI_ISL_484033, EPI_ISL_484034, EPI_ISL_484035, EPI_ISL_484036, EPI_ISL_484037, EPI_ISL_484038, EPI_ISL_484039, EPI_ISL_484040, EPI_ISL_484041, EPI_ISL_484042, EPI_ISL_484043, EPI_ISL_484044, EPI_ISL_484045, EPI_ISL_484046, EPI_ISL_484047, EPI_ISL_484048, EPI_ISL_484049, EPI_ISL_484050, EPI_ISL_484051, EPI_ISL_484052, EPI_ISL_484053, EPI_ISL_484054, EPI_ISL_484055, EPI_ISL_484056, EPI_ISL_484057, EPI_ISL_484058, EPI_ISL_484059, EPI_ISL_484060, EPI_ISL_484061, EPI_ISL_484062, EPI_ISL_484063, EPI_ISL_484064, EPI_ISL_484065, EPI_ISL_484066, EPI_ISL_484067, EPI_ISL_484068, EPI_ISL_484069, EPI_ISL_484070, EPI_ISL_484071, EPI_ISL_484072, EPI_ISL_484073, EPI_ISL_484074, EPI_ISL_484075, EPI_ISL_484076, EPI_ISL_484077, EPI_ISL_484078, EPI_ISL_484079, EPI_ISL_484080, EPI_ISL_484081, EPI_ISL_484082, EPI_ISL_484083, EPI_ISL_484084, EPI_ISL_484085, EPI_ISL_484086, EPI_ISL_484087, EPI_ISL_484088, EPI_ISL_484089, EPI_ISL_484090, EPI_ISL_484091, EPI_ISL_484092, EPI_ISL_484093, EPI_ISL_484094, EPI_ISL_484095, EPI_ISL_484096, EPI_ISL_484097, EPI_ISL_484098, EPI_ISL_484099, EPI_ISL_484100, EPI_ISL_484101, EPI_ISL_484102, EPI_ISL_484103, EPI_ISL_484104, EPI_ISL_484105, EPI_ISL_484106, EPI_ISL_484107, EPI_ISL_484108, EPI_ISL_484109, EPI_ISL_484110, EPI_ISL_484111, EPI_ISL_484112, EPI_ISL_484113, EPI_ISL_484114, EPI_ISL_484115, EPI_ISL_484116, EPI_ISL_484117, EPI_ISL_484118, EPI_ISL_484119, EPI_ISL_484120, EPI_ISL_484121, EPI_ISL_484122, EPI_ISL_484123, EPI_ISL_484124, EPI_ISL_484125, EPI_ISL_484126, EPI_ISL_484127, EPI_ISL_484128, EPI_ISL_484129, EPI_ISL_484130, EPI_ISL_484131, EPI_ISL_484132, EPI_ISL_484133, EPI_ISL_484134, EPI_ISL_484135, EPI_ISL_484136, EPI_ISL_484137, EPI_ISL_484138, EPI_ISL_484139, EPI_ISL_484140, EPI_ISL_484141, EPI_ISL_484142, EPI_ISL_484143, EPI_ISL_484144, EPI_ISL_484145, EPI_ISL_484146, EPI_ISL_484147, EPI_ISL_484148, EPI_ISL_484149, EPI_ISL_484150, EPI_ISL_484151, EPI_ISL_484152, EPI_ISL_484153, EPI_ISL_484154, EPI_ISL_484155, EPI_ISL_484156, EPI_ISL_484157, EPI_ISL_484158, EPI_ISL_484159, EPI_ISL_484160, EPI_ISL_484161, EPI_ISL_484162, EPI_ISL_484163, EPI_ISL_484164, EPI_ISL_484165, EPI_ISL_484166, EPI_ISL_484167, EPI_ISL_484168, EPI_ISL_484169, EPI_ISL_484170, EPI_ISL_484171, EPI_ISL_484172, EPI_ISL_484173, EPI_ISL_484174, EPI_ISL_484175, EPI_ISL_484176, EPI_ISL_484177, EPI_ISL_484178, EPI_ISL_484179, EPI_ISL_484180, EPI_ISL_484181, EPI_ISL_484182, EPI_ISL_484183, EPI_ISL_484184, EPI_ISL_484185, EPI_ISL_484186, EPI_ISL_484187, EPI_ISL_484188, EPI_ISL_484189, EPI_ISL_484190, EPI_ISL_484191, EPI_ISL_484192, EPI_ISL_484193, EPI_ISL_484194, EPI_ISL_484195, EPI_ISL_484196, EPI_ISL_484197, EPI_ISL_484198, EPI_ISL_484199, EPI_ISL_484200, EPI_ISL_484201, EPI_ISL_484202, EPI_ISL_484203, EPI_ISL_484204, EPI_ISL_484205, EPI_ISL_484206, EPI_ISL_484207, EPI_ISL_484208, EPI_ISL_484209, EPI_ISL_484210, EPI_ISL_484211, EPI_ISL_484212, EPI_ISL_484213, EPI_ISL_484214, EPI_ISL_484215, EPI_ISL_484216, EPI_ISL_484217 | | | |
| see above | Centre for Clinical Infection and Diagnostics Research and Genomics Innovation Unit, Guy's and St. Thomas' | COVID-19 Genomics UK (COG-UK) Consortium | Chloe Fisher, Luke Snell, Penny Cliff, Rahul Batra, Jonathan Edgeworth, Ali Raza Awan |

| | | | |
|---|---|---|---|
| | NHS Trust | | |
| EPI_ISL_484218, EPI_ISL_484219, EPI_ISL_484220 | University of Birmingham | COVID-19 Genomics UK (COG-UK) Consortium | Institute of Microbiology, University of Birmingham: Claire McMurray, Joanne Stockton, Samuel Nicholls, Radoslaw Poplawski, Will Rowe, Josh Quick, Nicholas Loman. University of Birmingham Testing Laboratory: Celina M Whalley, Andrew Bosworth, Charlotte Poxon, Kasun Wanigasooriya, Oliver Pickles, Mike Kidd, Alex Richter, Andrew D Beggs PHE Heartlands Lab: Husam Osman, Andrew Bosworth. Queen Elizabeth Hospital: Anna Casey |
| EPI_ISL_484221, EPI_ISL_484222, EPI_ISL_484223, EPI_ISL_484224, EPI_ISL_484225, EPI_ISL_484226, EPI_ISL_484227, EPI_ISL_484228, EPI_ISL_484229, EPI_ISL_484230, EPI_ISL_484231, EPI_ISL_484232, EPI_ISL_484233, EPI_ISL_484234, EPI_ISL_484235, EPI_ISL_484236, EPI_ISL_484237, EPI_ISL_484238, EPI_ISL_484239, EPI_ISL_484240, EPI_ISL_484241, EPI_ISL_484242, EPI_ISL_484243, EPI_ISL_484244, EPI_ISL_484245, EPI_ISL_484246, EPI_ISL_484247, EPI_ISL_484248, EPI_ISL_484249, EPI_ISL_484250, EPI_ISL_484251, EPI_ISL_484252, EPI_ISL_484253 | | | |
| see above | University Hospitals Of Leicester NHS Trust and DeepSeq Nottingham | COVID-19 Genomics UK (COG-UK) Consortium | Christopher Holmes, Paul Bird, Thomas Helmer, Karlie Fallon, Julian Tang, Jonathan Ball, Patrick McClure, Joeseph Chappell, Nadine Holmes, Matthew Carlisle, Christopher Moore, Fei Sang, Johnny Debebe, Victoria Wright, Matthew Loose |
| EPI_ISL_484254, EPI_ISL_484255, EPI_ISL_484256, EPI_ISL_484257, EPI_ISL_484258, EPI_ISL_484259, EPI_ISL_484260, EPI_ISL_484261, EPI_ISL_484262, EPI_ISL_484263 | Liverpool Clinical Laboratories | COVID-19 Genomics UK (COG-UK) Consortium | Sam Haldenby, Anita Lucaci, Steve Paterson, Julian Hiscox, Alistair Darby, M Almsaud, A Alrezaihi, Muhannad Alruwaili, Stuart D Armstrong, Jones Benjamin, Eleanor G Bentley, Anu Chawla, Jordan J Clark, Angela Cowell, Richard Eccles, Isabel García-Dorival, Matthew Gemmell, Alessandro Gerada, PKF Gilmore, Richard Gregory, Ximeng Han, Catherine Hartley, Margaret Hughes, Miren Iturriza-Gomara, James Johnson, L Luu, Jenifer Manson, Charlotte Nelson, Elaine O'Toole, Cassie Olateju, Rebekah Penrice-Randal , Lucille Rainbow, N.P Randle, Trevor Ian Robinson, Parul Sharma, Ghada T Shawli, James P Stewart, Neil Swainston, Ecaterina Vamos, Joanne Watts, Mark Whitehead |
| EPI_ISL_484264, EPI_ISL_484265, EPI_ISL_484266, EPI_ISL_484267, EPI_ISL_484268, EPI_ISL_484269, EPI_ISL_484270, EPI_ISL_484271, EPI_ISL_484272, EPI_ISL_484273, EPI_ISL_484274, EPI_ISL_484275, EPI_ISL_484276, EPI_ISL_484277, EPI_ISL_484278, EPI_ISL_484279, EPI_ISL_484280, EPI_ISL_484281, EPI_ISL_484282, EPI_ISL_484283, EPI_ISL_484284, EPI_ISL_484285, EPI_ISL_484286, EPI_ISL_484287, EPI_ISL_484288, EPI_ISL_484289, EPI_ISL_484290, EPI_ISL_484291, EPI_ISL_484292, EPI_ISL_484293, EPI_ISL_484294, EPI_ISL_484295, EPI_ISL_484296, EPI_ISL_484297, EPI_ISL_484298, EPI_ISL_484299, EPI_ISL_484300, EPI_ISL_484301, EPI_ISL_484302, EPI_ISL_484303, EPI_ISL_484304, EPI_ISL_484305, EPI_ISL_484306, EPI_ISL_484307, EPI_ISL_484308, EPI_ISL_484309, EPI_ISL_484310, EPI_ISL_484311, EPI_ISL_484312, EPI_ISL_484313, EPI_ISL_484314, EPI_ISL_484315, EPI_ISL_484316, EPI_ISL_484317, EPI_ISL_484318, EPI_ISL_484319, EPI_ISL_484320, EPI_ISL_484321, EPI_ISL_484322, EPI_ISL_484323, EPI_ISL_484324, EPI_ISL_484325, EPI_ISL_484326, EPI_ISL_484327 | | | |
| see above | Northumbria University / South Tees Hospitals NHS Foundation Trust / North Cumbria Integrated Care NHS Foundation Trust / North Tees and Hartlepool NHS Foundation Trust / Newcastle Hospitals NHS Foundation Trust | COVID-19 Genomics UK (COG-UK) Consortium | Darren L Smith,Andrew Nelson,Matthew Bashton,Greg R Young,Joshua Loh,John Allan,Mohammad A Tariq,Giles S Holt,Gary Black,Wen C Yew,Lynn Dover,Paul Baker,Steve Liggett,Sarah Essex,Jane Greenaway,Debra Padgett,Clive Graham,Garren Scott,Edward Barton,Emma Swindells,Brendan Payne,Jennifer Collins,Yusri Taha,Gary Eltringham |
| EPI_ISL_484328, EPI_ISL_484329, EPI_ISL_484330, EPI_ISL_484331, EPI_ISL_484332, EPI_ISL_484333, EPI_ISL_484334, EPI_ISL_484335, EPI_ISL_484336, EPI_ISL_484337, EPI_ISL_484338 | | | |
| see above | Quadram Institute Bioscience | COVID-19 Genomics UK (COG-UK) Consortium | Dave J. Baker, Gemma L. Kay, Alp Aydin, Thanh Le-Viet, Steven Rudder, Ana P. Tedim, Anastasia Kolyva, Maria Diaz, Leonardo de Oliveira Martins, Nabil-Fareed Alikhan, Lizzie Meadows, Rachael Stanley, Ngozi Elumogo, Muhammed Yasir, Nicholas M. Thomson, Alexander J Trotter, Rachel Gilroy, Samuel Bloomfield, Claire Stuart, Andrew Bell, Reenesh Prakash, Samir Dervisevic, Alison E. Mather, John Wain, Mark Webber, Andrew J. Page, Justin O'Grady |
| EPI_ISL_484339, EPI_ISL_484340, EPI_ISL_484341, EPI_ISL_484342, EPI_ISL_484343, EPI_ISL_484344, EPI_ISL_484345, EPI_ISL_484346, EPI_ISL_484347, EPI_ISL_484348, EPI_ISL_484349, EPI_ISL_484350, EPI_ISL_484351, EPI_ISL_484352, EPI_ISL_484353 | | | |
| see above | Queens Medical Centre, Clinical Microbiology Department / DeepSeq Nottingham | COVID-19 Genomics UK (COG-UK) Consortium | Gemma Clark, Wendy Smith, Manjinder Khakh, Vicki M Fleming, Michelle M Lister, Hannah Howson-Wells, Jonathan Ball, Patrick McClure, Joseph Chappell, Theocharis Tsoleridis, Nadine Holmes, Matthew Carlisle, Christopher Moore, Fei Sang, Johnny Debebe, Victoria Wright, Matthew Loose |
| EPI_ISL_484354, EPI_ISL_484355, EPI_ISL_484356, EPI_ISL_484357, EPI_ISL_484358, EPI_ISL_484359, EPI_ISL_484360, EPI_ISL_484361, EPI_ISL_484362, EPI_ISL_484363, EPI_ISL_484364, EPI_ISL_484365, EPI_ISL_484366, EPI_ISL_484367, EPI_ISL_484368, EPI_ISL_484369, EPI_ISL_484370, EPI_ISL_484371, EPI_ISL_484372, EPI_ISL_484373 | | | |
| see above | Lincolnshire Hospitals and DeepSeq Nottingham | COVID-19 Genomics UK (COG-UK) Consortium | Nichola Duckworth, Tim Sloan, Sarah Walsh, Jonathan Ball, Patrick McClure, Joeseph Chappell, Nadine Holmes, Matthew Carlisle, Christopher Moore, Fei Sang, Johnny Debebe, Victoria Wright, Matthew Loose |
| EPI_ISL_484374, EPI_ISL_484375 | Queens Medical Centre, Clinical Microbiology Department / DeepSeq Nottingham | COVID-19 Genomics UK (COG-UK) Consortium | Gemma Clark, Wendy Smith, Manjinder Khakh, Vicki M Fleming, Michelle M Lister, Hannah Howson-Wells, Jonathan Ball, Patrick McClure, Joseph Chappell, Theocharis Tsoleridis, Nadine Holmes, Matthew Carlisle, Christopher Moore, Fei Sang, Johnny Debebe, Victoria Wright, Matthew Loose |
| EPI_ISL_484376, EPI_ISL_484377, EPI_ISL_484378, EPI_ISL_484379, EPI_ISL_484380, EPI_ISL_484381, EPI_ISL_484382, EPI_ISL_484383, EPI_ISL_484384, EPI_ISL_484385, EPI_ISL_484386, EPI_ISL_484387, EPI_ISL_484388, EPI_ISL_484389 | | | |
| see above | University Hospitals Of Leicester NHS Trust and DeepSeq Nottingham | COVID-19 Genomics UK (COG-UK) Consortium | Christopher Holmes, Paul Bird, Thomas Helmer, Karlie Fallon, Julian Tang, Jonathan Ball, Patrick McClure, Joeseph Chappell, Nadine Holmes, Matthew Carlisle, Christopher Moore, Fei Sang, Johnny Debebe, Victoria Wright, Matthew Loose |
| EPI_ISL_484390, EPI_ISL_484391 | Queens Medical Centre, Clinical Microbiology Department / DeepSeq Nottingham | COVID-19 Genomics UK (COG-UK) Consortium | Gemma Clark, Wendy Smith, Manjinder Khakh, Vicki M Fleming, Michelle M Lister, Hannah Howson-Wells, Jonathan Ball, Patrick McClure, Joseph Chappell, Theocharis Tsoleridis, Nadine Holmes, Matthew Carlisle, Christopher Moore, Fei Sang, Johnny Debebe, Victoria Wright, Matthew Loose |
| EPI_ISL_484392, EPI_ISL_484393, EPI_ISL_484394, EPI_ISL_484395, EPI_ISL_484396, EPI_ISL_484397, EPI_ISL_484398, EPI_ISL_484399, EPI_ISL_484400, EPI_ISL_484401, EPI_ISL_484402, EPI_ISL_484403, EPI_ISL_484404, EPI_ISL_484405, EPI_ISL_484406 | | | |
| see above | Lincolnshire Hospitals and DeepSeq Nottingham | COVID-19 Genomics UK (COG-UK) Consortium | Nichola Duckworth, Tim Sloan, Sarah Walsh, Jonathan Ball, Patrick McClure, Joeseph Chappell, Nadine Holmes, Matthew Carlisle, Christopher Moore, Fei Sang, Johnny Debebe, Victoria Wright, Matthew Loose |
| EPI_ISL_484407, EPI_ISL_484408, EPI_ISL_484409, EPI_ISL_484410, EPI_ISL_484411, EPI_ISL_484412, EPI_ISL_484413, EPI_ISL_484414, EPI_ISL_484415, EPI_ISL_484416, EPI_ISL_484417, EPI_ISL_484418, EPI_ISL_484419, EPI_ISL_484420, EPI_ISL_484421, EPI_ISL_484422, EPI_ISL_484423, EPI_ISL_484424, EPI_ISL_484425, EPI_ISL_484426, EPI_ISL_484427, EPI_ISL_484428, EPI_ISL_484429, EPI_ISL_484430, EPI_ISL_484431, EPI_ISL_484432 | | | |
| see above | Centre for Enzyme Innovation, University of Portsmouth / Translational Research Laboratory, Portsmouth Hospitals NHS Trust | COVID-19 Genomics UK (COG-UK) Consortium | Angela Beckett,Yann Bourgeois,Garry Scarlett,Sharon Glaysher,Scott Elliott,Kelly Bicknell,Robert Impey,Allyson Lloyd,Sarah Wyllie,Ethan Butcher,Anoop Chauhan,Samuel Robson |
| EPI_ISL_484433, EPI_ISL_484434, EPI_ISL_484435, EPI_ISL_484436, EPI_ISL_484437, EPI_ISL_484438, EPI_ISL_484439, EPI_ISL_484440, EPI_ISL_484441, EPI_ISL_484442, EPI_ISL_484443, EPI_ISL_484444, EPI_ISL_484445, EPI_ISL_484446, EPI_ISL_484447, EPI_ISL_484448, EPI_ISL_484449, EPI_ISL_484450, EPI_ISL_484451, EPI_ISL_484452, EPI_ISL_484453, EPI_ISL_484454, EPI_ISL_484455, EPI_ISL_484456, EPI_ISL_484457, EPI_ISL_484458, EPI_ISL_484459, EPI_ISL_484460, EPI_ISL_484461, EPI_ISL_484462, EPI_ISL_484463, EPI_ISL_484464, EPI_ISL_484465, EPI_ISL_484466, EPI_ISL_484467, EPI_ISL_484468, EPI_ISL_484469, EPI_ISL_484470, EPI_ISL_484471, EPI_ISL_484472, EPI_ISL_484473, EPI_ISL_484474, EPI_ISL_484475, EPI_ISL_484476, EPI_ISL_484477, EPI_ISL_484478, EPI_ISL_484479, EPI_ISL_484480, EPI_ISL_484481, EPI_ISL_484482, EPI_ISL_484483, EPI_ISL_484484, EPI_ISL_484485, EPI_ISL_484486, EPI_ISL_484487, EPI_ISL_484488, EPI_ISL_484489, EPI_ISL_484490, EPI_ISL_484491, EPI_ISL_484492, EPI_ISL_484493, EPI_ISL_484494, EPI_ISL_484495, EPI_ISL_484496, EPI_ISL_484497, EPI_ISL_484498, EPI_ISL_484499, EPI_ISL_484500, EPI_ISL_484501, EPI_ISL_484502, EPI_ISL_484503, EPI_ISL_484504, EPI_ISL_484505, EPI_ISL_484506, EPI_ISL_484507, EPI_ISL_484508, EPI_ISL_484509, EPI_ISL_484510, EPI_ISL_484511, EPI_ISL_484512, EPI_ISL_484513, EPI_ISL_484514, EPI_ISL_484515, EPI_ISL_484516, EPI_ISL_484517, EPI_ISL_484518 | | | |
| see above | Virology Department, Sheffield Teaching Hospitals NHS Foundation Trust/Department of Infection, Immunity and Cardiovascular Disease, The Medical School, University of Sheffield | COVID-19 Genomics UK (COG-UK) Consortium | Thushan de Silva, Matthew Parker, Nikki Smith, Adri Angyal, Rebecca Brown, Luke Green, Rachel Tucker, Paul Parsons, Danielle Groves, Katie Johnson, Laura Carrilero, Alex Keeley, Dave Partridge, Matthew Wyles, Benjamin Lindsey, Mehmet Yavuz, Mohammad Raza, Cariad Evans |
| EPI_ISL_484519, EPI_ISL_484520, EPI_ISL_484521, EPI_ISL_484522, EPI_ISL_484523, EPI_ISL_484524, EPI_ISL_484525, EPI_ISL_484526, EPI_ISL_484527, EPI_ISL_484528, EPI_ISL_484529, EPI_ISL_484530, EPI_ISL_484531, EPI_ISL_484532, EPI_ISL_484533, EPI_ISL_484534, EPI_ISL_484535, EPI_ISL_484536, EPI_ISL_484537, EPI_ISL_484538, EPI_ISL_484539, EPI_ISL_484540, EPI_ISL_484541, EPI_ISL_484542, EPI_ISL_484543, EPI_ISL_484544, EPI_ISL_484545, EPI_ISL_484546, EPI_ISL_484547, EPI_ISL_484548, EPI_ISL_484549, EPI_ISL_484550, EPI_ISL_484551, EPI_ISL_484552, EPI_ISL_484553, EPI_ISL_484554, EPI_ISL_484555, EPI_ISL_484556, EPI_ISL_484557, EPI_ISL_484558, EPI_ISL_484559, EPI_ISL_484560, EPI_ISL_484561, EPI_ISL_484562, EPI_ISL_484563, EPI_ISL_484564, EPI_ISL_484565, EPI_ISL_484566, EPI_ISL_484567, EPI_ISL_484568, EPI_ISL_484569, EPI_ISL_484570, EPI_ISL_484571, EPI_ISL_484572, EPI_ISL_484573, EPI_ISL_484574, EPI_ISL_484575, EPI_ISL_484576, EPI_ISL_484577, EPI_ISL_484578, EPI_ISL_484579, EPI_ISL_484580, EPI_ISL_484581, EPI_ISL_484582, EPI_ISL_484583, EPI_ISL_484584, EPI_ISL_484585, EPI_ISL_484586, EPI_ISL_484587, EPI_ISL_484588, EPI_ISL_484589, EPI_ISL_484590, EPI_ISL_484591, EPI_ISL_484592, EPI_ISL_484593, EPI_ISL_484594, EPI_ISL_484595, EPI_ISL_484596, EPI_ISL_484597, EPI_ISL_484598, EPI_ISL_484599, EPI_ISL_484600, EPI_ISL_484601, EPI_ISL_484602, EPI_ISL_484603, EPI_ISL_484604, EPI_ISL_484605, EPI_ISL_484606, EPI_ISL_484607, EPI_ISL_484608, EPI_ISL_484609, EPI_ISL_484610, EPI_ISL_484611, EPI_ISL_484612, EPI_ISL_484613, EPI_ISL_484614, EPI_ISL_484615, EPI_ISL_484616, EPI_ISL_484617, EPI_ISL_484618, EPI_ISL_484619, EPI_ISL_484620, EPI_ISL_484621, EPI_ISL_484622, EPI_ISL_484623, EPI_ISL_484624, EPI_ISL_484625, EPI_ISL_484626, EPI_ISL_484627, EPI_ISL_484628, EPI_ISL_484629, EPI_ISL_484630, EPI_ISL_484631, EPI_ISL_484632, EPI_ISL_484633, EPI_ISL_484634, EPI_ISL_484635, EPI_ISL_484636, EPI_ISL_484637, EPI_ISL_484638, EPI_ISL_484639, EPI_ISL_484640, EPI_ISL_484641, EPI_ISL_484642, EPI_ISL_484643, EPI_ISL_484644, EPI_ISL_484645, EPI_ISL_484646, EPI_ISL_484647, EPI_ISL_484648, EPI_ISL_484649, EPI_ISL_484650, EPI_ISL_484651, EPI_ISL_484652, EPI_ISL_484653, EPI_ISL_484654, EPI_ISL_484655, EPI_ISL_484656, EPI_ISL_484657, EPI_ISL_484658, EPI_ISL_484659, EPI_ISL_484660, EPI_ISL_484661, EPI_ISL_484662, EPI_ISL_484663, EPI_ISL_484664, EPI_ISL_484665, EPI_ISL_484666, EPI_ISL_484667, EPI_ISL_484668, EPI_ISL_484669, EPI_ISL_484670, EPI_ISL_484671, EPI_ISL_484672, EPI_ISL_484673, EPI_ISL_484674, EPI_ISL_484675, EPI_ISL_484676, EPI_ISL_484677, EPI_ISL_484678, EPI_ISL_484679 | | | |
| see above | West of Scotland Specialist Virology Centre, NHSGGC / MRC-University of Glasgow Centre for Virus Research | COVID-19 Genomics UK (COG-UK) Consortium | Ana da Silva Filipe, Natasha Johnson, Kathy Smollett, Daniel Mair, Stephen Carmichael, Lily Tong, Jenna Nichols, Elihu Aranday-Cortes, Kirstyn Brunker, Yasmin Parr, Alice Broos, Kyriaki Nomikou; Sarah McDonald, Marc Niebel, Patawee Asamaphan; Richard Orton, Joseph Hughes, Sreenu Vattipally, David L Robertson; Alasdair MacLean, Rory Gunson; Kathy Li, Natasha Jesudason, Rajiv Shah, James Shepherd, Antonia Ho, Emma Thomson |
| EPI_ISL_484680, EPI_ISL_484681, EPI_ISL_484682, EPI_ISL_484683, EPI_ISL_484684 | Virology Department, Royal Infirmary of Edinburgh, NHS Lothian / School of Biological Sciences, University of Edinburgh / Institute of Genetics and Molecular Medicine, University of Edinburgh | COVID-19 Genomics UK (COG-UK) Consortium | McHugh M, Dewar R, Rooke S, Gallagher M, Balcaza C, O'Toole Á, Scher E, Hill V, McCrone JT, Colquhoun R, Yu X, Jackson B, Rambaut A, Williams TC, Templeton K |
| EPI_ISL_484685, EPI_ISL_484686, EPI_ISL_484687, EPI_ISL_484688, EPI_ISL_484689, EPI_ISL_484690, EPI_ISL_484691 | Originating lab: Wales Specialist Virology Centre Sequencing lab: Pathogen Genomics Unit | COVID-19 Genomics UK (COG-UK) Consortium | Catherine Moore, Johnathan Evans, Laura Gifford, Malorie Perry, Simon Cottrell, Angela Marchbank, Alec Birchley, Alexander Adams, Amy Gaskin, Bree Gatica-Wilcox, Jason Coombes, Joel Southgate, Lauren Gilbert, Lee Graham, Nicole Pacchiarini, Sara Kumziene-Summerhayes, Sarah Taylor, Sophie Jones, Sara Rey, Matthew Bull, Joanne Watkins, Sally Corden, Tom Connor |
| EPI_ISL_484692 | Minnesota Department of Health, Public Health Laboratory | Minnesota Department of Health, Public Health Laboratory | Matt Plumb, Jacob Garfin, and Xiong Wang |
| EPI_ISL_484693, EPI_ISL_484694, EPI_ISL_484695, EPI_ISL_484696, EPI_ISL_484697, EPI_ISL_484698, EPI_ISL_484699, EPI_ISL_484700, EPI_ISL_484701, EPI_ISL_484702, EPI_ISL_484703, EPI_ISL_484704, EPI_ISL_484705, EPI_ISL_484706, EPI_ISL_484707, EPI_ISL_484708 | | | |
| see above | Department of Clinical Microbiology | GIGA Medical Genomics | Keith Durkin, Maria Artesi, Sébastien Bontems, Raphaël Boreux, Cécile Meex, Axelle Chaslain, Céline Fombellida-Lopez, Pierrette Melin, Marie-Pierre Hayette, Vincent Bours. |
| EPI_ISL_484709, EPI_ISL_484710, EPI_ISL_484711, EPI_ISL_484712, EPI_ISL_484713, EPI_ISL_484714, EPI_ISL_484715, EPI_ISL_484716, EPI_ISL_484717, EPI_ISL_484718, EPI_ISL_484719, EPI_ISL_484720, EPI_ISL_484721, EPI_ISL_484722, EPI_ISL_484723, EPI_ISL_484724, EPI_ISL_484725, EPI_ISL_484726, EPI_ISL_484727, EPI_ISL_484728, EPI_ISL_484729, EPI_ISL_484730, EPI_ISL_484731, EPI_ISL_484732, EPI_ISL_484733, EPI_ISL_484734, EPI_ISL_484735, EPI_ISL_484736, EPI_ISL_484737, EPI_ISL_484738, EPI_ISL_484739, EPI_ISL_484740, EPI_ISL_484741, EPI_ISL_484742, EPI_ISL_484743, EPI_ISL_484744, EPI_ISL_484745, EPI_ISL_484746, EPI_ISL_484747, EPI_ISL_484748, EPI_ISL_484749, EPI_ISL_484750, EPI_ISL_484751, EPI_ISL_484752, EPI_ISL_484753, EPI_ISL_484754, EPI_ISL_484755, EPI_ISL_484756, EPI_ISL_484757, EPI_ISL_484758, EPI_ISL_484759, EPI_ISL_484760, EPI_ISL_484761, EPI_ISL_484762, EPI_ISL_484763, EPI_ISL_484764, EPI_ISL_484765, EPI_ISL_484766, EPI_ISL_484767, EPI_ISL_484768, EPI_ISL_484769, EPI_ISL_484770, EPI_ISL_484771, EPI_ISL_484772, EPI_ISL_484773, EPI_ISL_484774, EPI_ISL_484775, EPI_ISL_484776, EPI_ISL_484777, EPI_ISL_484778, EPI_ISL_484779, EPI_ISL_484780, EPI_ISL_484781, EPI_ISL_484782, EPI_ISL_484783, EPI_ISL_484784, EPI_ISL_484785, EPI_ISL_484786, EPI_ISL_484787, EPI_ISL_484788, EPI_ISL_484789, EPI_ISL_484790, EPI_ISL_484791, EPI_ISL_484792, EPI_ISL_484793, EPI_ISL_484794, EPI_ISL_484795, EPI_ISL_484796, EPI_ISL_484797, EPI_ISL_484798, EPI_ISL_484799, EPI_ISL_484800, EPI_ISL_484801, EPI_ISL_484802, EPI_ISL_484803, EPI_ISL_484804, EPI_ISL_484805, EPI_ISL_484806 | | | |
| see above | University of Michigan Clinical Microbiology Laboratory | Lauring Lab, University of Michigan, Department of Microbiology and Immunology | Valesano et al. |
| EPI_ISL_484807, EPI_ISL_484808, EPI_ISL_484809, EPI_ISL_484810, EPI_ISL_484811, EPI_ISL_484812, EPI_ISL_484813, EPI_ISL_484814, EPI_ISL_484815, EPI_ISL_484816, EPI_ISL_484817, EPI_ISL_484818, EPI_ISL_484819, EPI_ISL_484820, EPI_ISL_484821, EPI_ISL_484822, EPI_ISL_484823, EPI_ISL_484824, EPI_ISL_484825, EPI_ISL_484826, EPI_ISL_484827, EPI_ISL_484828, EPI_ISL_484829, EPI_ISL_484830, EPI_ISL_484831, EPI_ISL_484832, EPI_ISL_484833, EPI_ISL_484834, EPI_ISL_484835, EPI_ISL_484836, EPI_ISL_484837, EPI_ISL_484838, EPI_ISL_484839, EPI_ISL_484840, EPI_ISL_484841, EPI_ISL_484842, EPI_ISL_484843, EPI_ISL_484844, EPI_ISL_484845, EPI_ISL_484846, EPI_ISL_484847, EPI_ISL_484848, EPI_ISL_484849, EPI_ISL_484850, EPI_ISL_484851, EPI_ISL_484852, EPI_ISL_484853, EPI_ISL_484854, EPI_ISL_484855, EPI_ISL_484856, EPI_ISL_484857, EPI_ISL_484858, EPI_ISL_484859, EPI_ISL_484860, EPI_ISL_484861, EPI_ISL_484862, EPI_ISL_484863, EPI_ISL_484864, EPI_ISL_484865, EPI_ISL_484866, EPI_ISL_484867, EPI_ISL_484868, EPI_ISL_484869, EPI_ISL_484870, EPI_ISL_484871, EPI_ISL_484872, EPI_ISL_484873, EPI_ISL_484874, EPI_ISL_484875, EPI_ISL_484876, EPI_ISL_484877, EPI_ISL_484878, EPI_ISL_484879, EPI_ISL_484880, EPI_ISL_484881, EPI_ISL_484882, EPI_ISL_484883, EPI_ISL_484884, EPI_ISL_484885, EPI_ISL_484886, EPI_ISL_484887, EPI_ISL_484888, EPI_ISL_484889, EPI_ISL_484890, EPI_ISL_484891, EPI_ISL_484892, EPI_ISL_484893, EPI_ISL_484894, EPI_ISL_484895, EPI_ISL_484896, EPI_ISL_484897, EPI_ISL_484898, EPI_ISL_484899, EPI_ISL_484900, EPI_ISL_484901, EPI_ISL_484902, EPI_ISL_484903, EPI_ISL_484904, EPI_ISL_484905, EPI_ISL_484906, EPI_ISL_484907, EPI_ISL_484908, EPI_ISL_484909, EPI_ISL_484910, EPI_ISL_484911, EPI_ISL_484912, EPI_ISL_484913, EPI_ISL_484914, EPI_ISL_484915, EPI_ISL_484916, EPI_ISL_484917, EPI_ISL_484918, EPI_ISL_484919, EPI_ISL_484920, EPI_ISL_484921, EPI_ISL_484922, EPI_ISL_484923, EPI_ISL_484924, EPI_ISL_484925, EPI_ISL_484926, EPI_ISL_484927, EPI_ISL_484928, EPI_ISL_484929, EPI_ISL_484930, EPI_ISL_484931, EPI_ISL_484932, EPI_ISL_484933, EPI_ISL_484934, EPI_ISL_484935, EPI_ISL_484936, EPI_ISL_484937, EPI_ISL_484938, EPI_ISL_484939, EPI_ISL_484940, EPI_ISL_484941, EPI_ISL_484942, EPI_ISL_484943, EPI_ISL_484944, EPI_ISL_484945, EPI_ISL_484946, EPI_ISL_484947, EPI_ISL_484948, EPI_ISL_484949, EPI_ISL_484950, EPI_ISL_484951, EPI_ISL_484952, EPI_ISL_484953, EPI_ISL_484954, EPI_ISL_484955, EPI_ISL_484956, EPI_ISL_484957, EPI_ISL_484958, EPI_ISL_484959, EPI_ISL_484960, EPI_ISL_484961, EPI_ISL_484962, EPI_ISL_484963, EPI_ISL_484964, EPI_ISL_484965, EPI_ISL_484966, EPI_ISL_484967, EPI_ISL_484968, EPI_ISL_484969, EPI_ISL_484970, EPI_ISL_484971, EPI_ISL_484972, EPI_ISL_484973, EPI_ISL_484974, EPI_ISL_484975, EPI_ISL_484976, EPI_ISL_484977, EPI_ISL_484978, EPI_ISL_484979, EPI_ISL_484980, EPI_ISL_484981, EPI_ISL_484982, EPI_ISL_484983, EPI_ISL_484984, EPI_ISL_484985, EPI_ISL_484986, EPI_ISL_484987, EPI_ISL_484988, EPI_ISL_484989, EPI_ISL_484990, EPI_ISL_484991, EPI_ISL_484992, EPI_ISL_484993, EPI_ISL_484994, EPI_ISL_484995, EPI_ISL_484996, EPI_ISL_484997, EPI_ISL_484998, EPI_ISL_484999, EPI_ISL_485000 | | | |
| see above | University of Wisconsin-Madison AIDS Vaccine Research Laboratories | University of Wisconsin-Madison AIDS Vaccine Research Laboratories | Gage Moreno, Katarina Braun, et al. AIDS Vaccine Research Laboratories |
| EPI_ISL_485001, EPI_ISL_485002 | University of Ulsan College of Medicine and Asan Medical Center | University of Ulsan College of Medicine and Asan Medical Center | Kuenyoul Park, Jaewoong Lee, Kihyun Lee, Jiwon Jung, Sung-Han Kim, Jina Lee, Mauricio Chalita, Seok-Hwan Yoon, Jongsik Chun, Kyu-Hwa Hur, Heungsup Sung, Mi-Na Kim, and Hae Kyung Lee |
| EPI_ISL_485003, EPI_ISL_485004, EPI_ISL_485005, EPI_ISL_485006, EPI_ISL_485007, EPI_ISL_485008, EPI_ISL_485009, EPI_ISL_485010, EPI_ISL_485011, EPI_ISL_485012, EPI_ISL_485013, EPI_ISL_485014, EPI_ISL_485015, EPI_ISL_485016, EPI_ISL_485017, EPI_ISL_485018, EPI_ISL_485019, EPI_ISL_485020, EPI_ISL_485021, EPI_ISL_485022, EPI_ISL_485023, EPI_ISL_485024, EPI_ISL_485025, EPI_ISL_485026, EPI_ISL_485027, EPI_ISL_485028, EPI_ISL_485029, EPI_ISL_485030, EPI_ISL_485031, EPI_ISL_485032, EPI_ISL_485033, EPI_ISL_485034, EPI_ISL_485035, EPI_ISL_485036, EPI_ISL_485037, EPI_ISL_485038, EPI_ISL_485039, EPI_ISL_485040, EPI_ISL_485041, EPI_ISL_485042, EPI_ISL_485043, EPI_ISL_485044, EPI_ISL_485045, EPI_ISL_485046, EPI_ISL_485047, EPI_ISL_485048, EPI_ISL_485049, EPI_ISL_485050, EPI_ISL_485051, EPI_ISL_485052, EPI_ISL_485053, EPI_ISL_485054, EPI_ISL_485055, EPI_ISL_485056, EPI_ISL_485057, EPI_ISL_485058, EPI_ISL_485059, EPI_ISL_485060, EPI_ISL_485061, EPI_ISL_485062, EPI_ISL_485063, EPI_ISL_485064, EPI_ISL_485065, EPI_ISL_485066, EPI_ISL_485067, EPI_ISL_485068, EPI_ISL_485069, EPI_ISL_485070, EPI_ISL_485071, EPI_ISL_485072, EPI_ISL_485073, EPI_ISL_485074, | | | |

| | | | |
|---|---|---|---|
| EPI_ISL_485075, EPI_ISL_485076, EPI_ISL_485077, EPI_ISL_485078, EPI_ISL_485079, EPI_ISL_485080, EPI_ISL_485081, EPI_ISL_485082, EPI_ISL_485083, EPI_ISL_485084, EPI_ISL_485085, EPI_ISL_485086, EPI_ISL_485087, EPI_ISL_485088, EPI_ISL_485089, EPI_ISL_485090, EPI_ISL_485091, EPI_ISL_485092, EPI_ISL_485093, EPI_ISL_485094, EPI_ISL_485095, EPI_ISL_485096, EPI_ISL_485097, EPI_ISL_485098, EPI_ISL_485099, EPI_ISL_485100, EPI_ISL_485101, EPI_ISL_485102, EPI_ISL_485103, EPI_ISL_485104, EPI_ISL_485105, EPI_ISL_485106, EPI_ISL_485107, EPI_ISL_485108, EPI_ISL_485109, EPI_ISL_485110, EPI_ISL_485111, EPI_ISL_485112, EPI_ISL_485113, EPI_ISL_485114, EPI_ISL_485115, EPI_ISL_485116, EPI_ISL_485117, EPI_ISL_485118, EPI_ISL_485119, EPI_ISL_485120, EPI_ISL_485121, EPI_ISL_485122, EPI_ISL_485123, EPI_ISL_485124, EPI_ISL_485125, EPI_ISL_485126, EPI_ISL_485127, EPI_ISL_485128, EPI_ISL_485129, EPI_ISL_485130, EPI_ISL_485131, EPI_ISL_485132, EPI_ISL_485133, EPI_ISL_485134, EPI_ISL_485135, EPI_ISL_485136, EPI_ISL_485137, EPI_ISL_485138, EPI_ISL_485139, EPI_ISL_485140, EPI_ISL_485141, EPI_ISL_485142, EPI_ISL_485143, EPI_ISL_485144, EPI_ISL_485145, EPI_ISL_485146, EPI_ISL_485147, EPI_ISL_485148, EPI_ISL_485149, EPI_ISL_485150, EPI_ISL_485151, EPI_ISL_485152, EPI_ISL_485153, EPI_ISL_485154, EPI_ISL_485155, EPI_ISL_485156, EPI_ISL_485157, EPI_ISL_485158, EPI_ISL_485159, EPI_ISL_485160, EPI_ISL_485161, EPI_ISL_485162, EPI_ISL_485163, EPI_ISL_485164, EPI_ISL_485165, EPI_ISL_485166, EPI_ISL_485167, EPI_ISL_485168, EPI_ISL_485169, EPI_ISL_485170, EPI_ISL_485171, EPI_ISL_485172, EPI_ISL_485173, EPI_ISL_485174, EPI_ISL_485175, EPI_ISL_485176, EPI_ISL_485177, EPI_ISL_485178, EPI_ISL_485179, EPI_ISL_485180, EPI_ISL_485181, EPI_ISL_485182, EPI_ISL_485183, EPI_ISL_485184, EPI_ISL_485185, EPI_ISL_485186, EPI_ISL_485187, EPI_ISL_485188, EPI_ISL_485189, EPI_ISL_485190, EPI_ISL_485191, EPI_ISL_485192, EPI_ISL_485193, EPI_ISL_485194, EPI_ISL_485195, EPI_ISL_485196, EPI_ISL_485197, EPI_ISL_485198, EPI_ISL_485199, EPI_ISL_485200, EPI_ISL_485201, EPI_ISL_485202, EPI_ISL_485203, EPI_ISL_485204, EPI_ISL_485205, EPI_ISL_485206, EPI_ISL_485207, EPI_ISL_485208, EPI_ISL_485209, EPI_ISL_485210, EPI_ISL_485211, EPI_ISL_485212, EPI_ISL_485213, EPI_ISL_485214, EPI_ISL_485215, EPI_ISL_485216, EPI_ISL_485217, EPI_ISL_485218, EPI_ISL_485219, EPI_ISL_485220, EPI_ISL_485221, EPI_ISL_485222, EPI_ISL_485223, EPI_ISL_485224, EPI_ISL_485225, EPI_ISL_485226, EPI_ISL_485227, EPI_ISL_485228, EPI_ISL_485229, EPI_ISL_485230, EPI_ISL_485231, EPI_ISL_485232, EPI_ISL_485233, EPI_ISL_485234, EPI_ISL_485235, EPI_ISL_485236, EPI_ISL_485237, EPI_ISL_485238, EPI_ISL_485239, EPI_ISL_485240, EPI_ISL_485241, EPI_ISL_485242, EPI_ISL_485243, EPI_ISL_485244, EPI_ISL_485245, EPI_ISL_485246, EPI_ISL_485247, EPI_ISL_485248, EPI_ISL_485249, EPI_ISL_485250, EPI_ISL_485251, EPI_ISL_485252, EPI_ISL_485253, EPI_ISL_485254, EPI_ISL_485255, EPI_ISL_485256, EPI_ISL_485257, EPI_ISL_485258, EPI_ISL_485259, EPI_ISL_485260, EPI_ISL_485261, EPI_ISL_485262, EPI_ISL_485263, EPI_ISL_485264, EPI_ISL_485265, EPI_ISL_485266, EPI_ISL_485267, EPI_ISL_485268, EPI_ISL_485269, EPI_ISL_485270, EPI_ISL_485271, EPI_ISL_485272, EPI_ISL_485273, EPI_ISL_485274, EPI_ISL_485275, EPI_ISL_485276, EPI_ISL_485277, EPI_ISL_485278, EPI_ISL_485279, EPI_ISL_485280, EPI_ISL_485281, EPI_ISL_485282, EPI_ISL_485283, EPI_ISL_485284, EPI_ISL_485285, EPI_ISL_485286, EPI_ISL_485287, EPI_ISL_485288, EPI_ISL_485289, EPI_ISL_485290, EPI_ISL_485291, EPI_ISL_485292, EPI_ISL_485293, EPI_ISL_485294, EPI_ISL_485295, EPI_ISL_485296, EPI_ISL_485297, EPI_ISL_485298, EPI_ISL_485299, EPI_ISL_485300, EPI_ISL_485301, EPI_ISL_485302, EPI_ISL_485303, EPI_ISL_485304, EPI_ISL_485305, EPI_ISL_485306, EPI_ISL_485307, EPI_ISL_485308, EPI_ISL_485309, EPI_ISL_485310, EPI_ISL_485311, EPI_ISL_485312, EPI_ISL_485313, EPI_ISL_485314, EPI_ISL_485315, EPI_ISL_485316, EPI_ISL_485317, EPI_ISL_485318, EPI_ISL_485319, EPI_ISL_485320, EPI_ISL_485321, EPI_ISL_485322, EPI_ISL_485323, EPI_ISL_485324, EPI_ISL_485325, EPI_ISL_485326, EPI_ISL_485327, EPI_ISL_485328, EPI_ISL_485329, EPI_ISL_485330, EPI_ISL_485331, EPI_ISL_485332, EPI_ISL_485333, EPI_ISL_485334, EPI_ISL_485335, EPI_ISL_485336, EPI_ISL_485337, EPI_ISL_485338, EPI_ISL_485339, EPI_ISL_485340, EPI_ISL_485341, EPI_ISL_485342, EPI_ISL_485343, EPI_ISL_485344, EPI_ISL_485345, EPI_ISL_485346, EPI_ISL_485347, EPI_ISL_485348, EPI_ISL_485349, EPI_ISL_485350, EPI_ISL_485351, EPI_ISL_485352, EPI_ISL_485353, EPI_ISL_485354, EPI_ISL_485355, EPI_ISL_485356, EPI_ISL_485357, EPI_ISL_485358, EPI_ISL_485359, EPI_ISL_485360, EPI_ISL_485361, EPI_ISL_485362, EPI_ISL_485363, EPI_ISL_485364, EPI_ISL_485365, EPI_ISL_485366, EPI_ISL_485367, EPI_ISL_485368, EPI_ISL_485369, EPI_ISL_485370, EPI_ISL_485371, EPI_ISL_485372, EPI_ISL_485373, EPI_ISL_485374, EPI_ISL_485375, EPI_ISL_485376, EPI_ISL_485377, EPI_ISL_485378, EPI_ISL_485379, EPI_ISL_485380, EPI_ISL_485381, EPI_ISL_485382, EPI_ISL_485383, EPI_ISL_485384, EPI_ISL_485385, EPI_ISL_485386, EPI_ISL_485387 | | | |
| see above | River Road Testing Lab | Ginkgo Bioworks Clinical Laboratory | Rebecca C. Christofferson, Stephania A. Cormier, Luan V. Dinh, E. Handly Mayton, Hollis R. O'Neil, Thaya Stoufflet, Malaika Mckenzie-Bennett, James McGann, Jim Griffin, Keith Robison, Alex Plocik, Becky Schilling, Rebecca Littlefield, Michelle Spencer, Birgitte Simen |
| EPI_ISL_485388, EPI_ISL_485389, EPI_ISL_485390, EPI_ISL_485391, EPI_ISL_485392, EPI_ISL_485393, EPI_ISL_485394, EPI_ISL_485395, EPI_ISL_485396 | University of Ulsan College of Medicine and Asan Medical Center | University of Ulsan College of Medicine and Asan Medical Center | Kuenyoul Park, Jaewoong Lee, Kihyun Lee, Jiwon Jung, Sung-Han Kim, Jina Lee, Mauricio Chalita, Seok-Hwan Yoon, Jongsik Chun, Kyu-Hwa Hur, Heungsup Sung, Mi-Na Kim, and Hae Kyung Lee |
| EPI_ISL_485398 | Department of Internal Medicine, College of Medicine, Chosun University | Department of Internal Medicine, College of Medicine, Chosun University | Kim,D.-M. |
| EPI_ISL_485399 | Institute of Human Genetics, Polish Academy of Sciences | Institute of Human Genetics, Polish Academy of Sciences | Szymon Hryhorowicz, Adam Ustaszewski, Marta Kaczmarek-Ryś, Emilia Lis, Ewa Ziętkiewicz, Michał Witt, Andrzej Pławski |
| EPI_ISL_485400 | Institute of Human Genetics, Polish Academy of Sciences | Institute of Human Genetics, Polish Academy of Sciences, | Szymon Hryhorowicz, Adam Ustaszewski, Marta Kaczmarek-Ryś, Emilia Lis, Ewa Ziętkiewicz, Michał Witt, Andrzej Pławski |
| EPI_ISL_485401 | Communicable Disease Laboratory, Public Health Directorate | Communicable Disease Laboratory, Public Health Directorate | Zaed,A., Al-Wasti,H., Al-Taif,Z. and Shehab,F. |
| EPI_ISL_485575, EPI_ISL_485576, EPI_ISL_485577, EPI_ISL_485578, EPI_ISL_485579, EPI_ISL_485580, EPI_ISL_485581, EPI_ISL_485582, EPI_ISL_485583, EPI_ISL_485584, EPI_ISL_485585, EPI_ISL_485586, EPI_ISL_485587, EPI_ISL_485588, EPI_ISL_485589, EPI_ISL_485590, EPI_ISL_485591, EPI_ISL_485592, EPI_ISL_485593, EPI_ISL_485594, EPI_ISL_485595, EPI_ISL_485596, EPI_ISL_485597, EPI_ISL_485598, EPI_ISL_485599, EPI_ISL_485600, EPI_ISL_485601, EPI_ISL_485602 | | | |
| see above | Instituto de diagnóstico y Referencia Epidemiologicos (INDRE) | Instituto de diagnóstico y Referencia Epidemiologicos (INDRE) | Barrera-Badillo,G., Ramirez-Gonzalez,E. |
| EPI_ISL_485603 | Division of Infectious Disease | Steininger Lab | Jakob Thannesberger, Ingeborg Klymiuk, Nicolas Rascovan, Lorenz Schubert, Oliver Robak, Christoph Steininger |
| EPI_ISL_485604, EPI_ISL_485605, EPI_ISL_485606, EPI_ISL_485607, EPI_ISL_485608, EPI_ISL_485609, EPI_ISL_485610, EPI_ISL_485611 | Respiratory Virus Unit, Microbiology Services Colindale, Public Health England | Respiratory Virus Unit, Microbiology Services Colindale, Public Health England | PHE Covid Sequencing Team |
| EPI_ISL_485635, EPI_ISL_485708, EPI_ISL_485710, EPI_ISL_485711 | Institut Pasteur Dakar | Institut Pasteur de Dakar | Ndongo Dia, Moussa Moise Diagne, Mamadou diop, Marie Henriette Dior Ndione, Mamadou Malado Jallow, Safietou Sanke, Ousmane Faye, Amadou Alpha Sall. |
| EPI_ISL_485712 | Institut Pasteur | Institut Pasteur de Dakar | Ndongo Dia, Moussa Moise Diagne, Mamadou diop, Marie Henriette Dior Ndione, Mamadou Malado Jallow, Safietou Sanke, Ousmane Faye, Amadou Alpha Sall. |
| EPI_ISL_485713, EPI_ISL_485715, EPI_ISL_485716, EPI_ISL_485717 | Institut Pasteur Dakar | Institut Pasteur de Dakar | Ndongo Dia, Moussa Moise Diagne, Mamadou diop, Marie Henriette Dior Ndione, Mamadou Malado Jallow, Safietou Sanke, Ousmane Faye, Amadou Alpha Sall. |
| EPI_ISL_485809 | Institut für Virologie und Epidemiologie der Viruskrankheiten, Universitätsklinikum Tübingen | NGS Competence Center Tübingen, Institut für Medizinische Mikrobiologie und Hygiene, Universitätsklinikum Tübingen | Angelov at al. |
| EPI_ISL_485810, EPI_ISL_485811, EPI_ISL_485812, EPI_ISL_485813 | Institut für Virologie und Epidemiologie der Viruskrankheiten, Universitätsklinikum Tübingen | NGS Competence Center Tübingen, Institut für Medizinische Mikrobiologie und Hygiene, Universitätsklinikum Tübingen | Angelov et al. |
| EPI_ISL_485814, EPI_ISL_485815, EPI_ISL_485816, EPI_ISL_485817, EPI_ISL_485818, EPI_ISL_485819, EPI_ISL_485820, EPI_ISL_485821, EPI_ISL_485822, EPI_ISL_485823, EPI_ISL_485824, EPI_ISL_485825, EPI_ISL_485826, EPI_ISL_485827, EPI_ISL_485828, EPI_ISL_485829, EPI_ISL_485830, EPI_ISL_485831, EPI_ISL_485832, EPI_ISL_485833, EPI_ISL_485834, EPI_ISL_485835, EPI_ISL_485836, EPI_ISL_485837, EPI_ISL_485838, EPI_ISL_485839, EPI_ISL_485840, EPI_ISL_485841, EPI_ISL_485842, EPI_ISL_485843, EPI_ISL_485844, EPI_ISL_485845, EPI_ISL_485846, EPI_ISL_485847, EPI_ISL_485848, EPI_ISL_485849, EPI_ISL_485850, EPI_ISL_485851, EPI_ISL_485852, EPI_ISL_485853, EPI_ISL_485854, EPI_ISL_485855, EPI_ISL_485856, EPI_ISL_485857, EPI_ISL_485858, EPI_ISL_485859, EPI_ISL_485860, EPI_ISL_485861, EPI_ISL_485862, EPI_ISL_485863, EPI_ISL_485864, EPI_ISL_485865, EPI_ISL_485866, EPI_ISL_485867, EPI_ISL_485868, EPI_ISL_485869, EPI_ISL_485870, EPI_ISL_485871 | | | |
| see above | Virginia DCLS | Virginia DCLS | Virginia DCLS |
| EPI_ISL_485872 | New Mexico Department of Health Scientific Laboratory Division | Center for Global Health, University of New Mexico Health Sciences Center | Daryl Domman, Kurt Schwalm, Twila Kunde, Joseph Hicks, Michael Edwards, Darrell Dinwiddie |
| EPI_ISL_485873, EPI_ISL_485874, EPI_ISL_485875, EPI_ISL_485876, EPI_ISL_485877, EPI_ISL_485878, EPI_ISL_485879, EPI_ISL_485880, EPI_ISL_485881, EPI_ISL_485882, EPI_ISL_485883, EPI_ISL_485884, EPI_ISL_485885, EPI_ISL_485886, EPI_ISL_485887, EPI_ISL_485888, EPI_ISL_485889, EPI_ISL_485890, EPI_ISL_485891, EPI_ISL_485892, EPI_ISL_485893, EPI_ISL_485894, EPI_ISL_485895, EPI_ISL_485896, EPI_ISL_485897, EPI_ISL_485898, EPI_ISL_485899, EPI_ISL_485900, EPI_ISL_485901, EPI_ISL_485902, EPI_ISL_485903, EPI_ISL_485904, EPI_ISL_485905, EPI_ISL_485906 | | | |
| see above | River Road Testing Lab | Ginkgo Bioworks Clinical Laboratory | Rebecca C. Christofferson, Stephania A. Cormier, Luan V. Dinh, E. Handly Mayton, Hollis R. O'Neil, Thaya Stoufflet, Malaika Mckenzie-Bennett, James McGann, Jim Griffin, Keith Robison, Alex Plocik, Becky Schilling, Rebecca Littlefield, Michelle Spencer, Birgitte Simen |
| EPI_ISL_485917, EPI_ISL_485918, EPI_ISL_485919, EPI_ISL_485920, EPI_ISL_485921, EPI_ISL_485922, EPI_ISL_485923, EPI_ISL_485924, EPI_ISL_485925, EPI_ISL_485926, EPI_ISL_485927, EPI_ISL_485928, EPI_ISL_485929, EPI_ISL_485930, EPI_ISL_485931, EPI_ISL_485932, EPI_ISL_485933, EPI_ISL_485934, EPI_ISL_485935, EPI_ISL_485936, EPI_ISL_485937, EPI_ISL_485938, EPI_ISL_485939, EPI_ISL_485940, EPI_ISL_485941, EPI_ISL_485942, EPI_ISL_485944, EPI_ISL_485945, EPI_ISL_485946, EPI_ISL_485947, EPI_ISL_485948, EPI_ISL_485949, EPI_ISL_485950, EPI_ISL_485951, EPI_ISL_485952, EPI_ISL_485953, EPI_ISL_485954, EPI_ISL_485955, EPI_ISL_485956, EPI_ISL_485957, EPI_ISL_485958, EPI_ISL_485959, EPI_ISL_485960, EPI_ISL_485961, EPI_ISL_485963, EPI_ISL_485964, EPI_ISL_485967, EPI_ISL_485968, EPI_ISL_485969, EPI_ISL_485971, EPI_ISL_485972, EPI_ISL_485973, EPI_ISL_485974, EPI_ISL_485975, EPI_ISL_485976, EPI_ISL_485977, EPI_ISL_485978, EPI_ISL_485979, EPI_ISL_485980, EPI_ISL_485981, EPI_ISL_485982, EPI_ISL_485983, EPI_ISL_485984, EPI_ISL_485985, EPI_ISL_485986, EPI_ISL_485987, EPI_ISL_485988, EPI_ISL_485989, EPI_ISL_485990, EPI_ISL_485991, EPI_ISL_485992, EPI_ISL_485993, EPI_ISL_485994, EPI_ISL_485995, EPI_ISL_485996, EPI_ISL_485997, EPI_ISL_485998, EPI_ISL_485999, EPI_ISL_486000, EPI_ISL_486001, EPI_ISL_486002, EPI_ISL_486003, EPI_ISL_486004, EPI_ISL_486005, EPI_ISL_486006, EPI_ISL_486007, EPI_ISL_486008, EPI_ISL_486009, EPI_ISL_486010, EPI_ISL_486011, EPI_ISL_486012, EPI_ISL_486013, EPI_ISL_486014, EPI_ISL_486015, EPI_ISL_486016, EPI_ISL_486017, EPI_ISL_486018, EPI_ISL_486019, EPI_ISL_486020, EPI_ISL_486021, EPI_ISL_486022, EPI_ISL_486023, EPI_ISL_486024, EPI_ISL_486025, EPI_ISL_486026, EPI_ISL_486027, EPI_ISL_486028, EPI_ISL_486029, EPI_ISL_486030, EPI_ISL_486032, EPI_ISL_486033, EPI_ISL_486034, EPI_ISL_486035, EPI_ISL_486036, EPI_ISL_486037, EPI_ISL_486038, EPI_ISL_486039, EPI_ISL_486040, EPI_ISL_486041, EPI_ISL_486042, EPI_ISL_486043, EPI_ISL_486044, EPI_ISL_486045, EPI_ISL_486046, EPI_ISL_486047, EPI_ISL_486048, EPI_ISL_486049, EPI_ISL_486050, EPI_ISL_486051, EPI_ISL_486052, EPI_ISL_486053, EPI_ISL_486054, EPI_ISL_486055, EPI_ISL_486056, EPI_ISL_486057, EPI_ISL_486058, EPI_ISL_486059, EPI_ISL_486060, EPI_ISL_486061, EPI_ISL_486062, EPI_ISL_486063, EPI_ISL_486064, EPI_ISL_486065, EPI_ISL_486066, EPI_ISL_486067, EPI_ISL_486068, EPI_ISL_486069, EPI_ISL_486070, EPI_ISL_486071, EPI_ISL_486072, EPI_ISL_486073, EPI_ISL_486074, EPI_ISL_486075, EPI_ISL_486076, EPI_ISL_486077, EPI_ISL_486078, EPI_ISL_486079, EPI_ISL_486080, EPI_ISL_486081, EPI_ISL_486082, EPI_ISL_486083, EPI_ISL_486084, EPI_ISL_486085, EPI_ISL_486086, EPI_ISL_486087, EPI_ISL_486088, EPI_ISL_486089, EPI_ISL_486090, EPI_ISL_486093, EPI_ISL_486094, EPI_ISL_486095, EPI_ISL_486097, EPI_ISL_486098, EPI_ISL_486099, EPI_ISL_486102, EPI_ISL_486103, EPI_ISL_486105, EPI_ISL_486106, EPI_ISL_486108, EPI_ISL_486109, EPI_ISL_486110, EPI_ISL_486112, EPI_ISL_486113, EPI_ISL_486114 | | | |
| see above | UW Virology Lab | UW Virology Lab | Pavitra Roychoudhury, Hong Xie, Lasata Shrestha, Amin Addetia, Truong Nguyen, Victoria M Rachleff, Meei-Li Huang, Keith R Jerome, Alexander Greninger |
| EPI_ISL_486115, EPI_ISL_486116, EPI_ISL_486117, EPI_ISL_486118, EPI_ISL_486119 | County of Santa Clara Public Health Department | Chan-Zuckerberg Biohub | CZB Cliahub Consortium |
| EPI_ISL_486120, EPI_ISL_486121, EPI_ISL_486122, EPI_ISL_486123, EPI_ISL_486124, EPI_ISL_486125, EPI_ISL_486126, EPI_ISL_486127, EPI_ISL_486128, EPI_ISL_486129, EPI_ISL_486130, EPI_ISL_486131, EPI_ISL_486132, EPI_ISL_486133, EPI_ISL_486134, EPI_ISL_486135, EPI_ISL_486136, EPI_ISL_486137, EPI_ISL_486138, EPI_ISL_486139, EPI_ISL_486140, EPI_ISL_486141, EPI_ISL_486142, EPI_ISL_486143, EPI_ISL_486144, EPI_ISL_486145, EPI_ISL_486146, EPI_ISL_486147, EPI_ISL_486148, EPI_ISL_486149, EPI_ISL_486150, EPI_ISL_486151, EPI_ISL_486152, EPI_ISL_486153, EPI_ISL_486154, EPI_ISL_486155, EPI_ISL_486156, EPI_ISL_486157, EPI_ISL_486158, EPI_ISL_486159, EPI_ISL_486160, EPI_ISL_486161, EPI_ISL_486162, EPI_ISL_486163, EPI_ISL_486164, EPI_ISL_486165, EPI_ISL_486166, EPI_ISL_486167, EPI_ISL_486168, EPI_ISL_486169, EPI_ISL_486170, EPI_ISL_486171, EPI_ISL_486172, EPI_ISL_486173, EPI_ISL_486174, EPI_ISL_486175, EPI_ISL_486176, EPI_ISL_486177, EPI_ISL_486178, EPI_ISL_486179, EPI_ISL_486180, EPI_ISL_486181, EPI_ISL_486182, EPI_ISL_486183, EPI_ISL_486184, EPI_ISL_486185, EPI_ISL_486186, EPI_ISL_486187, EPI_ISL_486188, EPI_ISL_486189, EPI_ISL_486190, EPI_ISL_486191, EPI_ISL_486192, EPI_ISL_486193, EPI_ISL_486194, EPI_ISL_486195, EPI_ISL_486196, EPI_ISL_486197, EPI_ISL_486198, EPI_ISL_486199, EPI_ISL_486200, EPI_ISL_486201, EPI_ISL_486202, EPI_ISL_486203, EPI_ISL_486204, EPI_ISL_486205, EPI_ISL_486206, EPI_ISL_486207, EPI_ISL_486208, EPI_ISL_486209, EPI_ISL_486210, EPI_ISL_486211, EPI_ISL_486212, EPI_ISL_486213, EPI_ISL_486214, EPI_ISL_486215, EPI_ISL_486216, EPI_ISL_486217, EPI_ISL_486218, EPI_ISL_486219, EPI_ISL_486220, EPI_ISL_486221, EPI_ISL_486222, EPI_ISL_486223, EPI_ISL_486224, EPI_ISL_486225, EPI_ISL_486226, EPI_ISL_486227, EPI_ISL_486228, EPI_ISL_486229, EPI_ISL_486230, EPI_ISL_486231, EPI_ISL_486232, EPI_ISL_486233, EPI_ISL_486234, EPI_ISL_486235, EPI_ISL_486236, EPI_ISL_486237, EPI_ISL_486238, EPI_ISL_486239, EPI_ISL_486240, EPI_ISL_486241, EPI_ISL_486242, EPI_ISL_486243, EPI_ISL_486244, EPI_ISL_486245, EPI_ISL_486246, EPI_ISL_486247, EPI_ISL_486248, EPI_ISL_486249, EPI_ISL_486250, EPI_ISL_486251, EPI_ISL_486252, EPI_ISL_486253, EPI_ISL_486254, EPI_ISL_486255, EPI_ISL_486256, EPI_ISL_486257, EPI_ISL_486258, EPI_ISL_486259, EPI_ISL_486260, EPI_ISL_486261, EPI_ISL_486262, EPI_ISL_486263, EPI_ISL_486264, EPI_ISL_486265, EPI_ISL_486266, EPI_ISL_486267, EPI_ISL_486268, EPI_ISL_486269, EPI_ISL_486270, EPI_ISL_486271, EPI_ISL_486272, EPI_ISL_486273, EPI_ISL_486274, EPI_ISL_486275, EPI_ISL_486276, EPI_ISL_486277, EPI_ISL_486278, EPI_ISL_486279 | | | |
| see above | Orange County Public Health Laboratory | Chan-Zuckerberg Biohub | CZB Cliahub Consortium |
| EPI_ISL_486280, EPI_ISL_486281, EPI_ISL_486282, EPI_ISL_486283, EPI_ISL_486284, EPI_ISL_486285, EPI_ISL_486286 | Humboldt County Public Health Laboratory | Chan-Zuckerberg Biohub | CZB Cliahub Consortium |
| EPI_ISL_486287, EPI_ISL_486288, EPI_ISL_486289, EPI_ISL_486290, EPI_ISL_486291, EPI_ISL_486292, EPI_ISL_486293, EPI_ISL_486294, EPI_ISL_486295, EPI_ISL_486296, EPI_ISL_486297, EPI_ISL_486298, EPI_ISL_486299, EPI_ISL_486300, EPI_ISL_486301, EPI_ISL_486302, EPI_ISL_486303, EPI_ISL_486304, EPI_ISL_486305, EPI_ISL_486306, EPI_ISL_486307, EPI_ISL_486308, EPI_ISL_486309, EPI_ISL_486310, EPI_ISL_486311, EPI_ISL_486312, EPI_ISL_486313, EPI_ISL_486314, EPI_ISL_486315, EPI_ISL_486316, EPI_ISL_486317, EPI_ISL_486318, EPI_ISL_486319, EPI_ISL_486320, EPI_ISL_486321, EPI_ISL_486322, EPI_ISL_486323, EPI_ISL_486324, EPI_ISL_486325, EPI_ISL_486326, EPI_ISL_486327, EPI_ISL_486328, EPI_ISL_486329, EPI_ISL_486330, EPI_ISL_486331, EPI_ISL_486332, EPI_ISL_486333, EPI_ISL_486334, EPI_ISL_486335, EPI_ISL_486336, EPI_ISL_486337, EPI_ISL_486338 | | | |
| see above | San Joaquin County Public Health Lab | Chan-Zuckerberg Biohub | CZB Cliahub Consortium |
| EPI_ISL_486339, EPI_ISL_486340, EPI_ISL_486341, EPI_ISL_486342, EPI_ISL_486343, EPI_ISL_486344, EPI_ISL_486345, EPI_ISL_486346, EPI_ISL_486347, EPI_ISL_486348, EPI_ISL_486349, EPI_ISL_486350, EPI_ISL_486351, EPI_ISL_486352, EPI_ISL_486353, EPI_ISL_486354, EPI_ISL_486355, EPI_ISL_486356, EPI_ISL_486357, EPI_ISL_486358, EPI_ISL_486359, EPI_ISL_486360, EPI_ISL_486361, EPI_ISL_486362, EPI_ISL_486363, EPI_ISL_486364, EPI_ISL_486365 | | | |
| see above | UCSF Clinical Microbiology Laboratory | Chan-Zuckerberg Biohub | CZB Cliahub Consortium |
| EPI_ISL_486382 | District Surveillence Unit | Department of Neurovirology, National Institute of Mental Health and Neuroscience (NIMHANS) | Chitra Pattabiraman, Vijayalakshmi Reddy, Harsha PK, Risha Rasheed, Shafeeq S Hameed, Manjunatha Venkataswamy, Anita Desai, Ravi Vasanthapuram |
| EPI_ISL_486383 | CV Raman Hospital | Department of Neurovirology, National Institute of Mental Health and Neuroscience | Chitra Pattabiraman, Vijayalakshmi Reddy, Harsha PK, Risha Rasheed, Shafeeq S Hameed, Manjunatha Venkataswamy, Anita Desai, Ravi Vasanthapuram |

| | | | |
|---|---|---|---|
| | | (NIMHANS) | |
| EPI_ISL_486384 | DH | Department of Neurovirology, National Institute of Mental Health and Neuroscience (NIMHANS) | Chitra Pattabiraman, Vijayalakshmi Reddy, Harsha PK, Risha Rasheed, Shafeeq S Hameed, Manjunatha Venkataswamy, Anita Desai, Ravi Vasanthapuram |
| EPI_ISL_486385, EPI_ISL_486386 | Victoria Hospital | Department of Neurovirology, National Institute of Mental Health and Neuroscience (NIMHANS) | Chitra Pattabiraman, Vijayalakshmi Reddy, Harsha PK, Risha Rasheed, Shafeeq S Hameed, Manjunatha Venkataswamy, Anita Desai, Ravi Vasanthapuram |
| EPI_ISL_486387, EPI_ISL_486388, EPI_ISL_486389 | DH | Department of Neurovirology, National Institute of Mental Health and Neuroscience (NIMHANS) | Chitra Pattabiraman, Vijayalakshmi Reddy, Harsha PK, Risha Rasheed, Shafeeq S Hameed, Manjunatha Venkataswamy, Anita Desai, Ravi Vasanthapuram |
| EPI_ISL_486390, EPI_ISL_486391 | Centrālā laboratorija | Latvian Biomedical Research and Study Centre | Ivars Silamiķelis, Kaspars Megnis, Monta Ustinova, Ņikita Zrelovs, Vita Rovīte, Stella Lapiņa, Jana Osīte, Marta Priedīte, Uga Dumpis, Jānis Kloviņš |
| EPI_ISL_486392 | Victoria Hospital | Department of Neurovirology, National Institute of Mental Health and Neuroscience (NIMHANS) | Chitra Pattabiraman, Vijayalakshmi Reddy, Harsha PK, Risha Rasheed, Shafeeq S Hameed, Manjunatha Venkataswamy, Anita Desai, Ravi Vasanthapuram |
| EPI_ISL_486393 | SJMCH | Department of Neurovirology, National Institute of Mental Health and Neuroscience (NIMHANS) | Chitra Pattabiraman, Vijayalakshmi Reddy, Harsha PK, Risha Rasheed, Shafeeq S Hameed, Manjunatha Venkataswamy, Anita Desai, Ravi Vasanthapuram |
| EPI_ISL_486394 | MIMS | Department of Neurovirology, National Institute of Mental Health and Neuroscience (NIMHANS) | Chitra Pattabiraman, Vijayalakshmi Reddy, Harsha PK, Risha Rasheed, Shafeeq S Hameed, Manjunatha Venkataswamy, Anita Desai, Ravi Vasanthapuram |
| EPI_ISL_486395 | BIMS | Department of Neurovirology, National Institute of Mental Health and Neuroscience (NIMHANS) | Chitra Pattabiraman, Vijayalakshmi Reddy, Harsha PK, Risha Rasheed, Shafeeq S Hameed, Manjunatha Venkataswamy, Anita Desai, Ravi Vasanthapuram |
| EPI_ISL_486396 | Jayanagar General Hospital to Victoria Hospital | Department of Neurovirology, National Institute of Mental Health and Neuroscience (NIMHANS) | Chitra Pattabiraman, Vijayalakshmi Reddy, Harsha PK, Risha Rasheed, Shafeeq S Hameed, Manjunatha Venkataswamy, Anita Desai, Ravi Vasanthapuram |
| EPI_ISL_486397 | KC General Hospital | Department of Neurovirology, National Institute of Mental Health and Neuroscience (NIMHANS) | Chitra Pattabiraman, Vijayalakshmi Reddy, Harsha PK, Risha Rasheed, Shafeeq S Hameed, Manjunatha Venkataswamy, Anita Desai, Ravi Vasanthapuram |
| EPI_ISL_486398, EPI_ISL_486399 | MIMS | Department of Neurovirology, National Institute of Mental Health and Neuroscience (NIMHANS) | Chitra Pattabiraman, Vijayalakshmi Reddy, Harsha PK, Risha Rasheed, Shafeeq S Hameed, Manjunatha Venkataswamy, Anita Desai, Ravi Vasanthapuram |
| EPI_ISL_486400 | Victoria Hospital | Department of Neurovirology, National Institute of Mental Health and Neuroscience (NIMHANS) | Chitra Pattabiraman, Vijayalakshmi Reddy, Harsha PK, Risha Rasheed, Shafeeq S Hameed, Manjunatha Venkataswamy, Anita Desai, Ravi Vasanthapuram |
| EPI_ISL_486401, EPI_ISL_486402, EPI_ISL_486403 | DH | Department of Neurovirology, National Institute of Mental Health and Neuroscience (NIMHANS) | Chitra Pattabiraman, Vijayalakshmi Reddy, Harsha PK, Risha Rasheed, Shafeeq S Hameed, Manjunatha Venkataswamy, Anita Desai, Ravi Vasanthapuram |
| EPI_ISL_486404 | Victoria Hospital | Department of Neurovirology, National Institute of Mental Health and Neuroscience (NIMHANS) | Chitra Pattabiraman, Vijayalakshmi Reddy, Harsha PK, Risha Rasheed, Shafeeq S Hameed, Manjunatha Venkataswamy, Anita Desai, Ravi Vasanthapuram |
| EPI_ISL_486405, EPI_ISL_486406, EPI_ISL_486407, EPI_ISL_486408, EPI_ISL_486409 | DH | Department of Neurovirology, National Institute of Mental Health and Neuroscience (NIMHANS) | Chitra Pattabiraman, Vijayalakshmi Reddy, Harsha PK, Risha Rasheed, Shafeeq S Hameed, Manjunatha Venkataswamy, Anita Desai, Ravi Vasanthapuram |
| EPI_ISL_486410 | Centrala laboratorija | Latvian Biomedical Research and Study Centre | Ivars Silamiķelis, Kaspars Megnis, Monta Ustinova, Ņikita Zrelovs, Vita Rovīte, Stella Lapiņa, Jana Osīte, Marta Priedīte, Uga Dumpis, Jānis Kloviņš |
| EPI_ISL_486411 | Centrālā laboratorija | Latvian Biomedical Research and Study Centre | Ivars Silamiķelis, Kaspars Megnis, Monta Ustinova, Ņikita Zrelovs, Vita Rovīte, Stella Lapiņa, Jana Osīte, Marta Priedīte, Uga Dumpis, Jānis Kloviņš |
| EPI_ISL_486412 | Centrālā laboratorija | Latvian Biomedical Research and Study Centre | Ivars Silamiķelis, Kaspars Megnis, Monta Ustinova, Ņikita Zrelovs, Vita Rovīte, Stella Lapiņa, Jana Osīte, Marta Priedīte, Uga Dumpis, Jānis Kloviņš |
| EPI_ISL_486413, EPI_ISL_486414 | Centrālā laboratorija | Latvian Biomedical Research and Study Centre | Ivars Silamiķelis, Kaspars Megnis, Monta Ustinova, Ņikita Zrelovs, Vita Rovīte, Stella Lapiņa, Jana Osīte, Marta Priedīte, Uga Dumpis, Jānis Kloviņš |
| EPI_ISL_486415 | Centrālā laboratorija | Latvian Biomedical Research and Study Centre | Ivars Silamiķelis, Kaspars Megnis, Monta Ustinova, Ņikita Zrelovs, Vita Rovīte, Stella Lapiņa, Jana Osīte, Marta Priedīte, Uga Dumpis, Jānis Kloviņš |
| EPI_ISL_486416 | Centrālā laboratorija | Latvian Biomedical Research and Study Centre | Ivars Silamiķelis, Kaspars Megnis, Monta Ustinova, Ņikita Zrelovs, Vita Rovīte, Stella Lapiņa, Jana Osīte, Marta Priedīte, Uga Dumpis, Jānis Kloviņš |
| EPI_ISL_486417 | Centrala laboratorija | Latvian Biomedical Research and Study Centre | Ivars Silamiķelis, Kaspars Megnis, Monta Ustinova, Ņikita Zrelovs, Vita Rovīte, Stella Lapiņa, Jana Osīte, Marta Priedīte, Uga Dumpis, Jānis Kloviņš |
| EPI_ISL_486418 | Centrālā laboratorija | Latvian Biomedical Research and Study Centre | Ivars Silamiķelis, Kaspars Megnis, Monta Ustinova, Ņikita Zrelovs, Vita Rovīte, Stella Lapiņa, Jana Osīte, Marta Priedīte, Uga Dumpis, Jānis Kloviņš |
| EPI_ISL_486419 | Centrālā laboratorija | Latvian Biomedical Research and Study Centre | Ivars Silamiķelis, Kaspars Megnis, Monta Ustinova, Ņikita Zrelovs, Vita Rovīte, Stella Lapiņa, Jana Osīte, Marta Priedīte, Uga Dumpis, Jānis Kloviņš |
| EPI_ISL_486420, EPI_ISL_486421 | Centrālā laboratorija | Latvian Biomedical Research and Study Centre | Ivars Silamiķelis, Kaspars Megnis, Monta Ustinova, Ņikita Zrelovs, Vita Rovīte, Stella Lapiņa, Jana Osīte, Marta Priedīte, Uga Dumpis, Jānis Kloviņš |
| EPI_ISL_486422, EPI_ISL_486423, EPI_ISL_486424, EPI_ISL_486425, EPI_ISL_486426 | Latvijas Infektoloģijas centrs | Latvian Biomedical Research and Study Centre | Ivars Silamiķelis, Kaspars Megnis, Monta Ustinova, Ņikita Zrelovs, Vita Rovīte, Jeļena Storoženko, Tatjana Kolupajeva, Oksana Savicka, Uga Dumpis, Jānis Kloviņš |
| EPI_ISL_486427 | unknown | Clinical Laboratory, Hospital Israelita Albert Einstein | Amgarteb,D., Malta,F., Guedes,R.L., Santana,R.A., de Menezes,F.G., Mangueira,C.L. and Pinho,J.R. |
| EPI_ISL_486428 | Latvijas Infektoloģijas centrs | Latvian Biomedical Research and Study Centre | Ivars Silamiķelis, Kaspars Megnis, Monta Ustinova, Ņikita Zrelovs, Vita Rovīte, Jeļena Storoženko, Tatjana Kolupajeva, Oksana Savicka, Uga Dumpis, Jānis Kloviņš |
| EPI_ISL_486429 | unknown | Clinical Laboratory, Hospital Israelita Albert Einstein | Malta,F., Amgarten,D., Guedes,R.L., Santana,R.A., de Menezes,F.G., Mangueira,C.L. and Pinho,J.R. |
| EPI_ISL_486430, EPI_ISL_486431, EPI_ISL_486432, EPI_ISL_486433, EPI_ISL_486434, EPI_ISL_486435 | Latvijas Infektoloģijas centrs | Latvian Biomedical Research and Study Centre | Ivars Silamiķelis, Kaspars Megnis, Monta Ustinova, Ņikita Zrelovs, Vita Rovīte, Jeļena Storoženko, Tatjana Kolupajeva, Oksana Savicka, Uga Dumpis, Jānis Kloviņš |
| EPI_ISL_486436 | Latvijas Infektoloģijas centrs | Latvian Biomedical Research and Study Centre | Ivars Silamiķelis, Kaspars Megnis, Monta Ustinova, Ņikita Zrelovs, Vita Rovīte, Jeļena Storoženko, Tatjana Kolupajeva, Oksana Savicka, Uga Dumpis, Jānis Kloviņš |
| EPI_ISL_486437 | Centrālā laboratorija | Latvian Biomedical Research and Study Centre | Ivars Silamiķelis, Kaspars Megnis, Monta Ustinova, Ņikita Zrelovs, Vita Rovīte, Stella Lapiņa, Jana Osīte, Marta Priedīte, Uga Dumpis, Jānis Kloviņš |
| EPI_ISL_486438 | E. Gulbja laboratorija | Latvian Biomedical Research and Study Centre | Ivars Silamiķelis, Kaspars Megnis, Monta Ustinova, Ņikita Zrelovs, Vita Rovīte, Mikus Gavars, Dmitrijs Perminovs, Uga Dumpis, Jānis Kloviņš |
| EPI_ISL_486442, EPI_ISL_486443, EPI_ISL_486444, EPI_ISL_486445, EPI_ISL_486446, EPI_ISL_486447, EPI_ISL_486448, EPI_ISL_486449, EPI_ISL_486450, EPI_ISL_486451, EPI_ISL_486452, EPI_ISL_486453, EPI_ISL_486454, EPI_ISL_486455, EPI_ISL_486456, EPI_ISL_486457, EPI_ISL_486458, EPI_ISL_486459, EPI_ISL_486460, EPI_ISL_486461, EPI_ISL_486462, EPI_ISL_486463, EPI_ISL_486464, EPI_ISL_486465, EPI_ISL_486466, EPI_ISL_486467, EPI_ISL_486468, EPI_ISL_486469, EPI_ISL_486470, EPI_ISL_486471, EPI_ISL_486472, EPI_ISL_486473, EPI_ISL_486474, EPI_ISL_486475, EPI_ISL_486476, EPI_ISL_486477, EPI_ISL_486478, EPI_ISL_486479, EPI_ISL_486480, EPI_ISL_486481, EPI_ISL_486482, EPI_ISL_486483, EPI_ISL_486484, EPI_ISL_486485, EPI_ISL_486486, EPI_ISL_486487, EPI_ISL_486488, EPI_ISL_486489, EPI_ISL_486490, EPI_ISL_486491, EPI_ISL_486492, EPI_ISL_486493, EPI_ISL_486494, EPI_ISL_486495, EPI_ISL_486496, EPI_ISL_486497, EPI_ISL_486498, EPI_ISL_486499, EPI_ISL_486500, EPI_ISL_486501, EPI_ISL_486502, EPI_ISL_486503, EPI_ISL_486504, EPI_ISL_486505, EPI_ISL_486506, EPI_ISL_486507, EPI_ISL_486508, EPI_ISL_486509, EPI_ISL_486510, EPI_ISL_486511, EPI_ISL_486512, EPI_ISL_486513, EPI_ISL_486514, EPI_ISL_486515, EPI_ISL_486516, EPI_ISL_486517, EPI_ISL_486518, EPI_ISL_486519, EPI_ISL_486520, EPI_ISL_486521, EPI_ISL_486522, EPI_ISL_486523, EPI_ISL_486524, EPI_ISL_486525, EPI_ISL_486526, EPI_ISL_486527, EPI_ISL_486528, EPI_ISL_486529, EPI_ISL_486530, EPI_ISL_486531, EPI_ISL_486532, EPI_ISL_486533, EPI_ISL_486534, EPI_ISL_486535, EPI_ISL_486536, EPI_ISL_486537, EPI_ISL_486538 | | | |
| see above | Viollier AG | Department of Biosystems Science and Engineering, ETH Zürich | Christian Beisel, Sarah Nadeau, Ivan Topolsky, Pedro Ferreira, Philipp Jablonski, Susana Posada-Céspedes, Tobias Schär, Ina Nissen, Natascha Santacroce, Elodie Burcklen, Christiane Beckmann, Maurice Redondo, Olivier Kobel, Christoph Noppen, Sophie Seidel, Noemie Santamaria de Souza, Niko Beerenwinkel, Tanja Stadler |
| EPI_ISL_486646, EPI_ISL_486647 | Microbiology, Virology and Biemergency Laboratory-ASST FBF Sacco | Microbiology, Virology and Biemergency Laboratory-ASST FBF Sacco | Mancon A, Comandatore F, Romeri F, Micheli V, Rimoldi SG |
| EPI_ISL_486648 | Microbiology, Virology and Biemergency Laboratory-ASST FBF Sacco | Microbiology, Virology and Biemergency Laboratory-ASST FBF Sacco | Micheli V, Comandatore F, Romeri F, Mancon A, Rimoldi SG |

| | | | |
|---|---|---|---|
| EPI_ISL_486649 | Microbiology, Virology and Biemergency Laboratory-ASST FBF Sacco | Microbiology, Virology and Biemergency Laboratory-ASST FBF Sacco | Rimoldi SG, Comandatore F, Romeri F, Mancon A, Micheli V |
| EPI_ISL_486650 | Microbiology, Virology and Biemergency Laboratory-ASST FBF Sacco | Microbiology, Virology and Biemergency Laboratory-ASST FBF Sacco | Romeri F, Comandatore F, Mancon A, Micheli V, Rimoldi SG |
| EPI_ISL_486651 | Microbiology, Virology and Biemergency Laboratory-ASST FBF Sacco | Microbiology, Virology and Biemergency Laboratory-ASST FBF Sacco | Mancon A, Comandatore F, Romeri F, Micheli V, Rimoldi SG |
| EPI_ISL_486652 | Microbiology, Virology and Biemergency Laboratory-ASST FBF Sacco | Microbiology, Virology and Biemergency Laboratory-ASST FBF Sacco | Micheli V, Comandatore F, Romeri F, Mancon A, Rimoldi SG |
| EPI_ISL_486653 | Microbiology, Virology and Biemergency Laboratory-ASST FBF Sacco | Microbiology, Virology and Biemergency Laboratory-ASST FBF Sacco | Rimoldi SG, Comandatore F, Romeri F, Mancon A, Micheli V |
| EPI_ISL_486654 | Microbiology, Virology and Biemergency Laboratory-ASST FBF Sacco | Microbiology, Virology and Biemergency Laboratory-ASST FBF Sacco | Romeri F, Comandatore F, Mancon A, Micheli V, Rimoldi SG |
| EPI_ISL_486655 | Microbiology, Virology and Biemergency Laboratory-ASST FBF Sacco | Microbiology, Virology and Biemergency Laboratory-ASST FBF Sacco | Mancon A, Comandatore F, Romeri F, Micheli V, Rimoldi SG |
| EPI_ISL_486656 | Microbiology, Virology and Biemergency Laboratory-ASST FBF Sacco | Microbiology, Virology and Biemergency Laboratory-ASST FBF Sacco | Micheli V, Comandatore F, Romeri F, Mancon A, Rimoldi SG |
| EPI_ISL_486657 | Microbiology, Virology and Biemergency Laboratory-ASST FBF Sacco | Microbiology, Virology and Biemergency Laboratory-ASST FBF Sacco | Rimoldi SG, Comandatore F, Romeri F, Mancon A, Micheli V |
| EPI_ISL_486658 | Microbiology, Virology and Biemergency Laboratory-ASST FBF Sacco | Microbiology, Virology and Biemergency Laboratory-ASST FBF Sacco | Romeri F, Comandatore F, Mancon A, Micheli V, Rimoldi SG |
| EPI_ISL_486659 | Microbiology, Virology and Biemergency Laboratory-ASST FBF Sacco | Microbiology, Virology and Biemergency Laboratory-ASST FBF Sacco | Micheli V, Comandatore F, Romeri F, Mancon A, Rimoldi SG |
| EPI_ISL_486660 | Microbiology, Virology and Biemergency Laboratory-ASST FBF Sacco | Microbiology, Virology and Biemergency Laboratory-ASST FBF Sacco | Rimoldi SG, Comandatore F, Romeri F, Mancon A, Micheli V |
| EPI_ISL_486661 | Microbiology, Virology and Biemergency Laboratory-ASST FBF Sacco | Microbiology, Virology and Biemergency Laboratory-ASST FBF Sacco | Romeri F, Comandatore F, Mancon A, Micheli V, Rimoldi SG |
| EPI_ISL_486662 | Microbiology, Virology and Biemergency Laboratory-ASST FBF Sacco | Microbiology, Virology and Biemergency Laboratory-ASST FBF Sacco | Mancon A, Comandatore F, Romeri F, Micheli V, Rimoldi SG |
| EPI_ISL_486663 | Microbiology, Virology and Biemergency Laboratory-ASST FBF Sacco | Microbiology, Virology and Biemergency Laboratory-ASST FBF Sacco | Micheli V, Comandatore F, Romeri F, Mancon A, Rimoldi SG |
| EPI_ISL_486664 | Microbiology, Virology and Biemergency Laboratory-ASST FBF Sacco | Microbiology, Virology and Biemergency Laboratory-ASST FBF Sacco | Rimoldi SG, Comandatore F, Romeri F, Mancon A, Micheli V |
| EPI_ISL_486665 | Microbiology, Virology and Biemergency Laboratory-ASST FBF Sacco | Microbiology, Virology and Biemergency Laboratory-ASST FBF Sacco | Micheli V, Rimoldi SG, Comandatore F, Mancon A, Romeri F |
| EPI_ISL_486666, EPI_ISL_486667, EPI_ISL_486668, EPI_ISL_486669, EPI_ISL_486670, EPI_ISL_486671, EPI_ISL_486672 | Institute for Stem Cell Science and Regenerative Medicine | National Centre for Biological Sciences | Farhan Ali, Vanessa Molin Paynter, Srikar Krishna, Mohak Sharda, Shah-e-Jahan Gulzar, Awadhesh Pandit, Varadha Sundarmurthy, Uma Ramakrishnan, Dasaradhi Palakodeti, Aswin Seshasayee |
| EPI_ISL_486673, EPI_ISL_486674 | Institute for Stem Cell Science and Regenerative Medicine | National Centre for Biological Sciences | Farhan Ali, Vanessa Molin Paynter, Srikar Krishna, Mohak Sharda, Shah-e-Jahan Gulzar, Awadhesh Pandit, Varadha Sundarmurthy, Uma Ramakrishnan, Dasaradhi Palakodeti, Aswin Seshasayee |
| EPI_ISL_486815, EPI_ISL_486816, EPI_ISL_486817, EPI_ISL_486818, EPI_ISL_486819, EPI_ISL_486820, EPI_ISL_486821, EPI_ISL_486822, EPI_ISL_486823, EPI_ISL_486824, EPI_ISL_486825, EPI_ISL_486826, EPI_ISL_486827, EPI_ISL_486828, EPI_ISL_486829 | | | |
| see above | Molecular diagnostic laboratory of Federal Budget Institution of Science "Central Research Institute of Epidemiology" of The Federal Service on Customers' Rights Protection and Human Well-being Surveillance | Group of Genomics and Postgenomic Technologies of Central Research Institute of Epidemilology | Speranskaya AS, Kaptelova VV, Valdokhina AV, Bulanenko VP, Samoilov AE, Korneenko EV, Tivanova EV, Shipulina OY, Akimkin VG |
| EPI_ISL_486830, EPI_ISL_486831 | Providence St. Joseph Health Molecular Genomics Laboratory | Providence St. Joseph Health Molecular Genomics Laboratory | Alexa K Dowdell, Brian D Piening, Fred L Robinson, Carlo B Bifulco, Mary Campbell |
| EPI_ISL_486834 | Suceava County Energency Hospital "Sf. Ioan cel Nou" | SMU Metagenomics lab | Lobiuc Andrei, Antoniadis Panagiotis |
| EPI_ISL_486835, EPI_ISL_486836, EPI_ISL_486837, EPI_ISL_486838, EPI_ISL_486839, EPI_ISL_486840, EPI_ISL_486841 | Institute for Stem Cell Science and Regenerative Medicine | National Centre for Biological Sciences | Farhan Ali, Vanessa Molin Paynter, Srikar Krishna, Mohak Sharda, Shah-e-Jahan Gulzar, Awadhesh Pandit, Varadha Sundarmurthy, Uma Ramakrishnan, Dasaradhi Palakodeti, Aswin Seshasayee |
| EPI_ISL_486842, EPI_ISL_486843, EPI_ISL_486844 | Institute of Microbiology, Universidad San Francisco de Quito | Institute of Microbiology, Universidad San Francisco de Quito | Belén Prado-Vivar, Sully Márquez, Juan José Guadalupe, Monica Becerra-Wong, Carla Torres, Bernardo Gutiérrez, Fausto Maldonado, Geovanny Carzola, Verónica Barragán, Patricio Rojas-Silva, Gabriel Trueba, Michelle Grunauer, Paúl Cárdenas |
| EPI_ISL_486845, EPI_ISL_486846, EPI_ISL_486847, EPI_ISL_486848, EPI_ISL_486849, EPI_ISL_486850, EPI_ISL_486851 | Institute of Microbiology, Universidad San Francisco de Quito | Institute of Microbiology, Universidad San Francisco de Quito | Belén Prado-Vivar, Sully Márquez, Juan José Guadalupe, Monica Becerra-Wong, Carla Torres, Bernardo Gutiérrez, Jonathan Araujo, Verónica Barragán, Patricio Rojas-Silva, Gabriel Trueba, Michelle Grunauer, Paúl Cárdenas |
| EPI_ISL_486852 | CDRI/SGPGI | CSIR-CDRI/SGPGI | Saumya Sarkar, Dharam Veer Singh, Rahul Vishvkarma, Ujjala Ghoshal, Uday Ghoshal, Ravishankar Ramachandran, Tapas Kumar Kundu, Rajender Singh |
| EPI_ISL_486853 | CSIR-CDRI/SGPGI | CSIR-CDRI/SGPGI | Saumya Sarkar, Dharam Veer Singh, Rahul Vishvkarma, Ujjala Ghoshal, Uday Ghoshal, Ravishankar Ramachandran, Tapas Kumar Kundu, Rajender Singh |
| EPI_ISL_486854 | Emergency County Hospital Suceava | Stefan cel Mare, University Metagenomics lab | Lobiuc Andrei et al. |
| EPI_ISL_486855 | Emergency county Hospital Suceava | "Stefan cel Mare" University Metagenomics Lab | Lobiuc Andrei et al. |
| EPI_ISL_486856 | Emergency County Hospital | Stefan cel Mare, University Metagenomics lab | Lobiuc Andrei et al. |
| EPI_ISL_486857, EPI_ISL_486858, EPI_ISL_486859, EPI_ISL_486860, EPI_ISL_486861, EPI_ISL_486862, EPI_ISL_486863, EPI_ISL_486864, EPI_ISL_486865, EPI_ISL_486866, EPI_ISL_486867, EPI_ISL_486868, EPI_ISL_486869, EPI_ISL_486870, EPI_ISL_486871, EPI_ISL_486872, EPI_ISL_486873 | | | |
| see above | Institut Pasteur Dakar | Institut Pasteur de Dakar | Ndongo Dia, Moussa Moise Diagne, Mamadou Diop, Marie Henriette Dior Ndione, Mamadou Malado Jallow, Safietou Sanke, Ousmane Faye, Amadou Alpha Sall. |
| EPI_ISL_486876 | Clinical Microbiology Laboratory- Basurto University Hospital | Biocruces-Bizkaia | Mikel J. Urrutikoetxea-Gutierrez, Ana Belén Belén de la Hoz, Matxalen Vidal-García, Mº Carmen Nieto Toboso, Estibaliz Ugalde-Zarraga, José Luis Díaz de Tuesta del Arco |
| EPI_ISL_486881 | CV Raman Hospital | Department of Neurovirology, National Institute of Mental Health and Neuroscience (NIMHANS) | Chitra Pattabiraman, Vijayalakshmi Reddy, Harsha PK, Risha Rasheed, Shafeeq S Hameed, Manjunatha Venkataswamy, Anita Desai, Ravi Vasanthapuram |
| EPI_ISL_486884, EPI_ISL_486885 | Hamedan University of Medical Sciences | Hamedan University of Medical Sciences | Teimoori,A., Azizi jalilian,F., Ansari,N., Jamehdor,S., Zanjani,M., Nazari,A., Saadat,N. and Mazaheri,Z. |
| EPI_ISL_486887 | National Influenza Center, Bahrain | National Influenza Center, Bahrain | Zaed,A., Altaif,Z., Shehab,F., AlWasti,H. |
| EPI_ISL_486888 | National Influenza Center, Bahrain | National Influenza Center, Bahrain | AlWasti,H., Altaif,Z., Zaed,A., Shehab,F. |
| EPI_ISL_486889 | National Influenza Center, Bahrain | National Influenza Center, Bahrain | Altaif,Z., AlWasti,H., Shehab,F., Zaed,A. |
| EPI_ISL_486897, EPI_ISL_486898, EPI_ISL_486899, EPI_ISL_486900, EPI_ISL_486901, EPI_ISL_486902, EPI_ISL_486903, EPI_ISL_486904, EPI_ISL_486905, EPI_ISL_486906, EPI_ISL_486907, EPI_ISL_486908, EPI_ISL_486909, EPI_ISL_486910, EPI_ISL_486911 | | | |
| see above | Tokyo Metoropolitan Institute of Public Health | Tokyo Metoropolitan Institute of Public Health | Asakura,H., Yoshida,I., Kumagai,R., Chiba,T., Sadamasu,K., Nagashima,M. |
| EPI_ISL_486912, EPI_ISL_486913, EPI_ISL_486914, EPI_ISL_486915, EPI_ISL_486916, EPI_ISL_486917 | Maryland Department of Health | Maryland Department of Health | Keller,E. |
| EPI_ISL_487087, EPI_ISL_487089, EPI_ISL_487090, EPI_ISL_487091, EPI_ISL_487092 | Nigeria Centre for Disease Control (NCDC) | African Centre of Excellence for Genomics of Infectious Diseases (ACEGID), Redeemer's University, Ede, Osun State, Nigeria | Oluniyi P.E., Ajogbasile F.V., Kayode A., Oguzie J., Olawoye I., Uwanibe J., Olumade T., Folarin O.A., Ihekweazu C., Happi C.T. |
| EPI_ISL_487095 | Nigeria Centre for Disease Control (NCDC) | African Centre of Excellence for Genomics of Infectious Diseases (ACEGID), Redeemer's University, Ede, Osun State, Nigeria | Oluniyi P.E., Ajogbasile F.V., Kayode A., Oguzie J., Olawoye I., Uwanibe J., Olumade T., Folarin O.A., Ihekweazu C., Happi C.T. |
| EPI_ISL_487096, EPI_ISL_487097, EPI_ISL_487098, EPI_ISL_487099, EPI_ISL_487100, EPI_ISL_487101, EPI_ISL_487102, EPI_ISL_487103, EPI_ISL_487104, EPI_ISL_487105, EPI_ISL_487106, EPI_ISL_487107, EPI_ISL_487108, EPI_ISL_487109, EPI_ISL_487110, EPI_ISL_487111, EPI_ISL_487112 | | | |
| see above | Nigeria Centre for Disease Control (NCDC) | African Centre of Excellence for Genomics of Infectious Diseases (ACEGID), Redeemer's | Oluniyi P.E., Ajogbasile F.V., Kayode A., Oguzie J., Olawoye I., Uwanibe J., Olumade T., Folarin O.A., Ihekweazu C., Happi C.T. |

| | | | |
|---|---|---|---|
| | | University, Ede, Osun State, Nigeria | |
| EPI_ISL_487113 | Nigeria Centre for Disease Control (NCDC) | Redeemer's University, ACEGID | Oluniyi P.E., Ajogbasile F.V., Kayode A., Oguzie J., Olawoye I., Uwanibe J., Olumade T., Folarin O.A., Ihekweazu C., Happi C.T. |
| EPI_ISL_487192 | Viral Respiratory Lab, National institute for Biomedical Research (INRB) | Pathogen Sequencing Lab, National Institute for Biomedical Research (INRB) | Placide Mbala-Kingebeni, Edith Nkwembe, Eddy Kinganda-Lusamaki, Amuri Aziza, Francisca Muyembe-Mawete, Emmanuel Lokilo-Lofiko, Catherine Pratt, Matthias Pauthner, Josh Quick, Allison Black, James Hadfield, Trevor Bedford, Ian Goodfellow, Andrew Rambault, Nick Loman, Kristian Andersen, Michael Wiley, Steve Ahuka-Mundeke, Jean-Jacques Muyembe Tamfum. |
| EPI_ISL_487193, EPI_ISL_487194, EPI_ISL_487195, EPI_ISL_487196, EPI_ISL_487197, EPI_ISL_487198, EPI_ISL_487199, EPI_ISL_487200, EPI_ISL_487201, EPI_ISL_487202, EPI_ISL_487203, EPI_ISL_487204, EPI_ISL_487205, EPI_ISL_487206, EPI_ISL_487207, EPI_ISL_487208, EPI_ISL_487209, EPI_ISL_487210, EPI_ISL_487211, EPI_ISL_487212, EPI_ISL_487213, EPI_ISL_487214, EPI_ISL_487215, EPI_ISL_487216, EPI_ISL_487217, EPI_ISL_487218, EPI_ISL_487219, EPI_ISL_487220, EPI_ISL_487221, EPI_ISL_487222, EPI_ISL_487223, EPI_ISL_487224, EPI_ISL_487225, EPI_ISL_487226, EPI_ISL_487227, EPI_ISL_487228 | | | |
| see above | Utah Public Health Laboratory | Utah Public Health Laboratory | Heidi Butz, Erin Young, Kelly Oakeson |
| EPI_ISL_487229, EPI_ISL_487230 | University of Michigan Clinical Microbiology Laboratory | Lauring Lab, University of Michigan, Department of Microbiology and Immunology | Valesano et al. |
| EPI_ISL_487231 | Oklahoma State University Diagnostic Laboratory- Oklahoma Animal Disease Diagnostic Laboratory | Oklahoma Animal Disease Diagnostic Laboratory | Sai Narayanan, Teluguakula Narasaraju, John Ritchey, Girish Patil, Sunil More, Jerry Saliki, Anil Kaul, Akhilesh Ramachandran |
| EPI_ISL_487244, EPI_ISL_487245, EPI_ISL_487246, EPI_ISL_487247, EPI_ISL_487248, EPI_ISL_487249, EPI_ISL_487250, EPI_ISL_487251, EPI_ISL_487252, EPI_ISL_487253, EPI_ISL_487254, EPI_ISL_487255, EPI_ISL_487256, EPI_ISL_487257, EPI_ISL_487258, EPI_ISL_487259, EPI_ISL_487260, EPI_ISL_487261, EPI_ISL_487262, EPI_ISL_487263, EPI_ISL_487264, EPI_ISL_487265, EPI_ISL_487266, EPI_ISL_487267, EPI_ISL_487268 | | | |
| see above | Utah Public Health Laboratory | Utah Public Health Laboratory | Heidi Butz, Erin Young, Kelly Oakeson |
| EPI_ISL_487269 | Laboratorio de Referencia Nacional de Virus Respiratorio. Instituto Nacional de Salud. | Laboratorio de Referencia Nacional de Biotecnología y Biología Molecular. Instituto Nacional de Salud. | Carlos Padilla Rojas, Karolyn Chozo Vega, Priscila Lope Pari, Omar Caceres Rey, Marco Galarza Perez, Maribel Huaringa Nuñez, Johanna Balbuena Torrez, Henri Bailon Calderon, Nancy Rojas Serrano |
| EPI_ISL_487270 | unknown | Communicable Disease Laboratory, Public Health Directorate | AlWasti,H., AlTaif,Z., Zaed,A., Shehab,F. |
| EPI_ISL_487271 | unknown | MDU-PHL, The Peter Doherty Institute for Infection and Immunity | Caly,L., Seemann,T., Sait,M., Schultz,M.B., Sherry,N., Meumann,E., Baird,R., Leong,L., Lim,C.K., Turra,M., Bastian,I., Higgins,G., Soares da Silva,E., Dolores de Jesus da Costa,M., Salles de Sousa,A., Jayanti Pereira Tilman,A., Antonia da Costa,E., Baretto,I., Marr,I., Wapling,J., Francis,J., Ximenes,J., Canisia,D., Freeman,K., Dakh,F., Douglas,N. |
| EPI_ISL_487272 | unknown | Communicable Disease Laboratory, Public Health Directorate | Altaif,z., AlWasti,H., Shehab,F., Zaed,A. |
| EPI_ISL_487273 | unknown | Communicable Disease Laboratory, Public Health Directorate | Zaed,A., Shehab,F., AlWasti,H., Altaif,Z. |
| EPI_ISL_487274 | unknown | Communicable Disease Laboratory, Public Health Directorate | AlWasti,H., AlTaif,Z., Zaed,A., Shehab,F. |
| EPI_ISL_487275 | Department of Veterinary Pathology, University of Liege - FARAH | Department of Veterinary Pathology, University of Liege - FARAH | Garigliany,M., Vanlaere,A.-S., Clercx,C., Giet,D., Escriou,N., Huon,C., van der Werf,S., Eloit,M., Desmecht,D. |
| EPI_ISL_487276 | Department of Food Safety, Nutrition and Veterinary public health, Istituto Superiore di Sanita' | Department of Biomedical, Surgical and Dental Sciences and Department of Biomedical Sciences for Health | Delbue,S., Ferrante,P., Basilico,N., Parapini,S., Binda,S., D'Alessandro,S., Galli,C., Signorini,L., Primache,V., Anselmi,G., Pariani,E. |
| EPI_ISL_487277, EPI_ISL_487278, EPI_ISL_487279, EPI_ISL_487280, EPI_ISL_487281, EPI_ISL_487282, EPI_ISL_487283, EPI_ISL_487284, EPI_ISL_487285, EPI_ISL_487286, EPI_ISL_487287, EPI_ISL_487288, EPI_ISL_487289, EPI_ISL_487290, EPI_ISL_487291, EPI_ISL_487292, EPI_ISL_487293, EPI_ISL_487294, EPI_ISL_487295, EPI_ISL_487296, EPI_ISL_487297, EPI_ISL_487298, EPI_ISL_487299, EPI_ISL_487300, EPI_ISL_487301, EPI_ISL_487302, EPI_ISL_487303, EPI_ISL_487304, EPI_ISL_487305, EPI_ISL_487306, EPI_ISL_487307, EPI_ISL_487308, EPI_ISL_487309, EPI_ISL_487310, EPI_ISL_487311, EPI_ISL_487312, EPI_ISL_487313, EPI_ISL_487314, EPI_ISL_487315, EPI_ISL_487316, EPI_ISL_487317, EPI_ISL_487318, EPI_ISL_487319, EPI_ISL_487320, EPI_ISL_487321, EPI_ISL_487322, EPI_ISL_487323, EPI_ISL_487324, EPI_ISL_487325, EPI_ISL_487326, EPI_ISL_487327, EPI_ISL_487328 | | | |
| see above | NHLS-IALCH | KRISP, KZN Research Innovation and Sequencing Platform | Giandhari J, Pillay S, Lessells R, Chimukangara B, Mdlalose K, York D, Khan S, Tegally H, Wilkinson E, de Oliveira T |
| EPI_ISL_487329, EPI_ISL_487330, EPI_ISL_487331, EPI_ISL_487332, EPI_ISL_487333, EPI_ISL_487334, EPI_ISL_487335, EPI_ISL_487336, EPI_ISL_487337, EPI_ISL_487338, EPI_ISL_487339, EPI_ISL_487340, EPI_ISL_487341 | | | |
| see above | Molecular Diagnostics Services (MDS) | KRISP, KZN Research Innovation and Sequencing Platform | Giandhari J, Pillay S, Lessells R, Chimukangara B, Mdlalose K, York D, Khan S, Tegally H, Wilkinson E, de Oliveira T |
| EPI_ISL_487348 | NHLS-IALCH | KRISP, KZN Research Innovation and Sequencing Platform | Giandhari J, Pillay S, Lessells R, Chimukangara B, Mdlalose K, York D, Khan S, Tegally H, Wilkinson E, de Oliveira T |
| EPI_ISL_487362 | National Institute of Laboratory Medicine and Referral Center | Genomic Research Lab, BCSIR | Md. Ahasan Habib, Abu Sayeed Mohammad Mahmud, Mohammad Samir Uzzaman, Eshrar Osman, Shahina Akter, Tanjina Akhter Banu, Md. Murshed Hasan Sarkar, Barna Goswami, Iffat Jahan, Md. Saddam Hossain, Tasnim Nafisa, Md. Maruf Ahmed Molla, Mahmuda Yeasmin, Asish Kumar Ghosh, A. K. M. Shamsuzzaman, Sheikh Md. Selim Al Din, Utpal Chandra Ray, Salek Ahmed Sajib, Md. Salim Khan |
| EPI_ISL_487363, EPI_ISL_487364 | National Institute of Laboratory Medicine and Referral Center | Genomic Research Lab, BCSIR | Tanjina Akhter Banu, Abu Sayeed Mohammad Mahmud, Mohammad Samir Uzzaman, Eshrar Osman, Md. Ahasan Habib, Shahina Akter, Md. Murshed Hasan Sarkar, Barna Goswami, Iffat Jahan, Md. Saddam Hossain, Tasnim Nafisa, Md. Maruf Ahmed Molla, Mahmuda Yeasmin, Asish Kumar Ghosh, A. K. M. Shamsuzzaman, Sheikh Md. Selim Al Din, Utpal Chandra Ray, Salek Ahmed Sajib, Md. Salim Khan |
| EPI_ISL_487365 | Viral Respiratory Lab, National Institute for Biomedical Research (INRB) | Pathogen Sequencing Lab, National Institute for Biomedical Research (INRB) | Placide Mbala-Kingebeni, Edith Nkwembe, Eddy Kinganda-Lusamaki, Amuri Aziza, Francisca Muyembe-Mawete, Emmanuel Lokilo-Lofiko, Catherine Pratt, Matthias Pauthner, Josh Quick, Allison Black, James Hadfield, Trevor Bedford, Ian Goodfellow, Andrew Rambault, Nick Loman, Kristian Andersen, Michael Wiley, Steve Ahuka-Mundeke, Jean-Jacques Muyembe Tamfum |
| EPI_ISL_487366, EPI_ISL_487367 | National Institute of Laboratory Medicine and Referral Center | Genomic Research Lab, BCSIR | Shahina Akter, Abu Sayeed Mohammad Mahmud, Mohammad Samir Uzzaman, Eshrar Osman, Md. Ahasan Habib, Tanjina Akhter Banu, Md. Murshed Hasan Sarkar, Barna Goswami, Iffat Jahan, Md. Saddam Hossain, Tasnim Nafisa, Md. Maruf Ahmed Molla, Mahmuda Yeasmin, Asish Kumar Ghosh, A. K. M. Shamsuzzaman, Sheikh Md. Selim Al Din, Utpal Chandra Ray, Salek Ahmed Sajib, Md. Salim Khan |
| EPI_ISL_487368 | National Institute of Laboratory Medicine and Referral Center | Genomic Research Lab, BCSIR | Barna Goswami, Abu Sayeed Mohammad Mahmud, Mohammad Samir Uzzaman, Eshrar Osman, Md. Ahasan Habib, Shahina Akter, Tanjina Akhter Banu, Md. Murshed Hasan Sarkar, Iffat Jahan, Md. Saddam Hossain, Tasnim Nafisa, Md. Maruf Ahmed Molla, Mahmuda Yeasmin, Asish Kumar Ghosh, A. K. M. Shamsuzzaman, Sheikh Md. Selim Al Din, Utpal Chandra Ray, Salek Ahmed Sajib, Md. Salim Khan |
| EPI_ISL_487369 | Viral Respiratory Lab, National Institute for Biomedical Research (INRB) | Pathogen Sequencing Lab, National Institute for Biomedical Research (INRB) | Placide Mbala-Kingebeni, Edith Nkwembe, Eddy Kinganda-Lusamaki, Amuri Aziza, Francisca Muyembe-Mawete, Emmanuel Lokilo-Lofiko, Catherine Pratt, Matthias Pauthner, Josh Quick, Allison Black, James Hadfield, Trevor Bedford, Ian Goodfellow, Andrew Rambault, Nick Loman, Kristian Andersen, Michael Wiley, Steve Ahuka-Mundeke, Jean-Jacques Muyembe Tamfum |
| EPI_ISL_487370 | Hellenic Pasteur Institute, National Influenza Reference laboratory of Southern Greece & Unit of Bioinformatics and Applied Genomics | Hellenic Pasteur Institute, National Influenza Reference laboratory of Southern Greece & Unit of Bioinformatics and Applied Genomics | Vasiliki Pogka, Timokratis Karamitros, Athanasios Kossyvakis, Antonios Kalliaropoulos, Horefti Elina, Evangelidou Maria, Androniki Voulgari-Kokota, Aspasia Kontou, Andreas Mentis |
| EPI_ISL_487371 | National Institute of Laboratory Medicine and Referral Center | Genomic Research Lab, BCSIR | Barna Goswami, Abu Sayeed Mohammad Mahmud, Mohammad Samir Uzzaman, Eshrar Osman, Md. Ahasan Habib, Shahina Akter, Tanjina Akhter Banu, Md. Murshed Hasan Sarkar, Iffat Jahan, Md. Saddam Hossain, Tasnim Nafisa, Md. Maruf Ahmed Molla, Mahmuda Yeasmin, Asish Kumar Ghosh, A. K. M. Shamsuzzaman, Sheikh Md. Selim Al Din, Utpal Chandra Ray, Salek Ahmed Sajib, Md. Salim Khan |
| EPI_ISL_487372, EPI_ISL_487373 | National Institute of Laboratory Medicine and Referral Center | Genomic Research Lab, BCSIR | Iffat Jahan, Abu Sayeed Mohammad Mahmud, Mohammad Samir Uzzaman, Eshrar Osman, Md. Ahasan Habib, Shahina Akter, Tanjina Akhter Banu, Md. Murshed Hasan Sarkar, Barna Goswami, Md. Saddam Hossain, Tasnim Nafisa, Md. Maruf Ahmed Molla, Mahmuda Yeasmin, Asish Kumar Ghosh, A. K. M. Shamsuzzaman, Sheikh Md. Selim Al Din, Utpal Chandra Ray, Salek Ahmed Sajib, Md. Salim Khan |
| EPI_ISL_487375, EPI_ISL_487376 | National Institute of Laboratory Medicine and Referral Center | Genomic Research Lab, BCSIR | Md. Saddam Hossain, Abu Sayeed Mohammad Mahmud, Mohammad Samir Uzzaman, Eshrar Osman, Md. Ahasan Habib, Shahina Akter, Tanjina Akhter Banu, Md. Murshed Hasan Sarkar, Barna Goswami, Iffat Jahan, Tasnim Nafisa, Md. Maruf Ahmed Molla, Mahmuda Yeasmin, Asish Kumar Ghosh, A. K. M. Shamsuzzaman, Sheikh Md. Selim Al Din, Utpal Chandra Ray, Salek Ahmed Sajib, Md. Salim Khan |
| EPI_ISL_487377 | Hellenic Pasteur Institute, National Influenza Reference laboratory of Southern Greece & Unit of Bioinformatics and Applied Genomics | Hellenic Pasteur Institute, National Influenza Reference laboratory of Southern Greece & Unit of Bioinformatics and Applied Genomics | Vasiliki Pogka, Timokratis Karamitros, Athanasios Kossyvakis, Antonios Kalliaropoulos, Horefti Elina, Evangelidou Maria, Androniki Voulgari-Kokota, Aspasia Kontou, Andreas Mentis |
| EPI_ISL_487378 | National Institute of Laboratory Medicine and Referral Center | Genomic Research Lab, BCSIR | Md. Murshed Hasan Sarkar, Abu Sayeed Mohammad Mahmud, Mohammad Samir Uzzaman, Eshrar Osman, Md. Ahasan Habib, Shahina Akter, Tanjina Akhter Banu, Barna Goswami, Iffat Jahan, Md. Saddam Hossain, Tasnim Nafisa, Md. Maruf Ahmed Molla, Mahmuda Yeasmin, Asish Kumar Ghosh, A. K. M. Shamsuzzaman, Sheikh Md. Selim Al Din, Utpal Chandra Ray, Salek Ahmed Sajib, Md. Salim Khan |
| EPI_ISL_487379 | Hellenic Pasteur Institute, National Influenza Reference laboratory of Southern Greece & Unit of Bioinformatics and Applied Genomics | Hellenic Pasteur Institute, National Influenza Reference laboratory of Southern Greece & Unit of Bioinformatics and Applied Genomics | Vasiliki Pogka, Timokratis Karamitros, Athanasios Kossyvakis, Antonios Kalliaropoulos, Horefti Elina, Evangelidou Maria, Androniki Voulgari-Kokota, Aspasia Kontou, Andreas Mentis |
| EPI_ISL_487380 | National Institute of Laboratory Medicine and Referral Center | Genomic Research Lab, BCSIR | Md. Murshed Hasan Sarkar, Abu Sayeed Mohammad Mahmud, Mohammad Samir Uzzaman, Eshrar Osman, Md. Ahasan Habib, Shahina Akter, Tanjina Akhter Banu, Barna Goswami, Iffat Jahan, Md. Saddam Hossain, Tasnim Nafisa, Md. Maruf Ahmed Molla, Mahmuda Yeasmin, Asish Kumar Ghosh, A. K. M. Shamsuzzaman, Sheikh Md. Selim Al Din, Utpal Chandra Ray, Salek Ahmed Sajib, Md. Salim Khan |
| EPI_ISL_487381 | Hellenic Pasteur Institute, National Influenza Reference laboratory of Southern Greece & Unit of Bioinformatics and Applied Genomics | Hellenic Pasteur Institute, National Influenza Reference laboratory of Southern Greece & Unit of Bioinformatics and Applied Genomics | Vasiliki Pogka, Timokratis Karamitros, Athanasios Kossyvakis, Antonios Kalliaropoulos, Horefti Elina, Evangelidou Maria, Androniki Voulgari-Kokota, Aspasia Kontou, Andreas Mentis |
| EPI_ISL_487382, EPI_ISL_487383, EPI_ISL_487384, EPI_ISL_487385 | National Institute of Laboratory Medicine and Referral Center | Genomic Research Lab, BCSIR | Md. Murshed Hasan Sarkar, Abu Sayeed Mohammad Mahmud, Mohammad Samir Uzzaman, Eshrar Osman, Md. Ahasan Habib, Shahina Akter, Tanjina Akhter Banu, Barna Goswami, Iffat Jahan, Md. Saddam Hossain, Tasnim Nafisa, Md. Maruf Ahmed Molla, Mahmuda Yeasmin, Asish Kumar Ghosh, A. K. M. Shamsuzzaman, Sheikh Md. Selim Al Din, Utpal Chandra Ray, Salek Ahmed Sajib, Md. Salim Khan |
| EPI_ISL_487386, EPI_ISL_487392, EPI_ISL_487393, EPI_ISL_487394, EPI_ISL_487395, EPI_ISL_487396 | National Institute of Laboratory Medicine and Referral Center | Genomic Research Lab, BCSIR | Abu Sayeed Mohammad Mahmud, Mohammad Samir Uzzaman, Eshrar Osman, Md. Ahasan Habib, Shahina Akter, Tanjina Akhter Banu, Md. Murshed Hasan Sarkar, Barna Goswami, Iffat Jahan, Md. Saddam Hossain, Tasnim Nafisa, Md. Maruf Ahmed Molla, Mahmuda Yeasmin, Asish Kumar Ghosh, A. K. M. Shamsuzzaman, Sheikh Md. Selim Al Din, Utpal Chandra Ray, Salek Ahmed Sajib, Md. Salim Khan |
| EPI_ISL_487398, EPI_ISL_487399, EPI_ISL_487400, EPI_ISL_487401, EPI_ISL_487402, EPI_ISL_487403, EPI_ISL_487404, EPI_ISL_487405, EPI_ISL_487406, EPI_ISL_487407, EPI_ISL_487408, EPI_ISL_487409, EPI_ISL_487410, EPI_ISL_487411, EPI_ISL_487412, EPI_ISL_487413, EPI_ISL_487414, EPI_ISL_487415, EPI_ISL_487416, EPI_ISL_487417, EPI_ISL_487418, EPI_ISL_487419, EPI_ISL_487420, EPI_ISL_487421, EPI_ISL_487422, EPI_ISL_487423, EPI_ISL_487424, EPI_ISL_487425, EPI_ISL_487426, EPI_ISL_487427, EPI_ISL_487428, EPI_ISL_487429, EPI_ISL_487430, EPI_ISL_487431 | | | |
| see above | Labor Kneißler GmbH & Co. KG | Heinrich Pette Institute, Leibniz Institute for Experimental Virology | Thomas Günther, Adam Grundhoff, Manja Czech-Sioli, Nicole Fischer, Matthias Ottinger, Melanie M. Brinkmann |
| EPI_ISL_487432, EPI_ISL_487433, EPI_ISL_487434, EPI_ISL_487435, EPI_ISL_487436 | Queen Astrid Military Hospital | Institute of Tropical Medicine | Philippe Selhorst, Colin Anthony |
| EPI_ISL_487446, EPI_ISL_487447, EPI_ISL_487448, EPI_ISL_487449, EPI_ISL_487450, EPI_ISL_487451, EPI_ISL_487452, EPI_ISL_487453, EPI_ISL_487454, EPI_ISL_487455, EPI_ISL_487456, EPI_ISL_487457, EPI_ISL_487458, EPI_ISL_487459, EPI_ISL_487460, EPI_ISL_487461, EPI_ISL_487462, EPI_ISL_487463, EPI_ISL_487464, EPI_ISL_487465, EPI_ISL_487466 | | | |
| see above | CICM-Mali | Bundeswehr Institut of Microbiology | Kouriba, Dürr, Sangaré, Rehn, Traoré, Bestehorn-Willmann, Walter, Quedraogo, Zimmermann, Maiga, Heitzer, Sogodogo, Antwerpen, Wölfel |
| EPI_ISL_487524 | PHE South West Regional Laboratory, National | Wellcome Sanger Institute for the COVID-19 | Stephanie Hutchings, Hannah Pymont, Dr Peter Muir, Barry Vipond, Rich Hopes; and Alex Alderton, Roberto Amato, Sonia Goncalves, Ewan Harrison, David K. Jackson, Ian Johnston, Dominic Kwiatkowski, Cordelia Langford, John Sillitoe on behalf of the |

| | | | |
|---|---|---|---|
| | Infection Service | Genomics UK Consortium | Wellcome Sanger Institute COVID-19 Surveillance Team (http://www.sanger.ac.uk/covid-team) |
| EPI_ISL_487525 | University College London, Great Ormond Street Hospital for Children NHS Foundation Trust, Imperial College Healthcare NHS Trust | Wellcome Sanger Institute for the COVID-19 Genomics UK Consortium | Sergi Castellano, Rachel Williams, Mark Kristiansen, Paola Resende Silva, Sunando Roy, Tony Brooks, Helena Tutill, Paola Niola, Patricia Dyal, Charlotte Williams, Leysa Forrest, Yasmin Panchbhaya, Jacqueline Findlay, Sam Weeks, Julianne Brown, Kathryn Harris, Paul Randell, James Price, Alison Holmes, Judith Breuer and Alex Alderton, Roberto Amato, Sonia Goncalves, Ewan Harrison, David K. Jackson, Ian Johnston, Dominic Kwiatkowski, Cordelia Langford, John Sillitoe on behalf of the Wellcome Sanger Institute COVID-19 Surveillance Team (http://www.sanger.ac.uk/covid-team) |
| EPI_ISL_487526, EPI_ISL_487527, EPI_ISL_487528 | PHE South West Regional Laboratory, National Infection Service | Wellcome Sanger Institute for the COVID-19 Genomics UK Consortium | Stephanie Hutchings, Hannah Pymont, Dr Peter Muir, Barry Vipond, Rich Hopes; and Alex Alderton, Roberto Amato, Sonia Goncalves, Ewan Harrison, David K. Jackson, Ian Johnston, Dominic Kwiatkowski, Cordelia Langford, John Sillitoe on behalf of the Wellcome Sanger Institute COVID-19 Surveillance Team (http://www.sanger.ac.uk/covid-team) |
| EPI_ISL_487529 | University College London, Great Ormond Street Hospital for Children NHS Foundation Trust, Imperial College Healthcare NHS Trust | Wellcome Sanger Institute for the COVID-19 Genomics UK Consortium | Sergi Castellano, Rachel Williams, Mark Kristiansen, Paola Resende Silva, Sunando Roy, Tony Brooks, Helena Tutill, Paola Niola, Patricia Dyal, Charlotte Williams, Leysa Forrest, Yasmin Panchbhaya, Jacqueline Findlay, Sam Weeks, Julianne Brown, Kathryn Harris, Paul Randell, James Price, Alison Holmes, Judith Breuer and Alex Alderton, Roberto Amato, Sonia Goncalves, Ewan Harrison, David K. Jackson, Ian Johnston, Dominic Kwiatkowski, Cordelia Langford, John Sillitoe on behalf of the Wellcome Sanger Institute COVID-19 Surveillance Team (http://www.sanger.ac.uk/covid-team) |
| EPI_ISL_487530, EPI_ISL_487531, EPI_ISL_487532, EPI_ISL_487533 | PHE South West Regional Laboratory, National Infection Service | Wellcome Sanger Institute for the COVID-19 Genomics UK Consortium | Stephanie Hutchings, Hannah Pymont, Dr Peter Muir, Barry Vipond, Rich Hopes; and Alex Alderton, Roberto Amato, Sonia Goncalves, Ewan Harrison, David K. Jackson, Ian Johnston, Dominic Kwiatkowski, Cordelia Langford, John Sillitoe on behalf of the Wellcome Sanger Institute COVID-19 Surveillance Team (http://www.sanger.ac.uk/covid-team) |
| EPI_ISL_487534 | University College London, Great Ormond Street Hospital for Children NHS Foundation Trust, Imperial College Healthcare NHS Trust | Wellcome Sanger Institute for the COVID-19 Genomics UK Consortium | Sergi Castellano, Rachel Williams, Mark Kristiansen, Paola Resende Silva, Sunando Roy, Tony Brooks, Helena Tutill, Paola Niola, Patricia Dyal, Charlotte Williams, Leysa Forrest, Yasmin Panchbhaya, Jacqueline Findlay, Sam Weeks, Julianne Brown, Kathryn Harris, Paul Randell, James Price, Alison Holmes, Judith Breuer and Alex Alderton, Roberto Amato, Sonia Goncalves, Ewan Harrison, David K. Jackson, Ian Johnston, Dominic Kwiatkowski, Cordelia Langford, John Sillitoe on behalf of the Wellcome Sanger Institute COVID-19 Surveillance Team (http://www.sanger.ac.uk/covid-team) |
| EPI_ISL_487535, EPI_ISL_487536 | PHE South West Regional Laboratory, National Infection Service | Wellcome Sanger Institute for the COVID-19 Genomics UK Consortium | Stephanie Hutchings, Hannah Pymont, Dr Peter Muir, Barry Vipond, Rich Hopes; and Alex Alderton, Roberto Amato, Sonia Goncalves, Ewan Harrison, David K. Jackson, Ian Johnston, Dominic Kwiatkowski, Cordelia Langford, John Sillitoe on behalf of the Wellcome Sanger Institute COVID-19 Surveillance Team (http://www.sanger.ac.uk/covid-team) |
| EPI_ISL_487537 | University College London, Great Ormond Street Hospital for Children NHS Foundation Trust, Imperial College Healthcare NHS Trust | Wellcome Sanger Institute for the COVID-19 Genomics UK Consortium | Sergi Castellano, Rachel Williams, Mark Kristiansen, Paola Resende Silva, Sunando Roy, Tony Brooks, Helena Tutill, Paola Niola, Patricia Dyal, Charlotte Williams, Leysa Forrest, Yasmin Panchbhaya, Jacqueline Findlay, Sam Weeks, Julianne Brown, Kathryn Harris, Paul Randell, James Price, Alison Holmes, Judith Breuer and Alex Alderton, Roberto Amato, Sonia Goncalves, Ewan Harrison, David K. Jackson, Ian Johnston, Dominic Kwiatkowski, Cordelia Langford, John Sillitoe on behalf of the Wellcome Sanger Institute COVID-19 Surveillance Team (http://www.sanger.ac.uk/covid-team) |
| EPI_ISL_487538, EPI_ISL_487539 | PHE South West Regional Laboratory, National Infection Service | Wellcome Sanger Institute for the COVID-19 Genomics UK Consortium | Stephanie Hutchings, Hannah Pymont, Dr Peter Muir, Barry Vipond, Rich Hopes; and Alex Alderton, Roberto Amato, Sonia Goncalves, Ewan Harrison, David K. Jackson, Ian Johnston, Dominic Kwiatkowski, Cordelia Langford, John Sillitoe on behalf of the Wellcome Sanger Institute COVID-19 Surveillance Team (http://www.sanger.ac.uk/covid-team) |
| EPI_ISL_487540 | University College London, Great Ormond Street Hospital for Children NHS Foundation Trust, Imperial College Healthcare NHS Trust | Wellcome Sanger Institute for the COVID-19 Genomics UK Consortium | Sergi Castellano, Rachel Williams, Mark Kristiansen, Paola Resende Silva, Sunando Roy, Tony Brooks, Helena Tutill, Paola Niola, Patricia Dyal, Charlotte Williams, Leysa Forrest, Yasmin Panchbhaya, Jacqueline Findlay, Sam Weeks, Julianne Brown, Kathryn Harris, Paul Randell, James Price, Alison Holmes, Judith Breuer and Alex Alderton, Roberto Amato, Sonia Goncalves, Ewan Harrison, David K. Jackson, Ian Johnston, Dominic Kwiatkowski, Cordelia Langford, John Sillitoe on behalf of the Wellcome Sanger Institute COVID-19 Surveillance Team (http://www.sanger.ac.uk/covid-team) |
| EPI_ISL_487541, EPI_ISL_487542, EPI_ISL_487543, EPI_ISL_487544, EPI_ISL_487545 | PHE South West Regional Laboratory, National Infection Service | Wellcome Sanger Institute for the COVID-19 Genomics UK Consortium | Stephanie Hutchings, Hannah Pymont, Dr Peter Muir, Barry Vipond, Rich Hopes; and Alex Alderton, Roberto Amato, Sonia Goncalves, Ewan Harrison, David K. Jackson, Ian Johnston, Dominic Kwiatkowski, Cordelia Langford, John Sillitoe on behalf of the Wellcome Sanger Institute COVID-19 Surveillance Team (http://www.sanger.ac.uk/covid-team) |
| EPI_ISL_487546 | University College London, Great Ormond Street Hospital for Children NHS Foundation Trust, Imperial College Healthcare NHS Trust | Wellcome Sanger Institute for the COVID-19 Genomics UK Consortium | Sergi Castellano, Rachel Williams, Mark Kristiansen, Paola Resende Silva, Sunando Roy, Tony Brooks, Helena Tutill, Paola Niola, Patricia Dyal, Charlotte Williams, Leysa Forrest, Yasmin Panchbhaya, Jacqueline Findlay, Sam Weeks, Julianne Brown, Kathryn Harris, Paul Randell, James Price, Alison Holmes, Judith Breuer and Alex Alderton, Roberto Amato, Sonia Goncalves, Ewan Harrison, David K. Jackson, Ian Johnston, Dominic Kwiatkowski, Cordelia Langford, John Sillitoe on behalf of the Wellcome Sanger Institute COVID-19 Surveillance Team (http://www.sanger.ac.uk/covid-team) |
| EPI_ISL_487547, EPI_ISL_487548, EPI_ISL_487549 | PHE South West Regional Laboratory, National Infection Service | Wellcome Sanger Institute for the COVID-19 Genomics UK Consortium | Stephanie Hutchings, Hannah Pymont, Dr Peter Muir, Barry Vipond, Rich Hopes; and Alex Alderton, Roberto Amato, Sonia Goncalves, Ewan Harrison, David K. Jackson, Ian Johnston, Dominic Kwiatkowski, Cordelia Langford, John Sillitoe on behalf of the Wellcome Sanger Institute COVID-19 Surveillance Team (http://www.sanger.ac.uk/covid-team) |
| EPI_ISL_487550 | University College London, Great Ormond Street Hospital for Children NHS Foundation Trust, Imperial College Healthcare NHS Trust | Wellcome Sanger Institute for the COVID-19 Genomics UK Consortium | Sergi Castellano, Rachel Williams, Mark Kristiansen, Paola Resende Silva, Sunando Roy, Tony Brooks, Helena Tutill, Paola Niola, Patricia Dyal, Charlotte Williams, Leysa Forrest, Yasmin Panchbhaya, Jacqueline Findlay, Sam Weeks, Julianne Brown, Kathryn Harris, Paul Randell, James Price, Alison Holmes, Judith Breuer and Alex Alderton, Roberto Amato, Sonia Goncalves, Ewan Harrison, David K. Jackson, Ian Johnston, Dominic Kwiatkowski, Cordelia Langford, John Sillitoe on behalf of the Wellcome Sanger Institute COVID-19 Surveillance Team (http://www.sanger.ac.uk/covid-team) |
| EPI_ISL_487551, EPI_ISL_487552, EPI_ISL_487553, EPI_ISL_487554, EPI_ISL_487555 | PHE South West Regional Laboratory, National Infection Service | Wellcome Sanger Institute for the COVID-19 Genomics UK Consortium | Stephanie Hutchings, Hannah Pymont, Dr Peter Muir, Barry Vipond, Rich Hopes; and Alex Alderton, Roberto Amato, Sonia Goncalves, Ewan Harrison, David K. Jackson, Ian Johnston, Dominic Kwiatkowski, Cordelia Langford, John Sillitoe on behalf of the Wellcome Sanger Institute COVID-19 Surveillance Team (http://www.sanger.ac.uk/covid-team) |
| EPI_ISL_487556, EPI_ISL_487557, EPI_ISL_487558, EPI_ISL_487559 | University College London, Great Ormond Street Hospital for Children NHS Foundation Trust, Imperial College Healthcare NHS Trust | Wellcome Sanger Institute for the COVID-19 Genomics UK Consortium | Sergi Castellano, Rachel Williams, Mark Kristiansen, Paola Resende Silva, Sunando Roy, Tony Brooks, Helena Tutill, Paola Niola, Patricia Dyal, Charlotte Williams, Leysa Forrest, Yasmin Panchbhaya, Jacqueline Findlay, Sam Weeks, Julianne Brown, Kathryn Harris, Paul Randell, James Price, Alison Holmes, Judith Breuer and Alex Alderton, Roberto Amato, Sonia Goncalves, Ewan Harrison, David K. Jackson, Ian Johnston, Dominic Kwiatkowski, Cordelia Langford, John Sillitoe on behalf of the Wellcome Sanger Institute COVID-19 Surveillance Team (http://www.sanger.ac.uk/covid-team) |
| EPI_ISL_487560, EPI_ISL_487561 | PHE South West Regional Laboratory, National Infection Service | Wellcome Sanger Institute for the COVID-19 Genomics UK Consortium | Stephanie Hutchings, Hannah Pymont, Dr Peter Muir, Barry Vipond, Rich Hopes; and Alex Alderton, Roberto Amato, Sonia Goncalves, Ewan Harrison, David K. Jackson, Ian Johnston, Dominic Kwiatkowski, Cordelia Langford, John Sillitoe on behalf of the Wellcome Sanger Institute COVID-19 Surveillance Team (http://www.sanger.ac.uk/covid-team) |
| EPI_ISL_487562 | University College London, Great Ormond Street Hospital for Children NHS Foundation Trust, Imperial College Healthcare NHS Trust | Wellcome Sanger Institute for the COVID-19 Genomics UK Consortium | Sergi Castellano, Rachel Williams, Mark Kristiansen, Paola Resende Silva, Sunando Roy, Tony Brooks, Helena Tutill, Paola Niola, Patricia Dyal, Charlotte Williams, Leysa Forrest, Yasmin Panchbhaya, Jacqueline Findlay, Sam Weeks, Julianne Brown, Kathryn Harris, Paul Randell, James Price, Alison Holmes, Judith Breuer and Alex Alderton, Roberto Amato, Sonia Goncalves, Ewan Harrison, David K. Jackson, Ian Johnston, Dominic Kwiatkowski, Cordelia Langford, John Sillitoe on behalf of the Wellcome Sanger Institute COVID-19 Surveillance Team (http://www.sanger.ac.uk/covid-team) |
| EPI_ISL_487563 | PHE South West Regional Laboratory, National Infection Service | Wellcome Sanger Institute for the COVID-19 Genomics UK Consortium | Stephanie Hutchings, Hannah Pymont, Dr Peter Muir, Barry Vipond, Rich Hopes; and Alex Alderton, Roberto Amato, Sonia Goncalves, Ewan Harrison, David K. Jackson, Ian Johnston, Dominic Kwiatkowski, Cordelia Langford, John Sillitoe on behalf of the Wellcome Sanger Institute COVID-19 Surveillance Team (http://www.sanger.ac.uk/covid-team) |
| EPI_ISL_487564 | University College London, Great Ormond Street Hospital for Children NHS Foundation Trust, Imperial College Healthcare NHS Trust | Wellcome Sanger Institute for the COVID-19 Genomics UK Consortium | Sergi Castellano, Rachel Williams, Mark Kristiansen, Paola Resende Silva, Sunando Roy, Tony Brooks, Helena Tutill, Paola Niola, Patricia Dyal, Charlotte Williams, Leysa Forrest, Yasmin Panchbhaya, Jacqueline Findlay, Sam Weeks, Julianne Brown, Kathryn Harris, Paul Randell, James Price, Alison Holmes, Judith Breuer and Alex Alderton, Roberto Amato, Sonia Goncalves, Ewan Harrison, David K. Jackson, Ian Johnston, Dominic Kwiatkowski, Cordelia Langford, John Sillitoe on behalf of the Wellcome Sanger Institute COVID-19 Surveillance Team (http://www.sanger.ac.uk/covid-team) |
| EPI_ISL_487565, EPI_ISL_487566, EPI_ISL_487567 | PHE South West Regional Laboratory, National Infection Service | Wellcome Sanger Institute for the COVID-19 Genomics UK Consortium | Stephanie Hutchings, Hannah Pymont, Dr Peter Muir, Barry Vipond, Rich Hopes; and Alex Alderton, Roberto Amato, Sonia Goncalves, Ewan Harrison, David K. Jackson, Ian Johnston, Dominic Kwiatkowski, Cordelia Langford, John Sillitoe on behalf of the Wellcome Sanger Institute COVID-19 Surveillance Team (http://www.sanger.ac.uk/covid-team) |
| EPI_ISL_487568 | University College London, Great Ormond Street Hospital for Children NHS Foundation Trust, Imperial College Healthcare NHS Trust | Wellcome Sanger Institute for the COVID-19 Genomics UK Consortium | Sergi Castellano, Rachel Williams, Mark Kristiansen, Paola Resende Silva, Sunando Roy, Tony Brooks, Helena Tutill, Paola Niola, Patricia Dyal, Charlotte Williams, Leysa Forrest, Yasmin Panchbhaya, Jacqueline Findlay, Sam Weeks, Julianne Brown, Kathryn Harris, Paul Randell, James Price, Alison Holmes, Judith Breuer and Alex Alderton, Roberto Amato, Sonia Goncalves, Ewan Harrison, David K. Jackson, Ian Johnston, Dominic Kwiatkowski, Cordelia Langford, John Sillitoe on behalf of the Wellcome Sanger Institute COVID-19 Surveillance Team (http://www.sanger.ac.uk/covid-team) |
| EPI_ISL_487570, EPI_ISL_487571 | PHE South West Regional Laboratory, National Infection Service | Wellcome Sanger Institute for the COVID-19 Genomics UK Consortium | Stephanie Hutchings, Hannah Pymont, Dr Peter Muir, Barry Vipond, Rich Hopes; and Alex Alderton, Roberto Amato, Sonia Goncalves, Ewan Harrison, David K. Jackson, Ian Johnston, Dominic Kwiatkowski, Cordelia Langford, John Sillitoe on behalf of the Wellcome Sanger Institute COVID-19 Surveillance Team (http://www.sanger.ac.uk/covid-team) |
| EPI_ISL_487572 | University College London, Great Ormond Street Hospital for Children NHS Foundation Trust, Imperial College Healthcare NHS Trust | Wellcome Sanger Institute for the COVID-19 Genomics UK Consortium | Sergi Castellano, Rachel Williams, Mark Kristiansen, Paola Resende Silva, Sunando Roy, Tony Brooks, Helena Tutill, Paola Niola, Patricia Dyal, Charlotte Williams, Leysa Forrest, Yasmin Panchbhaya, Jacqueline Findlay, Sam Weeks, Julianne Brown, Kathryn Harris, Paul Randell, James Price, Alison Holmes, Judith Breuer and Alex Alderton, Roberto Amato, Sonia Goncalves, Ewan Harrison, David K. Jackson, Ian Johnston, Dominic Kwiatkowski, Cordelia Langford, John Sillitoe on behalf of the Wellcome Sanger Institute COVID-19 Surveillance Team (http://www.sanger.ac.uk/covid-team) |
| EPI_ISL_487573, EPI_ISL_487574, EPI_ISL_487575, EPI_ISL_487576, EPI_ISL_487577 | PHE South West Regional Laboratory, National Infection Service | Wellcome Sanger Institute for the COVID-19 Genomics UK Consortium | Stephanie Hutchings, Hannah Pymont, Dr Peter Muir, Barry Vipond, Rich Hopes; and Alex Alderton, Roberto Amato, Sonia Goncalves, Ewan Harrison, David K. Jackson, Ian Johnston, Dominic Kwiatkowski, Cordelia Langford, John Sillitoe on behalf of the Wellcome Sanger Institute COVID-19 Surveillance Team (http://www.sanger.ac.uk/covid-team) |
| EPI_ISL_487578 | University College London, Great Ormond Street Hospital for Children NHS Foundation Trust, Imperial College Healthcare NHS Trust | Wellcome Sanger Institute for the COVID-19 Genomics UK Consortium | Sergi Castellano, Rachel Williams, Mark Kristiansen, Paola Resende Silva, Sunando Roy, Tony Brooks, Helena Tutill, Paola Niola, Patricia Dyal, Charlotte Williams, Leysa Forrest, Yasmin Panchbhaya, Jacqueline Findlay, Sam Weeks, Julianne Brown, Kathryn Harris, Paul Randell, James Price, Alison Holmes, Judith Breuer and Alex Alderton, Roberto Amato, Sonia Goncalves, Ewan Harrison, David K. Jackson, Ian Johnston, Dominic Kwiatkowski, Cordelia Langford, John Sillitoe on behalf of the Wellcome Sanger Institute COVID-19 Surveillance Team (http://www.sanger.ac.uk/covid-team) |
| EPI_ISL_487579, EPI_ISL_487580 | PHE South West Regional Laboratory, National Infection Service | Wellcome Sanger Institute for the COVID-19 Genomics UK Consortium | Stephanie Hutchings, Hannah Pymont, Dr Peter Muir, Barry Vipond, Rich Hopes; and Alex Alderton, Roberto Amato, Sonia Goncalves, Ewan Harrison, David K. Jackson, Ian Johnston, Dominic Kwiatkowski, Cordelia Langford, John Sillitoe on behalf of the Wellcome Sanger Institute COVID-19 Surveillance Team (http://www.sanger.ac.uk/covid-team) |
| EPI_ISL_487581, EPI_ISL_487582 | University College London, Great Ormond Street Hospital for Children NHS Foundation Trust, Imperial College Healthcare NHS Trust | Wellcome Sanger Institute for the COVID-19 Genomics UK Consortium | Sergi Castellano, Rachel Williams, Mark Kristiansen, Paola Resende Silva, Sunando Roy, Tony Brooks, Helena Tutill, Paola Niola, Patricia Dyal, Charlotte Williams, Leysa Forrest, Yasmin Panchbhaya, Jacqueline Findlay, Sam Weeks, Julianne Brown, Kathryn Harris, Paul Randell, James Price, Alison Holmes, Judith Breuer and Alex Alderton, Roberto Amato, Sonia Goncalves, Ewan Harrison, David K. Jackson, Ian Johnston, Dominic Kwiatkowski, Cordelia Langford, John Sillitoe on behalf of the Wellcome Sanger Institute COVID-19 Surveillance Team (http://www.sanger.ac.uk/covid-team) |
| EPI_ISL_487583 | PHE South West Regional Laboratory, National Infection Service | Wellcome Sanger Institute for the COVID-19 Genomics UK Consortium | Stephanie Hutchings, Hannah Pymont, Dr Peter Muir, Barry Vipond, Rich Hopes; and Alex Alderton, Roberto Amato, Sonia Goncalves, Ewan Harrison, David K. Jackson, Ian Johnston, Dominic Kwiatkowski, Cordelia Langford, John Sillitoe on behalf of the Wellcome Sanger Institute COVID-19 Surveillance Team (http://www.sanger.ac.uk/covid-team) |
| EPI_ISL_487584, EPI_ISL_487585 | University College London, Great Ormond Street Hospital for Children NHS Foundation Trust, Imperial College Healthcare NHS Trust | Wellcome Sanger Institute for the COVID-19 Genomics UK Consortium | Sergi Castellano, Rachel Williams, Mark Kristiansen, Paola Resende Silva, Sunando Roy, Tony Brooks, Helena Tutill, Paola Niola, Patricia Dyal, Charlotte Williams, Leysa Forrest, Yasmin Panchbhaya, Jacqueline Findlay, Sam Weeks, Julianne Brown, Kathryn Harris, Paul Randell, James Price, Alison Holmes, Judith Breuer and Alex Alderton, Roberto Amato, Sonia Goncalves, Ewan Harrison, David K. Jackson, Ian Johnston, Dominic Kwiatkowski, Cordelia Langford, John Sillitoe on behalf of the Wellcome Sanger Institute COVID-19 Surveillance Team (http://www.sanger.ac.uk/covid-team) |
| EPI_ISL_487586, EPI_ISL_487587, EPI_ISL_487588 | PHE South West Regional Laboratory, National Infection Service | Wellcome Sanger Institute for the COVID-19 Genomics UK Consortium | Stephanie Hutchings, Hannah Pymont, Dr Peter Muir, Barry Vipond, Rich Hopes; and Alex Alderton, Roberto Amato, Sonia Goncalves, Ewan Harrison, David K. Jackson, Ian Johnston, Dominic Kwiatkowski, Cordelia Langford, John Sillitoe on behalf of the Wellcome Sanger Institute COVID-19 Surveillance Team (http://www.sanger.ac.uk/covid-team) |
| EPI_ISL_487589 | University College London, Great Ormond Street Hospital for Children NHS Foundation Trust, Imperial College Healthcare NHS Trust | Wellcome Sanger Institute for the COVID-19 Genomics UK Consortium | Sergi Castellano, Rachel Williams, Mark Kristiansen, Paola Resende Silva, Sunando Roy, Tony Brooks, Helena Tutill, Paola Niola, Patricia Dyal, Charlotte Williams, Leysa Forrest, Yasmin Panchbhaya, Jacqueline Findlay, Sam Weeks, Julianne Brown, Kathryn Harris, Paul Randell, James Price, Alison Holmes, Judith Breuer and Alex Alderton, Roberto Amato, Sonia Goncalves, Ewan Harrison, David K. Jackson, Ian Johnston, Dominic Kwiatkowski, Cordelia Langford, John Sillitoe on behalf of the Wellcome Sanger Institute COVID-19 Surveillance Team (http://www.sanger.ac.uk/covid-team) |
| EPI_ISL_487590, EPI_ISL_487591, EPI_ISL_487592, EPI_ISL_487594 | PHE South West Regional Laboratory, National Infection Service | Wellcome Sanger Institute for the COVID-19 Genomics UK Consortium | Stephanie Hutchings, Hannah Pymont, Dr Peter Muir, Barry Vipond, Rich Hopes; and Alex Alderton, Roberto Amato, Sonia Goncalves, Ewan Harrison, David K. Jackson, Ian Johnston, Dominic Kwiatkowski, Cordelia Langford, John Sillitoe on behalf of the Wellcome Sanger Institute COVID-19 Surveillance Team (http://www.sanger.ac.uk/covid-team) |
| EPI_ISL_487595 | University College London, Great Ormond Street Hospital for Children NHS Foundation Trust, Imperial College Healthcare NHS Trust | Wellcome Sanger Institute for the COVID-19 Genomics UK Consortium | Sergi Castellano, Rachel Williams, Mark Kristiansen, Paola Resende Silva, Sunando Roy, Tony Brooks, Helena Tutill, Paola Niola, Patricia Dyal, Charlotte Williams, Leysa Forrest, Yasmin Panchbhaya, Jacqueline Findlay, Sam Weeks, Julianne Brown, Kathryn Harris, Paul Randell, James Price, Alison Holmes, Judith Breuer and Alex Alderton, Roberto Amato, Sonia Goncalves, Ewan Harrison, David K. Jackson, Ian Johnston, Dominic Kwiatkowski, Cordelia Langford, John Sillitoe on behalf of the Wellcome Sanger Institute COVID-19 Surveillance Team (http://www.sanger.ac.uk/covid-team) |
| EPI_ISL_487596 | PHE South West Regional Laboratory, National Infection Service | Wellcome Sanger Institute for the COVID-19 Genomics UK Consortium | Stephanie Hutchings, Hannah Pymont, Dr Peter Muir, Barry Vipond, Rich Hopes; and Alex Alderton, Roberto Amato, Sonia Goncalves, Ewan Harrison, David K. Jackson, Ian Johnston, Dominic Kwiatkowski, Cordelia Langford, John Sillitoe on behalf of the Wellcome Sanger Institute COVID-19 Surveillance Team (http://www.sanger.ac.uk/covid-team) |
| EPI_ISL_487597 | University College London, Great Ormond Street Hospital for Children NHS Foundation Trust, Imperial College Healthcare NHS Trust | Wellcome Sanger Institute for the COVID-19 Genomics UK Consortium | Sergi Castellano, Rachel Williams, Mark Kristiansen, Paola Resende Silva, Sunando Roy, Tony Brooks, Helena Tutill, Paola Niola, Patricia Dyal, Charlotte Williams, Leysa Forrest, Yasmin Panchbhaya, Jacqueline Findlay, Sam Weeks, Julianne Brown, Kathryn Harris, Paul Randell, James Price, Alison Holmes, Judith Breuer and Alex Alderton, Roberto Amato, Sonia Goncalves, Ewan Harrison, David K. Jackson, Ian Johnston, Dominic Kwiatkowski, Cordelia Langford, John Sillitoe on behalf of the Wellcome Sanger Institute COVID-19 Surveillance Team (http://www.sanger.ac.uk/covid-team) |
| EPI_ISL_487598, EPI_ISL_487599 | PHE South West Regional Laboratory, National Infection Service | Wellcome Sanger Institute for the COVID-19 Genomics UK Consortium | Stephanie Hutchings, Hannah Pymont, Dr Peter Muir, Barry Vipond, Rich Hopes; and Alex Alderton, Roberto Amato, Sonia Goncalves, Ewan Harrison, David K. Jackson, Ian Johnston, Dominic Kwiatkowski, Cordelia Langford, John Sillitoe on behalf of the Wellcome Sanger Institute COVID-19 Surveillance Team (http://www.sanger.ac.uk/covid-team) |

| | | | |
|---|---|---|---|
| EPI_ISL_487600 | University College London, Great Ormond Street Hospital for Children NHS Foundation Trust, Imperial College Healthcare NHS Trust | Wellcome Sanger Institute for the COVID-19 Genomics UK Consortium | Sergi Castellano, Rachel Williams, Mark Kristiansen, Paola Resende Silva, Sunando Roy, Tony Brooks, Helena Tutill, Paola Niola, Patricia Dyal, Charlotte Williams, Leysa Forrest, Yasmin Panchbhaya, Jacqueline Findlay, Sam Weeks, Julianne Brown, Kathryn Harris, Paul Randell, James Price, Alison Holmes, Judith Breuer and Alex Alderton, Roberto Amato, Sonia Goncalves, Ewan Harrison, David K. Jackson, Ian Johnston, Dominic Kwiatkowski, Cordelia Langford, John Sillitoe on behalf of the Wellcome Sanger Institute COVID-19 Surveillance Team (http://www.sanger.ac.uk/covid-team) |
| EPI_ISL_487601, EPI_ISL_487602, EPI_ISL_487603, EPI_ISL_487604, EPI_ISL_487605 | PHE South West Regional Laboratory, National Infection Service | Wellcome Sanger Institute for the COVID-19 Genomics UK Consortium | Stephanie Hutchings, Hannah Pymont, Dr Peter Muir, Barry Vipond, Rich Hopes; and Alex Alderton, Roberto Amato, Sonia Goncalves, Ewan Harrison, David K. Jackson, Ian Johnston, Dominic Kwiatkowski, Cordelia Langford, John Sillitoe on behalf of the Wellcome Sanger Institute COVID-19 Surveillance Team (http://www.sanger.ac.uk/covid-team) |
| EPI_ISL_487606, EPI_ISL_487607 | University College London, Great Ormond Street Hospital for Children NHS Foundation Trust, Imperial College Healthcare NHS Trust | Wellcome Sanger Institute for the COVID-19 Genomics UK Consortium | Sergi Castellano, Rachel Williams, Mark Kristiansen, Paola Resende Silva, Sunando Roy, Tony Brooks, Helena Tutill, Paola Niola, Patricia Dyal, Charlotte Williams, Leysa Forrest, Yasmin Panchbhaya, Jacqueline Findlay, Sam Weeks, Julianne Brown, Kathryn Harris, Paul Randell, James Price, Alison Holmes, Judith Breuer and Alex Alderton, Roberto Amato, Sonia Goncalves, Ewan Harrison, David K. Jackson, Ian Johnston, Dominic Kwiatkowski, Cordelia Langford, John Sillitoe on behalf of the Wellcome Sanger Institute COVID-19 Surveillance Team (http://www.sanger.ac.uk/covid-team) |
| EPI_ISL_487608, EPI_ISL_487610, EPI_ISL_487611, EPI_ISL_487612, EPI_ISL_487613 | PHE South West Regional Laboratory, National Infection Service | Wellcome Sanger Institute for the COVID-19 Genomics UK Consortium | Stephanie Hutchings, Hannah Pymont, Dr Peter Muir, Barry Vipond, Rich Hopes; and Alex Alderton, Roberto Amato, Sonia Goncalves, Ewan Harrison, David K. Jackson, Ian Johnston, Dominic Kwiatkowski, Cordelia Langford, John Sillitoe on behalf of the Wellcome Sanger Institute COVID-19 Surveillance Team (http://www.sanger.ac.uk/covid-team) |
| EPI_ISL_487614, EPI_ISL_487615, EPI_ISL_487616 | University College London, Great Ormond Street Hospital for Children NHS Foundation Trust, Imperial College Healthcare NHS Trust | Wellcome Sanger Institute for the COVID-19 Genomics UK Consortium | Sergi Castellano, Rachel Williams, Mark Kristiansen, Paola Resende Silva, Sunando Roy, Tony Brooks, Helena Tutill, Paola Niola, Patricia Dyal, Charlotte Williams, Leysa Forrest, Yasmin Panchbhaya, Jacqueline Findlay, Sam Weeks, Julianne Brown, Kathryn Harris, Paul Randell, James Price, Alison Holmes, Judith Breuer and Alex Alderton, Roberto Amato, Sonia Goncalves, Ewan Harrison, David K. Jackson, Ian Johnston, Dominic Kwiatkowski, Cordelia Langford, John Sillitoe on behalf of the Wellcome Sanger Institute COVID-19 Surveillance Team (http://www.sanger.ac.uk/covid-team) |
| EPI_ISL_487617, EPI_ISL_487618, EPI_ISL_487619, EPI_ISL_487620 | PHE South West Regional Laboratory, National Infection Service | Wellcome Sanger Institute for the COVID-19 Genomics UK Consortium | Stephanie Hutchings, Hannah Pymont, Dr Peter Muir, Barry Vipond, Rich Hopes; and Alex Alderton, Roberto Amato, Sonia Goncalves, Ewan Harrison, David K. Jackson, Ian Johnston, Dominic Kwiatkowski, Cordelia Langford, John Sillitoe on behalf of the Wellcome Sanger Institute COVID-19 Surveillance Team (http://www.sanger.ac.uk/covid-team) |
| EPI_ISL_487621 | University College London, Great Ormond Street Hospital for Children NHS Foundation Trust, Imperial College Healthcare NHS Trust | Wellcome Sanger Institute for the COVID-19 Genomics UK Consortium | Sergi Castellano, Rachel Williams, Mark Kristiansen, Paola Resende Silva, Sunando Roy, Tony Brooks, Helena Tutill, Paola Niola, Patricia Dyal, Charlotte Williams, Leysa Forrest, Yasmin Panchbhaya, Jacqueline Findlay, Sam Weeks, Julianne Brown, Kathryn Harris, Paul Randell, James Price, Alison Holmes, Judith Breuer and Alex Alderton, Roberto Amato, Sonia Goncalves, Ewan Harrison, David K. Jackson, Ian Johnston, Dominic Kwiatkowski, Cordelia Langford, John Sillitoe on behalf of the Wellcome Sanger Institute COVID-19 Surveillance Team (http://www.sanger.ac.uk/covid-team) |
| EPI_ISL_487622, EPI_ISL_487623, EPI_ISL_487624 | PHE South West Regional Laboratory, National Infection Service | Wellcome Sanger Institute for the COVID-19 Genomics UK Consortium | Stephanie Hutchings, Hannah Pymont, Dr Peter Muir, Barry Vipond, Rich Hopes; and Alex Alderton, Roberto Amato, Sonia Goncalves, Ewan Harrison, David K. Jackson, Ian Johnston, Dominic Kwiatkowski, Cordelia Langford, John Sillitoe on behalf of the Wellcome Sanger Institute COVID-19 Surveillance Team (http://www.sanger.ac.uk/covid-team) |
| EPI_ISL_487625, EPI_ISL_487626, EPI_ISL_487627, EPI_ISL_487628 | University College London, Great Ormond Street Hospital for Children NHS Foundation Trust, Imperial College Healthcare NHS Trust | Wellcome Sanger Institute for the COVID-19 Genomics UK Consortium | Sergi Castellano, Rachel Williams, Mark Kristiansen, Paola Resende Silva, Sunando Roy, Tony Brooks, Helena Tutill, Paola Niola, Patricia Dyal, Charlotte Williams, Leysa Forrest, Yasmin Panchbhaya, Jacqueline Findlay, Sam Weeks, Julianne Brown, Kathryn Harris, Paul Randell, James Price, Alison Holmes, Judith Breuer and Alex Alderton, Roberto Amato, Sonia Goncalves, Ewan Harrison, David K. Jackson, Ian Johnston, Dominic Kwiatkowski, Cordelia Langford, John Sillitoe on behalf of the Wellcome Sanger Institute COVID-19 Surveillance Team (http://www.sanger.ac.uk/covid-team) |
| EPI_ISL_487629 | PHE South West Regional Laboratory, National Infection Service | Wellcome Sanger Institute for the COVID-19 Genomics UK Consortium | Stephanie Hutchings, Hannah Pymont, Dr Peter Muir, Barry Vipond, Rich Hopes; and Alex Alderton, Roberto Amato, Sonia Goncalves, Ewan Harrison, David K. Jackson, Ian Johnston, Dominic Kwiatkowski, Cordelia Langford, John Sillitoe on behalf of the Wellcome Sanger Institute COVID-19 Surveillance Team (http://www.sanger.ac.uk/covid-team) |
| EPI_ISL_487630, EPI_ISL_487631, EPI_ISL_487632, EPI_ISL_487633 | University College London, Great Ormond Street Hospital for Children NHS Foundation Trust, Imperial College Healthcare NHS Trust | Wellcome Sanger Institute for the COVID-19 Genomics UK Consortium | Sergi Castellano, Rachel Williams, Mark Kristiansen, Paola Resende Silva, Sunando Roy, Tony Brooks, Helena Tutill, Paola Niola, Patricia Dyal, Charlotte Williams, Leysa Forrest, Yasmin Panchbhaya, Jacqueline Findlay, Sam Weeks, Julianne Brown, Kathryn Harris, Paul Randell, James Price, Alison Holmes, Judith Breuer and Alex Alderton, Roberto Amato, Sonia Goncalves, Ewan Harrison, David K. Jackson, Ian Johnston, Dominic Kwiatkowski, Cordelia Langford, John Sillitoe on behalf of the Wellcome Sanger Institute COVID-19 Surveillance Team (http://www.sanger.ac.uk/covid-team) |
| EPI_ISL_487634 | PHE South West Regional Laboratory, National Infection Service | Wellcome Sanger Institute for the COVID-19 Genomics UK Consortium | Stephanie Hutchings, Hannah Pymont, Dr Peter Muir, Barry Vipond, Rich Hopes; and Alex Alderton, Roberto Amato, Sonia Goncalves, Ewan Harrison, David K. Jackson, Ian Johnston, Dominic Kwiatkowski, Cordelia Langford, John Sillitoe on behalf of the Wellcome Sanger Institute COVID-19 Surveillance Team (http://www.sanger.ac.uk/covid-team) |
| EPI_ISL_487635, EPI_ISL_487636, EPI_ISL_487637, EPI_ISL_487638, EPI_ISL_487639, EPI_ISL_487640, EPI_ISL_487641, EPI_ISL_487642, EPI_ISL_487643, EPI_ISL_487644, EPI_ISL_487645, EPI_ISL_487646 | | | |
| see above | University College London, Great Ormond Street Hospital for Children NHS Foundation Trust, Imperial College Healthcare NHS Trust | Wellcome Sanger Institute for the COVID-19 Genomics UK Consortium | Sergi Castellano, Rachel Williams, Mark Kristiansen, Paola Resende Silva, Sunando Roy, Tony Brooks, Helena Tutill, Paola Niola, Patricia Dyal, Charlotte Williams, Leysa Forrest, Yasmin Panchbhaya, Jacqueline Findlay, Sam Weeks, Julianne Brown, Kathryn Harris, Paul Randell, James Price, Alison Holmes, Judith Breuer and Alex Alderton, Roberto Amato, Sonia Goncalves, Ewan Harrison, David K. Jackson, Ian Johnston, Dominic Kwiatkowski, Cordelia Langford, John Sillitoe on behalf of the Wellcome Sanger Institute COVID-19 Surveillance Team (http://www.sanger.ac.uk/covid-team) |
| EPI_ISL_487647, EPI_ISL_487648, EPI_ISL_487649, EPI_ISL_487650, EPI_ISL_487651, EPI_ISL_487652, EPI_ISL_487653, EPI_ISL_487654, EPI_ISL_487655, EPI_ISL_487656, EPI_ISL_487657, EPI_ISL_487658, EPI_ISL_487659, EPI_ISL_487660, EPI_ISL_487661, EPI_ISL_487662, EPI_ISL_487663, EPI_ISL_487664, EPI_ISL_487665, EPI_ISL_487666, EPI_ISL_487667, EPI_ISL_487668, EPI_ISL_487669, EPI_ISL_487670, EPI_ISL_487671, EPI_ISL_487672, EPI_ISL_487673, EPI_ISL_487674, EPI_ISL_487675, EPI_ISL_487676, EPI_ISL_487677, EPI_ISL_487678, EPI_ISL_487679, EPI_ISL_487680, EPI_ISL_487681, EPI_ISL_487682, EPI_ISL_487683, EPI_ISL_487684, EPI_ISL_487685, EPI_ISL_487686, EPI_ISL_487687, EPI_ISL_487688, EPI_ISL_487689, EPI_ISL_487690, EPI_ISL_487691, EPI_ISL_487692, EPI_ISL_487693, EPI_ISL_487694, EPI_ISL_487695, EPI_ISL_487696, EPI_ISL_487697, EPI_ISL_487698, EPI_ISL_487699, EPI_ISL_487700, EPI_ISL_487701, EPI_ISL_487702, EPI_ISL_487703, EPI_ISL_487704, EPI_ISL_487705, EPI_ISL_487706, EPI_ISL_487707, EPI_ISL_487708, EPI_ISL_487709, EPI_ISL_487710, EPI_ISL_487711, EPI_ISL_487712, EPI_ISL_487713, EPI_ISL_487714, EPI_ISL_487715, EPI_ISL_487716, EPI_ISL_487717, EPI_ISL_487718, EPI_ISL_487719, EPI_ISL_487720, EPI_ISL_487721, EPI_ISL_487722, EPI_ISL_487723, EPI_ISL_487724, EPI_ISL_487725, EPI_ISL_487726, EPI_ISL_487727, EPI_ISL_487728, EPI_ISL_487729, EPI_ISL_487730, EPI_ISL_487731, EPI_ISL_487732, EPI_ISL_487733, EPI_ISL_487734, EPI_ISL_487735, EPI_ISL_487736, EPI_ISL_487737, EPI_ISL_487738, EPI_ISL_487739, EPI_ISL_487740, EPI_ISL_487741, EPI_ISL_487742, EPI_ISL_487743, EPI_ISL_487744, EPI_ISL_487745, EPI_ISL_487746, EPI_ISL_487747, EPI_ISL_487748, EPI_ISL_487749, EPI_ISL_487750, EPI_ISL_487751, EPI_ISL_487752, EPI_ISL_487753, EPI_ISL_487754, EPI_ISL_487755, EPI_ISL_487756, EPI_ISL_487757, EPI_ISL_487758, EPI_ISL_487759, EPI_ISL_487760, EPI_ISL_487761, EPI_ISL_487762, EPI_ISL_487763, EPI_ISL_487764, EPI_ISL_487765, EPI_ISL_487766, EPI_ISL_487768, EPI_ISL_487769, EPI_ISL_487770, EPI_ISL_487771, EPI_ISL_487772, EPI_ISL_487773, EPI_ISL_487774, EPI_ISL_487775, EPI_ISL_487776, EPI_ISL_487777, EPI_ISL_487778, EPI_ISL_487779, EPI_ISL_487780, EPI_ISL_487781, EPI_ISL_487782, EPI_ISL_487783, EPI_ISL_487784, EPI_ISL_487785, EPI_ISL_487786, EPI_ISL_487787, EPI_ISL_487788, EPI_ISL_487789, EPI_ISL_487790, EPI_ISL_487791, EPI_ISL_487792, EPI_ISL_487793, EPI_ISL_487794, EPI_ISL_487795, EPI_ISL_487796, EPI_ISL_487797, EPI_ISL_487798, EPI_ISL_487799, EPI_ISL_487800, EPI_ISL_487801, EPI_ISL_487802, EPI_ISL_487803, EPI_ISL_487804, EPI_ISL_487805, EPI_ISL_487806, EPI_ISL_487807, EPI_ISL_487808, EPI_ISL_487809, EPI_ISL_487810, EPI_ISL_487811, EPI_ISL_487812, EPI_ISL_487813, EPI_ISL_487814, EPI_ISL_487815, EPI_ISL_487816, EPI_ISL_487817, EPI_ISL_487818, EPI_ISL_487819, EPI_ISL_487820, EPI_ISL_487821, EPI_ISL_487822, EPI_ISL_487823, EPI_ISL_487824, EPI_ISL_487825, EPI_ISL_487826, EPI_ISL_487827, EPI_ISL_487828, EPI_ISL_487829, EPI_ISL_487830, EPI_ISL_487831, EPI_ISL_487832, EPI_ISL_487833, EPI_ISL_487834, EPI_ISL_487835, EPI_ISL_487836, EPI_ISL_487837, EPI_ISL_487838, EPI_ISL_487839, EPI_ISL_487840, EPI_ISL_487841, EPI_ISL_487842, EPI_ISL_487843, EPI_ISL_487844, EPI_ISL_487845, EPI_ISL_487846, EPI_ISL_487847, EPI_ISL_487848, EPI_ISL_487849, EPI_ISL_487850, EPI_ISL_487851, EPI_ISL_487852, EPI_ISL_487853, EPI_ISL_487854, EPI_ISL_487855, EPI_ISL_487856, EPI_ISL_487857, EPI_ISL_487858, EPI_ISL_487859, EPI_ISL_487860, EPI_ISL_487861, EPI_ISL_487862, EPI_ISL_487863, EPI_ISL_487864, EPI_ISL_487865, EPI_ISL_487866, EPI_ISL_487867, EPI_ISL_487868, EPI_ISL_487869, EPI_ISL_487870, EPI_ISL_487871, EPI_ISL_487872, EPI_ISL_487873, EPI_ISL_487874, EPI_ISL_487875, EPI_ISL_487876, EPI_ISL_487877, EPI_ISL_487878, EPI_ISL_487879, EPI_ISL_487880, EPI_ISL_487881, EPI_ISL_487882, EPI_ISL_487883, EPI_ISL_487884, EPI_ISL_487885, EPI_ISL_487886, EPI_ISL_487887, EPI_ISL_487888, EPI_ISL_487889, EPI_ISL_487890, EPI_ISL_487891, EPI_ISL_487892, EPI_ISL_487893, EPI_ISL_487894, EPI_ISL_487895, EPI_ISL_487896, EPI_ISL_487897, EPI_ISL_487898, EPI_ISL_487899, EPI_ISL_487900, EPI_ISL_487901, EPI_ISL_487902, EPI_ISL_487903, EPI_ISL_487904, EPI_ISL_487905, EPI_ISL_487906, EPI_ISL_487907, EPI_ISL_487908, EPI_ISL_487909, EPI_ISL_487910, EPI_ISL_487911, EPI_ISL_487912, EPI_ISL_487913, EPI_ISL_487914, EPI_ISL_487915, EPI_ISL_487916, EPI_ISL_487917, EPI_ISL_487918, EPI_ISL_487919, EPI_ISL_487920, EPI_ISL_487921, EPI_ISL_487922, EPI_ISL_487923, EPI_ISL_487924, EPI_ISL_487925, EPI_ISL_487926, EPI_ISL_487927, EPI_ISL_487928, EPI_ISL_487929, EPI_ISL_487930, EPI_ISL_487931, EPI_ISL_487932, EPI_ISL_487933, EPI_ISL_487934, EPI_ISL_487935, EPI_ISL_487936, EPI_ISL_487937, EPI_ISL_487938, EPI_ISL_487939, EPI_ISL_487940, EPI_ISL_487941, EPI_ISL_487942, EPI_ISL_487943, EPI_ISL_487944, EPI_ISL_487945, EPI_ISL_487946, EPI_ISL_487947, EPI_ISL_487948, EPI_ISL_487949, EPI_ISL_487950, EPI_ISL_487951, EPI_ISL_487952, EPI_ISL_487953, EPI_ISL_487954, EPI_ISL_487955, EPI_ISL_487956, EPI_ISL_487957, EPI_ISL_487958, EPI_ISL_487959, EPI_ISL_487960, EPI_ISL_487961, EPI_ISL_487962, EPI_ISL_487963, EPI_ISL_487964, EPI_ISL_487965, EPI_ISL_487966, EPI_ISL_487967, EPI_ISL_487968, EPI_ISL_487969, EPI_ISL_487970, EPI_ISL_487971, EPI_ISL_487972, EPI_ISL_487973, EPI_ISL_487974, EPI_ISL_487975, EPI_ISL_487976, EPI_ISL_487977, EPI_ISL_487978, EPI_ISL_487979, EPI_ISL_487980, EPI_ISL_487981, EPI_ISL_487982, EPI_ISL_487983, EPI_ISL_487984, EPI_ISL_487985, EPI_ISL_487986, EPI_ISL_487987, EPI_ISL_487988, EPI_ISL_487989, EPI_ISL_487990, EPI_ISL_487991, EPI_ISL_487992, EPI_ISL_487993, EPI_ISL_487994, EPI_ISL_487995, EPI_ISL_487996, EPI_ISL_487997, EPI_ISL_487998, EPI_ISL_487999, EPI_ISL_488000, EPI_ISL_488001, EPI_ISL_488002 | | | |
| see above | Virology Department, Royal Infirmary of Edinburgh, NHS Lothian / School of Biological Sciences, University of Edinburgh | Wellcome Sanger Institute for the COVID-19 Genomics UK Consortium | McHugh M, Dewar R, Rooke S, O'Toole Á, Scher E, Hill V, McCrone JT, Colquhoun R, Yu X, Jackson B, Rambaut A, Templeton K and Alex Alderton, Roberto Amato, Sonia Goncalves, Ewan Harrison, David K. Jackson, Ian Johnston, Dominic Kwiatkowski, Cordelia Langford, John Sillitoe on behalf of the Wellcome Sanger Institute COVID-19 Surveillance Team (http://www.sanger.ac.uk/covid-team) |
| EPI_ISL_488003, EPI_ISL_488004, EPI_ISL_488005, EPI_ISL_488006, EPI_ISL_488007, EPI_ISL_488008, EPI_ISL_488009, EPI_ISL_488010, EPI_ISL_488011, EPI_ISL_488012, EPI_ISL_488013, EPI_ISL_488014, EPI_ISL_488015, EPI_ISL_488016, EPI_ISL_488017, EPI_ISL_488018, EPI_ISL_488019, EPI_ISL_488020, EPI_ISL_488021, EPI_ISL_488022, EPI_ISL_488023, EPI_ISL_488024, EPI_ISL_488025, EPI_ISL_488026, EPI_ISL_488027, EPI_ISL_488028, EPI_ISL_488029, EPI_ISL_488030, EPI_ISL_488031, EPI_ISL_488032, EPI_ISL_488033, EPI_ISL_488034, EPI_ISL_488035, EPI_ISL_488036, EPI_ISL_488037, EPI_ISL_488038, EPI_ISL_488039, EPI_ISL_488040, EPI_ISL_488041, EPI_ISL_488042, EPI_ISL_488043, EPI_ISL_488044, EPI_ISL_488045, EPI_ISL_488046, EPI_ISL_488047, EPI_ISL_488048, EPI_ISL_488049, EPI_ISL_488050, EPI_ISL_488051, EPI_ISL_488052, EPI_ISL_488053, EPI_ISL_488054, EPI_ISL_488055, EPI_ISL_488056, EPI_ISL_488057, EPI_ISL_488058, EPI_ISL_488059, EPI_ISL_488060, EPI_ISL_488061, EPI_ISL_488062, EPI_ISL_488063, EPI_ISL_488064, EPI_ISL_488065, EPI_ISL_488066, EPI_ISL_488067, EPI_ISL_488068, EPI_ISL_488069, EPI_ISL_488070, EPI_ISL_488071, EPI_ISL_488072, EPI_ISL_488073, EPI_ISL_488074, EPI_ISL_488075, EPI_ISL_488076, EPI_ISL_488077, EPI_ISL_488078, EPI_ISL_488079, EPI_ISL_488080, EPI_ISL_488081, EPI_ISL_488082, EPI_ISL_488083, EPI_ISL_488084, EPI_ISL_488085, EPI_ISL_488086, EPI_ISL_488087, EPI_ISL_488088, EPI_ISL_488089, EPI_ISL_488090, EPI_ISL_488091, EPI_ISL_488092, EPI_ISL_488093, EPI_ISL_488094, EPI_ISL_488095, EPI_ISL_488096, EPI_ISL_488097, EPI_ISL_488098, EPI_ISL_488099, EPI_ISL_488100, EPI_ISL_488101, EPI_ISL_488102, EPI_ISL_488103, EPI_ISL_488104, EPI_ISL_488105, EPI_ISL_488106, EPI_ISL_488107, EPI_ISL_488108, EPI_ISL_488109, EPI_ISL_488110, EPI_ISL_488111, EPI_ISL_488112, EPI_ISL_488113, EPI_ISL_488114, EPI_ISL_488115, EPI_ISL_488116, EPI_ISL_488117, EPI_ISL_488118, EPI_ISL_488119, EPI_ISL_488120, EPI_ISL_488121, EPI_ISL_488122, EPI_ISL_488123, EPI_ISL_488124, EPI_ISL_488125, EPI_ISL_488126, EPI_ISL_488127, EPI_ISL_488128, EPI_ISL_488129, EPI_ISL_488130, EPI_ISL_488131, EPI_ISL_488132, EPI_ISL_488133, EPI_ISL_488134, EPI_ISL_488135, EPI_ISL_488136, EPI_ISL_488137, EPI_ISL_488138, EPI_ISL_488139, EPI_ISL_488140, EPI_ISL_488141, EPI_ISL_488142, EPI_ISL_488143, EPI_ISL_488144, EPI_ISL_488145, EPI_ISL_488146, EPI_ISL_488147, EPI_ISL_488148, EPI_ISL_488149, EPI_ISL_488150, EPI_ISL_488151, EPI_ISL_488152, EPI_ISL_488153, EPI_ISL_488154, EPI_ISL_488155, EPI_ISL_488156, EPI_ISL_488157, EPI_ISL_488158, EPI_ISL_488159, EPI_ISL_488160, EPI_ISL_488161, EPI_ISL_488162, EPI_ISL_488163, EPI_ISL_488164, EPI_ISL_488165, EPI_ISL_488166, EPI_ISL_488167, EPI_ISL_488168, EPI_ISL_488169, EPI_ISL_488170, EPI_ISL_488171, EPI_ISL_488172, EPI_ISL_488173, EPI_ISL_488174, EPI_ISL_488175, EPI_ISL_488176, EPI_ISL_488177, EPI_ISL_488178, EPI_ISL_488179, EPI_ISL_488180, EPI_ISL_488181, EPI_ISL_488182, EPI_ISL_488183, EPI_ISL_488184, EPI_ISL_488185, EPI_ISL_488186 | | | |
| see above | NU-OMICS DNA Sequencing research facility, Northumbria University | Wellcome Sanger Institute for the COVID-19 Genomics UK Consortium | Chris Duncan, Sheia Waugh, Shirelle Burton-Fanning, Gary Eltringham, Jennifer Collins, Brendan Payne, Yusri Taha, Emma Swindells, Jane Greenaway, Edward Barton, Garren Scott, Debra Padgett, Clive Graham, Sarah Essex, Steve Ligget, Paul Baker, Lynn Dover, Wen Yew, Gary Black, John Allan, Joshua Loh, Greg Young, Matthew Bashton, Andrew Nelson, Darren Smith and Alex Alderton, Roberto Amato, Sonia Goncalves, Ewan Harrison, David K. Jackson, Ian Johnston, Dominic Kwiatkowski, Cordelia Langford, John Sillitoe on behalf of the Wellcome Sanger Institute COVID-19 Surveillance Team (http://www.sanger.ac.uk/covid-team) |
| EPI_ISL_488187, EPI_ISL_488188, EPI_ISL_488189, EPI_ISL_488190, EPI_ISL_488191, EPI_ISL_488192, EPI_ISL_488193, EPI_ISL_488194, EPI_ISL_488195, EPI_ISL_488196, EPI_ISL_488197, EPI_ISL_488198, EPI_ISL_488199, EPI_ISL_488200, EPI_ISL_488201, EPI_ISL_488202, EPI_ISL_488203, EPI_ISL_488204, EPI_ISL_488205, EPI_ISL_488206, EPI_ISL_488207, EPI_ISL_488208, EPI_ISL_488209, EPI_ISL_488210, EPI_ISL_488211, EPI_ISL_488212, EPI_ISL_488213, EPI_ISL_488214, EPI_ISL_488215, EPI_ISL_488216, EPI_ISL_488217, EPI_ISL_488218, EPI_ISL_488219, EPI_ISL_488220, EPI_ISL_488221, EPI_ISL_488222, EPI_ISL_488223, EPI_ISL_488224, EPI_ISL_488225, EPI_ISL_488226, EPI_ISL_488227, EPI_ISL_488228, EPI_ISL_488229, EPI_ISL_488230, EPI_ISL_488231, EPI_ISL_488232, EPI_ISL_488233, EPI_ISL_488234, EPI_ISL_488235, EPI_ISL_488236, EPI_ISL_488237, EPI_ISL_488238, EPI_ISL_488239, EPI_ISL_488240, EPI_ISL_488241, EPI_ISL_488242, EPI_ISL_488243, EPI_ISL_488244, EPI_ISL_488245, EPI_ISL_488246, EPI_ISL_488247, EPI_ISL_488248, EPI_ISL_488249, EPI_ISL_488250, EPI_ISL_488251, EPI_ISL_488252, EPI_ISL_488253, EPI_ISL_488254, EPI_ISL_488255, EPI_ISL_488256, EPI_ISL_488257, EPI_ISL_488258, EPI_ISL_488259, EPI_ISL_488260, EPI_ISL_488261, EPI_ISL_488262, EPI_ISL_488263, EPI_ISL_488264, EPI_ISL_488265, EPI_ISL_488266, EPI_ISL_488267, EPI_ISL_488268, EPI_ISL_488269, EPI_ISL_488270, EPI_ISL_488271, EPI_ISL_488272, EPI_ISL_488273, EPI_ISL_488274, EPI_ISL_488275, EPI_ISL_488276, EPI_ISL_488277, EPI_ISL_488278, EPI_ISL_488279, EPI_ISL_488280, EPI_ISL_488281, EPI_ISL_488282, EPI_ISL_488283, EPI_ISL_488284, EPI_ISL_488285, EPI_ISL_488286, EPI_ISL_488287, EPI_ISL_488288, EPI_ISL_488289, EPI_ISL_488290, EPI_ISL_488291, EPI_ISL_488292, EPI_ISL_488293, EPI_ISL_488294, EPI_ISL_488295, EPI_ISL_488296, EPI_ISL_488297, EPI_ISL_488298, EPI_ISL_488299, EPI_ISL_488300, EPI_ISL_488301, EPI_ISL_488302, EPI_ISL_488303, EPI_ISL_488304, EPI_ISL_488305, EPI_ISL_488306, EPI_ISL_488307, EPI_ISL_488308, EPI_ISL_488309, EPI_ISL_488310, EPI_ISL_488311, EPI_ISL_488312, EPI_ISL_488313, EPI_ISL_488314, EPI_ISL_488315, EPI_ISL_488316, EPI_ISL_488317, EPI_ISL_488318, EPI_ISL_488319, EPI_ISL_488320, EPI_ISL_488321, EPI_ISL_488322, EPI_ISL_488323, EPI_ISL_488324, EPI_ISL_488325, EPI_ISL_488326, EPI_ISL_488327, EPI_ISL_488328, EPI_ISL_488329, EPI_ISL_488330, EPI_ISL_488331, EPI_ISL_488332, EPI_ISL_488333, EPI_ISL_488334, EPI_ISL_488335, EPI_ISL_488336, EPI_ISL_488337, EPI_ISL_488338, EPI_ISL_488339, EPI_ISL_488340, EPI_ISL_488341, EPI_ISL_488343, EPI_ISL_488344, EPI_ISL_488345, EPI_ISL_488346, EPI_ISL_488347, EPI_ISL_488348, EPI_ISL_488349, EPI_ISL_488350, EPI_ISL_488351, EPI_ISL_488352, EPI_ISL_488353, EPI_ISL_488354, EPI_ISL_488355, EPI_ISL_488356, EPI_ISL_488357, EPI_ISL_488358, EPI_ISL_488359, EPI_ISL_488360, EPI_ISL_488361, EPI_ISL_488362, EPI_ISL_488363, EPI_ISL_488364, EPI_ISL_488365, EPI_ISL_488366, EPI_ISL_488367, EPI_ISL_488368, EPI_ISL_488369, EPI_ISL_488370, EPI_ISL_488371, EPI_ISL_488372, EPI_ISL_488373, EPI_ISL_488374, EPI_ISL_488375, EPI_ISL_488376, EPI_ISL_488377, EPI_ISL_488378, EPI_ISL_488379, EPI_ISL_488380, EPI_ISL_488381, EPI_ISL_488382, EPI_ISL_488383, EPI_ISL_488384, EPI_ISL_488385, EPI_ISL_488386, EPI_ISL_488387, EPI_ISL_488388, EPI_ISL_488389, EPI_ISL_488390, EPI_ISL_488391, EPI_ISL_488392, EPI_ISL_488393, EPI_ISL_488394, EPI_ISL_488395, EPI_ISL_488396, EPI_ISL_488397, EPI_ISL_488398, EPI_ISL_488399, EPI_ISL_488400, EPI_ISL_488401, EPI_ISL_488402, EPI_ISL_488403, EPI_ISL_488404, EPI_ISL_488405, EPI_ISL_488406, EPI_ISL_488407, EPI_ISL_488408, EPI_ISL_488410, EPI_ISL_488412, EPI_ISL_488413, EPI_ISL_488414, EPI_ISL_488415, EPI_ISL_488416, EPI_ISL_488417, EPI_ISL_488418, EPI_ISL_488419, EPI_ISL_488420, EPI_ISL_488421, EPI_ISL_488422, EPI_ISL_488423, EPI_ISL_488424, EPI_ISL_488425, EPI_ISL_488426, EPI_ISL_488427, EPI_ISL_488428, EPI_ISL_488429, EPI_ISL_488430, EPI_ISL_488431, EPI_ISL_488432, EPI_ISL_488433, EPI_ISL_488434, EPI_ISL_488435, EPI_ISL_488436, EPI_ISL_488437, EPI_ISL_488438, EPI_ISL_488439, EPI_ISL_488440, EPI_ISL_488441, EPI_ISL_488442, EPI_ISL_488443, EPI_ISL_488444, EPI_ISL_488445, EPI_ISL_488446, EPI_ISL_488447, EPI_ISL_488448, EPI_ISL_488449, EPI_ISL_488450, EPI_ISL_488451, EPI_ISL_488452, EPI_ISL_488453, EPI_ISL_488454, EPI_ISL_488455 | | | |
| see above | PHE South West Regional Laboratory, National Infection Service | Wellcome Sanger Institute for the COVID-19 Genomics UK Consortium | Stephanie Hutchings, Hannah Pymont, Dr Peter Muir, Barry Vipond, Rich Hopes; and Alex Alderton, Roberto Amato, Sonia Goncalves, Ewan Harrison, David K. Jackson, Ian Johnston, Dominic Kwiatkowski, Cordelia Langford, John Sillitoe on behalf of the Wellcome Sanger Institute COVID-19 Surveillance Team (http://www.sanger.ac.uk/covid-team) |
| EPI_ISL_488456, EPI_ISL_488457, EPI_ISL_488458, EPI_ISL_488459, EPI_ISL_488460, EPI_ISL_488461, EPI_ISL_488462, EPI_ISL_488463, EPI_ISL_488464, EPI_ISL_488465, EPI_ISL_488466, EPI_ISL_488467, EPI_ISL_488468, EPI_ISL_488469, EPI_ISL_488470, EPI_ISL_488471, EPI_ISL_488472, EPI_ISL_488473, EPI_ISL_488474, EPI_ISL_488475, EPI_ISL_488476, EPI_ISL_488477, EPI_ISL_488478, EPI_ISL_488479, EPI_ISL_488480, EPI_ISL_488481, EPI_ISL_488482, EPI_ISL_488483, EPI_ISL_488484, EPI_ISL_488485, EPI_ISL_488486, EPI_ISL_488487, EPI_ISL_488488, EPI_ISL_488489, EPI_ISL_488490, EPI_ISL_488491, EPI_ISL_488492, EPI_ISL_488493, EPI_ISL_488494, EPI_ISL_488495, EPI_ISL_488496, EPI_ISL_488497, EPI_ISL_488498, EPI_ISL_488499, EPI_ISL_488500, EPI_ISL_488501, EPI_ISL_488502, EPI_ISL_488503, EPI_ISL_488504, EPI_ISL_488505, EPI_ISL_488506, EPI_ISL_488507, EPI_ISL_488508, EPI_ISL_488509, EPI_ISL_488510, EPI_ISL_488511, EPI_ISL_488512, EPI_ISL_488513, EPI_ISL_488514, EPI_ISL_488515, EPI_ISL_488516, EPI_ISL_488517, EPI_ISL_488518, EPI_ISL_488519, EPI_ISL_488520, EPI_ISL_488521, EPI_ISL_488522, EPI_ISL_488523, EPI_ISL_488524, EPI_ISL_488525, EPI_ISL_488526, EPI_ISL_488527, EPI_ISL_488528, EPI_ISL_488529, EPI_ISL_488530, EPI_ISL_488531, EPI_ISL_488532, EPI_ISL_488533, EPI_ISL_488534, EPI_ISL_488535, EPI_ISL_488536, EPI_ISL_488537, EPI_ISL_488538, EPI_ISL_488539, EPI_ISL_488540, EPI_ISL_488541, EPI_ISL_488542, EPI_ISL_488543, EPI_ISL_488544, EPI_ISL_488545, EPI_ISL_488546, EPI_ISL_488547, EPI_ISL_488548, EPI_ISL_488549, EPI_ISL_488550, EPI_ISL_488551, EPI_ISL_488552, EPI_ISL_488553, EPI_ISL_488554, EPI_ISL_488555, EPI_ISL_488556, EPI_ISL_488557, EPI_ISL_488558, EPI_ISL_488559, EPI_ISL_488560, EPI_ISL_488561, EPI_ISL_488562, EPI_ISL_488563, EPI_ISL_488564, EPI_ISL_488565, EPI_ISL_488566, EPI_ISL_488567, EPI_ISL_488568, EPI_ISL_488569, EPI_ISL_488570, EPI_ISL_488571, EPI_ISL_488572, EPI_ISL_488573, EPI_ISL_488574, EPI_ISL_488575, EPI_ISL_488576, EPI_ISL_488577, EPI_ISL_488578, EPI_ISL_488579, EPI_ISL_488580, EPI_ISL_488581, EPI_ISL_488582, EPI_ISL_488583, EPI_ISL_488584, EPI_ISL_488585, EPI_ISL_488586, EPI_ISL_488587, EPI_ISL_488588, EPI_ISL_488589, EPI_ISL_488590, EPI_ISL_488591, EPI_ISL_488592, EPI_ISL_488593, EPI_ISL_488594, EPI_ISL_488595, EPI_ISL_488596, EPI_ISL_488597, EPI_ISL_488598, EPI_ISL_488599, EPI_ISL_488600, EPI_ISL_488601, EPI_ISL_488602, EPI_ISL_488603, EPI_ISL_488604, EPI_ISL_488605, EPI_ISL_488606, EPI_ISL_488607, EPI_ISL_488608, EPI_ISL_488609, EPI_ISL_488610, EPI_ISL_488611, EPI_ISL_488612, EPI_ISL_488613, EPI_ISL_488614, EPI_ISL_488615, EPI_ISL_488616, EPI_ISL_488617, EPI_ISL_488618, EPI_ISL_488619, EPI_ISL_488620, EPI_ISL_488621, EPI_ISL_488622, EPI_ISL_488623, EPI_ISL_488624, EPI_ISL_488625, EPI_ISL_488626, EPI_ISL_488627, EPI_ISL_488628, EPI_ISL_488629, EPI_ISL_488630, EPI_ISL_488631, EPI_ISL_488632, EPI_ISL_488633, EPI_ISL_488634, EPI_ISL_488635, EPI_ISL_488636, EPI_ISL_488637, EPI_ISL_488638, EPI_ISL_488639, EPI_ISL_488640, EPI_ISL_488641, EPI_ISL_488642, EPI_ISL_488643, EPI_ISL_488644, EPI_ISL_488645, EPI_ISL_488646, EPI_ISL_488647, EPI_ISL_488648, EPI_ISL_488649, EPI_ISL_488650, EPI_ISL_488651, EPI_ISL_488652, EPI_ISL_488653, EPI_ISL_488654, EPI_ISL_488655, EPI_ISL_488656, EPI_ISL_488657, EPI_ISL_488658, EPI_ISL_488659, EPI_ISL_488660, EPI_ISL_488661, EPI_ISL_488662, EPI_ISL_488663, EPI_ISL_488664, EPI_ISL_488665, EPI_ISL_488666, EPI_ISL_488667, EPI_ISL_488668, EPI_ISL_488669, EPI_ISL_488670, EPI_ISL_488671, EPI_ISL_488672, EPI_ISL_488673, EPI_ISL_488674, EPI_ISL_488675, EPI_ISL_488676, EPI_ISL_488677, EPI_ISL_488678, EPI_ISL_488679, EPI_ISL_488680, EPI_ISL_488681, EPI_ISL_488682, EPI_ISL_488683, EPI_ISL_488684, EPI_ISL_488685, EPI_ISL_488686, EPI_ISL_488687, EPI_ISL_488688, EPI_ISL_488689, EPI_ISL_488690, EPI_ISL_488691, EPI_ISL_488692, EPI_ISL_488693, EPI_ISL_488694, EPI_ISL_488695, EPI_ISL_488696, EPI_ISL_488697, EPI_ISL_488698, EPI_ISL_488699, EPI_ISL_488700, EPI_ISL_488701, EPI_ISL_488702, EPI_ISL_488703, EPI_ISL_488704, EPI_ISL_488705, EPI_ISL_488706, EPI_ISL_488707, EPI_ISL_488708, EPI_ISL_488709, EPI_ISL_488710, EPI_ISL_488711, EPI_ISL_488712, EPI_ISL_488713, EPI_ISL_488714, EPI_ISL_488715, EPI_ISL_488716, EPI_ISL_488717, EPI_ISL_488718, EPI_ISL_488719, EPI_ISL_488720, EPI_ISL_488721, EPI_ISL_488722, EPI_ISL_488723, EPI_ISL_488724, EPI_ISL_488725, EPI_ISL_488726, EPI_ISL_488727, EPI_ISL_488728, EPI_ISL_488729, EPI_ISL_488730, EPI_ISL_488731, EPI_ISL_488732, EPI_ISL_488733, EPI_ISL_488734, EPI_ISL_488735, EPI_ISL_488736, EPI_ISL_488737, EPI_ISL_488738, EPI_ISL_488739, EPI_ISL_488740, EPI_ISL_488741, EPI_ISL_488742, EPI_ISL_488743, EPI_ISL_488744, EPI_ISL_488745, EPI_ISL_488746, EPI_ISL_488747, EPI_ISL_488748, EPI_ISL_488749, EPI_ISL_488750, EPI_ISL_488751, EPI_ISL_488752, EPI_ISL_488753, EPI_ISL_488754, EPI_ISL_488755, EPI_ISL_488756, EPI_ISL_488757, EPI_ISL_488758, EPI_ISL_488759, EPI_ISL_488760, EPI_ISL_488761, EPI_ISL_488762, EPI_ISL_488763, EPI_ISL_488764, EPI_ISL_488765, EPI_ISL_488766, EPI_ISL_488767, EPI_ISL_488768, EPI_ISL_488769, EPI_ISL_488770, EPI_ISL_488771, EPI_ISL_488772, EPI_ISL_488773, EPI_ISL_488774, EPI_ISL_488775, EPI_ISL_488776, EPI_ISL_488777, EPI_ISL_488778, EPI_ISL_488779, EPI_ISL_488780, EPI_ISL_488781, EPI_ISL_488782, EPI_ISL_488783, EPI_ISL_488784, EPI_ISL_488785, EPI_ISL_488786, EPI_ISL_488787, EPI_ISL_488788, EPI_ISL_488789, EPI_ISL_488790, EPI_ISL_488791, EPI_ISL_488792, EPI_ISL_488793, EPI_ISL_488794, EPI_ISL_488795, EPI_ISL_488796, EPI_ISL_488797, EPI_ISL_488798, EPI_ISL_488799, EPI_ISL_488800, EPI_ISL_488801, EPI_ISL_488802, EPI_ISL_488803, EPI_ISL_488804, EPI_ISL_488805, EPI_ISL_488806, EPI_ISL_488807, EPI_ISL_488808, EPI_ISL_488809, EPI_ISL_488810, EPI_ISL_488811, EPI_ISL_488812, EPI_ISL_488813, EPI_ISL_488814, EPI_ISL_488815, EPI_ISL_488816, EPI_ISL_488817, EPI_ISL_488818, EPI_ISL_488819, EPI_ISL_488820, EPI_ISL_488821, EPI_ISL_488822, EPI_ISL_488823, EPI_ISL_488824, EPI_ISL_488825, EPI_ISL_488826, EPI_ISL_488827, EPI_ISL_488828, EPI_ISL_488829, EPI_ISL_488830, EPI_ISL_488831, EPI_ISL_488832, EPI_ISL_488833, EPI_ISL_488834, EPI_ISL_488835, EPI_ISL_488836 | | | |
| see above | NU-OMICS DNA Sequencing research facility, Northumbria University | Wellcome Sanger Institute for the COVID-19 Genomics UK Consortium | Chris Duncan, Sheia Waugh, Shirelle Burton-Fanning, Gary Eltringham, Jennifer Collins, Brendan Payne, Yusri Taha, Emma Swindells, Jane Greenaway, Edward Barton, Garren Scott, Debra Padgett, Clive Graham, Sarah Essex, Steve Ligget, Paul Baker, Lynn Dover, Wen Yew, Gary Black, John Allan, Joshua Loh, Greg Young, Matthew Bashton, Andrew Nelson, Darren Smith and Alex Alderton, Roberto Amato, Sonia Goncalves, Ewan Harrison, David K. Jackson, Ian Johnston, Dominic Kwiatkowski, Cordelia Langford, John Sillitoe on behalf of the Wellcome Sanger Institute COVID-19 Surveillance Team (http://www.sanger.ac.uk/covid-team) |
| EPI_ISL_488837, EPI_ISL_488838, EPI_ISL_488839 | Microbiology Department, Hereford County Hospital | Wellcome Sanger Institute for the COVID-19 Genomics UK Consortium | Alison Johnson, Venkat Sivaprakasam, Fenella Halstead, Jane Thomas, Wendy Hogsden, Samantha Lamb and Alex Alderton, Roberto Amato, Sonia Goncalves, Ewan Harrison, David K. Jackson, Ian Johnston, Dominic Kwiatkowski, Cordelia Langford, John Sillitoe on behalf of the Wellcome Sanger Institute COVID-19 Surveillance Team (http://www.sanger.ac.uk/covid-team) |

| | | | |
|---|---|---|---|
| EPI_ISL_488840 | Department of Medical Microbiology, Western Sussex Hospitals NHS Foundation Trust, St Richard's Hospital | Wellcome Sanger Institute for the COVID-19 Genomics UK Consortium | Manasa Mutingwende, Sarah Lowdon, Olga Podplomyk, Michelle Erkiert, Jonathan Lewis, Paul Randell and Alex Alderton, Roberto Amato, Sonia Goncalves, Ewan Harrison, David K. Jackson, Ian Johnston, Dominic Kwiatkowski, Cordelia Langford, John Sillitoe on behalf of the Wellcome Sanger Institute COVID-19 Surveillance Team (http://www.sanger.ac.uk/covid-team) |
| EPI_ISL_488841, EPI_ISL_488842, EPI_ISL_488843, EPI_ISL_488844 | Microbiology Department, Hereford County Hospital | Wellcome Sanger Institute for the COVID-19 Genomics UK Consortium | Alison Johnson, Venkat Sivaprakasam, Fenella Halstead, Jane Thomas, Wendy Hogsden, Samantha Lamb and Alex Alderton, Roberto Amato, Sonia Goncalves, Ewan Harrison, David K. Jackson, Ian Johnston, Dominic Kwiatkowski, Cordelia Langford, John Sillitoe on behalf of the Wellcome Sanger Institute COVID-19 Surveillance Team (http://www.sanger.ac.uk/covid-team) |
| EPI_ISL_488845 | Department of Medical Microbiology, Western Sussex Hospitals NHS Foundation Trust, St Richard's Hospital | Wellcome Sanger Institute for the COVID-19 Genomics UK Consortium | Manasa Mutingwende, Sarah Lowdon, Olga Podplomyk, Michelle Erkiert, Jonathan Lewis, Paul Randell and Alex Alderton, Roberto Amato, Sonia Goncalves, Ewan Harrison, David K. Jackson, Ian Johnston, Dominic Kwiatkowski, Cordelia Langford, John Sillitoe on behalf of the Wellcome Sanger Institute COVID-19 Surveillance Team (http://www.sanger.ac.uk/covid-team) |
| EPI_ISL_488846 | Microbiology Department, Hereford County Hospital | Wellcome Sanger Institute for the COVID-19 Genomics UK Consortium | Alison Johnson, Venkat Sivaprakasam, Fenella Halstead, Jane Thomas, Wendy Hogsden, Samantha Lamb and Alex Alderton, Roberto Amato, Sonia Goncalves, Ewan Harrison, David K. Jackson, Ian Johnston, Dominic Kwiatkowski, Cordelia Langford, John Sillitoe on behalf of the Wellcome Sanger Institute COVID-19 Surveillance Team (http://www.sanger.ac.uk/covid-team) |
| EPI_ISL_488847 | Department of Medical Microbiology, Western Sussex Hospitals NHS Foundation Trust, St Richard's Hospital | Wellcome Sanger Institute for the COVID-19 Genomics UK Consortium | Manasa Mutingwende, Sarah Lowdon, Olga Podplomyk, Michelle Erkiert, Jonathan Lewis, Paul Randell and Alex Alderton, Roberto Amato, Sonia Goncalves, Ewan Harrison, David K. Jackson, Ian Johnston, Dominic Kwiatkowski, Cordelia Langford, John Sillitoe on behalf of the Wellcome Sanger Institute COVID-19 Surveillance Team (http://www.sanger.ac.uk/covid-team) |
| EPI_ISL_488848, EPI_ISL_488849, EPI_ISL_488850, EPI_ISL_488851, EPI_ISL_488852 | Microbiology Department, Hereford County Hospital | Wellcome Sanger Institute for the COVID-19 Genomics UK Consortium | Alison Johnson, Venkat Sivaprakasam, Fenella Halstead, Jane Thomas, Wendy Hogsden, Samantha Lamb and Alex Alderton, Roberto Amato, Sonia Goncalves, Ewan Harrison, David K. Jackson, Ian Johnston, Dominic Kwiatkowski, Cordelia Langford, John Sillitoe on behalf of the Wellcome Sanger Institute COVID-19 Surveillance Team (http://www.sanger.ac.uk/covid-team) |
| EPI_ISL_488853, EPI_ISL_488854, EPI_ISL_488855, EPI_ISL_488856 | Department of Medical Microbiology, Western Sussex Hospitals NHS Foundation Trust, St Richard's Hospital | Wellcome Sanger Institute for the COVID-19 Genomics UK Consortium | Manasa Mutingwende, Sarah Lowdon, Olga Podplomyk, Michelle Erkiert, Jonathan Lewis, Paul Randell and Alex Alderton, Roberto Amato, Sonia Goncalves, Ewan Harrison, David K. Jackson, Ian Johnston, Dominic Kwiatkowski, Cordelia Langford, John Sillitoe on behalf of the Wellcome Sanger Institute COVID-19 Surveillance Team (http://www.sanger.ac.uk/covid-team) |
| EPI_ISL_488857, EPI_ISL_488858, EPI_ISL_488859, EPI_ISL_488860 | Microbiology Department, Hereford County Hospital | Wellcome Sanger Institute for the COVID-19 Genomics UK Consortium | Alison Johnson, Venkat Sivaprakasam, Fenella Halstead, Jane Thomas, Wendy Hogsden, Samantha Lamb and Alex Alderton, Roberto Amato, Sonia Goncalves, Ewan Harrison, David K. Jackson, Ian Johnston, Dominic Kwiatkowski, Cordelia Langford, John Sillitoe on behalf of the Wellcome Sanger Institute COVID-19 Surveillance Team (http://www.sanger.ac.uk/covid-team) |
| EPI_ISL_488861 | Department of Medical Microbiology, Western Sussex Hospitals NHS Foundation Trust, St Richard's Hospital | Wellcome Sanger Institute for the COVID-19 Genomics UK Consortium | Manasa Mutingwende, Sarah Lowdon, Olga Podplomyk, Michelle Erkiert, Jonathan Lewis, Paul Randell and Alex Alderton, Roberto Amato, Sonia Goncalves, Ewan Harrison, David K. Jackson, Ian Johnston, Dominic Kwiatkowski, Cordelia Langford, John Sillitoe on behalf of the Wellcome Sanger Institute COVID-19 Surveillance Team (http://www.sanger.ac.uk/covid-team) |
| EPI_ISL_488862 | Microbiology Department, Hereford County Hospital | Wellcome Sanger Institute for the COVID-19 Genomics UK Consortium | Alison Johnson, Venkat Sivaprakasam, Fenella Halstead, Jane Thomas, Wendy Hogsden, Samantha Lamb and Alex Alderton, Roberto Amato, Sonia Goncalves, Ewan Harrison, David K. Jackson, Ian Johnston, Dominic Kwiatkowski, Cordelia Langford, John Sillitoe on behalf of the Wellcome Sanger Institute COVID-19 Surveillance Team (http://www.sanger.ac.uk/covid-team) |
| EPI_ISL_488863, EPI_ISL_488864 | Department of Medical Microbiology, Western Sussex Hospitals NHS Foundation Trust, St Richard's Hospital | Wellcome Sanger Institute for the COVID-19 Genomics UK Consortium | Manasa Mutingwende, Sarah Lowdon, Olga Podplomyk, Michelle Erkiert, Jonathan Lewis, Paul Randell and Alex Alderton, Roberto Amato, Sonia Goncalves, Ewan Harrison, David K. Jackson, Ian Johnston, Dominic Kwiatkowski, Cordelia Langford, John Sillitoe on behalf of the Wellcome Sanger Institute COVID-19 Surveillance Team (http://www.sanger.ac.uk/covid-team) |
| EPI_ISL_488865, EPI_ISL_488866, EPI_ISL_488867, EPI_ISL_488868, EPI_ISL_488869, EPI_ISL_488870, EPI_ISL_488871 | Microbiology Department, Hereford County Hospital | Wellcome Sanger Institute for the COVID-19 Genomics UK Consortium | Alison Johnson, Venkat Sivaprakasam, Fenella Halstead, Jane Thomas, Wendy Hogsden, Samantha Lamb and Alex Alderton, Roberto Amato, Sonia Goncalves, Ewan Harrison, David K. Jackson, Ian Johnston, Dominic Kwiatkowski, Cordelia Langford, John Sillitoe on behalf of the Wellcome Sanger Institute COVID-19 Surveillance Team (http://www.sanger.ac.uk/covid-team) |
| EPI_ISL_488872 | Department of Medical Microbiology, Western Sussex Hospitals NHS Foundation Trust, St Richard's Hospital | Wellcome Sanger Institute for the COVID-19 Genomics UK Consortium | Manasa Mutingwende, Sarah Lowdon, Olga Podplomyk, Michelle Erkiert, Jonathan Lewis, Paul Randell and Alex Alderton, Roberto Amato, Sonia Goncalves, Ewan Harrison, David K. Jackson, Ian Johnston, Dominic Kwiatkowski, Cordelia Langford, John Sillitoe on behalf of the Wellcome Sanger Institute COVID-19 Surveillance Team (http://www.sanger.ac.uk/covid-team) |
| EPI_ISL_488873 | Microbiology Department, Hereford County Hospital | Wellcome Sanger Institute for the COVID-19 Genomics UK Consortium | Alison Johnson, Venkat Sivaprakasam, Fenella Halstead, Jane Thomas, Wendy Hogsden, Samantha Lamb and Alex Alderton, Roberto Amato, Sonia Goncalves, Ewan Harrison, David K. Jackson, Ian Johnston, Dominic Kwiatkowski, Cordelia Langford, John Sillitoe on behalf of the Wellcome Sanger Institute COVID-19 Surveillance Team (http://www.sanger.ac.uk/covid-team) |
| EPI_ISL_488874, EPI_ISL_488875, EPI_ISL_488876 | Regional Virus Laboratory, Belfast Health and Social Care Trust | Wellcome Sanger Institute for the COVID-19 Genomics UK Consortium | Conall McCaughey, James McKenna, Tanya Curran, Susan Feeney, Alison Watt, Ciara Cox, Mairead Connor, Zoltan Molnar, David Simpson, Derek Fairley; and Alex Alderton, Roberto Amato, Sonia Goncalves, Ewan Harrison, David K. Jackson, Ian Johnston, Dominic Kwiatkowski, Cordelia Langford, John Sillitoe on behalf of the Wellcome Sanger Institute COVID-19 Surveillance Team (http://www.sanger.ac.uk/covid-team) |
| EPI_ISL_488877, EPI_ISL_488878 | PHE South West Regional Laboratory, National Infection Service | Wellcome Sanger Institute for the COVID-19 Genomics UK Consortium | Stephanie Hutchings, Hannah Pymont, Dr Peter Muir, Barry Vipond, Rich Hopes; and Alex Alderton, Roberto Amato, Sonia Goncalves, Ewan Harrison, David K. Jackson, Ian Johnston, Dominic Kwiatkowski, Cordelia Langford, John Sillitoe on behalf of the Wellcome Sanger Institute COVID-19 Surveillance Team (http://www.sanger.ac.uk/covid-team) |
| EPI_ISL_488879, EPI_ISL_488880, EPI_ISL_488881, EPI_ISL_488882, EPI_ISL_488883, EPI_ISL_488884, EPI_ISL_488885, EPI_ISL_488886, EPI_ISL_488887, EPI_ISL_488888, EPI_ISL_488889, EPI_ISL_488890, EPI_ISL_488891, EPI_ISL_488892, EPI_ISL_488893, EPI_ISL_488894, EPI_ISL_488895, EPI_ISL_488896, EPI_ISL_488897, EPI_ISL_488898, EPI_ISL_488899, EPI_ISL_488900, EPI_ISL_488901, EPI_ISL_488902, EPI_ISL_488903, EPI_ISL_488904, EPI_ISL_488905, EPI_ISL_488906, EPI_ISL_488907, EPI_ISL_488908, EPI_ISL_488909, EPI_ISL_488910, EPI_ISL_488911, EPI_ISL_488912, EPI_ISL_488913, EPI_ISL_488914, EPI_ISL_488915, EPI_ISL_488916, EPI_ISL_488917, EPI_ISL_488918, EPI_ISL_488919, EPI_ISL_488920, EPI_ISL_488921, EPI_ISL_488922, EPI_ISL_488923, EPI_ISL_488924, EPI_ISL_488925, EPI_ISL_488926, EPI_ISL_488927, EPI_ISL_488928, EPI_ISL_488929, EPI_ISL_488930, EPI_ISL_488931, EPI_ISL_488932, EPI_ISL_488933, EPI_ISL_488934, EPI_ISL_488935, EPI_ISL_488936, EPI_ISL_488937, EPI_ISL_488938, EPI_ISL_488939, EPI_ISL_488940, EPI_ISL_488941, EPI_ISL_488942, EPI_ISL_488943, EPI_ISL_488944, EPI_ISL_488945, EPI_ISL_488946, EPI_ISL_488947, EPI_ISL_488948, EPI_ISL_488949, EPI_ISL_488950, EPI_ISL_488951, EPI_ISL_488952, EPI_ISL_488953, EPI_ISL_488954, EPI_ISL_488955, EPI_ISL_488956, EPI_ISL_488957, EPI_ISL_488958, EPI_ISL_488959, EPI_ISL_488960, EPI_ISL_488961, EPI_ISL_488962, EPI_ISL_488963, EPI_ISL_488964, EPI_ISL_488965, EPI_ISL_488966, EPI_ISL_488967, EPI_ISL_488968, EPI_ISL_488969, EPI_ISL_488970, EPI_ISL_488971, EPI_ISL_488972, EPI_ISL_488973, EPI_ISL_488974, EPI_ISL_488975, EPI_ISL_488976, EPI_ISL_488977, EPI_ISL_488978, EPI_ISL_488979, EPI_ISL_488980, EPI_ISL_488981, EPI_ISL_488982, EPI_ISL_488983, EPI_ISL_488984, EPI_ISL_488985, EPI_ISL_488986, EPI_ISL_488987, EPI_ISL_488988, EPI_ISL_488989, EPI_ISL_488990, EPI_ISL_488991, EPI_ISL_488992, EPI_ISL_488993, EPI_ISL_488994, EPI_ISL_488995, EPI_ISL_488996, EPI_ISL_488997, EPI_ISL_488998, EPI_ISL_488999, EPI_ISL_489000, EPI_ISL_489001, EPI_ISL_489002, EPI_ISL_489003, EPI_ISL_489004, EPI_ISL_489005, EPI_ISL_489006, EPI_ISL_489007, EPI_ISL_489008, EPI_ISL_489009, EPI_ISL_489010, EPI_ISL_489011, EPI_ISL_489012, EPI_ISL_489013, EPI_ISL_489014, EPI_ISL_489015, EPI_ISL_489016, EPI_ISL_489017, EPI_ISL_489018, EPI_ISL_489019, EPI_ISL_489020, EPI_ISL_489021, EPI_ISL_489022, EPI_ISL_489023, EPI_ISL_489024, EPI_ISL_489025, EPI_ISL_489026, EPI_ISL_489027, EPI_ISL_489028, EPI_ISL_489029, EPI_ISL_489030, EPI_ISL_489031, EPI_ISL_489032, EPI_ISL_489033, EPI_ISL_489034, EPI_ISL_489035, EPI_ISL_489036, EPI_ISL_489037, EPI_ISL_489038, EPI_ISL_489039, EPI_ISL_489040, EPI_ISL_489041, EPI_ISL_489042, EPI_ISL_489044, EPI_ISL_489045, EPI_ISL_489046, EPI_ISL_489047, EPI_ISL_489048, EPI_ISL_489049, EPI_ISL_489050, EPI_ISL_489051, EPI_ISL_489052, EPI_ISL_489053, EPI_ISL_489054, EPI_ISL_489055, EPI_ISL_489056, EPI_ISL_489057, EPI_ISL_489058, EPI_ISL_489059, EPI_ISL_489060, EPI_ISL_489061, EPI_ISL_489062, EPI_ISL_489063 | | | |
| see above | Virology Department, Royal Infirmary of Edinburgh, NHS Lothian / School of Biological Sciences, University of Edinburgh | Wellcome Sanger Institute for the COVID-19 Genomics UK Consortium | McHugh M, Dewar R, Rooke S, O'Toole Á, Scher E, Hill V, McCrone JT, Colquhoun R, Yu X, Jackson B, Rambaut A, Templeton K and Alex Alderton, Roberto Amato, Sonia Goncalves, Ewan Harrison, David K. Jackson, Ian Johnston, Dominic Kwiatkowski, Cordelia Langford, John Sillitoe on behalf of the Wellcome Sanger Institute COVID-19 Surveillance Team (http://www.sanger.ac.uk/covid-team) |
| EPI_ISL_489064, EPI_ISL_489065, EPI_ISL_489066, EPI_ISL_489067, EPI_ISL_489068, EPI_ISL_489069, EPI_ISL_489070, EPI_ISL_489071, EPI_ISL_489072, EPI_ISL_489073, EPI_ISL_489074, EPI_ISL_489075, EPI_ISL_489076, EPI_ISL_489077, EPI_ISL_489078, EPI_ISL_489079, EPI_ISL_489080, EPI_ISL_489081, EPI_ISL_489082, EPI_ISL_489083, EPI_ISL_489084, EPI_ISL_489085, EPI_ISL_489086, EPI_ISL_489087, EPI_ISL_489088, EPI_ISL_489089, EPI_ISL_489090, EPI_ISL_489091, EPI_ISL_489092, EPI_ISL_489093, EPI_ISL_489094, EPI_ISL_489095, EPI_ISL_489096, EPI_ISL_489097, EPI_ISL_489098, EPI_ISL_489099, EPI_ISL_489100, EPI_ISL_489101, EPI_ISL_489102, EPI_ISL_489103, EPI_ISL_489104, EPI_ISL_489105, EPI_ISL_489106, EPI_ISL_489107, EPI_ISL_489108, EPI_ISL_489109, EPI_ISL_489110, EPI_ISL_489111, EPI_ISL_489112, EPI_ISL_489113, EPI_ISL_489114, EPI_ISL_489115, EPI_ISL_489116, EPI_ISL_489117, EPI_ISL_489118, EPI_ISL_489119 | | | |
| see above | NU-OMICS DNA Sequencing research facility, Northumbria University | Wellcome Sanger Institute for the COVID-19 Genomics UK Consortium | Chris Duncan, Sheia Waugh, Shirelle Burton-Fanning, Gary Eltringham, Jennifer Collins, Brendan Payne, Yusri Taha, Emma Swindells, Jane Greenaway, Edward Barton, Garren Scott, Debra Padgett, Clive Graham, Sarah Essex, Steve Ligget, Paul Baker, Lynn Dover, Wen Yew, Gary Black, John Allan, Joshua Loh, Greg Young, Matthew Bashton, Andrew Nelson, Darren Smith and Alex Alderton, Roberto Amato, Sonia Goncalves, Ewan Harrison, David K. Jackson, Ian Johnston, Dominic Kwiatkowski, Cordelia Langford, John Sillitoe on behalf of the Wellcome Sanger Institute COVID-19 Surveillance Team (http://www.sanger.ac.uk/covid-team) |
| EPI_ISL_489120 | PHE South West Regional Laboratory, National Infection Service | Wellcome Sanger Institute for the COVID-19 Genomics UK Consortium | Stephanie Hutchings, Hannah Pymont, Dr Peter Muir, Barry Vipond, Rich Hopes; and Alex Alderton, Roberto Amato, Sonia Goncalves, Ewan Harrison, David K. Jackson, Ian Johnston, Dominic Kwiatkowski, Cordelia Langford, John Sillitoe on behalf of the Wellcome Sanger Institute COVID-19 Surveillance Team (http://www.sanger.ac.uk/covid-team) |
| EPI_ISL_489121, EPI_ISL_489122, EPI_ISL_489123, EPI_ISL_489124, EPI_ISL_489125, EPI_ISL_489126, EPI_ISL_489127, EPI_ISL_489128, EPI_ISL_489129, EPI_ISL_489130, EPI_ISL_489131, EPI_ISL_489132, EPI_ISL_489133, EPI_ISL_489134, EPI_ISL_489135, EPI_ISL_489136, EPI_ISL_489137, EPI_ISL_489138, EPI_ISL_489139, EPI_ISL_489140, EPI_ISL_489141, EPI_ISL_489142, EPI_ISL_489143, EPI_ISL_489144, EPI_ISL_489145, EPI_ISL_489146, EPI_ISL_489147, EPI_ISL_489148, EPI_ISL_489149, EPI_ISL_489150, EPI_ISL_489151, EPI_ISL_489152, EPI_ISL_489153, EPI_ISL_489154, EPI_ISL_489155 | | | |
| see above | NU-OMICS DNA Sequencing research facility, Northumbria University | Wellcome Sanger Institute for the COVID-19 Genomics UK Consortium | Chris Duncan, Sheia Waugh, Shirelle Burton-Fanning, Gary Eltringham, Jennifer Collins, Brendan Payne, Yusri Taha, Emma Swindells, Jane Greenaway, Edward Barton, Garren Scott, Debra Padgett, Clive Graham, Sarah Essex, Steve Ligget, Paul Baker, Lynn Dover, Wen Yew, Gary Black, John Allan, Joshua Loh, Greg Young, Matthew Bashton, Andrew Nelson, Darren Smith and Alex Alderton, Roberto Amato, Sonia Goncalves, Ewan Harrison, David K. Jackson, Ian Johnston, Dominic Kwiatkowski, Cordelia Langford, John Sillitoe on behalf of the Wellcome Sanger Institute COVID-19 Surveillance Team (http://www.sanger.ac.uk/covid-team) |
| EPI_ISL_489156, EPI_ISL_489157, EPI_ISL_489158, EPI_ISL_489159, EPI_ISL_489160, EPI_ISL_489161, EPI_ISL_489162, EPI_ISL_489163, EPI_ISL_489164, EPI_ISL_489165, EPI_ISL_489166, EPI_ISL_489167, EPI_ISL_489168, EPI_ISL_489169, EPI_ISL_489170, EPI_ISL_489171, EPI_ISL_489172, EPI_ISL_489173, EPI_ISL_489174, EPI_ISL_489175, EPI_ISL_489176, EPI_ISL_489177, EPI_ISL_489178, EPI_ISL_489179, EPI_ISL_489180, EPI_ISL_489181, EPI_ISL_489182, EPI_ISL_489183, EPI_ISL_489184, EPI_ISL_489185, EPI_ISL_489186, EPI_ISL_489187, EPI_ISL_489188, EPI_ISL_489189, EPI_ISL_489190, EPI_ISL_489191, EPI_ISL_489192, EPI_ISL_489193, EPI_ISL_489194, EPI_ISL_489195, EPI_ISL_489196, EPI_ISL_489197, EPI_ISL_489198, EPI_ISL_489199, EPI_ISL_489200, EPI_ISL_489201, EPI_ISL_489202, EPI_ISL_489203, EPI_ISL_489204, EPI_ISL_489205, EPI_ISL_489206, EPI_ISL_489207, EPI_ISL_489208, EPI_ISL_489209, EPI_ISL_489210, EPI_ISL_489211, EPI_ISL_489212, EPI_ISL_489214, EPI_ISL_489215, EPI_ISL_489216, EPI_ISL_489217, EPI_ISL_489218, EPI_ISL_489219, EPI_ISL_489220, EPI_ISL_489221, EPI_ISL_489222, EPI_ISL_489223, EPI_ISL_489224, EPI_ISL_489225, EPI_ISL_489226, EPI_ISL_489227, EPI_ISL_489228, EPI_ISL_489229, EPI_ISL_489230, EPI_ISL_489231, EPI_ISL_489232, EPI_ISL_489233, EPI_ISL_489234, EPI_ISL_489235, EPI_ISL_489236, EPI_ISL_489237, EPI_ISL_489238, EPI_ISL_489239, EPI_ISL_489240, EPI_ISL_489241, EPI_ISL_489242, EPI_ISL_489243, EPI_ISL_489244, EPI_ISL_489245, EPI_ISL_489246, EPI_ISL_489247, EPI_ISL_489248, EPI_ISL_489249, EPI_ISL_489250, EPI_ISL_489251, EPI_ISL_489252, EPI_ISL_489253, EPI_ISL_489254, EPI_ISL_489255, EPI_ISL_489256, EPI_ISL_489257, EPI_ISL_489258, EPI_ISL_489259, EPI_ISL_489260, EPI_ISL_489261, EPI_ISL_489262, EPI_ISL_489263, EPI_ISL_489264, EPI_ISL_489265, EPI_ISL_489266, EPI_ISL_489267, EPI_ISL_489268, EPI_ISL_489269, EPI_ISL_489270, EPI_ISL_489271, EPI_ISL_489272, EPI_ISL_489273, EPI_ISL_489274, EPI_ISL_489275, EPI_ISL_489276, EPI_ISL_489277, EPI_ISL_489278, EPI_ISL_489279, EPI_ISL_489280, EPI_ISL_489281, EPI_ISL_489282, EPI_ISL_489283, EPI_ISL_489284, EPI_ISL_489285, EPI_ISL_489286, EPI_ISL_489287, EPI_ISL_489288, EPI_ISL_489289, EPI_ISL_489290, EPI_ISL_489291, EPI_ISL_489292, EPI_ISL_489293, EPI_ISL_489294, EPI_ISL_489295, EPI_ISL_489296, EPI_ISL_489297, EPI_ISL_489298, EPI_ISL_489299, EPI_ISL_489300, EPI_ISL_489301, EPI_ISL_489302, EPI_ISL_489303, EPI_ISL_489304, EPI_ISL_489305, EPI_ISL_489306, EPI_ISL_489307, EPI_ISL_489308, EPI_ISL_489309, EPI_ISL_489310, EPI_ISL_489311, EPI_ISL_489312, EPI_ISL_489313, EPI_ISL_489314, EPI_ISL_489315, EPI_ISL_489316, EPI_ISL_489317, EPI_ISL_489318, EPI_ISL_489319, EPI_ISL_489320, EPI_ISL_489321, EPI_ISL_489322, EPI_ISL_489323, EPI_ISL_489324, EPI_ISL_489325, EPI_ISL_489326, EPI_ISL_489327, EPI_ISL_489328, EPI_ISL_489329, EPI_ISL_489330, EPI_ISL_489331, EPI_ISL_489332, EPI_ISL_489333, EPI_ISL_489334, EPI_ISL_489335, EPI_ISL_489336, EPI_ISL_489337, EPI_ISL_489338, EPI_ISL_489339, EPI_ISL_489340, EPI_ISL_489341, EPI_ISL_489342, EPI_ISL_489343, EPI_ISL_489344, EPI_ISL_489345, EPI_ISL_489346, EPI_ISL_489347, EPI_ISL_489348, EPI_ISL_489349, EPI_ISL_489350, EPI_ISL_489351, EPI_ISL_489352, EPI_ISL_489353, EPI_ISL_489354, EPI_ISL_489355, EPI_ISL_489356, EPI_ISL_489359, EPI_ISL_489360, EPI_ISL_489361, EPI_ISL_489362, EPI_ISL_489363, EPI_ISL_489364, EPI_ISL_489365, EPI_ISL_489366, EPI_ISL_489367, EPI_ISL_489368, EPI_ISL_489369, EPI_ISL_489370, EPI_ISL_489371, EPI_ISL_489372, EPI_ISL_489373, EPI_ISL_489374, EPI_ISL_489375, EPI_ISL_489376, EPI_ISL_489377, EPI_ISL_489378, EPI_ISL_489379 | | | |
| see above | Regional Virus Laboratory, Belfast Health and Social Care Trust | Wellcome Sanger Institute for the COVID-19 Genomics UK Consortium | Conall McCaughey, James McKenna, Tanya Curran, Susan Feeney, Alison Watt, Ciara Cox, Mairead Connor, Zoltan Molnar, David Simpson, Derek Fairley; and Alex Alderton, Roberto Amato, Sonia Goncalves, Ewan Harrison, David K. Jackson, Ian Johnston, Dominic Kwiatkowski, Cordelia Langford, John Sillitoe on behalf of the Wellcome Sanger Institute COVID-19 Surveillance Team (http://www.sanger.ac.uk/covid-team) |
| EPI_ISL_489380 | NHSGGC West of Scotland Specialist Virology Centre / MRC-University of Glasgow Centre for Virus Research | Wellcome Sanger Institute for the COVID-19 Genomics UK Consortium | Ana da Silva Filipe, Natasha Johnson, Kathy Smollett, Daniel Mair, Stephen Carmichael, Lily Tong, Jenna Nichols, Elihu Aranday-Cortes, Kirstyn Brunker, Yasmin Parr, Kyriaki Nomikou; Sarah McDonald, Marc Niebel, Patawee Asamaphan; Richard Orton, Joseph Hughes, Sreenu Vattipally, David L Robertson; Alasdair MacLean, Rory Gunson; Kathy Li, Natasha Jesudason, Rajiv Shah, James Shepherd, Antonia Ho, Alice Broos, Emma Thomson and Alex Alderton, Roberto Amato, Sonia Goncalves, Ewan Harrison, David K. Jackson, Ian Johnston, Dominic Kwiatkowski, Cordelia Langford, John Sillitoe on behalf of the Wellcome Sanger Institute COVID-19 Surveillance Team (http://www.sanger.ac.uk/covid-team) |
| EPI_ISL_489381, EPI_ISL_489382, EPI_ISL_489383, EPI_ISL_489384, EPI_ISL_489385, EPI_ISL_489386, EPI_ISL_489387, EPI_ISL_489388, EPI_ISL_489389, EPI_ISL_489390 | Department of Pathology, University of Cambridge | Wellcome Sanger Institute for the COVID-19 Genomics UK Consortium | Luke W Meredith, M. Estée Török , Myra Hosmillo, William L. Hamilton, Martin D. Curran, Theresa Feltwell, Grant Hall, Anna Yakovleva, Fahad A Khokhar, Charlotte J. Houldcroft, Laura G Caller, Aminu S. Jahun, Sarah L. Caddy, Ian Goodfellow; and Alex Alderton, Roberto Amato, Sonia Goncalves, Ewan Harrison, David K. Jackson, Ian Johnston, Dominic Kwiatkowski, Cordelia Langford, John Sillitoe on behalf of the Wellcome Sanger Institute COVID-19 Surveillance Team (http://www.sanger.ac.uk/covid-team) |
| EPI_ISL_489391, EPI_ISL_489392, EPI_ISL_489393 | NHSGGC West of Scotland Specialist Virology Centre / MRC-University of Glasgow Centre for Virus Research | Wellcome Sanger Institute for the COVID-19 Genomics UK Consortium | Ana da Silva Filipe, Natasha Johnson, Kathy Smollett, Daniel Mair, Stephen Carmichael, Lily Tong, Jenna Nichols, Elihu Aranday-Cortes, Kirstyn Brunker, Yasmin Parr, Kyriaki Nomikou; Sarah McDonald, Marc Niebel, Patawee Asamaphan; Richard Orton, Joseph Hughes, Sreenu Vattipally, David L Robertson; Alasdair MacLean, Rory Gunson; Kathy Li, Natasha Jesudason, Rajiv Shah, James Shepherd, Antonia Ho, Alice Broos, Emma Thomson and Alex Alderton, Roberto Amato, Sonia Goncalves, Ewan Harrison, David K. Jackson, Ian Johnston, Dominic Kwiatkowski, Cordelia Langford, John Sillitoe on behalf of the Wellcome Sanger Institute COVID-19 Surveillance Team (http://www.sanger.ac.uk/covid-team) |
| EPI_ISL_489394, EPI_ISL_489395 | Department of Pathology, University of Cambridge | Wellcome Sanger Institute for the COVID-19 Genomics UK Consortium | Luke W Meredith, M. Estée Török , Myra Hosmillo, William L. Hamilton, Martin D. Curran, Theresa Feltwell, Grant Hall, Anna Yakovleva, Fahad A Khokhar, Charlotte J. Houldcroft, Laura G Caller, Aminu S. Jahun, Sarah L. Caddy, Ian Goodfellow; and Alex Alderton, Roberto Amato, Sonia Goncalves, Ewan Harrison, David K. Jackson, Ian Johnston, Dominic Kwiatkowski, Cordelia Langford, John Sillitoe on behalf of the Wellcome Sanger Institute COVID-19 Surveillance Team (http://www.sanger.ac.uk/covid-team) |
| EPI_ISL_489396 | NHSGGC West of Scotland Specialist Virology Centre / MRC-University of Glasgow Centre for Virus Research | Wellcome Sanger Institute for the COVID-19 Genomics UK Consortium | Ana da Silva Filipe, Natasha Johnson, Kathy Smollett, Daniel Mair, Stephen Carmichael, Lily Tong, Jenna Nichols, Elihu Aranday-Cortes, Kirstyn Brunker, Yasmin Parr, Kyriaki Nomikou; Sarah McDonald, Marc Niebel, Patawee Asamaphan; Richard Orton, Joseph Hughes, Sreenu Vattipally, David L Robertson; Alasdair MacLean, Rory Gunson; Kathy Li, Natasha Jesudason, Rajiv Shah, James Shepherd, Antonia Ho, Alice Broos, Emma Thomson and Alex Alderton, Roberto Amato, Sonia Goncalves, Ewan Harrison, David K. Jackson, Ian Johnston, Dominic Kwiatkowski, Cordelia Langford, John Sillitoe on behalf of the Wellcome Sanger Institute COVID-19 Surveillance Team (http://www.sanger.ac.uk/covid-team) |
| EPI_ISL_489397, EPI_ISL_489398, EPI_ISL_489399, EPI_ISL_489400 | Department of Pathology, University of Cambridge | Wellcome Sanger Institute for the COVID-19 Genomics UK Consortium | Luke W Meredith, M. Estée Török , Myra Hosmillo, William L. Hamilton, Martin D. Curran, Theresa Feltwell, Grant Hall, Anna Yakovleva, Fahad A Khokhar, Charlotte J. Houldcroft, Laura G Caller, Aminu S. Jahun, Sarah L. Caddy, Ian Goodfellow; and Alex Alderton, Roberto Amato, Sonia Goncalves, Ewan Harrison, David K. Jackson, Ian Johnston, Dominic Kwiatkowski, Cordelia Langford, John Sillitoe on behalf of the Wellcome Sanger Institute COVID-19 Surveillance Team (http://www.sanger.ac.uk/covid-team) |
| EPI_ISL_489401 | NHSGGC West of Scotland Specialist Virology Centre / MRC-University of Glasgow Centre for Virus Research | Wellcome Sanger Institute for the COVID-19 Genomics UK Consortium | Ana da Silva Filipe, Natasha Johnson, Kathy Smollett, Daniel Mair, Stephen Carmichael, Lily Tong, Jenna Nichols, Elihu Aranday-Cortes, Kirstyn Brunker, Yasmin Parr, Kyriaki Nomikou; Sarah McDonald, Marc Niebel, Patawee Asamaphan; Richard Orton, Joseph Hughes, Sreenu Vattipally, David L Robertson; Alasdair MacLean, Rory Gunson; Kathy Li, Natasha Jesudason, Rajiv Shah, James Shepherd, Antonia Ho, Alice Broos, Emma Thomson and Alex Alderton, Roberto Amato, Sonia Goncalves, Ewan Harrison, David K. Jackson, Ian Johnston, Dominic Kwiatkowski, Cordelia Langford, John Sillitoe on behalf of the Wellcome Sanger Institute COVID-19 Surveillance Team (http://www.sanger.ac.uk/covid-team) |
| EPI_ISL_489402, EPI_ISL_489403, EPI_ISL_489404, EPI_ISL_489405, EPI_ISL_489406 | Department of Pathology, University of Cambridge | Wellcome Sanger Institute for the COVID-19 Genomics UK Consortium | Luke W Meredith, M. Estée Török , Myra Hosmillo, William L. Hamilton, Martin D. Curran, Theresa Feltwell, Grant Hall, Anna Yakovleva, Fahad A Khokhar, Charlotte J. Houldcroft, Laura G Caller, Aminu S. Jahun, Sarah L. Caddy, Ian Goodfellow; and Alex Alderton, Roberto Amato, Sonia Goncalves, Ewan Harrison, David K. Jackson, Ian Johnston, Dominic Kwiatkowski, Cordelia Langford, John Sillitoe on behalf of the Wellcome Sanger Institute COVID-19 Surveillance Team (http://www.sanger.ac.uk/covid-team) |
| EPI_ISL_489407, EPI_ISL_489408 | NHSGGC West of Scotland Specialist Virology Centre / | Wellcome Sanger Institute for the COVID-19 | Ana da Silva Filipe, Natasha Johnson, Kathy Smollett, Daniel Mair, Stephen Carmichael, Lily Tong, Jenna Nichols, Elihu Aranday-Cortes, Kirstyn Brunker, Yasmin Parr, Kyriaki Nomikou; Sarah McDonald, Marc Niebel, Patawee Asamaphan; Richard Orton, Joseph |

| | | | |
|---|---|---|---|
| | MRC-University of Glasgow Centre for Virus Research | Genomics UK Consortium | Hughes, Sreenu Vattipally, David L Robertson; Alasdair MacLean, Rory Gunson; Kathy Li, Natasha Jesudason, Rajiv Shah, James Shepherd, Antonia Ho, Alice Broos, Emma Thomson and Alex Alderton, Roberto Amato, Sonia Goncalves, Ewan Harrison, David K. Jackson, Ian Johnston, Dominic Kwiatkowski, Cordelia Langford, John Sillitoe on behalf of the Wellcome Sanger Institute COVID-19 Surveillance Team (http://www.sanger.ac.uk/covid-team) |
| EPI_ISL_489409, EPI_ISL_489410, EPI_ISL_489411, EPI_ISL_489412 | Department of Pathology, University of Cambridge | Wellcome Sanger Institute for the COVID-19 Genomics UK Consortium | Luke W Meredith, M. Estée Török , Myra Hosmillo, William L. Hamilton, Martin D. Curran, Theresa Feltwell, Grant Hall, Anna Yakovleva, Fahad A Khokhar, Charlotte J. Houldcroft, Laura G Caller, Aminu S. Jahun, Sarah L. Caddy, Ian Goodfellow; and Alex Alderton, Roberto Amato, Sonia Goncalves, Ewan Harrison, David K. Jackson, Ian Johnston, Dominic Kwiatkowski, Cordelia Langford, John Sillitoe on behalf of the Wellcome Sanger Institute COVID-19 Surveillance Team (http://www.sanger.ac.uk/covid-team) |
| EPI_ISL_489413 | NHSGGC West of Scotland Specialist Virology Centre / MRC-University of Glasgow Centre for Virus Research | Wellcome Sanger Institute for the COVID-19 Genomics UK Consortium | Ana da Silva Filipe, Natasha Johnson, Kathy Smollett, Daniel Mair, Stephen Carmichael, Lily Tong, Jenna Nichols, Elihu Aranday-Cortes, Kirstyn Brunker, Yasmin Parr, Kyriaki Nomikou; Sarah McDonald, Marc Niebel, Patawee Asamaphan; Richard Orton, Joseph Hughes, Sreenu Vattipally, David L Robertson; Alasdair MacLean, Rory Gunson; Kathy Li, Natasha Jesudason, Rajiv Shah, James Shepherd, Antonia Ho, Alice Broos, Emma Thomson and Alex Alderton, Roberto Amato, Sonia Goncalves, Ewan Harrison, David K. Jackson, Ian Johnston, Dominic Kwiatkowski, Cordelia Langford, John Sillitoe on behalf of the Wellcome Sanger Institute COVID-19 Surveillance Team (http://www.sanger.ac.uk/covid-team) |
| EPI_ISL_489414, EPI_ISL_489415, EPI_ISL_489416, EPI_ISL_489417 | Department of Pathology, University of Cambridge | Wellcome Sanger Institute for the COVID-19 Genomics UK Consortium | Luke W Meredith, M. Estée Török , Myra Hosmillo, William L. Hamilton, Martin D. Curran, Theresa Feltwell, Grant Hall, Anna Yakovleva, Fahad A Khokhar, Charlotte J. Houldcroft, Laura G Caller, Aminu S. Jahun, Sarah L. Caddy, Ian Goodfellow; and Alex Alderton, Roberto Amato, Sonia Goncalves, Ewan Harrison, David K. Jackson, Ian Johnston, Dominic Kwiatkowski, Cordelia Langford, John Sillitoe on behalf of the Wellcome Sanger Institute COVID-19 Surveillance Team (http://www.sanger.ac.uk/covid-team) |
| EPI_ISL_489418 | NHSGGC West of Scotland Specialist Virology Centre / MRC-University of Glasgow Centre for Virus Research | Wellcome Sanger Institute for the COVID-19 Genomics UK Consortium | Ana da Silva Filipe, Natasha Johnson, Kathy Smollett, Daniel Mair, Stephen Carmichael, Lily Tong, Jenna Nichols, Elihu Aranday-Cortes, Kirstyn Brunker, Yasmin Parr, Kyriaki Nomikou; Sarah McDonald, Marc Niebel, Patawee Asamaphan; Richard Orton, Joseph Hughes, Sreenu Vattipally, David L Robertson; Alasdair MacLean, Rory Gunson; Kathy Li, Natasha Jesudason, Rajiv Shah, James Shepherd, Antonia Ho, Alice Broos, Emma Thomson and Alex Alderton, Roberto Amato, Sonia Goncalves, Ewan Harrison, David K. Jackson, Ian Johnston, Dominic Kwiatkowski, Cordelia Langford, John Sillitoe on behalf of the Wellcome Sanger Institute COVID-19 Surveillance Team (http://www.sanger.ac.uk/covid-team) |
| EPI_ISL_489419, EPI_ISL_489420, EPI_ISL_489421, EPI_ISL_489422, EPI_ISL_489423, EPI_ISL_489424, EPI_ISL_489425, EPI_ISL_489426, EPI_ISL_489427, EPI_ISL_489428 | Department of Pathology, University of Cambridge | Wellcome Sanger Institute for the COVID-19 Genomics UK Consortium | Luke W Meredith, M. Estée Török , Myra Hosmillo, William L. Hamilton, Martin D. Curran, Theresa Feltwell, Grant Hall, Anna Yakovleva, Fahad A Khokhar, Charlotte J. Houldcroft, Laura G Caller, Aminu S. Jahun, Sarah L. Caddy, Ian Goodfellow; and Alex Alderton, Roberto Amato, Sonia Goncalves, Ewan Harrison, David K. Jackson, Ian Johnston, Dominic Kwiatkowski, Cordelia Langford, John Sillitoe on behalf of the Wellcome Sanger Institute COVID-19 Surveillance Team (http://www.sanger.ac.uk/covid-team) |
| EPI_ISL_489429 | NHSGGC West of Scotland Specialist Virology Centre / MRC-University of Glasgow Centre for Virus Research | Wellcome Sanger Institute for the COVID-19 Genomics UK Consortium | Ana da Silva Filipe, Natasha Johnson, Kathy Smollett, Daniel Mair, Stephen Carmichael, Lily Tong, Jenna Nichols, Elihu Aranday-Cortes, Kirstyn Brunker, Yasmin Parr, Kyriaki Nomikou; Sarah McDonald, Marc Niebel, Patawee Asamaphan; Richard Orton, Joseph Hughes, Sreenu Vattipally, David L Robertson; Alasdair MacLean, Rory Gunson; Kathy Li, Natasha Jesudason, Rajiv Shah, James Shepherd, Antonia Ho, Alice Broos, Emma Thomson and Alex Alderton, Roberto Amato, Sonia Goncalves, Ewan Harrison, David K. Jackson, Ian Johnston, Dominic Kwiatkowski, Cordelia Langford, John Sillitoe on behalf of the Wellcome Sanger Institute COVID-19 Surveillance Team (http://www.sanger.ac.uk/covid-team) |
| EPI_ISL_489430, EPI_ISL_489431, EPI_ISL_489432, EPI_ISL_489433, EPI_ISL_489434, EPI_ISL_489435, EPI_ISL_489436, EPI_ISL_489437, EPI_ISL_489438, EPI_ISL_489439, EPI_ISL_489440, EPI_ISL_489441 | | | |
| see above | Department of Pathology, University of Cambridge | Wellcome Sanger Institute for the COVID-19 Genomics UK Consortium | Luke W Meredith, M. Estée Török , Myra Hosmillo, William L. Hamilton, Martin D. Curran, Theresa Feltwell, Grant Hall, Anna Yakovleva, Fahad A Khokhar, Charlotte J. Houldcroft, Laura G Caller, Aminu S. Jahun, Sarah L. Caddy, Ian Goodfellow; and Alex Alderton, Roberto Amato, Sonia Goncalves, Ewan Harrison, David K. Jackson, Ian Johnston, Dominic Kwiatkowski, Cordelia Langford, John Sillitoe on behalf of the Wellcome Sanger Institute COVID-19 Surveillance Team (http://www.sanger.ac.uk/covid-team) |
| EPI_ISL_489442 | NHSGGC West of Scotland Specialist Virology Centre / MRC-University of Glasgow Centre for Virus Research | Wellcome Sanger Institute for the COVID-19 Genomics UK Consortium | Ana da Silva Filipe, Natasha Johnson, Kathy Smollett, Daniel Mair, Stephen Carmichael, Lily Tong, Jenna Nichols, Elihu Aranday-Cortes, Kirstyn Brunker, Yasmin Parr, Kyriaki Nomikou; Sarah McDonald, Marc Niebel, Patawee Asamaphan; Richard Orton, Joseph Hughes, Sreenu Vattipally, David L Robertson; Alasdair MacLean, Rory Gunson; Kathy Li, Natasha Jesudason, Rajiv Shah, James Shepherd, Antonia Ho, Alice Broos, Emma Thomson and Alex Alderton, Roberto Amato, Sonia Goncalves, Ewan Harrison, David K. Jackson, Ian Johnston, Dominic Kwiatkowski, Cordelia Langford, John Sillitoe on behalf of the Wellcome Sanger Institute COVID-19 Surveillance Team (http://www.sanger.ac.uk/covid-team) |
| EPI_ISL_489443, EPI_ISL_489444, EPI_ISL_489445, EPI_ISL_489446, EPI_ISL_489447, EPI_ISL_489448 | Department of Pathology, University of Cambridge | Wellcome Sanger Institute for the COVID-19 Genomics UK Consortium | Luke W Meredith, M. Estée Török , Myra Hosmillo, William L. Hamilton, Martin D. Curran, Theresa Feltwell, Grant Hall, Anna Yakovleva, Fahad A Khokhar, Charlotte J. Houldcroft, Laura G Caller, Aminu S. Jahun, Sarah L. Caddy, Ian Goodfellow; and Alex Alderton, Roberto Amato, Sonia Goncalves, Ewan Harrison, David K. Jackson, Ian Johnston, Dominic Kwiatkowski, Cordelia Langford, John Sillitoe on behalf of the Wellcome Sanger Institute COVID-19 Surveillance Team (http://www.sanger.ac.uk/covid-team) |
| EPI_ISL_489449 | NHSGGC West of Scotland Specialist Virology Centre / MRC-University of Glasgow Centre for Virus Research | Wellcome Sanger Institute for the COVID-19 Genomics UK Consortium | Ana da Silva Filipe, Natasha Johnson, Kathy Smollett, Daniel Mair, Stephen Carmichael, Lily Tong, Jenna Nichols, Elihu Aranday-Cortes, Kirstyn Brunker, Yasmin Parr, Kyriaki Nomikou; Sarah McDonald, Marc Niebel, Patawee Asamaphan; Richard Orton, Joseph Hughes, Sreenu Vattipally, David L Robertson; Alasdair MacLean, Rory Gunson; Kathy Li, Natasha Jesudason, Rajiv Shah, James Shepherd, Antonia Ho, Alice Broos, Emma Thomson and Alex Alderton, Roberto Amato, Sonia Goncalves, Ewan Harrison, David K. Jackson, Ian Johnston, Dominic Kwiatkowski, Cordelia Langford, John Sillitoe on behalf of the Wellcome Sanger Institute COVID-19 Surveillance Team (http://www.sanger.ac.uk/covid-team) |
| EPI_ISL_489450, EPI_ISL_489451, EPI_ISL_489452, EPI_ISL_489453, EPI_ISL_489454 | Department of Pathology, University of Cambridge | Wellcome Sanger Institute for the COVID-19 Genomics UK Consortium | Luke W Meredith, M. Estée Török , Myra Hosmillo, William L. Hamilton, Martin D. Curran, Theresa Feltwell, Grant Hall, Anna Yakovleva, Fahad A Khokhar, Charlotte J. Houldcroft, Laura G Caller, Aminu S. Jahun, Sarah L. Caddy, Ian Goodfellow; and Alex Alderton, Roberto Amato, Sonia Goncalves, Ewan Harrison, David K. Jackson, Ian Johnston, Dominic Kwiatkowski, Cordelia Langford, John Sillitoe on behalf of the Wellcome Sanger Institute COVID-19 Surveillance Team (http://www.sanger.ac.uk/covid-team) |
| EPI_ISL_489455 | NHSGGC West of Scotland Specialist Virology Centre / MRC-University of Glasgow Centre for Virus Research | Wellcome Sanger Institute for the COVID-19 Genomics UK Consortium | Ana da Silva Filipe, Natasha Johnson, Kathy Smollett, Daniel Mair, Stephen Carmichael, Lily Tong, Jenna Nichols, Elihu Aranday-Cortes, Kirstyn Brunker, Yasmin Parr, Kyriaki Nomikou; Sarah McDonald, Marc Niebel, Patawee Asamaphan; Richard Orton, Joseph Hughes, Sreenu Vattipally, David L Robertson; Alasdair MacLean, Rory Gunson; Kathy Li, Natasha Jesudason, Rajiv Shah, James Shepherd, Antonia Ho, Alice Broos, Emma Thomson and Alex Alderton, Roberto Amato, Sonia Goncalves, Ewan Harrison, David K. Jackson, Ian Johnston, Dominic Kwiatkowski, Cordelia Langford, John Sillitoe on behalf of the Wellcome Sanger Institute COVID-19 Surveillance Team (http://www.sanger.ac.uk/covid-team) |
| EPI_ISL_489456, EPI_ISL_489457, EPI_ISL_489458, EPI_ISL_489459, EPI_ISL_489460, EPI_ISL_489461, EPI_ISL_489462, EPI_ISL_489463, EPI_ISL_489464, EPI_ISL_489465, EPI_ISL_489466, EPI_ISL_489467, EPI_ISL_489468, EPI_ISL_489469 | | | |
| see above | Department of Pathology, University of Cambridge | Wellcome Sanger Institute for the COVID-19 Genomics UK Consortium | Luke W Meredith, M. Estée Török , Myra Hosmillo, William L. Hamilton, Martin D. Curran, Theresa Feltwell, Grant Hall, Anna Yakovleva, Fahad A Khokhar, Charlotte J. Houldcroft, Laura G Caller, Aminu S. Jahun, Sarah L. Caddy, Ian Goodfellow; and Alex Alderton, Roberto Amato, Sonia Goncalves, Ewan Harrison, David K. Jackson, Ian Johnston, Dominic Kwiatkowski, Cordelia Langford, John Sillitoe on behalf of the Wellcome Sanger Institute COVID-19 Surveillance Team (http://www.sanger.ac.uk/covid-team) |
| EPI_ISL_489470 | NHSGGC West of Scotland Specialist Virology Centre / MRC-University of Glasgow Centre for Virus Research | Wellcome Sanger Institute for the COVID-19 Genomics UK Consortium | Ana da Silva Filipe, Natasha Johnson, Kathy Smollett, Daniel Mair, Stephen Carmichael, Lily Tong, Jenna Nichols, Elihu Aranday-Cortes, Kirstyn Brunker, Yasmin Parr, Kyriaki Nomikou; Sarah McDonald, Marc Niebel, Patawee Asamaphan; Richard Orton, Joseph Hughes, Sreenu Vattipally, David L Robertson; Alasdair MacLean, Rory Gunson; Kathy Li, Natasha Jesudason, Rajiv Shah, James Shepherd, Antonia Ho, Alice Broos, Emma Thomson and Alex Alderton, Roberto Amato, Sonia Goncalves, Ewan Harrison, David K. Jackson, Ian Johnston, Dominic Kwiatkowski, Cordelia Langford, John Sillitoe on behalf of the Wellcome Sanger Institute COVID-19 Surveillance Team (http://www.sanger.ac.uk/covid-team) |
| EPI_ISL_489471, EPI_ISL_489472, EPI_ISL_489473, EPI_ISL_489474, EPI_ISL_489475, EPI_ISL_489476, EPI_ISL_489477, EPI_ISL_489478, EPI_ISL_489479 | Department of Pathology, University of Cambridge | Wellcome Sanger Institute for the COVID-19 Genomics UK Consortium | Luke W Meredith, M. Estée Török , Myra Hosmillo, William L. Hamilton, Martin D. Curran, Theresa Feltwell, Grant Hall, Anna Yakovleva, Fahad A Khokhar, Charlotte J. Houldcroft, Laura G Caller, Aminu S. Jahun, Sarah L. Caddy, Ian Goodfellow; and Alex Alderton, Roberto Amato, Sonia Goncalves, Ewan Harrison, David K. Jackson, Ian Johnston, Dominic Kwiatkowski, Cordelia Langford, John Sillitoe on behalf of the Wellcome Sanger Institute COVID-19 Surveillance Team (http://www.sanger.ac.uk/covid-team) |
| EPI_ISL_489480 | NHSGGC West of Scotland Specialist Virology Centre / MRC-University of Glasgow Centre for Virus Research | Wellcome Sanger Institute for the COVID-19 Genomics UK Consortium | Ana da Silva Filipe, Natasha Johnson, Kathy Smollett, Daniel Mair, Stephen Carmichael, Lily Tong, Jenna Nichols, Elihu Aranday-Cortes, Kirstyn Brunker, Yasmin Parr, Kyriaki Nomikou; Sarah McDonald, Marc Niebel, Patawee Asamaphan; Richard Orton, Joseph Hughes, Sreenu Vattipally, David L Robertson; Alasdair MacLean, Rory Gunson; Kathy Li, Natasha Jesudason, Rajiv Shah, James Shepherd, Antonia Ho, Alice Broos, Emma Thomson and Alex Alderton, Roberto Amato, Sonia Goncalves, Ewan Harrison, David K. Jackson, Ian Johnston, Dominic Kwiatkowski, Cordelia Langford, John Sillitoe on behalf of the Wellcome Sanger Institute COVID-19 Surveillance Team (http://www.sanger.ac.uk/covid-team) |
| EPI_ISL_489481, EPI_ISL_489482, EPI_ISL_489483, EPI_ISL_489484, EPI_ISL_489485, EPI_ISL_489486, EPI_ISL_489487, EPI_ISL_489488 | Department of Pathology, University of Cambridge | Wellcome Sanger Institute for the COVID-19 Genomics UK Consortium | Luke W Meredith, M. Estée Török , Myra Hosmillo, William L. Hamilton, Martin D. Curran, Theresa Feltwell, Grant Hall, Anna Yakovleva, Fahad A Khokhar, Charlotte J. Houldcroft, Laura G Caller, Aminu S. Jahun, Sarah L. Caddy, Ian Goodfellow; and Alex Alderton, Roberto Amato, Sonia Goncalves, Ewan Harrison, David K. Jackson, Ian Johnston, Dominic Kwiatkowski, Cordelia Langford, John Sillitoe on behalf of the Wellcome Sanger Institute COVID-19 Surveillance Team (http://www.sanger.ac.uk/covid-team) |
| EPI_ISL_489489 | NHSGGC West of Scotland Specialist Virology Centre / MRC-University of Glasgow Centre for Virus Research | Wellcome Sanger Institute for the COVID-19 Genomics UK Consortium | Ana da Silva Filipe, Natasha Johnson, Kathy Smollett, Daniel Mair, Stephen Carmichael, Lily Tong, Jenna Nichols, Elihu Aranday-Cortes, Kirstyn Brunker, Yasmin Parr, Kyriaki Nomikou; Sarah McDonald, Marc Niebel, Patawee Asamaphan; Richard Orton, Joseph Hughes, Sreenu Vattipally, David L Robertson; Alasdair MacLean, Rory Gunson; Kathy Li, Natasha Jesudason, Rajiv Shah, James Shepherd, Antonia Ho, Alice Broos, Emma Thomson and Alex Alderton, Roberto Amato, Sonia Goncalves, Ewan Harrison, David K. Jackson, Ian Johnston, Dominic Kwiatkowski, Cordelia Langford, John Sillitoe on behalf of the Wellcome Sanger Institute COVID-19 Surveillance Team (http://www.sanger.ac.uk/covid-team) |
| EPI_ISL_489490, EPI_ISL_489491, EPI_ISL_489492, EPI_ISL_489493, EPI_ISL_489494, EPI_ISL_489495, EPI_ISL_489496, EPI_ISL_489497, EPI_ISL_489498, EPI_ISL_489499, EPI_ISL_489500, EPI_ISL_489501, EPI_ISL_489502, EPI_ISL_489503, EPI_ISL_489504, EPI_ISL_489505, EPI_ISL_489506, EPI_ISL_489507, EPI_ISL_489508, EPI_ISL_489509, EPI_ISL_489510, EPI_ISL_489511, EPI_ISL_489512, EPI_ISL_489513, EPI_ISL_489514, EPI_ISL_489515, EPI_ISL_489516, EPI_ISL_489517, EPI_ISL_489518, EPI_ISL_489519, EPI_ISL_489520, EPI_ISL_489521, EPI_ISL_489522 | | | |
| see above | Department of Pathology, University of Cambridge | Wellcome Sanger Institute for the COVID-19 Genomics UK Consortium | Luke W Meredith, M. Estée Török , Myra Hosmillo, William L. Hamilton, Martin D. Curran, Theresa Feltwell, Grant Hall, Anna Yakovleva, Fahad A Khokhar, Charlotte J. Houldcroft, Laura G Caller, Aminu S. Jahun, Sarah L. Caddy, Ian Goodfellow; and Alex Alderton, Roberto Amato, Sonia Goncalves, Ewan Harrison, David K. Jackson, Ian Johnston, Dominic Kwiatkowski, Cordelia Langford, John Sillitoe on behalf of the Wellcome Sanger Institute COVID-19 Surveillance Team (http://www.sanger.ac.uk/covid-team) |
| EPI_ISL_489523 | NHSGGC West of Scotland Specialist Virology Centre / MRC-University of Glasgow Centre for Virus Research | Wellcome Sanger Institute for the COVID-19 Genomics UK Consortium | Ana da Silva Filipe, Natasha Johnson, Kathy Smollett, Daniel Mair, Stephen Carmichael, Lily Tong, Jenna Nichols, Elihu Aranday-Cortes, Kirstyn Brunker, Yasmin Parr, Kyriaki Nomikou; Sarah McDonald, Marc Niebel, Patawee Asamaphan; Richard Orton, Joseph Hughes, Sreenu Vattipally, David L Robertson; Alasdair MacLean, Rory Gunson; Kathy Li, Natasha Jesudason, Rajiv Shah, James Shepherd, Antonia Ho, Alice Broos, Emma Thomson and Alex Alderton, Roberto Amato, Sonia Goncalves, Ewan Harrison, David K. Jackson, Ian Johnston, Dominic Kwiatkowski, Cordelia Langford, John Sillitoe on behalf of the Wellcome Sanger Institute COVID-19 Surveillance Team (http://www.sanger.ac.uk/covid-team) |
| EPI_ISL_489524, EPI_ISL_489525, EPI_ISL_489526, EPI_ISL_489527, EPI_ISL_489528, EPI_ISL_489529 | Department of Pathology, University of Cambridge | Wellcome Sanger Institute for the COVID-19 Genomics UK Consortium | Luke W Meredith, M. Estée Török , Myra Hosmillo, William L. Hamilton, Martin D. Curran, Theresa Feltwell, Grant Hall, Anna Yakovleva, Fahad A Khokhar, Charlotte J. Houldcroft, Laura G Caller, Aminu S. Jahun, Sarah L. Caddy, Ian Goodfellow; and Alex Alderton, Roberto Amato, Sonia Goncalves, Ewan Harrison, David K. Jackson, Ian Johnston, Dominic Kwiatkowski, Cordelia Langford, John Sillitoe on behalf of the Wellcome Sanger Institute COVID-19 Surveillance Team (http://www.sanger.ac.uk/covid-team) |
| EPI_ISL_489530 | NHSGGC West of Scotland Specialist Virology Centre / MRC-University of Glasgow Centre for Virus Research | Wellcome Sanger Institute for the COVID-19 Genomics UK Consortium | Ana da Silva Filipe, Natasha Johnson, Kathy Smollett, Daniel Mair, Stephen Carmichael, Lily Tong, Jenna Nichols, Elihu Aranday-Cortes, Kirstyn Brunker, Yasmin Parr, Kyriaki Nomikou; Sarah McDonald, Marc Niebel, Patawee Asamaphan; Richard Orton, Joseph Hughes, Sreenu Vattipally, David L Robertson; Alasdair MacLean, Rory Gunson; Kathy Li, Natasha Jesudason, Rajiv Shah, James Shepherd, Antonia Ho, Alice Broos, Emma Thomson and Alex Alderton, Roberto Amato, Sonia Goncalves, Ewan Harrison, David K. Jackson, Ian Johnston, Dominic Kwiatkowski, Cordelia Langford, John Sillitoe on behalf of the Wellcome Sanger Institute COVID-19 Surveillance Team (http://www.sanger.ac.uk/covid-team) |
| EPI_ISL_489531, EPI_ISL_489532, EPI_ISL_489533, EPI_ISL_489534, EPI_ISL_489535 | Department of Pathology, University of Cambridge | Wellcome Sanger Institute for the COVID-19 Genomics UK Consortium | Luke W Meredith, M. Estée Török , Myra Hosmillo, William L. Hamilton, Martin D. Curran, Theresa Feltwell, Grant Hall, Anna Yakovleva, Fahad A Khokhar, Charlotte J. Houldcroft, Laura G Caller, Aminu S. Jahun, Sarah L. Caddy, Ian Goodfellow; and Alex Alderton, Roberto Amato, Sonia Goncalves, Ewan Harrison, David K. Jackson, Ian Johnston, Dominic Kwiatkowski, Cordelia Langford, John Sillitoe on behalf of the Wellcome Sanger Institute COVID-19 Surveillance Team (http://www.sanger.ac.uk/covid-team) |
| EPI_ISL_489536, EPI_ISL_489537 | NHSGGC West of Scotland Specialist Virology Centre / MRC-University of Glasgow Centre for Virus Research | Wellcome Sanger Institute for the COVID-19 Genomics UK Consortium | Ana da Silva Filipe, Natasha Johnson, Kathy Smollett, Daniel Mair, Stephen Carmichael, Lily Tong, Jenna Nichols, Elihu Aranday-Cortes, Kirstyn Brunker, Yasmin Parr, Kyriaki Nomikou; Sarah McDonald, Marc Niebel, Patawee Asamaphan; Richard Orton, Joseph Hughes, Sreenu Vattipally, David L Robertson; Alasdair MacLean, Rory Gunson; Kathy Li, Natasha Jesudason, Rajiv Shah, James Shepherd, Antonia Ho, Alice Broos, Emma Thomson and Alex Alderton, Roberto Amato, Sonia Goncalves, Ewan Harrison, David K. Jackson, Ian Johnston, Dominic Kwiatkowski, Cordelia Langford, John Sillitoe on behalf of the Wellcome Sanger Institute COVID-19 Surveillance Team (http://www.sanger.ac.uk/covid-team) |
| EPI_ISL_489538, EPI_ISL_489539, EPI_ISL_489540, EPI_ISL_489541, EPI_ISL_489542 | Department of Pathology, University of Cambridge | Wellcome Sanger Institute for the COVID-19 Genomics UK Consortium | Luke W Meredith, M. Estée Török , Myra Hosmillo, William L. Hamilton, Martin D. Curran, Theresa Feltwell, Grant Hall, Anna Yakovleva, Fahad A Khokhar, Charlotte J. Houldcroft, Laura G Caller, Aminu S. Jahun, Sarah L. Caddy, Ian Goodfellow; and Alex Alderton, Roberto Amato, Sonia Goncalves, Ewan Harrison, David K. Jackson, Ian Johnston, Dominic Kwiatkowski, Cordelia Langford, John Sillitoe on behalf of the Wellcome Sanger Institute COVID-19 Surveillance Team (http://www.sanger.ac.uk/covid-team) |
| EPI_ISL_489543 | NHSGGC West of Scotland Specialist Virology Centre / MRC-University of Glasgow Centre for Virus Research | Wellcome Sanger Institute for the COVID-19 Genomics UK Consortium | Ana da Silva Filipe, Natasha Johnson, Kathy Smollett, Daniel Mair, Stephen Carmichael, Lily Tong, Jenna Nichols, Elihu Aranday-Cortes, Kirstyn Brunker, Yasmin Parr, Kyriaki Nomikou; Sarah McDonald, Marc Niebel, Patawee Asamaphan; Richard Orton, Joseph Hughes, Sreenu Vattipally, David L Robertson; Alasdair MacLean, Rory Gunson; Kathy Li, Natasha Jesudason, Rajiv Shah, James Shepherd, Antonia Ho, Alice Broos, Emma Thomson and Alex Alderton, Roberto Amato, Sonia Goncalves, Ewan Harrison, David K. Jackson, Ian Johnston, Dominic Kwiatkowski, Cordelia Langford, John Sillitoe on behalf of the Wellcome Sanger Institute COVID-19 Surveillance Team (http://www.sanger.ac.uk/covid-team) |
| EPI_ISL_489544, EPI_ISL_489545, EPI_ISL_489546 | Department of Pathology, University of Cambridge | Wellcome Sanger Institute for the COVID-19 Genomics UK Consortium | Luke W Meredith, M. Estée Török , Myra Hosmillo, William L. Hamilton, Martin D. Curran, Theresa Feltwell, Grant Hall, Anna Yakovleva, Fahad A Khokhar, Charlotte J. Houldcroft, Laura G Caller, Aminu S. Jahun, Sarah L. Caddy, Ian Goodfellow; and Alex Alderton, Roberto Amato, Sonia Goncalves, Ewan Harrison, David K. Jackson, Ian Johnston, Dominic Kwiatkowski, Cordelia Langford, John Sillitoe on behalf of the Wellcome Sanger Institute COVID-19 Surveillance Team (http://www.sanger.ac.uk/covid-team) |
| EPI_ISL_489547 | NHSGGC West of Scotland Specialist Virology Centre / MRC-University of Glasgow Centre for Virus Research | Wellcome Sanger Institute for the COVID-19 Genomics UK Consortium | Ana da Silva Filipe, Natasha Johnson, Kathy Smollett, Daniel Mair, Stephen Carmichael, Lily Tong, Jenna Nichols, Elihu Aranday-Cortes, Kirstyn Brunker, Yasmin Parr, Kyriaki Nomikou; Sarah McDonald, Marc Niebel, Patawee Asamaphan; Richard Orton, Joseph Hughes, Sreenu Vattipally, David L Robertson; Alasdair MacLean, Rory Gunson; Kathy Li, Natasha Jesudason, Rajiv Shah, James Shepherd, Antonia Ho, Alice Broos, Emma Thomson and Alex Alderton, Roberto Amato, Sonia Goncalves, Ewan Harrison, David K. Jackson, Ian Johnston, Dominic Kwiatkowski, Cordelia Langford, John Sillitoe on behalf of the Wellcome Sanger Institute COVID-19 Surveillance Team (http://www.sanger.ac.uk/covid-team) |
| EPI_ISL_489548, EPI_ISL_489549, EPI_ISL_489550, EPI_ISL_489551, EPI_ISL_489552, EPI_ISL_489553, EPI_ISL_489554, EPI_ISL_489555, EPI_ISL_489556, EPI_ISL_489557, EPI_ISL_489558, EPI_ISL_489559, EPI_ISL_489560, EPI_ISL_489561, EPI_ISL_489562, EPI_ISL_489563, EPI_ISL_489564, EPI_ISL_489565, EPI_ISL_489566 | | | |
| see above | Department of Pathology, University of Cambridge | Wellcome Sanger Institute for the COVID-19 Genomics UK Consortium | Luke W Meredith, M. Estée Török , Myra Hosmillo, William L. Hamilton, Martin D. Curran, Theresa Feltwell, Grant Hall, Anna Yakovleva, Fahad A Khokhar, Charlotte J. Houldcroft, Laura G Caller, Aminu S. Jahun, Sarah L. Caddy, Ian Goodfellow; and Alex Alderton, Roberto Amato, Sonia Goncalves, Ewan Harrison, David K. Jackson, Ian Johnston, Dominic Kwiatkowski, Cordelia Langford, John Sillitoe on behalf of the Wellcome Sanger Institute COVID-19 Surveillance Team (http://www.sanger.ac.uk/covid-team) |
| EPI_ISL_489567 | NHSGGC West of Scotland Specialist Virology Centre / MRC-University of Glasgow Centre for Virus Research | Wellcome Sanger Institute for the COVID-19 Genomics UK Consortium | Ana da Silva Filipe, Natasha Johnson, Kathy Smollett, Daniel Mair, Stephen Carmichael, Lily Tong, Jenna Nichols, Elihu Aranday-Cortes, Kirstyn Brunker, Yasmin Parr, Kyriaki Nomikou; Sarah McDonald, Marc Niebel, Patawee Asamaphan; Richard Orton, Joseph Hughes, Sreenu Vattipally, David L Robertson; Alasdair MacLean, Rory Gunson; Kathy Li, Natasha Jesudason, Rajiv Shah, James Shepherd, Antonia Ho, Alice Broos, Emma Thomson and Alex Alderton, Roberto Amato, Sonia Goncalves, Ewan Harrison, David K. Jackson, Ian Johnston, Dominic Kwiatkowski, Cordelia Langford, John Sillitoe on behalf of the Wellcome Sanger Institute COVID-19 Surveillance Team (http://www.sanger.ac.uk/covid-team) |

| | | | |
|---|---|---|---|
| EPI_ISL_489568, EPI_ISL_489569, EPI_ISL_489570, EPI_ISL_489571, EPI_ISL_489572, EPI_ISL_489573, EPI_ISL_489574, EPI_ISL_489575, EPI_ISL_489576, EPI_ISL_489577 | Department of Pathology, University of Cambridge | Wellcome Sanger Institute for the COVID-19 Genomics UK Consortium | Luke W Meredith, M. Estée Török , Myra Hosmillo, William L. Hamilton, Martin D. Curran, Theresa Feltwell, Grant Hall, Anna Yakovleva, Fahad A Khokhar, Charlotte J. Houldcroft, Laura G Caller, Aminu S. Jahun, Sarah L. Caddy, Ian Goodfellow; and Alex Alderton, Roberto Amato, Sonia Goncalves, Ewan Harrison, David K. Jackson, Ian Johnston, Dominic Kwiatkowski, Cordelia Langford, John Sillitoe on behalf of the Wellcome Sanger Institute COVID-19 Surveillance Team (http://www.sanger.ac.uk/covid-team) |
| EPI_ISL_489578, EPI_ISL_489579, EPI_ISL_489580, EPI_ISL_489582, EPI_ISL_489583, EPI_ISL_489584, EPI_ISL_489585, EPI_ISL_489586, EPI_ISL_489587, EPI_ISL_489588, EPI_ISL_489589, EPI_ISL_489590, EPI_ISL_489591, EPI_ISL_489592, EPI_ISL_489593, EPI_ISL_489594, EPI_ISL_489595, EPI_ISL_489596, EPI_ISL_489597, EPI_ISL_489598, EPI_ISL_489599, EPI_ISL_489600, EPI_ISL_489601, EPI_ISL_489602, EPI_ISL_489603, EPI_ISL_489604, EPI_ISL_489605, EPI_ISL_489606, EPI_ISL_489607, EPI_ISL_489608, EPI_ISL_489609, EPI_ISL_489610, EPI_ISL_489611, EPI_ISL_489612, EPI_ISL_489613, EPI_ISL_489614, EPI_ISL_489615, EPI_ISL_489616, EPI_ISL_489617, EPI_ISL_489618, EPI_ISL_489619, EPI_ISL_489620, EPI_ISL_489621, EPI_ISL_489622, EPI_ISL_489623, EPI_ISL_489624, EPI_ISL_489625, EPI_ISL_489626, EPI_ISL_489627, EPI_ISL_489628, EPI_ISL_489629, EPI_ISL_489630, EPI_ISL_489631, EPI_ISL_489632, EPI_ISL_489633, EPI_ISL_489634, EPI_ISL_489635, EPI_ISL_489636, EPI_ISL_489637, EPI_ISL_489638, EPI_ISL_489639, EPI_ISL_489640, EPI_ISL_489641, EPI_ISL_489642, EPI_ISL_489643, EPI_ISL_489644, EPI_ISL_489645, EPI_ISL_489646, EPI_ISL_489647, EPI_ISL_489648, EPI_ISL_489649, EPI_ISL_489650, EPI_ISL_489651, EPI_ISL_489652, EPI_ISL_489653, EPI_ISL_489654, EPI_ISL_489655, EPI_ISL_489656, EPI_ISL_489657, EPI_ISL_489658, EPI_ISL_489659, EPI_ISL_489660, EPI_ISL_489661, EPI_ISL_489662, EPI_ISL_489663, EPI_ISL_489664, EPI_ISL_489665, EPI_ISL_489666, EPI_ISL_489667, EPI_ISL_489668, EPI_ISL_489669, EPI_ISL_489670, EPI_ISL_489671, EPI_ISL_489672, EPI_ISL_489673, EPI_ISL_489674, EPI_ISL_489675, EPI_ISL_489676, EPI_ISL_489677, EPI_ISL_489678, EPI_ISL_489679, EPI_ISL_489680, EPI_ISL_489681, EPI_ISL_489682, EPI_ISL_489683, EPI_ISL_489684, EPI_ISL_489685, EPI_ISL_489686, EPI_ISL_489687, EPI_ISL_489688, EPI_ISL_489689, EPI_ISL_489690, EPI_ISL_489691, EPI_ISL_489692, EPI_ISL_489693, EPI_ISL_489694, EPI_ISL_489695, EPI_ISL_489696, EPI_ISL_489697, EPI_ISL_489698, EPI_ISL_489699, EPI_ISL_489700, EPI_ISL_489701, EPI_ISL_489702, EPI_ISL_489703, EPI_ISL_489704, EPI_ISL_489705, EPI_ISL_489706 | | | |
| see above | NHSGGC West of Scotland Specialist Virology Centre / MRC-University of Glasgow Centre for Virus Research | Wellcome Sanger Institute for the COVID-19 Genomics UK Consortium | Ana da Silva Filipe, Natasha Johnson, Kathy Smollett, Daniel Mair, Stephen Carmichael, Lily Tong, Jenna Nichols, Elihu Aranday-Cortes, Kirstyn Brunker, Yasmin Parr, Kyriaki Nomikou; Sarah McDonald, Marc Niebel, Patawee Asamaphan; Richard Orton, Joseph Hughes, Sreenu Vattipally, David L Robertson; Alasdair MacLean, Rory Gunson; Kathy Li, Natasha Jesudason, Rajiv Shah, James Shepherd, Antonia Ho, Alice Broos, Emma Thomson and Alex Alderton, Roberto Amato, Sonia Goncalves, Ewan Harrison, David K. Jackson, Ian Johnston, Dominic Kwiatkowski, Cordelia Langford, John Sillitoe on behalf of the Wellcome Sanger Institute COVID-19 Surveillance Team (http://www.sanger.ac.uk/covid-team) |
| EPI_ISL_489708 | The National Institute of Public Health | The National Institute of Public Health and State Veterinary Institute Prague | Nagy,A;Jirincova,H;Novakova,L;Trnka,D;Vecerova,J |
| EPI_ISL_489709 | The National Institute of Public Health | The National Institute of Public Health and State Veterinary Institute Prague | Nagy,A;Jirincova,H;Novakova,L;Trnka,D;Vecerova,J |
| EPI_ISL_489710, EPI_ISL_489711, EPI_ISL_489712, EPI_ISL_489713, EPI_ISL_489714, EPI_ISL_489715, EPI_ISL_489716, EPI_ISL_489717, EPI_ISL_489718, EPI_ISL_489719, EPI_ISL_489720, EPI_ISL_489721, EPI_ISL_489722, EPI_ISL_489723, EPI_ISL_489724, EPI_ISL_489725, EPI_ISL_489726, EPI_ISL_489727, EPI_ISL_489728, EPI_ISL_489729, EPI_ISL_489730, EPI_ISL_489731, EPI_ISL_489732, EPI_ISL_489733, EPI_ISL_489734, EPI_ISL_489735, EPI_ISL_489736, EPI_ISL_489737, EPI_ISL_489738, EPI_ISL_489739, EPI_ISL_489740, EPI_ISL_489741, EPI_ISL_489742, EPI_ISL_489743, EPI_ISL_489744, EPI_ISL_489745, EPI_ISL_489746, EPI_ISL_489747, EPI_ISL_489748, EPI_ISL_489749, EPI_ISL_489750, EPI_ISL_489751, EPI_ISL_489752, EPI_ISL_489753, EPI_ISL_489754, EPI_ISL_489755, EPI_ISL_489756, EPI_ISL_489757, EPI_ISL_489758, EPI_ISL_489759, EPI_ISL_489760, EPI_ISL_489761, EPI_ISL_489762, EPI_ISL_489763, EPI_ISL_489764, EPI_ISL_489765, EPI_ISL_489766, EPI_ISL_489767, EPI_ISL_489768, EPI_ISL_489769, EPI_ISL_489770, EPI_ISL_489771, EPI_ISL_489772, EPI_ISL_489773, EPI_ISL_489774, EPI_ISL_489775, EPI_ISL_489776, EPI_ISL_489777, EPI_ISL_489778, EPI_ISL_489779, EPI_ISL_489780, EPI_ISL_489781, EPI_ISL_489782, EPI_ISL_489783, EPI_ISL_489784, EPI_ISL_489785, EPI_ISL_489786, EPI_ISL_489787, EPI_ISL_489788, EPI_ISL_489789, EPI_ISL_489790, EPI_ISL_489791, EPI_ISL_489792, EPI_ISL_489793, EPI_ISL_489794, EPI_ISL_489795, EPI_ISL_489796, EPI_ISL_489797, EPI_ISL_489798, EPI_ISL_489799, EPI_ISL_489800, EPI_ISL_489801, EPI_ISL_489802, EPI_ISL_489803, EPI_ISL_489804, EPI_ISL_489805, EPI_ISL_489806, EPI_ISL_489807, EPI_ISL_489808, EPI_ISL_489809, EPI_ISL_489810, EPI_ISL_489811, EPI_ISL_489812, EPI_ISL_489813, EPI_ISL_489814, EPI_ISL_489815, EPI_ISL_489816, EPI_ISL_489817, EPI_ISL_489818, EPI_ISL_489819, EPI_ISL_489820, EPI_ISL_489821, EPI_ISL_489822, EPI_ISL_489823, EPI_ISL_489824, EPI_ISL_489825, EPI_ISL_489826, EPI_ISL_489827, EPI_ISL_489828, EPI_ISL_489829, EPI_ISL_489830, EPI_ISL_489831, EPI_ISL_489832 | | | |
| see above | Florida Bureau of Public Health Laboratories | Florida Bureau of Public Health Laboratories | Sarah Schmedes, Jason Blanton |
| EPI_ISL_489833 | Clinical Microbiology Laboratory- Basurto University Hospital | Biocruces-Bizkaia | Mikel J. Urrutikoetxea-Gutierrez, Ana Belén Belén de la Hoz, Matxalen Vidal-García, Mº Carmen Nieto Toboso, Estibaliz Ugalde-Zarraga, José Luis Díaz de Tuesta del Arco |
| EPI_ISL_489834 | Clinical Microbiology Laboratory- Basurto University Hospital | Biocruces-Bizkaia | Mikel J. Urrutikoetxea-Gutierrez, Ana Belén Belén de la Hoz, Matxalen Vidal-García, Mº Carmen Nieto Toboso, Estibaliz Ugalde-Zarraga, José Luis Díaz de Tuesta del Arco |
| EPI_ISL_489835 | Clinical Microbiology Laboratory- Basurto University Hospital | Biocruces-Bizkaia | Mikel J. Urrutikoetxea-Gutierrez, Ana Belén Belén de la Hoz, Matxalen Vidal-García, Mº Carmen Nieto Toboso, Estibaliz Ugalde-Zarraga, José Luis Díaz de Tuesta del Arco |
| EPI_ISL_489836 | Laboratorio de Referencia Nacional de Virus Respiratorio. Instituto Nacional de Salud Perú | Laboratorio de Referencia Nacional de Biotecnología y Biología Molecular. Instituto Nacional de Salud Perú | Carlos Padilla Rojas, Karolyn Chozo Vega, Priscila Lope Pari, Omar Caceres Rey, Marco Galarza Perez, Maribel Huaringa Nuñez, Johanna Balbuena Torres, Henri Bailon Calderon, Nancy Rojas Serrano |
| EPI_ISL_489837, EPI_ISL_489838 | Laboratorio de Referencia Nacional de Virus Respiratorio. Instituto Nacional de Salud Perú | Laboratorio de Referencia Nacional de Biotecnología y Biología Molecular. Instituto Nacional de Salud Perú | Carlos Padilla Rojas, Karolyn Chozo Vega, Priscila Lope Pari, Omar Caceres Rey, Marco Galarza Perez, Maribel Huaringa Nuñez, Johanna Balbuena Torrez, Henri Bailon Calderon, Nancy Rojas Serrano |
| EPI_ISL_489839 | Laboratorio de Referencia Nacional de Virus Respiratorio. Instituto Nacional de Salud Perú | Laboratorio de Referencia Nacional de Biotecnología y Biología Molecular. Instituto Nacional de Salud Perú | Carlos Padilla Rojas, Karolyn Chozo Vega, Priscila Lope Pari, Omar Caceres Rey, Marco Galarza Perez, Maribel Huaringa Nuñez, Johanna Balbuena Torres, Henri Bailon Calderon, Nancy Rojas Serrano. |
| EPI_ISL_489896, EPI_ISL_489897, EPI_ISL_489898, EPI_ISL_489899, EPI_ISL_489900, EPI_ISL_489901, EPI_ISL_489902, EPI_ISL_489903 | Gundersen Molecular Diagnostics Laboratory | Kabara Cancer Research Institute | Craig S. Richmond, Paraic A. Kenny |
| EPI_ISL_489904 | Gundersen Clinical Microbiology Laboratory | Kabara Cancer Research Institute | Craig S. Richmond, Paraic A. Kenny |
| EPI_ISL_489905, EPI_ISL_489906, EPI_ISL_489907, EPI_ISL_489908, EPI_ISL_489909, EPI_ISL_489910, EPI_ISL_489911, EPI_ISL_489912, EPI_ISL_489913, EPI_ISL_489914, EPI_ISL_489915, EPI_ISL_489916, EPI_ISL_489917, EPI_ISL_489918, EPI_ISL_489919, EPI_ISL_489920, EPI_ISL_489921, EPI_ISL_489922, EPI_ISL_489923, EPI_ISL_489924, EPI_ISL_489925, EPI_ISL_489926, EPI_ISL_489927, EPI_ISL_489928, EPI_ISL_489929, EPI_ISL_489930, EPI_ISL_489931, EPI_ISL_489932, EPI_ISL_489933, EPI_ISL_489934, EPI_ISL_489935, EPI_ISL_489936, EPI_ISL_489937, EPI_ISL_489938, EPI_ISL_489939, EPI_ISL_489940, EPI_ISL_489941, EPI_ISL_489942, EPI_ISL_489943, EPI_ISL_489944, EPI_ISL_489945, EPI_ISL_489946, EPI_ISL_489947, EPI_ISL_489948, EPI_ISL_489949, EPI_ISL_489950, EPI_ISL_489951, EPI_ISL_489952, EPI_ISL_489953, EPI_ISL_489954, EPI_ISL_489955 | | | |
| see above | Gundersen Molecular Diagnostics Laboratory | Kabara Cancer Research Institute | Craig S. Richmond, Paraic A. Kenny |
| EPI_ISL_489956, EPI_ISL_489957, EPI_ISL_489958 | Gundersen Clinical Microbiology Laboratory | Kabara Cancer Research Institute | Craig S. Richmond, Paraic A. Kenny |
| EPI_ISL_489959, EPI_ISL_489960, EPI_ISL_489961, EPI_ISL_489962, EPI_ISL_489963, EPI_ISL_489964, EPI_ISL_489965, EPI_ISL_489966, EPI_ISL_489967, EPI_ISL_489968, EPI_ISL_489969, EPI_ISL_489970, EPI_ISL_489971, EPI_ISL_489972, EPI_ISL_489973, EPI_ISL_489974, EPI_ISL_489975, EPI_ISL_489976, EPI_ISL_489977, EPI_ISL_489978, EPI_ISL_489979, EPI_ISL_489980, EPI_ISL_489981, EPI_ISL_489982, EPI_ISL_489983, EPI_ISL_489984, EPI_ISL_489985, EPI_ISL_489986 | | | |
| see above | Viollier AG | Department of Biosystems Science and Engineering, ETH Zürich | Christian Beisel, Sarah Nadeau, Ivan Topolsky, Pedro Ferreira, Philipp Jablonski, Susana Posada-Céspedes, Tobias Schär, Ina Nissen, Natascha Santacroce, Elodie Burcklen, Christiane Beckmann, Maurice Redondo, Olivier Kobel, Christoph Noppen, Sophie Seidel, Noemie Santamaria de Souza, Niko Beerenwinkel, Tanja Stadler |
| EPI_ISL_489987, EPI_ISL_489988, EPI_ISL_489989, EPI_ISL_489990 | Laboratorio de Referencia Nacional de Virus Respiratorio. Instituto Nacional de Salud Perú | Laboratorio de Referencia Nacional de Biotecnología y Biología Molecular. Instituto Nacional de Salud Perú | Carlos Padilla Rojas, Karolyn Chozo Vega, Priscila Lope Pari, Omar Caceres Rey, Marco Galarza Perez, Maribel Huaringa Nuñez, Johanna Balbuena Torres, Henri Bailon Calderon, Nancy Rojas Serrano. |
| EPI_ISL_489991 | National Institute of Health, Department of Medical Sciences, Ministry of Public Health, Thailand | National Institute of Health, Department of Medical Sciences, Ministry of Public Health, Thailand | Pilailuk,Okada; Siripaporn,Phuygun; Thanutsapa,Thanadachakul; Sittiporn,Parnmen;Warawan,Wongboot; Sunthareeya,Waicharoen; Malinee,Chittaganpitch |
| EPI_ISL_489992 | Institute for Medical Research, Infectious Disease Research Centre, National Institutes of Health, Ministry of Health Malaysia | Institute for Medical Research, Infectious Disease Research Centre, National Institutes of Health, Ministry of Health Malaysia | Suppiah J, Mohd-Zawawi Z, Kamel K, Kalyanasundram J, Thayan R |
| EPI_ISL_489993, EPI_ISL_489994 | Institute for Medical Research, Infectious Disease Research Centre, National Institutes of Health, Ministry of Health Malaysia | Institute for Medical Research, Infectious Disease Research Centre, National Institutes of Health, Ministry of Health Malaysia | Suppiah J, Mohd-Zawawi Z, Kamel K, Kalyanasundram J, Thayan R |
| EPI_ISL_489995 | CSIR-CDRI/SGPGI, Lucknow | CSIR-CDRI/SGPGI, Lucknow | Saumya Sarkar, Dharam Veer Singh, Rahul Vishvkarma, Ujjala Ghoshal, Uday Ghoshal, Ravishankar Ramachandran, Tapas Kumar Kundu, Rajender Singh |
| EPI_ISL_489996, EPI_ISL_489997, EPI_ISL_489998, EPI_ISL_489999, EPI_ISL_490000, EPI_ISL_490001, EPI_ISL_490002, EPI_ISL_490003, EPI_ISL_490004, EPI_ISL_490005, EPI_ISL_490006, EPI_ISL_490007, EPI_ISL_490008, EPI_ISL_490009, EPI_ISL_490010, EPI_ISL_490011, EPI_ISL_490012 | | | |
| see above | King Fahad Medical City | King Fahad Medical City | Alosaimi,B., Naeem,A., Alghoraibi,M., Enani,M. |
| EPI_ISL_490013 | CSIR-CDRI/SGPGI, Lucknow | CSIR-CDRI, Lucknow | Saumya Sarkar, Dharam Veer Singh, Rahul Vishvkarma, Ujjala Ghoshal, Uday Ghoshal, Ravishankar Ramachandran, Tapas Kumar Kundu, Rajender Singh |
| EPI_ISL_490014, EPI_ISL_490015 | Institute for Medical Research, Infectious Disease Research Centre, National Institutes of Health, Ministry of Health Malaysia | Institute for Medical Research, Infectious Disease Research Centre, National Institutes of Health, Ministry of Health Malaysia | Suppiah J, Mohd-Zawawi Z, Kamel K, Kalyanasundram J, Thayan R |
| EPI_ISL_490016 | Institute for Medical Research, Infectious Disease Research Centre, National Institutes of Health, Ministry of Health Malaysia | Institute for Medical Research, Infectious Disease Research Centre, National Institutes of Health, Ministry of Health Malaysia | Suppiah J, Mohd-Zawawi Z, Kamel K, Kalyanasundram J, Thayan R |
| EPI_ISL_490017, EPI_ISL_490018, EPI_ISL_490019 | South Eastern Area Laboratory Services (SEALS) | NSW Health Pathology - Institute of Clinical Pathology and Medical Research; Westmead Hospital; University of Sydney | CIDM-PH et al. |
| EPI_ISL_490020 | Sydney South West Pathology Service (SSWPS) - Liverpool Hospital - NSW Health Pathology | NSW Health Pathology - Institute of Clinical Pathology and Medical Research; Westmead Hospital; University of Sydney | CIDM-PH et al. |
| EPI_ISL_490021, EPI_ISL_490022, EPI_ISL_490023, EPI_ISL_490024, EPI_ISL_490025, EPI_ISL_490026, EPI_ISL_490027, EPI_ISL_490028, EPI_ISL_490029, EPI_ISL_490030, EPI_ISL_490031, EPI_ISL_490032, EPI_ISL_490033, EPI_ISL_490034, EPI_ISL_490035 | | | |
| see above | South Eastern Area Laboratory Services (SEALS) | NSW Health Pathology - Institute of Clinical Pathology and Medical Research; Westmead Hospital; University of Sydney | CIDM-PH et al. |
| EPI_ISL_490036 | Pathology West - NSW Health Pathology | NSW Health Pathology - Institute of Clinical Pathology and Medical Research; Westmead Hospital; University of Sydney | CIDM-PH et al. |
| EPI_ISL_490037 | Sydney South West Pathology Service (SSWPS) - Liverpool Hospital - NSW Health Pathology | NSW Health Pathology - Institute of Clinical Pathology and Medical Research; Westmead Hospital; University of Sydney | CIDM-PH et al. |
| EPI_ISL_490038, EPI_ISL_490039 | Pathology West - NSW Health Pathology | NSW Health Pathology - Institute of Clinical | CIDM-PH et al. |

| | | | |
|---|---|---|---|
| | | Pathology and Medical Research; Westmead Hospital; University of Sydney | |
| EPI_ISL_490040, EPI_ISL_490041, EPI_ISL_490042 | South Eastern Area Laboratory Services (SEALS) | NSW Health Pathology - Institute of Clinical Pathology and Medical Research; Westmead Hospital; University of Sydney | CIDM-PH et al. |
| EPI_ISL_490043 | Pathology North Hunter- NSW Health Pathology | NSW Health Pathology - Institute of Clinical Pathology and Medical Research; Westmead Hospital; University of Sydney | CIDM-PH et al. |
| EPI_ISL_490044 | Pathology West - NSW Health Pathology | NSW Health Pathology - Institute of Clinical Pathology and Medical Research; Westmead Hospital; University of Sydney | CIDM-PH et al. |
| EPI_ISL_490045 | South Eastern Area Laboratory Services (SEALS) | NSW Health Pathology - Institute of Clinical Pathology and Medical Research; Westmead Hospital; University of Sydney | CIDM-PH et al. |
| EPI_ISL_490046 | Pathology West - NSW Health Pathology | NSW Health Pathology - Institute of Clinical Pathology and Medical Research; Westmead Hospital; University of Sydney | CIDM-PH et al. |
| EPI_ISL_490047, EPI_ISL_490048 | Institute for Medical Research, Infectious Disease Research Centre, National Institutes of Health, Ministry of Health Malaysia | Institute for Medical Research, Infectious Disease Research Centre, National Institutes of Health, Ministry of Health Malaysia | Suppiah J, Mohd-Zawawi Z, Kamel K, Kalyanasundram J, Thayan R |
| EPI_ISL_490049, EPI_ISL_490050, EPI_ISL_490051, EPI_ISL_490052, EPI_ISL_490053, EPI_ISL_490054, EPI_ISL_490055, EPI_ISL_490056, EPI_ISL_490057, EPI_ISL_490058, EPI_ISL_490059, EPI_ISL_490060, EPI_ISL_490061, EPI_ISL_490062, EPI_ISL_490063, EPI_ISL_490064, EPI_ISL_490065, EPI_ISL_490066, EPI_ISL_490067, EPI_ISL_490068, EPI_ISL_490069, EPI_ISL_490070, EPI_ISL_490071, EPI_ISL_490072, EPI_ISL_490073, EPI_ISL_490074, EPI_ISL_490075, EPI_ISL_490076, EPI_ISL_490077, EPI_ISL_490078, EPI_ISL_490079 | | | |
| see above | National Public Health Laboratory, National Centre for Infectious Diseases | National Public Health Laboratory, National Centre for Infectious Diseases | Mak TM, Octavia S, Zhou Z, Chavatte JM, Cui L, Lin RTP |
| EPI_ISL_490089 | Institute for Medical Research, Infectious Disease Research Centre, National Institutes of Health, Ministry of Health Malaysia | Institute for Medical Research, Infectious Disease Research Centre, National Institutes of Health, Ministry of Health Malaysia | Suppiah J, Mohd-Zawawi Z, Kamel K, Kalyanasundram J, Thayan R |
| EPI_ISL_490090 | Institute for Medical Research, Infectious Disease Research Centre, National Institutes of Health, Ministry of Health Malaysia | Institute for Medical Research, Infectious Disease Research Centre, National Institutes of Health, Ministry of Health Malaysia | Suppiah J, Mohd-Zawawi Z, Kamel K, Kalyanasundram J, Thayan R |
| EPI_ISL_490091 | Institute for Medical Research, Infectious Disease Research Centre, National Institutes of Health, Ministry of Health Malaysia | Institute for Medical Research, Infectious Disease Research Centre, National Institutes of Health, Ministry of Health Malaysia | Suppiah J, Mohd-Zawawi Z, Kamel K, Kalyanasundram J, Thayan R |
| EPI_ISL_490092, EPI_ISL_490093 | Institute for Medical Research, Infectious Disease Research Centre, National Institutes of Health, Ministry of Health Malaysia | Institute for Medical Research, Infectious Disease Research Centre, National Institutes of Health, Ministry of Health Malaysia | Suppiah J, Mohd-Zawawi Z, Kamel K, Kalyanasundram J, Thayan R |
| EPI_ISL_490094 | Institute for Medical Research, Infectious Disease Research Centre, National Institutes of Health, Ministry of Health Malaysia | Institute for Medical Research, Infectious Disease Research Centre, National Institutes of Health, Ministry of Health Malaysia | Suppiah J, Mohd-Zawawi Z, Kamel K, Kalyanasundram J, Thayan R |
| EPI_ISL_490095, EPI_ISL_490096, EPI_ISL_490097 | Institute for Medical Research, Infectious Disease Research Centre, National Institutes of Health, Ministry of Health Malaysia | Institute for Medical Research, Infectious Disease Research Centre, National Institutes of Health, Ministry of Health Malaysia | Suppiah J, Mohd-Zawawi Z, Kamel K, Kalyanasundram J, Thayan R |
| EPI_ISL_490098 | Institute for Medical Research, Infectious Disease Research Centre, National Institutes of Health, Ministry of Health Malaysia | Institute for Medical Research, Infectious Disease Research Centre, National Institutes of Health, Ministry of Health Malaysia | Suppiah J, Mohd-Zawawi Z, Kamel K, Kalyanasundram J, Thayan R |
| EPI_ISL_490099 | Institute for Medical Research, Infectious Disease Research Centre, National Institutes of Health, Ministry of Health Malaysia | Institute for Medical Research, Infectious Disease Research Centre, National Institutes of Health, Ministry of Health Malaysia | Suppiah J, Mohd-Zawawi Z, Kamel K, Kalyanasundram J, Thayan R |
| EPI_ISL_490100 | Institute for Medical Research, Infectious Disease Research Centre, National Institutes of Health, Ministry of Health Malaysia | Institute for Medical Research, Infectious Disease Research Centre, National Institutes of Health, Ministry of Health Malaysia | Suppiah J, Mohd-Zawawi Z, Kamel K, Kalyanasundram J, Thayan R |
| EPI_ISL_490101 | Institute for Medical Research, Infectious Disease Research Centre, National Institutes of Health, Ministry of Health Malaysia | Institute for Medical Research, Infectious Disease Research Centre, National Institutes of Health, Ministry of Health Malaysia | Suppiah J, Mohd-Zawawi Z, Kamel K, Kalyanasundram J, Thayan R |
| EPI_ISL_490102 | Institute for Medical Research, Infectious Disease Research Centre, National Institutes of Health, Ministry of Health Malaysia | Institute for Medical Research, Infectious Disease Research Centre, National Institutes of Health, Ministry of Health Malaysia | Suppiah J, Mohd-Zawawi Z, Kamel K, Kalyanasundram J, Thayan R |
| EPI_ISL_490103 | Institute for Medical Research, Infectious Disease Research Centre, National Institutes of Health, Ministry of Health Malaysia | Institute for Medical Research, Infectious Disease Research Centre, National Institutes of Health, Ministry of Health Malaysia | Suppiah J, Mohd-Zawawi Z, Kamel K, Kalyanasundram J, Thayan R |
| EPI_ISL_490104 | CSIR-CDRI/SGPGI, Lucknow | CSIR-CDRI/SGPGI, Lucknow | Saumya Sarkar, Dharam Veer Singh, Rahul Vishvkarma, Ujjala Ghoshal, Uday Ghoshal, Ravishankar Ramachandran, Tapas Kumar Kundu, Rajender Singh |
| EPI_ISL_490105 | South Eastern Area Laboratory Services (SEALS) | NSW Health Pathology - Institute of Clinical Pathology and Medical Research; Westmead Hospital; University of Sydney | CIDM-PH et al. |
| EPI_ISL_490106 | CSIR-CDRI/SGPGI, Lucknow | CSIR-CDRI/SGPGI, Lucknow | Saumya Sarkar, Dharam Veer Singh, Rahul Vishvkarma, Ujjala Ghoshal, Uday Ghoshal, Ravishankar Ramachandran, Tapas Kumar Kundu, Rajender Singh |
| EPI_ISL_490109, EPI_ISL_490110, EPI_ISL_490111 | National Institute of Laboratory Medicine and Referral Center | Genomic Research Lab, BCSIR | Md. Murshed Hasan Sarkar, Abu Sayeed Mohammad Mahmud, Mohammad Samir Uzzaman, Eshrar Osman, Md. Ahasan Habib, Shahina Akter, Tanjina Akhter Banu, Barna Goswami, Iffat Jahan, Md. Saddam Hossain, Tasnim Nafisa, Md. Maruf Ahmed Molla, Mahmuda Yeasmin, Asish Kumar Ghosh, A. K. M. Shamsuzzaman, Sheikh Md. Selim Al Din, Utpal Chandra Ray, Salek Ahmed Sajib, Md. Salim Khan |
| EPI_ISL_490112 | The National Institute of Public Health | The National Institute of Public Health and State Veterinary Institute Prague | Nagy,A;Jirincova,H;Novakova,L;Trnka,D;Vecerova,J |
| EPI_ISL_490113 | National Institute of Laboratory Medicine and Referral Center | Genomic Research Lab, BCSIR | Shahina Akter, Abu Sayeed Mohammad Mahmud, Mohammad Samir Uzzaman, Eshrar Osman, Md. Ahasan Habib, Tanjina Akhter Banu, Md. Murshed Hasan Sarkar, Barna Goswami, Iffat Jahan, Md. Saddam Hossain, Tasnim Nafisa, Md. Maruf Ahmed Molla, Mahmuda Yeasmin, Asish Kumar Ghosh, A. K. M. Shamsuzzaman, Sheikh Md. Selim Al Din, Utpal Chandra Ray, Salek Ahmed Sajib, Md. Salim Khan |
| EPI_ISL_490114 | National Institute of Laboratory Medicine and Referral Center | Genomic Research Lab, BCSIR | Tanjina Akhter Banu, Abu Sayeed Mohammad Mahmud, Mohammad Samir Uzzaman, Eshrar Osman, Md. Ahasan Habib, Shahina Akter, Md. Murshed Hasan Sarkar, Barna Goswami, Iffat Jahan, Md. Saddam Hossain, Tasnim Nafisa, Md. Maruf Ahmed Molla, Mahmuda Yeasmin, Asish Kumar Ghosh, A. K. M. Shamsuzzaman, Sheikh Md. Selim Al Din, Utpal Chandra Ray, Salek Ahmed Sajib, Md. Salim Khan |
| EPI_ISL_490144 | National Institute of Laboratory Medicine and Referral Center | Genomic Research Lab, BCSIR | Barna Goswami, Abu Sayeed Mohammad Mahmud, Mohammad Samir Uzzaman, Eshrar Osman, Md. Ahasan Habib, Shahina Akter, Tanjina Akhter Banu, Md. Murshed Hasan Sarkar, Iffat Jahan, Md. Saddam Hossain, Tasnim Nafisa, Md. Maruf Ahmed Molla, Mahmuda Yeasmin, Asish Kumar Ghosh, A. K. M. Shamsuzzaman, Sheikh Md. Selim Al Din, Utpal Chandra Ray, Salek Ahmed Sajib, Md. Salim Khan |
| EPI_ISL_490164 | National Institute of Laboratory Medicine and Referral Center | Genomic Research Lab, BCSIR | Iffat Jahan, Abu Sayeed Mohammad Mahmud, Mohammad Samir Uzzaman, Eshrar Osman, Md. Ahasan Habib, Shahina Akter, Tanjina Akhter Banu, Md. Murshed Hasan Sarkar, Barna Goswami, Md. Saddam Hossain, Tasnim Nafisa, Md. Maruf Ahmed Molla, Mahmuda Yeasmin, Asish Kumar Ghosh, A. K. M. Shamsuzzaman, Sheikh Md. Selim Al Din, Utpal Chandra Ray, Salek Ahmed Sajib, Md. Salim Khan |
| EPI_ISL_490165 | National Institute of Laboratory Medicine and Referral Center | Genomic Research Lab, BCSIR | Md. Saddam Hossain, Abu Sayeed Mohammad Mahmud, Mohammad Samir Uzzaman, Eshrar Osman, Md. Ahasan Habib, Shahina Akter, Tanjina Akhter Banu, Md. Murshed Hasan Sarkar, Barna Goswami, Iffat Jahan, Tasnim Nafisa, Md. Maruf Ahmed Molla, Mahmuda Yeasmin, Asish Kumar Ghosh, A. K. M. Shamsuzzaman, Sheikh Md. Selim Al Din, Utpal Chandra Ray, Salek Ahmed Sajib, Md. Salim Khan |
| EPI_ISL_490167, EPI_ISL_490168 | National Institute of Laboratory Medicine and Referral Center | Genomic Research Lab, BCSIR | Abu Sayeed Mohammad Mahmud, Mohammad Samir Uzzaman, Eshrar Osman, Md. Ahasan Habib, Shahina Akter, Tanjina Akhter Banu, Md. Murshed Hasan Sarkar, Barna Goswami, Iffat Jahan, Md. Saddam Hossain, Tasnim Nafisa, Md. Maruf Ahmed Molla, Mahmuda Yeasmin, Asish Kumar Ghosh, A. K. M. Shamsuzzaman, Sheikh Md. Selim Al Din, Utpal Chandra Ray, Salek Ahmed Sajib, Md. Salim Khan |
| EPI_ISL_490201 | National Influenza Centre of INSPI-Ecuador | INSPI - Charité | Alfredo Bruno Caicedo, Domenica de Mora Coloma, Andres Moreira-Soto, Anna-Lena Sander, Nina Krause, Maritza Olmedo,Denisses Portugal, Manuel Gonzalez, Silvia Salgado, Alberto Orlando, Alexandra Usiña, Juan Carlos Zeballos,Jan Felix Drexler |
| EPI_ISL_490202 | Clinical Microbiology Laboratory- Basurto University Hospital | Biocruces-Bizkaia | Mikel J. Urrutikoetxea-Gutierrez, Ana Belén Belén de la Hoz, Matxalen Vidal-García, Mº Carmen Nieto Toboso, Estibaliz Ugalde-Zarraga, José Luis Díaz de Tuesta del Arco |
| EPI_ISL_490203 | Clinical Microbiology Laboratory- Basurto University Hospital | Biocruces-Bizkaia | Mikel J. Urrutikoetxea-Gutierrez, Ana Belén Belén de la Hoz, Matxalen Vidal-García, Mº Carmen Nieto Toboso, Estibaliz Ugalde-Zarraga, José Luis Díaz de Tuesta del Arco |
| EPI_ISL_490204 | Clinical Microbiology Laboratory- Basurto University Hospital | Biocruces-Bizkaia | Mikel J. Urrutikoetxea-Gutierrez, Ana Belén Belén de la Hoz, Matxalen Vidal-García, Mº Carmen Nieto Toboso, Estibaliz Ugalde-Zarraga, José Luis Díaz de Tuesta del Arco |
| EPI_ISL_490205, EPI_ISL_490206, EPI_ISL_490207, EPI_ISL_490208 | München Klinik Schwabing | MGZ Medical Genetics Center | Dieter A. Wolf, Elke Holinski-Feder |
| EPI_ISL_490209 | Laboratorio de Referencia Nacional de Virus Respiratorio. Instituto Nacional de Salud Perú | Laboratorio de Referencia Nacional de Biotecnología y Biología Molecular. Instituto Nacional de Salud Perú | Carlos Padilla Rojas, Karolyn Chozo Vega, Priscila Lope Pari, Omar Caceres Rey, Marco Galarza Perez, Maribel Huaringa Nuñez, Johanna Balbuena Torres, Henri Bailon Calderon, Nancy Rojas Serrano |

| | | | |
|---|---|---|---|
| EPI_ISL_490210, EPI_ISL_490211, EPI_ISL_490212, EPI_ISL_490213, EPI_ISL_490214, EPI_ISL_490215, EPI_ISL_490216, EPI_ISL_490217, EPI_ISL_490218, EPI_ISL_490219, EPI_ISL_490220, EPI_ISL_490221, EPI_ISL_490222, EPI_ISL_490223 | | | |
| see above | Quest Diagnostics | Q Squared Solutions - QRTP facility | Victor J Weigman |
| EPI_ISL_490224, EPI_ISL_490225, EPI_ISL_490226, EPI_ISL_490227, EPI_ISL_490228, EPI_ISL_490229, EPI_ISL_490230, EPI_ISL_490231, EPI_ISL_490232, EPI_ISL_490233, EPI_ISL_490234, EPI_ISL_490235, EPI_ISL_490236, EPI_ISL_490237, EPI_ISL_490238, EPI_ISL_490239, EPI_ISL_490240, EPI_ISL_490241, EPI_ISL_490242, EPI_ISL_490243, EPI_ISL_490244, EPI_ISL_490245, EPI_ISL_490246, EPI_ISL_490247, EPI_ISL_490248, EPI_ISL_490249, EPI_ISL_490250, EPI_ISL_490251, EPI_ISL_490252, EPI_ISL_490253, EPI_ISL_490254 | | | |
| see above | Respiratory Virus Unit, Microbiology Services Colindale, Public Health England | Respiratory Virus Unit, Microbiology Services Colindale, Public Health England | PHE Covid Sequencing Team |
| EPI_ISL_490255, EPI_ISL_490256, EPI_ISL_490257, EPI_ISL_490258, EPI_ISL_490259, EPI_ISL_490260, EPI_ISL_490261, EPI_ISL_490262, EPI_ISL_490263, EPI_ISL_490264, EPI_ISL_490265, EPI_ISL_490266, EPI_ISL_490267, EPI_ISL_490268, EPI_ISL_490269, EPI_ISL_490270, EPI_ISL_490271, EPI_ISL_490272, EPI_ISL_490273, EPI_ISL_490274, EPI_ISL_490275, EPI_ISL_490276, EPI_ISL_490277, EPI_ISL_490278, EPI_ISL_490279, EPI_ISL_490280, EPI_ISL_490281, EPI_ISL_490282, EPI_ISL_490283, EPI_ISL_490284, EPI_ISL_490285, EPI_ISL_490286, EPI_ISL_490287, EPI_ISL_490288, EPI_ISL_490289, EPI_ISL_490290, EPI_ISL_490291, EPI_ISL_490292, EPI_ISL_490293, EPI_ISL_490294, EPI_ISL_490295, EPI_ISL_490296, EPI_ISL_490297, EPI_ISL_490298, EPI_ISL_490299, EPI_ISL_490300, EPI_ISL_490301, EPI_ISL_490302, EPI_ISL_490303, EPI_ISL_490304, EPI_ISL_490305, EPI_ISL_490306, EPI_ISL_490307, EPI_ISL_490308, EPI_ISL_490309, EPI_ISL_490310, EPI_ISL_490311, EPI_ISL_490312, EPI_ISL_490313, EPI_ISL_490314 | | | |
| see above | National Institute for Communicable Diseases of the National Health Laboratory Service | National Institute for Communicable Diseases of the National Health Laboratory Service | Allam M, Ismail A, Khumalo Z, Kwenda S, Mtshali P, Mnyameni F, Mohale T, Subramoney K, Bhiman JN |
| EPI_ISL_490315, EPI_ISL_490316 | Laboratorio de Referencia Nacional de Virus Respiratorio. Instituto Nacional de Salud Perú | Laboratorio de Referencia Nacional de Biotecnología y Biología Molecular. Instituto Nacional de Salud Perú | Carlos Padilla Rojas, Karolyn Chozo Vega, Priscila Lope Pari, Omar Caceres Rey, Marco Galarza Perez, Maribel Huaringa Nuñez, Johanna Balbuena Torres, Henri Bailon Calderon, Nancy Rojas Serrano |
| EPI_ISL_490317, EPI_ISL_490318, EPI_ISL_490319, EPI_ISL_490320, EPI_ISL_490321, EPI_ISL_490322, EPI_ISL_490323, EPI_ISL_490324, EPI_ISL_490325, EPI_ISL_490326, EPI_ISL_490327, EPI_ISL_490328, EPI_ISL_490329 | | | |
| see above | Department of Pathology, University of Cambridge | COVID-19 Genomics UK (COG-UK) Consortium | Luke W Meredith, M. Estée Török, Myra Hosmillo, William L. Hamilton, Martin D. Curran, Theresa Feltwell, Grant Hall, Anna Yakovleva, Fahad A Khokhar, Charlotte J. Houldcroft, Laura G Caller, Aminu S. Jahun, Sarah L. Caddy, Yasmin Chaudhry, Malte Pinckert, Ian Goodfellow |
| EPI_ISL_490330 | West of Scotland Specialist Virology Centre, NHSGGC / MRC-University of Glasgow Centre for Virus Research | COVID-19 Genomics UK (COG-UK) Consortium | Ana da Silva Filipe, Natasha Johnson, Kathy Smollett, Daniel Mair, Stephen Carmichael, Lily Tong, Jenna Nichols, Elihu Aranday-Cortes, Kirstyn Brunker, Yasmin Parr, Alice Broos, Kyriaki Nomikou; Sarah McDonald, Marc Niebel, Patawee Asamaphan; Richard Orton, Joseph Hughes, Sreenu Vattipally, David L Robertson; Alasdair MacLean, Rory Gunson; Kathy Li, Natasha Jesudason, Rajiv Shah, James Shepherd, Antonia Ho, Emma Thomson |
| EPI_ISL_490331, EPI_ISL_490332, EPI_ISL_490333, EPI_ISL_490334, EPI_ISL_490335, EPI_ISL_490336, EPI_ISL_490337, EPI_ISL_490338, EPI_ISL_490339, EPI_ISL_490340, EPI_ISL_490341, EPI_ISL_490342, EPI_ISL_490343, EPI_ISL_490344, EPI_ISL_490345, EPI_ISL_490346, EPI_ISL_490347, EPI_ISL_490348, EPI_ISL_490349, EPI_ISL_490350, EPI_ISL_490351, EPI_ISL_490352, EPI_ISL_490353, EPI_ISL_490354, EPI_ISL_490355, EPI_ISL_490356, EPI_ISL_490357, EPI_ISL_490358, EPI_ISL_490359, EPI_ISL_490360, EPI_ISL_490361, EPI_ISL_490362, EPI_ISL_490363, EPI_ISL_490364, EPI_ISL_490365, EPI_ISL_490366, EPI_ISL_490367, EPI_ISL_490368, EPI_ISL_490369, EPI_ISL_490370, EPI_ISL_490371, EPI_ISL_490372, EPI_ISL_490373, EPI_ISL_490374, EPI_ISL_490375, EPI_ISL_490376, EPI_ISL_490377, EPI_ISL_490378, EPI_ISL_490379, EPI_ISL_490380, EPI_ISL_490381, EPI_ISL_490382, EPI_ISL_490383, EPI_ISL_490384, EPI_ISL_490385, EPI_ISL_490386, EPI_ISL_490387, EPI_ISL_490388, EPI_ISL_490389, EPI_ISL_490390, EPI_ISL_490391, EPI_ISL_490392, EPI_ISL_490393, EPI_ISL_490394, EPI_ISL_490395, EPI_ISL_490396, EPI_ISL_490397, EPI_ISL_490398, EPI_ISL_490399, EPI_ISL_490400, EPI_ISL_490401, EPI_ISL_490402, EPI_ISL_490403, EPI_ISL_490404, EPI_ISL_490405, EPI_ISL_490406, EPI_ISL_490407, EPI_ISL_490408, EPI_ISL_490409, EPI_ISL_490410, EPI_ISL_490411, EPI_ISL_490412, EPI_ISL_490413, EPI_ISL_490414, EPI_ISL_490415, EPI_ISL_490416, EPI_ISL_490417, EPI_ISL_490418, EPI_ISL_490419, EPI_ISL_490420 | | | |
| see above | Liverpool Clinical Laboratories | COVID-19 Genomics UK (COG-UK) Consortium | Sam Haldenby, Anita Lucaci, Steve Paterson, Julian Hiscox, Alistair Darby, M Almsaud, A Alrezaihi, Muhannad Alruwaili, Stuart D Armstrong, Jones Benjamin, Eleanor G Bentley, Anu Chawla, Jordan J Clark, Angela Cowell, Richard Eccles, Isabel García-Dorival, Matthew Gemmell, Alessandro Gerada, PKF Gilmore, Richard Gregory, Ximeng Han, Catherine Hartley, Margaret Hughes, Miren Iturriza-Gomara, James Johnson, L Luu, Jenifer Manson, Charlotte Nelson, Elaine O'Toole, Cassie Olateju, Rebekah Penrice-Randal , Lucille Rainbow, N.P Randle, Trevor Ian Robinson, Parul Sharma, Ghada T Shawli, James P Stewart, Neil Swainston, Ecaterina Vamos, Joanne Watts, Mark Whitehead |
| EPI_ISL_490421, EPI_ISL_490422, EPI_ISL_490423, EPI_ISL_490424, EPI_ISL_490425, EPI_ISL_490426, EPI_ISL_490427, EPI_ISL_490428, EPI_ISL_490429, EPI_ISL_490430, EPI_ISL_490431, EPI_ISL_490432, EPI_ISL_490433, EPI_ISL_490434, EPI_ISL_490435, EPI_ISL_490436, EPI_ISL_490437, EPI_ISL_490438, EPI_ISL_490439 | | | |
| see above | University College London Hospital | COVID-19 Genomics UK (COG-UK) Consortium | Judith Heaney, Matthew Byott, Dan Frampton, Moira Spyer and Eleni Nastouli |
| EPI_ISL_490440, EPI_ISL_490441, EPI_ISL_490442, EPI_ISL_490443, EPI_ISL_490444, EPI_ISL_490445, EPI_ISL_490446, EPI_ISL_490447, EPI_ISL_490448, EPI_ISL_490449, EPI_ISL_490450, EPI_ISL_490451, EPI_ISL_490452, EPI_ISL_490453, EPI_ISL_490454, EPI_ISL_490455, EPI_ISL_490456, EPI_ISL_490457, EPI_ISL_490458, EPI_ISL_490459, EPI_ISL_490460, EPI_ISL_490461, EPI_ISL_490462, EPI_ISL_490463, EPI_ISL_490464, EPI_ISL_490465, EPI_ISL_490466, EPI_ISL_490467, EPI_ISL_490468, EPI_ISL_490469, EPI_ISL_490470, EPI_ISL_490471, EPI_ISL_490472, EPI_ISL_490473, EPI_ISL_490474, EPI_ISL_490475, EPI_ISL_490476, EPI_ISL_490477, EPI_ISL_490478, EPI_ISL_490479, EPI_ISL_490480, EPI_ISL_490481, EPI_ISL_490482, EPI_ISL_490483, EPI_ISL_490484, EPI_ISL_490485, EPI_ISL_490486, EPI_ISL_490487, EPI_ISL_490488, EPI_ISL_490489, EPI_ISL_490490, EPI_ISL_490491, EPI_ISL_490492, EPI_ISL_490493, EPI_ISL_490494, EPI_ISL_490495 | | | |
| see above | Northumbria University / South Tees Hospitals NHS Foundation Trust / North Cumbria Integrated Care NHS Foundation Trust / North Tees and Hartlepool NHS Foundation Trust / Newcastle Hospitals NHS Foundation Trust | COVID-19 Genomics UK (COG-UK) Consortium | Darren L Smith,Andrew Nelson,Matthew Bashton,Greg R Young,Joshua Loh,John Allan,Mohammad A Tariq,Giles S Holt,Gary Black,Wen C Yew,Lynn Dover,Paul Baker,Steve Liggett,Sarah Essex,Jane Greenaway,Debra Padgett,Clive Graham,Garren Scott,Edward Barton,Emma Swindells,Brendan Payne,Jennifer Collins,Yusri Taha,Gary Eltringham |
| EPI_ISL_490496, EPI_ISL_490497, EPI_ISL_490498, EPI_ISL_490499, EPI_ISL_490500, EPI_ISL_490501, EPI_ISL_490502, EPI_ISL_490503, EPI_ISL_490504, EPI_ISL_490505, EPI_ISL_490506, EPI_ISL_490507, EPI_ISL_490508, EPI_ISL_490509, EPI_ISL_490510, EPI_ISL_490511, EPI_ISL_490512, EPI_ISL_490513, EPI_ISL_490514, EPI_ISL_490515, EPI_ISL_490516, EPI_ISL_490517, EPI_ISL_490518, EPI_ISL_490519, EPI_ISL_490520, EPI_ISL_490521, EPI_ISL_490522, EPI_ISL_490523, EPI_ISL_490524, EPI_ISL_490525, EPI_ISL_490526, EPI_ISL_490527, EPI_ISL_490528, EPI_ISL_490529, EPI_ISL_490530, EPI_ISL_490531, EPI_ISL_490532, EPI_ISL_490533, EPI_ISL_490534, EPI_ISL_490535, EPI_ISL_490536, EPI_ISL_490537, EPI_ISL_490538, EPI_ISL_490539, EPI_ISL_490540, EPI_ISL_490541, EPI_ISL_490542, EPI_ISL_490543, EPI_ISL_490544, EPI_ISL_490545, EPI_ISL_490546, EPI_ISL_490547, EPI_ISL_490548, EPI_ISL_490549, EPI_ISL_490550, EPI_ISL_490551, EPI_ISL_490552, EPI_ISL_490553, EPI_ISL_490554, EPI_ISL_490556, EPI_ISL_490557, EPI_ISL_490558, EPI_ISL_490559 | | | |
| see above | Quadram Institute Bioscience | COVID-19 Genomics UK (COG-UK) Consortium | Dave J. Baker, Gemma L. Kay, Alp Aydin, Thanh Le-Viet, Steven Rudder, Ana P. Tedim, Anastasia Kolyva, Maria Diaz, Leonardo de Oliveira Martins, Nabil-Fareed Alikhan, Lizzie Meadows, Rachael Stanley, Ngozi Elumogo, Muhammed Yasir, Nicholas M. Thomson, Alexander J Trotter, Rachel Gilroy, Samuel Bloomfield, Claire Stuart, Andrew Bell, Reenesh Prakash, Samir Dervisevic, Alison E. Mather, John Wain, Mark Webber, Andrew J. Page, Justin O'Grady |
| EPI_ISL_490560, EPI_ISL_490561, EPI_ISL_490562 | Queens Medical Centre, Clinical Microbiology Department / DeepSeq Nottingham | COVID-19 Genomics UK (COG-UK) Consortium | Gemma Clark, Wendy Smith, Manjinder Khakh, Vicki M Fleming, Michelle M Lister, Hannah Howson-Wells, Jonathan Ball, Patrick McClure, Joseph Chappell, Theocharis Tsoleridis, Nadine Holmes, Matthew Carlisle, Christopher Moore, Fei Sang, Johnny Debebe, Victoria Wright, Matthew Loose |
| EPI_ISL_490563, EPI_ISL_490564, EPI_ISL_490565, EPI_ISL_490566, EPI_ISL_490567, EPI_ISL_490568, EPI_ISL_490569, EPI_ISL_490570, EPI_ISL_490571, EPI_ISL_490572, EPI_ISL_490573, EPI_ISL_490574, EPI_ISL_490575, EPI_ISL_490576, EPI_ISL_490577, EPI_ISL_490578, EPI_ISL_490579, EPI_ISL_490580 | | | |
| see above | Lincolnshire Hospitals and DeepSeq Nottingham | COVID-19 Genomics UK (COG-UK) Consortium | Nichola Duckworth, Tim Sloan, Sarah Walsh, Jonathan Ball, Patrick McClure, Joeseph Chappell, Nadine Holmes, Matthew Carlisle, Christopher Moore, Fei Sang, Johnny Debebe, Victoria Wright, Matthew Loose |
| EPI_ISL_490581, EPI_ISL_490582, EPI_ISL_490583, EPI_ISL_490584, EPI_ISL_490585, EPI_ISL_490586, EPI_ISL_490587, EPI_ISL_490588, EPI_ISL_490589, EPI_ISL_490590, EPI_ISL_490591, EPI_ISL_490592, EPI_ISL_490593, EPI_ISL_490594, EPI_ISL_490595, EPI_ISL_490596, EPI_ISL_490597, EPI_ISL_490598, EPI_ISL_490599, EPI_ISL_490600, EPI_ISL_490601, EPI_ISL_490602, EPI_ISL_490603, EPI_ISL_490604, EPI_ISL_490605, EPI_ISL_490606, EPI_ISL_490607, EPI_ISL_490608, EPI_ISL_490609, EPI_ISL_490610, EPI_ISL_490611, EPI_ISL_490612, EPI_ISL_490613, EPI_ISL_490614, EPI_ISL_490615, EPI_ISL_490616, EPI_ISL_490617, EPI_ISL_490618, EPI_ISL_490619, EPI_ISL_490620, EPI_ISL_490621, EPI_ISL_490622, EPI_ISL_490623, EPI_ISL_490624, EPI_ISL_490625, EPI_ISL_490626, EPI_ISL_490627, EPI_ISL_490628, EPI_ISL_490629, EPI_ISL_490630, EPI_ISL_490631, EPI_ISL_490632, EPI_ISL_490633, EPI_ISL_490634, EPI_ISL_490635, EPI_ISL_490636, EPI_ISL_490637, EPI_ISL_490638, EPI_ISL_490639, EPI_ISL_490640, EPI_ISL_490641, EPI_ISL_490642, EPI_ISL_490643, EPI_ISL_490644, EPI_ISL_490645 | | | |
| see above | Virology Department, Sheffield Teaching Hospitals NHS Foundation Trust/Department of Infection, Immunity and Cardiovascular Disease, The Medical School, University of Sheffield | COVID-19 Genomics UK (COG-UK) Consortium | Thushan de Silva, Matthew Parker, Nikki Smith, Adri Angyal, Rebecca Brown, Luke Green, Rachel Tucker, Paul Parsons, Danielle Groves, Katie Johnson, Laura Carrilero, Alex Keeley, Dave Partridge, Matthew Wyles, Benjamin Lindsey, Mehmet Yavuz, Mohammad Raza, Cariad Evans |
| EPI_ISL_490646, EPI_ISL_490647, EPI_ISL_490648, EPI_ISL_490649, EPI_ISL_490650, EPI_ISL_490651, EPI_ISL_490652, EPI_ISL_490653, EPI_ISL_490654, EPI_ISL_490655, EPI_ISL_490656, EPI_ISL_490657, EPI_ISL_490658, EPI_ISL_490659, EPI_ISL_490660, EPI_ISL_490661, EPI_ISL_490662, EPI_ISL_490663, EPI_ISL_490664, EPI_ISL_490665, EPI_ISL_490666, EPI_ISL_490667, EPI_ISL_490668, EPI_ISL_490669, EPI_ISL_490670, EPI_ISL_490671, EPI_ISL_490672, EPI_ISL_490673, EPI_ISL_490674, EPI_ISL_490675, EPI_ISL_490676, EPI_ISL_490677, EPI_ISL_490678, EPI_ISL_490679, EPI_ISL_490680, EPI_ISL_490681, EPI_ISL_490682, EPI_ISL_490683, EPI_ISL_490684, EPI_ISL_490685, EPI_ISL_490686, EPI_ISL_490687, EPI_ISL_490688, EPI_ISL_490689, EPI_ISL_490690, EPI_ISL_490691, EPI_ISL_490692, EPI_ISL_490693, EPI_ISL_490694, EPI_ISL_490695, EPI_ISL_490696, EPI_ISL_490697, EPI_ISL_490698, EPI_ISL_490699, EPI_ISL_490700, EPI_ISL_490701, EPI_ISL_490702, EPI_ISL_490703, EPI_ISL_490704, EPI_ISL_490705, EPI_ISL_490706, EPI_ISL_490707, EPI_ISL_490708, EPI_ISL_490709 | | | |
| see above | West of Scotland Specialist Virology Centre, NHSGGC / MRC-University of Glasgow Centre for Virus Research | COVID-19 Genomics UK (COG-UK) Consortium | Ana da Silva Filipe, Natasha Johnson, Kathy Smollett, Daniel Mair, Stephen Carmichael, Lily Tong, Jenna Nichols, Elihu Aranday-Cortes, Kirstyn Brunker, Yasmin Parr, Alice Broos, Kyriaki Nomikou; Sarah McDonald, Marc Niebel, Patawee Asamaphan; Richard Orton, Joseph Hughes, Sreenu Vattipally, David L Robertson; Alasdair MacLean, Rory Gunson; Kathy Li, Natasha Jesudason, Rajiv Shah, James Shepherd, Antonia Ho, Emma Thomson |
| EPI_ISL_490710, EPI_ISL_490711, EPI_ISL_490712, EPI_ISL_490713, EPI_ISL_490714, EPI_ISL_490715, EPI_ISL_490716, EPI_ISL_490717, EPI_ISL_490718, EPI_ISL_490719, EPI_ISL_490720, EPI_ISL_490721, EPI_ISL_490722, EPI_ISL_490723, EPI_ISL_490724, EPI_ISL_490725, EPI_ISL_490726, EPI_ISL_490727, EPI_ISL_490728, EPI_ISL_490729, EPI_ISL_490730, EPI_ISL_490731, EPI_ISL_490732, EPI_ISL_490733, EPI_ISL_490734, EPI_ISL_490735, EPI_ISL_490736, EPI_ISL_490737, EPI_ISL_490738, EPI_ISL_490739, EPI_ISL_490740, EPI_ISL_490741, EPI_ISL_490742, EPI_ISL_490743, EPI_ISL_490744, EPI_ISL_490745, EPI_ISL_490746, EPI_ISL_490747, EPI_ISL_490748, EPI_ISL_490749, EPI_ISL_490750, EPI_ISL_490751, EPI_ISL_490752, EPI_ISL_490753, EPI_ISL_490754, EPI_ISL_490755, EPI_ISL_490756, EPI_ISL_490757, EPI_ISL_490758, EPI_ISL_490759, EPI_ISL_490760, EPI_ISL_490761, EPI_ISL_490762, EPI_ISL_490763, EPI_ISL_490764, EPI_ISL_490765, EPI_ISL_490766, EPI_ISL_490767, EPI_ISL_490768, EPI_ISL_490769, EPI_ISL_490770, EPI_ISL_490771, EPI_ISL_490772, EPI_ISL_490773, EPI_ISL_490774, EPI_ISL_490775, EPI_ISL_490776, EPI_ISL_490777, EPI_ISL_490778, EPI_ISL_490779, EPI_ISL_490780, EPI_ISL_490781, EPI_ISL_490782, EPI_ISL_490783, EPI_ISL_490784, EPI_ISL_490785, EPI_ISL_490786, EPI_ISL_490787, EPI_ISL_490788, EPI_ISL_490789, EPI_ISL_490790, EPI_ISL_490791, EPI_ISL_490792, EPI_ISL_490793, EPI_ISL_490794, EPI_ISL_490795, EPI_ISL_490796, EPI_ISL_490797, EPI_ISL_490798, EPI_ISL_490799, EPI_ISL_490800, EPI_ISL_490801, EPI_ISL_490802, EPI_ISL_490803, EPI_ISL_490804, EPI_ISL_490805, EPI_ISL_490806, EPI_ISL_490807, EPI_ISL_490808, EPI_ISL_490809, EPI_ISL_490810, EPI_ISL_490811, EPI_ISL_490812, EPI_ISL_490813, EPI_ISL_490814, EPI_ISL_490815, EPI_ISL_490816, EPI_ISL_490817, EPI_ISL_490818, EPI_ISL_490819, EPI_ISL_490820, EPI_ISL_490821, EPI_ISL_490822, EPI_ISL_490823, EPI_ISL_490824, EPI_ISL_490825, EPI_ISL_490826, EPI_ISL_490827, EPI_ISL_490828, EPI_ISL_490829, EPI_ISL_490830, EPI_ISL_490831, EPI_ISL_490832, EPI_ISL_490833, EPI_ISL_490834, EPI_ISL_490835, EPI_ISL_490836, EPI_ISL_490837, EPI_ISL_490838, EPI_ISL_490839, EPI_ISL_490840, EPI_ISL_490841, EPI_ISL_490842, EPI_ISL_490843, EPI_ISL_490844, EPI_ISL_490845, EPI_ISL_490846, EPI_ISL_490847, EPI_ISL_490848, EPI_ISL_490849, EPI_ISL_490850, EPI_ISL_490851, EPI_ISL_490852, EPI_ISL_490853, EPI_ISL_490854, EPI_ISL_490855, EPI_ISL_490856, EPI_ISL_490857, EPI_ISL_490858, EPI_ISL_490859, EPI_ISL_490860, EPI_ISL_490861, EPI_ISL_490862, EPI_ISL_490863, EPI_ISL_490864, EPI_ISL_490865, EPI_ISL_490866, EPI_ISL_490867 | | | |
| see above | Wales Specialist Virology Centre Sequencing lab: Pathogen Genomics Unit | COVID-19 Genomics UK (COG-UK) Consortium | Catherine Moore, Johnathan Evans, Laura Gifford, Malorie Perry, Simon Cottrell, Angela Marchbank, Alec Birchley, Alexander Adams, Amy Gaskin, Bree Gatica-Wilcox, Jason Coombes, Joel Southgate, Lauren Gilbert, Lee Graham, Nicole Pacchiarini, Sara Kumziene-Summerhayes, Sarah Taylor, Sophie Jones, Sara Rey, Matthew Bull, Joanne Watkins, Sally Corden, Tom Connor |
| EPI_ISL_490975, EPI_ISL_490976 | Laboratorio de Referencia Nacional de Virus Respiratorio. Instituto Nacional de Salud Perú | Laboratorio de Referencia Nacional de Biotecnología y Biología Molecular. Instituto Nacional de Salud Perú | Carlos Padilla Rojas, Karolyn Chozo Vega, Priscila Lope Pari, Omar Caceres Rey, Marco Galarza Perez, Maribel Huaringa Nuñez, Johanna Balbuena Torres, Henri Bailon Calderon, Nancy Rojas Serrano. |
| EPI_ISL_490977 | Clinical Microbiology Laboratory- Basurto University Hospital | Biocruces-Bizkaia | Mikel J. Urrutikoetxea-Gutierrez, Ana Belén Belén de la Hoz, Matxalen Vidal-García, Mº Carmen Nieto Toboso, Estibaliz Ugalde-Zarraga, José Luis Díaz de Tuesta del Arco |
| EPI_ISL_490978 | Mayo Clinic Laboratories | UW Virology Lab | Pavitra Roychoudhury, Hong Xie, Lasata Shrestha, Amin Addetia, Truong Nguyen, Victoria M Rachleff, Meei-Li Huang, Keith R Jerome, Alexander Greninger |
| EPI_ISL_490979, EPI_ISL_490980, EPI_ISL_490982, EPI_ISL_490983, EPI_ISL_490984, EPI_ISL_490985, EPI_ISL_490986, EPI_ISL_490987 | UW Virology Lab | UW Virology Lab | Pavitra Roychoudhury, Hong Xie, Lasata Shrestha, Amin Addetia, Truong Nguyen, Victoria M Rachleff, Meei-Li Huang, Keith R Jerome, Alexander Greninger |
| EPI_ISL_490988 | Mayo Clinic Laboratories | UW Virology Lab | Pavitra Roychoudhury, Hong Xie, Lasata Shrestha, Amin Addetia, Truong Nguyen, Victoria M Rachleff, Meei-Li Huang, Keith R Jerome, Alexander Greninger |
| EPI_ISL_490989, EPI_ISL_490990, EPI_ISL_490991, EPI_ISL_490992, EPI_ISL_490993, EPI_ISL_490994, EPI_ISL_490995, EPI_ISL_490996 | UW Virology Lab | UW Virology Lab | Pavitra Roychoudhury, Hong Xie, Lasata Shrestha, Amin Addetia, Truong Nguyen, Victoria M Rachleff, Meei-Li Huang, Keith R Jerome, Alexander Greninger |
| EPI_ISL_490997, EPI_ISL_490998, EPI_ISL_490999, EPI_ISL_491000 | Mayo Clinic Laboratories | UW Virology Lab | Pavitra Roychoudhury, Hong Xie, Lasata Shrestha, Amin Addetia, Truong Nguyen, Victoria M Rachleff, Meei-Li Huang, Keith R Jerome, Alexander Greninger |
| EPI_ISL_491001, EPI_ISL_491002, EPI_ISL_491003 | UW Virology Lab | UW Virology Lab | Pavitra Roychoudhury, Hong Xie, Lasata Shrestha, Amin Addetia, Truong Nguyen, Victoria M Rachleff, Meei-Li Huang, Keith R Jerome, Alexander Greninger |
| EPI_ISL_491004 | Mayo Clinic Laboratories | UW Virology Lab | Pavitra Roychoudhury, Hong Xie, Lasata Shrestha, Amin Addetia, Truong Nguyen, Victoria M Rachleff, Meei-Li Huang, Keith R Jerome, Alexander Greninger |
| EPI_ISL_491005, EPI_ISL_491006, EPI_ISL_491007, EPI_ISL_491009, EPI_ISL_491010, EPI_ISL_491011, EPI_ISL_491012, EPI_ISL_491013, EPI_ISL_491014, EPI_ISL_491015, EPI_ISL_491016, EPI_ISL_491017, EPI_ISL_491018, EPI_ISL_491019, EPI_ISL_491020, EPI_ISL_491021, EPI_ISL_491022, EPI_ISL_491023, EPI_ISL_491024, EPI_ISL_491025, EPI_ISL_491026, EPI_ISL_491027, EPI_ISL_491028, EPI_ISL_491029, EPI_ISL_491030, EPI_ISL_491031, EPI_ISL_491032, EPI_ISL_491033, EPI_ISL_491034 | | | |
| see above | UW Virology Lab | UW Virology Lab | Pavitra Roychoudhury, Hong Xie, Lasata Shrestha, Amin Addetia, Truong Nguyen, Victoria M Rachleff, Meei-Li Huang, Keith R Jerome, Alexander Greninger |
| EPI_ISL_491036, EPI_ISL_491037, EPI_ISL_491038 | Suceava County Emergency Hospital | "Stefan cel Mare" University Metagenomics Lab | Lobiuc Andrei, Antoniadis Panagiotis et al. |
| EPI_ISL_491039, EPI_ISL_491040, | Suceava County Emergency Hospital | "Stefan cel Mare" University Metagenomics | Lobiuc Andrei et al. |

| | | | |
|---|---|---|---|
| EPI_ISL_491041 | | Lab | |
| EPI_ISL_491042, EPI_ISL_491043 | Suceava County Emergency Hospital | "Stefan cel Mare" University Metagenomics Lab | Lobiuc Andrei, Antoniadis Panagiotis et al. |
| EPI_ISL_491044 | Suceava County Emergency Hospital | "Stefan cel Mare" University Metagenomics Lab | Lobiuc Andrei et al. |
| EPI_ISL_491045, EPI_ISL_491046, EPI_ISL_491047 | Suceava County Emergency Hospital | "Stefan cel Mare" University Metagenomics Lab | Lobiuc Andrei, Antoniadis Panagiotis et al. |
| EPI_ISL_491048 | Suceava County Emergency Hospital | "Stefan cel Mare" University Metagenomics Lab | Lobiuc Andrei et al. |
| EPI_ISL_491049 | Suceava County Emergency Hospital | "Stefan cel Mare" University Metagenomics Lab | Lobiuc Andrei, Antoniadis Panagiotis et al. |
| EPI_ISL_491050, EPI_ISL_491051, EPI_ISL_491052, EPI_ISL_491053, EPI_ISL_491054, EPI_ISL_491055, EPI_ISL_491056, EPI_ISL_491057 | Suceava County Emergency Hospital | "Stefan cel Mare" University Metagenomics Lab | Lobiuc Andrei et al. |
| EPI_ISL_491058, EPI_ISL_491059 | Suceava County Emergency Hospital | "Stefan cel Mare" University Metagenomics Lab | Lobiuc Andrei, Antoniadis Panagiotis et al. |
| EPI_ISL_491060, EPI_ISL_491061 | Suceava County Emergency Hospital | "Stefan cel Mare" University Metagenomics Lab | Lobiuc Andrei et al. |
| EPI_ISL_491062, EPI_ISL_491063, EPI_ISL_491064, EPI_ISL_491065 | Suceava County Emergency Hospital | "Stefan cel Mare" University Metagenomics Lab | Lobiuc Andrei, Antoniadis Panagiotis et al. |
| EPI_ISL_491066, EPI_ISL_491067, EPI_ISL_491068, EPI_ISL_491069, EPI_ISL_491070, EPI_ISL_491071, EPI_ISL_491072 | Suceava County Emergency Hospital | "Stefan cel Mare" University Metagenomics Lab | Lobiuc Andrei et al. |
| EPI_ISL_491073 | Suceava County Emergency Hospital | "Stefan cel Mare" University Metagenomics Lab | Lobiuc Andrei, Antoniadis Panagiotis et al. |
| EPI_ISL_491074 | Suceava County Emergency Hospital | "Stefan cel Mare" University Metagenomics Lab | Lobiuc Andrei et al. |
| EPI_ISL_491075 | Suceava County Emergency Hospital | "Stefan cel Mare" University Metagenomics Lab | Lobiuc Andrei, Antoniadis Panagiotis et al. |
| EPI_ISL_491076 | Suceava County Emergency Hospital | "Stefan cel Mare" University Metagenomics Lab | Lobiuc Andrei et al. |
| EPI_ISL_491077 | Suceava County Emergency Hospital | "Stefan cel Mare" University Metagenomics Lab | Lobiuc Andrei, Antoniadis Panagiotis et al. |
| EPI_ISL_491078 | Suceava County Emergency Hospital | "Stefan cel Mare" University Metagenomics Lab | Lobiuc Andrei et al. |
| EPI_ISL_491079, EPI_ISL_491080 | Suceava County Emergency Hospital | "Stefan cel Mare" University Metagenomics Lab | Lobiuc Andrei, Antoniadis Panagiotis et al. |
| EPI_ISL_491081 | Suceava County Emergency Hospital | "Stefan cel Mare" University Metagenomics Lab | Lobiuc Andrei et al. |
| EPI_ISL_491082 | Suceava County Emergency Hospital | "Stefan cel Mare" University Metagenomics Lab | Lobiuc Andrei, Antoniadis Panagiotis et al. |
| EPI_ISL_491083, EPI_ISL_491084 | Suceava County Emergency Hospital | "Stefan cel Mare" University Metagenomics Lab | Lobiuc Andrei et al. |
| EPI_ISL_491085 | Suceava County Emergency Hospital | "Stefan cel Mare" University Metagenomics Lab | Lobiuc Andrei, Antoniadis Panagiotis et al. |
| EPI_ISL_491086 | Suceava County Emergency Hospital | "Stefan cel Mare" University Metagenomics Lab | Lobiuc Andrei et al. |
| EPI_ISL_491087, EPI_ISL_491088 | Suceava County Emergency Hospital | "Stefan cel Mare" University Metagenomics Lab | Lobiuc Andrei, Antoniadis Panagiotis et al. |
| EPI_ISL_491089 | Suceava County Emergency Hospital | "Stefan cel Mare" University Metagenomics Lab | Lobiuc Andrei et al. |
| EPI_ISL_491090 | Suceava County Emergency Hospital | "Stefan cel Mare" University Metagenomics Lab | Lobiuc Andrei, Antoniadis Panagiotis et al. |
| EPI_ISL_491091 | Suceava County Emergency Hospital | "Stefan cel Mare" University Metagenomics Lab | Lobiuc Andrei et al. |
| EPI_ISL_491092 | The National Institute of Public Health | State Veterinary Institute Prague | Nagy,A;Jirincova,H;Novakova,L;Trnka,D;Vecerova,J |
| EPI_ISL_491093, EPI_ISL_491094, EPI_ISL_491095 | The National Institute of Public Health | The National Institute of Public Health and State Veterinary Institute Prague | Nagy,A; Jirincova,H; Novakova,L; Trnka,D; Vecerova,J |
| EPI_ISL_491096 | CSIR-CDRI/SGPGI, Lucknow | CSIR-CDRI/SGPGI, Lucknow | Saumya Sarkar, Dharam Veer Singh, Rahul Vishvkarma, Ujjala Ghoshal, Uday Ghoshal, Ravishankar Ramachandran, Tapas Kumar Kundu, Rajender Singh |
| EPI_ISL_491097, EPI_ISL_491098, EPI_ISL_491099, EPI_ISL_491100, EPI_ISL_491101, EPI_ISL_491102, EPI_ISL_491103, EPI_ISL_491104, EPI_ISL_491105, EPI_ISL_491106, EPI_ISL_491107, EPI_ISL_491108, EPI_ISL_491109, EPI_ISL_491110 | | | |
| see above | SC Department of Health and Environmental Control | SC Department of Health and Environmental Control | Flores,H. |
| EPI_ISL_491113, EPI_ISL_491114 | CSIR-CDRI/SGPGI, Lucknow | CSIR-CDRI/SGPGI, Lucknow | Saumya Sarkar, Dharam Veer Singh, Rahul Vishvkarma, Ujjala Ghoshal, Uday Ghoshal, Ravishankar Ramachandran, Tapas Kumar Kundu, Rajender Singh |
| EPI_ISL_491115 | Cicin-Sain Lab | Cicin-Sain Lab | M. Zeeshan Chaudhry, Kathrin Eschke, Yeonsu Kim, Luka Cicin-Sain |
| EPI_ISL_491116 | Oman-National Influenza Center | Biotechnology & OMICs Laboratory | Samira Al-Mahruqi, Abdul Latif Khan, Samiha Al-Kharusi, Adil Khan , Ahmed Al-Rawahi, Sajjad Asaf, Amina Al-Jardani, Hanan Al-Kindi, Intisar Al-Shukri, Ahlam Al-Amri, Aisha Al-Amri, Aisha Al-Busaidi, Adil Al-Wahaibi, Seif Al-Abri, Ahmed Al-Harrasi |
| EPI_ISL_491117 | The National Institute of Public Health | The National Institute of Public Health and State Veterinary Institute Prague | Nagy,A;Jirincova,H;Novakova,L;Trnka,D;Vecerova,J |
| EPI_ISL_491118 | The National Institute of Public Health | The National Institute of Public Health and State Veterinary Institute Prague | Nagy,A;Jirincova,H;Novakova,L;Trnka,D;Vecerova,J |
| EPI_ISL_491119 | The National Institute of Public Health | The National Institute of Public Health and State Veterinary Institute Prague | Nagy,A;Jirincova,H;Novakova,L;Trnka,D;Vecerova,J |
| EPI_ISL_491121, EPI_ISL_491122, EPI_ISL_491123, EPI_ISL_491124, EPI_ISL_491125, EPI_ISL_491126, EPI_ISL_491127, EPI_ISL_491128, EPI_ISL_491129, EPI_ISL_491130, EPI_ISL_491131, EPI_ISL_491132 | | | |
| see above | Oman-National Influenza Center | Biotechnology & OMICs Laboratory | Samira Al-Mahruqi, Abdul Latif Khan, Samiha Al-Kharusi, Adil Khan , Ahmed Al-Rawahi, Sajjad Asaf, Amina Al-Jardani, Hanan Al-Kindi, Intisar Al-Shukri, Ahlam Al-Amri, Aisha Al-Amri, Aisha Al-Busaidi, Adil Al-Wahaibi, Seif Al-Abri, Ahmed Al-Harrasi |
| EPI_ISL_491133, EPI_ISL_491134, EPI_ISL_491135, EPI_ISL_491136, EPI_ISL_491137, EPI_ISL_491138, EPI_ISL_491139, EPI_ISL_491140, EPI_ISL_491141, EPI_ISL_491142, EPI_ISL_491143, EPI_ISL_491144, EPI_ISL_491145 | | | |
| see above | Oman-National Influenza Center | Biotechnology & OMICs Laboratory | Samiha Al-Kharusi, Sajjad Asaf, Abdul Latif Khan, Samira Al-Mahruqi, Adil Khan, Ahmed Al-Rawahi, Amina Al-Jardani, Hanan Al-Kindi, Intisar Al-Shukri, Ahlam Al-Amri, Aisha Al-Amri, Aisha Al-Busaidi, Adil Al-Wahaibi, Seif Al-Abri, Ahmed Al-Harrasi |
| EPI_ISL_491146, EPI_ISL_491147, EPI_ISL_491148, EPI_ISL_491149, EPI_ISL_491150, EPI_ISL_491151, EPI_ISL_491152, EPI_ISL_491153, EPI_ISL_491154, EPI_ISL_491155, EPI_ISL_491156, EPI_ISL_491157, EPI_ISL_491158 | | | |
| see above | Oman-National Influenza Center | Biotechnology & OMICs Laboratory | Abdul Latif Khan, Samira Al-Mahruqi, Ahmed Al-Harrasi, Samiha Al-Kharusi, Adil Khan, Ahmed Al-Rawahi, Sajjad Asaf, Amina Al-Jardani, Hanan Al-Kindi, Intisar Al-Shukri, Ahlam Al-Amri, Aisha Al-Amri, Aisha Al-Busaidi, Adil Al-Wahaibi, Seif Al-Abri. |
| EPI_ISL_491159, EPI_ISL_491160, EPI_ISL_491161, EPI_ISL_491162, EPI_ISL_491163, EPI_ISL_491164, EPI_ISL_491165, EPI_ISL_491166, EPI_ISL_491167, EPI_ISL_491168, EPI_ISL_491169, EPI_ISL_491170, EPI_ISL_491171 | | | |
| see above | Oman-National Influenza Center | Biotechnology & OMICs Laboratory | Sajjad Asaf, Samiha Al-Kharusi, Ahmed Al-Harrasi, Samira Al-Mahruqi, Adil Khan, Ahmed Al-Rawahi, Abdul Latif Khan, Amina Al-Jardani, Hanan Al-Kindi, Intisar Al-Shukri, Ahlam Al-Amri, Aisha Al-Amri, Aisha Al-Busaidi, Adil Al-Wahaibi, Seif Al-Abri. |
| EPI_ISL_491172 | Laboratorio de Referencia Nacional de Virus Respiratorio. Instituto Nacional de Salud Perú | Laboratorio de Referencia Nacional de Biotecnología y Biología Molecular. Instituto Nacional de Salud Perú. | Carlos Padilla Rojas, Karolyn Chozo Vega, Priscila Lope Pari, Omar Caceres Rey, Marco Galarza Perez, Maribel Huaringa Nuñez, Johanna Balbuena Torres, Henri Bailon Calderon, Nancy Rojas Serrano |
| EPI_ISL_491174, EPI_ISL_491175, EPI_ISL_491176, EPI_ISL_491177, EPI_ISL_491178, EPI_ISL_491179, EPI_ISL_491180, EPI_ISL_491181, EPI_ISL_491182, EPI_ISL_491183, EPI_ISL_491184, EPI_ISL_491185, EPI_ISL_491186, EPI_ISL_491187, EPI_ISL_491188, EPI_ISL_491189, EPI_ISL_491190, EPI_ISL_491191, EPI_ISL_491192, EPI_ISL_491193, EPI_ISL_491194, EPI_ISL_491195, EPI_ISL_491196, EPI_ISL_491197, EPI_ISL_491198, EPI_ISL_491199, EPI_ISL_491200, EPI_ISL_491201 | | | |
| see above | Instituto Gulbenkian de Ciência | Instituto Gulbenkian de Ciência | João Costa, Cathy Paulino, Joao Sobral, Susana Ladeiro, Ricardo Leite |
| EPI_ISL_491202, EPI_ISL_491203, EPI_ISL_491204, EPI_ISL_491205, EPI_ISL_491206, EPI_ISL_491207, EPI_ISL_491208, EPI_ISL_491209, EPI_ISL_491210, EPI_ISL_491211, EPI_ISL_491212, EPI_ISL_491213, EPI_ISL_491214, EPI_ISL_491215, EPI_ISL_491216, EPI_ISL_491217, EPI_ISL_491218, EPI_ISL_491219, EPI_ISL_491220, EPI_ISL_491221, EPI_ISL_491222, EPI_ISL_491223, EPI_ISL_491224, EPI_ISL_491225, | | | |

| | | | |
|---|---|---|---|
| EPI_ISL_491226, EPI_ISL_491227, EPI_ISL_491228, EPI_ISL_491229, EPI_ISL_491230 | | | |
| see above | Instituto Gulbenkian de Ciência | Instituto Gulbenkian de Ciência | Cathy Paulino, Joao Sobral, Susana Ladeiro, João Costa, Ricardo Leite |
| EPI_ISL_491231, EPI_ISL_491232, EPI_ISL_491233, EPI_ISL_491234, EPI_ISL_491235, EPI_ISL_491236, EPI_ISL_491237, EPI_ISL_491238, EPI_ISL_491239, EPI_ISL_491240, EPI_ISL_491241, EPI_ISL_491242, EPI_ISL_491243, EPI_ISL_491244, EPI_ISL_491245, EPI_ISL_491246, EPI_ISL_491247, EPI_ISL_491248, EPI_ISL_491249, EPI_ISL_491250, EPI_ISL_491251, EPI_ISL_491252, EPI_ISL_491253, EPI_ISL_491254, EPI_ISL_491255, EPI_ISL_491256, EPI_ISL_491257, EPI_ISL_491258, EPI_ISL_491259, EPI_ISL_491260, EPI_ISL_491261, EPI_ISL_491262, EPI_ISL_491263, EPI_ISL_491264 | | | |
| see above | Instituto Gulbenkian de Ciência | Instituto Gulbenkian de Ciência | Joao Sobral, Susana Ladeiro, João Costa, Cathy Paulino, Ricardo Leite |
| EPI_ISL_491265, EPI_ISL_491266, EPI_ISL_491267, EPI_ISL_491268, EPI_ISL_491269, EPI_ISL_491270, EPI_ISL_491271, EPI_ISL_491272, EPI_ISL_491273, EPI_ISL_491274, EPI_ISL_491275, EPI_ISL_491276, EPI_ISL_491277, EPI_ISL_491278, EPI_ISL_491279, EPI_ISL_491280, EPI_ISL_491281, EPI_ISL_491282, EPI_ISL_491283, EPI_ISL_491284, EPI_ISL_491285, EPI_ISL_491286, EPI_ISL_491287, EPI_ISL_491288, EPI_ISL_491289, EPI_ISL_491290, EPI_ISL_491291, EPI_ISL_491292, EPI_ISL_491293, EPI_ISL_491294, EPI_ISL_491295, EPI_ISL_491296, EPI_ISL_491297 | | | |
| see above | Instituto Gulbenkian de Ciência | Instituto Gulbenkian de Ciência | Susana Ladeiro, João Costa, Cathy Paulino, Joao Sobral, Ricardo Leite |
| EPI_ISL_491298 | Rural Health Unit - Calauan, Laguna | Research Institute for Tropical Medicine | Tujan, M.A.A., Onza,O.J.T., Polotan,F.G.M., Medado,I.A.P., Bautista,C.T., Brunker, K., Mercado,E.S., Manalo, D.L., Demetria, C.S. |
| EPI_ISL_491299, EPI_ISL_491300, EPI_ISL_491301, EPI_ISL_491302, EPI_ISL_491303, EPI_ISL_491304, EPI_ISL_491305, EPI_ISL_491306, EPI_ISL_491307, EPI_ISL_491308, EPI_ISL_491309, EPI_ISL_491310, EPI_ISL_491311, EPI_ISL_491312, EPI_ISL_491313, EPI_ISL_491314, EPI_ISL_491315, EPI_ISL_491316, EPI_ISL_491317, EPI_ISL_491318, EPI_ISL_491319, EPI_ISL_491320, EPI_ISL_491321, EPI_ISL_491322, EPI_ISL_491323, EPI_ISL_491324, EPI_ISL_491325, EPI_ISL_491326, EPI_ISL_491327, EPI_ISL_491328, EPI_ISL_491329, EPI_ISL_491330, EPI_ISL_491331, EPI_ISL_491332, EPI_ISL_491333, EPI_ISL_491334, EPI_ISL_491335, EPI_ISL_491336, EPI_ISL_491337, EPI_ISL_491338, EPI_ISL_491339, EPI_ISL_491340, EPI_ISL_491341, EPI_ISL_491342, EPI_ISL_491343, EPI_ISL_491344, EPI_ISL_491345, EPI_ISL_491346, EPI_ISL_491347, EPI_ISL_491348, EPI_ISL_491349, EPI_ISL_491350, EPI_ISL_491351, EPI_ISL_491352, EPI_ISL_491353, EPI_ISL_491354, EPI_ISL_491355, EPI_ISL_491356, EPI_ISL_491357, EPI_ISL_491358, EPI_ISL_491359, EPI_ISL_491360, EPI_ISL_491361, EPI_ISL_491362, EPI_ISL_491363, EPI_ISL_491364, EPI_ISL_491365, EPI_ISL_491366, EPI_ISL_491367, EPI_ISL_491368, EPI_ISL_491369, EPI_ISL_491370, EPI_ISL_491371, EPI_ISL_491372, EPI_ISL_491373, EPI_ISL_491374, EPI_ISL_491375, EPI_ISL_491376, EPI_ISL_491377, EPI_ISL_491378, EPI_ISL_491379, EPI_ISL_491380, EPI_ISL_491381, EPI_ISL_491382, EPI_ISL_491383, EPI_ISL_491384, EPI_ISL_491385, EPI_ISL_491386, EPI_ISL_491387, EPI_ISL_491388, EPI_ISL_491389, EPI_ISL_491390, EPI_ISL_491391, EPI_ISL_491392, EPI_ISL_491393, EPI_ISL_491394, EPI_ISL_491395, EPI_ISL_491396, EPI_ISL_491397, EPI_ISL_491398, EPI_ISL_491399, EPI_ISL_491400, EPI_ISL_491401, EPI_ISL_491402, EPI_ISL_491403, EPI_ISL_491404, EPI_ISL_491405, EPI_ISL_491406, EPI_ISL_491407, EPI_ISL_491408, EPI_ISL_491409, EPI_ISL_491410, EPI_ISL_491411, EPI_ISL_491412, EPI_ISL_491413, EPI_ISL_491414, EPI_ISL_491415, EPI_ISL_491416, EPI_ISL_491417, EPI_ISL_491418, EPI_ISL_491419, EPI_ISL_491420, EPI_ISL_491421, EPI_ISL_491422, EPI_ISL_491423, EPI_ISL_491424, EPI_ISL_491425, EPI_ISL_491426 | | | |
| see above | University of Wisconsin-Madison AIDS Vaccine Research Laboratories | University of Wisconsin-Madison AIDS Vaccine Research Laboratories | Gage Moreno, Katarina Braun, et al. AIDS Vaccine Research Laboratories |
| EPI_ISL_491427 | Laboratorio de Referencia Nacional de Virus Respiratorio. Instituto Nacional de Salud Perú | Laboratorio de Referencia Nacional de Biotecnología y Biología Molecular. Instituto Nacional de Salud Perú | Carlos Padilla Rojas, Karolyn Chozo Vega, Priscila Lope Pari, Omar Caceres Rey, Marco Galarza Perez, Maribel Huaringa Nuñez, Johanna Balbuena Torres, Henri Bailon Calderon, Nancy Rojas Serrano |
| EPI_ISL_491428 | Laboratorio de Referencia Nacional de Virus Respiratorio. Instituto Nacional de Salud Perú | Laboratorio de Referencia Nacional de Biotecnología y Biología Molecular. Instituto Nacional de Salud Perú | Carlos Padilla Rojas, Karolyn Vega Chozo, Priscila Lope Pari, Omar Caceres Rey, Marco Galarza Perez, Maribel Huaringa Nuñez, Johanna Balbuena Torres, Henri Bailon Calderon, Nancy Rojas Serrano. |
| EPI_ISL_491429 | Laboratorio de Referencia Nacional de Virus Respiratorio. Instituto Nacional de Salud Perú | Laboratorio de Referencia Nacional de Biotecnología y Biología Molecular. Instituto Nacional de Salud Perú | Carlos Padilla Rojas, Karolyn Vega Chozo, Priscila Lope Pari, Omar Caceres Rey, Marco Galarza Perez, Maribel Huaringa Nuñez, Johanna Balbuena Torrez, Henri Bailon Calderon, Nancy Rojas Serrano |
| EPI_ISL_491430 | Laboratorio de Referencia Nacional de Virus Respiratorio. Instituto Nacional de Salud Perú | Laboratorio de Referencia Nacional de Biotecnología y Biología Molecular. Instituto Nacional de Salud Perú | Carlos Padilla Rojas, Karolyn Vega Chozo, Priscila Lope Pari, Omar Caceres Rey, Marco Galarza Perez, Maribel Huaringa Nuñez, Johanna Balbuena Torres, Henri Bailon Calderon, Nancy Rojas Serrano. |
| EPI_ISL_491431 | Laboratorio de Referencia Nacional de Virus Respiratorio. Instituto Nacional de Salud. Perú | Laboratorio de Referencia Nacional de Biotecnología y Biología Molecular. Instituto Nacional de Salud.Perú | Carlos Padilla Rojas, Karolyn Vega Chozo, Priscila Lope Pari, Omar Caceres Rey, Marco Galarza Perez, Maribel Huaringa Nuñez, Johanna Balbuena Torres, Henri Bailon Calderon, Nancy Rojas Serrano. |
| EPI_ISL_491432 | Laboratorio de Referencia Nacional de Virus Respiratorio. Instituto Nacional de Salud.Perú | Laboratorio de Referencia Nacional de Biotecnología y Biología Molecular. Instituto Nacional de Salud.Perú | Carlos Padilla Rojas, Karolyn Vega Chozo, Priscila Lope Pari, Omar Caceres Rey, Marco Galarza Perez, Maribel Huaringa Nuñez, Johanna Balbuena Torres, Henri Bailon Calderon, Nancy Rojas Serrano. |
| EPI_ISL_491433, EPI_ISL_491434, EPI_ISL_491435 | Laboratorio de Referencia Nacional de Virus Respiratorio. Instituto Nacional de Salud Perú | Laboratorio de Referencia Nacional de Biotecnología y Biología Molecular. Instituto Nacional de Salud Perú | Carlos Padilla Rojas, Karolyn Vega Chozo, Priscila Lope Pari, Omar Caceres Rey, Marco Galarza Perez, Maribel Huaringa Nuñez, Johanna Balbuena Torres, Henri Bailon Calderon, Nancy Rojas Serrano. |
| EPI_ISL_491436 | Laboratorio de Referencia Nacional de Virus Respiratorio. Instituto Nacional de Salud Perú | Laboratorio de Referencia Nacional de Biotecnología y Biología Molecular. Instituto Nacional de Salud Perú | Carlos Padilla Rojas, Karolyn Vega Chozo, Priscila Lope Pari, Omar Caceres Rey, Marco Galarza Perez, Maribel Huaringa Nuñez, Johanna Balbuena Torres, Henri Bailon Calderon, Nancy Rojas Serrano |
| EPI_ISL_492181, EPI_ISL_492182 | University of Arkansas for Medical Sciences (UAMS) | Department of Biomedical Informatics, University of Arkansas for Medical Sciences (UAMS) | Piroon Jenjaroenpun, David W Ussery, Thidathip Wongsurawat |
| EPI_ISL_512774 | Genomic Research Lab, BCSIR | Bangladesh Council of Scientific and Industrial Research | Abu Sayeed Mohammad Mahmud, Mohammad Samir Uzzaman, Eshrar Osman, Md. Ahasan Habib, Shahina Akter, Tanjina Akhter Banu, Md. Murshed Hasan Sarkar, Barna Goswami, Iffat Jahan, Md. Saddam Hossain, Tasnim Nafisa, Md. Maruf Ahmed Molla, Mahmuda Yeasmin, Asish Kumar Ghosh, A. K. M. Shamsuzzaman, Sheikh Md. Selim Al Din, Utpal Chandra Ray, Salek Ahmed Sajib, Md. Salim Khan |
| EPI_ISL_513298, EPI_ISL_513299, EPI_ISL_513300, EPI_ISL_513301, EPI_ISL_513302, EPI_ISL_513303, EPI_ISL_513304, EPI_ISL_513305, EPI_ISL_513306, EPI_ISL_513307 | Department of Infection Prevention and Infectious Diseases, University Hospital Regensburg | University Hospital Regensburg | Fritsch,J., Holzmann,T., Schneider-Brachert,W. |

We gratefully acknowledge the following Authors from the Originating laboratories responsible for obtaining the specimens, as well as the Submitting laboratories where the genome data were generated and shared via GISAID, on which this research is based.

All Submitters of data may be contacted directly via www.gisaid.org

| Accession ID | Originating Laboratory | Submitting Laboratory | Authors |
|---|---|---|---|
| EPI_ISL_508122, EPI_ISL_508123, EPI_ISL_508124, EPI_ISL_508125, EPI_ISL_508126, EPI_ISL_508127, EPI_ISL_508128, EPI_ISL_508129, EPI_ISL_508130, EPI_ISL_508131, EPI_ISL_508132, EPI_ISL_508133, EPI_ISL_508134, EPI_ISL_508135, EPI_ISL_508136, EPI_ISL_508137, EPI_ISL_508138, EPI_ISL_508139, EPI_ISL_508140, EPI_ISL_508141, EPI_ISL_508142, EPI_ISL_508143, EPI_ISL_508144, EPI_ISL_508145 | | | |
| see above | SA Pathology | SA Pathology | Lex Leong, Chuan Kok Lim, Mark Turra, Ivan Bastian, Geoff Higgins |
| EPI_ISL_508156, EPI_ISL_508157, EPI_ISL_508158, EPI_ISL_508159, EPI_ISL_508160, EPI_ISL_508161, EPI_ISL_508162, EPI_ISL_508163, EPI_ISL_508164, EPI_ISL_508165, EPI_ISL_508166, EPI_ISL_508167, EPI_ISL_508168, EPI_ISL_508169, EPI_ISL_508170, EPI_ISL_508171, EPI_ISL_508172, EPI_ISL_508173, EPI_ISL_508174, EPI_ISL_508175, EPI_ISL_508176, EPI_ISL_508177, EPI_ISL_508178, EPI_ISL_508179, EPI_ISL_508180, EPI_ISL_508181, EPI_ISL_508182, EPI_ISL_508183, EPI_ISL_508184, EPI_ISL_508185, EPI_ISL_508186, EPI_ISL_508187, EPI_ISL_508188, EPI_ISL_508189, EPI_ISL_508190, EPI_ISL_508191, EPI_ISL_508192, EPI_ISL_508193, EPI_ISL_508194, EPI_ISL_508195, EPI_ISL_508196, EPI_ISL_508197, EPI_ISL_508198, EPI_ISL_508199, EPI_ISL_508200, EPI_ISL_508201, EPI_ISL_508202, EPI_ISL_508203, EPI_ISL_508204, EPI_ISL_508205, EPI_ISL_508206 | | | |
| see above | All india institute of Medical Sciences Rishikesh | National Institute of Biomedical Genomics | Arindam Maitra, Deepjyoti Kalita, Amit Mangla, Ravi Kant, Saumitra Das |
| EPI_ISL_508207, EPI_ISL_508208, EPI_ISL_508209, EPI_ISL_508210, EPI_ISL_508211, EPI_ISL_508212, EPI_ISL_508213, EPI_ISL_508214, EPI_ISL_508215, EPI_ISL_508216, EPI_ISL_508217, EPI_ISL_508218, EPI_ISL_508219, EPI_ISL_508220, EPI_ISL_508221, EPI_ISL_508222, EPI_ISL_508223, EPI_ISL_508224, EPI_ISL_508225, EPI_ISL_508226, EPI_ISL_508227, EPI_ISL_508228, EPI_ISL_508229, EPI_ISL_508230, EPI_ISL_508231, EPI_ISL_508232, EPI_ISL_508233, EPI_ISL_508234, EPI_ISL_508235, EPI_ISL_508236, EPI_ISL_508237, EPI_ISL_508238, EPI_ISL_508239, EPI_ISL_508240, EPI_ISL_508241, EPI_ISL_508242, EPI_ISL_508243, EPI_ISL_508244, EPI_ISL_508245, EPI_ISL_508246, EPI_ISL_508247, EPI_ISL_508248, EPI_ISL_508249, EPI_ISL_508250, EPI_ISL_508251, EPI_ISL_508252, EPI_ISL_508253, EPI_ISL_508254, EPI_ISL_508255, EPI_ISL_508256, EPI_ISL_508257, EPI_ISL_508258, EPI_ISL_508259, EPI_ISL_508260, EPI_ISL_508261, EPI_ISL_508262, EPI_ISL_508263, EPI_ISL_508264, EPI_ISL_508265, EPI_ISL_508266, EPI_ISL_508267, EPI_ISL_508268, EPI_ISL_508269, EPI_ISL_508270, EPI_ISL_508271, EPI_ISL_508272, EPI_ISL_508273, EPI_ISL_508274, EPI_ISL_508275, EPI_ISL_508276, EPI_ISL_508277, EPI_ISL_508278, EPI_ISL_508279, EPI_ISL_508280, EPI_ISL_508281, EPI_ISL_508282, EPI_ISL_508283, EPI_ISL_508284, EPI_ISL_508285, EPI_ISL_508286 | | | |
| see above | Government Medical College | National Institute of Biomedical Genomics | Arindam Maitra, Jyoti Iravane, Dhaval Khatri, Maitrik Dave, Saumitra Das |
| EPI_ISL_508287, EPI_ISL_508288, EPI_ISL_508289, EPI_ISL_508290, EPI_ISL_508291, EPI_ISL_508292, EPI_ISL_508293, EPI_ISL_508294, EPI_ISL_508295, EPI_ISL_508296, EPI_ISL_508297, EPI_ISL_508298, EPI_ISL_508299, EPI_ISL_508300, EPI_ISL_508301, EPI_ISL_508302, EPI_ISL_508303, EPI_ISL_508304, EPI_ISL_508305, EPI_ISL_508306, EPI_ISL_508307, EPI_ISL_508308, EPI_ISL_508309, EPI_ISL_508310, EPI_ISL_508311, EPI_ISL_508312, EPI_ISL_508313, EPI_ISL_508314, EPI_ISL_508315, EPI_ISL_508316, EPI_ISL_508317, EPI_ISL_508318, EPI_ISL_508319, EPI_ISL_508320, EPI_ISL_508321, EPI_ISL_508322, EPI_ISL_508323, EPI_ISL_508324, EPI_ISL_508325, EPI_ISL_508326, EPI_ISL_508327, EPI_ISL_508328, EPI_ISL_508329, EPI_ISL_508330, EPI_ISL_508331, EPI_ISL_508332, EPI_ISL_508333, EPI_ISL_508334, EPI_ISL_508335, EPI_ISL_508336, EPI_ISL_508337 | | | |
| see above | Indian Institute of Science | National Institute of Biomedical Genomics | Arindam Maitra, Bharath K Sundararaj, Harsha Raheja, N. Srinivasan, Deepak K Saini, Amit Singh, Saumitra Das |
| EPI_ISL_508338, EPI_ISL_508339, EPI_ISL_508340, EPI_ISL_508341, EPI_ISL_508342, EPI_ISL_508343, EPI_ISL_508344, EPI_ISL_508345, EPI_ISL_508346, EPI_ISL_508347, EPI_ISL_508348, EPI_ISL_508349, EPI_ISL_508350, EPI_ISL_508351, EPI_ISL_508352, EPI_ISL_508353, EPI_ISL_508354, EPI_ISL_508355, EPI_ISL_508356, EPI_ISL_508357, EPI_ISL_508358, EPI_ISL_508359, EPI_ISL_508360, EPI_ISL_508361, EPI_ISL_508362, EPI_ISL_508363, EPI_ISL_508364, EPI_ISL_508365, EPI_ISL_508366, EPI_ISL_508367, EPI_ISL_508368, EPI_ISL_508369, EPI_ISL_508370, EPI_ISL_508371, EPI_ISL_508372, EPI_ISL_508373, EPI_ISL_508374, EPI_ISL_508375, EPI_ISL_508376, EPI_ISL_508377, EPI_ISL_508378, EPI_ISL_508379, EPI_ISL_508380, EPI_ISL_508381, EPI_ISL_508382, EPI_ISL_508383, EPI_ISL_508384, EPI_ISL_508385, EPI_ISL_508386, EPI_ISL_508387, EPI_ISL_508388, EPI_ISL_508389, EPI_ISL_508390, EPI_ISL_508391, EPI_ISL_508392, EPI_ISL_508393, EPI_ISL_508394, EPI_ISL_508395, EPI_ISL_508396, EPI_ISL_508397, EPI_ISL_508398, EPI_ISL_508399, EPI_ISL_508400, EPI_ISL_508401, EPI_ISL_508402, EPI_ISL_508403, EPI_ISL_508404, EPI_ISL_508405, EPI_ISL_508406, EPI_ISL_508407, EPI_ISL_508408, EPI_ISL_508409, EPI_ISL_508410, EPI_ISL_508411, EPI_ISL_508412, EPI_ISL_508413, EPI_ISL_508414 | | | |
| see above | Institute of Post Graduate Medical Education & Research | National Institute of Biomedical Genomics | Arindam Maitra, Aritra Biswas, Jayeeta Haldar, Raja Ray, Monimoy Banerjee, Saumitra Das |
| EPI_ISL_508415, EPI_ISL_508416, EPI_ISL_508417, EPI_ISL_508418, EPI_ISL_508419, EPI_ISL_508420, EPI_ISL_508421, EPI_ISL_508422 | Maulana Azad Medical College | National Institute of Biomedical Genomics | Arindam Maitra, Sonal Saxena, Vikas Manchanda, Oves Siddiqui, Saumitra Das |
| EPI_ISL_508423, EPI_ISL_508424, EPI_ISL_508425, EPI_ISL_508426, EPI_ISL_508427, EPI_ISL_508428, EPI_ISL_508429, EPI_ISL_508430, EPI_ISL_508431, EPI_ISL_508432, EPI_ISL_508433, EPI_ISL_508434, EPI_ISL_508435, EPI_ISL_508436, EPI_ISL_508437, EPI_ISL_508438, EPI_ISL_508439, EPI_ISL_508440 | | | |
| see above | Mahatma Gandhi Institute of Medical Sciences | National Institute of Biomedical Genomics | Arindam Maitra, Vijayshri Deotale, Rahul Narang, Deepashri Maraskolhe, Saumitra Das |
| EPI_ISL_508441, EPI_ISL_508442, EPI_ISL_508443, EPI_ISL_508444, EPI_ISL_508445, EPI_ISL_508446, EPI_ISL_508447, EPI_ISL_508448, EPI_ISL_508449, EPI_ISL_508450, EPI_ISL_508451, EPI_ISL_508452, EPI_ISL_508453, EPI_ISL_508454, EPI_ISL_508455, EPI_ISL_508456, EPI_ISL_508457, EPI_ISL_508458, EPI_ISL_508459, EPI_ISL_508460, EPI_ISL_508461, EPI_ISL_508462, EPI_ISL_508463, EPI_ISL_508464, EPI_ISL_508465, EPI_ISL_508466, EPI_ISL_508467, EPI_ISL_508468, EPI_ISL_508469, EPI_ISL_508470, EPI_ISL_508471, EPI_ISL_508472, EPI_ISL_508473, EPI_ISL_508474, EPI_ISL_508475, EPI_ISL_508476, EPI_ISL_508477, EPI_ISL_508478, EPI_ISL_508479, EPI_ISL_508480, EPI_ISL_508481, EPI_ISL_508482, EPI_ISL_508483, EPI_ISL_508484, EPI_ISL_508485, EPI_ISL_508486, EPI_ISL_508487, EPI_ISL_508488 | | | |
| see above | ICMR-National Institute of Cholera and Enteric Diseases | National Institute of Biomedical Genomics | Arindam Maitra, Mamta Chawla Sarkar, Sreedhar Chinnaswamy, Hasina Banu, Ananya Chatterjee, Shanta Dutta, Saumitra Das |
| EPI_ISL_508489, EPI_ISL_508490, EPI_ISL_508491, EPI_ISL_508492, EPI_ISL_508493, EPI_ISL_508494, EPI_ISL_508495, EPI_ISL_508496, EPI_ISL_508497, EPI_ISL_508498, EPI_ISL_508499, EPI_ISL_508500, EPI_ISL_508501, EPI_ISL_508502, EPI_ISL_508503, EPI_ISL_508504, EPI_ISL_508505, EPI_ISL_508506, EPI_ISL_508507, EPI_ISL_508508, EPI_ISL_508509 | | | |
| see above | Translational Health Science and Technology Institute | National Institute of Biomedical Genomics | Arindam Maitra, Guruprasad Medigeshi, Sharanabasava Patil, Anbalagan Ananthraj, Madhu Pareek, Imran Khan, Gagandeep Kang, Saumitra Das |
| EPI_ISL_508605, EPI_ISL_508606, EPI_ISL_508607, EPI_ISL_508608 | SA Pathology | SA Pathology | Lex Leong, Chuan Kok Lim, Mark Turra, Ivan Bastian, Geoff Higgins |
| EPI_ISL_508615, EPI_ISL_508616, EPI_ISL_508617, EPI_ISL_508618, EPI_ISL_508619, EPI_ISL_508620, EPI_ISL_508621, EPI_ISL_508622, EPI_ISL_508623, EPI_ISL_508624, EPI_ISL_508625, EPI_ISL_508626, EPI_ISL_508627, EPI_ISL_508628, EPI_ISL_508629, EPI_ISL_508630, EPI_ISL_508631, EPI_ISL_508632, EPI_ISL_508633, EPI_ISL_508634, EPI_ISL_508635, EPI_ISL_508636, EPI_ISL_508637, EPI_ISL_508638, EPI_ISL_508639, EPI_ISL_508640, EPI_ISL_508641, EPI_ISL_508642, EPI_ISL_508643, EPI_ISL_508644, EPI_ISL_508645, EPI_ISL_508646, EPI_ISL_508647, EPI_ISL_508648, EPI_ISL_508649, EPI_ISL_508650, EPI_ISL_508651, EPI_ISL_508652, EPI_ISL_508653, EPI_ISL_508654, EPI_ISL_508655, EPI_ISL_508656, EPI_ISL_508657, EPI_ISL_508658, EPI_ISL_508659, EPI_ISL_508660, EPI_ISL_508661, EPI_ISL_508662, EPI_ISL_508663, EPI_ISL_508664, EPI_ISL_508665, EPI_ISL_508666, EPI_ISL_508667, EPI_ISL_508668, EPI_ISL_508669, EPI_ISL_508670, EPI_ISL_508671, EPI_ISL_508672, EPI_ISL_508673, EPI_ISL_508674, EPI_ISL_508675, EPI_ISL_508676, EPI_ISL_508677, EPI_ISL_508678, EPI_ISL_508679, EPI_ISL_508680, EPI_ISL_508681, EPI_ISL_508682, EPI_ISL_508683, EPI_ISL_508684, EPI_ISL_508685, EPI_ISL_508686 | | | |
| see above | Departamento de Microbiología, CDB, Hospital Clínic, Barcelona | SeqCOVID-SPAIN consortium/IBV(CSIC) | Andrea Vergara, Mikel Martínez, Elisa Rubio, Jéssica Navero, Aida Peiró and SeqCOVID-SPAIN consortium |
| EPI_ISL_508687, EPI_ISL_508688, EPI_ISL_508689, EPI_ISL_508690, EPI_ISL_508691, EPI_ISL_508692, EPI_ISL_508693, EPI_ISL_508694, EPI_ISL_508695, EPI_ISL_508696, EPI_ISL_508697, EPI_ISL_508698, EPI_ISL_508699, EPI_ISL_508700, EPI_ISL_508701, EPI_ISL_508702, EPI_ISL_508703, EPI_ISL_508704 | | | |
| see above | Institut für Virologie und Epidemiologie der Viruskrankheiten, Universitätsklinikum Tübingen | NGS Competence Center Tübingen, Institut für Medizinische Mikrobiologie und Hygiene, Universitätsklinikum Tübingen | Angel Angelov |
| EPI_ISL_508708, EPI_ISL_508709, EPI_ISL_508710, EPI_ISL_508711, EPI_ISL_508712, EPI_ISL_508713, EPI_ISL_508714, EPI_ISL_508715, EPI_ISL_508716, EPI_ISL_508717, EPI_ISL_508718, EPI_ISL_508719, EPI_ISL_508720, EPI_ISL_508721, EPI_ISL_508722, EPI_ISL_508723, EPI_ISL_508724, EPI_ISL_508725, EPI_ISL_508726, EPI_ISL_508727, EPI_ISL_508728, EPI_ISL_508729, EPI_ISL_508730, EPI_ISL_508731, EPI_ISL_508732, EPI_ISL_508733, EPI_ISL_508734, EPI_ISL_508735, EPI_ISL_508736, EPI_ISL_508737, EPI_ISL_508738, EPI_ISL_508739, EPI_ISL_508740, EPI_ISL_508741, EPI_ISL_508742, EPI_ISL_508743, EPI_ISL_508744, EPI_ISL_508745, EPI_ISL_508746, EPI_ISL_508747, EPI_ISL_508748, EPI_ISL_508749, EPI_ISL_508750, EPI_ISL_508751, EPI_ISL_508752, EPI_ISL_508753, EPI_ISL_508754, EPI_ISL_508755, EPI_ISL_508756, EPI_ISL_508757, EPI_ISL_508758, EPI_ISL_508759, EPI_ISL_508760, EPI_ISL_508761, EPI_ISL_508762, EPI_ISL_508763, EPI_ISL_508765, EPI_ISL_508766, EPI_ISL_508767, EPI_ISL_508768, EPI_ISL_508769, EPI_ISL_508770, EPI_ISL_508771, EPI_ISL_508772, EPI_ISL_508773, EPI_ISL_508774, EPI_ISL_508775, EPI_ISL_508776, EPI_ISL_508777, EPI_ISL_508778, EPI_ISL_508779, EPI_ISL_508780, EPI_ISL_508781, EPI_ISL_508782, EPI_ISL_508783, EPI_ISL_508784, EPI_ISL_508785, EPI_ISL_508786, EPI_ISL_508787, EPI_ISL_508788, EPI_ISL_508789, EPI_ISL_508790, EPI_ISL_508791, EPI_ISL_508792, EPI_ISL_508793, EPI_ISL_508794, EPI_ISL_508795, EPI_ISL_508796, EPI_ISL_508797, EPI_ISL_508798, EPI_ISL_508799, EPI_ISL_508800, EPI_ISL_508801, EPI_ISL_508802, EPI_ISL_508803, EPI_ISL_508804, EPI_ISL_508805, EPI_ISL_508806, EPI_ISL_508807, EPI_ISL_508808, EPI_ISL_508809, EPI_ISL_508810, EPI_ISL_508811, EPI_ISL_508812, EPI_ISL_508813, EPI_ISL_508814, EPI_ISL_508815, EPI_ISL_508816, EPI_ISL_508817, EPI_ISL_508818, EPI_ISL_508819, EPI_ISL_508820, EPI_ISL_508821, EPI_ISL_508822, EPI_ISL_508823, EPI_ISL_508824, EPI_ISL_508825, EPI_ISL_508826, EPI_ISL_508827, EPI_ISL_508828, EPI_ISL_508829, EPI_ISL_508830, EPI_ISL_508831, EPI_ISL_508832, EPI_ISL_508833, EPI_ISL_508834, EPI_ISL_508835, EPI_ISL_508836, EPI_ISL_508837, EPI_ISL_508838, EPI_ISL_508839, EPI_ISL_508840, EPI_ISL_508841, EPI_ISL_508842, EPI_ISL_508843, EPI_ISL_508844, EPI_ISL_508845, EPI_ISL_508846, EPI_ISL_508847, EPI_ISL_508848, EPI_ISL_508849, EPI_ISL_508850, EPI_ISL_508851, EPI_ISL_508852, EPI_ISL_508853, EPI_ISL_508854, EPI_ISL_508855, EPI_ISL_508856, EPI_ISL_508857, EPI_ISL_508858, EPI_ISL_508859, EPI_ISL_508860, EPI_ISL_508861 | | | |
| see above | Florida Bureau of Public Health Laboratories | Florida Bureau of Public Health Laboratories | Sarah Schmedes, Jason Blanton |
| EPI_ISL_508862, EPI_ISL_508863 | Virology Unit, Institut Pasteur de Madagascar | Virology Unit, Institut Pasteur de Madagascar | Christian Ranaivoson, Cara Brook, Norosoa Razanajatovo, Vida Ahyong, Tsiry Randriambolamanantsoa, Michelle Tan, Vololoniaina Raharinosy, Helisoa Razafimanjato, Cristina M. Tato, Joseph L. DeRisi, Soa Fy Andriamandimby, Jean-Michel Heraud |
| EPI_ISL_508864 | Division of Infectious Diseases and Hospital Epidemiology, University Hospital Zürich | Institute of Medical Virology, University of Zurich | Verena Kufner, Maryam Zaheri, Dana Weissberg, Jürg Böni, Silvana K. Rampini, Peter W. Schreiber, Irene A. Abela, Hugo Sax, Aline Wolfensberger, Michael Huber |
| EPI_ISL_508865 | Division of Infectious Diseases and Hospital Epidemiology, University Hospital Zürich | Institute of Medical Virology, University of Zurich | Maryam Zaheri, Verena Kufner, Dana Weissberg, Jürg Böni, Silvana K. Rampini, Peter W. Schreiber, Irene A. Abela, Hugo Sax, Aline Wolfensberger, Michael Huber |
| EPI_ISL_508866 | Division of Infectious Diseases and Hospital Epidemiology, University Hospital Zürich | Institute of Medical Virology, University of Zurich | Verena Kufner, Maryam Zaheri, Dana Weissberg, Jürg Böni, Silvana K. Rampini, Peter W. Schreiber, Irene A. Abela, Hugo Sax, Aline Wolfensberger, Michael Huber |
| EPI_ISL_508867 | Division of Infectious Diseases and Hospital Epidemiology, University Hospital Zürich | Institute of Medical Virology, University of Zurich | Maryam Zaheri, Verena Kufner, Dana Weissberg, Jürg Böni, Silvana K. Rampini, Peter W. Schreiber, Irene A. Abela, Hugo Sax, Aline Wolfensberger, Michael Huber |
| EPI_ISL_508868 | Division of Infectious Diseases and Hospital Epidemiology, University Hospital Zürich | Institute of Medical Virology, University of Zurich | Verena Kufner, Maryam Zaheri, Dana Weissberg, Jürg Böni, Silvana K. Rampini, Peter W. Schreiber, Irene A. Abela, Hugo Sax, Aline Wolfensberger, Michael Huber |
| EPI_ISL_508869 | Division of Infectious Diseases and Hospital Epidemiology, University Hospital Zürich | Institute of Medical Virology, University of Zurich | Maryam Zaheri, Verena Kufner, Dana Weissberg, Jürg Böni, Silvana K. Rampini, Peter W. Schreiber, Irene A. Abela, Hugo Sax, Aline Wolfensberger, Michael Huber |
| EPI_ISL_508870 | Division of Infectious Diseases and Hospital Epidemiology, University Hospital Zürich | Institute of Medical Virology, University of Zurich | Verena Kufner, Maryam Zaheri, Dana Weissberg, Jürg Böni, Silvana K. Rampini, Peter W. Schreiber, Irene A. Abela, Hugo Sax, Aline Wolfensberger, Michael Huber |
| EPI_ISL_508871, EPI_ISL_508872, EPI_ISL_508873, EPI_ISL_508874 | Institut des Agents Infectieux (IAI), Hospices Civils de Lyon | CNR Virus des Infections Respiratoires - France SUD | Antonin Bal, Gregory Destras, Gwendolyne Burfin, Solenne Brun, Carine Moustaud, Raphaelle Lamy, Alexandre Gaymard, Maude Bouscambert-Duchamp, Florence Morfin-Sherpa, Martine Valette, Bruno Lina, Laurence Josset |
| EPI_ISL_508875, EPI_ISL_508876 | Centre Hospitalier de Macon | CNR Virus des Infections Respiratoires - France SUD | Antonin Bal, Gregory Destras, Gwendolyne Burfin, Solenne Brun, Carine Moustaud, Raphaelle Lamy, Alexandre Gaymard, Maude Bouscambert-Duchamp, Florence Morfin-Sherpa, Martine Valette, Bruno Lina, Laurence Josset |
| EPI_ISL_508877 | Institut des Agents Infectieux (IAI), Hospices Civils de Lyon | CNR Virus des Infections Respiratoires - France SUD | Antonin Bal, Gregory Destras, Gwendolyne Burfin, Solenne Brun, Carine Moustaud, Raphaelle Lamy, Alexandre Gaymard, Maude Bouscambert-Duchamp, Florence Morfin-Sherpa, Martine Valette, Bruno Lina, Laurence Josset |
| EPI_ISL_508878 | GH Les Portes du Sud | CNR Virus des Infections Respiratoires - France SUD | Antonin Bal, Gregory Destras, Gwendolyne Burfin, Solenne Brun, Carine Moustaud, Raphaelle Lamy, Alexandre Gaymard, Maude Bouscambert-Duchamp, Florence Morfin-Sherpa, Martine Valette, Bruno Lina, Laurence Josset |
| EPI_ISL_508879, EPI_ISL_508880 | Centre Hospitalier Saint Joseph Saint Luc | CNR Virus des Infections Respiratoires - France SUD | Antonin Bal, Gregory Destras, Gwendolyne Burfin, Solenne Brun, Carine Moustaud, Raphaelle Lamy, Alexandre Gaymard, Maude Bouscambert-Duchamp, Florence Morfin-Sherpa, Martine Valette, Bruno Lina, Laurence Josset |
| EPI_ISL_508881 | Centre Hospitalier de Valence | CNR Virus des Infections Respiratoires - France SUD | Antonin Bal, Gregory Destras, Gwendolyne Burfin, Solenne Brun, Carine Moustaud, Raphaelle Lamy, Alexandre Gaymard, Maude Bouscambert-Duchamp, Florence Morfin-Sherpa, Martine Valette, Bruno Lina, Laurence Josset |
| EPI_ISL_508882, EPI_ISL_508883, EPI_ISL_508884, EPI_ISL_508885, EPI_ISL_508886, EPI_ISL_508887, EPI_ISL_508888, EPI_ISL_508889, EPI_ISL_508890, EPI_ISL_508891, EPI_ISL_508892, EPI_ISL_508893, EPI_ISL_508894, EPI_ISL_508895, EPI_ISL_508896, EPI_ISL_508897, EPI_ISL_508898, EPI_ISL_508899, EPI_ISL_508900, EPI_ISL_508901, EPI_ISL_508902, EPI_ISL_508903, EPI_ISL_508904, EPI_ISL_508905, EPI_ISL_508906, EPI_ISL_508907, EPI_ISL_508908, EPI_ISL_508909, EPI_ISL_508910, EPI_ISL_508911 | | | |

| | | | |
|---|---|---|---|
| see above | Institut des Agents Infectieux (IAI), Hospices Civils de Lyon | CNR Virus des Infections Respiratoires - France SUD | Antonin Bal, Gregory Destras, Gwendolyne Burfin, Solenne Brun, Carine Moustaud, Raphaelle Lamy, Alexandre Gaymard, Maude Bouscambert-Duchamp, Florence Morfin-Sherpa, Martine Valette, Bruno Lina, Laurence Josset |
| EPI_ISL_508912, EPI_ISL_508913, EPI_ISL_508914, EPI_ISL_508915, EPI_ISL_508916, EPI_ISL_508917, EPI_ISL_508918, EPI_ISL_508919, EPI_ISL_508920, EPI_ISL_508921, EPI_ISL_508922, EPI_ISL_508923, EPI_ISL_508924, EPI_ISL_508925, EPI_ISL_508926, EPI_ISL_508927, EPI_ISL_508928, EPI_ISL_508929, EPI_ISL_508930 | | | |
| see above | CNR Virus des Infections Respiratoires - France SUD | CNR Virus des Infections Respiratoires - France SUD | Antonin Bal, Gregory Destras, Gwendolyne Burfin, Solenne Brun, Carine Moustaud, Raphaelle Lamy, Alexandre Gaymard, Maude Bouscambert-Duchamp, Florence Morfin-Sherpa, Martine Valette, Bruno Lina, Laurence Josset |
| EPI_ISL_508931 | L'Hôpital Nord-Ouest Tarare-Grandris | CNR Virus des Infections Respiratoires - France SUD | Antonin Bal, Gregory Destras, Gwendolyne Burfin, Solenne Brun, Carine Moustaud, Raphaelle Lamy, Alexandre Gaymard, Maude Bouscambert-Duchamp, Florence Morfin-Sherpa, Martine Valette, Bruno Lina, Laurence Josset |
| EPI_ISL_508932 | Centre Hospitalier de Villefranche | CNR Virus des Infections Respiratoires - France SUD | Antonin Bal, Gregory Destras, Gwendolyne Burfin, Solenne Brun, Carine Moustaud, Raphaelle Lamy, Alexandre Gaymard, Maude Bouscambert-Duchamp, Florence Morfin-Sherpa, Martine Valette, Bruno Lina, Laurence Josset |
| EPI_ISL_508933 | CNR Virus des Infections Respiratoires - France SUD | CNR Virus des Infections Respiratoires - France SUD | Antonin Bal, Gregory Destras, Gwendolyne Burfin, Solenne Brun, Carine Moustaud, Raphaelle Lamy, Alexandre Gaymard, Maude Bouscambert-Duchamp, Florence Morfin-Sherpa, Martine Valette, Bruno Lina, Laurence Josset |
| EPI_ISL_508934, EPI_ISL_508935, EPI_ISL_508936 | Centre hospitalier Métropole Savoie | CNR Virus des Infections Respiratoires - France SUD | Antonin Bal, Gregory Destras, Gwendolyne Burfin, Solenne Brun, Carine Moustaud, Raphaelle Lamy, Alexandre Gaymard, Maude Bouscambert-Duchamp, Florence Morfin-Sherpa, Martine Valette, Bruno Lina, Laurence Josset |
| EPI_ISL_508937 | CNR Virus des Infections Respiratoires - France SUD | CNR Virus des Infections Respiratoires - France SUD | Antonin Bal, Gregory Destras, Gwendolyne Burfin, Solenne Brun, Carine Moustaud, Raphaelle Lamy, Alexandre Gaymard, Maude Bouscambert-Duchamp, Florence Morfin-Sherpa, Martine Valette, Bruno Lina, Laurence Josset |
| EPI_ISL_508938 | Centre Hospitalier de Bourg en Bresse | CNR Virus des Infections Respiratoires - France SUD | Antonin Bal, Gregory Destras, Gwendolyne Burfin, Solenne Brun, Carine Moustaud, Raphaelle Lamy, Alexandre Gaymard, Maude Bouscambert-Duchamp, Florence Morfin-Sherpa, Martine Valette, Bruno Lina, Laurence Josset |
| EPI_ISL_508939, EPI_ISL_508940 | CNR Virus des Infections Respiratoires - France SUD | CNR Virus des Infections Respiratoires - France SUD | Antonin Bal, Gregory Destras, Gwendolyne Burfin, Solenne Brun, Carine Moustaud, Raphaelle Lamy, Alexandre Gaymard, Maude Bouscambert-Duchamp, Florence Morfin-Sherpa, Martine Valette, Bruno Lina, Laurence Josset |
| EPI_ISL_508941 | Centre Hospitalier de Macon | CNR Virus des Infections Respiratoires - France SUD | Antonin Bal, Gregory Destras, Gwendolyne Burfin, Solenne Brun, Carine Moustaud, Raphaelle Lamy, Alexandre Gaymard, Maude Bouscambert-Duchamp, Florence Morfin-Sherpa, Martine Valette, Bruno Lina, Laurence Josset |
| EPI_ISL_508942 | CNR Virus des Infections Respiratoires - France SUD | CNR Virus des Infections Respiratoires - France SUD | Antonin Bal, Gregory Destras, Gwendolyne Burfin, Solenne Brun, Carine Moustaud, Raphaelle Lamy, Alexandre Gaymard, Maude Bouscambert-Duchamp, Florence Morfin-Sherpa, Martine Valette, Bruno Lina, Laurence Josset |
| EPI_ISL_508943 | Centre Hospitalier de Macon | CNR Virus des Infections Respiratoires - France SUD | Antonin Bal, Gregory Destras, Gwendolyne Burfin, Solenne Brun, Carine Moustaud, Raphaelle Lamy, Alexandre Gaymard, Maude Bouscambert-Duchamp, Florence Morfin-Sherpa, Martine Valette, Bruno Lina, Laurence Josset |
| EPI_ISL_508944 | Centre Hospitalier de Bourg en Bresse | CNR Virus des Infections Respiratoires - France SUD | Antonin Bal, Gregory Destras, Gwendolyne Burfin, Solenne Brun, Carine Moustaud, Raphaelle Lamy, Alexandre Gaymard, Maude Bouscambert-Duchamp, Florence Morfin-Sherpa, Martine Valette, Bruno Lina, Laurence Josset |
| EPI_ISL_508945 | CNR Virus des Infections Respiratoires - France SUD | CNR Virus des Infections Respiratoires - France SUD | Antonin Bal, Gregory Destras, Gwendolyne Burfin, Solenne Brun, Carine Moustaud, Raphaelle Lamy, Alexandre Gaymard, Maude Bouscambert-Duchamp, Florence Morfin-Sherpa, Martine Valette, Bruno Lina, Laurence Josset |
| EPI_ISL_508946 | Centre Hospitalier de Macon | CNR Virus des Infections Respiratoires - France SUD | Antonin Bal, Gregory Destras, Gwendolyne Burfin, Solenne Brun, Carine Moustaud, Raphaelle Lamy, Alexandre Gaymard, Maude Bouscambert-Duchamp, Florence Morfin-Sherpa, Martine Valette, Bruno Lina, Laurence Josset |
| EPI_ISL_508947 | Centre Hospitalier Lucien Hussel | CNR Virus des Infections Respiratoires - France SUD | Antonin Bal, Gregory Destras, Gwendolyne Burfin, Solenne Brun, Carine Moustaud, Raphaelle Lamy, Alexandre Gaymard, Maude Bouscambert-Duchamp, Florence Morfin-Sherpa, Martine Valette, Bruno Lina, Laurence Josset |
| EPI_ISL_508948 | CNR Virus des Infections Respiratoires - France SUD | CNR Virus des Infections Respiratoires - France SUD | Antonin Bal, Gregory Destras, Gwendolyne Burfin, Solenne Brun, Carine Moustaud, Raphaelle Lamy, Alexandre Gaymard, Maude Bouscambert-Duchamp, Florence Morfin-Sherpa, Martine Valette, Bruno Lina, Laurence Josset |
| EPI_ISL_508949, EPI_ISL_508950 | Centre Hospitalier de Macon | CNR Virus des Infections Respiratoires - France SUD | Antonin Bal, Gregory Destras, Gwendolyne Burfin, Solenne Brun, Carine Moustaud, Raphaelle Lamy, Alexandre Gaymard, Maude Bouscambert-Duchamp, Florence Morfin-Sherpa, Martine Valette, Bruno Lina, Laurence Josset |
| EPI_ISL_508951 | CNR Virus des Infections Respiratoires - France SUD | CNR Virus des Infections Respiratoires - France SUD | Antonin Bal, Gregory Destras, Gwendolyne Burfin, Solenne Brun, Carine Moustaud, Raphaelle Lamy, Alexandre Gaymard, Maude Bouscambert-Duchamp, Florence Morfin-Sherpa, Martine Valette, Bruno Lina, Laurence Josset |
| EPI_ISL_508952 | Centre Hospitalier de Macon | CNR Virus des Infections Respiratoires - France SUD | Antonin Bal, Gregory Destras, Gwendolyne Burfin, Solenne Brun, Carine Moustaud, Raphaelle Lamy, Alexandre Gaymard, Maude Bouscambert-Duchamp, Florence Morfin-Sherpa, Martine Valette, Bruno Lina, Laurence Josset |
| EPI_ISL_508953, EPI_ISL_508954, EPI_ISL_508955, EPI_ISL_508956, EPI_ISL_508957 | CNR Virus des Infections Respiratoires - France SUD | CNR Virus des Infections Respiratoires - France SUD | Antonin Bal, Gregory Destras, Gwendolyne Burfin, Solenne Brun, Carine Moustaud, Raphaelle Lamy, Alexandre Gaymard, Maude Bouscambert-Duchamp, Florence Morfin-Sherpa, Martine Valette, Bruno Lina, Laurence Josset |
| EPI_ISL_508958 | Centre Hospitalier Alpes Leman | CNR Virus des Infections Respiratoires - France SUD | Antonin Bal, Gregory Destras, Gwendolyne Burfin, Solenne Brun, Carine Moustaud, Raphaelle Lamy, Alexandre Gaymard, Maude Bouscambert-Duchamp, Florence Morfin-Sherpa, Martine Valette, Bruno Lina, Laurence Josset |
| EPI_ISL_508959, EPI_ISL_508960 | Centre Hospitalier Saint Joseph Saint Luc | CNR Virus des Infections Respiratoires - France SUD | Antonin Bal, Gregory Destras, Gwendolyne Burfin, Solenne Brun, Carine Moustaud, Raphaelle Lamy, Alexandre Gaymard, Maude Bouscambert-Duchamp, Florence Morfin-Sherpa, Martine Valette, Bruno Lina, Laurence Josset |
| EPI_ISL_508961, EPI_ISL_508962, EPI_ISL_508963, EPI_ISL_508964, EPI_ISL_508965 | CNR Virus des Infections Respiratoires - France SUD | CNR Virus des Infections Respiratoires - France SUD | Antonin Bal, Gregory Destras, Gwendolyne Burfin, Solenne Brun, Carine Moustaud, Raphaelle Lamy, Alexandre Gaymard, Maude Bouscambert-Duchamp, Florence Morfin-Sherpa, Martine Valette, Bruno Lina, Laurence Josset |
| EPI_ISL_508966 | Centre Hospitalier Pierre Oudot | CNR Virus des Infections Respiratoires - France SUD | Antonin Bal, Gregory Destras, Gwendolyne Burfin, Solenne Brun, Carine Moustaud, Raphaelle Lamy, Alexandre Gaymard, Maude Bouscambert-Duchamp, Florence Morfin-Sherpa, Martine Valette, Bruno Lina, Laurence Josset |
| EPI_ISL_508967 | CNR Virus des Infections Respiratoires - France SUD | CNR Virus des Infections Respiratoires - France SUD | Antonin Bal, Gregory Destras, Gwendolyne Burfin, Solenne Brun, Carine Moustaud, Raphaelle Lamy, Alexandre Gaymard, Maude Bouscambert-Duchamp, Florence Morfin-Sherpa, Martine Valette, Bruno Lina, Laurence Josset |
| EPI_ISL_508968 | Centre Hospitalier de Bourg en Bresse | CNR Virus des Infections Respiratoires - France SUD | Antonin Bal, Gregory Destras, Gwendolyne Burfin, Solenne Brun, Carine Moustaud, Raphaelle Lamy, Alexandre Gaymard, Maude Bouscambert-Duchamp, Florence Morfin-Sherpa, Martine Valette, Bruno Lina, Laurence Josset |
| EPI_ISL_508969, EPI_ISL_508970, EPI_ISL_508971, EPI_ISL_508972, EPI_ISL_508973, EPI_ISL_508974 | CNR Virus des Infections Respiratoires - France SUD | CNR Virus des Infections Respiratoires - France SUD | Antonin Bal, Gregory Destras, Gwendolyne Burfin, Solenne Brun, Carine Moustaud, Raphaelle Lamy, Alexandre Gaymard, Maude Bouscambert-Duchamp, Florence Morfin-Sherpa, Martine Valette, Bruno Lina, Laurence Josset |
| EPI_ISL_508975 | Centre Hospitalier Pierre Oudot | CNR Virus des Infections Respiratoires - France SUD | Antonin Bal, Gregory Destras, Gwendolyne Burfin, Solenne Brun, Carine Moustaud, Raphaelle Lamy, Alexandre Gaymard, Maude Bouscambert-Duchamp, Florence Morfin-Sherpa, Martine Valette, Bruno Lina, Laurence Josset |
| EPI_ISL_508976, EPI_ISL_508977 | CNR Virus des Infections Respiratoires - France SUD | CNR Virus des Infections Respiratoires - France SUD | Antonin Bal, Gregory Destras, Gwendolyne Burfin, Solenne Brun, Carine Moustaud, Raphaelle Lamy, Alexandre Gaymard, Maude Bouscambert-Duchamp, Florence Morfin-Sherpa, Martine Valette, Bruno Lina, Laurence Josset |
| EPI_ISL_508978 | Centre Hospitalier de Villefranche | CNR Virus des Infections Respiratoires - France SUD | Antonin Bal, Gregory Destras, Gwendolyne Burfin, Solenne Brun, Carine Moustaud, Raphaelle Lamy, Alexandre Gaymard, Maude Bouscambert-Duchamp, Florence Morfin-Sherpa, Martine Valette, Bruno Lina, Laurence Josset |
| EPI_ISL_508979, EPI_ISL_508980, EPI_ISL_508981, EPI_ISL_508982, EPI_ISL_508983, EPI_ISL_508984, EPI_ISL_508985 | CNR Virus des Infections Respiratoires - France SUD | CNR Virus des Infections Respiratoires - France SUD | Antonin Bal, Gregory Destras, Gwendolyne Burfin, Solenne Brun, Carine Moustaud, Raphaelle Lamy, Alexandre Gaymard, Maude Bouscambert-Duchamp, Florence Morfin-Sherpa, Martine Valette, Bruno Lina, Laurence Josset |
| EPI_ISL_508986, EPI_ISL_508987, EPI_ISL_508988, EPI_ISL_508989 | CNR Virus des Infections Respiratoires - France SUD | CNR Virus des Infections Respiratoires - France SUD | Antonin Bal, Gregory Destras, Gwendolyne Burfin, Solenne Brun, Alexandre Gaymard, Maude Bouscambert-Duchamp, Florence Morfin-Sherpa, Martine Valette, Bruno Lina, Laurence Josset |
| EPI_ISL_508990, EPI_ISL_508991, EPI_ISL_508992, EPI_ISL_508993, EPI_ISL_508994, EPI_ISL_508995, EPI_ISL_508996, EPI_ISL_508997 | CNR Virus des Infections Respiratoires - France SUD | CNR Virus des Infections Respiratoires - France SUD | Antonin Bal, Gregory Destras, Gwendolyne Burfin, Solenne Brun, Carine Moustaud, Raphaelle Lamy, Alexandre Gaymard, Maude Bouscambert-Duchamp, Florence Morfin-Sherpa, Martine Valette, Bruno Lina, Laurence Josset |
| EPI_ISL_508998 | Centre Hospitalier Pierre Oudot | CNR Virus des Infections Respiratoires - France SUD | Antonin Bal, Gregory Destras, Gwendolyne Burfin, Solenne Brun, Carine Moustaud, Raphaelle Lamy, Alexandre Gaymard, Maude Bouscambert-Duchamp, Florence Morfin-Sherpa, Martine Valette, Bruno Lina, Laurence Josset |
| EPI_ISL_508999, EPI_ISL_509000, EPI_ISL_509001, EPI_ISL_509002 | CNR Virus des Infections Respiratoires - France SUD | CNR Virus des Infections Respiratoires - France SUD | Antonin Bal, Gregory Destras, Gwendolyne Burfin, Solenne Brun, Carine Moustaud, Raphaelle Lamy, Alexandre Gaymard, Maude Bouscambert-Duchamp, Florence Morfin-Sherpa, Martine Valette, Bruno Lina, Laurence Josset |
| EPI_ISL_509003, EPI_ISL_509004 | GH Les Portes du Sud | CNR Virus des Infections Respiratoires - France SUD | Antonin Bal, Gregory Destras, Gwendolyne Burfin, Solenne Brun, Carine Moustaud, Raphaelle Lamy, Alexandre Gaymard, Maude Bouscambert-Duchamp, Florence Morfin-Sherpa, Martine Valette, Bruno Lina, Laurence Josset |
| EPI_ISL_509005 | Centre Hospitalier de Macon | CNR Virus des Infections Respiratoires - France SUD | Antonin Bal, Gregory Destras, Gwendolyne Burfin, Solenne Brun, Carine Moustaud, Raphaelle Lamy, Alexandre Gaymard, Maude Bouscambert-Duchamp, Florence Morfin-Sherpa, Martine Valette, Bruno Lina, Laurence Josset |
| EPI_ISL_509006 | Centre Hospitalier de Villefranche | CNR Virus des Infections Respiratoires - France SUD | Antonin Bal, Gregory Destras, Gwendolyne Burfin, Solenne Brun, Carine Moustaud, Raphaelle Lamy, Alexandre Gaymard, Maude Bouscambert-Duchamp, Florence Morfin-Sherpa, Martine Valette, Bruno Lina, Laurence Josset |
| EPI_ISL_509007, EPI_ISL_509008, EPI_ISL_509009, EPI_ISL_509010, EPI_ISL_509011 | Institut des Agents Infectieux (IAI), Hospices Civils de Lyon | CNR Virus des Infections Respiratoires - France SUD | Antonin Bal, Gregory Destras, Gwendolyne Burfin, Solenne Brun, Carine Moustaud, Raphaelle Lamy, Alexandre Gaymard, Maude Bouscambert-Duchamp, Florence Morfin-Sherpa, Martine Valette, Bruno Lina, Laurence Josset |
| EPI_ISL_509012 | Centre Hospitalier Alpes Leman | CNR Virus des Infections Respiratoires - France SUD | Antonin Bal, Gregory Destras, Gwendolyne Burfin, Solenne Brun, Carine Moustaud, Raphaelle Lamy, Alexandre Gaymard, Maude Bouscambert-Duchamp, Florence Morfin-Sherpa, Martine Valette, Bruno Lina, Laurence Josset |
| EPI_ISL_509013, EPI_ISL_509014 | Institut des Agents Infectieux (IAI), Hospices Civils de | CNR Virus des Infections Respiratoires - France SUD | Antonin Bal, Gregory Destras, Gwendolyne Burfin, Solenne Brun, Carine Moustaud, Raphaelle Lamy, Alexandre Gaymard, Maude Bouscambert-Duchamp, Florence Morfin- |

| | | | |
|---|---|---|---|
| | Lyon | | Sherpa, Martine Valette, Bruno Lina, Laurence Josset |
| EPI_ISL_509015 | Centre Hospitalier de Villefranche | CNR Virus des Infections Respiratoires - France SUD | Antonin Bal, Gregory Destras, Gwendolyne Burfin, Solenne Brun, Carine Moustaud, Raphaelle Lamy, Alexandre Gaymard, Maude Bouscambert-Duchamp, Florence Morfin-Sherpa, Martine Valette, Bruno Lina, Laurence Josset |
| EPI_ISL_509016 | Centre Hospitalier du Haut-Bugey | CNR Virus des Infections Respiratoires - France SUD | Antonin Bal, Gregory Destras, Gwendolyne Burfin, Solenne Brun, Carine Moustaud, Raphaelle Lamy, Alexandre Gaymard, Maude Bouscambert-Duchamp, Florence Morfin-Sherpa, Martine Valette, Bruno Lina, Laurence Josset |
| EPI_ISL_509017, EPI_ISL_509018, EPI_ISL_509019, EPI_ISL_509020, EPI_ISL_509021, EPI_ISL_509022, EPI_ISL_509023, EPI_ISL_509024, EPI_ISL_509026, EPI_ISL_509028, EPI_ISL_509029, EPI_ISL_509031, EPI_ISL_509033, EPI_ISL_509035, EPI_ISL_509037, EPI_ISL_509038, EPI_ISL_509051 | | | |
| see above | genXone SA, Molecular Diagnostics Laboratory / NZOZ | genXone SA, Research & Development Laboratory | Maciej Sykulski, Grzegorz Nowicki, Monika Mańkowska-Woźniak, Jakub Grabowski, Natalia Drwęska-Matelska, Łukasz Krych, Michał Kaszuba |
| EPI_ISL_509059, EPI_ISL_509060, EPI_ISL_509061, EPI_ISL_509062, EPI_ISL_509063, EPI_ISL_509064, EPI_ISL_509065, EPI_ISL_509066, EPI_ISL_509067, EPI_ISL_509068, EPI_ISL_509069, EPI_ISL_509070, EPI_ISL_509071, EPI_ISL_509072, EPI_ISL_509073, EPI_ISL_509074, EPI_ISL_509075, EPI_ISL_509076, EPI_ISL_509077, EPI_ISL_509078, EPI_ISL_509079, EPI_ISL_509080, EPI_ISL_509081, EPI_ISL_509082, EPI_ISL_509083, EPI_ISL_509084, EPI_ISL_509085, EPI_ISL_509086, EPI_ISL_509087, EPI_ISL_509088, EPI_ISL_509089, EPI_ISL_509090, EPI_ISL_509091, EPI_ISL_509092, EPI_ISL_509093, EPI_ISL_509094, EPI_ISL_509095, EPI_ISL_509096, EPI_ISL_509097, EPI_ISL_509098, EPI_ISL_509099, EPI_ISL_509100, EPI_ISL_509101, EPI_ISL_509102, EPI_ISL_509103, EPI_ISL_509104, EPI_ISL_509105, EPI_ISL_509106, EPI_ISL_509107, EPI_ISL_509108, EPI_ISL_509109, EPI_ISL_509110, EPI_ISL_509111, EPI_ISL_509112, EPI_ISL_509113, EPI_ISL_509114, EPI_ISL_509115, EPI_ISL_509116, EPI_ISL_509117, EPI_ISL_509118, EPI_ISL_509119, EPI_ISL_509120, EPI_ISL_509121, EPI_ISL_509122, EPI_ISL_509123, EPI_ISL_509124, EPI_ISL_509125, EPI_ISL_509126, EPI_ISL_509127, EPI_ISL_509128, EPI_ISL_509129, EPI_ISL_509130, EPI_ISL_509131, EPI_ISL_509132, EPI_ISL_509133, EPI_ISL_509134, EPI_ISL_509135, EPI_ISL_509136, EPI_ISL_509137, EPI_ISL_509138, EPI_ISL_509139, EPI_ISL_509140, EPI_ISL_509141, EPI_ISL_509142, EPI_ISL_509143, EPI_ISL_509144, EPI_ISL_509145, EPI_ISL_509146, EPI_ISL_509147, EPI_ISL_509148, EPI_ISL_509149, EPI_ISL_509150, EPI_ISL_509151, EPI_ISL_509152, EPI_ISL_509153, EPI_ISL_509154, EPI_ISL_509155, EPI_ISL_509156, EPI_ISL_509157, EPI_ISL_509158, EPI_ISL_509159, EPI_ISL_509160, EPI_ISL_509161, EPI_ISL_509162, EPI_ISL_509163, EPI_ISL_509164, EPI_ISL_509165, EPI_ISL_509166, EPI_ISL_509167, EPI_ISL_509168, EPI_ISL_509169, EPI_ISL_509170, EPI_ISL_509171, EPI_ISL_509172, EPI_ISL_509173, EPI_ISL_509174, EPI_ISL_509175, EPI_ISL_509176, EPI_ISL_509177, EPI_ISL_509178, EPI_ISL_509179, EPI_ISL_509180, EPI_ISL_509181, EPI_ISL_509182, EPI_ISL_509183, EPI_ISL_509184, EPI_ISL_509185, EPI_ISL_509186, EPI_ISL_509187, EPI_ISL_509188, EPI_ISL_509189, EPI_ISL_509190, EPI_ISL_509191, EPI_ISL_509192, EPI_ISL_509193, EPI_ISL_509194, EPI_ISL_509195, EPI_ISL_509196, EPI_ISL_509197, EPI_ISL_509198, EPI_ISL_509199, EPI_ISL_509200, EPI_ISL_509201, EPI_ISL_509202, EPI_ISL_509203, EPI_ISL_509204, EPI_ISL_509205, EPI_ISL_509206, EPI_ISL_509207, EPI_ISL_509208, EPI_ISL_509209, EPI_ISL_509210, EPI_ISL_509211, EPI_ISL_509212, EPI_ISL_509213, EPI_ISL_509214 | | | |
| see above | OHSU Lab Services Molecular Microbiology Lab | Oregon SARS-CoV-2 Genome Sequencing Center | Brendan L. O'Connell, Ruth V. Nichols, Sally B. Grindstaff, Alec J. Hirsch, Guang Fan, Daniel N. Streblow, William B. Messer, Andrew C. Adey, Benjamin N. Bimber, Brian J. O'Roak |
| EPI_ISL_509222 | Division of Infectious Diseases and Hospital Epidemiology, University Hospital Zürich | Institute of Medical Virology, University of Zurich | Maryam Zaheri, Verena Kufner, Dana Weissberg, Jürg Böni, Silvana K. Rampini, Peter W. Schreiber, Irene A. Abela, Hugo Sax, Aline Wolfensberger, Michael Huber |
| EPI_ISL_509223, EPI_ISL_509224, EPI_ISL_509225, EPI_ISL_509226, EPI_ISL_509227, EPI_ISL_509228, EPI_ISL_509229, EPI_ISL_509230, EPI_ISL_509231, EPI_ISL_509232, EPI_ISL_509233, EPI_ISL_509234, EPI_ISL_509235, EPI_ISL_509236, EPI_ISL_509237, EPI_ISL_509238, EPI_ISL_509239, EPI_ISL_509240, EPI_ISL_509241, EPI_ISL_509242, EPI_ISL_509243, EPI_ISL_509244, EPI_ISL_509245, EPI_ISL_509246, EPI_ISL_509247, EPI_ISL_509248, EPI_ISL_509249, EPI_ISL_509250, EPI_ISL_509251, EPI_ISL_509252, EPI_ISL_509253, EPI_ISL_509254, EPI_ISL_509255, EPI_ISL_509256, EPI_ISL_509257, EPI_ISL_509258, EPI_ISL_509259, EPI_ISL_509260, EPI_ISL_509261, EPI_ISL_509262, EPI_ISL_509263, EPI_ISL_509264, EPI_ISL_509265, EPI_ISL_509266, EPI_ISL_509267, EPI_ISL_509268, EPI_ISL_509269, EPI_ISL_509270, EPI_ISL_509271, EPI_ISL_509272, EPI_ISL_509273, EPI_ISL_509274, EPI_ISL_509275, EPI_ISL_509276, EPI_ISL_509277, EPI_ISL_509278, EPI_ISL_509279, EPI_ISL_509280, EPI_ISL_509281, EPI_ISL_509282, EPI_ISL_509283, EPI_ISL_509284, EPI_ISL_509285, EPI_ISL_509286, EPI_ISL_509287, EPI_ISL_509288, EPI_ISL_509289, EPI_ISL_509290, EPI_ISL_509291, EPI_ISL_509292, EPI_ISL_509293, EPI_ISL_509294, EPI_ISL_509295, EPI_ISL_509296, EPI_ISL_509297, EPI_ISL_509298, EPI_ISL_509299, EPI_ISL_509300, EPI_ISL_509301, EPI_ISL_509302, EPI_ISL_509303, EPI_ISL_509304, EPI_ISL_509305, EPI_ISL_509306, EPI_ISL_509307, EPI_ISL_509308, EPI_ISL_509309, EPI_ISL_509310, EPI_ISL_509311, EPI_ISL_509312, EPI_ISL_509313, EPI_ISL_509314, EPI_ISL_509315, EPI_ISL_509316, EPI_ISL_509317, EPI_ISL_509318, EPI_ISL_509319, EPI_ISL_509320, EPI_ISL_509321, EPI_ISL_509322, EPI_ISL_509323, EPI_ISL_509324, EPI_ISL_509325, EPI_ISL_509326, EPI_ISL_509327, EPI_ISL_509328, EPI_ISL_509329, EPI_ISL_509330, EPI_ISL_509331, EPI_ISL_509332, EPI_ISL_509333, EPI_ISL_509334, EPI_ISL_509335, EPI_ISL_509336, EPI_ISL_509337, EPI_ISL_509338, EPI_ISL_509339, EPI_ISL_509340, EPI_ISL_509341, EPI_ISL_509342, EPI_ISL_509343, EPI_ISL_509344, EPI_ISL_509345, EPI_ISL_509346, EPI_ISL_509347, EPI_ISL_509348, EPI_ISL_509349, EPI_ISL_509350, EPI_ISL_509351, EPI_ISL_509352, EPI_ISL_509353, EPI_ISL_509354, EPI_ISL_509355, EPI_ISL_509356, EPI_ISL_509357, EPI_ISL_509358, EPI_ISL_509359, EPI_ISL_509360, EPI_ISL_509361, EPI_ISL_509362, EPI_ISL_509363, EPI_ISL_509364, EPI_ISL_509365, EPI_ISL_509366, EPI_ISL_509367, EPI_ISL_509368, EPI_ISL_509369, EPI_ISL_509370, EPI_ISL_509371 | | | |
| see above | NHLS-IALCH | KRISP, KZN Research Innovation and Sequencing Platform | Giandhari J, Pillay S, Lessells R, Mdlalose K, York D, Tegally H, Wilkinson E, de Oliveira T |
| EPI_ISL_509372, EPI_ISL_509373, EPI_ISL_509374, EPI_ISL_509375, EPI_ISL_509376, EPI_ISL_509377, EPI_ISL_509378, EPI_ISL_509379, EPI_ISL_509380, EPI_ISL_509381, EPI_ISL_509382, EPI_ISL_509383, EPI_ISL_509384, EPI_ISL_509385, EPI_ISL_509386, EPI_ISL_509387, EPI_ISL_509388, EPI_ISL_509389, EPI_ISL_509390, EPI_ISL_509391, EPI_ISL_509392, EPI_ISL_509393, EPI_ISL_509394, EPI_ISL_509395 | | | |
| see above | Singapore General Hospital | Department of Microbiology | Nurdyana Abdul Rahman, Kun Lee Lim, Chenhao Li, Kian Sing Chan, Lynette Oon, Kern Rei Chng, Niranjan Nagarajan, Karrie Ko |
| EPI_ISL_509412, EPI_ISL_509413, EPI_ISL_509414, EPI_ISL_509415, EPI_ISL_509416, EPI_ISL_509417 | Acibadem Labcell Cellular Therapy Laboratory | Acibadem Mehmet Ali Aydinlar University School of Medicine, Medical Genetics Department | Ozden Hatirnaz Ng, Sezer Akyoney, Ilayda Sahin, Gunseli Bayram Akcapinar, Ozkan Ozdemir, Derya Dilek Kancagi, Gozde Sir Karakus, Bulut Yurtsever, Cihan Tastan, Ercument Ovali, Ugur Ozbek |
| EPI_ISL_509418, EPI_ISL_509419, EPI_ISL_509420, EPI_ISL_509421, EPI_ISL_509422, EPI_ISL_509423 | Istituto Zooprofilattico Sperimentale del Mezzogiorno | Istituto Zooprofilattico Sperimentale del Mezzogiorno | Viscardi,M., Cozzolino,L., Fusco,G. |
| EPI_ISL_509424, EPI_ISL_509425, EPI_ISL_509426, EPI_ISL_509427, EPI_ISL_509428, EPI_ISL_509429 | Microbiology and Immunology, University of South Alabama | Microbiology and Immunology, University of South Alabama | Wood,R.R., Roberts,R.A., Houserova,D., Borchert,G.M., Fouty,B., Rayner,J.O. |
| EPI_ISL_509430, EPI_ISL_509431, EPI_ISL_509432, EPI_ISL_509433, EPI_ISL_509434, EPI_ISL_509435 | Centro de Desenvolvimento Tecnologico em Saude, Fundacao Oswaldo Cruz | Centro de Desenvolvimento Tecnologico em Saude, Fundacao Oswaldo Cruz | Souza,T.M., Fintelman-Rodrigues,N., De Paula,A.D., Saraiva,F.B., Ferreira,M.A., Sacramento,C.Q., Medeiros,M.A. |
| EPI_ISL_509436, EPI_ISL_509437, EPI_ISL_509438, EPI_ISL_509439, EPI_ISL_509440, EPI_ISL_509441, EPI_ISL_509442, EPI_ISL_509443 | The Princess Alexandra Hospital | Wellcome Sanger Institute for the COVID-19 Genomics UK Consortium | Nick Levene, Louise Lopez, Lynn Monaghan, Jessica Scott, Claudia McCrea and Alex Alderton, Roberto Amato, Sonia Goncalves, Ewan Harrison, David K. Jackson, Ian Johnston, Dominic Kwiatkowski, Cordelia Langford, John Sillitoe on behalf of the Wellcome Sanger Institute COVID-19 Surveillance Team (http://www.sanger.ac.uk/covid-team) |
| EPI_ISL_509444, EPI_ISL_509445, EPI_ISL_509446, EPI_ISL_509447, EPI_ISL_509448, EPI_ISL_509449, EPI_ISL_509450, EPI_ISL_509451, EPI_ISL_509452, EPI_ISL_509453, EPI_ISL_509454, EPI_ISL_509455, EPI_ISL_509456, EPI_ISL_509457, EPI_ISL_509458, EPI_ISL_509459, EPI_ISL_509460, EPI_ISL_509461, EPI_ISL_509462, EPI_ISL_509463, EPI_ISL_509464, EPI_ISL_509465, EPI_ISL_509466, EPI_ISL_509467, EPI_ISL_509468, EPI_ISL_509469, EPI_ISL_509470, EPI_ISL_509471, EPI_ISL_509472, EPI_ISL_509473, EPI_ISL_509474, EPI_ISL_509475, EPI_ISL_509476, EPI_ISL_509477, EPI_ISL_509478, EPI_ISL_509479, EPI_ISL_509480, EPI_ISL_509481, EPI_ISL_509482, EPI_ISL_509483, EPI_ISL_509484, EPI_ISL_509485, EPI_ISL_509486, EPI_ISL_509487, EPI_ISL_509488, EPI_ISL_509489, EPI_ISL_509490, EPI_ISL_509491 | | | |
| see above | Maryland Department of Health | Maryland Department of Health | Keller,E. |
| EPI_ISL_509492, EPI_ISL_509493, EPI_ISL_509494, EPI_ISL_509495, EPI_ISL_509496, EPI_ISL_509497, EPI_ISL_509498, EPI_ISL_509499, EPI_ISL_509500, EPI_ISL_509501, EPI_ISL_509502, EPI_ISL_509503, EPI_ISL_509504, EPI_ISL_509505, EPI_ISL_509506, EPI_ISL_509507, EPI_ISL_509508, EPI_ISL_509509, EPI_ISL_509510, EPI_ISL_509511, EPI_ISL_509512, EPI_ISL_509513, EPI_ISL_509514, EPI_ISL_509515, EPI_ISL_509516, EPI_ISL_509517, EPI_ISL_509518, EPI_ISL_509519, EPI_ISL_509520, EPI_ISL_509521, EPI_ISL_509522, EPI_ISL_509523 | | | |
| see above | Area of Virology, Serology and Virology Division (SAViD), New South Wales Health Pathology Randwick | Area of Virology, Serology and Virology Division (SAViD), New South Wales Health Pathology Randwick | Rawlinson, W. |
| EPI_ISL_509527, EPI_ISL_509528, EPI_ISL_509529, EPI_ISL_509530, EPI_ISL_509531, EPI_ISL_509532, EPI_ISL_509533, EPI_ISL_509534, EPI_ISL_509535, EPI_ISL_509536, EPI_ISL_509537, EPI_ISL_509538, EPI_ISL_509539, EPI_ISL_509540, EPI_ISL_509541, EPI_ISL_509542, EPI_ISL_509543, EPI_ISL_509544, EPI_ISL_509545, EPI_ISL_509546, EPI_ISL_509547, EPI_ISL_509548, EPI_ISL_509549, EPI_ISL_509550, EPI_ISL_509551, EPI_ISL_509552, EPI_ISL_509553, EPI_ISL_509554, EPI_ISL_509555, EPI_ISL_509556, EPI_ISL_509557, EPI_ISL_509558, EPI_ISL_509559, EPI_ISL_509560, EPI_ISL_509561, EPI_ISL_509562, EPI_ISL_509563, EPI_ISL_509567, EPI_ISL_509568, EPI_ISL_509569, EPI_ISL_509570, EPI_ISL_509571, EPI_ISL_509572, EPI_ISL_509573, EPI_ISL_509574, EPI_ISL_509575, EPI_ISL_509576, EPI_ISL_509577, EPI_ISL_509578, EPI_ISL_509579, EPI_ISL_509580, EPI_ISL_509581, EPI_ISL_509582, EPI_ISL_509583, EPI_ISL_509584, EPI_ISL_509585, EPI_ISL_509586, EPI_ISL_509587, EPI_ISL_509588, EPI_ISL_509589, EPI_ISL_509590, EPI_ISL_509591, EPI_ISL_509592, EPI_ISL_509593, EPI_ISL_509594, EPI_ISL_509595, EPI_ISL_509596, EPI_ISL_509597, EPI_ISL_509598, EPI_ISL_509599, EPI_ISL_509600 | | | |
| see above | Utah Public Health Laboratory | Utah Public Health Laboratory | Heidi Butz, Erin Young, Kelly Oakeson |
| EPI_ISL_509601 | Servicio de Microbiología. Hospital Universitario Donostia. OSI Donostialdea. Área de Enfermedades Infecciosas, Grupo de Infección Respiratoria y Resistencia Antimicrobiana. Instituto de Investigación Sanitaria Biodonostia | SeqCOVID-SPAIN consortium/IBV(CSIC) | Gustavo Cilla, Milagrosa Montes, Luis Piñeiro, Jose Maria Marimón and SeqCOVID-SPAIN consortium |
| EPI_ISL_509602, EPI_ISL_509603, EPI_ISL_509604, EPI_ISL_509605, EPI_ISL_509606, EPI_ISL_509607, EPI_ISL_509608, EPI_ISL_509609 | Servicio de Microbiología. HRU de Málaga. Servicio Andaluz de Salud | SeqCOVID-SPAIN consortium/IBV(CSIC) | Inmaculada de Toro Peinado. MªConcepción Mediavilla Gradolph. Begoña Palop Borrás and SeqCOVID-SPAIN consortium |
| EPI_ISL_509610 | Servicio de Microbiología. Hospital Universitario Donostia. OSI Donostialdea. Área de Enfermedades Infecciosas, Grupo de Infección Respiratoria y Resistencia Antimicrobiana. Instituto de Investigación Sanitaria Biodonostia | SeqCOVID-SPAIN consortium/IBV(CSIC) | Gustavo Cilla, Milagrosa Montes, Luis Piñeiro, Jose Maria Marimón and SeqCOVID-SPAIN consortium |
| EPI_ISL_509611 | Servicio de Microbiología. HRU de Málaga. Servicio Andaluz de Salud | SeqCOVID-SPAIN consortium/IBV(CSIC) | Inmaculada de Toro Peinado. MªConcepción Mediavilla Gradolph. Begoña Palop Borrás and SeqCOVID-SPAIN consortium |
| EPI_ISL_509612, EPI_ISL_509613, EPI_ISL_509614, EPI_ISL_509615 | Servicio de Microbiología. Hospital Universitario Donostia. OSI Donostialdea. Área de Enfermedades Infecciosas, Grupo de Infección Respiratoria y Resistencia Antimicrobiana. Instituto de Investigación Sanitaria Biodonostia | SeqCOVID-SPAIN consortium/IBV(CSIC) | Gustavo Cilla, Milagrosa Montes, Luis Piñeiro, Jose Maria Marimón and SeqCOVID-SPAIN consortium |
| EPI_ISL_509616, EPI_ISL_509617, EPI_ISL_509618 | Hospital Universitario Araba. Vitoria-Gasteiz | SeqCOVID-SPAIN consortium/IBV(CSIC) | Silvia Hernáez Crespo, Carmen Gómez González, Amaia Aguirre Quiñonero, Marina Fernández Torres, Mª Rosario Almela Ferrer, Mª Concepción Lecaroz Agara, Andrés Canut Blasco and SeqCOVID-SPAIN consortium |
| EPI_ISL_509619, EPI_ISL_509620, EPI_ISL_509621, EPI_ISL_509622, EPI_ISL_509623, EPI_ISL_509624, EPI_ISL_509625, EPI_ISL_509626, EPI_ISL_509627, EPI_ISL_509628, EPI_ISL_509629, EPI_ISL_509630, EPI_ISL_509631, EPI_ISL_509632 | | | |
| see above | Servicio de Microbiología. HRU de Málaga. Servicio Andaluz de Salud | SeqCOVID-SPAIN consortium/IBV(CSIC) | Inmaculada de Toro Peinado. MªConcepción Mediavilla Gradolph. Begoña Palop Borrás and SeqCOVID-SPAIN consortium |
| EPI_ISL_509633, EPI_ISL_509634, EPI_ISL_509635, EPI_ISL_509636, EPI_ISL_509637, EPI_ISL_509638, EPI_ISL_509639, EPI_ISL_509640, EPI_ISL_509641, EPI_ISL_509642, EPI_ISL_509643, EPI_ISL_509644, EPI_ISL_509645, EPI_ISL_509646, EPI_ISL_509647, EPI_ISL_509648, EPI_ISL_509649, EPI_ISL_509650, EPI_ISL_509651, EPI_ISL_509652, EPI_ISL_509653, EPI_ISL_509654 | | | |
| see above | Servicio de Microbiología. Hospital Universitario Donostia. OSI Donostialdea. Área de Enfermedades Infecciosas, Grupo de Infección Respiratoria y Resistencia Antimicrobiana. Instituto de Investigación Sanitaria Biodonostia | SeqCOVID-SPAIN consortium/IBV(CSIC) | Gustavo Cilla, Milagrosa Montes, Luis Piñeiro, Jose Maria Marimón and SeqCOVID-SPAIN consortium |
| EPI_ISL_509662 | Georgia Department of Health | Pathogen Discovery, Respiratory Viruses Branch, Division of Viral Diseases, Centers for Disease Control and Prevention | Yan Li, Anna Montmayeur, Jing Zhang, Krista Queen, Anna Uehara, Ying Tao, Rachel Marine, Clinton R. Paden, Haibin Wang, Suxiang Tong |
| EPI_ISL_509663, EPI_ISL_509664, EPI_ISL_509665, EPI_ISL_509666, EPI_ISL_509667, EPI_ISL_509668, EPI_ISL_509670, EPI_ISL_509671, EPI_ISL_509672, EPI_ISL_509673, EPI_ISL_509674, EPI_ISL_509675, EPI_ISL_509676, EPI_ISL_509677, EPI_ISL_509678, EPI_ISL_509679, EPI_ISL_509680, EPI_ISL_509681, EPI_ISL_509682 | | | |
| see above | AR Dept. of Health-Public Health Lab | Pathogen Discovery, Respiratory Viruses Branch, Division of Viral Diseases, Centers for Disease Control and Prevention | Yan Li, Anna Montmayeur, Jing Zhang, Krista Queen, Anna Uehara, Ying Tao, Rachel Marine, Clinton R. Paden, Haibin Wang, Suxiang Tong |
| EPI_ISL_509683, EPI_ISL_509684, EPI_ISL_509685 | AR Dept. of Health-Public Health Lab | Pathogen Discovery, Respiratory Viruses Branch, Division of Viral Diseases, Centers for Disease Control and Prevention | Jing Zhang, Yan Li, Anna Montmayeur, Krista Queen, Anna Uehara, Ying Tao, Rachel Marine, Clinton R. Paden, Haibin Wang, Suxiang Tong |
| EPI_ISL_509686 | M Health Fairview | Minnesota Department of Health, Public Health Laboratory | Matt Plumb, Jacob Garfin, and Xiong Wang |
| EPI_ISL_509688 | Alabama Department of Public Health Bureau of Clinical | Pathogen Discovery, Respiratory Viruses Branch, Division of Viral Diseases, Centers for Disease Control and Prevention | Ying Tao, Jing Zhang, Krista Queen, Anna Uehara, Yan Li, Clinton Paden, Haibin Wang, Suxiang Tong |

| | | | |
|---|---|---|---|
| | Laboratories | | |
| EPI_ISL_509689, EPI_ISL_509690, EPI_ISL_509691, EPI_ISL_509692, EPI_ISL_509693, EPI_ISL_509694 | Utah Public Health Laboratory | Pathogen Discovery, Respiratory Viruses Branch, Division of Viral Diseases, Centers for Disease Control and Prevention | Ying Tao, Jing Zhang, Krista Queen, Anna Uehara, Yan Li, Clinton Paden, Haibin Wang, Suxiang Tong |
| EPI_ISL_509695, EPI_ISL_509696, EPI_ISL_509697, EPI_ISL_509698, EPI_ISL_509699, EPI_ISL_509700, EPI_ISL_509701, EPI_ISL_509702, EPI_ISL_509703 | Guatemala Ministry of Public Health | Pathogen Discovery, Respiratory Viruses Branch, Division of Viral Diseases, Centers for Disease Control and Prevention | Ying Tao, Jing Zhang, Krista Queen, Anna Uehara, Yan Li, Clinton Paden, Haibin Wang, Suxiang Tong |
| EPI_ISL_509704, EPI_ISL_509705, EPI_ISL_509706 | Wisconsin Department of Health Services | Pathogen Discovery, Respiratory Viruses Branch, Division of Viral Diseases, Centers for Disease Control and Prevention | Ying Tao, Jing Zhang, Krista Queen, Anna Uehara, Yan Li, Clinton Paden, Haibin Wang, Suxiang Tong |
| EPI_ISL_509707, EPI_ISL_509708, EPI_ISL_509709 | Utah Public Health Laboratory | Pathogen Discovery, Respiratory Viruses Branch, Division of Viral Diseases, Centers for Disease Control and Prevention | Jing Zhang, Ying Tao, Krista Queen, Anna Uehara, Yan Li, Clinton Paden, Haibin Wang, Suxiang Tong |
| EPI_ISL_509710 | Guatemala Ministry of Public Health | Pathogen Discovery, Respiratory Viruses Branch, Division of Viral Diseases, Centers for Disease Control and Prevention | Jing Zhang, Ying Tao, Krista Queen, Anna Uehara, Yan Li, Clinton Paden, Haibin Wang, Suxiang Tong |
| EPI_ISL_509711, EPI_ISL_509712, EPI_ISL_509713, EPI_ISL_509714 | Belize Ministry of Health | Pathogen Discovery, Respiratory Viruses Branch, Division of Viral Diseases, Centers for Disease Control and Prevention | Jing Zhang, Ying Tao, Krista Queen, Anna Uehara, Yan Li, Clinton Paden, Haibin Wang, Suxiang Tong |
| EPI_ISL_509715 | Wisconsin Department of Health Services | Pathogen Discovery, Respiratory Viruses Branch, Division of Viral Diseases, Centers for Disease Control and Prevention | Jing Zhang, Ying Tao, Krista Queen, Anna Uehara, Yan Li, Clinton Paden, Haibin Wang, Suxiang Tong |
| EPI_ISL_509716, EPI_ISL_509717, EPI_ISL_509718, EPI_ISL_509719, EPI_ISL_509720, EPI_ISL_509721, EPI_ISL_509722, EPI_ISL_509723, EPI_ISL_509724, EPI_ISL_509725, EPI_ISL_509726, EPI_ISL_509727, EPI_ISL_509728, EPI_ISL_509729, EPI_ISL_509730, EPI_ISL_509731, EPI_ISL_509732, EPI_ISL_509733, EPI_ISL_509734, EPI_ISL_509735, EPI_ISL_509736, EPI_ISL_509737, EPI_ISL_509738, EPI_ISL_509739, EPI_ISL_509740, EPI_ISL_509741, EPI_ISL_509742, EPI_ISL_509743, EPI_ISL_509744, EPI_ISL_509745, EPI_ISL_509746, EPI_ISL_509747, EPI_ISL_509748, EPI_ISL_509749, EPI_ISL_509750, EPI_ISL_509751, EPI_ISL_509752, EPI_ISL_509753, EPI_ISL_509754, EPI_ISL_509755, EPI_ISL_509756, EPI_ISL_509757, EPI_ISL_509758, EPI_ISL_509759, EPI_ISL_509760, EPI_ISL_509761, EPI_ISL_509762, EPI_ISL_509763, EPI_ISL_509764, EPI_ISL_509765, EPI_ISL_509766, EPI_ISL_509767, EPI_ISL_509768, EPI_ISL_509769, EPI_ISL_509770, EPI_ISL_509771, EPI_ISL_509772, EPI_ISL_509773, EPI_ISL_509774, EPI_ISL_509775, EPI_ISL_509776, EPI_ISL_509777, EPI_ISL_509778, EPI_ISL_509779, EPI_ISL_509780, EPI_ISL_509781, EPI_ISL_509782, EPI_ISL_509783, EPI_ISL_509784, EPI_ISL_509785, EPI_ISL_509786, EPI_ISL_509787, EPI_ISL_509788, EPI_ISL_509789, EPI_ISL_509790, EPI_ISL_509791, EPI_ISL_509792, EPI_ISL_509793, EPI_ISL_509794, EPI_ISL_509795, EPI_ISL_509796, EPI_ISL_509797, EPI_ISL_509798 | | | |
| see above | Florida Bureau of Public Health Laboratories | Florida Bureau of Public Health Laboratories | Sarah Schmedes, Jason Blanton |
| EPI_ISL_509799, EPI_ISL_509800, EPI_ISL_509801, EPI_ISL_509802, EPI_ISL_509803, EPI_ISL_509804, EPI_ISL_509805, EPI_ISL_509806, EPI_ISL_509807, EPI_ISL_509808, EPI_ISL_509809, EPI_ISL_509810, EPI_ISL_509811, EPI_ISL_509812, EPI_ISL_509813, EPI_ISL_509814, EPI_ISL_509815, EPI_ISL_509816, EPI_ISL_509817, EPI_ISL_509818, EPI_ISL_509819, EPI_ISL_509820, EPI_ISL_509821, EPI_ISL_509822, EPI_ISL_509823, EPI_ISL_509824, EPI_ISL_509825, EPI_ISL_509826, EPI_ISL_509827, EPI_ISL_509828, EPI_ISL_509829, EPI_ISL_509830, EPI_ISL_509831, EPI_ISL_509832, EPI_ISL_509833, EPI_ISL_509834, EPI_ISL_509835, EPI_ISL_509836, EPI_ISL_509837, EPI_ISL_509838, EPI_ISL_509839, EPI_ISL_509840, EPI_ISL_509841, EPI_ISL_509842, EPI_ISL_509843, EPI_ISL_509844, EPI_ISL_509845, EPI_ISL_509846, EPI_ISL_509847, EPI_ISL_509848, EPI_ISL_509849, EPI_ISL_509850, EPI_ISL_509851, EPI_ISL_509852, EPI_ISL_509853, EPI_ISL_509854, EPI_ISL_509855, EPI_ISL_509856, EPI_ISL_509857, EPI_ISL_509858, EPI_ISL_509859, EPI_ISL_509860, EPI_ISL_509861, EPI_ISL_509862, EPI_ISL_509863, EPI_ISL_509864, EPI_ISL_509865, EPI_ISL_509866, EPI_ISL_509867, EPI_ISL_509868, EPI_ISL_509869, EPI_ISL_509870, EPI_ISL_509871, EPI_ISL_509872, EPI_ISL_509873, EPI_ISL_509874, EPI_ISL_509875, EPI_ISL_509876, EPI_ISL_509877, EPI_ISL_509878, EPI_ISL_509879, EPI_ISL_509880, EPI_ISL_509881, EPI_ISL_509882, EPI_ISL_509883, EPI_ISL_509884, EPI_ISL_509885, EPI_ISL_509886, EPI_ISL_509887, EPI_ISL_509888, EPI_ISL_509889, EPI_ISL_509890, EPI_ISL_509891, EPI_ISL_509892, EPI_ISL_509893, EPI_ISL_509894, EPI_ISL_509895, EPI_ISL_509896, EPI_ISL_509897, EPI_ISL_509898, EPI_ISL_509899, EPI_ISL_509900, EPI_ISL_509901, EPI_ISL_509902, EPI_ISL_509903, EPI_ISL_509904, EPI_ISL_509905, EPI_ISL_509906, EPI_ISL_509907, EPI_ISL_509908, EPI_ISL_509909, EPI_ISL_509910, EPI_ISL_509911, EPI_ISL_509912, EPI_ISL_509913, EPI_ISL_509914, EPI_ISL_509915, EPI_ISL_509916, EPI_ISL_509917, EPI_ISL_509918, EPI_ISL_509919, EPI_ISL_509920, EPI_ISL_509921, EPI_ISL_509922, EPI_ISL_509923, EPI_ISL_509924, EPI_ISL_509925, EPI_ISL_509926, EPI_ISL_509927, EPI_ISL_509928, EPI_ISL_509929, EPI_ISL_509930, EPI_ISL_509931, EPI_ISL_509932, EPI_ISL_509933, EPI_ISL_509934, EPI_ISL_509935, EPI_ISL_509936, EPI_ISL_509937, EPI_ISL_509938, EPI_ISL_509939, EPI_ISL_509940, EPI_ISL_509941, EPI_ISL_509942, EPI_ISL_509943, EPI_ISL_509944, EPI_ISL_509945, EPI_ISL_509946, EPI_ISL_509947, EPI_ISL_509948, EPI_ISL_509949, EPI_ISL_509950, EPI_ISL_509951, EPI_ISL_509952, EPI_ISL_509953, EPI_ISL_509954, EPI_ISL_509955, EPI_ISL_509956, EPI_ISL_509957, EPI_ISL_509958, EPI_ISL_509959, EPI_ISL_509960, EPI_ISL_509961, EPI_ISL_509962, EPI_ISL_509963, EPI_ISL_509964, EPI_ISL_509965, EPI_ISL_509966, EPI_ISL_509967, EPI_ISL_509968, EPI_ISL_509969, EPI_ISL_509970, EPI_ISL_509971, EPI_ISL_509972, EPI_ISL_509973, EPI_ISL_509974, EPI_ISL_509975, EPI_ISL_509976, EPI_ISL_509977, EPI_ISL_509978, EPI_ISL_509979, EPI_ISL_509980, EPI_ISL_509981, EPI_ISL_509982, EPI_ISL_509983, EPI_ISL_509984, EPI_ISL_509985, EPI_ISL_509986, EPI_ISL_509987, EPI_ISL_509988, EPI_ISL_509989, EPI_ISL_509990, EPI_ISL_509991, EPI_ISL_509992, EPI_ISL_509993, EPI_ISL_509994, EPI_ISL_509995, EPI_ISL_509996, EPI_ISL_509997, EPI_ISL_509998 | | | |
| see above | University of Wisconsin-Madison AIDS Vaccine Research Laboratories | University of Wisconsin-Madison AIDS Vaccine Research Laboratories | Gage Moreno, Katarina Braun, et al. AIDS Vaccine Research Laboratories |
| EPI_ISL_510044, EPI_ISL_510045, EPI_ISL_510046, EPI_ISL_510047, EPI_ISL_510048, EPI_ISL_510049, EPI_ISL_510050, EPI_ISL_510051, EPI_ISL_510052, EPI_ISL_510053, EPI_ISL_510054, EPI_ISL_510055, EPI_ISL_510056, EPI_ISL_510057, EPI_ISL_510058, EPI_ISL_510059, EPI_ISL_510060, EPI_ISL_510061, EPI_ISL_510062, EPI_ISL_510063 | | | |
| see above | Servicio de Microbiología. HRU de Málaga. Servicio Andaluz de Salud | SeqCOVID-SPAIN consortium/IBV(CSIC) | Inmaculada de Toro Peinado. MªConcepción Mediavilla Gradolph. Begoña Palop Borrás and SeqCOVID-SPAIN consortium |
| EPI_ISL_510064, EPI_ISL_510065, EPI_ISL_510066, EPI_ISL_510067, EPI_ISL_510068, EPI_ISL_510069, EPI_ISL_510070, EPI_ISL_510071, EPI_ISL_510072, EPI_ISL_510073, EPI_ISL_510074, EPI_ISL_510075, EPI_ISL_510076, EPI_ISL_510077 | | | |
| see above | Instituto de Investigaciones Biomédicas de Barcelona (CSIC), Hospital Clinic i Provincial de Barcelona, Instituto de Biomedicina de Valencia (CSIC), Hospital de Sant Pau | SeqCOVID-SPAIN consortium/IBV(CSIC) | Anna M. Planas, Mª Angeles Marcos, Miguel J. Martínez, Andrea Vergara, Alex Soriano, Jordi Pérez Tur, Israel Fernández Cadenas and SeqCOVID-SPAIN consortium |
| EPI_ISL_510081, EPI_ISL_510082 | Communicable Disease Branch | Hong Kong Department of Health | Mak Gannon C.K., Lam Edman T.K., Chan Rickjason C.W., Tsang Dominic N.C. |
| EPI_ISL_510083 | Princess Margaret Hospital | Hong Kong Department of Health | Mak Gannon C.K., Lam Edman T.K., Chan Rickjason C.W., Tsang Dominic N.C. |
| EPI_ISL_510084 | Prince of Wales Hospital | Hong Kong Department of Health | Mak Gannon C.K., Lam Edman T.K., Chan Rickjason C.W., Tsang Dominic N.C. |
| EPI_ISL_510085 | Yan Chai Hospital | Hong Kong Department of Health | Mak Gannon C.K., Lam Edman T.K., Chan Rickjason C.W., Tsang Dominic N.C. |
| EPI_ISL_510086 | Kwong Wah Hospital | Hong Kong Department of Health | Mak Gannon C.K., Lam Edman T.K., Chan Rickjason C.W., Tsang Dominic N.C. |
| EPI_ISL_510087 | Private medical practitioner | Hong Kong Department of Health | Mak Gannon C.K., Lam Edman T.K., Chan Rickjason C.W., Tsang Dominic N.C. |
| EPI_ISL_510088 | United Christian Hospital | Hong Kong Department of Health | Mak Gannon C.K., Lam Edman T.K., Chan Rickjason C.W., Tsang Dominic N.C. |
| EPI_ISL_510089 | Queen Elizabeth Hospital | Hong Kong Department of Health | Mak Gannon C.K., Lam Edman T.K., Chan Rickjason C.W., Tsang Dominic N.C. |
| EPI_ISL_510090 | Hong Kong Baptist Hospital | Hong Kong Department of Health | Mak Gannon C.K., Lam Edman T.K., Chan Rickjason C.W., Tsang Dominic N.C. |
| EPI_ISL_510091 | Kwong Wah Hospital | Hong Kong Department of Health | Mak Gannon C.K., Lam Edman T.K., Chan Rickjason C.W., Tsang Dominic N.C. |
| EPI_ISL_510092, EPI_ISL_510093, EPI_ISL_510094, EPI_ISL_510095, EPI_ISL_510096 | Queen Elizabeth Hospital | Hong Kong Department of Health | Mak Gannon C.K., Lam Edman T.K., Chan Rickjason C.W., Tsang Dominic N.C. |
| EPI_ISL_510097, EPI_ISL_510098 | United Christian Hospital | Hong Kong Department of Health | Mak Gannon C.K., Lam Edman T.K., Chan Rickjason C.W., Tsang Dominic N.C. |
| EPI_ISL_510099 | Queen Mary Hospital | Hong Kong Department of Health | Mak Gannon C.K., Lam Edman T.K., Chan Rickjason C.W., Tsang Dominic N.C. |
| EPI_ISL_510100 | Tuen Mun Hospital | Hong Kong Department of Health | Mak Gannon C.K., Lam Edman T.K., Chan Rickjason C.W., Tsang Dominic N.C. |
| EPI_ISL_510101 | Pamela Youde Nethersole Eastern Hospital | Hong Kong Department of Health | Mak Gannon C.K., Lam Edman T.K., Chan Rickjason C.W., Tsang Dominic N.C. |
| EPI_ISL_510103, EPI_ISL_510104, EPI_ISL_510105, EPI_ISL_510106, EPI_ISL_510107, EPI_ISL_510108, EPI_ISL_510109, EPI_ISL_510110, EPI_ISL_510111, EPI_ISL_510112, EPI_ISL_510113, EPI_ISL_510114, EPI_ISL_510115, EPI_ISL_510116, EPI_ISL_510117, EPI_ISL_510118, EPI_ISL_510119, EPI_ISL_510120, EPI_ISL_510121, EPI_ISL_510122, EPI_ISL_510123, EPI_ISL_510124, EPI_ISL_510125, EPI_ISL_510126, EPI_ISL_510127, EPI_ISL_510128, EPI_ISL_510129, EPI_ISL_510130, EPI_ISL_510131, EPI_ISL_510132, EPI_ISL_510133, EPI_ISL_510134, EPI_ISL_510135, EPI_ISL_510136, EPI_ISL_510137, EPI_ISL_510138, EPI_ISL_510139, EPI_ISL_510140, EPI_ISL_510141, EPI_ISL_510142, EPI_ISL_510143, EPI_ISL_510144, EPI_ISL_510145, EPI_ISL_510146, EPI_ISL_510147, EPI_ISL_510148, EPI_ISL_510149, EPI_ISL_510150, EPI_ISL_510151 | | | |
| see above | Hospital General Universitario Gregorio Marañón | SeqCOVID-SPAIN consortium/IBV(CSIC) | Laura Pérez-Lago, Marta Herranz, Jon Sicilia, Julia Suárez, Pilar Catalán, Patricia Muñoz, Darío García de Viedma and SeqCOVID-SPAIN consortium |
| EPI_ISL_510160 | Utah Public Health Laboratory | Pathogen Discovery, Respiratory Viruses Branch, Division of Viral Diseases, Centers for Disease Control and Prevention | Mary L. Killian, Kerrie Franzen, Krista Queen, Anna Uehara, Ying Tao, Yan Li, Jing Zhang, Clinton R. Paden, Haibin Wang, Suxiang Tong |
| EPI_ISL_510165, EPI_ISL_510166, EPI_ISL_510167, EPI_ISL_510168, EPI_ISL_510169, EPI_ISL_510170, EPI_ISL_510171, EPI_ISL_510172, EPI_ISL_510173, EPI_ISL_510174, EPI_ISL_510175, EPI_ISL_510176, EPI_ISL_510177, EPI_ISL_510178, EPI_ISL_510179, EPI_ISL_510180, EPI_ISL_510181, EPI_ISL_510182, EPI_ISL_510183, EPI_ISL_510184, EPI_ISL_510185, EPI_ISL_510186, EPI_ISL_510187, EPI_ISL_510188, EPI_ISL_510189, EPI_ISL_510190, EPI_ISL_510191, EPI_ISL_510192, EPI_ISL_510193, EPI_ISL_510194, EPI_ISL_510195, EPI_ISL_510196, EPI_ISL_510197, EPI_ISL_510198, EPI_ISL_510199, EPI_ISL_510200, EPI_ISL_510201, EPI_ISL_510202, EPI_ISL_510203, EPI_ISL_510204, EPI_ISL_510205, EPI_ISL_510206, EPI_ISL_510207, EPI_ISL_510208, EPI_ISL_510209, EPI_ISL_510210, EPI_ISL_510211, EPI_ISL_510212, EPI_ISL_510213, EPI_ISL_510214, EPI_ISL_510215, EPI_ISL_510216, EPI_ISL_510217, EPI_ISL_510218, EPI_ISL_510219, EPI_ISL_510220, EPI_ISL_510221, EPI_ISL_510222, EPI_ISL_510223, EPI_ISL_510224, EPI_ISL_510225, EPI_ISL_510226, EPI_ISL_510227, EPI_ISL_510228, EPI_ISL_510229, EPI_ISL_510230, EPI_ISL_510231, EPI_ISL_510232, EPI_ISL_510233, EPI_ISL_510234, EPI_ISL_510235, EPI_ISL_510236, EPI_ISL_510237, EPI_ISL_510238, EPI_ISL_510239, EPI_ISL_510240, EPI_ISL_510241, EPI_ISL_510242, EPI_ISL_510243, EPI_ISL_510244, EPI_ISL_510245 | | | |
| see above | Hospital General Universitario Gregorio Marañón | SeqCOVID-SPAIN consortium/IBV(CSIC) | Laura Pérez-Lago, Marta Herranz, Jon Sicilia, Julia Suárez, Pilar Catalán, Patricia Muñoz, Darío García de Viedma and SeqCOVID-SPAIN consortium |
| EPI_ISL_510246, EPI_ISL_510247, EPI_ISL_510248, EPI_ISL_510249, EPI_ISL_510250, EPI_ISL_510251, EPI_ISL_510252, EPI_ISL_510253, EPI_ISL_510254, EPI_ISL_510255, EPI_ISL_510256, EPI_ISL_510257, EPI_ISL_510258, EPI_ISL_510259, EPI_ISL_510260, EPI_ISL_510261, EPI_ISL_510262, EPI_ISL_510263, EPI_ISL_510264, EPI_ISL_510265, EPI_ISL_510266, EPI_ISL_510267 | | | |
| see above | Hospital de la Santa Creu i Sant Pau. Servicio de Microbiología | SeqCOVID-SPAIN consortium/IBV(CSIC) | Ferran Navarro, Núria Rabella, Elisenda Miró and SeqCOVID-SPAIN consortium |
| EPI_ISL_510268, EPI_ISL_510269, EPI_ISL_510270, EPI_ISL_510271, EPI_ISL_510272, EPI_ISL_510273, EPI_ISL_510274, EPI_ISL_510275, EPI_ISL_510276, EPI_ISL_510277, EPI_ISL_510278, EPI_ISL_510279, EPI_ISL_510280, EPI_ISL_510281, EPI_ISL_510282, EPI_ISL_510283, EPI_ISL_510284, EPI_ISL_510285, EPI_ISL_510286, EPI_ISL_510287, EPI_ISL_510288, EPI_ISL_510289, EPI_ISL_510290, EPI_ISL_510291, EPI_ISL_510292, EPI_ISL_510293, EPI_ISL_510294, EPI_ISL_510295, EPI_ISL_510296, EPI_ISL_510297, EPI_ISL_510298, EPI_ISL_510299, EPI_ISL_510300, EPI_ISL_510301, EPI_ISL_510302, EPI_ISL_510303, EPI_ISL_510304 | | | |
| see above | Hospital Clínico Universitario de Santiago de Compostela | SeqCOVID-SPAIN consortium/IBV(CSIC) | José Javier Costa Alcalde, Antonio Aguilera Guirao, Mª Luisa Pérez del Molino Bernal, Amparo Coira Nieto, Gema Barbeito Castiñeiras, Rocio Trastoy Pena and SeqCOVID-SPAIN consortium |
| EPI_ISL_510305, EPI_ISL_510306, EPI_ISL_510307, EPI_ISL_510308, EPI_ISL_510309, EPI_ISL_510310, EPI_ISL_510311, EPI_ISL_510312, EPI_ISL_510313, EPI_ISL_510314, EPI_ISL_510315, EPI_ISL_510316, EPI_ISL_510317, EPI_ISL_510318, EPI_ISL_510319, EPI_ISL_510320, EPI_ISL_510321, EPI_ISL_510322, EPI_ISL_510323, EPI_ISL_510324, EPI_ISL_510325, EPI_ISL_510326, EPI_ISL_510327, EPI_ISL_510328, EPI_ISL_510329, EPI_ISL_510330, EPI_ISL_510331, EPI_ISL_510332 | | | |
| see above | Hospital San Pedro de Alcántara (Cáceres) | SeqCOVID-SPAIN consortium/IBV(CSIC) | Cristina Muñoz Cuevas, Guadalupe Rodriguez Rodriguez and SeqCOVID-SPAIN consortium |
| EPI_ISL_510333, EPI_ISL_510334, EPI_ISL_510335, EPI_ISL_510336, EPI_ISL_510337, EPI_ISL_510338, EPI_ISL_510339, EPI_ISL_510340, EPI_ISL_510341, EPI_ISL_510342, EPI_ISL_510343, EPI_ISL_510344, EPI_ISL_510345, EPI_ISL_510346, EPI_ISL_510347, EPI_ISL_510348, EPI_ISL_510349, EPI_ISL_510350, EPI_ISL_510351, EPI_ISL_510352, EPI_ISL_510353, EPI_ISL_510354, EPI_ISL_510355, EPI_ISL_510356, EPI_ISL_510357, EPI_ISL_510358, EPI_ISL_510359, EPI_ISL_510360, EPI_ISL_510361, EPI_ISL_510362, EPI_ISL_510363, EPI_ISL_510364, EPI_ISL_510365, EPI_ISL_510366, EPI_ISL_510367, EPI_ISL_510368, EPI_ISL_510369, EPI_ISL_510370, EPI_ISL_510371, EPI_ISL_510372, EPI_ISL_510373, EPI_ISL_510374, EPI_ISL_510375, EPI_ISL_510376, EPI_ISL_510377, EPI_ISL_510378, EPI_ISL_510379, EPI_ISL_510380, EPI_ISL_510381, EPI_ISL_510382, EPI_ISL_510383, EPI_ISL_510384, EPI_ISL_510385, EPI_ISL_510386, EPI_ISL_510387, EPI_ISL_510388, EPI_ISL_510389, EPI_ISL_510390 | | | |
| see above | Servicio de Microbiología, Hospital Miguel Servet, Zaragoza | SeqCOVID-SPAIN consortium/IBV(CSIC) | Antonio Rezusta López, Alexander Tristancho Baró, Ana Milagro, Yolanda Gracia Grataloup, Nieves Martínez Cameo and SeqCOVID-SPAIN consortium |
| EPI_ISL_510391, EPI_ISL_510392, EPI_ISL_510393, EPI_ISL_510394, EPI_ISL_510395, EPI_ISL_510396, EPI_ISL_510397, EPI_ISL_510398, EPI_ISL_510399, EPI_ISL_510400, EPI_ISL_510401, EPI_ISL_510402, EPI_ISL_510403, EPI_ISL_510404, EPI_ISL_510405, EPI_ISL_510406, EPI_ISL_510407, EPI_ISL_510408, EPI_ISL_510409, EPI_ISL_510410, EPI_ISL_510411, EPI_ISL_510412, EPI_ISL_510413, EPI_ISL_510414, EPI_ISL_510415, EPI_ISL_510416, EPI_ISL_510417, EPI_ISL_510418, EPI_ISL_510419, EPI_ISL_510420, EPI_ISL_510421, EPI_ISL_510422, EPI_ISL_510423 | | | |
| see above | Servicio de Microbiología. Hospital Universitario Donostia. OSI Donostialdea. Área de Enfermedades Infecciosas, Grupo de Infección Respiratoria y Resistencia Antimicrobiana. Instituto de Investigación Sanitaria Biodonostia | SeqCOVID-SPAIN consortium/IBV(CSIC) | Gustavo Cilla, Milagrosa Montes, Luis Piñeiro, Jose Maria Marimón and SeqCOVID-SPAIN consortium |
| EPI_ISL_510424, EPI_ISL_510425, EPI_ISL_510426, EPI_ISL_510427, EPI_ISL_510428, EPI_ISL_510429, EPI_ISL_510430, EPI_ISL_510431, EPI_ISL_510432, EPI_ISL_510433, EPI_ISL_510434, EPI_ISL_510435, EPI_ISL_510436, EPI_ISL_510437, EPI_ISL_510438, EPI_ISL_510439, EPI_ISL_510440, EPI_ISL_510441, EPI_ISL_510442, EPI_ISL_510443, EPI_ISL_510444, EPI_ISL_510445, EPI_ISL_510446, EPI_ISL_510447, | | | |

| | | | |
|---|---|---|---|
| EPI_ISL_510448, EPI_ISL_510449, EPI_ISL_510450, EPI_ISL_510451, EPI_ISL_510452, EPI_ISL_510453, EPI_ISL_510454, EPI_ISL_510455, EPI_ISL_510456, EPI_ISL_510457, EPI_ISL_510458, EPI_ISL_510459, EPI_ISL_510460 | | | |
| see above | Hospital Universitario Virgen de las Nieves de Granada-SAS | SeqCOVID-SPAIN consortium/IBV(CSIC) | Mercedes Pérez Ruiz, Sara Sanbonmatsu Gámez, Irene Pedrosa Corral, José M. Navarro-Marí and SeqCOVID-SPAIN consortium |
| EPI_ISL_510461, EPI_ISL_510462, EPI_ISL_510463, EPI_ISL_510464, EPI_ISL_510465, EPI_ISL_510466 | Instituto de Investigaciones Biomédicas de Barcelona (CSIC), Hospital Clinic i Provincial de Barcelona, Instituto de Biomedicina de Valencia (CSIC), Hospital de Sant Pau | SeqCOVID-SPAIN consortium/IBV(CSIC) | Anna M. Planas, Mª Angeles Marcos, Miguel J. Martínez, Andrea Vergara, Alex Soriano, Jordi Pérez Tur, Israel Fernández Cadenas and SeqCOVID-SPAIN consortium |
| EPI_ISL_510467, EPI_ISL_510468, EPI_ISL_510469, EPI_ISL_510470, EPI_ISL_510471, EPI_ISL_510472, EPI_ISL_510473, EPI_ISL_510474, EPI_ISL_510475, EPI_ISL_510476, EPI_ISL_510477, EPI_ISL_510478, EPI_ISL_510479, EPI_ISL_510480, EPI_ISL_510481, EPI_ISL_510482, EPI_ISL_510483, EPI_ISL_510484, EPI_ISL_510485, EPI_ISL_510486, EPI_ISL_510487, EPI_ISL_510488, EPI_ISL_510489, EPI_ISL_510490, EPI_ISL_510491, EPI_ISL_510492, EPI_ISL_510493, EPI_ISL_510494, EPI_ISL_510495, EPI_ISL_510496, EPI_ISL_510497, EPI_ISL_510498, EPI_ISL_510499, EPI_ISL_510500, EPI_ISL_510501, EPI_ISL_510502, EPI_ISL_510503, EPI_ISL_510504, EPI_ISL_510505, EPI_ISL_510506, EPI_ISL_510507, EPI_ISL_510508, EPI_ISL_510509 | | | |
| see above | Servicio de Microbiología. Hospital Universitario Donostia. OSI Donostialdea. Área de Enfermedades Infecciosas, Grupo de Infección Respiratoria y Resistencia Antimicrobiana. Instituto de Investigación Sanitaria Biodonostia | SeqCOVID-SPAIN consortium/IBV(CSIC) | Gustavo Cilla, Milagrosa Montes, Luis Piñeiro, Jose Maria Marimón and SeqCOVID-SPAIN consortium |
| EPI_ISL_510510, EPI_ISL_510511, EPI_ISL_510512, EPI_ISL_510513, EPI_ISL_510514, EPI_ISL_510515, EPI_ISL_510516, EPI_ISL_510517, EPI_ISL_510518, EPI_ISL_510519, EPI_ISL_510520, EPI_ISL_510521, EPI_ISL_510522, EPI_ISL_510523 | | | |
| see above | Servicio de Microbiología, Laboratori Clínic Metropolitana Nord. Hospital Universitari Germans Trias i Pujol. Institut d'Investigació en Ciències de la Salut Germans Trias i Pujol (IGTP) | SeqCOVID-SPAIN consortium/IBV(CSIC) | Elisa Martró, Antoni E. Bordoy, Anna Not, Adrián Antuori, Anabel Fernández, Nona Romaní and SeqCOVID-SPAIN consortium |
| EPI_ISL_510524, EPI_ISL_510525 | University of Wisconsin-Madison AIDS Vaccine Research Laboratories | University of Wisconsin-Madison AIDS Vaccine Research Laboratories | Gage Moreno, Katarina Braun, et al. AIDS Vaccine Research Laboratories |
| EPI_ISL_510536 | Centro de Desenvolvimento Tecnologico em Saude, Fundacao Oswaldo Cruz | Centro de Desenvolvimento Tecnologico em Saude, Fundacao Oswaldo Cruz | Souza,T.M., Fintelman-Rodrigues,N., De Paula,A.D., Saraiva,F.B., Ferreira,M.A., Sacramento,C.Q. and Medeiros,M.A. |
| EPI_ISL_510537 | Microbiology Division, SC DHEC | Microbiology Division, SC DHEC | Flores,H. |
| EPI_ISL_510538, EPI_ISL_510539, EPI_ISL_510540 | Department of Microbiology, The University of Hong Kong | Department of Microbiology, The University of Hong Kong | Kelvin K.W. To, Kwok-Yung Yuen |
| EPI_ISL_510541 | Centro de Desenvolvimento Tecnologico em Saude, Fundacao Oswaldo Cruz | Centro de Desenvolvimento Tecnologico em Saude, Fundacao Oswaldo Cruz | Souza,T.M., Fintelman-Rodrigues,N., De Paula,A.D., Saraiva,F.B., Ferreira,M.A., Sacramento,C.Q. and Medeiros,M.A. |
| EPI_ISL_510542, EPI_ISL_510543, EPI_ISL_510544 | SA Pathology | SA Pathology | Lex Leong, Chuan Kok Lim, Mark Turra, Ivan Bastian, Geoff Higgins |
| EPI_ISL_510547, EPI_ISL_510548, EPI_ISL_510549, EPI_ISL_510550, EPI_ISL_510551, EPI_ISL_510552, EPI_ISL_510553, EPI_ISL_510554, EPI_ISL_510555, EPI_ISL_510556, EPI_ISL_510557, EPI_ISL_510558, EPI_ISL_510559, EPI_ISL_510560, EPI_ISL_510561, EPI_ISL_510562, EPI_ISL_510563, EPI_ISL_510564, EPI_ISL_510565, EPI_ISL_510566, EPI_ISL_510567, EPI_ISL_510568, EPI_ISL_510569, EPI_ISL_510570, EPI_ISL_510571, EPI_ISL_510572, EPI_ISL_510573, EPI_ISL_510574, EPI_ISL_510575, EPI_ISL_510576, EPI_ISL_510577, EPI_ISL_510578, EPI_ISL_510579, EPI_ISL_510580, EPI_ISL_510581, EPI_ISL_510582, EPI_ISL_510583, EPI_ISL_510584, EPI_ISL_510585, EPI_ISL_510586, EPI_ISL_510587, EPI_ISL_510588, EPI_ISL_510589, EPI_ISL_510590, EPI_ISL_510591, EPI_ISL_510592, EPI_ISL_510593, EPI_ISL_510594, EPI_ISL_510595, EPI_ISL_510596, EPI_ISL_510597, EPI_ISL_510598, EPI_ISL_510599, EPI_ISL_510600, EPI_ISL_510601, EPI_ISL_510602, EPI_ISL_510603, EPI_ISL_510604, EPI_ISL_510605, EPI_ISL_510606, EPI_ISL_510607, EPI_ISL_510608, EPI_ISL_510609, EPI_ISL_510610, EPI_ISL_510611, EPI_ISL_510612, EPI_ISL_510613, EPI_ISL_510614, EPI_ISL_510615, EPI_ISL_510616, EPI_ISL_510617, EPI_ISL_510618, EPI_ISL_510619, EPI_ISL_510620, EPI_ISL_510621, EPI_ISL_510622, EPI_ISL_510623, EPI_ISL_510624, EPI_ISL_510625, EPI_ISL_510626, EPI_ISL_510627, EPI_ISL_510628, EPI_ISL_510629, EPI_ISL_510630, EPI_ISL_510631, EPI_ISL_510632, EPI_ISL_510633, EPI_ISL_510634, EPI_ISL_510635, EPI_ISL_510636, EPI_ISL_510637, EPI_ISL_510638, EPI_ISL_510639, EPI_ISL_510640, EPI_ISL_510641, EPI_ISL_510642, EPI_ISL_510643, EPI_ISL_510644, EPI_ISL_510645, EPI_ISL_510646, EPI_ISL_510647, EPI_ISL_510648, EPI_ISL_510649, EPI_ISL_510650, EPI_ISL_510651, EPI_ISL_510652, EPI_ISL_510653, EPI_ISL_510654, EPI_ISL_510655, EPI_ISL_510656, EPI_ISL_510657, EPI_ISL_510658, EPI_ISL_510659, EPI_ISL_510660, EPI_ISL_510661, EPI_ISL_510662, EPI_ISL_510663, EPI_ISL_510664, EPI_ISL_510665, EPI_ISL_510666, EPI_ISL_510667, EPI_ISL_510668, EPI_ISL_510669, EPI_ISL_510670, EPI_ISL_510671, EPI_ISL_510672, EPI_ISL_510673, EPI_ISL_510674, EPI_ISL_510675, EPI_ISL_510676, EPI_ISL_510677, EPI_ISL_510678, EPI_ISL_510679, EPI_ISL_510680, EPI_ISL_510681, EPI_ISL_510682, EPI_ISL_510683, EPI_ISL_510684, EPI_ISL_510685, EPI_ISL_510686, EPI_ISL_510687, EPI_ISL_510688 | | | |
| see above | Division of Viral Diseases, Center for Laboratory Control of Infectious Diseases, Korea Centers for Diseases Control and Prevention | Division of Viral Diseases, Center for Laboratory Control of Infectious Diseases, Korea Centers for Diseases Control and Prevention | Jeong-Min Kim, Yoon-Seok Chung, Namjoo Lee, Sang Hee Woo, Hye-Jun Jo, Heui Man Kim, Jun-Sub Kim, Myung Guk Han |
| EPI_ISL_510689 | Hospital Universitari Germans Trias i Pujol(HUGTiP)/Fundació Lluita contra la SIDA (FLSida)/IRTA-CReSA | IrsiCaixa AIDS Research Lab | Pilar Armengol, Marc Noguera-Julian, Jordi Rodón, Julia Vergara, Lidia Ruiz, Nuria Izquierdo, Jorge Carrillo, Roger Paredes, Albert Bensaid, Julia Blanco, Joaquim Segalés, Bonaventura Clotet |
| EPI_ISL_510690, EPI_ISL_510691, EPI_ISL_510692, EPI_ISL_510693, EPI_ISL_510694, EPI_ISL_510695, EPI_ISL_510696, EPI_ISL_510697, EPI_ISL_510698, EPI_ISL_510699, EPI_ISL_510700, EPI_ISL_510701, EPI_ISL_510702, EPI_ISL_510703, EPI_ISL_510704, EPI_ISL_510705, EPI_ISL_510706, EPI_ISL_510707, EPI_ISL_510708, EPI_ISL_510709, EPI_ISL_510710, EPI_ISL_510711, EPI_ISL_510712, EPI_ISL_510713, EPI_ISL_510714, EPI_ISL_510715, EPI_ISL_510716, EPI_ISL_510717, EPI_ISL_510718, EPI_ISL_510719, EPI_ISL_510720, EPI_ISL_510721, EPI_ISL_510722, EPI_ISL_510723, EPI_ISL_510724, EPI_ISL_510725, EPI_ISL_510726, EPI_ISL_510727, EPI_ISL_510728, EPI_ISL_510729, EPI_ISL_510730, EPI_ISL_510731, EPI_ISL_510732, EPI_ISL_510733, EPI_ISL_510734, EPI_ISL_510735, EPI_ISL_510736, EPI_ISL_510737, EPI_ISL_510738, EPI_ISL_510739, EPI_ISL_510740, EPI_ISL_510741, EPI_ISL_510742, EPI_ISL_510743, EPI_ISL_510744, EPI_ISL_510745, EPI_ISL_510746, EPI_ISL_510747, EPI_ISL_510748, EPI_ISL_510749, EPI_ISL_510750, EPI_ISL_510751, EPI_ISL_510752, EPI_ISL_510753, EPI_ISL_510754, EPI_ISL_510755, EPI_ISL_510756, EPI_ISL_510757, EPI_ISL_510758, EPI_ISL_510759, EPI_ISL_510760, EPI_ISL_510761, EPI_ISL_510762, EPI_ISL_510763, EPI_ISL_510764, EPI_ISL_510765, EPI_ISL_510766, EPI_ISL_510767, EPI_ISL_510768, EPI_ISL_510769, EPI_ISL_510770, EPI_ISL_510771, EPI_ISL_510772, EPI_ISL_510773, EPI_ISL_510774, EPI_ISL_510775, EPI_ISL_510776, EPI_ISL_510777, EPI_ISL_510778, EPI_ISL_510779, EPI_ISL_510780, EPI_ISL_510781, EPI_ISL_510782, EPI_ISL_510783, EPI_ISL_510784, EPI_ISL_510785, EPI_ISL_510786, EPI_ISL_510787, EPI_ISL_510788, EPI_ISL_510789, EPI_ISL_510790, EPI_ISL_510791, EPI_ISL_510792, EPI_ISL_510793, EPI_ISL_510794, EPI_ISL_510795, EPI_ISL_510796, EPI_ISL_510797, EPI_ISL_510798, EPI_ISL_510799, EPI_ISL_510800, EPI_ISL_510801, EPI_ISL_510802, EPI_ISL_510803, EPI_ISL_510804, EPI_ISL_510805, EPI_ISL_510806, EPI_ISL_510807, EPI_ISL_510808, EPI_ISL_510809 | | | |
| see above | Viollier AG | Department of Biosystems Science and Engineering, ETH Zürich | Christian Beisel, Sarah Nadeau, Ivan Topolsky, Pedro Ferreira, Philipp Jablonski, Susana Posada-Céspedes, Tobias Schär, Ina Nissen, Natascha Santacroce, Elodie Burcklen, Christiane Beckmann, Maurice Redondo, Olivier Kobel, Christoph Noppen, Sophie Seidel, Noemie Santamaria de Souza, Niko Beerenwinkel, Tanja Stadler |
| EPI_ISL_510810, EPI_ISL_510811, EPI_ISL_510812, EPI_ISL_510813, EPI_ISL_510814, EPI_ISL_510815, EPI_ISL_510816 | NA | The Public Health Agency of Sweden | Oskar Karlsson Lindsjo, Maria Lind Karlberg, Mattias Haukland, Reza Advani, Olov Svartstrom, Anna-Malin Linde, Sandra Broddesson, Petra Edquist, Mia Brytting, Anna Risberg, Karin Tegmark-Wisell |
| EPI_ISL_510817 | Orebro klinisk mikrobiologi | The Public Health Agency of Sweden | Oskar Karlsson Lindsjo, Maria Lind Karlberg, Mattias Haukland, Reza Advani, Olov Svartstrom, Anna-Malin Linde, Sandra Broddesson, Petra Edquist, Mia Brytting, Anna Risberg, Karin Tegmark-Wisell |
| EPI_ISL_510818, EPI_ISL_510819 | Klinisk mikrobiologi Vasternorrland | The Public Health Agency of Sweden | Oskar Karlsson Lindsjo, Maria Lind Karlberg, Mattias Haukland, Reza Advani, Olov Svartstrom, Anna-Malin Linde, Sandra Broddesson, Petra Edquist, Mia Brytting, Anna Risberg, Karin Tegmark-Wisell |
| EPI_ISL_510820, EPI_ISL_510821, EPI_ISL_510822, EPI_ISL_510823, EPI_ISL_510824 | Klinisk mikrobiologi centralsjukhuset Karlstad | The Public Health Agency of Sweden | Oskar Karlsson Lindsjo, Maria Lind Karlberg, Mattias Haukland, Reza Advani, Olov Svartstrom, Anna-Malin Linde, Sandra Broddesson, Petra Edquist, Mia Brytting, Anna Risberg, Karin Tegmark-Wisell |
| EPI_ISL_510825, EPI_ISL_510826 | Kalmar klinisk mikrobiologi | The Public Health Agency of Sweden | Oskar Karlsson Lindsjo, Maria Lind Karlberg, Mattias Haukland, Reza Advani, Olov Svartstrom, Anna-Malin Linde, Sandra Broddesson, Petra Edquist, Mia Brytting, Anna Risberg, Karin Tegmark-Wisell |
| EPI_ISL_510827, EPI_ISL_510828, EPI_ISL_510829, EPI_ISL_510830 | Unilabs Eskilstuna | The Public Health Agency of Sweden | Oskar Karlsson Lindsjo, Maria Lind Karlberg, Mattias Haukland, Reza Advani, Olov Svartstrom, Anna-Malin Linde, Sandra Broddesson, Petra Edquist, Mia Brytting, Anna Risberg, Karin Tegmark-Wisell |
| EPI_ISL_510831 | Halmstad klinisk mikrobiologi | The Public Health Agency of Sweden | Oskar Karlsson Lindsjo, Maria Lind Karlberg, Mattias Haukland, Reza Advani, Olov Svartstrom, Anna-Malin Linde, Sandra Broddesson, Petra Edquist, Mia Brytting, Anna Risberg, Karin Tegmark-Wisell |
| EPI_ISL_510832, EPI_ISL_510833 | Kalmar klinisk mikrobiologi | The Public Health Agency of Sweden | Oskar Karlsson Lindsjo, Maria Lind Karlberg, Mattias Haukland, Reza Advani, Olov Svartstrom, Anna-Malin Linde, Sandra Broddesson, Petra Edquist, Mia Brytting, Anna Risberg, Karin Tegmark-Wisell |
| EPI_ISL_510834, EPI_ISL_510835 | Karolinska Universitetslaboratoriet | The Public Health Agency of Sweden | Oskar Karlsson Lindsjo, Maria Lind Karlberg, Mattias Haukland, Reza Advani, Olov Svartstrom, Anna-Malin Linde, Sandra Broddesson, Petra Edquist, Mia Brytting, Anna Risberg, Karin Tegmark-Wisell |
| EPI_ISL_510836, EPI_ISL_510837 | Kalmar klinisk mikrobiologi | The Public Health Agency of Sweden | Oskar Karlsson Lindsjo, Maria Lind Karlberg, Mattias Haukland, Reza Advani, Olov Svartstrom, Anna-Malin Linde, Sandra Broddesson, Petra Edquist, Mia Brytting, Anna Risberg, Karin Tegmark-Wisell |
| EPI_ISL_510838, EPI_ISL_510839, EPI_ISL_510840 | Karolinska Universitetslaboratoriet | The Public Health Agency of Sweden | Oskar Karlsson Lindsjo, Maria Lind Karlberg, Mattias Haukland, Reza Advani, Olov Svartstrom, Anna-Malin Linde, Sandra Broddesson, Petra Edquist, Mia Brytting, Anna Risberg, Karin Tegmark-Wisell |
| EPI_ISL_510841, EPI_ISL_510842, EPI_ISL_510844 | Klinisk mikrobiologi NAL Trollhattan | The Public Health Agency of Sweden | Oskar Karlsson Lindsjo, Maria Lind Karlberg, Mattias Haukland, Reza Advani, Olov Svartstrom, Anna-Malin Linde, Sandra Broddesson, Petra Edquist, Mia Brytting, Anna Risberg, Karin Tegmark-Wisell |
| EPI_ISL_510845, EPI_ISL_510846, EPI_ISL_510847, EPI_ISL_510848, EPI_ISL_510849, EPI_ISL_510850, EPI_ISL_510851 | Skovde/Unilabs | The Public Health Agency of Sweden | Oskar Karlsson Lindsjo, Maria Lind Karlberg, Mattias Haukland, Reza Advani, Olov Svartstrom, Anna-Malin Linde, Sandra Broddesson, Petra Edquist, Mia Brytting, Anna Risberg, Karin Tegmark-Wisell |
| EPI_ISL_510852 | Karolinska Universitetslaboratoriet | The Public Health Agency of Sweden | Oskar Karlsson Lindsjo, Maria Lind Karlberg, Mattias Haukland, Reza Advani, Olov Svartstrom, Anna-Malin Linde, Sandra Broddesson, Petra Edquist, Mia Brytting, Anna Risberg, Karin Tegmark-Wisell |
| EPI_ISL_510853, EPI_ISL_510854, EPI_ISL_510855, EPI_ISL_510856, EPI_ISL_510857, EPI_ISL_510858 | Klinisk mikrobiologi Vasternorrland | The Public Health Agency of Sweden | Oskar Karlsson Lindsjo, Maria Lind Karlberg, Mattias Haukland, Reza Advani, Olov Svartstrom, Anna-Malin Linde, Sandra Broddesson, Petra Edquist, Mia Brytting, Anna Risberg, Karin Tegmark-Wisell |
| EPI_ISL_510859 | Kalmar klinisk mikrobiologi | The Public Health Agency of Sweden | Oskar Karlsson Lindsjo, Maria Lind Karlberg, Mattias Haukland, Reza Advani, Olov Svartstrom, Anna-Malin Linde, Sandra Broddesson, Petra Edquist, Mia Brytting, Anna Risberg, Karin Tegmark-Wisell |
| EPI_ISL_510860, EPI_ISL_510861, EPI_ISL_510862, EPI_ISL_510863 | Karolinska Universitetslaboratoriet | The Public Health Agency of Sweden | Oskar Karlsson Lindsjo, Maria Lind Karlberg, Mattias Haukland, Reza Advani, Olov Svartstrom, Anna-Malin Linde, Sandra Broddesson, Petra Edquist, Mia Brytting, Anna Risberg, Karin Tegmark-Wisell |
| EPI_ISL_510864, EPI_ISL_510865 | Kalmar klinisk mikrobiologi | The Public Health Agency of Sweden | Oskar Karlsson Lindsjo, Maria Lind Karlberg, Mattias Haukland, Reza Advani, Olov Svartstrom, Anna-Malin Linde, Sandra Broddesson, Petra Edquist, Mia Brytting, Anna Risberg, Karin Tegmark-Wisell |
| EPI_ISL_510866 | Halmstad klinisk mikrobiologi | The Public Health Agency of Sweden | Oskar Karlsson Lindsjo, Maria Lind Karlberg, Mattias Haukland, Reza Advani, Olov Svartstrom, Anna-Malin Linde, Sandra Broddesson, Petra Edquist, Mia Brytting, Anna Risberg, Karin Tegmark-Wisell |
| EPI_ISL_510867 | Klinisk mikrobiologi centralsjukhuset Karlstad | The Public Health Agency of Sweden | Oskar Karlsson Lindsjo, Maria Lind Karlberg, Mattias Haukland, Reza Advani, Olov Svartstrom, Anna-Malin Linde, Sandra Broddesson, Petra Edquist, Mia Brytting, Anna Risberg, Karin Tegmark-Wisell |

| | | | |
|---|---|---|---|
| EPI_ISL_510868, EPI_ISL_510869, EPI_ISL_510870 | Klinisk mikrobiologi NAL Trollhattan | The Public Health Agency of Sweden | Oskar Karlsson Lindsjo, Maria Lind Karlberg, Mattias Haukland, Reza Advani, Olov Svartstrom, Anna-Malin Linde, Sandra Broddesson, Petra Edquist, Mia Brytting, Anna Risberg, Karin Tegmark-Wisell |
| EPI_ISL_510871, EPI_ISL_510872, EPI_ISL_510873, EPI_ISL_510874, EPI_ISL_510875, EPI_ISL_510876, EPI_ISL_510877 | Klinsisk mikrobiologi Linkoping | The Public Health Agency of Sweden | Oskar Karlsson Lindsjo, Maria Lind Karlberg, Mattias Haukland, Reza Advani, Olov Svartstrom, Anna-Malin Linde, Sandra Broddesson, Petra Edquist, Mia Brytting, Anna Risberg, Karin Tegmark-Wisell |
| EPI_ISL_510878, EPI_ISL_510879, EPI_ISL_510880, EPI_ISL_510881, EPI_ISL_510882, EPI_ISL_510883, EPI_ISL_510884, EPI_ISL_510885, EPI_ISL_510886, EPI_ISL_510887, EPI_ISL_510888 | | | |
| see above | New Mexico Department of Health Scientific Laboratory | New Mexico Department of Health Scientific Laboratory | Ellie Johnson, Anastacia Griego-Fisher |
| EPI_ISL_510889, EPI_ISL_510890, EPI_ISL_510891, EPI_ISL_510892, EPI_ISL_510893, EPI_ISL_510894, EPI_ISL_510895, EPI_ISL_510896, EPI_ISL_510897, EPI_ISL_510898, EPI_ISL_510899, EPI_ISL_510900, EPI_ISL_510901, EPI_ISL_510902, EPI_ISL_510903, EPI_ISL_510904, EPI_ISL_510905, EPI_ISL_510906, EPI_ISL_510907, EPI_ISL_510908, EPI_ISL_510909, EPI_ISL_510910, EPI_ISL_510911, EPI_ISL_510912, EPI_ISL_510913, EPI_ISL_510914, EPI_ISL_510915, EPI_ISL_510916, EPI_ISL_510917, EPI_ISL_510918, EPI_ISL_510919, EPI_ISL_510920, EPI_ISL_510921, EPI_ISL_510922, EPI_ISL_510923, EPI_ISL_510924, EPI_ISL_510925, EPI_ISL_510926, EPI_ISL_510927, EPI_ISL_510928, EPI_ISL_510929, EPI_ISL_510930, EPI_ISL_510931, EPI_ISL_510932, EPI_ISL_510933, EPI_ISL_510934, EPI_ISL_510935, EPI_ISL_510936, EPI_ISL_510937, EPI_ISL_510938, EPI_ISL_510939, EPI_ISL_510940, EPI_ISL_510941, EPI_ISL_510942, EPI_ISL_510943, EPI_ISL_510944, EPI_ISL_510945, EPI_ISL_510946, EPI_ISL_510947, EPI_ISL_510948, EPI_ISL_510949, EPI_ISL_510950, EPI_ISL_510951, EPI_ISL_510952, EPI_ISL_510953, EPI_ISL_510954, EPI_ISL_510955, EPI_ISL_510956, EPI_ISL_510957, EPI_ISL_510958, EPI_ISL_510959, EPI_ISL_510960, EPI_ISL_510961, EPI_ISL_510962, EPI_ISL_510963, EPI_ISL_510964, EPI_ISL_510965, EPI_ISL_510966, EPI_ISL_510967, EPI_ISL_510968, EPI_ISL_510969, EPI_ISL_510970, EPI_ISL_510971, EPI_ISL_510972, EPI_ISL_510973, EPI_ISL_510974, EPI_ISL_510975, EPI_ISL_510976, EPI_ISL_510977, EPI_ISL_510978, EPI_ISL_510979, EPI_ISL_510980, EPI_ISL_510981, EPI_ISL_510982, EPI_ISL_510983, EPI_ISL_510984, EPI_ISL_510985, EPI_ISL_510986, EPI_ISL_510987, EPI_ISL_510988, EPI_ISL_510989, EPI_ISL_510990, EPI_ISL_510991, EPI_ISL_510992, EPI_ISL_510993, EPI_ISL_510994, EPI_ISL_510995, EPI_ISL_510996, EPI_ISL_510997, EPI_ISL_510998, EPI_ISL_510999, EPI_ISL_511000, EPI_ISL_511001, EPI_ISL_511002, EPI_ISL_511003, EPI_ISL_511004, EPI_ISL_511005, EPI_ISL_511006, EPI_ISL_511007, EPI_ISL_511008, EPI_ISL_511009, EPI_ISL_511010, EPI_ISL_511011, EPI_ISL_511012, EPI_ISL_511013, EPI_ISL_511014, EPI_ISL_511015, EPI_ISL_511016, EPI_ISL_511017, EPI_ISL_511018, EPI_ISL_511019, EPI_ISL_511020, EPI_ISL_511021, EPI_ISL_511022, EPI_ISL_511023, EPI_ISL_511024, EPI_ISL_511025, EPI_ISL_511026, EPI_ISL_511027, EPI_ISL_511028, EPI_ISL_511029, EPI_ISL_511030, EPI_ISL_511031, EPI_ISL_511032, EPI_ISL_511033, EPI_ISL_511034, EPI_ISL_511035, EPI_ISL_511036, EPI_ISL_511037, EPI_ISL_511038, EPI_ISL_511039, EPI_ISL_511040, EPI_ISL_511041, EPI_ISL_511042, EPI_ISL_511043, EPI_ISL_511044, EPI_ISL_511045, EPI_ISL_511046, EPI_ISL_511047, EPI_ISL_511048, EPI_ISL_511049, EPI_ISL_511050, EPI_ISL_511051, EPI_ISL_511052, EPI_ISL_511053, EPI_ISL_511054, EPI_ISL_511055, EPI_ISL_511056, EPI_ISL_511057, EPI_ISL_511058, EPI_ISL_511059, EPI_ISL_511060, EPI_ISL_511061, EPI_ISL_511062, EPI_ISL_511063, EPI_ISL_511064, EPI_ISL_511065, EPI_ISL_511066, EPI_ISL_511067, EPI_ISL_511068, EPI_ISL_511069, EPI_ISL_511070, EPI_ISL_511071, EPI_ISL_511072, EPI_ISL_511073, EPI_ISL_511074, EPI_ISL_511075, EPI_ISL_511076, EPI_ISL_511077, EPI_ISL_511078, EPI_ISL_511079, EPI_ISL_511080, EPI_ISL_511081, EPI_ISL_511082, EPI_ISL_511083, EPI_ISL_511084, EPI_ISL_511085, EPI_ISL_511086, EPI_ISL_511087, EPI_ISL_511088, EPI_ISL_511089, EPI_ISL_511090, EPI_ISL_511091, EPI_ISL_511092, EPI_ISL_511093, EPI_ISL_511094, EPI_ISL_511095, EPI_ISL_511096, EPI_ISL_511097, EPI_ISL_511098, EPI_ISL_511099, EPI_ISL_511100, EPI_ISL_511101, EPI_ISL_511102, EPI_ISL_511103, EPI_ISL_511104, EPI_ISL_511105, EPI_ISL_511106, EPI_ISL_511107, EPI_ISL_511108, EPI_ISL_511109, EPI_ISL_511110, EPI_ISL_511111, EPI_ISL_511112, EPI_ISL_511113, EPI_ISL_511114, EPI_ISL_511115, EPI_ISL_511116, EPI_ISL_511117, EPI_ISL_511118, EPI_ISL_511119, EPI_ISL_511120, EPI_ISL_511121, EPI_ISL_511122, EPI_ISL_511123, EPI_ISL_511124, EPI_ISL_511125, EPI_ISL_511126, EPI_ISL_511127, EPI_ISL_511128, EPI_ISL_511129, EPI_ISL_511130, EPI_ISL_511131, EPI_ISL_511132, EPI_ISL_511133, EPI_ISL_511134, EPI_ISL_511135, EPI_ISL_511136, EPI_ISL_511137, EPI_ISL_511138, EPI_ISL_511139, EPI_ISL_511140, EPI_ISL_511141, EPI_ISL_511142, EPI_ISL_511143, EPI_ISL_511144, EPI_ISL_511145, EPI_ISL_511146, EPI_ISL_511147, EPI_ISL_511148, EPI_ISL_511149, EPI_ISL_511150, EPI_ISL_511151, EPI_ISL_511152, EPI_ISL_511153, EPI_ISL_511154, EPI_ISL_511155, EPI_ISL_511156, EPI_ISL_511157, EPI_ISL_511158, EPI_ISL_511159, EPI_ISL_511160, EPI_ISL_511161, EPI_ISL_511162, EPI_ISL_511163, EPI_ISL_511164, EPI_ISL_511165, EPI_ISL_511166, EPI_ISL_511167, EPI_ISL_511168, EPI_ISL_511169, EPI_ISL_511170, EPI_ISL_511171, EPI_ISL_511172, EPI_ISL_511173, EPI_ISL_511174, EPI_ISL_511175, EPI_ISL_511176, EPI_ISL_511177, EPI_ISL_511178, EPI_ISL_511179, EPI_ISL_511180, EPI_ISL_511181 | | | |
| see above | Instituto Nacional de Saude (INSA) | Instituto Nacional de Saude (INSA) | Borges et al |
| EPI_ISL_511182, EPI_ISL_511183, EPI_ISL_511184, EPI_ISL_511185, EPI_ISL_511186, EPI_ISL_511187, EPI_ISL_511188 | Instituto Nacional de Saude (INSA) and Instituto Gulbenkian de Ciencia (IGC) | Instituto Nacional de Saude (INSA) and Instituto Gulbenkian de Ciencia (IGC) | Borges et al |
| EPI_ISL_511189 | Instituto Nacional de Saude (INSA) | Instituto Nacional de Saude (INSA) | Borges et al |
| EPI_ISL_511190, EPI_ISL_511191, EPI_ISL_511192, EPI_ISL_511193, EPI_ISL_511194, EPI_ISL_511195, EPI_ISL_511196, EPI_ISL_511197, EPI_ISL_511198, EPI_ISL_511199, EPI_ISL_511200, EPI_ISL_511201, EPI_ISL_511202, EPI_ISL_511203, EPI_ISL_511204, EPI_ISL_511205, EPI_ISL_511206, EPI_ISL_511207, EPI_ISL_511208, EPI_ISL_511209, EPI_ISL_511210, EPI_ISL_511211, EPI_ISL_511212, EPI_ISL_511213, EPI_ISL_511214, EPI_ISL_511215, EPI_ISL_511216, EPI_ISL_511217, EPI_ISL_511218, EPI_ISL_511219, EPI_ISL_511220, EPI_ISL_511221, EPI_ISL_511222, EPI_ISL_511223, EPI_ISL_511224, EPI_ISL_511225, EPI_ISL_511226, EPI_ISL_511227, EPI_ISL_511228, EPI_ISL_511229, EPI_ISL_511230, EPI_ISL_511231, EPI_ISL_511232, EPI_ISL_511233, EPI_ISL_511234, EPI_ISL_511235, EPI_ISL_511236, EPI_ISL_511237, EPI_ISL_511238, EPI_ISL_511239, EPI_ISL_511240, EPI_ISL_511241, EPI_ISL_511242, EPI_ISL_511243, EPI_ISL_511244, EPI_ISL_511245, EPI_ISL_511246, EPI_ISL_511247, EPI_ISL_511248, EPI_ISL_511249, EPI_ISL_511250, EPI_ISL_511251, EPI_ISL_511252, EPI_ISL_511253, EPI_ISL_511254, EPI_ISL_511255, EPI_ISL_511256, EPI_ISL_511257, EPI_ISL_511258, EPI_ISL_511259, EPI_ISL_511260, EPI_ISL_511261, EPI_ISL_511262, EPI_ISL_511263, EPI_ISL_511264, EPI_ISL_511265, EPI_ISL_511266, EPI_ISL_511267, EPI_ISL_511268, EPI_ISL_511269, EPI_ISL_511270, EPI_ISL_511271, EPI_ISL_511272, EPI_ISL_511273, EPI_ISL_511274, EPI_ISL_511275, EPI_ISL_511276, EPI_ISL_511277, EPI_ISL_511278, EPI_ISL_511279, EPI_ISL_511280, EPI_ISL_511281, EPI_ISL_511282, EPI_ISL_511283, EPI_ISL_511284, EPI_ISL_511285, EPI_ISL_511286, EPI_ISL_511287, EPI_ISL_511288, EPI_ISL_511289, EPI_ISL_511290, EPI_ISL_511291, EPI_ISL_511292, EPI_ISL_511293, EPI_ISL_511294, EPI_ISL_511295, EPI_ISL_511296, EPI_ISL_511297, EPI_ISL_511298, EPI_ISL_511299, EPI_ISL_511300, EPI_ISL_511301, EPI_ISL_511302, EPI_ISL_511303, EPI_ISL_511304, EPI_ISL_511305, EPI_ISL_511306, EPI_ISL_511307, EPI_ISL_511308, EPI_ISL_511309, EPI_ISL_511310, EPI_ISL_511311, EPI_ISL_511312, EPI_ISL_511313, EPI_ISL_511314, EPI_ISL_511315 | | | |
| see above | Instituto Nacional de Saude (INSA) and Instituto Gulbenkian de Ciencia (IGC) | Instituto Nacional de Saude (INSA) and Instituto Gulbenkian de Ciencia (IGC) | Borges et al |
| EPI_ISL_511316, EPI_ISL_511317, EPI_ISL_511318, EPI_ISL_511319, EPI_ISL_511320, EPI_ISL_511321, EPI_ISL_511322, EPI_ISL_511323, EPI_ISL_511324, EPI_ISL_511325, EPI_ISL_511326, EPI_ISL_511327, EPI_ISL_511328, EPI_ISL_511329, EPI_ISL_511330, EPI_ISL_511331, EPI_ISL_511332, EPI_ISL_511333, EPI_ISL_511334, EPI_ISL_511335, EPI_ISL_511336, EPI_ISL_511337, EPI_ISL_511338, EPI_ISL_511339, EPI_ISL_511340, EPI_ISL_511341, EPI_ISL_511342, EPI_ISL_511343, EPI_ISL_511344, EPI_ISL_511345, EPI_ISL_511346, EPI_ISL_511347, EPI_ISL_511348, EPI_ISL_511349, EPI_ISL_511350, EPI_ISL_511351, EPI_ISL_511352, EPI_ISL_511353, EPI_ISL_511354, EPI_ISL_511355, EPI_ISL_511356, EPI_ISL_511357, EPI_ISL_511358, EPI_ISL_511359, EPI_ISL_511360, EPI_ISL_511361, EPI_ISL_511362, EPI_ISL_511363, EPI_ISL_511364, EPI_ISL_511365, EPI_ISL_511366, EPI_ISL_511367, EPI_ISL_511368, EPI_ISL_511369, EPI_ISL_511370, EPI_ISL_511371, EPI_ISL_511372, EPI_ISL_511373, EPI_ISL_511374, EPI_ISL_511375, EPI_ISL_511376, EPI_ISL_511377, EPI_ISL_511378, EPI_ISL_511379, EPI_ISL_511380, EPI_ISL_511381, EPI_ISL_511382, EPI_ISL_511383, EPI_ISL_511384, EPI_ISL_511385, EPI_ISL_511386, EPI_ISL_511387, EPI_ISL_511388, EPI_ISL_511389, EPI_ISL_511390, EPI_ISL_511391, EPI_ISL_511392, EPI_ISL_511393, EPI_ISL_511394, EPI_ISL_511395, EPI_ISL_511396, EPI_ISL_511397, EPI_ISL_511398, EPI_ISL_511399, EPI_ISL_511400, EPI_ISL_511401, EPI_ISL_511402, EPI_ISL_511403, EPI_ISL_511404, EPI_ISL_511405, EPI_ISL_511406, EPI_ISL_511407, EPI_ISL_511408, EPI_ISL_511409, EPI_ISL_511410, EPI_ISL_511411, EPI_ISL_511412, EPI_ISL_511413, EPI_ISL_511414, EPI_ISL_511415, EPI_ISL_511416, EPI_ISL_511417, EPI_ISL_511418, EPI_ISL_511419, EPI_ISL_511420, EPI_ISL_511421, EPI_ISL_511422, EPI_ISL_511423, EPI_ISL_511424, EPI_ISL_511425, EPI_ISL_511426, EPI_ISL_511427, EPI_ISL_511428, EPI_ISL_511429, EPI_ISL_511430, EPI_ISL_511431, EPI_ISL_511432, EPI_ISL_511433, EPI_ISL_511434, EPI_ISL_511435, EPI_ISL_511436, EPI_ISL_511437, EPI_ISL_511438, EPI_ISL_511439, EPI_ISL_511440, EPI_ISL_511441, EPI_ISL_511442, EPI_ISL_511443, EPI_ISL_511444, EPI_ISL_511445, EPI_ISL_511446, EPI_ISL_511447, EPI_ISL_511448, EPI_ISL_511449, EPI_ISL_511450, EPI_ISL_511451, EPI_ISL_511452, EPI_ISL_511453, EPI_ISL_511454, EPI_ISL_511455, EPI_ISL_511456, EPI_ISL_511457, EPI_ISL_511458, EPI_ISL_511459, EPI_ISL_511460, EPI_ISL_511461, EPI_ISL_511462, EPI_ISL_511463, EPI_ISL_511464, EPI_ISL_511465, EPI_ISL_511466, EPI_ISL_511467, EPI_ISL_511468, EPI_ISL_511469, EPI_ISL_511470, EPI_ISL_511471, EPI_ISL_511472, EPI_ISL_511473, EPI_ISL_511474, EPI_ISL_511475, EPI_ISL_511476, EPI_ISL_511477, EPI_ISL_511478, EPI_ISL_511479, EPI_ISL_511480, EPI_ISL_511481, EPI_ISL_511482, EPI_ISL_511483 | | | |
| see above | Instituto Nacional de Saude (INSA) | Instituto Nacional de Saude (INSA) | Borges et al |
| EPI_ISL_511484, EPI_ISL_511485, EPI_ISL_511486, EPI_ISL_511487, EPI_ISL_511488, EPI_ISL_511489, EPI_ISL_511490, EPI_ISL_511491, EPI_ISL_511492, EPI_ISL_511493, EPI_ISL_511494, EPI_ISL_511495, EPI_ISL_511496, EPI_ISL_511497, EPI_ISL_511498, EPI_ISL_511499, EPI_ISL_511500, EPI_ISL_511501, EPI_ISL_511502, EPI_ISL_511503, EPI_ISL_511504, EPI_ISL_511505, EPI_ISL_511506, EPI_ISL_511507, EPI_ISL_511508 | | | |
| see above | Instituto Nacional de Saude (INSA) | Instituto Nacional de Saude (INSA) and Instituto Gulbenkian de Ciencia (IGC) | Borges et al |
| EPI_ISL_511509, EPI_ISL_511510, EPI_ISL_511511, EPI_ISL_511512, EPI_ISL_511513, EPI_ISL_511514, EPI_ISL_511515, EPI_ISL_511516, EPI_ISL_511517, EPI_ISL_511518, EPI_ISL_511519, EPI_ISL_511520, EPI_ISL_511521, EPI_ISL_511522, EPI_ISL_511523, EPI_ISL_511524, EPI_ISL_511525, EPI_ISL_511526, EPI_ISL_511527, EPI_ISL_511528, EPI_ISL_511529, EPI_ISL_511530, EPI_ISL_511531, EPI_ISL_511532, EPI_ISL_511533, EPI_ISL_511534, EPI_ISL_511535, EPI_ISL_511536, EPI_ISL_511537, EPI_ISL_511538, EPI_ISL_511539, EPI_ISL_511540, EPI_ISL_511541, EPI_ISL_511542, EPI_ISL_511543, EPI_ISL_511544, EPI_ISL_511545, EPI_ISL_511546, EPI_ISL_511547, EPI_ISL_511548, EPI_ISL_511549, EPI_ISL_511550, EPI_ISL_511551, EPI_ISL_511552, EPI_ISL_511553, EPI_ISL_511554, EPI_ISL_511555, EPI_ISL_511556, EPI_ISL_511557, EPI_ISL_511558, EPI_ISL_511559, EPI_ISL_511560, EPI_ISL_511561, EPI_ISL_511562, EPI_ISL_511563, EPI_ISL_511564, EPI_ISL_511565, EPI_ISL_511566, EPI_ISL_511567, EPI_ISL_511568, EPI_ISL_511569, EPI_ISL_511570, EPI_ISL_511571, EPI_ISL_511572, EPI_ISL_511573, EPI_ISL_511574, EPI_ISL_511575, EPI_ISL_511576, EPI_ISL_511577, EPI_ISL_511578, EPI_ISL_511579, EPI_ISL_511580, EPI_ISL_511581, EPI_ISL_511582, EPI_ISL_511583, EPI_ISL_511584, EPI_ISL_511585, EPI_ISL_511586, EPI_ISL_511587, EPI_ISL_511588, EPI_ISL_511589, EPI_ISL_511590, EPI_ISL_511591, EPI_ISL_511592, EPI_ISL_511593, EPI_ISL_511594, EPI_ISL_511595, EPI_ISL_511596, EPI_ISL_511597, EPI_ISL_511598, EPI_ISL_511599, EPI_ISL_511600, EPI_ISL_511601, EPI_ISL_511602, EPI_ISL_511603, EPI_ISL_511604, EPI_ISL_511605, EPI_ISL_511606, EPI_ISL_511607, EPI_ISL_511608, EPI_ISL_511609, EPI_ISL_511610, EPI_ISL_511611, EPI_ISL_511612, EPI_ISL_511613, EPI_ISL_511614, EPI_ISL_511615, EPI_ISL_511616, EPI_ISL_511617, EPI_ISL_511618, EPI_ISL_511619, EPI_ISL_511620, EPI_ISL_511621, EPI_ISL_511622, EPI_ISL_511623, EPI_ISL_511624, EPI_ISL_511625, EPI_ISL_511626, EPI_ISL_511627, EPI_ISL_511628, EPI_ISL_511629, EPI_ISL_511630, EPI_ISL_511631, EPI_ISL_511632, EPI_ISL_511633, EPI_ISL_511634, EPI_ISL_511635, EPI_ISL_511636, EPI_ISL_511637, EPI_ISL_511638, EPI_ISL_511639, EPI_ISL_511640, EPI_ISL_511641, EPI_ISL_511642, EPI_ISL_511643, EPI_ISL_511644, EPI_ISL_511645, EPI_ISL_511646, EPI_ISL_511647, EPI_ISL_511648, EPI_ISL_511649, EPI_ISL_511650, EPI_ISL_511651, EPI_ISL_511652, EPI_ISL_511653, EPI_ISL_511654, EPI_ISL_511655, EPI_ISL_511656, EPI_ISL_511657, EPI_ISL_511658, EPI_ISL_511659, EPI_ISL_511660, EPI_ISL_511661, EPI_ISL_511662, EPI_ISL_511663, EPI_ISL_511664, EPI_ISL_511665, EPI_ISL_511666, EPI_ISL_511667, EPI_ISL_511668, EPI_ISL_511669, EPI_ISL_511670, EPI_ISL_511671, EPI_ISL_511672, EPI_ISL_511673, EPI_ISL_511674, EPI_ISL_511675, EPI_ISL_511676, EPI_ISL_511677, EPI_ISL_511678, EPI_ISL_511679, EPI_ISL_511680, EPI_ISL_511681, EPI_ISL_511682, EPI_ISL_511683, EPI_ISL_511684, EPI_ISL_511685, EPI_ISL_511686, EPI_ISL_511687, EPI_ISL_511688, EPI_ISL_511689, EPI_ISL_511690, EPI_ISL_511691, EPI_ISL_511692, EPI_ISL_511693, EPI_ISL_511694, EPI_ISL_511695, EPI_ISL_511696, EPI_ISL_511697, EPI_ISL_511698, EPI_ISL_511699, EPI_ISL_511700, EPI_ISL_511701, EPI_ISL_511702, EPI_ISL_511703, EPI_ISL_511704, EPI_ISL_511705, EPI_ISL_511706, EPI_ISL_511707, EPI_ISL_511708, EPI_ISL_511709, EPI_ISL_511710, EPI_ISL_511711, EPI_ISL_511712, EPI_ISL_511713, EPI_ISL_511714, EPI_ISL_511715, EPI_ISL_511716, EPI_ISL_511717, EPI_ISL_511718, EPI_ISL_511719, EPI_ISL_511720, EPI_ISL_511721, EPI_ISL_511722, EPI_ISL_511723, EPI_ISL_511724, EPI_ISL_511725, EPI_ISL_511726, EPI_ISL_511727, EPI_ISL_511728, EPI_ISL_511729, EPI_ISL_511730, EPI_ISL_511731, EPI_ISL_511732, EPI_ISL_511733, EPI_ISL_511734, EPI_ISL_511735, EPI_ISL_511736, EPI_ISL_511737, EPI_ISL_511738, EPI_ISL_511739, EPI_ISL_511740, EPI_ISL_511741, EPI_ISL_511742, EPI_ISL_511743, EPI_ISL_511744, EPI_ISL_511745, EPI_ISL_511746, EPI_ISL_511747, EPI_ISL_511748, EPI_ISL_511749, EPI_ISL_511750 | | | |
| see above | Instituto Nacional de Saude (INSA) | Instituto Nacional de Saude (INSA) | Borges et al |
| EPI_ISL_511751, EPI_ISL_511752, EPI_ISL_511753, EPI_ISL_511754, EPI_ISL_511755, EPI_ISL_511756, EPI_ISL_511757, EPI_ISL_511758, EPI_ISL_511759, EPI_ISL_511760, EPI_ISL_511761, EPI_ISL_511762, EPI_ISL_511763, EPI_ISL_511764, EPI_ISL_511765, EPI_ISL_511766, EPI_ISL_511767, EPI_ISL_511768, EPI_ISL_511769, EPI_ISL_511770, EPI_ISL_511771 | | | |
| see above | Instituto Nacional de Saude (INSA) | Instituto Nacional de Saude (INSA) and Instituto Gulbenkian de Ciencia (IGC) | Borges et al |
| EPI_ISL_511805, EPI_ISL_511806, EPI_ISL_511807, EPI_ISL_511808, EPI_ISL_511809, EPI_ISL_511810, EPI_ISL_511811, EPI_ISL_511812, EPI_ISL_511813, EPI_ISL_511814, EPI_ISL_511815, EPI_ISL_511816, EPI_ISL_511817, EPI_ISL_511818, EPI_ISL_511819, EPI_ISL_511820, EPI_ISL_511821, EPI_ISL_511822, EPI_ISL_511823, EPI_ISL_511824, EPI_ISL_511825, EPI_ISL_511826, EPI_ISL_511827, EPI_ISL_511828, EPI_ISL_511829, EPI_ISL_511830, EPI_ISL_511831, EPI_ISL_511832, EPI_ISL_511833, EPI_ISL_511834, EPI_ISL_511835, EPI_ISL_511836, EPI_ISL_511837, EPI_ISL_511838, EPI_ISL_511839, EPI_ISL_511840, EPI_ISL_511841, EPI_ISL_511842, EPI_ISL_511843, EPI_ISL_511844, EPI_ISL_511845, EPI_ISL_511846, EPI_ISL_511847, EPI_ISL_511848, EPI_ISL_511849, EPI_ISL_511850, EPI_ISL_511851 | | | |
| see above | Innovative Genomics Institute, UC Berkeley | Innovative Genomics Institute, UC Berkeley | Stacia Wyman, Haridha Shivram, Liana Lareau, Shana McDevitt, Justin Choi |
| EPI_ISL_511852, EPI_ISL_511853, EPI_ISL_511854, EPI_ISL_511855, EPI_ISL_511856, EPI_ISL_511857, EPI_ISL_511858, EPI_ISL_511859, EPI_ISL_511860, EPI_ISL_511861 | UW Virology Lab | UW Virology Lab | Pavitra Roychoudhury, Amin Addetia, Hong Xie, Lasata Shrestha, Truong Nguyen, Meei-Li Huang, Keith Jerome, Alexander Greninger |
| EPI_ISL_511862 | Innovative Genomics Institute, UC Berkeley | Innovative Genomics Institute, UC Berkeley | Stacia Wyman, Haridha Shivram, Liana Lareau, Shana McDevitt, Justin Choi |
| EPI_ISL_511863, EPI_ISL_511864, EPI_ISL_511865, EPI_ISL_511866, EPI_ISL_511867, EPI_ISL_511868, EPI_ISL_511869, EPI_ISL_511870, EPI_ISL_511871, EPI_ISL_511872, EPI_ISL_511873, EPI_ISL_511874, EPI_ISL_511875, EPI_ISL_511876, EPI_ISL_511877 | | | |
| see above | Johns Hopkins Hospital Department of Pathology | Johns Hopkins Hospital Department of Pathology | Peter M. Thielen, Thomas Mehoke, Shirlee Wohl, Srividya Ramakrishnan, Melanie Kirsche, Amanda Ernlund, Craig Howser, Kristina Zudock, Oluwaseun Falade-Nwulia, Norah Sadowski, Paul Morris, Mark Hopkins, Yunfan Fan, Nidia Trovao, Victoria Gniazdowski, Michael C. Schatz, Stuart C. Ray, Winston Timp, Heba H. Mostafa |
| EPI_ISL_511878 | Santosa Hospital Bandung Kopo | Laboratorium Kesehatan Provinsi Jawa Barat; Molecular Genetics Laboratory-Faculty of Medicine-Universitas Padjadjaran; School of Life Sciences and Technology & School of Pharmacy-Institut Teknologi Bandung | Ema Rahmawati, Marselina Irasonia Tan, Yunia Sribudiani, Catur Riani, Azzania Fibriani, Husna Nugrahapraja, Tarwadi, Hesti Lina Wiraswati, Lia Faridah, Savira Ekawardhani, Ryan Bayusantika Ristandi, Rifky Waluyajati Rachman, Cut Nur Cinthia Alamanda, Hammam Riza, Soni Solistia Wirawan, Agung Eru Wibowo |
| EPI_ISL_511879 | Laboratorium Kesehatan Provinsi Jawa Barat | Molecular Genetics Laboratory-Faculty of Medicine-Universitas Padjadjaran; School of Life Sciences and Technology & School of Pharmacy-Institut Teknologi Bandung; Laboratorium Kesehatan Provinsi Jawa Barat | Marselina Irasonia Tan, Yunia Sribudiani, Catur Riani, Azzania Fibriani, Husna Nugrahapraja, Tarwadi, Ema Rahmawati, Hesti Lina Wiraswati, Lia Faridah, Savira Ekawardhani, Ryan Bayusantika Ristandi, Rifky Waluyajati Rachman, Cut Nur Cinthia Alamanda, Hammam Riza, Soni Solistia Wirawan, Agung Eru Wibowo |
| EPI_ISL_511891, EPI_ISL_511892, EPI_ISL_511893, EPI_ISL_511894, EPI_ISL_511895, EPI_ISL_511896, EPI_ISL_511897, EPI_ISL_511898 | National Hospital of Tropical Diseases | Oxford University Clinical Research Unit, Hanoi, Vietnam | Nguyen Thi Tam, Van Dinh Trang, Nguyen Thi Hong Thuong, Vu Thi Ngoc Bich, Nguyen Thu Trang, Nguyen Thi Ngoc Diep, Le Nguyen Minh Hoa, Pham Ngoc Thach, H. Rogier van Doorn, on behalf of the OUCRU COVID-19 research group |
| EPI_ISL_511899 | ICMR-National Institute of Cholera and Enteric Diseases | National Institute of Biomedical Genomics - DBT's PAN-INDIA 1000 SARS--CoV-2 RNA Genome Sequencing Consortium | Arindam Maitra, Mamta Chawla Sarkar, Sreedhar Chinnaswamy, Hasina Banu, Ananya Chatterjee, Shanta Dutta, Saumitra Das |
| EPI_ISL_511900, EPI_ISL_511901, EPI_ISL_511902, EPI_ISL_511903, EPI_ISL_511904, EPI_ISL_511905, EPI_ISL_511906, EPI_ISL_511907 | Institute of Post Graduate Medical Education & Research | National Institute of Biomedical Genomics - DBT's PAN-INDIA 1000 SARS--CoV-2 RNA Genome Sequencing Consortium | Arindam Maitra, Aritra Biswas, Jayeeta Haldar, Raja Ray, Monimoy Banerjee, Saumitra Das |
| EPI_ISL_511908, EPI_ISL_511909, EPI_ISL_511910, EPI_ISL_511911, EPI_ISL_511912, EPI_ISL_511913, EPI_ISL_511914, EPI_ISL_511915, EPI_ISL_511916, EPI_ISL_511917, EPI_ISL_511918, EPI_ISL_511919, EPI_ISL_511920, EPI_ISL_511921, EPI_ISL_511922 | | | |
| see above | All india institute of Medical Sciences Rishikesh | National Institute of Biomedical Genomics - DBT's PAN-INDIA 1000 SARS--CoV-2 RNA Genome Sequencing Consortium | Arindam Maitra, Deepjyoti Kalita, Amit Mangla, Ravi Kant, Saumitra Das |
| EPI_ISL_511923, EPI_ISL_511924, EPI_ISL_511925, EPI_ISL_511926, EPI_ISL_511927, EPI_ISL_511928, EPI_ISL_511929 | Mahatma Gandhi Institute of Medical Sciences | National Institute of Biomedical Genomics - DBT's PAN-INDIA 1000 SARS--CoV-2 RNA Genome Sequencing Consortium | Arindam Maitra, Vijayshri Deotale, Rahul Narang, Deepashri Maraskolhe, Saumitra Das |
| EPI_ISL_511930, EPI_ISL_511931, EPI_ISL_511932, EPI_ISL_511933, EPI_ISL_511934, EPI_ISL_511935, EPI_ISL_511936, EPI_ISL_511937, EPI_ISL_511938, EPI_ISL_511939, EPI_ISL_511940, EPI_ISL_511941 | | | |
| see above | Government Medical College | National Institute of Biomedical Genomics - DBT's PAN-INDIA 1000 SARS--CoV-2 RNA Genome Sequencing Consortium | Arindam Maitra, Jyoti Iravane, Dhaval Khatri, Maitrik Dave, Saumitra Das |
| EPI_ISL_511942, EPI_ISL_511943, EPI_ISL_511944 | Translational Health Science and Technology Institute | National Institute of Biomedical Genomics - DBT's PAN-INDIA 1000 SARS--CoV-2 RNA Genome Sequencing Consortium | Arindam Maitra, Guruprasad Medigeshi, Sharanabasava Patil, Anbalagan Ananthraj, Madhu Pareek, Imran Khan, Gagandeep Kang, Saumitra Das |

| | | | |
|---|---|---|---|
| EPI_ISL_511945, EPI_ISL_511946, EPI_ISL_511947, EPI_ISL_511948, EPI_ISL_511949, EPI_ISL_511950, EPI_ISL_511951, EPI_ISL_511952, EPI_ISL_511953, EPI_ISL_511954 | Indian Institute of Science | National Institute of Biomedical Genomics - DBT's PAN-INDIA 1000 SARS--CoV-2 RNA Genome Sequencing Consortium | Arindam Maitra, Bharath K Sundararaj, Harsha Raheja, N. Srinivasan, Deepak K Saini, Amit Singh, Saumitra Das |
| EPI_ISL_511955, EPI_ISL_511956, EPI_ISL_511957, EPI_ISL_511958 | PHE South West Regional Laboratory, National Infection Service | Wellcome Sanger Institute for the COVID-19 Genomics UK Consortium | Stephanie Hutchings, Hannah Pymont, Dr Peter Muir, Barry Vipond, Rich Hopes; and Alex Alderton, Roberto Amato, Sonia Goncalves, Ewan Harrison, David K. Jackson, Ian Johnston, Dominic Kwiatkowski, Cordelia Langford, John Sillitoe on behalf of the Wellcome Sanger Institute COVID-19 Surveillance Team (http://www.sanger.ac.uk/covid-team) |
| EPI_ISL_511959 | Department of Pathology, University of Cambridge | Wellcome Sanger Institute for the COVID-19 Genomics UK Consortium | Luke W Meredith, M. Estée Török , Myra Hosmillo, William L. Hamilton, Martin D. Curran, Theresa Feltwell, Grant Hall, Anna Yakovleva, Fahad A Khokhar, Charlotte J. Houldcroft, Laura G Caller, Aminu S. Jahun, Sarah L. Caddy, Ian Goodfellow; and Alex Alderton, Roberto Amato, Sonia Goncalves, Ewan Harrison, David K. Jackson, Ian Johnston, Dominic Kwiatkowski, Cordelia Langford, John Sillitoe on behalf of the Wellcome Sanger Institute COVID-19 Surveillance Team (http://www.sanger.ac.uk/covid-team) |
| EPI_ISL_511960, EPI_ISL_511961, EPI_ISL_511962, EPI_ISL_511963, EPI_ISL_511964, EPI_ISL_511965, EPI_ISL_511966, EPI_ISL_511967, EPI_ISL_511968, EPI_ISL_511969 | PHE South West Regional Laboratory, National Infection Service | Wellcome Sanger Institute for the COVID-19 Genomics UK Consortium | Stephanie Hutchings, Hannah Pymont, Dr Peter Muir, Barry Vipond, Rich Hopes; and Alex Alderton, Roberto Amato, Sonia Goncalves, Ewan Harrison, David K. Jackson, Ian Johnston, Dominic Kwiatkowski, Cordelia Langford, John Sillitoe on behalf of the Wellcome Sanger Institute COVID-19 Surveillance Team (http://www.sanger.ac.uk/covid-team) |
| EPI_ISL_511970 | Department of Pathology, University of Cambridge | Wellcome Sanger Institute for the COVID-19 Genomics UK Consortium | Luke W Meredith, M. Estée Török , Myra Hosmillo, William L. Hamilton, Martin D. Curran, Theresa Feltwell, Grant Hall, Anna Yakovleva, Fahad A Khokhar, Charlotte J. Houldcroft, Laura G Caller, Aminu S. Jahun, Sarah L. Caddy, Ian Goodfellow; and Alex Alderton, Roberto Amato, Sonia Goncalves, Ewan Harrison, David K. Jackson, Ian Johnston, Dominic Kwiatkowski, Cordelia Langford, John Sillitoe on behalf of the Wellcome Sanger Institute COVID-19 Surveillance Team (http://www.sanger.ac.uk/covid-team) |
| EPI_ISL_511971, EPI_ISL_511972, EPI_ISL_511973, EPI_ISL_511974, EPI_ISL_511975, EPI_ISL_511976, EPI_ISL_511977, EPI_ISL_511978, EPI_ISL_511979, EPI_ISL_511980, EPI_ISL_511981 | | | |
| see above | PHE South West Regional Laboratory, National Infection Service | Wellcome Sanger Institute for the COVID-19 Genomics UK Consortium | Stephanie Hutchings, Hannah Pymont, Dr Peter Muir, Barry Vipond, Rich Hopes; and Alex Alderton, Roberto Amato, Sonia Goncalves, Ewan Harrison, David K. Jackson, Ian Johnston, Dominic Kwiatkowski, Cordelia Langford, John Sillitoe on behalf of the Wellcome Sanger Institute COVID-19 Surveillance Team (http://www.sanger.ac.uk/covid-team) |
| EPI_ISL_511982 | Department of Pathology, University of Cambridge | Wellcome Sanger Institute for the COVID-19 Genomics UK Consortium | Luke W Meredith, M. Estée Török , Myra Hosmillo, William L. Hamilton, Martin D. Curran, Theresa Feltwell, Grant Hall, Anna Yakovleva, Fahad A Khokhar, Charlotte J. Houldcroft, Laura G Caller, Aminu S. Jahun, Sarah L. Caddy, Ian Goodfellow; and Alex Alderton, Roberto Amato, Sonia Goncalves, Ewan Harrison, David K. Jackson, Ian Johnston, Dominic Kwiatkowski, Cordelia Langford, John Sillitoe on behalf of the Wellcome Sanger Institute COVID-19 Surveillance Team (http://www.sanger.ac.uk/covid-team) |
| EPI_ISL_511984, EPI_ISL_511985, EPI_ISL_511986, EPI_ISL_511987, EPI_ISL_511988, EPI_ISL_511989, EPI_ISL_511990, EPI_ISL_511991, EPI_ISL_511992, EPI_ISL_511993, EPI_ISL_511994, EPI_ISL_511995, EPI_ISL_511996, EPI_ISL_511997, EPI_ISL_511998, EPI_ISL_511999, EPI_ISL_512000, EPI_ISL_512001, EPI_ISL_512002, EPI_ISL_512003, EPI_ISL_512004, EPI_ISL_512005, EPI_ISL_512006, EPI_ISL_512007, EPI_ISL_512008, EPI_ISL_512009, EPI_ISL_512010, EPI_ISL_512011, EPI_ISL_512012, EPI_ISL_512013, EPI_ISL_512014, EPI_ISL_512015, EPI_ISL_512016, EPI_ISL_512017, EPI_ISL_512018, EPI_ISL_512019, EPI_ISL_512020, EPI_ISL_512021, EPI_ISL_512022, EPI_ISL_512023, EPI_ISL_512024, EPI_ISL_512025, EPI_ISL_512026, EPI_ISL_512027, EPI_ISL_512028, EPI_ISL_512029, EPI_ISL_512030, EPI_ISL_512031, EPI_ISL_512032, EPI_ISL_512033, EPI_ISL_512034, EPI_ISL_512035, EPI_ISL_512036, EPI_ISL_512037, EPI_ISL_512038, EPI_ISL_512039, EPI_ISL_512040, EPI_ISL_512041, EPI_ISL_512042, EPI_ISL_512043, EPI_ISL_512044, EPI_ISL_512045, EPI_ISL_512046, EPI_ISL_512047, EPI_ISL_512048, EPI_ISL_512049, EPI_ISL_512050, EPI_ISL_512051, EPI_ISL_512052, EPI_ISL_512053, EPI_ISL_512054 | | | |
| see above | Viollier AG | Department of Biosystems Science and Engineering, ETH Zürich | Christian Beisel, Sarah Nadeau, Ivan Topolsky, Pedro Ferreira, Philipp Jablonski, Susana Posada-Céspedes, Tobias Schär, Ina Nissen, Natascha Santacroce, Elodie Burcklen, Christiane Beckmann, Maurice Redondo, Olivier Kobel, Christoph Noppen, Sophie Seidel, Noemie Santamaria de Souza, Niko Beerenwinkel, Tanja Stadler |
| EPI_ISL_512058 | B.J. Medical College and Civil hospital, Ahmedabad | Gujarat Biotechnology Research Centre | Monika Gandhi, Pinal Trivedi, Maharshi Pandya, Nidhi Patel, Nitin Savaliya, Raghawendra Kumar, Dinesh Kumar, Zuber Saiyed, Komal Patel, Labdhi Pandya, Afzal Ansari, Nikha Trivedi, Pranay Shah, Kamlesh J Upadhyay, Sanjay Kapadia, Apurvasinh Puvar, Janvi Raval, Zarna Patel, R D Dixit, A M Kadri, Harsh Bakshi, Chaitanya Joshi, Madhvi Joshi |
| EPI_ISL_512059 | B.J. Medical College and Civil hospital, Ahmedabad | Gujarat Biotechnology Research Centre | Pinal Trivedi, Maharshi Pandya, Nidhi Patel, Nitin Savaliya, Raghawendra Kumar, Dinesh Kumar, Zuber Saiyed, Komal Patel, Labdhi Pandya, Afzal Ansari, Nikha Trivedi, Pranay Shah, Kamlesh J Upadhyay, Sanjay Kapadia, Apurvasinh Puvar, Janvi Raval, Zarna Patel, Monika Gandhi, R D Dixit, A M Kadri, Harsh Bakshi, Chaitanya Joshi, Madhvi Joshi |
| EPI_ISL_512060 | B.J. Medical College and Civil hospital, Ahmedabad | Gujarat Biotechnology Research Centre | Maharshi Pandya, Nidhi Patel, Nitin Savaliya, Raghawendra Kumar, Dinesh Kumar, Zuber Saiyed, Komal Patel, Labdhi Pandya, Afzal Ansari, Nikha Trivedi, Pranay Shah, Kamlesh J Upadhyay, Sanjay Kapadia, Apurvasinh Puvar, Janvi Raval, Zarna Patel, Monika Gandhi, Pinal Trivedi, R D Dixit, A M Kadri, Harsh Bakshi, Chaitanya Joshi, Madhvi Joshi |
| EPI_ISL_512061 | B.J. Medical College and Civil hospital, Ahmedabad | Gujarat Biotechnology Research Centre | Nidhi Patel, Nitin Savaliya, Raghawendra Kumar, Dinesh Kumar, Zuber Saiyed, Komal Patel, Labdhi Pandya, Afzal Ansari, Nikha Trivedi, Pranay Shah, Kamlesh J Upadhyay, Sanjay Kapadia, Apurvasinh Puvar, Janvi Raval, Zarna Patel, Monika Gandhi, Pinal Trivedi, Maharshi Pandya, R D Dixit, A M Kadri, Harsh Bakshi, Chaitanya Joshi, Madhvi Joshi |
| EPI_ISL_512062 | B.J. Medical College and Civil hospital, Ahmedabad | Gujarat Biotechnology Research Centre | Nitin Savaliya, Raghawendra Kumar, Dinesh Kumar, Zuber Saiyed, Komal Patel, Labdhi Pandya, Afzal Ansari, Nikha Trivedi, Pranay Shah, Kamlesh J Upadhyay, Sanjay Kapadia, Apurvasinh Puvar, Janvi Raval, Zarna Patel, Monika Gandhi, Pinal Trivedi, Maharshi Pandya, Nidhi Patel, R D Dixit, A M Kadri, Harsh Bakshi, Chaitanya Joshi, Madhvi Joshi |
| EPI_ISL_512063 | B.J. Medical College and Civil hospital, Ahmedabad | Gujarat Biotechnology Research Centre | Raghawendra Kumar, Dinesh Kumar, Zuber Saiyed, Komal Patel, Labdhi Pandya, Afzal Ansari, Nikha Trivedi, Pranay Shah, Kamlesh J Upadhyay, Sanjay Kapadia, Apurvasinh Puvar, Janvi Raval, Zarna Patel, Monika Gandhi, Pinal Trivedi, Maharshi Pandya, Nidhi Patel, Nitin Savaliya, R D Dixit, A M Kadri, Harsh Bakshi, Chaitanya Joshi, Madhvi Joshi |
| EPI_ISL_512064 | B.J. Medical College and Civil hospital, Ahmedabad | Gujarat Biotechnology Research Centre | Dinesh Kumar, Zuber Saiyed, Komal Patel, Labdhi Pandya, Afzal Ansari, Nikha Trivedi, Pranay Shah, Kamlesh J Upadhyay, Sanjay Kapadia, Apurvasinh Puvar, Janvi Raval, Zarna Patel, Monika Gandhi, Pinal Trivedi, Maharshi Pandya, Nidhi Patel, Nitin Savaliya, Raghawendra Kumar, R D Dixit, A M Kadri, Harsh Bakshi, Chaitanya Joshi, Madhvi Joshi |
| EPI_ISL_512065 | B.J. Medical College and Civil hospital, Ahmedabad | Gujarat Biotechnology Research Centre | Zuber Saiyed, Komal Patel, Labdhi Pandya, Afzal Ansari, Nikha Trivedi, Pranay Shah, Kamlesh J Upadhyay, Sanjay Kapadia, Apurvasinh Puvar, Janvi Raval, Zarna Patel, Monika Gandhi, Pinal Trivedi, Maharshi Pandya, Nidhi Patel, Nitin Savaliya, Raghawendra Kumar, Dinesh Kumar, R D Dixit, A M Kadri, Harsh Bakshi, Chaitanya Joshi, Madhvi Joshi |
| EPI_ISL_512066 | Sardar Vallabhbhai Patel Institute of Medical Sciences & Research | Gujarat Biotechnology Research Centre | Komal Patel, Labdhi Pandya, Afzal Ansari, Nikha Trivedi, Pranay Shah, Kamlesh J Upadhyay, Sanjay Kapadia, Apurvasinh Puvar, Janvi Raval, Zarna Patel, Monika Gandhi, Pinal Trivedi, Maharshi Pandya, Nidhi Patel, Nitin Savaliya, Raghawendra Kumar, Dinesh Kumar, Zuber Saiyed, R D Dixit, A M Kadri, Harsh Bakshi, Chaitanya Joshi, Madhvi Joshi |
| EPI_ISL_512067 | Sardar Vallabhbhai Patel Institute of Medical Sciences & Research | Gujarat Biotechnology Research Centre | Labdhi Pandya, Afzal Ansari, Nikha Trivedi, Pranay Shah, Kamlesh J Upadhyay, Sanjay Kapadia, Apurvasinh Puvar, Janvi Raval, Zarna Patel, Monika Gandhi, Pinal Trivedi, Maharshi Pandya, Nidhi Patel, Nitin Savaliya, Raghawendra Kumar, Dinesh Kumar, Zuber Saiyed, Komal Patel, R D Dixit, A M Kadri, Harsh Bakshi, Chaitanya Joshi, Madhvi Joshi |
| EPI_ISL_512068 | Sardar Vallabhbhai Patel Institute of Medical Sciences & Research | Gujarat Biotechnology Research Centre | Afzal Ansari, Nikha Trivedi, Pranay Shah, Kamlesh J Upadhyay, Sanjay Kapadia, Apurvasinh Puvar, Janvi Raval, Zarna Patel, Monika Gandhi, Pinal Trivedi, Maharshi Pandya, Nidhi Patel, Nitin Savaliya, Raghawendra Kumar, Dinesh Kumar, Zuber Saiyed, Komal Patel, Labdhi Pandya, R D Dixit, A M Kadri, Harsh Bakshi, Chaitanya Joshi, Madhvi Joshi |
| EPI_ISL_512069 | Sardar Vallabhbhai Patel Institute of Medical Sciences & Research | Gujarat Biotechnology Research Centre | Nikha Trivedi, Pranay Shah, Kamlesh J Upadhyay, Sanjay Kapadia, Apurvasinh Puvar, Janvi Raval, Zarna Patel, Monika Gandhi, Pinal Trivedi, Maharshi Pandya, Nidhi Patel, Nitin Savaliya, Raghawendra Kumar, Dinesh Kumar, Zuber Saiyed, Komal Patel, Labdhi Pandya, Afzal Ansari, R D Dixit, A M Kadri, Harsh Bakshi, Chaitanya Joshi, Madhvi Joshi |
| EPI_ISL_512070 | Department of MicroBiology, Government Medical College, Surat | Gujarat Biotechnology Research Centre | Naresh Chauhan, Summaiya Mullan, Amit gamit, Apurvasinh Puvar, Janvi Raval, Zarna Patel, Monika Gandhi, Pinal Trivedi, Maharshi Pandya, Nidhi Patel, Nitin Savaliya, Raghawendra Kumar, Dinesh Kumar, Zuber Saiyed, Komal Patel, Labdhi Pandya, Afzal Ansari, Nikha Trivedi, R D Dixit, A M Kadri, Harsh Bakshi, Chaitanya Joshi, Madhvi Joshi |
| EPI_ISL_512071 | Department of MicroBiology, Government Medical College, Surat | Gujarat Biotechnology Research Centre | Summaiya Mullan, Amit gamit, Apurvasinh Puvar, Janvi Raval, Zarna Patel, Monika Gandhi, Pinal Trivedi, Maharshi Pandya, Nidhi Patel, Nitin Savaliya, Raghawendra Kumar, Dinesh Kumar, Zuber Saiyed, Komal Patel, Labdhi Pandya, Afzal Ansari, Nikha Trivedi, Naresh Chauhan, R D Dixit, A M Kadri, Harsh Bakshi, Chaitanya Joshi, Madhvi Joshi |
| EPI_ISL_512072 | Saikrishna Hospital, Mehsana | Gujarat Biotechnology Research Centre | Harshadbhai Parmar, Apurvasinh Puvar, Janvi Raval, Zarna Patel, Monika Gandhi, Pinal Trivedi, Maharshi Pandya, Nidhi Patel, Nitin Savaliya, Raghawendra Kumar, Dinesh Kumar, Zuber Saiyed, Komal Patel, Labdhi Pandya, Afzal Ansari, Nikha Trivedi, R D Dixit, A M Kadri, Harsh Bakshi, Chaitanya Joshi, Madhvi Joshi |
| EPI_ISL_512073 | Saikrishna Hospital, Mehsana | Gujarat Biotechnology Research Centre | Apurvasinh Puvar, Janvi Raval, Zarna Patel, Monika Gandhi, Pinal Trivedi, Maharshi Pandya, Nidhi Patel, Nitin Savaliya, Raghawendra Kumar, Dinesh Kumar, Zuber Saiyed, Komal Patel, Labdhi Pandya, Afzal Ansari, Nikha Trivedi, Harshadbhai Parmar, R D Dixit, A M Kadri, Harsh Bakshi, Chaitanya Joshi, Madhvi Joshi |
| EPI_ISL_512074 | Saikrishna Hospital, Mehsana | Gujarat Biotechnology Research Centre | Janvi Raval, Zarna Patel, Monika Gandhi, Pinal Trivedi, Maharshi Pandya, Nidhi Patel, Nitin Savaliya, Raghawendra Kumar, Dinesh Kumar, Zuber Saiyed, Komal Patel, Labdhi Pandya, Afzal Ansari, Nikha Trivedi, Harshadbhai Parmar, Apurvasinh Puvar, R D Dixit, A M Kadri, Harsh Bakshi, Chaitanya Joshi, Madhvi Joshi |
| EPI_ISL_512075 | Dr. RSS Hospital, Modasa | Gujarat Biotechnology Research Centre | Monika Gandhi, Pinal Trivedi, Maharshi Pandya, Nidhi Patel, Nitin Savaliya, Raghawendra Kumar, Dinesh Kumar, Zuber Saiyed, Komal Patel, Labdhi Pandya, Afzal Ansari, Nikha Trivedi, Harsh Chaudhari, Apurvasinh Puvar, Janvi Raval, Zarna Patel, R D Dixit, A M Kadri, Harsh Bakshi, Chaitanya Joshi, Madhvi Joshi |
| EPI_ISL_512076 | Dr. RSS Hospital, Modasa | Gujarat Biotechnology Research Centre | Pinal Trivedi, Maharshi Pandya, Nidhi Patel, Nitin Savaliya, Raghawendra Kumar, Dinesh Kumar, Zuber Saiyed, Komal Patel, Labdhi Pandya, Afzal Ansari, Nikha Trivedi, Harsh Chaudhari, Apurvasinh Puvar, Janvi Raval, Zarna Patel, Monika Gandhi, R D Dixit, A M Kadri, Harsh Bakshi, Chaitanya Joshi, Madhvi Joshi |
| EPI_ISL_512077 | Dr. RSS Hospital, Modasa | Gujarat Biotechnology Research Centre | Maharshi Pandya, Nidhi Patel, Nitin Savaliya, Raghawendra Kumar, Dinesh Kumar, Zuber Saiyed, Komal Patel, Labdhi Pandya, Afzal Ansari, Nikha Trivedi, Harsh Chaudhari, Apurvasinh Puvar, Janvi Raval, Zarna Patel, Monika Gandhi, Pinal Trivedi, R D Dixit, A M Kadri, Harsh Bakshi, Chaitanya Joshi, Madhvi Joshi |
| EPI_ISL_512086 | UW Virology lab | UW Virology lab | Pavitra Roychoudhury, Amin Addetia, Hong Xie, Lasata Shrestha, Truong Nguyen, Meei-Li Huang, Keith Jerome, Alexander Greninger |
| EPI_ISL_512087, EPI_ISL_512088, EPI_ISL_512089, EPI_ISL_512090, EPI_ISL_512091, EPI_ISL_512092, EPI_ISL_512093, EPI_ISL_512094, EPI_ISL_512095, EPI_ISL_512096, EPI_ISL_512097, EPI_ISL_512098, EPI_ISL_512099, EPI_ISL_512100, EPI_ISL_512101, EPI_ISL_512102, EPI_ISL_512103, EPI_ISL_512104, EPI_ISL_512105, EPI_ISL_512106 | | | |
| see above | National Virus Reference Laboratory | National Virus Reference Laboratory | Michael Carr, Gabriel Gonzalez, Jonathan Dean, Aditi Chaturvedi, Suzie Coughlan, Cillian F De Gascun |
| EPI_ISL_512107 | Florida Bureau of Public Health Laboratories | Florida Bureau of Public Health Laboratories | Sarah Schmedes, Jason Blanton |
| EPI_ISL_512135, EPI_ISL_512136, EPI_ISL_512137, EPI_ISL_512138, EPI_ISL_512139, EPI_ISL_512140, EPI_ISL_512141, EPI_ISL_512142, EPI_ISL_512143, EPI_ISL_512144, EPI_ISL_512145, EPI_ISL_512146, EPI_ISL_512147, EPI_ISL_512148, EPI_ISL_512149, EPI_ISL_512150, EPI_ISL_512151, EPI_ISL_512152, EPI_ISL_512153, EPI_ISL_512154, EPI_ISL_512155, EPI_ISL_512156, EPI_ISL_512157 | | | |
| see above | Alaska State Virology Laboratory | Alaska State Virology Laboratory | Chen J et al with Pathogenomics group Dagdag R, Redlinger M, Milton E, George W, Kovalenko A, Drown DM, Bortz E |
| EPI_ISL_512158, EPI_ISL_512159, EPI_ISL_512160, EPI_ISL_512161, EPI_ISL_512162, EPI_ISL_512163, EPI_ISL_512164, EPI_ISL_512165, EPI_ISL_512166, EPI_ISL_512167, EPI_ISL_512168, EPI_ISL_512169, EPI_ISL_512170, EPI_ISL_512171, EPI_ISL_512172, EPI_ISL_512173, EPI_ISL_512174, EPI_ISL_512175, EPI_ISL_512176, EPI_ISL_512177, EPI_ISL_512178, EPI_ISL_512179, EPI_ISL_512180, EPI_ISL_512181, EPI_ISL_512182, EPI_ISL_512183, EPI_ISL_512184, EPI_ISL_512185, EPI_ISL_512186, EPI_ISL_512187, EPI_ISL_512188, EPI_ISL_512189, EPI_ISL_512190, EPI_ISL_512191, EPI_ISL_512192, EPI_ISL_512193, EPI_ISL_512194, EPI_ISL_512195, EPI_ISL_512196, EPI_ISL_512197, EPI_ISL_512198, EPI_ISL_512199, EPI_ISL_512200, EPI_ISL_512201, EPI_ISL_512202, EPI_ISL_512203, EPI_ISL_512204, EPI_ISL_512205, EPI_ISL_512206, EPI_ISL_512207, EPI_ISL_512208, EPI_ISL_512209, EPI_ISL_512210, EPI_ISL_512211, EPI_ISL_512212, EPI_ISL_512213, EPI_ISL_512214, EPI_ISL_512215, EPI_ISL_512216, EPI_ISL_512217, EPI_ISL_512218, EPI_ISL_512219, EPI_ISL_512220, EPI_ISL_512221, EPI_ISL_512222, EPI_ISL_512223, EPI_ISL_512224, EPI_ISL_512225, EPI_ISL_512226, EPI_ISL_512227, EPI_ISL_512228, EPI_ISL_512229, EPI_ISL_512230, EPI_ISL_512231, EPI_ISL_512232, EPI_ISL_512233, EPI_ISL_512234, EPI_ISL_512235, EPI_ISL_512236, EPI_ISL_512237, EPI_ISL_512238, EPI_ISL_512239, EPI_ISL_512240, EPI_ISL_512241, EPI_ISL_512242, EPI_ISL_512243, EPI_ISL_512244, EPI_ISL_512245, EPI_ISL_512246, EPI_ISL_512247, EPI_ISL_512248, EPI_ISL_512249, EPI_ISL_512250, EPI_ISL_512251, EPI_ISL_512252, EPI_ISL_512253, EPI_ISL_512254, EPI_ISL_512255, EPI_ISL_512256, EPI_ISL_512257, EPI_ISL_512258, EPI_ISL_512259, EPI_ISL_512260, EPI_ISL_512261, EPI_ISL_512262, EPI_ISL_512263, EPI_ISL_512264, EPI_ISL_512265, EPI_ISL_512266, EPI_ISL_512267, EPI_ISL_512268, EPI_ISL_512269, EPI_ISL_512270, EPI_ISL_512271, EPI_ISL_512272, EPI_ISL_512273 | | | |
| see above | San Diego County Public Health Laboratory | Andersen lab at Scripps Research | SEARCH Alliance San Diego with Tracy Basler, Jovan Shephard, Brett Austin |
| EPI_ISL_512296, EPI_ISL_512297, EPI_ISL_512298, EPI_ISL_512299, EPI_ISL_512300, EPI_ISL_512301, EPI_ISL_512302, EPI_ISL_512303 | Hematology Laboratory, Section of Molecular Diagnostics, University Clinical Centre, Medical University of Gdansk | Department of Virology, Faculty of Medicine, University of Helsinki, Helsinki, Finland | Maciej Grzybek, Marlena Robakowska, Aneta Szulc, Ewa Miłosz, Olli Vapalahti, Teemu Smura |
| EPI_ISL_512313, EPI_ISL_512314 | E. Gulbja Laboratorija | Latvian Biomedical Research and Study Centre | Ivars Silamiķelis, Kaspars Megnis, Monta Ustinova, Ņikita Zrelovs, Vita Rovīte, Mikus Gavars, Dmitrijs Perminovs, Uga Dumpis, Jānis Kloviņš |
| EPI_ISL_512330, EPI_ISL_512331, EPI_ISL_512332, EPI_ISL_512333, EPI_ISL_512334, EPI_ISL_512335 | Department of Pathology, University of Cambridge | COVID-19 Genomics UK (COG-UK) Consortium | Luke W Meredith, M. Estée Török, Myra Hosmillo, William L. Hamilton, Martin D. Curran, Theresa Feltwell, Grant Hall, Anna Yakovleva, Fahad A Khokhar, Charlotte J. Houldcroft, Laura G Caller, Aminu S. Jahun, Sarah L. Caddy, Yasmin Chaudhry, Malte Pinckert, Ian Goodfellow |
| EPI_ISL_512336, EPI_ISL_512337, EPI_ISL_512338, EPI_ISL_512339, | University of Exeter | COVID-19 Genomics UK (COG-UK) Consortium | Ben Temperton,Aaron Jeffries,Michelle Michelsen,Joanna Warwick-Dugdale,Audrey Farbos,Robyn Manley,Stephen Michell,Jane Masoli |

| | | | |
|---|---|---|---|
| EPI_ISL_512340, EPI_ISL_512341 | | | |
| EPI_ISL_512342, EPI_ISL_512343, EPI_ISL_512344, EPI_ISL_512345, EPI_ISL_512346, EPI_ISL_512347 | Liverpool Clinical Laboratories | COVID-19 Genomics UK (COG-UK) Consortium | Sam Haldenby, Anita Lucaci, Steve Paterson, Julian Hiscox, Alistair Darby, M Almsaud, A Alrezaihi, Muhannad Alruwaili, Stuart D Armstrong, Jones Benjamin, Eleanor G Bentley, Anu Chawla, Jordan J Clark, Angela Cowell, Richard Eccles, Isabel García-Dorival, Matthew Gemmell, Alessandro Gerada, PKF Gilmore, Richard Gregory, Ximeng Han, Catherine Hartley, Margaret Hughes, Miren Iturriza-Gomara, James Johnson, L Luu, Jenifer Manson, Charlotte Nelson, Elaine O'Toole, Cassie Olateju, Rebekah Penrice-Randal , Lucille Rainbow, N.P Randle, Trevor Ian Robinson, Parul Sharma, Ghada T Shawli, James P Stewart, Neil Swainston, Ecaterina Vamos, Joanne Watts, Mark Whitehead |
| EPI_ISL_512348, EPI_ISL_512349, EPI_ISL_512350, EPI_ISL_512351, EPI_ISL_512352, EPI_ISL_512353, EPI_ISL_512354, EPI_ISL_512355, EPI_ISL_512356, EPI_ISL_512357, EPI_ISL_512358, EPI_ISL_512359, EPI_ISL_512360, EPI_ISL_512361, EPI_ISL_512362, EPI_ISL_512363, EPI_ISL_512364, EPI_ISL_512365 | | | |
| see above | Northumbria University / South Tees Hospitals NHS Foundation Trust / North Cumbria Integrated Care NHS Foundation Trust / North Tees and Hartlepool NHS Foundation Trust / Newcastle Hospitals NHS Foundation Trust | COVID-19 Genomics UK (COG-UK) Consortium | Darren L Smith,Andrew Nelson,Matthew Bashton,Greg R Young,Joshua Loh,John Allan,Mohammad A Tariq,Giles S Holt,Gary Black,Wen C Yew,Lynn Dover,Paul Baker,Steve Liggett,Sarah Essex,Jane Greenaway,Debra Padgett,Clive Graham,Garren Scott,Edward Barton,Emma Swindells,Brendan Payne,Jennifer Collins,Yusri Taha,Gary Eltringham |
| EPI_ISL_512366, EPI_ISL_512367, EPI_ISL_512368, EPI_ISL_512369, EPI_ISL_512370, EPI_ISL_512371, EPI_ISL_512372, EPI_ISL_512373, EPI_ISL_512374, EPI_ISL_512375, EPI_ISL_512376, EPI_ISL_512377, EPI_ISL_512378 | | | |
| see above | Quadram Institute Bioscience | COVID-19 Genomics UK (COG-UK) Consortium | Dave J. Baker, Gemma L. Kay, Alp Aydin, Thanh Le-Viet, Steven Rudder, Ana P. Tedim, Anastasia Kolyva, Maria Diaz, Leonardo de Oliveira Martins, Nabil-Fareed Alikhan, Lizzie Meadows, Rachael Stanley, Ngozi Elumogo, Muhammed Yasir, Nicholas M. Thomson, Alexander J Trotter, Rachel Gilroy, Samuel Bloomfield, Claire Stuart, Andrew Bell, Reenesh Prakash, Samir Dervisevic, Alison E. Mather, John Wain, Mark Webber, Andrew J. Page, Justin O'Grady |
| EPI_ISL_512379, EPI_ISL_512380, EPI_ISL_512381, EPI_ISL_512382, EPI_ISL_512383 | Queens Medical Centre, Clinical Microbiology Department / DeepSeq Nottingham | COVID-19 Genomics UK (COG-UK) Consortium | Gemma Clark, Wendy Smith, Manjinder Khakh, Vicki M Fleming, Michelle M Lister, Hannah Howson-Wells, Jonathan Ball, Patrick McClure, Joseph Chappell, Theocharis Tsoleridis, Nadine Holmes, Matthew Carlisle, Christopher Moore, Fei Sang, Johnny Debebe, Victoria Wright, Matthew Loose |
| EPI_ISL_512384, EPI_ISL_512385, EPI_ISL_512386, EPI_ISL_512387, EPI_ISL_512388, EPI_ISL_512389, EPI_ISL_512390, EPI_ISL_512391, EPI_ISL_512392, EPI_ISL_512393, EPI_ISL_512394, EPI_ISL_512395, EPI_ISL_512396, EPI_ISL_512397, EPI_ISL_512398, EPI_ISL_512399, EPI_ISL_512400, EPI_ISL_512401, EPI_ISL_512402, EPI_ISL_512403, EPI_ISL_512404, EPI_ISL_512405, EPI_ISL_512406, EPI_ISL_512407, EPI_ISL_512408, EPI_ISL_512409, EPI_ISL_512410, EPI_ISL_512411, EPI_ISL_512412, EPI_ISL_512413, EPI_ISL_512414, EPI_ISL_512415, EPI_ISL_512416, EPI_ISL_512417, EPI_ISL_512418, EPI_ISL_512419, EPI_ISL_512420, EPI_ISL_512421, EPI_ISL_512422, EPI_ISL_512423, EPI_ISL_512424, EPI_ISL_512425, EPI_ISL_512426, EPI_ISL_512427, EPI_ISL_512428, EPI_ISL_512429, EPI_ISL_512430, EPI_ISL_512431, EPI_ISL_512432, EPI_ISL_512433, EPI_ISL_512434, EPI_ISL_512435 | | | |
| see above | Centre for Enzyme Innovation, University of Portsmouth / Translational Research Laboratory, Portsmouth Hospitals NHS Trust | COVID-19 Genomics UK (COG-UK) Consortium | Angela Beckett,Yann Bourgeois,Garry Scarlett,Sharon Glaysher,Scott Elliott,Kelly Bicknell,Robert Impey,Allyson Lloyd,Sarah Wyllie,Ethan Butcher,Anoop Chauhan,Samuel Robson |
| EPI_ISL_512436, EPI_ISL_512437, EPI_ISL_512438, EPI_ISL_512439, EPI_ISL_512440, EPI_ISL_512441, EPI_ISL_512442, EPI_ISL_512443, EPI_ISL_512444, EPI_ISL_512445, EPI_ISL_512446, EPI_ISL_512447, EPI_ISL_512448, EPI_ISL_512449, EPI_ISL_512450, EPI_ISL_512451, EPI_ISL_512452, EPI_ISL_512453, EPI_ISL_512454, EPI_ISL_512455, EPI_ISL_512456, EPI_ISL_512457, EPI_ISL_512458, EPI_ISL_512459, EPI_ISL_512460, EPI_ISL_512461, EPI_ISL_512462, EPI_ISL_512463, EPI_ISL_512464, EPI_ISL_512465, EPI_ISL_512466, EPI_ISL_512467, EPI_ISL_512468, EPI_ISL_512469, EPI_ISL_512470, EPI_ISL_512471, EPI_ISL_512472, EPI_ISL_512473, EPI_ISL_512474, EPI_ISL_512475, EPI_ISL_512476, EPI_ISL_512477, EPI_ISL_512478, EPI_ISL_512479, EPI_ISL_512480 | | | |
| see above | West of Scotland Specialist Virology Centre, NHSGGC / MRC-University of Glasgow Centre for Virus Research | COVID-19 Genomics UK (COG-UK) Consortium | Ana da Silva Filipe, Natasha Johnson, Kathy Smollett, Daniel Mair, Stephen Carmichael, Lily Tong, Jenna Nichols, Elihu Aranday-Cortes, Kirstyn Brunker, Yasmin Parr, Alice Broos, Kyriaki Nomikou; Sarah McDonald, Marc Niebel, Patawee Asamaphan; Richard Orton, Joseph Hughes, Sreenu Vattipally, David L Robertson; Alasdair MacLean, Rory Gunson; Kathy Li, Natasha Jesudason, Rajiv Shah, James Shepherd, Antonia Ho, Emma Thomson |
| EPI_ISL_512481, EPI_ISL_512482, EPI_ISL_512483, EPI_ISL_512484, EPI_ISL_512485, EPI_ISL_512486, EPI_ISL_512487, EPI_ISL_512488, EPI_ISL_512489, EPI_ISL_512490, EPI_ISL_512491, EPI_ISL_512492, EPI_ISL_512493, EPI_ISL_512494, EPI_ISL_512495, EPI_ISL_512496, EPI_ISL_512497, EPI_ISL_512498, EPI_ISL_512499, EPI_ISL_512500, EPI_ISL_512501, EPI_ISL_512502, EPI_ISL_512503, EPI_ISL_512504, EPI_ISL_512505, EPI_ISL_512506, EPI_ISL_512507, EPI_ISL_512508, EPI_ISL_512509, EPI_ISL_512510, EPI_ISL_512511, EPI_ISL_512512, EPI_ISL_512513, EPI_ISL_512514, EPI_ISL_512515, EPI_ISL_512516, EPI_ISL_512517, EPI_ISL_512518, EPI_ISL_512519, EPI_ISL_512520, EPI_ISL_512521, EPI_ISL_512522, EPI_ISL_512523, EPI_ISL_512524, EPI_ISL_512525, EPI_ISL_512526, EPI_ISL_512527, EPI_ISL_512528, EPI_ISL_512529, EPI_ISL_512530, EPI_ISL_512531, EPI_ISL_512532, EPI_ISL_512533, EPI_ISL_512534, EPI_ISL_512535, EPI_ISL_512536, EPI_ISL_512537, EPI_ISL_512538, EPI_ISL_512539, EPI_ISL_512540, EPI_ISL_512541, EPI_ISL_512542, EPI_ISL_512543, EPI_ISL_512544, EPI_ISL_512545, EPI_ISL_512546 | | | |
| see above | Wales Specialist Virology Centre Sequencing lab: Pathogen Genomics Unit | COVID-19 Genomics UK (COG-UK) Consortium | Catherine Moore, Johnathan Evans, Laura Gifford, Malorie Perry, Simon Cottrell, Angela Marchbank, Alec Birchley, Alexander Adams, Amy Gaskin, Bree Gatica-Wilcox, Jason Coombes, Joel Southgate, Lauren Gilbert, Lee Graham, Nicole Pacchiarini, Sara Kumziene-Summerhayes, Sarah Taylor, Sophie Jones, Sara Rey, Matthew Bull, Joanne Watkins, Sally Corden, Tom Connor |
| EPI_ISL_512547, EPI_ISL_512548, EPI_ISL_512549, EPI_ISL_512550, EPI_ISL_512551, EPI_ISL_512552, EPI_ISL_512553, EPI_ISL_512554, EPI_ISL_512555, EPI_ISL_512556, EPI_ISL_512557, EPI_ISL_512558, EPI_ISL_512559, EPI_ISL_512560, EPI_ISL_512561, EPI_ISL_512562, EPI_ISL_512563, EPI_ISL_512564, EPI_ISL_512565, EPI_ISL_512566, EPI_ISL_512567, EPI_ISL_512568, EPI_ISL_512569, EPI_ISL_512570, EPI_ISL_512571, EPI_ISL_512572, EPI_ISL_512573, EPI_ISL_512574, EPI_ISL_512575, EPI_ISL_512576, EPI_ISL_512577, EPI_ISL_512578, EPI_ISL_512579, EPI_ISL_512580, EPI_ISL_512581, EPI_ISL_512582, EPI_ISL_512583, EPI_ISL_512584, EPI_ISL_512585, EPI_ISL_512586, EPI_ISL_512587, EPI_ISL_512588, EPI_ISL_512589, EPI_ISL_512590, EPI_ISL_512591, EPI_ISL_512592, EPI_ISL_512593, EPI_ISL_512594, EPI_ISL_512595, EPI_ISL_512596 | | | |
| see above | Florida Bureau of Public Health Laboratories | Florida Bureau of Public Health Laboratories | Sarah Schmedes, Jason Blanton |
| EPI_ISL_512597, EPI_ISL_512598, EPI_ISL_512599, EPI_ISL_512600, EPI_ISL_512601, EPI_ISL_512602, EPI_ISL_512603, EPI_ISL_512604, EPI_ISL_512605, EPI_ISL_512606, EPI_ISL_512607, EPI_ISL_512608, EPI_ISL_512609, EPI_ISL_512610, EPI_ISL_512611, EPI_ISL_512612, EPI_ISL_512613, EPI_ISL_512614, EPI_ISL_512615, EPI_ISL_512616, EPI_ISL_512617, EPI_ISL_512618, EPI_ISL_512619, EPI_ISL_512620, EPI_ISL_512621, EPI_ISL_512622, EPI_ISL_512623, EPI_ISL_512624, EPI_ISL_512625, EPI_ISL_512626, EPI_ISL_512627, EPI_ISL_512628, EPI_ISL_512629, EPI_ISL_512630, EPI_ISL_512631, EPI_ISL_512632, EPI_ISL_512633, EPI_ISL_512634, EPI_ISL_512635, EPI_ISL_512636, EPI_ISL_512637, EPI_ISL_512638, EPI_ISL_512639, EPI_ISL_512640, EPI_ISL_512641, EPI_ISL_512642, EPI_ISL_512643, EPI_ISL_512644 | | | |
| see above | National Laboratory for Influenza/Virology reference laboratory, Public Health Center of the Ministry of Health of Ukraine | Respiratory Virus Unit, Microbiology Services Colindale, Public Health England | PHE Covid Sequencing Team, Dr. Iryna Demchyshyna |
| EPI_ISL_512645, EPI_ISL_512646 | E. Gulbja Laboratorija | Latvian Biomedical Research and Study Centre | Ivars Silamiķelis, Kaspars Megnis, Monta Ustinova, Ņikita Zrelovs, Vita Rovīte, Mikus Gavars, Dmitrijs Perminovs, Uga Dumpis, Jānis Kloviņš |
| EPI_ISL_512647, EPI_ISL_512648 | Latvijas Infektoloģijas centrs | Latvian Biomedical Research and Study Centre | Ivars Silamiķelis, Kaspars Megnis, Monta Ustinova, Ņikita Zrelovs, Vita Rovīte, Jeļena Storoženko, Tatjana Kolupajeva, Oksana Savicka, Uga Dumpis, Jānis Kloviņš |
| EPI_ISL_512649 | Latvijas Infektoloģijas centrs | Latvian Biomedical Research and Study Centre | Ivars Silamiķelis, Kaspars Megnis, Monta Ustinova, Ņikita Zrelovs, Vita Rovīte, Jeļena Storoženko, Tatjana Kolupajeva, Oksana Savicka, Uga Dumpis, Jānis Kloviņš |
| EPI_ISL_512650, EPI_ISL_512651, EPI_ISL_512652 | Latvijas Infektoloģijas centrs | Latvian Biomedical Research and Study Centre | Ivars Silamiķelis, Kaspars Megnis, Monta Ustinova, Ņikita Zrelovs, Vita Rovīte, Jeļena Storoženko, Tatjana Kolupajeva, Oksana Savicka, Uga Dumpis, Jānis Kloviņš |
| EPI_ISL_512653 | Area De Salud Desamparados 1 - Clinica Dr. Marcial Fallas [Grifo Alto/Desampara | Inciensa, Instituto Costarricense de Investigación y Enseñanza en Nutrición y Salud | Francisco Duarte, Hebleen Porras, Claudio Soto-Garita, Estela Cordero, Adriana Godinez & Melany Calderon |
| EPI_ISL_512654 | Hospital Dr. Rafael A. Calderon Guardia [San Jose/San Jose] | Inciensa, Instituto Costarricense de Investigación y Enseñanza en Nutrición y Salud | Francisco Duarte, Hebleen Porras, Claudio Soto-Garita, Estela Cordero, Adriana Godinez & Melany Calderon |
| EPI_ISL_512655 | Hospital De Niños Dr. Carlos Saenz Herrera [San Jose/San Jose] | Inciensa, Instituto Costarricense de Investigación y Enseñanza en Nutrición y Salud | Francisco Duarte, Hebleen Porras, Claudio Soto-Garita, Estela Cordero, Adriana Godinez & Melany Calderon |
| EPI_ISL_512656 | Area De Salud Pavas (Coopesalud) [Pavas/San Jose] | Inciensa, Instituto Costarricense de Investigación y Enseñanza en Nutrición y Salud | Francisco Duarte, Hebleen Porras, Claudio Soto-Garita, Estela Cordero, Adriana Godinez & Melany Calderon |
| EPI_ISL_512657 | Area De Salud Tibas-Uruca-Merced - Clinica Dr. Clorito Picado [Tibas/San Jose] | Inciensa, Instituto Costarricense de Investigación y Enseñanza en Nutrición y Salud | Francisco Duarte, Hebleen Porras, Claudio Soto-Garita, Estela Cordero, Adriana Godinez & Melany Calderon |
| EPI_ISL_512658 | Area De Salud Orotina-San Mateo [Orotina/Alajuela] | Inciensa, Instituto Costarricense de Investigación y Enseñanza en Nutrición y Salud | Francisco Duarte, Hebleen Porras, Claudio Soto-Garita, Estela Cordero, Adriana Godinez & Melany Calderon |
| EPI_ISL_512659 | Area De Salud Fortuna | Inciensa, Instituto Costarricense de Investigación y Enseñanza en Nutrición y Salud | Francisco Duarte, Hebleen Porras, Claudio Soto-Garita, Estela Cordero, Adriana Godinez & Melany Calderon |
| EPI_ISL_512660, EPI_ISL_512661 | Area De Salud Los Chiles | Inciensa, Instituto Costarricense de Investigación y Enseñanza en Nutrición y Salud | Francisco Duarte, Hebleen Porras, Claudio Soto-Garita, Estela Cordero, Adriana Godinez & Melany Calderon |
| EPI_ISL_512662 | Area De Salud La Cruz | Inciensa, Instituto Costarricense de Investigación y Enseñanza en Nutrición y Salud | Francisco Duarte, Hebleen Porras, Claudio Soto-Garita, Estela Cordero, Adriana Godinez & Melany Calderon |
| EPI_ISL_512663, EPI_ISL_512664 | Area De Salud Alajuela Norte - Clinica Dr. Marcial Rodriguez | Inciensa, Instituto Costarricense de Investigación y Enseñanza en Nutrición y Salud | Francisco Duarte, Hebleen Porras, Claudio Soto-Garita, Estela Cordero, Adriana Godinez & Melany Calderon |
| EPI_ISL_512665 | Hospital De Las Mujeres Dr. Adolfo Carit | Inciensa, Instituto Costarricense de Investigación y Enseñanza en Nutrición y Salud | Francisco Duarte, Hebleen Porras, Claudio Soto-Garita, Estela Cordero, Adriana Godinez & Melany Calderon |
| EPI_ISL_512666, EPI_ISL_512667 | Area De Salud La Cruz | Inciensa, Instituto Costarricense de Investigación y Enseñanza en Nutrición y Salud | Francisco Duarte, Hebleen Porras, Claudio Soto-Garita, Estela Cordero, Adriana Godinez & Melany Calderon |
| EPI_ISL_512668 | Area De Salud Corredores | Inciensa, Instituto Costarricense de Investigación y Enseñanza en Nutrición y Salud | Francisco Duarte, Hebleen Porras, Claudio Soto-Garita, Estela Cordero, Adriana Godinez & Melany Calderon |
| EPI_ISL_512669 | Area De Salud Alajuela Norte - Clinica Dr. Marcial Rodriguez | Inciensa, Instituto Costarricense de Investigación y Enseñanza en Nutrición y Salud | Francisco Duarte, Hebleen Porras, Claudio Soto-Garita, Estela Cordero, Adriana Godinez & Melany Calderon |
| EPI_ISL_512670 | Centro Nacional De Rehabilitacion Humberto Araya Rojas (Cenare) | Inciensa, Instituto Costarricense de Investigación y Enseñanza en Nutrición y Salud | Francisco Duarte, Hebleen Porras, Claudio Soto-Garita, Estela Cordero, Adriana Godinez & Melany Calderon |
| EPI_ISL_512671 | Area De Salud La Cruz | Inciensa, Instituto Costarricense de Investigación y Enseñanza en Nutrición y Salud | Francisco Duarte, Hebleen Porras, Claudio Soto-Garita, Estela Cordero, Adriana Godinez & Melany Calderon |
| EPI_ISL_512672, EPI_ISL_512673, EPI_ISL_512674, EPI_ISL_512675 | Hospital De Niños Dr. Carlos Saenz Herrera [San Jose/San Jose] | Inciensa, Instituto Costarricense de Investigación y Enseñanza en Nutrición y Salud | Francisco Duarte, Hebleen Porras, Claudio Soto-Garita, Estela Cordero, Adriana Godinez & Melany Calderon |
| EPI_ISL_512676, EPI_ISL_512677 | Florida Bureau of Public Health Laboratories | Florida Bureau of Public Health Laboratories | Sarah Schmedes, Jason Blanton |
| EPI_ISL_512712, EPI_ISL_512713, EPI_ISL_512714, EPI_ISL_512715, EPI_ISL_512716, EPI_ISL_512717, EPI_ISL_512718, EPI_ISL_512719, EPI_ISL_512720, EPI_ISL_512721, EPI_ISL_512722, EPI_ISL_512723, EPI_ISL_512724, EPI_ISL_512725, EPI_ISL_512726, EPI_ISL_512727, EPI_ISL_512728, EPI_ISL_512729, EPI_ISL_512730, EPI_ISL_512731, EPI_ISL_512732, EPI_ISL_512733, EPI_ISL_512734, EPI_ISL_512735, EPI_ISL_512736, EPI_ISL_512737, EPI_ISL_512738, EPI_ISL_512739, EPI_ISL_512740, EPI_ISL_512741, EPI_ISL_512742, EPI_ISL_512743, EPI_ISL_512744, EPI_ISL_512745, EPI_ISL_512746, EPI_ISL_512747, EPI_ISL_512748, EPI_ISL_512749, EPI_ISL_512750, EPI_ISL_512751, EPI_ISL_512752, EPI_ISL_512753, EPI_ISL_512754, EPI_ISL_512755, EPI_ISL_512756, EPI_ISL_512757, EPI_ISL_512758, EPI_ISL_512759, EPI_ISL_512760, EPI_ISL_512761, EPI_ISL_512762, EPI_ISL_512763, EPI_ISL_512764, EPI_ISL_512765 | | | |
| see above | PathWest Laboratory Medicine WA | PathWest Laboratory Medicine WA Microbial Surveillance Unit | PathWest Laboratory Medicine WA Microbial Surveillance Unit |
| EPI_ISL_512766, EPI_ISL_512767, EPI_ISL_512768, EPI_ISL_512769, EPI_ISL_512770, EPI_ISL_512771 | unknown | Center for Precision Medicine, Meizhou People's Hospital (Huangtang Hospital) | Guo,X., Wu,H., Yu,Z., Huang,Q. |
| EPI_ISL_512773 | Center for Precision Medicine, Meizhou People's Hospital (Huangtang Hospital) | Center for Precision Medicine, Meizhou People's Hospital (Huangtang Hospital) | Guo,X., Wu,H., Yu,Z. and Huang,Q. |
| EPI_ISL_512775, EPI_ISL_512776, EPI_ISL_512777, EPI_ISL_512778, EPI_ISL_512779, EPI_ISL_512780, EPI_ISL_512781, EPI_ISL_512782, EPI_ISL_512783, EPI_ISL_512784 | unknown | Utah Public Health Laboratory, Utah Public Health Laboratory Infectious Disease submission group | Butz,H.A., Young,E.L., Oakeson,K. |
| EPI_ISL_512785, EPI_ISL_512786, EPI_ISL_512787, EPI_ISL_512788, EPI_ISL_512789, EPI_ISL_512790, EPI_ISL_512791, EPI_ISL_512792, EPI_ISL_512793, EPI_ISL_512794, EPI_ISL_512795, EPI_ISL_512796, EPI_ISL_512797, EPI_ISL_512798, EPI_ISL_512799, EPI_ISL_512800, EPI_ISL_512801, EPI_ISL_512802, EPI_ISL_512803, EPI_ISL_512804, EPI_ISL_512805, EPI_ISL_512806, EPI_ISL_512807, EPI_ISL_512808, EPI_ISL_512809 | | | |

| | | | |
|---|---|---|---|
| see above | Public Health, United States Air Force School of Aerospace Medicine | Public Health, United States Air Force School of Aerospace Medicine | Fries,A.C., Purves,S.M., Meyer,J.R., Javorina,A.K., Connors,B.C., Macias,E.A., Lambert,A.W., Chapleau,R.R., Starr,C.R. |
| EPI_ISL_512810 | National Laboratory for Influenza/Virology reference laboratory, Public Health Center of the Ministry of Health of Ukraine | Respiratory Virus Unit, Microbiology Services Colindale, Public Health England | PHE Covid Sequencing Team, Dr. Iryna Demchyshyna |
| EPI_ISL_512811, EPI_ISL_512812, EPI_ISL_512813, EPI_ISL_512814, EPI_ISL_512815, EPI_ISL_512816, EPI_ISL_512817, EPI_ISL_512818, EPI_ISL_512819, EPI_ISL_512820, EPI_ISL_512821 | | | |
| see above | Kenema Government Hospital, Ministry of Health and Sanitation | Kenema Government Hospital, Ministry of Health and Sanitation | Goba,A., Momoh,M., Sandi,J., Tomkins-Tinch,C., Siddle,K., Mehta,S., Oluniyi,P., Jalloh,S., Park,D., Andersen,K., Garry,R., Happi,C., Grant,D., Olawoye,I. |
| EPI_ISL_512822, EPI_ISL_512823, EPI_ISL_512824, EPI_ISL_512825, EPI_ISL_512826, EPI_ISL_512827, EPI_ISL_512828, EPI_ISL_512829, EPI_ISL_512830, EPI_ISL_512831, EPI_ISL_512832, EPI_ISL_512833, EPI_ISL_512834, EPI_ISL_512835, EPI_ISL_512836, EPI_ISL_512837, EPI_ISL_512838, EPI_ISL_512839, EPI_ISL_512840, EPI_ISL_512841, EPI_ISL_512842 | | | |
| see above | National Public Health Laboratory, National Centre for Infectious Diseases | National Public Health Laboratory, National Centre for Infectious Diseases | Mak TM, Octavia S, Zhou Z, Chavatte JM, Cui L, Lin RTP |
| EPI_ISL_512844 | Department of Medical Research | DMR_Myanmar | Myat Htut Nyunt, Hnin Ohnmar Soe, Kay Thi Aye, Wah Wah Aung,Yi Yi Kyaw, Aung Kyaw Kyaw, Theingi Win Myat, Phyu Win Ei, Aung Zaw Latt, Nan Aye Thida Oo, Lai Lai San, Su Mon Win, Ni Ni Zaw, Htin Lin, Hlaing Myat Thu, Zaw Than Htun |
| EPI_ISL_512846 | O.I.J. MORGUE JUDICIAL | Inciensa, Instituto Costarricense de Investigación y Enseñanza en Nutrición y Salud | Francisco Duarte, Hebleen Porras, Claudio Soto-Garita, Estela Cordero, Adriana Godinez & Melany Calderon |
| EPI_ISL_512847, EPI_ISL_512848, EPI_ISL_512849, EPI_ISL_512850, EPI_ISL_512851, EPI_ISL_512852, EPI_ISL_512853, EPI_ISL_512854, EPI_ISL_512855, EPI_ISL_512856, EPI_ISL_512857, EPI_ISL_512858, EPI_ISL_512859, EPI_ISL_512860, EPI_ISL_512861, EPI_ISL_512862, EPI_ISL_512863, EPI_ISL_512864, EPI_ISL_512865, EPI_ISL_512866, EPI_ISL_512867, EPI_ISL_512868, EPI_ISL_512869, EPI_ISL_512870, EPI_ISL_512871, EPI_ISL_512872 | | | |
| see above | Ramathibodi Hospital | COVID-19 Network Investigations (CONI) Alliance | Elizabeth Batty, Wasun Chantratita, Thanat Chookajorn, Stefan Fernandez, Angkana Huang, Anthony R. Jones, Khajohn Joonsalak, Chonticha Klungtong, Theerarat Kochakarn, Namfon Kotanan, Krittikorn Kumpornsin, Wudtichai Manasatienkij, Bhakbhoom Panthan, Ekawat Pasomsub, Kingkan Rakmanee, Insee Sensorn, Janjira Thaipadungpanit, Arporn Wangwiwatsin,Treewat Watthanachockchai |
| EPI_ISL_512873 | Centre Pasteur of Cameroun | Virology Service, Centre Pasteur of Cameroun | Richard Njouom and Serge Alain SADEUH-Mba |
| EPI_ISL_512874, EPI_ISL_512875, EPI_ISL_512876, EPI_ISL_512877, EPI_ISL_512878, EPI_ISL_512879, EPI_ISL_512880, EPI_ISL_512881, EPI_ISL_512882, EPI_ISL_512883, EPI_ISL_512884, EPI_ISL_512885, EPI_ISL_512886, EPI_ISL_512887, EPI_ISL_512888, EPI_ISL_512889, EPI_ISL_512890, EPI_ISL_512891, EPI_ISL_512892, EPI_ISL_512893, EPI_ISL_512894, EPI_ISL_512895, EPI_ISL_512896, EPI_ISL_512897, EPI_ISL_512898, EPI_ISL_512899, EPI_ISL_512900, EPI_ISL_512901, EPI_ISL_512902, EPI_ISL_512903 | | | |
| see above | Pathogen Genomics Lab King Abdullah University of Science and Technology(KAUST) | Pathogen Genomics Lab King Abdullah University of Science and Technology(KAUST) | Raeece Naeem, Rahul P Salunke, Sharif Hala, Sara Mfarrej, Amit Kumar Subudhi, Fadwa Alofi, Fathia Ben Rached, Afrah Alsomali, Asim Khogeer, Ahmad Bakur Mahmoud, Anwar Hashem, Naif Almontashiri, Arnab Pain |
| EPI_ISL_512904, EPI_ISL_512905, EPI_ISL_512906, EPI_ISL_512907 | Pathogen Genomics Lab King Abdullah University of Science and Technology(KAUST) | Pathogen Genomics Lab King Abdullah University of Science and Technology(KAUST) | Fathia Ben Rached, Raeece Naeem, Sharif Hala, Fadwa Alofi, Rahul P Salunke, Sara Mfarrej, Amit Kumar Subudhi, Afrah Alsomali, Asim Khogeer, Ahmad Bakur Mahmoud, Anwar Hashem, Naif Almontashiri, Arnab Pain |
| EPI_ISL_512908, EPI_ISL_512909, EPI_ISL_512910, EPI_ISL_512911, EPI_ISL_512912, EPI_ISL_512913, EPI_ISL_512914, EPI_ISL_512915, EPI_ISL_512916, EPI_ISL_512917 | Pathogen Genomics Lab King Abdullah University of Science and Technology(KAUST) | Pathogen Genomics Lab King Abdullah University of Science and Technology(KAUST) | Sharif Hala, Fadwa Alofi, Sara Mfarrej, Amit Kumar Subudhi, Rahul P Salunke, Fathia Ben Rached, Amanda Ooi, Luke Esau, Afrah Alsomali, Asim Khogeer, Jumana Taha, Abdulaziz Alahmadi, Kahled Algithami, Raeece Naeem, Anwar Hashem, Naif Almontashiri, Arnab Pain |
| EPI_ISL_512918, EPI_ISL_512919, EPI_ISL_512920, EPI_ISL_512921, EPI_ISL_512922, EPI_ISL_512923, EPI_ISL_512924, EPI_ISL_512925, EPI_ISL_512926, EPI_ISL_512927, EPI_ISL_512928, EPI_ISL_512929, EPI_ISL_512930, EPI_ISL_512931, EPI_ISL_512932, EPI_ISL_512933, EPI_ISL_512934, EPI_ISL_512935, EPI_ISL_512936, EPI_ISL_512937, EPI_ISL_512938, EPI_ISL_512939, EPI_ISL_512940, EPI_ISL_512941, EPI_ISL_512942, EPI_ISL_512943, EPI_ISL_512944, EPI_ISL_512945 | | | |
| see above | Pathogen Genomics Lab King Abdullah University of Science and Technology(KAUST) | Pathogen Genomics Lab King Abdullah University of Science and Technology(KAUST) | Fadwa Alofi, Sharif Hala, Rahul P Salunke, Sara Mfarrej, Amit Kumar Subudhi, Fathia Ben Rached, Amanda, Luke, Afrah Alsomali, Asim Khogeer, Jumana Taha, Abdulaziz Alahmadi, Kahled Algithami, Raeece Naeem, Anwar Hashem, Naif Almontashiri, Arnab Pain |
| EPI_ISL_512946, EPI_ISL_512947, EPI_ISL_512948, EPI_ISL_512949, EPI_ISL_512950, EPI_ISL_512951, EPI_ISL_512952, EPI_ISL_512953, EPI_ISL_512954, EPI_ISL_512955, EPI_ISL_512956, EPI_ISL_512957, EPI_ISL_512958, EPI_ISL_512959, EPI_ISL_512960, EPI_ISL_512961, EPI_ISL_512962, EPI_ISL_512963, EPI_ISL_512964, EPI_ISL_512965, EPI_ISL_512966, EPI_ISL_512967, EPI_ISL_512968, EPI_ISL_512969, EPI_ISL_512970, EPI_ISL_512971, EPI_ISL_512972, EPI_ISL_512973, EPI_ISL_512974, EPI_ISL_512975, EPI_ISL_512976, EPI_ISL_512977, EPI_ISL_512978, EPI_ISL_512979, EPI_ISL_512980, EPI_ISL_512981, EPI_ISL_512982, EPI_ISL_512983, EPI_ISL_512984, EPI_ISL_512985, EPI_ISL_512986, EPI_ISL_512987, EPI_ISL_512988 | | | |
| see above | Pathogen Genomics Lab King Abdullah University of Science and Technology(KAUST) | Pathogen Genomics Lab King Abdullah University of Science and Technology(KAUST) | Sara Mfarrej, Raeece Naeem, Rahul P Salunke, Sharif Hala, Fadwa Alofi, Amit Kumar Subudhi, Fathia Ben Rached, Afrah Alsomali, Jumana Taha, Abdulaziz Alahmadi, Asim Khogeer, Nashwa Al-khotani, Anwar Hashem, Naif Almontashiri, Arnab Pain |
| EPI_ISL_512989, EPI_ISL_512990, EPI_ISL_512991, EPI_ISL_512992, EPI_ISL_512993, EPI_ISL_512994, EPI_ISL_512995, EPI_ISL_512996, EPI_ISL_512997, EPI_ISL_512998, EPI_ISL_512999, EPI_ISL_513000, EPI_ISL_513001 | | | |
| see above | Pathogen Genomics Lab King Abdullah University of Science and Technology(KAUST) | Pathogen Genomics Lab King Abdullah University of Science and Technology(KAUST) | Amit Kumar Subudhi, Rahul P Salunke, Sara Mfarrej, Sharif Hala, Fadwa Alofi, Fathia Ben Rached, Afrah Alsomali, Asim Khogeer, Nashwa Al-khotani, Raeece Naeem, Anwar Hashem, Naif Almontashiri, Arnab Pain |
| EPI_ISL_513002, EPI_ISL_513003, EPI_ISL_513004, EPI_ISL_513005, EPI_ISL_513006, EPI_ISL_513007, EPI_ISL_513008, EPI_ISL_513009, EPI_ISL_513010, EPI_ISL_513011, EPI_ISL_513012, EPI_ISL_513013, EPI_ISL_513014, EPI_ISL_513015, EPI_ISL_513016, EPI_ISL_513017, EPI_ISL_513018, EPI_ISL_513019, EPI_ISL_513020, EPI_ISL_513021, EPI_ISL_513022, EPI_ISL_513023, EPI_ISL_513024, EPI_ISL_513025, EPI_ISL_513026 | | | |
| see above | Pathogen Genomics Lab King Abdullah University of Science and Technology(KAUST) | Pathogen Genomics Lab King Abdullah University of Science and Technology(KAUST) | Afrah Alsomali, Fathia Ben Rached, Raeece Naeem, Sharif Hala,Rahul P Salunke, Amanda Ooi, Luke Esau, Sara Mfarrej, Amit Kumar Subudhi, Fadwa Alofi, Asim Khogeer, Kahled Algithami, Anwar Hashem, Naif Almontashiri, Arnab Pain |
| EPI_ISL_513027, EPI_ISL_513028, EPI_ISL_513029, EPI_ISL_513030, EPI_ISL_513031, EPI_ISL_513032, EPI_ISL_513033, EPI_ISL_513034, EPI_ISL_513035, EPI_ISL_513036, EPI_ISL_513037, EPI_ISL_513038, EPI_ISL_513039, EPI_ISL_513040, EPI_ISL_513041, EPI_ISL_513042, EPI_ISL_513043, EPI_ISL_513044, EPI_ISL_513045, EPI_ISL_513046, EPI_ISL_513047, EPI_ISL_513048, EPI_ISL_513049, EPI_ISL_513050, EPI_ISL_513051, EPI_ISL_513052, EPI_ISL_513053, EPI_ISL_513054, EPI_ISL_513055, EPI_ISL_513056, EPI_ISL_513057, EPI_ISL_513058, EPI_ISL_513059, EPI_ISL_513060, EPI_ISL_513061, EPI_ISL_513062, EPI_ISL_513063 | | | |
| see above | Pathogen Genomics Lab King Abdullah University of Science and Technology(KAUST) | Pathogen Genomics Lab King Abdullah University of Science and Technology(KAUST) | Rahul P Salunke, Sharif Hala, Raeece Naeem, Sara Mfarrej, Amit Kumar Subudhi, Amanda Ooi, Luke Esau, Fadwa Alofi, Fathia Ben Rached, Afrah Alsomali, Asim Khogeer, Ahmad Bakur Mahmoud, Anwar Hashem, Naif Almontashiri, Arnab Pain |
| EPI_ISL_513064, EPI_ISL_513065, EPI_ISL_513066, EPI_ISL_513067, EPI_ISL_513068, EPI_ISL_513069, EPI_ISL_513070, EPI_ISL_513071, EPI_ISL_513072, EPI_ISL_513073, EPI_ISL_513074, EPI_ISL_513075, EPI_ISL_513076 | | | |
| see above | Pathogen Genomics Lab King Abdullah University of Science and Technology(KAUST) | Pathogen Genomics Lab King Abdullah University of Science and Technology(KAUST) | Raeece Naeem, Rahul P Salunke, Sharif Hala, Sara Mfarrej, Amit Kumar Subudhi, Fadwa Alofi, Fathia Ben Rached, Afrah Alsomali, Asim Khogeer, Ahmad Bakur Mahmoud, Anwar Hashem, Naif Almontashiri, Arnab Pain |
| EPI_ISL_513077, EPI_ISL_513078, EPI_ISL_513079, EPI_ISL_513080, EPI_ISL_513081, EPI_ISL_513082, EPI_ISL_513083, EPI_ISL_513084, EPI_ISL_513085, EPI_ISL_513086, EPI_ISL_513087, EPI_ISL_513088, EPI_ISL_513089, EPI_ISL_513090, EPI_ISL_513091, EPI_ISL_513092, EPI_ISL_513093, EPI_ISL_513094, EPI_ISL_513095, EPI_ISL_513096, EPI_ISL_513097, EPI_ISL_513098, EPI_ISL_513099, EPI_ISL_513100, EPI_ISL_513101, EPI_ISL_513102, EPI_ISL_513103, EPI_ISL_513104, EPI_ISL_513105, EPI_ISL_513106, EPI_ISL_513107, EPI_ISL_513108, EPI_ISL_513109, EPI_ISL_513110, EPI_ISL_513111, EPI_ISL_513112, EPI_ISL_513113, EPI_ISL_513114, EPI_ISL_513115, EPI_ISL_513116, EPI_ISL_513117, EPI_ISL_513118, EPI_ISL_513119, EPI_ISL_513120 | | | |
| see above | Pathogen Genomics Lab King Abdullah University of Science and Technology(KAUST) | Pathogen Genomics Lab King Abdullah University of Science and Technology(KAUST) | Fathia Ben Rached, Raeece Naeem, Sharif Hala, Fadwa Alofi, Rahul P Salunke, Sara Mfarrej, Amit Kumar Subudhi, Afrah Alsomali, Asim Khogeer, Ahmad Bakur Mahmoud, Anwar Hashem, Naif Almontashiri, Arnab Pain |
| EPI_ISL_513121, EPI_ISL_513122, EPI_ISL_513123, EPI_ISL_513124, EPI_ISL_513125, EPI_ISL_513126, EPI_ISL_513127, EPI_ISL_513128, EPI_ISL_513129, EPI_ISL_513130, EPI_ISL_513131, EPI_ISL_513132, EPI_ISL_513133, EPI_ISL_513134, EPI_ISL_513135, EPI_ISL_513136, EPI_ISL_513137, EPI_ISL_513138, EPI_ISL_513139, EPI_ISL_513140, EPI_ISL_513141, EPI_ISL_513142, EPI_ISL_513143, EPI_ISL_513144, EPI_ISL_513145, EPI_ISL_513146 | | | |
| see above | Pathogen Genomics Lab King Abdullah University of Science and Technology(KAUST) | Pathogen Genomics Lab King Abdullah University of Science and Technology(KAUST) | Sharif Hala, Fadwa Alofi, Sara Mfarrej, Amit Kumar Subudhi, Rahul P Salunke, Fathia Ben Rached, Amanda Ooi, Luke Esau, Afrah Alsomali, Asim Khogeer, Jumana Taha, Abdulaziz Alahmadi, Kahled Algithami, Raeece Naeem, Anwar Hashem, Naif Almontashiri, Arnab Pain |
| EPI_ISL_513147, EPI_ISL_513148, EPI_ISL_513149, EPI_ISL_513150, EPI_ISL_513151, EPI_ISL_513152, EPI_ISL_513153, EPI_ISL_513154, EPI_ISL_513155, EPI_ISL_513156, EPI_ISL_513157, EPI_ISL_513158, EPI_ISL_513159, EPI_ISL_513160, EPI_ISL_513161, EPI_ISL_513162, EPI_ISL_513163, EPI_ISL_513164, EPI_ISL_513165, EPI_ISL_513166, EPI_ISL_513167 | | | |
| see above | Pathogen Genomics Lab King Abdullah University of Science and Technology(KAUST) | Pathogen Genomics Lab King Abdullah University of Science and Technology(KAUST) | Fadwa Alofi, Sharif Hala, Rahul P Salunke, Sara Mfarrej, Amit Kumar Subudhi, Fathia Ben Rached, Amanda, Luke, Afrah Alsomali, Asim Khogeer, Jumana Taha, Abdulaziz Alahmadi, Kahled Algithami, Raeece Naeem, Anwar Hashem, Naif Almontashiri, Arnab Pain |
| EPI_ISL_513168, EPI_ISL_513169, EPI_ISL_513170, EPI_ISL_513171, EPI_ISL_513172, EPI_ISL_513173, EPI_ISL_513174 | Pathogen Genomics Lab King Abdullah University of Science and Technology(KAUST) | Pathogen Genomics Lab King Abdullah University of Science and Technology(KAUST) | Sara Mfarrej, Raeece Naeem, Rahul P Salunke, Sharif Hala, Fadwa Alofi, Amit Kumar Subudhi, Fathia Ben Rached, Afrah Alsomali, Jumana Taha, Abdulaziz Alahmadi, Asim Khogeer, Nashwa Al-khotani, Anwar Hashem, Naif Almontashiri, Arnab Pain |
| EPI_ISL_513175, EPI_ISL_513176, EPI_ISL_513177, EPI_ISL_513178, EPI_ISL_513179, EPI_ISL_513180, EPI_ISL_513181, EPI_ISL_513182, EPI_ISL_513183, EPI_ISL_513184, EPI_ISL_513185, EPI_ISL_513186, EPI_ISL_513187, EPI_ISL_513188, EPI_ISL_513189 | | | |
| see above | Pathogen Genomics Lab King Abdullah University of Science and Technology(KAUST) | Pathogen Genomics Lab King Abdullah University of Science and Technology(KAUST) | Amit Kumar Subudhi, Rahul P Salunke, Sara Mfarrej, Sharif Hala, Fadwa Alofi, Fathia Ben Rached, Afrah Alsomali, Asim Khogeer, Nashwa Al-khotani, Raeece Naeem, Anwar Hashem, Naif Almontashiri, Arnab Pain |
| EPI_ISL_513190, EPI_ISL_513191, EPI_ISL_513192, EPI_ISL_513193, EPI_ISL_513194, EPI_ISL_513195, EPI_ISL_513196, EPI_ISL_513197 | Pathogen Genomics Lab King Abdullah University of Science and Technology(KAUST) | Pathogen Genomics Lab King Abdullah University of Science and Technology(KAUST) | Afrah Alsomali, Fathia Ben Rached, Raeece Naeem, Sharif Hala,Rahul P Salunke, Amanda Ooi, Luke Esau, Sara Mfarrej, Amit Kumar Subudhi, Fadwa Alofi, Asim Khogeer, Kahled Algithami, Anwar Hashem, Naif Almontashiri, Arnab Pain |
| EPI_ISL_513198, EPI_ISL_513199, EPI_ISL_513200, EPI_ISL_513201, EPI_ISL_513202, EPI_ISL_513203, EPI_ISL_513204, EPI_ISL_513205, EPI_ISL_513206, EPI_ISL_513207 | Pathogen Genomics Lab King Abdullah University of Science and Technology(KAUST) | Pathogen Genomics Lab King Abdullah University of Science and Technology(KAUST) | Rahul P Salunke, Sharif Hala, Raeece Naeem, Sara Mfarrej, Amit Kumar Subudhi, Amanda Ooi, Luke Esau, Fadwa Alofi, Fathia Ben Rached, Afrah Alsomali, Asim Khogeer, Ahmad Bakur Mahmoud, Anwar Hashem, Naif Almontashiri, Arnab Pain |
| EPI_ISL_513208, EPI_ISL_513209, EPI_ISL_513210, EPI_ISL_513211, EPI_ISL_513212, EPI_ISL_513213 | Pathogen Genomics Lab King Abdullah University of Science and Technology(KAUST) | Pathogen Genomics Lab King Abdullah University of Science and Technology(KAUST) | Raeece Naeem, Rahul P Salunke, Sharif Hala, Sara Mfarrej, Amit Kumar Subudhi, Fadwa Alofi, Fathia Ben Rached, Afrah Alsomali, Asim Khogeer, Ahmad Bakur Mahmoud, Anwar Hashem, Naif Almontashiri, Arnab Pain |
| EPI_ISL_513214, EPI_ISL_513215, EPI_ISL_513216, EPI_ISL_513217, EPI_ISL_513218, EPI_ISL_513219, EPI_ISL_513220, EPI_ISL_513221, EPI_ISL_513222, EPI_ISL_513223, EPI_ISL_513224, EPI_ISL_513225, EPI_ISL_513226 | | | |
| see above | Pathogen Genomics Lab King Abdullah University of Science and Technology(KAUST) | Pathogen Genomics Lab King Abdullah University of Science and Technology(KAUST) | Sharif Hala, Fadwa Alofi, Sara Mfarrej, Amit Kumar Subudhi, Rahul P Salunke, Fathia Ben Rached, Amanda Ooi, Luke Esau, Afrah Alsomali, Asim Khogeer, Jumana Taha, Abdulaziz Alahmadi, Kahled Algithami, Raeece Naeem, Anwar Hashem, Naif Almontashiri, Arnab Pain |
| EPI_ISL_513227, EPI_ISL_513228, EPI_ISL_513229, EPI_ISL_513230, EPI_ISL_513231, EPI_ISL_513232, EPI_ISL_513233, EPI_ISL_513234, EPI_ISL_513235, EPI_ISL_513236, EPI_ISL_513237, EPI_ISL_513238, EPI_ISL_513239, EPI_ISL_513240, EPI_ISL_513241, EPI_ISL_513242, EPI_ISL_513243, EPI_ISL_513244, EPI_ISL_513245, EPI_ISL_513246, EPI_ISL_513247, EPI_ISL_513248 | | | |
| see above | Pathogen Genomics Lab King Abdullah University of Science and Technology(KAUST) | Pathogen Genomics Lab King Abdullah University of Science and Technology(KAUST) | Amit Kumar Subudhi, Rahul P Salunke, Sara Mfarrej, Sharif Hala, Fadwa Alofi, Fathia Ben Rached, Afrah Alsomali, Asim Khogeer, Nashwa Al-khotani, Raeece Naeem, Anwar Hashem, Naif Almontashiri, Arnab Pain |
| EPI_ISL_513249, EPI_ISL_513250, EPI_ISL_513251, EPI_ISL_513252, EPI_ISL_513253, EPI_ISL_513254, EPI_ISL_513255, EPI_ISL_513256, EPI_ISL_513257, EPI_ISL_513258, EPI_ISL_513259, EPI_ISL_513260, EPI_ISL_513261, EPI_ISL_513262, EPI_ISL_513263, EPI_ISL_513264 | | | |

| | | | |
|---|---|---|---|
| see above | Pathogen Genomics Lab King Abdullah University of Science and Technology(KAUST) | Pathogen Genomics Lab King Abdullah University of Science and Technology(KAUST) | Afrah Alsomali, Fathia Ben Rached, Raeece Naeem, Sharif Hala,Rahul P Salunke, Amanda Ooi, Luke Esau, Sara Mfarrej, Amit Kumar Subudhi, Fadwa Alofi, Asim Khogeer, Kahled Algithami, Anwar Hashem, Naif Almontashiri, Arnab Pain |
| EPI_ISL_513265, EPI_ISL_513266, EPI_ISL_513267, EPI_ISL_513268, EPI_ISL_513269, EPI_ISL_513270, EPI_ISL_513271, EPI_ISL_513272, EPI_ISL_513273, EPI_ISL_513274, EPI_ISL_513275, EPI_ISL_513276, EPI_ISL_513277, EPI_ISL_513278, EPI_ISL_513279, EPI_ISL_513280, EPI_ISL_513281, EPI_ISL_513282, EPI_ISL_513283, EPI_ISL_513284, EPI_ISL_513285, EPI_ISL_513286, EPI_ISL_513287, EPI_ISL_513288, EPI_ISL_513289, EPI_ISL_513290, EPI_ISL_513291, EPI_ISL_513292, EPI_ISL_513293 | | | |
| see above | University of Miami Immunology and Histocompatibility Laboratory | University of Miami Immunology and Histocompatibility Laboratory | Emilio Margolles-Clark, PhD and Phillip Ruiz, MD, PhD |
| EPI_ISL_513294 | Baylor College of Medicine | Baylor College of Medicine: HGSC | Vasanthi Avadhanula, Erin Nicholson, David Henke, Pedro Piedra, Harsha Doddapaneni, Donna Muzny, Qingchang Meng, Hsu Chao, Zeineen Momin, Hua Shen, George Weissenberger, Kavya Kottapalli, Yimiti Meiheerguli, Sejal Salvi, Ginger Metcalf, Vipin Menon, Sara J.J. Cregeen, Matthew C. Ross, Tulin Ayvaz, Richard Sucgang, Kristi L. Hoffman, Matthew Wong, Joseph F. Petrosino |
| EPI_ISL_513296, EPI_ISL_513309 | Center for Precision Medicine, Meizhou People's Hospital (Huangtang Hospital) | Center for Precision Medicine, Meizhou People's Hospital (Huangtang Hospital) | Guo,X., Wu,H., Yu,Z. and Huang,Q. |
| EPI_ISL_513310 | Public Health, United States Air Force School of Aerospace Medicine | Public Health, United States Air Force School of Aerospace Medicine | Fries,A.C., Purves,S.M., Meyer,J.R., Javorina,A.K., Connors,B.C., Macias,E.A., Lambert,A.W., Chapleau,R.R. and Starr,C.R. |
| EPI_ISL_513311, EPI_ISL_513312, EPI_ISL_513313 | Public Health, United States Air Force School of Aerospace Medicine | Public Health, United States Air Force School of Aerospace Medicine | Fries,A.C., Purves,S.M., Meyer,J.R., Javorina,A.K., Connors,B.C., Macias,E.A., Lambert,A.W., Chapleau,R.R. and Starr,C.R. |
| EPI_ISL_513314 | The National Institute of Public Health | The National Institute of Public Health and State Veterinary Institute Prague | Nagy,A;Jirincova,H;Novakova,L;Trnka,D;Vecerova,J |
| EPI_ISL_513315, EPI_ISL_513316, EPI_ISL_513317 | South Eastern Area Laboratory Services (SEALS) | NSW Health Pathology - Institute of Clinical Pathology and Medical Research; Westmead Hospital; University of Sydney | CIDM-PH et al. |
| EPI_ISL_513318, EPI_ISL_513319, EPI_ISL_513320 | Histopath | NSW Health Pathology - Institute of Clinical Pathology and Medical Research; Westmead Hospital; University of Sydney | CIDM-PH et al. |
| EPI_ISL_513321, EPI_ISL_513322 | Pathology West - NSW Health Pathology | NSW Health Pathology - Institute of Clinical Pathology and Medical Research; Westmead Hospital; University of Sydney | CIDM-PH et al. |
| EPI_ISL_513323 | Sydney South West Pathology Service (SSWPS) - Liverpool Hospital - NSW Health Pathology | NSW Health Pathology - Institute of Clinical Pathology and Medical Research; Westmead Hospital; University of Sydney | CIDM-PH et al. |
| EPI_ISL_513324, EPI_ISL_513325, EPI_ISL_513326 | Pathology West - NSW Health Pathology | NSW Health Pathology - Institute of Clinical Pathology and Medical Research; Westmead Hospital; University of Sydney | CIDM-PH et al. |
| EPI_ISL_513327, EPI_ISL_513328, EPI_ISL_513329 | Sydney South West Pathology Service (SSWPS) - Liverpool Hospital - NSW Health Pathology | NSW Health Pathology - Institute of Clinical Pathology and Medical Research; Westmead Hospital; University of Sydney | CIDM-PH et al. |
| EPI_ISL_513330 | St Vincent's Pathology (SydPath) | NSW Health Pathology - Institute of Clinical Pathology and Medical Research; Westmead Hospital; University of Sydney | CIDM-PH et al. |
| EPI_ISL_513331 | Australian Clinical Labs | NSW Health Pathology - Institute of Clinical Pathology and Medical Research; Westmead Hospital; University of Sydney | CIDM-PH et al. |
| EPI_ISL_513332 | Pathology West - NSW Health Pathology | NSW Health Pathology - Institute of Clinical Pathology and Medical Research; Westmead Hospital; University of Sydney | CIDM-PH et al. |
| EPI_ISL_513333, EPI_ISL_513334 | Sydney South West Pathology Service (SSWPS) - Liverpool Hospital - NSW Health Pathology | NSW Health Pathology - Institute of Clinical Pathology and Medical Research; Westmead Hospital; University of Sydney | CIDM-PH et al. |
| EPI_ISL_513335 | Pathology North - Hunter - NSW Health Pathology | NSW Health Pathology - Institute of Clinical Pathology and Medical Research; Westmead Hospital; University of Sydney | CIDM-PH et al. |
| EPI_ISL_513336, EPI_ISL_513337 | Sydney South West Pathology Service (SSWPS) - Liverpool Hospital - NSW Health Pathology | NSW Health Pathology - Institute of Clinical Pathology and Medical Research; Westmead Hospital; University of Sydney | CIDM-PH et al. |
| EPI_ISL_513338, EPI_ISL_513339 | Pathology West - NSW Health Pathology | NSW Health Pathology - Institute of Clinical Pathology and Medical Research; Westmead Hospital; University of Sydney | CIDM-PH et al. |
| EPI_ISL_513340 | Austech Medical Laboratories | NSW Health Pathology - Institute of Clinical Pathology and Medical Research; Westmead Hospital; University of Sydney | CIDM-PH et al. |
| EPI_ISL_513341 | Medlab Pathology | NSW Health Pathology - Institute of Clinical Pathology and Medical Research; Westmead Hospital; University of Sydney | CIDM-PH et al. |
| EPI_ISL_513342 | Pathology West - NSW Health Pathology | NSW Health Pathology - Institute of Clinical Pathology and Medical Research; Westmead Hospital; University of Sydney | CIDM-PH et al. |
| EPI_ISL_513343, EPI_ISL_513344 | Children Westmead Hospital | NSW Health Pathology - Institute of Clinical Pathology and Medical Research; Westmead Hospital; University of Sydney | CIDM-PH et al. |
| EPI_ISL_513345, EPI_ISL_513346 | Pathology West - NSW Health Pathology | NSW Health Pathology - Institute of Clinical Pathology and Medical Research; Westmead Hospital; University of Sydney | CIDM-PH et al. |
| EPI_ISL_513347 | St Vincent's Pathology (SydPath) | NSW Health Pathology - Institute of Clinical Pathology and Medical Research; Westmead Hospital; University of Sydney | CIDM-PH et al. |
| EPI_ISL_513348, EPI_ISL_513349 | 4Cyte Pathology | NSW Health Pathology - Institute of Clinical Pathology and Medical Research; Westmead Hospital; University of Sydney | CIDM-PH et al. |
| EPI_ISL_513350, EPI_ISL_513351 | Pathology West - NSW Health Pathology | NSW Health Pathology - Institute of Clinical Pathology and Medical Research; Westmead Hospital; University of Sydney | CIDM-PH et al. |
| EPI_ISL_513352 | South Eastern Area Laboratory Services (SEALS) | NSW Health Pathology - Institute of Clinical Pathology and Medical Research; Westmead Hospital; University of Sydney | CIDM-PH et al. |
| EPI_ISL_513353, EPI_ISL_513354, EPI_ISL_513355, EPI_ISL_513356 | St Vincent's Pathology (SydPath) | NSW Health Pathology - Institute of Clinical Pathology and Medical Research; Westmead Hospital; University of Sydney | CIDM-PH et al. |
| EPI_ISL_513357 | Douglas Hanly Moir | NSW Health Pathology - Institute of Clinical Pathology and Medical Research; Westmead Hospital; University of Sydney | CIDM-PH et al. |
| EPI_ISL_513358, EPI_ISL_513359 | Pathology West - NSW Health Pathology | NSW Health Pathology - Institute of Clinical Pathology and Medical Research; Westmead Hospital; University of Sydney | CIDM-PH et al. |
| EPI_ISL_513360 | Pathology North - Hunter - NSW Health Pathology | NSW Health Pathology - Institute of Clinical Pathology and Medical Research; Westmead Hospital; University of Sydney | CIDM-PH et al. |
| EPI_ISL_513361, EPI_ISL_513362, EPI_ISL_513363, EPI_ISL_513364, EPI_ISL_513365 | St Vincent's Pathology (SydPath) | NSW Health Pathology - Institute of Clinical Pathology and Medical Research; Westmead Hospital; University of Sydney | CIDM-PH et al. |
| EPI_ISL_513366, EPI_ISL_513367 | Sydney South West Pathology Service (SSWPS) - Liverpool Hospital - NSW Health Pathology | NSW Health Pathology - Institute of Clinical Pathology and Medical Research; Westmead Hospital; University of Sydney | CIDM-PH et al. |
| EPI_ISL_513368 | Pathology North - Hunter - NSW Health Pathology | NSW Health Pathology - Institute of Clinical Pathology and Medical Research; Westmead Hospital; University of Sydney | CIDM-PH et al. |
| EPI_ISL_513369, EPI_ISL_513370, EPI_ISL_513371 | Pathology West - NSW Health Pathology | NSW Health Pathology - Institute of Clinical Pathology and Medical Research; Westmead Hospital; University of Sydney | CIDM-PH et al. |
| EPI_ISL_513372 | St Vincent's Pathology (SydPath) | NSW Health Pathology - Institute of Clinical Pathology and Medical Research; Westmead Hospital; University of Sydney | CIDM-PH et al. |
| EPI_ISL_513373, EPI_ISL_513374 | Sydney South West Pathology Service (SSWPS) - Liverpool Hospital - NSW Health Pathology | NSW Health Pathology - Institute of Clinical Pathology and Medical Research; Westmead Hospital; University of Sydney | CIDM-PH et al. |
| EPI_ISL_513375, EPI_ISL_513376 | Pathology North - Royal North Shore Hospital - NSW Health Pathology | NSW Health Pathology - Institute of Clinical Pathology and Medical Research; Westmead Hospital; University of Sydney | CIDM-PH et al. |
| EPI_ISL_513377 | South Eastern Area Laboratory Services (SEALS) | NSW Health Pathology - Institute of Clinical Pathology and Medical Research; Westmead Hospital; University of Sydney | CIDM-PH et al. |
| EPI_ISL_513378 | Pathology West - NSW Health Pathology | NSW Health Pathology - Institute of Clinical Pathology and Medical Research; Westmead Hospital; University of Sydney | CIDM-PH et al. |
| EPI_ISL_513379 | St Vincent's Pathology (SydPath) | NSW Health Pathology - Institute of Clinical Pathology and Medical Research; Westmead Hospital; University of Sydney | CIDM-PH et al. |
| EPI_ISL_513380 | Pathology West - NSW Health Pathology | NSW Health Pathology - Institute of Clinical Pathology and Medical Research; Westmead Hospital; University of Sydney | CIDM-PH et al. |
| EPI_ISL_513381, EPI_ISL_513382, EPI_ISL_513383 | Sydney South West Pathology Service (SSWPS) - Liverpool Hospital - NSW Health Pathology | NSW Health Pathology - Institute of Clinical Pathology and Medical Research; Westmead Hospital; University of Sydney | CIDM-PH et al. |
| EPI_ISL_513384, EPI_ISL_513385 | St Vincent's Pathology (SydPath) | NSW Health Pathology - Institute of Clinical Pathology and Medical Research; Westmead Hospital; University of Sydney | CIDM-PH et al. |
| EPI_ISL_513386, EPI_ISL_513387, EPI_ISL_513388 | Pathology North - NSW Health Pathology | NSW Health Pathology - Institute of Clinical Pathology and Medical Research; Westmead Hospital; University of Sydney | CIDM-PH et al. |
| EPI_ISL_513389 | Douglas Hanly Moir | NSW Health Pathology - Institute of Clinical Pathology and Medical Research; Westmead Hospital; University of Sydney | CIDM-PH et al. |
| EPI_ISL_513390, EPI_ISL_513391, EPI_ISL_513392, EPI_ISL_513393, EPI_ISL_513394, EPI_ISL_513395 | Pathology West - NSW Health Pathology | NSW Health Pathology - Institute of Clinical Pathology and Medical Research; Westmead Hospital; University of Sydney | CIDM-PH et al. |
| EPI_ISL_513396, EPI_ISL_513397, EPI_ISL_513398, EPI_ISL_513399 | St Vincent's Pathology (SydPath) | NSW Health Pathology - Institute of Clinical Pathology and Medical Research; Westmead Hospital; University of Sydney | CIDM-PH et al. |
| EPI_ISL_513400 | Sydney South West Pathology Service (SSWPS) - Royal Prince Alfred Hospital - NSW Health Pathology | NSW Health Pathology - Institute of Clinical Pathology and Medical Research; Westmead Hospital; University of Sydney | CIDM-PH et al. |
| EPI_ISL_513401 | Sydney South West Pathology Service (SSWPS) - Liverpool Hospital - NSW Health Pathology | NSW Health Pathology - Institute of Clinical Pathology and Medical Research; Westmead Hospital; University of Sydney | CIDM-PH et al. |
| EPI_ISL_513402 | 4Cyte Pathology | NSW Health Pathology - Institute of Clinical Pathology and Medical Research; Westmead Hospital; University of Sydney | CIDM-PH et al. |
| EPI_ISL_513403, EPI_ISL_513404, EPI_ISL_513405, EPI_ISL_513406, EPI_ISL_513407, EPI_ISL_513408 | Laverty Pathology | NSW Health Pathology - Institute of Clinical Pathology and Medical Research; Westmead Hospital; University of Sydney | CIDM-PH et al. |

| | | | |
|---|---|---|---|
| EPI_ISL_513409, EPI_ISL_513410, EPI_ISL_513411, EPI_ISL_513412 | Sydney South West Pathology Service (SSWPS) - Liverpool Hospital - NSW Health Pathology | NSW Health Pathology - Institute of Clinical Pathology and Medical Research; Westmead Hospital; University of Sydney | CIDM-PH et al. |
| EPI_ISL_513413 | 4Cyte Pathology | NSW Health Pathology - Institute of Clinical Pathology and Medical Research; Westmead Hospital; University of Sydney | CIDM-PH et al. |
| EPI_ISL_513414, EPI_ISL_513415, EPI_ISL_513416, EPI_ISL_513417, EPI_ISL_513418, EPI_ISL_513419, EPI_ISL_513420, EPI_ISL_513421, EPI_ISL_513422, EPI_ISL_513423, EPI_ISL_513424, EPI_ISL_513425, EPI_ISL_513426, EPI_ISL_513427, EPI_ISL_513428, EPI_ISL_513429, EPI_ISL_513430, EPI_ISL_513431, EPI_ISL_513432, EPI_ISL_513433, EPI_ISL_513434, EPI_ISL_513435, EPI_ISL_513436, EPI_ISL_513437, EPI_ISL_513438, EPI_ISL_513439, EPI_ISL_513440, EPI_ISL_513441, EPI_ISL_513442, EPI_ISL_513443, EPI_ISL_513444, EPI_ISL_513445, EPI_ISL_513446, EPI_ISL_513447, EPI_ISL_513448, EPI_ISL_513449, EPI_ISL_513450, EPI_ISL_513451, EPI_ISL_513452, EPI_ISL_513453, EPI_ISL_513454, EPI_ISL_513455, EPI_ISL_513456, EPI_ISL_513457, EPI_ISL_513458, EPI_ISL_513459, EPI_ISL_513460, EPI_ISL_513461, EPI_ISL_513462, EPI_ISL_513463, EPI_ISL_513464, EPI_ISL_513465, EPI_ISL_513466, EPI_ISL_513467, EPI_ISL_513468, EPI_ISL_513469, EPI_ISL_513470, EPI_ISL_513471, EPI_ISL_513472, EPI_ISL_513473, EPI_ISL_513474, EPI_ISL_513475, EPI_ISL_513476, EPI_ISL_513477, EPI_ISL_513478, EPI_ISL_513479, EPI_ISL_513480, EPI_ISL_513481, EPI_ISL_513482, EPI_ISL_513483, EPI_ISL_513484, EPI_ISL_513485, EPI_ISL_513486, EPI_ISL_513487, EPI_ISL_513488, EPI_ISL_513489, EPI_ISL_513490, EPI_ISL_513491, EPI_ISL_513492, EPI_ISL_513493, EPI_ISL_513494, EPI_ISL_513495, EPI_ISL_513496, EPI_ISL_513497, EPI_ISL_513498, EPI_ISL_513499, EPI_ISL_513500, EPI_ISL_513501, EPI_ISL_513502, EPI_ISL_513503, EPI_ISL_513504, EPI_ISL_513505, EPI_ISL_513506, EPI_ISL_513507, EPI_ISL_513508, EPI_ISL_513509, EPI_ISL_513510 | | | |
| see above | Maine HETL | Tewhey Lab, The Jackson Laboratory | Matluk,N., Dewey,H., Barter,M., Lynch,R., Munger,H. and Tewhey,R. |
| EPI_ISL_513511, EPI_ISL_513512 | The National Institute of Public Health | The National Institute of Public Health and State Veterinary Institute Prague | Nagy,A;Jirincova,H;Novakova,L;Trnka,D;Vecerova,J |
| EPI_ISL_513513, EPI_ISL_513514, EPI_ISL_513515, EPI_ISL_513516, EPI_ISL_513517, EPI_ISL_513518, EPI_ISL_513519, EPI_ISL_513520, EPI_ISL_513521, EPI_ISL_513522, EPI_ISL_513523, EPI_ISL_513524, EPI_ISL_513525, EPI_ISL_513526, EPI_ISL_513527, EPI_ISL_513528, EPI_ISL_513529, EPI_ISL_513530, EPI_ISL_513531, EPI_ISL_513532, EPI_ISL_513533, EPI_ISL_513534, EPI_ISL_513535, EPI_ISL_513536, EPI_ISL_513537, EPI_ISL_513538, EPI_ISL_513539, EPI_ISL_513540, EPI_ISL_513541, EPI_ISL_513542, EPI_ISL_513543, EPI_ISL_513544, EPI_ISL_513545, EPI_ISL_513546, EPI_ISL_513547, EPI_ISL_513548, EPI_ISL_513549, EPI_ISL_513550, EPI_ISL_513551, EPI_ISL_513552, EPI_ISL_513553, EPI_ISL_513554, EPI_ISL_513555, EPI_ISL_513556, EPI_ISL_513557, EPI_ISL_513558, EPI_ISL_513559, EPI_ISL_513560, EPI_ISL_513561, EPI_ISL_513562, EPI_ISL_513563, EPI_ISL_513564, EPI_ISL_513565, EPI_ISL_513566, EPI_ISL_513567, EPI_ISL_513568, EPI_ISL_513569, EPI_ISL_513570, EPI_ISL_513571, EPI_ISL_513572, EPI_ISL_513573, EPI_ISL_513574, EPI_ISL_513575, EPI_ISL_513576, EPI_ISL_513577, EPI_ISL_513578, EPI_ISL_513579, EPI_ISL_513580, EPI_ISL_513581, EPI_ISL_513582, EPI_ISL_513583 | | | |
| see above | Programa de Oncovirologia, Instituto Nacional de Câncer | Programa de Oncovirologia, Instituto Nacional de Câncer | Juliana D. Siqueira, Livia R. Goes, Brunna M. Alves, Claudia Cicala,James Arthos, João P.B. Viola, Andreia C. de Melo, Marcelo A. Soares |
| EPI_ISL_513584 | PHE South West Regional Laboratory, National Infection Service | Wellcome Sanger Institute for the COVID-19 Genomics UK Consortium | Stephanie Hutchings, Hannah Pymont, Dr Peter Muir, Barry Vipond, Rich Hopes; and Alex Alderton, Roberto Amato, Sonia Goncalves, Ewan Harrison, David K. Jackson, Ian Johnston, Dominic Kwiatkowski, Cordelia Langford, John Sillitoe on behalf of the Wellcome Sanger Institute COVID-19 Surveillance Team (http://www.sanger.ac.uk/covid-team) |
| EPI_ISL_513585, EPI_ISL_513586, EPI_ISL_513587, EPI_ISL_513588 | Microbiology, Pathology, Kettering General Hospital | Wellcome Sanger Institute for the COVID-19 Genomics UK Consortium | Sahar El-dirdiri, Anita Kenyon, Thomas Davis and Alex Alderton, Roberto Amato, Sonia Goncalves, Ewan Harrison, David K. Jackson, Ian Johnston, Dominic Kwiatkowski, Cordelia Langford, John Sillitoe on behalf of the Wellcome Sanger Institute COVID-19 Surveillance Team (http://www.sanger.ac.uk/covid-team) |
| EPI_ISL_513589, EPI_ISL_513590, EPI_ISL_513591, EPI_ISL_513592, EPI_ISL_513593, EPI_ISL_513594, EPI_ISL_513595, EPI_ISL_513596, EPI_ISL_513597, EPI_ISL_513598, EPI_ISL_513599, EPI_ISL_513600, EPI_ISL_513601, EPI_ISL_513602, EPI_ISL_513603, EPI_ISL_513604, EPI_ISL_513605, EPI_ISL_513606, EPI_ISL_513607, EPI_ISL_513608, EPI_ISL_513609, EPI_ISL_513610, EPI_ISL_513611, EPI_ISL_513612, EPI_ISL_513613, EPI_ISL_513614, EPI_ISL_513615, EPI_ISL_513616, EPI_ISL_513617, EPI_ISL_513618, EPI_ISL_513619, EPI_ISL_513620, EPI_ISL_513621, EPI_ISL_513622, EPI_ISL_513623, EPI_ISL_513624, EPI_ISL_513625, EPI_ISL_513626, EPI_ISL_513627, EPI_ISL_513628, EPI_ISL_513629, EPI_ISL_513630, EPI_ISL_513631 | | | |
| see above | Viral Respiratory Lab, National Institute for Biomedical Research (INRB) | Pathogen Sequencing Lab, National Institute for Biomedical Research (INRB) | Placide Mbala-Kingebeni, Edith Nkwembe, Eddy Kinganda-Lusamaki, Amuri Aziza, Francisca Muyembe Mawete, Emmanuel Lokilo Lofiko, Catherine Pratt, Matthias Pauthner, Josh Quick, Allison Black, James Hadfield, Trevor Bedford, Ian Goodfellow, Andrew Rambaut, Nick Loman, Kristian Andersen, Michael Wiley, Steve Ahuka-Mundeke, Jean-Jacques Muyembe Tamfum |
| EPI_ISL_513632, EPI_ISL_513633, EPI_ISL_513634, EPI_ISL_513635, EPI_ISL_513636, EPI_ISL_513637, EPI_ISL_513638 | University of Washington Virology Lab | University of Washington Virology Lab | Pavitra Roychoudhury, Hong Xie, Lasata Shrestha, Amin Addetia, Truong Nguyen, Victoria M Rachleff, Meei-Li Huang, Keith R Jerome, Alexander Greninger |
| EPI_ISL_513640, EPI_ISL_513641, EPI_ISL_513642, EPI_ISL_513643, EPI_ISL_513644, EPI_ISL_513645, EPI_ISL_513646, EPI_ISL_513647, EPI_ISL_513648, EPI_ISL_513649, EPI_ISL_513650, EPI_ISL_513651, EPI_ISL_513652, EPI_ISL_513653, EPI_ISL_513654, EPI_ISL_513655, EPI_ISL_513656, EPI_ISL_513657, EPI_ISL_513658, EPI_ISL_513659, EPI_ISL_513660, EPI_ISL_513661, EPI_ISL_513662, EPI_ISL_513663, EPI_ISL_513664, EPI_ISL_513665, EPI_ISL_513666, EPI_ISL_513667, EPI_ISL_513668, EPI_ISL_513669, EPI_ISL_513670, EPI_ISL_513671, EPI_ISL_513672, EPI_ISL_513673, EPI_ISL_513674, EPI_ISL_513675, EPI_ISL_513676, EPI_ISL_513677, EPI_ISL_513678, EPI_ISL_513679, EPI_ISL_513680, EPI_ISL_513681, EPI_ISL_513682, EPI_ISL_513683, EPI_ISL_513684, EPI_ISL_513685, EPI_ISL_513686, EPI_ISL_513687, EPI_ISL_513688, EPI_ISL_513689, EPI_ISL_513690, EPI_ISL_513691, EPI_ISL_513692, EPI_ISL_513693, EPI_ISL_513694, EPI_ISL_513695, EPI_ISL_513696, EPI_ISL_513697, EPI_ISL_513698, EPI_ISL_513699, EPI_ISL_513700, EPI_ISL_513701, EPI_ISL_513702, EPI_ISL_513703, EPI_ISL_513704, EPI_ISL_513705, EPI_ISL_513706 | | | |
| see above | Utah Public Health Laboratory | Utah Public Health Laboratory | Heidi Butz, Erin Young, Kelly Oakeson |
| EPI_ISL_513707, EPI_ISL_513708, EPI_ISL_513709, EPI_ISL_513710, EPI_ISL_513711, EPI_ISL_513712, EPI_ISL_513713, EPI_ISL_513714, EPI_ISL_513715, EPI_ISL_513716, EPI_ISL_513717, EPI_ISL_513718, EPI_ISL_513719, EPI_ISL_513720, EPI_ISL_513721, EPI_ISL_513722, EPI_ISL_513723, EPI_ISL_513724, EPI_ISL_513725, EPI_ISL_513726, EPI_ISL_513727, EPI_ISL_513728, EPI_ISL_513729, EPI_ISL_513730, EPI_ISL_513731, EPI_ISL_513732, EPI_ISL_513733, EPI_ISL_513734, EPI_ISL_513735, EPI_ISL_513736, EPI_ISL_513737, EPI_ISL_513738, EPI_ISL_513739, EPI_ISL_513740, EPI_ISL_513741, EPI_ISL_513742, EPI_ISL_513743, EPI_ISL_513744, EPI_ISL_513745, EPI_ISL_513746, EPI_ISL_513747, EPI_ISL_513748, EPI_ISL_513749, EPI_ISL_513750, EPI_ISL_513751, EPI_ISL_513752, EPI_ISL_513753, EPI_ISL_513754, EPI_ISL_513755, EPI_ISL_513756, EPI_ISL_513757, EPI_ISL_513758, EPI_ISL_513759, EPI_ISL_513760, EPI_ISL_513761, EPI_ISL_513762, EPI_ISL_513763, EPI_ISL_513764, EPI_ISL_513765, EPI_ISL_513766, EPI_ISL_513767, EPI_ISL_513768, EPI_ISL_513769, EPI_ISL_513770, EPI_ISL_513771, EPI_ISL_513772 | | | |
| see above | Orange County Public Health Lab | Chan-Zuckerberg Biohub | CZB Cliahub Consortium |
| EPI_ISL_513773, EPI_ISL_513774, EPI_ISL_513775, EPI_ISL_513776, EPI_ISL_513777, EPI_ISL_513778, EPI_ISL_513779, EPI_ISL_513780, EPI_ISL_513781, EPI_ISL_513782, EPI_ISL_513783, EPI_ISL_513784, EPI_ISL_513785, EPI_ISL_513786, EPI_ISL_513787, EPI_ISL_513788 | | | |
| see above | County of Santa Clara Public Health Department | Chan-Zuckerberg Biohub | CZB Cliahub Consortium |
| EPI_ISL_513789, EPI_ISL_513790, EPI_ISL_513791, EPI_ISL_513792, EPI_ISL_513793, EPI_ISL_513794, EPI_ISL_513795, EPI_ISL_513796, EPI_ISL_513797, EPI_ISL_513798, EPI_ISL_513799, EPI_ISL_513800, EPI_ISL_513801, EPI_ISL_513802, EPI_ISL_513803, EPI_ISL_513804, EPI_ISL_513805, EPI_ISL_513806, EPI_ISL_513807, EPI_ISL_513808, EPI_ISL_513809, EPI_ISL_513810, EPI_ISL_513811, EPI_ISL_513812, EPI_ISL_513813, EPI_ISL_513814, EPI_ISL_513815, EPI_ISL_513816, EPI_ISL_513817, EPI_ISL_513818, EPI_ISL_513819, EPI_ISL_513820, EPI_ISL_513821, EPI_ISL_513822, EPI_ISL_513823, EPI_ISL_513824, EPI_ISL_513825, EPI_ISL_513826, EPI_ISL_513827, EPI_ISL_513828, EPI_ISL_513829, EPI_ISL_513830, EPI_ISL_513831, EPI_ISL_513832, EPI_ISL_513833, EPI_ISL_513834, EPI_ISL_513835, EPI_ISL_513836, EPI_ISL_513837, EPI_ISL_513838, EPI_ISL_513839 | | | |
| see above | Orange County Public Health Laboratory | Chan-Zuckerberg Biohub | CZB Cliahub Consortium |
| EPI_ISL_513840, EPI_ISL_513841, EPI_ISL_513842, EPI_ISL_513843, EPI_ISL_513844, EPI_ISL_513845, EPI_ISL_513846, EPI_ISL_513847, EPI_ISL_513848, EPI_ISL_513849, EPI_ISL_513850, EPI_ISL_513851, EPI_ISL_513852, EPI_ISL_513853, EPI_ISL_513854, EPI_ISL_513855 | | | |
| see above | Humboldt County Public Health Laboratory | Chan-Zuckerberg Biohub | CZB Cliahub Consortium |
| EPI_ISL_513856, EPI_ISL_513857, EPI_ISL_513858, EPI_ISL_513859, EPI_ISL_513860, EPI_ISL_513861, EPI_ISL_513862, EPI_ISL_513863, EPI_ISL_513864, EPI_ISL_513865, EPI_ISL_513866, EPI_ISL_513867, EPI_ISL_513868, EPI_ISL_513869, EPI_ISL_513870, EPI_ISL_513871, EPI_ISL_513872, EPI_ISL_513873, EPI_ISL_513874, EPI_ISL_513875, EPI_ISL_513876, EPI_ISL_513877, EPI_ISL_513878, EPI_ISL_513879, EPI_ISL_513880, EPI_ISL_513881, EPI_ISL_513882 | | | |
| see above | San Francisco Public Health Laboratory | Chan-Zuckerberg Biohub | CZB Cliahub Consortium |
| EPI_ISL_513883, EPI_ISL_513884, EPI_ISL_513885, EPI_ISL_513886, EPI_ISL_513887, EPI_ISL_513888, EPI_ISL_513889, EPI_ISL_513890, EPI_ISL_513891, EPI_ISL_513892, EPI_ISL_513893, EPI_ISL_513894, EPI_ISL_513895, EPI_ISL_513896, EPI_ISL_513897, EPI_ISL_513898, EPI_ISL_513899, EPI_ISL_513900, EPI_ISL_513901, EPI_ISL_513902, EPI_ISL_513903, EPI_ISL_513904, EPI_ISL_513905, EPI_ISL_513906, EPI_ISL_513907, EPI_ISL_513908, EPI_ISL_513909, EPI_ISL_513910 | | | |
| see above | UCSF Clinical Microbiology Laboratory | Chan-Zuckerberg Biohub | CZB Cliahub Consortium |
| EPI_ISL_513911, EPI_ISL_513912, EPI_ISL_513913 | SA Pathology | SA Pathology | Lex Leong, Chuan Kok Lim, Mark Turra, Ivan Bastian, Geoff Higgins |
| EPI_ISL_513925 | Microbiology & Bioinformatics and Biostatistics, Kohat University of Science and Technology (Pakistan) & Shanghai Jiao Tong University (China) | Microbiology & Bioinformatics and Biostatistics, Kohat University of Science and Technology (Pakistan) & Shanghai Jiao Tong University (China) | Khan,M.T., Khan,T.A., Ali,S., Khan,A.S., Muhammad,N. and Wei,D.Q. |
| EPI_ISL_513941, EPI_ISL_513942 | Utah Public Health Laboratory, Utah Public Health Laboratory Infectious Disease submission group | Utah Public Health Laboratory, Utah Public Health Laboratory Infectious Disease submission group | Butz,H.A., Young,E.L. and Oakeson,K. |
| EPI_ISL_514062, EPI_ISL_514063, EPI_ISL_514064 | Navy and Marine Corps Public Health Center | Pathogen Discovery, Respiratory Viruses Branch, Division of Viral Diseases, Centers for Disease Control and Prevention | Yan Li, Anna Montmayeur, Krista Queen, Jing Zhang, Ying Tao, Anna Uehara, Rachel Marine, Clinton R. Paden, Haibin Wang, Suxiang Tong |
| EPI_ISL_514065 | Navy and Marine Corps Public Health Center | Pathogen Discovery, Respiratory Viruses Branch, Division of Viral Diseases, Centers for Disease Control and Prevention | Krista Queen, Yan Li, Anna Montmayeur, Jing Zhang, Ying Tao, Anna Uehara, Rachel Marine, Clinton R. Paden, Haibin Wang, Suxiang Tong |
| EPI_ISL_514066 | Navy and Marine Corps Public Health Center | Pathogen Discovery, Respiratory Viruses Branch, Division of Viral Diseases, Centers for Disease Control and Prevention | Yan Li, Anna Montmayeur, Krista Queen, Jing Zhang, Ying Tao, Anna Uehara, Rachel Marine, Clinton R. Paden, Haibin Wang, Suxiang Tong |
| EPI_ISL_514067, EPI_ISL_514068 | Navy and Marine Corps Public Health Center | Pathogen Discovery, Respiratory Viruses Branch, Division of Viral Diseases, Centers for Disease Control and Prevention | Krista Queen, Yan Li, Anna Montmayeur, Jing Zhang, Ying Tao, Anna Uehara, Rachel Marine, Clinton R. Paden, Haibin Wang, Suxiang Tong |
| EPI_ISL_514069, EPI_ISL_514070 | Navy and Marine Corps Public Health Center | Pathogen Discovery, Respiratory Viruses Branch, Division of Viral Diseases, Centers for Disease Control and Prevention | Yan Li, Anna Montmayeur, Krista Queen, Jing Zhang, Ying Tao, Anna Uehara, Rachel Marine, Clinton R. Paden, Haibin Wang, Suxiang Tong |
| EPI_ISL_514071, EPI_ISL_514072 | Navy and Marine Corps Public Health Center | Pathogen Discovery, Respiratory Viruses Branch, Division of Viral Diseases, Centers for Disease Control and Prevention | Krista Queen, Yan Li, Anna Montmayeur, Jing Zhang, Ying Tao, Anna Uehara, Rachel Marine, Clinton R. Paden, Haibin Wang, Suxiang Tong |
| EPI_ISL_514073, EPI_ISL_514074 | Navy and Marine Corps Public Health Center | Pathogen Discovery, Respiratory Viruses Branch, Division of Viral Diseases, Centers for Disease Control and Prevention | Yan Li, Anna Montmayeur, Krista Queen, Jing Zhang, Ying Tao, Anna Uehara, Rachel Marine, Clinton R. Paden, Haibin Wang, Suxiang Tong |
| EPI_ISL_514075, EPI_ISL_514076 | Navy and Marine Corps Public Health Center | Pathogen Discovery, Respiratory Viruses Branch, Division of Viral Diseases, Centers for Disease Control and Prevention | Krista Queen, Yan Li, Anna Montmayeur, Jing Zhang, Ying Tao, Anna Uehara, Rachel Marine, Clinton R. Paden, Haibin Wang, Suxiang Tong |
| EPI_ISL_514077, EPI_ISL_514078, EPI_ISL_514079 | Navy and Marine Corps Public Health Center | Pathogen Discovery, Respiratory Viruses Branch, Division of Viral Diseases, Centers for Disease Control and Prevention | Yan Li, Anna Montmayeur, Krista Queen, Jing Zhang, Ying Tao, Anna Uehara, Rachel Marine, Clinton R. Paden, Haibin Wang, Suxiang Tong |
| EPI_ISL_514080, EPI_ISL_514081, EPI_ISL_514082, EPI_ISL_514083 | Navy and Marine Corps Public Health Center | Pathogen Discovery, Respiratory Viruses Branch, Division of Viral Diseases, Centers for Disease Control and Prevention | Krista Queen, Yan Li, Anna Montmayeur, Jing Zhang, Ying Tao, Anna Uehara, Rachel Marine, Clinton R. Paden, Haibin Wang, Suxiang Tong |
| EPI_ISL_514084 | Navy and Marine Corps Public Health Center | Pathogen Discovery, Respiratory Viruses Branch, Division of Viral Diseases, Centers for Disease Control and Prevention | Yan Li, Anna Montmayeur, Krista Queen, Jing Zhang, Ying Tao, Anna Uehara, Rachel Marine, Clinton R. Paden, Haibin Wang, Suxiang Tong |
| EPI_ISL_514085, EPI_ISL_514086, EPI_ISL_514087 | Navy and Marine Corps Public Health Center | Pathogen Discovery, Respiratory Viruses Branch, Division of Viral Diseases, Centers for Disease Control and Prevention | Krista Queen, Yan Li, Anna Montmayeur, Jing Zhang, Ying Tao, Anna Uehara, Rachel Marine, Clinton R. Paden, Haibin Wang, Suxiang Tong |
| EPI_ISL_514088 | Navy and Marine Corps Public Health Center | Pathogen Discovery, Respiratory Viruses Branch, Division of Viral Diseases, Centers for Disease Control and Prevention | Yan Li, Anna Montmayeur, Krista Queen, Jing Zhang, Ying Tao, Anna Uehara, Rachel Marine, Clinton R. Paden, Haibin Wang, Suxiang Tong |
| EPI_ISL_514089 | Navy and Marine Corps Public Health Center | Pathogen Discovery, Respiratory Viruses Branch, Division of Viral Diseases, Centers for Disease Control and Prevention | Krista Queen, Yan Li, Anna Montmayeur, Jing Zhang, Ying Tao, Anna Uehara, Rachel Marine, Clinton R. Paden, Haibin Wang, Suxiang Tong |
| EPI_ISL_514090, EPI_ISL_514091, EPI_ISL_514092 | Navy and Marine Corps Public Health Center | Pathogen Discovery, Respiratory Viruses Branch, Division of Viral Diseases, Centers for Disease Control and Prevention | Yan Li, Anna Montmayeur, Krista Queen, Jing Zhang, Ying Tao, Anna Uehara, Rachel Marine, Clinton R. Paden, Haibin Wang, Suxiang Tong |
| EPI_ISL_514093 | National Scientific Center for Especially Dangerous Infections (NSCEDI) | Kazakh National Agrarian University (KazNAU) TreeGene LLP Genetic Laboratory | Beloussov Vyacheslav, Strochkov Vitaliy, Tabynov Kaissar, Sandybayev Nurlan, Tabynov Kairat, Turebekov Nurkeldy, Granica Joanna, Solomadin Maksim, Yerubayev Toktassyn, Yespolov Tlektes |
| EPI_ISL_514094, EPI_ISL_514095, EPI_ISL_514096, EPI_ISL_514097, EPI_ISL_514098, EPI_ISL_514099, EPI_ISL_514100, EPI_ISL_514101, EPI_ISL_514102 | Molecular Diagnostics, Central Public Health Laboratory | Molecular Diagnostics, Central Public Health Laboratory | Dler,H., Dlshad,H., Furat,S., Sharmeen,F.-A., Dalia,F., Mohsen,A., Hemdad,A., Fahmi,A., Hemn,M., Idrees,H. |
| EPI_ISL_514103, EPI_ISL_514104, EPI_ISL_514105, EPI_ISL_514106, EPI_ISL_514107, EPI_ISL_514108, EPI_ISL_514109, EPI_ISL_514110, EPI_ISL_514111, EPI_ISL_514112, EPI_ISL_514113, EPI_ISL_514114, EPI_ISL_514115, EPI_ISL_514116, EPI_ISL_514117, EPI_ISL_514118, EPI_ISL_514119, EPI_ISL_514120, EPI_ISL_514121, EPI_ISL_514122, EPI_ISL_514123, EPI_ISL_514124, EPI_ISL_514125 | | | |
| see above | Viral Respiratory Lab, National Institute for Biomedical Research (INRB) | Pathogen Sequencing Lab, National Institute for Biomedical Research (INRB) | Placide Mbala-Kingebeni, Edith Nkwembe, Eddy Kinganda-Lusamaki, Amuri Aziza, Francisca Muyembe Mawete, Emmanuel Lokilo Lofiko, Catherine Pratt, Matthias Pauthner, Josh Quick, Allison Black, James Hadfield, Trevor Bedford, Ian Goodfellow, Andrew Rambaut, Nick Loman, Kristian Andersen, Michael Wiley, Steve Ahuka-Mundeke, Jean-Jacques Muyembe Tamfum |
| EPI_ISL_514126 | National Scientific Center for Especially Dangerous Infections (NSCEDI) | Kazakh National Agrarian University (KazNAU) TreeGene LLP Genetic Laboratory | Tabynov Kaissar, Beloussov Vyacheslav, Strochkov Vitaliy, Sandybayev Nurlan, Tabynov Kairat, Turebekov Nurkeldy, Granica Joanna, Solomadin Maksim, Yerubayev Toktassyn, Yespolov Tlektes |

| | | | |
|---|---|---|---|
| EPI_ISL_514127 | National Scientific Center for Especially Dangerous Infections (NSCEDI) | Kazakh National Agrarian University (KazNAU) TreeGene LLP Genetic Laboratory | Tabynov Kairat, Beloussov Vyacheslav, Strochkov Vitaliy, Sandybayev Nurlan, Tabynov Kaissar, Turebekov Nurkeldy, Granica Joanna, Solomadin Maksim, Yerubayev Toktassyn, Yespolov Tlektes |
| EPI_ISL_514128 | Navy and Marine Corps Public Health Center | Centers for Disease Control and Prevention | Krista Queen, Yan Li, Anna Montmayeur, Jing Zhang, Ying Tao, Anna Uehara, Rachel Marine, Clinton R. Paden, Haibin Wang, Suxiang Tong |
| EPI_ISL_514129, EPI_ISL_514130 | National Institute of Laboratory Medicine and Referral Center | Genomic Research Lab, BCSIR | Md. Murshed Hasan Sarkar, Abu Sayeed Mohammad Mahmud, Mohammad Samir Uzzaman, Eshrar Osman, Md. Ahashan Habib, Shahina Akter, Tanjina Akhter Banu, Barna Goswami, Iffat Jahan, Md. Saddam Hossain, Tasnim Nafisa, Md. Maruf Ahmed Molla, Mahmuda Yeasmin, Asish Kumar Ghosh, A. K. M. Shamsuzzaman, Sheikh Md. Selim Al Din, Utpal Chandra Ray, Salek Ahmed Sajib, Md. Salim Khan |
| EPI_ISL_514131 | Rondônia Central Public Health Laboratory (LACEN/RO), vinculated to State Health Secretariat of Rondônia (SESAU/RO) | Molecular Virology Laboratory of Oswaldo Cruz Foundation of Rondônia | Luan Felipo Botelho-Souza, Felipe Souza Nogueira-Lima, Tárcio Peixoto Roca, Alcione de Oliveira dos Santos, Felipe Gomes Naveca, Adriana Cristina Salvador Maia, Cicileia Correia da Silva, Aline Linhares Ferreira de Melo Mendonça, Celina Aparecida Bertoni Lugtenburg, Camila Flávia Gomes Azzi, Juliana Loca Furtado, Suelen Cavalcante, Rita de Cássia Pontello Rampazzo, Caio Henrique Nemeth Santos, Alice Paula Di Sabatino Guimarães, Jansen Fernandes de Medeiros, Fernando Rodrigues Máximo, Juan Miguel Vilallobos-Salcedo and Deusilene Souza Vieira1 |
| EPI_ISL_514132 | Rondônia Central Public Health Laboratory (LACEN/RO), vinculated to State Health Secretariat of Rondônia (SESAU/RO) | Molecular Virology Laboratory of Oswaldo Cruz Foundation of Rondônia | Luan Felipo Botelho-Souza, Felipe Souza Nogueira-Lima, Tárcio Peixoto Roca, Alcione de Oliveira dos Santos, Felipe Gomes Naveca, Adriana Cristina Salvador Maia, Cicileia Correia da Silva, Aline Linhares Ferreira de Melo Mendonça, Celina Aparecida Bertoni Lugtenburg, Camila Flávia Gomes Azzi, Juliana Loca Furtado, Suelen Cavalcante, Rita de Cássia Pontello Rampazzo, Caio Henrique Nemeth Santos, Alice Paula Di Sabatino Guimarães, Jansen Fernandes de Medeiros, Fernando Rodrigues Máximo, Juan Miguel Vilallobos-Salcedo and Deusilene Souza Vieira. |
| EPI_ISL_514133, EPI_ISL_514134 | Rondônia Central Public Health Laboratory (LACEN/RO), vinculated to State Health Secretariat of Rondônia (SESAU/RO) | Molecular Virology Laboratory of Oswaldo Cruz Foundation of Rondônia | Luan Felipo Botelho-Souza, Felipe Souza Nogueira-Lima, Tárcio Peixoto Roca, Alcione de Oliveira dos Santos, Felipe Gomes Naveca, Adriana Cristina Salvador Maia, Cicileia Correia da Silva, Aline Linhares Ferreira de Melo Mendonça, Celina Aparecida Bertoni Lugtenburg, Camila Flávia Gomes Azzi, Juliana Loca Furtado, Suelen Cavalcante, Rita de Cássia Pontello Rampazzo, Caio Henrique Nemeth Santos, Alice Paula Di Sabatino Guimarães, Jansen Fernandes de Medeiros, Fernando Rodrigues Máximo, Juan Miguel Vilallobos-Salcedo and Deusilene Souza Vieira |
| EPI_ISL_514135 | Rondônia Central Public Health Laboratory (LACEN/RO), vinculated to State Health Secretariat of Rondônia (SESAU/RO) | Molecular Virology Laboratory of Oswaldo Cruz Foundation of Rondônia | Luan Felipo Botelho-Souza, Felipe Souza Nogueira-Lima, Tárcio Peixoto Roca, Alcione de Oliveira dos Santos, Felipe Gomes Naveca, Adriana Cristina Salvador Maia, Cicileia Correia da Silva, Aline Linhares Ferreira de Melo Mendonça, Celina Aparecida Bertoni Lugtenburg, Camila Flávia Gomes Azzi, Juliana Loca Furtado, Suelen Cavalcante, Rita de Cássia Pontello Rampazzo, Caio Henrique Nemeth Santos, Alice Paula Di Sabatino Guimarães, Jansen Fernandes de Medeiros, Fernando Rodrigues Máximo, Juan Miguel Vilallobos-Salcedo and Deusilene Souza Vieira |
| EPI_ISL_514136 | Rondônia Central Public Health Laboratory (LACEN/RO), vinculated to State Health Secretariat of Rondônia (SESAU/RO) | Molecular Virology Laboratory of Oswaldo Cruz Foundation of Rondônia | Luan Felipo Botelho-Souza, Felipe Souza Nogueira-Lima, Tárcio Peixoto Roca, Alcione de Oliveira dos Santos, Felipe Gomes Naveca, Adriana Cristina Salvador Maia, Cicileia Correia da Silva, Aline Linhares Ferreira de Melo Mendonça, Celina Aparecida Bertoni Lugtenburg, Camila Flávia Gomes Azzi, Juliana Loca Furtado, Suelen Cavalcante, Rita de Cássia Pontello Rampazzo, Caio Henrique Nemeth Santos, Alice Paula Di Sabatino Guimarães, Jansen Fernandes de Medeiros, Fernando Rodrigues Máximo, Juan Miguel Vilallobos-Salcedo and Deusilene Souza Vieira |
| EPI_ISL_514137 | Rondônia Central Public Health Laboratory (LACEN/RO), vinculated to State Health Secretariat of Rondônia (SESAU/RO) | Molecular Virology Laboratory of Oswaldo Cruz Foundation of Rondônia | Luan Felipo Botelho-Souza, Felipe Souza Nogueira-Lima, Tárcio Peixoto Roca, Alcione de Oliveira dos Santos, Felipe Gomes Naveca, Adriana Cristina Salvador Maia, Cicileia Correia da Silva, Aline Linhares Ferreira de Melo Mendonça, Celina Aparecida Bertoni Lugtenburg, Camila Flávia Gomes Azzi, Juliana Loca Furtado, Suelen Cavalcante, Rita de Cássia Pontello Rampazzo, Caio Henrique Nemeth Santos, Alice Paula Di Sabatino Guimarães, Jansen Fernandes de Medeiros, Fernando Rodrigues Máximo, Juan Miguel Vilallobos-Salcedo and Deusilene Souza Vieira |
| EPI_ISL_514138 | Rondônia Central Public Health Laboratory (LACEN/RO), vinculated to State Health Secretariat of Rondônia (SESAU/RO) | Molecular Virology Laboratory of Oswaldo Cruz Foundation of Rondônia | Luan Felipo Botelho-Souza, Felipe Souza Nogueira-Lima, Tárcio Peixoto Roca, Alcione de Oliveira dos Santos, Felipe Gomes Naveca, Adriana Cristina Salvador Maia, Cicileia Correia da Silva, Aline Linhares Ferreira de Melo Mendonça, Celina Aparecida Bertoni Lugtenburg, Camila Flávia Gomes Azzi, Juliana Loca Furtado, Suelen Cavalcante, Rita de Cássia Pontello Rampazzo, Caio Henrique Nemeth Santos, Alice Paula Di Sabatino Guimarães, Jansen Fernandes de Medeiros, Fernando Rodrigues Máximo, Juan Miguel Vilallobos-Salcedo and Deusilene Souza Vieira |
| EPI_ISL_514139, EPI_ISL_514140, EPI_ISL_514141, EPI_ISL_514142, EPI_ISL_514143, EPI_ISL_514144, EPI_ISL_514145, EPI_ISL_514146, EPI_ISL_514147, EPI_ISL_514148, EPI_ISL_514149, EPI_ISL_514150, EPI_ISL_514151, EPI_ISL_514152, EPI_ISL_514153, EPI_ISL_514154, EPI_ISL_514155, EPI_ISL_514156, EPI_ISL_514157, EPI_ISL_514158, EPI_ISL_514159, EPI_ISL_514160, EPI_ISL_514161, EPI_ISL_514162, EPI_ISL_514163, EPI_ISL_514164, EPI_ISL_514165, EPI_ISL_514166, EPI_ISL_514167, EPI_ISL_514168, EPI_ISL_514169, EPI_ISL_514170, EPI_ISL_514171, EPI_ISL_514172, EPI_ISL_514173, EPI_ISL_514174, EPI_ISL_514175, EPI_ISL_514176, EPI_ISL_514177, EPI_ISL_514178, EPI_ISL_514179, EPI_ISL_514180, EPI_ISL_514181, EPI_ISL_514182, EPI_ISL_514183, EPI_ISL_514184, EPI_ISL_514185, EPI_ISL_514186, EPI_ISL_514187, EPI_ISL_514188, EPI_ISL_514189, EPI_ISL_514190, EPI_ISL_514191, EPI_ISL_514192, EPI_ISL_514193, EPI_ISL_514194, EPI_ISL_514195, EPI_ISL_514196, EPI_ISL_514197, EPI_ISL_514198, EPI_ISL_514199, EPI_ISL_514200, EPI_ISL_514201, EPI_ISL_514202, EPI_ISL_514203, EPI_ISL_514204, EPI_ISL_514205, EPI_ISL_514206, EPI_ISL_514207, EPI_ISL_514208, EPI_ISL_514209, EPI_ISL_514210, EPI_ISL_514211, EPI_ISL_514212, EPI_ISL_514213, EPI_ISL_514214, EPI_ISL_514215, EPI_ISL_514216, EPI_ISL_514217, EPI_ISL_514218, EPI_ISL_514219, EPI_ISL_514220, EPI_ISL_514221, EPI_ISL_514222, EPI_ISL_514223, EPI_ISL_514224, EPI_ISL_514225 | | | |
| see above | Florida Bureau of Public Health Laboratories | Florida Bureau of Public Health Laboratories | Sarah Schmedes, Jason Blanton |
| EPI_ISL_514226 | Laboratorio de Referencia Nacional de Virus Respiratorio. Instituto Nacional de Salud Perú | Laboratorio de Referencia Nacional de Biotecnología y Biología Molecular. Instituto Nacional de Salud Perú | Carlos Padilla Rojas, Karolyn Vega Chozo, Priscila Lope Pari, Omar Caceres Rey, Marco Galarza Perez, Maribel Huaringa Nuñez, Johanna Balbuena Torrez, Henri Bailon Calderon, Nancy Rojas Serrano |
| EPI_ISL_514227 | Laboratorio de Referencia Nacional de Virus Respiratorio. Instituto Nacional de Salud. Peru | Laboratorio de Referencia Nacional de Biotecnologia y Biologia Molecular.Instituto Nacional de Salud.Peru | Carlos Padilla Rojas, Karolyn Vega Chozo, Priscila Lope Pari, Omar Caceres Rey, Marco Galarza Perez, Maribel Huaringa Nuñez, Johanna Balbuena Torrez, Henri Bailon Calderon, Nancy Rojas Serrano |
| EPI_ISL_514228, EPI_ISL_514229, EPI_ISL_514230 | National Institute of Laboratory Medicine and Referral Center | Genomic Research Lab, BCSIR | Barna Goswami, Abu Sayeed Mohammad Mahmud, Mohammad Samir Uzzaman, Eshrar Osman, Md. Ahashan Habib, Shahina Akter, Tanjina Akhter Banu, Md. Murshed Hasan Sarkar, Iffat Jahan, Md. Saddam Hossain, Tasnim Nafisa, Md. Maruf Ahmed Molla, Mahmuda Yeasmin, Asish Kumar Ghosh, A. K. M. Shamsuzzaman, Sheikh Md. Selim Al Din, Utpal Chandra Ray, Salek Ahmed Sajib, Md. Salim Khan |
| EPI_ISL_514231, EPI_ISL_514232 | National Institute of Laboratory Medicine and Referral Center | Genomic Research Lab, BCSIR | Shahina Akter, Abu Sayeed Mohammad Mahmud, Mohammad Samir Uzzaman, Eshrar Osman, Md. Ahashan Habib, Tanjina Akhter Banu, Md. Murshed Hasan Sarkar, Barna Goswami, Iffat Jahan, Md. Saddam Hossain, Tasnim Nafisa, Md. Maruf Ahmed Molla, Mahmuda Yeasmin, Asish Kumar Ghosh, A. K. M. Shamsuzzaman, Sheikh Md. Selim Al Din, Utpal Chandra Ray, Salek Ahmed Sajib, Md. Salim Khan |
| EPI_ISL_514233, EPI_ISL_514234, EPI_ISL_514235 | National Institute of Laboratory Medicine and Referral Center | Genomic Research Lab, BCSIR | Md. Murshed Hasan Sarkar, Abu Sayeed Mohammad Mahmud, Mohammad Samir Uzzaman, Eshrar Osman, Md. Ahashan Habib, Shahina Akter, Tanjina Akhter Banu, Barna Goswami, Iffat Jahan, Md. Saddam Hossain, Tasnim Nafisa, Md. Maruf Ahmed Molla, Mahmuda Yeasmin, Asish Kumar Ghosh, A. K. M. Shamsuzzaman, Sheikh Md. Selim Al Din, Utpal Chandra Ray, Salek Ahmed Sajib, Md. Salim Khan |
| EPI_ISL_514236 | National Institute of Laboratory Medicine and Referral Center | Genomic Research Lab, BCSIR | Md. Murshed Hasan Sarkar, Abu Sayeed Mohammad Mahmud, Mohammad Samir Uzzaman, Eshrar Osman, Md. Ahashan Habib, Shahina Akter, Tanjina Akhter Banu, Barna Goswami, Iffat Jahan, Md. Saddam Hossain, Tasnim Nafisa, Md. Maruf Ahmed Molla, Mahmuda Yeasmin, Asish Kumar Ghosh, A. K. M. Shamsuzzaman, Sheikh Md. Selim Al Din, Utpal Chandra Ray, Salek Ahmed Sajib, Md. Salim Khan |
| EPI_ISL_514237, EPI_ISL_514238, EPI_ISL_514239, EPI_ISL_514240, EPI_ISL_514241 | National Institute of Laboratory Medicine and Referral Center | Genomic Research Lab, BCSIR | Md. Saddam Hossain, Abu Sayeed Mohammad Mahmud, Mohammad Samir Uzzaman, Eshrar Osman, Md. Ahashan Habib, Shahina Akter, Tanjina Akhter Banu, Md. Murshed Hasan Sarkar, Barna Goswami, Iffat Jahan, Tasnim Nafisa, Md. Maruf Ahmed Molla, Mahmuda Yeasmin, Asish Kumar Ghosh, A. K. M. Shamsuzzaman, Sheikh Md. Selim Al Din, Utpal Chandra Ray, Salek Ahmed Sajib, Md. Salim Khan |
| EPI_ISL_514242, EPI_ISL_514243, EPI_ISL_514244 | National Institute of Laboratory Medicine and Referral Center | Genomic Research Lab, BCSIR | Tanjina Akhter Banu, Abu Sayeed Mohammad Mahmud, Mohammad Samir Uzzaman, Eshrar Osman, Md. Ahashan Habib, Shahina Akter, Md. Murshed Hasan Sarkar, Barna Goswami, Iffat Jahan, Md. Saddam Hossain, Tasnim Nafisa, Md. Maruf Ahmed Molla, Mahmuda Yeasmin, Asish Kumar Ghosh, A. K. M. Shamsuzzaman, Sheikh Md. Selim Al Din, Utpal Chandra Ray, Salek Ahmed Sajib, Md. Salim Khan |
| EPI_ISL_514245, EPI_ISL_514246, EPI_ISL_514247 | National Institute of Laboratory Medicine and Referral Center | Genomic Research Lab, BCSIR | Iffat Jahan, Abu Sayeed Mohammad Mahmud, Mohammad Samir Uzzaman, Eshrar Osman, Md. Ahashan Habib, Shahina Akter, Tanjina Akhter Banu, Md. Murshed Hasan Sarkar, Barna Goswami, Md. Saddam Hossain, Tasnim Nafisa, Md. Maruf Ahmed Molla, Mahmuda Yeasmin, Asish Kumar Ghosh, A. K. M. Shamsuzzaman, Sheikh Md. Selim Al Din, Utpal Chandra Ray, Salek Ahmed Sajib, Md. Salim Khan |
| EPI_ISL_514248, EPI_ISL_514249, EPI_ISL_514250, EPI_ISL_514251, EPI_ISL_514252 | National Institute of Laboratory Medicine and Referral Center | Genomic Research Lab, BCSIR | Abu Sayeed Mohammad Mahmud, Mohammad Samir Uzzaman, Eshrar Osman, Md. Ahashan Habib, Shahina Akter, Tanjina Akhter Banu, Md. Murshed Hasan Sarkar, Barna Goswami, Iffat Jahan, Md. Saddam Hossain, Tasnim Nafisa, Md. Maruf Ahmed Molla, Mahmuda Yeasmin, Asish Kumar Ghosh, A. K. M. Shamsuzzaman, Sheikh Md. Selim Al Din, Utpal Chandra Ray, Salek Ahmed Sajib, Md. Salim Khan |
| EPI_ISL_514253 | Advanced Biotechnology Laboratory | Genomic Research Lab, BCSIR | Abu Sayeed Mohammad Mahmud, Mohammad Samir Uzzaman, Eshrar Osman, Hossain Uddin Shekhar, M. Aftab Uddin, Md. Bayejid Hosen, Eunus Ali, Md. Ahashan Habib, Shahina Akter, Tanjina Akhter Banu, Md. Murshed Hasan Sarkar, Barna Goswami, Iffat Jahan, Md. Saddam Hossain, Utpal Chandra Ray, Salek Ahmed Sajib, Md. Salim Khan |
| EPI_ISL_514264, EPI_ISL_514265 | Laboratorio de Referencia Nacional de Virus Respiratorio. Instituto Nacional de Salud. Perú | Laboratorio de Referencia Nacional de Biotecnologia y Biologia Molecular.Instituto Nacional de Salud.Perú | Carlos Padilla Rojas, Karolyn Vega Chozo, Priscila Lope Pari, Omar Caceres Rey, Marco Galarza Perez, Maribel Huaringa Nuñez, Johanna Balbuena Torrez, Henri Bailon Calderon, Nancy Rojas Serrano |
| EPI_ISL_514266, EPI_ISL_514267, EPI_ISL_514268, EPI_ISL_514269, EPI_ISL_514270, EPI_ISL_514271, EPI_ISL_514272, EPI_ISL_514273, EPI_ISL_514274 | Israel Central Virology laboratory | Israel Central Virology laboratory | Neta Zuckerman, Efrat Dahan Bucris, Oran Erster, Ella Mendelson, Michal Mandelboim |
| EPI_ISL_514275 | Israeli Central Virology laboratory | Israel Central Virology laboratory | Neta Zuckerman, Efrat Dahan Bucris, Oran Erster, Ella Mendelson, Michal Mandelboim |
| EPI_ISL_514276, EPI_ISL_514277, EPI_ISL_514278, EPI_ISL_514279, EPI_ISL_514280, EPI_ISL_514281, EPI_ISL_514282, EPI_ISL_514283, EPI_ISL_514284, EPI_ISL_514285, EPI_ISL_514286, EPI_ISL_514287, EPI_ISL_514288, EPI_ISL_514289, EPI_ISL_514290, EPI_ISL_514291, EPI_ISL_514292, EPI_ISL_514293, EPI_ISL_514294, EPI_ISL_514295, EPI_ISL_514296, EPI_ISL_514297, EPI_ISL_514298, EPI_ISL_514299, EPI_ISL_514300, EPI_ISL_514301, EPI_ISL_514302, EPI_ISL_514303, EPI_ISL_514304, EPI_ISL_514305, EPI_ISL_514306, EPI_ISL_514307, EPI_ISL_514308, EPI_ISL_514309, EPI_ISL_514310, EPI_ISL_514311, EPI_ISL_514312, EPI_ISL_514313, EPI_ISL_514314, EPI_ISL_514315, EPI_ISL_514316 | | | |
| see above | Israel Central Virology laboratory | Israel Central Virology laboratory | Neta Zuckerman, Efrat Dahan Bucris, Oran Erster, Ella Mendelson, Michal Mandelboim |
| EPI_ISL_514317 | Laboratorio de Referencia Nacional de Virus Respiratorio. Instituto Nacional de Salud Perú | Laboratorio de Referencia Nacional de Biotecnología y Biología Molecular. Instituto Nacional de Salud Perú | Carlos Padilla Rojas, Karolyn Vega Chozo, Priscila Lope Pari, Omar Caceres Rey, Marco Galarza Perez, Maribel Huaringa Nuñez, Johanna Balbuena Torres, Henri Bailon Calderon, Nancy Rojas Serrano. |
| EPI_ISL_514338, EPI_ISL_514339, EPI_ISL_514340, EPI_ISL_514341, EPI_ISL_514342 | Laboratorio de Referencia Nacional de Virus Respiratorio. Instituto Nacional de Salud Perú | Laboratorio de Referencia Nacional de Biotecnología y Biología Molecular. Instituto Nacional de Salud Perú | Carlos Padilla Rojas, Karolyn Vega Chozo, Priscila Lope Pari, Omar Caceres Rey, Marco Galarza Perez, Maribel Huaringa Nuñez, Johanna Balbuena Torres, Henri Bailon Calderon, Nancy Rojas Serrano. |
| EPI_ISL_514343, EPI_ISL_514344, EPI_ISL_514345, EPI_ISL_514346, EPI_ISL_514347, EPI_ISL_514348, EPI_ISL_514349, EPI_ISL_514350, EPI_ISL_514351, EPI_ISL_514352, EPI_ISL_514353 | | | |
| see above | Respiratory Virus Unit, Microbiology Services Colindale, Public Health England | Respiratory Virus Unit, Microbiology Services Colindale, Public Health England | PHE Covid Sequencing Team |
| EPI_ISL_514354 | General Hospital - Prilep | Research Center for Genetic Engineering and Biotechnology "Georgi D. Efremov" , Macedonian Academy of Sciences and Arts | RCGEB - MASA |
| EPI_ISL_514355, EPI_ISL_514356, EPI_ISL_514357, EPI_ISL_514358, EPI_ISL_514359, EPI_ISL_514360, EPI_ISL_514361, EPI_ISL_514362, EPI_ISL_514363, EPI_ISL_514364, EPI_ISL_514365, EPI_ISL_514366, EPI_ISL_514367, EPI_ISL_514368, EPI_ISL_514369, EPI_ISL_514370, EPI_ISL_514371, EPI_ISL_514372, EPI_ISL_514373, EPI_ISL_514374, EPI_ISL_514375, EPI_ISL_514376, EPI_ISL_514377, EPI_ISL_514378, EPI_ISL_514379, EPI_ISL_514380, EPI_ISL_514381, EPI_ISL_514382, EPI_ISL_514383, EPI_ISL_514384, EPI_ISL_514385, EPI_ISL_514386, EPI_ISL_514387, EPI_ISL_514388, EPI_ISL_514389, EPI_ISL_514390, EPI_ISL_514391, EPI_ISL_514392, EPI_ISL_514393, EPI_ISL_514394, EPI_ISL_514395, EPI_ISL_514396, EPI_ISL_514397, EPI_ISL_514398, EPI_ISL_514399, EPI_ISL_514400, EPI_ISL_514401, EPI_ISL_514402, EPI_ISL_514403, EPI_ISL_514404, EPI_ISL_514405, EPI_ISL_514406, EPI_ISL_514407, EPI_ISL_514408, EPI_ISL_514409, EPI_ISL_514410, EPI_ISL_514411, EPI_ISL_514412, EPI_ISL_514413, EPI_ISL_514414, EPI_ISL_514415, EPI_ISL_514416, EPI_ISL_514417, EPI_ISL_514418, EPI_ISL_514419, EPI_ISL_514420, EPI_ISL_514421, EPI_ISL_514422, EPI_ISL_514423, EPI_ISL_514424 | | | |
| see above | National Institute for Communicable Diseases of the | National Institute for Communicable Diseases of the National Health Laboratory Service | Allam M, Ismail A, Khumalo Z, Kwenda S, Mtshali P, Mnyameni F, Mohale T, Bhiman JN |

| | | | |
|---|---|---|---|
| | National Health Laboratory Service | | |
| EPI_ISL_514425, EPI_ISL_514426, EPI_ISL_514427, EPI_ISL_514428, EPI_ISL_514429, EPI_ISL_514430, EPI_ISL_514431 | Laboratory Diagnostic, Veterinary Specialized Institute Kraljevo | Laboratory Diagnostic, Veterinary Specialized Institute Kraljevo | Vidanovic,D., Tesovic,B., Knezevic,A., Jankovic,M., Sekler,M., Banovic Djeri,B., Volkening,J., Afonso,C., Petrovic,T. |
| EPI_ISL_514432 | Prof. Massimo Zollo CEINGE TASK-FORCE COVID19 - Regione Campania | Prof. Massimo Zollo CEINGE TASK-FORCE COVID19 - Regione Campania | Veronica Ferrucci, Dae young Kong, Fatemeh asadzadeh, Laura Marrone, Roberto Siciliano, Rino Cerino, Giovanna Fusco, Marika Comegna, Angelo Boccia, Maurizio Viscardi, Giorgia Borriello, Sergio Brandi, Claudia Tiberio, Luigi Atripaldi, Giovanni Paolella, Giuseppe Castaldo, Stefano Pascarella, Martina Bianchi, Lorenzo Chiariotti, Jae Myun Lee, Jae Ho Jung, Kyong Seop Yun, Hong Yeoul Kim and Massimo Zollo |
| EPI_ISL_514434 | NSTU COVID-19 Diagnostic Center, | NSU Genome Research Institute (NGRI), North South University | Dr. Muhammad Maqsud Hossain, Aura Rahman, Prof. Firoz Ahmed, Tahrima Huq, Abdus Sadique, Jahidul Alam, Md Aminul Islam, Prof. Md. Didar-Ul-Alam, Prof. Kazi Nadim Hasan, Prof. Abdul Khaleque, Prof, Hasan Mahmud Reza |
| EPI_ISL_514435 | Dr. RSS Hospital, Modasa | Gujarat Biotechnology Research Centre | Harsh Chaudhari, Zarna Patel, Monika Gandhi, Pinal Trivedi, Maharshi Pandya, Nidhi Patel, Nitin Savaliya, Raghawendra Kumar, Dinesh Kumar, Zuber Saiyed, Komal Patel, Labdhi Pandya, Afzal Ansari, Nikha Trivedi, Apurvasinh Puvar, Janvi Raval, R D Dixit, A M Kadri, Harsh Bakshi, Chaitanya Joshi, Madhvi Joshi |
| EPI_ISL_514436 | Dr. RSS Hospital, Modasa | Gujarat Biotechnology Research Centre | Zarna Patel, Monika Gandhi, Pinal Trivedi, Maharshi Pandya, Nidhi Patel, Nitin Savaliya, Raghawendra Kumar, Dinesh Kumar, Zuber Saiyed, Komal Patel, Labdhi Pandya, Afzal Ansari, Nikha Trivedi, Harsh Chaudhari, Apurvasinh Puvar, Janvi Raval, R D Dixit, A M Kadri, Harsh Bakshi, Chaitanya Joshi, Madhvi Joshi |
| EPI_ISL_514437 | Dr. RSS Hospital, Modasa | Gujarat Biotechnology Research Centre | Nidhi Patel, Nitin Savaliya, Raghawendra Kumar, Dinesh Kumar, Zuber Saiyed, Komal Patel, Labdhi Pandya, Afzal Ansari, Nikha Trivedi, Harsh Chaudhari, Apurvasinh Puvar, Janvi Raval, Zarna Patel, Monika Gandhi, Pinal Trivedi, Maharshi Pandya, R D Dixit, A M Kadri, Harsh Bakshi, Chaitanya Joshi, Madhvi Joshi |
| EPI_ISL_514438, EPI_ISL_514439 | Tewhey Lab, The Jackson Laboratory | Tewhey Lab, The Jackson Laboratory | Matluk,N., Dewey,H., Barter,M., Lynch,R., Munger,H., Tewhey,R. |
| EPI_ISL_514440 | NSTU COVID-19 Diagnostic Center | NSU Genome Research Institute (NGRI), North South University | Dr. Muhammad Maqsud Hossain, Aura Rahman, Prof. Firoz Ahmed, Tahrima Huq, Abdus Sadique, Tamanna Afroze, Jahidul Alam, Md Aminul Islam, Prof. Md. Didar-Ul-Alam, Prof. Kazi Nadim Hasan, Prof. Abdul Khaleque, Prof, Hasan Mahmud Reza |
| EPI_ISL_514441 | NSTU COVID-19 Diagnostic Center | NSU Genome Research Institute (NGRI), North South University | Dr. Muhammad Maqsud Hossain, Aura Rahman, Prof. Firoz Ahmed, Tahrima Huq, Abdus Sadique, Jahidul Alam, Md Aminul Islam, Prof. Md. Didar-Ul-Alam, Prof. Kazi Nadim Hasan, Prof. Abdul Khaleque, Prof, Hasan Mahmud Reza |
| EPI_ISL_514442 | NSTU COVID-19 Diagnostic Center | NSU Genome Research Institute (NGRI), North South University | Dr. Muhammad Maqsud Hossain, Aura Rahman, Prof. Firoz Ahmed, Tahrima Huq, Abdus Sadique, Jahidul Alam, Tamanna Afroze, Md Aminul Islam, Prof. Md. Didar-Ul-Alam, Prof. Kazi Nadim Hasan, Prof. Abdul Khaleque, Prof, Hasan Mahmud Reza |
| EPI_ISL_514443, EPI_ISL_514444, EPI_ISL_514445, EPI_ISL_514446, EPI_ISL_514447, EPI_ISL_514448, EPI_ISL_514449, EPI_ISL_514450, EPI_ISL_514451 | Department of Pathology, University of Cambridge | COVID-19 Genomics UK (COG-UK) Consortium | Luke W Meredith, M. Estée Török, Myra Hosmillo, William L. Hamilton, Martin D. Curran, Theresa Feltwell, Grant Hall, Anna Yakovleva, Fahad A Khokhar, Charlotte J. Houldcroft, Laura G Caller, Aminu S. Jahun, Sarah L. Caddy, Yasmin Chaudhry, Malte Pinckert, Ian Goodfellow |
| EPI_ISL_514452, EPI_ISL_514453, EPI_ISL_514454 | Queens Medical Centre, Clinical Microbiology Department / DeepSeq Nottingham | COVID-19 Genomics UK (COG-UK) Consortium | Gemma Clark, Wendy Smith, Manjinder Khakh, Vicki M Fleming, Michelle M Lister, Hannah Howson-Wells, Jonathan Ball, Patrick McClure, Joseph Chappell, Theocharis Tsoleridis, Nadine Holmes, Matthew Carlisle, Christopher Moore, Fei Sang, Johnny Debebe, Victoria Wright, Matthew Loose |
| EPI_ISL_514455, EPI_ISL_514456, EPI_ISL_514457, EPI_ISL_514458, EPI_ISL_514459, EPI_ISL_514460, EPI_ISL_514461, EPI_ISL_514462, EPI_ISL_514463, EPI_ISL_514464, EPI_ISL_514465, EPI_ISL_514466, EPI_ISL_514467, EPI_ISL_514468, EPI_ISL_514469, EPI_ISL_514470, EPI_ISL_514471, EPI_ISL_514472, EPI_ISL_514473, EPI_ISL_514474, EPI_ISL_514475, EPI_ISL_514476, EPI_ISL_514477, EPI_ISL_514478, EPI_ISL_514479, EPI_ISL_514480, EPI_ISL_514481, EPI_ISL_514482, EPI_ISL_514483, EPI_ISL_514484, EPI_ISL_514485, EPI_ISL_514486, EPI_ISL_514487, EPI_ISL_514488, EPI_ISL_514489, EPI_ISL_514490, EPI_ISL_514491, EPI_ISL_514492, EPI_ISL_514493, EPI_ISL_514494, EPI_ISL_514495, EPI_ISL_514496, EPI_ISL_514497, EPI_ISL_514498, EPI_ISL_514499, EPI_ISL_514500, EPI_ISL_514501, EPI_ISL_514502, EPI_ISL_514503, EPI_ISL_514504, EPI_ISL_514505, EPI_ISL_514506, EPI_ISL_514507, EPI_ISL_514508, EPI_ISL_514509, EPI_ISL_514510, EPI_ISL_514511, EPI_ISL_514512, EPI_ISL_514513 | | | |
| see above | Centre for Enzyme Innovation, University of Portsmouth / Translational Research Laboratory, Portsmouth Hospitals NHS Trust | COVID-19 Genomics UK (COG-UK) Consortium | Angela Beckett,Yann Bourgeois,Garry Scarlett,Sharon Glaysher,Scott Elliott,Kelly Bicknell,Robert Impey,Allyson Lloyd,Sarah Wyllie,Ethan Butcher,Anoop Chauhan,Samuel Robson |
| EPI_ISL_514514, EPI_ISL_514515 | West of Scotland Specialist Virology Centre, NHSGGC / MRC-University of Glasgow Centre for Virus Research | COVID-19 Genomics UK (COG-UK) Consortium | Ana da Silva Filipe, Natasha Johnson, Kathy Smollett, Daniel Mair, Stephen Carmichael, Lily Tong, Jenna Nichols, Elihu Aranday-Cortes, Kirstyn Brunker, Yasmin Parr, Alice Broos, Kyriaki Nomikou; Sarah McDonald, Marc Niebel, Patawee Asamaphan; Richard Orton, Joseph Hughes, Sreenu Vattipally, David L Robertson; Alasdair MacLean, Rory Gunson; Kathy Li, Natasha Jesudason, Rajiv Shah, James Shepherd, Antonia Ho, Emma Thomson |
| EPI_ISL_514516, EPI_ISL_514517, EPI_ISL_514518, EPI_ISL_514519, EPI_ISL_514520, EPI_ISL_514521, EPI_ISL_514522, EPI_ISL_514523, EPI_ISL_514524, EPI_ISL_514525, EPI_ISL_514526, EPI_ISL_514527, EPI_ISL_514528, EPI_ISL_514529, EPI_ISL_514530, EPI_ISL_514531, EPI_ISL_514532, EPI_ISL_514533, EPI_ISL_514534, EPI_ISL_514535, EPI_ISL_514536, EPI_ISL_514537, EPI_ISL_514538, EPI_ISL_514539, EPI_ISL_514540, EPI_ISL_514541, EPI_ISL_514542, EPI_ISL_514543, EPI_ISL_514544, EPI_ISL_514545, EPI_ISL_514546, EPI_ISL_514547, EPI_ISL_514548, EPI_ISL_514549, EPI_ISL_514550, EPI_ISL_514551, EPI_ISL_514552, EPI_ISL_514553, EPI_ISL_514554, EPI_ISL_514555, EPI_ISL_514556, EPI_ISL_514557, EPI_ISL_514558, EPI_ISL_514559, EPI_ISL_514560, EPI_ISL_514561, EPI_ISL_514562 | | | |
| see above | Virology Department, Royal Infirmary of Edinburgh, NHS Lothian / School of Biological Sciences, University of Edinburgh / Institute of Genetics and Molecular Medicine, University of Edinburgh | COVID-19 Genomics UK (COG-UK) Consortium | McHugh M, Dewar R, Rooke S, Gallagher M, Balcaza C, O'Toole Á, Scher E, Hill V, McCrone JT, Colquhoun R, Yu X, Jackson B, Rambaut A, Williams TC, Templeton K |
| EPI_ISL_514563, EPI_ISL_514564, EPI_ISL_514565, EPI_ISL_514566, EPI_ISL_514567, EPI_ISL_514568, EPI_ISL_514569, EPI_ISL_514570, EPI_ISL_514571, EPI_ISL_514572, EPI_ISL_514573, EPI_ISL_514574, EPI_ISL_514575, EPI_ISL_514576, EPI_ISL_514577, EPI_ISL_514578, EPI_ISL_514579 | | | |
| see above | Wales Specialist Virology Centre Sequencing lab: Pathogen Genomics Unit | COVID-19 Genomics UK (COG-UK) Consortium | Catherine Moore, Johnathan Evans, Laura Gifford, Malorie Perry, Simon Cottrell, Angela Marchbank, Alec Birchley, Alexander Adams, Amy Gaskin, Bree Gatica-Wilcox, Jason Coombes, Joel Southgate, Lauren Gilbert, Lee Graham, Nicole Pacchiarini, Sara Kumziene-Summerhayes, Sarah Taylor, Sophie Jones, Sara Rey, Matthew Bull, Joanne Watkins, Sally Corden, Tom Connor |
| EPI_ISL_514580 | NSTU COVID-19 Diagnostic Center | NSU Genome Research Institute (NGRI), North South University | Dr. Muhammad Maqsud Hossain, Aura Rahman, Prof. Firoz Ahmed, Tahrima Huq, Abdus Sadique, Jahidul Alam, Tamanna Afroze, Md Aminul Islam, Prof. Md. Didar-Ul-Alam, Prof. Kazi Nadim Hasan, Prof. Abdul Khaleque, Prof, Hasan Mahmud Reza |
| EPI_ISL_514581 | B.J. Medical College and Civil hospital, Ahmedabad | Gujarat Biotechnology Research Centre | Nitin Savaliya, Raghawendra Kumar, Dinesh Kumar, Zuber Saiyed, Komal Patel, Labdhi Pandya, Afzal Ansari, Nikha Trivedi, Pranay Shah, Kamlesh J Upadhyay, Sanjay Kapadia, Apurvasinh Puvar, Janvi Raval, Zarna Patel, Monika Gandhi, Pinal Trivedi, Maharshi Pandya, Nidhi Patel, R D Dixit, A M Kadri, Harsh Bakshi, Chaitanya Joshi, Madhvi Joshi |
| EPI_ISL_514582 | B.J. Medical College and Civil hospital, Ahmedabad | Gujarat Biotechnology Research Centre | Raghawendra Kumar, Dinesh Kumar, Zuber Saiyed, Komal Patel, Labdhi Pandya, Afzal Ansari, Nikha Trivedi, Pranay Shah, Kamlesh J Upadhyay, Sanjay Kapadia, Apurvasinh Puvar, Janvi Raval, Zarna Patel, Monika Gandhi, Pinal Trivedi, Maharshi Pandya, Nidhi Patel, Nitin Savaliya, R D Dixit, A M Kadri, Harsh Bakshi, Chaitanya Joshi, Madhvi Joshi |
| EPI_ISL_514583 | B.J. Medical College and Civil hospital, Ahmedabad | Gujarat Biotechnology Research Centre | Dinesh Kumar, Zuber Saiyed, Komal Patel, Labdhi Pandya, Afzal Ansari, Nikha Trivedi, Pranay Shah, Kamlesh J Upadhyay, Sanjay Kapadia, Apurvasinh Puvar, Janvi Raval, Zarna Patel, Monika Gandhi, Pinal Trivedi, Maharshi Pandya, Nidhi Patel, Nitin Savaliya, Raghawendra Kumar, R D Dixit, A M Kadri, Harsh Bakshi, Chaitanya Joshi, Madhvi Joshi |
| EPI_ISL_514584 | B.J. Medical College and Civil hospital, Ahmedabad | Gujarat Biotechnology Research Centre | Zuber Saiyed, Komal Patel, Labdhi Pandya, Afzal Ansari, Nikha Trivedi, Pranay Shah, Kamlesh J Upadhyay, Sanjay Kapadia, Apurvasinh Puvar, Janvi Raval, Zarna Patel, Monika Gandhi, Pinal Trivedi, Maharshi Pandya, Nidhi Patel, Nitin Savaliya, Raghawendra Kumar, Dinesh Kumar, R D Dixit, A M Kadri, Harsh Bakshi, Chaitanya Joshi, Madhvi Joshi |
| EPI_ISL_514585 | B.J. Medical College and Civil hospital, Ahmedabad | Gujarat Biotechnology Research Centre | Komal Patel, Labdhi Pandya, Afzal Ansari, Nikha Trivedi, Pranay Shah, Kamlesh J Upadhyay, Sanjay Kapadia, Apurvasinh Puvar, Janvi Raval, Zarna Patel, Monika Gandhi, Pinal Trivedi, Maharshi Pandya, Nidhi Patel, Nitin Savaliya, Raghawendra Kumar, Dinesh Kumar, Zuber Saiyed, R D Dixit, A M Kadri, Harsh Bakshi, Chaitanya Joshi, Madhvi Joshi |
| EPI_ISL_514586 | B.J. Medical College and Civil hospital, Ahmedabad | Gujarat Biotechnology Research Centre | Labdhi Pandya, Afzal Ansari, Nikha Trivedi, Pranay Shah, Kamlesh J Upadhyay, Sanjay Kapadia, Apurvasinh Puvar, Janvi Raval, Zarna Patel, Monika Gandhi, Pinal Trivedi, Maharshi Pandya, Nidhi Patel, Nitin Savaliya, Raghawendra Kumar Dinesh Kumar Zuber Saiyed, Komal Patel, R D Dixit, A M Kadri, Harsh Bakshi, Chaitanya Joshi, Madhvi Joshi |
| EPI_ISL_514587 | B.J. Medical College and Civil hospital, Ahmedabad | Gujarat Biotechnology Research Centre | Afzal Ansari, Nikha Trivedi, Pranay Shah, Kamlesh J Upadhyay, Sanjay Kapadia, Apurvasinh Puvar, Janvi Raval, Zarna Patel, Monika Gandhi, Pinal Trivedi, Maharshi Pandya, Nidhi Patel, Nitin Savaliya, Raghawendra Kumar, Dinesh Kumar, Zuber Saiyed, Komal Patel, Labdhi Pandya, R D Dixit, A M Kadri, Harsh Bakshi, Chaitanya Joshi, Madhvi Joshi |
| EPI_ISL_514588 | B.J. Medical College and Civil hospital, Ahmedabad | Gujarat Biotechnology Research Centre | Nikha Trivedi, Pranay Shah, Kamlesh J Upadhyay, Sanjay Kapadia, Apurvasinh Puvar, Janvi Raval, Zarna Patel, Monika Gandhi, Pinal Trivedi, Maharshi Pandya, Nidhi Patel, Nitin Savaliya, Raghawendra Kumar, Dinesh Kumar, Zuber Saiyed, Komal Patel, Labdhi Pandya, Afzal Ansari, R D Dixit, A M Kadri, Harsh Bakshi, Chaitanya Joshi, Madhvi Joshi |
| EPI_ISL_514589 | B.J. Medical College and Civil hospital, Ahmedabad | Gujarat Biotechnology Research Centre | Pranay Shah, Kamlesh J Upadhyay, Sanjay Kapadia, Apurvasinh Puvar, Janvi Raval, Zarna Patel, Monika Gandhi, Pinal Trivedi, Maharshi Pandya, Nidhi Patel, Nitin Savaliya, Raghawendra Kumar, Dinesh Kumar, Zuber Saiyed, Komal Patel, Labdhi Pandya, Afzal Ansari, Nikha Trivedi, R D Dixit, A M Kadri, Harsh Bakshi, Chaitanya Joshi, Madhvi Joshi |
| EPI_ISL_514590 | B.J. Medical College and Civil hospital, Ahmedabad | Gujarat Biotechnology Research Centre | Kamlesh J Upadhyay, Sanjay Kapadia, Apurvasinh Puvar, Janvi Raval, Zarna Patel, Monika Gandhi, Pinal Trivedi, Maharshi Pandya, Nidhi Patel, Nitin Savaliya, Raghawendra Kumar, Dinesh Kumar, Zuber Saiyed, Komal Patel, Labdhi Pandya, Afzal Ansari, Nikha Trivedi, Pranay Shah, R D Dixit, A M Kadri, Harsh Bakshi, Chaitanya Joshi, Madhvi Joshi |
| EPI_ISL_514591 | B.J. Medical College and Civil hospital, Ahmedabad | Gujarat Biotechnology Research Centre | Sanjay Kapadia,Apurvasinh Puvar,Janvi Raval,Zarna Patel, Monika Gandhi, Pinal Trivedi, Maharshi Pandya, Nidhi Patel, Nitin Savaliya, Raghawendra Kumar, Dinesh Kumar, Zuber Saiyed, Komal Patel, Labdhi Pandya, Afzal Ansari, Nikha Trivedi, Pranay Shah, Kamlesh J Upadhyay, R D Dixit, A M Kadri, Harsh Bakshi, Chaitanya Joshi, Madhvi Joshi |
| EPI_ISL_514592 | B.J. Medical College and Civil hospital, Ahmedabad | Gujarat Biotechnology Research Centre | Apurvasinh Puvar, Janvi Raval, Zarna Patel, Monika Gandhi, Pinal Trivedi, Maharshi Pandya, Nidhi Patel, Nitin Savaliya, Raghawendra Kumar, Dinesh Kumar, Zuber Saiyed, Komal Patel, Labdhi Pandya, Afzal Ansari, Nikha Trivedi, Pranay Shah, Kamlesh J Upadhyay, Sanjay Kapadia, R D Dixit, A M Kadri, Harsh Bakshi, Chaitanya Joshi, Madhvi Joshi |
| EPI_ISL_514593 | B.J. Medical College and Civil hospital, Ahmedabad | Gujarat Biotechnology Research Centre | Janvi Raval, Zarna Patel, Monika Gandhi, Pinal Trivedi, Maharshi Pandya, Nidhi Patel, Nitin Savaliya, Raghawendra Kumar, Dinesh Kumar, Zuber Saiyed, Komal Patel, Labdhi Pandya, Afzal Ansari, Nikha Trivedi, Pranay Shah, Kamlesh J Upadhyay, Sanjay Kapadia, Apurvasinh Puvar, R D Dixit, A M Kadri, Harsh Bakshi, Chaitanya Joshi, Madhvi Joshi |
| EPI_ISL_514594 | B.J. Medical College and Civil hospital, Ahmedabad | Gujarat Biotechnology Research Centre | Zarna Patel, Monika Gandhi, Pinal Trivedi, Maharshi Pandya, Nidhi Patel, Nitin Savaliya, Raghawendra Kumar, Dinesh Kumar, Zuber Saiyed, Komal Patel, Labdhi Pandya, Afzal Ansari, Nikha Trivedi, Pranay Shah, Kamlesh J Upadhyay, Sanjay Kapadia, Apurvasinh Puvar, Janvi Raval, R D Dixit, A M Kadri, Harsh Bakshi, Chaitanya Joshi, Madhvi Joshi |
| EPI_ISL_514595 | B.J. Medical College and Civil hospital, Ahmedabad | Gujarat Biotechnology Research Centre | Monika Gandhi, Pinal Trivedi, Maharshi Pandya, Nidhi Patel, Nitin Savaliya, Raghawendra Kumar, Dinesh Kumar, Zuber Saiyed, Komal Patel, Labdhi Pandya, Afzal Ansari, Nikha Trivedi, Pranay Shah, Kamlesh J Upadhyay, Sanjay Kapadia, Apurvasinh Puvar, Janvi Raval, Zarna Patel, R D Dixit, A M Kadri, Harsh Bakshi, Chaitanya Joshi, Madhvi Joshi |
| EPI_ISL_514596 | B.J. Medical College and Civil hospital, Ahmedabad | Gujarat Biotechnology Research Centre | Pinal Trivedi, Maharshi Pandya, Nidhi Patel, Nitin Savaliya, Raghawendra Kumar, Dinesh Kumar, Zuber Saiyed, Komal Patel, Labdhi Pandya, Afzal Ansari, Nikha Trivedi, Pranay Shah, Kamlesh J Upadhyay, Sanjay Kapadia, Apurvasinh Puvar, Janvi Raval, Zarna Patel, Monika Gandhi, R D Dixit, A M Kadri, Harsh Bakshi, Chaitanya Joshi, Madhvi Joshi |
| EPI_ISL_514597 | B.J. Medical College and Civil hospital, Ahmedabad | Gujarat Biotechnology Research Centre | Maharshi Pandya, Nidhi Patel, Nitin Savaliya, Raghawendra Kumar, Dinesh Kumar, Zuber Saiyed, Komal Patel, Labdhi Pandya, Afzal Ansari, Nikha Trivedi, Pranay Shah, Kamlesh J Upadhyay, Sanjay Kapadia, Apurvasinh Puvar, Janvi Raval, Zarna Patel, Monika Gandhi, Pinal Trivedi, R D Dixit, A M Kadri, Harsh Bakshi, Chaitanya Joshi, Madhvi Joshi |
| EPI_ISL_514598 | B.J. Medical College and Civil hospital, Ahmedabad | Gujarat Biotechnology Research Centre | Nidhi Patel, Nitin Savaliya, Raghawendra Kumar, Dinesh Kumar, Zuber Saiyed, Komal Patel, Labdhi Pandya, Afzal Ansari, Nikha Trivedi, Pranay Shah, Kamlesh J Upadhyay, Sanjay Kapadia, Apurvasinh Puvar, Janvi Raval, Zarna Patel, Monika Gandhi, Pinal Trivedi, Maharshi Pandya, R D Dixit, A M Kadri, Harsh Bakshi, Chaitanya Joshi, Madhvi Joshi |
| EPI_ISL_514599 | B.J. Medical College and Civil hospital, Ahmedabad | Gujarat Biotechnology Research Centre | Nitin Savaliya, Raghawendra Kumar, Dinesh Kumar, Zuber Saiyed, Komal Patel, Labdhi Pandya, Afzal Ansari, Nikha Trivedi, Pranay Shah, Kamlesh J Upadhyay, Sanjay Kapadia, Apurvasinh Puvar, Janvi Raval, Zarna Patel, Monika Gandhi, Pinal Trivedi, Maharshi Pandya, Nidhi Patel, R D Dixit, A M Kadri, Harsh Bakshi, Chaitanya Joshi, Madhvi Joshi |
| EPI_ISL_514600 | B.J. Medical College and Civil hospital, Ahmedabad | Gujarat Biotechnology Research Centre | Raghawendra Kumar, Dinesh Kumar, Zuber Saiyed, Komal Patel, Labdhi Pandya, Afzal Ansari, Nikha Trivedi, Pranay Shah, Kamlesh J Upadhyay, Sanjay Kapadia, Apurvasinh Puvar, Janvi Raval, Zarna Patel, Monika Gandhi, Pinal Trivedi, Maharshi Pandya, Nidhi Patel, Nitin Savaliya, R D Dixit, A M Kadri, Harsh Bakshi, Chaitanya Joshi, Madhvi Joshi |
| EPI_ISL_514601 | B.J. Medical College and Civil hospital, Ahmedabad | Gujarat Biotechnology Research Centre | Dinesh Kumar, Zuber Saiyed, Komal Patel, Labdhi Pandya, Afzal Ansari, Nikha Trivedi, Pranay Shah, Kamlesh J Upadhyay, Sanjay Kapadia, Apurvasinh Puvar, Janvi Raval, Zarna |

| | | | |
|---|---|---|---|
| | | | Patel, Monika Gandhi, Pinal Trivedi, Maharshi Pandya, Nidhi Patel, Nitin Savaliya, Raghawendra Kumar, R D Dixit, A M Kadri, Harsh Bakshi, Chaitanya Joshi, Madhvi Joshi |
| EPI_ISL_514602 | B.J. Medical College and Civil hospital, Ahmedabad | Gujarat Biotechnology Research Centre | Zuber Saiyed, Komal Patel, Labdhi Pandya, Afzal Ansari, Nikha Trivedi, Pranay Shah, Kamlesh J Upadhyay, Sanjay Kapadia, Apurvasinh Puvar, Janvi Raval, Zarna Patel, Monika Gandhi, Pinal Trivedi, Maharshi Pandya, Nidhi Patel, Nitin Savaliya, Raghawendra Kumar, Dinesh Kumar, R D Dixit, A M Kadri, Harsh Bakshi, Chaitanya Joshi, Madhvi Joshi |
| EPI_ISL_514603 | B.J. Medical College and Civil hospital, Ahmedabad | Gujarat Biotechnology Research Centre | Komal Patel, Labdhi Pandya, Afzal Ansari, Nikha Trivedi, Pranay Shah, Kamlesh J Upadhyay, Sanjay Kapadia, Apurvasinh Puvar, Janvi Raval, Zarna Patel, Monika Gandhi, Pinal Trivedi, Maharshi Pandya, Nidhi Patel, Nitin Savaliya, Raghawendra Kumar, Dinesh Kumar, Zuber Saiyed, R D Dixit, A M Kadri, Harsh Bakshi, Chaitanya Joshi, Madhvi Joshi |
| EPI_ISL_514604 | B.J. Medical College and Civil hospital, Ahmedabad | Gujarat Biotechnology Research Centre | Labdhi Pandya, Afzal Ansari, Nikha Trivedi, Pranay Shah, Kamlesh J Upadhyay, Sanjay Kapadia, Apurvasinh Puvar, Janvi Raval, Zarna Patel, Monika Gandhi, Pinal Trivedi, Maharshi Pandya, Nidhi Patel, Nitin Savaliya, Raghawendra Kumar, Dinesh Kumar, Zuber Saiyed, Komal Patel, R D Dixit, A M Kadri, Harsh Bakshi, Chaitanya Joshi, Madhvi Joshi |
| EPI_ISL_514605 | B.J. Medical College and Civil hospital, Ahmedabad | Gujarat Biotechnology Research Centre | Afzal Ansari, Nikha Trivedi, Pranay Shah, Kamlesh J Upadhyay, Sanjay Kapadia, Apurvasinh Puvar, Janvi Raval, Zarna Patel, Monika Gandhi, Pinal Trivedi, Maharshi Pandya, Nidhi Patel, Nitin Savaliya, Raghawendra Kumar, Dinesh Kumar, Zuber Saiyed, Komal Patel, Labdhi Pandya, R D Dixit, A M Kadri, Harsh Bakshi, Chaitanya Joshi, Madhvi Joshi |
| EPI_ISL_514606 | B.J. Medical College and Civil hospital, Ahmedabad | Gujarat Biotechnology Research Centre | Nikha Trivedi, Pranay Shah, Kamlesh J Upadhyay, Sanjay Kapadia, Apurvasinh Puvar, Janvi Raval, Zarna Patel, Monika Gandhi, Pinal Trivedi, Maharshi Pandya, Nidhi Patel, Nitin Savaliya, Raghawendra Kumar, Dinesh Kumar, Zuber Saiyed, Komal Patel, Labdhi Pandya, Afzal Ansari, R D Dixit, A M Kadri, Harsh Bakshi, Chaitanya Joshi, Madhvi Joshi |
| EPI_ISL_514607 | B.J. Medical College and Civil hospital, Ahmedabad | Gujarat Biotechnology Research Centre | Pranay Shah, Kamlesh J Upadhyay, Sanjay Kapadia, Apurvasinh Puvar, Janvi Raval, Zarna Patel, Monika Gandhi, Pinal Trivedi, Maharshi Pandya, Nidhi Patel, Nitin Savaliya, Raghawendra Kumar, Dinesh Kumar, Zuber Saiyed, Komal Patel, Labdhi Pandya, Afzal Ansari, Nikha Trivedi, R D Dixit, A M Kadri, Harsh Bakshi, Chaitanya Joshi, Madhvi Joshi |
| EPI_ISL_514608 | B.J. Medical College and Civil hospital, Ahmedabad | Gujarat Biotechnology Research Centre | Kamlesh J Upadhyay, Sanjay Kapadia, Apurvasinh Puvar, Janvi Raval, Zarna Patel, Monika Gandhi, Pinal Trivedi, Maharshi Pandya, Nidhi Patel, Nitin Savaliya, Raghawendra Kumar, Dinesh Kumar, Zuber Saiyed, Komal Patel, Labdhi Pandya, Afzal Ansari, Nikha Trivedi, Pranay Shah, R D Dixit, A M Kadri, Harsh Bakshi, Chaitanya Joshi, Madhvi Joshi |
| EPI_ISL_514609 | B.J. Medical College and Civil hospital, Ahmedabad | Gujarat Biotechnology Research Centre | Sanjay Kapadia, Apurvasinh Puvar, Janvi Raval, Zarna Patel, Monika Gandhi, Pinal Trivedi, Maharshi Pandya, Nidhi Patel, Nitin Savaliya, Raghawendra Kumar, Dinesh Kumar, Zuber Saiyed, Komal Patel, Labdhi Pandya, Afzal Ansari, Nikha Trivedi, Pranay Shah, Kamlesh J Upadhyay, R D Dixit, A M Kadri, Harsh Bakshi, Chaitanya Joshi, Madhvi Joshi |
| EPI_ISL_514610 | GMERS Medical College & Hospital, Gotri, Vadodara | Gujarat Biotechnology Research Centre | Meenakshi Shah, Neena Doshi, Varsha Godbole, Apurvasinh Puvar, Janvi Raval, Zarna Patel, Monika Gandhi, Pinal Trivedi, Maharshi Pandya, Nidhi Patel, Nitin Savaliya, Raghawendra Kumar, Dinesh Kumar, Zuber Saiyed, Komal Patel, Labdhi Pandya, Afzal Ansari, Nikha Trivedi, Pranay Shah, Kamlesh J Upadhyay, Sanjay Kapadia, R D Dixit, A M Kadri, Harsh Bakshi, Chaitanya Joshi, Madhvi Joshi, |
| EPI_ISL_514611 | GMERS Medical College and Hospital, Gandhinagar | Gujarat Biotechnology Research Centre | Seema Bhatt, Gaurishankar Shrimali, Bhavesh Modi, Bharti Rajani, Apurvasinh Puvar, Janvi Raval, Zarna Patel, Monika Gandhi, Pinal Trivedi, Maharshi Pandya, Nidhi Patel, Nitin Savaliya, Raghawendra Kumar, Dinesh Kumar, Zuber Saiyed, Komal Patel, Labdhi Pandya, Afzal Ansari, Nikha Trivedi, Pranay Shah, Kamlesh J Upadhyay, Sanjay Kapadia, R D Dixit, A M Kadri, Harsh Bakshi, Chaitanya Joshi, Madhvi Joshi |
| EPI_ISL_514612 | GMERS Medical College and Hospital, Gandhinagar | Gujarat Biotechnology Research Centre | Gaurishankar Shrimali, Bhavesh Modi, Bharti Rajani, Apurvasinh Puvar, Janvi Raval, Zarna Patel, Monika Gandhi, Pinal Trivedi, Maharshi Pandya, Nidhi Patel, Nitin Savaliya, Raghawendra Kumar, Dinesh Kumar, Zuber Saiyed, Komal Patel, Labdhi Pandya, Afzal Ansari, Nikha Trivedi, Pranay Shah, Kamlesh J Upadhyay, Sanjay Kapadia, Seema Bhatt, R D Dixit, A M Kadri, Harsh Bakshi, Chaitanya Joshi, Madhvi Joshi |
| EPI_ISL_514613, EPI_ISL_514614 | NSTU COVID-19 Diagnostic Center | NSU Genome Research Institute (NGRI), North South University | Dr. Muhammad Maqsud Hossain, Aura Rahman, Prof. Firoz Ahmed, Tahrima Huq, Abdus Sadique, Jahidul Alam, Tamanna Afroze, Md Aminul Islam, Prof. Md. Didar-Ul-Alam, Prof. Kazi Nadim Hasan, Prof. Abdul Khaleque, Prof, Hasan Mahmud Reza |
| EPI_ISL_514615 | NSTU COVID-19 Diagnostic Center | NSU Genome Research Institute (NGRI), North South University | Dr. Muhammad Maqsud Hossain, Aura Rahman, Prof. Firoz Ahmed, Tahrima Huq, Abdus Sadique, Jahidul Alam, Tamanna Afroze, Md Aminul Islam, Prof. Md. Didar-Ul-Alam, Prof. Kazi Nadim Hasan, Prof. Abdul Khaleque, Prof, Hasan Mahmud Reza |
| EPI_ISL_514616, EPI_ISL_514617, EPI_ISL_514618 | Minnesota Department of Health, Public Health Laboratory | Minnesota Department of Health, Public Health Laboratory | Matt Plumb, Jacob Garfin, and Xiong Wang |
| EPI_ISL_514619, EPI_ISL_514620, EPI_ISL_514621, EPI_ISL_514622, EPI_ISL_514623, EPI_ISL_514624, EPI_ISL_514625, EPI_ISL_514626, EPI_ISL_514627, EPI_ISL_514628 | Mayo Clinic & Mayo Clinic Laboratories | Minnesota Department of Health, Public Health Laboratory | Matt Plumb, Jacob Garfin, and Xiong Wang |
| EPI_ISL_514629, EPI_ISL_514630, EPI_ISL_514631, EPI_ISL_514632, EPI_ISL_514633, EPI_ISL_514634, EPI_ISL_514635 | M Health Fairview | Minnesota Department of Health, Public Health Laboratory | Matt Plumb, Jacob Garfin, and Xiong Wang |
| EPI_ISL_514636 | Mayo Clinic & Mayo Clinic Laboratories | Minnesota Department of Health, Public Health Laboratory | Matt Plumb, Jacob Garfin, and Xiong Wang |
| EPI_ISL_514637 | M Health Fairview St. Joseph's Hospital | Minnesota Department of Health, Public Health Laboratory | Matt Plumb, Jacob Garfin, and Xiong Wang |
| EPI_ISL_514638, EPI_ISL_514639, EPI_ISL_514640 | Mayo Clinic & Mayo Clinic Laboratories | Minnesota Department of Health, Public Health Laboratory | Matt Plumb, Jacob Garfin, and Xiong Wang |
| EPI_ISL_514641 | St. Luke's Hospital | Minnesota Department of Health, Public Health Laboratory | Matt Plumb, Jacob Garfin, and Xiong Wang |
| EPI_ISL_514642 | Essentia Health-St. Mary's Medical Center | Minnesota Department of Health, Public Health Laboratory | Matt Plumb, Jacob Garfin, and Xiong Wang |
| EPI_ISL_514643 | M Health Fairview St. Joseph's Hospital | Minnesota Department of Health, Public Health Laboratory | Matt Plumb, Jacob Garfin, and Xiong Wang |
| EPI_ISL_514644 | Mayo Clinic & Mayo Clinic Laboratories | Minnesota Department of Health, Public Health Laboratory | Matt Plumb, Jacob Garfin, and Xiong Wang |
| EPI_ISL_514645 | Avera Mckennan Laboratory | Minnesota Department of Health, Public Health Laboratory | Matt Plumb, Jacob Garfin, and Xiong Wang |
| EPI_ISL_514646 | St. Luke's Hospital | Minnesota Department of Health, Public Health Laboratory | Matt Plumb, Jacob Garfin, and Xiong Wang |
| EPI_ISL_514647 | Allina Health Laboratory | Minnesota Department of Health, Public Health Laboratory | Matt Plumb, Jacob Garfin, and Xiong Wang |
| EPI_ISL_514648 | Essentia Health-St. Joseph's Medical Center | Minnesota Department of Health, Public Health Laboratory | Matt Plumb, Jacob Garfin, and Xiong Wang |
| EPI_ISL_514649 | Allina Health Laboratory | Minnesota Department of Health, Public Health Laboratory | Matt Plumb, Jacob Garfin, and Xiong Wang |
| EPI_ISL_514650 | M Health Fairview St. Joseph's Hospital | Minnesota Department of Health, Public Health Laboratory | Matt Plumb, Jacob Garfin, and Xiong Wang |
| EPI_ISL_514651 | Allina Health Laboratory | Minnesota Department of Health, Public Health Laboratory | Matt Plumb, Jacob Garfin, and Xiong Wang |
| EPI_ISL_514652 | M Health Fairview St. Joseph's Hospital | Minnesota Department of Health, Public Health Laboratory | Matt Plumb, Jacob Garfin, and Xiong Wang |
| EPI_ISL_514653, EPI_ISL_514654, EPI_ISL_514655, EPI_ISL_514656 | Mayo Clinic & Mayo Clinic Laboratories | Minnesota Department of Health, Public Health Laboratory | Matt Plumb, Jacob Garfin, and Xiong Wang |
| EPI_ISL_514657 | M Health Fairview St. Joseph's Hospital | Minnesota Department of Health, Public Health Laboratory | Matt Plumb, Jacob Garfin, and Xiong Wang |
| EPI_ISL_514658, EPI_ISL_514659, EPI_ISL_514660, EPI_ISL_514661, EPI_ISL_514662, EPI_ISL_514663, EPI_ISL_514664, EPI_ISL_514665, EPI_ISL_514666, EPI_ISL_514667, EPI_ISL_514668, EPI_ISL_514669, EPI_ISL_514670, EPI_ISL_514671, EPI_ISL_514672 | | | |
| see above | Minnesota Department of Health, Public Health Laboratory | Minnesota Department of Health, Public Health Laboratory | Matt Plumb, Jacob Garfin, and Xiong Wang |
| EPI_ISL_514673 | Nevada State Public Health Laboratory | Nevada State Public Health Laboratory | Richard Tillet, Joel R. Sevinsky, Paul Hartley, Heather Kerwin, David Jackson, Subhash C. Verma, Cyprian Rosetto, Andrew Gorzalski, Chris Laverdure, Natalie Crawford, Stephanie Van Hooser, and Mark Pandori |
| EPI_ISL_514674 | Nevada State Public Health Laboratory | Nevada State Public Health Laboratory | Richard Tillet, Joel R. Sevinsky, Paul Hartley, Heather Kerwin, David Jackson, Subhash C. Verma, Cyprian Rossetto, Andrew Gorzalski, Chris Laverdure, Natalie Crawford, Stephanie Van Hooser, and Mark Pandori |
| EPI_ISL_514750 | Pirogov Russian National Research Medical University | Pirogov Russian National Research Medical University, Research and Development | Blagodatskikh, K.A. |
| EPI_ISL_514751 | CoronaNet Lab- TaskForce Regione Campania, CEINGE Biotecnologie Avanzate, Via G. Salvatore | CoronaNet Lab- TaskForce Regione Campania, CEINGE Biotecnologie Avanzate, Via G. Salvatore | Zollo,M., Ferrucci,V., Kong,Dy., Asadzadeh,F., Marrone,L.,Siciliano,R., Cerino,R., Fusco,G., Comegna,M., Boccia,A.,Viscardi,M., Borriello,G., Brandi,S., Tiberio,C., Atripaldi,L.,Paolella,G., Castaldo,G., Pascarella,S., Bianchi,M., Chiariotti,L.,Lee,J.M., Jung,J.H., Yun,K.S. and Kim,H.Y. |
| EPI_ISL_514753 | Yaftabad Hospital, COVID Lab Center | University of Tabriz | Shahabzadeh,Z., Hosseinzadeh Gharajeh,N., Hashemian,S.M. and Barati,O. |
| EPI_ISL_514756, EPI_ISL_514757, EPI_ISL_514758, EPI_ISL_514759, EPI_ISL_514760, EPI_ISL_514761, EPI_ISL_514762, EPI_ISL_514763, EPI_ISL_514764, EPI_ISL_514765, EPI_ISL_514766, EPI_ISL_514767, EPI_ISL_514768, EPI_ISL_514769, EPI_ISL_514770, EPI_ISL_514771, EPI_ISL_514772, EPI_ISL_514773, EPI_ISL_514774, EPI_ISL_514775, EPI_ISL_514776, EPI_ISL_514777, EPI_ISL_514778, EPI_ISL_514779, EPI_ISL_514780, EPI_ISL_514781, EPI_ISL_514782, EPI_ISL_514783, EPI_ISL_514784, EPI_ISL_514785, EPI_ISL_514786, EPI_ISL_514787, EPI_ISL_514788, EPI_ISL_514789, EPI_ISL_514790, EPI_ISL_514791, EPI_ISL_514792, EPI_ISL_514793, EPI_ISL_514794, EPI_ISL_514795, EPI_ISL_514796, EPI_ISL_514797, EPI_ISL_514798, EPI_ISL_514799, EPI_ISL_514800, EPI_ISL_514801, EPI_ISL_514802, EPI_ISL_514803, EPI_ISL_514804, EPI_ISL_514805, EPI_ISL_514806, EPI_ISL_514807, EPI_ISL_514808, EPI_ISL_514809, EPI_ISL_514810, EPI_ISL_514811, EPI_ISL_514812, EPI_ISL_514813, EPI_ISL_514814, EPI_ISL_514815, EPI_ISL_514816, EPI_ISL_514817, EPI_ISL_514818, EPI_ISL_514819, EPI_ISL_514820, EPI_ISL_514821, EPI_ISL_514822, EPI_ISL_514823, EPI_ISL_514824, EPI_ISL_514825, EPI_ISL_514826, EPI_ISL_514827, EPI_ISL_514828, EPI_ISL_514829, EPI_ISL_514830, EPI_ISL_514831, EPI_ISL_514832, EPI_ISL_514833, EPI_ISL_514834, EPI_ISL_514835, EPI_ISL_514836, EPI_ISL_514837, EPI_ISL_514838, EPI_ISL_514839, EPI_ISL_514840, EPI_ISL_514841, EPI_ISL_514842, EPI_ISL_514843, EPI_ISL_514844, EPI_ISL_514845, EPI_ISL_514846, EPI_ISL_514847, EPI_ISL_514848, EPI_ISL_514849, EPI_ISL_514850, EPI_ISL_514851, EPI_ISL_514852, EPI_ISL_514853, EPI_ISL_514854, EPI_ISL_514855, EPI_ISL_514856, EPI_ISL_514857, EPI_ISL_514858, EPI_ISL_514859, EPI_ISL_514860, EPI_ISL_514861, EPI_ISL_514862, EPI_ISL_514863, EPI_ISL_514864, EPI_ISL_514865, EPI_ISL_514866, EPI_ISL_514867, EPI_ISL_514868, EPI_ISL_514869, EPI_ISL_514870, EPI_ISL_514871, EPI_ISL_514872, EPI_ISL_514873, EPI_ISL_514874, EPI_ISL_514875, EPI_ISL_514876, EPI_ISL_514877, EPI_ISL_514878, EPI_ISL_514879, EPI_ISL_514880, EPI_ISL_514881, EPI_ISL_514882, EPI_ISL_514883, EPI_ISL_514884, EPI_ISL_514885, EPI_ISL_514886, EPI_ISL_514887, EPI_ISL_514888, EPI_ISL_514889, EPI_ISL_514890, EPI_ISL_514891, EPI_ISL_514892, EPI_ISL_514893, EPI_ISL_514894, EPI_ISL_514895, EPI_ISL_514896, EPI_ISL_514897, EPI_ISL_514898, EPI_ISL_514899, EPI_ISL_514900, EPI_ISL_514901, EPI_ISL_514902, EPI_ISL_514903, EPI_ISL_514904, EPI_ISL_514905, EPI_ISL_514906, EPI_ISL_514907, EPI_ISL_514908, EPI_ISL_514909, EPI_ISL_514910, EPI_ISL_514911, EPI_ISL_514912, EPI_ISL_514913, EPI_ISL_514914, EPI_ISL_514915, EPI_ISL_514916, EPI_ISL_514917, EPI_ISL_514918, EPI_ISL_514919, EPI_ISL_514920, EPI_ISL_514921, EPI_ISL_514922, EPI_ISL_514923, EPI_ISL_514924, EPI_ISL_514925, EPI_ISL_514926, EPI_ISL_514927, EPI_ISL_514928, EPI_ISL_514929, EPI_ISL_514930, EPI_ISL_514931, EPI_ISL_514932, EPI_ISL_514933, EPI_ISL_514934, EPI_ISL_514935, EPI_ISL_514936, EPI_ISL_514937, EPI_ISL_514938, EPI_ISL_514939, EPI_ISL_514940, EPI_ISL_514941, EPI_ISL_514942, EPI_ISL_514943, EPI_ISL_514944, EPI_ISL_514945, EPI_ISL_514946, EPI_ISL_514947, EPI_ISL_514948, EPI_ISL_514949, EPI_ISL_514950, EPI_ISL_514951, EPI_ISL_514952, EPI_ISL_514953, EPI_ISL_514954, EPI_ISL_514955, EPI_ISL_514956, EPI_ISL_514957, EPI_ISL_514958, EPI_ISL_514959, EPI_ISL_514960, EPI_ISL_514961, EPI_ISL_514962, EPI_ISL_514963, EPI_ISL_514964, EPI_ISL_514965, EPI_ISL_514966, EPI_ISL_514967, EPI_ISL_514968, EPI_ISL_514969, EPI_ISL_514970, EPI_ISL_514971, EPI_ISL_514972, EPI_ISL_514973, EPI_ISL_514974, EPI_ISL_514975, EPI_ISL_514976, EPI_ISL_514977, EPI_ISL_514978, EPI_ISL_514979, EPI_ISL_514980, EPI_ISL_514981, EPI_ISL_514982, EPI_ISL_514983, EPI_ISL_514984, EPI_ISL_514985, EPI_ISL_514986, EPI_ISL_514987, EPI_ISL_514988, EPI_ISL_514989, EPI_ISL_514990, EPI_ISL_514991, EPI_ISL_514992, EPI_ISL_514993, EPI_ISL_514994, EPI_ISL_514995, EPI_ISL_514996, EPI_ISL_514997, EPI_ISL_514998, EPI_ISL_514999, EPI_ISL_515000, EPI_ISL_515001, EPI_ISL_515002, EPI_ISL_515003, EPI_ISL_515004, EPI_ISL_515005, EPI_ISL_515006, EPI_ISL_515007, EPI_ISL_515008, EPI_ISL_515009, EPI_ISL_515010, EPI_ISL_515011, EPI_ISL_515012, EPI_ISL_515013, EPI_ISL_515014, EPI_ISL_515015, EPI_ISL_515016, EPI_ISL_515017, EPI_ISL_515018, EPI_ISL_515019, EPI_ISL_515020, EPI_ISL_515021, EPI_ISL_515022, EPI_ISL_515023, EPI_ISL_515024, EPI_ISL_515025, EPI_ISL_515026, EPI_ISL_515027, EPI_ISL_515028, EPI_ISL_515029, EPI_ISL_515030, EPI_ISL_515031, EPI_ISL_515032, EPI_ISL_515033, EPI_ISL_515034, EPI_ISL_515035, EPI_ISL_515036, EPI_ISL_515037, EPI_ISL_515038, EPI_ISL_515039, EPI_ISL_515040, EPI_ISL_515041, EPI_ISL_515042, EPI_ISL_515043, EPI_ISL_515044, EPI_ISL_515045, EPI_ISL_515046, EPI_ISL_515047, EPI_ISL_515048, EPI_ISL_515049, EPI_ISL_515050, EPI_ISL_515051, EPI_ISL_515052, EPI_ISL_515053, EPI_ISL_515054 | | | |
| see above | Division of Viral Diseases, Center for Laboratory Control of Infectious Diseases, Korea Centers for Diseases Control and Prevention | Division of Viral Diseases, Center for Laboratory Control of Infectious Diseases, Korea Centers for Diseases Control and Prevention | Jeong-Min Kim, Yoon-Seok Chung, Namjoo Lee, Sang Hee Woo, Hye-Jun Jo, Heui Man Kim, Jun-Sub Kim, Myung Guk Han |
| EPI_ISL_515055, EPI_ISL_515056, EPI_ISL_515057, EPI_ISL_515058, EPI_ISL_515059, EPI_ISL_515060, EPI_ISL_515061, EPI_ISL_515062, EPI_ISL_515063, EPI_ISL_515064, EPI_ISL_515065, EPI_ISL_515066, EPI_ISL_515067, EPI_ISL_515068, EPI_ISL_515069, EPI_ISL_515070, EPI_ISL_515071, EPI_ISL_515072, EPI_ISL_515073, EPI_ISL_515074, EPI_ISL_515075, EPI_ISL_515076, EPI_ISL_515077, EPI_ISL_515078, EPI_ISL_515079, EPI_ISL_515080, EPI_ISL_515081 | | | |
| see above | Department of Clinical Microbiology | GIGA Medical Genomics | Keith Durkin, Maria Artesi, Sebastien Bontems, Raphael Boreux, Cecile Meex, Axelle Chaslain, Celine Fombellida-Lopez, Pierrette Melin, Marie-Pierre Hayette, Vincent Bours. |

| | | | |
|---|---|---|---|
| EPI_ISL_515082, EPI_ISL_515083, EPI_ISL_515084, EPI_ISL_515085, EPI_ISL_515086, EPI_ISL_515087, EPI_ISL_515088, EPI_ISL_515089, EPI_ISL_515090, EPI_ISL_515091, EPI_ISL_515092, EPI_ISL_515093, EPI_ISL_515094, EPI_ISL_515095, EPI_ISL_515096, EPI_ISL_515097, EPI_ISL_515098, EPI_ISL_515099, EPI_ISL_515100, EPI_ISL_515101, EPI_ISL_515102, EPI_ISL_515103, EPI_ISL_515104, EPI_ISL_515105, EPI_ISL_515106, EPI_ISL_515107, EPI_ISL_515108, EPI_ISL_515109, EPI_ISL_515110, EPI_ISL_515111, EPI_ISL_515112 | | | |
| see above | Department of Biochemistry, Cell and Molecular Biology | WACCBIP, University of Ghana | Ngoi,J.M., Quashie,P., Morang'a,C.M., Amuzu,D.S., Adu,B., Kumordjie,S., Eshun,M., Boatemaa,L., Magnussen,V., Kotey,E., Tei-Maya,F., Arjarquah,A., Mutungi,J.K., Bediako,Y., Asante,I., Bonney,E., Kyei,G.B., Bonney,K., Amenga-Etego,L.N., Anang,A.K., Awandare,G.A., Ampofo,W. |
| EPI_ISL_515133, EPI_ISL_515134, EPI_ISL_515135, EPI_ISL_515136, EPI_ISL_515137, EPI_ISL_515138, EPI_ISL_515139, EPI_ISL_515140, EPI_ISL_515141, EPI_ISL_515142, EPI_ISL_515143, EPI_ISL_515144, EPI_ISL_515145, EPI_ISL_515146, EPI_ISL_515147, EPI_ISL_515148, EPI_ISL_515149, EPI_ISL_515150, EPI_ISL_515151, EPI_ISL_515152, EPI_ISL_515153, EPI_ISL_515154, EPI_ISL_515155, EPI_ISL_515156, EPI_ISL_515157, EPI_ISL_515158, EPI_ISL_515159, EPI_ISL_515160, EPI_ISL_515161, EPI_ISL_515162, EPI_ISL_515163, EPI_ISL_515164, EPI_ISL_515165, EPI_ISL_515166, EPI_ISL_515167, EPI_ISL_515168, EPI_ISL_515169, EPI_ISL_515170, EPI_ISL_515171, EPI_ISL_515172, EPI_ISL_515173, EPI_ISL_515174, EPI_ISL_515175, EPI_ISL_515176, EPI_ISL_515177, EPI_ISL_515178, EPI_ISL_515179, EPI_ISL_515180 | | | |
| see above | National Institute for Communicable Diseases of the National Health Laboratory Service | National Institute for Communicable Diseases of the National Health Laboratory Service | Allam M, Ismail A, Khumalo Z, Kwenda S, Mtshali P, Mnyameni F, Mohale T, Bhiman JN |
| EPI_ISL_515181, EPI_ISL_515182, EPI_ISL_515183, EPI_ISL_515184 | Kumasi Centre for Collaborative Research in Tropical Medicine, Kumasi. | Institute of Virology, Charité – Universitätsmedizin Berlin | Augustina Sylverken, Philip El-Duah, Michael Owusu, Julia Schneider, Richmond Yeboah, Richmond Gorman, Eric Adu, Sherihane Aryeetey, Jesse Addo Asamoah,Jörn Beheim-Schwarzbach, Victor Max Corman, Christian Drosten, Richard Phillips. |
| EPI_ISL_515185, EPI_ISL_515186 | Latvijas Infektoloģijas centrs | Latvian Biomedical Research and Study Centre | Ivars Silamiķelis, Kaspars Megnis, Monta Ustinova, Ņikita Zrelovs, Vita Rovīte, Jeļena Storoženko, Tatjana Kolupajeva, Oksana Savicka, Uga Dumpis, Jānis Kloviņš |
| EPI_ISL_515187, EPI_ISL_515188, EPI_ISL_515189, EPI_ISL_515190, EPI_ISL_515191, EPI_ISL_515192, EPI_ISL_515193, EPI_ISL_515194, EPI_ISL_515195 | E. Gulbja Laboratorija | Latvian Biomedical Research and Study Centre | Ivars Silamiķelis, Kaspars Megnis, Monta Ustinova, Ņikita Zrelovs, Vita Rovīte, Mikus Gavars, Dmitrijs Perminovs, Uga Dumpis, Jānis Kloviņš |
| EPI_ISL_515196 | Centrālā laboratorija | Latvian Biomedical Research and Study Centre | Ivars Silamiķelis, Kaspars Megnis, Monta Ustinova, Ņikita Zrelovs, Vita Rovīte, Stella Lapiņa, Jana Osīte, Marta Priedīte, Uga Dumpis, Jānis Kloviņš |
| EPI_ISL_515197, EPI_ISL_515198, EPI_ISL_515199, EPI_ISL_515200, EPI_ISL_515201, EPI_ISL_515202, EPI_ISL_515203, EPI_ISL_515204, EPI_ISL_515205, EPI_ISL_515206, EPI_ISL_515207, EPI_ISL_515208, EPI_ISL_515209, EPI_ISL_515210, EPI_ISL_515211, EPI_ISL_515212, EPI_ISL_515213, EPI_ISL_515214, EPI_ISL_515215, EPI_ISL_515216, EPI_ISL_515217, EPI_ISL_515218, EPI_ISL_515219, EPI_ISL_515220, EPI_ISL_515221, EPI_ISL_515222, EPI_ISL_515223, EPI_ISL_515224, EPI_ISL_515225, EPI_ISL_515226, EPI_ISL_515227, EPI_ISL_515228, EPI_ISL_515229, EPI_ISL_515230, EPI_ISL_515231, EPI_ISL_515232, EPI_ISL_515233, EPI_ISL_515234, EPI_ISL_515235, EPI_ISL_515236, EPI_ISL_515237, EPI_ISL_515238, EPI_ISL_515239, EPI_ISL_515240, EPI_ISL_515241, EPI_ISL_515242, EPI_ISL_515243, EPI_ISL_515244, EPI_ISL_515245, EPI_ISL_515246 | | | |
| see above | Laboratoire de microbiologie, Hopital de Verdun | Smith Laboratory, Centre de Recherche CHU Sainte-Justine | Martin Smith, Marieke Rozendaal, Ivan Pavlov |
| EPI_ISL_515247, EPI_ISL_515248, EPI_ISL_515249 | Kumasi Centre for Collaborative Research in Tropical Medicine, Kumasi. | Institute of Virology, Charité – Universitätsmedizin Berlin | Augustina Sylverken, Philip El-Duah, Michael Owusu, Julia Schneider, Richmond Yeboah, Richmond Gorman, Eric Adu, Sherihane Aryeetey, Jesse Addo Asamoah,Jörn Beheim-Schwarzbach, Victor Max Corman, Christian Drosten, Richard Phillips. |
| EPI_ISL_515250, EPI_ISL_515251, EPI_ISL_515252, EPI_ISL_515253, EPI_ISL_515254, EPI_ISL_515255, EPI_ISL_515256, EPI_ISL_515257, EPI_ISL_515258, EPI_ISL_515259, EPI_ISL_515260, EPI_ISL_515261 | | | |
| see above | Texas Department of State Health Services | Texas Department of State Health Services | Rashmi Tuladhar, Bonnie Oh, Cara Akrout, Jenny Zhang, Maliha Rahman, Anita Pokharel, Myong Koag, Chun Wang, Rachel Lee, Grace Kubin |
| EPI_ISL_515262 | Minnesota Department of Health, Public Health Laboratory | Minnesota Department of Health, Public Health Laboratory | Matt Plumb, Jacob Garfin, and Xiong Wang |
| EPI_ISL_515263, EPI_ISL_515264, EPI_ISL_515265, EPI_ISL_515266 | Mayo Clinic & Mayo Clinic Laboratories | Minnesota Department of Health, Public Health Laboratory | Matt Plumb, Jacob Garfin, and Xiong Wang |
| EPI_ISL_515267, EPI_ISL_515268 | M Health Fairview | Minnesota Department of Health, Public Health Laboratory | Matt Plumb, Jacob Garfin, and Xiong Wang |
| EPI_ISL_515269 | Minnesota Department of Health, Public Health Laboratory | Minnesota Department of Health, Public Health Laboratory | Matt Plumb, Jacob Garfin, and Xiong Wang |
| EPI_ISL_515270, EPI_ISL_515271, EPI_ISL_515272, EPI_ISL_515273, EPI_ISL_515274, EPI_ISL_515275, EPI_ISL_515276, EPI_ISL_515277, EPI_ISL_515278, EPI_ISL_515279, EPI_ISL_515280, EPI_ISL_515281, EPI_ISL_515282, EPI_ISL_515283, EPI_ISL_515284, EPI_ISL_515285, EPI_ISL_515286 | | | |
| see above | University of Washington Virology Lab | University of Washington Virology Lab | Pavitra Roychoudhury, Hong Xie, Lasata Shrestha, Amin Addetia, Truong Nguyen, Victoria M Rachleff, Meei-Li Huang, Keith R Jerome, Alexander Greninger |
| EPI_ISL_515287, EPI_ISL_515288, EPI_ISL_515289, EPI_ISL_515290, EPI_ISL_515291 | National Institute of Health. Department of medical Sciences, Ministry of Public Health, Thailand | National Institute of Health. Department of medical Sciences, Ministry of Public Health, Thailand | Pilailuk,Okada; Siripaporn,Phuygun; Thanutsapa,Thanadachakul; Sittiporn,Parnmen;Warawan,Wongboot; Sunthareeya,Waicharoen; Malinee,Chittaganpitch |
| EPI_ISL_515292 | Ramkhamhaeng Hospital | National Institute of Health. Department of medical Sciences, Ministry of Public Health, Thailand | Pilailuk,Okada; Siripaporn,Phuygun; Thanutsapa,Thanadachakul; Sittiporn,Parnmen;Warawan,Wongboot; Sunthareeya,Waicharoen; Malinee,Chittaganpitch |
| EPI_ISL_515293, EPI_ISL_515294, EPI_ISL_515295, EPI_ISL_515296, EPI_ISL_515297, EPI_ISL_515298, EPI_ISL_515299, EPI_ISL_515300, EPI_ISL_515301, EPI_ISL_515302, EPI_ISL_515303, EPI_ISL_515304, EPI_ISL_515305, EPI_ISL_515306, EPI_ISL_515307, EPI_ISL_515308, EPI_ISL_515309, EPI_ISL_515310, EPI_ISL_515311, EPI_ISL_515312, EPI_ISL_515313, EPI_ISL_515314, EPI_ISL_515315, EPI_ISL_515316, EPI_ISL_515317, EPI_ISL_515318, EPI_ISL_515319, EPI_ISL_515320, EPI_ISL_515321, EPI_ISL_515322, EPI_ISL_515323, EPI_ISL_515324, EPI_ISL_515325, EPI_ISL_515326, EPI_ISL_515327, EPI_ISL_515328, EPI_ISL_515329, EPI_ISL_515330, EPI_ISL_515331, EPI_ISL_515332, EPI_ISL_515333, EPI_ISL_515334, EPI_ISL_515335, EPI_ISL_515336, EPI_ISL_515337, EPI_ISL_515338, EPI_ISL_515339, EPI_ISL_515340, EPI_ISL_515341, EPI_ISL_515342, EPI_ISL_515343, EPI_ISL_515344, EPI_ISL_515345, EPI_ISL_515346, EPI_ISL_515347, EPI_ISL_515348, EPI_ISL_515349, EPI_ISL_515350, EPI_ISL_515351, EPI_ISL_515352, EPI_ISL_515353, EPI_ISL_515354, EPI_ISL_515355, EPI_ISL_515356, EPI_ISL_515357, EPI_ISL_515358, EPI_ISL_515359, EPI_ISL_515360, EPI_ISL_515361, EPI_ISL_515362, EPI_ISL_515363, EPI_ISL_515364, EPI_ISL_515365, EPI_ISL_515366, EPI_ISL_515367, EPI_ISL_515368, EPI_ISL_515369, EPI_ISL_515370, EPI_ISL_515371, EPI_ISL_515372, EPI_ISL_515373, EPI_ISL_515374, EPI_ISL_515375, EPI_ISL_515376, EPI_ISL_515377, EPI_ISL_515378, EPI_ISL_515379, EPI_ISL_515380, EPI_ISL_515381, EPI_ISL_515382, EPI_ISL_515383, EPI_ISL_515384, EPI_ISL_515385, EPI_ISL_515386, EPI_ISL_515387, EPI_ISL_515388, EPI_ISL_515389, EPI_ISL_515390, EPI_ISL_515391, EPI_ISL_515392, EPI_ISL_515393, EPI_ISL_515394, EPI_ISL_515395, EPI_ISL_515396, EPI_ISL_515397, EPI_ISL_515398, EPI_ISL_515399, EPI_ISL_515400, EPI_ISL_515401, EPI_ISL_515402, EPI_ISL_515403, EPI_ISL_515404, EPI_ISL_515405, EPI_ISL_515406, EPI_ISL_515407, EPI_ISL_515408, EPI_ISL_515409, EPI_ISL_515410, EPI_ISL_515411, EPI_ISL_515412, EPI_ISL_515413, EPI_ISL_515414, EPI_ISL_515415, EPI_ISL_515416, EPI_ISL_515417, EPI_ISL_515418, EPI_ISL_515419, EPI_ISL_515420, EPI_ISL_515421, EPI_ISL_515422, EPI_ISL_515423, EPI_ISL_515424, EPI_ISL_515425, EPI_ISL_515426, EPI_ISL_515427, EPI_ISL_515428, EPI_ISL_515429, EPI_ISL_515430, EPI_ISL_515431, EPI_ISL_515432, EPI_ISL_515433, EPI_ISL_515434, EPI_ISL_515435, EPI_ISL_515436, EPI_ISL_515437, EPI_ISL_515438, EPI_ISL_515439, EPI_ISL_515440, EPI_ISL_515441, EPI_ISL_515442, EPI_ISL_515443, EPI_ISL_515444, EPI_ISL_515445, EPI_ISL_515446, EPI_ISL_515447, EPI_ISL_515448, EPI_ISL_515449, EPI_ISL_515450, EPI_ISL_515451, EPI_ISL_515452, EPI_ISL_515453, EPI_ISL_515454, EPI_ISL_515455, EPI_ISL_515456, EPI_ISL_515457, EPI_ISL_515458, EPI_ISL_515459, EPI_ISL_515460, EPI_ISL_515461, EPI_ISL_515462 | | | |
| see above | Nevada State Public Health Laboratory | Nevada State Public Health Laboratory | Richard Tillett, Joel R. Sevinsky, Paul Hartley, Heather Kerwin, David Jackson, Subhash C. Verma, Cyprian Rossetto, Andrew Gorzalski, Chris Laverdure, Natalie Crawford, Stephanie Van Hooser, and Mark Pandori |
| EPI_ISL_515463 | National Institute of Health. Department of medical Sciences, Ministry of Public Health, Thailand | National Institute of Health. Department of medical Sciences, Ministry of Public Health, Thailand | Pilailuk,Okada; Siripaporn,Phuygun; Thanutsapa,Thanadachakul; Sittiporn,Parnmen;Warawan,Wongboot; Sunthareeya,Waicharoen; Malinee,Chittaganpitch |
| EPI_ISL_515464 | Siriraj Hospital | National Institute of Health. Department of medical Sciences, Ministry of Public Health, Thailand | Pilailuk,Okada; Siripaporn,Phuygun; Thanutsapa,Thanadachakul; Sittiporn,Parnmen;Warawan,Wongboot; Sunthareeya,Waicharoen; Malinee,Chittaganpitch |
| EPI_ISL_515465 | Ramathibodi Hospital | National Institute of Health. Department of medical Sciences, Ministry of Public Health, Thailand | Pilailuk,Okada; Siripaporn,Phuygun; Thanutsapa,Thanadachakul; Sittiporn,Parnmen;Warawan,Wongboot; Sunthareeya,Waicharoen; Malinee,Chittaganpitch |
| EPI_ISL_515466 | Discovery DNA | Discovery DNA | Dustin Hittel, Marina Kerr, Leo Dimnik, Desmond Koo, Alice Li, Aneal Khan |
| EPI_ISL_515467, EPI_ISL_515468 | Ramathibodi Hospital | National Institute of Health. Department of medical Sciences, Ministry of Public Health, Thailand | Pilailuk,Okada; Siripaporn,Phuygun; Thanutsapa,Thanadachakul; Sittiporn,Parnmen;Warawan,Wongboot; Sunthareeya,Waicharoen; Malinee,Chittaganpitch |
| EPI_ISL_515469 | Bamrasnaradura hospital | National Institute of Health. Department of medical Sciences, Ministry of Public Health, Thailand | Pilailuk,Okada; Siripaporn,Phuygun; Thanutsapa,Thanadachakul; Sittiporn,Parnmen;Warawan,Wongboot; Sunthareeya,Waicharoen; Malinee,Chittaganpitch |
| EPI_ISL_515470 | National Institute of Health. Department of medical Sciences, Ministry of Public Health, Thailand | National Institute of Health. Department of medical Sciences, Ministry of Public Health, Thailand | Pilailuk,Okada; Siripaporn,Phuygun; Thanutsapa,Thanadachakul; Sittiporn,Parnmen;Warawan,Wongboot; Sunthareeya,Waicharoen; Malinee,Chittaganpitch |
| EPI_ISL_515471, EPI_ISL_515472, EPI_ISL_515473, EPI_ISL_515474, EPI_ISL_515475, EPI_ISL_515476, EPI_ISL_515477 | National Institute of Health, Department of Medical Sciences, Ministry of Public Health, Thailand | National Institute of Health, Department of Medical Sciences, Ministry of Public Health, Thailand | Pilailuk Okada; Siripaporn Phuygun; Thanutsapa Thanadachakul; Sittiporn Parnmen; Warawan Wongboot; Sunthareeya Waicharoen; Malinee Chittaganpitch |
| EPI_ISL_515520 | Hospital Municipal do Tatuape Carmino Caricchio | Instituto Adolfo Lutz, Interdiciplinary Procedures Center, Strategic Laboratory | Claudio Tavares Sacchi, Claudia Regina Gonçalves, Erica Valessa Ramos Gomes |
| EPI_ISL_515521 | Hospital Municipal Dr Waldemar Tebaldi | Instituto Adolfo Lutz, Interdiciplinary Procedures Center, Strategic Laboratory | Claudio Tavares Sacchi, Claudia Regina Gonçalves, Erica Valessa Ramos Gomes |
| EPI_ISL_515522 | UPA 24HS de Itatiba | Instituto Adolfo Lutz, Interdiciplinary Procedures Center, Strategic Laboratory | Claudio Tavares Sacchi, Claudia Regina Gonçalves, Erica Valessa Ramos Gomes |
| EPI_ISL_515523, EPI_ISL_515524 | PS Municipal Dr Lauro Ribas Braga | Instituto Adolfo Lutz, Interdiciplinary Procedures Center, Strategic Laboratory | Claudio Tavares Sacchi, Claudia Regina Gonçalves, Erica Valessa Ramos Gomes |
| EPI_ISL_515525 | National Influenza Center - Instituto Adolfo Lutz | Instituto Adolfo Lutz, Interdiciplinary Procedures Center, Strategic Laboratory | Claudio Tavares Sacchi, Claudia Regina Gonçalves, Erica Valessa Ramos Gomes |
| EPI_ISL_515526 | Hospital Municipal do Tatuape Carmino Caricchio | Instituto Adolfo Lutz, Interdiciplinary Procedures Center, Strategic Laboratory | Claudio Tavares Sacchi, Claudia Regina Gonçalves, Erica Valessa Ramos Gomes |
| EPI_ISL_515527 | Hospital Santa Clara | Instituto Adolfo Lutz, Interdiciplinary Procedures Center, Strategic Laboratory | Claudio Tavares Sacchi, Claudia Regina Gonçalves, Erica Valessa Ramos Gomes |
| EPI_ISL_515528 | Hospital Sao Paulo de Ensino da Unifesp | Instituto Adolfo Lutz, Interdiciplinary Procedures Center, Strategic Laboratory | Claudio Tavares Sacchi, Claudia Regina Gonçalves, Erica Valessa Ramos Gomes |
| EPI_ISL_515529 | Pronto Socorro Municipal Julio Tupy | Instituto Adolfo Lutz, Interdiciplinary Procedures Center, Strategic Laboratory | Claudio Tavares Sacchi, Claudia Regina Gonçalves, Erica Valessa Ramos Gomes |
| EPI_ISL_515541 | Hospital Montemagno | Instituto Adolfo Lutz, Interdiciplinary Procedures Center, Strategic Laboratory | Claudio Tavares Sacchi, Claudia Regina Gonçalves, Erica Valessa Ramos Gomes |
| EPI_ISL_515542 | Vigilância Epidemiológica de Leme | Instituto Adolfo Lutz, Interdiciplinary Procedures Center, Strategic Laboratory | Claudio Tavares Sacchi, Claudia Regina Gonçalves, Erica Valessa Ramos Gomes |
| EPI_ISL_515543 | Serviço de Vigilância Sanitária e Epidemiológica | Instituto Adolfo Lutz, Interdiciplinary Procedures Center, Strategic Laboratory | Claudio Tavares Sacchi, Claudia Regina Gonçalves, Erica Valessa Ramos Gomes |
| EPI_ISL_515544 | Ama Dr Jose Soares Hungria | Instituto Adolfo Lutz, Interdiciplinary Procedures Center, Strategic Laboratory | Claudio Tavares Sacchi, Claudia Regina Gonçalves, Erica Valessa Ramos Gomes |
| EPI_ISL_515545 | Hospital Sao Paulo de Ensino da Unifesp | Instituto Adolfo Lutz, Interdiciplinary Procedures Center, Strategic Laboratory | Claudio Tavares Sacchi, Claudia Regina Gonçalves, Erica Valessa Ramos Gomes |
| EPI_ISL_515546 | Hospital Municipal do Tatuape Carmino Caricchio | Instituto Adolfo Lutz, Interdiciplinary Procedures Center, Strategic Laboratory | Claudio Tavares Sacchi, Claudia Regina Gonçalves, Erica Valessa Ramos Gomes |
| EPI_ISL_515547 | Centro Medico da Policia Militar do Estado de Sao Paulo | Instituto Adolfo Lutz, Interdiciplinary Procedures Center, Strategic Laboratory | Claudio Tavares Sacchi, Claudia Regina Gonçalves, Erica Valessa Ramos Gomes |
| EPI_ISL_515548 | Hospital Municipal Dr. Jose Soares Hungria | Instituto Adolfo Lutz, Interdiciplinary Procedures Center, Strategic Laboratory | Claudio Tavares Sacchi, Claudia Regina Gonçalves, Erica Valessa Ramos Gomes |
| EPI_ISL_515550 | UPA Vila Santa Catarina | Instituto Adolfo Lutz, Interdiciplinary Procedures Center, Strategic Laboratory | Claudio Tavares Sacchi, Claudia Regina Gonçalves, Erica Valessa Ramos Gomes |
| EPI_ISL_515551, EPI_ISL_515552 | Hospital Municipal do Tatuape Carmino Caricchio | Instituto Adolfo Lutz, Interdiciplinary Procedures Center, Strategic Laboratory | Claudio Tavares Sacchi, Claudia Regina Gonçalves, Erica Valessa Ramos Gomes |
| EPI_ISL_515553 | Hospital Municipal Dr. Ignacio Proença de Gouvea | Instituto Adolfo Lutz, Interdiciplinary Procedures Center, Strategic Laboratory | Claudio Tavares Sacchi, Claudia Regina Gonçalves, Erica Valessa Ramos Gomes |
| EPI_ISL_515554 | Pronto Socorro Municipal de Perus | Instituto Adolfo Lutz, Interdiciplinary Procedures Center, Strategic Laboratory | Claudio Tavares Sacchi, Claudia Regina Gonçalves, Erica Valessa Ramos Gomes |
| EPI_ISL_515555 | Hospital Geral de Vila Nova Cachoeirinha | Instituto Adolfo Lutz, Interdiciplinary Procedures Center, Strategic Laboratory | Claudio Tavares Sacchi, Claudia Regina Gonçalves, Erica Valessa Ramos Gomes |
| EPI_ISL_515557 | Hospital Municipal Dr. Moysés Deutsch | Instituto Adolfo Lutz, Interdiciplinary Procedures Center, Strategic Laboratory | Claudio Tavares Sacchi, Claudia Regina Gonçalves, Erica Valessa Ramos Gomes |
| EPI_ISL_515559, EPI_ISL_515560 | Hospital Sao Paulo de Ensino da Unifesp | Instituto Adolfo Lutz, Interdiciplinary Procedures Center, Strategic Laboratory | Claudio Tavares Sacchi, Claudia Regina Gonçalves, Erica Valessa Ramos Gomes |

| | | | |
|---|---|---|---|
| EPI_ISL_515561 | Hospital Montemagno | Instituto Adolfo Lutz, Interdiciplinary Procedures Center, Strategic Laboratory | Claudio Tavares Sacchi, Claudia Regina Gonçalves, Erica Valessa Ramos Gomes |
| EPI_ISL_515562 | Hospital Municipal Doutor Alexandre Zaio | Instituto Adolfo Lutz, Interdiciplinary Procedures Center, Strategic Laboratory | Claudio Tavares Sacchi, Claudia Regina Gonçalves, Erica Valessa Ramos Gomes |
| EPI_ISL_515563 | Hospital Municipal Dr. Jose Soares Hungria | Instituto Adolfo Lutz, Interdiciplinary Procedures Center, Strategic Laboratory | Claudio Tavares Sacchi, Claudia Regina Gonçalves, Erica Valessa Ramos Gomes |
| EPI_ISL_515564 | Hosp. Municipal Prof. Dr. Alípio Corrêa Netto | Instituto Adolfo Lutz, Interdiciplinary Procedures Center, Strategic Laboratory | Claudio Tavares Sacchi, Claudia Regina Gonçalves, Erica Valessa Ramos Gomes |
| EPI_ISL_515565 | Hospital do Servidor Público Estadual Francisco Morato de Oliveira | Instituto Adolfo Lutz, Interdiciplinary Procedures Center, Strategic Laboratory | Claudio Tavares Sacchi, Claudia Regina Gonçalves, Erica Valessa Ramos Gomes |
| EPI_ISL_515566 | PS Municipal Dr Lauro Ribas Braga | Instituto Adolfo Lutz, Interdiciplinary Procedures Center, Strategic Laboratory | Claudio Tavares Sacchi, Claudia Regina Gonçalves, Erica Valessa Ramos Gomes |
| EPI_ISL_515568, EPI_ISL_515569, EPI_ISL_515570, EPI_ISL_515571, EPI_ISL_515572, EPI_ISL_515573, EPI_ISL_515574, EPI_ISL_515575, EPI_ISL_515576, EPI_ISL_515577, EPI_ISL_515578, EPI_ISL_515579, EPI_ISL_515580, EPI_ISL_515581, EPI_ISL_515582, EPI_ISL_515583, EPI_ISL_515584, EPI_ISL_515585, EPI_ISL_515586, EPI_ISL_515587, EPI_ISL_515588, EPI_ISL_515589, EPI_ISL_515590, EPI_ISL_515591, EPI_ISL_515592, EPI_ISL_515593, EPI_ISL_515594, EPI_ISL_515595, EPI_ISL_515596, EPI_ISL_515597, EPI_ISL_515598, EPI_ISL_515599, EPI_ISL_515600, EPI_ISL_515601, EPI_ISL_515602, EPI_ISL_515603, EPI_ISL_515604, EPI_ISL_515605, EPI_ISL_515606, EPI_ISL_515607, EPI_ISL_515608, EPI_ISL_515609, EPI_ISL_515610, EPI_ISL_515611, EPI_ISL_515612, EPI_ISL_515613, EPI_ISL_515614, EPI_ISL_515615, EPI_ISL_515616, EPI_ISL_515617, EPI_ISL_515618, EPI_ISL_515619, EPI_ISL_515620, EPI_ISL_515621, EPI_ISL_515622, EPI_ISL_515623, EPI_ISL_515624, EPI_ISL_515625, EPI_ISL_515626, EPI_ISL_515627, EPI_ISL_515628, EPI_ISL_515629, EPI_ISL_515630, EPI_ISL_515631, EPI_ISL_515632, EPI_ISL_515633, EPI_ISL_515634, EPI_ISL_515635, EPI_ISL_515636, EPI_ISL_515637, EPI_ISL_515638, EPI_ISL_515639, EPI_ISL_515640, EPI_ISL_515641, EPI_ISL_515642, EPI_ISL_515643, EPI_ISL_515644, EPI_ISL_515645, EPI_ISL_515646, EPI_ISL_515647, EPI_ISL_515648, EPI_ISL_515649, EPI_ISL_515650, EPI_ISL_515651, EPI_ISL_515652, EPI_ISL_515653, EPI_ISL_515654, EPI_ISL_515655, EPI_ISL_515656, EPI_ISL_515657, EPI_ISL_515658, EPI_ISL_515659, EPI_ISL_515660, EPI_ISL_515661, EPI_ISL_515662, EPI_ISL_515663, EPI_ISL_515664, EPI_ISL_515665, EPI_ISL_515666, EPI_ISL_515667, EPI_ISL_515668, EPI_ISL_515669, EPI_ISL_515670, EPI_ISL_515671, EPI_ISL_515672, EPI_ISL_515673, EPI_ISL_515674, EPI_ISL_515675, EPI_ISL_515676, EPI_ISL_515677, EPI_ISL_515678, EPI_ISL_515679, EPI_ISL_515680, EPI_ISL_515681, EPI_ISL_515682, EPI_ISL_515683, EPI_ISL_515684, EPI_ISL_515685, EPI_ISL_515686, EPI_ISL_515687, EPI_ISL_515688, EPI_ISL_515689, EPI_ISL_515690, EPI_ISL_515691, EPI_ISL_515692, EPI_ISL_515693, EPI_ISL_515694, EPI_ISL_515695, EPI_ISL_515696, EPI_ISL_515697, EPI_ISL_515698, EPI_ISL_515699, EPI_ISL_515700, EPI_ISL_515701, EPI_ISL_515702, EPI_ISL_515703, EPI_ISL_515704, EPI_ISL_515705, EPI_ISL_515706, EPI_ISL_515707, EPI_ISL_515708, EPI_ISL_515709, EPI_ISL_515710, EPI_ISL_515711, EPI_ISL_515712, EPI_ISL_515713, EPI_ISL_515714, EPI_ISL_515715, EPI_ISL_515716, EPI_ISL_515717, EPI_ISL_515718, EPI_ISL_515719, EPI_ISL_515720, EPI_ISL_515721, EPI_ISL_515722, EPI_ISL_515723, EPI_ISL_515724, EPI_ISL_515725, EPI_ISL_515726, EPI_ISL_515727, EPI_ISL_515728, EPI_ISL_515729, EPI_ISL_515730, EPI_ISL_515731, EPI_ISL_515732, EPI_ISL_515733, EPI_ISL_515734, EPI_ISL_515735, EPI_ISL_515736, EPI_ISL_515737, EPI_ISL_515738, EPI_ISL_515739, EPI_ISL_515740, EPI_ISL_515741, EPI_ISL_515742, EPI_ISL_515743, EPI_ISL_515744, EPI_ISL_515745, EPI_ISL_515746, EPI_ISL_515747, EPI_ISL_515748, EPI_ISL_515749, EPI_ISL_515750, EPI_ISL_515751, EPI_ISL_515752, EPI_ISL_515753, EPI_ISL_515754, EPI_ISL_515755, EPI_ISL_515756, EPI_ISL_515757, EPI_ISL_515758, EPI_ISL_515759, EPI_ISL_515760, EPI_ISL_515761, EPI_ISL_515762, EPI_ISL_515763, EPI_ISL_515764, EPI_ISL_515765, EPI_ISL_515766, EPI_ISL_515767, EPI_ISL_515768, EPI_ISL_515769, EPI_ISL_515770, EPI_ISL_515771, EPI_ISL_515772, EPI_ISL_515773, EPI_ISL_515774, EPI_ISL_515775, EPI_ISL_515776, EPI_ISL_515777, EPI_ISL_515778, EPI_ISL_515779, EPI_ISL_515780, EPI_ISL_515781, EPI_ISL_515782, EPI_ISL_515783, EPI_ISL_515784, EPI_ISL_515785, EPI_ISL_515786, EPI_ISL_515787, EPI_ISL_515788, EPI_ISL_515789, EPI_ISL_515790, EPI_ISL_515791, EPI_ISL_515792, EPI_ISL_515793, EPI_ISL_515794, EPI_ISL_515795, EPI_ISL_515796, EPI_ISL_515797, EPI_ISL_515798, EPI_ISL_515799 | | | |
| see above | NHLS-IALCH | KRISP, KZN Research Innovation and Sequencing Platform | Giandhari J, Pillay S, Lessells R, Mdlalose K, York D, Khan S, Tegally H, Wilkinson E, de Oliveira T |
| EPI_ISL_515800, EPI_ISL_515801, EPI_ISL_515802, EPI_ISL_515803, EPI_ISL_515804, EPI_ISL_515805, EPI_ISL_515806, EPI_ISL_515807, EPI_ISL_515808, EPI_ISL_515809, EPI_ISL_515810, EPI_ISL_515811, EPI_ISL_515812, EPI_ISL_515813, EPI_ISL_515814, EPI_ISL_515815, EPI_ISL_515816, EPI_ISL_515817, EPI_ISL_515818, EPI_ISL_515819, EPI_ISL_515820, EPI_ISL_515821, EPI_ISL_515822, EPI_ISL_515823, EPI_ISL_515824, EPI_ISL_515825, EPI_ISL_515826, EPI_ISL_515827, EPI_ISL_515828, EPI_ISL_515829, EPI_ISL_515830, EPI_ISL_515831, EPI_ISL_515832, EPI_ISL_515833, EPI_ISL_515834, EPI_ISL_515835, EPI_ISL_515836, EPI_ISL_515837, EPI_ISL_515838, EPI_ISL_515839, EPI_ISL_515840, EPI_ISL_515841, EPI_ISL_515842, EPI_ISL_515843, EPI_ISL_515844, EPI_ISL_515845, EPI_ISL_515846, EPI_ISL_515847, EPI_ISL_515848, EPI_ISL_515849, EPI_ISL_515850, EPI_ISL_515851, EPI_ISL_515852, EPI_ISL_515853, EPI_ISL_515854, EPI_ISL_515855, EPI_ISL_515856, EPI_ISL_515857, EPI_ISL_515858, EPI_ISL_515859, EPI_ISL_515860, EPI_ISL_515861, EPI_ISL_515862, EPI_ISL_515863, EPI_ISL_515864, EPI_ISL_515865, EPI_ISL_515866, EPI_ISL_515867, EPI_ISL_515868, EPI_ISL_515869, EPI_ISL_515870, EPI_ISL_515871, EPI_ISL_515872, EPI_ISL_515873, EPI_ISL_515874, EPI_ISL_515875, EPI_ISL_515876, EPI_ISL_515877, EPI_ISL_515878, EPI_ISL_515879, EPI_ISL_515880, EPI_ISL_515881, EPI_ISL_515882, EPI_ISL_515883, EPI_ISL_515884, EPI_ISL_515885, EPI_ISL_515886, EPI_ISL_515887, EPI_ISL_515888, EPI_ISL_515889, EPI_ISL_515890, EPI_ISL_515891, EPI_ISL_515892, EPI_ISL_515893 | | | |
| see above | Medical Disagnostics Services (MDS) | KRISP, KZN Research Innovation and Sequencing Platform | Giandhari J, Pillay S, Lessells R, ChimukangaraB, Mdlalose K, York D, Khan S, Tegally H, Wilkinson E, de Oliveira T |
| EPI_ISL_515894, EPI_ISL_515895, EPI_ISL_515896, EPI_ISL_515897, EPI_ISL_515898, EPI_ISL_515899, EPI_ISL_515900, EPI_ISL_515901, EPI_ISL_515902, EPI_ISL_515903, EPI_ISL_515904, EPI_ISL_515905, EPI_ISL_515906, EPI_ISL_515907, EPI_ISL_515908, EPI_ISL_515909, EPI_ISL_515910, EPI_ISL_515911, EPI_ISL_515912, EPI_ISL_515913, EPI_ISL_515914, EPI_ISL_515915, EPI_ISL_515916, EPI_ISL_515917, EPI_ISL_515918, EPI_ISL_515919, EPI_ISL_515920, EPI_ISL_515921, EPI_ISL_515922, EPI_ISL_515923, EPI_ISL_515924, EPI_ISL_515925, EPI_ISL_515926, EPI_ISL_515927, EPI_ISL_515928, EPI_ISL_515929 | | | |
| see above | California Department of Public Health | California Department of Public Health | CDPH IDLB COVIDNet |
| EPI_ISL_515932 | DH | Department of Neurovirology, National Institute of Mental Health and Neuroscience (NIMHANS) | Chitra Pattabiraman,Vijayalakshmi Reddy, Harsha PK, Risha Rasheed, Pramada Prasad, Shafeeq S Hameed, Manjunatha Venkataswamy, Anita Desai, Ravi Vasanthapuram |
| EPI_ISL_515933 | BIMS | Department of Neurovirology, National Institute of Mental Health and Neuroscience (NIMHANS) | Chitra Pattabiraman,Vijayalakshmi Reddy, Harsha PK, Risha Rasheed, Pramada Prasad, Shafeeq S Hameed, Manjunatha Venkataswamy, Anita Desai, Ravi Vasanthapuram |
| EPI_ISL_515934, EPI_ISL_515935, EPI_ISL_515936 | DH | Department of Neurovirology, National Institute of Mental Health and Neuroscience (NIMHANS) | Chitra Pattabiraman,Vijayalakshmi Reddy, Harsha PK, Risha Rasheed, Pramada Prasad, Shafeeq S Hameed, Manjunatha Venkataswamy, Anita Desai, Ravi Vasanthapuram |
| EPI_ISL_515937 | BIMS | Department of Neurovirology, National Institute of Mental Health and Neuroscience (NIMHANS) | Chitra Pattabiraman,Vijayalakshmi Reddy, Harsha PK, Risha Rasheed, Pramada Prasad, Shafeeq S Hameed, Manjunatha Venkataswamy, Anita Desai, Ravi Vasanthapuram |
| EPI_ISL_515938, EPI_ISL_515939, EPI_ISL_515940, EPI_ISL_515941, EPI_ISL_515942 | CV RAMAN HOSPITAL | Department of Neurovirology, National Institute of Mental Health and Neuroscience (NIMHANS) | Chitra Pattabiraman,Vijayalakshmi Reddy, Harsha PK, Risha Rasheed, Pramada Prasad, Shafeeq S Hameed, Manjunatha Venkataswamy, Anita Desai, Ravi Vasanthapuram |
| EPI_ISL_515943, EPI_ISL_515944 | ESIC | Department of Neurovirology, National Institute of Mental Health and Neuroscience (NIMHANS) | Chitra Pattabiraman,Vijayalakshmi Reddy, Harsha PK, Risha Rasheed, Pramada Prasad, Shafeeq S Hameed, Manjunatha Venkataswamy, Anita Desai, Ravi Vasanthapuram |
| EPI_ISL_515945, EPI_ISL_515946, EPI_ISL_515947, EPI_ISL_515948, EPI_ISL_515949 | DH | Department of Neurovirology, National Institute of Mental Health and Neuroscience (NIMHANS) | Chitra Pattabiraman,Vijayalakshmi Reddy, Harsha PK, Risha Rasheed, Pramada Prasad, Shafeeq S Hameed, Manjunatha Venkataswamy, Anita Desai, Ravi Vasanthapuram |
| EPI_ISL_515950, EPI_ISL_515951, EPI_ISL_515952, EPI_ISL_515953 | VICTORIA HOSPITAL | Department of Neurovirology, National Institute of Mental Health and Neuroscience (NIMHANS) | Chitra Pattabiraman,Vijayalakshmi Reddy, Harsha PK, Risha Rasheed, Pramada Prasad, Shafeeq S Hameed, Manjunatha Venkataswamy, Anita Desai, Ravi Vasanthapuram |
| EPI_ISL_515954, EPI_ISL_515955 | DH | Department of Neurovirology, National Institute of Mental Health and Neuroscience (NIMHANS) | Chitra Pattabiraman,Vijayalakshmi Reddy, Harsha PK, Risha Rasheed, Pramada Prasad, Shafeeq S Hameed, Manjunatha Venkataswamy, Anita Desai, Ravi Vasanthapuram |
| EPI_ISL_515956 | SHEKAR HOSPITAL | Department of Neurovirology, National Institute of Mental Health and Neuroscience (NIMHANS) | Chitra Pattabiraman,Vijayalakshmi Reddy, Harsha PK, Risha Rasheed, Pramada Prasad, Shafeeq S Hameed, Manjunatha Venkataswamy, Anita Desai, Ravi Vasanthapuram |
| EPI_ISL_515957, EPI_ISL_515958, EPI_ISL_515959 | DH | Department of Neurovirology, National Institute of Mental Health and Neuroscience (NIMHANS) | Chitra Pattabiraman,Vijayalakshmi Reddy, Harsha PK, Risha Rasheed, Pramada Prasad, Shafeeq S Hameed, Manjunatha Venkataswamy, Anita Desai, Ravi Vasanthapuram |
| EPI_ISL_515960 | JGH | Department of Neurovirology, National Institute of Mental Health and Neuroscience (NIMHANS) | Chitra Pattabiraman,Vijayalakshmi Reddy, Harsha PK, Risha Rasheed, Pramada Prasad, Shafeeq S Hameed, Manjunatha Venkataswamy, Anita Desai, Ravi Vasanthapuram |
| EPI_ISL_515961, EPI_ISL_515962 | DH | Department of Neurovirology, National Institute of Mental Health and Neuroscience (NIMHANS) | Chitra Pattabiraman,Vijayalakshmi Reddy, Harsha PK, Risha Rasheed, Pramada Prasad, Shafeeq S Hameed, Manjunatha Venkataswamy, Anita Desai, Ravi Vasanthapuram |
| EPI_ISL_515963, EPI_ISL_515964 | VICTORIA HOSPITAL | Department of Neurovirology, National Institute of Mental Health and Neuroscience (NIMHANS) | Chitra Pattabiraman,Vijayalakshmi Reddy, Harsha PK, Risha Rasheed, Pramada Prasad, Shafeeq S Hameed, Manjunatha Venkataswamy, Anita Desai, Ravi Vasanthapuram |
| EPI_ISL_515965 | DH | Department of Neurovirology, National Institute of Mental Health and Neuroscience (NIMHANS) | Chitra Pattabiraman,Vijayalakshmi Reddy, Harsha PK, Risha Rasheed, Pramada Prasad, Shafeeq S Hameed, Manjunatha Venkataswamy, Anita Desai, Ravi Vasanthapuram |
| EPI_ISL_515966, EPI_ISL_515967 | VICTORIA HOSPITAL | Department of Neurovirology, National Institute of Mental Health and Neuroscience (NIMHANS) | Chitra Pattabiraman,Vijayalakshmi Reddy, Harsha PK, Risha Rasheed, Pramada Prasad, Shafeeq S Hameed, Manjunatha Venkataswamy, Anita Desai, Ravi Vasanthapuram |
| EPI_ISL_515968, EPI_ISL_515969, EPI_ISL_515970, EPI_ISL_515971, EPI_ISL_515972, EPI_ISL_515973 | MIMS | Department of Neurovirology, National Institute of Mental Health and Neuroscience (NIMHANS) | Chitra Pattabiraman,Vijayalakshmi Reddy, Harsha PK, Risha Rasheed, Pramada Prasad, Shafeeq S Hameed, Manjunatha Venkataswamy, Anita Desai, Ravi Vasanthapuram |
| EPI_ISL_515974 | Department of Virus and Microbiological Special Diagnostics, Statens Serum Institut | Albertsen lab, Department of Chemistry and Bioscience, Aalborg University, Denmark | Danish Corona Danica Consortia |
| EPI_ISL_516075 | BIMS | Department of Neurovirology, National Institute of Mental Health and Neuroscience (NIMHANS) | Chitra Pattabiraman,Vijayalakshmi Reddy, Harsha PK, Risha Rasheed, Pramada Prasad, Shafeeq S Hameed, Manjunatha Venkataswamy, Anita Desai, Ravi Vasanthapuram |
| EPI_ISL_516076, EPI_ISL_516077, EPI_ISL_516078 | VICTORIA HOSPITAL | Department of Neurovirology, National Institute of Mental Health and Neuroscience (NIMHANS) | Chitra Pattabiraman,Vijayalakshmi Reddy, Harsha PK, Risha Rasheed, Pramada Prasad, Shafeeq S Hameed, Manjunatha Venkataswamy, Anita Desai, Ravi Vasanthapuram |
| EPI_ISL_516079, EPI_ISL_516080, EPI_ISL_516081, EPI_ISL_516082, EPI_ISL_516083, EPI_ISL_516084, EPI_ISL_516085, EPI_ISL_516086, EPI_ISL_516087, EPI_ISL_516088 | Biomedical Sciences and Public Health, Polytechnic University of Marche | Biomedical Sciences and Public Health, Polytechnic University of Marche | Bagnarelli,P., Caucci,S., Di Sante,L., Menzo,S., Alessandrini,F., Onofri,V., Turchi,C., Melchionda,F., Tagliabracci,A. |
| EPI_ISL_516193 | Hospital Universitari Germans Trias i Pujol. | IrsiCaixa AIDS Research Lab | Marc Noguera-Julian, Mariona Parera, Maria Pilar Armengol, Marta Massanella, Ester Ballana, Lidia Ruiz, Nuria Izquierdo, Jorge Carrillo, Roger Paredes, Julia Blanco, Joaquim Segalés, Bonaventura Clotet |
| EPI_ISL_516194, EPI_ISL_516195, EPI_ISL_516196, EPI_ISL_516197, EPI_ISL_516198, EPI_ISL_516199, EPI_ISL_516200, EPI_ISL_516201, EPI_ISL_516202, EPI_ISL_516203, EPI_ISL_516204, EPI_ISL_516205, EPI_ISL_516206, EPI_ISL_516207, EPI_ISL_516208, EPI_ISL_516209, EPI_ISL_516210, EPI_ISL_516211, EPI_ISL_516212, EPI_ISL_516213, EPI_ISL_516214, EPI_ISL_516215, EPI_ISL_516216, EPI_ISL_516217, EPI_ISL_516218, EPI_ISL_516219, EPI_ISL_516220, EPI_ISL_516221, EPI_ISL_516222, EPI_ISL_516223, EPI_ISL_516224, EPI_ISL_516225, EPI_ISL_516226, EPI_ISL_516227, EPI_ISL_516228, EPI_ISL_516229, EPI_ISL_516230, EPI_ISL_516231, EPI_ISL_516232, EPI_ISL_516233, EPI_ISL_516234, EPI_ISL_516235, EPI_ISL_516236, EPI_ISL_516237, EPI_ISL_516238, EPI_ISL_516239, EPI_ISL_516240, EPI_ISL_516241, EPI_ISL_516242, EPI_ISL_516243, EPI_ISL_516244, EPI_ISL_516245, EPI_ISL_516246, EPI_ISL_516247, EPI_ISL_516248, EPI_ISL_516249, EPI_ISL_516250, EPI_ISL_516251, EPI_ISL_516252, EPI_ISL_516253, EPI_ISL_516254, EPI_ISL_516255, EPI_ISL_516256, EPI_ISL_516257, EPI_ISL_516258, EPI_ISL_516259, EPI_ISL_516260, EPI_ISL_516261, EPI_ISL_516262, EPI_ISL_516263, EPI_ISL_516264, EPI_ISL_516265, EPI_ISL_516266, EPI_ISL_516267, EPI_ISL_516268, EPI_ISL_516269, EPI_ISL_516270, EPI_ISL_516271, EPI_ISL_516272, EPI_ISL_516273, EPI_ISL_516274, EPI_ISL_516275, EPI_ISL_516276, EPI_ISL_516277, EPI_ISL_516278, EPI_ISL_516279, EPI_ISL_516280, EPI_ISL_516281, EPI_ISL_516282, EPI_ISL_516283, EPI_ISL_516284, EPI_ISL_516285, EPI_ISL_516286, EPI_ISL_516287, EPI_ISL_516288, EPI_ISL_516289, EPI_ISL_516290, EPI_ISL_516291, EPI_ISL_516292, EPI_ISL_516293, EPI_ISL_516294, EPI_ISL_516295, EPI_ISL_516296, EPI_ISL_516297, EPI_ISL_516298, EPI_ISL_516299, EPI_ISL_516300, EPI_ISL_516301, EPI_ISL_516302, EPI_ISL_516303, EPI_ISL_516304, EPI_ISL_516305, EPI_ISL_516306, EPI_ISL_516307, EPI_ISL_516308, EPI_ISL_516309, EPI_ISL_516310, EPI_ISL_516311, EPI_ISL_516312, EPI_ISL_516313, EPI_ISL_516314, EPI_ISL_516315, EPI_ISL_516316, EPI_ISL_516317, EPI_ISL_516318, EPI_ISL_516319, EPI_ISL_516320, EPI_ISL_516321, EPI_ISL_516322, EPI_ISL_516323, EPI_ISL_516324, EPI_ISL_516325, EPI_ISL_516326, EPI_ISL_516327, EPI_ISL_516328, EPI_ISL_516329, EPI_ISL_516330, EPI_ISL_516331, EPI_ISL_516332, EPI_ISL_516333, EPI_ISL_516334, EPI_ISL_516335, EPI_ISL_516336, EPI_ISL_516337, EPI_ISL_516338, EPI_ISL_516339, EPI_ISL_516340, EPI_ISL_516341, EPI_ISL_516342, EPI_ISL_516343, EPI_ISL_516344, EPI_ISL_516345, EPI_ISL_516346, EPI_ISL_516347, EPI_ISL_516348, EPI_ISL_516349, EPI_ISL_516350, EPI_ISL_516351, EPI_ISL_516352, EPI_ISL_516353, EPI_ISL_516354, EPI_ISL_516355, EPI_ISL_516356, EPI_ISL_516357, EPI_ISL_516358, EPI_ISL_516359, EPI_ISL_516360, EPI_ISL_516361, EPI_ISL_516362, EPI_ISL_516363, EPI_ISL_516364, EPI_ISL_516365, EPI_ISL_516366, EPI_ISL_516367, EPI_ISL_516368, EPI_ISL_516369, EPI_ISL_516370, EPI_ISL_516371, EPI_ISL_516372, EPI_ISL_516373, EPI_ISL_516374, EPI_ISL_516375, EPI_ISL_516376, EPI_ISL_516377, EPI_ISL_516378, EPI_ISL_516379, EPI_ISL_516380, EPI_ISL_516381, EPI_ISL_516382, EPI_ISL_516383, EPI_ISL_516384, EPI_ISL_516385, EPI_ISL_516386, EPI_ISL_516387, EPI_ISL_516388, EPI_ISL_516389, EPI_ISL_516390, EPI_ISL_516391, EPI_ISL_516392, EPI_ISL_516393, EPI_ISL_516394, EPI_ISL_516395, EPI_ISL_516396, EPI_ISL_516397, EPI_ISL_516398, EPI_ISL_516399, EPI_ISL_516400, EPI_ISL_516401, EPI_ISL_516402, EPI_ISL_516403, EPI_ISL_516404, EPI_ISL_516405, EPI_ISL_516406, EPI_ISL_516407, EPI_ISL_516408, EPI_ISL_516409, EPI_ISL_516410, EPI_ISL_516411, EPI_ISL_516412 | | | |
| see above | Michigan Department of Health and Human Services, Bureau of Laboratories | Michigan Department of Health and Human Services, Bureau of Laboratories | Blankenship HM, Riner D, Soehnlen MK |
| EPI_ISL_516413 | General Hospital - Kumanovo | Research Center for Genetic Engineering and Biotechnology "Georgi D. Efremov" , Macedonian Academy of Sciences and Arts | RCGEB - MASA |
| EPI_ISL_516414, EPI_ISL_516415, EPI_ISL_516416, EPI_ISL_516417, EPI_ISL_516418, EPI_ISL_516419, EPI_ISL_516420, EPI_ISL_516421, EPI_ISL_516422, EPI_ISL_516423, EPI_ISL_516424, EPI_ISL_516425 | | | |
| see above | Center for public health - Skopje | Research Center for Genetic Engineering and Biotechnology "Georgi D. Efremov" , Macedonian Academy of Sciences and Arts | RCGEB - MASA |
| EPI_ISL_516426, EPI_ISL_516427, EPI_ISL_516428, EPI_ISL_516429, EPI_ISL_516430, EPI_ISL_516431 | Clinical Hospital - Shtip | Research Center for Genetic Engineering and Biotechnology "Georgi D. Efremov" , Macedonian Academy of Sciences and Arts | RCGEB - MASA |
| EPI_ISL_516432, EPI_ISL_516433, EPI_ISL_516434, EPI_ISL_516435, EPI_ISL_516436, EPI_ISL_516437, EPI_ISL_516438, EPI_ISL_516439, EPI_ISL_516440, EPI_ISL_516441, EPI_ISL_516442, EPI_ISL_516443, EPI_ISL_516444, EPI_ISL_516445, EPI_ISL_516446, EPI_ISL_516447, EPI_ISL_516448, EPI_ISL_516449, EPI_ISL_516450, EPI_ISL_516451, EPI_ISL_516452, EPI_ISL_516453, EPI_ISL_516454, EPI_ISL_516455, EPI_ISL_516456, EPI_ISL_516457, EPI_ISL_516458, EPI_ISL_516459, EPI_ISL_516460, EPI_ISL_516461, EPI_ISL_516462, EPI_ISL_516463, EPI_ISL_516464, EPI_ISL_516465, EPI_ISL_516466, EPI_ISL_516467, EPI_ISL_516468, EPI_ISL_516469, EPI_ISL_516470, EPI_ISL_516471, EPI_ISL_516472, EPI_ISL_516473, EPI_ISL_516474, EPI_ISL_516475, EPI_ISL_516476, EPI_ISL_516477, EPI_ISL_516478, EPI_ISL_516479, EPI_ISL_516480, EPI_ISL_516481, EPI_ISL_516482, EPI_ISL_516483, EPI_ISL_516484, EPI_ISL_516485, EPI_ISL_516486, EPI_ISL_516487, EPI_ISL_516488, EPI_ISL_516489, EPI_ISL_516490, EPI_ISL_516491, EPI_ISL_516492, EPI_ISL_516493, EPI_ISL_516494, EPI_ISL_516495, EPI_ISL_516496, EPI_ISL_516497, EPI_ISL_516498, EPI_ISL_516499, EPI_ISL_516500, EPI_ISL_516501, EPI_ISL_516502, EPI_ISL_516503, EPI_ISL_516504, EPI_ISL_516505, EPI_ISL_516506, EPI_ISL_516507, EPI_ISL_516508, EPI_ISL_516509, EPI_ISL_516510, EPI_ISL_516511, EPI_ISL_516512, EPI_ISL_516513, EPI_ISL_516514, EPI_ISL_516515, EPI_ISL_516516, EPI_ISL_516517, EPI_ISL_516518, EPI_ISL_516519, EPI_ISL_516520, EPI_ISL_516521, EPI_ISL_516522, EPI_ISL_516523, EPI_ISL_516524, EPI_ISL_516525, EPI_ISL_516526, EPI_ISL_516527, EPI_ISL_516528, EPI_ISL_516529, EPI_ISL_516530, EPI_ISL_516531, EPI_ISL_516532, EPI_ISL_516533, EPI_ISL_516534, EPI_ISL_516535, EPI_ISL_516536, EPI_ISL_516537, EPI_ISL_516538, EPI_ISL_516539, EPI_ISL_516540, EPI_ISL_516541, EPI_ISL_516542, EPI_ISL_516543, EPI_ISL_516544, EPI_ISL_516545, EPI_ISL_516546, EPI_ISL_516547, EPI_ISL_516548, EPI_ISL_516549 | | | |
| see above | University of Wisconsin-Madison AIDS Vaccine Research Laboratories | University of Wisconsin-Madison AIDS Vaccine Research Laboratories | Gage Moreno, Katarina Braun, et al. AIDS Vaccine Research Laboratories |
| EPI_ISL_516550, EPI_ISL_516551 | SA Pathology | SA Pathology | Lex Leong, Julien Soubrier, Luke Walters, Chuan Kok Lim, Song Gao, Mark Turra, Karin Kassahn, Ivan Bastian, Geoff Higgins |

| | | | |
|---|---|---|---|
| EPI_ISL_516552, EPI_ISL_516553, EPI_ISL_516554, EPI_ISL_516555, EPI_ISL_516556, EPI_ISL_516557, EPI_ISL_516558, EPI_ISL_516559, EPI_ISL_516560, EPI_ISL_516561, EPI_ISL_516562, EPI_ISL_516563, EPI_ISL_516564, EPI_ISL_516565, EPI_ISL_516566, EPI_ISL_516567, EPI_ISL_516568, EPI_ISL_516569, EPI_ISL_516570, EPI_ISL_516571, EPI_ISL_516572, EPI_ISL_516573, EPI_ISL_516574, EPI_ISL_516575, EPI_ISL_516576, EPI_ISL_516577, EPI_ISL_516578, EPI_ISL_516579, EPI_ISL_516580, EPI_ISL_516581, EPI_ISL_516582, EPI_ISL_516583, EPI_ISL_516584, EPI_ISL_516585, EPI_ISL_516586, EPI_ISL_516587, EPI_ISL_516588, EPI_ISL_516589, EPI_ISL_516590, EPI_ISL_516591, EPI_ISL_516592, EPI_ISL_516593, EPI_ISL_516594, EPI_ISL_516595, EPI_ISL_516596, EPI_ISL_516597, EPI_ISL_516598, EPI_ISL_516599, EPI_ISL_516600, EPI_ISL_516601, EPI_ISL_516602, EPI_ISL_516603, EPI_ISL_516604, EPI_ISL_516605, EPI_ISL_516606, EPI_ISL_516607 | | | |
| see above | Viollier AG | Department of Biosystems Science and Engineering, ETH Zürich | Christian Beisel, Sarah Nadeau, Ivan Topolsky, Pedro Ferreira, Philipp Jablonski, Susana Posada-Céspedes, Tobias Schär, Ina Nissen, Natascha Santacroce, Elodie Burcklen, Christiane Beckmann, Maurice Redondo, Olivier Kobel, Christoph Noppen, Sophie Seidel, Noemie Santamaria de Souza, Niko Beerenwinkel, Tanja Stadler |
| EPI_ISL_516608 | Instituto de Diagnostico y Referencia Epidemiologicos (INDRE) | Instituto de Diagnostico y Referencia Epidemiologicos (INDRE) | Ernesto Ramirez-Gonzalez, Abril Rodriguez-Maldonado, Claudia Wong-Arambula , Natividad Cruz-Ortiz, Tatiana Nunez-Garcia, Dayanira Arellano-Suarez, Adnan Araiza-Rodriguez, Edgar Mendieta-Condado, Lucia Hernandez-Rivas, Irma Lopez-Martinez, Gisela Barrera-Badillo. |
| EPI_ISL_516609, EPI_ISL_516610 | Instituto de Diagnostico y Referencia Epidemiologicos (INDRE) | Instituto de Diagnostico y Referencia Epidemiologicos (INDRE) | Ernesto Ramirez-Gonzalez, Abril Rodriguez-Maldonado, Claudia Wong-Arambula , Natividad Cruz-Ortiz, Tatiana Nunez-Garcia, Dayanira Arellano-Suarez, Adnan Araiza-Rodriguez, Fabiola Garces-Ayala, Lucia Hernandez-Rivas, Irma Lopez-Martinez, Gisela Barrera-Badillo. |
| EPI_ISL_516611 | Instituto de Diagnostico y Referencia Epidemiologicos (INDRE) | Instituto de Diagnostico y Referencia Epidemiologicos (INDRE) | Gisela Barrera-Badillo , Abril Rodriguez-Maldonado, Claudia Wong-Arambula , Natividad Cruz-Ortiz, Tatiana Nunez-Garcia, Dayanira Arellano-Suarez, Fabiola Garces-Ayala, Edgar Mendieta-Condado, Lucia Hernandez-Rivas, Irma Lopez-Martinez, Ernesto Ramirez-Gonzalez. |
| EPI_ISL_516612 | Instituto de Diagnostico y Referencia Epidemiologicos (INDRE) | Instituto de Diagnostico y Referencia Epidemiologicos (INDRE) | Ernesto Ramirez-Gonzalez, Abril Rodriguez-Maldonado, Claudia Wong-Arambula , Natividad Cruz-Ortiz, Tatiana Nunez-Garcia, Dayanira Arellano-Suarez, Adnan Araiza-Rodriguez, Fabiola Garces-Ayala, Lucia Hernandez-Rivas, Irma Lopez-Martinez, Gisela Barrera-Badillo. |
| EPI_ISL_516613 | Instituto de Diagnostico y Referencia Epidemiologicos (INDRE) | Instituto de Diagnostico y Referencia Epidemiologicos (INDRE) | Ernesto Ramirez-Gonzalez, Abril Rodriguez-Maldonado, Claudia Wong-Arambula , Natividad Cruz-Ortiz, Tatiana Nunez-Garcia, Dayanira Arellano-Suarez, Adnan Araiza-Rodriguez, Edgar Mendieta-Condado, Lucia Hernandez-Rivas, Irma Lopez-Martinez, Gisela Barrera-Badillo. |
| EPI_ISL_516614, EPI_ISL_516615, EPI_ISL_516616, EPI_ISL_516617 | Instituto de Diagnostico y Referencia Epidemiologicos (INDRE) | Instituto de Diagnostico y Referencia Epidemiologicos (INDRE) | Ernesto Ramirez-Gonzalez, Abril Rodriguez-Maldonado, Claudia Wong-Arambula , Natividad Cruz-Ortiz, Tatiana Nunez-Garcia, Dayanira Arellano-Suarez, Adnan Araiza-Rodriguez, Fabiola Garces-Ayala, Lucia Hernandez-Rivas, Irma Lopez-Martinez, Gisela Barrera-Badillo. |
| EPI_ISL_516618, EPI_ISL_516619 | Instituto de Diagnostico y Referencia Epidemiologicos (INDRE) | Instituto de Diagnostico y Referencia Epidemiologicos (INDRE) | Gisela Barrera-Badillo , Abril Rodriguez-Maldonado, Claudia Wong-Arambula , Natividad Cruz-Ortiz, Tatiana Nunez-Garcia, Dayanira Arellano-Suarez, Fabiola Garces-Ayala, Edgar Mendieta-Condado, Lucia Hernandez-Rivas, Irma Lopez-Martinez, Ernesto Ramirez-Gonzalez. |
| EPI_ISL_516620, EPI_ISL_516621 | Instituto de Diagnostico y Referencia Epidemiologicos (INDRE) | Instituto de Diagnostico y Referencia Epidemiologicos (INDRE) | Gisela Barrera-Badillo , Abril Rodriguez-Maldonado, Claudia Wong-Arambula , Natividad Cruz-Ortiz, Tatiana Nunez-Garcia, Dayanira Arellano-Suarez, Adnan Araiza-Rodriguez, Edgar Mendieta-Condado, Lucia Hernandez-Rivas, Irma Lopez-Martinez, Ernesto Ramirez-Gonzalez. |
| EPI_ISL_516622 | Instituto de Diagnostico y Referencia Epidemiologicos (INDRE) | Instituto de Diagnostico y Referencia Epidemiologicos (INDRE) | Gisela Barrera-Badillo , Abril Rodriguez-Maldonado, Claudia Wong-Arambula , Natividad Cruz-Ortiz, Tatiana Nunez-Garcia, Dayanira Arellano-Suarez, Fabiola Garces-Ayala, Edgar Mendieta-Condado, Lucia Hernandez-Rivas, Irma Lopez-Martinez, Ernesto Ramirez-Gonzalez. |
| EPI_ISL_516623 | Instituto de Diagnostico y Referencia Epidemiologicos (INDRE) | Instituto de Diagnostico y Referencia Epidemiologicos (INDRE) | Gisela Barrera-Badillo , Abril Rodriguez-Maldonado, Claudia Wong-Arambula , Natividad Cruz-Ortiz, Tatiana Nunez-Garcia, Dayanira Arellano-Suarez, Adnan Araiza-Rodriguez, Edgar Mendieta-Condado, Lucia Hernandez-Rivas, Irma Lopez-Martinez, Ernesto Ramirez-Gonzalez. |
| EPI_ISL_516624, EPI_ISL_516625 | Instituto de Diagnostico y Referencia Epidemiologicos (INDRE) | Instituto de Diagnostico y Referencia Epidemiologicos (INDRE) | Gisela Barrera-Badillo , Abril Rodriguez-Maldonado, Claudia Wong-Arambula , Natividad Cruz-Ortiz, Tatiana Nunez-Garcia, Dayanira Arellano-Suarez, Fabiola Garces-Ayala, Edgar Mendieta-Condado, Lucia Hernandez-Rivas, Irma Lopez-Martinez, Ernesto Ramirez-Gonzalez. |
| EPI_ISL_516626 | Laboratorio de Referencia Nacional de Virus Respiratorio. Centro Nacional de Salud Publica. Instituto Nacional de Salud Peru. | Laboratorio de Referencia Nacional de Biotecnología y Biología Molecular. Centro Nacional de Salud Publica. Instituto Nacional de Salud Peru. | Carlos Padilla Rojas, Karolyn Vega Chozo, Priscila Lope Pari, Omar Caceres Rey, Marco Galarza Perez, Maribel Huaringa Nuñez, Johanna Balbuena Torres, Henri Bailon Calderon, Nancy Rojas Serrano. |
| EPI_ISL_516627 | SYNLAB Eesti OÜ | Charite Universitatsmedizin Berlin, Institute of Virology | Victor M Corman, Jorn Beheim-Schwarzbach, Barbara Mühlemann, Talitha Veith, Julia Schneider, Paul Naaber, Terry Jones, Christian Drosten |
| EPI_ISL_516628 | Laboratorio de Referencia Nacional de Virus Respiratorio. Centro Nacional de Salud Publica. Instituto Nacional de Salud Peru. | Laboratorio de Referencia Nacional de Biotecnología y Biología Molecular. Centro Nacional de Salud Publica. Instituto Nacional de Salud Peru. | Carlos Padilla Rojas, Karolyn Vega Chozo, Priscila Lope Pari, Omar Caceres Rey, Marco Galarza Perez, Maribel Huaringa Nuñez, Johanna Balbuena Torres, Henri Bailon Calderon, Nancy Rojas Serrano. |
| EPI_ISL_516629, EPI_ISL_516630, EPI_ISL_516631, EPI_ISL_516632, EPI_ISL_516633, EPI_ISL_516634, EPI_ISL_516635, EPI_ISL_516636, EPI_ISL_516637, EPI_ISL_516638, EPI_ISL_516639, EPI_ISL_516640, EPI_ISL_516641, EPI_ISL_516642, EPI_ISL_516643, EPI_ISL_516644, EPI_ISL_516645 | | | |
| see above | Charité Universitätsmedizin Berlin, Institut für Virologie/Labor Berlin | Charité Universitätsmedizin Berlin, Institut für Virologie/Labor Berlin | Victor M Corman, Barbara Mühlemann, Jörn Beheim-Schwarzbach, Julia Schneider, Talitha Veith, Terry Jones, Christian Drosten |
| EPI_ISL_516646, EPI_ISL_516647 | Laboratorio de Referencia Nacional de Virus Respiratorio. Centro Nacional de Salud Publica. Instituto Nacional de Salud Peru. | Laboratorio de Referencia Nacional de Biotecnología y Biología Molecular. Centro Nacional de Salud Publica. Instituto Nacional de Salud Peru. | Carlos Padilla Rojas, Karolyn Vega Chozo, Priscila Lope Pari, Omar Caceres Rey, Marco Galarza Perez, Maribel Huaringa Nuñez, Johanna Balbuena Torres, Henri Bailon Calderon, Nancy Rojas Serrano. |
| EPI_ISL_516648 | Institute of Microbiology, Universidad San Francisco de Quito | Institute of Microbiology, Universidad San Francisco de Quito | Juan José Guadalupe, Monica Becerra-Wong, Prado-Vivar, Sully Márquez, Bernardo Gutiérrez, Nabih Dahik, Carlos Mena, Ligia Briceño, Verónica Barragán, Patricio Rojas-Silva, Gabriel Trueba, Michelle Grunauer, Paúl Cárdenas |
| EPI_ISL_516649 | Institute of Microbiology, Universidad San Francisco de Quito | Institute of Microbiology, Universidad San Francisco de Quito | Juan José Guadalupe, Monica Becerra-Wong, Bernardo Gutiérrez, Nabih Dahik, Carlos Mena, Ligia Briceño, Verónica Barragán, Patricio Rojas-Silva, Gabriel Trueba, Michelle Grunauer, Paúl Cárdenas |
| EPI_ISL_516650 | Institute of Microbiology, Universidad San Francisco de Quito | Institute of Microbiology, Universidad San Francisco de Quito | Prado-Vivar, Sully Márquez, Juan José Guadalupe, Monica Becerra-Wong, Bernardo Gutiérrez, Verónica Barragán, Patricio Rojas-Silva, Gabriel Trueba, Michelle Grunauer, Paúl Cárdenas |
| EPI_ISL_516651 | Institute of Microbiology, Universidad San Francisco de Quito | Institute of Microbiology, Universidad San Francisco de Quito | Prado-Vivar, Sully Márquez, Juan José Guadalupe, Monica Becerra-Wong, Bernardo Gutiérrez, Nabih Dahik, Carlos Mena, Eulalia Pazmiño, Carolina Pacheco, DamarisSandoya, Verónica Barragán, Patricio Rojas-Silva, Gabriel Trueba, Michelle Grunauer, Paúl Cárdenas |
| EPI_ISL_516652 | Institute of Microbiology, Universidad San Francisco de Quito | Institute of Microbiology, Universidad San Francisco de Quito | Prado-Vivar, Sully Márquez, Juan José Guadalupe, Monica Becerra-Wong, Bernardo Gutiérrez, Nabih Dahik, Carlos Mena, Edy Quizhpe, Yomara Napa, Verónica Barragán, Patricio Rojas-Silva, Gabriel Trueba, Michelle Grunauer, Paúl Cárdenas |
| EPI_ISL_516653, EPI_ISL_516654 | Laboratorio de Referencia Nacional de Virus Respiratorio. Centro Nacional de Salud Publica. Instituto Nacional de Salud Peru. | Laboratorio de Referencia Nacional de Biotecnología y Biología Molecular. Centro Nacional de Salud Publica. Instituto Nacional de Salud Peru. | Carlos Padilla Rojas, Karolyn Vega Chozo, Priscila Lope Pari, Omar Caceres Rey, Marco Galarza Perez, Maribel Huaringa Nuñez, Johanna Balbuena Torres, Henri Bailon Calderon, Nancy Rojas Serrano. |
| EPI_ISL_516655, EPI_ISL_516656, EPI_ISL_516657, EPI_ISL_516658, EPI_ISL_516659, EPI_ISL_516660, EPI_ISL_516661, EPI_ISL_516662, EPI_ISL_516663, EPI_ISL_516664, EPI_ISL_516665, EPI_ISL_516666, EPI_ISL_516667, EPI_ISL_516668, EPI_ISL_516669, EPI_ISL_516670, EPI_ISL_516671, EPI_ISL_516672, EPI_ISL_516673, EPI_ISL_516674, EPI_ISL_516675, EPI_ISL_516676, EPI_ISL_516677, EPI_ISL_516678, EPI_ISL_516679, EPI_ISL_516680, EPI_ISL_516681, EPI_ISL_516682, EPI_ISL_516683, EPI_ISL_516684 | | | |
| see above | Virginia DCLS | Virginia DCLS | Virginia DCLS |
| EPI_ISL_516685, EPI_ISL_516686 | Laboratorio de Referencia Nacional de Virus Respiratorio. Centro Nacional de Salud Publica. Instituto Nacional de Salud Peru. | Laboratorio de Referencia Nacional de Biotecnología y Biología Molecular. Centro Nacional de Salud Publica. Instituto Nacional de Salud Peru. | Carlos Padilla Rojas, Karolyn Vega Chozo, Priscila Lope Pari, Omar Caceres Rey, Marco Galarza Perez, Maribel Huaringa Nuñez, Johanna Balbuena Torres, Henri Bailon Calderon, Nancy Rojas Serrano. |
| EPI_ISL_516687, EPI_ISL_516688, EPI_ISL_516689, EPI_ISL_516690, EPI_ISL_516691, EPI_ISL_516692, EPI_ISL_516693, EPI_ISL_516694, EPI_ISL_516695, EPI_ISL_516696, EPI_ISL_516697, EPI_ISL_516698, EPI_ISL_516699, EPI_ISL_516700, EPI_ISL_516701, EPI_ISL_516702, EPI_ISL_516703, EPI_ISL_516704, EPI_ISL_516705, EPI_ISL_516706, EPI_ISL_516707, EPI_ISL_516708, EPI_ISL_516709, EPI_ISL_516710, EPI_ISL_516711, EPI_ISL_516712, EPI_ISL_516713, EPI_ISL_516714, EPI_ISL_516715, EPI_ISL_516716, EPI_ISL_516717, EPI_ISL_516718, EPI_ISL_516719, EPI_ISL_516720 | | | |
| see above | Virginia DCLS | Virginia DCLS | Virginia DCLS |
| EPI_ISL_516721, EPI_ISL_516722 | Laboratorio de Referencia Nacional de Virus Respiratorio. Centro Nacional de Salud Publica. Instituto Nacional de Salud Peru. | Laboratorio de Referencia Nacional de Biotecnología y Biología Molecular. Centro Nacional de Salud Publica. Instituto Nacional de Salud Peru. | Carlos Padilla Rojas, Karolyn Vega Chozo, Priscila Lope Pari, Omar Caceres Rey, Marco Galarza Perez, Maribel Huaringa Nuñez, Johanna Balbuena Torres, Henri Bailon Calderon, Nancy Rojas Serrano. |
| EPI_ISL_516723, EPI_ISL_516724, EPI_ISL_516725, EPI_ISL_516726, EPI_ISL_516727, EPI_ISL_516728, EPI_ISL_516729, EPI_ISL_516730, EPI_ISL_516731, EPI_ISL_516732, EPI_ISL_516733, EPI_ISL_516734, EPI_ISL_516735, EPI_ISL_516736, EPI_ISL_516737, EPI_ISL_516738, EPI_ISL_516739, EPI_ISL_516740, EPI_ISL_516741, EPI_ISL_516742, EPI_ISL_516743, EPI_ISL_516744, EPI_ISL_516745, EPI_ISL_516746, EPI_ISL_516747 | | | |
| see above | UCSF Clinical Microbiology Laboratory | Chan-Zuckerberg Biohub | CZB Cliahub Consortium |
| EPI_ISL_516748, EPI_ISL_516749, EPI_ISL_516750, EPI_ISL_516751, EPI_ISL_516752, EPI_ISL_516753, EPI_ISL_516754, EPI_ISL_516755, EPI_ISL_516756, EPI_ISL_516757, EPI_ISL_516758, EPI_ISL_516759, EPI_ISL_516760, EPI_ISL_516761, EPI_ISL_516762, EPI_ISL_516763, EPI_ISL_516764, EPI_ISL_516765, EPI_ISL_516766, EPI_ISL_516767, EPI_ISL_516768, EPI_ISL_516769, EPI_ISL_516770, EPI_ISL_516771, EPI_ISL_516772, EPI_ISL_516773, EPI_ISL_516774, EPI_ISL_516775, EPI_ISL_516776, EPI_ISL_516777, EPI_ISL_516778, EPI_ISL_516779, EPI_ISL_516780, EPI_ISL_516781, EPI_ISL_516782 | | | |
| see above | van Bakel Laboratory, Genetics and Genomics Sciences, Icahn School of Medicine at Mount Sinai | van Bakel Laboratory, Genetics and Genomics Sciences, Icahn School of Medicine at Mount Sinai | Andrew G. Letizia, Irene Ramos, Ajay Obla, Carl Goforth, Dawn Weir, Yongchao Ge, Marcas M. Bamman, Jayeeta Dutta, Ethan Ellis, Luis Estrella, Mary-Catherine George, Ana S. Gonzalez-Reiche, Darnell Graham, Adriana van de Guchte, Ramiro Gutierrez, Franca Jones, Aspasia Kalomoiri, Rhonda Lizewski, Stephen Lizewski, Jan Marayag, Nada Marjanovic, Eugene V. Millar, Venugopalan Nair, German Nudelman, Edgar Nunez, Brian Pike, James Regeimbal, Stas Rirak , Ernesto Santa Ana, Rachel S. Gelernter Sealfon, Robert Sebra, Mark Simons, Alessandra Soares-Schanoski, Michael Termini, Sindhu Vangeti, Carlos Williams, Harm van Bakel, Stuart C. Sealfon |
| EPI_ISL_516783, EPI_ISL_516784, EPI_ISL_516785 | Center for Laboratory Control of Infectious Diseases, Korea Centers for Diseases Control and Prevention | Center for Laboratory Control of Infectious Diseases, Korea Centers for Diseases Control and Prevention | Junyoung Kim, Ae Kyung Park, Eunkyung Shin, Jin Sun No, Jeong-Min Kim, Yoon-Seok Chung, Heui Man Kim, Myung Guk Han |
| EPI_ISL_516786 | Laboratorio de Referencia Nacional de Virus Respiratorio. Centro Nacional de Salud Publica. Instituto Nacional de Salud Peru. | Laboratorio de Referencia Nacional de Biotecnología y Biología Molecular. Centro Nacional de Salud Publica. Instituto Nacional de Salud Peru. | Carlos Padilla Rojas, Karolyn Vega Chozo, Priscila Lope Pari, Omar Caceres Rey, Marco Galarza Perez, Maribel Huaringa Nuñez, Johanna Balbuena Torres, Henri Bailon Calderon, Nancy Rojas Serrano. |
| EPI_ISL_516798, EPI_ISL_516799 | Department of Microbiology, The University of Hong Kong | Department of Microbiology, The University of Hong Kong | Kelvin K.W. To, Kwok-Yung Yuen |
| EPI_ISL_516800 | Rumah Sakit Akademik Universitas Gadjah Mada | Genetics Working Group (Pokja Genetik) Faculty of Medicine, Public Health and Nursing Universitas Gadjah Mada (FK-KMK UGM); Disease Investigation Center Wates Ministry of Agriculture Indonesia; Department of Microbiology FK-KMK UGM; Laboratorium Diagnostik Yayasan Tahija World Mosquito Program (WMP) Yogyakarta Center for Tropical Medicine FK-KMK UGM; Integrated Research center FK-KMK UGM; Department of Computer Science and Electronics FMIPA UGM | Gunadi, Hendra Wibawa, . Marcellus, Mohamad S. Hakim, Edwin W. Daniwijaya, Ludhang P. Rizki, Endah Supriyati, Eggi Arguni, Titik Nuryastuti, Tri Wibawa, Dwi AA Nugrahaningsih, Afiahayati, Siswanto, Alvin S. Kalim, Desyifa Mursalin |
| EPI_ISL_516801 | Department of Laboratory Medicine, Tan Tock Seng Hospital | Department of Laboratory Medicine, Tan Tock Seng Hospital | Chen YYC, Zair X, Li C, Tang WY, Maurer-Stroh S, Barkham TMS, Nagarajan N, Sessions OM |
| EPI_ISL_516802, EPI_ISL_516803 | Department of Laboratory Medicine, Tan Tock Seng Hospital | Department of Laboratory Medicine, Tan Tock Seng Hospital | Chen YYC, Zair X, Li C, Tang WY, Maurer-Stroh S, Barkham TMS, Nagarajan N, Sessions OM |
| EPI_ISL_516804 | Department of Laboratory Medicine, Tan Tock Seng Hospital | Department of Laboratory Medicine, Tan Tock Seng Hospital | Chen YYC, Zair X, Li C, Tang WY, Maurer-Stroh S, Barkham TMS, Nagarajan N, Sessions OM |
| EPI_ISL_516805 | Department of Laboratory Medicine, Tan Tock Seng | Department of Laboratory Medicine, Tan Tock Seng Hospital | Chen YYC, Zair X, Li C, Tang WY, Maurer-Stroh S, Barkham TMS, Nagarajan N, Sessions OM |

| | | | |
|---|---|---|---|
| | Hospital | | |
| EPI_ISL_516806 | Rumah Sakit PKU Gamping | Genetics Working Group (Pokja Genetik) Faculty of Medicine, Public Health and Nursing Universitas Gadjah Mada (FK-KMK UGM); Disease Investigation Center Wates Ministry of Agriculture Indonesia; Department of Microbiology FK-KMK UGM; Laboratorium Diagnostik Yayasan Tahija World Mosquito Program (WMP) Yogyakarta Center for Tropical Medicine FK-KMK UGM; Integrated Research center FK-KMK UGM; Department of Computer Science and Electronics FMIPA UGM | Gunadi, Hendra Wibawa, . Marcellus, Mohamad S. Hakim, Edwin W. Daniwijaya, Ludhang P. Rizki, Endah Supriyati, Eggi Arguni, Titik Nuryastuti, Tri Wibawa, Dwi AA Nugrahaningsih, . Afiahayati , . Siswanto, Ardorisye Saptaty Fornia, Kemala Athollah |
| EPI_ISL_516807, EPI_ISL_516808, EPI_ISL_516809, EPI_ISL_516810, EPI_ISL_516811, EPI_ISL_516812, EPI_ISL_516813, EPI_ISL_516814, EPI_ISL_516815, EPI_ISL_516816, EPI_ISL_516817, EPI_ISL_516818, EPI_ISL_516819, EPI_ISL_516820, EPI_ISL_516821, EPI_ISL_516822, EPI_ISL_516823, EPI_ISL_516824, EPI_ISL_516825, EPI_ISL_516826, EPI_ISL_516827, EPI_ISL_516828 | | | |
| see above | National Public Health Laboratory, National Centre for Infectious Diseases | National Public Health Laboratory, National Centre for Infectious Diseases | Mak TM, Octavia S, Zhou Z, Cui L, Lin RTP |
| EPI_ISL_516829 | RSUD Nyi Ageng Serang | Genetics Working Group (Pokja Genetik) Faculty of Medicine, Public Health and Nursing Universitas Gadjah Mada (FK-KMK UGM), Disease Investigation Center Wates Ministry of Agriculture Indonesia, Department of Microbiology FK-KMK UGM, Laboratorium Diagnostik Yayasan Tahija World Mosquito Program (WMP) Yogyakarta Center for Tropical Medicine FK-KMK UGM, Integrated Research Center FK-KMK UGM, Department of Computer Science and Electronics FMIPA UGM | Gunadi, Hendra Wibawa, . Marcellus, Mohamad S. Hakim, Edwin W. Daniwijaya, Ludhang P. Rizki, Endah Supriyati, Eggi Arguni, Titik Nuryastuti, Tri Wibawa, Dwi AA Nugrahaningsih, Afiahayati , . Siswanto, Beby Dewi Sartika, Dyah Ayu Puspitarani |
| EPI_ISL_516831, EPI_ISL_516832, EPI_ISL_516833, EPI_ISL_516834, EPI_ISL_516835, EPI_ISL_516836, EPI_ISL_516837, EPI_ISL_516838, EPI_ISL_516839, EPI_ISL_516840, EPI_ISL_516841, EPI_ISL_516842, EPI_ISL_516843, EPI_ISL_516844, EPI_ISL_516845, EPI_ISL_516846, EPI_ISL_516847, EPI_ISL_516848, EPI_ISL_516849, EPI_ISL_516850, EPI_ISL_516851, EPI_ISL_516852, EPI_ISL_516853, EPI_ISL_516854, EPI_ISL_516855, EPI_ISL_516856, EPI_ISL_516857, EPI_ISL_516858, EPI_ISL_516859, EPI_ISL_516860, EPI_ISL_516861, EPI_ISL_516862, EPI_ISL_516863, EPI_ISL_516864, EPI_ISL_516865, EPI_ISL_516866, EPI_ISL_516867, EPI_ISL_516868, EPI_ISL_516869, EPI_ISL_516870, EPI_ISL_516871, EPI_ISL_516872, EPI_ISL_516873, EPI_ISL_516874, EPI_ISL_516875, EPI_ISL_516876, EPI_ISL_516877, EPI_ISL_516878, EPI_ISL_516879, EPI_ISL_516880, EPI_ISL_516881, EPI_ISL_516882, EPI_ISL_516883 | | | |
| see above | North West London Pathology, Imperial College Healthcare NHS Trust | Wellcome Sanger Institute for the COVID-19 Genomics UK Consortium | Ling Li, Paul Randell, David Muir, Frankie Bolt, Alison Holmes, James Price, Aileen Rowan, Graham Taylor, Anjna Badhan, Carolina Herrera and Alex Alderton, Roberto Amato, Sonia Goncalves, Ewan Harrison, David K. Jackson, Ian Johnston, Dominic Kwiatkowski, Cordelia Langford, John Sillitoe on behalf of the Wellcome Sanger Institute COVID-19 Surveillance Team (http://www.sanger.ac.uk/covid-team) |
| EPI_ISL_516884, EPI_ISL_516885, EPI_ISL_516886, EPI_ISL_516887, EPI_ISL_516888, EPI_ISL_516889, EPI_ISL_516890, EPI_ISL_516891 | Israel Central Virology laboratory | Israel Central Virology laboratory | Neta Zuckerman, Efrat Dahan Bucris, Oran Erster, Ella Mendelson, Michal Mandelboim |
| EPI_ISL_516892 | Israeli Central Virology laboratory | Israel Central Virology laboratory | Neta Zuckerman, Efrat Dahan Bucris, Oran Erster, Ella Mendelson, Michal Mandelboim |
| EPI_ISL_516893, EPI_ISL_516894, EPI_ISL_516895, EPI_ISL_516896, EPI_ISL_516897, EPI_ISL_516898, EPI_ISL_516899, EPI_ISL_516900, EPI_ISL_516901, EPI_ISL_516902, EPI_ISL_516903, EPI_ISL_516904, EPI_ISL_516905, EPI_ISL_516906, EPI_ISL_516907, EPI_ISL_516908, EPI_ISL_516909, EPI_ISL_516910, EPI_ISL_516911, EPI_ISL_516912, EPI_ISL_516913, EPI_ISL_516914, EPI_ISL_516915, EPI_ISL_516916, EPI_ISL_516917, EPI_ISL_516918, EPI_ISL_516919, EPI_ISL_516920, EPI_ISL_516921 | | | |
| see above | Israel Central Virology laboratory | Israel Central Virology laboratory | Neta Zuckerman, Efrat Dahan Bucris, Oran Erster, Ella Mendelson, Michal Mandelboim |
| EPI_ISL_516922, EPI_ISL_516923, EPI_ISL_516924, EPI_ISL_516925, EPI_ISL_516926, EPI_ISL_516927, EPI_ISL_516928, EPI_ISL_516929, EPI_ISL_516930, EPI_ISL_516931, EPI_ISL_516932, EPI_ISL_516933 | | | |
| see above | Department for Molecular Diagnostics, Centre for Medical Microbiology, Institute of Public Health of Montenegro | Charité Universitätsmedizin Berlin, Institut für Virologie | Victor M Corman, Terry Jones, Jörn Beheim-Schwarzbach, Barbara Muehlemann, Talitha Veith, Julia Schneider, Marija Govedarica and Danijela Vujošević, Christian Drosten |
| EPI_ISL_516934, EPI_ISL_516935, EPI_ISL_516936, EPI_ISL_516938 | Nicolae Testemitanu State University of Medicine and Pharmacy | International Centre for Genetic Engineering and Biotechnology (ICGEB) and ARGO Open Lab Platform for Genome Sequencing | Ulinici M, Licastro D, Dal Monego S, Rajasekharan S, Marcello A |
| EPI_ISL_516940, EPI_ISL_516941, EPI_ISL_516942, EPI_ISL_516943, EPI_ISL_516944, EPI_ISL_516945, EPI_ISL_516946, EPI_ISL_516947, EPI_ISL_516948, EPI_ISL_516949, EPI_ISL_516950, EPI_ISL_516951, EPI_ISL_516952, EPI_ISL_516953, EPI_ISL_516954, EPI_ISL_516955, EPI_ISL_516956, EPI_ISL_516957, EPI_ISL_516958, EPI_ISL_516959, EPI_ISL_516960, EPI_ISL_516961, EPI_ISL_516962, EPI_ISL_516963, EPI_ISL_516964, EPI_ISL_516965, EPI_ISL_516966, EPI_ISL_516967, EPI_ISL_516968, EPI_ISL_516969, EPI_ISL_516970, EPI_ISL_516971, EPI_ISL_516972, EPI_ISL_516973, EPI_ISL_516974, EPI_ISL_516975, EPI_ISL_516976, EPI_ISL_516977, EPI_ISL_516978, EPI_ISL_516979, EPI_ISL_516980, EPI_ISL_516981, EPI_ISL_516982, EPI_ISL_516983, EPI_ISL_516984, EPI_ISL_516985, EPI_ISL_516986 | | | |
| see above | King Georges Medical University | CSIR-National Botanical Research Institute | Priti Prasad, Shantanu Prakash, Kishan Sahu, Babita Singh, Suruchi Shukla, Hricha Mishra, Danish Nasar Khan , Om Prakash, MLB Bhatt, SK Barik, Mehar H.Asif,Samir V. Sawant,Amita Jain, Sumit Kr. Bag |
| EPI_ISL_516987 | Laboratorio de Referencia Nacional de Virus Respiratorio. Centro Nacional de Salud Publica. Instituto Nacional de Salud Peru. | Laboratorio de Referencia Nacional de Biotecnología y Biología Molecular. Centro Nacional de Salud Publica. Instituto Nacional de Salud Peru. | Carlos Padilla Rojas, Karolyn Vega Chozo, Priscila Lope Pari, Omar Caceres Rey, Marco Galarza Perez, Maribel Huaringa Nuñez, Johanna Balbuena Torres, Henri Bailon Calderon, Nancy Rojas Serrano. |
| EPI_ISL_516988, EPI_ISL_516989 | Public Health Authority of the Slovak Republic, Department of Medical Microbiology | Charite Universitatsmedizin Berlin, Institute of Virology | Victor M Corman, Terry Jones, Jörn Beheim-Schwarzbach, Barbara Muehlemann, Talitha Veith, Julia Schneider, Mgr. Edita Staronova, Christian Drosten |
| EPI_ISL_516990, EPI_ISL_516991, EPI_ISL_516992, EPI_ISL_516993, EPI_ISL_516994, EPI_ISL_516995, EPI_ISL_516996, EPI_ISL_516997, EPI_ISL_516998, EPI_ISL_516999, EPI_ISL_517000 | | | |
| see above | Department of Pathology, University of Cambridge | COVID-19 Genomics UK (COG-UK) Consortium | Luke W Meredith, M. Estée Török, Myra Hosmillo, William L. Hamilton, Martin D. Curran, Theresa Feltwell, Grant Hall, Anna Yakovleva, Fahad A Khokhar, Charlotte J. Houldcroft, Laura G Caller, Aminu S. Jahun, Sarah L. Caddy, Yasmin Chaudhry, Malte Pinckert, Ian Goodfellow |
| EPI_ISL_517001, EPI_ISL_517002, EPI_ISL_517003, EPI_ISL_517004, EPI_ISL_517005, EPI_ISL_517006, EPI_ISL_517007, EPI_ISL_517008, EPI_ISL_517009, EPI_ISL_517010, EPI_ISL_517011, EPI_ISL_517012, EPI_ISL_517013, EPI_ISL_517014, EPI_ISL_517015, EPI_ISL_517016, EPI_ISL_517017, EPI_ISL_517018, EPI_ISL_517019, EPI_ISL_517020, EPI_ISL_517021, EPI_ISL_517022, EPI_ISL_517023, EPI_ISL_517024, EPI_ISL_517025, EPI_ISL_517026, EPI_ISL_517027, EPI_ISL_517028, EPI_ISL_517029, EPI_ISL_517030, EPI_ISL_517031, EPI_ISL_517032, EPI_ISL_517033, EPI_ISL_517034, EPI_ISL_517035, EPI_ISL_517036, EPI_ISL_517037, EPI_ISL_517038, EPI_ISL_517039, EPI_ISL_517040, EPI_ISL_517041, EPI_ISL_517042, EPI_ISL_517043, EPI_ISL_517044, EPI_ISL_517045, EPI_ISL_517046, EPI_ISL_517047, EPI_ISL_517048, EPI_ISL_517049, EPI_ISL_517050, EPI_ISL_517051, EPI_ISL_517052, EPI_ISL_517053, EPI_ISL_517054, EPI_ISL_517055, EPI_ISL_517056, EPI_ISL_517057, EPI_ISL_517058, EPI_ISL_517059, EPI_ISL_517060, EPI_ISL_517061, EPI_ISL_517062, EPI_ISL_517063, EPI_ISL_517064, EPI_ISL_517065, EPI_ISL_517066, EPI_ISL_517067, EPI_ISL_517068, EPI_ISL_517069, EPI_ISL_517070, EPI_ISL_517071, EPI_ISL_517072, EPI_ISL_517073, EPI_ISL_517074, EPI_ISL_517075, EPI_ISL_517076, EPI_ISL_517077, EPI_ISL_517078, EPI_ISL_517079, EPI_ISL_517080, EPI_ISL_517081, EPI_ISL_517082, EPI_ISL_517083, EPI_ISL_517084, EPI_ISL_517085, EPI_ISL_517086, EPI_ISL_517087, EPI_ISL_517088, EPI_ISL_517089, EPI_ISL_517090, EPI_ISL_517091, EPI_ISL_517092, EPI_ISL_517093, EPI_ISL_517094, EPI_ISL_517095, EPI_ISL_517096, EPI_ISL_517097, EPI_ISL_517098, EPI_ISL_517099, EPI_ISL_517100, EPI_ISL_517101, EPI_ISL_517102, EPI_ISL_517103, EPI_ISL_517104, EPI_ISL_517105, EPI_ISL_517106, EPI_ISL_517107, EPI_ISL_517108, EPI_ISL_517109, EPI_ISL_517110, EPI_ISL_517111, EPI_ISL_517112, EPI_ISL_517113, EPI_ISL_517114, EPI_ISL_517115, EPI_ISL_517116, EPI_ISL_517117, EPI_ISL_517118, EPI_ISL_517119, EPI_ISL_517120, EPI_ISL_517121, EPI_ISL_517122, EPI_ISL_517123, EPI_ISL_517124, EPI_ISL_517125, EPI_ISL_517126, EPI_ISL_517127, EPI_ISL_517128, EPI_ISL_517129, EPI_ISL_517130, EPI_ISL_517131, EPI_ISL_517132, EPI_ISL_517133, EPI_ISL_517134, EPI_ISL_517135, EPI_ISL_517136, EPI_ISL_517137, EPI_ISL_517138, EPI_ISL_517139, EPI_ISL_517140, EPI_ISL_517141, EPI_ISL_517142, EPI_ISL_517143, EPI_ISL_517144, EPI_ISL_517145, EPI_ISL_517146, EPI_ISL_517147, EPI_ISL_517148, EPI_ISL_517149, EPI_ISL_517150, EPI_ISL_517151, EPI_ISL_517152, EPI_ISL_517153, EPI_ISL_517154, EPI_ISL_517155, EPI_ISL_517156, EPI_ISL_517157, EPI_ISL_517158, EPI_ISL_517159, EPI_ISL_517160, EPI_ISL_517161, EPI_ISL_517162, EPI_ISL_517163, EPI_ISL_517164, EPI_ISL_517165, EPI_ISL_517166, EPI_ISL_517167, EPI_ISL_517168, EPI_ISL_517169, EPI_ISL_517170, EPI_ISL_517171, EPI_ISL_517172, EPI_ISL_517173, EPI_ISL_517174, EPI_ISL_517175, EPI_ISL_517176, EPI_ISL_517177, EPI_ISL_517178, EPI_ISL_517179, EPI_ISL_517180, EPI_ISL_517181, EPI_ISL_517182, EPI_ISL_517183, EPI_ISL_517184, EPI_ISL_517185, EPI_ISL_517186, EPI_ISL_517187, EPI_ISL_517188, EPI_ISL_517189, EPI_ISL_517190, EPI_ISL_517191, EPI_ISL_517192, EPI_ISL_517193, EPI_ISL_517194, EPI_ISL_517195, EPI_ISL_517196, EPI_ISL_517197, EPI_ISL_517198, EPI_ISL_517199, EPI_ISL_517200, EPI_ISL_517201, EPI_ISL_517202, EPI_ISL_517203, EPI_ISL_517204, EPI_ISL_517205, EPI_ISL_517206, EPI_ISL_517207, EPI_ISL_517208, EPI_ISL_517209, EPI_ISL_517210, EPI_ISL_517211, EPI_ISL_517212, EPI_ISL_517213, EPI_ISL_517214, EPI_ISL_517215, EPI_ISL_517216, EPI_ISL_517217, EPI_ISL_517218, EPI_ISL_517219, EPI_ISL_517220, EPI_ISL_517221, EPI_ISL_517222, EPI_ISL_517223, EPI_ISL_517224, EPI_ISL_517225, EPI_ISL_517226, EPI_ISL_517227, EPI_ISL_517228, EPI_ISL_517229, EPI_ISL_517230, EPI_ISL_517231, EPI_ISL_517232, EPI_ISL_517233, EPI_ISL_517234, EPI_ISL_517235, EPI_ISL_517236, EPI_ISL_517237, EPI_ISL_517238, EPI_ISL_517239, EPI_ISL_517240, EPI_ISL_517241, EPI_ISL_517242, EPI_ISL_517243, EPI_ISL_517244, EPI_ISL_517245, EPI_ISL_517246, EPI_ISL_517247, EPI_ISL_517248, EPI_ISL_517249, EPI_ISL_517250, EPI_ISL_517251, EPI_ISL_517252, EPI_ISL_517253, EPI_ISL_517254, EPI_ISL_517255, EPI_ISL_517256, EPI_ISL_517257, EPI_ISL_517258, EPI_ISL_517259, EPI_ISL_517260, EPI_ISL_517261, EPI_ISL_517262, EPI_ISL_517263, EPI_ISL_517264, EPI_ISL_517265, EPI_ISL_517266, EPI_ISL_517267, EPI_ISL_517268, EPI_ISL_517269, EPI_ISL_517270, EPI_ISL_517271, EPI_ISL_517272, EPI_ISL_517273, EPI_ISL_517274, EPI_ISL_517275, EPI_ISL_517276, EPI_ISL_517277, EPI_ISL_517278, EPI_ISL_517279, EPI_ISL_517280, EPI_ISL_517281, EPI_ISL_517282, EPI_ISL_517283, EPI_ISL_517284, EPI_ISL_517285, EPI_ISL_517286, EPI_ISL_517287, EPI_ISL_517288, EPI_ISL_517289, EPI_ISL_517290, EPI_ISL_517291, EPI_ISL_517292, EPI_ISL_517293, EPI_ISL_517294, EPI_ISL_517295, EPI_ISL_517296, EPI_ISL_517297, EPI_ISL_517298, EPI_ISL_517299, EPI_ISL_517300, EPI_ISL_517301, EPI_ISL_517302, EPI_ISL_517303, EPI_ISL_517304, EPI_ISL_517305, EPI_ISL_517306, EPI_ISL_517307, EPI_ISL_517308, EPI_ISL_517309, EPI_ISL_517310, EPI_ISL_517311, EPI_ISL_517312, EPI_ISL_517313, EPI_ISL_517314, EPI_ISL_517315, EPI_ISL_517316, EPI_ISL_517317, EPI_ISL_517318, EPI_ISL_517319, EPI_ISL_517320, EPI_ISL_517321, EPI_ISL_517322, EPI_ISL_517323, EPI_ISL_517324, EPI_ISL_517325, EPI_ISL_517326, EPI_ISL_517327, EPI_ISL_517328, EPI_ISL_517329, EPI_ISL_517330, EPI_ISL_517331, EPI_ISL_517332, EPI_ISL_517333, EPI_ISL_517334, EPI_ISL_517335, EPI_ISL_517336, EPI_ISL_517337, EPI_ISL_517338, EPI_ISL_517339, EPI_ISL_517340, EPI_ISL_517341, EPI_ISL_517342, EPI_ISL_517343, EPI_ISL_517344, EPI_ISL_517345, EPI_ISL_517346, EPI_ISL_517347, EPI_ISL_517348, EPI_ISL_517349, EPI_ISL_517350, EPI_ISL_517351, EPI_ISL_517352, EPI_ISL_517353, EPI_ISL_517354, EPI_ISL_517355, EPI_ISL_517356, EPI_ISL_517357, EPI_ISL_517358, EPI_ISL_517359, EPI_ISL_517360, EPI_ISL_517361, EPI_ISL_517362, EPI_ISL_517363, EPI_ISL_517364, EPI_ISL_517365, EPI_ISL_517366, EPI_ISL_517367, EPI_ISL_517368, EPI_ISL_517369, EPI_ISL_517370, EPI_ISL_517371, EPI_ISL_517372, EPI_ISL_517373, EPI_ISL_517374, EPI_ISL_517375, EPI_ISL_517376, EPI_ISL_517377, EPI_ISL_517378, EPI_ISL_517379, EPI_ISL_517380, EPI_ISL_517381, EPI_ISL_517382, EPI_ISL_517383, EPI_ISL_517384, EPI_ISL_517385, EPI_ISL_517386, EPI_ISL_517387, EPI_ISL_517388, EPI_ISL_517389, EPI_ISL_517390, EPI_ISL_517391, EPI_ISL_517392, EPI_ISL_517393, EPI_ISL_517394, EPI_ISL_517395, EPI_ISL_517396, EPI_ISL_517397, EPI_ISL_517398, EPI_ISL_517399, EPI_ISL_517400, EPI_ISL_517401, EPI_ISL_517402, EPI_ISL_517403, EPI_ISL_517404, EPI_ISL_517405, EPI_ISL_517406, EPI_ISL_517407, EPI_ISL_517408, EPI_ISL_517409, EPI_ISL_517410, EPI_ISL_517411, EPI_ISL_517412, EPI_ISL_517413, EPI_ISL_517414, EPI_ISL_517415, EPI_ISL_517416, EPI_ISL_517417, EPI_ISL_517418, EPI_ISL_517419, EPI_ISL_517420, EPI_ISL_517421, EPI_ISL_517422, EPI_ISL_517423, EPI_ISL_517424, EPI_ISL_517425, EPI_ISL_517426, EPI_ISL_517427, EPI_ISL_517428, EPI_ISL_517429, EPI_ISL_517430, EPI_ISL_517431, EPI_ISL_517432, EPI_ISL_517433, EPI_ISL_517434, EPI_ISL_517435, EPI_ISL_517436, EPI_ISL_517437, EPI_ISL_517438, EPI_ISL_517439, EPI_ISL_517440, EPI_ISL_517441, EPI_ISL_517442, EPI_ISL_517443, EPI_ISL_517444, EPI_ISL_517445, EPI_ISL_517446, EPI_ISL_517447, EPI_ISL_517448, EPI_ISL_517449, EPI_ISL_517450, EPI_ISL_517451, EPI_ISL_517452, EPI_ISL_517453, EPI_ISL_517454, EPI_ISL_517455, EPI_ISL_517456, EPI_ISL_517457, EPI_ISL_517458, EPI_ISL_517459, EPI_ISL_517460, EPI_ISL_517461, EPI_ISL_517462, EPI_ISL_517463, EPI_ISL_517464, EPI_ISL_517465, EPI_ISL_517466, EPI_ISL_517467, EPI_ISL_517468, EPI_ISL_517469, EPI_ISL_517470, EPI_ISL_517471, EPI_ISL_517472, EPI_ISL_517473, EPI_ISL_517474, EPI_ISL_517475, EPI_ISL_517476, EPI_ISL_517477, EPI_ISL_517478, EPI_ISL_517479, EPI_ISL_517480, EPI_ISL_517481, EPI_ISL_517482, EPI_ISL_517483, EPI_ISL_517484, EPI_ISL_517485, EPI_ISL_517486, EPI_ISL_517487, EPI_ISL_517488, EPI_ISL_517489, EPI_ISL_517490, EPI_ISL_517491, EPI_ISL_517492, EPI_ISL_517493, EPI_ISL_517494, EPI_ISL_517495, EPI_ISL_517496, EPI_ISL_517497, EPI_ISL_517498, EPI_ISL_517499, EPI_ISL_517500, EPI_ISL_517501, EPI_ISL_517502, EPI_ISL_517503, EPI_ISL_517504, EPI_ISL_517505, EPI_ISL_517506, EPI_ISL_517507, EPI_ISL_517508, EPI_ISL_517509 | | | |
| see above | Liverpool Clinical Laboratories | COVID-19 Genomics UK (COG-UK) Consortium | Sam Haldenby, Anita Lucaci, Steve Paterson, Julian Hiscox, Alistair Darby, M Almsaud, A Alrezaihi, Muhannad Alruwaili, Stuart D Armstrong, Jones Benjamin, Eleanor G Bentley, Anu Chawla, Jordan J Clark, Angela Cowell, Richard Eccles, Isabel García-Dorival, Matthew Gemmell, Alessandro Gerada, PKF Gilmore, Richard Gregory, Ximeng Han, Catherine Hartley, Margaret Hughes, Miren Iturriza-Gomara, James Johnson, L Luu, Jenifer Manson, Charlotte Nelson, Elaine O'Toole, Cassie Olateju, Rebekah Penrice-Randal , Lucille Rainbow, N.P Randle, Trevor Ian Robinson, Parul Sharma, Ghada T Shawli, James P Stewart, Neil Swainston, Ecaterina Vamos, Joanne Watts, Mark Whitehead |
| EPI_ISL_517510, EPI_ISL_517511, EPI_ISL_517512, EPI_ISL_517513, EPI_ISL_517514, EPI_ISL_517515, EPI_ISL_517516, EPI_ISL_517517, EPI_ISL_517518, EPI_ISL_517519, EPI_ISL_517520, EPI_ISL_517521, EPI_ISL_517522, EPI_ISL_517523, EPI_ISL_517524, EPI_ISL_517525, EPI_ISL_517526, EPI_ISL_517527, EPI_ISL_517528, EPI_ISL_517529, EPI_ISL_517530 | | | |
| see above | Centre for Enzyme Innovation, University of Portsmouth / Translational Research Laboratory, Portsmouth Hospitals NHS Trust | COVID-19 Genomics UK (COG-UK) Consortium | Angela Beckett,Yann Bourgeois,Garry Scarlett,Sharon Glaysher,Scott Elliott,Kelly Bicknell,Robert Impey,Allyson Lloyd,Sarah Wyllie,Ethan Butcher,Anoop Chauhan,Samuel Robson |
| EPI_ISL_517531 | Laboratorio de Referencia Nacional de Virus Respiratorio. Centro Nacional de Salud Publica. Instituto Nacional de Salud Peru. | Laboratorio de Referencia Nacional de Biotecnología y Biología Molecular. Centro Nacional de Salud Publica. Instituto Nacional de Salud Peru. | Carlos Padilla Rojas, Karolyn Vega Chozo, Priscila Lope Pari, Omar Caceres Rey, Marco Galarza Perez, Maribel Huaringa Nuñez, Johanna Balbuena Torres, Henri Bailon Calderon, Nancy Rojas Serrano. |
| EPI_ISL_517532, EPI_ISL_517533, EPI_ISL_517534, EPI_ISL_517535, EPI_ISL_517536, EPI_ISL_517537, EPI_ISL_517538, EPI_ISL_517539, EPI_ISL_517540, EPI_ISL_517541, EPI_ISL_517542 | | | |
| see above | Centre for Enzyme Innovation, University of Portsmouth / Translational Research Laboratory, Portsmouth Hospitals NHS Trust | COVID-19 Genomics UK (COG-UK) Consortium | Angela Beckett,Yann Bourgeois,Garry Scarlett,Sharon Glaysher,Scott Elliott,Kelly Bicknell,Robert Impey,Allyson Lloyd,Sarah Wyllie,Ethan Butcher,Anoop Chauhan,Samuel Robson |
| EPI_ISL_517543, EPI_ISL_517544, EPI_ISL_517545, EPI_ISL_517546, EPI_ISL_517547, EPI_ISL_517548, EPI_ISL_517549, EPI_ISL_517550, EPI_ISL_517551, EPI_ISL_517552, EPI_ISL_517553, EPI_ISL_517554, EPI_ISL_517555, EPI_ISL_517556, EPI_ISL_517557, EPI_ISL_517558, EPI_ISL_517559, EPI_ISL_517560, EPI_ISL_517561, EPI_ISL_517562, EPI_ISL_517563, EPI_ISL_517564, EPI_ISL_517565, EPI_ISL_517566, EPI_ISL_517567, EPI_ISL_517568, EPI_ISL_517569, EPI_ISL_517570, EPI_ISL_517571, EPI_ISL_517572, EPI_ISL_517573, EPI_ISL_517574, EPI_ISL_517575, EPI_ISL_517576, EPI_ISL_517577, EPI_ISL_517578, EPI_ISL_517579 | | | |
| see above | Virology Department, Sheffield Teaching Hospitals NHS Foundation Trust/Department of Infection, Immunity and Cardiovascular Disease, The Medical School, University of Sheffield | COVID-19 Genomics UK (COG-UK) Consortium | Thushan de Silva, Matthew Parker, Nikki Smith, Adri Angyal, Rebecca Brown, Luke Green, Rachel Tucker, Paul Parsons, Danielle Groves, Katie Johnson, Laura Carrilero, Alex Keeley, Dave Partridge, Matthew Wyles, Benjamin Lindsey, Mehmet Yavuz, Mohammad Raza, Cariad Evans |
| EPI_ISL_517580, EPI_ISL_517581, EPI_ISL_517582 | West of Scotland Specialist Virology Centre, NHSGGC / MRC-University of Glasgow Centre for Virus Research | COVID-19 Genomics UK (COG-UK) Consortium | Ana da Silva Filipe, Natasha Johnson, Kathy Smollett, Daniel Mair, Stephen Carmichael, Lily Tong, Jenna Nichols, Elihu Aranday-Cortes, Kirstyn Brunker, Yasmin Parr, Alice Broos, Kyriaki Nomikou; Sarah McDonald, Marc Niebel, Patawee Asamaphan; Richard Orton, Joseph Hughes, Sreenu Vattipally, David L Robertson; Alasdair MacLean, Rory Gunson; Kathy Li, Natasha Jesudason, Rajiv Shah, James Shepherd, Antonia Ho, Emma Thomson |
| EPI_ISL_517583, EPI_ISL_517584, EPI_ISL_517585, EPI_ISL_517586, EPI_ISL_517587, EPI_ISL_517588, EPI_ISL_517589 | Virology Department, Royal Infirmary of Edinburgh, NHS Lothian / School of Biological Sciences, University of Edinburgh / Institute of Genetics and Molecular Medicine, University of Edinburgh | COVID-19 Genomics UK (COG-UK) Consortium | McHugh M, Dewar R, Rooke S, Gallagher M, Balcaza C, O'Toole Á, Scher E, Hill V, McCrone JT, Colquhoun R, Yu X, Jackson B, Rambaut A, Williams TC, Templeton K |
| EPI_ISL_517590, EPI_ISL_517591, EPI_ISL_517592, EPI_ISL_517593, EPI_ISL_517594, EPI_ISL_517595, EPI_ISL_517596, EPI_ISL_517597, EPI_ISL_517598, EPI_ISL_517599, EPI_ISL_517600, EPI_ISL_517601, EPI_ISL_517602, EPI_ISL_517603, EPI_ISL_517604, EPI_ISL_517605, EPI_ISL_517606, EPI_ISL_517607, EPI_ISL_517608, EPI_ISL_517609, EPI_ISL_517610 | | | |
| see above | Wales Specialist Virology Centre Sequencing lab: Pathogen Genomics Unit | COVID-19 Genomics UK (COG-UK) Consortium | Catherine Moore, Johnathan Evans, Laura Gifford, Malorie Perry, Simon Cottrell, Angela Marchbank, Alec Birchley, Alexander Adams, Amy Gaskin, Bree Gatica-Wilcox, Jason Coombes, Joel Southgate, Lauren Gilbert, Lee Graham, Nicole Pacchiarini, Sara Kumziene-Summerhayes, Sarah Taylor, Sophie Jones, Sara Rey, Matthew Bull, Joanne Watkins, |

| | | | |
|---|---|---|---|
| | | | Sally Corden, Tom Connor |
| EPI_ISL_517611 | Institute of Medical Microbiology, University Medical Center Goettingen | Institute of Human Genetics, University Medical Center Goettingen | Arne Zibat, Gabriela Salinas-Riester, Andreas E Zautner, Maren Sitte, Antje Dickmanns, Maren Stegmann, Uwe Groß, Matthias Dobbelstein, Bernd Wollnik |
| EPI_ISL_517612, EPI_ISL_517613, EPI_ISL_517614, EPI_ISL_517615, EPI_ISL_517616, EPI_ISL_517617, EPI_ISL_517618, EPI_ISL_517619, EPI_ISL_517620, EPI_ISL_517621, EPI_ISL_517622, EPI_ISL_517623, EPI_ISL_517624, EPI_ISL_517625, EPI_ISL_517626, EPI_ISL_517627, EPI_ISL_517628, EPI_ISL_517629, EPI_ISL_517630, EPI_ISL_517631, EPI_ISL_517632, EPI_ISL_517633, EPI_ISL_517634, EPI_ISL_517635, EPI_ISL_517636, EPI_ISL_517637, EPI_ISL_517638, EPI_ISL_517639, EPI_ISL_517640, EPI_ISL_517641, EPI_ISL_517642, EPI_ISL_517643, EPI_ISL_517644, EPI_ISL_517645, EPI_ISL_517646, EPI_ISL_517647, EPI_ISL_517648, EPI_ISL_517649, EPI_ISL_517650, EPI_ISL_517651, EPI_ISL_517652, EPI_ISL_517653, EPI_ISL_517654, EPI_ISL_517655, EPI_ISL_517656, EPI_ISL_517657, EPI_ISL_517658, EPI_ISL_517659, EPI_ISL_517660, EPI_ISL_517661, EPI_ISL_517662 | | | |
| see above | Academic Hospital Paramaribo | Erasmus Medical Center | Bas Oude Munnink, Dion Gajadin, Ed Ijzerman, Emmanuelle Munger, Gary Gummels, Ingrid Krishnadath, Lycke Woittiez, Marion Koopmans, Mireille Van de Veer, Princes Wongsowidjojo, Radjesh Ori, Rohma Banwari, Stephen Vreden |
| EPI_ISL_517686, EPI_ISL_517687, EPI_ISL_517713 | Laboratorio de Referencia Nacional de Virus Respiratorio. Centro Nacional de Salud Publica. Instituto Nacional de Salud Peru. | Laboratorio de Referencia Nacional de Biotecnología y Biología Molecular. Centro Nacional de Salud Publica. Instituto Nacional de Salud Peru. | Carlos Padilla Rojas, Karolyn Vega Chozo, Priscila Lope Pari, Omar Caceres Rey, Marco Galarza Perez, Maribel Huaringa Nuñez, Johanna Balbuena Torres, Henri Bailon Calderon, Nancy Rojas Serrano. |
| EPI_ISL_517770 | Laboratorio de Referencia Nacional de Virus Respiratorio. Centro Nacional de Salud Publica. Instituto Nacional de Salud Peru | Laboratorio de Referencia Nacional de Biotecnología y Biología Molecular. Centro Nacional de Salud Publica. Instituto Nacional de Salud Peru. | Carlos Padilla Rojas, Karolyn Vega Chozo, Priscila Lope Pari, Omar Caceres Rey, Marco Galarza Perez, Maribel Huaringa Nuñez, Johanna Balbuena Torres, Henri Bailon Calderon, Nancy Rojas Serrano. |
| EPI_ISL_517771, EPI_ISL_517772, EPI_ISL_517773, EPI_ISL_517774, EPI_ISL_517775, EPI_ISL_517776, EPI_ISL_517777, EPI_ISL_517778, EPI_ISL_517779, EPI_ISL_517780, EPI_ISL_517781, EPI_ISL_517782, EPI_ISL_517783, EPI_ISL_517784, EPI_ISL_517785, EPI_ISL_517786, EPI_ISL_517787, EPI_ISL_517788, EPI_ISL_517789, EPI_ISL_517790, EPI_ISL_517791, EPI_ISL_517792, EPI_ISL_517793, EPI_ISL_517794, EPI_ISL_517795, EPI_ISL_517796, EPI_ISL_517797, EPI_ISL_517798, EPI_ISL_517799, EPI_ISL_517800, EPI_ISL_517801, EPI_ISL_517802, EPI_ISL_517803, EPI_ISL_517804, EPI_ISL_517805, EPI_ISL_517806, EPI_ISL_517807, EPI_ISL_517808, EPI_ISL_517809, EPI_ISL_517810, EPI_ISL_517811, EPI_ISL_517812, EPI_ISL_517813, EPI_ISL_517814, EPI_ISL_517815, EPI_ISL_517816, EPI_ISL_517817, EPI_ISL_517818, EPI_ISL_517819, EPI_ISL_517820, EPI_ISL_517821, EPI_ISL_517822, EPI_ISL_517823, EPI_ISL_517824, EPI_ISL_517825, EPI_ISL_517826, EPI_ISL_517827, EPI_ISL_517828, EPI_ISL_517829, EPI_ISL_517830, EPI_ISL_517831, EPI_ISL_517832, EPI_ISL_517833, EPI_ISL_517834, EPI_ISL_517835, EPI_ISL_517836, EPI_ISL_517837, EPI_ISL_517838, EPI_ISL_517839, EPI_ISL_517840, EPI_ISL_517841, EPI_ISL_517842, EPI_ISL_517843, EPI_ISL_517844, EPI_ISL_517845, EPI_ISL_517846, EPI_ISL_517847, EPI_ISL_517848, EPI_ISL_517849, EPI_ISL_517850, EPI_ISL_517851, EPI_ISL_517852, EPI_ISL_517853, EPI_ISL_517854, EPI_ISL_517855, EPI_ISL_517856, EPI_ISL_517857, EPI_ISL_517858, EPI_ISL_517859, EPI_ISL_517860, EPI_ISL_517861, EPI_ISL_517862, EPI_ISL_517863, EPI_ISL_517864, EPI_ISL_517865, EPI_ISL_517866, EPI_ISL_517867, EPI_ISL_517868, EPI_ISL_517869, EPI_ISL_517870, EPI_ISL_517871, EPI_ISL_517872, EPI_ISL_517873, EPI_ISL_517874, EPI_ISL_517875, EPI_ISL_517876, EPI_ISL_517877, EPI_ISL_517878, EPI_ISL_517879, EPI_ISL_517880, EPI_ISL_517881, EPI_ISL_517882, EPI_ISL_517883, EPI_ISL_517884, EPI_ISL_517885, EPI_ISL_517886, EPI_ISL_517887, EPI_ISL_517888, EPI_ISL_517889, EPI_ISL_517890, EPI_ISL_517891, EPI_ISL_517892, EPI_ISL_517893, EPI_ISL_517894, EPI_ISL_517895, EPI_ISL_517896, EPI_ISL_517897, EPI_ISL_517898, EPI_ISL_517899, EPI_ISL_517900, EPI_ISL_517901, EPI_ISL_517902, EPI_ISL_517903, EPI_ISL_517904, EPI_ISL_517905, EPI_ISL_517906, EPI_ISL_517907, EPI_ISL_517908, EPI_ISL_517909, EPI_ISL_517910, EPI_ISL_517911, EPI_ISL_517912, EPI_ISL_517913, EPI_ISL_517914, EPI_ISL_517915, EPI_ISL_517916, EPI_ISL_517917, EPI_ISL_517918, EPI_ISL_517919, EPI_ISL_517920, EPI_ISL_517921, EPI_ISL_517922, EPI_ISL_517923, EPI_ISL_517924, EPI_ISL_517925, EPI_ISL_517926, EPI_ISL_517927, EPI_ISL_517928, EPI_ISL_517929, EPI_ISL_517930, EPI_ISL_517931, EPI_ISL_517932, EPI_ISL_517933, EPI_ISL_517934, EPI_ISL_517935, EPI_ISL_517936, EPI_ISL_517937, EPI_ISL_517938, EPI_ISL_517939, EPI_ISL_517940, EPI_ISL_517941, EPI_ISL_517942, EPI_ISL_517943, EPI_ISL_517944, EPI_ISL_517945, EPI_ISL_517946, EPI_ISL_517947, EPI_ISL_517948, EPI_ISL_517949, EPI_ISL_517950, EPI_ISL_517951, EPI_ISL_517952, EPI_ISL_517953, EPI_ISL_517954, EPI_ISL_517955, EPI_ISL_517956, EPI_ISL_517957 | | | |
| see above | Florida Bureau of Public Health Laboratories | Florida Bureau of Public Health Laboratories | Sarah Schmedes, Jason Blanton |
| EPI_ISL_517958 | Laboratorio de Referencia Nacional de Virus Respiratorio. Centro Nacional de Salud Publica. Instituto Nacional de Salud Peru. | Laboratorio de Referencia Nacional de Biotecnología y Biología Molecular. Centro Nacional de Salud Publica. Instituto Nacional de Salud Peru. | Carlos Padilla Rojas, Karolyn Vega Chozo, Priscila Lope Pari, Omar Caceres Rey, Marco Galarza Perez, Maribel Huaringa Nuñez, Johanna Balbuena Torres, Henri Bailon Calderon, Nancy Rojas Serrano. |
| EPI_ISL_517959 | Laboratorio de Referencia Nacional de Virus Respiratorio. Centro Nacional de Salud Publica. Instituto Nacional de Salud Peru | Laboratorio de Referencia Nacional de Biotecnología y Biología Molecular. Centro Nacional de Salud Publica. Instituto Nacional de Salud Peru. | Carlos Padilla Rojas, Karolyn Vega Chozo, Priscila Lope Pari, Omar Caceres Rey, Marco Galarza Perez, Maribel Huaringa Nuñez, Johanna Balbuena Torres, Henri Bailon Calderon, Nancy Rojas Serrano. |
| EPI_ISL_517960, EPI_ISL_517961, EPI_ISL_517962, EPI_ISL_517963, EPI_ISL_517964, EPI_ISL_517965, EPI_ISL_517966, EPI_ISL_517967, EPI_ISL_517968, EPI_ISL_517969, EPI_ISL_517970, EPI_ISL_517971, EPI_ISL_517972, EPI_ISL_517973, EPI_ISL_517974, EPI_ISL_517975, EPI_ISL_517976, EPI_ISL_517977, EPI_ISL_517978, EPI_ISL_517979, EPI_ISL_517980, EPI_ISL_517981, EPI_ISL_517982, EPI_ISL_517983, EPI_ISL_517984, EPI_ISL_517985, EPI_ISL_517986, EPI_ISL_517987, EPI_ISL_517988, EPI_ISL_517989, EPI_ISL_517990, EPI_ISL_517991, EPI_ISL_517992, EPI_ISL_517993, EPI_ISL_517994, EPI_ISL_517995 | | | |
| see above | Texas Department of State Health Services | Texas Department of State Health Services | Rashmi Tuladhar, Bonnie Oh, Cara Akrout, Jenny Zhang, Maliha Rahman, Anita Pokharel, Myong Koag, Chun Wang, Rachel Lee, Grace Kubin |
| EPI_ISL_522407, EPI_ISL_522408, EPI_ISL_522409 | Department of Infection Prevention and Infectious Diseases, University Hospital Regensburg | Department of Infection Prevention and Infectious Diseases, University Hospital Regensburg | Fritsch,J., Holzmann,T., Schneider-Brachert,W. |
| EPI_ISL_525472, EPI_ISL_525473 | Institute of Clinical Microbiology and Hygiene, University Hospital Regensburg | Institute of Clinical Microbiology and Hygiene, University Hospital Regensburg | Hiergeist, A. |
| EPI_ISL_529139 | Centro de Desenvolvimento Tecnologico em Saude, Fundacao Oswaldo Cruz | Centro de Desenvolvimento Tecnologico em Saude, Fundacao Oswaldo Cruz | Souza,T.M., Fintelman-Rodrigues,N., De Paula,A.D., Saraiva,F.B., Ferreira,M.A., Sacramento,C.Q., Medeiros,M.A. |
| EPI_ISL_529140 | Centro de Desenvolvimento Tecnologico em Saude, Fundacao Oswaldo Cruz | Centro de Desenvolvimento Tecnologico em Saude, Fundacao Oswaldo Cruz | Souza,T.M., Fintelman-Rodrigues,N., De Paula,A.D., Saraiva,F.B., Ferreira,M.A., Sacramento,C.Q., Medeiros,M.A. |
| EPI_ISL_529167, EPI_ISL_529168 | Maryland Department of Health Laboratory | Maryland Department of Health Laboratory | Eric Keller, Maryland Department of Health Laboratories Administration |

We gratefully acknowledge the following Authors from the Originating laboratories responsible for obtaining the specimens, as well as the Submitting laboratories where the genome data were generated and shared via GISAID, on which this research is based.

All Submitters of data may be contacted directly via www.gisaid.org

| Accession ID | Originating Laboratory | Submitting Laboratory | Authors |
|---|---|---|---|
| EPI_ISL_517996, EPI_ISL_517997, EPI_ISL_517998, EPI_ISL_517999, EPI_ISL_518000, EPI_ISL_518001, EPI_ISL_518002, EPI_ISL_518003, EPI_ISL_518004, EPI_ISL_518005, EPI_ISL_518006, EPI_ISL_518007, EPI_ISL_518008, EPI_ISL_518009, EPI_ISL_518010, EPI_ISL_518011, EPI_ISL_518012, EPI_ISL_518013, EPI_ISL_518014, EPI_ISL_518015, EPI_ISL_518016, EPI_ISL_518017 | | | |
| see above | Singapore General Hospital | Department of Microbiology | Nurdyana Abdul Rahman, Kun Lee Lim, Chenhao Li, Kian Sing Chan, Lynette Oon, Kern Rei Chng, Niranjan Nagarajan, Karrie Ko |
| EPI_ISL_518031, EPI_ISL_518032 | Indian Institute of Science | National Institute of Biomedical Genomics - DBT's PAN-INDIA 1000 SARS--CoV-2 RNA Genome Sequencing Consortium | Arindam Maitra, Bharath K Sundararaj, Harsha Raheja, N. Srinivasan, Deepak K Saini, Amit Singh, Saumitra Das |
| EPI_ISL_518033, EPI_ISL_518034, EPI_ISL_518035, EPI_ISL_518036, EPI_ISL_518037, EPI_ISL_518038, EPI_ISL_518039, EPI_ISL_518040, EPI_ISL_518041, EPI_ISL_518042, EPI_ISL_518043, EPI_ISL_518044, EPI_ISL_518045, EPI_ISL_518046, EPI_ISL_518047, EPI_ISL_518048, EPI_ISL_518049, EPI_ISL_518050, EPI_ISL_518051, EPI_ISL_518052, EPI_ISL_518053 | | | |
| see above | NHLS-IALCH | KRISP, KZN Research Innovation and Sequencing Platform | Giandhari J, Pillay S, Lessells R, Mdlalose K, York D, Khan S, Tegally H, Wilkinson E, de Oliveira T |
| EPI_ISL_518063, EPI_ISL_518064, EPI_ISL_518065, EPI_ISL_518066, EPI_ISL_518067, EPI_ISL_518068, EPI_ISL_518069, EPI_ISL_518070, EPI_ISL_518071, EPI_ISL_518072, EPI_ISL_518073, EPI_ISL_518074, EPI_ISL_518075, EPI_ISL_518076, EPI_ISL_518077, EPI_ISL_518078, EPI_ISL_518079, EPI_ISL_518080 | | | |
| see above | Microbiological Diagnostic Unit - Public Health Laboratory (MDU-PHL) | MDU-PHL | Seemann T., Schultz M., Sait, M., Sherry, N. |
| EPI_ISL_518081, EPI_ISL_518082, EPI_ISL_518083, EPI_ISL_518084, EPI_ISL_518085, EPI_ISL_518086, EPI_ISL_518087, EPI_ISL_518088, EPI_ISL_518089, EPI_ISL_518090, EPI_ISL_518091, EPI_ISL_518092 | | | |
| see above | Victorian Infectious Diseases Reference Laboratory (VIDRL) | VIDRL and MDU-PHL | Caly L., Seemann T., Sait, M., Schultz M., Druce J., Sherry, N. |
| EPI_ISL_518093, EPI_ISL_518094, EPI_ISL_518095, EPI_ISL_518096, EPI_ISL_518097 | Microbiological Diagnostic Unit - Public Health Laboratory (MDU-PHL) | MDU-PHL | Seemann T., Schultz M., Sait, M., Sherry, N. |
| EPI_ISL_518098, EPI_ISL_518099, EPI_ISL_518100, EPI_ISL_518101, EPI_ISL_518102, EPI_ISL_518103, EPI_ISL_518104, EPI_ISL_518105, EPI_ISL_518106, EPI_ISL_518107, EPI_ISL_518108, EPI_ISL_518109, EPI_ISL_518110, EPI_ISL_518111, EPI_ISL_518112, EPI_ISL_518113, EPI_ISL_518114, EPI_ISL_518115, EPI_ISL_518116, EPI_ISL_518117, EPI_ISL_518118, EPI_ISL_518119, EPI_ISL_518120, EPI_ISL_518121, EPI_ISL_518122, EPI_ISL_518123, EPI_ISL_518124, EPI_ISL_518125, EPI_ISL_518126, EPI_ISL_518127, EPI_ISL_518128, EPI_ISL_518129, EPI_ISL_518130, EPI_ISL_518131, EPI_ISL_518132, EPI_ISL_518133, EPI_ISL_518134, EPI_ISL_518135, EPI_ISL_518136, EPI_ISL_518137, EPI_ISL_518138, EPI_ISL_518139, EPI_ISL_518140, EPI_ISL_518141, EPI_ISL_518142, EPI_ISL_518143, EPI_ISL_518144, EPI_ISL_518145, EPI_ISL_518146, EPI_ISL_518147, EPI_ISL_518148, EPI_ISL_518149, EPI_ISL_518150, EPI_ISL_518151, EPI_ISL_518152, EPI_ISL_518153, EPI_ISL_518154, EPI_ISL_518155, EPI_ISL_518156, EPI_ISL_518157, EPI_ISL_518158, EPI_ISL_518159, EPI_ISL_518160, EPI_ISL_518161, EPI_ISL_518162, EPI_ISL_518163, EPI_ISL_518164, EPI_ISL_518165, EPI_ISL_518166, EPI_ISL_518167, EPI_ISL_518168, EPI_ISL_518169, EPI_ISL_518170, EPI_ISL_518171, EPI_ISL_518172, EPI_ISL_518173, EPI_ISL_518174, EPI_ISL_518175, EPI_ISL_518176, EPI_ISL_518177, EPI_ISL_518178, EPI_ISL_518179, EPI_ISL_518180, EPI_ISL_518181, EPI_ISL_518182, EPI_ISL_518183, EPI_ISL_518184, EPI_ISL_518185, EPI_ISL_518186, EPI_ISL_518187, EPI_ISL_518188, EPI_ISL_518189, EPI_ISL_518190, EPI_ISL_518191, EPI_ISL_518192, EPI_ISL_518193, EPI_ISL_518194, EPI_ISL_518195, EPI_ISL_518196, EPI_ISL_518197, EPI_ISL_518198, EPI_ISL_518199, EPI_ISL_518200, EPI_ISL_518201, EPI_ISL_518202, EPI_ISL_518203, EPI_ISL_518204, EPI_ISL_518205, EPI_ISL_518206, EPI_ISL_518207, EPI_ISL_518208, EPI_ISL_518209, EPI_ISL_518210, EPI_ISL_518211, EPI_ISL_518212, EPI_ISL_518213, EPI_ISL_518214, EPI_ISL_518215, EPI_ISL_518216, EPI_ISL_518217, EPI_ISL_518218, EPI_ISL_518219, EPI_ISL_518220, EPI_ISL_518221, EPI_ISL_518222, EPI_ISL_518223, EPI_ISL_518224, EPI_ISL_518225, EPI_ISL_518226, EPI_ISL_518227, EPI_ISL_518228, EPI_ISL_518229, EPI_ISL_518230 | | | |
| see above | Victorian Infectious Diseases Reference Laboratory (VIDRL) | VIDRL and MDU-PHL | Caly L., Seemann T., Sait, M., Schultz M., Druce J., Sherry, N. |
| EPI_ISL_518231, EPI_ISL_518232, EPI_ISL_518233, EPI_ISL_518234, EPI_ISL_518235, EPI_ISL_518236, EPI_ISL_518237, EPI_ISL_518238 | Microbiological Diagnostic Unit - Public Health Laboratory (MDU-PHL) | MDU-PHL | Seemann T., Schultz M., Sait, M., Sherry, N. |
| EPI_ISL_518239, EPI_ISL_518240 | Victorian Infectious Diseases Reference Laboratory (VIDRL) | VIDRL and MDU-PHL | Caly L., Seemann T., Sait, M., Schultz M., Druce J., Sherry, N. |
| EPI_ISL_518241, EPI_ISL_518242, EPI_ISL_518243 | Microbiological Diagnostic Unit - Public Health Laboratory (MDU-PHL) | MDU-PHL | Seemann T., Schultz M., Sait, M., Sherry, N. |
| EPI_ISL_518244, EPI_ISL_518245, EPI_ISL_518246, EPI_ISL_518247, EPI_ISL_518248, EPI_ISL_518249 | Victorian Infectious Diseases Reference Laboratory (VIDRL) | VIDRL and MDU-PHL | Caly L., Seemann T., Sait, M., Schultz M., Druce J., Sherry, N. |
| EPI_ISL_518250, EPI_ISL_518251, EPI_ISL_518252, EPI_ISL_518253, EPI_ISL_518254, EPI_ISL_518255, EPI_ISL_518256, EPI_ISL_518257, EPI_ISL_518258, EPI_ISL_518259, EPI_ISL_518260, EPI_ISL_518261, EPI_ISL_518262, EPI_ISL_518263, EPI_ISL_518264, EPI_ISL_518265, EPI_ISL_518266, EPI_ISL_518267, EPI_ISL_518268, EPI_ISL_518269, EPI_ISL_518270, EPI_ISL_518271, EPI_ISL_518272, EPI_ISL_518273, EPI_ISL_518274, EPI_ISL_518275, EPI_ISL_518276, EPI_ISL_518277, EPI_ISL_518278, EPI_ISL_518279, EPI_ISL_518280, EPI_ISL_518281, EPI_ISL_518282, EPI_ISL_518283, EPI_ISL_518284, EPI_ISL_518285, EPI_ISL_518286, EPI_ISL_518287, EPI_ISL_518288, EPI_ISL_518289, EPI_ISL_518290, EPI_ISL_518292, EPI_ISL_518293, EPI_ISL_518294, EPI_ISL_518295, EPI_ISL_518296, EPI_ISL_518297, EPI_ISL_518298, EPI_ISL_518299, EPI_ISL_518300, EPI_ISL_518301, EPI_ISL_518302, EPI_ISL_518303, EPI_ISL_518304, EPI_ISL_518305, EPI_ISL_518306, EPI_ISL_518307, EPI_ISL_518308, EPI_ISL_518309, EPI_ISL_518310, EPI_ISL_518311, EPI_ISL_518312, EPI_ISL_518313, EPI_ISL_518314, EPI_ISL_518315, EPI_ISL_518316, EPI_ISL_518317, EPI_ISL_518318, EPI_ISL_518319, EPI_ISL_518320, EPI_ISL_518321, EPI_ISL_518322, EPI_ISL_518323, EPI_ISL_518324, EPI_ISL_518325, EPI_ISL_518326, EPI_ISL_518327, EPI_ISL_518328, EPI_ISL_518329, EPI_ISL_518330, EPI_ISL_518331, EPI_ISL_518332, EPI_ISL_518333, EPI_ISL_518334, EPI_ISL_518335, EPI_ISL_518336, EPI_ISL_518337, EPI_ISL_518339, EPI_ISL_518340, EPI_ISL_518341, EPI_ISL_518342, EPI_ISL_518343, EPI_ISL_518344, EPI_ISL_518345, EPI_ISL_518346, EPI_ISL_518347, EPI_ISL_518348, EPI_ISL_518349, EPI_ISL_518350, EPI_ISL_518351, EPI_ISL_518352, EPI_ISL_518353, EPI_ISL_518354, EPI_ISL_518355, EPI_ISL_518356, EPI_ISL_518357, EPI_ISL_518358, EPI_ISL_518359, EPI_ISL_518360, EPI_ISL_518361, EPI_ISL_518362, EPI_ISL_518363, EPI_ISL_518364, EPI_ISL_518365, EPI_ISL_518366, EPI_ISL_518367, EPI_ISL_518368, EPI_ISL_518369, EPI_ISL_518370, EPI_ISL_518371, EPI_ISL_518372, EPI_ISL_518373, EPI_ISL_518374, EPI_ISL_518375, EPI_ISL_518376, EPI_ISL_518377, EPI_ISL_518378, EPI_ISL_518379, EPI_ISL_518380, EPI_ISL_518381, EPI_ISL_518382, EPI_ISL_518383, EPI_ISL_518384, EPI_ISL_518385, EPI_ISL_518386, EPI_ISL_518387 | | | |
| see above | Microbiological Diagnostic Unit - Public Health Laboratory (MDU-PHL) | MDU-PHL | Seemann T., Schultz M., Sait, M., Sherry, N. |
| EPI_ISL_518388, EPI_ISL_518389, EPI_ISL_518390, EPI_ISL_518391, EPI_ISL_518392, EPI_ISL_518393, EPI_ISL_518394, EPI_ISL_518395, EPI_ISL_518396, EPI_ISL_518397, EPI_ISL_518398, EPI_ISL_518399, EPI_ISL_518400, EPI_ISL_518401 | | | |
| see above | Victorian Infectious Diseases Reference Laboratory (VIDRL) | VIDRL and MDU-PHL | Caly L., Seemann T., Sait, M., Schultz M., Druce J., Sherry, N. |
| EPI_ISL_518402, EPI_ISL_518403, EPI_ISL_518404, EPI_ISL_518405, EPI_ISL_518406, EPI_ISL_518407, EPI_ISL_518408, EPI_ISL_518409, EPI_ISL_518410, EPI_ISL_518411, EPI_ISL_518412, EPI_ISL_518413, EPI_ISL_518414, EPI_ISL_518415, EPI_ISL_518416, EPI_ISL_518417, EPI_ISL_518418, EPI_ISL_518419, EPI_ISL_518420, EPI_ISL_518421, EPI_ISL_518422, EPI_ISL_518423 | | | |
| see above | Microbiological Diagnostic Unit - Public Health Laboratory (MDU-PHL) | MDU-PHL | Seemann T., Schultz M., Sait, M., Sherry, N. |
| EPI_ISL_518424, EPI_ISL_518425, EPI_ISL_518426, EPI_ISL_518427, EPI_ISL_518428, EPI_ISL_518429, EPI_ISL_518430, EPI_ISL_518431, EPI_ISL_518432, EPI_ISL_518433, EPI_ISL_518434, EPI_ISL_518435, EPI_ISL_518436, EPI_ISL_518437, EPI_ISL_518438, EPI_ISL_518439, EPI_ISL_518440, EPI_ISL_518441, EPI_ISL_518442, EPI_ISL_518443, EPI_ISL_518444, EPI_ISL_518445, EPI_ISL_518446, EPI_ISL_518447, EPI_ISL_518448, EPI_ISL_518449, EPI_ISL_518450, EPI_ISL_518451, EPI_ISL_518452, EPI_ISL_518453, EPI_ISL_518454, EPI_ISL_518455, EPI_ISL_518456, EPI_ISL_518457, EPI_ISL_518458, EPI_ISL_518459, EPI_ISL_518460, EPI_ISL_518461, EPI_ISL_518462, EPI_ISL_518463, EPI_ISL_518464, EPI_ISL_518465, EPI_ISL_518466, EPI_ISL_518467, EPI_ISL_518468, EPI_ISL_518469, EPI_ISL_518470, EPI_ISL_518471 | | | |
| see above | Victorian Infectious Diseases Reference Laboratory (VIDRL) | VIDRL and MDU-PHL | Caly L., Seemann T., Sait, M., Schultz M., Druce J., Sherry, N. |
| EPI_ISL_518472, EPI_ISL_518473, EPI_ISL_518474, EPI_ISL_518475, EPI_ISL_518476, EPI_ISL_518477, EPI_ISL_518478, EPI_ISL_518479, EPI_ISL_518480, EPI_ISL_518481, EPI_ISL_518482, EPI_ISL_518483, EPI_ISL_518484, EPI_ISL_518485, EPI_ISL_518486, EPI_ISL_518487, EPI_ISL_518488, EPI_ISL_518489, EPI_ISL_518490, EPI_ISL_518491, EPI_ISL_518492, EPI_ISL_518493, EPI_ISL_518494, EPI_ISL_518495, EPI_ISL_518496, EPI_ISL_518497, EPI_ISL_518498, EPI_ISL_518499, EPI_ISL_518500, EPI_ISL_518501, EPI_ISL_518502, EPI_ISL_518503, EPI_ISL_518504, EPI_ISL_518505, EPI_ISL_518506, EPI_ISL_518507, EPI_ISL_518508, EPI_ISL_518509, EPI_ISL_518510, EPI_ISL_518511, EPI_ISL_518512, EPI_ISL_518513, EPI_ISL_518514, EPI_ISL_518515, EPI_ISL_518516, EPI_ISL_518517, EPI_ISL_518518, EPI_ISL_518519, EPI_ISL_518520, EPI_ISL_518521, EPI_ISL_518522, EPI_ISL_518523, EPI_ISL_518524, EPI_ISL_518525, EPI_ISL_518526, EPI_ISL_518527, EPI_ISL_518528, EPI_ISL_518529, EPI_ISL_518530, EPI_ISL_518531, EPI_ISL_518532, EPI_ISL_518533, EPI_ISL_518534, EPI_ISL_518535, EPI_ISL_518536, EPI_ISL_518537, EPI_ISL_518538, EPI_ISL_518539, EPI_ISL_518540, EPI_ISL_518541, EPI_ISL_518542, EPI_ISL_518543, EPI_ISL_518544, EPI_ISL_518545, EPI_ISL_518546, EPI_ISL_518547, EPI_ISL_518548, EPI_ISL_518549, EPI_ISL_518550, EPI_ISL_518551, EPI_ISL_518552, EPI_ISL_518553, EPI_ISL_518554, EPI_ISL_518555, EPI_ISL_518556, EPI_ISL_518557, EPI_ISL_518558, EPI_ISL_518559, EPI_ISL_518560, EPI_ISL_518561, EPI_ISL_518562, EPI_ISL_518563, EPI_ISL_518564, EPI_ISL_518565, EPI_ISL_518566, EPI_ISL_518567, EPI_ISL_518568, EPI_ISL_518569, EPI_ISL_518570, EPI_ISL_518571, EPI_ISL_518572, EPI_ISL_518573, EPI_ISL_518574, EPI_ISL_518575, EPI_ISL_518576, EPI_ISL_518577, EPI_ISL_518578, EPI_ISL_518579, EPI_ISL_518580, EPI_ISL_518581, EPI_ISL_518582, EPI_ISL_518583, EPI_ISL_518584, EPI_ISL_518585, EPI_ISL_518586, EPI_ISL_518587, EPI_ISL_518588, EPI_ISL_518589, EPI_ISL_518590, EPI_ISL_518591, EPI_ISL_518592, EPI_ISL_518593, EPI_ISL_518594, EPI_ISL_518595, EPI_ISL_518596, EPI_ISL_518597, EPI_ISL_518598, EPI_ISL_518599, EPI_ISL_518600, EPI_ISL_518601, EPI_ISL_518602, EPI_ISL_518603, EPI_ISL_518604, EPI_ISL_518605, EPI_ISL_518606, EPI_ISL_518607, EPI_ISL_518608, EPI_ISL_518609, EPI_ISL_518610, EPI_ISL_518611, EPI_ISL_518612, EPI_ISL_518613, EPI_ISL_518614, EPI_ISL_518615, EPI_ISL_518616, EPI_ISL_518617, EPI_ISL_518618, EPI_ISL_518619, EPI_ISL_518620, EPI_ISL_518621, EPI_ISL_518622, EPI_ISL_518623, EPI_ISL_518624, EPI_ISL_518625, EPI_ISL_518626, EPI_ISL_518627, EPI_ISL_518628, EPI_ISL_518629, EPI_ISL_518630, EPI_ISL_518631, EPI_ISL_518632, EPI_ISL_518633, EPI_ISL_518634, EPI_ISL_518635, EPI_ISL_518636, EPI_ISL_518637, EPI_ISL_518638, EPI_ISL_518639, EPI_ISL_518640, EPI_ISL_518641, EPI_ISL_518642, EPI_ISL_518643, EPI_ISL_518644, EPI_ISL_518645, EPI_ISL_518646, EPI_ISL_518647, EPI_ISL_518648, EPI_ISL_518649, EPI_ISL_518650 | | | |
| see above | Microbiological Diagnostic Unit - Public Health Laboratory (MDU-PHL) | MDU-PHL | Seemann T., Schultz M., Sait, M., Sherry, N. |
| EPI_ISL_518651 | Microbiological Diagnostic Unit - Public Health Laboratory (MDU-PHL) | Microbiological Diagnostic Unit - Public Health Laboratory (MDU-PHL) | Seemann T., Schultz M., Sait, M., Sherry, N. |
| EPI_ISL_518652, EPI_ISL_518653, EPI_ISL_518654, EPI_ISL_518655, EPI_ISL_518656, EPI_ISL_518657, EPI_ISL_518658, EPI_ISL_518659, EPI_ISL_518660, EPI_ISL_518661, EPI_ISL_518662, EPI_ISL_518663, EPI_ISL_518664, EPI_ISL_518665, EPI_ISL_518666, EPI_ISL_518667, EPI_ISL_518668, EPI_ISL_518669, EPI_ISL_518670, EPI_ISL_518671, EPI_ISL_518672, EPI_ISL_518673, EPI_ISL_518674, EPI_ISL_518675, EPI_ISL_518676, EPI_ISL_518677, EPI_ISL_518678, EPI_ISL_518679, EPI_ISL_518680, EPI_ISL_518681, EPI_ISL_518682, EPI_ISL_518683, EPI_ISL_518684, EPI_ISL_518685, EPI_ISL_518686, EPI_ISL_518687, EPI_ISL_518688, EPI_ISL_518689, EPI_ISL_518690, EPI_ISL_518691, EPI_ISL_518692, EPI_ISL_518693, EPI_ISL_518694, EPI_ISL_518695, EPI_ISL_518696, EPI_ISL_518697, EPI_ISL_518698, EPI_ISL_518699, EPI_ISL_518700, EPI_ISL_518701, EPI_ISL_518702, EPI_ISL_518703, EPI_ISL_518704, EPI_ISL_518705, EPI_ISL_518706, EPI_ISL_518707, EPI_ISL_518708, EPI_ISL_518709, EPI_ISL_518710, EPI_ISL_518711, EPI_ISL_518712, EPI_ISL_518713, EPI_ISL_518714, EPI_ISL_518715, EPI_ISL_518716, EPI_ISL_518717, EPI_ISL_518718, EPI_ISL_518719, EPI_ISL_518720, EPI_ISL_518721, EPI_ISL_518722 | | | |
| see above | Microbiological Diagnostic Unit - Public Health Laboratory (MDU-PHL) | MDU-PHL | Seemann T., Schultz M., Sait, M., Sherry, N. |
| EPI_ISL_518723 | Victorian Infectious Diseases Reference Laboratory (VIDRL) | VIDRL and MDU-PHL | Caly L., Seemann T., Sait, M., Schultz M., Druce J., Sherry, N. |
| EPI_ISL_518724, EPI_ISL_518725, EPI_ISL_518726, EPI_ISL_518727, EPI_ISL_518728, EPI_ISL_518729, EPI_ISL_518730, EPI_ISL_518731, EPI_ISL_518732, EPI_ISL_518733, EPI_ISL_518734, EPI_ISL_518735, EPI_ISL_518736, EPI_ISL_518737, EPI_ISL_518738, EPI_ISL_518739, EPI_ISL_518740, EPI_ISL_518741, EPI_ISL_518742, EPI_ISL_518743, EPI_ISL_518744, EPI_ISL_518745, EPI_ISL_518746, EPI_ISL_518747, EPI_ISL_518748, EPI_ISL_518749, EPI_ISL_518750, EPI_ISL_518751, EPI_ISL_518752, EPI_ISL_518753, EPI_ISL_518754, EPI_ISL_518755, EPI_ISL_518756, EPI_ISL_518757, EPI_ISL_518758, EPI_ISL_518759, EPI_ISL_518760, EPI_ISL_518761, EPI_ISL_518762, EPI_ISL_518763, EPI_ISL_518764, EPI_ISL_518765, EPI_ISL_518766, EPI_ISL_518767, EPI_ISL_518768, EPI_ISL_518769, EPI_ISL_518770, EPI_ISL_518771, EPI_ISL_518772, EPI_ISL_518773, EPI_ISL_518774, EPI_ISL_518775, EPI_ISL_518776, EPI_ISL_518777, EPI_ISL_518778, EPI_ISL_518779, EPI_ISL_518780, EPI_ISL_518781, EPI_ISL_518782, EPI_ISL_518783, EPI_ISL_518784, EPI_ISL_518785, EPI_ISL_518786, EPI_ISL_518787, EPI_ISL_518788, EPI_ISL_518789, EPI_ISL_518790, EPI_ISL_518791, EPI_ISL_518792, EPI_ISL_518793, EPI_ISL_518794, EPI_ISL_518795, EPI_ISL_518796 | | | |
| see above | Microbiological Diagnostic Unit - Public Health Laboratory (MDU-PHL) | MDU-PHL | Seemann T., Schultz M., Sait, M., Sherry, N. |
| EPI_ISL_518799, EPI_ISL_518800, EPI_ISL_518801, EPI_ISL_518802, EPI_ISL_518803, EPI_ISL_518804, EPI_ISL_518805, EPI_ISL_518806, EPI_ISL_518807, EPI_ISL_518808, EPI_ISL_518809, EPI_ISL_518810, EPI_ISL_518811, EPI_ISL_518812, EPI_ISL_518813, EPI_ISL_518814, EPI_ISL_518815, EPI_ISL_518816, EPI_ISL_518817, EPI_ISL_518818 | | | |
| see above | Academic Hospital Paramaribo | Erasmus Medical Center | Bas Oude Munnink, Dion Gajadin, Ed Ijzerman, Emmanuelle Munger, Gary Gummels, Ingrid Krishnadath, Lycke Woittiez, Marion Koopmans, Mireille Van de Veer, Princes Wongsowidjojo, Radjesh Ori, Rohma Banwari, Stephen Vreden |
| EPI_ISL_518819 | Qadr Hospital, Tangerang, Banten | Biosafety Level-3 Laboratory, Indonesian Institute of Sciences (LIPI) | Anik Budhi Dharmayanthi, Syam Budi Iryanto, Andri Wardiana, Anggia Prasetyoputri, Isa Nuryana, Ade Andriani, Ario Betha Juanssilfero, Asep M Ridwanuloh, Ahmad Fathoni, Rifki Sadikin, Ratih Asmana Ningrum, Wien Kusharyoto, Puspita Lisdiyanti |
| EPI_ISL_518820 | Klinik Apotek Dein, Jakarta, Indonesia | Biosafety Level-3 Laboratory, Indonesian Institute of Sciences (LIPI) | Anik Budhi Dharmayanthi, Syam Budi Iryanto, Andri Wardiana, Anggia Prasetyoputri, Isa Nuryana, Ade Andriani, Ario Betha Juanssilfero, Asep M Ridwanuloh, Ahmad Fathoni, Rifki Sadikin, Ratih Asmana Ningrum, Wien Kusharyoto, Puspita Lisdiyanti |
| EPI_ISL_518821, EPI_ISL_518822, EPI_ISL_518823 | Oman-National Influenza Center | Biotechnology & OMICs Laboratory, Natural & Medical Sciences Research Center, University of Nizwa | Samira Al-Mahruqi, Abdul Latif Khan, Samiha Al-Kharusi, Adil Khan , Ahmed Al-Rawahi, Sajjad Asaf, Amina Al-Jardani, Hanan Al-Kindi, Intisar Al-Shukri, Adil Al-Wahaibi, Seif Al-Abri, Ahmed Al-Harrasi |

| | | | |
|---|---|---|---|
| EPI_ISL_518824 | Oman-National Influenza Center | Biotechnology & OMICs Laboratory, Natural & Medical Sciences Research Center, University of Nizwa | Abdul Latif Khan, Samira Al-Mahruqi, Ahmed Al-Harrasi, Samiha Al-Kharusi, Adil Khan, Ahmed Al-Rawahi, Sajjad Asaf, Amina Al-Jardani, Hanan Al-Kindi, Intisar Al-Shukri, Ahlam Al-Amri, Aisha Al-Amri, Aisha Al-Busaidi, Adil Al-Wahaibi, Seif Al-Abri. |
| EPI_ISL_518825 | Oman-National Influenza Center | Biotechnology & OMICs Laboratory, Natural & Medical Sciences Research Center, University of Nizwa | Samira Al-Mahruqi, Abdul Latif Khan, Samiha Al-Kharusi, Adil Khan , Ahmed Al-Rawahi, Sajjad Asaf, Amina Al-Jardani, Hanan Al-Kindi, Intisar Al-Shukri, Adil Al-Wahaibi, Seif Al-Abri, Ahmed Al-Harrasi |
| EPI_ISL_518826, EPI_ISL_518827, EPI_ISL_518828, EPI_ISL_518829, EPI_ISL_518830, EPI_ISL_518831, EPI_ISL_518832 | Oman-National Influenza Center | Biotechnology & OMICs Laboratory, Natural & Medical Sciences Research Center, University of Nizwa | Sajjad Asaf, Samiha Al-Kharusi, Ahmed Al-Harrasi, Samira Al-Mahruqi, Adil Khan, Ahmed Al-Rawahi, Abdul Latif Khan, Amina Al-Jardani, Hanan Al-Kindi, Intisar Al-Shukri, Ahlam Al-Amri, Aisha Al-Amri, Aisha Al-Busaidi, Adil Al-Wahaibi, Seif Al-Abri. |
| EPI_ISL_518833, EPI_ISL_518834, EPI_ISL_518835, EPI_ISL_518836 | Oman-National Influenza Center | Biotechnology & OMICs Laboratory, Natural & Medical Sciences Research Center, University of Nizwa | Samira Al-Mahruqi, Abdul Latif Khan, Samiha Al-Kharusi, Adil Khan , Ahmed Al-Rawahi, Sajjad Asaf, Amina Al-Jardani, Hanan Al-Kindi, Intisar Al-Shukri, Adil Al-Wahaibi, Seif Al-Abri, Ahmed Al-Harrasi |
| EPI_ISL_518837, EPI_ISL_518838, EPI_ISL_518839, EPI_ISL_518840, EPI_ISL_518841, EPI_ISL_518842, EPI_ISL_518843, EPI_ISL_518844, EPI_ISL_518845 | Oman-National Influenza Center | Biotechnology & OMICs Laboratory, Natural & Medical Sciences Research Center, University of Nizwa | Samiha Al-Kharusi, Sajjad Asaf, Abdul Latif Khan, Samira Al-Mahruqi, Adil Khan, Ahmed Al-Rawahi, Amina Al-Jardani, Hanan Al-Kindi, Intisar Al-Shukri, Ahlam Al-Amri, Aisha Al-Amri, Aisha Al-Busaidi, Adil Al-Wahaibi, Seif Al-Abri, Ahmed Al-Harrasi |
| EPI_ISL_518846, EPI_ISL_518847, EPI_ISL_518848, EPI_ISL_518849, EPI_ISL_518850, EPI_ISL_518851, EPI_ISL_518852, EPI_ISL_518853, EPI_ISL_518854 | Oman-National Influenza Center | Biotechnology & OMICs Laboratory, Natural & Medical Sciences Research Center, University of Nizwa | Abdul Latif Khan, Samira Al-Mahruqi, Ahmed Al-Harrasi, Samiha Al-Kharusi, Adil Khan, Ahmed Al-Rawahi, Sajjad Asaf, Amina Al-Jardani, Hanan Al-Kindi, Intisar Al-Shukri, Ahlam Al-Amri, Aisha Al-Amri, Aisha Al-Busaidi, Adil Al-Wahaibi, Seif Al-Abri. |
| EPI_ISL_518855 | Minnesota Department of Health, Public Health Laboratory | Minnesota Department of Health, Public Health Laboratory | Matt Plumb, Jacob Garfin, and Xiong Wang |
| EPI_ISL_518856, EPI_ISL_518857, EPI_ISL_518858, EPI_ISL_518859, EPI_ISL_518860, EPI_ISL_518861, EPI_ISL_518862, EPI_ISL_518863, EPI_ISL_518864, EPI_ISL_518865, EPI_ISL_518866, EPI_ISL_518867, EPI_ISL_518868, EPI_ISL_518869, EPI_ISL_518870, EPI_ISL_518871, EPI_ISL_518872, EPI_ISL_518873, EPI_ISL_518874, EPI_ISL_518875, EPI_ISL_518876, EPI_ISL_518877, EPI_ISL_518878, EPI_ISL_518879, EPI_ISL_518880, EPI_ISL_518881, EPI_ISL_518882, EPI_ISL_518883, EPI_ISL_518884, EPI_ISL_518885, EPI_ISL_518886, EPI_ISL_518887, EPI_ISL_518888, EPI_ISL_518889, EPI_ISL_518890, EPI_ISL_518891, EPI_ISL_518892, EPI_ISL_518893, EPI_ISL_518894, EPI_ISL_518895, EPI_ISL_518896, EPI_ISL_518897, EPI_ISL_518898, EPI_ISL_518899, EPI_ISL_518900 | | | |
| see above | Mayo Clinic & Mayo Clinic Laboratories | Minnesota Department of Health, Public Health Laboratory | Matt Plumb, Jacob Garfin, and Xiong Wang |
| EPI_ISL_518901, EPI_ISL_518902, EPI_ISL_518903, EPI_ISL_518904, EPI_ISL_518905, EPI_ISL_518906, EPI_ISL_518907, EPI_ISL_518908, EPI_ISL_518909, EPI_ISL_518910, EPI_ISL_518911, EPI_ISL_518912, EPI_ISL_518913, EPI_ISL_518914, EPI_ISL_518915, EPI_ISL_518917, EPI_ISL_518918, EPI_ISL_518919, EPI_ISL_518920, EPI_ISL_518921, EPI_ISL_518922, EPI_ISL_518923, EPI_ISL_518924, EPI_ISL_518925, EPI_ISL_518926, EPI_ISL_518927, EPI_ISL_518928, EPI_ISL_518929, EPI_ISL_518930, EPI_ISL_518931, EPI_ISL_518932, EPI_ISL_518933, EPI_ISL_518934 | | | |
| see above | Microbiological Diagnostic Unit - Public Health Laboratory (MDU-PHL) | MDU-PHL | Seemann T., Schultz M., Sait, M., Sherry, N. |
| EPI_ISL_518935 | Microbiological Diagnostic Unit - Public Health Laboratory (MDU-PHL) | Microbiological Diagnostic Unit - Public Health Laboratory (MDU-PHL) | Seemann T., Schultz M., Sait, M., Sherry, N. |
| EPI_ISL_518936, EPI_ISL_518937, EPI_ISL_518938, EPI_ISL_518939, EPI_ISL_518940, EPI_ISL_518941, EPI_ISL_518942, EPI_ISL_518943, EPI_ISL_518944, EPI_ISL_518945, EPI_ISL_518946, EPI_ISL_518947, EPI_ISL_518948, EPI_ISL_518949, EPI_ISL_518950, EPI_ISL_518951, EPI_ISL_518952, EPI_ISL_518953, EPI_ISL_518954, EPI_ISL_518955, EPI_ISL_518956, EPI_ISL_518957, EPI_ISL_518958, EPI_ISL_518959, EPI_ISL_518960, EPI_ISL_518961, EPI_ISL_518962, EPI_ISL_518963, EPI_ISL_518964, EPI_ISL_518965, EPI_ISL_518966, EPI_ISL_518967, EPI_ISL_518968, EPI_ISL_518969, EPI_ISL_518970, EPI_ISL_518971, EPI_ISL_518972, EPI_ISL_518973, EPI_ISL_518975 | | | |
| see above | Microbiological Diagnostic Unit - Public Health Laboratory (MDU-PHL) | MDU-PHL | Seemann T., Schultz M., Sait, M., Sherry, N. |
| EPI_ISL_518976 | Victorian Infectious Diseases Reference Laboratory (VIDRL) | VIDRL and MDU-PHL | Caly L., Seemann T., Sait, M., Schultz M., Druce J., Sherry, N. |
| EPI_ISL_518977, EPI_ISL_518978, EPI_ISL_518979, EPI_ISL_518980 | Microbiological Diagnostic Unit - Public Health Laboratory (MDU-PHL) | MDU-PHL | Seemann T., Schultz M., Sait, M., Sherry, N. |
| EPI_ISL_518981, EPI_ISL_518982, EPI_ISL_518983, EPI_ISL_518984, EPI_ISL_518985, EPI_ISL_518986, EPI_ISL_518987, EPI_ISL_518988, EPI_ISL_518989, EPI_ISL_518990, EPI_ISL_518991, EPI_ISL_518992, EPI_ISL_518993, EPI_ISL_518994, EPI_ISL_518995, EPI_ISL_518996, EPI_ISL_518997, EPI_ISL_518998, EPI_ISL_518999, EPI_ISL_519000, EPI_ISL_519001, EPI_ISL_519002, EPI_ISL_519003, EPI_ISL_519004, EPI_ISL_519005, EPI_ISL_519006, EPI_ISL_519007, EPI_ISL_519008, EPI_ISL_519009, EPI_ISL_519010, EPI_ISL_519011, EPI_ISL_519012, EPI_ISL_519013, EPI_ISL_519014, EPI_ISL_519015, EPI_ISL_519016, EPI_ISL_519017, EPI_ISL_519018, EPI_ISL_519019, EPI_ISL_519020, EPI_ISL_519021, EPI_ISL_519022, EPI_ISL_519023, EPI_ISL_519025, EPI_ISL_519026, EPI_ISL_519027, EPI_ISL_519028, EPI_ISL_519029, EPI_ISL_519030, EPI_ISL_519031, EPI_ISL_519032, EPI_ISL_519033, EPI_ISL_519034, EPI_ISL_519035, EPI_ISL_519036, EPI_ISL_519037, EPI_ISL_519038, EPI_ISL_519039, EPI_ISL_519040, EPI_ISL_519041, EPI_ISL_519042, EPI_ISL_519043, EPI_ISL_519044, EPI_ISL_519045, EPI_ISL_519046, EPI_ISL_519047, EPI_ISL_519048, EPI_ISL_519049, EPI_ISL_519050, EPI_ISL_519051 | | | |
| see above | Victorian Infectious Diseases Reference Laboratory (VIDRL) | VIDRL and MDU-PHL | Caly L., Seemann T., Sait, M., Schultz M., Druce J., Sherry, N. |
| EPI_ISL_519052, EPI_ISL_519053, EPI_ISL_519054, EPI_ISL_519055, EPI_ISL_519056, EPI_ISL_519057, EPI_ISL_519058, EPI_ISL_519059, EPI_ISL_519060, EPI_ISL_519061, EPI_ISL_519062, EPI_ISL_519063, EPI_ISL_519064, EPI_ISL_519065, EPI_ISL_519066, EPI_ISL_519067, EPI_ISL_519068, EPI_ISL_519069, EPI_ISL_519070, EPI_ISL_519071, EPI_ISL_519072, EPI_ISL_519073, EPI_ISL_519074, EPI_ISL_519075, EPI_ISL_519076, EPI_ISL_519077, EPI_ISL_519078, EPI_ISL_519079, EPI_ISL_519080, EPI_ISL_519081, EPI_ISL_519082, EPI_ISL_519083, EPI_ISL_519084, EPI_ISL_519085, EPI_ISL_519086, EPI_ISL_519087, EPI_ISL_519088, EPI_ISL_519089, EPI_ISL_519090, EPI_ISL_519091, EPI_ISL_519092, EPI_ISL_519093, EPI_ISL_519094, EPI_ISL_519095, EPI_ISL_519096, EPI_ISL_519097, EPI_ISL_519098, EPI_ISL_519099, EPI_ISL_519100, EPI_ISL_519101, EPI_ISL_519102, EPI_ISL_519103, EPI_ISL_519104, EPI_ISL_519105, EPI_ISL_519106, EPI_ISL_519107, EPI_ISL_519108, EPI_ISL_519109, EPI_ISL_519110, EPI_ISL_519111, EPI_ISL_519112, EPI_ISL_519113, EPI_ISL_519114, EPI_ISL_519115, EPI_ISL_519116, EPI_ISL_519117, EPI_ISL_519118, EPI_ISL_519119, EPI_ISL_519120, EPI_ISL_519121, EPI_ISL_519122, EPI_ISL_519123, EPI_ISL_519124, EPI_ISL_519125, EPI_ISL_519126, EPI_ISL_519127, EPI_ISL_519128, EPI_ISL_519129, EPI_ISL_519130, EPI_ISL_519131, EPI_ISL_519132, EPI_ISL_519133, EPI_ISL_519134, EPI_ISL_519135, EPI_ISL_519136, EPI_ISL_519137, EPI_ISL_519138, EPI_ISL_519139, EPI_ISL_519140, EPI_ISL_519141, EPI_ISL_519142, EPI_ISL_519143, EPI_ISL_519144, EPI_ISL_519145, EPI_ISL_519146, EPI_ISL_519147, EPI_ISL_519148, EPI_ISL_519149, EPI_ISL_519150, EPI_ISL_519151, EPI_ISL_519152, EPI_ISL_519153, EPI_ISL_519154, EPI_ISL_519155, EPI_ISL_519156, EPI_ISL_519157, EPI_ISL_519158, EPI_ISL_519159, EPI_ISL_519160, EPI_ISL_519161, EPI_ISL_519162, EPI_ISL_519163, EPI_ISL_519164, EPI_ISL_519165, EPI_ISL_519166, EPI_ISL_519167, EPI_ISL_519168, EPI_ISL_519169, EPI_ISL_519170, EPI_ISL_519171, EPI_ISL_519172, EPI_ISL_519173, EPI_ISL_519174, EPI_ISL_519175, EPI_ISL_519176, EPI_ISL_519177, EPI_ISL_519178, EPI_ISL_519179, EPI_ISL_519180, EPI_ISL_519181, EPI_ISL_519182, EPI_ISL_519183, EPI_ISL_519184, EPI_ISL_519185, EPI_ISL_519186, EPI_ISL_519187, EPI_ISL_519188, EPI_ISL_519189, EPI_ISL_519190, EPI_ISL_519191, EPI_ISL_519192, EPI_ISL_519193, EPI_ISL_519194, EPI_ISL_519195, EPI_ISL_519196, EPI_ISL_519197, EPI_ISL_519198, EPI_ISL_519199, EPI_ISL_519200, EPI_ISL_519201, EPI_ISL_519202, EPI_ISL_519203, EPI_ISL_519204, EPI_ISL_519205, EPI_ISL_519206, EPI_ISL_519207, EPI_ISL_519208, EPI_ISL_519209, EPI_ISL_519210, EPI_ISL_519211, EPI_ISL_519212, EPI_ISL_519213, EPI_ISL_519214, EPI_ISL_519215, EPI_ISL_519216, EPI_ISL_519217, EPI_ISL_519218, EPI_ISL_519219, EPI_ISL_519220, EPI_ISL_519221, EPI_ISL_519222, EPI_ISL_519223, EPI_ISL_519224, EPI_ISL_519225, EPI_ISL_519226, EPI_ISL_519227, EPI_ISL_519228, EPI_ISL_519229, EPI_ISL_519230, EPI_ISL_519231, EPI_ISL_519232, EPI_ISL_519233, EPI_ISL_519234, EPI_ISL_519235, EPI_ISL_519236, EPI_ISL_519237, EPI_ISL_519238, EPI_ISL_519239, EPI_ISL_519240, EPI_ISL_519241, EPI_ISL_519242, EPI_ISL_519243, EPI_ISL_519244, EPI_ISL_519245, EPI_ISL_519246, EPI_ISL_519247, EPI_ISL_519248, EPI_ISL_519249, EPI_ISL_519250, EPI_ISL_519251, EPI_ISL_519252, EPI_ISL_519253, EPI_ISL_519254, EPI_ISL_519255 | | | |
| see above | Microbiological Diagnostic Unit - Public Health Laboratory (MDU-PHL) | MDU-PHL | Seemann T., Schultz M., Sait, M., Sherry, N. |
| EPI_ISL_519256, EPI_ISL_519257, EPI_ISL_519258, EPI_ISL_519259, EPI_ISL_519260, EPI_ISL_519261, EPI_ISL_519262, EPI_ISL_519263, EPI_ISL_519264, EPI_ISL_519265, EPI_ISL_519266, EPI_ISL_519267, EPI_ISL_519268, EPI_ISL_519269, EPI_ISL_519270, EPI_ISL_519271, EPI_ISL_519272, EPI_ISL_519273, EPI_ISL_519274, EPI_ISL_519275, EPI_ISL_519276, EPI_ISL_519277, EPI_ISL_519278, EPI_ISL_519279, EPI_ISL_519280, EPI_ISL_519281, EPI_ISL_519282, EPI_ISL_519283, EPI_ISL_519284, EPI_ISL_519285, EPI_ISL_519286, EPI_ISL_519287, EPI_ISL_519288, EPI_ISL_519289, EPI_ISL_519290, EPI_ISL_519291, EPI_ISL_519292, EPI_ISL_519293, EPI_ISL_519294, EPI_ISL_519295, EPI_ISL_519296, EPI_ISL_519297, EPI_ISL_519298, EPI_ISL_519299, EPI_ISL_519300, EPI_ISL_519301, EPI_ISL_519302, EPI_ISL_519303, EPI_ISL_519304, EPI_ISL_519305, EPI_ISL_519306, EPI_ISL_519307, EPI_ISL_519308, EPI_ISL_519309, EPI_ISL_519310, EPI_ISL_519311, EPI_ISL_519312, EPI_ISL_519313, EPI_ISL_519314, EPI_ISL_519315, EPI_ISL_519316, EPI_ISL_519317, EPI_ISL_519318, EPI_ISL_519319, EPI_ISL_519320, EPI_ISL_519321, EPI_ISL_519322, EPI_ISL_519323, EPI_ISL_519324, EPI_ISL_519325, EPI_ISL_519326 | | | |
| see above | Victorian Infectious Diseases Reference Laboratory (VIDRL) | VIDRL and MDU-PHL | Caly L., Seemann T., Sait, M., Schultz M., Druce J., Sherry, N. |
| EPI_ISL_519327, EPI_ISL_519328, EPI_ISL_519329, EPI_ISL_519330 | Microbiological Diagnostic Unit - Public Health Laboratory (MDU-PHL) | MDU-PHL | Seemann T., Schultz M., Sait, M., Sherry, N. |
| EPI_ISL_519331, EPI_ISL_519332 | Victorian Infectious Diseases Reference Laboratory (VIDRL) | VIDRL and MDU-PHL | Caly L., Seemann T., Sait, M., Schultz M., Druce J., Sherry, N. |
| EPI_ISL_519333, EPI_ISL_519334, EPI_ISL_519335, EPI_ISL_519336, EPI_ISL_519337, EPI_ISL_519338, EPI_ISL_519339, EPI_ISL_519340, EPI_ISL_519341, EPI_ISL_519342, EPI_ISL_519343, EPI_ISL_519344, EPI_ISL_519345, EPI_ISL_519346, EPI_ISL_519347, EPI_ISL_519348, EPI_ISL_519349, EPI_ISL_519350, EPI_ISL_519351, EPI_ISL_519352, EPI_ISL_519353, EPI_ISL_519354, EPI_ISL_519355, EPI_ISL_519356, EPI_ISL_519357, EPI_ISL_519358, EPI_ISL_519359, EPI_ISL_519360, EPI_ISL_519361, EPI_ISL_519362, EPI_ISL_519363, EPI_ISL_519364, EPI_ISL_519365, EPI_ISL_519366, EPI_ISL_519367, EPI_ISL_519368, EPI_ISL_519369, EPI_ISL_519371, EPI_ISL_519372, EPI_ISL_519373, EPI_ISL_519374, EPI_ISL_519375, EPI_ISL_519376, EPI_ISL_519377, EPI_ISL_519378, EPI_ISL_519379, EPI_ISL_519380, EPI_ISL_519381, EPI_ISL_519382, EPI_ISL_519383, EPI_ISL_519384 | | | |
| see above | Microbiological Diagnostic Unit - Public Health Laboratory (MDU-PHL) | MDU-PHL | Seemann T., Schultz M., Sait, M., Sherry, N. |
| EPI_ISL_519385 | Victorian Infectious Diseases Reference Laboratory (VIDRL) | VIDRL and MDU-PHL | Caly L., Seemann T., Sait, M., Schultz M., Druce J., Sherry, N. |
| EPI_ISL_519386, EPI_ISL_519387, EPI_ISL_519388, EPI_ISL_519389, EPI_ISL_519390, EPI_ISL_519391, EPI_ISL_519392, EPI_ISL_519393, EPI_ISL_519394, EPI_ISL_519395, EPI_ISL_519396, EPI_ISL_519397, EPI_ISL_519398, EPI_ISL_519399, EPI_ISL_519400, EPI_ISL_519401, EPI_ISL_519402, EPI_ISL_519403, EPI_ISL_519404, EPI_ISL_519405, EPI_ISL_519406, EPI_ISL_519407, EPI_ISL_519408, EPI_ISL_519409, EPI_ISL_519410, EPI_ISL_519411, EPI_ISL_519412, EPI_ISL_519413, EPI_ISL_519414, EPI_ISL_519415, EPI_ISL_519416, EPI_ISL_519417, EPI_ISL_519418, EPI_ISL_519419, EPI_ISL_519420, EPI_ISL_519421, EPI_ISL_519422, EPI_ISL_519423, EPI_ISL_519424, EPI_ISL_519425, EPI_ISL_519426, EPI_ISL_519427, EPI_ISL_519428, EPI_ISL_519429, EPI_ISL_519430, EPI_ISL_519431, EPI_ISL_519432, EPI_ISL_519433, EPI_ISL_519434, EPI_ISL_519435, EPI_ISL_519436, EPI_ISL_519437, EPI_ISL_519438, EPI_ISL_519439, EPI_ISL_519440, EPI_ISL_519441, EPI_ISL_519442, EPI_ISL_519443, EPI_ISL_519444, EPI_ISL_519445, EPI_ISL_519446, EPI_ISL_519447, EPI_ISL_519448, EPI_ISL_519449, EPI_ISL_519450, EPI_ISL_519451, EPI_ISL_519452, EPI_ISL_519453, EPI_ISL_519454, EPI_ISL_519455, EPI_ISL_519456, EPI_ISL_519457, EPI_ISL_519458, EPI_ISL_519459, EPI_ISL_519460, EPI_ISL_519461, EPI_ISL_519462, EPI_ISL_519463, EPI_ISL_519464, EPI_ISL_519465, EPI_ISL_519466, EPI_ISL_519467, EPI_ISL_519468, EPI_ISL_519469, EPI_ISL_519470, EPI_ISL_519471, EPI_ISL_519472, EPI_ISL_519473, EPI_ISL_519474, EPI_ISL_519475, EPI_ISL_519476, EPI_ISL_519477, EPI_ISL_519478, EPI_ISL_519479, EPI_ISL_519480, EPI_ISL_519481, EPI_ISL_519482, EPI_ISL_519483, EPI_ISL_519484, EPI_ISL_519485, EPI_ISL_519486, EPI_ISL_519487, EPI_ISL_519488, EPI_ISL_519489, EPI_ISL_519490, EPI_ISL_519491, EPI_ISL_519492, EPI_ISL_519493, EPI_ISL_519494, EPI_ISL_519495, EPI_ISL_519496, EPI_ISL_519497, EPI_ISL_519498, EPI_ISL_519499, EPI_ISL_519500, EPI_ISL_519501, EPI_ISL_519503, EPI_ISL_519504, EPI_ISL_519505, EPI_ISL_519506, EPI_ISL_519507, EPI_ISL_519508, EPI_ISL_519509, EPI_ISL_519510, EPI_ISL_519511, EPI_ISL_519512, EPI_ISL_519513, EPI_ISL_519514, EPI_ISL_519515, EPI_ISL_519516, EPI_ISL_519517, EPI_ISL_519518 | | | |
| see above | Microbiological Diagnostic Unit - Public Health Laboratory (MDU-PHL) | MDU-PHL | Seemann T., Schultz M., Sait, M., Sherry, N. |
| EPI_ISL_519519, EPI_ISL_519520, EPI_ISL_519521 | Victorian Infectious Diseases Reference Laboratory (VIDRL) | VIDRL and MDU-PHL | Caly L., Seemann T., Sait, M., Schultz M., Druce J., Sherry, N. |
| EPI_ISL_519522, EPI_ISL_519523, EPI_ISL_519524, EPI_ISL_519525, EPI_ISL_519526, EPI_ISL_519527, EPI_ISL_519528, EPI_ISL_519529, EPI_ISL_519530, EPI_ISL_519531, EPI_ISL_519532, EPI_ISL_519533, EPI_ISL_519534, EPI_ISL_519535, EPI_ISL_519536, EPI_ISL_519537, EPI_ISL_519538, EPI_ISL_519539, EPI_ISL_519540, EPI_ISL_519541, EPI_ISL_519542, EPI_ISL_519543, EPI_ISL_519544, EPI_ISL_519545, EPI_ISL_519546, EPI_ISL_519547, EPI_ISL_519548, EPI_ISL_519549, EPI_ISL_519550, EPI_ISL_519551, EPI_ISL_519552, EPI_ISL_519553, EPI_ISL_519554, EPI_ISL_519555, EPI_ISL_519556, EPI_ISL_519557, EPI_ISL_519558, EPI_ISL_519559, EPI_ISL_519560, EPI_ISL_519561, EPI_ISL_519562, EPI_ISL_519563, EPI_ISL_519564, EPI_ISL_519565, EPI_ISL_519566, EPI_ISL_519567, EPI_ISL_519568, EPI_ISL_519569, EPI_ISL_519570, EPI_ISL_519571, EPI_ISL_519572, EPI_ISL_519573, EPI_ISL_519574, EPI_ISL_519575, EPI_ISL_519576, EPI_ISL_519577, EPI_ISL_519578, EPI_ISL_519579, EPI_ISL_519580, EPI_ISL_519581, EPI_ISL_519582, EPI_ISL_519583, EPI_ISL_519584, EPI_ISL_519585, EPI_ISL_519586, EPI_ISL_519587, EPI_ISL_519588, EPI_ISL_519589, EPI_ISL_519590, EPI_ISL_519591, EPI_ISL_519592, EPI_ISL_519593, EPI_ISL_519594, EPI_ISL_519595, EPI_ISL_519596, EPI_ISL_519597, EPI_ISL_519598, EPI_ISL_519599, EPI_ISL_519600, EPI_ISL_519601, EPI_ISL_519602, EPI_ISL_519603, EPI_ISL_519604, EPI_ISL_519605, EPI_ISL_519606, EPI_ISL_519607, EPI_ISL_519608, EPI_ISL_519609, EPI_ISL_519610, EPI_ISL_519611, EPI_ISL_519612, EPI_ISL_519613, EPI_ISL_519614, EPI_ISL_519615, EPI_ISL_519616, EPI_ISL_519617, EPI_ISL_519618, EPI_ISL_519619, EPI_ISL_519620, EPI_ISL_519621, EPI_ISL_519622, EPI_ISL_519623, EPI_ISL_519624, EPI_ISL_519625, EPI_ISL_519626, EPI_ISL_519627, EPI_ISL_519628, EPI_ISL_519629, EPI_ISL_519630, EPI_ISL_519631, EPI_ISL_519632, EPI_ISL_519633, EPI_ISL_519634, EPI_ISL_519635, EPI_ISL_519636, EPI_ISL_519637, EPI_ISL_519638, EPI_ISL_519639, EPI_ISL_519640, EPI_ISL_519641, EPI_ISL_519642, EPI_ISL_519643, EPI_ISL_519645, EPI_ISL_519646, EPI_ISL_519647, EPI_ISL_519648, EPI_ISL_519649, EPI_ISL_519650, EPI_ISL_519651, EPI_ISL_519652, EPI_ISL_519653, EPI_ISL_519654, EPI_ISL_519655, EPI_ISL_519656, EPI_ISL_519657, EPI_ISL_519658, EPI_ISL_519659, EPI_ISL_519660, EPI_ISL_519661, EPI_ISL_519662, EPI_ISL_519663, EPI_ISL_519664, EPI_ISL_519665, EPI_ISL_519666, EPI_ISL_519667, EPI_ISL_519668, EPI_ISL_519669, EPI_ISL_519670, EPI_ISL_519671, EPI_ISL_519672, EPI_ISL_519673, EPI_ISL_519674, EPI_ISL_519675, EPI_ISL_519676, EPI_ISL_519677, EPI_ISL_519678, EPI_ISL_519679, EPI_ISL_519680, EPI_ISL_519681, EPI_ISL_519682, EPI_ISL_519683, EPI_ISL_519684, EPI_ISL_519685, EPI_ISL_519686, EPI_ISL_519687, EPI_ISL_519688, EPI_ISL_519689, EPI_ISL_519690, EPI_ISL_519691, EPI_ISL_519692, EPI_ISL_519693, EPI_ISL_519694, EPI_ISL_519695, EPI_ISL_519696, EPI_ISL_519697, EPI_ISL_519698, EPI_ISL_519699, EPI_ISL_519700, EPI_ISL_519701, EPI_ISL_519702, EPI_ISL_519703, EPI_ISL_519704, EPI_ISL_519705, EPI_ISL_519706, EPI_ISL_519707, EPI_ISL_519708, EPI_ISL_519709, EPI_ISL_519710, EPI_ISL_519711, EPI_ISL_519712, EPI_ISL_519713, EPI_ISL_519714, EPI_ISL_519715, EPI_ISL_519716, EPI_ISL_519717, EPI_ISL_519718, EPI_ISL_519719, EPI_ISL_519720, EPI_ISL_519721, EPI_ISL_519722, EPI_ISL_519723, EPI_ISL_519724, EPI_ISL_519725, EPI_ISL_519726, EPI_ISL_519727, EPI_ISL_519728, EPI_ISL_519729, EPI_ISL_519730, EPI_ISL_519731, EPI_ISL_519732, EPI_ISL_519733, EPI_ISL_519734, EPI_ISL_519735, EPI_ISL_519736, EPI_ISL_519737, EPI_ISL_519738, EPI_ISL_519739, EPI_ISL_519740, EPI_ISL_519741, EPI_ISL_519742, EPI_ISL_519743, EPI_ISL_519744, EPI_ISL_519745, EPI_ISL_519746, EPI_ISL_519747, EPI_ISL_519748, EPI_ISL_519749, EPI_ISL_519750, EPI_ISL_519751, EPI_ISL_519752, EPI_ISL_519753, EPI_ISL_519754, EPI_ISL_519755, EPI_ISL_519756, EPI_ISL_519757, EPI_ISL_519758, EPI_ISL_519759, EPI_ISL_519760, EPI_ISL_519761, EPI_ISL_519762, EPI_ISL_519763, EPI_ISL_519764, EPI_ISL_519765, EPI_ISL_519766, EPI_ISL_519767, EPI_ISL_519768, EPI_ISL_519769, EPI_ISL_519770, EPI_ISL_519771, EPI_ISL_519772, EPI_ISL_519773, EPI_ISL_519774, EPI_ISL_519775, EPI_ISL_519776, EPI_ISL_519777, EPI_ISL_519778, EPI_ISL_519779, EPI_ISL_519780, EPI_ISL_519781, EPI_ISL_519782, EPI_ISL_519783, EPI_ISL_519784, EPI_ISL_519785, EPI_ISL_519786, EPI_ISL_519787, EPI_ISL_519788, EPI_ISL_519789, EPI_ISL_519790, EPI_ISL_519791, EPI_ISL_519792, EPI_ISL_519793, EPI_ISL_519794, EPI_ISL_519795, EPI_ISL_519796, EPI_ISL_519797, EPI_ISL_519798, EPI_ISL_519799, EPI_ISL_519800, EPI_ISL_519801, EPI_ISL_519802, EPI_ISL_519803, EPI_ISL_519804, EPI_ISL_519805, EPI_ISL_519806, EPI_ISL_519807, EPI_ISL_519808, EPI_ISL_519809, EPI_ISL_519810, EPI_ISL_519811, EPI_ISL_519812, EPI_ISL_519813, EPI_ISL_519814, EPI_ISL_519815, EPI_ISL_519816, EPI_ISL_519817, EPI_ISL_519818, EPI_ISL_519819, EPI_ISL_519820, EPI_ISL_519821, EPI_ISL_519822, EPI_ISL_519823, EPI_ISL_519824, EPI_ISL_519825, EPI_ISL_519826, EPI_ISL_519827, EPI_ISL_519828, EPI_ISL_519829, EPI_ISL_519830, EPI_ISL_519831, EPI_ISL_519832, EPI_ISL_519833, EPI_ISL_519834, EPI_ISL_519835, EPI_ISL_519836, EPI_ISL_519837, EPI_ISL_519838, EPI_ISL_519839, EPI_ISL_519840, EPI_ISL_519841, EPI_ISL_519842, EPI_ISL_519843, EPI_ISL_519844, EPI_ISL_519845, EPI_ISL_519846, EPI_ISL_519847, EPI_ISL_519848, EPI_ISL_519849, EPI_ISL_519850, EPI_ISL_519851, EPI_ISL_519852, EPI_ISL_519853, EPI_ISL_519854, EPI_ISL_519855, EPI_ISL_519856, EPI_ISL_519857, EPI_ISL_519858, EPI_ISL_519859, EPI_ISL_519860, EPI_ISL_519861, EPI_ISL_519862, EPI_ISL_519863, EPI_ISL_519864, EPI_ISL_519865, EPI_ISL_519866, EPI_ISL_519867, EPI_ISL_519868, EPI_ISL_519869, EPI_ISL_519870, EPI_ISL_519871, EPI_ISL_519872, EPI_ISL_519873, EPI_ISL_519874, EPI_ISL_519875, EPI_ISL_519876, EPI_ISL_519877, EPI_ISL_519878, EPI_ISL_519879, EPI_ISL_519880, EPI_ISL_519881, EPI_ISL_519882, EPI_ISL_519883, EPI_ISL_519884, EPI_ISL_519885, EPI_ISL_519886, EPI_ISL_519887, EPI_ISL_519888, EPI_ISL_519889, EPI_ISL_519890, EPI_ISL_519891, EPI_ISL_519892, EPI_ISL_519893, EPI_ISL_519894 | | | |
| see above | Microbiological Diagnostic Unit - Public Health Laboratory (MDU-PHL) | MDU-PHL | Seemann T., Schultz M., Sait, M., Sherry, N. |
| EPI_ISL_519895 | Microbiological Diagnostic Unit - Public Health Laboratory (MDU-PHL) | Microbiological Diagnostic Unit - Public Health Laboratory (MDU-PHL) | Seemann T., Schultz M., Sait, M., Sherry, N. |
| EPI_ISL_519896, EPI_ISL_519897, EPI_ISL_519898, EPI_ISL_519899, EPI_ISL_519900, EPI_ISL_519901, EPI_ISL_519902, EPI_ISL_519903, EPI_ISL_519904, EPI_ISL_519905, EPI_ISL_519906, EPI_ISL_519907, EPI_ISL_519908, EPI_ISL_519909, EPI_ISL_519910, EPI_ISL_519911, EPI_ISL_519912, EPI_ISL_519913, EPI_ISL_519914, EPI_ISL_519915, EPI_ISL_519916, EPI_ISL_519917, EPI_ISL_519918, EPI_ISL_519919, | | | |

| | | | |
|---|---|---|---|
| EPI_ISL_519920, EPI_ISL_519921, EPI_ISL_519922, EPI_ISL_519923, EPI_ISL_519924, EPI_ISL_519925, EPI_ISL_519926, EPI_ISL_519927, EPI_ISL_519928, EPI_ISL_519929, EPI_ISL_519930, EPI_ISL_519931, EPI_ISL_519932, EPI_ISL_519933, EPI_ISL_519934, EPI_ISL_519935, EPI_ISL_519936, EPI_ISL_519937, EPI_ISL_519938, EPI_ISL_519939, EPI_ISL_519940, EPI_ISL_519941, EPI_ISL_519942, EPI_ISL_519943, EPI_ISL_519944, EPI_ISL_519945, EPI_ISL_519946, EPI_ISL_519947 | | | |
| see above | Microbiological Diagnostic Unit - Public Health Laboratory (MDU-PHL) | MDU-PHL | Seemann T., Schultz M., Sait, M., Sherry, N. |
| EPI_ISL_519948, EPI_ISL_519949, EPI_ISL_519950 | Victorian Infectious Diseases Reference Laboratory (VIDRL) | VIDRL and MDU-PHL | Caly L., Seemann T., Sait, M., Schultz M., Druce J., Sherry, N. |
| EPI_ISL_519951, EPI_ISL_519952, EPI_ISL_519953, EPI_ISL_519954, EPI_ISL_519955, EPI_ISL_519956, EPI_ISL_519957, EPI_ISL_519958, EPI_ISL_519959, EPI_ISL_519960, EPI_ISL_519961, EPI_ISL_519962, EPI_ISL_519963, EPI_ISL_519964, EPI_ISL_519965, EPI_ISL_519966, EPI_ISL_519967, EPI_ISL_519968, EPI_ISL_519969, EPI_ISL_519970, EPI_ISL_519971, EPI_ISL_519972, EPI_ISL_519973, EPI_ISL_519974, EPI_ISL_519975, EPI_ISL_519976, EPI_ISL_519977, EPI_ISL_519978, EPI_ISL_519979, EPI_ISL_519980, EPI_ISL_519981, EPI_ISL_519982, EPI_ISL_519983, EPI_ISL_519984, EPI_ISL_519985, EPI_ISL_519986, EPI_ISL_519987, EPI_ISL_519988, EPI_ISL_519989, EPI_ISL_519990, EPI_ISL_519991, EPI_ISL_519992, EPI_ISL_519993, EPI_ISL_519994, EPI_ISL_519995, EPI_ISL_519996, EPI_ISL_519997, EPI_ISL_519998, EPI_ISL_519999, EPI_ISL_520000, EPI_ISL_520001, EPI_ISL_520002, EPI_ISL_520003, EPI_ISL_520004, EPI_ISL_520005, EPI_ISL_520006, EPI_ISL_520007, EPI_ISL_520008, EPI_ISL_520009, EPI_ISL_520010, EPI_ISL_520011, EPI_ISL_520012, EPI_ISL_520013, EPI_ISL_520014, EPI_ISL_520015, EPI_ISL_520016, EPI_ISL_520017, EPI_ISL_520018, EPI_ISL_520019, EPI_ISL_520020, EPI_ISL_520021, EPI_ISL_520022, EPI_ISL_520023, EPI_ISL_520024, EPI_ISL_520025, EPI_ISL_520026, EPI_ISL_520027, EPI_ISL_520028, EPI_ISL_520029, EPI_ISL_520030, EPI_ISL_520031, EPI_ISL_520032, EPI_ISL_520033, EPI_ISL_520034, EPI_ISL_520035, EPI_ISL_520036, EPI_ISL_520037, EPI_ISL_520038, EPI_ISL_520039, EPI_ISL_520040, EPI_ISL_520041, EPI_ISL_520042, EPI_ISL_520043, EPI_ISL_520044, EPI_ISL_520045, EPI_ISL_520046, EPI_ISL_520047, EPI_ISL_520048, EPI_ISL_520049, EPI_ISL_520050, EPI_ISL_520051, EPI_ISL_520052, EPI_ISL_520053, EPI_ISL_520054, EPI_ISL_520055, EPI_ISL_520056, EPI_ISL_520057, EPI_ISL_520058, EPI_ISL_520059, EPI_ISL_520060, EPI_ISL_520061, EPI_ISL_520062, EPI_ISL_520063, EPI_ISL_520064, EPI_ISL_520065, EPI_ISL_520066, EPI_ISL_520067, EPI_ISL_520068, EPI_ISL_520069, EPI_ISL_520070, EPI_ISL_520071, EPI_ISL_520072, EPI_ISL_520073, EPI_ISL_520074, EPI_ISL_520075, EPI_ISL_520076, EPI_ISL_520077, EPI_ISL_520078, EPI_ISL_520079, EPI_ISL_520080, EPI_ISL_520081, EPI_ISL_520082, EPI_ISL_520083, EPI_ISL_520084, EPI_ISL_520085, EPI_ISL_520086, EPI_ISL_520087, EPI_ISL_520088, EPI_ISL_520089, EPI_ISL_520090, EPI_ISL_520091, EPI_ISL_520092, EPI_ISL_520093, EPI_ISL_520094, EPI_ISL_520095, EPI_ISL_520096, EPI_ISL_520097, EPI_ISL_520098, EPI_ISL_520099, EPI_ISL_520100, EPI_ISL_520101, EPI_ISL_520102, EPI_ISL_520103, EPI_ISL_520104, EPI_ISL_520105, EPI_ISL_520106, EPI_ISL_520107, EPI_ISL_520108, EPI_ISL_520109, EPI_ISL_520110, EPI_ISL_520111, EPI_ISL_520112, EPI_ISL_520113, EPI_ISL_520114, EPI_ISL_520115, EPI_ISL_520116, EPI_ISL_520117, EPI_ISL_520118, EPI_ISL_520119, EPI_ISL_520121, EPI_ISL_520122, EPI_ISL_520123, EPI_ISL_520124, EPI_ISL_520125, EPI_ISL_520126, EPI_ISL_520127, EPI_ISL_520128, EPI_ISL_520129, EPI_ISL_520130, EPI_ISL_520131, EPI_ISL_520132, EPI_ISL_520133, EPI_ISL_520134, EPI_ISL_520135, EPI_ISL_520136, EPI_ISL_520137, EPI_ISL_520138, EPI_ISL_520139, EPI_ISL_520140, EPI_ISL_520141, EPI_ISL_520142, EPI_ISL_520143, EPI_ISL_520144, EPI_ISL_520145, EPI_ISL_520146, EPI_ISL_520147, EPI_ISL_520148, EPI_ISL_520149, EPI_ISL_520150, EPI_ISL_520151, EPI_ISL_520152, EPI_ISL_520153, EPI_ISL_520154, EPI_ISL_520155, EPI_ISL_520156, EPI_ISL_520157, EPI_ISL_520158, EPI_ISL_520159, EPI_ISL_520160, EPI_ISL_520161, EPI_ISL_520162, EPI_ISL_520163, EPI_ISL_520164, EPI_ISL_520165, EPI_ISL_520166, EPI_ISL_520167, EPI_ISL_520168, EPI_ISL_520169, EPI_ISL_520170, EPI_ISL_520171, EPI_ISL_520172, EPI_ISL_520173, EPI_ISL_520174, EPI_ISL_520175, EPI_ISL_520176, EPI_ISL_520177, EPI_ISL_520178, EPI_ISL_520179, EPI_ISL_520180 | | | |
| see above | Microbiological Diagnostic Unit - Public Health Laboratory (MDU-PHL) | MDU-PHL | Seemann T., Schultz M., Sait, M., Sherry, N. |
| EPI_ISL_520181, EPI_ISL_520182, EPI_ISL_520183, EPI_ISL_520184, EPI_ISL_520185, EPI_ISL_520186, EPI_ISL_520187, EPI_ISL_520188, EPI_ISL_520189, EPI_ISL_520190, EPI_ISL_520191, EPI_ISL_520192, EPI_ISL_520193, EPI_ISL_520194, EPI_ISL_520195, EPI_ISL_520196, EPI_ISL_520197, EPI_ISL_520198, EPI_ISL_520199, EPI_ISL_520200, EPI_ISL_520202, EPI_ISL_520203, EPI_ISL_520204, EPI_ISL_520205, EPI_ISL_520206, EPI_ISL_520207, EPI_ISL_520208, EPI_ISL_520209, EPI_ISL_520210, EPI_ISL_520211, EPI_ISL_520212, EPI_ISL_520213, EPI_ISL_520214 | | | |
| see above | Victorian Infectious Diseases Reference Laboratory (VIDRL) | VIDRL and MDU-PHL | Caly L., Seemann T., Sait, M., Schultz M., Druce J., Sherry, N. |
| EPI_ISL_520215, EPI_ISL_520216, EPI_ISL_520217, EPI_ISL_520218, EPI_ISL_520219, EPI_ISL_520220, EPI_ISL_520221, EPI_ISL_520222, EPI_ISL_520223, EPI_ISL_520224, EPI_ISL_520225, EPI_ISL_520226, EPI_ISL_520227, EPI_ISL_520228, EPI_ISL_520229, EPI_ISL_520230, EPI_ISL_520231, EPI_ISL_520233, EPI_ISL_520234, EPI_ISL_520235, EPI_ISL_520236, EPI_ISL_520237, EPI_ISL_520238, EPI_ISL_520239, EPI_ISL_520240, EPI_ISL_520241, EPI_ISL_520242, EPI_ISL_520243, EPI_ISL_520244, EPI_ISL_520245, EPI_ISL_520246, EPI_ISL_520247, EPI_ISL_520248 | | | |
| see above | Microbiological Diagnostic Unit - Public Health Laboratory (MDU-PHL) | MDU-PHL | Seemann T., Schultz M., Sait, M., Sherry, N. |
| EPI_ISL_520249, EPI_ISL_520250, EPI_ISL_520251, EPI_ISL_520252, EPI_ISL_520253, EPI_ISL_520254, EPI_ISL_520255, EPI_ISL_520256, EPI_ISL_520257, EPI_ISL_520258, EPI_ISL_520259, EPI_ISL_520260, EPI_ISL_520261, EPI_ISL_520262, EPI_ISL_520263, EPI_ISL_520264, EPI_ISL_520265, EPI_ISL_520266, EPI_ISL_520267, EPI_ISL_520268, EPI_ISL_520269, EPI_ISL_520270, EPI_ISL_520271, EPI_ISL_520272, EPI_ISL_520273, EPI_ISL_520274, EPI_ISL_520275, EPI_ISL_520276, EPI_ISL_520277, EPI_ISL_520278, EPI_ISL_520279, EPI_ISL_520280, EPI_ISL_520281, EPI_ISL_520282, EPI_ISL_520283, EPI_ISL_520284, EPI_ISL_520285, EPI_ISL_520286, EPI_ISL_520287, EPI_ISL_520288, EPI_ISL_520289, EPI_ISL_520290, EPI_ISL_520291, EPI_ISL_520292, EPI_ISL_520293, EPI_ISL_520294, EPI_ISL_520295, EPI_ISL_520296, EPI_ISL_520297, EPI_ISL_520298, EPI_ISL_520299, EPI_ISL_520300, EPI_ISL_520301, EPI_ISL_520302, EPI_ISL_520303, EPI_ISL_520304, EPI_ISL_520305, EPI_ISL_520306, EPI_ISL_520307, EPI_ISL_520308, EPI_ISL_520309, EPI_ISL_520310, EPI_ISL_520311, EPI_ISL_520312, EPI_ISL_520313, EPI_ISL_520314, EPI_ISL_520315, EPI_ISL_520316, EPI_ISL_520317, EPI_ISL_520318, EPI_ISL_520319, EPI_ISL_520320, EPI_ISL_520321, EPI_ISL_520322, EPI_ISL_520323, EPI_ISL_520324, EPI_ISL_520325, EPI_ISL_520326, EPI_ISL_520327, EPI_ISL_520328, EPI_ISL_520329, EPI_ISL_520330, EPI_ISL_520331, EPI_ISL_520332, EPI_ISL_520333, EPI_ISL_520334, EPI_ISL_520335, EPI_ISL_520336, EPI_ISL_520337, EPI_ISL_520338, EPI_ISL_520339, EPI_ISL_520340, EPI_ISL_520341, EPI_ISL_520342, EPI_ISL_520343, EPI_ISL_520344, EPI_ISL_520345, EPI_ISL_520346, EPI_ISL_520347, EPI_ISL_520348, EPI_ISL_520349, EPI_ISL_520350, EPI_ISL_520351, EPI_ISL_520352, EPI_ISL_520353, EPI_ISL_520354, EPI_ISL_520355, EPI_ISL_520356, EPI_ISL_520357, EPI_ISL_520358, EPI_ISL_520359, EPI_ISL_520360, EPI_ISL_520361, EPI_ISL_520362, EPI_ISL_520363, EPI_ISL_520364, EPI_ISL_520365, EPI_ISL_520366, EPI_ISL_520367, EPI_ISL_520368, EPI_ISL_520369, EPI_ISL_520370, EPI_ISL_520371, EPI_ISL_520372, EPI_ISL_520373, EPI_ISL_520374, EPI_ISL_520375, EPI_ISL_520376, EPI_ISL_520377, EPI_ISL_520378, EPI_ISL_520379, EPI_ISL_520380, EPI_ISL_520381, EPI_ISL_520382, EPI_ISL_520383, EPI_ISL_520384, EPI_ISL_520385, EPI_ISL_520386, EPI_ISL_520387, EPI_ISL_520388, EPI_ISL_520389, EPI_ISL_520390, EPI_ISL_520391 | | | |
| see above | Victorian Infectious Diseases Reference Laboratory (VIDRL) | VIDRL and MDU-PHL | Caly L., Seemann T., Sait, M., Schultz M., Druce J., Sherry, N. |
| EPI_ISL_520392, EPI_ISL_520393, EPI_ISL_520394, EPI_ISL_520395, EPI_ISL_520396, EPI_ISL_520397, EPI_ISL_520398, EPI_ISL_520399, EPI_ISL_520400, EPI_ISL_520401, EPI_ISL_520402, EPI_ISL_520403, EPI_ISL_520404, EPI_ISL_520405, EPI_ISL_520406, EPI_ISL_520407, EPI_ISL_520408, EPI_ISL_520409, EPI_ISL_520410, EPI_ISL_520411, EPI_ISL_520412, EPI_ISL_520413, EPI_ISL_520414, EPI_ISL_520415, EPI_ISL_520416, EPI_ISL_520417, EPI_ISL_520418, EPI_ISL_520419, EPI_ISL_520420, EPI_ISL_520421, EPI_ISL_520422, EPI_ISL_520423, EPI_ISL_520424, EPI_ISL_520425, EPI_ISL_520426, EPI_ISL_520427, EPI_ISL_520428, EPI_ISL_520429, EPI_ISL_520430, EPI_ISL_520431, EPI_ISL_520432, EPI_ISL_520433, EPI_ISL_520434, EPI_ISL_520435, EPI_ISL_520436, EPI_ISL_520437, EPI_ISL_520438, EPI_ISL_520439, EPI_ISL_520440, EPI_ISL_520441, EPI_ISL_520442, EPI_ISL_520443, EPI_ISL_520444, EPI_ISL_520445, EPI_ISL_520446, EPI_ISL_520447, EPI_ISL_520448, EPI_ISL_520449, EPI_ISL_520450, EPI_ISL_520451, EPI_ISL_520452, EPI_ISL_520453, EPI_ISL_520454, EPI_ISL_520455, EPI_ISL_520456, EPI_ISL_520457, EPI_ISL_520458, EPI_ISL_520459, EPI_ISL_520460, EPI_ISL_520461, EPI_ISL_520462, EPI_ISL_520463, EPI_ISL_520464, EPI_ISL_520465, EPI_ISL_520466, EPI_ISL_520467, EPI_ISL_520468, EPI_ISL_520469, EPI_ISL_520470, EPI_ISL_520471, EPI_ISL_520472, EPI_ISL_520473, EPI_ISL_520474, EPI_ISL_520475, EPI_ISL_520476, EPI_ISL_520477, EPI_ISL_520478, EPI_ISL_520479, EPI_ISL_520480, EPI_ISL_520481, EPI_ISL_520482, EPI_ISL_520483, EPI_ISL_520484, EPI_ISL_520485, EPI_ISL_520486, EPI_ISL_520487, EPI_ISL_520488, EPI_ISL_520489, EPI_ISL_520490, EPI_ISL_520491, EPI_ISL_520492, EPI_ISL_520493, EPI_ISL_520494, EPI_ISL_520495, EPI_ISL_520496, EPI_ISL_520497, EPI_ISL_520498, EPI_ISL_520499, EPI_ISL_520500, EPI_ISL_520501, EPI_ISL_520502, EPI_ISL_520503, EPI_ISL_520504, EPI_ISL_520505, EPI_ISL_520506, EPI_ISL_520507, EPI_ISL_520508, EPI_ISL_520509, EPI_ISL_520510, EPI_ISL_520511, EPI_ISL_520512, EPI_ISL_520513, EPI_ISL_520514, EPI_ISL_520515, EPI_ISL_520516, EPI_ISL_520517, EPI_ISL_520518, EPI_ISL_520519, EPI_ISL_520520, EPI_ISL_520521, EPI_ISL_520522, EPI_ISL_520523, EPI_ISL_520524, EPI_ISL_520525, EPI_ISL_520526, EPI_ISL_520527, EPI_ISL_520528, EPI_ISL_520529, EPI_ISL_520530, EPI_ISL_520531, EPI_ISL_520532, EPI_ISL_520533, EPI_ISL_520534, EPI_ISL_520535, EPI_ISL_520536, EPI_ISL_520537, EPI_ISL_520538, EPI_ISL_520539, EPI_ISL_520540, EPI_ISL_520541, EPI_ISL_520542, EPI_ISL_520543, EPI_ISL_520544, EPI_ISL_520545, EPI_ISL_520546, EPI_ISL_520547, EPI_ISL_520548, EPI_ISL_520549, EPI_ISL_520550, EPI_ISL_520551, EPI_ISL_520552, EPI_ISL_520553, EPI_ISL_520554, EPI_ISL_520555, EPI_ISL_520556, EPI_ISL_520557, EPI_ISL_520558, EPI_ISL_520559, EPI_ISL_520560, EPI_ISL_520561, EPI_ISL_520562, EPI_ISL_520563, EPI_ISL_520564, EPI_ISL_520565, EPI_ISL_520566, EPI_ISL_520567, EPI_ISL_520568, EPI_ISL_520569, EPI_ISL_520570, EPI_ISL_520571, EPI_ISL_520572, EPI_ISL_520573, EPI_ISL_520574, EPI_ISL_520575, EPI_ISL_520576, EPI_ISL_520577, EPI_ISL_520578, EPI_ISL_520579, EPI_ISL_520580, EPI_ISL_520581, EPI_ISL_520582, EPI_ISL_520583, EPI_ISL_520584, EPI_ISL_520585, EPI_ISL_520586, EPI_ISL_520587, EPI_ISL_520588, EPI_ISL_520589, EPI_ISL_520590, EPI_ISL_520591, EPI_ISL_520592, EPI_ISL_520593, EPI_ISL_520594, EPI_ISL_520595, EPI_ISL_520596, EPI_ISL_520597, EPI_ISL_520598, EPI_ISL_520599, EPI_ISL_520600, EPI_ISL_520601, EPI_ISL_520602, EPI_ISL_520603 | | | |
| see above | Microbiological Diagnostic Unit - Public Health Laboratory (MDU-PHL) | MDU-PHL | Seemann T., Schultz M., Sait, M., Sherry, N. |
| EPI_ISL_520604, EPI_ISL_520605 | Microbiological Diagnostic Unit - Public Health Laboratory (MDU-PHL) | Microbiological Diagnostic Unit - Public Health Laboratory (MDU-PHL) | Seemann T., Schultz M., Sait, M., Sherry, N. |
| EPI_ISL_520606, EPI_ISL_520607, EPI_ISL_520608, EPI_ISL_520609, EPI_ISL_520610, EPI_ISL_520611, EPI_ISL_520612, EPI_ISL_520613, EPI_ISL_520614, EPI_ISL_520615, EPI_ISL_520616, EPI_ISL_520617, EPI_ISL_520618, EPI_ISL_520619, EPI_ISL_520620, EPI_ISL_520621, EPI_ISL_520622, EPI_ISL_520623, EPI_ISL_520624, EPI_ISL_520625, EPI_ISL_520626, EPI_ISL_520627, EPI_ISL_520628, EPI_ISL_520629, EPI_ISL_520630, EPI_ISL_520631, EPI_ISL_520632, EPI_ISL_520633, EPI_ISL_520634, EPI_ISL_520635, EPI_ISL_520636, EPI_ISL_520637, EPI_ISL_520638, EPI_ISL_520639, EPI_ISL_520640, EPI_ISL_520641, EPI_ISL_520642, EPI_ISL_520643 | | | |
| see above | Microbiological Diagnostic Unit - Public Health Laboratory (MDU-PHL) | MDU-PHL | Seemann T., Schultz M., Sait, M., Sherry, N. |
| EPI_ISL_520644 | Microbiological Diagnostic Unit - Public Health Laboratory (MDU-PHL) | Microbiological Diagnostic Unit - Public Health Laboratory (MDU-PHL) | Seemann T., Schultz M., Sait, M., Sherry, N. |
| EPI_ISL_520645, EPI_ISL_520646, EPI_ISL_520647, EPI_ISL_520648, EPI_ISL_520649, EPI_ISL_520650, EPI_ISL_520651, EPI_ISL_520652, EPI_ISL_520653, EPI_ISL_520654, EPI_ISL_520655, EPI_ISL_520656, EPI_ISL_520657, EPI_ISL_520658, EPI_ISL_520659, EPI_ISL_520660, EPI_ISL_520661, EPI_ISL_520662 | | | |
| see above | Microbiological Diagnostic Unit - Public Health Laboratory (MDU-PHL) | MDU-PHL | Seemann T., Schultz M., Sait, M., Sherry, N. |
| EPI_ISL_520663, EPI_ISL_520664, EPI_ISL_520665, EPI_ISL_520666, EPI_ISL_520667, EPI_ISL_520668, EPI_ISL_520669, EPI_ISL_520670, EPI_ISL_520671, EPI_ISL_520672, EPI_ISL_520673, EPI_ISL_520674, EPI_ISL_520675, EPI_ISL_520676, EPI_ISL_520677, EPI_ISL_520678, EPI_ISL_520679, EPI_ISL_520680, EPI_ISL_520681, EPI_ISL_520682, EPI_ISL_520683, EPI_ISL_520684, EPI_ISL_520685, EPI_ISL_520686, EPI_ISL_520687, EPI_ISL_520688, EPI_ISL_520689, EPI_ISL_520690, EPI_ISL_520691, EPI_ISL_520692, EPI_ISL_520693, EPI_ISL_520694, EPI_ISL_520695, EPI_ISL_520696, EPI_ISL_520697, EPI_ISL_520698, EPI_ISL_520699, EPI_ISL_520700, EPI_ISL_520701, EPI_ISL_520702, EPI_ISL_520703, EPI_ISL_520704, EPI_ISL_520705, EPI_ISL_520706, EPI_ISL_520707, EPI_ISL_520708, EPI_ISL_520709, EPI_ISL_520710, EPI_ISL_520711, EPI_ISL_520712, EPI_ISL_520713, EPI_ISL_520714, EPI_ISL_520715, EPI_ISL_520716, EPI_ISL_520717, EPI_ISL_520718, EPI_ISL_520719, EPI_ISL_520720, EPI_ISL_520721, EPI_ISL_520722, EPI_ISL_520723, EPI_ISL_520724, EPI_ISL_520725, EPI_ISL_520726, EPI_ISL_520727, EPI_ISL_520728, EPI_ISL_520729, EPI_ISL_520730, EPI_ISL_520731, EPI_ISL_520732, EPI_ISL_520733, EPI_ISL_520734, EPI_ISL_520735, EPI_ISL_520736, EPI_ISL_520737, EPI_ISL_520738, EPI_ISL_520739, EPI_ISL_520740, EPI_ISL_520741, EPI_ISL_520742, EPI_ISL_520743, EPI_ISL_520744 | | | |
| see above | Mohammed Bin Rashid University of Medicine and Health Sciences | Al Jalila Genomics Center | Ahmad Abou Tayoun, Tom Loney, Hamda Khansaheb, Sathishkumar Ramaswamy, Divinlal Harilal, Zulfa Omar Deesi, Rupa Murthy Varghese, Hanan Al Suwaidi, Abdulmajeed Alkhaja, Mohammed Uddin, Rifat Hamoudi, Rabih Halwani, Abiola Catherine Senok, Qutayba Hamid, Norbert Nowotny, Alawi Alsheikh-Ali |
| EPI_ISL_520745, EPI_ISL_520746, EPI_ISL_520747, EPI_ISL_520748, EPI_ISL_520749, EPI_ISL_520750, EPI_ISL_520751, EPI_ISL_520752, EPI_ISL_520753, EPI_ISL_520754, EPI_ISL_520755, EPI_ISL_520756, EPI_ISL_520757, EPI_ISL_520758, EPI_ISL_520759, EPI_ISL_520760, EPI_ISL_520761, EPI_ISL_520762, EPI_ISL_520763, EPI_ISL_520764 | | | |
| see above | Victorian Infectious Diseases Reference Laboratory (VIDRL) | VIDRL and MDU-PHL | Caly L., Seemann T., Sait, M., Schultz M., Druce J., Sherry, N. |
| EPI_ISL_520765, EPI_ISL_520766, EPI_ISL_520767, EPI_ISL_520768, EPI_ISL_520769, EPI_ISL_520770, EPI_ISL_520771, EPI_ISL_520772, EPI_ISL_520773, EPI_ISL_520774, EPI_ISL_520776, EPI_ISL_520777, EPI_ISL_520778, EPI_ISL_520779, EPI_ISL_520780, EPI_ISL_520781, EPI_ISL_520782, EPI_ISL_520783, EPI_ISL_520784, EPI_ISL_520785, EPI_ISL_520786, EPI_ISL_520787, EPI_ISL_520788, EPI_ISL_520789, EPI_ISL_520790 | | | |
| see above | Microbiological Diagnostic Unit - Public Health Laboratory (MDU-PHL) | MDU-PHL | Seemann T., Schultz M., Sait, M., Sherry, N. |
| EPI_ISL_520791, EPI_ISL_520792, EPI_ISL_520793, EPI_ISL_520794, EPI_ISL_520795, EPI_ISL_520796, EPI_ISL_520797, EPI_ISL_520798, EPI_ISL_520799, EPI_ISL_520800, EPI_ISL_520801, EPI_ISL_520802, EPI_ISL_520803, EPI_ISL_520804, EPI_ISL_520805, EPI_ISL_520806, EPI_ISL_520807, EPI_ISL_520808, EPI_ISL_520809, EPI_ISL_520810, EPI_ISL_520811, EPI_ISL_520812, EPI_ISL_520813, EPI_ISL_520814, EPI_ISL_520815, EPI_ISL_520816, EPI_ISL_520817, EPI_ISL_520818, EPI_ISL_520819, EPI_ISL_520820, EPI_ISL_520821, EPI_ISL_520822, EPI_ISL_520823, EPI_ISL_520824, EPI_ISL_520825, EPI_ISL_520826, EPI_ISL_520827, EPI_ISL_520828, EPI_ISL_520829, EPI_ISL_520830, EPI_ISL_520831, EPI_ISL_520832, EPI_ISL_520833, EPI_ISL_520834, EPI_ISL_520835, EPI_ISL_520836, EPI_ISL_520837, EPI_ISL_520838, EPI_ISL_520840, EPI_ISL_520841, EPI_ISL_520842, EPI_ISL_520843 | | | |
| see above | Victorian Infectious Diseases Reference Laboratory (VIDRL) | VIDRL and MDU-PHL | Caly L., Seemann T., Sait, M., Schultz M., Druce J., Sherry, N. |
| EPI_ISL_520844, EPI_ISL_520845, EPI_ISL_520846, EPI_ISL_520847, EPI_ISL_520848, EPI_ISL_520849, EPI_ISL_520850, EPI_ISL_520851, EPI_ISL_520852, EPI_ISL_520853, EPI_ISL_520854, EPI_ISL_520855, EPI_ISL_520856, EPI_ISL_520857, EPI_ISL_520858, EPI_ISL_520859, EPI_ISL_520860, EPI_ISL_520861, EPI_ISL_520862, EPI_ISL_520863 | | | |
| see above | Microbiological Diagnostic Unit - Public Health Laboratory (MDU-PHL) | MDU-PHL | Seemann T., Schultz M., Sait, M., Sherry, N. |
| EPI_ISL_520864, EPI_ISL_520865, EPI_ISL_520866, EPI_ISL_520867, EPI_ISL_520868, EPI_ISL_520869, EPI_ISL_520870, EPI_ISL_520871, EPI_ISL_520872, EPI_ISL_520873, EPI_ISL_520874, EPI_ISL_520875, EPI_ISL_520876, EPI_ISL_520877, EPI_ISL_520878, EPI_ISL_520879, EPI_ISL_520880, EPI_ISL_520881, EPI_ISL_520882, EPI_ISL_520883, EPI_ISL_520884, EPI_ISL_520885, EPI_ISL_520886, EPI_ISL_520887, EPI_ISL_520888, EPI_ISL_520889, EPI_ISL_520890, EPI_ISL_520891, EPI_ISL_520892, EPI_ISL_520893, EPI_ISL_520894, EPI_ISL_520895, EPI_ISL_520896, EPI_ISL_520897, EPI_ISL_520898, EPI_ISL_520899, EPI_ISL_520900, EPI_ISL_520901, EPI_ISL_520902, EPI_ISL_520903, EPI_ISL_520904, EPI_ISL_520905, EPI_ISL_520906, EPI_ISL_520907, EPI_ISL_520908, EPI_ISL_520909, EPI_ISL_520910, EPI_ISL_520911, EPI_ISL_520912, EPI_ISL_520913, EPI_ISL_520914, EPI_ISL_520915, EPI_ISL_520916, EPI_ISL_520917, EPI_ISL_520918, EPI_ISL_520919, EPI_ISL_520920, EPI_ISL_520921, EPI_ISL_520922, EPI_ISL_520923, EPI_ISL_520924, EPI_ISL_520925, EPI_ISL_520926, EPI_ISL_520927, EPI_ISL_520928 | | | |
| see above | Victorian Infectious Diseases Reference Laboratory (VIDRL) | VIDRL and MDU-PHL | Caly L., Seemann T., Sait, M., Schultz M., Druce J., Sherry, N. |
| EPI_ISL_520929, EPI_ISL_520930, EPI_ISL_520931, EPI_ISL_520932, EPI_ISL_520933, EPI_ISL_520934, EPI_ISL_520935, EPI_ISL_520936, EPI_ISL_520937, EPI_ISL_520938, EPI_ISL_520939, EPI_ISL_520940, EPI_ISL_520941, EPI_ISL_520942, EPI_ISL_520943, EPI_ISL_520944, EPI_ISL_520945, EPI_ISL_520946, EPI_ISL_520947, EPI_ISL_520948, EPI_ISL_520949, EPI_ISL_520950, EPI_ISL_520951, EPI_ISL_520952, EPI_ISL_520953, EPI_ISL_520954, EPI_ISL_520955, EPI_ISL_520956, EPI_ISL_520957, EPI_ISL_520958, EPI_ISL_520959, EPI_ISL_520960, EPI_ISL_520961, EPI_ISL_520962, EPI_ISL_520963, EPI_ISL_520964, EPI_ISL_520965, EPI_ISL_520966, EPI_ISL_520967, EPI_ISL_520968, EPI_ISL_520969, EPI_ISL_520970, EPI_ISL_520971, EPI_ISL_520972, EPI_ISL_520973, EPI_ISL_520974, EPI_ISL_520975, EPI_ISL_520976, EPI_ISL_520977, EPI_ISL_520978, EPI_ISL_520979, EPI_ISL_520980, EPI_ISL_520981, EPI_ISL_520982, EPI_ISL_520983, EPI_ISL_520984, EPI_ISL_520985, EPI_ISL_520986, EPI_ISL_520987, EPI_ISL_520988, EPI_ISL_520989, EPI_ISL_520990, EPI_ISL_520991, EPI_ISL_520992, EPI_ISL_520993, EPI_ISL_520994, EPI_ISL_520995, EPI_ISL_520996, EPI_ISL_520997, EPI_ISL_520998, EPI_ISL_520999, EPI_ISL_521000, EPI_ISL_521001, EPI_ISL_521002, EPI_ISL_521003, EPI_ISL_521004, EPI_ISL_521005, EPI_ISL_521006, EPI_ISL_521007, EPI_ISL_521008, EPI_ISL_521009, EPI_ISL_521010, EPI_ISL_521011, EPI_ISL_521012, EPI_ISL_521013, EPI_ISL_521014, EPI_ISL_521015, EPI_ISL_521016, EPI_ISL_521017, EPI_ISL_521018, EPI_ISL_521019, EPI_ISL_521020, EPI_ISL_521021, EPI_ISL_521022, EPI_ISL_521023, EPI_ISL_521024, EPI_ISL_521025, EPI_ISL_521026, EPI_ISL_521027, EPI_ISL_521028, EPI_ISL_521029, EPI_ISL_521030, EPI_ISL_521031, EPI_ISL_521032, EPI_ISL_521033, EPI_ISL_521034, EPI_ISL_521035, EPI_ISL_521036, EPI_ISL_521037, EPI_ISL_521038, EPI_ISL_521039, EPI_ISL_521040, EPI_ISL_521041, EPI_ISL_521042, EPI_ISL_521043, EPI_ISL_521044, EPI_ISL_521045, EPI_ISL_521046, EPI_ISL_521047, EPI_ISL_521048, EPI_ISL_521049, EPI_ISL_521050, EPI_ISL_521051, EPI_ISL_521052, EPI_ISL_521053, EPI_ISL_521054, EPI_ISL_521055, EPI_ISL_521056, EPI_ISL_521057, EPI_ISL_521058, EPI_ISL_521059, EPI_ISL_521060, EPI_ISL_521061, EPI_ISL_521062, EPI_ISL_521063, EPI_ISL_521064, EPI_ISL_521065, EPI_ISL_521066, EPI_ISL_521067, EPI_ISL_521068, EPI_ISL_521069, EPI_ISL_521070, EPI_ISL_521071, EPI_ISL_521072, EPI_ISL_521073, EPI_ISL_521074, EPI_ISL_521075, EPI_ISL_521076, EPI_ISL_521077, EPI_ISL_521078, EPI_ISL_521079, EPI_ISL_521080, EPI_ISL_521081, EPI_ISL_521082, EPI_ISL_521083, EPI_ISL_521084, EPI_ISL_521085, EPI_ISL_521086, EPI_ISL_521087, EPI_ISL_521088, EPI_ISL_521089, EPI_ISL_521090, EPI_ISL_521091, EPI_ISL_521092, EPI_ISL_521093, EPI_ISL_521094, EPI_ISL_521095, EPI_ISL_521096, EPI_ISL_521097, EPI_ISL_521098, EPI_ISL_521099, EPI_ISL_521100, EPI_ISL_521101 | | | |
| see above | Microbiological Diagnostic Unit - Public Health Laboratory (MDU-PHL) | MDU-PHL | Seemann T., Schultz M., Sait, M., Sherry, N. |
| EPI_ISL_521102, EPI_ISL_521103, EPI_ISL_521104, EPI_ISL_521105, EPI_ISL_521106, EPI_ISL_521107, EPI_ISL_521108, EPI_ISL_521109, EPI_ISL_521110, EPI_ISL_521111, EPI_ISL_521112, EPI_ISL_521113, EPI_ISL_521114, EPI_ISL_521115, EPI_ISL_521116, EPI_ISL_521117, EPI_ISL_521118, EPI_ISL_521119, EPI_ISL_521120, EPI_ISL_521121, EPI_ISL_521122, EPI_ISL_521123, EPI_ISL_521124, EPI_ISL_521125, EPI_ISL_521126, EPI_ISL_521127, EPI_ISL_521128, EPI_ISL_521129, EPI_ISL_521130, EPI_ISL_521131, EPI_ISL_521132, EPI_ISL_521133, EPI_ISL_521134, EPI_ISL_521135, EPI_ISL_521136, EPI_ISL_521137, EPI_ISL_521138, EPI_ISL_521139, EPI_ISL_521140, EPI_ISL_521141, EPI_ISL_521142, EPI_ISL_521143, EPI_ISL_521144, EPI_ISL_521145, EPI_ISL_521146, EPI_ISL_521147, EPI_ISL_521148, EPI_ISL_521149, EPI_ISL_521150, EPI_ISL_521151, EPI_ISL_521152 | | | |
| see above | Victorian Infectious Diseases Reference Laboratory (VIDRL) | VIDRL and MDU-PHL | Caly L., Seemann T., Sait, M., Schultz M., Druce J., Sherry, N. |
| EPI_ISL_521153 | Mayo Clinic & Mayo Clinic Laboratories | Minnesota Department of Health, Public Health Laboratory | Matt Plumb, Jacob Garfin, and Xiong Wang |
| EPI_ISL_521154, EPI_ISL_521156, EPI_ISL_521157, EPI_ISL_521158, EPI_ISL_521159, EPI_ISL_521160, EPI_ISL_521161, EPI_ISL_521162, EPI_ISL_521163, EPI_ISL_521164, EPI_ISL_521165, EPI_ISL_521166, EPI_ISL_521167, EPI_ISL_521168, EPI_ISL_521170, EPI_ISL_521171, EPI_ISL_521172, EPI_ISL_521173, EPI_ISL_521174, EPI_ISL_521175, EPI_ISL_521176, EPI_ISL_521177, EPI_ISL_521178, EPI_ISL_521179, EPI_ISL_521180, EPI_ISL_521181, EPI_ISL_521182, EPI_ISL_521183, EPI_ISL_521184, EPI_ISL_521185, EPI_ISL_521186, EPI_ISL_521187, EPI_ISL_521188, EPI_ISL_521189, EPI_ISL_521190, EPI_ISL_521191, EPI_ISL_521192, EPI_ISL_521193, EPI_ISL_521194, EPI_ISL_521195, EPI_ISL_521196, EPI_ISL_521197, EPI_ISL_521198, EPI_ISL_521199, EPI_ISL_521200, EPI_ISL_521201, EPI_ISL_521202, EPI_ISL_521203, | | | |

| | | | |
|---|---|---|---|
| EPI_ISL_521204, EPI_ISL_521205, EPI_ISL_521206, EPI_ISL_521207, EPI_ISL_521208, EPI_ISL_521209, EPI_ISL_521210, EPI_ISL_521211, EPI_ISL_521212, EPI_ISL_521213, EPI_ISL_521214, EPI_ISL_521215, EPI_ISL_521216, EPI_ISL_521217, EPI_ISL_521218, EPI_ISL_521219, EPI_ISL_521220, EPI_ISL_521221, EPI_ISL_521222, EPI_ISL_521223, EPI_ISL_521224, EPI_ISL_521226, EPI_ISL_521227, EPI_ISL_521228, EPI_ISL_521229, EPI_ISL_521230, EPI_ISL_521231, EPI_ISL_521232, EPI_ISL_521233, EPI_ISL_521234, EPI_ISL_521235, EPI_ISL_521236, EPI_ISL_521237, EPI_ISL_521238, EPI_ISL_521239, EPI_ISL_521240, EPI_ISL_521241, EPI_ISL_521242, EPI_ISL_521243, EPI_ISL_521244, EPI_ISL_521245, EPI_ISL_521246, EPI_ISL_521247, EPI_ISL_521248, EPI_ISL_521249, EPI_ISL_521250, EPI_ISL_521251, EPI_ISL_521252, EPI_ISL_521253, EPI_ISL_521254, EPI_ISL_521255, EPI_ISL_521256, EPI_ISL_521257, EPI_ISL_521258, EPI_ISL_521259, EPI_ISL_521260, EPI_ISL_521261, EPI_ISL_521262, EPI_ISL_521263, EPI_ISL_521264, EPI_ISL_521265, EPI_ISL_521266, EPI_ISL_521267, EPI_ISL_521268, EPI_ISL_521269 | | | |
| see above | Microbiological Diagnostic Unit - Public Health Laboratory (MDU-PHL) | MDU-PHL | Seemann T., Schultz M., Sait, M., Sherry, N. |
| EPI_ISL_521270, EPI_ISL_521271, EPI_ISL_521272, EPI_ISL_521273, EPI_ISL_521274, EPI_ISL_521275, EPI_ISL_521276, EPI_ISL_521277 | Victorian Infectious Diseases Reference Laboratory (VIDRL) | VIDRL and MDU-PHL | Caly L., Seemann T., Sait, M., Schultz M., Druce J., Sherry, N. |
| EPI_ISL_521278, EPI_ISL_521279, EPI_ISL_521280, EPI_ISL_521281, EPI_ISL_521282, EPI_ISL_521283, EPI_ISL_521284, EPI_ISL_521285, EPI_ISL_521286, EPI_ISL_521287, EPI_ISL_521288, EPI_ISL_521289, EPI_ISL_521290, EPI_ISL_521291, EPI_ISL_521292, EPI_ISL_521293, EPI_ISL_521294, EPI_ISL_521295, EPI_ISL_521296, EPI_ISL_521297, EPI_ISL_521298, EPI_ISL_521299, EPI_ISL_521300, EPI_ISL_521301, EPI_ISL_521302, EPI_ISL_521303, EPI_ISL_521304, EPI_ISL_521305, EPI_ISL_521306, EPI_ISL_521307, EPI_ISL_521308, EPI_ISL_521309, EPI_ISL_521310, EPI_ISL_521311, EPI_ISL_521312, EPI_ISL_521313, EPI_ISL_521314 | | | |
| see above | Microbiological Diagnostic Unit - Public Health Laboratory (MDU-PHL) | MDU-PHL | Seemann T., Schultz M., Sait, M., Sherry, N. |
| EPI_ISL_521315, EPI_ISL_521316, EPI_ISL_521317, EPI_ISL_521318, EPI_ISL_521319, EPI_ISL_521320, EPI_ISL_521321, EPI_ISL_521322, EPI_ISL_521323, EPI_ISL_521324, EPI_ISL_521325, EPI_ISL_521326, EPI_ISL_521327, EPI_ISL_521328, EPI_ISL_521329, EPI_ISL_521330, EPI_ISL_521331, EPI_ISL_521332, EPI_ISL_521333, EPI_ISL_521334, EPI_ISL_521335, EPI_ISL_521336, EPI_ISL_521337, EPI_ISL_521338, EPI_ISL_521339, EPI_ISL_521340, EPI_ISL_521341, EPI_ISL_521342, EPI_ISL_521343, EPI_ISL_521344, EPI_ISL_521345, EPI_ISL_521346, EPI_ISL_521347, EPI_ISL_521348, EPI_ISL_521349, EPI_ISL_521350, EPI_ISL_521351, EPI_ISL_521352, EPI_ISL_521353, EPI_ISL_521354, EPI_ISL_521355, EPI_ISL_521356, EPI_ISL_521357, EPI_ISL_521358, EPI_ISL_521359, EPI_ISL_521360, EPI_ISL_521361, EPI_ISL_521362, EPI_ISL_521363, EPI_ISL_521364, EPI_ISL_521365, EPI_ISL_521366, EPI_ISL_521367, EPI_ISL_521368, EPI_ISL_521369, EPI_ISL_521370, EPI_ISL_521371, EPI_ISL_521372, EPI_ISL_521373, EPI_ISL_521374, EPI_ISL_521375, EPI_ISL_521376, EPI_ISL_521377, EPI_ISL_521378, EPI_ISL_521379, EPI_ISL_521380, EPI_ISL_521381, EPI_ISL_521382, EPI_ISL_521383, EPI_ISL_521384, EPI_ISL_521385, EPI_ISL_521386, EPI_ISL_521387, EPI_ISL_521388, EPI_ISL_521389, EPI_ISL_521390, EPI_ISL_521391, EPI_ISL_521392, EPI_ISL_521393, EPI_ISL_521394, EPI_ISL_521395, EPI_ISL_521396, EPI_ISL_521397, EPI_ISL_521398, EPI_ISL_521399, EPI_ISL_521400, EPI_ISL_521401, EPI_ISL_521402, EPI_ISL_521403, EPI_ISL_521404, EPI_ISL_521405, EPI_ISL_521406, EPI_ISL_521407, EPI_ISL_521408, EPI_ISL_521409, EPI_ISL_521410, EPI_ISL_521411, EPI_ISL_521412, EPI_ISL_521413, EPI_ISL_521414, EPI_ISL_521415, EPI_ISL_521416, EPI_ISL_521417, EPI_ISL_521418, EPI_ISL_521419, EPI_ISL_521420, EPI_ISL_521421, EPI_ISL_521422, EPI_ISL_521423, EPI_ISL_521424, EPI_ISL_521425, EPI_ISL_521426, EPI_ISL_521427, EPI_ISL_521428, EPI_ISL_521429, EPI_ISL_521430, EPI_ISL_521431, EPI_ISL_521432, EPI_ISL_521433, EPI_ISL_521434, EPI_ISL_521435, EPI_ISL_521436, EPI_ISL_521437, EPI_ISL_521438, EPI_ISL_521439, EPI_ISL_521440, EPI_ISL_521441, EPI_ISL_521442, EPI_ISL_521443, EPI_ISL_521444, EPI_ISL_521445, EPI_ISL_521446, EPI_ISL_521447, EPI_ISL_521448, EPI_ISL_521449 | | | |
| see above | Victorian Infectious Diseases Reference Laboratory (VIDRL) | VIDRL and MDU-PHL | Caly L., Seemann T., Sait, M., Schultz M., Druce J., Sherry, N. |
| EPI_ISL_521450, EPI_ISL_521451, EPI_ISL_521452, EPI_ISL_521453, EPI_ISL_521454, EPI_ISL_521455, EPI_ISL_521456, EPI_ISL_521457, EPI_ISL_521458, EPI_ISL_521459, EPI_ISL_521460, EPI_ISL_521461 | | | |
| see above | Microbiological Diagnostic Unit - Public Health Laboratory (MDU-PHL) | MDU-PHL | Seemann T., Schultz M., Sait, M., Sherry, N. |
| EPI_ISL_521462, EPI_ISL_521463, EPI_ISL_521464, EPI_ISL_521465, EPI_ISL_521466, EPI_ISL_521467, EPI_ISL_521468, EPI_ISL_521469, EPI_ISL_521470, EPI_ISL_521471, EPI_ISL_521472, EPI_ISL_521473, EPI_ISL_521474, EPI_ISL_521475, EPI_ISL_521476, EPI_ISL_521477, EPI_ISL_521478, EPI_ISL_521479, EPI_ISL_521480, EPI_ISL_521481, EPI_ISL_521482, EPI_ISL_521483, EPI_ISL_521484, EPI_ISL_521485, EPI_ISL_521486, EPI_ISL_521487, EPI_ISL_521488, EPI_ISL_521489, EPI_ISL_521490, EPI_ISL_521491, EPI_ISL_521492, EPI_ISL_521493, EPI_ISL_521494, EPI_ISL_521495, EPI_ISL_521496, EPI_ISL_521497, EPI_ISL_521498, EPI_ISL_521499, EPI_ISL_521500, EPI_ISL_521501, EPI_ISL_521502, EPI_ISL_521503, EPI_ISL_521504, EPI_ISL_521505, EPI_ISL_521506, EPI_ISL_521507, EPI_ISL_521508, EPI_ISL_521509, EPI_ISL_521510, EPI_ISL_521511, EPI_ISL_521512, EPI_ISL_521513, EPI_ISL_521514, EPI_ISL_521515, EPI_ISL_521516, EPI_ISL_521517, EPI_ISL_521518, EPI_ISL_521519, EPI_ISL_521520, EPI_ISL_521521, EPI_ISL_521522, EPI_ISL_521523, EPI_ISL_521524, EPI_ISL_521525, EPI_ISL_521526, EPI_ISL_521527, EPI_ISL_521528, EPI_ISL_521529, EPI_ISL_521530, EPI_ISL_521531, EPI_ISL_521532, EPI_ISL_521533, EPI_ISL_521534, EPI_ISL_521535, EPI_ISL_521536, EPI_ISL_521537, EPI_ISL_521538, EPI_ISL_521539, EPI_ISL_521540, EPI_ISL_521541, EPI_ISL_521542, EPI_ISL_521543, EPI_ISL_521544, EPI_ISL_521545, EPI_ISL_521546, EPI_ISL_521547, EPI_ISL_521548, EPI_ISL_521549, EPI_ISL_521550, EPI_ISL_521551, EPI_ISL_521552, EPI_ISL_521553, EPI_ISL_521554, EPI_ISL_521555, EPI_ISL_521556, EPI_ISL_521557, EPI_ISL_521558, EPI_ISL_521559, EPI_ISL_521560, EPI_ISL_521561 | | | |
| see above | Victorian Infectious Diseases Reference Laboratory (VIDRL) | VIDRL and MDU-PHL | Caly L., Seemann T., Sait, M., Schultz M., Druce J., Sherry, N. |
| EPI_ISL_521562, EPI_ISL_521563, EPI_ISL_521564, EPI_ISL_521565, EPI_ISL_521566, EPI_ISL_521567, EPI_ISL_521568, EPI_ISL_521569, EPI_ISL_521570, EPI_ISL_521571, EPI_ISL_521572, EPI_ISL_521573, EPI_ISL_521574, EPI_ISL_521575, EPI_ISL_521576, EPI_ISL_521577, EPI_ISL_521578, EPI_ISL_521579, EPI_ISL_521580, EPI_ISL_521581, EPI_ISL_521582, EPI_ISL_521583, EPI_ISL_521584, EPI_ISL_521585, EPI_ISL_521586, EPI_ISL_521587, EPI_ISL_521588, EPI_ISL_521589, EPI_ISL_521590, EPI_ISL_521591, EPI_ISL_521592, EPI_ISL_521593, EPI_ISL_521594, EPI_ISL_521595, EPI_ISL_521596, EPI_ISL_521597, EPI_ISL_521598, EPI_ISL_521599, EPI_ISL_521600, EPI_ISL_521601, EPI_ISL_521602, EPI_ISL_521603, EPI_ISL_521604, EPI_ISL_521605, EPI_ISL_521606, EPI_ISL_521607, EPI_ISL_521608, EPI_ISL_521609, EPI_ISL_521610, EPI_ISL_521611, EPI_ISL_521612, EPI_ISL_521613, EPI_ISL_521614, EPI_ISL_521615, EPI_ISL_521616, EPI_ISL_521617, EPI_ISL_521618, EPI_ISL_521619, EPI_ISL_521620, EPI_ISL_521621, EPI_ISL_521622, EPI_ISL_521623, EPI_ISL_521624, EPI_ISL_521625, EPI_ISL_521626, EPI_ISL_521627, EPI_ISL_521628, EPI_ISL_521629, EPI_ISL_521630, EPI_ISL_521631, EPI_ISL_521632, EPI_ISL_521633, EPI_ISL_521634, EPI_ISL_521635, EPI_ISL_521636, EPI_ISL_521637, EPI_ISL_521638, EPI_ISL_521639, EPI_ISL_521640, EPI_ISL_521641, EPI_ISL_521642, EPI_ISL_521643, EPI_ISL_521644, EPI_ISL_521645, EPI_ISL_521646, EPI_ISL_521647, EPI_ISL_521648, EPI_ISL_521649, EPI_ISL_521650, EPI_ISL_521651, EPI_ISL_521652, EPI_ISL_521653, EPI_ISL_521654, EPI_ISL_521655, EPI_ISL_521656, EPI_ISL_521657, EPI_ISL_521658, EPI_ISL_521659, EPI_ISL_521660, EPI_ISL_521661, EPI_ISL_521662, EPI_ISL_521663, EPI_ISL_521664, EPI_ISL_521665, EPI_ISL_521666, EPI_ISL_521667, EPI_ISL_521668, EPI_ISL_521669, EPI_ISL_521670, EPI_ISL_521671, EPI_ISL_521672, EPI_ISL_521673, EPI_ISL_521674, EPI_ISL_521675, EPI_ISL_521676, EPI_ISL_521677, EPI_ISL_521678, EPI_ISL_521679, EPI_ISL_521680, EPI_ISL_521681, EPI_ISL_521682, EPI_ISL_521683, EPI_ISL_521684, EPI_ISL_521685, EPI_ISL_521686, EPI_ISL_521687, EPI_ISL_521688, EPI_ISL_521689, EPI_ISL_521690, EPI_ISL_521691, EPI_ISL_521692, EPI_ISL_521693, EPI_ISL_521694, EPI_ISL_521695, EPI_ISL_521696, EPI_ISL_521697, EPI_ISL_521698, EPI_ISL_521699, EPI_ISL_521700, EPI_ISL_521701, EPI_ISL_521702, EPI_ISL_521703, EPI_ISL_521704, EPI_ISL_521705, EPI_ISL_521706, EPI_ISL_521707, EPI_ISL_521708, EPI_ISL_521709, EPI_ISL_521710, EPI_ISL_521711, EPI_ISL_521712, EPI_ISL_521713, EPI_ISL_521714, EPI_ISL_521715, EPI_ISL_521716, EPI_ISL_521717, EPI_ISL_521718, EPI_ISL_521719, EPI_ISL_521720, EPI_ISL_521721, EPI_ISL_521722, EPI_ISL_521723, EPI_ISL_521724, EPI_ISL_521725, EPI_ISL_521726, EPI_ISL_521727, EPI_ISL_521728, EPI_ISL_521729, EPI_ISL_521730, EPI_ISL_521731, EPI_ISL_521732, EPI_ISL_521733, EPI_ISL_521734, EPI_ISL_521735, EPI_ISL_521736, EPI_ISL_521737, EPI_ISL_521738, EPI_ISL_521739, EPI_ISL_521740, EPI_ISL_521741, EPI_ISL_521742, EPI_ISL_521743, EPI_ISL_521744, EPI_ISL_521745, EPI_ISL_521746, EPI_ISL_521747, EPI_ISL_521748, EPI_ISL_521749, EPI_ISL_521750, EPI_ISL_521751, EPI_ISL_521752, EPI_ISL_521753, EPI_ISL_521754, EPI_ISL_521755, EPI_ISL_521756, EPI_ISL_521757, EPI_ISL_521758, EPI_ISL_521759, EPI_ISL_521760, EPI_ISL_521761, EPI_ISL_521762, EPI_ISL_521763, EPI_ISL_521764, EPI_ISL_521765, EPI_ISL_521766, EPI_ISL_521767, EPI_ISL_521768, EPI_ISL_521769, EPI_ISL_521770, EPI_ISL_521771, EPI_ISL_521772, EPI_ISL_521773, EPI_ISL_521774, EPI_ISL_521775, EPI_ISL_521776, EPI_ISL_521777, EPI_ISL_521778, EPI_ISL_521779, EPI_ISL_521780, EPI_ISL_521781, EPI_ISL_521782, EPI_ISL_521783, EPI_ISL_521784, EPI_ISL_521785, EPI_ISL_521786, EPI_ISL_521787, EPI_ISL_521788, EPI_ISL_521789, EPI_ISL_521790, EPI_ISL_521791, EPI_ISL_521792, EPI_ISL_521793, EPI_ISL_521794, EPI_ISL_521795, EPI_ISL_521796, EPI_ISL_521797, EPI_ISL_521798, EPI_ISL_521799, EPI_ISL_521800, EPI_ISL_521801, EPI_ISL_521802, EPI_ISL_521803, EPI_ISL_521804, EPI_ISL_521805, EPI_ISL_521806, EPI_ISL_521807, EPI_ISL_521808, EPI_ISL_521809, EPI_ISL_521810, EPI_ISL_521811, EPI_ISL_521812, EPI_ISL_521813, EPI_ISL_521814, EPI_ISL_521815, EPI_ISL_521816, EPI_ISL_521817, EPI_ISL_521818, EPI_ISL_521819, EPI_ISL_521820, EPI_ISL_521821, EPI_ISL_521822, EPI_ISL_521823, EPI_ISL_521824, EPI_ISL_521825, EPI_ISL_521826, EPI_ISL_521827, EPI_ISL_521828, EPI_ISL_521829, EPI_ISL_521830, EPI_ISL_521831, EPI_ISL_521832, EPI_ISL_521833, EPI_ISL_521834, EPI_ISL_521835, EPI_ISL_521836, EPI_ISL_521837, EPI_ISL_521838, EPI_ISL_521839 | | | |
| see above | Microbiological Diagnostic Unit - Public Health Laboratory (MDU-PHL) | MDU-PHL | Seemann T., Schultz M., Sait, M., Sherry, N. |
| EPI_ISL_521840, EPI_ISL_521841, EPI_ISL_521842, EPI_ISL_521843, EPI_ISL_521844, EPI_ISL_521845, EPI_ISL_521846, EPI_ISL_521847, EPI_ISL_521848, EPI_ISL_521849, EPI_ISL_521850, EPI_ISL_521851, EPI_ISL_521852, EPI_ISL_521853, EPI_ISL_521854, EPI_ISL_521855, EPI_ISL_521856, EPI_ISL_521857, EPI_ISL_521858, EPI_ISL_521859 | | | |
| see above | Victorian Infectious Diseases Reference Laboratory (VIDRL) | VIDRL and MDU-PHL | Caly L., Seemann T., Sait, M., Schultz M., Druce J., Sherry, N. |
| EPI_ISL_521860, EPI_ISL_521861, EPI_ISL_521862 | Royal Darwin Hospital Pathology | MDU-PHL | Meumann, E., Caly L., Seemann T., Sait, M., Schultz M., Druce J., Sherry, N. |
| EPI_ISL_521864, EPI_ISL_521865, EPI_ISL_521866, EPI_ISL_521867, EPI_ISL_521868, EPI_ISL_521870, EPI_ISL_521871, EPI_ISL_521872, EPI_ISL_521873, EPI_ISL_521874, EPI_ISL_521875, EPI_ISL_521876, EPI_ISL_521877, EPI_ISL_521878, EPI_ISL_521879, EPI_ISL_521880, EPI_ISL_521881, EPI_ISL_521882, EPI_ISL_521883, EPI_ISL_521884, EPI_ISL_521885, EPI_ISL_521886 | | | |
| see above | Victorian Infectious Diseases Reference Laboratory (VIDRL) | VIDRL and MDU-PHL | Caly L., Seemann T., Sait, M., Schultz M., Druce J., Sherry, N. |
| EPI_ISL_521887, EPI_ISL_521888 | Microbiological Diagnostic Unit - Public Health Laboratory (MDU-PHL) | MDU-PHL | Seemann T., Schultz M., Sait, M., Sherry, N. |
| EPI_ISL_521889, EPI_ISL_521890, EPI_ISL_521891, EPI_ISL_521892, EPI_ISL_521893, EPI_ISL_521894, EPI_ISL_521895, EPI_ISL_521896, EPI_ISL_521897, EPI_ISL_521898, EPI_ISL_521899, EPI_ISL_521900, EPI_ISL_521901, EPI_ISL_521902, EPI_ISL_521903, EPI_ISL_521904, EPI_ISL_521905 | | | |
| see above | Victorian Infectious Diseases Reference Laboratory (VIDRL) | VIDRL and MDU-PHL | Caly L., Seemann T., Sait, M., Schultz M., Druce J., Sherry, N. |
| EPI_ISL_521906, EPI_ISL_521907, EPI_ISL_521908, EPI_ISL_521909, EPI_ISL_521910 | Microbiological Diagnostic Unit - Public Health Laboratory (MDU-PHL) | MDU-PHL | Seemann T., Schultz M., Sait, M., Sherry, N. |
| EPI_ISL_521912, EPI_ISL_521913, EPI_ISL_521914, EPI_ISL_521915, EPI_ISL_521916, EPI_ISL_521917, EPI_ISL_521918, EPI_ISL_521919, EPI_ISL_521920, EPI_ISL_521921, EPI_ISL_521922 | | | |
| see above | Victorian Infectious Diseases Reference Laboratory (VIDRL) | VIDRL and MDU-PHL | Caly L., Seemann T., Sait, M., Schultz M., Druce J., Sherry, N. |
| EPI_ISL_521923 | Microbiological Diagnostic Unit - Public Health Laboratory (MDU-PHL) | MDU-PHL | Seemann T., Schultz M., Sait, M., Sherry, N. |
| EPI_ISL_521924, EPI_ISL_521925 | Victorian Infectious Diseases Reference Laboratory (VIDRL) | VIDRL and MDU-PHL | Caly L., Seemann T., Sait, M., Schultz M., Druce J., Sherry, N. |
| EPI_ISL_521926, EPI_ISL_521927, EPI_ISL_521928, EPI_ISL_521929, EPI_ISL_521930, EPI_ISL_521931, EPI_ISL_521932, EPI_ISL_521933, EPI_ISL_521934, EPI_ISL_521935, EPI_ISL_521936, EPI_ISL_521937, EPI_ISL_521938, EPI_ISL_521939, EPI_ISL_521940, EPI_ISL_521941, EPI_ISL_521942, EPI_ISL_521943, EPI_ISL_521944 | | | |
| see above | Microbiological Diagnostic Unit - Public Health Laboratory (MDU-PHL) | MDU-PHL | Seemann T., Schultz M., Sait, M., Sherry, N. |
| EPI_ISL_521945, EPI_ISL_521946, EPI_ISL_521947, EPI_ISL_521948, EPI_ISL_521949, EPI_ISL_521950, EPI_ISL_521951, EPI_ISL_521952, EPI_ISL_521953, EPI_ISL_521954, EPI_ISL_521955, EPI_ISL_521956, EPI_ISL_521957, EPI_ISL_521958, EPI_ISL_521959, EPI_ISL_521960, EPI_ISL_521961, EPI_ISL_521962, EPI_ISL_521963, EPI_ISL_521964, EPI_ISL_521965, EPI_ISL_521966, EPI_ISL_521967, EPI_ISL_521968, EPI_ISL_521969, EPI_ISL_521970, EPI_ISL_521971 | | | |
| see above | Victorian Infectious Diseases Reference Laboratory (VIDRL) | VIDRL and MDU-PHL | Caly L., Seemann T., Sait, M., Schultz M., Druce J., Sherry, N. |
| EPI_ISL_521972, EPI_ISL_521973, EPI_ISL_521974 | Microbiological Diagnostic Unit - Public Health Laboratory (MDU-PHL) | MDU-PHL | Seemann T., Schultz M., Sait, M., Sherry, N. |
| EPI_ISL_521975, EPI_ISL_521976, EPI_ISL_521977, EPI_ISL_521978, EPI_ISL_521980, EPI_ISL_521981, EPI_ISL_521982, EPI_ISL_521983, EPI_ISL_521984, EPI_ISL_521985, EPI_ISL_521986, EPI_ISL_521987, EPI_ISL_521988 | | | |
| see above | Victorian Infectious Diseases Reference Laboratory (VIDRL) | VIDRL and MDU-PHL | Caly L., Seemann T., Sait, M., Schultz M., Druce J., Sherry, N. |
| EPI_ISL_521989, EPI_ISL_521990, EPI_ISL_521991, EPI_ISL_521992, EPI_ISL_521993, EPI_ISL_521994, EPI_ISL_521995, EPI_ISL_521996, EPI_ISL_521997, EPI_ISL_521998, EPI_ISL_521999, EPI_ISL_522000, EPI_ISL_522001, EPI_ISL_522002, EPI_ISL_522003, EPI_ISL_522004, EPI_ISL_522005, EPI_ISL_522006, EPI_ISL_522007, EPI_ISL_522008, EPI_ISL_522009, EPI_ISL_522010, EPI_ISL_522011, EPI_ISL_522012, EPI_ISL_522013, EPI_ISL_522014, EPI_ISL_522015, EPI_ISL_522016, EPI_ISL_522017, EPI_ISL_522018, EPI_ISL_522019, EPI_ISL_522020, EPI_ISL_522021, EPI_ISL_522022, EPI_ISL_522023, EPI_ISL_522024, EPI_ISL_522025, EPI_ISL_522026, EPI_ISL_522027, EPI_ISL_522028, EPI_ISL_522029, EPI_ISL_522030, EPI_ISL_522031, EPI_ISL_522032, EPI_ISL_522033, EPI_ISL_522034, EPI_ISL_522035, EPI_ISL_522036, EPI_ISL_522037, EPI_ISL_522038, EPI_ISL_522039, EPI_ISL_522040, EPI_ISL_522041, EPI_ISL_522042, EPI_ISL_522043, EPI_ISL_522044, EPI_ISL_522045, EPI_ISL_522046, EPI_ISL_522047, EPI_ISL_522048, EPI_ISL_522049, EPI_ISL_522050, EPI_ISL_522051, EPI_ISL_522052, EPI_ISL_522053, EPI_ISL_522054 | | | |
| see above | Microbiological Diagnostic Unit - Public Health Laboratory (MDU-PHL) | MDU-PHL | Seemann T., Schultz M., Sait, M., Sherry, N. |
| EPI_ISL_522055, EPI_ISL_522056, EPI_ISL_522057, EPI_ISL_522058, EPI_ISL_522059, EPI_ISL_522060, EPI_ISL_522061, EPI_ISL_522062, EPI_ISL_522063, EPI_ISL_522064, EPI_ISL_522065, EPI_ISL_522066, EPI_ISL_522067, EPI_ISL_522068, EPI_ISL_522069, EPI_ISL_522070, EPI_ISL_522071, EPI_ISL_522072, EPI_ISL_522073, EPI_ISL_522074, EPI_ISL_522075, EPI_ISL_522076, EPI_ISL_522077, EPI_ISL_522078, EPI_ISL_522079, EPI_ISL_522080, EPI_ISL_522081, EPI_ISL_522082, EPI_ISL_522083, EPI_ISL_522084, EPI_ISL_522085, EPI_ISL_522086, EPI_ISL_522087, EPI_ISL_522088, EPI_ISL_522089, EPI_ISL_522090, EPI_ISL_522091, EPI_ISL_522092, EPI_ISL_522093, EPI_ISL_522094, EPI_ISL_522095, EPI_ISL_522096, EPI_ISL_522097, EPI_ISL_522098, EPI_ISL_522099, EPI_ISL_522100, EPI_ISL_522101, EPI_ISL_522102, EPI_ISL_522103, EPI_ISL_522104, EPI_ISL_522105, EPI_ISL_522106, EPI_ISL_522107, EPI_ISL_522108, EPI_ISL_522109, EPI_ISL_522110 | | | |
| see above | Victorian Infectious Diseases Reference Laboratory (VIDRL) | VIDRL and MDU-PHL | Caly L., Seemann T., Sait, M., Schultz M., Druce J., Sherry, N. |
| EPI_ISL_522111, EPI_ISL_522112, EPI_ISL_522113, EPI_ISL_522114, EPI_ISL_522115, EPI_ISL_522116, EPI_ISL_522117, EPI_ISL_522118 | Microbiological Diagnostic Unit - Public Health Laboratory (MDU-PHL) | MDU-PHL | Seemann T., Schultz M., Sait, M., Sherry, N. |
| EPI_ISL_522119, EPI_ISL_522120, EPI_ISL_522121, EPI_ISL_522122, EPI_ISL_522123, EPI_ISL_522124, EPI_ISL_522125, EPI_ISL_522126, EPI_ISL_522127, EPI_ISL_522128, EPI_ISL_522129, EPI_ISL_522130, EPI_ISL_522131, EPI_ISL_522132, EPI_ISL_522133, EPI_ISL_522134, EPI_ISL_522135, EPI_ISL_522136, EPI_ISL_522137, EPI_ISL_522138, EPI_ISL_522139, EPI_ISL_522140, EPI_ISL_522141, EPI_ISL_522142, EPI_ISL_522143, EPI_ISL_522144, EPI_ISL_522145, EPI_ISL_522146, EPI_ISL_522147, EPI_ISL_522148, EPI_ISL_522149, EPI_ISL_522150, EPI_ISL_522151, EPI_ISL_522152, EPI_ISL_522153, EPI_ISL_522154, EPI_ISL_522155, EPI_ISL_522156, EPI_ISL_522157, EPI_ISL_522158, EPI_ISL_522159, EPI_ISL_522160, EPI_ISL_522161, EPI_ISL_522162, EPI_ISL_522163, EPI_ISL_522164, EPI_ISL_522165, EPI_ISL_522166, EPI_ISL_522167, EPI_ISL_522168, EPI_ISL_522169, EPI_ISL_522170, EPI_ISL_522171, EPI_ISL_522172, EPI_ISL_522173, EPI_ISL_522174, EPI_ISL_522175, EPI_ISL_522176, EPI_ISL_522177, EPI_ISL_522178, EPI_ISL_522179, EPI_ISL_522180, EPI_ISL_522181, EPI_ISL_522182, EPI_ISL_522183, EPI_ISL_522184, EPI_ISL_522185, EPI_ISL_522186, EPI_ISL_522187 | | | |
| see above | Victorian Infectious Diseases Reference Laboratory (VIDRL) | VIDRL and MDU-PHL | Caly L., Seemann T., Sait, M., Schultz M., Druce J., Sherry, N. |
| EPI_ISL_522188, EPI_ISL_522189, EPI_ISL_522190, EPI_ISL_522191, EPI_ISL_522192, EPI_ISL_522193, EPI_ISL_522194, EPI_ISL_522195, EPI_ISL_522196, EPI_ISL_522197, EPI_ISL_522198, EPI_ISL_522199, EPI_ISL_522200, EPI_ISL_522201, EPI_ISL_522202, EPI_ISL_522203, EPI_ISL_522204, EPI_ISL_522205, EPI_ISL_522206, EPI_ISL_522207, EPI_ISL_522208, EPI_ISL_522209, EPI_ISL_522210, EPI_ISL_522211, EPI_ISL_522212, EPI_ISL_522213, EPI_ISL_522214, EPI_ISL_522215, EPI_ISL_522216, EPI_ISL_522217, EPI_ISL_522218, EPI_ISL_522219, EPI_ISL_522220, EPI_ISL_522221, EPI_ISL_522222, EPI_ISL_522223, EPI_ISL_522224, EPI_ISL_522225, EPI_ISL_522226, EPI_ISL_522227, EPI_ISL_522228, EPI_ISL_522229, EPI_ISL_522230, EPI_ISL_522231, EPI_ISL_522232 | | | |
| see above | Microbiological Diagnostic Unit - Public Health Laboratory (MDU- | MDU-PHL | Seemann T., Schultz M., Sait, M., Sherry, N. |

| | | | |
|---|---|---|---|
| | PHL) | | |
| EPI_ISL_522233, EPI_ISL_522234, EPI_ISL_522235, EPI_ISL_522236, EPI_ISL_522237, EPI_ISL_522238, EPI_ISL_522239, EPI_ISL_522240, EPI_ISL_522241, EPI_ISL_522242 | Victorian Infectious Diseases Reference Laboratory (VIDRL) | VIDRL and MDU-PHL | Caly L., Seemann T., Sait, M., Schultz M., Druce J., Sherry, N. |
| EPI_ISL_522243, EPI_ISL_522244, EPI_ISL_522245, EPI_ISL_522246, EPI_ISL_522247, EPI_ISL_522248, EPI_ISL_522249, EPI_ISL_522250, EPI_ISL_522251, EPI_ISL_522252, EPI_ISL_522253 | | | |
| see above | Microbiological Diagnostic Unit - Public Health Laboratory (MDU-PHL) | MDU-PHL | Seemann T., Schultz M., Sait, M., Sherry, N. |
| EPI_ISL_522254, EPI_ISL_522255, EPI_ISL_522256, EPI_ISL_522257, EPI_ISL_522258, EPI_ISL_522259, EPI_ISL_522260, EPI_ISL_522261, EPI_ISL_522262, EPI_ISL_522263, EPI_ISL_522264, EPI_ISL_522265, EPI_ISL_522266, EPI_ISL_522267, EPI_ISL_522268, EPI_ISL_522269, EPI_ISL_522270, EPI_ISL_522271, EPI_ISL_522272, EPI_ISL_522273, EPI_ISL_522274, EPI_ISL_522275, EPI_ISL_522276, EPI_ISL_522277, EPI_ISL_522278, EPI_ISL_522279, EPI_ISL_522280, EPI_ISL_522281, EPI_ISL_522282, EPI_ISL_522283, EPI_ISL_522284, EPI_ISL_522285, EPI_ISL_522286, EPI_ISL_522287, EPI_ISL_522288, EPI_ISL_522289, EPI_ISL_522290, EPI_ISL_522291, EPI_ISL_522292, EPI_ISL_522293, EPI_ISL_522294, EPI_ISL_522295, EPI_ISL_522296, EPI_ISL_522297, EPI_ISL_522298, EPI_ISL_522299, EPI_ISL_522300, EPI_ISL_522301, EPI_ISL_522302, EPI_ISL_522303, EPI_ISL_522304, EPI_ISL_522305, EPI_ISL_522306, EPI_ISL_522307, EPI_ISL_522308, EPI_ISL_522309, EPI_ISL_522310, EPI_ISL_522311, EPI_ISL_522312, EPI_ISL_522313, EPI_ISL_522314, EPI_ISL_522315, EPI_ISL_522316, EPI_ISL_522317, EPI_ISL_522318, EPI_ISL_522319, EPI_ISL_522320, EPI_ISL_522321, EPI_ISL_522322, EPI_ISL_522323, EPI_ISL_522324, EPI_ISL_522325, EPI_ISL_522326, EPI_ISL_522327, EPI_ISL_522328, EPI_ISL_522329, EPI_ISL_522330, EPI_ISL_522331, EPI_ISL_522332, EPI_ISL_522333, EPI_ISL_522334, EPI_ISL_522335, EPI_ISL_522336, EPI_ISL_522337, EPI_ISL_522338, EPI_ISL_522339, EPI_ISL_522340, EPI_ISL_522341, EPI_ISL_522342, EPI_ISL_522343, EPI_ISL_522344, EPI_ISL_522345, EPI_ISL_522346, EPI_ISL_522347, EPI_ISL_522348 | | | |
| see above | Utah Public Health Laboratory | Utah Public Health Laboratory | Erin Young, Kelly Oakeson |
| EPI_ISL_522349, EPI_ISL_522350 | KU Leuven, Rega Institute, Clinical and Epidemiological Virology | KU Leuven, Rega Institute, Clinical and Epidemiological Virology | Tony Wawina-Bokalanga, Joan Marti-Carerras, Bert Vanmechelen, Piet Maes |
| EPI_ISL_522351, EPI_ISL_522352, EPI_ISL_522353, EPI_ISL_522354, EPI_ISL_522355, EPI_ISL_522356, EPI_ISL_522357, EPI_ISL_522358, EPI_ISL_522359, EPI_ISL_522360, EPI_ISL_522361, EPI_ISL_522362, EPI_ISL_522363, EPI_ISL_522364, EPI_ISL_522365, EPI_ISL_522366, EPI_ISL_522367, EPI_ISL_522368, EPI_ISL_522369, EPI_ISL_522370, EPI_ISL_522371, EPI_ISL_522372, EPI_ISL_522373, EPI_ISL_522374, EPI_ISL_522375, EPI_ISL_522376, EPI_ISL_522377, EPI_ISL_522378, EPI_ISL_522379, EPI_ISL_522380, EPI_ISL_522381, EPI_ISL_522382, EPI_ISL_522383, EPI_ISL_522384, EPI_ISL_522385, EPI_ISL_522386, EPI_ISL_522387, EPI_ISL_522388, EPI_ISL_522389, EPI_ISL_522390, EPI_ISL_522391, EPI_ISL_522392, EPI_ISL_522393, EPI_ISL_522394, EPI_ISL_522395 | | | |
| see above | Texas Department of State Health Services | Texas Department of State Health Services | Rashmi Tuladhar, Bonnie Oh, Cara Akrout, Jenny Zhang, Maliha Rahman, Anita Pokharel, Myong Koag, Chun Wang, Rachel Lee, Grace Kubin |
| EPI_ISL_522396, EPI_ISL_522397, EPI_ISL_522398, EPI_ISL_522399, EPI_ISL_522400, EPI_ISL_522401, EPI_ISL_522402, EPI_ISL_522403, EPI_ISL_522404, EPI_ISL_522405 | Alaska State Virology Laboratory | Alaska State Virology Laboratory | Jack Chen, Ph.D. |
| EPI_ISL_522406 | Universidad Iberoamericana (UNIBE) | International Centre for Genetic Engineering and Biotechnology (ICGEB)and ARGO Open Lab Platform | Robert Paulino-Ramirez, Eileen Riego, Alejandro Vallejo, Victor Calderon, Leandro Tapia, Danilo Licastro, Simeone Dal Monego, Sreejith Rajasekharan, and Alessandro Marcello. |
| EPI_ISL_522437 | Mahatma Gandhi Institute of Medical Sciences | National Institute of Biomedical Genomics - DBT's PAN-INDIA 1000 SARS--CoV-2 RNA Genome Sequencing Consortium | Arindam Maitra, Vijayshri Deotale, Rahul Narang, Deepashri Maraskolhe, Saumitra Das |
| EPI_ISL_522439, EPI_ISL_522440, EPI_ISL_522441, EPI_ISL_522442, EPI_ISL_522443, EPI_ISL_522444, EPI_ISL_522445, EPI_ISL_522446, EPI_ISL_522447, EPI_ISL_522448, EPI_ISL_522449, EPI_ISL_522450, EPI_ISL_522451, EPI_ISL_522452, EPI_ISL_522453 | | | |
| see above | Center for Laboratory Control of Infectious Diseases, Korea Centers for Diseases Control and Prevention | Center for Laboratory Control of Infectious Diseases, Korea Centers for Diseases Control and Prevention | Junyoung Kim, Ae Kyung Park, Eunkyung Shin, Jin Sun No, Jeong-Min Kim, Yoon-Seok Chung, Heui Man Kim, Myung Guk Han |
| EPI_ISL_522454 | Division of Viral Diseases, Center for Laboratory Control of Infectious Diseases, Korea Centers for Diseases Control and Prevention | Division of Viral Diseases, Center for Laboratory Control of Infectious Diseases, Korea Centers for Diseases Control and Prevention | Jeong-Min Kim, Yoon-Seok Chung, Namjoo Lee, Sang Hee Woo, Hye-Jun Jo, Heui Man Kim, Jun-Sub Kim, Myung Guk Han |
| EPI_ISL_522455, EPI_ISL_522456, EPI_ISL_522457, EPI_ISL_522458, EPI_ISL_522459, EPI_ISL_522460, EPI_ISL_522461, EPI_ISL_522462, EPI_ISL_522463, EPI_ISL_522464 | Center for Laboratory Control of Infectious Diseases, Korea Centers for Diseases Control and Prevention | Center for Laboratory Control of Infectious Diseases, Korea Centers for Diseases Control and Prevention | Junyoung Kim, Ae Kyung Park, Eunkyung Shin, Jin Sun No, Jeong-Min Kim, Yoon-Seok Chung, Heui Man Kim, Myung Guk Han |
| EPI_ISL_522465, EPI_ISL_522466 | Division of Viral Diseases, Center for Laboratory Control of Infectious Diseases, Korea Centers for Diseases Control and Prevention | Division of Viral Diseases, Center for Laboratory Control of Infectious Diseases, Korea Centers for Diseases Control and Prevention | Jeong-Min Kim, Yoon-Seok Chung, Namjoo Lee, Sang Hee Woo, Hye-Jun Jo, Heui Man Kim, Jun-Sub Kim, Myung Guk Han |
| EPI_ISL_522467, EPI_ISL_522468, EPI_ISL_522469, EPI_ISL_522470, EPI_ISL_522471, EPI_ISL_522472, EPI_ISL_522473 | Center for Laboratory Control of Infectious Diseases, Korea Centers for Diseases Control and Prevention | Center for Laboratory Control of Infectious Diseases, Korea Centers for Diseases Control and Prevention | Junyoung Kim, Ae Kyung Park, Eunkyung Shin, Jin Sun No, Jeong-Min Kim, Yoon-Seok Chung, Heui Man Kim, Myung Guk Han |
| EPI_ISL_522474, EPI_ISL_522475 | Division of Viral Diseases, Center for Laboratory Control of Infectious Diseases, Korea Centers for Diseases Control and Prevention | Division of Viral Diseases, Center for Laboratory Control of Infectious Diseases, Korea Centers for Diseases Control and Prevention | Jeong-Min Kim, Yoon-Seok Chung, Namjoo Lee, Sang Hee Woo, Hye-Jun Jo, Heui Man Kim, Jun-Sub Kim, Myung Guk Han |
| EPI_ISL_522476 | Center for Laboratory Control of Infectious Diseases, Korea Centers for Diseases Control and Prevention | Center for Laboratory Control of Infectious Diseases, Korea Centers for Diseases Control and Prevention | Junyoung Kim, Ae Kyung Park, Eunkyung Shin, Jin Sun No, Jeong-Min Kim, Yoon-Seok Chung, Heui Man Kim, Myung Guk Han |
| EPI_ISL_522477, EPI_ISL_522478 | Division of Viral Diseases, Center for Laboratory Control of Infectious Diseases, Korea Centers for Diseases Control and Prevention | Division of Viral Diseases, Center for Laboratory Control of Infectious Diseases, Korea Centers for Diseases Control and Prevention | Jeong-Min Kim, Yoon-Seok Chung, Namjoo Lee, Sang Hee Woo, Hye-Jun Jo, Heui Man Kim, Jun-Sub Kim, Myung Guk Han |
| EPI_ISL_522479, EPI_ISL_522480, EPI_ISL_522481, EPI_ISL_522482, EPI_ISL_522483 | Center for Laboratory Control of Infectious Diseases, Korea Centers for Diseases Control and Prevention | Center for Laboratory Control of Infectious Diseases, Korea Centers for Diseases Control and Prevention | Junyoung Kim, Ae Kyung Park, Eunkyung Shin, Jin Sun No, Jeong-Min Kim, Yoon-Seok Chung, Heui Man Kim, Myung Guk Han |
| EPI_ISL_522484 | Division of Viral Diseases, Center for Laboratory Control of Infectious Diseases, Korea Centers for Diseases Control and Prevention | Division of Viral Diseases, Center for Laboratory Control of Infectious Diseases, Korea Centers for Diseases Control and Prevention | Jeong-Min Kim, Yoon-Seok Chung, Namjoo Lee, Sang Hee Woo, Hye-Jun Jo, Heui Man Kim, Jun-Sub Kim, Myung Guk Han |
| EPI_ISL_522485 | Center for Laboratory Control of Infectious Diseases, Korea Centers for Diseases Control and Prevention | Center for Laboratory Control of Infectious Diseases, Korea Centers for Diseases Control and Prevention | Junyoung Kim, Ae Kyung Park, Eunkyung Shin, Jin Sun No, Jeong-Min Kim, Yoon-Seok Chung, Heui Man Kim, Myung Guk Han |
| EPI_ISL_522486, EPI_ISL_522487, EPI_ISL_522488, EPI_ISL_522489, EPI_ISL_522490 | Division of Viral Diseases, Center for Laboratory Control of Infectious Diseases, Korea Centers for Diseases Control and Prevention | Division of Viral Diseases, Center for Laboratory Control of Infectious Diseases, Korea Centers for Diseases Control and Prevention | Jeong-Min Kim, Yoon-Seok Chung, Namjoo Lee, Sang Hee Woo, Hye-Jun Jo, Heui Man Kim, Jun-Sub Kim, Myung Guk Han |
| EPI_ISL_522491, EPI_ISL_522492, EPI_ISL_522493, EPI_ISL_522494, EPI_ISL_522495, EPI_ISL_522496, EPI_ISL_522497 | Center for Laboratory Control of Infectious Diseases, Korea Centers for Diseases Control and Prevention | Center for Laboratory Control of Infectious Diseases, Korea Centers for Diseases Control and Prevention | Junyoung Kim, Ae Kyung Park, Eunkyung Shin, Jin Sun No, Jeong-Min Kim, Yoon-Seok Chung, Heui Man Kim, Myung Guk Han |
| EPI_ISL_522498, EPI_ISL_522499, EPI_ISL_522500, EPI_ISL_522501 | Division of Viral Diseases, Center for Laboratory Control of Infectious Diseases, Korea Centers for Diseases Control and Prevention | Division of Viral Diseases, Center for Laboratory Control of Infectious Diseases, Korea Centers for Diseases Control and Prevention | Jeong-Min Kim, Yoon-Seok Chung, Namjoo Lee, Sang Hee Woo, Hye-Jun Jo, Heui Man Kim, Jun-Sub Kim, Myung Guk Han |
| EPI_ISL_522502 | Center for Laboratory Control of Infectious Diseases, Korea Centers for Diseases Control and Prevention | Center for Laboratory Control of Infectious Diseases, Korea Centers for Diseases Control and Prevention | Junyoung Kim, Ae Kyung Park, Eunkyung Shin, Jin Sun No, Jeong-Min Kim, Yoon-Seok Chung, Heui Man Kim, Myung Guk Han |
| EPI_ISL_522503, EPI_ISL_522504 | Division of Viral Diseases, Center for Laboratory Control of Infectious Diseases, Korea Centers for Diseases Control and Prevention | Division of Viral Diseases, Center for Laboratory Control of Infectious Diseases, Korea Centers for Diseases Control and Prevention | Jeong-Min Kim, Yoon-Seok Chung, Namjoo Lee, Sang Hee Woo, Hye-Jun Jo, Heui Man Kim, Jun-Sub Kim, Myung Guk Han |
| EPI_ISL_522505 | Center for Laboratory Control of Infectious Diseases, Korea Centers for Diseases Control and Prevention | Center for Laboratory Control of Infectious Diseases, Korea Centers for Diseases Control and Prevention | Junyoung Kim, Ae Kyung Park, Eunkyung Shin, Jin Sun No, Jeong-Min Kim, Yoon-Seok Chung, Heui Man Kim, Myung Guk Han |
| EPI_ISL_522506, EPI_ISL_522507 | Division of Viral Diseases, Center for Laboratory Control of Infectious Diseases, Korea Centers for Diseases Control and Prevention | Division of Viral Diseases, Center for Laboratory Control of Infectious Diseases, Korea Centers for Diseases Control and Prevention | Jeong-Min Kim, Yoon-Seok Chung, Namjoo Lee, Sang Hee Woo, Hye-Jun Jo, Heui Man Kim, Jun-Sub Kim, Myung Guk Han |
| EPI_ISL_522508 | Center for Laboratory Control of Infectious Diseases, Korea Centers for Diseases Control and Prevention | Center for Laboratory Control of Infectious Diseases, Korea Centers for Diseases Control and Prevention | Junyoung Kim, Ae Kyung Park, Eunkyung Shin, Jin Sun No, Jeong-Min Kim, Yoon-Seok Chung, Heui Man Kim, Myung Guk Han |
| EPI_ISL_522509 | Division of Viral Diseases, Center for Laboratory Control of Infectious Diseases, Korea Centers for Diseases Control and Prevention | Division of Viral Diseases, Center for Laboratory Control of Infectious Diseases, Korea Centers for Diseases Control and Prevention | Jeong-Min Kim, Yoon-Seok Chung, Namjoo Lee, Sang Hee Woo, Hye-Jun Jo, Heui Man Kim, Jun-Sub Kim, Myung Guk Han |
| EPI_ISL_522510 | Center for Laboratory Control of Infectious Diseases, Korea Centers for Diseases Control and Prevention | Center for Laboratory Control of Infectious Diseases, Korea Centers for Diseases Control and Prevention | Junyoung Kim, Ae Kyung Park, Eunkyung Shin, Jin Sun No, Jeong-Min Kim, Yoon-Seok Chung, Heui Man Kim, Myung Guk Han |
| EPI_ISL_522511, EPI_ISL_522512, EPI_ISL_522513, EPI_ISL_522514 | Division of Viral Diseases, Center for Laboratory Control of Infectious Diseases, Korea Centers for Diseases Control and Prevention | Division of Viral Diseases, Center for Laboratory Control of Infectious Diseases, Korea Centers for Diseases Control and Prevention | Jeong-Min Kim, Yoon-Seok Chung, Namjoo Lee, Sang Hee Woo, Hye-Jun Jo, Heui Man Kim, Jun-Sub Kim, Myung Guk Han |
| EPI_ISL_522515, EPI_ISL_522516 | Center for Laboratory Control of Infectious Diseases, Korea Centers for Diseases Control and Prevention | Center for Laboratory Control of Infectious Diseases, Korea Centers for Diseases Control and Prevention | Junyoung Kim, Ae Kyung Park, Eunkyung Shin, Jin Sun No, Jeong-Min Kim, Yoon-Seok Chung, Heui Man Kim, Myung Guk Han |
| EPI_ISL_522517 | Division of Viral Diseases, Center for Laboratory Control of | Division of Viral Diseases, Center for Laboratory Control of Infectious Diseases, Korea Centers for Diseases Control | Jeong-Min Kim, Yoon-Seok Chung, Namjoo Lee, Sang Hee Woo, Hye-Jun Jo, Heui Man Kim, Jun-Sub Kim, Myung Guk Han |

| | | | |
|---|---|---|---|
| | Infectious Diseases, Korea Centers for Diseases Control and Prevention | and Prevention | |
| EPI_ISL_522518 | Center for Laboratory Control of Infectious Diseases, Korea Centers for Diseases Control and Prevention | Center for Laboratory Control of Infectious Diseases, Korea Centers for Diseases Control and Prevention | Junyoung Kim, Ae Kyung Park, Eunkyung Shin, Jin Sun No, Jeong-Min Kim, Yoon-Seok Chung, Heui Man Kim, Myung Guk Han |
| EPI_ISL_522519, EPI_ISL_522520, EPI_ISL_522521, EPI_ISL_522522, EPI_ISL_522523, EPI_ISL_522524, EPI_ISL_522525, EPI_ISL_522526, EPI_ISL_522527, EPI_ISL_522528, EPI_ISL_522529, EPI_ISL_522530, EPI_ISL_522531, EPI_ISL_522532, EPI_ISL_522533, EPI_ISL_522534, EPI_ISL_522535, EPI_ISL_522536, EPI_ISL_522537, EPI_ISL_522538, EPI_ISL_522539, EPI_ISL_522540, EPI_ISL_522541, EPI_ISL_522542, EPI_ISL_522543, EPI_ISL_522544 | | | |
| see above | Division of Viral Diseases, Center for Laboratory Control of Infectious Diseases, Korea Centers for Diseases Control and Prevention | Division of Viral Diseases, Center for Laboratory Control of Infectious Diseases, Korea Centers for Diseases Control and Prevention | Jeong-Min Kim, Yoon-Seok Chung, Namjoo Lee, Sang Hee Woo, Hye-Jun Jo, Heui Man Kim, Jun-Sub Kim, Myung Guk Han |
| EPI_ISL_522547, EPI_ISL_522548 | Platforme CYROI | UMR PIMIT Université de La Réunion | David Wilkinson, Camille Lebarbenchon, Patrick Mavingui |
| EPI_ISL_522549, EPI_ISL_522550 | Félix Guyon Hospital | UMR PIMIT Université de La Réunion | David Wilkinson, Camille Lebarbenchon, Patrick Mavingui |
| EPI_ISL_522556, EPI_ISL_522557, EPI_ISL_522558, EPI_ISL_522559, EPI_ISL_522560, EPI_ISL_522561, EPI_ISL_522562, EPI_ISL_522563, EPI_ISL_522564, EPI_ISL_522565, EPI_ISL_522566, EPI_ISL_522567, EPI_ISL_522568, EPI_ISL_522569, EPI_ISL_522570, EPI_ISL_522571, EPI_ISL_522572, EPI_ISL_522573, EPI_ISL_522574, EPI_ISL_522575, EPI_ISL_522576, EPI_ISL_522577, EPI_ISL_522578, EPI_ISL_522579, EPI_ISL_522580, EPI_ISL_522581, EPI_ISL_522582, EPI_ISL_522583, EPI_ISL_522584, EPI_ISL_522585, EPI_ISL_522586, EPI_ISL_522587, EPI_ISL_522588, EPI_ISL_522589, EPI_ISL_522590, EPI_ISL_522591, EPI_ISL_522592, EPI_ISL_522593, EPI_ISL_522594, EPI_ISL_522595, EPI_ISL_522596, EPI_ISL_522597, EPI_ISL_522598, EPI_ISL_522599, EPI_ISL_522600, EPI_ISL_522601, EPI_ISL_522602, EPI_ISL_522603, EPI_ISL_522604, EPI_ISL_522605, EPI_ISL_522606, EPI_ISL_522607, EPI_ISL_522608, EPI_ISL_522609, EPI_ISL_522610, EPI_ISL_522611, EPI_ISL_522612, EPI_ISL_522613, EPI_ISL_522614, EPI_ISL_522615, EPI_ISL_522616, EPI_ISL_522617, EPI_ISL_522618, EPI_ISL_522619, EPI_ISL_522620, EPI_ISL_522621, EPI_ISL_522622, EPI_ISL_522623, EPI_ISL_522624, EPI_ISL_522625, EPI_ISL_522626, EPI_ISL_522627, EPI_ISL_522628, EPI_ISL_522629, EPI_ISL_522630, EPI_ISL_522631, EPI_ISL_522632, EPI_ISL_522633, EPI_ISL_522634, EPI_ISL_522635, EPI_ISL_522636, EPI_ISL_522637, EPI_ISL_522638, EPI_ISL_522639, EPI_ISL_522640, EPI_ISL_522641, EPI_ISL_522642, EPI_ISL_522643, EPI_ISL_522644, EPI_ISL_522645, EPI_ISL_522646, EPI_ISL_522648, EPI_ISL_522649, EPI_ISL_522650, EPI_ISL_522651, EPI_ISL_522652, EPI_ISL_522653, EPI_ISL_522654, EPI_ISL_522655, EPI_ISL_522656, EPI_ISL_522657, EPI_ISL_522658, EPI_ISL_522659, EPI_ISL_522660, EPI_ISL_522661, EPI_ISL_522662, EPI_ISL_522663, EPI_ISL_522664, EPI_ISL_522665, EPI_ISL_522666, EPI_ISL_522667, EPI_ISL_522668, EPI_ISL_522669, EPI_ISL_522670, EPI_ISL_522671, EPI_ISL_522672, EPI_ISL_522673, EPI_ISL_522674, EPI_ISL_522675, EPI_ISL_522676, EPI_ISL_522677, EPI_ISL_522678, EPI_ISL_522679, EPI_ISL_522680, EPI_ISL_522681, EPI_ISL_522682, EPI_ISL_522683, EPI_ISL_522684, EPI_ISL_522685, EPI_ISL_522686, EPI_ISL_522687, EPI_ISL_522688, EPI_ISL_522689, EPI_ISL_522690, EPI_ISL_522691, EPI_ISL_522692, EPI_ISL_522693, EPI_ISL_522694, EPI_ISL_522695, EPI_ISL_522696, EPI_ISL_522697, EPI_ISL_522698, EPI_ISL_522699, EPI_ISL_522700, EPI_ISL_522701, EPI_ISL_522702, EPI_ISL_522703, EPI_ISL_522704, EPI_ISL_522705, EPI_ISL_522706, EPI_ISL_522707, EPI_ISL_522708, EPI_ISL_522709, EPI_ISL_522710, EPI_ISL_522711, EPI_ISL_522712, EPI_ISL_522713, EPI_ISL_522714, EPI_ISL_522715, EPI_ISL_522716, EPI_ISL_522717, EPI_ISL_522718, EPI_ISL_522719, EPI_ISL_522720, EPI_ISL_522721, EPI_ISL_522722, EPI_ISL_522723, EPI_ISL_522724, EPI_ISL_522725, EPI_ISL_522726, EPI_ISL_522727, EPI_ISL_522728, EPI_ISL_522729, EPI_ISL_522730, EPI_ISL_522731, EPI_ISL_522732, EPI_ISL_522733, EPI_ISL_522734, EPI_ISL_522735, EPI_ISL_522736, EPI_ISL_522737, EPI_ISL_522738, EPI_ISL_522739, EPI_ISL_522740, EPI_ISL_522741, EPI_ISL_522742, EPI_ISL_522743, EPI_ISL_522744, EPI_ISL_522745, EPI_ISL_522746, EPI_ISL_522747, EPI_ISL_522748, EPI_ISL_522749, EPI_ISL_522750, EPI_ISL_522751, EPI_ISL_522752, EPI_ISL_522753, EPI_ISL_522754, EPI_ISL_522755, EPI_ISL_522756, EPI_ISL_522757, EPI_ISL_522758, EPI_ISL_522759, EPI_ISL_522760, EPI_ISL_522761, EPI_ISL_522762, EPI_ISL_522763, EPI_ISL_522764, EPI_ISL_522765, EPI_ISL_522766, EPI_ISL_522767, EPI_ISL_522768, EPI_ISL_522769 | | | |
| see above | Royal Hobart Hospital Microbiology Department | MDU-PHL | Cooley L., van Haeften R., Seemann T., Sait M., Schultz, M.B., Sherry N. |
| EPI_ISL_522777, EPI_ISL_522778, EPI_ISL_522779, EPI_ISL_522780, EPI_ISL_522781, EPI_ISL_522782, EPI_ISL_522783, EPI_ISL_522784, EPI_ISL_522785, EPI_ISL_522786, EPI_ISL_522787, EPI_ISL_522788, EPI_ISL_522789, EPI_ISL_522790, EPI_ISL_522791, EPI_ISL_522792, EPI_ISL_522793, EPI_ISL_522794, EPI_ISL_522795, EPI_ISL_522796, EPI_ISL_522797, EPI_ISL_522798, EPI_ISL_522799, EPI_ISL_522800, EPI_ISL_522801, EPI_ISL_522802, EPI_ISL_522803, EPI_ISL_522804, EPI_ISL_522805, EPI_ISL_522806, EPI_ISL_522807, EPI_ISL_522808, EPI_ISL_522809, EPI_ISL_522810, EPI_ISL_522811, EPI_ISL_522812, EPI_ISL_522813, EPI_ISL_522814, EPI_ISL_522815, EPI_ISL_522816, EPI_ISL_522817, EPI_ISL_522818, EPI_ISL_522819, EPI_ISL_522820, EPI_ISL_522821, EPI_ISL_522822, EPI_ISL_522823 | | | |
| see above | Virginia DCLS | Virginia DCLS | Virginia DCLS |
| EPI_ISL_522824, EPI_ISL_522825, EPI_ISL_522826, EPI_ISL_522827, EPI_ISL_522828, EPI_ISL_522829, EPI_ISL_522830, EPI_ISL_522831, EPI_ISL_522832, EPI_ISL_522833, EPI_ISL_522834, EPI_ISL_522835, EPI_ISL_522836, EPI_ISL_522837, EPI_ISL_522838, EPI_ISL_522839, EPI_ISL_522840, EPI_ISL_522841, EPI_ISL_522842, EPI_ISL_522843, EPI_ISL_522844, EPI_ISL_522845, EPI_ISL_522846, EPI_ISL_522847, EPI_ISL_522848, EPI_ISL_522849, EPI_ISL_522850, EPI_ISL_522851, EPI_ISL_522852, EPI_ISL_522853, EPI_ISL_522854 | | | |
| see above | Maryland Department of Health | Maryland Department of Health | Maryland Department of Health Laboratories Administration |
| EPI_ISL_522855 | ULSS9 Distretto di Bussolengo | Istituto Zooprofilattico Sperimentale delle Venezie | Adelaide Milani, Alessia Schivo, Annalisa Salviato, Erika Giorgia Quaranta, Gianpiero Zamperin, Ambra Pastori, Bianca Zecchin, Alice Fusaro, Calogero Terregino, Antonia Ricci |
| EPI_ISL_522856 | ULSS9 Distretto di San Bonifacio | Istituto Zooprofilattico Sperimentale delle Venezie | Adelaide Milani, Alessia Schivo, Annalisa Salviato, Erika Giorgia Quaranta, Gianpiero Zamperin, Ambra Pastori, Bianca Zecchin, Alice Fusaro, Calogero Terregino, Antonia Ricci |
| EPI_ISL_522857 | ULSS9 Scaligera | Istituto Zooprofilattico Sperimentale delle Venezie | Adelaide Milani, Alessia Schivo, Annalisa Salviato, Erika Giorgia Quaranta, Gianpiero Zamperin, Ambra Pastori, Bianca Zecchin, Alice Fusaro, Calogero Terregino, Antonia Ricci |
| EPI_ISL_522858 | ULSS9 Distretto di San Bonifacio | Istituto Zooprofilattico Sperimentale delle Venezie | Adelaide Milani, Alessia Schivo, Annalisa Salviato, Erika Giorgia Quaranta, Gianpiero Zamperin, Ambra Pastori, Bianca Zecchin, Alice Fusaro, Calogero Terregino, Antonia Ricci |
| EPI_ISL_522859 | ULSS9 Scaligera | Istituto Zooprofilattico Sperimentale delle Venezie | Adelaide Milani, Alessia Schivo, Annalisa Salviato, Erika Giorgia Quaranta, Gianpiero Zamperin, Ambra Pastori, Bianca Zecchin, Alice Fusaro, Calogero Terregino, Antonia Ricci |
| EPI_ISL_522860, EPI_ISL_522861, EPI_ISL_522862, EPI_ISL_522863, EPI_ISL_522864, EPI_ISL_522865, EPI_ISL_522866, EPI_ISL_522867, EPI_ISL_522868 | ULSS9 Distretto di Bussolengo | Istituto Zooprofilattico Sperimentale delle Venezie | Adelaide Milani, Alessia Schivo, Annalisa Salviato, Erika Giorgia Quaranta, Gianpiero Zamperin, Ambra Pastori, Bianca Zecchin, Alice Fusaro, Calogero Terregino, Antonia Ricci |
| EPI_ISL_522869, EPI_ISL_522870, EPI_ISL_522871 | Maryland Department of Health | Maryland Department of Health | Maryland Department of Health Laboratories Administration |
| EPI_ISL_522872 | Instituto Nacional de Medicina Genómica | Instituto Nacional de Medicina Genómica | Hidalgo-Miranda A, Mendoza-Vargas A, Reyes-Grajeda JP, Cisneros-Villanueva M, Cedro-Tanda A, Hurtado-Cordova E, Peñaloza-Figueroa F, Herrera-Montalvo LA |
| EPI_ISL_522873, EPI_ISL_522874, EPI_ISL_522875 | Instituto Nacional de Medicina Genomica | Instituto Nacional de Medicina Genomica | Hidalgo-Miranda A, Mendoza-Vargas A, Reyes-Grajeda JP, Cisneros-Villanueva M, Cedro-Tanda A, Hurtado-Cordova E, Peñaloza-Figueroa F, Herrera-Montalvo LA |
| EPI_ISL_522876 | Instituto Nacional de Medicina Genomica | Instituto Nacional de Medicina Genomica | Hidalgo-Miranda A, Mendoza-Vargas A, Reyes-Grajeda JP, Cisneros-Villanueva M, Cedro-Tanda A, Hurtado-Cordova E, Peñaloza-Figueroa F, Herrera-Montalvo LA |
| EPI_ISL_522877 | Instituto Nacional de Medicina Genomica | Instituto Nacional de Medicina Genomica | Hidalgo-Miranda A, Mendoza-Vargas A, Reyes-Grajeda JP, Cisneros-Villanueva M, Cedro-Tanda A, Hurtado-Cordova E, Peñaloza-Figueroa F, Herrera-Montalvo LA |
| EPI_ISL_522878 | Instituto Nacional de Medicina Genomica | Instituto Nacional de Medicina Genomica | Hidalgo-Miranda A, Mendoza-Vargas A, Reyes-Grajeda JP, Cisneros-Villanueva M, Hurtado-Cordova E, Cedro-Tanda A, Peñaloza-Figueroa F, Herrera-Montalvo LA |
| EPI_ISL_522879, EPI_ISL_522880 | Instituto Nacional de Medicina Genómica | Instituto Nacional de Medicina Genómica | Hidalgo-Miranda A, Mendoza-Vargas A, Reyes-Grajeda JP, Cisneros-Villanueva M, Cedro-Tanda A, Hurtado-Cordova E, Peñaloza-Figueroa F, Herrera-Montalvo LA |
| EPI_ISL_522881, EPI_ISL_522882, EPI_ISL_522883, EPI_ISL_522884, EPI_ISL_522885, EPI_ISL_522886, EPI_ISL_522887, EPI_ISL_522888, EPI_ISL_522889, EPI_ISL_522890, EPI_ISL_522891, EPI_ISL_522892, EPI_ISL_522893, EPI_ISL_522894, EPI_ISL_522895, EPI_ISL_522896, EPI_ISL_522897, EPI_ISL_522898, EPI_ISL_522899, EPI_ISL_522900, EPI_ISL_522901, EPI_ISL_522902, EPI_ISL_522903, EPI_ISL_522904, EPI_ISL_522905, EPI_ISL_522906, EPI_ISL_522907, EPI_ISL_522908, EPI_ISL_522909, EPI_ISL_522910, EPI_ISL_522911, EPI_ISL_522912, EPI_ISL_522913, EPI_ISL_522914, EPI_ISL_522915, EPI_ISL_522916, EPI_ISL_522917, EPI_ISL_522918, EPI_ISL_522919, EPI_ISL_522920, EPI_ISL_522921, EPI_ISL_522922, EPI_ISL_522923, EPI_ISL_522924, EPI_ISL_522925, EPI_ISL_522926, EPI_ISL_522927, EPI_ISL_522928, EPI_ISL_522929, EPI_ISL_522930, EPI_ISL_522931, EPI_ISL_522932, EPI_ISL_522933, EPI_ISL_522934, EPI_ISL_522935, EPI_ISL_522936, EPI_ISL_522937, EPI_ISL_522938, EPI_ISL_522939 | | | |
| see above | University of Wisconsin-Madison AIDS Vaccine Research Laboratories | University of Wisconsin-Madison AIDS Vaccine Research Laboratories | Gage Moreno, Katarina Braun, et al. AIDS Vaccine Research Laboratories |
| EPI_ISL_522940 | Instituto Nacional de Medicina Genómica | Instituto Nacional de Medicina Genómica | Hidalgo-Miranda A, Mendoza-Vargas A, Reyes-Grajeda JP, Cisneros-Villanueva M, Cedro-Tanda A, Hurtado-Cordova E, Peñaloza-Figueroa F, Herrera-Montalvo LA |
| EPI_ISL_522941 | Instituto Nacional de Medicina Genómica | Instituto Nacional de Medicina Genómica | Hidalgo-Miranda A, Mendoza-Vargas A, Reyes-Grajeda JP, Cisneros-Villanueva M, Cedro-Tanda A, Hurtado-Cordova E, Peñaloza-Figueroa F, Herrera-Montalvo LA |
| EPI_ISL_522942 | Instituto Nacional de Medicina Genómica | Instituto Nacional de Medicina Genómica | Hidalgo-Miranda A, Mendoza-Vargas A, Reyes-Grajeda JP, Cisneros-Villanueva M, Cedro-Tanda A, Hurtado-Cordova E, Peñaloza-Figueroa F, Herrera-Montalvo LA |
| EPI_ISL_522943, EPI_ISL_522944, EPI_ISL_522945, EPI_ISL_522946, EPI_ISL_522947, EPI_ISL_522948, EPI_ISL_522949, EPI_ISL_522950, EPI_ISL_522951, EPI_ISL_522952, EPI_ISL_522953, EPI_ISL_522954, EPI_ISL_522955, EPI_ISL_522956, EPI_ISL_522957, EPI_ISL_522958, EPI_ISL_522959, EPI_ISL_522960, EPI_ISL_522961, EPI_ISL_522962, EPI_ISL_522963, EPI_ISL_522964, EPI_ISL_522965, EPI_ISL_522966, EPI_ISL_522967, EPI_ISL_522968, EPI_ISL_522969, EPI_ISL_522970, EPI_ISL_522971, EPI_ISL_522972, EPI_ISL_522973, EPI_ISL_522974, EPI_ISL_522975, EPI_ISL_522976, EPI_ISL_522977 | | | |
| see above | Texas Department of State Health Services | Texas Department of State Health Services | Rashmi Tuladhar, Bonnie Oh, Cara Akrout, Jenny Zhang, Maliha Rahman, Anita Pokharel, Myong Koag, Chun Wang, Rachel Lee, Grace Kubin |
| EPI_ISL_522978 | Instituto Nacional de Medicina Genómica | Instituto Nacional de Medicina Genómica | Hidalgo-Miranda A, Mendoza-Vargas A, Reyes-Grajeda JP, Cisneros-Villanueva M, Cedro-Tanda A, Hurtado-Cordova E, Peñaloza-Figueroa F, Herrera-Montalvo LA |
| EPI_ISL_522979 | Instituto Nacional de Medicina Genómica | Instituto Nacional de Medicina Genómica | Hidalgo-Miranda A, Mendoza-Vargas A, Reyes-Grajeda JP, Cisneros-Villanueva M, Cedro-Tanda A, Hurtado-Cordova E, Peñaloza-Figueroa F, Herrera-Montalvo LA |
| EPI_ISL_522980, EPI_ISL_522981 | Instituto Nacional de Medicina Genómica | Instituto Nacional de Medicina Genómica | Hidalgo-Miranda A, Mendoza-Vargas A, Reyes-Grajeda JP, Cisneros-Villanueva M, Cedro-Tanda A, Hurtado-Cordova E, Peñaloza-Figueroa F, Herrera-Montalvo LA |
| EPI_ISL_522982 | Instituto Nacional de Medicina Genómica | Instituto Nacional de Medicina Genómica | Hidalgo-Miranda A, Mendoza-Vargas A, Reyes-Grajeda JP, Cisneros-Villanueva M, Cedro-Tanda A, Hurtado-Cordova E, Peñaloza-Figueroa F, Herrera-Montalvo LA |
| EPI_ISL_522983 | Instituto Nacional de Medicina Genómica | Instituto Nacional de Medicina Genómica | Hidalgo-Miranda A, Mendoza-Vargas A, Reyes-Grajeda JP, Cisneros-Villanueva M, Cedro-Tanda A, Hurtado-Cordova E, Peñaloza-Figueroa F, Herrera-Montalvo LA |
| EPI_ISL_522984 | Instituto Nacional de Medicina Genómica | Instituto Nacional de Medicina Genómica | Hidalgo-Miranda A, Mendoza-Vargas A, Reyes-Grajeda JP, Cisneros-Villanueva M, Cedro-Tanda A, Hurtado-Cordova E, Peñaloza-Figueroa F, Herrera-Montalvo LA |
| EPI_ISL_522985, EPI_ISL_522986 | Instituto Nacional de Medicina Genómica | Instituto Nacional de Medicina Genómica | Hidalgo-Miranda A, Mendoza-Vargas A, Reyes-Grajeda JP, Cisneros-Villanueva M, Cedro-Tanda A, Hurtado-Cordova E, Peñaloza-Figueroa F, Herrera-Montalvo LA |
| EPI_ISL_522987, EPI_ISL_522988, EPI_ISL_522989, EPI_ISL_522990, EPI_ISL_522991, EPI_ISL_522992, EPI_ISL_522993, EPI_ISL_522994, EPI_ISL_522995, EPI_ISL_522996, EPI_ISL_522997, EPI_ISL_522998, EPI_ISL_522999, EPI_ISL_523000, EPI_ISL_523001, EPI_ISL_523002, EPI_ISL_523003, EPI_ISL_523004, EPI_ISL_523005, EPI_ISL_523006, EPI_ISL_523007, EPI_ISL_523008, EPI_ISL_523009, EPI_ISL_523010, EPI_ISL_523011, EPI_ISL_523012, EPI_ISL_523013, EPI_ISL_523014, EPI_ISL_523015, EPI_ISL_523016, EPI_ISL_523017, EPI_ISL_523018, EPI_ISL_523019, EPI_ISL_523020, EPI_ISL_523021, EPI_ISL_523022, EPI_ISL_523023, EPI_ISL_523024, EPI_ISL_523025, EPI_ISL_523026, EPI_ISL_523027, EPI_ISL_523028, EPI_ISL_523029, EPI_ISL_523030, EPI_ISL_523031, EPI_ISL_523032, EPI_ISL_523033, EPI_ISL_523034, EPI_ISL_523035, EPI_ISL_523036, EPI_ISL_523037, EPI_ISL_523038, EPI_ISL_523039, EPI_ISL_523040, EPI_ISL_523041, EPI_ISL_523042, EPI_ISL_523043, EPI_ISL_523044, EPI_ISL_523045, EPI_ISL_523046, EPI_ISL_523047, EPI_ISL_523048, EPI_ISL_523049, EPI_ISL_523050, EPI_ISL_523051, EPI_ISL_523052, EPI_ISL_523053, EPI_ISL_523054, EPI_ISL_523055, EPI_ISL_523056, EPI_ISL_523057, EPI_ISL_523058, EPI_ISL_523059, EPI_ISL_523060, EPI_ISL_523061, EPI_ISL_523062, EPI_ISL_523063, EPI_ISL_523064, EPI_ISL_523065, EPI_ISL_523066, EPI_ISL_523067, EPI_ISL_523068, EPI_ISL_523069, EPI_ISL_523070, EPI_ISL_523071, EPI_ISL_523072, EPI_ISL_523073, EPI_ISL_523074, EPI_ISL_523075, EPI_ISL_523076, EPI_ISL_523077, EPI_ISL_523078, EPI_ISL_523079, EPI_ISL_523080, EPI_ISL_523081, EPI_ISL_523082, EPI_ISL_523083, EPI_ISL_523084, EPI_ISL_523085, EPI_ISL_523086, EPI_ISL_523087, EPI_ISL_523088, EPI_ISL_523089, EPI_ISL_523090, EPI_ISL_523091, EPI_ISL_523092, EPI_ISL_523093, EPI_ISL_523094, EPI_ISL_523095, EPI_ISL_523096, EPI_ISL_523097, EPI_ISL_523098, EPI_ISL_523099, EPI_ISL_523100, EPI_ISL_523101, EPI_ISL_523102, EPI_ISL_523103, EPI_ISL_523104, EPI_ISL_523105, EPI_ISL_523106, EPI_ISL_523107, EPI_ISL_523108, EPI_ISL_523109, EPI_ISL_523110, EPI_ISL_523111, EPI_ISL_523112, EPI_ISL_523113, EPI_ISL_523114, EPI_ISL_523115, EPI_ISL_523116, EPI_ISL_523117, EPI_ISL_523118, EPI_ISL_523119, EPI_ISL_523120 | | | |
| see above | Dutch COVID-19 response team | Erasmus Medical Center | OH consortium |
| EPI_ISL_523121, EPI_ISL_523122, EPI_ISL_523123, EPI_ISL_523124, EPI_ISL_523125, EPI_ISL_523126, EPI_ISL_523127, EPI_ISL_523128, EPI_ISL_523129, EPI_ISL_523130, EPI_ISL_523131, EPI_ISL_523132, EPI_ISL_523133, EPI_ISL_523134, EPI_ISL_523135, EPI_ISL_523136, EPI_ISL_523137, EPI_ISL_523138, EPI_ISL_523139, EPI_ISL_523140, EPI_ISL_523141, EPI_ISL_523142, EPI_ISL_523143, EPI_ISL_523144, EPI_ISL_523145, EPI_ISL_523146, EPI_ISL_523147, EPI_ISL_523148, EPI_ISL_523149, EPI_ISL_523150, EPI_ISL_523151, EPI_ISL_523152, EPI_ISL_523153, EPI_ISL_523154, EPI_ISL_523155, EPI_ISL_523156, EPI_ISL_523157, EPI_ISL_523158, EPI_ISL_523159, EPI_ISL_523160, EPI_ISL_523161, EPI_ISL_523162, EPI_ISL_523163, EPI_ISL_523164, EPI_ISL_523165, EPI_ISL_523166, EPI_ISL_523167, EPI_ISL_523168, EPI_ISL_523169, EPI_ISL_523170, EPI_ISL_523171, EPI_ISL_523172, EPI_ISL_523173, EPI_ISL_523174, EPI_ISL_523175, EPI_ISL_523176, EPI_ISL_523177, EPI_ISL_523178, EPI_ISL_523179, EPI_ISL_523180, EPI_ISL_523181, EPI_ISL_523182, EPI_ISL_523183, EPI_ISL_523184, EPI_ISL_523185, EPI_ISL_523186, EPI_ISL_523187, EPI_ISL_523188, EPI_ISL_523189, EPI_ISL_523190, EPI_ISL_523191, EPI_ISL_523192, EPI_ISL_523193, EPI_ISL_523194, EPI_ISL_523195, EPI_ISL_523196, EPI_ISL_523197, EPI_ISL_523198, EPI_ISL_523199, EPI_ISL_523200, EPI_ISL_523201, EPI_ISL_523202, EPI_ISL_523203, EPI_ISL_523204, EPI_ISL_523205, EPI_ISL_523206, EPI_ISL_523207, EPI_ISL_523208, EPI_ISL_523209, EPI_ISL_523210, EPI_ISL_523211, EPI_ISL_523212, EPI_ISL_523213, EPI_ISL_523214, EPI_ISL_523215, EPI_ISL_523216, EPI_ISL_523217, EPI_ISL_523218, EPI_ISL_523219, EPI_ISL_523220, EPI_ISL_523221, EPI_ISL_523222, EPI_ISL_523223, EPI_ISL_523224, EPI_ISL_523225, EPI_ISL_523226, EPI_ISL_523227, EPI_ISL_523228, EPI_ISL_523229, EPI_ISL_523230, EPI_ISL_523231, EPI_ISL_523232, EPI_ISL_523233, EPI_ISL_523234, EPI_ISL_523235, EPI_ISL_523236, EPI_ISL_523237, EPI_ISL_523238, EPI_ISL_523239, EPI_ISL_523240, EPI_ISL_523241, EPI_ISL_523242, EPI_ISL_523243, EPI_ISL_523244, EPI_ISL_523245, EPI_ISL_523246, EPI_ISL_523247, EPI_ISL_523248, EPI_ISL_523249, EPI_ISL_523250, EPI_ISL_523251, EPI_ISL_523252, EPI_ISL_523253, EPI_ISL_523254, EPI_ISL_523255, EPI_ISL_523256, EPI_ISL_523257, EPI_ISL_523258, EPI_ISL_523259, EPI_ISL_523260, EPI_ISL_523261, EPI_ISL_523262, EPI_ISL_523263, EPI_ISL_523264, EPI_ISL_523265, EPI_ISL_523266, EPI_ISL_523267, EPI_ISL_523268, EPI_ISL_523269, EPI_ISL_523270, EPI_ISL_523271, EPI_ISL_523272, EPI_ISL_523273, EPI_ISL_523274, EPI_ISL_523275, EPI_ISL_523276, EPI_ISL_523277, EPI_ISL_523278, EPI_ISL_523279, EPI_ISL_523280, EPI_ISL_523281, EPI_ISL_523282, EPI_ISL_523283, EPI_ISL_523284, EPI_ISL_523285, EPI_ISL_523286, EPI_ISL_523287, EPI_ISL_523288, EPI_ISL_523289, EPI_ISL_523290, EPI_ISL_523291, EPI_ISL_523292, EPI_ISL_523293, EPI_ISL_523294, EPI_ISL_523295, EPI_ISL_523296, EPI_ISL_523297, EPI_ISL_523298, EPI_ISL_523299, EPI_ISL_523300, EPI_ISL_523301, EPI_ISL_523302, EPI_ISL_523303, EPI_ISL_523304, EPI_ISL_523305, EPI_ISL_523306, EPI_ISL_523307, EPI_ISL_523308, EPI_ISL_523309, EPI_ISL_523310, EPI_ISL_523311, EPI_ISL_523312, EPI_ISL_523313, EPI_ISL_523314, EPI_ISL_523315, EPI_ISL_523316, EPI_ISL_523317, EPI_ISL_523318, EPI_ISL_523319, EPI_ISL_523320, EPI_ISL_523321, EPI_ISL_523322, EPI_ISL_523323, EPI_ISL_523324, EPI_ISL_523325, EPI_ISL_523326, EPI_ISL_523327, EPI_ISL_523328, EPI_ISL_523329, EPI_ISL_523330, EPI_ISL_523331, EPI_ISL_523332, EPI_ISL_523333, EPI_ISL_523334, EPI_ISL_523335, EPI_ISL_523336, EPI_ISL_523337, EPI_ISL_523338, EPI_ISL_523339, EPI_ISL_523340, EPI_ISL_523341, EPI_ISL_523342, EPI_ISL_523343, EPI_ISL_523344, EPI_ISL_523345, EPI_ISL_523346, EPI_ISL_523347, EPI_ISL_523348, EPI_ISL_523349, EPI_ISL_523350, EPI_ISL_523351, EPI_ISL_523352, EPI_ISL_523353, EPI_ISL_523354, EPI_ISL_523355, EPI_ISL_523356, EPI_ISL_523357, EPI_ISL_523358, EPI_ISL_523359, EPI_ISL_523360, EPI_ISL_523361, EPI_ISL_523362, EPI_ISL_523363, EPI_ISL_523364, EPI_ISL_523365, EPI_ISL_523366, EPI_ISL_523367, EPI_ISL_523368, EPI_ISL_523369, EPI_ISL_523370, EPI_ISL_523371, EPI_ISL_523372, EPI_ISL_523373, EPI_ISL_523374, EPI_ISL_523375, EPI_ISL_523376, EPI_ISL_523377, EPI_ISL_523378, EPI_ISL_523379, EPI_ISL_523380, EPI_ISL_523381, EPI_ISL_523382, EPI_ISL_523383, EPI_ISL_523384, EPI_ISL_523385, EPI_ISL_523386, EPI_ISL_523387, EPI_ISL_523388, EPI_ISL_523389, EPI_ISL_523390, EPI_ISL_523391, EPI_ISL_523392, EPI_ISL_523393, EPI_ISL_523394, EPI_ISL_523395, EPI_ISL_523396, EPI_ISL_523397, EPI_ISL_523398, EPI_ISL_523399, EPI_ISL_523400, EPI_ISL_523401, EPI_ISL_523402, EPI_ISL_523403, EPI_ISL_523404, EPI_ISL_523405, EPI_ISL_523406, EPI_ISL_523407, EPI_ISL_523408, EPI_ISL_523409, EPI_ISL_523410, EPI_ISL_523411, EPI_ISL_523412, EPI_ISL_523413, EPI_ISL_523414, EPI_ISL_523415, EPI_ISL_523416, EPI_ISL_523417, EPI_ISL_523418, EPI_ISL_523419, EPI_ISL_523420, EPI_ISL_523421, EPI_ISL_523422, EPI_ISL_523423, EPI_ISL_523424, EPI_ISL_523425, EPI_ISL_523426, EPI_ISL_523427, EPI_ISL_523428, EPI_ISL_523429, EPI_ISL_523430, EPI_ISL_523431, EPI_ISL_523432, EPI_ISL_523433, EPI_ISL_523434, EPI_ISL_523435, EPI_ISL_523436, EPI_ISL_523437, EPI_ISL_523438, EPI_ISL_523439, EPI_ISL_523440, EPI_ISL_523441, EPI_ISL_523442, EPI_ISL_523443, EPI_ISL_523444, EPI_ISL_523445, EPI_ISL_523446, EPI_ISL_523447, EPI_ISL_523448, EPI_ISL_523449, EPI_ISL_523450, EPI_ISL_523451, EPI_ISL_523452, EPI_ISL_523453, EPI_ISL_523454, EPI_ISL_523455, EPI_ISL_523456, EPI_ISL_523457, EPI_ISL_523458, EPI_ISL_523459, EPI_ISL_523460, EPI_ISL_523461, EPI_ISL_523462, EPI_ISL_523463, EPI_ISL_523464, EPI_ISL_523465, EPI_ISL_523466, EPI_ISL_523467, EPI_ISL_523468, EPI_ISL_523469, EPI_ISL_523470, EPI_ISL_523471, EPI_ISL_523472, EPI_ISL_523473, EPI_ISL_523474, EPI_ISL_523475, EPI_ISL_523476, EPI_ISL_523477, EPI_ISL_523478, EPI_ISL_523479, EPI_ISL_523480, EPI_ISL_523481, EPI_ISL_523482, EPI_ISL_523483, EPI_ISL_523484, EPI_ISL_523485, EPI_ISL_523486, EPI_ISL_523487, EPI_ISL_523488, EPI_ISL_523489, EPI_ISL_523490, EPI_ISL_523491, EPI_ISL_523492, EPI_ISL_523493, EPI_ISL_523494, EPI_ISL_523495, EPI_ISL_523496, EPI_ISL_523497, EPI_ISL_523498, EPI_ISL_523499 | | | |
| see above | Dutch COVID-19 response team | Erasmus Medical Center | Bas Oude Munnink, David Nieuwenhuijse, Reina Sikkema, Claudia Schapendonk, Irina Chestakova, Anne van der Linden, Theo Bestebroer, Stefan van Nieuwkoop, Mark Pronk, Pascal Lexmond, Corien Swaan, Manon Haverkate, Madelief Mollers, Mart Stein, Sandra Kengne Kamga Mobou, Jeroen van Kampen, Jolanda Voermans, Aura Timen, Corine GeurtsvanKessel, Annemiek van der Eijk, Richard Molenkamp, Marion Koopmans, on behalf of the Dutch national COVID-19 response team. |
| EPI_ISL_523500 | Instituto Nacional de Medicina Genómica | Instituto Nacional de Medicina Genómica | Hidalgo-Miranda A, Mendoza-Vargas A, Reyes-Grajeda JP, Cisneros-Villanueva M, Cedro-Tanda A, Hurtado-Cordova E, Peñaloza-Figueroa F, Herrera-Montalvo LA |

| | | | |
|---|---|---|---|
| EPI_ISL_523501, EPI_ISL_523502, EPI_ISL_523503, EPI_ISL_523504, EPI_ISL_523505, EPI_ISL_523506, EPI_ISL_523507, EPI_ISL_523508, EPI_ISL_523509, EPI_ISL_523510, EPI_ISL_523511, EPI_ISL_523512, EPI_ISL_523513, EPI_ISL_523514, EPI_ISL_523515, EPI_ISL_523516, EPI_ISL_523517, EPI_ISL_523518, EPI_ISL_523519, EPI_ISL_523520, EPI_ISL_523521, EPI_ISL_523522, EPI_ISL_523523, EPI_ISL_523524, EPI_ISL_523525, EPI_ISL_523526, EPI_ISL_523527, EPI_ISL_523528, EPI_ISL_523529, EPI_ISL_523530, EPI_ISL_523531, EPI_ISL_523532, EPI_ISL_523533, EPI_ISL_523534, EPI_ISL_523535, EPI_ISL_523536, EPI_ISL_523537, EPI_ISL_523538, EPI_ISL_523539, EPI_ISL_523540, EPI_ISL_523541, EPI_ISL_523542, EPI_ISL_523543, EPI_ISL_523544, EPI_ISL_523545, EPI_ISL_523546, EPI_ISL_523547, EPI_ISL_523548, EPI_ISL_523549, EPI_ISL_523550, EPI_ISL_523551, EPI_ISL_523552, EPI_ISL_523553, EPI_ISL_523554, EPI_ISL_523555, EPI_ISL_523556, EPI_ISL_523557, EPI_ISL_523558, EPI_ISL_523559, EPI_ISL_523560, EPI_ISL_523561, EPI_ISL_523562, EPI_ISL_523563, EPI_ISL_523564, EPI_ISL_523565, EPI_ISL_523566, EPI_ISL_523567, EPI_ISL_523568, EPI_ISL_523569, EPI_ISL_523570, EPI_ISL_523571, EPI_ISL_523572, EPI_ISL_523573, EPI_ISL_523574, EPI_ISL_523575, EPI_ISL_523576, EPI_ISL_523577, EPI_ISL_523578, EPI_ISL_523579, EPI_ISL_523580, EPI_ISL_523581, EPI_ISL_523582, EPI_ISL_523583, EPI_ISL_523584, EPI_ISL_523585, EPI_ISL_523586, EPI_ISL_523587, EPI_ISL_523588, EPI_ISL_523589, EPI_ISL_523590, EPI_ISL_523591, EPI_ISL_523592, EPI_ISL_523593, EPI_ISL_523594, EPI_ISL_523595, EPI_ISL_523596, EPI_ISL_523597, EPI_ISL_523598, EPI_ISL_523599, EPI_ISL_523600, EPI_ISL_523601, EPI_ISL_523602, EPI_ISL_523603, EPI_ISL_523604, EPI_ISL_523605, EPI_ISL_523606, EPI_ISL_523607, EPI_ISL_523608, EPI_ISL_523609, EPI_ISL_523610, EPI_ISL_523611, EPI_ISL_523612, EPI_ISL_523613, EPI_ISL_523614, EPI_ISL_523615, EPI_ISL_523616, EPI_ISL_523617, EPI_ISL_523618, EPI_ISL_523619, EPI_ISL_523620, EPI_ISL_523621, EPI_ISL_523622, EPI_ISL_523623, EPI_ISL_523624, EPI_ISL_523625, EPI_ISL_523626, EPI_ISL_523627, EPI_ISL_523628, EPI_ISL_523629, EPI_ISL_523630, EPI_ISL_523631, EPI_ISL_523632, EPI_ISL_523633, EPI_ISL_523634, EPI_ISL_523635, EPI_ISL_523636, EPI_ISL_523637, EPI_ISL_523638, EPI_ISL_523639, EPI_ISL_523640, EPI_ISL_523641, EPI_ISL_523642, EPI_ISL_523643, EPI_ISL_523644, EPI_ISL_523645, EPI_ISL_523646, EPI_ISL_523647, EPI_ISL_523648, EPI_ISL_523649, EPI_ISL_523650, EPI_ISL_523651, EPI_ISL_523652, EPI_ISL_523653, EPI_ISL_523654, EPI_ISL_523655, EPI_ISL_523656, EPI_ISL_523657, EPI_ISL_523658, EPI_ISL_523659, EPI_ISL_523660, EPI_ISL_523661, EPI_ISL_523662, EPI_ISL_523663, EPI_ISL_523664, EPI_ISL_523665, EPI_ISL_523666, EPI_ISL_523667, EPI_ISL_523668, EPI_ISL_523669, EPI_ISL_523670, EPI_ISL_523671, EPI_ISL_523672, EPI_ISL_523673, EPI_ISL_523674, EPI_ISL_523675, EPI_ISL_523676, EPI_ISL_523677, EPI_ISL_523678, EPI_ISL_523679, EPI_ISL_523680, EPI_ISL_523681, EPI_ISL_523682, EPI_ISL_523683, EPI_ISL_523684, EPI_ISL_523685, EPI_ISL_523686, EPI_ISL_523687, EPI_ISL_523688, EPI_ISL_523689, EPI_ISL_523690, EPI_ISL_523691, EPI_ISL_523692, EPI_ISL_523693, EPI_ISL_523694, EPI_ISL_523695, EPI_ISL_523696, EPI_ISL_523697, EPI_ISL_523698, EPI_ISL_523699, EPI_ISL_523700, EPI_ISL_523701, EPI_ISL_523702, EPI_ISL_523703, EPI_ISL_523704, EPI_ISL_523705, EPI_ISL_523706, EPI_ISL_523707, EPI_ISL_523708, EPI_ISL_523709, EPI_ISL_523710, EPI_ISL_523711, EPI_ISL_523712, EPI_ISL_523713, EPI_ISL_523714, EPI_ISL_523715, EPI_ISL_523716, EPI_ISL_523717, EPI_ISL_523718, EPI_ISL_523719, EPI_ISL_523720, EPI_ISL_523721, EPI_ISL_523722, EPI_ISL_523723, EPI_ISL_523724, EPI_ISL_523725, EPI_ISL_523726, EPI_ISL_523727, EPI_ISL_523728, EPI_ISL_523729, EPI_ISL_523730, EPI_ISL_523731, EPI_ISL_523732, EPI_ISL_523733, EPI_ISL_523734, EPI_ISL_523735, EPI_ISL_523736, EPI_ISL_523737, EPI_ISL_523738, EPI_ISL_523739, EPI_ISL_523740, EPI_ISL_523741, EPI_ISL_523742, EPI_ISL_523743, EPI_ISL_523744, EPI_ISL_523745, EPI_ISL_523746, EPI_ISL_523747, EPI_ISL_523748, EPI_ISL_523749, EPI_ISL_523750, EPI_ISL_523751, EPI_ISL_523752, EPI_ISL_523753, EPI_ISL_523754, EPI_ISL_523755, EPI_ISL_523756, EPI_ISL_523757, EPI_ISL_523758, EPI_ISL_523759, EPI_ISL_523760, EPI_ISL_523761, EPI_ISL_523762, EPI_ISL_523763, EPI_ISL_523764, EPI_ISL_523765, EPI_ISL_523766, EPI_ISL_523767, EPI_ISL_523768, EPI_ISL_523769, EPI_ISL_523770, EPI_ISL_523771, EPI_ISL_523772, EPI_ISL_523773, EPI_ISL_523774, EPI_ISL_523775, EPI_ISL_523776, EPI_ISL_523777, EPI_ISL_523778, EPI_ISL_523779, EPI_ISL_523780, EPI_ISL_523781, EPI_ISL_523782, EPI_ISL_523783, EPI_ISL_523784, EPI_ISL_523785, EPI_ISL_523786, EPI_ISL_523787, EPI_ISL_523788, EPI_ISL_523789, EPI_ISL_523790, EPI_ISL_523791, EPI_ISL_523792, EPI_ISL_523793, EPI_ISL_523794, EPI_ISL_523795, EPI_ISL_523796, EPI_ISL_523797, EPI_ISL_523798, EPI_ISL_523799, EPI_ISL_523800, EPI_ISL_523801, EPI_ISL_523802, EPI_ISL_523803, EPI_ISL_523804, EPI_ISL_523805, EPI_ISL_523806, EPI_ISL_523807, EPI_ISL_523808, EPI_ISL_523809 | | | |
| see above | Dutch COVID-19 response team | Erasmus Medical Center | Bas Oude Munnink, David Nieuwenhuijse, Reina Sikkema, Claudia Schapendonk, Irina Chestakova, Anne van der Linden, Theo Bestebroer, Stefan van Nieuwkoop, Mark Pronk, Pascal Lexmond, Corien Swaan, Manon Haverkate, Madelief Mollers, Mart Stein, Sandra Kengne Kamga Mobou, Jeroen van Kampen, Jolanda Voermans, Aura Timen, Corine GeurtsvanKessel, Annemiek van der Eijk, Richard Molenkamp, Marion Koopmans, on behalf of the Dutch national COVID-19 response team. |
| EPI_ISL_523810 | Laboratorio de Referencia Nacional de Virus Respiratorio. Centro Nacional de Salud Publica. Instituto Nacional de Salud Peru. | Laboratorio de Referencia Nacional de Biotecnología y Biología Molecular. Centro Nacional de Salud Publica. Instituto Nacional de Salud Peru. | Carlos Padilla Rojas, Karolyn Vega Chozo, Priscila Lope Pari, Omar Caceres Rey, Marco Galarza Perez, Maribel Huaringa Nuñez, Johanna Balbuena Torres, Henri Bailon Calderon, Nancy Rojas Serrano. |
| EPI_ISL_523811, EPI_ISL_523812 | Universidad Iberoamericana, Instituto de Medicina Tropical & Salud Global | International Centre for Genetic Engineering and Biotechnology (ICGEB) and ARGO Open Lab Platform | Robert Paulino-Ramirez, Eileen Riego, Alejandro Vallejo Degaudenzi, Victor Virgilio Calderon, Leandro Tapia, Danilo Licastro, Simeone Dal Monego, Sreejith Rajasekharan and Alessandro Marcello. |
| EPI_ISL_523824 | Communicable Disease Branch | Hong Kong Department of Health | Mak Gannon C.K., Lam Edman T.K., Chan Rickjason C.W., Tsang Dominic N.C. |
| EPI_ISL_523825 | United Christian Hospital | Hong Kong Department of Health | Mak Gannon C.K., Lam Edman T.K., Chan Rickjason C.W., Tsang Dominic N.C. |
| EPI_ISL_523826, EPI_ISL_523827 | Queen Mary Hospital | Hong Kong Department of Health | Mak Gannon C.K., Lam Edman T.K., Chan Rickjason C.W., Tsang Dominic N.C. |
| EPI_ISL_523828, EPI_ISL_523829 | Tuen Mun Hospital | Hong Kong Department of Health | Mak Gannon C.K., Lam Edman T.K., Chan Rickjason C.W., Tsang Dominic N.C. |
| EPI_ISL_523830, EPI_ISL_523831 | Our Lady of Maryknoll Hospital | Hong Kong Department of Health | Mak Gannon C.K., Lam Edman T.K., Chan Rickjason C.W., Tsang Dominic N.C. |
| EPI_ISL_523832, EPI_ISL_523833, EPI_ISL_523834 | Tuen Mun Hospital | Hong Kong Department of Health | Mak Gannon C.K., Lam Edman T.K., Chan Rickjason C.W., Tsang Dominic N.C. |
| EPI_ISL_523835, EPI_ISL_523836 | Queen Elizabeth Hospital | Hong Kong Department of Health | Mak Gannon C.K., Lam Edman T.K., Chan Rickjason C.W., Tsang Dominic N.C. |
| EPI_ISL_523837 | Yan Chai Hospital | Hong Kong Department of Health | Mak Gannon C.K., Lam Edman T.K., Chan Rickjason C.W., Tsang Dominic N.C. |
| EPI_ISL_523838 | Princess Margaret Hospital | Hong Kong Department of Health | Mak Gannon C.K., Lam Edman T.K., Chan Rickjason C.W., Tsang Dominic N.C. |
| EPI_ISL_523839 | Tseung Kwan O Hospital | Hong Kong Department of Health | Mak Gannon C.K., Lam Edman T.K., Chan Rickjason C.W., Tsang Dominic N.C. |
| EPI_ISL_523840 | Our Lady of Maryknoll Hospital | Hong Kong Department of Health | Mak Gannon C.K., Lam Edman T.K., Chan Rickjason C.W., Tsang Dominic N.C. |
| EPI_ISL_523841 | Queen Elizabeth Hospital | Hong Kong Department of Health | Mak Gannon C.K., Lam Edman T.K., Chan Rickjason C.W., Tsang Dominic N.C. |
| EPI_ISL_523842 | Kwong Wah Hospital | Hong Kong Department of Health | Mak Gannon C.K., Lam Edman T.K., Chan Rickjason C.W., Tsang Dominic N.C. |
| EPI_ISL_523843, EPI_ISL_523844, EPI_ISL_523845, EPI_ISL_523846, EPI_ISL_523847 | Queen Elizabeth Hospital | Hong Kong Department of Health | Mak Gannon C.K., Lam Edman T.K., Chan Rickjason C.W., Tsang Dominic N.C. |
| EPI_ISL_523848, EPI_ISL_523849 | United Christian Hospital | Hong Kong Department of Health | Mak Gannon C.K., Lam Edman T.K., Chan Rickjason C.W., Tsang Dominic N.C. |
| EPI_ISL_523850, EPI_ISL_523851, EPI_ISL_523852 | Tseung Kwan O Hospital | Hong Kong Department of Health | Mak Gannon C.K., Lam Edman T.K., Chan Rickjason C.W., Tsang Dominic N.C. |
| EPI_ISL_523853 | Hong Kong Baptist Hospital | Hong Kong Department of Health | Mak Gannon C.K., Lam Edman T.K., Chan Rickjason C.W., Tsang Dominic N.C. |
| EPI_ISL_523854 | Alice Ho Miu Ling Nethersole Hospital | Hong Kong Department of Health | Mak Gannon C.K., Lam Edman T.K., Chan Rickjason C.W., Tsang Dominic N.C. |
| EPI_ISL_523855 | Kwong Wah Hospital | Hong Kong Department of Health | Mak Gannon C.K., Lam Edman T.K., Chan Rickjason C.W., Tsang Dominic N.C. |
| EPI_ISL_523856, EPI_ISL_523857, EPI_ISL_523858, EPI_ISL_523859, EPI_ISL_523860, EPI_ISL_523861, EPI_ISL_523862, EPI_ISL_523863, EPI_ISL_523864, EPI_ISL_523865, EPI_ISL_523866, EPI_ISL_523867, EPI_ISL_523868, EPI_ISL_523869, EPI_ISL_523870, EPI_ISL_523871, EPI_ISL_523872, EPI_ISL_523873, EPI_ISL_523874, EPI_ISL_523875, EPI_ISL_523876, EPI_ISL_523877, EPI_ISL_523878, EPI_ISL_523879, EPI_ISL_523880, EPI_ISL_523881, EPI_ISL_523882, EPI_ISL_523883, EPI_ISL_523884, EPI_ISL_523885, EPI_ISL_523886, EPI_ISL_523887, EPI_ISL_523888, EPI_ISL_523889, EPI_ISL_523890, EPI_ISL_523891, EPI_ISL_523892, EPI_ISL_523893, EPI_ISL_523894, EPI_ISL_523895, EPI_ISL_523896, EPI_ISL_523897, EPI_ISL_523898, EPI_ISL_523899, EPI_ISL_523900, EPI_ISL_523901, EPI_ISL_523902, EPI_ISL_523903, EPI_ISL_523904, EPI_ISL_523905, EPI_ISL_523906, EPI_ISL_523907, EPI_ISL_523908, EPI_ISL_523909, EPI_ISL_523910, EPI_ISL_523911, EPI_ISL_523912, EPI_ISL_523913, EPI_ISL_523914, EPI_ISL_523915, EPI_ISL_523916, EPI_ISL_523917, EPI_ISL_523918, EPI_ISL_523919, EPI_ISL_523920, EPI_ISL_523921, EPI_ISL_523922, EPI_ISL_523923, EPI_ISL_523924, EPI_ISL_523925, EPI_ISL_523926 | | | |
| see above | Viollier AG | Department of Biosystems Science and Engineering, ETH Zürich | Christian Beisel, Sarah Nadeau, Ivan Topolsky, Pedro Ferreira, Philipp Jablonski, Susana Posada-Céspedes, Tobias Schär, Ina Nissen, Natascha Santacroce, Elodie Burcklen, Christiane Beckmann, Maurice Redondo, Olivier Kobel, Christoph Noppen, Sophie Seidel, Noemie Santamaria de Souza, Niko Beerenwinkel, Tanja Stadler |
| EPI_ISL_523927, EPI_ISL_523928, EPI_ISL_523929 | Center of Medical Microbiology, Virology, and Hospital Hygiene, University of Duesseldorf | Center of Medical Microbiology, Virology, and Hospital Hygiene, University of Duesseldorf | Maximilian Damagnez, Alexander Dilthey, Torsten Houwaart, Malte Kohns Vasconcelos, Marek Korencak, Jessica Nicolai, Klaus Pfeffer, Hendrik Streeck, Daniel Strelow, Jörg Timm, Andreas Walker, Tobias Wienemann |
| EPI_ISL_523930 | Center of Medical Microbiology, Virology, and Hospital Hygiene, University of Duesseldorf | Universitätstr.1 40225 Düsseldorf Germany | Maximilian Damagnez, Alexander Dilthey, Torsten Houwaart, Malte Kohns Vasconcelos, Marek Korencak, Jessica Nicolai, Klaus Pfeffer, Hendrik Streeck, Daniel Strelow, Jörg Timm, Andreas Walker, Tobias Wienemann |
| EPI_ISL_523931, EPI_ISL_523932, EPI_ISL_523933, EPI_ISL_523934, EPI_ISL_523935, EPI_ISL_523936, EPI_ISL_523937, EPI_ISL_523938, EPI_ISL_523939, EPI_ISL_523940, EPI_ISL_523941, EPI_ISL_523942, EPI_ISL_523943, EPI_ISL_523944, EPI_ISL_523945, EPI_ISL_523946, EPI_ISL_523947, EPI_ISL_523948, EPI_ISL_523949 | | | |
| see above | Center of Medical Microbiology, Virology, and Hospital Hygiene, University of Duesseldorf | Center of Medical Microbiology, Virology, and Hospital Hygiene, University of Duesseldorf | Maximilian Damagnez, Alexander Dilthey, Torsten Houwaart, Malte Kohns Vasconcelos, Marek Korencak, Jessica Nicolai, Klaus Pfeffer, Hendrik Streeck, Daniel Strelow, Jörg Timm, Andreas Walker, Tobias Wienemann |
| EPI_ISL_523950 | Center of Medical Microbiology, Virology, and Hospital Hygiene, University of Duesseldorf | Center of Medical Microbiology, Virology, and Hospital Hygiene, Heinrich Heine University Düsseldorf | Maximilian Damagnez, Alexander Dilthey, Torsten Houwaart, Malte Kohns Vasconcelos, Marek Korencak, Jessica Nicolai, Klaus Pfeffer, Hendrik Streeck, Daniel Strelow, Jörg Timm, Andreas Walker, Tobias Wienemann |
| EPI_ISL_523951 | Respiratory Virus Unit, Microbiology Services Colindale, Public Health England | Respiratory Virus Unit, Microbiology Services Colindale, Public Health England | PHE Covid Sequencing Team |
| EPI_ISL_523952, EPI_ISL_523953 | Mohammed Bin Rashid University of Medicine and Health Sciences | Al Jalila Genomics Center | Ahmad Abou Tayoun, Tom Loney, Hamda Khansaheb, Sathishkumar Ramaswamy, Divinlal Harilal, Zulfa Omar Deesi, Rupa Murthy Varghese, Hanan Al Suwaidi, Abdulmajeed Alkhaja, Mohammed Uddin, Rifat Hamoudi, Rabih Halwani, Abiola Catherine Senok, Qutayba Hamid, Norbert Nowotny, Alawi Alsheikh-Ali |
| EPI_ISL_523954 | Laboratorio de Referencia Nacional de Virus Respiratorio. Centro Nacional de Salud Publica. Instituto Nacional de Salud Peru. | Laboratorio de Referencia Nacional de Biotecnología y Biología Molecular. Centro Nacional de Salud Publica. Instituto Nacional de Salud Peru. | Carlos Padilla Rojas, Karolyn Vega Chozo, Priscila Lope Pari, Omar Caceres Rey, Marco Galarza Perez, Maribel Huaringa Nuñez, Johanna Balbuena Torres, Henri Bailon Calderon, Nancy Rojas Serrano. |
| EPI_ISL_523955 | Hospital Municipal do Tatuape Carmino Caricchio | Instituto Adolfo Lutz, Interdiciplinary Procedures Center, Strategic Laboratory | Claudio Tavares Sacchi, Claudia Regina Gonçalves, Erica Valessa Ramos Gomes |
| EPI_ISL_523956 | Hospital Regional de Assis | Instituto Adolfo Lutz, Interdiciplinary Procedures Center, Strategic Laboratory | Claudio Tavares Sacchi, Claudia Regina Gonçalves, Erica Valessa Ramos Gomes |
| EPI_ISL_523957 | Hospital Itamaraty | Instituto Adolfo Lutz, Interdiciplinary Procedures Center, Strategic Laboratory | Claudio Tavares Sacchi, Claudia Regina Gonçalves, Erica Valessa Ramos Gomes |
| EPI_ISL_523958, EPI_ISL_523959 | Pronto Socorro Municipal de Perus | Instituto Adolfo Lutz, Interdiciplinary Procedures Center, Strategic Laboratory | Claudio Tavares Sacchi, Claudia Regina Gonçalves, Erica Valessa Ramos Gomes |
| EPI_ISL_523960 | National Agency for Public Health, Republic of Moldova | Charite Universitatsmedizin Berlin, Institute of Virology | Victor M Corman, Jörn Beheim-Schwarzbach, Barbara Mühlemann, Talitha Veith, Julia Schneider, Ala Halacu, Mariana Apostol, Terry Jones, Christian Drosten |
| EPI_ISL_523961 | Pronto Socorro Municipal 21 de Junho | Instituto Adolfo Lutz, Interdiciplinary Procedures Center, Strategic Laboratory | Claudio Tavares Sacchi, Claudia Regina Gonçalves, Erica Valessa Ramos Gomes |
| EPI_ISL_523962 | National Agency for Public Health, Republic of Moldova | Charite Universitatsmedizin Berlin, Institute of Virology | Victor M Corman, Joern Beheim-Schwarzbach, Barbara Mühlemann, Talitha Veith, Julia Schneider, Elizabeta Jancheska, Maja Kuzmanovska, Golubinka Bosevska, Terry Jones, Christian Drosten |
| EPI_ISL_523963 | UBS Vila Silvia | Instituto Adolfo Lutz, Interdiciplinary Procedures Center, Strategic Laboratory | Claudio Tavares Sacchi, Claudia Regina Gonçalves, Erica Valessa Ramos Gomes |
| EPI_ISL_523964 | National Agency for Public Health, Republic of Moldova | Charite Universitatsmedizin Berlin, Institute of Virology | Victor M Corman, Jörn Beheim-Schwarzbach, Barbara Mühlemann, Talitha Veith, Julia Schneider, Ala Halacu, Mariana Apostol, Terry Jones, Christian Drosten |
| EPI_ISL_523965 | Hospital do Servidor Público Estadual Francisco Morato de Oliveira | Instituto Adolfo Lutz, Interdiciplinary Procedures Center, Strategic Laboratory | Claudio Tavares Sacchi, Claudia Regina Gonçalves, Erica Valessa Ramos Gomes |
| EPI_ISL_523966 | National Agency for Public Health, Republic of Moldova | Charite Universitatsmedizin Berlin, Institute of Virology | Victor M Corman, Jörn Beheim-Schwarzbach, Barbara Mühlemann, Talitha Veith, Julia Schneider, Ala Halacu, Mariana Apostol, Terry Jones, Christian Drosten |
| EPI_ISL_523967 | Hospital Sancta Maggiore | Instituto Adolfo Lutz, Interdiciplinary Procedures Center, Strategic Laboratory | Claudio Tavares Sacchi, Claudia Regina Gonçalves, Erica Valessa Ramos Gomes |
| EPI_ISL_523968 | National Agency for Public Health, Republic of Moldova | Charite Universitatsmedizin Berlin, Institute of Virology | Victor M Corman, Jörn Beheim-Schwarzbach, Barbara Mühlemann, Talitha Veith, Julia Schneider, Ala Halacu, Mariana Apostol, Terry Jones, Christian Drosten |
| EPI_ISL_523969 | Hospital Sao Paulo de Ensino da Unifesp | Instituto Adolfo Lutz, Interdiciplinary Procedures Center, Strategic Laboratory | Claudio Tavares Sacchi, Claudia Regina Gonçalves, Erica Valessa Ramos Gomes |
| EPI_ISL_523970 | Conjunto Hospitalar do Mandaqui | Instituto Adolfo Lutz, Interdiciplinary Procedures Center, Strategic Laboratory | Claudio Tavares Sacchi, Claudia Regina Gonçalves, Erica Valessa Ramos Gomes |
| EPI_ISL_523971 | Hospital Geral Santa Marcelina | Instituto Adolfo Lutz, Interdiciplinary Procedures Center, Strategic Laboratory | Claudio Tavares Sacchi, Claudia Regina Gonçalves, Erica Valessa Ramos Gomes |
| EPI_ISL_523972 | Hospital do Servidor Público Estadual Francisco Morato de Oliveira | Instituto Adolfo Lutz, Interdiciplinary Procedures Center, Strategic Laboratory | Claudio Tavares Sacchi, Claudia Regina Gonçalves, Erica Valessa Ramos Gomes |
| EPI_ISL_523973 | PS Municipal Dona Maria Antonieta Ferreira de Barros | Instituto Adolfo Lutz, Interdiciplinary Procedures Center, Strategic Laboratory | Claudio Tavares Sacchi, Claudia Regina Gonçalves, Erica Valessa Ramos Gomes |
| EPI_ISL_523974 | Hospital Municipal do Tatuape Carmino Caricchio | Instituto Adolfo Lutz, Interdiciplinary Procedures Center, Strategic Laboratory | Claudio Tavares Sacchi, Claudia Regina Gonçalves, Erica Valessa Ramos Gomes |
| EPI_ISL_523975 | UPA Tito Lopes | Instituto Adolfo Lutz, Interdiciplinary Procedures Center, Strategic Laboratory | Claudio Tavares Sacchi, Claudia Regina Gonçalves, Erica Valessa Ramos Gomes |
| EPI_ISL_523976 | Hospital Municipal do Tatuape Carmino Caricchio | Instituto Adolfo Lutz, Interdiciplinary Procedures Center, Strategic Laboratory | Claudio Tavares Sacchi, Claudia Regina Gonçalves, Erica Valessa Ramos Gomes |
| EPI_ISL_523977 | Hosp. Municipal Prof. Dr. Alípio Corrêa Netto | Instituto Adolfo Lutz, Interdiciplinary Procedures Center, Strategic Laboratory | Claudio Tavares Sacchi, Claudia Regina Gonçalves, Erica Valessa Ramos Gomes |

| | | | |
|---|---|---|---|
| EPI_ISL_523978 | Hospital do Servidor Público Estadual Francisco Morato de Oliveira | Instituto Adolfo Lutz, Interdiciplinary Procedures Center, Strategic Laboratory | Claudio Tavares Sacchi, Claudia Regina Gonçalves, Erica Valessa Ramos Gomes |
| EPI_ISL_523979 | Laboratorio de Referencia Nacional de Virus Respiratorio. Centro Nacional de Salud Publica. Instituto Nacional de Salud Peru. | Laboratorio de Referencia Nacional de Biotecnología y Biología Molecular. Centro Nacional de Salud Publica. Instituto Nacional de Salud Peru. | Carlos Padilla Rojas, Karolyn Vega Chozo, Priscila Lope Pari, Omar Caceres Rey, Marco Galarza Perez, Maribel Huaringa Nuñez, Johanna Balbuena Torres, Henri Bailon Calderon, Nancy Rojas Serrano. |
| EPI_ISL_523980 | UPA Tito Lopes | Instituto Adolfo Lutz, Interdiciplinary Procedures Center, Strategic Laboratory | Claudio Tavares Sacchi, Claudia Regina Gonçalves, Erica Valessa Ramos Gomes |
| EPI_ISL_523981 | Hospital Sao Paulo de Ensino da Unifesp | Instituto Adolfo Lutz, Interdiciplinary Procedures Center, Strategic Laboratory | Claudio Tavares Sacchi, Claudia Regina Gonçalves, Erica Valessa Ramos Gomes |
| EPI_ISL_523982 | Hospital do Servidor Público Estadual Francisco Morato de Oliveira | Instituto Adolfo Lutz, Interdiciplinary Procedures Center, Strategic Laboratory | Claudio Tavares Sacchi, Claudia Regina Gonçalves, Erica Valessa Ramos Gomes |
| EPI_ISL_523983 | UPA Campo Limpo | Instituto Adolfo Lutz, Interdiciplinary Procedures Center, Strategic Laboratory | Claudio Tavares Sacchi, Claudia Regina Gonçalves, Erica Valessa Ramos Gomes |
| EPI_ISL_523984 | Ama Dr Jose Soares Hungria | Instituto Adolfo Lutz, Interdiciplinary Procedures Center, Strategic Laboratory | Claudio Tavares Sacchi, Claudia Regina Gonçalves, Erica Valessa Ramos Gomes |
| EPI_ISL_523985 | Hospital Municipal Dr. Benedicto Montenegro | Instituto Adolfo Lutz, Interdiciplinary Procedures Center, Strategic Laboratory | Claudio Tavares Sacchi, Claudia Regina Gonçalves, Erica Valessa Ramos Gomes |
| EPI_ISL_523986 | Ama Dr Jose Soares Hungria | Instituto Adolfo Lutz, Interdiciplinary Procedures Center, Strategic Laboratory | Claudio Tavares Sacchi, Claudia Regina Gonçalves, Erica Valessa Ramos Gomes |
| EPI_ISL_523988 | Hospital Sao Paulo de Ensino da Unifesp | Instituto Adolfo Lutz, Interdiciplinary Procedures Center, Strategic Laboratory | Claudio Tavares Sacchi, Claudia Regina Gonçalves, Erica Valessa Ramos Gomes |
| EPI_ISL_523989 | AMA Jardim Joamar | Instituto Adolfo Lutz, Interdiciplinary Procedures Center, Strategic Laboratory | Claudio Tavares Sacchi, Claudia Regina Gonçalves, Erica Valessa Ramos Gomes |
| EPI_ISL_523990 | AMA Jardim Peri | Instituto Adolfo Lutz, Interdiciplinary Procedures Center, Strategic Laboratory | Claudio Tavares Sacchi, Claudia Regina Gonçalves, Erica Valessa Ramos Gomes |
| EPI_ISL_523991, EPI_ISL_523992 | Hospital Municipal Carmen Prudente | Instituto Adolfo Lutz, Interdiciplinary Procedures Center, Strategic Laboratory | Claudio Tavares Sacchi, Claudia Regina Gonçalves, Erica Valessa Ramos Gomes |
| EPI_ISL_523993 | UPA Campo Limpo | Instituto Adolfo Lutz, Interdiciplinary Procedures Center, Strategic Laboratory | Claudio Tavares Sacchi, Claudia Regina Gonçalves, Erica Valessa Ramos Gomes |
| EPI_ISL_523994 | Laboratorio de Referencia Nacional de Virus Respiratorio. Centro Nacional de Salud Publica. Instituto Nacional de Salud Peru. | Laboratorio de Referencia Nacional de Biotecnología y Biología Molecular. Centro Nacional de Salud Publica. Instituto Nacional de Salud Peru. | Carlos Padilla Rojas, Karolyn Vega Chozo, Priscila Lope Pari, Omar Caceres Rey, Marco Galarza Perez, Maribel Huaringa Nuñez, Johanna Balbuena Torres, Henri Bailon Calderon, Nancy Rojas Serrano. |
| EPI_ISL_523995, EPI_ISL_523996, EPI_ISL_523997, EPI_ISL_523998, EPI_ISL_523999, EPI_ISL_524000, EPI_ISL_524001, EPI_ISL_524002, EPI_ISL_524003, EPI_ISL_524004, EPI_ISL_524005, EPI_ISL_524006, EPI_ISL_524007, EPI_ISL_524008, EPI_ISL_524009, EPI_ISL_524010, EPI_ISL_524011, EPI_ISL_524012, EPI_ISL_524013, EPI_ISL_524014, EPI_ISL_524015, EPI_ISL_524016, EPI_ISL_524017, EPI_ISL_524018, EPI_ISL_524019, EPI_ISL_524020, EPI_ISL_524021, EPI_ISL_524022, EPI_ISL_524023, EPI_ISL_524024, EPI_ISL_524025, EPI_ISL_524026, EPI_ISL_524027, EPI_ISL_524028, EPI_ISL_524029, EPI_ISL_524030, EPI_ISL_524031, EPI_ISL_524032, EPI_ISL_524033, EPI_ISL_524034, EPI_ISL_524035, EPI_ISL_524036, EPI_ISL_524037, EPI_ISL_524038, EPI_ISL_524039, EPI_ISL_524040, EPI_ISL_524041, EPI_ISL_524042, EPI_ISL_524043, EPI_ISL_524044, EPI_ISL_524045, EPI_ISL_524046, EPI_ISL_524047, EPI_ISL_524048, EPI_ISL_524049, EPI_ISL_524050, EPI_ISL_524051, EPI_ISL_524052 | | | |
| see above | WHO National Influenza Centre Russian Federation | WHO National Influenza Centre Russian Federation | Andrey Komissarov, Artem Fadeev, Mariia Sergeeva, Anna Ivanova, Daria Danilenko |
| EPI_ISL_524053, EPI_ISL_524054, EPI_ISL_524055, EPI_ISL_524056, EPI_ISL_524057, EPI_ISL_524058, EPI_ISL_524059, EPI_ISL_524060, EPI_ISL_524061, EPI_ISL_524062, EPI_ISL_524063, EPI_ISL_524064, EPI_ISL_524065, EPI_ISL_524066, EPI_ISL_524067, EPI_ISL_524068, EPI_ISL_524069, EPI_ISL_524070, EPI_ISL_524071, EPI_ISL_524072, EPI_ISL_524073 | | | |
| see above | Texas Department of State Health Services | Texas Department of State Health Services | Rashmi Tuladhar, Bonnie Oh, Cara Akrout, Jenny Zhang, Maliha Rahman, Anita Pokharel, Myong Koag, Chun Wang, Rachel Lee, Grace Kubin |
| EPI_ISL_524074, EPI_ISL_524075, EPI_ISL_524076, EPI_ISL_524077, EPI_ISL_524078, EPI_ISL_524079, EPI_ISL_524080, EPI_ISL_524081, EPI_ISL_524082, EPI_ISL_524083, EPI_ISL_524084, EPI_ISL_524085, EPI_ISL_524086, EPI_ISL_524087, EPI_ISL_524088, EPI_ISL_524089, EPI_ISL_524090, EPI_ISL_524091, EPI_ISL_524092, EPI_ISL_524093, EPI_ISL_524094, EPI_ISL_524095, EPI_ISL_524096, EPI_ISL_524097, EPI_ISL_524098, EPI_ISL_524099, EPI_ISL_524100, EPI_ISL_524101, EPI_ISL_524102, EPI_ISL_524103, EPI_ISL_524104, EPI_ISL_524105, EPI_ISL_524106, EPI_ISL_524107, EPI_ISL_524108, EPI_ISL_524109, EPI_ISL_524110, EPI_ISL_524111, EPI_ISL_524112, EPI_ISL_524113, EPI_ISL_524114, EPI_ISL_524115, EPI_ISL_524116, EPI_ISL_524117, EPI_ISL_524118, EPI_ISL_524119, EPI_ISL_524120, EPI_ISL_524121, EPI_ISL_524122, EPI_ISL_524123, EPI_ISL_524124, EPI_ISL_524125, EPI_ISL_524126, EPI_ISL_524127, EPI_ISL_524128, EPI_ISL_524129, EPI_ISL_524130, EPI_ISL_524131, EPI_ISL_524132, EPI_ISL_524133, EPI_ISL_524134, EPI_ISL_524135, EPI_ISL_524136, EPI_ISL_524137, EPI_ISL_524138, EPI_ISL_524139, EPI_ISL_524140, EPI_ISL_524141, EPI_ISL_524142, EPI_ISL_524143, EPI_ISL_524144, EPI_ISL_524145, EPI_ISL_524146, EPI_ISL_524147, EPI_ISL_524148, EPI_ISL_524149, EPI_ISL_524150, EPI_ISL_524151, EPI_ISL_524152, EPI_ISL_524153, EPI_ISL_524154, EPI_ISL_524155, EPI_ISL_524156, EPI_ISL_524157, EPI_ISL_524158, EPI_ISL_524159, EPI_ISL_524160, EPI_ISL_524161, EPI_ISL_524162, EPI_ISL_524163, EPI_ISL_524164, EPI_ISL_524165, EPI_ISL_524166, EPI_ISL_524167, EPI_ISL_524168, EPI_ISL_524169, EPI_ISL_524170, EPI_ISL_524171, EPI_ISL_524172, EPI_ISL_524173, EPI_ISL_524174, EPI_ISL_524175, EPI_ISL_524176, EPI_ISL_524177, EPI_ISL_524178, EPI_ISL_524179, EPI_ISL_524180, EPI_ISL_524181, EPI_ISL_524182, EPI_ISL_524183, EPI_ISL_524184, EPI_ISL_524185, EPI_ISL_524186, EPI_ISL_524187, EPI_ISL_524188, EPI_ISL_524189, EPI_ISL_524190, EPI_ISL_524191, EPI_ISL_524192, EPI_ISL_524193, EPI_ISL_524194, EPI_ISL_524195, EPI_ISL_524196, EPI_ISL_524197, EPI_ISL_524198, EPI_ISL_524199, EPI_ISL_524200, EPI_ISL_524201, EPI_ISL_524202, EPI_ISL_524203, EPI_ISL_524204, EPI_ISL_524205, EPI_ISL_524206, EPI_ISL_524207, EPI_ISL_524208, EPI_ISL_524209, EPI_ISL_524210, EPI_ISL_524211, EPI_ISL_524212, EPI_ISL_524213, EPI_ISL_524214, EPI_ISL_524215, EPI_ISL_524216, EPI_ISL_524217, EPI_ISL_524218, EPI_ISL_524219, EPI_ISL_524220, EPI_ISL_524221, EPI_ISL_524222, EPI_ISL_524223, EPI_ISL_524224, EPI_ISL_524225, EPI_ISL_524226, EPI_ISL_524227, EPI_ISL_524228, EPI_ISL_524229, EPI_ISL_524230, EPI_ISL_524231, EPI_ISL_524232, EPI_ISL_524233, EPI_ISL_524234, EPI_ISL_524235, EPI_ISL_524236, EPI_ISL_524237, EPI_ISL_524238, EPI_ISL_524239, EPI_ISL_524240, EPI_ISL_524241, EPI_ISL_524242, EPI_ISL_524243, EPI_ISL_524244, EPI_ISL_524245, EPI_ISL_524246, EPI_ISL_524247, EPI_ISL_524248, EPI_ISL_524249, EPI_ISL_524250, EPI_ISL_524251, EPI_ISL_524252, EPI_ISL_524253, EPI_ISL_524254, EPI_ISL_524255, EPI_ISL_524256, EPI_ISL_524257, EPI_ISL_524258, EPI_ISL_524259, EPI_ISL_524260, EPI_ISL_524261, EPI_ISL_524262, EPI_ISL_524263, EPI_ISL_524264, EPI_ISL_524265, EPI_ISL_524266, EPI_ISL_524267, EPI_ISL_524268, EPI_ISL_524269, EPI_ISL_524270, EPI_ISL_524271, EPI_ISL_524272, EPI_ISL_524273, EPI_ISL_524274, EPI_ISL_524275, EPI_ISL_524276, EPI_ISL_524277, EPI_ISL_524278, EPI_ISL_524279, EPI_ISL_524280, EPI_ISL_524281, EPI_ISL_524282, EPI_ISL_524283, EPI_ISL_524284, EPI_ISL_524285, EPI_ISL_524286, EPI_ISL_524287, EPI_ISL_524288, EPI_ISL_524289, EPI_ISL_524290, EPI_ISL_524291, EPI_ISL_524292, EPI_ISL_524293, EPI_ISL_524294, EPI_ISL_524295, EPI_ISL_524296, EPI_ISL_524297, EPI_ISL_524298, EPI_ISL_524299, EPI_ISL_524300, EPI_ISL_524301, EPI_ISL_524302, EPI_ISL_524303, EPI_ISL_524304, EPI_ISL_524305, EPI_ISL_524306, EPI_ISL_524307, EPI_ISL_524308, EPI_ISL_524309, EPI_ISL_524310, EPI_ISL_524311, EPI_ISL_524312, EPI_ISL_524313, EPI_ISL_524314, EPI_ISL_524315, EPI_ISL_524316, EPI_ISL_524317, EPI_ISL_524318, EPI_ISL_524319, EPI_ISL_524320, EPI_ISL_524321, EPI_ISL_524322, EPI_ISL_524323, EPI_ISL_524324, EPI_ISL_524325, EPI_ISL_524326, EPI_ISL_524327, EPI_ISL_524328, EPI_ISL_524329, EPI_ISL_524330, EPI_ISL_524331, EPI_ISL_524332, EPI_ISL_524333, EPI_ISL_524334, EPI_ISL_524335, EPI_ISL_524336, EPI_ISL_524337, EPI_ISL_524338, EPI_ISL_524339, EPI_ISL_524340, EPI_ISL_524341, EPI_ISL_524342, EPI_ISL_524343, EPI_ISL_524344, EPI_ISL_524345, EPI_ISL_524346, EPI_ISL_524347, EPI_ISL_524348, EPI_ISL_524349, EPI_ISL_524350, EPI_ISL_524351, EPI_ISL_524352, EPI_ISL_524353, EPI_ISL_524354, EPI_ISL_524355, EPI_ISL_524356, EPI_ISL_524357, EPI_ISL_524358, EPI_ISL_524359, EPI_ISL_524360, EPI_ISL_524361, EPI_ISL_524362, EPI_ISL_524363, EPI_ISL_524364, EPI_ISL_524365, EPI_ISL_524366, EPI_ISL_524367, EPI_ISL_524368, EPI_ISL_524369, EPI_ISL_524370, EPI_ISL_524371, EPI_ISL_524372, EPI_ISL_524373, EPI_ISL_524374, EPI_ISL_524375, EPI_ISL_524376, EPI_ISL_524377, EPI_ISL_524378, EPI_ISL_524379, EPI_ISL_524380, EPI_ISL_524381, EPI_ISL_524382, EPI_ISL_524383, EPI_ISL_524384, EPI_ISL_524385, EPI_ISL_524386, EPI_ISL_524387, EPI_ISL_524388, EPI_ISL_524389, EPI_ISL_524390, EPI_ISL_524391, EPI_ISL_524392, EPI_ISL_524393, EPI_ISL_524394, EPI_ISL_524395, EPI_ISL_524396, EPI_ISL_524397, EPI_ISL_524398, EPI_ISL_524399, EPI_ISL_524400, EPI_ISL_524401, EPI_ISL_524402, EPI_ISL_524403, EPI_ISL_524404, EPI_ISL_524405, EPI_ISL_524406, EPI_ISL_524407, EPI_ISL_524408, EPI_ISL_524409, EPI_ISL_524410, EPI_ISL_524411, EPI_ISL_524412, EPI_ISL_524413, EPI_ISL_524414, EPI_ISL_524415, EPI_ISL_524416, EPI_ISL_524417, EPI_ISL_524418, EPI_ISL_524419, EPI_ISL_524420, EPI_ISL_524421, EPI_ISL_524422, EPI_ISL_524423, EPI_ISL_524424, EPI_ISL_524425 | | | |
| see above | Utah Public Health Laboratory | Utah Public Health Laboratory | Erin L. Young, Kelly Oakeson, Tara Gallagher, Michael T. Pyne, E. Susan Slechta, Melanie A. Mallory, Jeffrey B. Stevenson, Salika M. Shakir, David R. Hillyard |
| EPI_ISL_524428, EPI_ISL_524429, EPI_ISL_524430, EPI_ISL_524431, EPI_ISL_524432 | Dirk Dittmer | Dirk Dittmer | Bailey,A.G., Caro-Vegas,C.P., Dittmer,D., Eason,A.B., Juarez,A., Landis,J.T., McNamara,R.P., Miller,M.B., Moorad,R., Pluta,L.J.,Seltzer,T.A., Thompson,C., Vahrson,W. and Villamor,F. |
| EPI_ISL_524445, EPI_ISL_524446, EPI_ISL_524447, EPI_ISL_524448, EPI_ISL_524449 | Singapore General Hospital | Department of Microbiology | Nurdyana Abdul Rahman, Kun Lee Lim, Chenhao Li, Kian Sing Chan, Lynette Oon, Kern Rei Chng, Niranjan Nagarajan, Karrie Ko |
| EPI_ISL_524462 | Hospital Metropolitano | Instituto Adolfo Lutz, Interdiciplinary Procedures Center, Strategic Laboratory | Claudio Tavares Sacchi, Claudia Regina Gonçalves, Erica Valessa Ramos Gomes |
| EPI_ISL_524463 | Hospital Regional de Cotia | Instituto Adolfo Lutz, Interdiciplinary Procedures Center, Strategic Laboratory | Claudio Tavares Sacchi, Claudia Regina Gonçalves, Erica Valessa Ramos Gomes |
| EPI_ISL_524464 | Santa Casa de Santa Isabel | Instituto Adolfo Lutz, Interdiciplinary Procedures Center, Strategic Laboratory | Claudio Tavares Sacchi, Claudia Regina Gonçalves, Erica Valessa Ramos Gomes |
| EPI_ISL_524465 | PS Municipal Dr. Caetano Virgílio Neto | Instituto Adolfo Lutz, Interdiciplinary Procedures Center, Strategic Laboratory | Claudio Tavares Sacchi, Claudia Regina Gonçalves, Erica Valessa Ramos Gomes |
| EPI_ISL_524466 | PS Municipal Dr Lauro Ribas Braga | Instituto Adolfo Lutz, Interdiciplinary Procedures Center, Strategic Laboratory | Claudio Tavares Sacchi, Claudia Regina Gonçalves, Erica Valessa Ramos Gomes |
| EPI_ISL_524467 | Hospital Municipal Dr. Moysés Deutsch | Instituto Adolfo Lutz, Interdiciplinary Procedures Center, Strategic Laboratory | Claudio Tavares Sacchi, Claudia Regina Gonçalves, Erica Valessa Ramos Gomes |
| EPI_ISL_524468 | Hospital Municipal Vereador Jose Storopolli | Instituto Adolfo Lutz, Interdiciplinary Procedures Center, Strategic Laboratory | Claudio Tavares Sacchi, Claudia Regina Gonçalves, Erica Valessa Ramos Gomes |
| EPI_ISL_524469 | Santa Casa de Misericordia de Sao Paulo | Instituto Adolfo Lutz, Interdiciplinary Procedures Center, Strategic Laboratory | Claudio Tavares Sacchi, Claudia Regina Gonçalves, Erica Valessa Ramos Gomes |
| EPI_ISL_524470 | Hospital do Servidor Público Estadual Francisco Morato de Oliveira | Instituto Adolfo Lutz, Interdiciplinary Procedures Center, Strategic Laboratory | Claudio Tavares Sacchi, Claudia Regina Gonçalves, Erica Valessa Ramos Gomes |
| EPI_ISL_524471, EPI_ISL_524472, EPI_ISL_524473 | Laboratorio de Referencia Nacional de Virus Respiratorio. Centro Nacional de Salud Publica. Instituto Nacional de Salud Peru. | Laboratorio de Referencia Nacional de Biotecnología y Biología Molecular. Centro Nacional de Salud Publica. Instituto Nacional de Salud Peru. | Carlos Padilla Rojas, Karolyn Vega Chozo, Priscila Lope Pari, Omar Caceres Rey, Marco Galarza Perez, Maribel Huaringa Nuñez, Johanna Balbuena Torres, Henri Bailon Calderon, Nancy Rojas Serrano. |
| EPI_ISL_524474 | Department of Infectious Diseases, Cantonal Hospital Baden | Institute of Medical Virology, University of Zurich | Stefan Schmutz, Maryam Zaheri, Verena Kufner, Gabriela Ziltener, Patrick Redli, Fiona Steiner, Jon Huder, Riccarda Capaul, Andrea Zbinden, Jürg Böni, Michael Huber, Alexandra Trkola |
| EPI_ISL_524475 | Division of Infectious Diseases, University Hospital Zürich | Institute of Medical Virology, University of Zurich | Stefan Schmutz, Maryam Zaheri, Verena Kufner, Gabriela Ziltener, Patrick Redli, Fiona Steiner, Jon Huder, Riccarda Capaul, Andrea Zbinden, Jürg Böni, Michael Huber, Alexandra Trkola |
| EPI_ISL_524476 | Bülach Hospital | Institute of Medical Virology, University of Zurich | Stefan Schmutz, Maryam Zaheri, Verena Kufner, Gabriela Ziltener, Patrick Redli, Fiona Steiner, Jon Huder, Riccarda Capaul, Andrea Zbinden, Jürg Böni, Michael Huber, Alexandra Trkola |
| EPI_ISL_524477 | Division of Infectious Diseases, University Hospital Zürich | Institute of Medical Virology, University of Zurich | Stefan Schmutz, Maryam Zaheri, Verena Kufner, Gabriela Ziltener, Patrick Redli, Fiona Steiner, Jon Huder, Riccarda Capaul, Andrea Zbinden, Jürg Böni, Michael Huber, Alexandra Trkola |
| EPI_ISL_524478 | Cantonal Hospital Winterthur | Institute of Medical Virology, University of Zurich | Stefan Schmutz, Maryam Zaheri, Verena Kufner, Gabriela Ziltener, Patrick Redli, Fiona Steiner, Jon Huder, Riccarda Capaul, Andrea Zbinden, Jürg Böni, Michael Huber, Alexandra Trkola |
| EPI_ISL_524479 | Klinik Hirslanden Zürich | Institute of Medical Virology, University of Zurich | Stefan Schmutz, Maryam Zaheri, Verena Kufner, Gabriela Ziltener, Patrick Redli, Fiona Steiner, Jon Huder, Riccarda Capaul, Andrea Zbinden, Jürg Böni, Michael Huber, Alexandra Trkola |
| EPI_ISL_524480, EPI_ISL_524481 | Department of Infectious Diseases, Cantonal Hospital Baden | Institute of Medical Virology, University of Zurich | Stefan Schmutz, Maryam Zaheri, Verena Kufner, Gabriela Ziltener, Patrick Redli, Fiona Steiner, Jon Huder, Riccarda Capaul, Andrea Zbinden, Jürg Böni, Michael Huber, Alexandra Trkola |
| EPI_ISL_524482 | University Hospital Zürich | Institute of Medical Virology, University of Zurich | Verena Kufner, Maryam Zaheri, Gabriela Ziltener, Stefan Schmutz, Patrick Redli, Fiona Steiner, Jon Huder, Riccarda Capaul, Andrea Zbinden, Jürg Böni, Michael Huber, Alexandra Trkola |
| EPI_ISL_524483 | Division of Infectious Diseases, University Hospital Zürich | Institute of Medical Virology, University of Zurich | Verena Kufner, Maryam Zaheri, Gabriela Ziltener, Stefan Schmutz, Patrick Redli, Fiona Steiner, Jon Huder, Riccarda Capaul, Andrea Zbinden, Jürg Böni, Michael Huber, Alexandra Trkola |
| EPI_ISL_524484 | Hospital Schwyz | Institute of Medical Virology, University of Zurich | Verena Kufner, Maryam Zaheri, Gabriela Ziltener, Stefan Schmutz, Patrick Redli, Fiona Steiner, Jon Huder, Riccarda Capaul, Andrea Zbinden, Jürg Böni, Michael Huber, Alexandra Trkola |
| EPI_ISL_524485 | Division of Infectious Diseases, University Hospital Zürich | Institute of Medical Virology, University of Zurich | Verena Kufner, Maryam Zaheri, Gabriela Ziltener, Stefan Schmutz, Patrick Redli, Fiona Steiner, Jon Huder, Riccarda Capaul, Andrea Zbinden, Jürg Böni, Michael Huber, Alexandra Trkola |
| EPI_ISL_524486, EPI_ISL_524487, EPI_ISL_524488, EPI_ISL_524489, EPI_ISL_524490, EPI_ISL_524491, EPI_ISL_524492 | PHE South West Regional Laboratory, National Infection Service | Wellcome Sanger Institute for the COVID-19 Genomics UK Consortium | Stephanie Hutchings, Hannah Pymont, Dr Peter Muir, Barry Vipond, Rich Hopes; and Alex Alderton, Roberto Amato, Sonia Goncalves, Ewan Harrison, David K. Jackson, Ian Johnston, Dominic Kwiatkowski, Cordelia Langford, John Sillitoe on behalf of the Wellcome Sanger Institute COVID-19 Surveillance Team (http://www.sanger.ac.uk/covid-team) |
| EPI_ISL_524493 | Department of Pathology, University of Cambridge | Wellcome Sanger Institute for the COVID-19 Genomics UK Consortium | Luke W Meredith, M. Estée Török , Myra Hosmillo, William L. Hamilton, Martin D. Curran, Theresa Feltwell, Grant Hall, Anna Yakovleva, Fahad A Khokhar, Charlotte J. Houldcroft, Laura G Caller, Aminu S. Jahun, Sarah L. Caddy, Ian Goodfellow; and Alex Alderton, Roberto Amato, Sonia Goncalves, Ewan Harrison, David K. Jackson, Ian Johnston, Dominic Kwiatkowski, Cordelia Langford, John Sillitoe on behalf of the Wellcome Sanger Institute COVID-19 Surveillance Team (http://www.sanger.ac.uk/covid-team) |
| EPI_ISL_524494 | NHSGGC West of Scotland Specialist Virology Centre / MRC-University of Glasgow Centre for Virus Research | Wellcome Sanger Institute for the COVID-19 Genomics UK Consortium | Ana da Silva Filipe, Natasha Johnson, Kathy Smollett, Daniel Mair, Stephen Carmichael, Lily Tong, Jenna Nichols, Elihu Aranday-Cortes, Kirstyn Brunker, Yasmin Parr, Kyriaki Nomikou; Sarah McDonald, Marc Niebel, Patawee Asamaphan; Richard Orton, Joseph Hughes, Sreenu Vattipally, David L Robertson; Alasdair MacLean, Rory Gunson; Kathy Li, Natasha Jesudason, Rajiv Shah, James Shepherd, Antonia Ho, Alice Broos, Emma Thomson and Alex Alderton, Roberto Amato, Sonia Goncalves, Ewan Harrison, David K. Jackson, Ian Johnston, Dominic Kwiatkowski, Cordelia Langford, John Sillitoe on behalf of the Wellcome Sanger Institute COVID-19 Surveillance Team (http://www.sanger.ac.uk/covid-team) |

| | | | |
|---|---|---|---|
| EPI_ISL_524495 | Department of Pathology, University of Cambridge | Wellcome Sanger Institute for the COVID-19 Genomics UK Consortium | Luke W Meredith, M. Estée Török , Myra Hosmillo, William L. Hamilton, Martin D. Curran, Theresa Feltwell, Grant Hall, Anna Yakovleva, Fahad A Khokhar, Charlotte J. Houldcroft, Laura G Caller, Aminu S. Jahun, Sarah L. Caddy, Ian Goodfellow; and Alex Alderton, Roberto Amato, Sonia Goncalves, Ewan Harrison, David K. Jackson, Ian Johnston, Dominic Kwiatkowski, Cordelia Langford, John Sillitoe on behalf of the Wellcome Sanger Institute COVID-19 Surveillance Team (http://www.sanger.ac.uk/covid-team) |
| EPI_ISL_524496 | NHSGGC West of Scotland Specialist Virology Centre / MRC-University of Glasgow Centre for Virus Research | Wellcome Sanger Institute for the COVID-19 Genomics UK Consortium | Ana da Silva Filipe, Natasha Johnson, Kathy Smollett, Daniel Mair, Stephen Carmichael, Lily Tong, Jenna Nichols, Elihu Aranday-Cortes, Kirstyn Brunker, Yasmin Parr, Kyriaki Nomikou; Sarah McDonald, Marc Niebel, Patawee Asamaphan; Richard Orton, Joseph Hughes, Sreenu Vattipally, David L Robertson; Alasdair MacLean, Rory Gunson; Kathy Li, Natasha Jesudason, Rajiv Shah, James Shepherd, Antonia Ho, Alice Broos, Emma Thomson and Alex Alderton, Roberto Amato, Sonia Goncalves, Ewan Harrison, David K. Jackson, Ian Johnston, Dominic Kwiatkowski, Cordelia Langford, John Sillitoe on behalf of the Wellcome Sanger Institute COVID-19 Surveillance Team (http://www.sanger.ac.uk/covid-team) |
| EPI_ISL_524497, EPI_ISL_524498 | Department of Pathology, University of Cambridge | Wellcome Sanger Institute for the COVID-19 Genomics UK Consortium | Luke W Meredith, M. Estée Török , Myra Hosmillo, William L. Hamilton, Martin D. Curran, Theresa Feltwell, Grant Hall, Anna Yakovleva, Fahad A Khokhar, Charlotte J. Houldcroft, Laura G Caller, Aminu S. Jahun, Sarah L. Caddy, Ian Goodfellow; and Alex Alderton, Roberto Amato, Sonia Goncalves, Ewan Harrison, David K. Jackson, Ian Johnston, Dominic Kwiatkowski, Cordelia Langford, John Sillitoe on behalf of the Wellcome Sanger Institute COVID-19 Surveillance Team (http://www.sanger.ac.uk/covid-team) |
| EPI_ISL_524499, EPI_ISL_524500 | NHSGGC West of Scotland Specialist Virology Centre / MRC-University of Glasgow Centre for Virus Research | Wellcome Sanger Institute for the COVID-19 Genomics UK Consortium | Ana da Silva Filipe, Natasha Johnson, Kathy Smollett, Daniel Mair, Stephen Carmichael, Lily Tong, Jenna Nichols, Elihu Aranday-Cortes, Kirstyn Brunker, Yasmin Parr, Kyriaki Nomikou; Sarah McDonald, Marc Niebel, Patawee Asamaphan; Richard Orton, Joseph Hughes, Sreenu Vattipally, David L Robertson; Alasdair MacLean, Rory Gunson; Kathy Li, Natasha Jesudason, Rajiv Shah, James Shepherd, Antonia Ho, Alice Broos, Emma Thomson and Alex Alderton, Roberto Amato, Sonia Goncalves, Ewan Harrison, David K. Jackson, Ian Johnston, Dominic Kwiatkowski, Cordelia Langford, John Sillitoe on behalf of the Wellcome Sanger Institute COVID-19 Surveillance Team (http://www.sanger.ac.uk/covid-team) |
| EPI_ISL_524501, EPI_ISL_524502, EPI_ISL_524503 | Department of Pathology, University of Cambridge | Wellcome Sanger Institute for the COVID-19 Genomics UK Consortium | Luke W Meredith, M. Estée Török , Myra Hosmillo, William L. Hamilton, Martin D. Curran, Theresa Feltwell, Grant Hall, Anna Yakovleva, Fahad A Khokhar, Charlotte J. Houldcroft, Laura G Caller, Aminu S. Jahun, Sarah L. Caddy, Ian Goodfellow; and Alex Alderton, Roberto Amato, Sonia Goncalves, Ewan Harrison, David K. Jackson, Ian Johnston, Dominic Kwiatkowski, Cordelia Langford, John Sillitoe on behalf of the Wellcome Sanger Institute COVID-19 Surveillance Team (http://www.sanger.ac.uk/covid-team) |
| EPI_ISL_524504 | NHSGGC West of Scotland Specialist Virology Centre / MRC-University of Glasgow Centre for Virus Research | Wellcome Sanger Institute for the COVID-19 Genomics UK Consortium | Ana da Silva Filipe, Natasha Johnson, Kathy Smollett, Daniel Mair, Stephen Carmichael, Lily Tong, Jenna Nichols, Elihu Aranday-Cortes, Kirstyn Brunker, Yasmin Parr, Kyriaki Nomikou; Sarah McDonald, Marc Niebel, Patawee Asamaphan; Richard Orton, Joseph Hughes, Sreenu Vattipally, David L Robertson; Alasdair MacLean, Rory Gunson; Kathy Li, Natasha Jesudason, Rajiv Shah, James Shepherd, Antonia Ho, Alice Broos, Emma Thomson and Alex Alderton, Roberto Amato, Sonia Goncalves, Ewan Harrison, David K. Jackson, Ian Johnston, Dominic Kwiatkowski, Cordelia Langford, John Sillitoe on behalf of the Wellcome Sanger Institute COVID-19 Surveillance Team (http://www.sanger.ac.uk/covid-team) |
| EPI_ISL_524505, EPI_ISL_524506 | Department of Pathology, University of Cambridge | Wellcome Sanger Institute for the COVID-19 Genomics UK Consortium | Luke W Meredith, M. Estée Török , Myra Hosmillo, William L. Hamilton, Martin D. Curran, Theresa Feltwell, Grant Hall, Anna Yakovleva, Fahad A Khokhar, Charlotte J. Houldcroft, Laura G Caller, Aminu S. Jahun, Sarah L. Caddy, Ian Goodfellow; and Alex Alderton, Roberto Amato, Sonia Goncalves, Ewan Harrison, David K. Jackson, Ian Johnston, Dominic Kwiatkowski, Cordelia Langford, John Sillitoe on behalf of the Wellcome Sanger Institute COVID-19 Surveillance Team (http://www.sanger.ac.uk/covid-team) |
| EPI_ISL_524507 | NHSGGC West of Scotland Specialist Virology Centre / MRC-University of Glasgow Centre for Virus Research | Wellcome Sanger Institute for the COVID-19 Genomics UK Consortium | Ana da Silva Filipe, Natasha Johnson, Kathy Smollett, Daniel Mair, Stephen Carmichael, Lily Tong, Jenna Nichols, Elihu Aranday-Cortes, Kirstyn Brunker, Yasmin Parr, Kyriaki Nomikou; Sarah McDonald, Marc Niebel, Patawee Asamaphan; Richard Orton, Joseph Hughes, Sreenu Vattipally, David L Robertson; Alasdair MacLean, Rory Gunson; Kathy Li, Natasha Jesudason, Rajiv Shah, James Shepherd, Antonia Ho, Alice Broos, Emma Thomson and Alex Alderton, Roberto Amato, Sonia Goncalves, Ewan Harrison, David K. Jackson, Ian Johnston, Dominic Kwiatkowski, Cordelia Langford, John Sillitoe on behalf of the Wellcome Sanger Institute COVID-19 Surveillance Team (http://www.sanger.ac.uk/covid-team) |
| EPI_ISL_524508, EPI_ISL_524509 | Department of Pathology, University of Cambridge | Wellcome Sanger Institute for the COVID-19 Genomics UK Consortium | Luke W Meredith, M. Estée Török , Myra Hosmillo, William L. Hamilton, Martin D. Curran, Theresa Feltwell, Grant Hall, Anna Yakovleva, Fahad A Khokhar, Charlotte J. Houldcroft, Laura G Caller, Aminu S. Jahun, Sarah L. Caddy, Ian Goodfellow; and Alex Alderton, Roberto Amato, Sonia Goncalves, Ewan Harrison, David K. Jackson, Ian Johnston, Dominic Kwiatkowski, Cordelia Langford, John Sillitoe on behalf of the Wellcome Sanger Institute COVID-19 Surveillance Team (http://www.sanger.ac.uk/covid-team) |
| EPI_ISL_524510, EPI_ISL_524511, EPI_ISL_524512, EPI_ISL_524513, EPI_ISL_524514 | NHSGGC West of Scotland Specialist Virology Centre / MRC-University of Glasgow Centre for Virus Research | Wellcome Sanger Institute for the COVID-19 Genomics UK Consortium | Ana da Silva Filipe, Natasha Johnson, Kathy Smollett, Daniel Mair, Stephen Carmichael, Lily Tong, Jenna Nichols, Elihu Aranday-Cortes, Kirstyn Brunker, Yasmin Parr, Kyriaki Nomikou; Sarah McDonald, Marc Niebel, Patawee Asamaphan; Richard Orton, Joseph Hughes, Sreenu Vattipally, David L Robertson; Alasdair MacLean, Rory Gunson; Kathy Li, Natasha Jesudason, Rajiv Shah, James Shepherd, Antonia Ho, Alice Broos, Emma Thomson and Alex Alderton, Roberto Amato, Sonia Goncalves, Ewan Harrison, David K. Jackson, Ian Johnston, Dominic Kwiatkowski, Cordelia Langford, John Sillitoe on behalf of the Wellcome Sanger Institute COVID-19 Surveillance Team (http://www.sanger.ac.uk/covid-team) |
| EPI_ISL_524515 | Department of Pathology, University of Cambridge | Wellcome Sanger Institute for the COVID-19 Genomics UK Consortium | Luke W Meredith, M. Estée Török , Myra Hosmillo, William L. Hamilton, Martin D. Curran, Theresa Feltwell, Grant Hall, Anna Yakovleva, Fahad A Khokhar, Charlotte J. Houldcroft, Laura G Caller, Aminu S. Jahun, Sarah L. Caddy, Ian Goodfellow; and Alex Alderton, Roberto Amato, Sonia Goncalves, Ewan Harrison, David K. Jackson, Ian Johnston, Dominic Kwiatkowski, Cordelia Langford, John Sillitoe on behalf of the Wellcome Sanger Institute COVID-19 Surveillance Team (http://www.sanger.ac.uk/covid-team) |
| EPI_ISL_524516, EPI_ISL_524517, EPI_ISL_524518, EPI_ISL_524519, EPI_ISL_524520 | NHSGGC West of Scotland Specialist Virology Centre / MRC-University of Glasgow Centre for Virus Research | Wellcome Sanger Institute for the COVID-19 Genomics UK Consortium | Ana da Silva Filipe, Natasha Johnson, Kathy Smollett, Daniel Mair, Stephen Carmichael, Lily Tong, Jenna Nichols, Elihu Aranday-Cortes, Kirstyn Brunker, Yasmin Parr, Kyriaki Nomikou; Sarah McDonald, Marc Niebel, Patawee Asamaphan; Richard Orton, Joseph Hughes, Sreenu Vattipally, David L Robertson; Alasdair MacLean, Rory Gunson; Kathy Li, Natasha Jesudason, Rajiv Shah, James Shepherd, Antonia Ho, Alice Broos, Emma Thomson and Alex Alderton, Roberto Amato, Sonia Goncalves, Ewan Harrison, David K. Jackson, Ian Johnston, Dominic Kwiatkowski, Cordelia Langford, John Sillitoe on behalf of the Wellcome Sanger Institute COVID-19 Surveillance Team (http://www.sanger.ac.uk/covid-team) |
| EPI_ISL_524521, EPI_ISL_524522 | Department of Pathology, University of Cambridge | Wellcome Sanger Institute for the COVID-19 Genomics UK Consortium | Luke W Meredith, M. Estée Török , Myra Hosmillo, William L. Hamilton, Martin D. Curran, Theresa Feltwell, Grant Hall, Anna Yakovleva, Fahad A Khokhar, Charlotte J. Houldcroft, Laura G Caller, Aminu S. Jahun, Sarah L. Caddy, Ian Goodfellow; and Alex Alderton, Roberto Amato, Sonia Goncalves, Ewan Harrison, David K. Jackson, Ian Johnston, Dominic Kwiatkowski, Cordelia Langford, John Sillitoe on behalf of the Wellcome Sanger Institute COVID-19 Surveillance Team (http://www.sanger.ac.uk/covid-team) |
| EPI_ISL_524523, EPI_ISL_524524 | NHSGGC West of Scotland Specialist Virology Centre / MRC-University of Glasgow Centre for Virus Research | Wellcome Sanger Institute for the COVID-19 Genomics UK Consortium | Ana da Silva Filipe, Natasha Johnson, Kathy Smollett, Daniel Mair, Stephen Carmichael, Lily Tong, Jenna Nichols, Elihu Aranday-Cortes, Kirstyn Brunker, Yasmin Parr, Kyriaki Nomikou; Sarah McDonald, Marc Niebel, Patawee Asamaphan; Richard Orton, Joseph Hughes, Sreenu Vattipally, David L Robertson; Alasdair MacLean, Rory Gunson; Kathy Li, Natasha Jesudason, Rajiv Shah, James Shepherd, Antonia Ho, Alice Broos, Emma Thomson and Alex Alderton, Roberto Amato, Sonia Goncalves, Ewan Harrison, David K. Jackson, Ian Johnston, Dominic Kwiatkowski, Cordelia Langford, John Sillitoe on behalf of the Wellcome Sanger Institute COVID-19 Surveillance Team (http://www.sanger.ac.uk/covid-team) |
| EPI_ISL_524525 | Department of Pathology, University of Cambridge | Wellcome Sanger Institute for the COVID-19 Genomics UK Consortium | Luke W Meredith, M. Estée Török , Myra Hosmillo, William L. Hamilton, Martin D. Curran, Theresa Feltwell, Grant Hall, Anna Yakovleva, Fahad A Khokhar, Charlotte J. Houldcroft, Laura G Caller, Aminu S. Jahun, Sarah L. Caddy, Ian Goodfellow; and Alex Alderton, Roberto Amato, Sonia Goncalves, Ewan Harrison, David K. Jackson, Ian Johnston, Dominic Kwiatkowski, Cordelia Langford, John Sillitoe on behalf of the Wellcome Sanger Institute COVID-19 Surveillance Team (http://www.sanger.ac.uk/covid-team) |
| EPI_ISL_524526, EPI_ISL_524527 | NHSGGC West of Scotland Specialist Virology Centre / MRC-University of Glasgow Centre for Virus Research | Wellcome Sanger Institute for the COVID-19 Genomics UK Consortium | Ana da Silva Filipe, Natasha Johnson, Kathy Smollett, Daniel Mair, Stephen Carmichael, Lily Tong, Jenna Nichols, Elihu Aranday-Cortes, Kirstyn Brunker, Yasmin Parr, Kyriaki Nomikou; Sarah McDonald, Marc Niebel, Patawee Asamaphan; Richard Orton, Joseph Hughes, Sreenu Vattipally, David L Robertson; Alasdair MacLean, Rory Gunson; Kathy Li, Natasha Jesudason, Rajiv Shah, James Shepherd, Antonia Ho, Alice Broos, Emma Thomson and Alex Alderton, Roberto Amato, Sonia Goncalves, Ewan Harrison, David K. Jackson, Ian Johnston, Dominic Kwiatkowski, Cordelia Langford, John Sillitoe on behalf of the Wellcome Sanger Institute COVID-19 Surveillance Team (http://www.sanger.ac.uk/covid-team) |
| EPI_ISL_524528, EPI_ISL_524529, EPI_ISL_524530, EPI_ISL_524531 | Department of Pathology, University of Cambridge | Wellcome Sanger Institute for the COVID-19 Genomics UK Consortium | Luke W Meredith, M. Estée Török , Myra Hosmillo, William L. Hamilton, Martin D. Curran, Theresa Feltwell, Grant Hall, Anna Yakovleva, Fahad A Khokhar, Charlotte J. Houldcroft, Laura G Caller, Aminu S. Jahun, Sarah L. Caddy, Ian Goodfellow; and Alex Alderton, Roberto Amato, Sonia Goncalves, Ewan Harrison, David K. Jackson, Ian Johnston, Dominic Kwiatkowski, Cordelia Langford, John Sillitoe on behalf of the Wellcome Sanger Institute COVID-19 Surveillance Team (http://www.sanger.ac.uk/covid-team) |
| EPI_ISL_524532, EPI_ISL_524533, EPI_ISL_524534, EPI_ISL_524535, EPI_ISL_524536 | NHSGGC West of Scotland Specialist Virology Centre / MRC-University of Glasgow Centre for Virus Research | Wellcome Sanger Institute for the COVID-19 Genomics UK Consortium | Ana da Silva Filipe, Natasha Johnson, Kathy Smollett, Daniel Mair, Stephen Carmichael, Lily Tong, Jenna Nichols, Elihu Aranday-Cortes, Kirstyn Brunker, Yasmin Parr, Kyriaki Nomikou; Sarah McDonald, Marc Niebel, Patawee Asamaphan; Richard Orton, Joseph Hughes, Sreenu Vattipally, David L Robertson; Alasdair MacLean, Rory Gunson; Kathy Li, Natasha Jesudason, Rajiv Shah, James Shepherd, Antonia Ho, Alice Broos, Emma Thomson and Alex Alderton, Roberto Amato, Sonia Goncalves, Ewan Harrison, David K. Jackson, Ian Johnston, Dominic Kwiatkowski, Cordelia Langford, John Sillitoe on behalf of the Wellcome Sanger Institute COVID-19 Surveillance Team (http://www.sanger.ac.uk/covid-team) |
| EPI_ISL_524537, EPI_ISL_524538 | Department of Pathology, University of Cambridge | Wellcome Sanger Institute for the COVID-19 Genomics UK Consortium | Luke W Meredith, M. Estée Török , Myra Hosmillo, William L. Hamilton, Martin D. Curran, Theresa Feltwell, Grant Hall, Anna Yakovleva, Fahad A Khokhar, Charlotte J. Houldcroft, Laura G Caller, Aminu S. Jahun, Sarah L. Caddy, Ian Goodfellow; and Alex Alderton, Roberto Amato, Sonia Goncalves, Ewan Harrison, David K. Jackson, Ian Johnston, Dominic Kwiatkowski, Cordelia Langford, John Sillitoe on behalf of the Wellcome Sanger Institute COVID-19 Surveillance Team (http://www.sanger.ac.uk/covid-team) |
| EPI_ISL_524539, EPI_ISL_524540 | NHSGGC West of Scotland Specialist Virology Centre / MRC-University of Glasgow Centre for Virus Research | Wellcome Sanger Institute for the COVID-19 Genomics UK Consortium | Ana da Silva Filipe, Natasha Johnson, Kathy Smollett, Daniel Mair, Stephen Carmichael, Lily Tong, Jenna Nichols, Elihu Aranday-Cortes, Kirstyn Brunker, Yasmin Parr, Kyriaki Nomikou; Sarah McDonald, Marc Niebel, Patawee Asamaphan; Richard Orton, Joseph Hughes, Sreenu Vattipally, David L Robertson; Alasdair MacLean, Rory Gunson; Kathy Li, Natasha Jesudason, Rajiv Shah, James Shepherd, Antonia Ho, Alice Broos, Emma Thomson and Alex Alderton, Roberto Amato, Sonia Goncalves, Ewan Harrison, David K. Jackson, Ian Johnston, Dominic Kwiatkowski, Cordelia Langford, John Sillitoe on behalf of the Wellcome Sanger Institute COVID-19 Surveillance Team (http://www.sanger.ac.uk/covid-team) |
| EPI_ISL_524541 | Department of Pathology, University of Cambridge | Wellcome Sanger Institute for the COVID-19 Genomics UK Consortium | Luke W Meredith, M. Estée Török , Myra Hosmillo, William L. Hamilton, Martin D. Curran, Theresa Feltwell, Grant Hall, Anna Yakovleva, Fahad A Khokhar, Charlotte J. Houldcroft, Laura G Caller, Aminu S. Jahun, Sarah L. Caddy, Ian Goodfellow; and Alex Alderton, Roberto Amato, Sonia Goncalves, Ewan Harrison, David K. Jackson, Ian Johnston, Dominic Kwiatkowski, Cordelia Langford, John Sillitoe on behalf of the Wellcome Sanger Institute COVID-19 Surveillance Team (http://www.sanger.ac.uk/covid-team) |
| EPI_ISL_524542, EPI_ISL_524543 | NHSGGC West of Scotland Specialist Virology Centre / MRC-University of Glasgow Centre for Virus Research | Wellcome Sanger Institute for the COVID-19 Genomics UK Consortium | Ana da Silva Filipe, Natasha Johnson, Kathy Smollett, Daniel Mair, Stephen Carmichael, Lily Tong, Jenna Nichols, Elihu Aranday-Cortes, Kirstyn Brunker, Yasmin Parr, Kyriaki Nomikou; Sarah McDonald, Marc Niebel, Patawee Asamaphan; Richard Orton, Joseph Hughes, Sreenu Vattipally, David L Robertson; Alasdair MacLean, Rory Gunson; Kathy Li, Natasha Jesudason, Rajiv Shah, James Shepherd, Antonia Ho, Alice Broos, Emma Thomson and Alex Alderton, Roberto Amato, Sonia Goncalves, Ewan Harrison, David K. Jackson, Ian Johnston, Dominic Kwiatkowski, Cordelia Langford, John Sillitoe on behalf of the Wellcome Sanger Institute COVID-19 Surveillance Team (http://www.sanger.ac.uk/covid-team) |
| EPI_ISL_524544 | Department of Pathology, University of Cambridge | Wellcome Sanger Institute for the COVID-19 Genomics UK Consortium | Luke W Meredith, M. Estée Török , Myra Hosmillo, William L. Hamilton, Martin D. Curran, Theresa Feltwell, Grant Hall, Anna Yakovleva, Fahad A Khokhar, Charlotte J. Houldcroft, Laura G Caller, Aminu S. Jahun, Sarah L. Caddy, Ian Goodfellow; and Alex Alderton, Roberto Amato, Sonia Goncalves, Ewan Harrison, David K. Jackson, Ian Johnston, Dominic Kwiatkowski, Cordelia Langford, John Sillitoe on behalf of the Wellcome Sanger Institute COVID-19 Surveillance Team (http://www.sanger.ac.uk/covid-team) |
| EPI_ISL_524545 | NHSGGC West of Scotland Specialist Virology Centre / MRC-University of Glasgow Centre for Virus Research | Wellcome Sanger Institute for the COVID-19 Genomics UK Consortium | Ana da Silva Filipe, Natasha Johnson, Kathy Smollett, Daniel Mair, Stephen Carmichael, Lily Tong, Jenna Nichols, Elihu Aranday-Cortes, Kirstyn Brunker, Yasmin Parr, Kyriaki Nomikou; Sarah McDonald, Marc Niebel, Patawee Asamaphan; Richard Orton, Joseph Hughes, Sreenu Vattipally, David L Robertson; Alasdair MacLean, Rory Gunson; Kathy Li, Natasha Jesudason, Rajiv Shah, James Shepherd, Antonia Ho, Alice Broos, Emma Thomson and Alex Alderton, Roberto Amato, Sonia Goncalves, Ewan Harrison, David K. Jackson, Ian Johnston, Dominic Kwiatkowski, Cordelia Langford, John Sillitoe on behalf of the Wellcome Sanger Institute COVID-19 Surveillance Team (http://www.sanger.ac.uk/covid-team) |
| EPI_ISL_524546 | Department of Pathology, University of Cambridge | Wellcome Sanger Institute for the COVID-19 Genomics UK Consortium | Luke W Meredith, M. Estée Török , Myra Hosmillo, William L. Hamilton, Martin D. Curran, Theresa Feltwell, Grant Hall, Anna Yakovleva, Fahad A Khokhar, Charlotte J. Houldcroft, Laura G Caller, Aminu S. Jahun, Sarah L. Caddy, Ian Goodfellow; and Alex Alderton, Roberto Amato, Sonia Goncalves, Ewan Harrison, David K. Jackson, Ian Johnston, Dominic Kwiatkowski, Cordelia Langford, John Sillitoe on behalf of the Wellcome Sanger Institute COVID-19 Surveillance Team (http://www.sanger.ac.uk/covid-team) |

| | | | |
|---|---|---|---|
| EPI_ISL_524547 | NHSGGC West of Scotland Specialist Virology Centre / MRC-University of Glasgow Centre for Virus Research | Wellcome Sanger Institute for the COVID-19 Genomics UK Consortium | Ana da Silva Filipe, Natasha Johnson, Kathy Smollett, Daniel Mair, Stephen Carmichael, Lily Tong, Jenna Nichols, Elihu Aranday-Cortes, Kirstyn Brunker, Yasmin Parr, Kyriaki Nomikou; Sarah McDonald, Marc Niebel, Patawee Asamaphan; Richard Orton, Joseph Hughes, Sreenu Vattipally, David L Robertson; Alasdair MacLean, Rory Gunson; Kathy Li, Natasha Jesudason, Rajiv Shah, James Shepherd, Antonia Ho, Alice Broos, Emma Thomson and Alex Alderton, Roberto Amato, Sonia Goncalves, Ewan Harrison, David K. Jackson, Ian Johnston, Dominic Kwiatkowski, Cordelia Langford, John Sillitoe on behalf of the Wellcome Sanger Institute COVID-19 Surveillance Team (http://www.sanger.ac.uk/covid-team) |
| EPI_ISL_524548 | Department of Pathology, University of Cambridge | Wellcome Sanger Institute for the COVID-19 Genomics UK Consortium | Luke W Meredith, M. Estée Török , Myra Hosmillo, William L. Hamilton, Martin D. Curran, Theresa Feltwell, Grant Hall, Anna Yakovleva, Fahad A Khokhar, Charlotte J. Houldcroft, Laura G Caller, Aminu S. Jahun, Sarah L. Caddy, Ian Goodfellow; and Alex Alderton, Roberto Amato, Sonia Goncalves, Ewan Harrison, David K. Jackson, Ian Johnston, Dominic Kwiatkowski, Cordelia Langford, John Sillitoe on behalf of the Wellcome Sanger Institute COVID-19 Surveillance Team (http://www.sanger.ac.uk/covid-team) |
| EPI_ISL_524549, EPI_ISL_524550 | NHSGGC West of Scotland Specialist Virology Centre / MRC-University of Glasgow Centre for Virus Research | Wellcome Sanger Institute for the COVID-19 Genomics UK Consortium | Ana da Silva Filipe, Natasha Johnson, Kathy Smollett, Daniel Mair, Stephen Carmichael, Lily Tong, Jenna Nichols, Elihu Aranday-Cortes, Kirstyn Brunker, Yasmin Parr, Kyriaki Nomikou; Sarah McDonald, Marc Niebel, Patawee Asamaphan; Richard Orton, Joseph Hughes, Sreenu Vattipally, David L Robertson; Alasdair MacLean, Rory Gunson; Kathy Li, Natasha Jesudason, Rajiv Shah, James Shepherd, Antonia Ho, Alice Broos, Emma Thomson and Alex Alderton, Roberto Amato, Sonia Goncalves, Ewan Harrison, David K. Jackson, Ian Johnston, Dominic Kwiatkowski, Cordelia Langford, John Sillitoe on behalf of the Wellcome Sanger Institute COVID-19 Surveillance Team (http://www.sanger.ac.uk/covid-team) |
| EPI_ISL_524551 | Department of Pathology, University of Cambridge | Wellcome Sanger Institute for the COVID-19 Genomics UK Consortium | Luke W Meredith, M. Estée Török , Myra Hosmillo, William L. Hamilton, Martin D. Curran, Theresa Feltwell, Grant Hall, Anna Yakovleva, Fahad A Khokhar, Charlotte J. Houldcroft, Laura G Caller, Aminu S. Jahun, Sarah L. Caddy, Ian Goodfellow; and Alex Alderton, Roberto Amato, Sonia Goncalves, Ewan Harrison, David K. Jackson, Ian Johnston, Dominic Kwiatkowski, Cordelia Langford, John Sillitoe on behalf of the Wellcome Sanger Institute COVID-19 Surveillance Team (http://www.sanger.ac.uk/covid-team) |
| EPI_ISL_524552 | NHSGGC West of Scotland Specialist Virology Centre / MRC-University of Glasgow Centre for Virus Research | Wellcome Sanger Institute for the COVID-19 Genomics UK Consortium | Ana da Silva Filipe, Natasha Johnson, Kathy Smollett, Daniel Mair, Stephen Carmichael, Lily Tong, Jenna Nichols, Elihu Aranday-Cortes, Kirstyn Brunker, Yasmin Parr, Kyriaki Nomikou; Sarah McDonald, Marc Niebel, Patawee Asamaphan; Richard Orton, Joseph Hughes, Sreenu Vattipally, David L Robertson; Alasdair MacLean, Rory Gunson; Kathy Li, Natasha Jesudason, Rajiv Shah, James Shepherd, Antonia Ho, Alice Broos, Emma Thomson and Alex Alderton, Roberto Amato, Sonia Goncalves, Ewan Harrison, David K. Jackson, Ian Johnston, Dominic Kwiatkowski, Cordelia Langford, John Sillitoe on behalf of the Wellcome Sanger Institute COVID-19 Surveillance Team (http://www.sanger.ac.uk/covid-team) |
| EPI_ISL_524553 | Department of Pathology, University of Cambridge | Wellcome Sanger Institute for the COVID-19 Genomics UK Consortium | Luke W Meredith, M. Estée Török , Myra Hosmillo, William L. Hamilton, Martin D. Curran, Theresa Feltwell, Grant Hall, Anna Yakovleva, Fahad A Khokhar, Charlotte J. Houldcroft, Laura G Caller, Aminu S. Jahun, Sarah L. Caddy, Ian Goodfellow; and Alex Alderton, Roberto Amato, Sonia Goncalves, Ewan Harrison, David K. Jackson, Ian Johnston, Dominic Kwiatkowski, Cordelia Langford, John Sillitoe on behalf of the Wellcome Sanger Institute COVID-19 Surveillance Team (http://www.sanger.ac.uk/covid-team) |
| EPI_ISL_524554, EPI_ISL_524555, EPI_ISL_524556, EPI_ISL_524557 | NHSGGC West of Scotland Specialist Virology Centre / MRC-University of Glasgow Centre for Virus Research | Wellcome Sanger Institute for the COVID-19 Genomics UK Consortium | Ana da Silva Filipe, Natasha Johnson, Kathy Smollett, Daniel Mair, Stephen Carmichael, Lily Tong, Jenna Nichols, Elihu Aranday-Cortes, Kirstyn Brunker, Yasmin Parr, Kyriaki Nomikou; Sarah McDonald, Marc Niebel, Patawee Asamaphan; Richard Orton, Joseph Hughes, Sreenu Vattipally, David L Robertson; Alasdair MacLean, Rory Gunson; Kathy Li, Natasha Jesudason, Rajiv Shah, James Shepherd, Antonia Ho, Alice Broos, Emma Thomson and Alex Alderton, Roberto Amato, Sonia Goncalves, Ewan Harrison, David K. Jackson, Ian Johnston, Dominic Kwiatkowski, Cordelia Langford, John Sillitoe on behalf of the Wellcome Sanger Institute COVID-19 Surveillance Team (http://www.sanger.ac.uk/covid-team) |
| EPI_ISL_524558 | Department of Pathology, University of Cambridge | Wellcome Sanger Institute for the COVID-19 Genomics UK Consortium | Luke W Meredith, M. Estée Török , Myra Hosmillo, William L. Hamilton, Martin D. Curran, Theresa Feltwell, Grant Hall, Anna Yakovleva, Fahad A Khokhar, Charlotte J. Houldcroft, Laura G Caller, Aminu S. Jahun, Sarah L. Caddy, Ian Goodfellow; and Alex Alderton, Roberto Amato, Sonia Goncalves, Ewan Harrison, David K. Jackson, Ian Johnston, Dominic Kwiatkowski, Cordelia Langford, John Sillitoe on behalf of the Wellcome Sanger Institute COVID-19 Surveillance Team (http://www.sanger.ac.uk/covid-team) |
| EPI_ISL_524559 | NHSGGC West of Scotland Specialist Virology Centre / MRC-University of Glasgow Centre for Virus Research | Wellcome Sanger Institute for the COVID-19 Genomics UK Consortium | Ana da Silva Filipe, Natasha Johnson, Kathy Smollett, Daniel Mair, Stephen Carmichael, Lily Tong, Jenna Nichols, Elihu Aranday-Cortes, Kirstyn Brunker, Yasmin Parr, Kyriaki Nomikou; Sarah McDonald, Marc Niebel, Patawee Asamaphan; Richard Orton, Joseph Hughes, Sreenu Vattipally, David L Robertson; Alasdair MacLean, Rory Gunson; Kathy Li, Natasha Jesudason, Rajiv Shah, James Shepherd, Antonia Ho, Alice Broos, Emma Thomson and Alex Alderton, Roberto Amato, Sonia Goncalves, Ewan Harrison, David K. Jackson, Ian Johnston, Dominic Kwiatkowski, Cordelia Langford, John Sillitoe on behalf of the Wellcome Sanger Institute COVID-19 Surveillance Team (http://www.sanger.ac.uk/covid-team) |
| EPI_ISL_524560 | Department of Pathology, University of Cambridge | Wellcome Sanger Institute for the COVID-19 Genomics UK Consortium | Luke W Meredith, M. Estée Török , Myra Hosmillo, William L. Hamilton, Martin D. Curran, Theresa Feltwell, Grant Hall, Anna Yakovleva, Fahad A Khokhar, Charlotte J. Houldcroft, Laura G Caller, Aminu S. Jahun, Sarah L. Caddy, Ian Goodfellow; and Alex Alderton, Roberto Amato, Sonia Goncalves, Ewan Harrison, David K. Jackson, Ian Johnston, Dominic Kwiatkowski, Cordelia Langford, John Sillitoe on behalf of the Wellcome Sanger Institute COVID-19 Surveillance Team (http://www.sanger.ac.uk/covid-team) |
| EPI_ISL_524561, EPI_ISL_524562, EPI_ISL_524563 | NHSGGC West of Scotland Specialist Virology Centre / MRC-University of Glasgow Centre for Virus Research | Wellcome Sanger Institute for the COVID-19 Genomics UK Consortium | Ana da Silva Filipe, Natasha Johnson, Kathy Smollett, Daniel Mair, Stephen Carmichael, Lily Tong, Jenna Nichols, Elihu Aranday-Cortes, Kirstyn Brunker, Yasmin Parr, Kyriaki Nomikou; Sarah McDonald, Marc Niebel, Patawee Asamaphan; Richard Orton, Joseph Hughes, Sreenu Vattipally, David L Robertson; Alasdair MacLean, Rory Gunson; Kathy Li, Natasha Jesudason, Rajiv Shah, James Shepherd, Antonia Ho, Alice Broos, Emma Thomson and Alex Alderton, Roberto Amato, Sonia Goncalves, Ewan Harrison, David K. Jackson, Ian Johnston, Dominic Kwiatkowski, Cordelia Langford, John Sillitoe on behalf of the Wellcome Sanger Institute COVID-19 Surveillance Team (http://www.sanger.ac.uk/covid-team) |
| EPI_ISL_524564 | Department of Pathology, University of Cambridge | Wellcome Sanger Institute for the COVID-19 Genomics UK Consortium | Luke W Meredith, M. Estée Török , Myra Hosmillo, William L. Hamilton, Martin D. Curran, Theresa Feltwell, Grant Hall, Anna Yakovleva, Fahad A Khokhar, Charlotte J. Houldcroft, Laura G Caller, Aminu S. Jahun, Sarah L. Caddy, Ian Goodfellow; and Alex Alderton, Roberto Amato, Sonia Goncalves, Ewan Harrison, David K. Jackson, Ian Johnston, Dominic Kwiatkowski, Cordelia Langford, John Sillitoe on behalf of the Wellcome Sanger Institute COVID-19 Surveillance Team (http://www.sanger.ac.uk/covid-team) |
| EPI_ISL_524565, EPI_ISL_524566, EPI_ISL_524567 | NHSGGC West of Scotland Specialist Virology Centre / MRC-University of Glasgow Centre for Virus Research | Wellcome Sanger Institute for the COVID-19 Genomics UK Consortium | Ana da Silva Filipe, Natasha Johnson, Kathy Smollett, Daniel Mair, Stephen Carmichael, Lily Tong, Jenna Nichols, Elihu Aranday-Cortes, Kirstyn Brunker, Yasmin Parr, Kyriaki Nomikou; Sarah McDonald, Marc Niebel, Patawee Asamaphan; Richard Orton, Joseph Hughes, Sreenu Vattipally, David L Robertson; Alasdair MacLean, Rory Gunson; Kathy Li, Natasha Jesudason, Rajiv Shah, James Shepherd, Antonia Ho, Alice Broos, Emma Thomson and Alex Alderton, Roberto Amato, Sonia Goncalves, Ewan Harrison, David K. Jackson, Ian Johnston, Dominic Kwiatkowski, Cordelia Langford, John Sillitoe on behalf of the Wellcome Sanger Institute COVID-19 Surveillance Team (http://www.sanger.ac.uk/covid-team) |
| EPI_ISL_524568 | Department of Pathology, University of Cambridge | Wellcome Sanger Institute for the COVID-19 Genomics UK Consortium | Luke W Meredith, M. Estée Török , Myra Hosmillo, William L. Hamilton, Martin D. Curran, Theresa Feltwell, Grant Hall, Anna Yakovleva, Fahad A Khokhar, Charlotte J. Houldcroft, Laura G Caller, Aminu S. Jahun, Sarah L. Caddy, Ian Goodfellow; and Alex Alderton, Roberto Amato, Sonia Goncalves, Ewan Harrison, David K. Jackson, Ian Johnston, Dominic Kwiatkowski, Cordelia Langford, John Sillitoe on behalf of the Wellcome Sanger Institute COVID-19 Surveillance Team (http://www.sanger.ac.uk/covid-team) |
| EPI_ISL_524569 | NHSGGC West of Scotland Specialist Virology Centre / MRC-University of Glasgow Centre for Virus Research | Wellcome Sanger Institute for the COVID-19 Genomics UK Consortium | Ana da Silva Filipe, Natasha Johnson, Kathy Smollett, Daniel Mair, Stephen Carmichael, Lily Tong, Jenna Nichols, Elihu Aranday-Cortes, Kirstyn Brunker, Yasmin Parr, Kyriaki Nomikou; Sarah McDonald, Marc Niebel, Patawee Asamaphan; Richard Orton, Joseph Hughes, Sreenu Vattipally, David L Robertson; Alasdair MacLean, Rory Gunson; Kathy Li, Natasha Jesudason, Rajiv Shah, James Shepherd, Antonia Ho, Alice Broos, Emma Thomson and Alex Alderton, Roberto Amato, Sonia Goncalves, Ewan Harrison, David K. Jackson, Ian Johnston, Dominic Kwiatkowski, Cordelia Langford, John Sillitoe on behalf of the Wellcome Sanger Institute COVID-19 Surveillance Team (http://www.sanger.ac.uk/covid-team) |
| EPI_ISL_524570 | Department of Pathology, University of Cambridge | Wellcome Sanger Institute for the COVID-19 Genomics UK Consortium | Luke W Meredith, M. Estée Török , Myra Hosmillo, William L. Hamilton, Martin D. Curran, Theresa Feltwell, Grant Hall, Anna Yakovleva, Fahad A Khokhar, Charlotte J. Houldcroft, Laura G Caller, Aminu S. Jahun, Sarah L. Caddy, Ian Goodfellow; and Alex Alderton, Roberto Amato, Sonia Goncalves, Ewan Harrison, David K. Jackson, Ian Johnston, Dominic Kwiatkowski, Cordelia Langford, John Sillitoe on behalf of the Wellcome Sanger Institute COVID-19 Surveillance Team (http://www.sanger.ac.uk/covid-team) |
| EPI_ISL_524571, EPI_ISL_524572 | NHSGGC West of Scotland Specialist Virology Centre / MRC-University of Glasgow Centre for Virus Research | Wellcome Sanger Institute for the COVID-19 Genomics UK Consortium | Ana da Silva Filipe, Natasha Johnson, Kathy Smollett, Daniel Mair, Stephen Carmichael, Lily Tong, Jenna Nichols, Elihu Aranday-Cortes, Kirstyn Brunker, Yasmin Parr, Kyriaki Nomikou; Sarah McDonald, Marc Niebel, Patawee Asamaphan; Richard Orton, Joseph Hughes, Sreenu Vattipally, David L Robertson; Alasdair MacLean, Rory Gunson; Kathy Li, Natasha Jesudason, Rajiv Shah, James Shepherd, Antonia Ho, Alice Broos, Emma Thomson and Alex Alderton, Roberto Amato, Sonia Goncalves, Ewan Harrison, David K. Jackson, Ian Johnston, Dominic Kwiatkowski, Cordelia Langford, John Sillitoe on behalf of the Wellcome Sanger Institute COVID-19 Surveillance Team (http://www.sanger.ac.uk/covid-team) |
| EPI_ISL_524573, EPI_ISL_524574, EPI_ISL_524575, EPI_ISL_524576, EPI_ISL_524577, EPI_ISL_524578, EPI_ISL_524579, EPI_ISL_524580, EPI_ISL_524581, EPI_ISL_524582, EPI_ISL_524583, EPI_ISL_524584, EPI_ISL_524585, EPI_ISL_524586, EPI_ISL_524587, EPI_ISL_524588, EPI_ISL_524589, EPI_ISL_524591, EPI_ISL_524592, EPI_ISL_524593, EPI_ISL_524594, EPI_ISL_524596, EPI_ISL_524597, EPI_ISL_524598, EPI_ISL_524599, EPI_ISL_524600, EPI_ISL_524601, EPI_ISL_524602, EPI_ISL_524603, EPI_ISL_524604, EPI_ISL_524605, EPI_ISL_524606, EPI_ISL_524607, EPI_ISL_524608, EPI_ISL_524609, EPI_ISL_524610, EPI_ISL_524611, EPI_ISL_524612, EPI_ISL_524613, EPI_ISL_524614, EPI_ISL_524615, EPI_ISL_524616, EPI_ISL_524617, EPI_ISL_524618 | | | |
| see above | Department of Pathology, University of Cambridge | Wellcome Sanger Institute for the COVID-19 Genomics UK Consortium | Luke W Meredith, M. Estée Török , Myra Hosmillo, William L. Hamilton, Martin D. Curran, Theresa Feltwell, Grant Hall, Anna Yakovleva, Fahad A Khokhar, Charlotte J. Houldcroft, Laura G Caller, Aminu S. Jahun, Sarah L. Caddy, Ian Goodfellow; and Alex Alderton, Roberto Amato, Sonia Goncalves, Ewan Harrison, David K. Jackson, Ian Johnston, Dominic Kwiatkowski, Cordelia Langford, John Sillitoe on behalf of the Wellcome Sanger Institute COVID-19 Surveillance Team (http://www.sanger.ac.uk/covid-team) |
| EPI_ISL_524619, EPI_ISL_524620, EPI_ISL_524621, EPI_ISL_524622, EPI_ISL_524623, EPI_ISL_524624, EPI_ISL_524625, EPI_ISL_524626, EPI_ISL_524627, EPI_ISL_524628, EPI_ISL_524629, EPI_ISL_524630, EPI_ISL_524631, EPI_ISL_524632, EPI_ISL_524633, EPI_ISL_524634, EPI_ISL_524635, EPI_ISL_524636, EPI_ISL_524637, EPI_ISL_524638, EPI_ISL_524639, EPI_ISL_524640, EPI_ISL_524641, EPI_ISL_524642, EPI_ISL_524643, EPI_ISL_524644, EPI_ISL_524645, EPI_ISL_524646, EPI_ISL_524647, EPI_ISL_524648, EPI_ISL_524649, EPI_ISL_524650, EPI_ISL_524651, EPI_ISL_524652, EPI_ISL_524653, EPI_ISL_524654, EPI_ISL_524655, EPI_ISL_524656, EPI_ISL_524657, EPI_ISL_524658, EPI_ISL_524659, EPI_ISL_524660, EPI_ISL_524661, EPI_ISL_524662, EPI_ISL_524663, EPI_ISL_524664, EPI_ISL_524665, EPI_ISL_524666, EPI_ISL_524667, EPI_ISL_524668, EPI_ISL_524669, EPI_ISL_524670, EPI_ISL_524671, EPI_ISL_524672, EPI_ISL_524673, EPI_ISL_524674, EPI_ISL_524675, EPI_ISL_524676, EPI_ISL_524677, EPI_ISL_524678, EPI_ISL_524679, EPI_ISL_524680, EPI_ISL_524681, EPI_ISL_524682, EPI_ISL_524683, EPI_ISL_524684, EPI_ISL_524685, EPI_ISL_524686, EPI_ISL_524687, EPI_ISL_524688, EPI_ISL_524689, EPI_ISL_524690, EPI_ISL_524691, EPI_ISL_524692, EPI_ISL_524693, EPI_ISL_524694, EPI_ISL_524695, EPI_ISL_524696, EPI_ISL_524697, EPI_ISL_524698, EPI_ISL_524699, EPI_ISL_524700, EPI_ISL_524701, EPI_ISL_524702, EPI_ISL_524703, EPI_ISL_524704, EPI_ISL_524705, EPI_ISL_524706, EPI_ISL_524707, EPI_ISL_524708, EPI_ISL_524709, EPI_ISL_524710, EPI_ISL_524711, EPI_ISL_524712 | | | |
| see above | North West London Pathology, Imperial College Healthcare NHS Trust | Wellcome Sanger Institute for the COVID-19 Genomics UK Consortium | Ling Li, Paul Randell, David Muir, Frankie Bolt, Alison Holmes, James Price, Aileen Rowan, Graham Taylor, Anjna Badhan, Carolina Herrera and Alex Alderton, Roberto Amato, Sonia Goncalves, Ewan Harrison, David K. Jackson, Ian Johnston, Dominic Kwiatkowski, Cordelia Langford, John Sillitoe on behalf of the Wellcome Sanger Institute COVID-19 Surveillance Team (http://www.sanger.ac.uk/covid-team) |
| EPI_ISL_524713 | B.J. Medical College and Civil hospital, Ahmedabad | Gujarat Biotechnology Research Centre | Apurvasinh Puvar, Janvi Raval, Zarna Patel, Monika Gandhi, Pinal Trivedi, Maharshi Pandya, Nidhi Patel, Nitin Savaliya, Raghawendra Kumar, Dinesh Kumar, Zuber Saiyed, Komal Patel, Labdhi Pandya, Afzal Ansari, Nikha Trivedi, Pranay Shah, Kamlesh J Upadhyay, Sanjay Kapadia, R D Dixit, A M Kadri, Harsh Bakshi, Chaitanya Joshi, Madhvi Joshi |
| EPI_ISL_524714 | B.J. Medical College and Civil hospital, Ahmedabad | Gujarat Biotechnology Research Centre | Janvi Raval, Zarna Patel, Monika Gandhi, Pinal Trivedi, Maharshi Pandya, Nidhi Patel, Nitin Savaliya, Raghawendra Kumar, Dinesh Kumar, Zuber Saiyed, Komal Patel, Labdhi Pandya, Afzal Ansari, Nikha Trivedi, Pranay Shah, Kamlesh J Upadhyay, Sanjay Kapadia, Apurvasinh Puvar, R D Dixit, A M Kadri, Harsh Bakshi, Chaitanya Joshi, Madhvi Joshi |
| EPI_ISL_524715 | B.J. Medical College and Civil hospital, Ahmedabad | Gujarat Biotechnology Research Centre | Zarna Patel, Monika Gandhi, Pinal Trivedi, Maharshi Pandya, Nidhi Patel, Nitin Savaliya, Raghawendra Kumar, Dinesh Kumar, Zuber Saiyed, Komal Patel, Labdhi Pandya, Afzal Ansari, Nikha Trivedi, Pranay Shah, Kamlesh J Upadhyay, Sanjay Kapadia, Apurvasinh Puvar, Janvi Raval, R D Dixit, A M Kadri, Harsh Bakshi, Chaitanya Joshi, Madhvi Joshi |
| EPI_ISL_524716 | B.J. Medical College and Civil hospital, Ahmedabad | Gujarat Biotechnology Research Centre | Monika Gandhi, Pinal Trivedi, Maharshi Pandya, Nidhi Patel, Nitin Savaliya, Raghawendra Kumar, Dinesh Kumar, Zuber Saiyed, Komal Patel, Labdhi Pandya, Afzal Ansari, Nikha Trivedi, Pranay Shah, Kamlesh J Upadhyay, Sanjay Kapadia, Apurvasinh Puvar, Janvi Raval, Zarna Patel, R D Dixit, A M Kadri, Harsh Bakshi, Chaitanya Joshi, Madhvi Joshi |
| EPI_ISL_524717 | B.J. Medical College and Civil hospital, Ahmedabad | Gujarat Biotechnology Research Centre | Pinal Trivedi, Maharshi Pandya, Nidhi Patel, Nitin Savaliya, Raghawendra Kumar, Dinesh Kumar, Zuber Saiyed, Komal Patel, Labdhi Pandya, Afzal Ansari, Nikha Trivedi, Pranay Shah, Kamlesh J Upadhyay, Sanjay Kapadia, Apurvasinh Puvar, Janvi Raval, Zarna Patel, Monika Gandhi, R D Dixit, A M Kadri, Harsh Bakshi, Chaitanya Joshi, Madhvi Joshi |
| EPI_ISL_524718 | B.J. Medical College and Civil hospital, Ahmedabad | Gujarat Biotechnology Research Centre | Maharshi Pandya, Nidhi Patel, Nitin Savaliya, Raghawendra Kumar, Dinesh Kumar, Zuber Saiyed, Komal Patel, Labdhi Pandya, Afzal Ansari, Nikha Trivedi, Pranay Shah, Kamlesh J Upadhyay, Sanjay Kapadia, Apurvasinh Puvar, Janvi Raval, Zarna Patel, Monika Gandhi, Pinal Trivedi, R D Dixit, A M Kadri, Harsh Bakshi, Chaitanya Joshi, Madhvi Joshi |
| EPI_ISL_524719 | B.J. Medical College and Civil hospital, Ahmedabad | Gujarat Biotechnology Research Centre | Nidhi Patel, Nitin Savaliya, Raghawendra Kumar, Dinesh Kumar, Zuber Saiyed, Komal Patel, Labdhi Pandya, Afzal Ansari, Nikha Trivedi, Pranay Shah, Kamlesh J Upadhyay, Sanjay Kapadia, Apurvasinh Puvar, Janvi Raval, Zarna Patel, Monika Gandhi, Pinal Trivedi, Maharshi Pandya, R D Dixit, A M Kadri, Harsh Bakshi, Chaitanya Joshi, Madhvi Joshi |
| EPI_ISL_524720 | B.J. Medical College and Civil hospital, Ahmedabad | Gujarat Biotechnology Research Centre | Nitin Savaliya, Raghawendra Kumar, Dinesh Kumar, Zuber Saiyed, Komal Patel, Labdhi Pandya, Afzal Ansari, Nikha Trivedi, Pranay Shah, Kamlesh J Upadhyay, Sanjay |

| | | | |
|---|---|---|---|
| | | | Kapadia, Apurvasinh Puvar, Janvi Raval, Zarna Patel, Monika Gandhi, Pinal Trivedi, Maharshi Pandya, Nidhi Patel, R D Dixit, A M Kadri, Harsh Bakshi, Chaitanya Joshi, Madhvi Joshi |
| EPI_ISL_524721 | B.J. Medical College and Civil hospital, Ahmedabad | Gujarat Biotechnology Research Centre | Raghawendra Kumar, Dinesh Kumar, Zuber Saiyed, Komal Patel, Labdhi Pandya, Afzal Ansari, Nikha Trivedi, Pranay Shah, Kamlesh J Upadhyay, Sanjay Kapadia, Apurvasinh Puvar, Janvi Raval, Zarna Patel, Monika Gandhi, Pinal Trivedi, Maharshi Pandya, Nidhi Patel, Nitin Savaliya, R D Dixit, A M Kadri, Harsh Bakshi, Chaitanya Joshi, Madhvi Joshi |
| EPI_ISL_524722 | B.J. Medical College and Civil hospital, Ahmedabad | Gujarat Biotechnology Research Centre | Dinesh Kumar, Zuber Saiyed, Komal Patel, Labdhi Pandya, Afzal Ansari, Nikha Trivedi, Pranay Shah, Kamlesh J Upadhyay, Sanjay Kapadia, Apurvasinh Puvar, Janvi Raval, Zarna Patel, Monika Gandhi, Pinal Trivedi, Maharshi Pandya, Nidhi Patel, Nitin Savaliya, Raghawendra Kumar, R D Dixit, A M Kadri, Harsh Bakshi, Chaitanya Joshi, Madhvi Joshi |
| EPI_ISL_524723 | B.J. Medical College and Civil hospital, Ahmedabad | Gujarat Biotechnology Research Centre | Zuber Saiyed, Komal Patel, Labdhi Pandya, Afzal Ansari, Nikha Trivedi, Pranay Shah, Kamlesh J Upadhyay, Sanjay Kapadia, Apurvasinh Puvar, Janvi Raval, Zarna Patel, Monika Gandhi, Pinal Trivedi, Maharshi Pandya, Nidhi Patel, Nitin Savaliya, Raghawendra Kumar, Dinesh Kumar, R D Dixit, A M Kadri, Harsh Bakshi, Chaitanya Joshi, Madhvi Joshi |
| EPI_ISL_524724 | B.J. Medical College and Civil hospital, Ahmedabad | Gujarat Biotechnology Research Centre | Komal Patel, Labdhi Pandya, Afzal Ansari, Nikha Trivedi, Pranay Shah, Kamlesh J Upadhyay, Sanjay Kapadia, Apurvasinh Puvar, Janvi Raval, Zarna Patel, Monika Gandhi, Pinal Trivedi, Maharshi Pandya, Nidhi Patel, Nitin Savaliya, Raghawendra Kumar, Dinesh Kumar, Zuber Saiyed, R D Dixit, A M Kadri, Harsh Bakshi, Chaitanya Joshi, Madhvi Joshi |
| EPI_ISL_524725 | B.J. Medical College and Civil hospital, Ahmedabad | Gujarat Biotechnology Research Centre | Labdhi Pandya, Afzal Ansari, Nikha Trivedi, Pranay Shah, Kamlesh J Upadhyay, Sanjay Kapadia, Apurvasinh Puvar, Janvi Raval, Zarna Patel, Monika Gandhi, Pinal Trivedi, Maharshi Pandya, Nidhi Patel, Nitin Savaliya, Raghawendra Kumar, Dinesh Kumar, Zuber Saiyed, Komal Patel, R D Dixit, A M Kadri, Harsh Bakshi, Chaitanya Joshi, Madhvi Joshi |
| EPI_ISL_524726 | B.J. Medical College and Civil hospital, Ahmedabad | Gujarat Biotechnology Research Centre | Afzal Ansari, Nikha Trivedi, Pranay Shah, Kamlesh J Upadhyay, Sanjay Kapadia, Apurvasinh Puvar, Janvi Raval, Zarna Patel, Monika Gandhi, Pinal Trivedi, Maharshi Pandya, Nidhi Patel, Nitin Savaliya, Raghawendra Kumar, Dinesh Kumar, Zuber Saiyed, Komal Patel, Labdhi Pandya, R D Dixit, A M Kadri, Harsh Bakshi, Chaitanya Joshi, Madhvi Joshi |
| EPI_ISL_524727 | B.J. Medical College and Civil hospital, Ahmedabad | Gujarat Biotechnology Research Centre | Nikha Trivedi, Pranay Shah, Kamlesh J Upadhyay, Sanjay Kapadia, Apurvasinh Puvar, Janvi Raval, Zarna Patel, Monika Gandhi, Pinal Trivedi, Maharshi Pandya, Nidhi Patel, Nitin Savaliya, Raghawendra Kumar, Dinesh Kumar, Zuber Saiyed, Komal Patel, Labdhi Pandya, Afzal Ansari, R D Dixit, A M Kadri, Harsh Bakshi, Chaitanya Joshi, Madhvi Joshi |
| EPI_ISL_524728 | B.J. Medical College and Civil hospital, Ahmedabad | Gujarat Biotechnology Research Centre | Pranay Shah, Kamlesh J Upadhyay, Sanjay Kapadia, Apurvasinh Puvar, Janvi Raval, Zarna Patel, Monika Gandhi, Pinal Trivedi, Maharshi Pandya, Nidhi Patel, Nitin Savaliya, Raghawendra Kumar, Dinesh Kumar, Zuber Saiyed, Komal Patel, Labdhi Pandya, Afzal Ansari, Nikha Trivedi, R D Dixit, A M Kadri, Harsh Bakshi, Chaitanya Joshi, Madhvi Joshi |
| EPI_ISL_524729 | B.J. Medical College and Civil hospital, Ahmedabad | Gujarat Biotechnology Research Centre | Kamlesh J Upadhyay, Sanjay Kapadia, Apurvasinh Puvar, Janvi Raval, Zarna Patel, Monika Gandhi, Pinal Trivedi, Maharshi Pandya, Nidhi Patel, Nitin Savaliya, Raghawendra Kumar, Dinesh Kumar, Zuber Saiyed, Komal Patel, Labdhi Pandya, Afzal Ansari, Nikha Trivedi, Pranay Shah, R D Dixit, A M Kadri, Harsh Bakshi, Chaitanya Joshi, Madhvi Joshi |
| EPI_ISL_524730 | GMERS Medical College & Hospital, Gotri, Vadodara | Gujarat Biotechnology Research Centre | Meenakshi Shah, Neena Doshi, Varsha Godbole, Apurvasinh Puvar, Janvi Raval, Zarna Patel, Monika Gandhi, Pinal Trivedi, Maharshi Pandya, Nidhi Patel, Nitin Savaliya, Raghawendra Kumar, Dinesh Kumar, Zuber Saiyed, Komal Patel, Labdhi Pandya, Afzal Ansari, Nikha Trivedi, R D Dixit, A M Kadri, Harsh Bakshi, Chaitanya Joshi, Madhvi Joshi |
| EPI_ISL_524731 | GMERS Medical College & Hospital, Gotri, Vadodara | Gujarat Biotechnology Research Centre | Neena Doshi, Varsha Godbole, Apurvasinh Puvar, Janvi Raval, Zarna Patel, Monika Gandhi, Pinal Trivedi, Maharshi Pandya, Nidhi Patel, Nitin Savaliya, Raghawendra Kumar, Dinesh Kumar, Zuber Saiyed, Komal Patel, Labdhi Pandya, Afzal Ansari, Nikha Trivedi, Meenakshi Shah, R D Dixit, A M Kadri, Harsh Bakshi, Chaitanya Joshi, Madhvi Joshi |
| EPI_ISL_524732 | Dr. RSS Hospital, Modasa | Gujarat Biotechnology Research Centre | Harsh Chaudhari, Apurvasinh Puvar, Janvi Raval, Zarna Patel, Monika Gandhi, Pinal Trivedi, Maharshi Pandya, Nidhi Patel, Nitin Savaliya, Raghawendra Kumar, Dinesh Kumar, Zuber Saiyed, Komal Patel, Labdhi Pandya, Afzal Ansari, Nikha Trivedi, R D Dixit, A M Kadri, Harsh Bakshi, Chaitanya Joshi, Madhvi Joshi, Neena Doshi, Varsha Godbole |
| EPI_ISL_524733 | GMERS Medical College and Hospital, Dharpur, Patan | Gujarat Biotechnology Research Centre | A N Parmar, Apurvasinh Puvar, Janvi Raval, Zarna Patel, Monika Gandhi, Pinal Trivedi, Maharshi Pandya, Nidhi Patel, Nitin Savaliya, Raghawendra Kumar, Dinesh Kumar, Zuber Saiyed, Komal Patel, Labdhi Pandya, Afzal Ansari, Nikha Trivedi, R D Dixit, A M Kadri, Harsh Bakshi, Chaitanya Joshi, Madhvi Joshi |
| EPI_ISL_524734 | GMERS Medical College and Hospital, Dharpur, Patan | Gujarat Biotechnology Research Centre | Apurvasinh Puvar, Janvi Raval, Zarna Patel, Monika Gandhi, Pinal Trivedi, Maharshi Pandya, Nidhi Patel, Nitin Savaliya, Raghawendra Kumar, Dinesh Kumar, Zuber Saiyed, Komal Patel, Labdhi Pandya, Afzal Ansari, Nikha Trivedi, A N Parmar, R D Dixit, A M Kadri, Harsh Bakshi, Chaitanya Joshi, Madhvi Joshi |
| EPI_ISL_524735 | Government General Hospital, Jam Khambhaliya, Devbhoomi Dwarka | Gujarat Biotechnology Research Centre | Harish Matani, Janvi Raval, Zarna Patel, Monika Gandhi, Pinal Trivedi, Maharshi Pandya, Nidhi Patel, Nitin Savaliya, Raghawendra Kumar, Dinesh Kumar, Zuber Saiyed, Komal Patel, Labdhi Pandya, Afzal Ansari, Nikha Trivedi, Apurvasinh Puvar, R D Dixit, A M Kadri, Harsh Bakshi, Chaitanya Joshi, Madhvi Joshi |
| EPI_ISL_524736 | Government General Hospital, Jam Khambhaliya, Devbhoomi Dwarka | Gujarat Biotechnology Research Centre | Janvi Raval, Zarna Patel, Monika Gandhi, Pinal Trivedi, Maharshi Pandya, Nidhi Patel, Nitin Savaliya, Raghawendra Kumar, Dinesh Kumar, Zuber Saiyed, Komal Patel, Labdhi Pandya, Afzal Ansari, Nikha Trivedi, Harish Matani, Apurvasinh Puvar, R D Dixit, A M Kadri, Harsh Bakshi, Chaitanya Joshi, Madhvi Joshi |
| EPI_ISL_524737 | GMERS Medical College and Hospital, Gandhinagar | Gujarat Biotechnology Research Centre | Zarna Patel, Monika Gandhi, Pinal Trivedi, Maharshi Pandya, Nidhi Patel, Nitin Savaliya, Raghawendra Kumar, Dinesh Kumar, Zuber Saiyed, Komal Patel, Labdhi Pandya, Afzal Ansari, Nikha Trivedi, Seema Bhatt, Gaurishankar Shrimali, Bhavesh Modi, Bharti Rajani, Apurvasinh Puvar, Janvi Raval, R D Dixit, A M Kadri, Harsh Bakshi, Chaitanya Joshi, Madhvi Joshi |
| EPI_ISL_524738 | GMERS Medical College and Hospital, Gandhinagar | Gujarat Biotechnology Research Centre | Monika Gandhi, Pinal Trivedi, Maharshi Pandya, Nidhi Patel, Nitin Savaliya, Raghawendra Kumar, Dinesh Kumar, Zuber Saiyed, Komal Patel, Labdhi Pandya, Afzal Ansari, Nikha Trivedi, Seema Bhatt, Gaurishankar Shrimali, Bhavesh Modi, Bharti Rajani, Apurvasinh Puvar, Janvi Raval, Zarna Patel, R D Dixit, A M Kadri, Harsh Bakshi, Chaitanya Joshi, Madhvi Joshi |
| EPI_ISL_524739 | GMERS Medical College and Hospital, Gandhinagar | Gujarat Biotechnology Research Centre | Pinal Trivedi, Maharshi Pandya, Nidhi Patel, Nitin Savaliya, Raghawendra Kumar, Dinesh Kumar, Zuber Saiyed, Komal Patel, Labdhi Pandya, Afzal Ansari, Nikha Trivedi, Seema Bhatt, Gaurishankar Shrimali, Bhavesh Modi, Bharti Rajani, Apurvasinh Puvar, Janvi Raval, Zarna Patel, Monika Gandhi, R D Dixit, A M Kadri, Harsh Bakshi, Chaitanya Joshi, Madhvi Joshi |
| EPI_ISL_524740 | GMERS Medical College and Hospital, Gandhinagar | Gujarat Biotechnology Research Centre | Maharshi Pandya, Nidhi Patel, Nitin Savaliya, Raghawendra Kumar, Dinesh Kumar, Zuber Saiyed, Komal Patel, Labdhi Pandya, Afzal Ansari, Nikha Trivedi, Seema Bhatt, Gaurishankar Shrimali, Bhavesh Modi, Bharti Rajani, Apurvasinh Puvar, Janvi Raval, Zarna Patel, Monika Gandhi, Pinal Trivedi, R D Dixit, A M Kadri, Harsh Bakshi, Chaitanya Joshi, Madhvi Joshi |
| EPI_ISL_524741 | GMERS Medical College and Hospital, Gandhinagar | Gujarat Biotechnology Research Centre | Nidhi Patel, Nitin Savaliya, Raghawendra Kumar, Dinesh Kumar, Zuber Saiyed, Komal Patel, Labdhi Pandya, Afzal Ansari, Nikha Trivedi, Seema Bhatt, Gaurishankar Shrimali, Bhavesh Modi, Bharti Rajani, Apurvasinh Puvar, Janvi Raval, Zarna Patel, Monika Gandhi, Pinal Trivedi, Maharshi Pandya, R D Dixit, A M Kadri, Harsh Bakshi, Chaitanya Joshi, Madhvi Joshi |
| EPI_ISL_524742 | GMERS Medical College and Hospital, Gandhinagar | Gujarat Biotechnology Research Centre | Dinesh Kumar, Zuber Saiyed, Komal Patel, Labdhi Pandya, Afzal Ansari, Nikha Trivedi, Seema Bhatt, Gaurishankar Shrimali, Bhavesh Modi, Bharti Rajani, Apurvasinh Puvar, Janvi Raval, Zarna Patel, Monika Gandhi, Pinal Trivedi, Maharshi Pandya, Nidhi Patel, Nitin Savaliya, Raghawendra Kumar, R D Dixit, A M Kadri, Harsh Bakshi, Chaitanya Joshi, Madhvi Joshi |
| EPI_ISL_524743 | GMERS Medical College and Hospital, Gandhinagar | Gujarat Biotechnology Research Centre | Zuber Saiyed, Komal Patel, Labdhi Pandya, Afzal Ansari, Nikha Trivedi, Seema Bhatt, Gaurishankar Shrimali, Bhavesh Modi, Bharti Rajani, Apurvasinh Puvar, Janvi Raval, Zarna Patel, Monika Gandhi, Pinal Trivedi, Maharshi Pandya, Nidhi Patel, Nitin Savaliya, Raghawendra Kumar, Dinesh Kumar, R D Dixit, A M Kadri, Harsh Bakshi, Chaitanya Joshi, Madhvi Joshi |
| EPI_ISL_524744 | GMERS Medical College and Hospital, Gandhinagar | Gujarat Biotechnology Research Centre | Komal Patel, Labdhi Pandya, Afzal Ansari, Nikha Trivedi, Seema Bhatt, Gaurishankar Shrimali, Bhavesh Modi, Bharti Rajani, Apurvasinh Puvar, Janvi Raval, Zarna Patel, Monika Gandhi, Pinal Trivedi, Maharshi Pandya, Nidhi Patel, Nitin Savaliya, Raghawendra Kumar, Dinesh Kumar, Zuber Saiyed, R D Dixit, A M Kadri, Harsh Bakshi, Chaitanya Joshi, Madhvi Joshi |
| EPI_ISL_524745 | GMERS Medical College and Hospital, Gandhinagar | Gujarat Biotechnology Research Centre | Labdhi Pandya, Afzal Ansari, Nikha Trivedi, Seema Bhatt, Gaurishankar Shrimali, Bhavesh Modi, Bharti Rajani, Apurvasinh Puvar, Janvi Raval, Zarna Patel, Monika Gandhi, Pinal Trivedi, Maharshi Pandya, Nidhi Patel, Nitin Savaliya, Raghawendra Kumar, Dinesh Kumar, Zuber Saiyed, Komal Patel, R D Dixit, A M Kadri, Harsh Bakshi, Chaitanya Joshi, Madhvi Joshi |
| EPI_ISL_524746 | Sir M P Shah Government Medical College, Jamnagar | Gujarat Biotechnology Research Centre | Binita Aring, Afzal Ansari, Nikha Trivedi, Apurvasinh Puvar, Janvi Raval, Zarna Patel, Monika Gandhi, Pinal Trivedi, Maharshi Pandya, Nidhi Patel, Nitin Savaliya, Raghawendra Kumar, Dinesh Kumar, Zuber Saiyed, Komal Patel, Labdhi Pandya, R D Dixit, A M Kadri, Harsh Bakshi, Chaitanya Joshi, Madhvi Joshi |
| EPI_ISL_524747 | Sir M P Shah Government Medical College, Jamnagar | Gujarat Biotechnology Research Centre | Afzal Ansari, Nikha Trivedi, Binita Aring, Apurvasinh Puvar, Janvi Raval, Zarna Patel, Monika Gandhi, Pinal Trivedi, Maharshi Pandya, Nidhi Patel, Nitin Savaliya, Raghawendra Kumar, Dinesh Kumar, Zuber Saiyed, Komal Patel, Labdhi Pandya, R D Dixit, A M Kadri, Harsh Bakshi, Chaitanya Joshi, Madhvi Joshi |
| EPI_ISL_524748 | Sir M P Shah Government Medical College, Jamnagar | Gujarat Biotechnology Research Centre | Nikha Trivedi, Apurvasinh Puvar, Janvi Raval, Binita Aring, Zarna Patel, Monika Gandhi, Pinal Trivedi, Maharshi Pandya, Nidhi Patel, Nitin Savaliya, Raghawendra Kumar, Dinesh Kumar, Zuber Saiyed, Komal Patel, Labdhi Pandya, Afzal Ansari, R D Dixit, A M Kadri, Harsh Bakshi, Chaitanya Joshi, Madhvi Joshi |
| EPI_ISL_524749 | Sir M P Shah Government Medical College, Jamnagar | Gujarat Biotechnology Research Centre | Apurvasinh Puvar, Janvi Raval, Zarna Patel, Monika Gandhi, Pinal Trivedi, Maharshi Pandya, Nidhi Patel, Nitin Savaliya, Raghawendra Kumar, Dinesh Kumar, Zuber Saiyed, Komal Patel, Labdhi Pandya, Afzal Ansari, Nikha Trivedi, Binita Aring, R D Dixit, A M Kadri, Harsh Bakshi, Chaitanya Joshi, Madhvi Joshi |
| EPI_ISL_524750 | Sir M P Shah Government Medical College, Jamnagar | Gujarat Biotechnology Research Centre | Janvi Raval, Zarna Patel, Monika Gandhi, Pinal Trivedi, Maharshi Pandya, Nidhi Patel, Nitin Savaliya, Raghawendra Kumar, Dinesh Kumar, Zuber Saiyed, Komal Patel, Labdhi Pandya, Afzal Ansari, Nikha Trivedi, Binita Aring, Apurvasinh Puvar, R D Dixit, A M Kadri, Harsh Bakshi, Chaitanya Joshi, Madhvi Joshi |
| EPI_ISL_524751 | Sir M P Shah Government Medical College, Jamnagar | Gujarat Biotechnology Research Centre | Zarna Patel, Monika Gandhi, Pinal Trivedi, Maharshi Pandya, Nidhi Patel, Nitin Savaliya, Raghawendra Kumar, Dinesh Kumar, Zuber Saiyed, Komal Patel, Labdhi Pandya, Afzal Ansari, Nikha Trivedi, Binita Aring, Apurvasinh Puvar, Janvi Raval, R D Dixit, A M Kadri, Harsh Bakshi, Chaitanya Joshi, Madhvi Joshi |
| EPI_ISL_524752 | GMERS Medical College and Hospital, Dharpur, Patan | Gujarat Biotechnology Research Centre | Monika Gandhi, Pinal Trivedi, Maharshi Pandya, Nidhi Patel, Nitin Savaliya, Raghawendra Kumar, Dinesh Kumar, Zuber Saiyed, Komal Patel, Labdhi Pandya, Afzal Ansari, Nikha Trivedi, A N Parmar, Apurvasinh Puvar, Janvi Raval, Zarna Patel, R D Dixit, A M Kadri, Harsh Bakshi, Chaitanya Joshi, Madhvi Joshi |
| EPI_ISL_524753 | GMERS Medical College and Hospital, Dharpur, Patan | Gujarat Biotechnology Research Centre | Pinal Trivedi, Maharshi Pandya, Nidhi Patel, Nitin Savaliya, Raghawendra Kumar, Dinesh Kumar, Zuber Saiyed, Komal Patel, Labdhi Pandya, Afzal Ansari, Nikha Trivedi, A N Parmar, Apurvasinh Puvar, Janvi Raval, Zarna Patel, Monika Gandhi, R D Dixit, A M Kadri, Harsh Bakshi, Chaitanya Joshi, Madhvi Joshi |
| EPI_ISL_524754 | GMERS Medical College and Hospital, Dharpur, Patan | Gujarat Biotechnology Research Centre | Maharshi Pandya, Nidhi Patel, Nitin Savaliya, Raghawendra Kumar, Dinesh Kumar, Zuber Saiyed, Komal Patel, Labdhi Pandya, Afzal Ansari, Nikha Trivedi, A N Parmar, Apurvasinh Puvar, Janvi Raval, Zarna Patel, Monika Gandhi, Pinal Trivedi, R D Dixit, A M Kadri, Harsh Bakshi, Chaitanya Joshi, Madhvi Joshi |
| EPI_ISL_524755 | GMERS Medical College and Hospital, Dharpur, Patan | Gujarat Biotechnology Research Centre | Nidhi Patel, Nitin Savaliya, Raghawendra Kumar, Dinesh Kumar, Zuber Saiyed, Komal Patel, Labdhi Pandya, Afzal Ansari, Nikha Trivedi, A N Parmar, Apurvasinh Puvar, Janvi Raval, Zarna Patel, Monika Gandhi, Pinal Trivedi, Maharshi Pandya, R D Dixit, A M Kadri, Harsh Bakshi, Chaitanya Joshi, Madhvi Joshi |
| EPI_ISL_524756 | GMERS Medical College and Hospital, Dharpur, Patan | Gujarat Biotechnology Research Centre | Nitin Savaliya, Raghawendra Kumar, Dinesh Kumar, Zuber Saiyed, Komal Patel, Labdhi Pandya, Afzal Ansari, Nikha Trivedi, A N Parmar, Apurvasinh Puvar, Janvi Raval, Zarna Patel, Monika Gandhi, Pinal Trivedi, Maharshi Pandya, Nidhi Patel, R D Dixit, A M Kadri, Harsh Bakshi, Chaitanya Joshi, Madhvi Joshi |
| EPI_ISL_524757 | GMERS Medical College and Hospital, Dharpur, Patan | Gujarat Biotechnology Research Centre | Raghawendra Kumar, Dinesh Kumar, Zuber Saiyed, Komal Patel, Labdhi Pandya, Afzal Ansari, Nikha Trivedi, A N Parmar, Apurvasinh Puvar, Janvi Raval, Zarna Patel, Monika Gandhi, Pinal Trivedi, Maharshi Pandya, Nidhi Patel, Nitin Savaliya, R D Dixit, A M Kadri, Harsh Bakshi, Chaitanya Joshi, Madhvi Joshi |
| EPI_ISL_524758 | GMERS Medical College and Hospital, Dharpur, Patan | Gujarat Biotechnology Research Centre | Dinesh Kumar, Zuber Saiyed, Komal Patel, Labdhi Pandya, Afzal Ansari, Nikha Trivedi, A N Parmar, Apurvasinh Puvar, Janvi Raval, Zarna Patel, Monika Gandhi, Pinal Trivedi, Maharshi Pandya, Nidhi Patel, Nitin Savaliya, Raghawendra Kumar, R D Dixit, A M Kadri, Harsh Bakshi, Chaitanya Joshi, Madhvi Joshi |
| EPI_ISL_524759 | GMERS Medical College and Hospital, Dharpur, Patan | Gujarat Biotechnology Research Centre | Zuber Saiyed, Komal Patel, Labdhi Pandya, Afzal Ansari, Nikha Trivedi, A N Parmar, Apurvasinh Puvar, Janvi Raval, Zarna Patel, Monika Gandhi, Pinal Trivedi, Maharshi Pandya, Nidhi Patel, Nitin Savaliya, Raghawendra Kumar, Dinesh Kumar, R D Dixit, A M Kadri, Harsh Bakshi, Chaitanya Joshi, Madhvi Joshi |
| EPI_ISL_524760 | GMERS Medical College and Hospital, Dharpur, Patan | Gujarat Biotechnology Research Centre | Komal Patel, Labdhi Pandya, Afzal Ansari, Nikha Trivedi, A N Parmar, Apurvasinh Puvar, Janvi Raval, Zarna Patel, Monika Gandhi, Pinal Trivedi, Maharshi Pandya, Nidhi Patel, Nitin Savaliya, Raghawendra Kumar, Dinesh Kumar, Zuber Saiyed, R D Dixit, A M Kadri, Harsh Bakshi, Chaitanya Joshi, Madhvi Joshi |
| EPI_ISL_524761 | GMERS Medical College and Hospital, Dharpur, Patan | Gujarat Biotechnology Research Centre | Labdhi Pandya, Afzal Ansari, Nikha Trivedi, A N Parmar, Apurvasinh Puvar, Janvi Raval, Zarna Patel, Monika Gandhi, Pinal Trivedi, Maharshi Pandya, Nidhi Patel, Nitin Savaliya, |

| | | | |
|---|---|---|---|
| | | | Raghawendra Kumar, Dinesh Kumar, Zuber Saiyed, Komal Patel, R D Dixit, A M Kadri, Harsh Bakshi, Chaitanya Joshi, Madhvi Joshi |
| EPI_ISL_524762 | GMERS Medical College and Hospital, Dharpur, Patan | Gujarat Biotechnology Research Centre | Afzal Ansari, Nikha Trivedi, A N Parmar, Apurvasinh Puvar, Janvi Raval, Zarna Patel, Monika Gandhi, Pinal Trivedi, Maharshi Pandya, Nidhi Patel, Nitin Savaliya, Raghawendra Kumar, Dinesh Kumar, Zuber Saiyed, Komal Patel, Labdhi Pandya, R D Dixit, A M Kadri, Harsh Bakshi, Chaitanya Joshi, Madhvi Joshi |
| EPI_ISL_524763 | GMERS Medical College and Hospital, Dharpur, Patan | Gujarat Biotechnology Research Centre | Nikha Trivedi, A N Parmar, Apurvasinh Puvar, Janvi Raval, Zarna Patel, Monika Gandhi, Pinal Trivedi, Maharshi Pandya, Nidhi Patel, Nitin Savaliya, Raghawendra Kumar, Dinesh Kumar, Zuber Saiyed, Komal Patel, Labdhi Pandya, Afzal Ansari, R D Dixit, A M Kadri, Harsh Bakshi, Chaitanya Joshi, Madhvi Joshi |
| EPI_ISL_524764 | GMERS Medical College and Hospital, Dharpur, Patan | Gujarat Biotechnology Research Centre | A N Parmar, Apurvasinh Puvar, Janvi Raval, Zarna Patel, Monika Gandhi, Pinal Trivedi, Maharshi Pandya, Nidhi Patel, Nitin Savaliya, Raghawendra Kumar, Dinesh Kumar, Zuber Saiyed, Komal Patel, Labdhi Pandya, Afzal Ansari, Nikha Trivedi, R D Dixit, A M Kadri, Harsh Bakshi, Chaitanya Joshi, Madhvi Joshi |
| EPI_ISL_524765 | GMERS Medical College and Hospital, Dharpur, Patan | Gujarat Biotechnology Research Centre | Apurvasinh Puvar, Janvi Raval, Zarna Patel, Monika Gandhi, Pinal Trivedi, Maharshi Pandya, Nidhi Patel, Nitin Savaliya, Raghawendra Kumar, Dinesh Kumar, Zuber Saiyed, Komal Patel, Labdhi Pandya, Afzal Ansari, Nikha Trivedi, A N Parmar, R D Dixit, A M Kadri, Harsh Bakshi, Chaitanya Joshi, Madhvi Joshi |
| EPI_ISL_524766 | GMERS Medical College and Hospital, Dharpur, Patan | Gujarat Biotechnology Research Centre | Janvi Raval, Zarna Patel, Monika Gandhi, Pinal Trivedi, Maharshi Pandya, Nidhi Patel, Nitin Savaliya, Raghawendra Kumar, Dinesh Kumar, Zuber Saiyed, Komal Patel, Labdhi Pandya, Afzal Ansari, Nikha Trivedi, Apurvasinh Puvar, A N Parmar, R D Dixit, A M Kadri, Harsh Bakshi, Chaitanya Joshi, Madhvi Joshi |
| EPI_ISL_524767, EPI_ISL_524768, EPI_ISL_524769, EPI_ISL_524770, EPI_ISL_524771, EPI_ISL_524772, EPI_ISL_524773, EPI_ISL_524774, EPI_ISL_524775, EPI_ISL_524776, EPI_ISL_524777, EPI_ISL_524778, EPI_ISL_524779, EPI_ISL_524780, EPI_ISL_524781 | | | |
| see above | Utah Public Health Laboratory | Utah Public Health Laboratory | Erin L. Young, Kelly Oakeson, Tara Gallagher, Michael T. Pyne, E. Susan Slechta, Melanie A. Mallory, Jeffrey B. Stevenson, Salika M. Shakir, David R. Hillyard |
| EPI_ISL_524782 | University Hospital Zürich | Institute of Medical Virology, University of Zurich | Stefan Schmutz, Maryam Zaheri, Verena Kufner, Gabriela Ziltener, Patrick Redli, Fiona Steiner, Jon Huder, Riccarda Capaul, Andrea Zbinden, Jürg Böni, Michael Huber, Alexandra Trkola |
| EPI_ISL_524783, EPI_ISL_524784, EPI_ISL_524785, EPI_ISL_524786, EPI_ISL_524787, EPI_ISL_524788, EPI_ISL_524789, EPI_ISL_524790, EPI_ISL_524791, EPI_ISL_524792, EPI_ISL_524793, EPI_ISL_524794, EPI_ISL_524795, EPI_ISL_524796, EPI_ISL_524797, EPI_ISL_524798, EPI_ISL_524799, EPI_ISL_524800, EPI_ISL_524801 | | | |
| see above | Evandro Chagas Institute | Evandro Chagas Institute | Santos, M.C.; Silva, A.M.; Junior, W.D.C.; Barbagelata, L.S.; Ferreira, J.A.; Sousa, E.M.A.; da Silva, P.S.; Resque, H.R; Martins, L.C.; Sousa Junior, E.C.;Viana, G.M.R |
| EPI_ISL_524802, EPI_ISL_524803, EPI_ISL_524804, EPI_ISL_524805, EPI_ISL_524806, EPI_ISL_524807, EPI_ISL_524808, EPI_ISL_524809, EPI_ISL_524810, EPI_ISL_524811, EPI_ISL_524812, EPI_ISL_524813, EPI_ISL_524814, EPI_ISL_524815, EPI_ISL_524816, EPI_ISL_524817, EPI_ISL_524818, EPI_ISL_524819, EPI_ISL_524820, EPI_ISL_524821, EPI_ISL_524822, EPI_ISL_524823, EPI_ISL_524824, EPI_ISL_524825, EPI_ISL_524826, EPI_ISL_524827, EPI_ISL_524828, EPI_ISL_524829, EPI_ISL_524830, EPI_ISL_524831, EPI_ISL_524832, EPI_ISL_524833, EPI_ISL_524834, EPI_ISL_524835, EPI_ISL_524836, EPI_ISL_524837, EPI_ISL_524838, EPI_ISL_524839, EPI_ISL_524840, EPI_ISL_524841, EPI_ISL_524842, EPI_ISL_524843, EPI_ISL_524844, EPI_ISL_524845, EPI_ISL_524846, EPI_ISL_524847, EPI_ISL_524848, EPI_ISL_524849, EPI_ISL_524850, EPI_ISL_524851, EPI_ISL_524852, EPI_ISL_524853, EPI_ISL_524854, EPI_ISL_524855, EPI_ISL_524856, EPI_ISL_524857, EPI_ISL_524858, EPI_ISL_524859, EPI_ISL_524860, EPI_ISL_524861, EPI_ISL_524862, EPI_ISL_524863, EPI_ISL_524864, EPI_ISL_524865, EPI_ISL_524866, EPI_ISL_524867, EPI_ISL_524868, EPI_ISL_524869, EPI_ISL_524870, EPI_ISL_524871, EPI_ISL_524872 | | | |
| see above | Utah Public Health Laboratory | Utah Public Health Laboratory | Erin L. Young, Kelly Oakeson, Tara Gallagher, Michael T. Pyne, E. Susan Slechta, Melanie A. Mallory, Jeffrey B. Stevenson, Salika M. Shakir, David R. Hillyard |
| EPI_ISL_524873, EPI_ISL_524874, EPI_ISL_524875, EPI_ISL_524876, EPI_ISL_524877, EPI_ISL_524878, EPI_ISL_524879, EPI_ISL_524880, EPI_ISL_524881, EPI_ISL_524882, EPI_ISL_524883, EPI_ISL_524884, EPI_ISL_524885, EPI_ISL_524886, EPI_ISL_524887, EPI_ISL_524888, EPI_ISL_524889, EPI_ISL_524890, EPI_ISL_524891, EPI_ISL_524892, EPI_ISL_524893, EPI_ISL_524894, EPI_ISL_524895, EPI_ISL_524896, EPI_ISL_524897, EPI_ISL_524898, EPI_ISL_524899, EPI_ISL_524900, EPI_ISL_524901, EPI_ISL_524902, EPI_ISL_524903 | | | |
| see above | MD PHL | MD PHL | Maryland Department of Health Laboratories Administration |
| EPI_ISL_524904, EPI_ISL_524905, EPI_ISL_524906, EPI_ISL_524907, EPI_ISL_524908, EPI_ISL_524909, EPI_ISL_524910, EPI_ISL_524911, EPI_ISL_524912, EPI_ISL_524913, EPI_ISL_524914, EPI_ISL_524915, EPI_ISL_524916, EPI_ISL_524917, EPI_ISL_524918, EPI_ISL_524919, EPI_ISL_524920, EPI_ISL_524921, EPI_ISL_524922, EPI_ISL_524923, EPI_ISL_524924, EPI_ISL_524925, EPI_ISL_524926, EPI_ISL_524927, EPI_ISL_524928, EPI_ISL_524929, EPI_ISL_524930, EPI_ISL_524931, EPI_ISL_524932, EPI_ISL_524933, EPI_ISL_524934, EPI_ISL_524935, EPI_ISL_524936, EPI_ISL_524937, EPI_ISL_524938, EPI_ISL_524939, EPI_ISL_524940, EPI_ISL_524941, EPI_ISL_524942, EPI_ISL_524943, EPI_ISL_524944, EPI_ISL_524945, EPI_ISL_524946, EPI_ISL_524947, EPI_ISL_524948, EPI_ISL_524949, EPI_ISL_524950, EPI_ISL_524951, EPI_ISL_524952, EPI_ISL_524953, EPI_ISL_524954, EPI_ISL_524955, EPI_ISL_524956, EPI_ISL_524957, EPI_ISL_524958, EPI_ISL_524959, EPI_ISL_524960, EPI_ISL_524961, EPI_ISL_524962, EPI_ISL_524963, EPI_ISL_524964, EPI_ISL_524965, EPI_ISL_524966, EPI_ISL_524967, EPI_ISL_524968, EPI_ISL_524969, EPI_ISL_524970, EPI_ISL_524971, EPI_ISL_524972, EPI_ISL_524973, EPI_ISL_524974, EPI_ISL_524975, EPI_ISL_524976, EPI_ISL_524977, EPI_ISL_524978, EPI_ISL_524979, EPI_ISL_524980, EPI_ISL_524981, EPI_ISL_524982, EPI_ISL_524983, EPI_ISL_524984, EPI_ISL_524985, EPI_ISL_524986, EPI_ISL_524987, EPI_ISL_524988, EPI_ISL_524989, EPI_ISL_524990, EPI_ISL_524991, EPI_ISL_524992, EPI_ISL_524993, EPI_ISL_524994, EPI_ISL_524995, EPI_ISL_524996, EPI_ISL_524997, EPI_ISL_524998, EPI_ISL_524999, EPI_ISL_525000, EPI_ISL_525001, EPI_ISL_525002, EPI_ISL_525003, EPI_ISL_525004, EPI_ISL_525005, EPI_ISL_525006, EPI_ISL_525007, EPI_ISL_525008, EPI_ISL_525009, EPI_ISL_525010, EPI_ISL_525011, EPI_ISL_525012, EPI_ISL_525013, EPI_ISL_525014, EPI_ISL_525015, EPI_ISL_525016, EPI_ISL_525017, EPI_ISL_525018, EPI_ISL_525019, EPI_ISL_525020, EPI_ISL_525021, EPI_ISL_525022, EPI_ISL_525023, EPI_ISL_525024, EPI_ISL_525025, EPI_ISL_525026, EPI_ISL_525027, EPI_ISL_525028, EPI_ISL_525029, EPI_ISL_525030, EPI_ISL_525031, EPI_ISL_525032, EPI_ISL_525033, EPI_ISL_525034, EPI_ISL_525035, EPI_ISL_525036, EPI_ISL_525037, EPI_ISL_525038, EPI_ISL_525039, EPI_ISL_525040, EPI_ISL_525041, EPI_ISL_525042, EPI_ISL_525043, EPI_ISL_525044, EPI_ISL_525045, EPI_ISL_525046, EPI_ISL_525047, EPI_ISL_525048, EPI_ISL_525049, EPI_ISL_525050, EPI_ISL_525051, EPI_ISL_525052, EPI_ISL_525053, EPI_ISL_525054, EPI_ISL_525055, EPI_ISL_525056, EPI_ISL_525057, EPI_ISL_525058, EPI_ISL_525059, EPI_ISL_525060, EPI_ISL_525061, EPI_ISL_525062, EPI_ISL_525063, EPI_ISL_525064, EPI_ISL_525065, EPI_ISL_525066, EPI_ISL_525067, EPI_ISL_525068, EPI_ISL_525069, EPI_ISL_525070, EPI_ISL_525071, EPI_ISL_525072, EPI_ISL_525073, EPI_ISL_525074, EPI_ISL_525075, EPI_ISL_525076, EPI_ISL_525077, EPI_ISL_525078, EPI_ISL_525079, EPI_ISL_525080, EPI_ISL_525081, EPI_ISL_525082, EPI_ISL_525083, EPI_ISL_525084, EPI_ISL_525085, EPI_ISL_525086, EPI_ISL_525087, EPI_ISL_525088, EPI_ISL_525089, EPI_ISL_525090, EPI_ISL_525091, EPI_ISL_525092, EPI_ISL_525093, EPI_ISL_525094, EPI_ISL_525095, EPI_ISL_525096, EPI_ISL_525097, EPI_ISL_525098, EPI_ISL_525099, EPI_ISL_525100, EPI_ISL_525101, EPI_ISL_525102, EPI_ISL_525103, EPI_ISL_525104, EPI_ISL_525105, EPI_ISL_525106, EPI_ISL_525107, EPI_ISL_525108, EPI_ISL_525109, EPI_ISL_525110, EPI_ISL_525111, EPI_ISL_525112, EPI_ISL_525113, EPI_ISL_525114, EPI_ISL_525115, EPI_ISL_525116, EPI_ISL_525117 | | | |
| see above | Utah Public Health Laboratory | Utah Public Health Laboratory | Erin L. Young, Kelly Oakeson, Tara Gallagher, Michael T. Pyne, E. Susan Slechta, Melanie A. Mallory, Jeffrey B. Stevenson, Salika M. Shakir, David R. Hillyard |
| EPI_ISL_525118 | Maryland Department of Health Laboratory | Maryland Department of Health Laboratory | Maryland Department of Health Laboratories Administration |
| EPI_ISL_525119, EPI_ISL_525120, EPI_ISL_525121, EPI_ISL_525122, EPI_ISL_525123, EPI_ISL_525124, EPI_ISL_525125, EPI_ISL_525126, EPI_ISL_525127, EPI_ISL_525128, EPI_ISL_525129, EPI_ISL_525130, EPI_ISL_525131, EPI_ISL_525132, EPI_ISL_525133, EPI_ISL_525134, EPI_ISL_525135, EPI_ISL_525136, EPI_ISL_525137, EPI_ISL_525138, EPI_ISL_525139, EPI_ISL_525140, EPI_ISL_525141, EPI_ISL_525142, EPI_ISL_525143, EPI_ISL_525144, EPI_ISL_525145, EPI_ISL_525146, EPI_ISL_525147, EPI_ISL_525148, EPI_ISL_525149, EPI_ISL_525150, EPI_ISL_525151, EPI_ISL_525152, EPI_ISL_525153, EPI_ISL_525154, EPI_ISL_525155, EPI_ISL_525156, EPI_ISL_525157, EPI_ISL_525158, EPI_ISL_525159, EPI_ISL_525160, EPI_ISL_525161, EPI_ISL_525162, EPI_ISL_525163, EPI_ISL_525164, EPI_ISL_525165, EPI_ISL_525166, EPI_ISL_525167, EPI_ISL_525168, EPI_ISL_525169, EPI_ISL_525170, EPI_ISL_525171, EPI_ISL_525172, EPI_ISL_525173, EPI_ISL_525174, EPI_ISL_525175, EPI_ISL_525176, EPI_ISL_525177, EPI_ISL_525178, EPI_ISL_525179, EPI_ISL_525180, EPI_ISL_525181, EPI_ISL_525182, EPI_ISL_525183, EPI_ISL_525184, EPI_ISL_525185, EPI_ISL_525186, EPI_ISL_525187, EPI_ISL_525188 | | | |
| see above | Utah Public Health Laboratory | Utah Public Health Laboratory | Erin L. Young, Kelly Oakeson, Tara Gallagher, Michael T. Pyne, E. Susan Slechta, Melanie A. Mallory, Jeffrey B. Stevenson, Salika M. Shakir, David R. Hillyard |
| EPI_ISL_525189, EPI_ISL_525190, EPI_ISL_525191, EPI_ISL_525192, EPI_ISL_525193, EPI_ISL_525194, EPI_ISL_525195, EPI_ISL_525196, EPI_ISL_525197, EPI_ISL_525198, EPI_ISL_525199, EPI_ISL_525200, EPI_ISL_525201, EPI_ISL_525202, EPI_ISL_525203, EPI_ISL_525204, EPI_ISL_525205 | | | |
| see above | Virginia DCLS | Virginia DCLS | Virginia DCLS |
| EPI_ISL_525206 | Laboratorio de Referencia Nacional de Virus Respiratorio. Centro Nacional de Salud Publica. Instituto Nacional de Salud Peru. | Laboratorio de Referencia Nacional de Biotecnología y Biología Molecular. Centro Nacional de Salud Publica. Instituto Nacional de Salud Peru. | Carlos Padilla Rojas, Karolyn Vega Chozo, Priscila Lope Pari, Omar Caceres Rey, Marco Galarza Perez, Maribel Huaringa Nuñez, Johanna Balbuena Torres, Henri Bailon Calderon, Nancy Rojas Serrano. |
| EPI_ISL_525207, EPI_ISL_525208, EPI_ISL_525209, EPI_ISL_525210, EPI_ISL_525211, EPI_ISL_525212, EPI_ISL_525213, EPI_ISL_525214, EPI_ISL_525215, EPI_ISL_525216, EPI_ISL_525217, EPI_ISL_525218, EPI_ISL_525219, EPI_ISL_525220, EPI_ISL_525221, EPI_ISL_525222, EPI_ISL_525223, EPI_ISL_525224, EPI_ISL_525225, EPI_ISL_525226, EPI_ISL_525227, EPI_ISL_525228, EPI_ISL_525229, EPI_ISL_525230, EPI_ISL_525231, EPI_ISL_525232, EPI_ISL_525233, EPI_ISL_525234, EPI_ISL_525235, EPI_ISL_525236, EPI_ISL_525237, EPI_ISL_525238, EPI_ISL_525239, EPI_ISL_525240, EPI_ISL_525241, EPI_ISL_525242, EPI_ISL_525243, EPI_ISL_525244, EPI_ISL_525245, EPI_ISL_525246, EPI_ISL_525247, EPI_ISL_525248, EPI_ISL_525249, EPI_ISL_525250, EPI_ISL_525251, EPI_ISL_525252, EPI_ISL_525253, EPI_ISL_525254, EPI_ISL_525255, EPI_ISL_525256, EPI_ISL_525257, EPI_ISL_525258, EPI_ISL_525259, EPI_ISL_525260, EPI_ISL_525261, EPI_ISL_525262, EPI_ISL_525263, EPI_ISL_525264, EPI_ISL_525265, EPI_ISL_525266, EPI_ISL_525267, EPI_ISL_525268, EPI_ISL_525269, EPI_ISL_525270, EPI_ISL_525271, EPI_ISL_525272, EPI_ISL_525273, EPI_ISL_525274, EPI_ISL_525275, EPI_ISL_525276, EPI_ISL_525277, EPI_ISL_525278, EPI_ISL_525279, EPI_ISL_525280, EPI_ISL_525281, EPI_ISL_525282, EPI_ISL_525283, EPI_ISL_525284, EPI_ISL_525285, EPI_ISL_525286, EPI_ISL_525287, EPI_ISL_525288, EPI_ISL_525289, EPI_ISL_525290, EPI_ISL_525291, EPI_ISL_525292, EPI_ISL_525293, EPI_ISL_525294, EPI_ISL_525295, EPI_ISL_525296, EPI_ISL_525297, EPI_ISL_525298, EPI_ISL_525299, EPI_ISL_525300, EPI_ISL_525301, EPI_ISL_525302, EPI_ISL_525303, EPI_ISL_525304, EPI_ISL_525305, EPI_ISL_525306, EPI_ISL_525307, EPI_ISL_525308, EPI_ISL_525309, EPI_ISL_525310, EPI_ISL_525311, EPI_ISL_525312, EPI_ISL_525313, EPI_ISL_525314, EPI_ISL_525315, EPI_ISL_525316, EPI_ISL_525317, EPI_ISL_525318, EPI_ISL_525319, EPI_ISL_525320, EPI_ISL_525321, EPI_ISL_525322, EPI_ISL_525323, EPI_ISL_525324, EPI_ISL_525325, EPI_ISL_525326, EPI_ISL_525327, EPI_ISL_525328, EPI_ISL_525329, EPI_ISL_525330, EPI_ISL_525331, EPI_ISL_525332, EPI_ISL_525333, EPI_ISL_525334, EPI_ISL_525335, EPI_ISL_525336 | | | |
| see above | Utah Public Health Laboratory | Utah Public Health Laboratory | Erin L. Young, Kelly Oakeson, Tara Gallagher, Michael T. Pyne, E. Susan Slechta, Melanie A. Mallory, Jeffrey B. Stevenson, Salika M. Shakir, David R. Hillyard |
| EPI_ISL_525337, EPI_ISL_525338, EPI_ISL_525339, EPI_ISL_525340, EPI_ISL_525341, EPI_ISL_525342, EPI_ISL_525343, EPI_ISL_525344, EPI_ISL_525345, EPI_ISL_525346, EPI_ISL_525347, EPI_ISL_525348, EPI_ISL_525349, EPI_ISL_525350, EPI_ISL_525351 | | | |
| see above | Utah Public Health Laboratory | Utah Public Health Laboratory | Erin Young, Kelly Oakeson |
| EPI_ISL_525352, EPI_ISL_525353, EPI_ISL_525354, EPI_ISL_525355, EPI_ISL_525356, EPI_ISL_525357, EPI_ISL_525358, EPI_ISL_525359, EPI_ISL_525360, EPI_ISL_525361, EPI_ISL_525362, EPI_ISL_525363, EPI_ISL_525364, EPI_ISL_525365, EPI_ISL_525366, EPI_ISL_525367, EPI_ISL_525368, EPI_ISL_525369, EPI_ISL_525370, EPI_ISL_525371, EPI_ISL_525372, EPI_ISL_525373, EPI_ISL_525374, EPI_ISL_525375, EPI_ISL_525376, EPI_ISL_525377, EPI_ISL_525378, EPI_ISL_525379, EPI_ISL_525380, EPI_ISL_525381, EPI_ISL_525382, EPI_ISL_525383, EPI_ISL_525384, EPI_ISL_525385, EPI_ISL_525386, EPI_ISL_525387, EPI_ISL_525388, EPI_ISL_525389, EPI_ISL_525390, EPI_ISL_525391, EPI_ISL_525392, EPI_ISL_525393, EPI_ISL_525394, EPI_ISL_525395, EPI_ISL_525396, EPI_ISL_525397, EPI_ISL_525398, EPI_ISL_525399, EPI_ISL_525400, EPI_ISL_525401, EPI_ISL_525402, EPI_ISL_525403, EPI_ISL_525404, EPI_ISL_525405, EPI_ISL_525406, EPI_ISL_525407, EPI_ISL_525408, EPI_ISL_525409, EPI_ISL_525410, EPI_ISL_525411, EPI_ISL_525412, EPI_ISL_525413, EPI_ISL_525414, EPI_ISL_525415, EPI_ISL_525416, EPI_ISL_525417, EPI_ISL_525418 | | | |
| see above | National Virus Reference Laboratory | National Virus Reference Laboratory | Michael Carr, Gabriel Gonzalez, Jonathan Dean, Aditi Chaturvedi, Suzie Coughlan, Cillian F De Gascun |
| EPI_ISL_525419 | GMERS Medical College and Hospital, Gandhinagar | Gujarat Biotechnology Research Centre | Nitin Savaliya, Raghawendra Kumar, Dinesh Kumar, Zuber Saiyed, Komal Patel, Labdhi Pandya, Afzal Ansari, Nikha Trivedi, Seema Bhatt, Gaurishankar Shrimali, Bhavesh Modi, Bharti Rajani, Apurvasinh Puvar, Janvi Raval, Zarna Patel, Monika Gandhi, Pinal Trivedi, Maharshi Pandya, Nidhi Patel, R D Dixit, A M Kadri, Harsh Bakshi, Chaitanya Joshi, Madhvi Joshi |
| EPI_ISL_525420 | GMERS Medical College and Hospital, Gandhinagar | Gujarat Biotechnology Research Centre | Raghawendra Kumar, Dinesh Kumar, Zuber Saiyed, Komal Patel, Labdhi Pandya, Afzal Ansari, Nikha Trivedi, Seema Bhatt, Gaurishankar Shrimali, Bhavesh Modi, Bharti Rajani, Apurvasinh Puvar, Janvi Raval, Zarna Patel, Monika Gandhi, Pinal Trivedi, Maharshi Pandya, Nidhi Patel, Nitin Savaliya, R D Dixit, A M Kadri, Harsh Bakshi, Chaitanya Joshi, Madhvi Joshi |
| EPI_ISL_525421 | B.J. Medical College and Civil hospital, Ahmedabad | Gujarat Biotechnology Research Centre | Zarna Patel, Monika Gandhi, Pinal Trivedi, Maharshi Pandya, Nidhi Patel, Nitin Savaliya, Raghawendra Kumar, Dinesh Kumar, Zuber Saiyed, Komal Patel, Labdhi Pandya, Afzal Ansari, Nikha Trivedi, Pranay Shah, Kamlesh J Upadhyay, Sanjay Kapadia, Apurvasinh Puvar, Janvi Raval, R D Dixit, A M Kadri, Harsh Bakshi, Chaitanya Joshi, Madhvi Joshi |
| EPI_ISL_525422 | B.J. Medical College and Civil hospital, Ahmedabad | Gujarat Biotechnology Research Centre | Monika Gandhi, Pinal Trivedi, Maharshi Pandya, Nidhi Patel, Nitin Savaliya, Raghawendra Kumar, Dinesh Kumar, Zuber Saiyed, Komal Patel, Labdhi Pandya, Afzal Ansari, Nikha Trivedi, Pranay Shah, Kamlesh J Upadhyay, Sanjay Kapadia, Apurvasinh Puvar, Janvi Raval, Zarna Patel, R D Dixit, A M Kadri, Harsh Bakshi, Chaitanya Joshi, Madhvi Joshi |
| EPI_ISL_525423 | Oman-National Influenza Center | Biotechnology & OMICs Laboratory | Sajjad Asaf, Samiha Al-Kharusi, Ahmed Al-Harrasi, Samira Al-Mahruqi, Adil Khan, Ahmed Al-Rawahi, Abdul Latif Khan, Amina Al-Jardani, Hanan Al-Kindi, Intisar Al-Shukri, Ahlam Al-Amri, Aisha Al-Amri, Aisha Al-Busaidi, Adil Al-Wahaibi, Seif Al-Abri. |
| EPI_ISL_525424 | Oman-National Influenza Center | Biotechnology & OMICs Laboratory | Samira Al-Mahruqi, Abdul Latif Khan, Samiha Al-Kharusi, Adil Khan , Ahmed Al-Rawahi, Sajjad Asaf, Amina Al-Jardani, Hanan Al-Kindi, Intisar Al-Shukri, Ahlam Al-Amri, Aisha Al-Amri, Aisha Al-Busaidi, Adil Al-Wahaibi, Seif Al-Abri, Ahmed Al-Harrasi |
| EPI_ISL_525425 | Oman-National Influenza Center | Biotechnology & OMICs Laboratory | Samira Al-Mahruqi, Abdul Latif Khan, Samiha Al-Kharusi, Adil Khan , Ahmed Al-Rawahi, Sajjad Asaf, Amina Al-Jardani, Hanan Al-Kindi, Intisar Al-Shukri, Adil Al-Wahaibi, Seif Al-Abri, Ahmed Al-Harrasi |
| EPI_ISL_525426, EPI_ISL_525427, EPI_ISL_525428 | Oman-National Influenza Center | Biotechnology & OMICs Laboratory | Sajjad Asaf, Samiha Al-Kharusi, Ahmed Al-Harrasi, Samira Al-Mahruqi, Adil Khan, Ahmed Al-Rawahi, Abdul Latif Khan, Amina Al-Jardani, Hanan Al-Kindi, Intisar Al-Shukri, Ahlam Al-Amri, Aisha Al-Amri, Aisha Al-Busaidi, Adil Al-Wahaibi, Seif Al-Abri. |
| EPI_ISL_525429 | Oman-National Influenza Center | Biotechnology & OMICs Laboratory | Samira Al-Mahruqi, Abdul Latif Khan, Samiha Al-Kharusi, Adil Khan , Ahmed Al-Rawahi, Sajjad Asaf, Amina Al-Jardani, Hanan Al-Kindi, Intisar Al-Shukri, Adil Al-Wahaibi, Seif Al-Abri, Ahmed Al-Harrasi |
| EPI_ISL_525430 | Institute of Microbiology, Universidad San Francisco de Quito | Institute of Microbiology, Universidad San Francisco de Quito | Juan José Guadalupe, Monica Becerra-Wong, Belén Prado-Vivar, Sully Márquez, Bernardo Gutiérrez, Verónica Barragán, Patricio Rojas-Silva, Gabriel Trueba, Michelle Grunauer, Paúl Cárdenas |
| EPI_ISL_525431, EPI_ISL_525432 | Institute of Microbiology, Universidad San Francisco de Quito | Institute of Microbiology, Universidad San Francisco de Quito | Juan José Guadalupe, Monica Becerra-Wong, Belén Prado-Vivar, Sully Márquez, Ligia Briceño, Carlos Mena, Nabih Dahik, Bernardo Gutiérrez, Verónica Barragán, Patricio Rojas-Silva, Gabriel Trueba, Michelle Grunauer, Paúl Cárdenas |
| EPI_ISL_525433, EPI_ISL_525434 | Institute of Microbiology, Universidad San Francisco de Quito | Institute of Microbiology, Universidad San Francisco de Quito | Juan José Guadalupe, Monica Becerra-Wong, Belén Prado-Vivar, Sully Márquez, Bernardo Gutiérrez, Eulalia Pazmiño, Carolina Pacheco, Damaris Zandoya, Carlos Mena, Nabih Dahik, Verónica Barragán, Patricio Rojas-Silva, Gabriel Trueba, Michelle Grunauer, Paúl Cárdenas |
| EPI_ISL_525435 | Institute of Microbiology, Universidad San Francisco de Quito | Institute of Microbiology, Universidad San Francisco de Quito | Juan José Guadalupe, Monica Becerra-Wong, Belén Prado-Vivar, Sully Márquez, Bernardo Gutiérrez, Yomara Napa, Edy Quizhpe, Carlos Mena, Nabih Dahik, Verónica Barragán, Patricio Rojas-Silva, Gabriel Trueba, Michelle Grunauer, Paúl Cárdenas |
| EPI_ISL_525436, EPI_ISL_525437 | Institute of Microbiology, Universidad San Francisco de Quito | Institute of Microbiology, Universidad San Francisco de Quito | Juan José Guadalupe, Monica Becerra-Wong, Belén Prado-Vivar, Sully Márquez, Bernardo Gutiérrez, Dayron Brossard, Carlos Mena, Nabih Dahik, Verónica Barragán, Patricio Rojas-Silva, Gabriel Trueba, Michelle Grunauer, Paúl Cárdenas |
| EPI_ISL_525438 | Institute of Microbiology, Universidad San Francisco de Quito | Institute of Microbiology, Universidad San Francisco de Quito | Diego Egas, Juan José Guadalupe, Monica Becerra-Wong, Belén Prado-Vivar, Sully Márquez, Bernardo Gutiérrez, Stalin Castillo, Eddy Chavez, Francisco Rodriguez, Verónica Barragán, Patricio Rojas-Silva, Gabriel Trueba, Michelle Grunauer, Paúl Cárdenas |
| EPI_ISL_525466 | Oman-National Influenza Center | Biotechnology & OMICs Laboratory | Samiha Al-Kharusi, Sajjad Asaf, Abdul Latif Khan, Samira Al-Mahruqi, Adil Khan, Ahmed Al-Rawahi, Amina Al-Jardani, Hanan Al-Kindi, Intisar Al-Shukri, Aisha Al-Busaidi, Adil Al-Wahaibi, Seif Al-Abri, Ahmed Al-Harrasi |

| | | | |
|---|---|---|---|
| EPI_ISL_525467 | Universidad Iberoamericana | International Centre for Genetic Engineering and Biotechnology (ICGEB) and ARGO Open Lab Platform | Robert Paulino-Ramirez, Eileen Riego, Alejandro Vallejo Degaudenzi, Victor Virgilio Calderon, Leandro Tapia, Patricia Leon, Danilo Licastro, Simeone Dal Monego, Sreejith Rajasekharan and Alessandro Marcello. |
| EPI_ISL_525468 | Universidad Iberoamericana | International Centre for Genetic Engineering and Biotechnology (ICGEB) and ARGO Open Lab Platform | Robert Paulino-Ramirez, Eileen Riego, Alejandro Vallejo Degaudenzi, Victor Virgilio Calderon, Leandro Tapia,Patricia Leon,Danilo Licastro, Simeone Dal Monego, Sreejith Rajasekharan and Alessandro Marcello. |
| EPI_ISL_525469, EPI_ISL_525470, EPI_ISL_525471 | Universidad Iberoamericana | International Centre for Genetic Engineering and Biotechnology (ICGEB) and ARGO Open Lab Platform | Robert Paulino-Ramirez, Eileen Riego, Alejandro Vallejo Degaudenzi, Victor Virgilio Calderon, Leandro Tapia, Patricia Leon, Danilo Licastro, Simeone Dal Monego, Sreejith Rajasekharan and Alessandro Marcello. |
| EPI_ISL_525474 | Centre for Dengue Research | Centre for Dengue Research, USJ, SL | Chandima Jeewandara, Deshni Jayathilaka, Dinuka Ariyaratne, Laksiri Gomes, Diyanath Ranasinghe, Dinuka Guruge, Ruwan Wijayamuni, Gathsaurie Neelika Malavige |
| EPI_ISL_525476 | Centre for Dengue Research | Centre for Dengue Research | Chandima Jeewandara, Deshni Jayathilaka, Dinuka Ariyaratne, Laksiri Gomes, Diyanath Ranasinghe, Dinuka Guruge, Ruwan Wijayamuni, Gathsaurie Neelika Malavige |
| EPI_ISL_525478, EPI_ISL_525479, EPI_ISL_525481 | Centre for Dengue Research | Centre for Dengue Research | Chandima Jeewandara, Deshni Jayathilaka, Dinuka Ariyaratne, Laksiri Gomes, Diyanath Ranasinghe, Ananda Wijewickrama, Eranga Narangoda, Damayanthi Idampitiya, Gathsaurie Neelika Malavige |
| EPI_ISL_525486, EPI_ISL_525488, EPI_ISL_525489 | Centre for Dengue Research | Centre for Dengue Research | Chandima Jeewandara, Deshni Jayathilaka, Dinuka Ariyaratne, Laksiri Gomes, Diyanath Ranasinghe, Ananda Wijewickrama, Malika Karunaratne, Eranga Narangoda, Damayanthi Idampitiya, Gathsaurie Neelika Malavige |
| EPI_ISL_525492 | RSUP dr. SOERADJI TIRTONEGORO | Genetics Working Group (Pokja Genetik) Faculty of Medicine, Public Health and Nursing Universitas Gadjah Mada (FK-KMK UGM), Disease Investigation Center Wates Ministry of Agriculture Indonesia, Department of Microbiology FK-KMK UGM, Laboratorium Diagnostik Yayasan Tahija World Mosquito Program (WMP) Yogyakarta Center for Tropical Medicine FK-KMK UGM, Integrated Research Center FK-KMK UGM, Department of Computer Science and Electronics FMIPA UGM | Gunadi, Hendra Wibawa, . Marcellus, Mohamad S. Hakim, Edwin W. Daniwijaya, Ludhang P. Rizki, Endah Supriyati, Eggi Arguni, Titik Nuryastuti, Tri Wibawa, Dwi AA Nugrahaningsih, Afiahayati , . Siswanto, Kurniyanto, Indah Juliana, Alvin S. Kalim, Dwiki Afandy |
| EPI_ISL_525495, EPI_ISL_525496 | Laboratory of Molecular Virology of the International Centre for Genetic Engineering and Biotechnology (ICGEB) | ARGO Open Lab Platform for Genome Sequencing | Licastro D, Rajasekharan S, Dal Monego S, Segat L, D'Agaro P, Marcello A |
| EPI_ISL_525498, EPI_ISL_525499, EPI_ISL_525500, EPI_ISL_525501, EPI_ISL_525502, EPI_ISL_525503, EPI_ISL_525504, EPI_ISL_525505, EPI_ISL_525506, EPI_ISL_525507, EPI_ISL_525508, EPI_ISL_525509, EPI_ISL_525510, EPI_ISL_525511, EPI_ISL_525512, EPI_ISL_525513, EPI_ISL_525514, EPI_ISL_525515, EPI_ISL_525516, EPI_ISL_525517, EPI_ISL_525518, EPI_ISL_525519, EPI_ISL_525520, EPI_ISL_525521, EPI_ISL_525522, EPI_ISL_525523, EPI_ISL_525524, EPI_ISL_525525, EPI_ISL_525526, EPI_ISL_525527, EPI_ISL_525528, EPI_ISL_525529, EPI_ISL_525530, EPI_ISL_525531, EPI_ISL_525532, EPI_ISL_525533, EPI_ISL_525534, EPI_ISL_525535 | | | |
| see above | Clinical Laboratory Sciences, University of Babylon | Clinical Laboratory Sciences, University of Babylon | Hashim,H.O., Mohammed,M.K., Mousa,M.J., Abdulameer,H.H., Alhassnawi,A.T., Hassan,S.A., Al-Shuhaib,M.B.S. |
| EPI_ISL_525536, EPI_ISL_525537, EPI_ISL_525538 | CNR Virus des Infections Respiratoires - France SUD | CNR Virus des Infections Respiratoires - France SUD | Antonin Bal, Gregory Destras, Gwendolyne Burfin, Solenne Brun, Alexandre Gaymard, Maude Bouscambert-Duchamp, Florence Morfin-Sherpa, Martine Valette, Bruno Lina, Laurence Josset |
| EPI_ISL_525539 | Centre Hospitalier de Bourg en Bresse | CNR Virus des Infections Respiratoires - France SUD | Antonin Bal, Gregory Destras, Gwendolyne Burfin, Solenne Brun, Alexandre Gaymard, Maude Bouscambert-Duchamp, Florence Morfin-Sherpa, Martine Valette, Bruno Lina, Laurence Josset |
| EPI_ISL_525540, EPI_ISL_525541, EPI_ISL_525542, EPI_ISL_525543 | CNR Virus des Infections Respiratoires - France SUD | CNR Virus des Infections Respiratoires - France SUD | Antonin Bal, Gregory Destras, Gwendolyne Burfin, Solenne Brun, Alexandre Gaymard, Maude Bouscambert-Duchamp, Florence Morfin-Sherpa, Martine Valette, Bruno Lina, Laurence Josset |
| EPI_ISL_525553, EPI_ISL_525554, EPI_ISL_525555, EPI_ISL_525556, EPI_ISL_525557, EPI_ISL_525558, EPI_ISL_525559, EPI_ISL_525560, EPI_ISL_525561, EPI_ISL_525562, EPI_ISL_525563, EPI_ISL_525564, EPI_ISL_525565, EPI_ISL_525566, EPI_ISL_525567, EPI_ISL_525568, EPI_ISL_525569, EPI_ISL_525570, EPI_ISL_525571, EPI_ISL_525572, EPI_ISL_525573, EPI_ISL_525574 | | | |
| see above | Istituto Zooprofilattico Sperimentale Puglia e Basilicata; Dipartimento di Bioscienze, Biotecnologie e Biofarmaceutica dell'Università degli Studi di Bari "A.Moro"; Istituto di Biomembrane. Bioenergetica e Biotecnologie Molecolari del Consiglio Nazionale delle Ricerche di Bari | Beaconlab (Bioinformatics, Evolution and Comparative Genomics lab), Dept of Biosciences, University on Milan | Parisi A.,Pesole G., Manzari C., Chiara M |
| EPI_ISL_525575, EPI_ISL_525576, EPI_ISL_525577, EPI_ISL_525578, EPI_ISL_525579, EPI_ISL_525580, EPI_ISL_525581, EPI_ISL_525582, EPI_ISL_525583, EPI_ISL_525584, EPI_ISL_525585, EPI_ISL_525586, EPI_ISL_525587, EPI_ISL_525588, EPI_ISL_525589, EPI_ISL_525590, EPI_ISL_525591, EPI_ISL_525592, EPI_ISL_525593, EPI_ISL_525594, EPI_ISL_525595, EPI_ISL_525596, EPI_ISL_525597, EPI_ISL_525598, EPI_ISL_525599, EPI_ISL_525600, EPI_ISL_525601, EPI_ISL_525602, EPI_ISL_525603, EPI_ISL_525604, EPI_ISL_525605, EPI_ISL_525606, EPI_ISL_525607, EPI_ISL_525608, EPI_ISL_525609, EPI_ISL_525610, EPI_ISL_525611, EPI_ISL_525612, EPI_ISL_525613, EPI_ISL_525614, EPI_ISL_525615, EPI_ISL_525616, EPI_ISL_525617, EPI_ISL_525618, EPI_ISL_525619, EPI_ISL_525620, EPI_ISL_525621, EPI_ISL_525622, EPI_ISL_525623, EPI_ISL_525624, EPI_ISL_525625, EPI_ISL_525626, EPI_ISL_525627, EPI_ISL_525628, EPI_ISL_525629, EPI_ISL_525630, EPI_ISL_525631, EPI_ISL_525632, EPI_ISL_525633, EPI_ISL_525634, EPI_ISL_525635, EPI_ISL_525636, EPI_ISL_525637, EPI_ISL_525638, EPI_ISL_525639, EPI_ISL_525640, EPI_ISL_525641, EPI_ISL_525642, EPI_ISL_525643, EPI_ISL_525644, EPI_ISL_525645, EPI_ISL_525646, EPI_ISL_525647, EPI_ISL_525648, EPI_ISL_525649, EPI_ISL_525650, EPI_ISL_525651, EPI_ISL_525652, EPI_ISL_525653, EPI_ISL_525654, EPI_ISL_525655, EPI_ISL_525656, EPI_ISL_525657, EPI_ISL_525658, EPI_ISL_525659, EPI_ISL_525660, EPI_ISL_525661, EPI_ISL_525662, EPI_ISL_525663, EPI_ISL_525664, EPI_ISL_525665, EPI_ISL_525666, EPI_ISL_525667, EPI_ISL_525668, EPI_ISL_525669, EPI_ISL_525670, EPI_ISL_525671, EPI_ISL_525672, EPI_ISL_525673, EPI_ISL_525674, EPI_ISL_525675, EPI_ISL_525676, EPI_ISL_525677, EPI_ISL_525678, EPI_ISL_525679, EPI_ISL_525680, EPI_ISL_525681, EPI_ISL_525682, EPI_ISL_525683, EPI_ISL_525684, EPI_ISL_525685, EPI_ISL_525686 | | | |
| see above | Wadsworth Center, New York State Department of Health | Wadsworth Center, New York State Department of Health | Kirsten St. George, Daryl M. Lamson, Sara Griesemer, Jonathan Plitnick, Navjot Singh, Matthew D. Shudt, Erica Lasek-Nesselquist |
| EPI_ISL_525699 | Seattle Flu Study | Seattle Flu Study | Deborah A. Nickerson, Chris D. Frazar, Jover Lee, Benjamin Pelle, Matthew Richardson, Amanda Adler, Elisabeth Brandstetter, Peter D. Han, Kairsten Fay, Misja Ilcisin, Kirsten Lacombe, Thomas R. Sibley, Melissa Truong, Caitlin R. Wolf, Karen Cowgill, Stephanie Schrag, Jeff Duchin, Michael Boeckh, Janet A. Englund, Michael Famulare, Barry R. Lutz, Mark J. Rieder, Lea M. Starita, Matthew Thompson, Helen Y. Chu, Trevor Bedford, Jay Shendure |
| EPI_ISL_525700, EPI_ISL_525701 | Seattle Flu Study | Seattle Flu Study | Deborah A. Nickerson, Chris D. Frazar, Jover Lee, Benjamin Pelle, Matthew Richardson, Amanda Adler, Elisabeth Brandstetter, Peter D. Han, Kairsten Fay, Misja Ilcisin, Kirsten Lacombe, Thomas R. Sibley, Melissa Truong, Caitlin R. Wolf, Michael Boeckh, Janet A. Englund, Michael Famulare, Barry R. Lutz, Mark J. Rieder, Lea M. Starita, Matthew Thompson, Jay Shendure, Trevor Bedford, Helen Y. Chu |
| EPI_ISL_525702, EPI_ISL_525703, EPI_ISL_525704, EPI_ISL_525705 | Seattle Flu Study | Seattle Flu Study | Deborah A. Nickerson, Chris D. Frazar, Jover Lee, Benjamin Pelle, Matthew Richardson, Amanda Adler, Elisabeth Brandstetter, Peter D. Han, Kairsten Fay, Misja Ilcisin, Kirsten Lacombe, Thomas R. Sibley, Melissa Truong, Caitlin R. Wolf, Karen Cowgill, Stephanie Schrag, Jeff Duchin, Michael Boeckh, Janet A. Englund, Michael Famulare, Barry R. Lutz, Mark J. Rieder, Lea M. Starita, Matthew Thompson, Helen Y. Chu, Trevor Bedford, Jay Shendure |
| EPI_ISL_525706 | Seattle Flu Study | Seattle Flu Study | Deborah A. Nickerson, Chris D. Frazar, Jover Lee, Benjamin Pelle, Matthew Richardson, Amanda Adler, Elisabeth Brandstetter, Peter D. Han, Kairsten Fay, Misja Ilcisin, Kirsten Lacombe, Thomas R. Sibley, Melissa Truong, Caitlin R. Wolf, Michael Boeckh, Janet A. Englund, Michael Famulare, Barry R. Lutz, Mark J. Rieder, Lea M. Starita, Matthew Thompson, Jay Shendure, Trevor Bedford, Helen Y. Chu |
| EPI_ISL_525707, EPI_ISL_525708, EPI_ISL_525709, EPI_ISL_525710, EPI_ISL_525711, EPI_ISL_525712, EPI_ISL_525713, EPI_ISL_525714, EPI_ISL_525715, EPI_ISL_525716, EPI_ISL_525717, EPI_ISL_525718, EPI_ISL_525719, EPI_ISL_525720, EPI_ISL_525721, EPI_ISL_525722, EPI_ISL_525723, EPI_ISL_525724, EPI_ISL_525725, EPI_ISL_525726, EPI_ISL_525727 | | | |
| see above | Seattle Flu Study | Seattle Flu Study | Deborah A. Nickerson, Chris D. Frazar, Jover Lee, Benjamin Pelle, Matthew Richardson, Amanda Adler, Elisabeth Brandstetter, Peter D. Han, Kairsten Fay, Misja Ilcisin, Kirsten Lacombe, Thomas R. Sibley, Melissa Truong, Caitlin R. Wolf, Karen Cowgill, Stephanie Schrag, Jeff Duchin, Michael Boeckh, Janet A. Englund, Michael Famulare, Barry R. Lutz, Mark J. Rieder, Lea M. Starita, Matthew Thompson, Helen Y. Chu, Trevor Bedford, Jay Shendure |
| EPI_ISL_525728 | Seattle Flu Study | Seattle Flu Study | Deborah A. Nickerson, Chris D. Frazar, Jover Lee, Benjamin Pelle, Matthew Richardson, Amanda Adler, Elisabeth Brandstetter, Peter D. Han, Kairsten Fay, Misja Ilcisin, Kirsten Lacombe, Thomas R. Sibley, Melissa Truong, Caitlin R. Wolf, Michael Boeckh, Janet A. Englund, Michael Famulare, Barry R. Lutz, Mark J. Rieder, Lea M. Starita, Matthew Thompson, Jay Shendure, Trevor Bedford, Helen Y. Chu |
| EPI_ISL_525729, EPI_ISL_525730, EPI_ISL_525731, EPI_ISL_525732, EPI_ISL_525733, EPI_ISL_525734, EPI_ISL_525735, EPI_ISL_525736, EPI_ISL_525737, EPI_ISL_525738, EPI_ISL_525739, EPI_ISL_525740, EPI_ISL_525741 | | | |
| see above | Seattle Flu Study | Seattle Flu Study | Deborah A. Nickerson, Chris D. Frazar, Jover Lee, Benjamin Pelle, Matthew Richardson, Amanda Adler, Elisabeth Brandstetter, Peter D. Han, Kairsten Fay, Misja Ilcisin, Kirsten Lacombe, Thomas R. Sibley, Melissa Truong, Caitlin R. Wolf, Karen Cowgill, Stephanie Schrag, Jeff Duchin, Michael Boeckh, Janet A. Englund, Michael Famulare, Barry R. Lutz, Mark J. Rieder, Lea M. Starita, Matthew Thompson, Helen Y. Chu, Trevor Bedford, Jay Shendure |
| EPI_ISL_525742 | Seattle Flu Study | Seattle Flu Study | Deborah A. Nickerson, Chris D. Frazar, Jover Lee, Benjamin Pelle, Matthew Richardson, Amanda Adler, Elisabeth Brandstetter, Peter D. Han, Kairsten Fay, Misja Ilcisin, Kirsten Lacombe, Thomas R. Sibley, Melissa Truong, Caitlin R. Wolf, Michael Boeckh, Janet A. Englund, Michael Famulare, Barry R. Lutz, Mark J. Rieder, Lea M. Starita, Matthew Thompson, Jay Shendure, Trevor Bedford, Helen Y. Chu |
| EPI_ISL_525743, EPI_ISL_525744, EPI_ISL_525745, EPI_ISL_525746, EPI_ISL_525747, EPI_ISL_525748, EPI_ISL_525749, EPI_ISL_525750, EPI_ISL_525751, EPI_ISL_525752, EPI_ISL_525753, EPI_ISL_525754, EPI_ISL_525755, EPI_ISL_525756 | | | |
| see above | Seattle Flu Study | Seattle Flu Study | Deborah A. Nickerson, Chris D. Frazar, Jover Lee, Benjamin Pelle, Matthew Richardson, Amanda Adler, Elisabeth Brandstetter, Peter D. Han, Kairsten Fay, Misja Ilcisin, Kirsten Lacombe, Thomas R. Sibley, Melissa Truong, Caitlin R. Wolf, Karen Cowgill, Stephanie Schrag, Jeff Duchin, Michael Boeckh, Janet A. Englund, Michael Famulare, Barry R. Lutz, Mark J. Rieder, Lea M. Starita, Matthew Thompson, Helen Y. Chu, Trevor Bedford, Jay Shendure |
| EPI_ISL_525758, EPI_ISL_525759, EPI_ISL_525760 | Alaska State Virology Laboratory | Alaska State Virology Laboratory | Jack Chen, Ph.D. |
| EPI_ISL_525763, EPI_ISL_525764, EPI_ISL_525765, EPI_ISL_525766, EPI_ISL_525767, EPI_ISL_525768, EPI_ISL_525769, EPI_ISL_525770, EPI_ISL_525771, EPI_ISL_525772, EPI_ISL_525773, EPI_ISL_525774, EPI_ISL_525775, EPI_ISL_525776, EPI_ISL_525777, EPI_ISL_525778, EPI_ISL_525779, EPI_ISL_525780, EPI_ISL_525781, EPI_ISL_525782, EPI_ISL_525783, EPI_ISL_525784, EPI_ISL_525785, EPI_ISL_525786, EPI_ISL_525787, EPI_ISL_525788, EPI_ISL_525789 | | | |
| see above | Texas Department of State Health Services | Texas Department of State Health Services | Jenny Zhang, Rashmi Tuladhar, Bonnie Oh, Maliha Rahman, Anita Pokharel, Myong Koag, Chun Wang, Rachel Lee, Grace Kubin |
| EPI_ISL_525802 | OHSU Lab Services Molecular Microbiology Lab | Oregon SARS-CoV-2 Genome Sequencing Center | Brendan L. O'Connell, Ruth V. Nichols, Alec J. Hirsch, Guang Fan, Daniel N. Streblow, William B. Messer, Andrew C. Adey, Benjamin N. Bimber, Brian J. O'Roak |
| EPI_ISL_525803 | OHSU Lab Services Molecular Microbiology Lab | Ginkgo Bioworks Clinical Laboratory | Brendan L. O'Connell, Ruth V. Nichols, Alec J. Hirsch, Guang Fan, Daniel N. Streblow, Malaika Mckenzie-Bennett, James McGann, Jim Griffin, Keith Robison, Alex Plocik, Becky Schilling, Rebecca Littlefield, Michelle Spencer, Birgitte Simen, William B. Messer, Andrew C. Adey, Benjamin N. Bimber, Brian J. O'Roak |
| EPI_ISL_525804, EPI_ISL_525805, EPI_ISL_525806, EPI_ISL_525807 | OHSU Lab Services Molecular Microbiology Lab | Oregon SARS-CoV-2 Genome Sequencing Center | Brendan L. O'Connell, Ruth V. Nichols, Alec J. Hirsch, Guang Fan, Daniel N. Streblow, William B. Messer, Andrew C. Adey, Benjamin N. Bimber, Brian J. O'Roak |
| EPI_ISL_525808, EPI_ISL_525809, EPI_ISL_525810, EPI_ISL_525811, EPI_ISL_525812, EPI_ISL_525813, EPI_ISL_525814, EPI_ISL_525815, EPI_ISL_525816, EPI_ISL_525817, EPI_ISL_525818, EPI_ISL_525819, EPI_ISL_525820, EPI_ISL_525821, EPI_ISL_525822, EPI_ISL_525823, EPI_ISL_525824 | | | |
| see above | OHSU Lab Services Molecular Microbiology Lab | Ginkgo Bioworks Clinical Laboratory | Brendan L. O'Connell, Ruth V. Nichols, Alec J. Hirsch, Guang Fan, Daniel N. Streblow, Malaika Mckenzie-Bennett, James McGann, Jim Griffin, Keith Robison, Alex Plocik, Becky Schilling, Rebecca Littlefield, Michelle Spencer, Birgitte Simen, William B. Messer, Andrew C. Adey, Benjamin N. Bimber, Brian J. O'Roak |
| EPI_ISL_525825 | OHSU Lab Services Molecular Microbiology Lab | Oregon SARS-CoV-2 Genome Sequencing Center | Brendan L. O'Connell, Ruth V. Nichols, Alec J. Hirsch, Guang Fan, Daniel N. Streblow, William B. Messer, Andrew C. Adey, Benjamin N. Bimber, Brian J. O'Roak |
| EPI_ISL_525826 | OHSU Lab Services Molecular Microbiology Lab | Ginkgo Bioworks Clinical Laboratory | Brendan L. O'Connell, Ruth V. Nichols, Alec J. Hirsch, Guang Fan, Daniel N. Streblow, Malaika Mckenzie-Bennett, James McGann, Jim Griffin, Keith Robison, Alex Plocik, Becky Schilling, Rebecca Littlefield, Michelle Spencer, Birgitte Simen, William B. Messer, Andrew C. Adey, Benjamin N. Bimber, Brian J. O'Roak |
| EPI_ISL_525827 | OHSU Lab Services Molecular Microbiology Lab | Oregon SARS-CoV-2 Genome Sequencing Center | Brendan L. O'Connell, Ruth V. Nichols, Alec J. Hirsch, Guang Fan, Daniel N. Streblow, William B. Messer, Andrew C. Adey, Benjamin N. Bimber, Brian J. O'Roak |
| EPI_ISL_525828, EPI_ISL_525829, EPI_ISL_525830, EPI_ISL_525831, EPI_ISL_525832 | OHSU Lab Services Molecular Microbiology Lab | Ginkgo Bioworks Clinical Laboratory | Brendan L. O'Connell, Ruth V. Nichols, Alec J. Hirsch, Guang Fan, Daniel N. Streblow, Malaika Mckenzie-Bennett, James McGann, Jim Griffin, Keith Robison, Alex Plocik, Becky Schilling, Rebecca Littlefield, Michelle Spencer, Birgitte Simen, William B. Messer, Andrew C. Adey, Benjamin N. Bimber, Brian J. O'Roak |
| EPI_ISL_525833 | OHSU Lab Services Molecular Microbiology Lab | Oregon SARS-CoV-2 Genome Sequencing Center | Brendan L. O'Connell, Ruth V. Nichols, Alec J. Hirsch, Guang Fan, Daniel N. Streblow, William B. Messer, Andrew C. Adey, Benjamin N. Bimber, Brian J. O'Roak |

| | | | |
|---|---|---|---|
| EPI_ISL_525834, EPI_ISL_525835, EPI_ISL_525836, EPI_ISL_525837, EPI_ISL_525838, EPI_ISL_525839, EPI_ISL_525840 | OHSU Lab Services Molecular Microbiology Lab | Ginkgo Bioworks Clinical Laboratory | Brendan L. O'Connell, Ruth V. Nichols, Alec J. Hirsch, Guang Fan, Daniel N. Streblow, Malaika Mckenzie-Bennett, James McGann, Jim Griffin, Keith Robison, Alex Plocik, Becky Schilling, Rebecca Littlefield, Michelle Spencer, Birgitte Simen, William B. Messer, Andrew C. Adey, Benjamin N. Bimber, Brian J. O’Roak |
| EPI_ISL_525841 | OHSU Lab Services Molecular Microbiology Lab | Oregon SARS-CoV-2 Genome Sequencing Center | Brendan L. O'Connell, Ruth V. Nichols, Alec J. Hirsch, Guang Fan, Daniel N. Streblow, William B. Messer, Andrew C. Adey, Benjamin N. Bimber, Brian J. O’Roak |
| EPI_ISL_525842, EPI_ISL_525843, EPI_ISL_525844, EPI_ISL_525845, EPI_ISL_525846 | OHSU Lab Services Molecular Microbiology Lab | Ginkgo Bioworks Clinical Laboratory | Brendan L. O'Connell, Ruth V. Nichols, Alec J. Hirsch, Guang Fan, Daniel N. Streblow, Malaika Mckenzie-Bennett, James McGann, Jim Griffin, Keith Robison, Alex Plocik, Becky Schilling, Rebecca Littlefield, Michelle Spencer, Birgitte Simen, William B. Messer, Andrew C. Adey, Benjamin N. Bimber, Brian J. O’Roak |
| EPI_ISL_525847 | OHSU Lab Services Molecular Microbiology Lab | Oregon SARS-CoV-2 Genome Sequencing Center | Brendan L. O'Connell, Ruth V. Nichols, Alec J. Hirsch, Guang Fan, Daniel N. Streblow, William B. Messer, Andrew C. Adey, Benjamin N. Bimber, Brian J. O’Roak |
| EPI_ISL_525848 | OHSU Lab Services Molecular Microbiology Lab | Ginkgo Bioworks Clinical Laboratory | Brendan L. O'Connell, Ruth V. Nichols, Alec J. Hirsch, Guang Fan, Daniel N. Streblow, Malaika Mckenzie-Bennett, James McGann, Jim Griffin, Keith Robison, Alex Plocik, Becky Schilling, Rebecca Littlefield, Michelle Spencer, Birgitte Simen, William B. Messer, Andrew C. Adey, Benjamin N. Bimber, Brian J. O’Roak |
| EPI_ISL_525849 | OHSU Lab Services Molecular Microbiology Lab | Oregon SARS-CoV-2 Genome Sequencing Center | Brendan L. O'Connell, Ruth V. Nichols, Alec J. Hirsch, Guang Fan, Daniel N. Streblow, William B. Messer, Andrew C. Adey, Benjamin N. Bimber, Brian J. O’Roak |
| EPI_ISL_525850 | OHSU Lab Services Molecular Microbiology Lab | Ginkgo Bioworks Clinical Laboratory | Brendan L. O'Connell, Ruth V. Nichols, Alec J. Hirsch, Guang Fan, Daniel N. Streblow, Malaika Mckenzie-Bennett, James McGann, Jim Griffin, Keith Robison, Alex Plocik, Becky Schilling, Rebecca Littlefield, Michelle Spencer, Birgitte Simen, William B. Messer, Andrew C. Adey, Benjamin N. Bimber, Brian J. O’Roak |
| EPI_ISL_525851 | OHSU Lab Services Molecular Microbiology Lab | Oregon SARS-CoV-2 Genome Sequencing Center | Brendan L. O'Connell, Ruth V. Nichols, Alec J. Hirsch, Guang Fan, Daniel N. Streblow, William B. Messer, Andrew C. Adey, Benjamin N. Bimber, Brian J. O’Roak |
| EPI_ISL_525852, EPI_ISL_525853, EPI_ISL_525854, EPI_ISL_525855 | OHSU Lab Services Molecular Microbiology Lab | Ginkgo Bioworks Clinical Laboratory | Brendan L. O'Connell, Ruth V. Nichols, Alec J. Hirsch, Guang Fan, Daniel N. Streblow, Malaika Mckenzie-Bennett, James McGann, Jim Griffin, Keith Robison, Alex Plocik, Becky Schilling, Rebecca Littlefield, Michelle Spencer, Birgitte Simen, William B. Messer, Andrew C. Adey, Benjamin N. Bimber, Brian J. O’Roak |
| EPI_ISL_525856, EPI_ISL_525857, EPI_ISL_525858 | OHSU Lab Services Molecular Microbiology Lab | Oregon SARS-CoV-2 Genome Sequencing Center | Brendan L. O'Connell, Ruth V. Nichols, Alec J. Hirsch, Guang Fan, Daniel N. Streblow, William B. Messer, Andrew C. Adey, Benjamin N. Bimber, Brian J. O’Roak |
| EPI_ISL_525859 | OHSU Lab Services Molecular Microbiology Lab | Ginkgo Bioworks Clinical Laboratory | Brendan L. O'Connell, Ruth V. Nichols, Alec J. Hirsch, Guang Fan, Daniel N. Streblow, Malaika Mckenzie-Bennett, James McGann, Jim Griffin, Keith Robison, Alex Plocik, Becky Schilling, Rebecca Littlefield, Michelle Spencer, Birgitte Simen, William B. Messer, Andrew C. Adey, Benjamin N. Bimber, Brian J. O’Roak |
| EPI_ISL_525860 | OHSU Lab Services Molecular Microbiology Lab | Oregon SARS-CoV-2 Genome Sequencing Center | Brendan L. O'Connell, Ruth V. Nichols, Alec J. Hirsch, Guang Fan, Daniel N. Streblow, William B. Messer, Andrew C. Adey, Benjamin N. Bimber, Brian J. O’Roak |
| EPI_ISL_525861 | OHSU Lab Services Molecular Microbiology Lab | Ginkgo Bioworks Clinical Laboratory | Brendan L. O'Connell, Ruth V. Nichols, Alec J. Hirsch, Guang Fan, Daniel N. Streblow, Malaika Mckenzie-Bennett, James McGann, Jim Griffin, Keith Robison, Alex Plocik, Becky Schilling, Rebecca Littlefield, Michelle Spencer, Birgitte Simen, William B. Messer, Andrew C. Adey, Benjamin N. Bimber, Brian J. O’Roak |
| EPI_ISL_525862 | OHSU Lab Services Molecular Microbiology Lab | Oregon SARS-CoV-2 Genome Sequencing Center | Brendan L. O'Connell, Ruth V. Nichols, Alec J. Hirsch, Guang Fan, Daniel N. Streblow, William B. Messer, Andrew C. Adey, Benjamin N. Bimber, Brian J. O’Roak |
| EPI_ISL_525863, EPI_ISL_525864 | OHSU Lab Services Molecular Microbiology Lab | Ginkgo Bioworks Clinical Laboratory | Brendan L. O'Connell, Ruth V. Nichols, Alec J. Hirsch, Guang Fan, Daniel N. Streblow, Malaika Mckenzie-Bennett, James McGann, Jim Griffin, Keith Robison, Alex Plocik, Becky Schilling, Rebecca Littlefield, Michelle Spencer, Birgitte Simen, William B. Messer, Andrew C. Adey, Benjamin N. Bimber, Brian J. O’Roak |
| EPI_ISL_525865 | OHSU Lab Services Molecular Microbiology Lab | Oregon SARS-CoV-2 Genome Sequencing Center | Brendan L. O'Connell, Ruth V. Nichols, Alec J. Hirsch, Guang Fan, Daniel N. Streblow, William B. Messer, Andrew C. Adey, Benjamin N. Bimber, Brian J. O’Roak |
| EPI_ISL_525866, EPI_ISL_525867 | OHSU Lab Services Molecular Microbiology Lab | Ginkgo Bioworks Clinical Laboratory | Brendan L. O'Connell, Ruth V. Nichols, Alec J. Hirsch, Guang Fan, Daniel N. Streblow, Malaika Mckenzie-Bennett, James McGann, Jim Griffin, Keith Robison, Alex Plocik, Becky Schilling, Rebecca Littlefield, Michelle Spencer, Birgitte Simen, William B. Messer, Andrew C. Adey, Benjamin N. Bimber, Brian J. O’Roak |
| EPI_ISL_525868 | OHSU Lab Services Molecular Microbiology Lab | Oregon SARS-CoV-2 Genome Sequencing Center | Brendan L. O'Connell, Ruth V. Nichols, Alec J. Hirsch, Guang Fan, Daniel N. Streblow, William B. Messer, Andrew C. Adey, Benjamin N. Bimber, Brian J. O’Roak |
| EPI_ISL_525869 | OHSU Lab Services Molecular Microbiology Lab | Ginkgo Bioworks Clinical Laboratory | Brendan L. O'Connell, Ruth V. Nichols, Alec J. Hirsch, Guang Fan, Daniel N. Streblow, Malaika Mckenzie-Bennett, James McGann, Jim Griffin, Keith Robison, Alex Plocik, Becky Schilling, Rebecca Littlefield, Michelle Spencer, Birgitte Simen, William B. Messer, Andrew C. Adey, Benjamin N. Bimber, Brian J. O’Roak |
| EPI_ISL_525870 | OHSU Lab Services Molecular Microbiology Lab | Oregon SARS-CoV-2 Genome Sequencing Center | Brendan L. O'Connell, Ruth V. Nichols, Alec J. Hirsch, Guang Fan, Daniel N. Streblow, William B. Messer, Andrew C. Adey, Benjamin N. Bimber, Brian J. O’Roak |
| EPI_ISL_525871, EPI_ISL_525872, EPI_ISL_525873 | OHSU Lab Services Molecular Microbiology Lab | Ginkgo Bioworks Clinical Laboratory | Brendan L. O'Connell, Ruth V. Nichols, Alec J. Hirsch, Guang Fan, Daniel N. Streblow, Malaika Mckenzie-Bennett, James McGann, Jim Griffin, Keith Robison, Alex Plocik, Becky Schilling, Rebecca Littlefield, Michelle Spencer, Birgitte Simen, William B. Messer, Andrew C. Adey, Benjamin N. Bimber, Brian J. O’Roak |
| EPI_ISL_525874 | OHSU Lab Services Molecular Microbiology Lab | Oregon SARS-CoV-2 Genome Sequencing Center | Brendan L. O'Connell, Ruth V. Nichols, Alec J. Hirsch, Guang Fan, Daniel N. Streblow, William B. Messer, Andrew C. Adey, Benjamin N. Bimber, Brian J. O’Roak |
| EPI_ISL_525875 | OHSU Lab Services Molecular Microbiology Lab | Ginkgo Bioworks Clinical Laboratory | Brendan L. O'Connell, Ruth V. Nichols, Alec J. Hirsch, Guang Fan, Daniel N. Streblow, Malaika Mckenzie-Bennett, James McGann, Jim Griffin, Keith Robison, Alex Plocik, Becky Schilling, Rebecca Littlefield, Michelle Spencer, Birgitte Simen, William B. Messer, Andrew C. Adey, Benjamin N. Bimber, Brian J. O’Roak |
| EPI_ISL_525876 | OHSU Lab Services Molecular Microbiology Lab | Oregon SARS-CoV-2 Genome Sequencing Center | Brendan L. O'Connell, Ruth V. Nichols, Alec J. Hirsch, Guang Fan, Daniel N. Streblow, William B. Messer, Andrew C. Adey, Benjamin N. Bimber, Brian J. O’Roak |
| EPI_ISL_525877 | OHSU Lab Services Molecular Microbiology Lab | Ginkgo Bioworks Clinical Laboratory | Brendan L. O'Connell, Ruth V. Nichols, Alec J. Hirsch, Guang Fan, Daniel N. Streblow, Malaika Mckenzie-Bennett, James McGann, Jim Griffin, Keith Robison, Alex Plocik, Becky Schilling, Rebecca Littlefield, Michelle Spencer, Birgitte Simen, William B. Messer, Andrew C. Adey, Benjamin N. Bimber, Brian J. O’Roak |
| EPI_ISL_525878 | OHSU Lab Services Molecular Microbiology Lab | Oregon SARS-CoV-2 Genome Sequencing Center | Brendan L. O'Connell, Ruth V. Nichols, Alec J. Hirsch, Guang Fan, Daniel N. Streblow, William B. Messer, Andrew C. Adey, Benjamin N. Bimber, Brian J. O’Roak |
| EPI_ISL_525879 | OHSU Lab Services Molecular Microbiology Lab | Ginkgo Bioworks Clinical Laboratory | Brendan L. O'Connell, Ruth V. Nichols, Alec J. Hirsch, Guang Fan, Daniel N. Streblow, Malaika Mckenzie-Bennett, James McGann, Jim Griffin, Keith Robison, Alex Plocik, Becky Schilling, Rebecca Littlefield, Michelle Spencer, Birgitte Simen, William B. Messer, Andrew C. Adey, Benjamin N. Bimber, Brian J. O’Roak |
| EPI_ISL_525880, EPI_ISL_525881 | OHSU Lab Services Molecular Microbiology Lab | Oregon SARS-CoV-2 Genome Sequencing Center | Brendan L. O'Connell, Ruth V. Nichols, Alec J. Hirsch, Guang Fan, Daniel N. Streblow, William B. Messer, Andrew C. Adey, Benjamin N. Bimber, Brian J. O’Roak |
| EPI_ISL_525882 | OHSU Lab Services Molecular Microbiology Lab | Ginkgo Bioworks Clinical Laboratory | Brendan L. O'Connell, Ruth V. Nichols, Alec J. Hirsch, Guang Fan, Daniel N. Streblow, Malaika Mckenzie-Bennett, James McGann, Jim Griffin, Keith Robison, Alex Plocik, Becky Schilling, Rebecca Littlefield, Michelle Spencer, Birgitte Simen, William B. Messer, Andrew C. Adey, Benjamin N. Bimber, Brian J. O’Roak |
| EPI_ISL_525883 | OHSU Lab Services Molecular Microbiology Lab | Oregon SARS-CoV-2 Genome Sequencing Center | Brendan L. O'Connell, Ruth V. Nichols, Alec J. Hirsch, Guang Fan, Daniel N. Streblow, William B. Messer, Andrew C. Adey, Benjamin N. Bimber, Brian J. O’Roak |
| EPI_ISL_525884 | OHSU Lab Services Molecular Microbiology Lab | Ginkgo Bioworks Clinical Laboratory | Brendan L. O'Connell, Ruth V. Nichols, Alec J. Hirsch, Guang Fan, Daniel N. Streblow, Malaika Mckenzie-Bennett, James McGann, Jim Griffin, Keith Robison, Alex Plocik, Becky Schilling, Rebecca Littlefield, Michelle Spencer, Birgitte Simen, William B. Messer, Andrew C. Adey, Benjamin N. Bimber, Brian J. O’Roak |
| EPI_ISL_525885 | OHSU Lab Services Molecular Microbiology Lab | Oregon SARS-CoV-2 Genome Sequencing Center | Brendan L. O'Connell, Ruth V. Nichols, Alec J. Hirsch, Guang Fan, Daniel N. Streblow, William B. Messer, Andrew C. Adey, Benjamin N. Bimber, Brian J. O’Roak |
| EPI_ISL_525886, EPI_ISL_525887, EPI_ISL_525888, EPI_ISL_525889, EPI_ISL_525890 | OHSU Lab Services Molecular Microbiology Lab | Ginkgo Bioworks Clinical Laboratory | Brendan L. O'Connell, Ruth V. Nichols, Alec J. Hirsch, Guang Fan, Daniel N. Streblow, Malaika Mckenzie-Bennett, James McGann, Jim Griffin, Keith Robison, Alex Plocik, Becky Schilling, Rebecca Littlefield, Michelle Spencer, Birgitte Simen, William B. Messer, Andrew C. Adey, Benjamin N. Bimber, Brian J. O’Roak |
| EPI_ISL_525891, EPI_ISL_525892, EPI_ISL_525893, EPI_ISL_525894, EPI_ISL_525895, EPI_ISL_525896, EPI_ISL_525897, EPI_ISL_525898, EPI_ISL_525899, EPI_ISL_525900, EPI_ISL_525901, EPI_ISL_525902, EPI_ISL_525903, EPI_ISL_525904, EPI_ISL_525905, EPI_ISL_525906, EPI_ISL_525907, EPI_ISL_525908, EPI_ISL_525909, EPI_ISL_525910, EPI_ISL_525911, EPI_ISL_525912, EPI_ISL_525913, EPI_ISL_525914, EPI_ISL_525915, EPI_ISL_525916, EPI_ISL_525917, EPI_ISL_525918, EPI_ISL_525919, EPI_ISL_525920, EPI_ISL_525921, EPI_ISL_525922, EPI_ISL_525923, EPI_ISL_525924, EPI_ISL_525925, EPI_ISL_525926, EPI_ISL_525927, EPI_ISL_525928, EPI_ISL_525929, EPI_ISL_525930, EPI_ISL_525931, EPI_ISL_525932, EPI_ISL_525933, EPI_ISL_525934, EPI_ISL_525935, EPI_ISL_525936, EPI_ISL_525937, EPI_ISL_525938, EPI_ISL_525939, EPI_ISL_525940, EPI_ISL_525941, EPI_ISL_525942, EPI_ISL_525943, EPI_ISL_525944, EPI_ISL_525945, EPI_ISL_525946, EPI_ISL_525947, EPI_ISL_525948, EPI_ISL_525949, EPI_ISL_525950, EPI_ISL_525951, EPI_ISL_525952, EPI_ISL_525953, EPI_ISL_525954, EPI_ISL_525955, EPI_ISL_525956, EPI_ISL_525957, EPI_ISL_525958, EPI_ISL_525959, EPI_ISL_525960, EPI_ISL_525961, EPI_ISL_525962, EPI_ISL_525963, EPI_ISL_525964, EPI_ISL_525965, EPI_ISL_525966, EPI_ISL_525967, EPI_ISL_525968, EPI_ISL_525969, EPI_ISL_525970, EPI_ISL_525971, EPI_ISL_525972, EPI_ISL_525973, EPI_ISL_525974, EPI_ISL_525975, EPI_ISL_525976, EPI_ISL_525977, EPI_ISL_525978, EPI_ISL_525979, EPI_ISL_525980, EPI_ISL_525981, EPI_ISL_525982, EPI_ISL_525983, EPI_ISL_525984, EPI_ISL_525985, EPI_ISL_525986, EPI_ISL_525987, EPI_ISL_525988, EPI_ISL_525989, EPI_ISL_525990, EPI_ISL_525991, EPI_ISL_525992, EPI_ISL_525993, EPI_ISL_525994, EPI_ISL_525995, EPI_ISL_525996, EPI_ISL_525997, EPI_ISL_525998, EPI_ISL_525999, EPI_ISL_526000, EPI_ISL_526001, EPI_ISL_526002, EPI_ISL_526003, EPI_ISL_526004, EPI_ISL_526005, EPI_ISL_526006, EPI_ISL_526007, EPI_ISL_526008, EPI_ISL_526009, EPI_ISL_526010, EPI_ISL_526011, EPI_ISL_526012, EPI_ISL_526013, EPI_ISL_526014, EPI_ISL_526015, EPI_ISL_526016, EPI_ISL_526017, EPI_ISL_526018, EPI_ISL_526019, EPI_ISL_526020, EPI_ISL_526021, EPI_ISL_526022, EPI_ISL_526023, EPI_ISL_526024, EPI_ISL_526025, EPI_ISL_526026, EPI_ISL_526027, EPI_ISL_526028, EPI_ISL_526029, EPI_ISL_526030, EPI_ISL_526031, EPI_ISL_526032, EPI_ISL_526033, EPI_ISL_526034, EPI_ISL_526035, EPI_ISL_526036, EPI_ISL_526037, EPI_ISL_526038, EPI_ISL_526039, EPI_ISL_526040, EPI_ISL_526041, EPI_ISL_526042, EPI_ISL_526043, EPI_ISL_526044, EPI_ISL_526045, EPI_ISL_526046, EPI_ISL_526047, EPI_ISL_526048, EPI_ISL_526049, EPI_ISL_526050, EPI_ISL_526051, EPI_ISL_526052, EPI_ISL_526053, EPI_ISL_526054, EPI_ISL_526055, EPI_ISL_526056, EPI_ISL_526057, EPI_ISL_526058, EPI_ISL_526059, EPI_ISL_526060, EPI_ISL_526061, EPI_ISL_526062, EPI_ISL_526063, EPI_ISL_526064, EPI_ISL_526065, EPI_ISL_526066, EPI_ISL_526067, EPI_ISL_526068, EPI_ISL_526069, EPI_ISL_526070, EPI_ISL_526071, EPI_ISL_526072, EPI_ISL_526073, EPI_ISL_526074, EPI_ISL_526075, EPI_ISL_526076, EPI_ISL_526077, EPI_ISL_526078, EPI_ISL_526079, EPI_ISL_526080, EPI_ISL_526081, EPI_ISL_526082, EPI_ISL_526083, EPI_ISL_526084, EPI_ISL_526085, EPI_ISL_526086 | | | |
| see above | OHSU Lab Services Molecular Microbiology Lab | Oregon SARS-CoV-2 Genome Sequencing Center | Brendan L. O'Connell, Ruth V. Nichols, Alec J. Hirsch, Guang Fan, Daniel N. Streblow, William B. Messer, Andrew C. Adey, Benjamin N. Bimber, Brian J. O’Roak |
| EPI_ISL_526087, EPI_ISL_526088, EPI_ISL_526089, EPI_ISL_526090 | OHSU Lab Services Molecular Microbiology Lab | Ginkgo Bioworks Clinical Laboratory | Brendan L. O'Connell, Ruth V. Nichols, Alec J. Hirsch, Guang Fan, Daniel N. Streblow, Malaika Mckenzie-Bennett, James McGann, Jim Griffin, Keith Robison, Alex Plocik, Becky Schilling, Rebecca Littlefield, Michelle Spencer, Birgitte Simen, William B. Messer, Andrew C. Adey, Benjamin N. Bimber, Brian J. O’Roak |
| EPI_ISL_526091 | OHSU Lab Services Molecular Microbiology Lab | Oregon SARS-CoV-2 Genome Sequencing Center | Brendan L. O'Connell, Ruth V. Nichols, Alec J. Hirsch, Guang Fan, Daniel N. Streblow, William B. Messer, Andrew C. Adey, Benjamin N. Bimber, Brian J. O’Roak |
| EPI_ISL_526092, EPI_ISL_526093 | OHSU Lab Services Molecular Microbiology Lab | Ginkgo Bioworks Clinical Laboratory | Brendan L. O'Connell, Ruth V. Nichols, Alec J. Hirsch, Guang Fan, Daniel N. Streblow, Malaika Mckenzie-Bennett, James McGann, Jim Griffin, Keith Robison, Alex Plocik, Becky Schilling, Rebecca Littlefield, Michelle Spencer, Birgitte Simen, William B. Messer, Andrew C. Adey, Benjamin N. Bimber, Brian J. O’Roak |
| EPI_ISL_526094, EPI_ISL_526095, EPI_ISL_526096 | OHSU Lab Services Molecular Microbiology Lab | Oregon SARS-CoV-2 Genome Sequencing Center | Brendan L. O'Connell, Ruth V. Nichols, Alec J. Hirsch, Guang Fan, Daniel N. Streblow, William B. Messer, Andrew C. Adey, Benjamin N. Bimber, Brian J. O’Roak |
| EPI_ISL_526097 | OHSU Lab Services Molecular Microbiology Lab | Ginkgo Bioworks Clinical Laboratory | Brendan L. O'Connell, Ruth V. Nichols, Alec J. Hirsch, Guang Fan, Daniel N. Streblow, Malaika Mckenzie-Bennett, James McGann, Jim Griffin, Keith Robison, Alex Plocik, Becky Schilling, Rebecca Littlefield, Michelle Spencer, Birgitte Simen, William B. Messer, Andrew C. Adey, Benjamin N. Bimber, Brian J. O’Roak |
| EPI_ISL_526098, EPI_ISL_526099, EPI_ISL_526100, EPI_ISL_526101, EPI_ISL_526102, EPI_ISL_526103, EPI_ISL_526104, EPI_ISL_526105, EPI_ISL_526106, EPI_ISL_526107, EPI_ISL_526108, EPI_ISL_526109, EPI_ISL_526110, EPI_ISL_526111, EPI_ISL_526112, EPI_ISL_526113, EPI_ISL_526114 | | | |
| see above | OHSU Lab Services Molecular Microbiology Lab | Oregon SARS-CoV-2 Genome Sequencing Center | Brendan L. O'Connell, Ruth V. Nichols, Alec J. Hirsch, Guang Fan, Daniel N. Streblow, William B. Messer, Andrew C. Adey, Benjamin N. Bimber, Brian J. O’Roak |
| EPI_ISL_526115, EPI_ISL_526116, EPI_ISL_526117, EPI_ISL_526118 | Pathology West - NSW Health Pathology | NSW Health Pathology - Institute of Clinical Pathology and Medical Research; Westmead Hospital; University of Sydney | CIDM-PH et al. |
| EPI_ISL_526119, EPI_ISL_526120 | Sydney South West Pathology Service (SSWPS) - Liverpool Hospital - NSW Health Pathology | NSW Health Pathology - Institute of Clinical Pathology and Medical Research; Westmead Hospital; University of Sydney | CIDM-PH et al. |
| EPI_ISL_526121, EPI_ISL_526122, EPI_ISL_526123 | St Vincent's Pathology (SydPath) | NSW Health Pathology - Institute of Clinical Pathology and Medical Research; Westmead Hospital; University of Sydney | CIDM-PH et al. |
| EPI_ISL_526124 | Sydney South West Pathology Service (SSWPS) - Liverpool Hospital - NSW Health Pathology | NSW Health Pathology - Institute of Clinical Pathology and Medical Research; Westmead Hospital; University of Sydney | CIDM-PH et al. |
| EPI_ISL_526125 | Sydney South West Pathology Service (SSWPS) - Concord Repatriation General Hospital - NSW Health Pathology | NSW Health Pathology - Institute of Clinical Pathology and Medical Research; Westmead Hospital; University of Sydney | CIDM-PH et al. |
| EPI_ISL_526126, EPI_ISL_526127 | Sydney South West Pathology Service (SSWPS) - Liverpool Hospital - NSW Health Pathology | NSW Health Pathology - Institute of Clinical Pathology and Medical Research; Westmead Hospital; University of Sydney | CIDM-PH et al. |
| EPI_ISL_526128 | Pathology North - Royal North Shore Hospital - NSW Health | NSW Health Pathology - Institute of Clinical Pathology and Medical Research; Westmead Hospital; University of | CIDM-PH et al. |

| | | | |
|---|---|---|---|
| | Pathology | Sydney | |
| EPI_ISL_526129 | The Children's Hospital at Westmead | NSW Health Pathology - Institute of Clinical Pathology and Medical Research; Westmead Hospital; University of Sydney | CIDM-PH et al. |
| EPI_ISL_526130 | Pathology North - Royal North Shore Hospital - NSW Health Pathology | NSW Health Pathology - Institute of Clinical Pathology and Medical Research; Westmead Hospital; University of Sydney | CIDM-PH et al. |
| EPI_ISL_526131 | Sydney South West Pathology Service (SSWPS) - Liverpool Hospital - NSW Health Pathology | NSW Health Pathology - Institute of Clinical Pathology and Medical Research; Westmead Hospital; University of Sydney | CIDM-PH et al. |
| EPI_ISL_526132, EPI_ISL_526133 | Pathology North - Royal North Shore Hospital - NSW Health Pathology | NSW Health Pathology - Institute of Clinical Pathology and Medical Research; Westmead Hospital; University of Sydney | CIDM-PH et al. |
| EPI_ISL_526134, EPI_ISL_526135, EPI_ISL_526136, EPI_ISL_526137 | Sydney South West Pathology Service (SSWPS) - Royal Prince Alfred Hospital - NSW Health Pathology | NSW Health Pathology - Institute of Clinical Pathology and Medical Research; Westmead Hospital; University of Sydney | CIDM-PH et al. |
| EPI_ISL_526138, EPI_ISL_526139, EPI_ISL_526140, EPI_ISL_526141 | Pathology West - NSW Health Pathology | NSW Health Pathology - Institute of Clinical Pathology and Medical Research; Westmead Hospital; University of Sydney | CIDM-PH et al. |
| EPI_ISL_526142 | South Eastern Area Laboratory Services (SEALS) | NSW Health Pathology - Institute of Clinical Pathology and Medical Research; Westmead Hospital; University of Sydney | CIDM-PH et al. |
| EPI_ISL_526143 | Laverty Pathology | NSW Health Pathology - Institute of Clinical Pathology and Medical Research; Westmead Hospital; University of Sydney | CIDM-PH et al. |
| EPI_ISL_526144, EPI_ISL_526145, EPI_ISL_526146, EPI_ISL_526147, EPI_ISL_526148, EPI_ISL_526149 | South Eastern Area Laboratory Services (SEALS) | NSW Health Pathology - Institute of Clinical Pathology and Medical Research; Westmead Hospital; University of Sydney | CIDM-PH et al. |
| EPI_ISL_526150, EPI_ISL_526151 | Pathology West - NSW Health Pathology | NSW Health Pathology - Institute of Clinical Pathology and Medical Research; Westmead Hospital; University of Sydney | CIDM-PH et al. |
| EPI_ISL_526152 | Pathology North - Hunter - NSW Health Pathology | NSW Health Pathology - Institute of Clinical Pathology and Medical Research; Westmead Hospital; University of Sydney | CIDM-PH et al. |
| EPI_ISL_526153 | Douglass Hanly Moir Pathology | NSW Health Pathology - Institute of Clinical Pathology and Medical Research; Westmead Hospital; University of Sydney | CIDM-PH et al. |
| EPI_ISL_526154, EPI_ISL_526155 | Pathology North - Royal North Shore Hospital - NSW Health Pathology | NSW Health Pathology - Institute of Clinical Pathology and Medical Research; Westmead Hospital; University of Sydney | CIDM-PH et al. |
| EPI_ISL_526156 | Pathology West - NSW Health Pathology | NSW Health Pathology - Institute of Clinical Pathology and Medical Research; Westmead Hospital; University of Sydney | CIDM-PH et al. |
| EPI_ISL_526157, EPI_ISL_526158, EPI_ISL_526159, EPI_ISL_526160 | Sydney South West Pathology Service (SSWPS) - Liverpool Hospital - NSW Health Pathology | NSW Health Pathology - Institute of Clinical Pathology and Medical Research; Westmead Hospital; University of Sydney | CIDM-PH et al. |
| EPI_ISL_526161, EPI_ISL_526162, EPI_ISL_526163, EPI_ISL_526164, EPI_ISL_526165 | Pathology West - NSW Health Pathology | NSW Health Pathology - Institute of Clinical Pathology and Medical Research; Westmead Hospital; University of Sydney | CIDM-PH et al. |
| EPI_ISL_526166, EPI_ISL_526167, EPI_ISL_526168, EPI_ISL_526169 | Austech Medical Laboratories | NSW Health Pathology - Institute of Clinical Pathology and Medical Research; Westmead Hospital; University of Sydney | CIDM-PH et al. |
| EPI_ISL_526170 | Pathology North - Royal North Shore Hospital - NSW Health Pathology | NSW Health Pathology - Institute of Clinical Pathology and Medical Research; Westmead Hospital; University of Sydney | CIDM-PH et al. |
| EPI_ISL_526171 | St Vincent's Pathology (SydPath) | NSW Health Pathology - Institute of Clinical Pathology and Medical Research; Westmead Hospital; University of Sydney | CIDM-PH et al. |
| EPI_ISL_526172, EPI_ISL_526173, EPI_ISL_526174, EPI_ISL_526175, EPI_ISL_526176 | Sydney South West Pathology Service (SSWPS) - Liverpool Hospital - NSW Health Pathology | NSW Health Pathology - Institute of Clinical Pathology and Medical Research; Westmead Hospital; University of Sydney | CIDM-PH et al. |
| EPI_ISL_526177, EPI_ISL_526178, EPI_ISL_526179, EPI_ISL_526180, EPI_ISL_526181, EPI_ISL_526182, EPI_ISL_526183, EPI_ISL_526184, EPI_ISL_526185 | St Vincent's Pathology (SydPath) | NSW Health Pathology - Institute of Clinical Pathology and Medical Research; Westmead Hospital; University of Sydney | CIDM-PH et al. |
| EPI_ISL_526186 | 4Cyte Pathology | NSW Health Pathology - Institute of Clinical Pathology and Medical Research; Westmead Hospital; University of Sydney | CIDM-PH et al. |
| EPI_ISL_526187, EPI_ISL_526188, EPI_ISL_526189, EPI_ISL_526190 | St Vincent's Pathology (SydPath) | NSW Health Pathology - Institute of Clinical Pathology and Medical Research; Westmead Hospital; University of Sydney | CIDM-PH et al. |
| EPI_ISL_526191 | South Eastern Area Laboratory Services (SEALS) | NSW Health Pathology - Institute of Clinical Pathology and Medical Research; Westmead Hospital; University of Sydney | CIDM-PH et al. |
| EPI_ISL_526192 | Medlab Pathology | NSW Health Pathology - Institute of Clinical Pathology and Medical Research; Westmead Hospital; University of Sydney | CIDM-PH et al. |
| EPI_ISL_526193 | Sydney South West Pathology Service (SSWPS) - Liverpool Hospital - NSW Health Pathology | NSW Health Pathology - Institute of Clinical Pathology and Medical Research; Westmead Hospital; University of Sydney | CIDM-PH et al. |
| EPI_ISL_526194, EPI_ISL_526195 | St Vincent's Pathology (SydPath) | NSW Health Pathology - Institute of Clinical Pathology and Medical Research; Westmead Hospital; University of Sydney | CIDM-PH et al. |
| EPI_ISL_526196 | Sydney South West Pathology Service (SSWPS) - Royal Prince Alfred Hospital - NSW Health Pathology | NSW Health Pathology - Institute of Clinical Pathology and Medical Research; Westmead Hospital; University of Sydney | CIDM-PH et al. |
| EPI_ISL_526197, EPI_ISL_526198 | Sydney South West Pathology Service (SSWPS) - Liverpool Hospital - NSW Health Pathology | NSW Health Pathology - Institute of Clinical Pathology and Medical Research; Westmead Hospital; University of Sydney | CIDM-PH et al. |
| EPI_ISL_526199 | Pathology West - NSW Health Pathology | NSW Health Pathology - Institute of Clinical Pathology and Medical Research; Westmead Hospital; University of Sydney | CIDM-PH et al. |
| EPI_ISL_526200 | St Vincent's Pathology (SydPath) | NSW Health Pathology - Institute of Clinical Pathology and Medical Research; Westmead Hospital; University of Sydney | CIDM-PH et al. |
| EPI_ISL_526201 | Sydney South West Pathology Service (SSWPS) - Royal Prince Alfred Hospital - NSW Health Pathology | NSW Health Pathology - Institute of Clinical Pathology and Medical Research; Westmead Hospital; University of Sydney | CIDM-PH et al. |
| EPI_ISL_526202, EPI_ISL_526203, EPI_ISL_526204 | Laverty Pathology | NSW Health Pathology - Institute of Clinical Pathology and Medical Research; Westmead Hospital; University of Sydney | CIDM-PH et al. |
| EPI_ISL_526205 | The Children's Hospital at Westmead | NSW Health Pathology - Institute of Clinical Pathology and Medical Research; Westmead Hospital; University of Sydney | CIDM-PH et al. |
| EPI_ISL_526206 | Sydney South West Pathology Service (SSWPS) - Liverpool Hospital - NSW Health Pathology | NSW Health Pathology - Institute of Clinical Pathology and Medical Research; Westmead Hospital; University of Sydney | CIDM-PH et al. |
| EPI_ISL_526207, EPI_ISL_526208, EPI_ISL_526209 | Sydney South West Pathology Service (SSWPS) - Royal Prince Alfred Hospital - NSW Health Pathology | NSW Health Pathology - Institute of Clinical Pathology and Medical Research; Westmead Hospital; University of Sydney | CIDM-PH et al. |
| EPI_ISL_526210, EPI_ISL_526211 | Sydney South West Pathology Service (SSWPS) - Liverpool Hospital - NSW Health Pathology | NSW Health Pathology - Institute of Clinical Pathology and Medical Research; Westmead Hospital; University of Sydney | CIDM-PH et al. |
| EPI_ISL_526212 | Australian Clinical Labs (formerly Healthscope Pathology) | NSW Health Pathology - Institute of Clinical Pathology and Medical Research; Westmead Hospital; University of Sydney | CIDM-PH et al. |
| EPI_ISL_526213, EPI_ISL_526214 | Douglass Hanly Moir Pathology | NSW Health Pathology - Institute of Clinical Pathology and Medical Research; Westmead Hospital; University of Sydney | CIDM-PH et al. |
| EPI_ISL_526215, EPI_ISL_526216, EPI_ISL_526217, EPI_ISL_526218, EPI_ISL_526219, EPI_ISL_526220, EPI_ISL_526221, EPI_ISL_526222, EPI_ISL_526223, EPI_ISL_526224, EPI_ISL_526225, EPI_ISL_526226, EPI_ISL_526227, EPI_ISL_526228, EPI_ISL_526229, EPI_ISL_526230, EPI_ISL_526231, EPI_ISL_526232, EPI_ISL_526233, EPI_ISL_526234, EPI_ISL_526235, EPI_ISL_526236, EPI_ISL_526237, EPI_ISL_526238 | | | |
| see above | Hungarian Defence Forces Military Medical Centre | National Laboratory of Virology, Szentágothai Research Centre | Endre Gábor Tóth, Balázs Somogyi, Bálint Eszenyi, Ferenc Jakab, Gábor Kemenesi |
| EPI_ISL_526253, EPI_ISL_526254, EPI_ISL_526255, EPI_ISL_526256, EPI_ISL_526257, EPI_ISL_526258, EPI_ISL_526259, EPI_ISL_526260, EPI_ISL_526261, EPI_ISL_526262, EPI_ISL_526263, EPI_ISL_526264, EPI_ISL_526265, EPI_ISL_526266, EPI_ISL_526267, EPI_ISL_526268, EPI_ISL_526269, EPI_ISL_526270, EPI_ISL_526271, EPI_ISL_526272, EPI_ISL_526273, EPI_ISL_526274, EPI_ISL_526275, EPI_ISL_526276, EPI_ISL_526277, EPI_ISL_526278, EPI_ISL_526279, EPI_ISL_526280, EPI_ISL_526281, EPI_ISL_526282, EPI_ISL_526283, EPI_ISL_526284, EPI_ISL_526285, EPI_ISL_526286 | | | |
| see above | Unity Health Toronto | Ontario Institute for Cancer Research | Ramzi Fattouh, Larissa M. Matukas, Mark Downing, Annette Gower, Karel Boissinot, Samira Mubareka, TIBDN, Ilinca Lungu, Bernard Lam, Jeremy Johns, Paul Krzyzanowski, |

| | | | |
|---|---|---|---|
| | | | Richard de Borja, Felicia Vincelli, Philip Zuzarte, Jared Simpson |
| EPI_ISL_526287, EPI_ISL_526288, EPI_ISL_526289, EPI_ISL_526290, EPI_ISL_526291, EPI_ISL_526292, EPI_ISL_526293, EPI_ISL_526294, EPI_ISL_526295, EPI_ISL_526296, EPI_ISL_526297, EPI_ISL_526298, EPI_ISL_526299, EPI_ISL_526300, EPI_ISL_526301, EPI_ISL_526302, EPI_ISL_526303, EPI_ISL_526304, EPI_ISL_526305, EPI_ISL_526306, EPI_ISL_526307, EPI_ISL_526308, EPI_ISL_526309, EPI_ISL_526310, EPI_ISL_526311, EPI_ISL_526312, EPI_ISL_526313, EPI_ISL_526314, EPI_ISL_526315, EPI_ISL_526316, EPI_ISL_526317, EPI_ISL_526318, EPI_ISL_526319, EPI_ISL_526320, EPI_ISL_526321, EPI_ISL_526322, EPI_ISL_526323, EPI_ISL_526324, EPI_ISL_526325, EPI_ISL_526326, EPI_ISL_526327, EPI_ISL_526328, EPI_ISL_526329, EPI_ISL_526330, EPI_ISL_526331, EPI_ISL_526332, EPI_ISL_526333, EPI_ISL_526334, EPI_ISL_526335, EPI_ISL_526336 | | | |
| see above | University of Birmingham | COVID-19 Genomics UK (COG-UK) Consortium | Institute of Microbiology, University of Birmingham: Claire McMurray, Joanne Stockton, Samuel Nicholls, Radoslaw Poplawski, Will Rowe, Josh Quick, Nicholas Loman. University of Birmingham Testing Laboratory: Celina M Whalley, Andrew Bosworth, Charlotte Poxon, Kasun Wanigasooriya, Oliver Pickles, Mike Kidd, Alex Richter, Andrew D Beggs PHE Heartlands Lab: Husam Osman, Andrew Bosworth. Queen Elizabeth Hospital: Anna Casey |
| EPI_ISL_526337, EPI_ISL_526338, EPI_ISL_526339, EPI_ISL_526340, EPI_ISL_526341, EPI_ISL_526342, EPI_ISL_526343, EPI_ISL_526344, EPI_ISL_526345, EPI_ISL_526346, EPI_ISL_526347, EPI_ISL_526348, EPI_ISL_526349, EPI_ISL_526350, EPI_ISL_526351, EPI_ISL_526352, EPI_ISL_526353, EPI_ISL_526354, EPI_ISL_526355, EPI_ISL_526356, EPI_ISL_526357, EPI_ISL_526358, EPI_ISL_526359, EPI_ISL_526360, EPI_ISL_526361, EPI_ISL_526362, EPI_ISL_526363, EPI_ISL_526364, EPI_ISL_526365, EPI_ISL_526366 | | | |
| see above | Queens Medical Centre, Clinical Microbiology Department / DeepSeq Nottingham | COVID-19 Genomics UK (COG-UK) Consortium | Gemma Clark, Wendy Smith, Manjinder Khakh, Vicki M Fleming, Michelle M Lister, Hannah Howson-Wells, Jonathan Ball, Patrick McClure, Joseph Chappell, Theocharis Tsoleridis, Nadine Holmes, Matthew Carlisle, Christopher Moore, Fei Sang, Johnny Debebe, Victoria Wright, Matthew Loose |
| EPI_ISL_526367, EPI_ISL_526368, EPI_ISL_526369, EPI_ISL_526370, EPI_ISL_526371, EPI_ISL_526372, EPI_ISL_526373, EPI_ISL_526374, EPI_ISL_526375, EPI_ISL_526376, EPI_ISL_526377, EPI_ISL_526378, EPI_ISL_526379, EPI_ISL_526380, EPI_ISL_526381, EPI_ISL_526382, EPI_ISL_526383, EPI_ISL_526384, EPI_ISL_526385, EPI_ISL_526386, EPI_ISL_526387, EPI_ISL_526388, EPI_ISL_526389, EPI_ISL_526390, EPI_ISL_526391, EPI_ISL_526392, EPI_ISL_526393, EPI_ISL_526394 | | | |
| see above | Liverpool Clinical Laboratories | COVID-19 Genomics UK (COG-UK) Consortium | Sam Haldenby, Anita Lucaci, Steve Paterson, Julian Hiscox, Alistair Darby, M Almsaud, A Alrezaihi, Muhannad Alruwaili, Stuart D Armstrong, Jones Benjamin, Eleanor G Bentley, Anu Chawla, Jordan J Clark, Angela Cowell, Richard Eccles, Isabel García-Dorival, Matthew Gemmell, Alessandro Gerada, PKF Gilmore, Richard Gregory, Ximeng Han, Catherine Hartley, Margaret Hughes, Miren Iturriza-Gomara, James Johnson, L Luu, Jenifer Manson, Charlotte Nelson, Elaine O'Toole, Cassie Olateju, Rebekah Penrice-Randal , Lucille Rainbow, N.P Randle, Trevor Ian Robinson, Parul Sharma, Ghada T Shawli, James P Stewart, Neil Swainston, Ecaterina Vamos, Joanne Watts, Mark Whitehead |
| EPI_ISL_526395, EPI_ISL_526396, EPI_ISL_526397, EPI_ISL_526398, EPI_ISL_526399, EPI_ISL_526400, EPI_ISL_526401, EPI_ISL_526402, EPI_ISL_526403, EPI_ISL_526404, EPI_ISL_526405, EPI_ISL_526406, EPI_ISL_526407, EPI_ISL_526408, EPI_ISL_526409, EPI_ISL_526410, EPI_ISL_526411, EPI_ISL_526412, EPI_ISL_526413, EPI_ISL_526414, EPI_ISL_526415, EPI_ISL_526416, EPI_ISL_526417, EPI_ISL_526418, EPI_ISL_526419, EPI_ISL_526420, EPI_ISL_526421, EPI_ISL_526422, EPI_ISL_526423, EPI_ISL_526424, EPI_ISL_526425, EPI_ISL_526426, EPI_ISL_526427, EPI_ISL_526428, EPI_ISL_526429, EPI_ISL_526430 | | | |
| see above | Queens Medical Centre, Clinical Microbiology Department / DeepSeq Nottingham | COVID-19 Genomics UK (COG-UK) Consortium | Gemma Clark, Wendy Smith, Manjinder Khakh, Vicki M Fleming, Michelle M Lister, Hannah Howson-Wells, Jonathan Ball, Patrick McClure, Joseph Chappell, Theocharis Tsoleridis, Nadine Holmes, Matthew Carlisle, Christopher Moore, Fei Sang, Johnny Debebe, Victoria Wright, Matthew Loose |
| EPI_ISL_526431, EPI_ISL_526432 | Lincolnshire Hospitals and DeepSeq Nottingham | COVID-19 Genomics UK (COG-UK) Consortium | Nichola Duckworth, Tim Sloan, Sarah Walsh, Jonathan Ball, Patrick McClure, Joeseph Chappell, Nadine Holmes, Matthew Carlisle, Christopher Moore, Fei Sang, Johnny Debebe, Victoria Wright, Matthew Loose |
| EPI_ISL_526433, EPI_ISL_526434, EPI_ISL_526435, EPI_ISL_526436 | Centre for Enzyme Innovation, University of Portsmouth / Translational Research Laboratory, Portsmouth Hospitals NHS Trust | COVID-19 Genomics UK (COG-UK) Consortium | Angela Beckett,Yann Bourgeois,Garry Scarlett,Sharon Glaysher,Scott Elliott,Kelly Bicknell,Robert Impey,Allyson Lloyd,Sarah Wyllie,Ethan Butcher,Anoop Chauhan,Samuel Robson |
| EPI_ISL_526437, EPI_ISL_526438, EPI_ISL_526439, EPI_ISL_526440, EPI_ISL_526441, EPI_ISL_526442, EPI_ISL_526443, EPI_ISL_526444, EPI_ISL_526445, EPI_ISL_526446, EPI_ISL_526447, EPI_ISL_526448, EPI_ISL_526449, EPI_ISL_526450 | | | |
| see above | Virology Department, Sheffield Teaching Hospitals NHS Foundation Trust/Department of Infection, Immunity and Cardiovascular Disease, The Medical School, University of Sheffield | COVID-19 Genomics UK (COG-UK) Consortium | Thushan de Silva, Matthew Parker, Nikki Smith, Adri Angyal, Rebecca Brown, Luke Green, Rachel Tucker, Paul Parsons, Danielle Groves, Katie Johnson, Laura Carrilero, Alex Keeley, Dave Partridge, Matthew Wyles, Benjamin Lindsey, Mehmet Yavuz, Mohammad Raza, Cariad Evans |
| EPI_ISL_526451, EPI_ISL_526452, EPI_ISL_526453, EPI_ISL_526454, EPI_ISL_526455, EPI_ISL_526456, EPI_ISL_526457, EPI_ISL_526458 | West of Scotland Specialist Virology Centre, NHSGGC / MRC-University of Glasgow Centre for Virus Research | COVID-19 Genomics UK (COG-UK) Consortium | Ana da Silva Filipe, Natasha Johnson, Kathy Smollett, Daniel Mair, Stephen Carmichael, Lily Tong, Jenna Nichols, Elihu Aranday-Cortes, Kirstyn Brunker, Yasmin Parr, Alice Broos, Kyriaki Nomikou; Sarah McDonald, Marc Niebel, Patawee Asamaphan; Richard Orton, Joseph Hughes, Sreenu Vattipally, David L Robertson; Alasdair MacLean, Rory Gunson; Kathy Li, Natasha Jesudason, Rajiv Shah, James Shepherd, Antonia Ho, Emma Thomson |
| EPI_ISL_526459, EPI_ISL_526460, EPI_ISL_526461, EPI_ISL_526462, EPI_ISL_526463, EPI_ISL_526464, EPI_ISL_526465, EPI_ISL_526466, EPI_ISL_526467, EPI_ISL_526468, EPI_ISL_526469, EPI_ISL_526470, EPI_ISL_526471, EPI_ISL_526472, EPI_ISL_526473, EPI_ISL_526474, EPI_ISL_526475, EPI_ISL_526476, EPI_ISL_526477, EPI_ISL_526478, EPI_ISL_526479, EPI_ISL_526480, EPI_ISL_526481, EPI_ISL_526482, EPI_ISL_526483, EPI_ISL_526484, EPI_ISL_526485, EPI_ISL_526486, EPI_ISL_526487, EPI_ISL_526488, EPI_ISL_526489, EPI_ISL_526490, EPI_ISL_526491, EPI_ISL_526492, EPI_ISL_526493, EPI_ISL_526494, EPI_ISL_526495, EPI_ISL_526496, EPI_ISL_526497, EPI_ISL_526498, EPI_ISL_526499, EPI_ISL_526500, EPI_ISL_526501, EPI_ISL_526502, EPI_ISL_526503, EPI_ISL_526504, EPI_ISL_526505, EPI_ISL_526506, EPI_ISL_526507, EPI_ISL_526508, EPI_ISL_526509, EPI_ISL_526510, EPI_ISL_526511, EPI_ISL_526512, EPI_ISL_526513, EPI_ISL_526514, EPI_ISL_526515, EPI_ISL_526516, EPI_ISL_526517, EPI_ISL_526518, EPI_ISL_526519, EPI_ISL_526520, EPI_ISL_526521, EPI_ISL_526522, EPI_ISL_526523, EPI_ISL_526524, EPI_ISL_526525, EPI_ISL_526526, EPI_ISL_526527, EPI_ISL_526528, EPI_ISL_526529, EPI_ISL_526530, EPI_ISL_526531, EPI_ISL_526532, EPI_ISL_526533, EPI_ISL_526534, EPI_ISL_526535, EPI_ISL_526536 | | | |
| see above | Virology Department, Royal Infirmary of Edinburgh, NHS Lothian / School of Biological Sciences, University of Edinburgh / Institute of Genetics and Molecular Medicine, University of Edinburgh | COVID-19 Genomics UK (COG-UK) Consortium | McHugh M, Dewar R, Rooke S, Gallagher M, Balcaza C, O'Toole Á, Scher E, Hill V, McCrone JT, Colquhoun R, Yu X, Jackson B, Rambaut A, Williams TC, Templeton K |
| EPI_ISL_526537, EPI_ISL_526538, EPI_ISL_526539, EPI_ISL_526540, EPI_ISL_526541, EPI_ISL_526542, EPI_ISL_526543, EPI_ISL_526544, EPI_ISL_526545, EPI_ISL_526546, EPI_ISL_526547, EPI_ISL_526548, EPI_ISL_526549, EPI_ISL_526550 | | | |
| see above | Respiratory Virus Unit, Microbiology Services Colindale, Public Health England | Respiratory Virus Unit, Microbiology Services Colindale, Public Health England | PHE Covid Sequencing Team |
| EPI_ISL_526551, EPI_ISL_526552 | OHSU Lab Services Molecular Microbiology Lab | Oregon SARS-CoV-2 Genome Sequencing Center | Brendan L. O'Connell, Ruth V. Nichols, Alec J. Hirsch, Guang Fan, Daniel N. Streblow, William B. Messer, Andrew C. Adey, Benjamin N. Bimber, Brian J. O'Roak |
| EPI_ISL_526553, EPI_ISL_526554, EPI_ISL_526555, EPI_ISL_526556, EPI_ISL_526557, EPI_ISL_526558, EPI_ISL_526559, EPI_ISL_526560, EPI_ISL_526561, EPI_ISL_526562, EPI_ISL_526563, EPI_ISL_526564, EPI_ISL_526565, EPI_ISL_526566, EPI_ISL_526567, EPI_ISL_526568, EPI_ISL_526569, EPI_ISL_526570, EPI_ISL_526571, EPI_ISL_526572, EPI_ISL_526573, EPI_ISL_526574, EPI_ISL_526575, EPI_ISL_526576, EPI_ISL_526577, EPI_ISL_526578, EPI_ISL_526579, EPI_ISL_526580, EPI_ISL_526581, EPI_ISL_526582, EPI_ISL_526583, EPI_ISL_526584, EPI_ISL_526585, EPI_ISL_526586, EPI_ISL_526587, EPI_ISL_526588, EPI_ISL_526589, EPI_ISL_526590, EPI_ISL_526591, EPI_ISL_526592, EPI_ISL_526593, EPI_ISL_526594, EPI_ISL_526595, EPI_ISL_526596, EPI_ISL_526597, EPI_ISL_526598, EPI_ISL_526599, EPI_ISL_526600, EPI_ISL_526601, EPI_ISL_526602, EPI_ISL_526603, EPI_ISL_526604, EPI_ISL_526605, EPI_ISL_526606, EPI_ISL_526607, EPI_ISL_526608, EPI_ISL_526609, EPI_ISL_526610, EPI_ISL_526611, EPI_ISL_526612, EPI_ISL_526613, EPI_ISL_526614, EPI_ISL_526615, EPI_ISL_526616, EPI_ISL_526617, EPI_ISL_526618, EPI_ISL_526619, EPI_ISL_526620, EPI_ISL_526621, EPI_ISL_526622, EPI_ISL_526623, EPI_ISL_526624, EPI_ISL_526625, EPI_ISL_526626, EPI_ISL_526627, EPI_ISL_526628, EPI_ISL_526629, EPI_ISL_526630, EPI_ISL_526631, EPI_ISL_526632, EPI_ISL_526633, EPI_ISL_526634, EPI_ISL_526635, EPI_ISL_526636, EPI_ISL_526637, EPI_ISL_526638, EPI_ISL_526639, EPI_ISL_526640, EPI_ISL_526641, EPI_ISL_526642, EPI_ISL_526643, EPI_ISL_526644, EPI_ISL_526645, EPI_ISL_526646, EPI_ISL_526647, EPI_ISL_526648, EPI_ISL_526649, EPI_ISL_526650, EPI_ISL_526651, EPI_ISL_526652, EPI_ISL_526653, EPI_ISL_526654, EPI_ISL_526655, EPI_ISL_526656, EPI_ISL_526657, EPI_ISL_526658, EPI_ISL_526659, EPI_ISL_526660, EPI_ISL_526661, EPI_ISL_526662, EPI_ISL_526663, EPI_ISL_526664, EPI_ISL_526665, EPI_ISL_526666, EPI_ISL_526667, EPI_ISL_526668, EPI_ISL_526669, EPI_ISL_526670, EPI_ISL_526671, EPI_ISL_526672, EPI_ISL_526673, EPI_ISL_526674, EPI_ISL_526675, EPI_ISL_526676, EPI_ISL_526677, EPI_ISL_526678, EPI_ISL_526679, EPI_ISL_526680, EPI_ISL_526681, EPI_ISL_526682, EPI_ISL_526683, EPI_ISL_526684, EPI_ISL_526685 | | | |
| see above | Florida Bureau of Public Health Laboratories | Florida Bureau of Public Health Laboratories | Sarah Schmedes, Jason Blanton |
| EPI_ISL_526686, EPI_ISL_526687, EPI_ISL_526688 | Faith Laboratory, Immunology Institute, Icahn School of Medicine at Mount Sinai | van Bakel Laboratory, Genetics and Genomics Sciences, Icahn School of Medicine at Mount Sinai | Graham J. Britton, Alice Chen-Liaw, Francesca Cossarini, Alexandra Livanos, Matthew P. Spindler, Tamar Plitt, Joseph Eggers, Ilaria Mogno, Ana S. Gonzalez-Reiche, Sophia Sui, Michael Tankelevich, Lauren Tal Grinspan, Rebekah E. Dixon, Divya Jha, Gustavo Martinez-Delgado, Fatima Amanat, Daisy Hoagland, Benjamin R. tenOever, Marla C. Dubinsky, Miriam Merad, Harm Van Bakel, Florian Krammer, Gerold Bongers, Saurabh Mehandru and Jeremiah J. Faith |
| EPI_ISL_526689 | Pathology West - NSW Health Pathology | NSW Health Pathology - Institute of Clinical Pathology and Medical Research; Westmead Hospital; University of Sydney | CIDM-PH et al. |
| EPI_ISL_526697 | South Eastern Area Laboratory Services (SEALS) | NSW Health Pathology - Institute of Clinical Pathology and Medical Research; Westmead Hospital; University of Sydney | CIDM-PH et al. |
| EPI_ISL_526700, EPI_ISL_526701, EPI_ISL_526702, EPI_ISL_526703, EPI_ISL_526704, EPI_ISL_526705, EPI_ISL_526706, EPI_ISL_526707, EPI_ISL_526708, EPI_ISL_526709, EPI_ISL_526710, EPI_ISL_526711, EPI_ISL_526712, EPI_ISL_526713, EPI_ISL_526714, EPI_ISL_526715, EPI_ISL_526716, EPI_ISL_526717, EPI_ISL_526718, EPI_ISL_526719, EPI_ISL_526720, EPI_ISL_526721, EPI_ISL_526722, EPI_ISL_526723, EPI_ISL_526724, EPI_ISL_526725, EPI_ISL_526726, EPI_ISL_526727, EPI_ISL_526728, EPI_ISL_526729 | | | |
| see above | Division of Viral Diseases, Center for Laboratory Control of Infectious Diseases, Korea Centers for Diseases Control and Prevention | Division of Viral Diseases, Center for Laboratory Control of Infectious Diseases, Korea Centers for Diseases Control and Prevention | Jeong-Min Kim, Yoon-Seok Chung, Namjoo Lee, Sang Hee Woo, Hye-Jun Jo, Heui Man Kim, Jun-Sub Kim, Myung Guk Han |
| EPI_ISL_526730, EPI_ISL_526731 | Center for Laboratory Control of Infectious Diseases, Korea Centers for Diseases Control and Prevention | Center for Laboratory Control of Infectious Diseases, Korea Centers for Diseases Control and Prevention | Junyoung Kim, Ae Kyung Park, Eunkyung Shin, Jin Sun No, Jeong-Min Kim, Yoon-Seok Chung, Heui Man Kim, Myung Guk Han |
| EPI_ISL_526732, EPI_ISL_526733 | Division of Viral Diseases, Center for Laboratory Control of Infectious Diseases, Korea Centers for Diseases Control and Prevention | Division of Viral Diseases, Center for Laboratory Control of Infectious Diseases, Korea Centers for Diseases Control and Prevention | Jeong-Min Kim, Yoon-Seok Chung, Namjoo Lee, Sang Hee Woo, Hye-Jun Jo, Heui Man Kim, Jun-Sub Kim, Myung Guk Han |
| EPI_ISL_526734, EPI_ISL_526735, EPI_ISL_526736, EPI_ISL_526737, EPI_ISL_526738, EPI_ISL_526739, EPI_ISL_526740, EPI_ISL_526741, EPI_ISL_526742, EPI_ISL_526743, EPI_ISL_526744, EPI_ISL_526745, EPI_ISL_526746 | | | |
| see above | Center for Laboratory Control of Infectious Diseases, Korea Centers for Diseases Control and Prevention | Center for Laboratory Control of Infectious Diseases, Korea Centers for Diseases Control and Prevention | Junyoung Kim, Ae Kyung Park, Eunkyung Shin, Jin Sun No, Jeong-Min Kim, Yoon-Seok Chung, Heui Man Kim, Myung Guk Han |
| EPI_ISL_526754, EPI_ISL_526755, EPI_ISL_526756, EPI_ISL_526757, EPI_ISL_526758, EPI_ISL_526759, EPI_ISL_526760, EPI_ISL_526761, EPI_ISL_526762, EPI_ISL_526763, EPI_ISL_526764, EPI_ISL_526765, EPI_ISL_526766, EPI_ISL_526767, EPI_ISL_526768, EPI_ISL_526769, EPI_ISL_526770, EPI_ISL_526771, EPI_ISL_526772, EPI_ISL_526773 | | | |
| see above | Respiratory Virus Unit, Microbiology Services Colindale, Public Health England | Respiratory Virus Unit, Microbiology Services Colindale, Public Health England | PHE Covid Sequencing Team |
| EPI_ISL_526775, EPI_ISL_526776, EPI_ISL_526777, EPI_ISL_526778, EPI_ISL_526779, EPI_ISL_526780, EPI_ISL_526781, EPI_ISL_526782, EPI_ISL_526783, EPI_ISL_526784, EPI_ISL_526785, EPI_ISL_526786, EPI_ISL_526787, EPI_ISL_526788, EPI_ISL_526789, EPI_ISL_526790, EPI_ISL_526791, EPI_ISL_526792, EPI_ISL_526793, EPI_ISL_526794, EPI_ISL_526795, EPI_ISL_526796, EPI_ISL_526797, EPI_ISL_526798, EPI_ISL_526799, EPI_ISL_526800, EPI_ISL_526801, EPI_ISL_526802, EPI_ISL_526803, EPI_ISL_526804, EPI_ISL_526805, EPI_ISL_526806, EPI_ISL_526807, EPI_ISL_526808, EPI_ISL_526809, EPI_ISL_526810, EPI_ISL_526811, EPI_ISL_526812, EPI_ISL_526813, EPI_ISL_526814, EPI_ISL_526815, EPI_ISL_526816, EPI_ISL_526817, EPI_ISL_526818, EPI_ISL_526819, EPI_ISL_526820, EPI_ISL_526821, EPI_ISL_526822, EPI_ISL_526823, EPI_ISL_526824, EPI_ISL_526825, EPI_ISL_526826, EPI_ISL_526827, EPI_ISL_526828, EPI_ISL_526829, EPI_ISL_526830, EPI_ISL_526831, EPI_ISL_526832, EPI_ISL_526833, EPI_ISL_526834, EPI_ISL_526835, EPI_ISL_526836, EPI_ISL_526837, EPI_ISL_526838, EPI_ISL_526839, EPI_ISL_526840, EPI_ISL_526841, EPI_ISL_526842, EPI_ISL_526843, EPI_ISL_526844, EPI_ISL_526845, EPI_ISL_526846, EPI_ISL_526847, EPI_ISL_526848, EPI_ISL_526849, EPI_ISL_526850, EPI_ISL_526851, EPI_ISL_526852, EPI_ISL_526853, EPI_ISL_526854, EPI_ISL_526855, EPI_ISL_526856, EPI_ISL_526857, EPI_ISL_526858, EPI_ISL_526859, EPI_ISL_526860, EPI_ISL_526861, EPI_ISL_526862, EPI_ISL_526863, EPI_ISL_526864, EPI_ISL_526865, EPI_ISL_526866, EPI_ISL_526867, EPI_ISL_526868, EPI_ISL_526869, EPI_ISL_526870, EPI_ISL_526871, EPI_ISL_526872, EPI_ISL_526873, EPI_ISL_526874, EPI_ISL_526875, EPI_ISL_526876, EPI_ISL_526877, EPI_ISL_526878, EPI_ISL_526879, EPI_ISL_526880, EPI_ISL_526881, EPI_ISL_526882, EPI_ISL_526883, EPI_ISL_526884, EPI_ISL_526885, EPI_ISL_526886, EPI_ISL_526887, EPI_ISL_526888, EPI_ISL_526889, EPI_ISL_526890, EPI_ISL_526891, EPI_ISL_526892, EPI_ISL_526893, EPI_ISL_526894, EPI_ISL_526895, EPI_ISL_526896, EPI_ISL_526897, EPI_ISL_526898, EPI_ISL_526899, EPI_ISL_526900, EPI_ISL_526901, EPI_ISL_526902, EPI_ISL_526903, EPI_ISL_526904, EPI_ISL_526905, EPI_ISL_526906, EPI_ISL_526907, EPI_ISL_526908, EPI_ISL_526909, EPI_ISL_526910, EPI_ISL_526911, EPI_ISL_526912, EPI_ISL_526913, EPI_ISL_526914, EPI_ISL_526915, EPI_ISL_526916, EPI_ISL_526917, EPI_ISL_526918, EPI_ISL_526919, EPI_ISL_526920, EPI_ISL_526921, EPI_ISL_526922, EPI_ISL_526923, EPI_ISL_526924, EPI_ISL_526925, EPI_ISL_526926, EPI_ISL_526927, EPI_ISL_526928, EPI_ISL_526929, EPI_ISL_526930, EPI_ISL_526931 | | | |
| see above | Virginia DCLS | Virginia DCLS | Virginia DCLS |
| EPI_ISL_526932, EPI_ISL_526933, EPI_ISL_526934 | Instituto Nacional de Salud, Bogotá, Colombia | Instituto Nacional de Salud, Bogotá, Colombia | Katherine Laiton-Donato, Diego A. Álvarez-Díaz, Carlos Franco-Muñoz, Mauricio Pacheco-Montealegre, Jonathan Reales, Diego Andrés Prada, Jose A. Usme-Ciro, Zulma M. Cucunubá, Christian Julian Villabona-Arenas, Liz Villabona-Arenas, Sussy Echeverria, Astrid C. Flórez, Carolina Ferro, Diana Marcela Walteros-Acero, Franklin Prieto, Carlos Andrés Durán, Martha Lucia Ospina Martinez, Marcela Mercado-Reyes |
| EPI_ISL_526935, EPI_ISL_526936, EPI_ISL_526937, EPI_ISL_526938, EPI_ISL_526939, EPI_ISL_526940, EPI_ISL_526941, EPI_ISL_526942, EPI_ISL_526943, EPI_ISL_526944, EPI_ISL_526945, EPI_ISL_526946, EPI_ISL_526947, EPI_ISL_526948 | | | |
| see above | Faroese National Reference Laboratory for Fish and Animal Diseases | Faroese National Reference Laboratory for Fish and Animal Diseases | Maria Marjunardóttir Dahl, Petra Elisabeth Petersen, Debes Hammershaimb Christiansen |
| EPI_ISL_526949, EPI_ISL_526950 | Instituto Nacional de Salud, Bogotá, Colombia | Instituto Nacional de Salud, Bogotá, Colombia | Katherine Laiton-Donato, Diego A. Álvarez-Díaz, Carlos Franco-Muñoz, Mauricio Pacheco-Montealegre, Jonathan Reales, Diego Andrés Prada, Jose A. Usme-Ciro, Zulma M. Cucunubá, Christian Julian Villabona-Arenas, Liz Villabona-Arenas, Sussy Echeverria, Astrid C. Flórez, Carolina Ferro, Diana Marcela Walteros-Acero, Franklin Prieto, Carlos Andrés Durán, Martha Lucia Ospina Martinez, Marcela Mercado-Reyes |
| EPI_ISL_526951 | Instituto Nacional de Salud, Bogotá, Colombia | Instituto Nacional de Salud, Bogotá, Colombia | Katherine Laiton-Donato, Diego A. Álvarez-Díaz, Carlos Franco-Muñoz, Mauricio Pacheco-Montealegre, Jonathan Reales, Diego Andrés Prada, Jose A. Usme-Ciro, Zulma M. Cucunubá, Christian Julian Villabona-Arenas, Liz Villabona-Arenas, Sussy Echeverria, Astrid C. Flórez, Carolina Ferro, Diana Marcela Walteros-Acero, Franklin Prieto, Carlos |

| | | | |
|---|---|---|---|
| | | | Andrés Durán, Martha Lucia Ospina Martinez, Marcela Mercado-Reyes |
| EPI_ISL_526953, EPI_ISL_526954, EPI_ISL_526955, EPI_ISL_526956, EPI_ISL_526957, EPI_ISL_526958, EPI_ISL_526960, EPI_ISL_526961 | Instituto Nacional de Salud, Bogotá, Colombia | Instituto Nacional de Salud, Bogotá, Colombia | Katherine Laiton-Donato, Diego A. Álvarez-Díaz, Carlos Franco-Muñoz, Mauricio Pacheco-Montealegre, Jonathan Reales, Diego Andrés Prada, Jose A. Usme-Ciro, Zulma M. Cucunubá, Christian Julian Villabona-Arenas, Liz Villabona-Arenas, Sussy Echeverria, Astrid C. Flórez, Carolina Ferro, Diana Marcela Walteros-Acero, Franklin Prieto, Carlos Andrés Durán, Martha Lucia Ospina Martinez, Marcela Mercado-Reyes |
| EPI_ISL_526962, EPI_ISL_526963, EPI_ISL_526964, EPI_ISL_526965 | Instituto Nacional de Salud, Bogotá, Colombia | Instituto Nacional de Salud, Bogotá, Colombia | Katherine Laiton-Donato, Diego A. Álvarez-Díaz, Carlos Franco-Muñoz, Jonathan Reales, Diego Andrés Prada, Jeadran Malagón-Rojas, Felix Betzler, Wendy K. Jo, Edmilson F. de Oliveira-Filho, Carolina Ferro, Diana Marcela Walteros-Acero, Franklin Prieto, Carlos Andrés Durán, Martha Lucia Ospina Martinez, Marcela Mercado-Reyes |
| EPI_ISL_526967, EPI_ISL_526968, EPI_ISL_526969, EPI_ISL_526970 | Instituto Nacional de Salud, Bogotá, Colombia | Instituto Nacional de Salud, Bogotá, Colombia | Katherine Laiton-Donato, Diego A. Álvarez-Díaz, Carlos Franco-Muñoz, Mauricio Pacheco-Montealegre, Jonathan Reales, Diego Andrés Prada, Jose A. Usme-Ciro, Zulma M. Cucunubá, Christian Julian Villabona-Arenas, Liz Villabona-Arenas, Sussy Echeverria, Astrid C. Flórez, Carolina Ferro, Diana Marcela Walteros-Acero, Franklin Prieto, Carlos Andrés Durán, Martha Lucia Ospina Martinez, Marcela Mercado-Reyes |
| EPI_ISL_526971, EPI_ISL_526973 | Instituto Nacional de Salud, Bogotá, Colombia | Instituto Nacional de Salud, Bogotá, Colombia | Katherine Laiton-Donato, Diego A. Álvarez-Díaz, Carlos Franco-Muñoz, Mauricio Pacheco-Montealegre, Jonathan Reales, Diego Andrés Prada, Jose A. Usme-Ciro, Zulma M. Cucunubá, Christian Julian Villabona-Arenas, Liz Villabona-Arenas, Sussy Echeverria, Astrid C. Flórez, Carolina Ferro, Diana Marcela Walteros-Acero, Franklin Prieto, Carlos Andrés Durán, Martha Lucia Ospina Martinez, Marcela Mercado-Reyes |
| EPI_ISL_526974 | Instituto Nacional de Salud, Bogotá, Colombia | Instituto Nacional de Salud, Bogotá, Colombia | Katherine Laiton-Donato, Diego A. Álvarez-Díaz, Carlos Franco-Muñoz, Mauricio Pacheco-Montealegre, Jonathan Reales, Diego Andrés Prada, Jose A. Usme-Ciro, Zulma M. Cucunubá, Christian Julian Villabona-Arenas, Liz Villabona-Arenas, Sussy Echeverria, Astrid C. Flórez, Carolina Ferro, Diana Marcela Walteros-Acero, Franklin Prieto, Carlos Andrés Durán, Martha Lucia Ospina Martinez, Marcela Mercado-Reyes |
| EPI_ISL_526975, EPI_ISL_526976, EPI_ISL_526977, EPI_ISL_526978, EPI_ISL_526979, EPI_ISL_526980, EPI_ISL_526981, EPI_ISL_526982, EPI_ISL_526983, EPI_ISL_526984, EPI_ISL_526985, EPI_ISL_526986, EPI_ISL_526987, EPI_ISL_526988, EPI_ISL_526989, EPI_ISL_526990, EPI_ISL_526991, EPI_ISL_526992, EPI_ISL_526993, EPI_ISL_526994, EPI_ISL_526995, EPI_ISL_526996 | | | |
| see above | Biological prevention, army | Biological prevention, army | Seadawy, M.G., Gad, A.F., Harty, B.E., Elhosieny, M.F., Shamel, M.D. |
| EPI_ISL_526997, EPI_ISL_526998 | Biological prevention, army | Biological prevention, army | Seadawy, M.G., Harty,B.E., Gad,A.F., Elhoseiny,M.F., Shamel,M.D., Shabaan,A.E., Ageez,A.M. |
| EPI_ISL_526999, EPI_ISL_527000, EPI_ISL_527001 | Biological prevention, army | Biological prevention, army | Seadawy,M.G., Harty,B.E., Gad,A.F., Elhoseiny,M.F., Shamel,M.D., Shabaan,A.E., Ageez,A.M. |
| EPI_ISL_527002 | Biological prevention, army | Biological prevention, army | Seadawy, M.G., Harty,B.E., Gad,A.F., Elhoseiny,M.F., Shamel,M.D., Shabaan,A.E., Ageez,A.M. |
| EPI_ISL_527003, EPI_ISL_527004, EPI_ISL_527005, EPI_ISL_527006 | Biological prevention, army | Biological prevention, army | Seadawy,M.G., Harty,B.E., Gad,A.F., Elhoseiny,M.F., Shamel,M.D., Shabaan,A.E., Ageez,A.M. |
| EPI_ISL_527007 | Biological Prevention, Army | Biological Prevention, Army | Seadawy, M.G., Harty,B.E., Gad,A.F., Elhoseiny,M.F., Shamel,M.D., Shabaan,A.E., Ageez,A.M. |
| EPI_ISL_527008, EPI_ISL_527009, EPI_ISL_527010, EPI_ISL_527011, EPI_ISL_527012, EPI_ISL_527013, EPI_ISL_527014, EPI_ISL_527015, EPI_ISL_527016, EPI_ISL_527017, EPI_ISL_527018, EPI_ISL_527019, EPI_ISL_527020, EPI_ISL_527021, EPI_ISL_527022, EPI_ISL_527023, EPI_ISL_527024, EPI_ISL_527025, EPI_ISL_527026, EPI_ISL_527027, EPI_ISL_527028, EPI_ISL_527029, EPI_ISL_527030, EPI_ISL_527031, EPI_ISL_527032, EPI_ISL_527033, EPI_ISL_527034, EPI_ISL_527035, EPI_ISL_527036, EPI_ISL_527037, EPI_ISL_527038, EPI_ISL_527039, EPI_ISL_527040, EPI_ISL_527041, EPI_ISL_527042, EPI_ISL_527043, EPI_ISL_527044, EPI_ISL_527045, EPI_ISL_527046, EPI_ISL_527047, EPI_ISL_527048, EPI_ISL_527049, EPI_ISL_527050, EPI_ISL_527051, EPI_ISL_527052, EPI_ISL_527053, EPI_ISL_527054, EPI_ISL_527055, EPI_ISL_527056, EPI_ISL_527057, EPI_ISL_527058, EPI_ISL_527059, EPI_ISL_527060, EPI_ISL_527061, EPI_ISL_527062, EPI_ISL_527063, EPI_ISL_527064, EPI_ISL_527179, EPI_ISL_527180 | | | |
| see above | Area of Virology, Serology and Virology Division (SAViD), New South Wales Health Pathology Randwick | Area of Virology, Serology and Virology Division (SAViD), New South Wales Health Pathology Randwick | Rawlinson, W. |
| EPI_ISL_527257, EPI_ISL_527258, EPI_ISL_527329, EPI_ISL_527330, EPI_ISL_527331, EPI_ISL_527332, EPI_ISL_527333, EPI_ISL_527334, EPI_ISL_527335, EPI_ISL_527336, EPI_ISL_527337, EPI_ISL_527338, EPI_ISL_527339, EPI_ISL_527340, EPI_ISL_527341, EPI_ISL_527342, EPI_ISL_527343, EPI_ISL_527344, EPI_ISL_527345, EPI_ISL_527346, EPI_ISL_527347, EPI_ISL_527348, EPI_ISL_527349, EPI_ISL_527350, EPI_ISL_527351, EPI_ISL_527352, EPI_ISL_527353, EPI_ISL_527354, EPI_ISL_527355, EPI_ISL_527356, EPI_ISL_527357, EPI_ISL_527358 | | | |
| see above | Respiratory Virus Unit, Microbiology Services Colindale, Public Health England | Respiratory Virus Unit, Microbiology Services Colindale, Public Health England | PHE Covid Sequencing Team |
| EPI_ISL_527359, EPI_ISL_527360, EPI_ISL_527361, EPI_ISL_527362, EPI_ISL_527363, EPI_ISL_527364, EPI_ISL_527365, EPI_ISL_527366, EPI_ISL_527367, EPI_ISL_527368, EPI_ISL_527369, EPI_ISL_527370, EPI_ISL_527371, EPI_ISL_527372, EPI_ISL_527373, EPI_ISL_527374, EPI_ISL_527375, EPI_ISL_527376, EPI_ISL_527377, EPI_ISL_527378, EPI_ISL_527379 | | | |
| see above | National Public Health Laboratory, National Centre for Infectious Diseases | National Public Health Laboratory, National Centre for Infectious Diseases | Mak TM, Octavia S, Zhou Z, Cui L, Lin RTP |
| EPI_ISL_527380 | Istituto Zooprofilattico Sperimentale Puglia e Basilicata; Dipartimento di Bioscienze, Biotecnologie e Biofarmaceutica dell'Università degli Studi di Bari "A.Moro"; Istituto di Biomembrane. Bioenergetica e Biotecnologie Molecolari del Consiglio Nazionale delle Ricerche di Bari | Beaconlab (Bioinformatics, Evolution and Comparative Genomics lab), Dept of Biosciences, University on Milan | Parisi A.,Pesole G., Manzari C., Chiara M |
| EPI_ISL_527381, EPI_ISL_527382, EPI_ISL_527383, EPI_ISL_527384, EPI_ISL_527385, EPI_ISL_527386, EPI_ISL_527387, EPI_ISL_527388, EPI_ISL_527389, EPI_ISL_527390, EPI_ISL_527391, EPI_ISL_527392, EPI_ISL_527393, EPI_ISL_527394, EPI_ISL_527395, EPI_ISL_527396, EPI_ISL_527397, EPI_ISL_527398, EPI_ISL_527399 | | | |
| see above | University of Miami Immunology and Histocompatibility Laboratory | University of Miami Immunology and Histocompatibility Laboratory | Emilio Margolles-Clark, PhD and Phillip Ruiz, MD, PhD |
| EPI_ISL_527400 | Area of Virology, Serology and Virology Division (SAViD), New South Wales Health Pathology Randwick | Area of Virology, Serology and Virology Division (SAViD), New South Wales Health Pathology Randwick | Rawlinson, W., Deveson, I., Bull, R. |
| EPI_ISL_527401, EPI_ISL_527402, EPI_ISL_527403, EPI_ISL_527404, EPI_ISL_527405, EPI_ISL_527406, EPI_ISL_527407, EPI_ISL_527408, EPI_ISL_527409, EPI_ISL_527410, EPI_ISL_527411, EPI_ISL_527412, EPI_ISL_527413, EPI_ISL_527414, EPI_ISL_527415, EPI_ISL_527416, EPI_ISL_527417, EPI_ISL_527418, EPI_ISL_527419, EPI_ISL_527420, EPI_ISL_527421, EPI_ISL_527422, EPI_ISL_527423, EPI_ISL_527424, EPI_ISL_527425, EPI_ISL_527426, EPI_ISL_527427, EPI_ISL_527428, EPI_ISL_527429, EPI_ISL_527430, EPI_ISL_527431, EPI_ISL_527432, EPI_ISL_527433, EPI_ISL_527434, EPI_ISL_527435, EPI_ISL_527436, EPI_ISL_527437, EPI_ISL_527438, EPI_ISL_527439, EPI_ISL_527440, EPI_ISL_527441, EPI_ISL_527442, EPI_ISL_527443, EPI_ISL_527444, EPI_ISL_527445, EPI_ISL_527446, EPI_ISL_527447, EPI_ISL_527448, EPI_ISL_527449, EPI_ISL_527450, EPI_ISL_527451, EPI_ISL_527452, EPI_ISL_527453, EPI_ISL_527454, EPI_ISL_527455, EPI_ISL_527456, EPI_ISL_527457, EPI_ISL_527458, EPI_ISL_527459, EPI_ISL_527460, EPI_ISL_527461, EPI_ISL_527462, EPI_ISL_527463, EPI_ISL_527464, EPI_ISL_527465, EPI_ISL_527466, EPI_ISL_527467, EPI_ISL_527468, EPI_ISL_527469, EPI_ISL_527470, EPI_ISL_527471, EPI_ISL_527472, EPI_ISL_527473, EPI_ISL_527474, EPI_ISL_527475, EPI_ISL_527476, EPI_ISL_527477, EPI_ISL_527478, EPI_ISL_527479, EPI_ISL_527480, EPI_ISL_527481, EPI_ISL_527482, EPI_ISL_527483, EPI_ISL_527484, EPI_ISL_527485, EPI_ISL_527486, EPI_ISL_527487, EPI_ISL_527488 | | | |
| see above | Colorado State University - Ebel Lab | Colorado State University - Ebel Lab | Greg Ebel et al. |
| EPI_ISL_527489, EPI_ISL_527490, EPI_ISL_527491, EPI_ISL_527492, EPI_ISL_527493, EPI_ISL_527494, EPI_ISL_527495, EPI_ISL_527496, EPI_ISL_527497, EPI_ISL_527498, EPI_ISL_527499, EPI_ISL_527500, EPI_ISL_527501, EPI_ISL_527502, EPI_ISL_527503, EPI_ISL_527504, EPI_ISL_527505, EPI_ISL_527506, EPI_ISL_527507, EPI_ISL_527508, EPI_ISL_527509, EPI_ISL_527510, EPI_ISL_527511, EPI_ISL_527512, EPI_ISL_527513, EPI_ISL_527514, EPI_ISL_527515, EPI_ISL_527516, EPI_ISL_527517, EPI_ISL_527518, EPI_ISL_527519, EPI_ISL_527520, EPI_ISL_527521, EPI_ISL_527522, EPI_ISL_527523, EPI_ISL_527524, EPI_ISL_527525, EPI_ISL_527526, EPI_ISL_527527, EPI_ISL_527528, EPI_ISL_527529, EPI_ISL_527530, EPI_ISL_527531, EPI_ISL_527532, EPI_ISL_527533, EPI_ISL_527534, EPI_ISL_527535, EPI_ISL_527536, EPI_ISL_527537, EPI_ISL_527538, EPI_ISL_527539, EPI_ISL_527540, EPI_ISL_527541, EPI_ISL_527542, EPI_ISL_527543, EPI_ISL_527544, EPI_ISL_527545, EPI_ISL_527546, EPI_ISL_527547, EPI_ISL_527548, EPI_ISL_527549, EPI_ISL_527550, EPI_ISL_527551, EPI_ISL_527552, EPI_ISL_527553, EPI_ISL_527554, EPI_ISL_527555, EPI_ISL_527556, EPI_ISL_527557, EPI_ISL_527558, EPI_ISL_527559, EPI_ISL_527560, EPI_ISL_527561, EPI_ISL_527562, EPI_ISL_527563, EPI_ISL_527564, EPI_ISL_527565, EPI_ISL_527566, EPI_ISL_527567, EPI_ISL_527568, EPI_ISL_527569, EPI_ISL_527570, EPI_ISL_527571, EPI_ISL_527572 | | | |
| see above | Viral Respiratory Lab, National Institute for Biomedical Research (INRB) | Pathogen Sequencing Lab, National Institute for Biomedical Research (INRB) | Placide Mbala-Kingebeni, Edith Nkwembe, Eddy Kinganda-Lusamaki, Amuri Aziza, Francisca Muyembe Mawete, Emmanuel Lokilo Lofiko, Catherine Pratt, Matthias Pauthner, Josh Quick, Allison Black, James Hadfield, Trevor Bedford, Ian Goodfellow, Andrew Rambaut, Nick Loman, Kristian Andersen, Michael Wiley, Steve Ahuka-Mundeke, Jean-Jacques Muyembe Tamfum |
| EPI_ISL_527573, EPI_ISL_527574, EPI_ISL_527575, EPI_ISL_527576, EPI_ISL_527577, EPI_ISL_527578, EPI_ISL_527579, EPI_ISL_527580, EPI_ISL_527581, EPI_ISL_527582, EPI_ISL_527583, EPI_ISL_527584, EPI_ISL_527585, EPI_ISL_527586, EPI_ISL_527587, EPI_ISL_527588, EPI_ISL_527589, EPI_ISL_527590, EPI_ISL_527591, EPI_ISL_527592, EPI_ISL_527593, EPI_ISL_527594, EPI_ISL_527595, EPI_ISL_527596, EPI_ISL_527597, EPI_ISL_527598, EPI_ISL_527599, EPI_ISL_527600, EPI_ISL_527601, EPI_ISL_527602, EPI_ISL_527603, EPI_ISL_527604, EPI_ISL_527605, EPI_ISL_527606, EPI_ISL_527607, EPI_ISL_527608, EPI_ISL_527609, EPI_ISL_527610, EPI_ISL_527611, EPI_ISL_527612, EPI_ISL_527613, EPI_ISL_527614, EPI_ISL_527615, EPI_ISL_527616, EPI_ISL_527617, EPI_ISL_527618, EPI_ISL_527619, EPI_ISL_527620, EPI_ISL_527621, EPI_ISL_527622, EPI_ISL_527623, EPI_ISL_527624, EPI_ISL_527625, EPI_ISL_527626, EPI_ISL_527627, EPI_ISL_527628, EPI_ISL_527629 | | | |
| see above | Minnesota Department of Health, Public Health Laboratory | Minnesota Department of Health, Public Health Laboratory | Matt Plumb, Jacob Garfin, and Xiong Wang |
| EPI_ISL_527630, EPI_ISL_527631 | Mayo Clinic & Mayo Clinic Laboratories | Minnesota Department of Health, Public Health Laboratory | Matt Plumb, Jacob Garfin, and Xiong Wang |
| EPI_ISL_527632, EPI_ISL_527633, EPI_ISL_527634, EPI_ISL_527635, EPI_ISL_527636, EPI_ISL_527637, EPI_ISL_527638, EPI_ISL_527639, EPI_ISL_527640, EPI_ISL_527641, EPI_ISL_527642, EPI_ISL_527643, EPI_ISL_527644, EPI_ISL_527645, EPI_ISL_527646, EPI_ISL_527647, EPI_ISL_527648, EPI_ISL_527649, EPI_ISL_527650, EPI_ISL_527651, EPI_ISL_527652, EPI_ISL_527653, EPI_ISL_527654, EPI_ISL_527655, EPI_ISL_527656, EPI_ISL_527657 | | | |
| see above | AR Dept. of Health-Public Health Lab | Pathogen Discovery, Respiratory Viruses Branch, Division of Viral Diseases, Centers for Disease Control and Prevention | Ying Tao, Jing Zhang, Yan Li, Krista Queen, Anna Uehara, Clinton Paden, Haibin Wang, Suxiang Tong |
| EPI_ISL_527658, EPI_ISL_527659, EPI_ISL_527660, EPI_ISL_527661, EPI_ISL_527662, EPI_ISL_527663, EPI_ISL_527664, EPI_ISL_527665 | GA Department of Public Health Laboratory | Pathogen Discovery, Respiratory Viruses Branch, Division of Viral Diseases, Centers for Disease Control and Prevention | Yan Li, Anna Montmayeur, Jing Zhang, Krista Queen, Ying Tao, Anna Uehara, Rachel Marine, Clinton R. Paden, Haibin Wang, Suxiang Tong |
| EPI_ISL_527666 | IN State Department of Health Laboratory Services | Pathogen Discovery, Respiratory Viruses Branch, Division of Viral Diseases, Centers for Disease Control and Prevention | Krista Queen, Brian Lynch, Yan Li, Anna Montmayeur, Jing Zhang, Ying Tao, Anna Uehara, Rachel Marine, Clinton R. Paden, Haibin Wang, Suxiang Tong |
| EPI_ISL_527667, EPI_ISL_527668, EPI_ISL_527669, EPI_ISL_527670, EPI_ISL_527671, EPI_ISL_527672, EPI_ISL_527673, EPI_ISL_527674, EPI_ISL_527675, EPI_ISL_527676, EPI_ISL_527677, EPI_ISL_527678, EPI_ISL_527679, EPI_ISL_527680, EPI_ISL_527681, EPI_ISL_527682, EPI_ISL_527683, EPI_ISL_527684, EPI_ISL_527685, EPI_ISL_527686, EPI_ISL_527687, EPI_ISL_527688, EPI_ISL_527689, EPI_ISL_527690, EPI_ISL_527691 | | | |
| see above | MN PHL Division, Minnesota Department of Health | Pathogen Discovery, Respiratory Viruses Branch, Division of Viral Diseases, Centers for Disease Control and Prevention | Yan Li, Anna Montmayeur, Jing Zhang, Krista Queen, Ying Tao, Anna Uehara, Rachel Marine, Clinton R. Paden, Haibin Wang, Suxiang Tong |
| EPI_ISL_527692 | MN PHL Division, Minnesota Department of Health | Pathogen Discovery, Respiratory Viruses Branch, Division of Viral Diseases, Centers for Disease Control and Prevention | Krista Queen, Brian Lynch, Yan Li, Anna Montmayeur, Jing Zhang, Ying Tao, Anna Uehara, Rachel Marine, Clinton R. Paden, Haibin Wang, Suxiang Tong |
| EPI_ISL_527693, EPI_ISL_527694, EPI_ISL_527695, EPI_ISL_527696, EPI_ISL_527697, EPI_ISL_527698 | MN PHL Division, Minnesota Department of Health | Pathogen Discovery, Respiratory Viruses Branch, Division of Viral Diseases, Centers for Disease Control and Prevention | Yan Li, Anna Montmayeur, Jing Zhang, Krista Queen, Ying Tao, Anna Uehara, Rachel Marine, Clinton R. Paden, Haibin Wang, Suxiang Tong |
| EPI_ISL_527699 | MN PHL Division, Minnesota Department of Health | Pathogen Discovery, Respiratory Viruses Branch, Division of Viral Diseases, Centers for Disease Control and Prevention | Krista Queen, Brian Lynch, Yan Li, Anna Montmayeur, Jing Zhang, Ying Tao, Anna Uehara, Rachel Marine, Clinton R. Paden, Haibin Wang, Suxiang Tong |
| EPI_ISL_527700 | MN PHL Division, Minnesota Department of Health | Pathogen Discovery, Respiratory Viruses Branch, Division of Viral Diseases, Centers for Disease Control and Prevention | Yan Li, Anna Montmayeur, Jing Zhang, Krista Queen, Ying Tao, Anna Uehara, Rachel Marine, Clinton R. Paden, Haibin Wang, Suxiang Tong |
| EPI_ISL_527701 | MN PHL Division, Minnesota Department of Health | Pathogen Discovery, Respiratory Viruses Branch, Division of Viral Diseases, Centers for Disease Control and Prevention | Krista Queen, Brian Lynch, Yan Li, Anna Montmayeur, Jing Zhang, Ying Tao, Anna Uehara, Rachel Marine, Clinton R. Paden, Haibin Wang, Suxiang Tong |
| EPI_ISL_527702, EPI_ISL_527703, EPI_ISL_527704, EPI_ISL_527705, EPI_ISL_527706, EPI_ISL_527707, EPI_ISL_527708, EPI_ISL_527709, EPI_ISL_527710, EPI_ISL_527711, EPI_ISL_527712, EPI_ISL_527713 | | | |
| see above | MN PHL Division, Minnesota Department of Health | Pathogen Discovery, Respiratory Viruses Branch, Division of Viral Diseases, Centers for Disease Control and Prevention | Yan Li, Anna Montmayeur, Jing Zhang, Krista Queen, Ying Tao, Anna Uehara, Rachel Marine, Clinton R. Paden, Haibin Wang, Suxiang Tong |

| | | | |
|---|---|---|---|
| EPI_ISL_527714 | MN PHL Division, Minnesota Department of Health | Pathogen Discovery, Respiratory Viruses Branch, Division of Viral Diseases, Centers for Disease Control and Prevention | Krista Queen, Brian Lynch, Yan Li, Anna Montmayeur, Jing Zhang, Ying Tao, Anna Uehara, Rachel Marine, Clinton R. Paden, Haibin Wang, Suxiang Tong |
| EPI_ISL_527715 | MN PHL Division, Minnesota Department of Health | Pathogen Discovery, Respiratory Viruses Branch, Division of Viral Diseases, Centers for Disease Control and Prevention | Yan Li, Anna Montmayeur, Jing Zhang, Krista Queen, Ying Tao, Anna Uehara, Rachel Marine, Clinton R. Paden, Haibin Wang, Suxiang Tong |
| EPI_ISL_527716, EPI_ISL_527717 | MN PHL Division, Minnesota Department of Health | Pathogen Discovery, Respiratory Viruses Branch, Division of Viral Diseases, Centers for Disease Control and Prevention | Krista Queen, Brian Lynch, Yan Li, Anna Montmayeur, Jing Zhang, Ying Tao, Anna Uehara, Rachel Marine, Clinton R. Paden, Haibin Wang, Suxiang Tong |
| EPI_ISL_527718, EPI_ISL_527719 | MN PHL Division, Minnesota Department of Health | Pathogen Discovery, Respiratory Viruses Branch, Division of Viral Diseases, Centers for Disease Control and Prevention | Yan Li, Anna Montmayeur, Jing Zhang, Krista Queen, Ying Tao, Anna Uehara, Rachel Marine, Clinton R. Paden, Haibin Wang, Suxiang Tong |
| EPI_ISL_527720 | MN PHL Division, Minnesota Department of Health | Pathogen Discovery, Respiratory Viruses Branch, Division of Viral Diseases, Centers for Disease Control and Prevention | Krista Queen, Brian Lynch, Yan Li, Anna Montmayeur, Jing Zhang, Ying Tao, Anna Uehara, Rachel Marine, Clinton R. Paden, Haibin Wang, Suxiang Tong |
| EPI_ISL_527721, EPI_ISL_527722, EPI_ISL_527723 | MN PHL Division, Minnesota Department of Health | Pathogen Discovery, Respiratory Viruses Branch, Division of Viral Diseases, Centers for Disease Control and Prevention | Yan Li, Anna Montmayeur, Jing Zhang, Krista Queen, Ying Tao, Anna Uehara, Rachel Marine, Clinton R. Paden, Haibin Wang, Suxiang Tong |
| EPI_ISL_527724 | MN PHL Division, Minnesota Department of Health | Pathogen Discovery, Respiratory Viruses Branch, Division of Viral Diseases, Centers for Disease Control and Prevention | Krista Queen, Brian Lynch, Yan Li, Anna Montmayeur, Jing Zhang, Ying Tao, Anna Uehara, Rachel Marine, Clinton R. Paden, Haibin Wang, Suxiang Tong |
| EPI_ISL_527725, EPI_ISL_527726, EPI_ISL_527727, EPI_ISL_527728, EPI_ISL_527729, EPI_ISL_527730 | MN PHL Division, Minnesota Department of Health | Pathogen Discovery, Respiratory Viruses Branch, Division of Viral Diseases, Centers for Disease Control and Prevention | Yan Li, Anna Montmayeur, Jing Zhang, Krista Queen, Ying Tao, Anna Uehara, Rachel Marine, Clinton R. Paden, Haibin Wang, Suxiang Tong |
| EPI_ISL_527731 | MN PHL Division, Minnesota Department of Health | Pathogen Discovery, Respiratory Viruses Branch, Division of Viral Diseases, Centers for Disease Control and Prevention | Krista Queen, Brian Lynch, Yan Li, Anna Montmayeur, Jing Zhang, Ying Tao, Anna Uehara, Rachel Marine, Clinton R. Paden, Haibin Wang, Suxiang Tong |
| EPI_ISL_527732, EPI_ISL_527733 | MN PHL Division, Minnesota Department of Health | Pathogen Discovery, Respiratory Viruses Branch, Division of Viral Diseases, Centers for Disease Control and Prevention | Yan Li, Anna Montmayeur, Jing Zhang, Krista Queen, Ying Tao, Anna Uehara, Rachel Marine, Clinton R. Paden, Haibin Wang, Suxiang Tong |
| EPI_ISL_527734, EPI_ISL_527735 | MN PHL Division, Minnesota Department of Health | Pathogen Discovery, Respiratory Viruses Branch, Division of Viral Diseases, Centers for Disease Control and Prevention | Krista Queen, Brian Lynch, Yan Li, Anna Montmayeur, Jing Zhang, Ying Tao, Anna Uehara, Rachel Marine, Clinton R. Paden, Haibin Wang, Suxiang Tong |
| EPI_ISL_527736 | MN PHL Division, Minnesota Department of Health | Pathogen Discovery, Respiratory Viruses Branch, Division of Viral Diseases, Centers for Disease Control and Prevention | Yan Li, Anna Montmayeur, Jing Zhang, Krista Queen, Ying Tao, Anna Uehara, Rachel Marine, Clinton R. Paden, Haibin Wang, Suxiang Tong |
| EPI_ISL_527737 | WI State Laboratory of Hygiene | Pathogen Discovery, Respiratory Viruses Branch, Division of Viral Diseases, Centers for Disease Control and Prevention | Ying Tao, Jing Zhang, Yan Li, Krista Queen, Anna Uehara, Clinton Paden, Haibin Wang, Suxiang Tong |
| EPI_ISL_527738 | Connecticut State Department of Public Health | Grubaugh Lab - Yale School of Public Health | Joseph Fauver, Mary Petrone, Anderson Brito, Tara Alpert, Nathan Grubaugh |
| EPI_ISL_527739 | Hospital Mexico [San Jose/San Jose] | Inciensa, Instituto Costarricense de Investigación y Enseñanza en Nutrición y Salud | Francisco Duarte, Hebleen Porras, Claudio Soto-Garita, Estela Cordero, Adriana Godinez & Melany Calderon |
| EPI_ISL_527740 | Area De Salud Alajuela Norte - Clinica Dr. Marcial Rodriguez | Inciensa, Instituto Costarricense de Investigación y Enseñanza en Nutrición y Salud | Francisco Duarte, Hebleen Porras, Claudio Soto-Garita, Estela Cordero, Adriana Godinez & Melany Calderon |
| EPI_ISL_527741 | Hospital Metropolitano | Inciensa, Instituto Costarricense de Investigación y Enseñanza en Nutrición y Salud | Francisco Duarte, Hebleen Porras, Claudio Soto-Garita, Estela Cordero, Adriana Godinez & Melany Calderon |
| EPI_ISL_527742 | Centro Nacional De Rehabilitacion Humberto Araya Rojas (Cenare) | Inciensa, Instituto Costarricense de Investigación y Enseñanza en Nutrición y Salud | Francisco Duarte, Hebleen Porras, Claudio Soto-Garita, Estela Cordero, Adriana Godinez & Melany Calderon |
| EPI_ISL_527743, EPI_ISL_527744 | Hospital Cima | Inciensa, Instituto Costarricense de Investigación y Enseñanza en Nutrición y Salud | Francisco Duarte, Hebleen Porras, Claudio Soto-Garita, Estela Cordero, Adriana Godinez & Melany Calderon |
| EPI_ISL_527745 | Area De Salud Corredores | Inciensa, Instituto Costarricense de Investigación y Enseñanza en Nutrición y Salud | Francisco Duarte, Hebleen Porras, Claudio Soto-Garita, Estela Cordero, Adriana Godinez & Melany Calderon |
| EPI_ISL_527746, EPI_ISL_527747 | Area De Salud La Cruz | Inciensa, Instituto Costarricense de Investigación y Enseñanza en Nutrición y Salud | Francisco Duarte, Hebleen Porras, Claudio Soto-Garita, Estela Cordero, Adriana Godinez & Melany Calderon |
| EPI_ISL_527748 | Area De Salud Corredores | Inciensa, Instituto Costarricense de Investigación y Enseñanza en Nutrición y Salud | Francisco Duarte, Hebleen Porras, Claudio Soto-Garita, Estela Cordero, Adriana Godinez & Melany Calderon |
| EPI_ISL_527749 | Area De Salud Alajuela Norte - Clinica Dr. Marcial Rodriguez | Inciensa, Instituto Costarricense de Investigación y Enseñanza en Nutrición y Salud | Francisco Duarte, Hebleen Porras, Claudio Soto-Garita, Estela Cordero, Adriana Godinez & Melany Calderon |
| EPI_ISL_527750 | Hospital De Niños Dr. Carlos Saenz Herrera [San Jose/San Jose] | Inciensa, Instituto Costarricense de Investigación y Enseñanza en Nutrición y Salud | Francisco Duarte, Hebleen Porras, Claudio Soto-Garita, Estela Cordero, Adriana Godinez & Melany Calderon |
| EPI_ISL_527751 | Area De Salud Corredores | Inciensa, Instituto Costarricense de Investigación y Enseñanza en Nutrición y Salud | Francisco Duarte, Hebleen Porras, Claudio Soto-Garita, Estela Cordero, Adriana Godinez & Melany Calderon |
| EPI_ISL_527752 | Hospital Dr. Rafael A. Calderon Guardia | Inciensa, Instituto Costarricense de Investigación y Enseñanza en Nutrición y Salud | Francisco Duarte, Hebleen Porras, Claudio Soto-Garita, Estela Cordero, Adriana Godinez & Melany Calderon |
| EPI_ISL_527753, EPI_ISL_527754 | Hospital San Vicente De Paul | Inciensa, Instituto Costarricense de Investigación y Enseñanza en Nutrición y Salud | Francisco Duarte, Hebleen Porras, Claudio Soto-Garita, Estela Cordero, Adriana Godinez & Melany Calderon |
| EPI_ISL_527755 | Centro Nacional De Rehabilitacion Humberto Araya Rojas (Cenare) | Inciensa, Instituto Costarricense de Investigación y Enseñanza en Nutrición y Salud | Francisco Duarte, Hebleen Porras, Claudio Soto-Garita, Estela Cordero, Adriana Godinez & Melany Calderon |
| EPI_ISL_527756 | Area De Salud Aserri | Inciensa, Instituto Costarricense de Investigación y Enseñanza en Nutrición y Salud | Francisco Duarte, Hebleen Porras, Claudio Soto-Garita, Estela Cordero, Adriana Godinez & Melany Calderon |
| EPI_ISL_527757 | Area De Salud Goicoechea 1 | Inciensa, Instituto Costarricense de Investigación y Enseñanza en Nutrición y Salud | Francisco Duarte, Hebleen Porras, Claudio Soto-Garita, Estela Cordero, Adriana Godinez & Melany Calderon |
| EPI_ISL_527758 | Hospital Mexico [San Jose/San Jose] | Inciensa, Instituto Costarricense de Investigación y Enseñanza en Nutrición y Salud | Francisco Duarte, Hebleen Porras, Claudio Soto-Garita, Estela Cordero, Adriana Godinez & Melany Calderon |
| EPI_ISL_527759 | Clinica Biblica | Inciensa, Instituto Costarricense de Investigación y Enseñanza en Nutrición y Salud | Francisco Duarte, Hebleen Porras, Claudio Soto-Garita, Estela Cordero, Adriana Godinez & Melany Calderon |
| EPI_ISL_527760 | Area De Salud La Cruz | Inciensa, Instituto Costarricense de Investigación y Enseñanza en Nutrición y Salud | Francisco Duarte, Hebleen Porras, Claudio Soto-Garita, Estela Cordero, Adriana Godinez & Melany Calderon |
| EPI_ISL_527761, EPI_ISL_527762, EPI_ISL_527763, EPI_ISL_527764, EPI_ISL_527765, EPI_ISL_527766, EPI_ISL_527767, EPI_ISL_527768, EPI_ISL_527769, EPI_ISL_527770, EPI_ISL_527771, EPI_ISL_527772, EPI_ISL_527773, EPI_ISL_527774, EPI_ISL_527775, EPI_ISL_527776, EPI_ISL_527777, EPI_ISL_527778, EPI_ISL_527779, EPI_ISL_527780, EPI_ISL_527781, EPI_ISL_527782, EPI_ISL_527783, EPI_ISL_527784, EPI_ISL_527785 | | | |
| see above | Connecticut State Department of Public Health | Grubaugh Lab - Yale School of Public Health | Joseph Fauver, Mary Petrone, Chantal Vogels, Anderson Brito, Tara Alpert, Anthony Muyombwe, Jafar Razeq, Albert Ko, Nathan Grubaugh |
| EPI_ISL_527787, EPI_ISL_527789 | Laboratorio de Referencia Nacional de Virus Respiratorio. Centro Nacional de Salud Publica. Instituto Nacional de Salud Peru. | Laboratorio de Referencia Nacional de Biotecnología y Biología Molecular. Centro Nacional de Salud Publica. Instituto Nacional de Salud Peru. | Carlos Padilla Rojas, Karolyn Vega Chozo, Priscila Lope Pari, Omar Caceres Rey, Marco Galarza Perez, Maribel Huaringa Nuñez, Johanna Balbuena Torres, Henri Bailon Calderon, Nancy Rojas Serrano. |
| EPI_ISL_527794, EPI_ISL_527795, EPI_ISL_527796, EPI_ISL_527797, EPI_ISL_527798, EPI_ISL_527799, EPI_ISL_527800, EPI_ISL_527801, EPI_ISL_527802, EPI_ISL_527803, EPI_ISL_527804, EPI_ISL_527805, EPI_ISL_527806, EPI_ISL_527807, EPI_ISL_527808 | | | |
| see above | University of Wisconsin-Madison AIDS Vaccine Research Laboratories | University of Wisconsin-Madison AIDS Vaccine Research Laboratories | Gage Moreno, Katarina Braun, et al. AIDS Vaccine Research Laboratories |
| EPI_ISL_527809, EPI_ISL_527810, EPI_ISL_527811 | Institute of Microbiology, Universidad San Francisco de Quito | Institute of Microbiology, Universidad San Francisco de Quito | Belén Prado-Vivar, Sully Márquez, Juan José Guadalupe, Monica Becerra-Wong, Bernardo Gutiérrez, Stephanie Arregui, Rene Bracho, Karina Barragan, Anita Garcia, Carlos Tobar, Verónica Barragán, Patricio Rojas-Silva, Gabriel Trueba, Michelle Grunauer, Paúl Cárdenas |
| EPI_ISL_527812, EPI_ISL_527813, EPI_ISL_527814, EPI_ISL_527815, EPI_ISL_527816, EPI_ISL_527817 | Institute of Microbiology, Universidad San Francisco de Quito | Institute of Microbiology, Universidad San Francisco de Quito | Belén Prado-Vivar, Sully Márquez, Juan José Guadalupe, Monica Becerra-Wong, Bernardo Gutiérrez, Khurram Mahbbob, Verónica Barragán, Patricio Rojas-Silva, Gabriel Trueba, Michelle Grunauer, Paúl Cárdenas |
| EPI_ISL_527818, EPI_ISL_527819 | Centro de Investigaciones, Universidad de Especialidades Espíritu Santo | Institute of Microbiology, Universidad San Francisco de Quito | Derly Andrade, Juan Carlos Fernandez, Belén Prado-Vivar, Sully Márquez, Juan José Guadalupe, Monica Becerra-Wong, Bernardo Gutiérrez, Gabriel Morey, Ruben Armas, Jose Pedro Barberan, Fernando Espinoza, Edith Lopez, Verónica Barragán, Patricio Rojas-Silva, Gabriel Trueba, Michelle Grunauer, Paúl Cárdenas |
| EPI_ISL_527820, EPI_ISL_527821, EPI_ISL_527822, EPI_ISL_527823, EPI_ISL_527824, EPI_ISL_527825, EPI_ISL_527826, EPI_ISL_527827, EPI_ISL_527828, EPI_ISL_527829, EPI_ISL_527830, EPI_ISL_527831, EPI_ISL_527832, EPI_ISL_527833, EPI_ISL_527834, EPI_ISL_527835, EPI_ISL_527836, EPI_ISL_527837, EPI_ISL_527838, EPI_ISL_527839, EPI_ISL_527840, EPI_ISL_527841, EPI_ISL_527842, EPI_ISL_527843, EPI_ISL_527844, EPI_ISL_527845, EPI_ISL_527846, EPI_ISL_527847, EPI_ISL_527848, EPI_ISL_527849, EPI_ISL_527850, EPI_ISL_527851, EPI_ISL_527852, EPI_ISL_527853, EPI_ISL_527854, EPI_ISL_527855 | | | |
| see above | Texas Department of State Health Services | Texas Department of State Health Services | Bonnie Oh, Rashmi Tuladhar, Jenny Zhang, Maliha Rahman, Anita Pokharel, Myong Koag, Chun Wang, Rachel Lee, Grace Kubin |
| EPI_ISL_527856 | Hospital Municipal Prof. Waldomiro de Paula | Instituto Adolfo Lutz, Interdiciplinary Procedures Center, Strategic Laboratory | Claudio Tavares Sacchi, Claudia Regina Gonçalves, Erica Valessa Ramos Gomes |
| EPI_ISL_527857 | Hospital Regional Vale do Ribeira | Instituto Adolfo Lutz, Interdiciplinary Procedures Center, Strategic Laboratory | Claudio Tavares Sacchi, Claudia Regina Gonçalves, Erica Valessa Ramos Gomes |
| EPI_ISL_527858 | Pronto Atendimento Sancta Maggiore Jardim Paulista | Instituto Adolfo Lutz, Interdiciplinary Procedures Center, Strategic Laboratory | Claudio Tavares Sacchi, Claudia Regina Gonçalves, Erica Valessa Ramos Gomes |
| EPI_ISL_527859 | Hospital Municipal Vereador Jose Storopolli | Instituto Adolfo Lutz, Interdiciplinary Procedures Center, Strategic Laboratory | Claudio Tavares Sacchi, Claudia Regina Gonçalves, Erica Valessa Ramos Gomes |
| EPI_ISL_527860 | Hospital Municipal de Parelheiros Josanias Castanha Braga | Instituto Adolfo Lutz, Interdiciplinary Procedures Center, Strategic Laboratory | Claudio Tavares Sacchi, Claudia Regina Gonçalves, Erica Valessa Ramos Gomes |
| EPI_ISL_527861 | Hospital e Maternidade Celso Pierro | Instituto Adolfo Lutz, Interdiciplinary Procedures Center, Strategic Laboratory | Av. Dr. Arnaldo, 355 - Brazil, Cerqueira Cesar, São Paulo - SP, 01246-1301 |
| EPI_ISL_527862 | Hospital Municipal de Urgência | Instituto Adolfo Lutz, Interdiciplinary Procedures Center, Strategic Laboratory | Claudio Tavares Sacchi, Claudia Regina Gonçalves, Erica Valessa Ramos Gomes |
| EPI_ISL_527863 | Hospital Municipal do Tatuape Carmino Caricchio | Instituto Adolfo Lutz, Interdiciplinary Procedures Center, Strategic Laboratory | Claudio Tavares Sacchi, Claudia Regina Gonçalves, Erica Valessa Ramos Gomes |
| EPI_ISL_527864 | Hospital e Pronto Socorro Comunitário Vila Iolanda | Instituto Adolfo Lutz, Interdiciplinary Procedures Center, Strategic Laboratory | Claudio Tavares Sacchi, Claudia Regina Gonçalves, Erica Valessa Ramos Gomes |
| EPI_ISL_527865 | Hospital e Maternidade São Cristóvão | Instituto Adolfo Lutz, Interdiciplinary Procedures Center, Strategic Laboratory | Claudio Tavares Sacchi, Claudia Regina Gonçalves, Erica Valessa Ramos Gomes |
| EPI_ISL_527866 | PS Municipal Dr Lauro Ribas Braga | Instituto Adolfo Lutz, Interdiciplinary Procedures Center, Strategic Laboratory | Av. Dr. Arnaldo, 355 - Brazil, Cerqueira Cesar, São Paulo - SP, 01246-1301 |
| EPI_ISL_527867 | Pronto Socorro Municipal - Balneario São José | Instituto Adolfo Lutz, Interdiciplinary Procedures Center, Strategic Laboratory | Claudio Tavares Sacchi, Claudia Regina Gonçalves, Erica Valessa Ramos Gomes |
| EPI_ISL_527868 | Hospital e Maternidade do Braz | Instituto Adolfo Lutz, Interdiciplinary Procedures Center, Strategic Laboratory | Claudio Tavares Sacchi, Claudia Regina Gonçalves, Erica Valessa Ramos Gomes |
| EPI_ISL_527869 | Hospital Municipal Carmen Prudente | Instituto Adolfo Lutz, Interdiciplinary Procedures Center, Strategic Laboratory | Claudio Tavares Sacchi, Claudia Regina Gonçalves, Erica Valessa Ramos Gomes |
| EPI_ISL_527870 | Hospital Municipal Mário Gatti | Instituto Adolfo Lutz, Interdiciplinary Procedures Center, Strategic Laboratory | Claudio Tavares Sacchi, Claudia Regina Gonçalves, Erica Valessa Ramos Gomes |
| EPI_ISL_527872, EPI_ISL_527873, EPI_ISL_527874, EPI_ISL_527875, EPI_ISL_527876, EPI_ISL_527877, EPI_ISL_527878, EPI_ISL_527879, EPI_ISL_527880, EPI_ISL_527881, EPI_ISL_527882, EPI_ISL_527883, EPI_ISL_527884, EPI_ISL_527885, EPI_ISL_527886, EPI_ISL_527887, EPI_ISL_527888, EPI_ISL_527889, EPI_ISL_527890, EPI_ISL_527891, EPI_ISL_527892, EPI_ISL_527893, EPI_ISL_527894, EPI_ISL_527895, EPI_ISL_527896, EPI_ISL_527897, EPI_ISL_527898, EPI_ISL_527899, EPI_ISL_527900, EPI_ISL_527901, EPI_ISL_527902, EPI_ISL_527903, EPI_ISL_527904, EPI_ISL_527905, EPI_ISL_527906, EPI_ISL_527907, EPI_ISL_527908, EPI_ISL_527909, EPI_ISL_527910, EPI_ISL_527911, EPI_ISL_527912, EPI_ISL_527913, EPI_ISL_527914, EPI_ISL_527915 | | | |
| see above | Nigeria Centre for Disease Control (NCDC) | African Centre of Excellence for Genomics of Infectious Diseases (ACEGID), Redeemer's University, Ede, Osun State, Nigeria | Oluniyi P.E. et al |

| | | | |
|---|---|---|---|
| EPI_ISL_527916, EPI_ISL_527917, EPI_ISL_527918, EPI_ISL_527919, EPI_ISL_527920, EPI_ISL_527921, EPI_ISL_527922, EPI_ISL_527923, EPI_ISL_527924, EPI_ISL_527925, EPI_ISL_527926, EPI_ISL_527927, EPI_ISL_527928, EPI_ISL_527929, EPI_ISL_527930, EPI_ISL_527931, EPI_ISL_527932, EPI_ISL_527933, EPI_ISL_527934, EPI_ISL_527935, EPI_ISL_527936, EPI_ISL_527937, EPI_ISL_527938, EPI_ISL_527939, EPI_ISL_527940, EPI_ISL_527941, EPI_ISL_527942, EPI_ISL_527943, EPI_ISL_527944, EPI_ISL_527945, EPI_ISL_527946, EPI_ISL_527947, EPI_ISL_527948, EPI_ISL_527949, EPI_ISL_527950, EPI_ISL_527951, EPI_ISL_527952, EPI_ISL_527953, EPI_ISL_527954, EPI_ISL_527955, EPI_ISL_527956, EPI_ISL_527957, EPI_ISL_527958, EPI_ISL_527959, EPI_ISL_527960, EPI_ISL_527961, EPI_ISL_527962, EPI_ISL_527963, EPI_ISL_527964, EPI_ISL_527965, EPI_ISL_527966, EPI_ISL_527967, EPI_ISL_527968, EPI_ISL_527969, EPI_ISL_527970, EPI_ISL_527971, EPI_ISL_527972, EPI_ISL_527973, EPI_ISL_527974, EPI_ISL_527975, EPI_ISL_527976, EPI_ISL_527977, EPI_ISL_527978, EPI_ISL_527979, EPI_ISL_527980, EPI_ISL_527981, EPI_ISL_527982, EPI_ISL_527983, EPI_ISL_527984, EPI_ISL_527985, EPI_ISL_527986, EPI_ISL_527987, EPI_ISL_527988, EPI_ISL_527989, EPI_ISL_527990, EPI_ISL_527991, EPI_ISL_527992, EPI_ISL_527993, EPI_ISL_527994, EPI_ISL_527995, EPI_ISL_527996, EPI_ISL_527997, EPI_ISL_527998, EPI_ISL_527999, EPI_ISL_528000, EPI_ISL_528001, EPI_ISL_528002, EPI_ISL_528003, EPI_ISL_528004, EPI_ISL_528005, EPI_ISL_528006, EPI_ISL_528007, EPI_ISL_528008, EPI_ISL_528009, EPI_ISL_528010, EPI_ISL_528011, EPI_ISL_528012, EPI_ISL_528013, EPI_ISL_528014, EPI_ISL_528015, EPI_ISL_528016, EPI_ISL_528017, EPI_ISL_528018, EPI_ISL_528019, EPI_ISL_528020, EPI_ISL_528021, EPI_ISL_528022, EPI_ISL_528023, EPI_ISL_528024, EPI_ISL_528025, EPI_ISL_528026, EPI_ISL_528027, EPI_ISL_528028, EPI_ISL_528029, EPI_ISL_528030, EPI_ISL_528031, EPI_ISL_528032, EPI_ISL_528033, EPI_ISL_528034, EPI_ISL_528035, EPI_ISL_528036, EPI_ISL_528037, EPI_ISL_528038, EPI_ISL_528039, EPI_ISL_528040, EPI_ISL_528041, EPI_ISL_528042, EPI_ISL_528043, EPI_ISL_528044, EPI_ISL_528045, EPI_ISL_528046, EPI_ISL_528047, EPI_ISL_528048, EPI_ISL_528049, EPI_ISL_528050, EPI_ISL_528051, EPI_ISL_528052, EPI_ISL_528053, EPI_ISL_528054, EPI_ISL_528055, EPI_ISL_528056, EPI_ISL_528057, EPI_ISL_528058, EPI_ISL_528059, EPI_ISL_528060, EPI_ISL_528061, EPI_ISL_528062, EPI_ISL_528063, EPI_ISL_528064, EPI_ISL_528065, EPI_ISL_528066, EPI_ISL_528067, EPI_ISL_528068, EPI_ISL_528069, EPI_ISL_528070, EPI_ISL_528071, EPI_ISL_528072, EPI_ISL_528073, EPI_ISL_528074, EPI_ISL_528075, EPI_ISL_528076, EPI_ISL_528077, EPI_ISL_528078, EPI_ISL_528079, EPI_ISL_528080, EPI_ISL_528081, EPI_ISL_528082, EPI_ISL_528083, EPI_ISL_528084, EPI_ISL_528085, EPI_ISL_528086, EPI_ISL_528087, EPI_ISL_528088, EPI_ISL_528089, EPI_ISL_528090, EPI_ISL_528091, EPI_ISL_528092, EPI_ISL_528093, EPI_ISL_528094, EPI_ISL_528095, EPI_ISL_528096, EPI_ISL_528097, EPI_ISL_528098, EPI_ISL_528099, EPI_ISL_528100, EPI_ISL_528101, EPI_ISL_528102, EPI_ISL_528103, EPI_ISL_528104, EPI_ISL_528105, EPI_ISL_528106, EPI_ISL_528107, EPI_ISL_528108, EPI_ISL_528109, EPI_ISL_528110, EPI_ISL_528111, EPI_ISL_528112, EPI_ISL_528113, EPI_ISL_528114, EPI_ISL_528115, EPI_ISL_528116, EPI_ISL_528117, EPI_ISL_528118, EPI_ISL_528119, EPI_ISL_528120, EPI_ISL_528121, EPI_ISL_528122, EPI_ISL_528123, EPI_ISL_528124, EPI_ISL_528125, EPI_ISL_528126, EPI_ISL_528127, EPI_ISL_528128, EPI_ISL_528129, EPI_ISL_528130, EPI_ISL_528131, EPI_ISL_528132, EPI_ISL_528133, EPI_ISL_528134, EPI_ISL_528135, EPI_ISL_528136, EPI_ISL_528137, EPI_ISL_528138, EPI_ISL_528139, EPI_ISL_528140, EPI_ISL_528141, EPI_ISL_528142, EPI_ISL_528143, EPI_ISL_528144, EPI_ISL_528145, EPI_ISL_528146, EPI_ISL_528147, EPI_ISL_528148, EPI_ISL_528149, EPI_ISL_528150, EPI_ISL_528151, EPI_ISL_528152, EPI_ISL_528153, EPI_ISL_528154, EPI_ISL_528155, EPI_ISL_528156, EPI_ISL_528157, EPI_ISL_528158, EPI_ISL_528159, EPI_ISL_528160, EPI_ISL_528161, EPI_ISL_528162, EPI_ISL_528163, EPI_ISL_528164, EPI_ISL_528165, EPI_ISL_528166, EPI_ISL_528167, EPI_ISL_528168, EPI_ISL_528169, EPI_ISL_528170, EPI_ISL_528171, EPI_ISL_528172, EPI_ISL_528173, EPI_ISL_528174, EPI_ISL_528175, EPI_ISL_528176, EPI_ISL_528177, EPI_ISL_528178, EPI_ISL_528179, EPI_ISL_528180, EPI_ISL_528181, EPI_ISL_528182, EPI_ISL_528183, EPI_ISL_528184, EPI_ISL_528185, EPI_ISL_528186, EPI_ISL_528187, EPI_ISL_528188, EPI_ISL_528189, EPI_ISL_528190, EPI_ISL_528191, EPI_ISL_528192, EPI_ISL_528193, EPI_ISL_528194, EPI_ISL_528195, EPI_ISL_528196, EPI_ISL_528197, EPI_ISL_528198, EPI_ISL_528199, EPI_ISL_528200, EPI_ISL_528201, EPI_ISL_528202, EPI_ISL_528203, EPI_ISL_528204, EPI_ISL_528205, EPI_ISL_528206, EPI_ISL_528207, EPI_ISL_528208, EPI_ISL_528209, EPI_ISL_528210, EPI_ISL_528211, EPI_ISL_528212, EPI_ISL_528213, EPI_ISL_528214, EPI_ISL_528215, EPI_ISL_528216, EPI_ISL_528217, EPI_ISL_528218, EPI_ISL_528219, EPI_ISL_528220, EPI_ISL_528221, EPI_ISL_528222, EPI_ISL_528223, EPI_ISL_528224, EPI_ISL_528225, EPI_ISL_528226, EPI_ISL_528227, EPI_ISL_528228, EPI_ISL_528229, EPI_ISL_528230, EPI_ISL_528231, EPI_ISL_528232, EPI_ISL_528233, EPI_ISL_528234, EPI_ISL_528235, EPI_ISL_528236, EPI_ISL_528237, EPI_ISL_528238, EPI_ISL_528239, EPI_ISL_528240, EPI_ISL_528241, EPI_ISL_528242, EPI_ISL_528243, EPI_ISL_528244, EPI_ISL_528245, EPI_ISL_528246, EPI_ISL_528247, EPI_ISL_528248, EPI_ISL_528249, EPI_ISL_528250, EPI_ISL_528251, EPI_ISL_528252, EPI_ISL_528253, EPI_ISL_528254, EPI_ISL_528255, EPI_ISL_528256, EPI_ISL_528257, EPI_ISL_528258, EPI_ISL_528259, EPI_ISL_528260, EPI_ISL_528261, EPI_ISL_528262, EPI_ISL_528263, EPI_ISL_528264, EPI_ISL_528265, EPI_ISL_528266, EPI_ISL_528267, EPI_ISL_528268, EPI_ISL_528269, EPI_ISL_528270, EPI_ISL_528271, EPI_ISL_528272, EPI_ISL_528273, EPI_ISL_528274, EPI_ISL_528275, EPI_ISL_528276, EPI_ISL_528277, EPI_ISL_528278, EPI_ISL_528279, EPI_ISL_528280, EPI_ISL_528281, EPI_ISL_528282, EPI_ISL_528283, EPI_ISL_528284, EPI_ISL_528285, EPI_ISL_528286, EPI_ISL_528287, EPI_ISL_528288, EPI_ISL_528289, EPI_ISL_528290, EPI_ISL_528291, EPI_ISL_528292, EPI_ISL_528293, EPI_ISL_528294, EPI_ISL_528295, EPI_ISL_528296, EPI_ISL_528297, EPI_ISL_528298, EPI_ISL_528299, EPI_ISL_528300, EPI_ISL_528301, EPI_ISL_528302, EPI_ISL_528303, EPI_ISL_528304, EPI_ISL_528305, EPI_ISL_528306, EPI_ISL_528307, EPI_ISL_528308, EPI_ISL_528309, EPI_ISL_528310, EPI_ISL_528311, EPI_ISL_528312, EPI_ISL_528313, EPI_ISL_528314, EPI_ISL_528315, EPI_ISL_528316, EPI_ISL_528317, EPI_ISL_528318, EPI_ISL_528319, EPI_ISL_528320, EPI_ISL_528321, EPI_ISL_528322, EPI_ISL_528323, EPI_ISL_528324, EPI_ISL_528325, EPI_ISL_528326, EPI_ISL_528327, EPI_ISL_528328, EPI_ISL_528329, EPI_ISL_528330, EPI_ISL_528331, EPI_ISL_528332, EPI_ISL_528333, EPI_ISL_528334, EPI_ISL_528335, EPI_ISL_528336, EPI_ISL_528337, EPI_ISL_528338, EPI_ISL_528339, EPI_ISL_528340, EPI_ISL_528341, EPI_ISL_528342, EPI_ISL_528343, EPI_ISL_528344, EPI_ISL_528345, EPI_ISL_528346, EPI_ISL_528347, EPI_ISL_528348, EPI_ISL_528349, EPI_ISL_528350, EPI_ISL_528351, EPI_ISL_528352, EPI_ISL_528353, EPI_ISL_528354, EPI_ISL_528355, EPI_ISL_528356, EPI_ISL_528357, EPI_ISL_528358, EPI_ISL_528359, EPI_ISL_528360, EPI_ISL_528361, EPI_ISL_528362, EPI_ISL_528363, EPI_ISL_528364, EPI_ISL_528365, EPI_ISL_528366, EPI_ISL_528367, EPI_ISL_528368, EPI_ISL_528369, EPI_ISL_528370, EPI_ISL_528371, EPI_ISL_528372, EPI_ISL_528373, EPI_ISL_528374, EPI_ISL_528375, EPI_ISL_528376, EPI_ISL_528377, EPI_ISL_528378, EPI_ISL_528379, EPI_ISL_528380, EPI_ISL_528381 | | | |
| see above | University Hospital Basel, Clinical Virology | University Hospital Basel, Clinical Bacteriology | Madlen Stange, Alfredo Mari, Tim Roloff, Helena MB Seth-Smith, Michael Schweitzer, Myrta Brunner, Karoline Leuzinger, Kirstine K. Soegaard, Alexander Gensch, Sarah Tschudin-Sutter, Simon Fuchs, Julia Bielicki, Hans Pargger, Martin Siegemund, Christian Nickel, Roland Bingisser, Michael Osthoff, Stefano Bassetti, Rita Schneider-Sliwa, Manuel Battegay, Hans Hirsch, Adrian Egli |
| EPI_ISL_528386 | Viral vaccines, VSVRI- Veterinary serum and vaccine research institute | Viral vaccines, VSVRI- Veterinary serum and vaccine research institute | Saleh,A.A., Saad,M.A. |

We gratefully acknowledge the following Authors from the Originating laboratories responsible for obtaining the specimens, as well as the Submitting laboratories where the genome data were generated and shared via GISAID, on which this research is based.

All Submitters of data may be contacted directly via www.gisaid.org

| Accession ID | Originating Laboratory | Submitting Laboratory | Authors |
|---|---|---|---|
| EPI_ISL_528382, EPI_ISL_528383, EPI_ISL_528384, EPI_ISL_528385 | Translational Health Science and Technology Institute -ESIC medical college and hospital, Faridabad | THSTI Bioassay laboratory | Saurabh Kumar, Jigme Wangchuk, Anil Kumar Pandey, Asim Das, Guruprasad R. Medigeshi |
| EPI_ISL_528388, EPI_ISL_528389, EPI_ISL_528390, EPI_ISL_528391, EPI_ISL_528392, EPI_ISL_528393, EPI_ISL_528394, EPI_ISL_528395, EPI_ISL_528396, EPI_ISL_528397, EPI_ISL_528398, EPI_ISL_528399, EPI_ISL_528400, EPI_ISL_528401, EPI_ISL_528402, EPI_ISL_528403, EPI_ISL_528404, EPI_ISL_528405, EPI_ISL_528406, EPI_ISL_528407, EPI_ISL_528408, EPI_ISL_528409, EPI_ISL_528410, EPI_ISL_528411, EPI_ISL_528412, EPI_ISL_528413, EPI_ISL_528414, EPI_ISL_528415, EPI_ISL_528416, EPI_ISL_528417, EPI_ISL_528418 | | | |
| see above | Respiratory Virus Unit, Microbiology Services Colindale, Public Health England | Respiratory Virus Unit, Microbiology Services Colindale, Public Health England | PHE Covid Sequencing Team |
| EPI_ISL_528419, EPI_ISL_528420, EPI_ISL_528421, EPI_ISL_528422, EPI_ISL_528423, EPI_ISL_528424 | TNMC & BYL NAIR CH. HOSPITAL | Institute of Genomics and Integrative Biology - Council of Scientific and Industrial Research | Rajesh Pandey, Jayanthi Shastri, Akshay Kanakan, Vivekanand A, Janani Srinivasa Vasudevan, Ranjeet Maurya, Sachee Agrawal, Nirjhar Chatterjee, Swapneil Parikh, Manish Pathak, Subrat Thanapati, Jasmina Savak, Suresh Poojari, Mahesh Sangar, Amol Borse, Shweta Kawankar, Vasil Nachan, Mayuresh Vishwanathan, Shruthi Sachidanandan, Shrutika Pophale, Utkarsha Yelve |
| EPI_ISL_528425, EPI_ISL_528426 | P. D. Hinduja Hospital and Medical Research Centre | Institute of Genomics and Integrative Biology - Council of Scientific and Industrial Research | Rajesh Pandey, Jayanthi Shastri, Akshay Kanakan, Vivekanand A, Janani Srinivasa Vasudevan, Ranjeet Maurya, Sachee Agrawal, Nirjhar Chatterjee, Swapneil Parikh, Manish Pathak, Subrat Thanapati, Jasmina Savak, Suresh Poojari, Mahesh Sangar, Amol Borse, Shweta Kawankar, Vasil Nachan, Mayuresh Vishwanathan, Shruthi Sachidanandan, Shrutika Pophale, Utkarsha Yelve |
| EPI_ISL_528427 | National Genomics Core-Center for DNA Fingerprinting and Diagnostics | National Genomics Core- Center for DNA Fingerprinting and Diagnostics (NGC-CDFD)- DBT's PAN-INDIA-1000 Genome consortium | Bala Pratyusha, Heena Shah, G Shashikanth, Vinay Donipadi, Edurugatla Dinesh, Guru Raja, Hilal Ahmad Reshi, J. Mallikarjun, K. Viswakalyan, Kaisar Ahmad Lone,Kausika Kumar Malik, N. Sudheer, R Harinarayanan, Rashna Bhandari, Murali Dharan Bashyam, Debashish Mitra, Divya Vashisht, Ashwin Dalal |
| EPI_ISL_528428, EPI_ISL_528429 | National Genomics Core-Center for DNA Fingerprinting and Diagnostics | National Genomics Core- Center for DNA Fingerprinting and Diagnostics (NGC-CDFD)- DBT's PAN-INDIA-1000 Genome consortium | Ashwin Dalal, Bala Pratyusha, Heena Shah, G Shashikanth, Vinay Donipadi, Neeraj Kumar, Niteen Pathak, Pradipta Hore, Rahul Baroi, Sayantan Goswami, Shaffiqu T S, Shalini Aricthota, Sobhan Babu, R Harinarayanan, Rashna Bhandari, Murali Dharan Bashyam, Debashish Mitra, Divya Vashisht |
| EPI_ISL_528430, EPI_ISL_528431, EPI_ISL_528432, EPI_ISL_528433, EPI_ISL_528434, EPI_ISL_528435, EPI_ISL_528436, EPI_ISL_528437, EPI_ISL_528438, EPI_ISL_528439 | Respiratory Virus Unit, Microbiology Services Colindale, Public Health England | Respiratory Virus Unit, Microbiology Services Colindale, Public Health England | PHE Covid Sequencing Team |
| EPI_ISL_528440, EPI_ISL_528441, EPI_ISL_528442, EPI_ISL_528443, EPI_ISL_528444, EPI_ISL_528445, EPI_ISL_528446, EPI_ISL_528447, EPI_ISL_528448, EPI_ISL_528449, EPI_ISL_528450, EPI_ISL_528451, EPI_ISL_528452, EPI_ISL_528453, EPI_ISL_528454, EPI_ISL_528455, EPI_ISL_528456, EPI_ISL_528457, EPI_ISL_528458, EPI_ISL_528459, EPI_ISL_528460, EPI_ISL_528461, EPI_ISL_528462, EPI_ISL_528463, EPI_ISL_528464, EPI_ISL_528465, EPI_ISL_528466, EPI_ISL_528467, EPI_ISL_528468, EPI_ISL_528469, EPI_ISL_528470, EPI_ISL_528471, EPI_ISL_528472, EPI_ISL_528473, EPI_ISL_528474, EPI_ISL_528475, EPI_ISL_528476, EPI_ISL_528477, EPI_ISL_528478, EPI_ISL_528479, EPI_ISL_528480, EPI_ISL_528481, EPI_ISL_528482, EPI_ISL_528483, EPI_ISL_528484 | | | |
| see above | National Virus Reference Laboratory | National Virus Reference Laboratory | Michael Carr, Gabriel Gonzalez, Jonathan Dean, Suzie Coughlan, Cillian F De Gascun |
| EPI_ISL_528485, EPI_ISL_528486, EPI_ISL_528487, EPI_ISL_528488, EPI_ISL_528489, EPI_ISL_528490, EPI_ISL_528491, EPI_ISL_528492, EPI_ISL_528493, EPI_ISL_528494, EPI_ISL_528495, EPI_ISL_528496, EPI_ISL_528497, EPI_ISL_528498, EPI_ISL_528499, EPI_ISL_528500, EPI_ISL_528501, EPI_ISL_528502, EPI_ISL_528503, EPI_ISL_528504, EPI_ISL_528505, EPI_ISL_528506, EPI_ISL_528507, EPI_ISL_528508, EPI_ISL_528509, EPI_ISL_528510, EPI_ISL_528511, EPI_ISL_528512, EPI_ISL_528513, EPI_ISL_528514, EPI_ISL_528515, EPI_ISL_528516, EPI_ISL_528517, EPI_ISL_528518, EPI_ISL_528519, EPI_ISL_528520, EPI_ISL_528521, EPI_ISL_528522, EPI_ISL_528523, EPI_ISL_528524, EPI_ISL_528525, EPI_ISL_528526, EPI_ISL_528527, EPI_ISL_528528, EPI_ISL_528529, EPI_ISL_528530, EPI_ISL_528531, EPI_ISL_528532, EPI_ISL_528533, EPI_ISL_528534, EPI_ISL_528535, EPI_ISL_528536, EPI_ISL_528537 | | | |
| see above | Alaska State Virology Laboratory | Alaska State Virology Laboratory | Chen J et al with Pathogenomics group Dagdag R, Redlinger M, Milton E, George W, Kovalenko A, Drown DM, Bortz E |
| EPI_ISL_528538 | Alsafar | Alsafar | Andreas Henschel, Gihan Elsir Ahmed Daw Elbait, Samuel Feng, Rifat, Ernesto Damiani, Guan Tay, Habiba Alsafar |
| EPI_ISL_528539 | LVM/UFRJ | LNCC | Gustavo M. Romário M. de Souza; Bruno B. Bezerra; Lucio A. Caldas; Fabio Limonte; Elena Cobos; Sharton V. A. Coelho; Luiz Almeida; Luiza Higga; Isadora A. Correa; Diana Marianni; Luciana B. Arruda; Marcelo Bozza; Orlando Ferreira; Wanderley de Souza; Ana Teresa R. Vasconcelos; Terezinha M. Castineiras; Amilcar Tanuri; Luciana J. Costa. |
| EPI_ISL_528540, EPI_ISL_528541 | National Genomics Core-Center for DNA Fingerprinting and Diagnostics | National Genomics Core- Center for DNA Fingerprinting and Diagnostics (NGC-CDFD)- DBT's PAN-INDIA-1000 Genome consortium | G Shashikanth, Heena Shah, Bala Pratyusha, Vinay Donipadi, K.Manohar, Madhumohan Rao, Shubhra Ganguli, Suchitra Upreti, Swathi Chodisetty, Vani Singh, R Harinarayanan, Rashna Bhandari, Murali Dharan Bashyam, Debashish Mitra, Divya Vashisht, Ashwin Dalal |
| EPI_ISL_528542, EPI_ISL_528543, EPI_ISL_528544, EPI_ISL_528545, EPI_ISL_528546, EPI_ISL_528547, EPI_ISL_528548, EPI_ISL_528549 | National Genomics Core-Center for DNA Fingerprinting and Diagnostics | National Genomics Core- Center for DNA Fingerprinting and Diagnostics (NGC-CDFD)- DBT's PAN-INDIA-1000 Genome consortium | G Shashikanth, Heena Shah, Bala Pratyusha, Vinay Donipadi, S Vasantha Rani, M Sri Lalitha, R. Angalena, Usha Rani Dutta, Nimmala Naresh, Nanci Rani K, Ch Venkateshwar Goud, Devinder Singh Negi, R Harinarayanan, Rashna Bhandari, Murali Dharan Bashyam, Debashish Mitra, Divya Vashisht, Ashwin Dalal |
| EPI_ISL_528550, EPI_ISL_528551, EPI_ISL_528552, EPI_ISL_528553, EPI_ISL_528554, EPI_ISL_528555, EPI_ISL_528556 | National Genomics Core-Center for DNA Fingerprinting and Diagnostics | National Genomics Core- Center for DNA Fingerprinting and Diagnostics (NGC-CDFD)- DBT's PAN-INDIA-1000 Genome consortium | Heena Shah, G Shashikanth, Bala Pratyusha, Vinay Donipadi, Shruti Dasgupta, Kandali Sreethi Sreenivasulu Reddy, Chandra Shekhar Singh, Sunke Vijayakumar, R Lakshmi Vaishna, Jenige Aravindh Kumar, Muthulakshmi, V Naga Sailaja, R Harinarayanan, Rashna Bhandari, Murali Dharan Bashyam, Debashish Mitra, Divya Vashisht, Ashwin Dalal |
| EPI_ISL_528557, EPI_ISL_528558, EPI_ISL_528559, EPI_ISL_528560, EPI_ISL_528561, EPI_ISL_528562, EPI_ISL_528563, EPI_ISL_528564 | National Genomics Core-Center for DNA Fingerprinting and Diagnostics | National Genomics Core- Center for DNA Fingerprinting and Diagnostics (NGC-CDFD)- DBT's PAN-INDIA-1000 Genome consortium | Heena Shah, G Shashikanth, Bala Pratyusha, Vinay Donipadi, Binod Bihari Pradhan, Jamal Md Nurul Jain, Srinivas G, C Bala Maddeleti, R. Manorama, T. Navaneetha, Surya Vamshi, Chendra Shekar P, R Harinarayanan, Rashna Bhandari, Murali Dharan Bashyam, Debashish Mitra, Divya Vashisht, Ashwin Dalal |
| EPI_ISL_528565, EPI_ISL_528566, EPI_ISL_528567, EPI_ISL_528568, EPI_ISL_528569, EPI_ISL_528570, EPI_ISL_528571, EPI_ISL_528572 | National Genomics Core-Center for DNA Fingerprinting and Diagnostics | National Genomics Core- Center for DNA Fingerprinting and Diagnostics (NGC-CDFD)- DBT's PAN-INDIA-1000 Genome consortium | G Shashikanth, Heena Shah, Bala Pratyusha, Vinay Donipadi, Rajitha Ponnala, Seyed Khaja Ali, B. Krishna Murthy, Akruti Shah, Jayashree Ladke, Shivangi Wagh, Asodu Sandeep Sarma, Sunu Joseph, R Harinarayanan, Rashna Bhandari, Murali Dharan Bashyam, Debashish Mitra, Divya Vashisht, Ashwin Dalal |
| EPI_ISL_528573, EPI_ISL_528574, EPI_ISL_528575, EPI_ISL_528576, EPI_ISL_528577, EPI_ISL_528578, EPI_ISL_528579, EPI_ISL_528580 | National Genomics Core-Center for DNA Fingerprinting and Diagnostics | National Genomics Core- Center for DNA Fingerprinting and Diagnostics (NGC-CDFD)- DBT's PAN-INDIA-1000 Genome consortium | Vinay Donipadi, G Shashikanth, Heena Shah, Bala Pratyusha, Parveen Kumar, Sandip Patra, Mugdha Singh, Reelina Basu, Dhanraj Adey, Bharath Kumar, Bhavani Sontam, Shaik Nasar Vali, R Harinarayanan, Rashna Bhandari, Murali Dharan Bashyam, Debashish Mitra, Divya Vashisht, Ashwin Dalal |
| EPI_ISL_528581, EPI_ISL_528582, EPI_ISL_528583, EPI_ISL_528584, EPI_ISL_528585, EPI_ISL_528586, EPI_ISL_528587 | National Genomics Core-Center for DNA Fingerprinting and Diagnostics | National Genomics Core- Center for DNA Fingerprinting and Diagnostics (NGC-CDFD)- DBT's PAN-INDIA-1000 Genome consortium | G Shashikanth, Heena Shah, Bala Pratyusha, Vinay Donipadi, Bathula Siddardha, Vineesha Oddi, Lavanya Banda, Surya Chodisetty, Abhijeeth Singh Thakur, Mohammad Mudassir, Nalini Raghunathan, Rajeshree Sanyal, R Harinarayanan, Rashna Bhandari, Murali Dharan Bashyam, Debashish Mitra, Divya Vashisht, Ashwin Dalal |
| EPI_ISL_528588, EPI_ISL_528589, EPI_ISL_528590, EPI_ISL_528591, EPI_ISL_528592, EPI_ISL_528593, EPI_ISL_528594, EPI_ISL_528595 | National Genomics Core-Center for DNA Fingerprinting and Diagnostics | National Genomics Core- Center for DNA Fingerprinting and Diagnostics (NGC-CDFD)- DBT's PAN-INDIA-1000 Genome consortium | Heena Shah, G Shashikanth, Bala Pratyusha, Vinay Donipadi, Raju Kumar, Ajay Kumar Chaudhary, Akash Chinchole, Brahmaji Sontyana, C. Arun Kumar, Chandra Shekhar V, Chilakala Gangi Reddy, Chinthakindi KrishnaPrasad, R Harinarayanan, Rashna Bhandari, Murali Dharan Bashyam, Debashish Mitra, Divya Vashisht, Ashwin Dalal |
| EPI_ISL_528596, EPI_ISL_528597, EPI_ISL_528598, EPI_ISL_528599, EPI_ISL_528600, EPI_ISL_528601, EPI_ISL_528602 | National Genomics Core-Center for DNA Fingerprinting and Diagnostics | National Genomics Core- Center for DNA Fingerprinting and Diagnostics (NGC-CDFD)- DBT's PAN-INDIA-1000 Genome consortium | Bala Pratyusha, Heena Shah, G Shashikanth, Vinay Donipadi, Edurugatla Dinesh, Guru Raja, Hilal Ahmad Reshi, J. Mallikarjun, K. Viswakalyan, Kaisar Ahmad Lone,Kausika Kumar Malik, N. Sudheer, R Harinarayanan, Rashna Bhandari, Murali Dharan Bashyam, Debashish Mitra, Divya Vashisht, Ashwin Dalal |
| EPI_ISL_528603, EPI_ISL_528604 | National Genomics Core-Center for DNA Fingerprinting and Diagnostics | National Genomics Core- Center for DNA Fingerprinting and Diagnostics (NGC-CDFD)- DBT's PAN-INDIA-1000 Genome consortium | Ashwin Dalal, Bala Pratyusha, Heena Shah, G Shashikanth, Vinay Donipadi, Neeraj Kumar, Niteen Pathak, Pradipta Hore, Rahul Baroi, Sayantan Goswami, Shaffiqu T S, Shalini Aricthota, Sobhan Babu, R Harinarayanan, Rashna Bhandari, Murali Dharan Bashyam, Debashish Mitra, Divya Vashisht |
| EPI_ISL_528605, EPI_ISL_528606, EPI_ISL_528607, EPI_ISL_528608 | National Genomics Core-Center for DNA Fingerprinting and Diagnostics | National Genomics Core- Center for DNA Fingerprinting and Diagnostics (NGC-CDFD)- DBT's PAN-INDIA-1000 Genome consortium | Ashwin Dalal, Bala Pratyusha, Heena Shah, G Shashikanth, Vinay Donipadi, K.Manohar, Madhumohan Rao,Neeraj Kumar, Niteen Pathak, Pradipta Hore, Rahul Baroi, Sayantan Goswami, Shaffiqu T S, Shalini Aricthota, Sobhan Babu, R Harinarayanan, Rashna Bhandari, Murali Dharan Bashyam, Debashish Mitra, Divya Vashisht |
| EPI_ISL_528609, EPI_ISL_528610, EPI_ISL_528611, EPI_ISL_528612, EPI_ISL_528613, EPI_ISL_528614, EPI_ISL_528615, EPI_ISL_528616 | National Genomics Core-Center for DNA Fingerprinting and Diagnostics | National Genomics Core- Center for DNA Fingerprinting and Diagnostics (NGC-CDFD)- DBT's PAN-INDIA-1000 Genome consortium | Divya Vashisht, Bala Pratyusha, Heena Shah, G Shashikanth, Vinay Donipadi, K.Manohar, Madhumohan Rao, SPR Prasad, Yogesh Patidar, Arjita Jaiswal, Arpita Singh, Devanshi Gupta, Romila Moirangthem, Sanjana Sarkar, Shivani Yadav, R Harinarayanan, Rashna Bhandari, Murali Dharan Bashyam, Debashish Mitra, Ashwin Dalal |
| EPI_ISL_528617, EPI_ISL_528618, EPI_ISL_528619, EPI_ISL_528620, EPI_ISL_528621, EPI_ISL_528622, EPI_ISL_528623, EPI_ISL_528624 | National Genomics Core-Center for DNA Fingerprinting and Diagnostics | National Genomics Core- Center for DNA Fingerprinting and Diagnostics (NGC-CDFD)- DBT's PAN-INDIA-1000 Genome consortium | Heena Shah, G Shashikanth, Bala Pratyusha, Vinay Donipadi, K.Manohar, Madhumohan Rao, Shubhra Ganguli, Suchitra Upreti, Swathi Chodisetty, Vani Singh, R Harinarayanan, Rashna Bhandari, Murali Dharan Bashyam, Debashish Mitra, Divya Vashisht, Ashwin Dalal |
| EPI_ISL_528625, EPI_ISL_528626, EPI_ISL_528627, EPI_ISL_528628, EPI_ISL_528629, EPI_ISL_528630 | National Genomics Core-Center for DNA Fingerprinting and Diagnostics | National Genomics Core- Center for DNA Fingerprinting and Diagnostics (NGC-CDFD)- DBT's PAN-INDIA-1000 Genome consortium | G Shashikanth, Heena Shah, Bala Pratyusha, Vinay Donipadi, K.Manohar, Madhumohan Rao, Shubhra Ganguli, Suchitra Upreti, Swathi Chodisetty, Vani Singh, R Harinarayanan, Rashna Bhandari, Murali Dharan Bashyam, Debashish Mitra, Divya Vashisht, Ashwin Dalal |
| EPI_ISL_528631, EPI_ISL_528632, EPI_ISL_528633, EPI_ISL_528634, EPI_ISL_528635, EPI_ISL_528636 | National Genomics Core-Center for DNA Fingerprinting and Diagnostics | National Genomics Core- Center for DNA Fingerprinting and Diagnostics (NGC-CDFD)- DBT's PAN-INDIA-1000 Genome consortium | Heena Shah, G Shashikanth, Bala Pratyusha, Vinay Donipadi, K.Manohar, Madhumohan Rao, Shruti Dasgupta, Kandali Sreethi Sreenivasulu Reddy, Chandra Shekhar Singh, Sunke Vijayakumar, R Lakshmi Vaishna, Jenige Aravindh Kumar, Muthulakshmi, V Naga Sailaja, R Harinarayanan, Rashna Bhandari, Murali Dharan Bashyam, Debashish Mitra, Divya Vashisht, Ashwin Dalal |
| EPI_ISL_528637, EPI_ISL_528638 | LVM/UFRJ | Bioinformatics Laboratory / LNCC | Gustavo D. P. Silva; M. Romário M. de Souza; Bruno B. Bezerra; Lucio A. Caldas; Fabio Limonte; Elena Cobos; Sharton V. A. Coelho; Luiz Almeida; Luiza Higga; Isadora A. Correa; Diana Marianni; Luciana B. Arruda; Marcelo Bozza; Orlando Ferreira; Wanderley de Souza; Ana Teresa R. Vasconcelos; Terezinha M. Castineiras; Amilcar Tanuri; Luciana J. Costa |
| EPI_ISL_528639, EPI_ISL_528640, EPI_ISL_528641, EPI_ISL_528642, EPI_ISL_528643, EPI_ISL_528644, EPI_ISL_528645, EPI_ISL_528646, EPI_ISL_528647, EPI_ISL_528648, EPI_ISL_528649, EPI_ISL_528650, EPI_ISL_528651, EPI_ISL_528652, EPI_ISL_528653, EPI_ISL_528654, EPI_ISL_528655, EPI_ISL_528656, EPI_ISL_528657, EPI_ISL_528658, EPI_ISL_528659, EPI_ISL_528660, EPI_ISL_528661, EPI_ISL_528662 | | | |
| see above | Virginia DCLS | Virginia DCLS | Virginia DCLS |
| EPI_ISL_528663, EPI_ISL_528664, EPI_ISL_528665, EPI_ISL_528666, EPI_ISL_528667, EPI_ISL_528668, EPI_ISL_528669, EPI_ISL_528670, EPI_ISL_528671, EPI_ISL_528672, EPI_ISL_528673, EPI_ISL_528674, EPI_ISL_528675, EPI_ISL_528676, EPI_ISL_528677, EPI_ISL_528678, EPI_ISL_528679, EPI_ISL_528680, EPI_ISL_528681, EPI_ISL_528682 | | | |
| see above | Virginia Division of Consolidated Laboratory Services (DCLS) | Virginia Division of Consolidated Laboratory Services (DCLS) | Virginia DCLS |

| | | | |
|---|---|---|---|
| EPI_ISL_528683, EPI_ISL_528684 | NCDC Institute of Genomics and Integrative Biology | NCDC Institute of Genomics and Integrative Biology | Vivekanand A, Mahesh S. Dhar, Bharathram Uppili, Akshay Kanakan, Simmi Tiwari, RadhaKrishnan VS, Robin Marwal, Azka Khan, Ajit Shewale, Pooja Sharma, Tushar Nale, Rajesh Pandey, Sandhya Kabra, Mohammed Faruq, Sujeet Singh, Anurag Agrawal, Partha Rakshit |
| EPI_ISL_528685 | IIP Institute of Genomics and Integrative Biology | IIP Institute of Genomics and Integrative Biology | Vivekanand A, Mahesh S. Dhar, Bharathram Uppili, Akshay Kanakan, Simmi Tiwari, RadhaKrishnan VS, Robin Marwal, Azka Khan, Ajit Shewale, Pooja Sharma, Tushar Nale, Rajesh Pandey, Sandhya Kabra, Mohammed Faruq, Sujeet Singh, Anurag Agrawal, Partha Rakshit |
| EPI_ISL_528686, EPI_ISL_528687, EPI_ISL_528688, EPI_ISL_528689, EPI_ISL_528690, EPI_ISL_528691, EPI_ISL_528692, EPI_ISL_528693, EPI_ISL_528694, EPI_ISL_528695, EPI_ISL_528696, EPI_ISL_528697, EPI_ISL_528698, EPI_ISL_528699, EPI_ISL_528700, EPI_ISL_528701, EPI_ISL_528702, EPI_ISL_528703, EPI_ISL_528704, EPI_ISL_528705, EPI_ISL_528706, EPI_ISL_528707, EPI_ISL_528708, EPI_ISL_528709, EPI_ISL_528710, EPI_ISL_528711, EPI_ISL_528712, EPI_ISL_528713, EPI_ISL_528714, EPI_ISL_528715, EPI_ISL_528716, EPI_ISL_528717, EPI_ISL_528718, EPI_ISL_528719, EPI_ISL_528720, EPI_ISL_528721 | | | |
| see above | Alsafar - Khalifa University Abu Dhabi | Alsafar - Khalifa University Abu Dhabi | Andreas Henschel, Gihan Daw Elbait, Samuel Feng, Rifat Hamoudi, Ernesto Damiani, Guan Tay, Habiba Alsafar |
| EPI_ISL_528738, EPI_ISL_528739, EPI_ISL_528740, EPI_ISL_528741 | Malaysia Genome Institute | Malaysia Genome Institute | Mohd Noor Mat Isa, Irni Suhayu Sapian, Yusuf Muhammad Noor, Nurhezreen Md Iqbal, Mohd Faizal Abu Bakar, Enizza Kasim, Shamsidar Sopie, Siti Noraini Othman, Azrin Ahmad, Nor Azfa Johari, Shahrul Hisham Zainal Ariffin |
| EPI_ISL_528742 | Malaysia Genome Institute | Malaysia Genome Institute | Mohd Noor Mat Isa, Irni Suhayu Sapian, Gan Han Ming, Yusuf Muhammad Noor, Tan Ju Lin, Nurhezreen Md Iqbal, Mohd Faizal Abu Bakar, Enizza Kasim, Shamsidar Sopie, Siti Noraini Othman, Azrin Ahmad, Nor Azfa Johari, Shahrul Hisham Zainal Ariffin |
| EPI_ISL_528743, EPI_ISL_528744 | Malaysia Genome Institute | Malaysia Genome Institute | Mohd Noor Mat Isa, Irni Suhayu Sapian, Yusuf Muhammad Noor, Nurhezreen Md Iqbal, Mohd Faizal Abu Bakar, Enizza Kasim, Shamsidar Sopie, Siti Noraini Othman, Azrin Ahmad, Nor Azfa Johari, Shahrul Hisham Zainal Ariffin |
| EPI_ISL_528745 | Laboratorium Kesehatan Provinsi Jawa Barat | School of Life Sciences and Technology & School of Pharmacy-Institut Teknologi Bandung; Molecular Genetics Laboratory-Faculty of Medicine-Universitas Padjadjaran; Laboratorium Kesehatan Provinsi Jawa Barat | Marselina Irasonia Tan, Yunia Sribudiani, Catur Riani, Azzania Fibriani, Husna Nugrahapraja, Tarwadi, Ema Rahmawati, Savira Ekawardhani, Hesti Lina Wiraswati, Ryan Bayusantika Ristandi, Rifky Waluyajati Rachman, Cut Nur Cinthia Alamanda, Lia Faridah, Tri Hanggono Achmad, Mas Rizky A.A. Syamsunarno, Fensi Amalina, Hammam Riza, Sony Solistia Wirawan, Agung Eru Wibowo, Irvan Faizal |
| EPI_ISL_528746 | Immanuel Hospital | Molecular Genetics Laboratory-Faculty of Medicine-Universitas Padjadjaran; School of Life Sciences and Technology & School of Pharmacy-Institut Teknologi Bandung; Laboratorium Kesehatan Provinsi Jawa Barat | Yunia Sribudiani,Tri Hanggono Achmad, Mas Rizky A.A. Syamsunarno, Fensi Amalina, Catur Riani, Azzania Fibriani, Husna Nugrahapraja, Marselina Irasonia Tan, Tarwadi, Ema Rahmawati, Savira Ekawardhani, Hesti Lina Wiraswati, Ryan Bayusantika Ristandi, Rifky Waluyajati Rachman, Cut Nur Cinthia Alamanda, Lia Faridah, Hammam Riza, Sony Solistia Wirawan, Agung Eru Wibowo, Irvan Faizal |
| EPI_ISL_528747 | Santo Borromeus Hospital | School of Pharmacy & School of Life Sciences and Technology - Institut Teknologi Bandung; Molecular Genetics Laboratory-Faculty of Medicine-Universitas Padjadjaran; Laboratorium Kesehatan Provinsi Jawa Barat | Catur Riani, Marselina Irasonia Tan, Yunia Sribudiani, Azzania Fibriani, Husna Nugrahapraja, Tarwadi, Ema Rahmawati, Savira Ekawardhani, Hesti Lina Wiraswati, Ryan Bayusantika Ristandi, Rifky Waluyajati Rachman, Cut Nur Cinthia Alamanda, Lia Faridah, Miftahul Faridl, Karimatu Khoirunnisa, Hammam Riza, Sony Solistia Wirawan, Agung Eru Wibowo, Irvan Faizal |
| EPI_ISL_528748 | Dinkes Provinsi Jawa Barat | School of Life Sciences and Technology & School of Pharmacy-Institut Teknologi Bandung; Molecular Genetics Laboratory-Faculty of Medicine-Universitas Padjadjaran; Laboratorium Kesehatan Provinsi Jawa Barat | Azzania Fibriani, Catur Riani, Marselina Irasonia Tan, Yunia Sribudiani, Husna Nugrahapraja, Tarwadi, Ema Rahmawati, Savira Ekawardhani, Hesti Lina Wiraswati, Ryan Bayusantika Ristandi, Rifky Waluyajati Rachman, Cut Nur Cinthia Alamanda, Lia Faridah, Gusti Ayu Prani Pradani, Adelina Khristiani Rahayu, Hammam Riza, Sony Solistia Wirawan, Agung Eru Wibowo, Irvan Faizal |
| EPI_ISL_528749 | Santosa Hospital Bandung Central | School of Life Sciences and Technology & School of Pharmacy-Institut Teknologi Bandung; Molecular Genetics Laboratory-Faculty of Medicine-Universitas Padjadjaran; Laboratorium Kesehatan Provinsi Jawa Barat | Husna Nugrahapraja, Azzania Fibriani, Catur Riani, Marselina Irasonia Tan, Yunia Sribudiani, Tarwadi, Ema Rahmawati, Savira Ekawardhani, Hesti Lina Wiraswati, Ryan Bayusantika Ristandi, Rifky Waluyajati Rachman, Cut Nur Cinthia Alamanda, Lia Faridah, Davin H. E. Setiamarga, Rizki Mardian, Hammam Riza, Sony Solistia Wirawan, Agung Eru Wibowo, Irvan Faizal |
| EPI_ISL_528750 | Santo Borromeus Hospital | School of Life Sciences and Technology & School of Pharmacy-Institut Teknologi Bandung; Molecular Genetics Laboratory-Faculty of Medicine-Universitas Padjadjaran; Laboratorium Kesehatan Provinsi Jawa Barat | Marselina Irasonia Tan, Yunia Sribudiani, Catur Riani, Azzania Fibriani, Husna Nugrahapraja, Tarwadi, Ema Rahmawati, Savira Ekawardhani, Hesti Lina Wiraswati, Ryan Bayusantika Ristandi, Rifky Waluyajati Rachman, Cut Nur Cinthia Alamanda, Lia Faridah, Miftahul Faridl, Karimatu Khoirunnisa, Hammam Riza, Sony Solistia Wirawan, Agung Eru Wibowo, Irvan Faizal |
| EPI_ISL_528751 | Santo Borromeus Hospital | Molecular Genetics Laboratory-Faculty of Medicine-Universitas Padjadjaran; School of Life Sciences and Technology & School of Pharmacy-Institut Teknologi Bandung; Laboratorium Kesehatan Provinsi Jawa Barat | Yunia Sribudiani, Tri Hanggono Achmad, Mas Rizky A.A. Syamsunarno, Fensi Amalina, Catur Riani, Azzania Fibriani, Husna Nugrahapraja, Marselina Irasonia Tan, Tarwadi, Ema Rahmawati, Savira Ekawardhani, Hesti Lina Wiraswati, Ryan Bayusantika Ristandi, Rifky Waluyajati Rachman, Cut Nur Cinthia Alamanda, Lia Faridah, Hammam Riza, Sony Solistia Wirawan, Agung Eru Wibowo, Irvan Faizal |
| EPI_ISL_528752 | Dr. H. A. Rotinsulu Lung Hospital | School of Pharmacy & School of Life Sciences and Technology - Institut Teknologi Bandung; Molecular Genetics Laboratory-Faculty of Medicine-Universitas Padjadjaran; Laboratorium Kesehatan Provinsi Jawa Barat | Catur Riani, Marselina Irasonia Tan, Yunia Sribudiani, Azzania Fibriani, Husna Nugrahapraja, Tarwadi, Ema Rahmawati, Savira Ekawardhani, Hesti Lina Wiraswati, Ryan Bayusantika Ristandi, Rifky Waluyajati Rachman, Cut Nur Cinthia Alamanda, Lia Faridah, Gusti Ayu Prani Pradani, Adelina Khristiani Rahayu, Hammam Riza, Sony Solistia Wirawan, Agung Eru Wibowo, Irvan Faizal |
| EPI_ISL_528753 | Dinkes Kota Bogor | School of Life Sciences and Technology & School of Pharmacy-Institut Teknologi Bandung; Molecular Genetics Laboratory-Faculty of Medicine-Universitas Padjadjaran; Laboratorium Kesehatan Provinsi Jawa Barat | Azzania Fibriani, Catur Riani, Marselina Irasonia Tan, Yunia Sribudiani, Husna Nugrahapraja, Tarwadi, Ema Rahmawati, Savira Ekawardhani, Hesti Lina Wiraswati, Ryan Bayusantika Ristandi, Rifky Waluyajati Rachman, Cut Nur Cinthia Alamanda, Lia Faridah, Davin H. E. Setiamarga, Rizki Mardian , Hammam Riza, Sony Solistia Wirawan, Agung Eru Wibowo, Irvan Faizal |
| EPI_ISL_528759 | Santo Borromeus Hospital | School of Life Sciences and Technology & School of Pharmacy-Institut Teknologi Bandung; Molecular Genetics Laboratory-Faculty of Medicine-Universitas Padjadjaran; Laboratorium Kesehatan Provinsi Jawa Barat | Husna Nugrahapraja, Azzania Fibriani, Catur Riani, Marselina Irasonia Tan, Yunia Sribudiani, Tarwadi, Ema Rahmawati, Savira Ekawardhani, Hesti Lina Wiraswati, Ryan Bayusantika Ristandi, Rifky Waluyajati Rachman, Cut Nur Cinthia Alamanda, Lia Faridah, Tri Hanggono Achmad, Mas Rizky A.A. Syamsunarno, Fensi Amalina, Hammam Riza, Sony Solistia Wirawan, Agung Eru Wibowo, Irvan Faizal |
| EPI_ISL_528788, EPI_ISL_528789, EPI_ISL_528790, EPI_ISL_528791, EPI_ISL_528792, EPI_ISL_528793, EPI_ISL_528794, EPI_ISL_528795, EPI_ISL_528796, EPI_ISL_528797, EPI_ISL_528798, EPI_ISL_528799, EPI_ISL_528800, EPI_ISL_528801, EPI_ISL_528802, EPI_ISL_528803, EPI_ISL_528804, EPI_ISL_528805, EPI_ISL_528806, EPI_ISL_528807, EPI_ISL_528808 | | | |
| see above | Microbiology Department, Barking Havering and Redbridge University Hospitals NHS trust | Wellcome Sanger Institute for the COVID-19 Genomics UK Consortium | Amy Ash, Fatima Ali, Cherian Koshy and Alex Alderton, Roberto Amato, Sonia Goncalves, Ewan Harrison, David K. Jackson, Ian Johnston, Dominic Kwiatkowski, Cordelia Langford, John Sillitoe on behalf of the Wellcome Sanger Institute COVID-19 Surveillance Team (http://www.sanger.ac.uk/covid-team) |
| EPI_ISL_528809 | Department of Medicine, Gandhi hospital, Hyderabad | CSIR-Centre for Cellular and Molecular Biology | Rajarao Mesipogu , Thrilok Chander Bingi ,Vinayasekhar Aedula,Tulasi Nagabandi, Namami Gaur, Sakshi Shambhavi, Lamuk Zaveri, Shagufta Khan, Nikhil Hajirnis, M Soujanya Reddy, Pratheusa Maccha, Purushotham Vodnala, Payel Mukherjee, Sofia Banu, Priya Singh, Onkar Kulkarni, Dhiviya Vedagiri, Divya Gupta, Vishal Sah, Santosh Kumar Kuncha, Krishnan Harinivas Harshan, Archana Bharadwaj Siva, Karthik Bharadwaj Tallapaka,G. Aditya Kumar, Koushick Sivakumar, Pooja Ramesh Gupta, Rajan Kumar Jha, Shraddha Vijay Lahoti, Rakesh K Mishra, Divya Tej Sowpati |
| EPI_ISL_528810, EPI_ISL_528811, EPI_ISL_528812 | Department of Medicine, Gandhi hospital, Hyderabad | CSIR-Centre for Cellular and Molecular Biology | Thrilok Chander Bingi,Rajarao Mesipogu ,Vinayasekhar Aedula,Lamuk Zaveri, Shagufta Khan, Namami Gaur, Sakshi Shambhavi, Nikhil Hajirnis, M Soujanya Reddy, Pratheusa Maccha, Tulasi Nagabandi, Purushotham Vodnala, Payel Mukherjee, Sofia Banu, Priya Singh, Onkar Kulkarni, Dhiviya Vedagiri, Divya Gupta, Vishal Sah, Santosh Kumar Kuncha, Krishnan Harinivas Harshan, Archana Bharadwaj Siva, Karthik Bharadwaj Tallapaka, Renu Sudhakar, Somesh Gorde, Gangumala Srinivas Reddy, Sujoy Deb, Swati Bayyana, Rakesh K Mishra, Divya Tej Sowpati |
| EPI_ISL_528813 | Department of Medicine, Gandhi hospital, Hyderabad | CSIR-Centre for Cellular and Molecular Biology | Vinayasekhar Aedula,Thrilok Chander Bingi, Rajarao Mesipogu, Shagufta Khan, Lamuk Zaveri, Namami Gaur, Sakshi Shambhavi, Nikhil Hajirnis, M Soujanya Reddy, Pratheusa Maccha,Tulasi Nagabandi, Purushotham Vodnala, Payel Mukherjee, Sofia Banu, Priya Singh, Onkar Kulkarni, Dhiviya Vedagiri, Divya Gupta, Vishal Sah, Santosh Kumar Kuncha, Krishnan Harinivas Harshan, Archana Bharadwaj Siva, Karthik Bharadwaj Tallapaka,Umesh Kumar, Unis Ahmad Bhat, Ajay Sarawagi, Priyanka Pant, Rajkanwar Nathawat, Rakesh K Mishra, Divya Tej Sowpati |
| EPI_ISL_528814 | Department of Medicine, Gandhi hospital, Hyderabad | CSIR-Centre for Cellular and Molecular Biology | Rajarao Mesipogu , Thrilok Chander Bingi ,Vinayasekhar Aedula,Tulasi Nagabandi, Namami Gaur, Sakshi Shambhavi, Lamuk Zaveri, Shagufta Khan, Nikhil Hajirnis, M Soujanya Reddy, Pratheusa Maccha, Purushotham Vodnala, Payel Mukherjee, Sofia Banu, Priya Singh, Onkar Kulkarni, Dhiviya Vedagiri, Divya Gupta, Vishal Sah, Santosh Kumar Kuncha, Krishnan Harinivas Harshan, Archana Bharadwaj Siva, Karthik Bharadwaj Tallapaka,G. Aditya Kumar, Koushick Sivakumar, Pooja Ramesh Gupta, Rajan Kumar Jha, Shraddha Vijay Lahoti, Rakesh K Mishra, Divya Tej Sowpati |
| EPI_ISL_528815 | Department of Medicine, Gandhi hospital, Hyderabad | CSIR-Centre for Cellular and Molecular Biology | Vinayasekhar Aedula,Thrilok Chander Bingi, Rajarao Mesipogu, Shagufta Khan, Lamuk Zaveri, Namami Gaur, Sakshi Shambhavi, Nikhil Hajirnis, M Soujanya Reddy, Pratheusa Maccha,Tulasi Nagabandi, Purushotham Vodnala, Payel Mukherjee, Sofia Banu, Priya Singh, Onkar Kulkarni, Dhiviya Vedagiri, Divya Gupta, Vishal Sah, Santosh Kumar Kuncha, Krishnan Harinivas Harshan, Archana Bharadwaj Siva, Karthik Bharadwaj Tallapaka,Umesh Kumar, Unis Ahmad Bhat, Ajay Sarawagi, Priyanka Pant, Rajkanwar Nathawat, Rakesh K Mishra, Divya Tej Sowpati |
| EPI_ISL_528816 | Department of Medicine, Gandhi hospital, Hyderabad | CSIR-Centre for Cellular and Molecular Biology | Rajarao Mesipogu , Thrilok Chander Bingi ,Vinayasekhar Aedula,Tulasi Nagabandi, Namami Gaur, Sakshi Shambhavi, Lamuk Zaveri, Shagufta Khan, Nikhil Hajirnis, M Soujanya Reddy, Pratheusa Maccha, Purushotham Vodnala, Payel Mukherjee, Sofia Banu, Priya Singh, Onkar Kulkarni, Dhiviya Vedagiri, Divya Gupta, Vishal Sah, Santosh Kumar Kuncha, Krishnan Harinivas Harshan, Archana Bharadwaj Siva, Karthik Bharadwaj Tallapaka,G. Aditya Kumar, Koushick Sivakumar, Pooja Ramesh Gupta, Rajan Kumar Jha, Shraddha Vijay Lahoti, Rakesh K Mishra, Divya Tej Sowpati |
| EPI_ISL_528817 | Department of Medicine, Gandhi hospital, Hyderabad | CSIR-Centre for Cellular and Molecular Biology | Thrilok Chander Bingi,Rajarao Mesipogu ,Vinayasekhar Aedula,Lamuk Zaveri, Shagufta Khan, Namami Gaur, Sakshi Shambhavi, Nikhil Hajirnis, M Soujanya Reddy, Pratheusa Maccha, Tulasi Nagabandi, Purushotham Vodnala, Payel Mukherjee, Sofia Banu, Priya Singh, Onkar Kulkarni, Dhiviya Vedagiri, Divya Gupta, Vishal Sah, Santosh Kumar Kuncha, Krishnan Harinivas Harshan, Archana Bharadwaj Siva, Karthik Bharadwaj Tallapaka, Renu Sudhakar, Somesh Gorde, Gangumala Srinivas Reddy, Sujoy Deb, Swati Bayyana, Rakesh K Mishra, Divya Tej Sowpati |
| EPI_ISL_528818 | Department of Medicine, Gandhi hospital, Hyderabad | CSIR-Centre for Cellular and Molecular Biology | Vinayasekhar Aedula,Thrilok Chander Bingi, Rajarao Mesipogu, Shagufta Khan, Lamuk Zaveri, Namami Gaur, Sakshi Shambhavi, Nikhil Hajirnis, M Soujanya Reddy, Pratheusa Maccha,Tulasi Nagabandi, Purushotham Vodnala, Payel Mukherjee, Sofia Banu, Priya Singh, Onkar Kulkarni, Dhiviya Vedagiri, Divya Gupta, Vishal Sah, Santosh Kumar Kuncha, Krishnan Harinivas Harshan, Archana Bharadwaj Siva, Karthik Bharadwaj Tallapaka,Umesh Kumar, Unis Ahmad Bhat, Ajay Sarawagi, Priyanka Pant, Rajkanwar Nathawat, Rakesh K Mishra, Divya Tej Sowpati |
| EPI_ISL_528819, EPI_ISL_528820 | Department of Medicine, Gandhi hospital, Hyderabad | CSIR-Centre for Cellular and Molecular Biology | Rajarao Mesipogu , Thrilok Chander Bingi ,Vinayasekhar Aedula,Tulasi Nagabandi, Namami Gaur, Sakshi Shambhavi, Lamuk Zaveri, Shagufta Khan, Nikhil Hajirnis, M Soujanya Reddy, Pratheusa Maccha, Purushotham Vodnala, Payel Mukherjee, Sofia Banu, Priya Singh, Onkar Kulkarni, Dhiviya Vedagiri, Divya Gupta, Vishal Sah, Santosh Kumar Kuncha, Krishnan Harinivas Harshan, Archana Bharadwaj Siva, Karthik Bharadwaj Tallapaka,G. Aditya Kumar, Koushick Sivakumar, Pooja Ramesh Gupta, Rajan Kumar Jha, Shraddha Vijay Lahoti, Rakesh K Mishra, Divya Tej Sowpati |
| EPI_ISL_528821 | Department of Medicine, Gandhi hospital, Hyderabad | CSIR-Centre for Cellular and Molecular Biology | Thrilok Chander Bingi,Rajarao Mesipogu ,Vinayasekhar Aedula,Lamuk Zaveri, Shagufta Khan, Namami Gaur, Sakshi Shambhavi, Nikhil Hajirnis, M Soujanya Reddy, Pratheusa Maccha, Tulasi Nagabandi, Purushotham Vodnala, Payel Mukherjee, Sofia Banu, Priya Singh, Onkar Kulkarni, Dhiviya Vedagiri, Divya Gupta, Vishal Sah, Santosh Kumar Kuncha, Krishnan Harinivas Harshan, Archana Bharadwaj Siva, Karthik Bharadwaj Tallapaka, Renu Sudhakar, Somesh Gorde, Gangumala Srinivas Reddy, Sujoy Deb, Swati Bayyana, Rakesh K Mishra, Divya Tej Sowpati |
| EPI_ISL_528822 | Department of Medicine, Gandhi hospital, Hyderabad | CSIR-Centre for Cellular and Molecular Biology | Vinayasekhar Aedula,Thrilok Chander Bingi, Rajarao Mesipogu, Shagufta Khan, Lamuk Zaveri, Namami Gaur, Sakshi Shambhavi, Nikhil Hajirnis, M Soujanya Reddy, Pratheusa Maccha,Tulasi Nagabandi, Purushotham Vodnala, Payel Mukherjee, Sofia Banu, Priya Singh, Onkar Kulkarni, Dhiviya Vedagiri, Divya Gupta, Vishal Sah, Santosh Kumar Kuncha, Krishnan Harinivas Harshan, Archana Bharadwaj Siva, Karthik Bharadwaj Tallapaka,Umesh Kumar, Unis Ahmad Bhat, Ajay Sarawagi, Priyanka Pant, Rajkanwar Nathawat, Rakesh K Mishra, Divya Tej Sowpati |
| EPI_ISL_528823 | CSIR-Centre for Cellular and Molecular Biology | CSIR-Centre for Cellular and Molecular Biology | Shagufta Khan, Lamuk Zaveri, Namami Gaur, Sakshi Shambhavi, Nikhil Hajirnis, M Soujanya Reddy, Pratheusa Maccha, Tulasi Nagabandi, Purushotham Vodnala, Payel Mukherjee, Sofia Banu, Priya Singh, Onkar Kulkarni, Dhiviya Vedagiri, Divya Gupta, Vishal Sah, Santosh Kumar Kuncha, Krishnan Harinivas Harshan, Archana Bharadwaj Siva, Karthik Bharadwaj Tallapaka, Renu Sudhakar, Somesh Gorde, Gangumala Srinivas Reddy, Sujoy Deb, Swati Bayyana, Rakesh K Mishra, Divya Tej Sowpati |
| EPI_ISL_528824 | CSIR-Centre for Cellular and Molecular Biology | CSIR-Centre for Cellular and Molecular Biology | Tulasi Nagabandi, Namami Gaur, Sakshi Shambhavi, Lamuk Zaveri, Shagufta Khan, Nikhil Hajirnis, M Soujanya Reddy, Pratheusa Maccha, Purushotham Vodnala, Payel Mukherjee, Sofia Banu, Priya Singh, Onkar Kulkarni, Dhiviya Vedagiri, Divya Gupta, Vishal Sah, Santosh Kumar Kuncha, Krishnan Harinivas Harshan, Archana Bharadwaj Siva, Karthik Bharadwaj Tallapaka,G. Aditya Kumar, Koushick Sivakumar, Pooja Ramesh Gupta, Rajan Kumar Jha, Shraddha Vijay Lahoti, Rakesh K Mishra, Divya Tej Sowpati |
| EPI_ISL_528825, EPI_ISL_528826 | CSIR-Centre for Cellular and Molecular Biology | CSIR-Centre for Cellular and Molecular Biology | Shagufta Khan, Lamuk Zaveri, Namami Gaur, Sakshi Shambhavi, Nikhil Hajirnis, M Soujanya Reddy, Pratheusa Maccha, Tulasi Nagabandi, Purushotham Vodnala, Payel Mukherjee, Sofia Banu, Priya Singh, Onkar |

| | | | |
|---|---|---|---|
| | | | Kulkarni, Dhiviya Vedagiri, Divya Gupta, Vishal Sah, Santosh Kumar Kuncha, Krishnan Harinivas Harshan, Archana Bharadwaj Siva, Karthik Bharadwaj Tallapaka, Renu Sudhakar, Somesh Gorde, Gangumala Srinivas Reddy, Sujoy Deb, Swati Bayyana, Rakesh K Mishra, Divya Tej Sowpati |
| EPI_ISL_528827 | CSIR-Centre for Cellular and Molecular Biology | CSIR-Centre for Cellular and Molecular Biology | Namami Gaur, Sakshi Shambhavi, Lamuk Zaveri, Shagufta Khan, Nikhil Hajirnis, M Soujanya Reddy, Pratheusa Maccha, Tulasi Nagabandi, Purushotham Vodnala, Payel Mukherjee, Sofia Banu, Priya Singh, Onkar Kulkarni, Dhiviya Vedagiri, Divya Gupta, Vishal Sah, Santosh Kumar Kuncha, Krishnan Harinivas Harshan, Archana Bharadwaj Siva, Karthik Bharadwaj Tallapaka, Zeba Rizvi, Zuberwasim Sayyad, Kakade Aishwarya Arun, Amrutha H C, Ananga Ghosh, Rakesh K Mishra, Divya Tej Sowpati |
| EPI_ISL_528828 | CSIR-Centre for Cellular and Molecular Biology | CSIR-Centre for Cellular and Molecular Biology | Sofia Banu, Payel Mukherjee, Priya Singh,Onkar Kulkarni, Dhiviya Vedagiri, Divya Gupta, Vishal Sah, Santosh Kumar Kuncha, Krishnan Harinivas Harshan, Archana Bharadwaj Siva, Karthik Bharadwaj Tallapaka, Shagufta Khan, Lamuk Zaveri, Namami Gaur, Sakshi Shambhavi, Nikhil Hajirnis, M Soujanya Reddy, Pratheusa Maccha,Tulasi Nagabandi, Purushotham Vodnala, Deepak Kumar, Devi Prasad Vijayashankar, Disha Nanda, Divya Das, Jotin Gogoi, Manish Bhattacharjee, Rakesh K Mishra, Divya Tej Sowpati |
| EPI_ISL_528829 | CSIR-Centre for Cellular and Molecular Biology | CSIR-Centre for Cellular and Molecular Biology | Sakshi Shambhavi, Lamuk Zaveri, Shagufta Khan, Namami Gaur, Nikhil Hajirnis, M Soujanya Reddy, Pratheusa Maccha, Tulasi Nagabandi, Purushotham Vodnala, Payel Mukherjee, Sofia Banu, Priya Singh, Onkar Kulkarni, Dhiviya Vedagiri, Divya Gupta, Vishal Sah, Santosh Kumar Kuncha, Krishnan Harinivas Harshan, Archana Bharadwaj Siva, Karthik Bharadwaj Tallapaka, Deepak Kumar, Devi Prasad Vijayashankar, Disha Nanda, Divya Das, Jotin Gogoi, Manish Bhattacharjee, Rakesh K Mishra, Divya Tej Sowpati |
| EPI_ISL_528830 | CSIR-Centre for Cellular and Molecular Biology | CSIR-Centre for Cellular and Molecular Biology | Tulasi Nagabandi, Namami Gaur, Sakshi Shambhavi, Lamuk Zaveri, Shagufta Khan, Nikhil Hajirnis, M Soujanya Reddy, Pratheusa Maccha, Purushotham Vodnala, Payel Mukherjee, Sofia Banu, Priya Singh, Onkar Kulkarni, Dhiviya Vedagiri, Divya Gupta, Vishal Sah, Santosh Kumar Kuncha, Krishnan Harinivas Harshan, Archana Bharadwaj Siva, Karthik Bharadwaj Tallapaka,G. Aditya Kumar, Koushick Sivakumar, Pooja Ramesh Gupta, Rajan Kumar Jha, Shraddha Vijay Lahoti, Rakesh K Mishra, Divya Tej Sowpati |
| EPI_ISL_528831 | CSIR-Centre for Cellular and Molecular Biology | CSIR-Centre for Cellular and Molecular Biology | Lamuk Zaveri, Shagufta Khan, Namami Gaur, Sakshi Shambhavi, Nikhil Hajirnis, M Soujanya Reddy, Pratheusa Maccha, Tulasi Nagabandi, Purushotham Vodnala, Payel Mukherjee, Sofia Banu, Priya Singh, Onkar Kulkarni, Dhiviya Vedagiri, Divya Gupta, Vishal Sah, Santosh Kumar Kuncha, Krishnan Harinivas Harshan, Archana Bharadwaj Siva, Karthik Bharadwaj Tallapaka, Renu Sudhakar, Somesh Gorde, Gangumala Srinivas Reddy, Sujoy Deb, Swati Bayyana, Rakesh K Mishra, Divya Tej Sowpati |
| EPI_ISL_528832 | CSIR-Centre for Cellular and Molecular Biology | CSIR-Centre for Cellular and Molecular Biology | Sofia Banu, Payel Mukherjee, Priya Singh,Onkar Kulkarni, Dhiviya Vedagiri, Divya Gupta, Vishal Sah, Santosh Kumar Kuncha, Krishnan Harinivas Harshan, Archana Bharadwaj Siva, Karthik Bharadwaj Tallapaka, Shagufta Khan, Lamuk Zaveri, Namami Gaur, Sakshi Shambhavi, Nikhil Hajirnis, M Soujanya Reddy, Pratheusa Maccha,Tulasi Nagabandi, Purushotham Vodnala, Deepak Kumar, Devi Prasad Vijayashankar, Disha Nanda, Divya Das, Jotin Gogoi, Manish Bhattacharjee, Rakesh K Mishra, Divya Tej Sowpati |
| EPI_ISL_528833 | CSIR-Centre for Cellular and Molecular Biology | CSIR-Centre for Cellular and Molecular Biology | Lamuk Zaveri, Shagufta Khan, Namami Gaur, Sakshi Shambhavi, Nikhil Hajirnis, M Soujanya Reddy, Pratheusa Maccha, Tulasi Nagabandi, Purushotham Vodnala, Payel Mukherjee, Sofia Banu, Priya Singh, Onkar Kulkarni, Dhiviya Vedagiri, Divya Gupta, Vishal Sah, Santosh Kumar Kuncha, Krishnan Harinivas Harshan, Archana Bharadwaj Siva, Karthik Bharadwaj Tallapaka, Renu Sudhakar, Somesh Gorde, Gangumala Srinivas Reddy, Sujoy Deb, Swati Bayyana, Rakesh K Mishra, Divya Tej Sowpati |
| EPI_ISL_528834 | CSIR-Centre for Cellular and Molecular Biology | CSIR-Centre for Cellular and Molecular Biology | Pratheusa Maccha, Sakshi Shambhavi, Lamuk Zaveri, Shagufta Khan, Namami Gaur, Nikhil Hajirnis, M Soujanya Reddy, Tulasi Nagabandi, Purushotham Vodnala, Payel Mukherjee, Sofia Banu, Priya Singh,Onkar Kulkarni, Dhiviya Vedagiri, Divya Gupta, Vishal Sah, Santosh Kumar Kuncha, Krishnan Harinivas Harshan, Archana Bharadwaj Siva, Karthik Bharadwaj Tallapaka, G. Aditya Kumar, Koushick Sivakumar,Disha Nanda, Divya Das, Jotin Gogoi, Manish Bhattacharjee, Ravi Prasad Mukku, Rakesh K Mishra, Divya Tej Sowpati |
| EPI_ISL_528835 | CSIR-Centre for Cellular and Molecular Biology | CSIR-Centre for Cellular and Molecular Biology | M Soujanya Reddy, Nikhil Hajirnis, Pratheusa Maccha, Namami Gaur, Sakshi Shambhavi, Lamuk Zaveri, Shagufta Khan, Tulasi Nagabandi, Purushotham Vodnala, Payel Mukherjee, Sofia Banu, Priya Singh, Onkar Kulkarni, Dhiviya Vedagiri, Divya Gupta, Vishal Sah, Santosh Kumar Kuncha, Krishnan Harinivas Harshan, Archana Bharadwaj Siva, Karthik Bharadwaj Tallapaka, Zeba Rizvi, Zuberwasim Sayyad, Kakade Aishwarya Arun, Amrutha H C, Ananga Ghosh, Rakesh K Mishra, Divya Tej Sowpati |
| EPI_ISL_528836 | CSIR-Centre for Cellular and Molecular Biology | CSIR-Centre for Cellular and Molecular Biology | Tulasi Nagabandi, Namami Gaur, Sakshi Shambhavi, Lamuk Zaveri, Shagufta Khan, Nikhil Hajirnis, M Soujanya Reddy, Pratheusa Maccha, Purushotham Vodnala, Payel Mukherjee, Sofia Banu, Priya Singh, Onkar Kulkarni, Dhiviya Vedagiri, Divya Gupta, Vishal Sah, Santosh Kumar Kuncha, Krishnan Harinivas Harshan, Archana Bharadwaj Siva, Karthik Bharadwaj Tallapaka,G. Aditya Kumar, Koushick Sivakumar, Pooja Ramesh Gupta, Rajan Kumar Jha, Shraddha Vijay Lahoti, Rakesh K Mishra, Divya Tej Sowpati |
| EPI_ISL_528837 | CSIR-Centre for Cellular and Molecular Biology | CSIR-Centre for Cellular and Molecular Biology | Sakshi Shambhavi, Lamuk Zaveri, Shagufta Khan, Namami Gaur, Nikhil Hajirnis, M Soujanya Reddy, Pratheusa Maccha, Tulasi Nagabandi, Purushotham Vodnala, Payel Mukherjee, Sofia Banu, Priya Singh, Onkar Kulkarni, Dhiviya Vedagiri, Divya Gupta, Vishal Sah, Santosh Kumar Kuncha, Krishnan Harinivas Harshan, Archana Bharadwaj Siva, Karthik Bharadwaj Tallapaka, Deepak Kumar, Devi Prasad Vijayashankar, Disha Nanda, Divya Das, Jotin Gogoi, Manish Bhattacharjee, Rakesh K Mishra, Divya Tej Sowpati |
| EPI_ISL_528838 | CSIR-Centre for Cellular and Molecular Biology | CSIR-Centre for Cellular and Molecular Biology | Payel Mukherjee, Sofia Banu, Priya Singh, Onkar Kulkarni, Dhiviya Vedagiri, Divya Gupta, Vishal Sah, Santosh Kumar Kuncha, Krishnan Harinivas Harshan, Archana Bharadwaj Siva, Karthik Bharadwaj Tallapaka, Shagufta Khan, Lamuk Zaveri, Nikhil Hajirnis, M Soujanya Reddy, Pratheusa Maccha, Namami Gaur, Sakshi Shambhavi, Tulasi Nagabandi, Purushotham Vodnala, Rakesh K Mishra, Sonu Uday, Sudipta Mondal, Annapoorna P Karthyayani, Debabrata Jana, Debrya Saha, Divya Tej Sowpati |
| EPI_ISL_528839 | CSIR-Centre for Cellular and Molecular Biology | CSIR-Centre for Cellular and Molecular Biology | Pratheusa Maccha, Sakshi Shambhavi, Lamuk Zaveri, Shagufta Khan, Namami Gaur, Nikhil Hajirnis, M Soujanya Reddy, Tulasi Nagabandi, Purushotham Vodnala, Payel Mukherjee, Sofia Banu, Priya Singh,Onkar Kulkarni, Dhiviya Vedagiri, Divya Gupta, Vishal Sah, Santosh Kumar Kuncha, Krishnan Harinivas Harshan, Archana Bharadwaj Siva, Karthik Bharadwaj Tallapaka, G. Aditya Kumar, Koushick Sivakumar,Disha Nanda, Divya Das, Jotin Gogoi, Manish Bhattacharjee, Ravi Prasad Mukku, Rakesh K Mishra, Divya Tej Sowpati |
| EPI_ISL_528840 | CSIR-Centre for Cellular and Molecular Biology | CSIR-Centre for Cellular and Molecular Biology | Tulasi Nagabandi, Namami Gaur, Sakshi Shambhavi, Lamuk Zaveri, Shagufta Khan, Nikhil Hajirnis, M Soujanya Reddy, Pratheusa Maccha, Purushotham Vodnala, Payel Mukherjee, Sofia Banu, Priya Singh, Onkar Kulkarni, Dhiviya Vedagiri, Divya Gupta, Vishal Sah, Santosh Kumar Kuncha, Krishnan Harinivas Harshan, Archana Bharadwaj Siva, Karthik Bharadwaj Tallapaka,G. Aditya Kumar, Koushick Sivakumar, Pooja Ramesh Gupta, Rajan Kumar Jha, Shraddha Vijay Lahoti, Rakesh K Mishra, Divya Tej Sowpati |
| EPI_ISL_528841, EPI_ISL_528842 | CSIR-Centre for Cellular and Molecular Biology | CSIR-Centre for Cellular and Molecular Biology | Payel Mukherjee, Sofia Banu, Priya Singh, Onkar Kulkarni, Dhiviya Vedagiri, Divya Gupta, Vishal Sah, Santosh Kumar Kuncha, Krishnan Harinivas Harshan, Archana Bharadwaj Siva, Karthik Bharadwaj Tallapaka, Shagufta Khan, Lamuk Zaveri, Nikhil Hajirnis, M Soujanya Reddy, Pratheusa Maccha, Namami Gaur, Sakshi Shambhavi, Tulasi Nagabandi, Purushotham Vodnala, Rakesh K Mishra, Sonu Uday, Sudipta Mondal, Annapoorna P Karthyayani, Debabrata Jana, Debrya Saha, Divya Tej Sowpati |
| EPI_ISL_528843, EPI_ISL_528844 | CSIR-Centre for Cellular and Molecular Biology | CSIR-Centre for Cellular and Molecular Biology | Namami Gaur, Sakshi Shambhavi, Lamuk Zaveri, Shagufta Khan, Nikhil Hajirnis, M Soujanya Reddy, Pratheusa Maccha, Tulasi Nagabandi, Purushotham Vodnala, Payel Mukherjee, Sofia Banu, Priya Singh, Onkar Kulkarni, Dhiviya Vedagiri, Divya Gupta, Vishal Sah, Santosh Kumar Kuncha, Krishnan Harinivas Harshan, Archana Bharadwaj Siva, Karthik Bharadwaj Tallapaka, Zeba Rizvi, Zuberwasim Sayyad, Kakade Aishwarya Arun, Amrutha H C, Ananga Ghosh, Rakesh K Mishra, Divya Tej Sowpati |
| EPI_ISL_528845 | CSIR-Centre for Cellular and Molecular Biology | CSIR-Centre for Cellular and Molecular Biology | Payel Mukherjee, Sofia Banu, Priya Singh, Onkar Kulkarni, Dhiviya Vedagiri, Divya Gupta, Vishal Sah, Santosh Kumar Kuncha, Krishnan Harinivas Harshan, Archana Bharadwaj Siva, Karthik Bharadwaj Tallapaka, Shagufta Khan, Lamuk Zaveri, Nikhil Hajirnis, M Soujanya Reddy, Pratheusa Maccha, Namami Gaur, Sakshi Shambhavi, Tulasi Nagabandi, Purushotham Vodnala, Rakesh K Mishra, Sonu Uday, Sudipta Mondal, Annapoorna P Karthyayani, Debabrata Jana, Debrya Saha, Divya Tej Sowpati |
| EPI_ISL_528846 | CSIR-Centre for Cellular and Molecular Biology | CSIR-Centre for Cellular and Molecular Biology | Namami Gaur, Sakshi Shambhavi, Lamuk Zaveri, Shagufta Khan, Nikhil Hajirnis, M Soujanya Reddy, Pratheusa Maccha, Tulasi Nagabandi, Purushotham Vodnala, Payel Mukherjee, Sofia Banu, Priya Singh, Onkar Kulkarni, Dhiviya Vedagiri, Divya Gupta, Vishal Sah, Santosh Kumar Kuncha, Krishnan Harinivas Harshan, Archana Bharadwaj Siva, Karthik Bharadwaj Tallapaka, Zeba Rizvi, Zuberwasim Sayyad, Kakade Aishwarya Arun, Amrutha H C, Ananga Ghosh, Rakesh K Mishra, Divya Tej Sowpati |
| EPI_ISL_528847 | CSIR-Centre for Cellular and Molecular Biology | CSIR-Centre for Cellular and Molecular Biology | Shagufta Khan, Lamuk Zaveri, Namami Gaur, Sakshi Shambhavi, Nikhil Hajirnis, M Soujanya Reddy, Pratheusa Maccha, Tulasi Nagabandi, Purushotham Vodnala, Payel Mukherjee, Sofia Banu, Priya Singh, Onkar Kulkarni, Dhiviya Vedagiri, Divya Gupta, Vishal Sah, Santosh Kumar Kuncha, Krishnan Harinivas Harshan, Archana Bharadwaj Siva, Karthik Bharadwaj Tallapaka, Renu Sudhakar, Somesh Gorde, Gangumala Srinivas Reddy, Sujoy Deb, Swati Bayyana, Rakesh K Mishra, Divya Tej Sowpati |
| EPI_ISL_528848 | CSIR-Centre for Cellular and Molecular Biology | CSIR-Centre for Cellular and Molecular Biology | Sofia Banu, Payel Mukherjee, Priya Singh,Onkar Kulkarni, Dhiviya Vedagiri, Divya Gupta, Vishal Sah, Santosh Kumar Kuncha, Krishnan Harinivas Harshan, Archana Bharadwaj Siva, Karthik Bharadwaj Tallapaka, Shagufta Khan, Lamuk Zaveri, Namami Gaur, Sakshi Shambhavi, Nikhil Hajirnis, M Soujanya Reddy, Pratheusa Maccha,Tulasi Nagabandi, Purushotham Vodnala, Deepak Kumar, Devi Prasad Vijayashankar, Disha Nanda, Divya Das, Jotin Gogoi, Manish Bhattacharjee, Rakesh K Mishra, Divya Tej Sowpati |
| EPI_ISL_528849 | CSIR-Centre for Cellular and Molecular Biology | CSIR-Centre for Cellular and Molecular Biology | Nikhil Hajirnis, M Soujanya Reddy, Pratheusa Maccha, Lamuk Zaveri, Shagufta Khan, Namami Gaur, Sakshi Shambhavi, Tulasi Nagabandi, Purushotham Vodnala, Payel Mukherjee, Sofia Banu, Priya Singh, Onkar Kulkarni, Dhiviya Vedagiri, Divya Gupta, Vishal Sah, Santosh Kumar Kuncha, Krishnan Harinivas Harshan, Archana Bharadwaj Siva, Karthik Bharadwaj Tallapaka,Zeba Rizvi, Zuberwasim Sayyad, Kakade Aishwarya Arun, Amrutha H C, Ananga Ghosh, Rakesh K Mishra, Divya Tej Sowpati |
| EPI_ISL_528850 | CSIR-Centre for Cellular and Molecular Biology | CSIR-Centre for Cellular and Molecular Biology | Lamuk Zaveri, Shagufta Khan, Namami Gaur, Sakshi Shambhavi, Nikhil Hajirnis, M Soujanya Reddy, Pratheusa Maccha, Tulasi Nagabandi, Purushotham Vodnala, Payel Mukherjee, Sofia Banu, Priya Singh, Onkar Kulkarni, Dhiviya Vedagiri, Divya Gupta, Vishal Sah, Santosh Kumar Kuncha, Krishnan Harinivas Harshan, Archana Bharadwaj Siva, Karthik Bharadwaj Tallapaka, Renu Sudhakar, Somesh Gorde, Gangumala Srinivas Reddy, Sujoy Deb, Swati Bayyana, Rakesh K Mishra, Divya Tej Sowpati |
| EPI_ISL_528851 | CSIR-Centre for Cellular and Molecular Biology | CSIR-Centre for Cellular and Molecular Biology | Payel Mukherjee, Sofia Banu, Priya Singh, Onkar Kulkarni, Dhiviya Vedagiri, Divya Gupta, Vishal Sah, Santosh Kumar Kuncha, Krishnan Harinivas Harshan, Archana Bharadwaj Siva, Karthik Bharadwaj Tallapaka, Shagufta Khan, Lamuk Zaveri, Nikhil Hajirnis, M Soujanya Reddy, Pratheusa Maccha, Namami Gaur, Sakshi Shambhavi, Tulasi Nagabandi, Purushotham Vodnala, Rakesh K Mishra, Sonu Uday, Sudipta Mondal, Annapoorna P Karthyayani, Debabrata Jana, Debrya Saha, Divya Tej Sowpati |
| EPI_ISL_528852 | CSIR-Centre for Cellular and Molecular Biology | CSIR-Centre for Cellular and Molecular Biology | Namami Gaur, Sakshi Shambhavi, Lamuk Zaveri, Shagufta Khan, Nikhil Hajirnis, M Soujanya Reddy, Pratheusa Maccha, Tulasi Nagabandi, Purushotham Vodnala, Payel Mukherjee, Sofia Banu, Priya Singh, Onkar Kulkarni, Dhiviya Vedagiri, Divya Gupta, Vishal Sah, Santosh Kumar Kuncha, Krishnan Harinivas Harshan, Archana Bharadwaj Siva, Karthik Bharadwaj Tallapaka, Zeba Rizvi, Zuberwasim Sayyad, Kakade Aishwarya Arun, Amrutha H C, Ananga Ghosh, Rakesh K Mishra, Divya Tej Sowpati |
| EPI_ISL_528853 | CSIR-Centre for Cellular and Molecular Biology | CSIR-Centre for Cellular and Molecular Biology | Tulasi Nagabandi, Namami Gaur, Sakshi Shambhavi, Lamuk Zaveri, Shagufta Khan, Nikhil Hajirnis, M Soujanya Reddy, Pratheusa Maccha, Purushotham Vodnala, Payel Mukherjee, Sofia Banu, Priya Singh, Onkar Kulkarni, Dhiviya Vedagiri, Divya Gupta, Vishal Sah, Santosh Kumar Kuncha, Krishnan Harinivas Harshan, Archana Bharadwaj Siva, Karthik Bharadwaj Tallapaka,G. Aditya Kumar, Koushick Sivakumar, Pooja Ramesh Gupta, Rajan Kumar Jha, Shraddha Vijay Lahoti, Rakesh K Mishra, Divya Tej Sowpati |
| EPI_ISL_528854 | CSIR-Centre for Cellular and Molecular Biology | CSIR-Centre for Cellular and Molecular Biology | Payel Mukherjee, Sofia Banu, Priya Singh, Onkar Kulkarni, Dhiviya Vedagiri, Divya Gupta, Vishal Sah, Santosh Kumar Kuncha, Krishnan Harinivas Harshan, Archana Bharadwaj Siva, Karthik Bharadwaj Tallapaka, Shagufta Khan, Lamuk Zaveri, Nikhil Hajirnis, M Soujanya Reddy, Pratheusa Maccha, Namami Gaur, Sakshi Shambhavi, Tulasi Nagabandi, Purushotham Vodnala, Rakesh K Mishra, Sonu Uday, Sudipta Mondal, Annapoorna P Karthyayani, Debabrata Jana, Debrya Saha, Divya Tej Sowpati |
| EPI_ISL_528855 | CSIR-Centre for Cellular and Molecular Biology | CSIR-Centre for Cellular and Molecular Biology | Nikhil Hajirnis, M Soujanya Reddy, Pratheusa Maccha, Lamuk Zaveri, Shagufta Khan, Namami Gaur, Sakshi Shambhavi, Tulasi Nagabandi, Purushotham Vodnala, Payel Mukherjee, Sofia Banu, Priya Singh, Onkar Kulkarni, Dhiviya Vedagiri, Divya Gupta, Vishal Sah, Santosh Kumar Kuncha, Krishnan Harinivas Harshan, Archana Bharadwaj Siva, Karthik Bharadwaj Tallapaka,Zeba Rizvi, Zuberwasim Sayyad, Kakade Aishwarya Arun, Amrutha H C, Ananga Ghosh, Rakesh K Mishra, Divya Tej Sowpati |
| EPI_ISL_528856 | CSIR-Centre for Cellular and Molecular Biology | CSIR-Centre for Cellular and Molecular Biology | Pratheusa Maccha, Sakshi Shambhavi, Lamuk Zaveri, Shagufta Khan, Namami Gaur, Nikhil Hajirnis, M Soujanya Reddy, Tulasi Nagabandi, Purushotham Vodnala, Payel Mukherjee, Sofia Banu, Priya Singh,Onkar Kulkarni, Dhiviya Vedagiri, Divya Gupta, Vishal Sah, Santosh Kumar Kuncha, Krishnan Harinivas Harshan, Archana Bharadwaj Siva, Karthik Bharadwaj Tallapaka, G. Aditya Kumar, Koushick Sivakumar,Disha Nanda, Divya Das, Jotin Gogoi, Manish Bhattacharjee, Ravi Prasad Mukku, Rakesh K Mishra, Divya Tej Sowpati |
| EPI_ISL_528857 | CSIR-Centre for Cellular and Molecular Biology | CSIR-Centre for Cellular and Molecular Biology | Nikhil Hajirnis, M Soujanya Reddy, Pratheusa Maccha, Lamuk Zaveri, Shagufta Khan, Namami Gaur, Sakshi Shambhavi, Tulasi Nagabandi, Purushotham Vodnala, Payel Mukherjee, Sofia Banu, Priya Singh, Onkar Kulkarni, Dhiviya Vedagiri, Divya Gupta, Vishal Sah, Santosh Kumar Kuncha, Krishnan Harinivas Harshan, Archana Bharadwaj Siva, Karthik Bharadwaj Tallapaka,Zeba Rizvi, Zuberwasim Sayyad, Kakade Aishwarya Arun, Amrutha H C, Ananga Ghosh, Rakesh K Mishra, Divya Tej Sowpati |
| EPI_ISL_528858, EPI_ISL_528859 | CSIR-Centre for Cellular and Molecular Biology | CSIR-Centre for Cellular and Molecular Biology | Pratheusa Maccha, Sakshi Shambhavi, Lamuk Zaveri, Shagufta Khan, Namami Gaur, Nikhil Hajirnis, M Soujanya Reddy, Tulasi Nagabandi, Purushotham Vodnala, Payel Mukherjee, Sofia Banu, Priya Singh,Onkar Kulkarni, Dhiviya Vedagiri, Divya Gupta, Vishal Sah, Santosh Kumar Kuncha, Krishnan Harinivas Harshan, Archana Bharadwaj Siva, Karthik Bharadwaj Tallapaka, G. Aditya Kumar, Koushick Sivakumar,Disha Nanda, Divya Das, Jotin Gogoi, Manish Bhattacharjee, Ravi Prasad Mukku, Rakesh K Mishra, Divya Tej Sowpati |
| EPI_ISL_528860 | CSIR-Centre for Cellular and Molecular Biology | CSIR-Centre for Cellular and Molecular Biology | Tulasi Nagabandi, Namami Gaur, Sakshi Shambhavi, Lamuk Zaveri, Shagufta Khan, Nikhil Hajirnis, M Soujanya Reddy, Pratheusa Maccha, Purushotham Vodnala, Payel Mukherjee, Sofia Banu, Priya Singh, Onkar Kulkarni, Dhiviya Vedagiri, Divya Gupta, Vishal Sah, Santosh Kumar Kuncha, Krishnan Harinivas Harshan, Archana Bharadwaj Siva, Karthik Bharadwaj Tallapaka,G. Aditya Kumar, Koushick Sivakumar, Pooja Ramesh Gupta, Rajan Kumar Jha, Shraddha Vijay Lahoti, Rakesh K Mishra, Divya Tej Sowpati |

| | | | |
|---|---|---|---|
| EPI_ISL_528861 | CSIR-Centre for Cellular and Molecular Biology | CSIR-Centre for Cellular and Molecular Biology | Sakshi Shambhavi, Lamuk Zaveri, Shagufta Khan, Namami Gaur, Nikhil Hajirnis, M Soujanya Reddy, Pratheusa Maccha, Tulasi Nagabandi, Purushotham Vodnala, Payel Mukherjee, Sofia Banu, Priya Singh, Onkar Kulkarni, Dhiviya Vedagiri, Divya Gupta, Vishal Sah, Santosh Kumar Kuncha, Krishnan Harinivas Harshan, Archana Bharadwaj Siva, Karthik Bharadwaj Tallapaka, Deepak Kumar, Devi Prasad Vijayashankar, Disha Nanda, Divya Das, Jotin Gogoi, Manish Bhattacharjee, Rakesh K Mishra, Divya Tej Sowpati |
| EPI_ISL_528862 | CSIR-Centre for Cellular and Molecular Biology | CSIR-Centre for Cellular and Molecular Biology | Lamuk Zaveri, Shagufta Khan, Namami Gaur, Sakshi Shambhavi, Nikhil Hajirnis, M Soujanya Reddy, Pratheusa Maccha, Tulasi Nagabandi, Purushotham Vodnala, Payel Mukherjee, Sofia Banu, Priya Singh, Onkar Kulkarni, Dhiviya Vedagiri, Divya Gupta, Vishal Sah, Santosh Kumar Kuncha, Krishnan Harinivas Harshan, Archana Bharadwaj Siva, Karthik Bharadwaj Tallapaka, Renu Sudhakar, Somesh Gorde, Gangumala Srinivas Reddy, Sujoy Deb, Swati Bayyana, Rakesh K Mishra, Divya Tej Sowpati |
| EPI_ISL_528863 | CSIR-Centre for Cellular and Molecular Biology | CSIR-Centre for Cellular and Molecular Biology | Payel Mukherjee, Sofia Banu, Priya Singh, Onkar Kulkarni, Dhiviya Vedagiri, Divya Gupta, Vishal Sah, Santosh Kumar Kuncha, Krishnan Harinivas Harshan, Archana Bharadwaj Siva, Karthik Bharadwaj Tallapaka, Shagufta Khan, Lamuk Zaveri, Nikhil Hajirnis, M Soujanya Reddy, Pratheusa Maccha, Namami Gaur, Sakshi Shambhavi, Tulasi Nagabandi, Purushotham Vodnala, Rakesh K Mishra, Sonu Uday, Sudipta Mondal, Annapoorna P Karthyayani, Debabrata Jana, Debrya Saha, Divya Tej Sowpati |
| EPI_ISL_528864 | CSIR-Centre for Cellular and Molecular Biology | CSIR-Centre for Cellular and Molecular Biology | Sofia Banu, Payel Mukherjee, Priya Singh,Onkar Kulkarni, Dhiviya Vedagiri, Divya Gupta, Vishal Sah, Santosh Kumar Kuncha, Krishnan Harinivas Harshan, Archana Bharadwaj Siva, Karthik Bharadwaj Tallapaka, Shagufta Khan, Lamuk Zaveri, Namami Gaur, Sakshi Shambhavi, Nikhil Hajirnis, M Soujanya Reddy, Pratheusa Maccha,Tulasi Nagabandi, Purushotham Vodnala, Deepak Kumar, Devi Prasad Vijayashankar, Disha Nanda, Divya Das, Jotin Gogoi, Manish Bhattacharjee, Rakesh K Mishra, Divya Tej Sowpati |
| EPI_ISL_528865 | CSIR-Centre for Cellular and Molecular Biology | CSIR-Centre for Cellular and Molecular Biology | Shagufta Khan, Lamuk Zaveri, Namami Gaur, Sakshi Shambhavi, Nikhil Hajirnis, M Soujanya Reddy, Pratheusa Maccha, Tulasi Nagabandi, Purushotham Vodnala, Payel Mukherjee, Sofia Banu, Priya Singh, Onkar Kulkarni, Dhiviya Vedagiri, Divya Gupta, Vishal Sah, Santosh Kumar Kuncha, Krishnan Harinivas Harshan, Archana Bharadwaj Siva, Karthik Bharadwaj Tallapaka, Renu Sudhakar, Somesh Gorde, Gangumala Srinivas Reddy, Sujoy Deb, Swati Bayyana, Rakesh K Mishra, Divya Tej Sowpati |
| EPI_ISL_528866 | CSIR-Centre for Cellular and Molecular Biology | CSIR-Centre for Cellular and Molecular Biology | Sakshi Shambhavi, Lamuk Zaveri, Shagufta Khan, Namami Gaur, Nikhil Hajirnis, M Soujanya Reddy, Pratheusa Maccha, Tulasi Nagabandi, Purushotham Vodnala, Payel Mukherjee, Sofia Banu, Priya Singh, Onkar Kulkarni, Dhiviya Vedagiri, Divya Gupta, Vishal Sah, Santosh Kumar Kuncha, Krishnan Harinivas Harshan, Archana Bharadwaj Siva, Karthik Bharadwaj Tallapaka, Deepak Kumar, Devi Prasad Vijayashankar, Disha Nanda, Divya Das, Jotin Gogoi, Manish Bhattacharjee, Rakesh K Mishra, Divya Tej Sowpati |
| EPI_ISL_528867 | CSIR-Centre for Cellular and Molecular Biology | CSIR-Centre for Cellular and Molecular Biology | M Soujanya Reddy, Nikhil Hajirnis, Pratheusa Maccha, Namami Gaur, Sakshi Shambhavi, Lamuk Zaveri, Shagufta Khan, Tulasi Nagabandi, Purushotham Vodnala, Payel Mukherjee, Sofia Banu, Priya Singh, Onkar Kulkarni, Dhiviya Vedagiri, Divya Gupta, Vishal Sah, Santosh Kumar Kuncha, Krishnan Harinivas Harshan, Archana Bharadwaj Siva, Karthik Bharadwaj Tallapaka, Zeba Rizvi, Zuberwasim Sayyad, Kakade Aishwarya Arun, Amrutha H C, Ananga Ghosh, Rakesh K Mishra, Divya Tej Sowpati |
| EPI_ISL_528868, EPI_ISL_528869 | CSIR-Centre for Cellular and Molecular Biology | CSIR-Centre for Cellular and Molecular Biology | Shagufta Khan, Lamuk Zaveri, Namami Gaur, Sakshi Shambhavi, Nikhil Hajirnis, M Soujanya Reddy, Pratheusa Maccha, Tulasi Nagabandi, Purushotham Vodnala, Payel Mukherjee, Sofia Banu, Priya Singh, Onkar Kulkarni, Dhiviya Vedagiri, Divya Gupta, Vishal Sah, Santosh Kumar Kuncha, Krishnan Harinivas Harshan, Archana Bharadwaj Siva, Karthik Bharadwaj Tallapaka, Renu Sudhakar, Somesh Gorde, Gangumala Srinivas Reddy, Sujoy Deb, Swati Bayyana, Rakesh K Mishra, Divya Tej Sowpati |
| EPI_ISL_528880 | National Genomics Core-Center for DNA Fingerprinting and Diagnostics | National Genomics Core- Center for DNA Fingerprinting and Diagnostics (NGC-CDFD)- DBT's PAN-INDIA-1000 Genome consortium | G Shashikanth, Heena Shah, Bala Pratyusha, Vinay Donipadi, Bathula Siddardha, Vineesha Oddi, Lavanya Banda, Surya Chodisetty, Abhijeeth Singh Thakur, Mohammad Mudassir, Nalini Raghunathan, Rajeshree Sanyal, R Harinarayanan, Rashna Bhandari, Murali Dharan Bashyam, Debashish Mitra, Divya Vashisht, Ashwin Dalal |
| EPI_ISL_528919 | Ospedale Civile S. Liberatore-Atri | Istituto Zooprofilattico Sperimentale dell'Abruzzo e Molise "G.Caporale" | Lorusso A, Marcacci M, Di Domenico M, Curini V, Ancora M, Cammà C, Rinaldi A, Mangone I, Di Pasquale A, Puglia I, Savini G. |
| EPI_ISL_528920, EPI_ISL_528921 | Presidio Ospedaliero "Santo Spirito"-Pescara | Istituto Zooprofilattico Sperimentale dell'Abruzzo e Molise "G.Caporale" | Lorusso A, Marcacci M, Di Domenico M, Curini V, Ancora M, Cammà C, Rinaldi A, Mangone I, Di Pasquale A, Puglia I, Savini G. |
| EPI_ISL_528922 | Ospedale "Giuseppe Mazzini"-Teramo | Istituto Zooprofilattico Sperimentale dell'Abruzzo e Molise "G.Caporale" | Lorusso A, Marcacci M, Di Domenico M, Curini V, Ancora M, Cammà C, Rinaldi A, Mangone I, Di Pasquale A, Puglia I, Savini G. |
| EPI_ISL_528923 | Ospedale Civile S. Liberatore-Atri | Istituto Zooprofilattico Sperimentale dell'Abruzzo e Molise "G.Caporale" | Lorusso A, Marcacci M, Di Domenico M, Curini V, Ancora M, Cammà C, Rinaldi A, Mangone I, Di Pasquale A, Puglia I, Savini G. |
| EPI_ISL_528924 | Ospedale "Giuseppe Mazzini"-Teramo | Istituto Zooprofilattico Sperimentale dell'Abruzzo e Molise "G.Caporale" | Lorusso A, Marcacci M, Di Domenico M, Curini V, Ancora M, Cammà C, Rinaldi A, Mangone I, Di Pasquale A, Puglia I, Savini G. |
| EPI_ISL_528925 | Ospedale Regionale San Salvatore-L'Aquila | Istituto Zooprofilattico Sperimentale dell'Abruzzo e Molise "G.Caporale" | Lorusso A, Marcacci M, Di Domenico M, Curini V, Ancora M, Cammà C, Rinaldi A, Mangone I, Di Pasquale A, Puglia I, Savini G. |
| EPI_ISL_528926, EPI_ISL_528927, EPI_ISL_528928 | Ospedale "Giuseppe Mazzini"-Teramo | Istituto Zooprofilattico Sperimentale dell'Abruzzo e Molise "G.Caporale" | Lorusso A, Marcacci M, Di Domenico M, Curini V, Ancora M, Cammà C, Rinaldi A, Mangone I, Di Pasquale A, Puglia I, Savini G. |
| EPI_ISL_528929 | Ospedale Civile S. Liberatore-Atri | Istituto Zooprofilattico Sperimentale dell'Abruzzo e Molise "G.Caporale" | Lorusso A, Marcacci M, Di Domenico M, Curini V, Ancora M, Cammà C, Rinaldi A, Mangone I, Di Pasquale A, Puglia I, Savini G. |
| EPI_ISL_528930, EPI_ISL_528931, EPI_ISL_528932, EPI_ISL_528933 | Respiratory Virus Unit, Microbiology Services Colindale, Public Health England | Respiratory Virus Unit, Microbiology Services Colindale, Public Health England | PHE Covid Sequencing Team |
| EPI_ISL_528934, EPI_ISL_528935, EPI_ISL_528936, EPI_ISL_528937, EPI_ISL_528938, EPI_ISL_528939, EPI_ISL_528940, EPI_ISL_528941, EPI_ISL_528942, EPI_ISL_528943, EPI_ISL_528944, EPI_ISL_528945, EPI_ISL_528946, EPI_ISL_528947, EPI_ISL_528948, EPI_ISL_528949 | | | |
| see above | Agenzia di Tutela della Salute di Bergamo | Istituto Zooprofilattico Sperimentale dell'Abruzzo e Molise "G.Caporale" | Lorusso A, Marcacci M, Di Domenico M, Curini V, Ancora M, Cammà C, Rinaldi A, Mangone I, Di Pasquale A, Puglia I, Savini G. |
| EPI_ISL_528950 | Respiratory Virus Unit, Microbiology Services Colindale, Public Health England | Respiratory Virus Unit, Microbiology Services Colindale, Public Health England | PHE Covid Sequencing Team |
| EPI_ISL_528990 | Ospedale Civile Maria SS. dello Splendore | Istituto Zooprofilattico Sperimentale dell'Abruzzo e Molise "G.Caporale" | Lorusso A, Marcacci M, Di Domenico M, Curini V, Ancora M, Cammà C, Rinaldi A, Mangone I, Di Pasquale A, Puglia I, Savini G. |
| EPI_ISL_528991, EPI_ISL_528992 | Ospedale SS Annunziata-Sulmona | Istituto Zooprofilattico Sperimentale dell'Abruzzo e Molise "G.Caporale" | Lorusso A, Marcacci M, Di Domenico M, Curini V, Ancora M, Cammà C, Rinaldi A, Mangone I, Di Pasquale A, Puglia I, Savini G. |
| EPI_ISL_528993 | Ospedale Civile S. Liberatore-Atri | Istituto Zooprofilattico Sperimentale dell'Abruzzo e Molise "G.Caporale" | Lorusso A, Marcacci M, Di Domenico M, Curini V, Ancora M, Cammà C, Rinaldi A, Mangone I, Di Pasquale A, Puglia I, Savini G. |
| EPI_ISL_528994, EPI_ISL_528995, EPI_ISL_528996, EPI_ISL_528997, EPI_ISL_528998, EPI_ISL_528999, EPI_ISL_529000, EPI_ISL_529001, EPI_ISL_529002, EPI_ISL_529003, EPI_ISL_529004, EPI_ISL_529005 | | | |
| see above | Servizio di igiene epidemiologia e sanità pubblica (SIESP)-Chieti | Istituto Zooprofilattico Sperimentale dell'Abruzzo e Molise "G.Caporale" | Lorusso A, Marcacci M, Di Domenico M, Curini V, Ancora M, Cammà C, Rinaldi A, Mangone I, Di Pasquale A, Puglia I, Savini G. |
| EPI_ISL_529006 | Servizio di igiene e sanità pubblica (SIESP)-Teramo | Istituto Zooprofilattico Sperimentale dell'Abruzzo e Molise "G.Caporale" | Lorusso A, Marcacci M, Di Domenico M, Curini V, Ancora M, Cammà C, Rinaldi A, Mangone I, Di Pasquale A, Puglia I, Savini G. |
| EPI_ISL_529007 | Ospedale Civile S. Liberatore-Atri | Istituto Zooprofilattico Sperimentale dell'Abruzzo e Molise "G.Caporale" | Lorusso A, Marcacci M, Di Domenico M, Curini V, Ancora M, Cammà C, Rinaldi A, Mangone I, Di Pasquale A, Puglia I, Savini G. |
| EPI_ISL_529008 | Servizio di igiene e sanità pubblica (SIESP)-Teramo | Istituto Zooprofilattico Sperimentale dell'Abruzzo e Molise "G.Caporale" | Lorusso A, Marcacci M, Di Domenico M, Curini V, Ancora M, Cammà C, Rinaldi A, Mangone I, Di Pasquale A, Puglia I, Savini G. |
| EPI_ISL_529009 | Ospedale Civile S. Liberatore-Atri | Istituto Zooprofilattico Sperimentale dell'Abruzzo e Molise "G.Caporale" | Lorusso A, Marcacci M, Di Domenico M, Curini V, Ancora M, Cammà C, Rinaldi A, Mangone I, Di Pasquale A, Puglia I, Savini G. |
| EPI_ISL_529010, EPI_ISL_529011, EPI_ISL_529012 | Servizio Igiene Epidemiologia e Sanità Pubblica (SIESP)-L'Aquila | Istituto Zooprofilattico Sperimentale dell'Abruzzo e Molise "G.Caporale" | Lorusso A, Marcacci M, Di Domenico M, Curini V, Ancora M, Cammà C, Rinaldi A, Mangone I, Di Pasquale A, Puglia I, Savini G. |
| EPI_ISL_529013 | Presidio Ospedaliero "S.Filippo e Nicola"-Avezzano | Istituto Zooprofilattico Sperimentale dell'Abruzzo e Molise "G.Caporale" | Lorusso A, Marcacci M, Di Domenico M, Curini V, Ancora M, Cammà C, Rinaldi A, Mangone I, Di Pasquale A, Puglia I, Savini G. |
| EPI_ISL_529014, EPI_ISL_529015 | Ospedale "Ss. Annunziata" | Istituto Zooprofilattico Sperimentale dell'Abruzzo e Molise "G.Caporale" | Lorusso A, Marcacci M, Di Domenico M, Curini V, Ancora M, Cammà C, Rinaldi A, Mangone I, Di Pasquale A, Puglia I, Savini G. |
| EPI_ISL_529016 | Ospedale SS Annunziata-Sulmona | Istituto Zooprofilattico Sperimentale dell'Abruzzo e Molise "G.Caporale" | Lorusso A, Marcacci M, Di Domenico M, Curini V, Ancora M, Cammà C, Rinaldi A, Mangone I, Di Pasquale A, Puglia I, Savini G. |
| EPI_ISL_529017 | RSA San Raffaele Sulmona | Istituto Zooprofilattico Sperimentale dell'Abruzzo e Molise "G.Caporale" | Lorusso A, Marcacci M, Di Domenico M, Curini V, Ancora M, Cammà C, Rinaldi A, Mangone I, Di Pasquale A, Puglia I, Savini G. |
| EPI_ISL_529018 | Ospedale "Giuseppe Mazzini"-Teramo | Istituto Zooprofilattico Sperimentale dell'Abruzzo e Molise "G.Caporale" | Lorusso A, Marcacci M, Di Domenico M, Curini V, Ancora M, Cammà C, Rinaldi A, Mangone I, Di Pasquale A, Puglia I, Savini G. |
| EPI_ISL_529019 | RSA/RP Villa San Giovanni - Gruppo Edos | Istituto Zooprofilattico Sperimentale dell'Abruzzo e Molise "G.Caporale" | Lorusso A, Marcacci M, Di Domenico M, Curini V, Ancora M, Cammà C, Rinaldi A, Mangone I, Di Pasquale A, Puglia I, Savini G. |
| EPI_ISL_529020, EPI_ISL_529021 | Ospedale Civile S. Liberatore-Atri | Istituto Zooprofilattico Sperimentale dell'Abruzzo e Molise "G.Caporale" | Lorusso A, Marcacci M, Di Domenico M, Curini V, Ancora M, Cammà C, Rinaldi A, Mangone I, Di Pasquale A, Puglia I, Savini G. |
| EPI_ISL_529022 | Ospedale "Ss. Annunziata" | Istituto Zooprofilattico Sperimentale dell'Abruzzo e Molise "G.Caporale" | Lorusso A, Marcacci M, Di Domenico M, Curini V, Ancora M, Cammà C, Rinaldi A, Mangone I, Di Pasquale A, Puglia I, Savini G. |
| EPI_ISL_529023, EPI_ISL_529024, EPI_ISL_529025 | Servizio di igiene epidemiologia e sanità pubblica (SIESP)-Chieti | Istituto Zooprofilattico Sperimentale dell'Abruzzo e Molise "G.Caporale" | Lorusso A, Marcacci M, Di Domenico M, Curini V, Ancora M, Cammà C, Rinaldi A, Mangone I, Di Pasquale A, Puglia I, Savini G. |
| EPI_ISL_529026 | Ospedale "Giuseppe Mazzini"-Teramo | Istituto Zooprofilattico Sperimentale dell'Abruzzo e Molise "G.Caporale" | Lorusso A, Marcacci M, Di Domenico M, Curini V, Ancora M, Cammà C, Rinaldi A, Mangone I, Di Pasquale A, Puglia I, Savini G. |
| EPI_ISL_529027, EPI_ISL_529028 | Respiratory Virus Unit, Microbiology Services Colindale, Public Health England | Respiratory Virus Unit, Microbiology Services Colindale, Public Health England | PHE Covid Sequencing Team |

| | | | |
|---|---|---|---|
| EPI_ISL_529029, EPI_ISL_529030 | Kingston Health Sciences Center | Queen's Genomics Lab at Ongwanada (Q-GLO) | Sjaarda CP, Rustom N, Huang D, Perez-Patrigeon S, Hudson ML, Wong H,Guan H, Ayub M, Soares CN, Colautti R, Evans GA, Sheth P |
| EPI_ISL_529031 | Central Molecular Microbiology Laboratory, Clinical and Chemical Pathology Department, Faculty of Medicine, CAIRO UNIVERSITY | Next Generation Sequencing Reference Laboratory, Faculty of Medicine, Cairo University and The Center for Genome and Microbiome Research, Faculty of Pharmacy, CAIRO UNIVERSITY | May Sherif Soliman, May Abdelfattah, Ramy Karam Aziz |
| EPI_ISL_529032 | Central Molecular Microbiology Laboratory and Next Generation Sequencing Reference Laboratory, Clinical and Chemical Pathology Department, Faculty of Medicine, CAIRO UNIVERSITY | Next Generation Sequencing Reference Laboratory, Faculty of Medicine, CAIRO UNIVERSITY and The Center for Genome and Microbiome Research, Faculty of Pharmacy, CAIRO UNIVERSITY | May Sherif Soliman, May Abdelfattah, Ramy Karam Aziz |
| EPI_ISL_529033, EPI_ISL_529034, EPI_ISL_529035, EPI_ISL_529036 | Wadsworth Center, New York State Department of Health | Wadsworth Center, New York State Department of Health | Kirsten St. George, Daryl M. Lamson, Sara Griesemer, Jonathan Plitnick, Navjot Singh, Matthew D. Shudt, Erica Lasek-Nesselquist |
| EPI_ISL_529065, EPI_ISL_529066, EPI_ISL_529067, EPI_ISL_529068, EPI_ISL_529069, EPI_ISL_529070, EPI_ISL_529071, EPI_ISL_529072, EPI_ISL_529073, EPI_ISL_529074, EPI_ISL_529075, EPI_ISL_529076, EPI_ISL_529077, EPI_ISL_529078, EPI_ISL_529079 | | | |
| see above | Laboratorio de Referencia Nacional de Virus Respiratorios, Instituto Nacional de Salud Peru | Laboratorio de Genómica Microbiana, Universidad Peruana Cayetano Heredia | Pablo Tsukayama, Alejandra Dávila-Barclay, Luis González, Pedro E. Romero, Brenda Ayzanoa, Janet Huancachoque, Pool Marcos, Maribel Huaringa |
| EPI_ISL_529080, EPI_ISL_529081, EPI_ISL_529082, EPI_ISL_529083, EPI_ISL_529084, EPI_ISL_529085, EPI_ISL_529086, EPI_ISL_529087, EPI_ISL_529088, EPI_ISL_529089, EPI_ISL_529090, EPI_ISL_529091, EPI_ISL_529092, EPI_ISL_529093, EPI_ISL_529094, EPI_ISL_529095, EPI_ISL_529096, EPI_ISL_529097, EPI_ISL_529098, EPI_ISL_529099, EPI_ISL_529100, EPI_ISL_529101, EPI_ISL_529102, EPI_ISL_529103, EPI_ISL_529104, EPI_ISL_529105, EPI_ISL_529106, EPI_ISL_529107, EPI_ISL_529108 | | | |
| see above | Microbiology Division, SC DHEC | Microbiology Division, SC DHEC | Flores,H. |
| EPI_ISL_529135 | Department of Virology III, National Institute of Infectious Diseases | Pathogen Genomics Center, National Institute of Infectious Diseases | Tsuyoshi Sekizuka, Shutoku Matsuyama, Kentaro Itokawa, Rina Tanaka, Masanori Hashino, Makoto Takeda, Takaji Wakita, Makoto Kuroda |
| EPI_ISL_529138 | RSAL Dr. Ramelan Surabaya | Institute of Tropical Disease, Universitas Airlangga | Jezzy R Dewantari, Rima R Prasetya, Krisnoadi Rahardjo, Aldise M Nastri, Radito Soesanto, Gatot Soegiarto, Laksmi Wulandari, Retno A Setyoningrum, Resti Yudhawati, Yohko K Shimizu, Mitsuhiro Nishimura, Yasuko Mori, Soetjipto, Kazufumi Shimizu, Maria I Lusida |
| EPI_ISL_529218, EPI_ISL_529219, EPI_ISL_529220, EPI_ISL_529221, EPI_ISL_529222, EPI_ISL_529223 | University of Birmingham | COVID-19 Genomics UK (COG-UK) Consortium | Institute of Microbiology, University of Birmingham: Claire McMurray, Joanne Stockton, Samuel Nicholls, Radoslaw Poplawski, Will Rowe, Josh Quick, Nicholas Loman. University of Birmingham Testing Laboratory: Celina M Whalley, Andrew Bosworth, Charlotte Poxon, Kasun Wanigasooriya, Oliver Pickles, Mike Kidd, Alex Richter, Andrew D Beggs PHE Heartlands Lab: Husam Osman, Andrew Bosworth. Queen Elizabeth Hospital: Anna Casey |
| EPI_ISL_529224, EPI_ISL_529225 | Department of Pathology, University of Cambridge | COVID-19 Genomics UK (COG-UK) Consortium | Luke W Meredith, M. Estée Török, Myra Hosmillo, William L. Hamilton, Martin D. Curran, Theresa Feltwell, Grant Hall, Anna Yakovleva, Fahad A Khokhar, Charlotte J. Houldcroft, Laura G Caller, Aminu S. Jahun, Sarah L. Caddy, Yasmin Chaudhry, Malte Pinckert, Ian Goodfellow |
| EPI_ISL_529226, EPI_ISL_529227, EPI_ISL_529228, EPI_ISL_529229, EPI_ISL_529230, EPI_ISL_529231, EPI_ISL_529232, EPI_ISL_529233 | Wales Specialist Virology Centre Sequencing lab: Pathogen Genomics Unit | COVID-19 Genomics UK (COG-UK) Consortium | Catherine Moore, Johnathan Evans, Laura Gifford, Malorie Perry, Simon Cottrell, Angela Marchbank, Alec Birchley, Alexander Adams, Amy Gaskin, Bree Gatica-Wilcox, Jason Coombes, Joel Southgate, Lauren Gilbert, Lee Graham, Nicole Pacchiarini, Sara Kumziene-Summerhayes, Sarah Taylor, Sophie Jones, Sara Rey, Matthew Bull, Joanne Watkins, Sally Corden, Tom Connor |
| EPI_ISL_529234, EPI_ISL_529235 | University of Birmingham | COVID-19 Genomics UK (COG-UK) Consortium | Institute of Microbiology, University of Birmingham: Claire McMurray, Joanne Stockton, Samuel Nicholls, Radoslaw Poplawski, Will Rowe, Josh Quick, Nicholas Loman. University of Birmingham Testing Laboratory: Celina M Whalley, Andrew Bosworth, Charlotte Poxon, Kasun Wanigasooriya, Oliver Pickles, Mike Kidd, Alex Richter, Andrew D Beggs PHE Heartlands Lab: Husam Osman, Andrew Bosworth. Queen Elizabeth Hospital: Anna Casey |
| EPI_ISL_529236 | Department of Pathology, University of Cambridge | COVID-19 Genomics UK (COG-UK) Consortium | Luke W Meredith, M. Estée Török, Myra Hosmillo, William L. Hamilton, Martin D. Curran, Theresa Feltwell, Grant Hall, Anna Yakovleva, Fahad A Khokhar, Charlotte J. Houldcroft, Laura G Caller, Aminu S. Jahun, Sarah L. Caddy, Yasmin Chaudhry, Malte Pinckert, Ian Goodfellow |
| EPI_ISL_529237 | University of Birmingham | COVID-19 Genomics UK (COG-UK) Consortium | Institute of Microbiology, University of Birmingham: Claire McMurray, Joanne Stockton, Samuel Nicholls, Radoslaw Poplawski, Will Rowe, Josh Quick, Nicholas Loman. University of Birmingham Testing Laboratory: Celina M Whalley, Andrew Bosworth, Charlotte Poxon, Kasun Wanigasooriya, Oliver Pickles, Mike Kidd, Alex Richter, Andrew D Beggs PHE Heartlands Lab: Husam Osman, Andrew Bosworth. Queen Elizabeth Hospital: Anna Casey |
| EPI_ISL_529238 | Quadram Institute Bioscience | COVID-19 Genomics UK (COG-UK) Consortium | Dave J. Baker, Gemma L. Kay, Alp Aydin, Thanh Le-Viet, Steven Rudder, Ana P. Tedim, Anastasia Kolyva, Maria Diaz, Leonardo de Oliveira Martins, Nabil-Fareed Alikhan, Lizzie Meadows, Rachael Stanley, Ngozi Elumogo, Muhammed Yasir, Nicholas M. Thomson, Alexander J Trotter, Rachel Gilroy, Samuel Bloomfield, Claire Stuart, Andrew Bell, Reenesh Prakash, Samir Dervisevic, Alison E. Mather, John Wain, Mark Webber, Andrew J. Page, Justin O'Grady |
| EPI_ISL_529239 | University of Birmingham | COVID-19 Genomics UK (COG-UK) Consortium | Institute of Microbiology, University of Birmingham: Claire McMurray, Joanne Stockton, Samuel Nicholls, Radoslaw Poplawski, Will Rowe, Josh Quick, Nicholas Loman. University of Birmingham Testing Laboratory: Celina M Whalley, Andrew Bosworth, Charlotte Poxon, Kasun Wanigasooriya, Oliver Pickles, Mike Kidd, Alex Richter, Andrew D Beggs PHE Heartlands Lab: Husam Osman, Andrew Bosworth. Queen Elizabeth Hospital: Anna Casey |
| EPI_ISL_529240, EPI_ISL_529241, EPI_ISL_529242, EPI_ISL_529243, EPI_ISL_529244, EPI_ISL_529245, EPI_ISL_529246, EPI_ISL_529247, EPI_ISL_529248, EPI_ISL_529249, EPI_ISL_529250 | | | |
| see above | Quadram Institute Bioscience | COVID-19 Genomics UK (COG-UK) Consortium | Dave J. Baker, Gemma L. Kay, Alp Aydin, Thanh Le-Viet, Steven Rudder, Ana P. Tedim, Anastasia Kolyva, Maria Diaz, Leonardo de Oliveira Martins, Nabil-Fareed Alikhan, Lizzie Meadows, Rachael Stanley, Ngozi Elumogo, Muhammed Yasir, Nicholas M. Thomson, Alexander J Trotter, Rachel Gilroy, Samuel Bloomfield, Claire Stuart, Andrew Bell, Reenesh Prakash, Samir Dervisevic, Alison E. Mather, John Wain, Mark Webber, Andrew J. Page, Justin O'Grady |
| EPI_ISL_529251, EPI_ISL_529252 | University of Birmingham | COVID-19 Genomics UK (COG-UK) Consortium | Institute of Microbiology, University of Birmingham: Claire McMurray, Joanne Stockton, Samuel Nicholls, Radoslaw Poplawski, Will Rowe, Josh Quick, Nicholas Loman. University of Birmingham Testing Laboratory: Celina M Whalley, Andrew Bosworth, Charlotte Poxon, Kasun Wanigasooriya, Oliver Pickles, Mike Kidd, Alex Richter, Andrew D Beggs PHE Heartlands Lab: Husam Osman, Andrew Bosworth. Queen Elizabeth Hospital: Anna Casey |
| EPI_ISL_529253 | Wales Specialist Virology Centre Sequencing lab: Pathogen Genomics Unit | COVID-19 Genomics UK (COG-UK) Consortium | Catherine Moore, Johnathan Evans, Laura Gifford, Malorie Perry, Simon Cottrell, Angela Marchbank, Alec Birchley, Alexander Adams, Amy Gaskin, Bree Gatica-Wilcox, Jason Coombes, Joel Southgate, Lauren Gilbert, Lee Graham, Nicole Pacchiarini, Sara Kumziene-Summerhayes, Sarah Taylor, Sophie Jones, Sara Rey, Matthew Bull, Joanne Watkins, Sally Corden, Tom Connor |
| EPI_ISL_529254 | Liverpool Clinical Laboratories | COVID-19 Genomics UK (COG-UK) Consortium | Sam Haldenby, Anita Lucaci, Steve Paterson, Julian Hiscox, Alistair Darby, M Almsaud, A Alrezaihi, Muhannad Alruwaili, Stuart D Armstrong, Jones Benjamin, Eleanor G Bentley, Anu Chawla, Jordan J Clark, Angela Cowell, Richard Eccles, Isabel García-Dorival, Matthew Gemmell, Alessandro Gerada, PKF Gilmore, Richard Gregory, Ximeng Han, Catherine Hartley, Margaret Hughes, Miren Iturriza-Gomara, James Johnson, L Luu, Jenifer Manson, Charlotte Nelson, Elaine O'Toole, Cassie Olateju, Rebekah Penrice-Randal , Lucille Rainbow, N.P Randle, Trevor Ian Robinson, Parul Sharma, Ghada T Shawli, James P Stewart, Neil Swainston, Ecaterina Vamos, Joanne Watts, Mark Whitehead |
| EPI_ISL_529255, EPI_ISL_529256 | University of Birmingham | COVID-19 Genomics UK (COG-UK) Consortium | Institute of Microbiology, University of Birmingham: Claire McMurray, Joanne Stockton, Samuel Nicholls, Radoslaw Poplawski, Will Rowe, Josh Quick, Nicholas Loman. University of Birmingham Testing Laboratory: Celina M Whalley, Andrew Bosworth, Charlotte Poxon, Kasun Wanigasooriya, Oliver Pickles, Mike Kidd, Alex Richter, Andrew D Beggs PHE Heartlands Lab: Husam Osman, Andrew Bosworth. Queen Elizabeth Hospital: Anna Casey |
| EPI_ISL_529257 | Department of Pathology, University of Cambridge | COVID-19 Genomics UK (COG-UK) Consortium | Luke W Meredith, M. Estée Török, Myra Hosmillo, William L. Hamilton, Martin D. Curran, Theresa Feltwell, Grant Hall, Anna Yakovleva, Fahad A Khokhar, Charlotte J. Houldcroft, Laura G Caller, Aminu S. Jahun, Sarah L. Caddy, Yasmin Chaudhry, Malte Pinckert, Ian Goodfellow |
| EPI_ISL_529258, EPI_ISL_529259, EPI_ISL_529260 | Liverpool Clinical Laboratories | COVID-19 Genomics UK (COG-UK) Consortium | Sam Haldenby, Anita Lucaci, Steve Paterson, Julian Hiscox, Alistair Darby, M Almsaud, A Alrezaihi, Muhannad Alruwaili, Stuart D Armstrong, Jones Benjamin, Eleanor G Bentley, Anu Chawla, Jordan J Clark, Angela Cowell, Richard Eccles, Isabel García-Dorival, Matthew Gemmell, Alessandro Gerada, PKF Gilmore, Richard Gregory, Ximeng Han, Catherine Hartley, Margaret Hughes, Miren Iturriza-Gomara, James Johnson, L Luu, Jenifer Manson, Charlotte Nelson, Elaine O'Toole, Cassie Olateju, Rebekah Penrice-Randal , Lucille Rainbow, N.P Randle, Trevor Ian Robinson, Parul Sharma, Ghada T Shawli, James P Stewart, Neil Swainston, Ecaterina Vamos, Joanne Watts, Mark Whitehead |
| EPI_ISL_529261 | Department of Pathology, University of Cambridge | COVID-19 Genomics UK (COG-UK) Consortium | Luke W Meredith, M. Estée Török, Myra Hosmillo, William L. Hamilton, Martin D. Curran, Theresa Feltwell, Grant Hall, Anna Yakovleva, Fahad A Khokhar, Charlotte J. Houldcroft, Laura G Caller, Aminu S. Jahun, Sarah L. Caddy, Yasmin Chaudhry, Malte Pinckert, Ian Goodfellow |
| EPI_ISL_529262 | University of Birmingham | COVID-19 Genomics UK (COG-UK) Consortium | Institute of Microbiology, University of Birmingham: Claire McMurray, Joanne Stockton, Samuel Nicholls, Radoslaw Poplawski, Will Rowe, Josh Quick, Nicholas Loman. University of Birmingham Testing Laboratory: Celina M Whalley, Andrew Bosworth, Charlotte Poxon, Kasun Wanigasooriya, Oliver Pickles, Mike Kidd, Alex Richter, Andrew D Beggs PHE Heartlands Lab: Husam Osman, Andrew Bosworth. Queen Elizabeth Hospital: Anna Casey |
| EPI_ISL_529263 | Liverpool Clinical Laboratories | COVID-19 Genomics UK (COG-UK) Consortium | Sam Haldenby, Anita Lucaci, Steve Paterson, Julian Hiscox, Alistair Darby, M Almsaud, A Alrezaihi, Muhannad Alruwaili, Stuart D Armstrong, Jones Benjamin, Eleanor G Bentley, Anu Chawla, Jordan J Clark, Angela Cowell, Richard Eccles, Isabel García-Dorival, Matthew Gemmell, Alessandro Gerada, PKF Gilmore, Richard Gregory, Ximeng Han, Catherine Hartley, Margaret Hughes, Miren Iturriza-Gomara, James Johnson, L Luu, Jenifer Manson, Charlotte Nelson, Elaine O'Toole, Cassie Olateju, Rebekah Penrice-Randal , Lucille Rainbow, N.P Randle, Trevor Ian Robinson, Parul Sharma, Ghada T Shawli, James P Stewart, Neil Swainston, Ecaterina Vamos, Joanne Watts, Mark Whitehead |
| EPI_ISL_529264 | Queens Medical Centre, Clinical Microbiology Department / DeepSeq Nottingham | COVID-19 Genomics UK (COG-UK) Consortium | Gemma Clark, Wendy Smith, Manjinder Khakh, Vicki M Fleming, Michelle M Lister, Hannah Howson-Wells, Jonathan Ball, Patrick McClure, Joseph Chappell, Theocharis Tsoleridis, Nadine Holmes, Matthew Carlisle, Christopher Moore, Fei Sang, Johnny Debebe, Victoria Wright, Matthew Loose |
| EPI_ISL_529265 | Department of Pathology, University of Cambridge | COVID-19 Genomics UK (COG-UK) Consortium | Luke W Meredith, M. Estée Török, Myra Hosmillo, William L. Hamilton, Martin D. Curran, Theresa Feltwell, Grant Hall, Anna Yakovleva, Fahad A Khokhar, Charlotte J. Houldcroft, Laura G Caller, Aminu S. Jahun, Sarah L. Caddy, Yasmin Chaudhry, Malte Pinckert, Ian Goodfellow |
| EPI_ISL_529266 | Queens Medical Centre, Clinical Microbiology Department / DeepSeq Nottingham | COVID-19 Genomics UK (COG-UK) Consortium | Gemma Clark, Wendy Smith, Manjinder Khakh, Vicki M Fleming, Michelle M Lister, Hannah Howson-Wells, Jonathan Ball, Patrick McClure, Joseph Chappell, Theocharis Tsoleridis, Nadine Holmes, Matthew Carlisle, Christopher Moore, Fei Sang, Johnny Debebe, Victoria Wright, Matthew Loose |
| EPI_ISL_529267 | Quadram Institute Bioscience | COVID-19 Genomics UK (COG-UK) Consortium | Dave J. Baker, Gemma L. Kay, Alp Aydin, Thanh Le-Viet, Steven Rudder, Ana P. Tedim, Anastasia Kolyva, Maria Diaz, Leonardo de Oliveira Martins, Nabil-Fareed Alikhan, Lizzie Meadows, Rachael Stanley, Ngozi Elumogo, Muhammed Yasir, Nicholas M. Thomson, Alexander J Trotter, Rachel Gilroy, Samuel Bloomfield, Claire Stuart, Andrew Bell, Reenesh Prakash, Samir Dervisevic, Alison E. Mather, John Wain, Mark Webber, Andrew J. Page, Justin O'Grady |
| EPI_ISL_529268 | University of Birmingham | COVID-19 Genomics UK (COG-UK) Consortium | Institute of Microbiology, University of Birmingham: Claire McMurray, Joanne Stockton, Samuel Nicholls, Radoslaw Poplawski, Will Rowe, Josh Quick, Nicholas Loman. University of Birmingham Testing Laboratory: Celina M Whalley, Andrew Bosworth, Charlotte Poxon, Kasun Wanigasooriya, Oliver Pickles, Mike Kidd, Alex Richter, Andrew D Beggs PHE Heartlands Lab: Husam Osman, Andrew Bosworth. Queen Elizabeth Hospital: Anna Casey |
| EPI_ISL_529269 | Department of Pathology, University of Cambridge | COVID-19 Genomics UK (COG-UK) Consortium | Luke W Meredith, M. Estée Török, Myra Hosmillo, William L. Hamilton, Martin D. Curran, Theresa Feltwell, Grant Hall, Anna Yakovleva, Fahad A Khokhar, Charlotte J. Houldcroft, Laura G Caller, Aminu S. Jahun, Sarah L. Caddy, Yasmin Chaudhry, Malte Pinckert, Ian Goodfellow |
| EPI_ISL_529270, EPI_ISL_529271 | Liverpool Clinical Laboratories | COVID-19 Genomics UK (COG-UK) Consortium | Sam Haldenby, Anita Lucaci, Steve Paterson, Julian Hiscox, Alistair Darby, M Almsaud, A Alrezaihi, Muhannad Alruwaili, Stuart D Armstrong, Jones Benjamin, Eleanor G Bentley, Anu Chawla, Jordan J Clark, Angela Cowell, Richard Eccles, Isabel García-Dorival, Matthew Gemmell, Alessandro Gerada, PKF Gilmore, Richard Gregory, Ximeng Han, Catherine Hartley, Margaret Hughes, Miren Iturriza-Gomara, James Johnson, L Luu, Jenifer Manson, Charlotte Nelson, Elaine O'Toole, Cassie Olateju, Rebekah Penrice-Randal , Lucille Rainbow, N.P Randle, Trevor Ian Robinson, Parul Sharma, Ghada T Shawli, James P Stewart, Neil Swainston, Ecaterina Vamos, Joanne Watts, Mark Whitehead |
| EPI_ISL_529272 | Centre for Enzyme Innovation, University of Portsmouth / Translational Research Laboratory, Portsmouth Hospitals NHS Trust | COVID-19 Genomics UK (COG-UK) Consortium | Angela Beckett,Yann Bourgeois,Garry Scarlett,Sharon Glaysher,Scott Elliott,Kelly Bicknell,Robert Impey,Allyson Lloyd,Sarah Wyllie,Ethan Butcher,Anoop Chauhan,Samuel Robson |
| EPI_ISL_529273 | Quadram Institute Bioscience | COVID-19 Genomics UK (COG-UK) Consortium | Dave J. Baker, Gemma L. Kay, Alp Aydin, Thanh Le-Viet, Steven Rudder, Ana P. Tedim, Anastasia Kolyva, Maria Diaz, Leonardo de Oliveira Martins, Nabil-Fareed Alikhan, Lizzie Meadows, Rachael Stanley, Ngozi Elumogo, |

| | | | |
|---|---|---|---|
| | | | Muhammed Yasir, Nicholas M. Thomson, Alexander J Trotter, Rachel Gilroy, Samuel Bloomfield, Claire Stuart, Andrew Bell, Reenesh Prakash, Samir Dervisevic, Alison E. Mather, John Wain, Mark Webber, Andrew J. Page, Justin O’Grady |
| EPI_ISL_529274 | Liverpool Clinical Laboratories | COVID-19 Genomics UK (COG-UK) Consortium | Sam Haldenby, Anita Lucaci, Steve Paterson, Julian Hiscox, Alistair Darby, M Almsaud, A Alrezaihi, Muhannad Alruwaili, Stuart D Armstrong, Jones Benjamin, Eleanor G Bentley, Anu Chawla, Jordan J Clark, Angela Cowell, Richard Eccles, Isabel García-Dorival, Matthew Gemmell, Alessandro Gerada, PKF Gilmore, Richard Gregory, Ximeng Han, Catherine Hartley, Margaret Hughes, Miren Iturriza-Gomara, James Johnson, L Luu, Jenifer Manson, Charlotte Nelson, Elaine O’Toole, Cassie Olateju, Rebekah Penrice-Randal , Lucille Rainbow, N.P Randle, Trevor Ian Robinson, Parul Sharma, Ghada T Shawli, James P Stewart, Neil Swainston, Ecaterina Vamos, Joanne Watts, Mark Whitehead |
| EPI_ISL_529275 | University of Birmingham | COVID-19 Genomics UK (COG-UK) Consortium | Institute of Microbiology, University of Birmingham: Claire McMurray, Joanne Stockton, Samuel Nicholls, Radoslaw Poplawski, Will Rowe, Josh Quick, Nicholas Loman. University of Birmingham Testing Laboratory: Celina M Whalley, Andrew Bosworth, Charlotte Poxon, Kasun Wanigasooriya, Oliver Pickles, Mike Kidd, Alex Richter, Andrew D Beggs PHE Heartlands Lab: Husam Osman, Andrew Bosworth. Queen Elizabeth Hospital: Anna Casey |
| EPI_ISL_529276 | Queens Medical Centre, Clinical Microbiology Department / DeepSeq Nottingham | COVID-19 Genomics UK (COG-UK) Consortium | Gemma Clark, Wendy Smith, Manjinder Khakh, Vicki M Fleming, Michelle M Lister, Hannah Howson-Wells, Jonathan Ball, Patrick McClure, Joseph Chappell, Theocharis Tsoleridis, Nadine Holmes, Matthew Carlisle, Christopher Moore, Fei Sang, Johnny Debebe, Victoria Wright, Matthew Loose |
| EPI_ISL_529277, EPI_ISL_529278 | University of Birmingham | COVID-19 Genomics UK (COG-UK) Consortium | Institute of Microbiology, University of Birmingham: Claire McMurray, Joanne Stockton, Samuel Nicholls, Radoslaw Poplawski, Will Rowe, Josh Quick, Nicholas Loman. University of Birmingham Testing Laboratory: Celina M Whalley, Andrew Bosworth, Charlotte Poxon, Kasun Wanigasooriya, Oliver Pickles, Mike Kidd, Alex Richter, Andrew D Beggs PHE Heartlands Lab: Husam Osman, Andrew Bosworth. Queen Elizabeth Hospital: Anna Casey |
| EPI_ISL_529279 | Queens Medical Centre, Clinical Microbiology Department / DeepSeq Nottingham | COVID-19 Genomics UK (COG-UK) Consortium | Gemma Clark, Wendy Smith, Manjinder Khakh, Vicki M Fleming, Michelle M Lister, Hannah Howson-Wells, Jonathan Ball, Patrick McClure, Joseph Chappell, Theocharis Tsoleridis, Nadine Holmes, Matthew Carlisle, Christopher Moore, Fei Sang, Johnny Debebe, Victoria Wright, Matthew Loose |
| EPI_ISL_529280 | Liverpool Clinical Laboratories | COVID-19 Genomics UK (COG-UK) Consortium | Sam Haldenby, Anita Lucaci, Steve Paterson, Julian Hiscox, Alistair Darby, M Almsaud, A Alrezaihi, Muhannad Alruwaili, Stuart D Armstrong, Jones Benjamin, Eleanor G Bentley, Anu Chawla, Jordan J Clark, Angela Cowell, Richard Eccles, Isabel García-Dorival, Matthew Gemmell, Alessandro Gerada, PKF Gilmore, Richard Gregory, Ximeng Han, Catherine Hartley, Margaret Hughes, Miren Iturriza-Gomara, James Johnson, L Luu, Jenifer Manson, Charlotte Nelson, Elaine O’Toole, Cassie Olateju, Rebekah Penrice-Randal , Lucille Rainbow, N.P Randle, Trevor Ian Robinson, Parul Sharma, Ghada T Shawli, James P Stewart, Neil Swainston, Ecaterina Vamos, Joanne Watts, Mark Whitehead |
| EPI_ISL_529281, EPI_ISL_529282 | Quadram Institute Bioscience | COVID-19 Genomics UK (COG-UK) Consortium | Dave J. Baker, Gemma L. Kay, Alp Aydin, Thanh Le-Viet, Steven Rudder, Ana P. Tedim, Anastasia Kolyva, Maria Diaz, Leonardo de Oliveira Martins, Nabil-Fareed Alikhan, Lizzie Meadows, Rachael Stanley, Ngozi Elumogo, Muhammed Yasir, Nicholas M. Thomson, Alexander J Trotter, Rachel Gilroy, Samuel Bloomfield, Claire Stuart, Andrew Bell, Reenesh Prakash, Samir Dervisevic, Alison E. Mather, John Wain, Mark Webber, Andrew J. Page, Justin O’Grady |
| EPI_ISL_529283 | Department of Pathology, University of Cambridge | COVID-19 Genomics UK (COG-UK) Consortium | Luke W Meredith, M. Estée Török, Myra Hosmillo, William L. Hamilton, Martin D. Curran, Theresa Feltwell, Grant Hall, Anna Yakovleva, Fahad A Khokhar, Charlotte J. Houldcroft, Laura G Caller, Aminu S. Jahun, Sarah L. Caddy, Yasmin Chaudhry, Malte Pinckert, Ian Goodfellow |
| EPI_ISL_529284 | Liverpool Clinical Laboratories | COVID-19 Genomics UK (COG-UK) Consortium | Sam Haldenby, Anita Lucaci, Steve Paterson, Julian Hiscox, Alistair Darby, M Almsaud, A Alrezaihi, Muhannad Alruwaili, Stuart D Armstrong, Jones Benjamin, Eleanor G Bentley, Anu Chawla, Jordan J Clark, Angela Cowell, Richard Eccles, Isabel García-Dorival, Matthew Gemmell, Alessandro Gerada, PKF Gilmore, Richard Gregory, Ximeng Han, Catherine Hartley, Margaret Hughes, Miren Iturriza-Gomara, James Johnson, L Luu, Jenifer Manson, Charlotte Nelson, Elaine O’Toole, Cassie Olateju, Rebekah Penrice-Randal , Lucille Rainbow, N.P Randle, Trevor Ian Robinson, Parul Sharma, Ghada T Shawli, James P Stewart, Neil Swainston, Ecaterina Vamos, Joanne Watts, Mark Whitehead |
| EPI_ISL_529285 | Virology Department, Royal Infirmary of Edinburgh, NHS Lothian / School of Biological Sciences, University of Edinburgh / Institute of Genetics and Molecular Medicine, University of Edinburgh | COVID-19 Genomics UK (COG-UK) Consortium | McHugh M, Dewar R, Rooke S, Gallagher M, Balcaza C, O’Toole Á, Scher E, Hill V, McCrone JT, Colquhoun R, Yu X, Jackson B, Rambaut A, Williams TC, Templeton K |
| EPI_ISL_529286 | Quadram Institute Bioscience | COVID-19 Genomics UK (COG-UK) Consortium | Dave J. Baker, Gemma L. Kay, Alp Aydin, Thanh Le-Viet, Steven Rudder, Ana P. Tedim, Anastasia Kolyva, Maria Diaz, Leonardo de Oliveira Martins, Nabil-Fareed Alikhan, Lizzie Meadows, Rachael Stanley, Ngozi Elumogo, Muhammed Yasir, Nicholas M. Thomson, Alexander J Trotter, Rachel Gilroy, Samuel Bloomfield, Claire Stuart, Andrew Bell, Reenesh Prakash, Samir Dervisevic, Alison E. Mather, John Wain, Mark Webber, Andrew J. Page, Justin O’Grady |
| EPI_ISL_529287 | Liverpool Clinical Laboratories | COVID-19 Genomics UK (COG-UK) Consortium | Sam Haldenby, Anita Lucaci, Steve Paterson, Julian Hiscox, Alistair Darby, M Almsaud, A Alrezaihi, Muhannad Alruwaili, Stuart D Armstrong, Jones Benjamin, Eleanor G Bentley, Anu Chawla, Jordan J Clark, Angela Cowell, Richard Eccles, Isabel García-Dorival, Matthew Gemmell, Alessandro Gerada, PKF Gilmore, Richard Gregory, Ximeng Han, Catherine Hartley, Margaret Hughes, Miren Iturriza-Gomara, James Johnson, L Luu, Jenifer Manson, Charlotte Nelson, Elaine O’Toole, Cassie Olateju, Rebekah Penrice-Randal , Lucille Rainbow, N.P Randle, Trevor Ian Robinson, Parul Sharma, Ghada T Shawli, James P Stewart, Neil Swainston, Ecaterina Vamos, Joanne Watts, Mark Whitehead |
| EPI_ISL_529288 | Queens Medical Centre, Clinical Microbiology Department / DeepSeq Nottingham | COVID-19 Genomics UK (COG-UK) Consortium | Gemma Clark, Wendy Smith, Manjinder Khakh, Vicki M Fleming, Michelle M Lister, Hannah Howson-Wells, Jonathan Ball, Patrick McClure, Joseph Chappell, Theocharis Tsoleridis, Nadine Holmes, Matthew Carlisle, Christopher Moore, Fei Sang, Johnny Debebe, Victoria Wright, Matthew Loose |
| EPI_ISL_529289 | University of Birmingham | COVID-19 Genomics UK (COG-UK) Consortium | Institute of Microbiology, University of Birmingham: Claire McMurray, Joanne Stockton, Samuel Nicholls, Radoslaw Poplawski, Will Rowe, Josh Quick, Nicholas Loman. University of Birmingham Testing Laboratory: Celina M Whalley, Andrew Bosworth, Charlotte Poxon, Kasun Wanigasooriya, Oliver Pickles, Mike Kidd, Alex Richter, Andrew D Beggs PHE Heartlands Lab: Husam Osman, Andrew Bosworth. Queen Elizabeth Hospital: Anna Casey |
| EPI_ISL_529290 | Quadram Institute Bioscience | COVID-19 Genomics UK (COG-UK) Consortium | Dave J. Baker, Gemma L. Kay, Alp Aydin, Thanh Le-Viet, Steven Rudder, Ana P. Tedim, Anastasia Kolyva, Maria Diaz, Leonardo de Oliveira Martins, Nabil-Fareed Alikhan, Lizzie Meadows, Rachael Stanley, Ngozi Elumogo, Muhammed Yasir, Nicholas M. Thomson, Alexander J Trotter, Rachel Gilroy, Samuel Bloomfield, Claire Stuart, Andrew Bell, Reenesh Prakash, Samir Dervisevic, Alison E. Mather, John Wain, Mark Webber, Andrew J. Page, Justin O’Grady |
| EPI_ISL_529291 | University of Birmingham | COVID-19 Genomics UK (COG-UK) Consortium | Institute of Microbiology, University of Birmingham: Claire McMurray, Joanne Stockton, Samuel Nicholls, Radoslaw Poplawski, Will Rowe, Josh Quick, Nicholas Loman. University of Birmingham Testing Laboratory: Celina M Whalley, Andrew Bosworth, Charlotte Poxon, Kasun Wanigasooriya, Oliver Pickles, Mike Kidd, Alex Richter, Andrew D Beggs PHE Heartlands Lab: Husam Osman, Andrew Bosworth. Queen Elizabeth Hospital: Anna Casey |
| EPI_ISL_529292 | Quadram Institute Bioscience | COVID-19 Genomics UK (COG-UK) Consortium | Dave J. Baker, Gemma L. Kay, Alp Aydin, Thanh Le-Viet, Steven Rudder, Ana P. Tedim, Anastasia Kolyva, Maria Diaz, Leonardo de Oliveira Martins, Nabil-Fareed Alikhan, Lizzie Meadows, Rachael Stanley, Ngozi Elumogo, Muhammed Yasir, Nicholas M. Thomson, Alexander J Trotter, Rachel Gilroy, Samuel Bloomfield, Claire Stuart, Andrew Bell, Reenesh Prakash, Samir Dervisevic, Alison E. Mather, John Wain, Mark Webber, Andrew J. Page, Justin O’Grady |
| EPI_ISL_529293 | Liverpool Clinical Laboratories | COVID-19 Genomics UK (COG-UK) Consortium | Sam Haldenby, Anita Lucaci, Steve Paterson, Julian Hiscox, Alistair Darby, M Almsaud, A Alrezaihi, Muhannad Alruwaili, Stuart D Armstrong, Jones Benjamin, Eleanor G Bentley, Anu Chawla, Jordan J Clark, Angela Cowell, Richard Eccles, Isabel García-Dorival, Matthew Gemmell, Alessandro Gerada, PKF Gilmore, Richard Gregory, Ximeng Han, Catherine Hartley, Margaret Hughes, Miren Iturriza-Gomara, James Johnson, L Luu, Jenifer Manson, Charlotte Nelson, Elaine O’Toole, Cassie Olateju, Rebekah Penrice-Randal , Lucille Rainbow, N.P Randle, Trevor Ian Robinson, Parul Sharma, Ghada T Shawli, James P Stewart, Neil Swainston, Ecaterina Vamos, Joanne Watts, Mark Whitehead |
| EPI_ISL_529294 | Quadram Institute Bioscience | COVID-19 Genomics UK (COG-UK) Consortium | Dave J. Baker, Gemma L. Kay, Alp Aydin, Thanh Le-Viet, Steven Rudder, Ana P. Tedim, Anastasia Kolyva, Maria Diaz, Leonardo de Oliveira Martins, Nabil-Fareed Alikhan, Lizzie Meadows, Rachael Stanley, Ngozi Elumogo, Muhammed Yasir, Nicholas M. Thomson, Alexander J Trotter, Rachel Gilroy, Samuel Bloomfield, Claire Stuart, Andrew Bell, Reenesh Prakash, Samir Dervisevic, Alison E. Mather, John Wain, Mark Webber, Andrew J. Page, Justin O’Grady |
| EPI_ISL_529295, EPI_ISL_529296 | Liverpool Clinical Laboratories | COVID-19 Genomics UK (COG-UK) Consortium | Sam Haldenby, Anita Lucaci, Steve Paterson, Julian Hiscox, Alistair Darby, M Almsaud, A Alrezaihi, Muhannad Alruwaili, Stuart D Armstrong, Jones Benjamin, Eleanor G Bentley, Anu Chawla, Jordan J Clark, Angela Cowell, Richard Eccles, Isabel García-Dorival, Matthew Gemmell, Alessandro Gerada, PKF Gilmore, Richard Gregory, Ximeng Han, Catherine Hartley, Margaret Hughes, Miren Iturriza-Gomara, James Johnson, L Luu, Jenifer Manson, Charlotte Nelson, Elaine O’Toole, Cassie Olateju, Rebekah Penrice-Randal , Lucille Rainbow, N.P Randle, Trevor Ian Robinson, Parul Sharma, Ghada T Shawli, James P Stewart, Neil Swainston, Ecaterina Vamos, Joanne Watts, Mark Whitehead |
| EPI_ISL_529297 | Wales Specialist Virology Centre Sequencing lab: Pathogen Genomics Unit | COVID-19 Genomics UK (COG-UK) Consortium | Catherine Moore, Johnathan Evans, Laura Gifford, Malorie Perry, Simon Cottrell, Angela Marchbank, Alec Birchley, Alexander Adams, Amy Gaskin, Bree Gatica-Wilcox, Jason Coombes, Joel Southgate, Lauren Gilbert, Lee Graham, Nicole Pacchiarini, Sara Kumziene-Summerhayes, Sarah Taylor, Sophie Jones, Sara Rey, Matthew Bull, Joanne Watkins, Sally Corden, Tom Connor |
| EPI_ISL_529298 | University of Birmingham | COVID-19 Genomics UK (COG-UK) Consortium | Institute of Microbiology, University of Birmingham: Claire McMurray, Joanne Stockton, Samuel Nicholls, Radoslaw Poplawski, Will Rowe, Josh Quick, Nicholas Loman. University of Birmingham Testing Laboratory: Celina M Whalley, Andrew Bosworth, Charlotte Poxon, Kasun Wanigasooriya, Oliver Pickles, Mike Kidd, Alex Richter, Andrew D Beggs PHE Heartlands Lab: Husam Osman, Andrew Bosworth. Queen Elizabeth Hospital: Anna Casey |
| EPI_ISL_529299 | Department of Pathology, University of Cambridge | COVID-19 Genomics UK (COG-UK) Consortium | Luke W Meredith, M. Estée Török, Myra Hosmillo, William L. Hamilton, Martin D. Curran, Theresa Feltwell, Grant Hall, Anna Yakovleva, Fahad A Khokhar, Charlotte J. Houldcroft, Laura G Caller, Aminu S. Jahun, Sarah L. Caddy, Yasmin Chaudhry, Malte Pinckert, Ian Goodfellow |
| EPI_ISL_529300 | University of Birmingham | COVID-19 Genomics UK (COG-UK) Consortium | Institute of Microbiology, University of Birmingham: Claire McMurray, Joanne Stockton, Samuel Nicholls, Radoslaw Poplawski, Will Rowe, Josh Quick, Nicholas Loman. University of Birmingham Testing Laboratory: Celina M Whalley, Andrew Bosworth, Charlotte Poxon, Kasun Wanigasooriya, Oliver Pickles, Mike Kidd, Alex Richter, Andrew D Beggs PHE Heartlands Lab: Husam Osman, Andrew Bosworth. Queen Elizabeth Hospital: Anna Casey |
| EPI_ISL_529301 | Virology Department, Sheffield Teaching Hospitals NHS Foundation Trust/Department of Infection, Immunity and Cardiovascular Disease, The Medical School, University of Sheffield | COVID-19 Genomics UK (COG-UK) Consortium | Thushan de Silva, Matthew Parker, Nikki Smith, Adri Angyal, Rebecca Brown, Luke Green, Rachel Tucker, Paul Parsons, Danielle Groves, Katie Johnson, Laura Carrilero, Alex Keeley, Dave Partridge, Matthew Wyles, Benjamin Lindsey, Mehmet Yavuz, Mohammad Raza, Cariad Evans |
| EPI_ISL_529302 | Liverpool Clinical Laboratories | COVID-19 Genomics UK (COG-UK) Consortium | Sam Haldenby, Anita Lucaci, Steve Paterson, Julian Hiscox, Alistair Darby, M Almsaud, A Alrezaihi, Muhannad Alruwaili, Stuart D Armstrong, Jones Benjamin, Eleanor G Bentley, Anu Chawla, Jordan J Clark, Angela Cowell, Richard Eccles, Isabel García-Dorival, Matthew Gemmell, Alessandro Gerada, PKF Gilmore, Richard Gregory, Ximeng Han, Catherine Hartley, Margaret Hughes, Miren Iturriza-Gomara, James Johnson, L Luu, Jenifer Manson, Charlotte Nelson, Elaine O’Toole, Cassie Olateju, Rebekah Penrice-Randal , Lucille Rainbow, N.P Randle, Trevor Ian Robinson, Parul Sharma, Ghada T Shawli, James P Stewart, Neil Swainston, Ecaterina Vamos, Joanne Watts, Mark Whitehead |
| EPI_ISL_529303 | Centre for Enzyme Innovation, University of Portsmouth / Translational Research Laboratory, Portsmouth Hospitals NHS Trust | COVID-19 Genomics UK (COG-UK) Consortium | Angela Beckett,Yann Bourgeois,Garry Scarlett,Sharon Glaysher,Scott Elliott,Kelly Bicknell,Robert Impey,Allyson Lloyd,Sarah Wyllie,Ethan Butcher,Anoop Chauhan,Samuel Robson |
| EPI_ISL_529304, EPI_ISL_529305 | Liverpool Clinical Laboratories | COVID-19 Genomics UK (COG-UK) Consortium | Sam Haldenby, Anita Lucaci, Steve Paterson, Julian Hiscox, Alistair Darby, M Almsaud, A Alrezaihi, Muhannad Alruwaili, Stuart D Armstrong, Jones Benjamin, Eleanor G Bentley, Anu Chawla, Jordan J Clark, Angela Cowell, Richard Eccles, Isabel García-Dorival, Matthew Gemmell, Alessandro Gerada, PKF Gilmore, Richard Gregory, Ximeng Han, Catherine Hartley, Margaret Hughes, Miren Iturriza-Gomara, James Johnson, L Luu, Jenifer Manson, Charlotte Nelson, Elaine O’Toole, Cassie Olateju, Rebekah Penrice-Randal , Lucille Rainbow, N.P Randle, Trevor Ian Robinson, Parul Sharma, Ghada T Shawli, James P Stewart, Neil Swainston, Ecaterina Vamos, Joanne Watts, Mark Whitehead |
| EPI_ISL_529306, EPI_ISL_529307, EPI_ISL_529308, EPI_ISL_529309, EPI_ISL_529310, EPI_ISL_529311 | Queens Medical Centre, Clinical Microbiology Department / DeepSeq Nottingham | COVID-19 Genomics UK (COG-UK) Consortium | Gemma Clark, Wendy Smith, Manjinder Khakh, Vicki M Fleming, Michelle M Lister, Hannah Howson-Wells, Jonathan Ball, Patrick McClure, Joseph Chappell, Theocharis Tsoleridis, Nadine Holmes, Matthew Carlisle, Christopher Moore, Fei Sang, Johnny Debebe, Victoria Wright, Matthew Loose |
| EPI_ISL_529312, EPI_ISL_529313, EPI_ISL_529314 | University of Birmingham | COVID-19 Genomics UK (COG-UK) Consortium | Institute of Microbiology, University of Birmingham: Claire McMurray, Joanne Stockton, Samuel Nicholls, Radoslaw Poplawski, Will Rowe, Josh Quick, Nicholas Loman. University of Birmingham Testing Laboratory: Celina M Whalley, Andrew Bosworth, Charlotte Poxon, Kasun Wanigasooriya, Oliver Pickles, Mike Kidd, Alex Richter, Andrew D Beggs PHE Heartlands Lab: Husam Osman, Andrew Bosworth. Queen Elizabeth Hospital: Anna Casey |
| EPI_ISL_529315, EPI_ISL_529316 | West of Scotland Specialist Virology Centre, NHSGGC / MRC-University of Glasgow Centre for Virus Research | COVID-19 Genomics UK (COG-UK) Consortium | Ana da Silva Filipe, Natasha Johnson, Kathy Smollett, Daniel Mair, Stephen Carmichael, Lily Tong, Jenna Nichols, Elihu Aranday-Cortes, Kirstyn Brunker, Yasmin Parr, Alice Broos, Kyriaki Nomikou; Sarah McDonald, Marc Niebel, Patawee Asamaphan; Richard Orton, Joseph Hughes, Sreenu Vattipally, David L Robertson; Alasdair MacLean, Rory Gunson; Kathy Li, Natasha Jesudason, Rajiv Shah, James Shepherd, Antonia Ho, Emma |

| | | | |
|---|---|---|---|
| | | | Thomson |
| EPI_ISL_529317, EPI_ISL_529318 | Virology Department, Royal Infirmary of Edinburgh, NHS Lothian / School of Biological Sciences, University of Edinburgh / Institute of Genetics and Molecular Medicine, University of Edinburgh | COVID-19 Genomics UK (COG-UK) Consortium | McHugh M, Dewar R, Rooke S, Gallagher M, Balcaza C, O'Toole Á, Scher E, Hill V, McCrone JT, Colquhoun R, Yu X, Jackson B, Rambaut A, Williams TC, Templeton K |
| EPI_ISL_529319 | West of Scotland Specialist Virology Centre, NHSGGC / MRC-University of Glasgow Centre for Virus Research | COVID-19 Genomics UK (COG-UK) Consortium | Ana da Silva Filipe, Natasha Johnson, Kathy Smollett, Daniel Mair, Stephen Carmichael, Lily Tong, Jenna Nichols, Elihu Aranday-Cortes, Kirstyn Brunker, Yasmin Parr, Alice Broos, Kyriaki Nomikou; Sarah McDonald, Marc Niebel, Patawee Asamaphan; Richard Orton, Joseph Hughes, Sreenu Vattipally, David L Robertson; Alasdair MacLean, Rory Gunson; Kathy Li, Natasha Jesudason, Rajiv Shah, James Shepherd, Antonia Ho, Emma Thomson |
| EPI_ISL_529320, EPI_ISL_529321, EPI_ISL_529322, EPI_ISL_529323, EPI_ISL_529324, EPI_ISL_529325, EPI_ISL_529326, EPI_ISL_529327, EPI_ISL_529328, EPI_ISL_529329, EPI_ISL_529330, EPI_ISL_529331, EPI_ISL_529332, EPI_ISL_529333, EPI_ISL_529334, EPI_ISL_529335, EPI_ISL_529336, EPI_ISL_529337, EPI_ISL_529338, EPI_ISL_529339, EPI_ISL_529340 | | | |
| see above | University of Birmingham | COVID-19 Genomics UK (COG-UK) Consortium | Institute of Microbiology, University of Birmingham: Claire McMurray, Joanne Stockton, Samuel Nicholls, Radoslaw Poplawski, Will Rowe, Josh Quick, Nicholas Loman. University of Birmingham Testing Laboratory: Celina M Whalley, Andrew Bosworth, Charlotte Poxon, Kasun Wanigasooriya, Oliver Pickles, Mike Kidd, Alex Richter, Andrew D Beggs PHE Heartlands Lab: Husam Osman, Andrew Bosworth. Queen Elizabeth Hospital: Anna Casey |
| EPI_ISL_529341, EPI_ISL_529342, EPI_ISL_529343, EPI_ISL_529344, EPI_ISL_529345, EPI_ISL_529346, EPI_ISL_529347, EPI_ISL_529348, EPI_ISL_529349, EPI_ISL_529350, EPI_ISL_529351, EPI_ISL_529352 | | | |
| see above | Centre for Enzyme Innovation, University of Portsmouth / Translational Research Laboratory, Portsmouth Hospitals NHS Trust | COVID-19 Genomics UK (COG-UK) Consortium | Angela Beckett,Yann Bourgeois,Garry Scarlett,Sharon Glaysher,Scott Elliott,Kelly Bicknell,Robert Impey,Allyson Lloyd,Sarah Wyllie,Ethan Butcher,Anoop Chauhan,Samuel Robson |
| EPI_ISL_529353, EPI_ISL_529354, EPI_ISL_529355, EPI_ISL_529356 | Virology Department, Sheffield Teaching Hospitals NHS Foundation Trust/Department of Infection, Immunity and Cardiovascular Disease, The Medical School, University of Sheffield | COVID-19 Genomics UK (COG-UK) Consortium | Thushan de Silva, Matthew Parker, Nikki Smith, Adri Angyal, Rebecca Brown, Luke Green, Rachel Tucker, Paul Parsons, Danielle Groves, Katie Johnson, Laura Carrilero, Alex Keeley, Dave Partridge, Matthew Wyles, Benjamin Lindsey, Mehmet Yavuz, Mohammad Raza, Cariad Evans |
| EPI_ISL_529357, EPI_ISL_529358, EPI_ISL_529359, EPI_ISL_529360, EPI_ISL_529361, EPI_ISL_529362, EPI_ISL_529363, EPI_ISL_529364, EPI_ISL_529365, EPI_ISL_529366, EPI_ISL_529367, EPI_ISL_529368, EPI_ISL_529369, EPI_ISL_529370, EPI_ISL_529371, EPI_ISL_529372 | | | |
| see above | University of Birmingham | COVID-19 Genomics UK (COG-UK) Consortium | Institute of Microbiology, University of Birmingham: Claire McMurray, Joanne Stockton, Samuel Nicholls, Radoslaw Poplawski, Will Rowe, Josh Quick, Nicholas Loman. University of Birmingham Testing Laboratory: Celina M Whalley, Andrew Bosworth, Charlotte Poxon, Kasun Wanigasooriya, Oliver Pickles, Mike Kidd, Alex Richter, Andrew D Beggs PHE Heartlands Lab: Husam Osman, Andrew Bosworth. Queen Elizabeth Hospital: Anna Casey |
| EPI_ISL_529373, EPI_ISL_529374, EPI_ISL_529375, EPI_ISL_529376, EPI_ISL_529377 | Centre for Enzyme Innovation, University of Portsmouth / Translational Research Laboratory, Portsmouth Hospitals NHS Trust | COVID-19 Genomics UK (COG-UK) Consortium | Angela Beckett,Yann Bourgeois,Garry Scarlett,Sharon Glaysher,Scott Elliott,Kelly Bicknell,Robert Impey,Allyson Lloyd,Sarah Wyllie,Ethan Butcher,Anoop Chauhan,Samuel Robson |
| EPI_ISL_529378, EPI_ISL_529379, EPI_ISL_529380, EPI_ISL_529381 | Quadram Institute Bioscience | COVID-19 Genomics UK (COG-UK) Consortium | Dave J. Baker, Gemma L. Kay, Alp Aydin, Thanh Le-Viet, Steven Rudder, Ana P. Tedim, Anastasia Kolyva, Maria Diaz, Leonardo de Oliveira Martins, Nabil-Fareed Alikhan, Lizzie Meadows, Rachael Stanley, Ngozi Elumogo, Muhammed Yasir, Nicholas M. Thomson, Alexander J Trotter, Rachel Gilroy, Samuel Bloomfield, Claire Stuart, Andrew Bell, Reenesh Prakash, Samir Dervisevic, Alison E. Mather, John Wain, Mark Webber, Andrew J. Page, Justin O'Grady |
| EPI_ISL_529382 | Virology Department, Sheffield Teaching Hospitals NHS Foundation Trust/Department of Infection, Immunity and Cardiovascular Disease, The Medical School, University of Sheffield | COVID-19 Genomics UK (COG-UK) Consortium | Thushan de Silva, Matthew Parker, Nikki Smith, Adri Angyal, Rebecca Brown, Luke Green, Rachel Tucker, Paul Parsons, Danielle Groves, Katie Johnson, Laura Carrilero, Alex Keeley, Dave Partridge, Matthew Wyles, Benjamin Lindsey, Mehmet Yavuz, Mohammad Raza, Cariad Evans |
| EPI_ISL_529383 | Queens Medical Centre, Clinical Microbiology Department / DeepSeq Nottingham | COVID-19 Genomics UK (COG-UK) Consortium | Gemma Clark, Wendy Smith, Manjinder Khakh, Vicki M Fleming, Michelle M Lister, Hannah Howson-Wells, Jonathan Ball, Patrick McClure, Joseph Chappell, Theocharis Tsoleridis, Nadine Holmes, Matthew Carlisle, Christopher Moore, Fei Sang, Johnny Debebe, Victoria Wright, Matthew Loose |
| EPI_ISL_529384 | University of Birmingham | COVID-19 Genomics UK (COG-UK) Consortium | Institute of Microbiology, University of Birmingham: Claire McMurray, Joanne Stockton, Samuel Nicholls, Radoslaw Poplawski, Will Rowe, Josh Quick, Nicholas Loman. University of Birmingham Testing Laboratory: Celina M Whalley, Andrew Bosworth, Charlotte Poxon, Kasun Wanigasooriya, Oliver Pickles, Mike Kidd, Alex Richter, Andrew D Beggs PHE Heartlands Lab: Husam Osman, Andrew Bosworth. Queen Elizabeth Hospital: Anna Casey |
| EPI_ISL_529385 | Queens Medical Centre, Clinical Microbiology Department / DeepSeq Nottingham | COVID-19 Genomics UK (COG-UK) Consortium | Gemma Clark, Wendy Smith, Manjinder Khakh, Vicki M Fleming, Michelle M Lister, Hannah Howson-Wells, Jonathan Ball, Patrick McClure, Joseph Chappell, Theocharis Tsoleridis, Nadine Holmes, Matthew Carlisle, Christopher Moore, Fei Sang, Johnny Debebe, Victoria Wright, Matthew Loose |
| EPI_ISL_529386 | University of Birmingham | COVID-19 Genomics UK (COG-UK) Consortium | Institute of Microbiology, University of Birmingham: Claire McMurray, Joanne Stockton, Samuel Nicholls, Radoslaw Poplawski, Will Rowe, Josh Quick, Nicholas Loman. University of Birmingham Testing Laboratory: Celina M Whalley, Andrew Bosworth, Charlotte Poxon, Kasun Wanigasooriya, Oliver Pickles, Mike Kidd, Alex Richter, Andrew D Beggs PHE Heartlands Lab: Husam Osman, Andrew Bosworth. Queen Elizabeth Hospital: Anna Casey |
| EPI_ISL_529387 | Queens Medical Centre, Clinical Microbiology Department / DeepSeq Nottingham | COVID-19 Genomics UK (COG-UK) Consortium | Gemma Clark, Wendy Smith, Manjinder Khakh, Vicki M Fleming, Michelle M Lister, Hannah Howson-Wells, Jonathan Ball, Patrick McClure, Joseph Chappell, Theocharis Tsoleridis, Nadine Holmes, Matthew Carlisle, Christopher Moore, Fei Sang, Johnny Debebe, Victoria Wright, Matthew Loose |
| EPI_ISL_529388, EPI_ISL_529389 | Quadram Institute Bioscience | COVID-19 Genomics UK (COG-UK) Consortium | Dave J. Baker, Gemma L. Kay, Alp Aydin, Thanh Le-Viet, Steven Rudder, Ana P. Tedim, Anastasia Kolyva, Maria Diaz, Leonardo de Oliveira Martins, Nabil-Fareed Alikhan, Lizzie Meadows, Rachael Stanley, Ngozi Elumogo, Muhammed Yasir, Nicholas M. Thomson, Alexander J Trotter, Rachel Gilroy, Samuel Bloomfield, Claire Stuart, Andrew Bell, Reenesh Prakash, Samir Dervisevic, Alison E. Mather, John Wain, Mark Webber, Andrew J. Page, Justin O'Grady |
| EPI_ISL_529390, EPI_ISL_529391, EPI_ISL_529392, EPI_ISL_529393, EPI_ISL_529394, EPI_ISL_529395, EPI_ISL_529396, EPI_ISL_529397, EPI_ISL_529398, EPI_ISL_529399, EPI_ISL_529400, EPI_ISL_529401, EPI_ISL_529402, EPI_ISL_529403, EPI_ISL_529404, EPI_ISL_529405, EPI_ISL_529406 | | | |
| see above | University of Birmingham | COVID-19 Genomics UK (COG-UK) Consortium | Institute of Microbiology, University of Birmingham: Claire McMurray, Joanne Stockton, Samuel Nicholls, Radoslaw Poplawski, Will Rowe, Josh Quick, Nicholas Loman. University of Birmingham Testing Laboratory: Celina M Whalley, Andrew Bosworth, Charlotte Poxon, Kasun Wanigasooriya, Oliver Pickles, Mike Kidd, Alex Richter, Andrew D Beggs PHE Heartlands Lab: Husam Osman, Andrew Bosworth. Queen Elizabeth Hospital: Anna Casey |
| EPI_ISL_529407 | Virology Department, Sheffield Teaching Hospitals NHS Foundation Trust/Department of Infection, Immunity and Cardiovascular Disease, The Medical School, University of Sheffield | COVID-19 Genomics UK (COG-UK) Consortium | Thushan de Silva, Matthew Parker, Nikki Smith, Adri Angyal, Rebecca Brown, Luke Green, Rachel Tucker, Paul Parsons, Danielle Groves, Katie Johnson, Laura Carrilero, Alex Keeley, Dave Partridge, Matthew Wyles, Benjamin Lindsey, Mehmet Yavuz, Mohammad Raza, Cariad Evans |
| EPI_ISL_529408 | University of Birmingham | COVID-19 Genomics UK (COG-UK) Consortium | Institute of Microbiology, University of Birmingham: Claire McMurray, Joanne Stockton, Samuel Nicholls, Radoslaw Poplawski, Will Rowe, Josh Quick, Nicholas Loman. University of Birmingham Testing Laboratory: Celina M Whalley, Andrew Bosworth, Charlotte Poxon, Kasun Wanigasooriya, Oliver Pickles, Mike Kidd, Alex Richter, Andrew D Beggs PHE Heartlands Lab: Husam Osman, Andrew Bosworth. Queen Elizabeth Hospital: Anna Casey |
| EPI_ISL_529409 | West of Scotland Specialist Virology Centre, NHSGGC / MRC-University of Glasgow Centre for Virus Research | COVID-19 Genomics UK (COG-UK) Consortium | Ana da Silva Filipe, Natasha Johnson, Kathy Smollett, Daniel Mair, Stephen Carmichael, Lily Tong, Jenna Nichols, Elihu Aranday-Cortes, Kirstyn Brunker, Yasmin Parr, Alice Broos, Kyriaki Nomikou; Sarah McDonald, Marc Niebel, Patawee Asamaphan; Richard Orton, Joseph Hughes, Sreenu Vattipally, David L Robertson; Alasdair MacLean, Rory Gunson; Kathy Li, Natasha Jesudason, Rajiv Shah, James Shepherd, Antonia Ho, Emma Thomson |
| EPI_ISL_529410, EPI_ISL_529411, EPI_ISL_529412 | Queens Medical Centre, Clinical Microbiology Department / DeepSeq Nottingham | COVID-19 Genomics UK (COG-UK) Consortium | Gemma Clark, Wendy Smith, Manjinder Khakh, Vicki M Fleming, Michelle M Lister, Hannah Howson-Wells, Jonathan Ball, Patrick McClure, Joseph Chappell, Theocharis Tsoleridis, Nadine Holmes, Matthew Carlisle, Christopher Moore, Fei Sang, Johnny Debebe, Victoria Wright, Matthew Loose |
| EPI_ISL_529413 | Quadram Institute Bioscience | COVID-19 Genomics UK (COG-UK) Consortium | Dave J. Baker, Gemma L. Kay, Alp Aydin, Thanh Le-Viet, Steven Rudder, Ana P. Tedim, Anastasia Kolyva, Maria Diaz, Leonardo de Oliveira Martins, Nabil-Fareed Alikhan, Lizzie Meadows, Rachael Stanley, Ngozi Elumogo, Muhammed Yasir, Nicholas M. Thomson, Alexander J Trotter, Rachel Gilroy, Samuel Bloomfield, Claire Stuart, Andrew Bell, Reenesh Prakash, Samir Dervisevic, Alison E. Mather, John Wain, Mark Webber, Andrew J. Page, Justin O'Grady |
| EPI_ISL_529414 | Queens Medical Centre, Clinical Microbiology Department / DeepSeq Nottingham | COVID-19 Genomics UK (COG-UK) Consortium | Gemma Clark, Wendy Smith, Manjinder Khakh, Vicki M Fleming, Michelle M Lister, Hannah Howson-Wells, Jonathan Ball, Patrick McClure, Joseph Chappell, Theocharis Tsoleridis, Nadine Holmes, Matthew Carlisle, Christopher Moore, Fei Sang, Johnny Debebe, Victoria Wright, Matthew Loose |
| EPI_ISL_529415, EPI_ISL_529416, EPI_ISL_529417 | Quadram Institute Bioscience | COVID-19 Genomics UK (COG-UK) Consortium | Dave J. Baker, Gemma L. Kay, Alp Aydin, Thanh Le-Viet, Steven Rudder, Ana P. Tedim, Anastasia Kolyva, Maria Diaz, Leonardo de Oliveira Martins, Nabil-Fareed Alikhan, Lizzie Meadows, Rachael Stanley, Ngozi Elumogo, Muhammed Yasir, Nicholas M. Thomson, Alexander J Trotter, Rachel Gilroy, Samuel Bloomfield, Claire Stuart, Andrew Bell, Reenesh Prakash, Samir Dervisevic, Alison E. Mather, John Wain, Mark Webber, Andrew J. Page, Justin O'Grady |
| EPI_ISL_529418 | Wales Specialist Virology Centre Sequencing lab: Pathogen Genomics Unit | COVID-19 Genomics UK (COG-UK) Consortium | Catherine Moore, Johnathan Evans, Laura Gifford, Malorie Perry, Simon Cottrell, Angela Marchbank, Alec Birchley, Alexander Adams, Amy Gaskin, Bree Gatica-Wilcox, Jason Coombes, Joel Southgate, Lauren Gilbert, Lee Graham, Nicole Pacchiarini, Sara Kumziene-Summerhayes, Sarah Taylor, Sophie Jones, Sara Rey, Matthew Bull, Joanne Watkins, Sally Corden, Tom Connor |
| EPI_ISL_529419 | Virology Department, Sheffield Teaching Hospitals NHS Foundation Trust/Department of Infection, Immunity and Cardiovascular Disease, The Medical School, University of Sheffield | COVID-19 Genomics UK (COG-UK) Consortium | Thushan de Silva, Matthew Parker, Nikki Smith, Adri Angyal, Rebecca Brown, Luke Green, Rachel Tucker, Paul Parsons, Danielle Groves, Katie Johnson, Laura Carrilero, Alex Keeley, Dave Partridge, Matthew Wyles, Benjamin Lindsey, Mehmet Yavuz, Mohammad Raza, Cariad Evans |
| EPI_ISL_529420, EPI_ISL_529421 | University of Birmingham | COVID-19 Genomics UK (COG-UK) Consortium | Institute of Microbiology, University of Birmingham: Claire McMurray, Joanne Stockton, Samuel Nicholls, Radoslaw Poplawski, Will Rowe, Josh Quick, Nicholas Loman. University of Birmingham Testing Laboratory: Celina M Whalley, Andrew Bosworth, Charlotte Poxon, Kasun Wanigasooriya, Oliver Pickles, Mike Kidd, Alex Richter, Andrew D Beggs PHE Heartlands Lab: Husam Osman, Andrew Bosworth. Queen Elizabeth Hospital: Anna Casey |
| EPI_ISL_529422 | Quadram Institute Bioscience | COVID-19 Genomics UK (COG-UK) Consortium | Dave J. Baker, Gemma L. Kay, Alp Aydin, Thanh Le-Viet, Steven Rudder, Ana P. Tedim, Anastasia Kolyva, Maria Diaz, Leonardo de Oliveira Martins, Nabil-Fareed Alikhan, Lizzie Meadows, Rachael Stanley, Ngozi Elumogo, Muhammed Yasir, Nicholas M. Thomson, Alexander J Trotter, Rachel Gilroy, Samuel Bloomfield, Claire Stuart, Andrew Bell, Reenesh Prakash, Samir Dervisevic, Alison E. Mather, John Wain, Mark Webber, Andrew J. Page, Justin O'Grady |
| EPI_ISL_529423 | Queens Medical Centre, Clinical Microbiology Department / DeepSeq Nottingham | COVID-19 Genomics UK (COG-UK) Consortium | Gemma Clark, Wendy Smith, Manjinder Khakh, Vicki M Fleming, Michelle M Lister, Hannah Howson-Wells, Jonathan Ball, Patrick McClure, Joseph Chappell, Theocharis Tsoleridis, Nadine Holmes, Matthew Carlisle, Christopher Moore, Fei Sang, Johnny Debebe, Victoria Wright, Matthew Loose |
| EPI_ISL_529424, EPI_ISL_529425, EPI_ISL_529426, EPI_ISL_529427, EPI_ISL_529428, EPI_ISL_529429, EPI_ISL_529430 | University of Birmingham | COVID-19 Genomics UK (COG-UK) Consortium | Institute of Microbiology, University of Birmingham: Claire McMurray, Joanne Stockton, Samuel Nicholls, Radoslaw Poplawski, Will Rowe, Josh Quick, Nicholas Loman. University of Birmingham Testing Laboratory: Celina M Whalley, Andrew Bosworth, Charlotte Poxon, Kasun Wanigasooriya, Oliver Pickles, Mike Kidd, Alex Richter, Andrew D Beggs PHE Heartlands Lab: Husam Osman, Andrew Bosworth. Queen Elizabeth Hospital: Anna Casey |
| EPI_ISL_529431, EPI_ISL_529432 | Centre for Enzyme Innovation, University of Portsmouth / Translational Research Laboratory, Portsmouth Hospitals NHS Trust | COVID-19 Genomics UK (COG-UK) Consortium | Angela Beckett,Yann Bourgeois,Garry Scarlett,Sharon Glaysher,Scott Elliott,Kelly Bicknell,Robert Impey,Allyson Lloyd,Sarah Wyllie,Ethan Butcher,Anoop Chauhan,Samuel Robson |
| EPI_ISL_529433 | Virology Department, Sheffield Teaching Hospitals NHS Foundation Trust/Department of Infection, Immunity and | COVID-19 Genomics UK (COG-UK) Consortium | Thushan de Silva, Matthew Parker, Nikki Smith, Adri Angyal, Rebecca Brown, Luke Green, Rachel Tucker, Paul Parsons, Danielle Groves, Katie Johnson, Laura Carrilero, Alex Keeley, Dave Partridge, Matthew Wyles, Benjamin Lindsey, Mehmet Yavuz, Mohammad Raza, Cariad Evans |

| | | | |
|---|---|---|---|
| | Cardiovascular Disease, The Medical School, University of Sheffield | | |
| EPI_ISL_529434, EPI_ISL_529435 | University of Birmingham | COVID-19 Genomics UK (COG-UK) Consortium | Institute of Microbiology, University of Birmingham: Claire McMurray, Joanne Stockton, Samuel Nicholls, Radoslaw Poplawski, Will Rowe, Josh Quick, Nicholas Loman. University of Birmingham Testing Laboratory: Celina M Whalley, Andrew Bosworth, Charlotte Poxon, Kasun Wanigasooriya, Oliver Pickles, Mike Kidd, Alex Richter, Andrew D Beggs PHE Heartlands Lab: Husam Osman, Andrew Bosworth. Queen Elizabeth Hospital: Anna Casey |
| EPI_ISL_529436 | Queens Medical Centre, Clinical Microbiology Department / DeepSeq Nottingham | COVID-19 Genomics UK (COG-UK) Consortium | Gemma Clark, Wendy Smith, Manjinder Khakh, Vicki M Fleming, Michelle M Lister, Hannah Howson-Wells, Jonathan Ball, Patrick McClure, Joseph Chappell, Theocharis Tsoleridis, Nadine Holmes, Matthew Carlisle, Christopher Moore, Fei Sang, Johnny Debebe, Victoria Wright, Matthew Loose |
| EPI_ISL_529437 | Quadram Institute Bioscience | COVID-19 Genomics UK (COG-UK) Consortium | Dave J. Baker, Gemma L. Kay, Alp Aydin, Thanh Le-Viet, Steven Rudder, Ana P. Tedim, Anastasia Kolyva, Maria Diaz, Leonardo de Oliveira Martins, Nabil-Fareed Alikhan, Lizzie Meadows, Rachael Stanley, Ngozi Elumogo, Muhammed Yasir, Nicholas M. Thomson, Alexander J Trotter, Rachel Gilroy, Samuel Bloomfield, Claire Stuart, Andrew Bell, Reenesh Prakash, Samir Dervisevic, Alison E. Mather, John Wain, Mark Webber, Andrew J. Page, Justin O’Grady |
| EPI_ISL_529438, EPI_ISL_529439, EPI_ISL_529440, EPI_ISL_529441, EPI_ISL_529442, EPI_ISL_529443, EPI_ISL_529444 | University of Birmingham | COVID-19 Genomics UK (COG-UK) Consortium | Institute of Microbiology, University of Birmingham: Claire McMurray, Joanne Stockton, Samuel Nicholls, Radoslaw Poplawski, Will Rowe, Josh Quick, Nicholas Loman. University of Birmingham Testing Laboratory: Celina M Whalley, Andrew Bosworth, Charlotte Poxon, Kasun Wanigasooriya, Oliver Pickles, Mike Kidd, Alex Richter, Andrew D Beggs PHE Heartlands Lab: Husam Osman, Andrew Bosworth. Queen Elizabeth Hospital: Anna Casey |
| EPI_ISL_529445 | Department of Pathology, University of Cambridge | COVID-19 Genomics UK (COG-UK) Consortium | Luke W Meredith, M. Estée Török, Myra Hosmillo, William L. Hamilton, Martin D. Curran, Theresa Feltwell, Grant Hall, Anna Yakovleva, Fahad A Khokhar, Charlotte J. Houldcroft, Laura G Caller, Aminu S. Jahun, Sarah L. Caddy, Yasmin Chaudhry, Malte Pinckert, Ian Goodfellow |
| EPI_ISL_529446 | Virology Department, Sheffield Teaching Hospitals NHS Foundation Trust/Department of Infection, Immunity and Cardiovascular Disease, The Medical School, University of Sheffield | COVID-19 Genomics UK (COG-UK) Consortium | Thushan de Silva, Matthew Parker, Nikki Smith, Adri Angyal, Rebecca Brown, Luke Green, Rachel Tucker, Paul Parsons, Danielle Groves, Katie Johnson, Laura Carrilero, Alex Keeley, Dave Partridge, Matthew Wyles, Benjamin Lindsey, Mehmet Yavuz, Mohammad Raza, Cariad Evans |
| EPI_ISL_529447 | Wales Specialist Virology Centre Sequencing lab: Pathogen Genomics Unit | COVID-19 Genomics UK (COG-UK) Consortium | Catherine Moore, Johnathan Evans, Laura Gifford, Malorie Perry, Simon Cottrell, Angela Marchbank, Alec Birchley, Alexander Adams, Amy Gaskin, Bree Gatica-Wilcox, Jason Coombes, Joel Southgate, Lauren Gilbert, Lee Graham, Nicole Pacchiarini, Sara Kumziene-Summerhayes, Sarah Taylor, Sophie Jones, Sara Rey, Matthew Bull, Joanne Watkins, Sally Corden, Tom Connor |
| EPI_ISL_529448 | University of Birmingham | COVID-19 Genomics UK (COG-UK) Consortium | Institute of Microbiology, University of Birmingham: Claire McMurray, Joanne Stockton, Samuel Nicholls, Radoslaw Poplawski, Will Rowe, Josh Quick, Nicholas Loman. University of Birmingham Testing Laboratory: Celina M Whalley, Andrew Bosworth, Charlotte Poxon, Kasun Wanigasooriya, Oliver Pickles, Mike Kidd, Alex Richter, Andrew D Beggs PHE Heartlands Lab: Husam Osman, Andrew Bosworth. Queen Elizabeth Hospital: Anna Casey |
| EPI_ISL_529449 | Virology Department, Sheffield Teaching Hospitals NHS Foundation Trust/Department of Infection, Immunity and Cardiovascular Disease, The Medical School, University of Sheffield | COVID-19 Genomics UK (COG-UK) Consortium | Thushan de Silva, Matthew Parker, Nikki Smith, Adri Angyal, Rebecca Brown, Luke Green, Rachel Tucker, Paul Parsons, Danielle Groves, Katie Johnson, Laura Carrilero, Alex Keeley, Dave Partridge, Matthew Wyles, Benjamin Lindsey, Mehmet Yavuz, Mohammad Raza, Cariad Evans |
| EPI_ISL_529450 | Queens Medical Centre, Clinical Microbiology Department / DeepSeq Nottingham | COVID-19 Genomics UK (COG-UK) Consortium | Gemma Clark, Wendy Smith, Manjinder Khakh, Vicki M Fleming, Michelle M Lister, Hannah Howson-Wells, Jonathan Ball, Patrick McClure, Joseph Chappell, Theocharis Tsoleridis, Nadine Holmes, Matthew Carlisle, Christopher Moore, Fei Sang, Johnny Debebe, Victoria Wright, Matthew Loose |
| EPI_ISL_529451, EPI_ISL_529452 | Quadram Institute Bioscience | COVID-19 Genomics UK (COG-UK) Consortium | Dave J. Baker, Gemma L. Kay, Alp Aydin, Thanh Le-Viet, Steven Rudder, Ana P. Tedim, Anastasia Kolyva, Maria Diaz, Leonardo de Oliveira Martins, Nabil-Fareed Alikhan, Lizzie Meadows, Rachael Stanley, Ngozi Elumogo, Muhammed Yasir, Nicholas M. Thomson, Alexander J Trotter, Rachel Gilroy, Samuel Bloomfield, Claire Stuart, Andrew Bell, Reenesh Prakash, Samir Dervisevic, Alison E. Mather, John Wain, Mark Webber, Andrew J. Page, Justin O’Grady |
| EPI_ISL_529453 | University of Birmingham | COVID-19 Genomics UK (COG-UK) Consortium | Institute of Microbiology, University of Birmingham: Claire McMurray, Joanne Stockton, Samuel Nicholls, Radoslaw Poplawski, Will Rowe, Josh Quick, Nicholas Loman. University of Birmingham Testing Laboratory: Celina M Whalley, Andrew Bosworth, Charlotte Poxon, Kasun Wanigasooriya, Oliver Pickles, Mike Kidd, Alex Richter, Andrew D Beggs PHE Heartlands Lab: Husam Osman, Andrew Bosworth. Queen Elizabeth Hospital: Anna Casey |
| EPI_ISL_529454, EPI_ISL_529455, EPI_ISL_529456 | Queens Medical Centre, Clinical Microbiology Department / DeepSeq Nottingham | COVID-19 Genomics UK (COG-UK) Consortium | Gemma Clark, Wendy Smith, Manjinder Khakh, Vicki M Fleming, Michelle M Lister, Hannah Howson-Wells, Jonathan Ball, Patrick McClure, Joseph Chappell, Theocharis Tsoleridis, Nadine Holmes, Matthew Carlisle, Christopher Moore, Fei Sang, Johnny Debebe, Victoria Wright, Matthew Loose |
| EPI_ISL_529457, EPI_ISL_529458, EPI_ISL_529459, EPI_ISL_529460 | Quadram Institute Bioscience | COVID-19 Genomics UK (COG-UK) Consortium | Dave J. Baker, Gemma L. Kay, Alp Aydin, Thanh Le-Viet, Steven Rudder, Ana P. Tedim, Anastasia Kolyva, Maria Diaz, Leonardo de Oliveira Martins, Nabil-Fareed Alikhan, Lizzie Meadows, Rachael Stanley, Ngozi Elumogo, Muhammed Yasir, Nicholas M. Thomson, Alexander J Trotter, Rachel Gilroy, Samuel Bloomfield, Claire Stuart, Andrew Bell, Reenesh Prakash, Samir Dervisevic, Alison E. Mather, John Wain, Mark Webber, Andrew J. Page, Justin O’Grady |
| EPI_ISL_529461, EPI_ISL_529462 | University of Birmingham | COVID-19 Genomics UK (COG-UK) Consortium | Institute of Microbiology, University of Birmingham: Claire McMurray, Joanne Stockton, Samuel Nicholls, Radoslaw Poplawski, Will Rowe, Josh Quick, Nicholas Loman. University of Birmingham Testing Laboratory: Celina M Whalley, Andrew Bosworth, Charlotte Poxon, Kasun Wanigasooriya, Oliver Pickles, Mike Kidd, Alex Richter, Andrew D Beggs PHE Heartlands Lab: Husam Osman, Andrew Bosworth. Queen Elizabeth Hospital: Anna Casey |
| EPI_ISL_529463, EPI_ISL_529464 | Queens Medical Centre, Clinical Microbiology Department / DeepSeq Nottingham | COVID-19 Genomics UK (COG-UK) Consortium | Gemma Clark, Wendy Smith, Manjinder Khakh, Vicki M Fleming, Michelle M Lister, Hannah Howson-Wells, Jonathan Ball, Patrick McClure, Joseph Chappell, Theocharis Tsoleridis, Nadine Holmes, Matthew Carlisle, Christopher Moore, Fei Sang, Johnny Debebe, Victoria Wright, Matthew Loose |
| EPI_ISL_529465 | Centre for Enzyme Innovation, University of Portsmouth / Translational Research Laboratory, Portsmouth Hospitals NHS Trust | COVID-19 Genomics UK (COG-UK) Consortium | Angela Beckett,Yann Bourgeois,Garry Scarlett,Sharon Glaysher,Scott Elliott,Kelly Bicknell,Robert Impey,Allyson Lloyd,Sarah Wyllie,Ethan Butcher,Anoop Chauhan,Samuel Robson |
| EPI_ISL_529466 | Liverpool Clinical Laboratories | COVID-19 Genomics UK (COG-UK) Consortium | Sam Haldenby, Anita Lucaci, Steve Paterson, Julian Hiscox, Alistair Darby, M Almsaud, A Alrezaihi, Muhannad Alruwaili, Stuart D Armstrong, Jones Benjamin, Eleanor G Bentley, Anu Chawla, Jordan J Clark, Angela Cowell, Richard Eccles, Isabel García-Dorival, Matthew Gemmell, Alessandro Gerada, PKF Gilmore, Richard Gregory, Ximeng Han, Catherine Hartley, Margaret Hughes, Miren Iturriza-Gomara, James Johnson, L Luu, Jenifer Manson, Charlotte Nelson, Elaine O’Toole, Cassie Olateju, Rebekah Penrice-Randal , Lucille Rainbow, N.P Randle, Trevor Ian Robinson, Parul Sharma, Ghada T Shawli, James P Stewart, Neil Swainston, Ecaterina Vamos, Joanne Watts, Mark Whitehead |
| EPI_ISL_529467 | Queens Medical Centre, Clinical Microbiology Department / DeepSeq Nottingham | COVID-19 Genomics UK (COG-UK) Consortium | Gemma Clark, Wendy Smith, Manjinder Khakh, Vicki M Fleming, Michelle M Lister, Hannah Howson-Wells, Jonathan Ball, Patrick McClure, Joseph Chappell, Theocharis Tsoleridis, Nadine Holmes, Matthew Carlisle, Christopher Moore, Fei Sang, Johnny Debebe, Victoria Wright, Matthew Loose |
| EPI_ISL_529468 | University of Birmingham | COVID-19 Genomics UK (COG-UK) Consortium | Institute of Microbiology, University of Birmingham: Claire McMurray, Joanne Stockton, Samuel Nicholls, Radoslaw Poplawski, Will Rowe, Josh Quick, Nicholas Loman. University of Birmingham Testing Laboratory: Celina M Whalley, Andrew Bosworth, Charlotte Poxon, Kasun Wanigasooriya, Oliver Pickles, Mike Kidd, Alex Richter, Andrew D Beggs PHE Heartlands Lab: Husam Osman, Andrew Bosworth. Queen Elizabeth Hospital: Anna Casey |
| EPI_ISL_529469 | Quadram Institute Bioscience | COVID-19 Genomics UK (COG-UK) Consortium | Dave J. Baker, Gemma L. Kay, Alp Aydin, Thanh Le-Viet, Steven Rudder, Ana P. Tedim, Anastasia Kolyva, Maria Diaz, Leonardo de Oliveira Martins, Nabil-Fareed Alikhan, Lizzie Meadows, Rachael Stanley, Ngozi Elumogo, Muhammed Yasir, Nicholas M. Thomson, Alexander J Trotter, Rachel Gilroy, Samuel Bloomfield, Claire Stuart, Andrew Bell, Reenesh Prakash, Samir Dervisevic, Alison E. Mather, John Wain, Mark Webber, Andrew J. Page, Justin O’Grady |
| EPI_ISL_529470, EPI_ISL_529471, EPI_ISL_529472, EPI_ISL_529473, EPI_ISL_529474, EPI_ISL_529475 | University of Birmingham | COVID-19 Genomics UK (COG-UK) Consortium | Institute of Microbiology, University of Birmingham: Claire McMurray, Joanne Stockton, Samuel Nicholls, Radoslaw Poplawski, Will Rowe, Josh Quick, Nicholas Loman. University of Birmingham Testing Laboratory: Celina M Whalley, Andrew Bosworth, Charlotte Poxon, Kasun Wanigasooriya, Oliver Pickles, Mike Kidd, Alex Richter, Andrew D Beggs PHE Heartlands Lab: Husam Osman, Andrew Bosworth. Queen Elizabeth Hospital: Anna Casey |
| EPI_ISL_529476, EPI_ISL_529477 | Quadram Institute Bioscience | COVID-19 Genomics UK (COG-UK) Consortium | Dave J. Baker, Gemma L. Kay, Alp Aydin, Thanh Le-Viet, Steven Rudder, Ana P. Tedim, Anastasia Kolyva, Maria Diaz, Leonardo de Oliveira Martins, Nabil-Fareed Alikhan, Lizzie Meadows, Rachael Stanley, Ngozi Elumogo, Muhammed Yasir, Nicholas M. Thomson, Alexander J Trotter, Rachel Gilroy, Samuel Bloomfield, Claire Stuart, Andrew Bell, Reenesh Prakash, Samir Dervisevic, Alison E. Mather, John Wain, Mark Webber, Andrew J. Page, Justin O’Grady |
| EPI_ISL_529478 | Virology Department, Royal Infirmary of Edinburgh, NHS Lothian / School of Biological Sciences, University of Edinburgh / Institute of Genetics and Molecular Medicine, University of Edinburgh | COVID-19 Genomics UK (COG-UK) Consortium | McHugh M, Dewar R, Rooke S, Gallagher M, Balcaza C, O’Toole Á, Scher E, Hill V, McCrone JT, Colquhoun R, Yu X, Jackson B, Rambaut A, Williams TC, Templeton K |
| EPI_ISL_529479 | Queens Medical Centre, Clinical Microbiology Department / DeepSeq Nottingham | COVID-19 Genomics UK (COG-UK) Consortium | Gemma Clark, Wendy Smith, Manjinder Khakh, Vicki M Fleming, Michelle M Lister, Hannah Howson-Wells, Jonathan Ball, Patrick McClure, Joseph Chappell, Theocharis Tsoleridis, Nadine Holmes, Matthew Carlisle, Christopher Moore, Fei Sang, Johnny Debebe, Victoria Wright, Matthew Loose |
| EPI_ISL_529480 | University of Birmingham | COVID-19 Genomics UK (COG-UK) Consortium | Institute of Microbiology, University of Birmingham: Claire McMurray, Joanne Stockton, Samuel Nicholls, Radoslaw Poplawski, Will Rowe, Josh Quick, Nicholas Loman. University of Birmingham Testing Laboratory: Celina M Whalley, Andrew Bosworth, Charlotte Poxon, Kasun Wanigasooriya, Oliver Pickles, Mike Kidd, Alex Richter, Andrew D Beggs PHE Heartlands Lab: Husam Osman, Andrew Bosworth. Queen Elizabeth Hospital: Anna Casey |
| EPI_ISL_529481 | Centre for Enzyme Innovation, University of Portsmouth / Translational Research Laboratory, Portsmouth Hospitals NHS Trust | COVID-19 Genomics UK (COG-UK) Consortium | Angela Beckett,Yann Bourgeois,Garry Scarlett,Sharon Glaysher,Scott Elliott,Kelly Bicknell,Robert Impey,Allyson Lloyd,Sarah Wyllie,Ethan Butcher,Anoop Chauhan,Samuel Robson |
| EPI_ISL_529482 | University of Birmingham | COVID-19 Genomics UK (COG-UK) Consortium | Institute of Microbiology, University of Birmingham: Claire McMurray, Joanne Stockton, Samuel Nicholls, Radoslaw Poplawski, Will Rowe, Josh Quick, Nicholas Loman. University of Birmingham Testing Laboratory: Celina M Whalley, Andrew Bosworth, Charlotte Poxon, Kasun Wanigasooriya, Oliver Pickles, Mike Kidd, Alex Richter, Andrew D Beggs PHE Heartlands Lab: Husam Osman, Andrew Bosworth. Queen Elizabeth Hospital: Anna Casey |
| EPI_ISL_529483 | Queens Medical Centre, Clinical Microbiology Department / DeepSeq Nottingham | COVID-19 Genomics UK (COG-UK) Consortium | Gemma Clark, Wendy Smith, Manjinder Khakh, Vicki M Fleming, Michelle M Lister, Hannah Howson-Wells, Jonathan Ball, Patrick McClure, Joseph Chappell, Theocharis Tsoleridis, Nadine Holmes, Matthew Carlisle, Christopher Moore, Fei Sang, Johnny Debebe, Victoria Wright, Matthew Loose |
| EPI_ISL_529484 | Liverpool Clinical Laboratories | COVID-19 Genomics UK (COG-UK) Consortium | Sam Haldenby, Anita Lucaci, Steve Paterson, Julian Hiscox, Alistair Darby, M Almsaud, A Alrezaihi, Muhannad Alruwaili, Stuart D Armstrong, Jones Benjamin, Eleanor G Bentley, Anu Chawla, Jordan J Clark, Angela Cowell, Richard Eccles, Isabel García-Dorival, Matthew Gemmell, Alessandro Gerada, PKF Gilmore, Richard Gregory, Ximeng Han, Catherine Hartley, Margaret Hughes, Miren Iturriza-Gomara, James Johnson, L Luu, Jenifer Manson, Charlotte Nelson, Elaine O’Toole, Cassie Olateju, Rebekah Penrice-Randal , Lucille Rainbow, N.P Randle, Trevor Ian Robinson, Parul Sharma, Ghada T Shawli, James P Stewart, Neil Swainston, Ecaterina Vamos, Joanne Watts, Mark Whitehead |
| EPI_ISL_529485 | University of Birmingham | COVID-19 Genomics UK (COG-UK) Consortium | Institute of Microbiology, University of Birmingham: Claire McMurray, Joanne Stockton, Samuel Nicholls, Radoslaw Poplawski, Will Rowe, Josh Quick, Nicholas Loman. University of Birmingham Testing Laboratory: Celina M Whalley, Andrew Bosworth, Charlotte Poxon, Kasun Wanigasooriya, Oliver Pickles, Mike Kidd, Alex Richter, Andrew D Beggs PHE Heartlands Lab: Husam Osman, Andrew Bosworth. Queen Elizabeth Hospital: Anna Casey |
| EPI_ISL_529486 | Queens Medical Centre, Clinical Microbiology Department / DeepSeq Nottingham | COVID-19 Genomics UK (COG-UK) Consortium | Gemma Clark, Wendy Smith, Manjinder Khakh, Vicki M Fleming, Michelle M Lister, Hannah Howson-Wells, Jonathan Ball, Patrick McClure, Joseph Chappell, Theocharis Tsoleridis, Nadine Holmes, Matthew Carlisle, Christopher Moore, Fei Sang, Johnny Debebe, Victoria Wright, Matthew Loose |
| EPI_ISL_529487, EPI_ISL_529488, EPI_ISL_529489 | Quadram Institute Bioscience | COVID-19 Genomics UK (COG-UK) Consortium | Dave J. Baker, Gemma L. Kay, Alp Aydin, Thanh Le-Viet, Steven Rudder, Ana P. Tedim, Anastasia Kolyva, Maria Diaz, Leonardo de Oliveira Martins, Nabil-Fareed Alikhan, Lizzie Meadows, Rachael Stanley, Ngozi Elumogo, Muhammed Yasir, Nicholas M. Thomson, Alexander J Trotter, Rachel Gilroy, Samuel Bloomfield, Claire Stuart, Andrew Bell, Reenesh Prakash, Samir Dervisevic, Alison E. Mather, John Wain, Mark Webber, Andrew J. Page, Justin O’Grady |

| | | | |
|---|---|---|---|
| EPI_ISL_529490 | Liverpool Clinical Laboratories | COVID-19 Genomics UK (COG-UK) Consortium | Sam Haldenby, Anita Lucaci, Steve Paterson, Julian Hiscox, Alistair Darby, M Almsaud, A Alrezaihi, Muhannad Alruwaili, Stuart D Armstrong, Jones Benjamin, Eleanor G Bentley, Anu Chawla, Jordan J Clark, Angela Cowell, Richard Eccles, Isabel García-Dorival, Matthew Gemmell, Alessandro Gerada, PKF Gilmore, Richard Gregory, Ximeng Han, Catherine Hartley, Margaret Hughes, Miren Iturriza-Gomara, James Johnson, L Luu, Jenifer Manson, Charlotte Nelson, Elaine O'Toole, Cassie Olateju, Rebekah Penrice-Randal , Lucille Rainbow, N.P Randle, Trevor Ian Robinson, Parul Sharma, Ghada T Shawli, James P Stewart, Neil Swainston, Ecaterina Vamos, Joanne Watts, Mark Whitehead |
| EPI_ISL_529491 | Centre for Enzyme Innovation, University of Portsmouth / Translational Research Laboratory, Portsmouth Hospitals NHS Trust | COVID-19 Genomics UK (COG-UK) Consortium | Angela Beckett,Yann Bourgeois,Garry Scarlett,Sharon Glaysher,Scott Elliott,Kelly Bicknell,Robert Impey,Allyson Lloyd,Sarah Wyllie,Ethan Butcher,Anoop Chauhan,Samuel Robson |
| EPI_ISL_529492 | Quadram Institute Bioscience | COVID-19 Genomics UK (COG-UK) Consortium | Dave J. Baker, Gemma L. Kay, Alp Aydin, Thanh Le-Viet, Steven Rudder, Ana P. Tedim, Anastasia Kolyva, Maria Diaz, Leonardo de Oliveira Martins, Nabil-Fareed Alikhan, Lizzie Meadows, Rachael Stanley, Ngozi Elumogo, Muhammed Yasir, Nicholas M. Thomson, Alexander J Trotter, Rachel Gilroy, Samuel Bloomfield, Claire Stuart, Andrew Bell, Reenesh Prakash, Samir Dervisevic, Alison E. Mather, John Wain, Mark Webber, Andrew J. Page, Justin O'Grady |
| EPI_ISL_529493, EPI_ISL_529494, EPI_ISL_529495, EPI_ISL_529496, EPI_ISL_529497 | University of Birmingham | COVID-19 Genomics UK (COG-UK) Consortium | Institute of Microbiology, University of Birmingham: Claire McMurray, Joanne Stockton, Samuel Nicholls, Radoslaw Poplawski, Will Rowe, Josh Quick, Nicholas Loman. University of Birmingham Testing Laboratory: Celina M Whalley, Andrew Bosworth, Charlotte Poxon, Kasun Wanigasooriya, Oliver Pickles, Mike Kidd, Alex Richter, Andrew D Beggs PHE Heartlands Lab: Husam Osman, Andrew Bosworth. Queen Elizabeth Hospital: Anna Casey |
| EPI_ISL_529498 | Liverpool Clinical Laboratories | COVID-19 Genomics UK (COG-UK) Consortium | Sam Haldenby, Anita Lucaci, Steve Paterson, Julian Hiscox, Alistair Darby, M Almsaud, A Alrezaihi, Muhannad Alruwaili, Stuart D Armstrong, Jones Benjamin, Eleanor G Bentley, Anu Chawla, Jordan J Clark, Angela Cowell, Richard Eccles, Isabel García-Dorival, Matthew Gemmell, Alessandro Gerada, PKF Gilmore, Richard Gregory, Ximeng Han, Catherine Hartley, Margaret Hughes, Miren Iturriza-Gomara, James Johnson, L Luu, Jenifer Manson, Charlotte Nelson, Elaine O'Toole, Cassie Olateju, Rebekah Penrice-Randal , Lucille Rainbow, N.P Randle, Trevor Ian Robinson, Parul Sharma, Ghada T Shawli, James P Stewart, Neil Swainston, Ecaterina Vamos, Joanne Watts, Mark Whitehead |
| EPI_ISL_529499 | University of Birmingham | COVID-19 Genomics UK (COG-UK) Consortium | Institute of Microbiology, University of Birmingham: Claire McMurray, Joanne Stockton, Samuel Nicholls, Radoslaw Poplawski, Will Rowe, Josh Quick, Nicholas Loman. University of Birmingham Testing Laboratory: Celina M Whalley, Andrew Bosworth, Charlotte Poxon, Kasun Wanigasooriya, Oliver Pickles, Mike Kidd, Alex Richter, Andrew D Beggs PHE Heartlands Lab: Husam Osman, Andrew Bosworth. Queen Elizabeth Hospital: Anna Casey |
| EPI_ISL_529500, EPI_ISL_529501, EPI_ISL_529502, EPI_ISL_529503 | Virology Department, Sheffield Teaching Hospitals NHS Foundation Trust/Department of Infection, Immunity and Cardiovascular Disease, The Medical School, University of Sheffield | COVID-19 Genomics UK (COG-UK) Consortium | Thushan de Silva, Matthew Parker, Nikki Smith, Adri Angyal, Rebecca Brown, Luke Green, Rachel Tucker, Paul Parsons, Danielle Groves, Katie Johnson, Laura Carrilero, Alex Keeley, Dave Partridge, Matthew Wyles, Benjamin Lindsey, Mehmet Yavuz, Mohammad Raza, Cariad Evans |
| EPI_ISL_529504 | Queens Medical Centre, Clinical Microbiology Department / DeepSeq Nottingham | COVID-19 Genomics UK (COG-UK) Consortium | Gemma Clark, Wendy Smith, Manjinder Khakh, Vicki M Fleming, Michelle M Lister, Hannah Howson-Wells, Jonathan Ball, Patrick McClure, Joseph Chappell, Theocharis Tsoleridis, Nadine Holmes, Matthew Carlisle, Christopher Moore, Fei Sang, Johnny Debebe, Victoria Wright, Matthew Loose |
| EPI_ISL_529505 | Wales Specialist Virology Centre Sequencing lab: Pathogen Genomics Unit | COVID-19 Genomics UK (COG-UK) Consortium | Catherine Moore, Johnathan Evans, Laura Gifford, Malorie Perry, Simon Cottrell, Angela Marchbank, Alec Birchley, Alexander Adams, Amy Gaskin, Bree Gatica-Wilcox, Jason Coombes, Joel Southgate, Lauren Gilbert, Lee Graham, Nicole Pacchiarini, Sara Kumziene-Summerhayes, Sarah Taylor, Sophie Jones, Sara Rey, Matthew Bull, Joanne Watkins, Sally Corden, Tom Connor |
| EPI_ISL_529506, EPI_ISL_529507 | Quadram Institute Bioscience | COVID-19 Genomics UK (COG-UK) Consortium | Dave J. Baker, Gemma L. Kay, Alp Aydin, Thanh Le-Viet, Steven Rudder, Ana P. Tedim, Anastasia Kolyva, Maria Diaz, Leonardo de Oliveira Martins, Nabil-Fareed Alikhan, Lizzie Meadows, Rachael Stanley, Ngozi Elumogo, Muhammed Yasir, Nicholas M. Thomson, Alexander J Trotter, Rachel Gilroy, Samuel Bloomfield, Claire Stuart, Andrew Bell, Reenesh Prakash, Samir Dervisevic, Alison E. Mather, John Wain, Mark Webber, Andrew J. Page, Justin O'Grady |
| EPI_ISL_529508 | University of Birmingham | COVID-19 Genomics UK (COG-UK) Consortium | Institute of Microbiology, University of Birmingham: Claire McMurray, Joanne Stockton, Samuel Nicholls, Radoslaw Poplawski, Will Rowe, Josh Quick, Nicholas Loman. University of Birmingham Testing Laboratory: Celina M Whalley, Andrew Bosworth, Charlotte Poxon, Kasun Wanigasooriya, Oliver Pickles, Mike Kidd, Alex Richter, Andrew D Beggs PHE Heartlands Lab: Husam Osman, Andrew Bosworth. Queen Elizabeth Hospital: Anna Casey |
| EPI_ISL_529509, EPI_ISL_529510 | Liverpool Clinical Laboratories | COVID-19 Genomics UK (COG-UK) Consortium | Sam Haldenby, Anita Lucaci, Steve Paterson, Julian Hiscox, Alistair Darby, M Almsaud, A Alrezaihi, Muhannad Alruwaili, Stuart D Armstrong, Jones Benjamin, Eleanor G Bentley, Anu Chawla, Jordan J Clark, Angela Cowell, Richard Eccles, Isabel García-Dorival, Matthew Gemmell, Alessandro Gerada, PKF Gilmore, Richard Gregory, Ximeng Han, Catherine Hartley, Margaret Hughes, Miren Iturriza-Gomara, James Johnson, L Luu, Jenifer Manson, Charlotte Nelson, Elaine O'Toole, Cassie Olateju, Rebekah Penrice-Randal , Lucille Rainbow, N.P Randle, Trevor Ian Robinson, Parul Sharma, Ghada T Shawli, James P Stewart, Neil Swainston, Ecaterina Vamos, Joanne Watts, Mark Whitehead |
| EPI_ISL_529511 | Centre for Enzyme Innovation, University of Portsmouth / Translational Research Laboratory, Portsmouth Hospitals NHS Trust | COVID-19 Genomics UK (COG-UK) Consortium | Angela Beckett,Yann Bourgeois,Garry Scarlett,Sharon Glaysher,Scott Elliott,Kelly Bicknell,Robert Impey,Allyson Lloyd,Sarah Wyllie,Ethan Butcher,Anoop Chauhan,Samuel Robson |
| EPI_ISL_529512 | University of Birmingham | COVID-19 Genomics UK (COG-UK) Consortium | Institute of Microbiology, University of Birmingham: Claire McMurray, Joanne Stockton, Samuel Nicholls, Radoslaw Poplawski, Will Rowe, Josh Quick, Nicholas Loman. University of Birmingham Testing Laboratory: Celina M Whalley, Andrew Bosworth, Charlotte Poxon, Kasun Wanigasooriya, Oliver Pickles, Mike Kidd, Alex Richter, Andrew D Beggs PHE Heartlands Lab: Husam Osman, Andrew Bosworth. Queen Elizabeth Hospital: Anna Casey |
| EPI_ISL_529513 | Liverpool Clinical Laboratories | COVID-19 Genomics UK (COG-UK) Consortium | Sam Haldenby, Anita Lucaci, Steve Paterson, Julian Hiscox, Alistair Darby, M Almsaud, A Alrezaihi, Muhannad Alruwaili, Stuart D Armstrong, Jones Benjamin, Eleanor G Bentley, Anu Chawla, Jordan J Clark, Angela Cowell, Richard Eccles, Isabel García-Dorival, Matthew Gemmell, Alessandro Gerada, PKF Gilmore, Richard Gregory, Ximeng Han, Catherine Hartley, Margaret Hughes, Miren Iturriza-Gomara, James Johnson, L Luu, Jenifer Manson, Charlotte Nelson, Elaine O'Toole, Cassie Olateju, Rebekah Penrice-Randal , Lucille Rainbow, N.P Randle, Trevor Ian Robinson, Parul Sharma, Ghada T Shawli, James P Stewart, Neil Swainston, Ecaterina Vamos, Joanne Watts, Mark Whitehead |
| EPI_ISL_529514, EPI_ISL_529515 | University of Birmingham | COVID-19 Genomics UK (COG-UK) Consortium | Institute of Microbiology, University of Birmingham: Claire McMurray, Joanne Stockton, Samuel Nicholls, Radoslaw Poplawski, Will Rowe, Josh Quick, Nicholas Loman. University of Birmingham Testing Laboratory: Celina M Whalley, Andrew Bosworth, Charlotte Poxon, Kasun Wanigasooriya, Oliver Pickles, Mike Kidd, Alex Richter, Andrew D Beggs PHE Heartlands Lab: Husam Osman, Andrew Bosworth. Queen Elizabeth Hospital: Anna Casey |
| EPI_ISL_529516 | Liverpool Clinical Laboratories | COVID-19 Genomics UK (COG-UK) Consortium | Sam Haldenby, Anita Lucaci, Steve Paterson, Julian Hiscox, Alistair Darby, M Almsaud, A Alrezaihi, Muhannad Alruwaili, Stuart D Armstrong, Jones Benjamin, Eleanor G Bentley, Anu Chawla, Jordan J Clark, Angela Cowell, Richard Eccles, Isabel García-Dorival, Matthew Gemmell, Alessandro Gerada, PKF Gilmore, Richard Gregory, Ximeng Han, Catherine Hartley, Margaret Hughes, Miren Iturriza-Gomara, James Johnson, L Luu, Jenifer Manson, Charlotte Nelson, Elaine O'Toole, Cassie Olateju, Rebekah Penrice-Randal , Lucille Rainbow, N.P Randle, Trevor Ian Robinson, Parul Sharma, Ghada T Shawli, James P Stewart, Neil Swainston, Ecaterina Vamos, Joanne Watts, Mark Whitehead |
| EPI_ISL_529517, EPI_ISL_529518 | University of Birmingham | COVID-19 Genomics UK (COG-UK) Consortium | Institute of Microbiology, University of Birmingham: Claire McMurray, Joanne Stockton, Samuel Nicholls, Radoslaw Poplawski, Will Rowe, Josh Quick, Nicholas Loman. University of Birmingham Testing Laboratory: Celina M Whalley, Andrew Bosworth, Charlotte Poxon, Kasun Wanigasooriya, Oliver Pickles, Mike Kidd, Alex Richter, Andrew D Beggs PHE Heartlands Lab: Husam Osman, Andrew Bosworth. Queen Elizabeth Hospital: Anna Casey |
| EPI_ISL_529519 | Queens Medical Centre, Clinical Microbiology Department / DeepSeq Nottingham | COVID-19 Genomics UK (COG-UK) Consortium | Gemma Clark, Wendy Smith, Manjinder Khakh, Vicki M Fleming, Michelle M Lister, Hannah Howson-Wells, Jonathan Ball, Patrick McClure, Joseph Chappell, Theocharis Tsoleridis, Nadine Holmes, Matthew Carlisle, Christopher Moore, Fei Sang, Johnny Debebe, Victoria Wright, Matthew Loose |
| EPI_ISL_529520 | University of Birmingham | COVID-19 Genomics UK (COG-UK) Consortium | Institute of Microbiology, University of Birmingham: Claire McMurray, Joanne Stockton, Samuel Nicholls, Radoslaw Poplawski, Will Rowe, Josh Quick, Nicholas Loman. University of Birmingham Testing Laboratory: Celina M Whalley, Andrew Bosworth, Charlotte Poxon, Kasun Wanigasooriya, Oliver Pickles, Mike Kidd, Alex Richter, Andrew D Beggs PHE Heartlands Lab: Husam Osman, Andrew Bosworth. Queen Elizabeth Hospital: Anna Casey |
| EPI_ISL_529521 | Virology Department, Sheffield Teaching Hospitals NHS Foundation Trust/Department of Infection, Immunity and Cardiovascular Disease, The Medical School, University of Sheffield | COVID-19 Genomics UK (COG-UK) Consortium | Thushan de Silva, Matthew Parker, Nikki Smith, Adri Angyal, Rebecca Brown, Luke Green, Rachel Tucker, Paul Parsons, Danielle Groves, Katie Johnson, Laura Carrilero, Alex Keeley, Dave Partridge, Matthew Wyles, Benjamin Lindsey, Mehmet Yavuz, Mohammad Raza, Cariad Evans |
| EPI_ISL_529522 | University of Birmingham | COVID-19 Genomics UK (COG-UK) Consortium | Institute of Microbiology, University of Birmingham: Claire McMurray, Joanne Stockton, Samuel Nicholls, Radoslaw Poplawski, Will Rowe, Josh Quick, Nicholas Loman. University of Birmingham Testing Laboratory: Celina M Whalley, Andrew Bosworth, Charlotte Poxon, Kasun Wanigasooriya, Oliver Pickles, Mike Kidd, Alex Richter, Andrew D Beggs PHE Heartlands Lab: Husam Osman, Andrew Bosworth. Queen Elizabeth Hospital: Anna Casey |
| EPI_ISL_529523 | Quadram Institute Bioscience | COVID-19 Genomics UK (COG-UK) Consortium | Dave J. Baker, Gemma L. Kay, Alp Aydin, Thanh Le-Viet, Steven Rudder, Ana P. Tedim, Anastasia Kolyva, Maria Diaz, Leonardo de Oliveira Martins, Nabil-Fareed Alikhan, Lizzie Meadows, Rachael Stanley, Ngozi Elumogo, Muhammed Yasir, Nicholas M. Thomson, Alexander J Trotter, Rachel Gilroy, Samuel Bloomfield, Claire Stuart, Andrew Bell, Reenesh Prakash, Samir Dervisevic, Alison E. Mather, John Wain, Mark Webber, Andrew J. Page, Justin O'Grady |
| EPI_ISL_529524 | Department of Pathology, University of Cambridge | COVID-19 Genomics UK (COG-UK) Consortium | Luke W Meredith, M. Estée Török, Myra Hosmillo, William L. Hamilton, Martin D. Curran, Theresa Feltwell, Grant Hall, Anna Yakovleva, Fahad A Khokhar, Charlotte J. Houldcroft, Laura G Caller, Aminu S. Jahun, Sarah L. Caddy, Yasmin Chaudhry, Malte Pinckert, Ian Goodfellow |
| EPI_ISL_529525 | Centre for Enzyme Innovation, University of Portsmouth / Translational Research Laboratory, Portsmouth Hospitals NHS Trust | COVID-19 Genomics UK (COG-UK) Consortium | Angela Beckett,Yann Bourgeois,Garry Scarlett,Sharon Glaysher,Scott Elliott,Kelly Bicknell,Robert Impey,Allyson Lloyd,Sarah Wyllie,Ethan Butcher,Anoop Chauhan,Samuel Robson |
| EPI_ISL_529526 | Quadram Institute Bioscience | COVID-19 Genomics UK (COG-UK) Consortium | Dave J. Baker, Gemma L. Kay, Alp Aydin, Thanh Le-Viet, Steven Rudder, Ana P. Tedim, Anastasia Kolyva, Maria Diaz, Leonardo de Oliveira Martins, Nabil-Fareed Alikhan, Lizzie Meadows, Rachael Stanley, Ngozi Elumogo, Muhammed Yasir, Nicholas M. Thomson, Alexander J Trotter, Rachel Gilroy, Samuel Bloomfield, Claire Stuart, Andrew Bell, Reenesh Prakash, Samir Dervisevic, Alison E. Mather, John Wain, Mark Webber, Andrew J. Page, Justin O'Grady |
| EPI_ISL_529527 | Wales Specialist Virology Centre Sequencing lab: Pathogen Genomics Unit | COVID-19 Genomics UK (COG-UK) Consortium | Catherine Moore, Johnathan Evans, Laura Gifford, Malorie Perry, Simon Cottrell, Angela Marchbank, Alec Birchley, Alexander Adams, Amy Gaskin, Bree Gatica-Wilcox, Jason Coombes, Joel Southgate, Lauren Gilbert, Lee Graham, Nicole Pacchiarini, Sara Kumziene-Summerhayes, Sarah Taylor, Sophie Jones, Sara Rey, Matthew Bull, Joanne Watkins, Sally Corden, Tom Connor |
| EPI_ISL_529528, EPI_ISL_529529, EPI_ISL_529530, EPI_ISL_529531, EPI_ISL_529532, EPI_ISL_529533, EPI_ISL_529534, EPI_ISL_529535, EPI_ISL_529536, EPI_ISL_529537, EPI_ISL_529538, EPI_ISL_529539, EPI_ISL_529540, EPI_ISL_529541, EPI_ISL_529542, EPI_ISL_529543, EPI_ISL_529544, EPI_ISL_529545, EPI_ISL_529546, EPI_ISL_529547, EPI_ISL_529548, EPI_ISL_529549, EPI_ISL_529550, EPI_ISL_529551, EPI_ISL_529552, EPI_ISL_529553, EPI_ISL_529554, EPI_ISL_529555, EPI_ISL_529556 | | | |
| see above | Queens Medical Centre, Clinical Microbiology Department / DeepSeq Nottingham | COVID-19 Genomics UK (COG-UK) Consortium | Gemma Clark, Wendy Smith, Manjinder Khakh, Vicki M Fleming, Michelle M Lister, Hannah Howson-Wells, Jonathan Ball, Patrick McClure, Joseph Chappell, Theocharis Tsoleridis, Nadine Holmes, Matthew Carlisle, Christopher Moore, Fei Sang, Johnny Debebe, Victoria Wright, Matthew Loose |
| EPI_ISL_529557, EPI_ISL_529558, EPI_ISL_529559, EPI_ISL_529560, EPI_ISL_529561, EPI_ISL_529562, EPI_ISL_529563, EPI_ISL_529564, EPI_ISL_529565, EPI_ISL_529566, EPI_ISL_529567, EPI_ISL_529568, EPI_ISL_529569, EPI_ISL_529570, EPI_ISL_529571, EPI_ISL_529572, EPI_ISL_529573, EPI_ISL_529574, EPI_ISL_529575, EPI_ISL_529576, EPI_ISL_529577, EPI_ISL_529578, EPI_ISL_529579, EPI_ISL_529580, EPI_ISL_529581, EPI_ISL_529582, EPI_ISL_529583, EPI_ISL_529584, EPI_ISL_529585, EPI_ISL_529586, EPI_ISL_529587, EPI_ISL_529588, EPI_ISL_529589 | | | |
| see above | Quadram Institute Bioscience | COVID-19 Genomics UK (COG-UK) Consortium | Dave J. Baker, Gemma L. Kay, Alp Aydin, Thanh Le-Viet, Steven Rudder, Ana P. Tedim, Anastasia Kolyva, Maria Diaz, Leonardo de Oliveira Martins, Nabil-Fareed Alikhan, Lizzie Meadows, Rachael Stanley, Ngozi Elumogo, Muhammed Yasir, Nicholas M. Thomson, Alexander J Trotter, Rachel Gilroy, Samuel Bloomfield, Claire Stuart, Andrew Bell, Reenesh Prakash, Samir Dervisevic, Alison E. Mather, John Wain, Mark Webber, Andrew J. Page, Justin O'Grady |
| EPI_ISL_529590, EPI_ISL_529591, EPI_ISL_529592, EPI_ISL_529593, EPI_ISL_529594, EPI_ISL_529595, EPI_ISL_529596, EPI_ISL_529597, EPI_ISL_529598, EPI_ISL_529599, EPI_ISL_529600, EPI_ISL_529601, EPI_ISL_529602, EPI_ISL_529603, EPI_ISL_529604, EPI_ISL_529605, EPI_ISL_529606, EPI_ISL_529607, EPI_ISL_529608, EPI_ISL_529609, EPI_ISL_529610, EPI_ISL_529611, EPI_ISL_529612, EPI_ISL_529613, EPI_ISL_529614, EPI_ISL_529615, EPI_ISL_529616, EPI_ISL_529617, EPI_ISL_529618, EPI_ISL_529619, EPI_ISL_529620, EPI_ISL_529621, EPI_ISL_529622, EPI_ISL_529623, EPI_ISL_529624, EPI_ISL_529625, EPI_ISL_529626, EPI_ISL_529627, EPI_ISL_529628, EPI_ISL_529629, EPI_ISL_529630, EPI_ISL_529631, EPI_ISL_529632, EPI_ISL_529633, EPI_ISL_529634, EPI_ISL_529635, EPI_ISL_529636, EPI_ISL_529637, EPI_ISL_529638, EPI_ISL_529639, EPI_ISL_529640, EPI_ISL_529641, EPI_ISL_529642, EPI_ISL_529643, EPI_ISL_529644, EPI_ISL_529645, EPI_ISL_529646, EPI_ISL_529647, EPI_ISL_529648, EPI_ISL_529649, EPI_ISL_529650, EPI_ISL_529651, EPI_ISL_529652, EPI_ISL_529653, EPI_ISL_529654, EPI_ISL_529655 | | | |

| | | | |
|---|---|---|---|
| see above | University of Birmingham | COVID-19 Genomics UK (COG-UK) Consortium | Institute of Microbiology, University of Birmingham: Claire McMurray, Joanne Stockton, Samuel Nicholls, Radoslaw Poplawski, Will Rowe, Josh Quick, Nicholas Loman. University of Birmingham Testing Laboratory: Celina M Whalley, Andrew Bosworth, Charlotte Poxon, Kasun Wanigasooriya, Oliver Pickles, Mike Kidd, Alex Richter, Andrew D Beggs PHE Heartlands Lab: Husam Osman, Andrew Bosworth. Queen Elizabeth Hospital: Anna Casey |
| EPI_ISL_529656, EPI_ISL_529657, EPI_ISL_529658, EPI_ISL_529659, EPI_ISL_529660 | Department of Pathology, University of Cambridge | COVID-19 Genomics UK (COG-UK) Consortium | Luke W Meredith, M. Estée Török, Myra Hosmillo, William L. Hamilton, Martin D. Curran, Theresa Feltwell, Grant Hall, Anna Yakovleva, Fahad A Khokhar, Charlotte J. Houldcroft, Laura G Caller, Aminu S. Jahun, Sarah L. Caddy, Yasmin Chaudhry, Malte Pinckert, Ian Goodfellow |
| EPI_ISL_529661, EPI_ISL_529662, EPI_ISL_529663, EPI_ISL_529664, EPI_ISL_529665, EPI_ISL_529666 | University of Birmingham | COVID-19 Genomics UK (COG-UK) Consortium | Institute of Microbiology, University of Birmingham: Claire McMurray, Joanne Stockton, Samuel Nicholls, Radoslaw Poplawski, Will Rowe, Josh Quick, Nicholas Loman. University of Birmingham Testing Laboratory: Celina M Whalley, Andrew Bosworth, Charlotte Poxon, Kasun Wanigasooriya, Oliver Pickles, Mike Kidd, Alex Richter, Andrew D Beggs PHE Heartlands Lab: Husam Osman, Andrew Bosworth. Queen Elizabeth Hospital: Anna Casey |
| EPI_ISL_529667 | Queens Medical Centre, Clinical Microbiology Department / DeepSeq Nottingham | COVID-19 Genomics UK (COG-UK) Consortium | Gemma Clark, Wendy Smith, Manjinder Khakh, Vicki M Fleming, Michelle M Lister, Hannah Howson-Wells, Jonathan Ball, Patrick McClure, Joseph Chappell, Theocharis Tsoleridis, Nadine Holmes, Matthew Carlisle, Christopher Moore, Fei Sang, Johnny Debebe, Victoria Wright, Matthew Loose |
| EPI_ISL_529668, EPI_ISL_529669, EPI_ISL_529670, EPI_ISL_529671, EPI_ISL_529672, EPI_ISL_529673, EPI_ISL_529674 | Centre for Enzyme Innovation, University of Portsmouth / Translational Research Laboratory, Portsmouth Hospitals NHS Trust | COVID-19 Genomics UK (COG-UK) Consortium | Angela Beckett,Yann Bourgeois,Garry Scarlett,Sharon Glaysher,Scott Elliott,Kelly Bicknell,Robert Impey,Allyson Lloyd,Sarah Wyllie,Ethan Butcher,Anoop Chauhan,Samuel Robson |
| EPI_ISL_529675, EPI_ISL_529676, EPI_ISL_529677, EPI_ISL_529678, EPI_ISL_529679, EPI_ISL_529680, EPI_ISL_529681, EPI_ISL_529682, EPI_ISL_529683 | Virology Department, Sheffield Teaching Hospitals NHS Foundation Trust/Department of Infection, Immunity and Cardiovascular Disease, The Medical School, University of Sheffield | COVID-19 Genomics UK (COG-UK) Consortium | Thushan de Silva, Matthew Parker, Nikki Smith, Adri Angyal, Rebecca Brown, Luke Green, Rachel Tucker, Paul Parsons, Danielle Groves, Katie Johnson, Laura Carrilero, Alex Keeley, Dave Partridge, Matthew Wyles, Benjamin Lindsey, Mehmet Yavuz, Mohammad Raza, Cariad Evans |
| EPI_ISL_529684, EPI_ISL_529685, EPI_ISL_529686, EPI_ISL_529687, EPI_ISL_529688 | West of Scotland Specialist Virology Centre, NHSGGC / MRC-University of Glasgow Centre for Virus Research | COVID-19 Genomics UK (COG-UK) Consortium | Ana da Silva Filipe, Natasha Johnson, Kathy Smollett, Daniel Mair, Stephen Carmichael, Lily Tong, Jenna Nichols, Elihu Aranday-Cortes, Kirstyn Brunker, Yasmin Parr, Alice Broos, Kyriaki Nomikou; Sarah McDonald, Marc Niebel, Patawee Asamaphan; Richard Orton, Joseph Hughes, Sreenu Vattipally, David L Robertson; Alasdair MacLean, Rory Gunson; Kathy Li, Natasha Jesudason, Rajiv Shah, James Shepherd, Antonia Ho, Emma Thomson |
| EPI_ISL_529689, EPI_ISL_529690, EPI_ISL_529691, EPI_ISL_529692, EPI_ISL_529693, EPI_ISL_529694, EPI_ISL_529695, EPI_ISL_529696 | Virology Department, Royal Infirmary of Edinburgh, NHS Lothian / School of Biological Sciences, University of Edinburgh / Institute of Genetics and Molecular Medicine, University of Edinburgh | COVID-19 Genomics UK (COG-UK) Consortium | McHugh M, Dewar R, Rooke S, Gallagher M, Balcaza C, O'Toole Á, Scher E, Hill V, McCrone JT, Colquhoun R, Yu X, Jackson B, Rambaut A, Williams TC, Templeton K |
| EPI_ISL_529697, EPI_ISL_529698, EPI_ISL_529699, EPI_ISL_529700, EPI_ISL_529701, EPI_ISL_529702, EPI_ISL_529703, EPI_ISL_529704, EPI_ISL_529705, EPI_ISL_529706, EPI_ISL_529707, EPI_ISL_529708, EPI_ISL_529709, EPI_ISL_529710, EPI_ISL_529711, EPI_ISL_529712, EPI_ISL_529713, EPI_ISL_529714, EPI_ISL_529715, EPI_ISL_529716, EPI_ISL_529717 | | | |
| see above | Wales Specialist Virology Centre Sequencing lab: Pathogen Genomics Unit | COVID-19 Genomics UK (COG-UK) Consortium | Catherine Moore, Johnathan Evans, Laura Gifford, Malorie Perry, Simon Cottrell, Angela Marchbank, Alec Birchley, Alexander Adams, Amy Gaskin, Bree Gatica-Wilcox, Jason Coombes, Joel Southgate, Lauren Gilbert, Lee Graham, Nicole Pacchiarini, Sara Kumziene-Summerhayes, Sarah Taylor, Sophie Jones, Sara Rey, Matthew Bull, Joanne Watkins, Sally Corden, Tom Connor |
| EPI_ISL_529718 | Santo Borromeus Hospital | Laboratorium Kesehatan Provinsi Jawa Barat; School of Life Sciences and Technology & School of Pharmacy-Institut Teknologi Bandung; Molecular Genetics Laboratory-Faculty of Medicine-Universitas Padjadjaran | Ema Rahmawati, Marselina Irasonia Tan, Yunia Sribudiani, Catur Riani, Azzania Fibriani, Husna Nugrahapraja, Tarwadi, Savira Ekawardhani, Hesti Lina Wiraswati, Ryan Bayusantika Ristandi, Rifky Waluyajati Rachman, Cut Nur Cinthia Alamanda, Lia Faridah, Gusti Ayu Prani Pradani, Adelina Khristiani Rahayu, Hammam Riza, Sony Solistia Wirawan, Agung Eru Wibowo, Irvan Faizal |
| EPI_ISL_529719 | Sumedang Regional General Hospital | Laboratorium Kesehatan Provinsi Jawa Barat; School of Life Sciences and Technology & School of Pharmacy-Institut Teknologi Bandung; Molecular Genetics Laboratory-Faculty of Medicine-Universitas Padjadjaran | Ema Rahmawati, Marselina Irasonia Tan, Yunia Sribudiani, Catur Riani, Azzania Fibriani, Husna Nugrahapraja, Tarwadi, Savira Ekawardhani, Hesti Lina Wiraswati, Ryan Bayusantika Ristandi, Rifky Waluyajati Rachman, Cut Nur Cinthia Alamanda, Lia Faridah, Gusti Ayu Prani Pradani, Adelina Khristiani Rahayu, Hammam Riza, Sony Solistia Wirawan, Agung Eru Wibowo, Irvan Faizal |
| EPI_ISL_529720, EPI_ISL_529721, EPI_ISL_529722, EPI_ISL_529723, EPI_ISL_529724, EPI_ISL_529725, EPI_ISL_529726, EPI_ISL_529727, EPI_ISL_529728, EPI_ISL_529729, EPI_ISL_529730, EPI_ISL_529731, EPI_ISL_529732, EPI_ISL_529733, EPI_ISL_529734, EPI_ISL_529735, EPI_ISL_529736, EPI_ISL_529737, EPI_ISL_529738, EPI_ISL_529739, EPI_ISL_529740, EPI_ISL_529741, EPI_ISL_529742, EPI_ISL_529743, EPI_ISL_529744, EPI_ISL_529745, EPI_ISL_529746, EPI_ISL_529747, EPI_ISL_529748, EPI_ISL_529749, EPI_ISL_529750, EPI_ISL_529751, EPI_ISL_529752, EPI_ISL_529753, EPI_ISL_529754, EPI_ISL_529755, EPI_ISL_529756, EPI_ISL_529757, EPI_ISL_529758, EPI_ISL_529759, EPI_ISL_529760, EPI_ISL_529761, EPI_ISL_529762, EPI_ISL_529763, EPI_ISL_529764, EPI_ISL_529765, EPI_ISL_529766, EPI_ISL_529767, EPI_ISL_529768, EPI_ISL_529769, EPI_ISL_529770, EPI_ISL_529771, EPI_ISL_529772, EPI_ISL_529773, EPI_ISL_529774, EPI_ISL_529775, EPI_ISL_529776, EPI_ISL_529777, EPI_ISL_529778, EPI_ISL_529779, EPI_ISL_529780, EPI_ISL_529781, EPI_ISL_529782, EPI_ISL_529783, EPI_ISL_529784, EPI_ISL_529785, EPI_ISL_529786, EPI_ISL_529787, EPI_ISL_529788, EPI_ISL_529789, EPI_ISL_529790, EPI_ISL_529791, EPI_ISL_529792, EPI_ISL_529793, EPI_ISL_529794, EPI_ISL_529795, EPI_ISL_529796, EPI_ISL_529797, EPI_ISL_529798, EPI_ISL_529799, EPI_ISL_529800, EPI_ISL_529801, EPI_ISL_529802, EPI_ISL_529803 | | | |
| see above | NHLS-IALCH | KRISP, KZN Research Innovation and Sequencing Platform | Giandhari J, Pillay S, Lessells R, Mdlalose K, York D, Khan S, Tegally H, Wilkinson E, de Oliveira T |
| EPI_ISL_529804, EPI_ISL_529805, EPI_ISL_529806, EPI_ISL_529807, EPI_ISL_529808, EPI_ISL_529809, EPI_ISL_529810, EPI_ISL_529811, EPI_ISL_529812, EPI_ISL_529813, EPI_ISL_529814, EPI_ISL_529815, EPI_ISL_529816, EPI_ISL_529817, EPI_ISL_529818, EPI_ISL_529819, EPI_ISL_529820, EPI_ISL_529821, EPI_ISL_529822, EPI_ISL_529823, EPI_ISL_529824, EPI_ISL_529825, EPI_ISL_529826, EPI_ISL_529827, EPI_ISL_529828, EPI_ISL_529829, EPI_ISL_529830, EPI_ISL_529831, EPI_ISL_529832, EPI_ISL_529833, EPI_ISL_529834, EPI_ISL_529835, EPI_ISL_529836, EPI_ISL_529837, EPI_ISL_529838, EPI_ISL_529839, EPI_ISL_529840, EPI_ISL_529841, EPI_ISL_529842, EPI_ISL_529843, EPI_ISL_529844, EPI_ISL_529845, EPI_ISL_529846, EPI_ISL_529847, EPI_ISL_529848, EPI_ISL_529849, EPI_ISL_529850, EPI_ISL_529851, EPI_ISL_529852, EPI_ISL_529853, EPI_ISL_529854, EPI_ISL_529855, EPI_ISL_529856, EPI_ISL_529857, EPI_ISL_529858, EPI_ISL_529859, EPI_ISL_529860, EPI_ISL_529861, EPI_ISL_529862, EPI_ISL_529863, EPI_ISL_529864, EPI_ISL_529865, EPI_ISL_529866, EPI_ISL_529867, EPI_ISL_529868, EPI_ISL_529869, EPI_ISL_529870, EPI_ISL_529871, EPI_ISL_529872, EPI_ISL_529873, EPI_ISL_529874, EPI_ISL_529875, EPI_ISL_529876, EPI_ISL_529877, EPI_ISL_529878, EPI_ISL_529879, EPI_ISL_529880, EPI_ISL_529881, EPI_ISL_529882, EPI_ISL_529883, EPI_ISL_529884, EPI_ISL_529885, EPI_ISL_529886, EPI_ISL_529887, EPI_ISL_529888, EPI_ISL_529889, EPI_ISL_529890, EPI_ISL_529891, EPI_ISL_529892, EPI_ISL_529893, EPI_ISL_529894, EPI_ISL_529895, EPI_ISL_529896, EPI_ISL_529897, EPI_ISL_529898, EPI_ISL_529899, EPI_ISL_529900, EPI_ISL_529901, EPI_ISL_529902, EPI_ISL_529903, EPI_ISL_529904, EPI_ISL_529905, EPI_ISL_529906, EPI_ISL_529907 | | | |
| see above | Michigan Department of Health and Human Services, Bureau of Laboratories | Michigan Department of Health and Human Services, Bureau of Laboratories | Blankenship HM, Riner D, Soehnlen MK |
| EPI_ISL_529908, EPI_ISL_529909, EPI_ISL_529910, EPI_ISL_529911, EPI_ISL_529912, EPI_ISL_529913, EPI_ISL_529914, EPI_ISL_529915, EPI_ISL_529916, EPI_ISL_529917, EPI_ISL_529918, EPI_ISL_529919, EPI_ISL_529920, EPI_ISL_529921, EPI_ISL_529922, EPI_ISL_529923, EPI_ISL_529924, EPI_ISL_529925, EPI_ISL_529926, EPI_ISL_529927, EPI_ISL_529928, EPI_ISL_529929, EPI_ISL_529930, EPI_ISL_529931, EPI_ISL_529932, EPI_ISL_529933, EPI_ISL_529934, EPI_ISL_529935, EPI_ISL_529936 | | | |
| see above | Virginia Division of Consolidated Laboratory Services | Virginia Division of Consolidated Laboratory Services | Virginia DCLS |
| EPI_ISL_529937, EPI_ISL_529938, EPI_ISL_529939, EPI_ISL_529940, EPI_ISL_529941, EPI_ISL_529942, EPI_ISL_529943, EPI_ISL_529944, EPI_ISL_529945, EPI_ISL_529946, EPI_ISL_529947, EPI_ISL_529948, EPI_ISL_529949, EPI_ISL_529950, EPI_ISL_529951, EPI_ISL_529952, EPI_ISL_529953, EPI_ISL_529954, EPI_ISL_529955, EPI_ISL_529956, EPI_ISL_529957, EPI_ISL_529958, EPI_ISL_529959, EPI_ISL_529960 | | | |
| see above | Virginia DCLS | Virginia DCLS | Virginia DCLS |
| EPI_ISL_529961 | Husada Utama Hospital | Institute of Tropical Disease, Universitas Airlangga | Krisnoadi Rahardjo, Aldise M Nastri, Jezzy R Dewantari, Rima R Prasetya, Didi Dewanto, Gatot Soegiarto, Laksmi Wulandari, Retno A Setyoningrum, Resti Yudhawati, Yohko K Shimizu, Mitsuhiro Nishimura, Yasuko Mori, Soetjipto, Kazufumi Shimizu, Maria I Lusida |
| EPI_ISL_529962 | Universitas Airlangga Hospital | Institute of Tropical Disease, Universitas Airlangga | Jezzy R Dewantari, Rima R Prasetya, Krisnoadi Rahardjo, Aldise M Nastri, Nasronudin, Gatot Soegiarto, Laksmi Wulandari, Retno A Setyoningrum, Resti Yudhawati, Yohko K Shimizu, Mitsuhiro Nishimura, Yasuko Mori, Soetjipto, Kazufumi Shimizu, Maria I Lusida |
| EPI_ISL_529963 | Universitas Airlangga Hospital | Institute of Tropical Disease, Universitas Airlangga | Rima R Prasetya, Krisnoadi Rahardjo, Aldise M Nastri, Jezzy R Dewantari, Nasronudin, Gatot Soegiarto, Laksmi Wulandari, Retno A Setyoningrum, Resti Yudhawati, Yohko K Shimizu, Mitsuhiro Nishimura, Yasuko Mori, Soetjipto, Kazufumi Shimizu, Maria I Lusida |
| EPI_ISL_529964 | RSUD Bangil Pasuruan | Institute of Tropical Disease, Universitas Airlangga | Aldise M Nastri, Jezzy R Dewantari, Rima R Prasetya, Krisnoadi Rahardjo, Arma Roosalina, Gatot Soegiarto, Laksmi Wulandari, Retno A Setyoningrum, Resti Yudhawati, Yohko K Shimizu, Mitsuhiro Nishimura, Yasuko Mori, Soetjipto, Kazufumi Shimizu, Maria I Lusida |
| EPI_ISL_529965 | RSUD Dr. Soetomo | Institute of Tropical Disease, Universitas Airlangga | Krisnoadi Rahardjo, Aldise M Nastri, Jezzy R Dewantari, Rima R Prasetya, Joni Wahyuhadi, Gatot Soegiarto, Laksmi Wulandari, Retno A Setyoningrum, Resti Yudhawati, Yohko K Shimizu, Mitsuhiro Nishimura, Yasuko Mori, Soetjipto, Kazufumi Shimizu, Maria I Lusida |
| EPI_ISL_529966 | RSUD Sidoarjo | Institute of Tropical Disease, Universitas Airlangga | Kazufumi Shimizu, Krisnoadi Rahardjo, Aldise M Nastri, Jezzy R Dewantari, Rima R Prasetya, Atok Irawan, Gatot Soegiarto, Laksmi Wulandari, Retno A Setyoningrum, Resti Yudhawati, Yohko K Shimizu, Mitsuhiro Nishimura, Yasuko Mori, Soetjipto, Maria I Lusida |
| EPI_ISL_529967 | RSUD Sidoarjo | Institute of Tropical Disease, Universitas Airlangga | Aldise M Nastri, Jezzy R Dewantari, Rima R Prasetya, Krisnoadi Rahardjo, Atok Irawan, Gatot Soegiarto, Laksmi Wulandari, Retno A Setyoningrum, Resti Yudhawati, Yohko K Shimizu, Mitsuhiro Nishimura, Yasuko Mori, Soetjipto, Kazufumi Shimizu, Maria I Lusida |
| EPI_ISL_529972, EPI_ISL_529973, EPI_ISL_529974, EPI_ISL_529975, EPI_ISL_529976, EPI_ISL_529977, EPI_ISL_529978, EPI_ISL_529979, EPI_ISL_529980, EPI_ISL_529981, EPI_ISL_529982, EPI_ISL_529983, EPI_ISL_529984, EPI_ISL_529985, EPI_ISL_529986, EPI_ISL_529987, EPI_ISL_529988 | | | |
| see above | Hospital Universitario 12 de Octubre | Hospital Universitario 12 de Octubre | Raúl Recio, Sara González, Esther Viedma, Elias Dahdouh, Fernando Lázaro, Natalia Stella, Julio García, Juan Carlos Galán, Rafael Cantón, Mª Dolores Folgueira, Rafael Delgado, Jesús Mingorance |
| EPI_ISL_529989, EPI_ISL_529990, EPI_ISL_529991, EPI_ISL_529992, EPI_ISL_529993, EPI_ISL_529994, EPI_ISL_529995, EPI_ISL_529996, EPI_ISL_529997, EPI_ISL_529998, EPI_ISL_529999, EPI_ISL_530000, EPI_ISL_530001, EPI_ISL_530002, EPI_ISL_530003, EPI_ISL_530004, EPI_ISL_530005, EPI_ISL_530006 | | | |
| see above | Hospital Universitario 12 de Octubre | Hospital Universitario 12 de Octubre | Sara González, Esther Viedma, Raúl Recio, Elias Dahdouh, Fernando Lázaro, Natalia Stella, Julio García, Juan Carlos Galán, Rafael Cantón, Mª Dolores Folgueira, Rafael Delgado, Jesús Mingorance |
| EPI_ISL_530007, EPI_ISL_530008, EPI_ISL_530009, EPI_ISL_530010, EPI_ISL_530011, EPI_ISL_530012, EPI_ISL_530013, EPI_ISL_530014, EPI_ISL_530015, EPI_ISL_530016, EPI_ISL_530017, EPI_ISL_530018, EPI_ISL_530019, EPI_ISL_530020, EPI_ISL_530021, EPI_ISL_530022, EPI_ISL_530023, EPI_ISL_530024 | | | |
| see above | Hospital Universitario 12 de Octubre | Hospital Universitario 12 de Octubre | Esther Viedma, Raúl Recio, Sara González, Elias Dahdouh, Fernando Lázaro, Natalia Stella, Julio García, Juan Carlos Galán, Rafael Cantón, Mª Dolores Folgueira, Rafael Delgado, Jesús Mingorance |
| EPI_ISL_530025, EPI_ISL_530026, EPI_ISL_530027, EPI_ISL_530028, EPI_ISL_530029, EPI_ISL_530030, EPI_ISL_530031, EPI_ISL_530032, EPI_ISL_530033, EPI_ISL_530034, EPI_ISL_530035, EPI_ISL_530036, EPI_ISL_530037, EPI_ISL_530038, EPI_ISL_530039, EPI_ISL_530040, EPI_ISL_530041, EPI_ISL_530042, EPI_ISL_530043 | | | |
| see above | Hospital Universitario La Paz | Hospital Universitario La Paz | María Rodríguez, Elias Dahdouh, Sara González, Raúl Recio, Fernando Lázaro, Esther Viedma, Natalia Stella, Julio García, Juan Carlos Galán, Rafael Cantón, Mª Dolores Folgueira, Rafael Delgado, Jesús Mingorance |
| EPI_ISL_530044, EPI_ISL_530045, EPI_ISL_530046, EPI_ISL_530047, EPI_ISL_530048, EPI_ISL_530049, EPI_ISL_530050, EPI_ISL_530051, EPI_ISL_530052, EPI_ISL_530053, EPI_ISL_530054, EPI_ISL_530055, EPI_ISL_530056, EPI_ISL_530057, EPI_ISL_530058, EPI_ISL_530059, EPI_ISL_530060, EPI_ISL_530061, EPI_ISL_530062, EPI_ISL_530063, EPI_ISL_530064, EPI_ISL_530065, EPI_ISL_530066, EPI_ISL_530067, EPI_ISL_530068, EPI_ISL_530069 | | | |
| see above | Hospital Universitario La Paz | Hospital Universitario La Paz | Elias Dahdouh, Sara González, Raúl Recio, Fernando Lázaro, Esther Viedma, Natalia Stella, Julio García, Juan Carlos Galán, Rafael Cantón, Mª Dolores Folgueira, Rafael Delgado, Jesús Mingorance |
| EPI_ISL_530070, EPI_ISL_530071, EPI_ISL_530072, EPI_ISL_530073, EPI_ISL_530074, EPI_ISL_530075, EPI_ISL_530076, EPI_ISL_530077, EPI_ISL_530078, EPI_ISL_530079, EPI_ISL_530080, EPI_ISL_530081, EPI_ISL_530082, EPI_ISL_530083, EPI_ISL_530084, EPI_ISL_530085, EPI_ISL_530086, EPI_ISL_530087, EPI_ISL_530088, EPI_ISL_530089, EPI_ISL_530090, EPI_ISL_530091, EPI_ISL_530092, EPI_ISL_530093 | | | |
| see above | Hospital Universitario La Paz | Hospital Universitario La Paz | María Rodríguez, Elias Dahdouh, Sara González, Raúl Recio, Fernando Lázaro, Esther Viedma, Natalia Stella, Julio García, Juan Carlos Galán, Rafael Cantón, Mª Dolores Folgueira, Rafael Delgado, Jesús Mingorance |
| EPI_ISL_530094, EPI_ISL_530095, EPI_ISL_530096, EPI_ISL_530097, EPI_ISL_530098, EPI_ISL_530099 | Hospital Universitario Ramón y Cajal | Hospital Universitario 12 de Octubre | Raúl Recio, Sara González, Elias Dahdouh, Fernando Lázaro, Esther Viedma, Natalia Stella, Julio García, Juan Carlos Galán, Rafael Cantón, Mª Dolores Folgueira, Rafael Delgado, Jesús Mingorance |
| EPI_ISL_530100, EPI_ISL_530101, EPI_ISL_530102, EPI_ISL_530103, EPI_ISL_530104, EPI_ISL_530105, EPI_ISL_530106, EPI_ISL_530107, EPI_ISL_530108, EPI_ISL_530109, EPI_ISL_530110, EPI_ISL_530111, EPI_ISL_530112, EPI_ISL_530113 | | | |
| see above | Hospital Universitario Ramón y Cajal | Hospital Universitario La Paz | Raúl Recio, Sara González, Elias Dahdouh, Fernando Lázaro, Esther Viedma, Natalia Stella, Julio García, Juan Carlos Galán, Rafael Cantón, Mª Dolores Folgueira, Rafael Delgado, Jesús Mingorance |
| EPI_ISL_530114, EPI_ISL_530115, EPI_ISL_530116, EPI_ISL_530117, EPI_ISL_530118, EPI_ISL_530119 | Hospital Universitario Ramón y Cajal | Hospital Universitario La Paz | María Rodríguez, Elias Dahdouh, Sara González, Raúl Recio, Fernando Lázaro, Esther Viedma, Natalia Stella, Julio García, Juan Carlos Galán, Rafael Cantón, Mª Dolores Folgueira, Rafael Delgado, Jesús Mingorance |
| EPI_ISL_530122, EPI_ISL_530123 | University Hospital Basel, Clinical Virology | University Hospital Basel, Clinical Bacteriology | Madlen Stange, Alfredo Mari, Tim Roloff, Helena MB Seth-Smith, Michael Schweitzer, Myrta Brunner, Karoline Leuzinger, Kirstine K. Soegaard, Alexander Gensch, Sarah Tschudin-Sutter, Simon Fuchs, Julia Bielicki, Hans Pargger, Martin Siegemund, Christian Nickel, Roland Bingisser, Michael Osthoff, Stefano Bassetti, Rita Schneider-Sliwa, Manuel Battegay, Hans Hirsch, Adrian Egli |
| EPI_ISL_530124, EPI_ISL_530125 | Seattle Flu Study | Seattle Flu Study | Deborah A. Nickerson, Chris D. Frazar, Jover Lee, Benjamin Pelle, Matthew Richardson, Amanda Adler, Elisabeth Brandstetter, Peter D. Han, Kairsten Fay, Misja Ilcisin, Kirsten Lacombe, Thomas R. Sibley, Melissa Truong, Caitlin R. Wolf, Michael Boeckh, Janet A. Englund, Michael Famulare, Barry R. Lutz, Mark J. Rieder, Lea M. Starita, Matthew Thompson, Jay Shendure, Trevor Bedford, Helen Y. Chu |
| EPI_ISL_530126, EPI_ISL_530127, EPI_ISL_530128, | Seattle Flu Study | Seattle Flu Study | Deborah A. Nickerson, Chris D. Frazar, Jover Lee, Benjamin Pelle, Matthew Richardson, Amanda Adler, Elisabeth Brandstetter, Peter D. Han, Kairsten Fay, Misja Ilcisin, Kirsten Lacombe, Thomas R. Sibley, Melissa Truong, |

| | | | |
|---|---|---|---|
| EPI_ISL_530129, EPI_ISL_530130, EPI_ISL_530131 | | | Caitlin R. Wolf, Karen Cowgill, Stephanie Schrag, Jeff Duchin, Michael Boeckh, Janet A. Englund, Michael Famulare, Barry R. Lutz, Mark J. Rieder, Lea M. Starita, Matthew Thompson, Helen Y. Chu, Trevor Bedford, Jay Shendure |
| EPI_ISL_530132 | Seattle Flu Study | Seattle Flu Study | Deborah A. Nickerson, Chris D. Frazar, Jover Lee, Benjamin Pelle, Matthew Richardson, Amanda Adler, Elisabeth Brandstetter, Peter D. Han, Kairsten Fay, Misja Ilcisin, Kirsten Lacombe, Thomas R. Sibley, Melissa Truong, Caitlin R. Wolf, Michael Boeckh, Janet A. Englund, Michael Famulare, Barry R. Lutz, Mark J. Rieder, Lea M. Starita, Matthew Thompson, Jay Shendure, Trevor Bedford, Helen Y. Chu |
| EPI_ISL_530133, EPI_ISL_530134, EPI_ISL_530135, EPI_ISL_530136, EPI_ISL_530137, EPI_ISL_530138, EPI_ISL_530139, EPI_ISL_530140, EPI_ISL_530141, EPI_ISL_530142, EPI_ISL_530143, EPI_ISL_530144, EPI_ISL_530145, EPI_ISL_530146, EPI_ISL_530147, EPI_ISL_530148, EPI_ISL_530149, EPI_ISL_530150, EPI_ISL_530151, EPI_ISL_530152, EPI_ISL_530153, EPI_ISL_530154, EPI_ISL_530155, EPI_ISL_530156, EPI_ISL_530157, EPI_ISL_530158, EPI_ISL_530159, EPI_ISL_530160, EPI_ISL_530161, EPI_ISL_530162 | | | |
| see above | Seattle Flu Study | Seattle Flu Study | Deborah A. Nickerson, Chris D. Frazar, Jover Lee, Benjamin Pelle, Matthew Richardson, Amanda Adler, Elisabeth Brandstetter, Peter D. Han, Kairsten Fay, Misja Ilcisin, Kirsten Lacombe, Thomas R. Sibley, Melissa Truong, Caitlin R. Wolf, Karen Cowgill, Stephanie Schrag, Jeff Duchin, Michael Boeckh, Janet A. Englund, Michael Famulare, Barry R. Lutz, Mark J. Rieder, Lea M. Starita, Matthew Thompson, Helen Y. Chu, Trevor Bedford, Jay Shendure |
| EPI_ISL_530163, EPI_ISL_530164, EPI_ISL_530165, EPI_ISL_530166, EPI_ISL_530167, EPI_ISL_530168 | M Health Fairview | Minnesota Department of Health, Public Health Laboratory | Matt Plumb, Jacob Garfin, and Xiong Wang |
| EPI_ISL_530169, EPI_ISL_530170, EPI_ISL_530171 | Hennepin County Medical Center | Minnesota Department of Health, Public Health Laboratory | Matt Plumb, Jacob Garfin, and Xiong Wang |
| EPI_ISL_530172, EPI_ISL_530173, EPI_ISL_530174, EPI_ISL_530175, EPI_ISL_530176, EPI_ISL_530177, EPI_ISL_530178, EPI_ISL_530179, EPI_ISL_530180, EPI_ISL_530181, EPI_ISL_530182, EPI_ISL_530183, EPI_ISL_530184, EPI_ISL_530185, EPI_ISL_530186, EPI_ISL_530187, EPI_ISL_530188, EPI_ISL_530189, EPI_ISL_530190, EPI_ISL_530191, EPI_ISL_530192, EPI_ISL_530193, EPI_ISL_530194, EPI_ISL_530195, EPI_ISL_530196, EPI_ISL_530197, EPI_ISL_530198, EPI_ISL_530199, EPI_ISL_530200, EPI_ISL_530201, EPI_ISL_530202, EPI_ISL_530203, EPI_ISL_530204, EPI_ISL_530205, EPI_ISL_530206, EPI_ISL_530207, EPI_ISL_530208, EPI_ISL_530209, EPI_ISL_530210, EPI_ISL_530211, EPI_ISL_530212, EPI_ISL_530213, EPI_ISL_530214, EPI_ISL_530215, EPI_ISL_530216, EPI_ISL_530217, EPI_ISL_530218, EPI_ISL_530219, EPI_ISL_530220, EPI_ISL_530221, EPI_ISL_530222, EPI_ISL_530223, EPI_ISL_530224 | | | |
| see above | Minnesota Department of Health, Public Health Laboratory | Minnesota Department of Health, Public Health Laboratory | Matt Plumb, Jacob Garfin, and Xiong Wang |
| EPI_ISL_530225, EPI_ISL_530226, EPI_ISL_530227, EPI_ISL_530228, EPI_ISL_530229, EPI_ISL_530230, EPI_ISL_530231, EPI_ISL_530232, EPI_ISL_530233, EPI_ISL_530234, EPI_ISL_530235, EPI_ISL_530236, EPI_ISL_530237, EPI_ISL_530238, EPI_ISL_530239, EPI_ISL_530240, EPI_ISL_530241, EPI_ISL_530242, EPI_ISL_530243, EPI_ISL_530244, EPI_ISL_530245, EPI_ISL_530246, EPI_ISL_530247, EPI_ISL_530248, EPI_ISL_530249, EPI_ISL_530250, EPI_ISL_530251, EPI_ISL_530252, EPI_ISL_530253, EPI_ISL_530254, EPI_ISL_530255, EPI_ISL_530256, EPI_ISL_530257, EPI_ISL_530258, EPI_ISL_530259, EPI_ISL_530260, EPI_ISL_530261, EPI_ISL_530262, EPI_ISL_530263, EPI_ISL_530264, EPI_ISL_530265, EPI_ISL_530266, EPI_ISL_530267, EPI_ISL_530268, EPI_ISL_530269, EPI_ISL_530270, EPI_ISL_530271, EPI_ISL_530272, EPI_ISL_530273, EPI_ISL_530274, EPI_ISL_530275, EPI_ISL_530276, EPI_ISL_530277, EPI_ISL_530278, EPI_ISL_530279, EPI_ISL_530280, EPI_ISL_530281, EPI_ISL_530282, EPI_ISL_530283 | | | |
| see above | Queensland Health Forensic and Scientific Services, Public Health Virology | Public Health Virology Laboratory, Forensic and Scientific Services, Queensland Health | Son Nguyen et al |
| EPI_ISL_530334, EPI_ISL_530335, EPI_ISL_530336, EPI_ISL_530337, EPI_ISL_530338, EPI_ISL_530339, EPI_ISL_530340 | Area of Virology, Serology and Virology Division (SAViD), New South Wales Health Pathology Randwick | Area of Virology, Serology and Virology Division (SAViD), New South Wales Health Pathology Randwick | Rawlinson, W., Deveson, I., Bull, R., Van Hal, S. |
| EPI_ISL_530347 | The National Institute of Public Health | State Veterinary Institute Prague | Nagy,A;Jirincova,H;Novakova,L;Trnka,D;Vecerova,J |
| EPI_ISL_530348, EPI_ISL_530349, EPI_ISL_530350 | The National Institute of Public Health | State Veterinary Institute Prague | Nagy,A;Jirincova,H;Novakova,L;Trnka,D;Vecerova,J |
| EPI_ISL_530351, EPI_ISL_530352, EPI_ISL_530353, EPI_ISL_530354, EPI_ISL_530355, EPI_ISL_530356, EPI_ISL_530357, EPI_ISL_530358, EPI_ISL_530359, EPI_ISL_530360 | Lighthouse Lab in Glasgow | Wellcome Sanger Institute for the COVID-19 Genomics UK Consortium | Harper VanSteenhouse, Yumi Kasai, David Gray, Carol Clugston, Anna Dominiczak and Alex Alderton, Roberto Amato, Sonia Goncalves, Ewan Harrison, David K. Jackson, Ian Johnston, Dominic Kwiatkowski, Cordelia Langford, John Sillitoe |
| EPI_ISL_530361 | NHSGGC West of Scotland Specialist Virology Centre / MRC-University of Glasgow Centre for Virus Research | Wellcome Sanger Institute for the COVID-19 Genomics UK Consortium | Ana da Silva Filipe, Natasha Johnson, Kathy Smollett, Daniel Mair, Stephen Carmichael, Lily Tong, Jenna Nichols, Elihu Aranday-Cortes, Kirstyn Brunker, Yasmin Parr, Kyriaki Nomikou; Sarah McDonald, Marc Niebel, Patawee Asamaphan; Richard Orton, Joseph Hughes, Sreenu Vattipally, David L Robertson; Alasdair MacLean, Rory Gunson; Kathy Li, Natasha Jesudason, Rajiv Shah, James Shepherd, Antonia Ho, Alice Broos, Emma Thomson and Alex Alderton, Roberto Amato, Sonia Goncalves, Ewan Harrison, David K. Jackson, Ian Johnston, Dominic Kwiatkowski, Cordelia Langford, John Sillitoe |
| EPI_ISL_530362, EPI_ISL_530363, EPI_ISL_530364, EPI_ISL_530365 | Lighthouse Lab in Glasgow | Wellcome Sanger Institute for the COVID-19 Genomics UK Consortium | Harper VanSteenhouse, Yumi Kasai, David Gray, Carol Clugston, Anna Dominiczak and Alex Alderton, Roberto Amato, Sonia Goncalves, Ewan Harrison, David K. Jackson, Ian Johnston, Dominic Kwiatkowski, Cordelia Langford, John Sillitoe |
| EPI_ISL_530366 | NHSGGC West of Scotland Specialist Virology Centre / MRC-University of Glasgow Centre for Virus Research | Wellcome Sanger Institute for the COVID-19 Genomics UK Consortium | Ana da Silva Filipe, Natasha Johnson, Kathy Smollett, Daniel Mair, Stephen Carmichael, Lily Tong, Jenna Nichols, Elihu Aranday-Cortes, Kirstyn Brunker, Yasmin Parr, Kyriaki Nomikou; Sarah McDonald, Marc Niebel, Patawee Asamaphan; Richard Orton, Joseph Hughes, Sreenu Vattipally, David L Robertson; Alasdair MacLean, Rory Gunson; Kathy Li, Natasha Jesudason, Rajiv Shah, James Shepherd, Antonia Ho, Alice Broos, Emma Thomson and Alex Alderton, Roberto Amato, Sonia Goncalves, Ewan Harrison, David K. Jackson, Ian Johnston, Dominic Kwiatkowski, Cordelia Langford, John Sillitoe |
| EPI_ISL_530367, EPI_ISL_530368, EPI_ISL_530369 | Lighthouse Lab in Glasgow | Wellcome Sanger Institute for the COVID-19 Genomics UK Consortium | Harper VanSteenhouse, Yumi Kasai, David Gray, Carol Clugston, Anna Dominiczak and Alex Alderton, Roberto Amato, Sonia Goncalves, Ewan Harrison, David K. Jackson, Ian Johnston, Dominic Kwiatkowski, Cordelia Langford, John Sillitoe |
| EPI_ISL_530370 | NHSGGC West of Scotland Specialist Virology Centre / MRC-University of Glasgow Centre for Virus Research | Wellcome Sanger Institute for the COVID-19 Genomics UK Consortium | Ana da Silva Filipe, Natasha Johnson, Kathy Smollett, Daniel Mair, Stephen Carmichael, Lily Tong, Jenna Nichols, Elihu Aranday-Cortes, Kirstyn Brunker, Yasmin Parr, Kyriaki Nomikou; Sarah McDonald, Marc Niebel, Patawee Asamaphan; Richard Orton, Joseph Hughes, Sreenu Vattipally, David L Robertson; Alasdair MacLean, Rory Gunson; Kathy Li, Natasha Jesudason, Rajiv Shah, James Shepherd, Antonia Ho, Alice Broos, Emma Thomson and Alex Alderton, Roberto Amato, Sonia Goncalves, Ewan Harrison, David K. Jackson, Ian Johnston, Dominic Kwiatkowski, Cordelia Langford, John Sillitoe |
| EPI_ISL_530371, EPI_ISL_530372, EPI_ISL_530373, EPI_ISL_530374, EPI_ISL_530375, EPI_ISL_530376 | Lighthouse Lab in Glasgow | Wellcome Sanger Institute for the COVID-19 Genomics UK Consortium | Harper VanSteenhouse, Yumi Kasai, David Gray, Carol Clugston, Anna Dominiczak and Alex Alderton, Roberto Amato, Sonia Goncalves, Ewan Harrison, David K. Jackson, Ian Johnston, Dominic Kwiatkowski, Cordelia Langford, John Sillitoe |
| EPI_ISL_530377 | NHSGGC West of Scotland Specialist Virology Centre / MRC-University of Glasgow Centre for Virus Research | Wellcome Sanger Institute for the COVID-19 Genomics UK Consortium | Ana da Silva Filipe, Natasha Johnson, Kathy Smollett, Daniel Mair, Stephen Carmichael, Lily Tong, Jenna Nichols, Elihu Aranday-Cortes, Kirstyn Brunker, Yasmin Parr, Kyriaki Nomikou; Sarah McDonald, Marc Niebel, Patawee Asamaphan; Richard Orton, Joseph Hughes, Sreenu Vattipally, David L Robertson; Alasdair MacLean, Rory Gunson; Kathy Li, Natasha Jesudason, Rajiv Shah, James Shepherd, Antonia Ho, Alice Broos, Emma Thomson and Alex Alderton, Roberto Amato, Sonia Goncalves, Ewan Harrison, David K. Jackson, Ian Johnston, Dominic Kwiatkowski, Cordelia Langford, John Sillitoe |
| EPI_ISL_530378, EPI_ISL_530379, EPI_ISL_530380, EPI_ISL_530381, EPI_ISL_530382, EPI_ISL_530383, EPI_ISL_530384, EPI_ISL_530385 | Lighthouse Lab in Glasgow | Wellcome Sanger Institute for the COVID-19 Genomics UK Consortium | Harper VanSteenhouse, Yumi Kasai, David Gray, Carol Clugston, Anna Dominiczak and Alex Alderton, Roberto Amato, Sonia Goncalves, Ewan Harrison, David K. Jackson, Ian Johnston, Dominic Kwiatkowski, Cordelia Langford, John Sillitoe |
| EPI_ISL_530386 | NHSGGC West of Scotland Specialist Virology Centre / MRC-University of Glasgow Centre for Virus Research | Wellcome Sanger Institute for the COVID-19 Genomics UK Consortium | Ana da Silva Filipe, Natasha Johnson, Kathy Smollett, Daniel Mair, Stephen Carmichael, Lily Tong, Jenna Nichols, Elihu Aranday-Cortes, Kirstyn Brunker, Yasmin Parr, Kyriaki Nomikou; Sarah McDonald, Marc Niebel, Patawee Asamaphan; Richard Orton, Joseph Hughes, Sreenu Vattipally, David L Robertson; Alasdair MacLean, Rory Gunson; Kathy Li, Natasha Jesudason, Rajiv Shah, James Shepherd, Antonia Ho, Alice Broos, Emma Thomson and Alex Alderton, Roberto Amato, Sonia Goncalves, Ewan Harrison, David K. Jackson, Ian Johnston, Dominic Kwiatkowski, Cordelia Langford, John Sillitoe |
| EPI_ISL_530387 | Lighthouse Lab in Glasgow | Wellcome Sanger Institute for the COVID-19 Genomics UK Consortium | Harper VanSteenhouse, Yumi Kasai, David Gray, Carol Clugston, Anna Dominiczak and Alex Alderton, Roberto Amato, Sonia Goncalves, Ewan Harrison, David K. Jackson, Ian Johnston, Dominic Kwiatkowski, Cordelia Langford, John Sillitoe |
| EPI_ISL_530388 | NHSGGC West of Scotland Specialist Virology Centre / MRC-University of Glasgow Centre for Virus Research | Wellcome Sanger Institute for the COVID-19 Genomics UK Consortium | Ana da Silva Filipe, Natasha Johnson, Kathy Smollett, Daniel Mair, Stephen Carmichael, Lily Tong, Jenna Nichols, Elihu Aranday-Cortes, Kirstyn Brunker, Yasmin Parr, Kyriaki Nomikou; Sarah McDonald, Marc Niebel, Patawee Asamaphan; Richard Orton, Joseph Hughes, Sreenu Vattipally, David L Robertson; Alasdair MacLean, Rory Gunson; Kathy Li, Natasha Jesudason, Rajiv Shah, James Shepherd, Antonia Ho, Alice Broos, Emma Thomson and Alex Alderton, Roberto Amato, Sonia Goncalves, Ewan Harrison, David K. Jackson, Ian Johnston, Dominic Kwiatkowski, Cordelia Langford, John Sillitoe |
| EPI_ISL_530389, EPI_ISL_530390 | Lighthouse Lab in Glasgow | Wellcome Sanger Institute for the COVID-19 Genomics UK Consortium | Harper VanSteenhouse, Yumi Kasai, David Gray, Carol Clugston, Anna Dominiczak and Alex Alderton, Roberto Amato, Sonia Goncalves, Ewan Harrison, David K. Jackson, Ian Johnston, Dominic Kwiatkowski, Cordelia Langford, John Sillitoe |
| EPI_ISL_530391 | NHSGGC West of Scotland Specialist Virology Centre / MRC-University of Glasgow Centre for Virus Research | Wellcome Sanger Institute for the COVID-19 Genomics UK Consortium | Ana da Silva Filipe, Natasha Johnson, Kathy Smollett, Daniel Mair, Stephen Carmichael, Lily Tong, Jenna Nichols, Elihu Aranday-Cortes, Kirstyn Brunker, Yasmin Parr, Kyriaki Nomikou; Sarah McDonald, Marc Niebel, Patawee Asamaphan; Richard Orton, Joseph Hughes, Sreenu Vattipally, David L Robertson; Alasdair MacLean, Rory Gunson; Kathy Li, Natasha Jesudason, Rajiv Shah, James Shepherd, Antonia Ho, Alice Broos, Emma Thomson and Alex Alderton, Roberto Amato, Sonia Goncalves, Ewan Harrison, David K. Jackson, Ian Johnston, Dominic Kwiatkowski, Cordelia Langford, John Sillitoe |
| EPI_ISL_530392, EPI_ISL_530394, EPI_ISL_530395, EPI_ISL_530396, EPI_ISL_530397 | Lighthouse Lab in Glasgow | Wellcome Sanger Institute for the COVID-19 Genomics UK Consortium | Harper VanSteenhouse, Yumi Kasai, David Gray, Carol Clugston, Anna Dominiczak and Alex Alderton, Roberto Amato, Sonia Goncalves, Ewan Harrison, David K. Jackson, Ian Johnston, Dominic Kwiatkowski, Cordelia Langford, John Sillitoe |
| EPI_ISL_530398 | NHSGGC West of Scotland Specialist Virology Centre / MRC-University of Glasgow Centre for Virus Research | Wellcome Sanger Institute for the COVID-19 Genomics UK Consortium | Ana da Silva Filipe, Natasha Johnson, Kathy Smollett, Daniel Mair, Stephen Carmichael, Lily Tong, Jenna Nichols, Elihu Aranday-Cortes, Kirstyn Brunker, Yasmin Parr, Kyriaki Nomikou; Sarah McDonald, Marc Niebel, Patawee Asamaphan; Richard Orton, Joseph Hughes, Sreenu Vattipally, David L Robertson; Alasdair MacLean, Rory Gunson; Kathy Li, Natasha Jesudason, Rajiv Shah, James Shepherd, Antonia Ho, Alice Broos, Emma Thomson and Alex Alderton, Roberto Amato, Sonia Goncalves, Ewan Harrison, David K. Jackson, Ian Johnston, Dominic Kwiatkowski, Cordelia Langford, John Sillitoe |
| EPI_ISL_530399, EPI_ISL_530400, EPI_ISL_530401, EPI_ISL_530402, EPI_ISL_530403, EPI_ISL_530404, EPI_ISL_530405, EPI_ISL_530406, EPI_ISL_530407, EPI_ISL_530408, EPI_ISL_530409 | | | |
| see above | Lighthouse Lab in Glasgow | Wellcome Sanger Institute for the COVID-19 Genomics UK Consortium | Harper VanSteenhouse, Yumi Kasai, David Gray, Carol Clugston, Anna Dominiczak and Alex Alderton, Roberto Amato, Sonia Goncalves, Ewan Harrison, David K. Jackson, Ian Johnston, Dominic Kwiatkowski, Cordelia Langford, John Sillitoe |
| EPI_ISL_530410 | NHSGGC West of Scotland Specialist Virology Centre / MRC-University of Glasgow Centre for Virus Research | Wellcome Sanger Institute for the COVID-19 Genomics UK Consortium | Ana da Silva Filipe, Natasha Johnson, Kathy Smollett, Daniel Mair, Stephen Carmichael, Lily Tong, Jenna Nichols, Elihu Aranday-Cortes, Kirstyn Brunker, Yasmin Parr, Kyriaki Nomikou; Sarah McDonald, Marc Niebel, Patawee Asamaphan; Richard Orton, Joseph Hughes, Sreenu Vattipally, David L Robertson; Alasdair MacLean, Rory Gunson; Kathy Li, Natasha Jesudason, Rajiv Shah, James Shepherd, Antonia Ho, Alice Broos, Emma Thomson and Alex Alderton, Roberto Amato, Sonia Goncalves, Ewan Harrison, David K. Jackson, Ian Johnston, Dominic Kwiatkowski, Cordelia Langford, John Sillitoe |
| EPI_ISL_530411, EPI_ISL_530412, EPI_ISL_530413, EPI_ISL_530414, EPI_ISL_530415, EPI_ISL_530416, EPI_ISL_530417, EPI_ISL_530418, EPI_ISL_530419, EPI_ISL_530420, EPI_ISL_530421, EPI_ISL_530422, EPI_ISL_530423, EPI_ISL_530424, EPI_ISL_530425, EPI_ISL_530426, EPI_ISL_530427, EPI_ISL_530428 | | | |
| see above | Lighthouse Lab in Glasgow | Wellcome Sanger Institute for the COVID-19 Genomics UK Consortium | Harper VanSteenhouse, Yumi Kasai, David Gray, Carol Clugston, Anna Dominiczak and Alex Alderton, Roberto Amato, Sonia Goncalves, Ewan Harrison, David K. Jackson, Ian Johnston, Dominic Kwiatkowski, Cordelia Langford, John Sillitoe |
| EPI_ISL_530429 | NHSGGC West of Scotland Specialist Virology Centre / MRC-University of Glasgow Centre for Virus Research | Wellcome Sanger Institute for the COVID-19 Genomics UK Consortium | Ana da Silva Filipe, Natasha Johnson, Kathy Smollett, Daniel Mair, Stephen Carmichael, Lily Tong, Jenna Nichols, Elihu Aranday-Cortes, Kirstyn Brunker, Yasmin Parr, Kyriaki Nomikou; Sarah McDonald, Marc Niebel, Patawee Asamaphan; Richard Orton, Joseph Hughes, Sreenu Vattipally, David L Robertson; Alasdair MacLean, Rory Gunson; Kathy Li, Natasha Jesudason, Rajiv Shah, James Shepherd, Antonia Ho, Alice Broos, Emma Thomson and Alex Alderton, Roberto Amato, Sonia Goncalves, Ewan Harrison, David K. Jackson, Ian Johnston, Dominic Kwiatkowski, Cordelia Langford, John Sillitoe |
| EPI_ISL_530430, EPI_ISL_530431, EPI_ISL_530432, EPI_ISL_530433, EPI_ISL_530434 | Lighthouse Lab in Glasgow | Wellcome Sanger Institute for the COVID-19 Genomics UK Consortium | Harper VanSteenhouse, Yumi Kasai, David Gray, Carol Clugston, Anna Dominiczak and Alex Alderton, Roberto Amato, Sonia Goncalves, Ewan Harrison, David K. Jackson, Ian Johnston, Dominic Kwiatkowski, Cordelia Langford, John Sillitoe |
| EPI_ISL_530435 | NHSGGC West of Scotland Specialist Virology Centre / MRC-University of Glasgow Centre for Virus Research | Wellcome Sanger Institute for the COVID-19 Genomics UK Consortium | Ana da Silva Filipe, Natasha Johnson, Kathy Smollett, Daniel Mair, Stephen Carmichael, Lily Tong, Jenna Nichols, Elihu Aranday-Cortes, Kirstyn Brunker, Yasmin Parr, Kyriaki Nomikou; Sarah McDonald, Marc Niebel, Patawee Asamaphan; Richard Orton, Joseph Hughes, Sreenu Vattipally, David L Robertson; Alasdair MacLean, Rory Gunson; Kathy Li, Natasha Jesudason, Rajiv Shah, James Shepherd, Antonia Ho, Alice Broos, Emma Thomson and Alex Alderton, Roberto Amato, Sonia Goncalves, Ewan Harrison, David K. Jackson, Ian Johnston, Dominic Kwiatkowski, Cordelia Langford, John Sillitoe |
| EPI_ISL_530436, EPI_ISL_530437, EPI_ISL_530438, EPI_ISL_530439 | Lighthouse Lab in Glasgow | Wellcome Sanger Institute for the COVID-19 Genomics UK Consortium | Harper VanSteenhouse, Yumi Kasai, David Gray, Carol Clugston, Anna Dominiczak and Alex Alderton, Roberto Amato, Sonia Goncalves, Ewan Harrison, David K. Jackson, Ian Johnston, Dominic Kwiatkowski, Cordelia Langford, John Sillitoe |
| EPI_ISL_530440 | NHSGGC West of Scotland Specialist Virology Centre / MRC-University of Glasgow Centre for Virus Research | Wellcome Sanger Institute for the COVID-19 Genomics UK Consortium | Ana da Silva Filipe, Natasha Johnson, Kathy Smollett, Daniel Mair, Stephen Carmichael, Lily Tong, Jenna Nichols, Elihu Aranday-Cortes, Kirstyn Brunker, Yasmin Parr, Kyriaki Nomikou; Sarah McDonald, Marc Niebel, Patawee Asamaphan; Richard Orton, Joseph Hughes, Sreenu Vattipally, David L Robertson; Alasdair MacLean, Rory Gunson; Kathy Li, Natasha Jesudason, Rajiv Shah, James Shepherd, Antonia Ho, Alice Broos, Emma Thomson and Alex Alderton, Roberto Amato, Sonia Goncalves, Ewan Harrison, David K. Jackson, Ian Johnston, Dominic Kwiatkowski, Cordelia Langford, John Sillitoe |
| EPI_ISL_530441, EPI_ISL_530442, EPI_ISL_530443, EPI_ISL_530444, EPI_ISL_530445, EPI_ISL_530446, EPI_ISL_530447, EPI_ISL_530448, EPI_ISL_530449, EPI_ISL_530450, EPI_ISL_530451, EPI_ISL_530452, EPI_ISL_530453, EPI_ISL_530454, EPI_ISL_530455, EPI_ISL_530456, EPI_ISL_530457, EPI_ISL_530458, EPI_ISL_530459, EPI_ISL_530460, EPI_ISL_530461, EPI_ISL_530462, EPI_ISL_530463, EPI_ISL_530464, EPI_ISL_530465, EPI_ISL_530466, EPI_ISL_530467 | | | |

| | | | |
|---|---|---|---|
| see above | Lighthouse Lab in Glasgow | Wellcome Sanger Institute for the COVID-19 Genomics UK Consortium | Harper VanSteenhouse, Yumi Kasai, David Gray, Carol Clugston, Anna Dominiczak and Alex Alderton, Roberto Amato, Sonia Goncalves, Ewan Harrison, David K. Jackson, Ian Johnston, Dominic Kwiatkowski, Cordelia Langford, John Sillitoe |
| EPI_ISL_530468 | NHSGGC West of Scotland Specialist Virology Centre / MRC-University of Glasgow Centre for Virus Research | Wellcome Sanger Institute for the COVID-19 Genomics UK Consortium | Ana da Silva Filipe, Natasha Johnson, Kathy Smollett, Daniel Mair, Stephen Carmichael, Lily Tong, Jenna Nichols, Elihu Aranday-Cortes, Kirstyn Brunker, Yasmin Parr, Kyriaki Nomikou; Sarah McDonald, Marc Niebel, Patawee Asamaphan; Richard Orton, Joseph Hughes, Sreenu Vattipally, David L Robertson; Alasdair MacLean, Rory Gunson; Kathy Li, Natasha Jesudason, Rajiv Shah, James Shepherd, Antonia Ho, Alice Broos, Emma Thomson and Alex Alderton, Roberto Amato, Sonia Goncalves, Ewan Harrison, David K. Jackson, Ian Johnston, Dominic Kwiatkowski, Cordelia Langford, John Sillitoe |
| EPI_ISL_530470, EPI_ISL_530471, EPI_ISL_530472, EPI_ISL_530473, EPI_ISL_530474, EPI_ISL_530475, EPI_ISL_530476, EPI_ISL_530477, EPI_ISL_530478, EPI_ISL_530479 | Lighthouse Lab in Glasgow | Wellcome Sanger Institute for the COVID-19 Genomics UK Consortium | Harper VanSteenhouse, Yumi Kasai, David Gray, Carol Clugston, Anna Dominiczak and Alex Alderton, Roberto Amato, Sonia Goncalves, Ewan Harrison, David K. Jackson, Ian Johnston, Dominic Kwiatkowski, Cordelia Langford, John Sillitoe |
| EPI_ISL_530480 | NHSGGC West of Scotland Specialist Virology Centre / MRC-University of Glasgow Centre for Virus Research | Wellcome Sanger Institute for the COVID-19 Genomics UK Consortium | Ana da Silva Filipe, Natasha Johnson, Kathy Smollett, Daniel Mair, Stephen Carmichael, Lily Tong, Jenna Nichols, Elihu Aranday-Cortes, Kirstyn Brunker, Yasmin Parr, Kyriaki Nomikou; Sarah McDonald, Marc Niebel, Patawee Asamaphan; Richard Orton, Joseph Hughes, Sreenu Vattipally, David L Robertson; Alasdair MacLean, Rory Gunson; Kathy Li, Natasha Jesudason, Rajiv Shah, James Shepherd, Antonia Ho, Alice Broos, Emma Thomson and Alex Alderton, Roberto Amato, Sonia Goncalves, Ewan Harrison, David K. Jackson, Ian Johnston, Dominic Kwiatkowski, Cordelia Langford, John Sillitoe |
| EPI_ISL_530481, EPI_ISL_530483, EPI_ISL_530484, EPI_ISL_530485, EPI_ISL_530486, EPI_ISL_530487 | Lighthouse Lab in Glasgow | Wellcome Sanger Institute for the COVID-19 Genomics UK Consortium | Harper VanSteenhouse, Yumi Kasai, David Gray, Carol Clugston, Anna Dominiczak and Alex Alderton, Roberto Amato, Sonia Goncalves, Ewan Harrison, David K. Jackson, Ian Johnston, Dominic Kwiatkowski, Cordelia Langford, John Sillitoe |
| EPI_ISL_530488, EPI_ISL_530489 | NHSGGC West of Scotland Specialist Virology Centre / MRC-University of Glasgow Centre for Virus Research | Wellcome Sanger Institute for the COVID-19 Genomics UK Consortium | Ana da Silva Filipe, Natasha Johnson, Kathy Smollett, Daniel Mair, Stephen Carmichael, Lily Tong, Jenna Nichols, Elihu Aranday-Cortes, Kirstyn Brunker, Yasmin Parr, Kyriaki Nomikou; Sarah McDonald, Marc Niebel, Patawee Asamaphan; Richard Orton, Joseph Hughes, Sreenu Vattipally, David L Robertson; Alasdair MacLean, Rory Gunson; Kathy Li, Natasha Jesudason, Rajiv Shah, James Shepherd, Antonia Ho, Alice Broos, Emma Thomson and Alex Alderton, Roberto Amato, Sonia Goncalves, Ewan Harrison, David K. Jackson, Ian Johnston, Dominic Kwiatkowski, Cordelia Langford, John Sillitoe |
| EPI_ISL_530490, EPI_ISL_530491, EPI_ISL_530492, EPI_ISL_530493, EPI_ISL_530494, EPI_ISL_530495, EPI_ISL_530496, EPI_ISL_530498, EPI_ISL_530499, EPI_ISL_530500, EPI_ISL_530501, EPI_ISL_530502, EPI_ISL_530503, EPI_ISL_530504, EPI_ISL_530505, EPI_ISL_530506, EPI_ISL_530507, EPI_ISL_530508, EPI_ISL_530509, EPI_ISL_530510, EPI_ISL_530511, EPI_ISL_530512, EPI_ISL_530513, EPI_ISL_530514, EPI_ISL_530515, EPI_ISL_530516, EPI_ISL_530517, EPI_ISL_530518, EPI_ISL_530519, EPI_ISL_530520, EPI_ISL_530521, EPI_ISL_530522 | | | |
| see above | Lighthouse Lab in Glasgow | Wellcome Sanger Institute for the COVID-19 Genomics UK Consortium | Harper VanSteenhouse, Yumi Kasai, David Gray, Carol Clugston, Anna Dominiczak and Alex Alderton, Roberto Amato, Sonia Goncalves, Ewan Harrison, David K. Jackson, Ian Johnston, Dominic Kwiatkowski, Cordelia Langford, John Sillitoe |
| EPI_ISL_530523 | NHSGGC West of Scotland Specialist Virology Centre / MRC-University of Glasgow Centre for Virus Research | Wellcome Sanger Institute for the COVID-19 Genomics UK Consortium | Ana da Silva Filipe, Natasha Johnson, Kathy Smollett, Daniel Mair, Stephen Carmichael, Lily Tong, Jenna Nichols, Elihu Aranday-Cortes, Kirstyn Brunker, Yasmin Parr, Kyriaki Nomikou; Sarah McDonald, Marc Niebel, Patawee Asamaphan; Richard Orton, Joseph Hughes, Sreenu Vattipally, David L Robertson; Alasdair MacLean, Rory Gunson; Kathy Li, Natasha Jesudason, Rajiv Shah, James Shepherd, Antonia Ho, Alice Broos, Emma Thomson and Alex Alderton, Roberto Amato, Sonia Goncalves, Ewan Harrison, David K. Jackson, Ian Johnston, Dominic Kwiatkowski, Cordelia Langford, John Sillitoe |
| EPI_ISL_530524, EPI_ISL_530525 | Lighthouse Lab in Glasgow | Wellcome Sanger Institute for the COVID-19 Genomics UK Consortium | Harper VanSteenhouse, Yumi Kasai, David Gray, Carol Clugston, Anna Dominiczak and Alex Alderton, Roberto Amato, Sonia Goncalves, Ewan Harrison, David K. Jackson, Ian Johnston, Dominic Kwiatkowski, Cordelia Langford, John Sillitoe |
| EPI_ISL_530528 | NHSGGC West of Scotland Specialist Virology Centre / MRC-University of Glasgow Centre for Virus Research | Wellcome Sanger Institute for the COVID-19 Genomics UK Consortium | Ana da Silva Filipe, Natasha Johnson, Kathy Smollett, Daniel Mair, Stephen Carmichael, Lily Tong, Jenna Nichols, Elihu Aranday-Cortes, Kirstyn Brunker, Yasmin Parr, Kyriaki Nomikou; Sarah McDonald, Marc Niebel, Patawee Asamaphan; Richard Orton, Joseph Hughes, Sreenu Vattipally, David L Robertson; Alasdair MacLean, Rory Gunson; Kathy Li, Natasha Jesudason, Rajiv Shah, James Shepherd, Antonia Ho, Alice Broos, Emma Thomson and Alex Alderton, Roberto Amato, Sonia Goncalves, Ewan Harrison, David K. Jackson, Ian Johnston, Dominic Kwiatkowski, Cordelia Langford, John Sillitoe |
| EPI_ISL_530529, EPI_ISL_530530, EPI_ISL_530531, EPI_ISL_530532 | Lighthouse Lab in Glasgow | Wellcome Sanger Institute for the COVID-19 Genomics UK Consortium | Harper VanSteenhouse, Yumi Kasai, David Gray, Carol Clugston, Anna Dominiczak and Alex Alderton, Roberto Amato, Sonia Goncalves, Ewan Harrison, David K. Jackson, Ian Johnston, Dominic Kwiatkowski, Cordelia Langford, John Sillitoe |
| EPI_ISL_530533 | NHSGGC West of Scotland Specialist Virology Centre / MRC-University of Glasgow Centre for Virus Research | Wellcome Sanger Institute for the COVID-19 Genomics UK Consortium | Ana da Silva Filipe, Natasha Johnson, Kathy Smollett, Daniel Mair, Stephen Carmichael, Lily Tong, Jenna Nichols, Elihu Aranday-Cortes, Kirstyn Brunker, Yasmin Parr, Kyriaki Nomikou; Sarah McDonald, Marc Niebel, Patawee Asamaphan; Richard Orton, Joseph Hughes, Sreenu Vattipally, David L Robertson; Alasdair MacLean, Rory Gunson; Kathy Li, Natasha Jesudason, Rajiv Shah, James Shepherd, Antonia Ho, Alice Broos, Emma Thomson and Alex Alderton, Roberto Amato, Sonia Goncalves, Ewan Harrison, David K. Jackson, Ian Johnston, Dominic Kwiatkowski, Cordelia Langford, John Sillitoe |
| EPI_ISL_530534 | Lighthouse Lab in Glasgow | Wellcome Sanger Institute for the COVID-19 Genomics UK Consortium | Harper VanSteenhouse, Yumi Kasai, David Gray, Carol Clugston, Anna Dominiczak and Alex Alderton, Roberto Amato, Sonia Goncalves, Ewan Harrison, David K. Jackson, Ian Johnston, Dominic Kwiatkowski, Cordelia Langford, John Sillitoe |
| EPI_ISL_530535 | NHSGGC West of Scotland Specialist Virology Centre / MRC-University of Glasgow Centre for Virus Research | Wellcome Sanger Institute for the COVID-19 Genomics UK Consortium | Ana da Silva Filipe, Natasha Johnson, Kathy Smollett, Daniel Mair, Stephen Carmichael, Lily Tong, Jenna Nichols, Elihu Aranday-Cortes, Kirstyn Brunker, Yasmin Parr, Kyriaki Nomikou; Sarah McDonald, Marc Niebel, Patawee Asamaphan; Richard Orton, Joseph Hughes, Sreenu Vattipally, David L Robertson; Alasdair MacLean, Rory Gunson; Kathy Li, Natasha Jesudason, Rajiv Shah, James Shepherd, Antonia Ho, Alice Broos, Emma Thomson and Alex Alderton, Roberto Amato, Sonia Goncalves, Ewan Harrison, David K. Jackson, Ian Johnston, Dominic Kwiatkowski, Cordelia Langford, John Sillitoe |
| EPI_ISL_530536, EPI_ISL_530537, EPI_ISL_530538, EPI_ISL_530539, EPI_ISL_530540, EPI_ISL_530541 | Lighthouse Lab in Glasgow | Wellcome Sanger Institute for the COVID-19 Genomics UK Consortium | Harper VanSteenhouse, Yumi Kasai, David Gray, Carol Clugston, Anna Dominiczak and Alex Alderton, Roberto Amato, Sonia Goncalves, Ewan Harrison, David K. Jackson, Ian Johnston, Dominic Kwiatkowski, Cordelia Langford, John Sillitoe |
| EPI_ISL_530542 | NHSGGC West of Scotland Specialist Virology Centre / MRC-University of Glasgow Centre for Virus Research | Wellcome Sanger Institute for the COVID-19 Genomics UK Consortium | Ana da Silva Filipe, Natasha Johnson, Kathy Smollett, Daniel Mair, Stephen Carmichael, Lily Tong, Jenna Nichols, Elihu Aranday-Cortes, Kirstyn Brunker, Yasmin Parr, Kyriaki Nomikou; Sarah McDonald, Marc Niebel, Patawee Asamaphan; Richard Orton, Joseph Hughes, Sreenu Vattipally, David L Robertson; Alasdair MacLean, Rory Gunson; Kathy Li, Natasha Jesudason, Rajiv Shah, James Shepherd, Antonia Ho, Alice Broos, Emma Thomson and Alex Alderton, Roberto Amato, Sonia Goncalves, Ewan Harrison, David K. Jackson, Ian Johnston, Dominic Kwiatkowski, Cordelia Langford, John Sillitoe |
| EPI_ISL_530543, EPI_ISL_530544, EPI_ISL_530545, EPI_ISL_530546, EPI_ISL_530547, EPI_ISL_530548, EPI_ISL_530549, EPI_ISL_530550, EPI_ISL_530551, EPI_ISL_530552 | Lighthouse Lab in Glasgow | Wellcome Sanger Institute for the COVID-19 Genomics UK Consortium | Harper VanSteenhouse, Yumi Kasai, David Gray, Carol Clugston, Anna Dominiczak and Alex Alderton, Roberto Amato, Sonia Goncalves, Ewan Harrison, David K. Jackson, Ian Johnston, Dominic Kwiatkowski, Cordelia Langford, John Sillitoe |
| EPI_ISL_530553 | NHSGGC West of Scotland Specialist Virology Centre / MRC-University of Glasgow Centre for Virus Research | Wellcome Sanger Institute for the COVID-19 Genomics UK Consortium | Ana da Silva Filipe, Natasha Johnson, Kathy Smollett, Daniel Mair, Stephen Carmichael, Lily Tong, Jenna Nichols, Elihu Aranday-Cortes, Kirstyn Brunker, Yasmin Parr, Kyriaki Nomikou; Sarah McDonald, Marc Niebel, Patawee Asamaphan; Richard Orton, Joseph Hughes, Sreenu Vattipally, David L Robertson; Alasdair MacLean, Rory Gunson; Kathy Li, Natasha Jesudason, Rajiv Shah, James Shepherd, Antonia Ho, Alice Broos, Emma Thomson and Alex Alderton, Roberto Amato, Sonia Goncalves, Ewan Harrison, David K. Jackson, Ian Johnston, Dominic Kwiatkowski, Cordelia Langford, John Sillitoe |
| EPI_ISL_530554, EPI_ISL_530555, EPI_ISL_530556, EPI_ISL_530557, EPI_ISL_530558, EPI_ISL_530559, EPI_ISL_530560, EPI_ISL_530561 | Lighthouse Lab in Glasgow | Wellcome Sanger Institute for the COVID-19 Genomics UK Consortium | Harper VanSteenhouse, Yumi Kasai, David Gray, Carol Clugston, Anna Dominiczak and Alex Alderton, Roberto Amato, Sonia Goncalves, Ewan Harrison, David K. Jackson, Ian Johnston, Dominic Kwiatkowski, Cordelia Langford, John Sillitoe |
| EPI_ISL_530562 | NHSGGC West of Scotland Specialist Virology Centre / MRC-University of Glasgow Centre for Virus Research | Wellcome Sanger Institute for the COVID-19 Genomics UK Consortium | Ana da Silva Filipe, Natasha Johnson, Kathy Smollett, Daniel Mair, Stephen Carmichael, Lily Tong, Jenna Nichols, Elihu Aranday-Cortes, Kirstyn Brunker, Yasmin Parr, Kyriaki Nomikou; Sarah McDonald, Marc Niebel, Patawee Asamaphan; Richard Orton, Joseph Hughes, Sreenu Vattipally, David L Robertson; Alasdair MacLean, Rory Gunson; Kathy Li, Natasha Jesudason, Rajiv Shah, James Shepherd, Antonia Ho, Alice Broos, Emma Thomson and Alex Alderton, Roberto Amato, Sonia Goncalves, Ewan Harrison, David K. Jackson, Ian Johnston, Dominic Kwiatkowski, Cordelia Langford, John Sillitoe |
| EPI_ISL_530563, EPI_ISL_530564, EPI_ISL_530565, EPI_ISL_530566, EPI_ISL_530567, EPI_ISL_530568, EPI_ISL_530569, EPI_ISL_530570, EPI_ISL_530571, EPI_ISL_530572, EPI_ISL_530573, EPI_ISL_530574, EPI_ISL_530575, EPI_ISL_530576, EPI_ISL_530578, EPI_ISL_530579, EPI_ISL_530580, EPI_ISL_530581, EPI_ISL_530582, EPI_ISL_530583, EPI_ISL_530584, EPI_ISL_530585, EPI_ISL_530586, EPI_ISL_530587, EPI_ISL_530588, EPI_ISL_530589, EPI_ISL_530590, EPI_ISL_530591, EPI_ISL_530592, EPI_ISL_530593, EPI_ISL_530594, EPI_ISL_530595 | | | |
| see above | Lighthouse Lab in Glasgow | Wellcome Sanger Institute for the COVID-19 Genomics UK Consortium | Harper VanSteenhouse, Yumi Kasai, David Gray, Carol Clugston, Anna Dominiczak and Alex Alderton, Roberto Amato, Sonia Goncalves, Ewan Harrison, David K. Jackson, Ian Johnston, Dominic Kwiatkowski, Cordelia Langford, John Sillitoe |
| EPI_ISL_530596 | NHSGGC West of Scotland Specialist Virology Centre / MRC-University of Glasgow Centre for Virus Research | Wellcome Sanger Institute for the COVID-19 Genomics UK Consortium | Ana da Silva Filipe, Natasha Johnson, Kathy Smollett, Daniel Mair, Stephen Carmichael, Lily Tong, Jenna Nichols, Elihu Aranday-Cortes, Kirstyn Brunker, Yasmin Parr, Kyriaki Nomikou; Sarah McDonald, Marc Niebel, Patawee Asamaphan; Richard Orton, Joseph Hughes, Sreenu Vattipally, David L Robertson; Alasdair MacLean, Rory Gunson; Kathy Li, Natasha Jesudason, Rajiv Shah, James Shepherd, Antonia Ho, Alice Broos, Emma Thomson and Alex Alderton, Roberto Amato, Sonia Goncalves, Ewan Harrison, David K. Jackson, Ian Johnston, Dominic Kwiatkowski, Cordelia Langford, John Sillitoe |
| EPI_ISL_530597, EPI_ISL_530598, EPI_ISL_530599, EPI_ISL_530600, EPI_ISL_530601, EPI_ISL_530602, EPI_ISL_530603, EPI_ISL_530604, EPI_ISL_530605, EPI_ISL_530606, EPI_ISL_530607, EPI_ISL_530608, EPI_ISL_530609, EPI_ISL_530610, EPI_ISL_530611, EPI_ISL_530612, EPI_ISL_530613, EPI_ISL_530614, EPI_ISL_530615, EPI_ISL_530616, EPI_ISL_530617, EPI_ISL_530618, EPI_ISL_530619, EPI_ISL_530620, EPI_ISL_530621, EPI_ISL_530622, EPI_ISL_530623, EPI_ISL_530624, EPI_ISL_530625, EPI_ISL_530626, EPI_ISL_530627, EPI_ISL_530628, EPI_ISL_530629, EPI_ISL_530630, EPI_ISL_530631, EPI_ISL_530632, EPI_ISL_530633, EPI_ISL_530634, EPI_ISL_530635, EPI_ISL_530636, EPI_ISL_530637, EPI_ISL_530638, EPI_ISL_530639, EPI_ISL_530640, EPI_ISL_530641 | | | |
| see above | Lighthouse Lab in Glasgow | Wellcome Sanger Institute for the COVID-19 Genomics UK Consortium | Harper VanSteenhouse, Yumi Kasai, David Gray, Carol Clugston, Anna Dominiczak and Alex Alderton, Roberto Amato, Sonia Goncalves, Ewan Harrison, David K. Jackson, Ian Johnston, Dominic Kwiatkowski, Cordelia Langford, John Sillitoe |
| EPI_ISL_530642 | NHSGGC West of Scotland Specialist Virology Centre / MRC-University of Glasgow Centre for Virus Research | Wellcome Sanger Institute for the COVID-19 Genomics UK Consortium | Ana da Silva Filipe, Natasha Johnson, Kathy Smollett, Daniel Mair, Stephen Carmichael, Lily Tong, Jenna Nichols, Elihu Aranday-Cortes, Kirstyn Brunker, Yasmin Parr, Kyriaki Nomikou; Sarah McDonald, Marc Niebel, Patawee Asamaphan; Richard Orton, Joseph Hughes, Sreenu Vattipally, David L Robertson; Alasdair MacLean, Rory Gunson; Kathy Li, Natasha Jesudason, Rajiv Shah, James Shepherd, Antonia Ho, Alice Broos, Emma Thomson and Alex Alderton, Roberto Amato, Sonia Goncalves, Ewan Harrison, David K. Jackson, Ian Johnston, Dominic Kwiatkowski, Cordelia Langford, John Sillitoe |
| EPI_ISL_530643, EPI_ISL_530644, EPI_ISL_530645, EPI_ISL_530646, EPI_ISL_530647, EPI_ISL_530648, EPI_ISL_530649, EPI_ISL_530650, EPI_ISL_530651, EPI_ISL_530652, EPI_ISL_530653, EPI_ISL_530654, EPI_ISL_530655, EPI_ISL_530656, EPI_ISL_530657, EPI_ISL_530658 | | | |
| see above | Lighthouse Lab in Glasgow | Wellcome Sanger Institute for the COVID-19 Genomics UK Consortium | Harper VanSteenhouse, Yumi Kasai, David Gray, Carol Clugston, Anna Dominiczak and Alex Alderton, Roberto Amato, Sonia Goncalves, Ewan Harrison, David K. Jackson, Ian Johnston, Dominic Kwiatkowski, Cordelia Langford, John Sillitoe |
| EPI_ISL_530659 | NHSGGC West of Scotland Specialist Virology Centre / MRC-University of Glasgow Centre for Virus Research | Wellcome Sanger Institute for the COVID-19 Genomics UK Consortium | Ana da Silva Filipe, Natasha Johnson, Kathy Smollett, Daniel Mair, Stephen Carmichael, Lily Tong, Jenna Nichols, Elihu Aranday-Cortes, Kirstyn Brunker, Yasmin Parr, Kyriaki Nomikou; Sarah McDonald, Marc Niebel, Patawee Asamaphan; Richard Orton, Joseph Hughes, Sreenu Vattipally, David L Robertson; Alasdair MacLean, Rory Gunson; Kathy Li, Natasha Jesudason, Rajiv Shah, James Shepherd, Antonia Ho, Alice Broos, Emma Thomson and Alex Alderton, Roberto Amato, Sonia Goncalves, Ewan Harrison, David K. Jackson, Ian Johnston, Dominic Kwiatkowski, Cordelia Langford, John Sillitoe |
| EPI_ISL_530660, EPI_ISL_530661, EPI_ISL_530662, EPI_ISL_530663, EPI_ISL_530664, EPI_ISL_530665, EPI_ISL_530666, EPI_ISL_530667, EPI_ISL_530668, EPI_ISL_530669, EPI_ISL_530670, EPI_ISL_530671, EPI_ISL_530672 | | | |
| see above | Lighthouse Lab in Glasgow | Wellcome Sanger Institute for the COVID-19 Genomics UK Consortium | Harper VanSteenhouse, Yumi Kasai, David Gray, Carol Clugston, Anna Dominiczak and Alex Alderton, Roberto Amato, Sonia Goncalves, Ewan Harrison, David K. Jackson, Ian Johnston, Dominic Kwiatkowski, Cordelia Langford, John Sillitoe |
| EPI_ISL_530673 | NHSGGC West of Scotland Specialist Virology Centre / MRC-University of Glasgow Centre for Virus Research | Wellcome Sanger Institute for the COVID-19 Genomics UK Consortium | Ana da Silva Filipe, Natasha Johnson, Kathy Smollett, Daniel Mair, Stephen Carmichael, Lily Tong, Jenna Nichols, Elihu Aranday-Cortes, Kirstyn Brunker, Yasmin Parr, Kyriaki Nomikou; Sarah McDonald, Marc Niebel, Patawee Asamaphan; Richard Orton, Joseph Hughes, Sreenu Vattipally, David L Robertson; Alasdair MacLean, Rory Gunson; Kathy Li, Natasha Jesudason, Rajiv Shah, James Shepherd, Antonia Ho, Alice Broos, Emma Thomson and Alex Alderton, Roberto Amato, Sonia Goncalves, Ewan Harrison, David K. Jackson, Ian Johnston, Dominic Kwiatkowski, Cordelia Langford, John Sillitoe |
| EPI_ISL_530674, EPI_ISL_530675, EPI_ISL_530676, EPI_ISL_530677, EPI_ISL_530678, EPI_ISL_530679, EPI_ISL_530680, EPI_ISL_530681 | Lighthouse Lab in Glasgow | Wellcome Sanger Institute for the COVID-19 Genomics UK Consortium | Harper VanSteenhouse, Yumi Kasai, David Gray, Carol Clugston, Anna Dominiczak and Alex Alderton, Roberto Amato, Sonia Goncalves, Ewan Harrison, David K. Jackson, Ian Johnston, Dominic Kwiatkowski, Cordelia Langford, John Sillitoe |
| EPI_ISL_530682 | NHSGGC West of Scotland Specialist Virology Centre / MRC-University of Glasgow Centre for Virus Research | Wellcome Sanger Institute for the COVID-19 Genomics UK Consortium | Ana da Silva Filipe, Natasha Johnson, Kathy Smollett, Daniel Mair, Stephen Carmichael, Lily Tong, Jenna Nichols, Elihu Aranday-Cortes, Kirstyn Brunker, Yasmin Parr, Kyriaki Nomikou; Sarah McDonald, Marc Niebel, Patawee Asamaphan; Richard Orton, Joseph Hughes, Sreenu Vattipally, David L Robertson; Alasdair MacLean, Rory Gunson; Kathy Li, Natasha Jesudason, Rajiv Shah, James Shepherd, Antonia Ho, Alice Broos, Emma Thomson and Alex Alderton, Roberto Amato, Sonia Goncalves, Ewan Harrison, David K. Jackson, Ian Johnston, Dominic Kwiatkowski, Cordelia Langford, John Sillitoe |
| EPI_ISL_530683, EPI_ISL_530684, EPI_ISL_530685, EPI_ISL_530687, EPI_ISL_530688, EPI_ISL_530690, EPI_ISL_530691, EPI_ISL_530692, EPI_ISL_530693, EPI_ISL_530694, EPI_ISL_530695, EPI_ISL_530696, EPI_ISL_530697, EPI_ISL_530698, EPI_ISL_530699, EPI_ISL_530700, EPI_ISL_530701, EPI_ISL_530702, EPI_ISL_530703, EPI_ISL_530704, EPI_ISL_530705, EPI_ISL_530706, EPI_ISL_530707 | | | |
| see above | Lighthouse Lab in Glasgow | Wellcome Sanger Institute for the COVID-19 Genomics UK Consortium | Harper VanSteenhouse, Yumi Kasai, David Gray, Carol Clugston, Anna Dominiczak and Alex Alderton, Roberto Amato, Sonia Goncalves, Ewan Harrison, David K. Jackson, Ian Johnston, Dominic Kwiatkowski, Cordelia Langford, John Sillitoe |
| EPI_ISL_530708 | NHSGGC West of Scotland Specialist Virology Centre / MRC- | Wellcome Sanger Institute for the COVID-19 Genomics UK | Ana da Silva Filipe, Natasha Johnson, Kathy Smollett, Daniel Mair, Stephen Carmichael, Lily Tong, Jenna Nichols, Elihu Aranday-Cortes, Kirstyn Brunker, Yasmin Parr, Kyriaki Nomikou; Sarah McDonald, Marc Niebel, |

| | | | |
|---|---|---|---|
| | University of Glasgow Centre for Virus Research | Consortium | Patawee Asamaphan; Richard Orton, Joseph Hughes, Sreenu Vattipally, David L Robertson; Alasdair MacLean, Rory Gunson; Kathy Li, Natasha Jesudason, Rajiv Shah, James Shepherd, Antonia Ho, Alice Broos, Emma Thomson and Alex Alderton, Roberto Amato, Sonia Goncalves, Ewan Harrison, David K. Jackson, Ian Johnston, Dominic Kwiatkowski, Cordelia Langford, John Sillitoe |
| EPI_ISL_530709, EPI_ISL_530710, EPI_ISL_530711, EPI_ISL_530712, EPI_ISL_530713, EPI_ISL_530714, EPI_ISL_530715, EPI_ISL_530716, EPI_ISL_530717, EPI_ISL_530718, EPI_ISL_530719, EPI_ISL_530720, EPI_ISL_530721 | | | |
| see above | Lighthouse Lab in Glasgow | Wellcome Sanger Institute for the COVID-19 Genomics UK Consortium | Harper VanSteenhouse, Yumi Kasai, David Gray, Carol Clugston, Anna Dominiczak and Alex Alderton, Roberto Amato, Sonia Goncalves, Ewan Harrison, David K. Jackson, Ian Johnston, Dominic Kwiatkowski, Cordelia Langford, John Sillitoe |
| EPI_ISL_530722 | NHSGGC West of Scotland Specialist Virology Centre / MRC-University of Glasgow Centre for Virus Research | Wellcome Sanger Institute for the COVID-19 Genomics UK Consortium | Ana da Silva Filipe, Natasha Johnson, Kathy Smollett, Daniel Mair, Stephen Carmichael, Lily Tong, Jenna Nichols, Elihu Aranday-Cortes, Kirstyn Brunker, Yasmin Parr, Kyriaki Nomikou; Sarah McDonald, Marc Niebel, Patawee Asamaphan; Richard Orton, Joseph Hughes, Sreenu Vattipally, David L Robertson; Alasdair MacLean, Rory Gunson; Kathy Li, Natasha Jesudason, Rajiv Shah, James Shepherd, Antonia Ho, Alice Broos, Emma Thomson and Alex Alderton, Roberto Amato, Sonia Goncalves, Ewan Harrison, David K. Jackson, Ian Johnston, Dominic Kwiatkowski, Cordelia Langford, John Sillitoe |
| EPI_ISL_530723, EPI_ISL_530724, EPI_ISL_530725, EPI_ISL_530726, EPI_ISL_530727, EPI_ISL_530728, EPI_ISL_530729, EPI_ISL_530730, EPI_ISL_530731, EPI_ISL_530732, EPI_ISL_530733, EPI_ISL_530734, EPI_ISL_530735, EPI_ISL_530736, EPI_ISL_530737, EPI_ISL_530738, EPI_ISL_530739 | | | |
| see above | Lighthouse Lab in Glasgow | Wellcome Sanger Institute for the COVID-19 Genomics UK Consortium | Harper VanSteenhouse, Yumi Kasai, David Gray, Carol Clugston, Anna Dominiczak and Alex Alderton, Roberto Amato, Sonia Goncalves, Ewan Harrison, David K. Jackson, Ian Johnston, Dominic Kwiatkowski, Cordelia Langford, John Sillitoe |
| EPI_ISL_530740 | NHSGGC West of Scotland Specialist Virology Centre / MRC-University of Glasgow Centre for Virus Research | Wellcome Sanger Institute for the COVID-19 Genomics UK Consortium | Ana da Silva Filipe, Natasha Johnson, Kathy Smollett, Daniel Mair, Stephen Carmichael, Lily Tong, Jenna Nichols, Elihu Aranday-Cortes, Kirstyn Brunker, Yasmin Parr, Kyriaki Nomikou; Sarah McDonald, Marc Niebel, Patawee Asamaphan; Richard Orton, Joseph Hughes, Sreenu Vattipally, David L Robertson; Alasdair MacLean, Rory Gunson; Kathy Li, Natasha Jesudason, Rajiv Shah, James Shepherd, Antonia Ho, Alice Broos, Emma Thomson and Alex Alderton, Roberto Amato, Sonia Goncalves, Ewan Harrison, David K. Jackson, Ian Johnston, Dominic Kwiatkowski, Cordelia Langford, John Sillitoe |
| EPI_ISL_530741, EPI_ISL_530742 | Lighthouse Lab in Glasgow | Wellcome Sanger Institute for the COVID-19 Genomics UK Consortium | Harper VanSteenhouse, Yumi Kasai, David Gray, Carol Clugston, Anna Dominiczak and Alex Alderton, Roberto Amato, Sonia Goncalves, Ewan Harrison, David K. Jackson, Ian Johnston, Dominic Kwiatkowski, Cordelia Langford, John Sillitoe |
| EPI_ISL_530743 | NHSGGC West of Scotland Specialist Virology Centre / MRC-University of Glasgow Centre for Virus Research | Wellcome Sanger Institute for the COVID-19 Genomics UK Consortium | Ana da Silva Filipe, Natasha Johnson, Kathy Smollett, Daniel Mair, Stephen Carmichael, Lily Tong, Jenna Nichols, Elihu Aranday-Cortes, Kirstyn Brunker, Yasmin Parr, Kyriaki Nomikou; Sarah McDonald, Marc Niebel, Patawee Asamaphan; Richard Orton, Joseph Hughes, Sreenu Vattipally, David L Robertson; Alasdair MacLean, Rory Gunson; Kathy Li, Natasha Jesudason, Rajiv Shah, James Shepherd, Antonia Ho, Alice Broos, Emma Thomson and Alex Alderton, Roberto Amato, Sonia Goncalves, Ewan Harrison, David K. Jackson, Ian Johnston, Dominic Kwiatkowski, Cordelia Langford, John Sillitoe |
| EPI_ISL_530744, EPI_ISL_530745, EPI_ISL_530747, EPI_ISL_530748, EPI_ISL_530749, EPI_ISL_530750, EPI_ISL_530751, EPI_ISL_530752, EPI_ISL_530753 | Lighthouse Lab in Glasgow | Wellcome Sanger Institute for the COVID-19 Genomics UK Consortium | Harper VanSteenhouse, Yumi Kasai, David Gray, Carol Clugston, Anna Dominiczak and Alex Alderton, Roberto Amato, Sonia Goncalves, Ewan Harrison, David K. Jackson, Ian Johnston, Dominic Kwiatkowski, Cordelia Langford, John Sillitoe |
| EPI_ISL_530754 | NHSGGC West of Scotland Specialist Virology Centre / MRC-University of Glasgow Centre for Virus Research | Wellcome Sanger Institute for the COVID-19 Genomics UK Consortium | Ana da Silva Filipe, Natasha Johnson, Kathy Smollett, Daniel Mair, Stephen Carmichael, Lily Tong, Jenna Nichols, Elihu Aranday-Cortes, Kirstyn Brunker, Yasmin Parr, Kyriaki Nomikou; Sarah McDonald, Marc Niebel, Patawee Asamaphan; Richard Orton, Joseph Hughes, Sreenu Vattipally, David L Robertson; Alasdair MacLean, Rory Gunson; Kathy Li, Natasha Jesudason, Rajiv Shah, James Shepherd, Antonia Ho, Alice Broos, Emma Thomson and Alex Alderton, Roberto Amato, Sonia Goncalves, Ewan Harrison, David K. Jackson, Ian Johnston, Dominic Kwiatkowski, Cordelia Langford, John Sillitoe |
| EPI_ISL_530755, EPI_ISL_530756, EPI_ISL_530757, EPI_ISL_530758, EPI_ISL_530759, EPI_ISL_530760, EPI_ISL_530761, EPI_ISL_530762, EPI_ISL_530763 | Lighthouse Lab in Glasgow | Wellcome Sanger Institute for the COVID-19 Genomics UK Consortium | Harper VanSteenhouse, Yumi Kasai, David Gray, Carol Clugston, Anna Dominiczak and Alex Alderton, Roberto Amato, Sonia Goncalves, Ewan Harrison, David K. Jackson, Ian Johnston, Dominic Kwiatkowski, Cordelia Langford, John Sillitoe |
| EPI_ISL_530764 | NHSGGC West of Scotland Specialist Virology Centre / MRC-University of Glasgow Centre for Virus Research | Wellcome Sanger Institute for the COVID-19 Genomics UK Consortium | Ana da Silva Filipe, Natasha Johnson, Kathy Smollett, Daniel Mair, Stephen Carmichael, Lily Tong, Jenna Nichols, Elihu Aranday-Cortes, Kirstyn Brunker, Yasmin Parr, Kyriaki Nomikou; Sarah McDonald, Marc Niebel, Patawee Asamaphan; Richard Orton, Joseph Hughes, Sreenu Vattipally, David L Robertson; Alasdair MacLean, Rory Gunson; Kathy Li, Natasha Jesudason, Rajiv Shah, James Shepherd, Antonia Ho, Alice Broos, Emma Thomson and Alex Alderton, Roberto Amato, Sonia Goncalves, Ewan Harrison, David K. Jackson, Ian Johnston, Dominic Kwiatkowski, Cordelia Langford, John Sillitoe |
| EPI_ISL_530765 | Lighthouse Lab in Glasgow | Wellcome Sanger Institute for the COVID-19 Genomics UK Consortium | Harper VanSteenhouse, Yumi Kasai, David Gray, Carol Clugston, Anna Dominiczak and Alex Alderton, Roberto Amato, Sonia Goncalves, Ewan Harrison, David K. Jackson, Ian Johnston, Dominic Kwiatkowski, Cordelia Langford, John Sillitoe |
| EPI_ISL_530766, EPI_ISL_530767, EPI_ISL_530768 | NHSGGC West of Scotland Specialist Virology Centre / MRC-University of Glasgow Centre for Virus Research | Wellcome Sanger Institute for the COVID-19 Genomics UK Consortium | Ana da Silva Filipe, Natasha Johnson, Kathy Smollett, Daniel Mair, Stephen Carmichael, Lily Tong, Jenna Nichols, Elihu Aranday-Cortes, Kirstyn Brunker, Yasmin Parr, Kyriaki Nomikou; Sarah McDonald, Marc Niebel, Patawee Asamaphan; Richard Orton, Joseph Hughes, Sreenu Vattipally, David L Robertson; Alasdair MacLean, Rory Gunson; Kathy Li, Natasha Jesudason, Rajiv Shah, James Shepherd, Antonia Ho, Alice Broos, Emma Thomson and Alex Alderton, Roberto Amato, Sonia Goncalves, Ewan Harrison, David K. Jackson, Ian Johnston, Dominic Kwiatkowski, Cordelia Langford, John Sillitoe |
| EPI_ISL_530769, EPI_ISL_530770, EPI_ISL_530771, EPI_ISL_530772, EPI_ISL_530773, EPI_ISL_530774, EPI_ISL_530775, EPI_ISL_530776, EPI_ISL_530777, EPI_ISL_530778, EPI_ISL_530779, EPI_ISL_530780, EPI_ISL_530781, EPI_ISL_530782, EPI_ISL_530783, EPI_ISL_530784, EPI_ISL_530785, EPI_ISL_530786, EPI_ISL_530787, EPI_ISL_530788, EPI_ISL_530789, EPI_ISL_530790, EPI_ISL_530791, EPI_ISL_530792, EPI_ISL_530793, EPI_ISL_530794, EPI_ISL_530796, EPI_ISL_530797, EPI_ISL_530798, EPI_ISL_530799, EPI_ISL_530801, EPI_ISL_530802, EPI_ISL_530803, EPI_ISL_530804, EPI_ISL_530805, EPI_ISL_530806, EPI_ISL_530807, EPI_ISL_530808, EPI_ISL_530809, EPI_ISL_530810, EPI_ISL_530811, EPI_ISL_530812, EPI_ISL_530813, EPI_ISL_530814, EPI_ISL_530815, EPI_ISL_530816, EPI_ISL_530817, EPI_ISL_530818, EPI_ISL_530819, EPI_ISL_530820, EPI_ISL_530821, EPI_ISL_530822, EPI_ISL_530823, EPI_ISL_530824, EPI_ISL_530825 | | | |
| see above | Lighthouse Lab in Glasgow | Wellcome Sanger Institute for the COVID-19 Genomics UK Consortium | Harper VanSteenhouse, Yumi Kasai, David Gray, Carol Clugston, Anna Dominiczak and Alex Alderton, Roberto Amato, Sonia Goncalves, Ewan Harrison, David K. Jackson, Ian Johnston, Dominic Kwiatkowski, Cordelia Langford, John Sillitoe |
| EPI_ISL_530826 | NHSGGC West of Scotland Specialist Virology Centre / MRC-University of Glasgow Centre for Virus Research | Wellcome Sanger Institute for the COVID-19 Genomics UK Consortium | Ana da Silva Filipe, Natasha Johnson, Kathy Smollett, Daniel Mair, Stephen Carmichael, Lily Tong, Jenna Nichols, Elihu Aranday-Cortes, Kirstyn Brunker, Yasmin Parr, Kyriaki Nomikou; Sarah McDonald, Marc Niebel, Patawee Asamaphan; Richard Orton, Joseph Hughes, Sreenu Vattipally, David L Robertson; Alasdair MacLean, Rory Gunson; Kathy Li, Natasha Jesudason, Rajiv Shah, James Shepherd, Antonia Ho, Alice Broos, Emma Thomson and Alex Alderton, Roberto Amato, Sonia Goncalves, Ewan Harrison, David K. Jackson, Ian Johnston, Dominic Kwiatkowski, Cordelia Langford, John Sillitoe |
| EPI_ISL_530827, EPI_ISL_530828, EPI_ISL_530829, EPI_ISL_530830, EPI_ISL_530831, EPI_ISL_530832, EPI_ISL_530833, EPI_ISL_530834, EPI_ISL_530835 | Lighthouse Lab in Glasgow | Wellcome Sanger Institute for the COVID-19 Genomics UK Consortium | Harper VanSteenhouse, Yumi Kasai, David Gray, Carol Clugston, Anna Dominiczak and Alex Alderton, Roberto Amato, Sonia Goncalves, Ewan Harrison, David K. Jackson, Ian Johnston, Dominic Kwiatkowski, Cordelia Langford, John Sillitoe |
| EPI_ISL_530836 | NHSGGC West of Scotland Specialist Virology Centre / MRC-University of Glasgow Centre for Virus Research | Wellcome Sanger Institute for the COVID-19 Genomics UK Consortium | Ana da Silva Filipe, Natasha Johnson, Kathy Smollett, Daniel Mair, Stephen Carmichael, Lily Tong, Jenna Nichols, Elihu Aranday-Cortes, Kirstyn Brunker, Yasmin Parr, Kyriaki Nomikou; Sarah McDonald, Marc Niebel, Patawee Asamaphan; Richard Orton, Joseph Hughes, Sreenu Vattipally, David L Robertson; Alasdair MacLean, Rory Gunson; Kathy Li, Natasha Jesudason, Rajiv Shah, James Shepherd, Antonia Ho, Alice Broos, Emma Thomson and Alex Alderton, Roberto Amato, Sonia Goncalves, Ewan Harrison, David K. Jackson, Ian Johnston, Dominic Kwiatkowski, Cordelia Langford, John Sillitoe |
| EPI_ISL_530837, EPI_ISL_530838, EPI_ISL_530839, EPI_ISL_530840, EPI_ISL_530841, EPI_ISL_530842, EPI_ISL_530843, EPI_ISL_530844 | Lighthouse Lab in Glasgow | Wellcome Sanger Institute for the COVID-19 Genomics UK Consortium | Harper VanSteenhouse, Yumi Kasai, David Gray, Carol Clugston, Anna Dominiczak and Alex Alderton, Roberto Amato, Sonia Goncalves, Ewan Harrison, David K. Jackson, Ian Johnston, Dominic Kwiatkowski, Cordelia Langford, John Sillitoe |
| EPI_ISL_530845 | NHSGGC West of Scotland Specialist Virology Centre / MRC-University of Glasgow Centre for Virus Research | Wellcome Sanger Institute for the COVID-19 Genomics UK Consortium | Ana da Silva Filipe, Natasha Johnson, Kathy Smollett, Daniel Mair, Stephen Carmichael, Lily Tong, Jenna Nichols, Elihu Aranday-Cortes, Kirstyn Brunker, Yasmin Parr, Kyriaki Nomikou; Sarah McDonald, Marc Niebel, Patawee Asamaphan; Richard Orton, Joseph Hughes, Sreenu Vattipally, David L Robertson; Alasdair MacLean, Rory Gunson; Kathy Li, Natasha Jesudason, Rajiv Shah, James Shepherd, Antonia Ho, Alice Broos, Emma Thomson and Alex Alderton, Roberto Amato, Sonia Goncalves, Ewan Harrison, David K. Jackson, Ian Johnston, Dominic Kwiatkowski, Cordelia Langford, John Sillitoe |
| EPI_ISL_530846, EPI_ISL_530847, EPI_ISL_530848, EPI_ISL_530849, EPI_ISL_530850, EPI_ISL_530851, EPI_ISL_530852, EPI_ISL_530853, EPI_ISL_530854, EPI_ISL_530855, EPI_ISL_530856, EPI_ISL_530857, EPI_ISL_530858, EPI_ISL_530859, EPI_ISL_530860, EPI_ISL_530861 | | | |
| see above | Lighthouse Lab in Glasgow | Wellcome Sanger Institute for the COVID-19 Genomics UK Consortium | Harper VanSteenhouse, Yumi Kasai, David Gray, Carol Clugston, Anna Dominiczak and Alex Alderton, Roberto Amato, Sonia Goncalves, Ewan Harrison, David K. Jackson, Ian Johnston, Dominic Kwiatkowski, Cordelia Langford, John Sillitoe |
| EPI_ISL_530862 | NHSGGC West of Scotland Specialist Virology Centre / MRC-University of Glasgow Centre for Virus Research | Wellcome Sanger Institute for the COVID-19 Genomics UK Consortium | Ana da Silva Filipe, Natasha Johnson, Kathy Smollett, Daniel Mair, Stephen Carmichael, Lily Tong, Jenna Nichols, Elihu Aranday-Cortes, Kirstyn Brunker, Yasmin Parr, Kyriaki Nomikou; Sarah McDonald, Marc Niebel, Patawee Asamaphan; Richard Orton, Joseph Hughes, Sreenu Vattipally, David L Robertson; Alasdair MacLean, Rory Gunson; Kathy Li, Natasha Jesudason, Rajiv Shah, James Shepherd, Antonia Ho, Alice Broos, Emma Thomson and Alex Alderton, Roberto Amato, Sonia Goncalves, Ewan Harrison, David K. Jackson, Ian Johnston, Dominic Kwiatkowski, Cordelia Langford, John Sillitoe |
| EPI_ISL_530863, EPI_ISL_530864, EPI_ISL_530865, EPI_ISL_530866, EPI_ISL_530867, EPI_ISL_530869, EPI_ISL_530870 | Lighthouse Lab in Glasgow | Wellcome Sanger Institute for the COVID-19 Genomics UK Consortium | Harper VanSteenhouse, Yumi Kasai, David Gray, Carol Clugston, Anna Dominiczak and Alex Alderton, Roberto Amato, Sonia Goncalves, Ewan Harrison, David K. Jackson, Ian Johnston, Dominic Kwiatkowski, Cordelia Langford, John Sillitoe |
| EPI_ISL_530871 | NHSGGC West of Scotland Specialist Virology Centre / MRC-University of Glasgow Centre for Virus Research | Wellcome Sanger Institute for the COVID-19 Genomics UK Consortium | Ana da Silva Filipe, Natasha Johnson, Kathy Smollett, Daniel Mair, Stephen Carmichael, Lily Tong, Jenna Nichols, Elihu Aranday-Cortes, Kirstyn Brunker, Yasmin Parr, Kyriaki Nomikou; Sarah McDonald, Marc Niebel, Patawee Asamaphan; Richard Orton, Joseph Hughes, Sreenu Vattipally, David L Robertson; Alasdair MacLean, Rory Gunson; Kathy Li, Natasha Jesudason, Rajiv Shah, James Shepherd, Antonia Ho, Alice Broos, Emma Thomson and Alex Alderton, Roberto Amato, Sonia Goncalves, Ewan Harrison, David K. Jackson, Ian Johnston, Dominic Kwiatkowski, Cordelia Langford, John Sillitoe |
| EPI_ISL_530872 | Lighthouse Lab in Glasgow | Wellcome Sanger Institute for the COVID-19 Genomics UK Consortium | Harper VanSteenhouse, Yumi Kasai, David Gray, Carol Clugston, Anna Dominiczak and Alex Alderton, Roberto Amato, Sonia Goncalves, Ewan Harrison, David K. Jackson, Ian Johnston, Dominic Kwiatkowski, Cordelia Langford, John Sillitoe |
| EPI_ISL_530873 | NHSGGC West of Scotland Specialist Virology Centre / MRC-University of Glasgow Centre for Virus Research | Wellcome Sanger Institute for the COVID-19 Genomics UK Consortium | Ana da Silva Filipe, Natasha Johnson, Kathy Smollett, Daniel Mair, Stephen Carmichael, Lily Tong, Jenna Nichols, Elihu Aranday-Cortes, Kirstyn Brunker, Yasmin Parr, Kyriaki Nomikou; Sarah McDonald, Marc Niebel, Patawee Asamaphan; Richard Orton, Joseph Hughes, Sreenu Vattipally, David L Robertson; Alasdair MacLean, Rory Gunson; Kathy Li, Natasha Jesudason, Rajiv Shah, James Shepherd, Antonia Ho, Alice Broos, Emma Thomson and Alex Alderton, Roberto Amato, Sonia Goncalves, Ewan Harrison, David K. Jackson, Ian Johnston, Dominic Kwiatkowski, Cordelia Langford, John Sillitoe |
| EPI_ISL_530875, EPI_ISL_530876, EPI_ISL_530877, EPI_ISL_530878, EPI_ISL_530879, EPI_ISL_530880, EPI_ISL_530881, EPI_ISL_530882, EPI_ISL_530883, EPI_ISL_530884, EPI_ISL_530885, EPI_ISL_530886, EPI_ISL_530887, EPI_ISL_530888, EPI_ISL_530889, EPI_ISL_530890, EPI_ISL_530891, EPI_ISL_530892, EPI_ISL_530893, EPI_ISL_530894, EPI_ISL_530895, EPI_ISL_530896, EPI_ISL_530897, EPI_ISL_530900, EPI_ISL_530901, EPI_ISL_530902, EPI_ISL_530903 | | | |
| see above | Lighthouse Lab in Glasgow | Wellcome Sanger Institute for the COVID-19 Genomics UK Consortium | Harper VanSteenhouse, Yumi Kasai, David Gray, Carol Clugston, Anna Dominiczak and Alex Alderton, Roberto Amato, Sonia Goncalves, Ewan Harrison, David K. Jackson, Ian Johnston, Dominic Kwiatkowski, Cordelia Langford, John Sillitoe |
| EPI_ISL_530904 | NHSGGC West of Scotland Specialist Virology Centre / MRC-University of Glasgow Centre for Virus Research | Wellcome Sanger Institute for the COVID-19 Genomics UK Consortium | Ana da Silva Filipe, Natasha Johnson, Kathy Smollett, Daniel Mair, Stephen Carmichael, Lily Tong, Jenna Nichols, Elihu Aranday-Cortes, Kirstyn Brunker, Yasmin Parr, Kyriaki Nomikou; Sarah McDonald, Marc Niebel, Patawee Asamaphan; Richard Orton, Joseph Hughes, Sreenu Vattipally, David L Robertson; Alasdair MacLean, Rory Gunson; Kathy Li, Natasha Jesudason, Rajiv Shah, James Shepherd, Antonia Ho, Alice Broos, Emma Thomson and Alex Alderton, Roberto Amato, Sonia Goncalves, Ewan Harrison, David K. Jackson, Ian Johnston, Dominic Kwiatkowski, Cordelia Langford, John Sillitoe |
| EPI_ISL_530905 | Lighthouse Lab in Glasgow | Wellcome Sanger Institute for the COVID-19 Genomics UK Consortium | Harper VanSteenhouse, Yumi Kasai, David Gray, Carol Clugston, Anna Dominiczak and Alex Alderton, Roberto Amato, Sonia Goncalves, Ewan Harrison, David K. Jackson, Ian Johnston, Dominic Kwiatkowski, Cordelia Langford, John Sillitoe |
| EPI_ISL_530906 | NHSGGC West of Scotland Specialist Virology Centre / MRC-University of Glasgow Centre for Virus Research | Wellcome Sanger Institute for the COVID-19 Genomics UK Consortium | Ana da Silva Filipe, Natasha Johnson, Kathy Smollett, Daniel Mair, Stephen Carmichael, Lily Tong, Jenna Nichols, Elihu Aranday-Cortes, Kirstyn Brunker, Yasmin Parr, Kyriaki Nomikou; Sarah McDonald, Marc Niebel, Patawee Asamaphan; Richard Orton, Joseph Hughes, Sreenu Vattipally, David L Robertson; Alasdair MacLean, Rory Gunson; Kathy Li, Natasha Jesudason, Rajiv Shah, James Shepherd, Antonia Ho, Alice Broos, Emma Thomson and Alex Alderton, Roberto Amato, Sonia Goncalves, Ewan Harrison, David K. Jackson, Ian Johnston, Dominic Kwiatkowski, Cordelia Langford, John Sillitoe |
| EPI_ISL_530907, EPI_ISL_530908, EPI_ISL_530909, EPI_ISL_530910, EPI_ISL_530911, EPI_ISL_530912, EPI_ISL_530913, EPI_ISL_530914, EPI_ISL_530915, EPI_ISL_530916, EPI_ISL_530917, EPI_ISL_530918, EPI_ISL_530919, EPI_ISL_530920, EPI_ISL_530921, EPI_ISL_530922 | | | |
| see above | Lighthouse Lab in Glasgow | Wellcome Sanger Institute for the COVID-19 Genomics UK Consortium | Harper VanSteenhouse, Yumi Kasai, David Gray, Carol Clugston, Anna Dominiczak and Alex Alderton, Roberto Amato, Sonia Goncalves, Ewan Harrison, David K. Jackson, Ian Johnston, Dominic Kwiatkowski, Cordelia Langford, John Sillitoe |
| EPI_ISL_530923 | NHSGGC West of Scotland Specialist Virology Centre / MRC-University of Glasgow Centre for Virus Research | Wellcome Sanger Institute for the COVID-19 Genomics UK Consortium | Ana da Silva Filipe, Natasha Johnson, Kathy Smollett, Daniel Mair, Stephen Carmichael, Lily Tong, Jenna Nichols, Elihu Aranday-Cortes, Kirstyn Brunker, Yasmin Parr, Kyriaki Nomikou; Sarah McDonald, Marc Niebel, Patawee Asamaphan; Richard Orton, Joseph Hughes, Sreenu Vattipally, David L Robertson; Alasdair MacLean, Rory Gunson; Kathy Li, Natasha Jesudason, Rajiv Shah, James Shepherd, Antonia Ho, Alice Broos, Emma Thomson and Alex Alderton, Roberto Amato, Sonia Goncalves, Ewan Harrison, David K. Jackson, Ian Johnston, Dominic Kwiatkowski, Cordelia Langford, John Sillitoe |
| EPI_ISL_530924, EPI_ISL_530925, EPI_ISL_530926, EPI_ISL_530927, EPI_ISL_530928 | Lighthouse Lab in Glasgow | Wellcome Sanger Institute for the COVID-19 Genomics UK Consortium | Harper VanSteenhouse, Yumi Kasai, David Gray, Carol Clugston, Anna Dominiczak and Alex Alderton, Roberto Amato, Sonia Goncalves, Ewan Harrison, David K. Jackson, Ian Johnston, Dominic Kwiatkowski, Cordelia Langford, John Sillitoe |

| | | | |
|---|---|---|---|
| EPI_ISL_530929 | NHSGGC West of Scotland Specialist Virology Centre / MRC-University of Glasgow Centre for Virus Research | Wellcome Sanger Institute for the COVID-19 Genomics UK Consortium | Ana da Silva Filipe, Natasha Johnson, Kathy Smollett, Daniel Mair, Stephen Carmichael, Lily Tong, Jenna Nichols, Elihu Aranday-Cortes, Kirstyn Brunker, Yasmin Parr, Kyriaki Nomikou; Sarah McDonald, Marc Niebel, Patawee Asamaphan; Richard Orton, Joseph Hughes, Sreenu Vattipally, David L Robertson; Alasdair MacLean, Rory Gunson; Kathy Li, Natasha Jesudason, Rajiv Shah, James Shepherd, Antonia Ho, Alice Broos, Emma Thomson and Alex Alderton, Roberto Amato, Sonia Goncalves, Ewan Harrison, David K. Jackson, Ian Johnston, Dominic Kwiatkowski, Cordelia Langford, John Sillitoe |
| EPI_ISL_530930, EPI_ISL_530931, EPI_ISL_530932, EPI_ISL_530933, EPI_ISL_530934, EPI_ISL_530935, EPI_ISL_530936, EPI_ISL_530937, EPI_ISL_530938, EPI_ISL_530939, EPI_ISL_530940, EPI_ISL_530941, EPI_ISL_530942, EPI_ISL_530943, EPI_ISL_530944, EPI_ISL_530945, EPI_ISL_530946, EPI_ISL_530947, EPI_ISL_530948, EPI_ISL_530949, EPI_ISL_530950, EPI_ISL_530951, EPI_ISL_530952 | | | |
| see above | Lighthouse Lab in Glasgow | Wellcome Sanger Institute for the COVID-19 Genomics UK Consortium | Harper VanSteenhouse, Yumi Kasai, David Gray, Carol Clugston, Anna Dominiczak and Alex Alderton, Roberto Amato, Sonia Goncalves, Ewan Harrison, David K. Jackson, Ian Johnston, Dominic Kwiatkowski, Cordelia Langford, John Sillitoe |
| EPI_ISL_530953 | NHSGGC West of Scotland Specialist Virology Centre / MRC-University of Glasgow Centre for Virus Research | Wellcome Sanger Institute for the COVID-19 Genomics UK Consortium | Ana da Silva Filipe, Natasha Johnson, Kathy Smollett, Daniel Mair, Stephen Carmichael, Lily Tong, Jenna Nichols, Elihu Aranday-Cortes, Kirstyn Brunker, Yasmin Parr, Kyriaki Nomikou; Sarah McDonald, Marc Niebel, Patawee Asamaphan; Richard Orton, Joseph Hughes, Sreenu Vattipally, David L Robertson; Alasdair MacLean, Rory Gunson; Kathy Li, Natasha Jesudason, Rajiv Shah, James Shepherd, Antonia Ho, Alice Broos, Emma Thomson and Alex Alderton, Roberto Amato, Sonia Goncalves, Ewan Harrison, David K. Jackson, Ian Johnston, Dominic Kwiatkowski, Cordelia Langford, John Sillitoe |
| EPI_ISL_530954, EPI_ISL_530955, EPI_ISL_530956, EPI_ISL_530957, EPI_ISL_530958 | Lighthouse Lab in Glasgow | Wellcome Sanger Institute for the COVID-19 Genomics UK Consortium | Harper VanSteenhouse, Yumi Kasai, David Gray, Carol Clugston, Anna Dominiczak and Alex Alderton, Roberto Amato, Sonia Goncalves, Ewan Harrison, David K. Jackson, Ian Johnston, Dominic Kwiatkowski, Cordelia Langford, John Sillitoe |
| EPI_ISL_530959 | NHSGGC West of Scotland Specialist Virology Centre / MRC-University of Glasgow Centre for Virus Research | Wellcome Sanger Institute for the COVID-19 Genomics UK Consortium | Ana da Silva Filipe, Natasha Johnson, Kathy Smollett, Daniel Mair, Stephen Carmichael, Lily Tong, Jenna Nichols, Elihu Aranday-Cortes, Kirstyn Brunker, Yasmin Parr, Kyriaki Nomikou; Sarah McDonald, Marc Niebel, Patawee Asamaphan; Richard Orton, Joseph Hughes, Sreenu Vattipally, David L Robertson; Alasdair MacLean, Rory Gunson; Kathy Li, Natasha Jesudason, Rajiv Shah, James Shepherd, Antonia Ho, Alice Broos, Emma Thomson and Alex Alderton, Roberto Amato, Sonia Goncalves, Ewan Harrison, David K. Jackson, Ian Johnston, Dominic Kwiatkowski, Cordelia Langford, John Sillitoe |
| EPI_ISL_530960 | Lighthouse Lab in Glasgow | Wellcome Sanger Institute for the COVID-19 Genomics UK Consortium | Harper VanSteenhouse, Yumi Kasai, David Gray, Carol Clugston, Anna Dominiczak and Alex Alderton, Roberto Amato, Sonia Goncalves, Ewan Harrison, David K. Jackson, Ian Johnston, Dominic Kwiatkowski, Cordelia Langford, John Sillitoe |
| EPI_ISL_530961, EPI_ISL_530962 | NHSGGC West of Scotland Specialist Virology Centre / MRC-University of Glasgow Centre for Virus Research | Wellcome Sanger Institute for the COVID-19 Genomics UK Consortium | Ana da Silva Filipe, Natasha Johnson, Kathy Smollett, Daniel Mair, Stephen Carmichael, Lily Tong, Jenna Nichols, Elihu Aranday-Cortes, Kirstyn Brunker, Yasmin Parr, Kyriaki Nomikou; Sarah McDonald, Marc Niebel, Patawee Asamaphan; Richard Orton, Joseph Hughes, Sreenu Vattipally, David L Robertson; Alasdair MacLean, Rory Gunson; Kathy Li, Natasha Jesudason, Rajiv Shah, James Shepherd, Antonia Ho, Alice Broos, Emma Thomson and Alex Alderton, Roberto Amato, Sonia Goncalves, Ewan Harrison, David K. Jackson, Ian Johnston, Dominic Kwiatkowski, Cordelia Langford, John Sillitoe |
| EPI_ISL_530964, EPI_ISL_530965, EPI_ISL_530966, EPI_ISL_530967, EPI_ISL_530968, EPI_ISL_530969, EPI_ISL_530970, EPI_ISL_530971, EPI_ISL_530972 | Lighthouse Lab in Glasgow | Wellcome Sanger Institute for the COVID-19 Genomics UK Consortium | Harper VanSteenhouse, Yumi Kasai, David Gray, Carol Clugston, Anna Dominiczak and Alex Alderton, Roberto Amato, Sonia Goncalves, Ewan Harrison, David K. Jackson, Ian Johnston, Dominic Kwiatkowski, Cordelia Langford, John Sillitoe |
| EPI_ISL_530973 | NHSGGC West of Scotland Specialist Virology Centre / MRC-University of Glasgow Centre for Virus Research | Wellcome Sanger Institute for the COVID-19 Genomics UK Consortium | Ana da Silva Filipe, Natasha Johnson, Kathy Smollett, Daniel Mair, Stephen Carmichael, Lily Tong, Jenna Nichols, Elihu Aranday-Cortes, Kirstyn Brunker, Yasmin Parr, Kyriaki Nomikou; Sarah McDonald, Marc Niebel, Patawee Asamaphan; Richard Orton, Joseph Hughes, Sreenu Vattipally, David L Robertson; Alasdair MacLean, Rory Gunson; Kathy Li, Natasha Jesudason, Rajiv Shah, James Shepherd, Antonia Ho, Alice Broos, Emma Thomson and Alex Alderton, Roberto Amato, Sonia Goncalves, Ewan Harrison, David K. Jackson, Ian Johnston, Dominic Kwiatkowski, Cordelia Langford, John Sillitoe |
| EPI_ISL_530974, EPI_ISL_530975 | Lighthouse Lab in Glasgow | Wellcome Sanger Institute for the COVID-19 Genomics UK Consortium | Harper VanSteenhouse, Yumi Kasai, David Gray, Carol Clugston, Anna Dominiczak and Alex Alderton, Roberto Amato, Sonia Goncalves, Ewan Harrison, David K. Jackson, Ian Johnston, Dominic Kwiatkowski, Cordelia Langford, John Sillitoe |
| EPI_ISL_530976 | NHSGGC West of Scotland Specialist Virology Centre / MRC-University of Glasgow Centre for Virus Research | Wellcome Sanger Institute for the COVID-19 Genomics UK Consortium | Ana da Silva Filipe, Natasha Johnson, Kathy Smollett, Daniel Mair, Stephen Carmichael, Lily Tong, Jenna Nichols, Elihu Aranday-Cortes, Kirstyn Brunker, Yasmin Parr, Kyriaki Nomikou; Sarah McDonald, Marc Niebel, Patawee Asamaphan; Richard Orton, Joseph Hughes, Sreenu Vattipally, David L Robertson; Alasdair MacLean, Rory Gunson; Kathy Li, Natasha Jesudason, Rajiv Shah, James Shepherd, Antonia Ho, Alice Broos, Emma Thomson and Alex Alderton, Roberto Amato, Sonia Goncalves, Ewan Harrison, David K. Jackson, Ian Johnston, Dominic Kwiatkowski, Cordelia Langford, John Sillitoe |
| EPI_ISL_530977, EPI_ISL_530978 | Lighthouse Lab in Glasgow | Wellcome Sanger Institute for the COVID-19 Genomics UK Consortium | Harper VanSteenhouse, Yumi Kasai, David Gray, Carol Clugston, Anna Dominiczak and Alex Alderton, Roberto Amato, Sonia Goncalves, Ewan Harrison, David K. Jackson, Ian Johnston, Dominic Kwiatkowski, Cordelia Langford, John Sillitoe |
| EPI_ISL_530979 | NHSGGC West of Scotland Specialist Virology Centre / MRC-University of Glasgow Centre for Virus Research | Wellcome Sanger Institute for the COVID-19 Genomics UK Consortium | Ana da Silva Filipe, Natasha Johnson, Kathy Smollett, Daniel Mair, Stephen Carmichael, Lily Tong, Jenna Nichols, Elihu Aranday-Cortes, Kirstyn Brunker, Yasmin Parr, Kyriaki Nomikou; Sarah McDonald, Marc Niebel, Patawee Asamaphan; Richard Orton, Joseph Hughes, Sreenu Vattipally, David L Robertson; Alasdair MacLean, Rory Gunson; Kathy Li, Natasha Jesudason, Rajiv Shah, James Shepherd, Antonia Ho, Alice Broos, Emma Thomson and Alex Alderton, Roberto Amato, Sonia Goncalves, Ewan Harrison, David K. Jackson, Ian Johnston, Dominic Kwiatkowski, Cordelia Langford, John Sillitoe |
| EPI_ISL_530980, EPI_ISL_530981, EPI_ISL_530982 | Lighthouse Lab in Glasgow | Wellcome Sanger Institute for the COVID-19 Genomics UK Consortium | Harper VanSteenhouse, Yumi Kasai, David Gray, Carol Clugston, Anna Dominiczak and Alex Alderton, Roberto Amato, Sonia Goncalves, Ewan Harrison, David K. Jackson, Ian Johnston, Dominic Kwiatkowski, Cordelia Langford, John Sillitoe |
| EPI_ISL_530983 | NHSGGC West of Scotland Specialist Virology Centre / MRC-University of Glasgow Centre for Virus Research | Wellcome Sanger Institute for the COVID-19 Genomics UK Consortium | Ana da Silva Filipe, Natasha Johnson, Kathy Smollett, Daniel Mair, Stephen Carmichael, Lily Tong, Jenna Nichols, Elihu Aranday-Cortes, Kirstyn Brunker, Yasmin Parr, Kyriaki Nomikou; Sarah McDonald, Marc Niebel, Patawee Asamaphan; Richard Orton, Joseph Hughes, Sreenu Vattipally, David L Robertson; Alasdair MacLean, Rory Gunson; Kathy Li, Natasha Jesudason, Rajiv Shah, James Shepherd, Antonia Ho, Alice Broos, Emma Thomson and Alex Alderton, Roberto Amato, Sonia Goncalves, Ewan Harrison, David K. Jackson, Ian Johnston, Dominic Kwiatkowski, Cordelia Langford, John Sillitoe |
| EPI_ISL_530984 | Lighthouse Lab in Glasgow | Wellcome Sanger Institute for the COVID-19 Genomics UK Consortium | Harper VanSteenhouse, Yumi Kasai, David Gray, Carol Clugston, Anna Dominiczak and Alex Alderton, Roberto Amato, Sonia Goncalves, Ewan Harrison, David K. Jackson, Ian Johnston, Dominic Kwiatkowski, Cordelia Langford, John Sillitoe |
| EPI_ISL_530985, EPI_ISL_530986 | NHSGGC West of Scotland Specialist Virology Centre / MRC-University of Glasgow Centre for Virus Research | Wellcome Sanger Institute for the COVID-19 Genomics UK Consortium | Ana da Silva Filipe, Natasha Johnson, Kathy Smollett, Daniel Mair, Stephen Carmichael, Lily Tong, Jenna Nichols, Elihu Aranday-Cortes, Kirstyn Brunker, Yasmin Parr, Kyriaki Nomikou; Sarah McDonald, Marc Niebel, Patawee Asamaphan; Richard Orton, Joseph Hughes, Sreenu Vattipally, David L Robertson; Alasdair MacLean, Rory Gunson; Kathy Li, Natasha Jesudason, Rajiv Shah, James Shepherd, Antonia Ho, Alice Broos, Emma Thomson and Alex Alderton, Roberto Amato, Sonia Goncalves, Ewan Harrison, David K. Jackson, Ian Johnston, Dominic Kwiatkowski, Cordelia Langford, John Sillitoe |
| EPI_ISL_530987, EPI_ISL_530988, EPI_ISL_530989 | Lighthouse Lab in Glasgow | Wellcome Sanger Institute for the COVID-19 Genomics UK Consortium | Harper VanSteenhouse, Yumi Kasai, David Gray, Carol Clugston, Anna Dominiczak and Alex Alderton, Roberto Amato, Sonia Goncalves, Ewan Harrison, David K. Jackson, Ian Johnston, Dominic Kwiatkowski, Cordelia Langford, John Sillitoe |
| EPI_ISL_530990 | NHSGGC West of Scotland Specialist Virology Centre / MRC-University of Glasgow Centre for Virus Research | Wellcome Sanger Institute for the COVID-19 Genomics UK Consortium | Ana da Silva Filipe, Natasha Johnson, Kathy Smollett, Daniel Mair, Stephen Carmichael, Lily Tong, Jenna Nichols, Elihu Aranday-Cortes, Kirstyn Brunker, Yasmin Parr, Kyriaki Nomikou; Sarah McDonald, Marc Niebel, Patawee Asamaphan; Richard Orton, Joseph Hughes, Sreenu Vattipally, David L Robertson; Alasdair MacLean, Rory Gunson; Kathy Li, Natasha Jesudason, Rajiv Shah, James Shepherd, Antonia Ho, Alice Broos, Emma Thomson and Alex Alderton, Roberto Amato, Sonia Goncalves, Ewan Harrison, David K. Jackson, Ian Johnston, Dominic Kwiatkowski, Cordelia Langford, John Sillitoe |
| EPI_ISL_530991, EPI_ISL_530992, EPI_ISL_530993, EPI_ISL_530994, EPI_ISL_530995 | Lighthouse Lab in Glasgow | Wellcome Sanger Institute for the COVID-19 Genomics UK Consortium | Harper VanSteenhouse, Yumi Kasai, David Gray, Carol Clugston, Anna Dominiczak and Alex Alderton, Roberto Amato, Sonia Goncalves, Ewan Harrison, David K. Jackson, Ian Johnston, Dominic Kwiatkowski, Cordelia Langford, John Sillitoe |
| EPI_ISL_530996, EPI_ISL_530997 | NHSGGC West of Scotland Specialist Virology Centre / MRC-University of Glasgow Centre for Virus Research | Wellcome Sanger Institute for the COVID-19 Genomics UK Consortium | Ana da Silva Filipe, Natasha Johnson, Kathy Smollett, Daniel Mair, Stephen Carmichael, Lily Tong, Jenna Nichols, Elihu Aranday-Cortes, Kirstyn Brunker, Yasmin Parr, Kyriaki Nomikou; Sarah McDonald, Marc Niebel, Patawee Asamaphan; Richard Orton, Joseph Hughes, Sreenu Vattipally, David L Robertson; Alasdair MacLean, Rory Gunson; Kathy Li, Natasha Jesudason, Rajiv Shah, James Shepherd, Antonia Ho, Alice Broos, Emma Thomson and Alex Alderton, Roberto Amato, Sonia Goncalves, Ewan Harrison, David K. Jackson, Ian Johnston, Dominic Kwiatkowski, Cordelia Langford, John Sillitoe |
| EPI_ISL_530998, EPI_ISL_530999, EPI_ISL_531000, EPI_ISL_531001 | Lighthouse Lab in Glasgow | Wellcome Sanger Institute for the COVID-19 Genomics UK Consortium | Harper VanSteenhouse, Yumi Kasai, David Gray, Carol Clugston, Anna Dominiczak and Alex Alderton, Roberto Amato, Sonia Goncalves, Ewan Harrison, David K. Jackson, Ian Johnston, Dominic Kwiatkowski, Cordelia Langford, John Sillitoe |
| EPI_ISL_531002 | NHSGGC West of Scotland Specialist Virology Centre / MRC-University of Glasgow Centre for Virus Research | Wellcome Sanger Institute for the COVID-19 Genomics UK Consortium | Ana da Silva Filipe, Natasha Johnson, Kathy Smollett, Daniel Mair, Stephen Carmichael, Lily Tong, Jenna Nichols, Elihu Aranday-Cortes, Kirstyn Brunker, Yasmin Parr, Kyriaki Nomikou; Sarah McDonald, Marc Niebel, Patawee Asamaphan; Richard Orton, Joseph Hughes, Sreenu Vattipally, David L Robertson; Alasdair MacLean, Rory Gunson; Kathy Li, Natasha Jesudason, Rajiv Shah, James Shepherd, Antonia Ho, Alice Broos, Emma Thomson and Alex Alderton, Roberto Amato, Sonia Goncalves, Ewan Harrison, David K. Jackson, Ian Johnston, Dominic Kwiatkowski, Cordelia Langford, John Sillitoe |
| EPI_ISL_531003 | Lighthouse Lab in Glasgow | Wellcome Sanger Institute for the COVID-19 Genomics UK Consortium | Harper VanSteenhouse, Yumi Kasai, David Gray, Carol Clugston, Anna Dominiczak and Alex Alderton, Roberto Amato, Sonia Goncalves, Ewan Harrison, David K. Jackson, Ian Johnston, Dominic Kwiatkowski, Cordelia Langford, John Sillitoe |
| EPI_ISL_531004 | NHSGGC West of Scotland Specialist Virology Centre / MRC-University of Glasgow Centre for Virus Research | Wellcome Sanger Institute for the COVID-19 Genomics UK Consortium | Ana da Silva Filipe, Natasha Johnson, Kathy Smollett, Daniel Mair, Stephen Carmichael, Lily Tong, Jenna Nichols, Elihu Aranday-Cortes, Kirstyn Brunker, Yasmin Parr, Kyriaki Nomikou; Sarah McDonald, Marc Niebel, Patawee Asamaphan; Richard Orton, Joseph Hughes, Sreenu Vattipally, David L Robertson; Alasdair MacLean, Rory Gunson; Kathy Li, Natasha Jesudason, Rajiv Shah, James Shepherd, Antonia Ho, Alice Broos, Emma Thomson and Alex Alderton, Roberto Amato, Sonia Goncalves, Ewan Harrison, David K. Jackson, Ian Johnston, Dominic Kwiatkowski, Cordelia Langford, John Sillitoe |
| EPI_ISL_531005 | Lighthouse Lab in Glasgow | Wellcome Sanger Institute for the COVID-19 Genomics UK Consortium | Harper VanSteenhouse, Yumi Kasai, David Gray, Carol Clugston, Anna Dominiczak and Alex Alderton, Roberto Amato, Sonia Goncalves, Ewan Harrison, David K. Jackson, Ian Johnston, Dominic Kwiatkowski, Cordelia Langford, John Sillitoe |
| EPI_ISL_531006 | NHSGGC West of Scotland Specialist Virology Centre / MRC-University of Glasgow Centre for Virus Research | Wellcome Sanger Institute for the COVID-19 Genomics UK Consortium | Ana da Silva Filipe, Natasha Johnson, Kathy Smollett, Daniel Mair, Stephen Carmichael, Lily Tong, Jenna Nichols, Elihu Aranday-Cortes, Kirstyn Brunker, Yasmin Parr, Kyriaki Nomikou; Sarah McDonald, Marc Niebel, Patawee Asamaphan; Richard Orton, Joseph Hughes, Sreenu Vattipally, David L Robertson; Alasdair MacLean, Rory Gunson; Kathy Li, Natasha Jesudason, Rajiv Shah, James Shepherd, Antonia Ho, Alice Broos, Emma Thomson and Alex Alderton, Roberto Amato, Sonia Goncalves, Ewan Harrison, David K. Jackson, Ian Johnston, Dominic Kwiatkowski, Cordelia Langford, John Sillitoe |
| EPI_ISL_531007, EPI_ISL_531008, EPI_ISL_531009, EPI_ISL_531010, EPI_ISL_531011, EPI_ISL_531012, EPI_ISL_531013, EPI_ISL_531014, EPI_ISL_531015, EPI_ISL_531016 | Lighthouse Lab in Glasgow | Wellcome Sanger Institute for the COVID-19 Genomics UK Consortium | Harper VanSteenhouse, Yumi Kasai, David Gray, Carol Clugston, Anna Dominiczak and Alex Alderton, Roberto Amato, Sonia Goncalves, Ewan Harrison, David K. Jackson, Ian Johnston, Dominic Kwiatkowski, Cordelia Langford, John Sillitoe |
| EPI_ISL_531017 | NHSGGC West of Scotland Specialist Virology Centre / MRC-University of Glasgow Centre for Virus Research | Wellcome Sanger Institute for the COVID-19 Genomics UK Consortium | Ana da Silva Filipe, Natasha Johnson, Kathy Smollett, Daniel Mair, Stephen Carmichael, Lily Tong, Jenna Nichols, Elihu Aranday-Cortes, Kirstyn Brunker, Yasmin Parr, Kyriaki Nomikou; Sarah McDonald, Marc Niebel, Patawee Asamaphan; Richard Orton, Joseph Hughes, Sreenu Vattipally, David L Robertson; Alasdair MacLean, Rory Gunson; Kathy Li, Natasha Jesudason, Rajiv Shah, James Shepherd, Antonia Ho, Alice Broos, Emma Thomson and Alex Alderton, Roberto Amato, Sonia Goncalves, Ewan Harrison, David K. Jackson, Ian Johnston, Dominic Kwiatkowski, Cordelia Langford, John Sillitoe |
| EPI_ISL_531018, EPI_ISL_531019 | Lighthouse Lab in Glasgow | Wellcome Sanger Institute for the COVID-19 Genomics UK Consortium | Harper VanSteenhouse, Yumi Kasai, David Gray, Carol Clugston, Anna Dominiczak and Alex Alderton, Roberto Amato, Sonia Goncalves, Ewan Harrison, David K. Jackson, Ian Johnston, Dominic Kwiatkowski, Cordelia Langford, John Sillitoe |
| EPI_ISL_531020 | NHSGGC West of Scotland Specialist Virology Centre / MRC-University of Glasgow Centre for Virus Research | Wellcome Sanger Institute for the COVID-19 Genomics UK Consortium | Ana da Silva Filipe, Natasha Johnson, Kathy Smollett, Daniel Mair, Stephen Carmichael, Lily Tong, Jenna Nichols, Elihu Aranday-Cortes, Kirstyn Brunker, Yasmin Parr, Kyriaki Nomikou; Sarah McDonald, Marc Niebel, Patawee Asamaphan; Richard Orton, Joseph Hughes, Sreenu Vattipally, David L Robertson; Alasdair MacLean, Rory Gunson; Kathy Li, Natasha Jesudason, Rajiv Shah, James Shepherd, Antonia Ho, Alice Broos, Emma Thomson and Alex Alderton, Roberto Amato, Sonia Goncalves, Ewan Harrison, David K. Jackson, Ian Johnston, Dominic Kwiatkowski, Cordelia Langford, John Sillitoe |
| EPI_ISL_531021, EPI_ISL_531022, EPI_ISL_531023, EPI_ISL_531024, EPI_ISL_531026, EPI_ISL_531027, EPI_ISL_531028, EPI_ISL_531029, EPI_ISL_531030, EPI_ISL_531031 | Lighthouse Lab in Glasgow | Wellcome Sanger Institute for the COVID-19 Genomics UK Consortium | Harper VanSteenhouse, Yumi Kasai, David Gray, Carol Clugston, Anna Dominiczak and Alex Alderton, Roberto Amato, Sonia Goncalves, Ewan Harrison, David K. Jackson, Ian Johnston, Dominic Kwiatkowski, Cordelia Langford, John Sillitoe |
| EPI_ISL_531032, EPI_ISL_531033 | NHSGGC West of Scotland Specialist Virology Centre / MRC-University of Glasgow Centre for Virus Research | Wellcome Sanger Institute for the COVID-19 Genomics UK Consortium | Ana da Silva Filipe, Natasha Johnson, Kathy Smollett, Daniel Mair, Stephen Carmichael, Lily Tong, Jenna Nichols, Elihu Aranday-Cortes, Kirstyn Brunker, Yasmin Parr, Kyriaki Nomikou; Sarah McDonald, Marc Niebel, Patawee Asamaphan; Richard Orton, Joseph Hughes, Sreenu Vattipally, David L Robertson; Alasdair MacLean, Rory Gunson; Kathy Li, Natasha Jesudason, Rajiv Shah, James Shepherd, Antonia Ho, Alice Broos, Emma Thomson and Alex Alderton, Roberto Amato, Sonia Goncalves, Ewan Harrison, David K. Jackson, Ian Johnston, Dominic Kwiatkowski, Cordelia Langford, John Sillitoe |
| EPI_ISL_531034, EPI_ISL_531035, EPI_ISL_531036, EPI_ISL_531037 | Lighthouse Lab in Glasgow | Wellcome Sanger Institute for the COVID-19 Genomics UK Consortium | Harper VanSteenhouse, Yumi Kasai, David Gray, Carol Clugston, Anna Dominiczak and Alex Alderton, Roberto Amato, Sonia Goncalves, Ewan Harrison, David K. Jackson, Ian Johnston, Dominic Kwiatkowski, Cordelia Langford, John Sillitoe |
| EPI_ISL_531038 | NHSGGC West of Scotland Specialist Virology Centre / MRC-University of Glasgow Centre for Virus Research | Wellcome Sanger Institute for the COVID-19 Genomics UK Consortium | Ana da Silva Filipe, Natasha Johnson, Kathy Smollett, Daniel Mair, Stephen Carmichael, Lily Tong, Jenna Nichols, Elihu Aranday-Cortes, Kirstyn Brunker, Yasmin Parr, Kyriaki Nomikou; Sarah McDonald, Marc Niebel, Patawee Asamaphan; Richard Orton, Joseph Hughes, Sreenu Vattipally, David L Robertson; Alasdair MacLean, Rory Gunson; Kathy Li, Natasha Jesudason, Rajiv Shah, James Shepherd, Antonia Ho, Alice Broos, Emma Thomson and Alex Alderton, Roberto Amato, Sonia Goncalves, Ewan Harrison, David K. Jackson, Ian Johnston, Dominic Kwiatkowski, Cordelia Langford, John Sillitoe |

| | | | |
|---|---|---|---|
| EPI_ISL_531039 | Lighthouse Lab in Glasgow | Wellcome Sanger Institute for the COVID-19 Genomics UK Consortium | Harper VanSteenhouse, Yumi Kasai, David Gray, Carol Clugston, Anna Dominiczak and Alex Alderton, Roberto Amato, Sonia Goncalves, Ewan Harrison, David K. Jackson, Ian Johnston, Dominic Kwiatkowski, Cordelia Langford, John Sillitoe |
| EPI_ISL_531040 | NHSGGC West of Scotland Specialist Virology Centre / MRC-University of Glasgow Centre for Virus Research | Wellcome Sanger Institute for the COVID-19 Genomics UK Consortium | Ana da Silva Filipe, Natasha Johnson, Kathy Smollett, Daniel Mair, Stephen Carmichael, Lily Tong, Jenna Nichols, Elihu Aranday-Cortes, Kirstyn Brunker, Yasmin Parr, Kyriaki Nomikou; Sarah McDonald, Marc Niebel, Patawee Asamaphan; Richard Orton, Joseph Hughes, Sreenu Vattipally, David L Robertson; Alasdair MacLean, Rory Gunson; Kathy Li, Natasha Jesudason, Rajiv Shah, James Shepherd, Antonia Ho, Alice Broos, Emma Thomson and Alex Alderton, Roberto Amato, Sonia Goncalves, Ewan Harrison, David K. Jackson, Ian Johnston, Dominic Kwiatkowski, Cordelia Langford, John Sillitoe |
| EPI_ISL_531041, EPI_ISL_531042, EPI_ISL_531043, EPI_ISL_531044 | Lighthouse Lab in Glasgow | Wellcome Sanger Institute for the COVID-19 Genomics UK Consortium | Harper VanSteenhouse, Yumi Kasai, David Gray, Carol Clugston, Anna Dominiczak and Alex Alderton, Roberto Amato, Sonia Goncalves, Ewan Harrison, David K. Jackson, Ian Johnston, Dominic Kwiatkowski, Cordelia Langford, John Sillitoe |
| EPI_ISL_531045 | NHSGGC West of Scotland Specialist Virology Centre / MRC-University of Glasgow Centre for Virus Research | Wellcome Sanger Institute for the COVID-19 Genomics UK Consortium | Ana da Silva Filipe, Natasha Johnson, Kathy Smollett, Daniel Mair, Stephen Carmichael, Lily Tong, Jenna Nichols, Elihu Aranday-Cortes, Kirstyn Brunker, Yasmin Parr, Kyriaki Nomikou; Sarah McDonald, Marc Niebel, Patawee Asamaphan; Richard Orton, Joseph Hughes, Sreenu Vattipally, David L Robertson; Alasdair MacLean, Rory Gunson; Kathy Li, Natasha Jesudason, Rajiv Shah, James Shepherd, Antonia Ho, Alice Broos, Emma Thomson and Alex Alderton, Roberto Amato, Sonia Goncalves, Ewan Harrison, David K. Jackson, Ian Johnston, Dominic Kwiatkowski, Cordelia Langford, John Sillitoe |
| EPI_ISL_531046, EPI_ISL_531047 | Lighthouse Lab in Glasgow | Wellcome Sanger Institute for the COVID-19 Genomics UK Consortium | Harper VanSteenhouse, Yumi Kasai, David Gray, Carol Clugston, Anna Dominiczak and Alex Alderton, Roberto Amato, Sonia Goncalves, Ewan Harrison, David K. Jackson, Ian Johnston, Dominic Kwiatkowski, Cordelia Langford, John Sillitoe |
| EPI_ISL_531048, EPI_ISL_531049 | NHSGGC West of Scotland Specialist Virology Centre / MRC-University of Glasgow Centre for Virus Research | Wellcome Sanger Institute for the COVID-19 Genomics UK Consortium | Ana da Silva Filipe, Natasha Johnson, Kathy Smollett, Daniel Mair, Stephen Carmichael, Lily Tong, Jenna Nichols, Elihu Aranday-Cortes, Kirstyn Brunker, Yasmin Parr, Kyriaki Nomikou; Sarah McDonald, Marc Niebel, Patawee Asamaphan; Richard Orton, Joseph Hughes, Sreenu Vattipally, David L Robertson; Alasdair MacLean, Rory Gunson; Kathy Li, Natasha Jesudason, Rajiv Shah, James Shepherd, Antonia Ho, Alice Broos, Emma Thomson and Alex Alderton, Roberto Amato, Sonia Goncalves, Ewan Harrison, David K. Jackson, Ian Johnston, Dominic Kwiatkowski, Cordelia Langford, John Sillitoe |
| EPI_ISL_531050 | Lighthouse Lab in Glasgow | Wellcome Sanger Institute for the COVID-19 Genomics UK Consortium | Harper VanSteenhouse, Yumi Kasai, David Gray, Carol Clugston, Anna Dominiczak and Alex Alderton, Roberto Amato, Sonia Goncalves, Ewan Harrison, David K. Jackson, Ian Johnston, Dominic Kwiatkowski, Cordelia Langford, John Sillitoe |
| EPI_ISL_531052 | NHSGGC West of Scotland Specialist Virology Centre / MRC-University of Glasgow Centre for Virus Research | Wellcome Sanger Institute for the COVID-19 Genomics UK Consortium | Ana da Silva Filipe, Natasha Johnson, Kathy Smollett, Daniel Mair, Stephen Carmichael, Lily Tong, Jenna Nichols, Elihu Aranday-Cortes, Kirstyn Brunker, Yasmin Parr, Kyriaki Nomikou; Sarah McDonald, Marc Niebel, Patawee Asamaphan; Richard Orton, Joseph Hughes, Sreenu Vattipally, David L Robertson; Alasdair MacLean, Rory Gunson; Kathy Li, Natasha Jesudason, Rajiv Shah, James Shepherd, Antonia Ho, Alice Broos, Emma Thomson and Alex Alderton, Roberto Amato, Sonia Goncalves, Ewan Harrison, David K. Jackson, Ian Johnston, Dominic Kwiatkowski, Cordelia Langford, John Sillitoe |
| EPI_ISL_531053 | Lighthouse Lab in Glasgow | Wellcome Sanger Institute for the COVID-19 Genomics UK Consortium | Harper VanSteenhouse, Yumi Kasai, David Gray, Carol Clugston, Anna Dominiczak and Alex Alderton, Roberto Amato, Sonia Goncalves, Ewan Harrison, David K. Jackson, Ian Johnston, Dominic Kwiatkowski, Cordelia Langford, John Sillitoe |
| EPI_ISL_531054 | NHSGGC West of Scotland Specialist Virology Centre / MRC-University of Glasgow Centre for Virus Research | Wellcome Sanger Institute for the COVID-19 Genomics UK Consortium | Ana da Silva Filipe, Natasha Johnson, Kathy Smollett, Daniel Mair, Stephen Carmichael, Lily Tong, Jenna Nichols, Elihu Aranday-Cortes, Kirstyn Brunker, Yasmin Parr, Kyriaki Nomikou; Sarah McDonald, Marc Niebel, Patawee Asamaphan; Richard Orton, Joseph Hughes, Sreenu Vattipally, David L Robertson; Alasdair MacLean, Rory Gunson; Kathy Li, Natasha Jesudason, Rajiv Shah, James Shepherd, Antonia Ho, Alice Broos, Emma Thomson and Alex Alderton, Roberto Amato, Sonia Goncalves, Ewan Harrison, David K. Jackson, Ian Johnston, Dominic Kwiatkowski, Cordelia Langford, John Sillitoe |
| EPI_ISL_531055 | Lighthouse Lab in Glasgow | Wellcome Sanger Institute for the COVID-19 Genomics UK Consortium | Harper VanSteenhouse, Yumi Kasai, David Gray, Carol Clugston, Anna Dominiczak and Alex Alderton, Roberto Amato, Sonia Goncalves, Ewan Harrison, David K. Jackson, Ian Johnston, Dominic Kwiatkowski, Cordelia Langford, John Sillitoe |
| EPI_ISL_531056, EPI_ISL_531057 | NHSGGC West of Scotland Specialist Virology Centre / MRC-University of Glasgow Centre for Virus Research | Wellcome Sanger Institute for the COVID-19 Genomics UK Consortium | Ana da Silva Filipe, Natasha Johnson, Kathy Smollett, Daniel Mair, Stephen Carmichael, Lily Tong, Jenna Nichols, Elihu Aranday-Cortes, Kirstyn Brunker, Yasmin Parr, Kyriaki Nomikou; Sarah McDonald, Marc Niebel, Patawee Asamaphan; Richard Orton, Joseph Hughes, Sreenu Vattipally, David L Robertson; Alasdair MacLean, Rory Gunson; Kathy Li, Natasha Jesudason, Rajiv Shah, James Shepherd, Antonia Ho, Alice Broos, Emma Thomson and Alex Alderton, Roberto Amato, Sonia Goncalves, Ewan Harrison, David K. Jackson, Ian Johnston, Dominic Kwiatkowski, Cordelia Langford, John Sillitoe |
| EPI_ISL_531058, EPI_ISL_531059, EPI_ISL_531060, EPI_ISL_531061, EPI_ISL_531062, EPI_ISL_531063, EPI_ISL_531064 | Lighthouse Lab in Glasgow | Wellcome Sanger Institute for the COVID-19 Genomics UK Consortium | Harper VanSteenhouse, Yumi Kasai, David Gray, Carol Clugston, Anna Dominiczak and Alex Alderton, Roberto Amato, Sonia Goncalves, Ewan Harrison, David K. Jackson, Ian Johnston, Dominic Kwiatkowski, Cordelia Langford, John Sillitoe |
| EPI_ISL_531065 | NHSGGC West of Scotland Specialist Virology Centre / MRC-University of Glasgow Centre for Virus Research | Wellcome Sanger Institute for the COVID-19 Genomics UK Consortium | Ana da Silva Filipe, Natasha Johnson, Kathy Smollett, Daniel Mair, Stephen Carmichael, Lily Tong, Jenna Nichols, Elihu Aranday-Cortes, Kirstyn Brunker, Yasmin Parr, Kyriaki Nomikou; Sarah McDonald, Marc Niebel, Patawee Asamaphan; Richard Orton, Joseph Hughes, Sreenu Vattipally, David L Robertson; Alasdair MacLean, Rory Gunson; Kathy Li, Natasha Jesudason, Rajiv Shah, James Shepherd, Antonia Ho, Alice Broos, Emma Thomson and Alex Alderton, Roberto Amato, Sonia Goncalves, Ewan Harrison, David K. Jackson, Ian Johnston, Dominic Kwiatkowski, Cordelia Langford, John Sillitoe |
| EPI_ISL_531066, EPI_ISL_531067, EPI_ISL_531068, EPI_ISL_531069, EPI_ISL_531070, EPI_ISL_531071, EPI_ISL_531072, EPI_ISL_531073, EPI_ISL_531074, EPI_ISL_531075, EPI_ISL_531076, EPI_ISL_531077, EPI_ISL_531078 | | | |
| see above | Lighthouse Lab in Glasgow | Wellcome Sanger Institute for the COVID-19 Genomics UK Consortium | Harper VanSteenhouse, Yumi Kasai, David Gray, Carol Clugston, Anna Dominiczak and Alex Alderton, Roberto Amato, Sonia Goncalves, Ewan Harrison, David K. Jackson, Ian Johnston, Dominic Kwiatkowski, Cordelia Langford, John Sillitoe |
| EPI_ISL_531079 | NHSGGC West of Scotland Specialist Virology Centre / MRC-University of Glasgow Centre for Virus Research | Wellcome Sanger Institute for the COVID-19 Genomics UK Consortium | Ana da Silva Filipe, Natasha Johnson, Kathy Smollett, Daniel Mair, Stephen Carmichael, Lily Tong, Jenna Nichols, Elihu Aranday-Cortes, Kirstyn Brunker, Yasmin Parr, Kyriaki Nomikou; Sarah McDonald, Marc Niebel, Patawee Asamaphan; Richard Orton, Joseph Hughes, Sreenu Vattipally, David L Robertson; Alasdair MacLean, Rory Gunson; Kathy Li, Natasha Jesudason, Rajiv Shah, James Shepherd, Antonia Ho, Alice Broos, Emma Thomson and Alex Alderton, Roberto Amato, Sonia Goncalves, Ewan Harrison, David K. Jackson, Ian Johnston, Dominic Kwiatkowski, Cordelia Langford, John Sillitoe |
| EPI_ISL_531080, EPI_ISL_531081, EPI_ISL_531082 | Lighthouse Lab in Glasgow | Wellcome Sanger Institute for the COVID-19 Genomics UK Consortium | Harper VanSteenhouse, Yumi Kasai, David Gray, Carol Clugston, Anna Dominiczak and Alex Alderton, Roberto Amato, Sonia Goncalves, Ewan Harrison, David K. Jackson, Ian Johnston, Dominic Kwiatkowski, Cordelia Langford, John Sillitoe |
| EPI_ISL_531083 | NHSGGC West of Scotland Specialist Virology Centre / MRC-University of Glasgow Centre for Virus Research | Wellcome Sanger Institute for the COVID-19 Genomics UK Consortium | Ana da Silva Filipe, Natasha Johnson, Kathy Smollett, Daniel Mair, Stephen Carmichael, Lily Tong, Jenna Nichols, Elihu Aranday-Cortes, Kirstyn Brunker, Yasmin Parr, Kyriaki Nomikou; Sarah McDonald, Marc Niebel, Patawee Asamaphan; Richard Orton, Joseph Hughes, Sreenu Vattipally, David L Robertson; Alasdair MacLean, Rory Gunson; Kathy Li, Natasha Jesudason, Rajiv Shah, James Shepherd, Antonia Ho, Alice Broos, Emma Thomson and Alex Alderton, Roberto Amato, Sonia Goncalves, Ewan Harrison, David K. Jackson, Ian Johnston, Dominic Kwiatkowski, Cordelia Langford, John Sillitoe |
| EPI_ISL_531084, EPI_ISL_531085, EPI_ISL_531086 | Lighthouse Lab in Glasgow | Wellcome Sanger Institute for the COVID-19 Genomics UK Consortium | Harper VanSteenhouse, Yumi Kasai, David Gray, Carol Clugston, Anna Dominiczak and Alex Alderton, Roberto Amato, Sonia Goncalves, Ewan Harrison, David K. Jackson, Ian Johnston, Dominic Kwiatkowski, Cordelia Langford, John Sillitoe |
| EPI_ISL_531087 | NHSGGC West of Scotland Specialist Virology Centre / MRC-University of Glasgow Centre for Virus Research | Wellcome Sanger Institute for the COVID-19 Genomics UK Consortium | Ana da Silva Filipe, Natasha Johnson, Kathy Smollett, Daniel Mair, Stephen Carmichael, Lily Tong, Jenna Nichols, Elihu Aranday-Cortes, Kirstyn Brunker, Yasmin Parr, Kyriaki Nomikou; Sarah McDonald, Marc Niebel, Patawee Asamaphan; Richard Orton, Joseph Hughes, Sreenu Vattipally, David L Robertson; Alasdair MacLean, Rory Gunson; Kathy Li, Natasha Jesudason, Rajiv Shah, James Shepherd, Antonia Ho, Alice Broos, Emma Thomson and Alex Alderton, Roberto Amato, Sonia Goncalves, Ewan Harrison, David K. Jackson, Ian Johnston, Dominic Kwiatkowski, Cordelia Langford, John Sillitoe |
| EPI_ISL_531088, EPI_ISL_531089, EPI_ISL_531090 | Lighthouse Lab in Glasgow | Wellcome Sanger Institute for the COVID-19 Genomics UK Consortium | Harper VanSteenhouse, Yumi Kasai, David Gray, Carol Clugston, Anna Dominiczak and Alex Alderton, Roberto Amato, Sonia Goncalves, Ewan Harrison, David K. Jackson, Ian Johnston, Dominic Kwiatkowski, Cordelia Langford, John Sillitoe |
| EPI_ISL_531091 | NHSGGC West of Scotland Specialist Virology Centre / MRC-University of Glasgow Centre for Virus Research | Wellcome Sanger Institute for the COVID-19 Genomics UK Consortium | Ana da Silva Filipe, Natasha Johnson, Kathy Smollett, Daniel Mair, Stephen Carmichael, Lily Tong, Jenna Nichols, Elihu Aranday-Cortes, Kirstyn Brunker, Yasmin Parr, Kyriaki Nomikou; Sarah McDonald, Marc Niebel, Patawee Asamaphan; Richard Orton, Joseph Hughes, Sreenu Vattipally, David L Robertson; Alasdair MacLean, Rory Gunson; Kathy Li, Natasha Jesudason, Rajiv Shah, James Shepherd, Antonia Ho, Alice Broos, Emma Thomson and Alex Alderton, Roberto Amato, Sonia Goncalves, Ewan Harrison, David K. Jackson, Ian Johnston, Dominic Kwiatkowski, Cordelia Langford, John Sillitoe |
| EPI_ISL_531092, EPI_ISL_531093, EPI_ISL_531094, EPI_ISL_531095 | Lighthouse Lab in Glasgow | Wellcome Sanger Institute for the COVID-19 Genomics UK Consortium | Harper VanSteenhouse, Yumi Kasai, David Gray, Carol Clugston, Anna Dominiczak and Alex Alderton, Roberto Amato, Sonia Goncalves, Ewan Harrison, David K. Jackson, Ian Johnston, Dominic Kwiatkowski, Cordelia Langford, John Sillitoe |
| EPI_ISL_531096 | NHSGGC West of Scotland Specialist Virology Centre / MRC-University of Glasgow Centre for Virus Research | Wellcome Sanger Institute for the COVID-19 Genomics UK Consortium | Ana da Silva Filipe, Natasha Johnson, Kathy Smollett, Daniel Mair, Stephen Carmichael, Lily Tong, Jenna Nichols, Elihu Aranday-Cortes, Kirstyn Brunker, Yasmin Parr, Kyriaki Nomikou; Sarah McDonald, Marc Niebel, Patawee Asamaphan; Richard Orton, Joseph Hughes, Sreenu Vattipally, David L Robertson; Alasdair MacLean, Rory Gunson; Kathy Li, Natasha Jesudason, Rajiv Shah, James Shepherd, Antonia Ho, Alice Broos, Emma Thomson and Alex Alderton, Roberto Amato, Sonia Goncalves, Ewan Harrison, David K. Jackson, Ian Johnston, Dominic Kwiatkowski, Cordelia Langford, John Sillitoe |
| EPI_ISL_531097, EPI_ISL_531098 | Lighthouse Lab in Glasgow | Wellcome Sanger Institute for the COVID-19 Genomics UK Consortium | Harper VanSteenhouse, Yumi Kasai, David Gray, Carol Clugston, Anna Dominiczak and Alex Alderton, Roberto Amato, Sonia Goncalves, Ewan Harrison, David K. Jackson, Ian Johnston, Dominic Kwiatkowski, Cordelia Langford, John Sillitoe |
| EPI_ISL_531099, EPI_ISL_531100 | NHSGGC West of Scotland Specialist Virology Centre / MRC-University of Glasgow Centre for Virus Research | Wellcome Sanger Institute for the COVID-19 Genomics UK Consortium | Ana da Silva Filipe, Natasha Johnson, Kathy Smollett, Daniel Mair, Stephen Carmichael, Lily Tong, Jenna Nichols, Elihu Aranday-Cortes, Kirstyn Brunker, Yasmin Parr, Kyriaki Nomikou; Sarah McDonald, Marc Niebel, Patawee Asamaphan; Richard Orton, Joseph Hughes, Sreenu Vattipally, David L Robertson; Alasdair MacLean, Rory Gunson; Kathy Li, Natasha Jesudason, Rajiv Shah, James Shepherd, Antonia Ho, Alice Broos, Emma Thomson and Alex Alderton, Roberto Amato, Sonia Goncalves, Ewan Harrison, David K. Jackson, Ian Johnston, Dominic Kwiatkowski, Cordelia Langford, John Sillitoe |
| EPI_ISL_531101, EPI_ISL_531102 | Lighthouse Lab in Glasgow | Wellcome Sanger Institute for the COVID-19 Genomics UK Consortium | Harper VanSteenhouse, Yumi Kasai, David Gray, Carol Clugston, Anna Dominiczak and Alex Alderton, Roberto Amato, Sonia Goncalves, Ewan Harrison, David K. Jackson, Ian Johnston, Dominic Kwiatkowski, Cordelia Langford, John Sillitoe |
| EPI_ISL_531103, EPI_ISL_531104 | NHSGGC West of Scotland Specialist Virology Centre / MRC-University of Glasgow Centre for Virus Research | Wellcome Sanger Institute for the COVID-19 Genomics UK Consortium | Ana da Silva Filipe, Natasha Johnson, Kathy Smollett, Daniel Mair, Stephen Carmichael, Lily Tong, Jenna Nichols, Elihu Aranday-Cortes, Kirstyn Brunker, Yasmin Parr, Kyriaki Nomikou; Sarah McDonald, Marc Niebel, Patawee Asamaphan; Richard Orton, Joseph Hughes, Sreenu Vattipally, David L Robertson; Alasdair MacLean, Rory Gunson; Kathy Li, Natasha Jesudason, Rajiv Shah, James Shepherd, Antonia Ho, Alice Broos, Emma Thomson and Alex Alderton, Roberto Amato, Sonia Goncalves, Ewan Harrison, David K. Jackson, Ian Johnston, Dominic Kwiatkowski, Cordelia Langford, John Sillitoe |
| EPI_ISL_531105, EPI_ISL_531106, EPI_ISL_531107, EPI_ISL_531108, EPI_ISL_531109, EPI_ISL_531110, EPI_ISL_531111, EPI_ISL_531112, EPI_ISL_531113 | Lighthouse Lab in Glasgow | Wellcome Sanger Institute for the COVID-19 Genomics UK Consortium | Harper VanSteenhouse, Yumi Kasai, David Gray, Carol Clugston, Anna Dominiczak and Alex Alderton, Roberto Amato, Sonia Goncalves, Ewan Harrison, David K. Jackson, Ian Johnston, Dominic Kwiatkowski, Cordelia Langford, John Sillitoe |
| EPI_ISL_531114 | NHSGGC West of Scotland Specialist Virology Centre / MRC-University of Glasgow Centre for Virus Research | Wellcome Sanger Institute for the COVID-19 Genomics UK Consortium | Ana da Silva Filipe, Natasha Johnson, Kathy Smollett, Daniel Mair, Stephen Carmichael, Lily Tong, Jenna Nichols, Elihu Aranday-Cortes, Kirstyn Brunker, Yasmin Parr, Kyriaki Nomikou; Sarah McDonald, Marc Niebel, Patawee Asamaphan; Richard Orton, Joseph Hughes, Sreenu Vattipally, David L Robertson; Alasdair MacLean, Rory Gunson; Kathy Li, Natasha Jesudason, Rajiv Shah, James Shepherd, Antonia Ho, Alice Broos, Emma Thomson and Alex Alderton, Roberto Amato, Sonia Goncalves, Ewan Harrison, David K. Jackson, Ian Johnston, Dominic Kwiatkowski, Cordelia Langford, John Sillitoe |
| EPI_ISL_531115 | Lighthouse Lab in Glasgow | Wellcome Sanger Institute for the COVID-19 Genomics UK Consortium | Harper VanSteenhouse, Yumi Kasai, David Gray, Carol Clugston, Anna Dominiczak and Alex Alderton, Roberto Amato, Sonia Goncalves, Ewan Harrison, David K. Jackson, Ian Johnston, Dominic Kwiatkowski, Cordelia Langford, John Sillitoe |
| EPI_ISL_531116 | NHSGGC West of Scotland Specialist Virology Centre / MRC-University of Glasgow Centre for Virus Research | Wellcome Sanger Institute for the COVID-19 Genomics UK Consortium | Ana da Silva Filipe, Natasha Johnson, Kathy Smollett, Daniel Mair, Stephen Carmichael, Lily Tong, Jenna Nichols, Elihu Aranday-Cortes, Kirstyn Brunker, Yasmin Parr, Kyriaki Nomikou; Sarah McDonald, Marc Niebel, Patawee Asamaphan; Richard Orton, Joseph Hughes, Sreenu Vattipally, David L Robertson; Alasdair MacLean, Rory Gunson; Kathy Li, Natasha Jesudason, Rajiv Shah, James Shepherd, Antonia Ho, Alice Broos, Emma Thomson and Alex Alderton, Roberto Amato, Sonia Goncalves, Ewan Harrison, David K. Jackson, Ian Johnston, Dominic Kwiatkowski, Cordelia Langford, John Sillitoe |
| EPI_ISL_531117 | Lighthouse Lab in Glasgow | Wellcome Sanger Institute for the COVID-19 Genomics UK Consortium | Harper VanSteenhouse, Yumi Kasai, David Gray, Carol Clugston, Anna Dominiczak and Alex Alderton, Roberto Amato, Sonia Goncalves, Ewan Harrison, David K. Jackson, Ian Johnston, Dominic Kwiatkowski, Cordelia Langford, John Sillitoe |
| EPI_ISL_531118 | NHSGGC West of Scotland Specialist Virology Centre / MRC-University of Glasgow Centre for Virus Research | Wellcome Sanger Institute for the COVID-19 Genomics UK Consortium | Ana da Silva Filipe, Natasha Johnson, Kathy Smollett, Daniel Mair, Stephen Carmichael, Lily Tong, Jenna Nichols, Elihu Aranday-Cortes, Kirstyn Brunker, Yasmin Parr, Kyriaki Nomikou; Sarah McDonald, Marc Niebel, Patawee Asamaphan; Richard Orton, Joseph Hughes, Sreenu Vattipally, David L Robertson; Alasdair MacLean, Rory Gunson; Kathy Li, Natasha Jesudason, Rajiv Shah, James Shepherd, Antonia Ho, Alice Broos, Emma Thomson and Alex Alderton, Roberto Amato, Sonia Goncalves, Ewan Harrison, David K. Jackson, Ian Johnston, Dominic Kwiatkowski, Cordelia Langford, John Sillitoe |
| EPI_ISL_531119, EPI_ISL_531120, EPI_ISL_531121, EPI_ISL_531122 | Lighthouse Lab in Glasgow | Wellcome Sanger Institute for the COVID-19 Genomics UK Consortium | Harper VanSteenhouse, Yumi Kasai, David Gray, Carol Clugston, Anna Dominiczak and Alex Alderton, Roberto Amato, Sonia Goncalves, Ewan Harrison, David K. Jackson, Ian Johnston, Dominic Kwiatkowski, Cordelia Langford, John Sillitoe |
| EPI_ISL_531123 | NHSGGC West of Scotland Specialist Virology Centre / MRC-University of Glasgow Centre for Virus Research | Wellcome Sanger Institute for the COVID-19 Genomics UK Consortium | Ana da Silva Filipe, Natasha Johnson, Kathy Smollett, Daniel Mair, Stephen Carmichael, Lily Tong, Jenna Nichols, Elihu Aranday-Cortes, Kirstyn Brunker, Yasmin Parr, Kyriaki Nomikou; Sarah McDonald, Marc Niebel, Patawee Asamaphan; Richard Orton, Joseph Hughes, Sreenu Vattipally, David L Robertson; Alasdair MacLean, Rory Gunson; Kathy Li, Natasha Jesudason, Rajiv Shah, James Shepherd, Antonia Ho, Alice Broos, Emma Thomson and Alex Alderton, Roberto Amato, Sonia Goncalves, Ewan Harrison, David K. Jackson, Ian Johnston, Dominic Kwiatkowski, Cordelia Langford, John Sillitoe |

| | | | |
|---|---|---|---|
| EPI_ISL_531124, EPI_ISL_531125 | Lighthouse Lab in Glasgow | Wellcome Sanger Institute for the COVID-19 Genomics UK Consortium | Harper VanSteenhouse, Yumi Kasai, David Gray, Carol Clugston, Anna Dominiczak and Alex Alderton, Roberto Amato, Sonia Goncalves, Ewan Harrison, David K. Jackson, Ian Johnston, Dominic Kwiatkowski, Cordelia Langford, John Sillitoe |
| EPI_ISL_531126 | NHSGGC West of Scotland Specialist Virology Centre / MRC-University of Glasgow Centre for Virus Research | Wellcome Sanger Institute for the COVID-19 Genomics UK Consortium | Ana da Silva Filipe, Natasha Johnson, Kathy Smollett, Daniel Mair, Stephen Carmichael, Lily Tong, Jenna Nichols, Elihu Aranday-Cortes, Kirstyn Brunker, Yasmin Parr, Kyriaki Nomikou; Sarah McDonald, Marc Niebel, Patawee Asamaphan; Richard Orton, Joseph Hughes, Sreenu Vattipally, David L Robertson; Alasdair MacLean, Rory Gunson; Kathy Li, Natasha Jesudason, Rajiv Shah, James Shepherd, Antonia Ho, Alice Broos, Emma Thomson and Alex Alderton, Roberto Amato, Sonia Goncalves, Ewan Harrison, David K. Jackson, Ian Johnston, Dominic Kwiatkowski, Cordelia Langford, John Sillitoe |
| EPI_ISL_531127, EPI_ISL_531128, EPI_ISL_531129 | Lighthouse Lab in Glasgow | Wellcome Sanger Institute for the COVID-19 Genomics UK Consortium | Harper VanSteenhouse, Yumi Kasai, David Gray, Carol Clugston, Anna Dominiczak and Alex Alderton, Roberto Amato, Sonia Goncalves, Ewan Harrison, David K. Jackson, Ian Johnston, Dominic Kwiatkowski, Cordelia Langford, John Sillitoe |
| EPI_ISL_531130, EPI_ISL_531131 | NHSGGC West of Scotland Specialist Virology Centre / MRC-University of Glasgow Centre for Virus Research | Wellcome Sanger Institute for the COVID-19 Genomics UK Consortium | Ana da Silva Filipe, Natasha Johnson, Kathy Smollett, Daniel Mair, Stephen Carmichael, Lily Tong, Jenna Nichols, Elihu Aranday-Cortes, Kirstyn Brunker, Yasmin Parr, Kyriaki Nomikou; Sarah McDonald, Marc Niebel, Patawee Asamaphan; Richard Orton, Joseph Hughes, Sreenu Vattipally, David L Robertson; Alasdair MacLean, Rory Gunson; Kathy Li, Natasha Jesudason, Rajiv Shah, James Shepherd, Antonia Ho, Alice Broos, Emma Thomson and Alex Alderton, Roberto Amato, Sonia Goncalves, Ewan Harrison, David K. Jackson, Ian Johnston, Dominic Kwiatkowski, Cordelia Langford, John Sillitoe |
| EPI_ISL_531132, EPI_ISL_531133, EPI_ISL_531134, EPI_ISL_531135, EPI_ISL_531136, EPI_ISL_531137, EPI_ISL_531138, EPI_ISL_531139, EPI_ISL_531140, EPI_ISL_531141 | Lighthouse Lab in Glasgow | Wellcome Sanger Institute for the COVID-19 Genomics UK Consortium | Harper VanSteenhouse, Yumi Kasai, David Gray, Carol Clugston, Anna Dominiczak and Alex Alderton, Roberto Amato, Sonia Goncalves, Ewan Harrison, David K. Jackson, Ian Johnston, Dominic Kwiatkowski, Cordelia Langford, John Sillitoe |
| EPI_ISL_531142 | NHSGGC West of Scotland Specialist Virology Centre / MRC-University of Glasgow Centre for Virus Research | Wellcome Sanger Institute for the COVID-19 Genomics UK Consortium | Ana da Silva Filipe, Natasha Johnson, Kathy Smollett, Daniel Mair, Stephen Carmichael, Lily Tong, Jenna Nichols, Elihu Aranday-Cortes, Kirstyn Brunker, Yasmin Parr, Kyriaki Nomikou; Sarah McDonald, Marc Niebel, Patawee Asamaphan; Richard Orton, Joseph Hughes, Sreenu Vattipally, David L Robertson; Alasdair MacLean, Rory Gunson; Kathy Li, Natasha Jesudason, Rajiv Shah, James Shepherd, Antonia Ho, Alice Broos, Emma Thomson and Alex Alderton, Roberto Amato, Sonia Goncalves, Ewan Harrison, David K. Jackson, Ian Johnston, Dominic Kwiatkowski, Cordelia Langford, John Sillitoe |
| EPI_ISL_531143, EPI_ISL_531144, EPI_ISL_531145, EPI_ISL_531146 | Lighthouse Lab in Glasgow | Wellcome Sanger Institute for the COVID-19 Genomics UK Consortium | Harper VanSteenhouse, Yumi Kasai, David Gray, Carol Clugston, Anna Dominiczak and Alex Alderton, Roberto Amato, Sonia Goncalves, Ewan Harrison, David K. Jackson, Ian Johnston, Dominic Kwiatkowski, Cordelia Langford, John Sillitoe |
| EPI_ISL_531147 | NHSGGC West of Scotland Specialist Virology Centre / MRC-University of Glasgow Centre for Virus Research | Wellcome Sanger Institute for the COVID-19 Genomics UK Consortium | Ana da Silva Filipe, Natasha Johnson, Kathy Smollett, Daniel Mair, Stephen Carmichael, Lily Tong, Jenna Nichols, Elihu Aranday-Cortes, Kirstyn Brunker, Yasmin Parr, Kyriaki Nomikou; Sarah McDonald, Marc Niebel, Patawee Asamaphan; Richard Orton, Joseph Hughes, Sreenu Vattipally, David L Robertson; Alasdair MacLean, Rory Gunson; Kathy Li, Natasha Jesudason, Rajiv Shah, James Shepherd, Antonia Ho, Alice Broos, Emma Thomson and Alex Alderton, Roberto Amato, Sonia Goncalves, Ewan Harrison, David K. Jackson, Ian Johnston, Dominic Kwiatkowski, Cordelia Langford, John Sillitoe |
| EPI_ISL_531148, EPI_ISL_531149, EPI_ISL_531150, EPI_ISL_531151, EPI_ISL_531152, EPI_ISL_531153 | Lighthouse Lab in Glasgow | Wellcome Sanger Institute for the COVID-19 Genomics UK Consortium | Harper VanSteenhouse, Yumi Kasai, David Gray, Carol Clugston, Anna Dominiczak and Alex Alderton, Roberto Amato, Sonia Goncalves, Ewan Harrison, David K. Jackson, Ian Johnston, Dominic Kwiatkowski, Cordelia Langford, John Sillitoe |
| EPI_ISL_531154, EPI_ISL_531155, EPI_ISL_531156, EPI_ISL_531157 | NHSGGC West of Scotland Specialist Virology Centre / MRC-University of Glasgow Centre for Virus Research | Wellcome Sanger Institute for the COVID-19 Genomics UK Consortium | Ana da Silva Filipe, Natasha Johnson, Kathy Smollett, Daniel Mair, Stephen Carmichael, Lily Tong, Jenna Nichols, Elihu Aranday-Cortes, Kirstyn Brunker, Yasmin Parr, Kyriaki Nomikou; Sarah McDonald, Marc Niebel, Patawee Asamaphan; Richard Orton, Joseph Hughes, Sreenu Vattipally, David L Robertson; Alasdair MacLean, Rory Gunson; Kathy Li, Natasha Jesudason, Rajiv Shah, James Shepherd, Antonia Ho, Alice Broos, Emma Thomson and Alex Alderton, Roberto Amato, Sonia Goncalves, Ewan Harrison, David K. Jackson, Ian Johnston, Dominic Kwiatkowski, Cordelia Langford, John Sillitoe |
| EPI_ISL_531158 | Lighthouse Lab in Glasgow | Wellcome Sanger Institute for the COVID-19 Genomics UK Consortium | Harper VanSteenhouse, Yumi Kasai, David Gray, Carol Clugston, Anna Dominiczak and Alex Alderton, Roberto Amato, Sonia Goncalves, Ewan Harrison, David K. Jackson, Ian Johnston, Dominic Kwiatkowski, Cordelia Langford, John Sillitoe |
| EPI_ISL_531159 | NHSGGC West of Scotland Specialist Virology Centre / MRC-University of Glasgow Centre for Virus Research | Wellcome Sanger Institute for the COVID-19 Genomics UK Consortium | Ana da Silva Filipe, Natasha Johnson, Kathy Smollett, Daniel Mair, Stephen Carmichael, Lily Tong, Jenna Nichols, Elihu Aranday-Cortes, Kirstyn Brunker, Yasmin Parr, Kyriaki Nomikou; Sarah McDonald, Marc Niebel, Patawee Asamaphan; Richard Orton, Joseph Hughes, Sreenu Vattipally, David L Robertson; Alasdair MacLean, Rory Gunson; Kathy Li, Natasha Jesudason, Rajiv Shah, James Shepherd, Antonia Ho, Alice Broos, Emma Thomson and Alex Alderton, Roberto Amato, Sonia Goncalves, Ewan Harrison, David K. Jackson, Ian Johnston, Dominic Kwiatkowski, Cordelia Langford, John Sillitoe |
| EPI_ISL_531160, EPI_ISL_531161, EPI_ISL_531162, EPI_ISL_531163, EPI_ISL_531164, EPI_ISL_531165 | Lighthouse Lab in Glasgow | Wellcome Sanger Institute for the COVID-19 Genomics UK Consortium | Harper VanSteenhouse, Yumi Kasai, David Gray, Carol Clugston, Anna Dominiczak and Alex Alderton, Roberto Amato, Sonia Goncalves, Ewan Harrison, David K. Jackson, Ian Johnston, Dominic Kwiatkowski, Cordelia Langford, John Sillitoe |
| EPI_ISL_531166 | NHSGGC West of Scotland Specialist Virology Centre / MRC-University of Glasgow Centre for Virus Research | Wellcome Sanger Institute for the COVID-19 Genomics UK Consortium | Ana da Silva Filipe, Natasha Johnson, Kathy Smollett, Daniel Mair, Stephen Carmichael, Lily Tong, Jenna Nichols, Elihu Aranday-Cortes, Kirstyn Brunker, Yasmin Parr, Kyriaki Nomikou; Sarah McDonald, Marc Niebel, Patawee Asamaphan; Richard Orton, Joseph Hughes, Sreenu Vattipally, David L Robertson; Alasdair MacLean, Rory Gunson; Kathy Li, Natasha Jesudason, Rajiv Shah, James Shepherd, Antonia Ho, Alice Broos, Emma Thomson and Alex Alderton, Roberto Amato, Sonia Goncalves, Ewan Harrison, David K. Jackson, Ian Johnston, Dominic Kwiatkowski, Cordelia Langford, John Sillitoe |
| EPI_ISL_531167, EPI_ISL_531168, EPI_ISL_531169, EPI_ISL_531170, EPI_ISL_531171, EPI_ISL_531172, EPI_ISL_531173 | Lighthouse Lab in Glasgow | Wellcome Sanger Institute for the COVID-19 Genomics UK Consortium | Harper VanSteenhouse, Yumi Kasai, David Gray, Carol Clugston, Anna Dominiczak and Alex Alderton, Roberto Amato, Sonia Goncalves, Ewan Harrison, David K. Jackson, Ian Johnston, Dominic Kwiatkowski, Cordelia Langford, John Sillitoe |
| EPI_ISL_531174, EPI_ISL_531175 | NHSGGC West of Scotland Specialist Virology Centre / MRC-University of Glasgow Centre for Virus Research | Wellcome Sanger Institute for the COVID-19 Genomics UK Consortium | Ana da Silva Filipe, Natasha Johnson, Kathy Smollett, Daniel Mair, Stephen Carmichael, Lily Tong, Jenna Nichols, Elihu Aranday-Cortes, Kirstyn Brunker, Yasmin Parr, Kyriaki Nomikou; Sarah McDonald, Marc Niebel, Patawee Asamaphan; Richard Orton, Joseph Hughes, Sreenu Vattipally, David L Robertson; Alasdair MacLean, Rory Gunson; Kathy Li, Natasha Jesudason, Rajiv Shah, James Shepherd, Antonia Ho, Alice Broos, Emma Thomson and Alex Alderton, Roberto Amato, Sonia Goncalves, Ewan Harrison, David K. Jackson, Ian Johnston, Dominic Kwiatkowski, Cordelia Langford, John Sillitoe |
| EPI_ISL_531176, EPI_ISL_531177, EPI_ISL_531178, EPI_ISL_531179, EPI_ISL_531180, EPI_ISL_531181, EPI_ISL_531182, EPI_ISL_531183, EPI_ISL_531184, EPI_ISL_531185, EPI_ISL_531186, EPI_ISL_531187 | | | |
| see above | Lighthouse Lab in Glasgow | Wellcome Sanger Institute for the COVID-19 Genomics UK Consortium | Harper VanSteenhouse, Yumi Kasai, David Gray, Carol Clugston, Anna Dominiczak and Alex Alderton, Roberto Amato, Sonia Goncalves, Ewan Harrison, David K. Jackson, Ian Johnston, Dominic Kwiatkowski, Cordelia Langford, John Sillitoe |
| EPI_ISL_531188, EPI_ISL_531189 | NHSGGC West of Scotland Specialist Virology Centre / MRC-University of Glasgow Centre for Virus Research | Wellcome Sanger Institute for the COVID-19 Genomics UK Consortium | Ana da Silva Filipe, Natasha Johnson, Kathy Smollett, Daniel Mair, Stephen Carmichael, Lily Tong, Jenna Nichols, Elihu Aranday-Cortes, Kirstyn Brunker, Yasmin Parr, Kyriaki Nomikou; Sarah McDonald, Marc Niebel, Patawee Asamaphan; Richard Orton, Joseph Hughes, Sreenu Vattipally, David L Robertson; Alasdair MacLean, Rory Gunson; Kathy Li, Natasha Jesudason, Rajiv Shah, James Shepherd, Antonia Ho, Alice Broos, Emma Thomson and Alex Alderton, Roberto Amato, Sonia Goncalves, Ewan Harrison, David K. Jackson, Ian Johnston, Dominic Kwiatkowski, Cordelia Langford, John Sillitoe |
| EPI_ISL_531190, EPI_ISL_531191, EPI_ISL_531192, EPI_ISL_531193, EPI_ISL_531194, EPI_ISL_531195, EPI_ISL_531196, EPI_ISL_531197 | Lighthouse Lab in Glasgow | Wellcome Sanger Institute for the COVID-19 Genomics UK Consortium | Harper VanSteenhouse, Yumi Kasai, David Gray, Carol Clugston, Anna Dominiczak and Alex Alderton, Roberto Amato, Sonia Goncalves, Ewan Harrison, David K. Jackson, Ian Johnston, Dominic Kwiatkowski, Cordelia Langford, John Sillitoe |
| EPI_ISL_531198 | NHSGGC West of Scotland Specialist Virology Centre / MRC-University of Glasgow Centre for Virus Research | Wellcome Sanger Institute for the COVID-19 Genomics UK Consortium | Ana da Silva Filipe, Natasha Johnson, Kathy Smollett, Daniel Mair, Stephen Carmichael, Lily Tong, Jenna Nichols, Elihu Aranday-Cortes, Kirstyn Brunker, Yasmin Parr, Kyriaki Nomikou; Sarah McDonald, Marc Niebel, Patawee Asamaphan; Richard Orton, Joseph Hughes, Sreenu Vattipally, David L Robertson; Alasdair MacLean, Rory Gunson; Kathy Li, Natasha Jesudason, Rajiv Shah, James Shepherd, Antonia Ho, Alice Broos, Emma Thomson and Alex Alderton, Roberto Amato, Sonia Goncalves, Ewan Harrison, David K. Jackson, Ian Johnston, Dominic Kwiatkowski, Cordelia Langford, John Sillitoe |
| EPI_ISL_531199, EPI_ISL_531200, EPI_ISL_531201, EPI_ISL_531202, EPI_ISL_531203, EPI_ISL_531204, EPI_ISL_531205, EPI_ISL_531206 | Lighthouse Lab in Glasgow | Wellcome Sanger Institute for the COVID-19 Genomics UK Consortium | Harper VanSteenhouse, Yumi Kasai, David Gray, Carol Clugston, Anna Dominiczak and Alex Alderton, Roberto Amato, Sonia Goncalves, Ewan Harrison, David K. Jackson, Ian Johnston, Dominic Kwiatkowski, Cordelia Langford, John Sillitoe |
| EPI_ISL_531207 | NHSGGC West of Scotland Specialist Virology Centre / MRC-University of Glasgow Centre for Virus Research | Wellcome Sanger Institute for the COVID-19 Genomics UK Consortium | Ana da Silva Filipe, Natasha Johnson, Kathy Smollett, Daniel Mair, Stephen Carmichael, Lily Tong, Jenna Nichols, Elihu Aranday-Cortes, Kirstyn Brunker, Yasmin Parr, Kyriaki Nomikou; Sarah McDonald, Marc Niebel, Patawee Asamaphan; Richard Orton, Joseph Hughes, Sreenu Vattipally, David L Robertson; Alasdair MacLean, Rory Gunson; Kathy Li, Natasha Jesudason, Rajiv Shah, James Shepherd, Antonia Ho, Alice Broos, Emma Thomson and Alex Alderton, Roberto Amato, Sonia Goncalves, Ewan Harrison, David K. Jackson, Ian Johnston, Dominic Kwiatkowski, Cordelia Langford, John Sillitoe |
| EPI_ISL_531208, EPI_ISL_531209, EPI_ISL_531210 | Lighthouse Lab in Glasgow | Wellcome Sanger Institute for the COVID-19 Genomics UK Consortium | Harper VanSteenhouse, Yumi Kasai, David Gray, Carol Clugston, Anna Dominiczak and Alex Alderton, Roberto Amato, Sonia Goncalves, Ewan Harrison, David K. Jackson, Ian Johnston, Dominic Kwiatkowski, Cordelia Langford, John Sillitoe |
| EPI_ISL_531212 | NHSGGC West of Scotland Specialist Virology Centre / MRC-University of Glasgow Centre for Virus Research | Wellcome Sanger Institute for the COVID-19 Genomics UK Consortium | Ana da Silva Filipe, Natasha Johnson, Kathy Smollett, Daniel Mair, Stephen Carmichael, Lily Tong, Jenna Nichols, Elihu Aranday-Cortes, Kirstyn Brunker, Yasmin Parr, Kyriaki Nomikou; Sarah McDonald, Marc Niebel, Patawee Asamaphan; Richard Orton, Joseph Hughes, Sreenu Vattipally, David L Robertson; Alasdair MacLean, Rory Gunson; Kathy Li, Natasha Jesudason, Rajiv Shah, James Shepherd, Antonia Ho, Alice Broos, Emma Thomson and Alex Alderton, Roberto Amato, Sonia Goncalves, Ewan Harrison, David K. Jackson, Ian Johnston, Dominic Kwiatkowski, Cordelia Langford, John Sillitoe |
| EPI_ISL_531213 | Lighthouse Lab in Glasgow | Wellcome Sanger Institute for the COVID-19 Genomics UK Consortium | Harper VanSteenhouse, Yumi Kasai, David Gray, Carol Clugston, Anna Dominiczak and Alex Alderton, Roberto Amato, Sonia Goncalves, Ewan Harrison, David K. Jackson, Ian Johnston, Dominic Kwiatkowski, Cordelia Langford, John Sillitoe |
| EPI_ISL_531214 | NHSGGC West of Scotland Specialist Virology Centre / MRC-University of Glasgow Centre for Virus Research | Wellcome Sanger Institute for the COVID-19 Genomics UK Consortium | Ana da Silva Filipe, Natasha Johnson, Kathy Smollett, Daniel Mair, Stephen Carmichael, Lily Tong, Jenna Nichols, Elihu Aranday-Cortes, Kirstyn Brunker, Yasmin Parr, Kyriaki Nomikou; Sarah McDonald, Marc Niebel, Patawee Asamaphan; Richard Orton, Joseph Hughes, Sreenu Vattipally, David L Robertson; Alasdair MacLean, Rory Gunson; Kathy Li, Natasha Jesudason, Rajiv Shah, James Shepherd, Antonia Ho, Alice Broos, Emma Thomson and Alex Alderton, Roberto Amato, Sonia Goncalves, Ewan Harrison, David K. Jackson, Ian Johnston, Dominic Kwiatkowski, Cordelia Langford, John Sillitoe |
| EPI_ISL_531215, EPI_ISL_531216, EPI_ISL_531217, EPI_ISL_531218, EPI_ISL_531219, EPI_ISL_531220, EPI_ISL_531221, EPI_ISL_531222 | Lighthouse Lab in Glasgow | Wellcome Sanger Institute for the COVID-19 Genomics UK Consortium | Harper VanSteenhouse, Yumi Kasai, David Gray, Carol Clugston, Anna Dominiczak and Alex Alderton, Roberto Amato, Sonia Goncalves, Ewan Harrison, David K. Jackson, Ian Johnston, Dominic Kwiatkowski, Cordelia Langford, John Sillitoe |
| EPI_ISL_531223 | NHSGGC West of Scotland Specialist Virology Centre / MRC-University of Glasgow Centre for Virus Research | Wellcome Sanger Institute for the COVID-19 Genomics UK Consortium | Ana da Silva Filipe, Natasha Johnson, Kathy Smollett, Daniel Mair, Stephen Carmichael, Lily Tong, Jenna Nichols, Elihu Aranday-Cortes, Kirstyn Brunker, Yasmin Parr, Kyriaki Nomikou; Sarah McDonald, Marc Niebel, Patawee Asamaphan; Richard Orton, Joseph Hughes, Sreenu Vattipally, David L Robertson; Alasdair MacLean, Rory Gunson; Kathy Li, Natasha Jesudason, Rajiv Shah, James Shepherd, Antonia Ho, Alice Broos, Emma Thomson and Alex Alderton, Roberto Amato, Sonia Goncalves, Ewan Harrison, David K. Jackson, Ian Johnston, Dominic Kwiatkowski, Cordelia Langford, John Sillitoe |
| EPI_ISL_531224, EPI_ISL_531226, EPI_ISL_531227, EPI_ISL_531228, EPI_ISL_531229, EPI_ISL_531230, EPI_ISL_531232, EPI_ISL_531233, EPI_ISL_531234, EPI_ISL_531235, EPI_ISL_531236, EPI_ISL_531237, EPI_ISL_531238, EPI_ISL_531239, EPI_ISL_531240, EPI_ISL_531241, EPI_ISL_531242, EPI_ISL_531243, EPI_ISL_531244, EPI_ISL_531245, EPI_ISL_531246, EPI_ISL_531247, EPI_ISL_531248 | | | |
| see above | Lighthouse Lab in Glasgow | Wellcome Sanger Institute for the COVID-19 Genomics UK Consortium | Harper VanSteenhouse, Yumi Kasai, David Gray, Carol Clugston, Anna Dominiczak and Alex Alderton, Roberto Amato, Sonia Goncalves, Ewan Harrison, David K. Jackson, Ian Johnston, Dominic Kwiatkowski, Cordelia Langford, John Sillitoe |
| EPI_ISL_531249 | NHSGGC West of Scotland Specialist Virology Centre / MRC-University of Glasgow Centre for Virus Research | Wellcome Sanger Institute for the COVID-19 Genomics UK Consortium | Ana da Silva Filipe, Natasha Johnson, Kathy Smollett, Daniel Mair, Stephen Carmichael, Lily Tong, Jenna Nichols, Elihu Aranday-Cortes, Kirstyn Brunker, Yasmin Parr, Kyriaki Nomikou; Sarah McDonald, Marc Niebel, Patawee Asamaphan; Richard Orton, Joseph Hughes, Sreenu Vattipally, David L Robertson; Alasdair MacLean, Rory Gunson; Kathy Li, Natasha Jesudason, Rajiv Shah, James Shepherd, Antonia Ho, Alice Broos, Emma Thomson and Alex Alderton, Roberto Amato, Sonia Goncalves, Ewan Harrison, David K. Jackson, Ian Johnston, Dominic Kwiatkowski, Cordelia Langford, John Sillitoe |
| EPI_ISL_531250, EPI_ISL_531251, EPI_ISL_531252, EPI_ISL_531253, EPI_ISL_531254, EPI_ISL_531255, EPI_ISL_531256, EPI_ISL_531257, EPI_ISL_531258, EPI_ISL_531259, EPI_ISL_531260, EPI_ISL_531261, EPI_ISL_531262, EPI_ISL_531263, EPI_ISL_531264, EPI_ISL_531265 | | | |
| see above | Lighthouse Lab in Glasgow | Wellcome Sanger Institute for the COVID-19 Genomics UK Consortium | Harper VanSteenhouse, Yumi Kasai, David Gray, Carol Clugston, Anna Dominiczak and Alex Alderton, Roberto Amato, Sonia Goncalves, Ewan Harrison, David K. Jackson, Ian Johnston, Dominic Kwiatkowski, Cordelia Langford, John Sillitoe |
| EPI_ISL_531266 | NHSGGC West of Scotland Specialist Virology Centre / MRC-University of Glasgow Centre for Virus Research | Wellcome Sanger Institute for the COVID-19 Genomics UK Consortium | Ana da Silva Filipe, Natasha Johnson, Kathy Smollett, Daniel Mair, Stephen Carmichael, Lily Tong, Jenna Nichols, Elihu Aranday-Cortes, Kirstyn Brunker, Yasmin Parr, Kyriaki Nomikou; Sarah McDonald, Marc Niebel, Patawee Asamaphan; Richard Orton, Joseph Hughes, Sreenu Vattipally, David L Robertson; Alasdair MacLean, Rory Gunson; Kathy Li, Natasha Jesudason, Rajiv Shah, James Shepherd, Antonia Ho, Alice Broos, Emma Thomson and Alex Alderton, Roberto Amato, Sonia Goncalves, Ewan Harrison, David K. Jackson, Ian Johnston, Dominic Kwiatkowski, Cordelia Langford, John Sillitoe |
| EPI_ISL_531267, EPI_ISL_531268, EPI_ISL_531269, EPI_ISL_531270, EPI_ISL_531271, EPI_ISL_531272, EPI_ISL_531273, EPI_ISL_531274, EPI_ISL_531275, EPI_ISL_531276, EPI_ISL_531277, EPI_ISL_531278, EPI_ISL_531279, EPI_ISL_531280, EPI_ISL_531281, EPI_ISL_531282, EPI_ISL_531283, EPI_ISL_531284, EPI_ISL_531285, EPI_ISL_531286, EPI_ISL_531287, EPI_ISL_531288, EPI_ISL_531289, EPI_ISL_531290, EPI_ISL_531291, EPI_ISL_531292, EPI_ISL_531293, EPI_ISL_531294, EPI_ISL_531295, EPI_ISL_531296, EPI_ISL_531297, EPI_ISL_531298, EPI_ISL_531299, EPI_ISL_531300, EPI_ISL_531301, EPI_ISL_531302, EPI_ISL_531303, EPI_ISL_531304, EPI_ISL_531305, EPI_ISL_531306, EPI_ISL_531307, EPI_ISL_531308, EPI_ISL_531309, EPI_ISL_531310, EPI_ISL_531311, EPI_ISL_531312, EPI_ISL_531313, EPI_ISL_531314, EPI_ISL_531315, EPI_ISL_531316, EPI_ISL_531317, EPI_ISL_531318, EPI_ISL_531319, EPI_ISL_531320 | | | |
| see above | Lighthouse Lab in Glasgow | Wellcome Sanger Institute for the COVID-19 Genomics UK Consortium | Harper VanSteenhouse, Yumi Kasai, David Gray, Carol Clugston, Anna Dominiczak and Alex Alderton, Roberto Amato, Sonia Goncalves, Ewan Harrison, David K. Jackson, Ian Johnston, Dominic Kwiatkowski, Cordelia Langford, John Sillitoe |

| | | | |
|---|---|---|---|
| EPI_ISL_531321 | NHSGGC West of Scotland Specialist Virology Centre / MRC-University of Glasgow Centre for Virus Research | Wellcome Sanger Institute for the COVID-19 Genomics UK Consortium | Ana da Silva Filipe, Natasha Johnson, Kathy Smollett, Daniel Mair, Stephen Carmichael, Lily Tong, Jenna Nichols, Elihu Aranday-Cortes, Kirstyn Brunker, Yasmin Parr, Kyriaki Nomikou; Sarah McDonald, Marc Niebel, Patawee Asamaphan; Richard Orton, Joseph Hughes, Sreenu Vattipally, David L Robertson; Alasdair MacLean, Rory Gunson; Kathy Li, Natasha Jesudason, Rajiv Shah, James Shepherd, Antonia Ho, Alice Broos, Emma Thomson and Alex Alderton, Roberto Amato, Sonia Goncalves, Ewan Harrison, David K. Jackson, Ian Johnston, Dominic Kwiatkowski, Cordelia Langford, John Sillitoe |
| EPI_ISL_531322, EPI_ISL_531323, EPI_ISL_531324, EPI_ISL_531325, EPI_ISL_531326, EPI_ISL_531327, EPI_ISL_531328, EPI_ISL_531329 | Lighthouse Lab in Glasgow | Wellcome Sanger Institute for the COVID-19 Genomics UK Consortium | Harper VanSteenhouse, Yumi Kasai, David Gray, Carol Clugston, Anna Dominiczak and Alex Alderton, Roberto Amato, Sonia Goncalves, Ewan Harrison, David K. Jackson, Ian Johnston, Dominic Kwiatkowski, Cordelia Langford, John Sillitoe |
| EPI_ISL_531330 | NHSGGC West of Scotland Specialist Virology Centre / MRC-University of Glasgow Centre for Virus Research | Wellcome Sanger Institute for the COVID-19 Genomics UK Consortium | Ana da Silva Filipe, Natasha Johnson, Kathy Smollett, Daniel Mair, Stephen Carmichael, Lily Tong, Jenna Nichols, Elihu Aranday-Cortes, Kirstyn Brunker, Yasmin Parr, Kyriaki Nomikou; Sarah McDonald, Marc Niebel, Patawee Asamaphan; Richard Orton, Joseph Hughes, Sreenu Vattipally, David L Robertson; Alasdair MacLean, Rory Gunson; Kathy Li, Natasha Jesudason, Rajiv Shah, James Shepherd, Antonia Ho, Alice Broos, Emma Thomson and Alex Alderton, Roberto Amato, Sonia Goncalves, Ewan Harrison, David K. Jackson, Ian Johnston, Dominic Kwiatkowski, Cordelia Langford, John Sillitoe |
| EPI_ISL_531331, EPI_ISL_531332, EPI_ISL_531333, EPI_ISL_531334, EPI_ISL_531335, EPI_ISL_531336, EPI_ISL_531337, EPI_ISL_531338, EPI_ISL_531339, EPI_ISL_531340, EPI_ISL_531341, EPI_ISL_531342, EPI_ISL_531343, EPI_ISL_531344, EPI_ISL_531345, EPI_ISL_531346, EPI_ISL_531347, EPI_ISL_531348, EPI_ISL_531349, EPI_ISL_531350, EPI_ISL_531351, EPI_ISL_531352, EPI_ISL_531353, EPI_ISL_531354, EPI_ISL_531355, EPI_ISL_531356, EPI_ISL_531357, EPI_ISL_531358, EPI_ISL_531359, EPI_ISL_531360, EPI_ISL_531361, EPI_ISL_531362, EPI_ISL_531363, EPI_ISL_531364 | | | |
| see above | Lighthouse Lab in Glasgow | Wellcome Sanger Institute for the COVID-19 Genomics UK Consortium | Harper VanSteenhouse, Yumi Kasai, David Gray, Carol Clugston, Anna Dominiczak and Alex Alderton, Roberto Amato, Sonia Goncalves, Ewan Harrison, David K. Jackson, Ian Johnston, Dominic Kwiatkowski, Cordelia Langford, John Sillitoe |
| EPI_ISL_531365 | NHSGGC West of Scotland Specialist Virology Centre / MRC-University of Glasgow Centre for Virus Research | Wellcome Sanger Institute for the COVID-19 Genomics UK Consortium | Ana da Silva Filipe, Natasha Johnson, Kathy Smollett, Daniel Mair, Stephen Carmichael, Lily Tong, Jenna Nichols, Elihu Aranday-Cortes, Kirstyn Brunker, Yasmin Parr, Kyriaki Nomikou; Sarah McDonald, Marc Niebel, Patawee Asamaphan; Richard Orton, Joseph Hughes, Sreenu Vattipally, David L Robertson; Alasdair MacLean, Rory Gunson; Kathy Li, Natasha Jesudason, Rajiv Shah, James Shepherd, Antonia Ho, Alice Broos, Emma Thomson and Alex Alderton, Roberto Amato, Sonia Goncalves, Ewan Harrison, David K. Jackson, Ian Johnston, Dominic Kwiatkowski, Cordelia Langford, John Sillitoe |
| EPI_ISL_531366, EPI_ISL_531367, EPI_ISL_531368, EPI_ISL_531369, EPI_ISL_531370, EPI_ISL_531371, EPI_ISL_531372, EPI_ISL_531373 | Lighthouse Lab in Glasgow | Wellcome Sanger Institute for the COVID-19 Genomics UK Consortium | Harper VanSteenhouse, Yumi Kasai, David Gray, Carol Clugston, Anna Dominiczak and Alex Alderton, Roberto Amato, Sonia Goncalves, Ewan Harrison, David K. Jackson, Ian Johnston, Dominic Kwiatkowski, Cordelia Langford, John Sillitoe |
| EPI_ISL_531374 | NHSGGC West of Scotland Specialist Virology Centre / MRC-University of Glasgow Centre for Virus Research | Wellcome Sanger Institute for the COVID-19 Genomics UK Consortium | Ana da Silva Filipe, Natasha Johnson, Kathy Smollett, Daniel Mair, Stephen Carmichael, Lily Tong, Jenna Nichols, Elihu Aranday-Cortes, Kirstyn Brunker, Yasmin Parr, Kyriaki Nomikou; Sarah McDonald, Marc Niebel, Patawee Asamaphan; Richard Orton, Joseph Hughes, Sreenu Vattipally, David L Robertson; Alasdair MacLean, Rory Gunson; Kathy Li, Natasha Jesudason, Rajiv Shah, James Shepherd, Antonia Ho, Alice Broos, Emma Thomson and Alex Alderton, Roberto Amato, Sonia Goncalves, Ewan Harrison, David K. Jackson, Ian Johnston, Dominic Kwiatkowski, Cordelia Langford, John Sillitoe |
| EPI_ISL_531375, EPI_ISL_531376, EPI_ISL_531377, EPI_ISL_531378, EPI_ISL_531379, EPI_ISL_531380 | Lighthouse Lab in Glasgow | Wellcome Sanger Institute for the COVID-19 Genomics UK Consortium | Harper VanSteenhouse, Yumi Kasai, David Gray, Carol Clugston, Anna Dominiczak and Alex Alderton, Roberto Amato, Sonia Goncalves, Ewan Harrison, David K. Jackson, Ian Johnston, Dominic Kwiatkowski, Cordelia Langford, John Sillitoe |
| EPI_ISL_531382 | NHSGGC West of Scotland Specialist Virology Centre / MRC-University of Glasgow Centre for Virus Research | Wellcome Sanger Institute for the COVID-19 Genomics UK Consortium | Ana da Silva Filipe, Natasha Johnson, Kathy Smollett, Daniel Mair, Stephen Carmichael, Lily Tong, Jenna Nichols, Elihu Aranday-Cortes, Kirstyn Brunker, Yasmin Parr, Kyriaki Nomikou; Sarah McDonald, Marc Niebel, Patawee Asamaphan; Richard Orton, Joseph Hughes, Sreenu Vattipally, David L Robertson; Alasdair MacLean, Rory Gunson; Kathy Li, Natasha Jesudason, Rajiv Shah, James Shepherd, Antonia Ho, Alice Broos, Emma Thomson and Alex Alderton, Roberto Amato, Sonia Goncalves, Ewan Harrison, David K. Jackson, Ian Johnston, Dominic Kwiatkowski, Cordelia Langford, John Sillitoe |
| EPI_ISL_531383, EPI_ISL_531384, EPI_ISL_531385, EPI_ISL_531386, EPI_ISL_531387, EPI_ISL_531388, EPI_ISL_531389, EPI_ISL_531390, EPI_ISL_531391, EPI_ISL_531392, EPI_ISL_531393, EPI_ISL_531394, EPI_ISL_531395, EPI_ISL_531396, EPI_ISL_531397, EPI_ISL_531398, EPI_ISL_531399, EPI_ISL_531400, EPI_ISL_531401, EPI_ISL_531402, EPI_ISL_531403, EPI_ISL_531404, EPI_ISL_531405, EPI_ISL_531406, EPI_ISL_531407, EPI_ISL_531408, EPI_ISL_531409, EPI_ISL_531410, EPI_ISL_531411 | | | |
| see above | Lighthouse Lab in Glasgow | Wellcome Sanger Institute for the COVID-19 Genomics UK Consortium | Harper VanSteenhouse, Yumi Kasai, David Gray, Carol Clugston, Anna Dominiczak and Alex Alderton, Roberto Amato, Sonia Goncalves, Ewan Harrison, David K. Jackson, Ian Johnston, Dominic Kwiatkowski, Cordelia Langford, John Sillitoe |
| EPI_ISL_531412 | NHSGGC West of Scotland Specialist Virology Centre / MRC-University of Glasgow Centre for Virus Research | Wellcome Sanger Institute for the COVID-19 Genomics UK Consortium | Ana da Silva Filipe, Natasha Johnson, Kathy Smollett, Daniel Mair, Stephen Carmichael, Lily Tong, Jenna Nichols, Elihu Aranday-Cortes, Kirstyn Brunker, Yasmin Parr, Kyriaki Nomikou; Sarah McDonald, Marc Niebel, Patawee Asamaphan; Richard Orton, Joseph Hughes, Sreenu Vattipally, David L Robertson; Alasdair MacLean, Rory Gunson; Kathy Li, Natasha Jesudason, Rajiv Shah, James Shepherd, Antonia Ho, Alice Broos, Emma Thomson and Alex Alderton, Roberto Amato, Sonia Goncalves, Ewan Harrison, David K. Jackson, Ian Johnston, Dominic Kwiatkowski, Cordelia Langford, John Sillitoe |
| EPI_ISL_531413, EPI_ISL_531414, EPI_ISL_531415 | Lighthouse Lab in Glasgow | Wellcome Sanger Institute for the COVID-19 Genomics UK Consortium | Harper VanSteenhouse, Yumi Kasai, David Gray, Carol Clugston, Anna Dominiczak and Alex Alderton, Roberto Amato, Sonia Goncalves, Ewan Harrison, David K. Jackson, Ian Johnston, Dominic Kwiatkowski, Cordelia Langford, John Sillitoe |
| EPI_ISL_531416 | NHSGGC West of Scotland Specialist Virology Centre / MRC-University of Glasgow Centre for Virus Research | Wellcome Sanger Institute for the COVID-19 Genomics UK Consortium | Ana da Silva Filipe, Natasha Johnson, Kathy Smollett, Daniel Mair, Stephen Carmichael, Lily Tong, Jenna Nichols, Elihu Aranday-Cortes, Kirstyn Brunker, Yasmin Parr, Kyriaki Nomikou; Sarah McDonald, Marc Niebel, Patawee Asamaphan; Richard Orton, Joseph Hughes, Sreenu Vattipally, David L Robertson; Alasdair MacLean, Rory Gunson; Kathy Li, Natasha Jesudason, Rajiv Shah, James Shepherd, Antonia Ho, Alice Broos, Emma Thomson and Alex Alderton, Roberto Amato, Sonia Goncalves, Ewan Harrison, David K. Jackson, Ian Johnston, Dominic Kwiatkowski, Cordelia Langford, John Sillitoe |
| EPI_ISL_531417, EPI_ISL_531418, EPI_ISL_531419 | Lighthouse Lab in Glasgow | Wellcome Sanger Institute for the COVID-19 Genomics UK Consortium | Harper VanSteenhouse, Yumi Kasai, David Gray, Carol Clugston, Anna Dominiczak and Alex Alderton, Roberto Amato, Sonia Goncalves, Ewan Harrison, David K. Jackson, Ian Johnston, Dominic Kwiatkowski, Cordelia Langford, John Sillitoe |
| EPI_ISL_531420 | NHSGGC West of Scotland Specialist Virology Centre / MRC-University of Glasgow Centre for Virus Research | Wellcome Sanger Institute for the COVID-19 Genomics UK Consortium | Ana da Silva Filipe, Natasha Johnson, Kathy Smollett, Daniel Mair, Stephen Carmichael, Lily Tong, Jenna Nichols, Elihu Aranday-Cortes, Kirstyn Brunker, Yasmin Parr, Kyriaki Nomikou; Sarah McDonald, Marc Niebel, Patawee Asamaphan; Richard Orton, Joseph Hughes, Sreenu Vattipally, David L Robertson; Alasdair MacLean, Rory Gunson; Kathy Li, Natasha Jesudason, Rajiv Shah, James Shepherd, Antonia Ho, Alice Broos, Emma Thomson and Alex Alderton, Roberto Amato, Sonia Goncalves, Ewan Harrison, David K. Jackson, Ian Johnston, Dominic Kwiatkowski, Cordelia Langford, John Sillitoe |
| EPI_ISL_531421 | Lighthouse Lab in Glasgow | Wellcome Sanger Institute for the COVID-19 Genomics UK Consortium | Harper VanSteenhouse, Yumi Kasai, David Gray, Carol Clugston, Anna Dominiczak and Alex Alderton, Roberto Amato, Sonia Goncalves, Ewan Harrison, David K. Jackson, Ian Johnston, Dominic Kwiatkowski, Cordelia Langford, John Sillitoe |
| EPI_ISL_531422 | NHSGGC West of Scotland Specialist Virology Centre / MRC-University of Glasgow Centre for Virus Research | Wellcome Sanger Institute for the COVID-19 Genomics UK Consortium | Ana da Silva Filipe, Natasha Johnson, Kathy Smollett, Daniel Mair, Stephen Carmichael, Lily Tong, Jenna Nichols, Elihu Aranday-Cortes, Kirstyn Brunker, Yasmin Parr, Kyriaki Nomikou; Sarah McDonald, Marc Niebel, Patawee Asamaphan; Richard Orton, Joseph Hughes, Sreenu Vattipally, David L Robertson; Alasdair MacLean, Rory Gunson; Kathy Li, Natasha Jesudason, Rajiv Shah, James Shepherd, Antonia Ho, Alice Broos, Emma Thomson and Alex Alderton, Roberto Amato, Sonia Goncalves, Ewan Harrison, David K. Jackson, Ian Johnston, Dominic Kwiatkowski, Cordelia Langford, John Sillitoe |
| EPI_ISL_531423, EPI_ISL_531424, EPI_ISL_531425, EPI_ISL_531426, EPI_ISL_531427, EPI_ISL_531428, EPI_ISL_531429, EPI_ISL_531430, EPI_ISL_531431 | Lighthouse Lab in Glasgow | Wellcome Sanger Institute for the COVID-19 Genomics UK Consortium | Harper VanSteenhouse, Yumi Kasai, David Gray, Carol Clugston, Anna Dominiczak and Alex Alderton, Roberto Amato, Sonia Goncalves, Ewan Harrison, David K. Jackson, Ian Johnston, Dominic Kwiatkowski, Cordelia Langford, John Sillitoe |
| EPI_ISL_531432 | NHSGGC West of Scotland Specialist Virology Centre / MRC-University of Glasgow Centre for Virus Research | Wellcome Sanger Institute for the COVID-19 Genomics UK Consortium | Ana da Silva Filipe, Natasha Johnson, Kathy Smollett, Daniel Mair, Stephen Carmichael, Lily Tong, Jenna Nichols, Elihu Aranday-Cortes, Kirstyn Brunker, Yasmin Parr, Kyriaki Nomikou; Sarah McDonald, Marc Niebel, Patawee Asamaphan; Richard Orton, Joseph Hughes, Sreenu Vattipally, David L Robertson; Alasdair MacLean, Rory Gunson; Kathy Li, Natasha Jesudason, Rajiv Shah, James Shepherd, Antonia Ho, Alice Broos, Emma Thomson and Alex Alderton, Roberto Amato, Sonia Goncalves, Ewan Harrison, David K. Jackson, Ian Johnston, Dominic Kwiatkowski, Cordelia Langford, John Sillitoe |
| EPI_ISL_531433, EPI_ISL_531434, EPI_ISL_531435, EPI_ISL_531436, EPI_ISL_531437, EPI_ISL_531438, EPI_ISL_531439, EPI_ISL_531440, EPI_ISL_531441 | Lighthouse Lab in Glasgow | Wellcome Sanger Institute for the COVID-19 Genomics UK Consortium | Harper VanSteenhouse, Yumi Kasai, David Gray, Carol Clugston, Anna Dominiczak and Alex Alderton, Roberto Amato, Sonia Goncalves, Ewan Harrison, David K. Jackson, Ian Johnston, Dominic Kwiatkowski, Cordelia Langford, John Sillitoe |
| EPI_ISL_531442 | NHSGGC West of Scotland Specialist Virology Centre / MRC-University of Glasgow Centre for Virus Research | Wellcome Sanger Institute for the COVID-19 Genomics UK Consortium | Ana da Silva Filipe, Natasha Johnson, Kathy Smollett, Daniel Mair, Stephen Carmichael, Lily Tong, Jenna Nichols, Elihu Aranday-Cortes, Kirstyn Brunker, Yasmin Parr, Kyriaki Nomikou; Sarah McDonald, Marc Niebel, Patawee Asamaphan; Richard Orton, Joseph Hughes, Sreenu Vattipally, David L Robertson; Alasdair MacLean, Rory Gunson; Kathy Li, Natasha Jesudason, Rajiv Shah, James Shepherd, Antonia Ho, Alice Broos, Emma Thomson and Alex Alderton, Roberto Amato, Sonia Goncalves, Ewan Harrison, David K. Jackson, Ian Johnston, Dominic Kwiatkowski, Cordelia Langford, John Sillitoe |
| EPI_ISL_531443, EPI_ISL_531444, EPI_ISL_531445, EPI_ISL_531446, EPI_ISL_531447, EPI_ISL_531448, EPI_ISL_531449, EPI_ISL_531450, EPI_ISL_531451, EPI_ISL_531452, EPI_ISL_531453, EPI_ISL_531454, EPI_ISL_531455, EPI_ISL_531456, EPI_ISL_531457, EPI_ISL_531458, EPI_ISL_531459, EPI_ISL_531460, EPI_ISL_531461, EPI_ISL_531462, EPI_ISL_531463 | | | |
| see above | Lighthouse Lab in Glasgow | Wellcome Sanger Institute for the COVID-19 Genomics UK Consortium | Harper VanSteenhouse, Yumi Kasai, David Gray, Carol Clugston, Anna Dominiczak and Alex Alderton, Roberto Amato, Sonia Goncalves, Ewan Harrison, David K. Jackson, Ian Johnston, Dominic Kwiatkowski, Cordelia Langford, John Sillitoe |
| EPI_ISL_531464 | NHSGGC West of Scotland Specialist Virology Centre / MRC-University of Glasgow Centre for Virus Research | Wellcome Sanger Institute for the COVID-19 Genomics UK Consortium | Ana da Silva Filipe, Natasha Johnson, Kathy Smollett, Daniel Mair, Stephen Carmichael, Lily Tong, Jenna Nichols, Elihu Aranday-Cortes, Kirstyn Brunker, Yasmin Parr, Kyriaki Nomikou; Sarah McDonald, Marc Niebel, Patawee Asamaphan; Richard Orton, Joseph Hughes, Sreenu Vattipally, David L Robertson; Alasdair MacLean, Rory Gunson; Kathy Li, Natasha Jesudason, Rajiv Shah, James Shepherd, Antonia Ho, Alice Broos, Emma Thomson and Alex Alderton, Roberto Amato, Sonia Goncalves, Ewan Harrison, David K. Jackson, Ian Johnston, Dominic Kwiatkowski, Cordelia Langford, John Sillitoe |
| EPI_ISL_531465, EPI_ISL_531466, EPI_ISL_531467, EPI_ISL_531468, EPI_ISL_531469 | Lighthouse Lab in Glasgow | Wellcome Sanger Institute for the COVID-19 Genomics UK Consortium | Harper VanSteenhouse, Yumi Kasai, David Gray, Carol Clugston, Anna Dominiczak and Alex Alderton, Roberto Amato, Sonia Goncalves, Ewan Harrison, David K. Jackson, Ian Johnston, Dominic Kwiatkowski, Cordelia Langford, John Sillitoe |
| EPI_ISL_531470 | NHSGGC West of Scotland Specialist Virology Centre / MRC-University of Glasgow Centre for Virus Research | Wellcome Sanger Institute for the COVID-19 Genomics UK Consortium | Ana da Silva Filipe, Natasha Johnson, Kathy Smollett, Daniel Mair, Stephen Carmichael, Lily Tong, Jenna Nichols, Elihu Aranday-Cortes, Kirstyn Brunker, Yasmin Parr, Kyriaki Nomikou; Sarah McDonald, Marc Niebel, Patawee Asamaphan; Richard Orton, Joseph Hughes, Sreenu Vattipally, David L Robertson; Alasdair MacLean, Rory Gunson; Kathy Li, Natasha Jesudason, Rajiv Shah, James Shepherd, Antonia Ho, Alice Broos, Emma Thomson and Alex Alderton, Roberto Amato, Sonia Goncalves, Ewan Harrison, David K. Jackson, Ian Johnston, Dominic Kwiatkowski, Cordelia Langford, John Sillitoe |
| EPI_ISL_531471, EPI_ISL_531472 | Lighthouse Lab in Glasgow | Wellcome Sanger Institute for the COVID-19 Genomics UK Consortium | Harper VanSteenhouse, Yumi Kasai, David Gray, Carol Clugston, Anna Dominiczak and Alex Alderton, Roberto Amato, Sonia Goncalves, Ewan Harrison, David K. Jackson, Ian Johnston, Dominic Kwiatkowski, Cordelia Langford, John Sillitoe |
| EPI_ISL_531473 | NHSGGC West of Scotland Specialist Virology Centre / MRC-University of Glasgow Centre for Virus Research | Wellcome Sanger Institute for the COVID-19 Genomics UK Consortium | Ana da Silva Filipe, Natasha Johnson, Kathy Smollett, Daniel Mair, Stephen Carmichael, Lily Tong, Jenna Nichols, Elihu Aranday-Cortes, Kirstyn Brunker, Yasmin Parr, Kyriaki Nomikou; Sarah McDonald, Marc Niebel, Patawee Asamaphan; Richard Orton, Joseph Hughes, Sreenu Vattipally, David L Robertson; Alasdair MacLean, Rory Gunson; Kathy Li, Natasha Jesudason, Rajiv Shah, James Shepherd, Antonia Ho, Alice Broos, Emma Thomson and Alex Alderton, Roberto Amato, Sonia Goncalves, Ewan Harrison, David K. Jackson, Ian Johnston, Dominic Kwiatkowski, Cordelia Langford, John Sillitoe |
| EPI_ISL_531474, EPI_ISL_531475 | Lighthouse Lab in Glasgow | Wellcome Sanger Institute for the COVID-19 Genomics UK Consortium | Harper VanSteenhouse, Yumi Kasai, David Gray, Carol Clugston, Anna Dominiczak and Alex Alderton, Roberto Amato, Sonia Goncalves, Ewan Harrison, David K. Jackson, Ian Johnston, Dominic Kwiatkowski, Cordelia Langford, John Sillitoe |
| EPI_ISL_531476, EPI_ISL_531477 | NHSGGC West of Scotland Specialist Virology Centre / MRC-University of Glasgow Centre for Virus Research | Wellcome Sanger Institute for the COVID-19 Genomics UK Consortium | Ana da Silva Filipe, Natasha Johnson, Kathy Smollett, Daniel Mair, Stephen Carmichael, Lily Tong, Jenna Nichols, Elihu Aranday-Cortes, Kirstyn Brunker, Yasmin Parr, Kyriaki Nomikou; Sarah McDonald, Marc Niebel, Patawee Asamaphan; Richard Orton, Joseph Hughes, Sreenu Vattipally, David L Robertson; Alasdair MacLean, Rory Gunson; Kathy Li, Natasha Jesudason, Rajiv Shah, James Shepherd, Antonia Ho, Alice Broos, Emma Thomson and Alex Alderton, Roberto Amato, Sonia Goncalves, Ewan Harrison, David K. Jackson, Ian Johnston, Dominic Kwiatkowski, Cordelia Langford, John Sillitoe |
| EPI_ISL_531478, EPI_ISL_531479 | Lighthouse Lab in Glasgow | Wellcome Sanger Institute for the COVID-19 Genomics UK Consortium | Harper VanSteenhouse, Yumi Kasai, David Gray, Carol Clugston, Anna Dominiczak and Alex Alderton, Roberto Amato, Sonia Goncalves, Ewan Harrison, David K. Jackson, Ian Johnston, Dominic Kwiatkowski, Cordelia Langford, John Sillitoe |
| EPI_ISL_531480 | NHSGGC West of Scotland Specialist Virology Centre / MRC-University of Glasgow Centre for Virus Research | Wellcome Sanger Institute for the COVID-19 Genomics UK Consortium | Ana da Silva Filipe, Natasha Johnson, Kathy Smollett, Daniel Mair, Stephen Carmichael, Lily Tong, Jenna Nichols, Elihu Aranday-Cortes, Kirstyn Brunker, Yasmin Parr, Kyriaki Nomikou; Sarah McDonald, Marc Niebel, Patawee Asamaphan; Richard Orton, Joseph Hughes, Sreenu Vattipally, David L Robertson; Alasdair MacLean, Rory Gunson; Kathy Li, Natasha Jesudason, Rajiv Shah, James Shepherd, Antonia Ho, Alice Broos, Emma Thomson and Alex Alderton, Roberto Amato, Sonia Goncalves, Ewan Harrison, David K. Jackson, Ian Johnston, Dominic Kwiatkowski, Cordelia Langford, John Sillitoe |
| EPI_ISL_531481, EPI_ISL_531482 | Lighthouse Lab in Glasgow | Wellcome Sanger Institute for the COVID-19 Genomics UK Consortium | Harper VanSteenhouse, Yumi Kasai, David Gray, Carol Clugston, Anna Dominiczak and Alex Alderton, Roberto Amato, Sonia Goncalves, Ewan Harrison, David K. Jackson, Ian Johnston, Dominic Kwiatkowski, Cordelia Langford, John Sillitoe |
| EPI_ISL_531483, EPI_ISL_531484, EPI_ISL_531485 | NHSGGC West of Scotland Specialist Virology Centre / MRC-University of Glasgow Centre for Virus Research | Wellcome Sanger Institute for the COVID-19 Genomics UK Consortium | Ana da Silva Filipe, Natasha Johnson, Kathy Smollett, Daniel Mair, Stephen Carmichael, Lily Tong, Jenna Nichols, Elihu Aranday-Cortes, Kirstyn Brunker, Yasmin Parr, Kyriaki Nomikou; Sarah McDonald, Marc Niebel, Patawee Asamaphan; Richard Orton, Joseph Hughes, Sreenu Vattipally, David L Robertson; Alasdair MacLean, Rory Gunson; Kathy Li, Natasha Jesudason, Rajiv Shah, James Shepherd, Antonia Ho, Alice Broos, Emma Thomson and Alex Alderton, Roberto Amato, Sonia Goncalves, Ewan Harrison, David K. Jackson, Ian Johnston, Dominic Kwiatkowski, Cordelia Langford, John Sillitoe |
| EPI_ISL_531486, EPI_ISL_531487, EPI_ISL_531488, EPI_ISL_531489, EPI_ISL_531490, EPI_ISL_531491 | Lighthouse Lab in Glasgow | Wellcome Sanger Institute for the COVID-19 Genomics UK Consortium | Harper VanSteenhouse, Yumi Kasai, David Gray, Carol Clugston, Anna Dominiczak and Alex Alderton, Roberto Amato, Sonia Goncalves, Ewan Harrison, David K. Jackson, Ian Johnston, Dominic Kwiatkowski, Cordelia Langford, John Sillitoe |

| | | | |
|---|---|---|---|
| EPI_ISL_531492 | NHSGGC West of Scotland Specialist Virology Centre / MRC-University of Glasgow Centre for Virus Research | Wellcome Sanger Institute for the COVID-19 Genomics UK Consortium | Ana da Silva Filipe, Natasha Johnson, Kathy Smollett, Daniel Mair, Stephen Carmichael, Lily Tong, Jenna Nichols, Elihu Aranday-Cortes, Kirstyn Brunker, Yasmin Parr, Kyriaki Nomikou; Sarah McDonald, Marc Niebel, Patawee Asamaphan; Richard Orton, Joseph Hughes, Sreenu Vattipally, David L Robertson; Alasdair MacLean, Rory Gunson; Kathy Li, Natasha Jesudason, Rajiv Shah, James Shepherd, Antonia Ho, Alice Broos, Emma Thomson and Alex Alderton, Roberto Amato, Sonia Goncalves, Ewan Harrison, David K. Jackson, Ian Johnston, Dominic Kwiatkowski, Cordelia Langford, John Sillitoe |
| EPI_ISL_531493, EPI_ISL_531494 | Lighthouse Lab in Glasgow | Wellcome Sanger Institute for the COVID-19 Genomics UK Consortium | Harper VanSteenhouse, Yumi Kasai, David Gray, Carol Clugston, Anna Dominiczak and Alex Alderton, Roberto Amato, Sonia Goncalves, Ewan Harrison, David K. Jackson, Ian Johnston, Dominic Kwiatkowski, Cordelia Langford, John Sillitoe |
| EPI_ISL_531495 | NHSGGC West of Scotland Specialist Virology Centre / MRC-University of Glasgow Centre for Virus Research | Wellcome Sanger Institute for the COVID-19 Genomics UK Consortium | Ana da Silva Filipe, Natasha Johnson, Kathy Smollett, Daniel Mair, Stephen Carmichael, Lily Tong, Jenna Nichols, Elihu Aranday-Cortes, Kirstyn Brunker, Yasmin Parr, Kyriaki Nomikou; Sarah McDonald, Marc Niebel, Patawee Asamaphan; Richard Orton, Joseph Hughes, Sreenu Vattipally, David L Robertson; Alasdair MacLean, Rory Gunson; Kathy Li, Natasha Jesudason, Rajiv Shah, James Shepherd, Antonia Ho, Alice Broos, Emma Thomson and Alex Alderton, Roberto Amato, Sonia Goncalves, Ewan Harrison, David K. Jackson, Ian Johnston, Dominic Kwiatkowski, Cordelia Langford, John Sillitoe |
| EPI_ISL_531496, EPI_ISL_531497, EPI_ISL_531498, EPI_ISL_531499 | Lighthouse Lab in Glasgow | Wellcome Sanger Institute for the COVID-19 Genomics UK Consortium | Harper VanSteenhouse, Yumi Kasai, David Gray, Carol Clugston, Anna Dominiczak and Alex Alderton, Roberto Amato, Sonia Goncalves, Ewan Harrison, David K. Jackson, Ian Johnston, Dominic Kwiatkowski, Cordelia Langford, John Sillitoe |
| EPI_ISL_531500 | NHSGGC West of Scotland Specialist Virology Centre / MRC-University of Glasgow Centre for Virus Research | Wellcome Sanger Institute for the COVID-19 Genomics UK Consortium | Ana da Silva Filipe, Natasha Johnson, Kathy Smollett, Daniel Mair, Stephen Carmichael, Lily Tong, Jenna Nichols, Elihu Aranday-Cortes, Kirstyn Brunker, Yasmin Parr, Kyriaki Nomikou; Sarah McDonald, Marc Niebel, Patawee Asamaphan; Richard Orton, Joseph Hughes, Sreenu Vattipally, David L Robertson; Alasdair MacLean, Rory Gunson; Kathy Li, Natasha Jesudason, Rajiv Shah, James Shepherd, Antonia Ho, Alice Broos, Emma Thomson and Alex Alderton, Roberto Amato, Sonia Goncalves, Ewan Harrison, David K. Jackson, Ian Johnston, Dominic Kwiatkowski, Cordelia Langford, John Sillitoe |
| EPI_ISL_531501, EPI_ISL_531502, EPI_ISL_531503 | Lighthouse Lab in Glasgow | Wellcome Sanger Institute for the COVID-19 Genomics UK Consortium | Harper VanSteenhouse, Yumi Kasai, David Gray, Carol Clugston, Anna Dominiczak and Alex Alderton, Roberto Amato, Sonia Goncalves, Ewan Harrison, David K. Jackson, Ian Johnston, Dominic Kwiatkowski, Cordelia Langford, John Sillitoe |
| EPI_ISL_531504 | NHSGGC West of Scotland Specialist Virology Centre / MRC-University of Glasgow Centre for Virus Research | Wellcome Sanger Institute for the COVID-19 Genomics UK Consortium | Ana da Silva Filipe, Natasha Johnson, Kathy Smollett, Daniel Mair, Stephen Carmichael, Lily Tong, Jenna Nichols, Elihu Aranday-Cortes, Kirstyn Brunker, Yasmin Parr, Kyriaki Nomikou; Sarah McDonald, Marc Niebel, Patawee Asamaphan; Richard Orton, Joseph Hughes, Sreenu Vattipally, David L Robertson; Alasdair MacLean, Rory Gunson; Kathy Li, Natasha Jesudason, Rajiv Shah, James Shepherd, Antonia Ho, Alice Broos, Emma Thomson and Alex Alderton, Roberto Amato, Sonia Goncalves, Ewan Harrison, David K. Jackson, Ian Johnston, Dominic Kwiatkowski, Cordelia Langford, John Sillitoe |
| EPI_ISL_531505, EPI_ISL_531506, EPI_ISL_531507, EPI_ISL_531508, EPI_ISL_531509, EPI_ISL_531510, EPI_ISL_531511, EPI_ISL_531512, EPI_ISL_531513, EPI_ISL_531514, EPI_ISL_531515, EPI_ISL_531516, EPI_ISL_531517, EPI_ISL_531518, EPI_ISL_531519, EPI_ISL_531520, EPI_ISL_531521, EPI_ISL_531522, EPI_ISL_531523 | | | |
| see above | Lighthouse Lab in Glasgow | Wellcome Sanger Institute for the COVID-19 Genomics UK Consortium | Harper VanSteenhouse, Yumi Kasai, David Gray, Carol Clugston, Anna Dominiczak and Alex Alderton, Roberto Amato, Sonia Goncalves, Ewan Harrison, David K. Jackson, Ian Johnston, Dominic Kwiatkowski, Cordelia Langford, John Sillitoe |
| EPI_ISL_531524 | NHSGGC West of Scotland Specialist Virology Centre / MRC-University of Glasgow Centre for Virus Research | Wellcome Sanger Institute for the COVID-19 Genomics UK Consortium | Ana da Silva Filipe, Natasha Johnson, Kathy Smollett, Daniel Mair, Stephen Carmichael, Lily Tong, Jenna Nichols, Elihu Aranday-Cortes, Kirstyn Brunker, Yasmin Parr, Kyriaki Nomikou; Sarah McDonald, Marc Niebel, Patawee Asamaphan; Richard Orton, Joseph Hughes, Sreenu Vattipally, David L Robertson; Alasdair MacLean, Rory Gunson; Kathy Li, Natasha Jesudason, Rajiv Shah, James Shepherd, Antonia Ho, Alice Broos, Emma Thomson and Alex Alderton, Roberto Amato, Sonia Goncalves, Ewan Harrison, David K. Jackson, Ian Johnston, Dominic Kwiatkowski, Cordelia Langford, John Sillitoe |
| EPI_ISL_531525, EPI_ISL_531526, EPI_ISL_531527, EPI_ISL_531528, EPI_ISL_531529, EPI_ISL_531530, EPI_ISL_531531, EPI_ISL_531532, EPI_ISL_531533, EPI_ISL_531534, EPI_ISL_531535, EPI_ISL_531536, EPI_ISL_531537, EPI_ISL_531538, EPI_ISL_531539, EPI_ISL_531540, EPI_ISL_531541, EPI_ISL_531542, EPI_ISL_531543, EPI_ISL_531544, EPI_ISL_531545 | | | |
| see above | Lighthouse Lab in Glasgow | Wellcome Sanger Institute for the COVID-19 Genomics UK Consortium | Harper VanSteenhouse, Yumi Kasai, David Gray, Carol Clugston, Anna Dominiczak and Alex Alderton, Roberto Amato, Sonia Goncalves, Ewan Harrison, David K. Jackson, Ian Johnston, Dominic Kwiatkowski, Cordelia Langford, John Sillitoe |
| EPI_ISL_531546 | NHSGGC West of Scotland Specialist Virology Centre / MRC-University of Glasgow Centre for Virus Research | Wellcome Sanger Institute for the COVID-19 Genomics UK Consortium | Ana da Silva Filipe, Natasha Johnson, Kathy Smollett, Daniel Mair, Stephen Carmichael, Lily Tong, Jenna Nichols, Elihu Aranday-Cortes, Kirstyn Brunker, Yasmin Parr, Kyriaki Nomikou; Sarah McDonald, Marc Niebel, Patawee Asamaphan; Richard Orton, Joseph Hughes, Sreenu Vattipally, David L Robertson; Alasdair MacLean, Rory Gunson; Kathy Li, Natasha Jesudason, Rajiv Shah, James Shepherd, Antonia Ho, Alice Broos, Emma Thomson and Alex Alderton, Roberto Amato, Sonia Goncalves, Ewan Harrison, David K. Jackson, Ian Johnston, Dominic Kwiatkowski, Cordelia Langford, John Sillitoe |
| EPI_ISL_531547 | Lighthouse Lab in Glasgow | Wellcome Sanger Institute for the COVID-19 Genomics UK Consortium | Harper VanSteenhouse, Yumi Kasai, David Gray, Carol Clugston, Anna Dominiczak and Alex Alderton, Roberto Amato, Sonia Goncalves, Ewan Harrison, David K. Jackson, Ian Johnston, Dominic Kwiatkowski, Cordelia Langford, John Sillitoe |
| EPI_ISL_531548 | NHSGGC West of Scotland Specialist Virology Centre / MRC-University of Glasgow Centre for Virus Research | Wellcome Sanger Institute for the COVID-19 Genomics UK Consortium | Ana da Silva Filipe, Natasha Johnson, Kathy Smollett, Daniel Mair, Stephen Carmichael, Lily Tong, Jenna Nichols, Elihu Aranday-Cortes, Kirstyn Brunker, Yasmin Parr, Kyriaki Nomikou; Sarah McDonald, Marc Niebel, Patawee Asamaphan; Richard Orton, Joseph Hughes, Sreenu Vattipally, David L Robertson; Alasdair MacLean, Rory Gunson; Kathy Li, Natasha Jesudason, Rajiv Shah, James Shepherd, Antonia Ho, Alice Broos, Emma Thomson and Alex Alderton, Roberto Amato, Sonia Goncalves, Ewan Harrison, David K. Jackson, Ian Johnston, Dominic Kwiatkowski, Cordelia Langford, John Sillitoe |
| EPI_ISL_531549, EPI_ISL_531550, EPI_ISL_531551, EPI_ISL_531552, EPI_ISL_531553, EPI_ISL_531554, EPI_ISL_531555, EPI_ISL_531556 | Lighthouse Lab in Glasgow | Wellcome Sanger Institute for the COVID-19 Genomics UK Consortium | Harper VanSteenhouse, Yumi Kasai, David Gray, Carol Clugston, Anna Dominiczak and Alex Alderton, Roberto Amato, Sonia Goncalves, Ewan Harrison, David K. Jackson, Ian Johnston, Dominic Kwiatkowski, Cordelia Langford, John Sillitoe |
| EPI_ISL_531557 | NHSGGC West of Scotland Specialist Virology Centre / MRC-University of Glasgow Centre for Virus Research | Wellcome Sanger Institute for the COVID-19 Genomics UK Consortium | Ana da Silva Filipe, Natasha Johnson, Kathy Smollett, Daniel Mair, Stephen Carmichael, Lily Tong, Jenna Nichols, Elihu Aranday-Cortes, Kirstyn Brunker, Yasmin Parr, Kyriaki Nomikou; Sarah McDonald, Marc Niebel, Patawee Asamaphan; Richard Orton, Joseph Hughes, Sreenu Vattipally, David L Robertson; Alasdair MacLean, Rory Gunson; Kathy Li, Natasha Jesudason, Rajiv Shah, James Shepherd, Antonia Ho, Alice Broos, Emma Thomson and Alex Alderton, Roberto Amato, Sonia Goncalves, Ewan Harrison, David K. Jackson, Ian Johnston, Dominic Kwiatkowski, Cordelia Langford, John Sillitoe |
| EPI_ISL_531558, EPI_ISL_531559, EPI_ISL_531560, EPI_ISL_531561, EPI_ISL_531562 | Lighthouse Lab in Glasgow | Wellcome Sanger Institute for the COVID-19 Genomics UK Consortium | Harper VanSteenhouse, Yumi Kasai, David Gray, Carol Clugston, Anna Dominiczak and Alex Alderton, Roberto Amato, Sonia Goncalves, Ewan Harrison, David K. Jackson, Ian Johnston, Dominic Kwiatkowski, Cordelia Langford, John Sillitoe |
| EPI_ISL_531563, EPI_ISL_531564, EPI_ISL_531565 | NHSGGC West of Scotland Specialist Virology Centre / MRC-University of Glasgow Centre for Virus Research | Wellcome Sanger Institute for the COVID-19 Genomics UK Consortium | Ana da Silva Filipe, Natasha Johnson, Kathy Smollett, Daniel Mair, Stephen Carmichael, Lily Tong, Jenna Nichols, Elihu Aranday-Cortes, Kirstyn Brunker, Yasmin Parr, Kyriaki Nomikou; Sarah McDonald, Marc Niebel, Patawee Asamaphan; Richard Orton, Joseph Hughes, Sreenu Vattipally, David L Robertson; Alasdair MacLean, Rory Gunson; Kathy Li, Natasha Jesudason, Rajiv Shah, James Shepherd, Antonia Ho, Alice Broos, Emma Thomson and Alex Alderton, Roberto Amato, Sonia Goncalves, Ewan Harrison, David K. Jackson, Ian Johnston, Dominic Kwiatkowski, Cordelia Langford, John Sillitoe |
| EPI_ISL_531566, EPI_ISL_531567, EPI_ISL_531568 | Lighthouse Lab in Glasgow | Wellcome Sanger Institute for the COVID-19 Genomics UK Consortium | Harper VanSteenhouse, Yumi Kasai, David Gray, Carol Clugston, Anna Dominiczak and Alex Alderton, Roberto Amato, Sonia Goncalves, Ewan Harrison, David K. Jackson, Ian Johnston, Dominic Kwiatkowski, Cordelia Langford, John Sillitoe |
| EPI_ISL_531569, EPI_ISL_531570 | NHSGGC West of Scotland Specialist Virology Centre / MRC-University of Glasgow Centre for Virus Research | Wellcome Sanger Institute for the COVID-19 Genomics UK Consortium | Ana da Silva Filipe, Natasha Johnson, Kathy Smollett, Daniel Mair, Stephen Carmichael, Lily Tong, Jenna Nichols, Elihu Aranday-Cortes, Kirstyn Brunker, Yasmin Parr, Kyriaki Nomikou; Sarah McDonald, Marc Niebel, Patawee Asamaphan; Richard Orton, Joseph Hughes, Sreenu Vattipally, David L Robertson; Alasdair MacLean, Rory Gunson; Kathy Li, Natasha Jesudason, Rajiv Shah, James Shepherd, Antonia Ho, Alice Broos, Emma Thomson and Alex Alderton, Roberto Amato, Sonia Goncalves, Ewan Harrison, David K. Jackson, Ian Johnston, Dominic Kwiatkowski, Cordelia Langford, John Sillitoe |
| EPI_ISL_531571, EPI_ISL_531572, EPI_ISL_531573, EPI_ISL_531574 | Lighthouse Lab in Glasgow | Wellcome Sanger Institute for the COVID-19 Genomics UK Consortium | Harper VanSteenhouse, Yumi Kasai, David Gray, Carol Clugston, Anna Dominiczak and Alex Alderton, Roberto Amato, Sonia Goncalves, Ewan Harrison, David K. Jackson, Ian Johnston, Dominic Kwiatkowski, Cordelia Langford, John Sillitoe |
| EPI_ISL_531575 | NHSGGC West of Scotland Specialist Virology Centre / MRC-University of Glasgow Centre for Virus Research | Wellcome Sanger Institute for the COVID-19 Genomics UK Consortium | Ana da Silva Filipe, Natasha Johnson, Kathy Smollett, Daniel Mair, Stephen Carmichael, Lily Tong, Jenna Nichols, Elihu Aranday-Cortes, Kirstyn Brunker, Yasmin Parr, Kyriaki Nomikou; Sarah McDonald, Marc Niebel, Patawee Asamaphan; Richard Orton, Joseph Hughes, Sreenu Vattipally, David L Robertson; Alasdair MacLean, Rory Gunson; Kathy Li, Natasha Jesudason, Rajiv Shah, James Shepherd, Antonia Ho, Alice Broos, Emma Thomson and Alex Alderton, Roberto Amato, Sonia Goncalves, Ewan Harrison, David K. Jackson, Ian Johnston, Dominic Kwiatkowski, Cordelia Langford, John Sillitoe |
| EPI_ISL_531576 | Lighthouse Lab in Glasgow | Wellcome Sanger Institute for the COVID-19 Genomics UK Consortium | Harper VanSteenhouse, Yumi Kasai, David Gray, Carol Clugston, Anna Dominiczak and Alex Alderton, Roberto Amato, Sonia Goncalves, Ewan Harrison, David K. Jackson, Ian Johnston, Dominic Kwiatkowski, Cordelia Langford, John Sillitoe |
| EPI_ISL_531577 | NHSGGC West of Scotland Specialist Virology Centre / MRC-University of Glasgow Centre for Virus Research | Wellcome Sanger Institute for the COVID-19 Genomics UK Consortium | Ana da Silva Filipe, Natasha Johnson, Kathy Smollett, Daniel Mair, Stephen Carmichael, Lily Tong, Jenna Nichols, Elihu Aranday-Cortes, Kirstyn Brunker, Yasmin Parr, Kyriaki Nomikou; Sarah McDonald, Marc Niebel, Patawee Asamaphan; Richard Orton, Joseph Hughes, Sreenu Vattipally, David L Robertson; Alasdair MacLean, Rory Gunson; Kathy Li, Natasha Jesudason, Rajiv Shah, James Shepherd, Antonia Ho, Alice Broos, Emma Thomson and Alex Alderton, Roberto Amato, Sonia Goncalves, Ewan Harrison, David K. Jackson, Ian Johnston, Dominic Kwiatkowski, Cordelia Langford, John Sillitoe |
| EPI_ISL_531578, EPI_ISL_531579, EPI_ISL_531580, EPI_ISL_531581, EPI_ISL_531582 | Lighthouse Lab in Glasgow | Wellcome Sanger Institute for the COVID-19 Genomics UK Consortium | Harper VanSteenhouse, Yumi Kasai, David Gray, Carol Clugston, Anna Dominiczak and Alex Alderton, Roberto Amato, Sonia Goncalves, Ewan Harrison, David K. Jackson, Ian Johnston, Dominic Kwiatkowski, Cordelia Langford, John Sillitoe |
| EPI_ISL_531583 | NHSGGC West of Scotland Specialist Virology Centre / MRC-University of Glasgow Centre for Virus Research | Wellcome Sanger Institute for the COVID-19 Genomics UK Consortium | Ana da Silva Filipe, Natasha Johnson, Kathy Smollett, Daniel Mair, Stephen Carmichael, Lily Tong, Jenna Nichols, Elihu Aranday-Cortes, Kirstyn Brunker, Yasmin Parr, Kyriaki Nomikou; Sarah McDonald, Marc Niebel, Patawee Asamaphan; Richard Orton, Joseph Hughes, Sreenu Vattipally, David L Robertson; Alasdair MacLean, Rory Gunson; Kathy Li, Natasha Jesudason, Rajiv Shah, James Shepherd, Antonia Ho, Alice Broos, Emma Thomson and Alex Alderton, Roberto Amato, Sonia Goncalves, Ewan Harrison, David K. Jackson, Ian Johnston, Dominic Kwiatkowski, Cordelia Langford, John Sillitoe |
| EPI_ISL_531584, EPI_ISL_531586, EPI_ISL_531587, EPI_ISL_531588, EPI_ISL_531589 | Lighthouse Lab in Glasgow | Wellcome Sanger Institute for the COVID-19 Genomics UK Consortium | Harper VanSteenhouse, Yumi Kasai, David Gray, Carol Clugston, Anna Dominiczak and Alex Alderton, Roberto Amato, Sonia Goncalves, Ewan Harrison, David K. Jackson, Ian Johnston, Dominic Kwiatkowski, Cordelia Langford, John Sillitoe |
| EPI_ISL_531590 | NHSGGC West of Scotland Specialist Virology Centre / MRC-University of Glasgow Centre for Virus Research | Wellcome Sanger Institute for the COVID-19 Genomics UK Consortium | Ana da Silva Filipe, Natasha Johnson, Kathy Smollett, Daniel Mair, Stephen Carmichael, Lily Tong, Jenna Nichols, Elihu Aranday-Cortes, Kirstyn Brunker, Yasmin Parr, Kyriaki Nomikou; Sarah McDonald, Marc Niebel, Patawee Asamaphan; Richard Orton, Joseph Hughes, Sreenu Vattipally, David L Robertson; Alasdair MacLean, Rory Gunson; Kathy Li, Natasha Jesudason, Rajiv Shah, James Shepherd, Antonia Ho, Alice Broos, Emma Thomson and Alex Alderton, Roberto Amato, Sonia Goncalves, Ewan Harrison, David K. Jackson, Ian Johnston, Dominic Kwiatkowski, Cordelia Langford, John Sillitoe |
| EPI_ISL_531591, EPI_ISL_531592, EPI_ISL_531593, EPI_ISL_531594 | Lighthouse Lab in Glasgow | Wellcome Sanger Institute for the COVID-19 Genomics UK Consortium | Harper VanSteenhouse, Yumi Kasai, David Gray, Carol Clugston, Anna Dominiczak and Alex Alderton, Roberto Amato, Sonia Goncalves, Ewan Harrison, David K. Jackson, Ian Johnston, Dominic Kwiatkowski, Cordelia Langford, John Sillitoe |
| EPI_ISL_531595 | NHSGGC West of Scotland Specialist Virology Centre / MRC-University of Glasgow Centre for Virus Research | Wellcome Sanger Institute for the COVID-19 Genomics UK Consortium | Ana da Silva Filipe, Natasha Johnson, Kathy Smollett, Daniel Mair, Stephen Carmichael, Lily Tong, Jenna Nichols, Elihu Aranday-Cortes, Kirstyn Brunker, Yasmin Parr, Kyriaki Nomikou; Sarah McDonald, Marc Niebel, Patawee Asamaphan; Richard Orton, Joseph Hughes, Sreenu Vattipally, David L Robertson; Alasdair MacLean, Rory Gunson; Kathy Li, Natasha Jesudason, Rajiv Shah, James Shepherd, Antonia Ho, Alice Broos, Emma Thomson and Alex Alderton, Roberto Amato, Sonia Goncalves, Ewan Harrison, David K. Jackson, Ian Johnston, Dominic Kwiatkowski, Cordelia Langford, John Sillitoe |
| EPI_ISL_531596, EPI_ISL_531597 | Lighthouse Lab in Glasgow | Wellcome Sanger Institute for the COVID-19 Genomics UK Consortium | Harper VanSteenhouse, Yumi Kasai, David Gray, Carol Clugston, Anna Dominiczak and Alex Alderton, Roberto Amato, Sonia Goncalves, Ewan Harrison, David K. Jackson, Ian Johnston, Dominic Kwiatkowski, Cordelia Langford, John Sillitoe |
| EPI_ISL_531598, EPI_ISL_531599 | NHSGGC West of Scotland Specialist Virology Centre / MRC-University of Glasgow Centre for Virus Research | Wellcome Sanger Institute for the COVID-19 Genomics UK Consortium | Ana da Silva Filipe, Natasha Johnson, Kathy Smollett, Daniel Mair, Stephen Carmichael, Lily Tong, Jenna Nichols, Elihu Aranday-Cortes, Kirstyn Brunker, Yasmin Parr, Kyriaki Nomikou; Sarah McDonald, Marc Niebel, Patawee Asamaphan; Richard Orton, Joseph Hughes, Sreenu Vattipally, David L Robertson; Alasdair MacLean, Rory Gunson; Kathy Li, Natasha Jesudason, Rajiv Shah, James Shepherd, Antonia Ho, Alice Broos, Emma Thomson and Alex Alderton, Roberto Amato, Sonia Goncalves, Ewan Harrison, David K. Jackson, Ian Johnston, Dominic Kwiatkowski, Cordelia Langford, John Sillitoe |
| EPI_ISL_531600, EPI_ISL_531601, EPI_ISL_531602, EPI_ISL_531603, EPI_ISL_531604, EPI_ISL_531605, EPI_ISL_531606 | Lighthouse Lab in Glasgow | Wellcome Sanger Institute for the COVID-19 Genomics UK Consortium | Harper VanSteenhouse, Yumi Kasai, David Gray, Carol Clugston, Anna Dominiczak and Alex Alderton, Roberto Amato, Sonia Goncalves, Ewan Harrison, David K. Jackson, Ian Johnston, Dominic Kwiatkowski, Cordelia Langford, John Sillitoe |
| EPI_ISL_531607 | NHSGGC West of Scotland Specialist Virology Centre / MRC-University of Glasgow Centre for Virus Research | Wellcome Sanger Institute for the COVID-19 Genomics UK Consortium | Ana da Silva Filipe, Natasha Johnson, Kathy Smollett, Daniel Mair, Stephen Carmichael, Lily Tong, Jenna Nichols, Elihu Aranday-Cortes, Kirstyn Brunker, Yasmin Parr, Kyriaki Nomikou; Sarah McDonald, Marc Niebel, Patawee Asamaphan; Richard Orton, Joseph Hughes, Sreenu Vattipally, David L Robertson; Alasdair MacLean, Rory Gunson; Kathy Li, Natasha Jesudason, Rajiv Shah, James Shepherd, Antonia Ho, Alice Broos, Emma Thomson and Alex Alderton, Roberto Amato, Sonia Goncalves, Ewan Harrison, David K. Jackson, Ian Johnston, Dominic Kwiatkowski, Cordelia Langford, John Sillitoe |
| EPI_ISL_531608 | Lighthouse Lab in Glasgow | Wellcome Sanger Institute for the COVID-19 Genomics UK Consortium | Harper VanSteenhouse, Yumi Kasai, David Gray, Carol Clugston, Anna Dominiczak and Alex Alderton, Roberto Amato, Sonia Goncalves, Ewan Harrison, David K. Jackson, Ian Johnston, Dominic Kwiatkowski, Cordelia Langford, John Sillitoe |
| EPI_ISL_531609 | NHSGGC West of Scotland Specialist Virology Centre / MRC-University of Glasgow Centre for Virus Research | Wellcome Sanger Institute for the COVID-19 Genomics UK Consortium | Ana da Silva Filipe, Natasha Johnson, Kathy Smollett, Daniel Mair, Stephen Carmichael, Lily Tong, Jenna Nichols, Elihu Aranday-Cortes, Kirstyn Brunker, Yasmin Parr, Kyriaki Nomikou; Sarah McDonald, Marc Niebel, Patawee Asamaphan; Richard Orton, Joseph Hughes, Sreenu Vattipally, David L Robertson; Alasdair MacLean, Rory Gunson; Kathy Li, Natasha Jesudason, Rajiv Shah, James Shepherd, Antonia Ho, Alice Broos, Emma Thomson and Alex Alderton, Roberto Amato, Sonia Goncalves, Ewan Harrison, David K. Jackson, Ian Johnston, Dominic Kwiatkowski, Cordelia Langford, John Sillitoe |
| EPI_ISL_531610, EPI_ISL_531611, EPI_ISL_531612, | Lighthouse Lab in Glasgow | Wellcome Sanger Institute for the COVID-19 Genomics UK | Harper VanSteenhouse, Yumi Kasai, David Gray, Carol Clugston, Anna Dominiczak and Alex Alderton, Roberto Amato, Sonia Goncalves, Ewan Harrison, David K. Jackson, Ian Johnston, Dominic Kwiatkowski, Cordelia |

| | | | |
|---|---|---|---|
| EPI_ISL_531613, EPI_ISL_531614, EPI_ISL_531615, EPI_ISL_531616 | | Consortium | Langford, John Sillitoe |
| EPI_ISL_531617, EPI_ISL_531618 | NHSGGC West of Scotland Specialist Virology Centre / MRC-University of Glasgow Centre for Virus Research | Wellcome Sanger Institute for the COVID-19 Genomics UK Consortium | Ana da Silva Filipe, Natasha Johnson, Kathy Smollett, Daniel Mair, Stephen Carmichael, Lily Tong, Jenna Nichols, Elihu Aranday-Cortes, Kirstyn Brunker, Yasmin Parr, Kyriaki Nomikou; Sarah McDonald, Marc Niebel, Patawee Asamaphan; Richard Orton, Joseph Hughes, Sreenu Vattipally, David L Robertson; Alasdair MacLean, Rory Gunson; Kathy Li, Natasha Jesudason, Rajiv Shah, James Shepherd, Antonia Ho, Alice Broos, Emma Thomson and Alex Alderton, Roberto Amato, Sonia Goncalves, Ewan Harrison, David K. Jackson, Ian Johnston, Dominic Kwiatkowski, Cordelia Langford, John Sillitoe |
| EPI_ISL_531619, EPI_ISL_531620, EPI_ISL_531621, EPI_ISL_531622, EPI_ISL_531623, EPI_ISL_531624 | Lighthouse Lab in Glasgow | Wellcome Sanger Institute for the COVID-19 Genomics UK Consortium | Harper VanSteenhouse, Yumi Kasai, David Gray, Carol Clugston, Anna Dominiczak and Alex Alderton, Roberto Amato, Sonia Goncalves, Ewan Harrison, David K. Jackson, Ian Johnston, Dominic Kwiatkowski, Cordelia Langford, John Sillitoe |
| EPI_ISL_531625 | NHSGGC West of Scotland Specialist Virology Centre / MRC-University of Glasgow Centre for Virus Research | Wellcome Sanger Institute for the COVID-19 Genomics UK Consortium | Ana da Silva Filipe, Natasha Johnson, Kathy Smollett, Daniel Mair, Stephen Carmichael, Lily Tong, Jenna Nichols, Elihu Aranday-Cortes, Kirstyn Brunker, Yasmin Parr, Kyriaki Nomikou; Sarah McDonald, Marc Niebel, Patawee Asamaphan; Richard Orton, Joseph Hughes, Sreenu Vattipally, David L Robertson; Alasdair MacLean, Rory Gunson; Kathy Li, Natasha Jesudason, Rajiv Shah, James Shepherd, Antonia Ho, Alice Broos, Emma Thomson and Alex Alderton, Roberto Amato, Sonia Goncalves, Ewan Harrison, David K. Jackson, Ian Johnston, Dominic Kwiatkowski, Cordelia Langford, John Sillitoe |
| EPI_ISL_531626, EPI_ISL_531627, EPI_ISL_531628, EPI_ISL_531629, EPI_ISL_531630 | Lighthouse Lab in Glasgow | Wellcome Sanger Institute for the COVID-19 Genomics UK Consortium | Harper VanSteenhouse, Yumi Kasai, David Gray, Carol Clugston, Anna Dominiczak and Alex Alderton, Roberto Amato, Sonia Goncalves, Ewan Harrison, David K. Jackson, Ian Johnston, Dominic Kwiatkowski, Cordelia Langford, John Sillitoe |
| EPI_ISL_531631 | NHSGGC West of Scotland Specialist Virology Centre / MRC-University of Glasgow Centre for Virus Research | Wellcome Sanger Institute for the COVID-19 Genomics UK Consortium | Ana da Silva Filipe, Natasha Johnson, Kathy Smollett, Daniel Mair, Stephen Carmichael, Lily Tong, Jenna Nichols, Elihu Aranday-Cortes, Kirstyn Brunker, Yasmin Parr, Kyriaki Nomikou; Sarah McDonald, Marc Niebel, Patawee Asamaphan; Richard Orton, Joseph Hughes, Sreenu Vattipally, David L Robertson; Alasdair MacLean, Rory Gunson; Kathy Li, Natasha Jesudason, Rajiv Shah, James Shepherd, Antonia Ho, Alice Broos, Emma Thomson and Alex Alderton, Roberto Amato, Sonia Goncalves, Ewan Harrison, David K. Jackson, Ian Johnston, Dominic Kwiatkowski, Cordelia Langford, John Sillitoe |
| EPI_ISL_531632, EPI_ISL_531633 | Lighthouse Lab in Glasgow | Wellcome Sanger Institute for the COVID-19 Genomics UK Consortium | Harper VanSteenhouse, Yumi Kasai, David Gray, Carol Clugston, Anna Dominiczak and Alex Alderton, Roberto Amato, Sonia Goncalves, Ewan Harrison, David K. Jackson, Ian Johnston, Dominic Kwiatkowski, Cordelia Langford, John Sillitoe |
| EPI_ISL_531634 | NHSGGC West of Scotland Specialist Virology Centre / MRC-University of Glasgow Centre for Virus Research | Wellcome Sanger Institute for the COVID-19 Genomics UK Consortium | Ana da Silva Filipe, Natasha Johnson, Kathy Smollett, Daniel Mair, Stephen Carmichael, Lily Tong, Jenna Nichols, Elihu Aranday-Cortes, Kirstyn Brunker, Yasmin Parr, Kyriaki Nomikou; Sarah McDonald, Marc Niebel, Patawee Asamaphan; Richard Orton, Joseph Hughes, Sreenu Vattipally, David L Robertson; Alasdair MacLean, Rory Gunson; Kathy Li, Natasha Jesudason, Rajiv Shah, James Shepherd, Antonia Ho, Alice Broos, Emma Thomson and Alex Alderton, Roberto Amato, Sonia Goncalves, Ewan Harrison, David K. Jackson, Ian Johnston, Dominic Kwiatkowski, Cordelia Langford, John Sillitoe |
| EPI_ISL_531635, EPI_ISL_531636, EPI_ISL_531637 | Lighthouse Lab in Glasgow | Wellcome Sanger Institute for the COVID-19 Genomics UK Consortium | Harper VanSteenhouse, Yumi Kasai, David Gray, Carol Clugston, Anna Dominiczak and Alex Alderton, Roberto Amato, Sonia Goncalves, Ewan Harrison, David K. Jackson, Ian Johnston, Dominic Kwiatkowski, Cordelia Langford, John Sillitoe |
| EPI_ISL_531638, EPI_ISL_531639 | NHSGGC West of Scotland Specialist Virology Centre / MRC-University of Glasgow Centre for Virus Research | Wellcome Sanger Institute for the COVID-19 Genomics UK Consortium | Ana da Silva Filipe, Natasha Johnson, Kathy Smollett, Daniel Mair, Stephen Carmichael, Lily Tong, Jenna Nichols, Elihu Aranday-Cortes, Kirstyn Brunker, Yasmin Parr, Kyriaki Nomikou; Sarah McDonald, Marc Niebel, Patawee Asamaphan; Richard Orton, Joseph Hughes, Sreenu Vattipally, David L Robertson; Alasdair MacLean, Rory Gunson; Kathy Li, Natasha Jesudason, Rajiv Shah, James Shepherd, Antonia Ho, Alice Broos, Emma Thomson and Alex Alderton, Roberto Amato, Sonia Goncalves, Ewan Harrison, David K. Jackson, Ian Johnston, Dominic Kwiatkowski, Cordelia Langford, John Sillitoe |
| EPI_ISL_531640, EPI_ISL_531641, EPI_ISL_531642, EPI_ISL_531643, EPI_ISL_531644, EPI_ISL_531645, EPI_ISL_531646, EPI_ISL_531647, EPI_ISL_531648 | Lighthouse Lab in Glasgow | Wellcome Sanger Institute for the COVID-19 Genomics UK Consortium | Harper VanSteenhouse, Yumi Kasai, David Gray, Carol Clugston, Anna Dominiczak and Alex Alderton, Roberto Amato, Sonia Goncalves, Ewan Harrison, David K. Jackson, Ian Johnston, Dominic Kwiatkowski, Cordelia Langford, John Sillitoe |
| EPI_ISL_531649 | NHSGGC West of Scotland Specialist Virology Centre / MRC-University of Glasgow Centre for Virus Research | Wellcome Sanger Institute for the COVID-19 Genomics UK Consortium | Ana da Silva Filipe, Natasha Johnson, Kathy Smollett, Daniel Mair, Stephen Carmichael, Lily Tong, Jenna Nichols, Elihu Aranday-Cortes, Kirstyn Brunker, Yasmin Parr, Kyriaki Nomikou; Sarah McDonald, Marc Niebel, Patawee Asamaphan; Richard Orton, Joseph Hughes, Sreenu Vattipally, David L Robertson; Alasdair MacLean, Rory Gunson; Kathy Li, Natasha Jesudason, Rajiv Shah, James Shepherd, Antonia Ho, Alice Broos, Emma Thomson and Alex Alderton, Roberto Amato, Sonia Goncalves, Ewan Harrison, David K. Jackson, Ian Johnston, Dominic Kwiatkowski, Cordelia Langford, John Sillitoe |
| EPI_ISL_531650, EPI_ISL_531651, EPI_ISL_531652, EPI_ISL_531653, EPI_ISL_531654, EPI_ISL_531655, EPI_ISL_531656, EPI_ISL_531657, EPI_ISL_531658, EPI_ISL_531659 | Lighthouse Lab in Glasgow | Wellcome Sanger Institute for the COVID-19 Genomics UK Consortium | Harper VanSteenhouse, Yumi Kasai, David Gray, Carol Clugston, Anna Dominiczak and Alex Alderton, Roberto Amato, Sonia Goncalves, Ewan Harrison, David K. Jackson, Ian Johnston, Dominic Kwiatkowski, Cordelia Langford, John Sillitoe |
| EPI_ISL_531660 | NHSGGC West of Scotland Specialist Virology Centre / MRC-University of Glasgow Centre for Virus Research | Wellcome Sanger Institute for the COVID-19 Genomics UK Consortium | Ana da Silva Filipe, Natasha Johnson, Kathy Smollett, Daniel Mair, Stephen Carmichael, Lily Tong, Jenna Nichols, Elihu Aranday-Cortes, Kirstyn Brunker, Yasmin Parr, Kyriaki Nomikou; Sarah McDonald, Marc Niebel, Patawee Asamaphan; Richard Orton, Joseph Hughes, Sreenu Vattipally, David L Robertson; Alasdair MacLean, Rory Gunson; Kathy Li, Natasha Jesudason, Rajiv Shah, James Shepherd, Antonia Ho, Alice Broos, Emma Thomson and Alex Alderton, Roberto Amato, Sonia Goncalves, Ewan Harrison, David K. Jackson, Ian Johnston, Dominic Kwiatkowski, Cordelia Langford, John Sillitoe |
| EPI_ISL_531661, EPI_ISL_531662, EPI_ISL_531663, EPI_ISL_531664, EPI_ISL_531665, EPI_ISL_531666, EPI_ISL_531667 | Lighthouse Lab in Glasgow | Wellcome Sanger Institute for the COVID-19 Genomics UK Consortium | Harper VanSteenhouse, Yumi Kasai, David Gray, Carol Clugston, Anna Dominiczak and Alex Alderton, Roberto Amato, Sonia Goncalves, Ewan Harrison, David K. Jackson, Ian Johnston, Dominic Kwiatkowski, Cordelia Langford, John Sillitoe |
| EPI_ISL_531668 | NHSGGC West of Scotland Specialist Virology Centre / MRC-University of Glasgow Centre for Virus Research | Wellcome Sanger Institute for the COVID-19 Genomics UK Consortium | Ana da Silva Filipe, Natasha Johnson, Kathy Smollett, Daniel Mair, Stephen Carmichael, Lily Tong, Jenna Nichols, Elihu Aranday-Cortes, Kirstyn Brunker, Yasmin Parr, Kyriaki Nomikou; Sarah McDonald, Marc Niebel, Patawee Asamaphan; Richard Orton, Joseph Hughes, Sreenu Vattipally, David L Robertson; Alasdair MacLean, Rory Gunson; Kathy Li, Natasha Jesudason, Rajiv Shah, James Shepherd, Antonia Ho, Alice Broos, Emma Thomson and Alex Alderton, Roberto Amato, Sonia Goncalves, Ewan Harrison, David K. Jackson, Ian Johnston, Dominic Kwiatkowski, Cordelia Langford, John Sillitoe |
| EPI_ISL_531669, EPI_ISL_531670, EPI_ISL_531671 | Lighthouse Lab in Glasgow | Wellcome Sanger Institute for the COVID-19 Genomics UK Consortium | Harper VanSteenhouse, Yumi Kasai, David Gray, Carol Clugston, Anna Dominiczak and Alex Alderton, Roberto Amato, Sonia Goncalves, Ewan Harrison, David K. Jackson, Ian Johnston, Dominic Kwiatkowski, Cordelia Langford, John Sillitoe |
| EPI_ISL_531672 | NHSGGC West of Scotland Specialist Virology Centre / MRC-University of Glasgow Centre for Virus Research | Wellcome Sanger Institute for the COVID-19 Genomics UK Consortium | Ana da Silva Filipe, Natasha Johnson, Kathy Smollett, Daniel Mair, Stephen Carmichael, Lily Tong, Jenna Nichols, Elihu Aranday-Cortes, Kirstyn Brunker, Yasmin Parr, Kyriaki Nomikou; Sarah McDonald, Marc Niebel, Patawee Asamaphan; Richard Orton, Joseph Hughes, Sreenu Vattipally, David L Robertson; Alasdair MacLean, Rory Gunson; Kathy Li, Natasha Jesudason, Rajiv Shah, James Shepherd, Antonia Ho, Alice Broos, Emma Thomson and Alex Alderton, Roberto Amato, Sonia Goncalves, Ewan Harrison, David K. Jackson, Ian Johnston, Dominic Kwiatkowski, Cordelia Langford, John Sillitoe |
| EPI_ISL_531673, EPI_ISL_531674, EPI_ISL_531675, EPI_ISL_531676 | Lighthouse Lab in Glasgow | Wellcome Sanger Institute for the COVID-19 Genomics UK Consortium | Harper VanSteenhouse, Yumi Kasai, David Gray, Carol Clugston, Anna Dominiczak and Alex Alderton, Roberto Amato, Sonia Goncalves, Ewan Harrison, David K. Jackson, Ian Johnston, Dominic Kwiatkowski, Cordelia Langford, John Sillitoe |
| EPI_ISL_531677, EPI_ISL_531678 | NHSGGC West of Scotland Specialist Virology Centre / MRC-University of Glasgow Centre for Virus Research | Wellcome Sanger Institute for the COVID-19 Genomics UK Consortium | Ana da Silva Filipe, Natasha Johnson, Kathy Smollett, Daniel Mair, Stephen Carmichael, Lily Tong, Jenna Nichols, Elihu Aranday-Cortes, Kirstyn Brunker, Yasmin Parr, Kyriaki Nomikou; Sarah McDonald, Marc Niebel, Patawee Asamaphan; Richard Orton, Joseph Hughes, Sreenu Vattipally, David L Robertson; Alasdair MacLean, Rory Gunson; Kathy Li, Natasha Jesudason, Rajiv Shah, James Shepherd, Antonia Ho, Alice Broos, Emma Thomson and Alex Alderton, Roberto Amato, Sonia Goncalves, Ewan Harrison, David K. Jackson, Ian Johnston, Dominic Kwiatkowski, Cordelia Langford, John Sillitoe |
| EPI_ISL_531679, EPI_ISL_531680, EPI_ISL_531681, EPI_ISL_531682, EPI_ISL_531683, EPI_ISL_531684, EPI_ISL_531685, EPI_ISL_531686 | Lighthouse Lab in Glasgow | Wellcome Sanger Institute for the COVID-19 Genomics UK Consortium | Harper VanSteenhouse, Yumi Kasai, David Gray, Carol Clugston, Anna Dominiczak and Alex Alderton, Roberto Amato, Sonia Goncalves, Ewan Harrison, David K. Jackson, Ian Johnston, Dominic Kwiatkowski, Cordelia Langford, John Sillitoe |
| EPI_ISL_531687, EPI_ISL_531689, EPI_ISL_531690, EPI_ISL_531691 | NHSGGC West of Scotland Specialist Virology Centre / MRC-University of Glasgow Centre for Virus Research | Wellcome Sanger Institute for the COVID-19 Genomics UK Consortium | Ana da Silva Filipe, Natasha Johnson, Kathy Smollett, Daniel Mair, Stephen Carmichael, Lily Tong, Jenna Nichols, Elihu Aranday-Cortes, Kirstyn Brunker, Yasmin Parr, Kyriaki Nomikou; Sarah McDonald, Marc Niebel, Patawee Asamaphan; Richard Orton, Joseph Hughes, Sreenu Vattipally, David L Robertson; Alasdair MacLean, Rory Gunson; Kathy Li, Natasha Jesudason, Rajiv Shah, James Shepherd, Antonia Ho, Alice Broos, Emma Thomson and Alex Alderton, Roberto Amato, Sonia Goncalves, Ewan Harrison, David K. Jackson, Ian Johnston, Dominic Kwiatkowski, Cordelia Langford, John Sillitoe |
| EPI_ISL_531692, EPI_ISL_531693, EPI_ISL_531694, EPI_ISL_531695, EPI_ISL_531696, EPI_ISL_531697, EPI_ISL_531698, EPI_ISL_531699, EPI_ISL_531700, EPI_ISL_531701 | Lighthouse Lab in Glasgow | Wellcome Sanger Institute for the COVID-19 Genomics UK Consortium | Harper VanSteenhouse, Yumi Kasai, David Gray, Carol Clugston, Anna Dominiczak and Alex Alderton, Roberto Amato, Sonia Goncalves, Ewan Harrison, David K. Jackson, Ian Johnston, Dominic Kwiatkowski, Cordelia Langford, John Sillitoe |
| EPI_ISL_531702, EPI_ISL_531703 | NHSGGC West of Scotland Specialist Virology Centre / MRC-University of Glasgow Centre for Virus Research | Wellcome Sanger Institute for the COVID-19 Genomics UK Consortium | Ana da Silva Filipe, Natasha Johnson, Kathy Smollett, Daniel Mair, Stephen Carmichael, Lily Tong, Jenna Nichols, Elihu Aranday-Cortes, Kirstyn Brunker, Yasmin Parr, Kyriaki Nomikou; Sarah McDonald, Marc Niebel, Patawee Asamaphan; Richard Orton, Joseph Hughes, Sreenu Vattipally, David L Robertson; Alasdair MacLean, Rory Gunson; Kathy Li, Natasha Jesudason, Rajiv Shah, James Shepherd, Antonia Ho, Alice Broos, Emma Thomson and Alex Alderton, Roberto Amato, Sonia Goncalves, Ewan Harrison, David K. Jackson, Ian Johnston, Dominic Kwiatkowski, Cordelia Langford, John Sillitoe |
| EPI_ISL_531704, EPI_ISL_531705, EPI_ISL_531706, EPI_ISL_531707, EPI_ISL_531708, EPI_ISL_531709, EPI_ISL_531710, EPI_ISL_531711, EPI_ISL_531712, EPI_ISL_531713 | Lighthouse Lab in Glasgow | Wellcome Sanger Institute for the COVID-19 Genomics UK Consortium | Harper VanSteenhouse, Yumi Kasai, David Gray, Carol Clugston, Anna Dominiczak and Alex Alderton, Roberto Amato, Sonia Goncalves, Ewan Harrison, David K. Jackson, Ian Johnston, Dominic Kwiatkowski, Cordelia Langford, John Sillitoe |
| EPI_ISL_531714 | NHSGGC West of Scotland Specialist Virology Centre / MRC-University of Glasgow Centre for Virus Research | Wellcome Sanger Institute for the COVID-19 Genomics UK Consortium | Ana da Silva Filipe, Natasha Johnson, Kathy Smollett, Daniel Mair, Stephen Carmichael, Lily Tong, Jenna Nichols, Elihu Aranday-Cortes, Kirstyn Brunker, Yasmin Parr, Kyriaki Nomikou; Sarah McDonald, Marc Niebel, Patawee Asamaphan; Richard Orton, Joseph Hughes, Sreenu Vattipally, David L Robertson; Alasdair MacLean, Rory Gunson; Kathy Li, Natasha Jesudason, Rajiv Shah, James Shepherd, Antonia Ho, Alice Broos, Emma Thomson and Alex Alderton, Roberto Amato, Sonia Goncalves, Ewan Harrison, David K. Jackson, Ian Johnston, Dominic Kwiatkowski, Cordelia Langford, John Sillitoe |
| EPI_ISL_531715, EPI_ISL_531716, EPI_ISL_531717, EPI_ISL_531718, EPI_ISL_531719, EPI_ISL_531720, EPI_ISL_531721, EPI_ISL_531722, EPI_ISL_531723, EPI_ISL_531724, EPI_ISL_531725 | | | |
| see above | Lighthouse Lab in Glasgow | Wellcome Sanger Institute for the COVID-19 Genomics UK Consortium | Harper VanSteenhouse, Yumi Kasai, David Gray, Carol Clugston, Anna Dominiczak and Alex Alderton, Roberto Amato, Sonia Goncalves, Ewan Harrison, David K. Jackson, Ian Johnston, Dominic Kwiatkowski, Cordelia Langford, John Sillitoe |
| EPI_ISL_531727 | NHSGGC West of Scotland Specialist Virology Centre / MRC-University of Glasgow Centre for Virus Research | Wellcome Sanger Institute for the COVID-19 Genomics UK Consortium | Ana da Silva Filipe, Natasha Johnson, Kathy Smollett, Daniel Mair, Stephen Carmichael, Lily Tong, Jenna Nichols, Elihu Aranday-Cortes, Kirstyn Brunker, Yasmin Parr, Kyriaki Nomikou; Sarah McDonald, Marc Niebel, Patawee Asamaphan; Richard Orton, Joseph Hughes, Sreenu Vattipally, David L Robertson; Alasdair MacLean, Rory Gunson; Kathy Li, Natasha Jesudason, Rajiv Shah, James Shepherd, Antonia Ho, Alice Broos, Emma Thomson and Alex Alderton, Roberto Amato, Sonia Goncalves, Ewan Harrison, David K. Jackson, Ian Johnston, Dominic Kwiatkowski, Cordelia Langford, John Sillitoe |
| EPI_ISL_531728, EPI_ISL_531729 | Lighthouse Lab in Glasgow | Wellcome Sanger Institute for the COVID-19 Genomics UK Consortium | Harper VanSteenhouse, Yumi Kasai, David Gray, Carol Clugston, Anna Dominiczak and Alex Alderton, Roberto Amato, Sonia Goncalves, Ewan Harrison, David K. Jackson, Ian Johnston, Dominic Kwiatkowski, Cordelia Langford, John Sillitoe |
| EPI_ISL_531730 | NHSGGC West of Scotland Specialist Virology Centre / MRC-University of Glasgow Centre for Virus Research | Wellcome Sanger Institute for the COVID-19 Genomics UK Consortium | Ana da Silva Filipe, Natasha Johnson, Kathy Smollett, Daniel Mair, Stephen Carmichael, Lily Tong, Jenna Nichols, Elihu Aranday-Cortes, Kirstyn Brunker, Yasmin Parr, Kyriaki Nomikou; Sarah McDonald, Marc Niebel, Patawee Asamaphan; Richard Orton, Joseph Hughes, Sreenu Vattipally, David L Robertson; Alasdair MacLean, Rory Gunson; Kathy Li, Natasha Jesudason, Rajiv Shah, James Shepherd, Antonia Ho, Alice Broos, Emma Thomson and Alex Alderton, Roberto Amato, Sonia Goncalves, Ewan Harrison, David K. Jackson, Ian Johnston, Dominic Kwiatkowski, Cordelia Langford, John Sillitoe |
| EPI_ISL_531731 | Lighthouse Lab in Glasgow | Wellcome Sanger Institute for the COVID-19 Genomics UK Consortium | Harper VanSteenhouse, Yumi Kasai, David Gray, Carol Clugston, Anna Dominiczak and Alex Alderton, Roberto Amato, Sonia Goncalves, Ewan Harrison, David K. Jackson, Ian Johnston, Dominic Kwiatkowski, Cordelia Langford, John Sillitoe |
| EPI_ISL_531732 | NHSGGC West of Scotland Specialist Virology Centre / MRC-University of Glasgow Centre for Virus Research | Wellcome Sanger Institute for the COVID-19 Genomics UK Consortium | Ana da Silva Filipe, Natasha Johnson, Kathy Smollett, Daniel Mair, Stephen Carmichael, Lily Tong, Jenna Nichols, Elihu Aranday-Cortes, Kirstyn Brunker, Yasmin Parr, Kyriaki Nomikou; Sarah McDonald, Marc Niebel, Patawee Asamaphan; Richard Orton, Joseph Hughes, Sreenu Vattipally, David L Robertson; Alasdair MacLean, Rory Gunson; Kathy Li, Natasha Jesudason, Rajiv Shah, James Shepherd, Antonia Ho, Alice Broos, Emma Thomson and Alex Alderton, Roberto Amato, Sonia Goncalves, Ewan Harrison, David K. Jackson, Ian Johnston, Dominic Kwiatkowski, Cordelia Langford, John Sillitoe |
| EPI_ISL_531733, EPI_ISL_531735, EPI_ISL_531736, EPI_ISL_531737, EPI_ISL_531738, EPI_ISL_531739, EPI_ISL_531740, EPI_ISL_531741, EPI_ISL_531742 | Lighthouse Lab in Glasgow | Wellcome Sanger Institute for the COVID-19 Genomics UK Consortium | Harper VanSteenhouse, Yumi Kasai, David Gray, Carol Clugston, Anna Dominiczak and Alex Alderton, Roberto Amato, Sonia Goncalves, Ewan Harrison, David K. Jackson, Ian Johnston, Dominic Kwiatkowski, Cordelia Langford, John Sillitoe |
| EPI_ISL_531743 | NHSGGC West of Scotland Specialist Virology Centre / MRC- | Wellcome Sanger Institute for the COVID-19 Genomics UK | Ana da Silva Filipe, Natasha Johnson, Kathy Smollett, Daniel Mair, Stephen Carmichael, Lily Tong, Jenna Nichols, Elihu Aranday-Cortes, Kirstyn Brunker, Yasmin Parr, Kyriaki Nomikou; Sarah McDonald, Marc Niebel, |

| | | | |
|---|---|---|---|
| | University of Glasgow Centre for Virus Research | Consortium | Patawee Asamaphan; Richard Orton, Joseph Hughes, Sreenu Vattipally, David L Robertson; Alasdair MacLean, Rory Gunson; Kathy Li, Natasha Jesudason, Rajiv Shah, James Shepherd, Antonia Ho, Alice Broos, Emma Thomson and Alex Alderton, Roberto Amato, Sonia Goncalves, Ewan Harrison, David K. Jackson, Ian Johnston, Dominic Kwiatkowski, Cordelia Langford, John Sillitoe |
| EPI_ISL_531744, EPI_ISL_531745, EPI_ISL_531746, EPI_ISL_531747, EPI_ISL_531748, EPI_ISL_531749, EPI_ISL_531750, EPI_ISL_531751, EPI_ISL_531752 | Lighthouse Lab in Glasgow | Wellcome Sanger Institute for the COVID-19 Genomics UK Consortium | Harper VanSteenhouse, Yumi Kasai, David Gray, Carol Clugston, Anna Dominiczak and Alex Alderton, Roberto Amato, Sonia Goncalves, Ewan Harrison, David K. Jackson, Ian Johnston, Dominic Kwiatkowski, Cordelia Langford, John Sillitoe |
| EPI_ISL_531753, EPI_ISL_531754, EPI_ISL_531755 | NHSGGC West of Scotland Specialist Virology Centre / MRC-University of Glasgow Centre for Virus Research | Wellcome Sanger Institute for the COVID-19 Genomics UK Consortium | Ana da Silva Filipe, Natasha Johnson, Kathy Smollett, Daniel Mair, Stephen Carmichael, Lily Tong, Jenna Nichols, Elihu Aranday-Cortes, Kirstyn Brunker, Yasmin Parr, Kyriaki Nomikou; Sarah McDonald, Marc Niebel, Patawee Asamaphan; Richard Orton, Joseph Hughes, Sreenu Vattipally, David L Robertson; Alasdair MacLean, Rory Gunson; Kathy Li, Natasha Jesudason, Rajiv Shah, James Shepherd, Antonia Ho, Alice Broos, Emma Thomson and Alex Alderton, Roberto Amato, Sonia Goncalves, Ewan Harrison, David K. Jackson, Ian Johnston, Dominic Kwiatkowski, Cordelia Langford, John Sillitoe |
| EPI_ISL_531756, EPI_ISL_531757 | Lighthouse Lab in Glasgow | Wellcome Sanger Institute for the COVID-19 Genomics UK Consortium | Harper VanSteenhouse, Yumi Kasai, David Gray, Carol Clugston, Anna Dominiczak and Alex Alderton, Roberto Amato, Sonia Goncalves, Ewan Harrison, David K. Jackson, Ian Johnston, Dominic Kwiatkowski, Cordelia Langford, John Sillitoe |
| EPI_ISL_531758 | NHSGGC West of Scotland Specialist Virology Centre / MRC-University of Glasgow Centre for Virus Research | Wellcome Sanger Institute for the COVID-19 Genomics UK Consortium | Ana da Silva Filipe, Natasha Johnson, Kathy Smollett, Daniel Mair, Stephen Carmichael, Lily Tong, Jenna Nichols, Elihu Aranday-Cortes, Kirstyn Brunker, Yasmin Parr, Kyriaki Nomikou; Sarah McDonald, Marc Niebel, Patawee Asamaphan; Richard Orton, Joseph Hughes, Sreenu Vattipally, David L Robertson; Alasdair MacLean, Rory Gunson; Kathy Li, Natasha Jesudason, Rajiv Shah, James Shepherd, Antonia Ho, Alice Broos, Emma Thomson and Alex Alderton, Roberto Amato, Sonia Goncalves, Ewan Harrison, David K. Jackson, Ian Johnston, Dominic Kwiatkowski, Cordelia Langford, John Sillitoe |
| EPI_ISL_531759, EPI_ISL_531760, EPI_ISL_531761, EPI_ISL_531762, EPI_ISL_531763, EPI_ISL_531764, EPI_ISL_531765, EPI_ISL_531766, EPI_ISL_531767 | Lighthouse Lab in Glasgow | Wellcome Sanger Institute for the COVID-19 Genomics UK Consortium | Harper VanSteenhouse, Yumi Kasai, David Gray, Carol Clugston, Anna Dominiczak and Alex Alderton, Roberto Amato, Sonia Goncalves, Ewan Harrison, David K. Jackson, Ian Johnston, Dominic Kwiatkowski, Cordelia Langford, John Sillitoe |
| EPI_ISL_531768 | NHSGGC West of Scotland Specialist Virology Centre / MRC-University of Glasgow Centre for Virus Research | Wellcome Sanger Institute for the COVID-19 Genomics UK Consortium | Ana da Silva Filipe, Natasha Johnson, Kathy Smollett, Daniel Mair, Stephen Carmichael, Lily Tong, Jenna Nichols, Elihu Aranday-Cortes, Kirstyn Brunker, Yasmin Parr, Kyriaki Nomikou; Sarah McDonald, Marc Niebel, Patawee Asamaphan; Richard Orton, Joseph Hughes, Sreenu Vattipally, David L Robertson; Alasdair MacLean, Rory Gunson; Kathy Li, Natasha Jesudason, Rajiv Shah, James Shepherd, Antonia Ho, Alice Broos, Emma Thomson and Alex Alderton, Roberto Amato, Sonia Goncalves, Ewan Harrison, David K. Jackson, Ian Johnston, Dominic Kwiatkowski, Cordelia Langford, John Sillitoe |
| EPI_ISL_531769, EPI_ISL_531770, EPI_ISL_531771, EPI_ISL_531772, EPI_ISL_531773, EPI_ISL_531774, EPI_ISL_531775, EPI_ISL_531776, EPI_ISL_531777, EPI_ISL_531778 | Lighthouse Lab in Glasgow | Wellcome Sanger Institute for the COVID-19 Genomics UK Consortium | Harper VanSteenhouse, Yumi Kasai, David Gray, Carol Clugston, Anna Dominiczak and Alex Alderton, Roberto Amato, Sonia Goncalves, Ewan Harrison, David K. Jackson, Ian Johnston, Dominic Kwiatkowski, Cordelia Langford, John Sillitoe |
| EPI_ISL_531779 | NHSGGC West of Scotland Specialist Virology Centre / MRC-University of Glasgow Centre for Virus Research | Wellcome Sanger Institute for the COVID-19 Genomics UK Consortium | Ana da Silva Filipe, Natasha Johnson, Kathy Smollett, Daniel Mair, Stephen Carmichael, Lily Tong, Jenna Nichols, Elihu Aranday-Cortes, Kirstyn Brunker, Yasmin Parr, Kyriaki Nomikou; Sarah McDonald, Marc Niebel, Patawee Asamaphan; Richard Orton, Joseph Hughes, Sreenu Vattipally, David L Robertson; Alasdair MacLean, Rory Gunson; Kathy Li, Natasha Jesudason, Rajiv Shah, James Shepherd, Antonia Ho, Alice Broos, Emma Thomson and Alex Alderton, Roberto Amato, Sonia Goncalves, Ewan Harrison, David K. Jackson, Ian Johnston, Dominic Kwiatkowski, Cordelia Langford, John Sillitoe |
| EPI_ISL_531780, EPI_ISL_531781, EPI_ISL_531782, EPI_ISL_531783, EPI_ISL_531784, EPI_ISL_531785, EPI_ISL_531786, EPI_ISL_531787 | Lighthouse Lab in Glasgow | Wellcome Sanger Institute for the COVID-19 Genomics UK Consortium | Harper VanSteenhouse, Yumi Kasai, David Gray, Carol Clugston, Anna Dominiczak and Alex Alderton, Roberto Amato, Sonia Goncalves, Ewan Harrison, David K. Jackson, Ian Johnston, Dominic Kwiatkowski, Cordelia Langford, John Sillitoe |
| EPI_ISL_531788 | NHSGGC West of Scotland Specialist Virology Centre / MRC-University of Glasgow Centre for Virus Research | Wellcome Sanger Institute for the COVID-19 Genomics UK Consortium | Ana da Silva Filipe, Natasha Johnson, Kathy Smollett, Daniel Mair, Stephen Carmichael, Lily Tong, Jenna Nichols, Elihu Aranday-Cortes, Kirstyn Brunker, Yasmin Parr, Kyriaki Nomikou; Sarah McDonald, Marc Niebel, Patawee Asamaphan; Richard Orton, Joseph Hughes, Sreenu Vattipally, David L Robertson; Alasdair MacLean, Rory Gunson; Kathy Li, Natasha Jesudason, Rajiv Shah, James Shepherd, Antonia Ho, Alice Broos, Emma Thomson and Alex Alderton, Roberto Amato, Sonia Goncalves, Ewan Harrison, David K. Jackson, Ian Johnston, Dominic Kwiatkowski, Cordelia Langford, John Sillitoe |
| EPI_ISL_531789, EPI_ISL_531790, EPI_ISL_531791 | Lighthouse Lab in Glasgow | Wellcome Sanger Institute for the COVID-19 Genomics UK Consortium | Harper VanSteenhouse, Yumi Kasai, David Gray, Carol Clugston, Anna Dominiczak and Alex Alderton, Roberto Amato, Sonia Goncalves, Ewan Harrison, David K. Jackson, Ian Johnston, Dominic Kwiatkowski, Cordelia Langford, John Sillitoe |
| EPI_ISL_531792 | NHSGGC West of Scotland Specialist Virology Centre / MRC-University of Glasgow Centre for Virus Research | Wellcome Sanger Institute for the COVID-19 Genomics UK Consortium | Ana da Silva Filipe, Natasha Johnson, Kathy Smollett, Daniel Mair, Stephen Carmichael, Lily Tong, Jenna Nichols, Elihu Aranday-Cortes, Kirstyn Brunker, Yasmin Parr, Kyriaki Nomikou; Sarah McDonald, Marc Niebel, Patawee Asamaphan; Richard Orton, Joseph Hughes, Sreenu Vattipally, David L Robertson; Alasdair MacLean, Rory Gunson; Kathy Li, Natasha Jesudason, Rajiv Shah, James Shepherd, Antonia Ho, Alice Broos, Emma Thomson and Alex Alderton, Roberto Amato, Sonia Goncalves, Ewan Harrison, David K. Jackson, Ian Johnston, Dominic Kwiatkowski, Cordelia Langford, John Sillitoe |
| EPI_ISL_531793, EPI_ISL_531794, EPI_ISL_531795, EPI_ISL_531796, EPI_ISL_531797, EPI_ISL_531798, EPI_ISL_531799, EPI_ISL_531800 | Lighthouse Lab in Glasgow | Wellcome Sanger Institute for the COVID-19 Genomics UK Consortium | Harper VanSteenhouse, Yumi Kasai, David Gray, Carol Clugston, Anna Dominiczak and Alex Alderton, Roberto Amato, Sonia Goncalves, Ewan Harrison, David K. Jackson, Ian Johnston, Dominic Kwiatkowski, Cordelia Langford, John Sillitoe |
| EPI_ISL_531801, EPI_ISL_531802 | NHSGGC West of Scotland Specialist Virology Centre / MRC-University of Glasgow Centre for Virus Research | Wellcome Sanger Institute for the COVID-19 Genomics UK Consortium | Ana da Silva Filipe, Natasha Johnson, Kathy Smollett, Daniel Mair, Stephen Carmichael, Lily Tong, Jenna Nichols, Elihu Aranday-Cortes, Kirstyn Brunker, Yasmin Parr, Kyriaki Nomikou; Sarah McDonald, Marc Niebel, Patawee Asamaphan; Richard Orton, Joseph Hughes, Sreenu Vattipally, David L Robertson; Alasdair MacLean, Rory Gunson; Kathy Li, Natasha Jesudason, Rajiv Shah, James Shepherd, Antonia Ho, Alice Broos, Emma Thomson and Alex Alderton, Roberto Amato, Sonia Goncalves, Ewan Harrison, David K. Jackson, Ian Johnston, Dominic Kwiatkowski, Cordelia Langford, John Sillitoe |
| EPI_ISL_531803, EPI_ISL_531804 | Lighthouse Lab in Glasgow | Wellcome Sanger Institute for the COVID-19 Genomics UK Consortium | Harper VanSteenhouse, Yumi Kasai, David Gray, Carol Clugston, Anna Dominiczak and Alex Alderton, Roberto Amato, Sonia Goncalves, Ewan Harrison, David K. Jackson, Ian Johnston, Dominic Kwiatkowski, Cordelia Langford, John Sillitoe |
| EPI_ISL_531805 | NHSGGC West of Scotland Specialist Virology Centre / MRC-University of Glasgow Centre for Virus Research | Wellcome Sanger Institute for the COVID-19 Genomics UK Consortium | Ana da Silva Filipe, Natasha Johnson, Kathy Smollett, Daniel Mair, Stephen Carmichael, Lily Tong, Jenna Nichols, Elihu Aranday-Cortes, Kirstyn Brunker, Yasmin Parr, Kyriaki Nomikou; Sarah McDonald, Marc Niebel, Patawee Asamaphan; Richard Orton, Joseph Hughes, Sreenu Vattipally, David L Robertson; Alasdair MacLean, Rory Gunson; Kathy Li, Natasha Jesudason, Rajiv Shah, James Shepherd, Antonia Ho, Alice Broos, Emma Thomson and Alex Alderton, Roberto Amato, Sonia Goncalves, Ewan Harrison, David K. Jackson, Ian Johnston, Dominic Kwiatkowski, Cordelia Langford, John Sillitoe |
| EPI_ISL_531806, EPI_ISL_531807, EPI_ISL_531808, EPI_ISL_531809, EPI_ISL_531810, EPI_ISL_531811, EPI_ISL_531812, EPI_ISL_531813 | Lighthouse Lab in Glasgow | Wellcome Sanger Institute for the COVID-19 Genomics UK Consortium | Harper VanSteenhouse, Yumi Kasai, David Gray, Carol Clugston, Anna Dominiczak and Alex Alderton, Roberto Amato, Sonia Goncalves, Ewan Harrison, David K. Jackson, Ian Johnston, Dominic Kwiatkowski, Cordelia Langford, John Sillitoe |
| EPI_ISL_531814, EPI_ISL_531815 | NHSGGC West of Scotland Specialist Virology Centre / MRC-University of Glasgow Centre for Virus Research | Wellcome Sanger Institute for the COVID-19 Genomics UK Consortium | Ana da Silva Filipe, Natasha Johnson, Kathy Smollett, Daniel Mair, Stephen Carmichael, Lily Tong, Jenna Nichols, Elihu Aranday-Cortes, Kirstyn Brunker, Yasmin Parr, Kyriaki Nomikou; Sarah McDonald, Marc Niebel, Patawee Asamaphan; Richard Orton, Joseph Hughes, Sreenu Vattipally, David L Robertson; Alasdair MacLean, Rory Gunson; Kathy Li, Natasha Jesudason, Rajiv Shah, James Shepherd, Antonia Ho, Alice Broos, Emma Thomson and Alex Alderton, Roberto Amato, Sonia Goncalves, Ewan Harrison, David K. Jackson, Ian Johnston, Dominic Kwiatkowski, Cordelia Langford, John Sillitoe |
| EPI_ISL_531816, EPI_ISL_531817, EPI_ISL_531818, EPI_ISL_531819, EPI_ISL_531820 | Lighthouse Lab in Glasgow | Wellcome Sanger Institute for the COVID-19 Genomics UK Consortium | Harper VanSteenhouse, Yumi Kasai, David Gray, Carol Clugston, Anna Dominiczak and Alex Alderton, Roberto Amato, Sonia Goncalves, Ewan Harrison, David K. Jackson, Ian Johnston, Dominic Kwiatkowski, Cordelia Langford, John Sillitoe |
| EPI_ISL_531821 | NHSGGC West of Scotland Specialist Virology Centre / MRC-University of Glasgow Centre for Virus Research | Wellcome Sanger Institute for the COVID-19 Genomics UK Consortium | Ana da Silva Filipe, Natasha Johnson, Kathy Smollett, Daniel Mair, Stephen Carmichael, Lily Tong, Jenna Nichols, Elihu Aranday-Cortes, Kirstyn Brunker, Yasmin Parr, Kyriaki Nomikou; Sarah McDonald, Marc Niebel, Patawee Asamaphan; Richard Orton, Joseph Hughes, Sreenu Vattipally, David L Robertson; Alasdair MacLean, Rory Gunson; Kathy Li, Natasha Jesudason, Rajiv Shah, James Shepherd, Antonia Ho, Alice Broos, Emma Thomson and Alex Alderton, Roberto Amato, Sonia Goncalves, Ewan Harrison, David K. Jackson, Ian Johnston, Dominic Kwiatkowski, Cordelia Langford, John Sillitoe |
| EPI_ISL_531822, EPI_ISL_531823 | Lighthouse Lab in Glasgow | Wellcome Sanger Institute for the COVID-19 Genomics UK Consortium | Harper VanSteenhouse, Yumi Kasai, David Gray, Carol Clugston, Anna Dominiczak and Alex Alderton, Roberto Amato, Sonia Goncalves, Ewan Harrison, David K. Jackson, Ian Johnston, Dominic Kwiatkowski, Cordelia Langford, John Sillitoe |
| EPI_ISL_531824 | NHSGGC West of Scotland Specialist Virology Centre / MRC-University of Glasgow Centre for Virus Research | Wellcome Sanger Institute for the COVID-19 Genomics UK Consortium | Ana da Silva Filipe, Natasha Johnson, Kathy Smollett, Daniel Mair, Stephen Carmichael, Lily Tong, Jenna Nichols, Elihu Aranday-Cortes, Kirstyn Brunker, Yasmin Parr, Kyriaki Nomikou; Sarah McDonald, Marc Niebel, Patawee Asamaphan; Richard Orton, Joseph Hughes, Sreenu Vattipally, David L Robertson; Alasdair MacLean, Rory Gunson; Kathy Li, Natasha Jesudason, Rajiv Shah, James Shepherd, Antonia Ho, Alice Broos, Emma Thomson and Alex Alderton, Roberto Amato, Sonia Goncalves, Ewan Harrison, David K. Jackson, Ian Johnston, Dominic Kwiatkowski, Cordelia Langford, John Sillitoe |
| EPI_ISL_531825, EPI_ISL_531827, EPI_ISL_531828, EPI_ISL_531829, EPI_ISL_531830, EPI_ISL_531831 | Lighthouse Lab in Glasgow | Wellcome Sanger Institute for the COVID-19 Genomics UK Consortium | Harper VanSteenhouse, Yumi Kasai, David Gray, Carol Clugston, Anna Dominiczak and Alex Alderton, Roberto Amato, Sonia Goncalves, Ewan Harrison, David K. Jackson, Ian Johnston, Dominic Kwiatkowski, Cordelia Langford, John Sillitoe |
| EPI_ISL_531832 | NHSGGC West of Scotland Specialist Virology Centre / MRC-University of Glasgow Centre for Virus Research | Wellcome Sanger Institute for the COVID-19 Genomics UK Consortium | Ana da Silva Filipe, Natasha Johnson, Kathy Smollett, Daniel Mair, Stephen Carmichael, Lily Tong, Jenna Nichols, Elihu Aranday-Cortes, Kirstyn Brunker, Yasmin Parr, Kyriaki Nomikou; Sarah McDonald, Marc Niebel, Patawee Asamaphan; Richard Orton, Joseph Hughes, Sreenu Vattipally, David L Robertson; Alasdair MacLean, Rory Gunson; Kathy Li, Natasha Jesudason, Rajiv Shah, James Shepherd, Antonia Ho, Alice Broos, Emma Thomson and Alex Alderton, Roberto Amato, Sonia Goncalves, Ewan Harrison, David K. Jackson, Ian Johnston, Dominic Kwiatkowski, Cordelia Langford, John Sillitoe |
| EPI_ISL_531833, EPI_ISL_531834, EPI_ISL_531835, EPI_ISL_531836, EPI_ISL_531837, EPI_ISL_531838, EPI_ISL_531839, EPI_ISL_531840, EPI_ISL_531841 | Lighthouse Lab in Glasgow | Wellcome Sanger Institute for the COVID-19 Genomics UK Consortium | Harper VanSteenhouse, Yumi Kasai, David Gray, Carol Clugston, Anna Dominiczak and Alex Alderton, Roberto Amato, Sonia Goncalves, Ewan Harrison, David K. Jackson, Ian Johnston, Dominic Kwiatkowski, Cordelia Langford, John Sillitoe |
| EPI_ISL_531842 | NHSGGC West of Scotland Specialist Virology Centre / MRC-University of Glasgow Centre for Virus Research | Wellcome Sanger Institute for the COVID-19 Genomics UK Consortium | Ana da Silva Filipe, Natasha Johnson, Kathy Smollett, Daniel Mair, Stephen Carmichael, Lily Tong, Jenna Nichols, Elihu Aranday-Cortes, Kirstyn Brunker, Yasmin Parr, Kyriaki Nomikou; Sarah McDonald, Marc Niebel, Patawee Asamaphan; Richard Orton, Joseph Hughes, Sreenu Vattipally, David L Robertson; Alasdair MacLean, Rory Gunson; Kathy Li, Natasha Jesudason, Rajiv Shah, James Shepherd, Antonia Ho, Alice Broos, Emma Thomson and Alex Alderton, Roberto Amato, Sonia Goncalves, Ewan Harrison, David K. Jackson, Ian Johnston, Dominic Kwiatkowski, Cordelia Langford, John Sillitoe |
| EPI_ISL_531843, EPI_ISL_531844, EPI_ISL_531845, EPI_ISL_531846, EPI_ISL_531847, EPI_ISL_531848, EPI_ISL_531849, EPI_ISL_531850 | Lighthouse Lab in Glasgow | Wellcome Sanger Institute for the COVID-19 Genomics UK Consortium | Harper VanSteenhouse, Yumi Kasai, David Gray, Carol Clugston, Anna Dominiczak and Alex Alderton, Roberto Amato, Sonia Goncalves, Ewan Harrison, David K. Jackson, Ian Johnston, Dominic Kwiatkowski, Cordelia Langford, John Sillitoe |
| EPI_ISL_531851 | NHSGGC West of Scotland Specialist Virology Centre / MRC-University of Glasgow Centre for Virus Research | Wellcome Sanger Institute for the COVID-19 Genomics UK Consortium | Ana da Silva Filipe, Natasha Johnson, Kathy Smollett, Daniel Mair, Stephen Carmichael, Lily Tong, Jenna Nichols, Elihu Aranday-Cortes, Kirstyn Brunker, Yasmin Parr, Kyriaki Nomikou; Sarah McDonald, Marc Niebel, Patawee Asamaphan; Richard Orton, Joseph Hughes, Sreenu Vattipally, David L Robertson; Alasdair MacLean, Rory Gunson; Kathy Li, Natasha Jesudason, Rajiv Shah, James Shepherd, Antonia Ho, Alice Broos, Emma Thomson and Alex Alderton, Roberto Amato, Sonia Goncalves, Ewan Harrison, David K. Jackson, Ian Johnston, Dominic Kwiatkowski, Cordelia Langford, John Sillitoe |
| EPI_ISL_531852, EPI_ISL_531853, EPI_ISL_531854, EPI_ISL_531855 | Lighthouse Lab in Glasgow | Wellcome Sanger Institute for the COVID-19 Genomics UK Consortium | Harper VanSteenhouse, Yumi Kasai, David Gray, Carol Clugston, Anna Dominiczak and Alex Alderton, Roberto Amato, Sonia Goncalves, Ewan Harrison, David K. Jackson, Ian Johnston, Dominic Kwiatkowski, Cordelia Langford, John Sillitoe |
| EPI_ISL_531856, EPI_ISL_531857, EPI_ISL_531858 | NHSGGC West of Scotland Specialist Virology Centre / MRC-University of Glasgow Centre for Virus Research | Wellcome Sanger Institute for the COVID-19 Genomics UK Consortium | Ana da Silva Filipe, Natasha Johnson, Kathy Smollett, Daniel Mair, Stephen Carmichael, Lily Tong, Jenna Nichols, Elihu Aranday-Cortes, Kirstyn Brunker, Yasmin Parr, Kyriaki Nomikou; Sarah McDonald, Marc Niebel, Patawee Asamaphan; Richard Orton, Joseph Hughes, Sreenu Vattipally, David L Robertson; Alasdair MacLean, Rory Gunson; Kathy Li, Natasha Jesudason, Rajiv Shah, James Shepherd, Antonia Ho, Alice Broos, Emma Thomson and Alex Alderton, Roberto Amato, Sonia Goncalves, Ewan Harrison, David K. Jackson, Ian Johnston, Dominic Kwiatkowski, Cordelia Langford, John Sillitoe |
| EPI_ISL_531859, EPI_ISL_531860, EPI_ISL_531861 | Lighthouse Lab in Glasgow | Wellcome Sanger Institute for the COVID-19 Genomics UK Consortium | Harper VanSteenhouse, Yumi Kasai, David Gray, Carol Clugston, Anna Dominiczak and Alex Alderton, Roberto Amato, Sonia Goncalves, Ewan Harrison, David K. Jackson, Ian Johnston, Dominic Kwiatkowski, Cordelia Langford, John Sillitoe |
| EPI_ISL_531862, EPI_ISL_531863 | NHSGGC West of Scotland Specialist Virology Centre / MRC-University of Glasgow Centre for Virus Research | Wellcome Sanger Institute for the COVID-19 Genomics UK Consortium | Ana da Silva Filipe, Natasha Johnson, Kathy Smollett, Daniel Mair, Stephen Carmichael, Lily Tong, Jenna Nichols, Elihu Aranday-Cortes, Kirstyn Brunker, Yasmin Parr, Kyriaki Nomikou; Sarah McDonald, Marc Niebel, Patawee Asamaphan; Richard Orton, Joseph Hughes, Sreenu Vattipally, David L Robertson; Alasdair MacLean, Rory Gunson; Kathy Li, Natasha Jesudason, Rajiv Shah, James Shepherd, Antonia Ho, Alice Broos, Emma Thomson and Alex Alderton, Roberto Amato, Sonia Goncalves, Ewan Harrison, David K. Jackson, Ian Johnston, Dominic Kwiatkowski, Cordelia Langford, John Sillitoe |
| EPI_ISL_531864 | Lighthouse Lab in Glasgow | Wellcome Sanger Institute for the COVID-19 Genomics UK Consortium | Harper VanSteenhouse, Yumi Kasai, David Gray, Carol Clugston, Anna Dominiczak and Alex Alderton, Roberto Amato, Sonia Goncalves, Ewan Harrison, David K. Jackson, Ian Johnston, Dominic Kwiatkowski, Cordelia Langford, John Sillitoe |
| EPI_ISL_531865 | NHSGGC West of Scotland Specialist Virology Centre / MRC-University of Glasgow Centre for Virus Research | Wellcome Sanger Institute for the COVID-19 Genomics UK Consortium | Ana da Silva Filipe, Natasha Johnson, Kathy Smollett, Daniel Mair, Stephen Carmichael, Lily Tong, Jenna Nichols, Elihu Aranday-Cortes, Kirstyn Brunker, Yasmin Parr, Kyriaki Nomikou; Sarah McDonald, Marc Niebel, Patawee Asamaphan; Richard Orton, Joseph Hughes, Sreenu Vattipally, David L Robertson; Alasdair MacLean, Rory Gunson; Kathy Li, Natasha Jesudason, Rajiv Shah, James Shepherd, Antonia Ho, Alice Broos, Emma |

| | | | |
|---|---|---|---|
| | | | Thomson and Alex Alderton, Roberto Amato, Sonia Goncalves, Ewan Harrison, David K. Jackson, Ian Johnston, Dominic Kwiatkowski, Cordelia Langford, John Sillitoe |
| EPI_ISL_531866, EPI_ISL_531867, EPI_ISL_531868, EPI_ISL_531869, EPI_ISL_531870, EPI_ISL_531871 | Lighthouse Lab in Glasgow | Wellcome Sanger Institute for the COVID-19 Genomics UK Consortium | Harper VanSteenhouse, Yumi Kasai, David Gray, Carol Clugston, Anna Dominiczak and Alex Alderton, Roberto Amato, Sonia Goncalves, Ewan Harrison, David K. Jackson, Ian Johnston, Dominic Kwiatkowski, Cordelia Langford, John Sillitoe |
| EPI_ISL_531873 | NHSGGC West of Scotland Specialist Virology Centre / MRC-University of Glasgow Centre for Virus Research | Wellcome Sanger Institute for the COVID-19 Genomics UK Consortium | Ana da Silva Filipe, Natasha Johnson, Kathy Smollett, Daniel Mair, Stephen Carmichael, Lily Tong, Jenna Nichols, Elihu Aranday-Cortes, Kirstyn Brunker, Yasmin Parr, Kyriaki Nomikou; Sarah McDonald, Marc Niebel, Patawee Asamaphan; Richard Orton, Joseph Hughes, Sreenu Vattipally, David L Robertson; Alasdair MacLean, Rory Gunson; Kathy Li, Natasha Jesudason, Rajiv Shah, James Shepherd, Antonia Ho, Alice Broos, Emma Thomson and Alex Alderton, Roberto Amato, Sonia Goncalves, Ewan Harrison, David K. Jackson, Ian Johnston, Dominic Kwiatkowski, Cordelia Langford, John Sillitoe |
| EPI_ISL_531874, EPI_ISL_531875, EPI_ISL_531876 | Lighthouse Lab in Glasgow | Wellcome Sanger Institute for the COVID-19 Genomics UK Consortium | Harper VanSteenhouse, Yumi Kasai, David Gray, Carol Clugston, Anna Dominiczak and Alex Alderton, Roberto Amato, Sonia Goncalves, Ewan Harrison, David K. Jackson, Ian Johnston, Dominic Kwiatkowski, Cordelia Langford, John Sillitoe |
| EPI_ISL_531877 | NHSGGC West of Scotland Specialist Virology Centre / MRC-University of Glasgow Centre for Virus Research | Wellcome Sanger Institute for the COVID-19 Genomics UK Consortium | Ana da Silva Filipe, Natasha Johnson, Kathy Smollett, Daniel Mair, Stephen Carmichael, Lily Tong, Jenna Nichols, Elihu Aranday-Cortes, Kirstyn Brunker, Yasmin Parr, Kyriaki Nomikou; Sarah McDonald, Marc Niebel, Patawee Asamaphan; Richard Orton, Joseph Hughes, Sreenu Vattipally, David L Robertson; Alasdair MacLean, Rory Gunson; Kathy Li, Natasha Jesudason, Rajiv Shah, James Shepherd, Antonia Ho, Alice Broos, Emma Thomson and Alex Alderton, Roberto Amato, Sonia Goncalves, Ewan Harrison, David K. Jackson, Ian Johnston, Dominic Kwiatkowski, Cordelia Langford, John Sillitoe |
| EPI_ISL_531878, EPI_ISL_531879 | Lighthouse Lab in Glasgow | Wellcome Sanger Institute for the COVID-19 Genomics UK Consortium | Harper VanSteenhouse, Yumi Kasai, David Gray, Carol Clugston, Anna Dominiczak and Alex Alderton, Roberto Amato, Sonia Goncalves, Ewan Harrison, David K. Jackson, Ian Johnston, Dominic Kwiatkowski, Cordelia Langford, John Sillitoe |
| EPI_ISL_531880, EPI_ISL_531881 | NHSGGC West of Scotland Specialist Virology Centre / MRC-University of Glasgow Centre for Virus Research | Wellcome Sanger Institute for the COVID-19 Genomics UK Consortium | Ana da Silva Filipe, Natasha Johnson, Kathy Smollett, Daniel Mair, Stephen Carmichael, Lily Tong, Jenna Nichols, Elihu Aranday-Cortes, Kirstyn Brunker, Yasmin Parr, Kyriaki Nomikou; Sarah McDonald, Marc Niebel, Patawee Asamaphan; Richard Orton, Joseph Hughes, Sreenu Vattipally, David L Robertson; Alasdair MacLean, Rory Gunson; Kathy Li, Natasha Jesudason, Rajiv Shah, James Shepherd, Antonia Ho, Alice Broos, Emma Thomson and Alex Alderton, Roberto Amato, Sonia Goncalves, Ewan Harrison, David K. Jackson, Ian Johnston, Dominic Kwiatkowski, Cordelia Langford, John Sillitoe |
| EPI_ISL_531882, EPI_ISL_531883 | Lighthouse Lab in Glasgow | Wellcome Sanger Institute for the COVID-19 Genomics UK Consortium | Harper VanSteenhouse, Yumi Kasai, David Gray, Carol Clugston, Anna Dominiczak and Alex Alderton, Roberto Amato, Sonia Goncalves, Ewan Harrison, David K. Jackson, Ian Johnston, Dominic Kwiatkowski, Cordelia Langford, John Sillitoe |
| EPI_ISL_531884 | NHSGGC West of Scotland Specialist Virology Centre / MRC-University of Glasgow Centre for Virus Research | Wellcome Sanger Institute for the COVID-19 Genomics UK Consortium | Ana da Silva Filipe, Natasha Johnson, Kathy Smollett, Daniel Mair, Stephen Carmichael, Lily Tong, Jenna Nichols, Elihu Aranday-Cortes, Kirstyn Brunker, Yasmin Parr, Kyriaki Nomikou; Sarah McDonald, Marc Niebel, Patawee Asamaphan; Richard Orton, Joseph Hughes, Sreenu Vattipally, David L Robertson; Alasdair MacLean, Rory Gunson; Kathy Li, Natasha Jesudason, Rajiv Shah, James Shepherd, Antonia Ho, Alice Broos, Emma Thomson and Alex Alderton, Roberto Amato, Sonia Goncalves, Ewan Harrison, David K. Jackson, Ian Johnston, Dominic Kwiatkowski, Cordelia Langford, John Sillitoe |
| EPI_ISL_531885, EPI_ISL_531886 | Lighthouse Lab in Glasgow | Wellcome Sanger Institute for the COVID-19 Genomics UK Consortium | Harper VanSteenhouse, Yumi Kasai, David Gray, Carol Clugston, Anna Dominiczak and Alex Alderton, Roberto Amato, Sonia Goncalves, Ewan Harrison, David K. Jackson, Ian Johnston, Dominic Kwiatkowski, Cordelia Langford, John Sillitoe |
| EPI_ISL_531887 | NHSGGC West of Scotland Specialist Virology Centre / MRC-University of Glasgow Centre for Virus Research | Wellcome Sanger Institute for the COVID-19 Genomics UK Consortium | Ana da Silva Filipe, Natasha Johnson, Kathy Smollett, Daniel Mair, Stephen Carmichael, Lily Tong, Jenna Nichols, Elihu Aranday-Cortes, Kirstyn Brunker, Yasmin Parr, Kyriaki Nomikou; Sarah McDonald, Marc Niebel, Patawee Asamaphan; Richard Orton, Joseph Hughes, Sreenu Vattipally, David L Robertson; Alasdair MacLean, Rory Gunson; Kathy Li, Natasha Jesudason, Rajiv Shah, James Shepherd, Antonia Ho, Alice Broos, Emma Thomson and Alex Alderton, Roberto Amato, Sonia Goncalves, Ewan Harrison, David K. Jackson, Ian Johnston, Dominic Kwiatkowski, Cordelia Langford, John Sillitoe |
| EPI_ISL_531888 | Lighthouse Lab in Glasgow | Wellcome Sanger Institute for the COVID-19 Genomics UK Consortium | Harper VanSteenhouse, Yumi Kasai, David Gray, Carol Clugston, Anna Dominiczak and Alex Alderton, Roberto Amato, Sonia Goncalves, Ewan Harrison, David K. Jackson, Ian Johnston, Dominic Kwiatkowski, Cordelia Langford, John Sillitoe |
| EPI_ISL_531889, EPI_ISL_531890 | NHSGGC West of Scotland Specialist Virology Centre / MRC-University of Glasgow Centre for Virus Research | Wellcome Sanger Institute for the COVID-19 Genomics UK Consortium | Ana da Silva Filipe, Natasha Johnson, Kathy Smollett, Daniel Mair, Stephen Carmichael, Lily Tong, Jenna Nichols, Elihu Aranday-Cortes, Kirstyn Brunker, Yasmin Parr, Kyriaki Nomikou; Sarah McDonald, Marc Niebel, Patawee Asamaphan; Richard Orton, Joseph Hughes, Sreenu Vattipally, David L Robertson; Alasdair MacLean, Rory Gunson; Kathy Li, Natasha Jesudason, Rajiv Shah, James Shepherd, Antonia Ho, Alice Broos, Emma Thomson and Alex Alderton, Roberto Amato, Sonia Goncalves, Ewan Harrison, David K. Jackson, Ian Johnston, Dominic Kwiatkowski, Cordelia Langford, John Sillitoe |
| EPI_ISL_531891 | Lighthouse Lab in Glasgow | Wellcome Sanger Institute for the COVID-19 Genomics UK Consortium | Harper VanSteenhouse, Yumi Kasai, David Gray, Carol Clugston, Anna Dominiczak and Alex Alderton, Roberto Amato, Sonia Goncalves, Ewan Harrison, David K. Jackson, Ian Johnston, Dominic Kwiatkowski, Cordelia Langford, John Sillitoe |
| EPI_ISL_531892 | NHSGGC West of Scotland Specialist Virology Centre / MRC-University of Glasgow Centre for Virus Research | Wellcome Sanger Institute for the COVID-19 Genomics UK Consortium | Ana da Silva Filipe, Natasha Johnson, Kathy Smollett, Daniel Mair, Stephen Carmichael, Lily Tong, Jenna Nichols, Elihu Aranday-Cortes, Kirstyn Brunker, Yasmin Parr, Kyriaki Nomikou; Sarah McDonald, Marc Niebel, Patawee Asamaphan; Richard Orton, Joseph Hughes, Sreenu Vattipally, David L Robertson; Alasdair MacLean, Rory Gunson; Kathy Li, Natasha Jesudason, Rajiv Shah, James Shepherd, Antonia Ho, Alice Broos, Emma Thomson and Alex Alderton, Roberto Amato, Sonia Goncalves, Ewan Harrison, David K. Jackson, Ian Johnston, Dominic Kwiatkowski, Cordelia Langford, John Sillitoe |
| EPI_ISL_531893 | Lighthouse Lab in Glasgow | Wellcome Sanger Institute for the COVID-19 Genomics UK Consortium | Harper VanSteenhouse, Yumi Kasai, David Gray, Carol Clugston, Anna Dominiczak and Alex Alderton, Roberto Amato, Sonia Goncalves, Ewan Harrison, David K. Jackson, Ian Johnston, Dominic Kwiatkowski, Cordelia Langford, John Sillitoe |
| EPI_ISL_531894 | NHSGGC West of Scotland Specialist Virology Centre / MRC-University of Glasgow Centre for Virus Research | Wellcome Sanger Institute for the COVID-19 Genomics UK Consortium | Ana da Silva Filipe, Natasha Johnson, Kathy Smollett, Daniel Mair, Stephen Carmichael, Lily Tong, Jenna Nichols, Elihu Aranday-Cortes, Kirstyn Brunker, Yasmin Parr, Kyriaki Nomikou; Sarah McDonald, Marc Niebel, Patawee Asamaphan; Richard Orton, Joseph Hughes, Sreenu Vattipally, David L Robertson; Alasdair MacLean, Rory Gunson; Kathy Li, Natasha Jesudason, Rajiv Shah, James Shepherd, Antonia Ho, Alice Broos, Emma Thomson and Alex Alderton, Roberto Amato, Sonia Goncalves, Ewan Harrison, David K. Jackson, Ian Johnston, Dominic Kwiatkowski, Cordelia Langford, John Sillitoe |
| EPI_ISL_531895, EPI_ISL_531896, EPI_ISL_531897, EPI_ISL_531898, EPI_ISL_531899, EPI_ISL_531900, EPI_ISL_531901, EPI_ISL_531902, EPI_ISL_531903, EPI_ISL_531904 | Lighthouse Lab in Glasgow | Wellcome Sanger Institute for the COVID-19 Genomics UK Consortium | Harper VanSteenhouse, Yumi Kasai, David Gray, Carol Clugston, Anna Dominiczak and Alex Alderton, Roberto Amato, Sonia Goncalves, Ewan Harrison, David K. Jackson, Ian Johnston, Dominic Kwiatkowski, Cordelia Langford, John Sillitoe |
| EPI_ISL_531905 | NHSGGC West of Scotland Specialist Virology Centre / MRC-University of Glasgow Centre for Virus Research | Wellcome Sanger Institute for the COVID-19 Genomics UK Consortium | Ana da Silva Filipe, Natasha Johnson, Kathy Smollett, Daniel Mair, Stephen Carmichael, Lily Tong, Jenna Nichols, Elihu Aranday-Cortes, Kirstyn Brunker, Yasmin Parr, Kyriaki Nomikou; Sarah McDonald, Marc Niebel, Patawee Asamaphan; Richard Orton, Joseph Hughes, Sreenu Vattipally, David L Robertson; Alasdair MacLean, Rory Gunson; Kathy Li, Natasha Jesudason, Rajiv Shah, James Shepherd, Antonia Ho, Alice Broos, Emma Thomson and Alex Alderton, Roberto Amato, Sonia Goncalves, Ewan Harrison, David K. Jackson, Ian Johnston, Dominic Kwiatkowski, Cordelia Langford, John Sillitoe |
| EPI_ISL_531906, EPI_ISL_531907, EPI_ISL_531908, EPI_ISL_531909, EPI_ISL_531910, EPI_ISL_531911 | Lighthouse Lab in Glasgow | Wellcome Sanger Institute for the COVID-19 Genomics UK Consortium | Harper VanSteenhouse, Yumi Kasai, David Gray, Carol Clugston, Anna Dominiczak and Alex Alderton, Roberto Amato, Sonia Goncalves, Ewan Harrison, David K. Jackson, Ian Johnston, Dominic Kwiatkowski, Cordelia Langford, John Sillitoe |
| EPI_ISL_531912, EPI_ISL_531913 | NHSGGC West of Scotland Specialist Virology Centre / MRC-University of Glasgow Centre for Virus Research | Wellcome Sanger Institute for the COVID-19 Genomics UK Consortium | Ana da Silva Filipe, Natasha Johnson, Kathy Smollett, Daniel Mair, Stephen Carmichael, Lily Tong, Jenna Nichols, Elihu Aranday-Cortes, Kirstyn Brunker, Yasmin Parr, Kyriaki Nomikou; Sarah McDonald, Marc Niebel, Patawee Asamaphan; Richard Orton, Joseph Hughes, Sreenu Vattipally, David L Robertson; Alasdair MacLean, Rory Gunson; Kathy Li, Natasha Jesudason, Rajiv Shah, James Shepherd, Antonia Ho, Alice Broos, Emma Thomson and Alex Alderton, Roberto Amato, Sonia Goncalves, Ewan Harrison, David K. Jackson, Ian Johnston, Dominic Kwiatkowski, Cordelia Langford, John Sillitoe |
| EPI_ISL_531914, EPI_ISL_531915, EPI_ISL_531916, EPI_ISL_531917 | Lighthouse Lab in Glasgow | Wellcome Sanger Institute for the COVID-19 Genomics UK Consortium | Harper VanSteenhouse, Yumi Kasai, David Gray, Carol Clugston, Anna Dominiczak and Alex Alderton, Roberto Amato, Sonia Goncalves, Ewan Harrison, David K. Jackson, Ian Johnston, Dominic Kwiatkowski, Cordelia Langford, John Sillitoe |
| EPI_ISL_531918, EPI_ISL_531919 | NHSGGC West of Scotland Specialist Virology Centre / MRC-University of Glasgow Centre for Virus Research | Wellcome Sanger Institute for the COVID-19 Genomics UK Consortium | Ana da Silva Filipe, Natasha Johnson, Kathy Smollett, Daniel Mair, Stephen Carmichael, Lily Tong, Jenna Nichols, Elihu Aranday-Cortes, Kirstyn Brunker, Yasmin Parr, Kyriaki Nomikou; Sarah McDonald, Marc Niebel, Patawee Asamaphan; Richard Orton, Joseph Hughes, Sreenu Vattipally, David L Robertson; Alasdair MacLean, Rory Gunson; Kathy Li, Natasha Jesudason, Rajiv Shah, James Shepherd, Antonia Ho, Alice Broos, Emma Thomson and Alex Alderton, Roberto Amato, Sonia Goncalves, Ewan Harrison, David K. Jackson, Ian Johnston, Dominic Kwiatkowski, Cordelia Langford, John Sillitoe |
| EPI_ISL_531920, EPI_ISL_531921, EPI_ISL_531922, EPI_ISL_531923, EPI_ISL_531924 | Lighthouse Lab in Glasgow | Wellcome Sanger Institute for the COVID-19 Genomics UK Consortium | Harper VanSteenhouse, Yumi Kasai, David Gray, Carol Clugston, Anna Dominiczak and Alex Alderton, Roberto Amato, Sonia Goncalves, Ewan Harrison, David K. Jackson, Ian Johnston, Dominic Kwiatkowski, Cordelia Langford, John Sillitoe |
| EPI_ISL_531925 | NHSGGC West of Scotland Specialist Virology Centre / MRC-University of Glasgow Centre for Virus Research | Wellcome Sanger Institute for the COVID-19 Genomics UK Consortium | Ana da Silva Filipe, Natasha Johnson, Kathy Smollett, Daniel Mair, Stephen Carmichael, Lily Tong, Jenna Nichols, Elihu Aranday-Cortes, Kirstyn Brunker, Yasmin Parr, Kyriaki Nomikou; Sarah McDonald, Marc Niebel, Patawee Asamaphan; Richard Orton, Joseph Hughes, Sreenu Vattipally, David L Robertson; Alasdair MacLean, Rory Gunson; Kathy Li, Natasha Jesudason, Rajiv Shah, James Shepherd, Antonia Ho, Alice Broos, Emma Thomson and Alex Alderton, Roberto Amato, Sonia Goncalves, Ewan Harrison, David K. Jackson, Ian Johnston, Dominic Kwiatkowski, Cordelia Langford, John Sillitoe |
| EPI_ISL_531926, EPI_ISL_531927, EPI_ISL_531928 | Lighthouse Lab in Glasgow | Wellcome Sanger Institute for the COVID-19 Genomics UK Consortium | Harper VanSteenhouse, Yumi Kasai, David Gray, Carol Clugston, Anna Dominiczak and Alex Alderton, Roberto Amato, Sonia Goncalves, Ewan Harrison, David K. Jackson, Ian Johnston, Dominic Kwiatkowski, Cordelia Langford, John Sillitoe |
| EPI_ISL_531929 | NHSGGC West of Scotland Specialist Virology Centre / MRC-University of Glasgow Centre for Virus Research | Wellcome Sanger Institute for the COVID-19 Genomics UK Consortium | Ana da Silva Filipe, Natasha Johnson, Kathy Smollett, Daniel Mair, Stephen Carmichael, Lily Tong, Jenna Nichols, Elihu Aranday-Cortes, Kirstyn Brunker, Yasmin Parr, Kyriaki Nomikou; Sarah McDonald, Marc Niebel, Patawee Asamaphan; Richard Orton, Joseph Hughes, Sreenu Vattipally, David L Robertson; Alasdair MacLean, Rory Gunson; Kathy Li, Natasha Jesudason, Rajiv Shah, James Shepherd, Antonia Ho, Alice Broos, Emma Thomson and Alex Alderton, Roberto Amato, Sonia Goncalves, Ewan Harrison, David K. Jackson, Ian Johnston, Dominic Kwiatkowski, Cordelia Langford, John Sillitoe |
| EPI_ISL_531930, EPI_ISL_531931 | Lighthouse Lab in Glasgow | Wellcome Sanger Institute for the COVID-19 Genomics UK Consortium | Harper VanSteenhouse, Yumi Kasai, David Gray, Carol Clugston, Anna Dominiczak and Alex Alderton, Roberto Amato, Sonia Goncalves, Ewan Harrison, David K. Jackson, Ian Johnston, Dominic Kwiatkowski, Cordelia Langford, John Sillitoe |
| EPI_ISL_531932, EPI_ISL_531933 | NHSGGC West of Scotland Specialist Virology Centre / MRC-University of Glasgow Centre for Virus Research | Wellcome Sanger Institute for the COVID-19 Genomics UK Consortium | Ana da Silva Filipe, Natasha Johnson, Kathy Smollett, Daniel Mair, Stephen Carmichael, Lily Tong, Jenna Nichols, Elihu Aranday-Cortes, Kirstyn Brunker, Yasmin Parr, Kyriaki Nomikou; Sarah McDonald, Marc Niebel, Patawee Asamaphan; Richard Orton, Joseph Hughes, Sreenu Vattipally, David L Robertson; Alasdair MacLean, Rory Gunson; Kathy Li, Natasha Jesudason, Rajiv Shah, James Shepherd, Antonia Ho, Alice Broos, Emma Thomson and Alex Alderton, Roberto Amato, Sonia Goncalves, Ewan Harrison, David K. Jackson, Ian Johnston, Dominic Kwiatkowski, Cordelia Langford, John Sillitoe |
| EPI_ISL_531934 | Lighthouse Lab in Glasgow | Wellcome Sanger Institute for the COVID-19 Genomics UK Consortium | Harper VanSteenhouse, Yumi Kasai, David Gray, Carol Clugston, Anna Dominiczak and Alex Alderton, Roberto Amato, Sonia Goncalves, Ewan Harrison, David K. Jackson, Ian Johnston, Dominic Kwiatkowski, Cordelia Langford, John Sillitoe |
| EPI_ISL_531935, EPI_ISL_531936 | NHSGGC West of Scotland Specialist Virology Centre / MRC-University of Glasgow Centre for Virus Research | Wellcome Sanger Institute for the COVID-19 Genomics UK Consortium | Ana da Silva Filipe, Natasha Johnson, Kathy Smollett, Daniel Mair, Stephen Carmichael, Lily Tong, Jenna Nichols, Elihu Aranday-Cortes, Kirstyn Brunker, Yasmin Parr, Kyriaki Nomikou; Sarah McDonald, Marc Niebel, Patawee Asamaphan; Richard Orton, Joseph Hughes, Sreenu Vattipally, David L Robertson; Alasdair MacLean, Rory Gunson; Kathy Li, Natasha Jesudason, Rajiv Shah, James Shepherd, Antonia Ho, Alice Broos, Emma Thomson and Alex Alderton, Roberto Amato, Sonia Goncalves, Ewan Harrison, David K. Jackson, Ian Johnston, Dominic Kwiatkowski, Cordelia Langford, John Sillitoe |
| EPI_ISL_531937, EPI_ISL_531938, EPI_ISL_531939, EPI_ISL_531940, EPI_ISL_531941, EPI_ISL_531942 | Lighthouse Lab in Glasgow | Wellcome Sanger Institute for the COVID-19 Genomics UK Consortium | Harper VanSteenhouse, Yumi Kasai, David Gray, Carol Clugston, Anna Dominiczak and Alex Alderton, Roberto Amato, Sonia Goncalves, Ewan Harrison, David K. Jackson, Ian Johnston, Dominic Kwiatkowski, Cordelia Langford, John Sillitoe |
| EPI_ISL_531943 | NHSGGC West of Scotland Specialist Virology Centre / MRC-University of Glasgow Centre for Virus Research | Wellcome Sanger Institute for the COVID-19 Genomics UK Consortium | Ana da Silva Filipe, Natasha Johnson, Kathy Smollett, Daniel Mair, Stephen Carmichael, Lily Tong, Jenna Nichols, Elihu Aranday-Cortes, Kirstyn Brunker, Yasmin Parr, Kyriaki Nomikou; Sarah McDonald, Marc Niebel, Patawee Asamaphan; Richard Orton, Joseph Hughes, Sreenu Vattipally, David L Robertson; Alasdair MacLean, Rory Gunson; Kathy Li, Natasha Jesudason, Rajiv Shah, James Shepherd, Antonia Ho, Alice Broos, Emma Thomson and Alex Alderton, Roberto Amato, Sonia Goncalves, Ewan Harrison, David K. Jackson, Ian Johnston, Dominic Kwiatkowski, Cordelia Langford, John Sillitoe |
| EPI_ISL_531944 | Lighthouse Lab in Glasgow | Wellcome Sanger Institute for the COVID-19 Genomics UK Consortium | Harper VanSteenhouse, Yumi Kasai, David Gray, Carol Clugston, Anna Dominiczak and Alex Alderton, Roberto Amato, Sonia Goncalves, Ewan Harrison, David K. Jackson, Ian Johnston, Dominic Kwiatkowski, Cordelia Langford, John Sillitoe |
| EPI_ISL_531945 | NHSGGC West of Scotland Specialist Virology Centre / MRC-University of Glasgow Centre for Virus Research | Wellcome Sanger Institute for the COVID-19 Genomics UK Consortium | Ana da Silva Filipe, Natasha Johnson, Kathy Smollett, Daniel Mair, Stephen Carmichael, Lily Tong, Jenna Nichols, Elihu Aranday-Cortes, Kirstyn Brunker, Yasmin Parr, Kyriaki Nomikou; Sarah McDonald, Marc Niebel, Patawee Asamaphan; Richard Orton, Joseph Hughes, Sreenu Vattipally, David L Robertson; Alasdair MacLean, Rory Gunson; Kathy Li, Natasha Jesudason, Rajiv Shah, James Shepherd, Antonia Ho, Alice Broos, Emma Thomson and Alex Alderton, Roberto Amato, Sonia Goncalves, Ewan Harrison, David K. Jackson, Ian Johnston, Dominic Kwiatkowski, Cordelia Langford, John Sillitoe |
| EPI_ISL_531946, EPI_ISL_531947, EPI_ISL_531948, EPI_ISL_531949, EPI_ISL_531950 | Lighthouse Lab in Glasgow | Wellcome Sanger Institute for the COVID-19 Genomics UK Consortium | Harper VanSteenhouse, Yumi Kasai, David Gray, Carol Clugston, Anna Dominiczak and Alex Alderton, Roberto Amato, Sonia Goncalves, Ewan Harrison, David K. Jackson, Ian Johnston, Dominic Kwiatkowski, Cordelia Langford, John Sillitoe |
| EPI_ISL_531951 | NHSGGC West of Scotland Specialist Virology Centre / MRC-University of Glasgow Centre for Virus Research | Wellcome Sanger Institute for the COVID-19 Genomics UK Consortium | Ana da Silva Filipe, Natasha Johnson, Kathy Smollett, Daniel Mair, Stephen Carmichael, Lily Tong, Jenna Nichols, Elihu Aranday-Cortes, Kirstyn Brunker, Yasmin Parr, Kyriaki Nomikou; Sarah McDonald, Marc Niebel, Patawee Asamaphan; Richard Orton, Joseph Hughes, Sreenu Vattipally, David L Robertson; Alasdair MacLean, Rory Gunson; Kathy Li, Natasha Jesudason, Rajiv Shah, James Shepherd, Antonia Ho, Alice Broos, Emma Thomson and Alex Alderton, Roberto Amato, Sonia Goncalves, Ewan Harrison, David K. Jackson, Ian Johnston, Dominic Kwiatkowski, Cordelia Langford, John Sillitoe |
| EPI_ISL_531952, EPI_ISL_531953 | Lighthouse Lab in Glasgow | Wellcome Sanger Institute for the COVID-19 Genomics UK | Harper VanSteenhouse, Yumi Kasai, David Gray, Carol Clugston, Anna Dominiczak and Alex Alderton, Roberto Amato, Sonia Goncalves, Ewan Harrison, David K. Jackson, Ian Johnston, Dominic Kwiatkowski, Cordelia |

| | | | |
|---|---|---|---|
| | | Consortium | Langford, John Sillitoe |
| EPI_ISL_531954 | NHSGGC West of Scotland Specialist Virology Centre / MRC-University of Glasgow Centre for Virus Research | Wellcome Sanger Institute for the COVID-19 Genomics UK Consortium | Ana da Silva Filipe, Natasha Johnson, Kathy Smollett, Daniel Mair, Stephen Carmichael, Lily Tong, Jenna Nichols, Elihu Aranday-Cortes, Kirstyn Brunker, Yasmin Parr, Kyriaki Nomikou; Sarah McDonald, Marc Niebel, Patawee Asamaphan; Richard Orton, Joseph Hughes, Sreenu Vattipally, David L Robertson; Alasdair MacLean, Rory Gunson; Kathy Li, Natasha Jesudason, Rajiv Shah, James Shepherd, Antonia Ho, Alice Broos, Emma Thomson and Alex Alderton, Roberto Amato, Sonia Goncalves, Ewan Harrison, David K. Jackson, Ian Johnston, Dominic Kwiatkowski, Cordelia Langford, John Sillitoe |
| EPI_ISL_531955, EPI_ISL_531956, EPI_ISL_531957, EPI_ISL_531958, EPI_ISL_531959, EPI_ISL_531960, EPI_ISL_531961, EPI_ISL_531962 | Lighthouse Lab in Glasgow | Wellcome Sanger Institute for the COVID-19 Genomics UK Consortium | Harper VanSteenhouse, Yumi Kasai, David Gray, Carol Clugston, Anna Dominiczak and Alex Alderton, Roberto Amato, Sonia Goncalves, Ewan Harrison, David K. Jackson, Ian Johnston, Dominic Kwiatkowski, Cordelia Langford, John Sillitoe |
| EPI_ISL_531963 | NHSGGC West of Scotland Specialist Virology Centre / MRC-University of Glasgow Centre for Virus Research | Wellcome Sanger Institute for the COVID-19 Genomics UK Consortium | Ana da Silva Filipe, Natasha Johnson, Kathy Smollett, Daniel Mair, Stephen Carmichael, Lily Tong, Jenna Nichols, Elihu Aranday-Cortes, Kirstyn Brunker, Yasmin Parr, Kyriaki Nomikou; Sarah McDonald, Marc Niebel, Patawee Asamaphan; Richard Orton, Joseph Hughes, Sreenu Vattipally, David L Robertson; Alasdair MacLean, Rory Gunson; Kathy Li, Natasha Jesudason, Rajiv Shah, James Shepherd, Antonia Ho, Alice Broos, Emma Thomson and Alex Alderton, Roberto Amato, Sonia Goncalves, Ewan Harrison, David K. Jackson, Ian Johnston, Dominic Kwiatkowski, Cordelia Langford, John Sillitoe |
| EPI_ISL_531964, EPI_ISL_531965, EPI_ISL_531966, EPI_ISL_531967, EPI_ISL_531968, EPI_ISL_531969, EPI_ISL_531970, EPI_ISL_531971, EPI_ISL_531972 | Lighthouse Lab in Glasgow | Wellcome Sanger Institute for the COVID-19 Genomics UK Consortium | Harper VanSteenhouse, Yumi Kasai, David Gray, Carol Clugston, Anna Dominiczak and Alex Alderton, Roberto Amato, Sonia Goncalves, Ewan Harrison, David K. Jackson, Ian Johnston, Dominic Kwiatkowski, Cordelia Langford, John Sillitoe |
| EPI_ISL_531973 | NHSGGC West of Scotland Specialist Virology Centre / MRC-University of Glasgow Centre for Virus Research | Wellcome Sanger Institute for the COVID-19 Genomics UK Consortium | Ana da Silva Filipe, Natasha Johnson, Kathy Smollett, Daniel Mair, Stephen Carmichael, Lily Tong, Jenna Nichols, Elihu Aranday-Cortes, Kirstyn Brunker, Yasmin Parr, Kyriaki Nomikou; Sarah McDonald, Marc Niebel, Patawee Asamaphan; Richard Orton, Joseph Hughes, Sreenu Vattipally, David L Robertson; Alasdair MacLean, Rory Gunson; Kathy Li, Natasha Jesudason, Rajiv Shah, James Shepherd, Antonia Ho, Alice Broos, Emma Thomson and Alex Alderton, Roberto Amato, Sonia Goncalves, Ewan Harrison, David K. Jackson, Ian Johnston, Dominic Kwiatkowski, Cordelia Langford, John Sillitoe |
| EPI_ISL_531974, EPI_ISL_531975, EPI_ISL_531976, EPI_ISL_531977 | Lighthouse Lab in Glasgow | Wellcome Sanger Institute for the COVID-19 Genomics UK Consortium | Harper VanSteenhouse, Yumi Kasai, David Gray, Carol Clugston, Anna Dominiczak and Alex Alderton, Roberto Amato, Sonia Goncalves, Ewan Harrison, David K. Jackson, Ian Johnston, Dominic Kwiatkowski, Cordelia Langford, John Sillitoe |
| EPI_ISL_531978, EPI_ISL_531979 | NHSGGC West of Scotland Specialist Virology Centre / MRC-University of Glasgow Centre for Virus Research | Wellcome Sanger Institute for the COVID-19 Genomics UK Consortium | Ana da Silva Filipe, Natasha Johnson, Kathy Smollett, Daniel Mair, Stephen Carmichael, Lily Tong, Jenna Nichols, Elihu Aranday-Cortes, Kirstyn Brunker, Yasmin Parr, Kyriaki Nomikou; Sarah McDonald, Marc Niebel, Patawee Asamaphan; Richard Orton, Joseph Hughes, Sreenu Vattipally, David L Robertson; Alasdair MacLean, Rory Gunson; Kathy Li, Natasha Jesudason, Rajiv Shah, James Shepherd, Antonia Ho, Alice Broos, Emma Thomson and Alex Alderton, Roberto Amato, Sonia Goncalves, Ewan Harrison, David K. Jackson, Ian Johnston, Dominic Kwiatkowski, Cordelia Langford, John Sillitoe |
| EPI_ISL_531980, EPI_ISL_531981 | Lighthouse Lab in Glasgow | Wellcome Sanger Institute for the COVID-19 Genomics UK Consortium | Harper VanSteenhouse, Yumi Kasai, David Gray, Carol Clugston, Anna Dominiczak and Alex Alderton, Roberto Amato, Sonia Goncalves, Ewan Harrison, David K. Jackson, Ian Johnston, Dominic Kwiatkowski, Cordelia Langford, John Sillitoe |
| EPI_ISL_531982 | NHSGGC West of Scotland Specialist Virology Centre / MRC-University of Glasgow Centre for Virus Research | Wellcome Sanger Institute for the COVID-19 Genomics UK Consortium | Ana da Silva Filipe, Natasha Johnson, Kathy Smollett, Daniel Mair, Stephen Carmichael, Lily Tong, Jenna Nichols, Elihu Aranday-Cortes, Kirstyn Brunker, Yasmin Parr, Kyriaki Nomikou; Sarah McDonald, Marc Niebel, Patawee Asamaphan; Richard Orton, Joseph Hughes, Sreenu Vattipally, David L Robertson; Alasdair MacLean, Rory Gunson; Kathy Li, Natasha Jesudason, Rajiv Shah, James Shepherd, Antonia Ho, Alice Broos, Emma Thomson and Alex Alderton, Roberto Amato, Sonia Goncalves, Ewan Harrison, David K. Jackson, Ian Johnston, Dominic Kwiatkowski, Cordelia Langford, John Sillitoe |
| EPI_ISL_531983 | Lighthouse Lab in Glasgow | Wellcome Sanger Institute for the COVID-19 Genomics UK Consortium | Harper VanSteenhouse, Yumi Kasai, David Gray, Carol Clugston, Anna Dominiczak and Alex Alderton, Roberto Amato, Sonia Goncalves, Ewan Harrison, David K. Jackson, Ian Johnston, Dominic Kwiatkowski, Cordelia Langford, John Sillitoe |
| EPI_ISL_531984 | NHSGGC West of Scotland Specialist Virology Centre / MRC-University of Glasgow Centre for Virus Research | Wellcome Sanger Institute for the COVID-19 Genomics UK Consortium | Ana da Silva Filipe, Natasha Johnson, Kathy Smollett, Daniel Mair, Stephen Carmichael, Lily Tong, Jenna Nichols, Elihu Aranday-Cortes, Kirstyn Brunker, Yasmin Parr, Kyriaki Nomikou; Sarah McDonald, Marc Niebel, Patawee Asamaphan; Richard Orton, Joseph Hughes, Sreenu Vattipally, David L Robertson; Alasdair MacLean, Rory Gunson; Kathy Li, Natasha Jesudason, Rajiv Shah, James Shepherd, Antonia Ho, Alice Broos, Emma Thomson and Alex Alderton, Roberto Amato, Sonia Goncalves, Ewan Harrison, David K. Jackson, Ian Johnston, Dominic Kwiatkowski, Cordelia Langford, John Sillitoe |
| EPI_ISL_531985, EPI_ISL_531986 | Lighthouse Lab in Glasgow | Wellcome Sanger Institute for the COVID-19 Genomics UK Consortium | Harper VanSteenhouse, Yumi Kasai, David Gray, Carol Clugston, Anna Dominiczak and Alex Alderton, Roberto Amato, Sonia Goncalves, Ewan Harrison, David K. Jackson, Ian Johnston, Dominic Kwiatkowski, Cordelia Langford, John Sillitoe |
| EPI_ISL_531987 | NHSGGC West of Scotland Specialist Virology Centre / MRC-University of Glasgow Centre for Virus Research | Wellcome Sanger Institute for the COVID-19 Genomics UK Consortium | Ana da Silva Filipe, Natasha Johnson, Kathy Smollett, Daniel Mair, Stephen Carmichael, Lily Tong, Jenna Nichols, Elihu Aranday-Cortes, Kirstyn Brunker, Yasmin Parr, Kyriaki Nomikou; Sarah McDonald, Marc Niebel, Patawee Asamaphan; Richard Orton, Joseph Hughes, Sreenu Vattipally, David L Robertson; Alasdair MacLean, Rory Gunson; Kathy Li, Natasha Jesudason, Rajiv Shah, James Shepherd, Antonia Ho, Alice Broos, Emma Thomson and Alex Alderton, Roberto Amato, Sonia Goncalves, Ewan Harrison, David K. Jackson, Ian Johnston, Dominic Kwiatkowski, Cordelia Langford, John Sillitoe |
| EPI_ISL_531988, EPI_ISL_531989 | Lighthouse Lab in Glasgow | Wellcome Sanger Institute for the COVID-19 Genomics UK Consortium | Harper VanSteenhouse, Yumi Kasai, David Gray, Carol Clugston, Anna Dominiczak and Alex Alderton, Roberto Amato, Sonia Goncalves, Ewan Harrison, David K. Jackson, Ian Johnston, Dominic Kwiatkowski, Cordelia Langford, John Sillitoe |
| EPI_ISL_531990 | NHSGGC West of Scotland Specialist Virology Centre / MRC-University of Glasgow Centre for Virus Research | Wellcome Sanger Institute for the COVID-19 Genomics UK Consortium | Ana da Silva Filipe, Natasha Johnson, Kathy Smollett, Daniel Mair, Stephen Carmichael, Lily Tong, Jenna Nichols, Elihu Aranday-Cortes, Kirstyn Brunker, Yasmin Parr, Kyriaki Nomikou; Sarah McDonald, Marc Niebel, Patawee Asamaphan; Richard Orton, Joseph Hughes, Sreenu Vattipally, David L Robertson; Alasdair MacLean, Rory Gunson; Kathy Li, Natasha Jesudason, Rajiv Shah, James Shepherd, Antonia Ho, Alice Broos, Emma Thomson and Alex Alderton, Roberto Amato, Sonia Goncalves, Ewan Harrison, David K. Jackson, Ian Johnston, Dominic Kwiatkowski, Cordelia Langford, John Sillitoe |
| EPI_ISL_531991, EPI_ISL_531992, EPI_ISL_531993, EPI_ISL_531994 | Lighthouse Lab in Glasgow | Wellcome Sanger Institute for the COVID-19 Genomics UK Consortium | Harper VanSteenhouse, Yumi Kasai, David Gray, Carol Clugston, Anna Dominiczak and Alex Alderton, Roberto Amato, Sonia Goncalves, Ewan Harrison, David K. Jackson, Ian Johnston, Dominic Kwiatkowski, Cordelia Langford, John Sillitoe |
| EPI_ISL_531995 | NHSGGC West of Scotland Specialist Virology Centre / MRC-University of Glasgow Centre for Virus Research | Wellcome Sanger Institute for the COVID-19 Genomics UK Consortium | Ana da Silva Filipe, Natasha Johnson, Kathy Smollett, Daniel Mair, Stephen Carmichael, Lily Tong, Jenna Nichols, Elihu Aranday-Cortes, Kirstyn Brunker, Yasmin Parr, Kyriaki Nomikou; Sarah McDonald, Marc Niebel, Patawee Asamaphan; Richard Orton, Joseph Hughes, Sreenu Vattipally, David L Robertson; Alasdair MacLean, Rory Gunson; Kathy Li, Natasha Jesudason, Rajiv Shah, James Shepherd, Antonia Ho, Alice Broos, Emma Thomson and Alex Alderton, Roberto Amato, Sonia Goncalves, Ewan Harrison, David K. Jackson, Ian Johnston, Dominic Kwiatkowski, Cordelia Langford, John Sillitoe |
| EPI_ISL_531996, EPI_ISL_531997, EPI_ISL_531998, EPI_ISL_531999, EPI_ISL_532000, EPI_ISL_532001, EPI_ISL_532002 | Lighthouse Lab in Glasgow | Wellcome Sanger Institute for the COVID-19 Genomics UK Consortium | Harper VanSteenhouse, Yumi Kasai, David Gray, Carol Clugston, Anna Dominiczak and Alex Alderton, Roberto Amato, Sonia Goncalves, Ewan Harrison, David K. Jackson, Ian Johnston, Dominic Kwiatkowski, Cordelia Langford, John Sillitoe |
| EPI_ISL_532003, EPI_ISL_532004, EPI_ISL_532005 | NHSGGC West of Scotland Specialist Virology Centre / MRC-University of Glasgow Centre for Virus Research | Wellcome Sanger Institute for the COVID-19 Genomics UK Consortium | Ana da Silva Filipe, Natasha Johnson, Kathy Smollett, Daniel Mair, Stephen Carmichael, Lily Tong, Jenna Nichols, Elihu Aranday-Cortes, Kirstyn Brunker, Yasmin Parr, Kyriaki Nomikou; Sarah McDonald, Marc Niebel, Patawee Asamaphan; Richard Orton, Joseph Hughes, Sreenu Vattipally, David L Robertson; Alasdair MacLean, Rory Gunson; Kathy Li, Natasha Jesudason, Rajiv Shah, James Shepherd, Antonia Ho, Alice Broos, Emma Thomson and Alex Alderton, Roberto Amato, Sonia Goncalves, Ewan Harrison, David K. Jackson, Ian Johnston, Dominic Kwiatkowski, Cordelia Langford, John Sillitoe |
| EPI_ISL_532006 | Lighthouse Lab in Glasgow | Wellcome Sanger Institute for the COVID-19 Genomics UK Consortium | Harper VanSteenhouse, Yumi Kasai, David Gray, Carol Clugston, Anna Dominiczak and Alex Alderton, Roberto Amato, Sonia Goncalves, Ewan Harrison, David K. Jackson, Ian Johnston, Dominic Kwiatkowski, Cordelia Langford, John Sillitoe |
| EPI_ISL_532007 | NHSGGC West of Scotland Specialist Virology Centre / MRC-University of Glasgow Centre for Virus Research | Wellcome Sanger Institute for the COVID-19 Genomics UK Consortium | Ana da Silva Filipe, Natasha Johnson, Kathy Smollett, Daniel Mair, Stephen Carmichael, Lily Tong, Jenna Nichols, Elihu Aranday-Cortes, Kirstyn Brunker, Yasmin Parr, Kyriaki Nomikou; Sarah McDonald, Marc Niebel, Patawee Asamaphan; Richard Orton, Joseph Hughes, Sreenu Vattipally, David L Robertson; Alasdair MacLean, Rory Gunson; Kathy Li, Natasha Jesudason, Rajiv Shah, James Shepherd, Antonia Ho, Alice Broos, Emma Thomson and Alex Alderton, Roberto Amato, Sonia Goncalves, Ewan Harrison, David K. Jackson, Ian Johnston, Dominic Kwiatkowski, Cordelia Langford, John Sillitoe |
| EPI_ISL_532008, EPI_ISL_532009, EPI_ISL_532010, EPI_ISL_532011, EPI_ISL_532013, EPI_ISL_532014, EPI_ISL_532015, EPI_ISL_532016, EPI_ISL_532017 | Lighthouse Lab in Glasgow | Wellcome Sanger Institute for the COVID-19 Genomics UK Consortium | Harper VanSteenhouse, Yumi Kasai, David Gray, Carol Clugston, Anna Dominiczak and Alex Alderton, Roberto Amato, Sonia Goncalves, Ewan Harrison, David K. Jackson, Ian Johnston, Dominic Kwiatkowski, Cordelia Langford, John Sillitoe |
| EPI_ISL_532018 | NHSGGC West of Scotland Specialist Virology Centre / MRC-University of Glasgow Centre for Virus Research | Wellcome Sanger Institute for the COVID-19 Genomics UK Consortium | Ana da Silva Filipe, Natasha Johnson, Kathy Smollett, Daniel Mair, Stephen Carmichael, Lily Tong, Jenna Nichols, Elihu Aranday-Cortes, Kirstyn Brunker, Yasmin Parr, Kyriaki Nomikou; Sarah McDonald, Marc Niebel, Patawee Asamaphan; Richard Orton, Joseph Hughes, Sreenu Vattipally, David L Robertson; Alasdair MacLean, Rory Gunson; Kathy Li, Natasha Jesudason, Rajiv Shah, James Shepherd, Antonia Ho, Alice Broos, Emma Thomson and Alex Alderton, Roberto Amato, Sonia Goncalves, Ewan Harrison, David K. Jackson, Ian Johnston, Dominic Kwiatkowski, Cordelia Langford, John Sillitoe |
| EPI_ISL_532019, EPI_ISL_532020, EPI_ISL_532021, EPI_ISL_532022, EPI_ISL_532023 | Lighthouse Lab in Glasgow | Wellcome Sanger Institute for the COVID-19 Genomics UK Consortium | Harper VanSteenhouse, Yumi Kasai, David Gray, Carol Clugston, Anna Dominiczak and Alex Alderton, Roberto Amato, Sonia Goncalves, Ewan Harrison, David K. Jackson, Ian Johnston, Dominic Kwiatkowski, Cordelia Langford, John Sillitoe |
| EPI_ISL_532024 | NHSGGC West of Scotland Specialist Virology Centre / MRC-University of Glasgow Centre for Virus Research | Wellcome Sanger Institute for the COVID-19 Genomics UK Consortium | Ana da Silva Filipe, Natasha Johnson, Kathy Smollett, Daniel Mair, Stephen Carmichael, Lily Tong, Jenna Nichols, Elihu Aranday-Cortes, Kirstyn Brunker, Yasmin Parr, Kyriaki Nomikou; Sarah McDonald, Marc Niebel, Patawee Asamaphan; Richard Orton, Joseph Hughes, Sreenu Vattipally, David L Robertson; Alasdair MacLean, Rory Gunson; Kathy Li, Natasha Jesudason, Rajiv Shah, James Shepherd, Antonia Ho, Alice Broos, Emma Thomson and Alex Alderton, Roberto Amato, Sonia Goncalves, Ewan Harrison, David K. Jackson, Ian Johnston, Dominic Kwiatkowski, Cordelia Langford, John Sillitoe |
| EPI_ISL_532025, EPI_ISL_532026, EPI_ISL_532027, EPI_ISL_532028, EPI_ISL_532029, EPI_ISL_532030, EPI_ISL_532031 | Lighthouse Lab in Glasgow | Wellcome Sanger Institute for the COVID-19 Genomics UK Consortium | Harper VanSteenhouse, Yumi Kasai, David Gray, Carol Clugston, Anna Dominiczak and Alex Alderton, Roberto Amato, Sonia Goncalves, Ewan Harrison, David K. Jackson, Ian Johnston, Dominic Kwiatkowski, Cordelia Langford, John Sillitoe |
| EPI_ISL_532032 | NHSGGC West of Scotland Specialist Virology Centre / MRC-University of Glasgow Centre for Virus Research | Wellcome Sanger Institute for the COVID-19 Genomics UK Consortium | Ana da Silva Filipe, Natasha Johnson, Kathy Smollett, Daniel Mair, Stephen Carmichael, Lily Tong, Jenna Nichols, Elihu Aranday-Cortes, Kirstyn Brunker, Yasmin Parr, Kyriaki Nomikou; Sarah McDonald, Marc Niebel, Patawee Asamaphan; Richard Orton, Joseph Hughes, Sreenu Vattipally, David L Robertson; Alasdair MacLean, Rory Gunson; Kathy Li, Natasha Jesudason, Rajiv Shah, James Shepherd, Antonia Ho, Alice Broos, Emma Thomson and Alex Alderton, Roberto Amato, Sonia Goncalves, Ewan Harrison, David K. Jackson, Ian Johnston, Dominic Kwiatkowski, Cordelia Langford, John Sillitoe |
| EPI_ISL_532033, EPI_ISL_532034, EPI_ISL_532035, EPI_ISL_532036, EPI_ISL_532037 | Lighthouse Lab in Glasgow | Wellcome Sanger Institute for the COVID-19 Genomics UK Consortium | Harper VanSteenhouse, Yumi Kasai, David Gray, Carol Clugston, Anna Dominiczak and Alex Alderton, Roberto Amato, Sonia Goncalves, Ewan Harrison, David K. Jackson, Ian Johnston, Dominic Kwiatkowski, Cordelia Langford, John Sillitoe |
| EPI_ISL_532038 | NHSGGC West of Scotland Specialist Virology Centre / MRC-University of Glasgow Centre for Virus Research | Wellcome Sanger Institute for the COVID-19 Genomics UK Consortium | Ana da Silva Filipe, Natasha Johnson, Kathy Smollett, Daniel Mair, Stephen Carmichael, Lily Tong, Jenna Nichols, Elihu Aranday-Cortes, Kirstyn Brunker, Yasmin Parr, Kyriaki Nomikou; Sarah McDonald, Marc Niebel, Patawee Asamaphan; Richard Orton, Joseph Hughes, Sreenu Vattipally, David L Robertson; Alasdair MacLean, Rory Gunson; Kathy Li, Natasha Jesudason, Rajiv Shah, James Shepherd, Antonia Ho, Alice Broos, Emma Thomson and Alex Alderton, Roberto Amato, Sonia Goncalves, Ewan Harrison, David K. Jackson, Ian Johnston, Dominic Kwiatkowski, Cordelia Langford, John Sillitoe |
| EPI_ISL_532039, EPI_ISL_532040, EPI_ISL_532041, EPI_ISL_532042, EPI_ISL_532043, EPI_ISL_532044, EPI_ISL_532045, EPI_ISL_532046, EPI_ISL_532047, EPI_ISL_532048 | Lighthouse Lab in Glasgow | Wellcome Sanger Institute for the COVID-19 Genomics UK Consortium | Harper VanSteenhouse, Yumi Kasai, David Gray, Carol Clugston, Anna Dominiczak and Alex Alderton, Roberto Amato, Sonia Goncalves, Ewan Harrison, David K. Jackson, Ian Johnston, Dominic Kwiatkowski, Cordelia Langford, John Sillitoe |
| EPI_ISL_532049 | NHSGGC West of Scotland Specialist Virology Centre / MRC-University of Glasgow Centre for Virus Research | Wellcome Sanger Institute for the COVID-19 Genomics UK Consortium | Ana da Silva Filipe, Natasha Johnson, Kathy Smollett, Daniel Mair, Stephen Carmichael, Lily Tong, Jenna Nichols, Elihu Aranday-Cortes, Kirstyn Brunker, Yasmin Parr, Kyriaki Nomikou; Sarah McDonald, Marc Niebel, Patawee Asamaphan; Richard Orton, Joseph Hughes, Sreenu Vattipally, David L Robertson; Alasdair MacLean, Rory Gunson; Kathy Li, Natasha Jesudason, Rajiv Shah, James Shepherd, Antonia Ho, Alice Broos, Emma Thomson and Alex Alderton, Roberto Amato, Sonia Goncalves, Ewan Harrison, David K. Jackson, Ian Johnston, Dominic Kwiatkowski, Cordelia Langford, John Sillitoe |
| EPI_ISL_532050, EPI_ISL_532051, EPI_ISL_532052 | Lighthouse Lab in Glasgow | Wellcome Sanger Institute for the COVID-19 Genomics UK Consortium | Harper VanSteenhouse, Yumi Kasai, David Gray, Carol Clugston, Anna Dominiczak and Alex Alderton, Roberto Amato, Sonia Goncalves, Ewan Harrison, David K. Jackson, Ian Johnston, Dominic Kwiatkowski, Cordelia Langford, John Sillitoe |
| EPI_ISL_532053 | NHSGGC West of Scotland Specialist Virology Centre / MRC-University of Glasgow Centre for Virus Research | Wellcome Sanger Institute for the COVID-19 Genomics UK Consortium | Ana da Silva Filipe, Natasha Johnson, Kathy Smollett, Daniel Mair, Stephen Carmichael, Lily Tong, Jenna Nichols, Elihu Aranday-Cortes, Kirstyn Brunker, Yasmin Parr, Kyriaki Nomikou; Sarah McDonald, Marc Niebel, Patawee Asamaphan; Richard Orton, Joseph Hughes, Sreenu Vattipally, David L Robertson; Alasdair MacLean, Rory Gunson; Kathy Li, Natasha Jesudason, Rajiv Shah, James Shepherd, Antonia Ho, Alice Broos, Emma Thomson and Alex Alderton, Roberto Amato, Sonia Goncalves, Ewan Harrison, David K. Jackson, Ian Johnston, Dominic Kwiatkowski, Cordelia Langford, John Sillitoe |
| EPI_ISL_532054 | Lighthouse Lab in Glasgow | Wellcome Sanger Institute for the COVID-19 Genomics UK Consortium | Harper VanSteenhouse, Yumi Kasai, David Gray, Carol Clugston, Anna Dominiczak and Alex Alderton, Roberto Amato, Sonia Goncalves, Ewan Harrison, David K. Jackson, Ian Johnston, Dominic Kwiatkowski, Cordelia Langford, John Sillitoe |
| EPI_ISL_532055 | NHSGGC West of Scotland Specialist Virology Centre / MRC- | Wellcome Sanger Institute for the COVID-19 Genomics UK | Ana da Silva Filipe, Natasha Johnson, Kathy Smollett, Daniel Mair, Stephen Carmichael, Lily Tong, Jenna Nichols, Elihu Aranday-Cortes, Kirstyn Brunker, Yasmin Parr, Kyriaki Nomikou; Sarah McDonald, Marc Niebel, |

| | | | |
|---|---|---|---|
| | University of Glasgow Centre for Virus Research | Consortium | Patawee Asamaphan; Richard Orton, Joseph Hughes, Sreenu Vattipally, David L Robertson; Alasdair MacLean, Rory Gunson; Kathy Li, Natasha Jesudason, Rajiv Shah, James Shepherd, Antonia Ho, Alice Broos, Emma Thomson and Alex Alderton, Roberto Amato, Sonia Goncalves, Ewan Harrison, David K. Jackson, Ian Johnston, Dominic Kwiatkowski, Cordelia Langford, John Sillitoe |
| EPI_ISL_532056 | Lighthouse Lab in Glasgow | Wellcome Sanger Institute for the COVID-19 Genomics UK Consortium | Harper VanSteenhouse, Yumi Kasai, David Gray, Carol Clugston, Anna Dominiczak and Alex Alderton, Roberto Amato, Sonia Goncalves, Ewan Harrison, David K. Jackson, Ian Johnston, Dominic Kwiatkowski, Cordelia Langford, John Sillitoe |
| EPI_ISL_532057 | NHSGGC West of Scotland Specialist Virology Centre / MRC-University of Glasgow Centre for Virus Research | Wellcome Sanger Institute for the COVID-19 Genomics UK Consortium | Ana da Silva Filipe, Natasha Johnson, Kathy Smollett, Daniel Mair, Stephen Carmichael, Lily Tong, Jenna Nichols, Elihu Aranday-Cortes, Kirstyn Brunker, Yasmin Parr, Kyriaki Nomikou; Sarah McDonald, Marc Niebel, Patawee Asamaphan; Richard Orton, Joseph Hughes, Sreenu Vattipally, David L Robertson; Alasdair MacLean, Rory Gunson; Kathy Li, Natasha Jesudason, Rajiv Shah, James Shepherd, Antonia Ho, Alice Broos, Emma Thomson and Alex Alderton, Roberto Amato, Sonia Goncalves, Ewan Harrison, David K. Jackson, Ian Johnston, Dominic Kwiatkowski, Cordelia Langford, John Sillitoe |
| EPI_ISL_532058, EPI_ISL_532059, EPI_ISL_532060, EPI_ISL_532061 | Lighthouse Lab in Glasgow | Wellcome Sanger Institute for the COVID-19 Genomics UK Consortium | Harper VanSteenhouse, Yumi Kasai, David Gray, Carol Clugston, Anna Dominiczak and Alex Alderton, Roberto Amato, Sonia Goncalves, Ewan Harrison, David K. Jackson, Ian Johnston, Dominic Kwiatkowski, Cordelia Langford, John Sillitoe |
| EPI_ISL_532062 | NHSGGC West of Scotland Specialist Virology Centre / MRC-University of Glasgow Centre for Virus Research | Wellcome Sanger Institute for the COVID-19 Genomics UK Consortium | Ana da Silva Filipe, Natasha Johnson, Kathy Smollett, Daniel Mair, Stephen Carmichael, Lily Tong, Jenna Nichols, Elihu Aranday-Cortes, Kirstyn Brunker, Yasmin Parr, Kyriaki Nomikou; Sarah McDonald, Marc Niebel, Patawee Asamaphan; Richard Orton, Joseph Hughes, Sreenu Vattipally, David L Robertson; Alasdair MacLean, Rory Gunson; Kathy Li, Natasha Jesudason, Rajiv Shah, James Shepherd, Antonia Ho, Alice Broos, Emma Thomson and Alex Alderton, Roberto Amato, Sonia Goncalves, Ewan Harrison, David K. Jackson, Ian Johnston, Dominic Kwiatkowski, Cordelia Langford, John Sillitoe |
| EPI_ISL_532063, EPI_ISL_532064, EPI_ISL_532065, EPI_ISL_532066, EPI_ISL_532067, EPI_ISL_532068, EPI_ISL_532069 | Lighthouse Lab in Glasgow | Wellcome Sanger Institute for the COVID-19 Genomics UK Consortium | Harper VanSteenhouse, Yumi Kasai, David Gray, Carol Clugston, Anna Dominiczak and Alex Alderton, Roberto Amato, Sonia Goncalves, Ewan Harrison, David K. Jackson, Ian Johnston, Dominic Kwiatkowski, Cordelia Langford, John Sillitoe |
| EPI_ISL_532070, EPI_ISL_532071 | NHSGGC West of Scotland Specialist Virology Centre / MRC-University of Glasgow Centre for Virus Research | Wellcome Sanger Institute for the COVID-19 Genomics UK Consortium | Ana da Silva Filipe, Natasha Johnson, Kathy Smollett, Daniel Mair, Stephen Carmichael, Lily Tong, Jenna Nichols, Elihu Aranday-Cortes, Kirstyn Brunker, Yasmin Parr, Kyriaki Nomikou; Sarah McDonald, Marc Niebel, Patawee Asamaphan; Richard Orton, Joseph Hughes, Sreenu Vattipally, David L Robertson; Alasdair MacLean, Rory Gunson; Kathy Li, Natasha Jesudason, Rajiv Shah, James Shepherd, Antonia Ho, Alice Broos, Emma Thomson and Alex Alderton, Roberto Amato, Sonia Goncalves, Ewan Harrison, David K. Jackson, Ian Johnston, Dominic Kwiatkowski, Cordelia Langford, John Sillitoe |
| EPI_ISL_532072, EPI_ISL_532073, EPI_ISL_532074, EPI_ISL_532075 | Lighthouse Lab in Glasgow | Wellcome Sanger Institute for the COVID-19 Genomics UK Consortium | Harper VanSteenhouse, Yumi Kasai, David Gray, Carol Clugston, Anna Dominiczak and Alex Alderton, Roberto Amato, Sonia Goncalves, Ewan Harrison, David K. Jackson, Ian Johnston, Dominic Kwiatkowski, Cordelia Langford, John Sillitoe |
| EPI_ISL_532076, EPI_ISL_532077, EPI_ISL_532078 | NHSGGC West of Scotland Specialist Virology Centre / MRC-University of Glasgow Centre for Virus Research | Wellcome Sanger Institute for the COVID-19 Genomics UK Consortium | Ana da Silva Filipe, Natasha Johnson, Kathy Smollett, Daniel Mair, Stephen Carmichael, Lily Tong, Jenna Nichols, Elihu Aranday-Cortes, Kirstyn Brunker, Yasmin Parr, Kyriaki Nomikou; Sarah McDonald, Marc Niebel, Patawee Asamaphan; Richard Orton, Joseph Hughes, Sreenu Vattipally, David L Robertson; Alasdair MacLean, Rory Gunson; Kathy Li, Natasha Jesudason, Rajiv Shah, James Shepherd, Antonia Ho, Alice Broos, Emma Thomson and Alex Alderton, Roberto Amato, Sonia Goncalves, Ewan Harrison, David K. Jackson, Ian Johnston, Dominic Kwiatkowski, Cordelia Langford, John Sillitoe |
| EPI_ISL_532079 | Lighthouse Lab in Glasgow | Wellcome Sanger Institute for the COVID-19 Genomics UK Consortium | Harper VanSteenhouse, Yumi Kasai, David Gray, Carol Clugston, Anna Dominiczak and Alex Alderton, Roberto Amato, Sonia Goncalves, Ewan Harrison, David K. Jackson, Ian Johnston, Dominic Kwiatkowski, Cordelia Langford, John Sillitoe |
| EPI_ISL_532080, EPI_ISL_532081 | NHSGGC West of Scotland Specialist Virology Centre / MRC-University of Glasgow Centre for Virus Research | Wellcome Sanger Institute for the COVID-19 Genomics UK Consortium | Ana da Silva Filipe, Natasha Johnson, Kathy Smollett, Daniel Mair, Stephen Carmichael, Lily Tong, Jenna Nichols, Elihu Aranday-Cortes, Kirstyn Brunker, Yasmin Parr, Kyriaki Nomikou; Sarah McDonald, Marc Niebel, Patawee Asamaphan; Richard Orton, Joseph Hughes, Sreenu Vattipally, David L Robertson; Alasdair MacLean, Rory Gunson; Kathy Li, Natasha Jesudason, Rajiv Shah, James Shepherd, Antonia Ho, Alice Broos, Emma Thomson and Alex Alderton, Roberto Amato, Sonia Goncalves, Ewan Harrison, David K. Jackson, Ian Johnston, Dominic Kwiatkowski, Cordelia Langford, John Sillitoe |
| EPI_ISL_532082, EPI_ISL_532083, EPI_ISL_532084 | Lighthouse Lab in Glasgow | Wellcome Sanger Institute for the COVID-19 Genomics UK Consortium | Harper VanSteenhouse, Yumi Kasai, David Gray, Carol Clugston, Anna Dominiczak and Alex Alderton, Roberto Amato, Sonia Goncalves, Ewan Harrison, David K. Jackson, Ian Johnston, Dominic Kwiatkowski, Cordelia Langford, John Sillitoe |
| EPI_ISL_532085 | NHSGGC West of Scotland Specialist Virology Centre / MRC-University of Glasgow Centre for Virus Research | Wellcome Sanger Institute for the COVID-19 Genomics UK Consortium | Ana da Silva Filipe, Natasha Johnson, Kathy Smollett, Daniel Mair, Stephen Carmichael, Lily Tong, Jenna Nichols, Elihu Aranday-Cortes, Kirstyn Brunker, Yasmin Parr, Kyriaki Nomikou; Sarah McDonald, Marc Niebel, Patawee Asamaphan; Richard Orton, Joseph Hughes, Sreenu Vattipally, David L Robertson; Alasdair MacLean, Rory Gunson; Kathy Li, Natasha Jesudason, Rajiv Shah, James Shepherd, Antonia Ho, Alice Broos, Emma Thomson and Alex Alderton, Roberto Amato, Sonia Goncalves, Ewan Harrison, David K. Jackson, Ian Johnston, Dominic Kwiatkowski, Cordelia Langford, John Sillitoe |
| EPI_ISL_532086, EPI_ISL_532087, EPI_ISL_532088, EPI_ISL_532089, EPI_ISL_532090, EPI_ISL_532091, EPI_ISL_532092 | Lighthouse Lab in Glasgow | Wellcome Sanger Institute for the COVID-19 Genomics UK Consortium | Harper VanSteenhouse, Yumi Kasai, David Gray, Carol Clugston, Anna Dominiczak and Alex Alderton, Roberto Amato, Sonia Goncalves, Ewan Harrison, David K. Jackson, Ian Johnston, Dominic Kwiatkowski, Cordelia Langford, John Sillitoe |
| EPI_ISL_532093, EPI_ISL_532094 | NHSGGC West of Scotland Specialist Virology Centre / MRC-University of Glasgow Centre for Virus Research | Wellcome Sanger Institute for the COVID-19 Genomics UK Consortium | Ana da Silva Filipe, Natasha Johnson, Kathy Smollett, Daniel Mair, Stephen Carmichael, Lily Tong, Jenna Nichols, Elihu Aranday-Cortes, Kirstyn Brunker, Yasmin Parr, Kyriaki Nomikou; Sarah McDonald, Marc Niebel, Patawee Asamaphan; Richard Orton, Joseph Hughes, Sreenu Vattipally, David L Robertson; Alasdair MacLean, Rory Gunson; Kathy Li, Natasha Jesudason, Rajiv Shah, James Shepherd, Antonia Ho, Alice Broos, Emma Thomson and Alex Alderton, Roberto Amato, Sonia Goncalves, Ewan Harrison, David K. Jackson, Ian Johnston, Dominic Kwiatkowski, Cordelia Langford, John Sillitoe |
| EPI_ISL_532095, EPI_ISL_532096, EPI_ISL_532097 | Lighthouse Lab in Glasgow | Wellcome Sanger Institute for the COVID-19 Genomics UK Consortium | Harper VanSteenhouse, Yumi Kasai, David Gray, Carol Clugston, Anna Dominiczak and Alex Alderton, Roberto Amato, Sonia Goncalves, Ewan Harrison, David K. Jackson, Ian Johnston, Dominic Kwiatkowski, Cordelia Langford, John Sillitoe |
| EPI_ISL_532098 | NHSGGC West of Scotland Specialist Virology Centre / MRC-University of Glasgow Centre for Virus Research | Wellcome Sanger Institute for the COVID-19 Genomics UK Consortium | Ana da Silva Filipe, Natasha Johnson, Kathy Smollett, Daniel Mair, Stephen Carmichael, Lily Tong, Jenna Nichols, Elihu Aranday-Cortes, Kirstyn Brunker, Yasmin Parr, Kyriaki Nomikou; Sarah McDonald, Marc Niebel, Patawee Asamaphan; Richard Orton, Joseph Hughes, Sreenu Vattipally, David L Robertson; Alasdair MacLean, Rory Gunson; Kathy Li, Natasha Jesudason, Rajiv Shah, James Shepherd, Antonia Ho, Alice Broos, Emma Thomson and Alex Alderton, Roberto Amato, Sonia Goncalves, Ewan Harrison, David K. Jackson, Ian Johnston, Dominic Kwiatkowski, Cordelia Langford, John Sillitoe |
| EPI_ISL_532099, EPI_ISL_532100, EPI_ISL_532101 | Lighthouse Lab in Glasgow | Wellcome Sanger Institute for the COVID-19 Genomics UK Consortium | Harper VanSteenhouse, Yumi Kasai, David Gray, Carol Clugston, Anna Dominiczak and Alex Alderton, Roberto Amato, Sonia Goncalves, Ewan Harrison, David K. Jackson, Ian Johnston, Dominic Kwiatkowski, Cordelia Langford, John Sillitoe |
| EPI_ISL_532102 | NHSGGC West of Scotland Specialist Virology Centre / MRC-University of Glasgow Centre for Virus Research | Wellcome Sanger Institute for the COVID-19 Genomics UK Consortium | Ana da Silva Filipe, Natasha Johnson, Kathy Smollett, Daniel Mair, Stephen Carmichael, Lily Tong, Jenna Nichols, Elihu Aranday-Cortes, Kirstyn Brunker, Yasmin Parr, Kyriaki Nomikou; Sarah McDonald, Marc Niebel, Patawee Asamaphan; Richard Orton, Joseph Hughes, Sreenu Vattipally, David L Robertson; Alasdair MacLean, Rory Gunson; Kathy Li, Natasha Jesudason, Rajiv Shah, James Shepherd, Antonia Ho, Alice Broos, Emma Thomson and Alex Alderton, Roberto Amato, Sonia Goncalves, Ewan Harrison, David K. Jackson, Ian Johnston, Dominic Kwiatkowski, Cordelia Langford, John Sillitoe |
| EPI_ISL_532103, EPI_ISL_532104 | Lighthouse Lab in Glasgow | Wellcome Sanger Institute for the COVID-19 Genomics UK Consortium | Harper VanSteenhouse, Yumi Kasai, David Gray, Carol Clugston, Anna Dominiczak and Alex Alderton, Roberto Amato, Sonia Goncalves, Ewan Harrison, David K. Jackson, Ian Johnston, Dominic Kwiatkowski, Cordelia Langford, John Sillitoe |
| EPI_ISL_532105 | NHSGGC West of Scotland Specialist Virology Centre / MRC-University of Glasgow Centre for Virus Research | Wellcome Sanger Institute for the COVID-19 Genomics UK Consortium | Ana da Silva Filipe, Natasha Johnson, Kathy Smollett, Daniel Mair, Stephen Carmichael, Lily Tong, Jenna Nichols, Elihu Aranday-Cortes, Kirstyn Brunker, Yasmin Parr, Kyriaki Nomikou; Sarah McDonald, Marc Niebel, Patawee Asamaphan; Richard Orton, Joseph Hughes, Sreenu Vattipally, David L Robertson; Alasdair MacLean, Rory Gunson; Kathy Li, Natasha Jesudason, Rajiv Shah, James Shepherd, Antonia Ho, Alice Broos, Emma Thomson and Alex Alderton, Roberto Amato, Sonia Goncalves, Ewan Harrison, David K. Jackson, Ian Johnston, Dominic Kwiatkowski, Cordelia Langford, John Sillitoe |
| EPI_ISL_532106, EPI_ISL_532107, EPI_ISL_532108, EPI_ISL_532109, EPI_ISL_532110, EPI_ISL_532111, EPI_ISL_532112, EPI_ISL_532113, EPI_ISL_532114, EPI_ISL_532115, EPI_ISL_532116, EPI_ISL_532117 | | | |
| see above | Lighthouse Lab in Glasgow | Wellcome Sanger Institute for the COVID-19 Genomics UK Consortium | Harper VanSteenhouse, Yumi Kasai, David Gray, Carol Clugston, Anna Dominiczak and Alex Alderton, Roberto Amato, Sonia Goncalves, Ewan Harrison, David K. Jackson, Ian Johnston, Dominic Kwiatkowski, Cordelia Langford, John Sillitoe |
| EPI_ISL_532118 | NHSGGC West of Scotland Specialist Virology Centre / MRC-University of Glasgow Centre for Virus Research | Wellcome Sanger Institute for the COVID-19 Genomics UK Consortium | Ana da Silva Filipe, Natasha Johnson, Kathy Smollett, Daniel Mair, Stephen Carmichael, Lily Tong, Jenna Nichols, Elihu Aranday-Cortes, Kirstyn Brunker, Yasmin Parr, Kyriaki Nomikou; Sarah McDonald, Marc Niebel, Patawee Asamaphan; Richard Orton, Joseph Hughes, Sreenu Vattipally, David L Robertson; Alasdair MacLean, Rory Gunson; Kathy Li, Natasha Jesudason, Rajiv Shah, James Shepherd, Antonia Ho, Alice Broos, Emma Thomson and Alex Alderton, Roberto Amato, Sonia Goncalves, Ewan Harrison, David K. Jackson, Ian Johnston, Dominic Kwiatkowski, Cordelia Langford, John Sillitoe |
| EPI_ISL_532119, EPI_ISL_532120, EPI_ISL_532121, EPI_ISL_532122 | Lighthouse Lab in Glasgow | Wellcome Sanger Institute for the COVID-19 Genomics UK Consortium | Harper VanSteenhouse, Yumi Kasai, David Gray, Carol Clugston, Anna Dominiczak and Alex Alderton, Roberto Amato, Sonia Goncalves, Ewan Harrison, David K. Jackson, Ian Johnston, Dominic Kwiatkowski, Cordelia Langford, John Sillitoe |
| EPI_ISL_532123 | NHSGGC West of Scotland Specialist Virology Centre / MRC-University of Glasgow Centre for Virus Research | Wellcome Sanger Institute for the COVID-19 Genomics UK Consortium | Ana da Silva Filipe, Natasha Johnson, Kathy Smollett, Daniel Mair, Stephen Carmichael, Lily Tong, Jenna Nichols, Elihu Aranday-Cortes, Kirstyn Brunker, Yasmin Parr, Kyriaki Nomikou; Sarah McDonald, Marc Niebel, Patawee Asamaphan; Richard Orton, Joseph Hughes, Sreenu Vattipally, David L Robertson; Alasdair MacLean, Rory Gunson; Kathy Li, Natasha Jesudason, Rajiv Shah, James Shepherd, Antonia Ho, Alice Broos, Emma Thomson and Alex Alderton, Roberto Amato, Sonia Goncalves, Ewan Harrison, David K. Jackson, Ian Johnston, Dominic Kwiatkowski, Cordelia Langford, John Sillitoe |
| EPI_ISL_532124 | Lighthouse Lab in Glasgow | Wellcome Sanger Institute for the COVID-19 Genomics UK Consortium | Harper VanSteenhouse, Yumi Kasai, David Gray, Carol Clugston, Anna Dominiczak and Alex Alderton, Roberto Amato, Sonia Goncalves, Ewan Harrison, David K. Jackson, Ian Johnston, Dominic Kwiatkowski, Cordelia Langford, John Sillitoe |
| EPI_ISL_532125 | NHSGGC West of Scotland Specialist Virology Centre / MRC-University of Glasgow Centre for Virus Research | Wellcome Sanger Institute for the COVID-19 Genomics UK Consortium | Ana da Silva Filipe, Natasha Johnson, Kathy Smollett, Daniel Mair, Stephen Carmichael, Lily Tong, Jenna Nichols, Elihu Aranday-Cortes, Kirstyn Brunker, Yasmin Parr, Kyriaki Nomikou; Sarah McDonald, Marc Niebel, Patawee Asamaphan; Richard Orton, Joseph Hughes, Sreenu Vattipally, David L Robertson; Alasdair MacLean, Rory Gunson; Kathy Li, Natasha Jesudason, Rajiv Shah, James Shepherd, Antonia Ho, Alice Broos, Emma Thomson and Alex Alderton, Roberto Amato, Sonia Goncalves, Ewan Harrison, David K. Jackson, Ian Johnston, Dominic Kwiatkowski, Cordelia Langford, John Sillitoe |
| EPI_ISL_532126, EPI_ISL_532127, EPI_ISL_532128 | Lighthouse Lab in Glasgow | Wellcome Sanger Institute for the COVID-19 Genomics UK Consortium | Harper VanSteenhouse, Yumi Kasai, David Gray, Carol Clugston, Anna Dominiczak and Alex Alderton, Roberto Amato, Sonia Goncalves, Ewan Harrison, David K. Jackson, Ian Johnston, Dominic Kwiatkowski, Cordelia Langford, John Sillitoe |
| EPI_ISL_532129, EPI_ISL_532130, EPI_ISL_532131 | NHSGGC West of Scotland Specialist Virology Centre / MRC-University of Glasgow Centre for Virus Research | Wellcome Sanger Institute for the COVID-19 Genomics UK Consortium | Ana da Silva Filipe, Natasha Johnson, Kathy Smollett, Daniel Mair, Stephen Carmichael, Lily Tong, Jenna Nichols, Elihu Aranday-Cortes, Kirstyn Brunker, Yasmin Parr, Kyriaki Nomikou; Sarah McDonald, Marc Niebel, Patawee Asamaphan; Richard Orton, Joseph Hughes, Sreenu Vattipally, David L Robertson; Alasdair MacLean, Rory Gunson; Kathy Li, Natasha Jesudason, Rajiv Shah, James Shepherd, Antonia Ho, Alice Broos, Emma Thomson and Alex Alderton, Roberto Amato, Sonia Goncalves, Ewan Harrison, David K. Jackson, Ian Johnston, Dominic Kwiatkowski, Cordelia Langford, John Sillitoe |
| EPI_ISL_532132, EPI_ISL_532133, EPI_ISL_532134, EPI_ISL_532135, EPI_ISL_532136, EPI_ISL_532137 | Lighthouse Lab in Glasgow | Wellcome Sanger Institute for the COVID-19 Genomics UK Consortium | Harper VanSteenhouse, Yumi Kasai, David Gray, Carol Clugston, Anna Dominiczak and Alex Alderton, Roberto Amato, Sonia Goncalves, Ewan Harrison, David K. Jackson, Ian Johnston, Dominic Kwiatkowski, Cordelia Langford, John Sillitoe |
| EPI_ISL_532138 | NHSGGC West of Scotland Specialist Virology Centre / MRC-University of Glasgow Centre for Virus Research | Wellcome Sanger Institute for the COVID-19 Genomics UK Consortium | Ana da Silva Filipe, Natasha Johnson, Kathy Smollett, Daniel Mair, Stephen Carmichael, Lily Tong, Jenna Nichols, Elihu Aranday-Cortes, Kirstyn Brunker, Yasmin Parr, Kyriaki Nomikou; Sarah McDonald, Marc Niebel, Patawee Asamaphan; Richard Orton, Joseph Hughes, Sreenu Vattipally, David L Robertson; Alasdair MacLean, Rory Gunson; Kathy Li, Natasha Jesudason, Rajiv Shah, James Shepherd, Antonia Ho, Alice Broos, Emma Thomson and Alex Alderton, Roberto Amato, Sonia Goncalves, Ewan Harrison, David K. Jackson, Ian Johnston, Dominic Kwiatkowski, Cordelia Langford, John Sillitoe |
| EPI_ISL_532139, EPI_ISL_532140, EPI_ISL_532141, EPI_ISL_532142, EPI_ISL_532143, EPI_ISL_532144, EPI_ISL_532145, EPI_ISL_532146, EPI_ISL_532147, EPI_ISL_532148, EPI_ISL_532149, EPI_ISL_532150, EPI_ISL_532151, EPI_ISL_532152 | | | |
| see above | Lighthouse Lab in Glasgow | Wellcome Sanger Institute for the COVID-19 Genomics UK Consortium | Harper VanSteenhouse, Yumi Kasai, David Gray, Carol Clugston, Anna Dominiczak and Alex Alderton, Roberto Amato, Sonia Goncalves, Ewan Harrison, David K. Jackson, Ian Johnston, Dominic Kwiatkowski, Cordelia Langford, John Sillitoe |
| EPI_ISL_532153 | NHSGGC West of Scotland Specialist Virology Centre / MRC-University of Glasgow Centre for Virus Research | Wellcome Sanger Institute for the COVID-19 Genomics UK Consortium | Ana da Silva Filipe, Natasha Johnson, Kathy Smollett, Daniel Mair, Stephen Carmichael, Lily Tong, Jenna Nichols, Elihu Aranday-Cortes, Kirstyn Brunker, Yasmin Parr, Kyriaki Nomikou; Sarah McDonald, Marc Niebel, Patawee Asamaphan; Richard Orton, Joseph Hughes, Sreenu Vattipally, David L Robertson; Alasdair MacLean, Rory Gunson; Kathy Li, Natasha Jesudason, Rajiv Shah, James Shepherd, Antonia Ho, Alice Broos, Emma Thomson and Alex Alderton, Roberto Amato, Sonia Goncalves, Ewan Harrison, David K. Jackson, Ian Johnston, Dominic Kwiatkowski, Cordelia Langford, John Sillitoe |
| EPI_ISL_532154, EPI_ISL_532155, EPI_ISL_532156, EPI_ISL_532157, EPI_ISL_532158, EPI_ISL_532159, EPI_ISL_532160, EPI_ISL_532161, EPI_ISL_532162, EPI_ISL_532163, EPI_ISL_532164, EPI_ISL_532165, EPI_ISL_532166, EPI_ISL_532167 | | | |
| see above | Lighthouse Lab in Glasgow | Wellcome Sanger Institute for the COVID-19 Genomics UK Consortium | Harper VanSteenhouse, Yumi Kasai, David Gray, Carol Clugston, Anna Dominiczak and Alex Alderton, Roberto Amato, Sonia Goncalves, Ewan Harrison, David K. Jackson, Ian Johnston, Dominic Kwiatkowski, Cordelia Langford, John Sillitoe |
| EPI_ISL_532168 | NHSGGC West of Scotland Specialist Virology Centre / MRC-University of Glasgow Centre for Virus Research | Wellcome Sanger Institute for the COVID-19 Genomics UK Consortium | Ana da Silva Filipe, Natasha Johnson, Kathy Smollett, Daniel Mair, Stephen Carmichael, Lily Tong, Jenna Nichols, Elihu Aranday-Cortes, Kirstyn Brunker, Yasmin Parr, Kyriaki Nomikou; Sarah McDonald, Marc Niebel, Patawee Asamaphan; Richard Orton, Joseph Hughes, Sreenu Vattipally, David L Robertson; Alasdair MacLean, Rory Gunson; Kathy Li, Natasha Jesudason, Rajiv Shah, James Shepherd, Antonia Ho, Alice Broos, Emma Thomson and Alex Alderton, Roberto Amato, Sonia Goncalves, Ewan Harrison, David K. Jackson, Ian Johnston, Dominic Kwiatkowski, Cordelia Langford, John Sillitoe |
| EPI_ISL_532169, EPI_ISL_532170, EPI_ISL_532171, | Lighthouse Lab in Glasgow | Wellcome Sanger Institute for the COVID-19 Genomics UK | Harper VanSteenhouse, Yumi Kasai, David Gray, Carol Clugston, Anna Dominiczak and Alex Alderton, Roberto Amato, Sonia Goncalves, Ewan Harrison, David K. Jackson, Ian Johnston, Dominic Kwiatkowski, Cordelia |

| | | | |
|---|---|---|---|
| EPI_ISL_532172, EPI_ISL_532173, EPI_ISL_532174, EPI_ISL_532175, EPI_ISL_532176, EPI_ISL_532177 | | Consortium | Langford, John Sillitoe |
| EPI_ISL_532178 | NHSGGC West of Scotland Specialist Virology Centre / MRC-University of Glasgow Centre for Virus Research | Wellcome Sanger Institute for the COVID-19 Genomics UK Consortium | Ana da Silva Filipe, Natasha Johnson, Kathy Smollett, Daniel Mair, Stephen Carmichael, Lily Tong, Jenna Nichols, Elihu Aranday-Cortes, Kirstyn Brunker, Yasmin Parr, Kyriaki Nomikou; Sarah McDonald, Marc Niebel, Patawee Asamaphan; Richard Orton, Joseph Hughes, Sreenu Vattipally, David L Robertson; Alasdair MacLean, Rory Gunson; Kathy Li, Natasha Jesudason, Rajiv Shah, James Shepherd, Antonia Ho, Alice Broos, Emma Thomson and Alex Alderton, Roberto Amato, Sonia Goncalves, Ewan Harrison, David K. Jackson, Ian Johnston, Dominic Kwiatkowski, Cordelia Langford, John Sillitoe |
| EPI_ISL_532179, EPI_ISL_532180, EPI_ISL_532181, EPI_ISL_532182, EPI_ISL_532183, EPI_ISL_532184, EPI_ISL_532185 | Lighthouse Lab in Glasgow | Wellcome Sanger Institute for the COVID-19 Genomics UK Consortium | Harper VanSteenhouse, Yumi Kasai, David Gray, Carol Clugston, Anna Dominiczak and Alex Alderton, Roberto Amato, Sonia Goncalves, Ewan Harrison, David K. Jackson, Ian Johnston, Dominic Kwiatkowski, Cordelia Langford, John Sillitoe |
| EPI_ISL_532186 | NHSGGC West of Scotland Specialist Virology Centre / MRC-University of Glasgow Centre for Virus Research | Wellcome Sanger Institute for the COVID-19 Genomics UK Consortium | Ana da Silva Filipe, Natasha Johnson, Kathy Smollett, Daniel Mair, Stephen Carmichael, Lily Tong, Jenna Nichols, Elihu Aranday-Cortes, Kirstyn Brunker, Yasmin Parr, Kyriaki Nomikou; Sarah McDonald, Marc Niebel, Patawee Asamaphan; Richard Orton, Joseph Hughes, Sreenu Vattipally, David L Robertson; Alasdair MacLean, Rory Gunson; Kathy Li, Natasha Jesudason, Rajiv Shah, James Shepherd, Antonia Ho, Alice Broos, Emma Thomson and Alex Alderton, Roberto Amato, Sonia Goncalves, Ewan Harrison, David K. Jackson, Ian Johnston, Dominic Kwiatkowski, Cordelia Langford, John Sillitoe |
| EPI_ISL_532187, EPI_ISL_532188, EPI_ISL_532189, EPI_ISL_532190, EPI_ISL_532191 | Lighthouse Lab in Glasgow | Wellcome Sanger Institute for the COVID-19 Genomics UK Consortium | Harper VanSteenhouse, Yumi Kasai, David Gray, Carol Clugston, Anna Dominiczak and Alex Alderton, Roberto Amato, Sonia Goncalves, Ewan Harrison, David K. Jackson, Ian Johnston, Dominic Kwiatkowski, Cordelia Langford, John Sillitoe |
| EPI_ISL_532192 | NHSGGC West of Scotland Specialist Virology Centre / MRC-University of Glasgow Centre for Virus Research | Wellcome Sanger Institute for the COVID-19 Genomics UK Consortium | Ana da Silva Filipe, Natasha Johnson, Kathy Smollett, Daniel Mair, Stephen Carmichael, Lily Tong, Jenna Nichols, Elihu Aranday-Cortes, Kirstyn Brunker, Yasmin Parr, Kyriaki Nomikou; Sarah McDonald, Marc Niebel, Patawee Asamaphan; Richard Orton, Joseph Hughes, Sreenu Vattipally, David L Robertson; Alasdair MacLean, Rory Gunson; Kathy Li, Natasha Jesudason, Rajiv Shah, James Shepherd, Antonia Ho, Alice Broos, Emma Thomson and Alex Alderton, Roberto Amato, Sonia Goncalves, Ewan Harrison, David K. Jackson, Ian Johnston, Dominic Kwiatkowski, Cordelia Langford, John Sillitoe |
| EPI_ISL_532193 | Lighthouse Lab in Glasgow | Wellcome Sanger Institute for the COVID-19 Genomics UK Consortium | Harper VanSteenhouse, Yumi Kasai, David Gray, Carol Clugston, Anna Dominiczak and Alex Alderton, Roberto Amato, Sonia Goncalves, Ewan Harrison, David K. Jackson, Ian Johnston, Dominic Kwiatkowski, Cordelia Langford, John Sillitoe |
| EPI_ISL_532194, EPI_ISL_532195, EPI_ISL_532196, EPI_ISL_532197 | NHSGGC West of Scotland Specialist Virology Centre / MRC-University of Glasgow Centre for Virus Research | Wellcome Sanger Institute for the COVID-19 Genomics UK Consortium | Ana da Silva Filipe, Natasha Johnson, Kathy Smollett, Daniel Mair, Stephen Carmichael, Lily Tong, Jenna Nichols, Elihu Aranday-Cortes, Kirstyn Brunker, Yasmin Parr, Kyriaki Nomikou; Sarah McDonald, Marc Niebel, Patawee Asamaphan; Richard Orton, Joseph Hughes, Sreenu Vattipally, David L Robertson; Alasdair MacLean, Rory Gunson; Kathy Li, Natasha Jesudason, Rajiv Shah, James Shepherd, Antonia Ho, Alice Broos, Emma Thomson and Alex Alderton, Roberto Amato, Sonia Goncalves, Ewan Harrison, David K. Jackson, Ian Johnston, Dominic Kwiatkowski, Cordelia Langford, John Sillitoe |
| EPI_ISL_532198, EPI_ISL_532199 | Lighthouse Lab in Glasgow | Wellcome Sanger Institute for the COVID-19 Genomics UK Consortium | Harper VanSteenhouse, Yumi Kasai, David Gray, Carol Clugston, Anna Dominiczak and Alex Alderton, Roberto Amato, Sonia Goncalves, Ewan Harrison, David K. Jackson, Ian Johnston, Dominic Kwiatkowski, Cordelia Langford, John Sillitoe |
| EPI_ISL_532200 | NHSGGC West of Scotland Specialist Virology Centre / MRC-University of Glasgow Centre for Virus Research | Wellcome Sanger Institute for the COVID-19 Genomics UK Consortium | Ana da Silva Filipe, Natasha Johnson, Kathy Smollett, Daniel Mair, Stephen Carmichael, Lily Tong, Jenna Nichols, Elihu Aranday-Cortes, Kirstyn Brunker, Yasmin Parr, Kyriaki Nomikou; Sarah McDonald, Marc Niebel, Patawee Asamaphan; Richard Orton, Joseph Hughes, Sreenu Vattipally, David L Robertson; Alasdair MacLean, Rory Gunson; Kathy Li, Natasha Jesudason, Rajiv Shah, James Shepherd, Antonia Ho, Alice Broos, Emma Thomson and Alex Alderton, Roberto Amato, Sonia Goncalves, Ewan Harrison, David K. Jackson, Ian Johnston, Dominic Kwiatkowski, Cordelia Langford, John Sillitoe |
| EPI_ISL_532201 | Lighthouse Lab in Glasgow | Wellcome Sanger Institute for the COVID-19 Genomics UK Consortium | Harper VanSteenhouse, Yumi Kasai, David Gray, Carol Clugston, Anna Dominiczak and Alex Alderton, Roberto Amato, Sonia Goncalves, Ewan Harrison, David K. Jackson, Ian Johnston, Dominic Kwiatkowski, Cordelia Langford, John Sillitoe |
| EPI_ISL_532202, EPI_ISL_532203 | NHSGGC West of Scotland Specialist Virology Centre / MRC-University of Glasgow Centre for Virus Research | Wellcome Sanger Institute for the COVID-19 Genomics UK Consortium | Ana da Silva Filipe, Natasha Johnson, Kathy Smollett, Daniel Mair, Stephen Carmichael, Lily Tong, Jenna Nichols, Elihu Aranday-Cortes, Kirstyn Brunker, Yasmin Parr, Kyriaki Nomikou; Sarah McDonald, Marc Niebel, Patawee Asamaphan; Richard Orton, Joseph Hughes, Sreenu Vattipally, David L Robertson; Alasdair MacLean, Rory Gunson; Kathy Li, Natasha Jesudason, Rajiv Shah, James Shepherd, Antonia Ho, Alice Broos, Emma Thomson and Alex Alderton, Roberto Amato, Sonia Goncalves, Ewan Harrison, David K. Jackson, Ian Johnston, Dominic Kwiatkowski, Cordelia Langford, John Sillitoe |
| EPI_ISL_532204, EPI_ISL_532205, EPI_ISL_532206, EPI_ISL_532207, EPI_ISL_532208, EPI_ISL_532209 | Lighthouse Lab in Glasgow | Wellcome Sanger Institute for the COVID-19 Genomics UK Consortium | Harper VanSteenhouse, Yumi Kasai, David Gray, Carol Clugston, Anna Dominiczak and Alex Alderton, Roberto Amato, Sonia Goncalves, Ewan Harrison, David K. Jackson, Ian Johnston, Dominic Kwiatkowski, Cordelia Langford, John Sillitoe |
| EPI_ISL_532210 | NHSGGC West of Scotland Specialist Virology Centre / MRC-University of Glasgow Centre for Virus Research | Wellcome Sanger Institute for the COVID-19 Genomics UK Consortium | Ana da Silva Filipe, Natasha Johnson, Kathy Smollett, Daniel Mair, Stephen Carmichael, Lily Tong, Jenna Nichols, Elihu Aranday-Cortes, Kirstyn Brunker, Yasmin Parr, Kyriaki Nomikou; Sarah McDonald, Marc Niebel, Patawee Asamaphan; Richard Orton, Joseph Hughes, Sreenu Vattipally, David L Robertson; Alasdair MacLean, Rory Gunson; Kathy Li, Natasha Jesudason, Rajiv Shah, James Shepherd, Antonia Ho, Alice Broos, Emma Thomson and Alex Alderton, Roberto Amato, Sonia Goncalves, Ewan Harrison, David K. Jackson, Ian Johnston, Dominic Kwiatkowski, Cordelia Langford, John Sillitoe |
| EPI_ISL_532211, EPI_ISL_532212, EPI_ISL_532213 | Lighthouse Lab in Glasgow | Wellcome Sanger Institute for the COVID-19 Genomics UK Consortium | Harper VanSteenhouse, Yumi Kasai, David Gray, Carol Clugston, Anna Dominiczak and Alex Alderton, Roberto Amato, Sonia Goncalves, Ewan Harrison, David K. Jackson, Ian Johnston, Dominic Kwiatkowski, Cordelia Langford, John Sillitoe |
| EPI_ISL_532214 | NHSGGC West of Scotland Specialist Virology Centre / MRC-University of Glasgow Centre for Virus Research | Wellcome Sanger Institute for the COVID-19 Genomics UK Consortium | Ana da Silva Filipe, Natasha Johnson, Kathy Smollett, Daniel Mair, Stephen Carmichael, Lily Tong, Jenna Nichols, Elihu Aranday-Cortes, Kirstyn Brunker, Yasmin Parr, Kyriaki Nomikou; Sarah McDonald, Marc Niebel, Patawee Asamaphan; Richard Orton, Joseph Hughes, Sreenu Vattipally, David L Robertson; Alasdair MacLean, Rory Gunson; Kathy Li, Natasha Jesudason, Rajiv Shah, James Shepherd, Antonia Ho, Alice Broos, Emma Thomson and Alex Alderton, Roberto Amato, Sonia Goncalves, Ewan Harrison, David K. Jackson, Ian Johnston, Dominic Kwiatkowski, Cordelia Langford, John Sillitoe |
| EPI_ISL_532215, EPI_ISL_532216, EPI_ISL_532217, EPI_ISL_532218 | Lighthouse Lab in Glasgow | Wellcome Sanger Institute for the COVID-19 Genomics UK Consortium | Harper VanSteenhouse, Yumi Kasai, David Gray, Carol Clugston, Anna Dominiczak and Alex Alderton, Roberto Amato, Sonia Goncalves, Ewan Harrison, David K. Jackson, Ian Johnston, Dominic Kwiatkowski, Cordelia Langford, John Sillitoe |
| EPI_ISL_532219 | NHSGGC West of Scotland Specialist Virology Centre / MRC-University of Glasgow Centre for Virus Research | Wellcome Sanger Institute for the COVID-19 Genomics UK Consortium | Ana da Silva Filipe, Natasha Johnson, Kathy Smollett, Daniel Mair, Stephen Carmichael, Lily Tong, Jenna Nichols, Elihu Aranday-Cortes, Kirstyn Brunker, Yasmin Parr, Kyriaki Nomikou; Sarah McDonald, Marc Niebel, Patawee Asamaphan; Richard Orton, Joseph Hughes, Sreenu Vattipally, David L Robertson; Alasdair MacLean, Rory Gunson; Kathy Li, Natasha Jesudason, Rajiv Shah, James Shepherd, Antonia Ho, Alice Broos, Emma Thomson and Alex Alderton, Roberto Amato, Sonia Goncalves, Ewan Harrison, David K. Jackson, Ian Johnston, Dominic Kwiatkowski, Cordelia Langford, John Sillitoe |
| EPI_ISL_532220 | Lighthouse Lab in Glasgow | Wellcome Sanger Institute for the COVID-19 Genomics UK Consortium | Harper VanSteenhouse, Yumi Kasai, David Gray, Carol Clugston, Anna Dominiczak and Alex Alderton, Roberto Amato, Sonia Goncalves, Ewan Harrison, David K. Jackson, Ian Johnston, Dominic Kwiatkowski, Cordelia Langford, John Sillitoe |
| EPI_ISL_532221 | NHSGGC West of Scotland Specialist Virology Centre / MRC-University of Glasgow Centre for Virus Research | Wellcome Sanger Institute for the COVID-19 Genomics UK Consortium | Ana da Silva Filipe, Natasha Johnson, Kathy Smollett, Daniel Mair, Stephen Carmichael, Lily Tong, Jenna Nichols, Elihu Aranday-Cortes, Kirstyn Brunker, Yasmin Parr, Kyriaki Nomikou; Sarah McDonald, Marc Niebel, Patawee Asamaphan; Richard Orton, Joseph Hughes, Sreenu Vattipally, David L Robertson; Alasdair MacLean, Rory Gunson; Kathy Li, Natasha Jesudason, Rajiv Shah, James Shepherd, Antonia Ho, Alice Broos, Emma Thomson and Alex Alderton, Roberto Amato, Sonia Goncalves, Ewan Harrison, David K. Jackson, Ian Johnston, Dominic Kwiatkowski, Cordelia Langford, John Sillitoe |
| EPI_ISL_532222, EPI_ISL_532223, EPI_ISL_532224, EPI_ISL_532225, EPI_ISL_532226, EPI_ISL_532227, EPI_ISL_532228, EPI_ISL_532229, EPI_ISL_532230, EPI_ISL_532231, EPI_ISL_532232, EPI_ISL_532233, EPI_ISL_532234 | | | |
| see above | Lighthouse Lab in Glasgow | Wellcome Sanger Institute for the COVID-19 Genomics UK Consortium | Harper VanSteenhouse, Yumi Kasai, David Gray, Carol Clugston, Anna Dominiczak and Alex Alderton, Roberto Amato, Sonia Goncalves, Ewan Harrison, David K. Jackson, Ian Johnston, Dominic Kwiatkowski, Cordelia Langford, John Sillitoe |
| EPI_ISL_532235 | NHSGGC West of Scotland Specialist Virology Centre / MRC-University of Glasgow Centre for Virus Research | Wellcome Sanger Institute for the COVID-19 Genomics UK Consortium | Ana da Silva Filipe, Natasha Johnson, Kathy Smollett, Daniel Mair, Stephen Carmichael, Lily Tong, Jenna Nichols, Elihu Aranday-Cortes, Kirstyn Brunker, Yasmin Parr, Kyriaki Nomikou; Sarah McDonald, Marc Niebel, Patawee Asamaphan; Richard Orton, Joseph Hughes, Sreenu Vattipally, David L Robertson; Alasdair MacLean, Rory Gunson; Kathy Li, Natasha Jesudason, Rajiv Shah, James Shepherd, Antonia Ho, Alice Broos, Emma Thomson and Alex Alderton, Roberto Amato, Sonia Goncalves, Ewan Harrison, David K. Jackson, Ian Johnston, Dominic Kwiatkowski, Cordelia Langford, John Sillitoe |
| EPI_ISL_532236, EPI_ISL_532237, EPI_ISL_532238, EPI_ISL_532239, EPI_ISL_532240 | Lighthouse Lab in Glasgow | Wellcome Sanger Institute for the COVID-19 Genomics UK Consortium | Harper VanSteenhouse, Yumi Kasai, David Gray, Carol Clugston, Anna Dominiczak and Alex Alderton, Roberto Amato, Sonia Goncalves, Ewan Harrison, David K. Jackson, Ian Johnston, Dominic Kwiatkowski, Cordelia Langford, John Sillitoe |
| EPI_ISL_532241 | NHSGGC West of Scotland Specialist Virology Centre / MRC-University of Glasgow Centre for Virus Research | Wellcome Sanger Institute for the COVID-19 Genomics UK Consortium | Ana da Silva Filipe, Natasha Johnson, Kathy Smollett, Daniel Mair, Stephen Carmichael, Lily Tong, Jenna Nichols, Elihu Aranday-Cortes, Kirstyn Brunker, Yasmin Parr, Kyriaki Nomikou; Sarah McDonald, Marc Niebel, Patawee Asamaphan; Richard Orton, Joseph Hughes, Sreenu Vattipally, David L Robertson; Alasdair MacLean, Rory Gunson; Kathy Li, Natasha Jesudason, Rajiv Shah, James Shepherd, Antonia Ho, Alice Broos, Emma Thomson and Alex Alderton, Roberto Amato, Sonia Goncalves, Ewan Harrison, David K. Jackson, Ian Johnston, Dominic Kwiatkowski, Cordelia Langford, John Sillitoe |
| EPI_ISL_532242, EPI_ISL_532243 | Lighthouse Lab in Glasgow | Wellcome Sanger Institute for the COVID-19 Genomics UK Consortium | Harper VanSteenhouse, Yumi Kasai, David Gray, Carol Clugston, Anna Dominiczak and Alex Alderton, Roberto Amato, Sonia Goncalves, Ewan Harrison, David K. Jackson, Ian Johnston, Dominic Kwiatkowski, Cordelia Langford, John Sillitoe |
| EPI_ISL_532244 | NHSGGC West of Scotland Specialist Virology Centre / MRC-University of Glasgow Centre for Virus Research | Wellcome Sanger Institute for the COVID-19 Genomics UK Consortium | Ana da Silva Filipe, Natasha Johnson, Kathy Smollett, Daniel Mair, Stephen Carmichael, Lily Tong, Jenna Nichols, Elihu Aranday-Cortes, Kirstyn Brunker, Yasmin Parr, Kyriaki Nomikou; Sarah McDonald, Marc Niebel, Patawee Asamaphan; Richard Orton, Joseph Hughes, Sreenu Vattipally, David L Robertson; Alasdair MacLean, Rory Gunson; Kathy Li, Natasha Jesudason, Rajiv Shah, James Shepherd, Antonia Ho, Alice Broos, Emma Thomson and Alex Alderton, Roberto Amato, Sonia Goncalves, Ewan Harrison, David K. Jackson, Ian Johnston, Dominic Kwiatkowski, Cordelia Langford, John Sillitoe |
| EPI_ISL_532245, EPI_ISL_532246, EPI_ISL_532247, EPI_ISL_532248 | Lighthouse Lab in Glasgow | Wellcome Sanger Institute for the COVID-19 Genomics UK Consortium | Harper VanSteenhouse, Yumi Kasai, David Gray, Carol Clugston, Anna Dominiczak and Alex Alderton, Roberto Amato, Sonia Goncalves, Ewan Harrison, David K. Jackson, Ian Johnston, Dominic Kwiatkowski, Cordelia Langford, John Sillitoe |
| EPI_ISL_532249 | NHSGGC West of Scotland Specialist Virology Centre / MRC-University of Glasgow Centre for Virus Research | Wellcome Sanger Institute for the COVID-19 Genomics UK Consortium | Ana da Silva Filipe, Natasha Johnson, Kathy Smollett, Daniel Mair, Stephen Carmichael, Lily Tong, Jenna Nichols, Elihu Aranday-Cortes, Kirstyn Brunker, Yasmin Parr, Kyriaki Nomikou; Sarah McDonald, Marc Niebel, Patawee Asamaphan; Richard Orton, Joseph Hughes, Sreenu Vattipally, David L Robertson; Alasdair MacLean, Rory Gunson; Kathy Li, Natasha Jesudason, Rajiv Shah, James Shepherd, Antonia Ho, Alice Broos, Emma Thomson and Alex Alderton, Roberto Amato, Sonia Goncalves, Ewan Harrison, David K. Jackson, Ian Johnston, Dominic Kwiatkowski, Cordelia Langford, John Sillitoe |
| EPI_ISL_532250, EPI_ISL_532251, EPI_ISL_532252, EPI_ISL_532253, EPI_ISL_532254, EPI_ISL_532255, EPI_ISL_532256, EPI_ISL_532257 | Lighthouse Lab in Glasgow | Wellcome Sanger Institute for the COVID-19 Genomics UK Consortium | Harper VanSteenhouse, Yumi Kasai, David Gray, Carol Clugston, Anna Dominiczak and Alex Alderton, Roberto Amato, Sonia Goncalves, Ewan Harrison, David K. Jackson, Ian Johnston, Dominic Kwiatkowski, Cordelia Langford, John Sillitoe |
| EPI_ISL_532258 | NHSGGC West of Scotland Specialist Virology Centre / MRC-University of Glasgow Centre for Virus Research | Wellcome Sanger Institute for the COVID-19 Genomics UK Consortium | Ana da Silva Filipe, Natasha Johnson, Kathy Smollett, Daniel Mair, Stephen Carmichael, Lily Tong, Jenna Nichols, Elihu Aranday-Cortes, Kirstyn Brunker, Yasmin Parr, Kyriaki Nomikou; Sarah McDonald, Marc Niebel, Patawee Asamaphan; Richard Orton, Joseph Hughes, Sreenu Vattipally, David L Robertson; Alasdair MacLean, Rory Gunson; Kathy Li, Natasha Jesudason, Rajiv Shah, James Shepherd, Antonia Ho, Alice Broos, Emma Thomson and Alex Alderton, Roberto Amato, Sonia Goncalves, Ewan Harrison, David K. Jackson, Ian Johnston, Dominic Kwiatkowski, Cordelia Langford, John Sillitoe |
| EPI_ISL_532259, EPI_ISL_532260, EPI_ISL_532261, EPI_ISL_532262 | Lighthouse Lab in Glasgow | Wellcome Sanger Institute for the COVID-19 Genomics UK Consortium | Harper VanSteenhouse, Yumi Kasai, David Gray, Carol Clugston, Anna Dominiczak and Alex Alderton, Roberto Amato, Sonia Goncalves, Ewan Harrison, David K. Jackson, Ian Johnston, Dominic Kwiatkowski, Cordelia Langford, John Sillitoe |
| EPI_ISL_532263 | NHSGGC West of Scotland Specialist Virology Centre / MRC-University of Glasgow Centre for Virus Research | Wellcome Sanger Institute for the COVID-19 Genomics UK Consortium | Ana da Silva Filipe, Natasha Johnson, Kathy Smollett, Daniel Mair, Stephen Carmichael, Lily Tong, Jenna Nichols, Elihu Aranday-Cortes, Kirstyn Brunker, Yasmin Parr, Kyriaki Nomikou; Sarah McDonald, Marc Niebel, Patawee Asamaphan; Richard Orton, Joseph Hughes, Sreenu Vattipally, David L Robertson; Alasdair MacLean, Rory Gunson; Kathy Li, Natasha Jesudason, Rajiv Shah, James Shepherd, Antonia Ho, Alice Broos, Emma Thomson and Alex Alderton, Roberto Amato, Sonia Goncalves, Ewan Harrison, David K. Jackson, Ian Johnston, Dominic Kwiatkowski, Cordelia Langford, John Sillitoe |
| EPI_ISL_532264, EPI_ISL_532265, EPI_ISL_532266, EPI_ISL_532267, EPI_ISL_532268, EPI_ISL_532269, EPI_ISL_532270 | Lighthouse Lab in Glasgow | Wellcome Sanger Institute for the COVID-19 Genomics UK Consortium | Harper VanSteenhouse, Yumi Kasai, David Gray, Carol Clugston, Anna Dominiczak and Alex Alderton, Roberto Amato, Sonia Goncalves, Ewan Harrison, David K. Jackson, Ian Johnston, Dominic Kwiatkowski, Cordelia Langford, John Sillitoe |
| EPI_ISL_532271, EPI_ISL_532272 | NHSGGC West of Scotland Specialist Virology Centre / MRC-University of Glasgow Centre for Virus Research | Wellcome Sanger Institute for the COVID-19 Genomics UK Consortium | Ana da Silva Filipe, Natasha Johnson, Kathy Smollett, Daniel Mair, Stephen Carmichael, Lily Tong, Jenna Nichols, Elihu Aranday-Cortes, Kirstyn Brunker, Yasmin Parr, Kyriaki Nomikou; Sarah McDonald, Marc Niebel, Patawee Asamaphan; Richard Orton, Joseph Hughes, Sreenu Vattipally, David L Robertson; Alasdair MacLean, Rory Gunson; Kathy Li, Natasha Jesudason, Rajiv Shah, James Shepherd, Antonia Ho, Alice Broos, Emma Thomson and Alex Alderton, Roberto Amato, Sonia Goncalves, Ewan Harrison, David K. Jackson, Ian Johnston, Dominic Kwiatkowski, Cordelia Langford, John Sillitoe |
| EPI_ISL_532273, EPI_ISL_532274 | Lighthouse Lab in Glasgow | Wellcome Sanger Institute for the COVID-19 Genomics UK Consortium | Harper VanSteenhouse, Yumi Kasai, David Gray, Carol Clugston, Anna Dominiczak and Alex Alderton, Roberto Amato, Sonia Goncalves, Ewan Harrison, David K. Jackson, Ian Johnston, Dominic Kwiatkowski, Cordelia Langford, John Sillitoe |
| EPI_ISL_532275, EPI_ISL_532276 | NHSGGC West of Scotland Specialist Virology Centre / MRC-University of Glasgow Centre for Virus Research | Wellcome Sanger Institute for the COVID-19 Genomics UK Consortium | Ana da Silva Filipe, Natasha Johnson, Kathy Smollett, Daniel Mair, Stephen Carmichael, Lily Tong, Jenna Nichols, Elihu Aranday-Cortes, Kirstyn Brunker, Yasmin Parr, Kyriaki Nomikou; Sarah McDonald, Marc Niebel, Patawee Asamaphan; Richard Orton, Joseph Hughes, Sreenu Vattipally, David L Robertson; Alasdair MacLean, Rory Gunson; Kathy Li, Natasha Jesudason, Rajiv Shah, James Shepherd, Antonia Ho, Alice Broos, Emma Thomson and Alex Alderton, Roberto Amato, Sonia Goncalves, Ewan Harrison, David K. Jackson, Ian Johnston, Dominic Kwiatkowski, Cordelia Langford, John Sillitoe |

| | | | |
|---|---|---|---|
| EPI_ISL_532277 | Lighthouse Lab in Glasgow | Wellcome Sanger Institute for the COVID-19 Genomics UK Consortium | Harper VanSteenhouse, Yumi Kasai, David Gray, Carol Clugston, Anna Dominiczak and Alex Alderton, Roberto Amato, Sonia Goncalves, Ewan Harrison, David K. Jackson, Ian Johnston, Dominic Kwiatkowski, Cordelia Langford, John Sillitoe |
| EPI_ISL_532278 | NHSGGC West of Scotland Specialist Virology Centre / MRC-University of Glasgow Centre for Virus Research | Wellcome Sanger Institute for the COVID-19 Genomics UK Consortium | Ana da Silva Filipe, Natasha Johnson, Kathy Smollett, Daniel Mair, Stephen Carmichael, Lily Tong, Jenna Nichols, Elihu Aranday-Cortes, Kirstyn Brunker, Yasmin Parr, Kyriaki Nomikou; Sarah McDonald, Marc Niebel, Patawee Asamaphan; Richard Orton, Joseph Hughes, Sreenu Vattipally, David L Robertson; Alasdair MacLean, Rory Gunson; Kathy Li, Natasha Jesudason, Rajiv Shah, James Shepherd, Antonia Ho, Alice Broos, Emma Thomson and Alex Alderton, Roberto Amato, Sonia Goncalves, Ewan Harrison, David K. Jackson, Ian Johnston, Dominic Kwiatkowski, Cordelia Langford, John Sillitoe |
| EPI_ISL_532279, EPI_ISL_532280, EPI_ISL_532281, EPI_ISL_532282 | Lighthouse Lab in Glasgow | Wellcome Sanger Institute for the COVID-19 Genomics UK Consortium | Harper VanSteenhouse, Yumi Kasai, David Gray, Carol Clugston, Anna Dominiczak and Alex Alderton, Roberto Amato, Sonia Goncalves, Ewan Harrison, David K. Jackson, Ian Johnston, Dominic Kwiatkowski, Cordelia Langford, John Sillitoe |
| EPI_ISL_532283 | NHSGGC West of Scotland Specialist Virology Centre / MRC-University of Glasgow Centre for Virus Research | Wellcome Sanger Institute for the COVID-19 Genomics UK Consortium | Ana da Silva Filipe, Natasha Johnson, Kathy Smollett, Daniel Mair, Stephen Carmichael, Lily Tong, Jenna Nichols, Elihu Aranday-Cortes, Kirstyn Brunker, Yasmin Parr, Kyriaki Nomikou; Sarah McDonald, Marc Niebel, Patawee Asamaphan; Richard Orton, Joseph Hughes, Sreenu Vattipally, David L Robertson; Alasdair MacLean, Rory Gunson; Kathy Li, Natasha Jesudason, Rajiv Shah, James Shepherd, Antonia Ho, Alice Broos, Emma Thomson and Alex Alderton, Roberto Amato, Sonia Goncalves, Ewan Harrison, David K. Jackson, Ian Johnston, Dominic Kwiatkowski, Cordelia Langford, John Sillitoe |
| EPI_ISL_532284, EPI_ISL_532285, EPI_ISL_532286, EPI_ISL_532287 | Lighthouse Lab in Glasgow | Wellcome Sanger Institute for the COVID-19 Genomics UK Consortium | Harper VanSteenhouse, Yumi Kasai, David Gray, Carol Clugston, Anna Dominiczak and Alex Alderton, Roberto Amato, Sonia Goncalves, Ewan Harrison, David K. Jackson, Ian Johnston, Dominic Kwiatkowski, Cordelia Langford, John Sillitoe |
| EPI_ISL_532288 | NHSGGC West of Scotland Specialist Virology Centre / MRC-University of Glasgow Centre for Virus Research | Wellcome Sanger Institute for the COVID-19 Genomics UK Consortium | Ana da Silva Filipe, Natasha Johnson, Kathy Smollett, Daniel Mair, Stephen Carmichael, Lily Tong, Jenna Nichols, Elihu Aranday-Cortes, Kirstyn Brunker, Yasmin Parr, Kyriaki Nomikou; Sarah McDonald, Marc Niebel, Patawee Asamaphan; Richard Orton, Joseph Hughes, Sreenu Vattipally, David L Robertson; Alasdair MacLean, Rory Gunson; Kathy Li, Natasha Jesudason, Rajiv Shah, James Shepherd, Antonia Ho, Alice Broos, Emma Thomson and Alex Alderton, Roberto Amato, Sonia Goncalves, Ewan Harrison, David K. Jackson, Ian Johnston, Dominic Kwiatkowski, Cordelia Langford, John Sillitoe |
| EPI_ISL_532289, EPI_ISL_532290, EPI_ISL_532291, EPI_ISL_532292 | Lighthouse Lab in Glasgow | Wellcome Sanger Institute for the COVID-19 Genomics UK Consortium | Harper VanSteenhouse, Yumi Kasai, David Gray, Carol Clugston, Anna Dominiczak and Alex Alderton, Roberto Amato, Sonia Goncalves, Ewan Harrison, David K. Jackson, Ian Johnston, Dominic Kwiatkowski, Cordelia Langford, John Sillitoe |
| EPI_ISL_532293 | NHSGGC West of Scotland Specialist Virology Centre / MRC-University of Glasgow Centre for Virus Research | Wellcome Sanger Institute for the COVID-19 Genomics UK Consortium | Ana da Silva Filipe, Natasha Johnson, Kathy Smollett, Daniel Mair, Stephen Carmichael, Lily Tong, Jenna Nichols, Elihu Aranday-Cortes, Kirstyn Brunker, Yasmin Parr, Kyriaki Nomikou; Sarah McDonald, Marc Niebel, Patawee Asamaphan; Richard Orton, Joseph Hughes, Sreenu Vattipally, David L Robertson; Alasdair MacLean, Rory Gunson; Kathy Li, Natasha Jesudason, Rajiv Shah, James Shepherd, Antonia Ho, Alice Broos, Emma Thomson and Alex Alderton, Roberto Amato, Sonia Goncalves, Ewan Harrison, David K. Jackson, Ian Johnston, Dominic Kwiatkowski, Cordelia Langford, John Sillitoe |
| EPI_ISL_532295, EPI_ISL_532296, EPI_ISL_532297, EPI_ISL_532298, EPI_ISL_532299, EPI_ISL_532300 | Lighthouse Lab in Glasgow | Wellcome Sanger Institute for the COVID-19 Genomics UK Consortium | Harper VanSteenhouse, Yumi Kasai, David Gray, Carol Clugston, Anna Dominiczak and Alex Alderton, Roberto Amato, Sonia Goncalves, Ewan Harrison, David K. Jackson, Ian Johnston, Dominic Kwiatkowski, Cordelia Langford, John Sillitoe |
| EPI_ISL_532301 | NHSGGC West of Scotland Specialist Virology Centre / MRC-University of Glasgow Centre for Virus Research | Wellcome Sanger Institute for the COVID-19 Genomics UK Consortium | Ana da Silva Filipe, Natasha Johnson, Kathy Smollett, Daniel Mair, Stephen Carmichael, Lily Tong, Jenna Nichols, Elihu Aranday-Cortes, Kirstyn Brunker, Yasmin Parr, Kyriaki Nomikou; Sarah McDonald, Marc Niebel, Patawee Asamaphan; Richard Orton, Joseph Hughes, Sreenu Vattipally, David L Robertson; Alasdair MacLean, Rory Gunson; Kathy Li, Natasha Jesudason, Rajiv Shah, James Shepherd, Antonia Ho, Alice Broos, Emma Thomson and Alex Alderton, Roberto Amato, Sonia Goncalves, Ewan Harrison, David K. Jackson, Ian Johnston, Dominic Kwiatkowski, Cordelia Langford, John Sillitoe |
| EPI_ISL_532302, EPI_ISL_532303, EPI_ISL_532304, EPI_ISL_532305, EPI_ISL_532306, EPI_ISL_532307, EPI_ISL_532308, EPI_ISL_532309, EPI_ISL_532310 | Lighthouse Lab in Glasgow | Wellcome Sanger Institute for the COVID-19 Genomics UK Consortium | Harper VanSteenhouse, Yumi Kasai, David Gray, Carol Clugston, Anna Dominiczak and Alex Alderton, Roberto Amato, Sonia Goncalves, Ewan Harrison, David K. Jackson, Ian Johnston, Dominic Kwiatkowski, Cordelia Langford, John Sillitoe |
| EPI_ISL_532311 | NHSGGC West of Scotland Specialist Virology Centre / MRC-University of Glasgow Centre for Virus Research | Wellcome Sanger Institute for the COVID-19 Genomics UK Consortium | Ana da Silva Filipe, Natasha Johnson, Kathy Smollett, Daniel Mair, Stephen Carmichael, Lily Tong, Jenna Nichols, Elihu Aranday-Cortes, Kirstyn Brunker, Yasmin Parr, Kyriaki Nomikou; Sarah McDonald, Marc Niebel, Patawee Asamaphan; Richard Orton, Joseph Hughes, Sreenu Vattipally, David L Robertson; Alasdair MacLean, Rory Gunson; Kathy Li, Natasha Jesudason, Rajiv Shah, James Shepherd, Antonia Ho, Alice Broos, Emma Thomson and Alex Alderton, Roberto Amato, Sonia Goncalves, Ewan Harrison, David K. Jackson, Ian Johnston, Dominic Kwiatkowski, Cordelia Langford, John Sillitoe |
| EPI_ISL_532312, EPI_ISL_532313, EPI_ISL_532314, EPI_ISL_532315 | Lighthouse Lab in Glasgow | Wellcome Sanger Institute for the COVID-19 Genomics UK Consortium | Harper VanSteenhouse, Yumi Kasai, David Gray, Carol Clugston, Anna Dominiczak and Alex Alderton, Roberto Amato, Sonia Goncalves, Ewan Harrison, David K. Jackson, Ian Johnston, Dominic Kwiatkowski, Cordelia Langford, John Sillitoe |
| EPI_ISL_532316, EPI_ISL_532317 | NHSGGC West of Scotland Specialist Virology Centre / MRC-University of Glasgow Centre for Virus Research | Wellcome Sanger Institute for the COVID-19 Genomics UK Consortium | Ana da Silva Filipe, Natasha Johnson, Kathy Smollett, Daniel Mair, Stephen Carmichael, Lily Tong, Jenna Nichols, Elihu Aranday-Cortes, Kirstyn Brunker, Yasmin Parr, Kyriaki Nomikou; Sarah McDonald, Marc Niebel, Patawee Asamaphan; Richard Orton, Joseph Hughes, Sreenu Vattipally, David L Robertson; Alasdair MacLean, Rory Gunson; Kathy Li, Natasha Jesudason, Rajiv Shah, James Shepherd, Antonia Ho, Alice Broos, Emma Thomson and Alex Alderton, Roberto Amato, Sonia Goncalves, Ewan Harrison, David K. Jackson, Ian Johnston, Dominic Kwiatkowski, Cordelia Langford, John Sillitoe |
| EPI_ISL_532318, EPI_ISL_532319, EPI_ISL_532320 | Lighthouse Lab in Glasgow | Wellcome Sanger Institute for the COVID-19 Genomics UK Consortium | Harper VanSteenhouse, Yumi Kasai, David Gray, Carol Clugston, Anna Dominiczak and Alex Alderton, Roberto Amato, Sonia Goncalves, Ewan Harrison, David K. Jackson, Ian Johnston, Dominic Kwiatkowski, Cordelia Langford, John Sillitoe |
| EPI_ISL_532321 | NHSGGC West of Scotland Specialist Virology Centre / MRC-University of Glasgow Centre for Virus Research | Wellcome Sanger Institute for the COVID-19 Genomics UK Consortium | Ana da Silva Filipe, Natasha Johnson, Kathy Smollett, Daniel Mair, Stephen Carmichael, Lily Tong, Jenna Nichols, Elihu Aranday-Cortes, Kirstyn Brunker, Yasmin Parr, Kyriaki Nomikou; Sarah McDonald, Marc Niebel, Patawee Asamaphan; Richard Orton, Joseph Hughes, Sreenu Vattipally, David L Robertson; Alasdair MacLean, Rory Gunson; Kathy Li, Natasha Jesudason, Rajiv Shah, James Shepherd, Antonia Ho, Alice Broos, Emma Thomson and Alex Alderton, Roberto Amato, Sonia Goncalves, Ewan Harrison, David K. Jackson, Ian Johnston, Dominic Kwiatkowski, Cordelia Langford, John Sillitoe |
| EPI_ISL_532322, EPI_ISL_532323, EPI_ISL_532324, EPI_ISL_532325, EPI_ISL_532326 | Lighthouse Lab in Glasgow | Wellcome Sanger Institute for the COVID-19 Genomics UK Consortium | Harper VanSteenhouse, Yumi Kasai, David Gray, Carol Clugston, Anna Dominiczak and Alex Alderton, Roberto Amato, Sonia Goncalves, Ewan Harrison, David K. Jackson, Ian Johnston, Dominic Kwiatkowski, Cordelia Langford, John Sillitoe |
| EPI_ISL_532327, EPI_ISL_532328 | NHSGGC West of Scotland Specialist Virology Centre / MRC-University of Glasgow Centre for Virus Research | Wellcome Sanger Institute for the COVID-19 Genomics UK Consortium | Ana da Silva Filipe, Natasha Johnson, Kathy Smollett, Daniel Mair, Stephen Carmichael, Lily Tong, Jenna Nichols, Elihu Aranday-Cortes, Kirstyn Brunker, Yasmin Parr, Kyriaki Nomikou; Sarah McDonald, Marc Niebel, Patawee Asamaphan; Richard Orton, Joseph Hughes, Sreenu Vattipally, David L Robertson; Alasdair MacLean, Rory Gunson; Kathy Li, Natasha Jesudason, Rajiv Shah, James Shepherd, Antonia Ho, Alice Broos, Emma Thomson and Alex Alderton, Roberto Amato, Sonia Goncalves, Ewan Harrison, David K. Jackson, Ian Johnston, Dominic Kwiatkowski, Cordelia Langford, John Sillitoe |
| EPI_ISL_532329, EPI_ISL_532330, EPI_ISL_532331, EPI_ISL_532332, EPI_ISL_532333 | Lighthouse Lab in Glasgow | Wellcome Sanger Institute for the COVID-19 Genomics UK Consortium | Harper VanSteenhouse, Yumi Kasai, David Gray, Carol Clugston, Anna Dominiczak and Alex Alderton, Roberto Amato, Sonia Goncalves, Ewan Harrison, David K. Jackson, Ian Johnston, Dominic Kwiatkowski, Cordelia Langford, John Sillitoe |
| EPI_ISL_532334, EPI_ISL_532335 | NHSGGC West of Scotland Specialist Virology Centre / MRC-University of Glasgow Centre for Virus Research | Wellcome Sanger Institute for the COVID-19 Genomics UK Consortium | Ana da Silva Filipe, Natasha Johnson, Kathy Smollett, Daniel Mair, Stephen Carmichael, Lily Tong, Jenna Nichols, Elihu Aranday-Cortes, Kirstyn Brunker, Yasmin Parr, Kyriaki Nomikou; Sarah McDonald, Marc Niebel, Patawee Asamaphan; Richard Orton, Joseph Hughes, Sreenu Vattipally, David L Robertson; Alasdair MacLean, Rory Gunson; Kathy Li, Natasha Jesudason, Rajiv Shah, James Shepherd, Antonia Ho, Alice Broos, Emma Thomson and Alex Alderton, Roberto Amato, Sonia Goncalves, Ewan Harrison, David K. Jackson, Ian Johnston, Dominic Kwiatkowski, Cordelia Langford, John Sillitoe |
| EPI_ISL_532336, EPI_ISL_532337 | Lighthouse Lab in Glasgow | Wellcome Sanger Institute for the COVID-19 Genomics UK Consortium | Harper VanSteenhouse, Yumi Kasai, David Gray, Carol Clugston, Anna Dominiczak and Alex Alderton, Roberto Amato, Sonia Goncalves, Ewan Harrison, David K. Jackson, Ian Johnston, Dominic Kwiatkowski, Cordelia Langford, John Sillitoe |
| EPI_ISL_532338 | NHSGGC West of Scotland Specialist Virology Centre / MRC-University of Glasgow Centre for Virus Research | Wellcome Sanger Institute for the COVID-19 Genomics UK Consortium | Ana da Silva Filipe, Natasha Johnson, Kathy Smollett, Daniel Mair, Stephen Carmichael, Lily Tong, Jenna Nichols, Elihu Aranday-Cortes, Kirstyn Brunker, Yasmin Parr, Kyriaki Nomikou; Sarah McDonald, Marc Niebel, Patawee Asamaphan; Richard Orton, Joseph Hughes, Sreenu Vattipally, David L Robertson; Alasdair MacLean, Rory Gunson; Kathy Li, Natasha Jesudason, Rajiv Shah, James Shepherd, Antonia Ho, Alice Broos, Emma Thomson and Alex Alderton, Roberto Amato, Sonia Goncalves, Ewan Harrison, David K. Jackson, Ian Johnston, Dominic Kwiatkowski, Cordelia Langford, John Sillitoe |
| EPI_ISL_532339 | Lighthouse Lab in Glasgow | Wellcome Sanger Institute for the COVID-19 Genomics UK Consortium | Harper VanSteenhouse, Yumi Kasai, David Gray, Carol Clugston, Anna Dominiczak and Alex Alderton, Roberto Amato, Sonia Goncalves, Ewan Harrison, David K. Jackson, Ian Johnston, Dominic Kwiatkowski, Cordelia Langford, John Sillitoe |
| EPI_ISL_532340 | NHSGGC West of Scotland Specialist Virology Centre / MRC-University of Glasgow Centre for Virus Research | Wellcome Sanger Institute for the COVID-19 Genomics UK Consortium | Ana da Silva Filipe, Natasha Johnson, Kathy Smollett, Daniel Mair, Stephen Carmichael, Lily Tong, Jenna Nichols, Elihu Aranday-Cortes, Kirstyn Brunker, Yasmin Parr, Kyriaki Nomikou; Sarah McDonald, Marc Niebel, Patawee Asamaphan; Richard Orton, Joseph Hughes, Sreenu Vattipally, David L Robertson; Alasdair MacLean, Rory Gunson; Kathy Li, Natasha Jesudason, Rajiv Shah, James Shepherd, Antonia Ho, Alice Broos, Emma Thomson and Alex Alderton, Roberto Amato, Sonia Goncalves, Ewan Harrison, David K. Jackson, Ian Johnston, Dominic Kwiatkowski, Cordelia Langford, John Sillitoe |
| EPI_ISL_532341 | Lighthouse Lab in Glasgow | Wellcome Sanger Institute for the COVID-19 Genomics UK Consortium | Harper VanSteenhouse, Yumi Kasai, David Gray, Carol Clugston, Anna Dominiczak and Alex Alderton, Roberto Amato, Sonia Goncalves, Ewan Harrison, David K. Jackson, Ian Johnston, Dominic Kwiatkowski, Cordelia Langford, John Sillitoe |
| EPI_ISL_532342 | NHSGGC West of Scotland Specialist Virology Centre / MRC-University of Glasgow Centre for Virus Research | Wellcome Sanger Institute for the COVID-19 Genomics UK Consortium | Ana da Silva Filipe, Natasha Johnson, Kathy Smollett, Daniel Mair, Stephen Carmichael, Lily Tong, Jenna Nichols, Elihu Aranday-Cortes, Kirstyn Brunker, Yasmin Parr, Kyriaki Nomikou; Sarah McDonald, Marc Niebel, Patawee Asamaphan; Richard Orton, Joseph Hughes, Sreenu Vattipally, David L Robertson; Alasdair MacLean, Rory Gunson; Kathy Li, Natasha Jesudason, Rajiv Shah, James Shepherd, Antonia Ho, Alice Broos, Emma Thomson and Alex Alderton, Roberto Amato, Sonia Goncalves, Ewan Harrison, David K. Jackson, Ian Johnston, Dominic Kwiatkowski, Cordelia Langford, John Sillitoe |
| EPI_ISL_532343, EPI_ISL_532344, EPI_ISL_532345, EPI_ISL_532346 | Lighthouse Lab in Glasgow | Wellcome Sanger Institute for the COVID-19 Genomics UK Consortium | Harper VanSteenhouse, Yumi Kasai, David Gray, Carol Clugston, Anna Dominiczak and Alex Alderton, Roberto Amato, Sonia Goncalves, Ewan Harrison, David K. Jackson, Ian Johnston, Dominic Kwiatkowski, Cordelia Langford, John Sillitoe |
| EPI_ISL_532347 | NHSGGC West of Scotland Specialist Virology Centre / MRC-University of Glasgow Centre for Virus Research | Wellcome Sanger Institute for the COVID-19 Genomics UK Consortium | Ana da Silva Filipe, Natasha Johnson, Kathy Smollett, Daniel Mair, Stephen Carmichael, Lily Tong, Jenna Nichols, Elihu Aranday-Cortes, Kirstyn Brunker, Yasmin Parr, Kyriaki Nomikou; Sarah McDonald, Marc Niebel, Patawee Asamaphan; Richard Orton, Joseph Hughes, Sreenu Vattipally, David L Robertson; Alasdair MacLean, Rory Gunson; Kathy Li, Natasha Jesudason, Rajiv Shah, James Shepherd, Antonia Ho, Alice Broos, Emma Thomson and Alex Alderton, Roberto Amato, Sonia Goncalves, Ewan Harrison, David K. Jackson, Ian Johnston, Dominic Kwiatkowski, Cordelia Langford, John Sillitoe |
| EPI_ISL_532348 | Lighthouse Lab in Glasgow | Wellcome Sanger Institute for the COVID-19 Genomics UK Consortium | Harper VanSteenhouse, Yumi Kasai, David Gray, Carol Clugston, Anna Dominiczak and Alex Alderton, Roberto Amato, Sonia Goncalves, Ewan Harrison, David K. Jackson, Ian Johnston, Dominic Kwiatkowski, Cordelia Langford, John Sillitoe |
| EPI_ISL_532349 | NHSGGC West of Scotland Specialist Virology Centre / MRC-University of Glasgow Centre for Virus Research | Wellcome Sanger Institute for the COVID-19 Genomics UK Consortium | Ana da Silva Filipe, Natasha Johnson, Kathy Smollett, Daniel Mair, Stephen Carmichael, Lily Tong, Jenna Nichols, Elihu Aranday-Cortes, Kirstyn Brunker, Yasmin Parr, Kyriaki Nomikou; Sarah McDonald, Marc Niebel, Patawee Asamaphan; Richard Orton, Joseph Hughes, Sreenu Vattipally, David L Robertson; Alasdair MacLean, Rory Gunson; Kathy Li, Natasha Jesudason, Rajiv Shah, James Shepherd, Antonia Ho, Alice Broos, Emma Thomson and Alex Alderton, Roberto Amato, Sonia Goncalves, Ewan Harrison, David K. Jackson, Ian Johnston, Dominic Kwiatkowski, Cordelia Langford, John Sillitoe |
| EPI_ISL_532350 | Lighthouse Lab in Glasgow | Wellcome Sanger Institute for the COVID-19 Genomics UK Consortium | Harper VanSteenhouse, Yumi Kasai, David Gray, Carol Clugston, Anna Dominiczak and Alex Alderton, Roberto Amato, Sonia Goncalves, Ewan Harrison, David K. Jackson, Ian Johnston, Dominic Kwiatkowski, Cordelia Langford, John Sillitoe |
| EPI_ISL_532351 | NHSGGC West of Scotland Specialist Virology Centre / MRC-University of Glasgow Centre for Virus Research | Wellcome Sanger Institute for the COVID-19 Genomics UK Consortium | Ana da Silva Filipe, Natasha Johnson, Kathy Smollett, Daniel Mair, Stephen Carmichael, Lily Tong, Jenna Nichols, Elihu Aranday-Cortes, Kirstyn Brunker, Yasmin Parr, Kyriaki Nomikou; Sarah McDonald, Marc Niebel, Patawee Asamaphan; Richard Orton, Joseph Hughes, Sreenu Vattipally, David L Robertson; Alasdair MacLean, Rory Gunson; Kathy Li, Natasha Jesudason, Rajiv Shah, James Shepherd, Antonia Ho, Alice Broos, Emma Thomson and Alex Alderton, Roberto Amato, Sonia Goncalves, Ewan Harrison, David K. Jackson, Ian Johnston, Dominic Kwiatkowski, Cordelia Langford, John Sillitoe |
| EPI_ISL_532352 | Lighthouse Lab in Glasgow | Wellcome Sanger Institute for the COVID-19 Genomics UK Consortium | Harper VanSteenhouse, Yumi Kasai, David Gray, Carol Clugston, Anna Dominiczak and Alex Alderton, Roberto Amato, Sonia Goncalves, Ewan Harrison, David K. Jackson, Ian Johnston, Dominic Kwiatkowski, Cordelia Langford, John Sillitoe |
| EPI_ISL_532353 | NHSGGC West of Scotland Specialist Virology Centre / MRC-University of Glasgow Centre for Virus Research | Wellcome Sanger Institute for the COVID-19 Genomics UK Consortium | Ana da Silva Filipe, Natasha Johnson, Kathy Smollett, Daniel Mair, Stephen Carmichael, Lily Tong, Jenna Nichols, Elihu Aranday-Cortes, Kirstyn Brunker, Yasmin Parr, Kyriaki Nomikou; Sarah McDonald, Marc Niebel, Patawee Asamaphan; Richard Orton, Joseph Hughes, Sreenu Vattipally, David L Robertson; Alasdair MacLean, Rory Gunson; Kathy Li, Natasha Jesudason, Rajiv Shah, James Shepherd, Antonia Ho, Alice Broos, Emma Thomson and Alex Alderton, Roberto Amato, Sonia Goncalves, Ewan Harrison, David K. Jackson, Ian Johnston, Dominic Kwiatkowski, Cordelia Langford, John Sillitoe |
| EPI_ISL_532354 | Lighthouse Lab in Glasgow | Wellcome Sanger Institute for the COVID-19 Genomics UK Consortium | Harper VanSteenhouse, Yumi Kasai, David Gray, Carol Clugston, Anna Dominiczak and Alex Alderton, Roberto Amato, Sonia Goncalves, Ewan Harrison, David K. Jackson, Ian Johnston, Dominic Kwiatkowski, Cordelia Langford, John Sillitoe |
| EPI_ISL_532355, EPI_ISL_532356 | NHSGGC West of Scotland Specialist Virology Centre / MRC-University of Glasgow Centre for Virus Research | Wellcome Sanger Institute for the COVID-19 Genomics UK Consortium | Ana da Silva Filipe, Natasha Johnson, Kathy Smollett, Daniel Mair, Stephen Carmichael, Lily Tong, Jenna Nichols, Elihu Aranday-Cortes, Kirstyn Brunker, Yasmin Parr, Kyriaki Nomikou; Sarah McDonald, Marc Niebel, Patawee Asamaphan; Richard Orton, Joseph Hughes, Sreenu Vattipally, David L Robertson; Alasdair MacLean, Rory Gunson; Kathy Li, Natasha Jesudason, Rajiv Shah, James Shepherd, Antonia Ho, Alice Broos, Emma Thomson and Alex Alderton, Roberto Amato, Sonia Goncalves, Ewan Harrison, David K. Jackson, Ian Johnston, Dominic Kwiatkowski, Cordelia Langford, John Sillitoe |
| EPI_ISL_532357, EPI_ISL_532358, EPI_ISL_532359, EPI_ISL_532360, EPI_ISL_532361, EPI_ISL_532362 | Lighthouse Lab in Glasgow | Wellcome Sanger Institute for the COVID-19 Genomics UK Consortium | Harper VanSteenhouse, Yumi Kasai, David Gray, Carol Clugston, Anna Dominiczak and Alex Alderton, Roberto Amato, Sonia Goncalves, Ewan Harrison, David K. Jackson, Ian Johnston, Dominic Kwiatkowski, Cordelia Langford, John Sillitoe |

| | | | |
|---|---|---|---|
| EPI_ISL_532363, EPI_ISL_532364 | NHSGGC West of Scotland Specialist Virology Centre / MRC-University of Glasgow Centre for Virus Research | Wellcome Sanger Institute for the COVID-19 Genomics UK Consortium | Ana da Silva Filipe, Natasha Johnson, Kathy Smollett, Daniel Mair, Stephen Carmichael, Lily Tong, Jenna Nichols, Elihu Aranday-Cortes, Kirstyn Brunker, Yasmin Parr, Kyriaki Nomikou; Sarah McDonald, Marc Niebel, Patawee Asamaphan; Richard Orton, Joseph Hughes, Sreenu Vattipally, David L Robertson; Alasdair MacLean, Rory Gunson; Kathy Li, Natasha Jesudason, Rajiv Shah, James Shepherd, Antonia Ho, Alice Broos, Emma Thomson and Alex Alderton, Roberto Amato, Sonia Goncalves, Ewan Harrison, David K. Jackson, Ian Johnston, Dominic Kwiatkowski, Cordelia Langford, John Sillitoe |
| EPI_ISL_532366, EPI_ISL_532367, EPI_ISL_532368, EPI_ISL_532369, EPI_ISL_532370, EPI_ISL_532371, EPI_ISL_532372, EPI_ISL_532373, EPI_ISL_532374, EPI_ISL_532375, EPI_ISL_532376, EPI_ISL_532377, EPI_ISL_532378 | | | |
| see above | Lighthouse Lab in Glasgow | Wellcome Sanger Institute for the COVID-19 Genomics UK Consortium | Harper VanSteenhouse, Yumi Kasai, David Gray, Carol Clugston, Anna Dominiczak and Alex Alderton, Roberto Amato, Sonia Goncalves, Ewan Harrison, David K. Jackson, Ian Johnston, Dominic Kwiatkowski, Cordelia Langford, John Sillitoe |
| EPI_ISL_532379 | NHSGGC West of Scotland Specialist Virology Centre / MRC-University of Glasgow Centre for Virus Research | Wellcome Sanger Institute for the COVID-19 Genomics UK Consortium | Ana da Silva Filipe, Natasha Johnson, Kathy Smollett, Daniel Mair, Stephen Carmichael, Lily Tong, Jenna Nichols, Elihu Aranday-Cortes, Kirstyn Brunker, Yasmin Parr, Kyriaki Nomikou; Sarah McDonald, Marc Niebel, Patawee Asamaphan; Richard Orton, Joseph Hughes, Sreenu Vattipally, David L Robertson; Alasdair MacLean, Rory Gunson; Kathy Li, Natasha Jesudason, Rajiv Shah, James Shepherd, Antonia Ho, Alice Broos, Emma Thomson and Alex Alderton, Roberto Amato, Sonia Goncalves, Ewan Harrison, David K. Jackson, Ian Johnston, Dominic Kwiatkowski, Cordelia Langford, John Sillitoe |
| EPI_ISL_532380, EPI_ISL_532381, EPI_ISL_532382, EPI_ISL_532383 | Lighthouse Lab in Glasgow | Wellcome Sanger Institute for the COVID-19 Genomics UK Consortium | Harper VanSteenhouse, Yumi Kasai, David Gray, Carol Clugston, Anna Dominiczak and Alex Alderton, Roberto Amato, Sonia Goncalves, Ewan Harrison, David K. Jackson, Ian Johnston, Dominic Kwiatkowski, Cordelia Langford, John Sillitoe |
| EPI_ISL_532384 | NHSGGC West of Scotland Specialist Virology Centre / MRC-University of Glasgow Centre for Virus Research | Wellcome Sanger Institute for the COVID-19 Genomics UK Consortium | Ana da Silva Filipe, Natasha Johnson, Kathy Smollett, Daniel Mair, Stephen Carmichael, Lily Tong, Jenna Nichols, Elihu Aranday-Cortes, Kirstyn Brunker, Yasmin Parr, Kyriaki Nomikou; Sarah McDonald, Marc Niebel, Patawee Asamaphan; Richard Orton, Joseph Hughes, Sreenu Vattipally, David L Robertson; Alasdair MacLean, Rory Gunson; Kathy Li, Natasha Jesudason, Rajiv Shah, James Shepherd, Antonia Ho, Alice Broos, Emma Thomson and Alex Alderton, Roberto Amato, Sonia Goncalves, Ewan Harrison, David K. Jackson, Ian Johnston, Dominic Kwiatkowski, Cordelia Langford, John Sillitoe |
| EPI_ISL_532385, EPI_ISL_532386, EPI_ISL_532387 | Lighthouse Lab in Glasgow | Wellcome Sanger Institute for the COVID-19 Genomics UK Consortium | Harper VanSteenhouse, Yumi Kasai, David Gray, Carol Clugston, Anna Dominiczak and Alex Alderton, Roberto Amato, Sonia Goncalves, Ewan Harrison, David K. Jackson, Ian Johnston, Dominic Kwiatkowski, Cordelia Langford, John Sillitoe |
| EPI_ISL_532388 | NHSGGC West of Scotland Specialist Virology Centre / MRC-University of Glasgow Centre for Virus Research | Wellcome Sanger Institute for the COVID-19 Genomics UK Consortium | Ana da Silva Filipe, Natasha Johnson, Kathy Smollett, Daniel Mair, Stephen Carmichael, Lily Tong, Jenna Nichols, Elihu Aranday-Cortes, Kirstyn Brunker, Yasmin Parr, Kyriaki Nomikou; Sarah McDonald, Marc Niebel, Patawee Asamaphan; Richard Orton, Joseph Hughes, Sreenu Vattipally, David L Robertson; Alasdair MacLean, Rory Gunson; Kathy Li, Natasha Jesudason, Rajiv Shah, James Shepherd, Antonia Ho, Alice Broos, Emma Thomson and Alex Alderton, Roberto Amato, Sonia Goncalves, Ewan Harrison, David K. Jackson, Ian Johnston, Dominic Kwiatkowski, Cordelia Langford, John Sillitoe |
| EPI_ISL_532389 | Lighthouse Lab in Glasgow | Wellcome Sanger Institute for the COVID-19 Genomics UK Consortium | Harper VanSteenhouse, Yumi Kasai, David Gray, Carol Clugston, Anna Dominiczak and Alex Alderton, Roberto Amato, Sonia Goncalves, Ewan Harrison, David K. Jackson, Ian Johnston, Dominic Kwiatkowski, Cordelia Langford, John Sillitoe |
| EPI_ISL_532390 | NHSGGC West of Scotland Specialist Virology Centre / MRC-University of Glasgow Centre for Virus Research | Wellcome Sanger Institute for the COVID-19 Genomics UK Consortium | Ana da Silva Filipe, Natasha Johnson, Kathy Smollett, Daniel Mair, Stephen Carmichael, Lily Tong, Jenna Nichols, Elihu Aranday-Cortes, Kirstyn Brunker, Yasmin Parr, Kyriaki Nomikou; Sarah McDonald, Marc Niebel, Patawee Asamaphan; Richard Orton, Joseph Hughes, Sreenu Vattipally, David L Robertson; Alasdair MacLean, Rory Gunson; Kathy Li, Natasha Jesudason, Rajiv Shah, James Shepherd, Antonia Ho, Alice Broos, Emma Thomson and Alex Alderton, Roberto Amato, Sonia Goncalves, Ewan Harrison, David K. Jackson, Ian Johnston, Dominic Kwiatkowski, Cordelia Langford, John Sillitoe |
| EPI_ISL_532391, EPI_ISL_532392, EPI_ISL_532393 | Lighthouse Lab in Glasgow | Wellcome Sanger Institute for the COVID-19 Genomics UK Consortium | Harper VanSteenhouse, Yumi Kasai, David Gray, Carol Clugston, Anna Dominiczak and Alex Alderton, Roberto Amato, Sonia Goncalves, Ewan Harrison, David K. Jackson, Ian Johnston, Dominic Kwiatkowski, Cordelia Langford, John Sillitoe |
| EPI_ISL_532394 | NHSGGC West of Scotland Specialist Virology Centre / MRC-University of Glasgow Centre for Virus Research | Wellcome Sanger Institute for the COVID-19 Genomics UK Consortium | Ana da Silva Filipe, Natasha Johnson, Kathy Smollett, Daniel Mair, Stephen Carmichael, Lily Tong, Jenna Nichols, Elihu Aranday-Cortes, Kirstyn Brunker, Yasmin Parr, Kyriaki Nomikou; Sarah McDonald, Marc Niebel, Patawee Asamaphan; Richard Orton, Joseph Hughes, Sreenu Vattipally, David L Robertson; Alasdair MacLean, Rory Gunson; Kathy Li, Natasha Jesudason, Rajiv Shah, James Shepherd, Antonia Ho, Alice Broos, Emma Thomson and Alex Alderton, Roberto Amato, Sonia Goncalves, Ewan Harrison, David K. Jackson, Ian Johnston, Dominic Kwiatkowski, Cordelia Langford, John Sillitoe |
| EPI_ISL_532395, EPI_ISL_532396, EPI_ISL_532397, EPI_ISL_532398 | Lighthouse Lab in Glasgow | Wellcome Sanger Institute for the COVID-19 Genomics UK Consortium | Harper VanSteenhouse, Yumi Kasai, David Gray, Carol Clugston, Anna Dominiczak and Alex Alderton, Roberto Amato, Sonia Goncalves, Ewan Harrison, David K. Jackson, Ian Johnston, Dominic Kwiatkowski, Cordelia Langford, John Sillitoe |
| EPI_ISL_532399 | NHSGGC West of Scotland Specialist Virology Centre / MRC-University of Glasgow Centre for Virus Research | Wellcome Sanger Institute for the COVID-19 Genomics UK Consortium | Ana da Silva Filipe, Natasha Johnson, Kathy Smollett, Daniel Mair, Stephen Carmichael, Lily Tong, Jenna Nichols, Elihu Aranday-Cortes, Kirstyn Brunker, Yasmin Parr, Kyriaki Nomikou; Sarah McDonald, Marc Niebel, Patawee Asamaphan; Richard Orton, Joseph Hughes, Sreenu Vattipally, David L Robertson; Alasdair MacLean, Rory Gunson; Kathy Li, Natasha Jesudason, Rajiv Shah, James Shepherd, Antonia Ho, Alice Broos, Emma Thomson and Alex Alderton, Roberto Amato, Sonia Goncalves, Ewan Harrison, David K. Jackson, Ian Johnston, Dominic Kwiatkowski, Cordelia Langford, John Sillitoe |
| EPI_ISL_532400, EPI_ISL_532401 | Lighthouse Lab in Glasgow | Wellcome Sanger Institute for the COVID-19 Genomics UK Consortium | Harper VanSteenhouse, Yumi Kasai, David Gray, Carol Clugston, Anna Dominiczak and Alex Alderton, Roberto Amato, Sonia Goncalves, Ewan Harrison, David K. Jackson, Ian Johnston, Dominic Kwiatkowski, Cordelia Langford, John Sillitoe |
| EPI_ISL_532402 | NHSGGC West of Scotland Specialist Virology Centre / MRC-University of Glasgow Centre for Virus Research | Wellcome Sanger Institute for the COVID-19 Genomics UK Consortium | Ana da Silva Filipe, Natasha Johnson, Kathy Smollett, Daniel Mair, Stephen Carmichael, Lily Tong, Jenna Nichols, Elihu Aranday-Cortes, Kirstyn Brunker, Yasmin Parr, Kyriaki Nomikou; Sarah McDonald, Marc Niebel, Patawee Asamaphan; Richard Orton, Joseph Hughes, Sreenu Vattipally, David L Robertson; Alasdair MacLean, Rory Gunson; Kathy Li, Natasha Jesudason, Rajiv Shah, James Shepherd, Antonia Ho, Alice Broos, Emma Thomson and Alex Alderton, Roberto Amato, Sonia Goncalves, Ewan Harrison, David K. Jackson, Ian Johnston, Dominic Kwiatkowski, Cordelia Langford, John Sillitoe |
| EPI_ISL_532403 | Lighthouse Lab in Glasgow | Wellcome Sanger Institute for the COVID-19 Genomics UK Consortium | Harper VanSteenhouse, Yumi Kasai, David Gray, Carol Clugston, Anna Dominiczak and Alex Alderton, Roberto Amato, Sonia Goncalves, Ewan Harrison, David K. Jackson, Ian Johnston, Dominic Kwiatkowski, Cordelia Langford, John Sillitoe |
| EPI_ISL_532404 | NHSGGC West of Scotland Specialist Virology Centre / MRC-University of Glasgow Centre for Virus Research | Wellcome Sanger Institute for the COVID-19 Genomics UK Consortium | Ana da Silva Filipe, Natasha Johnson, Kathy Smollett, Daniel Mair, Stephen Carmichael, Lily Tong, Jenna Nichols, Elihu Aranday-Cortes, Kirstyn Brunker, Yasmin Parr, Kyriaki Nomikou; Sarah McDonald, Marc Niebel, Patawee Asamaphan; Richard Orton, Joseph Hughes, Sreenu Vattipally, David L Robertson; Alasdair MacLean, Rory Gunson; Kathy Li, Natasha Jesudason, Rajiv Shah, James Shepherd, Antonia Ho, Alice Broos, Emma Thomson and Alex Alderton, Roberto Amato, Sonia Goncalves, Ewan Harrison, David K. Jackson, Ian Johnston, Dominic Kwiatkowski, Cordelia Langford, John Sillitoe |
| EPI_ISL_532405, EPI_ISL_532406, EPI_ISL_532407, EPI_ISL_532408, EPI_ISL_532409, EPI_ISL_532410, EPI_ISL_532411, EPI_ISL_532412, EPI_ISL_532413 | Lighthouse Lab in Glasgow | Wellcome Sanger Institute for the COVID-19 Genomics UK Consortium | Harper VanSteenhouse, Yumi Kasai, David Gray, Carol Clugston, Anna Dominiczak and Alex Alderton, Roberto Amato, Sonia Goncalves, Ewan Harrison, David K. Jackson, Ian Johnston, Dominic Kwiatkowski, Cordelia Langford, John Sillitoe |
| EPI_ISL_532414 | NHSGGC West of Scotland Specialist Virology Centre / MRC-University of Glasgow Centre for Virus Research | Wellcome Sanger Institute for the COVID-19 Genomics UK Consortium | Ana da Silva Filipe, Natasha Johnson, Kathy Smollett, Daniel Mair, Stephen Carmichael, Lily Tong, Jenna Nichols, Elihu Aranday-Cortes, Kirstyn Brunker, Yasmin Parr, Kyriaki Nomikou; Sarah McDonald, Marc Niebel, Patawee Asamaphan; Richard Orton, Joseph Hughes, Sreenu Vattipally, David L Robertson; Alasdair MacLean, Rory Gunson; Kathy Li, Natasha Jesudason, Rajiv Shah, James Shepherd, Antonia Ho, Alice Broos, Emma Thomson and Alex Alderton, Roberto Amato, Sonia Goncalves, Ewan Harrison, David K. Jackson, Ian Johnston, Dominic Kwiatkowski, Cordelia Langford, John Sillitoe |
| EPI_ISL_532415, EPI_ISL_532416, EPI_ISL_532417, EPI_ISL_532418, EPI_ISL_532419, EPI_ISL_532420, EPI_ISL_532421, EPI_ISL_532422, EPI_ISL_532423, EPI_ISL_532424, EPI_ISL_532425, EPI_ISL_532426, EPI_ISL_532427, EPI_ISL_532428, EPI_ISL_532429 | | | |
| see above | Lighthouse Lab in Glasgow | Wellcome Sanger Institute for the COVID-19 Genomics UK Consortium | Harper VanSteenhouse, Yumi Kasai, David Gray, Carol Clugston, Anna Dominiczak and Alex Alderton, Roberto Amato, Sonia Goncalves, Ewan Harrison, David K. Jackson, Ian Johnston, Dominic Kwiatkowski, Cordelia Langford, John Sillitoe |
| EPI_ISL_532430, EPI_ISL_532431, EPI_ISL_532432 | NHSGGC West of Scotland Specialist Virology Centre / MRC-University of Glasgow Centre for Virus Research | Wellcome Sanger Institute for the COVID-19 Genomics UK Consortium | Ana da Silva Filipe, Natasha Johnson, Kathy Smollett, Daniel Mair, Stephen Carmichael, Lily Tong, Jenna Nichols, Elihu Aranday-Cortes, Kirstyn Brunker, Yasmin Parr, Kyriaki Nomikou; Sarah McDonald, Marc Niebel, Patawee Asamaphan; Richard Orton, Joseph Hughes, Sreenu Vattipally, David L Robertson; Alasdair MacLean, Rory Gunson; Kathy Li, Natasha Jesudason, Rajiv Shah, James Shepherd, Antonia Ho, Alice Broos, Emma Thomson and Alex Alderton, Roberto Amato, Sonia Goncalves, Ewan Harrison, David K. Jackson, Ian Johnston, Dominic Kwiatkowski, Cordelia Langford, John Sillitoe |
| EPI_ISL_532433, EPI_ISL_532434, EPI_ISL_532435, EPI_ISL_532436, EPI_ISL_532437 | Lighthouse Lab in Glasgow | Wellcome Sanger Institute for the COVID-19 Genomics UK Consortium | Harper VanSteenhouse, Yumi Kasai, David Gray, Carol Clugston, Anna Dominiczak and Alex Alderton, Roberto Amato, Sonia Goncalves, Ewan Harrison, David K. Jackson, Ian Johnston, Dominic Kwiatkowski, Cordelia Langford, John Sillitoe |
| EPI_ISL_532438 | NHSGGC West of Scotland Specialist Virology Centre / MRC-University of Glasgow Centre for Virus Research | Wellcome Sanger Institute for the COVID-19 Genomics UK Consortium | Ana da Silva Filipe, Natasha Johnson, Kathy Smollett, Daniel Mair, Stephen Carmichael, Lily Tong, Jenna Nichols, Elihu Aranday-Cortes, Kirstyn Brunker, Yasmin Parr, Kyriaki Nomikou; Sarah McDonald, Marc Niebel, Patawee Asamaphan; Richard Orton, Joseph Hughes, Sreenu Vattipally, David L Robertson; Alasdair MacLean, Rory Gunson; Kathy Li, Natasha Jesudason, Rajiv Shah, James Shepherd, Antonia Ho, Alice Broos, Emma Thomson and Alex Alderton, Roberto Amato, Sonia Goncalves, Ewan Harrison, David K. Jackson, Ian Johnston, Dominic Kwiatkowski, Cordelia Langford, John Sillitoe |
| EPI_ISL_532439 | Lighthouse Lab in Glasgow | Wellcome Sanger Institute for the COVID-19 Genomics UK Consortium | Harper VanSteenhouse, Yumi Kasai, David Gray, Carol Clugston, Anna Dominiczak and Alex Alderton, Roberto Amato, Sonia Goncalves, Ewan Harrison, David K. Jackson, Ian Johnston, Dominic Kwiatkowski, Cordelia Langford, John Sillitoe |
| EPI_ISL_532440 | NHSGGC West of Scotland Specialist Virology Centre / MRC-University of Glasgow Centre for Virus Research | Wellcome Sanger Institute for the COVID-19 Genomics UK Consortium | Ana da Silva Filipe, Natasha Johnson, Kathy Smollett, Daniel Mair, Stephen Carmichael, Lily Tong, Jenna Nichols, Elihu Aranday-Cortes, Kirstyn Brunker, Yasmin Parr, Kyriaki Nomikou; Sarah McDonald, Marc Niebel, Patawee Asamaphan; Richard Orton, Joseph Hughes, Sreenu Vattipally, David L Robertson; Alasdair MacLean, Rory Gunson; Kathy Li, Natasha Jesudason, Rajiv Shah, James Shepherd, Antonia Ho, Alice Broos, Emma Thomson and Alex Alderton, Roberto Amato, Sonia Goncalves, Ewan Harrison, David K. Jackson, Ian Johnston, Dominic Kwiatkowski, Cordelia Langford, John Sillitoe |
| EPI_ISL_532441, EPI_ISL_532442, EPI_ISL_532443 | Lighthouse Lab in Glasgow | Wellcome Sanger Institute for the COVID-19 Genomics UK Consortium | Harper VanSteenhouse, Yumi Kasai, David Gray, Carol Clugston, Anna Dominiczak and Alex Alderton, Roberto Amato, Sonia Goncalves, Ewan Harrison, David K. Jackson, Ian Johnston, Dominic Kwiatkowski, Cordelia Langford, John Sillitoe |
| EPI_ISL_532444 | NHSGGC West of Scotland Specialist Virology Centre / MRC-University of Glasgow Centre for Virus Research | Wellcome Sanger Institute for the COVID-19 Genomics UK Consortium | Ana da Silva Filipe, Natasha Johnson, Kathy Smollett, Daniel Mair, Stephen Carmichael, Lily Tong, Jenna Nichols, Elihu Aranday-Cortes, Kirstyn Brunker, Yasmin Parr, Kyriaki Nomikou; Sarah McDonald, Marc Niebel, Patawee Asamaphan; Richard Orton, Joseph Hughes, Sreenu Vattipally, David L Robertson; Alasdair MacLean, Rory Gunson; Kathy Li, Natasha Jesudason, Rajiv Shah, James Shepherd, Antonia Ho, Alice Broos, Emma Thomson and Alex Alderton, Roberto Amato, Sonia Goncalves, Ewan Harrison, David K. Jackson, Ian Johnston, Dominic Kwiatkowski, Cordelia Langford, John Sillitoe |
| EPI_ISL_532445 | Lighthouse Lab in Glasgow | Wellcome Sanger Institute for the COVID-19 Genomics UK Consortium | Harper VanSteenhouse, Yumi Kasai, David Gray, Carol Clugston, Anna Dominiczak and Alex Alderton, Roberto Amato, Sonia Goncalves, Ewan Harrison, David K. Jackson, Ian Johnston, Dominic Kwiatkowski, Cordelia Langford, John Sillitoe |
| EPI_ISL_532446 | NHSGGC West of Scotland Specialist Virology Centre / MRC-University of Glasgow Centre for Virus Research | Wellcome Sanger Institute for the COVID-19 Genomics UK Consortium | Ana da Silva Filipe, Natasha Johnson, Kathy Smollett, Daniel Mair, Stephen Carmichael, Lily Tong, Jenna Nichols, Elihu Aranday-Cortes, Kirstyn Brunker, Yasmin Parr, Kyriaki Nomikou; Sarah McDonald, Marc Niebel, Patawee Asamaphan; Richard Orton, Joseph Hughes, Sreenu Vattipally, David L Robertson; Alasdair MacLean, Rory Gunson; Kathy Li, Natasha Jesudason, Rajiv Shah, James Shepherd, Antonia Ho, Alice Broos, Emma Thomson and Alex Alderton, Roberto Amato, Sonia Goncalves, Ewan Harrison, David K. Jackson, Ian Johnston, Dominic Kwiatkowski, Cordelia Langford, John Sillitoe |
| EPI_ISL_532447, EPI_ISL_532448, EPI_ISL_532449, EPI_ISL_532450, EPI_ISL_532451 | Lighthouse Lab in Glasgow | Wellcome Sanger Institute for the COVID-19 Genomics UK Consortium | Harper VanSteenhouse, Yumi Kasai, David Gray, Carol Clugston, Anna Dominiczak and Alex Alderton, Roberto Amato, Sonia Goncalves, Ewan Harrison, David K. Jackson, Ian Johnston, Dominic Kwiatkowski, Cordelia Langford, John Sillitoe |
| EPI_ISL_532452 | NHSGGC West of Scotland Specialist Virology Centre / MRC-University of Glasgow Centre for Virus Research | Wellcome Sanger Institute for the COVID-19 Genomics UK Consortium | Ana da Silva Filipe, Natasha Johnson, Kathy Smollett, Daniel Mair, Stephen Carmichael, Lily Tong, Jenna Nichols, Elihu Aranday-Cortes, Kirstyn Brunker, Yasmin Parr, Kyriaki Nomikou; Sarah McDonald, Marc Niebel, Patawee Asamaphan; Richard Orton, Joseph Hughes, Sreenu Vattipally, David L Robertson; Alasdair MacLean, Rory Gunson; Kathy Li, Natasha Jesudason, Rajiv Shah, James Shepherd, Antonia Ho, Alice Broos, Emma Thomson and Alex Alderton, Roberto Amato, Sonia Goncalves, Ewan Harrison, David K. Jackson, Ian Johnston, Dominic Kwiatkowski, Cordelia Langford, John Sillitoe |
| EPI_ISL_532453 | Lighthouse Lab in Glasgow | Wellcome Sanger Institute for the COVID-19 Genomics UK Consortium | Harper VanSteenhouse, Yumi Kasai, David Gray, Carol Clugston, Anna Dominiczak and Alex Alderton, Roberto Amato, Sonia Goncalves, Ewan Harrison, David K. Jackson, Ian Johnston, Dominic Kwiatkowski, Cordelia Langford, John Sillitoe |
| EPI_ISL_532454 | NHSGGC West of Scotland Specialist Virology Centre / MRC-University of Glasgow Centre for Virus Research | Wellcome Sanger Institute for the COVID-19 Genomics UK Consortium | Ana da Silva Filipe, Natasha Johnson, Kathy Smollett, Daniel Mair, Stephen Carmichael, Lily Tong, Jenna Nichols, Elihu Aranday-Cortes, Kirstyn Brunker, Yasmin Parr, Kyriaki Nomikou; Sarah McDonald, Marc Niebel, Patawee Asamaphan; Richard Orton, Joseph Hughes, Sreenu Vattipally, David L Robertson; Alasdair MacLean, Rory Gunson; Kathy Li, Natasha Jesudason, Rajiv Shah, James Shepherd, Antonia Ho, Alice Broos, Emma Thomson and Alex Alderton, Roberto Amato, Sonia Goncalves, Ewan Harrison, David K. Jackson, Ian Johnston, Dominic Kwiatkowski, Cordelia Langford, John Sillitoe |
| EPI_ISL_532455 | Lighthouse Lab in Glasgow | Wellcome Sanger Institute for the COVID-19 Genomics UK Consortium | Harper VanSteenhouse, Yumi Kasai, David Gray, Carol Clugston, Anna Dominiczak and Alex Alderton, Roberto Amato, Sonia Goncalves, Ewan Harrison, David K. Jackson, Ian Johnston, Dominic Kwiatkowski, Cordelia Langford, John Sillitoe |
| EPI_ISL_532456 | NHSGGC West of Scotland Specialist Virology Centre / MRC-University of Glasgow Centre for Virus Research | Wellcome Sanger Institute for the COVID-19 Genomics UK Consortium | Ana da Silva Filipe, Natasha Johnson, Kathy Smollett, Daniel Mair, Stephen Carmichael, Lily Tong, Jenna Nichols, Elihu Aranday-Cortes, Kirstyn Brunker, Yasmin Parr, Kyriaki Nomikou; Sarah McDonald, Marc Niebel, Patawee Asamaphan; Richard Orton, Joseph Hughes, Sreenu Vattipally, David L Robertson; Alasdair MacLean, Rory Gunson; Kathy Li, Natasha Jesudason, Rajiv Shah, James Shepherd, Antonia Ho, Alice Broos, Emma Thomson and Alex Alderton, Roberto Amato, Sonia Goncalves, Ewan Harrison, David K. Jackson, Ian Johnston, Dominic Kwiatkowski, Cordelia Langford, John Sillitoe |
| EPI_ISL_532457, EPI_ISL_532458, EPI_ISL_532459, EPI_ISL_532460, EPI_ISL_532461, EPI_ISL_532462 | Lighthouse Lab in Glasgow | Wellcome Sanger Institute for the COVID-19 Genomics UK Consortium | Harper VanSteenhouse, Yumi Kasai, David Gray, Carol Clugston, Anna Dominiczak and Alex Alderton, Roberto Amato, Sonia Goncalves, Ewan Harrison, David K. Jackson, Ian Johnston, Dominic Kwiatkowski, Cordelia Langford, John Sillitoe |

| | | | |
|---|---|---|---|
| EPI_ISL_532463, EPI_ISL_532464 | NHSGGC West of Scotland Specialist Virology Centre / MRC-University of Glasgow Centre for Virus Research | Wellcome Sanger Institute for the COVID-19 Genomics UK Consortium | Ana da Silva Filipe, Natasha Johnson, Kathy Smollett, Daniel Mair, Stephen Carmichael, Lily Tong, Jenna Nichols, Elihu Aranday-Cortes, Kirstyn Brunker, Yasmin Parr, Kyriaki Nomikou; Sarah McDonald, Marc Niebel, Patawee Asamaphan; Richard Orton, Joseph Hughes, Sreenu Vattipally, David L Robertson; Alasdair MacLean, Rory Gunson; Kathy Li, Natasha Jesudason, Rajiv Shah, James Shepherd, Antonia Ho, Alice Broos, Emma Thomson and Alex Alderton, Roberto Amato, Sonia Goncalves, Ewan Harrison, David K. Jackson, Ian Johnston, Dominic Kwiatkowski, Cordelia Langford, John Sillitoe |
| EPI_ISL_532465, EPI_ISL_532466, EPI_ISL_532467, EPI_ISL_532468 | Lighthouse Lab in Glasgow | Wellcome Sanger Institute for the COVID-19 Genomics UK Consortium | Harper VanSteenhouse, Yumi Kasai, David Gray, Carol Clugston, Anna Dominiczak and Alex Alderton, Roberto Amato, Sonia Goncalves, Ewan Harrison, David K. Jackson, Ian Johnston, Dominic Kwiatkowski, Cordelia Langford, John Sillitoe |
| EPI_ISL_532469 | NHSGGC West of Scotland Specialist Virology Centre / MRC-University of Glasgow Centre for Virus Research | Wellcome Sanger Institute for the COVID-19 Genomics UK Consortium | Ana da Silva Filipe, Natasha Johnson, Kathy Smollett, Daniel Mair, Stephen Carmichael, Lily Tong, Jenna Nichols, Elihu Aranday-Cortes, Kirstyn Brunker, Yasmin Parr, Kyriaki Nomikou; Sarah McDonald, Marc Niebel, Patawee Asamaphan; Richard Orton, Joseph Hughes, Sreenu Vattipally, David L Robertson; Alasdair MacLean, Rory Gunson; Kathy Li, Natasha Jesudason, Rajiv Shah, James Shepherd, Antonia Ho, Alice Broos, Emma Thomson and Alex Alderton, Roberto Amato, Sonia Goncalves, Ewan Harrison, David K. Jackson, Ian Johnston, Dominic Kwiatkowski, Cordelia Langford, John Sillitoe |
| EPI_ISL_532470, EPI_ISL_532471 | Lighthouse Lab in Glasgow | Wellcome Sanger Institute for the COVID-19 Genomics UK Consortium | Harper VanSteenhouse, Yumi Kasai, David Gray, Carol Clugston, Anna Dominiczak and Alex Alderton, Roberto Amato, Sonia Goncalves, Ewan Harrison, David K. Jackson, Ian Johnston, Dominic Kwiatkowski, Cordelia Langford, John Sillitoe |
| EPI_ISL_532472, EPI_ISL_532473 | NHSGGC West of Scotland Specialist Virology Centre / MRC-University of Glasgow Centre for Virus Research | Wellcome Sanger Institute for the COVID-19 Genomics UK Consortium | Ana da Silva Filipe, Natasha Johnson, Kathy Smollett, Daniel Mair, Stephen Carmichael, Lily Tong, Jenna Nichols, Elihu Aranday-Cortes, Kirstyn Brunker, Yasmin Parr, Kyriaki Nomikou; Sarah McDonald, Marc Niebel, Patawee Asamaphan; Richard Orton, Joseph Hughes, Sreenu Vattipally, David L Robertson; Alasdair MacLean, Rory Gunson; Kathy Li, Natasha Jesudason, Rajiv Shah, James Shepherd, Antonia Ho, Alice Broos, Emma Thomson and Alex Alderton, Roberto Amato, Sonia Goncalves, Ewan Harrison, David K. Jackson, Ian Johnston, Dominic Kwiatkowski, Cordelia Langford, John Sillitoe |
| EPI_ISL_532474, EPI_ISL_532475, EPI_ISL_532476, EPI_ISL_532477, EPI_ISL_532478, EPI_ISL_532479 | Lighthouse Lab in Glasgow | Wellcome Sanger Institute for the COVID-19 Genomics UK Consortium | Harper VanSteenhouse, Yumi Kasai, David Gray, Carol Clugston, Anna Dominiczak and Alex Alderton, Roberto Amato, Sonia Goncalves, Ewan Harrison, David K. Jackson, Ian Johnston, Dominic Kwiatkowski, Cordelia Langford, John Sillitoe |
| EPI_ISL_532480 | NHSGGC West of Scotland Specialist Virology Centre / MRC-University of Glasgow Centre for Virus Research | Wellcome Sanger Institute for the COVID-19 Genomics UK Consortium | Ana da Silva Filipe, Natasha Johnson, Kathy Smollett, Daniel Mair, Stephen Carmichael, Lily Tong, Jenna Nichols, Elihu Aranday-Cortes, Kirstyn Brunker, Yasmin Parr, Kyriaki Nomikou; Sarah McDonald, Marc Niebel, Patawee Asamaphan; Richard Orton, Joseph Hughes, Sreenu Vattipally, David L Robertson; Alasdair MacLean, Rory Gunson; Kathy Li, Natasha Jesudason, Rajiv Shah, James Shepherd, Antonia Ho, Alice Broos, Emma Thomson and Alex Alderton, Roberto Amato, Sonia Goncalves, Ewan Harrison, David K. Jackson, Ian Johnston, Dominic Kwiatkowski, Cordelia Langford, John Sillitoe |
| EPI_ISL_532481, EPI_ISL_532482, EPI_ISL_532483 | Lighthouse Lab in Glasgow | Wellcome Sanger Institute for the COVID-19 Genomics UK Consortium | Harper VanSteenhouse, Yumi Kasai, David Gray, Carol Clugston, Anna Dominiczak and Alex Alderton, Roberto Amato, Sonia Goncalves, Ewan Harrison, David K. Jackson, Ian Johnston, Dominic Kwiatkowski, Cordelia Langford, John Sillitoe |
| EPI_ISL_532484 | NHSGGC West of Scotland Specialist Virology Centre / MRC-University of Glasgow Centre for Virus Research | Wellcome Sanger Institute for the COVID-19 Genomics UK Consortium | Ana da Silva Filipe, Natasha Johnson, Kathy Smollett, Daniel Mair, Stephen Carmichael, Lily Tong, Jenna Nichols, Elihu Aranday-Cortes, Kirstyn Brunker, Yasmin Parr, Kyriaki Nomikou; Sarah McDonald, Marc Niebel, Patawee Asamaphan; Richard Orton, Joseph Hughes, Sreenu Vattipally, David L Robertson; Alasdair MacLean, Rory Gunson; Kathy Li, Natasha Jesudason, Rajiv Shah, James Shepherd, Antonia Ho, Alice Broos, Emma Thomson and Alex Alderton, Roberto Amato, Sonia Goncalves, Ewan Harrison, David K. Jackson, Ian Johnston, Dominic Kwiatkowski, Cordelia Langford, John Sillitoe |
| EPI_ISL_532485, EPI_ISL_532486, EPI_ISL_532487, EPI_ISL_532488, EPI_ISL_532489 | Lighthouse Lab in Glasgow | Wellcome Sanger Institute for the COVID-19 Genomics UK Consortium | Harper VanSteenhouse, Yumi Kasai, David Gray, Carol Clugston, Anna Dominiczak and Alex Alderton, Roberto Amato, Sonia Goncalves, Ewan Harrison, David K. Jackson, Ian Johnston, Dominic Kwiatkowski, Cordelia Langford, John Sillitoe |
| EPI_ISL_532490 | NHSGGC West of Scotland Specialist Virology Centre / MRC-University of Glasgow Centre for Virus Research | Wellcome Sanger Institute for the COVID-19 Genomics UK Consortium | Ana da Silva Filipe, Natasha Johnson, Kathy Smollett, Daniel Mair, Stephen Carmichael, Lily Tong, Jenna Nichols, Elihu Aranday-Cortes, Kirstyn Brunker, Yasmin Parr, Kyriaki Nomikou; Sarah McDonald, Marc Niebel, Patawee Asamaphan; Richard Orton, Joseph Hughes, Sreenu Vattipally, David L Robertson; Alasdair MacLean, Rory Gunson; Kathy Li, Natasha Jesudason, Rajiv Shah, James Shepherd, Antonia Ho, Alice Broos, Emma Thomson and Alex Alderton, Roberto Amato, Sonia Goncalves, Ewan Harrison, David K. Jackson, Ian Johnston, Dominic Kwiatkowski, Cordelia Langford, John Sillitoe |
| EPI_ISL_532491 | Lighthouse Lab in Glasgow | Wellcome Sanger Institute for the COVID-19 Genomics UK Consortium | Harper VanSteenhouse, Yumi Kasai, David Gray, Carol Clugston, Anna Dominiczak and Alex Alderton, Roberto Amato, Sonia Goncalves, Ewan Harrison, David K. Jackson, Ian Johnston, Dominic Kwiatkowski, Cordelia Langford, John Sillitoe |
| EPI_ISL_532492, EPI_ISL_532493 | NHSGGC West of Scotland Specialist Virology Centre / MRC-University of Glasgow Centre for Virus Research | Wellcome Sanger Institute for the COVID-19 Genomics UK Consortium | Ana da Silva Filipe, Natasha Johnson, Kathy Smollett, Daniel Mair, Stephen Carmichael, Lily Tong, Jenna Nichols, Elihu Aranday-Cortes, Kirstyn Brunker, Yasmin Parr, Kyriaki Nomikou; Sarah McDonald, Marc Niebel, Patawee Asamaphan; Richard Orton, Joseph Hughes, Sreenu Vattipally, David L Robertson; Alasdair MacLean, Rory Gunson; Kathy Li, Natasha Jesudason, Rajiv Shah, James Shepherd, Antonia Ho, Alice Broos, Emma Thomson and Alex Alderton, Roberto Amato, Sonia Goncalves, Ewan Harrison, David K. Jackson, Ian Johnston, Dominic Kwiatkowski, Cordelia Langford, John Sillitoe |
| EPI_ISL_532494, EPI_ISL_532495, EPI_ISL_532496, EPI_ISL_532497, EPI_ISL_532498, EPI_ISL_532499, EPI_ISL_532500, EPI_ISL_532501, EPI_ISL_532502, EPI_ISL_532503, EPI_ISL_532504, EPI_ISL_532505 | | | |
| see above | Lighthouse Lab in Glasgow | Wellcome Sanger Institute for the COVID-19 Genomics UK Consortium | Harper VanSteenhouse, Yumi Kasai, David Gray, Carol Clugston, Anna Dominiczak and Alex Alderton, Roberto Amato, Sonia Goncalves, Ewan Harrison, David K. Jackson, Ian Johnston, Dominic Kwiatkowski, Cordelia Langford, John Sillitoe |
| EPI_ISL_532506 | NHSGGC West of Scotland Specialist Virology Centre / MRC-University of Glasgow Centre for Virus Research | Wellcome Sanger Institute for the COVID-19 Genomics UK Consortium | Ana da Silva Filipe, Natasha Johnson, Kathy Smollett, Daniel Mair, Stephen Carmichael, Lily Tong, Jenna Nichols, Elihu Aranday-Cortes, Kirstyn Brunker, Yasmin Parr, Kyriaki Nomikou; Sarah McDonald, Marc Niebel, Patawee Asamaphan; Richard Orton, Joseph Hughes, Sreenu Vattipally, David L Robertson; Alasdair MacLean, Rory Gunson; Kathy Li, Natasha Jesudason, Rajiv Shah, James Shepherd, Antonia Ho, Alice Broos, Emma Thomson and Alex Alderton, Roberto Amato, Sonia Goncalves, Ewan Harrison, David K. Jackson, Ian Johnston, Dominic Kwiatkowski, Cordelia Langford, John Sillitoe |
| EPI_ISL_532507, EPI_ISL_532508, EPI_ISL_532509 | Lighthouse Lab in Glasgow | Wellcome Sanger Institute for the COVID-19 Genomics UK Consortium | Harper VanSteenhouse, Yumi Kasai, David Gray, Carol Clugston, Anna Dominiczak and Alex Alderton, Roberto Amato, Sonia Goncalves, Ewan Harrison, David K. Jackson, Ian Johnston, Dominic Kwiatkowski, Cordelia Langford, John Sillitoe |
| EPI_ISL_532510, EPI_ISL_532512 | NHSGGC West of Scotland Specialist Virology Centre / MRC-University of Glasgow Centre for Virus Research | Wellcome Sanger Institute for the COVID-19 Genomics UK Consortium | Ana da Silva Filipe, Natasha Johnson, Kathy Smollett, Daniel Mair, Stephen Carmichael, Lily Tong, Jenna Nichols, Elihu Aranday-Cortes, Kirstyn Brunker, Yasmin Parr, Kyriaki Nomikou; Sarah McDonald, Marc Niebel, Patawee Asamaphan; Richard Orton, Joseph Hughes, Sreenu Vattipally, David L Robertson; Alasdair MacLean, Rory Gunson; Kathy Li, Natasha Jesudason, Rajiv Shah, James Shepherd, Antonia Ho, Alice Broos, Emma Thomson and Alex Alderton, Roberto Amato, Sonia Goncalves, Ewan Harrison, David K. Jackson, Ian Johnston, Dominic Kwiatkowski, Cordelia Langford, John Sillitoe |
| EPI_ISL_532513 | Lighthouse Lab in Glasgow | Wellcome Sanger Institute for the COVID-19 Genomics UK Consortium | Harper VanSteenhouse, Yumi Kasai, David Gray, Carol Clugston, Anna Dominiczak and Alex Alderton, Roberto Amato, Sonia Goncalves, Ewan Harrison, David K. Jackson, Ian Johnston, Dominic Kwiatkowski, Cordelia Langford, John Sillitoe |
| EPI_ISL_532514, EPI_ISL_532515 | NHSGGC West of Scotland Specialist Virology Centre / MRC-University of Glasgow Centre for Virus Research | Wellcome Sanger Institute for the COVID-19 Genomics UK Consortium | Ana da Silva Filipe, Natasha Johnson, Kathy Smollett, Daniel Mair, Stephen Carmichael, Lily Tong, Jenna Nichols, Elihu Aranday-Cortes, Kirstyn Brunker, Yasmin Parr, Kyriaki Nomikou; Sarah McDonald, Marc Niebel, Patawee Asamaphan; Richard Orton, Joseph Hughes, Sreenu Vattipally, David L Robertson; Alasdair MacLean, Rory Gunson; Kathy Li, Natasha Jesudason, Rajiv Shah, James Shepherd, Antonia Ho, Alice Broos, Emma Thomson and Alex Alderton, Roberto Amato, Sonia Goncalves, Ewan Harrison, David K. Jackson, Ian Johnston, Dominic Kwiatkowski, Cordelia Langford, John Sillitoe |
| EPI_ISL_532516, EPI_ISL_532517 | Lighthouse Lab in Glasgow | Wellcome Sanger Institute for the COVID-19 Genomics UK Consortium | Harper VanSteenhouse, Yumi Kasai, David Gray, Carol Clugston, Anna Dominiczak and Alex Alderton, Roberto Amato, Sonia Goncalves, Ewan Harrison, David K. Jackson, Ian Johnston, Dominic Kwiatkowski, Cordelia Langford, John Sillitoe |
| EPI_ISL_532518 | NHSGGC West of Scotland Specialist Virology Centre / MRC-University of Glasgow Centre for Virus Research | Wellcome Sanger Institute for the COVID-19 Genomics UK Consortium | Ana da Silva Filipe, Natasha Johnson, Kathy Smollett, Daniel Mair, Stephen Carmichael, Lily Tong, Jenna Nichols, Elihu Aranday-Cortes, Kirstyn Brunker, Yasmin Parr, Kyriaki Nomikou; Sarah McDonald, Marc Niebel, Patawee Asamaphan; Richard Orton, Joseph Hughes, Sreenu Vattipally, David L Robertson; Alasdair MacLean, Rory Gunson; Kathy Li, Natasha Jesudason, Rajiv Shah, James Shepherd, Antonia Ho, Alice Broos, Emma Thomson and Alex Alderton, Roberto Amato, Sonia Goncalves, Ewan Harrison, David K. Jackson, Ian Johnston, Dominic Kwiatkowski, Cordelia Langford, John Sillitoe |
| EPI_ISL_532519, EPI_ISL_532520, EPI_ISL_532521, EPI_ISL_532522 | Lighthouse Lab in Glasgow | Wellcome Sanger Institute for the COVID-19 Genomics UK Consortium | Harper VanSteenhouse, Yumi Kasai, David Gray, Carol Clugston, Anna Dominiczak and Alex Alderton, Roberto Amato, Sonia Goncalves, Ewan Harrison, David K. Jackson, Ian Johnston, Dominic Kwiatkowski, Cordelia Langford, John Sillitoe |
| EPI_ISL_532523 | NHSGGC West of Scotland Specialist Virology Centre / MRC-University of Glasgow Centre for Virus Research | Wellcome Sanger Institute for the COVID-19 Genomics UK Consortium | Ana da Silva Filipe, Natasha Johnson, Kathy Smollett, Daniel Mair, Stephen Carmichael, Lily Tong, Jenna Nichols, Elihu Aranday-Cortes, Kirstyn Brunker, Yasmin Parr, Kyriaki Nomikou; Sarah McDonald, Marc Niebel, Patawee Asamaphan; Richard Orton, Joseph Hughes, Sreenu Vattipally, David L Robertson; Alasdair MacLean, Rory Gunson; Kathy Li, Natasha Jesudason, Rajiv Shah, James Shepherd, Antonia Ho, Alice Broos, Emma Thomson and Alex Alderton, Roberto Amato, Sonia Goncalves, Ewan Harrison, David K. Jackson, Ian Johnston, Dominic Kwiatkowski, Cordelia Langford, John Sillitoe |
| EPI_ISL_532524, EPI_ISL_532525, EPI_ISL_532526, EPI_ISL_532527, EPI_ISL_532528, EPI_ISL_532529, EPI_ISL_532530 | Lighthouse Lab in Glasgow | Wellcome Sanger Institute for the COVID-19 Genomics UK Consortium | Harper VanSteenhouse, Yumi Kasai, David Gray, Carol Clugston, Anna Dominiczak and Alex Alderton, Roberto Amato, Sonia Goncalves, Ewan Harrison, David K. Jackson, Ian Johnston, Dominic Kwiatkowski, Cordelia Langford, John Sillitoe |
| EPI_ISL_532531 | NHSGGC West of Scotland Specialist Virology Centre / MRC-University of Glasgow Centre for Virus Research | Wellcome Sanger Institute for the COVID-19 Genomics UK Consortium | Ana da Silva Filipe, Natasha Johnson, Kathy Smollett, Daniel Mair, Stephen Carmichael, Lily Tong, Jenna Nichols, Elihu Aranday-Cortes, Kirstyn Brunker, Yasmin Parr, Kyriaki Nomikou; Sarah McDonald, Marc Niebel, Patawee Asamaphan; Richard Orton, Joseph Hughes, Sreenu Vattipally, David L Robertson; Alasdair MacLean, Rory Gunson; Kathy Li, Natasha Jesudason, Rajiv Shah, James Shepherd, Antonia Ho, Alice Broos, Emma Thomson and Alex Alderton, Roberto Amato, Sonia Goncalves, Ewan Harrison, David K. Jackson, Ian Johnston, Dominic Kwiatkowski, Cordelia Langford, John Sillitoe |
| EPI_ISL_532532, EPI_ISL_532533, EPI_ISL_532534, EPI_ISL_532535, EPI_ISL_532536, EPI_ISL_532537 | Lighthouse Lab in Glasgow | Wellcome Sanger Institute for the COVID-19 Genomics UK Consortium | Harper VanSteenhouse, Yumi Kasai, David Gray, Carol Clugston, Anna Dominiczak and Alex Alderton, Roberto Amato, Sonia Goncalves, Ewan Harrison, David K. Jackson, Ian Johnston, Dominic Kwiatkowski, Cordelia Langford, John Sillitoe |
| EPI_ISL_532538 | NHSGGC West of Scotland Specialist Virology Centre / MRC-University of Glasgow Centre for Virus Research | Wellcome Sanger Institute for the COVID-19 Genomics UK Consortium | Ana da Silva Filipe, Natasha Johnson, Kathy Smollett, Daniel Mair, Stephen Carmichael, Lily Tong, Jenna Nichols, Elihu Aranday-Cortes, Kirstyn Brunker, Yasmin Parr, Kyriaki Nomikou; Sarah McDonald, Marc Niebel, Patawee Asamaphan; Richard Orton, Joseph Hughes, Sreenu Vattipally, David L Robertson; Alasdair MacLean, Rory Gunson; Kathy Li, Natasha Jesudason, Rajiv Shah, James Shepherd, Antonia Ho, Alice Broos, Emma Thomson and Alex Alderton, Roberto Amato, Sonia Goncalves, Ewan Harrison, David K. Jackson, Ian Johnston, Dominic Kwiatkowski, Cordelia Langford, John Sillitoe |
| EPI_ISL_532539 | Lighthouse Lab in Glasgow | Wellcome Sanger Institute for the COVID-19 Genomics UK Consortium | Harper VanSteenhouse, Yumi Kasai, David Gray, Carol Clugston, Anna Dominiczak and Alex Alderton, Roberto Amato, Sonia Goncalves, Ewan Harrison, David K. Jackson, Ian Johnston, Dominic Kwiatkowski, Cordelia Langford, John Sillitoe |
| EPI_ISL_532540 | NHSGGC West of Scotland Specialist Virology Centre / MRC-University of Glasgow Centre for Virus Research | Wellcome Sanger Institute for the COVID-19 Genomics UK Consortium | Ana da Silva Filipe, Natasha Johnson, Kathy Smollett, Daniel Mair, Stephen Carmichael, Lily Tong, Jenna Nichols, Elihu Aranday-Cortes, Kirstyn Brunker, Yasmin Parr, Kyriaki Nomikou; Sarah McDonald, Marc Niebel, Patawee Asamaphan; Richard Orton, Joseph Hughes, Sreenu Vattipally, David L Robertson; Alasdair MacLean, Rory Gunson; Kathy Li, Natasha Jesudason, Rajiv Shah, James Shepherd, Antonia Ho, Alice Broos, Emma Thomson and Alex Alderton, Roberto Amato, Sonia Goncalves, Ewan Harrison, David K. Jackson, Ian Johnston, Dominic Kwiatkowski, Cordelia Langford, John Sillitoe |
| EPI_ISL_532541, EPI_ISL_532542, EPI_ISL_532543, EPI_ISL_532544, EPI_ISL_532545, EPI_ISL_532546, EPI_ISL_532547, EPI_ISL_532548, EPI_ISL_532549, EPI_ISL_532550, EPI_ISL_532551, EPI_ISL_532552, EPI_ISL_532553, EPI_ISL_532554, EPI_ISL_532555, EPI_ISL_532556, EPI_ISL_532557, EPI_ISL_532558, EPI_ISL_532559, EPI_ISL_532560, EPI_ISL_532561, EPI_ISL_532562 | | | |
| see above | Lighthouse Lab in Glasgow | Wellcome Sanger Institute for the COVID-19 Genomics UK Consortium | Harper VanSteenhouse, Yumi Kasai, David Gray, Carol Clugston, Anna Dominiczak and Alex Alderton, Roberto Amato, Sonia Goncalves, Ewan Harrison, David K. Jackson, Ian Johnston, Dominic Kwiatkowski, Cordelia Langford, John Sillitoe |
| EPI_ISL_532563, EPI_ISL_532564 | NHSGGC West of Scotland Specialist Virology Centre / MRC-University of Glasgow Centre for Virus Research | Wellcome Sanger Institute for the COVID-19 Genomics UK Consortium | Ana da Silva Filipe, Natasha Johnson, Kathy Smollett, Daniel Mair, Stephen Carmichael, Lily Tong, Jenna Nichols, Elihu Aranday-Cortes, Kirstyn Brunker, Yasmin Parr, Kyriaki Nomikou; Sarah McDonald, Marc Niebel, Patawee Asamaphan; Richard Orton, Joseph Hughes, Sreenu Vattipally, David L Robertson; Alasdair MacLean, Rory Gunson; Kathy Li, Natasha Jesudason, Rajiv Shah, James Shepherd, Antonia Ho, Alice Broos, Emma Thomson and Alex Alderton, Roberto Amato, Sonia Goncalves, Ewan Harrison, David K. Jackson, Ian Johnston, Dominic Kwiatkowski, Cordelia Langford, John Sillitoe |
| EPI_ISL_532565, EPI_ISL_532566, EPI_ISL_532567, EPI_ISL_532568, EPI_ISL_532569, EPI_ISL_532570, EPI_ISL_532571 | Lighthouse Lab in Glasgow | Wellcome Sanger Institute for the COVID-19 Genomics UK Consortium | Harper VanSteenhouse, Yumi Kasai, David Gray, Carol Clugston, Anna Dominiczak and Alex Alderton, Roberto Amato, Sonia Goncalves, Ewan Harrison, David K. Jackson, Ian Johnston, Dominic Kwiatkowski, Cordelia Langford, John Sillitoe |
| EPI_ISL_532572 | NHSGGC West of Scotland Specialist Virology Centre / MRC-University of Glasgow Centre for Virus Research | Wellcome Sanger Institute for the COVID-19 Genomics UK Consortium | Ana da Silva Filipe, Natasha Johnson, Kathy Smollett, Daniel Mair, Stephen Carmichael, Lily Tong, Jenna Nichols, Elihu Aranday-Cortes, Kirstyn Brunker, Yasmin Parr, Kyriaki Nomikou; Sarah McDonald, Marc Niebel, Patawee Asamaphan; Richard Orton, Joseph Hughes, Sreenu Vattipally, David L Robertson; Alasdair MacLean, Rory Gunson; Kathy Li, Natasha Jesudason, Rajiv Shah, James Shepherd, Antonia Ho, Alice Broos, Emma Thomson and Alex Alderton, Roberto Amato, Sonia Goncalves, Ewan Harrison, David K. Jackson, Ian Johnston, Dominic Kwiatkowski, Cordelia Langford, John Sillitoe |
| EPI_ISL_532573, EPI_ISL_532574, EPI_ISL_532575, EPI_ISL_532576, EPI_ISL_532577, EPI_ISL_532578, EPI_ISL_532579, EPI_ISL_532580, EPI_ISL_532581 | Lighthouse Lab in Glasgow | Wellcome Sanger Institute for the COVID-19 Genomics UK Consortium | Harper VanSteenhouse, Yumi Kasai, David Gray, Carol Clugston, Anna Dominiczak and Alex Alderton, Roberto Amato, Sonia Goncalves, Ewan Harrison, David K. Jackson, Ian Johnston, Dominic Kwiatkowski, Cordelia Langford, John Sillitoe |
| EPI_ISL_532582 | NHSGGC West of Scotland Specialist Virology Centre / MRC-University of Glasgow Centre for Virus Research | Wellcome Sanger Institute for the COVID-19 Genomics UK Consortium | Ana da Silva Filipe, Natasha Johnson, Kathy Smollett, Daniel Mair, Stephen Carmichael, Lily Tong, Jenna Nichols, Elihu Aranday-Cortes, Kirstyn Brunker, Yasmin Parr, Kyriaki Nomikou; Sarah McDonald, Marc Niebel, Patawee Asamaphan; Richard Orton, Joseph Hughes, Sreenu Vattipally, David L Robertson; Alasdair MacLean, Rory Gunson; Kathy Li, Natasha Jesudason, Rajiv Shah, James Shepherd, Antonia Ho, Alice Broos, Emma Thomson and Alex Alderton, Roberto Amato, Sonia Goncalves, Ewan Harrison, David K. Jackson, Ian Johnston, Dominic Kwiatkowski, Cordelia Langford, John Sillitoe |
| EPI_ISL_532584, EPI_ISL_532585, EPI_ISL_532586 | Lighthouse Lab in Glasgow | Wellcome Sanger Institute for the COVID-19 Genomics UK | Harper VanSteenhouse, Yumi Kasai, David Gray, Carol Clugston, Anna Dominiczak and Alex Alderton, Roberto Amato, Sonia Goncalves, Ewan Harrison, David K. Jackson, Ian Johnston, Dominic Kwiatkowski, Cordelia |

| | | | |
|---|---|---|---|
| | | Consortium | Langford, John Sillitoe |
| EPI_ISL_532587 | NHSGGC West of Scotland Specialist Virology Centre / MRC-University of Glasgow Centre for Virus Research | Wellcome Sanger Institute for the COVID-19 Genomics UK Consortium | Ana da Silva Filipe, Natasha Johnson, Kathy Smollett, Daniel Mair, Stephen Carmichael, Lily Tong, Jenna Nichols, Elihu Aranday-Cortes, Kirstyn Brunker, Yasmin Parr, Kyriaki Nomikou; Sarah McDonald, Marc Niebel, Patawee Asamaphan; Richard Orton, Joseph Hughes, Sreenu Vattipally, David L Robertson; Alasdair MacLean, Rory Gunson; Kathy Li, Natasha Jesudason, Rajiv Shah, James Shepherd, Antonia Ho, Alice Broos, Emma Thomson and Alex Alderton, Roberto Amato, Sonia Goncalves, Ewan Harrison, David K. Jackson, Ian Johnston, Dominic Kwiatkowski, Cordelia Langford, John Sillitoe |
| EPI_ISL_532588, EPI_ISL_532589, EPI_ISL_532590, EPI_ISL_532591, EPI_ISL_532592, EPI_ISL_532593 | Lighthouse Lab in Glasgow | Wellcome Sanger Institute for the COVID-19 Genomics UK Consortium | Harper VanSteenhouse, Yumi Kasai, David Gray, Carol Clugston, Anna Dominiczak and Alex Alderton, Roberto Amato, Sonia Goncalves, Ewan Harrison, David K. Jackson, Ian Johnston, Dominic Kwiatkowski, Cordelia Langford, John Sillitoe |
| EPI_ISL_532594 | NHSGGC West of Scotland Specialist Virology Centre / MRC-University of Glasgow Centre for Virus Research | Wellcome Sanger Institute for the COVID-19 Genomics UK Consortium | Ana da Silva Filipe, Natasha Johnson, Kathy Smollett, Daniel Mair, Stephen Carmichael, Lily Tong, Jenna Nichols, Elihu Aranday-Cortes, Kirstyn Brunker, Yasmin Parr, Kyriaki Nomikou; Sarah McDonald, Marc Niebel, Patawee Asamaphan; Richard Orton, Joseph Hughes, Sreenu Vattipally, David L Robertson; Alasdair MacLean, Rory Gunson; Kathy Li, Natasha Jesudason, Rajiv Shah, James Shepherd, Antonia Ho, Alice Broos, Emma Thomson and Alex Alderton, Roberto Amato, Sonia Goncalves, Ewan Harrison, David K. Jackson, Ian Johnston, Dominic Kwiatkowski, Cordelia Langford, John Sillitoe |
| EPI_ISL_532595, EPI_ISL_532596, EPI_ISL_532597 | Lighthouse Lab in Glasgow | Wellcome Sanger Institute for the COVID-19 Genomics UK Consortium | Harper VanSteenhouse, Yumi Kasai, David Gray, Carol Clugston, Anna Dominiczak and Alex Alderton, Roberto Amato, Sonia Goncalves, Ewan Harrison, David K. Jackson, Ian Johnston, Dominic Kwiatkowski, Cordelia Langford, John Sillitoe |
| EPI_ISL_532598 | NHSGGC West of Scotland Specialist Virology Centre / MRC-University of Glasgow Centre for Virus Research | Wellcome Sanger Institute for the COVID-19 Genomics UK Consortium | Ana da Silva Filipe, Natasha Johnson, Kathy Smollett, Daniel Mair, Stephen Carmichael, Lily Tong, Jenna Nichols, Elihu Aranday-Cortes, Kirstyn Brunker, Yasmin Parr, Kyriaki Nomikou; Sarah McDonald, Marc Niebel, Patawee Asamaphan; Richard Orton, Joseph Hughes, Sreenu Vattipally, David L Robertson; Alasdair MacLean, Rory Gunson; Kathy Li, Natasha Jesudason, Rajiv Shah, James Shepherd, Antonia Ho, Alice Broos, Emma Thomson and Alex Alderton, Roberto Amato, Sonia Goncalves, Ewan Harrison, David K. Jackson, Ian Johnston, Dominic Kwiatkowski, Cordelia Langford, John Sillitoe |
| EPI_ISL_532599, EPI_ISL_532600, EPI_ISL_532601, EPI_ISL_532602, EPI_ISL_532603, EPI_ISL_532604, EPI_ISL_532605, EPI_ISL_532606, EPI_ISL_532607, EPI_ISL_532608, EPI_ISL_532609, EPI_ISL_532610, EPI_ISL_532611, EPI_ISL_532612, EPI_ISL_532613, EPI_ISL_532614, EPI_ISL_532615, EPI_ISL_532616, EPI_ISL_532617, EPI_ISL_532618, EPI_ISL_532619 | | | |
| see above | Lighthouse Lab in Glasgow | Wellcome Sanger Institute for the COVID-19 Genomics UK Consortium | Harper VanSteenhouse, Yumi Kasai, David Gray, Carol Clugston, Anna Dominiczak and Alex Alderton, Roberto Amato, Sonia Goncalves, Ewan Harrison, David K. Jackson, Ian Johnston, Dominic Kwiatkowski, Cordelia Langford, John Sillitoe |
| EPI_ISL_532620, EPI_ISL_532621 | NHSGGC West of Scotland Specialist Virology Centre / MRC-University of Glasgow Centre for Virus Research | Wellcome Sanger Institute for the COVID-19 Genomics UK Consortium | Ana da Silva Filipe, Natasha Johnson, Kathy Smollett, Daniel Mair, Stephen Carmichael, Lily Tong, Jenna Nichols, Elihu Aranday-Cortes, Kirstyn Brunker, Yasmin Parr, Kyriaki Nomikou; Sarah McDonald, Marc Niebel, Patawee Asamaphan; Richard Orton, Joseph Hughes, Sreenu Vattipally, David L Robertson; Alasdair MacLean, Rory Gunson; Kathy Li, Natasha Jesudason, Rajiv Shah, James Shepherd, Antonia Ho, Alice Broos, Emma Thomson and Alex Alderton, Roberto Amato, Sonia Goncalves, Ewan Harrison, David K. Jackson, Ian Johnston, Dominic Kwiatkowski, Cordelia Langford, John Sillitoe |
| EPI_ISL_532622, EPI_ISL_532623, EPI_ISL_532624 | Lighthouse Lab in Glasgow | Wellcome Sanger Institute for the COVID-19 Genomics UK Consortium | Harper VanSteenhouse, Yumi Kasai, David Gray, Carol Clugston, Anna Dominiczak and Alex Alderton, Roberto Amato, Sonia Goncalves, Ewan Harrison, David K. Jackson, Ian Johnston, Dominic Kwiatkowski, Cordelia Langford, John Sillitoe |
| EPI_ISL_532625, EPI_ISL_532626 | NHSGGC West of Scotland Specialist Virology Centre / MRC-University of Glasgow Centre for Virus Research | Wellcome Sanger Institute for the COVID-19 Genomics UK Consortium | Ana da Silva Filipe, Natasha Johnson, Kathy Smollett, Daniel Mair, Stephen Carmichael, Lily Tong, Jenna Nichols, Elihu Aranday-Cortes, Kirstyn Brunker, Yasmin Parr, Kyriaki Nomikou; Sarah McDonald, Marc Niebel, Patawee Asamaphan; Richard Orton, Joseph Hughes, Sreenu Vattipally, David L Robertson; Alasdair MacLean, Rory Gunson; Kathy Li, Natasha Jesudason, Rajiv Shah, James Shepherd, Antonia Ho, Alice Broos, Emma Thomson and Alex Alderton, Roberto Amato, Sonia Goncalves, Ewan Harrison, David K. Jackson, Ian Johnston, Dominic Kwiatkowski, Cordelia Langford, John Sillitoe |
| EPI_ISL_532627, EPI_ISL_532628, EPI_ISL_532629, EPI_ISL_532630, EPI_ISL_532631, EPI_ISL_532632 | Lighthouse Lab in Glasgow | Wellcome Sanger Institute for the COVID-19 Genomics UK Consortium | Harper VanSteenhouse, Yumi Kasai, David Gray, Carol Clugston, Anna Dominiczak and Alex Alderton, Roberto Amato, Sonia Goncalves, Ewan Harrison, David K. Jackson, Ian Johnston, Dominic Kwiatkowski, Cordelia Langford, John Sillitoe |
| EPI_ISL_532633 | NHSGGC West of Scotland Specialist Virology Centre / MRC-University of Glasgow Centre for Virus Research | Wellcome Sanger Institute for the COVID-19 Genomics UK Consortium | Ana da Silva Filipe, Natasha Johnson, Kathy Smollett, Daniel Mair, Stephen Carmichael, Lily Tong, Jenna Nichols, Elihu Aranday-Cortes, Kirstyn Brunker, Yasmin Parr, Kyriaki Nomikou; Sarah McDonald, Marc Niebel, Patawee Asamaphan; Richard Orton, Joseph Hughes, Sreenu Vattipally, David L Robertson; Alasdair MacLean, Rory Gunson; Kathy Li, Natasha Jesudason, Rajiv Shah, James Shepherd, Antonia Ho, Alice Broos, Emma Thomson and Alex Alderton, Roberto Amato, Sonia Goncalves, Ewan Harrison, David K. Jackson, Ian Johnston, Dominic Kwiatkowski, Cordelia Langford, John Sillitoe |
| EPI_ISL_532634, EPI_ISL_532635, EPI_ISL_532636, EPI_ISL_532637, EPI_ISL_532638 | Lighthouse Lab in Glasgow | Wellcome Sanger Institute for the COVID-19 Genomics UK Consortium | Harper VanSteenhouse, Yumi Kasai, David Gray, Carol Clugston, Anna Dominiczak and Alex Alderton, Roberto Amato, Sonia Goncalves, Ewan Harrison, David K. Jackson, Ian Johnston, Dominic Kwiatkowski, Cordelia Langford, John Sillitoe |
| EPI_ISL_532639 | NHSGGC West of Scotland Specialist Virology Centre / MRC-University of Glasgow Centre for Virus Research | Wellcome Sanger Institute for the COVID-19 Genomics UK Consortium | Ana da Silva Filipe, Natasha Johnson, Kathy Smollett, Daniel Mair, Stephen Carmichael, Lily Tong, Jenna Nichols, Elihu Aranday-Cortes, Kirstyn Brunker, Yasmin Parr, Kyriaki Nomikou; Sarah McDonald, Marc Niebel, Patawee Asamaphan; Richard Orton, Joseph Hughes, Sreenu Vattipally, David L Robertson; Alasdair MacLean, Rory Gunson; Kathy Li, Natasha Jesudason, Rajiv Shah, James Shepherd, Antonia Ho, Alice Broos, Emma Thomson and Alex Alderton, Roberto Amato, Sonia Goncalves, Ewan Harrison, David K. Jackson, Ian Johnston, Dominic Kwiatkowski, Cordelia Langford, John Sillitoe |
| EPI_ISL_532640, EPI_ISL_532641, EPI_ISL_532642, EPI_ISL_532643, EPI_ISL_532644, EPI_ISL_532645, EPI_ISL_532646, EPI_ISL_532647 | Lighthouse Lab in Glasgow | Wellcome Sanger Institute for the COVID-19 Genomics UK Consortium | Harper VanSteenhouse, Yumi Kasai, David Gray, Carol Clugston, Anna Dominiczak and Alex Alderton, Roberto Amato, Sonia Goncalves, Ewan Harrison, David K. Jackson, Ian Johnston, Dominic Kwiatkowski, Cordelia Langford, John Sillitoe |
| EPI_ISL_532648 | NHSGGC West of Scotland Specialist Virology Centre / MRC-University of Glasgow Centre for Virus Research | Wellcome Sanger Institute for the COVID-19 Genomics UK Consortium | Ana da Silva Filipe, Natasha Johnson, Kathy Smollett, Daniel Mair, Stephen Carmichael, Lily Tong, Jenna Nichols, Elihu Aranday-Cortes, Kirstyn Brunker, Yasmin Parr, Kyriaki Nomikou; Sarah McDonald, Marc Niebel, Patawee Asamaphan; Richard Orton, Joseph Hughes, Sreenu Vattipally, David L Robertson; Alasdair MacLean, Rory Gunson; Kathy Li, Natasha Jesudason, Rajiv Shah, James Shepherd, Antonia Ho, Alice Broos, Emma Thomson and Alex Alderton, Roberto Amato, Sonia Goncalves, Ewan Harrison, David K. Jackson, Ian Johnston, Dominic Kwiatkowski, Cordelia Langford, John Sillitoe |
| EPI_ISL_532649, EPI_ISL_532650, EPI_ISL_532651, EPI_ISL_532652, EPI_ISL_532653, EPI_ISL_532654, EPI_ISL_532655, EPI_ISL_532656, EPI_ISL_532657, EPI_ISL_532658 | Lighthouse Lab in Glasgow | Wellcome Sanger Institute for the COVID-19 Genomics UK Consortium | Harper VanSteenhouse, Yumi Kasai, David Gray, Carol Clugston, Anna Dominiczak and Alex Alderton, Roberto Amato, Sonia Goncalves, Ewan Harrison, David K. Jackson, Ian Johnston, Dominic Kwiatkowski, Cordelia Langford, John Sillitoe |
| EPI_ISL_532659 | Virology Department, Royal Infirmary of Edinburgh, NHS Lothian / School of Biological Sciences, University of Edinburgh | Wellcome Sanger Institute for the COVID-19 Genomics UK Consortium | McHugh M, Dewar R, Rooke S, O'Toole Á, Scher E, Hill V, McCrone JT, Colquhoun R, Yu X, Jackson B, Rambaut A, Templeton K and Alex Alderton, Roberto Amato, Sonia Goncalves, Ewan Harrison, David K. Jackson, Ian Johnston, Dominic Kwiatkowski, Cordelia Langford, John Sillitoe |
| EPI_ISL_532660, EPI_ISL_532661, EPI_ISL_532662, EPI_ISL_532663, EPI_ISL_532664, EPI_ISL_532665, EPI_ISL_532666, EPI_ISL_532667, EPI_ISL_532668, EPI_ISL_532669 | Lighthouse Lab in Glasgow | Wellcome Sanger Institute for the COVID-19 Genomics UK Consortium | Harper VanSteenhouse, Yumi Kasai, David Gray, Carol Clugston, Anna Dominiczak and Alex Alderton, Roberto Amato, Sonia Goncalves, Ewan Harrison, David K. Jackson, Ian Johnston, Dominic Kwiatkowski, Cordelia Langford, John Sillitoe |
| EPI_ISL_532670 | NHSGGC West of Scotland Specialist Virology Centre / MRC-University of Glasgow Centre for Virus Research | Wellcome Sanger Institute for the COVID-19 Genomics UK Consortium | Ana da Silva Filipe, Natasha Johnson, Kathy Smollett, Daniel Mair, Stephen Carmichael, Lily Tong, Jenna Nichols, Elihu Aranday-Cortes, Kirstyn Brunker, Yasmin Parr, Kyriaki Nomikou; Sarah McDonald, Marc Niebel, Patawee Asamaphan; Richard Orton, Joseph Hughes, Sreenu Vattipally, David L Robertson; Alasdair MacLean, Rory Gunson; Kathy Li, Natasha Jesudason, Rajiv Shah, James Shepherd, Antonia Ho, Alice Broos, Emma Thomson and Alex Alderton, Roberto Amato, Sonia Goncalves, Ewan Harrison, David K. Jackson, Ian Johnston, Dominic Kwiatkowski, Cordelia Langford, John Sillitoe |
| EPI_ISL_532671, EPI_ISL_532672, EPI_ISL_532673, EPI_ISL_532674, EPI_ISL_532675, EPI_ISL_532676, EPI_ISL_532677, EPI_ISL_532678, EPI_ISL_532679, EPI_ISL_532680, EPI_ISL_532681, EPI_ISL_532682 | | | |
| see above | Lighthouse Lab in Glasgow | Wellcome Sanger Institute for the COVID-19 Genomics UK Consortium | Harper VanSteenhouse, Yumi Kasai, David Gray, Carol Clugston, Anna Dominiczak and Alex Alderton, Roberto Amato, Sonia Goncalves, Ewan Harrison, David K. Jackson, Ian Johnston, Dominic Kwiatkowski, Cordelia Langford, John Sillitoe |
| EPI_ISL_532683 | Virology Department, Royal Infirmary of Edinburgh, NHS Lothian / School of Biological Sciences, University of Edinburgh | Wellcome Sanger Institute for the COVID-19 Genomics UK Consortium | McHugh M, Dewar R, Rooke S, O'Toole Á, Scher E, Hill V, McCrone JT, Colquhoun R, Yu X, Jackson B, Rambaut A, Templeton K and Alex Alderton, Roberto Amato, Sonia Goncalves, Ewan Harrison, David K. Jackson, Ian Johnston, Dominic Kwiatkowski, Cordelia Langford, John Sillitoe |
| EPI_ISL_532684, EPI_ISL_532685, EPI_ISL_532686, EPI_ISL_532687, EPI_ISL_532688 | Lighthouse Lab in Glasgow | Wellcome Sanger Institute for the COVID-19 Genomics UK Consortium | Harper VanSteenhouse, Yumi Kasai, David Gray, Carol Clugston, Anna Dominiczak and Alex Alderton, Roberto Amato, Sonia Goncalves, Ewan Harrison, David K. Jackson, Ian Johnston, Dominic Kwiatkowski, Cordelia Langford, John Sillitoe |
| EPI_ISL_532689 | Virology Department, Royal Infirmary of Edinburgh, NHS Lothian / School of Biological Sciences, University of Edinburgh | Wellcome Sanger Institute for the COVID-19 Genomics UK Consortium | McHugh M, Dewar R, Rooke S, O'Toole Á, Scher E, Hill V, McCrone JT, Colquhoun R, Yu X, Jackson B, Rambaut A, Templeton K and Alex Alderton, Roberto Amato, Sonia Goncalves, Ewan Harrison, David K. Jackson, Ian Johnston, Dominic Kwiatkowski, Cordelia Langford, John Sillitoe |
| EPI_ISL_532690, EPI_ISL_532691, EPI_ISL_532692, EPI_ISL_532693, EPI_ISL_532694, EPI_ISL_532695, EPI_ISL_532696 | Lighthouse Lab in Glasgow | Wellcome Sanger Institute for the COVID-19 Genomics UK Consortium | Harper VanSteenhouse, Yumi Kasai, David Gray, Carol Clugston, Anna Dominiczak and Alex Alderton, Roberto Amato, Sonia Goncalves, Ewan Harrison, David K. Jackson, Ian Johnston, Dominic Kwiatkowski, Cordelia Langford, John Sillitoe |
| EPI_ISL_532697 | NHSGGC West of Scotland Specialist Virology Centre / MRC-University of Glasgow Centre for Virus Research | Wellcome Sanger Institute for the COVID-19 Genomics UK Consortium | Ana da Silva Filipe, Natasha Johnson, Kathy Smollett, Daniel Mair, Stephen Carmichael, Lily Tong, Jenna Nichols, Elihu Aranday-Cortes, Kirstyn Brunker, Yasmin Parr, Kyriaki Nomikou; Sarah McDonald, Marc Niebel, Patawee Asamaphan; Richard Orton, Joseph Hughes, Sreenu Vattipally, David L Robertson; Alasdair MacLean, Rory Gunson; Kathy Li, Natasha Jesudason, Rajiv Shah, James Shepherd, Antonia Ho, Alice Broos, Emma Thomson and Alex Alderton, Roberto Amato, Sonia Goncalves, Ewan Harrison, David K. Jackson, Ian Johnston, Dominic Kwiatkowski, Cordelia Langford, John Sillitoe |
| EPI_ISL_532698, EPI_ISL_532699, EPI_ISL_532700, EPI_ISL_532701, EPI_ISL_532702, EPI_ISL_532703, EPI_ISL_532704 | Lighthouse Lab in Glasgow | Wellcome Sanger Institute for the COVID-19 Genomics UK Consortium | Harper VanSteenhouse, Yumi Kasai, David Gray, Carol Clugston, Anna Dominiczak and Alex Alderton, Roberto Amato, Sonia Goncalves, Ewan Harrison, David K. Jackson, Ian Johnston, Dominic Kwiatkowski, Cordelia Langford, John Sillitoe |
| EPI_ISL_532705 | Virology Department, Royal Infirmary of Edinburgh, NHS Lothian / School of Biological Sciences, University of Edinburgh | Wellcome Sanger Institute for the COVID-19 Genomics UK Consortium | McHugh M, Dewar R, Rooke S, O'Toole Á, Scher E, Hill V, McCrone JT, Colquhoun R, Yu X, Jackson B, Rambaut A, Templeton K and Alex Alderton, Roberto Amato, Sonia Goncalves, Ewan Harrison, David K. Jackson, Ian Johnston, Dominic Kwiatkowski, Cordelia Langford, John Sillitoe |
| EPI_ISL_532706, EPI_ISL_532707, EPI_ISL_532708, EPI_ISL_532709, EPI_ISL_532710, EPI_ISL_532711, EPI_ISL_532712, EPI_ISL_532713, EPI_ISL_532714 | Lighthouse Lab in Glasgow | Wellcome Sanger Institute for the COVID-19 Genomics UK Consortium | Harper VanSteenhouse, Yumi Kasai, David Gray, Carol Clugston, Anna Dominiczak and Alex Alderton, Roberto Amato, Sonia Goncalves, Ewan Harrison, David K. Jackson, Ian Johnston, Dominic Kwiatkowski, Cordelia Langford, John Sillitoe |
| EPI_ISL_532715 | NHSGGC West of Scotland Specialist Virology Centre / MRC-University of Glasgow Centre for Virus Research | Wellcome Sanger Institute for the COVID-19 Genomics UK Consortium | Ana da Silva Filipe, Natasha Johnson, Kathy Smollett, Daniel Mair, Stephen Carmichael, Lily Tong, Jenna Nichols, Elihu Aranday-Cortes, Kirstyn Brunker, Yasmin Parr, Kyriaki Nomikou; Sarah McDonald, Marc Niebel, Patawee Asamaphan; Richard Orton, Joseph Hughes, Sreenu Vattipally, David L Robertson; Alasdair MacLean, Rory Gunson; Kathy Li, Natasha Jesudason, Rajiv Shah, James Shepherd, Antonia Ho, Alice Broos, Emma Thomson and Alex Alderton, Roberto Amato, Sonia Goncalves, Ewan Harrison, David K. Jackson, Ian Johnston, Dominic Kwiatkowski, Cordelia Langford, John Sillitoe |
| EPI_ISL_532716, EPI_ISL_532717, EPI_ISL_532718, EPI_ISL_532719, EPI_ISL_532720, EPI_ISL_532721, EPI_ISL_532722, EPI_ISL_532723, EPI_ISL_532724, EPI_ISL_532725, EPI_ISL_532726, EPI_ISL_532727, EPI_ISL_532728, EPI_ISL_532729, EPI_ISL_532730, EPI_ISL_532731, EPI_ISL_532732 | | | |
| see above | Lighthouse Lab in Glasgow | Wellcome Sanger Institute for the COVID-19 Genomics UK Consortium | Harper VanSteenhouse, Yumi Kasai, David Gray, Carol Clugston, Anna Dominiczak and Alex Alderton, Roberto Amato, Sonia Goncalves, Ewan Harrison, David K. Jackson, Ian Johnston, Dominic Kwiatkowski, Cordelia Langford, John Sillitoe |
| EPI_ISL_532733 | NHSGGC West of Scotland Specialist Virology Centre / MRC-University of Glasgow Centre for Virus Research | Wellcome Sanger Institute for the COVID-19 Genomics UK Consortium | Ana da Silva Filipe, Natasha Johnson, Kathy Smollett, Daniel Mair, Stephen Carmichael, Lily Tong, Jenna Nichols, Elihu Aranday-Cortes, Kirstyn Brunker, Yasmin Parr, Kyriaki Nomikou; Sarah McDonald, Marc Niebel, Patawee Asamaphan; Richard Orton, Joseph Hughes, Sreenu Vattipally, David L Robertson; Alasdair MacLean, Rory Gunson; Kathy Li, Natasha Jesudason, Rajiv Shah, James Shepherd, Antonia Ho, Alice Broos, Emma Thomson and Alex Alderton, Roberto Amato, Sonia Goncalves, Ewan Harrison, David K. Jackson, Ian Johnston, Dominic Kwiatkowski, Cordelia Langford, John Sillitoe |
| EPI_ISL_532734, EPI_ISL_532735, EPI_ISL_532736, EPI_ISL_532737 | Lighthouse Lab in Glasgow | Wellcome Sanger Institute for the COVID-19 Genomics UK Consortium | Harper VanSteenhouse, Yumi Kasai, David Gray, Carol Clugston, Anna Dominiczak and Alex Alderton, Roberto Amato, Sonia Goncalves, Ewan Harrison, David K. Jackson, Ian Johnston, Dominic Kwiatkowski, Cordelia Langford, John Sillitoe |
| EPI_ISL_532738 | NHSGGC West of Scotland Specialist Virology Centre / MRC-University of Glasgow Centre for Virus Research | Wellcome Sanger Institute for the COVID-19 Genomics UK Consortium | Ana da Silva Filipe, Natasha Johnson, Kathy Smollett, Daniel Mair, Stephen Carmichael, Lily Tong, Jenna Nichols, Elihu Aranday-Cortes, Kirstyn Brunker, Yasmin Parr, Kyriaki Nomikou; Sarah McDonald, Marc Niebel, Patawee Asamaphan; Richard Orton, Joseph Hughes, Sreenu Vattipally, David L Robertson; Alasdair MacLean, Rory Gunson; Kathy Li, Natasha Jesudason, Rajiv Shah, James Shepherd, Antonia Ho, Alice Broos, Emma Thomson and Alex Alderton, Roberto Amato, Sonia Goncalves, Ewan Harrison, David K. Jackson, Ian Johnston, Dominic Kwiatkowski, Cordelia Langford, John Sillitoe |
| EPI_ISL_532739, EPI_ISL_532740, EPI_ISL_532741, EPI_ISL_532742, EPI_ISL_532743 | Lighthouse Lab in Glasgow | Wellcome Sanger Institute for the COVID-19 Genomics UK Consortium | Harper VanSteenhouse, Yumi Kasai, David Gray, Carol Clugston, Anna Dominiczak and Alex Alderton, Roberto Amato, Sonia Goncalves, Ewan Harrison, David K. Jackson, Ian Johnston, Dominic Kwiatkowski, Cordelia Langford, John Sillitoe |

| | | | |
|---|---|---|---|
| EPI_ISL_532744 | NHSGGC West of Scotland Specialist Virology Centre / MRC-University of Glasgow Centre for Virus Research | Wellcome Sanger Institute for the COVID-19 Genomics UK Consortium | Ana da Silva Filipe, Natasha Johnson, Kathy Smollett, Daniel Mair, Stephen Carmichael, Lily Tong, Jenna Nichols, Elihu Aranday-Cortes, Kirstyn Brunker, Yasmin Parr, Kyriaki Nomikou; Sarah McDonald, Marc Niebel, Patawee Asamaphan; Richard Orton, Joseph Hughes, Sreenu Vattipally, David L Robertson; Alasdair MacLean, Rory Gunson; Kathy Li, Natasha Jesudason, Rajiv Shah, James Shepherd, Antonia Ho, Alice Broos, Emma Thomson and Alex Alderton, Roberto Amato, Sonia Goncalves, Ewan Harrison, David K. Jackson, Ian Johnston, Dominic Kwiatkowski, Cordelia Langford, John Sillitoe |
| EPI_ISL_532745 | Lighthouse Lab in Glasgow | Wellcome Sanger Institute for the COVID-19 Genomics UK Consortium | Harper VanSteenhouse, Yumi Kasai, David Gray, Carol Clugston, Anna Dominiczak and Alex Alderton, Roberto Amato, Sonia Goncalves, Ewan Harrison, David K. Jackson, Ian Johnston, Dominic Kwiatkowski, Cordelia Langford, John Sillitoe |
| EPI_ISL_532746 | NHSGGC West of Scotland Specialist Virology Centre / MRC-University of Glasgow Centre for Virus Research | Wellcome Sanger Institute for the COVID-19 Genomics UK Consortium | Ana da Silva Filipe, Natasha Johnson, Kathy Smollett, Daniel Mair, Stephen Carmichael, Lily Tong, Jenna Nichols, Elihu Aranday-Cortes, Kirstyn Brunker, Yasmin Parr, Kyriaki Nomikou; Sarah McDonald, Marc Niebel, Patawee Asamaphan; Richard Orton, Joseph Hughes, Sreenu Vattipally, David L Robertson; Alasdair MacLean, Rory Gunson; Kathy Li, Natasha Jesudason, Rajiv Shah, James Shepherd, Antonia Ho, Alice Broos, Emma Thomson and Alex Alderton, Roberto Amato, Sonia Goncalves, Ewan Harrison, David K. Jackson, Ian Johnston, Dominic Kwiatkowski, Cordelia Langford, John Sillitoe |
| EPI_ISL_532747, EPI_ISL_532748, EPI_ISL_532749 | Lighthouse Lab in Glasgow | Wellcome Sanger Institute for the COVID-19 Genomics UK Consortium | Harper VanSteenhouse, Yumi Kasai, David Gray, Carol Clugston, Anna Dominiczak and Alex Alderton, Roberto Amato, Sonia Goncalves, Ewan Harrison, David K. Jackson, Ian Johnston, Dominic Kwiatkowski, Cordelia Langford, John Sillitoe |
| EPI_ISL_532750 | NHSGGC West of Scotland Specialist Virology Centre / MRC-University of Glasgow Centre for Virus Research | Wellcome Sanger Institute for the COVID-19 Genomics UK Consortium | Ana da Silva Filipe, Natasha Johnson, Kathy Smollett, Daniel Mair, Stephen Carmichael, Lily Tong, Jenna Nichols, Elihu Aranday-Cortes, Kirstyn Brunker, Yasmin Parr, Kyriaki Nomikou; Sarah McDonald, Marc Niebel, Patawee Asamaphan; Richard Orton, Joseph Hughes, Sreenu Vattipally, David L Robertson; Alasdair MacLean, Rory Gunson; Kathy Li, Natasha Jesudason, Rajiv Shah, James Shepherd, Antonia Ho, Alice Broos, Emma Thomson and Alex Alderton, Roberto Amato, Sonia Goncalves, Ewan Harrison, David K. Jackson, Ian Johnston, Dominic Kwiatkowski, Cordelia Langford, John Sillitoe |
| EPI_ISL_532751, EPI_ISL_532752 | Lighthouse Lab in Glasgow | Wellcome Sanger Institute for the COVID-19 Genomics UK Consortium | Harper VanSteenhouse, Yumi Kasai, David Gray, Carol Clugston, Anna Dominiczak and Alex Alderton, Roberto Amato, Sonia Goncalves, Ewan Harrison, David K. Jackson, Ian Johnston, Dominic Kwiatkowski, Cordelia Langford, John Sillitoe |
| EPI_ISL_532753 | NHSGGC West of Scotland Specialist Virology Centre / MRC-University of Glasgow Centre for Virus Research | Wellcome Sanger Institute for the COVID-19 Genomics UK Consortium | Ana da Silva Filipe, Natasha Johnson, Kathy Smollett, Daniel Mair, Stephen Carmichael, Lily Tong, Jenna Nichols, Elihu Aranday-Cortes, Kirstyn Brunker, Yasmin Parr, Kyriaki Nomikou; Sarah McDonald, Marc Niebel, Patawee Asamaphan; Richard Orton, Joseph Hughes, Sreenu Vattipally, David L Robertson; Alasdair MacLean, Rory Gunson; Kathy Li, Natasha Jesudason, Rajiv Shah, James Shepherd, Antonia Ho, Alice Broos, Emma Thomson and Alex Alderton, Roberto Amato, Sonia Goncalves, Ewan Harrison, David K. Jackson, Ian Johnston, Dominic Kwiatkowski, Cordelia Langford, John Sillitoe |
| EPI_ISL_532754 | Lighthouse Lab in Glasgow | Wellcome Sanger Institute for the COVID-19 Genomics UK Consortium | Harper VanSteenhouse, Yumi Kasai, David Gray, Carol Clugston, Anna Dominiczak and Alex Alderton, Roberto Amato, Sonia Goncalves, Ewan Harrison, David K. Jackson, Ian Johnston, Dominic Kwiatkowski, Cordelia Langford, John Sillitoe |
| EPI_ISL_532755 | NHSGGC West of Scotland Specialist Virology Centre / MRC-University of Glasgow Centre for Virus Research | Wellcome Sanger Institute for the COVID-19 Genomics UK Consortium | Ana da Silva Filipe, Natasha Johnson, Kathy Smollett, Daniel Mair, Stephen Carmichael, Lily Tong, Jenna Nichols, Elihu Aranday-Cortes, Kirstyn Brunker, Yasmin Parr, Kyriaki Nomikou; Sarah McDonald, Marc Niebel, Patawee Asamaphan; Richard Orton, Joseph Hughes, Sreenu Vattipally, David L Robertson; Alasdair MacLean, Rory Gunson; Kathy Li, Natasha Jesudason, Rajiv Shah, James Shepherd, Antonia Ho, Alice Broos, Emma Thomson and Alex Alderton, Roberto Amato, Sonia Goncalves, Ewan Harrison, David K. Jackson, Ian Johnston, Dominic Kwiatkowski, Cordelia Langford, John Sillitoe |
| EPI_ISL_532756, EPI_ISL_532757, EPI_ISL_532758, EPI_ISL_532759, EPI_ISL_532760, EPI_ISL_532761, EPI_ISL_532762, EPI_ISL_532763, EPI_ISL_532764 | Lighthouse Lab in Glasgow | Wellcome Sanger Institute for the COVID-19 Genomics UK Consortium | Harper VanSteenhouse, Yumi Kasai, David Gray, Carol Clugston, Anna Dominiczak and Alex Alderton, Roberto Amato, Sonia Goncalves, Ewan Harrison, David K. Jackson, Ian Johnston, Dominic Kwiatkowski, Cordelia Langford, John Sillitoe |
| EPI_ISL_532765 | NHSGGC West of Scotland Specialist Virology Centre / MRC-University of Glasgow Centre for Virus Research | Wellcome Sanger Institute for the COVID-19 Genomics UK Consortium | Ana da Silva Filipe, Natasha Johnson, Kathy Smollett, Daniel Mair, Stephen Carmichael, Lily Tong, Jenna Nichols, Elihu Aranday-Cortes, Kirstyn Brunker, Yasmin Parr, Kyriaki Nomikou; Sarah McDonald, Marc Niebel, Patawee Asamaphan; Richard Orton, Joseph Hughes, Sreenu Vattipally, David L Robertson; Alasdair MacLean, Rory Gunson; Kathy Li, Natasha Jesudason, Rajiv Shah, James Shepherd, Antonia Ho, Alice Broos, Emma Thomson and Alex Alderton, Roberto Amato, Sonia Goncalves, Ewan Harrison, David K. Jackson, Ian Johnston, Dominic Kwiatkowski, Cordelia Langford, John Sillitoe |
| EPI_ISL_532766 | Lighthouse Lab in Glasgow | Wellcome Sanger Institute for the COVID-19 Genomics UK Consortium | Harper VanSteenhouse, Yumi Kasai, David Gray, Carol Clugston, Anna Dominiczak and Alex Alderton, Roberto Amato, Sonia Goncalves, Ewan Harrison, David K. Jackson, Ian Johnston, Dominic Kwiatkowski, Cordelia Langford, John Sillitoe |
| EPI_ISL_532767 | NHSGGC West of Scotland Specialist Virology Centre / MRC-University of Glasgow Centre for Virus Research | Wellcome Sanger Institute for the COVID-19 Genomics UK Consortium | Ana da Silva Filipe, Natasha Johnson, Kathy Smollett, Daniel Mair, Stephen Carmichael, Lily Tong, Jenna Nichols, Elihu Aranday-Cortes, Kirstyn Brunker, Yasmin Parr, Kyriaki Nomikou; Sarah McDonald, Marc Niebel, Patawee Asamaphan; Richard Orton, Joseph Hughes, Sreenu Vattipally, David L Robertson; Alasdair MacLean, Rory Gunson; Kathy Li, Natasha Jesudason, Rajiv Shah, James Shepherd, Antonia Ho, Alice Broos, Emma Thomson and Alex Alderton, Roberto Amato, Sonia Goncalves, Ewan Harrison, David K. Jackson, Ian Johnston, Dominic Kwiatkowski, Cordelia Langford, John Sillitoe |
| EPI_ISL_532768, EPI_ISL_532769, EPI_ISL_532770, EPI_ISL_532771, EPI_ISL_532772, EPI_ISL_532773, EPI_ISL_532774, EPI_ISL_532775, EPI_ISL_532776, EPI_ISL_532777, EPI_ISL_532778, EPI_ISL_532779, EPI_ISL_532780, EPI_ISL_532781, EPI_ISL_532782, EPI_ISL_532783 | | | |
| see above | Lighthouse Lab in Glasgow | Wellcome Sanger Institute for the COVID-19 Genomics UK Consortium | Harper VanSteenhouse, Yumi Kasai, David Gray, Carol Clugston, Anna Dominiczak and Alex Alderton, Roberto Amato, Sonia Goncalves, Ewan Harrison, David K. Jackson, Ian Johnston, Dominic Kwiatkowski, Cordelia Langford, John Sillitoe |
| EPI_ISL_532784 | NHSGGC West of Scotland Specialist Virology Centre / MRC-University of Glasgow Centre for Virus Research | Wellcome Sanger Institute for the COVID-19 Genomics UK Consortium | Ana da Silva Filipe, Natasha Johnson, Kathy Smollett, Daniel Mair, Stephen Carmichael, Lily Tong, Jenna Nichols, Elihu Aranday-Cortes, Kirstyn Brunker, Yasmin Parr, Kyriaki Nomikou; Sarah McDonald, Marc Niebel, Patawee Asamaphan; Richard Orton, Joseph Hughes, Sreenu Vattipally, David L Robertson; Alasdair MacLean, Rory Gunson; Kathy Li, Natasha Jesudason, Rajiv Shah, James Shepherd, Antonia Ho, Alice Broos, Emma Thomson and Alex Alderton, Roberto Amato, Sonia Goncalves, Ewan Harrison, David K. Jackson, Ian Johnston, Dominic Kwiatkowski, Cordelia Langford, John Sillitoe |
| EPI_ISL_532785 | Lighthouse Lab in Glasgow | Wellcome Sanger Institute for the COVID-19 Genomics UK Consortium | Harper VanSteenhouse, Yumi Kasai, David Gray, Carol Clugston, Anna Dominiczak and Alex Alderton, Roberto Amato, Sonia Goncalves, Ewan Harrison, David K. Jackson, Ian Johnston, Dominic Kwiatkowski, Cordelia Langford, John Sillitoe |
| EPI_ISL_532786 | NHSGGC West of Scotland Specialist Virology Centre / MRC-University of Glasgow Centre for Virus Research | Wellcome Sanger Institute for the COVID-19 Genomics UK Consortium | Ana da Silva Filipe, Natasha Johnson, Kathy Smollett, Daniel Mair, Stephen Carmichael, Lily Tong, Jenna Nichols, Elihu Aranday-Cortes, Kirstyn Brunker, Yasmin Parr, Kyriaki Nomikou; Sarah McDonald, Marc Niebel, Patawee Asamaphan; Richard Orton, Joseph Hughes, Sreenu Vattipally, David L Robertson; Alasdair MacLean, Rory Gunson; Kathy Li, Natasha Jesudason, Rajiv Shah, James Shepherd, Antonia Ho, Alice Broos, Emma Thomson and Alex Alderton, Roberto Amato, Sonia Goncalves, Ewan Harrison, David K. Jackson, Ian Johnston, Dominic Kwiatkowski, Cordelia Langford, John Sillitoe |
| EPI_ISL_532787, EPI_ISL_532788, EPI_ISL_532789, EPI_ISL_532790 | Lighthouse Lab in Glasgow | Wellcome Sanger Institute for the COVID-19 Genomics UK Consortium | Harper VanSteenhouse, Yumi Kasai, David Gray, Carol Clugston, Anna Dominiczak and Alex Alderton, Roberto Amato, Sonia Goncalves, Ewan Harrison, David K. Jackson, Ian Johnston, Dominic Kwiatkowski, Cordelia Langford, John Sillitoe |
| EPI_ISL_532791 | NHSGGC West of Scotland Specialist Virology Centre / MRC-University of Glasgow Centre for Virus Research | Wellcome Sanger Institute for the COVID-19 Genomics UK Consortium | Ana da Silva Filipe, Natasha Johnson, Kathy Smollett, Daniel Mair, Stephen Carmichael, Lily Tong, Jenna Nichols, Elihu Aranday-Cortes, Kirstyn Brunker, Yasmin Parr, Kyriaki Nomikou; Sarah McDonald, Marc Niebel, Patawee Asamaphan; Richard Orton, Joseph Hughes, Sreenu Vattipally, David L Robertson; Alasdair MacLean, Rory Gunson; Kathy Li, Natasha Jesudason, Rajiv Shah, James Shepherd, Antonia Ho, Alice Broos, Emma Thomson and Alex Alderton, Roberto Amato, Sonia Goncalves, Ewan Harrison, David K. Jackson, Ian Johnston, Dominic Kwiatkowski, Cordelia Langford, John Sillitoe |
| EPI_ISL_532792, EPI_ISL_532793, EPI_ISL_532794, EPI_ISL_532795, EPI_ISL_532796, EPI_ISL_532797, EPI_ISL_532798, EPI_ISL_532799, EPI_ISL_532800, EPI_ISL_532801, EPI_ISL_532802, EPI_ISL_532803, EPI_ISL_532804, EPI_ISL_532805, EPI_ISL_532806, EPI_ISL_532807, EPI_ISL_532808, EPI_ISL_532809, EPI_ISL_532810 | | | |
| see above | Lighthouse Lab in Glasgow | Wellcome Sanger Institute for the COVID-19 Genomics UK Consortium | Harper VanSteenhouse, Yumi Kasai, David Gray, Carol Clugston, Anna Dominiczak and Alex Alderton, Roberto Amato, Sonia Goncalves, Ewan Harrison, David K. Jackson, Ian Johnston, Dominic Kwiatkowski, Cordelia Langford, John Sillitoe |
| EPI_ISL_532811 | NHSGGC West of Scotland Specialist Virology Centre / MRC-University of Glasgow Centre for Virus Research | Wellcome Sanger Institute for the COVID-19 Genomics UK Consortium | Ana da Silva Filipe, Natasha Johnson, Kathy Smollett, Daniel Mair, Stephen Carmichael, Lily Tong, Jenna Nichols, Elihu Aranday-Cortes, Kirstyn Brunker, Yasmin Parr, Kyriaki Nomikou; Sarah McDonald, Marc Niebel, Patawee Asamaphan; Richard Orton, Joseph Hughes, Sreenu Vattipally, David L Robertson; Alasdair MacLean, Rory Gunson; Kathy Li, Natasha Jesudason, Rajiv Shah, James Shepherd, Antonia Ho, Alice Broos, Emma Thomson and Alex Alderton, Roberto Amato, Sonia Goncalves, Ewan Harrison, David K. Jackson, Ian Johnston, Dominic Kwiatkowski, Cordelia Langford, John Sillitoe |
| EPI_ISL_532812, EPI_ISL_532813, EPI_ISL_532814, EPI_ISL_532815, EPI_ISL_532816, EPI_ISL_532817, EPI_ISL_532818, EPI_ISL_532819, EPI_ISL_532820, EPI_ISL_532821, EPI_ISL_532822, EPI_ISL_532823, EPI_ISL_532824, EPI_ISL_532825, EPI_ISL_532826, EPI_ISL_532827, EPI_ISL_532828, EPI_ISL_532829, EPI_ISL_532830, EPI_ISL_532831, EPI_ISL_532832, EPI_ISL_532833, EPI_ISL_532834 | | | |
| see above | Lighthouse Lab in Glasgow | Wellcome Sanger Institute for the COVID-19 Genomics UK Consortium | Harper VanSteenhouse, Yumi Kasai, David Gray, Carol Clugston, Anna Dominiczak and Alex Alderton, Roberto Amato, Sonia Goncalves, Ewan Harrison, David K. Jackson, Ian Johnston, Dominic Kwiatkowski, Cordelia Langford, John Sillitoe |
| EPI_ISL_532835 | NHSGGC West of Scotland Specialist Virology Centre / MRC-University of Glasgow Centre for Virus Research | Wellcome Sanger Institute for the COVID-19 Genomics UK Consortium | Ana da Silva Filipe, Natasha Johnson, Kathy Smollett, Daniel Mair, Stephen Carmichael, Lily Tong, Jenna Nichols, Elihu Aranday-Cortes, Kirstyn Brunker, Yasmin Parr, Kyriaki Nomikou; Sarah McDonald, Marc Niebel, Patawee Asamaphan; Richard Orton, Joseph Hughes, Sreenu Vattipally, David L Robertson; Alasdair MacLean, Rory Gunson; Kathy Li, Natasha Jesudason, Rajiv Shah, James Shepherd, Antonia Ho, Alice Broos, Emma Thomson and Alex Alderton, Roberto Amato, Sonia Goncalves, Ewan Harrison, David K. Jackson, Ian Johnston, Dominic Kwiatkowski, Cordelia Langford, John Sillitoe |
| EPI_ISL_532836, EPI_ISL_532837 | Lighthouse Lab in Glasgow | Wellcome Sanger Institute for the COVID-19 Genomics UK Consortium | Harper VanSteenhouse, Yumi Kasai, David Gray, Carol Clugston, Anna Dominiczak and Alex Alderton, Roberto Amato, Sonia Goncalves, Ewan Harrison, David K. Jackson, Ian Johnston, Dominic Kwiatkowski, Cordelia Langford, John Sillitoe |
| EPI_ISL_532838 | NHSGGC West of Scotland Specialist Virology Centre / MRC-University of Glasgow Centre for Virus Research | Wellcome Sanger Institute for the COVID-19 Genomics UK Consortium | Ana da Silva Filipe, Natasha Johnson, Kathy Smollett, Daniel Mair, Stephen Carmichael, Lily Tong, Jenna Nichols, Elihu Aranday-Cortes, Kirstyn Brunker, Yasmin Parr, Kyriaki Nomikou; Sarah McDonald, Marc Niebel, Patawee Asamaphan; Richard Orton, Joseph Hughes, Sreenu Vattipally, David L Robertson; Alasdair MacLean, Rory Gunson; Kathy Li, Natasha Jesudason, Rajiv Shah, James Shepherd, Antonia Ho, Alice Broos, Emma Thomson and Alex Alderton, Roberto Amato, Sonia Goncalves, Ewan Harrison, David K. Jackson, Ian Johnston, Dominic Kwiatkowski, Cordelia Langford, John Sillitoe |
| EPI_ISL_532839, EPI_ISL_532840, EPI_ISL_532841, EPI_ISL_532842, EPI_ISL_532843 | Lighthouse Lab in Glasgow | Wellcome Sanger Institute for the COVID-19 Genomics UK Consortium | Harper VanSteenhouse, Yumi Kasai, David Gray, Carol Clugston, Anna Dominiczak and Alex Alderton, Roberto Amato, Sonia Goncalves, Ewan Harrison, David K. Jackson, Ian Johnston, Dominic Kwiatkowski, Cordelia Langford, John Sillitoe |
| EPI_ISL_532844 | NHSGGC West of Scotland Specialist Virology Centre / MRC-University of Glasgow Centre for Virus Research | Wellcome Sanger Institute for the COVID-19 Genomics UK Consortium | Ana da Silva Filipe, Natasha Johnson, Kathy Smollett, Daniel Mair, Stephen Carmichael, Lily Tong, Jenna Nichols, Elihu Aranday-Cortes, Kirstyn Brunker, Yasmin Parr, Kyriaki Nomikou; Sarah McDonald, Marc Niebel, Patawee Asamaphan; Richard Orton, Joseph Hughes, Sreenu Vattipally, David L Robertson; Alasdair MacLean, Rory Gunson; Kathy Li, Natasha Jesudason, Rajiv Shah, James Shepherd, Antonia Ho, Alice Broos, Emma Thomson and Alex Alderton, Roberto Amato, Sonia Goncalves, Ewan Harrison, David K. Jackson, Ian Johnston, Dominic Kwiatkowski, Cordelia Langford, John Sillitoe |
| EPI_ISL_532845, EPI_ISL_532846, EPI_ISL_532847, EPI_ISL_532848, EPI_ISL_532849, EPI_ISL_532850, EPI_ISL_532852, EPI_ISL_532853, EPI_ISL_532854, EPI_ISL_532855, EPI_ISL_532856, EPI_ISL_532857 | | | |
| see above | Lighthouse Lab in Glasgow | Wellcome Sanger Institute for the COVID-19 Genomics UK Consortium | Harper VanSteenhouse, Yumi Kasai, David Gray, Carol Clugston, Anna Dominiczak and Alex Alderton, Roberto Amato, Sonia Goncalves, Ewan Harrison, David K. Jackson, Ian Johnston, Dominic Kwiatkowski, Cordelia Langford, John Sillitoe |
| EPI_ISL_532858 | NHSGGC West of Scotland Specialist Virology Centre / MRC-University of Glasgow Centre for Virus Research | Wellcome Sanger Institute for the COVID-19 Genomics UK Consortium | Ana da Silva Filipe, Natasha Johnson, Kathy Smollett, Daniel Mair, Stephen Carmichael, Lily Tong, Jenna Nichols, Elihu Aranday-Cortes, Kirstyn Brunker, Yasmin Parr, Kyriaki Nomikou; Sarah McDonald, Marc Niebel, Patawee Asamaphan; Richard Orton, Joseph Hughes, Sreenu Vattipally, David L Robertson; Alasdair MacLean, Rory Gunson; Kathy Li, Natasha Jesudason, Rajiv Shah, James Shepherd, Antonia Ho, Alice Broos, Emma Thomson and Alex Alderton, Roberto Amato, Sonia Goncalves, Ewan Harrison, David K. Jackson, Ian Johnston, Dominic Kwiatkowski, Cordelia Langford, John Sillitoe |
| EPI_ISL_532859, EPI_ISL_532860, EPI_ISL_532861, EPI_ISL_532862, EPI_ISL_532863 | Lighthouse Lab in Glasgow | Wellcome Sanger Institute for the COVID-19 Genomics UK Consortium | Harper VanSteenhouse, Yumi Kasai, David Gray, Carol Clugston, Anna Dominiczak and Alex Alderton, Roberto Amato, Sonia Goncalves, Ewan Harrison, David K. Jackson, Ian Johnston, Dominic Kwiatkowski, Cordelia Langford, John Sillitoe |
| EPI_ISL_532864, EPI_ISL_532865 | NHSGGC West of Scotland Specialist Virology Centre / MRC-University of Glasgow Centre for Virus Research | Wellcome Sanger Institute for the COVID-19 Genomics UK Consortium | Ana da Silva Filipe, Natasha Johnson, Kathy Smollett, Daniel Mair, Stephen Carmichael, Lily Tong, Jenna Nichols, Elihu Aranday-Cortes, Kirstyn Brunker, Yasmin Parr, Kyriaki Nomikou; Sarah McDonald, Marc Niebel, Patawee Asamaphan; Richard Orton, Joseph Hughes, Sreenu Vattipally, David L Robertson; Alasdair MacLean, Rory Gunson; Kathy Li, Natasha Jesudason, Rajiv Shah, James Shepherd, Antonia Ho, Alice Broos, Emma Thomson and Alex Alderton, Roberto Amato, Sonia Goncalves, Ewan Harrison, David K. Jackson, Ian Johnston, Dominic Kwiatkowski, Cordelia Langford, John Sillitoe |
| EPI_ISL_532866, EPI_ISL_532867, EPI_ISL_532868, EPI_ISL_532869, EPI_ISL_532870, EPI_ISL_532871, EPI_ISL_532872, EPI_ISL_532873 | Lighthouse Lab in Glasgow | Wellcome Sanger Institute for the COVID-19 Genomics UK Consortium | Harper VanSteenhouse, Yumi Kasai, David Gray, Carol Clugston, Anna Dominiczak and Alex Alderton, Roberto Amato, Sonia Goncalves, Ewan Harrison, David K. Jackson, Ian Johnston, Dominic Kwiatkowski, Cordelia Langford, John Sillitoe |
| EPI_ISL_532874, EPI_ISL_532875, EPI_ISL_532876 | NHSGGC West of Scotland Specialist Virology Centre / MRC-University of Glasgow Centre for Virus Research | Wellcome Sanger Institute for the COVID-19 Genomics UK Consortium | Ana da Silva Filipe, Natasha Johnson, Kathy Smollett, Daniel Mair, Stephen Carmichael, Lily Tong, Jenna Nichols, Elihu Aranday-Cortes, Kirstyn Brunker, Yasmin Parr, Kyriaki Nomikou; Sarah McDonald, Marc Niebel, Patawee Asamaphan; Richard Orton, Joseph Hughes, Sreenu Vattipally, David L Robertson; Alasdair MacLean, Rory Gunson; Kathy Li, Natasha Jesudason, Rajiv Shah, James Shepherd, Antonia Ho, Alice Broos, Emma Thomson and Alex Alderton, Roberto Amato, Sonia Goncalves, Ewan Harrison, David K. Jackson, Ian Johnston, Dominic Kwiatkowski, Cordelia Langford, John Sillitoe |
| EPI_ISL_532877 | Lighthouse Lab in Glasgow | Wellcome Sanger Institute for the COVID-19 Genomics UK Consortium | Harper VanSteenhouse, Yumi Kasai, David Gray, Carol Clugston, Anna Dominiczak and Alex Alderton, Roberto Amato, Sonia Goncalves, Ewan Harrison, David K. Jackson, Ian Johnston, Dominic Kwiatkowski, Cordelia Langford, John Sillitoe |
| EPI_ISL_532878 | NHSGGC West of Scotland Specialist Virology Centre / MRC-University of Glasgow Centre for Virus Research | Wellcome Sanger Institute for the COVID-19 Genomics UK Consortium | Ana da Silva Filipe, Natasha Johnson, Kathy Smollett, Daniel Mair, Stephen Carmichael, Lily Tong, Jenna Nichols, Elihu Aranday-Cortes, Kirstyn Brunker, Yasmin Parr, Kyriaki Nomikou; Sarah McDonald, Marc Niebel, Patawee Asamaphan; Richard Orton, Joseph Hughes, Sreenu Vattipally, David L Robertson; Alasdair MacLean, Rory Gunson; Kathy Li, Natasha Jesudason, Rajiv Shah, James Shepherd, Antonia Ho, Alice Broos, Emma |

| | | | |
|---|---|---|---|
| | | | Thomson and Alex Alderton, Roberto Amato, Sonia Goncalves, Ewan Harrison, David K. Jackson, Ian Johnston, Dominic Kwiatkowski, Cordelia Langford, John Sillitoe |
| EPI_ISL_532879, EPI_ISL_532880, EPI_ISL_532881, EPI_ISL_532882, EPI_ISL_532883, EPI_ISL_532884 | Lighthouse Lab in Glasgow | Wellcome Sanger Institute for the COVID-19 Genomics UK Consortium | Harper VanSteenhouse, Yumi Kasai, David Gray, Carol Clugston, Anna Dominiczak and Alex Alderton, Roberto Amato, Sonia Goncalves, Ewan Harrison, David K. Jackson, Ian Johnston, Dominic Kwiatkowski, Cordelia Langford, John Sillitoe |
| EPI_ISL_532885, EPI_ISL_532886 | NHSGGC West of Scotland Specialist Virology Centre / MRC-University of Glasgow Centre for Virus Research | Wellcome Sanger Institute for the COVID-19 Genomics UK Consortium | Ana da Silva Filipe, Natasha Johnson, Kathy Smollett, Daniel Mair, Stephen Carmichael, Lily Tong, Jenna Nichols, Elihu Aranday-Cortes, Kirstyn Brunker, Yasmin Parr, Kyriaki Nomikou; Sarah McDonald, Marc Niebel, Patawee Asamaphan; Richard Orton, Joseph Hughes, Sreenu Vattipally, David L Robertson; Alasdair MacLean, Rory Gunson; Kathy Li, Natasha Jesudason, Rajiv Shah, James Shepherd, Antonia Ho, Alice Broos, Emma Thomson and Alex Alderton, Roberto Amato, Sonia Goncalves, Ewan Harrison, David K. Jackson, Ian Johnston, Dominic Kwiatkowski, Cordelia Langford, John Sillitoe |
| EPI_ISL_532887, EPI_ISL_532888, EPI_ISL_532889, EPI_ISL_532890 | Lighthouse Lab in Glasgow | Wellcome Sanger Institute for the COVID-19 Genomics UK Consortium | Harper VanSteenhouse, Yumi Kasai, David Gray, Carol Clugston, Anna Dominiczak and Alex Alderton, Roberto Amato, Sonia Goncalves, Ewan Harrison, David K. Jackson, Ian Johnston, Dominic Kwiatkowski, Cordelia Langford, John Sillitoe |
| EPI_ISL_532891 | NHSGGC West of Scotland Specialist Virology Centre / MRC-University of Glasgow Centre for Virus Research | Wellcome Sanger Institute for the COVID-19 Genomics UK Consortium | Ana da Silva Filipe, Natasha Johnson, Kathy Smollett, Daniel Mair, Stephen Carmichael, Lily Tong, Jenna Nichols, Elihu Aranday-Cortes, Kirstyn Brunker, Yasmin Parr, Kyriaki Nomikou; Sarah McDonald, Marc Niebel, Patawee Asamaphan; Richard Orton, Joseph Hughes, Sreenu Vattipally, David L Robertson; Alasdair MacLean, Rory Gunson; Kathy Li, Natasha Jesudason, Rajiv Shah, James Shepherd, Antonia Ho, Alice Broos, Emma Thomson and Alex Alderton, Roberto Amato, Sonia Goncalves, Ewan Harrison, David K. Jackson, Ian Johnston, Dominic Kwiatkowski, Cordelia Langford, John Sillitoe |
| EPI_ISL_532892, EPI_ISL_532893 | Lighthouse Lab in Glasgow | Wellcome Sanger Institute for the COVID-19 Genomics UK Consortium | Harper VanSteenhouse, Yumi Kasai, David Gray, Carol Clugston, Anna Dominiczak and Alex Alderton, Roberto Amato, Sonia Goncalves, Ewan Harrison, David K. Jackson, Ian Johnston, Dominic Kwiatkowski, Cordelia Langford, John Sillitoe |
| EPI_ISL_532894 | NHSGGC West of Scotland Specialist Virology Centre / MRC-University of Glasgow Centre for Virus Research | Wellcome Sanger Institute for the COVID-19 Genomics UK Consortium | Ana da Silva Filipe, Natasha Johnson, Kathy Smollett, Daniel Mair, Stephen Carmichael, Lily Tong, Jenna Nichols, Elihu Aranday-Cortes, Kirstyn Brunker, Yasmin Parr, Kyriaki Nomikou; Sarah McDonald, Marc Niebel, Patawee Asamaphan; Richard Orton, Joseph Hughes, Sreenu Vattipally, David L Robertson; Alasdair MacLean, Rory Gunson; Kathy Li, Natasha Jesudason, Rajiv Shah, James Shepherd, Antonia Ho, Alice Broos, Emma Thomson and Alex Alderton, Roberto Amato, Sonia Goncalves, Ewan Harrison, David K. Jackson, Ian Johnston, Dominic Kwiatkowski, Cordelia Langford, John Sillitoe |
| EPI_ISL_532895 | Lighthouse Lab in Glasgow | Wellcome Sanger Institute for the COVID-19 Genomics UK Consortium | Harper VanSteenhouse, Yumi Kasai, David Gray, Carol Clugston, Anna Dominiczak and Alex Alderton, Roberto Amato, Sonia Goncalves, Ewan Harrison, David K. Jackson, Ian Johnston, Dominic Kwiatkowski, Cordelia Langford, John Sillitoe |
| EPI_ISL_532896 | NHSGGC West of Scotland Specialist Virology Centre / MRC-University of Glasgow Centre for Virus Research | Wellcome Sanger Institute for the COVID-19 Genomics UK Consortium | Ana da Silva Filipe, Natasha Johnson, Kathy Smollett, Daniel Mair, Stephen Carmichael, Lily Tong, Jenna Nichols, Elihu Aranday-Cortes, Kirstyn Brunker, Yasmin Parr, Kyriaki Nomikou; Sarah McDonald, Marc Niebel, Patawee Asamaphan; Richard Orton, Joseph Hughes, Sreenu Vattipally, David L Robertson; Alasdair MacLean, Rory Gunson; Kathy Li, Natasha Jesudason, Rajiv Shah, James Shepherd, Antonia Ho, Alice Broos, Emma Thomson and Alex Alderton, Roberto Amato, Sonia Goncalves, Ewan Harrison, David K. Jackson, Ian Johnston, Dominic Kwiatkowski, Cordelia Langford, John Sillitoe |
| EPI_ISL_532897, EPI_ISL_532898, EPI_ISL_532899, EPI_ISL_532900, EPI_ISL_532901, EPI_ISL_532902, EPI_ISL_532903, EPI_ISL_532904 | Lighthouse Lab in Glasgow | Wellcome Sanger Institute for the COVID-19 Genomics UK Consortium | Harper VanSteenhouse, Yumi Kasai, David Gray, Carol Clugston, Anna Dominiczak and Alex Alderton, Roberto Amato, Sonia Goncalves, Ewan Harrison, David K. Jackson, Ian Johnston, Dominic Kwiatkowski, Cordelia Langford, John Sillitoe |
| EPI_ISL_532905, EPI_ISL_532906 | NHSGGC West of Scotland Specialist Virology Centre / MRC-University of Glasgow Centre for Virus Research | Wellcome Sanger Institute for the COVID-19 Genomics UK Consortium | Ana da Silva Filipe, Natasha Johnson, Kathy Smollett, Daniel Mair, Stephen Carmichael, Lily Tong, Jenna Nichols, Elihu Aranday-Cortes, Kirstyn Brunker, Yasmin Parr, Kyriaki Nomikou; Sarah McDonald, Marc Niebel, Patawee Asamaphan; Richard Orton, Joseph Hughes, Sreenu Vattipally, David L Robertson; Alasdair MacLean, Rory Gunson; Kathy Li, Natasha Jesudason, Rajiv Shah, James Shepherd, Antonia Ho, Alice Broos, Emma Thomson and Alex Alderton, Roberto Amato, Sonia Goncalves, Ewan Harrison, David K. Jackson, Ian Johnston, Dominic Kwiatkowski, Cordelia Langford, John Sillitoe |
| EPI_ISL_532907, EPI_ISL_532908, EPI_ISL_532909, EPI_ISL_532910, EPI_ISL_532911, EPI_ISL_532912, EPI_ISL_532913, EPI_ISL_532914 | Lighthouse Lab in Glasgow | Wellcome Sanger Institute for the COVID-19 Genomics UK Consortium | Harper VanSteenhouse, Yumi Kasai, David Gray, Carol Clugston, Anna Dominiczak and Alex Alderton, Roberto Amato, Sonia Goncalves, Ewan Harrison, David K. Jackson, Ian Johnston, Dominic Kwiatkowski, Cordelia Langford, John Sillitoe |
| EPI_ISL_532915 | NHSGGC West of Scotland Specialist Virology Centre / MRC-University of Glasgow Centre for Virus Research | Wellcome Sanger Institute for the COVID-19 Genomics UK Consortium | Ana da Silva Filipe, Natasha Johnson, Kathy Smollett, Daniel Mair, Stephen Carmichael, Lily Tong, Jenna Nichols, Elihu Aranday-Cortes, Kirstyn Brunker, Yasmin Parr, Kyriaki Nomikou; Sarah McDonald, Marc Niebel, Patawee Asamaphan; Richard Orton, Joseph Hughes, Sreenu Vattipally, David L Robertson; Alasdair MacLean, Rory Gunson; Kathy Li, Natasha Jesudason, Rajiv Shah, James Shepherd, Antonia Ho, Alice Broos, Emma Thomson and Alex Alderton, Roberto Amato, Sonia Goncalves, Ewan Harrison, David K. Jackson, Ian Johnston, Dominic Kwiatkowski, Cordelia Langford, John Sillitoe |
| EPI_ISL_532916, EPI_ISL_532917, EPI_ISL_532918, EPI_ISL_532919, EPI_ISL_532920, EPI_ISL_532921, EPI_ISL_532922, EPI_ISL_532923, EPI_ISL_532924, EPI_ISL_532925 | Lighthouse Lab in Glasgow | Wellcome Sanger Institute for the COVID-19 Genomics UK Consortium | Harper VanSteenhouse, Yumi Kasai, David Gray, Carol Clugston, Anna Dominiczak and Alex Alderton, Roberto Amato, Sonia Goncalves, Ewan Harrison, David K. Jackson, Ian Johnston, Dominic Kwiatkowski, Cordelia Langford, John Sillitoe |
| EPI_ISL_532926 | Virology Department, Royal Infirmary of Edinburgh, NHS Lothian / School of Biological Sciences, University of Edinburgh | Wellcome Sanger Institute for the COVID-19 Genomics UK Consortium | McHugh M, Dewar R, Rooke S, O'Toole Á, Scher E, Hill V, McCrone JT, Colquhoun R, Yu X, Jackson B, Rambaut A, Templeton K and Alex Alderton, Roberto Amato, Sonia Goncalves, Ewan Harrison, David K. Jackson, Ian Johnston, Dominic Kwiatkowski, Cordelia Langford, John Sillitoe |
| EPI_ISL_532927, EPI_ISL_532928, EPI_ISL_532929, EPI_ISL_532930, EPI_ISL_532931 | Lighthouse Lab in Glasgow | Wellcome Sanger Institute for the COVID-19 Genomics UK Consortium | Harper VanSteenhouse, Yumi Kasai, David Gray, Carol Clugston, Anna Dominiczak and Alex Alderton, Roberto Amato, Sonia Goncalves, Ewan Harrison, David K. Jackson, Ian Johnston, Dominic Kwiatkowski, Cordelia Langford, John Sillitoe |
| EPI_ISL_532932, EPI_ISL_532933 | NHSGGC West of Scotland Specialist Virology Centre / MRC-University of Glasgow Centre for Virus Research | Wellcome Sanger Institute for the COVID-19 Genomics UK Consortium | Ana da Silva Filipe, Natasha Johnson, Kathy Smollett, Daniel Mair, Stephen Carmichael, Lily Tong, Jenna Nichols, Elihu Aranday-Cortes, Kirstyn Brunker, Yasmin Parr, Kyriaki Nomikou; Sarah McDonald, Marc Niebel, Patawee Asamaphan; Richard Orton, Joseph Hughes, Sreenu Vattipally, David L Robertson; Alasdair MacLean, Rory Gunson; Kathy Li, Natasha Jesudason, Rajiv Shah, James Shepherd, Antonia Ho, Alice Broos, Emma Thomson and Alex Alderton, Roberto Amato, Sonia Goncalves, Ewan Harrison, David K. Jackson, Ian Johnston, Dominic Kwiatkowski, Cordelia Langford, John Sillitoe |
| EPI_ISL_532934, EPI_ISL_532935, EPI_ISL_532936, EPI_ISL_532937, EPI_ISL_532938, EPI_ISL_532939, EPI_ISL_532940, EPI_ISL_532941, EPI_ISL_532942, EPI_ISL_532943, EPI_ISL_532944, EPI_ISL_532945, EPI_ISL_532946, EPI_ISL_532947 | | | |
| see above | Lighthouse Lab in Glasgow | Wellcome Sanger Institute for the COVID-19 Genomics UK Consortium | Harper VanSteenhouse, Yumi Kasai, David Gray, Carol Clugston, Anna Dominiczak and Alex Alderton, Roberto Amato, Sonia Goncalves, Ewan Harrison, David K. Jackson, Ian Johnston, Dominic Kwiatkowski, Cordelia Langford, John Sillitoe |
| EPI_ISL_532948 | NHSGGC West of Scotland Specialist Virology Centre / MRC-University of Glasgow Centre for Virus Research | Wellcome Sanger Institute for the COVID-19 Genomics UK Consortium | Ana da Silva Filipe, Natasha Johnson, Kathy Smollett, Daniel Mair, Stephen Carmichael, Lily Tong, Jenna Nichols, Elihu Aranday-Cortes, Kirstyn Brunker, Yasmin Parr, Kyriaki Nomikou; Sarah McDonald, Marc Niebel, Patawee Asamaphan; Richard Orton, Joseph Hughes, Sreenu Vattipally, David L Robertson; Alasdair MacLean, Rory Gunson; Kathy Li, Natasha Jesudason, Rajiv Shah, James Shepherd, Antonia Ho, Alice Broos, Emma Thomson and Alex Alderton, Roberto Amato, Sonia Goncalves, Ewan Harrison, David K. Jackson, Ian Johnston, Dominic Kwiatkowski, Cordelia Langford, John Sillitoe |
| EPI_ISL_532949, EPI_ISL_532950, EPI_ISL_532951, EPI_ISL_532952, EPI_ISL_532953, EPI_ISL_532954, EPI_ISL_532955, EPI_ISL_532956, EPI_ISL_532957, EPI_ISL_532958, EPI_ISL_532959, EPI_ISL_532960, EPI_ISL_532961 | | | |
| see above | Lighthouse Lab in Glasgow | Wellcome Sanger Institute for the COVID-19 Genomics UK Consortium | Harper VanSteenhouse, Yumi Kasai, David Gray, Carol Clugston, Anna Dominiczak and Alex Alderton, Roberto Amato, Sonia Goncalves, Ewan Harrison, David K. Jackson, Ian Johnston, Dominic Kwiatkowski, Cordelia Langford, John Sillitoe |
| EPI_ISL_532962 | NHSGGC West of Scotland Specialist Virology Centre / MRC-University of Glasgow Centre for Virus Research | Wellcome Sanger Institute for the COVID-19 Genomics UK Consortium | Ana da Silva Filipe, Natasha Johnson, Kathy Smollett, Daniel Mair, Stephen Carmichael, Lily Tong, Jenna Nichols, Elihu Aranday-Cortes, Kirstyn Brunker, Yasmin Parr, Kyriaki Nomikou; Sarah McDonald, Marc Niebel, Patawee Asamaphan; Richard Orton, Joseph Hughes, Sreenu Vattipally, David L Robertson; Alasdair MacLean, Rory Gunson; Kathy Li, Natasha Jesudason, Rajiv Shah, James Shepherd, Antonia Ho, Alice Broos, Emma Thomson and Alex Alderton, Roberto Amato, Sonia Goncalves, Ewan Harrison, David K. Jackson, Ian Johnston, Dominic Kwiatkowski, Cordelia Langford, John Sillitoe |
| EPI_ISL_532963, EPI_ISL_532964, EPI_ISL_532965, EPI_ISL_532966, EPI_ISL_532967, EPI_ISL_532968, EPI_ISL_532969, EPI_ISL_532970, EPI_ISL_532971, EPI_ISL_532972, EPI_ISL_532973, EPI_ISL_532974, EPI_ISL_532975, EPI_ISL_532976, EPI_ISL_532977 | | | |
| see above | Lighthouse Lab in Glasgow | Wellcome Sanger Institute for the COVID-19 Genomics UK Consortium | Harper VanSteenhouse, Yumi Kasai, David Gray, Carol Clugston, Anna Dominiczak and Alex Alderton, Roberto Amato, Sonia Goncalves, Ewan Harrison, David K. Jackson, Ian Johnston, Dominic Kwiatkowski, Cordelia Langford, John Sillitoe |
| EPI_ISL_532978 | Virology Department, Royal Infirmary of Edinburgh, NHS Lothian / School of Biological Sciences, University of Edinburgh | Wellcome Sanger Institute for the COVID-19 Genomics UK Consortium | McHugh M, Dewar R, Rooke S, O'Toole Á, Scher E, Hill V, McCrone JT, Colquhoun R, Yu X, Jackson B, Rambaut A, Templeton K and Alex Alderton, Roberto Amato, Sonia Goncalves, Ewan Harrison, David K. Jackson, Ian Johnston, Dominic Kwiatkowski, Cordelia Langford, John Sillitoe |
| EPI_ISL_532979, EPI_ISL_532980, EPI_ISL_532981, EPI_ISL_532982 | Lighthouse Lab in Glasgow | Wellcome Sanger Institute for the COVID-19 Genomics UK Consortium | Harper VanSteenhouse, Yumi Kasai, David Gray, Carol Clugston, Anna Dominiczak and Alex Alderton, Roberto Amato, Sonia Goncalves, Ewan Harrison, David K. Jackson, Ian Johnston, Dominic Kwiatkowski, Cordelia Langford, John Sillitoe |
| EPI_ISL_532983 | Virology Department, Royal Infirmary of Edinburgh, NHS Lothian / School of Biological Sciences, University of Edinburgh | Wellcome Sanger Institute for the COVID-19 Genomics UK Consortium | McHugh M, Dewar R, Rooke S, O'Toole Á, Scher E, Hill V, McCrone JT, Colquhoun R, Yu X, Jackson B, Rambaut A, Templeton K and Alex Alderton, Roberto Amato, Sonia Goncalves, Ewan Harrison, David K. Jackson, Ian Johnston, Dominic Kwiatkowski, Cordelia Langford, John Sillitoe |
| EPI_ISL_532984, EPI_ISL_532985, EPI_ISL_532986, EPI_ISL_532987 | Lighthouse Lab in Glasgow | Wellcome Sanger Institute for the COVID-19 Genomics UK Consortium | Harper VanSteenhouse, Yumi Kasai, David Gray, Carol Clugston, Anna Dominiczak and Alex Alderton, Roberto Amato, Sonia Goncalves, Ewan Harrison, David K. Jackson, Ian Johnston, Dominic Kwiatkowski, Cordelia Langford, John Sillitoe |
| EPI_ISL_532988 | NHSGGC West of Scotland Specialist Virology Centre / MRC-University of Glasgow Centre for Virus Research | Wellcome Sanger Institute for the COVID-19 Genomics UK Consortium | Ana da Silva Filipe, Natasha Johnson, Kathy Smollett, Daniel Mair, Stephen Carmichael, Lily Tong, Jenna Nichols, Elihu Aranday-Cortes, Kirstyn Brunker, Yasmin Parr, Kyriaki Nomikou; Sarah McDonald, Marc Niebel, Patawee Asamaphan; Richard Orton, Joseph Hughes, Sreenu Vattipally, David L Robertson; Alasdair MacLean, Rory Gunson; Kathy Li, Natasha Jesudason, Rajiv Shah, James Shepherd, Antonia Ho, Alice Broos, Emma Thomson and Alex Alderton, Roberto Amato, Sonia Goncalves, Ewan Harrison, David K. Jackson, Ian Johnston, Dominic Kwiatkowski, Cordelia Langford, John Sillitoe |
| EPI_ISL_532989 | Lighthouse Lab in Glasgow | Wellcome Sanger Institute for the COVID-19 Genomics UK Consortium | Harper VanSteenhouse, Yumi Kasai, David Gray, Carol Clugston, Anna Dominiczak and Alex Alderton, Roberto Amato, Sonia Goncalves, Ewan Harrison, David K. Jackson, Ian Johnston, Dominic Kwiatkowski, Cordelia Langford, John Sillitoe |
| EPI_ISL_532990 | Virology Department, Royal Infirmary of Edinburgh, NHS Lothian / School of Biological Sciences, University of Edinburgh | Wellcome Sanger Institute for the COVID-19 Genomics UK Consortium | McHugh M, Dewar R, Rooke S, O'Toole Á, Scher E, Hill V, McCrone JT, Colquhoun R, Yu X, Jackson B, Rambaut A, Templeton K and Alex Alderton, Roberto Amato, Sonia Goncalves, Ewan Harrison, David K. Jackson, Ian Johnston, Dominic Kwiatkowski, Cordelia Langford, John Sillitoe |
| EPI_ISL_532991 | Lighthouse Lab in Glasgow | Wellcome Sanger Institute for the COVID-19 Genomics UK Consortium | Harper VanSteenhouse, Yumi Kasai, David Gray, Carol Clugston, Anna Dominiczak and Alex Alderton, Roberto Amato, Sonia Goncalves, Ewan Harrison, David K. Jackson, Ian Johnston, Dominic Kwiatkowski, Cordelia Langford, John Sillitoe |
| EPI_ISL_532992 | Virology Department, Royal Infirmary of Edinburgh, NHS Lothian / School of Biological Sciences, University of Edinburgh | Wellcome Sanger Institute for the COVID-19 Genomics UK Consortium | McHugh M, Dewar R, Rooke S, O'Toole Á, Scher E, Hill V, McCrone JT, Colquhoun R, Yu X, Jackson B, Rambaut A, Templeton K and Alex Alderton, Roberto Amato, Sonia Goncalves, Ewan Harrison, David K. Jackson, Ian Johnston, Dominic Kwiatkowski, Cordelia Langford, John Sillitoe |
| EPI_ISL_532993 | Lighthouse Lab in Glasgow | Wellcome Sanger Institute for the COVID-19 Genomics UK Consortium | Harper VanSteenhouse, Yumi Kasai, David Gray, Carol Clugston, Anna Dominiczak and Alex Alderton, Roberto Amato, Sonia Goncalves, Ewan Harrison, David K. Jackson, Ian Johnston, Dominic Kwiatkowski, Cordelia Langford, John Sillitoe |
| EPI_ISL_532994 | Virology Department, Royal Infirmary of Edinburgh, NHS Lothian / School of Biological Sciences, University of Edinburgh | Wellcome Sanger Institute for the COVID-19 Genomics UK Consortium | McHugh M, Dewar R, Rooke S, O'Toole Á, Scher E, Hill V, McCrone JT, Colquhoun R, Yu X, Jackson B, Rambaut A, Templeton K and Alex Alderton, Roberto Amato, Sonia Goncalves, Ewan Harrison, David K. Jackson, Ian Johnston, Dominic Kwiatkowski, Cordelia Langford, John Sillitoe |
| EPI_ISL_532995, EPI_ISL_532996, EPI_ISL_532997, EPI_ISL_532998, EPI_ISL_532999 | Lighthouse Lab in Glasgow | Wellcome Sanger Institute for the COVID-19 Genomics UK Consortium | Harper VanSteenhouse, Yumi Kasai, David Gray, Carol Clugston, Anna Dominiczak and Alex Alderton, Roberto Amato, Sonia Goncalves, Ewan Harrison, David K. Jackson, Ian Johnston, Dominic Kwiatkowski, Cordelia Langford, John Sillitoe |
| EPI_ISL_533000 | NHSGGC West of Scotland Specialist Virology Centre / MRC-University of Glasgow Centre for Virus Research | Wellcome Sanger Institute for the COVID-19 Genomics UK Consortium | Ana da Silva Filipe, Natasha Johnson, Kathy Smollett, Daniel Mair, Stephen Carmichael, Lily Tong, Jenna Nichols, Elihu Aranday-Cortes, Kirstyn Brunker, Yasmin Parr, Kyriaki Nomikou; Sarah McDonald, Marc Niebel, Patawee Asamaphan; Richard Orton, Joseph Hughes, Sreenu Vattipally, David L Robertson; Alasdair MacLean, Rory Gunson; Kathy Li, Natasha Jesudason, Rajiv Shah, James Shepherd, Antonia Ho, Alice Broos, Emma Thomson and Alex Alderton, Roberto Amato, Sonia Goncalves, Ewan Harrison, David K. Jackson, Ian Johnston, Dominic Kwiatkowski, Cordelia Langford, John Sillitoe |
| EPI_ISL_533001, EPI_ISL_533002, EPI_ISL_533003, EPI_ISL_533004, EPI_ISL_533005, EPI_ISL_533006, EPI_ISL_533007, EPI_ISL_533008, EPI_ISL_533009, EPI_ISL_533010, EPI_ISL_533011, EPI_ISL_533012, EPI_ISL_533013, EPI_ISL_533014, EPI_ISL_533015 | | | |
| see above | Lighthouse Lab in Glasgow | Wellcome Sanger Institute for the COVID-19 Genomics UK Consortium | Harper VanSteenhouse, Yumi Kasai, David Gray, Carol Clugston, Anna Dominiczak and Alex Alderton, Roberto Amato, Sonia Goncalves, Ewan Harrison, David K. Jackson, Ian Johnston, Dominic Kwiatkowski, Cordelia Langford, John Sillitoe |
| EPI_ISL_533016 | NHSGGC West of Scotland Specialist Virology Centre / MRC-University of Glasgow Centre for Virus Research | Wellcome Sanger Institute for the COVID-19 Genomics UK Consortium | Ana da Silva Filipe, Natasha Johnson, Kathy Smollett, Daniel Mair, Stephen Carmichael, Lily Tong, Jenna Nichols, Elihu Aranday-Cortes, Kirstyn Brunker, Yasmin Parr, Kyriaki Nomikou; Sarah McDonald, Marc Niebel, Patawee Asamaphan; Richard Orton, Joseph Hughes, Sreenu Vattipally, David L Robertson; Alasdair MacLean, Rory Gunson; Kathy Li, Natasha Jesudason, Rajiv Shah, James Shepherd, Antonia Ho, Alice Broos, Emma |

| | | | |
|---|---|---|---|
| | | | Thomson and Alex Alderton, Roberto Amato, Sonia Goncalves, Ewan Harrison, David K. Jackson, Ian Johnston, Dominic Kwiatkowski, Cordelia Langford, John Sillitoe |
| EPI_ISL_533017, EPI_ISL_533018, EPI_ISL_533019, EPI_ISL_533020, EPI_ISL_533021, EPI_ISL_533022, EPI_ISL_533023, EPI_ISL_533024, EPI_ISL_533025, EPI_ISL_533026 | Lighthouse Lab in Glasgow | Wellcome Sanger Institute for the COVID-19 Genomics UK Consortium | Harper VanSteenhouse, Yumi Kasai, David Gray, Carol Clugston, Anna Dominiczak and Alex Alderton, Roberto Amato, Sonia Goncalves, Ewan Harrison, David K. Jackson, Ian Johnston, Dominic Kwiatkowski, Cordelia Langford, John Sillitoe |
| EPI_ISL_533027 | NHSGGC West of Scotland Specialist Virology Centre / MRC-University of Glasgow Centre for Virus Research | Wellcome Sanger Institute for the COVID-19 Genomics UK Consortium | Ana da Silva Filipe, Natasha Johnson, Kathy Smollett, Daniel Mair, Stephen Carmichael, Lily Tong, Jenna Nichols, Elihu Aranday-Cortes, Kirstyn Brunker, Yasmin Parr, Kyriaki Nomikou; Sarah McDonald, Marc Niebel, Patawee Asamaphan; Richard Orton, Joseph Hughes, Sreenu Vattipally, David L Robertson; Alasdair MacLean, Rory Gunson; Kathy Li, Natasha Jesudason, Rajiv Shah, James Shepherd, Antonia Ho, Alice Broos, Emma Thomson and Alex Alderton, Roberto Amato, Sonia Goncalves, Ewan Harrison, David K. Jackson, Ian Johnston, Dominic Kwiatkowski, Cordelia Langford, John Sillitoe |
| EPI_ISL_533028, EPI_ISL_533029, EPI_ISL_533030, EPI_ISL_533031, EPI_ISL_533032, EPI_ISL_533033, EPI_ISL_533034, EPI_ISL_533035, EPI_ISL_533036, EPI_ISL_533037, EPI_ISL_533038, EPI_ISL_533039, EPI_ISL_533040, EPI_ISL_533041, EPI_ISL_533042, EPI_ISL_533043, EPI_ISL_533044, EPI_ISL_533045, EPI_ISL_533046, EPI_ISL_533047, EPI_ISL_533048, EPI_ISL_533049, EPI_ISL_533050, EPI_ISL_533051, EPI_ISL_533052, EPI_ISL_533053, EPI_ISL_533054, EPI_ISL_533055, EPI_ISL_533056, EPI_ISL_533057, EPI_ISL_533058, EPI_ISL_533059, EPI_ISL_533060, EPI_ISL_533061, EPI_ISL_533062, EPI_ISL_533063, EPI_ISL_533064, EPI_ISL_533065, EPI_ISL_533066, EPI_ISL_533067, EPI_ISL_533068, EPI_ISL_533069, EPI_ISL_533070, EPI_ISL_533071 | | | |
| see above | Lighthouse Lab in Glasgow | Wellcome Sanger Institute for the COVID-19 Genomics UK Consortium | Harper VanSteenhouse, Yumi Kasai, David Gray, Carol Clugston, Anna Dominiczak and Alex Alderton, Roberto Amato, Sonia Goncalves, Ewan Harrison, David K. Jackson, Ian Johnston, Dominic Kwiatkowski, Cordelia Langford, John Sillitoe |
| EPI_ISL_533072 | NHSGGC West of Scotland Specialist Virology Centre / MRC-University of Glasgow Centre for Virus Research | Wellcome Sanger Institute for the COVID-19 Genomics UK Consortium | Ana da Silva Filipe, Natasha Johnson, Kathy Smollett, Daniel Mair, Stephen Carmichael, Lily Tong, Jenna Nichols, Elihu Aranday-Cortes, Kirstyn Brunker, Yasmin Parr, Kyriaki Nomikou; Sarah McDonald, Marc Niebel, Patawee Asamaphan; Richard Orton, Joseph Hughes, Sreenu Vattipally, David L Robertson; Alasdair MacLean, Rory Gunson; Kathy Li, Natasha Jesudason, Rajiv Shah, James Shepherd, Antonia Ho, Alice Broos, Emma Thomson and Alex Alderton, Roberto Amato, Sonia Goncalves, Ewan Harrison, David K. Jackson, Ian Johnston, Dominic Kwiatkowski, Cordelia Langford, John Sillitoe |
| EPI_ISL_533073, EPI_ISL_533074, EPI_ISL_533075, EPI_ISL_533076, EPI_ISL_533077, EPI_ISL_533078, EPI_ISL_533079, EPI_ISL_533080, EPI_ISL_533081, EPI_ISL_533082, EPI_ISL_533083, EPI_ISL_533084, EPI_ISL_533085, EPI_ISL_533087, EPI_ISL_533088, EPI_ISL_533089, EPI_ISL_533090, EPI_ISL_533091, EPI_ISL_533092, EPI_ISL_533093, EPI_ISL_533094, EPI_ISL_533095, EPI_ISL_533096, EPI_ISL_533097, EPI_ISL_533098, EPI_ISL_533099, EPI_ISL_533100, EPI_ISL_533101, EPI_ISL_533102, EPI_ISL_533103, EPI_ISL_533104, EPI_ISL_533105, EPI_ISL_533106, EPI_ISL_533107, EPI_ISL_533108, EPI_ISL_533109, EPI_ISL_533110, EPI_ISL_533111, EPI_ISL_533112, EPI_ISL_533113, EPI_ISL_533114, EPI_ISL_533115, EPI_ISL_533116, EPI_ISL_533117, EPI_ISL_533118, EPI_ISL_533119, EPI_ISL_533120, EPI_ISL_533121, EPI_ISL_533122, EPI_ISL_533123, EPI_ISL_533124, EPI_ISL_533125, EPI_ISL_533126, EPI_ISL_533127, EPI_ISL_533128, EPI_ISL_533129, EPI_ISL_533130, EPI_ISL_533131, EPI_ISL_533132, EPI_ISL_533133, EPI_ISL_533134, EPI_ISL_533135, EPI_ISL_533136, EPI_ISL_533137, EPI_ISL_533138, EPI_ISL_533139, EPI_ISL_533140, EPI_ISL_533141, EPI_ISL_533142, EPI_ISL_533143, EPI_ISL_533144, EPI_ISL_533145, EPI_ISL_533146, EPI_ISL_533147, EPI_ISL_533148, EPI_ISL_533149, EPI_ISL_533150, EPI_ISL_533151, EPI_ISL_533152, EPI_ISL_533153, EPI_ISL_533154, EPI_ISL_533155, EPI_ISL_533156, EPI_ISL_533157, EPI_ISL_533158, EPI_ISL_533159, EPI_ISL_533160, EPI_ISL_533161, EPI_ISL_533162, EPI_ISL_533163, EPI_ISL_533164, EPI_ISL_533165, EPI_ISL_533166, EPI_ISL_533167, EPI_ISL_533168, EPI_ISL_533169, EPI_ISL_533170, EPI_ISL_533171, EPI_ISL_533172, EPI_ISL_533173, EPI_ISL_533174, EPI_ISL_533175, EPI_ISL_533176, EPI_ISL_533177, EPI_ISL_533178, EPI_ISL_533179, EPI_ISL_533180, EPI_ISL_533181, EPI_ISL_533182, EPI_ISL_533183, EPI_ISL_533184, EPI_ISL_533185, EPI_ISL_533186, EPI_ISL_533187, EPI_ISL_533188, EPI_ISL_533189, EPI_ISL_533190, EPI_ISL_533191, EPI_ISL_533192, EPI_ISL_533193, EPI_ISL_533194, EPI_ISL_533195, EPI_ISL_533196, EPI_ISL_533197, EPI_ISL_533198, EPI_ISL_533199, EPI_ISL_533200, EPI_ISL_533201, EPI_ISL_533202, EPI_ISL_533203, EPI_ISL_533204, EPI_ISL_533205, EPI_ISL_533206, EPI_ISL_533207, EPI_ISL_533208, EPI_ISL_533209, EPI_ISL_533210, EPI_ISL_533211, EPI_ISL_533212, EPI_ISL_533213, EPI_ISL_533214, EPI_ISL_533215, EPI_ISL_533216, EPI_ISL_533217, EPI_ISL_533218, EPI_ISL_533219, EPI_ISL_533220, EPI_ISL_533221, EPI_ISL_533222, EPI_ISL_533223, EPI_ISL_533224, EPI_ISL_533225, EPI_ISL_533226, EPI_ISL_533227 | | | |
| see above | Lighthouse Lab in Glasgow | Wellcome Sanger Institute for the COVID-19 Genomics UK Consortium | Harper VanSteenhouse, Yumi Kasai, David Gray, Carol Clugston, Anna Dominiczak and Alex Alderton, Roberto Amato, Sonia Goncalves, Ewan Harrison, David K. Jackson, Ian Johnston, Dominic Kwiatkowski, Cordelia Langford, John Sillitoe |
| EPI_ISL_533228 | NHSGGC West of Scotland Specialist Virology Centre / MRC-University of Glasgow Centre for Virus Research | Wellcome Sanger Institute for the COVID-19 Genomics UK Consortium | Ana da Silva Filipe, Natasha Johnson, Kathy Smollett, Daniel Mair, Stephen Carmichael, Lily Tong, Jenna Nichols, Elihu Aranday-Cortes, Kirstyn Brunker, Yasmin Parr, Kyriaki Nomikou; Sarah McDonald, Marc Niebel, Patawee Asamaphan; Richard Orton, Joseph Hughes, Sreenu Vattipally, David L Robertson; Alasdair MacLean, Rory Gunson; Kathy Li, Natasha Jesudason, Rajiv Shah, James Shepherd, Antonia Ho, Alice Broos, Emma Thomson and Alex Alderton, Roberto Amato, Sonia Goncalves, Ewan Harrison, David K. Jackson, Ian Johnston, Dominic Kwiatkowski, Cordelia Langford, John Sillitoe |
| EPI_ISL_533229 | Lighthouse Lab in Glasgow | Wellcome Sanger Institute for the COVID-19 Genomics UK Consortium | Harper VanSteenhouse, Yumi Kasai, David Gray, Carol Clugston, Anna Dominiczak and Alex Alderton, Roberto Amato, Sonia Goncalves, Ewan Harrison, David K. Jackson, Ian Johnston, Dominic Kwiatkowski, Cordelia Langford, John Sillitoe |
| EPI_ISL_533230, EPI_ISL_533231, EPI_ISL_533232 | NHSGGC West of Scotland Specialist Virology Centre / MRC-University of Glasgow Centre for Virus Research | Wellcome Sanger Institute for the COVID-19 Genomics UK Consortium | Ana da Silva Filipe, Natasha Johnson, Kathy Smollett, Daniel Mair, Stephen Carmichael, Lily Tong, Jenna Nichols, Elihu Aranday-Cortes, Kirstyn Brunker, Yasmin Parr, Kyriaki Nomikou; Sarah McDonald, Marc Niebel, Patawee Asamaphan; Richard Orton, Joseph Hughes, Sreenu Vattipally, David L Robertson; Alasdair MacLean, Rory Gunson; Kathy Li, Natasha Jesudason, Rajiv Shah, James Shepherd, Antonia Ho, Alice Broos, Emma Thomson and Alex Alderton, Roberto Amato, Sonia Goncalves, Ewan Harrison, David K. Jackson, Ian Johnston, Dominic Kwiatkowski, Cordelia Langford, John Sillitoe |
| EPI_ISL_533233, EPI_ISL_533234, EPI_ISL_533235 | Lighthouse Lab in Glasgow | Wellcome Sanger Institute for the COVID-19 Genomics UK Consortium | Harper VanSteenhouse, Yumi Kasai, David Gray, Carol Clugston, Anna Dominiczak and Alex Alderton, Roberto Amato, Sonia Goncalves, Ewan Harrison, David K. Jackson, Ian Johnston, Dominic Kwiatkowski, Cordelia Langford, John Sillitoe |
| EPI_ISL_533236, EPI_ISL_533237 | NHSGGC West of Scotland Specialist Virology Centre / MRC-University of Glasgow Centre for Virus Research | Wellcome Sanger Institute for the COVID-19 Genomics UK Consortium | Ana da Silva Filipe, Natasha Johnson, Kathy Smollett, Daniel Mair, Stephen Carmichael, Lily Tong, Jenna Nichols, Elihu Aranday-Cortes, Kirstyn Brunker, Yasmin Parr, Kyriaki Nomikou; Sarah McDonald, Marc Niebel, Patawee Asamaphan; Richard Orton, Joseph Hughes, Sreenu Vattipally, David L Robertson; Alasdair MacLean, Rory Gunson; Kathy Li, Natasha Jesudason, Rajiv Shah, James Shepherd, Antonia Ho, Alice Broos, Emma Thomson and Alex Alderton, Roberto Amato, Sonia Goncalves, Ewan Harrison, David K. Jackson, Ian Johnston, Dominic Kwiatkowski, Cordelia Langford, John Sillitoe |
| EPI_ISL_533238, EPI_ISL_533239, EPI_ISL_533240 | Lighthouse Lab in Glasgow | Wellcome Sanger Institute for the COVID-19 Genomics UK Consortium | Harper VanSteenhouse, Yumi Kasai, David Gray, Carol Clugston, Anna Dominiczak and Alex Alderton, Roberto Amato, Sonia Goncalves, Ewan Harrison, David K. Jackson, Ian Johnston, Dominic Kwiatkowski, Cordelia Langford, John Sillitoe |
| EPI_ISL_533241 | NHSGGC West of Scotland Specialist Virology Centre / MRC-University of Glasgow Centre for Virus Research | Wellcome Sanger Institute for the COVID-19 Genomics UK Consortium | Ana da Silva Filipe, Natasha Johnson, Kathy Smollett, Daniel Mair, Stephen Carmichael, Lily Tong, Jenna Nichols, Elihu Aranday-Cortes, Kirstyn Brunker, Yasmin Parr, Kyriaki Nomikou; Sarah McDonald, Marc Niebel, Patawee Asamaphan; Richard Orton, Joseph Hughes, Sreenu Vattipally, David L Robertson; Alasdair MacLean, Rory Gunson; Kathy Li, Natasha Jesudason, Rajiv Shah, James Shepherd, Antonia Ho, Alice Broos, Emma Thomson and Alex Alderton, Roberto Amato, Sonia Goncalves, Ewan Harrison, David K. Jackson, Ian Johnston, Dominic Kwiatkowski, Cordelia Langford, John Sillitoe |
| EPI_ISL_533242, EPI_ISL_533243, EPI_ISL_533244, EPI_ISL_533245, EPI_ISL_533246, EPI_ISL_533247, EPI_ISL_533248, EPI_ISL_533249, EPI_ISL_533250, EPI_ISL_533251 | Lighthouse Lab in Glasgow | Wellcome Sanger Institute for the COVID-19 Genomics UK Consortium | Harper VanSteenhouse, Yumi Kasai, David Gray, Carol Clugston, Anna Dominiczak and Alex Alderton, Roberto Amato, Sonia Goncalves, Ewan Harrison, David K. Jackson, Ian Johnston, Dominic Kwiatkowski, Cordelia Langford, John Sillitoe |
| EPI_ISL_533252 | NHSGGC West of Scotland Specialist Virology Centre / MRC-University of Glasgow Centre for Virus Research | Wellcome Sanger Institute for the COVID-19 Genomics UK Consortium | Ana da Silva Filipe, Natasha Johnson, Kathy Smollett, Daniel Mair, Stephen Carmichael, Lily Tong, Jenna Nichols, Elihu Aranday-Cortes, Kirstyn Brunker, Yasmin Parr, Kyriaki Nomikou; Sarah McDonald, Marc Niebel, Patawee Asamaphan; Richard Orton, Joseph Hughes, Sreenu Vattipally, David L Robertson; Alasdair MacLean, Rory Gunson; Kathy Li, Natasha Jesudason, Rajiv Shah, James Shepherd, Antonia Ho, Alice Broos, Emma Thomson and Alex Alderton, Roberto Amato, Sonia Goncalves, Ewan Harrison, David K. Jackson, Ian Johnston, Dominic Kwiatkowski, Cordelia Langford, John Sillitoe |
| EPI_ISL_533253, EPI_ISL_533254, EPI_ISL_533255 | Lighthouse Lab in Glasgow | Wellcome Sanger Institute for the COVID-19 Genomics UK Consortium | Harper VanSteenhouse, Yumi Kasai, David Gray, Carol Clugston, Anna Dominiczak and Alex Alderton, Roberto Amato, Sonia Goncalves, Ewan Harrison, David K. Jackson, Ian Johnston, Dominic Kwiatkowski, Cordelia Langford, John Sillitoe |
| EPI_ISL_533256 | NHSGGC West of Scotland Specialist Virology Centre / MRC-University of Glasgow Centre for Virus Research | Wellcome Sanger Institute for the COVID-19 Genomics UK Consortium | Ana da Silva Filipe, Natasha Johnson, Kathy Smollett, Daniel Mair, Stephen Carmichael, Lily Tong, Jenna Nichols, Elihu Aranday-Cortes, Kirstyn Brunker, Yasmin Parr, Kyriaki Nomikou; Sarah McDonald, Marc Niebel, Patawee Asamaphan; Richard Orton, Joseph Hughes, Sreenu Vattipally, David L Robertson; Alasdair MacLean, Rory Gunson; Kathy Li, Natasha Jesudason, Rajiv Shah, James Shepherd, Antonia Ho, Alice Broos, Emma Thomson and Alex Alderton, Roberto Amato, Sonia Goncalves, Ewan Harrison, David K. Jackson, Ian Johnston, Dominic Kwiatkowski, Cordelia Langford, John Sillitoe |
| EPI_ISL_533257, EPI_ISL_533258, EPI_ISL_533259, EPI_ISL_533260, EPI_ISL_533261 | Lighthouse Lab in Glasgow | Wellcome Sanger Institute for the COVID-19 Genomics UK Consortium | Harper VanSteenhouse, Yumi Kasai, David Gray, Carol Clugston, Anna Dominiczak and Alex Alderton, Roberto Amato, Sonia Goncalves, Ewan Harrison, David K. Jackson, Ian Johnston, Dominic Kwiatkowski, Cordelia Langford, John Sillitoe |
| EPI_ISL_533262 | NHSGGC West of Scotland Specialist Virology Centre / MRC-University of Glasgow Centre for Virus Research | Wellcome Sanger Institute for the COVID-19 Genomics UK Consortium | Ana da Silva Filipe, Natasha Johnson, Kathy Smollett, Daniel Mair, Stephen Carmichael, Lily Tong, Jenna Nichols, Elihu Aranday-Cortes, Kirstyn Brunker, Yasmin Parr, Kyriaki Nomikou; Sarah McDonald, Marc Niebel, Patawee Asamaphan; Richard Orton, Joseph Hughes, Sreenu Vattipally, David L Robertson; Alasdair MacLean, Rory Gunson; Kathy Li, Natasha Jesudason, Rajiv Shah, James Shepherd, Antonia Ho, Alice Broos, Emma Thomson and Alex Alderton, Roberto Amato, Sonia Goncalves, Ewan Harrison, David K. Jackson, Ian Johnston, Dominic Kwiatkowski, Cordelia Langford, John Sillitoe |
| EPI_ISL_533263, EPI_ISL_533264, EPI_ISL_533265, EPI_ISL_533266, EPI_ISL_533267, EPI_ISL_533268, EPI_ISL_533269 | Lighthouse Lab in Glasgow | Wellcome Sanger Institute for the COVID-19 Genomics UK Consortium | Harper VanSteenhouse, Yumi Kasai, David Gray, Carol Clugston, Anna Dominiczak and Alex Alderton, Roberto Amato, Sonia Goncalves, Ewan Harrison, David K. Jackson, Ian Johnston, Dominic Kwiatkowski, Cordelia Langford, John Sillitoe |
| EPI_ISL_533270, EPI_ISL_533271 | NHSGGC West of Scotland Specialist Virology Centre / MRC-University of Glasgow Centre for Virus Research | Wellcome Sanger Institute for the COVID-19 Genomics UK Consortium | Ana da Silva Filipe, Natasha Johnson, Kathy Smollett, Daniel Mair, Stephen Carmichael, Lily Tong, Jenna Nichols, Elihu Aranday-Cortes, Kirstyn Brunker, Yasmin Parr, Kyriaki Nomikou; Sarah McDonald, Marc Niebel, Patawee Asamaphan; Richard Orton, Joseph Hughes, Sreenu Vattipally, David L Robertson; Alasdair MacLean, Rory Gunson; Kathy Li, Natasha Jesudason, Rajiv Shah, James Shepherd, Antonia Ho, Alice Broos, Emma Thomson and Alex Alderton, Roberto Amato, Sonia Goncalves, Ewan Harrison, David K. Jackson, Ian Johnston, Dominic Kwiatkowski, Cordelia Langford, John Sillitoe |
| EPI_ISL_533272, EPI_ISL_533273, EPI_ISL_533274, EPI_ISL_533275, EPI_ISL_533276, EPI_ISL_533277, EPI_ISL_533278, EPI_ISL_533279, EPI_ISL_533280, EPI_ISL_533281, EPI_ISL_533282 | | | |
| see above | Lighthouse Lab in Glasgow | Wellcome Sanger Institute for the COVID-19 Genomics UK Consortium | Harper VanSteenhouse, Yumi Kasai, David Gray, Carol Clugston, Anna Dominiczak and Alex Alderton, Roberto Amato, Sonia Goncalves, Ewan Harrison, David K. Jackson, Ian Johnston, Dominic Kwiatkowski, Cordelia Langford, John Sillitoe |
| EPI_ISL_533283 | NHSGGC West of Scotland Specialist Virology Centre / MRC-University of Glasgow Centre for Virus Research | Wellcome Sanger Institute for the COVID-19 Genomics UK Consortium | Ana da Silva Filipe, Natasha Johnson, Kathy Smollett, Daniel Mair, Stephen Carmichael, Lily Tong, Jenna Nichols, Elihu Aranday-Cortes, Kirstyn Brunker, Yasmin Parr, Kyriaki Nomikou; Sarah McDonald, Marc Niebel, Patawee Asamaphan; Richard Orton, Joseph Hughes, Sreenu Vattipally, David L Robertson; Alasdair MacLean, Rory Gunson; Kathy Li, Natasha Jesudason, Rajiv Shah, James Shepherd, Antonia Ho, Alice Broos, Emma Thomson and Alex Alderton, Roberto Amato, Sonia Goncalves, Ewan Harrison, David K. Jackson, Ian Johnston, Dominic Kwiatkowski, Cordelia Langford, John Sillitoe |
| EPI_ISL_533284, EPI_ISL_533285, EPI_ISL_533286, EPI_ISL_533287, EPI_ISL_533288 | Lighthouse Lab in Glasgow | Wellcome Sanger Institute for the COVID-19 Genomics UK Consortium | Harper VanSteenhouse, Yumi Kasai, David Gray, Carol Clugston, Anna Dominiczak and Alex Alderton, Roberto Amato, Sonia Goncalves, Ewan Harrison, David K. Jackson, Ian Johnston, Dominic Kwiatkowski, Cordelia Langford, John Sillitoe |
| EPI_ISL_533289 | NHSGGC West of Scotland Specialist Virology Centre / MRC-University of Glasgow Centre for Virus Research | Wellcome Sanger Institute for the COVID-19 Genomics UK Consortium | Ana da Silva Filipe, Natasha Johnson, Kathy Smollett, Daniel Mair, Stephen Carmichael, Lily Tong, Jenna Nichols, Elihu Aranday-Cortes, Kirstyn Brunker, Yasmin Parr, Kyriaki Nomikou; Sarah McDonald, Marc Niebel, Patawee Asamaphan; Richard Orton, Joseph Hughes, Sreenu Vattipally, David L Robertson; Alasdair MacLean, Rory Gunson; Kathy Li, Natasha Jesudason, Rajiv Shah, James Shepherd, Antonia Ho, Alice Broos, Emma Thomson and Alex Alderton, Roberto Amato, Sonia Goncalves, Ewan Harrison, David K. Jackson, Ian Johnston, Dominic Kwiatkowski, Cordelia Langford, John Sillitoe |
| EPI_ISL_533290, EPI_ISL_533291, EPI_ISL_533292, EPI_ISL_533293, EPI_ISL_533294, EPI_ISL_533295, EPI_ISL_533296, EPI_ISL_533297 | Lighthouse Lab in Glasgow | Wellcome Sanger Institute for the COVID-19 Genomics UK Consortium | Harper VanSteenhouse, Yumi Kasai, David Gray, Carol Clugston, Anna Dominiczak and Alex Alderton, Roberto Amato, Sonia Goncalves, Ewan Harrison, David K. Jackson, Ian Johnston, Dominic Kwiatkowski, Cordelia Langford, John Sillitoe |
| EPI_ISL_533298 | NHSGGC West of Scotland Specialist Virology Centre / MRC-University of Glasgow Centre for Virus Research | Wellcome Sanger Institute for the COVID-19 Genomics UK Consortium | Ana da Silva Filipe, Natasha Johnson, Kathy Smollett, Daniel Mair, Stephen Carmichael, Lily Tong, Jenna Nichols, Elihu Aranday-Cortes, Kirstyn Brunker, Yasmin Parr, Kyriaki Nomikou; Sarah McDonald, Marc Niebel, Patawee Asamaphan; Richard Orton, Joseph Hughes, Sreenu Vattipally, David L Robertson; Alasdair MacLean, Rory Gunson; Kathy Li, Natasha Jesudason, Rajiv Shah, James Shepherd, Antonia Ho, Alice Broos, Emma Thomson and Alex Alderton, Roberto Amato, Sonia Goncalves, Ewan Harrison, David K. Jackson, Ian Johnston, Dominic Kwiatkowski, Cordelia Langford, John Sillitoe |
| EPI_ISL_533299, EPI_ISL_533300, EPI_ISL_533301, EPI_ISL_533302, EPI_ISL_533303, EPI_ISL_533304, EPI_ISL_533305, EPI_ISL_533306, EPI_ISL_533307, EPI_ISL_533308, EPI_ISL_533309, EPI_ISL_533310 | | | |
| see above | Lighthouse Lab in Glasgow | Wellcome Sanger Institute for the COVID-19 Genomics UK Consortium | Harper VanSteenhouse, Yumi Kasai, David Gray, Carol Clugston, Anna Dominiczak and Alex Alderton, Roberto Amato, Sonia Goncalves, Ewan Harrison, David K. Jackson, Ian Johnston, Dominic Kwiatkowski, Cordelia Langford, John Sillitoe |
| EPI_ISL_533311 | NHSGGC West of Scotland Specialist Virology Centre / MRC-University of Glasgow Centre for Virus Research | Wellcome Sanger Institute for the COVID-19 Genomics UK Consortium | Ana da Silva Filipe, Natasha Johnson, Kathy Smollett, Daniel Mair, Stephen Carmichael, Lily Tong, Jenna Nichols, Elihu Aranday-Cortes, Kirstyn Brunker, Yasmin Parr, Kyriaki Nomikou; Sarah McDonald, Marc Niebel, Patawee Asamaphan; Richard Orton, Joseph Hughes, Sreenu Vattipally, David L Robertson; Alasdair MacLean, Rory Gunson; Kathy Li, Natasha Jesudason, Rajiv Shah, James Shepherd, Antonia Ho, Alice Broos, Emma Thomson and Alex Alderton, Roberto Amato, Sonia Goncalves, Ewan Harrison, David K. Jackson, Ian Johnston, Dominic Kwiatkowski, Cordelia Langford, John Sillitoe |
| EPI_ISL_533312, EPI_ISL_533313, EPI_ISL_533314, EPI_ISL_533315, EPI_ISL_533316, EPI_ISL_533317, EPI_ISL_533318, EPI_ISL_533319, EPI_ISL_533320, EPI_ISL_533321, EPI_ISL_533322, EPI_ISL_533323, EPI_ISL_533324, EPI_ISL_533325, EPI_ISL_533326, EPI_ISL_533327, EPI_ISL_533328, EPI_ISL_533329, EPI_ISL_533330, EPI_ISL_533331, EPI_ISL_533332, EPI_ISL_533333, EPI_ISL_533334, EPI_ISL_533335, EPI_ISL_533336, EPI_ISL_533337, EPI_ISL_533338, EPI_ISL_533339, EPI_ISL_533340, EPI_ISL_533341, EPI_ISL_533342 | | | |
| see above | Lighthouse Lab in Glasgow | Wellcome Sanger Institute for the COVID-19 Genomics UK Consortium | Harper VanSteenhouse, Yumi Kasai, David Gray, Carol Clugston, Anna Dominiczak and Alex Alderton, Roberto Amato, Sonia Goncalves, Ewan Harrison, David K. Jackson, Ian Johnston, Dominic Kwiatkowski, Cordelia Langford, John Sillitoe |
| EPI_ISL_533343 | NHSGGC West of Scotland Specialist Virology Centre / MRC-University of Glasgow Centre for Virus Research | Wellcome Sanger Institute for the COVID-19 Genomics UK Consortium | Ana da Silva Filipe, Natasha Johnson, Kathy Smollett, Daniel Mair, Stephen Carmichael, Lily Tong, Jenna Nichols, Elihu Aranday-Cortes, Kirstyn Brunker, Yasmin Parr, Kyriaki Nomikou; Sarah McDonald, Marc Niebel, Patawee Asamaphan; Richard Orton, Joseph Hughes, Sreenu Vattipally, David L Robertson; Alasdair MacLean, Rory Gunson; Kathy Li, Natasha Jesudason, Rajiv Shah, James Shepherd, Antonia Ho, Alice Broos, Emma |

| | | | |
|---|---|---|---|
| | | | Thomson and Alex Alderton, Roberto Amato, Sonia Goncalves, Ewan Harrison, David K. Jackson, Ian Johnston, Dominic Kwiatkowski, Cordelia Langford, John Sillitoe |
| EPI_ISL_533344, EPI_ISL_533345 | Lighthouse Lab in Glasgow | Wellcome Sanger Institute for the COVID-19 Genomics UK Consortium | Harper VanSteenhouse, Yumi Kasai, David Gray, Carol Clugston, Anna Dominiczak and Alex Alderton, Roberto Amato, Sonia Goncalves, Ewan Harrison, David K. Jackson, Ian Johnston, Dominic Kwiatkowski, Cordelia Langford, John Sillitoe |
| EPI_ISL_533346 | NHSGGC West of Scotland Specialist Virology Centre / MRC-University of Glasgow Centre for Virus Research | Wellcome Sanger Institute for the COVID-19 Genomics UK Consortium | Ana da Silva Filipe, Natasha Johnson, Kathy Smollett, Daniel Mair, Stephen Carmichael, Lily Tong, Jenna Nichols, Elihu Aranday-Cortes, Kirstyn Brunker, Yasmin Parr, Kyriaki Nomikou; Sarah McDonald, Marc Niebel, Patawee Asamaphan; Richard Orton, Joseph Hughes, Sreenu Vattipally, David L Robertson; Alasdair MacLean, Rory Gunson; Kathy Li, Natasha Jesudason, Rajiv Shah, James Shepherd, Antonia Ho, Alice Broos, Emma Thomson and Alex Alderton, Roberto Amato, Sonia Goncalves, Ewan Harrison, David K. Jackson, Ian Johnston, Dominic Kwiatkowski, Cordelia Langford, John Sillitoe |
| EPI_ISL_533347, EPI_ISL_533348, EPI_ISL_533349 | Lighthouse Lab in Glasgow | Wellcome Sanger Institute for the COVID-19 Genomics UK Consortium | Harper VanSteenhouse, Yumi Kasai, David Gray, Carol Clugston, Anna Dominiczak and Alex Alderton, Roberto Amato, Sonia Goncalves, Ewan Harrison, David K. Jackson, Ian Johnston, Dominic Kwiatkowski, Cordelia Langford, John Sillitoe |
| EPI_ISL_533350, EPI_ISL_533351 | NHSGGC West of Scotland Specialist Virology Centre / MRC-University of Glasgow Centre for Virus Research | Wellcome Sanger Institute for the COVID-19 Genomics UK Consortium | Ana da Silva Filipe, Natasha Johnson, Kathy Smollett, Daniel Mair, Stephen Carmichael, Lily Tong, Jenna Nichols, Elihu Aranday-Cortes, Kirstyn Brunker, Yasmin Parr, Kyriaki Nomikou; Sarah McDonald, Marc Niebel, Patawee Asamaphan; Richard Orton, Joseph Hughes, Sreenu Vattipally, David L Robertson; Alasdair MacLean, Rory Gunson; Kathy Li, Natasha Jesudason, Rajiv Shah, James Shepherd, Antonia Ho, Alice Broos, Emma Thomson and Alex Alderton, Roberto Amato, Sonia Goncalves, Ewan Harrison, David K. Jackson, Ian Johnston, Dominic Kwiatkowski, Cordelia Langford, John Sillitoe |
| EPI_ISL_533352, EPI_ISL_533353, EPI_ISL_533354 | Lighthouse Lab in Glasgow | Wellcome Sanger Institute for the COVID-19 Genomics UK Consortium | Harper VanSteenhouse, Yumi Kasai, David Gray, Carol Clugston, Anna Dominiczak and Alex Alderton, Roberto Amato, Sonia Goncalves, Ewan Harrison, David K. Jackson, Ian Johnston, Dominic Kwiatkowski, Cordelia Langford, John Sillitoe |
| EPI_ISL_533355 | NHSGGC West of Scotland Specialist Virology Centre / MRC-University of Glasgow Centre for Virus Research | Wellcome Sanger Institute for the COVID-19 Genomics UK Consortium | Ana da Silva Filipe, Natasha Johnson, Kathy Smollett, Daniel Mair, Stephen Carmichael, Lily Tong, Jenna Nichols, Elihu Aranday-Cortes, Kirstyn Brunker, Yasmin Parr, Kyriaki Nomikou; Sarah McDonald, Marc Niebel, Patawee Asamaphan; Richard Orton, Joseph Hughes, Sreenu Vattipally, David L Robertson; Alasdair MacLean, Rory Gunson; Kathy Li, Natasha Jesudason, Rajiv Shah, James Shepherd, Antonia Ho, Alice Broos, Emma Thomson and Alex Alderton, Roberto Amato, Sonia Goncalves, Ewan Harrison, David K. Jackson, Ian Johnston, Dominic Kwiatkowski, Cordelia Langford, John Sillitoe |
| EPI_ISL_533356, EPI_ISL_533357 | Lighthouse Lab in Glasgow | Wellcome Sanger Institute for the COVID-19 Genomics UK Consortium | Harper VanSteenhouse, Yumi Kasai, David Gray, Carol Clugston, Anna Dominiczak and Alex Alderton, Roberto Amato, Sonia Goncalves, Ewan Harrison, David K. Jackson, Ian Johnston, Dominic Kwiatkowski, Cordelia Langford, John Sillitoe |
| EPI_ISL_533358 | NHSGGC West of Scotland Specialist Virology Centre / MRC-University of Glasgow Centre for Virus Research | Wellcome Sanger Institute for the COVID-19 Genomics UK Consortium | Ana da Silva Filipe, Natasha Johnson, Kathy Smollett, Daniel Mair, Stephen Carmichael, Lily Tong, Jenna Nichols, Elihu Aranday-Cortes, Kirstyn Brunker, Yasmin Parr, Kyriaki Nomikou; Sarah McDonald, Marc Niebel, Patawee Asamaphan; Richard Orton, Joseph Hughes, Sreenu Vattipally, David L Robertson; Alasdair MacLean, Rory Gunson; Kathy Li, Natasha Jesudason, Rajiv Shah, James Shepherd, Antonia Ho, Alice Broos, Emma Thomson and Alex Alderton, Roberto Amato, Sonia Goncalves, Ewan Harrison, David K. Jackson, Ian Johnston, Dominic Kwiatkowski, Cordelia Langford, John Sillitoe |
| EPI_ISL_533359, EPI_ISL_533360, EPI_ISL_533361, EPI_ISL_533362, EPI_ISL_533363, EPI_ISL_533364 | Lighthouse Lab in Glasgow | Wellcome Sanger Institute for the COVID-19 Genomics UK Consortium | Harper VanSteenhouse, Yumi Kasai, David Gray, Carol Clugston, Anna Dominiczak and Alex Alderton, Roberto Amato, Sonia Goncalves, Ewan Harrison, David K. Jackson, Ian Johnston, Dominic Kwiatkowski, Cordelia Langford, John Sillitoe |
| EPI_ISL_533365 | NHSGGC West of Scotland Specialist Virology Centre / MRC-University of Glasgow Centre for Virus Research | Wellcome Sanger Institute for the COVID-19 Genomics UK Consortium | Ana da Silva Filipe, Natasha Johnson, Kathy Smollett, Daniel Mair, Stephen Carmichael, Lily Tong, Jenna Nichols, Elihu Aranday-Cortes, Kirstyn Brunker, Yasmin Parr, Kyriaki Nomikou; Sarah McDonald, Marc Niebel, Patawee Asamaphan; Richard Orton, Joseph Hughes, Sreenu Vattipally, David L Robertson; Alasdair MacLean, Rory Gunson; Kathy Li, Natasha Jesudason, Rajiv Shah, James Shepherd, Antonia Ho, Alice Broos, Emma Thomson and Alex Alderton, Roberto Amato, Sonia Goncalves, Ewan Harrison, David K. Jackson, Ian Johnston, Dominic Kwiatkowski, Cordelia Langford, John Sillitoe |
| EPI_ISL_533366, EPI_ISL_533367, EPI_ISL_533368, EPI_ISL_533369, EPI_ISL_533370, EPI_ISL_533371, EPI_ISL_533372, EPI_ISL_533373, EPI_ISL_533374, EPI_ISL_533375 | Lighthouse Lab in Glasgow | Wellcome Sanger Institute for the COVID-19 Genomics UK Consortium | Harper VanSteenhouse, Yumi Kasai, David Gray, Carol Clugston, Anna Dominiczak and Alex Alderton, Roberto Amato, Sonia Goncalves, Ewan Harrison, David K. Jackson, Ian Johnston, Dominic Kwiatkowski, Cordelia Langford, John Sillitoe |
| EPI_ISL_533376 | NHSGGC West of Scotland Specialist Virology Centre / MRC-University of Glasgow Centre for Virus Research | Wellcome Sanger Institute for the COVID-19 Genomics UK Consortium | Ana da Silva Filipe, Natasha Johnson, Kathy Smollett, Daniel Mair, Stephen Carmichael, Lily Tong, Jenna Nichols, Elihu Aranday-Cortes, Kirstyn Brunker, Yasmin Parr, Kyriaki Nomikou; Sarah McDonald, Marc Niebel, Patawee Asamaphan; Richard Orton, Joseph Hughes, Sreenu Vattipally, David L Robertson; Alasdair MacLean, Rory Gunson; Kathy Li, Natasha Jesudason, Rajiv Shah, James Shepherd, Antonia Ho, Alice Broos, Emma Thomson and Alex Alderton, Roberto Amato, Sonia Goncalves, Ewan Harrison, David K. Jackson, Ian Johnston, Dominic Kwiatkowski, Cordelia Langford, John Sillitoe |
| EPI_ISL_533377, EPI_ISL_533378, EPI_ISL_533379, EPI_ISL_533380, EPI_ISL_533381, EPI_ISL_533382, EPI_ISL_533383, EPI_ISL_533384, EPI_ISL_533385, EPI_ISL_533386, EPI_ISL_533387, EPI_ISL_533388, EPI_ISL_533389, EPI_ISL_533390, EPI_ISL_533391, EPI_ISL_533392, EPI_ISL_533393 | | | |
| see above | Lighthouse Lab in Glasgow | Wellcome Sanger Institute for the COVID-19 Genomics UK Consortium | Harper VanSteenhouse, Yumi Kasai, David Gray, Carol Clugston, Anna Dominiczak and Alex Alderton, Roberto Amato, Sonia Goncalves, Ewan Harrison, David K. Jackson, Ian Johnston, Dominic Kwiatkowski, Cordelia Langford, John Sillitoe |
| EPI_ISL_533394 | NHSGGC West of Scotland Specialist Virology Centre / MRC-University of Glasgow Centre for Virus Research | Wellcome Sanger Institute for the COVID-19 Genomics UK Consortium | Ana da Silva Filipe, Natasha Johnson, Kathy Smollett, Daniel Mair, Stephen Carmichael, Lily Tong, Jenna Nichols, Elihu Aranday-Cortes, Kirstyn Brunker, Yasmin Parr, Kyriaki Nomikou; Sarah McDonald, Marc Niebel, Patawee Asamaphan; Richard Orton, Joseph Hughes, Sreenu Vattipally, David L Robertson; Alasdair MacLean, Rory Gunson; Kathy Li, Natasha Jesudason, Rajiv Shah, James Shepherd, Antonia Ho, Alice Broos, Emma Thomson and Alex Alderton, Roberto Amato, Sonia Goncalves, Ewan Harrison, David K. Jackson, Ian Johnston, Dominic Kwiatkowski, Cordelia Langford, John Sillitoe |
| EPI_ISL_533395 | Lighthouse Lab in Glasgow | Wellcome Sanger Institute for the COVID-19 Genomics UK Consortium | Harper VanSteenhouse, Yumi Kasai, David Gray, Carol Clugston, Anna Dominiczak and Alex Alderton, Roberto Amato, Sonia Goncalves, Ewan Harrison, David K. Jackson, Ian Johnston, Dominic Kwiatkowski, Cordelia Langford, John Sillitoe |
| EPI_ISL_533396 | NHSGGC West of Scotland Specialist Virology Centre / MRC-University of Glasgow Centre for Virus Research | Wellcome Sanger Institute for the COVID-19 Genomics UK Consortium | Ana da Silva Filipe, Natasha Johnson, Kathy Smollett, Daniel Mair, Stephen Carmichael, Lily Tong, Jenna Nichols, Elihu Aranday-Cortes, Kirstyn Brunker, Yasmin Parr, Kyriaki Nomikou; Sarah McDonald, Marc Niebel, Patawee Asamaphan; Richard Orton, Joseph Hughes, Sreenu Vattipally, David L Robertson; Alasdair MacLean, Rory Gunson; Kathy Li, Natasha Jesudason, Rajiv Shah, James Shepherd, Antonia Ho, Alice Broos, Emma Thomson and Alex Alderton, Roberto Amato, Sonia Goncalves, Ewan Harrison, David K. Jackson, Ian Johnston, Dominic Kwiatkowski, Cordelia Langford, John Sillitoe |
| EPI_ISL_533397, EPI_ISL_533398, EPI_ISL_533399 | Lighthouse Lab in Glasgow | Wellcome Sanger Institute for the COVID-19 Genomics UK Consortium | Harper VanSteenhouse, Yumi Kasai, David Gray, Carol Clugston, Anna Dominiczak and Alex Alderton, Roberto Amato, Sonia Goncalves, Ewan Harrison, David K. Jackson, Ian Johnston, Dominic Kwiatkowski, Cordelia Langford, John Sillitoe |
| EPI_ISL_533400 | Lighthouse Lab in Glasgow | Wellcome Sanger Institute for the COVID-19 Genomics UK Consortium | Harper VanSteenhouse, Yumi Kasai, David Gray, Carol Clugston, Anna Dominiczak and Alex Alderton, Roberto Amato, Sonia Goncalves, Ewan Harrison, David K. Jackson, Ian Johnston, Dominic Kwiatkowski, Cordelia Langford, John Sillitoe on behalf of the Wellcome Sanger Institute COVID-19 Surveillance Team |
| EPI_ISL_533401, EPI_ISL_533402, EPI_ISL_533404, EPI_ISL_533405 | Lighthouse Lab in Glasgow | Wellcome Sanger Institute for the COVID-19 Genomics UK Consortium | Harper VanSteenhouse, Yumi Kasai, David Gray, Carol Clugston, Anna Dominiczak and Alex Alderton, Roberto Amato, Sonia Goncalves, Ewan Harrison, David K. Jackson, Ian Johnston, Dominic Kwiatkowski, Cordelia Langford, John Sillitoe |
| EPI_ISL_533406 | NHSGGC West of Scotland Specialist Virology Centre / MRC-University of Glasgow Centre for Virus Research | Wellcome Sanger Institute for the COVID-19 Genomics UK Consortium | Ana da Silva Filipe, Natasha Johnson, Kathy Smollett, Daniel Mair, Stephen Carmichael, Lily Tong, Jenna Nichols, Elihu Aranday-Cortes, Kirstyn Brunker, Yasmin Parr, Kyriaki Nomikou; Sarah McDonald, Marc Niebel, Patawee Asamaphan; Richard Orton, Joseph Hughes, Sreenu Vattipally, David L Robertson; Alasdair MacLean, Rory Gunson; Kathy Li, Natasha Jesudason, Rajiv Shah, James Shepherd, Antonia Ho, Alice Broos, Emma Thomson and Alex Alderton, Roberto Amato, Sonia Goncalves, Ewan Harrison, David K. Jackson, Ian Johnston, Dominic Kwiatkowski, Cordelia Langford, John Sillitoe |
| EPI_ISL_533407, EPI_ISL_533408, EPI_ISL_533409, EPI_ISL_533410, EPI_ISL_533411, EPI_ISL_533412, EPI_ISL_533413, EPI_ISL_533414, EPI_ISL_533415, EPI_ISL_533416, EPI_ISL_533417, EPI_ISL_533418 | | | |
| see above | Lighthouse Lab in Glasgow | Wellcome Sanger Institute for the COVID-19 Genomics UK Consortium | Harper VanSteenhouse, Yumi Kasai, David Gray, Carol Clugston, Anna Dominiczak and Alex Alderton, Roberto Amato, Sonia Goncalves, Ewan Harrison, David K. Jackson, Ian Johnston, Dominic Kwiatkowski, Cordelia Langford, John Sillitoe |
| EPI_ISL_533419 | NHSGGC West of Scotland Specialist Virology Centre / MRC-University of Glasgow Centre for Virus Research | Wellcome Sanger Institute for the COVID-19 Genomics UK Consortium | Ana da Silva Filipe, Natasha Johnson, Kathy Smollett, Daniel Mair, Stephen Carmichael, Lily Tong, Jenna Nichols, Elihu Aranday-Cortes, Kirstyn Brunker, Yasmin Parr, Kyriaki Nomikou; Sarah McDonald, Marc Niebel, Patawee Asamaphan; Richard Orton, Joseph Hughes, Sreenu Vattipally, David L Robertson; Alasdair MacLean, Rory Gunson; Kathy Li, Natasha Jesudason, Rajiv Shah, James Shepherd, Antonia Ho, Alice Broos, Emma Thomson and Alex Alderton, Roberto Amato, Sonia Goncalves, Ewan Harrison, David K. Jackson, Ian Johnston, Dominic Kwiatkowski, Cordelia Langford, John Sillitoe |
| EPI_ISL_533420 | Lighthouse Lab in Glasgow | Wellcome Sanger Institute for the COVID-19 Genomics UK Consortium | Harper VanSteenhouse, Yumi Kasai, David Gray, Carol Clugston, Anna Dominiczak and Alex Alderton, Roberto Amato, Sonia Goncalves, Ewan Harrison, David K. Jackson, Ian Johnston, Dominic Kwiatkowski, Cordelia Langford, John Sillitoe |
| EPI_ISL_533421 | NHSGGC West of Scotland Specialist Virology Centre / MRC-University of Glasgow Centre for Virus Research | Wellcome Sanger Institute for the COVID-19 Genomics UK Consortium | Ana da Silva Filipe, Natasha Johnson, Kathy Smollett, Daniel Mair, Stephen Carmichael, Lily Tong, Jenna Nichols, Elihu Aranday-Cortes, Kirstyn Brunker, Yasmin Parr, Kyriaki Nomikou; Sarah McDonald, Marc Niebel, Patawee Asamaphan; Richard Orton, Joseph Hughes, Sreenu Vattipally, David L Robertson; Alasdair MacLean, Rory Gunson; Kathy Li, Natasha Jesudason, Rajiv Shah, James Shepherd, Antonia Ho, Alice Broos, Emma Thomson and Alex Alderton, Roberto Amato, Sonia Goncalves, Ewan Harrison, David K. Jackson, Ian Johnston, Dominic Kwiatkowski, Cordelia Langford, John Sillitoe |
| EPI_ISL_533422, EPI_ISL_533423, EPI_ISL_533424, EPI_ISL_533425, EPI_ISL_533426, EPI_ISL_533427, EPI_ISL_533428, EPI_ISL_533429, EPI_ISL_533430, EPI_ISL_533431 | Lighthouse Lab in Glasgow | Wellcome Sanger Institute for the COVID-19 Genomics UK Consortium | Harper VanSteenhouse, Yumi Kasai, David Gray, Carol Clugston, Anna Dominiczak and Alex Alderton, Roberto Amato, Sonia Goncalves, Ewan Harrison, David K. Jackson, Ian Johnston, Dominic Kwiatkowski, Cordelia Langford, John Sillitoe |
| EPI_ISL_533432, EPI_ISL_533433, EPI_ISL_533434 | Lighthouse Lab in Glasgow | Wellcome Sanger Institute for the COVID-19 Genomics UK Consortium | Harper VanSteenhouse, Yumi Kasai, David Gray, Carol Clugston, Anna Dominiczak and Alex Alderton, Roberto Amato, Sonia Goncalves, Ewan Harrison, David K. Jackson, Ian Johnston, Dominic Kwiatkowski, Cordelia Langford, John Sillitoe on behalf of the Wellcome Sanger Institute COVID-19 Surveillance Team |
| EPI_ISL_534199 | E. Gulbja Laboratorija | Latvian Biomedical Research and Study Centre | Ivars Silamiķelis, Jānis Pjalkovskis, Kaspars Megnis, Monta Ustinova, Ņikita Zrelovs, Vita Rovīte, Mikus Gavars, Dmitrijs Perminovs, Uga Dumpis, Jānis Kloviņš |
| EPI_ISL_534200, EPI_ISL_534201, EPI_ISL_534202 | Centrālā laboratorija | Latvian Biomedical Research and Study Centre | Ivars Silamiķelis, Jānis Pjalkovskis, Kaspars Megnis, Monta Ustinova, Ņikita Zrelovs, Vita Rovīte, Stella Lapiņa, Jana Osīte, Marta Priedīte, Uga Dumpis, Jānis Kloviņš |
| EPI_ISL_534203 | E. Gulbja Laboratorija | Latvian Biomedical Research and Study Centre | Ivars Silamiķelis, Jānis Pjalkovskis, Kaspars Megnis, Monta Ustinova, Ņikita Zrelovs, Vita Rovīte, Mikus Gavars, Dmitrijs Perminovs, Uga Dumpis, Jānis Kloviņš |
| EPI_ISL_534204, EPI_ISL_534205 | Latvijas Infektoloģijas centrs | Latvian Biomedical Research and Study Centre | Ivars Silamiķelis, Jānis Pjalkovskis, Kaspars Megnis, Monta Ustinova, Ņikita Zrelovs, Vita Rovīte, Jeļena Storoženko, Tatjana Kolupajeva, Oksana Savicka, Uga Dumpis, Jānis Kloviņš |
| EPI_ISL_534206, EPI_ISL_534207, EPI_ISL_534208 | Centrālā laboratorija | Latvian Biomedical Research and Study Centre | Ivars Silamiķelis, Jānis Pjalkovskis, Kaspars Megnis, Monta Ustinova, Ņikita Zrelovs, Vita Rovīte, Stella Lapiņa, Jana Osīte, Marta Priedīte, Uga Dumpis, Jānis Kloviņš |
| EPI_ISL_534209 | Latvijas Infektoloģijas centrs | Latvian Biomedical Research and Study Centre | Ivars Silamiķelis, Jānis Pjalkovskis, Kaspars Megnis, Monta Ustinova, Ņikita Zrelovs, Vita Rovīte, Jeļena Storoženko, Tatjana Kolupajeva, Oksana Savicka, Uga Dumpis, Jānis Kloviņš |
| EPI_ISL_534210, EPI_ISL_534211 | Centrālā laboratorija | Latvian Biomedical Research and Study Centre | Ivars Silamiķelis, Jānis Pjalkovskis, Kaspars Megnis, Monta Ustinova, Ņikita Zrelovs, Vita Rovīte, Stella Lapiņa, Jana Osīte, Marta Priedīte, Uga Dumpis, Jānis Kloviņš |
| EPI_ISL_534212, EPI_ISL_534213, EPI_ISL_534214, EPI_ISL_534215, EPI_ISL_534216 | E. Gulbja Laboratorija | Latvian Biomedical Research and Study Centre | Ivars Silamiķelis, Jānis Pjalkovskis, Kaspars Megnis, Monta Ustinova, Ņikita Zrelovs, Vita Rovīte, Mikus Gavars, Dmitrijs Perminovs, Uga Dumpis, Jānis Kloviņš |
| EPI_ISL_534217 | M Health Fairview | Minnesota Department of Health, Public Health Laboratory | Matt Plumb, Jacob Garfin, and Xiong Wang |
| EPI_ISL_534218, EPI_ISL_534219 | E. Gulbja Laboratorija | Latvian Biomedical Research and Study Centre | Ivars Silamiķelis, Jānis Pjalkovskis, Kaspars Megnis, Monta Ustinova, Ņikita Zrelovs, Vita Rovīte, Mikus Gavars, Dmitrijs Perminovs, Uga Dumpis, Jānis Kloviņš |
| EPI_ISL_534220 | Centrālā laboratorija | Latvian Biomedical Research and Study Centre | Ivars Silamiķelis, Jānis Pjalkovskis, Kaspars Megnis, Monta Ustinova, Ņikita Zrelovs, Vita Rovīte, Stella Lapiņa, Jana Osīte, Marta Priedīte, Uga Dumpis, Jānis Kloviņš |
| EPI_ISL_534221 | E. Gulbja Laboratorija | Latvian Biomedical Research and Study Centre | Ivars Silamiķelis, Jānis Pjalkovskis, Kaspars Megnis, Monta Ustinova, Ņikita Zrelovs, Vita Rovīte, Mikus Gavars, Dmitrijs Perminovs, Uga Dumpis, Jānis Kloviņš |
| EPI_ISL_534222, EPI_ISL_534223 | Centrālā laboratorija | Latvian Biomedical Research and Study Centre | Ivars Silamiķelis, Jānis Pjalkovskis, Kaspars Megnis, Monta Ustinova, Ņikita Zrelovs, Vita Rovīte, Stella Lapiņa, Jana Osīte, Marta Priedīte, Uga Dumpis, Jānis Kloviņš |
| EPI_ISL_534224 | Universitetssjukhuset i Linkoping | The Public Health Agency of Sweden | Anna-Malin Linde, Maria Lind Karlberg, Mattias Haukland, Reza Advani, Olov Svartstrom, Oskar Karlsson Lindsjo, Sandra Broddesson, Petra Edquist, Mia Brytting, Anna Risberg, Karin Tegmark-Wisell |
| EPI_ISL_534225, EPI_ISL_534226, EPI_ISL_534227, EPI_ISL_534228, EPI_ISL_534229 | Skanes universitetssjukhus Lund | The Public Health Agency of Sweden | Anna-Malin Linde, Maria Lind Karlberg, Mattias Haukland, Reza Advani, Olov Svartstrom, Oskar Karlsson Lindsjo, Sandra Broddesson, Petra Edquist, Mia Brytting, Anna Risberg, Karin Tegmark-Wisell |
| EPI_ISL_534230, EPI_ISL_534231, EPI_ISL_534232, EPI_ISL_534233 | Capio S:t Gorans sjukhus | The Public Health Agency of Sweden | Anna-Malin Linde, Maria Lind Karlberg, Mattias Haukland, Reza Advani, Olov Svartstrom, Oskar Karlsson Lindsjo, Sandra Broddesson, Petra Edquist, Mia Brytting, Anna Risberg, Karin Tegmark-Wisell |
| EPI_ISL_534234, EPI_ISL_534235, EPI_ISL_534236 | Karolinska universitetslaboratoriet SOLNA | The Public Health Agency of Sweden | Anna-Malin Linde, Maria Lind Karlberg, Mattias Haukland, Reza Advani, Olov Svartstrom, Oskar Karlsson Lindsjo, Sandra Broddesson, Petra Edquist, Mia Brytting, Anna Risberg, Karin Tegmark-Wisell |

| | | | |
|---|---|---|---|
| EPI_ISL_534237, EPI_ISL_534238 | Lanssjukhuset Kalmar | The Public Health Agency of Sweden | Anna-Malin Linde, Maria Lind Karlberg, Mattias Haukland, Reza Advani, Olov Svartstrom, Oskar Karlsson Lindsjo, Sandra Broddesson, Petra Edquist, Mia Brytting, Anna Risberg, Karin Tegmark-Wisell |
| EPI_ISL_534239, EPI_ISL_534240, EPI_ISL_534241, EPI_ISL_534242, EPI_ISL_534243 | Norra Alvsborgs lanssjukhus | The Public Health Agency of Sweden | Anna-Malin Linde, Maria Lind Karlberg, Mattias Haukland, Reza Advani, Olov Svartstrom, Oskar Karlsson Lindsjo, Sandra Broddesson, Petra Edquist, Mia Brytting, Anna Risberg, Karin Tegmark-Wisell |
| EPI_ISL_534244 | Sundsvalls sjukhus | The Public Health Agency of Sweden | Anna-Malin Linde, Maria Lind Karlberg, Mattias Haukland, Reza Advani, Olov Svartstrom, Oskar Karlsson Lindsjo, Sandra Broddesson, Petra Edquist, Mia Brytting, Anna Risberg, Karin Tegmark-Wisell |
| EPI_ISL_534245 | Kliniskt mikrobiologiska laboratoriet | The Public Health Agency of Sweden | Anna-Malin Linde, Maria Lind Karlberg, Mattias Haukland, Reza Advani, Olov Svartstrom, Oskar Karlsson Lindsjo, Sandra Broddesson, Petra Edquist, Mia Brytting, Anna Risberg, Karin Tegmark-Wisell |
| EPI_ISL_534246, EPI_ISL_534247 | Universitetssjukhuset i Linkoping | The Public Health Agency of Sweden | Anna-Malin Linde, Maria Lind Karlberg, Mattias Haukland, Reza Advani, Olov Svartstrom, Oskar Karlsson Lindsjo, Sandra Broddesson, Petra Edquist, Mia Brytting, Anna Risberg, Karin Tegmark-Wisell |
| EPI_ISL_534248 | Laboratoriemedicin Vasternorrland | The Public Health Agency of Sweden | Anna-Malin Linde, Maria Lind Karlberg, Mattias Haukland, Reza Advani, Olov Svartstrom, Oskar Karlsson Lindsjo, Sandra Broddesson, Petra Edquist, Mia Brytting, Anna Risberg, Karin Tegmark-Wisell |
| EPI_ISL_534249, EPI_ISL_534250, EPI_ISL_534251 | Gavle Sjukhus | The Public Health Agency of Sweden | Anna-Malin Linde, Maria Lind Karlberg, Mattias Haukland, Reza Advani, Olov Svartstrom, Oskar Karlsson Lindsjo, Sandra Broddesson, Petra Edquist, Mia Brytting, Anna Risberg, Karin Tegmark-Wisell |
| EPI_ISL_534252, EPI_ISL_534253, EPI_ISL_534254, EPI_ISL_534255 | Laboratoriemedicin Vasternorrland | The Public Health Agency of Sweden | Anna-Malin Linde, Maria Lind Karlberg, Mattias Haukland, Reza Advani, Olov Svartstrom, Oskar Karlsson Lindsjo, Sandra Broddesson, Petra Edquist, Mia Brytting, Anna Risberg, Karin Tegmark-Wisell |
| EPI_ISL_534256 | Klinisk mikrobiologi, Laboratoriemedicin Gavleborg | The Public Health Agency of Sweden | Anna-Malin Linde, Maria Lind Karlberg, Mattias Haukland, Reza Advani, Olov Svartstrom, Oskar Karlsson Lindsjo, Sandra Broddesson, Petra Edquist, Mia Brytting, Anna Risberg, Karin Tegmark-Wisell |
| EPI_ISL_534257, EPI_ISL_534258 | Laboratoriemedicin Vasternorrland | The Public Health Agency of Sweden | Anna-Malin Linde, Maria Lind Karlberg, Mattias Haukland, Reza Advani, Olov Svartstrom, Oskar Karlsson Lindsjo, Sandra Broddesson, Petra Edquist, Mia Brytting, Anna Risberg, Karin Tegmark-Wisell |
| EPI_ISL_534259 | Karolinska universitetslaboratoriet | The Public Health Agency of Sweden | Anna-Malin Linde, Maria Lind Karlberg, Mattias Haukland, Reza Advani, Olov Svartstrom, Oskar Karlsson Lindsjo, Sandra Broddesson, Petra Edquist, Mia Brytting, Anna Risberg, Karin Tegmark-Wisell |
| EPI_ISL_534260, EPI_ISL_534261, EPI_ISL_534262, EPI_ISL_534263, EPI_ISL_534264, EPI_ISL_534265, EPI_ISL_534266, EPI_ISL_534267, EPI_ISL_534268, EPI_ISL_534269, EPI_ISL_534270, EPI_ISL_534271, EPI_ISL_534272, EPI_ISL_534273, EPI_ISL_534274, EPI_ISL_534275, EPI_ISL_534276, EPI_ISL_534277, EPI_ISL_534278, EPI_ISL_534279, EPI_ISL_534280, EPI_ISL_534281, EPI_ISL_534282, EPI_ISL_534283, EPI_ISL_534284, EPI_ISL_534285, EPI_ISL_534286, EPI_ISL_534287, EPI_ISL_534288, EPI_ISL_534289, EPI_ISL_534290, EPI_ISL_534291, EPI_ISL_534292, EPI_ISL_534293, EPI_ISL_534294, EPI_ISL_534295, EPI_ISL_534296, EPI_ISL_534297, EPI_ISL_534298, EPI_ISL_534299, EPI_ISL_534300, EPI_ISL_534301, EPI_ISL_534302 | | | |
| see above | Texas Department of State Health Services | Texas Department of State Health Services | Rashmi Tuladhar, Bonnie Oh, Jenny Zhang, Maliha Rahman, Anita Pokharel, Myong Koag, Chun Wang, Rachel Lee, Grace Kubin |
| EPI_ISL_534311 | UPA III 26 de Agosto | Instituto Adolfo Lutz, Interdiciplinary Procedures Center, Strategic Laboratory | Claudio Tavares Sacchi, Claudia Regina Gonçalves, Erica Valessa Ramos Gomes |
| EPI_ISL_534312 | Distrito Sanitario Sul | Instituto Adolfo Lutz, Interdiciplinary Procedures Center, Strategic Laboratory | Claudio Tavares Sacchi, Claudia Regina Gonçalves, Erica Valessa Ramos Gomes |
| EPI_ISL_534313 | Hospital da Sta Casa de Sto Amaro | Instituto Adolfo Lutz, Interdiciplinary Procedures Center, Strategic Laboratory | Claudio Tavares Sacchi, Claudia Regina Gonçalves, Erica Valessa Ramos Gomes |
| EPI_ISL_534314 | Hospital Universitario da USP de SP | Instituto Adolfo Lutz, Interdiciplinary Procedures Center, Strategic Laboratory | Claudio Tavares Sacchi, Claudia Regina Gonçalves, Erica Valessa Ramos Gomes |
| EPI_ISL_534315 | Serviço de Verificação de Óbitos SVO Guaruulhos | Instituto Adolfo Lutz, Interdiciplinary Procedures Center, Strategic Laboratory | Claudio Tavares Sacchi, Claudia Regina Gonçalves, Erica Valessa Ramos Gomes |
| EPI_ISL_534316 | OS Mun Santana Lauro Ribas Braga | Instituto Adolfo Lutz, Interdiciplinary Procedures Center, Strategic Laboratory | Claudio Tavares Sacchi, Claudia Regina Gonçalves, Erica Valessa Ramos Gomes |
| EPI_ISL_534317 | Hospital Geral de Itapevi | Instituto Adolfo Lutz, Interdiciplinary Procedures Center, Strategic Laboratory | Claudio Tavares Sacchi, Claudia Regina Gonçalves, Erica Valessa Ramos Gomes |
| EPI_ISL_534318 | Hospital Municipal Antonio Giglio | Instituto Adolfo Lutz, Interdiciplinary Procedures Center, Strategic Laboratory | Claudio Tavares Sacchi, Claudia Regina Gonçalves, Erica Valessa Ramos Gomes |
| EPI_ISL_534319, EPI_ISL_534320 | Hospital do Serv Pub ESTAFCO Morato de Oliveira | Instituto Adolfo Lutz, Interdiciplinary Procedures Center, Strategic Laboratory | Claudio Tavares Sacchi, Claudia Regina Gonçalves, Erica Valessa Ramos Gomes |
| EPI_ISL_534321 | PS e Maternidade Nair Fonseca Leitao Arantes | Instituto Adolfo Lutz, Interdiciplinary Procedures Center, Strategic Laboratory | Claudio Tavares Sacchi, Claudia Regina Gonçalves, Erica Valessa Ramos Gomes |
| EPI_ISL_534322 | PS Mun Julio Tupy | Instituto Adolfo Lutz, Interdiciplinary Procedures Center, Strategic Laboratory | Claudio Tavares Sacchi, Claudia Regina Gonçalves, Erica Valessa Ramos Gomes |
| EPI_ISL_534323 | Hospital e Pronto Socorro Comunitario Vila Yolanda | Instituto Adolfo Lutz, Interdiciplinary Procedures Center, Strategic Laboratory | Claudio Tavares Sacchi, Claudia Regina Gonçalves, Erica Valessa Ramos Gomes |
| EPI_ISL_534324 | Hospital Mun Ver Jose Storopolli | Instituto Adolfo Lutz, Interdiciplinary Procedures Center, Strategic Laboratory | Claudio Tavares Sacchi, Claudia Regina Gonçalves, Erica Valessa Ramos Gomes |
| EPI_ISL_534325 | Unidade de Vigilancia em Saude de Guarulhos | Instituto Adolfo Lutz, Interdiciplinary Procedures Center, Strategic Laboratory | Claudio Tavares Sacchi, Claudia Regina Gonçalves, Erica Valessa Ramos Gomes |
| EPI_ISL_534326 | Notre Dame Intermedica Saude AS | Instituto Adolfo Lutz, Interdiciplinary Procedures Center, Strategic Laboratory | Claudio Tavares Sacchi, Claudia Regina Gonçalves, Erica Valessa Ramos Gomes |
| EPI_ISL_534327, EPI_ISL_534328 | Hospital Universitario 12 de Octubre | Hospital Universitario 12 de Octubre | Raúl Recio, Sara González, Esther Viedma, Elias Dahdouh, Fernando Lázaro, Natalia Stella, Julio García, Juan Carlos Galán, Rafael Cantón, Ma Dolores Folgueira, Rafael Delgado, Jesús Mingorance |
| EPI_ISL_534329 | Hospital Universitario 12 de Octubre | Hospital Universitario 12 de Octubre | Esther Viedma, Raúl Recio, Sara González, Elias Dahdouh, Fernando Lázaro, Natalia Stella, Julio García, Juan Carlos Galán, Rafael Cantón, Ma Dolores Folgueira, Rafael Delgado, Jesús Mingorance |
| EPI_ISL_534330, EPI_ISL_534331, EPI_ISL_534332, EPI_ISL_534333 | Hospital Universitario La Paz | Hospital Universitario La Paz | María Rodríguez, Elias Dahdouh, Sara González, Raúl Recio, Fernando Lázaro, Esther Viedma, Natalia Stella, Julio García, Juan Carlos Galán, Rafael Cantón, Ma Dolores Folgueira, Rafael Delgado, Jesús Mingorance |
| EPI_ISL_534334 | Hospital Universitario Ramón y Cajal | Hospital Universitario La Paz | María Rodríguez, Elias Dahdouh, Sara González, Raúl Recio, Fernando Lázaro, Esther Viedma, Natalia Stella, Julio García, Juan Carlos Galán, Rafael Cantón, Ma Dolores Folgueira, Rafael Delgado, Jesús Mingorance |
| EPI_ISL_534335 | LabPLUS | Institute of Environmental Science and Research (ESR) | Matt Storey, Xiaoyun Ren, Anja Werno, Antje van der Linden, Arlo Upton, Chris Mansell, David Hammer, Dragana Drinkovic, Gary McAuliffe, Hana Sofia Andersson, James Ussher, Jill Sherwood, Josh Freeman, Julia Howard, Juliet Elvy, Mary DeAlmeida, Matt Blakiston, Matthew Rogers, Max Bloomfield, Michael Addidle, Michelle Balm, Sally Roberts, Sarah Jefferies, Sharmini Muttaiyah, Susan Morpeth, Susan Taylor, Timothy Blackmore, Vani Sathyendran, Veronica Playle, Virginia Hope, Erasmus Smit, Lauren Jelly, Joep de Ligt |
| EPI_ISL_534336 | Department of Laboratory Medicine, National Taiwan University Hospital | Microbial Genomics Core Lab, National Taiwan University Centers of Genomic and Precision Medicine | Shiou-Hwei Yeh, You-Yu Lin, Ya-Yun Lai, Chiao-Ling Li, Shan-Chwen Chang, Pei-Jer Chen, Sui-Yuan Chang |
| EPI_ISL_534337, EPI_ISL_534338, EPI_ISL_534339, EPI_ISL_534340, EPI_ISL_534341, EPI_ISL_534342, EPI_ISL_534343, EPI_ISL_534344, EPI_ISL_534345 | Molecular diagnostic laboratory of Federal Budget Institution of Science "Central Research Institute of Epidemiology" of The Federal Service on Customers' Rights Protection and Human Well-being Surveillance | Group of Genomics and Postgenomic Technologies of Central Research Institute of Epidemilology | Speranskaya AS, Kaptelova VV, Samoilov AE, Valdokhina AV, Bulanenko VP, Bukharina A.Y., Tivanova EV, Shipulina OY, Akimkin VG |
| EPI_ISL_534346, EPI_ISL_534347, EPI_ISL_534348, EPI_ISL_534349 | Molecular diagnostic laboratory of Federal Budget Institution of Science "Central Research Institute of Epidemiology" of The Federal Service on Customers' Rights Protection and Human Well-being Surveillance | Group of Genomics and Postgenomic Technologies of Central Research Institute of Epidemilology | Speranskaya AS, Kaptelova VV, Valdokhina AV, Bulanenko VP, Samoilov AE, Korneenko EV, Tivanova EV, Shipulina OY, Akimkin VG |
| EPI_ISL_534365, EPI_ISL_534366, EPI_ISL_534367, EPI_ISL_534368, EPI_ISL_534369, EPI_ISL_534370, EPI_ISL_534371, EPI_ISL_534372, EPI_ISL_534373, EPI_ISL_534374, EPI_ISL_534375, EPI_ISL_534376, EPI_ISL_534377, EPI_ISL_534378, EPI_ISL_534379, EPI_ISL_534380, EPI_ISL_534381, EPI_ISL_534382, EPI_ISL_534383, EPI_ISL_534384, EPI_ISL_534385, EPI_ISL_534386, EPI_ISL_534387, EPI_ISL_534388, EPI_ISL_534389, EPI_ISL_534390, EPI_ISL_534391, EPI_ISL_534392, EPI_ISL_534393, EPI_ISL_534394, EPI_ISL_534396, EPI_ISL_534397, EPI_ISL_534398, EPI_ISL_534399, EPI_ISL_534401, EPI_ISL_534402, EPI_ISL_534403, EPI_ISL_534404, EPI_ISL_534405, EPI_ISL_534406, EPI_ISL_534407, EPI_ISL_534408, EPI_ISL_534409, EPI_ISL_534410, EPI_ISL_534411, EPI_ISL_534412, EPI_ISL_534413, EPI_ISL_534414, EPI_ISL_534415, EPI_ISL_534416, EPI_ISL_534417, EPI_ISL_534418, EPI_ISL_534419, EPI_ISL_534420, EPI_ISL_534421, EPI_ISL_534422, EPI_ISL_534423, EPI_ISL_534424, EPI_ISL_534425, EPI_ISL_534426, EPI_ISL_534427, EPI_ISL_534428, EPI_ISL_534429, EPI_ISL_534430, EPI_ISL_534431, EPI_ISL_534432, EPI_ISL_534433, EPI_ISL_534434, EPI_ISL_534435, EPI_ISL_534436, EPI_ISL_534437, EPI_ISL_534438, EPI_ISL_534439, EPI_ISL_534440, EPI_ISL_534441, EPI_ISL_534442, EPI_ISL_534443, EPI_ISL_534444, EPI_ISL_534445, EPI_ISL_534446, EPI_ISL_534447, EPI_ISL_534448, EPI_ISL_534449, EPI_ISL_534450, EPI_ISL_534451, EPI_ISL_534452, EPI_ISL_534453, EPI_ISL_534454, EPI_ISL_534455, EPI_ISL_534456, EPI_ISL_534457, EPI_ISL_534458, EPI_ISL_534459, EPI_ISL_534460, EPI_ISL_534461, EPI_ISL_534462, EPI_ISL_534463, EPI_ISL_534464, EPI_ISL_534465, EPI_ISL_534466, EPI_ISL_534467, EPI_ISL_534468, EPI_ISL_534469, EPI_ISL_534470, EPI_ISL_534471, EPI_ISL_534472, EPI_ISL_534473, EPI_ISL_534474, EPI_ISL_534475, EPI_ISL_534476, EPI_ISL_534477, EPI_ISL_534478, EPI_ISL_534479, EPI_ISL_534480, EPI_ISL_534481, EPI_ISL_534482, EPI_ISL_534483, EPI_ISL_534484, EPI_ISL_534485, EPI_ISL_534486, EPI_ISL_534487, EPI_ISL_534488, EPI_ISL_534489, EPI_ISL_534490, EPI_ISL_534491, EPI_ISL_534492, EPI_ISL_534493, EPI_ISL_534494, EPI_ISL_534495, EPI_ISL_534496, EPI_ISL_534497, EPI_ISL_534498, EPI_ISL_534499, EPI_ISL_534500, EPI_ISL_534501, EPI_ISL_534502, EPI_ISL_534503, EPI_ISL_534504, EPI_ISL_534505, EPI_ISL_534506, EPI_ISL_534507, EPI_ISL_534508, EPI_ISL_534509, EPI_ISL_534510, EPI_ISL_534511, EPI_ISL_534512, EPI_ISL_534513, EPI_ISL_534514, EPI_ISL_534515, EPI_ISL_534516, EPI_ISL_534517, EPI_ISL_534518, EPI_ISL_534519, EPI_ISL_534520, EPI_ISL_534521, EPI_ISL_534522, EPI_ISL_534523, EPI_ISL_534524, EPI_ISL_534525, EPI_ISL_534526, EPI_ISL_534527, EPI_ISL_534528, EPI_ISL_534529, EPI_ISL_534530, EPI_ISL_534531, EPI_ISL_534532, EPI_ISL_534533, EPI_ISL_534534, EPI_ISL_534535, EPI_ISL_534536, EPI_ISL_534537, EPI_ISL_534538, EPI_ISL_534539, EPI_ISL_534540, EPI_ISL_534541, EPI_ISL_534542, EPI_ISL_534543, EPI_ISL_534544, EPI_ISL_534545, EPI_ISL_534546, EPI_ISL_534547, EPI_ISL_534548, EPI_ISL_534549, EPI_ISL_534550, EPI_ISL_534551, EPI_ISL_534552, EPI_ISL_534553, EPI_ISL_534554, EPI_ISL_534555, EPI_ISL_534556, EPI_ISL_534557, EPI_ISL_534558, EPI_ISL_534559, EPI_ISL_534560, EPI_ISL_534561, EPI_ISL_534562, EPI_ISL_534563, EPI_ISL_534564, EPI_ISL_534565, EPI_ISL_534566, EPI_ISL_534567, EPI_ISL_534568, EPI_ISL_534569, EPI_ISL_534570, EPI_ISL_534571, EPI_ISL_534572, EPI_ISL_534573, EPI_ISL_534574, EPI_ISL_534575, EPI_ISL_534576, EPI_ISL_534577, EPI_ISL_534578, EPI_ISL_534579, EPI_ISL_534580, EPI_ISL_534581, EPI_ISL_534582, EPI_ISL_534583, EPI_ISL_534584, EPI_ISL_534585, EPI_ISL_534586, EPI_ISL_534587, EPI_ISL_534588, EPI_ISL_534589, EPI_ISL_534590, EPI_ISL_534591, EPI_ISL_534592, EPI_ISL_534593, EPI_ISL_534594, EPI_ISL_534595, EPI_ISL_534596, EPI_ISL_534597, EPI_ISL_534598, EPI_ISL_534599, EPI_ISL_534600, EPI_ISL_534601, EPI_ISL_534602, EPI_ISL_534603, EPI_ISL_534604, EPI_ISL_534605, EPI_ISL_534606, EPI_ISL_534607, EPI_ISL_534608, EPI_ISL_534609, EPI_ISL_534610, EPI_ISL_534611, EPI_ISL_534612, EPI_ISL_534613, EPI_ISL_534614, EPI_ISL_534615, EPI_ISL_534616, EPI_ISL_534617, EPI_ISL_534618, EPI_ISL_534619, EPI_ISL_534620, EPI_ISL_534621, EPI_ISL_534622, EPI_ISL_534623, EPI_ISL_534624, EPI_ISL_534625, EPI_ISL_534626, EPI_ISL_534627, EPI_ISL_534628, EPI_ISL_534629, EPI_ISL_534630, EPI_ISL_534631, EPI_ISL_534632, EPI_ISL_534633, EPI_ISL_534635, EPI_ISL_534636, EPI_ISL_534637, EPI_ISL_534638, EPI_ISL_534639, EPI_ISL_534640, EPI_ISL_534641, EPI_ISL_534642, EPI_ISL_534643, EPI_ISL_534644, EPI_ISL_534645, EPI_ISL_534646, EPI_ISL_534647, EPI_ISL_534648, EPI_ISL_534649, EPI_ISL_534650, EPI_ISL_534651, EPI_ISL_534652, EPI_ISL_534653, EPI_ISL_534654, EPI_ISL_534655, EPI_ISL_534656, EPI_ISL_534657, EPI_ISL_534658, EPI_ISL_534659, EPI_ISL_534660, EPI_ISL_534661, EPI_ISL_534662, EPI_ISL_534663, EPI_ISL_534664, EPI_ISL_534665, EPI_ISL_534666, EPI_ISL_534667, EPI_ISL_534668, EPI_ISL_534669, EPI_ISL_534670, EPI_ISL_534671, EPI_ISL_534672, EPI_ISL_534673, EPI_ISL_534674, EPI_ISL_534675, EPI_ISL_534676, EPI_ISL_534677, EPI_ISL_534678, EPI_ISL_534679, EPI_ISL_534680, EPI_ISL_534681, EPI_ISL_534682, EPI_ISL_534683, EPI_ISL_534684, EPI_ISL_534685, EPI_ISL_534686, EPI_ISL_534687, EPI_ISL_534688, EPI_ISL_534689, EPI_ISL_534690 | | | |
| see above | NHSGGC West of Scotland Specialist Virology Centre / MRC-University of Glasgow Centre for Virus Research | Wellcome Sanger Institute for the COVID-19 Genomics UK Consortium | Ana da Silva Filipe, Natasha Johnson, Kathy Smollett, Daniel Mair, Stephen Carmichael, Lily Tong, Jenna Nichols, Elihu Aranday-Cortes, Kirstyn Brunker, Yasmin Parr, Kyriaki Nomikou; Sarah McDonald, Marc Niebel, Patawee Asamaphan; Richard Orton, Joseph Hughes, Sreenu Vattipally, David L Robertson; Alasdair MacLean, Rory Gunson; Kathy Li, Natasha Jesudason, Rajiv Shah, James Shepherd, Antonia Ho, Alice Broos, Emma Thomson and Alex Alderton, Roberto Amato, Sonia Goncalves, Ewan Harrison, David K. Jackson, Ian Johnston, Dominic Kwiatkowski, Cordelia Langford, John Sillitoe on behalf of the Wellcome Sanger Institute COVID-19 Surveillance Team (http://www.sanger.ac.uk/covid-team) |
| EPI_ISL_534693, EPI_ISL_534694, EPI_ISL_534695, EPI_ISL_534696, EPI_ISL_534697, EPI_ISL_534698 | University of Miami Immunology and Histocompatibility Laboratory | University of Miami Immunology and Histocompatibility Laboratory | Emilio Margolles-Clark, PhD and Phillip Ruiz, MD, PhD |
| EPI_ISL_534699, EPI_ISL_534700, EPI_ISL_534701, EPI_ISL_534702, EPI_ISL_534703, EPI_ISL_534704, EPI_ISL_534705, EPI_ISL_534706, EPI_ISL_534707, EPI_ISL_534708, EPI_ISL_534709, EPI_ISL_534710, EPI_ISL_534711, EPI_ISL_534712, EPI_ISL_534713, EPI_ISL_534714, EPI_ISL_534715, EPI_ISL_534716 | | | |
| see above | MD PHL | MD PHL | Maryland Department of Health Laboratories Administration |
| EPI_ISL_534717, EPI_ISL_534718, EPI_ISL_534719, EPI_ISL_534720, EPI_ISL_534721, EPI_ISL_534722, EPI_ISL_534723, EPI_ISL_534724, EPI_ISL_534725, EPI_ISL_534726, EPI_ISL_534727, EPI_ISL_534728, EPI_ISL_534729, EPI_ISL_534730 | | | |
| see above | Respiratory Virus Unit, Microbiology Services Colindale, Public Health England | Respiratory Virus Unit, Microbiology Services Colindale, Public Health England | PHE Covid Sequencing Team |
| EPI_ISL_534731 | Wadsworth Center, New York State Department of Health | Wadsworth Center, New York State Department of Health | Kirsten St. George, Daryl M. Lamson, Sara Griesemer, Jonathan Plitnick, Navjot Singh, Matthew D. Shudt, Erica Lasek-Nesselquist |
| EPI_ISL_534732, EPI_ISL_534733, EPI_ISL_534734, EPI_ISL_534735, EPI_ISL_534736, EPI_ISL_534737, EPI_ISL_534738, EPI_ISL_534739, EPI_ISL_534740, EPI_ISL_534741, EPI_ISL_534742, EPI_ISL_534743, EPI_ISL_534744, EPI_ISL_534745, EPI_ISL_534746, EPI_ISL_534747, EPI_ISL_534748, EPI_ISL_534749, EPI_ISL_534750, EPI_ISL_534751, EPI_ISL_534752, EPI_ISL_534753, EPI_ISL_534754, EPI_ISL_534755 | | | |

| | | | |
|---|---|---|---|
| see above | Liverpool Clinical Laboratories | COVID-19 Genomics UK (COG-UK) Consortium | Sam Haldenby, Anita Lucaci, Steve Paterson, Julian Hiscox, Alistair Darby, M Almsaud, A Alrezaihi, Muhannad Alruwaili, Stuart D Armstrong, Jones Benjamin, Eleanor G Bentley, Anu Chawla, Jordan J Clark, Angela Cowell, Richard Eccles, Isabel García-Dorival, Matthew Gemmell, Alessandro Gerada, PKF Gilmore, Richard Gregory, Ximeng Han, Catherine Hartley, Margaret Hughes, Miren Iturriza-Gomara, James Johnson, L Luu, Jenifer Manson, Charlotte Nelson, Elaine O'Toole, Cassie Olateju, Rebekah Penrice-Randal , Lucille Rainbow, N.P Randle, Trevor Ian Robinson, Parul Sharma, Ghada T Shawli, James P Stewart, Neil Swainston, Ecaterina Vamos, Joanne Watts, Mark Whitehead |
| EPI_ISL_534756, EPI_ISL_534757, EPI_ISL_534758, EPI_ISL_534759, EPI_ISL_534760, EPI_ISL_534761, EPI_ISL_534762 | Queens Medical Centre, Clinical Microbiology Department / DeepSeq Nottingham | COVID-19 Genomics UK (COG-UK) Consortium | Gemma Clark, Wendy Smith, Manjinder Khakh, Vicki M Fleming, Michelle M Lister, Hannah Howson-Wells, Jonathan Ball, Patrick McClure, Joseph Chappell, Theocharis Tsoleridis, Nadine Holmes, Matthew Carlisle, Christopher Moore, Fei Sang, Johnny Debebe, Victoria Wright, Matthew Loose |
| EPI_ISL_534763, EPI_ISL_534764, EPI_ISL_534765, EPI_ISL_534766, EPI_ISL_534767, EPI_ISL_534768, EPI_ISL_534769, EPI_ISL_534770, EPI_ISL_534771, EPI_ISL_534772, EPI_ISL_534773, EPI_ISL_534774, EPI_ISL_534775, EPI_ISL_534776, EPI_ISL_534777, EPI_ISL_534778, EPI_ISL_534779, EPI_ISL_534780, EPI_ISL_534781, EPI_ISL_534782, EPI_ISL_534783, EPI_ISL_534784, EPI_ISL_534785, EPI_ISL_534786, EPI_ISL_534787, EPI_ISL_534788, EPI_ISL_534789, EPI_ISL_534790, EPI_ISL_534791, EPI_ISL_534792, EPI_ISL_534793, EPI_ISL_534794, EPI_ISL_534795, EPI_ISL_534796, EPI_ISL_534797, EPI_ISL_534798, EPI_ISL_534799, EPI_ISL_534800, EPI_ISL_534801, EPI_ISL_534802, EPI_ISL_534803, EPI_ISL_534804, EPI_ISL_534805, EPI_ISL_534806, EPI_ISL_534807, EPI_ISL_534808, EPI_ISL_534809, EPI_ISL_534810, EPI_ISL_534811, EPI_ISL_534812, EPI_ISL_534813, EPI_ISL_534814, EPI_ISL_534815, EPI_ISL_534816, EPI_ISL_534817, EPI_ISL_534818, EPI_ISL_534819, EPI_ISL_534820, EPI_ISL_534821, EPI_ISL_534822, EPI_ISL_534823, EPI_ISL_534824, EPI_ISL_534825, EPI_ISL_534826, EPI_ISL_534827, EPI_ISL_534828, EPI_ISL_534829, EPI_ISL_534830, EPI_ISL_534831, EPI_ISL_534832, EPI_ISL_534833, EPI_ISL_534834, EPI_ISL_534835, EPI_ISL_534836, EPI_ISL_534837, EPI_ISL_534838, EPI_ISL_534839, EPI_ISL_534840, EPI_ISL_534841, EPI_ISL_534842, EPI_ISL_534843, EPI_ISL_534844, EPI_ISL_534845, EPI_ISL_534846, EPI_ISL_534847, EPI_ISL_534848, EPI_ISL_534849, EPI_ISL_534850, EPI_ISL_534851, EPI_ISL_534852, EPI_ISL_534853, EPI_ISL_534854, EPI_ISL_534855, EPI_ISL_534856, EPI_ISL_534857, EPI_ISL_534858, EPI_ISL_534859, EPI_ISL_534860, EPI_ISL_534861, EPI_ISL_534862, EPI_ISL_534863, EPI_ISL_534864, EPI_ISL_534865, EPI_ISL_534866, EPI_ISL_534867, EPI_ISL_534868, EPI_ISL_534869, EPI_ISL_534870, EPI_ISL_534871, EPI_ISL_534872, EPI_ISL_534873, EPI_ISL_534874, EPI_ISL_534875, EPI_ISL_534876, EPI_ISL_534877, EPI_ISL_534878, EPI_ISL_534879, EPI_ISL_534880, EPI_ISL_534881, EPI_ISL_534882, EPI_ISL_534883, EPI_ISL_534884, EPI_ISL_534885, EPI_ISL_534886, EPI_ISL_534887, EPI_ISL_534888, EPI_ISL_534889, EPI_ISL_534890, EPI_ISL_534891, EPI_ISL_534892, EPI_ISL_534893, EPI_ISL_534894, EPI_ISL_534895, EPI_ISL_534896, EPI_ISL_534897, EPI_ISL_534898, EPI_ISL_534899, EPI_ISL_534900, EPI_ISL_534901, EPI_ISL_534902, EPI_ISL_534903, EPI_ISL_534904, EPI_ISL_534905, EPI_ISL_534906, EPI_ISL_534907, EPI_ISL_534908, EPI_ISL_534909, EPI_ISL_534910, EPI_ISL_534911, EPI_ISL_534912, EPI_ISL_534913, EPI_ISL_534914, EPI_ISL_534915, EPI_ISL_534916, EPI_ISL_534917, EPI_ISL_534918, EPI_ISL_534919, EPI_ISL_534920, EPI_ISL_534921, EPI_ISL_534922, EPI_ISL_534923, EPI_ISL_534924, EPI_ISL_534925, EPI_ISL_534926, EPI_ISL_534927, EPI_ISL_534928, EPI_ISL_534929, EPI_ISL_534930, EPI_ISL_534931, EPI_ISL_534932, EPI_ISL_534933, EPI_ISL_534934, EPI_ISL_534935, EPI_ISL_534936, EPI_ISL_534937, EPI_ISL_534938, EPI_ISL_534939, EPI_ISL_534940, EPI_ISL_534941, EPI_ISL_534942, EPI_ISL_534943, EPI_ISL_534944, EPI_ISL_534945, EPI_ISL_534946, EPI_ISL_534947, EPI_ISL_534948, EPI_ISL_534949, EPI_ISL_534950, EPI_ISL_534951, EPI_ISL_534952, EPI_ISL_534953, EPI_ISL_534954, EPI_ISL_534955, EPI_ISL_534956, EPI_ISL_534957, EPI_ISL_534958, EPI_ISL_534959, EPI_ISL_534960, EPI_ISL_534961, EPI_ISL_534962, EPI_ISL_534963, EPI_ISL_534964, EPI_ISL_534965, EPI_ISL_534966, EPI_ISL_534967, EPI_ISL_534968, EPI_ISL_534969, EPI_ISL_534970, EPI_ISL_534971, EPI_ISL_534972, EPI_ISL_534973, EPI_ISL_534974, EPI_ISL_534975, EPI_ISL_534976, EPI_ISL_534977, EPI_ISL_534978, EPI_ISL_534979, EPI_ISL_534980, EPI_ISL_534981, EPI_ISL_534982, EPI_ISL_534983, EPI_ISL_534984, EPI_ISL_534985, EPI_ISL_534986, EPI_ISL_534987, EPI_ISL_534988, EPI_ISL_534989, EPI_ISL_534990, EPI_ISL_534991, EPI_ISL_534992, EPI_ISL_534993, EPI_ISL_534994, EPI_ISL_534995, EPI_ISL_534996, EPI_ISL_534997, EPI_ISL_534998, EPI_ISL_534999, EPI_ISL_535000, EPI_ISL_535001, EPI_ISL_535002, EPI_ISL_535003, EPI_ISL_535004, EPI_ISL_535005, EPI_ISL_535006, EPI_ISL_535007, EPI_ISL_535008, EPI_ISL_535009, EPI_ISL_535010, EPI_ISL_535011, EPI_ISL_535012, EPI_ISL_535013, EPI_ISL_535014, EPI_ISL_535015, EPI_ISL_535016, EPI_ISL_535017, EPI_ISL_535018, EPI_ISL_535019, EPI_ISL_535020, EPI_ISL_535021, EPI_ISL_535022, EPI_ISL_535023 | | | |
| see above | Oxford Viromics, NDM, University of Oxford; Oxford University Hospitals; Basingstoke and North Hampshire Hospital | COVID-19 Genomics UK (COG-UK) Consortium | Tanya Golubchik, David Bonsall, George Macintyre, Amy Trebes, Mariateresa de Cesare, Catrin Moore, Alex Mobbs, Anita Justice, Robert Shaw, Monique Andersson, Timothy Peto, Emma Wise, Nathan Moore, Jessica Lynch, Nick Cortes, Matilde Mori, Stephen Kidd, David Buck, John Todd, Christophe Fraser |
| EPI_ISL_535024, EPI_ISL_535025, EPI_ISL_535026, EPI_ISL_535027, EPI_ISL_535028, EPI_ISL_535029, EPI_ISL_535030, EPI_ISL_535031, EPI_ISL_535032, EPI_ISL_535033, EPI_ISL_535034, EPI_ISL_535035, EPI_ISL_535036, EPI_ISL_535037, EPI_ISL_535038, EPI_ISL_535039, EPI_ISL_535040, EPI_ISL_535041, EPI_ISL_535042 | | | |
| see above | Centre for Enzyme Innovation, University of Portsmouth / Translational Research Laboratory, Portsmouth Hospitals NHS Trust | COVID-19 Genomics UK (COG-UK) Consortium | Angela Beckett,Yann Bourgeois,Garry Scarlett,Sharon Glaysher,Scott Elliott,Kelly Bicknell,Robert Impey,Allyson Lloyd,Sarah Wyllie,Ethan Butcher,Anoop Chauhan,Samuel Robson |
| EPI_ISL_535043, EPI_ISL_535044, EPI_ISL_535045, EPI_ISL_535046, EPI_ISL_535047, EPI_ISL_535048, EPI_ISL_535049, EPI_ISL_535050, EPI_ISL_535051, EPI_ISL_535052, EPI_ISL_535053, EPI_ISL_535054, EPI_ISL_535055, EPI_ISL_535056, EPI_ISL_535057, EPI_ISL_535058, EPI_ISL_535059, EPI_ISL_535060, EPI_ISL_535061, EPI_ISL_535062, EPI_ISL_535063, EPI_ISL_535064, EPI_ISL_535065, EPI_ISL_535066, EPI_ISL_535067, EPI_ISL_535068, EPI_ISL_535069, EPI_ISL_535070, EPI_ISL_535071, EPI_ISL_535072, EPI_ISL_535073, EPI_ISL_535074, EPI_ISL_535075, EPI_ISL_535076, EPI_ISL_535077, EPI_ISL_535078, EPI_ISL_535079, EPI_ISL_535080, EPI_ISL_535081, EPI_ISL_535082, EPI_ISL_535083, EPI_ISL_535084, EPI_ISL_535085, EPI_ISL_535086, EPI_ISL_535087, EPI_ISL_535088, EPI_ISL_535089, EPI_ISL_535090, EPI_ISL_535091, EPI_ISL_535092, EPI_ISL_535093, EPI_ISL_535094, EPI_ISL_535095, EPI_ISL_535096, EPI_ISL_535097, EPI_ISL_535098, EPI_ISL_535099, EPI_ISL_535100, EPI_ISL_535101 | | | |
| see above | Virology Department, Sheffield Teaching Hospitals NHS Foundation Trust/Department of Infection, Immunity and Cardiovascular Disease, The Medical School, University of Sheffield | COVID-19 Genomics UK (COG-UK) Consortium | Thushan de Silva, Matthew Parker, Nikki Smith, Adri Angyal, Rebecca Brown, Luke Green, Rachel Tucker, Paul Parsons, Danielle Groves, Katie Johnson, Laura Carrilero, Alex Keeley, Dave Partridge, Matthew Wyles, Benjamin Lindsey, Mehmet Yavuz, Mohammad Raza, Cariad Evans |
| EPI_ISL_535102, EPI_ISL_535103, EPI_ISL_535104, EPI_ISL_535105, EPI_ISL_535106, EPI_ISL_535107, EPI_ISL_535108, EPI_ISL_535109, EPI_ISL_535110, EPI_ISL_535111, EPI_ISL_535112, EPI_ISL_535113, EPI_ISL_535114, EPI_ISL_535115, EPI_ISL_535116, EPI_ISL_535117, EPI_ISL_535118, EPI_ISL_535119, EPI_ISL_535120, EPI_ISL_535121, EPI_ISL_535122, EPI_ISL_535123, EPI_ISL_535124, EPI_ISL_535125, EPI_ISL_535126, EPI_ISL_535127, EPI_ISL_535128, EPI_ISL_535129, EPI_ISL_535130, EPI_ISL_535131, EPI_ISL_535132, EPI_ISL_535133, EPI_ISL_535134, EPI_ISL_535135, EPI_ISL_535136, EPI_ISL_535137, EPI_ISL_535138, EPI_ISL_535139, EPI_ISL_535140, EPI_ISL_535141, EPI_ISL_535142, EPI_ISL_535143, EPI_ISL_535144, EPI_ISL_535145, EPI_ISL_535146, EPI_ISL_535147, EPI_ISL_535148, EPI_ISL_535149, EPI_ISL_535150, EPI_ISL_535151, EPI_ISL_535152, EPI_ISL_535153, EPI_ISL_535154, EPI_ISL_535155, EPI_ISL_535156, EPI_ISL_535157, EPI_ISL_535158, EPI_ISL_535159, EPI_ISL_535160, EPI_ISL_535161, EPI_ISL_535162, EPI_ISL_535163, EPI_ISL_535164, EPI_ISL_535165, EPI_ISL_535166, EPI_ISL_535167, EPI_ISL_535168, EPI_ISL_535169, EPI_ISL_535170, EPI_ISL_535171, EPI_ISL_535172, EPI_ISL_535173, EPI_ISL_535174, EPI_ISL_535175, EPI_ISL_535176, EPI_ISL_535177, EPI_ISL_535178, EPI_ISL_535179, EPI_ISL_535180, EPI_ISL_535181, EPI_ISL_535182 | | | |
| see above | West of Scotland Specialist Virology Centre, NHSGGC / MRC-University of Glasgow Centre for Virus Research | COVID-19 Genomics UK (COG-UK) Consortium | Ana da Silva Filipe, Natasha Johnson, Kathy Smollett, Daniel Mair, Stephen Carmichael, Lily Tong, Jenna Nichols, Elihu Aranday-Cortes, Yasmin Parr, Alice Broos, Kyriaki Nomikou; Sarah McDonald, Marc Niebel, Patawee Asamaphan; Richard Orton, Joseph Hughes, Sreenu Vattipally, David L Robertson; Alasdair MacLean, Rory Gunson; Kathy Li, Natasha Jesudason, Rajiv Shah, James Shepherd, Antonia Ho, Emma Thomson |
| EPI_ISL_535183 | Queens Medical Centre, Clinical Microbiology Department / DeepSeq Nottingham | COVID-19 Genomics UK (COG-UK) Consortium | Gemma Clark, Wendy Smith, Manjinder Khakh, Vicki M Fleming, Michelle M Lister, Hannah Howson-Wells, Jonathan Ball, Patrick McClure, Joseph Chappell, Theocharis Tsoleridis, Nadine Holmes, Matthew Carlisle, Christopher Moore, Fei Sang, Johnny Debebe, Victoria Wright, Matthew Loose |
| EPI_ISL_535184, EPI_ISL_535185, EPI_ISL_535186, EPI_ISL_535187, EPI_ISL_535188, EPI_ISL_535189, EPI_ISL_535190, EPI_ISL_535191, EPI_ISL_535192, EPI_ISL_535193, EPI_ISL_535194, EPI_ISL_535195, EPI_ISL_535196, EPI_ISL_535197, EPI_ISL_535198, EPI_ISL_535199, EPI_ISL_535200, EPI_ISL_535201, EPI_ISL_535202, EPI_ISL_535203, EPI_ISL_535204, EPI_ISL_535205, EPI_ISL_535206, EPI_ISL_535207, EPI_ISL_535208, EPI_ISL_535209, EPI_ISL_535210, EPI_ISL_535211, EPI_ISL_535212, EPI_ISL_535213, EPI_ISL_535214, EPI_ISL_535215, EPI_ISL_535216, EPI_ISL_535217, EPI_ISL_535218, EPI_ISL_535219, EPI_ISL_535220, EPI_ISL_535221, EPI_ISL_535222, EPI_ISL_535223, EPI_ISL_535224, EPI_ISL_535225, EPI_ISL_535226, EPI_ISL_535227, EPI_ISL_535228, EPI_ISL_535229, EPI_ISL_535230, EPI_ISL_535231, EPI_ISL_535232, EPI_ISL_535233, EPI_ISL_535234, EPI_ISL_535235, EPI_ISL_535236, EPI_ISL_535237, EPI_ISL_535238, EPI_ISL_535239, EPI_ISL_535240, EPI_ISL_535241, EPI_ISL_535242, EPI_ISL_535243, EPI_ISL_535244, EPI_ISL_535245, EPI_ISL_535246, EPI_ISL_535247, EPI_ISL_535248, EPI_ISL_535249, EPI_ISL_535250, EPI_ISL_535251, EPI_ISL_535252, EPI_ISL_535253, EPI_ISL_535254, EPI_ISL_535255, EPI_ISL_535256, EPI_ISL_535257, EPI_ISL_535258, EPI_ISL_535259, EPI_ISL_535260, EPI_ISL_535261, EPI_ISL_535262 | | | |
| see above | Wales Specialist Virology Centre Sequencing lab: Pathogen Genomics Unit | COVID-19 Genomics UK (COG-UK) Consortium | Catherine Moore, Johnathan Evans, Laura Gifford, Malorie Perry, Simon Cottrell, Angela Marchbank, Alec Birchley, Alexander Adams, Amy Gaskin, Bree Gatica-Wilcox, Jason Coombes, Joel Southgate, Lauren Gilbert, Lee Graham, Nicole Pacchiarini, Sara Kumziene-Summerhayes, Sarah Taylor, Sophie Jones, Sara Rey, Matthew Bull, Joanne Watkins, Sally Corden, Tom Connor |
| EPI_ISL_535263, EPI_ISL_535264, EPI_ISL_535265, EPI_ISL_535266, EPI_ISL_535267, EPI_ISL_535268, EPI_ISL_535269, EPI_ISL_535270, EPI_ISL_535271, EPI_ISL_535272, EPI_ISL_535273, EPI_ISL_535274, EPI_ISL_535275, EPI_ISL_535276, EPI_ISL_535277, EPI_ISL_535278, EPI_ISL_535279, EPI_ISL_535280, EPI_ISL_535281, EPI_ISL_535282, EPI_ISL_535283, EPI_ISL_535284, EPI_ISL_535285, EPI_ISL_535286, EPI_ISL_535287, EPI_ISL_535288, EPI_ISL_535289, EPI_ISL_535290, EPI_ISL_535291, EPI_ISL_535292, EPI_ISL_535293, EPI_ISL_535294, EPI_ISL_535295, EPI_ISL_535296, EPI_ISL_535297, EPI_ISL_535298, EPI_ISL_535299, EPI_ISL_535300, EPI_ISL_535301, EPI_ISL_535302, EPI_ISL_535303 | | | |
| see above | New Mexico Department of Health Scientific Laboratory | New Mexico Department of Health Scientific Laboratory | Ellie Johnson, Anastacia Griego-Fisher, D'Eldra Malone |
| EPI_ISL_535305, EPI_ISL_535306, EPI_ISL_535307, EPI_ISL_535308, EPI_ISL_535309, EPI_ISL_535310, EPI_ISL_535311, EPI_ISL_535312, EPI_ISL_535313, EPI_ISL_535314, EPI_ISL_535315, EPI_ISL_535316, EPI_ISL_535317, EPI_ISL_535318, EPI_ISL_535319, EPI_ISL_535320, EPI_ISL_535321, EPI_ISL_535322, EPI_ISL_535323, EPI_ISL_535324, EPI_ISL_535325, EPI_ISL_535326, EPI_ISL_535327, EPI_ISL_535328, EPI_ISL_535329, EPI_ISL_535330, EPI_ISL_535331, EPI_ISL_535332, EPI_ISL_535333, EPI_ISL_535334, EPI_ISL_535335, EPI_ISL_535336, EPI_ISL_535337, EPI_ISL_535338, EPI_ISL_535339, EPI_ISL_535340 | | | |
| see above | LA Office of Public Health Laboratories | Pathogen Discovery, Respiratory Viruses Branch, Division of Viral Diseases, Centers for Disease Control and Prevention | Ying Tao, Jing Zhang, Yan Li, Krista Queen, Anna Uehara, Clinton Paden, Haibin Wang, Suxiang Tong |
| EPI_ISL_535341, EPI_ISL_535342 | LA Office of Public Health Laboratories | Pathogen Discovery, Respiratory Viruses Branch, Division of Viral Diseases, Centers for Disease Control and Prevention | Jing Zhang, Ying Tao, Yan Li, Krista Queen, Anna Uehara, Clinton Paden, Haibin Wang, Suxiang Tong |
| EPI_ISL_535343 | LA Office of Public Health Laboratories | Pathogen Discovery, Respiratory Viruses Branch, Division of Viral Diseases, Centers for Disease Control and Prevention | Ying Tao, Jing Zhang, Yan Li, Krista Queen, Anna Uehara, Clinton Paden, Haibin Wang, Suxiang Tong |
| EPI_ISL_535344, EPI_ISL_535345, EPI_ISL_535346 | LA Office of Public Health Laboratories | Pathogen Discovery, Respiratory Viruses Branch, Division of Viral Diseases, Centers for Disease Control and Prevention | Jing Zhang, Ying Tao, Yan Li, Krista Queen, Anna Uehara, Clinton Paden, Haibin Wang, Suxiang Tong |
| EPI_ISL_535347 | LA Office of Public Health Laboratories | Pathogen Discovery, Respiratory Viruses Branch, Division of Viral Diseases, Centers for Disease Control and Prevention | Ying Tao, Jing Zhang, Yan Li, Krista Queen, Anna Uehara, Clinton Paden, Haibin Wang, Suxiang Tong |
| EPI_ISL_535348 | LA Office of Public Health Laboratories | Pathogen Discovery, Respiratory Viruses Branch, Division of Viral Diseases, Centers for Disease Control and Prevention | Jing Zhang, Ying Tao, Yan Li, Krista Queen, Anna Uehara, Clinton Paden, Haibin Wang, Suxiang Tong |
| EPI_ISL_535349, EPI_ISL_535350 | LA Office of Public Health Laboratories | Pathogen Discovery, Respiratory Viruses Branch, Division of Viral Diseases, Centers for Disease Control and Prevention | Ying Tao, Jing Zhang, Yan Li, Krista Queen, Anna Uehara, Clinton Paden, Haibin Wang, Suxiang Tong |
| EPI_ISL_535351 | LA Office of Public Health Laboratories | Pathogen Discovery, Respiratory Viruses Branch, Division of Viral Diseases, Centers for Disease Control and Prevention | Jing Zhang, Ying Tao, Yan Li, Krista Queen, Anna Uehara, Clinton Paden, Haibin Wang, Suxiang Tong |
| EPI_ISL_535352, EPI_ISL_535353, EPI_ISL_535354, EPI_ISL_535355, EPI_ISL_535356 | LA Office of Public Health Laboratories | Pathogen Discovery, Respiratory Viruses Branch, Division of Viral Diseases, Centers for Disease Control and Prevention | Ying Tao, Jing Zhang, Yan Li, Krista Queen, Anna Uehara, Clinton Paden, Haibin Wang, Suxiang Tong |
| EPI_ISL_535357 | LA Office of Public Health Laboratories | Pathogen Discovery, Respiratory Viruses Branch, Division of Viral Diseases, Centers for Disease Control and Prevention | Jing Zhang, Ying Tao, Yan Li, Krista Queen, Anna Uehara, Clinton Paden, Haibin Wang, Suxiang Tong |
| EPI_ISL_535358 | LA Office of Public Health Laboratories | Pathogen Discovery, Respiratory Viruses Branch, Division of Viral Diseases, Centers for Disease Control and Prevention | Ying Tao, Jing Zhang, Yan Li, Krista Queen, Anna Uehara, Clinton Paden, Haibin Wang, Suxiang Tong |
| EPI_ISL_535359, EPI_ISL_535360 | RI State Health Laboratories | Pathogen Discovery, Respiratory Viruses Branch, Division of Viral Diseases, Centers for Disease Control and Prevention | Jing Zhang, Ying Tao, Yan Li, Krista Queen, Anna Uehara, Clinton Paden, Haibin Wang, Suxiang Tong |
| EPI_ISL_535361, EPI_ISL_535362, EPI_ISL_535363, EPI_ISL_535364 | Oklahoma Animal Disease Diagnostic Laboratory | Oklahoma Animal Disease Diagnostic Laboratory | Sai Narayanan, John C Ritchey, Girish Patil, Teluguakula Narasaraju, Sunil More, Jerry Malayer, Jeremiah Saliki, Anil Kaul, Akhilesh Ramachandran |
| EPI_ISL_535390, EPI_ISL_535391, EPI_ISL_535392, EPI_ISL_535393, EPI_ISL_535394, EPI_ISL_535395, EPI_ISL_535396, EPI_ISL_535397, EPI_ISL_535398, EPI_ISL_535399, EPI_ISL_535400, EPI_ISL_535401, EPI_ISL_535402, EPI_ISL_535403, EPI_ISL_535404, EPI_ISL_535405, EPI_ISL_535406, EPI_ISL_535407, EPI_ISL_535408, EPI_ISL_535409, EPI_ISL_535410, EPI_ISL_535411, EPI_ISL_535412, EPI_ISL_535413, EPI_ISL_535414, EPI_ISL_535415, EPI_ISL_535416, EPI_ISL_535417, EPI_ISL_535418, EPI_ISL_535419, EPI_ISL_535420, EPI_ISL_535421, EPI_ISL_535422, EPI_ISL_535423, EPI_ISL_535424, EPI_ISL_535425, EPI_ISL_535426, EPI_ISL_535427, EPI_ISL_535428, EPI_ISL_535429, EPI_ISL_535430, EPI_ISL_535431, EPI_ISL_535432, EPI_ISL_535433, EPI_ISL_535434, EPI_ISL_535435, EPI_ISL_535436, EPI_ISL_535437, EPI_ISL_535438, EPI_ISL_535439, EPI_ISL_535440, EPI_ISL_535441, EPI_ISL_535442, EPI_ISL_535443, EPI_ISL_535444, EPI_ISL_535445, EPI_ISL_535446, EPI_ISL_535447, EPI_ISL_535448, EPI_ISL_535449, EPI_ISL_535450, EPI_ISL_535451, EPI_ISL_535452, EPI_ISL_535453, EPI_ISL_535454, EPI_ISL_535455, EPI_ISL_535456, EPI_ISL_535457, EPI_ISL_535458, EPI_ISL_535459, EPI_ISL_535460, EPI_ISL_535461, EPI_ISL_535462, EPI_ISL_535463, EPI_ISL_535464, EPI_ISL_535465, EPI_ISL_535466, EPI_ISL_535467, EPI_ISL_535468, EPI_ISL_535469, EPI_ISL_535470, EPI_ISL_535471, EPI_ISL_535472, EPI_ISL_535473, EPI_ISL_535474, EPI_ISL_535475, EPI_ISL_535476, EPI_ISL_535477, EPI_ISL_535478, EPI_ISL_535479, EPI_ISL_535480, EPI_ISL_535481, EPI_ISL_535482, EPI_ISL_535483, EPI_ISL_535484, EPI_ISL_535485, EPI_ISL_535486, EPI_ISL_535487, EPI_ISL_535488, EPI_ISL_535496, EPI_ISL_535497, EPI_ISL_535498, EPI_ISL_535499, EPI_ISL_535500, EPI_ISL_535501, EPI_ISL_535502, EPI_ISL_535503, EPI_ISL_535504, EPI_ISL_535505, EPI_ISL_535506, EPI_ISL_535507, EPI_ISL_535508, EPI_ISL_535509, EPI_ISL_535510, EPI_ISL_535511, EPI_ISL_535512, EPI_ISL_535513, EPI_ISL_535514, EPI_ISL_535515, EPI_ISL_535516, EPI_ISL_535517, EPI_ISL_535518, EPI_ISL_535519, EPI_ISL_535520, EPI_ISL_535521, EPI_ISL_535522, EPI_ISL_535523, EPI_ISL_535524, EPI_ISL_535525, EPI_ISL_535526, EPI_ISL_535527, EPI_ISL_535528, EPI_ISL_535529, EPI_ISL_535530, EPI_ISL_535531, EPI_ISL_535532, EPI_ISL_535533, EPI_ISL_535534, EPI_ISL_535535, EPI_ISL_535536, EPI_ISL_535537, EPI_ISL_535538, EPI_ISL_535539, EPI_ISL_535540, EPI_ISL_535541, EPI_ISL_535542, EPI_ISL_535543, EPI_ISL_535544, EPI_ISL_535545, EPI_ISL_535546, EPI_ISL_535547, EPI_ISL_535548, EPI_ISL_535549, EPI_ISL_535550, EPI_ISL_535551, EPI_ISL_535552, EPI_ISL_535553, EPI_ISL_535554, EPI_ISL_535555, EPI_ISL_535556, EPI_ISL_535557, EPI_ISL_535558, EPI_ISL_535559, EPI_ISL_535560, EPI_ISL_535561, EPI_ISL_535562, EPI_ISL_535563, EPI_ISL_535564, EPI_ISL_535565, EPI_ISL_535566, EPI_ISL_535567, EPI_ISL_535568, EPI_ISL_535569, EPI_ISL_535570, EPI_ISL_535571, EPI_ISL_535572 | | | |
| see above | NHLS-IALCH | KRISP, KZN Research Innovation and Sequencing Platform | Giandhari J, Pillay S, Lessells R, Mdlalose K, York D, Khan S, Tegally H, Wilkinson E, de Oliveira T |
| EPI_ISL_535575 | Hospital Universitari Germans Trias i Pujol(HUGTiP)/Fundació Lluita contra la SIDA (FLSida) | IrsiCaixa AIDS Research Lab | Marc Noguera-Julian, Mariona Parera, Maria Pilar Armengol, Marta Massanella, Ester Ballana, Lidia Ruiz, Nuria Izquierdo, Jorge Carrillo, Roger Paredes, Julia Blanco, Joaquim Segalés, Bonaventura Clotet |
| EPI_ISL_535585, EPI_ISL_535586, EPI_ISL_535587, EPI_ISL_535588, EPI_ISL_535589, EPI_ISL_535590, EPI_ISL_535591, EPI_ISL_535592, EPI_ISL_535593, EPI_ISL_535594, EPI_ISL_535595, EPI_ISL_535596, EPI_ISL_535597, EPI_ISL_535598, EPI_ISL_535599, EPI_ISL_535600, EPI_ISL_535601, EPI_ISL_535602, EPI_ISL_535603, EPI_ISL_535604, EPI_ISL_535605, EPI_ISL_535606, EPI_ISL_535607, EPI_ISL_535608, EPI_ISL_535609, EPI_ISL_535610, EPI_ISL_535611, EPI_ISL_535612, EPI_ISL_535613, EPI_ISL_535614, EPI_ISL_535615, EPI_ISL_535616, EPI_ISL_535617, EPI_ISL_535618, EPI_ISL_535619, EPI_ISL_535620, EPI_ISL_535621, EPI_ISL_535622, EPI_ISL_535623, EPI_ISL_535624, EPI_ISL_535625, EPI_ISL_535626, EPI_ISL_535627, EPI_ISL_535628, EPI_ISL_535629, EPI_ISL_535630, EPI_ISL_535631, EPI_ISL_535632, EPI_ISL_535633, EPI_ISL_535634, EPI_ISL_535635, EPI_ISL_535636, EPI_ISL_535637, EPI_ISL_535638, EPI_ISL_535639, EPI_ISL_535640, EPI_ISL_535641, EPI_ISL_535642, EPI_ISL_535643, EPI_ISL_535644, EPI_ISL_535645, EPI_ISL_535646, EPI_ISL_535647, EPI_ISL_535648, EPI_ISL_535649 | | | |
| see above | Viollier AG | Department of Biosystems Science and Engineering, ETH Zürich | Christian Beisel, Sarah Nadeau, Ivan Topolsky, Pedro Ferreira, Philipp Jablonski, Susana Posada-Céspedes, Tobias Schär, Ina Nissen, Natascha Santacroce, Elodie Burcklen, Christiane Beckmann, Maurice Redondo, Olivier Kobel, Christoph Noppen, Sophie Seidel, Noemie Santamaria de Souza, Niko Beerenwinkel, Tanja Stadler |
| EPI_ISL_535650 | AR Dept. of Health-Public Health Lab | Pathogen Discovery, Respiratory Viruses Branch, Division of Viral | Brian Lynch, Yan Li, Jing Zhang, Ying Tao, Krista Queen, Anna Uehara, Clinton R. Paden, Rachel Marine, Haibin Wang, Suxiang Tong |

| | | | |
|---|---|---|---|
| | | Diseases, Centers for Disease Control and Prevention | |
| EPI_ISL_535651 | AR Dept. of Health-Public Health Lab | Pathogen Discovery, Respiratory Viruses Branch, Division of Viral Diseases, Centers for Disease Control and Prevention | Yan Li, Jing Zhang, Ying Tao, Krista Queen, Brian Lynch, Anna Uehara, Clinton R. Paden, Rachel Marine, Haibin Wang, Suxiang Tong |
| EPI_ISL_535652 | AR Dept. of Health-Public Health Lab | Pathogen Discovery, Respiratory Viruses Branch, Division of Viral Diseases, Centers for Disease Control and Prevention | Brian Lynch, Yan Li, Jing Zhang, Ying Tao, Krista Queen, Anna Uehara, Clinton R. Paden, Rachel Marine, Haibin Wang, Suxiang Tong |
| EPI_ISL_535653 | AR Dept. of Health-Public Health Lab | Pathogen Discovery, Respiratory Viruses Branch, Division of Viral Diseases, Centers for Disease Control and Prevention | Yan Li, Jing Zhang, Ying Tao, Krista Queen, Brian Lynch, Anna Uehara, Clinton R. Paden, Rachel Marine, Haibin Wang, Suxiang Tong |
| EPI_ISL_535654 | AR Dept. of Health-Public Health Lab | Pathogen Discovery, Respiratory Viruses Branch, Division of Viral Diseases, Centers for Disease Control and Prevention | Brian Lynch, Yan Li, Jing Zhang, Ying Tao, Krista Queen, Anna Uehara, Clinton R. Paden, Rachel Marine, Haibin Wang, Suxiang Tong |
| EPI_ISL_535655, EPI_ISL_535656, EPI_ISL_535657 | AR Dept. of Health-Public Health Lab | Pathogen Discovery, Respiratory Viruses Branch, Division of Viral Diseases, Centers for Disease Control and Prevention | Yan Li, Jing Zhang, Ying Tao, Krista Queen, Brian Lynch, Anna Uehara, Clinton R. Paden, Rachel Marine, Haibin Wang, Suxiang Tong |
| EPI_ISL_535658, EPI_ISL_535659 | AR Dept. of Health-Public Health Lab | Pathogen Discovery, Respiratory Viruses Branch, Division of Viral Diseases, Centers for Disease Control and Prevention | Brian Lynch, Yan Li, Jing Zhang, Ying Tao, Krista Queen, Anna Uehara, Clinton R. Paden, Rachel Marine, Haibin Wang, Suxiang Tong |
| EPI_ISL_535660 | AR Dept. of Health-Public Health Lab | Pathogen Discovery, Respiratory Viruses Branch, Division of Viral Diseases, Centers for Disease Control and Prevention | Yan Li, Jing Zhang, Ying Tao, Krista Queen, Brian Lynch, Anna Uehara, Clinton R. Paden, Rachel Marine, Haibin Wang, Suxiang Tong |
| EPI_ISL_535661 | AR Dept. of Health-Public Health Lab | Pathogen Discovery, Respiratory Viruses Branch, Division of Viral Diseases, Centers for Disease Control and Prevention | Brian Lynch, Yan Li, Jing Zhang, Ying Tao, Krista Queen, Anna Uehara, Clinton R. Paden, Rachel Marine, Haibin Wang, Suxiang Tong |
| EPI_ISL_535662, EPI_ISL_535663, EPI_ISL_535664, EPI_ISL_535665, EPI_ISL_535666, EPI_ISL_535667, EPI_ISL_535668, EPI_ISL_535669, EPI_ISL_535670, EPI_ISL_535671, EPI_ISL_535672, EPI_ISL_535673, EPI_ISL_535674 | | | |
| see above | CDPH, Microbial Diseases Laboratory | Pathogen Discovery, Respiratory Viruses Branch, Division of Viral Diseases, Centers for Disease Control and Prevention | Yan Li, Jing Zhang, Ying Tao, Krista Queen, Brian Lynch, Anna Uehara, Clinton R. Paden, Rachel Marine, Haibin Wang, Suxiang Tong |
| EPI_ISL_535675 | CDPH, Microbial Diseases Laboratory | Pathogen Discovery, Respiratory Viruses Branch, Division of Viral Diseases, Centers for Disease Control and Prevention | Brian Lynch, Yan Li, Jing Zhang, Ying Tao, Krista Queen, Anna Uehara, Clinton R. Paden, Rachel Marine, Haibin Wang, Suxiang Tong |
| EPI_ISL_535676, EPI_ISL_535677, EPI_ISL_535678, EPI_ISL_535679, EPI_ISL_535680, EPI_ISL_535681, EPI_ISL_535682, EPI_ISL_535683, EPI_ISL_535684, EPI_ISL_535685, EPI_ISL_535686, EPI_ISL_535687, EPI_ISL_535688, EPI_ISL_535689, EPI_ISL_535690, EPI_ISL_535691, EPI_ISL_535692, EPI_ISL_535693, EPI_ISL_535694, EPI_ISL_535695, EPI_ISL_535696, EPI_ISL_535697, EPI_ISL_535698, EPI_ISL_535699, EPI_ISL_535700 | | | |
| see above | CDPH, Microbial Diseases Laboratory | Pathogen Discovery, Respiratory Viruses Branch, Division of Viral Diseases, Centers for Disease Control and Prevention | Yan Li, Jing Zhang, Ying Tao, Krista Queen, Brian Lynch, Anna Uehara, Clinton R. Paden, Rachel Marine, Haibin Wang, Suxiang Tong |
| EPI_ISL_535701 | CDPH, Microbial Diseases Laboratory | Pathogen Discovery, Respiratory Viruses Branch, Division of Viral Diseases, Centers for Disease Control and Prevention | Brian Lynch, Yan Li, Jing Zhang, Ying Tao, Krista Queen, Anna Uehara, Clinton R. Paden, Rachel Marine, Haibin Wang, Suxiang Tong |
| EPI_ISL_535702 | CDPH, Microbial Diseases Laboratory | Pathogen Discovery, Respiratory Viruses Branch, Division of Viral Diseases, Centers for Disease Control and Prevention | Yan Li, Jing Zhang, Ying Tao, Krista Queen, Brian Lynch, Anna Uehara, Clinton R. Paden, Rachel Marine, Haibin Wang, Suxiang Tong |
| EPI_ISL_535703 | CDPH, Microbial Diseases Laboratory | Pathogen Discovery, Respiratory Viruses Branch, Division of Viral Diseases, Centers for Disease Control and Prevention | Ying Tao, Jing Zhang, Yan Li, Krista Queen, Anna Uehara, Clinton R. Paden, Haibin Wang, Suxiang Tong |
| EPI_ISL_535704, EPI_ISL_535705 | CDPH, Microbial Diseases Laboratory | Pathogen Discovery, Respiratory Viruses Branch, Division of Viral Diseases, Centers for Disease Control and Prevention | Yan Li, Jing Zhang, Ying Tao, Krista Queen, Brian Lynch, Anna Uehara, Clinton R. Paden, Rachel Marine, Haibin Wang, Suxiang Tong |
| EPI_ISL_535706 | CDPH, Microbial Diseases Laboratory | Pathogen Discovery, Respiratory Viruses Branch, Division of Viral Diseases, Centers for Disease Control and Prevention | Ying Tao, Jing Zhang, Yan Li, Krista Queen, Anna Uehara, Clinton R. Paden, Haibin Wang, Suxiang Tong |
| EPI_ISL_535707 | CDPH, Microbial Diseases Laboratory | Pathogen Discovery, Respiratory Viruses Branch, Division of Viral Diseases, Centers for Disease Control and Prevention | Yan Li, Jing Zhang, Ying Tao, Krista Queen, Brian Lynch, Anna Uehara, Clinton R. Paden, Rachel Marine, Haibin Wang, Suxiang Tong |
| EPI_ISL_535708 | CDPH, Microbial Diseases Laboratory | Pathogen Discovery, Respiratory Viruses Branch, Division of Viral Diseases, Centers for Disease Control and Prevention | Brian Lynch, Yan Li, Jing Zhang, Ying Tao, Krista Queen, Anna Uehara, Clinton R. Paden, Rachel Marine, Haibin Wang, Suxiang Tong |
| EPI_ISL_535709, EPI_ISL_535710, EPI_ISL_535711, EPI_ISL_535712, EPI_ISL_535713, EPI_ISL_535714, EPI_ISL_535715 | CDPH, Microbial Diseases Laboratory | Pathogen Discovery, Respiratory Viruses Branch, Division of Viral Diseases, Centers for Disease Control and Prevention | Yan Li, Jing Zhang, Ying Tao, Krista Queen, Brian Lynch, Anna Uehara, Clinton R. Paden, Rachel Marine, Haibin Wang, Suxiang Tong |
| EPI_ISL_535716 | Hôpital de Verdun | Laboratoire de santé publique du Québec | Sandrine Moreira, Ioannis Ragoussis, Guillaume Bourque, Jesse Shapiro, Mark Lathrop and Michel Roger |
| EPI_ISL_535717 | Hôpital Honoré-Mercier | Laboratoire de santé publique du Québec | Sandrine Moreira, Ioannis Ragoussis, Guillaume Bourque, Jesse Shapiro, Mark Lathrop and Michel Roger |
| EPI_ISL_535718 | Hôpital Pierre-Boucher | Laboratoire de santé publique du Québec | Sandrine Moreira, Ioannis Ragoussis, Guillaume Bourque, Jesse Shapiro, Mark Lathrop and Michel Roger |
| EPI_ISL_535719, EPI_ISL_535720 | Hôpital général Juif | Laboratoire de santé publique du Québec | Sandrine Moreira, Ioannis Ragoussis, Guillaume Bourque, Jesse Shapiro, Mark Lathrop and Michel Roger |
| EPI_ISL_535721 | Hôpital Notre-Dame | Laboratoire de santé publique du Québec | Sandrine Moreira, Ioannis Ragoussis, Guillaume Bourque, Jesse Shapiro, Mark Lathrop and Michel Roger |
| EPI_ISL_535722 | Centre de SSS de Trois-Rivières | Laboratoire de santé publique du Québec | Sandrine Moreira, Ioannis Ragoussis, Guillaume Bourque, Jesse Shapiro, Mark Lathrop and Michel Roger |
| EPI_ISL_535723 | Hôpital de Verdun | Laboratoire de santé publique du Québec | Sandrine Moreira, Ioannis Ragoussis, Guillaume Bourque, Jesse Shapiro, Mark Lathrop and Michel Roger |
| EPI_ISL_535724 | Centre de SSS La Pommeraie | Laboratoire de santé publique du Québec | Sandrine Moreira, Ioannis Ragoussis, Guillaume Bourque, Jesse Shapiro, Mark Lathrop and Michel Roger |
| EPI_ISL_535725 | Hôtel-Dieu de Lévis | Laboratoire de santé publique du Québec | Sandrine Moreira, Ioannis Ragoussis, Guillaume Bourque, Jesse Shapiro, Mark Lathrop and Michel Roger |
| EPI_ISL_535726 | CHUM - Microbiologie - Hôpital Saint-Luc | Laboratoire de santé publique du Québec | Sandrine Moreira, Ioannis Ragoussis, Guillaume Bourque, Jesse Shapiro, Mark Lathrop and Michel Roger |
| EPI_ISL_535727 | Centre de SSS de la Haute-Yamaska | Laboratoire de santé publique du Québec | Sandrine Moreira, Ioannis Ragoussis, Guillaume Bourque, Jesse Shapiro, Mark Lathrop and Michel Roger |
| EPI_ISL_535728 | Hôpital de Verdun | Laboratoire de santé publique du Québec | Sandrine Moreira, Ioannis Ragoussis, Guillaume Bourque, Jesse Shapiro, Mark Lathrop and Michel Roger |
| EPI_ISL_535729, EPI_ISL_535730 | CHUM - Microbiologie - Hôpital Saint-Luc | Laboratoire de santé publique du Québec | Sandrine Moreira, Ioannis Ragoussis, Guillaume Bourque, Jesse Shapiro, Mark Lathrop and Michel Roger |
| EPI_ISL_535731 | Hôpital Pierre-Le Gardeur | Laboratoire de santé publique du Québec | Sandrine Moreira, Ioannis Ragoussis, Guillaume Bourque, Jesse Shapiro, Mark Lathrop and Michel Roger |
| EPI_ISL_535732 | CHU Sainte-Justine | Laboratoire de santé publique du Québec | Sandrine Moreira, Ioannis Ragoussis, Guillaume Bourque, Jesse Shapiro, Mark Lathrop and Michel Roger |
| EPI_ISL_535733, EPI_ISL_535734 | Hôpital Pierre-Boucher | Laboratoire de santé publique du Québec | Sandrine Moreira, Ioannis Ragoussis, Guillaume Bourque, Jesse Shapiro, Mark Lathrop and Michel Roger |
| EPI_ISL_535735 | CHUM - Microbiologie - Hôpital Saint-Luc | Laboratoire de santé publique du Québec | Sandrine Moreira, Ioannis Ragoussis, Guillaume Bourque, Jesse Shapiro, Mark Lathrop and Michel Roger |
| EPI_ISL_535736, EPI_ISL_535737 | CHUL-LABO MULTI / MICRO | Laboratoire de santé publique du Québec | Sandrine Moreira, Ioannis Ragoussis, Guillaume Bourque, Jesse Shapiro, Mark Lathrop and Michel Roger |
| EPI_ISL_535738 | Hôpital Charles-LeMoyne | Laboratoire de santé publique du Québec | Sandrine Moreira, Ioannis Ragoussis, Guillaume Bourque, Jesse Shapiro, Mark Lathrop and Michel Roger |
| EPI_ISL_535739 | Hôpital de l'Enfant-Jésus | Laboratoire de santé publique du Québec | Sandrine Moreira, Ioannis Ragoussis, Guillaume Bourque, Jesse Shapiro, Mark Lathrop and Michel Roger |
| EPI_ISL_535740 | CHUM - Microbiologie - Hôpital Saint-Luc | Laboratoire de santé publique du Québec | Sandrine Moreira, Ioannis Ragoussis, Guillaume Bourque, Jesse Shapiro, Mark Lathrop and Michel Roger |
| EPI_ISL_535741 | CHUL-LABO MULTI / MICRO | Laboratoire de santé publique du Québec | Sandrine Moreira, Ioannis Ragoussis, Guillaume Bourque, Jesse Shapiro, Mark Lathrop and Michel Roger |
| EPI_ISL_535742, EPI_ISL_535743 | Centre de SSS La Pommeraie | Laboratoire de santé publique du Québec | Sandrine Moreira, Ioannis Ragoussis, Guillaume Bourque, Jesse Shapiro, Mark Lathrop and Michel Roger |
| EPI_ISL_535744 | Hôpital de Lasalle | Laboratoire de santé publique du Québec | Sandrine Moreira, Ioannis Ragoussis, Guillaume Bourque, Jesse Shapiro, Mark Lathrop and Michel Roger |
| EPI_ISL_535745 | Hôpital Laurentien | Laboratoire de santé publique du Québec | Sandrine Moreira, Ioannis Ragoussis, Guillaume Bourque, Jesse Shapiro, Mark Lathrop and Michel Roger |
| EPI_ISL_535746 | Centre hospitalier de St-Mary | Laboratoire de santé publique du Québec | Sandrine Moreira, Ioannis Ragoussis, Guillaume Bourque, Jesse Shapiro, Mark Lathrop and Michel Roger |
| EPI_ISL_535747, EPI_ISL_535748 | Centre de SSS de Trois-Rivières | Laboratoire de santé publique du Québec | Sandrine Moreira, Ioannis Ragoussis, Guillaume Bourque, Jesse Shapiro, Mark Lathrop and Michel Roger |
| EPI_ISL_535749 | CUSM-Site Glen-LAB Microbiologie | Laboratoire de santé publique du Québec | Sandrine Moreira, Ioannis Ragoussis, Guillaume Bourque, Jesse Shapiro, Mark Lathrop and Michel Roger |
| EPI_ISL_535750, EPI_ISL_535751 | Hôtel-Dieu de Lévis | Laboratoire de santé publique du Québec | Sandrine Moreira, Ioannis Ragoussis, Guillaume Bourque, Jesse Shapiro, Mark Lathrop and Michel Roger |
| EPI_ISL_535752 | Hôpital Pierre-Le Gardeur | Laboratoire de santé publique du Québec | Sandrine Moreira, Ioannis Ragoussis, Guillaume Bourque, Jesse Shapiro, Mark Lathrop and Michel Roger |
| EPI_ISL_535753, EPI_ISL_535754 | CHUM - Microbiologie - Hôpital Saint-Luc | Laboratoire de santé publique du Québec | Sandrine Moreira, Ioannis Ragoussis, Guillaume Bourque, Jesse Shapiro, Mark Lathrop and Michel Roger |
| EPI_ISL_535755 | Hôpital Maisonneuve-Rosemont | Laboratoire de santé publique du Québec | Sandrine Moreira, Ioannis Ragoussis, Guillaume Bourque, Jesse Shapiro, Mark Lathrop and Michel Roger |
| EPI_ISL_535756, EPI_ISL_535757 | Hôpital général Juif | Laboratoire de santé publique du Québec | Sandrine Moreira, Ioannis Ragoussis, Guillaume Bourque, Jesse Shapiro, Mark Lathrop and Michel Roger |
| EPI_ISL_535758, EPI_ISL_535759 | Hôtel-Dieu de Lévis | Laboratoire de santé publique du Québec | Sandrine Moreira, Ioannis Ragoussis, Guillaume Bourque, Jesse Shapiro, Mark Lathrop and Michel Roger |
| EPI_ISL_535760 | CHUM - Microbiologie - Hôpital Saint-Luc | Laboratoire de santé publique du Québec | Sandrine Moreira, Ioannis Ragoussis, Guillaume Bourque, Jesse Shapiro, Mark Lathrop and Michel Roger |
| EPI_ISL_535761 | CUSM-Site Glen-LAB Microbiologie | Laboratoire de santé publique du Québec | Sandrine Moreira, Ioannis Ragoussis, Guillaume Bourque, Jesse Shapiro, Mark Lathrop and Michel Roger |
| EPI_ISL_535762 | Hôpital Pierre-Le Gardeur | Laboratoire de santé publique du Québec | Sandrine Moreira, Ioannis Ragoussis, Guillaume Bourque, Jesse Shapiro, Mark Lathrop and Michel Roger |
| EPI_ISL_535763 | Hôpital Pierre-Boucher | Laboratoire de santé publique du Québec | Sandrine Moreira, Ioannis Ragoussis, Guillaume Bourque, Jesse Shapiro, Mark Lathrop and Michel Roger |
| EPI_ISL_535764, EPI_ISL_535765 | CUSM-Site Glen-LAB Microbiologie | Laboratoire de santé publique du Québec | Sandrine Moreira, Ioannis Ragoussis, Guillaume Bourque, Jesse Shapiro, Mark Lathrop and Michel Roger |
| EPI_ISL_535766 | Hôpital Pierre-Le Gardeur | Laboratoire de santé publique du Québec | Sandrine Moreira, Ioannis Ragoussis, Guillaume Bourque, Jesse Shapiro, Mark Lathrop and Michel Roger |

| | | | |
|---|---|---|---|
| EPI_ISL_535767, EPI_ISL_535768 | Centre hospitalier de St-Mary | Laboratoire de santé publique du Québec | Sandrine Moreira, Ioannis Ragoussis, Guillaume Bourque, Jesse Shapiro, Mark Lathrop and Michel Roger |
| EPI_ISL_535769 | Hôpital général Juif | Laboratoire de santé publique du Québec | Sandrine Moreira, Ioannis Ragoussis, Guillaume Bourque, Jesse Shapiro, Mark Lathrop and Michel Roger |
| EPI_ISL_535770, EPI_ISL_535771 | Hôpital Pierre-Le Gardeur | Laboratoire de santé publique du Québec | Sandrine Moreira, Ioannis Ragoussis, Guillaume Bourque, Jesse Shapiro, Mark Lathrop and Michel Roger |
| EPI_ISL_535772, EPI_ISL_535773, EPI_ISL_535774, EPI_ISL_535775 | CHUL-LABO MULTI / MICRO | Laboratoire de santé publique du Québec | Sandrine Moreira, Ioannis Ragoussis, Guillaume Bourque, Jesse Shapiro, Mark Lathrop and Michel Roger |
| EPI_ISL_535776, EPI_ISL_535777, EPI_ISL_535778, EPI_ISL_535779, EPI_ISL_535780 | Centre de SSS de la Haute-Yamaska | Laboratoire de santé publique du Québec | Sandrine Moreira, Ioannis Ragoussis, Guillaume Bourque, Jesse Shapiro, Mark Lathrop and Michel Roger |
| EPI_ISL_535781, EPI_ISL_535782 | CHUL-LABO MULTI / MICRO | Laboratoire de santé publique du Québec | Sandrine Moreira, Ioannis Ragoussis, Guillaume Bourque, Jesse Shapiro, Mark Lathrop and Michel Roger |
| EPI_ISL_535783 | CHUM - Microbiologie - Hôpital Saint-Luc | Laboratoire de santé publique du Québec | Sandrine Moreira, Ioannis Ragoussis, Guillaume Bourque, Jesse Shapiro, Mark Lathrop and Michel Roger |
| EPI_ISL_535784, EPI_ISL_535785 | CHUL-LABO MULTI / MICRO | Laboratoire de santé publique du Québec | Sandrine Moreira, Ioannis Ragoussis, Guillaume Bourque, Jesse Shapiro, Mark Lathrop and Michel Roger |
| EPI_ISL_535786 | Hôpital Pierre-Boucher | Laboratoire de santé publique du Québec | Sandrine Moreira, Ioannis Ragoussis, Guillaume Bourque, Jesse Shapiro, Mark Lathrop and Michel Roger |
| EPI_ISL_535787, EPI_ISL_535788 | Centre Hospitalier Régional de Lanaudière | Laboratoire de santé publique du Québec | Sandrine Moreira, Ioannis Ragoussis, Guillaume Bourque, Jesse Shapiro, Mark Lathrop and Michel Roger |
| EPI_ISL_535789 | CHUM - Microbiologie - Hôpital Saint-Luc | Laboratoire de santé publique du Québec | Sandrine Moreira, Ioannis Ragoussis, Guillaume Bourque, Jesse Shapiro, Mark Lathrop and Michel Roger |
| EPI_ISL_535790 | Centre Hospitalier Régional de Lanaudière | Laboratoire de santé publique du Québec | Sandrine Moreira, Ioannis Ragoussis, Guillaume Bourque, Jesse Shapiro, Mark Lathrop and Michel Roger |
| EPI_ISL_535791 | Hôpital régional de Saint-Jérôme | Laboratoire de santé publique du Québec | Sandrine Moreira, Ioannis Ragoussis, Guillaume Bourque, Jesse Shapiro, Mark Lathrop and Michel Roger |
| EPI_ISL_535792 | Hôpital Fleurimont | Laboratoire de santé publique du Québec | Sandrine Moreira, Ioannis Ragoussis, Guillaume Bourque, Jesse Shapiro, Mark Lathrop and Michel Roger |
| EPI_ISL_535793 | Hôpital de Saint-Eustache | Laboratoire de santé publique du Québec | Sandrine Moreira, Ioannis Ragoussis, Guillaume Bourque, Jesse Shapiro, Mark Lathrop and Michel Roger |
| EPI_ISL_535794 | Hôpital de Hull | Laboratoire de santé publique du Québec | Sandrine Moreira, Ioannis Ragoussis, Guillaume Bourque, Jesse Shapiro, Mark Lathrop and Michel Roger |
| EPI_ISL_535795 | Hôpital Notre-Dame | Laboratoire de santé publique du Québec | Sandrine Moreira, Ioannis Ragoussis, Guillaume Bourque, Jesse Shapiro, Mark Lathrop and Michel Roger |
| EPI_ISL_535796 | Hôpital Charles-LeMoyne | Laboratoire de santé publique du Québec | Sandrine Moreira, Ioannis Ragoussis, Guillaume Bourque, Jesse Shapiro, Mark Lathrop and Michel Roger |
| EPI_ISL_535797 | Hôpital Honoré-Mercier | Laboratoire de santé publique du Québec | Sandrine Moreira, Ioannis Ragoussis, Guillaume Bourque, Jesse Shapiro, Mark Lathrop and Michel Roger |
| EPI_ISL_535798 | Hôpital régional de Saint-Jérôme | Laboratoire de santé publique du Québec | Sandrine Moreira, Ioannis Ragoussis, Guillaume Bourque, Jesse Shapiro, Mark Lathrop and Michel Roger |
| EPI_ISL_535799, EPI_ISL_535800 | CSSS Haut-Richelieu/Rouville (Hôpital) | Laboratoire de santé publique du Québec | Sandrine Moreira, Ioannis Ragoussis, Guillaume Bourque, Jesse Shapiro, Mark Lathrop and Michel Roger |
| EPI_ISL_535801 | CHU Sainte-Justine | Laboratoire de santé publique du Québec | Sandrine Moreira, Ioannis Ragoussis, Guillaume Bourque, Jesse Shapiro, Mark Lathrop and Michel Roger |
| EPI_ISL_535802 | CHUM - Microbiologie - Hôpital Saint-Luc | Laboratoire de santé publique du Québec | Sandrine Moreira, Ioannis Ragoussis, Guillaume Bourque, Jesse Shapiro, Mark Lathrop and Michel Roger |
| EPI_ISL_535803, EPI_ISL_535804, EPI_ISL_535805 | Centre de SSS de la Haute-Yamaska | Laboratoire de santé publique du Québec | Sandrine Moreira, Ioannis Ragoussis, Guillaume Bourque, Jesse Shapiro, Mark Lathrop and Michel Roger |
| EPI_ISL_535806 | CHUM - Microbiologie - Hôpital Saint-Luc | Laboratoire de santé publique du Québec | Sandrine Moreira, Ioannis Ragoussis, Guillaume Bourque, Jesse Shapiro, Mark Lathrop and Michel Roger |
| EPI_ISL_535807 | Centre hospitalier régional du Grand Portage | Laboratoire de santé publique du Québec | Sandrine Moreira, Ioannis Ragoussis, Guillaume Bourque, Jesse Shapiro, Mark Lathrop and Michel Roger |
| EPI_ISL_535808 | Centre de santé Chibougamau | Laboratoire de santé publique du Québec | Sandrine Moreira, Ioannis Ragoussis, Guillaume Bourque, Jesse Shapiro, Mark Lathrop and Michel Roger |
| EPI_ISL_535809 | Hôpital de Hull | Laboratoire de santé publique du Québec | Sandrine Moreira, Ioannis Ragoussis, Guillaume Bourque, Jesse Shapiro, Mark Lathrop and Michel Roger |
| EPI_ISL_535810 | Hôpital Sainte-Croix | Laboratoire de santé publique du Québec | Sandrine Moreira, Ioannis Ragoussis, Guillaume Bourque, Jesse Shapiro, Mark Lathrop and Michel Roger |
| EPI_ISL_535811 | CUSM-Site Glen-LAB Microbiologie | Laboratoire de santé publique du Québec | Sandrine Moreira, Ioannis Ragoussis, Guillaume Bourque, Jesse Shapiro, Mark Lathrop and Michel Roger |
| EPI_ISL_535812 | Hôpital du Sacré-Coeur de Montréal | Laboratoire de santé publique du Québec | Sandrine Moreira, Ioannis Ragoussis, Guillaume Bourque, Jesse Shapiro, Mark Lathrop and Michel Roger |
| EPI_ISL_535813 | CSSS Haut-Richelieu/Rouville (Hôpital) | Laboratoire de santé publique du Québec | Sandrine Moreira, Ioannis Ragoussis, Guillaume Bourque, Jesse Shapiro, Mark Lathrop and Michel Roger |
| EPI_ISL_535814 | CHUM - Microbiologie - Hôpital Saint-Luc | Laboratoire de santé publique du Québec | Sandrine Moreira, Ioannis Ragoussis, Guillaume Bourque, Jesse Shapiro, Mark Lathrop and Michel Roger |
| EPI_ISL_535815 | CSSS Haut-Richelieu/Rouville (Hôpital) | Laboratoire de santé publique du Québec | Sandrine Moreira, Ioannis Ragoussis, Guillaume Bourque, Jesse Shapiro, Mark Lathrop and Michel Roger |
| EPI_ISL_535816 | Centre de SSS de la Haute-Yamaska | Laboratoire de santé publique du Québec | Sandrine Moreira, Ioannis Ragoussis, Guillaume Bourque, Jesse Shapiro, Mark Lathrop and Michel Roger |
| EPI_ISL_535817 | Hôpital Maisonneuve-Rosemont | Laboratoire de santé publique du Québec | Sandrine Moreira, Ioannis Ragoussis, Guillaume Bourque, Jesse Shapiro, Mark Lathrop and Michel Roger |
| EPI_ISL_535818 | Hôpital Pierre-Boucher | Laboratoire de santé publique du Québec | Sandrine Moreira, Ioannis Ragoussis, Guillaume Bourque, Jesse Shapiro, Mark Lathrop and Michel Roger |
| EPI_ISL_535819 | Hôpital Charles-LeMoyne | Laboratoire de santé publique du Québec | Sandrine Moreira, Ioannis Ragoussis, Guillaume Bourque, Jesse Shapiro, Mark Lathrop and Michel Roger |
| EPI_ISL_535820, EPI_ISL_535821 | Hôtel-Dieu de Lévis | Laboratoire de santé publique du Québec | Sandrine Moreira, Ioannis Ragoussis, Guillaume Bourque, Jesse Shapiro, Mark Lathrop and Michel Roger |
| EPI_ISL_535822, EPI_ISL_535823, EPI_ISL_535824, EPI_ISL_535825 | CHUL-LABO MULTI / MICRO | Laboratoire de santé publique du Québec | Sandrine Moreira, Ioannis Ragoussis, Guillaume Bourque, Jesse Shapiro, Mark Lathrop and Michel Roger |
| EPI_ISL_535826, EPI_ISL_535827 | CHUM - Microbiologie - Hôpital Saint-Luc | Laboratoire de santé publique du Québec | Sandrine Moreira, Ioannis Ragoussis, Guillaume Bourque, Jesse Shapiro, Mark Lathrop and Michel Roger |
| EPI_ISL_535828 | CHUL-LABO MULTI / MICRO | Laboratoire de santé publique du Québec | Sandrine Moreira, Ioannis Ragoussis, Guillaume Bourque, Jesse Shapiro, Mark Lathrop and Michel Roger |
| EPI_ISL_535829 | Hôpital de Chicoutimi | Laboratoire de santé publique du Québec | Sandrine Moreira, Ioannis Ragoussis, Guillaume Bourque, Jesse Shapiro, Mark Lathrop and Michel Roger |
| EPI_ISL_535830 | CHUM - Microbiologie - Hôpital Saint-Luc | Laboratoire de santé publique du Québec | Sandrine Moreira, Ioannis Ragoussis, Guillaume Bourque, Jesse Shapiro, Mark Lathrop and Michel Roger |
| EPI_ISL_535831 | Hôpital Maisonneuve-Rosemont | Laboratoire de santé publique du Québec | Sandrine Moreira, Ioannis Ragoussis, Guillaume Bourque, Jesse Shapiro, Mark Lathrop and Michel Roger |
| EPI_ISL_535832 | CHUM - Microbiologie - Hôpital Saint-Luc | Laboratoire de santé publique du Québec | Sandrine Moreira, Ioannis Ragoussis, Guillaume Bourque, Jesse Shapiro, Mark Lathrop and Michel Roger |
| EPI_ISL_535833, EPI_ISL_535834, EPI_ISL_535835, EPI_ISL_535836 | Hôtel-Dieu de Lévis | Laboratoire de santé publique du Québec | Sandrine Moreira, Ioannis Ragoussis, Guillaume Bourque, Jesse Shapiro, Mark Lathrop and Michel Roger |
| EPI_ISL_535837, EPI_ISL_535838 | Hôpital régional de Saint-Jérôme | Laboratoire de santé publique du Québec | Sandrine Moreira, Ioannis Ragoussis, Guillaume Bourque, Jesse Shapiro, Mark Lathrop and Michel Roger |
| EPI_ISL_535839 | Hôpital Cité de la Santé | Laboratoire de santé publique du Québec | Sandrine Moreira, Ioannis Ragoussis, Guillaume Bourque, Jesse Shapiro, Mark Lathrop and Michel Roger |
| EPI_ISL_535840 | Hôpital Pierre-Boucher | Laboratoire de santé publique du Québec | Sandrine Moreira, Ioannis Ragoussis, Guillaume Bourque, Jesse Shapiro, Mark Lathrop and Michel Roger |
| EPI_ISL_535841 | CSSS Haut-Richelieu/Rouville (Hôpital) | Laboratoire de santé publique du Québec | Sandrine Moreira, Ioannis Ragoussis, Guillaume Bourque, Jesse Shapiro, Mark Lathrop and Michel Roger |
| EPI_ISL_535842, EPI_ISL_535843, EPI_ISL_535844 | Centre Hospitalier Régional de Lanaudière | Laboratoire de santé publique du Québec | Sandrine Moreira, Ioannis Ragoussis, Guillaume Bourque, Jesse Shapiro, Mark Lathrop and Michel Roger |
| EPI_ISL_535845 | Centre de SSS D'Arthabaska-et-de-l'érable - Hôtel-Dieu | Laboratoire de santé publique du Québec | Sandrine Moreira, Ioannis Ragoussis, Guillaume Bourque, Jesse Shapiro, Mark Lathrop and Michel Roger |
| EPI_ISL_535846 | CSSS Haut-Richelieu/Rouville (Hôpital) | Laboratoire de santé publique du Québec | Sandrine Moreira, Ioannis Ragoussis, Guillaume Bourque, Jesse Shapiro, Mark Lathrop and Michel Roger |
| EPI_ISL_535847 | Hôpital du Suroît | Laboratoire de santé publique du Québec | Sandrine Moreira, Ioannis Ragoussis, Guillaume Bourque, Jesse Shapiro, Mark Lathrop and Michel Roger |
| EPI_ISL_535848 | Hôpital Charles-LeMoyne | Laboratoire de santé publique du Québec | Sandrine Moreira, Ioannis Ragoussis, Guillaume Bourque, Jesse Shapiro, Mark Lathrop and Michel Roger |
| EPI_ISL_535849 | Centre de SSS de Trois-Rivières | Laboratoire de santé publique du Québec | Sandrine Moreira, Ioannis Ragoussis, Guillaume Bourque, Jesse Shapiro, Mark Lathrop and Michel Roger |
| EPI_ISL_535850 | Hôpital Pierre-Le Gardeur | Laboratoire de santé publique du Québec | Sandrine Moreira, Ioannis Ragoussis, Guillaume Bourque, Jesse Shapiro, Mark Lathrop and Michel Roger |
| EPI_ISL_535851 | Hôpital Charles-LeMoyne | Laboratoire de santé publique du Québec | Sandrine Moreira, Ioannis Ragoussis, Guillaume Bourque, Jesse Shapiro, Mark Lathrop and Michel Roger |
| EPI_ISL_535852 | Hôpital général du Lakeshore | Laboratoire de santé publique du Québec | Sandrine Moreira, Ioannis Ragoussis, Guillaume Bourque, Jesse Shapiro, Mark Lathrop and Michel Roger |
| EPI_ISL_535853 | Hôpital Charles-LeMoyne | Laboratoire de santé publique du Québec | Sandrine Moreira, Ioannis Ragoussis, Guillaume Bourque, Jesse Shapiro, Mark Lathrop and Michel Roger |
| EPI_ISL_535854, EPI_ISL_535855 | Centre de SSS de la Haute-Yamaska | Laboratoire de santé publique du Québec | Sandrine Moreira, Ioannis Ragoussis, Guillaume Bourque, Jesse Shapiro, Mark Lathrop and Michel Roger |
| EPI_ISL_535856 | Hôpital de Verdun | Laboratoire de santé publique du Québec | Sandrine Moreira, Ioannis Ragoussis, Guillaume Bourque, Jesse Shapiro, Mark Lathrop and Michel Roger |
| EPI_ISL_535857 | Hôpital Pierre-Boucher | Laboratoire de santé publique du Québec | Sandrine Moreira, Ioannis Ragoussis, Guillaume Bourque, Jesse Shapiro, Mark Lathrop and Michel Roger |
| EPI_ISL_535858 | Hôpital de Gatineau | Laboratoire de santé publique du Québec | Sandrine Moreira, Ioannis Ragoussis, Guillaume Bourque, Jesse Shapiro, Mark Lathrop and Michel Roger |
| EPI_ISL_535859, EPI_ISL_535860 | Hôpital Sainte-Croix | Laboratoire de santé publique du Québec | Sandrine Moreira, Ioannis Ragoussis, Guillaume Bourque, Jesse Shapiro, Mark Lathrop and Michel Roger |
| EPI_ISL_535861, EPI_ISL_535862, EPI_ISL_535863 | Hôpital Pierre-Le Gardeur | Laboratoire de santé publique du Québec | Sandrine Moreira, Ioannis Ragoussis, Guillaume Bourque, Jesse Shapiro, Mark Lathrop and Michel Roger |
| EPI_ISL_535864 | CSSS Haut-Richelieu/Rouville (Hôpital) | Laboratoire de santé publique du Québec | Sandrine Moreira, Ioannis Ragoussis, Guillaume Bourque, Jesse Shapiro, Mark Lathrop and Michel Roger |
| EPI_ISL_535865 | Hôpital de Saint-Eustache | Laboratoire de santé publique du Québec | Sandrine Moreira, Ioannis Ragoussis, Guillaume Bourque, Jesse Shapiro, Mark Lathrop and Michel Roger |
| EPI_ISL_535866 | Hôpital Charles-LeMoyne | Laboratoire de santé publique du Québec | Sandrine Moreira, Ioannis Ragoussis, Guillaume Bourque, Jesse Shapiro, Mark Lathrop and Michel Roger |
| EPI_ISL_535867 | Hôpital de Saint-Eustache | Laboratoire de santé publique du Québec | Sandrine Moreira, Ioannis Ragoussis, Guillaume Bourque, Jesse Shapiro, Mark Lathrop and Michel Roger |
| EPI_ISL_535868 | Hôpital du Suroît | Laboratoire de santé publique du Québec | Sandrine Moreira, Ioannis Ragoussis, Guillaume Bourque, Jesse Shapiro, Mark Lathrop and Michel Roger |
| EPI_ISL_535869 | Hôpital Charles-LeMoyne | Laboratoire de santé publique du Québec | Sandrine Moreira, Ioannis Ragoussis, Guillaume Bourque, Jesse Shapiro, Mark Lathrop and Michel Roger |
| EPI_ISL_535870, EPI_ISL_535871 | Centre Hospitalier Régional de Lanaudière | Laboratoire de santé publique du Québec | Sandrine Moreira, Ioannis Ragoussis, Guillaume Bourque, Jesse Shapiro, Mark Lathrop and Michel Roger |

| | | | |
|---|---|---|---|
| EPI_ISL_535872, EPI_ISL_535873 | Centre hospitalier régional du Grand Portage | Laboratoire de santé publique du Québec | Sandrine Moreira, Ioannis Ragoussis, Guillaume Bourque, Jesse Shapiro, Mark Lathrop and Michel Roger |
| EPI_ISL_535874 | Hôpital Charles-LeMoyne | Laboratoire de santé publique du Québec | Sandrine Moreira, Ioannis Ragoussis, Guillaume Bourque, Jesse Shapiro, Mark Lathrop and Michel Roger |
| EPI_ISL_535875 | Hôpital général du Lakeshore | Laboratoire de santé publique du Québec | Sandrine Moreira, Ioannis Ragoussis, Guillaume Bourque, Jesse Shapiro, Mark Lathrop and Michel Roger |
| EPI_ISL_535876, EPI_ISL_535877 | CSSS Haut-Richelieu/Rouville (Hôpital) | Laboratoire de santé publique du Québec | Sandrine Moreira, Ioannis Ragoussis, Guillaume Bourque, Jesse Shapiro, Mark Lathrop and Michel Roger |
| EPI_ISL_535878 | Hôpital de Saint-Eustache | Laboratoire de santé publique du Québec | Sandrine Moreira, Ioannis Ragoussis, Guillaume Bourque, Jesse Shapiro, Mark Lathrop and Michel Roger |
| EPI_ISL_535879 | Hôpital Pierre-Boucher | Laboratoire de santé publique du Québec | Sandrine Moreira, Ioannis Ragoussis, Guillaume Bourque, Jesse Shapiro, Mark Lathrop and Michel Roger |
| EPI_ISL_535880 | Hôpital du Suroît | Laboratoire de santé publique du Québec | Sandrine Moreira, Ioannis Ragoussis, Guillaume Bourque, Jesse Shapiro, Mark Lathrop and Michel Roger |
| EPI_ISL_535881, EPI_ISL_535882, EPI_ISL_535883 | Hôpital Pierre-Le Gardeur | Laboratoire de santé publique du Québec | Sandrine Moreira, Ioannis Ragoussis, Guillaume Bourque, Jesse Shapiro, Mark Lathrop and Michel Roger |
| EPI_ISL_535884 | CSSS Haut-Richelieu/Rouville (Hôpital) | Laboratoire de santé publique du Québec | Sandrine Moreira, Ioannis Ragoussis, Guillaume Bourque, Jesse Shapiro, Mark Lathrop and Michel Roger |
| EPI_ISL_535885 | Hôpital de Maria | Laboratoire de santé publique du Québec | Sandrine Moreira, Ioannis Ragoussis, Guillaume Bourque, Jesse Shapiro, Mark Lathrop and Michel Roger |
| EPI_ISL_535886, EPI_ISL_535887 | Hôpital de Hull | Laboratoire de santé publique du Québec | Sandrine Moreira, Ioannis Ragoussis, Guillaume Bourque, Jesse Shapiro, Mark Lathrop and Michel Roger |
| EPI_ISL_535888 | Hôpital Charles-LeMoyne | Laboratoire de santé publique du Québec | Sandrine Moreira, Ioannis Ragoussis, Guillaume Bourque, Jesse Shapiro, Mark Lathrop and Michel Roger |
| EPI_ISL_535889 | Hôpital de Saint-Eustache | Laboratoire de santé publique du Québec | Sandrine Moreira, Ioannis Ragoussis, Guillaume Bourque, Jesse Shapiro, Mark Lathrop and Michel Roger |
| EPI_ISL_535890, EPI_ISL_535891 | Hôpital Charles-LeMoyne | Laboratoire de santé publique du Québec | Sandrine Moreira, Ioannis Ragoussis, Guillaume Bourque, Jesse Shapiro, Mark Lathrop and Michel Roger |
| EPI_ISL_535892, EPI_ISL_535893 | Hôpital Pierre-Le Gardeur | Laboratoire de santé publique du Québec | Sandrine Moreira, Ioannis Ragoussis, Guillaume Bourque, Jesse Shapiro, Mark Lathrop and Michel Roger |
| EPI_ISL_535894, EPI_ISL_535895, EPI_ISL_535896 | Centre Hospitalier Régional de Lanaudière | Laboratoire de santé publique du Québec | Sandrine Moreira, Ioannis Ragoussis, Guillaume Bourque, Jesse Shapiro, Mark Lathrop and Michel Roger |
| EPI_ISL_535897, EPI_ISL_535898, EPI_ISL_535899, EPI_ISL_535900, EPI_ISL_535901 | Hôpital Sainte-Croix | Laboratoire de santé publique du Québec | Sandrine Moreira, Ioannis Ragoussis, Guillaume Bourque, Jesse Shapiro, Mark Lathrop and Michel Roger |
| EPI_ISL_535902 | CSSS Haut-Richelieu/Rouville (Hôpital) | Laboratoire de santé publique du Québec | Sandrine Moreira, Ioannis Ragoussis, Guillaume Bourque, Jesse Shapiro, Mark Lathrop and Michel Roger |
| EPI_ISL_535903, EPI_ISL_535904 | Hôtel-Dieu de Sorel | Laboratoire de santé publique du Québec | Sandrine Moreira, Ioannis Ragoussis, Guillaume Bourque, Jesse Shapiro, Mark Lathrop and Michel Roger |
| EPI_ISL_535905, EPI_ISL_535906 | Hôpital de Chicoutimi | Laboratoire de santé publique du Québec | Sandrine Moreira, Ioannis Ragoussis, Guillaume Bourque, Jesse Shapiro, Mark Lathrop and Michel Roger |
| EPI_ISL_535907 | Centre Hospitalier Régional de Lanaudière | Laboratoire de santé publique du Québec | Sandrine Moreira, Ioannis Ragoussis, Guillaume Bourque, Jesse Shapiro, Mark Lathrop and Michel Roger |
| EPI_ISL_535908, EPI_ISL_535909 | Hôpital Pierre-Boucher | Laboratoire de santé publique du Québec | Sandrine Moreira, Ioannis Ragoussis, Guillaume Bourque, Jesse Shapiro, Mark Lathrop and Michel Roger |
| EPI_ISL_535910 | CSSS Haut-Richelieu/Rouville (Hôpital) | Laboratoire de santé publique du Québec | Sandrine Moreira, Ioannis Ragoussis, Guillaume Bourque, Jesse Shapiro, Mark Lathrop and Michel Roger |
| EPI_ISL_535911 | Hôpital de Hull | Laboratoire de santé publique du Québec | Sandrine Moreira, Ioannis Ragoussis, Guillaume Bourque, Jesse Shapiro, Mark Lathrop and Michel Roger |
| EPI_ISL_535912 | Hôpital Honoré-Mercier | Laboratoire de santé publique du Québec | Sandrine Moreira, Ioannis Ragoussis, Guillaume Bourque, Jesse Shapiro, Mark Lathrop and Michel Roger |
| EPI_ISL_535913 | Hôpital du Suroît | Laboratoire de santé publique du Québec | Sandrine Moreira, Ioannis Ragoussis, Guillaume Bourque, Jesse Shapiro, Mark Lathrop and Michel Roger |
| EPI_ISL_535914 | Hôpital du Centre-de-la-Mauricie | Laboratoire de santé publique du Québec | Sandrine Moreira, Ioannis Ragoussis, Guillaume Bourque, Jesse Shapiro, Mark Lathrop and Michel Roger |
| EPI_ISL_535915 | Hôpital Honoré-Mercier | Laboratoire de santé publique du Québec | Sandrine Moreira, Ioannis Ragoussis, Guillaume Bourque, Jesse Shapiro, Mark Lathrop and Michel Roger |
| EPI_ISL_535916 | Centre de SSS de Trois-Rivières | Laboratoire de santé publique du Québec | Sandrine Moreira, Ioannis Ragoussis, Guillaume Bourque, Jesse Shapiro, Mark Lathrop and Michel Roger |
| EPI_ISL_535917, EPI_ISL_535918, EPI_ISL_535919 | Hôpital du Suroît | Laboratoire de santé publique du Québec | Sandrine Moreira, Ioannis Ragoussis, Guillaume Bourque, Jesse Shapiro, Mark Lathrop and Michel Roger |
| EPI_ISL_535920, EPI_ISL_535921, EPI_ISL_535922, EPI_ISL_535923 | Hôpital Charles-LeMoyne | Laboratoire de santé publique du Québec | Sandrine Moreira, Ioannis Ragoussis, Guillaume Bourque, Jesse Shapiro, Mark Lathrop and Michel Roger |
| EPI_ISL_535924 | Hôpital de Saint-Eustache | Laboratoire de santé publique du Québec | Sandrine Moreira, Ioannis Ragoussis, Guillaume Bourque, Jesse Shapiro, Mark Lathrop and Michel Roger |
| EPI_ISL_535925 | CSSS Haut-Richelieu/Rouville (Hôpital) | Laboratoire de santé publique du Québec | Sandrine Moreira, Ioannis Ragoussis, Guillaume Bourque, Jesse Shapiro, Mark Lathrop and Michel Roger |
| EPI_ISL_535926 | Hôpital de Saint-Eustache | Laboratoire de santé publique du Québec | Sandrine Moreira, Ioannis Ragoussis, Guillaume Bourque, Jesse Shapiro, Mark Lathrop and Michel Roger |
| EPI_ISL_535927 | Hôpital Charles-LeMoyne | Laboratoire de santé publique du Québec | Sandrine Moreira, Ioannis Ragoussis, Guillaume Bourque, Jesse Shapiro, Mark Lathrop and Michel Roger |
| EPI_ISL_535928 | CSSS Haut-Richelieu/Rouville (Hôpital) | Laboratoire de santé publique du Québec | Sandrine Moreira, Ioannis Ragoussis, Guillaume Bourque, Jesse Shapiro, Mark Lathrop and Michel Roger |
| EPI_ISL_535929, EPI_ISL_535930 | Hôpital de Saint-Eustache | Laboratoire de santé publique du Québec | Sandrine Moreira, Ioannis Ragoussis, Guillaume Bourque, Jesse Shapiro, Mark Lathrop and Michel Roger |
| EPI_ISL_535931, EPI_ISL_535932, EPI_ISL_535933, EPI_ISL_535934, EPI_ISL_535935 | Centre Hospitalier Régional de Lanaudière | Laboratoire de santé publique du Québec | Sandrine Moreira, Ioannis Ragoussis, Guillaume Bourque, Jesse Shapiro, Mark Lathrop and Michel Roger |
| EPI_ISL_535936 | Hôpital Pierre-Le Gardeur | Laboratoire de santé publique du Québec | Sandrine Moreira, Ioannis Ragoussis, Guillaume Bourque, Jesse Shapiro, Mark Lathrop and Michel Roger |
| EPI_ISL_535937 | Hôpital du Centre-de-la-Mauricie | Laboratoire de santé publique du Québec | Sandrine Moreira, Ioannis Ragoussis, Guillaume Bourque, Jesse Shapiro, Mark Lathrop and Michel Roger |
| EPI_ISL_535938 | Hôpital Pierre-Le Gardeur | Laboratoire de santé publique du Québec | Sandrine Moreira, Ioannis Ragoussis, Guillaume Bourque, Jesse Shapiro, Mark Lathrop and Michel Roger |
| EPI_ISL_535939, EPI_ISL_535940 | Centre de SSS D'Arthabaska-et-de-l'érable - Hôtel-Dieu | Laboratoire de santé publique du Québec | Sandrine Moreira, Ioannis Ragoussis, Guillaume Bourque, Jesse Shapiro, Mark Lathrop and Michel Roger |
| EPI_ISL_535941, EPI_ISL_535942, EPI_ISL_535943 | Hôpital de Saint-Eustache | Laboratoire de santé publique du Québec | Sandrine Moreira, Ioannis Ragoussis, Guillaume Bourque, Jesse Shapiro, Mark Lathrop and Michel Roger |
| EPI_ISL_535944 | Hôpital Charles-LeMoyne | Laboratoire de santé publique du Québec | Sandrine Moreira, Ioannis Ragoussis, Guillaume Bourque, Jesse Shapiro, Mark Lathrop and Michel Roger |
| EPI_ISL_535945 | Hôpital de Mont-Laurier | Laboratoire de santé publique du Québec | Sandrine Moreira, Ioannis Ragoussis, Guillaume Bourque, Jesse Shapiro, Mark Lathrop and Michel Roger |
| EPI_ISL_535946 | CSSS Haut-Richelieu/Rouville (Hôpital) | Laboratoire de santé publique du Québec | Sandrine Moreira, Ioannis Ragoussis, Guillaume Bourque, Jesse Shapiro, Mark Lathrop and Michel Roger |
| EPI_ISL_535947 | Hôpital de Saint-Eustache | Laboratoire de santé publique du Québec | Sandrine Moreira, Ioannis Ragoussis, Guillaume Bourque, Jesse Shapiro, Mark Lathrop and Michel Roger |
| EPI_ISL_535948 | Hôpital Pierre-Boucher | Laboratoire de santé publique du Québec | Sandrine Moreira, Ioannis Ragoussis, Guillaume Bourque, Jesse Shapiro, Mark Lathrop and Michel Roger |
| EPI_ISL_535949, EPI_ISL_535950 | Hôpital Charles-LeMoyne | Laboratoire de santé publique du Québec | Sandrine Moreira, Ioannis Ragoussis, Guillaume Bourque, Jesse Shapiro, Mark Lathrop and Michel Roger |
| EPI_ISL_535951 | Hôpital de Saint-Eustache | Laboratoire de santé publique du Québec | Sandrine Moreira, Ioannis Ragoussis, Guillaume Bourque, Jesse Shapiro, Mark Lathrop and Michel Roger |
| EPI_ISL_535952, EPI_ISL_535953 | Hôpital Charles-LeMoyne | Laboratoire de santé publique du Québec | Sandrine Moreira, Ioannis Ragoussis, Guillaume Bourque, Jesse Shapiro, Mark Lathrop and Michel Roger |
| EPI_ISL_535954 | Hôpital Pierre-Le Gardeur | Laboratoire de santé publique du Québec | Sandrine Moreira, Ioannis Ragoussis, Guillaume Bourque, Jesse Shapiro, Mark Lathrop and Michel Roger |
| EPI_ISL_535955 | Centre Hospitalier Régional de Lanaudière | Laboratoire de santé publique du Québec | Sandrine Moreira, Ioannis Ragoussis, Guillaume Bourque, Jesse Shapiro, Mark Lathrop and Michel Roger |
| EPI_ISL_535956 | Hôpital Pierre-Le Gardeur | Laboratoire de santé publique du Québec | Sandrine Moreira, Ioannis Ragoussis, Guillaume Bourque, Jesse Shapiro, Mark Lathrop and Michel Roger |
| EPI_ISL_535957 | Hôpital Charles-LeMoyne | Laboratoire de santé publique du Québec | Sandrine Moreira, Ioannis Ragoussis, Guillaume Bourque, Jesse Shapiro, Mark Lathrop and Michel Roger |
| EPI_ISL_535958, EPI_ISL_535959 | Centre hospitalier régional du Grand Portage | Laboratoire de santé publique du Québec | Sandrine Moreira, Ioannis Ragoussis, Guillaume Bourque, Jesse Shapiro, Mark Lathrop and Michel Roger |
| EPI_ISL_535960 | Centre hospitalier Anna-Laberge | Laboratoire de santé publique du Québec | Sandrine Moreira, Ioannis Ragoussis, Guillaume Bourque, Jesse Shapiro, Mark Lathrop and Michel Roger |
| EPI_ISL_535961, EPI_ISL_535962 | Hôpital Charles-LeMoyne | Laboratoire de santé publique du Québec | Sandrine Moreira, Ioannis Ragoussis, Guillaume Bourque, Jesse Shapiro, Mark Lathrop and Michel Roger |
| EPI_ISL_535963 | Hôpital d'Alma | Laboratoire de santé publique du Québec | Sandrine Moreira, Ioannis Ragoussis, Guillaume Bourque, Jesse Shapiro, Mark Lathrop and Michel Roger |
| EPI_ISL_535964, EPI_ISL_535965 | Hôpital de Chicoutimi | Laboratoire de santé publique du Québec | Sandrine Moreira, Ioannis Ragoussis, Guillaume Bourque, Jesse Shapiro, Mark Lathrop and Michel Roger |
| EPI_ISL_535966 | Hôpital Charles-LeMoyne | Laboratoire de santé publique du Québec | Sandrine Moreira, Ioannis Ragoussis, Guillaume Bourque, Jesse Shapiro, Mark Lathrop and Michel Roger |
| EPI_ISL_535967 | CSSS Haut-Richelieu/Rouville (Hôpital) | Laboratoire de santé publique du Québec | Sandrine Moreira, Ioannis Ragoussis, Guillaume Bourque, Jesse Shapiro, Mark Lathrop and Michel Roger |
| EPI_ISL_535968 | Hôpital du Centre-de-la-Mauricie | Laboratoire de santé publique du Québec | Sandrine Moreira, Ioannis Ragoussis, Guillaume Bourque, Jesse Shapiro, Mark Lathrop and Michel Roger |
| EPI_ISL_535969 | Hôpital Sainte-Croix | Laboratoire de santé publique du Québec | Sandrine Moreira, Ioannis Ragoussis, Guillaume Bourque, Jesse Shapiro, Mark Lathrop and Michel Roger |
| EPI_ISL_535970 | CSSS de la Minganie | Laboratoire de santé publique du Québec | Sandrine Moreira, Ioannis Ragoussis, Guillaume Bourque, Jesse Shapiro, Mark Lathrop and Michel Roger |
| EPI_ISL_535971 | Hôpital du Centre-de-la-Mauricie | Laboratoire de santé publique du Québec | Sandrine Moreira, Ioannis Ragoussis, Guillaume Bourque, Jesse Shapiro, Mark Lathrop and Michel Roger |
| EPI_ISL_535972, EPI_ISL_535973 | Hôpital Charles-LeMoyne | Laboratoire de santé publique du Québec | Sandrine Moreira, Ioannis Ragoussis, Guillaume Bourque, Jesse Shapiro, Mark Lathrop and Michel Roger |
| EPI_ISL_535974, EPI_ISL_535975 | Hôpital du Centre-de-la-Mauricie | Laboratoire de santé publique du Québec | Sandrine Moreira, Ioannis Ragoussis, Guillaume Bourque, Jesse Shapiro, Mark Lathrop and Michel Roger |
| EPI_ISL_535976, EPI_ISL_535977 | Centre de SSS de Trois-Rivières | Laboratoire de santé publique du Québec | Sandrine Moreira, Ioannis Ragoussis, Guillaume Bourque, Jesse Shapiro, Mark Lathrop and Michel Roger |
| EPI_ISL_535978 | Hôpital du Centre-de-la-Mauricie | Laboratoire de santé publique du Québec | Sandrine Moreira, Ioannis Ragoussis, Guillaume Bourque, Jesse Shapiro, Mark Lathrop and Michel Roger |
| EPI_ISL_535979 | Centre de SSS de Trois-Rivières | Laboratoire de santé publique du Québec | Sandrine Moreira, Ioannis Ragoussis, Guillaume Bourque, Jesse Shapiro, Mark Lathrop and Michel Roger |
| EPI_ISL_535980, EPI_ISL_535981 | Centre hospitalier Anna-Laberge | Laboratoire de santé publique du Québec | Sandrine Moreira, Ioannis Ragoussis, Guillaume Bourque, Jesse Shapiro, Mark Lathrop and Michel Roger |
| EPI_ISL_535982 | Hôpital de Saint-Eustache | Laboratoire de santé publique du Québec | Sandrine Moreira, Ioannis Ragoussis, Guillaume Bourque, Jesse Shapiro, Mark Lathrop and Michel Roger |
| EPI_ISL_535983 | Hôpital Charles-LeMoyne | Laboratoire de santé publique du Québec | Sandrine Moreira, Ioannis Ragoussis, Guillaume Bourque, Jesse Shapiro, Mark Lathrop and Michel Roger |

| | | | |
|---|---|---|---|
| EPI_ISL_535984 | CSSS Haut-Richelieu/Rouville (Hôpital) | Laboratoire de santé publique du Québec | Sandrine Moreira, Ioannis Ragoussis, Guillaume Bourque, Jesse Shapiro, Mark Lathrop and Michel Roger |
| EPI_ISL_535985, EPI_ISL_535986, EPI_ISL_535987 | Hôpital de Rouyn-Noranda | Laboratoire de santé publique du Québec | Sandrine Moreira, Ioannis Ragoussis, Guillaume Bourque, Jesse Shapiro, Mark Lathrop and Michel Roger |
| EPI_ISL_535988 | Hôpital du Suroît | Laboratoire de santé publique du Québec | Sandrine Moreira, Ioannis Ragoussis, Guillaume Bourque, Jesse Shapiro, Mark Lathrop and Michel Roger |
| EPI_ISL_535989 | Hôpital de Hull | Laboratoire de santé publique du Québec | Sandrine Moreira, Ioannis Ragoussis, Guillaume Bourque, Jesse Shapiro, Mark Lathrop and Michel Roger |
| EPI_ISL_535990 | Hôtel-Dieu de Sorel | Laboratoire de santé publique du Québec | Sandrine Moreira, Ioannis Ragoussis, Guillaume Bourque, Jesse Shapiro, Mark Lathrop and Michel Roger |
| EPI_ISL_535991 | Hôpital et Centre d'Hébergement de Sept-Iles | Laboratoire de santé publique du Québec | Sandrine Moreira, Ioannis Ragoussis, Guillaume Bourque, Jesse Shapiro, Mark Lathrop and Michel Roger |
| EPI_ISL_535992 | Hôtel-Dieu de Sorel | Laboratoire de santé publique du Québec | Sandrine Moreira, Ioannis Ragoussis, Guillaume Bourque, Jesse Shapiro, Mark Lathrop and Michel Roger |
| EPI_ISL_535993 | Hôpital de Hull | Laboratoire de santé publique du Québec | Sandrine Moreira, Ioannis Ragoussis, Guillaume Bourque, Jesse Shapiro, Mark Lathrop and Michel Roger |
| EPI_ISL_535994 | Hôpital Laurentien | Laboratoire de santé publique du Québec | Sandrine Moreira, Ioannis Ragoussis, Guillaume Bourque, Jesse Shapiro, Mark Lathrop and Michel Roger |
| EPI_ISL_535995 | Hôpital de Hull | Laboratoire de santé publique du Québec | Sandrine Moreira, Ioannis Ragoussis, Guillaume Bourque, Jesse Shapiro, Mark Lathrop and Michel Roger |
| EPI_ISL_535996, EPI_ISL_535997, EPI_ISL_535998, EPI_ISL_535999 | Hôpital du Centre-de-la-Mauricie | Laboratoire de santé publique du Québec | Sandrine Moreira, Ioannis Ragoussis, Guillaume Bourque, Jesse Shapiro, Mark Lathrop and Michel Roger |
| EPI_ISL_536000 | Centre hospitalier Anna-Laberge | Laboratoire de santé publique du Québec | Sandrine Moreira, Ioannis Ragoussis, Guillaume Bourque, Jesse Shapiro, Mark Lathrop and Michel Roger |
| EPI_ISL_536001 | Hôpital du Centre-de-la-Mauricie | Laboratoire de santé publique du Québec | Sandrine Moreira, Ioannis Ragoussis, Guillaume Bourque, Jesse Shapiro, Mark Lathrop and Michel Roger |
| EPI_ISL_536002 | Centre Hospitalier Régional de Lanaudière | Laboratoire de santé publique du Québec | Sandrine Moreira, Ioannis Ragoussis, Guillaume Bourque, Jesse Shapiro, Mark Lathrop and Michel Roger |
| EPI_ISL_536003 | Hôpital Sainte-Croix | Laboratoire de santé publique du Québec | Sandrine Moreira, Ioannis Ragoussis, Guillaume Bourque, Jesse Shapiro, Mark Lathrop and Michel Roger |
| EPI_ISL_536004 | Centre hospitalier Anna-Laberge | Laboratoire de santé publique du Québec | Sandrine Moreira, Ioannis Ragoussis, Guillaume Bourque, Jesse Shapiro, Mark Lathrop and Michel Roger |
| EPI_ISL_536005, EPI_ISL_536006 | Centre Hospitalier Régional de Lanaudière | Laboratoire de santé publique du Québec | Sandrine Moreira, Ioannis Ragoussis, Guillaume Bourque, Jesse Shapiro, Mark Lathrop and Michel Roger |
| EPI_ISL_536007 | Centre de SSS de Trois-Rivières | Laboratoire de santé publique du Québec | Sandrine Moreira, Ioannis Ragoussis, Guillaume Bourque, Jesse Shapiro, Mark Lathrop and Michel Roger |
| EPI_ISL_536008 | Hôpital de Saint-Eustache | Laboratoire de santé publique du Québec | Sandrine Moreira, Ioannis Ragoussis, Guillaume Bourque, Jesse Shapiro, Mark Lathrop and Michel Roger |
| EPI_ISL_536009 | Centre Hospitalier Régional de Lanaudière | Laboratoire de santé publique du Québec | Sandrine Moreira, Ioannis Ragoussis, Guillaume Bourque, Jesse Shapiro, Mark Lathrop and Michel Roger |
| EPI_ISL_536010, EPI_ISL_536011 | Hôpital Sainte-Croix | Laboratoire de santé publique du Québec | Sandrine Moreira, Ioannis Ragoussis, Guillaume Bourque, Jesse Shapiro, Mark Lathrop and Michel Roger |
| EPI_ISL_536012 | Centre Hospitalier Régional de Lanaudière | Laboratoire de santé publique du Québec | Sandrine Moreira, Ioannis Ragoussis, Guillaume Bourque, Jesse Shapiro, Mark Lathrop and Michel Roger |
| EPI_ISL_536013 | Hôpital Honoré-Mercier | Laboratoire de santé publique du Québec | Sandrine Moreira, Ioannis Ragoussis, Guillaume Bourque, Jesse Shapiro, Mark Lathrop and Michel Roger |
| EPI_ISL_536014, EPI_ISL_536015, EPI_ISL_536016 | Centre Hospitalier Régional de Lanaudière | Laboratoire de santé publique du Québec | Sandrine Moreira, Ioannis Ragoussis, Guillaume Bourque, Jesse Shapiro, Mark Lathrop and Michel Roger |
| EPI_ISL_536017, EPI_ISL_536018 | Hôpital Charles-LeMoyne | Laboratoire de santé publique du Québec | Sandrine Moreira, Ioannis Ragoussis, Guillaume Bourque, Jesse Shapiro, Mark Lathrop and Michel Roger |
| EPI_ISL_536019 | Hôpital de Chicoutimi | Laboratoire de santé publique du Québec | Sandrine Moreira, Ioannis Ragoussis, Guillaume Bourque, Jesse Shapiro, Mark Lathrop and Michel Roger |
| EPI_ISL_536020 | Hôpital Charles-LeMoyne | Laboratoire de santé publique du Québec | Sandrine Moreira, Ioannis Ragoussis, Guillaume Bourque, Jesse Shapiro, Mark Lathrop and Michel Roger |
| EPI_ISL_536021 | Hôpital Pierre-Boucher | Laboratoire de santé publique du Québec | Sandrine Moreira, Ioannis Ragoussis, Guillaume Bourque, Jesse Shapiro, Mark Lathrop and Michel Roger |
| EPI_ISL_536022 | Hôpital régional de Saint-Jérôme | Laboratoire de santé publique du Québec | Sandrine Moreira, Ioannis Ragoussis, Guillaume Bourque, Jesse Shapiro, Mark Lathrop and Michel Roger |
| EPI_ISL_536023 | Centre hospitalier Anna-Laberge | Laboratoire de santé publique du Québec | Sandrine Moreira, Ioannis Ragoussis, Guillaume Bourque, Jesse Shapiro, Mark Lathrop and Michel Roger |
| EPI_ISL_536024, EPI_ISL_536025 | Hôpital Charles-LeMoyne | Laboratoire de santé publique du Québec | Sandrine Moreira, Ioannis Ragoussis, Guillaume Bourque, Jesse Shapiro, Mark Lathrop and Michel Roger |
| EPI_ISL_536026 | Hôpital du Suroît | Laboratoire de santé publique du Québec | Sandrine Moreira, Ioannis Ragoussis, Guillaume Bourque, Jesse Shapiro, Mark Lathrop and Michel Roger |
| EPI_ISL_536027, EPI_ISL_536028 | CSSS Haut-Richelieu/Rouville (Hôpital) | Laboratoire de santé publique du Québec | Sandrine Moreira, Ioannis Ragoussis, Guillaume Bourque, Jesse Shapiro, Mark Lathrop and Michel Roger |
| EPI_ISL_536029 | Hôpital Honoré-Mercier | Laboratoire de santé publique du Québec | Sandrine Moreira, Ioannis Ragoussis, Guillaume Bourque, Jesse Shapiro, Mark Lathrop and Michel Roger |
| EPI_ISL_536030 | Hôpital du Centre-de-la-Mauricie | Laboratoire de santé publique du Québec | Sandrine Moreira, Ioannis Ragoussis, Guillaume Bourque, Jesse Shapiro, Mark Lathrop and Michel Roger |
| EPI_ISL_536031 | Centre de SSS de Trois-Rivières | Laboratoire de santé publique du Québec | Sandrine Moreira, Ioannis Ragoussis, Guillaume Bourque, Jesse Shapiro, Mark Lathrop and Michel Roger |
| EPI_ISL_536032, EPI_ISL_536033 | Hôpital du Centre-de-la-Mauricie | Laboratoire de santé publique du Québec | Sandrine Moreira, Ioannis Ragoussis, Guillaume Bourque, Jesse Shapiro, Mark Lathrop and Michel Roger |
| EPI_ISL_536034, EPI_ISL_536035, EPI_ISL_536036 | Centre de SSS de Trois-Rivières | Laboratoire de santé publique du Québec | Sandrine Moreira, Ioannis Ragoussis, Guillaume Bourque, Jesse Shapiro, Mark Lathrop and Michel Roger |
| EPI_ISL_536037 | Hôpital du Centre-de-la-Mauricie | Laboratoire de santé publique du Québec | Sandrine Moreira, Ioannis Ragoussis, Guillaume Bourque, Jesse Shapiro, Mark Lathrop and Michel Roger |
| EPI_ISL_536038 | Centre de SSS de Trois-Rivières | Laboratoire de santé publique du Québec | Sandrine Moreira, Ioannis Ragoussis, Guillaume Bourque, Jesse Shapiro, Mark Lathrop and Michel Roger |
| EPI_ISL_536039 | Hôpital Pierre-Boucher | Laboratoire de santé publique du Québec | Sandrine Moreira, Ioannis Ragoussis, Guillaume Bourque, Jesse Shapiro, Mark Lathrop and Michel Roger |
| EPI_ISL_536040 | CSSS Haut-Richelieu/Rouville (Hôpital) | Laboratoire de santé publique du Québec | Sandrine Moreira, Ioannis Ragoussis, Guillaume Bourque, Jesse Shapiro, Mark Lathrop and Michel Roger |
| EPI_ISL_536041 | Hôpital Pierre-Boucher | Laboratoire de santé publique du Québec | Sandrine Moreira, Ioannis Ragoussis, Guillaume Bourque, Jesse Shapiro, Mark Lathrop and Michel Roger |
| EPI_ISL_536042 | CSSS Haut-Richelieu/Rouville (Hôpital) | Laboratoire de santé publique du Québec | Sandrine Moreira, Ioannis Ragoussis, Guillaume Bourque, Jesse Shapiro, Mark Lathrop and Michel Roger |
| EPI_ISL_536043 | Centre hospitalier Anna-Laberge | Laboratoire de santé publique du Québec | Sandrine Moreira, Ioannis Ragoussis, Guillaume Bourque, Jesse Shapiro, Mark Lathrop and Michel Roger |
| EPI_ISL_536044 | CSSS Haut-Richelieu/Rouville (Hôpital) | Laboratoire de santé publique du Québec | Sandrine Moreira, Ioannis Ragoussis, Guillaume Bourque, Jesse Shapiro, Mark Lathrop and Michel Roger |
| EPI_ISL_536045 | Centre hospitalier Anna-Laberge | Laboratoire de santé publique du Québec | Sandrine Moreira, Ioannis Ragoussis, Guillaume Bourque, Jesse Shapiro, Mark Lathrop and Michel Roger |
| EPI_ISL_536046 | Hôpital d'Alma | Laboratoire de santé publique du Québec | Sandrine Moreira, Ioannis Ragoussis, Guillaume Bourque, Jesse Shapiro, Mark Lathrop and Michel Roger |
| EPI_ISL_536047, EPI_ISL_536048 | Centre Hospitalier Régional de Lanaudière | Laboratoire de santé publique du Québec | Sandrine Moreira, Ioannis Ragoussis, Guillaume Bourque, Jesse Shapiro, Mark Lathrop and Michel Roger |
| EPI_ISL_536049 | Hôpital de Saint-Eustache | Laboratoire de santé publique du Québec | Sandrine Moreira, Ioannis Ragoussis, Guillaume Bourque, Jesse Shapiro, Mark Lathrop and Michel Roger |
| EPI_ISL_536050 | Centre Hospitalier Régional de Lanaudière | Laboratoire de santé publique du Québec | Sandrine Moreira, Ioannis Ragoussis, Guillaume Bourque, Jesse Shapiro, Mark Lathrop and Michel Roger |
| EPI_ISL_536051 | Hôpital de Hull | Laboratoire de santé publique du Québec | Sandrine Moreira, Ioannis Ragoussis, Guillaume Bourque, Jesse Shapiro, Mark Lathrop and Michel Roger |
| EPI_ISL_536052, EPI_ISL_536053, EPI_ISL_536054 | Centre Hospitalier Régional de Lanaudière | Laboratoire de santé publique du Québec | Sandrine Moreira, Ioannis Ragoussis, Guillaume Bourque, Jesse Shapiro, Mark Lathrop and Michel Roger |
| EPI_ISL_536055 | Hôpital Honoré-Mercier | Laboratoire de santé publique du Québec | Sandrine Moreira, Ioannis Ragoussis, Guillaume Bourque, Jesse Shapiro, Mark Lathrop and Michel Roger |
| EPI_ISL_536056, EPI_ISL_536057 | Hôpital Sainte-Croix | Laboratoire de santé publique du Québec | Sandrine Moreira, Ioannis Ragoussis, Guillaume Bourque, Jesse Shapiro, Mark Lathrop and Michel Roger |
| EPI_ISL_536058 | Hôpital de Saint-Eustache | Laboratoire de santé publique du Québec | Sandrine Moreira, Ioannis Ragoussis, Guillaume Bourque, Jesse Shapiro, Mark Lathrop and Michel Roger |
| EPI_ISL_536059, EPI_ISL_536060 | Centre Hospitalier Régional de Lanaudière | Laboratoire de santé publique du Québec | Sandrine Moreira, Ioannis Ragoussis, Guillaume Bourque, Jesse Shapiro, Mark Lathrop and Michel Roger |
| EPI_ISL_536061, EPI_ISL_536062 | Hôpital Pierre-Le Gardeur | Laboratoire de santé publique du Québec | Sandrine Moreira, Ioannis Ragoussis, Guillaume Bourque, Jesse Shapiro, Mark Lathrop and Michel Roger |
| EPI_ISL_536063 | Centre Hospitalier Régional de Lanaudière | Laboratoire de santé publique du Québec | Sandrine Moreira, Ioannis Ragoussis, Guillaume Bourque, Jesse Shapiro, Mark Lathrop and Michel Roger |
| EPI_ISL_536064, EPI_ISL_536065, EPI_ISL_536066, EPI_ISL_536067, EPI_ISL_536068 | Hôpital Pierre-Le Gardeur | Laboratoire de santé publique du Québec | Sandrine Moreira, Ioannis Ragoussis, Guillaume Bourque, Jesse Shapiro, Mark Lathrop and Michel Roger |
| EPI_ISL_536069 | Hôpital de Saint-Eustache | Laboratoire de santé publique du Québec | Sandrine Moreira, Ioannis Ragoussis, Guillaume Bourque, Jesse Shapiro, Mark Lathrop and Michel Roger |
| EPI_ISL_536070 | Hôpital Pierre-Le Gardeur | Laboratoire de santé publique du Québec | Sandrine Moreira, Ioannis Ragoussis, Guillaume Bourque, Jesse Shapiro, Mark Lathrop and Michel Roger |
| EPI_ISL_536071, EPI_ISL_536072 | Hôpital de Saint-Eustache | Laboratoire de santé publique du Québec | Sandrine Moreira, Ioannis Ragoussis, Guillaume Bourque, Jesse Shapiro, Mark Lathrop and Michel Roger |
| EPI_ISL_536073, EPI_ISL_536074 | Centre Hospitalier Régional de Lanaudière | Laboratoire de santé publique du Québec | Sandrine Moreira, Ioannis Ragoussis, Guillaume Bourque, Jesse Shapiro, Mark Lathrop and Michel Roger |
| EPI_ISL_536075 | Hôpital Charles-LeMoyne | Laboratoire de santé publique du Québec | Sandrine Moreira, Ioannis Ragoussis, Guillaume Bourque, Jesse Shapiro, Mark Lathrop and Michel Roger |
| EPI_ISL_536076 | Centre Hospitalier Régional de Lanaudière | Laboratoire de santé publique du Québec | Sandrine Moreira, Ioannis Ragoussis, Guillaume Bourque, Jesse Shapiro, Mark Lathrop and Michel Roger |
| EPI_ISL_536077, EPI_ISL_536078 | Hôpital Charles-LeMoyne | Laboratoire de santé publique du Québec | Sandrine Moreira, Ioannis Ragoussis, Guillaume Bourque, Jesse Shapiro, Mark Lathrop and Michel Roger |
| EPI_ISL_536079 | Centre Hospitalier Régional de Lanaudière | Laboratoire de santé publique du Québec | Sandrine Moreira, Ioannis Ragoussis, Guillaume Bourque, Jesse Shapiro, Mark Lathrop and Michel Roger |
| EPI_ISL_536080 | Centre hospitalier Anna-Laberge | Laboratoire de santé publique du Québec | Sandrine Moreira, Ioannis Ragoussis, Guillaume Bourque, Jesse Shapiro, Mark Lathrop and Michel Roger |
| EPI_ISL_536081, EPI_ISL_536082 | Hôpital Charles-LeMoyne | Laboratoire de santé publique du Québec | Sandrine Moreira, Ioannis Ragoussis, Guillaume Bourque, Jesse Shapiro, Mark Lathrop and Michel Roger |
| EPI_ISL_536083, EPI_ISL_536084, EPI_ISL_536085, EPI_ISL_536086 | Hôpital Pierre-Le Gardeur | Laboratoire de santé publique du Québec | Sandrine Moreira, Ioannis Ragoussis, Guillaume Bourque, Jesse Shapiro, Mark Lathrop and Michel Roger |
| EPI_ISL_536087, EPI_ISL_536088, EPI_ISL_536089, EPI_ISL_536090, EPI_ISL_536091 | Hôpital Charles-LeMoyne | Laboratoire de santé publique du Québec | Sandrine Moreira, Ioannis Ragoussis, Guillaume Bourque, Jesse Shapiro, Mark Lathrop and Michel Roger |
| EPI_ISL_536092 | Hôpital Pierre-Boucher | Laboratoire de santé publique du Québec | Sandrine Moreira, Ioannis Ragoussis, Guillaume Bourque, Jesse Shapiro, Mark Lathrop and Michel Roger |
| EPI_ISL_536093, EPI_ISL_536094 | Hôpital Charles-LeMoyne | Laboratoire de santé publique du Québec | Sandrine Moreira, Ioannis Ragoussis, Guillaume Bourque, Jesse Shapiro, Mark Lathrop and Michel Roger |

| | | | |
|---|---|---|---|
| EPI_ISL_536095 | Hôpital du Suroît | Laboratoire de santé publique du Québec | Sandrine Moreira, Ioannis Ragoussis, Guillaume Bourque, Jesse Shapiro, Mark Lathrop and Michel Roger |
| EPI_ISL_536096 | Hôpital Pierre-Boucher | Laboratoire de santé publique du Québec | Sandrine Moreira, Ioannis Ragoussis, Guillaume Bourque, Jesse Shapiro, Mark Lathrop and Michel Roger |
| EPI_ISL_536097, EPI_ISL_536098 | Hôpital du Suroît | Laboratoire de santé publique du Québec | Sandrine Moreira, Ioannis Ragoussis, Guillaume Bourque, Jesse Shapiro, Mark Lathrop and Michel Roger |
| EPI_ISL_536099 | Hôpital Pierre-Boucher | Laboratoire de santé publique du Québec | Sandrine Moreira, Ioannis Ragoussis, Guillaume Bourque, Jesse Shapiro, Mark Lathrop and Michel Roger |
| EPI_ISL_536100 | Hôpital Charles-LeMoyne | Laboratoire de santé publique du Québec | Sandrine Moreira, Ioannis Ragoussis, Guillaume Bourque, Jesse Shapiro, Mark Lathrop and Michel Roger |
| EPI_ISL_536101 | Hôpital, CLSC et Centre d'Hébergement de Roberval | Laboratoire de santé publique du Québec | Sandrine Moreira, Ioannis Ragoussis, Guillaume Bourque, Jesse Shapiro, Mark Lathrop and Michel Roger |
| EPI_ISL_536102 | Hôpital du Suroît | Laboratoire de santé publique du Québec | Sandrine Moreira, Ioannis Ragoussis, Guillaume Bourque, Jesse Shapiro, Mark Lathrop and Michel Roger |
| EPI_ISL_536103 | Hôpital Pierre-Boucher | Laboratoire de santé publique du Québec | Sandrine Moreira, Ioannis Ragoussis, Guillaume Bourque, Jesse Shapiro, Mark Lathrop and Michel Roger |
| EPI_ISL_536104 | Centre hospitalier Anna-Laberge | Laboratoire de santé publique du Québec | Sandrine Moreira, Ioannis Ragoussis, Guillaume Bourque, Jesse Shapiro, Mark Lathrop and Michel Roger |
| EPI_ISL_536105 | Hôpital du Suroît | Laboratoire de santé publique du Québec | Sandrine Moreira, Ioannis Ragoussis, Guillaume Bourque, Jesse Shapiro, Mark Lathrop and Michel Roger |
| EPI_ISL_536106 | Hôpital régional de Rimouski | Laboratoire de santé publique du Québec | Sandrine Moreira, Ioannis Ragoussis, Guillaume Bourque, Jesse Shapiro, Mark Lathrop and Michel Roger |
| EPI_ISL_536107 | Hôpital de Chicoutimi | Laboratoire de santé publique du Québec | Sandrine Moreira, Ioannis Ragoussis, Guillaume Bourque, Jesse Shapiro, Mark Lathrop and Michel Roger |
| EPI_ISL_536108 | Hôpital Le Royer | Laboratoire de santé publique du Québec | Sandrine Moreira, Ioannis Ragoussis, Guillaume Bourque, Jesse Shapiro, Mark Lathrop and Michel Roger |
| EPI_ISL_536109 | CSSS de Port-Cartier | Laboratoire de santé publique du Québec | Sandrine Moreira, Ioannis Ragoussis, Guillaume Bourque, Jesse Shapiro, Mark Lathrop and Michel Roger |
| EPI_ISL_536110 | CSSS Haut-Richelieu/Rouville (Hôpital) | Laboratoire de santé publique du Québec | Sandrine Moreira, Ioannis Ragoussis, Guillaume Bourque, Jesse Shapiro, Mark Lathrop and Michel Roger |
| EPI_ISL_536111, EPI_ISL_536112 | Hôpital de Maria | Laboratoire de santé publique du Québec | Sandrine Moreira, Ioannis Ragoussis, Guillaume Bourque, Jesse Shapiro, Mark Lathrop and Michel Roger |
| EPI_ISL_536113, EPI_ISL_536114 | Hôpital et Centre d'Hébergement de Sept-Iles | Laboratoire de santé publique du Québec | Sandrine Moreira, Ioannis Ragoussis, Guillaume Bourque, Jesse Shapiro, Mark Lathrop and Michel Roger |
| EPI_ISL_536115 | Centre hospitalier régional du Grand Portage | Laboratoire de santé publique du Québec | Sandrine Moreira, Ioannis Ragoussis, Guillaume Bourque, Jesse Shapiro, Mark Lathrop and Michel Roger |
| EPI_ISL_536116 | Hôtel-Dieu de Sorel | Laboratoire de santé publique du Québec | Sandrine Moreira, Ioannis Ragoussis, Guillaume Bourque, Jesse Shapiro, Mark Lathrop and Michel Roger |
| EPI_ISL_536117, EPI_ISL_536118 | Hôpital de Hull | Laboratoire de santé publique du Québec | Sandrine Moreira, Ioannis Ragoussis, Guillaume Bourque, Jesse Shapiro, Mark Lathrop and Michel Roger |
| EPI_ISL_536119, EPI_ISL_536120 | Centre Hospitalier Régional de Lanaudière | Laboratoire de santé publique du Québec | Sandrine Moreira, Ioannis Ragoussis, Guillaume Bourque, Jesse Shapiro, Mark Lathrop and Michel Roger |
| EPI_ISL_536121 | Hôtel-Dieu de Sorel | Laboratoire de santé publique du Québec | Sandrine Moreira, Ioannis Ragoussis, Guillaume Bourque, Jesse Shapiro, Mark Lathrop and Michel Roger |
| EPI_ISL_536122 | Hôpital de Hull | Laboratoire de santé publique du Québec | Sandrine Moreira, Ioannis Ragoussis, Guillaume Bourque, Jesse Shapiro, Mark Lathrop and Michel Roger |
| EPI_ISL_536123 | Centre Hospitalier Régional de Lanaudière | Laboratoire de santé publique du Québec | Sandrine Moreira, Ioannis Ragoussis, Guillaume Bourque, Jesse Shapiro, Mark Lathrop and Michel Roger |
| EPI_ISL_536124 | Hôpital de Hull | Laboratoire de santé publique du Québec | Sandrine Moreira, Ioannis Ragoussis, Guillaume Bourque, Jesse Shapiro, Mark Lathrop and Michel Roger |
| EPI_ISL_536125 | Hôpital Pierre-Le Gardeur | Laboratoire de santé publique du Québec | Sandrine Moreira, Ioannis Ragoussis, Guillaume Bourque, Jesse Shapiro, Mark Lathrop and Michel Roger |
| EPI_ISL_536126 | Centre Hospitalier Régional de Lanaudière | Laboratoire de santé publique du Québec | Sandrine Moreira, Ioannis Ragoussis, Guillaume Bourque, Jesse Shapiro, Mark Lathrop and Michel Roger |
| EPI_ISL_536127 | Hôpital de Hull | Laboratoire de santé publique du Québec | Sandrine Moreira, Ioannis Ragoussis, Guillaume Bourque, Jesse Shapiro, Mark Lathrop and Michel Roger |
| EPI_ISL_536128 | Hôpital Pierre-Le Gardeur | Laboratoire de santé publique du Québec | Sandrine Moreira, Ioannis Ragoussis, Guillaume Bourque, Jesse Shapiro, Mark Lathrop and Michel Roger |
| EPI_ISL_536129, EPI_ISL_536130 | Centre Hospitalier Régional de Lanaudière | Laboratoire de santé publique du Québec | Sandrine Moreira, Ioannis Ragoussis, Guillaume Bourque, Jesse Shapiro, Mark Lathrop and Michel Roger |
| EPI_ISL_536131 | Hôpital Pierre-Le Gardeur | Laboratoire de santé publique du Québec | Sandrine Moreira, Ioannis Ragoussis, Guillaume Bourque, Jesse Shapiro, Mark Lathrop and Michel Roger |
| EPI_ISL_536132 | Hôpital de Hull | Laboratoire de santé publique du Québec | Sandrine Moreira, Ioannis Ragoussis, Guillaume Bourque, Jesse Shapiro, Mark Lathrop and Michel Roger |
| EPI_ISL_536133, EPI_ISL_536134 | Centre Hospitalier Régional de Lanaudière | Laboratoire de santé publique du Québec | Sandrine Moreira, Ioannis Ragoussis, Guillaume Bourque, Jesse Shapiro, Mark Lathrop and Michel Roger |
| EPI_ISL_536135, EPI_ISL_536136 | Hôpital de Hull | Laboratoire de santé publique du Québec | Sandrine Moreira, Ioannis Ragoussis, Guillaume Bourque, Jesse Shapiro, Mark Lathrop and Michel Roger |
| EPI_ISL_536137 | Hôpital Pierre-Le Gardeur | Laboratoire de santé publique du Québec | Sandrine Moreira, Ioannis Ragoussis, Guillaume Bourque, Jesse Shapiro, Mark Lathrop and Michel Roger |
| EPI_ISL_536138 | Centre Hospitalier Régional de Lanaudière | Laboratoire de santé publique du Québec | Sandrine Moreira, Ioannis Ragoussis, Guillaume Bourque, Jesse Shapiro, Mark Lathrop and Michel Roger |
| EPI_ISL_536139, EPI_ISL_536140 | Hôpital Pierre-Le Gardeur | Laboratoire de santé publique du Québec | Sandrine Moreira, Ioannis Ragoussis, Guillaume Bourque, Jesse Shapiro, Mark Lathrop and Michel Roger |
| EPI_ISL_536141, EPI_ISL_536142 | Centre hospitalier Anna-Laberge | Laboratoire de santé publique du Québec | Sandrine Moreira, Ioannis Ragoussis, Guillaume Bourque, Jesse Shapiro, Mark Lathrop and Michel Roger |
| EPI_ISL_536143, EPI_ISL_536144 | Hôpital Pierre-Le Gardeur | Laboratoire de santé publique du Québec | Sandrine Moreira, Ioannis Ragoussis, Guillaume Bourque, Jesse Shapiro, Mark Lathrop and Michel Roger |
| EPI_ISL_536145 | Centre hospitalier Anna-Laberge | Laboratoire de santé publique du Québec | Sandrine Moreira, Ioannis Ragoussis, Guillaume Bourque, Jesse Shapiro, Mark Lathrop and Michel Roger |
| EPI_ISL_536146, EPI_ISL_536147, EPI_ISL_536148, EPI_ISL_536149 | Hôpital Charles-LeMoyne | Laboratoire de santé publique du Québec | Sandrine Moreira, Ioannis Ragoussis, Guillaume Bourque, Jesse Shapiro, Mark Lathrop and Michel Roger |
| EPI_ISL_536150 | CSSS Haut-Richelieu/Rouville (Hôpital) | Laboratoire de santé publique du Québec | Sandrine Moreira, Ioannis Ragoussis, Guillaume Bourque, Jesse Shapiro, Mark Lathrop and Michel Roger |
| EPI_ISL_536151 | Centre Hospitalier Régional de Lanaudière | Laboratoire de santé publique du Québec | Sandrine Moreira, Ioannis Ragoussis, Guillaume Bourque, Jesse Shapiro, Mark Lathrop and Michel Roger |
| EPI_ISL_536152 | Hôpital Pierre-Le Gardeur | Laboratoire de santé publique du Québec | Sandrine Moreira, Ioannis Ragoussis, Guillaume Bourque, Jesse Shapiro, Mark Lathrop and Michel Roger |
| EPI_ISL_536153, EPI_ISL_536154 | Centre Hospitalier Régional de Lanaudière | Laboratoire de santé publique du Québec | Sandrine Moreira, Ioannis Ragoussis, Guillaume Bourque, Jesse Shapiro, Mark Lathrop and Michel Roger |
| EPI_ISL_536155, EPI_ISL_536156 | Centre hospitalier Anna-Laberge | Laboratoire de santé publique du Québec | Sandrine Moreira, Ioannis Ragoussis, Guillaume Bourque, Jesse Shapiro, Mark Lathrop and Michel Roger |
| EPI_ISL_536157 | Hôpital Pierre-Le Gardeur | Laboratoire de santé publique du Québec | Sandrine Moreira, Ioannis Ragoussis, Guillaume Bourque, Jesse Shapiro, Mark Lathrop and Michel Roger |
| EPI_ISL_536158 | CSSS Haut-Richelieu/Rouville (Hôpital) | Laboratoire de santé publique du Québec | Sandrine Moreira, Ioannis Ragoussis, Guillaume Bourque, Jesse Shapiro, Mark Lathrop and Michel Roger |
| EPI_ISL_536159 | Centre hospitalier Anna-Laberge | Laboratoire de santé publique du Québec | Sandrine Moreira, Ioannis Ragoussis, Guillaume Bourque, Jesse Shapiro, Mark Lathrop and Michel Roger |
| EPI_ISL_536160, EPI_ISL_536161 | CSSS Haut-Richelieu/Rouville (Hôpital) | Laboratoire de santé publique du Québec | Sandrine Moreira, Ioannis Ragoussis, Guillaume Bourque, Jesse Shapiro, Mark Lathrop and Michel Roger |
| EPI_ISL_536162 | Centre hospitalier Anna-Laberge | Laboratoire de santé publique du Québec | Sandrine Moreira, Ioannis Ragoussis, Guillaume Bourque, Jesse Shapiro, Mark Lathrop and Michel Roger |
| EPI_ISL_536163 | Hôpital du Suroît | Laboratoire de santé publique du Québec | Sandrine Moreira, Ioannis Ragoussis, Guillaume Bourque, Jesse Shapiro, Mark Lathrop and Michel Roger |
| EPI_ISL_536164 | CSSS Haut-Richelieu/Rouville (Hôpital) | Laboratoire de santé publique du Québec | Sandrine Moreira, Ioannis Ragoussis, Guillaume Bourque, Jesse Shapiro, Mark Lathrop and Michel Roger |
| EPI_ISL_536165 | Hôpital du Suroît | Laboratoire de santé publique du Québec | Sandrine Moreira, Ioannis Ragoussis, Guillaume Bourque, Jesse Shapiro, Mark Lathrop and Michel Roger |
| EPI_ISL_536166 | CSSS Haut-Richelieu/Rouville (Hôpital) | Laboratoire de santé publique du Québec | Sandrine Moreira, Ioannis Ragoussis, Guillaume Bourque, Jesse Shapiro, Mark Lathrop and Michel Roger |
| EPI_ISL_536167 | Hôpital du Suroît | Laboratoire de santé publique du Québec | Sandrine Moreira, Ioannis Ragoussis, Guillaume Bourque, Jesse Shapiro, Mark Lathrop and Michel Roger |
| EPI_ISL_536168, EPI_ISL_536169 | Hôpital de Hull | Laboratoire de santé publique du Québec | Sandrine Moreira, Ioannis Ragoussis, Guillaume Bourque, Jesse Shapiro, Mark Lathrop and Michel Roger |
| EPI_ISL_536170 | Hôpital du Suroît | Laboratoire de santé publique du Québec | Sandrine Moreira, Ioannis Ragoussis, Guillaume Bourque, Jesse Shapiro, Mark Lathrop and Michel Roger |
| EPI_ISL_536171 | Hôpital de Papineau | Laboratoire de santé publique du Québec | Sandrine Moreira, Ioannis Ragoussis, Guillaume Bourque, Jesse Shapiro, Mark Lathrop and Michel Roger |
| EPI_ISL_536172 | CSSS Haut-Richelieu/Rouville (Hôpital) | Laboratoire de santé publique du Québec | Sandrine Moreira, Ioannis Ragoussis, Guillaume Bourque, Jesse Shapiro, Mark Lathrop and Michel Roger |
| EPI_ISL_536173, EPI_ISL_536174, EPI_ISL_536175, EPI_ISL_536176 | Hôpital Pierre-Boucher | Laboratoire de santé publique du Québec | Sandrine Moreira, Ioannis Ragoussis, Guillaume Bourque, Jesse Shapiro, Mark Lathrop and Michel Roger |
| EPI_ISL_536177 | Hôpital Honoré-Mercier | Laboratoire de santé publique du Québec | Sandrine Moreira, Ioannis Ragoussis, Guillaume Bourque, Jesse Shapiro, Mark Lathrop and Michel Roger |
| EPI_ISL_536178, EPI_ISL_536179 | Hôpital Pierre-Boucher | Laboratoire de santé publique du Québec | Sandrine Moreira, Ioannis Ragoussis, Guillaume Bourque, Jesse Shapiro, Mark Lathrop and Michel Roger |
| EPI_ISL_536180 | Hôpital Honoré-Mercier | Laboratoire de santé publique du Québec | Sandrine Moreira, Ioannis Ragoussis, Guillaume Bourque, Jesse Shapiro, Mark Lathrop and Michel Roger |
| EPI_ISL_536181 | CSSS Haut-Richelieu/Rouville (Hôpital) | Laboratoire de santé publique du Québec | Sandrine Moreira, Ioannis Ragoussis, Guillaume Bourque, Jesse Shapiro, Mark Lathrop and Michel Roger |
| EPI_ISL_536182, EPI_ISL_536183, EPI_ISL_536184 | Centre hospitalier Anna-Laberge | Laboratoire de santé publique du Québec | Sandrine Moreira, Ioannis Ragoussis, Guillaume Bourque, Jesse Shapiro, Mark Lathrop and Michel Roger |
| EPI_ISL_536185, EPI_ISL_536186, EPI_ISL_536187 | Hôpital Charles-LeMoyne | Laboratoire de santé publique du Québec | Sandrine Moreira, Ioannis Ragoussis, Guillaume Bourque, Jesse Shapiro, Mark Lathrop and Michel Roger |
| EPI_ISL_536188, EPI_ISL_536189 | Hôpital Pierre-Boucher | Laboratoire de santé publique du Québec | Sandrine Moreira, Ioannis Ragoussis, Guillaume Bourque, Jesse Shapiro, Mark Lathrop and Michel Roger |
| EPI_ISL_536190 | Hôpital du Suroît | Laboratoire de santé publique du Québec | Sandrine Moreira, Ioannis Ragoussis, Guillaume Bourque, Jesse Shapiro, Mark Lathrop and Michel Roger |
| EPI_ISL_536191 | Hôpital Charles-LeMoyne | Laboratoire de santé publique du Québec | Sandrine Moreira, Ioannis Ragoussis, Guillaume Bourque, Jesse Shapiro, Mark Lathrop and Michel Roger |
| EPI_ISL_536192, EPI_ISL_536193 | Hôpital du Suroît | Laboratoire de santé publique du Québec | Sandrine Moreira, Ioannis Ragoussis, Guillaume Bourque, Jesse Shapiro, Mark Lathrop and Michel Roger |
| EPI_ISL_536194 | Hôpital Charles-LeMoyne | Laboratoire de santé publique du Québec | Sandrine Moreira, Ioannis Ragoussis, Guillaume Bourque, Jesse Shapiro, Mark Lathrop and Michel Roger |
| EPI_ISL_536195, EPI_ISL_536196 | Hôpital Pierre-Boucher | Laboratoire de santé publique du Québec | Sandrine Moreira, Ioannis Ragoussis, Guillaume Bourque, Jesse Shapiro, Mark Lathrop and Michel Roger |
| EPI_ISL_536197 | CSSS Haut-Richelieu/Rouville (Hôpital) | Laboratoire de santé publique du Québec | Sandrine Moreira, Ioannis Ragoussis, Guillaume Bourque, Jesse Shapiro, Mark Lathrop and Michel Roger |
| EPI_ISL_536198 | Hôpital du Suroît | Laboratoire de santé publique du Québec | Sandrine Moreira, Ioannis Ragoussis, Guillaume Bourque, Jesse Shapiro, Mark Lathrop and Michel Roger |

| | | | |
|---|---|---|---|
| EPI_ISL_536199 | Centre hospitalier régional du Grand Portage | Laboratoire de santé publique du Québec | Sandrine Moreira, Ioannis Ragoussis, Guillaume Bourque, Jesse Shapiro, Mark Lathrop and Michel Roger |
| EPI_ISL_536200, EPI_ISL_536201, EPI_ISL_536202 | Hôpital du Suroît | Laboratoire de santé publique du Québec | Sandrine Moreira, Ioannis Ragoussis, Guillaume Bourque, Jesse Shapiro, Mark Lathrop and Michel Roger |
| EPI_ISL_536203 | Hôpital Charles-LeMoyne | Laboratoire de santé publique du Québec | Sandrine Moreira, Ioannis Ragoussis, Guillaume Bourque, Jesse Shapiro, Mark Lathrop and Michel Roger |
| EPI_ISL_536204 | Centre hospitalier régional du Grand Portage | Laboratoire de santé publique du Québec | Sandrine Moreira, Ioannis Ragoussis, Guillaume Bourque, Jesse Shapiro, Mark Lathrop and Michel Roger |
| EPI_ISL_536205 | Hôpital régional de Rimouski | Laboratoire de santé publique du Québec | Sandrine Moreira, Ioannis Ragoussis, Guillaume Bourque, Jesse Shapiro, Mark Lathrop and Michel Roger |
| EPI_ISL_536206 | CSSS de la Minganie | Laboratoire de santé publique du Québec | Sandrine Moreira, Ioannis Ragoussis, Guillaume Bourque, Jesse Shapiro, Mark Lathrop and Michel Roger |
| EPI_ISL_536207, EPI_ISL_536208, EPI_ISL_536209, EPI_ISL_536210 | Hôpital de Hull | Laboratoire de santé publique du Québec | Sandrine Moreira, Ioannis Ragoussis, Guillaume Bourque, Jesse Shapiro, Mark Lathrop and Michel Roger |
| EPI_ISL_536211, EPI_ISL_536212 | Centre hospitalier Anna-Laberge | Laboratoire de santé publique du Québec | Sandrine Moreira, Ioannis Ragoussis, Guillaume Bourque, Jesse Shapiro, Mark Lathrop and Michel Roger |
| EPI_ISL_536213 | CSSS Haut-Richelieu/Rouville (Hôpital) | Laboratoire de santé publique du Québec | Sandrine Moreira, Ioannis Ragoussis, Guillaume Bourque, Jesse Shapiro, Mark Lathrop and Michel Roger |
| EPI_ISL_536214 | Hôpital Charles-LeMoyne | Laboratoire de santé publique du Québec | Sandrine Moreira, Ioannis Ragoussis, Guillaume Bourque, Jesse Shapiro, Mark Lathrop and Michel Roger |
| EPI_ISL_536215, EPI_ISL_536216, EPI_ISL_536217 | CSSS Haut-Richelieu/Rouville (Hôpital) | Laboratoire de santé publique du Québec | Sandrine Moreira, Ioannis Ragoussis, Guillaume Bourque, Jesse Shapiro, Mark Lathrop and Michel Roger |
| EPI_ISL_536218, EPI_ISL_536219, EPI_ISL_536220, EPI_ISL_536221 | Centre hospitalier Anna-Laberge | Laboratoire de santé publique du Québec | Sandrine Moreira, Ioannis Ragoussis, Guillaume Bourque, Jesse Shapiro, Mark Lathrop and Michel Roger |
| EPI_ISL_536222 | Conseil Cri de la SSS de la Baie-James | Laboratoire de santé publique du Québec | Sandrine Moreira, Ioannis Ragoussis, Guillaume Bourque, Jesse Shapiro, Mark Lathrop and Michel Roger |
| EPI_ISL_536223 | Hôpital Charles-LeMoyne | Laboratoire de santé publique du Québec | Sandrine Moreira, Ioannis Ragoussis, Guillaume Bourque, Jesse Shapiro, Mark Lathrop and Michel Roger |
| EPI_ISL_536224 | Centre hospitalier Anna-Laberge | Laboratoire de santé publique du Québec | Sandrine Moreira, Ioannis Ragoussis, Guillaume Bourque, Jesse Shapiro, Mark Lathrop and Michel Roger |
| EPI_ISL_536225 | Hôpital Charles-LeMoyne | Laboratoire de santé publique du Québec | Sandrine Moreira, Ioannis Ragoussis, Guillaume Bourque, Jesse Shapiro, Mark Lathrop and Michel Roger |
| EPI_ISL_536226, EPI_ISL_536227, EPI_ISL_536228 | Hôpital du Suroît | Laboratoire de santé publique du Québec | Sandrine Moreira, Ioannis Ragoussis, Guillaume Bourque, Jesse Shapiro, Mark Lathrop and Michel Roger |
| EPI_ISL_536229 | Hôpital Pierre-Boucher | Laboratoire de santé publique du Québec | Sandrine Moreira, Ioannis Ragoussis, Guillaume Bourque, Jesse Shapiro, Mark Lathrop and Michel Roger |
| EPI_ISL_536230 | Hôpital de Hull | Laboratoire de santé publique du Québec | Sandrine Moreira, Ioannis Ragoussis, Guillaume Bourque, Jesse Shapiro, Mark Lathrop and Michel Roger |
| EPI_ISL_536231 | Hôpital Pierre-Boucher | Laboratoire de santé publique du Québec | Sandrine Moreira, Ioannis Ragoussis, Guillaume Bourque, Jesse Shapiro, Mark Lathrop and Michel Roger |
| EPI_ISL_536232 | Centre hospitalier Anna-Laberge | Laboratoire de santé publique du Québec | Sandrine Moreira, Ioannis Ragoussis, Guillaume Bourque, Jesse Shapiro, Mark Lathrop and Michel Roger |
| EPI_ISL_536233 | Hôpital Charles-LeMoyne | Laboratoire de santé publique du Québec | Sandrine Moreira, Ioannis Ragoussis, Guillaume Bourque, Jesse Shapiro, Mark Lathrop and Michel Roger |
| EPI_ISL_536234 | Hôpital Pierre-Boucher | Laboratoire de santé publique du Québec | Sandrine Moreira, Ioannis Ragoussis, Guillaume Bourque, Jesse Shapiro, Mark Lathrop and Michel Roger |
| EPI_ISL_536235 | CSSS Haut-Richelieu/Rouville (Hôpital) | Laboratoire de santé publique du Québec | Sandrine Moreira, Ioannis Ragoussis, Guillaume Bourque, Jesse Shapiro, Mark Lathrop and Michel Roger |
| EPI_ISL_536236 | Hôpital Pierre-Boucher | Laboratoire de santé publique du Québec | Sandrine Moreira, Ioannis Ragoussis, Guillaume Bourque, Jesse Shapiro, Mark Lathrop and Michel Roger |
| EPI_ISL_536237 | CSSS Haut-Richelieu/Rouville (Hôpital) | Laboratoire de santé publique du Québec | Sandrine Moreira, Ioannis Ragoussis, Guillaume Bourque, Jesse Shapiro, Mark Lathrop and Michel Roger |
| EPI_ISL_536238, EPI_ISL_536239 | Hôpital de Hull | Laboratoire de santé publique du Québec | Sandrine Moreira, Ioannis Ragoussis, Guillaume Bourque, Jesse Shapiro, Mark Lathrop and Michel Roger |
| EPI_ISL_536240, EPI_ISL_536241 | Centre hospitalier Anna-Laberge | Laboratoire de santé publique du Québec | Sandrine Moreira, Ioannis Ragoussis, Guillaume Bourque, Jesse Shapiro, Mark Lathrop and Michel Roger |
| EPI_ISL_536242 | Hôpital de Hull | Laboratoire de santé publique du Québec | Sandrine Moreira, Ioannis Ragoussis, Guillaume Bourque, Jesse Shapiro, Mark Lathrop and Michel Roger |
| EPI_ISL_536243 | CSSS Haut-Richelieu/Rouville (Hôpital) | Laboratoire de santé publique du Québec | Sandrine Moreira, Ioannis Ragoussis, Guillaume Bourque, Jesse Shapiro, Mark Lathrop and Michel Roger |
| EPI_ISL_536244 | Hôpital de Hull | Laboratoire de santé publique du Québec | Sandrine Moreira, Ioannis Ragoussis, Guillaume Bourque, Jesse Shapiro, Mark Lathrop and Michel Roger |
| EPI_ISL_536245 | Hôpital Pierre-Boucher | Laboratoire de santé publique du Québec | Sandrine Moreira, Ioannis Ragoussis, Guillaume Bourque, Jesse Shapiro, Mark Lathrop and Michel Roger |
| EPI_ISL_536246 | Hôpital du Centre-de-la-Mauricie | Laboratoire de santé publique du Québec | Sandrine Moreira, Ioannis Ragoussis, Guillaume Bourque, Jesse Shapiro, Mark Lathrop and Michel Roger |
| EPI_ISL_536247 | Hôpital Charles-LeMoyne | Laboratoire de santé publique du Québec | Sandrine Moreira, Ioannis Ragoussis, Guillaume Bourque, Jesse Shapiro, Mark Lathrop and Michel Roger |
| EPI_ISL_536248, EPI_ISL_536249, EPI_ISL_536250, EPI_ISL_536251 | Hôpital du Centre-de-la-Mauricie | Laboratoire de santé publique du Québec | Sandrine Moreira, Ioannis Ragoussis, Guillaume Bourque, Jesse Shapiro, Mark Lathrop and Michel Roger |
| EPI_ISL_536252 | Hôpital Sainte-Croix | Laboratoire de santé publique du Québec | Sandrine Moreira, Ioannis Ragoussis, Guillaume Bourque, Jesse Shapiro, Mark Lathrop and Michel Roger |
| EPI_ISL_536253 | Hôpital Pierre-Boucher | Laboratoire de santé publique du Québec | Sandrine Moreira, Ioannis Ragoussis, Guillaume Bourque, Jesse Shapiro, Mark Lathrop and Michel Roger |
| EPI_ISL_536254 | Hôpital du Suroît | Laboratoire de santé publique du Québec | Sandrine Moreira, Ioannis Ragoussis, Guillaume Bourque, Jesse Shapiro, Mark Lathrop and Michel Roger |
| EPI_ISL_536255 | Centre hospitalier Anna-Laberge | Laboratoire de santé publique du Québec | Sandrine Moreira, Ioannis Ragoussis, Guillaume Bourque, Jesse Shapiro, Mark Lathrop and Michel Roger |
| EPI_ISL_536256 | CSSS Haut-Richelieu/Rouville (Hôpital) | Laboratoire de santé publique du Québec | Sandrine Moreira, Ioannis Ragoussis, Guillaume Bourque, Jesse Shapiro, Mark Lathrop and Michel Roger |
| EPI_ISL_536257 | Hôpital du Suroît | Laboratoire de santé publique du Québec | Sandrine Moreira, Ioannis Ragoussis, Guillaume Bourque, Jesse Shapiro, Mark Lathrop and Michel Roger |
| EPI_ISL_536258 | CSSS Haut-Richelieu/Rouville (Hôpital) | Laboratoire de santé publique du Québec | Sandrine Moreira, Ioannis Ragoussis, Guillaume Bourque, Jesse Shapiro, Mark Lathrop and Michel Roger |
| EPI_ISL_536259 | Hôpital du Suroît | Laboratoire de santé publique du Québec | Sandrine Moreira, Ioannis Ragoussis, Guillaume Bourque, Jesse Shapiro, Mark Lathrop and Michel Roger |
| EPI_ISL_536260 | CSSS de la Minganie | Laboratoire de santé publique du Québec | Sandrine Moreira, Ioannis Ragoussis, Guillaume Bourque, Jesse Shapiro, Mark Lathrop and Michel Roger |
| EPI_ISL_536261 | Hôpital du Suroît | Laboratoire de santé publique du Québec | Sandrine Moreira, Ioannis Ragoussis, Guillaume Bourque, Jesse Shapiro, Mark Lathrop and Michel Roger |
| EPI_ISL_536262 | Hôpital de Maria | Laboratoire de santé publique du Québec | Sandrine Moreira, Ioannis Ragoussis, Guillaume Bourque, Jesse Shapiro, Mark Lathrop and Michel Roger |
| EPI_ISL_536263, EPI_ISL_536264 | Point de Service de Témiscaming-et-Kipawa | Laboratoire de santé publique du Québec | Sandrine Moreira, Ioannis Ragoussis, Guillaume Bourque, Jesse Shapiro, Mark Lathrop and Michel Roger |
| EPI_ISL_536265 | Hôpital de Chandler | Laboratoire de santé publique du Québec | Sandrine Moreira, Ioannis Ragoussis, Guillaume Bourque, Jesse Shapiro, Mark Lathrop and Michel Roger |
| EPI_ISL_536266 | Hôpital de Gatineau | Laboratoire de santé publique du Québec | Sandrine Moreira, Ioannis Ragoussis, Guillaume Bourque, Jesse Shapiro, Mark Lathrop and Michel Roger |
| EPI_ISL_536267 | Hôpital de Hull | Laboratoire de santé publique du Québec | Sandrine Moreira, Ioannis Ragoussis, Guillaume Bourque, Jesse Shapiro, Mark Lathrop and Michel Roger |
| EPI_ISL_536268, EPI_ISL_536269, EPI_ISL_536270 | Hôpital Charles-LeMoyne | Laboratoire de santé publique du Québec | Sandrine Moreira, Ioannis Ragoussis, Guillaume Bourque, Jesse Shapiro, Mark Lathrop and Michel Roger |
| EPI_ISL_536271 | Hôpital Honoré-Mercier | Laboratoire de santé publique du Québec | Sandrine Moreira, Ioannis Ragoussis, Guillaume Bourque, Jesse Shapiro, Mark Lathrop and Michel Roger |
| EPI_ISL_536272 | CSSS Haut-Richelieu/Rouville (Hôpital) | Laboratoire de santé publique du Québec | Sandrine Moreira, Ioannis Ragoussis, Guillaume Bourque, Jesse Shapiro, Mark Lathrop and Michel Roger |
| EPI_ISL_536273, EPI_ISL_536274 | Hôpital du Suroît | Laboratoire de santé publique du Québec | Sandrine Moreira, Ioannis Ragoussis, Guillaume Bourque, Jesse Shapiro, Mark Lathrop and Michel Roger |
| EPI_ISL_536275 | Hôpital de Hull | Laboratoire de santé publique du Québec | Sandrine Moreira, Ioannis Ragoussis, Guillaume Bourque, Jesse Shapiro, Mark Lathrop and Michel Roger |
| EPI_ISL_536276, EPI_ISL_536277 | Hôpital Pierre-Boucher | Laboratoire de santé publique du Québec | Sandrine Moreira, Ioannis Ragoussis, Guillaume Bourque, Jesse Shapiro, Mark Lathrop and Michel Roger |
| EPI_ISL_536278 | CSSS Haut-Richelieu/Rouville (Hôpital) | Laboratoire de santé publique du Québec | Sandrine Moreira, Ioannis Ragoussis, Guillaume Bourque, Jesse Shapiro, Mark Lathrop and Michel Roger |
| EPI_ISL_536279, EPI_ISL_536280, EPI_ISL_536281, EPI_ISL_536282 | Hôpital Charles-LeMoyne | Laboratoire de santé publique du Québec | Sandrine Moreira, Ioannis Ragoussis, Guillaume Bourque, Jesse Shapiro, Mark Lathrop and Michel Roger |
| EPI_ISL_536283 | Centre hospitalier Anna-Laberge | Laboratoire de santé publique du Québec | Sandrine Moreira, Ioannis Ragoussis, Guillaume Bourque, Jesse Shapiro, Mark Lathrop and Michel Roger |
| EPI_ISL_536284, EPI_ISL_536285 | Hôpital Charles-LeMoyne | Laboratoire de santé publique du Québec | Sandrine Moreira, Ioannis Ragoussis, Guillaume Bourque, Jesse Shapiro, Mark Lathrop and Michel Roger |
| EPI_ISL_536286 | CSSS Haut-Richelieu/Rouville (Hôpital) | Laboratoire de santé publique du Québec | Sandrine Moreira, Ioannis Ragoussis, Guillaume Bourque, Jesse Shapiro, Mark Lathrop and Michel Roger |
| EPI_ISL_536287 | Hôpital du Suroît | Laboratoire de santé publique du Québec | Sandrine Moreira, Ioannis Ragoussis, Guillaume Bourque, Jesse Shapiro, Mark Lathrop and Michel Roger |
| EPI_ISL_536288 | Conseil Cri de la SSS de la Baie-James | Laboratoire de santé publique du Québec | Sandrine Moreira, Ioannis Ragoussis, Guillaume Bourque, Jesse Shapiro, Mark Lathrop and Michel Roger |
| EPI_ISL_536289 | Hôpital de Maria | Laboratoire de santé publique du Québec | Sandrine Moreira, Ioannis Ragoussis, Guillaume Bourque, Jesse Shapiro, Mark Lathrop and Michel Roger |
| EPI_ISL_536290 | Hôpital Charles-LeMoyne | Laboratoire de santé publique du Québec | Sandrine Moreira, Ioannis Ragoussis, Guillaume Bourque, Jesse Shapiro, Mark Lathrop and Michel Roger |
| EPI_ISL_536291 | Hôpital de Hull | Laboratoire de santé publique du Québec | Sandrine Moreira, Ioannis Ragoussis, Guillaume Bourque, Jesse Shapiro, Mark Lathrop and Michel Roger |
| EPI_ISL_536292, EPI_ISL_536293, EPI_ISL_536294 | Hôpital Charles-LeMoyne | Laboratoire de santé publique du Québec | Sandrine Moreira, Ioannis Ragoussis, Guillaume Bourque, Jesse Shapiro, Mark Lathrop and Michel Roger |
| EPI_ISL_536295 | Hôpital Pierre-Boucher | Laboratoire de santé publique du Québec | Sandrine Moreira, Ioannis Ragoussis, Guillaume Bourque, Jesse Shapiro, Mark Lathrop and Michel Roger |
| EPI_ISL_536296, EPI_ISL_536297, EPI_ISL_536298 | Hôpital Charles-LeMoyne | Laboratoire de santé publique du Québec | Sandrine Moreira, Ioannis Ragoussis, Guillaume Bourque, Jesse Shapiro, Mark Lathrop and Michel Roger |
| EPI_ISL_536299 | CSSS Haut-Richelieu/Rouville (Hôpital) | Laboratoire de santé publique du Québec | Sandrine Moreira, Ioannis Ragoussis, Guillaume Bourque, Jesse Shapiro, Mark Lathrop and Michel Roger |
| EPI_ISL_536300 | Hôpital Pierre-Boucher | Laboratoire de santé publique du Québec | Sandrine Moreira, Ioannis Ragoussis, Guillaume Bourque, Jesse Shapiro, Mark Lathrop and Michel Roger |
| EPI_ISL_536301, EPI_ISL_536302 | CSSS Haut-Richelieu/Rouville (Hôpital) | Laboratoire de santé publique du Québec | Sandrine Moreira, Ioannis Ragoussis, Guillaume Bourque, Jesse Shapiro, Mark Lathrop and Michel Roger |
| EPI_ISL_536303 | Hôpital Pierre-Boucher | Laboratoire de santé publique du Québec | Sandrine Moreira, Ioannis Ragoussis, Guillaume Bourque, Jesse Shapiro, Mark Lathrop and Michel Roger |
| EPI_ISL_536304 | CSSS Haut-Richelieu/Rouville (Hôpital) | Laboratoire de santé publique du Québec | Sandrine Moreira, Ioannis Ragoussis, Guillaume Bourque, Jesse Shapiro, Mark Lathrop and Michel Roger |

| | | | |
|---|---|---|---|
| EPI_ISL_536305, EPI_ISL_536306, EPI_ISL_536307, EPI_ISL_536308, EPI_ISL_536309 | Hôpital Pierre-Boucher | Laboratoire de santé publique du Québec | Sandrine Moreira, Ioannis Ragoussis, Guillaume Bourque, Jesse Shapiro, Mark Lathrop and Michel Roger |
| EPI_ISL_536310 | CLSC et Centre d'Hébergement la Petite-Nation | Laboratoire de santé publique du Québec | Sandrine Moreira, Ioannis Ragoussis, Guillaume Bourque, Jesse Shapiro, Mark Lathrop and Michel Roger |
| EPI_ISL_536311 | Hôpital de Gatineau | Laboratoire de santé publique du Québec | Sandrine Moreira, Ioannis Ragoussis, Guillaume Bourque, Jesse Shapiro, Mark Lathrop and Michel Roger |
| EPI_ISL_536312 | Hôpital de Hull | Laboratoire de santé publique du Québec | Sandrine Moreira, Ioannis Ragoussis, Guillaume Bourque, Jesse Shapiro, Mark Lathrop and Michel Roger |
| EPI_ISL_536313, EPI_ISL_536314 | Hôpital Charles-LeMoyne | Laboratoire de santé publique du Québec | Sandrine Moreira, Ioannis Ragoussis, Guillaume Bourque, Jesse Shapiro, Mark Lathrop and Michel Roger |
| EPI_ISL_536315 | Centre hospitalier Anna-Laberge | Laboratoire de santé publique du Québec | Sandrine Moreira, Ioannis Ragoussis, Guillaume Bourque, Jesse Shapiro, Mark Lathrop and Michel Roger |
| EPI_ISL_536316, EPI_ISL_536317 | Hôpital Charles-LeMoyne | Laboratoire de santé publique du Québec | Sandrine Moreira, Ioannis Ragoussis, Guillaume Bourque, Jesse Shapiro, Mark Lathrop and Michel Roger |
| EPI_ISL_536318, EPI_ISL_536319, EPI_ISL_536320 | Hôpital du Suroît | Laboratoire de santé publique du Québec | Sandrine Moreira, Ioannis Ragoussis, Guillaume Bourque, Jesse Shapiro, Mark Lathrop and Michel Roger |
| EPI_ISL_536321 | CSSS Haut-Richelieu/Rouville (Hôpital) | Laboratoire de santé publique du Québec | Sandrine Moreira, Ioannis Ragoussis, Guillaume Bourque, Jesse Shapiro, Mark Lathrop and Michel Roger |
| EPI_ISL_536322, EPI_ISL_536323, EPI_ISL_536324 | Hôpital Pierre-Boucher | Laboratoire de santé publique du Québec | Sandrine Moreira, Ioannis Ragoussis, Guillaume Bourque, Jesse Shapiro, Mark Lathrop and Michel Roger |
| EPI_ISL_536325 | Centre hospitalier Anna-Laberge | Laboratoire de santé publique du Québec | Sandrine Moreira, Ioannis Ragoussis, Guillaume Bourque, Jesse Shapiro, Mark Lathrop and Michel Roger |
| EPI_ISL_536326, EPI_ISL_536327, EPI_ISL_536328 | Hôpital Charles-LeMoyne | Laboratoire de santé publique du Québec | Sandrine Moreira, Ioannis Ragoussis, Guillaume Bourque, Jesse Shapiro, Mark Lathrop and Michel Roger |
| EPI_ISL_536329 | Centre hospitalier Anna-Laberge | Laboratoire de santé publique du Québec | Sandrine Moreira, Ioannis Ragoussis, Guillaume Bourque, Jesse Shapiro, Mark Lathrop and Michel Roger |
| EPI_ISL_536330 | Hôpital Honoré-Mercier | Laboratoire de santé publique du Québec | Sandrine Moreira, Ioannis Ragoussis, Guillaume Bourque, Jesse Shapiro, Mark Lathrop and Michel Roger |
| EPI_ISL_536331 | Hôpital Charles-LeMoyne | Laboratoire de santé publique du Québec | Sandrine Moreira, Ioannis Ragoussis, Guillaume Bourque, Jesse Shapiro, Mark Lathrop and Michel Roger |
| EPI_ISL_536332, EPI_ISL_536333, EPI_ISL_536334, EPI_ISL_536335 | Hôpital Pierre-Boucher | Laboratoire de santé publique du Québec | Sandrine Moreira, Ioannis Ragoussis, Guillaume Bourque, Jesse Shapiro, Mark Lathrop and Michel Roger |
| EPI_ISL_536336 | Hôpital Charles-LeMoyne | Laboratoire de santé publique du Québec | Sandrine Moreira, Ioannis Ragoussis, Guillaume Bourque, Jesse Shapiro, Mark Lathrop and Michel Roger |
| EPI_ISL_536337 | CUSM-Site Glen-LAB Microbiologie | Laboratoire de santé publique du Québec | Sandrine Moreira, Ioannis Ragoussis, Guillaume Bourque, Jesse Shapiro, Mark Lathrop and Michel Roger |
| EPI_ISL_536338, EPI_ISL_536339, EPI_ISL_536340, EPI_ISL_536341, EPI_ISL_536342, EPI_ISL_536343 | Hôpital Charles-LeMoyne | Laboratoire de santé publique du Québec | Sandrine Moreira, Ioannis Ragoussis, Guillaume Bourque, Jesse Shapiro, Mark Lathrop and Michel Roger |
| EPI_ISL_536344, EPI_ISL_536345 | CSSS Haut-Richelieu/Rouville (Hôpital) | Laboratoire de santé publique du Québec | Sandrine Moreira, Ioannis Ragoussis, Guillaume Bourque, Jesse Shapiro, Mark Lathrop and Michel Roger |
| EPI_ISL_536346, EPI_ISL_536347 | Hôpital Pierre-Boucher | Laboratoire de santé publique du Québec | Sandrine Moreira, Ioannis Ragoussis, Guillaume Bourque, Jesse Shapiro, Mark Lathrop and Michel Roger |
| EPI_ISL_536348 | Centre hospitalier Anna-Laberge | Laboratoire de santé publique du Québec | Sandrine Moreira, Ioannis Ragoussis, Guillaume Bourque, Jesse Shapiro, Mark Lathrop and Michel Roger |
| EPI_ISL_536349 | Hôpital du Suroît | Laboratoire de santé publique du Québec | Sandrine Moreira, Ioannis Ragoussis, Guillaume Bourque, Jesse Shapiro, Mark Lathrop and Michel Roger |
| EPI_ISL_536350 | Hôpital Charles-LeMoyne | Laboratoire de santé publique du Québec | Sandrine Moreira, Ioannis Ragoussis, Guillaume Bourque, Jesse Shapiro, Mark Lathrop and Michel Roger |
| EPI_ISL_536351, EPI_ISL_536352 | Hôpital Pierre-Boucher | Laboratoire de santé publique du Québec | Sandrine Moreira, Ioannis Ragoussis, Guillaume Bourque, Jesse Shapiro, Mark Lathrop and Michel Roger |
| EPI_ISL_536353 | Centre de santé Inuulitsivik | Laboratoire de santé publique du Québec | Sandrine Moreira, Ioannis Ragoussis, Guillaume Bourque, Jesse Shapiro, Mark Lathrop and Michel Roger |
| EPI_ISL_536354, EPI_ISL_536355, EPI_ISL_536356, EPI_ISL_536357, EPI_ISL_536358, EPI_ISL_536359 | Hôpital du Suroît | Laboratoire de santé publique du Québec | Sandrine Moreira, Ioannis Ragoussis, Guillaume Bourque, Jesse Shapiro, Mark Lathrop and Michel Roger |
| EPI_ISL_536360 | Hôpital Pierre-Boucher | Laboratoire de santé publique du Québec | Sandrine Moreira, Ioannis Ragoussis, Guillaume Bourque, Jesse Shapiro, Mark Lathrop and Michel Roger |
| EPI_ISL_536361 | CSSS Haut-Richelieu/Rouville (Hôpital) | Laboratoire de santé publique du Québec | Sandrine Moreira, Ioannis Ragoussis, Guillaume Bourque, Jesse Shapiro, Mark Lathrop and Michel Roger |
| EPI_ISL_536362 | Hôpital du Suroît | Laboratoire de santé publique du Québec | Sandrine Moreira, Ioannis Ragoussis, Guillaume Bourque, Jesse Shapiro, Mark Lathrop and Michel Roger |
| EPI_ISL_536363, EPI_ISL_536364 | Hôpital de Hull | Laboratoire de santé publique du Québec | Sandrine Moreira, Ioannis Ragoussis, Guillaume Bourque, Jesse Shapiro, Mark Lathrop and Michel Roger |
| EPI_ISL_536365, EPI_ISL_536366 | CUSM-Site Glen-LAB Microbiologie | Laboratoire de santé publique du Québec | Sandrine Moreira, Ioannis Ragoussis, Guillaume Bourque, Jesse Shapiro, Mark Lathrop and Michel Roger |
| EPI_ISL_536367 | Hôpital de Hull | Laboratoire de santé publique du Québec | Sandrine Moreira, Ioannis Ragoussis, Guillaume Bourque, Jesse Shapiro, Mark Lathrop and Michel Roger |
| EPI_ISL_536368, EPI_ISL_536369, EPI_ISL_536370 | Centre hospitalier Anna-Laberge | Laboratoire de santé publique du Québec | Sandrine Moreira, Ioannis Ragoussis, Guillaume Bourque, Jesse Shapiro, Mark Lathrop and Michel Roger |
| EPI_ISL_536371, EPI_ISL_536372, EPI_ISL_536373 | Hôpital Charles-LeMoyne | Laboratoire de santé publique du Québec | Sandrine Moreira, Ioannis Ragoussis, Guillaume Bourque, Jesse Shapiro, Mark Lathrop and Michel Roger |
| EPI_ISL_536374, EPI_ISL_536375 | Hôpital Pierre-Boucher | Laboratoire de santé publique du Québec | Sandrine Moreira, Ioannis Ragoussis, Guillaume Bourque, Jesse Shapiro, Mark Lathrop and Michel Roger |
| EPI_ISL_536376, EPI_ISL_536377, EPI_ISL_536378 | Centre hospitalier Anna-Laberge | Laboratoire de santé publique du Québec | Sandrine Moreira, Ioannis Ragoussis, Guillaume Bourque, Jesse Shapiro, Mark Lathrop and Michel Roger |
| EPI_ISL_536379 | Hôpital Honoré-Mercier | Laboratoire de santé publique du Québec | Sandrine Moreira, Ioannis Ragoussis, Guillaume Bourque, Jesse Shapiro, Mark Lathrop and Michel Roger |
| EPI_ISL_536380 | Hôpital Pierre-Boucher | Laboratoire de santé publique du Québec | Sandrine Moreira, Ioannis Ragoussis, Guillaume Bourque, Jesse Shapiro, Mark Lathrop and Michel Roger |
| EPI_ISL_536381 | Centre Hospitalier Régional de Lanaudière | Laboratoire de santé publique du Québec | Sandrine Moreira, Ioannis Ragoussis, Guillaume Bourque, Jesse Shapiro, Mark Lathrop and Michel Roger |
| EPI_ISL_536382, EPI_ISL_536383 | CSSS Haut-Richelieu/Rouville (Hôpital) | Laboratoire de santé publique du Québec | Sandrine Moreira, Ioannis Ragoussis, Guillaume Bourque, Jesse Shapiro, Mark Lathrop and Michel Roger |
| EPI_ISL_536384 | Hôpital de Gatineau | Laboratoire de santé publique du Québec | Sandrine Moreira, Ioannis Ragoussis, Guillaume Bourque, Jesse Shapiro, Mark Lathrop and Michel Roger |
| EPI_ISL_536385, EPI_ISL_536386, EPI_ISL_536387 | Hôpital de Hull | Laboratoire de santé publique du Québec | Sandrine Moreira, Ioannis Ragoussis, Guillaume Bourque, Jesse Shapiro, Mark Lathrop and Michel Roger |
| EPI_ISL_536388, EPI_ISL_536389 | Hôpital Pierre-Boucher | Laboratoire de santé publique du Québec | Sandrine Moreira, Ioannis Ragoussis, Guillaume Bourque, Jesse Shapiro, Mark Lathrop and Michel Roger |
| EPI_ISL_536390 | Hôpital du Suroît | Laboratoire de santé publique du Québec | Sandrine Moreira, Ioannis Ragoussis, Guillaume Bourque, Jesse Shapiro, Mark Lathrop and Michel Roger |
| EPI_ISL_536391 | Hôpital Pierre-Boucher | Laboratoire de santé publique du Québec | Sandrine Moreira, Ioannis Ragoussis, Guillaume Bourque, Jesse Shapiro, Mark Lathrop and Michel Roger |
| EPI_ISL_536392, EPI_ISL_536393, EPI_ISL_536394 | Hôpital de Hull | Laboratoire de santé publique du Québec | Sandrine Moreira, Ioannis Ragoussis, Guillaume Bourque, Jesse Shapiro, Mark Lathrop and Michel Roger |
| EPI_ISL_536395 | Hôpital Pierre-Boucher | Laboratoire de santé publique du Québec | Sandrine Moreira, Ioannis Ragoussis, Guillaume Bourque, Jesse Shapiro, Mark Lathrop and Michel Roger |
| EPI_ISL_536396 | Hôpital de Hull | Laboratoire de santé publique du Québec | Sandrine Moreira, Ioannis Ragoussis, Guillaume Bourque, Jesse Shapiro, Mark Lathrop and Michel Roger |
| EPI_ISL_536397 | Hôpital Pierre-Boucher | Laboratoire de santé publique du Québec | Sandrine Moreira, Ioannis Ragoussis, Guillaume Bourque, Jesse Shapiro, Mark Lathrop and Michel Roger |
| EPI_ISL_536398 | Lithuanian University of Health Sciences Hospital, Department of Laboratory Medicine | Lithuanian University of Health Sciences, Molecular cardiology lab. | Lukas Zemaitis, Arnoldas Pautienius, Kamile Tamusauskaite, Dovydas Gecys, Vaiva Lesauskaite, Astra Vitkauskiene |
| EPI_ISL_536399 | Laboratory of Immunovirology. Universidad de Antioquia | Instituto Nacional de Salud - Unidad de Secuenciación y Genómica | Wbeimar Aguilar-Jimenez, Lizdany Flórez, Francisco J. Díaz, Katherine Laiton-Donato, Carlos Franco-Muñoz, Diego Álvarez-Diaz and Marcela Mercado-Reyes |
| EPI_ISL_536400 | Fareham Creek Veterinary Surgery | MRC-University of Glasgow Centre for Virus Research | Margaret J Hosie, Ilaria Epifano, Vanessa Herder, Richard J Orton, Andrew Stevenson, Natasha Johnson, Daniel Goldfarb, Emma MacDonald, Lynn Stevenson, Frazer Bell, Dawn Dunbar, Michael McDonald, Fiona Howie, Bryn Tennant, Darcy Herrity, Ana Da Silva Filipe, Daniel G Streicker, Brian J Willett, Pablo R Murcia, Ruth F Jarrett, David L Robertson and William Weir |
| EPI_ISL_538390, EPI_ISL_538391, EPI_ISL_538392, EPI_ISL_538393, EPI_ISL_538394, EPI_ISL_538395, EPI_ISL_538396, EPI_ISL_538397, EPI_ISL_538398, EPI_ISL_538399, EPI_ISL_538400, EPI_ISL_538401, EPI_ISL_538402, EPI_ISL_538403, EPI_ISL_538404, EPI_ISL_538405, EPI_ISL_538406, EPI_ISL_538407, EPI_ISL_538408, EPI_ISL_538409, EPI_ISL_538410, EPI_ISL_538411, EPI_ISL_538412, EPI_ISL_538413, EPI_ISL_538414, EPI_ISL_538415, EPI_ISL_538416, EPI_ISL_538417, EPI_ISL_538418, EPI_ISL_538419, EPI_ISL_538420, EPI_ISL_538421, EPI_ISL_538422, EPI_ISL_538423, EPI_ISL_538424, EPI_ISL_538425, EPI_ISL_538426, EPI_ISL_538427, EPI_ISL_538428, EPI_ISL_538429, EPI_ISL_538430, EPI_ISL_538431, EPI_ISL_538432, EPI_ISL_538433, EPI_ISL_538434 | | | |
| see above | Microbiology Division, South Carolina Department of Health and Environmental Control | Microbiology Division, South Carolina Department of Health and Environmental Control | Flores,H. |
| EPI_ISL_538514, EPI_ISL_538515, EPI_ISL_538516, EPI_ISL_538517, EPI_ISL_538518, EPI_ISL_538519, EPI_ISL_538520, EPI_ISL_538521 | Infectious Diseases, North Carolina State Laboratory of Public Health COVID-19 Response Team | Infectious Diseases, North Carolina State Laboratory of Public Health COVID-19 Response Team | Chase,K. |
| EPI_ISL_538522, EPI_ISL_538523 | Infectious Diseases, North Carolina State Laboratory of Public Health COVID-19 Response Team | North Carolina State Laboratory of Public Health | Chase,K. |
